# Adropin protects against cardiac remodeling and metabolic dysfunction in a mouse model of HFpEF

**DOI:** 10.1101/2025.05.03.652038

**Authors:** Bellina A.S. Mushala, Michael W. Stoner, Maryam Sharifi-Sanjani, Janet R. Manning, Paramesha Bugga, Nisha Bhattarai, Brenda McMahon, Amber Vandevender, Steven J. Mullet, Brett A. Kaufman, Sruti S. Shiva, Cheng Zhang, Eric Goetzman, Stephen Y. Chan, Stacy L. Gelhaus, Michael J. Jurczak, Iain Scott

## Abstract

Cardiometabolic heart failure with preserved ejection fraction (HFpEF) is a heterogenous metabolic disease, which in the heart presents as left ventricle diastolic dysfunction, ventricular stiffness, and myocardial structural remodeling. Deleterious changes in cardiac metabolism are central to HFpEF pathophysiology, and proposed treatments for the disease have focused on repairing these defects. In this study, we used a preclinical mouse model that recapitulates cardiometabolic HFpEF to elucidate the molecular mechanisms driving cardiac dysfunction, and tested whether recombinant Adropin (a liver- and brain-derived endogenous peptide hormone) could reverse observed defects. We show that long-term treatment with Adropin reversed multiple markers of HFpEF-related cardiac dysfunction (including fibrosis, diastolic dysfunction, and cardiomyocyte hypertrophy). Using untargeted metabolomics, we found that Adropin treatment reduced hexosamine biosynthesis pathway activity, leading to a reduction in the O-GlcNAcylation of the cardiac fatty acid oxidation enzyme long chain acyl-CoA dehydrogenase (LCAD). Reducing LCAD O-GlcNAcylation increased LCAD activity *in vitro*, and reduced the accumulation of long-chain acylcarnitines in HFpEF mouse hearts *in vivo.* Our results suggest that Adropin may restore cardiac metabolic function in HFpEF, and that targeting this pathway may be a novel therapeutic avenue for this disease.

**CLINICAL PERSPECTIVE:** - Adropin is a circulating liver- and brain-derived peptide that regulates energy metabolism in the heart and other high metabolic-demand tissues. The plasma abundance of Adropin is decreased in diabetic, hypertensive, and aged individuals; all comorbid risk factors for the development of heart failure with preserved ejection (HFpEF). We therefore examined the potential role of Adropin in HFpEF pathophysiology.

- Patients with HFpEF display significant reductions in circulating Adropin levels, matching those seen in comorbid diseases. In a mouse model of HFpEF, treatment with recombinant Adropin reduced diastolic dysfunction, cardiac fibrosis, and cardiomyocyte hypertrophy.

- These data suggest that targeting the Adropin pathway may represent a new therapeutic approach in HFpEF.

## INTRODUCTION

Heart failure is a complex clinical syndrome that results from any structural or functional impairments of ventricular filling and/or ejection of blood, and remains the leading cause of morbidity and mortality worldwide (Heidenreich et al., 2023). Heart failure with preserved ejection fraction (HFpEF), a subtype of heart failure, is primarily characterized by cardiac diastolic dysfunction. This presentation of the disease currently accounts for >50% of all heart failure cases, with a global prevalence of approximately 32 million (Shah et al., 2020; Redfield and Borlaug, 2023). HFpEF is recognized as a heterogenous syndrome, whose underlying pathophysiological mechanisms include several extracardiac abnormalities such as metabolic derangements, arterial hypertension, microvascular endothelial dysfunction, inflammation, and renal insufficiency (Shah et al., 2016, Shah et al., 2020).

Partly due to a lack of specifically-designed drugs for the disease, treatment for HFpEF currently focuses on the management of comorbidities (e.g. obesity, type 2 diabetes, hypertension), with new guidelines recommending the use of glycemia-reducing therapeutics (e.g. SGLT2 inhibitors) that improve cardiovascular outcomes in heart failure (Kittleson et al., 2023). The recent STEP-HFpEF trial in obesity-related HFpEF demonstrated that therapy-mediated reductions in body weight may also be a key tool in disease regression (Kosiborod et al., 2023). These developments suggest that approaches targeting metabolic defects can offer substantial benefits in this disease. On this basis, we examined whether treatment with recombinant Adropin, a brain- and liver-derived endogenous peptide hormone (Kumar et al., 2008), may have beneficial effects in a preclinical model of cardiometabolic HFpEF.

Previous studies have shown that Adropin improves physical activity (Wong et al, 2014, Ganesh- Kumar et al., 2012), attenuates hepatic steatosis and injury (Kumar et al., 2008, Chen et al., 2019), and reduces vascular stiffness in diet-induced obesity (Jurrissen et al., 2022). In the heart, we recently demonstrated that short-term Adropin treatment (≤ 3 days) can improve cardiac fuel substrate flexibility by restoring glucose oxidation in the diabetic heart (Thapa et al., 2019b). Combined, these beneficial features raised the question of whether this peptide can be used as a therapeutic intervention in HFpEF. Here, we show that long-term treatment (4 weeks) with recombinant Adropin restored cardiac diastolic function, reduced cardiac hypertrophy and fibrosis, and improved whole-body glucose tolerance. These changes were linked to a reversal of metabolic remodeling and improved fatty acid oxidation enzyme activity, suggesting that Adropin treatment mitigates the impact of HFpEF stimuli via improved cardiometabolic function.

## MATERIALS AND METHODS

### Animal Care and Use

Male C57BL/6J 8-week-old mice were obtained from the Jackson Laboratory and maintained on chow for 4 weeks to acclimate to their new environment. Animals were housed in the University of Pittsburgh animal facility under standard conditions, with *ad libitum* access to water and food, and maintained on a constant 12-hour light/dark cycle. At 12 weeks of age, mice were exposed to normal chow (Chow; 60% carbohydrate, 26% protein, 14% fat; ProLab IsoPro RMH 3000), or a combination of high-fat diet (HFD; 20% carbohydrate, 20% protein, 60% fat; Research Diets D12492) plus N-ω- nitro-L-arginine methyl ester (L-NAME: 0.5 g/L, pH 7.4; ThermoFisher) supplemented in drinking water to induce HFpEF (Schiattarella et al., 2019). After 8 weeks of diet, mice were randomly assigned to a control or treatment group receiving daily intraperitoneal injections of either vehicle or recombinant Adropin (Adr.) at 450 nmol/kg animal weight, prepared in PBS with 0.1% BSA, for the remaining 4 weeks of the study. The Adropin dose used was based on prior studies by our group and others (see, e.g., Gao et al., 2015; Thapa et al., 2019a; Thapa et al., 2019b). This dose has become standard in the field for *in vivo* studies using this peptide. L-NAME supplemented drinking water was changed weekly, and Adropin was made fresh and sterile filtered before each use. Animals were euthanized by isofluorane anesthesia followed by cervical dislocation. Experiments were conducted in compliance with National Institutes of Health guidelines, and followed procedures approved by the University of Pittsburgh Institutional Animal Care and Use Committee.

### Echocardiography

Mice were anesthetized using isofluorane (1.5%–2.0% v/v by inhalation) and monitored for cardiac functional parameters in the supine position using a Visual Sonics Vevo 3100. Core temperature was maintained at 37 °C by imaging mice while on a heating pad, and heart rates were kept consistent between experimental groups (∼400–500 beats per min). Short axis M-mode was used to assess LV dimensions and motion patterns, pulsed-wave (PW) doppler mode to evaluate blood flow velocities across the mitral valve, and tissue doppler mode to measure mitral annular plane velocity. Doppler profiles were acquired in the parasternal long axis apical 4-chamber view. The left atrial area was quantified in the apical 4-chamber view by tracing the border of the left atrium. Markers of systolic and diastolic function were calculated using standard echocardiography equations. At the end of the procedure, all mice recovered from anesthesia without difficulties. Image analysis was performed independently by a blinded sonographer, and all parameters were measured at least 3 times with averages presented.

### Intraperitoneal Glucose Tolerance Test (IPGTT)

Glucose tolerance tests were performed as previously described with minor modifications (Jurczak et al., 2011). After an overnight fast (∼16 h), mice were set-up in restrainers with tails snipped < 5 mm to prime for blood collection and left to acclimate under a heated bulb for 2 h prior to GTT start time at 9 a.m. Basal plasma samples (t = 0) were collected, and D-glucose was diluted in water (20% final concentration) and sterile filtered for intraperitoneal administration at 1.5 g/kg body weight. Blood glucose was measured by tail bleed at set time points using a Bayer Contour Next EZ handheld glucometer (t = 15, 30, 45, 60, and 120 min).

### Histology

Preparation and staining of all histological samples were conducted by the Pitt Biospecimen Core at the University of Pittsburgh. All mice were sacrificed, and tissue harvested for analysis within one hour of completion of the IPGTT study. The apex of the heart was excised from each mouse following euthanasia, fixed in 10% buffered formalin phosphate overnight at shaking incubation, washed 3 times for 5 min with 1X phosphate-buffered saline (PBS), and then transferred to 70% ethanol.

Samples were then embedded in paraffin, sectioned into 4 μm slides and stained with hematoxylin and eosin (H&E), wheat germ agglutinin (WGA) or Masson’s Trichrome for analysis. Histology sections were visualized with the Evos FL Auto 2 Microscope, observed changes in structure were quantified using ImageJ software, and representative images are shown. All histological analysis was carried out under blinded conditions.

### Transcriptomics

Total RNA was isolated from the left ventricle of the heart using the RNeasy Plus Mini Kit (Qiagen REF 74134). Approximately 2 μg RNA was used for bulk RNA sequencing performed by Azenta/GENEWIZ, based on company recommendations to identify differential gene expression patterns between the experimental groups.

### TGFβ Assay

Human adult ventricular fibroblasts were purchased from LONZA (CC-2904). Cells were plated and expanded according to manufacturer’s recommended protocol using fibroblast medium supplemented with FGM-3 Bullet Kit (LONZA). Cells were treated with TGF-β (10 ng/mL; Prospec) and/or Adropin (0.5µg/ml; AbClonal) for 48 h, prior to mRNA isolation using RNeasy Mini Kit RNA (Qiagen), followed by cDNA production using Maxima First Strand cDNA Synthesis Kit (Thermo Scientific). Quantitative PCR was performed using PowerUp SYBR Green Master Mix (Thermo Fisher Scientific) and validated gene-specific primers (Qiagen) with GAPDH as the internal control for transcript level measurement.

### Quantitative Metabolomics

Metabolic quenching and polar metabolite pool extraction was performed by adding ice cold 80% methanol (aqueous) at a ratio of 1:15 wt input tissue:vol. (13C1)-creatinine, (D3)-taurine, (D3)-lactate and (D3)-alanine (Sigma-Aldrich) were added to the sample lysates as an internal standard for a final concentration of 10 μM. Samples are homogenized using an MP Bio FastPrep system using Matrix D (ceramic sphere) for 60 s at 60 hz. The supernatant was then cleared of protein by centrifugation at 16,000 g. Cleared supernatant (2 μL) was subjected to online LC-MS analysis. Analyses were performed by untargeted liquid chromatography-high-resolution mass spectrometry (LC-HRMS).

Briefly, samples were injected via a Thermo Vanquish UHPLC and separated over a reversed phase. Thermo HyperCarb porous graphite column (2.1×100 mm, 3 μm particle size) maintained at 55 °C. For the 20 min LC gradient, the mobile phase consisted of the following: solvent A (water/0.1% FA) and solvent B (ACN/0.1% FA). The gradient used was: 0-1 min 1% B, increase to 15% B over 5 min, continue increasing to 98% B over 5 min, hold at 98% B for 5 min, re-equillibrate at 1% B for 5 min. The Thermo IDX tribrid mass spectrometer was operated in both positive and negative ion mode, scanning in ddMS2 mode (2 μscans) from 70 to 800 m/z at 120,000 resolution with an AGC target of 2e^5^ for full scan, 2e^4^ for ms2 scans using HCD fragmentation at stepped 15, 35, 50 collision energies. Source ionization setting was 3.0 and 2.4 kV spray voltage, respectively, for positive and negative mode. Source gas parameters were 35 sheath gas, 12 auxiliary gas at 320 °C, and 8 sweep gas.

Calibration was performed prior to analysis using the PierceTM FlexMix Ion Calibration Solutions (Thermo Fisher Scientific). Integrated peak areas were then extracted manually using Quan Browser (Thermo Fisher Xcalibur ver. 2.7). Untargeted differential comparisons were performed using Compound Discoverer 3.0 (Thermo Fisher) to generate a ranked list of significant compounds with tentative identifications from BioCyc, KEGG, and internal compound databases. Purified standards were then purchased and compared in retention time, m/z, along with ms2 fragmentation patterns to validate the identity of significant hits.

### Protein Isolation and Immunoblotting

Heart tissues were rapidly harvested following euthanasia, weighed, RV discarded, and LV flash-frozen in liquid nitrogen. For protein isolation, tissues were minced and lysed in CHAPS buffer (1% CHAPS, 150 mM NaCl, 10 mM HEPES, pH 7.4) using a VWR 4-Place Mini Bead Mill, then incubated on ice for ∼ 2.5 h. Homogenates were spun at 10,000 g at 4 °C for 10 min, and the supernatants collected for immunoblotting. For immunoblotting, protein lysates were quantitated using a BioDrop μLITE Analyzer, prepared in LDS sample buffer, separated using Bolt SDS-PAGE 4–12% or 12% Bis– Tris Plus gels, and transferred to nitrocellulose membranes (all Invitrogen). Membranes were blocked using SuperBlock (PBS) Blocking Buffer and incubated overnight in the following primary antibodies: mouse O-GlcNAc (9875), rabbit GFAT (D12F4), rabbit OGT (D1D8Q), rabbit OGA (E9C5U), and rabbit GAPDH (2118S) from Cell Signaling Technologies; rabbit LCAD (17526-1-AP) from Protein Tech. Protein loading was confirmed using GAPDH as a loading control. Fluorescent anti-goat or anti-rabbit secondary antibodies (red, 700 nm; green, 800 nm) from LiCor were used to detect expression levels. Images were obtained using Licor Odyssey CLx System and protein densitometry was measured using the LiCor Image Studio Lite Ver. 5.2 Software.

### Co-Immunoprecipitation

For co-immunoprecipitation experiments, protein lysates were harvested in CHAPS lysis buffer (1% CHAPS, 150 mM NaCl, 10 mM HEPES, pH 7.4), and 500 μg of protein was incubated overnight at 4 °C with 5 μL mouse O-GlcNAc antibody (Cell Signaling). Immunocaptured proteins were isolated using Protein-G and Protein-A agarose beads (Cell Signaling Technology, catalog number 9007), washed 5 times with 500 μL 1% CHAPS buffer, and then eluted in LDS sample buffer (Life Technologies) at 95 °C. Samples were separated on 4-12% Bis–Tris Bolt gels, transferred to nitrocellulose membranes, and probed with appropriate antibodies. Images were obtained using LiCor Odyssey CLx System and protein densitometry was measured using the LiCor Image Studio Lite Ver. 5.2 Software.

### Biochemical Assays

To assess the activity of long chain acyl-CoA dehydrogenase enzymes (LCAD), homogenized protein samples were incubated with palmitoyl-CoA as described previously (Thapa et al., 2017). Briefly, ∼10 μg of protein was incubated with 0.1 M potassium phosphate, 50 μM 2,6- dichlorophenolindophenol, 2 mM phenazine ethosulfate, 0.2 mM N-ethylmaleimide, 0.4 mM potassium cyanide, and 0.1% Triton X-100 at 37 °C for 4 min. The reaction was initiated with 60 μM palmitoyl-CoA, and the rate of absorbance change was measured/observed at 600 nm over 45 min. Activities were converted to moles of substrate oxidized/min/mass of protein. *In vitro* LCAD activity was measured after incubating a reaction mixture of 4 μg of recombinant LCAD, 0.8 mM UDP-GlcNAc (Sigma, Catalog # U4375), 1 μg of recombinant human O-GlcNAc Transferase (rhOGT) (Catalog # 8446-GT) in assay buffer (25 mM Tris, 10 mM CaCl2, pH 7.5 and 10 mM MgCl2) at 37 °C for 30 min.

### Statistics

Means ± SEM were calculated for all data sets. Data sets containing three or more groups were analyzed using one-way ANOVA with Tukey’s post-hoc testing to determine differences between genotypes and feeding/treatment groups. Data sets containing two groups were analyzed using Student’s T-test. Time course data was analyzed using one-way ANOVA with Sidak’s post- hoc testing. *P* ≤ 0.05 was considered statistically significant. Statistical analyses were performed using GraphPad Prism 9.5 Software.

## RESULTS

### Circulating Adropin levels are diminished in HFpEF patients

Circulating Adropin levels are known to decrease in patients with type 2 diabetes (Soltani et al., 2023), aging (Yang et al., 2018), and hypertension (Gulen et al, 2016); all comorbidities for HFpEF. We therefore measured plasma levels of Adropin in deidentified age- and gender-matched control and HFpEF patient samples (n = 19-20) obtained from a University of Pittsburgh biobank **(Sup. Table 1)**. Consistent with the studies in comorbid disease states, there was a significant decrease in plasma Adropin levels in HFpEF patients relative to non-affected controls **(Fig. 1A)**. These data suggest that a reduction in circulating Adropin levels may be linked to HFpEF disease development or progression.

**Figure 1:**
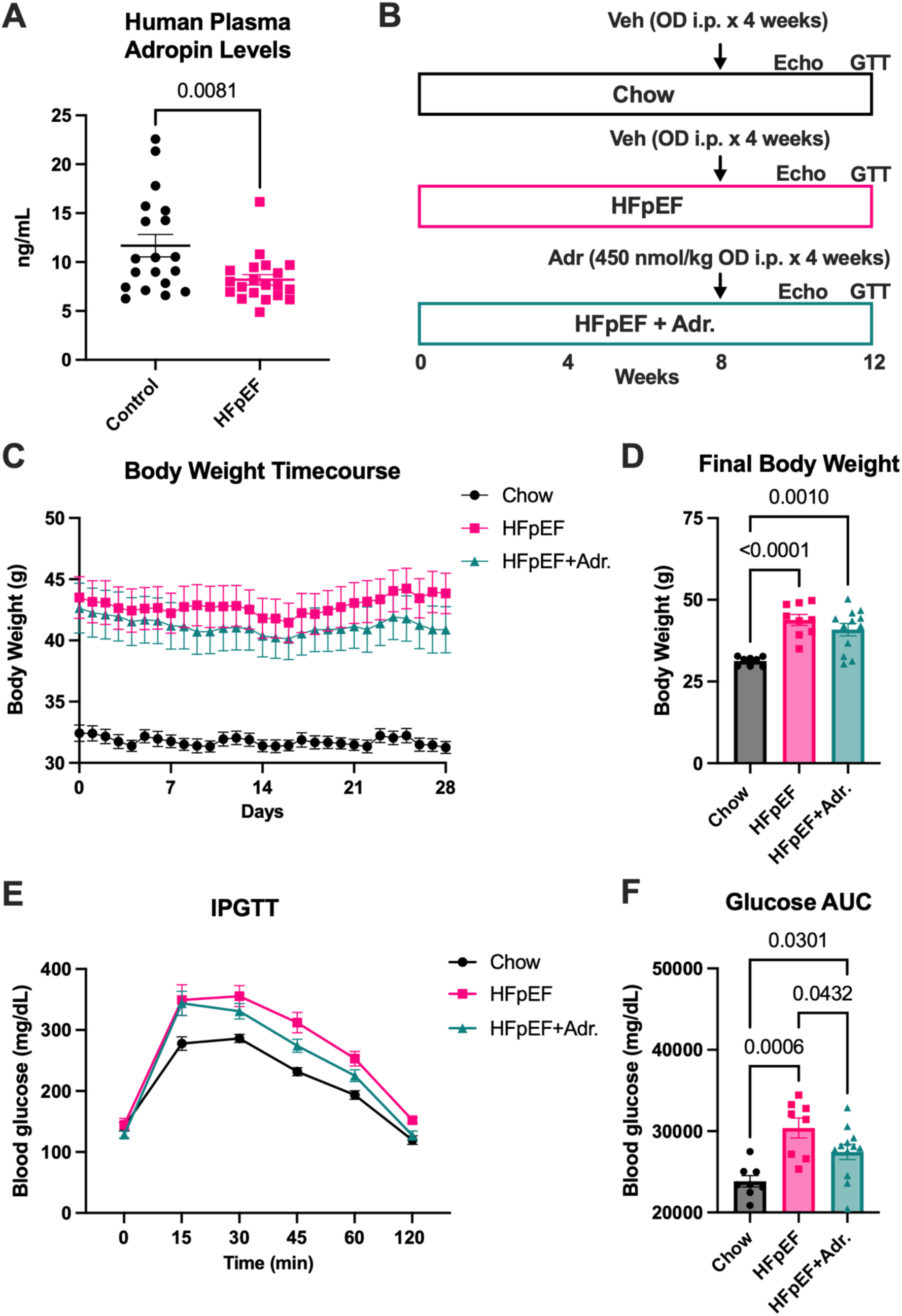
Recombinant Adropin treatment improves glucose tolerance in HFpEF without changes in body weight. A. Total circulating Adropin levels in age- and gender-matched control and HFpEF patients. N = 19-20**. B.** Timeline and treatment protocol used in mouse HFpEF studies. **C,D.** Daily and final body weights of mice. N = 8-12. **E,F.** IPGTT time-course and area under-the-curve (AUC). N = 8-12.

### Recombinant Adropin treatment improves glucose tolerance in HFpEF without changes in body weight

The potential link between reduced circulating Adropin and HFpEF in patients supported the premise that therapeutic restoration of Adropin levels may be protective against the disease. We therefore used a recently developed two-hit mouse model of HFpEF (Schiattarella et al, 2019) to test this hypothesis. Male C57BL/6J mice were placed on either a chow (control) or HFpEF (60% high fat diet plus L-NAME-supplemented drinking water) diet for 8 weeks. After 8 weeks, HFpEF mice were randomly given either vehicle (PBS) or Adropin (450 nmol/kg/day; Thapa et al., 2019b) by intraperitoneal (I.P.) injection once daily for 4 weeks **(Fig. 1B)**. Daily Adropin treatment significantly increased circulating Adropin levels in HFpEF mice (15.9 vs. 2.2 ng/mL, *P* = 0.0067), demonstrating that the administration of recombinant Adropin could restore plasma concentrations to those observed in healthy humans **(as seen in Fig. 1A)**.

The recent STEP-HFpEF Phase 3 clinical trial showed that the GLP-1 receptor agonist semaglutide improved patient-reported outcomes in HFpEF, which was at least partially related to weight loss (Kosiborod et al, 2023). We therefore first examined whether Adropin treatment promoted weight loss in mice, and found that there was no significant difference in body weight between vehicle- and Adropin-treated HFpEF animals **(Fig. 1C,D)**. Our group and others have previously shown that short-term Adropin treatment (≤ 3 days) leads to improved whole-body glucose tolerance in diet-induced obese mice (Gao et al., 2015; Thapa et al., 2019a). We next tested whether long-term Adropin treatment (4 weeks) in HFpEF mice had the same effect, and found that there was a significant decrease in insulin resistance in HFpEF mice treated with Adropin relative to non-treated HFpEF controls **(Fig. 1E,F)**. Combined, these data suggest that Adropin treatment may have a positive effect on whole-body physiology by reducing glucose intolerance. However, unlike semaglutide in humans, this improvement appears to be unrelated to the beneficial effects of weight loss.

### Recombinant Adropin treatment limits cardiac diastolic dysfunction in a mouse model of HFpEF

We next examined the effect of Adropin treatment on cardiac functional parameters and tissue structure **(Fig. 2A)**. Mice with HFpEF displayed a significant increase in cardiac weight and a reduction in cardiac output, both of which were absent in Adropin-treated HFpEF mice **(Fig. 2B,E)**. As expected from a HFpEF model, all mouse groups displayed normal systolic function, as measured by left ventricular ejection fraction **(Fig. 2F)**. Both control and Adropin-treated HFpEF groups displayed a non-significant ∼25% increase in E/e’ ratio, and a significant increase in mitral annulus tissue velocity during early diastole (e’), suggestive of mild-to-moderate diastolic dysfunction **(Fig. 2H,I)**. Adropin-treated mice displayed a significant decrease in isovolumetric relaxation time (IVRT), a significant decrease in myocardial performance index (MPI; also known as Tei index), and a trend towards a decreased E/A ratio relative to untreated HFpEF mice **(Fig. 2J-L)**. Combined, these data indicate that Adropin treatment may reduce mild-to-moderate diastolic dysfunction in HFpEF by improving ventricular relaxation capacity (IVRT and MPI), whilst having a more limited impact on echocardiography measures of left ventricle filling pressures (such as E/e’ ratio and e’).

**Figure 2:**
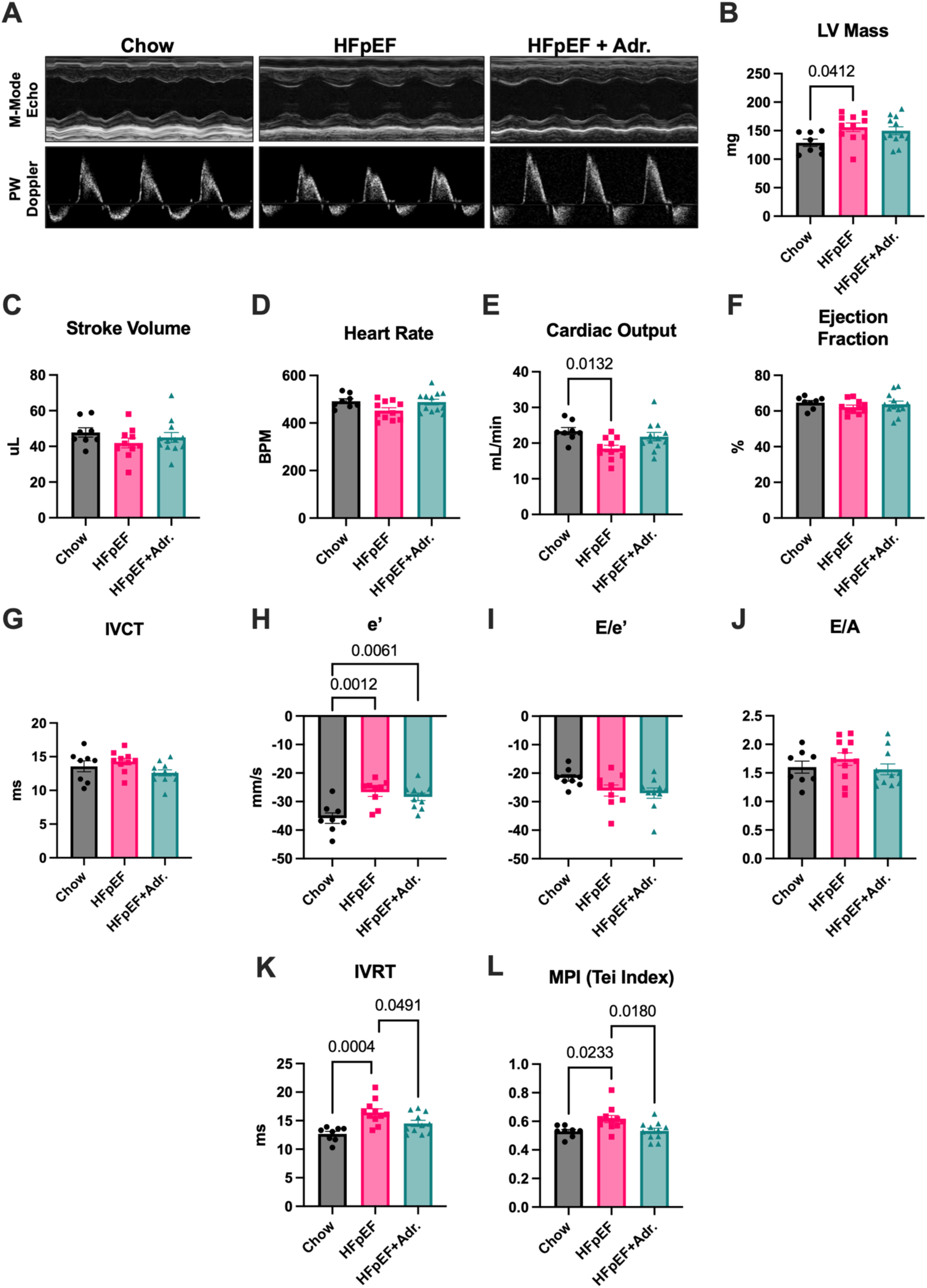
Recombinant Adropin treatment limits cardiac diastolic dysfunction in a mouse model of HFpEF. **A.** Representative echocardiography (M-mode and Doppler) images. **B-L.** Echocardiography measurements of heart rate, stroke volume, cardiac output, calculated left ventricle (LV) mass, ejection fraction, E/e’ ratio, E/A ratio, mitral annulus tissue velocity during early diastole (e’), isovolumetric relaxation time (IVRT), and myocardial performance index/Tei index) (MPI/Tei). N = 8-12.

### Long-term Adropin treatment prevents cardiomyocyte hypertrophy and reduces cardiac fibrosis

Impairment of cardiac relaxation (such as IVRT) often correlates with cardiac remodeling and increased myocardial fibrosis (e.g. Brilla et al., 2000) in disease states. Histological analysis of cardiac tissue sections demonstrated that untreated HFpEF mice displayed a significant increase in myocardial interstitial fibrosis and cardiomyocyte cross-sectional area, which was completely reversed following long-term Adropin treatment **(Fig. 3A-C)**. The positive impact of Adropin on reducing myocardial fibrosis was evident in bulk RNA-seq analysis of untreated vs. Adropin-treated HFpEF hearts **(Sup. Table 2)**, with extracellular matrix processes being identified as the top hit in pathway analyses **(Fig. 3D)**. Closer examination demonstrated that multiple genes related to extracellular matrix formation and fibrosis (e.g. *Postn, Fn1, Col1a1*) were significantly downregulated in HFpEF hearts after Adropin treatment **(Fig. 3E, Sup. Fig. 1)**. To test whether these changes were the result of a direct effect of Adropin on cardiac fibrosis/extracellular matrix composition, we treated human primary cardiac fibroblasts with TGFβ in the presence or absence of Adropin, and measured related gene expression after 48 h. TGFβ treatment led to a significant increase in *Col1a1* and *Mmp2* gene expression, which was attenuated by co-treatment with Adropin **(Fig. 3F)**. Combined with the earlier studies, these data demonstrate that long-term Adropin treatment in HFpEF promoted whole-body metabolic homeostasis, improved myocardial relaxation capacity, prevented maladaptive cardiac tissue remodeling, and limited cardiomyocyte hypertrophy.

**Figure 3:**
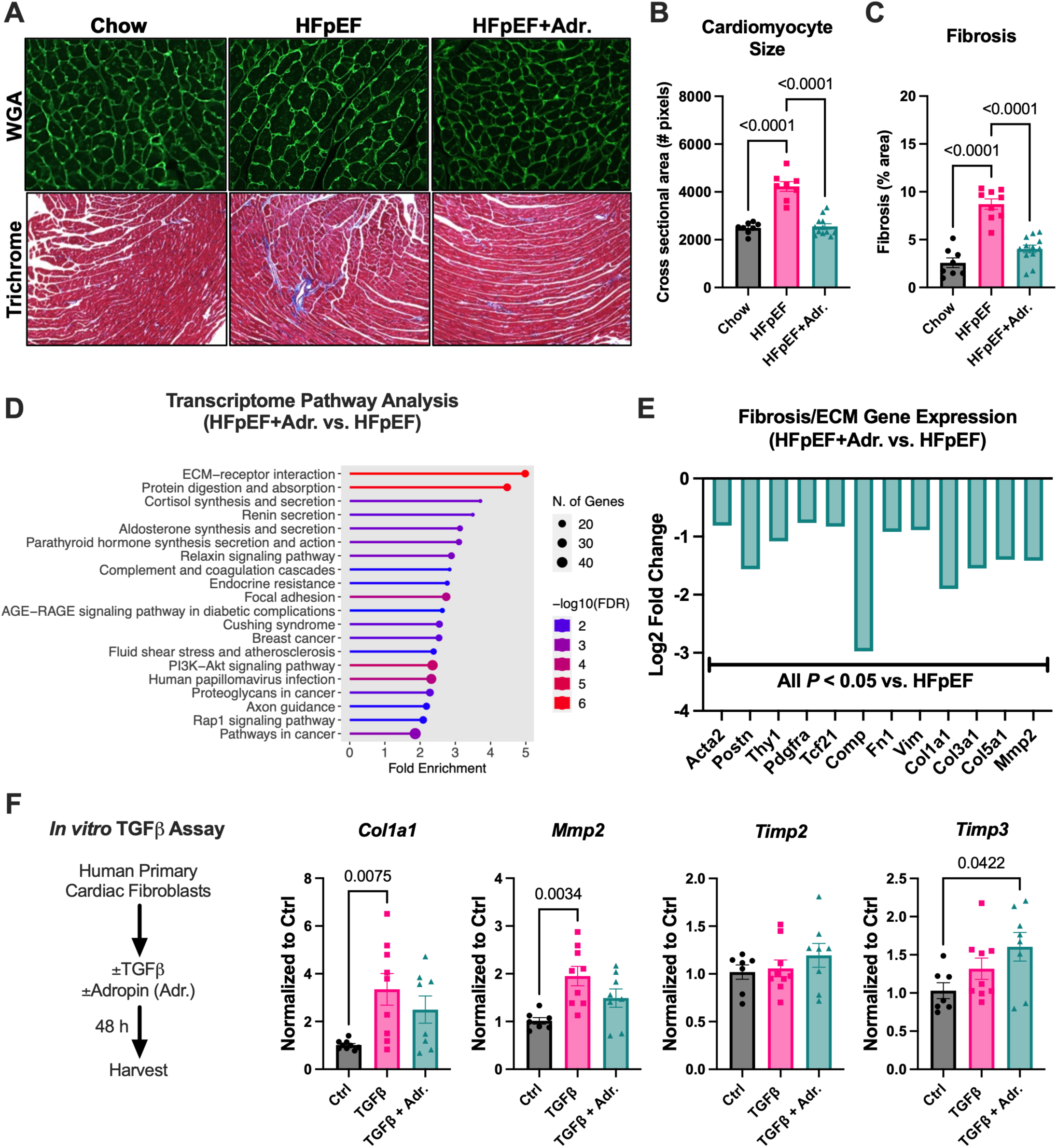
Long-term Adropin treatment prevents cardiomyocyte hypertrophy and reduces cardiac fibrosis. **A.** Representative wheat germ agglutinin (WGA) and Masson’s trichrome images. **B.** Cardiomyocyte cross-sectional area. **C.** Cardiac interstitial fibrosis. **D**. Bulk RNA-seq pathway analysis of mouse hearts using ShinyGO in HFpEF vs. HFpEF+Adr. mice. N = 4. **E**. Fibrosis/ECM gene expression (fold change) in HFpEF vs. HFpEF+Adr. mice. N = 4. **F.** *In vitro* assay of fibrosis/ECM gene expression in primary human cardiac fibroblasts following TGFβ or TGFβ + Adr. treatment. N = 8.

### Adropin treatment limits cardiac protein O-GlcNAcylation in HFpEF mice

The decrease in cardiac fibrosis in Adropin-treated HFpEF mice is likely to account for at least some of the functional improvements observed, particularly in terms of myocardial relaxation. However, it is unlikely to explain all of the improvements that were found to result from Adropin treatment, such as improved metabolic homeostasis. To better understand the underlying processes involved, we next performed untargeted metabolomic analyses of hearts from untreated and Adropin-treated HFpEF mice. Our analysis demonstrated that metabolites related to the hexosamine biosynthesis pathway (HBP) were specifically enriched in Adropin-treated vs. untreated HFpEF hearts **(Fig. 4A; Sup. Table 3)**. As the HBP provides substrates for the generation of UDP-GlcNAc, the co-factor for protein O-GlcNAcylation (**Fig. 4B)**, we next examined the metabolites and enzymes that regulate this pathway.

**Figure 4:**
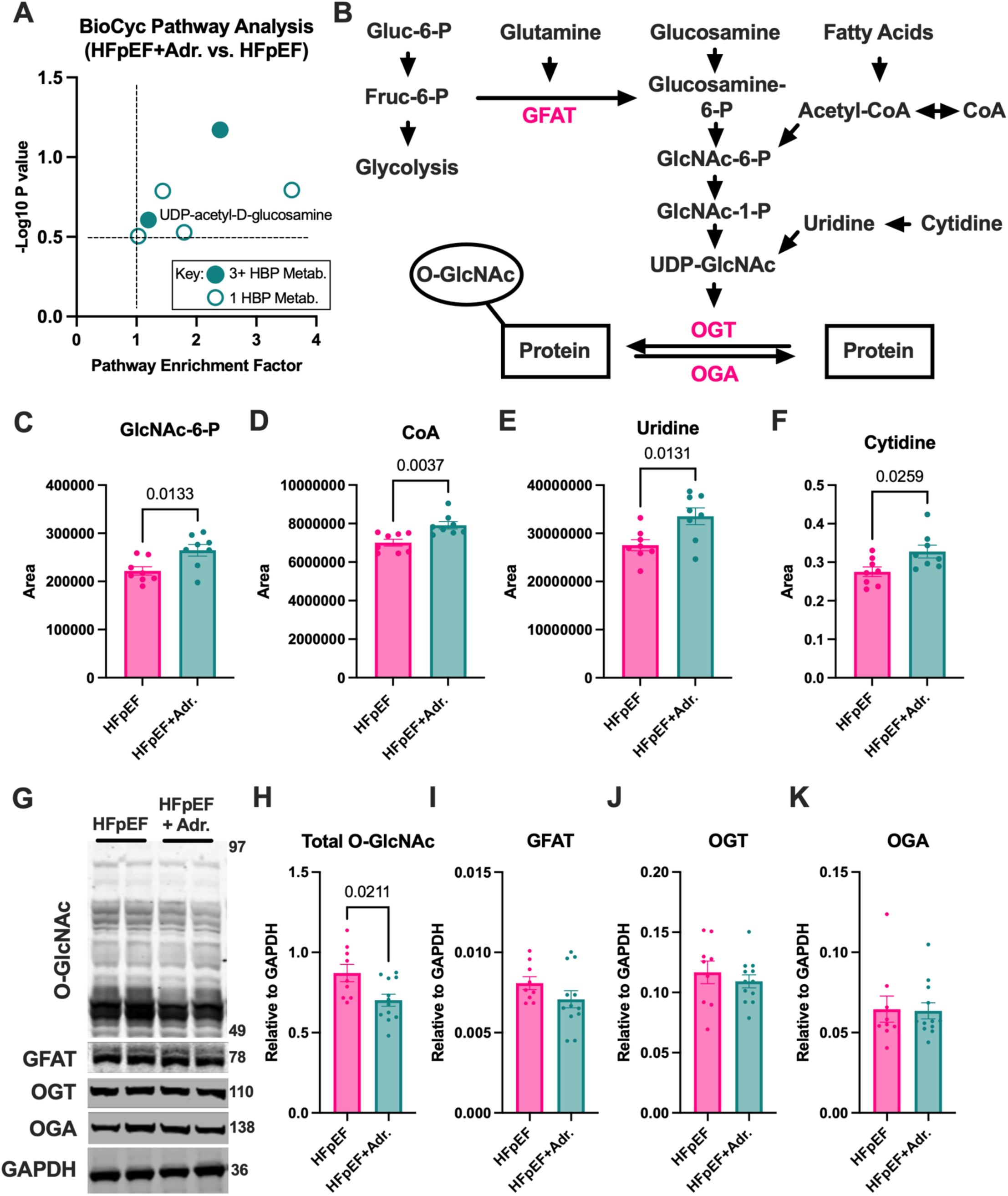
Adropin treatment limits cardiac protein O-GlcNAcylation in HFpEF mice. **A.** BioCyc pathway analysis of untargeted metabolites in HFpEF vs. HFpEF+Adr. hearts. HBP = hexosamine biosynthesis pathway. N = 8. **B.** Schematic of hexosamine biosynthesis pathway (HBP) for the generation of the O-GlcNAcylation substrate, UDP-GlcNAc. **C-F.** Quantification of HBP metabolites GlcNAc-6-P, CoA, Uridine, and Cytidine in untreated and Adropin-treated HFpEF mice. N = 8. **G.** Representative immunoblots of total O-GlcNAcylation and HBP/O-GlcNAcylation regulatory proteins. **H-K.** Quantification of immunoblots. N = 8-12.

Detailed analysis of our metabolomic data revealed an accumulation of GlcNAc-6-P, CoA, Uridine, and Cytidine in Adropin-treated HFpEF hearts relative to control and untreated HFpEF mice **(Fig. 4C- F)**. As these metabolites feed into the HBP at different points, their general accumulation in Adropin-treated mice suggests a possible overall reduction in metabolites entering this pathway to generate UDP-GlcNAc, the co-factor for protein O-GlcNAcylation. Corroborating our metabolomic data, immunoblotting using an O-GlcNAc-specific antibody demonstrated that there was a significant decrease in cardiac protein O-GlcNAcylation in Adropin-treated HFpEF hearts relative to untreated HFpEF controls **(Fig. 4G)**.

O-GlcNAcylation is controlled at the protein level by the rate-limiting enzyme GFAT, and by the opposing activities of the O-GlcNAc transferase (OGT) and O-GlcNAcase (OGA) which, respectively, add and remove this modification to/from other proteins (e.g., Umapathy et al., 2021, Tran et al., 2020, Prakoso et al., 2022). Unexpectedly, despite the difference in O-GlcNAcylation levels observed in untreated and Adropin-treated HFpEF hearts, there were no significant changes in the abundance of any of these regulatory enzymes **(Fig. 4H-K)**. Combined, these data suggest that part of the function of Adropin in protecting from HFpEF is mediated by its modulatory effect on cardiac protein O-GlcNAcylation. By comparing our metabolomic and biochemical data, it would appear that this modulation occurs at the level of HBP metabolite abundance, rather than as an effect on HBP regulatory enzyme activity.

### Adropin-mediated reductions in LCAD O-GlcNAcylation improve its enzymatic activity

Studies in humans and mice have demonstrated that pathways related to cardiac energy metabolism, particularly those in fatty acid utilization, are negatively impacted by HFpEF progression (Tong et al., 2021; Hahn et al., 2023; O’Sullivan et al., 2024). We therefore examined whether specific fatty acid oxidation enzymes were impacted by the increased protein O-GlcNAcylation observed above. Using O-GlcNAc-specific immunoprecipitation, we observed that the beta oxidation enzyme long-chain acyl-CoA dehydrogenase (LCAD) displayed elevated O-GlcNAcylation in HFpEF hearts, which was reversed by Adropin treatment **(Fig. 5A)**. We examined the effect of O-GlcNAcylation on LCAD enzymatic activity *ex vivo*, and found that the ability of LCAD to oxidize C16 palmitoyl-CoA was reduced in vehicle-treated HFpEF hearts, but not those treated with Adropin **(Fig. 5B)**. Combined, there was a significant negative correlation between LCAD O-GlcNAcylation abundance and its enzymatic activity across all cardiac tissues **(Fig. 5C)**. To determine if these enzymatic changes had an impact on the fatty acid composition of the heart, we examined the acylcarnitine profile from our untargeted metabolomics analysis. Long-chain acylcarnitines (C16 and C14) that are metabolized by LCAD were significantly accumulated in untreated HFpEF cardiac tissues relative to Adropin-treated hearts. In contrast, there were no significant differences in medium- and short-chain acylcarnitines (C12 and below) that are metabolized by MCAD and SCAD **(Fig. 5D)**. These findings suggest a specific negative impact of O-GlcNAcylation on LCAD under HFpEF conditions, without impairment of other cardiac acyl-CoA dehydrogenases. We next investigated potential outcomes of the fatty acid accumulation in HFpEF hearts. PLIN5 (a lipid droplet membrane protein that has been shown to block fatty acid oxidation; Wang et al., 2019), was significantly elevated at the protein level in HFpEF mice relative to Adropin-treated animals **(Fig. 5E)**. Combined with the accumulation of long-chain acylcarnitines in our metabolomics analysis, these data suggest that there was an increase in non-metabolized, lipid droplet-encapsulated fatty acids in response to a HFpEF diet.

**Figure 5:**
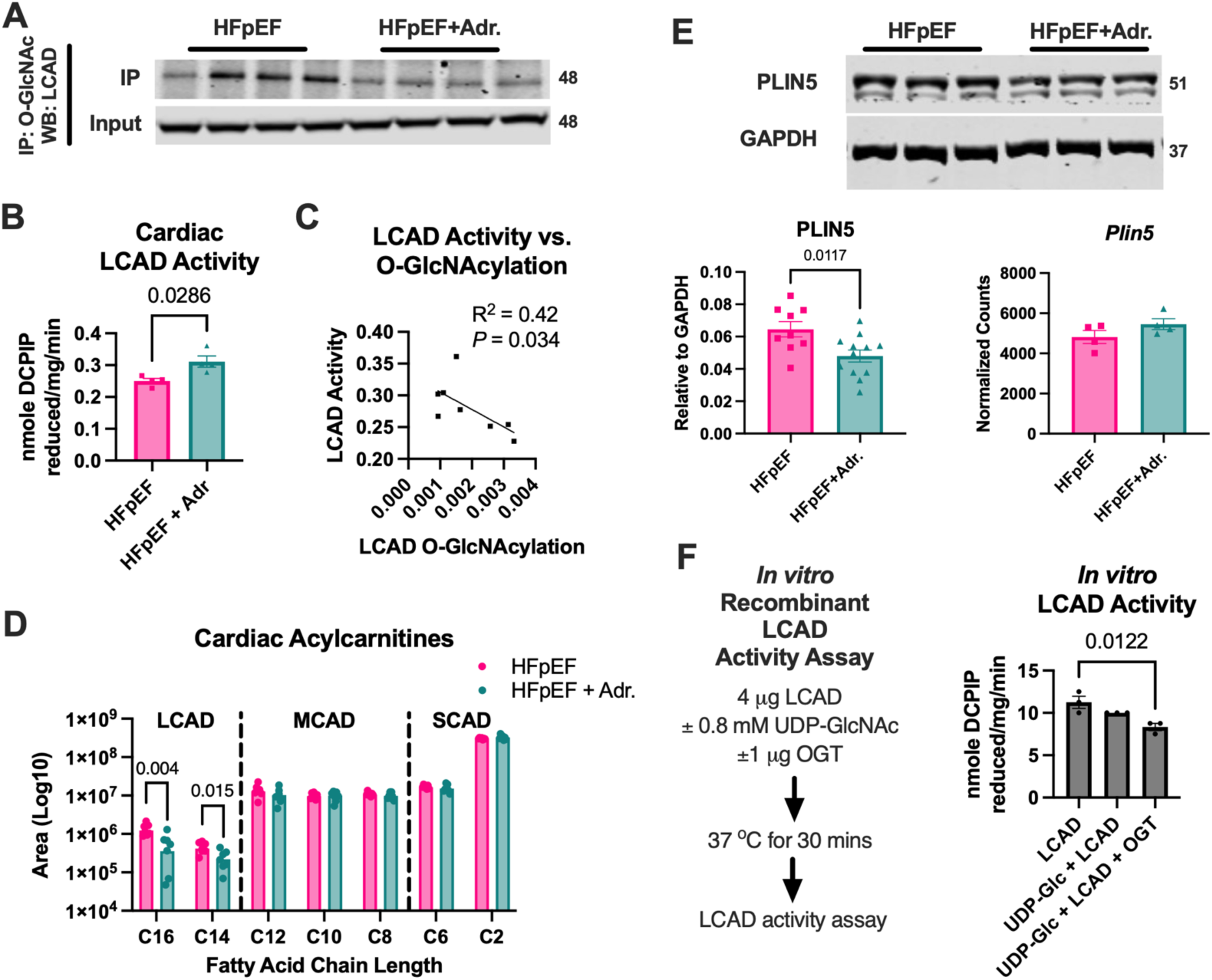
Adropin-mediated reductions in LCAD O-GlcNAcylation improve its enzymatic activity. **A.** Immunoprecipitation analysis of LCAD O-GlcNAcylation status, N = 4. **B.** LCAD enzymatic activity from bulk cardiac tissue, N = 4. **C.** Linear regression analysis of the relationship between LCAD O-GlcNAcylation and LCAD activity. **D.** Profile of long-, medium-, and short-chain cardiac acylcarnitines. N = 8. **E.** Protein abundance and gene expression of cardiac lipid droplet protein PLIN5. N = 4-12. **F.** *In vitro* enzymatic activity of recombinant LCAD after exposure to O- GlcNAcylation substrate (UDP-GlcNAc) and/or recombinant O-GlcNAc transferase (OGT), N = 3.

Finally, to confirm the regulatory effect of increased O-GlcNAcylation on LCAD activity, we performed a series of *in vitro* assays. Recombinant LCAD activity was significantly decreased in the presence of the O-GlcNAcylation substrate (UDP-GlcNAc) and transferase enzyme (OGT), but not in the presence of the substrate alone **(Fig. 5F)**, demonstrating the regulatory effect of this modification. Combined, our data suggest that increased O-GlcNAcylation of fatty acid enzymes like LCAD in HFpEF may decrease their enzymatic activity, which can be reversed by Adropin treatment *in vivo*.

## DISCUSSION

Previous studies on Adropin signaling in cardiometabolic disease have focused on its effects on fuel substrate metabolism, with limited exploration of its role on myocardial structure and function (reviewed in Mushala et al., 2021). Abnormalities of left ventricle passive elasticity are often caused by structural impediments, such as increased cardiomyocyte diameter and extracellular matrix deposition, which contribute to diastolic dysfunction (Aurigemma et al., 2006). Our data shows that Adropin treatment attenuates left ventricular hypertrophy and fibrosis **(Fig. 3)**, indicative of improved structural remodeling. Transcriptomic analyses from our mouse models demonstrated similar increases in pro-fibrotic pathways as seen in HFpEF patients (Ye et al., 2023), which was attenuated upon Adropin treatment **(Fig. 3; Sup. Fig. 1)**, providing further evidence of restored myocardial structure and function. These data match similar findings in a model of systemic sclerosis, where Adropin treatment significantly reduced fibroblast activation and extracellular matrix deposition (Liang et al., 2024). Collectively, these findings highlight the therapeutic potential of prolonged Adropin treatment in preventing cardiac remodeling and functional decline in HFpEF.

Unlike previous acute studies from our group and others, the ability of Adropin treatment to modulate cardiac substrate utilization was independent of the inhibitory effects of PDK4 or GCN5L1 on pyruvate dehydrogenase activity (Thapa et al., 2019b, Altamimi et al., 2019). Instead, we found that Adropin treatment may reduce metabolite entry into and/or flux through the HBP, which inhibits cardiac protein O-GlcNAcylation to promote fatty acid oxidation **(Figs. 4,5)**. The HBP is a non-oxidative branch of glycolysis where the rate-limiting enzyme, GFAT, uses fructose-6-phosphate and glutamine to catalyze its conversion into glucosamine-6-phosphate, which is subsequently metabolized through a series of reactions utilizing substrates from major metabolic pathways to generate the end-product UDP-GlcNAc (Umapathi et al., 2021). UDP-GlcNAc serves as the co-factor for the attachment of an O-linked β-N-acetyl glucosamine moiety (O-GlcNAc) to serine and threonine residues of nuclear, cytoplasmic, and mitochondrial proteins, using 2-3% of the total cellular glucose found in the heart (Tran et al., 2020).

The dynamic on- and off-cycling of the O-GlcNAc modification is regulated by the activity of two enzymes, O-GlcNAc transferase (OGT) and O-GlcNAcase (OGA), respectively (Prakoso et al., 2022). Several studies have implicated aberrant regulation of the HBP, and consequent maladaptive protein O-GlcNAcylation, in heart failure (e.g., Umapathy et al., 2021, Tran et al., 2020, Prakoso et al., 2022). These studies have identified changes in OGT and OGA enzyme expression as the predominant mechanism that triggers excessive protein O-GlcNAcylation in diabetic hearts. However, here we found that changes in HBP metabolite abundance following Adropin treatment appeared to decrease UDP-GlcNAc availability for cardiac protein O-GlcNAcylation levels **(Fig. 4)**, contrasting with the idea that OGT/OGA abundance is the only regulatory component of the pathway.

In this study, we identified the fatty acid oxidation enzyme LCAD as a target of O-GlcNAcylation in HFpEF mice, and demonstrated that Adropin treatment could limit this deleterious modification **(Fig. 5)**. LCAD O-GlcNAcylation was associated with decreased cardiac enzyme activity in HFpEF mouse hearts, which was restored upon Adropin treatment *ex vivo* **(Fig. 5)**. The inhibitory effect of O-GlcNAcylation on LCAD enzyme activity was confirmed *in vitro* using recombinant protein assays, suggesting that Adropin treatment may augment cardiac fatty acid oxidation activity via this mechanism. Further studies need to be conducted to determine whether Adropin-mediated regulation of cardiac LCAD O-GlcNAcylation regulates fatty acid utilization in the heart *in vivo*.

Connections between the metabolic processes (i.e. O-GlcNAcylation and fatty acid oxidation) and functional processes (i.e. cardiac dysfunction and structural remodeling) described in this study are complex, and are likely to include several areas of crosstalk/overlap. For example, a decrease in cardiac fatty acid oxidation has been shown to result in cardiac hypertrophy by driving glucose towards anabolic pathways that support abnormal cellular growth (Ritterhoff et al., 2020). As such, the promotion of fatty acid oxidation enzyme activity observed here following Adropin treatment may help to block glucose-mediated cardiomyocyte hypertrophy. Similarly, there may be a direct relationship between O-GlcNAcylation levels and cardiac fibrosis. One key report, which used cardiac-specific overexpression of a dominant-negative OGA enzyme, found that sustained increases in O-GlcNAcylation (via reduced removal of O-GlcNAc modifications) led to cardiac diastolic dysfunction and mitochondrial impairment (Ha et al., 2023). In a related diabetic cardiomyopathy model, Prakoso et al. (2022) found that AAV-mediated cardiac overexpression of OGT led to diastolic dysfunction.

Crucially, in both of these studies, the overall increase in cardiac O-GlcNAcylation correlated with a significant increase in cardiac fibrosis. As such, increased myocardial O-GlcNAcylation may exacerbate diastolic dysfunction by directly driving increased fibrosis in the heart. Further work will be required to separate these pathways to understand the relative contribution of each process to HFpEF pathogenesis.

Our study has three notable limitations. Firstly, the exclusive use of male animals prohibits the investigation into possible sex differences in Adropin-mediated effects observed in the heart. Previous studies using the current HFpEF model have demonstrated that female sex is protective against cardiac dysfunction (Tong et al., 2019). On this basis, our study exclusively examined male mice, and it is unknown whether the findings are relevant for female mice. Epidemiological data show that women display a higher prevalence of HFpEF compared to men, particularly as age increases (Gori et al., 2014, Beale et al., 2018, Duca et al., 2018). Sexual dimorphism in metabolic homeostasis influences the development of cardiac dysfunction (Lindman et al., 2014, Lagou et al., 2021), suggesting a potential role of reproductive hormones in cardiometabolic HFpEF. Interestingly, hepatic Adropin production has been shown to be regulated by estrogen (Stokar et al., 2022), suggesting a potential difference in basal circulating Adropin levels between male and female mice. We did not detect any sex differences in circulating Adropin levels between male and female patients, regardless of HFpEF status (data not shown). However, the members of our female patient cohort were most likely to be in a post-menopausal state (mean age 67, range 56-75; **Sup. Table 1**), which may limit our ability to extrapolate to these findings to the general population, and more work in this area is required.

Secondly, our metabolomics data relied on a cross-sectional analysis of HFpEF hearts, which precluded flux measurements of metabolites related to the HBP. While cross-sectional metabolomics data has allowed us to make major advances of our understanding of HFpEF (e.g., Hahn et al, 2023), it does limit our understanding of how metabolite abundance changes in response to disease progression. Future work will likely include the use of stably-labeled isotope tracers for specific metabolites, or the measurement of metabolites entering and exiting the heart (e.g., O’Sullivan et al, 2024), to achieve a better understanding of HBP metabolite flux. In addition, the direct measurement of some metabolites (e.g. UDP-GlcNAc) using a newly-developed enzymatic assay (Upadhyay et al., 2024) will provide further information on the modification of the HBP pathway in HFpEF.

Finally, we acknowledge that despite our use of a well-established “two-hit” HFpEF mouse model (HFD+L-NAME; Schiattarella et al., 2019), we did not observe a significant change in E/e’ or E/A ratios during our echocardiography analysis **(Fig. 2)**. While these ratios have become the *de facto* norm for detecting diastolic dysfunction in mice (as discussed in Sun et al., 2024), changes in age, heart rate, and anesthesia levels can affect the outcomes observed (Rottman et al., 2007; Sun et al., 2024). While changes in E/e’ and E/A ratio were not observed, we did detect a significant increase in both isovolumetric relaxation time (IVRT) and myocardial performance index (MPI; also known as Tei index) in untreated HFpEF animals, which was absent following Adropin treatment **(Fig. 2)**. Both IVRT and MPI have been well validated as measures of diastolic dysfunction (e.g. Tei et al., 1996; Schnelle et al., 2018), and as such we believe that the improvements observed indicate that Adropin can mitigate aspects of mild-to-moderate diastolic dysfunction in HFpEF. In future studies, the use of invasive *in vivo* hemodynamic measurements, such as left ventricle end diastolic pressure (e.g. EDPVR; Wearing et al., 2025), would be merited to determine load-independent measured of diastolic dysfunction.

In summary, we found that HFpEF-driven disruptions to normal cardiac metabolism led to a potential increase in metabolites entering the HBP, resulting in increased in cardiac protein O-GlcNAcylation, and decreased fatty acid oxidation enzyme activity. This occurred in concert with an increase in diastolic dysfunction, fibrosis, and cardiac hypertrophy in HFpEF animals. Long-term Adropin treatment reversed the metabolic and structural remodeling observed in HFpEF hearts, leading to improved myocardial relaxation capacity **(Fig. 6)**. These findings significantly increase our understanding of the protective effects of Adropin in HFpEF, and highlight this pathway as a potential target for future therapeutic interventions.

**Figure 6:**
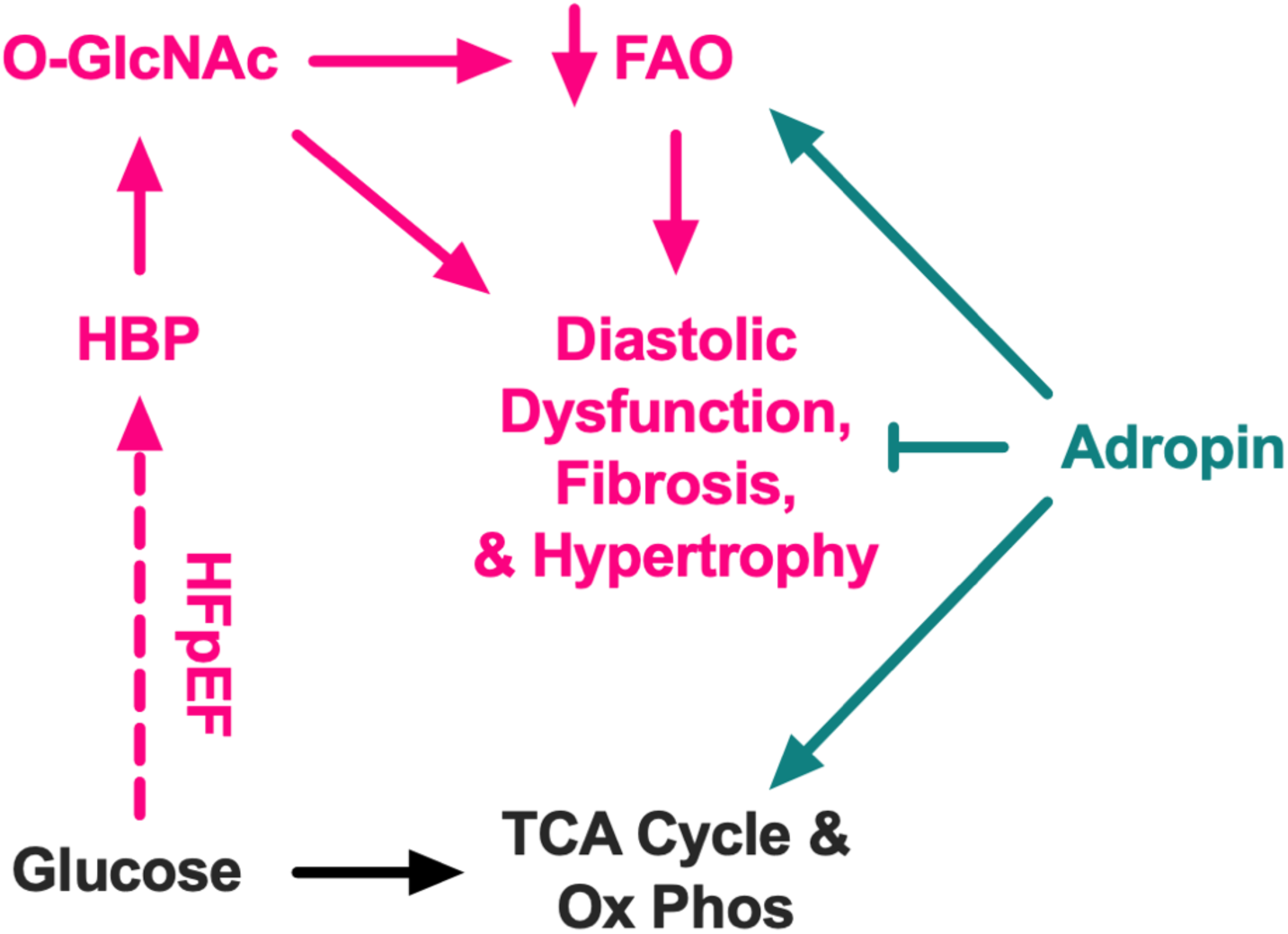
Summary. Adropin treatment prevents deleterious changes in cardiac metabolism, leading to a reduction in diastolic dysfunction, cardiac fibrosis, and cardiomyocyte hypertrophy.

## DATA AVAILABILITY

All data reported herein can be found in the results or supplementary information sections of this article.

## ACKNOWLEDGEMENTS

This work was supported by: National Institute of Health Fellowships (F31DK134089 and T32HL110849) to B.A.S.M; National Institute of Health Shared Instrumentation Grants (S10OD023402, S10OD032141) to S.L.G.; National Institutes of Health Research Grants (R01HL124021, R01HL122596, R01HL151228) to S.Y.C.; National Institute of Health Research Grants (R01HL147861, R0HL156874) and American Heart Association Established Investigator Award (23EIA1037834) to I.S. The University of Pittsburgh Center for Metabolism and Mitochondrial Medicine is supported by the Pittsburgh Foundation (MR2020 109502) grant to M.J.J. This project used the University of Pittsburgh Medical Center (UPMC) Hillman Cancer Center and Tissue and Research Pathology/Pitt Biospecimen Core shared resource, which is supported in part by award P30CA047904. Echocardiography was carried out by the University of Pittsburgh Rodent Ultrasonography Core, which received funding from the NIH Shared Instrumentation Grant Program (S10OD023684).

## AUTHOR CONTRIBUTIONS

B.A.S.M., M.J.J., and I.S. conceived the study and designed experiments. E.S.G., Y.A.A. and S.Y.C. provided essential biological samples. B.A.S.M, M.W.S., M.S.S., B.M., A.M.V., and S.J.M. performed experiments. B.A.S.M., B.M., S.J.M. and I.S. analyzed data. B.A.S.M. and I.S. prepared figures. J.R.M., P.B., N.B., M.S.S., S.S.S., B.A.K., C.Z., E.S.G, S.Y.C., and S.L.G provided critical input and expertise. B.A.S.M and I.S. drafted the manuscript. B.A.S.M. and I.S. edited and revised the manuscript. All authors approved the final submission.

## DISCLOSURES

S.Y.C. has served as a consultant for Merck, Janssen, and United Therapeutics. S.Y.C. is a director, officer, and shareholder in Synhale Therapeutics. S.Y.C. has held grants from Bayer and United Therapeutics. I.S. is listed as the inventor on patent applications for the use of Adropin in the detection and treatment of cardiometabolic disease.

## Notes

### Competing Interest Statement

The authors have declared no competing interest.

### Summary of Updates

Addition of new data to Figures 2, 3, and 5. Changes to text based on new data. Increased discussion.

