## Supplementary figures and images for "Adropin protects against cardiac remodeling and metabolic dysfunction in a mouse model of HFpEF"

### Supplemental Figure 1

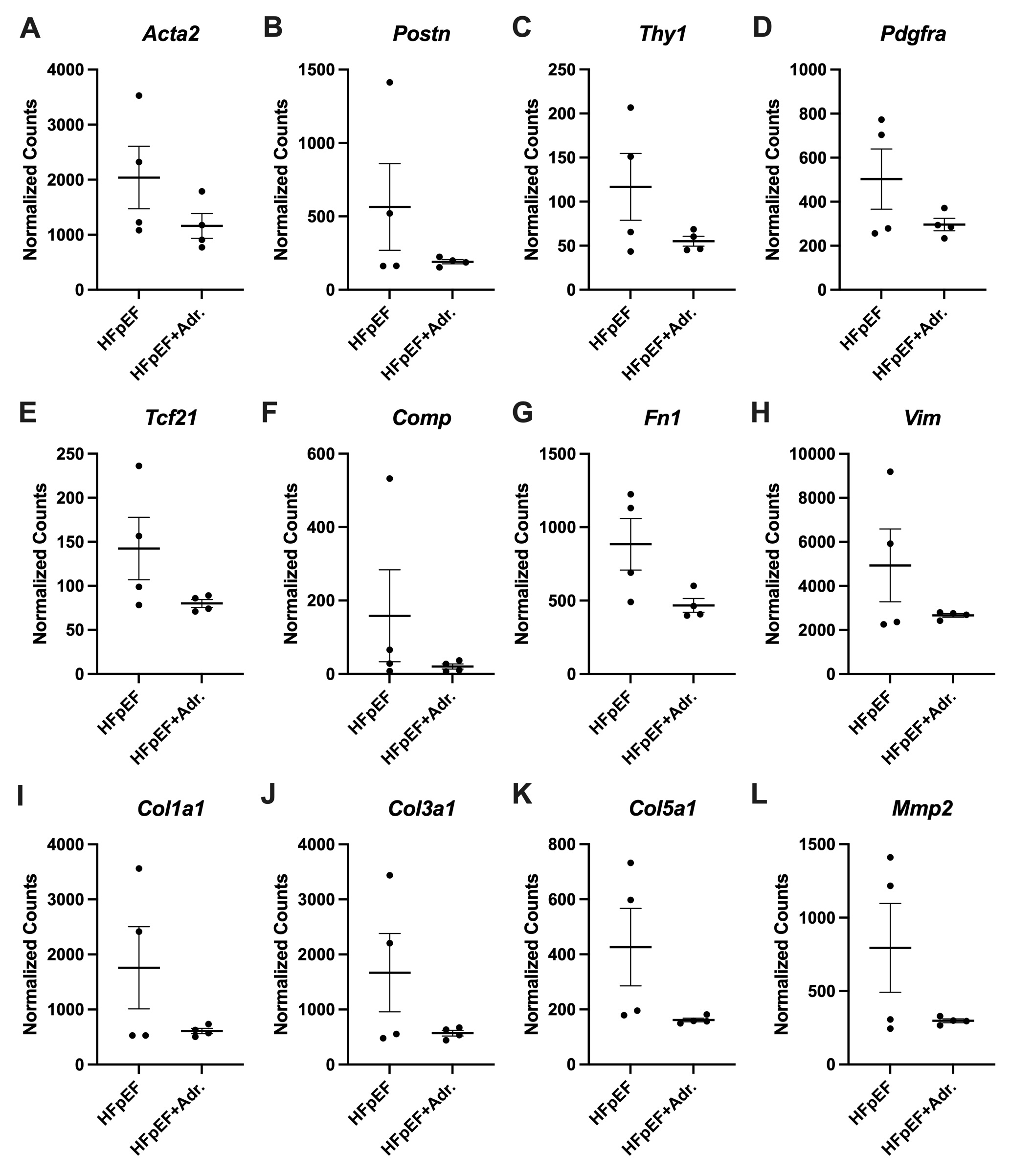
