## Supplemental Table 1 for "Adropin protects against cardiac remodeling and metabolic dysfunction in a mouse model of HFpEF"

| Control | | | | | HFpEF | | | | |
|---|---|---|---|---|---|---|---|---|---|
| **ID** | **Age** | **Gender** | **Race** | **Ethnicity** | **ID** | **Age** | **Gender** | **Race** | **Ethnicity** |
| 758 | 78 | M | W | NH/NL | 166 | 77 | M | W | NH/NL |
| 843 | 61 | F | W | NH/NL | 434 | 58 | F | W | NH/NL |
| 854 | 77 | M | W | NH/NL | 442 | 76 | M | W | NH/NL |
| 872 | 75 | F | W | NH/NL | 445 | 72 | F | W | NH/NL |
| 884 | 73 | F | W | NH/NL | 459 | 72 | F | W | NH/NL |
| 909 | 75 | F | W | NH/NL | 462 | 74 | F | W | NH/NL |
| 933 | 56 | F | W | NH/NL | 466 | 58 | F | W | NH/NL |
| 935 | 72 | M | W | NH/NL | 470 | 74 | M | W | NH/NL |
| 896 | 62 | F | W | NH/NL | 476 | 59 | F | W | NH/NL |
| 958 | 65 | F | W | NH/NL | 482 | 65 | F | W | NH/NL |
| 930 | 63 | F | W | NH/NL | 492 | 68 | F | W | NH/NL |
| 946 | 45 | M | W | NH/NL | 499 | 46 | M | W | NH/NL |
| 947 | 72 | F | W | NH/NL | 514 | 74 | F | W | NH/NL |
| 974 | 66 | M | W | NH/NL | 562 | 69 | M | W | NH/NL |
| 1137 | 59 | F | W | NH/NL | 576 | 70 | F | W | NH/NL |
| 992 | 74 | M | W | NH/NL | 583 | 63 | F | W | NH/NL |
| 1014 | 73 | F | W | NH/NL | 633 | 71 | M | W | NH/NL |
| 1122 | 50 | M | W | NH/NL | 636 | 74 | F | W | NH/NL |
| 1052 | 72 | M | W | NH/NL | 663 | 48 | M | W | NH/NL |
| | | | | | 675 | 72 | M | W | NH/NL |

**Supplemental Table 1 – Control and HFpEF Patient Characteristics.** M = male, F = female, W = white, NH/NL = non-Hispanic/non-Latino. N = 19-20.
