## Supplemental Table 2 for "Adropin protects against cardiac remodeling and metabolic dysfunction in a mouse model of HFpEF"

Sup Table 2

| ID | log2FoldChange | pvalue | padj | A1HRT | A2HRT | A3HRT | A4HRT | H1HRT | H2HRT | H3HRT | H4HRT | Gene.name |
|---|---|---|---|---|---|---|---|---|---|---|---|---|
| **ENSMUSG00000000001** | -0.402589154594001 | 0.0136551284267496 | 0.635862253526407 | 488.800920663853 | 442.22774080048 | 388.046504345863 | 399.556771753977 | 681.063997799478 | 616.433444965225 | 463.279750297481 | 512.368066523536 | Gnai3 |
| **ENSMUSG00000000028** | 0.166812155280671 | 0.449734552858307 | 0.89108031640712 | 54.2108229941427 | 51.3081909216027 | 59.6266579848522 | 62.8410885152968 | 62.4126171616885 | 50.1463715150282 | 44.7663578939139 | 45.5920737160773 | Cdc45 |
| **ENSMUSG00000000031** | -0.570059660688536 | 0.294334844622863 | 0.851785577030345 | 316.229800799166 | 247.582381589956 | 171.308334845369 | 186.647412157374 | 652.595084708182 | 439.086521314516 | 221.749633288457 | 55.3618037980939 | H19 |
| **ENSMUSG00000000049** | -0.362123377580488 | 0.230557150302713 | 0.832048466795696 | 362.309000344187 | 701.211942595237 | 377.63550057073 | 432.384206053012 | 411.70428162798 | 513.694537471021 | 548.648153723084 | 934.637511179585 | Apoh |
| **ENSMUSG00000000056** | 0.0455122212369412 | 0.866655806387048 | 0.974868041734905 | 2344.61809449667 | 2668.84034365225 | 2516.62354891622 | 2130.03152266028 | 2030.05249504861 | 1231.64380769838 | 3126.35750594008 | 2971.08347049771 | Narf |
| **ENSMUSG00000000058** | -0.181718868419708 | 0.449947055449652 | 0.89108031640712 | 1251.36649744813 | 802.199492980614 | 933.204520207369 | 1502.55856420157 | 1481.47843894324 | 1501.9449809867 | 928.641656776074 | 1179.96628879467 | Cav2 |
| **ENSMUSG00000000078** | -0.321513553301027 | 0.181327021323265 | 0.803056849297055 | 994.768601942519 | 920.289773673191 | 847.07712534036 | 1018.58838996436 | 1570.17005280458 | 1486.04491196974 | 794.342583094332 | 874.933605122817 | Klf6 |
| **ENSMUSG00000000085** | 0.219854914460097 | 0.323756368551158 | 0.855977678491063 | 889.961010820509 | 1052.22512175731 | 1042.99328729059 | 814.120370616084 | 732.527033002976 | 524.702277559686 | 1015.05113829223 | 988.91378941301 | Scmh1 |
| **ENSMUSG00000000088** | 0.0719125743232897 | 0.552843159730834 | 0.914977712327965 | 17107.1287095183 | 14692.0597496145 | 14784.5718155774 | 18088.8542254629 | 16315.9720759004 | 16860.1885691382 | 13839.0510577627 | 14513.4767996179 | Cox5a |
| **ENSMUSG00000000093** | -0.721197208121327 | 0.0874998102174972 | 0.74812051555524 | 85.8338030740593 | 96.9154717408051 | 65.3053873167428 | 70.3445020693621 | 163.148771177045 | 233.608706326107 | 75.998700610598 | 53.1907526687569 | Tbx2 |
| **ENSMUSG00000000126** | 0.258414100512365 | 0.293903063688154 | 0.851785577030345 | 151.7903043836 | 146.594831204579 | 260.275094378323 | 229.79204009325 | 151.104231023035 | 154.108361241306 | 141.586620315635 | 211.677485110359 | Wnt9a |
| **ENSMUSG00000000127** | 0.237093248274642 | 0.100218027173709 | 0.759800953500741 | 480.669297214732 | 503.308920469055 | 526.228918088536 | 450.204813243917 | 404.039574257247 | 337.570696052385 | 492.429936833053 | 427.697072479392 | Fer |
| **ENSMUSG00000000131** | -0.242558346935931 | 0.214473255851375 | 0.824248286808591 | 99.3865088225949 | 97.7298874697194 | 130.610774633486 | 127.55803041911 | 144.534481848121 | 161.446854633749 | 114.518589961175 | 118.322286548867 | Xpo6 |
| **ENSMUSG00000000134** | -0.219045247229553 | 0.216136861849557 | 0.825444325292919 | 170.764092431549 | 252.468875963442 | 243.238906382651 | 211.971432902344 | 269.359716171498 | 293.539735697726 | 243.612273190136 | 217.105112933702 | Tfe3 |
| **ENSMUSG00000000142** | -0.0585161164790679 | 0.870680122099639 | 0.975613767803763 | 52.4037955610046 | 65.9676740420606 | 106.002947528626 | 74.0962088463947 | 51.4630352034976 | 45.2540425867328 | 116.600746142287 | 96.6117752554972 | Axin2 |
| **ENSMUSG00000000148** | -0.281471292618107 | 0.0458580701078179 | 0.697906909182231 | 144.562194651047 | 157.99665140938 | 147.646962629158 | 157.571684635371 | 197.092475247437 | 193.24699266767 | 158.243869764533 | 191.052499381657 | Brat1 |
| **ENSMUSG00000000149** | 0.135288481187496 | 0.326850379750869 | 0.855977678491063 | 5707.49614756666 | 5954.19339409266 | 6117.88440022356 | 5328.36155008061 | 5000.67408030582 | 4223.30294735104 | 5819.6265262088 | 5994.2721680995 | Gna12 |
| **ENSMUSG00000000154** | -0.508017303599758 | 0.199726895229401 | 0.812921911423739 | 11.7456783153976 | 15.4738988493722 | 19.8755526616174 | 26.2619474392285 | 25.1840385038392 | 22.0154801773295 | 17.6983275394543 | 39.0789203280663 | Slc22a18 |
| **ENSMUSG00000000168** | 0.173397198632263 | 0.333483239485804 | 0.85715806744223 | 12808.2104460828 | 14188.7508291454 | 13747.2572576187 | 12493.1835675187 | 10946.2970836035 | 8365.8824673852 | 14186.7711400085 | 13708.0168306339 | Dlat |
| **ENSMUSG00000000171** | 0.105626214474712 | 0.46398356733553 | 0.895328112556867 | 9251.07694395045 | 7818.39099757756 | 7745.78680869889 | 8220.92747517278 | 6965.02908360528 | 6371.03534687273 | 8260.95464856294 | 9105.38843643944 | Sdhd |
| **ENSMUSG00000000183** | 0.986467918285268 | 0.0569228110663832 | 0.707207375413293 | 37.0440623793308 | 32.5766291565731 | 14.1968233297267 | 49.7101147956825 | 14.2344565456483 | 4.89232892829544 | 22.903717992235 | 24.9670879873757 | Fgf6 |
| **ENSMUSG00000000184** | 0.0875937094219768 | 0.628563379492852 | 0.931825800800958 | 2470.20650109977 | 2226.61260285177 | 1695.10070556937 | 2372.95453647315 | 1738.79361496073 | 1786.92314105991 | 2152.94949127009 | 2569.43901157036 | Ccnd2 |
| **ENSMUSG00000000194** | -0.162444003646614 | 0.0493706904988041 | 0.703396849069134 | 800.513152880174 | 803.013908709528 | 836.666121565227 | 807.554883756277 | 966.848086908263 | 906.30393396673 | 868.259127523818 | 893.387539722182 | Gpr107 |
| **ENSMUSG00000000197** | -0.828399071603439 | 0.20760395926164 | 0.820372930586885 | 8.13162344912141 | 10.5874044758863 | 9.46454888648447 | 4.68963347129081 | 14.2344565456483 | 33.0232202659942 | 4.16431236222455 | 7.59867895267955 | Nalcn |
| **ENSMUSG00000000253** | 0.0618084089684906 | 0.6050878263094 | 0.926448151609826 | 2032.00234856378 | 1838.95071588855 | 1849.37285241907 | 2088.76274811293 | 2059.61636633572 | 2060.89356104445 | 1671.97141343316 | 1690.16330418887 | Gmpr |
| **ENSMUSG00000000266** | 0.171324544777461 | 0.263688061746211 | 0.842510961779757 | 374.958192376154 | 462.588134023339 | 422.118880337207 | 332.026049767389 | 341.626957095558 | 306.993640250539 | 370.623800237985 | 394.045779974668 | Mid2 |
| **ENSMUSG00000000275** | -0.149340877337089 | 0.350357340111358 | 0.861774532843376 | 460.791995450213 | 560.318021493058 | 542.31865119556 | 460.522006880757 | 705.153078107498 | 527.148442023833 | 482.019155927491 | 530.8220011229 | Trim25 |
| **ENSMUSG00000000276** | -0.0250233788868591 | 0.891912305950071 | 0.982421229269187 | 242.141676040504 | 266.313943354985 | 326.526936583714 | 349.846656958294 | 329.582416941548 | 231.162541861959 | 317.528817619622 | 325.657669400552 | Dgke |
| **ENSMUSG00000000278** | -0.336739272201012 | 0.205087680447036 | 0.819289008170643 | 262.018977805023 | 280.973426475443 | 275.418372596698 | 298.260688774095 | 440.173194719277 | 491.679057293692 | 211.338852382896 | 268.124814473121 | Scpep1 |
| **ENSMUSG00000000282** | -0.407265865004227 | 0.0140274527958795 | 0.635862253526407 | 277.378710986697 | 246.767965861042 | 294.347470369667 | 293.571055302804 | 428.128654565267 | 311.885969178834 | 424.759860946904 | 308.289260365856 | Mnt |
| **ENSMUSG00000000290** | -0.465190963152806 | 0.185975327138078 | 0.80598979530356 | 62.3424464432641 | 79.8127414336042 | 108.842312194571 | 72.2203554578784 | 118.255485148462 | 187.131581507301 | 64.5468416144805 | 77.072315091464 | Itgb2 |
| **ENSMUSG00000000295** | -0.0690491560937394 | 0.827378176619722 | 0.967319591893432 | 26.2018977805023 | 23.6180561385155 | 28.3936466594534 | 32.8274342990356 | 28.4689130912965 | 42.8078781225851 | 22.903717992235 | 22.7960368580387 | Hddc2 |
| **ENSMUSG00000000296** | 0.330985958748896 | 0.0875201438877519 | 0.74812051555524 | 179.79922959724 | 141.708336831093 | 188.344522841041 | 211.971432902344 | 141.249607260664 | 111.300483118721 | 165.531416398426 | 154.144630182928 | Tpd52l1 |
| **ENSMUSG00000000301** | -0.17036720244366 | 0.586707134488618 | 0.923058967588378 | 94.8689402397497 | 113.203786319092 | 92.7525790875478 | 108.799496533947 | 90.881530252985 | 113.746647582869 | 65.5879197050366 | 191.052499381657 | Pemt |
| **ENSMUSG00000000305** | 0.0712543162254655 | 0.747686869591056 | 0.956119556124081 | 444.52874855197 | 578.235167529173 | 420.22597055991 | 450.204813243917 | 407.324448844704 | 298.432064626022 | 553.853544175865 | 541.677256769585 | Cdh4 |
| **ENSMUSG00000000317** | -0.442937405291965 | 0.0251459218209995 | 0.655303666217042 | 268.343573821006 | 193.83094348161 | 211.059440168604 | 299.198615468353 | 394.184950494875 | 261.739597663806 | 325.857442344071 | 338.683976176574 | Bcl6b |
| **ENSMUSG00000000318** | -0.593264876130152 | 0.292012544979799 | 0.851785577030345 | 10.8421645988285 | 10.5874044758863 | 36.9117406572894 | 21.5723139679377 | 30.6588294829347 | 50.1463715150282 | 9.36970281500523 | 30.3947158107182 | Clec10a |
| **ENSMUSG00000000320** | 0.0560030877541427 | 0.871411727068953 | 0.975613767803763 | 103.90407740544 | 109.13170767452 | 267.84673348751 | 175.392291826276 | 171.908436743598 | 216.485555077073 | 106.189965236726 | 136.776221148232 | Alox12 |
| **ENSMUSG00000000325** | 0.197032383519597 | 0.158309471509994 | 0.789230661864335 | 684.863397159336 | 655.604661776035 | 565.033568523123 | 674.369293171618 | 526.674892188986 | 516.140701935169 | 524.703357640293 | 682.795580176491 | Arvcf |
| **ENSMUSG00000000326** | 0.402515321185326 | 0.0177375333379287 | 0.642273034804458 | 8142.46561372023 | 7162.78633580152 | 10789.5857305923 | 11150.072541341 | 6878.52738613557 | 7846.07251875381 | 6880.48510048551 | 6571.77176850314 | Comt |
| **ENSMUSG00000000339** | -0.0128964111160674 | 0.905069861218969 | 0.98504921011344 | 519.520387027201 | 472.361122770311 | 506.353365426919 | 523.363095396054 | 480.686647964584 | 574.848649074714 | 495.553171104721 | 489.572029665497 | Rtca |
| **ENSMUSG00000000340** | 0.144607084541612 | 0.519303135390326 | 0.905962179970576 | 1536.87683188395 | 1613.35755897928 | 1696.99361534667 | 1419.0830884126 | 1096.05315401492 | 978.465785659088 | 1884.35134390661 | 1708.61723878823 | Dbt |
| **ENSMUSG00000000346** | 0.100585809203883 | 0.133309436538931 | 0.77303252008843 | 5005.46598979251 | 4779.80591299819 | 5222.53807556213 | 5157.65889172563 | 4618.53366996495 | 4628.14316616749 | 4822.27371545602 | 4736.1480386487 | Dazap2 |
| **ENSMUSG00000000355** | 0.0338317469845541 | 0.778284255322293 | 0.960153504434377 | 369.537110076739 | 351.82759489099 | 322.74111702912 | 413.625672167849 | 365.716037403578 | 342.463024980681 | 360.213019332423 | 354.966859646602 | Mcts1 |
| **ENSMUSG00000000359** | -0.551046875326622 | 0.100371032033235 | 0.759800953500741 | 34.3335212296237 | 27.6901347830872 | 37.8581955459379 | 57.2135283497478 | 67.887408140784 | 79.5003450848009 | 42.6842017128016 | 40.1644458927348 | Rem1 |
| **ENSMUSG00000000374** | 0.207789113529012 | 0.439412055886205 | 0.889583654898613 | 1406.770856698 | 1669.55224427437 | 1377.09186298349 | 1177.09800129399 | 970.13296149572 | 705.718447906617 | 1611.5888841809 | 1587.03837554536 | Trappc10 |
| **ENSMUSG00000000378** | 0.0291925691793944 | 0.759075335790819 | 0.957357521291862 | 624.32797814921 | 623.842448348376 | 664.41133183121 | 687.500266891232 | 675.589206820383 | 649.456665231219 | 596.537745888666 | 626.348250813729 | Ccm2 |
| **ENSMUSG00000000384** | 0.102688442984196 | 0.464745022925501 | 0.895328112556867 | 1684.1495676847 | 1815.33265975004 | 1609.91976559101 | 1530.69636502932 | 1457.38935863522 | 1256.10545233985 | 1781.28461294155 | 1687.99225305953 | Tbrg4 |
| **ENSMUSG00000000392** | -0.309545546165296 | 0.39047358450777 | 0.877797322358001 | 26.2018977805023 | 24.4324718674299 | 32.1794662140472 | 28.1378008277448 | 49.2731188118594 | 26.9078091056249 | 17.6983275394543 | 43.4210225867403 | Fap |
| **ENSMUSG00000000399** | 0.0738737691223948 | 0.449565939263221 | 0.89108031640712 | 21392.4942672053 | 18243.7267434099 | 18685.8588665863 | 21206.522557177 | 19410.3239372851 | 20086.679497349 | 17719.1491012654 | 18343.2109917684 | Ndufa9 |
| **ENSMUSG00000000420** | 0.105441731162372 | 0.547484896198743 | 0.914699131434573 | 2528.03137896019 | 2812.17751194118 | 2776.89864329454 | 2488.3195198669 | 2123.12394169323 | 1867.64656837678 | 3007.67460361668 | 2858.18881177218 | Galnt1 |
| **ENSMUSG00000000439** | 0.164541294256464 | 0.252187156908082 | 0.83723604018117 | 871.890736489128 | 965.082638763479 | 942.669069093853 | 950.119741283517 | 823.408563255961 | 643.34125407085 | 920.313032051625 | 938.979613438259 | Mkrn2 |
| **ENSMUSG00000000440** | -0.181412659244692 | 0.406550331602211 | 0.881296485258502 | 287.317361868956 | 297.26174105373 | 349.241853911277 | 294.508981997063 | 416.084114411257 | 443.978850242811 | 234.242570375131 | 299.605055848508 | Pparg |
| **ENSMUSG00000000441** | 0.121396061593639 | 0.662745944759779 | 0.939724497010345 | 1207.99783905281 | 1474.09246933493 | 1491.61290450995 | 1054.22960434617 | 1065.39432453198 | 634.779678446333 | 1542.87773020419 | 1562.07128755798 | Raf1 |
| **ENSMUSG00000000486** | 0.337396241415532 | 0.555730106015913 | 0.916239900758826 | 18.9737880479499 | 13.0306516626293 | 39.7511053232348 | 27.1998741334867 | 29.5638712871156 | 31.8001380339204 | 5.20539045278068 | 11.9407812113536 | Sept1 |
| **ENSMUSG00000000489** | -0.821699274931657 | 0.0221206778164367 | 0.648762042745232 | 614.389327266951 | 574.977504613516 | 723.091534927413 | 638.728078789808 | 1784.78185918513 | 1547.19902357343 | 614.236073428121 | 563.387768062955 | Pdgfb |
| **ENSMUSG00000000530** | -0.57209642798522 | 0.0891178413770747 | 0.751206603737452 | 793.285043147622 | 648.274920215806 | 690.912068713366 | 668.741733006069 | 1388.40699229861 | 1614.46854633749 | 536.15521663641 | 626.348250813729 | Acvrl1 |
| **ENSMUSG00000000531** | -0.484511907187782 | 0.106479252839134 | 0.762094734041179 | 206.904641094311 | 186.501201921381 | 172.254789734017 | 233.543746870282 | 365.716037403578 | 398.724807656078 | 188.435134390661 | 166.085411394282 | Grasp |
| **ENSMUSG00000000532** | 0.0113370599725247 | 0.952728591827342 | 0.992000114323903 | 337.010616280254 | 339.611358957275 | 343.563124579386 | 275.750448111899 | 336.152166116463 | 225.04713070159 | 397.691830592444 | 325.657669400552 | Acvr1b |
| **ENSMUSG00000000538** | 0.18633771069192 | 0.387414009171441 | 0.875951433019341 | 2827.09441914454 | 3919.78290326466 | 3835.03520880351 | 2945.08981997063 | 2676.07783058187 | 2049.88582095579 | 3664.5948787576 | 3496.47784379726 | Tom1l2 |
| **ENSMUSG00000000552** | -0.652091579399422 | 0.0319817136805306 | 0.671606027118506 | 174.378147297826 | 131.935348084121 | 165.629605513478 | 140.689004138724 | 272.644590758955 | 379.155491942897 | 146.792010768415 | 164.999885829613 | Zfp385a |
| **ENSMUSG00000000555** | 0.172613854436702 | 0.297009067989342 | 0.851785577030345 | 759.855035634567 | 934.949256793649 | 1146.15687015327 | 1163.02910088012 | 914.290093508946 | 916.088591823321 | 782.890724098215 | 938.979613438259 | Itga5 |
| **ENSMUSG00000000561** | 0.195189013153328 | 0.108968678813955 | 0.763154315676794 | 505.064167562096 | 523.669313691913 | 478.906173656114 | 590.893817382642 | 468.642107810573 | 402.3940543523 | 485.14239019916 | 475.460197324806 | Wdr77 |
| **ENSMUSG00000000563** | 0.0999189024615332 | 0.281758221998522 | 0.850048154552447 | 30023.7608015894 | 25394.2968432777 | 27192.5954057585 | 30673.0166823246 | 26962.2506138494 | 25480.4721407947 | 26193.5247583924 | 27065.4089038799 | Atp5f1 |
| **ENSMUSG00000000567** | -1.37932385741228 | 0.02034068287451 | 0.645050422664255 | 12.6491920319666 | 11.4018202048006 | 9.46454888648447 | 33.7653609932938 | 50.3680770076785 | 85.6157562451702 | 22.903717992235 | 16.2828834700276 | Sox9 |
| **ENSMUSG00000000568** | 0.0867616248418989 | 0.417083406959236 | 0.882316253832438 | 1194.44513330428 | 1233.83982930521 | 1372.35958854025 | 1459.4139362657 | 1305.19016941636 | 1245.09771225119 | 1255.5401772107 | 1146.31499628994 | Hnrnpd |
| **ENSMUSG00000000579** | -0.105002625377968 | 0.666539514691611 | 0.942024383071524 | 37.0440623793308 | 44.7928650902881 | 35.0188308799925 | 40.3308478531009 | 51.4630352034976 | 39.1386314263635 | 37.4788112600209 | 41.2499714574033 | Dynlt1c |
| **ENSMUSG00000000581** | 0.109923464437055 | 0.450897426676045 | 0.89108031640712 | 973.987786461431 | 710.984931342209 | 769.467824471187 | 858.202925246218 | 763.185862485911 | 654.348994159515 | 801.630129728225 | 848.880991570773 | C1d |
| **ENSMUSG00000000594** | -0.242801849870083 | 0.296951700642291 | 0.851785577030345 | 393.931980424104 | 356.714089264476 | 374.796135904785 | 406.122258613784 | 500.395895489327 | 647.010500767072 | 314.405583347953 | 351.710282952596 | Gm2a |
| **ENSMUSG00000000600** | 0.290646552847191 | 0.314083647005861 | 0.853228028313558 | 463.50253659992 | 570.905425968944 | 530.961192531779 | 402.370551836751 | 360.241246424483 | 209.14706168463 | 595.49666779811 | 442.894430384751 | Krit1 |
| **ENSMUSG00000000605** | -0.0464371063379174 | 0.477971217927073 | 0.897674307047416 | 2013.02856051583 | 2015.67892906296 | 2135.2022287909 | 2047.49397356557 | 2088.08527942702 | 2122.04767264815 | 2117.55283619118 | 2151.51166917298 | Clcn4 |
| **ENSMUSG00000000628** | -0.0961273361449198 | 0.546137875083727 | 0.914211323944814 | 5928.85700812607 | 6192.00278693564 | 5474.29507594262 | 5271.14802173087 | 5692.68766006349 | 4662.38946866555 | 7047.05759497449 | 7038.5477613106 | Hk2 |
| **ENSMUSG00000000631** | 0.264852248715615 | 0.227289423752081 | 0.832048466795696 | 7419.654640465 | 8519.60294017279 | 6968.74734511851 | 6725.87232452527 | 5333.54137183482 | 4042.28677700411 | 7331.27191369631 | 7955.81686345549 | Myo18a |
| **ENSMUSG00000000673** | -0.405381594499019 | 0.183304061583094 | 0.805963874499992 | 78.6056933415069 | 93.6578088251478 | 68.1447519826882 | 93.7926694258161 | 82.1218646864323 | 108.854318654574 | 72.8754663389296 | 179.111718170304 | Haao |
| **ENSMUSG00000000682** | 0.745525751323186 | 0.114518870164311 | 0.767518180080285 | 29.8159526467785 | 40.7207864457164 | 28.3936466594534 | 22.5102406621959 | 12.0445401540101 | 37.9155491942896 | 14.5750932677859 | 8.68420451734806 | Cd52 |
| **ENSMUSG00000000686** | -0.320493607942191 | 0.257283547456825 | 0.837517407767513 | 20.780815481088 | 30.1333819698302 | 31.2330113253987 | 37.5170677703264 | 33.943704070392 | 34.2463024980681 | 36.4377331694648 | 44.5065481514088 | Abhd15 |
| **ENSMUSG00000000693** | -0.962801758573124 | 0.064357251479353 | 0.716991331547963 | 19.877301764519 | 29.3189662409158 | 35.0188308799925 | 27.1998741334867 | 62.4126171616885 | 108.854318654574 | 30.191264626128 | 16.2828834700276 | Loxl3 |
| **ENSMUSG00000000708** | 0.293015254722631 | 0.248313450444858 | 0.834621660998471 | 2452.13622676839 | 2933.52545554941 | 3018.2446398999 | 2310.11344795785 | 1763.97765346457 | 1325.82113956806 | 2912.93649737607 | 2740.95205078798 | Kat2b |
| **ENSMUSG00000000711** | 0.144089653984355 | 0.410771575229393 | 0.882192126163777 | 2085.30965784136 | 2146.79986141817 | 2196.72179655304 | 1814.88815338954 | 1640.24737733701 | 1413.88306027738 | 2271.63239359349 | 2133.05773457362 | Rab5b |
| **ENSMUSG00000000732** | -0.539439550943633 | 0.0643562820264902 | 0.716991331547963 | 69.5705561758165 | 51.3081909216027 | 53.001473764313 | 72.2203554578784 | 130.300025302473 | 99.0696607979826 | 68.711153976705 | 59.7039060567679 | Icosl |
| **ENSMUSG00000000738** | 0.162269432612541 | 0.139386191482925 | 0.779711473022872 | 4033.28523076422 | 3850.55756630695 | 3992.14672031915 | 4438.26911722962 | 3455.68806600507 | 3216.70627035425 | 3907.16607385718 | 3996.90512910944 | Spg7 |
| **ENSMUSG00000000740** | 0.00786604964396656 | 0.967203644859675 | 0.994284908685797 | 10830.4189205131 | 7714.96020000543 | 8878.69331041108 | 11443.6435966438 | 10042.9565720528 | 12279.7456100216 | 7972.57601747889 | 8361.80342464151 | Rpl13 |
| **ENSMUSG00000000743** | 0.107432070751218 | 0.229173824224555 | 0.832048466795696 | 1640.78090928939 | 1553.90521076854 | 1703.6187995672 | 1740.79194454315 | 1461.76919141849 | 1511.72963884329 | 1675.09464770482 | 1512.13711158323 | Chmp1a |
| **ENSMUSG00000000751** | -0.0101946341635252 | 0.916207497094766 | 0.986713918026868 | 1253.17352488127 | 1237.09749222087 | 1396.96741564511 | 1219.30470253561 | 1258.10696699614 | 1181.49743618335 | 1339.86750254575 | 1361.24905809431 | Rpa1 |

| | | | | | | | | | | | | |
|---|---|---|---|---|---|---|---|---|---|---|---|---|
| ENSMUSG00000000753 | -1.05135195581965 | 0.0133651859284921 | 0.635862253526407 | 309.905204783182 | 322.508628650074 | 285.829376371831 | 351.72251034681 | 715.00770186987 | 1294.02100153414 | 235.283648465687 | 388.618152151326 | Serpinf1 |
| ENSMUSG00000000759 | -0.0348723611495578 | 0.803719199261766 | 0.963461896828986 | 206.001127377742 | 228.036404096012 | 243.238906382651 | 229.79204009325 | 211.326931793086 | 205.477814988408 | 256.10521027681 | 255.098507697099 | Tubgcp3 |
| ENSMUSG00000000776 | 0.146624135734288 | 0.253940346720961 | 0.837406774112203 | 371.344137509877 | 403.135785812593 | 336.937940358847 | 446.453106466885 | 349.291664466292 | 332.67836712409 | 357.089785060755 | 367.993166422624 | Polr3d |
| ENSMUSG00000000782 | -0.800704519266159 | 0.0890660279069941 | 0.751206603737452 | 16.2632468982428 | 8.95857301805762 | 8.51809399783602 | 6.56548685980713 | 21.8991639163819 | 23.2385624094033 | 15.616171358342 | 9.76973008201657 | Tcf7 |
| ENSMUSG00000000787 | 2.74405545377147E-0 | 0.999841052747827 | 0.999913040727721 | 2932.80552398312 | 3840.78457755997 | 4074.48829563156 | 3302.43989048299 | 3735.99736413476 | 3133.53667857323 | 3572.98000678866 | 3707.06980334295 | Ddx3x |
| ENSMUSG00000000794 | -0.821544026613749 | 0.1398757749611 | 0.779711473022872 | 9.93865088225949 | 12.2162359337149 | 10.4110037751329 | 2.81378008277448 | 20.8042057205628 | 23.2385624094033 | 11.4518589961175 | 7.59867895267955 | Kcnn3 |
| ENSMUSG00000000804 | 0.159315290257674 | 0.595806065531073 | 0.924461858177432 | 580.055806037327 | 780.210268299927 | 800.700835796586 | 500.852854733858 | 482.876564356222 | 321.670627035425 | 817.246301086567 | 760.953420832624 | Usp32 |
| ENSMUSG00000000805 | -0.442167458138576 | 0.191610852979127 | 0.808824684419004 | 857.434517024024 | 561.946852950887 | 819.629933569555 | 978.257542111262 | 1535.13139053837 | 1564.32217482247 | 667.331056046484 | 604.637739520359 | Car4 |
| ENSMUSG00000000811 | 0.128899645428732 | 0.345570544322224 | 0.861266047293794 | 525.844983043184 | 585.564909089402 | 516.764369202052 | 456.770300103725 | 445.647985698372 | 415.847958905112 | 520.539045278068 | 524.308847734889 | Txnrd3 |
| ENSMUSG00000000823 | 0.0743783165433801 | 0.775580170524582 | 0.9595799687189 | 202.387072511466 | 208.490426602068 | 223.363353721033 | 219.47484645641 | 212.421889988905 | 105.185071958352 | 220.708555197901 | 271.381391167127 | Zfp512b |
| ENSMUSG00000000826 | 0.123567400692974 | 0.419981432611499 | 0.882568531013176 | 1612.77198407575 | 1821.84798558135 | 1959.16161950228 | 1547.57904552597 | 1495.71289548889 | 1284.23634367755 | 1846.87253264659 | 1743.35405685762 | Dnajc5 |
| ENSMUSG00000000827 | 0.0300179633423394 | 0.756159956333076 | 0.956566265296272 | 911.645340018166 | 791.612088504727 | 853.702309560899 | 949.181814589259 | 850.782518151438 | 884.2884537894 | 854.725112346588 | 843.45336374743 | Tpd52l2 |
| ENSMUSG00000000838 | 0.0162324715881638 | 0.926062343050852 | 0.986713918026868 | 121.974351736821 | 180.800291818981 | 177.933519065908 | 172.578511743502 | 168.623562156141 | 144.323703384715 | 166.572494488982 | 166.085411394282 | Fmr1 |
| ENSMUSG00000000861 | -0.184787818885593 | 0.581357317761871 | 0.922450244597442 | 36.1405486627618 | 48.0505280059454 | 28.3936466594534 | 47.8342614071662 | 26.2789966996583 | 41.5847958905112 | 66.6289977955927 | 47.7631248454143 | Bcl11a |
| ENSMUSG00000000869 | -0.456433592953908 | 0.311961826022251 | 0.851808485049943 | 10.8421645988285 | 10.5874044758863 | 6.62518422053913 | 10.3171936368398 | 15.3294147414674 | 7.33849339244316 | 13.5340151772298 | 16.2828834700276 | Il4 |
| ENSMUSG00000000876 | 0.120030387836319 | 0.208237224673639 | 0.820372930586885 | 1075.18132271716 | 1015.57641395617 | 1112.08449416193 | 1085.18118525669 | 983.272459845549 | 885.511536021474 | 1016.09221638279 | 1058.38742555179 | Pxmp4 |
| ENSMUSG00000000881 | -0.119182704929212 | 0.719247441387902 | 0.951674101666722 | 45.1756858284523 | 52.9370223794314 | 46.3762895437739 | 61.9031618210386 | 61.3176589658694 | 86.838838477244 | 29.1501865355718 | 47.7631248454143 | Dlg3 |
| ENSMUSG00000000884 | -0.328725038226576 | 0.165875533692809 | 0.796071270505327 | 71.3775836089546 | 47.2361122770311 | 44.483379766477 | 49.7101147956825 | 77.7420319031559 | 57.4848649074714 | 68.711153976705 | 62.9604827507734 | Gnb1l |
| ENSMUSG00000000889 | -0.460823937537321 | 0.13655370711937 | 0.776456211338551 | 59.631905293557 | 44.7928650902881 | 27.447191770805 | 47.8342614071662 | 54.7479097909548 | 88.0619207093179 | 46.8485140750261 | 58.6183804920994 | Dbh |
| ENSMUSG00000000901 | -0.566528378984021 | 0.0773605945803657 | 0.739855948845396 | 55.1143367107117 | 34.2054606144018 | 40.6975602118832 | 77.8479156234274 | 88.6916138613469 | 92.9542496376133 | 75.998700610598 | 49.9341759747513 | Mmp11 |
| ENSMUSG00000000902 | -0.227871641067671 | 0.307802065834446 | 0.851785577030345 | 251.176813206195 | 239.438224300813 | 206.327165725361 | 284.191788360223 | 331.772333333186 | 374.263163014601 | 199.886993386778 | 244.243252050414 | Smarcb1 |
| ENSMUSG00000000915 | 0.0835217238885045 | 0.604039692130041 | 0.926448151609826 | 495.125516679837 | 587.193740547231 | 505.406910538271 | 412.687745473591 | 547.479097909548 | 451.317343635254 | 494.512093014165 | 395.131305539337 | Hip1r |
| ENSMUSG00000000916 | 0.180980415508808 | 0.251508381813344 | 0.83723604018117 | 148.176249517323 | 131.120932355207 | 158.057966404291 | 135.999370667433 | 116.065568756824 | 125.977469903608 | 145.750932677859 | 117.236760984199 | Nsun5 |
| ENSMUSG00000000934 | 0.0318585207673685 | 0.83531475787531 | 0.968814461162998 | 177.088688447533 | 156.367819951551 | 172.254789734017 | 151.006197775564 | 180.668102310151 | 133.315963296051 | 166.572494488982 | 160.657783570939 | Top1mt |
| ENSMUSG00000000957 | -1.11505044856836 | 0.00344580946431842 | 0.635862253526407 | 124.684892886528 | 99.3587189275481 | 173.201244622666 | 147.254490998531 | 435.793361936001 | 430.524945689999 | 166.572494488982 | 146.545951230249 | Mmp14 |
| ENSMUSG00000000958 | -0.643397935138043 | 0.0533405562140316 | 0.703396849069134 | 43.3686583953142 | 42.3496179035451 | 35.0188308799925 | 53.4618215727152 | 59.1277425742312 | 114.969729814943 | 45.80743598447 | 53.1907526687569 | Slc7a7 |
| ENSMUSG00000000959 | 0.0402466020558296 | 0.625851472616694 | 0.931698409533928 | 2303.95997725106 | 2132.95479402663 | 2396.42377805787 | 2357.00978267076 | 2277.51304730372 | 2095.13986354252 | 2325.76845430241 | 2236.18266321713 | Oxa1l |
| ENSMUSG00000000976 | 0.0618151052649683 | 0.730222887885119 | 0.952921973914044 | 176.185174730964 | 246.767965861042 | 231.881447718869 | 207.281799431054 | 218.991639163819 | 162.669936865823 | 239.447960827911 | 204.078806157679 | Heatr6 |
| ENSMUSG00000001016 | -0.0615770851272529 | 0.588699994040447 | 0.92352104528132 | 560.178504272808 | 469.103459854653 | 515.817914313404 | 480.218467460179 | 578.137927392483 | 496.571386221987 | 501.799639648058 | 536.249628946243 | Ilf2 |
| ENSMUSG00000001017 | -0.0846142296346599 | 0.299578656676415 | 0.851785577030345 | 1844.07149551742 | 1832.43539005724 | 1696.04716045802 | 1793.3158394216 | 1774.92723542276 | 1867.64656837678 | 1971.80190351332 | 1984.34073221403 | Chtop |
| ENSMUSG00000001018 | 0.204889344894324 | 0.14082673848839 | 0.779711473022872 | 1666.07929335332 | 1480.60779516625 | 1485.93417517806 | 1590.72367346184 | 1171.60526952643 | 1164.37428493431 | 1587.64408809811 | 1474.14371681983 | Snapin |
| ENSMUSG00000001020 | -0.594491608510927 | 0.161639434140882 | 0.792786631432503 | 28.0089252136404 | 14.6594831204579 | 17.036187995672 | 33.7653609932938 | 40.5134532453066 | 55.0387004433237 | 19.7804837205666 | 26.0526135520442 | S100a4 |
| ENSMUSG00000001025 | -0.501558191891452 | 0.163352970933794 | 0.793968226938511 | 270.150601254144 | 245.953550132127 | 206.327165725361 | 325.460562907582 | 376.665619361769 | 661.687487551958 | 176.983275394543 | 269.21034003779 | S100a6 |
| ENSMUSG00000001027 | 0.292922433334557 | 0.420509054795888 | 0.882568531013176 | 770.697200233395 | 1018.01966114291 | 947.401343537095 | 602.14893771374 | 486.161438943679 | 292.316653465652 | 877.628830338823 | 1068.15715563381 | Scn4a |
| ENSMUSG00000001029 | -0.672476025166841 | 0.0916474122556814 | 0.753823363185192 | 343.335212296237 | 175.913797445495 | 206.327165725361 | 295.446908691321 | 594.56230032977 | 606.648787108634 | 181.147587756768 | 245.328777615083 | Icam2 |
| ENSMUSG00000001034 | -0.341669693394171 | 0.1693443707752 | 0.79907358295209 | 70.4740698923855 | 68.4109212288036 | 86.1273948670087 | 87.227182566009 | 109.49581958191 | 134.539045528125 | 68.711153976705 | 83.5854684794751 | Mapk7 |
| ENSMUSG00000001036 | -0.0967737028978492 | 0.363841349656838 | 0.86924083181804 | 536.687147642013 | 598.595560752032 | 586.802030962037 | 589.955890688383 | 656.974917491458 | 665.35673424818 | 561.141090809758 | 590.525907179668 | Epn2 |
| ENSMUSG00000001039 | 0.137942505811136 | 0.727580454127853 | 0.952237115597464 | 108.421645988285 | 59.452348210746 | 24.6078271048596 | 61.9031618210386 | 65.6974917491458 | 66.0464405319884 | 54.1360607089191 | 45.5920737160773 | B9d1 |
| ENSMUSG00000001052 | -0.00531967597327152 | 0.964648436533728 | 0.994284908685797 | 490.607948096991 | 559.503605764144 | 565.033568523123 | 483.970174237211 | 568.283303630111 | 461.102001491845 | 538.237372817523 | 538.42068007558 | Sec24b |
| ENSMUSG00000001053 | -0.736372992476758 | 0.0399569076490133 | 0.679566075569698 | 205.097613661173 | 177.542628903324 | 212.952349945901 | 236.357526953057 | 502.585811880965 | 507.579126310652 | 160.326025945645 | 216.019587369033 | N4bp3 |
| ENSMUSG00000001054 | -0.13744955160606 | 0.446230105884298 | 0.89108031640712 | 262.018977805023 | 252.468875963442 | 198.755526616174 | 250.426427366929 | 308.778211220985 | 304.547475786391 | 239.447960827911 | 208.420908416353 | Rmnd5b |
| ENSMUSG00000001056 | 0.0998967011942873 | 0.614117268022184 | 0.928075610917191 | 799.609639163605 | 654.79024604712 | 651.160963390131 | 929.485354009838 | 791.654775577207 | 879.396124861105 | 579.880496439768 | 581.84170266232 | Nhp2 |
| ENSMUSG00000001062 | -0.0569019683362219 | 0.696519421906241 | 0.944933386973863 | 728.23205555465 | 790.797672775813 | 664.41133183121 | 699.693313916588 | 639.455586358353 | 659.24132308781 | 810.99983254323 | 889.045437463508 | Vps9d1 |
| ENSMUSG00000001065 | 0.165373447586269 | 0.230082004752687 | 0.832048466795696 | 191.544907912638 | 218.26341534904 | 212.005895057252 | 172.578511743502 | 175.193311331055 | 179.793088114857 | 173.860041122875 | 180.197243734972 | Zfp276 |
| ENSMUSG00000001082 | -1.12494931235141 | 0.00550069216926635 | 0.635862253526407 | 33.4300075130547 | 52.122606650517 | 29.3401015481019 | 50.6480414899407 | 105.115986798633 | 154.108361241306 | 55.1771387994752 | 47.7631248454143 | Mfsd10 |
| ENSMUSG00000001089 | 0.0592668005961196 | 0.698955325948195 | 0.945141109561958 | 1045.36537007039 | 1067.69902060668 | 1059.08302039761 | 860.078778634734 | 925.239675467137 | 770.541806206532 | 1125.40541589118 | 1047.53216990511 | Luzp1 |
| ENSMUSG00000001098 | -0.0595780884419279 | 0.66836556174016 | 0.942024383071524 | 1125.77809084503 | 1233.02541357629 | 1293.80383278243 | 918.23023367874 | 1074.15399009853 | 1174.15894279091 | 1265.95095811626 | 1249.43992493345 | Kctd10 |
| ENSMUSG00000001100 | 0.173147974332823 | 0.146310380397341 | 0.780607990324772 | 3400.82562916589 | 3233.23044378988 | 3387.36204647279 | 3178.63356684091 | 2701.26186908571 | 2509.76474021556 | 3187.78111328289 | 3306.51086998027 | Poldip2 |
| ENSMUSG00000001105 | -0.0581956333282322 | 0.743193299331552 | 0.954383574136061 | 781.539364832224 | 583.121661902659 | 547.997380527451 | 767.224035903176 | 650.405168316544 | 858.60372691585 | 635.057635239243 | 646.973236542431 | Ift20 |
| ENSMUSG00000001119 | -1.27247255153706 | 0.00465995546103714 | 0.635862253526407 | 381.282788392137 | 442.22774080048 | 383.314229902621 | 418.31530563914 | 1446.43977667703 | 1692.74580919022 | 317.528817619622 | 470.032569501464 | Col6a1 |
| ENSMUSG00000001120 | -0.64160753878824 | 0.118640627018534 | 0.769868040577702 | 28.9124389302094 | 35.0198763433161 | 18.9290977729689 | 24.3860940507122 | 37.2285786578493 | 77.0541806206532 | 26.0269522639034 | 28.2236646813812 | Pcbp3 |
| ENSMUSG00000001123 | -0.723501757140583 | 0.00644363862141524 | 0.635862253526407 | 315.326287082597 | 306.220314071788 | 283.936466594534 | 299.198615468353 | 617.556422441971 | 703.272283442469 | 310.241270985729 | 359.308961905276 | Lgals9 |
| ENSMUSG00000001127 | 0.246611757787204 | 0.112452232737687 | 0.765080324297948 | 1535.97331816738 | 1337.27062687733 | 1523.792370724 | 1619.79940098384 | 1181.45989328881 | 1006.59667699679 | 1544.95988638531 | 1336.28197010693 | Araf |
| ENSMUSG00000001128 | -0.825194258021607 | 0.0249926030642551 | 0.655303666217042 | 42.4651446787451 | 52.9370223794314 | 57.7337482075553 | 42.2067012416173 | 104.021028602814 | 140.654456688494 | 43.7252798033577 | 58.6183804920994 | Cfp |
| ENSMUSG00000001131 | -1.32007675762674 | 0.0841849736795368 | 0.742565114108635 | 27.1054114970714 | 23.6180561385155 | 9.46454888648447 | 19.6964605794214 | 41.6084114411257 | 139.43137445642 | 10.4107809055614 | 8.68420451734806 | Timp1 |
| ENSMUSG00000001134 | 0.0912322353220032 | 0.627092281025249 | 0.931825800800958 | 459.888481733644 | 351.013179162076 | 362.492222352355 | 484.908100931469 | 339.43704070392 | 503.90987961443 | 358.130863151311 | 356.05238521127 | Uxt |
| ENSMUSG00000001138 | -0.150405380826945 | 0.237190634709449 | 0.832048466795696 | 235.817080024521 | 225.593156909269 | 222.416898832385 | 199.778385876988 | 223.371471947096 | 242.170281950624 | 266.515991182371 | 248.585354309088 | Cnnm3 |
| ENSMUSG00000001143 | 0.0711371607181598 | 0.7202200205887 | 0.951674101666722 | 660.468526811972 | 773.694942468612 | 878.310136665759 | 685.624413502716 | 595.657258525589 | 536.933099880424 | 850.560799984364 | 869.505977299475 | Lman2l |
| ENSMUSG00000001150 | 0.106241939745887 | 0.473261646017752 | 0.89728597737823 | 422.844419354313 | 428.382673408937 | 433.476339000989 | 381.736164563072 | 397.469825082332 | 300.878229090169 | 447.663578939139 | 400.558933362679 | Mcm3ap |
| ENSMUSG00000001151 | 0.204733574962425 | 0.368797405247184 | 0.871495599696183 | 2032.90586228035 | 2576.81136628494 | 2689.82479353889 | 1806.44681314122 | 1849.38439273845 | 1317.25956394355 | 2576.66827412644 | 2156.93929699632 | Pcnt |
| ENSMUSG00000001155 | -0.225184920608975 | 0.554429516062527 | 0.915945640531442 | 24.3948703473642 | 45.6072808192024 | 24.6078271048596 | 31.8895076047775 | 26.2789966996583 | 33.0232202659942 | 27.0680303544596 | 61.8749571861049 | Ftcd |
| ENSMUSG00000001156 | -0.284860079420879 | 0.125505203619209 | 0.771949105062466 | 113.8427282877 | 131.935348084121 | 92.7525790875478 | 97.5443762028488 | 154.389105610493 | 116.192812047017 | 135.340151772298 | 125.920965501547 | Mxd1 |
| ENSMUSG00000001157 | 0.183368784800682 | 0.262493786583644 | 0.841952818014037 | 118.360296870545 | 126.234437981721 | 100.324218196735 | 117.24083678227 | 112.780694169367 | 100.292743030056 | 98.902418602833 | 95.5262496908287 | Gmcl1 |
| ENSMUSG00000001158 | -0.0677327225923702 | 0.725877910971242 | 0.951802528404659 | 941.461292664945 | 686.552459474779 | 670.0900611631 | 904.161333264867 | 815.743855885227 | 1081.20469315329 | 680.865071223713 | 779.407355431988 | Snrnp27 |
| ENSMUSG00000001173 | 0.159351364260834 | 0.605245839712083 | 0.926448151609826 | 232.203025158245 | 333.09603312596 | 308.544293699394 | 183.8336320746 | 192.712642464161 | 138.208292224346 | 355.007628879643 | 260.526135520442 | Ocrl |
| ENSMUSG00000001175 | -0.305895068565083 | 0.0705963072111535 | 0.72606739345142 | 5665.93451660448 | 4653.57147501647 | 4056.50565274724 | 4734.6539526152 | 6455.8735225494 | 7123.23091959816 | 4876.40977616494 | 5170.3582645161 | Calm1 |
| ENSMUSG00000001211 | 0.0757326377254424 | 0.499603264589643 | 0.900927059989498 | 5486.13528700724 | 6078.79900061655 | 6967.80089022987 | 5874.23488613886 | 5512.01955775333 | 5486.74689308333 | 6390.13731983357 | 5768.48285064845 | Agpat3 |
| ENSMUSG00000001227 | -0.525164771250645 | 0.146239904295259 | 0.780607990324772 | 312.61574593289 | 267.1283590839 | 287.722286149128 | 236.357526953057 | 482.876564356222 | 676.364474336844 | 194.681602933998 | 235.559047533066 | Sema6b |
| ENSMUSG00000001228 | -0.189393348747622 | 0.659058220606188 | 0.938989601151841 | 29.8159526467785 | 8.14415728914329 | 22.7149173275627 | 17.8206071909051 | 25.1840385038392 | 17.123151249034 | 27.0680303544596 | 19.5394601640331 | Uhrf1 |
| ENSMUSG00000001229 | 0.134496230374087 | 0.422591813926945 | 0.883739881879745 | 1700.41281458294 | 1715.15952509358 | 1670.49287846451 | 1569.1513594939 | 1468.33894059341 | 1090.98935100988 | 1805.22940902434 | 1696.67645757688 | Dpp9 |
| ENSMUSG00000001240 | -0.79252343294261 | 0.0577047414679565 | 0.710862211775517 | 566.503100288791 | 482.948527246197 | 431.583429223692 | 443.63932638411 | 1185.83972607208 | 1413.88306027738 | 383.116737324658 | 351.710282952596 | Ramp2 |
| ENSMUSG00000001247 | -0.663074956386536 | 0.0699066158999661 | 0.724916894583216 | 26.2018977805023 | 26.0613033252585 | 24.6078271048596 | 20.6343872736795 | 35.0386622662111 | 66.0464405319884 | 23.9447960827911 | 30.3947158107182 | Lsr |
| ENSMUSG00000001248 | 0.0207163469786147 | 0.850976170004094 | 0.972692784625411 | 534.880120208875 | 485.39177443294 | 557.461929413935 | 536.494069115668 | 562.808512651016 | 474.555906044658 | 549.68923181364 | 494.999657488839 | Gramd1a |
| ENSMUSG00000001249 | 0.268671963225131 | 0.237432459183281 | 0.832048466795696 | 156.307872966445 | 153.110157035894 | 146.700507740509 | 149.130344387048 | 117.160526952643 | 92.9542496376133 | 107.231043327282 | 184.539345993646 | Hpn |
| ENSMUSG00000001270 | -0.531333128866672 | 0.051173501938002 | 0.703396849069134 | 912.548853734736 | 717.500257173524 | 710.787621374984 | 755.03098887782 | 1228.54309570903 | 1726.99211168829 | 755.822693743755 | 764.209997526629 | Ckb |
| ENSMUSG00000001280 | -0.277181530481791 | 0.0494898708593534 | 0.703396849069134 | 382.186302108706 | 456.072808192024 | 473.227444324223 | 462.397860269274 | 620.841297029428 | 453.763508099402 | 553.853544175865 | 519.966745476215 | Sp1 |
| ENSMUSG00000001281 | -0.675138458816184 | 0.164290699413541 | 0.794844240631867 | 10.8421645988285 | 12.2162359337149 | 13.2503684410783 | 7.50341355406529 | 14.2344565456483 | 33.0232202659942 | 11.4518589961175 | 11.9407812113536 | Itgb7 |
| ENSMUSG00000001285 | -0.17042846056559 | 0.393792765289186 | 0.878942305112115 | 333.396561413978 | 315.99330281876 | 266.900278598862 | 335.777756544422 | 373.380744774312 | 470.886659348436 | 286.296474902938 | 280.065595684475 | Myg1 |
| ENSMUSG00000001288 | -0.514703863802631 | 0.0593116749933524 | 0.712406493939829 | 275.571683553559 | 242.69588721647 | 314.223023031284 | 362.977630677908 | 552.953888888644 | 550.387004433237 | 246.735507461804 | 358.223436340608 | Rarg |
| ENSMUSG00000001289 | 0.0741855026535404 | 0.690340677592642 | 0.94469184542218 | 3348.42183360488 | 2098.74933341223 | 2465.5149849292 | 3159.87503295575 | 2543.58788888776 | 3161.66756991093 | 2274.75562786516 | 2537.95877019497 | Pfdn5 |
| ENSMUSG00000001300 | -0.525406828609401 | 0.00421318438272385 | 0.635862253526407 | 494.222002963268 | 470.732291312482 | 422.118880337207 | 464.27371365779 | 794.939650164664 | 804.7881087046 | 549.68923181364 | 516.71016878221 | Efnb2 |
| ENSMUSG00000001305 | 0.0709703162506376 | 0.668004404565157 | 0.942024383071524 | 210.518695960587 | 178.357044632238 | 148.593417517806 | 185.709485463116 | 163.148771177045 | 157.777607937528 | 167.613572579538 | 199.736703899005 | Rrp15 |
| ENSMUSG00000001313 | -0.515776848372766 | 0.0353530954469572 | 0.679566075569698 | 110.228673421423 | 115.647033505835 | 78.5557557578211 | 85.3513291774927 | 156.579022002131 | 184.685417043153 | 113.477511870619 | 104.210454208177 | Rnd2 |
| ENSMUSG00000001323 | 0.157205622095895 | 0.435817136279405 | 0.888687115594624 | 468.923618899334 | 498.422426095569 | 481.745538322059 | 433.32213274727 | 326.297542354091 | 321.670627035425 | 500.758561557502 | 538.42068007558 | Srr |
| ENSMUSG00000001333 | 0.10146364813817 | 0.743117970539228 | 0.954383574136061 | 324.361424248287 | 431.640336324594 | 329.36630124966 | 290.75727522003 | 244.175677667659 | 165.116101329971 | 432.047407580797 | 440.723379255414 | Sync |
| ENSMUSG00000001334 | -0.0363548440606343 | 0.705041809081219 | 0.947468143202218 | 11748.3888565473 | 10330.0491055493 | 10322.0370156 | 11305.7683725879 | 10440.4263971351 | 12329.8919815366 | 11215.5342695613 | 10836.8017120857 | Fndc5 |
| ENSMUSG00000001348 | -0.0642599090713924 | 0.842982821846889 | 0.971386364129692 | 27.1054114970714 | 38.2775392589734 | 40.6975602118832 | 29.075727522003 | 21.8991639163819 | 44.0309603546589 | 29.1501865355718 | 46.6775992807458 | Acp5 |

| | | | | | | | | | | | | |
|---|---|---|---|---|---|---|---|---|---|---|---|---|
| **ENSMUSG00000001349** | -0.720309953121475 | 0.185736558320872 | 0.805963874499992 | 33.4300075130547 | 93.6578088251478 | 39.7511053232348 | 30.9515809105193 | 106.210944994452 | 151.662196777159 | 34.3555769883525 | 34.7368180693922 | Cnn1 |
| **ENSMUSG00000001366** | 0.041467302111654 | 0.825372369926215 | 0.966806404871388 | 306.291149916906 | 343.683437601847 | 404.136237452887 | 394.867138282686 | 285.784089108784 | 291.093571233579 | 396.650752501888 | 433.124700302735 | Fbxo9 |
| **ENSMUSG00000001376** | 0.223419377581487 | 0.344930489340541 | 0.861266047293794 | 460.791995450213 | 491.09268453534 | 535.693466975021 | 438.949692912819 | 344.911831683016 | 259.293433199658 | 549.68923181364 | 494.999657488839 | Vps50 |
| **ENSMUSG00000001380** | 0.0024384854815402 | 0.983650595720294 | 0.996940244108351 | 856.531003307455 | 845.363526613073 | 873.577862222516 | 871.333898965832 | 933.99934103369 | 972.350374498718 | 787.055036460439 | 749.01263962127 | Hars |
| **ENSMUSG00000001383** | 0.0589200428083131 | 0.661977159487119 | 0.939611397743053 | 1398.63923324888 | 1177.64514401012 | 1300.42901700297 | 1490.36551717622 | 1310.66496039546 | 1505.61422768292 | 1203.48627268289 | 1133.28868951392 | Zmat2 |
| **ENSMUSG00000001403** | 0.204343870104849 | 0.71144792283313 | 0.950020479830613 | 37.0440623793308 | 23.6180561385155 | 75.7163910918757 | 18.7585338851632 | 29.5638712871156 | 67.2695227640623 | 21.8626399016789 | 16.2828834700276 | Ube2c |
| **ENSMUSG00000001415** | 0.115633439119833 | 0.558787903965808 | 0.916574868225118 | 1364.30571201926 | 1422.78427841333 | 1485.93417517806 | 1165.84288096289 | 954.803546754252 | 1006.59667699679 | 1480.41304477083 | 1577.26864546334 | Smg5 |
| **ENSMUSG00000001416** | -0.0143101995119184 | 0.905800440272566 | 0.98504921011344 | 3148.74530224312 | 2726.66386040517 | 2631.14459044268 | 3203.01966089162 | 3170.9989350921 | 3214.2601058901 | 2640.17403765036 | 2801.74148240942 | Cct3 |
| **ENSMUSG00000001418** | -0.405641086357368 | 0.0818520908952353 | 0.739855948845396 | 486.993893230715 | 532.627886709971 | 530.01473764313 | 547.749189446766 | 822.313605060142 | 957.673387713832 | 512.210420553619 | 487.40097853616 | Glmp |
| **ENSMUSG00000001419** | 0.184093718813305 | 0.368847533464503 | 0.871495599696183 | 3132.48205534488 | 4283.82673408937 | 4023.37973164455 | 3442.19096792745 | 3235.60146864543 | 2190.54027764428 | 3982.12369637722 | 3689.70139430826 | Mef2d |
| **ENSMUSG00000001420** | 0.502778740964419 | 0.0441525628412409 | 0.691978859031708 | 51.5002818444356 | 56.1946852950887 | 62.4660226507975 | 51.5859681841989 | 26.2789966996583 | 47.7002070508805 | 43.7252798033577 | 39.0789203280663 | Tmem79 |
| **ENSMUSG00000001424** | 0.0100674795359736 | 0.92677814288227 | 0.986790505471895 | 2209.09103701132 | 2378.90834415875 | 2131.4164092363 | 2015.60446596079 | 2349.78028822778 | 2311.62541861959 | 2096.73127438006 | 1918.12367276925 | Snd1 |
| **ENSMUSG00000001435** | -0.836522946116842 | 0.0194455496148595 | 0.642273034804458 | 170.764092431549 | 333.09603312596 | 170.36187995672 | 185.709485463116 | 454.407651264925 | 590.748718091674 | 193.640524843441 | 298.51953028384 | Col18a1 |
| **ENSMUSG00000001436** | -0.0620526493699238 | 0.7935139464351 | 0.961804768755116 | 45.1756858284523 | 72.4829998733752 | 58.6802030962037 | 48.7721881014244 | 60.2227007700503 | 68.4926049961361 | 54.1360607089191 | 53.1907526687569 | Slc19a1 |
| **ENSMUSG00000001440** | 0.230541718129199 | 0.120068737070781 | 0.769868040577702 | 3437.86969154522 | 3642.06713970488 | 4005.39708876023 | 3309.00537734279 | 2930.1081320119 | 2416.81049057795 | 3491.77591572528 | 3428.08973322315 | Kpnb1 |
| **ENSMUSG00000001441** | 0.0484395271959126 | 0.526877184231655 | 0.908217276872524 | 3909.50385159426 | 3704.77715083128 | 3659.94105440354 | 3852.06493331827 | 3445.8334422427 | 3911.4169781722 | 3631.2803798598 | 3639.76721833351 | Npepps |
| **ENSMUSG00000001445** | 0.0237597760789452 | 0.818280694407551 | 0.966315773308182 | 1375.14787661809 | 1167.87215526315 | 1122.49549793706 | 1313.09737196143 | 1208.83384818428 | 1188.83592957579 | 1219.10244404124 | 1279.83464074417 | Mrpl10 |
| **ENSMUSG00000001467** | 0.165550083758715 | 0.310000926809868 | 0.851785577030345 | 383.993329541844 | 411.279943101736 | 397.511053232348 | 342.343243404229 | 362.431162816121 | 253.178022039289 | 363.336253604092 | 388.618152151326 | Cyp51 |
| **ENSMUSG00000001472** | 0.143254965031552 | 0.0799198934463849 | 0.739855948845396 | 3529.12457691869 | 3661.61311719882 | 4112.3464911775 | 3745.14129017284 | 3330.86283168169 | 3457.6534700728 | 3486.5705252725 | 3349.93189256701 | Tcf25 |
| **ENSMUSG00000001473** | 0.0447902278309937 | 0.736510425998909 | 0.954383574136061 | 697.512589191303 | 610.811796685746 | 785.557557578211 | 831.940977806989 | 770.850569856644 | 715.503105763208 | 668.37213413704 | 680.624529047154 | Tubb6 |
| **ENSMUSG00000001482** | 0.0729617996744333 | 0.670593977099422 | 0.942863684776898 | 1803.41337827181 | 1727.37576102729 | 1490.6664496213 | 1514.75161122693 | 1411.40111441082 | 1176.60510725505 | 1852.07792309937 | 1772.66324710367 | Def8 |
| **ENSMUSG00000001493** | -0.480198434615472 | 0.188953659000928 | 0.808677608936066 | 202.387072511466 | 131.935348084121 | 100.324218196735 | 227.916186704733 | 290.163921892061 | 343.686107212755 | 127.011527047849 | 163.914360264945 | Meox1 |
| **ENSMUSG00000001504** | 0.328507553332282 | 0.492119685324036 | 0.899236955296196 | 197.869503928621 | 52.122606650517 | 242.292451494002 | 286.067641748739 | 212.421889988905 | 124.754387671534 | 73.9165444294857 | 208.420908416353 | Irx2 |
| **ENSMUSG00000001506** | -1.90195456758139 | 0.00474928394952846 | 0.635862253526407 | 367.730082643601 | 436.52683069808 | 369.117406572894 | 427.694572581722 | 2164.73235313435 | 2942.73585036971 | 471.60837502193 | 404.901035621353 | Col1a1 |
| **ENSMUSG00000001507** | -0.0439152822414039 | 0.800207585556558 | 0.963461896828986 | 167.150037565273 | 203.603932228582 | 175.094154399963 | 154.757904552597 | 222.276513751277 | 178.570005882784 | 154.079557402308 | 168.256462523619 | Itga3 |
| **ENSMUSG00000001508** | 0.0481474700228799 | 0.657416613978335 | 0.938389079274206 | 4034.18874448079 | 3725.13754405414 | 3920.21614878187 | 4195.34610341675 | 4083.09911220941 | 4277.11856556229 | 3568.81569442644 | 3425.91868209381 | Sgca |
| **ENSMUSG00000001518** | 0.13603361373954 | 0.442729284370695 | 0.89108031640712 | 271.054114970713 | 292.375246680244 | 363.438677241004 | 230.729966787508 | 257.315176017488 | 232.385624094033 | 260.269522639034 | 302.861632542514 | Itfg2 |
| **ENSMUSG00000001520** | -0.190671999658926 | 0.611631588416873 | 0.928055903343287 | 55.1143367107117 | 35.0198763433161 | 35.965285768641 | 34.703287687552 | 59.1277425742312 | 69.71568722821 | 26.0269522639034 | 29.3091902460497 | Nrip2 |
| **ENSMUSG00000001521** | -0.336238037890144 | 0.0483481721158175 | 0.703396849069134 | 124.684892886528 | 98.5443031986338 | 111.681676860517 | 113.489130005238 | 134.679858085749 | 147.992950080937 | 163.449260217313 | 119.407812113536 | Tulp3 |
| **ENSMUSG00000001524** | -0.290824380225827 | 0.107272282040547 | 0.762287160109383 | 300.870067617492 | 225.593156909269 | 207.27362061401 | 258.867767615252 | 352.576539053749 | 336.347613820311 | 266.515991182371 | 259.440609955773 | Gtf2h4 |
| **ENSMUSG00000001525** | -0.468574475994 | 0.0732162225276646 | 0.730864975478592 | 1679.63199910185 | 1356.81660437127 | 1537.98919405373 | 1731.41267760057 | 2682.64757975679 | 3089.50571821857 | 1434.60560878636 | 1519.73579053591 | Tubb5 |
| **ENSMUSG00000001542** | 0.452201402159771 | 0.0280330606597389 | 0.665956123973759 | 998.382656808795 | 1140.99643620897 | 875.470771999813 | 741.900015158206 | 623.031213421066 | 506.356044078578 | 770.397787011541 | 845.624414876767 | Ell2 |
| **ENSMUSG00000001552** | -0.097409102393953 | 0.48355331348056 | 0.89863290032536 | 4619.66563281753 | 4978.52335085329 | 5589.76257235773 | 5403.39568562127 | 6545.66009460656 | 5797.40978003009 | 4915.97074360608 | 4770.88485671809 | Jup |
| **ENSMUSG00000001555** | 0.220786947340648 | 0.274554212985994 | 0.845953420225256 | 599.029594085277 | 665.377650523007 | 959.705257089525 | 743.775868546722 | 686.538788778574 | 781.549546295196 | 503.88179582917 | 575.328549274309 | Fkbp10 |
| **ENSMUSG00000001558** | 0.0179434227299472 | 0.970407509105307 | 0.9951105526746 | 12.6491920319666 | 10.5874044758863 | 15.1432782183751 | 7.50341355406529 | 8.75966556655277 | 12.2308223207386 | 17.6983275394543 | 6.51315338801105 | Klhl10 |
| **ENSMUSG00000001569** | 0.119585809435252 | 0.419622028478587 | 0.882568531013176 | 638.784197614315 | 649.08933594472 | 724.98444470471 | 611.528204656321 | 582.51776017576 | 475.778988276731 | 666.289977955927 | 689.308733564502 | Nom1 |
| **ENSMUSG00000001576** | 0.112137888165243 | 0.272833516042237 | 0.844738346035895 | 868.276681622852 | 841.291447968501 | 760.003275584703 | 734.39660160414 | 721.577451044785 | 716.726187995282 | 789.137192641552 | 737.071858409917 | Ergic1 |
| **ENSMUSG00000001604** | 0.176785102636153 | 0.118328136376302 | 0.769868040577702 | 3084.59582836672 | 2624.86189429088 | 2720.11134997564 | 3136.42686559929 | 2627.89966996583 | 2838.77386064343 | 2343.46678184186 | 2422.89306034011 | Tcea3 |
| **ENSMUSG00000001627** | 0.429801402684065 | 0.00735756224430992 | 0.635862253526407 | 688.477452025612 | 800.570661522785 | 1004.188636856 | 805.679030367761 | 699.678287128403 | 536.933099880424 | 555.935700356977 | 655.657441059779 | Ifrd1 |
| **ENSMUSG00000001630** | -0.0824283337129882 | 0.450257496511866 | 0.89108031640712 | 316.229800799166 | 330.652785939217 | 292.45456059237 | 293.571055302804 | 325.202584158272 | 311.885969178834 | 328.980676615739 | 339.769501741243 | Stk38l |
| **ENSMUSG00000001632** | 0.143649612218238 | 0.407442203370054 | 0.881573350402261 | 452.660372001092 | 532.627886709971 | 468.495169880981 | 333.901903155905 | 444.553027502553 | 346.132271676902 | 421.636626675235 | 405.986561186022 | Brpf1 |
| **ENSMUSG00000001642** | -0.0114968130167094 | 0.900288273412635 | 0.984802943363903 | 5957.76944705628 | 5003.77023844964 | 4929.13706008111 | 5349.93386404855 | 5289.74304400206 | 5397.46189014194 | 5278.26591911961 | 5444.99623237723 | Akr1b3 |
| **ENSMUSG00000001663** | -0.221671633226485 | 0.244709108895698 | 0.832048466795696 | 235.817080024521 | 256.540954608014 | 236.613722162112 | 289.819348525772 | 348.196706270473 | 333.901449356164 | 207.174540020671 | 299.605055848508 | Gstt1 |
| **ENSMUSG00000001665** | -0.601697546068252 | 0.092011326989423 | 0.754058637722175 | 56.0178504272808 | 57.009101024003 | 42.5904699891801 | 78.7858423176855 | 145.62944004394 | 92.9542496376133 | 47.8895921655823 | 69.4736361387845 | Gstt3 |
| **ENSMUSG00000001666** | 0.0813684218872257 | 0.532648054602742 | 0.909082872831377 | 740.881247586617 | 583.936077631574 | 697.537252933905 | 753.155135489303 | 701.868203520041 | 708.164612370765 | 594.455589707554 | 618.749571861049 | Ddt |
| **ENSMUSG00000001670** | -0.148271573455877 | 0.754844082885981 | 0.956497786093635 | 73.1846110420926 | 294.818493866987 | 66.2518422053913 | 110.675349922463 | 82.1218646864323 | 140.654456688494 | 187.394056300105 | 194.309076075663 | Tat |
| **ENSMUSG00000001674** | 0.0386368846329051 | 0.775863506803253 | 0.959602406551077 | 428.265501653727 | 392.548381336706 | 443.887342776122 | 471.777127211855 | 486.161438943679 | 442.755768010737 | 362.295175513536 | 399.473407798011 | Ddx18 |
| **ENSMUSG00000001687** | 0.0244589980177205 | 0.870247771184569 | 0.975613767803763 | 2254.26672283977 | 2200.55129952652 | 2315.0286576341 | 2036.23885323447 | 2037.71720241934 | 1680.51498686948 | 2491.29987070084 | 2446.77462276282 | Ubl3 |
| **ENSMUSG00000001700** | -0.00746417736329833 | 0.961312510072543 | 0.993822507448007 | 318.036828232304 | 341.240190415104 | 324.634026806417 | 334.839829850164 | 321.917709570815 | 254.401104271363 | 386.239971596327 | 361.480013034613 | Gramd3 |
| **ENSMUSG00000001707** | 0.035598429833942 | 0.80339693512549 | 0.963461896828986 | 385.800356974982 | 329.838370210303 | 283.936466594534 | 317.019222659259 | 294.543754675337 | 308.216722482613 | 360.213019332423 | 321.315567141878 | Eef1e1 |
| **ENSMUSG00000001729** | 0.0899082283937512 | 0.279807811744235 | 0.848943557485723 | 2866.84902267358 | 2904.2064893085 | 3174.40969652689 | 3184.26112700646 | 2957.48208690738 | 2718.91180190019 | 2767.18556469821 | 2951.54401033367 | Akt1 |
| **ENSMUSG00000001739** | -0.851308753739294 | 0.0678933172747794 | 0.722112580225498 | 86.7373167906283 | 72.4829998733752 | 68.1447519826882 | 91.9168160372998 | 246.365594059297 | 212.816308380852 | 49.9717483466946 | 67.3025850094475 | Cldn15 |
| **ENSMUSG00000001741** | -0.278053149021188 | 0.410242958236568 | 0.882192126163777 | 50.5967681278665 | 69.2253369577179 | 47.3227444324223 | 100.358156285623 | 109.49581958191 | 86.838838477244 | 45.80743598447 | 82.4999429148066 | Il16 |
| **ENSMUSG00000001750** | -0.220441590115458 | 0.379939258384718 | 0.87432013076003 | 85.8338030740593 | 62.7100111264033 | 54.8943835416099 | 81.59962240046 | 90.881530252985 | 103.961989726278 | 79.1219348822664 | 58.6183804920994 | Tcirg1 |
| **ENSMUSG00000001751** | -0.0903192171514747 | 0.578360905071807 | 0.922450244597442 | 157.211386683014 | 170.212887343095 | 182.66579350915 | 143.502784221499 | 201.472308030714 | 181.016170346931 | 145.750932677859 | 168.256462523619 | Naglu |
| **ENSMUSG00000001755** | 0.087560047514322 | 0.46604762717473 | 0.895328112556867 | 421.037391921175 | 393.362797065621 | 456.191256328551 | 366.729337454941 | 401.849657865609 | 392.609396495709 | 395.609674411332 | 350.624757387928 | Ppat |
| **ENSMUSG00000001761** | -0.121061218547658 | 0.520364226168869 | 0.905962179970576 | 176.185174730964 | 266.313943354985 | 218.631079277791 | 188.52326554589 | 251.840385038392 | 261.739597663806 | 188.435134390661 | 223.618266321713 | Smo |
| **ENSMUSG00000001763** | 0.500543197917521 | 0.10670811520554 | 0.762287160109383 | 70.4740698923855 | 71.6685841444609 | 126.824955078892 | 51.5859681841989 | 45.9882442244021 | 66.0464405319884 | 63.5057635239243 | 51.0197015394199 | Tspan33 |
| **ENSMUSG00000001767** | -0.0687861387939352 | 0.526426378886606 | 0.908217276872524 | 430.072529086865 | 500.865673282312 | 469.44162476963 | 444.577253078368 | 492.731188118594 | 465.99433042014 | 522.621201459181 | 453.749686031436 | Crnkl1 |
| **ENSMUSG00000001768** | -0.29444487732162 | 0.0363339337650815 | 0.679566075569698 | 389.414411841258 | 392.548381336706 | 405.082692341535 | 450.204813243917 | 512.440435643337 | 594.417964787896 | 483.060234018047 | 419.012867962044 | Rin2 |
| **ENSMUSG00000001774** | 0.0960960572901519 | 0.535192712117455 | 0.910813988293389 | 1126.6816045616 | 1193.11904285949 | 1277.7140996754 | 1098.31215897631 | 1066.4892827278 | 834.142082274372 | 1228.47214685624 | 1262.46623170947 | Chordc1 |
| **ENSMUSG00000001783** | 0.193474469572645 | 0.0777580900692078 | 0.739855948845396 | 1835.9398720683 | 1748.55056997906 | 2088.82593924712 | 2056.87324050815 | 1871.28355665484 | 1620.58395749786 | 1695.91620951595 | 1570.75549207533 | Rtcb |
| **ENSMUSG00000001785** | 0.25942231392027 | 0.0141310511933107 | 0.635862253526407 | 436.397125102849 | 476.433201414882 | 487.42426765395 | 486.783954319986 | 389.805117711598 | 364.47850515801 | 395.609674411332 | 425.526021350055 | Pwp1 |
| **ENSMUSG00000001786** | -0.168353879029524 | 0.41739799333101 | 0.882436023575407 | 266.536546387868 | 253.283291692356 | 294.347470369667 | 298.260688774095 | 390.900075907418 | 377.932409710823 | 229.03717992235 | 252.927456567762 | Fbxo7 |
| **ENSMUSG00000001794** | -0.116069185425052 | 0.45112873254917 | 0.891094723422636 | 8198.48346414752 | 7552.89146995148 | 7677.6420567162 | 9007.84797165538 | 8834.12272386847 | 11077.4557758929 | 7903.86486350219 | 7340.32386828845 | Capns1 |
| **ENSMUSG00000001802** | 0.173382251449706 | 0.25237944347341 | 0.83723604018117 | 252.983840639333 | 290.746415222415 | 300.972654590206 | 262.619474392285 | 270.454674367317 | 194.470074899744 | 247.776585552361 | 268.124814473121 | Lrp3 |
| **ENSMUSG00000001829** | 0.121809081258256 | 0.166091759677785 | 0.796071270505327 | 3180.36828232304 | 3080.12028675399 | 3059.88865500043 | 3263.9848960184 | 2707.83161826063 | 2701.78865065116 | 3074.30360141227 | 3079.63602696456 | Clpb |
| **ENSMUSG00000001833** | 0.0512996191561472 | 0.701595515563578 | 0.945991110533699 | 2650.00573069701 | 2677.79891667031 | 2629.25168066539 | 2512.70561391761 | 2275.32313091208 | 2102.47835693496 | 2804.66437595823 | 2920.06376895829 | Sept7 |
| **ENSMUSG00000001844** | -0.0515233050611174 | 0.717813695618378 | 0.951674101666722 | 267.440060104437 | 261.427448981499 | 251.757000380487 | 328.274342990356 | 283.594172717146 | 305.770558018465 | 254.023054095697 | 306.118209236519 | Zdhhc4 |
| **ENSMUSG00000001847** | -0.031589704281001 | 0.765887752114908 | 0.958345041873658 | 3537.25620036781 | 3994.70915032478 | 4202.2597055991 | 3654.1624008298 | 4234.20334323245 | 4131.5717799455 | 3813.46904570713 | 3550.75412203069 | Rac1 |
| **ENSMUSG00000001855** | -0.0101105674727249 | 0.948042212978908 | 0.991192530276814 | 365.019541493894 | 433.269167782423 | 385.207139679918 | 336.71568323868 | 445.647985698372 | 338.793778284459 | 417.472314313011 | 328.914246094558 | Nup214 |
| **ENSMUSG00000001864** | -0.293144699673342 | 0.474246904607553 | 0.897549741595057 | 34.3335212296237 | 27.6901347830872 | 23.6613722162112 | 28.1378008277448 | 30.6588294829347 | 66.0464405319884 | 19.7804837205666 | 23.8815624227072 | Aif1l |
| **ENSMUSG00000001870** | -0.2074784767932 | 0.157489359629377 | 0.789197677076549 | 629.749060448624 | 799.756245793871 | 680.501064938233 | 550.562969529541 | 837.643019801609 | 792.557286383861 | 689.193695948162 | 754.440267444613 | Ltbp1 |
| **ENSMUSG00000001891** | 0.264923997650139 | 0.191345136115298 | 0.808824684419004 | 7071.80185958592 | 6726.25950510344 | 6386.67758859972 | 7296.13175463424 | 5476.98089548712 | 3672.9159429178 | 6751.39141725655 | 6967.98859960715 | Ugp2 |
| **ENSMUSG00000001909** | 0.186093500710748 | 0.123711689166919 | 0.770326056866107 | 927.908586916409 | 868.167167022674 | 945.508433759798 | 1003.58156285623 | 950.423713970976 | 842.703657898889 | 771.438865102097 | 727.3021283279 | Trmt1 |
| **ENSMUSG00000001910** | -0.0674054213783121 | 0.568044189656669 | 0.920432752255281 | 949.592916114066 | 998.473683648967 | 918.061241988993 | 794.423910036663 | 904.435469746574 | 912.419345127099 | 1073.35151136338 | 945.49276682627 | Nacc1 |
| **ENSMUSG00000001911** | 0.148476164437985 | 0.429603654068191 | 0.885743677088682 | 4655.80618148029 | 5271.71301326245 | 4353.69248778286 | 4190.65646994546 | 3612.2670880072 | 3080.94414259405 | 4760.85010811321 | 5210.52271040884 | Nfix |
| **ENSMUSG00000001918** | -0.838357792408267 | 0.0368173332136658 | 0.679566075569698 | 93.9654265231807 | 107.502876216691 | 132.503684410783 | 154.757904552597 | 279.21433993387 | 374.263163014601 | 114.518589961175 | 106.381505337514 | Slc1a5 |
| **ENSMUSG00000001924** | 0.00221398101986202 | 0.97564851461869 | 0.99575536949131 | 4807.59648586389 | 5160.9524741301 | 5097.60603026053 | 4707.45407848171 | 5081.70098679643 | 4719.87433357302 | 5007.58561557502 | 4933.71369141837 | Uba1 |
| **ENSMUSG00000001930** | -0.676087421991683 | 0.291148218524043 | 0.851785577030345 | 1175.47134525633 | 910.516784926219 | 1227.55199057704 | 504.604561510891 | 2575.34167656652 | 2377.67185915158 | 725.631429117627 | 422.269444656049 | Vwf |
| **ENSMUSG00000001942** | -0.25247968819996 | 0.147459721202238 | 0.780607990324772 | 252.983840639333 | 341.240190415104 | 323.687571917769 | 262.619474392285 | 315.3479603959 | 304.547475786391 | 431.006329490241 | 354.966859646602 | Siae |
| **ENSMUSG00000001943** | 0.594423955115124 | 0.215858712924621 | 0.825162496944895 | 13.5527057485357 | 13.0306516626293 | 22.7149173275627 | 9.37926694258161 | 14.2344565456483 | 8.56157562451702 | 6.24646854333682 | 9.76973008201657 | Vsig2 |
| **ENSMUSG00000001946** | -0.722252321082332 | 0.0645756771610153 | 0.716991331547963 | 1059.82158953549 | 770.437279552955 | 732.556083813898 | 930.423280704096 | 2180.06176787582 | 2195.43260657258 | 667.331056046484 | 720.788974939889 | Esam |
| **ENSMUSG00000001948** | 0.0529548790720569 | 0.829674335719424 | 0.967523998969336 | 136.430571201926 | 80.6271571625185 | 103.163582862681 | 90.9788893430416 | 104.021028602814 | 121.085140975312 | 102.025652874501 | 69.4736361387845 | Spa17 |
| **ENSMUSG00000001962** | 0.107400815953636 | 0.46135532314872 | 0.895328112556867 | 674.924746277077 | 495.979178908826 | 519.603733867997 | 528.990655561603 | 533.2446413639 | 576.071731306788 | 464.320828388037 | 487.40097853616 | Fam50a |
| **ENSMUSG00000001964** | 0.0690689678144159 | 0.535209774834459 | 0.910813988293389 | 867.373167906283 | 758.22104361924 | 907.65023821386 | 876.961459131381 | 878.156473046916 | 806.011190936674 | 838.06786289769 | 727.3021283279 | Emd |
| **ENSMUSG00000001983** | -0.0506287814230246 | 0.900260645032405 | 0.984802943363903 | 609.871758684105 | 851.064436715473 | 637.910594949053 | 408.936038696558 | 460.97740043984 | 236.054870790255 | 960.915077583314 | 938.979613438259 | Taco1 |

| | | | | | | | | | | | | |
|---|---|---|---|---|---|---|---|---|---|---|---|---|
| **ENSMUSG00000001986** | -0.352068992891678 | 0.443904616935982 | 0.89108031640712 | 18.9737880479499 | 18.7315617650296 | 17.036187995672 | 16.8826804966469 | 30.6588294829347 | 37.9155491942896 | 9.36970281500523 | 14.1118323406906 | Gria3 |
| **ENSMUSG00000001988** | -0.202406520412728 | 0.691835408849668 | 0.94469184542218 | 10.8421645988285 | 16.2883145782866 | 3.78581955459379 | 12.1930470253561 | 7.66470737073368 | 14.6769867848863 | 13.5340151772298 | 14.1118323406906 | Npas1 |
| **ENSMUSG00000001995** | -0.0918766610454438 | 0.49119732724893 | 0.899236955296196 | 866.469654189714 | 1042.45213301034 | 976.741445085197 | 784.106716399823 | 894.580845984202 | 917.311674055395 | 1016.09221638279 | 1083.35451353917 | Sipa1l2 |
| **ENSMUSG00000001998** | 0.522314402817869 | 0.0791638338610957 | 0.739855948845396 | 84.0267756409212 | 122.162359337149 | 103.163582862681 | 69.4065753751039 | 48.1781606160403 | 45.2540425867328 | 89.5327157878278 | 80.3288917854696 | Ap4e1 |
| **ENSMUSG00000001999** | 0.0463787282376281 | 0.827730099416364 | 0.967319591893432 | 337.914129996823 | 230.479651282755 | 249.86409060319 | 345.157023487003 | 329.582416941548 | 327.786038195794 | 213.421008564008 | 256.184033261768 | Blvra |
| **ENSMUSG00000002006** | -0.211493297490735 | 0.393137695859899 | 0.878770935869648 | 46.0791995450213 | 30.9477976987445 | 44.483379766477 | 42.2067012416173 | 44.893286028583 | 57.4848649074714 | 42.6842017128016 | 44.5065481514088 | Pdzd4 |
| **ENSMUSG00000002007** | -0.0381649284199959 | 0.825180166142859 | 0.96672635991852 | 430.072529086865 | 386.847471234306 | 265.007368821565 | 392.99128489417 | 386.520243124141 | 341.239942748607 | 388.322127777439 | 398.387882233342 | Srpk3 |
| **ENSMUSG00000002010** | 0.067808269290817 | 0.410576075235824 | 0.882192126163777 | 17512.8063682578 | 15925.0851631908 | 17173.4239545261 | 17769.0212227209 | 15744.4038976828 | 15330.1126968138 | 16885.24555073 | 17278.3104128286 | Idh3g |
| **ENSMUSG00000002012** | -0.0164630683208763 | 0.921561958763457 | 0.986713918026868 | 220.457346842847 | 171.841718800923 | 204.434255948065 | 166.950951577953 | 181.76306050597 | 176.123841418636 | 231.119336103462 | 182.368294864309 | Pnck |
| **ENSMUSG00000002014** | 0.0473605401152093 | 0.764694933956167 | 0.958110466593105 | 1131.19917314444 | 935.763672522564 | 1054.35074595437 | 1193.98068179064 | 1078.53382288181 | 1286.6825081417 | 943.21675004386 | 868.420451734806 | Ssr4 |
| **ENSMUSG00000002015** | -0.139375330158283 | 0.405653915946965 | 0.881158207349486 | 1387.79706865005 | 1255.01463825698 | 1242.69526879541 | 1368.43504692266 | 1466.14902420177 | 1890.88513078619 | 1196.198726049 | 1235.32809259276 | Bcap31 |
| **ENSMUSG00000002017** | 0.0957238041585367 | 0.598667232930057 | 0.924848033480823 | 932.426155499254 | 1022.9061555164 | 901.97150888197 | 925.733647232805 | 829.978312430875 | 621.325773893521 | 1065.02288663893 | 1022.56508191773 | Fam98a |
| **ENSMUSG00000002020** | -1.32098651282157 | 0.0209670736359888 | 0.648762042745232 | 34.3335212296237 | 36.6487078011448 | 22.7149173275627 | 28.1378008277448 | 63.5075753575076 | 184.685417043153 | 34.3555769883525 | 22.7960368580387 | Ltbp2 |
| **ENSMUSG00000002028** | 0.251600394781076 | 0.162044989843115 | 0.793654763371249 | 698.416102907872 | 906.444706281648 | 801.647290685234 | 644.355638955357 | 600.037091308865 | 494.125221757839 | 733.960053842076 | 733.815281715911 | Kmt2a |
| **ENSMUSG00000002031** | -0.283304924813244 | 0.212356150780856 | 0.823370267313127 | 175.281661014395 | 143.337168288922 | 142.914688185915 | 199.778385876988 | 241.98576127602 | 249.508775343067 | 147.833088858971 | 166.085411394282 | Ift46 |
| **ENSMUSG00000002032** | -0.165333366084464 | 0.483384736747303 | 0.89863290032536 | 123.781379169959 | 120.533527879321 | 87.0738497556571 | 100.358156285623 | 93.0714466446232 | 94.1773318696872 | 146.792010768415 | 149.802527924254 | Tmem25 |
| **ENSMUSG00000002043** | 0.0868400952429251 | 0.735127703068046 | 0.954383574136061 | 316.229800799166 | 237.809392842984 | 282.990011705886 | 406.122258613784 | 278.119381738051 | 429.301863457925 | 247.776585552361 | 216.019587369033 | Trappc6a |
| **ENSMUSG00000002052** | 0.134958895891747 | 0.353282203536837 | 0.862728204353539 | 1388.70058236662 | 1735.51991831643 | 1675.22515290775 | 1336.54553931788 | 1382.93220131952 | 1207.1821630569 | 1626.16397744869 | 1371.01878817632 | Supt6 |
| **ENSMUSG00000002058** | -0.0954222481991361 | 0.633734150606861 | 0.9325328237529 | 111.132187137993 | 106.688460487777 | 123.039135524298 | 99.4202295913651 | 112.780694169367 | 140.654456688494 | 85.3684034256032 | 132.434118889558 | Unc119 |
| **ENSMUSG00000002059** | -0.43803545148386 | 0.227427296379983 | 0.832048466795696 | 60.535419010126 | 95.2866402829764 | 70.9841166486335 | 111.613276616721 | 155.484063806312 | 166.339183562045 | 61.4236073428121 | 75.9867895267955 | Rab34 |
| **ENSMUSG00000002064** | -0.00181096306451868 | 0.991085492012672 | 0.998628446692557 | 1087.83051474913 | 817.673391829986 | 912.382512657103 | 1037.34692384953 | 998.601874587016 | 1180.27435395127 | 850.560799984364 | 831.512582536077 | Sdf2 |
| **ENSMUSG00000002068** | 0.706182712865421 | 0.128649956760313 | 0.77303252008843 | 26.2018977805023 | 22.8036404096012 | 21.7684624389143 | 13.1309737196143 | 17.5193311331055 | 7.33849339244316 | 19.7804837205666 | 6.51315338801105 | Ccne1 |
| **ENSMUSG00000002076** | 0.643923542414435 | 0.134730204857961 | 0.773946462149327 | 24.3948703473642 | 15.4738988493722 | 10.4110037751329 | 24.3860940507122 | 10.949581958191 | 14.6769867848863 | 12.4929370866736 | 9.76973008201657 | Hsf2bp |
| **ENSMUSG00000002083** | -0.568461488448999 | 0.0750459697259727 | 0.735399943172214 | 61.4389327266951 | 48.0505280059454 | 50.1621090983677 | 77.8479156234274 | 123.730276127558 | 108.854318654574 | 67.6700758861489 | 52.1052271040884 | Bbc3 |
| **ENSMUSG00000002100** | 0.220014250235365 | 0.311106591406198 | 0.851785577030345 | 92921.8681805434 | 93866.2992517499 | 75640.6747007839 | 96583.0013412342 | 73564.766386106 | 48316.6404958457 | 95433.5464050999 | 90916.0226176815 | Mybpc3 |
| **ENSMUSG00000002102** | 0.0726073388125435 | 0.615910699597093 | 0.928422072203769 | 7149.50403921085 | 5693.58036084007 | 6454.82234058241 | 7746.33656787815 | 6802.97527062405 | 7378.8551061016 | 5653.05403171982 | 5882.46303493864 | Psmc3 |
| **ENSMUSG00000002103** | -0.252836180977401 | 0.0707949816925987 | 0.726501344493452 | 373.151164943016 | 448.743066631795 | 405.082692341535 | 444.577253078368 | 471.926982398031 | 577.294813538862 | 520.539045278068 | 423.354970220718 | Acp2 |
| **ENSMUSG00000002104** | -0.0622295008193182 | 0.697986872402849 | 0.944933386973863 | 390.317925557827 | 391.733965607792 | 275.418372596698 | 393.929211588428 | 422.653863586171 | 384.047820871192 | 338.350379430744 | 371.24974311663 | Rapsn |
| **ENSMUSG00000002105** | -0.0333423517588213 | 0.785204421115365 | 0.961141387070613 | 370.440623793308 | 361.600583637962 | 343.563124579386 | 335.777756544422 | 329.582416941548 | 419.517205601334 | 349.802238426862 | 347.368180693922 | Slc39a13 |
| **ENSMUSG00000002107** | -0.187913954860672 | 0.327948373743372 | 0.855977678491063 | 1171.85729039005 | 1040.0088858236 | 762.842640250648 | 1190.22897501361 | 1214.30863916338 | 1476.26025411315 | 1037.95485628447 | 1017.13745409439 | Celf2 |
| **ENSMUSG00000002108** | -0.129366082453118 | 0.398404330448848 | 0.880459952732461 | 588.187429486448 | 496.79359463774 | 576.391027186904 | 591.8317440769 | 644.930377337448 | 747.303243797128 | 569.469715534207 | 503.683862006187 | Nr1h3 |
| **ENSMUSG00000002109** | 0.908273046351927 | 0.0202411763640291 | 0.644586421602923 | 18.9737880479499 | 16.2883145782866 | 24.6078271048596 | 21.5723139679377 | 8.75966556655277 | 13.4539045528125 | 12.4929370866736 | 8.68420451734806 | Ddb2 |
| **ENSMUSG00000002111** | -0.400905328374594 | 0.116682266010394 | 0.769868040577702 | 61.4389327266951 | 56.1946852950887 | 87.0738497556571 | 80.6616957062019 | 86.5016974697087 | 132.092881063977 | 75.998700610598 | 82.4999429148066 | Spi1 |
| **ENSMUSG00000002129** | -0.163278241193209 | 0.153364203612049 | 0.784037095388796 | 805.030721463019 | 819.302223287815 | 740.127722923085 | 684.686486808458 | 944.948922991881 | 856.157562451702 | 771.438865102097 | 843.45336374743 | Sf3a1 |
| **ENSMUSG00000002147** | 0.0453590588979263 | 0.746195422791389 | 0.956085046483146 | 787.863960848207 | 964.268223034565 | 1121.54904304841 | 929.485354009838 | 1022.69095489504 | 841.480575666815 | 888.039611244385 | 932.466460050248 | Stat6 |
| **ENSMUSG00000002205** | 0.0364405332365453 | 0.728411701055718 | 0.952401994188708 | 468.923618899334 | 449.557482360709 | 435.369248778286 | 408.936038696558 | 404.039574257247 | 458.655837027697 | 439.33495421469 | 417.927342397375 | Vrk3 |
| **ENSMUSG00000002210** | -0.269008213467202 | 0.172817495634128 | 0.799889021902745 | 91.2548853734735 | 112.389370590177 | 123.039135524298 | 122.868396947819 | 150.009272827216 | 163.893019097897 | 113.477511870619 | 115.065709854862 | Smg9 |
| **ENSMUSG00000002221** | 0.105618851607866 | 0.59128032098029 | 0.923935529355527 | 349.65980831222 | 298.890572511559 | 327.473391472363 | 286.067641748739 | 259.505092409126 | 215.262472844999 | 384.157815415214 | 312.63136262453 | Paxip1 |
| **ENSMUSG00000002222** | 0.282700254595538 | 0.252679283259311 | 0.837241231982784 | 2888.53335187124 | 3831.82600454192 | 4133.16849872777 | 2787.51813533526 | 2292.84246204519 | 1848.0772526636 | 3517.80286798919 | 3554.01069872469 | Rmnd5a |
| **ENSMUSG00000002227** | -0.308594243813808 | 0.305520034786904 | 0.851785577030345 | 65.9565013095403 | 31.7622134276588 | 66.2518422053913 | 54.3997482669734 | 84.3117810780705 | 66.0464405319884 | 46.8485140750261 | 72.73021283279 | Mov10 |
| **ENSMUSG00000002228** | 0.112263548058217 | 0.430863750315534 | 0.886096756275211 | 477.055242348456 | 355.899673535562 | 369.117406572894 | 442.701399689852 | 402.944616061428 | 403.617136584374 | 361.254097422979 | 353.881334081933 | Ppm1j |
| **ENSMUSG00000002233** | -0.286229168710866 | 0.249410994499433 | 0.835314899646652 | 1142.94485145984 | 876.311324311818 | 848.023580229008 | 1207.11165551025 | 1342.41874807421 | 1784.47697659576 | 981.736639394437 | 860.821772782126 | Rhoc |
| **ENSMUSG00000002249** | 0.190181509307639 | 0.232258732676732 | 0.832048466795696 | 160.82544154929 | 197.088606397268 | 168.468970179424 | 186.647412157374 | 135.774816281568 | 152.885279009232 | 185.311900118992 | 150.888053488923 | Tead3 |
| **ENSMUSG00000002250** | 0.0922473614034891 | 0.532187451274905 | 0.909082872831377 | 758.048008201429 | 842.105863697416 | 831.933847121985 | 723.141481273042 | 684.348872386936 | 599.310293716191 | 868.259127523818 | 806.545494548701 | Ppard |
| **ENSMUSG00000002257** | -0.770670897956788 | 0.0375263208708845 | 0.679566075569698 | 44.2721721118832 | 39.0919549878878 | 31.2330113253987 | 62.8410885152968 | 93.0714466446232 | 113.746647582869 | 56.2182168900314 | 40.1644458927348 | Def6 |
| **ENSMUSG00000002265** | -0.176127967394643 | 0.522835444058475 | 0.90648515894888 | 84.9302893574902 | 87.9568987227475 | 110.735221971868 | 90.0409626487835 | 123.730276127558 | 151.662196777159 | 77.0397787011541 | 70.559161703453 | Peg3 |
| **ENSMUSG00000002266** | -0.165385224435195 | 0.692675416488634 | 0.94469184542218 | 21.6843291976571 | 30.9477976987445 | 26.5007368821565 | 16.8826804966469 | 19.7092475247437 | 12.2308223207386 | 39.5609674411332 | 35.8223436340608 | Zim1 |
| **ENSMUSG00000002274** | -0.257913065920297 | 0.18335172733865 | 0.805963874499992 | 98.4829951060259 | 101.801966114291 | 79.5022106464695 | 82.5375490947182 | 120.445401540101 | 110.077400886647 | 84.3273253350471 | 119.407812113536 | Metrn |
| **ENSMUSG00000002279** | -0.239791346418912 | 0.156536245073337 | 0.787275540343856 | 728.23205555465 | 704.469605510894 | 538.532831640966 | 597.459304242449 | 757.711071506815 | 936.880989768576 | 712.097413940397 | 628.519301943066 | Lmf1 |
| **ENSMUSG00000002280** | -0.0493168427492037 | 0.656133352348518 | 0.938034442101846 | 893.575065686785 | 811.158065998671 | 856.541674226844 | 754.093062183562 | 770.850569856644 | 923.427085215764 | 828.698160082685 | 908.584897627541 | Narfl |
| **ENSMUSG00000002289** | 0.501014809876389 | 0.271774372740105 | 0.844129257012535 | 582.766347187034 | 459.330471107681 | 2040.55673992605 | 1943.38411050291 | 1264.67671617106 | 829.249753346077 | 625.687932424238 | 831.512582536077 | Angptl4 |
| **ENSMUSG00000002297** | 0.0212211393053538 | 0.954472117373009 | 0.992371505557225 | 17.1667606148119 | 19.5459774939439 | 24.6078271048596 | 26.2619474392285 | 20.8042057205628 | 15.9000690169602 | 15.616171358342 | 33.6512925047237 | Dbf4 |
| **ENSMUSG00000002307** | 0.00359769264101776 | 0.977639372809866 | 0.996215309832663 | 458.984968017075 | 478.876448601625 | 508.246275204216 | 472.715053906113 | 551.858930692825 | 408.509465512669 | 502.840717738614 | 449.407583772762 | Daxx |
| **ENSMUSG00000002308** | -0.0402845479337413 | 0.802591665850579 | 0.963461896828986 | 464.406050316489 | 355.085257806647 | 401.296872786941 | 412.687745473591 | 474.116898789669 | 484.340563901248 | 353.966550789086 | 367.993166422624 | Cd320 |
| **ENSMUSG00000002319** | 0.10088225587881 | 0.387154544130393 | 0.875798327171006 | 1187.21702357173 | 1268.85970564852 | 1384.66350209268 | 1557.89623916281 | 1338.03891529094 | 1242.65154778704 | 1239.92400585236 | 1212.53205573472 | Ipo4 |
| **ENSMUSG00000002320** | -0.0715523110647813 | 0.690251892051788 | 0.94469184542218 | 556.564449406532 | 668.635313438664 | 662.518422053913 | 472.715053906113 | 664.639624862192 | 469.663577116362 | 648.591650416473 | 696.907412517182 | Tm9sf1 |
| **ENSMUSG00000002325** | -0.309218444437543 | 0.196283754888766 | 0.811906300001476 | 192.448421629207 | 200.346269312925 | 196.862616838877 | 201.654239265505 | 319.727793179176 | 306.993640250539 | 158.243869764533 | 196.480127205 | Irf9 |
| **ENSMUSG00000002326** | -0.113119491097842 | 0.584034306147141 | 0.922450244597442 | 170.764092431549 | 172.656134529838 | 154.272146849697 | 188.52326554589 | 199.282391639076 | 233.608706326107 | 124.929370866736 | 185.624871558315 | Gmpr2 |
| **ENSMUSG00000002329** | 0.0621615939393668 | 0.67162926717362 | 0.943173644832837 | 1094.15511076511 | 841.291447968501 | 878.310136665759 | 1027.96765690694 | 774.135444444101 | 998.035101372269 | 880.752064610492 | 1026.90718417641 | Mdp1 |
| **ENSMUSG00000002332** | -0.0537797340653218 | 0.685455865046631 | 0.944452327903597 | 725.521514404943 | 715.871425715695 | 585.855576073389 | 654.672832592197 | 664.639624862192 | 599.310293716191 | 748.535147109862 | 770.72315091464 | Dhrs1 |
| **ENSMUSG00000002342** | -0.0476085894321454 | 0.603938966309733 | 0.926448151609826 | 1328.1651633565 | 1338.08504260624 | 1392.23514120187 | 1306.53188510162 | 1378.55236853624 | 1539.86053018099 | 1336.74426827408 | 1291.77542195552 | Tmem161a |
| **ENSMUSG00000002343** | -0.0838602327460763 | 0.624895226378673 | 0.931529302402539 | 129.202461469373 | 119.719112150406 | 106.949402417274 | 98.4823028971069 | 117.160526952643 | 139.43137445642 | 118.6829023234 | 107.467030902182 | Armc6 |
| **ENSMUSG00000002345** | 0.138398902553352 | 0.402468277029596 | 0.881158207349486 | 252.983840639333 | 282.602257933272 | 286.775831260479 | 200.716312571247 | 213.516848184724 | 218.931719541221 | 229.03717992235 | 268.124814473121 | Borcs8 |
| **ENSMUSG00000002346** | 0.134650896794066 | 0.612557203960074 | 0.928055903343287 | 758.951521917998 | 1065.25577341994 | 940.776159316556 | 756.906842266336 | 603.321965896322 | 505.132961846504 | 1028.58515346946 | 1070.32820676315 | Slc25a42 |
| **ENSMUSG00000002365** | 0.0388327879708207 | 0.794635275194212 | 0.961804768755116 | 686.670424592474 | 819.302223287815 | 760.003275584703 | 648.107345732389 | 674.494248624564 | 591.971800323748 | 846.396487622139 | 722.960026069226 | Snx9 |
| **ENSMUSG00000002372** | -0.141506002864296 | 0.24235760215827 | 0.832048466795696 | 835.750187826367 | 890.970807432275 | 813.951204237664 | 916.354380290224 | 1009.55145654521 | 1088.54318654574 | 868.259127523818 | 848.880991570773 | Ranbp3 |
| **ENSMUSG00000002379** | 0.0185802966947191 | 0.915808390467637 | 0.986713918026868 | 7746.72660586299 | 5417.49342873811 | 5545.27919259125 | 7156.38067718977 | 6374.84661605878 | 7843.62635428966 | 5578.09640919978 | 5739.1736604024 | Ndufa11 |
| **ENSMUSG00000002393** | -0.239474453554909 | 0.0263897251495114 | 0.664353746352133 | 1147.46242004269 | 1060.36927904646 | 968.223351087361 | 1127.38788649831 | 1315.04479317874 | 1372.29826438687 | 1253.45802102959 | 1140.8873684666 | Nr2f6 |
| **ENSMUSG00000002395** | 0.0827823157924945 | 0.672610281423837 | 0.943216235131458 | 3497.50159683877 | 2321.89924313475 | 2503.37318047514 | 3495.65278950017 | 2858.93584928366 | 3449.09189444828 | 2347.63109420409 | 2504.30747769025 | Use1 |
| **ENSMUSG00000002396** | -0.138666297668465 | 0.487371520490154 | 0.899236955296196 | 199.676531361759 | 228.850819824926 | 221.470443943737 | 258.867767615252 | 238.700886688563 | 338.793778284459 | 232.160414194018 | 192.138024946326 | Ocel1 |
| **ENSMUSG00000002409** | -0.0229932056400269 | 0.843516815211425 | 0.971654415184575 | 360.501972911049 | 443.856572258309 | 446.726707442067 | 420.191159027656 | 446.742943894192 | 409.732547744743 | 426.842017128016 | 414.67076570337 | Dyrk1b |
| **ENSMUSG00000002413** | 0.213746278079306 | 0.368864145909517 | 0.871495599696183 | 246.659244623349 | 341.240190415104 | 377.63550057073 | 303.888248939644 | 254.03030143003 | 174.900759186562 | 323.775286162958 | 340.855027305911 | Braf |
| **ENSMUSG00000002416** | 0.0362030555347886 | 0.854670644068933 | 0.973268281968411 | 6283.03438502114 | 4173.06619495702 | 4574.21647683794 | 6110.59241309192 | 5388.28928162578 | 6481.11274775938 | 4347.54210616243 | 4400.72063916613 | Ndufb2 |
| **ENSMUSG00000002428** | 0.102634609851121 | 0.411634132783231 | 0.882192126163777 | 275.571683553559 | 265.499527626071 | 242.292451494002 | 287.005568442997 | 273.739548954774 | 239.724117486476 | 251.940897914585 | 231.216945274392 | Hltf |
| **ENSMUSG00000002455** | 0.0334144303789449 | 0.817937990673208 | 0.966315773308182 | 1258.59460718068 | 1407.31037956396 | 1433.8791563024 | 1347.80065964898 | 1617.25325522481 | 1407.76764911701 | 1181.62363278122 | 1117.00580604389 | Prpf6 |
| **ENSMUSG00000002458** | -0.367638192996275 | 0.244693492109674 | 0.832048466795696 | 49.6932544112975 | 35.0198763433161 | 40.6975602118832 | 51.5859681841989 | 56.937826182593 | 84.3926740130963 | 32.2734208072402 | 55.3618037980939 | Rgs19 |
| **ENSMUSG00000002475** | -0.0938636441103862 | 0.772174374840628 | 0.959045638244974 | 59.631905293557 | 123.791190794978 | 102.217127974032 | 60.9652351267805 | 63.5075753575076 | 74.6080161565054 | 112.436433780063 | 119.407812113536 | Abhd3 |
| **ENSMUSG00000002477** | -0.0758723175442565 | 0.711618062631258 | 0.950020479830613 | 294.545471601509 | 220.706662535783 | 192.130342395635 | 293.571055302804 | 304.398378437709 | 300.878229090169 | 236.324726556243 | 213.848536239696 | Snrpd1 |
| **ENSMUSG00000002486** | 0.346127809851786 | 0.0153748585193423 | 0.635862253526407 | 234.913566307952 | 242.69588721647 | 253.649910157784 | 270.12288794635 | 189.427767876704 | 174.900759186562 | 234.242570375131 | 187.795922687652 | Tchp |
| **ENSMUSG00000002489** | 0.21371858637677 | 0.547819819047029 | 0.914699131434573 | 50.5967681278665 | 95.2866402829764 | 53.001473764313 | 39.3929211588428 | 44.893286028583 | 37.9155491942896 | 44.7663578939139 | 78.1578406561325 | Tiam1 |
| **ENSMUSG00000002496** | 0.205079996096204 | 0.212737460895671 | 0.823370267313127 | 3422.50995836354 | 3950.73070096341 | 4111.40003628885 | 3338.0811048648 | 2948.72242134083 | 2496.31083566275 | 3603.17127141479 | 3809.10920642179 | Tsc2 |
| **ENSMUSG00000002500** | 0.0506800176461136 | 0.676760000668177 | 0.94409858171788 | 8203.90454644693 | 7054.46904385591 | 8977.12461883052 | 8593.28437279327 | 8418.03860945722 | 8719.35323245454 | 7240.69811981793 | 7317.52783143041 | Rpl3l |
| **ENSMUSG00000002504** | -0.571658107608638 | 0.0644321692323239 | 0.716991331547963 | 2620.18977805023 | 2288.50819824926 | 2633.03750021998 | 2472.37476606451 | 4574.73534213219 | 5865.90238502623 | 2199.79800534512 | 2243.78134216981 | Slc9a3r2 |
| **ENSMUSG00000002524** | -0.0922382000197096 | 0.549879338832009 | 0.914699131434573 | 1429.35869961223 | 1313.65257073881 | 1394.12805097916 | 1471.60698329106 | 1631.48771177045 | 1828.50793695042 | 1302.38869128573 | 1217.95968355807 | Puf60 |
| **ENSMUSG00000002546** | 0.295529698496899 | 0.0729983253811531 | 0.730383296285387 | 2730.41845147165 | 3158.30419672977 | 3379.7904073636 | 2651.51876466782 | 2291.74750384937 | 1873.76197953715 | 2761.98017424543 | 2783.28754781005 | Golga2 |
| **ENSMUSG00000002550** | -0.0378108799600068 | 0.766692705999011 | 0.958624690460551 | 334.300075130547 | 322.508628650074 | 274.47191770805 | 292.633128608546 | 306.588294829347 | 347.355353908976 | 287.337552993494 | 316.973464883204 | Uck1 |
| **ENSMUSG00000002565** | 0.344210939371823 | 0.268634595624934 | 0.843793744782845 | 38.8510898124689 | 45.6072808192024 | 33.1259211026956 | 51.5859681841989 | 36.1336204620302 | 22.0154801773295 | 27.0680303544596 | 47.7631248454143 | Scin |

| | | | | | | | | | | | | |
|---|---|---|---|---|---|---|---|---|---|---|---|---|
| **ENSMUSG00000002580** | 0.0203232218244536 | 0.915692064438528 | 0.986713918026868 | 995.672115659088 | 737.046234667467 | 736.341903368492 | 1033.59521707249 | 780.705193619016 | 1120.34332457966 | 759.98700610598 | 793.519187772679 | Mien1 |
| **ENSMUSG00000002588** | -0.383804559213888 | 0.159049982038838 | 0.789393312599749 | 187.027339329792 | 213.376920975554 | 165.629605513478 | 200.716312571247 | 186.142893289246 | 218.931719541221 | 188.435134390661 | 407.07208675069 | Pon1 |
| **ENSMUSG00000002602** | -1.02147302867274 | 0.0102055814833394 | 0.635862253526407 | 559.274990556239 | 579.049583258088 | 651.160963390131 | 672.493439783102 | 1751.93311331055 | 2086.578287918 | 596.537745888666 | 563.387768062955 | Axl |
| **ENSMUSG00000002603** | -0.410575499062161 | 0.132355308706552 | 0.77303252008843 | 446.335775985108 | 390.919549878878 | 465.655805215036 | 476.466760683146 | 800.41444114376 | 785.218792991418 | 363.336253604092 | 416.841816832707 | Tgfb1 |
| **ENSMUSG00000002608** | 0.168974440965404 | 0.253400724887915 | 0.837406774112203 | 1033.61969175499 | 1161.35682943183 | 1341.12657721485 | 1086.11911195095 | 917.574968096403 | 885.511536021474 | 1167.04853951343 | 1139.80184290193 | Ccdc97 |
| **ENSMUSG00000002617** | -0.320248142650652 | 0.479246591702016 | 0.897732059433541 | 6.32459601598331 | 25.2468875963442 | 14.1968233297267 | 15.0068271081306 | 22.994122112201 | 15.9000690169602 | 15.616171358342 | 21.7105112933701 | Zfp40 |
| **ENSMUSG00000002625** | 0.0370683726034929 | 0.779891274679617 | 0.960643510321816 | 865.566140473145 | 708.541684155466 | 875.470771999813 | 995.140222607909 | 882.536305830192 | 859.826809147923 | 845.355409531583 | 768.552099785303 | Akap8l |
| **ENSMUSG00000002635** | -0.0583033129093746 | 0.678525206254765 | 0.944452327903597 | 337.914129996823 | 310.292392716359 | 302.865564367503 | 370.481044231974 | 398.564783278151 | 352.247682837272 | 300.871568170723 | 324.572143835884 | Pdcd2l |
| **ENSMUSG00000002658** | -0.0984523423197143 | 0.568500858686892 | 0.92059532906293 | 554.757421973394 | 520.411650776256 | 531.907647420427 | 586.204183911351 | 610.986673267056 | 750.97249049335 | 542.401685179747 | 445.065481514088 | Gtf2f1 |
| **ENSMUSG00000002660** | 0.00559308239595908 | 0.975962768692111 | 0.99575536949131 | 2032.00234856378 | 1571.00794107574 | 1690.36843112613 | 2174.11407729042 | 2013.62812211132 | 2331.19473433278 | 1582.43869764533 | 1512.13711158323 | Clpp |
| **ENSMUSG00000002661** | -0.114607919492028 | 0.463684193090122 | 0.895328112556867 | 1350.75300627072 | 926.805099504506 | 945.508433759798 | 1065.48472467727 | 1161.75064576406 | 1311.14415278318 | 1073.35151136338 | 1097.46634587986 | Alkbh7 |
| **ENSMUSG00000002679** | -0.136781823473185 | 0.447535425357506 | 0.89108031640712 | 181.606257030378 | 154.738988493722 | 173.201244622666 | 156.633757941113 | 187.237851485066 | 233.608706326107 | 147.833088858971 | 164.999885829613 | Med6 |
| **ENSMUSG00000002688** | -0.514953446965053 | 0.0108112485567283 | 0.635862253526407 | 139.141112351633 | 114.018202048006 | 168.468970179424 | 113.489130005238 | 237.605928492744 | 187.131581507301 | 161.367104036201 | 178.026192605635 | Prkd1 |
| **ENSMUSG00000002699** | -0.311404077608734 | 0.370895272182177 | 0.873323500696802 | 24.3948703473642 | 13.8450673915436 | 35.965285768641 | 27.1998741334867 | 28.4689130912965 | 30.5770558018465 | 27.0680303544596 | 39.0789203280663 | Lcp2 |
| **ENSMUSG00000002718** | 0.0702094938796001 | 0.702364909479592 | 0.946434671388391 | 923.391018333564 | 952.866402829765 | 967.276896198713 | 707.196727470654 | 760.995946094272 | 650.679747463293 | 957.791843311646 | 1011.70982627105 | Cse1l |
| **ENSMUSG00000002728** | 0.0853024559047552 | 0.495721156551824 | 0.900927059989498 | 1373.34084918495 | 1057.92603185971 | 1135.74586637814 | 1375.00053378246 | 1228.54309570903 | 1108.11250225892 | 1181.62363278122 | 1138.71631733726 | Naa20 |
| **ENSMUSG00000002731** | 0.067052454179137 | 0.617603048748111 | 0.929314954729851 | 332.493047697409 | 300.519403969387 | 274.47191770805 | 299.198615468353 | 284.689130912965 | 254.401104271363 | 281.091084450157 | 331.085297223895 | Prkra |
| **ENSMUSG00000002732** | -0.237423269348385 | 0.250016646964242 | 0.835795722342557 | 301.773581334061 | 244.324718674299 | 256.489274823729 | 329.212269684615 | 388.710159515779 | 417.071041137186 | 267.557069272927 | 261.61166108511 | Fkbp7 |
| **ENSMUSG00000002733** | 0.0811745058747901 | 0.424549869851919 | 0.884903615282587 | 477.055242348456 | 477.247617143797 | 480.799083433411 | 511.170048370698 | 457.692525852383 | 485.563646133322 | 479.936999746379 | 416.841816832707 | Plekha3 |
| **ENSMUSG00000002741** | 0.0689963261074255 | 0.631239218340871 | 0.931825800800958 | 1574.82440797985 | 1720.04601946706 | 1564.48993093588 | 1502.55856420157 | 1297.52546204563 | 1296.46716599829 | 1714.65561514596 | 1755.29483806898 | Ykt6 |
| **ENSMUSG00000002748** | 0.234184467702346 | 0.192222512184328 | 0.808824684419004 | 1870.27339329792 | 2159.01609735189 | 2333.95775540707 | 1737.04023776611 | 1549.36584708402 | 1319.70572840769 | 2011.36287095446 | 2004.96571794273 | Baz1b |
| **ENSMUSG00000002763** | -0.0513410722684907 | 0.641131952406359 | 0.934544499347759 | 1822.38716631976 | 1898.4030640993 | 2087.87948435847 | 2015.60446596079 | 2210.72059735876 | 2195.43260657258 | 1838.54390792214 | 1863.84739453583 | Pex6 |
| **ENSMUSG00000002767** | 0.043228412354612 | 0.773937973225492 | 0.959203117135841 | 2440.39054845299 | 1991.24645719553 | 2030.14573615092 | 2402.96819068941 | 2270.94329812881 | 2580.70350967584 | 1857.28331355215 | 1895.32763591121 | Mrpl2 |
| **ENSMUSG00000002768** | 0.0225536590597283 | 0.88460068610229 | 0.979520370692028 | 539.39768879172 | 412.09435883065 | 458.084166105848 | 506.480414899407 | 475.211856985488 | 561.394744521902 | 391.445362049107 | 459.177313854779 | Mea1 |
| **ENSMUSG00000002769** | -0.154558676080611 | 0.585563676589726 | 0.922964550512322 | 538.494175075151 | 708.541684155466 | 572.60520763231 | 672.493439783102 | 427.033696369448 | 611.54111603693 | 593.414511616998 | 1141.97289403127 | Gnmt |
| **ENSMUSG00000002771** | 0.456203289721638 | 0.372952001129223 | 0.873700364743909 | 22.5878429142261 | 33.3910448854875 | 19.8755526616174 | 5.62756016554897 | 19.7092475247437 | 12.2308223207386 | 13.5340151772298 | 14.1118323406906 | Grin2d |
| **ENSMUSG00000002778** | -0.0451723326823791 | 0.65665017393464 | 0.938361205344872 | 1704.93038316579 | 1682.582895937 | 1738.6376304472 | 1699.52316999579 | 1699.37511991124 | 2033.98575193883 | 1661.56063252759 | 1649.99885829613 | Kdelr1 |
| **ENSMUSG00000002781** | 0.159453148253408 | 0.441235310081695 | 0.89108031640712 | 3123.44691817919 | 3370.86670197641 | 3384.52268180685 | 2972.28969410411 | 2656.36858305713 | 1855.41574605605 | 3500.10454044973 | 3493.22126710326 | Tmem143 |
| **ENSMUSG00000002797** | 0.0210487207988193 | 0.898490215423459 | 0.984462340635154 | 462.599022883351 | 370.55915665602 | 353.974128354519 | 463.335786963532 | 383.235368536684 | 500.240632918209 | 368.541644056872 | 375.591845375304 | Ggct |
| **ENSMUSG00000002799** | -0.458447278789096 | 0.0413039264109697 | 0.683543358336265 | 560.178504272808 | 562.761268679801 | 642.642869392295 | 506.480414899407 | 1062.10944994452 | 897.742358342213 | 554.894622266421 | 607.894316214364 | Jag2 |
| **ENSMUSG00000002803** | 0.0246399376201028 | 0.799683865114565 | 0.96336590515536 | 500.546598979251 | 455.25839246311 | 493.102996985841 | 478.342614071662 | 503.680770076785 | 452.540425867328 | 474.731609293598 | 462.433890548784 | Btbd6 |
| **ENSMUSG00000002804** | -0.131337136892145 | 0.535368701687793 | 0.910861252626585 | 163.535982698997 | 130.306516626293 | 120.199770858353 | 166.013024883695 | 171.908436743598 | 203.031650524261 | 135.340151772298 | 125.920965501547 | Nudt14 |
| **ENSMUSG00000002808** | 0.214574245304312 | 0.20222777619256 | 0.81455046200345 | 674.924746277077 | 846.992358070902 | 803.540200462531 | 637.79015209555 | 618.65138063779 | 517.363784167243 | 782.890724098215 | 633.946929766408 | Epdr1 |
| **ENSMUSG00000002812** | 0.0866770675419664 | 0.452933752804263 | 0.891364384592125 | 3231.86856416747 | 3726.76637551197 | 3837.87457346945 | 3230.21953502511 | 3471.01748074654 | 2873.0201631415 | 3488.65268145361 | 3374.89898055439 | Flii |
| **ENSMUSG00000002814** | 0.0613517735661573 | 0.698150809391614 | 0.944933386973863 | 127.395434036235 | 105.059629029948 | 120.199770858353 | 112.551203310979 | 112.780694169367 | 100.292743030056 | 112.436433780063 | 119.407812113536 | Top3a |
| **ENSMUSG00000002820** | -0.0262804003201297 | 0.797049571683554 | 0.962278883153809 | 1648.00901902194 | 1437.44376153379 | 1434.82561119105 | 1355.30407320304 | 1485.85827172651 | 1386.97525117176 | 1584.52085382644 | 1525.16341835925 | Atg4d |
| **ENSMUSG00000002825** | 0.283224452188447 | 0.176420176080829 | 0.799889021902745 | 112.939214571131 | 103.43079757212 | 95.5919437534931 | 80.6616957062019 | 78.836990098975 | 102.738907494204 | 77.0397787011541 | 65.1315338801105 | Qtrt1 |
| **ENSMUSG00000002831** | 0.174304457694172 | 0.502502496604752 | 0.900927059989498 | 4004.37279183401 | 5489.16201288258 | 5494.17062860423 | 3613.8315529767 | 3627.59650274867 | 2572.14193405133 | 5804.01035485046 | 4479.96400538693 | Plin4 |
| **ENSMUSG00000002833** | 0.129804988631003 | 0.282830579742592 | 0.850587941933635 | 1016.45293114018 | 920.289773673191 | 1019.33191507438 | 1187.41519493083 | 1057.72961716125 | 924.650167447838 | 901.573626421614 | 902.07174423953 | Hdgfl2 |
| **ENSMUSG00000002835** | 0.229960758842114 | 0.290711718659331 | 0.851785577030345 | 65.0529875929712 | 66.7820897709749 | 68.1447519826882 | 81.59962240046 | 67.887408140784 | 50.1463715150282 | 49.9717483466946 | 71.6446872681215 | Chaf1a |
| **ENSMUSG00000002844** | -0.153828755121127 | 0.285687713194575 | 0.851362869828931 | 830.329105526952 | 835.590537866101 | 769.467824471187 | 854.451218469185 | 1001.88674917447 | 1088.54318654574 | 799.547973547113 | 771.808676479309 | Adprh |
| **ENSMUSG00000002845** | -0.0179923268647636 | 0.914503622345854 | 0.986675105979797 | 121.974351736821 | 177.542628903324 | 163.736695736181 | 138.813150750208 | 159.863896589588 | 151.662196777159 | 147.833088858971 | 150.888053488923 | Tmem39a |
| **ENSMUSG00000002846** | 0.0150187745010618 | 0.901001730123819 | 0.984802943363903 | 1188.12053728829 | 981.370953341766 | 940.776159316556 | 1148.02227377199 | 1054.44474257379 | 962.565716642128 | 1069.18719900115 | 1126.77553612591 | Timmdc1 |
| **ENSMUSG00000002847** | -0.845801513562197 | 0.0389165221758636 | 0.679566075569698 | 115.649755720838 | 118.090280692578 | 104.110037751329 | 108.799496533947 | 270.454674367317 | 344.909189444828 | 90.5737938783839 | 97.6973008201657 | Pla1a |
| **ENSMUSG00000002870** | -0.130275782108466 | 0.329589209831114 | 0.855977678491063 | 152.693818100169 | 182.42912327681 | 157.111511515642 | 159.447538023887 | 175.193311331055 | 178.570005882784 | 179.065431575656 | 181.282769299641 | Mcm2 |
| **ENSMUSG00000002871** | -0.00989612744565605 | 0.944240088062244 | 0.990505437194639 | 410.195227322346 | 345.312269059675 | 358.706402797761 | 432.384206053012 | 430.318570956905 | 341.239942748607 | 423.718782856348 | 360.394487469945 | Tpra1 |
| **ENSMUSG00000002881** | -0.0786068258250056 | 0.462964031597314 | 0.895328112556867 | 701.126644057579 | 676.779470727807 | 641.696414503647 | 604.024791102256 | 656.974917491458 | 643.34125407085 | 711.056335849841 | 758.782369703287 | Nab1 |
| **ENSMUSG00000002885** | -0.35549311847227 | 0.102195168666503 | 0.759800953500741 | 472.537673765611 | 515.52515640277 | 565.980023411771 | 436.135912830045 | 848.5926017598 | 728.95701031602 | 505.963952010282 | 463.519416113453 | Adgre5 |
| **ENSMUSG00000002897** | -0.705805075159206 | 0.0545931375019865 | 0.703396849069134 | 46.9827132615903 | 43.9784493613737 | 67.1982970940397 | 50.6480414899407 | 117.160526952643 | 121.085140975312 | 63.5057635239243 | 39.0789203280663 | Il17ra |
| **ENSMUSG00000002900** | -0.363734410157888 | 0.0911255020174416 | 0.753823363185192 | 1041.75131520411 | 1186.60371702818 | 1255.94563723649 | 1148.02227377199 | 1952.31046314545 | 1784.47697659576 | 1080.63905799727 | 1144.14394516061 | Lamb1 |
| **ENSMUSG00000002908** | 0.258165018735128 | 0.0384799380743042 | 0.679566075569698 | 543.011743657996 | 502.494504740141 | 543.265106084208 | 560.88016316638 | 442.363111110915 | 375.486245246675 | 499.717483466946 | 477.631248454143 | Kcnn1 |
| **ENSMUSG00000002910** | 0.229688100594013 | 0.59249200832668 | 0.923935529355527 | 406.58117245607 | 342.054606144018 | 275.418372596698 | 92.854742731558 | 308.778211220985 | 327.786038195794 | 188.435134390661 | 128.092016630884 | Arrdc2 |
| **ENSMUSG00000002944** | 0.0122371929594565 | 0.884426096646646 | 0.979450877350433 | 54274.9724680191 | 54617.1620281816 | 60726.4385654616 | 55579.6600483501 | 60028.8931693904 | 56595.6841247537 | 52678.5513821405 | 53992.9560610469 | Cd36 |
| **ENSMUSG00000002948** | 0.226077798926598 | 0.0679715106001447 | 0.722112580225498 | 1254.07703859783 | 1284.3336044979 | 1280.55346434135 | 1188.35312162509 | 965.753128712444 | 925.873249679912 | 1200.36303841123 | 1187.56496774735 | Map2k7 |
| **ENSMUSG00000002949** | 0.12195031233672 | 0.301812105257225 | 0.851785577030345 | 2394.31134890797 | 2226.61260285177 | 2135.2022287909 | 2401.09233730089 | 2065.09115731482 | 1772.24615427502 | 2258.09837841626 | 2317.59708056726 | Timm44 |
| **ENSMUSG00000002957** | 0.0259528797984223 | 0.797037843013749 | 0.962278883153809 | 1828.71176233575 | 2068.61595144239 | 2046.23546925794 | 1982.77703166175 | 1978.58945984511 | 1713.53820713548 | 2008.23963668279 | 2083.12355859887 | Ap2a2 |
| **ENSMUSG00000002963** | -0.0172195348076445 | 0.893269838561084 | 0.983433576156181 | 445.432262268539 | 437.341246426994 | 382.367775013973 | 407.060185308042 | 465.357233223116 | 458.655837027697 | 404.979377226337 | 364.736589728619 | Pnkp |
| **ENSMUSG00000002968** | 0.113579657031484 | 0.358993252271318 | 0.866754558180356 | 1727.51822608001 | 1941.56709773176 | 2155.07778145251 | 1831.77083388619 | 1682.95074697395 | 1572.88375044698 | 2009.28071477334 | 1809.5711163024 | Med25 |
| **ENSMUSG00000002980** | -0.119068263373549 | 0.508456969143816 | 0.902394714365738 | 6211.65680141218 | 6354.88593271851 | 7122.07303707956 | 5940.82768143119 | 8082.98140153657 | 8604.3835026396 | 5721.76518569653 | 5426.54229777787 | Bcam |
| **ENSMUSG00000002981** | 0.094279138333952 | 0.294254463460605 | 0.851785577030345 | 3321.31642210781 | 3206.35472473571 | 3613.56476485977 | 3336.20525147628 | 3354.95191198971 | 3320.66826008053 | 2976.44226089999 | 2973.25452162704 | Clptm1 |
| **ENSMUSG00000002983** | -0.220078052626651 | 0.324014235174429 | 0.855977678491063 | 146.369222084185 | 132.749763813036 | 108.842312194571 | 149.130344387048 | 183.952976897608 | 195.693157131818 | 112.436433780063 | 134.605170018895 | Relb |
| **ENSMUSG00000002984** | 0.0231957862279758 | 0.803490624122532 | 0.963461896828986 | 1836.84338578487 | 1663.03691844306 | 1571.11511515642 | 1732.35060429482 | 1721.27428382762 | 1775.91540097124 | 1576.19222910199 | 1622.86071917942 | Tomm40 |
| **ENSMUSG00000002985** | -0.483145866462658 | 0.053688564875605 | 0.703396849069134 | 7839.7885186696 | 10891.1815427713 | 8448.05633607604 | 10884.639286866 | 13312.5017447686 | 17952.4010023801 | 7710.22433865875 | 14231.2401528041 | Apoe |
| **ENSMUSG00000002996** | 0.0269401552835791 | 0.751661299605874 | 0.956197192463153 | 1829.61527605232 | 1825.10564849701 | 1742.42345000179 | 1716.40585049244 | 1812.15581408061 | 1582.66840830357 | 1797.94186239045 | 1787.86060500903 | Hbp1 |
| **ENSMUSG00000002997** | 0.162929898750297 | 0.818893377974689 | 0.966315773308182 | 28.0089252136404 | 30.1333819698302 | 57.7337482075553 | 22.5102406621959 | 41.6084114411257 | 56.2617826753975 | 1.04107809055614 | 24.9670879873757 | Prkar2b |
| **ENSMUSG00000003031** | -0.0982000351995187 | 0.452434216846931 | 0.891110666112841 | 702.933671490717 | 763.107537992726 | 813.951204237664 | 706.258800776395 | 843.117810780705 | 648.233582999146 | 856.8072685277 | 846.709940441436 | Cdkn1b |
| **ENSMUSG00000003032** | -0.786102856409024 | 0.0325236751658631 | 0.671606027118506 | 973.987786461431 | 851.878852444388 | 889.66759532954 | 761.596475737627 | 2322.4063333323 | 2139.17082389718 | 860.971580889925 | 674.111375659143 | Klf4 |
| **ENSMUSG00000003033** | 0.0497365859391293 | 0.638189406824097 | 0.933198322011444 | 835.750187826367 | 875.496908582903 | 963.491076644119 | 866.644265494541 | 866.111932892906 | 754.641737189571 | 892.203923606609 | 906.413846498204 | Ap1m1 |
| **ENSMUSG00000003037** | -0.157574033011344 | 0.238741373308771 | 0.832048466795696 | 526.748496759753 | 430.82592059568 | 431.583429223692 | 505.542488205149 | 543.099265126272 | 595.64104701997 | 466.402984569149 | 509.11148982953 | Rab8a |
| **ENSMUSG00000003038** | 0.240669824772762 | 0.0463904221491274 | 0.69805115405055 | 1950.68611407257 | 1535.98806473242 | 1974.30489772066 | 1906.80496942684 | 1588.78434213351 | 1428.56004706227 | 1683.42327242927 | 1532.76209731193 | Hmgn2 |
| **ENSMUSG00000003039** | -0.134465859138417 | 0.40100277302794 | 0.880899654809123 | 674.924746277077 | 518.782819318427 | 661.571967165264 | 800.98939689647 | 751.1413223319 | 813.349684329117 | 648.591650416473 | 702.335040340524 | Fam32a |
| **ENSMUSG00000003053** | -0.0289248986918427 | 0.922395155955194 | 0.986713918026868 | 178.895715880671 | 254.097707421271 | 106.949402417274 | 205.405946042537 | 170.813478547779 | 194.470074899744 | 134.299073681742 | 261.61166108511 | Cyp2c29 |
| **ENSMUSG00000003062** | 0.191811812440519 | 0.269748857308369 | 0.843793744782845 | 284.606820719249 | 228.036404096012 | 282.043556817237 | 315.143369270742 | 280.309298129689 | 212.816308380852 | 204.051305749003 | 273.552442296464 | Stard3nl |
| **ENSMUSG00000003068** | 0.0907907482648518 | 0.300338186388052 | 0.851785577030345 | 4323.31313378288 | 4030.54344239701 | 4282.70837113422 | 4721.52297889558 | 4064.48482288049 | 4186.61048038882 | 4236.14675047292 | 3811.28025755113 | Stk11 |
| **ENSMUSG00000003070** | 0.0295953752239647 | 0.930162889160782 | 0.987529878729394 | 22.5878429142261 | 26.0613033252585 | 14.1968233297267 | 30.9515809105193 | 21.8991639163819 | 24.4616446414772 | 20.8215618111227 | 24.9670879873757 | Efna2 |
| **ENSMUSG00000003072** | -0.159575014981585 | 0.336438138459468 | 0.858570960161243 | 15308.2328998293 | 12849.8513708103 | 12433.5778721746 | 15078.1095368942 | 16356.4855291457 | 19539.961739612 | 13201.9112663424 | 13083.8396309495 | Atp5d |
| **ENSMUSG00000003099** | 0.152686531246756 | 0.167264506666198 | 0.797837911261499 | 2170.23994719885 | 2193.22155796629 | 2125.73767990441 | 2115.02469555215 | 1830.77010340953 | 1698.86122035059 | 2235.19466042403 | 1973.48547656735 | Ppp5c |
| **ENSMUSG00000003119** | 0.105780223625528 | 0.696357288207438 | 0.944933386973863 | 222.264374275985 | 324.951875836817 | 362.492222352355 | 212.909359596603 | 249.650468646754 | 162.669936865823 | 343.555769883525 | 286.578749072486 | Cdk12 |
| **ENSMUSG00000003123** | -0.170862292661744 | 0.491488432473529 | 0.899236955296196 | 1296.54218327658 | 1526.21507598545 | 2132.36286412495 | 1744.54365132018 | 2360.72987018597 | 2409.4719971855 | 1350.27828345131 | 1422.03848971574 | Lipe |
| **ENSMUSG00000003131** | 0.0704700967946547 | 0.373932918102503 | 0.873700364743909 | 774.311255099672 | 810.343650269757 | 796.915016241992 | 769.099889291692 | 742.381656765348 | 732.626257012242 | 758.945928015424 | 766.381048655966 | Pafah1b2 |
| **ENSMUSG00000003134** | -0.107680359887805 | 0.590988977615397 | 0.923935529355527 | 70.4740698923855 | 112.389370590177 | 91.8061241988993 | 95.6685228143324 | 105.115986798633 | 88.0619207093179 | 110.35427759895 | 95.5262496908287 | Tbc1d8 |
| **ENSMUSG00000003135** | 0.00963577792760335 | 0.935053858310057 | 0.98818087308472 | 404.774145022932 | 394.177212794535 | 359.65285768641 | 378.922384480297 | 393.089992299056 | 346.132271676902 | 427.883095218572 | 359.308961905276 | Cnot11 |
| **ENSMUSG00000003153** | 0.0188724747778867 | 0.96642982553281 | 0.994284908685797 | 41.5616309621761 | 46.4216965481167 | 95.5919437534931 | 29.075727522003 | 59.1277425742312 | 84.3926740130963 | 39.5609674411332 | 27.1381391167127 | Slc2a3 |
| **ENSMUSG00000003154** | 0.0734530626300537 | 0.668230434608011 | 0.942024383071524 | 884.539928521095 | 1058.74044758863 | 1202.94416347218 | 984.823028971069 | 1062.10944994452 | 727.733928083946 | 1028.58515346946 | 1106.15055039721 | Foxj2 |
| **ENSMUSG00000003161** | -0.0837400315607887 | 0.491985627126825 | 0.899236955296196 | 1123.06754969532 | 950.423155643021 | 1077.06566328193 | 1105.81557253037 | 1240.58763586304 | 1232.86688993045 | 1012.96898211112 | 1024.73613304707 | Sri |
| **ENSMUSG00000003166** | -0.217340140614334 | 0.0393613032198792 | 0.679566075569698 | 673.117718843939 | 749.262470601182 | 723.091534927413 | 670.617586394585 | 891.295971396745 | 839.034411202668 | 813.081988724343 | 731.644230586574 | Dgcr2 |
| **ENSMUSG00000003184** | -0.201111816532116 | 0.373663184345602 | 0.873700364743909 | 337.914129996823 | 276.901347830872 | 277.311282373995 | 300.136542162612 | 428.128654565267 | 423.186452297555 | 222.790711379013 | 297.434004719171 | Irf3 |

| | | | | | | | | | | | | |
|---|---|---|---|---|---|---|---|---|---|---|---|---|
| **ENSMUSG00000003190** | -0.258053437373391 | 0.401807150432223 | 0.881158207349486 | 150.886790667031 | 123.791190794978 | 101.270673085384 | 86.2892558717508 | 158.768938393769 | 204.254732756335 | 106.189965236726 | 84.6709940441436 | Bcl2l12 |
| **ENSMUSG00000003198** | -0.101264965755925 | 0.671465463764989 | 0.943173644832837 | 63.2459601598332 | 53.7514381083457 | 39.7511053232348 | 45.9584080186499 | 45.9882442244021 | 53.8156182112498 | 62.4646854333682 | 55.3618037980939 | Zfp959 |
| **ENSMUSG00000003199** | -0.13621186032472 | 0.318223376497073 | 0.854351922245337 | 1028.19860945557 | 960.196144389993 | 862.220403558735 | 1103.93971914186 | 1082.91365566509 | 1274.45168582096 | 953.627530949421 | 1036.67691425842 | Mpnd |
| **ENSMUSG00000003200** | -0.328598359629184 | 0.132834971464264 | 0.77303252008843 | 177.088688447533 | 193.83094348161 | 240.399541716705 | 206.343872736795 | 317.537876787538 | 305.770558018465 | 226.955023741238 | 176.940667040967 | Sh3gl1 |
| **ENSMUSG00000003208** | -0.158394806991008 | 0.497541516743531 | 0.900927059989498 | 79.509207058076 | 86.3280672649188 | 87.0738497556571 | 88.1651092602672 | 106.210944994452 | 125.977469903608 | 86.4094815161593 | 62.9604827507734 | Ccdc94 |
| **ENSMUSG00000003226** | 0.262401855576043 | 0.306663518379895 | 0.851785577030345 | 1415.80599386369 | 2071.87361435805 | 1992.28754060498 | 1334.66968592936 | 1154.08593839333 | 945.442565393093 | 1908.2961399894 | 1672.79489515417 | Ranbp2 |
| **ENSMUSG00000003227** | 0.308836286722425 | 0.42388425534271 | 0.884584010063004 | 18.9737880479499 | 20.3603932228582 | 23.6613722162112 | 12.1930470253561 | 18.6142893289246 | 13.4539045528125 | 17.6983275394543 | 10.8552556466851 | Edar |
| **ENSMUSG00000003228** | 0.0101776144477897 | 0.948938616003695 | 0.991267322147698 | 970.373731595154 | 1187.41813275709 | 1118.70967838246 | 1054.22960434617 | 931.809424642051 | 1040.84297949485 | 1376.30523571521 | 950.920394649613 | Grk5 |
| **ENSMUSG00000003233** | 0.185534962075998 | 0.630239278287434 | 0.931825800800958 | 606.257703817829 | 994.401605004395 | 995.670542858166 | 513.045901759214 | 444.553027502553 | 315.555215875056 | 1068.1461209106 | 905.328320933535 | Dvl3 |
| **ENSMUSG00000003234** | -0.00860690031653543 | 0.936916372213795 | 0.988852035581444 | 1299.25272442629 | 1417.08336831093 | 1441.45079541158 | 1466.91734981976 | 1570.17005280458 | 1399.2060734925 | 1453.34501441637 | 1235.32809259276 | Abcf3 |
| **ENSMUSG00000003235** | 0.0676280732308828 | 0.504737478319369 | 0.90122958035533 | 1756.43066501022 | 1504.22585130477 | 1820.97920575961 | 1761.42633181683 | 1743.173447744 | 1660.9456711563 | 1564.74037010587 | 1559.90023642865 | Eif2b5 |
| **ENSMUSG00000003269** | 0.05493585065555 | 0.708430754093752 | 0.949174314815223 | 976.698327611138 | 880.383402956389 | 982.420174417088 | 1040.1607039323 | 1045.68507700724 | 1086.09702208159 | 828.698160082685 | 775.065253173314 | Cyth2 |
| **ENSMUSG00000003273** | -0.0614028802614654 | 0.79655531240354 | 0.962278883153809 | 78.6056933415069 | 64.338842584232 | 85.1809399783602 | 107.861569839689 | 72.2672409240604 | 110.077400886647 | 73.9165444294857 | 94.4407241261602 | Car11 |
| **ENSMUSG00000003279** | 0.0257283785694808 | 0.970742688589255 | 0.995122272737203 | 28.0089252136404 | 31.7622134276588 | 6.62518422053913 | 2.81378008277448 | 25.1840385038392 | 22.0154801773295 | 12.4929370866736 | 8.68420451734806 | Dlgap1 |
| **ENSMUSG00000003282** | -0.19409333749443 | 0.761431580784966 | 0.957479700684068 | 7.22810973255236 | 7.32974156022896 | 19.8755526616174 | 7.50341355406529 | 6.56974917491458 | 3.66924669622158 | 16.6572494488982 | 20.6249857287016 | Plag1 |
| **ENSMUSG00000003283** | -0.508121933187969 | 0.297620604599674 | 0.851785577030345 | 7.22810973255236 | 19.5459774939439 | 19.8755526616174 | 15.9447538023887 | 25.1840385038392 | 35.4693847301419 | 9.36970281500523 | 19.5394601640331 | Hck |
| **ENSMUSG00000003299** | -0.0406249343824494 | 0.650028040283573 | 0.936463929813731 | 2172.95048834855 | 1898.4030640993 | 1909.94596529257 | 2146.91420315693 | 2104.5096523643 | 2181.97870201977 | 2031.14335467502 | 2042.95911270613 | Mrpl4 |
| **ENSMUSG00000003308** | -0.0902674914104052 | 0.292270008135371 | 0.851785577030345 | 1850.3960915334 | 1916.32021013542 | 1765.13836732935 | 1793.3158394216 | 1973.11466886601 | 1783.25389436369 | 2075.90971256894 | 1964.80127205 | Keap1 |
| **ENSMUSG00000003316** | 0.0735905191653443 | 0.502697718560509 | 0.900927059989498 | 4148.03147276849 | 5134.07675507593 | 5147.7681393589 | 4402.62790284781 | 4291.14116941504 | 4328.48801930939 | 4512.0324444703 | 4764.37170333008 | Glg1 |
| **ENSMUSG00000003344** | 0.0111331009194312 | 0.904762031385853 | 0.98504921011344 | 1036.33023290469 | 1012.31875104051 | 1019.33191507438 | 938.864620952419 | 1077.43886468599 | 1011.48900592508 | 960.915077583314 | 927.038832226905 | Btbd2 |
| **ENSMUSG00000003345** | 0.0609739821899681 | 0.459875144025609 | 0.894774044236998 | 2818.96279569542 | 2880.58843316998 | 2695.50352287078 | 2838.1661768252 | 2552.34755445431 | 2811.8660515378 | 2867.1290613916 | 2537.95877019497 | Csnk1g2 |
| **ENSMUSG00000003346** | -0.193648923825255 | 0.366429541858524 | 0.869754677013001 | 1053.49699351951 | 842.92027942633 | 910.489602879806 | 1095.49837889353 | 1129.99685808531 | 1575.32991491113 | 857.848346618257 | 900.986218674861 | Abhd17a |
| **ENSMUSG00000003348** | -0.591216923277669 | 0.149803434270812 | 0.782641594375979 | 32.5264937964856 | 30.9477976987445 | 44.483379766477 | 11.2551203310979 | 40.5134532453066 | 61.154111603693 | 48.9306702561384 | 29.3091902460497 | Mob3a |
| **ENSMUSG00000003352** | -0.967957018657422 | 0.0120762794699633 | 0.635862253526407 | 34.3335212296237 | 46.4216965481167 | 35.0188308799925 | 44.0825546301336 | 84.3117810780705 | 138.208292224346 | 48.9306702561384 | 42.3354970220718 | Cacnb3 |
| **ENSMUSG00000003355** | 0.366421792752027 | 0.130130468466875 | 0.77303252008843 | 80.412720774645 | 49.679359463774 | 58.6802030962037 | 76.9099889291692 | 51.4630352034976 | 51.3694537471021 | 58.3003730711437 | 44.5065481514088 | Fkbp11 |
| **ENSMUSG00000003360** | 0.226435023432843 | 0.120006485976572 | 0.769868040577702 | 2100.66939102303 | 2240.45767024332 | 2294.20665008384 | 2345.75466233966 | 1776.02219361858 | 1520.29121446781 | 2227.90711379013 | 2150.42614360831 | Ddx23 |
| **ENSMUSG00000003363** | -0.237870548279067 | 0.0979275616720837 | 0.759800953500741 | 655.950958229127 | 588.008156276145 | 604.784673846357 | 599.335157630965 | 736.906865786252 | 881.842289325253 | 669.413212227596 | 600.295637261685 | Pld3 |
| **ENSMUSG00000003378** | -0.188485340686899 | 0.414141822697254 | 0.882192126163777 | 98.4829951060259 | 73.2974156022896 | 74.7699362032273 | 117.24083678227 | 108.400861386091 | 125.977469903608 | 97.8613405122768 | 82.4999429148066 | Grik5 |
| **ENSMUSG00000003380** | -0.128520155798613 | 0.58429165844405 | 0.922450244597442 | 6533.30768451076 | 5116.97402476873 | 5976.86262181494 | 7273.62151397204 | 7445.71573156986 | 9661.12655115142 | 5009.66777175613 | 5105.22673063599 | Rabac1 |
| **ENSMUSG00000003382** | 0.0873992376143457 | 0.658472114274868 | 0.938989601151841 | 518.616873310632 | 672.707392083235 | 800.700835796586 | 620.907471598903 | 520.105143014071 | 494.125221757839 | 704.809867306505 | 739.242909539254 | Etv3 |
| **ENSMUSG00000003402** | -0.125924509518555 | 0.313042394303577 | 0.852747373382367 | 1171.85729039005 | 1036.75122290794 | 1145.21041526462 | 1051.4158242634 | 1283.29100549998 | 1371.0751821548 | 1116.03571307618 | 1037.76243982309 | Prkcsh |
| **ENSMUSG00000003418** | -0.66037145249796 | 0.0702370471864776 | 0.726006819419135 | 94.8689402397497 | 106.688460487777 | 74.7699362032273 | 81.59962240046 | 188.332809680885 | 221.377884005369 | 83.2862472444909 | 73.8157383974585 | St8sia6 |
| **ENSMUSG00000003420** | -0.340916075052618 | 0.114989584791006 | 0.767518180080285 | 1836.84338578487 | 1707.82978353335 | 1639.25986713911 | 1899.30155587278 | 2670.60303960278 | 2974.53598840363 | 1696.9572876065 | 1630.4593981321 | Fcgrt |
| **ENSMUSG00000003421** | -0.230627901527914 | 0.244165727441174 | 0.832048466795696 | 782.442878548793 | 684.92362801695 | 712.68053115228 | 904.161333264867 | 998.601874587016 | 1164.37428493431 | 790.178270732108 | 666.512696706464 | Nosip |
| **ENSMUSG00000003423** | -0.113438291327536 | 0.556999979986143 | 0.916239900758826 | 1062.5321306852 | 757.406627890326 | 823.415753124149 | 1026.09180351843 | 1039.11532783232 | 1270.78243912474 | 794.342583094332 | 866.249400605469 | Pih1d1 |
| **ENSMUSG00000003429** | 0.0753773251308539 | 0.69720485571849 | 0.944933386973863 | 11277.6582102148 | 7305.30908836153 | 8562.5773776025 | 11001.8801236482 | 8690.68320021617 | 11492.080652566 | 7655.04719985927 | 8368.31657802953 | Rps11 |
| **ENSMUSG00000003435** | 0.248504141005002 | 0.018835615703456 | 0.642273034804458 | 2749.3922395196 | 2830.09465797729 | 3043.7989218934 | 2618.69133036879 | 2339.92566446541 | 2096.3629457746 | 2618.31139774868 | 2406.61017687008 | Supt5 |
| **ENSMUSG00000003437** | -0.00119694655797321 | 0.990970808312212 | 0.998628446692557 | 1413.09545271399 | 1325.86880667253 | 1351.53758098998 | 1488.4896637877 | 1490.23810450979 | 1505.61422768292 | 1360.68906435687 | 1227.72941364008 | Paf1 |
| **ENSMUSG00000003438** | 0.00451575237948381 | 0.966393279801786 | 0.994284908685797 | 1637.16685442311 | 1426.04194132899 | 1405.48550964294 | 1595.41330693313 | 1631.48771177045 | 1594.89923062431 | 1416.9072812469 | 1402.49902955171 | Timm50 |
| **ENSMUSG00000003444** | 0.0270403091921549 | 0.895198806605834 | 0.983733637008786 | 438.204152535987 | 325.766291565731 | 365.3315870183 | 405.184331919526 | 390.900075907418 | 503.90987961443 | 333.144988977964 | 278.980070119806 | Med29 |
| **ENSMUSG00000003452** | 0.0853428587767733 | 0.750361739481243 | 0.956197192463153 | 76.7986659083688 | 78.1839099757755 | 55.8408384302584 | 61.9031618210386 | 65.6974917491458 | 37.9155491942896 | 81.2040910633787 | 71.6446872681215 | Bicd1 |
| **ENSMUSG00000003458** | -0.160053348448011 | 0.096973628320246 | 0.757809915356977 | 842.07478384235 | 842.105863697416 | 841.398396008469 | 791.610129953888 | 996.411958195378 | 952.781058785537 | 928.641656776074 | 829.34153140674 | Ncstn |
| **ENSMUSG00000003464** | 0.173621765011496 | 0.291353356293249 | 0.851785577030345 | 3603.21270167735 | 3809.83677986123 | 3459.29261801007 | 3260.23318924137 | 2745.06019691848 | 2433.93364182698 | 3663.55380066704 | 3686.44481761425 | Pex19 |
| **ENSMUSG00000003469** | -0.374337015662213 | 0.0386745215022045 | 0.679566075569698 | 358.694945477911 | 252.468875963442 | 220.523989055088 | 241.985087118606 | 354.766455445387 | 310.66288694676 | 360.213019332423 | 365.822115293287 | Phyhip |
| **ENSMUSG00000003476** | 0.0591353810284388 | 0.694961914187538 | 0.944799940106186 | 419.230364488037 | 457.701639649853 | 351.134763688574 | 327.336416296098 | 331.772333333186 | 384.047820871192 | 381.034581143546 | 397.302356668674 | Crhr2 |
| **ENSMUSG00000003477** | -0.3241597636493 | 0.330029193491664 | 0.855977678491063 | 224.974915425692 | 166.140808698523 | 321.794662140472 | 248.550573978413 | 321.917709570815 | 492.902139525765 | 186.352978209548 | 202.993280593011 | Inmt |
| **ENSMUSG00000003484** | -0.630076151815703 | 0.212453855669062 | 0.823370267313127 | 5.42108229941427 | 8.14415728914329 | 19.8755526616174 | 11.2551203310979 | 26.2789966996583 | 17.123151249034 | 13.5340151772298 | 11.9407812113536 | Cyp4f18 |
| **ENSMUSG00000003500** | -0.253708265005769 | 0.126372926518298 | 0.771949105062466 | 534.880120208875 | 516.339572131684 | 523.389553422591 | 570.259430108962 | 753.331238723539 | 748.526326029202 | 555.935700356977 | 500.427285312182 | Impdh1 |
| **ENSMUSG00000003518** | -0.125312108890308 | 0.335674793908021 | 0.858570960161243 | 3277.04424999593 | 3456.38035351241 | 2590.4470302308 | 2879.43495137256 | 3199.4678481834 | 2996.55146858096 | 3601.08911523368 | 3513.84625283196 | Dusp3 |
| **ENSMUSG00000003526** | -0.0241403864163762 | 0.864313512985393 | 0.974868041734905 | 396.642521573811 | 397.434875710192 | 375.742590793433 | 372.35689762049 | 344.911831683016 | 335.124531588238 | 454.951125573032 | 432.039174738066 | Prodh |
| **ENSMUSG00000003527** | -0.0744309017833602 | 0.608545889379727 | 0.927932112008768 | 211.422209677157 | 179.985876090067 | 196.862616838877 | 204.468019348279 | 244.175677667659 | 198.139321595965 | 208.215618111227 | 183.453820428978 | Dgcr14 |
| **ENSMUSG00000003528** | -0.116412254374467 | 0.61133688221553 | 0.927932112008768 | 122.87786545339 | 179.985876090067 | 146.700507740509 | 199.778385876988 | 189.427767876704 | 222.600966237442 | 130.134761319517 | 162.828834700276 | Slc25a1 |
| **ENSMUSG00000003531** | -0.0374198897326389 | 0.833867202043428 | 0.96835936700622 | 983.022923627121 | 769.622863824041 | 953.080072868986 | 1079.55362509114 | 963.563212320805 | 1245.09771225119 | 873.464517976599 | 803.288917854696 | Dgcr6 |
| **ENSMUSG00000003534** | -0.0705570594486365 | 0.597882502594599 | 0.924496535139492 | 589.090943203017 | 685.738043745865 | 728.770264259304 | 590.893817382642 | 681.063997799478 | 731.403174780168 | 731.877897660964 | 580.756177097652 | Ddr1 |
| **ENSMUSG00000003541** | -0.487587009125249 | 0.0271890487710192 | 0.664353746352133 | 630.652574165193 | 492.721515993169 | 486.477812765302 | 695.941607139556 | 988.747250824645 | 984.581196819457 | 666.289977955927 | 593.782483873674 | Ier3 |
| **ENSMUSG00000003545** | 0.807136851143999 | 0.387638799745241 | 0.875951433019341 | 65.9565013095403 | 7.32974156022896 | 4.73227444324223 | 5.62756016554897 | 19.7092475247437 | 20.7923979452556 | 3.12323427166841 | 4.34210225867403 | Fosb |
| **ENSMUSG00000003546** | -0.0399317440460758 | 0.690213701104261 | 0.94469184542218 | 833.03964667666 | 798.127414336042 | 746.752907143624 | 841.320244749571 | 771.945528052463 | 809.680437632895 | 911.984407327176 | 815.229699066049 | Klc4 |
| **ENSMUSG00000003549** | -0.161268760744586 | 0.482065309672876 | 0.897732059433541 | 416.51982333833 | 329.838370210303 | 408.868511896129 | 427.694572581722 | 477.401773377126 | 626.218102821816 | 334.18606706852 | 333.256348353232 | Ercc1 |
| **ENSMUSG00000003559** | 0.185945415244605 | 0.184629194906823 | 0.805963874499992 | 1591.08765487809 | 1423.59869414225 | 1527.57819027859 | 1350.61443973175 | 1138.75652365186 | 1166.82044939846 | 1285.73144183683 | 1588.12390111003 | As3mt |
| **ENSMUSG00000003573** | -0.387041726517673 | 0.0335165796265247 | 0.671782526713461 | 99.3865088225949 | 117.275864963663 | 115.467496415111 | 113.489130005238 | 171.908436743598 | 165.116101329971 | 120.765058504512 | 125.920965501547 | Homer3 |
| **ENSMUSG00000003575** | 0.176256659837772 | 0.384853717057939 | 0.875126835008975 | 207.80815481088 | 238.623808571898 | 269.739643264807 | 151.006197775564 | 198.187433443257 | 170.008430258266 | 186.352978209548 | 212.763010675027 | Crtc1 |
| **ENSMUSG00000003581** | -0.249420751299319 | 0.284599660843017 | 0.851125503777472 | 156.307872966445 | 124.605606523892 | 142.914688185915 | 189.461192240149 | 215.706764576362 | 231.162541861959 | 144.709854587303 | 137.8617467129 | Rnf215 |
| **ENSMUSG00000003585** | -0.364355287234198 | 0.365844919606768 | 0.869253398153585 | 48.7897406947284 | 112.389370590177 | 52.0550188756646 | 77.8479156234274 | 67.887408140784 | 72.1618516923577 | 63.5057635239243 | 171.513039217624 | Sec14l2 |
| **ENSMUSG00000003604** | 0.0543460587174686 | 0.733284231045503 | 0.953733898884637 | 547.529312240841 | 452.000729547452 | 506.353365426919 | 608.714424573547 | 535.434557755538 | 609.094951572782 | 451.827891301363 | 440.723379255414 | Aven |
| **ENSMUSG00000003617** | 0.0969623607297199 | 0.530061927655316 | 0.90873366762927 | 1315.51597132453 | 1543.31780629265 | 1861.67676597149 | 1798.0054728929 | 1354.46328822822 | 1731.88444061659 | 1614.71211845257 | 1393.81482503436 | Cp |
| **ENSMUSG00000003623** | 0.0468586582892129 | 0.755947564247361 | 0.956566265296272 | 932.426155499254 | 1038.38005436577 | 992.831178192221 | 928.54742731558 | 796.034608360484 | 818.242013257412 | 985.900951756661 | 1166.93998201865 | Crot |
| **ENSMUSG00000003644** | -0.433895699395102 | 0.0317078127111126 | 0.671606027118506 | 128.298947752804 | 97.7298874697194 | 117.360406192407 | 144.440710915757 | 170.813478547779 | 206.700897220482 | 137.42230795341 | 144.374900100912 | Rps6ka1 |
| **ENSMUSG00000003660** | 0.155492985950842 | 0.2091771596189 | 0.821566820583092 | 2425.93432898789 | 2769.01347830872 | 2822.32847794967 | 2344.8167356454 | 2349.78028822778 | 1987.50862712002 | 2578.75043030755 | 2385.98519114138 | Snrnp200 |
| **ENSMUSG00000003662** | -0.0901822547527273 | 0.597282847646765 | 0.924496535139492 | 913.452367451305 | 851.064436715473 | 747.699362032273 | 857.264998551959 | 1022.69095489504 | 1067.75078860048 | 821.410613448792 | 676.28242678848 | Ciao1 |
| **ENSMUSG00000003680** | -0.189316311933118 | 0.288758067960395 | 0.851785577030345 | 264.72951895473 | 228.850819824926 | 225.25626349833 | 188.52326554589 | 282.499214521327 | 309.439804714686 | 229.03717992235 | 214.934061804364 | Taf6l |
| **ENSMUSG00000003721** | 0.134640206168192 | 0.397582175656234 | 0.88020150974241 | 1308.28786159198 | 1397.53739081699 | 1378.98477276079 | 1304.6560317131 | 1020.5010385034 | 1011.48900592508 | 1385.67493853022 | 1490.42660028986 | Insig2 |
| **ENSMUSG00000003731** | 0.129281208802114 | 0.581506049091823 | 0.922450244597442 | 1322.74408105708 | 1623.94496345517 | 1563.54347604723 | 1176.16007459973 | 992.032125412102 | 914.865509591247 | 1601.17810327534 | 1690.16330418887 | Kpna6 |
| **ENSMUSG00000003746** | -0.168965147254841 | 0.287016937876375 | 0.851785577030345 | 838.460728976074 | 1153.21267214269 | 887.774685552243 | 773.789522762983 | 1069.77415731526 | 983.358114587383 | 1142.06266534008 | 912.926999886215 | Man1a |
| **ENSMUSG00000003752** | 0.278400062634884 | 0.306169637774873 | 0.851785577030345 | 55.1143367107117 | 111.574954861263 | 110.735221971868 | 77.8479156234274 | 73.3621991198795 | 58.7079471395453 | 75.998700610598 | 84.6709940441436 | Itpkc |
| **ENSMUSG00000003762** | 0.176936231169937 | 0.209672686634075 | 0.821566820583092 | 250.273299489625 | 261.427448981499 | 225.25626349833 | 237.295453647315 | 234.321053905287 | 238.501035254403 | 182.188665847324 | 208.420908416353 | Coq8b |
| **ENSMUSG00000003778** | 0.0412542969527342 | 0.782740195014235 | 0.960786612650028 | 346.949267162513 | 388.476302692135 | 438.208613444231 | 415.501525556365 | 409.514365236342 | 336.347613820311 | 452.868969391919 | 344.111603999917 | Brd8 |
| **ENSMUSG00000003779** | 0.218202993447923 | 0.512444141908778 | 0.902707562601512 | 25.2983840639333 | 35.8342920722305 | 17.9826428843205 | 29.075727522003 | 25.1840385038392 | 28.1308913376988 | 21.8626399016789 | 18.4539345993646 | Kif20a |
| **ENSMUSG00000003808** | -0.106673962221657 | 0.38720500555487 | 0.875798327171006 | 369.537110076739 | 341.240190415104 | 334.098575692902 | 339.529463321454 | 433.603445544362 | 360.809258461789 | 343.555769883525 | 352.795808517265 | Farsa |
| **ENSMUSG00000003809** | 0.0917873550293351 | 0.375409422498843 | 0.873821423608136 | 2060.01127377742 | 1803.11642381632 | 1820.03275087096 | 2173.17615059616 | 1865.80876567574 | 1761.23841418636 | 1782.32569103211 | 1961.54469535599 | Gcdh |
| **ENSMUSG00000003810** | 0.0956491167021398 | 0.591257498835063 | 0.923935529355527 | 3190.3069332053 | 3624.96440939768 | 3396.82659535928 | 2913.20031236585 | 2609.28538063691 | 2407.02583272136 | 3903.00176149496 | 3362.95819934304 | Mast2 |
| **ENSMUSG00000003812** | 0.17802084731228 | 0.361215268651708 | 0.868539734006824 | 155.404359249876 | 121.347943608235 | 122.09268063565 | 139.751077444466 | 131.394983498292 | 145.546785616789 | 94.7381062406084 | 105.295979772845 | Dnase2a |
| **ENSMUSG00000003813** | 0.118507593942562 | 0.162261343358474 | 0.793874624603454 | 2236.19644850839 | 2263.26131065292 | 2390.74504872598 | 2347.63051572818 | 1964.35500329946 | 2059.67047881238 | 2251.85190987292 | 2231.84056095845 | Rad23a |
| **ENSMUSG00000003814** | -0.14893502401666 | 0.296261875757758 | 0.851785577030345 | 6930.85371980114 | 6073.09809051415 | 5393.8464104075 | 6328.19140615981 | 7247.5282981266 | 8034.42718249318 | 5779.02448067711 | 6355.75218113411 | Calr |
| **ENSMUSG00000003824** | -0.2027968276216 | 0.686646238686351 | 0.944452327903597 | 23.4913566307952 | 11.4018202048006 | 16.0897331070236 | 17.8206071909051 | 18.6142893289246 | 19.5693157131818 | 6.24646854333682 | 34.7368180693922 | Syce2 |
| **ENSMUSG00000003847** | -0.0504377395275429 | 0.694066255041998 | 0.94469184542218 | 348.756294595651 | 430.011504866766 | 406.975602118832 | 357.350070512359 | 421.558905390352 | 360.809258461789 | 429.965251399684 | 385.36157545732 | Nfat5 |
| **ENSMUSG00000003848** | 0.17443059913379 | 0.198563462785928 | 0.812921911423739 | 526.748496759753 | 529.370223794314 | 618.981497176084 | 633.100518624259 | 571.568178217569 | 516.140701935169 | 432.047407580797 | 525.394373299558 | Nob1 |
| **ENSMUSG00000003849** | -0.0213959933651434 | 0.89198543268453 | 0.982421229269187 | 2037.4234308632 | 2128.06829965314 | 1378.98477276079 | 1883.35680207039 | 1781.49698459767 | 1772.24615427502 | 2045.71844794281 | 1939.83418406262 | Nqo1 |

| | | | | | | | | | | | | |
|---|---|---|---|---|---|---|---|---|---|---|---|---|
| ENSMUSG00000003865 | 0.209863327377712 | 0.374325796133345 | 0.873700364743909 | 2602.11950371885 | 3260.92057857297 | 4244.85017558828 | 2656.20839813911 | 2542.49293069194 | 1887.21588408997 | 3564.65138206421 | 3040.55710663649 | Gys1 |
| ENSMUSG00000003868 | -0.144810430128172 | 0.30237782878785 | 0.851785577030345 | 575.538237454482 | 475.618785685968 | 430.636974335043 | 534.618215727152 | 616.461464246152 | 587.079471395453 | 494.512093014165 | 531.907526687569 | Ruvbl2 |
| ENSMUSG00000003872 | 0.0799059735835345 | 0.851648489157078 | 0.972732497553372 | 21.6843291976571 | 13.0306516626293 | 14.1968233297267 | 16.8826804966469 | 6.56974917491458 | 19.5693157131818 | 17.6983275394543 | 18.4539345993646 | Lin7b |
| ENSMUSG00000003873 | -0.273937459583166 | 0.264392602605785 | 0.842510961779757 | 400.256576440087 | 354.270842077733 | 387.100049457215 | 513.045901759214 | 490.541271726955 | 746.080161565054 | 388.322127777439 | 376.677370939972 | Bax |
| ENSMUSG00000003923 | 0.136277180477509 | 0.448091290233436 | 0.89108031640712 | 1030.90915060528 | 1039.19447009468 | 930.365155541423 | 980.133395499779 | 808.079148514494 | 660.464405319884 | 1029.62623156002 | 1122.43343386724 | Tfam |
| ENSMUSG00000003929 | -0.135375534242341 | 0.566833421934954 | 0.919931558693298 | 68.6670424592474 | 64.338842584232 | 82.3415753124149 | 83.4754757889764 | 75.5521155115177 | 86.838838477244 | 58.3003730711437 | 107.467030902182 | Zfp81 |
| ENSMUSG00000003934 | 0.0200843453490879 | 0.924940332359115 | 0.986713918026868 | 2611.15464088454 | 3543.52283650624 | 2264.86654853573 | 2031.54921976318 | 2138.4533564347 | 2188.09411318014 | 3020.16754070335 | 2960.22821485102 | Efnb3 |
| ENSMUSG00000003948 | -0.146963646720471 | 0.476494000663189 | 0.897553735056661 | 418.326850771468 | 443.856572258309 | 590.587850516631 | 681.872706725683 | 564.998429042654 | 480.671317205027 | 591.332355435886 | 725.131077198563 | Mmd |
| ENSMUSG00000003949 | 0.00243386594063083 | 0.990300973063105 | 0.998628446692557 | 449.046317134815 | 457.701639649853 | 300.026199701558 | 391.115431505653 | 419.368988998714 | 289.870489001505 | 438.293876124134 | 447.236532643425 | Hlf |
| ENSMUSG00000003955 | 0.0166673240223441 | 0.920341764840181 | 0.986713918026868 | 2912.02470850203 | 2192.40714223737 | 2195.7753416644 | 2997.61371484908 | 2618.04504620346 | 3016.12078429414 | 2214.3730986129 | 2331.70891290795 | Fam162a |
| ENSMUSG00000003970 | -0.0440827171663896 | 0.78212772202788 | 0.960742105865155 | 8680.95978879538 | 6894.84356098871 | 8126.26167393556 | 9421.47364382323 | 8972.08745654168 | 10339.9371899524 | 7406.22953621636 | 7433.67906684994 | Rpl8 |
| ENSMUSG00000003992 | 0.0786787175601927 | 0.54946413952498 | 0.914699131434573 | 446.335775985108 | 513.896324944941 | 507.299820315568 | 414.563598862107 | 420.463947194533 | 415.847958905112 | 509.087186281951 | 436.38127699674 | Ssbp2 |
| ENSMUSG00000004018 | 0.0326233111646586 | 0.874199168387017 | 0.97671366425653 | 117.456783153976 | 81.4415728914329 | 102.217127974032 | 98.4823028971069 | 117.160526952643 | 78.277262852727 | 106.189965236726 | 87.9275707381491 | Fancl |
| ENSMUSG00000004032 | -0.0844976882501474 | 0.613739020793985 | 0.928075610917191 | 1145.65539260955 | 886.08431305879 | 994.724087969518 | 1161.1532474916 | 1171.60526952643 | 1367.40593545857 | 935.929203409967 | 966.117752554972 | Gstm5 |
| ENSMUSG00000004035 | -0.0719520218235896 | 0.447730470936262 | 0.89108031640712 | 1010.12833512419 | 900.743796179247 | 1010.81382107654 | 884.464872685446 | 974.512794278996 | 976.01962119494 | 1054.61210573337 | 994.341417236353 | Gstm7 |
| ENSMUSG00000004040 | -0.107890779424465 | 0.537469587184379 | 0.912099403004057 | 1590.18414116152 | 2335.74431052629 | 1914.67823973581 | 1538.19977858338 | 1986.25416721584 | 1718.43053606377 | 2349.7132503852 | 1897.49868704055 | Stat3 |
| ENSMUSG00000004043 | 0.19068129339757 | 0.40520660533543 | 0.881158207349486 | 288.220875585525 | 340.425774686189 | 315.169477919933 | 252.302280755445 | 260.600050604945 | 162.669936865823 | 340.432535611857 | 283.322172378481 | Stat5a |
| ENSMUSG00000004044 | -0.269082357814405 | 0.118324585417338 | 0.769868040577702 | 4621.47266025067 | 4462.99819445052 | 4944.28033829949 | 5698.84259431259 | 7401.91740373709 | 6465.21267874242 | 4723.37129685319 | 5182.29904572746 | Cavin1 |
| ENSMUSG00000004054 | -0.51456158385584 | 0.030941597185064 | 0.667184591665619 | 669.503663977662 | 705.284021239809 | 756.217456030109 | 739.086235075431 | 1352.27337183658 | 1297.69024823037 | 772.479943192653 | 678.453477917817 | Map3k11 |
| ENSMUSG00000004056 | 0.128538608190013 | 0.303498028990753 | 0.851785577030345 | 4854.57919912548 | 5341.75276594908 | 6341.24775394459 | 4989.77001345342 | 4686.42107810573 | 4500.9426140318 | 5365.71647872633 | 5137.79249757605 | Akt2 |
| ENSMUSG00000004069 | 0.18514411378672 | 0.311767866621242 | 0.851785577030345 | 4138.09282188623 | 3957.24602679472 | 3815.15965614189 | 3853.94078670678 | 3276.11492189074 | 2383.78727031195 | 4094.56013015729 | 4109.79978783497 | Dnaja3 |
| ENSMUSG00000004070 | -0.0250696729577321 | 0.812345040940651 | 0.965224347191244 | 1389.60409608319 | 1250.12814388349 | 1421.57524274997 | 1461.28978965422 | 1517.61205940527 | 1283.01326144548 | 1487.70059140472 | 1328.68329115425 | Hmox2 |
| ENSMUSG00000004071 | 0.111362630964769 | 0.328846597901066 | 0.855977678491063 | 1527.84169471826 | 1603.58457023231 | 1548.40019782886 | 1501.62063750732 | 1324.89941694111 | 1245.09771225119 | 1589.72624427922 | 1560.98576199331 | Cdip1 |
| ENSMUSG00000004085 | 0.308402364752373 | 0.319459623467555 | 0.854670142081901 | 4099.24173207376 | 5629.24151825584 | 4943.33388341084 | 3907.4026082795 | 2635.56437733657 | 1994.84712051247 | 5236.62279549737 | 5135.62144644671 | Map3k20 |
| ENSMUSG00000004096 | 0.0383229774846633 | 0.762754178055692 | 0.957842155335615 | 1590.18414116152 | 1308.76607636533 | 1465.1121676278 | 1498.80685742454 | 1481.47843894324 | 1651.16101329971 | 1310.71731601018 | 1266.80833396815 | Cwc15 |
| ENSMUSG00000004098 | -1.05007796865674 | 0.00741670897499749 | 0.635862253526407 | 143.658680934478 | 151.481325578065 | 127.77140996754 | 137.87522405595 | 465.357233223116 | 413.401794440965 | 130.134761319517 | 153.05910461826 | Col5a3 |
| ENSMUSG00000004099 | -0.184510844849521 | 0.19383913995595 | 0.809162446871198 | 271.957628687283 | 311.921224174188 | 262.16800415562 | 260.743621003769 | 372.285786578493 | 286.201242305283 | 289.419709174606 | 310.460311495193 | Dnmt1 |
| ENSMUSG00000004100 | -0.0955874608892033 | 0.67304735030127 | 0.94344814771265 | 96.6759676728878 | 99.3587189275481 | 77.6093008691726 | 131.309737196143 | 113.875652365186 | 133.315963296051 | 86.4094815161593 | 99.8683519495027 | Ppan |
| ENSMUSG00000004105 | -0.486077469156709 | 0.0093611047250085 | 0.635862253526407 | 622.520950716072 | 765.550785179469 | 574.498117409607 | 570.259430108962 | 855.162350934715 | 1169.26661386261 | 802.671207818781 | 722.960026069226 | Angptl2 |
| ENSMUSG00000004127 | 0.188475245309178 | 0.443176493409222 | 0.89108031640712 | 50.5967681278665 | 39.0919549878878 | 51.1085639870161 | 50.6480414899407 | 38.3235368536684 | 33.0232202659942 | 44.7663578939139 | 51.0197015394199 | Trmt10a |
| ENSMUSG00000004151 | -0.191760658990513 | 0.399159412190189 | 0.880459952732461 | 48.7897406947284 | 66.7820897709749 | 84.2344850897118 | 80.6616957062019 | 84.3117810780705 | 89.2850029413918 | 71.8343882483734 | 74.901263962127 | Etv1 |
| ENSMUSG00000004187 | -0.0198792672245871 | 0.951857452725552 | 0.991915011884503 | 38.8510898124689 | 52.122606650517 | 35.965285768641 | 40.3308478531009 | 27.3739548954774 | 28.1308913376988 | 55.1771387994752 | 58.6183804920994 | Kifc2 |
| ENSMUSG00000004207 | 0.00928840797771121 | 0.939415029657633 | 0.989269716906553 | 38355.0607820725 | 44941.9031686794 | 48708.3543894037 | 36771.4160483912 | 42732.9335082319 | 39081.146561456 | 44347.8445015103 | 41531.1225786524 | Psap |
| ENSMUSG00000004221 | 0.0721007723455927 | 0.700331002954862 | 0.94561797945347 | 846.592352425195 | 972.412380323708 | 944.56197887115 | 787.858423176855 | 752.23628052772 | 613.987280501078 | 1003.59927929612 | 1007.36772401237 | Ikbkg |
| ENSMUSG00000004233 | 0.174796723173301 | 0.388654560683336 | 0.876405748044371 | 140.948139784771 | 144.151584017836 | 144.807597963212 | 145.378637610015 | 105.115986798633 | 94.1773318696872 | 157.202791673977 | 151.973579053591 | Wars2 |
| ENSMUSG00000004263 | 0.0148575839178005 | 0.914050831659639 | 0.986339811354287 | 250.273299489625 | 225.593156909269 | 258.382184601026 | 246.674720589896 | 277.024423542232 | 244.616446414772 | 242.57119509958 | 206.249857287016 | Atn1 |
| ENSMUSG00000004264 | 0.159258111949883 | 0.105495181788531 | 0.762094734041179 | 8927.61903341873 | 7280.06220076518 | 8472.6641631809 | 8674.88399519373 | 7859.60992958948 | 7611.24073019563 | 7294.83418052685 | 7102.59376962605 | Phb2 |
| ENSMUSG00000004266 | -0.0379185929515236 | 0.850868849220338 | 0.972692784625411 | 83.1232619243521 | 89.5857301805761 | 105.056492639978 | 83.4754757889764 | 100.736154015357 | 113.746647582869 | 82.2451691539348 | 74.901263962127 | Ptpn6 |
| ENSMUSG00000004268 | 0.017798722943534 | 0.888711323791571 | 0.981177989624427 | 1423.03410359625 | 1149.14059349812 | 1177.38988147867 | 1413.45552824705 | 1285.48092189162 | 1424.89080036605 | 1150.39129006453 | 1239.67019485144 | Emg1 |
| ENSMUSG00000004270 | 0.156429726659656 | 0.415247074870427 | 0.882192126163777 | 883.636414804526 | 1170.31540244989 | 1502.02390828509 | 951.995594672034 | 1061.0144917487 | 884.2884537894 | 1135.81619679675 | 962.861175860966 | Lpcat3 |
| ENSMUSG00000004285 | -0.0454227653180892 | 0.821857128882756 | 0.966637987288264 | 1666.98280706989 | 1172.75864963663 | 1311.78647566675 | 1622.61318106662 | 1469.43389878923 | 2002.18561390491 | 1217.02028786012 | 1271.15043622682 | Atp6v1f |
| ENSMUSG00000004317 | -0.274755636574976 | 0.488759242198245 | 0.899236955296196 | 18.0702743313809 | 13.0306516626293 | 23.6613722162112 | 17.8206071909051 | 35.0386622662111 | 18.3462334811079 | 14.5750932677859 | 19.5394601640331 | Clcn5 |
| ENSMUSG00000004319 | -0.00456797309392702 | 0.984214442754208 | 0.996940244108351 | 623.424464432641 | 746.004807685525 | 567.872933189068 | 565.569796637671 | 536.529515951357 | 407.286383280595 | 791.219348822664 | 775.065253173314 | Clcn3 |
| ENSMUSG00000004328 | 0.452652505400582 | 0.256985131402018 | 0.837517407767513 | 40.658117245607 | 49.679359463774 | 51.1085639870161 | 16.8826804966469 | 39.4184950494875 | 33.0232202659942 | 20.8215618111227 | 22.7960368580387 | Hif3a |
| ENSMUSG00000004347 | 0.121697344119285 | 0.682171086873752 | 0.944452327903597 | 677.635287426784 | 853.507683902216 | 686.179794270124 | 481.156394154437 | 506.965644664242 | 340.016860516533 | 839.108940988246 | 793.519187772679 | Pde1c |
| ENSMUSG00000004356 | 0.105941064824621 | 0.54924260320976 | 0.914699131434573 | 314.422773366028 | 338.796943228361 | 239.453086828057 | 268.247034557834 | 292.353838283699 | 215.262472844999 | 309.200192895173 | 261.61166108511 | Utp20 |
| ENSMUSG00000004364 | 0.25571647910913 | 0.150295029435203 | 0.782641594375979 | 4014.31144271627 | 4239.84828472799 | 4582.73457083578 | 3774.21701769484 | 3174.28380967956 | 2534.22638485704 | 4075.82072452727 | 4127.16819686967 | Cul3 |
| ENSMUSG00000004383 | 0.212544933152559 | 0.409097756026445 | 0.882192126163777 | 1699.50930086637 | 1913.06254721976 | 2177.79269878008 | 1798.94339958715 | 1386.21707590698 | 931.988660840281 | 2162.3191940851 | 2067.92620069351 | Large1 |
| ENSMUSG00000004393 | 0.113439575578051 | 0.284603339625448 | 0.851125503777472 | 509.581736144941 | 477.247617143797 | 528.121827865833 | 550.562969529541 | 511.345477447518 | 490.455975061618 | 466.402984569149 | 440.723379255414 | Ddx56 |
| ENSMUSG00000004394 | 0.0471574655739914 | 0.529420893847505 | 0.908560090446144 | 1954.30016893884 | 2058.02854696651 | 2048.12837903524 | 1934.94277025459 | 1912.89196809596 | 1898.22362417863 | 2051.96491648615 | 1874.70265018251 | Tmed4 |
| ENSMUSG00000004446 | -0.128323834108201 | 0.730139491643786 | 0.952921973914044 | 10.8421645988285 | 22.8036404096012 | 15.1432782183751 | 19.6964605794214 | 18.6142893289246 | 15.9000690169602 | 18.7394056300105 | 21.7105112933701 | Bid |
| ENSMUSG00000004451 | -0.127197704073786 | 0.294842853814686 | 0.851785577030345 | 544.818771091134 | 483.762942975111 | 475.12035410152 | 550.562969529541 | 598.942133113046 | 596.864129252044 | 485.14239019916 | 563.387768062955 | Ralb |
| ENSMUSG00000004455 | 0.0842406357700336 | 0.464385029043326 | 0.895328112556867 | 1562.17521594788 | 1540.060143377 | 1676.1716077964 | 1525.06880486377 | 1384.02715951534 | 1307.47490608696 | 1701.12159996873 | 1551.2160319113 | Ppp1cc |
| ENSMUSG00000004460 | -0.00441067977166699 | 0.974344862261927 | 0.995510898988495 | 1365.20922573583 | 1184.16046984143 | 1131.01359193489 | 1387.19358080782 | 1343.51370627003 | 1453.02169170375 | 1079.59797990671 | 1208.18995347605 | Dnajb11 |
| ENSMUSG00000004462 | 0.0113323769742829 | 0.948609840888125 | 0.991233613135933 | 156.307872966445 | 184.057954734638 | 168.468970179424 | 127.55803041911 | 146.724398239759 | 141.877538920568 | 185.311900118992 | 157.401206876934 | Tbccd1 |
| ENSMUSG00000004473 | -0.733314611194345 | 0.164559637579949 | 0.795216960550936 | 13.5527057485357 | 18.7315617650296 | 25.5542819935081 | 9.37926694258161 | 24.0890803080201 | 57.4848649074714 | 15.616171358342 | 15.1973579053591 | Clec11a |
| ENSMUSG00000004500 | -0.408983753727001 | 0.0483900621063775 | 0.703396849069134 | 42.4651446787451 | 44.7928650902881 | 48.2691993210708 | 44.0825546301336 | 58.0327843784121 | 63.6002760678407 | 55.1771387994752 | 61.8749571861049 | Zfp324 |
| ENSMUSG00000004508 | 0.153302717738564 | 0.365348875544593 | 0.86924083181804 | 683.959883442767 | 707.727268426552 | 755.271001141461 | 721.265627884526 | 579.232885588302 | 483.117481669175 | 771.438865102097 | 743.585011797928 | Gab2 |
| ENSMUSG00000004530 | -0.470985088261864 | 0.0118478993688263 | 0.635862253526407 | 415.616309621761 | 397.434875710192 | 398.457508120996 | 386.425798034362 | 687.633746974393 | 651.902829695367 | 422.677704765791 | 453.749686031436 | Coro1c |
| ENSMUSG00000004535 | 0.155911449819554 | 0.292964282838078 | 0.851785577030345 | 3069.23609518505 | 3360.27929750052 | 3864.37531035161 | 2914.13823906011 | 2746.1551551143 | 2499.98008235897 | 3383.50379430744 | 3224.01092706547 | Tax1bp1 |
| ENSMUSG00000004552 | 1.05795996856752 | 0.295896668741171 | 0.851785577030345 | 10.8421645988285 | 1.62883145782866 | 39.7511053232348 | 2.81378008277448 | 7.66470737073368 | 14.6769867848863 | 3.12323427166841 | 1.08552556466851 | Ctse |
| ENSMUSG00000004558 | 0.180634307592702 | 0.160149145475913 | 0.791063880035713 | 50968.1122653764 | 51699.9248872105 | 51227.8173029858 | 51499.6789283271 | 41042.3180538872 | 37953.4647434839 | 52610.8813062544 | 49615.0314587388 | Ndrg2 |
| ENSMUSG00000004561 | -0.0232622491056012 | 0.867867683778036 | 0.974868041734905 | 215.939778260002 | 245.139134403213 | 229.042083052924 | 211.971432902344 | 263.884925192402 | 209.14706168463 | 237.365804646799 | 206.249857287016 | Mettl17 |
| ENSMUSG00000004562 | -0.354139098308812 | 0.131520029532811 | 0.77303252008843 | 257.501409222178 | 287.488752306758 | 282.990011705886 | 339.529463321454 | 443.458069306734 | 499.017550686135 | 294.625099627387 | 256.184033261768 | Arhgef40 |
| ENSMUSG00000004565 | -0.273325512502876 | 0.113652004669902 | 0.767518180080285 | 116.553269437407 | 138.450673915436 | 148.593417517806 | 130.371810501884 | 187.237851485066 | 177.34692365071 | 146.792010768415 | 134.605170018895 | Pnpla6 |
| ENSMUSG00000004567 | 0.0411530042827325 | 0.741868412970556 | 0.954383574136061 | 543.011743657996 | 567.647763053287 | 701.323072488499 | 634.038445318517 | 579.232885588302 | 629.887349518038 | 620.482541971457 | 547.104884592928 | Mcoln1 |
| ENSMUSG00000004568 | 0.107464798322105 | 0.497784238039267 | 0.900927059989498 | 352.370349461928 | 452.000729547452 | 408.868511896129 | 408.936038696558 | 428.128654565267 | 291.093571233579 | 388.322127777439 | 397.302356668674 | Arhgef18 |
| ENSMUSG00000004591 | 0.108053276779934 | 0.503685988013671 | 0.900927059989498 | 952.303457263773 | 1136.9243575644 | 1151.83559948516 | 849.761584997894 | 858.447225522172 | 801.118862008378 | 1065.02288663893 | 1070.32820676315 | Pkn2 |
| ENSMUSG00000004609 | 0.0531270935476162 | 0.865491849730002 | 0.974868041734905 | 37.9475760958999 | 44.7928650902881 | 75.7163910918757 | 41.2687745473591 | 49.2731188118594 | 63.6002760678407 | 34.3555769883525 | 45.5920737160773 | Cd33 |
| ENSMUSG00000004610 | -0.0614297103846671 | 0.776190138164343 | 0.959731259983925 | 9000.80364446083 | 6595.95298847715 | 7590.56820696054 | 9498.3836327524 | 9031.21519911591 | 11636.4043559507 | 6512.98453451919 | 6927.82415371441 | Etfb |
| ENSMUSG00000004626 | 0.0222420488085971 | 0.943188576867525 | 0.990505437194639 | 19.877301764519 | 25.2468875963442 | 28.3936466594534 | 19.6964605794214 | 19.7092475247437 | 23.2385624094033 | 26.0269522639034 | 22.7960368580387 | Stxbp2 |
| ENSMUSG00000004631 | -0.564180331435309 | 0.0719090240488324 | 0.728531250210272 | 200.580045078328 | 177.542628903324 | 175.094154399963 | 204.468019348279 | 363.52612101194 | 417.071041137186 | 186.352978209548 | 154.144630182928 | Sgce |
| ENSMUSG00000004633 | -0.661961385473675 | 0.0488137124881874 | 0.703396849069134 | 28.0089252136404 | 33.3910448854875 | 39.7511053232348 | 31.8895076047775 | 58.0327843784121 | 80.7234273168747 | 30.191264626128 | 42.3354970220718 | Chn2 |
| ENSMUSG00000004637 | 0.283012050065988 | 0.155966207486262 | 0.787275540343856 | 147.272735800754 | 152.295741306979 | 110.735221971868 | 99.4202295913651 | 95.2613630362614 | 108.854318654574 | 113.477511870619 | 102.03940307884 | Wwox |
| ENSMUSG00000004642 | -0.358723048160286 | 0.114649453984719 | 0.767518180080285 | 119.263810587114 | 91.2145616384048 | 106.949402417274 | 145.378637610015 | 174.098353135236 | 168.785348026193 | 105.14888714617 | 145.46042566558 | Slbp |
| ENSMUSG00000004655 | -0.678270334580437 | 0.0517854662790855 | 0.703396849069134 | 7133.24079231261 | 4613.66510429967 | 5257.55690644212 | 6905.95424982284 | 17005.7957392664 | 10061.0744410396 | 5668.67020307816 | 5526.41064972737 | Aqp1 |
| ENSMUSG00000004661 | 0.355149665291884 | 0.301077312144844 | 0.851785577030345 | 50.5967681278665 | 42.3496179035451 | 45.4298346551254 | 38.4549944645846 | 26.2789966996583 | 19.5693157131818 | 34.3555769883525 | 57.5328549274309 | Arid3b |
| ENSMUSG00000004665 | -0.391714729573925 | 0.0083631015739748 | 0.635862253526407 | 625.231491865779 | 612.440628143575 | 654.946782944725 | 636.852225401292 | 918.669926292222 | 985.804279051531 | 743.329756657082 | 671.940324529806 | Cnn2 |
| ENSMUSG00000004667 | 0.148373548833303 | 0.29332231434802 | 0.851785577030345 | 3180.36828232304 | 2545.86356858619 | 2524.19518802541 | 3170.19222659259 | 2840.32155995474 | 2825.31995609062 | 2291.41287731406 | 2347.99179637798 | Polr2e |
| ENSMUSG00000004677 | 0.348960770949047 | 0.036477771587406 | 0.679566075569698 | 385.800356974982 | 500.865673282312 | 445.780252553418 | 412.687745473591 | 294.543754675337 | 289.870489001505 | 371.664878328541 | 413.585240138701 | Myo9b |
| ENSMUSG00000004698 | -0.113588541762285 | 0.674141910219209 | 0.943983909911292 | 136.430571201926 | 174.284965987666 | 142.914688185915 | 202.592165959763 | 163.148771177045 | 101.51582526213 | 219.667477107345 | 224.703791886381 | Hdac9 |
| ENSMUSG00000004730 | -0.338797452876684 | 0.141836886168882 | 0.779934421569979 | 86.7373167906283 | 114.018202048006 | 124.932045301595 | 116.302910088012 | 151.104231023035 | 176.123841418636 | 93.6970281500523 | 138.947272277569 | Adgre1 |
| ENSMUSG00000004748 | -0.119677162252812 | 0.492475215981112 | 0.899236955296196 | 4504.91939081326 | 3756.8997574818 | 2819.48911328372 | 3618.52118644799 | 3699.86374367273 | 3406.2840163257 | 4212.20195439013 | 4652.56257016922 | Mtfp1 |
| ENSMUSG00000004768 | -0.352828235536704 | 0.0238082102625027 | 0.654124840968651 | 94.8689402397497 | 119.719112150406 | 109.78876708322 | 111.613276616721 | 125.920192519196 | 144.323703384715 | 139.504464134522 | 147.631476794917 | Rab23 |
| ENSMUSG00000004771 | 0.0365342130298939 | 0.627035680843632 | 0.931825800800958 | 1937.13340832403 | 1790.08577215369 | 1925.08924351094 | 1918.9980164522 | 1934.79113201234 | 1846.85417043153 | 1779.20245676044 | 1820.42637194909 | Rab11a |
| ENSMUSG00000004788 | -0.109307942255298 | 0.500307801323488 | 0.900927059989498 | 692.091506891888 | 561.132437221972 | 586.802030962037 | 625.597105070194 | 748.951405940262 | 790.111121919713 | 550.730309904196 | 570.986447015635 | Eif2b2 |
| ENSMUSG00000004789 | 0.150387848524455 | 0.386356586817368 | 0.875126835008975 | 14934.1782211697 | 16212.5739154975 | 15242.6559816832 | 14703.8767858852 | 12974.1596622605 | 9790.77326775125 | 16069.040327734 | 16210.1532571948 | Dlst |

| | | | | | | | | | | | | |
|---|---|---|---|---|---|---|---|---|---|---|---|---|
| **ENSMUSG00000004791** | -0.804332418230627 | 0.0652298977984607 | 0.718339708336358 | 58.7283915769879 | 60.2667639396603 | 53.001473764313 | 63.779015209555 | 152.199189218854 | 167.562265794119 | 41.6431236222455 | 51.0197015394199 | Pgf |
| **ENSMUSG00000004798** | 0.183337423855413 | 0.456687648361946 | 0.892805623137304 | 1960.62476495483 | 2416.37146768881 | 2331.11839074112 | 1857.09485463116 | 1459.57927502686 | 1215.74373868142 | 2430.91734144858 | 2435.91936711613 | Ulk2 |
| **ENSMUSG00000004814** | -0.17476884422309 | 0.734666368415898 | 0.954383574136061 | 20.780815481088 | 4.88649437348597 | 19.8755526616174 | 15.9447538023887 | 21.8991639163819 | 18.3462334811079 | 8.32862472444909 | 20.6249857287016 | Ccl24 |
| **ENSMUSG00000004815** | -0.0655359845944583 | 0.868596136671839 | 0.975147618296101 | 18.0702743313809 | 25.2468875963442 | 35.965285768641 | 15.9447538023887 | 25.1840385038392 | 14.6769867848863 | 35.3966550789086 | 23.8815624227072 | Dgkq |
| **ENSMUSG00000004837** | -0.481507861874813 | 0.0620677013461724 | 0.715615581523459 | 170.764092431549 | 134.378595270864 | 115.467496415111 | 162.261318106662 | 278.119381738051 | 238.501035254403 | 144.709854587303 | 153.05910461826 | Grap |
| **ENSMUSG00000004843** | 0.147217248492297 | 0.25705141640077 | 0.837517407767513 | 966.759676728878 | 1040.82330155251 | 1036.36810307005 | 1105.81557253037 | 903.340511550755 | 769.318723974458 | 1033.79054392224 | 1038.84796538776 | Chmp2b |
| **ENSMUSG00000004846** | -0.54937922156585 | 0.0386251894864653 | 0.679566075569698 | 724.618000688374 | 723.201167275924 | 724.98444470471 | 783.168789705565 | 1361.03303740314 | 1534.96820125269 | 705.850945397061 | 725.131077198563 | Plod3 |
| **ENSMUSG00000004849** | -0.13065618840151 | 0.278386150295294 | 0.848075053184273 | 700.22313034101 | 601.853223667689 | 619.927952064733 | 659.362466063487 | 646.025335533267 | 821.911259953634 | 679.823993133157 | 679.539003482486 | Ap1s1 |
| **ENSMUSG00000004865** | -0.00134370548454808 | 0.99285383830795 | 0.999111773824344 | 569.213641438498 | 558.689190035229 | 655.893237833374 | 603.086864407998 | 556.238763476101 | 477.002070508805 | 682.947227404826 | 670.854798965138 | Srpk1 |
| **ENSMUSG00000004880** | -0.263761564797081 | 0.037207562879768 | 0.679566075569698 | 383.089815825275 | 383.589808318649 | 375.742590793433 | 357.350070512359 | 531.054724972262 | 441.532685778663 | 398.732908683 | 429.868123608729 | Lbr |
| **ENSMUSG00000004891** | -0.342849671490485 | 0.00198994940234112 | 0.635862253526407 | 931.522641782685 | 866.538335564846 | 790.289832021453 | 990.450589136618 | 1226.35317931739 | 1094.6585977061 | 1072.31043327282 | 1145.22947072528 | Nes |
| **ENSMUSG00000004895** | -0.0648885123907089 | 0.438996000748673 | 0.889583654898613 | 1250.46298373156 | 1246.05606523892 | 1346.80530654674 | 1242.75286989206 | 1432.20532013138 | 1298.91333046244 | 1288.8546761085 | 1299.3741009082 | Prcc |
| **ENSMUSG00000004896** | 0.206350250055173 | 0.36754989137458 | 0.870908903308243 | 49.6932544112975 | 61.0811796685746 | 60.5731128735006 | 78.7858423176855 | 55.8428679867739 | 62.3771938357668 | 48.9306702561384 | 49.9341759747513 | Rrnad1 |
| **ENSMUSG00000004897** | 0.0818008760902921 | 0.330808295782898 | 0.856303993370193 | 3947.45142769016 | 3879.87653254786 | 3710.10316350191 | 4055.59502597229 | 3366.99645214372 | 3647.23121604425 | 3876.97480923105 | 3841.67497336185 | Hdgf |
| **ENSMUSG00000004929** | -0.0867185275141098 | 0.637898489786661 | 0.933164350662676 | 442.721721118832 | 453.629561005281 | 447.673162330715 | 545.87333605825 | 590.182467546493 | 594.417964787896 | 450.786813210807 | 372.335268681298 | Thop1 |
| **ENSMUSG00000004931** | -0.00041445628541971 | 0.996995940151399 | 0.999262355603273 | 1741.97444554512 | 1527.84390744328 | 1428.20042697051 | 1777.37108561922 | 1526.37172497182 | 1689.076562494 | 1682.38219433872 | 1579.43969659268 | Apba3 |
| **ENSMUSG00000004933** | -0.55889788536012 | 0.330375652673455 | 0.85614138351386 | 2.71054114970714 | 8.14415728914329 | 14.1968233297267 | 12.1930470253561 | 17.5193311331055 | 7.33849339244316 | 19.7804837205666 | 9.76973008201657 | Matk |
| **ENSMUSG00000004934** | -0.141461274973499 | 0.112946852510253 | 0.767518180080285 | 513.195791011218 | 493.535931722083 | 497.835271429083 | 489.59773440276 | 548.574056105368 | 541.82542880872 | 577.798340258656 | 530.8220011229 | Pias4 |
| **ENSMUSG00000004936** | -0.0414750388163872 | 0.675028722212706 | 0.944023556644042 | 869.180195339421 | 917.032110757534 | 864.113313336032 | 868.520118883057 | 935.094299229509 | 797.449615312156 | 972.366936579432 | 915.098051015552 | Map2k1 |
| **ENSMUSG00000004937** | 0.017153027130683 | 0.857990023459065 | 0.974350331019239 | 2682.53222449349 | 2392.7534115503 | 2607.48321822647 | 2771.57338153287 | 2674.98287238605 | 2783.7351602001 | 2477.76585552361 | 2394.66939565873 | Sgta |
| **ENSMUSG00000004939** | -0.169083431645845 | 0.590924194152342 | 0.923935529355527 | 41.5616309621761 | 34.2054606144018 | 35.965285768641 | 64.7169419038131 | 43.7983278327639 | 68.4926049961361 | 35.3966550789086 | 51.0197015394199 | Nmrk2 |
| **ENSMUSG00000004945** | 0.0182202339837852 | 0.892029988735996 | 0.982421229269187 | 2884.91929700496 | 2118.29531090617 | 2085.04011969253 | 2451.74037879083 | 2223.86009570859 | 2468.17994432505 | 2293.49503349517 | 2434.83384155146 | Tmem242 |
| **ENSMUSG00000004947** | -0.201274517262881 | 0.264516409100866 | 0.842510961779757 | 173.474633581257 | 160.439898596123 | 178.879973954556 | 181.019851991825 | 227.751304730372 | 240.94719971855 | 155.120635492864 | 174.76961591163 | Dtx2 |
| **ENSMUSG00000004951** | 0.0637304984497178 | 0.797779972199514 | 0.962575035949553 | 9824.8081539718 | 8423.5018841609 | 9024.44736326294 | 14822.993476056 | 11645.9753707319 | 13053.9566629243 | 7221.95871418792 | 8355.2902712535 | Hspb1 |
| **ENSMUSG00000004952** | -0.252816149526169 | 0.44977280541988 | 0.89108031640712 | 71.3775836089546 | 52.122606650517 | 64.3589324280944 | 88.1651092602672 | 85.4067392738896 | 134.539045528125 | 52.0539045278068 | 57.5328549274309 | Rasa4 |
| **ENSMUSG00000004980** | 0.11526624802703 | 0.224237382884138 | 0.832048466795696 | 13162.3878229778 | 12312.3369897268 | 11802.2924614461 | 13370.1450266501 | 11365.6660726022 | 10607.7921987766 | 12508.553258032 | 12274.0375597068 | Hnrnpa2b1 |
| **ENSMUSG00000004994** | -0.193583314420606 | 0.406274856538234 | 0.881192654171779 | 83.1232619243521 | 88.7713144516618 | 64.3589324280944 | 104.109863062656 | 107.305903190271 | 122.308223207386 | 77.0397787011541 | 83.5854684794751 | Ccdc130 |
| **ENSMUSG00000004996** | 0.0103771341760287 | 0.957358166881453 | 0.992403427270738 | 317.133314515735 | 258.984201794756 | 245.131816159948 | 285.129715054481 | 281.404256325508 | 364.47850515801 | 223.831789469569 | 230.131419709724 | Mri1 |
| **ENSMUSG00000005034** | -0.0505393650034146 | 0.765247395983173 | 0.958197631858494 | 1104.99727536394 | 1400.79505373265 | 1264.46373123432 | 962.312788308873 | 1121.23719251876 | 1017.60441708545 | 1385.67493853022 | 1376.44641599967 | Prkacb |
| **ENSMUSG00000005043** | -0.36939207607998 | 0.132400420450577 | 0.77303252008843 | 83.1232619243521 | 82.2559886203472 | 81.3951204237664 | 75.0341355406529 | 141.249607260664 | 121.085140975312 | 77.0397787011541 | 77.072315091464 | Sgsh |
| **ENSMUSG00000005054** | 0.159773493891673 | 0.404536973240132 | 0.881158207349486 | 334.300075130547 | 258.984201794756 | 201.594891282119 | 259.805694309511 | 258.410134213307 | 262.96267989588 | 196.76375911511 | 226.874843015718 | Cstb |
| **ENSMUSG00000005069** | 0.0921777085798912 | 0.477180568211518 | 0.897674307047416 | 506.871194995234 | 692.253369577179 | 621.82086184203 | 614.341984739096 | 564.998429042654 | 528.371524255907 | 604.866370613115 | 586.183804920994 | Pex5 |
| **ENSMUSG00000005078** | -0.298597660672846 | 0.0516695489825724 | 0.703396849069134 | 415.616309621761 | 361.600583637962 | 350.188308799925 | 390.177504811395 | 527.769850384805 | 538.156182112498 | 390.404283958551 | 411.414189009364 | Jkamp |
| **ENSMUSG00000005087** | 0.23821908345573 | 0.386944451172744 | 0.875691730512848 | 137.334084918495 | 170.212887343095 | 201.594891282119 | 295.446908691321 | 197.092475247437 | 210.370143916704 | 147.833088858971 | 127.006491066215 | Cd44 |
| **ENSMUSG00000005089** | -0.391389287415775 | 0.118174124211553 | 0.769868040577702 | 64.1494738764022 | 111.574954861263 | 67.1982970940397 | 82.5375490947182 | 85.4067392738896 | 130.869798831903 | 92.6559500594962 | 119.407812113536 | Slc1a2 |
| **ENSMUSG00000005102** | -0.167909535205633 | 0.430761223215311 | 0.886017087288711 | 57.8248778604189 | 94.4722245540621 | 90.8596693102509 | 77.8479156234274 | 96.3563212320805 | 78.277262852727 | 96.8202624217207 | 89.0130963028176 | Eif2ak4 |
| **ENSMUSG00000005103** | 0.0249094566272263 | 0.845834356683518 | 0.972180580805179 | 1953.39665522228 | 2067.80153571348 | 2279.06337186546 | 2145.97627646267 | 1961.070128712 | 1760.01533195428 | 2161.27811599454 | 2417.46543251677 | Wdr1 |
| **ENSMUSG00000005107** | -0.097983233459545 | 0.775972226809337 | 0.959602406551077 | 20.780815481088 | 17.9171460361152 | 26.5007368821565 | 30.0136542162612 | 29.5638712871156 | 33.0232202659942 | 16.6572494488982 | 22.7960368580387 | Slc2a9 |
| **ENSMUSG00000005125** | -0.725488963984536 | 0.0680569765648639 | 0.722112580225498 | 992.058060792811 | 1048.96745884166 | 986.205993971682 | 812.244517227568 | 2208.53068096712 | 2660.20385476064 | 687.11153976705 | 793.519187772679 | Ndrg1 |
| **ENSMUSG00000005142** | -0.032892333879111 | 0.845034138627403 | 0.972136081650911 | 703.837185207286 | 895.857301805761 | 1087.47666705707 | 884.464872685446 | 1013.93128932848 | 1017.60441708545 | 812.040910633787 | 810.887596807375 | Man2b1 |
| **ENSMUSG00000005150** | -0.128439565132203 | 0.290530402801238 | 0.851785577030345 | 1015.54941742361 | 862.466256920274 | 812.058294460367 | 914.478526901707 | 1017.21616391594 | 1092.21243324196 | 951.545374768309 | 880.36123294616 | Wdr83 |
| **ENSMUSG00000005161** | 0.0568816225021663 | 0.651865535024551 | 0.936720465895231 | 11643.5812654253 | 8989.52081575636 | 9643.42886043902 | 11411.754089039 | 10014.4876589615 | 11097.0250916061 | 9196.88385197291 | 9768.6445564519 | Prdx2 |
| **ENSMUSG00000005198** | -0.169784324440324 | 0.168490956185787 | 0.79907358295209 | 648.722848496574 | 813.601313185414 | 811.111839571719 | 691.251973668265 | 905.530427942393 | 764.426395046162 | 821.410613448792 | 843.45336374743 | Polr2a |
| **ENSMUSG00000005204** | 0.130349570218238 | 0.214841180480779 | 0.824338622350122 | 816.776399778417 | 766.365200908383 | 822.4692982355 | 862.892558717508 | 769.755611660825 | 807.234273168747 | 751.658381381531 | 657.828492189116 | Senp3 |
| **ENSMUSG00000005220** | 0.252422440999644 | 0.306733469228375 | 0.851785577030345 | 4038.70631306363 | 5848.31934933379 | 6896.81677358123 | 4019.01588489622 | 4173.9806424624 | 2983.09756402814 | 5573.93209683756 | 4731.80593639002 | Corin |
| **ENSMUSG00000005225** | 0.254356960939081 | 0.0933972003252002 | 0.754786063592529 | 223.167887992554 | 195.459774939439 | 230.934992830221 | 227.916186704733 | 199.282391639076 | 146.769867848863 | 197.804837205666 | 189.966973816989 | Plekha8 |
| **ENSMUSG00000005233** | 0.348437182364502 | 0.35478519831194 | 0.863949229409685 | 22.5878429142261 | 12.2162359337149 | 18.9290977729689 | 22.5102406621959 | 16.4243729372865 | 12.2308223207386 | 15.616171358342 | 15.1973579053591 | Spc25 |
| **ENSMUSG00000005237** | 0.00615996986354965 | 0.98884038905718 | 0.998437672973644 | 22.5878429142261 | 9.77298874697194 | 15.1432782183751 | 17.8206071909051 | 24.0890803080201 | 8.56157562451702 | 15.616171358342 | 16.2828834700276 | Dnah2 |
| **ENSMUSG00000005262** | 0.0976769153057701 | 0.365746811678633 | 0.869253398153585 | 887.250469670802 | 976.48445896828 | 923.739971320884 | 934.174987481129 | 776.32536083574 | 817.018931025338 | 926.559500594962 | 957.433548037624 | Ufd1 |
| **ENSMUSG00000005267** | -0.318250066254714 | 0.335463766849205 | 0.858570960161243 | 28.0089252136404 | 34.2054606144018 | 30.2865564367503 | 23.448167356454 | 52.5579933993167 | 41.5847958905112 | 24.9858741733473 | 26.0526135520442 | Zfp287 |
| **ENSMUSG00000005268** | -0.0433204009851897 | 0.808011065203626 | 0.964026716319739 | 175.281661014395 | 190.573280565953 | 136.289503965376 | 157.571684635371 | 142.344565456483 | 203.031650524261 | 162.408182126757 | 173.684090346961 | Prlr |
| **ENSMUSG00000005299** | 0.0289182940470103 | 0.782271761859265 | 0.960742105865155 | 3183.98233718931 | 3105.36717435034 | 2647.23432354971 | 3073.58577708399 | 2892.87955335405 | 2668.76543038516 | 3123.23427166841 | 3086.14918035257 | Letm1 |
| **ENSMUSG00000005312** | 0.199144712457877 | 0.111433598277924 | 0.764116514278315 | 1511.57844782001 | 1709.45861499118 | 1761.35254777476 | 1528.8205116408 | 1471.62381518087 | 1179.0512717192 | 1456.46824868804 | 1563.15681312265 | Ubqln1 |
| **ENSMUSG00000005338** | -1.40264626315088 | 0.0133167697452517 | 0.635862253526407 | 16.2632468982428 | 28.5045505120015 | 9.46454888648447 | 16.8826804966469 | 58.0327843784121 | 86.838838477244 | 32.2734208072402 | 11.9407812113536 | Cadm3 |
| **ENSMUSG00000005354** | -0.0386690726149954 | 0.731032966087567 | 0.952984317124008 | 5511.43367107118 | 4669.85978959476 | 4666.02260103684 | 5399.64397884423 | 5021.47828602638 | 5873.24087841867 | 5032.57148974836 | 4870.75320866759 | Txn2 |
| **ENSMUSG00000005360** | -0.758783002026475 | 0.137663508266264 | 0.779425998172399 | 7.22810973255236 | 13.8450673915436 | 9.46454888648447 | 5.62756016554897 | 21.8991639163819 | 20.7923979452556 | 8.32862472444909 | 10.8552556466851 | Slc1a3 |
| **ENSMUSG00000005362** | 0.223111873340276 | 0.378263754038799 | 0.87422797717331 | 745.398816169462 | 953.680818558679 | 995.670542858166 | 802.865250284986 | 710.627869086594 | 418.29412336926 | 930.723812957186 | 935.723036744253 | Crbn |
| **ENSMUSG00000005370** | 0.0199173355520309 | 0.911102331952377 | 0.986222832825632 | 163.535982698997 | 186.501201921381 | 210.112985279955 | 180.081925297567 | 188.332809680885 | 135.762127760198 | 212.379930473452 | 192.138024946326 | Msh6 |
| **ENSMUSG00000005371** | 0.0965663026373029 | 0.635098391606593 | 0.932543559775713 | 636.977170181177 | 747.633639143354 | 822.4692982355 | 655.610759286455 | 596.752216721408 | 468.440494884288 | 841.191097169358 | 769.637625349972 | Fbxo11 |
| **ENSMUSG00000005373** | 0.00607457046643761 | 0.978365662592974 | 0.996589839646261 | 157.211386683014 | 232.922898469498 | 179.826428843205 | 200.716312571247 | 162.053812981226 | 139.43137445642 | 236.324726556243 | 229.045894145055 | Mlxipl |
| **ENSMUSG00000005374** | -0.195110272777469 | 0.255100768024665 | 0.837406774112203 | 210.518695960587 | 245.139134403213 | 274.47191770805 | 199.778385876988 | 285.784089108784 | 242.170281950624 | 310.241270985729 | 225.78931745105 | Tbl2 |
| **ENSMUSG00000005378** | -0.0991449432065515 | 0.619687943657309 | 0.929594515086012 | 324.361424248287 | 223.96432545144 | 213.898804834549 | 274.812521417641 | 267.16979977986 | 354.693847301419 | 247.776585552361 | 242.072200921077 | Bud23 |
| **ENSMUSG00000005397** | -0.671913101974483 | 0.0344005907633299 | 0.678229434609245 | 949.592916114066 | 1157.28475078726 | 1279.6070094527 | 1078.61569839689 | 2586.29125852471 | 2437.6028885232 | 1097.29630744617 | 993.255891671684 | Nid1 |
| **ENSMUSG00000005410** | -0.265379845503633 | 0.4250096156109 | 0.885066977635295 | 48.7897406947284 | 30.1333819698302 | 29.3401015481019 | 43.1446279358754 | 48.1781606160403 | 68.4926049961361 | 32.2734208072402 | 33.6512925047237 | Mcm5 |
| **ENSMUSG00000005413** | 0.245707973101652 | 0.491584760910365 | 0.899236955296196 | 375.861706092723 | 421.867347577622 | 713.626986040929 | 209.15765281957 | 372.285786578493 | 522.256113095538 | 249.858741733473 | 307.203734801188 | Hmox1 |
| **ENSMUSG00000005417** | 0.0298936769932152 | 0.741838833371 | 0.954383574136061 | 4098.33821835719 | 4269.98166669782 | 4657.50450703901 | 4004.00905778809 | 4222.15880307844 | 3884.50916906658 | 4428.74619722581 | 4143.45108033969 | Mprip |
| **ENSMUSG00000005442** | 0.0123599217531827 | 0.938819590774126 | 0.989269716906553 | 769.793686516826 | 930.877178149078 | 1037.3145579587 | 708.134654164912 | 877.061514851097 | 726.510845851873 | 872.423439886043 | 940.065139002928 | Cic |
| **ENSMUSG00000005447** | -0.883961587460428 | 0.0221437103861993 | 0.648762042745232 | 17.1667606148119 | 21.1748089517725 | 22.7149173275627 | 15.0068271081306 | 33.943704070392 | 57.4848649074714 | 30.191264626128 | 19.5394601640331 | Pafah1b3 |
| **ENSMUSG00000005465** | -0.597809225451932 | 0.214124823898409 | 0.824248286808591 | 26.2018977805023 | 21.1748089517725 | 18.9290977729689 | 10.3171936368398 | 51.4630352034976 | 33.0232202659942 | 16.6572494488982 | 15.1973579053591 | Il27ra |
| **ENSMUSG00000005469** | 0.137128481404313 | 0.421304775753702 | 0.882568531013176 | 5294.5903790946 | 5589.33514753904 | 5055.01556027135 | 4955.06672576587 | 4202.44955555369 | 3566.50778872737 | 5707.19009242874 | 5522.0685474687 | Prkaca |
| **ENSMUSG00000005470** | -0.399558187014005 | 0.48714473700122 | 0.899236955296196 | 19.877301764519 | 16.2883145782866 | 8.51809399783602 | 5.62756016554897 | 18.6142893289246 | 29.3539735697726 | 12.4929370866736 | 6.51315338801105 | Asf1b |
| **ENSMUSG00000005481** | -0.0467191592109824 | 0.719893504257456 | 0.951674101666722 | 353.273863178497 | 269.571606270643 | 313.276568142636 | 329.212269684615 | 346.006789878835 | 331.455284892016 | 324.816364253515 | 303.947158107182 | Ddx39 |
| **ENSMUSG00000005483** | -0.0873397849592461 | 0.731169259709576 | 0.952992645818943 | 403.870631306363 | 445.485403716138 | 314.223023031284 | 285.129715054481 | 524.484975797347 | 442.755768010737 | 309.200192895173 | 263.782712214447 | Dnajb1 |
| **ENSMUSG00000005501** | 0.362396066460547 | 0.264742951050235 | 0.842510961779757 | 353.273863178497 | 438.970077884823 | 571.658752743662 | 406.122258613784 | 250.745426842573 | 173.677676954488 | 469.526218840818 | 481.973350712817 | Usp40 |
| **ENSMUSG00000005505** | 0.0561577141721179 | 0.78521769408273 | 0.961141387070613 | 125.588406603097 | 135.193010999779 | 160.897331070236 | 116.302910088012 | 107.305903190271 | 103.961989726278 | 146.792010768415 | 158.486732441602 | Kbtbd4 |
| **ENSMUSG00000005506** | 0.144139672039201 | 0.484146822127139 | 0.899157555735522 | 1722.0971437806 | 2145.98544568926 | 2045.28901436929 | 1693.89560983024 | 1472.71877337669 | 1214.52065644934 | 2029.06119849391 | 2166.70902707834 | Celf1 |
| **ENSMUSG00000005510** | -0.00720680335318625 | 0.953124528161825 | 0.992049433957868 | 10674.1110475467 | 8662.94010846171 | 9649.10758977092 | 10827.4257585162 | 10213.7700506005 | 11331.8568801643 | 9104.22790191341 | 9363.74352083055 | Ndufs3 |
| **ENSMUSG00000005514** | -0.111451113244377 | 0.216432384882593 | 0.825447777181429 | 1824.1941937529 | 1921.2067045089 | 1929.82151795418 | 1958.39093761104 | 2265.46850714971 | 2101.25527470289 | 1961.39112260776 | 1919.20919833392 | Por |
| **ENSMUSG00000005533** | 0.0848603274959906 | 0.678527139869337 | 0.944452327903597 | 623.424464432641 | 720.757920089181 | 895.346324661431 | 705.320874082137 | 937.284215621147 | 671.472145408549 | 643.386259963692 | 524.308847734889 | Igf1r |
| **ENSMUSG00000005534** | 0.478390701636649 | 0.0615308174930612 | 0.71282992338528 | 3372.81670395225 | 4705.69408166699 | 4708.61307102602 | 3747.95507025561 | 2925.72829922863 | 1642.59943767519 | 3659.38948830482 | 3639.76721833351 | Insr |
| **ENSMUSG00000005547** | -0.233591632548812 | 0.525330154603723 | 0.908006595346551 | 145.465708367616 | 210.933673788811 | 152.3792370724 | 187.585338851632 | 104.021028602814 | 194.470074899744 | 140.545542225078 | 379.933947633978 | Cyp2a5 |
| **ENSMUSG00000005566** | -0.0919646950194284 | 0.519390182830221 | 0.905962179970576 | 1132.10268686101 | 1311.20932355207 | 1294.75028767108 | 1237.12530972651 | 1537.32130693001 | 1440.79086938301 | 1263.86880193515 | 1061.6440022458 | Trim28 |
| **ENSMUSG00000005575** | 0.0112678623347303 | 0.955399139864322 | 0.992371505557225 | 3783.91544499116 | 3701.51948791562 | 4110.4535814002 | 4912.86002452425 | 4398.44707260531 | 5404.80038353439 | 3592.76049050923 | 2985.1953028384 | Ube2m |
| **ENSMUSG00000005580** | 0.261424851507772 | 0.609721581580497 | 0.927932112008768 | 234.913566307952 | 506.566583384712 | 472.280989435575 | 159.447538023887 | 186.142893289246 | 80.7234273168747 | 452.868969391919 | 425.526021350055 | Adcy9 |
| **ENSMUSG00000005583** | -0.0489340773740092 | 0.705810084005645 | 0.947767021156073 | 1007.41779397449 | 1100.27564976326 | 808.272474905774 | 958.561081531841 | 938.379173816966 | 966.234963338349 | 1107.70708835173 | 996.51246836569 | Mef2c |
| **ENSMUSG00000005609** | -0.0447933423637705 | 0.723129763853075 | 0.951674101666722 | 344.238726012806 | 330.652785939217 | 386.153594568566 | 321.708856130549 | 377.760577557588 | 315.555215875056 | 387.281049686883 | 344.111603999917 | Ctr9 |
| **ENSMUSG00000005610** | 0.145033835536274 | 0.484380717407388 | 0.899236955296196 | 19928.8020463634 | 22960.0082295528 | 22083.6319168342 | 17664.9113596582 | 15551.6912552186 | 13264.326806841 | 22523.724489182 | 23393.0759186063 | Eif4g2 |

| | | | | | | | | | | | | |
|---|---|---|---|---|---|---|---|---|---|---|---|---|
| **ENSMUSG00000005611** | -0.319195490689133 | 0.192904540292713 | 0.808824684419004 | 184.316798180085 | 259.798617523671 | 160.897331070236 | 157.571684635371 | 245.270635863478 | 297.208982393948 | 250.899819824029 | 159.572258006271 | Mrvi1 |
| **ENSMUSG00000005615** | 0.155140768291109 | 0.527337169708378 | 0.908217276872524 | 2434.06595243701 | 3251.96200555491 | 2971.86835035612 | 2176.92785737319 | 1951.21550494963 | 1616.91471080164 | 2951.45638672665 | 3209.89909472478 | Pcyt1a |
| **ENSMUSG00000005621** | 0.127398333682852 | 0.478659566225837 | 0.897729054128663 | 511.388763578079 | 640.945178655577 | 531.907647420427 | 389.239578117137 | 460.97740043984 | 408.509465512669 | 486.183468289716 | 542.762782334254 | Zfp592 |
| **ENSMUSG00000005625** | 0.117242754894034 | 0.528814510279855 | 0.908389011598712 | 3486.65943223994 | 2899.31999493501 | 3596.5285768641 | 4294.76633300812 | 3671.39483058143 | 4030.05595468337 | 2729.70675343819 | 2732.26784627063 | Psmd4 |
| **ENSMUSG00000005628** | 0.0614155443361072 | 0.704448464103065 | 0.947128948445608 | 738.17070643691 | 542.400875456943 | 574.498117409607 | 763.472329126143 | 608.796756875418 | 711.833859066986 | 548.648153723084 | 640.46008315442 | Tmod4 |
| **ENSMUSG00000005656** | 0.0811110599835986 | 0.381773916378936 | 0.875126835008975 | 578.248778604189 | 541.586459728029 | 573.551662520959 | 560.88016316638 | 514.630352034976 | 511.248373006873 | 534.073060455298 | 569.900921450967 | Snx6 |
| **ENSMUSG00000005667** | 0.0992827998597913 | 0.659388810377596 | 0.93918279082699 | 306.291149916906 | 341.240190415104 | 298.133289924261 | 216.661066373635 | 209.137015401448 | 374.263163014601 | 240.489038918468 | 262.697186649779 | Mthfd2 |
| **ENSMUSG00000005672** | -0.664443287063478 | 0.0651412349351175 | 0.718339708336358 | 16.2632468982428 | 13.8450673915436 | 15.1432782183751 | 19.6964605794214 | 38.3235368536684 | 19.5693157131818 | 21.8626399016789 | 22.7960368580387 | Kit |
| **ENSMUSG00000005674** | -0.0558934363341423 | 0.755005049285565 | 0.956497786093635 | 782.442878548793 | 778.581436842098 | 669.143606274452 | 677.183073254392 | 601.132049504684 | 621.325773893521 | 881.793142701048 | 917.269102144889 | Tomm40l |
| **ENSMUSG00000005677** | -0.00205039544566211 | 0.994550111364551 | 0.999262355603273 | 63.2459601598332 | 46.4216965481167 | 31.2330113253987 | 45.9584080186499 | 41.6084114411257 | 57.4848649074714 | 33.3144988977964 | 55.3618037980939 | Nr1i3 |
| **ENSMUSG00000005681** | -0.117120407356225 | 0.687486140427225 | 0.944452327903597 | 1188.12053728829 | 2223.35493993612 | 1369.5202238743 | 1514.75161122693 | 1234.01788668812 | 1689.076562494 | 1245.12939630514 | 2660.62315900251 | Apoa2 |
| **ENSMUSG00000005682** | 0.173377342218649 | 0.380330686281379 | 0.87432013076003 | 93.9654265231807 | 88.7713144516618 | 121.146225747001 | 120.054616865045 | 107.305903190271 | 77.0541806206532 | 91.61487196894 | 98.7828263848342 | Pan2 |
| **ENSMUSG00000005683** | 0.215993206123737 | 0.23107164642133 | 0.832048466795696 | 27798.4065176798 | 30630.1755644679 | 32888.3609256449 | 27752.3129564047 | 23641.2424059301 | 18363.3566323569 | 30364.0835891603 | 30142.8738797151 | Cs |
| **ENSMUSG00000005686** | 0.16088503176565 | 0.501022622817133 | 0.900927059989498 | 749.012871035738 | 946.35107699845 | 1027.85000907221 | 716.575994413235 | 733.621991198795 | 483.117481669175 | 1029.62623156002 | 829.34153140674 | Ampd3 |
| **ENSMUSG00000005687** | 0.105129255985694 | 0.32142656973408 | 0.855704427283138 | 1143.84836517641 | 982.999784799595 | 1126.28131749165 | 1137.70508013515 | 997.506916391197 | 928.319414144059 | 1113.95355689507 | 1039.93349095243 | Bcas2 |
| **ENSMUSG00000005698** | 0.0184544003089306 | 0.897686666486485 | 0.984332015045905 | 931.522641782685 | 972.412380323708 | 1078.95857305923 | 815.9962240046 | 892.390929592564 | 787.664957455566 | 1036.91377819391 | 1032.33481199975 | Ctcf |
| **ENSMUSG00000005699** | -0.322375362657113 | 0.14683531633877 | 0.780607990324772 | 138.237598635064 | 78.1839099757755 | 86.1273948670087 | 105.047789756914 | 132.489941694111 | 145.546785616789 | 115.559668051731 | 116.15123541953 | Pard6a |
| **ENSMUSG00000005718** | -0.20737434953664 | 0.627483555713468 | 0.931825800800958 | 40.658117245607 | 23.6180561385155 | 49.2156542097192 | 64.7169419038131 | 77.7420319031559 | 68.4926049961361 | 30.191264626128 | 29.3091902460497 | Tfap4 |
| **ENSMUSG00000005732** | -0.0876167752307584 | 0.596423657966382 | 0.92448662081833 | 782.442878548793 | 636.058684282091 | 508.246275204216 | 751.279282100787 | 770.850569856644 | 750.97249049335 | 632.975479058131 | 691.479784693839 | Ranbp1 |
| **ENSMUSG00000005774** | 0.303582049038503 | 0.18295629258219 | 0.805716353538343 | 100.290022539164 | 98.5443031986338 | 126.824955078892 | 124.744250336335 | 68.9823663366031 | 74.6080161565054 | 100.984574783945 | 119.407812113536 | Rfx5 |
| **ENSMUSG00000005779** | -0.109317191959075 | 0.541702455589594 | 0.913837378403038 | 5160.87034904238 | 3962.13252116821 | 4046.09464897211 | 5333.0511835519 | 5123.30939823755 | 6302.5427418766 | 4237.18782856348 | 4296.51018495795 | Psmb4 |
| **ENSMUSG00000005800** | -0.00061765902125917 | 0.999349600997691 | 0.999709467616704 | 17.1667606148119 | 9.77298874697194 | 35.965285768641 | 5.62756016554897 | 21.8991639163819 | 34.2463024980681 | 10.4107809055614 | 2.17105112933702 | Mmp8 |
| **ENSMUSG00000005802** | -0.0119931443809052 | 0.93445632866081 | 0.98818087308472 | 246.659244623349 | 333.910448854875 | 293.401015481019 | 320.770929436291 | 304.398378437709 | 271.524255520397 | 334.18606706852 | 294.177428025166 | Slc30a4 |
| **ENSMUSG00000005803** | 0.0832138690086253 | 0.361326483130404 | 0.868564573435346 | 1321.84056734051 | 1320.16789657013 | 1380.87768253808 | 1342.17309948343 | 1396.07169966935 | 1280.56709698133 | 1170.1717737851 | 1217.95968355807 | Sqor |
| **ENSMUSG00000005804** | 0.0329396235838546 | 0.803384829046441 | 0.963461896828986 | 600.836621518415 | 575.79192034243 | 488.370722542599 | 571.19735680322 | 504.775728272604 | 480.671317205027 | 613.194995337564 | 586.183804920994 | Bloc1s6 |
| **ENSMUSG00000005813** | 0.152171687387074 | 0.496078015971783 | 0.900927059989498 | 895.382093119924 | 1056.29720040188 | 1089.36957683436 | 952.933521366292 | 775.230402639921 | 577.294813538862 | 1147.26805579286 | 1093.12424362119 | Metap1 |
| **ENSMUSG00000005823** | -0.0583880705611693 | 0.680198937571402 | 0.944452327903597 | 555.660935689963 | 573.348673155687 | 725.930899593359 | 594.645524159674 | 591.277425742312 | 733.849339244316 | 652.755962778698 | 573.157498144972 | Gpr108 |
| **ENSMUSG00000005836** | -0.0201907740267553 | 0.866745195200935 | 0.974868041734905 | 1415.80599386369 | 1494.45286255779 | 1355.32340054458 | 1172.4083678227 | 1409.21119801918 | 1264.66702796437 | 1530.38479311752 | 1310.22935655489 | Gata6 |
| **ENSMUSG00000005846** | 0.228029175483522 | 0.0218926109578174 | 0.648762042745232 | 1303.77029300913 | 1235.46866076304 | 1236.07008457487 | 1463.16564304273 | 1177.08006050553 | 1059.18921297596 | 1092.09091699339 | 1143.05841959594 | Rsl1d1 |
| **ENSMUSG00000005871** | 0.206878932503213 | 0.449181798546026 | 0.89108031640712 | 1034.52320547156 | 1295.7354247027 | 1173.60406192407 | 912.602673513191 | 791.654775577207 | 533.263853184203 | 1279.48497329349 | 1221.21626025207 | Apc |
| **ENSMUSG00000005873** | -0.0233769114814034 | 0.808323918345333 | 0.964026716319739 | 6645.34338536533 | 6661.10624679029 | 6592.05829943643 | 6456.68736327318 | 6058.40369746706 | 6251.1732881295 | 7081.41317196284 | 7393.5146209572 | Reep5 |
| **ENSMUSG00000005881** | -0.0583296660388006 | 0.698823610280422 | 0.945141109561958 | 1217.93648993507 | 1066.07018914886 | 1177.38988147867 | 1241.81494319781 | 1244.96746864631 | 1528.85279009232 | 1031.70838774113 | 1093.12424362119 | Ergic3 |
| **ENSMUSG00000005882** | 0.0730664652563952 | 0.669671412825456 | 0.942388853397102 | 4436.25234835401 | 3997.96681324044 | 3451.72097890089 | 3844.5615197642 | 3247.64600879944 | 2945.18201483385 | 4229.90028192958 | 4529.89818136168 | Uqcc1 |
| **ENSMUSG00000005886** | 0.343211350375598 | 0.326597489848893 | 0.855977678491063 | 739.074220153479 | 1295.7354247027 | 1268.24955078892 | 620.907471598903 | 615.366506050332 | 402.3940543523 | 1133.73404061563 | 941.150664567596 | Ncoa2 |
| **ENSMUSG00000005893** | 0.0127230878873501 | 0.971847778330764 | 0.995123114262614 | 97.5794813894569 | 175.099381716581 | 216.738169500494 | 96.6064495085906 | 118.255485148462 | 80.7234273168747 | 195.722681024554 | 185.624871558315 | Nr2c2 |
| **ENSMUSG00000005897** | -0.0261722119230227 | 0.864376902925641 | 0.974868041734905 | 153.597331816738 | 150.666909849151 | 188.344522841041 | 159.447538023887 | 190.522726072523 | 162.669936865823 | 157.202791673977 | 153.05910461826 | Nr2c1 |
| **ENSMUSG00000005899** | 0.194673849817087 | 0.294144366848472 | 0.851785577030345 | 290.027903018663 | 320.879797192245 | 276.364827485346 | 265.43325447506 | 237.605928492744 | 179.793088114857 | 294.625099627387 | 294.177428025166 | Smpd4 |
| **ENSMUSG00000005907** | 0.00070894111993544 | 0.997256638321773 | 0.999262355603273 | 139.141112351633 | 176.728213174409 | 144.807597963212 | 151.006197775564 | 139.059690869025 | 108.854318654574 | 172.818963032319 | 189.966973816989 | Pex1 |
| **ENSMUSG00000005936** | 0.113232519538055 | 0.308629218114216 | 0.851785577030345 | 1035.42671918813 | 1171.1298181788 | 1202.94416347218 | 1107.69142591889 | 1041.30524422396 | 907.527016198804 | 1108.74816644229 | 1117.00580604389 | Kctd20 |
| **ENSMUSG00000005949** | -0.383908405830036 | 0.133861553268994 | 0.77303252008843 | 79.509207058076 | 86.3280672649188 | 128.717864856189 | 164.137171495178 | 165.338687568684 | 147.992950080937 | 155.120635492864 | 129.177542195552 | Ctns |
| **ENSMUSG00000005950** | 0.267555135675437 | 0.0442075020779623 | 0.691978859031708 | 619.810409566365 | 578.235167529173 | 645.482234058241 | 679.996853337167 | 453.312693069106 | 483.117481669175 | 610.071761065896 | 548.190410157596 | P2rx5 |
| **ENSMUSG00000005951** | -0.432225594039252 | 0.175942156821601 | 0.799889021902745 | 19.877301764519 | 21.9892246806869 | 22.7149173275627 | 30.0136542162612 | 44.893286028583 | 30.5770558018465 | 27.0680303544596 | 24.9670879873757 | Shpk |
| **ENSMUSG00000005958** | -0.0403715245613021 | 0.81362697866127 | 0.966168994922206 | 263.826005238161 | 244.324718674299 | 208.220075502658 | 222.288626539184 | 187.237851485066 | 256.847268735511 | 235.283648465687 | 286.578749072486 | Ephb3 |
| **ENSMUSG00000005973** | 0.0146181364910757 | 0.881702129483256 | 0.978854883720278 | 663.179067961679 | 689.810122390436 | 674.822335606343 | 714.700141024719 | 739.09678217789 | 703.272283442469 | 634.016557148687 | 639.374557589751 | Rcn1 |
| **ENSMUSG00000005980** | 0.268307802195832 | 0.602069569095846 | 0.926448151609826 | 17.1667606148119 | 30.9477976987445 | 16.0897331070236 | 23.448167356454 | 6.56974917491458 | 9.78465785659088 | 31.2323427166841 | 24.9670879873757 | Dnase1 |
| **ENSMUSG00000005981** | 0.305238021056774 | 0.0864690717957957 | 0.746460787597018 | 3353.8429159043 | 3397.74242103058 | 3876.67922390404 | 3673.85886140922 | 2511.83410120901 | 2164.85555077073 | 3558.40491352088 | 3337.99111135566 | Trap1 |
| **ENSMUSG00000005982** | -0.0019004162757248 | 0.983800272077461 | 0.996940244108351 | 728.23205555465 | 840.477032239587 | 759.056820696054 | 740.962088463947 | 738.001823982071 | 759.534066117867 | 790.178270732108 | 785.920508819999 | Naa60 |
| **ENSMUSG00000005983** | 0.403888687700471 | 0.27831333643973 | 0.848075053184273 | 15.3597331816738 | 28.5045505120015 | 36.9117406572894 | 30.0136542162612 | 18.6142893289246 | 29.3539735697726 | 19.7804837205666 | 16.2828834700276 | 1700037C18 |
| **ENSMUSG00000006005** | 0.379834940876083 | 0.0428144970668295 | 0.687730553581215 | 4148.93498648505 | 4609.5930256551 | 4364.10349155799 | 3934.60248241299 | 3024.27453685235 | 2281.04836281775 | 4054.99916271615 | 3747.23424923569 | Tpr |
| **ENSMUSG00000006010** | 0.255275820060906 | 0.0305792181449962 | 0.667184591665619 | 753.530439618584 | 749.262470601182 | 856.541674226844 | 882.58901929693 | 625.221129812704 | 644.564336302924 | 681.906149314269 | 763.124471961961 | BC003331 |
| **ENSMUSG00000006014** | -0.69599985911366 | 0.125087025102288 | 0.771949105062466 | 117.456783153976 | 142.522752560008 | 137.235958854025 | 203.530092654021 | 330.677375137367 | 434.19419238622 | 89.5327157878278 | 119.407812113536 | Prg4 |
| **ENSMUSG00000006019** | 0.169708762436717 | 0.268042480716352 | 0.843793744782845 | 213.229237110295 | 236.99497711407 | 264.060913932917 | 205.405946042537 | 190.522726072523 | 199.362403828039 | 239.447960827911 | 187.795922687652 | Dhx34 |
| **ENSMUSG00000006021** | -0.03632911469246 | 0.750780294285195 | 0.956197192463153 | 378.57224724243 | 372.187988113848 | 364.385132129652 | 350.784583652552 | 399.65974147397 | 412.178712208891 | 343.555769883525 | 349.539231823259 | Kptn |
| **ENSMUSG00000006024** | 0.0365129217578878 | 0.690841253308716 | 0.94469184542218 | 2307.57403211734 | 2169.60350182777 | 2320.70738696599 | 2479.87817961858 | 2288.46262926191 | 2448.61062861187 | 2216.45525479401 | 2092.89328868088 | Napa |
| **ENSMUSG00000006050** | -0.03419149836338 | 0.845128608974097 | 0.972136081650911 | 1251.36649744813 | 974.855627510451 | 1216.19453191325 | 1367.4971202284 | 1155.18089658915 | 1571.66066821491 | 1019.21545065446 | 1179.96628879467 | Sra1 |
| **ENSMUSG00000006057** | 0.00013903900145242 | 0.999302255591537 | 0.999709467616704 | 42797.6377264425 | 32507.4038196154 | 34777.4848833872 | 41738.6758211824 | 39070.298343217 | 46225.1698789994 | 32957.4091127356 | 33554.6807294682 | Atp5g1 |
| **ENSMUSG00000006058** | -0.078829160138961 | 0.622397509067141 | 0.93078489223231 | 789.670988281345 | 652.346998860377 | 648.321598724186 | 758.782695654852 | 767.565695269187 | 934.434825304429 | 624.646854333682 | 683.88110574116 | Snf8 |
| **ENSMUSG00000006095** | -0.037215598136439 | 0.859743748903774 | 0.974868041734905 | 1174.56783153976 | 822.559886203472 | 1020.27836996303 | 1266.20103724852 | 1204.45401540101 | 1417.5523069736 | 855.766190437144 | 918.354627709557 | Tbcb |
| **ENSMUSG00000006127** | 0.134028035514232 | 0.267589613297955 | 0.843793744782845 | 758.951521917998 | 840.477032239587 | 988.098903748979 | 872.27182566009 | 842.022852584886 | 825.580506649855 | 727.71358529874 | 757.696844138618 | Inpp5k |
| **ENSMUSG00000006134** | 0.00847352022721141 | 0.975821791760835 | 0.99575536949131 | 402.967117589794 | 583.936077631574 | 585.855576073389 | 313.267515882226 | 440.173194719277 | 288.647406769431 | 575.716184077544 | 569.900921450967 | Crkl |
| **ENSMUSG00000006154** | 0.271588261186165 | 0.629103650667681 | 0.931825800800958 | 87.6408305071974 | 112.389370590177 | 151.432782183751 | 201.654239265505 | 232.131137513649 | 114.969729814943 | 93.6970281500523 | 17.3684090346961 | Eps8l1 |
| **ENSMUSG00000006169** | 0.320689614881141 | 0.121246542475873 | 0.769868040577702 | 1587.47360001181 | 2062.10062561108 | 2053.80710836713 | 1681.70256280488 | 1410.306156215 | 989.473525747752 | 1689.66974097261 | 1822.59742307842 | Clint1 |
| **ENSMUSG00000006191** | 0.17217165854824 | 0.40863087966799 | 0.882113132676277 | 260.211950371885 | 233.737314198412 | 199.701981504822 | 212.909359596603 | 178.478185918513 | 150.439114545085 | 272.762459725708 | 201.907755028342 | Cdkal1 |
| **ENSMUSG00000006205** | -0.230799552146315 | 0.100748387729795 | 0.759800953500741 | 1669.6933482196 | 1632.90353647323 | 1584.3654835975 | 1564.46172602261 | 2126.40881628069 | 2207.66342889332 | 1648.02661735036 | 1590.29495223936 | Htra1 |
| **ENSMUSG00000006215** | 0.0304303040282351 | 0.726367717799945 | 0.952038256048006 | 814.06585862871 | 745.190391956611 | 733.502538702546 | 764.410255820401 | 751.1413223319 | 764.426395046162 | 714.17957012151 | 764.209997526629 | Zbtb17 |
| **ENSMUSG00000006216** | -0.123130062516969 | 0.780359499538644 | 0.960742105865155 | 35.2370349461928 | 19.5459774939439 | 15.1432782183751 | 46.8963347129081 | 27.3739548954774 | 51.3694537471021 | 27.0680303544596 | 21.7105112933701 | Clcnkb |
| **ENSMUSG00000006218** | -0.0305511654915922 | 0.931602512360143 | 0.987783121884152 | 15.3597331816738 | 24.4324718674299 | 14.1968233297267 | 20.6343872736795 | 18.6142893289246 | 22.0154801773295 | 20.8215618111227 | 15.1973579053591 | Fam131c |
| **ENSMUSG00000006219** | -0.0832785317438686 | 0.736942675774272 | 0.954383574136061 | 2765.65548641785 | 3490.58581412681 | 2913.18814725992 | 2302.61003440379 | 2385.91390868981 | 1999.73944944076 | 3979.00046210555 | 3788.48422069309 | Fblim1 |
| **ENSMUSG00000006221** | 0.169082804966354 | 0.236494784777371 | 0.832048466795696 | 89839.0793796099 | 80756.6492634159 | 69379.8756123744 | 84821.4005952368 | 60642.069759049 | 68744.5599359434 | 85073.7783259758 | 74416.0340347202 | Hspb7 |
| **ENSMUSG00000006235** | 0.0279607917816275 | 0.894341851882822 | 0.983733637008786 | 55.1143367107117 | 52.122606650517 | 56.7872933189068 | 63.779015209555 | 63.5075753575076 | 46.4771248188067 | 55.1771387994752 | 57.5328549274309 | Epor |
| **ENSMUSG00000006241** | 0.011026101259232 | 0.951868715566414 | 0.991915011884503 | 70.4740698923855 | 74.9262470601182 | 71.930571537282 | 73.1582821521366 | 66.7924499449649 | 78.277262852727 | 65.5879197050366 | 78.1578406561325 | Ccdc159 |
| **ENSMUSG00000006262** | 0.239579290955605 | 0.29376104222624 | 0.851785577030345 | 382.186302108706 | 544.029706914772 | 542.31865119556 | 486.783954319986 | 360.241246424483 | 280.085831144914 | 539.278450908079 | 475.460197324806 | Mob1b |
| **ENSMUSG00000006273** | 0.137672996236959 | 0.376821014164388 | 0.87422797717331 | 1454.65708367616 | 1558.79170514202 | 1837.06893886664 | 1490.36551717622 | 1438.77506930629 | 1113.00483118721 | 1620.9585869959 | 1589.2094266747 | Atp6v1b2 |
| **ENSMUSG00000006276** | 0.0845452818849201 | 0.581420293083873 | 0.922450244597442 | 654.143930795989 | 706.098436968723 | 671.036516051749 | 580.576623745802 | 589.087509350674 | 481.894399437101 | 700.64555494428 | 690.394259129171 | Eps15l1 |
| **ENSMUSG00000006281** | 0.143750856763776 | 0.46876398150262 | 0.895328112556867 | 166.246523848704 | 267.942774812814 | 242.292451494002 | 244.79886720138 | 215.706764576362 | 162.669936865823 | 226.955023741238 | 227.960368580387 | Tep1 |
| **ENSMUSG00000006288** | 0.00363039942298095 | 0.982693545797609 | 0.996940244108351 | 176.185174730964 | 159.625482867208 | 163.736695736181 | 161.323391412404 | 194.902558855799 | 184.685417043153 | 144.709854587303 | 135.690695583563 | Ttc5 |
| **ENSMUSG00000006289** | 0.0495360695213164 | 0.725109035013305 | 0.951674101666722 | 1015.54941742361 | 816.858976101072 | 936.043884873314 | 1125.51203310979 | 998.601874587016 | 1018.82749931753 | 931.764891047742 | 813.058647936712 | Osgep |
| **ENSMUSG00000006299** | 0.0238806588914 | 0.829361444989093 | 0.967483872587428 | 2706.92709484086 | 2308.05417574321 | 2461.72916537461 | 2750.00106756493 | 2569.86688558742 | 2788.6274891284 | 2426.75302908635 | 2274.17605798052 | Aamp |
| **ENSMUSG00000006301** | 0.12845696928325 | 0.385208292660729 | 0.875126835008975 | 1307.38434787541 | 1167.05773953423 | 1491.61290450995 | 1476.29661676235 | 1423.44565456483 | 1327.04422180014 | 1210.77381931679 | 1017.13745409439 | Tmbim1 |
| **ENSMUSG00000006304** | -0.0394779632650138 | 0.627592941452988 | 0.931825800800958 | 3414.37833491442 | 3133.87172486234 | 3144.12314009014 | 3161.75088634426 | 3360.42670296881 | 3535.93073292553 | 3166.95955147177 | 3149.10966310334 | Arpc2 |
| **ENSMUSG00000006307** | 0.14565214051317 | 0.38116303797061 | 0.875099933456492 | 395.739007857242 | 470.732291312482 | 478.906173656114 | 374.232751009006 | 414.989156215438 | 295.985900161874 | 431.006329490241 | 411.414189009364 | Kmt2b |
| **ENSMUSG00000006315** | 0.0805864376974259 | 0.639668861084373 | 0.933896249399425 | 2296.73186751851 | 1696.42796332855 | 1702.67234467856 | 2411.40953093773 | 1911.79700990014 | 2283.4945272819 | 1714.65561514596 | 1757.46588919831 | Tmem147 |
| **ENSMUSG00000006333** | 0.0778541680604404 | 0.686808358651425 | 0.944452327903597 | 12523.6036253635 | 8612.44633326902 | 11043.2356407501 | 13079.3877514301 | 10854.3205951547 | 13751.1135352064 | 8712.78253986431 | 9563.48022472955 | Rps9 |
| **ENSMUSG00000006335** | 0.00135628074448572 | 0.993060174482923 | 0.999133450854041 | 402.063603873225 | 340.425774686189 | 377.63550057073 | 387.363724728621 | 397.469825082332 | 458.655837027697 | 309.200192895173 | 341.94055287058 | Tfpt |
| **ENSMUSG00000006342** | -1.41051221089708 | 0.0098206364109752 | 0.635862253526407 | 25.2983840639333 | 25.2468875963442 | 19.8755526616174 | 23.448167356454 | 100.736154015357 | 103.961989726278 | 30.191264626128 | 15.1973579053591 | Susd2 |
| **ENSMUSG00000006344** | -1.01955683200707 | 0.0239475903677735 | 0.654124840968651 | 80.412720774645 | 71.6685841444609 | 111.681676860517 | 123.806323642077 | 297.828629262794 | 314.332133642982 | 100.984574783945 | 72.73021283279 | Ggt5 |
| **ENSMUSG00000006356** | -0.232643306516609 | 0.075131815124241 | 0.735399943172214 | 30647.185266022 | 27063.8490875521 | 25791.8421705588 | 26985.0889205016 | 31804.1557557615 | 39193.6701268068 | 29969.5149928395 | 28856.5260855829 | Crip2 |

| | | | | | | | | | | | | |
|---|---|---|---|---|---|---|---|---|---|---|---|---|
| **ENSMUSG00000006360** | -0.361785042445995 | 0.263381155198446 | 0.842510961779757 | 411.098741038915 | 345.312269059675 | 260.275094378323 | 361.101777289392 | 564.998429042654 | 671.472145408549 | 257.146288367366 | 277.894544555138 | Crip1 |
| **ENSMUSG00000006362** | -0.0807424677598184 | 0.670136376483891 | 0.94249140110542 | 625.231491865779 | 599.409976480946 | 651.160963390131 | 665.927952923294 | 884.72622222183 | 725.287763619799 | 494.512093014165 | 584.012753791657 | Cbfa2t3 |
| **ENSMUSG00000006369** | -0.940733483050523 | 0.00835212095054444 | 0.635862253526407 | 1227.87514081733 | 930.877178149078 | 1524.73882561265 | 1036.40899715527 | 2313.64666776575 | 4100.99472414365 | 1413.78404697523 | 1232.07151589876 | Fbln1 |
| **ENSMUSG00000006373** | -0.269353760518357 | 0.208928654608277 | 0.821566820583092 | 1074.27780900059 | 1084.80175091389 | 990.938268414924 | 1273.70445080258 | 1223.06830472993 | 1649.93793106764 | 865.13589325215 | 1594.63705449804 | Pgrmc1 |
| **ENSMUSG00000006378** | -0.155564264511204 | 0.327481866280356 | 0.855977678491063 | 300.870067617492 | 230.479651282755 | 217.684624389143 | 260.743621003769 | 320.822751374995 | 283.755077841135 | 274.84461590682 | 245.328777615083 | Gcat |
| **ENSMUSG00000006386** | -0.472950873901453 | 0.0948169541382973 | 0.757085157577544 | 717.389890955822 | 691.438953848265 | 821.522843346852 | 910.726820124675 | 1423.44565456483 | 1530.0758723244 | 688.152617857606 | 718.617923810552 | Tek |
| **ENSMUSG00000006390** | -0.629103921625353 | 0.106039195522106 | 0.762094734041179 | 219.553833126278 | 196.274190668353 | 218.631079277791 | 289.819348525772 | 568.283303630111 | 507.579126310652 | 174.901119213431 | 179.111718170304 | Elovl1 |
| **ENSMUSG00000006392** | 0.133933640002951 | 0.283249100449029 | 0.850587941933635 | 280.089252136404 | 285.859920848929 | 339.777305024792 | 328.274342990356 | 293.448796479518 | 281.308913376988 | 286.296474902938 | 262.697186649779 | Med8 |
| **ENSMUSG00000006395** | -0.0848299391158951 | 0.817970373062912 | 0.966315773308182 | 49.6932544112975 | 29.3189662409158 | 36.9117406572894 | 54.3997482669734 | 54.7479097909548 | 57.4848649074714 | 19.7804837205666 | 48.8486504100828 | Hyi |
| **ENSMUSG00000006398** | -0.380996821601451 | 0.297530546404157 | 0.851785577030345 | 19.877301764519 | 19.5459774939439 | 27.447191770805 | 28.1378008277448 | 36.1336204620302 | 46.4771248188067 | 19.7804837205666 | 21.7105112933701 | Cdc20 |
| **ENSMUSG00000006403** | -0.223837225759724 | 0.474959849159887 | 0.897549741595057 | 22.5878429142261 | 30.9477976987445 | 26.5007368821565 | 31.8895076047775 | 47.0832024202212 | 31.8001380339204 | 28.1091084450157 | 23.8815624227072 | Adamts4 |
| **ENSMUSG00000006412** | 0.0821499461520424 | 0.541629165685621 | 0.913837378403038 | 1397.73571953231 | 1079.9152565404 | 1104.51285505274 | 1390.94528758485 | 1168.32039493898 | 1291.57483707 | 1148.30913388342 | 1089.86766692718 | Pfdn2 |
| **ENSMUSG00000006418** | 0.308488818128457 | 0.036111758231006 | 0.679566075569698 | 2725.90088288881 | 2456.27783840562 | 3512.29409177439 | 3186.13698039497 | 2682.64757975679 | 2140.39390612925 | 2337.22031329853 | 2431.57726485746 | Rnf114 |
| **ENSMUSG00000006423** | 0.0343089064460693 | 0.867722814261167 | 0.974868041734905 | 331.58953398084 | 412.908774559565 | 415.493696116668 | 369.543117537716 | 279.21433993387 | 299.655146858096 | 461.197594116369 | 452.664160466768 | C330007P06 |
| **ENSMUSG00000006435** | 0.140214512112482 | 0.549113966989336 | 0.914699131434573 | 72.2810973255236 | 104.245213301034 | 79.5022106464695 | 56.2756016554897 | 67.887408140784 | 66.0464405319884 | 72.8754663389296 | 77.072315091464 | Neurl1a |
| **ENSMUSG00000006442** | -0.0557088979467915 | 0.724824914275884 | 0.951674101666722 | 234.010052591383 | 269.571606270643 | 216.738169500494 | 287.005568442997 | 298.923587458613 | 280.085831144914 | 227.996101831794 | 240.986675356409 | Srm |
| **ENSMUSG00000006445** | -0.833774515846912 | 0.0248294331478715 | 0.655303666217042 | 123.781379169959 | 172.656134529838 | 101.270673085384 | 76.9099889291692 | 331.772333333186 | 260.516515431732 | 133.257995591185 | 121.578863242873 | Epha2 |
| **ENSMUSG00000006456** | -0.258157828635055 | 0.20100332419065 | 0.812921911423739 | 192.448421629207 | 230.479651282755 | 173.201244622666 | 172.578511743502 | 281.404256325508 | 266.631926592101 | 190.517290571773 | 182.368294864309 | Rbm14 |
| **ENSMUSG00000006462** | 0.471424844923389 | 0.0347832395433218 | 0.679566075569698 | 77.7021796249379 | 64.338842584232 | 69.0912068713366 | 66.5927952923295 | 51.4630352034976 | 63.6002760678407 | 47.8895921655823 | 37.9933947633978 | A530013C23 |
| **ENSMUSG00000006463** | -0.33294317924987 | 0.195743775913836 | 0.811193545163524 | 43.3686583953142 | 39.9063707168021 | 38.8046504345863 | 49.7101147956825 | 64.6025335533267 | 67.2695227640623 | 41.6431236222455 | 43.4210225867403 | Zdhhc24 |
| **ENSMUSG00000006464** | -0.34242346815533 | 0.374959177052253 | 0.873700364743909 | 32.5264937964856 | 22.8036404096012 | 46.3762895437739 | 47.8342614071662 | 42.7033696369448 | 23.2385624094033 | 51.0128264372507 | 71.6446872681215 | Bbs1 |
| **ENSMUSG00000006471** | 0.0659108838591746 | 0.664747702227385 | 0.941513723457524 | 141.85165350134 | 130.306516626293 | 144.807597963212 | 136.937297361692 | 134.679858085749 | 118.638976511164 | 151.997401221196 | 122.664388807541 | Ndor1 |
| **ENSMUSG00000006476** | -0.509604277753479 | 0.0402162213188115 | 0.679566075569698 | 80.412720774645 | 85.5136515360045 | 68.1447519826882 | 86.2892558717508 | 116.065568756824 | 165.116101329971 | 90.5737938783839 | 85.7565196088121 | Nsmf |
| **ENSMUSG00000006494** | 0.385851047879391 | 0.148375398421095 | 0.781357001591751 | 6431.21063453846 | 8110.7662442578 | 9086.91338591374 | 6603.00392757746 | 4494.80339383739 | 3487.00744364257 | 7589.45928015424 | 7565.02766017483 | Pdk1 |
| **ENSMUSG00000006498** | -0.456995484566164 | 0.0638178339107091 | 0.716991331547963 | 321.65088309858 | 357.52850499339 | 422.118880337207 | 468.963347129081 | 631.790878987619 | 736.295503708463 | 408.102611498006 | 379.933947633978 | Ptbp1 |
| **ENSMUSG00000006517** | -0.0690507115302487 | 0.720792682485352 | 0.951674101666722 | 103.000563688871 | 74.9262470601182 | 82.3415753124149 | 102.23400967414 | 102.926070406995 | 105.185071958352 | 85.3684034256032 | 86.8420451734806 | Mvd |
| **ENSMUSG00000006519** | -0.254175023857363 | 0.31696957338166 | 0.854283473529811 | 553.853908256825 | 394.991628523449 | 569.765842966365 | 690.314046974007 | 705.153078107498 | 903.857769502582 | 431.006329490241 | 594.868009438342 | Cyba |
| **ENSMUSG00000006522** | -0.440273529738873 | 0.313923395176547 | 0.853228028313558 | 59.631905293557 | 202.789516499668 | 88.0203046443056 | 96.6064495085906 | 68.9823663366031 | 123.53130543946 | 182.188665847324 | 232.302470839061 | Itih3 |
| **ENSMUSG00000006526** | -0.310583430535239 | 0.152944402703384 | 0.783412422888014 | 186.123825613223 | 178.357044632238 | 130.610774633486 | 201.654239265505 | 245.270635863478 | 270.301173288323 | 173.860041122875 | 175.855141476298 | Tmem110 |
| **ENSMUSG00000006527** | 0.0411130669345219 | 0.793210439527835 | 0.961804768755116 | 319.843855665442 | 336.353696041618 | 281.097101928589 | 282.315934971707 | 307.683253025166 | 231.162541861959 | 306.076958623504 | 339.769501741243 | Sfmbt1 |
| **ENSMUSG00000006529** | -0.495993579208583 | 0.139319635668811 | 0.779711473022872 | 55.1143367107117 | 103.43079757212 | 72.8770264259304 | 69.4065753751039 | 71.1722827282413 | 79.5003450848009 | 96.8202624217207 | 176.940667040967 | Itih1 |
| **ENSMUSG00000006542** | 0.18826498918713 | 0.499560325101462 | 0.900927059989498 | 103.90407740544 | 84.6992358070902 | 48.2691993210708 | 61.9031618210386 | 75.5521155115177 | 53.8156182112498 | 63.5057635239243 | 69.4736361387845 | Prkag3 |
| **ENSMUSG00000006574** | 1.24083395109323 | 0.33296841286868 | 0.856918526232803 | 65.9565013095403 | 41.5352021746308 | 494.995906763138 | 6.56548685980713 | 39.4184950494875 | 181.016170346931 | 13.5340151772298 | 23.8815624227072 | Slc4a1 |
| **ENSMUSG00000006575** | 0.134195160844304 | 0.566110961816984 | 0.919931558693298 | 56.0178504272808 | 52.122606650517 | 53.001473764313 | 51.5859681841989 | 39.4184950494875 | 64.8233582999146 | 42.6842017128016 | 47.7631248454143 | Rundc3a |
| **ENSMUSG00000006576** | 0.0441013802587022 | 0.699232678821309 | 0.945240088450412 | 5510.53015735461 | 6280.7741013873 | 6958.33634134338 | 5502.81591521263 | 5732.10615511297 | 5753.37881967544 | 6382.84977319967 | 5653.41714079359 | Slc4a3 |
| **ENSMUSG00000006585** | -0.105694074843579 | 0.80134502367592 | 0.963461896828986 | 19.877301764519 | 13.0306516626293 | 28.3936466594534 | 23.448167356454 | 25.1840385038392 | 35.4693847301419 | 18.7394056300105 | 11.9407812113536 | Cdt1 |
| **ENSMUSG00000006586** | -0.500767083504489 | 0.131017698872288 | 0.77303252008843 | 64.1494738764022 | 62.7100111264033 | 38.8046504345863 | 39.3929211588428 | 108.400861386091 | 73.3849339244316 | 66.6289977955927 | 42.3354970220718 | Runx1t1 |
| **ENSMUSG00000006587** | -0.107438786459023 | 0.820404726087843 | 0.966398681144067 | 21.6843291976571 | 14.6594831204579 | 16.0897331070236 | 36.5791410760683 | 19.7092475247437 | 11.0077400886647 | 40.6020455316893 | 23.8815624227072 | Snai3 |
| **ENSMUSG00000006589** | -0.0653136025304792 | 0.75573492659431 | 0.956566265296272 | 369.537110076739 | 250.025628776699 | 260.275094378323 | 341.405316709971 | 390.900075907418 | 359.586176229715 | 280.050006359601 | 247.49982874442 | Aprt |
| **ENSMUSG00000006599** | 0.112236492050475 | 0.412675438129315 | 0.882192126163777 | 592.704998069294 | 609.997380956832 | 551.783200082044 | 524.301022090312 | 491.636229922775 | 440.309603546589 | 593.414511616998 | 581.84170266232 | Gtf2h1 |
| **ENSMUSG00000006611** | 0.00606488261553297 | 0.944439087462601 | 0.990505437194639 | 931.522641782685 | 962.639391576736 | 1006.0815466333 | 967.940348474422 | 1005.17162376193 | 1012.71208815716 | 900.532548331058 | 934.637511179585 | Hfe |
| **ENSMUSG00000006638** | 0.972630768633071 | 0.0517056603662044 | 0.703396849069134 | 17.1667606148119 | 13.0306516626293 | 33.1259211026956 | 15.0068271081306 | 9.85462376237187 | 7.33849339244316 | 6.24646854333682 | 16.2828834700276 | Abhd1 |
| **ENSMUSG00000006641** | -0.0394735848119553 | 0.840298174492986 | 0.969957774070128 | 265.633032671299 | 290.746415222415 | 318.955297474527 | 195.088752405698 | 257.315176017488 | 226.270212933664 | 320.65205189129 | 295.262953589834 | Slc5a6 |
| **ENSMUSG00000006642** | -0.0801083558974184 | 0.834426476123608 | 0.968516656512387 | 160.82544154929 | 216.634583891211 | 282.990011705886 | 156.633757941113 | 300.018545654433 | 329.009120427868 | 132.216917500629 | 103.124928643508 | Tcf23 |
| **ENSMUSG00000006651** | -0.459567066279106 | 0.174837119377559 | 0.799889021902745 | 51.5002818444356 | 61.895595397489 | 58.6802030962037 | 39.3929211588428 | 95.2613630362614 | 99.0696607979826 | 61.4236073428121 | 35.8223436340608 | Aplp1 |
| **ENSMUSG00000006673** | 0.0706877848605666 | 0.548098287052908 | 0.914699131434573 | 819.486940928124 | 867.35275129376 | 784.611102689562 | 741.900015158206 | 819.028730472685 | 649.456665231219 | 784.972880279327 | 805.459968984033 | Qrich1 |
| **ENSMUSG00000006675** | -0.0747845578596679 | 0.698006559643908 | 0.944933386973863 | 274.66816983699 | 272.014853457386 | 331.259211026956 | 310.453735799451 | 359.146288228664 | 377.932409710823 | 293.584021536831 | 221.447215192376 | P4htm |
| **ENSMUSG00000006676** | 0.118790326842356 | 0.530994106249497 | 0.909082872831377 | 2415.99567810563 | 3036.95625312153 | 3103.42557987826 | 2442.36111184825 | 2446.13660945986 | 1833.40026587872 | 3182.57572283011 | 2666.05078682585 | Usp19 |
| **ENSMUSG00000006678** | -0.00390403703087145 | 0.985579149430734 | 0.997573007767463 | 75.8951521917998 | 80.6271571625185 | 53.001473764313 | 75.9720622349111 | 68.9823663366031 | 59.9310293716191 | 80.1630129728225 | 77.072315091464 | Pola1 |
| **ENSMUSG00000006699** | -0.032527218562065 | 0.782496432228402 | 0.960742105865155 | 3999.85522325116 | 3664.87078011448 | 3619.24349419166 | 3809.85823207665 | 4210.11426292443 | 4296.68788127547 | 3289.80676615739 | 3643.02379502751 | Cdc42 |
| **ENSMUSG00000006705** | 0.00446495228259532 | 0.980134384973915 | 0.996872525500296 | 184.316798180085 | 201.160685041839 | 273.525462819401 | 195.088752405698 | 204.757182618171 | 185.908499275227 | 245.694429371248 | 213.848536239696 | Pknox1 |
| **ENSMUSG00000006715** | 0.118448496921711 | 0.384390405248351 | 0.875126835008975 | 1385.99004121691 | 1081.54408799823 | 1015.54609551978 | 1241.81494319781 | 1196.78930803027 | 988.250443515679 | 1069.18719900115 | 1097.46634587986 | Gmnn |
| **ENSMUSG00000006717** | 0.16108466775846 | 0.244590361952456 | 0.832048466795696 | 6261.35005582348 | 4558.2848347335 | 4885.60013520328 | 6007.42047672352 | 5053.23207370513 | 5236.01503550819 | 4620.30456588813 | 4509.27319563298 | Acot13 |
| **ENSMUSG00000006728** | 0.0302222654022108 | 0.8331751245447 | 0.967868401766821 | 849.302893574902 | 732.974156022896 | 683.340429604179 | 822.561710864407 | 840.927894389066 | 853.711397987554 | 666.289977955927 | 664.341645577127 | Cdk4 |
| **ENSMUSG00000006731** | -0.647138386771038 | 0.0182760365898338 | 0.642273034804458 | 23.4913566307952 | 34.2054606144018 | 36.9117406572894 | 46.8963347129081 | 52.5579933993167 | 67.2695227640623 | 55.1771387994752 | 46.6775992807458 | B4galnt1 |
| **ENSMUSG00000006732** | 0.161534213915466 | 0.410209162728764 | 0.882192126163777 | 299.966553900923 | 264.685111897157 | 299.079744812909 | 374.232751009006 | 307.683253025166 | 346.132271676902 | 230.078258012906 | 223.618266321713 | Mettl1 |
| **ENSMUSG00000006736** | -0.171316674082986 | 0.0962685926736506 | 0.757809915356977 | 949.592916114066 | 848.62118952873 | 797.861471130641 | 854.451218469185 | 1020.5010385034 | 1005.37359476471 | 883.87529888216 | 976.973008201657 | Tspan31 |
| **ENSMUSG00000006740** | 0.0614664778596804 | 0.574762344520996 | 0.922450244597442 | 7886.77123193119 | 7500.76886330097 | 6446.30424658457 | 7638.47499803846 | 6386.89115621279 | 6819.90652604384 | 7646.71857513482 | 7389.17251869853 | Kif5b |
| **ENSMUSG00000006763** | -0.248049492825743 | 0.254260493297417 | 0.837406774112203 | 81.316234491214 | 96.9154717408051 | 84.2344850897118 | 107.861569839689 | 143.439523652302 | 116.192812047017 | 91.61487196894 | 89.0130963028176 | Saal1 |
| **ENSMUSG00000006782** | -0.601178190460783 | 0.0759235318763737 | 0.735399943172214 | 121.974351736821 | 107.502876216691 | 89.9132144216024 | 120.992543559303 | 208.042057205628 | 260.516515431732 | 85.3684034256032 | 115.065709854862 | Cnp |
| **ENSMUSG00000006800** | -0.170272923381776 | 0.200262766132528 | 0.812921911423739 | 1035.42671918813 | 1008.24667239594 | 855.595219338196 | 907.9130400419 | 955.898504950072 | 1260.99778126815 | 1086.88552654061 | 982.400636024999 | Sulf2 |
| **ENSMUSG00000006818** | 0.0313862353112678 | 0.727412235230424 | 0.952237115597464 | 16264.1504119594 | 16353.4678365997 | 15453.7154218518 | 15828.4508923007 | 15052.3903179251 | 14085.0149845626 | 16571.8810454726 | 16813.7054711505 | Sod2 |
| **ENSMUSG00000006850** | -0.177935965561078 | 0.429268577690033 | 0.885743677088682 | 62.3424464432641 | 44.7928650902881 | 48.2691993210708 | 45.0204813243917 | 59.1277425742312 | 53.8156182112498 | 63.5057635239243 | 49.9341759747513 | Tmco6 |
| **ENSMUSG00000006906** | 0.0394659715537989 | 0.81115084742929 | 0.964955219787201 | 259.308436655316 | 247.582381589956 | 251.757000380487 | 299.198615468353 | 273.739548954774 | 315.555215875056 | 219.667477107345 | 221.447215192376 | Stambp |
| **ENSMUSG00000006920** | 0.266642912006152 | 0.186462491327274 | 0.807093800104656 | 892.671551970216 | 1294.92100897378 | 1662.92123935532 | 1150.83605385476 | 1179.26997689717 | 894.073111645991 | 1087.92660463116 | 995.426942801021 | Ezh1 |
| **ENSMUSG00000006930** | -0.449510885538599 | 0.340007919635504 | 0.858570960161243 | 9.03513716569045 | 15.4738988493722 | 16.0897331070236 | 11.2551203310979 | 21.8991639163819 | 28.1308913376988 | 8.32862472444909 | 13.0263067760221 | Hap1 |
| **ENSMUSG00000006931** | 0.0582493423208684 | 0.669275680865887 | 0.942250071683273 | 995.672115659088 | 1031.05031280554 | 1325.98329899647 | 1179.91178137677 | 1101.52794499401 | 1226.75147877008 | 1026.50299728835 | 998.683519495027 | P3h4 |
| **ENSMUSG00000006932** | 0.0803738840605131 | 0.539129935442219 | 0.913567744698356 | 8751.43385868777 | 9889.45019620669 | 10428.9864180172 | 9396.14962307826 | 8524.24955445167 | 7617.356141356 | 10252.5370357968 | 9985.7496693856 | Ctnnb1 |
| **ENSMUSG00000006941** | 0.023962191833838 | 0.872562791754054 | 0.976162349063696 | 1185.40999613859 | 1006.61784093811 | 927.525790875478 | 1049.53997087488 | 1167.22543674316 | 1155.8127093098 | 877.628830338823 | 900.986218674861 | Eif1b |
| **ENSMUSG00000006958** | -1.16770766036523 | 0.0192717654260465 | 0.642273034804458 | 13.5527057485357 | 6.51532583131463 | 12.3039135524298 | 14.0689004138724 | 44.893286028583 | 30.5770558018465 | 15.616171358342 | 13.0263067760221 | Chrd |
| **ENSMUSG00000006998** | 0.183021495710072 | 0.0430912250957128 | 0.687730553581215 | 8418.94081099036 | 8399.06941229347 | 8956.30261128025 | 9170.10928976204 | 7959.25112540902 | 6848.03741738154 | 7967.37062702611 | 8003.57998830091 | Psmd2 |
| **ENSMUSG00000007029** | 0.0362224207520389 | 0.749497643369905 | 0.956197192463153 | 1104.09376164737 | 1157.28475078726 | 1154.67496415111 | 1120.8223996385 | 1215.4035973592 | 1223.08223207386 | 991.106342209442 | 996.51246836569 | Vars |
| **ENSMUSG00000007030** | 0.150256700404416 | 0.465678349095293 | 0.895328112556867 | 515.002818444356 | 650.718167402549 | 518.657278979349 | 519.611388619021 | 387.61520131996 | 382.824738639118 | 630.893322877019 | 584.012753791657 | Vwa7 |
| **ENSMUSG00000007033** | 0.25767093955353 | 0.114400479403111 | 0.767518180080285 | 296.352499034647 | 291.56083095133 | 247.024725937245 | 334.839829850164 | 218.991639163819 | 250.731857575141 | 293.584021536831 | 214.934061804364 | Hspa1l |
| **ENSMUSG00000007036** | 0.0407977707059027 | 0.692412342888345 | 0.94469184542218 | 1381.47247263407 | 1309.58049209424 | 1356.26985543322 | 1512.87575783841 | 1432.20532013138 | 1413.88306027738 | 1362.77122053798 | 1196.2491722647 | Abhd16a |
| **ENSMUSG00000007038** | -0.2948068654917 | 0.0241588224202245 | 0.654124840968651 | 440.011179969125 | 377.888898216248 | 440.101523221528 | 438.949692912819 | 589.087509350674 | 536.933099880424 | 512.210420553619 | 442.894430384751 | Neu1 |
| **ENSMUSG00000007039** | -0.588045665003891 | 0.0755158455145013 | 0.735399943172214 | 309.905204783182 | 245.139134403213 | 226.202718386979 | 326.39848960184 | 504.775728272604 | 661.687487551958 | 226.955023741238 | 272.466916731795 | Ddah2 |
| **ENSMUSG00000007041** | -0.517378738699858 | 0.0953784086344418 | 0.757085157577544 | 571.924182588206 | 365.672662282534 | 466.602260103684 | 600.273084325223 | 832.168228822514 | 1114.22791341929 | 443.499266576914 | 479.80229958348 | Clic1 |
| **ENSMUSG00000007050** | 0.00303073406039665 | 0.989828774685572 | 0.998628446692557 | 214.132750826864 | 160.439898596123 | 111.681676860517 | 172.578511743502 | 205.85214081399 | 170.008430258266 | 131.175839410073 | 150.888053488923 | Lsm2 |
| **ENSMUSG00000007080** | 0.0187081116523674 | 0.935185982150934 | 0.98818087308472 | 40.658117245607 | 43.1640336324594 | 43.5369248778286 | 45.9584080186499 | 39.4184950494875 | 51.3694537471021 | 40.6020455316893 | 40.1644458927348 | Pole |
| **ENSMUSG00000007097** | 0.185601323943639 | 0.0240296633401609 | 0.654124840968651 | 28535.6737104001 | 29471.2619822228 | 32945.1482189638 | 31858.556023867 | 27045.4674367317 | 25832.719823632 | 27091.9751505423 | 28013.0727218355 | Atp1a2 |
| **ENSMUSG00000007122** | -0.607750140079434 | 0.430246317092585 | 0.886017087288711 | 231.299511441676 | 188.944449108124 | 24.6078271048596 | 171.640585049243 | 98.5462376237187 | 479.448234972953 | 245.694429371248 | 116.15123541953 | Casq1 |
| **ENSMUSG00000007216** | -0.00692575822211132 | 0.950073396778315 | 0.991548537174153 | 585.476888336741 | 536.699965354543 | 504.460455649622 | 543.997482669733 | 498.205979097689 | 558.948580057754 | 600.702058250891 | 523.223322170221 | Zfp775 |
| **ENSMUSG00000007279** | -0.054297294031402 | 0.835568296841431 | 0.968865818776815 | 517.713359594063 | 502.494504740141 | 318.008842585878 | 433.32213274727 | 329.582416941548 | 333.901449356164 | 559.058934628645 | 616.578520731712 | Scube2 |
| **ENSMUSG00000007338** | 0.00021929230757973 | 0.998471003125833 | 0.999262355603273 | 803.223694029881 | 699.583111137408 | 688.072704047421 | 745.651721935238 | 685.443830582755 | 687.372214425509 | 729.795741479852 | 832.598108100745 | Mrpl49 |
| **ENSMUSG00000007379** | 0.0479501145328875 | 0.878563759245065 | 0.978059679082628 | 56.9213641438498 | 27.6901347830872 | 38.8046504345863 | 45.0204813243917 | 55.8428679867739 | 34.2463024980681 | 32.2734208072402 | 40.1644458927348 | Dennd2c |
| **ENSMUSG00000007411** | 0.059877191634199 | 0.670955012102691 | 0.943084913793401 | 934.233182932393 | 1018.01966114291 | 953.080072868986 | 882.58901929693 | 920.859842683861 | 711.833859066986 | 1035.87270010336 | 963.946701425635 | Mark3 |

| | | | | | | | | | | | | |
|---|---|---|---|---|---|---|---|---|---|---|---|---|
| **ENSMUSG00000007415** | 0.067429253808325 | 0.626583361561965 | 0.931698409533928 | 2236.19644850839 | 2403.34081602618 | 2432.38906382651 | 2034.36299984595 | 1999.39366556567 | 1811.38478570139 | 2413.21901390912 | 2465.22855736218 | Gatad1 |
| **ENSMUSG00000007458** | 0.0128486924381951 | 0.891044325077969 | 0.982142775127356 | 1001.99671167507 | 971.597964594794 | 1084.63730239112 | 1068.29850476005 | 1067.58424092362 | 1028.61215717412 | 933.847047228855 | 1059.47295111646 | M6pr |
| **ENSMUSG00000007476** | -0.401162792204038 | 0.19234025330179 | 0.808824684419004 | 486.993893230715 | 473.175538499225 | 335.04503058155 | 367.667264149199 | 752.23628052772 | 753.418654957498 | 383.116737324658 | 308.289260365856 | Lrrc8a |
| **ENSMUSG00000007564** | -0.134748701796598 | 0.0517073225901499 | 0.703396849069134 | 3503.82619285476 | 3558.99673535562 | 3509.45472710844 | 3349.33622519589 | 3861.91755665395 | 3976.24033647212 | 3827.00306088436 | 3621.31328373414 | Ppp2r1a |
| **ENSMUSG00000007570** | 0.0181951662490146 | 0.928973950445018 | 0.986954885397071 | 178.895715880671 | 207.676010873154 | 159.004421292939 | 203.530092654021 | 197.092475247437 | 243.393364182698 | 148.874166949528 | 151.973579053591 | Fance |
| **ENSMUSG00000007589** | -0.0503383387101738 | 0.747524424315354 | 0.956119556124081 | 181.606257030378 | 162.068730053951 | 170.36187995672 | 202.592165959763 | 221.181555555458 | 176.123841418636 | 176.983275394543 | 167.17093695895 | Tinf2 |
| **ENSMUSG00000007603** | 0.00904746888193437 | 0.945349955429295 | 0.990505437194639 | 280.089252136404 | 315.99330281876 | 342.616669690738 | 340.467390015713 | 316.442918591719 | 326.562955963721 | 348.761160336306 | 278.980070119806 | Dus3l |
| **ENSMUSG00000007610** | 0.104740312766755 | 0.375323846583863 | 0.873821423608136 | 267.440060104437 | 267.1283590839 | 299.079744812909 | 278.564228194674 | 264.979883388221 | 237.277953022329 | 252.981976005141 | 277.894544555138 | Gtpbp3 |
| **ENSMUSG00000007613** | 0.0253300771849757 | 0.867899094167226 | 0.974868041734905 | 383.993329541844 | 509.82424630037 | 468.495169880981 | 355.474217123843 | 440.173194719277 | 402.3940543523 | 397.691830592444 | 448.322058208094 | Tgfbr1 |
| **ENSMUSG00000007617** | -0.0701801740172003 | 0.742091375701726 | 0.954383574136061 | 1067.04969926804 | 1160.54241370292 | 982.420174417088 | 816.934150698859 | 855.162350934715 | 796.226533080083 | 1405.45542225078 | 1170.19655871265 | Homer1 |
| **ENSMUSG00000007646** | 0.350455115189885 | 0.262552987655794 | 0.841952818014037 | 24.3948703473642 | 26.0613033252585 | 27.447191770805 | 35.6412143818101 | 26.2789966996583 | 25.6847268735511 | 16.6572494488982 | 20.6249857287016 | Rad51c |
| **ENSMUSG00000007655** | -0.646820969666729 | 0.0607416641082121 | 0.71282992338528 | 3092.72745181584 | 2767.38464685089 | 3143.17668520149 | 3788.28591810871 | 6886.1920935063 | 7691.9641575125 | 2593.32552357534 | 2857.10328620751 | Cav1 |
| **ENSMUSG00000007656** | 0.024081468229915 | 0.849775726925994 | 0.972640876584422 | 2436.77649358671 | 2289.32261397818 | 2354.77976295734 | 2710.60814640609 | 2180.06176787582 | 2836.32769617928 | 2454.86213753137 | 2159.11034812566 | Arpp19 |
| **ENSMUSG00000007659** | 0.0650718954554191 | 0.785652374259555 | 0.961141387070613 | 2341.0040396304 | 2455.4634226767 | 2742.8262673032 | 1370.31090031117 | 2634.46941914075 | 2273.7098694253 | 1992.62346532445 | 1616.34756579141 | Bcl2l1 |
| **ENSMUSG00000007670** | 0.155238501450536 | 0.10755815259765 | 0.762287160109383 | 721.003945822098 | 803.828324438442 | 816.79056890361 | 771.913669374467 | 742.381656765348 | 662.910569784032 | 668.37213413704 | 721.874500504558 | Khsrp |
| **ENSMUSG00000007682** | -0.472845473343984 | 0.354328787516558 | 0.863443308527191 | 27.1054114970714 | 22.8036404096012 | 14.1968233297267 | 26.2619474392285 | 15.3294147414674 | 61.154111603693 | 36.4377331694648 | 13.0263067760221 | Dio2 |
| **ENSMUSG00000007721** | -0.0638974593263034 | 0.768005571770952 | 0.95879680569409 | 2167.52940604914 | 1612.54314325037 | 1694.15425068072 | 2223.8241920861 | 2051.95165896499 | 2782.51207796803 | 1677.17680388594 | 1536.01867400594 | Ccdc124 |
| **ENSMUSG00000007739** | 0.172018497977476 | 0.118306314822261 | 0.769868040577702 | 4309.76042803434 | 3626.59324085551 | 4037.57655497427 | 4737.46773269797 | 3778.7007337717 | 3802.56265951763 | 3486.5705252725 | 3764.60265827038 | Cct4 |
| **ENSMUSG00000007777** | 0.0010171292504529 | 0.992670129269971 | 0.999071668397934 | 455.370913150799 | 454.443976734195 | 387.100049457215 | 472.715053906113 | 425.938738173629 | 431.748027922072 | 458.0743598447 | 452.664160466768 | 0610009B22I |
| **ENSMUSG00000007783** | -1.46690746912368 | 0.00993872819514987 | 0.635862253526407 | 8.13162344912141 | 3.25766291565731 | 11.3574586637814 | 10.3171936368398 | 22.994122112201 | 42.8078781225851 | 14.5750932677859 | 10.8552556466851 | Cpt1c |
| **ENSMUSG00000007812** | 0.25954762802629 | 0.0534448077320285 | 0.703396849069134 | 644.205279913729 | 570.905425968944 | 752.431636475515 | 592.769670771158 | 549.669014301187 | 469.663577116362 | 570.510793624763 | 547.104884592928 | Zfp655 |
| **ENSMUSG00000007815** | -0.0317040846492647 | 0.778850248118023 | 0.960254744040417 | 5479.81069099126 | 5029.83154177489 | 5688.19388077717 | 5725.10454175182 | 6407.69536193336 | 5776.61738208484 | 5033.61256783892 | 5192.06877580947 | Rhoa |
| **ENSMUSG00000007817** | -0.201440853364369 | 0.219236674782664 | 0.827535829146756 | 780.635851115655 | 798.941830064956 | 969.16980597601 | 713.762214330461 | 1158.4657711766 | 868.38838477244 | 936.970281500523 | 787.006034384668 | Zmiz1 |
| **ENSMUSG00000007827** | -0.301498726159133 | 0.241923305373159 | 0.832048466795696 | 69.5705561758165 | 59.452348210746 | 45.4298346551254 | 64.7169419038131 | 89.7865720571659 | 48.9232892829544 | 68.711153976705 | 86.8420451734806 | Ankrd26 |
| **ENSMUSG00000007833** | -0.0439743643074365 | 0.795474329038184 | 0.962043367573119 | 330.68602026427 | 329.838370210303 | 368.170951684246 | 332.963976461647 | 348.196706270473 | 458.655837027697 | 296.707255808499 | 301.776106977845 | Aldh16a1 |
| **ENSMUSG00000007836** | -0.0910519871867718 | 0.633785352700743 | 0.9325328237529 | 1672.4038893693 | 2078.38894018937 | 2120.05895057252 | 2109.3971353866 | 2607.09546424527 | 2517.103233608 | 1732.35394268541 | 1644.57123047279 | Hnrnpa0 |
| **ENSMUSG00000007837** | -0.169901592139842 | 0.480624435704161 | 0.897732059433541 | 165.343010132135 | 136.007426728693 | 108.842312194571 | 152.88205116408 | 173.003394939417 | 215.262472844999 | 130.134761319517 | 116.15123541953 | Prrg2 |
| **ENSMUSG00000007850** | -0.0458241855038157 | 0.748103436077777 | 0.956197192463153 | 1615.48252522545 | 1652.44951396717 | 1602.34812648182 | 1629.17866792643 | 1578.92971837114 | 1319.70572840769 | 1965.55543496999 | 1843.22240880713 | Hnrnph1 |
| **ENSMUSG00000007867** | -0.0210785946556764 | 0.924066004447117 | 0.986713918026868 | 609.871758684105 | 413.723190288479 | 425.904699891801 | 625.597105070194 | 494.921104510232 | 706.941530138691 | 424.759860946904 | 479.80229958348 | Ift43 |
| **ENSMUSG00000007872** | -0.678877478290531 | 0.0506638092757691 | 0.703396849069134 | 713.775836089546 | 475.618785685968 | 590.587850516631 | 841.320244749571 | 1523.08685038436 | 1473.814089649 | 577.798340258656 | 622.006148555055 | Id3 |
| **ENSMUSG00000007877** | -0.241089320445354 | 0.306922793341462 | 0.851785577030345 | 46981.8097478738 | 32659.6995609224 | 31735.5788712711 | 36633.5408243353 | 46674.7830131806 | 59226.5340059446 | 40875.8490695056 | 28155.2765708071 | Tcap |
| **ENSMUSG00000007880** | -0.0300999095047257 | 0.859251945121192 | 0.974868041734905 | 757.14449448486 | 1063.62694196211 | 969.16980597601 | 785.044643094081 | 942.759006600242 | 722.841599155651 | 1020.25652874501 | 963.946701425635 | Arid1a |
| **ENSMUSG00000007888** | -2.20864627577956 | 0.00718698472853088 | 0.635862253526407 | 9.93865088225949 | 4.88649437348597 | 1.89290977729689 | 11.2551203310979 | 32.8487458745729 | 81.9465095489486 | 9.36970281500523 | 5.42762782334254 | Crlf1 |
| **ENSMUSG00000007891** | -0.111204102128434 | 0.348269430351383 | 0.861266047293794 | 23891.6132072353 | 21961.5345459038 | 22450.8564136298 | 24160.9916440902 | 25349.3771914079 | 29426.135421465 | 22244.715560913 | 22855.7407640954 | Ctsd |
| **ENSMUSG00000007892** | 0.0380950378336224 | 0.84824503605128 | 0.972640876584422 | 12323.9270940018 | 8761.48441166035 | 10283.2323651654 | 12687.3343932301 | 11129.1551023053 | 14064.2225866173 | 8527.47063974532 | 9187.88837935425 | Rplp1 |
| **ENSMUSG00000007950** | -0.467795130545143 | 0.0159038855105849 | 0.63777231095375 | 276.475197270128 | 262.241864710414 | 224.309808609682 | 279.502154888932 | 370.095870186855 | 480.671317205027 | 306.076958623504 | 286.578749072486 | Abhd8 |
| **ENSMUSG00000007987** | 0.0349262726419962 | 0.831272163970661 | 0.967583158900775 | 274.66816983699 | 228.850819824926 | 212.952349945901 | 242.923013812864 | 199.282391639076 | 269.078091056249 | 262.351678820146 | 206.249857287016 | Ift22 |
| **ENSMUSG00000007989** | 0.271834653769116 | 0.521542845088248 | 0.90609507420585 | 15.3597331816738 | 15.4738988493722 | 12.3039135524298 | 10.3171936368398 | 7.66470737073368 | 13.4539045528125 | 10.4107809055614 | 13.0263067760221 | Fzd3 |
| **ENSMUSG00000008035** | -0.157395441188113 | 0.258908779453679 | 0.838606660810377 | 2900.27903018663 | 2709.56113009797 | 2384.11986450544 | 2045.61812017705 | 2779.00390098887 | 3089.50571821857 | 2768.22664278877 | 2561.84033261768 | Mid1ip1 |
| **ENSMUSG00000008036** | -0.0360196362365044 | 0.851398544846732 | 0.972692784625411 | 898.9961479862 | 818.4878075589 | 770.414279359836 | 1018.58838996436 | 916.480009900584 | 1202.2898341286 | 729.795741479852 | 747.927114056602 | Ap2s1 |
| **ENSMUSG00000008090** | 0.182394720103058 | 0.148834623653359 | 0.78151662300109 | 580.055806037327 | 595.337897836374 | 526.228918088536 | 576.824916968769 | 592.372383938131 | 445.201932474885 | 497.635327285833 | 472.203620630801 | Fgfrl1 |
| **ENSMUSG00000008136** | -0.121851374851216 | 0.31825245776759 | 0.854351922245337 | 19330.6759659947 | 17786.8395194889 | 15575.8081024875 | 17279.4234883181 | 16587.5217084635 | 18062.4784032668 | 20280.2012040335 | 21208.9984824933 | Fhl2 |
| **ENSMUSG00000008140** | -0.229068017223343 | 0.0636855308809096 | 0.716991331547963 | 2004.89693706671 | 2156.57285016514 | 2185.36433788926 | 2070.00421422776 | 2633.37446094493 | 2862.01242305283 | 2203.96231770734 | 2167.79455264301 | Emc10 |
| **ENSMUSG00000008153** | -0.0279023000932358 | 0.957669440755128 | 0.992540556043032 | 8.13162344912141 | 33.3910448854875 | 18.9290977729689 | 13.1309737196143 | 10.949581958191 | 18.3462334811079 | 14.5750932677859 | 31.4802413753867 | Clstn3 |
| **ENSMUSG00000008167** | -0.118978209940504 | 0.639854170086805 | 0.933921420069286 | 40.658117245607 | 31.7622134276588 | 51.1085639870161 | 37.5170677703264 | 39.4184950494875 | 51.3694537471021 | 41.6431236222455 | 42.3354970220718 | Fbxw9 |
| **ENSMUSG00000008200** | 0.0465669676725716 | 0.880489258129254 | 0.978433072023387 | 95.7724539563188 | 124.605606523892 | 184.558703286447 | 137.87522405595 | 119.350443344282 | 73.3849339244316 | 188.435134390661 | 143.289374536243 | Fnbp4 |
| **ENSMUSG00000008206** | -0.125757923459598 | 0.446560274432 | 0.89108031640712 | 358.694945477911 | 414.537606017393 | 344.509579468035 | 456.770300103725 | 492.731188118594 | 478.225152740879 | 351.884394607974 | 396.216831104005 | Cers4 |
| **ENSMUSG00000008226** | 0.221788640589169 | 0.298551648975128 | 0.851785577030345 | 786.960447131638 | 900.743796179247 | 917.114787100345 | 823.499637558666 | 643.835419141629 | 484.340563901248 | 949.463218587197 | 860.821772782126 | Scrn3 |
| **ENSMUSG00000008301** | -0.0321486850614103 | 0.863799452219806 | 0.974868041734905 | 667.696636544524 | 572.534257426773 | 592.480760293928 | 714.700141024719 | 811.364023101951 | 747.303243797128 | 526.785513821405 | 519.966745476215 | Phax |
| **ENSMUSG00000008305** | -0.242505894433137 | 0.14136547669289 | 0.779725819159865 | 203.290586228035 | 212.56250524664 | 212.005895057252 | 197.902532488472 | 274.834507150593 | 292.316653465652 | 208.215618111227 | 202.993280593011 | Tle1 |
| **ENSMUSG00000008318** | 0.0861073622037777 | 0.765189295411822 | 0.958197631858494 | 52.4037955610046 | 50.4937751926884 | 33.1259211026956 | 51.5859681841989 | 35.0386622662111 | 31.8001380339204 | 54.1360607089191 | 55.3618037980939 | Relt |
| **ENSMUSG00000008333** | -0.117984354248532 | 0.483700518862352 | 0.898689166265124 | 507.774708711803 | 437.341246426994 | 403.189782564238 | 530.866508950119 | 546.384139713729 | 617.656527197299 | 449.745735120251 | 426.611546914723 | Snrpb2 |
| **ENSMUSG00000008348** | -0.2945638846973 | 0.184903670784238 | 0.805963874499992 | 3483.94889109024 | 2663.95384927877 | 2792.04192151292 | 3500.34242297146 | 4077.62432123032 | 5309.39996943262 | 3088.87869468006 | 2783.28754781005 | Ubc |
| **ENSMUSG00000008373** | 0.090637098634363 | 0.423482578327559 | 0.884482395183227 | 475.248214915318 | 440.598909342652 | 493.102996985841 | 449.266886549659 | 455.502609460744 | 468.440494884288 | 439.33495421469 | 382.104998763315 | Prpf31 |
| **ENSMUSG00000008384** | -0.293966768813706 | 0.378291874303216 | 0.87422797717331 | 182.509770746947 | 131.120932355207 | 152.3792370724 | 188.52326554589 | 303.30342024189 | 261.739597663806 | 125.970448957293 | 111.809133160856 | Sertad1 |
| **ENSMUSG00000008393** | -0.272025365397809 | 0.254611834321388 | 0.837406774112203 | 626.135005582348 | 535.071133896714 | 709.841166486335 | 615.279911433354 | 849.687559955619 | 1061.63537744011 | 539.278450908079 | 552.53251241627 | Carhsp1 |
| **ENSMUSG00000008398** | -0.312641276013135 | 0.16541249776969 | 0.795216960550936 | 653.24041707942 | 550.545032746086 | 679.554610049585 | 808.492810450535 | 1057.72961716125 | 1022.49674601375 | 642.345181873136 | 620.920622990386 | Elk3 |
| **ENSMUSG00000008429** | 0.245246752577854 | 0.043499662658693 | 0.688841110980169 | 1153.78701605867 | 1306.32282917858 | 1455.64761874131 | 1299.96639824181 | 1166.13047854734 | 947.888729857241 | 1126.44649398174 | 1158.2557775013 | Herpud2 |
| **ENSMUSG00000008435** | 0.149721256358886 | 0.386398419697298 | 0.875126835008975 | 638.784197614315 | 737.860650396382 | 650.214508501483 | 547.749189446766 | 556.238763476101 | 441.532685778663 | 631.934400967575 | 690.394259129171 | Rdh13 |
| **ENSMUSG00000008450** | 0.114263262690789 | 0.308037350061572 | 0.851785577030345 | 806.837748896157 | 727.273245920495 | 683.340429604179 | 781.292936317048 | 636.170711770895 | 689.818378889657 | 671.495368408708 | 772.894202043977 | Nutf2 |
| **ENSMUSG00000008475** | -0.153186112054708 | 0.295903942931963 | 0.851785577030345 | 750.819898468876 | 710.984931342209 | 655.893237833374 | 715.638067718977 | 898.960678767479 | 879.396124861105 | 620.482541971457 | 753.354741879944 | Arpc5 |
| **ENSMUSG00000008496** | 0.513764735498237 | 0.291936165925106 | 0.851785577030345 | 10.8421645988285 | 14.6594831204579 | 31.2330113253987 | 30.9515809105193 | 15.3294147414674 | 7.33849339244316 | 20.8215618111227 | 17.3684090346961 | Pou2f2 |
| **ENSMUSG00000008540** | -0.0534504429866002 | 0.776341643412293 | 0.959733396167384 | 2116.0291242047 | 2431.84536653819 | 2828.00720728156 | 2838.1661768252 | 2342.11558085705 | 3040.58242893562 | 2000.95209004889 | 3216.41224811279 | Mgst1 |
| **ENSMUSG00000008575** | -0.203486896564154 | 0.328467068312551 | 0.855977678491063 | 5204.2390074377 | 3912.45316170443 | 3418.59505779819 | 4968.19769948548 | 5390.47919801741 | 6508.02055686501 | 3969.63075929055 | 4287.82598044061 | Nfib |
| **ENSMUSG00000008604** | 0.282935911411383 | 0.147792594666542 | 0.780607990324772 | 2937.32309256597 | 3229.97278087423 | 2697.39643264807 | 2717.17363326589 | 2094.65502860193 | 1665.8380000846 | 2895.23816983662 | 2862.53091403085 | Ubqln4 |
| **ENSMUSG00000008658** | 0.102170694196427 | 0.596422518617951 | 0.92448662081833 | 805.934235179588 | 1197.19112150406 | 1062.86883995221 | 766.286109208918 | 810.269064906132 | 743.633997100907 | 1034.8316220128 | 981.315110460331 | Rbfox1 |
| **ENSMUSG00000008668** | 0.109925241998841 | 0.523335714067104 | 0.906635460329092 | 6816.10747779688 | 4562.35691337807 | 5523.51073015234 | 6594.56258732913 | 5637.93975027253 | 6437.08178740472 | 4600.52408216757 | 5097.62805168331 | Rps18 |
| **ENSMUSG00000008682** | 0.0327189052733969 | 0.85913265124749 | 0.974868041734905 | 8406.2916189584 | 6276.70202274273 | 7201.57524772603 | 8616.73254014973 | 7639.52333222984 | 9630.54949534957 | 6036.17076904448 | 6512.06786244638 | Rpl10 |
| **ENSMUSG00000008683** | 0.138042890317681 | 0.374340597500994 | 0.873700364743909 | 4560.03372752397 | 3185.99433151285 | 3841.66039302405 | 4571.45470781428 | 3652.78054125251 | 4230.64144074348 | 3289.80676615739 | 3511.67520170262 | Rps15a |
| **ENSMUSG00000008690** | 0.00020669383194504 | 0.997682677211348 | 0.999262355603273 | 1889.24718134587 | 1748.55056997906 | 1863.56967574879 | 1905.86704273258 | 1829.67514521371 | 1884.76971962582 | 1825.00989274491 | 1866.01844566516 | Ncaph2 |
| **ENSMUSG00000008730** | 0.125914791729663 | 0.515787808990251 | 0.904239892322932 | 2378.9516157263 | 2722.5917817606 | 2527.98100758 | 1550.39282560874 | 2393.57861606055 | 1784.47697659576 | 2109.22421146673 | 2125.45905562094 | Hipk1 |
| **ENSMUSG00000008734** | -0.694218744413039 | 0.0362490942112007 | 0.679566075569698 | 177.088688447533 | 178.357044632238 | 171.308334845369 | 225.102406621959 | 389.805117711598 | 470.886659348436 | 190.517290571773 | 166.085411394282 | Gprc5b |
| **ENSMUSG00000008763** | 0.0378387086752151 | 0.845918410175626 | 0.972180580805179 | 564.696072855653 | 764.736369450555 | 627.49959117392 | 526.176875478828 | 551.858930692825 | 457.432754795624 | 737.083288113745 | 671.940324529806 | Man1a2 |
| **ENSMUSG00000008822** | 0.160004488055714 | 0.39547048565797 | 0.879215782741314 | 290.027903018663 | 223.96432545144 | 196.862616838877 | 299.198615468353 | 191.617684268342 | 233.608706326107 | 259.228444548478 | 219.276164063039 | Acyp1 |
| **ENSMUSG00000008845** | -0.306340560815297 | 0.290212946294317 | 0.851785577030345 | 226.78194285883 | 381.146561131906 | 527.175372977185 | 374.232751009006 | 618.65138063779 | 546.717757737015 | 356.048706970199 | 345.197129564585 | Cd163 |
| **ENSMUSG00000008855** | 0.075101888607361 | 0.413453971791849 | 0.882192126163777 | 2837.93658374337 | 3089.07885977205 | 3244.44735828688 | 3148.61991262465 | 2733.01565676447 | 2891.3663966226 | 3191.94542564511 | 2877.72827193621 | Hdac5 |
| **ENSMUSG00000008859** | 0.0686630681843468 | 0.414870921189419 | 0.882192126163777 | 1202.5767567534 | 1075.84317789583 | 1085.58375727977 | 1133.95337335812 | 1024.88087128667 | 1099.5509266344 | 1097.29630744617 | 1067.07163006914 | Rala |
| **ENSMUSG00000008892** | 0.0108760197938831 | 0.895720910577142 | 0.983911700112021 | 8775.82872903513 | 7880.28659297504 | 7457.11806766111 | 7636.59914464995 | 7690.98636743334 | 7954.92683740838 | 7931.9739719472 | 7934.10635216212 | Vdac3 |
| **ENSMUSG00000008958** | -0.100579431424368 | 0.517578094450932 | 0.905207722478398 | 571.924182588206 | 539.143212541285 | 539.479286529615 | 610.590277962063 | 580.327843784121 | 774.211052902753 | 560.100012719201 | 511.282540958867 | Vps72 |
| **ENSMUSG00000008976** | 0.217741263753158 | 0.276294583725598 | 0.846688575024234 | 854.723975874317 | 1141.81085193789 | 1060.02947528626 | 865.706338800283 | 826.693437843418 | 580.964060235083 | 1006.72251356778 | 957.433548037624 | Gabpa |
| **ENSMUSG00000008999** | -0.164622232811173 | 0.722670468652825 | 0.951674101666722 | 35.2370349461928 | 37.4631235300591 | 14.1968233297267 | 17.8206071909051 | 20.8042057205628 | 14.6769867848863 | 40.6020455316893 | 41.2499714574033 | Bmp7 |
| **ENSMUSG00000009013** | -0.382933849516283 | 0.161339434899681 | 0.792786631432503 | 607.161217534398 | 394.991628523449 | 420.22597055991 | 526.176875478828 | 670.114415841287 | 963.788798874201 | 473.690531203042 | 434.210225867403 | Dynll1 |
| **ENSMUSG00000009030** | -0.070835911129278 | 0.566545943970108 | 0.919931558693298 | 239.431134890797 | 277.715763559786 | 273.525462819401 | 289.819348525772 | 296.733671066975 | 256.847268735511 | 288.37863108405 | 292.006376895829 | Pdcl |
| **ENSMUSG00000009035** | -0.631216179640582 | 0.00760708208058734 | 0.635862253526407 | 651.433389646282 | 633.615437095348 | 493.102996985841 | 592.769670771158 | 1125.61702530203 | 1216.96682091349 | 679.823993133157 | 651.315338801105 | Tmem184b |
| **ENSMUSG00000009073** | 0.175057287362003 | 0.542270810057508 | 0.913837378403038 | 3004.18310759207 | 3504.43088151836 | 3345.71803137226 | 2493.94708003245 | 1806.68102310151 | 1700.08430258266 | 3906.12499576662 | 3523.61598291398 | Nf2 |
| **ENSMUSG00000009076** | 0.00613448763485161 | 0.977453094299915 | 0.996179425433073 | 367.730082643601 | 267.1283590839 | 257.435729712378 | 364.853484066425 | 273.739548954774 | 429.301863457925 | 268.598147363483 | 281.151121249143 | Zmat5 |
| **ENSMUSG00000009079** | -0.0142938064161577 | 0.876324648755826 | 0.977469019294424 | 1273.05082664578 | 1378.80582905196 | 1264.46373123432 | 1445.34503585183 | 1384.02715951534 | 1258.551616804 | 1396.08571943578 | 1375.360890435 | Ewsr1 |

| | | | | | | | | | | | | |
|---|---|---|---|---|---|---|---|---|---|---|---|---|
| ENSMUSG00000009090 | -0.113265821403807 | 0.383195578518198 | 0.875126835008975 | 1342.6213828216 | 1500.15377266019 | 1278.66055456405 | 1154.5877606318 | 1332.56412431184 | 1294.02100153414 | 1450.2217801447 | 1630.4593981321 | Ap1b1 |
| ENSMUSG00000009097 | 0.249300153535769 | 0.523670766055623 | 0.906842904938611 | 21.6843291976571 | 11.4018202048006 | 27.447191770805 | 25.3240207449704 | 20.8042057205628 | 14.6769867848863 | 14.5750932677859 | 21.7105112933701 | Tbx1 |
| ENSMUSG00000009112 | 0.144172873696483 | 0.568612607510587 | 0.92059532906293 | 1872.08042073106 | 2344.70288354435 | 2408.7276916103 | 1688.26804966469 | 1520.89693399273 | 1185.16668287957 | 2371.57589028688 | 2444.60357163348 | Bcl2l13 |
| ENSMUSG00000009207 | 0.00349861366913759 | 0.984611951281968 | 0.99711600049115 | 174.378147297826 | 232.108482740584 | 212.952349945901 | 192.274972322923 | 203.662224422352 | 151.662196777159 | 233.201492284575 | 220.361689627707 | Lnpk |
| ENSMUSG00000009210 | -0.399520294612611 | 0.271048798428403 | 0.844129257012535 | 28.9124389302094 | 13.8450673915436 | 30.2865564367503 | 23.448167356454 | 41.6084114411257 | 33.0232202659942 | 19.7804837205666 | 32.5657669400552 | Prr29 |
| ENSMUSG00000009281 | -0.578976293930004 | 0.0644949659350027 | 0.716991331547963 | 273.764656120421 | 271.200437728471 | 229.042083052924 | 299.198615468353 | 443.458069306734 | 613.987280501078 | 185.311900118992 | 361.480013034613 | Rarres2 |
| ENSMUSG00000009291 | -0.111112304397702 | 0.452612364969481 | 0.891110666112841 | 2609.3476134514 | 2564.59513035122 | 2772.1663688513 | 2462.05757242767 | 3278.30483828238 | 3210.59085919388 | 2410.09577963746 | 2343.64969411931 | Pttg1ip |
| ENSMUSG00000009293 | 0.0573441441259364 | 0.536139143088898 | 0.911291453479697 | 3185.78936462245 | 2934.33987127833 | 3371.27231336577 | 3095.15809105193 | 2898.35434433315 | 2974.53598840363 | 3326.24449932686 | 2895.09668097091 | Ube2g2 |
| ENSMUSG00000009376 | -0.160059213175722 | 0.724934997172968 | 0.951674101666722 | 23.4913566307952 | 65.9676740420606 | 31.2330113253987 | 34.703287687552 | 27.3739548954774 | 31.8001380339204 | 33.3144988977964 | 81.4144173501381 | Met |
| ENSMUSG00000009378 | -0.28609545029924 | 0.436747708421528 | 0.888701139843674 | 14.4562194651047 | 28.5045505120015 | 23.6613722162112 | 34.703287687552 | 26.2789966996583 | 35.4693847301419 | 20.8215618111227 | 41.2499714574033 | Slc16a12 |
| ENSMUSG00000009394 | -0.109197679771576 | 0.683416148153234 | 0.944452327903597 | 136.430571201926 | 79.8127414336042 | 168.468970179424 | 105.047789756914 | 146.724398239759 | 149.216032313011 | 136.381229862854 | 95.5262496908287 | Syn2 |
| ENSMUSG00000009406 | -0.0164140624834354 | 0.922473243608098 | 0.986713918026868 | 185.220311896654 | 204.418347957496 | 169.415425068072 | 181.957778686083 | 164.243729372865 | 154.108361241306 | 213.421008564008 | 217.105112933702 | Elk1 |
| ENSMUSG00000009418 | -0.484975986581332 | 0.0244291878991057 | 0.655303666217042 | 326.168451681425 | 248.39679731887 | 278.257737262643 | 306.702029022419 | 499.300937293508 | 502.686797382356 | 318.569895710178 | 302.861632542514 | Nav1 |
| ENSMUSG00000009470 | 0.147743623063886 | 0.410362465829183 | 0.882192126163777 | 1174.56783153976 | 1538.43131191917 | 1520.00655116941 | 1239.00116311503 | 1202.26409900937 | 917.311674055395 | 1459.5914829597 | 1359.07800696497 | Tnpo1 |
| ENSMUSG00000009535 | 0.220192031075726 | 0.169223675405159 | 0.79907358295209 | 269.247087537575 | 249.211213047785 | 310.437203476691 | 261.681547698027 | 255.12525962585 | 179.793088114857 | 249.858741733473 | 249.670879873757 | Rnmt |
| ENSMUSG00000009545 | -0.100434031671191 | 0.565030739823954 | 0.919648110634488 | 546.625798524272 | 573.348673155687 | 396.564598343699 | 437.073839524303 | 425.938738173629 | 539.379264344572 | 529.908748093074 | 600.295637261685 | Kcnq1 |
| ENSMUSG00000009549 | -0.0283800886829678 | 0.867775550523791 | 0.974868041734905 | 1755.52715129365 | 1288.40568314247 | 1439.55788563429 | 1752.9849915685 | 1449.72465126448 | 1997.29328497661 | 1383.59278234911 | 1530.5910461826 | Srp14 |
| ENSMUSG00000009555 | -0.0886743162272487 | 0.644668282746748 | 0.934857223264556 | 599.933107801846 | 507.380999113627 | 547.997380527451 | 636.852225401292 | 705.153078107498 | 760.757148349941 | 514.292576734731 | 458.09178829011 | Cdk9 |
| ENSMUSG00000009563 | -0.194618278519851 | 0.450729108913651 | 0.89108031640712 | 148.176249517323 | 124.605606523892 | 120.199770858353 | 115.364983393754 | 179.573144114332 | 193.24699266767 | 118.6829023234 | 91.1841474321546 | Tor2a |
| ENSMUSG00000009566 | -0.163014461605269 | 0.37778111603397 | 0.87422797717331 | 81.316234491214 | 96.9154717408051 | 81.3951204237664 | 88.1651092602672 | 109.49581958191 | 105.185071958352 | 78.0808567917102 | 97.6973008201657 | Fpgs |
| ENSMUSG00000009569 | 0.0881978926398785 | 0.593357375789151 | 0.923935529355527 | 643.30176619716 | 877.940155769646 | 689.019158936069 | 559.942236472122 | 712.817785478232 | 623.771938357668 | 683.988305495382 | 586.183804920994 | Mkl2 |
| ENSMUSG00000009575 | -0.00730771868048211 | 0.969250346192119 | 0.995039712387919 | 898.092634269631 | 996.030436462224 | 830.040937344688 | 905.099259959126 | 712.817785478232 | 721.618516923577 | 1088.96768272172 | 1123.51895943191 | Cbx5 |
| ENSMUSG00000009585 | -0.197725551147439 | 0.560173078211881 | 0.916574868225118 | 88.5443442237664 | 69.2253369577179 | 77.6093008691726 | 68.4686486808458 | 118.255485148462 | 125.977469903608 | 60.3825292522559 | 44.5065481514088 | Apobec3 |
| ENSMUSG00000009614 | -0.233593688541709 | 0.411642642245354 | 0.882192126163777 | 229.492484008537 | 353.456426348819 | 202.541346170768 | 295.446908691321 | 279.21433993387 | 269.078091056249 | 226.955023741238 | 496.085183053508 | Sardh |
| ENSMUSG00000009621 | -0.0037780260280494 | 0.983233846203721 | 0.996940244108351 | 293.64195788494 | 317.622134276588 | 259.328639489674 | 271.060814640609 | 256.220217821669 | 217.708637309147 | 325.857442344071 | 344.111603999917 | Vav2 |
| ENSMUSG00000009630 | 0.149995741227038 | 0.18435415234362 | 0.805963874499992 | 1088.7340284657 | 1066.07018914886 | 1154.67496415111 | 1290.58713129923 | 1089.48340484 | 1125.23565350795 | 956.75076522109 | 974.80195707232 | Ppp2cb |
| ENSMUSG00000009633 | -0.183560137972319 | 0.384975308129595 | 0.875126835008975 | 1054.40050723608 | 622.213616890547 | 1208.62289280407 | 1062.6709445945 | 1215.4035973592 | 1234.08997216252 | 987.983107937774 | 1045.36111877577 | G0s2 |
| ENSMUSG00000009640 | -0.0854769572063258 | 0.630063221986558 | 0.931825800800958 | 337.914129996823 | 312.735639903102 | 309.490748588042 | 358.287997206618 | 383.235368536684 | 415.847958905112 | 350.843316517418 | 249.670879873757 | Dmap1 |
| ENSMUSG00000009646 | -0.24044804315696 | 0.542694261655774 | 0.913837378403038 | 14.4562194651047 | 17.1027303072009 | 18.9290977729689 | 24.3860940507122 | 20.8042057205628 | 15.9000690169602 | 15.616171358342 | 35.8223436340608 | Pla2g12b |
| ENSMUSG00000009647 | 0.0395457821789689 | 0.841823503959509 | 0.970689728540393 | 506.871194995234 | 404.764617270421 | 461.869985660442 | 452.080666632434 | 385.425284928322 | 316.77829810713 | 535.114138545854 | 537.335154510911 | Mcu |
| ENSMUSG00000009681 | -0.231970399992395 | 0.185911321374944 | 0.805963874499992 | 220.457346842847 | 302.96265115613 | 257.435729712378 | 213.847286290861 | 327.39250054991 | 329.009120427868 | 251.940897914585 | 261.61166108511 | Bcr |
| ENSMUSG00000009687 | -0.684870430135663 | 0.0864652970680264 | 0.746460787597018 | 212.325723393726 | 160.439898596123 | 173.201244622666 | 213.847286290861 | 381.045452145046 | 541.82542880872 | 160.326025945645 | 138.947272277569 | Fxyd5 |
| ENSMUSG00000009733 | 0.211192188932845 | 0.517999969674696 | 0.905473752527583 | 107.518132271716 | 174.284965987666 | 196.862616838877 | 135.999370667433 | 107.305903190271 | 72.1618516923577 | 201.96914956789 | 148.717002359586 | Tfcp2 |
| ENSMUSG00000009739 | 0.117992854941694 | 0.542863636035821 | 0.913837378403038 | 204.194099944604 | 294.818493866987 | 227.149173275627 | 248.550573978413 | 190.522726072523 | 183.462334811079 | 266.515991182371 | 257.269558826436 | Pou6f1 |
| ENSMUSG00000009741 | 0.259553181420566 | 0.219550951704867 | 0.828235393585171 | 513.195791011218 | 608.368549499004 | 641.696414503647 | 510.23212167644 | 432.508487348543 | 316.77829810713 | 580.921574530324 | 567.72987032163 | Ubp1 |
| ENSMUSG00000009772 | -0.259749486207057 | 0.33310500892029 | 0.856918526232803 | 49.6932544112975 | 36.6487078011448 | 52.0550188756646 | 34.703287687552 | 65.6974917491458 | 50.1463715150282 | 53.094982618363 | 37.9933947633978 | Nuak2 |
| ENSMUSG00000009828 | 0.139089384449139 | 0.296560117709208 | 0.851785577030345 | 281.896279569542 | 323.323044378988 | 290.561650815073 | 347.03287687552 | 314.253002200081 | 287.424324537357 | 256.10521027681 | 271.381391167127 | Ick |
| ENSMUSG00000009863 | 0.0291073072416289 | 0.778568005744599 | 0.960247700212438 | 32536.4324473679 | 26475.026515547 | 27097.003462005 | 31085.7044277982 | 29557.3015379407 | 29890.9066696531 | 27204.4115843224 | 28200.8686445232 | Sdhb |
| ENSMUSG00000009876 | -0.395582627083964 | 0.326599328668968 | 0.855977678491063 | 110.228673421423 | 70.0397526866323 | 83.2880302010633 | 97.5443762028488 | 150.009272827216 | 203.031650524261 | 52.0539045278068 | 70.559161703453 | Cox4i2 |
| ENSMUSG00000009894 | -0.0689141554042804 | 0.431779308736958 | 0.886559861988535 | 791.478015714483 | 796.498582878213 | 781.771738023617 | 739.086235075431 | 866.111932892906 | 825.580506649855 | 801.630129728225 | 768.552099785303 | Snap47 |
| ENSMUSG00000009905 | -0.236020263512754 | 0.260740545045182 | 0.840316047847749 | 138.237598635064 | 111.574954861263 | 109.78876708322 | 135.999370667433 | 182.858018701789 | 165.116101329971 | 111.395355689507 | 124.835439936878 | Kdsr |
| ENSMUSG00000009907 | -0.0171708590736645 | 0.849757501553458 | 0.972640876584422 | 711.968808656408 | 783.467931215584 | 779.87882824632 | 755.968915572078 | 809.174106710313 | 717.949270227356 | 780.808567917102 | 758.782369703287 | Vps4b |
| ENSMUSG00000009927 | 0.215233757529944 | 0.201035764272177 | 0.812921911423739 | 6879.35343795671 | 4746.41486811271 | 5833.94793362903 | 7333.64882240456 | 5178.05730802851 | 6276.85801500305 | 4797.28784128268 | 5105.22673063599 | Rps25 |
| ENSMUSG00000009995 | 0.125545072864647 | 0.376225887969033 | 0.8741697248747 | 997.479143092226 | 919.475357944277 | 837.612576453875 | 970.754128557197 | 839.832936193247 | 680.033721033066 | 961.95615567387 | 931.380934485579 | Taz |
| ENSMUSG00000010025 | -0.148587947358567 | 0.28386131641421 | 0.851125503777472 | 531.266065342598 | 627.914526992947 | 583.016211407443 | 528.052728867345 | 719.387534653147 | 598.087211484117 | 526.785513821405 | 673.025850094475 | Aldh3a2 |
| ENSMUSG00000010045 | -0.339313360261403 | 0.149912232154869 | 0.782641594375979 | 506.871194995234 | 517.153987860599 | 506.353365426919 | 543.059555975475 | 842.022852584886 | 850.042151291332 | 490.34778065194 | 441.808904820083 | Tmem115 |
| ENSMUSG00000010047 | -0.169653028125342 | 0.403088584393155 | 0.881158207349486 | 1018.25995857331 | 885.269897329875 | 866.952678001977 | 1012.02290310456 | 1243.87251045049 | 1358.84435983406 | 899.491470240502 | 753.354741879944 | Hyal2 |
| ENSMUSG00000010048 | 0.14978690944829 | 0.156043049146288 | 0.787275540343856 | 1457.36762482587 | 1378.80582905196 | 1349.64467121269 | 1567.27550610539 | 1140.9464400435 | 1327.04422180014 | 1365.89445480965 | 1351.47932801229 | Ifrd2 |
| ENSMUSG00000010051 | -0.223685562774379 | 0.378869782565418 | 0.87422797717331 | 75.8951521917998 | 79.8127414336042 | 90.8596693102509 | 77.8479156234274 | 127.015150715015 | 111.300483118721 | 80.1630129728225 | 60.7894316214364 | Hyal1 |
| ENSMUSG00000010054 | 0.0767866609651616 | 0.417908141367776 | 0.882468496126562 | 822.197482077831 | 751.705717787925 | 873.577862222516 | 850.699511692152 | 810.269064906132 | 807.234273168747 | 756.863771834311 | 752.269216315276 | Tusc2 |
| ENSMUSG00000010057 | -0.20053772394654 | 0.234197775858163 | 0.832048466795696 | 309.905204783182 | 236.180561385155 | 197.809071727525 | 270.12288794635 | 289.068963696242 | 303.324393554317 | 309.200192895173 | 263.782712214447 | Nprl2 |
| ENSMUSG00000010064 | -0.0595874598115689 | 0.787502449148321 | 0.961224689823763 | 1409.48139784771 | 1777.05512049107 | 1809.62174709583 | 1700.46109669005 | 1631.48771177045 | 1122.7894890438 | 2348.67217229464 | 1874.70265018251 | Slc38a3 |
| ENSMUSG00000010066 | -0.809690745875664 | NA | NA | 50.5967681278665 | 54.56585383726 | 50.1621090983677 | 46.8963347129081 | 21.8991639163819 | 264.185762127954 | 34.3555769883525 | 34.7368180693922 | Cacna2d2 |
| ENSMUSG00000010067 | -0.157267535488489 | 0.430325906082668 | 0.886017087288711 | 130.105975185942 | 121.347943608235 | 143.861143074564 | 151.944124469822 | 191.617684268342 | 170.008430258266 | 119.723980413956 | 129.177542195552 | Rassf1 |
| ENSMUSG00000010080 | 0.402897924654534 | 0.0667755851740406 | 0.719725956118125 | 323.457910531718 | 218.26341534904 | 356.813493020464 | 398.618845059719 | 240.890803080201 | 203.031650524261 | 234.242570375131 | 301.776106977845 | Epn3 |
| ENSMUSG00000010095 | -0.358624226941287 | 0.138458989434584 | 0.779711473022872 | 838.460728976074 | 868.167167022674 | 821.522843346852 | 760.658549043369 | 1221.97334653411 | 1504.39114545085 | 776.644255554878 | 715.361347116547 | Slc3a2 |
| ENSMUSG00000010097 | -0.0864503079788814 | 0.599186899725508 | 0.924950659834108 | 2103.37993217274 | 1569.37910961791 | 1647.77796113695 | 1949.01167066846 | 1921.65163366251 | 2399.68733932891 | 1717.77884941763 | 1680.39357410685 | Nxf1 |
| ENSMUSG00000010110 | -0.217291161298415 | 0.317518252327314 | 0.854351922245337 | 271.054114970713 | 329.838370210303 | 318.008842585878 | 322.646782824807 | 456.597567656563 | 443.978850242811 | 280.050006359601 | 263.782712214447 | Stx5a |
| ENSMUSG00000010122 | 0.128739185192134 | 0.363368519831689 | 0.86924083181804 | 1890.15069506244 | 2185.89181640606 | 1739.58408533585 | 1809.26059322399 | 1645.7221683161 | 1495.82956982633 | 2087.36157156505 | 1744.43958242229 | Slc47a1 |
| ENSMUSG00000010175 | 0.0671429430848732 | 0.845452834851223 | 0.972136081650911 | 626.135005582348 | 737.860650396382 | 1069.49402417274 | 989.51266244236 | 549.669014301187 | 414.624876673038 | 1255.5401772107 | 1046.44664434044 | Prox1 |
| ENSMUSG00000010277 | 0.225213097749523 | 0.31566382903973 | 0.853277176422981 | 10314.5125883522 | 12226.8233381908 | 10659.9214108475 | 9241.39171852566 | 7550.83171836849 | 6006.55684171472 | 11348.7922651524 | 11401.2750057133 | 2610507B11 |
| ENSMUSG00000010290 | 0.129924865458967 | 0.589934853033966 | 0.923810046070101 | 810.451803762433 | 844.549110884159 | 766.628459805242 | 666.865879617553 | 529.959766776443 | 473.332823812584 | 939.052437681635 | 879.275707381491 | AI597479 |
| ENSMUSG00000010307 | -0.44263808301513 | 0.0376127473351601 | 0.679566075569698 | 298.159526467785 | 280.159010746529 | 256.489274823729 | 336.71568323868 | 476.306815181307 | 503.90987961443 | 313.364505257397 | 299.605055848508 | Tmem86a |
| ENSMUSG00000010358 | -0.154010195300203 | 0.465420311020879 | 0.895328112556867 | 226.78194285883 | 214.191336704468 | 216.738169500494 | 275.750448111899 | 305.493336633528 | 329.009120427868 | 201.96914956789 | 202.993280593011 | Ifi35 |
| ENSMUSG00000010362 | 0.10701890578567 | 0.295970119952224 | 0.851785577030345 | 944.171833814652 | 821.745470474558 | 847.07712534036 | 985.760955665328 | 824.50352145178 | 798.67269754423 | 850.560799984364 | 866.249400605469 | Rdm1 |
| ENSMUSG00000010376 | -0.114614846721665 | 0.581372757376762 | 0.922450244597442 | 2016.64261538211 | 1611.72872752146 | 1585.31193848615 | 2096.26616166699 | 1942.45583938308 | 2743.37344654167 | 1696.9572876065 | 1532.76209731193 | Nedd8 |
| ENSMUSG00000010392 | 0.0479845752999842 | 0.748412561409308 | 0.956197192463153 | 377.668733525861 | 459.330471107681 | 439.155068332879 | 392.053358199911 | 351.48158085793 | 363.255422925936 | 412.26692386023 | 486.315452971491 | Gosr1 |
| ENSMUSG00000010406 | 0.111540240073984 | 0.528526966601945 | 0.908232038364594 | 642.398252480591 | 456.887223920938 | 453.351891662606 | 587.142110605609 | 479.591689768764 | 599.310293716191 | 421.636626675235 | 480.887825148149 | Mrpl52 |
| ENSMUSG00000010453 | 0.189129134976833 | 0.39559460474889 | 0.879215782741314 | 1344.42841025474 | 1593.81158148534 | 1790.69264932286 | 1488.4896637877 | 1084.00861386091 | 949.111812089315 | 1708.40914660262 | 1710.78828991757 | Kansl3 |
| ENSMUSG00000010476 | -0.17969935081922 | 0.5004388916361 | 0.900927059989498 | 136.430571201926 | 108.317291945606 | 178.879973954556 | 135.061443973175 | 211.326931793086 | 190.800828203522 | 111.395355689507 | 119.407812113536 | Ebf3 |
| ENSMUSG00000010492 | -0.035714367922501 | 0.910789528602432 | 0.986166948757759 | 38.8510898124689 | 58.6379324818317 | 60.5731128735006 | 30.0136542162612 | 66.7924499449649 | 46.4771248188067 | 36.4377331694648 | 43.4210225867403 | Uckl1os |
| ENSMUSG00000010517 | 0.303389177704142 | 0.0400177878692278 | 0.679566075569698 | 2208.18752329475 | 2211.1387040024 | 2382.22695472814 | 2406.71989746644 | 1846.099518151 | 1421.22155366982 | 2158.15488172287 | 2034.27490818878 | Faf1 |
| ENSMUSG00000010538 | 0.135607797155788 | 0.662530887652661 | 0.939611397743053 | 60.535419010126 | 42.3496179035451 | 41.6440151005317 | 41.2687745473591 | 22.994122112201 | 53.8156182112498 | 41.6431236222455 | 51.0197015394199 | Tsacc |
| ENSMUSG00000010554 | 0.174735228576267 | 0.282663933507385 | 0.850587941933635 | 618.003382133227 | 649.903751673634 | 518.657278979349 | 567.445650026188 | 473.02194059385 | 417.071041137186 | 570.510793624763 | 624.177199684392 | Mettl16 |
| ENSMUSG00000010607 | 0.00084024836932158 | 0.996432262694611 | 0.999262355603273 | 563.792559139084 | 369.744740927105 | 411.707876562074 | 516.797608536247 | 525.579933993166 | 527.148442023833 | 386.239971596327 | 422.269444656049 | Pigyl |
| ENSMUSG00000010608 | -0.102710580202332 | 0.466572441602195 | 0.895328112556867 | 1704.93038316579 | 1435.00051434705 | 1389.39577653592 | 1631.9924480092 | 1650.10200109938 | 1993.62403828039 | 1515.80969984973 | 1457.86083334981 | Rbm25 |
| ENSMUSG00000010609 | -0.243560685063453 | 0.0801160903346623 | 0.739855948845396 | 359.59845919448 | 368.930325198191 | 340.723759913441 | 319.833002742033 | 451.122776677468 | 469.663577116362 | 368.541644056872 | 357.137910775939 | Psen2 |
| ENSMUSG00000010651 | -0.324594549721973 | 0.529977428600184 | 0.908700960777257 | 127.395434036235 | 335.539280312703 | 143.861143074564 | 191.337045628665 | 111.685735973548 | 152.885279009232 | 146.792010768415 | 588.354856050331 | Acaa1b |
| ENSMUSG00000010660 | -0.138474900236214 | 0.324351550938467 | 0.855977678491063 | 207.80815481088 | 212.56250524664 | 235.667267273463 | 256.053987532478 | 273.739548954774 | 232.385624094033 | 269.639225454039 | 226.874843015718 | Plcd1 |
| ENSMUSG00000010663 | -0.0330516930718465 | 0.823798197522813 | 0.966637987288264 | 412.905768472054 | 500.865673282312 | 529.068282754482 | 476.466760683146 | 450.027818481649 | 584.633306931305 | 425.80093903746 | 504.769387570856 | Fads1 |
| ENSMUSG00000010721 | 0.124927659515323 | 0.37402233360207 | 0.873700364743909 | 312.61574593289 | 330.652785939217 | 360.599312575058 | 303.888248939644 | 301.113503850252 | 245.839528646846 | 314.405583347953 | 336.512925047237 | Lmbr1 |
| ENSMUSG00000010751 | -0.739996399400104 | 0.0187087826720245 | 0.642273034804458 | 26.2018977805023 | 17.1027303072009 | 29.3401015481019 | 25.3240207449704 | 45.9882442244021 | 52.592535979176 | 28.1091084450157 | 36.9078691987293 | Tnfrsf22 |
| ENSMUSG00000010755 | 0.110322378125329 | 0.3917170359604 | 0.878285927324016 | 558.37147683967 | 516.339572131684 | 632.231865617162 | 682.810633419941 | 602.227007700503 | 546.717757737015 | 535.114138545854 | 528.650949993563 | Cars |
| ENSMUSG00000010830 | -0.446741537630513 | 0.226795224127658 | 0.832048466795696 | 39.754603529038 | 25.2468875963442 | 29.3401015481019 | 40.3308478531009 | 56.937826182593 | 69.71568722821 | 30.191264626128 | 27.1381391167127 | Kdelr3 |
| ENSMUSG00000010911 | 0.15649380941621 | 0.243204750768419 | 0.832048466795696 | 1621.80712124144 | 1290.84893032921 | 1435.77206607969 | 1759.55047842831 | 1450.8196094603 | 1455.46785616789 | 1221.18460022235 | 1352.56485357696 | Apip |
| ENSMUSG00000010914 | 0.183267125099406 | 0.265893378463168 | 0.843277879501733 | 3397.21157429961 | 3304.89902793435 | 3327.73538848794 | 3121.42003849116 | 2757.10473707249 | 2109.81685032741 | 3385.58595048856 | 3328.22138127364 | Pdhx |
| ENSMUSG00000010936 | 0.0599657563092634 | 0.532906641983517 | 0.909082872831377 | 477.055242348456 | 487.020605890769 | 444.83379766477 | 490.535661097018 | 447.837902090011 | 435.417274618294 | 469.526218840818 | 468.947043936795 | Vac14 |

| | | | | | | | | | | | | |
|---|---|---|---|---|---|---|---|---|---|---|---|---|
| **ENSMUSG00000011034** | 0.301290959572157 | 0.468151750559181 | 0.895328112556867 | 39.754603529038 | 20.3603932228582 | 37.8581955459379 | 28.1378008277448 | 21.8991639163819 | 23.2385624094033 | 43.7252798033577 | 13.0263067760221 | Slc5a1 |
| **ENSMUSG00000011096** | -0.044150980124594 | 0.705467295863817 | 0.947760509100828 | 2134.09939853608 | 1940.75268200285 | 2088.82593924712 | 2107.52128199809 | 2137.35839823888 | 2490.19542450238 | 2043.6362917617 | 1858.41976671248 | Akt1s1 |
| **ENSMUSG00000011114** | -0.0990318659408698 | 0.571286022327574 | 0.921130164032017 | 708.354753790131 | 638.501931468834 | 702.269527377147 | 815.9962240046 | 872.68168206782 | 933.211743072355 | 634.016557148687 | 629.604827507734 | Tbrg1 |
| **ENSMUSG00000011148** | -0.0413313340908861 | 0.674715012827678 | 0.944023556644042 | 4981.97463316171 | 4463.81261017944 | 4253.36826958612 | 4750.59870641759 | 5103.60015071281 | 4312.58795029243 | 4896.19025988551 | 4672.10203033326 | Adssl1 |
| **ENSMUSG00000011158** | 0.0834227032557366 | 0.484537906654214 | 0.899236955296196 | 555.660935689963 | 598.595560752032 | 630.338955839866 | 575.886990274511 | 525.579933993166 | 502.686797382356 | 644.427338054249 | 553.618037980939 | Brf1 |
| **ENSMUSG00000011171** | -0.272921913613777 | 0.647007185150326 | 0.935063240511793 | 10.8421645988285 | 5.7009101024003 | 10.4110037751329 | 18.7585338851632 | 24.0890803080201 | 18.3462334811079 | 5.20539045278068 | 7.59867895267955 | Vipr2 |
| **ENSMUSG00000011179** | 0.24280377630386 | 0.103232049768056 | 0.759800953500741 | 2524.41732409391 | 2173.67558047234 | 2581.92893623296 | 2562.4157287133 | 2083.70544664374 | 1602.23772401676 | 2368.45265601521 | 2261.1497512045 | Odc1 |
| **ENSMUSG00000011254** | 0.0434135827971946 | 0.760486000380926 | 0.957357521291862 | 176.185174730964 | 148.223662662408 | 151.432782183751 | 156.633757941113 | 155.484063806312 | 139.43137445642 | 153.038479311752 | 164.999885829613 | Thg1l |
| **ENSMUSG00000011256** | 0.241570354175507 | 0.561343891117228 | 0.917031459819748 | 184.316798180085 | 280.973426475443 | 439.155068332879 | 513.983828453472 | 165.338687568684 | 223.824048469516 | 245.694429371248 | 564.473293627624 | Adam19 |
| **ENSMUSG00000011257** | 0.179774569847368 | 0.10643410582143 | 0.762094734041179 | 13809.3036440413 | 13210.6375387193 | 13827.7059231538 | 14973.9996738315 | 11281.3542915242 | 10990.6169374157 | 13587.1101598481 | 13420.3525559968 | Pabpc4 |
| **ENSMUSG00000011263** | -0.827746743151923 | 0.0519941878609798 | 0.703396849069134 | 11.7456783153976 | 9.77298874697194 | 12.3039135524298 | 18.7585338851632 | 25.1840385038392 | 35.4693847301419 | 19.7804837205666 | 13.0263067760221 | Exoc3l2 |
| **ENSMUSG00000011267** | 0.326942973064709 | 0.464029894285624 | 0.895328112556867 | 12.6491920319666 | 13.8450673915436 | 20.8220075502658 | 8.44134024832345 | 10.949581958191 | 13.4539045528125 | 11.4518589961175 | 8.68420451734806 | Zfp296 |
| **ENSMUSG00000011305** | 0.00498323490874067 | 0.971014024946633 | 0.995122272737203 | 5403.01202508289 | 4985.85309241352 | 6227.67316730678 | 5212.99656668686 | 6022.27007700503 | 4542.52740992231 | 6053.86909658393 | 5133.45039531737 | Plin5 |
| **ENSMUSG00000011306** | 0.14756440457315 | 0.226376675375179 | 0.832048466795696 | 797.802611730467 | 729.716493107238 | 802.593745573883 | 891.968286239511 | 760.995946094272 | 817.018931025338 | 672.536446499264 | 658.914017753784 | Sugp1 |
| **ENSMUSG00000011382** | 0.346462412161879 | 0.0880827790244211 | 0.750135871546083 | 1291.12110097717 | 1452.91766038316 | 1250.2669079046 | 1163.96702757438 | 880.346389438554 | 708.164612370765 | 1243.04724012403 | 1224.47283694608 | Dhdh |
| **ENSMUSG00000011427** | 0.0382969155229247 | 0.86760466315851 | 0.974868041734905 | 180.702743313809 | 171.841718800923 | 226.202718386979 | 147.254490998531 | 166.433645764503 | 118.638976511164 | 215.50316474512 | 205.164331722348 | Zfp790 |
| **ENSMUSG00000011658** | -0.107484450839845 | 0.582235789309535 | 0.922450244597442 | 228.588970291968 | 181.614707547895 | 212.952349945901 | 194.150825711439 | 228.846262926191 | 288.647406769431 | 194.681602933998 | 169.341988088287 | Fuz |
| **ENSMUSG00000011751** | 0.998517073288869 | 0.0614372184303055 | 0.71282992338528 | 7.22810973255236 | 15.4738988493722 | 16.0897331070236 | 15.9447538023887 | 8.75966556655277 | 2.44616446414772 | 9.36970281500523 | 6.51315338801105 | Sptbn4 |
| **ENSMUSG00000011752** | -0.11119636444815 | 0.35781567553163 | 0.865834559821921 | 939.654265231807 | 940.65016689605 | 977.687899973846 | 786.920496482597 | 1036.92541144068 | 1067.75078860048 | 964.038311854983 | 869.505977299475 | Pgam1 |
| **ENSMUSG00000011831** | 0.00030017455534842 | 0.998058559979339 | 0.999262355603273 | 325.264937964856 | 336.353696041618 | 361.545767463707 | 312.329589187968 | 338.342082508101 | 309.439804714686 | 309.200192895173 | 377.762896504641 | Evi5 |
| **ENSMUSG00000011832** | -0.104416143923733 | 0.634899972503882 | 0.932543559775713 | 463.50253659992 | 588.008156276145 | 479.852628544763 | 399.556771753977 | 433.603445544362 | 374.263163014601 | 671.495368408708 | 595.953535003011 | Evi5l |
| **ENSMUSG00000011837** | -0.193855668506191 | 0.276689916570455 | 0.847094277204072 | 333.396561413978 | 267.942774812814 | 327.473391472363 | 318.895076047775 | 350.386622662111 | 461.102001491845 | 333.144988977964 | 283.322172378481 | Snapc2 |
| **ENSMUSG00000011877** | 0.0719520759007768 | 0.352899874259367 | 0.862728204353539 | 2226.25779762613 | 2393.56782727921 | 2525.14164291406 | 2326.05820176024 | 2226.05001210022 | 2238.24048469516 | 2334.09707902686 | 2211.21557522975 | Git1 |
| **ENSMUSG00000011884** | -0.338252689234438 | 0.165982730243312 | 0.796071270505327 | 289.124389302094 | 252.468875963442 | 274.47191770805 | 315.143369270742 | 366.810995599397 | 534.486935416277 | 275.885693997376 | 254.012982132431 | Gltp |
| **ENSMUSG00000011958** | 0.130974360483946 | 0.335384673208416 | 0.858570960161243 | 3028.57797793944 | 2959.58675887467 | 2619.7871317789 | 2882.24873145533 | 2310.36179317829 | 2248.02514255175 | 2949.37423054554 | 2984.10977727373 | Bnip2 |
| **ENSMUSG00000011960** | 0.0184598257518979 | 0.868054034625032 | 0.974898560893217 | 493.318489246699 | 534.2567181678 | 498.781726317731 | 473.652980600371 | 489.446313531136 | 459.878919259771 | 468.485140750262 | 556.874614674944 | Ccnt1 |
| **ENSMUSG00000012017** | -1.0084114496866 | 0.029358588712873 | 0.665956123973759 | 56.9213641438498 | 53.7514381083457 | 73.8234813145789 | 63.779015209555 | 193.80760065998 | 203.031650524261 | 54.1360607089191 | 48.8486504100828 | Scarf2 |
| **ENSMUSG00000012076** | 0.117718022836172 | 0.532926817758316 | 0.909082872831377 | 329.782506547701 | 292.375246680244 | 325.580481695066 | 308.577882410935 | 263.884925192402 | 210.370143916704 | 375.829190690765 | 306.118209236519 | Brms1l |
| **ENSMUSG00000012114** | 0.0536004234193875 | 0.683271459072131 | 0.944452327903597 | 478.862269781594 | 610.811796685746 | 628.446046062569 | 517.735535230505 | 544.194223322091 | 549.163922201163 | 567.387559353095 | 493.914131924171 | Med15 |
| **ENSMUSG00000012117** | -0.0248941302963022 | 0.86415458114463 | 0.974868041734905 | 1104.09376164737 | 1123.07929017286 | 820.576388458203 | 907.9130400419 | 949.328755775157 | 908.750098430878 | 1067.10504282004 | 1099.6373970092 | Dhdds |
| **ENSMUSG00000012126** | 0.0016139146490887 | 0.996950004657097 | 0.999262355603273 | 15.3597331816738 | 20.3603932228582 | 12.3039135524298 | 11.2551203310979 | 14.2344565456483 | 13.4539045528125 | 21.8626399016789 | 9.76973008201657 | Ubxn11 |
| **ENSMUSG00000012296** | 0.239846775678591 | 0.094884501829616 | 0.757085157577544 | 430.976042803434 | 422.681763306537 | 551.783200082044 | 542.121629281217 | 399.65974147397 | 381.601656407044 | 455.992203663588 | 410.328663444696 | Tjap1 |
| **ENSMUSG00000012405** | 0.0903897313836626 | 0.474641717685679 | 0.897549741595057 | 3314.99182609183 | 2659.06735490528 | 2881.00868104587 | 3280.86757651505 | 2811.85264686344 | 3285.19887535039 | 2620.3935539298 | 2682.33367029588 | Rpl15 |
| **ENSMUSG00000012422** | 0.187667493234785 | 0.210441024447712 | 0.822692324677376 | 402.967117589794 | 335.539280312703 | 309.490748588042 | 368.605190843457 | 266.074841584041 | 359.586176229715 | 296.707255808499 | 322.401092706547 | Tmem167 |
| **ENSMUSG00000012428** | -0.689213670523073 | 0.024938678500874 | 0.655303666217042 | 309.905204783182 | 367.301493740362 | 300.026199701558 | 304.826175633902 | 823.408563255961 | 602.979540412413 | 302.953724351836 | 338.683976176574 | Steap4 |
| **ENSMUSG00000012429** | -0.111037091726718 | 0.667923816350715 | 0.942024383071524 | 113.8427282877 | 92.8433930962335 | 77.6093008691726 | 120.054616865045 | 98.5462376237187 | 157.777607937528 | 102.025652874501 | 79.2433662208011 | Mplkip |
| **ENSMUSG00000012443** | -0.586079123061615 | 0.311814195388303 | 0.851785577030345 | 2.71054114970714 | 13.0306516626293 | 6.62518422053913 | 15.9447538023887 | 22.994122112201 | 12.2308223207386 | 9.36970281500523 | 13.0263067760221 | Kif11 |
| **ENSMUSG00000012483** | 0.0306199916944808 | 0.873582779226194 | 0.976584692430731 | 163.535982698997 | 107.502876216691 | 109.78876708322 | 117.24083678227 | 127.015150715015 | 125.977469903608 | 124.929370866736 | 109.638082031519 | Rpa3 |
| **ENSMUSG00000012519** | -0.253726762995696 | 0.32164426504184 | 0.855704427283138 | 32.5264937964856 | 35.0198763433161 | 34.0723759913441 | 41.2687745473591 | 45.9882442244021 | 51.3694537471021 | 32.2734208072402 | 41.2499714574033 | Mlkl |
| **ENSMUSG00000012535** | 0.15223242615475 | 0.373093169434523 | 0.873700364743909 | 1337.20030052219 | 1613.35755897928 | 1537.04273916508 | 1277.45615757962 | 1154.08593839333 | 1017.60441708545 | 1564.74037010587 | 1450.26215439713 | Tnpo3 |
| **ENSMUSG00000012609** | 0.539657721870185 | 0.0329379235686381 | 0.671606027118506 | 412.905768472054 | 413.723190288479 | 477.013263878817 | 313.267515882226 | 264.979883388221 | 162.669936865823 | 363.336253604092 | 320.23004157721 | Ttll5 |
| **ENSMUSG00000012640** | 0.0891988315821587 | 0.537746156658235 | 0.912224489006215 | 190.641394196068 | 197.088606397268 | 197.809071727525 | 209.15765281957 | 167.528603960322 | 162.669936865823 | 206.133461930115 | 209.506433981022 | Zfp715 |
| **ENSMUSG00000012819** | -0.518295827149585 | 0.0461833079644921 | 0.69805115405055 | 48.7897406947284 | 51.3081909216027 | 46.3762895437739 | 59.0893817382642 | 104.021028602814 | 74.6080161565054 | 53.094982618363 | 62.9604827507734 | Cdh23 |
| **ENSMUSG00000012848** | 0.0737695278836129 | 0.696537599977121 | 0.944933386973863 | 7314.84704934299 | 5020.05855302792 | 6142.49222732842 | 7739.77108101835 | 6479.96260285742 | 7680.95641742384 | 5035.69472402003 | 5714.20657241502 | Rps5 |
| **ENSMUSG00000013033** | 0.0710213229275229 | 0.67207188197924 | 0.943216235131458 | 320.747369382011 | 325.766291565731 | 287.722286149128 | 285.129715054481 | 285.784089108784 | 215.262472844999 | 338.350379430744 | 320.23004157721 | Adgrl1 |
| **ENSMUSG00000013076** | 0.162441963972587 | 0.417670030686254 | 0.882468496126562 | 2556.9438178904 | 2975.87507345296 | 3088.28230165988 | 2202.25187811816 | 2045.38190979007 | 1822.39252579005 | 3002.4692131639 | 2799.57043128008 | Amotl1 |
| **ENSMUSG00000013089** | 0.234539401520449 | 0.333130810530234 | 0.856918526232803 | 314.422773366028 | 305.405898342873 | 447.673162330715 | 384.549944645846 | 328.487458745729 | 187.131581507301 | 393.52751823022 | 323.486618271215 | Etv5 |
| **ENSMUSG00000013150** | -0.263920532875398 | 0.100910096848279 | 0.759800953500741 | 102.097049972302 | 107.502876216691 | 100.324218196735 | 113.489130005238 | 140.154649064844 | 139.43137445642 | 113.477511870619 | 116.15123541953 | Gfod2 |
| **ENSMUSG00000013155** | -0.0755796765979226 | 0.846547593490381 | 0.972262781013841 | 28.0089252136404 | 11.4018202048006 | 15.1432782183751 | 21.5723139679377 | 24.0890803080201 | 22.0154801773295 | 16.6572494488982 | 17.3684090346961 | Enkd1 |
| **ENSMUSG00000013160** | -0.0715878192339486 | 0.545079289625978 | 0.914211323944814 | 892.671551970216 | 937.392503980392 | 871.68495244522 | 908.850966736158 | 1023.78591309086 | 1073.86619976085 | 841.191097169358 | 857.565196088121 | Atp6v0d1 |
| **ENSMUSG00000013236** | 0.0223572511577676 | 0.92186108780798 | 0.986713918026868 | 993.86508822595 | 1157.28475078726 | 841.398396008469 | 763.472329126143 | 821.218646864323 | 624.995020589742 | 1166.00746142287 | 1085.52556466851 | Ptprs |
| **ENSMUSG00000013275** | 0.112298251925282 | 0.690348936349889 | 0.94469184542218 | 1197.15567445398 | 1397.53739081699 | 1590.04421292939 | 1072.05021153708 | 1121.23719251876 | 653.125911927441 | 1794.81862811878 | 1292.86094752019 | Slc41a1 |
| **ENSMUSG00000013419** | 0.139807874889736 | 0.373531202475003 | 0.873700364743909 | 4310.66394175091 | 4251.2501049328 | 3532.169644436 | 3980.56089043164 | 2947.62746314501 | 3220.37551705047 | 4198.6679392129 | 4222.69444656049 | Zfp651 |
| **ENSMUSG00000013465** | -0.0445435356647082 | 0.610171724122732 | 0.927932112008768 | 741.784761303186 | 725.644414462667 | 701.323072488499 | 758.782695654852 | 800.41444114376 | 759.534066117867 | 716.261726302622 | 743.585011797928 | Nelfb |
| **ENSMUSG00000013495** | -0.41271247334351 | 0.00408313985406273 | 0.635862253526407 | 112.939214571131 | 117.275864963663 | 108.842312194571 | 102.23400967414 | 144.534481848121 | 146.769867848863 | 142.627698406191 | 154.144630182928 | Tmem175 |
| **ENSMUSG00000013539** | 0.17819781387021 | 0.20136968482533 | 0.813395633194566 | 3997.14468210146 | 3547.59491515082 | 4335.70984489854 | 4270.38023895741 | 3350.57207920644 | 2946.40509706593 | 4028.97221045225 | 3945.88542757003 | Tango2 |
| **ENSMUSG00000013584** | -0.458534593255959 | 0.291487890198225 | 0.851785577030345 | 34.3335212296237 | 45.6072808192024 | 28.3936466594534 | 59.0893817382642 | 50.3680770076785 | 111.300483118721 | 30.191264626128 | 39.0789203280663 | Aldh1a2 |
| **ENSMUSG00000013593** | 0.0418837330058051 | 0.657480382973115 | 0.938389079274206 | 37573.5214172403 | 32356.7369097663 | 34962.0435866736 | 39293.5009292514 | 35728.4859295771 | 36787.8673763175 | 33589.3435137032 | 33954.1541372662 | Ndufs2 |
| **ENSMUSG00000013622** | 0.0234078135567896 | 0.86049003804774 | 0.974868041734905 | 1094.15511076511 | 920.289773673191 | 1060.02947528626 | 1145.20849368921 | 1121.23719251876 | 1155.8127093098 | 877.628830338823 | 997.597993930358 | Atraid |
| **ENSMUSG00000013629** | 0.0796720250365335 | 0.575747493408674 | 0.922450244597442 | 287.317361868956 | 311.106808445274 | 314.223023031284 | 269.184961252092 | 321.917709570815 | 253.178022039289 | 243.612273190136 | 299.605055848508 | Cad |
| **ENSMUSG00000013646** | 0.1125984125014 | 0.478457833600982 | 0.897674307047416 | 608.064731250967 | 687.366875203693 | 646.428688946889 | 533.680289032894 | 534.339599559719 | 459.878919259771 | 618.400385790345 | 676.28242678848 | Sh3bp5l |
| **ENSMUSG00000013662** | 0.20168711935718 | 0.2398565349647 | 0.832048466795696 | 2316.60916928303 | 2475.82381589956 | 2479.71180825893 | 2099.07994174977 | 1909.6070935085 | 1492.16032313011 | 2401.76715491301 | 2343.64969411931 | Atad1 |
| **ENSMUSG00000013663** | 0.0652616213595858 | 0.63947382615468 | 0.933801135937988 | 2459.36433650094 | 2564.59513035122 | 2484.44408270217 | 2100.01786844402 | 2157.06764576362 | 1894.55437748241 | 2555.84671231532 | 2574.8666393937 | Pten |
| **ENSMUSG00000013698** | -0.321324655038886 | 0.13414214856391 | 0.77303252008843 | 1353.46354742043 | 1208.59294170886 | 1016.49255040843 | 1357.17992659156 | 1744.26840593982 | 2048.66273872371 | 1134.77511870619 | 1240.7557204161 | Pea15a |
| **ENSMUSG00000013701** | 0.208629646030377 | 0.062473123809223 | 0.715615581523459 | 6030.05054438181 | 5114.53077758198 | 5424.13296684425 | 5591.91895116716 | 4629.48325192314 | 4181.71815146053 | 5239.74602976904 | 5123.68066523536 | Timm23 |
| **ENSMUSG00000013707** | -0.463151764254034 | 0.289393662686122 | 0.851785577030345 | 32.5264937964856 | 12.2162359337149 | 44.483379766477 | 33.7653609932938 | 44.893286028583 | 64.8233582999146 | 23.9447960827911 | 35.8223436340608 | Tnfaip8l2 |
| **ENSMUSG00000013736** | 0.226457078578209 | 0.0527965293352328 | 0.703396849069134 | 710.161781223269 | 780.210268299927 | 758.110365807406 | 768.161962597434 | 649.310210120724 | 540.602346576646 | 699.604476853724 | 687.137682435165 | Trnt1 |
| **ENSMUSG00000013787** | 0.149706198715434 | 0.188180391554177 | 0.808677608936066 | 1790.76418623985 | 1642.6765252202 | 1797.3178335434 | 1934.94277025459 | 1772.73731903112 | 1729.43827615244 | 1538.71341784197 | 1418.78191302174 | Ehmt2 |
| **ENSMUSG00000013822** | 0.0216859878441201 | 0.865196942758625 | 0.974868041734905 | 569.213641438498 | 457.701639649853 | 482.691993210708 | 507.418341593665 | 478.496731572945 | 571.179402378492 | 475.772687384154 | 462.433890548784 | Elof1 |
| **ENSMUSG00000013833** | 0.141240007737917 | 0.34841381903573 | 0.861266047293794 | 1438.39383677792 | 1464.31948058796 | 1325.98329899647 | 1340.29724609491 | 1184.74476787626 | 978.465785659088 | 1430.44129642413 | 1454.6042566558 | Med16 |
| **ENSMUSG00000013846** | -0.248665710972853 | 0.322973056863503 | 0.855977678491063 | 530.362551626029 | 581.492830444831 | 801.647290685234 | 440.825546301336 | 877.061514851097 | 853.711397987554 | 510.128264372507 | 556.874614674944 | St3gal1 |
| **ENSMUSG00000013858** | -0.00409471182551403 | 0.955544919686054 | 0.992371505557225 | 3372.81670395225 | 3178.66458995262 | 3212.26789207283 | 3452.50816156429 | 3266.26029812837 | 3420.96100311059 | 3404.32535611857 | 3162.13596987936 | Tmem259 |
| **ENSMUSG00000013878** | 0.166751337447673 | 0.461588432366193 | 0.895328112556867 | 391.221439274397 | 378.703313945163 | 307.597838810745 | 302.012395551128 | 254.03030143003 | 211.593226148778 | 368.541644056872 | 394.045779974668 | Rnf170 |
| **ENSMUSG00000013921** | -0.735116013282927 | 0.0191409073287928 | 0.642273034804458 | 91.2548853734735 | 80.6271571625185 | 84.2344850897118 | 90.0409626487835 | 173.003394939417 | 222.600966237442 | 97.8613405122768 | 83.5854684794751 | Clip3 |
| **ENSMUSG00000013936** | 0.121853045463887 | 0.454811721977835 | 0.892775943845882 | 228516.689194643 | 154993.086201144 | 162520.501204268 | 224559.347065983 | 182943.425441063 | 189808.90851331 | 163376.384750975 | 172048.203321006 | Myl2 |
| **ENSMUSG00000013973** | 0.170375977265723 | 0.214904948003595 | 0.824338622350122 | 363.212514060756 | 351.82759489099 | 434.422793889637 | 361.101777289392 | 328.487458745729 | 306.993640250539 | 389.363205867995 | 315.887939318536 | Dedd |
| **ENSMUSG00000013997** | 0.175916999161465 | 0.114940836128486 | 0.767518180080285 | 703.837185207286 | 635.244268553176 | 586.802030962037 | 649.983199120906 | 540.909348734634 | 529.594606487981 | 624.646854333682 | 584.012753791657 | Nit1 |
| **ENSMUSG00000014039** | 0.153437505158903 | 0.58384286491913 | 0.922450244597442 | 60.535419010126 | 108.317291945606 | 87.0738497556571 | 55.3376749612315 | 63.5075753575076 | 68.4926049961361 | 58.3003730711437 | 90.0986218674861 | Prdm15 |
| **ENSMUSG00000014074** | 0.0206822883529018 | 0.846428555231749 | 0.972262781013841 | 338.817643713392 | 337.982527499446 | 377.63550057073 | 342.343243404229 | 325.202584158272 | 338.793778284459 | 359.171941241867 | 352.795808517265 | Rnf168 |
| **ENSMUSG00000014075** | -0.0458374468501151 | 0.825811676704263 | 0.967079014368284 | 122.87786545339 | 100.987550385377 | 126.824955078892 | 104.109863062656 | 104.021028602814 | 156.554525705454 | 93.6970281500523 | 116.15123541953 | Tctex1d2 |
| **ENSMUSG00000014077** | 0.130682190279632 | 0.312675138043033 | 0.852613848149207 | 2402.44297235709 | 2577.62578201385 | 2757.96954552157 | 2351.38222250521 | 2158.16260395944 | 1944.70074899744 | 2503.79280778751 | 2607.43240633376 | Chp1 |
| **ENSMUSG00000014158** | -0.548757066578647 | 0.140407373123844 | 0.779711473022872 | 206.904641094311 | 183.243539005724 | 234.720812384815 | 139.751077444466 | 429.223612761086 | 390.163232031561 | 150.95632313064 | 148.717002359586 | Trpv4 |
| **ENSMUSG00000014164** | 0.425190232538995 | 0.430679663055929 | 0.886017087288711 | 19.877301764519 | 45.6072808192024 | 24.6078271048596 | 13.1309737196143 | 25.1840385038392 | 4.89232892829544 | 26.0269522639034 | 20.6249857287016 | Klhl3 |
| **ENSMUSG00000014177** | 0.156623301356511 | 0.233017531205487 | 0.832048466795696 | 512.292277294649 | 540.772043999114 | 565.033568523123 | 506.480414899407 | 539.814390538815 | 467.217412652214 | 393.52751823022 | 505.854913135525 | Tvp23b |
| **ENSMUSG00000014195** | 0.216714579695294 | 0.0262025667835703 | 0.664353746352133 | 2135.00291225265 | 1845.46604171987 | 2264.86654853573 | 2184.43127092726 | 1769.45244444366 | 1767.35382534673 | 1842.70822028436 | 1872.53159905318 | Dnajc7 |
| **ENSMUSG00000014226** | -0.0313387142291099 | 0.810481606154047 | 0.964412705343932 | 1519.71007126913 | 1293.29217751595 | 1225.65908079974 | 1448.1588159346 | 1276.72125632507 | 1257.32853457193 | 1474.16657622749 | 1597.89363119204 | Cacybp |
| **ENSMUSG00000014232** | 0.133689596789942 | 0.291169263328229 | 0.851785577030345 | 309.905204783182 | 293.189662409158 | 289.615195926425 | 327.336416296098 | 258.410134213307 | 320.447544803351 | 275.885693997376 | 258.355084391105 | Cluap1 |

| | | | | | | | | | | | | |
|---|---|---|---|---|---|---|---|---|---|---|---|---|
| **ENSMUSG00000014243** | 0.144880409013095 | 0.408627444732412 | 0.882113132676277 | 337.010616280254 | 300.519403969387 | 303.812019256151 | 401.432625142493 | 339.43704070392 | 347.355353908976 | 236.324726556243 | 292.006376895829 | Zswim7 |
| **ENSMUSG00000014245** | 0.37886858751492 | 0.0727704343996086 | 0.729806017191742 | 85.8338030740593 | 94.4722245540621 | 75.7163910918757 | 73.1582821521366 | 74.4571573156986 | 48.9232892829544 | 69.7522320672611 | 59.7039060567679 | Pigl |
| **ENSMUSG00000014294** | 0.0648670329152334 | 0.761979569988027 | 0.957666649379949 | 5625.27639935887 | 3886.39185837918 | 4354.6389426715 | 5424.03007289495 | 4680.94628712664 | 6328.22746875015 | 3704.15584619873 | 3729.86584020099 | Ndufa2 |
| **ENSMUSG00000014301** | 0.0909467895868523 | 0.631596821884413 | 0.931825800800958 | 1009.22482140762 | 688.181290932608 | 821.522843346852 | 967.002421780164 | 862.827058305448 | 1024.94291047789 | 721.467116755403 | 664.341645577127 | Pam16 |
| **ENSMUSG00000014303** | -0.56777246762152 | 0.0629052703259348 | 0.715615581523459 | 62.3424464432641 | 65.9676740420606 | 79.5022106464695 | 87.227182566009 | 136.869774477387 | 156.554525705454 | 74.9576225200418 | 69.4736361387845 | Glis2 |
| **ENSMUSG00000014313** | 0.0822179152796126 | 0.651239793903617 | 0.936463929813731 | 23072.1262663071 | 17185.8007115502 | 18911.1151300846 | 23239.9476303287 | 19546.0987535667 | 25178.3708294725 | 16298.0775076563 | 16822.3896756679 | Cox6c |
| **ENSMUSG00000014329** | -0.513453037139946 | 0.0726569212106807 | 0.729721356194038 | 189.737880479499 | 216.634583891211 | 193.076797284283 | 204.468019348279 | 375.57066116595 | 403.617136584374 | 194.681602933998 | 174.76961591163 | Bicc1 |
| **ENSMUSG00000014349** | 0.101283532264746 | 0.302020304671927 | 0.851785577030345 | 1494.4116872052 | 1702.94328915986 | 1728.22662667206 | 1532.57221841784 | 1508.85239383872 | 1382.08292224346 | 1580.35654146422 | 1547.95945521729 | Ube2z |
| **ENSMUSG00000014353** | 0.0365709161092202 | 0.805127216333216 | 0.963475137341661 | 402.063603873225 | 450.371898089624 | 447.673162330715 | 320.770929436291 | 399.65974147397 | 354.693847301419 | 413.308001950786 | 412.499714574033 | Tmem87b |
| **ENSMUSG00000014355** | 0.0625177071732505 | 0.72539073903339 | 0.951698749807259 | 834.846674109798 | 1061.18369477537 | 1038.26101284735 | 811.30659053331 | 837.643019801609 | 686.149132193435 | 1027.54407537891 | 1034.50586312909 | Anapc1 |
| **ENSMUSG00000014361** | -0.445661975804247 | 0.155920538035713 | 0.787275540343856 | 116.553269437407 | 193.83094348161 | 165.629605513478 | 131.309737196143 | 280.309298129689 | 281.308913376988 | 137.42230795341 | 129.177542195552 | Mertk |
| **ENSMUSG00000014402** | 0.147412861044427 | 0.0600543836872537 | 0.71282992338528 | 1348.04246512102 | 1314.46698646773 | 1305.16129144621 | 1416.26930832982 | 1197.88426622609 | 1163.15120270224 | 1259.70448957293 | 1238.58466928677 | Tsg101 |
| **ENSMUSG00000014418** | -0.144573562302524 | 0.487604053924575 | 0.899236955296196 | 95.7724539563188 | 162.883145782866 | 119.253315969704 | 121.930470253561 | 134.679858085749 | 114.969729814943 | 147.833088858971 | 155.230155747597 | Hps5 |
| **ENSMUSG00000014426** | 0.137466294911794 | 0.360948248504905 | 0.868539734006824 | 1965.14233353767 | 2144.35661423143 | 1822.87211553691 | 1763.30218520534 | 1608.49358965825 | 1408.99073134909 | 1944.73387315886 | 2033.18938262411 | Map3k4 |
| **ENSMUSG00000014444** | -0.114305202665471 | 0.569871858569479 | 0.920872458148558 | 394.835494140673 | 473.175538499225 | 480.799083433411 | 376.108604397523 | 650.405168316544 | 414.624876673038 | 407.061533407449 | 395.131305539337 | Piezo1 |
| **ENSMUSG00000014453** | 0.281378968952778 | 0.593618325788138 | 0.923935529355527 | 10.8421645988285 | 18.7315617650296 | 4.73227444324223 | 14.0689004138724 | 6.56974917491458 | 13.4539045528125 | 11.4518589961175 | 8.68420451734806 | Blk |
| **ENSMUSG00000014470** | 0.117534970298686 | 0.366499217030748 | 0.869754677013001 | 904.417230285614 | 754.148964974668 | 883.042411109001 | 941.678401035194 | 686.538788778574 | 857.380644683776 | 782.890724098215 | 883.617809640165 | Rnf166 |
| **ENSMUSG00000014496** | 0.238845262785435 | 0.411157827528278 | 0.882192126163777 | 551.143367107117 | 769.622863824041 | 572.60520763231 | 489.59773440276 | 365.716037403578 | 300.878229090169 | 694.399086400943 | 657.828492189116 | Ankrd28 |
| **ENSMUSG00000014498** | 0.277684499280926 | 0.571400623482509 | 0.921130164032017 | 778.828823682517 | 1444.77350309402 | 1319.35811477593 | 610.590277962063 | 517.915226622433 | 229.939459629886 | 1474.16657622749 | 1203.84785121737 | Ankrd52 |
| **ENSMUSG00000014503** | 0.304518608477498 | 0.151216623957223 | 0.783412422888014 | 178.895715880671 | 243.510302945384 | 227.149173275627 | 168.826804966469 | 135.774816281568 | 134.539045528125 | 204.051305749003 | 187.795922687652 | Pkd2l2 |
| **ENSMUSG00000014504** | 0.00800472914626667 | 0.961536026348676 | 0.993822507448007 | 774.311255099672 | 585.564909089402 | 636.017685171756 | 804.741103673502 | 619.746338833609 | 846.372904595111 | 698.563398763168 | 620.920622990386 | Srp19 |
| **ENSMUSG00000014542** | -0.10039242956197 | 0.762817640375063 | 0.957842155335615 | 37.9475760958999 | 30.1333819698302 | 27.447191770805 | 25.3240207449704 | 19.7092475247437 | 47.7002070508805 | 28.1091084450157 | 34.7368180693922 | Clec4f |
| **ENSMUSG00000014547** | -0.488792569598203 | 0.213150288132403 | 0.824236498373909 | 14.4562194651047 | 26.0613033252585 | 21.7684624389143 | 21.5723139679377 | 25.1840385038392 | 15.9000690169602 | 46.8485140750261 | 29.3091902460497 | Wdfy2 |
| **ENSMUSG00000014550** | 0.163259451391191 | 0.355449481243469 | 0.864479425231231 | 545.722284807703 | 614.069459601404 | 663.464876942561 | 537.431995809926 | 483.971522552041 | 399.947889888152 | 568.428637443651 | 654.57191549511 | Rbsn |
| **ENSMUSG00000014551** | -0.0243097559467915 | 0.865715413975842 | 0.974868041734905 | 962.242108146033 | 754.963380703583 | 740.127722923085 | 958.561081531841 | 866.111932892906 | 985.804279051531 | 792.26042691322 | 830.427056971408 | Mrps25 |
| **ENSMUSG00000014554** | -0.044696415796493 | 0.765767753265937 | 0.958345041873658 | 424.651446787451 | 384.404224047563 | 298.133289924261 | 419.253232333398 | 400.754699669789 | 393.832478727783 | 358.130863151311 | 422.269444656049 | Dguok |
| **ENSMUSG00000014592** | 0.0986668558434197 | 0.324434809284532 | 0.855977678491063 | 3293.30749689417 | 2897.69116347718 | 2975.65416991072 | 3372.78439255235 | 2817.32743784254 | 2734.81187091715 | 3197.1508160979 | 2959.14268928635 | Camta1 |
| **ENSMUSG00000014599** | -0.298930413536314 | 0.169011126268395 | 0.79907358295209 | 443.625234835401 | 324.137460107903 | 492.156542097192 | 604.962717796514 | 625.221129812704 | 678.810638800992 | 460.156516025812 | 529.736475558232 | Csf1 |
| **ENSMUSG00000014601** | -0.179895773199137 | 0.140517142692396 | 0.779711473022872 | 453.563885717661 | 463.402549752253 | 417.386605893965 | 371.418970926232 | 510.250519251699 | 447.648096939033 | 513.251498644175 | 461.348364984116 | Strip1 |
| **ENSMUSG00000014602** | -0.324736685676316 | 0.606162385946484 | 0.926859923029135 | 19.877301764519 | 16.2883145782866 | 11.3574586637814 | 20.6343872736795 | 17.5193311331055 | 52.592535979176 | 10.4107809055614 | 5.42762782334254 | Kif1a |
| **ENSMUSG00000014606** | 0.0997576729518623 | 0.370253815725041 | 0.872997029438264 | 15381.4175108714 | 14437.1476264643 | 14098.3920213073 | 14468.4571856264 | 13811.8026820621 | 11374.6647582869 | 14646.9276560343 | 14649.1674952015 | Slc25a11 |
| **ENSMUSG00000014633** | -0.169058822719355 | 0.155221865702983 | 0.787275540343856 | 806.837748896157 | 724.015583004838 | 604.784673846357 | 697.817460528072 | 750.046364136081 | 806.011190936674 | 815.164144905455 | 815.229699066049 | Cmc2 |
| **ENSMUSG00000014668** | 0.130193088995095 | 0.463163848800305 | 0.895328112556867 | 237.624107457659 | 248.39679731887 | 298.133289924261 | 252.302280755445 | 223.371471947096 | 178.570005882784 | 261.31060072959 | 282.236646813812 | Chfr |
| **ENSMUSG00000014748** | -0.0217206767156362 | 0.796599908340528 | 0.962278883153809 | 2068.14289722654 | 2144.35661423143 | 2022.57409704173 | 1943.38411050291 | 1921.65163366251 | 2200.32493550087 | 2056.12922884837 | 2126.54458118561 | Tex261 |
| **ENSMUSG00000014763** | -0.0154386577850307 | 0.912932202590064 | 0.986226496507248 | 529.45903790946 | 515.52515640277 | 524.33600831124 | 511.170048370698 | 505.870686468423 | 431.748027922072 | 641.30410378258 | 522.137796605552 | Fam120b |
| **ENSMUSG00000014767** | -0.128129901626179 | 0.4240440449449 | 0.884586455610141 | 137.334084918495 | 120.533527879321 | 143.861143074564 | 146.316564304273 | 127.015150715015 | 144.323703384715 | 156.16171358342 | 170.427513652956 | Tbp |
| **ENSMUSG00000014769** | -0.0107642541523761 | 0.947635026076946 | 0.991041028853253 | 3383.65886855107 | 2634.63488303785 | 2701.18225220267 | 3668.23130124367 | 3176.4737260712 | 3693.70834086306 | 2646.4205061937 | 2964.57031710969 | Psmb1 |
| **ENSMUSG00000014773** | -0.472910846720991 | 0.198290461917671 | 0.812921911423739 | 74.0881247586617 | 52.9370223794314 | 83.2880302010633 | 61.9031618210386 | 119.350443344282 | 150.439114545085 | 51.0128264372507 | 57.5328549274309 | Dll1 |
| **ENSMUSG00000014776** | -0.117523389243711 | 0.391587716201094 | 0.878285927324016 | 1851.29960524997 | 1603.58457023231 | 1493.50581428725 | 1732.35060429482 | 1804.49110670987 | 2163.63246853866 | 1711.53238087429 | 1569.66996651066 | Nol3 |
| **ENSMUSG00000014778** | -0.291496435621958 | 0.206923241284263 | 0.819960256425221 | 141.85165350134 | 122.976775066064 | 160.897331070236 | 179.143998603309 | 240.890803080201 | 203.031650524261 | 127.011527047849 | 169.341988088287 | Fhod1 |
| **ENSMUSG00000014786** | -0.368875565472466 | 0.421462878068115 | 0.882568531013176 | 6.32459601598331 | 11.4018202048006 | 11.3574586637814 | 15.9447538023887 | 9.85462376237187 | 20.7923979452556 | 11.4518589961175 | 16.2828834700276 | Slc9a5 |
| **ENSMUSG00000014791** | 0.471160719027794 | 0.113252255596295 | 0.767518180080285 | 31.6229800799166 | 28.5045505120015 | 26.5007368821565 | 27.1998741334867 | 20.8042057205628 | 24.4616446414772 | 18.7394056300105 | 18.4539345993646 | Elmo3 |
| **ENSMUSG00000014813** | -0.692326559407232 | 0.203864243024536 | 0.816074141171399 | 69.5705561758165 | 43.1640336324594 | 48.2691993210708 | 22.5102406621959 | 128.110108910834 | 112.523565350795 | 28.1091084450157 | 28.2236646813812 | Stc1 |
| **ENSMUSG00000014837** | 0.158662931438051 | 0.479552640320471 | 0.897732059433541 | 119.263810587114 | 100.173134656462 | 66.2518422053913 | 114.427056699496 | 83.2168228822514 | 100.292743030056 | 85.3684034256032 | 90.0986218674861 | 4931428F04[illegible] |
| **ENSMUSG00000014846** | -0.714952449161725 | 0.0803328774557111 | 0.739855948845396 | 263.826005238161 | 236.180561385155 | 180.772883731853 | 296.384835385579 | 447.837902090011 | 759.534066117867 | 198.845915296222 | 198.651178334337 | Tppp3 |
| **ENSMUSG00000014850** | 0.178553285938063 | 0.294175065347167 | 0.851785577030345 | 265.633032671299 | 247.582381589956 | 292.45456059237 | 261.681547698027 | 200.377349834895 | 196.916239363891 | 295.666177717943 | 248.585354309088 | Msh3 |
| **ENSMUSG00000014856** | 0.105253569617997 | 0.51270089044365 | 0.902972980953514 | 318.036828232304 | 222.335493993612 | 285.829376371831 | 308.577882410935 | 282.499214521327 | 287.424324537357 | 238.406882737355 | 246.414303179751 | Tmem208 |
| **ENSMUSG00000014859** | 0.0629141683245981 | 0.656151377930996 | 0.938034442101846 | 490.607948096991 | 566.018931595458 | 539.479286529615 | 538.369922504185 | 574.853052805026 | 401.170972120226 | 531.990904274186 | 534.078577816906 | E2f4 |
| **ENSMUSG00000014867** | -0.0785548272493774 | 0.335318462627178 | 0.858570960161243 | 2865.04199524044 | 3001.1219610493 | 3008.78009101341 | 2767.82167475583 | 3077.92748844748 | 3314.55284892016 | 2920.22404400996 | 2984.10977727373 | Surf4 |
| **ENSMUSG00000014873** | 0.148778081619361 | 0.316212960272719 | 0.853356198600137 | 228.588970291968 | 219.892246806869 | 253.649910157784 | 245.736793895638 | 175.193311331055 | 206.700897220482 | 229.03717992235 | 243.157726485746 | Surf2 |
| **ENSMUSG00000014905** | 0.150532457836619 | 0.259917014688892 | 0.840316047847749 | 1236.91027798302 | 1381.2490762387 | 1639.25986713911 | 1389.06943419634 | 1334.75404070348 | 1076.312364225 | 1334.66211209297 | 1339.53854680094 | Dnajb9 |
| **ENSMUSG00000014907** | 0.0184937331529348 | 0.902366620721003 | 0.984902747274457 | 131.913002619081 | 158.811067138294 | 170.36187995672 | 144.440710915757 | 150.009272827216 | 138.208292224346 | 160.326025945645 | 148.717002359586 | Naf1 |
| **ENSMUSG00000014932** | 0.0589463227189231 | 0.764225895081973 | 0.958041307101859 | 252.983840639333 | 225.593156909269 | 303.812019256151 | 371.418970926232 | 260.600050604945 | 234.831788558181 | 289.419709174606 | 321.315567141878 | Yes1 |
| **ENSMUSG00000014956** | 0.155285257708719 | 0.45621855302729 | 0.892805623137304 | 9045.07581657271 | 9983.10800503184 | 10155.4609551978 | 8533.25706436075 | 7015.39716061295 | 5867.1254672583 | 10550.2853696959 | 10434.0717275937 | Ppp1cb |
| **ENSMUSG00000014959** | -0.0861208205343153 | 0.266243495433157 | 0.843513595393324 | 1678.72848538529 | 1605.21340169014 | 1752.83445377692 | 1705.15073016134 | 1792.44656655586 | 1890.88513078619 | 1756.2987387682 | 1717.30144330558 | Gorasp2 |
| **ENSMUSG00000014980** | 0.103208596804917 | 0.443504538601188 | 0.89108031640712 | 234.010052591383 | 210.933673788811 | 247.971180825893 | 241.047160424347 | 245.270635863478 | 198.139321595965 | 221.749633288457 | 202.993280593011 | Tsen15 |
| **ENSMUSG00000015002** | 0.0538942802905351 | 0.740280069292363 | 0.954383574136061 | 1306.48083415884 | 1586.48183992511 | 1528.52464516724 | 1267.13896394278 | 1101.52794499401 | 1234.08997216252 | 1621.99966508646 | 1521.90684166525 | Efr3a |
| **ENSMUSG00000015013** | 0.05854798370354 | 0.747602325506474 | 0.956119556124081 | 859.241544457162 | 613.255043872489 | 622.767316730678 | 805.679030367761 | 732.527033002976 | 850.042151291332 | 585.085886892549 | 618.749571861049 | Trappc2l |
| **ENSMUSG00000015016** | 0.135602514692688 | 0.210003992517143 | 0.821677593257218 | 871.890736489128 | 831.518459221529 | 800.700835796586 | 872.27182566009 | 742.381656765348 | 680.033721033066 | 857.848346618257 | 791.348136643342 | Acsf3 |
| **ENSMUSG00000015023** | 0.166697961142717 | 0.267672505637336 | 0.843793744782845 | 525.844983043184 | 609.997380956832 | 607.624038512303 | 490.535661097018 | 492.731188118594 | 404.840218816448 | 514.292576734731 | 577.499600403646 | Ddx19a |
| **ENSMUSG00000015027** | 0.0735645097314549 | 0.780779277945444 | 0.960742105865155 | 122.87786545339 | 124.605606523892 | 113.574586637814 | 112.551203310979 | 114.970610561005 | 147.992950080937 | 128.052605138405 | 59.7039060567679 | Galns |
| **ENSMUSG00000015053** | -0.571818062016993 | 0.101086032346667 | 0.759800953500741 | 354.177376895066 | 280.973426475443 | 263.114459044268 | 249.488500672671 | 568.283303630111 | 664.133652016106 | 250.899819824029 | 223.618266321713 | Gata2 |
| **ENSMUSG00000015083** | -0.282103375778975 | 0.330363566842769 | 0.85614138351386 | 184.316798180085 | 181.614707547895 | 173.201244622666 | 176.330218520534 | 166.433645764503 | 226.270212933664 | 128.052605138405 | 349.539231823259 | C8g |
| **ENSMUSG00000015085** | -0.857876904017363 | 0.0215656573458354 | 0.648762042745232 | 201.483558794897 | 191.387696294867 | 159.950876181588 | 154.757904552597 | 400.754699669789 | 545.494675504941 | 160.326025945645 | 176.940667040967 | Entpd2 |
| **ENSMUSG00000015087** | 0.0174212653126699 | 0.894626404156686 | 0.983733637008786 | 2548.81219444128 | 2508.40044505613 | 2250.66972520601 | 2236.95516580571 | 1950.12054675381 | 2243.13281362346 | 2614.14708538646 | 2622.62976423911 | Rabl6 |
| **ENSMUSG00000015090** | 0.0859317836175903 | 0.550023269592654 | 0.914699131434573 | 11065.3324868211 | 7749.98007634875 | 8560.6844678252 | 10372.531311801 | 9551.32034212998 | 8121.26602097043 | 9277.04686494573 | 8614.73088120928 | Ptgds |
| **ENSMUSG00000015092** | -0.0260984654043673 | 0.8851529591962 | 0.979866999597423 | 3688.14299103484 | 2816.24959058575 | 3054.20992566854 | 3877.38895406324 | 3426.12419471795 | 4399.42678876967 | 2918.14188782885 | 2938.51770355765 | Edf1 |
| **ENSMUSG00000015094** | -0.652062242469424 | 0.0432035401027977 | 0.687730553581215 | 317.133314515735 | 250.840044505613 | 267.84673348751 | 327.336416296098 | 516.820268426614 | 757.087901653719 | 277.967850178489 | 276.809018990469 | Npdc1 |
| **ENSMUSG00000015095** | 0.108572245294441 | 0.399111003401126 | 0.880459952732461 | 1800.70283712211 | 1863.38318775598 | 1790.69264932286 | 1712.6541437154 | 1494.61793729307 | 1413.88306027738 | 1843.74929837492 | 1894.24211034655 | Fbxw5 |
| **ENSMUSG00000015112** | 0.0683575155243258 | 0.773166756303865 | 0.959121750920843 | 2596.69842141944 | 3045.10041041067 | 2664.27051154538 | 2562.4157287133 | 2580.81646754561 | 1457.91402063204 | 3246.08148635403 | 3079.63602696456 | Slc25a13 |
| **ENSMUSG00000015120** | -0.00922236027168136 | 0.91257830404861 | 0.986226496507248 | 807.741262612726 | 794.869751420385 | 844.237760674415 | 785.982569788339 | 856.257309130534 | 810.903519864969 | 810.99983254323 | 775.065253173314 | Ube2i |
| **ENSMUSG00000015126** | 0.00284833015007718 | 0.988419363546998 | 0.998437672973644 | 146.369222084185 | 114.018202048006 | 113.574586637814 | 124.744250336335 | 124.825234323377 | 160.223772401676 | 103.066730965058 | 110.723607596188 | Tsr3 |
| **ENSMUSG00000015127** | -0.0742276899003035 | 0.743147770204227 | 0.954383574136061 | 59.631905293557 | 65.9676740420606 | 50.1621090983677 | 56.2756016554897 | 74.4571573156986 | 45.2540425867328 | 62.4646854333682 | 61.8749571861049 | Unkl |
| **ENSMUSG00000015133** | -0.416943756231096 | 0.0359606987304966 | 0.679566075569698 | 177.992202164102 | 232.108482740584 | 246.078271048596 | 202.592165959763 | 354.766455445387 | 319.224462571277 | 255.064132186253 | 218.19063849837 | Lrrk1 |
| **ENSMUSG00000015134** | -0.916382486664213 | 0.102008697586832 | 0.759800953500741 | 12.6491920319666 | 4.07207864457164 | 14.1968233297267 | 9.37926694258161 | 25.1840385038392 | 30.5770558018465 | 8.32862472444909 | 11.9407812113536 | Aldh1a3 |
| **ENSMUSG00000015143** | -0.272045337118181 | 0.271990828066657 | 0.84423521828958 | 505.967681278665 | 568.462178782201 | 453.351891662606 | 459.584080186499 | 771.945528052463 | 787.664957455566 | 425.80093903746 | 415.756291268038 | Actn1 |
| **ENSMUSG00000015149** | 0.120277735121414 | 0.193295836284149 | 0.809022817984525 | 2386.17972545885 | 2132.14037829771 | 2411.56705627624 | 2427.35428474012 | 1977.49450164929 | 2230.90199130272 | 2268.50915932182 | 2130.88668344428 | Sirt2 |
| **ENSMUSG00000015165** | 0.0358030925444698 | 0.733233475661053 | 0.953733898884637 | 1793.47472738955 | 1856.05344619575 | 1775.54937110449 | 1960.26679099956 | 1896.46759515868 | 1560.65292812624 | 1955.14465406442 | 1790.03165613837 | Hnrnpl |
| **ENSMUSG00000015176** | 0.045661239420678 | 0.671930693481525 | 0.943216235131458 | 536.687147642013 | 552.173864203915 | 566.92647830042 | 556.19052969509 | 592.372383938131 | 513.694537471021 | 477.854843565267 | 559.045665804281 | Nolc1 |
| **ENSMUSG00000015189** | 0.127279074799806 | 0.466787990922723 | 0.895328112556867 | 558.37147683967 | 736.231818938553 | 708.894711597687 | 571.19735680322 | 512.440435643337 | 490.455975061618 | 712.097413940397 | 641.545608719088 | Casd1 |
| **ENSMUSG00000015202** | 0.33398422612068 | 0.359204943814476 | 0.866960324862394 | 489.704434380422 | 658.862324691692 | 1298.53610722567 | 983.885102276811 | 497.11102090187 | 385.270903103266 | 843.273253350471 | 995.426942801021 | Magi1 |
| **ENSMUSG00000015214** | 0.225474843994249 | 0.32659545885371 | 0.855977678491063 | 423.747933070882 | 574.163088884602 | 579.230391852849 | 446.453106466885 | 381.045452145046 | 286.201242305283 | 537.196294726966 | 525.394373299558 | Mtmr1 |
| **ENSMUSG00000015217** | 0.0826019815196614 | 0.653482903406265 | 0.936991241819063 | 1233.29622311675 | 1135.29552610657 | 1110.19158438463 | 1139.58093352367 | 912.100177117308 | 814.572766561191 | 1326.33348736852 | 1306.97277986088 | Hmgb3 |
| **ENSMUSG00000015222** | -0.29795657756069 | 0.272545279658719 | 0.844738346035895 | 41.5616309621761 | 37.4631235300591 | 27.447191770805 | 38.4549944645846 | 48.1781606160403 | 57.4848649074714 | 35.3966550789086 | 37.9933947633978 | Map2 |
| **ENSMUSG00000015224** | -0.735877354659952 | 0.176272637774728 | 0.799889021902745 | 30.7194663633475 | 17.1027303072009 | 8.51809399783602 | 30.9515809105193 | 58.0327843784121 | 52.592535979176 | 18.7394056300105 | 16.2828834700276 | Cyp2j9 |
| **ENSMUSG00000015243** | -0.225857911379712 | 0.321654942551424 | 0.855704427283138 | 455.370913150799 | 637.687515739919 | 742.967087589031 | 394.867138282686 | 631.790878987619 | 541.82542880872 | 792.26042691322 | 642.631134283757 | Abca1 |
| **ENSMUSG00000015247** | -0.128575129142058 | 0.365594308097983 | 0.86924083181804 | 145.465708367616 | 134.378595270864 | 132.503684410783 | 126.620103724852 | 150.009272827216 | 146.769867848863 | 135.340151772298 | 157.401206876934 | Nipsnap3b |

| | | | | | | | | | | | | |
|---|---|---|---|---|---|---|---|---|---|---|---|---|
| ENSMUSG00000015289 | 0.112059304715308 | 0.580591043479663 | 0.922450244597442 | 418.326850771468 | 298.890572511559 | 285.829376371831 | 406.122258613784 | 292.353838283699 | 419.517205601334 | 275.885693997376 | 316.973464883204 | Lage3 |
| ENSMUSG00000015290 | 0.10842628350471 | 0.44629410723172 | 0.89108031640712 | 2533.4524612596 | 2479.89589454413 | 2447.53234204488 | 2369.20282969612 | 2010.34324752386 | 1873.76197953715 | 2632.88649101647 | 2599.83372738108 | Ubl4a |
| ENSMUSG00000015291 | -0.105785245596175 | 0.356971243126173 | 0.865163598066054 | 727.328541838081 | 876.311324311818 | 839.505486231172 | 814.120370616084 | 833.263187018333 | 976.01962119494 | 899.491470240502 | 797.861290031353 | Gdi1 |
| ENSMUSG00000015305 | -0.134851404222369 | 0.437328957590578 | 0.888722970093854 | 558.37147683967 | 638.501931468834 | 617.088587398787 | 467.087493740564 | 747.856447744443 | 689.818378889657 | 561.141090809758 | 506.940438700193 | Sash1 |
| ENSMUSG00000015312 | -0.00614102933132966 | 0.976112678977227 | 0.99575536949131 | 156.307872966445 | 141.708336831093 | 149.539872406455 | 207.281799431054 | 176.288269526875 | 204.254732756335 | 132.216917500629 | 145.46042566558 | Gadd45b |
| ENSMUSG00000015335 | -0.213745503277381 | 0.508808707773627 | 0.902394714365738 | 46.9827132615903 | 39.9063707168021 | 43.5369248778286 | 58.151455044006 | 52.5579933993167 | 83.1695917810225 | 54.1360607089191 | 29.3091902460497 | Zdhhc12 |
| ENSMUSG00000015337 | -0.174498074747754 | 0.308490111134428 | 0.851785577030345 | 1610.06144292604 | 1173.57306536555 | 1218.08744169055 | 1513.81368453267 | 1618.34821342063 | 1900.66978864278 | 1366.93553290021 | 1339.53854680094 | Endog |
| ENSMUSG00000015340 | -0.202259372183122 | 0.559509618540027 | 0.916574868225118 | 38.8510898124689 | 64.338842584232 | 118.306861081056 | 120.054616865045 | 91.9764884488041 | 99.0696607979826 | 87.4505596067155 | 113.980184290193 | Cybb |
| ENSMUSG00000015341 | 0.00131324084129505 | 0.990806610746259 | 0.998628446692557 | 737.267192720341 | 747.633639143354 | 758.110365807406 | 732.520748215624 | 672.304332232926 | 684.926049961361 | 751.658381381531 | 862.992823911464 | Golga7 |
| ENSMUSG00000015342 | 0.295939905715339 | 0.159152962354254 | 0.789393312599749 | 325.264937964856 | 359.971752180133 | 283.936466594534 | 250.426427366929 | 213.516848184724 | 184.685417043153 | 311.282349076285 | 283.322172378481 | Xk |
| ENSMUSG00000015354 | -0.218273719710951 | 0.442380925140333 | 0.89108031640712 | 292.738444168371 | 208.490426602068 | 220.523989055088 | 340.467390015713 | 328.487458745729 | 465.99433042014 | 224.872867560126 | 217.105112933702 | Pcolce2 |
| ENSMUSG00000015355 | 0.0483145634694117 | 0.918201884692788 | 0.986713918026868 | 22.5878429142261 | 13.8450673915436 | 21.7684624389143 | 9.37926694258161 | 21.8991639163819 | 12.2308223207386 | 8.32862472444909 | 22.7960368580387 | Cd48 |
| ENSMUSG00000015357 | 0.332921169602882 | 0.133869496992681 | 0.77303252008843 | 3068.33258146848 | 3939.32888075861 | 4714.29180035791 | 3562.2455847925 | 3407.50990538903 | 1931.24684444462 | 3580.26755342255 | 3214.24119698345 | Clpx |
| ENSMUSG00000015363 | 0.220128504410207 | 0.00499634046411002 | 0.635862253526407 | 1081.50591873315 | 1064.44135769103 | 1090.31603172301 | 1148.96020046625 | 914.290093508946 | 920.980920751616 | 960.915077583314 | 967.20327811964 | Trabd |
| ENSMUSG00000015365 | -0.197386010466604 | 0.0796351900537288 | 0.739855948845396 | 985.733464776828 | 949.608739914107 | 847.07712534036 | 877.899385825639 | 967.943045104082 | 963.788798874201 | 1108.74816644229 | 1156.08472637196 | Mov10l1 |
| ENSMUSG00000015377 | 0.0794589882744582 | 0.556140521901376 | 0.916239900758826 | 279.185738419835 | 236.180561385155 | 288.668741037776 | 279.502154888932 | 234.321053905287 | 239.724117486476 | 263.392756910703 | 286.578749072486 | Dennd6b |
| ENSMUSG00000015396 | -0.0847429126913334 | 0.819176978452401 | 0.966315773308182 | 21.6843291976571 | 34.2054606144018 | 37.8581955459379 | 70.3445020693621 | 47.0832024202212 | 50.1463715150282 | 33.3144988977964 | 43.4210225867403 | Cd83 |
| ENSMUSG00000015405 | -0.122205850490172 | 0.783285003573261 | 0.961051227381536 | 23.4913566307952 | 10.5874044758863 | 26.5007368821565 | 24.3860940507122 | 37.2285786578493 | 23.2385624094033 | 19.7804837205666 | 11.9407812113536 | Ace2 |
| ENSMUSG00000015461 | -0.209580291378888 | 0.203053219507149 | 0.81561862896307 | 356.887918044773 | 400.69253862585 | 427.797609669098 | 367.667264149199 | 492.731188118594 | 544.271593272868 | 398.732908683 | 361.480013034613 | Atf6b |
| ENSMUSG00000015468 | -0.729874261751458 | 0.0302102293111324 | 0.665956123973759 | 791.478015714483 | 683.294796559122 | 723.091534927413 | 616.217838127612 | 1819.82052145134 | 1547.19902357343 | 653.797040869254 | 646.973236542431 | Notch4 |
| ENSMUSG00000015474 | -0.0313501735858436 | 0.786174011768802 | 0.961141387070613 | 720.100432105529 | 693.067785306094 | 713.626986040929 | 686.562340196974 | 794.939650164664 | 786.441875223492 | 661.084587503147 | 633.946929766408 | Ppt2 |
| ENSMUSG00000015476 | -0.971803992131492 | 0.105659105180954 | 0.762094734041179 | 15.3597331816738 | 4.88649437348597 | 13.2503684410783 | 22.5102406621959 | 32.8487458745729 | 51.3694537471021 | 9.36970281500523 | 16.2828834700276 | Prrt1 |
| ENSMUSG00000015478 | -0.051008026109656 | 0.660612793885029 | 0.939611397743053 | 927.908586916409 | 840.477032239587 | 756.217456030109 | 969.816201862939 | 865.016974697087 | 963.788798874201 | 868.259127523818 | 923.7822555329 | Rnf5 |
| ENSMUSG00000015488 | -0.00737337291815675 | 0.944558356726719 | 0.990505437194639 | 1538.68385931708 | 1549.01871639505 | 1694.15425068072 | 1776.43315892496 | 1629.29779537882 | 1464.02943179241 | 1765.66844158321 | 1730.3277500816 | Cacfd1 |
| ENSMUSG00000015501 | 0.0803056776403815 | 0.678430178405416 | 0.944452327903597 | 354.177376895066 | 521.22606650517 | 376.689045682082 | 319.833002742033 | 361.336204620302 | 306.993640250539 | 428.924173309128 | 389.703677715994 | Hivep2 |
| ENSMUSG00000015522 | 0.0737184447319625 | 0.697536707904018 | 0.944933386973863 | 729.135569271219 | 900.743796179247 | 866.952678001977 | 746.589648629496 | 767.565695269187 | 529.594606487981 | 916.1487196894 | 867.334926170138 | Arnt |
| ENSMUSG00000015536 | -0.0317864286843064 | 0.838312860821635 | 0.969135716755098 | 447.239289701677 | 350.198763433161 | 401.296872786941 | 469.901273823339 | 363.52612101194 | 496.571386221987 | 398.732908683 | 447.236532643425 | Mocs2 |
| ENSMUSG00000015542 | 0.128513694137309 | 0.375998496674477 | 0.874078513608115 | 474.344701198749 | 451.186313818538 | 478.906173656114 | 499.9149280396 | 497.11102090187 | 494.125221757839 | 395.609674411332 | 356.05238521127 | Nat9 |
| ENSMUSG00000015568 | 0.0876991857314347 | 0.416020137335297 | 0.882192126163777 | 154190.940328524 | 137527.126478847 | 173458.680352378 | 166114.320966674 | 153911.703837116 | 158516.3496057 | 141275.337966558 | 140355.198934944 | Lpl |
| ENSMUSG00000015575 | -0.143270643519226 | 0.447706177182181 | 0.89108031640712 | 868.276681622852 | 695.511032492837 | 809.218929794422 | 872.27182566009 | 951.518672166795 | 1176.60510725505 | 705.850945397061 | 751.183690750607 | Atp6v0e |
| ENSMUSG00000015579 | -0.271653419023282 | 0.0820575889986792 | 0.739855948845396 | 2051.8796503283 | 1831.62097432833 | 1809.62174709583 | 2043.74226678853 | 2287.36767106609 | 2967.19749501118 | 2191.46938062067 | 1895.32763591121 | Nkx2-5 |
| ENSMUSG00000015597 | 0.414206813400703 | 0.132566234296113 | 0.77303252008843 | 333.396561413978 | 480.505280059454 | 392.778778789105 | 319.833002742033 | 217.896680968 | 177.34692365071 | 392.486440139664 | 357.137910775939 | Zfp318 |
| ENSMUSG00000015605 | 0.0997620157198901 | 0.574350477745787 | 0.922280709351328 | 2805.41008994688 | 3264.17824148863 | 3694.95988528354 | 3294.93647692892 | 2567.67696919578 | 2413.14124388173 | 3609.41773995813 | 3595.2606701821 | Srf |
| ENSMUSG00000015619 | -0.660615126160872 | 0.275346055593161 | 0.846286635233373 | 8.13162344912141 | 5.7009101024003 | 17.036187995672 | 13.1309737196143 | 35.0386622662111 | 18.3462334811079 | 7.28754663389296 | 8.68420451734806 | Gata3 |
| ENSMUSG00000015647 | 0.106344943937043 | 0.582264915521679 | 0.922450244597442 | 2461.17136393408 | 3182.7366685972 | 3732.81808082947 | 2190.05883109281 | 2911.49384268298 | 2298.17151406678 | 2985.811963715 | 2548.81402584166 | Lama5 |
| ENSMUSG00000015653 | -0.474238728801014 | 0.306560748380022 | 0.851785577030345 | 10.8421645988285 | 5.7009101024003 | 11.3574586637814 | 7.50341355406529 | 9.85462376237187 | 15.9000690169602 | 12.4929370866736 | 10.8552556466851 | Steap2 |
| ENSMUSG00000015656 | 0.178680529273713 | 0.0935590304707572 | 0.755231402633518 | 70072.9098022289 | 56674.3761594192 | 57365.577255871 | 65954.0672135396 | 52771.5102475014 | 58624.7775477642 | 53332.3484230098 | 56208.5137385353 | Hspa8 |
| ENSMUSG00000015659 | -0.0363035125667966 | 0.927011909742561 | 0.986790505471895 | 108.421645988285 | 150.666909849151 | 135.343049076728 | 93.7926694258161 | 89.7865720571659 | 46.4771248188067 | 172.818963032319 | 191.052499381657 | Serac1 |
| ENSMUSG00000015668 | 0.00113103630089878 | 0.992616107119271 | 0.999071668397934 | 267.440060104437 | 259.798617523671 | 260.275094378323 | 288.881421831514 | 244.175677667659 | 292.316653465652 | 284.214318721825 | 255.098507697099 | Pdzd11 |
| ENSMUSG00000015671 | -0.0146294571190656 | 0.910784511985622 | 0.986166948757759 | 2532.54894754303 | 2005.09152458708 | 1966.73325861147 | 2535.21585457981 | 2325.69120791976 | 2468.17994432505 | 2151.90841317953 | 2186.24848724237 | Psma2 |
| ENSMUSG00000015672 | 0.222569905570474 | 0.0775868600275855 | 0.739855948845396 | 559.274990556239 | 479.69086433054 | 453.351891662606 | 518.673461924763 | 409.514365236342 | 408.509465512669 | 413.308001950786 | 491.743080794834 | Mrpl32 |
| ENSMUSG00000015697 | 0.044369400341225 | 0.789484593902155 | 0.961659329450003 | 242.141676040504 | 266.313943354985 | 257.435729712378 | 242.923013812864 | 238.700886688563 | 185.908499275227 | 252.981976005141 | 299.605055848508 | Setdb1 |
| ENSMUSG00000015711 | 0.140190450338944 | 0.322264140686401 | 0.855977678491063 | 658.661499378834 | 803.828324438442 | 879.256591554407 | 741.900015158206 | 695.298454345127 | 590.748718091674 | 771.438865102097 | 739.242909539254 | Prune1 |
| ENSMUSG00000015714 | -0.00483500537087654 | 0.967808420445113 | 0.994406701590602 | 553.853908256825 | 556.245942848486 | 559.354839191232 | 580.576623745802 | 579.232885588302 | 627.44118505389 | 470.567296931374 | 581.84170266232 | Cers2 |
| ENSMUSG00000015733 | 0.102160582351195 | 0.482110487635276 | 0.897732059433541 | 3261.68451681425 | 3329.33149980178 | 3350.4503058155 | 3248.04014221601 | 2731.92069856865 | 2477.96460218164 | 3617.74636468257 | 3458.48444903387 | Capza2 |
| ENSMUSG00000015745 | -0.00243486333541989 | 0.989538223558211 | 0.998596768760793 | 1494.4116872052 | 1235.46866076304 | 1205.78352813812 | 1829.89498049767 | 1589.87930032933 | 1695.19197365437 | 1233.67753730902 | 1257.03860388613 | Plekho1 |
| ENSMUSG00000015747 | -0.102389975979038 | 0.54402151362112 | 0.913837378403038 | 230.395997725106 | 189.758864837039 | 160.897331070236 | 172.578511743502 | 208.042057205628 | 231.162541861959 | 181.147587756768 | 189.966973816989 | Vps45 |
| ENSMUSG00000015748 | 0.0897358879419465 | 0.459538737471046 | 0.894774044236998 | 367.730082643601 | 328.209538752474 | 332.205665915605 | 401.432625142493 | 338.342082508101 | 316.77829810713 | 328.980676615739 | 358.223436340608 | Prpf3 |
| ENSMUSG00000015749 | 0.116237159377224 | 0.444773394074608 | 0.89108031640712 | 2550.61922187441 | 2645.22228751374 | 2501.48027069784 | 2297.92040093249 | 1980.77937623675 | 1906.78519980315 | 2755.73370570209 | 2577.03769052304 | Anp32e |
| ENSMUSG00000015750 | -0.0932809085980901 | 0.299177923713361 | 0.851785577030345 | 1118.54998111248 | 1092.94590820303 | 1090.31603172301 | 1064.54679798301 | 1179.26997689717 | 1276.89785028511 | 1082.72121417838 | 1121.34790830257 | Aph1a |
| ENSMUSG00000015755 | 0.140120678896192 | 0.455170550696164 | 0.892775943845882 | 1723.00065749717 | 2003.46269312925 | 1961.05452927958 | 1643.2475683403 | 1401.54649064844 | 1252.43620564363 | 1949.93926361164 | 2047.30121496481 | Tab2 |
| ENSMUSG00000015757 | 0.0719412472481793 | 0.602186106871008 | 0.926448151609826 | 400.256576440087 | 399.063707168021 | 511.085639870161 | 418.31530563914 | 369.000911991036 | 428.078781225851 | 454.951125573032 | 391.874728845331 | Ppil4 |
| ENSMUSG00000015759 | 0.0207632860395721 | 0.878362167263848 | 0.977992024312213 | 1316.4194850411 | 1005.8034252092 | 947.401343537095 | 1232.43567625522 | 1103.71786138565 | 1083.65085761744 | 1103.5427759895 | 1146.31499628994 | Cnih1 |
| ENSMUSG00000015766 | -0.463939451286569 | 0.145507446663442 | 0.780607990324772 | 217.74680569314 | 225.593156909269 | 273.525462819401 | 171.640585049243 | 431.413529152724 | 418.29412336926 | 181.147587756768 | 195.394601640331 | Eps8 |
| ENSMUSG00000015776 | -0.0289637365733118 | 0.781225018751014 | 0.960742105865155 | 351.466835745359 | 333.09603312596 | 377.63550057073 | 374.232751009006 | 364.621079207759 | 366.924669622158 | 377.911346871878 | 354.966859646602 | Med22 |
| ENSMUSG00000015790 | -0.0408146994532629 | 0.747343742308349 | 0.956119556124081 | 851.10992100804 | 718.314672902438 | 818.683478680906 | 852.575365080669 | 802.604357535398 | 968.681127802497 | 733.960053842076 | 829.34153140674 | Surf1 |
| ENSMUSG00000015804 | -0.0661012397767211 | 0.605820617193987 | 0.926859923029135 | 809.548290045864 | 673.52180781215 | 707.948256709038 | 840.382318055312 | 842.022852584886 | 851.265233523406 | 689.193695948162 | 791.348136643342 | Med28 |
| ENSMUSG00000015806 | -0.0102863017241087 | 0.919781934810284 | 0.986713918026868 | 1844.07149551742 | 1735.51991831643 | 1605.18749114777 | 1875.85338851632 | 1726.74907480672 | 1936.13917337292 | 1626.16397744869 | 1823.68294864309 | Qdpr |
| ENSMUSG00000015837 | 0.0702000744404064 | 0.618246680341953 | 0.929568738467614 | 23916.9115912992 | 25259.9182480068 | 27665.8228500827 | 26401.698516673 | 28188.6037931668 | 27527.9117972864 | 22760.0492157383 | 19865.1178334337 | Sqstm1 |
| ENSMUSG00000015839 | -0.132190350737073 | 0.472634593356651 | 0.89728597737823 | 1227.87514081733 | 1435.81493007596 | 937.936794650611 | 1008.27119632752 | 1496.80785368471 | 1347.83661974539 | 1166.00746142287 | 1043.19006764644 | Nfe2l2 |
| ENSMUSG00000015843 | -0.0561245509907224 | 0.66483715794641 | 0.941513723457524 | 1874.79096188077 | 1600.32690731666 | 1396.02096075646 | 1422.83479518963 | 1647.91208470774 | 1538.63744794892 | 1581.39761955477 | 1775.91982379768 | Rxrg |
| ENSMUSG00000015846 | 0.0100440850054918 | 0.949272583391226 | 0.991267322147698 | 1971.46692955366 | 2397.63990592378 | 2490.12281203406 | 1913.37045628665 | 2079.32561386047 | 1758.79224972221 | 2397.60284255078 | 2474.9982874442 | Rxra |
| ENSMUSG00000015850 | 0.0422503763186773 | 0.739925527359529 | 0.954383574136061 | 832.13613296009 | 945.536661269536 | 889.66759532954 | 772.851596068725 | 884.72622222183 | 856.157562451702 | 899.491470240502 | 701.249514775856 | Adamtsl4 |
| ENSMUSG00000015852 | -1.00676099418311 | 0.00351985908760649 | 0.635862253526407 | 47.8862269781594 | 64.338842584232 | 41.6440151005317 | 45.9584080186499 | 110.590777777729 | 159.000690169602 | 52.0539045278068 | 81.4144173501381 | Fcrls |
| ENSMUSG00000015854 | -0.600889821519902 | 0.351801404214676 | 0.862728204353539 | 7.22810973255236 | 17.1027303072009 | 3.78581955459379 | 8.44134024832345 | 6.56974917491458 | 7.33849339244316 | 13.5340151772298 | 28.2236646813812 | Cd5l |
| ENSMUSG00000015869 | -0.0564204061688734 | 0.682657912056335 | 0.944452327903597 | 433.686583953142 | 379.517729674077 | 442.940887887473 | 459.584080186499 | 482.876564356222 | 492.902139525765 | 439.33495421469 | 369.078691987293 | Prpsap1 |
| ENSMUSG00000015882 | 0.176947262523359 | 0.360870301793263 | 0.868539734006824 | 104.807591122009 | 83.8848200781758 | 99.3777633080869 | 82.5375490947182 | 73.3621991198795 | 69.71568722821 | 92.6559500594962 | 91.1841474321546 | Lcorl |
| ENSMUSG00000015889 | -0.211699614265898 | 0.17980608954637 | 0.802731406993634 | 497.836057829544 | 455.25839246311 | 489.317177431247 | 549.625042835282 | 635.075753575076 | 689.818378889657 | 469.526218840818 | 513.453592088204 | Lta4h |
| ENSMUSG00000015890 | -0.462055078827065 | 0.446792226693161 | 0.89108031640712 | 15.3597331816738 | 34.2054606144018 | 9.46454888648447 | 15.9447538023887 | 7.66470737073368 | 20.7923979452556 | 19.7804837205666 | 55.3618037980939 | Amdhd1 |
| ENSMUSG00000015932 | 0.0395613414997149 | 0.558598758810121 | 0.916574868225118 | 2495.5048851637 | 2726.66386040517 | 2591.39348511945 | 2598.99486978936 | 2494.3147700759 | 2537.89563155326 | 2517.32682296474 | 2582.46531834638 | Dstn |
| ENSMUSG00000015937 | -0.191375184315877 | 0.10701209913275 | 0.762287160109383 | 699.319616624441 | 687.366875203693 | 641.696414503647 | 717.513921107493 | 785.085026402292 | 905.080851734656 | 763.110240377648 | 683.88110574116 | H2afy |
| ENSMUSG00000015942 | 0.0842463569246994 | 0.494203284549261 | 0.900141680336539 | 217.74680569314 | 232.922898469498 | 250.810545491838 | 225.102406621959 | 224.466430142915 | 233.608706326107 | 210.29777429234 | 206.249857287016 | Gtf2ird2 |
| ENSMUSG00000015943 | -0.0400037781956348 | 0.881599761765468 | 0.978854883720278 | 541.204716224858 | 375.445651029505 | 443.887342776122 | 522.425168701796 | 549.669014301187 | 713.05694129906 | 332.103910887408 | 341.94055287058 | Bola1 |
| ENSMUSG00000015944 | -0.0800443385394821 | 0.593352904342733 | 0.923935529355527 | 570.117155155067 | 758.22104361924 | 558.408384302584 | 588.080037299867 | 705.153078107498 | 671.472145408549 | 686.070461676494 | 554.703563545607 | Gatsl2 |
| ENSMUSG00000015947 | -0.0874092044948616 | 0.776631206032569 | 0.959738279197945 | 20.780815481088 | 20.3603932228582 | 24.6078271048596 | 24.3860940507122 | 25.1840385038392 | 29.3539735697726 | 21.8626399016789 | 19.5394601640331 | Fcgr1 |
| ENSMUSG00000015950 | -0.241004199483731 | 0.457532257670091 | 0.892966448175628 | 40.658117245607 | 70.8541684155466 | 82.3415753124149 | 64.7169419038131 | 87.5966556655277 | 111.300483118721 | 52.0539045278068 | 55.3618037980939 | Ncf1 |
| ENSMUSG00000015957 | -0.366456768235932 | 0.21321203560042 | 0.824245804199788 | 85.8338030740593 | 109.946123403434 | 76.6628459805242 | 91.9168160372998 | 173.003394939417 | 133.315963296051 | 95.7791843311646 | 68.388110574116 | Wnt11 |
| ENSMUSG00000015961 | -0.270708847047558 | 0.267501314888405 | 0.843793744782845 | 149.983276950461 | 121.347943608235 | 147.646962629158 | 144.440710915757 | 197.092475247437 | 234.831788558181 | 127.011527047849 | 121.578863242873 | Adss |
| ENSMUSG00000015968 | -1.26377798693134 | 0.0277361471753887 | 0.665956123973759 | 8.13162344912141 | 13.0306516626293 | 2.83936466594534 | 4.68963347129081 | 9.85462376237187 | 29.3539735697726 | 16.6572494488982 | 14.1118323406906 | Cacna1d |
| ENSMUSG00000015970 | -0.349043444021247 | 0.505860029756981 | 0.901494649536949 | 16.2632468982428 | 42.3496179035451 | 17.036187995672 | 12.1930470253561 | 25.1840385038392 | 18.3462334811079 | 16.6572494488982 | 52.1052271040884 | Chdh |
| ENSMUSG00000015971 | 0.193001262347382 | 0.170034279833265 | 0.79907358295209 | 850.206407291471 | 876.311324311818 | 1045.83265195653 | 1048.60204418062 | 816.838814081046 | 699.603036746248 | 934.888125319411 | 889.045437463508 | Actr8 |
| ENSMUSG00000015980 | 0.0711192595894951 | 0.72298310068596 | 0.951674101666722 | 215.036264543433 | 136.007426728693 | 153.325691961048 | 168.826804966469 | 143.439523652302 | 194.470074899744 | 139.504464134522 | 163.914360264945 | Lrrc27 |
| ENSMUSG00000015981 | -0.368305706894252 | 0.325288225055078 | 0.855977678491063 | 13.5527057485357 | 25.2468875963442 | 10.4110037751329 | 23.448167356454 | 25.1840385038392 | 24.4616446414772 | 22.903717992235 | 21.7105112933701 | Stk32c |
| ENSMUSG00000015994 | 0.108209985306881 | 0.23764302810258 | 0.832048466795696 | 1907.31745567725 | 1611.72872752146 | 1773.65646132719 | 1809.26059322399 | 1727.84403300253 | 1570.43758598284 | 1608.46564990923 | 1680.39357410685 | Fnta |
| ENSMUSG00000016018 | 0.0681836496693764 | 0.633140848275069 | 0.932281134354785 | 694.802048041596 | 846.177942341987 | 848.970035117657 | 748.465502018013 | 755.521155115177 | 606.648787108634 | 849.519721893808 | 780.492880996657 | Skiv2l2 |
| ENSMUSG00000016024 | -1.03781505349431 | 0.0104182614790082 | 0.635862253526407 | 91.2548853734735 | 112.389370590177 | 106.002947528626 | 90.9788893430416 | 238.700886688563 | 379.155491942897 | 114.518589961175 | 91.1841474321546 | Lbp |
| ENSMUSG00000016028 | 0.12900554546925 | 0.732442578145327 | 0.953658362433314 | 103.90407740544 | 124.605606523892 | 92.7525790875478 | 84.4134024832345 | 60.2227007700503 | 40.3617136584374 | 137.42230795341 | 132.434118889558 | Celsr1 |

| | | | | | | | | | | | | |
|---|---|---|---|---|---|---|---|---|---|---|---|---|
| **ENSMUSG00000016087** | -0.308700233753476 | 0.191064360214259 | 0.808824684419004 | 446.335775985108 | 306.220314071788 | 400.350417898293 | 488.659807708502 | 675.589206820383 | 576.071731306788 | 409.143689588562 | 372.335268681298 | Fli1 |
| **ENSMUSG00000016128** | -0.0770339893581594 | 0.677424405749807 | 0.944214159784677 | 545.722284807703 | 636.873100011005 | 485.531357876653 | 394.867138282686 | 581.42280197994 | 453.763508099402 | 630.893322877019 | 510.197015394199 | Stard13 |
| **ENSMUSG00000016181** | 0.0583072739243817 | 0.725111675911084 | 0.951674101666722 | 118.360296870545 | 140.079505373265 | 109.78876708322 | 144.440710915757 | 127.015150715015 | 112.523565350795 | 133.257995591185 | 119.407812113536 | Diexf |
| **ENSMUSG00000016194** | -0.303013087650811 | 0.150343018505397 | 0.782641594375979 | 438.204152535987 | 400.69253862585 | 461.869985660442 | 486.783954319986 | 623.031213421066 | 687.372214425509 | 351.884394607974 | 543.848307898922 | Hsd11b1 |
| **ENSMUSG00000016206** | -0.250595365933115 | 0.527952483299795 | 0.908232038364594 | 62.3424464432641 | 35.8342920722305 | 46.3762895437739 | 67.5307219865876 | 99.6411958195378 | 80.7234273168747 | 40.6020455316893 | 31.4802413753867 | H2-M3 |
| **ENSMUSG00000016239** | 0.136054194420685 | 0.763277314147561 | 0.95785116771606 | 22.5878429142261 | 25.2468875963442 | 40.6975602118832 | 21.5723139679377 | 14.2344565456483 | 12.2308223207386 | 41.6431236222455 | 31.4802413753867 | Lonrf3 |
| **ENSMUSG00000016252** | 0.0862362573611261 | 0.650025599733225 | 0.936463929813731 | 13890.6198785325 | 9245.24735463546 | 10107.1917558768 | 13039.056903577 | 10465.6104356389 | 13896.6603208232 | 9312.44352002464 | 9922.78918663483 | Atp5e |
| **ENSMUSG00000016253** | -0.0646463564258122 | 0.623761767422856 | 0.931145988862144 | 552.046880823686 | 557.060358577401 | 421.172425448559 | 481.156394154437 | 535.434557755538 | 525.92535979176 | 558.017856538089 | 485.229927406823 | Nelfcd |
| **ENSMUSG00000016255** | 0.809964320153378 | 0.310304618433285 | 0.851785577030345 | 34.3335212296237 | 48.0505280059454 | 124.932045301595 | 2.81378008277448 | 28.4689130912965 | 55.0387004433237 | 8.32862472444909 | 28.2236646813812 | Tubb1 |
| **ENSMUSG00000016256** | -0.584892304392388 | 0.119901408112279 | 0.769868040577702 | 473.44118748218 | 478.062032872711 | 399.403963009645 | 624.659178375935 | 900.055636963298 | 1289.12867260585 | 385.198893505771 | 389.703677715994 | Ctsz |
| **ENSMUSG00000016257** | 0.0584787374241651 | 0.536041845797232 | 0.911291453479697 | 1616.38603894202 | 1513.99884005174 | 1370.46667876295 | 1592.59952685036 | 1401.54649064844 | 1449.35244500752 | 1443.97531160136 | 1556.64365973464 | Prelid3b |
| **ENSMUSG00000016262** | -0.636181255962591 | 0.0970981993088581 | 0.757809915356977 | 31.6229800799166 | 50.4937751926884 | 41.6440151005317 | 74.0962088463947 | 93.0714466446232 | 117.415894279091 | 54.1360607089191 | 43.4210225867403 | Sertad4 |
| **ENSMUSG00000016308** | 0.0638452386356506 | 0.582117226726153 | 0.922450244597442 | 926.101559483271 | 760.664290805983 | 752.431636475515 | 891.968286239511 | 758.806029702634 | 853.711397987554 | 769.356708920985 | 805.459968984033 | Ube2a |
| **ENSMUSG00000016319** | -0.0117198528764251 | 0.910766977904451 | 0.986166948757759 | 7015.78400915863 | 6266.92903399576 | 6463.34043458024 | 6475.44589715835 | 7292.42158415519 | 6936.09933809086 | 6416.16427209747 | 5791.27888750649 | Slc25a5 |
| **ENSMUSG00000016344** | -0.117458892379098 | 0.501292695602322 | 0.900927059989498 | 901.706689135907 | 744.375976227696 | 752.431636475515 | 962.312788308873 | 930.714466446232 | 1157.03579154187 | 804.753363999894 | 754.440267444613 | Ppdpf |
| **ENSMUSG00000016349** | 0.0715929382482036 | 0.368166986859154 | 0.871495599696183 | 63509.7861650713 | 62461.6143290844 | 64704.3884624511 | 70425.1637650683 | 64241.1973487064 | 63978.2084775515 | 62060.7471342324 | 58179.8281639733 | Eef1a2 |
| **ENSMUSG00000016382** | -0.413896853192473 | 0.0439620874935535 | 0.691603097094229 | 901.706689135907 | 954.495234287593 | 798.807926019289 | 915.416453595965 | 1371.98261936133 | 1566.76833928661 | 905.737938783839 | 914.012525450883 | Pls3 |
| **ENSMUSG00000016386** | 0.107918780981395 | 0.38784389633863 | 0.875951433019341 | 299.063040184354 | 316.807718547674 | 355.867038131816 | 315.143369270742 | 286.879047304603 | 267.855008824175 | 311.282349076285 | 326.743194965221 | Mpped2 |
| **ENSMUSG00000016409** | -0.111780464678631 | 0.382245176789365 | 0.875126835008975 | 288.220875585525 | 316.807718547674 | 250.810545491838 | 265.43325447506 | 287.974005500422 | 305.770558018465 | 325.857442344071 | 293.091902460497 | Nkap |
| **ENSMUSG00000016427** | 0.00598587328930926 | 0.979583774727635 | 0.996872525500296 | 5846.63725991829 | 4265.09517232434 | 4338.54920956448 | 5557.21566347961 | 5026.95307700547 | 7243.0929783414 | 3874.89265304994 | 3780.88554174041 | Ndufa1 |
| **ENSMUSG00000016458** | -0.70423494264615 | 0.0427698509791865 | 0.687730553581215 | 106.614618555147 | 67.5965054998893 | 93.6990339761962 | 47.8342614071662 | 188.332809680885 | 150.439114545085 | 100.984574783945 | 74.901263962127 | Wt1 |
| **ENSMUSG00000016477** | 0.133314008620674 | 0.573640825007798 | 0.922207298536842 | 234.010052591383 | 246.767965861042 | 245.131816159948 | 231.667893481766 | 176.288269526875 | 139.43137445642 | 268.598147363483 | 287.664274637155 | E2f3 |
| **ENSMUSG00000016481** | -0.0556383553337958 | 0.476821259566438 | 0.897622440754504 | 1322.74408105708 | 1291.66334605813 | 1323.14393433053 | 1238.06323642077 | 1273.43638173761 | 1410.21381358116 | 1345.07289299853 | 1351.47932801229 | Cr1l |
| **ENSMUSG00000016487** | 0.180328324559739 | 0.338681473040859 | 0.858570960161243 | 2433.16243872044 | 2841.49647818209 | 2723.89716953023 | 2160.04517687655 | 2074.94578107719 | 1631.59169758653 | 2819.23946922602 | 2438.09041824547 | Ppfibp1 |
| **ENSMUSG00000016494** | -0.854014592875892 | 0.0163783356587675 | 0.641523953292557 | 1076.08483643373 | 980.556537612852 | 781.771738023617 | 1157.40154071457 | 2494.3147700759 | 2784.95824243218 | 891.162845516053 | 1052.95979772845 | Cd34 |
| **ENSMUSG00000016495** | 0.303426830283197 | 0.0513253786938484 | 0.703396849069134 | 245.75573090678 | 215.005752433383 | 212.005895057252 | 241.985087118606 | 221.181555555458 | 193.24699266767 | 168.654650670094 | 158.486732441602 | Plgrkt |
| **ENSMUSG00000016496** | -0.00626020514156747 | 0.977722208356375 | 0.996226633954669 | 93.0619128066116 | 91.2145616384048 | 121.146225747001 | 107.861569839689 | 125.920192519196 | 107.6312364225 | 69.7522320672611 | 111.809133160856 | Cd274 |
| **ENSMUSG00000016503** | 0.240997782286621 | 0.0518268888761188 | 0.703396849069134 | 419.230364488037 | 352.642010619904 | 365.3315870183 | 431.446279358754 | 323.012667766634 | 340.016860516533 | 310.241270985729 | 353.881334081933 | Gtf3a |
| **ENSMUSG00000016510** | -0.0921279927986796 | 0.500409898318899 | 0.900927059989498 | 378.57224724243 | 375.445651029505 | 355.867038131816 | 432.384206053012 | 355.861413641207 | 380.37857417497 | 451.827891301363 | 454.835211596105 | Mtif3 |
| **ENSMUSG00000016520** | 0.159574326787659 | 0.58762276663011 | 0.923207807769736 | 505.967681278665 | 757.406627890326 | 375.742590793433 | 264.495327780801 | 431.413529152724 | 491.679057293692 | 389.363205867995 | 392.96025441 | Lnx2 |
| **ENSMUSG00000016526** | 0.439232265061319 | 0.428225659754192 | 0.885378691567738 | 8.13162344912141 | 13.8450673915436 | 10.4110037751329 | 13.1309737196143 | 2.18991639163819 | 14.6769867848863 | 7.28754663389296 | 9.76973008201657 | Dyrk3 |
| **ENSMUSG00000016528** | 0.23171731643951 | 0.156385055796696 | 0.787275540343856 | 10054.3006379803 | 11154.2378232106 | 13390.4437645982 | 11223.2308234932 | 9752.7926501607 | 7443.67846440151 | 11434.160668578 | 10390.650705007 | Mapkapk2 |
| **ENSMUSG00000016534** | 0.0600640023511846 | 0.650755072520698 | 0.936463929813731 | 3034.90257395542 | 3829.38275735517 | 4296.90519446395 | 3356.83963874996 | 3369.18636853536 | 3319.44517784845 | 3566.73353824532 | 3670.16193414422 | Lamp2 |
| **ENSMUSG00000016541** | 0.067476937412149 | 0.442763455357194 | 0.89108031640712 | 1805.22040570495 | 1690.72705322615 | 1871.14131485798 | 1814.88815338954 | 1797.92135753496 | 1782.03081213161 | 1709.45022469318 | 1564.24233868732 | Atxn10 |
| **ENSMUSG00000016554** | -0.147409180424161 | 0.378669640983994 | 0.87422797717331 | 1436.58680934478 | 1340.52828979298 | 1367.62731409701 | 1480.98625023364 | 1816.53564686388 | 1876.2081440013 | 1262.82772384459 | 1276.57806405016 | Eif3d |
| **ENSMUSG00000016559** | -0.235194403521244 | 0.189311870525202 | 0.808677608936066 | 2105.18695960588 | 1926.09319888239 | 2414.40642094219 | 2705.9185129348 | 3007.85016391506 | 3248.50640838817 | 2335.13815711741 | 2180.82085941903 | H3f3b |
| **ENSMUSG00000016619** | 0.0149641948699416 | 0.920996659844358 | 0.986713918026868 | 369.537110076739 | 482.948527246197 | 443.887342776122 | 368.605190843457 | 443.458069306734 | 349.801518373124 | 404.979377226337 | 449.407583772762 | Nup50 |
| **ENSMUSG00000016624** | -0.290815524109977 | 0.598709899074259 | 0.924848033480823 | 9.93865088225949 | 5.7009101024003 | 15.1432782183751 | 11.2551203310979 | 12.0445401540101 | 6.1154111603693 | 23.9447960827911 | 8.68420451734806 | Phf21b |
| **ENSMUSG00000016637** | -0.219728439390626 | 0.434969124521588 | 0.888379878975956 | 160.82544154929 | 115.647033505835 | 132.503684410783 | 141.626930832982 | 146.724398239759 | 258.070350967584 | 108.272121417838 | 129.177542195552 | Ift27 |
| **ENSMUSG00000016664** | 0.135749898382029 | 0.502531723289266 | 0.900927059989498 | 4412.76099172322 | 5393.06095687068 | 4956.58425185192 | 4196.28403011101 | 3603.50742244065 | 3137.20592526945 | 4953.4495548661 | 5561.14746779676 | Pacsin2 |
| **ENSMUSG00000016756** | -0.374193383054535 | 0.328838019392835 | 0.855977678491063 | 18.0702743313809 | 35.8342920722305 | 34.0723759913441 | 18.7585338851632 | 45.9882442244021 | 20.7923979452556 | 28.1091084450157 | 43.4210225867403 | Cmah |
| **ENSMUSG00000016757** | -0.173385933501263 | 0.182292669474614 | 0.804505478598291 | 170.764092431549 | 190.573280565953 | 176.040609288611 | 166.950951577953 | 216.801722772181 | 194.470074899744 | 193.640524843441 | 189.966973816989 | Ttll12 |
| **ENSMUSG00000016758** | -0.100456339789205 | 0.756239703635262 | 0.956566265296272 | 64.1494738764022 | 48.0505280059454 | 84.2344850897118 | 135.999370667433 | 99.6411958195378 | 92.9542496376133 | 92.6559500594962 | 70.559161703453 | Bik |
| **ENSMUSG00000016763** | 0.00592354661976584 | 0.986378710678201 | 0.997945975039712 | 22.5878429142261 | 39.9063707168021 | 42.5904699891801 | 30.0136542162612 | 31.7537876787538 | 51.3694537471021 | 22.903717992235 | 29.3091902460497 | Scube1 |
| **ENSMUSG00000016831** | 0.0825071898985385 | 0.519792548490271 | 0.905962179970576 | 845.688838708626 | 982.18536907068 | 986.205993971682 | 900.409626487835 | 912.100177117308 | 720.395434691503 | 957.791843311646 | 916.18357658022 | Tox4 |
| **ENSMUSG00000016833** | 0.0748383546984571 | 0.688046932266267 | 0.944453919646345 | 959.531566996326 | 609.182965227918 | 683.340429604179 | 869.458045577315 | 728.147200219699 | 891.626947181844 | 701.686633034836 | 642.631134283757 | Mrps18c |
| **ENSMUSG00000016918** | -0.856627434140843 | 0.0588738337090833 | 0.711712402279519 | 196.965990212052 | 177.542628903324 | 176.98706417726 | 163.19924480092 | 456.597567656563 | 578.517895770936 | 144.709854587303 | 115.065709854862 | Sulf1 |
| **ENSMUSG00000016921** | 0.0525989057014929 | 0.679775153099615 | 0.944452327903597 | 969.470217878585 | 1044.89538019708 | 1067.60111439545 | 1197.73238856767 | 1042.40020241978 | 909.973180662952 | 1192.03441368678 | 980.229584895662 | Srsf6 |
| **ENSMUSG00000016933** | 0.0202637257812293 | 0.847368162774441 | 0.972401171590961 | 1295.63866956001 | 1397.53739081699 | 1445.23661496618 | 1291.52505799349 | 1509.94735203453 | 1227.97456100216 | 1336.74426827408 | 1278.7491151795 | Plcg1 |
| **ENSMUSG00000016940** | -0.0555364766141494 | 0.743291334180086 | 0.954383574136061 | 568.310127721929 | 574.977504613516 | 608.570493400951 | 708.134654164912 | 713.912743674051 | 772.987970670679 | 554.894622266421 | 515.624643217541 | Kctd2 |
| **ENSMUSG00000016942** | -0.278144799898029 | 0.557529477481602 | 0.916341990969984 | 39.754603529038 | 113.203786319092 | 49.2156542097192 | 65.6548685980713 | 51.4630352034976 | 50.1463715150282 | 56.2182168900314 | 167.17093695895 | Tmprss6 |
| **ENSMUSG00000016946** | -0.12639574808803 | 0.45165852033413 | 0.891110666112841 | 177.088688447533 | 136.007426728693 | 168.468970179424 | 167.888878272211 | 208.042057205628 | 184.685417043153 | 149.915245040084 | 166.085411394282 | Kctd5 |
| **ENSMUSG00000016984** | 0.0246681836894856 | 0.913052813166438 | 0.986226496507248 | 55.1143367107117 | 56.1946852950887 | 72.8770264259304 | 57.2135283497478 | 52.5579933993167 | 47.7002070508805 | 65.5879197050366 | 70.559161703453 | Etaa1 |
| **ENSMUSG00000017009** | -0.424119826575583 | 0.0111997535945449 | 0.635862253526407 | 311.712232216321 | 286.674336577844 | 282.990011705886 | 352.660437041069 | 477.401773377126 | 477.002070508805 | 363.336253604092 | 338.683976176574 | Sdc4 |
| **ENSMUSG00000017057** | 0.193207644176351 | 0.246124201152526 | 0.832048466795696 | 784.249905981931 | 829.889627763701 | 585.855576073389 | 753.155135489303 | 695.298454345127 | 507.579126310652 | 716.261726302622 | 663.256120012458 | Il13ra1 |
| **ENSMUSG00000017119** | 0.0608489363478317 | 0.732882703474351 | 0.953733898884637 | 2214.51211931073 | 2896.06233201935 | 2932.11724503289 | 2228.51382555739 | 2470.22568976788 | 1872.53889730508 | 2933.75805918719 | 2569.43901157036 | Nbr1 |
| **ENSMUSG00000017132** | -0.00974125665400307 | 0.940652984955395 | 0.989523626251926 | 1157.40107092495 | 1218.36593045584 | 1238.90944924082 | 933.23706078687 | 1228.54309570903 | 1149.69729814943 | 1018.1743725639 | 1183.22286548867 | Cyth1 |
| **ENSMUSG00000017144** | 0.0694185726252056 | 0.772109865945623 | 0.959045638244974 | 167.150037565273 | 294.004078138073 | 239.453086828057 | 296.384835385579 | 199.282391639076 | 248.285693110993 | 193.640524843441 | 309.374785930525 | Rnd3 |
| **ENSMUSG00000017167** | 0.424558273088362 | 0.152767395208839 | 0.783412422888014 | 46.9827132615903 | 43.9784493613737 | 52.0550188756646 | 36.5791410760683 | 29.5638712871156 | 20.7923979452556 | 46.8485140750261 | 35.8223436340608 | Cntnap1 |
| **ENSMUSG00000017176** | 0.18155216086829 | 0.119306334041907 | 0.769868040577702 | 489.704434380422 | 479.69086433054 | 512.03209475881 | 535.55614242141 | 429.223612761086 | 497.794468454061 | 460.156516025812 | 391.874728845331 | Nt5c3b |
| **ENSMUSG00000017188** | -0.0124851784417696 | 0.936985466346755 | 0.988852035581444 | 1871.17690701449 | 1323.42555948578 | 1277.7140996754 | 1660.13024883695 | 1461.76919141849 | 1734.33060508073 | 1435.64668687691 | 1554.4726086053 | Coa3 |
| **ENSMUSG00000017204** | 0.681778417608294 | 0.194253396749397 | 0.809162446871198 | 15.3597331816738 | 12.2162359337149 | 5.67872933189068 | 21.5723139679377 | 6.56974917491458 | 7.33849339244316 | 11.4518589961175 | 8.68420451734806 | Gsdma |
| **ENSMUSG00000017210** | 0.0231911233385543 | 0.818399696941789 | 0.966315773308182 | 1158.30458464152 | 1149.95500922703 | 1020.27836996303 | 1078.61569839689 | 1019.40608030758 | 1013.93517038923 | 1181.62363278122 | 1121.34790830257 | Med24 |
| **ENSMUSG00000017221** | 0.0396241823827774 | 0.626632580407916 | 0.931698409533928 | 3319.50939467467 | 3333.40357844635 | 3416.70214802089 | 3423.43243404229 | 3478.68218811727 | 3455.20730560865 | 3160.71308292843 | 3034.04395324848 | Psmd3 |
| **ENSMUSG00000017264** | 0.119463300709806 | 0.233500858738509 | 0.832048466795696 | 936.9437240821 | 937.392503980392 | 1044.88619706789 | 898.533773099319 | 939.474132012785 | 900.188522806361 | 868.259127523818 | 806.545494548701 | Exosc10 |
| **ENSMUSG00000017286** | 0.0493955193493094 | 0.559308191694526 | 0.916574868225118 | 915.259394884443 | 867.35275129376 | 942.669069093853 | 919.168160372998 | 824.50352145178 | 883.065371557327 | 908.861173055507 | 904.242795368867 | Glod4 |
| **ENSMUSG00000017288** | 0.100611665709265 | 0.294942340827561 | 0.851785577030345 | 714.679349806115 | 818.4878075589 | 850.862944894954 | 770.975742680209 | 753.331238723539 | 728.95701031602 | 733.960053842076 | 726.216602763232 | Vps53 |
| **ENSMUSG00000017291** | 0.173789745746889 | 0.33165982434182 | 0.856333478432716 | 1367.91976688553 | 1565.30703097334 | 1304.21483655756 | 1156.46361402031 | 1039.11532783232 | 939.327154232724 | 1454.38609250692 | 1348.22275131829 | Taok1 |
| **ENSMUSG00000017299** | 0.0966021099072582 | 0.52335515629364 | 0.906635460329092 | 549.336339673979 | 491.09268453534 | 526.228918088536 | 637.79015209555 | 558.428679867739 | 596.864129252044 | 469.526218840818 | 437.466802561409 | Dnttip1 |
| **ENSMUSG00000017307** | -0.124360102957334 | 0.527384912053636 | 0.908217276872524 | 196.062476495483 | 149.852494120236 | 164.68315062483 | 207.281799431054 | 214.611806380543 | 237.277953022329 | 149.915245040084 | 181.282769299641 | Acot8 |
| **ENSMUSG00000017309** | -0.580830371788438 | 0.0920095629631647 | 0.754058637722175 | 2071.75695209282 | 1904.91838993061 | 2311.24283807951 | 2451.74037879083 | 4633.86308470642 | 4908.2289973124 | 1745.88795786264 | 1784.60402831503 | Cd300lg |
| **ENSMUSG00000017314** | -0.427634244255819 | 0.173892354969333 | 0.799889021902745 | 47.8862269781594 | 78.1839099757755 | 47.3227444324223 | 62.8410885152968 | 88.6916138613469 | 99.0696607979826 | 89.5327157878278 | 41.2499714574033 | Mpp2 |
| **ENSMUSG00000017344** | -0.718578656348784 | 0.00902196383338625 | 0.635862253526407 | 464.406050316489 | 784.282346944498 | 406.029147230184 | 538.369922504185 | 981.082543453911 | 1136.24339359662 | 540.319528998635 | 953.09144577895 | Vtn |
| **ENSMUSG00000017376** | 0.0309411333715621 | 0.848936825456157 | 0.972640876584422 | 352.370349461928 | 326.580707294646 | 279.204192151292 | 282.315934971707 | 292.353838283699 | 242.170281950624 | 348.761160336306 | 329.999771659226 | Nlk |
| **ENSMUSG00000017386** | -0.0482072927935741 | 0.823152875007343 | 0.966637987288264 | 74.9916384752307 | 70.0397526866323 | 77.6093008691726 | 91.9168160372998 | 107.305903190271 | 68.4926049961361 | 74.9576225200418 | 73.8157383974585 | Traf4 |
| **ENSMUSG00000017404** | 0.086576213356696 | 0.637201218120767 | 0.932931968211785 | 8343.94917251513 | 5507.07915891869 | 6660.20305141912 | 8267.82380988569 | 6866.48284598156 | 8277.82054667588 | 5674.9166716215 | 6284.10749386599 | Rpl19 |
| **ENSMUSG00000017405** | 0.11646846196315 | 0.680043145889758 | 0.944452327903597 | 38.8510898124689 | 35.8342920722305 | 23.6613722162112 | 37.5170677703264 | 32.8487458745729 | 36.6924669622158 | 29.1501865355718 | 27.1381391167127 | Nek8 |
| **ENSMUSG00000017417** | -0.95461638796652 | 0.0256266450108013 | 0.663332857280604 | 32.5264937964856 | 38.2775392589734 | 31.2330113253987 | 29.075727522003 | 93.0714466446232 | 99.0696607979826 | 29.1501865355718 | 33.6512925047237 | Plxdc1 |
| **ENSMUSG00000017418** | -0.00986974017514731 | 0.939099074319417 | 0.989269716906553 | 321.65088309858 | 294.004078138073 | 298.133289924261 | 271.998741334867 | 325.202584158272 | 258.070350967584 | 317.528817619622 | 292.006376895829 | Arl5b |
| **ENSMUSG00000017421** | -0.0136741896174994 | 0.877220213117018 | 0.977569405470617 | 1069.76024041775 | 1090.50266101629 | 954.972982646283 | 969.816201862939 | 1031.45062046159 | 1048.1814728873 | 1019.21545065446 | 1026.90718417641 | Zfp207 |
| **ENSMUSG00000017428** | 0.253561926666906 | 0.0747046761069529 | 0.735399943172214 | 2071.75695209282 | 2260.00364773726 | 2540.28492113243 | 2458.30586565064 | 1892.0877623754 | 1581.4453260715 | 2234.15358233347 | 2116.77485110359 | Psmd11 |
| **ENSMUSG00000017446** | -0.333563154259191 | 0.198465230113246 | 0.812921911423739 | 140.948139784771 | 197.903022126182 | 135.343049076728 | 119.116690170786 | 229.94122112201 | 234.831788558181 | 125.970448957293 | 158.486732441602 | C1qtnf1 |
| **ENSMUSG00000017453** | 0.180485773037833 | 0.682908178330012 | 0.944452327903597 | 30.7194663633475 | 70.0397526866323 | 25.5542819935081 | 48.7721881014244 | 22.994122112201 | 34.2463024980681 | 29.1501865355718 | 68.388110574116 | Pipox |
| **ENSMUSG00000017466** | -0.934012413001275 | 0.00502140302442564 | 0.635862253526407 | 816.776399778417 | 877.125740040732 | 912.382512657103 | 870.395972271574 | 2172.39706050509 | 2618.61905887013 | 954.668609039977 | 897.729641980856 | Timp2 |
| **ENSMUSG00000017478** | 0.171268388181911 | 0.24718557453267 | 0.833120697990749 | 944.171833814652 | 988.700694901995 | 1073.27984372734 | 912.602673513191 | 838.737977997428 | 681.25680326514 | 962.997233764426 | 995.426942801021 | Zc3h18 |
| **ENSMUSG00000017485** | 0.151516082718554 | 0.348568244782621 | 0.861266047293794 | 1331.77921822277 | 1549.83313212397 | 1525.6852805013 | 1328.10419906956 | 1170.51031133061 | 1005.37359476471 | 1496.02921612917 | 1490.42660028986 | Top2b |
| **ENSMUSG00000017491** | 0.15439047704939 | 0.569648250052938 | 0.920872458148558 | 184.316798180085 | 258.169786065842 | 265.007368821565 | 248.550573978413 | 162.053812981226 | 136.985209992272 | 306.076958623504 | 252.927456567762 | Rarb |

| | | | | | | | | | | | | |
|---|---|---|---|---|---|---|---|---|---|---|---|---|
| ENSMUSG00000017493 | -0.73800967234166 | 0.0157737093773737 | 0.63777231095375 | 1306.48083415884 | 1532.73040181677 | 1271.08891545486 | 1415.33138163557 | 2506.35931022991 | 3822.13197523081 | 1311.75839410073 | 1577.26864546334 | Igfbp4 |
| ENSMUSG00000017548 | -0.0730196483383494 | 0.695849733104181 | 0.944933386973863 | 534.880120208875 | 609.997380956832 | 603.838218957709 | 524.301022090312 | 555.143805280282 | 423.186452297555 | 678.782915042601 | 732.729756151243 | Suz12 |
| ENSMUSG00000017550 | -0.19713772524241 | 0.693925758530923 | 0.94469184542218 | 17.1667606148119 | 8.14415728914329 | 6.62518422053913 | 15.0068271081306 | 10.949581958191 | 9.78465785659088 | 21.8626399016789 | 10.8552556466851 | Atad5 |
| ENSMUSG00000017561 | -0.058786571158461 | 0.760518288889032 | 0.957357521291862 | 132.81651633565 | 100.987550385377 | 127.77140996754 | 105.047789756914 | 147.819356435578 | 124.754387671534 | 114.518589961175 | 98.7828263848342 | Crlf3 |
| ENSMUSG00000017615 | -0.254253816228788 | 0.028318968654374 | 0.665956123973759 | 1263.11217576352 | 1245.24164951001 | 1149.94268970786 | 1188.35312162509 | 1698.28016171542 | 1424.89080036605 | 1326.33348736852 | 1331.93986784826 | Tnfaip1 |
| ENSMUSG00000017631 | -0.232496536930064 | 0.143680298041638 | 0.780607990324772 | 510.48524986151 | 598.595560752032 | 586.802030962037 | 448.328959855401 | 682.158955995297 | 705.718447906617 | 626.729010514794 | 505.854913135525 | Abr |
| ENSMUSG00000017652 | -0.531256912270004 | 0.257455810813454 | 0.837877509887272 | 18.0702743313809 | 18.7315617650296 | 11.3574586637814 | 32.8274342990356 | 36.1336204620302 | 41.5847958905112 | 11.4518589961175 | 28.2236646813812 | Cd40 |
| ENSMUSG00000017664 | -0.032798537710249 | 0.771489504703546 | 0.959045638244974 | 389.414411841258 | 383.589808318649 | 441.047978110176 | 395.805064976944 | 421.558905390352 | 418.29412336926 | 437.252798033577 | 369.078691987293 | Slc35c2 |
| ENSMUSG00000017667 | -0.194246188268382 | 0.339819911650983 | 0.858570960161243 | 77.7021796249379 | 65.1532583131463 | 67.1982970940397 | 86.2892558717508 | 91.9764884488041 | 96.6234963338349 | 83.2862472444909 | 67.3025850094475 | Zfp334 |
| ENSMUSG00000017670 | -0.214852917421126 | 0.144478430650949 | 0.780607990324772 | 252.983840639333 | 306.220314071788 | 344.509579468035 | 255.11606083822 | 335.057207920644 | 351.024600605198 | 351.884394607974 | 307.203734801188 | Elmo2 |
| ENSMUSG00000017677 | -0.352997250241242 | 0.0398876119402406 | 0.679566075569698 | 140.948139784771 | 126.234437981721 | 104.110037751329 | 140.689004138724 | 175.193311331055 | 140.654456688494 | 183.22974393788 | 154.144630182928 | Wsb1 |
| ENSMUSG00000017679 | 0.00853209662836465 | 0.960953080184084 | 0.993822507448007 | 199.676531361759 | 272.8292691863 | 217.684624389143 | 193.212899017181 | 192.712642464161 | 214.039390612925 | 261.31060072959 | 210.59195954569 | Ttpal |
| ENSMUSG00000017686 | 0.166560377603852 | 0.510561113810834 | 0.902394714365738 | 1160.11161207465 | 1399.16622227482 | 1234.17717479757 | 1012.96082979881 | 863.922016501267 | 654.348994159515 | 1410.66081270357 | 1352.56485357696 | Rhot1 |
| ENSMUSG00000017692 | 0.146941523016786 | 0.532619496591406 | 0.909082872831377 | 568.310127721929 | 550.545032746086 | 538.532831640966 | 486.783954319986 | 361.336204620302 | 336.347613820311 | 679.823993133157 | 557.960140239613 | Rhbdl3 |
| ENSMUSG00000017697 | -0.228132009728574 | 0.653498523629854 | 0.936991241819063 | 12.6491920319666 | 5.7009101024003 | 14.1968233297267 | 6.56548685980713 | 12.0445401540101 | 14.6769867848863 | 12.4929370866736 | 6.51315338801105 | Ada |
| ENSMUSG00000017707 | -0.246590071388247 | 0.189672379143474 | 0.808824684419004 | 9274.56830058125 | 10015.6846341884 | 11983.065345178 | 9162.60587620798 | 14489.5818052741 | 14181.6384808964 | 10044.3214176856 | 9258.4475410577 | Serinc3 |
| ENSMUSG00000017713 | 0.278415387948439 | 0.0409722752476901 | 0.683226273624098 | 337.914129996823 | 334.724864583789 | 302.865564367503 | 369.543117537716 | 273.739548954774 | 234.831788558181 | 309.200192895173 | 289.835325766492 | Tha1 |
| ENSMUSG00000017715 | -0.092580424035751 | 0.380195014949741 | 0.87432013076003 | 486.993893230715 | 499.236841824483 | 518.657278979349 | 528.052728867345 | 502.585811880965 | 514.917619703095 | 545.524919451416 | 603.55221395569 | Pgs1 |
| ENSMUSG00000017718 | 0.0993681887958671 | 0.754445789930392 | 0.956497786093635 | 75.8951521917998 | 57.009101024003 | 53.9479286529615 | 38.4549944645846 | 38.3235368536684 | 36.6924669622158 | 70.7933101578173 | 64.046008315442 | Afmid |
| ENSMUSG00000017721 | -0.0866981483972298 | 0.499699240761695 | 0.900927059989498 | 765.276117933981 | 878.754571498561 | 873.577862222516 | 816.934150698859 | 935.094299229509 | 1024.94291047789 | 808.917676362118 | 773.979727608646 | Pigt |
| ENSMUSG00000017734 | -0.327645177923466 | 0.0223981460285515 | 0.64949947460664 | 183.413284463516 | 153.110157035894 | 203.487801059416 | 173.51643843776 | 216.801722772181 | 237.277953022329 | 233.201492284575 | 207.335382851685 | Dbndd2 |
| ENSMUSG00000017737 | 0.0659105453427368 | 0.91403398136973 | 0.986339811354287 | 21.6843291976571 | 4.88649437348597 | 29.3401015481019 | 15.9447538023887 | 29.5638712871156 | 22.0154801773295 | 8.32862472444909 | 8.68420451734806 | Mmp9 |
| ENSMUSG00000017740 | -0.157037827184427 | 0.668257347329589 | 0.942024383071524 | 23.4913566307952 | 26.0613033252585 | 21.7684624389143 | 39.3929211588428 | 31.7537876787538 | 47.7002070508805 | 24.9858741733473 | 19.5394601640331 | Slc12a5 |
| ENSMUSG00000017747 | -0.089692833400105 | 0.550860924006494 | 0.914699131434573 | 346.949267162513 | 250.840044505613 | 339.777305024792 | 329.212269684615 | 316.442918591719 | 377.932409710823 | 336.268223249632 | 316.973464883204 | Ghdc |
| ENSMUSG00000017754 | -0.791731351888943 | 0.0409899098404889 | 0.683226273624098 | 389.414411841258 | 373.816819571677 | 461.869985660442 | 456.770300103725 | 1013.93128932848 | 1190.05901180787 | 320.65205189129 | 387.532626586657 | Pltp |
| ENSMUSG00000017756 | -0.0834512457842062 | 0.680096128345088 | 0.944452327903597 | 3491.17700082279 | 4086.7381276921 | 3415.75569313224 | 2876.62117128978 | 3260.78550714927 | 2613.72672994184 | 4421.45865059191 | 4399.63511360146 | Slc12a7 |
| ENSMUSG00000017760 | -0.165688965255809 | 0.24846424391453 | 0.834621660998471 | 2527.12786524362 | 2332.48664761064 | 2359.51203740058 | 2575.54670243291 | 2819.51735423417 | 3384.26853614837 | 2503.79280778751 | 2280.68921136853 | Ctsa |
| ENSMUSG00000017764 | -0.262812274996285 | 0.224943101891799 | 0.832048466795696 | 104.807591122009 | 122.976775066064 | 105.056492639978 | 97.5443762028488 | 165.338687568684 | 106.408154190426 | 146.792010768415 | 97.6973008201657 | Zswim1 |
| ENSMUSG00000017765 | 0.114661260984759 | 0.396909195236122 | 0.879613019351548 | 1363.40219830269 | 1365.77517738933 | 1449.96888940942 | 1541.01355866616 | 1566.88517821713 | 1396.75990902835 | 1192.03441368678 | 1127.86106169058 | Slc12a4 |
| ENSMUSG00000017774 | -0.0412703425158135 | 0.710387212664476 | 0.949886733585872 | 2464.78541880035 | 2808.10543329661 | 3086.38939188259 | 2558.66402193626 | 3084.4972376224 | 2674.88084154553 | 2730.74783152875 | 2744.20862748199 | Myo1c |
| ENSMUSG00000017776 | 0.0458802147049862 | 0.775511543332143 | 0.9595799687189 | 2103.37993217274 | 2362.62002958047 | 2548.80301513027 | 2134.72115613157 | 2083.70544664374 | 1697.63813811852 | 2543.35377522864 | 2536.8732446303 | Crk |
| ENSMUSG00000017778 | 0.158780252791684 | 0.341233011576453 | 0.859283395807012 | 14505.0092057994 | 10525.5088804888 | 12288.7702742114 | 14412.1815839709 | 11938.3292090156 | 14190.2000565209 | 10020.3766216028 | 10191.9995266726 | Cox7c |
| ENSMUSG00000017781 | 0.0680168024732682 | 0.290246441455775 | 0.851785577030345 | 2249.74915425692 | 2284.43611960469 | 2198.61470633034 | 2272.59638018752 | 2118.74410890995 | 2096.3629457746 | 2206.04447388845 | 2168.88007820768 | Pitpna |
| ENSMUSG00000017801 | -0.0556718954744695 | 0.691662260134873 | 0.94469184542218 | 314.422773366028 | 226.407572638183 | 274.47191770805 | 284.191788360223 | 284.689130912965 | 292.316653465652 | 295.666177717943 | 269.21034003779 | Mlx |
| ENSMUSG00000017802 | 0.0197120269832025 | 0.839725385932117 | 0.969879194284716 | 1101.38322049767 | 1018.83407687183 | 1094.1018512776 | 1031.71936368398 | 1032.54557865741 | 952.781058785537 | 1045.24240291836 | 1156.08472637196 | Retreg3 |
| ENSMUSG00000017817 | 0.0357750235057142 | 0.863461843965086 | 0.974868041734905 | 15816.0076085411 | 17667.9348230674 | 15002.2564399665 | 15219.7364677272 | 11971.1779548902 | 11415.0264719453 | 19708.6493323182 | 19049.8881343676 | Jph2 |
| ENSMUSG00000017830 | -0.565568077706078 | 0.194525865161336 | 0.809698611654466 | 9.93865088225949 | 17.1027303072009 | 23.6613722162112 | 21.5723139679377 | 31.7537876787538 | 41.5847958905112 | 18.7394056300105 | 15.1973579053591 | Dhx58 |
| ENSMUSG00000017831 | 0.156823382776365 | 0.363030734695903 | 0.86924083181804 | 1094.15511076511 | 1311.20932355207 | 1471.73735184833 | 1105.81557253037 | 1006.26658195775 | 896.519276110139 | 1282.60820756516 | 1283.09121743818 | Rab5a |
| ENSMUSG00000017837 | 0.0295744248004354 | 0.833876629860432 | 0.96835936700622 | 1346.23543768788 | 1274.56061575092 | 1381.82413742673 | 1235.249456338 | 1133.28173267277 | 1076.312364225 | 1415.86620315635 | 1504.53843263055 | Nkiras2 |
| ENSMUSG00000017843 | 0.121364974640547 | 0.556143126725445 | 0.916239900758826 | 2318.41619671617 | 2699.788141351 | 2519.46291358217 | 2195.68639125836 | 1919.46171727088 | 1536.19128348477 | 2728.66567534763 | 2762.66256208135 | Ppp2r5c |
| ENSMUSG00000017858 | 0.136469288967062 | 0.245094595224493 | 0.832048466795696 | 540.301202508289 | 535.885549625628 | 535.693466975021 | 588.080037299867 | 486.161438943679 | 430.524945689999 | 548.648153723084 | 534.078577816906 | Ift52 |
| ENSMUSG00000017897 | -0.337265096697858 | 0.33209657107389 | 0.856655692540538 | 37.9475760958999 | 21.1748089517725 | 45.4298346551254 | 25.3240207449704 | 55.8428679867739 | 45.2540425867328 | 29.1501865355718 | 33.6512925047237 | Eya2 |
| ENSMUSG00000017929 | 0.0064320440290133 | 0.970082988671063 | 0.9951105526746 | 189.737880479499 | 233.737314198412 | 183.612248397799 | 210.095579513828 | 252.935343234211 | 190.800828203522 | 171.777884941763 | 198.651178334337 | B4galt5 |
| ENSMUSG00000017950 | -0.179221458314945 | 0.670193489468555 | 0.94249140110542 | 57.8248778604189 | 182.42912327681 | 86.1273948670087 | 81.59962240046 | 90.881530252985 | 97.8465785659088 | 71.8343882483734 | 201.907755028342 | Hnf4a |
| ENSMUSG00000017969 | -1.12415563228002 | 0.0268816014493505 | 0.664353746352133 | 168.053551281842 | 153.924572764808 | 145.754052851861 | 121.930470253561 | 390.900075907418 | 671.472145408549 | 123.88829277618 | 99.8683519495027 | Ptgis |
| ENSMUSG00000017978 | -0.146011143361992 | 0.651779353660637 | 0.936720465895231 | 21.6843291976571 | 19.5459774939439 | 26.5007368821565 | 19.6964605794214 | 27.3739548954774 | 28.1308913376988 | 24.9858741733473 | 16.2828834700276 | Cadps2 |
| ENSMUSG00000017999 | 0.178767675730293 | 0.187287824997534 | 0.807540361382744 | 412.905768472054 | 399.063707168021 | 475.12035410152 | 485.846027625728 | 443.458069306734 | 403.617136584374 | 375.829190690765 | 343.026078435248 | Ddx27 |
| ENSMUSG00000018001 | -0.277700794738674 | 0.127827299691053 | 0.772950109316303 | 502.353626412389 | 702.026358324151 | 603.838218957709 | 548.687116141024 | 823.408563255961 | 843.926740130963 | 618.400385790345 | 573.157498144972 | Cyth3 |
| ENSMUSG00000018008 | -0.33869881019851 | 0.196910664004578 | 0.812065429583558 | 37.9475760958999 | 61.0811796685746 | 81.3951204237664 | 71.2824287636203 | 73.3621991198795 | 94.1773318696872 | 79.1219348822664 | 71.6446872681215 | Cyth4 |
| ENSMUSG00000018012 | -1.15499842740837 | 0.0260135664930519 | 0.664353746352133 | 5.42108229941427 | 9.77298874697194 | 9.46454888648447 | 5.62756016554897 | 12.0445401540101 | 29.3539735697726 | 9.36970281500523 | 17.3684090346961 | Rac3 |
| ENSMUSG00000018040 | 0.044363800215328 | 0.768828108853484 | 0.95879680569409 | 857.434517024024 | 842.92027942633 | 1054.35074595437 | 1003.58156285623 | 956.993463145891 | 1072.64311752877 | 863.053737071037 | 752.269216315276 | Rrp7a |
| ENSMUSG00000018042 | -0.130021505183115 | 0.287246532426 | 0.851785577030345 | 3453.22942472689 | 3146.90237652497 | 2979.43998946531 | 3356.83963874996 | 3496.20151925038 | 4187.8335626209 | 3299.1764689724 | 3175.16227665538 | Cyb5r3 |
| ENSMUSG00000018068 | -0.142373430562504 | 0.576044662062774 | 0.922450244597442 | 42.4651446787451 | 41.5352021746308 | 42.5904699891801 | 31.8895076047775 | 43.7983278327639 | 31.8001380339204 | 51.0128264372507 | 47.7631248454143 | Ints2 |
| ENSMUSG00000018076 | 0.170431237691093 | 0.328229657117774 | 0.855977678491063 | 482.47632464787 | 645.831673029063 | 653.053873167428 | 534.618215727152 | 544.194223322091 | 388.940149799487 | 552.812466085309 | 570.986447015635 | Med13l |
| ENSMUSG00000018102 | 0.0479369315639969 | 0.686677742368697 | 0.944452327903597 | 1396.83220581574 | 1225.69567201606 | 1395.07450586781 | 1338.4213927064 | 1255.9170506045 | 1525.1835433961 | 1194.11656986789 | 1207.10442791138 | Hist1h2bc |
| ENSMUSG00000018143 | 0.264905217445256 | 0.0325728518521938 | 0.671606027118506 | 2101.5729047396 | 2373.20743405635 | 2244.04454098547 | 2328.87198184301 | 1859.23901650083 | 1560.65292812624 | 2145.6619446362 | 1962.63022092066 | Mafk |
| ENSMUSG00000018160 | 0.163314013291082 | 0.271037782513702 | 0.844129257012535 | 640.591225047453 | 673.52180781215 | 708.894711597687 | 605.900644490772 | 557.33372167192 | 463.548165955993 | 669.413212227596 | 655.657441059779 | Med1 |
| ENSMUSG00000018166 | 0.269042506084755 | 0.389054565313744 | 0.876536136761342 | 173.474633581257 | 237.809392842984 | 245.131816159948 | 208.219726125312 | 155.484063806312 | 86.838838477244 | 201.96914956789 | 272.466916731795 | Erbb3 |
| ENSMUSG00000018167 | -0.151401405266315 | 0.218870253704884 | 0.827378912571506 | 389.414411841258 | 390.105134149963 | 423.065335225856 | 384.549944645846 | 494.921104510232 | 463.548165955993 | 424.759860946904 | 379.933947633978 | Stard3 |
| ENSMUSG00000018169 | -0.327646712587014 | 0.351075925005299 | 0.86217195868522 | 112.035700854562 | 77.3694942468612 | 93.6990339761962 | 166.013024883695 | 194.902558855799 | 179.793088114857 | 71.8343882483734 | 117.236760984199 | Mfng |
| ENSMUSG00000018171 | 0.0430381494948122 | 0.669103640199754 | 0.942250071683273 | 1722.0971437806 | 1588.11067138294 | 1770.81709666124 | 1482.86210362215 | 1493.52297909725 | 1536.19128348477 | 1678.21788197649 | 1661.93963950749 | Vmp1 |
| ENSMUSG00000018189 | 0.122301582328988 | 0.209777098386811 | 0.821566820583092 | 864.662626756576 | 756.592212161411 | 783.664647800914 | 855.389145163443 | 742.381656765348 | 799.895779776304 | 710.015257759285 | 743.585011797928 | Uchl5 |
| ENSMUSG00000018196 | 0.230174642900926 | 0.0720365376651408 | 0.728760020516246 | 833.943160393229 | 746.819223414439 | 674.822335606343 | 886.340726073962 | 666.82954125383 | 721.618516923577 | 604.866370613115 | 686.052156870497 | Glrx2 |
| ENSMUSG00000018199 | 0.08429119084432 | 0.388548548421904 | 0.876405748044371 | 533.073092775737 | 560.318021493058 | 550.836745193396 | 535.55614242141 | 562.808512651016 | 516.140701935169 | 489.306702561384 | 488.486504100828 | Trove2 |
| ENSMUSG00000018209 | 0.104682071301534 | 0.723884356013029 | 0.951674101666722 | 322.554396815149 | 443.042156529395 | 466.602260103684 | 257.929840920994 | 267.16979977986 | 214.039390612925 | 437.252798033577 | 466.775992807458 | Stk4 |
| ENSMUSG00000018217 | -0.818452328213243 | 0.0399340602615843 | 0.679566075569698 | 585.476888336741 | 453.629561005281 | 407.922057007481 | 454.894446715208 | 1114.66744334384 | 1442.01395161508 | 397.691830592444 | 400.558933362679 | Pmp22 |
| ENSMUSG00000018239 | 0.0243845972195354 | 0.881941372210985 | 0.978854883720278 | 112.939214571131 | 110.760539132349 | 103.163582862681 | 93.7926694258161 | 105.115986798633 | 113.746647582869 | 92.6559500594962 | 103.124928643508 | Zcchc10 |
| ENSMUSG00000018263 | -0.348868274558228 | 0.0361622672854835 | 0.679566075569698 | 624.32797814921 | 592.080234920717 | 747.699362032273 | 657.486612674971 | 739.09678217789 | 843.926740130963 | 1058.77641809559 | 695.821886952513 | Tbx5 |
| ENSMUSG00000018286 | -0.0248646858017101 | 0.901475796463233 | 0.984802943363903 | 4211.27743292832 | 3137.94380350691 | 3382.62977202955 | 4117.49818779333 | 3892.57638613689 | 5089.24516765933 | 2998.30490080167 | 3128.48467737464 | Psmb6 |
| ENSMUSG00000018287 | 0.0231611764157188 | 0.866439303146999 | 0.974868041734905 | 2317.5126829996 | 1850.35253609335 | 1833.28311931204 | 2339.18917547985 | 2187.72647524656 | 2245.57897808761 | 1953.06249788331 | 1821.51189751376 | Spag7 |
| ENSMUSG00000018293 | -0.118609488229282 | 0.463464016315823 | 0.895328112556867 | 4735.31538853837 | 3816.35210569254 | 3446.04224956899 | 4668.99908401713 | 4974.39508360616 | 5199.32256854598 | 3918.6179328533 | 4003.41828249746 | Pfn1 |
| ENSMUSG00000018322 | 0.122159525997635 | 0.396355810867753 | 0.879359239049715 | 369.537110076739 | 358.342920722305 | 454.298346551254 | 429.570425970238 | 424.84377997781 | 381.601656407044 | 330.021754706295 | 344.111603999917 | Tomm34 |
| ENSMUSG00000018326 | -0.141011132217223 | 0.356630184145561 | 0.864965376583892 | 2039.23045829633 | 2141.09895131577 | 2134.25577390225 | 2039.05263331724 | 2555.63242904177 | 2802.08139368121 | 1914.54260853274 | 1940.91970962729 | Ywhab |
| ENSMUSG00000018334 | -0.61768623803065 | 0.0295735378307134 | 0.665956123973759 | 114.746242004269 | 105.874044758863 | 98.4313084194385 | 110.675349922463 | 241.98576127602 | 193.24699266767 | 128.052605138405 | 96.6117752554972 | Ksr1 |
| ENSMUSG00000018339 | -0.0779617637069641 | 0.834784584190841 | 0.968603949077836 | 5272.90604989695 | 5674.84879907504 | 8829.47765620136 | 9594.05215556673 | 6741.65761165818 | 14717.3484985448 | 5086.70755045728 | 4457.16796852889 | Gpx3 |
| ENSMUSG00000018340 | -0.0044786588392981 | 0.949600938816522 | 0.991267322147698 | 4704.59592217502 | 4677.18953115499 | 4710.50598080332 | 4350.10400796935 | 4587.87484048202 | 4831.17481669175 | 4445.4034466747 | 4637.36521226386 | Anxa6 |
| ENSMUSG00000018347 | -0.0995160322529267 | 0.623191657163324 | 0.931059568526633 | 77.7021796249379 | 71.6685841444609 | 100.324218196735 | 85.3513291774927 | 81.0269064906132 | 74.6080161565054 | 98.902418602833 | 103.124928643508 | Zkscan6 |
| ENSMUSG00000018362 | -0.0225663743289834 | 0.878240502461202 | 0.977934951033036 | 846.592352425195 | 892.599638890104 | 728.770264259304 | 712.824287636203 | 943.853964796062 | 671.472145408549 | 835.985706716578 | 779.407355431988 | Kpna2 |
| ENSMUSG00000018363 | 0.19741472050576 | 0.332921816242199 | 0.856918526232803 | 1098.67267934796 | 1418.71219976876 | 1347.75176143539 | 1004.51948955049 | 889.106055005107 | 809.680437632895 | 1227.43106876569 | 1319.99908663691 | Smurf2 |
| ENSMUSG00000018372 | -0.0610321178658699 | 0.776980341942469 | 0.959738279197945 | 97.5794813894569 | 112.389370590177 | 68.1447519826882 | 89.1030359545253 | 77.7420319031559 | 108.854318654574 | 102.025652874501 | 95.5262496908287 | Cep95 |
| ENSMUSG00000018377 | 0.06236808974255 | 0.681866690416275 | 0.944452327903597 | 1283.89299124461 | 1323.42555948578 | 1612.75913025695 | 1483.80003031641 | 1234.01788668812 | 1132.57414690039 | 1597.01379091311 | 1496.93975367787 | Vezf1 |
| ENSMUSG00000018378 | 0.193861160659501 | 0.0532910810813232 | 0.703396849069134 | 1594.70170974436 | 1548.20430066614 | 1648.72441602559 | 1695.77146321876 | 1288.76579647908 | 1353.95203090576 | 1585.561931917 | 1441.57794987978 | Cuedc1 |
| ENSMUSG00000018379 | 0.0465411109288503 | 0.714246730228906 | 0.951176382858794 | 1340.81435538846 | 1386.9499863411 | 1567.32929560183 | 1269.95274402555 | 1231.82797029648 | 1194.95134073616 | 1477.28981049916 | 1482.82792133718 | Srsf1 |
| ENSMUSG00000018381 | -0.496732521659614 | 0.199431404490445 | 0.812921911423739 | 74.9916384752307 | 92.0289773673191 | 85.1809399783602 | 86.2892558717508 | 185.047935093427 | 171.23151249034 | 66.6289977955927 | 55.3618037980939 | Abi3 |
| ENSMUSG00000018387 | -0.44414735248263 | 0.312888854684832 | 0.852615634676169 | 13.5527057485357 | 9.77298874697194 | 11.3574586637814 | 20.6343872736795 | 20.8042057205628 | 25.6847268735511 | 9.36970281500523 | 19.5394601640331 | Shroom1 |
| ENSMUSG00000018395 | -0.00141739813076747 | 0.99255550859894 | 0.999071668397934 | 296.352499034647 | 288.303168035672 | 282.990011705886 | 349.846656958294 | 283.594172717146 | 373.040080782527 | 284.214318721825 | 278.980070119806 | Kif3a |
| ENSMUSG00000018398 | -0.325964630106732 | 0.0101788150668361 | 0.635862253526407 | 612.582299833812 | 631.172189908605 | 610.463403178248 | 684.686486808458 | 777.420319031559 | 950.334894321389 | 715.220648212066 | 741.413960668591 | Sept8 |

| | | | | | | | | | | | | |
|---|---|---|---|---|---|---|---|---|---|---|---|---|
| **ENSMUSG00000018401** | 0.078851532363163 | 0.72194302575555 | 0.951674101666722 | 351.466835745359 | 370.55915665602 | 448.619617219364 | 318.895076047775 | 358.051330032845 | 225.04713070159 | 383.116737324658 | 442.894430384751 | Mtmr4 |
| **ENSMUSG00000018405** | -0.197468192273407 | 0.204634954503827 | 0.818186389769189 | 326.168451681425 | 260.613033252585 | 227.149173275627 | 267.309107863576 | 313.158044004262 | 267.855008824175 | 322.734208072402 | 335.427399482569 | Mrm1 |
| **ENSMUSG00000018411** | 0.13787335577716 | 0.463117327999806 | 0.895328112556867 | 3499.30862427191 | 3404.25774686189 | 2795.82774106751 | 3652.28654744128 | 2567.67696919578 | 2305.51000745923 | 3676.04673775372 | 3584.40541453541 | Mapt |
| **ENSMUSG00000018412** | -0.00433652052587437 | 0.984567986099669 | 0.99711600049115 | 444.52874855197 | 588.82257200506 | 562.194203857177 | 373.294824314748 | 459.882442244021 | 344.909189444828 | 578.839418349212 | 590.525907179668 | Kansl1 |
| **ENSMUSG00000018415** | 0.244017387321939 | 0.275393491091062 | 0.846286635233373 | 2079.88857554194 | 2374.02184978527 | 2393.58441339192 | 1994.97007868711 | 1490.23810450979 | 1264.66702796437 | 2277.87886213683 | 2432.66279042213 | Gid4 |
| **ENSMUSG00000018417** | -0.388750548039096 | 0.0121441433431722 | 0.635862253526407 | 260.211950371885 | 308.663561258531 | 254.596365046432 | 288.881421831514 | 427.033696369448 | 390.163232031561 | 299.830490080167 | 340.855027305911 | Myo1b |
| **ENSMUSG00000018425** | 0.268109213104325 | 0.0292334699197751 | 0.665956123973759 | 592.704998069294 | 654.79024604712 | 779.87882824632 | 674.369293171618 | 602.227007700503 | 522.256113095538 | 563.22324699087 | 554.703563545607 | Dhx40 |
| **ENSMUSG00000018427** | 0.008208133649177 | 0.970411043136156 | 0.9951105526746 | 284.606820719249 | 282.602257933272 | 315.169477919933 | 301.07446885687 | 239.795844884382 | 203.031650524261 | 331.062832796851 | 401.644458927348 | Ypel2 |
| **ENSMUSG00000018428** | 0.161442119266751 | 0.55300903480831 | 0.914984573375513 | 4596.17427618673 | 4996.44049688941 | 4462.53479997743 | 4153.13940217514 | 3221.36701209978 | 2257.80980040834 | 4968.02464813388 | 5832.52885896389 | Akap1 |
| **ENSMUSG00000018433** | -0.00408773457990153 | 0.984191333428868 | 0.996940244108351 | 374.958192376154 | 455.25839246311 | 388.046504345863 | 336.71568323868 | 384.330326732503 | 259.293433199658 | 461.197594116369 | 453.749686031436 | Nol11 |
| **ENSMUSG00000018442** | 0.0886457832447951 | 0.444681053671582 | 0.89108031640712 | 934.233182932393 | 825.817549119129 | 865.05976822468 | 957.623154837583 | 782.895110010654 | 962.565716642128 | 849.519721893808 | 775.065253173314 | Derl2 |
| **ENSMUSG00000018446** | 0.167804607301835 | 0.0996895904623397 | 0.759800953500741 | 3505.63322028789 | 3162.37627537434 | 3057.04929033448 | 3791.09969819149 | 3144.71993839245 | 3030.79777107902 | 2927.51159064386 | 2928.74797347563 | C1qbp |
| **ENSMUSG00000018449** | -0.0250587131020304 | 0.902663464478506 | 0.985027469232262 | 163.535982698997 | 139.26508964435 | 121.146225747001 | 185.709485463116 | 187.237851485066 | 165.116101329971 | 122.847214685624 | 145.46042566558 | Rpain |
| **ENSMUSG00000018474** | 0.0359664156115986 | 0.786448797197722 | 0.961141387070613 | 595.415539219001 | 634.429852824262 | 717.412805595523 | 512.107975064956 | 573.758094609207 | 585.856389163379 | 598.619902069779 | 640.46008315442 | Chd3 |
| **ENSMUSG00000018476** | 0.0879067990906084 | 0.576127543079705 | 0.922450244597442 | 658.661499378834 | 732.159740293981 | 721.198625150116 | 656.548685980713 | 833.263187018333 | 547.940839969089 | 641.30410378258 | 581.84170266232 | Kdm6b |
| **ENSMUSG00000018481** | -0.0145060554678498 | 0.935672389353051 | 0.98818087308472 | 486.090379514146 | 548.916201288258 | 645.482234058241 | 496.163221262567 | 546.384139713729 | 421.963370065482 | 687.11153976705 | 541.677256769585 | Appbp2 |
| **ENSMUSG00000018486** | -1.1988169518656 | 0.0558046861205686 | 0.703396849069134 | 16.2632468982428 | 8.95857301805762 | 3.78581955459379 | 7.50341355406529 | 27.3739548954774 | 30.5770558018465 | 21.8626399016789 | 4.34210225867403 | Wnt9b |
| **ENSMUSG00000018501** | 0.221572614336004 | 0.243319350891533 | 0.832048466795696 | 3155.97341197567 | 3754.45651029506 | 3708.21025372461 | 2845.66959037926 | 2470.22568976788 | 2192.98644210843 | 3434.51662074469 | 3448.71471895185 | Ncor1 |
| **ENSMUSG00000018507** | -0.233325463081636 | 0.418575992359838 | 0.882468496126562 | 90.3513716569045 | 69.2253369577179 | 118.306861081056 | 84.4134024832345 | 135.774816281568 | 135.762127760198 | 65.5879197050366 | 89.0130963028176 | Trpv2 |
| **ENSMUSG00000018509** | 0.0739403467073224 | 0.64124002681753 | 0.934544499347759 | 233.106538874814 | 201.975100770753 | 229.988537941573 | 246.674720589896 | 247.460552255116 | 226.270212933664 | 221.749633288457 | 170.427513652956 | Cenpv |
| **ENSMUSG00000018537** | -0.436691093343461 | 0.21688471041855 | 0.825692108906787 | 39.754603529038 | 34.2054606144018 | 58.6802030962037 | 39.3929211588428 | 82.1218646864323 | 74.6080161565054 | 45.80743598447 | 30.3947158107182 | Pcgf2 |
| **ENSMUSG00000018541** | -0.19568457701808 | 0.152203089333621 | 0.783412422888014 | 184.316798180085 | 184.057954734638 | 194.96970706158 | 176.330218520534 | 213.516848184724 | 239.724117486476 | 211.338852382896 | 183.453820428978 | Cwc25 |
| **ENSMUSG00000018547** | 0.0634681406143745 | 0.780865816909992 | 0.960742105865155 | 620.713923282934 | 767.179616637298 | 730.663174036601 | 538.369922504185 | 500.395895489327 | 458.655837027697 | 758.945928015424 | 823.913903583397 | Pip4k2b |
| **ENSMUSG00000018548** | 0.199641514527622 | 0.290196665458739 | 0.851785577030345 | 234.010052591383 | 329.838370210303 | 357.759947909113 | 251.364354061187 | 231.036179317829 | 238.501035254403 | 303.994802442392 | 247.49982874442 | Trim37 |
| **ENSMUSG00000018554** | -0.114609410030351 | 0.506112048208506 | 0.901494649536949 | 271.957628687283 | 218.26341534904 | 205.380710836713 | 246.674720589896 | 244.175677667659 | 320.447544803351 | 234.242570375131 | 222.532740757044 | Ybx2 |
| **ENSMUSG00000018559** | -0.00780715605331665 | 0.926772296469479 | 0.986790505471895 | 2112.41506933843 | 1924.46436742456 | 1817.19338620502 | 1986.52873843879 | 1940.26592299144 | 2069.45513666897 | 1921.83015516663 | 1952.86049083865 | Ctdnep1 |
| **ENSMUSG00000018565** | 0.0111638091714098 | 0.916647705104757 | 0.986713918026868 | 691.187993175319 | 636.873100011005 | 689.019158936069 | 732.520748215624 | 681.063997799478 | 715.503105763208 | 725.631429117627 | 605.723265085027 | Elp5 |
| **ENSMUSG00000018566** | -0.175482472401534 | 0.136752965675872 | 0.776456211338551 | 6593.84310352089 | 6403.75087645337 | 5536.76109859341 | 5421.21629281217 | 7339.50478657541 | 6418.73555392362 | 7205.30146473902 | 6090.883943355 | Slc2a4 |
| **ENSMUSG00000018567** | -0.00284455530874122 | 0.982124106921653 | 0.996940244108351 | 5642.44315997369 | 4761.07435123317 | 5189.41215445943 | 5757.93197605085 | 5641.22462485999 | 6139.87280501078 | 4768.13765474711 | 4844.70059511555 | Gabarap |
| **ENSMUSG00000018572** | -0.240550976397616 | 0.215884846212941 | 0.825162496944895 | 458.984968017075 | 410.465527372822 | 499.72818120638 | 467.087493740564 | 623.031213421066 | 689.818378889657 | 408.102611498006 | 449.407583772762 | Phf23 |
| **ENSMUSG00000018574** | 0.162132451459322 | 0.0910680694677167 | 0.753823363185192 | 30870.3531540146 | 31151.4016309731 | 35277.2130645936 | 33524.3138328695 | 29188.3006259497 | 26359.8682656558 | 29885.1876675045 | 31481.3269009514 | Acadvl |
| **ENSMUSG00000018583** | 0.0395579465919657 | 0.639377366007862 | 0.933801135937988 | 2428.64487013759 | 2591.47084940539 | 2481.60471803623 | 2493.94708003245 | 2492.12485368426 | 2195.43260657258 | 2561.0521027681 | 2474.9982874442 | G3bp1 |
| **ENSMUSG00000018585** | -0.0897075514397538 | 0.729243638009887 | 0.952797867741259 | 609.871758684105 | 487.020605890769 | 507.299820315568 | 625.597105070194 | 648.215251924905 | 889.180782717696 | 416.431236222455 | 420.098393526712 | Atox1 |
| **ENSMUSG00000018593** | -0.669490715458912 | 0.0488081893441843 | 0.703396849069134 | 5255.73928928214 | 4580.27405941418 | 3906.96578034079 | 5616.30504521787 | 9742.93802639832 | 12312.7688302875 | 3862.39971596327 | 4874.0097853616 | Sparc |
| **ENSMUSG00000018599** | 0.0618753648449927 | 0.577874964127577 | 0.922450244597442 | 756.240980768291 | 752.52013351684 | 775.146553803078 | 749.403428712271 | 735.811907590433 | 612.764198269004 | 768.315630830429 | 787.006034384668 | Mief2 |
| **ENSMUSG00000018604** | -0.567693150477228 | 0.0823833669650807 | 0.739855948845396 | 282.799793286111 | 241.067055758641 | 360.599312575058 | 348.908730264036 | 628.506004400162 | 654.348994159515 | 291.501865355718 | 254.012982132431 | Tbx3 |
| **ENSMUSG00000018651** | -0.102584940125994 | 0.567708580492276 | 0.920429362097163 | 123.781379169959 | 115.647033505835 | 115.467496415111 | 121.930470253561 | 124.825234323377 | 105.185071958352 | 162.408182126757 | 118.322286548867 | Tada2a |
| **ENSMUSG00000018654** | -0.00211597764603344 | 0.99556215083139 | 0.999262355603273 | 13.5527057485357 | 13.8450673915436 | 27.447191770805 | 24.3860940507122 | 19.7092475247437 | 24.4616446414772 | 18.7394056300105 | 16.2828834700276 | Ikzf1 |
| **ENSMUSG00000018659** | 0.168328152342117 | 0.126218278273251 | 0.771949105062466 | 533.073092775737 | 505.752167655798 | 600.052399403115 | 513.983828453472 | 464.262275027297 | 468.440494884288 | 461.197594116369 | 521.052271040884 | Pnpo |
| **ENSMUSG00000018661** | -0.0510356460986187 | 0.630921696525103 | 0.931825800800958 | 466.213077749627 | 517.153987860599 | 501.621090983677 | 527.114802173087 | 573.758094609207 | 502.686797382356 | 522.621201459181 | 485.229927406823 | Cog1 |
| **ENSMUSG00000018666** | -0.0312729833596532 | 0.759354202601925 | 0.957357521291862 | 800.513152880174 | 747.633639143354 | 717.412805595523 | 780.35500962279 | 741.286698569529 | 766.87255951031 | 735.001131932632 | 869.505977299475 | Cbx1 |
| **ENSMUSG00000018669** | -0.00877290924230321 | 0.961463879309423 | 0.993822507448007 | 637.880683897746 | 580.678414715916 | 638.857049837702 | 758.782695654852 | 689.823663366031 | 850.042151291332 | 554.894622266421 | 538.42068007558 | Cdk5rap3 |
| **ENSMUSG00000018672** | -0.129021722788054 | 0.45773370129665 | 0.892966448175628 | 1056.20753466921 | 842.92027942633 | 809.218929794422 | 1106.75349922463 | 1001.88674917447 | 1311.14415278318 | 955.709687130533 | 904.242795368867 | Copz2 |
| **ENSMUSG00000018677** | 0.0844258956086418 | 0.497121689512685 | 0.900927059989498 | 4354.03260014623 | 4135.60307142696 | 4875.18913142815 | 4739.34358608649 | 4432.3907766757 | 4952.25995766706 | 3800.97610862045 | 3890.52362377193 | Slc25a39 |
| **ENSMUSG00000018678** | -0.175357360545705 | 0.214046683550264 | 0.824248286808591 | 175.281661014395 | 193.016527752696 | 232.827902607518 | 192.274972322923 | 228.846262926191 | 211.593226148778 | 237.365804646799 | 217.105112933702 | Sp2 |
| **ENSMUSG00000018697** | 0.326920187762235 | 0.0150903734890371 | 0.635862253526407 | 291.834930451802 | 289.117583764587 | 286.775831260479 | 288.881421831514 | 274.834507150593 | 204.254732756335 | 213.421008564008 | 229.045894145055 | Aatf |
| **ENSMUSG00000018707** | 0.236714396517099 | 0.0359173009875509 | 0.679566075569698 | 8536.39759414434 | 10398.4600267781 | 9440.88751426826 | 9114.77161480081 | 9089.24798349432 | 7395.97825735063 | 7774.77118027323 | 7557.42898122215 | Dync1h1 |
| **ENSMUSG00000018733** | 0.345185891154614 | 0.00372588371203083 | 0.635862253526407 | 332.493047697409 | 324.951875836817 | 351.134763688574 | 332.026049767389 | 267.16979977986 | 228.716377397812 | 286.296474902938 | 271.381391167127 | Pex12 |
| **ENSMUSG00000018736** | 0.163137147836578 | 0.0825041529171262 | 0.739855948845396 | 1527.84169471826 | 1563.67819951551 | 1488.77353984401 | 1427.52442866092 | 1215.4035973592 | 1312.36723501525 | 1452.30393632581 | 1385.13062051702 | Ndel1 |
| **ENSMUSG00000018740** | -0.12036601603639 | 0.570255790435962 | 0.920872458148558 | 113.8427282877 | 101.801966114291 | 72.8770264259304 | 91.9168160372998 | 98.5462376237187 | 85.6157562451702 | 127.011527047849 | 102.03940307884 | Slc25a35 |
| **ENSMUSG00000018750** | -0.0731908509191038 | 0.493664207876029 | 0.900117970535229 | 1138.427282877 | 1239.54073940761 | 1353.43049076728 | 1145.20849368921 | 1270.15150715015 | 1243.87463001912 | 1413.78404697523 | 1201.67680008804 | Zbtb4 |
| **ENSMUSG00000018752** | -0.420674690961071 | 0.185521481965128 | 0.805963874499992 | 54.2108229941427 | 33.3910448854875 | 34.0723759913441 | 59.0893817382642 | 70.0773245324222 | 81.9465095489486 | 47.8895921655823 | 42.3354970220718 | Tnfsfm13 |
| **ENSMUSG00000018761** | -0.123219399158797 | 0.508838683970306 | 0.902394714365738 | 162.632468982428 | 158.811067138294 | 194.023252172932 | 239.171307035831 | 208.042057205628 | 200.585486060113 | 175.942197303987 | 236.644573097735 | Mpdu1 |
| **ENSMUSG00000018765** | 0.233359855571696 | 0.0334678954359453 | 0.671782526713461 | 4661.2272637797 | 4861.24748588963 | 5159.12559802268 | 4960.69428593141 | 3975.79320901914 | 3670.46977845365 | 4770.21981092822 | 4289.99703156994 | Fxr2 |
| **ENSMUSG00000018770** | -0.00772625007987374 | 0.958854327795057 | 0.993101678702062 | 41659.2104435655 | 33452.1260651561 | 34931.7570302369 | 41849.3511711049 | 40982.0953531172 | 45055.9032651368 | 32996.9700801768 | 33674.0885415818 | Atp5g3 |
| **ENSMUSG00000018774** | -0.559660611116595 | 0.127313839561933 | 0.772837268577874 | 43.3686583953142 | 45.6072808192024 | 54.8943835416099 | 48.7721881014244 | 85.4067392738896 | 117.415894279091 | 40.6020455316893 | 41.2499714574033 | Cd68 |
| **ENSMUSG00000018796** | 0.203639042459098 | 0.38266782935793 | 0.875126835008975 | 18729.8393444763 | 23867.2673515633 | 25833.4861856594 | 18766.9752254115 | 17543.4202134136 | 11822.3128552259 | 23547.1042521987 | 22804.721062556 | Acsl1 |
| **ENSMUSG00000018800** | 0.213215734525829 | 0.452009918848468 | 0.891110666112841 | 134.623543768788 | 226.407572638183 | 217.684624389143 | 160.385464718146 | 132.489941694111 | 101.51582526213 | 225.913945650682 | 176.940667040967 | Abca5 |
| **ENSMUSG00000018809** | -0.0648982345456056 | 0.637313979248334 | 0.932931968211785 | 149.079763233892 | 149.038078391322 | 150.486327295103 | 150.068271081306 | 139.059690869025 | 154.108361241306 | 165.531416398426 | 167.17093695895 | Smyd4 |
| **ENSMUSG00000018819** | -0.398813704373144 | 0.0172683776206934 | 0.642273034804458 | 258.404922938747 | 239.438224300813 | 313.276568142636 | 278.564228194674 | 340.531998899739 | 450.09426140318 | 341.473613702413 | 305.032683671851 | Lsp1 |
| **ENSMUSG00000018820** | 0.0891319167541913 | 0.569835758134911 | 0.920872458148558 | 381.282788392137 | 459.330471107681 | 562.194203857177 | 403.308478531009 | 421.558905390352 | 384.047820871192 | 452.868969391919 | 438.552328126077 | Zfyve27 |
| **ENSMUSG00000018821** | -0.0168710830398287 | 0.93750203035319 | 0.988970340396118 | 281.896279569542 | 218.26341534904 | 238.506631939409 | 325.460562907582 | 312.063085808443 | 322.893709267499 | 250.899819824029 | 191.052499381657 | Avpi1 |
| **ENSMUSG00000018822** | -2.02606290151533 | NA | NA | 22.5878429142261 | 30.1333819698302 | 21.7684624389143 | 33.7653609932938 | 30.6588294829347 | 315.555215875056 | 52.0539045278068 | 43.4210225867403 | Sfrp5 |
| **ENSMUSG00000018830** | -0.495775260678741 | 0.186680544843413 | 0.807540361382744 | 750.819898468876 | 1245.24164951001 | 708.894711597687 | 542.121629281217 | 1653.38687568684 | 1607.13005294505 | 752.699459472087 | 566.644344756961 | Myh11 |
| **ENSMUSG00000018841** | 0.153822626285652 | 0.425173224991124 | 0.885066977635295 | 156.307872966445 | 170.212887343095 | 180.772883731853 | 158.509611329629 | 122.635317931739 | 196.916239363891 | 122.847214685624 | 157.401206876934 | Rad51d |
| **ENSMUSG00000018845** | 0.109602685564778 | 0.329670161200194 | 0.855977678491063 | 5934.27809042549 | 6086.12874217678 | 5780.94645986471 | 6676.16220972959 | 5621.51537733524 | 5015.8602337349 | 5545.82298839254 | 6502.29813236436 | Unc45b |
| **ENSMUSG00000018846** | -0.00205410476615993 | 0.993666629956639 | 0.999133450854041 | 524.941469326615 | 697.139863950665 | 692.804978490663 | 470.839200517597 | 478.496731572945 | 384.047820871192 | 746.45299092875 | 779.407355431988 | Pank3 |
| **ENSMUSG00000018848** | -0.0252441406595205 | 0.836144385778203 | 0.968924788429433 | 889.05749710394 | 853.507683902216 | 816.79056890361 | 898.533773099319 | 908.81530252985 | 996.812019140196 | 880.752064610492 | 733.815281715911 | Rars |
| **ENSMUSG00000018849** | -0.319387930398032 | 0.567334080929785 | 0.9203643243484 | 6.32459601598331 | 17.9171460361152 | 7.57163910918757 | 12.1930470253561 | 5.47479097909548 | 12.2308223207386 | 13.5340151772298 | 23.8815624227072 | Wwc1 |
| **ENSMUSG00000018858** | 0.0507177412928349 | 0.70453321444536 | 0.947151214541817 | 1759.14120615993 | 1319.35348084121 | 1416.84296830672 | 1655.44061536565 | 1574.54988558786 | 1610.79929964127 | 1365.89445480965 | 1387.30167164635 | Mrpl58 |
| **ENSMUSG00000018861** | -0.031615774240007 | 0.807962407273248 | 0.964026716319739 | 211.422209677157 | 214.191336704468 | 216.738169500494 | 250.426427366929 | 246.365594059297 | 229.939459629886 | 208.215618111227 | 227.960368580387 | Fdxr |
| **ENSMUSG00000018882** | -0.0958178069009234 | 0.417804071244171 | 0.882468496126562 | 1400.44626068202 | 1147.51176204029 | 1057.19011062032 | 1254.00799022316 | 1255.9170506045 | 1382.08292224346 | 1294.06006656128 | 1261.38070614481 | Mrpl45 |
| **ENSMUSG00000018893** | -0.098296253316358 | 0.519438775081277 | 0.905962179970576 | 464516.278989911 | 354793.696975696 | 347130.313054702 | 437133.866832736 | 425709.891910703 | 512371.162495917 | 387683.946907928 | 390876.045325836 | Mb |
| **ENSMUSG00000018899** | -0.535054326523679 | 0.130047253925142 | 0.77303252008843 | 328.878992831132 | 221.521078264697 | 271.632553042104 | 316.081295965 | 605.511882287961 | 588.302553627526 | 207.174540020671 | 248.585354309088 | Irf1 |
| **ENSMUSG00000018900** | 0.28848656297445 | 0.169942263709679 | 0.79907358295209 | 589.090943203017 | 698.768695408494 | 772.307189137133 | 590.893817382642 | 455.502609460744 | 388.940149799487 | 671.495368408708 | 653.486389930442 | Slc22a5 |
| **ENSMUSG00000018906** | -0.162098027502846 | 0.211383267988659 | 0.823363318105013 | 1532.3592633011 | 1269.67412137744 | 1193.47961458569 | 1425.64857527241 | 1534.03643234255 | 1720.87670052792 | 1357.5658300852 | 1454.6042566558 | P4ha2 |
| **ENSMUSG00000018909** | -0.114498376762287 | 0.609576017640262 | 0.927932112008768 | 349.65980831222 | 494.350347450997 | 545.158015861505 | 353.598363735327 | 535.434557755538 | 588.302553627526 | 385.198893505771 | 378.848422069309 | Arrb1 |
| **ENSMUSG00000018920** | -0.36057996549165 | 0.270017967407316 | 0.843793744782845 | 115.649755720838 | 99.3587189275481 | 92.7525790875478 | 118.178763476528 | 190.522726072523 | 190.800828203522 | 71.8343882483734 | 94.4407241261602 | Cxcl16 |
| **ENSMUSG00000018921** | 0.0876215902805142 | 0.377680498983148 | 0.87422797717331 | 746.302329886031 | 688.995706661522 | 750.538726698218 | 665.927952923294 | 711.722827282413 | 648.233582999146 | 695.440164491499 | 627.433776378397 | Pelp1 |
| **ENSMUSG00000018923** | -0.192348526383646 | 0.387926622163225 | 0.875951433019341 | 80.412720774645 | 65.9676740420606 | 63.4124775394459 | 91.9168160372998 | 89.7865720571659 | 107.6312364225 | 69.7522320672611 | 78.1578406561325 | Med11 |
| **ENSMUSG00000018927** | -0.240608643439078 | 0.405929102440156 | 0.881158207349486 | 324.361424248287 | 292.375246680244 | 463.762895437739 | 376.108604397523 | 506.965644664242 | 624.995020589742 | 250.899819824029 | 338.683976176574 | Ccl6 |
| **ENSMUSG00000018931** | -0.100558272001886 | 0.543859542869206 | 0.913837378403038 | 160.82544154929 | 156.367819951551 | 194.96970706158 | 161.323391412404 | 199.282391639076 | 141.877538920568 | 192.599446752885 | 186.710397122983 | Natd1 |
| **ENSMUSG00000018932** | 0.126538758043152 | 0.240235250233432 | 0.832048466795696 | 1664.27226592018 | 1717.60277228032 | 1943.07188639526 | 1722.97133735224 | 1702.6599944987 | 1410.21381358116 | 1709.45022469318 | 1631.54492369677 | Map2k3 |
| **ENSMUSG00000018965** | -0.173255579673851 | 0.0972768235479834 | 0.757809915356977 | 1890.15069506244 | 1781.12719913564 | 1643.99214158235 | 1694.8335365245 | 2108.88948514758 | 2137.94774166511 | 1792.73647193767 | 1867.10397122983 | Ywhah |
| **ENSMUSG00000018974** | 0.010871021242019 | 0.920313425502173 | 0.986713918026868 | 377.668733525861 | 375.445651029505 | 395.618143455051 | 381.736164563072 | 388.710159515779 | 349.801518373124 | 418.513392403567 | 360.394487469945 | Sart3 |
| **ENSMUSG00000018983** | 0.498579511844318 | 0.601005122611891 | 0.925969784079517 | 12.6491920319666 | 11.4018202048006 | 74.7699362032273 | 1.87585338851632 | 8.75966556655277 | 48.9232892829544 | 7.28754663389296 | 6.51315338801105 | E2f2 |
| **ENSMUSG00000018986** | -0.309298669300913 | 0.293346659933291 | 0.851785577030345 | 61.4389327266951 | 55.3802695661743 | 55.8408384302584 | 45.0204813243917 | 106.210944994452 | 57.4848649074714 | 59.3414511616998 | 46.6775992807458 | Slfn3 |

| | | | | | | | | | | | | |
|---|---|---|---|---|---|---|---|---|---|---|---|---|
| ENSMUSG00000018995 | 0.156780695120518 | 0.57735871086264 | 0.922450244597442 | 294.545471601509 | 320.065381463331 | 350.188308799925 | 254.178134143962 | 235.416012101106 | 144.323703384715 | 367.500565966316 | 345.197129564585 | Nars2 |
| ENSMUSG00000018999 | -0.317930074021183 | 0.0400833754016036 | 0.679566075569698 | 112.939214571131 | 114.018202048006 | 113.574586637814 | 135.999370667433 | 159.863896589588 | 159.000690169602 | 134.299073681742 | 141.118323406906 | Slc35b4 |
| ENSMUSG00000019039 | 0.169733550146106 | 0.203354045701973 | 0.81605904225138 | 426.458474220589 | 413.723190288479 | 443.887342776122 | 507.418341593665 | 366.810995599397 | 399.947889888152 | 460.156516025812 | 364.736589728619 | Dalrd3 |
| ENSMUSG00000019054 | -0.125372555138157 | 0.528694806733876 | 0.908295716206993 | 2180.17859808111 | 1684.21172739483 | 1778.38873577043 | 2323.24442167747 | 2186.63151705074 | 2910.93571233579 | 1737.55933313819 | 1855.16319001848 | Fis1 |
| ENSMUSG00000019055 | -0.384916088949844 | 0.0604115496802326 | 0.71282992338528 | 1748.2990415611 | 1812.8894125633 | 1784.06746510232 | 1710.77829032689 | 2819.51735423417 | 2930.50502804897 | 1729.23070841374 | 1735.75537790494 | Plod1 |
| ENSMUSG00000019066 | -0.254816539444939 | 0.0807157715936933 | 0.739855948845396 | 469.827132615903 | 545.6585383726 | 548.943835416099 | 500.852854733858 | 717.197618261509 | 666.579816480253 | 578.839418349212 | 502.598336441519 | Rab3d |
| ENSMUSG00000019080 | -0.186224436827435 | 0.468896070071417 | 0.895328112556867 | 137.334084918495 | 107.502876216691 | 79.5022106464695 | 101.296082979881 | 130.300025302473 | 159.000690169602 | 114.518589961175 | 81.4144173501381 | Mfsd3 |
| ENSMUSG00000019082 | 0.126350572798801 | 0.575936617960654 | 0.922450244597442 | 636.073656464608 | 726.458830191581 | 902.917963770618 | 810.368663839051 | 690.91862156185 | 441.532685778663 | 883.87529888216 | 800.03234116069 | Slc25a22 |
| ENSMUSG00000019087 | -0.159424923600709 | 0.108251483895076 | 0.762287160109383 | 1958.81773752169 | 2086.53309747851 | 1993.23399549363 | 1909.61874950962 | 2351.97020461942 | 2382.56418807988 | 2194.59261489234 | 1949.60391414464 | Atp6ap1 |
| ENSMUSG00000019088 | 0.219490881197709 | 0.138703876563263 | 0.779711473022872 | 297.256012751216 | 269.571606270643 | 313.276568142636 | 381.736164563072 | 293.448796479518 | 264.185762127954 | 257.146288367366 | 268.124814473121 | Dnase1l1 |
| ENSMUSG00000019122 | -0.274154803244821 | 0.253728256675131 | 0.837406774112203 | 234.913566307952 | 220.706662535783 | 282.990011705886 | 293.571055302804 | 335.057207920644 | 432.971110154146 | 201.96914956789 | 278.980070119806 | Ccl9 |
| ENSMUSG00000019124 | -0.330839608906453 | 0.243795400415693 | 0.832048466795696 | 88.5443442237664 | 118.090280692578 | 135.343049076728 | 129.433883807626 | 140.154649064844 | 233.608706326107 | 106.189965236726 | 113.980184290193 | Scrn1 |
| ENSMUSG00000019132 | 0.219633259717538 | 0.146490709780837 | 0.780607990324772 | 4286.26907140355 | 4837.62942975111 | 5403.31095929398 | 4663.37152385158 | 3846.58814191249 | 3291.31428651076 | 4729.61776539653 | 4611.31259871182 | BC005537 |
| ENSMUSG00000019139 | 0.121959417664376 | 0.691449429519648 | 0.94469184542218 | 351.466835745359 | 394.991628523449 | 728.770264259304 | 458.646153492241 | 512.440435643337 | 629.887349518038 | 352.92547269853 | 282.236646813812 | Isyna1 |
| ENSMUSG00000019143 | 0.14929223927789 | 0.456563164054079 | 0.892805623137304 | 410.195227322346 | 438.155662155909 | 376.689045682082 | 418.31530563914 | 348.196706270473 | 245.839528646846 | 439.33495421469 | 447.236532643425 | Hars2 |
| ENSMUSG00000019158 | -0.316122420886556 | 0.241336887740081 | 0.832048466795696 | 1735.64984952914 | 1254.20022252807 | 1166.97887770353 | 1754.86084495702 | 1869.0936402632 | 2811.8660515378 | 1300.30653510461 | 1379.70299269367 | Tmem160 |
| ENSMUSG00000019173 | -0.318290211410029 | 0.11748737453671 | 0.769868040577702 | 826.715050660676 | 831.518459221529 | 891.560505106837 | 950.119741283517 | 1253.72713421287 | 1429.78312929434 | 787.055036460439 | 894.47306528685 | Rab5c |
| ENSMUSG00000019179 | 0.00467827078739334 | 0.966045911507612 | 0.994284908685797 | 40793.6443030924 | 35498.7527919178 | 36717.7174051165 | 42127.9153992996 | 41556.9484059222 | 41658.1808244357 | 35057.2636213873 | 36364.0208908303 | Mdh2 |
| ENSMUSG00000019188 | -0.0917013657481564 | 0.659071534317298 | 0.938989601151841 | 675.828259993646 | 644.202841571234 | 807.326020017125 | 708.134654164912 | 838.737977997428 | 1004.15051253264 | 644.427338054249 | 535.164103381574 | H13 |
| ENSMUSG00000019189 | -0.375839335283775 | 0.0650630013521946 | 0.718339708336358 | 185.220311896654 | 184.057954734638 | 163.736695736181 | 181.957778686083 | 274.834507150593 | 291.093571233579 | 190.517290571773 | 172.598564782293 | Rnf145 |
| ENSMUSG00000019194 | -0.206389777713078 | 0.374194490640516 | 0.873700364743909 | 454.46739943423 | 416.166437475222 | 343.563124579386 | 523.363095396054 | 416.084114411257 | 735.07242147639 | 417.472314313011 | 437.466802561409 | Scn1b |
| ENSMUSG00000019210 | 0.0754917970713582 | 0.272119968325043 | 0.844447354788841 | 2697.89195767517 | 2571.11045618254 | 2624.51940622214 | 2622.44303714582 | 2487.74502090099 | 2375.22569468744 | 2601.65414829979 | 2512.9916822076 | Atp6v1e1 |
| ENSMUSG00000019214 | 0.0282036039222333 | 0.93799943086656 | 0.989054284880932 | 28.9124389302094 | 19.5459774939439 | 27.447191770805 | 18.7585338851632 | 26.2789966996583 | 33.0232202659942 | 19.7804837205666 | 14.1118323406906 | Chtf18 |
| ENSMUSG00000019232 | -0.0207372042143457 | 0.951736815064016 | 0.991915011884503 | 18.9737880479499 | 21.9892246806869 | 22.7149173275627 | 21.5723139679377 | 12.0445401540101 | 25.6847268735511 | 24.9858741733473 | 23.8815624227072 | Etnppl |
| ENSMUSG00000019235 | -0.415694785556717 | 0.107985309691752 | 0.762287160109383 | 43.3686583953142 | 29.3189662409158 | 42.5904699891801 | 37.5170677703264 | 62.4126171616885 | 55.0387004433237 | 40.6020455316893 | 45.5920737160773 | Rps6kl1 |
| ENSMUSG00000019254 | 0.144215400024767 | 0.473783835206156 | 0.897549741595057 | 1702.21984201608 | 1828.36331141267 | 2591.39348511945 | 2223.8241920861 | 2119.83906710577 | 2223.56349791028 | 1835.42067365047 | 1373.18983930566 | Ppp1r12c |
| ENSMUSG00000019256 | -0.212325136187388 | 0.375119329915463 | 0.873700364743909 | 125.588406603097 | 135.193010999779 | 188.344522841041 | 190.399118934407 | 174.098353135236 | 250.731857575141 | 136.381229862854 | 180.197243734972 | Ahr |
| ENSMUSG00000019261 | -0.27395184474495 | 0.0936278591956553 | 0.755231402633518 | 265.633032671299 | 261.427448981499 | 245.131816159948 | 259.805694309511 | 363.52612101194 | 362.032340693862 | 272.762459725708 | 250.756405438425 | Map1s |
| ENSMUSG00000019278 | -0.747358308352849 | 0.0642174837104451 | 0.716991331547963 | 726.425028121512 | 507.380999113627 | 784.611102689562 | 763.472329126143 | 1734.41378217745 | 1843.18492373531 | 568.428637443651 | 524.308847734889 | Dpep1 |
| ENSMUSG00000019295 | -0.0746005378884311 | 0.461854507810167 | 0.895328112556867 | 393.931980424104 | 380.332145402991 | 389.93941412316 | 391.115431505653 | 414.989156215438 | 428.078781225851 | 422.677704765791 | 372.335268681298 | Tmem129 |
| ENSMUSG00000019297 | -0.0696483051934417 | 0.659291093268204 | 0.939139912367485 | 318.036828232304 | 351.013179162076 | 335.991485470199 | 365.791410760683 | 412.7992398238 | 418.29412336926 | 309.200192895173 | 299.605055848508 | Nop9 |
| ENSMUSG00000019302 | 0.0951177922889513 | 0.489082672470568 | 0.899236955296196 | 2001.28288220043 | 2330.85781615281 | 2360.45849228923 | 2129.09359596603 | 1964.35500329946 | 1691.52272695815 | 2368.45265601521 | 2232.92608652312 | Atp6v0a1 |
| ENSMUSG00000019303 | 0.289116893239787 | 0.16620101855046 | 0.79616206680466 | 60.535419010126 | 69.2253369577179 | 59.6266579848522 | 60.0273084325223 | 47.0832024202212 | 47.7002070508805 | 60.3825292522559 | 48.8486504100828 | Psmc3ip |
| ENSMUSG00000019312 | 0.059793208015438 | 0.894745389650108 | 0.983733637008786 | 6.32459601598331 | 12.2162359337149 | 13.2503684410783 | 12.1930470253561 | 9.85462376237187 | 12.2308223207386 | 9.36970281500523 | 10.8552556466851 | Grb7 |
| ENSMUSG00000019326 | -1.14708354407482 | 0.0140171809868576 | 0.635862253526407 | 215.939778260002 | 185.686786192467 | 119.253315969704 | 145.378637610015 | 555.143805280282 | 624.995020589742 | 154.079557402308 | 142.203848971575 | Aoc3 |
| ENSMUSG00000019338 | 0.286519524142966 | 0.116749671249465 | 0.769868040577702 | 241.238162323935 | 271.200437728471 | 281.097101928589 | 230.729966787508 | 206.947099009809 | 155.33144347338 | 262.351678820146 | 213.848536239696 | Zfp687 |
| ENSMUSG00000019362 | -0.098093742960368 | 0.633904689861337 | 0.9325328237529 | 2019.35315653182 | 1618.24405335277 | 1736.7447206699 | 2281.97564713011 | 2005.96341474059 | 2824.09687385854 | 1709.45022469318 | 1656.51201168414 | D8Ertd738e |
| ENSMUSG00000019368 | -0.467689955052163 | 0.362920762909756 | 0.86924083181804 | 31.6229800799166 | 49.679359463774 | 26.5007368821565 | 25.3240207449704 | 24.0890803080201 | 34.2463024980681 | 22.903717992235 | 103.124928643508 | Sec14l4 |
| ENSMUSG00000019370 | -0.102681706903812 | 0.278136369700806 | 0.848075053184273 | 5998.42756430189 | 5753.03270905082 | 5407.09677884858 | 4939.12197196348 | 5646.69941583908 | 6026.12615742791 | 5838.36593183881 | 6217.89043442121 | Calm3 |
| ENSMUSG00000019373 | 0.0900321924953135 | 0.478482309009302 | 0.897674307047416 | 1675.11443051901 | 1474.09246933493 | 1453.75470896401 | 1605.73050056997 | 1403.73640704008 | 1218.18990314556 | 1591.80840046033 | 1617.43309135608 | Cops3 |
| ENSMUSG00000019428 | -0.0980252957667131 | 0.501391181602587 | 0.900927059989498 | 2818.96279569542 | 2405.78406321293 | 2613.16194755836 | 2893.50385178643 | 2831.56189438818 | 3568.95395319152 | 2518.36790105529 | 2568.35348600569 | Fkbp8 |
| ENSMUSG00000019432 | -0.110943486896304 | 0.352782276862512 | 0.862728204353539 | 1287.50704611089 | 1170.31540244989 | 1201.05125369488 | 1328.10419906956 | 1462.86414961431 | 1501.9449809867 | 1193.07549177733 | 1228.81493920475 | Ddx39b |
| ENSMUSG00000019433 | -0.0579407337408687 | 0.671033253060952 | 0.943099452091129 | 539.39768879172 | 450.371898089624 | 464.709350326387 | 521.487242007538 | 491.636229922775 | 606.648787108634 | 509.087186281951 | 450.493109337431 | Gipc1 |
| ENSMUSG00000019437 | 0.254512052770423 | 0.109147138691795 | 0.763154315676794 | 146.369222084185 | 129.492100897378 | 173.201244622666 | 144.440710915757 | 119.350443344282 | 124.754387671534 | 134.299073681742 | 118.322286548867 | Tlcd1 |
| ENSMUSG00000019461 | -0.142181402690231 | 0.369526831277134 | 0.872319457246667 | 346.949267162513 | 351.013179162076 | 343.563124579386 | 347.970803569778 | 458.787484048202 | 432.971110154146 | 326.898520434627 | 315.887939318536 | Plscr3 |
| ENSMUSG00000019464 | 0.199028874162205 | 0.57389833990622 | 0.922280709351328 | 37.0440623793308 | 34.2054606144018 | 25.5542819935081 | 37.5170677703264 | 37.2285786578493 | 12.2308223207386 | 33.3144988977964 | 33.6512925047237 | Ptger1 |
| ENSMUSG00000019467 | -0.24975801941557 | 0.370739085923195 | 0.873323500696802 | 194.255449062345 | 150.666909849151 | 214.845259723197 | 252.302280755445 | 271.549632563136 | 352.247682837272 | 170.736806851206 | 171.513039217624 | Arhgef25 |
| ENSMUSG00000019470 | -0.189479197412972 | 0.274907688419173 | 0.846286635233373 | 367.730082643601 | 381.96097686082 | 377.63550057073 | 471.777127211855 | 531.054724972262 | 528.371524255907 | 392.486440139664 | 372.335268681298 | Xab2 |
| ENSMUSG00000019471 | 0.0569411164545152 | 0.578139624613789 | 0.922450244597442 | 4687.42916156021 | 4129.90216132456 | 4478.62453308445 | 4850.01893600895 | 4499.18322662067 | 4792.03618526538 | 4149.73726895676 | 4003.41828249746 | Cdc37 |
| ENSMUSG00000019478 | -0.191767436361681 | 0.290005540863188 | 0.851785577030345 | 132.81651633565 | 125.420022252807 | 109.78876708322 | 112.551203310979 | 122.635317931739 | 168.785348026193 | 144.709854587303 | 113.980184290193 | Rab4a |
| ENSMUSG00000019487 | 0.164286252768655 | 0.0934108886476284 | 0.754786063592529 | 3448.71185614404 | 3387.15501655469 | 3281.35909894416 | 3621.33496653076 | 3327.57795709424 | 2712.79639073982 | 3211.72590936568 | 3005.8202885671 | Trip10 |
| ENSMUSG00000019494 | 0.0246620522573122 | 0.850914913264737 | 0.972692784625411 | 3848.06491886756 | 3147.71679225388 | 3622.08285885761 | 4146.57391531533 | 3829.06881077938 | 4047.1791059324 | 3197.1508160979 | 3441.11603999917 | Cops6 |
| ENSMUSG00000019505 | -0.156933509624526 | 0.504542890963437 | 0.901187590673496 | 15958.762775759 | 12643.804191395 | 15289.032271227 | 16714.7916183747 | 18383.2531496068 | 24356.4595695188 | 12565.8125530126 | 12266.4388807541 | Ubb |
| ENSMUSG00000019518 | 0.0710593202751862 | 0.583607010347045 | 0.922450244597442 | 204.194099944604 | 190.573280565953 | 192.130342395635 | 194.150825711439 | 177.383227722694 | 170.008430258266 | 205.092383839559 | 189.966973816989 | Ap4m1 |
| ENSMUSG00000019528 | 0.136616268719245 | 0.153149524423368 | 0.783412422888014 | 4522.98966514464 | 4071.26422884273 | 4150.20468672344 | 4379.17973549136 | 3617.7418789863 | 3626.43881809899 | 4219.48950102402 | 4110.88531339964 | Gyg |
| ENSMUSG00000019539 | -1.1830074862935 | 0.0155501368227277 | 0.635862253526407 | 241.238162323935 | 267.1283590839 | 181.719338620502 | 302.950322245386 | 712.817785478232 | 1124.01257127588 | 185.311900118992 | 233.387996403729 | Rcn3 |
| ENSMUSG00000019558 | 0.24165885394858 | 0.364538003822189 | 0.86924083181804 | 4644.96401688146 | 6461.57439320628 | 6130.18831377599 | 4832.19832881805 | 3850.96797469576 | 2706.68097957945 | 6257.92040233294 | 5848.81174243392 | Slc6a8 |
| ENSMUSG00000019564 | -0.129972659679398 | 0.667599045042278 | 0.942024383071524 | 33.4300075130547 | 30.9477976987445 | 59.6266579848522 | 38.4549944645846 | 39.4184950494875 | 37.9155491942896 | 58.3003730711437 | 41.2499714574033 | Arid3a |
| ENSMUSG00000019577 | 0.684229353768732 | 0.0194243925162892 | 0.642273034804458 | 56757.8281611508 | 67461.3124888895 | 108109.756110757 | 84479.0573518326 | 57653.9288426587 | 25485.364469723 | 57235.3501845047 | 56786.013338939 | Pdk4 |
| ENSMUSG00000019578 | 0.0742748013556454 | 0.600192064587838 | 0.925516139867954 | 2409.67108208964 | 1947.26800783416 | 2306.51056363627 | 2618.69133036879 | 2346.49541364032 | 2510.98782244763 | 1923.91231134774 | 2035.36043375345 | Ubxn6 |
| ENSMUSG00000019579 | -0.147923942080958 | 0.469198352854685 | 0.895328112556867 | 354.177376895066 | 281.787842204358 | 352.081218577222 | 365.791410760683 | 394.184950494875 | 502.686797382356 | 321.693129981846 | 282.236646813812 | Mydgf |
| ENSMUSG00000019590 | -0.544790964421324 | 0.0450204657617091 | 0.693563614452513 | 209.615182244018 | 221.521078264697 | 162.790240847533 | 151.006197775564 | 320.822751374995 | 384.047820871192 | 197.804837205666 | 185.624871558315 | Cyb561 |
| ENSMUSG00000019647 | -0.267404770204027 | 0.444941017354425 | 0.89108031640712 | 136.430571201926 | 127.86326943955 | 153.325691961048 | 307.639955716677 | 292.353838283699 | 277.639666680766 | 149.915245040084 | 153.05910461826 | Sema6a |
| ENSMUSG00000019659 | -0.109992711506798 | 0.593545651411069 | 0.923935529355527 | 863.759113040007 | 694.696616763922 | 712.68053115228 | 947.305961200743 | 958.08842134171 | 1124.01257127588 | 751.658381381531 | 640.46008315442 | Ccdc12 |
| ENSMUSG00000019689 | 0.15180795748086 | 0.365592684335221 | 0.86924083181804 | 1957.91422380512 | 1387.76440207002 | 1612.75913025695 | 2044.68019348279 | 1679.66587238649 | 1799.15396338065 | 1333.62103400241 | 1491.51212585453 | Fmc1 |
| ENSMUSG00000019699 | 0.0392422300805324 | 0.78700012318828 | 0.961141387070613 | 581.862833470465 | 528.555808065399 | 557.461929413935 | 423.942865804689 | 493.826146314413 | 453.763508099402 | 511.169342463063 | 576.414074838978 | Akt3 |
| ENSMUSG00000019710 | 0.0603821137790814 | 0.681103916575089 | 0.944452327903597 | 1624.51766239114 | 1298.17867188944 | 1327.87620877377 | 1555.08245908003 | 1539.51122332165 | 1583.89149053565 | 1260.74556766348 | 1184.30839105334 | Mrpl24 |
| ENSMUSG00000019715 | 0.0522175196108542 | 0.590847242006751 | 0.923935529355527 | 751.723412185445 | 764.736369450555 | 727.823809370656 | 649.045272426648 | 698.583328932584 | 669.025980944401 | 713.138492030954 | 709.933719293204 | Gle1 |
| ENSMUSG00000019718 | -0.0860274992209356 | 0.689801339354973 | 0.94469184542218 | 174.378147297826 | 138.450673915436 | 120.199770858353 | 193.212899017181 | 175.193311331055 | 154.108361241306 | 132.216917500629 | 202.993280593011 | L3hypdh |
| ENSMUSG00000019726 | 0.149976074365479 | 0.549503277571997 | 0.914699131434573 | 65.9565013095403 | 116.461449234749 | 116.413951303759 | 70.3445020693621 | 86.5016974697087 | 90.5080851734656 | 77.0397787011541 | 79.2433662208011 | Lyst |
| ENSMUSG00000019731 | 0.120721915820022 | 0.653594683367169 | 0.936991241819063 | 1464.59573455842 | 1994.50412011119 | 1963.89389394553 | 1248.38043005761 | 1235.11284488394 | 967.458045570423 | 2092.56696201783 | 1839.96583211312 | Slc35e1 |
| ENSMUSG00000019732 | 0.0240267710896238 | 0.857047229737633 | 0.974012929230545 | 632.459601598332 | 672.707392083235 | 777.985918469023 | 763.472329126143 | 624.126171616885 | 787.664957455566 | 748.535147109862 | 639.374557589751 | Calr3 |
| ENSMUSG00000019738 | 0.111122428695932 | 0.502292987061465 | 0.900927059989498 | 469.827132615903 | 373.816819571677 | 388.046504345863 | 425.818719193205 | 387.61520131996 | 483.117481669175 | 336.268223249632 | 328.914246094558 | Polr2i |
| ENSMUSG00000019761 | -0.233425227294093 | 0.40776517652054 | 0.881573350402261 | 61.4389327266951 | 40.7207864457164 | 30.2865564367503 | 45.0204813243917 | 55.8428679867739 | 53.8156182112498 | 39.5609674411332 | 59.7039060567679 | Krt10 |
| ENSMUSG00000019762 | -0.348385087839802 | 0.330964898716832 | 0.856333478432716 | 15.3597331816738 | 26.0613033252585 | 27.447191770805 | 23.448167356454 | 19.7092475247437 | 30.5770558018465 | 23.9447960827911 | 43.4210225867403 | Iyd |
| ENSMUSG00000019763 | 0.268238614223465 | 0.066589959683822 | 0.719725956118125 | 1273.95434036235 | 1091.3170767452 | 1201.05125369488 | 1330.91797915233 | 962.468254124986 | 821.911259953634 | 1189.95225750566 | 1089.86766692718 | Rmnd1 |
| ENSMUSG00000019773 | 0.0961203403432597 | 0.829548047529912 | 0.967523998969336 | 12.6491920319666 | 10.5874044758863 | 12.3039135524298 | 9.37926694258161 | 13.1394983498292 | 9.78465785659088 | 12.4929370866736 | 6.51315338801105 | Fbxo5 |
| ENSMUSG00000019774 | 0.0154307746878028 | 0.910279967447867 | 0.98603671070003 | 271.054114970713 | 289.931999493501 | 265.007368821565 | 233.543746870282 | 267.16979977986 | 225.04713070159 | 286.296474902938 | 269.21034003779 | Mtrf1l |
| ENSMUSG00000019775 | -0.757951718661754 | 0.182738013935449 | 0.805531898941092 | 21.6843291976571 | 15.4738988493722 | 23.6613722162112 | 20.6343872736795 | 55.8428679867739 | 58.7079471395453 | 9.36970281500523 | 14.1118323406906 | Rgs17 |
| ENSMUSG00000019777 | 0.103932829644542 | 0.284333787280329 | 0.851125503777472 | 1061.62861696863 | 1154.84150360052 | 1119.65613327111 | 1112.38105939018 | 1056.63465896543 | 916.088591823321 | 1110.8303226234 | 1054.04532329312 | Hdac2 |
| ENSMUSG00000019779 | -1.05921426034222 | 0.00388613498477151 | 0.635862253526407 | 10.8421645988285 | 13.0306516626293 | 16.0897331070236 | 19.6964605794214 | 36.1336204620302 | 41.5847958905112 | 24.9858741733473 | 21.7105112933701 | Frk |
| ENSMUSG00000019782 | 0.132903947604756 | 0.300637844920063 | 0.851785577030345 | 2054.59019147801 | 1581.59534555163 | 1841.80121330988 | 2155.35554340525 | 1841.71968536772 | 1822.39252579005 | 1646.98553925981 | 1649.99885829613 | Rwdd1 |
| ENSMUSG00000019787 | -0.0651502088679616 | 0.720788671947468 | 0.951674101666722 | 10301.8633963203 | 8399.06941229347 | 7610.44375962216 | 9980.47795360109 | 8403.80415291157 | 12572.0622634872 | 8136.02527769621 | 8857.88860769502 | Trdn |
| ENSMUSG00000019789 | -0.251381577610778 | 0.276262898959482 | 0.846688575024234 | 58.7283915769879 | 50.4937751926884 | 67.1982970940397 | 66.5927952923295 | 91.9764884488041 | 53.8156182112498 | 73.9165444294857 | 68.388110574116 | Hey2 |
| ENSMUSG00000019790 | 0.166523746598391 | 0.316704149473603 | 0.854012936553746 | 374.958192376154 | 437.341246426994 | 421.172425448559 | 327.336416296098 | 360.241246424483 | 272.747337752471 | 402.897221045225 | 353.881334081933 | Stxbp5 |
| ENSMUSG00000019791 | 0.199322014628512 | 0.179608558178125 | 0.802391035126198 | 1011.93536255733 | 765.550785179469 | 904.810873547915 | 1081.42947847966 | 769.755611660825 | 824.357424417781 | 745.411912838194 | 937.894087873591 | Hint3 |
| ENSMUSG00000019792 | -0.245216768587304 | 0.217797527123964 | 0.826333693458579 | 81.316234491214 | 77.3694942468612 | 68.1447519826882 | 92.854742731558 | 98.5462376237187 | 73.3849339244316 | 108.272121417838 | 97.6973008201657 | Trmt11 |

| | | | | | | | | | | | | |
|---|---|---|---|---|---|---|---|---|---|---|---|---|
| ENSMUSG00000019794 | 0.00354483881881908 | 0.980111191206694 | 0.996872525500296 | 386.703870691551 | 286.674336577844 | 293.401015481019 | 333.901903155905 | 350.386622662111 | 316.77829810713 | 325.857442344071 | 303.947158107182 | Katna1 |
| ENSMUSG00000019795 | 0.105616943006966 | 0.411096490732656 | 0.882192126163777 | 2468.39947366663 | 2506.7716135983 | 2496.7479962546 | 2347.63051572818 | 1974.20962706183 | 2005.85486060113 | 2551.68239995309 | 2593.32057399306 | Pcmt1 |
| ENSMUSG00000019796 | 0.45072212110681 | 0.0879006469370096 | 0.749502753809124 | 183.413284463516 | 231.294067011669 | 183.612248397799 | 169.764731660727 | 116.065568756824 | 90.5080851734656 | 211.338852382896 | 143.289374536243 | Lrp11 |
| ENSMUSG00000019797 | -0.028276333838786 | 0.859697016936936 | 0.974868041734905 | 1282.98947752804 | 1037.56563863685 | 984.313084194385 | 1308.40773849013 | 1189.12460065954 | 1419.99847143775 | 1059.81749618615 | 1036.67691425842 | 1700021F05F |
| ENSMUSG00000019802 | 0.186102210478166 | 0.0127755242174826 | 0.635862253526407 | 1723.00065749717 | 1671.99549146112 | 1721.60144245152 | 1702.33695007856 | 1408.11623982336 | 1481.15258304144 | 1554.32958920031 | 1549.04498078196 | Sec63 |
| ENSMUSG00000019804 | 0.0651514592507418 | 0.494757285571969 | 0.900200915538898 | 3221.02639956865 | 2785.301792887 | 2948.20697813991 | 3184.26112700646 | 2924.63334103281 | 3145.76750089397 | 2714.09058207985 | 2819.10989144411 | Snx3 |
| ENSMUSG00000019806 | 0.152087620680964 | 0.130962474144824 | 0.77303252008843 | 1032.71617803842 | 899.114964721419 | 1052.45783617707 | 1087.05703864521 | 964.658170516624 | 923.427085215764 | 889.080689334941 | 885.788860769502 | Aig1 |
| ENSMUSG00000019808 | 0.0884342915593239 | 0.674850431831009 | 0.944023556644042 | 73.1846110420926 | 55.3802695661743 | 55.8408384302584 | 59.0893817382642 | 61.3176589658694 | 51.3694537471021 | 54.1360607089191 | 61.8749571861049 | Adat2 |
| ENSMUSG00000019809 | 0.179276207659366 | 0.306426668177082 | 0.851785577030345 | 300.870067617492 | 310.292392716359 | 305.704929033448 | 264.495327780801 | 235.416012101106 | 198.139321595965 | 315.446661438509 | 293.091902460497 | Pex3 |
| ENSMUSG00000019810 | 0.076828201575767 | 0.469422155992228 | 0.895491044377672 | 5187.07224682289 | 4458.92611580595 | 4289.33355535476 | 5078.87304940794 | 4599.91938063603 | 4118.11787539269 | 4778.54843565267 | 4529.89818136168 | Fuca2 |
| ENSMUSG00000019813 | 0.00147661002713647 | 0.995791315031266 | 0.999262355603273 | 37.0440623793308 | 35.8342920722305 | 41.6440151005317 | 45.9584080186499 | 55.8428679867739 | 33.0232202659942 | 41.6431236222455 | 29.3091902460497 | Cep57l1 |
| ENSMUSG00000019814 | 0.229655274457842 | 0.048601216069587 | 0.703396849069134 | 498.739571546113 | 458.516055378767 | 493.102996985841 | 505.542488205149 | 473.02194059385 | 392.609396495709 | 425.80093903746 | 375.591845375304 | Ltv1 |
| ENSMUSG00000019817 | -0.146386307691464 | 0.73988045569999 | 0.954383574136061 | 18.0702743313809 | 11.4018202048006 | 12.3039135524298 | 7.50341355406529 | 13.1394983498292 | 17.123151249034 | 9.36970281500523 | 15.1973579053591 | Plagl1 |
| ENSMUSG00000019818 | 0.055077982930071 | 0.695747440402446 | 0.944933386973863 | 2491.89083029743 | 2890.36142191695 | 3016.3517301226 | 2424.54050465735 | 2378.24920131908 | 2234.57123799894 | 2961.86716763221 | 2841.90592830215 | Cd164 |
| ENSMUSG00000019820 | 0.19302102960449 | 0.318431905861288 | 0.854351922245337 | 1633.55279955683 | 2102.8214120568 | 1873.98067952392 | 1377.81431386524 | 1507.7574356429 | 1125.23565350795 | 1772.9559882171 | 1706.44618765889 | Utrn |
| ENSMUSG00000019822 | 0.0025605172450233 | 0.985230976213857 | 0.99735301701069 | 750.819898468876 | 666.192066251921 | 700.376617599851 | 746.589648629496 | 838.737977997428 | 770.541806206532 | 644.427338054249 | 605.723265085027 | Smpd2 |
| ENSMUSG00000019823 | -0.095589562102402 | 0.706573127662303 | 0.948424888213123 | 141.85165350134 | 167.769640156352 | 151.432782183751 | 211.033506208086 | 228.846262926191 | 227.493295165738 | 131.175839410073 | 131.348593324889 | Mical1 |
| ENSMUSG00000019826 | 0.119071300707956 | 0.445363810184716 | 0.89108031640712 | 169.86057871498 | 140.079505373265 | 190.237432618338 | 164.137171495178 | 158.768938393769 | 138.208292224346 | 163.449260217313 | 149.802527924254 | Zbtb24 |
| ENSMUSG00000019828 | 0.16959832634682 | 0.490719760634976 | 0.899236955296196 | 661.372040528541 | 852.693268173302 | 929.418700652775 | 659.362466063487 | 572.663136413388 | 451.317343635254 | 938.011359591079 | 795.690238902016 | Grm1 |
| ENSMUSG00000019832 | -0.355297236952845 | 0.288022015565218 | 0.851785577030345 | 21.6843291976571 | 29.3189662409158 | 17.036187995672 | 42.2067012416173 | 35.0386622662111 | 39.1386314263635 | 30.191264626128 | 36.9078691987293 | Rab32 |
| ENSMUSG00000019836 | 0.029664699833692 | 0.952455531352131 | 0.992000114323903 | 18.0702743313809 | 18.7315617650296 | 11.3574586637814 | 11.2551203310979 | 6.56974917491458 | 15.9000690169602 | 26.0269522639034 | 9.76973008201657 | Amd-ps4 |
| ENSMUSG00000019837 | 0.00341483677761879 | 0.980333431015266 | 0.996888334226108 | 964.049135579171 | 803.828324438442 | 714.573440929577 | 772.851596068725 | 705.153078107498 | 801.118862008378 | 888.039611244385 | 853.223093829447 | Gtf3c6 |
| ENSMUSG00000019838 | -0.149035096800281 | 0.3773636324815 | 0.87422797717331 | 402.967117589794 | 452.000729547452 | 345.456034356683 | 289.819348525772 | 410.609323432161 | 373.040080782527 | 462.238672206925 | 407.07208675069 | Slc16a10 |
| ENSMUSG00000019841 | 0.223413785436957 | 0.468885718793835 | 0.895328112556867 | 123.781379169959 | 162.883145782866 | 159.950876181588 | 140.689004138724 | 74.4571573156986 | 84.3926740130963 | 192.599446752885 | 150.888053488923 | Rev3l |
| ENSMUSG00000019842 | -0.573835391610672 | 0.0149512253240597 | 0.635862253526407 | 54.2108229941427 | 48.0505280059454 | 70.0376617599851 | 78.7858423176855 | 110.590777777729 | 99.0696607979826 | 89.5327157878278 | 73.8157383974585 | Traf3ip2 |
| ENSMUSG00000019843 | 0.0361285811529209 | 0.702655728215881 | 0.946451777392559 | 1096.86565191482 | 992.772773546567 | 921.847061543587 | 967.002421780164 | 986.557334433006 | 952.781058785537 | 935.929203409967 | 1005.19667288304 | Fyn |
| ENSMUSG00000019845 | 0.00783802320916906 | 0.98873460972316 | 0.998437672973644 | 17.1667606148119 | 10.5874044758863 | 8.51809399783602 | 6.56548685980713 | 16.4243729372865 | 4.89232892829544 | 14.5750932677859 | 6.51315338801105 | Tube1 |
| ENSMUSG00000019846 | -0.541629147033704 | 0.0181099233431884 | 0.642273034804458 | 468.923618899334 | 614.883875330318 | 519.603733867997 | 625.597105070194 | 1040.21028602814 | 1000.48126583642 | 581.96265262088 | 623.091674119723 | Lama4 |
| ENSMUSG00000019848 | 0.193401940870137 | 0.0788482526777237 | 0.739855948845396 | 1603.73684691005 | 1377.17699759413 | 1304.21483655756 | 1576.65477304797 | 1312.8548767871 | 1190.05901180787 | 1276.36173902182 | 1346.05170018895 | Popdc3 |
| ENSMUSG00000019849 | 0.123521601970425 | 0.302990847330487 | 0.851785577030345 | 1217.93648993507 | 1300.62191907618 | 1339.23366743755 | 1189.29104831935 | 1170.51031133061 | 965.011881106275 | 1217.02028786012 | 1278.7491151795 | Prep |
| ENSMUSG00000019850 | -0.482973082409794 | 0.0999393278278828 | 0.759800953500741 | 37.9475760958999 | 28.5045505120015 | 25.5542819935081 | 30.0136542162612 | 47.0832024202212 | 51.3694537471021 | 27.0680303544596 | 45.5920737160773 | Tnfaip3 |
| ENSMUSG00000019851 | 0.108191537596188 | 0.303654134329621 | 0.851785577030345 | 3541.77376895066 | 3186.80874724177 | 2987.0116285745 | 3342.77073833609 | 2835.94172717146 | 2813.08913376988 | 3351.2303735002 | 3113.28731946928 | Perp |
| ENSMUSG00000019853 | -0.954468171358919 | 0.0452881557615378 | 0.695534138838186 | 10.8421645988285 | 10.5874044758863 | 10.4110037751329 | 9.37926694258161 | 21.8991639163819 | 34.2463024980681 | 15.616171358342 | 8.68420451734806 | Hebp2 |
| ENSMUSG00000019854 | 0.121484145257541 | 0.323600967817187 | 0.855977678491063 | 661.372040528541 | 686.552459474779 | 695.644343156608 | 638.728078789808 | 612.081631462875 | 519.80994863139 | 710.015257759285 | 622.006148555055 | Reps1 |
| ENSMUSG00000019856 | -0.0206921657038999 | 0.966047585682188 | 0.994284908685797 | 9.93865088225949 | 17.9171460361152 | 8.51809399783602 | 12.1930470253561 | 10.949581958191 | 7.33849339244316 | 10.4107809055614 | 20.6249857287016 | Fam184a |
| ENSMUSG00000019857 | 0.130340115017735 | 0.436074470451298 | 0.888687115594624 | 496.029030396406 | 460.144886836596 | 406.029147230184 | 406.122258613784 | 436.88832013182 | 297.208982393948 | 431.006329490241 | 449.407583772762 | Asf1a |
| ENSMUSG00000019861 | -0.0885035569156216 | 0.429376401102978 | 0.885743677088682 | 443.625234835401 | 448.743066631795 | 427.797609669098 | 414.563598862107 | 483.971522552041 | 397.501725424004 | 482.019155927491 | 479.80229958348 | Gopc |
| ENSMUSG00000019863 | -0.00048442566367410 | 0.997830514828799 | 0.999262355603273 | 529.45903790946 | 445.485403716138 | 402.24332767559 | 419.253232333398 | 390.900075907418 | 357.140011765567 | 522.621201459181 | 525.394373299558 | Qrsl1 |
| ENSMUSG00000019864 | 0.0404891895352668 | 0.850819278860014 | 0.972692784625411 | 1759.14120615993 | 1787.64252496695 | 1273.92828012081 | 1508.18612436712 | 1394.97674147353 | 1042.06606172693 | 1877.06379727271 | 1838.88030654845 | Rtn4ip1 |
| ENSMUSG00000019866 | -0.0473016224935653 | 0.814445748154223 | 0.966177091475091 | 212.325723393726 | 273.643684915214 | 233.774357496166 | 224.164479927701 | 290.163921892061 | 295.985900161874 | 179.065431575656 | 211.677485110359 | Crybg1 |
| ENSMUSG00000019868 | -0.0150018798595453 | 0.885054870385871 | 0.979866999597423 | 425.55496050402 | 412.09435883065 | 443.887342776122 | 417.377378944882 | 430.318570956905 | 391.386314263635 | 464.320828388037 | 428.78259804406 | Vta1 |
| ENSMUSG00000019872 | 0.132644128732499 | 0.207821899465481 | 0.820372930586885 | 1704.93038316579 | 1637.79003084671 | 1972.41198794336 | 1896.48777579 | 1742.07848954818 | 1715.98437159963 | 1526.2204807553 | 1593.55152893337 | Smpdl3a |
| ENSMUSG00000019873 | -0.278894382594793 | 0.131439400709785 | 0.77303252008843 | 526.748496759753 | 548.101785559343 | 614.249222732842 | 589.017963994125 | 849.687559955619 | 806.011190936674 | 516.374732915844 | 592.696958309005 | Reep3 |
| ENSMUSG00000019876 | 0.245875927015684 | 0.673365748975502 | 0.943631400280823 | 12.6491920319666 | 10.5874044758863 | 7.57163910918757 | 22.5102406621959 | 18.6142893289246 | 6.1154111603693 | 15.616171358342 | 4.34210225867403 | Pkib |
| ENSMUSG00000019877 | 0.0454770283045415 | 0.65862767100224 | 0.938989601151841 | 5150.93169816013 | 5955.00780982157 | 6476.59080302132 | 5421.21629281217 | 5668.59857975546 | 5492.8623042437 | 5788.39418349212 | 5339.70025260439 | Serinc1 |
| ENSMUSG00000019878 | 0.0279755388146463 | 0.862268389180553 | 0.974868041734905 | 271.957628687283 | 297.26174105373 | 239.453086828057 | 312.329589187968 | 278.119381738051 | 234.831788558181 | 254.023054095697 | 332.170822788563 | Hsf2 |
| ENSMUSG00000019883 | 0.129869238359864 | 0.736090043668663 | 0.954383574136061 | 173.474633581257 | 228.850819824926 | 212.952349945901 | 151.944124469822 | 128.110108910834 | 64.8233582999146 | 264.433835001259 | 243.157726485746 | Echdc1 |
| ENSMUSG00000019889 | 0.07728216816369 | 0.542713911974418 | 0.913837378403038 | 386.703870691551 | 418.609684661965 | 445.780252553418 | 384.549944645846 | 414.989156215438 | 387.717067567413 | 417.472314313011 | 329.999771659226 | Ptprk |
| ENSMUSG00000019890 | -1.14604434668344 | 0.0150332480105993 | 0.635862253526407 | 9.03513716569045 | 8.95857301805762 | 8.51809399783602 | 13.1309737196143 | 24.0890803080201 | 35.4693847301419 | 18.7394056300105 | 9.76973008201657 | Nts |
| ENSMUSG00000019891 | -0.343596648716949 | 0.291567515044055 | 0.851785577030345 | 114.746242004269 | 81.4415728914329 | 67.1982970940397 | 69.4065753751039 | 146.724398239759 | 138.208292224346 | 70.7933101578173 | 67.3025850094475 | Dcbld1 |
| ENSMUSG00000019897 | 0.0904963114591367 | 0.500129430799144 | 0.900927059989498 | 285.510334435818 | 276.086932101957 | 302.865564367503 | 324.522636213324 | 298.923587458613 | 310.66288694676 | 265.474913091815 | 242.072200921077 | Ccdc59 |
| ENSMUSG00000019899 | 0.02639851334918 | 0.780904868424574 | 0.960742105865155 | 1407.67437041457 | 1612.54314325037 | 1431.03979163645 | 1435.02784221499 | 1567.98013641295 | 1424.89080036605 | 1362.77122053798 | 1425.29506640975 | Lama2 |
| ENSMUSG00000019906 | -0.980716609728002 | 0.0168849433151122 | 0.642273034804458 | 24.3948703473642 | 45.6072808192024 | 23.6613722162112 | 22.5102406621959 | 83.2168228822514 | 78.277262852727 | 36.4377331694648 | 32.5657669400552 | Lin7a |
| ENSMUSG00000019907 | 0.0846528075113117 | 0.62415359269118 | 0.931145988862144 | 622.520950716072 | 785.096762673413 | 717.412805595523 | 541.183702586959 | 580.327843784121 | 516.140701935169 | 693.358008310387 | 724.045551633895 | Ppp1r12a |
| ENSMUSG00000019916 | -0.214359274860495 | 0.18737702400393 | 0.807540361382744 | 2047.36208174546 | 1914.69137867759 | 1484.04126540076 | 1876.79131521058 | 2093.56007040611 | 2614.94981217391 | 1782.32569103211 | 2007.13676907207 | P4ha1 |
| ENSMUSG00000019917 | 0.109564191505732 | 0.380919302721325 | 0.874829549404631 | 327.975479114563 | 376.26006675842 | 336.937940358847 | 308.577882410935 | 324.107625962453 | 277.639666680766 | 334.18606706852 | 314.802413753867 | Sept10 |
| ENSMUSG00000019920 | 0.10232426599788 | 0.418985446270798 | 0.882550938324998 | 2148.55561800119 | 2590.65643367648 | 2193.8824318871 | 2241.64479927701 | 1934.79113201234 | 1899.4467064107 | 2322.64522003074 | 2389.24176783538 | Lims1 |
| ENSMUSG00000019923 | 0.200259585925023 | 0.320847771631347 | 0.855228644171892 | 1077.89186386687 | 1338.08504260624 | 1533.25691961048 | 1024.21595012991 | 952.613630362614 | 835.365164506446 | 1328.41564354963 | 1211.44653017005 | Zwint |
| ENSMUSG00000019927 | 0.0940187558579706 | 0.449445713975342 | 0.89108031640712 | 877.311818788543 | 908.887953468391 | 821.522843346852 | 880.713165908413 | 730.337116611338 | 724.064681387725 | 942.175671953304 | 870.591502864143 | Ube2d1 |
| ENSMUSG00000019929 | -0.841383533076178 | 0.0529675360927206 | 0.703396849069134 | 4543.77048062573 | 3509.31737589184 | 3619.24349419166 | 5011.34232742136 | 9434.15981517734 | 14099.6919713475 | 3101.37163176673 | 3258.74774513486 | Dcn |
| ENSMUSG00000019933 | 0.42348857050987 | 0.3074494231488 | 0.851785577030345 | 12.6491920319666 | 17.9171460361152 | 15.1432782183751 | 21.5723139679377 | 15.3294147414674 | 17.123151249034 | 9.36970281500523 | 8.68420451734806 | Mrln |
| ENSMUSG00000019942 | -1.19979066217246 | 0.0367171132124802 | 0.679566075569698 | 11.7456783153976 | 7.32974156022896 | 2.83936466594534 | 12.1930470253561 | 30.6588294829347 | 23.2385624094033 | 7.28754663389296 | 17.3684090346961 | Cdk1 |
| ENSMUSG00000019943 | -0.0405352183001649 | 0.758440271224394 | 0.956833366694536 | 1543.20142789993 | 1674.43873864786 | 1568.27575049048 | 1330.91797915233 | 1463.95910781013 | 1360.06744206613 | 1746.9290359532 | 1719.47249443492 | Atp2b1 |
| ENSMUSG00000019944 | 0.515587707665609 | 0.0581409897019778 | 0.710862211775517 | 3646.58136007267 | 5358.85549625628 | 4758.77518012439 | 3022.93773559405 | 2637.7542937282 | 1764.90766088258 | 3912.37146430996 | 3427.00420765848 | Rhobtb1 |
| ENSMUSG00000019945 | 0.110820435918137 | 0.346827569115375 | 0.861266047293794 | 271.957628687283 | 290.746415222415 | 303.812019256151 | 288.881421831514 | 270.454674367317 | 253.178022039289 | 294.625099627387 | 250.756405438425 | Cabcoco1 |
| ENSMUSG00000019947 | 0.051650797573094 | 0.822978444147381 | 0.966637987288264 | 133.720030052219 | 229.665235553841 | 275.418372596698 | 171.640585049243 | 187.237851485066 | 200.585486060113 | 199.886993386778 | 194.309076075663 | Arid5b |
| ENSMUSG00000019948 | 0.0127839148294744 | 0.936608263214431 | 0.988788384589834 | 276.475197270128 | 205.232763686411 | 228.095628164276 | 239.171307035831 | 243.08071947184 | 190.800828203522 | 243.612273190136 | 261.61166108511 | Actr6 |
| ENSMUSG00000019951 | 0.0997341413020825 | 0.676368644465945 | 0.94409858171788 | 561.985531705946 | 717.500257173524 | 820.576388458203 | 791.610129953888 | 669.019457645468 | 407.286383280595 | 860.971580889925 | 759.867895267955 | Uhrf1bp1l |
| ENSMUSG00000019952 | -0.0740774459198505 | 0.559624113285076 | 0.916574868225118 | 177.088688447533 | 177.542628903324 | 198.755526616174 | 183.8336320746 | 203.662224422352 | 195.693157131818 | 181.147587756768 | 195.394601640331 | Poc1b |
| ENSMUSG00000019960 | -0.35818197213408 | 0.145082797829742 | 0.780607990324772 | 206.001127377742 | 190.573280565953 | 223.363353721033 | 234.48167356454 | 413.894198019619 | 269.078091056249 | 212.379930473452 | 199.736703899005 | Dusp6 |
| ENSMUSG00000019961 | -0.247829534798578 | 0.156181492474421 | 0.787275540343856 | 478.862269781594 | 477.247617143797 | 594.373670071225 | 536.494069115668 | 764.28082068173 | 678.810638800992 | 544.483841360859 | 490.657555230165 | Tmpo |
| ENSMUSG00000019966 | -0.0871518929736205 | 0.514556747502645 | 0.904135765061573 | 1542.29791418336 | 1897.58864837039 | 1559.75765649264 | 1738.91609115463 | 1906.32221892105 | 2046.21657425957 | 1570.98683864921 | 1636.97255152011 | Kitl |
| ENSMUSG00000019969 | 0.00073492553270665 | 0.99519762288321 | 0.999262355603273 | 766.17963165055 | 759.035459348154 | 896.292779550079 | 724.079407967301 | 846.402685368162 | 850.042151291332 | 737.083288113745 | 711.019244857872 | Psen1 |
| ENSMUSG00000019970 | -0.380636098764451 | 0.337430670480217 | 0.858570960161243 | 1448.33248766018 | 1200.44878441972 | 1328.82266366242 | 778.479156234274 | 2416.57273817275 | 2244.35589585553 | 812.040910633787 | 719.703449375221 | Sgk1 |
| ENSMUSG00000019971 | 0.403373595919505 | 0.089950104897565 | 0.753823363185192 | 159.018414116152 | 221.521078264697 | 223.363353721033 | 186.647412157374 | 127.015150715015 | 101.51582526213 | 197.804837205666 | 170.427513652956 | Cep290 |
| ENSMUSG00000019975 | -0.410355647950364 | 0.0496523360422403 | 0.703396849069134 | 224.071401709123 | 210.119258059897 | 215.791714611846 | 205.405946042537 | 290.163921892061 | 398.724807656078 | 227.996101831794 | 221.447215192376 | Ikbip |
| ENSMUSG00000019977 | 0.22659143063183 | 0.176121420898278 | 0.799889021902745 | 1941.65097690688 | 2215.21078264697 | 2245.93745076276 | 2023.10787951485 | 1587.68938393769 | 1384.52908670761 | 2148.78517890787 | 2078.78145634019 | Hbs1l |
| ENSMUSG00000019978 | -0.100563420609036 | 0.400532223366247 | 0.880487015130533 | 578.248778604189 | 574.977504613516 | 570.712297855013 | 494.287367874051 | 672.304332232926 | 604.202622644487 | 563.22324699087 | 539.506205640248 | Epb41l2 |
| ENSMUSG00000019979 | -0.320359179017273 | 0.170647823472724 | 0.799531027639052 | 61.4389327266951 | 77.3694942468612 | 74.7699362032273 | 48.7721881014244 | 102.926070406995 | 78.277262852727 | 67.6700758861489 | 79.2433662208011 | Apaf1 |
| ENSMUSG00000019984 | -0.078928811646621 | 0.632841502072367 | 0.932149359892384 | 193.351935345776 | 241.881471487556 | 208.220075502658 | 199.778385876988 | 224.466430142915 | 178.570005882784 | 265.474913091815 | 221.447215192376 | Med23 |
| ENSMUSG00000019986 | 0.0676882022025289 | 0.860970401840166 | 0.974868041734905 | 17.1667606148119 | 38.2775392589734 | 36.9117406572894 | 20.6343872736795 | 20.8042057205628 | 36.6924669622158 | 31.2323427166841 | 19.5394601640331 | Ahi1 |
| ENSMUSG00000019987 | -0.282973435262096 | 0.293116033407512 | 0.851785577030345 | 352.370349461928 | 538.328796812371 | 348.295399022628 | 372.35689762049 | 304.398378437709 | 454.986590331476 | 484.101312108604 | 717.532398245884 | Arg1 |
| ENSMUSG00000019988 | 0.0227910920223327 | 0.920211397872956 | 0.986713918026868 | 529.45903790946 | 648.274920215806 | 574.498117409607 | 498.039074651084 | 452.217734873287 | 371.816998550453 | 696.481242582055 | 693.650835823176 | Nedd1 |
| ENSMUSG00000019989 | -0.57132080638667 | 0.167413315125448 | 0.797999638672775 | 28.0089252136404 | 44.7928650902881 | 35.0188308799925 | 46.8963347129081 | 94.1664048404423 | 74.6080161565054 | 31.2323427166841 | 30.3947158107182 | Enpp3 |
| ENSMUSG00000019990 | 0.424511171918136 | 0.103381519386958 | 0.759800953500741 | 495.125516679837 | 597.781145023117 | 612.356312955545 | 475.528833988888 | 390.900075907418 | 218.931719541221 | 478.895921655823 | 535.164103381574 | Pde7b |
| ENSMUSG00000019996 | 0.560309256821371 | 0.05026673923814 | 0.703396849069134 | 768.890172800257 | 897.48613326359 | 1419.68233297267 | 892.90621293377 | 596.752216721408 | 396.278643191931 | 917.189797779956 | 787.006034384668 | Map7 |
| ENSMUSG00000019997 | 0.213933830466284 | 0.595872981026172 | 0.924461858177432 | 530.362551626029 | 902.372627637076 | 1444.29016007753 | 1874.91546182206 | 1044.59011881142 | 1684.1842335657 | 742.288678566525 | 626.348250813729 | Ctgf |

| | | | | | | | | | | | | |
|---|---|---|---|---|---|---|---|---|---|---|---|---|
| ENSMUSG00000019998 | 0.0630204912614167 | 0.637947140106276 | 0.933164350662676 | 1093.25159704854 | 1108.4198070524 | 1213.35516724731 | 1042.97448401508 | 1045.68507700724 | 864.719138076219 | 1206.60950695456 | 1148.48604741928 | Stx7 |
| ENSMUSG00000020000 | -1.20151567768163 | 0.0484916933148074 | 0.703396849069134 | 5.42108229941427 | 17.1027303072009 | 1.89290977729689 | 9.37926694258161 | 26.2789966996583 | 26.9078091056249 | 13.5340151772298 | 11.9407812113536 | Moxd1 |
| ENSMUSG00000020003 | 0.123456054099969 | 0.501749579452322 | 0.900927059989498 | 740.881247586617 | 704.469605510894 | 707.00180182039 | 772.851596068725 | 569.37826182593 | 499.017550686135 | 783.931802188771 | 831.512582536077 | Pex7 |
| ENSMUSG00000020009 | -0.477493904225372 | 0.106082660581276 | 0.762094734041179 | 889.05749710394 | 715.871425715695 | 649.268053612834 | 769.099889291692 | 1369.79270296969 | 1512.95272107536 | 609.03068297534 | 718.617923810552 | Ifngr1 |
| ENSMUSG00000020010 | -0.281130101451324 | 0.486573202600774 | 0.899236955296196 | 15.3597331816738 | 26.0613033252585 | 15.1432782183751 | 16.8826804966469 | 14.2344565456483 | 20.7923979452556 | 18.7394056300105 | 35.8223436340608 | Vnn3 |
| ENSMUSG00000020015 | -0.264427160984147 | 0.061319466665915 | 0.71282992338528 | 220.457346842847 | 226.407572638183 | 269.739643264807 | 274.812521417641 | 331.772333333186 | 304.547475786391 | 270.680303544596 | 283.322172378481 | Cdk17 |
| ENSMUSG00000020017 | -0.291252053053343 | 0.384546634642578 | 0.875126835008975 | 36.1405486627618 | 59.452348210746 | 45.4298346551254 | 39.3929211588428 | 33.943704070392 | 42.8078781225851 | 58.3003730711437 | 85.7565196088121 | Hal |
| ENSMUSG00000020018 | -0.012911877505807 | 0.951217926354001 | 0.991915011884503 | 309.905204783182 | 237.809392842984 | 240.399541716705 | 322.646782824807 | 290.163921892061 | 370.593916318379 | 212.379930473452 | 248.585354309088 | Snrpf |
| ENSMUSG00000020019 | -0.536249144250448 | 0.0510461859756079 | 0.703396849069134 | 155.404359249876 | 179.985876090067 | 145.754052851861 | 125.682177030594 | 296.733671066975 | 280.085831144914 | 167.613572579538 | 136.776221148232 | Ntn4 |
| ENSMUSG00000020021 | 0.0249978034739682 | 0.935765605839216 | 0.98818087308472 | 107.518132271716 | 175.099381716581 | 151.432782183751 | 84.4134024832345 | 116.065568756824 | 78.277262852727 | 176.983275394543 | 137.8617467129 | Fgd6 |
| ENSMUSG00000020022 | 0.0545928015196089 | 0.740065703611793 | 0.954383574136061 | 10287.4071768551 | 8192.20781714923 | 8918.44441573432 | 10833.0533186818 | 9462.62872826864 | 11461.5035967641 | 7970.49386129778 | 7917.82346869209 | Ndufa12 |
| ENSMUSG00000020023 | -0.197420872108401 | 0.429532067735655 | 0.885743677088682 | 1165.53269437407 | 1334.82737969058 | 1206.72998302677 | 1098.31215897631 | 1853.76422552173 | 1761.23841418636 | 944.257828134416 | 952.005920214281 | Tmcc3 |
| ENSMUSG00000020024 | 0.200366496799448 | 0.362962280838596 | 0.86924083181804 | 458.081454300506 | 485.39177443294 | 493.102996985841 | 496.163221262567 | 351.48158085793 | 275.193502216618 | 531.990904274186 | 522.137796605552 | Cep83 |
| ENSMUSG00000020027 | -0.133527182011458 | 0.582223353413386 | 0.922450244597442 | 488.800920663853 | 293.189662409158 | 420.22597055991 | 606.83857118503 | 545.28918151791 | 607.871869340708 | 413.308001950786 | 417.927342397375 | Socs2 |
| ENSMUSG00000020029 | -0.0307901346498558 | 0.83449167727921 | 0.968516656512387 | 6524.27254734507 | 6595.95298847715 | 6006.20272336304 | 5322.73398991506 | 4986.43962376017 | 6328.22746875015 | 7355.21670977911 | 6306.90353072403 | Nudt4 |
| ENSMUSG00000020032 | 0.538664358209592 | 0.0112867965180326 | 0.635862253526407 | 1592.89468231123 | 2193.22155796629 | 1745.26281466774 | 1605.73050056997 | 931.809424642051 | 1000.48126583642 | 1327.37456545907 | 1653.25543499014 | Nuak1 |
| ENSMUSG00000020034 | 0.153353115762473 | 0.490612744765632 | 0.899236955296196 | 1600.12279204378 | 2350.40379364675 | 3010.67300079071 | 2275.4101602703 | 2425.3324037393 | 1689.076562494 | 2446.53351280692 | 1743.35405685762 | Tcp11l2 |
| ENSMUSG00000020038 | -0.0278715712783476 | 0.844187349011293 | 0.971721411930626 | 244.852217190211 | 246.767965861042 | 225.25626349833 | 244.79886720138 | 248.555510450935 | 292.316653465652 | 212.379930473452 | 229.045894145055 | Cry1 |
| ENSMUSG00000020042 | 0.106184893033081 | 0.76087979305925 | 0.957468401713057 | 39.754603529038 | 69.2253369577179 | 46.3762895437739 | 31.8895076047775 | 30.6588294829347 | 30.5770558018465 | 55.1771387994752 | 57.5328549274309 | Btbd11 |
| ENSMUSG00000020044 | -0.431246199002027 | 0.158262062054363 | 0.789230661864335 | 3457.74699330974 | 3450.67944341001 | 3408.18405402306 | 2462.99549912193 | 6519.3810979069 | 5373.00024550047 | 2856.71828048604 | 2483.68249196155 | Timp3 |
| ENSMUSG00000020048 | 0.027537962788511 | 0.718077149086775 | 0.951674101666722 | 9030.6195971076 | 8278.53588441415 | 8222.80007257771 | 8860.59348065685 | 8395.04448734502 | 8812.30748209216 | 7978.82248602223 | 8557.19802628185 | Hsp90b1 |
| ENSMUSG00000020051 | -0.196419004451046 | 0.549533492716096 | 0.914699131434573 | 102.097049972302 | 229.665235553841 | 105.056492639978 | 141.626930832982 | 148.914314631397 | 127.200552135681 | 135.340151772298 | 251.841931003094 | Pah |
| ENSMUSG00000020053 | -0.513991044268701 | 0.092981834851414 | 0.754058637722175 | 137.334084918495 | 215.820168162297 | 142.914688185915 | 252.302280755445 | 282.499214521327 | 392.609396495709 | 163.449260217313 | 231.216945274392 | Igf1 |
| ENSMUSG00000020056 | 0.0212009042591045 | 0.904079687254904 | 0.985039825636533 | 485.186865797577 | 386.033055505392 | 374.796135904785 | 584.328330522834 | 475.211856985488 | 477.002070508805 | 399.773986773556 | 451.578634902099 | Washc3 |
| ENSMUSG00000020057 | -0.105625638924462 | 0.598935304523888 | 0.924848033480823 | 94.8689402397497 | 78.9983257046899 | 77.6093008691726 | 89.1030359545253 | 109.49581958191 | 100.292743030056 | 86.4094815161593 | 70.559161703453 | Dram1 |
| ENSMUSG00000020061 | 1.41585134191973 | 0.00551473824120012 | 0.635862253526407 | 14.4562194651047 | 36.6487078011448 | 16.0897331070236 | 20.6343872736795 | 12.0445401540101 | 6.1154111603693 | 10.4107809055614 | 4.34210225867403 | Mybpc1 |
| ENSMUSG00000020063 | 0.194458312476733 | 0.157498629291303 | 0.789197677076549 | 251.176813206195 | 317.622134276588 | 323.687571917769 | 267.309107863576 | 246.365594059297 | 240.94719971855 | 263.392756910703 | 262.697186649779 | Sirt1 |
| ENSMUSG00000020064 | 0.292612929673829 | 0.174917094080435 | 0.799889021902745 | 607.161217534398 | 710.170515613295 | 715.519895818226 | 637.79015209555 | 455.502609460744 | 368.147751854232 | 667.331056046484 | 688.223207999834 | Herc4 |
| ENSMUSG00000020067 | 0.32969623213702 | 0.171532052134937 | 0.799531027639052 | 4005.27630555058 | 5315.69146262382 | 5567.04765503016 | 4339.78681433251 | 3216.98717931651 | 2436.37980629113 | 5027.36609929558 | 4617.82575209983 | Mypn |
| ENSMUSG00000020069 | 0.0682226414297702 | 0.5330906596119 | 0.909101198527844 | 1384.18301378378 | 1191.49021140166 | 1247.42754323865 | 1454.72430279441 | 1304.09521122054 | 1203.51291636068 | 1349.23720536075 | 1175.62418653599 | Hnrnph3 |
| ENSMUSG00000020070 | -0.287376679913951 | 0.234495570458654 | 0.832048466795696 | 57.8248778604189 | 55.3802695661743 | 47.3227444324223 | 45.9584080186499 | 77.7420319031559 | 44.0309603546589 | 59.3414511616998 | 70.559161703453 | Rufy2 |
| ENSMUSG00000020072 | 0.0913320030003924 | 0.696485993639177 | 0.944933386973863 | 146.369222084185 | 198.717437855096 | 232.827902607518 | 190.399118934407 | 141.249607260664 | 214.039390612925 | 138.463386043966 | 227.960368580387 | Pbld2 |
| ENSMUSG00000020074 | 0.0510277673447864 | 0.662854880573716 | 0.939783024514536 | 1941.65097690688 | 1744.47849133449 | 1989.44817593904 | 1794.25376611586 | 1557.03055445476 | 2037.65499863505 | 1742.76472359097 | 1873.61712461784 | Ccar1 |
| ENSMUSG00000020075 | 0.302157988273795 | 0.0675484282953782 | 0.722038584319139 | 1486.28006375608 | 1796.60109798501 | 1881.55231863311 | 1604.79257387571 | 1353.3683300324 | 1031.05832163826 | 1507.48107512529 | 1596.80810562737 | Ddx21 |
| ENSMUSG00000020076 | 0.290130173597573 | 0.00819973437540317 | 0.635862253526407 | 1182.69945498888 | 1201.26320014863 | 1300.42901700297 | 1195.85653517916 | 964.658170516624 | 887.957700485622 | 1139.98050915897 | 996.51246836569 | Ddx50 |
| ENSMUSG00000020077 | -0.372568767366485 | 0.307138883879161 | 0.851785577030345 | 657.757985662265 | 498.422426095569 | 745.806452254976 | 723.141481273042 | 1108.09769416893 | 1362.51360653028 | 370.623800237985 | 557.960140239613 | Srgn |
| ENSMUSG00000020078 | 0.19016189840574 | 0.217949521905186 | 0.826684560148288 | 2174.75751578169 | 2252.67390617703 | 2380.33404495084 | 2191.93468448132 | 1825.29531243043 | 1512.95272107536 | 2290.3717992235 | 2257.8931745105 | Vps26a |
| ENSMUSG00000020079 | -0.00062782294439681 | 0.997116531408716 | 0.999262355603273 | 739.074220153479 | 664.563234794092 | 555.569019636638 | 556.19052969509 | 623.031213421066 | 479.448234972953 | 716.261726302622 | 696.907412517182 | Supv3l1 |
| ENSMUSG00000020083 | -0.138980326067614 | 0.629442721349785 | 0.931825800800958 | 70.4740698923855 | 57.8235167529173 | 34.0723759913441 | 72.2203554578784 | 50.3680770076785 | 64.8233582999146 | 81.2040910633787 | 61.8749571861049 | Fam241b |
| ENSMUSG00000020085 | 0.182364202935256 | 0.377009946295919 | 0.87422797717331 | 770.697200233395 | 788.35442558907 | 1004.188636856 | 707.196727470654 | 628.506004400162 | 519.80994863139 | 866.176971342706 | 866.249400605469 | Aifm2 |
| ENSMUSG00000020086 | -0.528291353157104 | 0.019702070333789 | 0.642273034804458 | 50.5967681278665 | 70.0397526866323 | 77.6093008691726 | 64.7169419038131 | 90.881530252985 | 123.53130543946 | 84.3273253350471 | 81.4144173501381 | H2afy2 |
| ENSMUSG00000020087 | -0.210889894348547 | 0.11749935849513 | 0.769868040577702 | 654.143930795989 | 605.110886583346 | 588.694940739334 | 619.969544904645 | 779.610235423197 | 826.803588881929 | 610.071761065896 | 641.545608719088 | Tysnd1 |
| ENSMUSG00000020088 | 0.143582588528201 | 0.234792516745697 | 0.832048466795696 | 5872.83915769879 | 6025.86197823712 | 6679.13214919209 | 5852.66257217093 | 5229.52034323201 | 4722.32049803717 | 6229.81129388792 | 5932.39721091339 | Sar1a |
| ENSMUSG00000020089 | 0.0181950650936947 | 0.862069386921549 | 0.974868041734905 | 1296.54218327658 | 1149.14059349812 | 1048.67201662248 | 1266.20103724852 | 1119.04727612712 | 1168.04353163054 | 1210.77381931679 | 1202.76232565271 | Ppa1 |
| ENSMUSG00000020091 | 0.250046009007411 | 0.503937770342471 | 0.900927059989498 | 414.712795905192 | 675.965054998893 | 755.271001141461 | 353.598363735327 | 372.285786578493 | 204.254732756335 | 631.934400967575 | 640.46008315442 | Eif4ebp2 |
| ENSMUSG00000020092 | -0.489356669955774 | 0.0642796566185227 | 0.716991331547963 | 254.790868072471 | 238.623808571898 | 157.111511515642 | 209.15765281957 | 447.837902090011 | 305.770558018465 | 223.831789469569 | 230.131419709724 | Pald1 |
| ENSMUSG00000020097 | 0.14725885326778 | 0.341733727749311 | 0.859283395807012 | 418.326850771468 | 484.577358704026 | 515.817914313404 | 392.99128489417 | 406.229490648885 | 335.124531588238 | 426.842017128016 | 466.775992807458 | Sgpl1 |
| ENSMUSG00000020098 | -0.185514156788757 | 0.457102303063553 | 0.892805623137304 | 56.9213641438498 | 87.9568987227475 | 67.1982970940397 | 70.3445020693621 | 62.4126171616885 | 83.1695917810225 | 67.6700758861489 | 108.552556466851 | Pcbd1 |
| ENSMUSG00000020099 | -0.747325775266521 | 0.00057978083427366 | 0.525706734317154 | 233.106538874814 | 209.304842330982 | 221.470443943737 | 294.508981997063 | 457.692525852383 | 519.80994863139 | 301.91264626128 | 329.999771659226 | Unc5b |
| ENSMUSG00000020100 | -0.252895414569013 | 0.200244677974756 | 0.812921911423739 | 54.2108229941427 | 57.009101024003 | 60.5731128735006 | 59.0893817382642 | 74.4571573156986 | 74.6080161565054 | 68.711153976705 | 57.5328549274309 | Slc29a3 |
| ENSMUSG00000020101 | -0.44355669413366 | 0.156455514849026 | 0.787275540343856 | 322.554396815149 | 320.065381463331 | 480.799083433411 | 558.066383083606 | 743.476614961167 | 779.103381831049 | 326.898520434627 | 437.466802561409 | Vsir |
| ENSMUSG00000020102 | -0.0842552150978543 | 0.730262923372172 | 0.952921973914044 | 84.9302893574902 | 97.7298874697194 | 131.557229522134 | 78.7858423176855 | 106.210944994452 | 86.838838477244 | 86.4094815161593 | 136.776221148232 | Slc16a7 |
| ENSMUSG00000020105 | -0.476508714758895 | 0.120615157301489 | 0.769868040577702 | 18.9737880479499 | 17.9171460361152 | 21.7684624389143 | 19.6964605794214 | 31.7537876787538 | 30.5770558018465 | 22.903717992235 | 23.8815624227072 | Lrig3 |
| ENSMUSG00000020107 | -0.0193891883021942 | 0.888356829482668 | 0.980942551992548 | 1108.61133023022 | 876.311324311818 | 1122.49549793706 | 1109.5672793074 | 895.675804180021 | 1164.37428493431 | 1119.15894734785 | 1094.20976918586 | Anapc16 |
| ENSMUSG00000020108 | 0.455365745504902 | 0.563983879279694 | 0.918590066040684 | 2753.00629438588 | 3403.44333113298 | 1201.99770858353 | 360.163850595134 | 1766.1675698562 | 2623.51138779843 | 676.700758861489 | 563.387768062955 | Ddit4 |
| ENSMUSG00000020109 | 0.143481556351276 | 0.208682067428161 | 0.821131421126672 | 1042.65482892068 | 1212.66502035344 | 1328.82266366242 | 1114.2569127787 | 1061.0144917487 | 1007.81975922886 | 1149.35021197397 | 1034.50586312909 | Dnajb12 |
| ENSMUSG00000020111 | -0.0137802415119806 | 0.89106980770881 | 0.982142775127356 | 412.905768472054 | 377.074482487334 | 390.885869011809 | 394.867138282686 | 382.140410340865 | 412.178712208891 | 381.034581143546 | 415.756291268038 | Micu1 |
| ENSMUSG00000020114 | 0.0172834101649667 | 0.879599767979787 | 0.978393501240173 | 623.424464432641 | 732.974156022896 | 789.343377132805 | 649.045272426648 | 684.348872386936 | 682.479885497214 | 721.467116755403 | 673.025850094475 | Cand1 |
| ENSMUSG00000020115 | 0.137658279971954 | 0.40348138581297 | 0.881158207349486 | 397.54603529038 | 406.39344872825 | 423.065335225856 | 361.101777289392 | 356.956371837026 | 266.631926592101 | 406.020455316893 | 412.499714574033 | Tbk1 |
| ENSMUSG00000020116 | 0.106598090592573 | 0.482708278629461 | 0.898407877551013 | 806.837748896157 | 764.736369450555 | 748.645816920921 | 856.327071857701 | 654.78500109982 | 600.533375948265 | 813.081988724343 | 880.36123294616 | Pno1 |
| ENSMUSG00000020120 | 0.434876713580387 | 0.321050905861748 | 0.855469551206808 | 86.7373167906283 | 146.594831204579 | 323.687571917769 | 82.5375490947182 | 123.730276127558 | 165.116101329971 | 77.0397787011541 | 107.467030902182 | Plek |
| ENSMUSG00000020121 | 0.0804372415828793 | 0.737569537918812 | 0.954383574136061 | 41.5616309621761 | 48.8649437348597 | 51.1085639870161 | 47.8342614071662 | 52.5579933993167 | 36.6924669622158 | 37.4788112600209 | 52.1052271040884 | Srgap1 |
| ENSMUSG00000020122 | -0.875786099237589 | 0.00286920180110781 | 0.635862253526407 | 55.1143367107117 | 115.647033505835 | 66.2518422053913 | 99.4202295913651 | 168.623562156141 | 167.562265794119 | 99.9434966933891 | 182.368294864309 | Egfr |
| ENSMUSG00000020123 | -0.147306593558136 | 0.73577616278659 | 0.954383574136061 | 12.6491920319666 | 23.6180561385155 | 22.7149173275627 | 25.3240207449704 | 29.5638712871156 | 35.4693847301419 | 10.4107809055614 | 18.4539345993646 | Avpr1a |
| ENSMUSG00000020124 | 0.317207436201919 | 0.143497762564262 | 0.780607990324772 | 2200.05589984562 | 2368.32093968287 | 2527.98100758 | 2061.56287397944 | 1574.54988558786 | 1209.62832752105 | 2282.04317449905 | 2282.86026249787 | Usp15 |
| ENSMUSG00000020128 | 0.240685467383069 | 0.187084323409373 | 0.807540361382744 | 658.661499378834 | 721.572335818095 | 693.751433379311 | 624.659178375935 | 476.306815181307 | 439.086521314516 | 720.426038664847 | 646.973236542431 | Vps54 |
| ENSMUSG00000020130 | 0.0780131152647035 | 0.581257067213343 | 0.922450244597442 | 677.635287426784 | 792.426504233642 | 806.379565128477 | 664.052099534778 | 688.728705170212 | 572.402484610566 | 779.767489826546 | 743.585011797928 | Tbc1d15 |
| ENSMUSG00000020131 | 0.432915383655747 | 0.171420307498305 | 0.799531027639052 | 39.754603529038 | 30.1333819698302 | 36.9117406572894 | 61.9031618210386 | 27.3739548954774 | 25.6847268735511 | 32.2734208072402 | 39.0789203280663 | Pcsk4 |
| ENSMUSG00000020132 | 0.0797615063241126 | 0.668357754823692 | 0.942024383071524 | 1726.61471236345 | 1963.55632241245 | 1802.99656287529 | 1592.59952685036 | 1542.79609790911 | 1190.05901180787 | 1909.33721807995 | 2061.4130473055 | Rab21 |
| ENSMUSG00000020133 | -0.0570117092257847 | 0.682229081557047 | 0.944452327903597 | 649.626362213143 | 536.699965354543 | 585.855576073389 | 603.086864407998 | 695.298454345127 | 689.818378889657 | 533.031982364742 | 553.618037980939 | 2310011J03R |
| ENSMUSG00000020134 | -0.148778149053777 | 0.398634620062143 | 0.880459952732461 | 284.606820719249 | 315.99330281876 | 269.739643264807 | 198.84045918273 | 324.107625962453 | 293.539735697726 | 312.323427166841 | 256.184033261768 | Peli1 |
| ENSMUSG00000020135 | -0.325797013508825 | 0.161636394052812 | 0.792786631432503 | 78.6056933415069 | 93.6578088251478 | 61.519567762149 | 83.4754757889764 | 106.210944994452 | 128.423634367755 | 89.5327157878278 | 74.901263962127 | Apc2 |
| ENSMUSG00000020137 | 0.436268139060889 | 0.0830513098102144 | 0.741377052226143 | 97.5794813894569 | 127.86326943955 | 98.4313084194385 | 91.9168160372998 | 62.4126171616885 | 53.8156182112498 | 85.3684034256032 | 105.295979772845 | Thap2 |
| ENSMUSG00000020142 | -0.487132099637838 | 0.167095185811913 | 0.797814084166983 | 36.1405486627618 | 22.8036404096012 | 22.7149173275627 | 21.5723139679377 | 27.3739548954774 | 57.4848649074714 | 31.2323427166841 | 29.3091902460497 | Slc1a4 |
| ENSMUSG00000020143 | -0.416090931149203 | 0.170967196564992 | 0.799531027639052 | 34.3335212296237 | 29.3189662409158 | 23.6613722162112 | 18.7585338851632 | 32.8487458745729 | 35.4693847301419 | 29.1501865355718 | 44.5065481514088 | Dock2 |
| ENSMUSG00000020149 | -0.0388860117140756 | 0.603833421603869 | 0.926448151609826 | 3572.493235314 | 3330.9603312596 | 3300.28819671713 | 3259.29526254711 | 3432.69394389287 | 3277.86038195794 | 3585.47294387533 | 3533.38571299599 | Rab1 |
| ENSMUSG00000020150 | -0.174601736699708 | 0.524201688159125 | 0.907437243979788 | 121.070838020252 | 87.9568987227475 | 85.1809399783602 | 116.302910088012 | 113.875652365186 | 171.23151249034 | 74.9576225200418 | 104.210454208177 | Gamt |
| ENSMUSG00000020151 | -0.274979129287583 | 0.465519940815646 | 0.895328112556867 | 58.7283915769879 | 53.7514381083457 | 55.8408384302584 | 53.4618215727152 | 94.1664048404423 | 97.8465785659088 | 27.0680303544596 | 49.9341759747513 | Ptprr |
| ENSMUSG00000020152 | 0.0497765594553582 | 0.548869622609764 | 0.914699131434573 | 2013.02856051583 | 2175.30441193017 | 1960.10807439093 | 1893.67399570723 | 1959.97517051618 | 1936.13917337292 | 1863.52978209548 | 2011.47887133074 | Actr2 |
| ENSMUSG00000020153 | -0.0621081891726189 | 0.721762483356006 | 0.951674101666722 | 15553.0851170195 | 12029.7347317935 | 12971.164248927 | 16045.1119586744 | 14968.0785368471 | 18807.3354825997 | 12992.6545701406 | 12321.8006845522 | Ndufs7 |
| ENSMUSG00000020154 | -0.216607056172117 | 0.26831925241865 | 0.843793744782845 | 2435.87297987015 | 2515.73018661636 | 2823.27493283832 | 2489.25744656116 | 4087.47894499269 | 3115.19044509212 | 2406.97254536579 | 2317.59708056726 | Ptprb |
| ENSMUSG00000020155 | -0.582007882198413 | 0.254727406562356 | 0.837406774112203 | 30.7194663633475 | 30.1333819698302 | 15.1432782183751 | 14.0689004138724 | 41.6084114411257 | 61.154111603693 | 16.6572494488982 | 16.2828834700276 | Kcnmb1 |
| ENSMUSG00000020156 | -0.405614615726699 | 0.0562616198211525 | 0.703396849069134 | 190.641394196068 | 184.057954734638 | 186.451613063744 | 233.543746870282 | 329.582416941548 | 314.332133642982 | 199.886993386778 | 209.506433981022 | Mum1 |
| ENSMUSG00000020160 | -0.0575918973669961 | 0.726940479961167 | 0.95219229341535 | 309.905204783182 | 273.643684915214 | 262.16800415562 | 296.384835385579 | 234.321053905287 | 272.747337752471 | 325.857442344071 | 354.966859646602 | Meis1 |
| ENSMUSG00000020163 | 0.044643551597874 | 0.83382434061757 | 0.96835936700622 | 13121.7297057322 | 8868.17287214812 | 9542.15818735364 | 11842.2624417035 | 10642.9936633616 | 14325.9621842811 | 8576.40131000145 | 8508.34937587176 | Uqcr11 |
| ENSMUSG00000020166 | 0.0119403562121241 | 0.898774193373528 | 0.984540500469898 | 659.565013095403 | 725.644414462667 | 662.518422053913 | 697.817460528072 | 685.443830582755 | 627.44118505389 | 703.768789215948 | 705.59161703453 | Cnot2 |
| ENSMUSG00000020167 | -0.312274376254927 | 0.12291940863264 | 0.770251097497048 | 352.370349461928 | 314.364471360931 | 374.796135904785 | 419.253232333398 | 502.585811880965 | 590.748718091674 | 372.705956419097 | 348.453706258591 | Tcf3 |

| | | | | | | | | | | | | |
|---|---|---|---|---|---|---|---|---|---|---|---|---|
| **ENSMUSG00000020169** | 0.698891143781181 | 0.15665730911343 | 0.787538915521367 | 78.6056933415069 | 171.027303072009 | 194.023252172932 | 64.7169419038131 | 40.5134532453066 | 34.2463024980681 | 119.723980413956 | 118.322286548867 | Best3 |
| **ENSMUSG00000020170** | 0.107361086750549 | 0.698596396754248 | 0.945023758367404 | 300.870067617492 | 434.083583511337 | 446.726707442067 | 271.060814640609 | 319.727793179176 | 195.693157131818 | 437.252798033577 | 395.131305539337 | Frs2 |
| **ENSMUSG00000020171** | 0.0433234613169806 | 0.698076411607727 | 0.944933386973863 | 590.897970636155 | 518.782819318427 | 478.906173656114 | 585.266257217093 | 520.105143014071 | 523.479195327612 | 542.401685179747 | 523.223322170221 | Yeats4 |
| **ENSMUSG00000020173** | 0.230575187505887 | 0.221678143084201 | 0.831293036565752 | 234.010052591383 | 281.787842204358 | 173.201244622666 | 228.854113398991 | 163.148771177045 | 217.708637309147 | 208.215618111227 | 194.309076075663 | Cobl |
| **ENSMUSG00000020176** | 0.00245393088405536 | 0.987307080136678 | 0.998437672973644 | 1261.30514833039 | 1358.4454358291 | 1176.44342659002 | 1009.20912302178 | 1025.97582948249 | 1068.97387083255 | 1314.8816283724 | 1387.30167164635 | Grb10 |
| **ENSMUSG00000020178** | -0.537926853309797 | 0.164694336982093 | 0.795216960550936 | 247.562758339918 | 210.933673788811 | 444.83379766477 | 317.957149353517 | 626.316088008523 | 666.579816480253 | 239.447960827911 | 240.986675356409 | Adora2a |
| **ENSMUSG00000020180** | -0.0456663737975173 | 0.7824889037611 | 0.960742105865155 | 1597.41225089407 | 1245.24164951001 | 1194.42606947434 | 1498.80685742454 | 1518.70701760109 | 1709.86896043926 | 1311.75839410073 | 1174.53866097133 | Snrpd3 |
| **ENSMUSG00000020181** | 0.0492835339604952 | 0.880497328731443 | 0.978433072023387 | 23.4913566307952 | 38.2775392589734 | 28.3936466594534 | 36.5791410760683 | 43.7983278327639 | 23.2385624094033 | 22.903717992235 | 32.5657669400552 | Nav3 |
| **ENSMUSG00000020182** | -0.559123653635929 | 0.171347053670105 | 0.799531027639052 | 15.3597331816738 | 58.6379324818317 | 35.0188308799925 | 31.8895076047775 | 35.0386622662111 | 74.6080161565054 | 47.8895921655823 | 51.0197015394199 | Ddc |
| **ENSMUSG00000020183** | -0.659687593322276 | 0.0439459015198072 | 0.691603097094229 | 108.421645988285 | 96.9154717408051 | 90.8596693102509 | 75.0341355406529 | 181.76306050597 | 221.377884005369 | 107.231043327282 | 77.072315091464 | Cpm |
| **ENSMUSG00000020184** | 0.0883057562659958 | 0.618340273378811 | 0.929568738467614 | 1611.86847035918 | 1790.90018788261 | 1571.11511515642 | 1372.18675369969 | 1347.89353905331 | 1125.23565350795 | 1705.28591233095 | 1790.03165613837 | Mdm2 |
| **ENSMUSG00000020185** | -0.469415354445537 | 0.377391994385146 | 0.87422797717331 | 28.0089252136404 | 6.51532583131463 | 11.3574586637814 | 12.1930470253561 | 32.8487458745729 | 12.2308223207386 | 19.7804837205666 | 15.1973579053591 | E2f7 |
| **ENSMUSG00000020186** | -0.382967938445244 | 0.249571535172076 | 0.835314899646652 | 181.606257030378 | 206.047179415325 | 103.163582862681 | 140.689004138724 | 241.98576127602 | 309.439804714686 | 156.16171358342 | 117.236760984199 | Csrp2 |
| **ENSMUSG00000020189** | 0.214739394423762 | 0.351158603200932 | 0.862222555853093 | 1501.63979693775 | 1909.8048843041 | 1480.25544584617 | 1313.09737196143 | 1172.70022772225 | 875.726878164883 | 1555.37066729087 | 1742.26853129295 | Osbpl8 |
| **ENSMUSG00000020190** | 0.450929096223128 | 0.0106863036996619 | 0.635862253526407 | 8978.2158015466 | 9930.17098265241 | 11928.1709616364 | 10947.4803753813 | 6682.52986908395 | 5950.29505903933 | 9086.52957437396 | 8847.03335204834 | Mknk2 |
| **ENSMUSG00000020193** | 0.833781701834297 | 0.0168776034507167 | 0.642273034804458 | 43.3686583953142 | 48.8649437348597 | 40.6975602118832 | 45.9584080186499 | 27.3739548954774 | 11.0077400886647 | 39.5609674411332 | 21.7105112933701 | Zpbp |
| **ENSMUSG00000020196** | -0.0442947824682974 | 0.765917084941647 | 0.958345041873658 | 271.054114970713 | 357.52850499339 | 369.117406572894 | 333.901903155905 | 365.716037403578 | 297.208982393948 | 370.623800237985 | 338.683976176574 | Cabin1 |
| **ENSMUSG00000020198** | 0.0488867294101996 | 0.709123820189283 | 0.949280056132338 | 1530.55223586796 | 1743.66407560558 | 1835.17602908934 | 1576.65477304797 | 1571.2650110004 | 1353.95203090576 | 1835.42067365047 | 1701.01855983555 | Ap3d1 |
| **ENSMUSG00000020205** | 0.0219162759403054 | 0.933787858473388 | 0.98818087308472 | 221.360860559416 | 157.182235680465 | 261.221549266971 | 314.205442576484 | 301.113503850252 | 253.178022039289 | 160.326025945645 | 224.703791886381 | Phlda1 |
| **ENSMUSG00000020211** | -0.326034825512185 | 0.195506449681647 | 0.811094945724481 | 628.845546732055 | 540.772043999114 | 579.230391852849 | 588.080037299867 | 872.68168206782 | 1042.06606172693 | 540.319528998635 | 475.460197324806 | Sf3a2 |
| **ENSMUSG00000020212** | 0.410339618302417 | 0.0949962231605099 | 0.757085157577544 | 56.9213641438498 | 79.8127414336042 | 83.2880302010633 | 52.523894878457 | 45.9882442244021 | 45.2540425867328 | 58.3003730711437 | 55.3618037980939 | Mdm1 |
| **ENSMUSG00000020216** | -0.525400372208421 | 0.180362812447161 | 0.802808628271642 | 18.9737880479499 | 18.7315617650296 | 11.3574586637814 | 22.5102406621959 | 25.1840385038392 | 40.3617136584374 | 21.8626399016789 | 16.2828834700276 | Jsrp1 |
| **ENSMUSG00000020218** | -0.600832353912937 | 0.0808133703491795 | 0.739855948845396 | 92.1583990900426 | 95.2866402829764 | 35.0188308799925 | 75.0341355406529 | 141.249607260664 | 138.208292224346 | 71.8343882483734 | 100.953877514171 | Wif1 |
| **ENSMUSG00000020219** | -0.0231367859903805 | 0.90486830172132 | 0.98504921011344 | 1697.70227343324 | 1204.52086306429 | 1245.53463346136 | 1472.54490998531 | 1400.45153245262 | 1894.55437748241 | 1198.28088223011 | 1219.04520912273 | Timm13 |
| **ENSMUSG00000020220** | 0.293164200559794 | 0.222154997139377 | 0.831327997735572 | 2734.9360200545 | 3401.00008394624 | 3315.43147493551 | 2457.36793895638 | 2086.9903212312 | 1528.85279009232 | 3146.13798966064 | 2955.88611259235 | Vps13d |
| **ENSMUSG00000020224** | 0.074357293463426 | 0.724803016713631 | 0.951674101666722 | 193.351935345776 | 122.976775066064 | 165.629605513478 | 213.847286290861 | 147.819356435578 | 194.470074899744 | 171.777884941763 | 146.545951230249 | Llph |
| **ENSMUSG00000020225** | -0.0364013144591476 | 0.771181494538559 | 0.959045638244974 | 1685.05308140127 | 1506.66909849151 | 1627.90240847533 | 1838.336320746 | 1649.00704290356 | 2014.41643622565 | 1567.86360437754 | 1597.89363119204 | Tmbim4 |
| **ENSMUSG00000020227** | -0.563262919576343 | 0.0970980385977458 | 0.757809915356977 | 85.8338030740593 | 78.9983257046899 | 123.039135524298 | 86.2892558717508 | 182.858018701789 | 201.808568292187 | 93.6970281500523 | 74.901263962127 | Irak3 |
| **ENSMUSG00000020228** | -0.183665854190196 | 0.400278849058364 | 0.880487015130533 | 148.176249517323 | 114.83261777692 | 132.503684410783 | 157.571684635371 | 141.249607260664 | 110.077400886647 | 184.270822028436 | 191.052499381657 | Helb |
| **ENSMUSG00000020230** | 0.0519057404168 | 0.672250046275808 | 0.943216235131458 | 423.747933070882 | 429.197089137851 | 401.296872786941 | 521.487242007538 | 424.84377997781 | 445.201932474885 | 416.431236222455 | 426.611546914723 | Prmt2 |
| **ENSMUSG00000020231** | -0.257837244037663 | 0.101893490773911 | 0.759800953500741 | 168.957064998411 | 167.769640156352 | 164.68315062483 | 174.454365132018 | 243.08071947184 | 214.039390612925 | 181.147587756768 | 170.427513652956 | Dip2a |
| **ENSMUSG00000020232** | -0.0558361452456997 | 0.603347188839264 | 0.926448151609826 | 829.425591810383 | 690.624538119351 | 679.554610049585 | 749.403428712271 | 763.185862485911 | 812.126602097043 | 759.98700610598 | 730.558705021906 | Hmg20b |
| **ENSMUSG00000020235** | -0.120678558698663 | 0.275585012634093 | 0.846289746011912 | 789.670988281345 | 724.015583004838 | 720.252170261468 | 666.865879617553 | 799.319482947941 | 703.272283442469 | 865.13589325215 | 784.834983255331 | Fzr1 |
| **ENSMUSG00000020238** | 0.0753939426981864 | 0.456812831234972 | 0.892805623137304 | 1324.55110849022 | 1350.30127853996 | 1542.72146849697 | 1475.35869006809 | 1420.16077997737 | 1427.33696483019 | 1315.92270646296 | 1239.67019485144 | Ncln |
| **ENSMUSG00000020241** | -1.41916564568192 | 0.00347459071090153 | 0.635862253526407 | 402.063603873225 | 523.669313691913 | 358.706402797761 | 502.728708122374 | 1738.79361496073 | 2183.20178425184 | 363.336253604092 | 494.999657488839 | Col6a2 |
| **ENSMUSG00000020246** | 0.118611976660405 | 0.543790600101761 | 0.913837378403038 | 424.651446787451 | 437.341246426994 | 503.514000760974 | 370.481044231974 | 402.944616061428 | 278.86274891284 | 493.471014923609 | 422.269444656049 | Hcfc2 |
| **ENSMUSG00000020248** | 0.0113812550163709 | 0.925332033154934 | 0.986713918026868 | 276.475197270128 | 309.477976987445 | 283.936466594534 | 276.688374806158 | 286.879047304603 | 249.508775343067 | 299.830490080167 | 300.690581413177 | Nfyb |
| **ENSMUSG00000020250** | 0.0771723058552852 | 0.789506440630855 | 0.961659329450003 | 1911.8350242601 | 3017.41027562759 | 2409.67414649895 | 2175.98993067893 | 1889.89784598376 | 1212.07449198519 | 3191.94542564511 | 2724.66916731795 | Txnrd1 |
| **ENSMUSG00000020251** | -1.13463298080437 | 0.0144341872776729 | 0.635862253526407 | 9.03513716569045 | 10.5874044758863 | 5.67872933189068 | 4.68963347129081 | 22.994122112201 | 14.6769867848863 | 13.5340151772298 | 15.1973579053591 | Glt8d2 |
| **ENSMUSG00000020253** | -0.313925364542544 | 0.184847415459598 | 0.805963874499992 | 102.097049972302 | 100.987550385377 | 141.021778408619 | 124.744250336335 | 174.098353135236 | 184.685417043153 | 108.272121417838 | 116.15123541953 | Ppm1m |
| **ENSMUSG00000020255** | 0.0460563275770858 | 0.740308173928369 | 0.954383574136061 | 215.036264543433 | 254.097707421271 | 199.701981504822 | 250.426427366929 | 232.131137513649 | 214.039390612925 | 211.338852382896 | 233.387996403729 | D10Wsu102e |
| **ENSMUSG00000020256** | 0.306052448532755 | 0.549014940229909 | 0.914699131434573 | 62.3424464432641 | 124.605606523892 | 125.878500190243 | 48.7721881014244 | 36.1336204620302 | 26.9078091056249 | 99.9434966933891 | 129.177542195552 | Aldh1l2 |
| **ENSMUSG00000020257** | -0.0588497010530323 | 0.615074271452111 | 0.928325723184812 | 760.758549351136 | 820.931054745643 | 797.861471130641 | 767.224035903176 | 869.396807480363 | 678.810638800992 | 868.259127523818 | 859.736247217458 | Wdr82 |
| **ENSMUSG00000020258** | 0.117180172717409 | 0.719159564306331 | 0.951674101666722 | 40.658117245607 | 54.56585383726 | 38.8046504345863 | 43.1446279358754 | 38.3235368536684 | 23.2385624094033 | 37.4788112600209 | 64.046008315442 | Glyctk |
| **ENSMUSG00000020260** | -0.137099621626937 | 0.165615135580401 | 0.795216960550936 | 1024.5845545893 | 998.473683648967 | 926.579335986829 | 966.064495085906 | 1019.40608030758 | 1132.57414690039 | 1166.00746142287 | 988.91378941301 | Pofut2 |
| **ENSMUSG00000020261** | 0.0277552992515911 | 0.875126824272463 | 0.976986380159756 | 128.298947752804 | 141.708336831093 | 137.235958854025 | 133.185590584659 | 139.059690869025 | 96.6234963338349 | 148.874166949528 | 144.374900100912 | Slc36a1 |
| **ENSMUSG00000020262** | -0.0981047911583856 | 0.45209175344136 | 0.891110666112841 | 389.414411841258 | 534.2567181678 | 512.978549647458 | 486.783954319986 | 526.674892188986 | 502.686797382356 | 520.539045278068 | 509.11148982953 | Adarb1 |
| **ENSMUSG00000020263** | 0.341611322924617 | 0.131554066567481 | 0.77303252008843 | 521.327414460339 | 595.337897836374 | 671.036516051749 | 490.535661097018 | 428.128654565267 | 281.308913376988 | 544.483841360859 | 542.762782334254 | Appl2 |
| **ENSMUSG00000020264** | -0.0174926024132464 | 0.90613918597694 | 0.98504921011344 | 348.756294595651 | 296.447325324816 | 361.545767463707 | 306.702029022419 | 339.43704070392 | 267.855008824175 | 369.582722147429 | 350.624757387928 | Slc36a2 |
| **ENSMUSG00000020265** | -0.109102074243177 | 0.364751570348619 | 0.86924083181804 | 1297.44569699315 | 1285.14802022681 | 1035.4216481814 | 1076.73984500837 | 1262.48679977942 | 1379.63675777931 | 1200.36303841123 | 1223.38731138141 | Sumo3 |
| **ENSMUSG00000020267** | 0.106969598215149 | 0.485566738352084 | 0.899236955296196 | 3829.09113081961 | 2896.87674774827 | 3308.80629071497 | 4022.76759167325 | 3200.56280637922 | 3897.96307361939 | 2927.51159064386 | 3027.53079986047 | Hint1 |
| **ENSMUSG00000020268** | 0.0920545205350196 | 0.71222015475784 | 0.950233477782287 | 402.063603873225 | 412.09435883065 | 307.597838810745 | 309.515809105193 | 262.789966996583 | 225.04713070159 | 458.0743598447 | 396.216831104005 | Lyrm7 |
| **ENSMUSG00000020271** | 0.217171304679229 | 0.309275825572788 | 0.851785577030345 | 1021.87401343959 | 1403.23830091939 | 1480.25544584617 | 1096.43630558779 | 1000.79179097865 | 754.641737189571 | 1355.48367390409 | 1190.82154444135 | Fbxw11 |
| **ENSMUSG00000020272** | -0.33565692034134 | 0.103717430745065 | 0.759800953500741 | 179.79922959724 | 208.490426602068 | 207.27362061401 | 135.061443973175 | 277.024423542232 | 256.847268735511 | 188.435134390661 | 200.822229463674 | Stk10 |
| **ENSMUSG00000020273** | -0.0995169744797828 | 0.506887307059414 | 0.901698316485955 | 128.298947752804 | 145.780415475665 | 141.968233297267 | 137.87522405595 | 146.724398239759 | 125.977469903608 | 160.326025945645 | 159.572258006271 | Papolg |
| **ENSMUSG00000020275** | 0.116344787933478 | 0.628287561917185 | 0.931825800800958 | 110.228673421423 | 146.594831204579 | 160.897331070236 | 128.495957113368 | 108.400861386091 | 86.838838477244 | 167.613572579538 | 140.032797842237 | Rel |
| **ENSMUSG00000020277** | -0.186710759789008 | 0.0676150258559562 | 0.722038584319139 | 4934.08840618355 | 4698.36434010676 | 4538.2511910693 | 3922.40943538763 | 5044.47240813858 | 5039.0987961443 | 5475.02967823472 | 5034.66756893254 | Pfkl |
| **ENSMUSG00000020280** | 0.108172852165111 | 0.642905615052413 | 0.934616627390525 | 142.755167217909 | 164.511977240694 | 135.343049076728 | 120.054616865045 | 144.534481848121 | 79.5003450848009 | 157.202791673977 | 140.032797842237 | Pus10 |
| **ENSMUSG00000020282** | -0.0142251217671933 | 0.876603512853191 | 0.977469019294424 | 539.39768879172 | 508.195414842541 | 503.514000760974 | 508.356268287923 | 526.674892188986 | 508.802208542726 | 544.483841360859 | 499.341759747513 | Rhbdf1 |
| **ENSMUSG00000020283** | 0.196996241154387 | 0.120922337774986 | 0.769868040577702 | 816.776399778417 | 767.179616637298 | 848.970035117657 | 764.410255820401 | 646.025335533267 | 594.417964787896 | 789.137192641552 | 757.696844138618 | Pex13 |
| **ENSMUSG00000020284** | 0.102545043222301 | 0.504233762732783 | 0.900927059989498 | 423.747933070882 | 334.724864583789 | 401.296872786941 | 323.584709519066 | 286.879047304603 | 352.247682837272 | 366.45948787576 | 375.591845375304 | 1810043G02 |
| **ENSMUSG00000020287** | -0.0472234210381003 | 0.830359929772747 | 0.967583158900775 | 69.5705561758165 | 49.679359463774 | 61.519567762149 | 72.2203554578784 | 77.7420319031559 | 56.2617826753975 | 60.3825292522559 | 66.217059444779 | Mpg |
| **ENSMUSG00000020288** | 0.220958268174743 | 0.212306175603897 | 0.823370267313127 | 720.100432105529 | 683.294796559122 | 721.198625150116 | 755.03098887782 | 578.137927392483 | 442.755768010737 | 765.19239655876 | 682.795580176491 | Ahsa2 |
| **ENSMUSG00000020289** | 0.146841195403243 | 0.240999702560306 | 0.832048466795696 | 275.571683553559 | 298.076156782644 | 327.473391472363 | 337.653609932938 | 292.353838283699 | 258.070350967584 | 293.584021536831 | 273.552442296464 | Nprl3 |
| **ENSMUSG00000020290** | -0.113369542773004 | 0.493571754099719 | 0.900117970535229 | 684.863397159336 | 691.438953848265 | 604.784673846357 | 698.75538722233 | 720.482492848966 | 552.833168897385 | 909.902251146063 | 714.275821551878 | Xpo1 |
| **ENSMUSG00000020300** | 0.198218370682021 | 0.43026897187825 | 0.886017087288711 | 1257.69109346411 | 1522.14299734088 | 1281.49991923 | 1035.47107046101 | 1017.21616391594 | 644.564336302924 | 1375.26415762466 | 1404.67008068105 | Cpeb4 |
| **ENSMUSG00000020303** | -0.291270471059731 | 0.145052945096478 | 0.780607990324772 | 209.615182244018 | 124.605606523892 | 192.130342395635 | 214.785212985119 | 216.801722772181 | 255.624186503437 | 211.338852382896 | 222.532740757044 | Stc2 |
| **ENSMUSG00000020305** | -0.318295859243642 | 0.0525773441426734 | 0.703396849069134 | 131.009488902512 | 163.69756151178 | 129.664319744837 | 166.013024883695 | 200.377349834895 | 198.139321595965 | 180.106509666212 | 158.486732441602 | Asb3 |
| **ENSMUSG00000020307** | 0.0470819301941912 | 0.696896103887673 | 0.944933386973863 | 3250.84235221542 | 2975.06065772404 | 2965.24316613558 | 3569.74899834656 | 2948.72242134083 | 3575.06936435189 | 3019.1264626128 | 2809.3401613621 | Cdc34 |
| **ENSMUSG00000020308** | -0.364355467692584 | 0.176854344887943 | 0.799889021902745 | 388.510898124689 | 337.168111770532 | 342.616669690738 | 442.701399689852 | 542.004306930453 | 726.510845851873 | 377.911346871878 | 299.605055848508 | Tpgs1 |
| **ENSMUSG00000020309** | -0.0799390434254615 | 0.598760829158806 | 0.924848033480823 | 387.60738440812 | 315.178887089845 | 296.240380146964 | 359.225923900876 | 313.158044004262 | 326.562955963721 | 398.732908683 | 396.216831104005 | Chac2 |
| **ENSMUSG00000020311** | 0.131221542126725 | 0.264312923867635 | 0.842510961779757 | 765.276117933981 | 888.527560245533 | 911.436057768454 | 933.23706078687 | 765.375778877549 | 717.949270227356 | 870.34128370493 | 839.111261488756 | Erlec1 |
| **ENSMUSG00000020312** | 0.160558981376609 | 0.514385607364775 | 0.90408556099417 | 51.5002818444356 | 48.8649437348597 | 52.0550188756646 | 72.2203554578784 | 60.2227007700503 | 37.9155491942896 | 51.0128264372507 | 51.0197015394199 | Shc2 |
| **ENSMUSG00000020315** | 0.210330088580121 | 0.233136365901146 | 0.832048466795696 | 24720.1352853291 | 27206.3718401121 | 25568.4788168378 | 21625.7757895104 | 21422.8571012006 | 15149.0965264668 | 24502.8139393292 | 24598.0092953884 | Sptbn1 |
| **ENSMUSG00000020319** | 0.0243226088837946 | 0.913567810086698 | 0.986234232499696 | 59.631905293557 | 53.7514381083457 | 72.8770264259304 | 62.8410885152968 | 47.0832024202212 | 74.6080161565054 | 59.3414511616998 | 64.046008315442 | Wdpcp |
| **ENSMUSG00000020321** | 0.0191180641822316 | 0.7854860131938 | 0.961141387070613 | 48210.5884024077 | 44104.6837993555 | 42851.6915384471 | 45533.527226151 | 45307.1802266026 | 43448.7732121918 | 45403.4976853342 | 44161.3510218442 | Mdh1 |
| **ENSMUSG00000020325** | -0.0767506711523357 | 0.773595839089434 | 0.959203117135841 | 103.000563688871 | 83.8848200781758 | 72.8770264259304 | 120.992543559303 | 93.0714466446232 | 146.769867848863 | 82.2451691539348 | 80.3288917854696 | Fstl3 |
| **ENSMUSG00000020326** | 0.102103312252878 | 0.690163281020829 | 0.94469184542218 | 20861.2282018627 | 21631.6961756935 | 18340.4028322296 | 16280.5315589332 | 12986.2042024145 | 11695.1123030902 | 23338.8886340875 | 23824.0295677797 | Ccng1 |
| **ENSMUSG00000020328** | 0.142853582761175 | 0.366950748386115 | 0.870096026116028 | 334.300075130547 | 276.901347830872 | 277.311282373995 | 336.71568323868 | 251.840385038392 | 277.639666680766 | 243.612273190136 | 336.512925047237 | Nudcd2 |
| **ENSMUSG00000020329** | -0.0473862098788636 | 0.680971789927727 | 0.944452327903597 | 620.713923282934 | 668.635313438664 | 583.962666296092 | 575.886990274511 | 664.639624862192 | 562.617826753975 | 610.071761065896 | 693.650835823176 | Polrmt |
| **ENSMUSG00000020331** | 0.169950601860909 | 0.131360217081595 | 0.77303252008843 | 643.30176619716 | 618.95595397489 | 572.60520763231 | 552.438822918057 | 592.372383938131 | 501.463715150282 | 497.635327285833 | 530.8220011229 | Hcn2 |
| **ENSMUSG00000020333** | 0.127235620738381 | 0.59938713995894 | 0.924951380294376 | 439.107666252556 | 402.321370083678 | 422.118880337207 | 306.702029022419 | 324.107625962453 | 222.600966237442 | 462.238672206925 | 427.697072479392 | Acsl6 |
| **ENSMUSG00000020334** | 0.145663080357534 | 0.608715827263238 | 0.927932112008768 | 72.2810973255236 | 61.0811796685746 | 65.3053873167428 | 67.5307219865876 | 47.0832024202212 | 36.6924669622158 | 87.4505596067155 | 68.388110574116 | Slc22a4 |
| **ENSMUSG00000020335** | 0.165001199350535 | 0.496929162556632 | 0.900927059989498 | 53.3073092775737 | 40.7207864457164 | 44.483379766477 | 45.9584080186499 | 43.7983278327639 | 45.2540425867328 | 31.2323427166841 | 44.5065481514088 | Zfp354b |
| **ENSMUSG00000020340** | 0.340284433994105 | 0.155207270008212 | 0.787275540343856 | 4632.31482484949 | 5011.91439573878 | 5402.36450440533 | 3879.26480745176 | 3209.32247194577 | 2326.30240540448 | 4551.59341191143 | 4860.98347858558 | Cyfip2 |
| **ENSMUSG00000020346** | -0.210329796578849 | 0.0459063505206788 | 0.697906909182231 | 1607.35090177633 | 1483.05104235299 | 1366.68085920836 | 1560.71001924558 | 1823.1053960388 | 1898.22362417863 | 1624.08182126757 | 1618.51861692074 | Mgat1 |
| **ENSMUSG00000020349** | 0.146711103133287 | 0.197839086836454 | 0.812921911423739 | 5716.53128473235 | 5892.29779869517 | 6962.12216089797 | 6139.66814061392 | 5888.6851771151 | 4992.6216713255 | 6110.08731347397 | 5327.75947139303 | Ppp2ca |

| | | | | | | | | | | | | |
|---|---|---|---|---|---|---|---|---|---|---|---|---|
| **ENSMUSG00000020354** | 0.236488965750544 | 0.537244969864963 | 0.912099403004057 | 243.045189757073 | 429.197089137851 | 456.191256328551 | 219.47484645641 | 173.003394939417 | 155.33144347338 | 431.006329490241 | 384.276049892652 | Sgcd |
| **ENSMUSG00000020357** | -0.60467482660404 | 0.0174265574592826 | 0.642273034804458 | 269.247087537575 | 297.26174105373 | 291.508105703722 | 242.923013812864 | 544.194223322091 | 550.387004433237 | 280.050006359601 | 300.690581413177 | Flt4 |
| **ENSMUSG00000020358** | -0.00993518390022278 | 0.917092583128598 | 0.986713918026868 | 5281.94118706264 | 4796.09422757648 | 4834.49157121627 | 5519.69859570928 | 5399.23886358397 | 5442.71593272868 | 4979.47650713 | 4752.43092211873 | Hnrnpab |
| **ENSMUSG00000020359** | 0.150877085733985 | 0.320116856932154 | 0.855228644171892 | 212.325723393726 | 215.005752433383 | 200.648436393471 | 235.419600258798 | 154.389105610493 | 209.14706168463 | 212.379930473452 | 201.907755028342 | Phykpl |
| **ENSMUSG00000020361** | 0.16205400161499 | 0.260158802365868 | 0.840316047847749 | 5767.12805286021 | 6021.78989959255 | 5403.31095929398 | 5090.12816973904 | 4564.88071836982 | 4043.50985923618 | 5653.05403171982 | 5652.33161522892 | Hspa4 |
| **ENSMUSG00000020362** | -0.113500150487497 | 0.348634846312368 | 0.861266047293794 | 402.967117589794 | 381.146561131906 | 371.010316350191 | 331.088123073131 | 445.647985698372 | 407.286383280595 | 374.788112600209 | 381.019473198646 | Cnot6 |
| **ENSMUSG00000020363** | -1.05941320855235 | 0.00777777808243474 | 0.635862253526407 | 140.948139784771 | 117.275864963663 | 130.610774633486 | 145.378637610015 | 385.425284928322 | 456.20967256355 | 157.202791673977 | 115.065709854862 | Gfpt2 |
| **ENSMUSG00000020364** | -0.0688856708675146 | 0.840236092260955 | 0.969957774070128 | 22.5878429142261 | 43.9784493613737 | 24.6078271048596 | 42.2067012416173 | 32.8487458745729 | 28.1308913376988 | 30.191264626128 | 48.8486504100828 | Zfp354a |
| **ENSMUSG00000020366** | 0.0631056144035218 | 0.752947119666952 | 0.956497786093635 | 888.153983387371 | 1113.30630142589 | 1023.11773462897 | 832.878904501247 | 815.743855885227 | 661.687487551958 | 1076.47474563505 | 1137.6307917726 | Mapk9 |
| **ENSMUSG00000020368** | 0.217372410056474 | 0.200381574651261 | 0.812921911423739 | 7940.98205492534 | 8745.19609708206 | 8853.13902841757 | 7625.34402431885 | 6674.86516171321 | 5271.48442023833 | 7853.89311515549 | 8724.36896324079 | Canx |
| **ENSMUSG00000020372** | 0.088169599818718 | 0.553758961515421 | 0.915567635607008 | 9755.23759779598 | 6810.14432518162 | 8082.72474905774 | 9361.44633539071 | 7768.72839933649 | 9136.42427359173 | 7204.26038664847 | 7884.17217618737 | Rack1 |
| **ENSMUSG00000020375** | 0.0882490214555966 | 0.403450162786391 | 0.881158207349486 | 1023.68104087273 | 1017.205245414 | 1016.49255040843 | 952.933521366292 | 903.340511550755 | 839.034411202668 | 1055.65318382392 | 972.630905942983 | Rufy1 |
| **ENSMUSG00000020376** | -0.0899788333992386 | 0.520555022500672 | 0.905962179970576 | 925.198045766702 | 919.475357944277 | 1005.13509174465 | 1063.60887128875 | 1167.22543674316 | 1187.61284734372 | 927.600578685518 | 883.617809640165 | Rnf130 |
| **ENSMUSG00000020377** | -0.385183650168695 | 0.437917102561849 | 0.888759286175347 | 20.780815481088 | 10.5874044758863 | 15.1432782183751 | 15.0068271081306 | 26.2789966996583 | 34.2463024980681 | 10.4107809055614 | 9.76973008201657 | Ltc4s |
| **ENSMUSG00000020380** | 0.124732702530216 | 0.425645769459154 | 0.885066977635295 | 294.545471601509 | 320.879797192245 | 303.812019256151 | 250.426427366929 | 264.979883388221 | 215.262472844999 | 303.994802442392 | 287.664274637155 | Rad50 |
| **ENSMUSG00000020381** | -0.367363669478563 | 0.322852223998243 | 0.855977678491063 | 28.0089252136404 | 21.9892246806869 | 14.1968233297267 | 30.0136542162612 | 41.6084114411257 | 35.4693847301419 | 18.7394056300105 | 26.0526135520442 | Mrnip |
| **ENSMUSG00000020385** | 0.292158632023855 | 0.0148094070135872 | 0.635862253526407 | 357.791431761342 | 314.364471360931 | 330.312756138308 | 339.529463321454 | 250.745426842573 | 259.293433199658 | 283.173240631269 | 301.776106977845 | Clk4 |
| **ENSMUSG00000020386** | 0.0755867565640852 | 0.566528879685873 | 0.919931558693298 | 3849.8719463007 | 3311.41435376566 | 3473.4894413398 | 3397.17048660306 | 3035.22411881054 | 2827.76612055476 | 3611.49989613924 | 3839.50392223251 | Sar1b |
| **ENSMUSG00000020387** | -0.00103547326861479 | 0.995562393212618 | 0.999262355603273 | 495.125516679837 | 552.173864203915 | 575.444572298256 | 348.908730264036 | 568.283303630111 | 454.986590331476 | 507.005030100839 | 442.894430384751 | Jade2 |
| **ENSMUSG00000020388** | -0.349815903913574 | 0.131243665003384 | 0.77303252008843 | 212.325723393726 | 162.883145782866 | 185.505158175096 | 171.640585049243 | 234.321053905287 | 336.347613820311 | 173.860041122875 | 189.966973816989 | Pdlim4 |
| **ENSMUSG00000020389** | -0.0252695716512901 | 0.898871532680183 | 0.984569481028921 | 154.500845533307 | 149.852494120236 | 158.057966404291 | 156.633757941113 | 140.154649064844 | 111.300483118721 | 193.640524843441 | 183.453820428978 | Cdkl3 |
| **ENSMUSG00000020390** | 0.192871385331172 | 0.162787610888849 | 0.793968226938511 | 7454.89167541119 | 7280.06220076518 | 8382.75094875929 | 7859.82569788339 | 6091.25244334164 | 5653.08607664538 | 7834.11263143493 | 7520.52111202342 | Ube2b |
| **ENSMUSG00000020392** | 0.0307337290203468 | 0.822628703778739 | 0.966637987288264 | 277.378710986697 | 252.468875963442 | 267.84673348751 | 287.943495137256 | 271.549632563136 | 305.770558018465 | 224.872867560126 | 261.61166108511 | Cdkn2aipnl |
| **ENSMUSG00000020393** | 0.183515246121347 | 0.389166099732849 | 0.876536136761342 | 3937.5127768079 | 4879.97904765466 | 4515.53627374174 | 3998.38149762254 | 3324.29308250678 | 2545.2341249457 | 4836.84880872381 | 4553.77974378439 | Kremen1 |
| **ENSMUSG00000020397** | 0.0599822369931235 | 0.657176517528298 | 0.938389079274206 | 236.72059374109 | 237.809392842984 | 243.238906382651 | 284.191788360223 | 229.94122112201 | 247.06261087892 | 266.515991182371 | 217.105112933702 | Med7 |
| **ENSMUSG00000020400** | 0.12035541165521 | 0.421064850698599 | 0.882568531013176 | 1009.22482140762 | 904.815874823819 | 839.505486231172 | 1238.06323642077 | 835.453103409971 | 978.465785659088 | 948.422140496641 | 909.670423192209 | Tnip1 |
| **ENSMUSG00000020402** | 0.164361259507679 | 0.246299004046774 | 0.832123990973914 | 26779.2430453899 | 28164.1247373153 | 28528.0432536415 | 26518.9393534553 | 22076.5471441046 | 19886.0940112889 | 27886.3177336367 | 28296.394894214 | Vdac1 |
| **ENSMUSG00000020407** | 0.0817612942758783 | 0.765220179552954 | 0.958197631858494 | 126.491920319666 | 60.2667639396603 | 87.0738497556571 | 128.495957113368 | 107.305903190271 | 102.738907494204 | 91.61487196894 | 78.1578406561325 | Upp1 |
| **ENSMUSG00000020409** | 0.064943721635923 | 0.453058948145712 | 0.891468041670908 | 983.92643734369 | 1020.46290832965 | 1095.04830616625 | 1009.20912302178 | 956.993463145891 | 1049.40455511937 | 969.243702307763 | 953.09144577895 | Slu7 |
| **ENSMUSG00000020412** | 0.163269283202092 | 0.289126474495285 | 0.851785577030345 | 529.45903790946 | 495.164763179912 | 540.425741418263 | 512.107975064956 | 425.938738173629 | 362.032340693862 | 528.867670002517 | 536.249628946243 | Ascc2 |
| **ENSMUSG00000020413** | 0.011539022048892 | 0.94600392649563 | 0.990505437194639 | 175.281661014395 | 162.883145782866 | 170.36187995672 | 131.309737196143 | 156.579022002131 | 129.646716599829 | 169.69572876065 | 178.026192605635 | Hus1 |
| **ENSMUSG00000020415** | 0.0977135506339034 | 0.509610825520703 | 0.902394714365738 | 1498.92925578805 | 1268.85970564852 | 1439.55788563429 | 1554.14453238577 | 1294.24058745817 | 1681.73806910156 | 1231.59538112791 | 1177.79523766533 | Pttg1 |
| **ENSMUSG00000020420** | -0.19785942862393 | 0.501137463540665 | 0.900927059989498 | 36.1405486627618 | 32.5766291565731 | 31.2330113253987 | 38.4549944645846 | 29.5638712871156 | 30.5770558018465 | 40.6020455316893 | 57.5328549274309 | Zfp607a |
| **ENSMUSG00000020422** | 0.138160943524023 | 0.430177550346868 | 0.886017087288711 | 454.46739943423 | 596.966729294203 | 463.762895437739 | 370.481044231974 | 388.710159515779 | 388.940149799487 | 471.60837502193 | 464.604941678121 | Tns3 |
| **ENSMUSG00000020423** | -0.0643991840599672 | 0.761434180975969 | 0.957479700684068 | 529.45903790946 | 702.840774053066 | 566.92647830042 | 460.522006880757 | 754.426196919358 | 618.879609429373 | 572.592949805875 | 417.927342397375 | Btg2 |
| **ENSMUSG00000020424** | -0.782284556912513 | 0.0926112467727111 | 0.754058637722175 | 82.2197482077831 | 76.5550785179469 | 53.9479286529615 | 66.5927952923295 | 145.62944004394 | 231.162541861959 | 43.7252798033577 | 60.7894316214364 | Gatsl3 |
| **ENSMUSG00000020427** | -0.922359334848642 | 0.0121071729054552 | 0.635862253526407 | 309.905204783182 | 237.809392842984 | 201.594891282119 | 231.667893481766 | 669.019457645468 | 698.379954514174 | 234.242570375131 | 258.355084391105 | Igfbp3 |
| **ENSMUSG00000020430** | 0.0964736840393928 | 0.363026136948548 | 0.86924083181804 | 1067.04969926804 | 1114.1207171548 | 1087.47666705707 | 1245.56664997484 | 1173.79518591807 | 1000.48126583642 | 990.065264118886 | 1057.30189998713 | Pes1 |
| **ENSMUSG00000020431** | -0.39639647644646 | 0.244681515644835 | 0.832048466795696 | 30.7194663633475 | 17.9171460361152 | 22.7149173275627 | 23.448167356454 | 19.7092475247437 | 26.9078091056249 | 37.4788112600209 | 40.1644458927348 | Adcy1 |
| **ENSMUSG00000020432** | -0.471087208084826 | 0.0871269502730187 | 0.747269672026173 | 954.110484696911 | 804.642740167357 | 908.596693102509 | 952.933521366292 | 1661.05158305757 | 1731.88444061659 | 840.150019078802 | 785.920508819999 | Tcn2 |
| **ENSMUSG00000020434** | 0.168870545192118 | 0.643497108760905 | 0.934616627390525 | 18.0702743313809 | 29.3189662409158 | 18.9290977729689 | 18.7585338851632 | 14.2344565456483 | 15.9000690169602 | 26.0269522639034 | 19.5394601640331 | 4921536K21I |
| **ENSMUSG00000020435** | 0.263785089336989 | 0.155591067997635 | 0.787275540343856 | 668.600150261093 | 955.309650016507 | 842.344850897118 | 658.424539369229 | 652.595084708182 | 499.017550686135 | 735.001131932632 | 715.361347116547 | Osbp2 |
| **ENSMUSG00000020437** | -0.228052398376751 | 0.595398257547932 | 0.924461858177432 | 10.8421645988285 | 18.7315617650296 | 16.0897331070236 | 24.3860940507122 | 21.8991639163819 | 33.0232202659942 | 13.5340151772298 | 14.1118323406906 | Myo1g |
| **ENSMUSG00000020439** | -0.423397671010169 | 0.0224755194146025 | 0.650119841310877 | 1282.08596381147 | 1193.11904285949 | 814.897659126313 | 952.933521366292 | 1550.46080527984 | 1653.60717776386 | 1245.12939630514 | 1242.92677154544 | Smtn |
| **ENSMUSG00000020440** | -0.194473383044366 | 0.307291293715589 | 0.851785577030345 | 2546.10165329157 | 1946.45359210525 | 1864.51613063744 | 2468.62305928748 | 2595.05092409126 | 3276.63729972587 | 2095.6901962895 | 2133.05773457362 | Arf5 |
| **ENSMUSG00000020441** | -0.113759298791198 | 0.597792221720611 | 0.924496535139492 | 242.141676040504 | 216.634583891211 | 181.719338620502 | 251.364354061187 | 243.08071947184 | 331.455284892016 | 205.092383839559 | 186.710397122983 | 2310033P09I |
| **ENSMUSG00000020444** | 0.136419390266142 | 0.428486803901848 | 0.885378691567738 | 3118.92934959634 | 1995.31853584011 | 2156.02423634116 | 2878.4970246783 | 2281.892880087 | 2524.44172700045 | 2084.23833729339 | 2342.56416855464 | Guk1 |
| **ENSMUSG00000020448** | -0.121692954117515 | 0.285554912634412 | 0.851332418221073 | 376.765219809292 | 406.39344872825 | 433.476339000989 | 371.418970926232 | 408.419407040523 | 403.617136584374 | 464.320828388037 | 450.493109337431 | Rnf185 |
| **ENSMUSG00000020451** | 0.139023943351441 | 0.325961937231602 | 0.855977678491063 | 566.503100288791 | 495.164763179912 | 533.800557197724 | 465.211640352048 | 434.698403740182 | 406.063301048521 | 558.017856538089 | 471.118095066132 | Limk2 |
| **ENSMUSG00000020453** | -0.0757365369662188 | 0.687861522667649 | 0.944452327903597 | 250.273299489625 | 288.303168035672 | 375.742590793433 | 318.895076047775 | 353.671497249568 | 276.416584448692 | 391.445362049107 | 276.809018990469 | Patz1 |
| **ENSMUSG00000020454** | 0.213700289243731 | 0.36212420847824 | 0.86924083181804 | 1159.20809835808 | 1215.92268326909 | 1167.92533259218 | 958.561081531841 | 812.45898129777 | 612.764198269004 | 1243.04724012403 | 1212.53205573472 | Eif4enif1 |
| **ENSMUSG00000020455** | -0.0827903214373631 | 0.545711504505818 | 0.914211323944814 | 234.010052591383 | 217.448999620126 | 213.898804834549 | 234.48167356454 | 240.890803080201 | 278.86274891284 | 221.749633288457 | 212.763010675027 | Trim11 |
| **ENSMUSG00000020456** | 0.205428189516124 | 0.26677252637566 | 0.843513595393324 | 66176.0551426666 | 77358.0924266564 | 74708.4166354652 | 63182.4938320068 | 57899.1994785222 | 42780.9703134795 | 70817.2549539001 | 72576.0682026071 | Ogdh |
| **ENSMUSG00000020457** | 0.105033981961148 | 0.298387635316482 | 0.851785577030345 | 1338.10381423876 | 1105.16214413674 | 1266.35664101162 | 1352.49029312027 | 1154.08593839333 | 1144.80496922113 | 1210.77381931679 | 1195.16364670003 | Drg1 |
| **ENSMUSG00000020458** | -0.144114059550156 | 0.397266384690826 | 0.8799282544021 | 1910.93151054353 | 2409.8561418575 | 1966.73325861147 | 2146.91420315693 | 2303.79204400338 | 2978.20523509985 | 2207.08555197901 | 1833.45267872511 | Rtn4 |
| **ENSMUSG00000020459** | -0.0132758801377958 | 0.940059286613616 | 0.989269716906553 | 894.478579403355 | 920.289773673191 | 845.184215563063 | 731.582821521366 | 747.856447744443 | 664.133652016106 | 1053.57102764281 | 956.348022472955 | Mtif2 |
| **ENSMUSG00000020460** | 0.0564715590362474 | 0.756600915957029 | 0.956566265296272 | 4776.87701950054 | 3231.60161233206 | 3790.55182903703 | 4848.14308262044 | 3968.12850164841 | 4942.47529981047 | 3487.61160336306 | 3610.45802808746 | Rps27a |
| **ENSMUSG00000020462** | 0.153274502321484 | 0.221960963013816 | 0.831327997735572 | 458.081454300506 | 395.806044252364 | 414.54724122802 | 472.715053906113 | 350.386622662111 | 424.409534529629 | 420.595548584679 | 370.164217551961 | Cfap36 |
| **ENSMUSG00000020463** | 0.058018462495329 | 0.783578484528142 | 0.961051227381536 | 525.844983043184 | 611.626212414661 | 668.197151385803 | 493.349441179793 | 459.882442244021 | 403.617136584374 | 714.17957012151 | 629.604827507734 | Ppp4r3b |
| **ENSMUSG00000020464** | 0.0496797904534503 | 0.789330968336243 | 0.961659329450003 | 440.011179969125 | 414.537606017393 | 403.189782564238 | 355.474217123843 | 342.721915291377 | 295.985900161874 | 417.472314313011 | 501.51281087685 | Pnpt1 |
| **ENSMUSG00000020467** | -1.02808259303955 | 0.00853400068431455 | 0.635862253526407 | 58.7283915769879 | 57.009101024003 | 64.3589324280944 | 75.0341355406529 | 160.958854785407 | 222.600966237442 | 62.4646854333682 | 74.901263962127 | Efemp1 |
| **ENSMUSG00000020469** | -5.60404746133055 | NA | NA | 70.4740698923855 | 2.44324718674299 | 0.946454888648447 | 40.3308478531009 | 4.37983278327639 | 5536.89326459836 | 7.28754663389296 | 5.42762782334254 | Myl7 |
| **ENSMUSG00000020471** | 0.0797655560401775 | 0.62014823859776 | 0.929720349068849 | 586.38040205331 | 651.532583131463 | 830.040937344688 | 725.017334661559 | 747.856447744443 | 617.656527197299 | 721.467116755403 | 554.703563545607 | Pold2 |
| **ENSMUSG00000020472** | 0.0830426433248131 | 0.546893496668897 | 0.914451747770672 | 426.458474220589 | 464.216965481167 | 498.781726317731 | 498.039074651084 | 494.921104510232 | 501.463715150282 | 393.52751823022 | 392.96025441 | Zkscan17 |
| **ENSMUSG00000020473** | -1.29486371028735 | 0.0223260083824493 | 0.648762042745232 | 100.290022539164 | 109.13170767452 | 91.8061241988993 | 94.7305961200743 | 347.101748074654 | 491.679057293692 | 79.1219348822664 | 54.2762782334254 | Aebp1 |
| **ENSMUSG00000020474** | -0.0557226225450596 | 0.909938142636968 | 0.985944245354498 | 9.93865088225949 | 11.4018202048006 | 18.9290977729689 | 39.3929211588428 | 16.4243729372865 | 25.6847268735511 | 16.6572494488982 | 23.8815624227072 | Polm |
| **ENSMUSG00000020475** | -0.0760140912674362 | 0.678921943051322 | 0.944452327903597 | 19251.1667589366 | 14310.9131884826 | 12823.5172862978 | 17806.5382904912 | 18173.0211760096 | 20443.8195091146 | 14710.4334195582 | 14338.7071837063 | Pgam2 |
| **ENSMUSG00000020476** | -0.121776006263975 | 0.301241713668113 | 0.851785577030345 | 526.748496759753 | 522.854897962999 | 524.33600831124 | 606.83857118503 | 675.589206820383 | 554.056251129458 | 578.839418349212 | 563.387768062955 | Dbnl |
| **ENSMUSG00000020477** | 0.0599990986771005 | 0.69804306943664 | 0.944933386973863 | 1902.79988709441 | 1505.85468276259 | 1563.54347604723 | 2130.96944935454 | 1713.60957645689 | 1965.49314694269 | 1573.06899483032 | 1562.07128755798 | Mrps24 |
| **ENSMUSG00000020482** | 0.171929647896714 | 0.364359086527008 | 0.86924083181804 | 767.083145367119 | 827.446380576958 | 723.091534927413 | 607.776497879289 | 650.405168316544 | 456.20967256355 | 766.233474649317 | 722.960026069226 | Ccdc117 |
| **ENSMUSG00000020483** | 0.14576616872165 | 0.348465463418508 | 0.861266047293794 | 25073.4091485076 | 24614.9009907067 | 21480.7401527651 | 22115.3735239132 | 18174.1161342054 | 17257.6902945622 | 24759.9602276966 | 24126.8912003223 | Dynll2 |
| **ENSMUSG00000020484** | -0.167708961287649 | 0.306975529652269 | 0.851785577030345 | 1135.71674172729 | 1158.91358224509 | 1383.71704720403 | 1249.31835675187 | 1538.41626512583 | 1681.73806910156 | 1183.70578896233 | 1132.20316394925 | Xbp1 |
| **ENSMUSG00000020485** | 0.0675607775645483 | 0.720823804830742 | 0.951674101666722 | 1082.40943244972 | 775.323773926441 | 972.955625530603 | 1100.18801236482 | 1017.21616391594 | 1169.26661386261 | 740.206522385413 | 824.999429148066 | Supt4a |
| **ENSMUSG00000020486** | -0.569035884327822 | 0.0281077429601186 | 0.665956123973759 | 255.69438178904 | 225.593156909269 | 212.952349945901 | 384.549944645846 | 448.93286028583 | 527.148442023833 | 291.501865355718 | 333.256348353232 | Sept4 |
| **ENSMUSG00000020491** | 0.140619470257182 | 0.76686001654674 | 0.958655025595455 | 19.877301764519 | 39.0919549878878 | 31.2330113253987 | 32.8274342990356 | 10.949581958191 | 17.123151249034 | 31.2323427166841 | 52.1052271040884 | 2810021J22F |
| **ENSMUSG00000020492** | -0.0152246833907934 | 0.942483483888562 | 0.990324199350338 | 202.387072511466 | 193.016527752696 | 205.380710836713 | 207.281799431054 | 163.148771177045 | 147.992950080937 | 240.489038918468 | 263.782712214447 | Ska2 |
| **ENSMUSG00000020495** | -0.121802623556653 | 0.500434512168103 | 0.900927059989498 | 148.176249517323 | 181.614707547895 | 180.772883731853 | 151.006197775564 | 208.042057205628 | 177.34692365071 | 197.804837205666 | 136.776221148232 | Smg8 |
| **ENSMUSG00000020496** | -0.0532601658799699 | 0.509067417203844 | 0.902394714365738 | 4219.40905637744 | 4328.61959917966 | 4227.81398759261 | 4248.80792498947 | 4241.86805060318 | 4825.05940553138 | 4459.97853994249 | 4140.19450364569 | Rnf187 |
| **ENSMUSG00000020513** | -0.0728677995098789 | 0.678376953224602 | 0.944452327903597 | 108.421645988285 | 92.8433930962335 | 88.966759532954 | 108.799496533947 | 122.635317931739 | 97.8465785659088 | 94.7381062406084 | 104.210454208177 | Tubd1 |
| **ENSMUSG00000020514** | 0.0141116514643327 | 0.919796948307966 | 0.986713918026868 | 663.179067961679 | 530.184639523228 | 523.389553422591 | 597.459304242449 | 615.366506050332 | 639.672007374629 | 542.401685179747 | 494.999657488839 | Mrpl22 |
| **ENSMUSG00000020515** | -0.0368315706420705 | 0.722048411851851 | 0.951674101666722 | 898.9961479862 | 886.08431305879 | 812.058294460367 | 911.664746818933 | 997.506916391197 | 892.850029413917 | 895.327157878277 | 814.144173501381 | Cnot8 |
| **ENSMUSG00000020516** | 0.215966287752578 | 0.0366912343655582 | 0.679566075569698 | 1035.42671918813 | 994.401605004395 | 997.563452635463 | 962.312788308873 | 819.028730472685 | 764.426395046162 | 940.093515772191 | 909.670423192209 | Rps6kb1 |
| **ENSMUSG00000020519** | 0.00853484967806842 | 0.96573673733501 | 0.994284908685797 | 793.285043147622 | 867.35275129376 | 890.614050218188 | 900.409626487835 | 793.844691968845 | 1191.28209403994 | 826.616003901573 | 620.920622990386 | Sap30l |
| **ENSMUSG00000020520** | -0.54257399213819 | 0.0399392086005967 | 0.679566075569698 | 266.536546387868 | 215.005752433383 | 224.309808609682 | 227.916186704733 | 475.211856985488 | 420.740287833408 | 223.831789469569 | 240.986675356409 | Galnt10 |
| **ENSMUSG00000020521** | 0.18284891854431 | 0.147390039751324 | 0.780607990324772 | 658.661499378834 | 658.047908962778 | 724.037989816062 | 698.75538722233 | 632.885837183438 | 489.232892829544 | 648.591650416473 | 640.46008315442 | Rnft1 |
| **ENSMUSG00000020522** | -0.325524028418573 | 0.0184004887114203 | 0.642273034804458 | 208.711668527449 | 181.614707547895 | 161.843785958884 | 173.51643843776 | 243.08071947184 | 229.939459629886 | 213.421008564008 | 223.618266321713 | Mfap3 |
| **ENSMUSG00000020523** | 0.183370426822697 | 0.0788520554802108 | 0.739855948845396 | 1295.63866956001 | 1127.96578454635 | 1309.89356588945 | 1437.84162229776 | 1157.37081298079 | 1110.55866672306 | 1157.67883669842 | 1126.77553612591 | Fam114a2 |

| | | | | | | | | | | | | |
|---|---|---|---|---|---|---|---|---|---|---|---|---|
| **ENSMUSG00000020525** | -0.259711894143423 | 0.192154023234831 | 0.808824684419004 | 204.194099944604 | 140.893921102179 | 182.66579350915 | 184.771558768858 | 257.315176017488 | 228.716377397812 | 208.215618111227 | 158.486732441602 | Ppm1d |
| **ENSMUSG00000020526** | 0.139441340077003 | 0.362447772661885 | 0.86924083181804 | 177.088688447533 | 149.038078391322 | 156.165056626994 | 171.640585049243 | 130.300025302473 | 146.769867848863 | 144.709854587303 | 171.513039217624 | Znhit3 |
| **ENSMUSG00000020527** | 0.317298236762819 | 0.374444637260081 | 0.873700364743909 | 18.0702743313809 | 19.5459774939439 | 19.8755526616174 | 16.8826804966469 | 16.4243729372865 | 14.6769867848863 | 17.6983275394543 | 10.8552556466851 | Myo19 |
| **ENSMUSG00000020528** | -0.143301260615447 | 0.463108606395697 | 0.895328112556867 | 127.395434036235 | 122.162359337149 | 82.3415753124149 | 117.24083678227 | 135.774816281568 | 103.961989726278 | 124.929370866736 | 131.348593324889 | Prpsap2 |
| **ENSMUSG00000020530** | 0.0731836640713151 | 0.392461618086363 | 0.878537584780694 | 1800.70283712211 | 1602.7701545034 | 1667.65351379856 | 1792.37791272735 | 1688.42553795305 | 1578.99916160735 | 1690.71081906317 | 1564.24233868732 | Ggnbp2 |
| **ENSMUSG00000020532** | -0.0622434780917779 | 0.795322024528096 | 0.962032824235414 | 187.930853046361 | 297.26174105373 | 306.651383922097 | 166.950951577953 | 243.08071947184 | 203.031650524261 | 277.967850178489 | 276.809018990469 | Acaca |
| **ENSMUSG00000020534** | 0.244598493172998 | 0.209464067582344 | 0.821566820583092 | 286.413848152387 | 241.881471487556 | 241.345996605354 | 293.571055302804 | 240.890803080201 | 157.777607937528 | 279.008928269045 | 218.19063849837 | Shmt1 |
| **ENSMUSG00000020536** | -0.0812223975452311 | 0.593250766068891 | 0.923935529355527 | 179.79922959724 | 221.521078264697 | 248.917635714542 | 204.468019348279 | 248.555510450935 | 209.14706168463 | 234.242570375131 | 211.677485110359 | Llgl1 |
| **ENSMUSG00000020537** | 0.00433355492592262 | 0.975834707178406 | 0.99575536949131 | 464.406050316489 | 447.114235173966 | 481.745538322059 | 439.887619607078 | 466.452191418935 | 533.263853184203 | 468.485140750262 | 360.394487469945 | Drg2 |
| **ENSMUSG00000020538** | -0.364906223968246 | 0.0442886397177242 | 0.691978859031708 | 638.784197614315 | 568.462178782201 | 734.448993591195 | 501.790781428116 | 946.0438811877 | 669.025980944401 | 865.13589325215 | 665.427171141795 | Srebf1 |
| **ENSMUSG00000020541** | 0.168176396538272 | 0.400823630043563 | 0.880784720978499 | 327.975479114563 | 301.333819698302 | 337.884395247495 | 321.708856130549 | 245.270635863478 | 201.808568292187 | 343.555769883525 | 354.966859646602 | Tom1l1 |
| **ENSMUSG00000020542** | 0.120350452525846 | 0.674440669519663 | 0.944023556644042 | 463.50253659992 | 602.667639396603 | 386.153594568566 | 390.177504811395 | 351.48158085793 | 238.501035254403 | 571.551871715319 | 532.993052252237 | Myocd |
| **ENSMUSG00000020544** | -0.00784162427209365 | 0.950385719692421 | 0.991696503889296 | 337.010616280254 | 345.312269059675 | 306.651383922097 | 343.281170098487 | 343.816873487196 | 289.870489001505 | 376.870268781321 | 327.828720529889 | Cox11 |
| **ENSMUSG00000020546** | -0.168017354451317 | 0.462406432937277 | 0.895328112556867 | 60.535419010126 | 63.5244268553176 | 76.6628459805242 | 45.0204813243917 | 63.5075753575076 | 72.1618516923577 | 61.4236073428121 | 79.2433662208011 | Stxbp4 |
| **ENSMUSG00000020547** | 0.0679769698743517 | 0.487007498438249 | 0.899236955296196 | 2646.39167583073 | 2836.60998380861 | 3047.584741448 | 2959.1587203845 | 2628.99462816165 | 2656.53460806442 | 3055.56419578226 | 2618.28766198044 | Bzw2 |
| **ENSMUSG00000020549** | 0.237339773790908 | 0.168434641389546 | 0.79907358295209 | 693.898534325027 | 741.932729040953 | 865.05976822468 | 760.658549043369 | 575.948011000845 | 500.240632918209 | 746.45299092875 | 772.894202043977 | Elac2 |
| **ENSMUSG00000020553** | -0.312127859975353 | 0.0969220770787659 | 0.757809915356977 | 138.237598635064 | 150.666909849151 | 111.681676860517 | 103.171936368398 | 133.58489988993 | 147.992950080937 | 167.613572579538 | 176.940667040967 | Pctp |
| **ENSMUSG00000020561** | 0.113123541923636 | 0.526062995813521 | 0.90806673740897 | 231.299511441676 | 217.448999620126 | 177.933519065908 | 248.550573978413 | 182.858018701789 | 254.401104271363 | 181.147587756768 | 192.138024946326 | Twistnb |
| **ENSMUSG00000020564** | 0.0713611647058068 | 0.609734853772631 | 0.927932112008768 | 245.75573090678 | 272.014853457386 | 255.542819935081 | 223.226553233442 | 209.137015401448 | 221.377884005369 | 254.023054095697 | 263.782712214447 | Atxn7l1 |
| **ENSMUSG00000020570** | -0.0272552593399414 | 0.765591737785404 | 0.958345041873658 | 1889.24718134587 | 1846.28045744878 | 1935.50024728607 | 1874.91546182206 | 2085.89536303538 | 2026.64725854639 | 1794.81862811878 | 1783.51850275036 | Sypl |
| **ENSMUSG00000020571** | -0.21900889690405 | 0.208253056389443 | 0.820372930586885 | 2129.58182995324 | 2109.33673788811 | 1816.24693131637 | 2079.38348117034 | 2568.7719273916 | 2997.77455081303 | 1807.31156520545 | 2096.14986537489 | Pdia6 |
| **ENSMUSG00000020572** | 0.175164361789464 | 0.257243830901498 | 0.837517407767513 | 7276.89947324709 | 8302.96835628158 | 7964.41788797668 | 6600.19014749468 | 6333.23820461766 | 5240.90736443649 | 7520.74812617753 | 7601.93552937356 | Nampt |
| **ENSMUSG00000020573** | -0.141694621781897 | 0.652906554380456 | 0.936991241819063 | 19.877301764519 | 21.9892246806869 | 28.3936466594534 | 24.3860940507122 | 33.943704070392 | 26.9078091056249 | 23.9447960827911 | 19.5394601640331 | Pik3cg |
| **ENSMUSG00000020576** | 0.0634585825481776 | 0.612690614374222 | 0.928055903343287 | 672.214205127369 | 742.747144769868 | 819.629933569555 | 701.569167305105 | 731.432074807157 | 593.194882555822 | 747.494069019306 | 735.986332845248 | Nbas |
| **ENSMUSG00000020577** | -0.490904938726483 | 0.148452204982047 | 0.781357001591751 | 741.784761303186 | 549.730617017172 | 624.660226507975 | 1164.90495426864 | 1382.93220131952 | 1581.4453260715 | 654.83811895981 | 711.019244857872 | Tspan13 |
| **ENSMUSG00000020580** | 0.254742313480115 | 0.31984795437008 | 0.855057089594033 | 3105.37664384781 | 4120.12917257759 | 3298.39528693984 | 2817.53178955152 | 2181.15672607164 | 1771.02307204295 | 3775.99023444711 | 3453.05682121052 | Rock2 |
| **ENSMUSG00000020585** | -0.0746409509427918 | 0.421134086212252 | 0.882568531013176 | 10527.7418254625 | 9220.81488276803 | 8901.40822773864 | 9341.74987481129 | 10088.9448162772 | 10736.2158331443 | 9512.33051341142 | 9673.11830676107 | Laptm4a |
| **ENSMUSG00000020589** | -0.452991684323463 | 0.0893448378720373 | 0.751593644682828 | 121.070838020252 | 108.317291945606 | 88.0203046443056 | 116.302910088012 | 144.534481848121 | 227.493295165738 | 116.600746142287 | 106.381505337514 | Fam49a |
| **ENSMUSG00000020590** | 0.313875376324382 | 0.22516810366894 | 0.832048466795696 | 1995.86179990102 | 2358.5479509359 | 2304.61765385897 | 1798.0054728929 | 1376.3624521446 | 1002.92743030056 | 2218.53741097513 | 2204.70242184174 | Snx13 |
| **ENSMUSG00000020591** | 0.0217988799669867 | 0.925407288118826 | 0.986713918026868 | 72.2810973255236 | 73.2974156022896 | 60.5731128735006 | 72.2203554578784 | 83.2168228822514 | 56.2617826753975 | 83.2862472444909 | 51.0197015394199 | Ntsr2 |
| **ENSMUSG00000020592** | -0.697015451920034 | 0.028124266847549 | 0.665956123973759 | 99.3865088225949 | 98.5443031986338 | 71.930571537282 | 74.0962088463947 | 150.009272827216 | 222.600966237442 | 82.2451691539348 | 104.210454208177 | Sdc1 |
| **ENSMUSG00000020593** | -0.19436140829699 | 0.277664035523317 | 0.848075053184273 | 3511.95781630388 | 4026.47136375244 | 4060.29147230184 | 2903.82104542327 | 3926.52009020728 | 3275.4142174938 | 5178.32242242622 | 4212.92471647848 | Lpin1 |
| **ENSMUSG00000020594** | 0.126128835295363 | 0.418498500660568 | 0.882468496126562 | 3415.28184863099 | 3782.96106080706 | 3535.00900910195 | 3117.66833171413 | 2884.1198877875 | 2506.09549351934 | 3560.48706970199 | 3739.63557028301 | Pum2 |
| **ENSMUSG00000020599** | 0.251931540636103 | 0.497966728542223 | 0.900927059989498 | 34.3335212296237 | 21.9892246806869 | 27.447191770805 | 50.6480414899407 | 37.2285786578493 | 33.0232202659942 | 17.6983275394543 | 24.9670879873757 | Rgs9 |
| **ENSMUSG00000020601** | -0.282985583052505 | 0.0333627184097746 | 0.671606027118506 | 574.634723737913 | 605.92530231226 | 646.428688946889 | 686.562340196974 | 914.290093508946 | 739.964750404685 | 714.17957012151 | 689.308733564502 | Trib2 |
| **ENSMUSG00000020604** | 0.239167094783951 | 0.433865593992437 | 0.887868762931002 | 25.2983840639333 | 44.7928650902881 | 34.0723759913441 | 34.703287687552 | 26.2789966996583 | 25.6847268735511 | 39.5609674411332 | 26.0526135520442 | Arsg |
| **ENSMUSG00000020605** | -0.257459324848275 | 0.0530881265039122 | 0.703396849069134 | 178.895715880671 | 193.83094348161 | 183.612248397799 | 207.281799431054 | 257.315176017488 | 226.270212933664 | 214.462086654564 | 214.934061804364 | Hs1bp3 |
| **ENSMUSG00000020607** | -0.287860847799608 | 0.612197359291901 | 0.928055903343287 | 11.7456783153976 | 5.7009101024003 | 12.3039135524298 | 10.3171936368398 | 9.85462376237187 | 23.2385624094033 | 4.16431236222455 | 11.9407812113536 | Fam84a |
| **ENSMUSG00000020608** | 0.122719745446221 | 0.183646589750527 | 0.805963874499992 | 1059.82158953549 | 1162.98566088966 | 1085.58375727977 | 1062.6709445945 | 1039.11532783232 | 907.527016198804 | 1036.91377819391 | 1030.16376087041 | Smc6 |
| **ENSMUSG00000020609** | -0.562835517105575 | 0.229453192826085 | 0.832048466795696 | 282.799793286111 | 753.334549245754 | 271.632553042104 | 438.949692912819 | 354.766455445387 | 385.270903103266 | 470.567296931374 | 1369.93326261166 | Apob |
| **ENSMUSG00000020610** | 0.156277892888435 | 0.104383965487566 | 0.761198544536855 | 1236.91027798302 | 1188.23254848601 | 1212.40871235866 | 1276.51823088536 | 1054.44474257379 | 1013.93517038923 | 1113.95355689507 | 1225.55836251075 | Amz2 |
| **ENSMUSG00000020611** | -0.0163603616919325 | 0.912485693345926 | 0.986226496507248 | 479.765783498163 | 603.482055125517 | 575.444572298256 | 550.562969529541 | 535.434557755538 | 452.540425867328 | 630.893322877019 | 614.407469602375 | Gna13 |
| **ENSMUSG00000020612** | -0.0417951624525325 | 0.799297278105657 | 0.963317934306948 | 8165.05345663446 | 9412.2025790629 | 8699.81333645652 | 7825.12241019584 | 7854.13513861038 | 6779.5448123854 | 10117.1968840245 | 10352.6573102436 | Prkar1a |
| **ENSMUSG00000020614** | -0.741999932177621 | 0.0628045425686914 | 0.715615581523459 | 29.8159526467785 | 35.0198763433161 | 57.7337482075553 | 55.3376749612315 | 86.5016974697087 | 114.969729814943 | 32.2734208072402 | 64.046008315442 | Fam20a |
| **ENSMUSG00000020620** | -0.218650156433489 | 0.379016398078709 | 0.87422797717331 | 157.211386683014 | 196.274190668353 | 131.557229522134 | 147.254490998531 | 212.421889988905 | 245.839528646846 | 122.847214685624 | 156.315681312265 | Abca8b |
| **ENSMUSG00000020621** | -0.0235712655026213 | 0.831488124999491 | 0.967583158900775 | 1260.40163461382 | 1057.92603185971 | 1101.67349038679 | 1199.60824195619 | 1191.31451705118 | 1192.50517627201 | 1269.07419238793 | 1042.10454208177 | Rdh14 |
| **ENSMUSG00000020623** | 0.0285627663988428 | 0.924759387785529 | 0.986713918026868 | 27.1054114970714 | 35.8342920722305 | 34.0723759913441 | 20.6343872736795 | 27.3739548954774 | 34.2463024980681 | 23.9447960827911 | 30.3947158107182 | Map2k6 |
| **ENSMUSG00000020628** | 0.0794461620571058 | 0.582783269493382 | 0.922450244597442 | 521.327414460339 | 654.79024604712 | 609.5169482896 | 604.024791102256 | 615.366506050332 | 453.763508099402 | 623.605776243126 | 567.72987032163 | Trappc12 |
| **ENSMUSG00000020629** | 0.0570019692759787 | 0.723829658051579 | 0.951674101666722 | 1087.83051474913 | 1199.63436869081 | 1105.45930994139 | 1118.00861955573 | 924.144717271318 | 891.626947181844 | 1143.10374343064 | 1376.44641599967 | Adi1 |
| **ENSMUSG00000020630** | 0.199525712330403 | 0.23038112941916 | 0.832048466795696 | 152.693818100169 | 124.605606523892 | 115.467496415111 | 147.254490998531 | 107.305903190271 | 116.192812047017 | 127.011527047849 | 119.407812113536 | Rnaseh1 |
| **ENSMUSG00000020634** | 0.173726834091836 | 0.508390753509536 | 0.902394714365738 | 1605.54387434319 | 1999.39061448468 | 1964.84034883418 | 1431.27613543795 | 1088.38844664418 | 1040.84297949485 | 2018.65041758835 | 2058.15647061149 | Ubxn2a |
| **ENSMUSG00000020638** | -0.42587518872557 | 0.0996932773915537 | 0.759800953500741 | 246.659244623349 | 294.004078138073 | 210.112985279955 | 242.923013812864 | 355.861413641207 | 492.902139525765 | 216.544242835676 | 271.381391167127 | Cmpk2 |
| **ENSMUSG00000020639** | 0.13111736082337 | 0.570243764773547 | 0.920872458148558 | 43.3686583953142 | 42.3496179035451 | 53.9479286529615 | 58.151455044006 | 43.7983278327639 | 41.5847958905112 | 47.8895921655823 | 46.6775992807458 | Pfn4 |
| **ENSMUSG00000020640** | 0.268809343232282 | 0.231939679375078 | 0.832048466795696 | 529.45903790946 | 728.08766164941 | 651.160963390131 | 501.790781428116 | 413.894198019619 | 358.363093997641 | 635.057635239243 | 592.696958309005 | Itsn2 |
| **ENSMUSG00000020641** | 0.130116610009349 | 0.747301184758793 | 0.956119556124081 | 452.660372001092 | 416.166437475222 | 1108.29867460733 | 549.625042835282 | 602.227007700503 | 996.812019140196 | 323.775286162958 | 386.447101021989 | Rsad2 |
| **ENSMUSG00000020642** | -0.033929930685123 | 0.854914411166898 | 0.973268281968411 | 328.878992831132 | 364.043830824705 | 443.887342776122 | 520.54931531328 | 463.167316831478 | 452.540425867328 | 345.637926064637 | 435.295751432072 | Rnf144a |
| **ENSMUSG00000020644** | -0.559421336425438 | 0.00924701076905134 | 0.635862253526407 | 120.167324303683 | 114.018202048006 | 107.895857305923 | 137.87522405595 | 226.656346534553 | 194.470074899744 | 124.929370866736 | 161.743309135608 | Id2 |
| **ENSMUSG00000020646** | -0.0323241208552278 | 0.881094443091037 | 0.978524171626649 | 134.623543768788 | 128.677685168464 | 100.324218196735 | 115.364983393754 | 87.5966556655277 | 113.746647582869 | 133.257995591185 | 155.230155747597 | Mboat2 |
| **ENSMUSG00000020647** | 0.0187985300315521 | 0.915831352407646 | 0.986713918026868 | 446.335775985108 | 579.863998987002 | 559.354839191232 | 422.067012416173 | 473.02194059385 | 388.940149799487 | 564.264325081426 | 554.703563545607 | Ncoa1 |
| **ENSMUSG00000020648** | 0.108740385315634 | 0.61810005452853 | 0.929568738467614 | 136.430571201926 | 97.7298874697194 | 108.842312194571 | 125.682177030594 | 89.7865720571659 | 84.3926740130963 | 130.134761319517 | 129.177542195552 | Dus4l |
| **ENSMUSG00000020649** | 0.349639061119422 | 0.187288006274734 | 0.807540361382744 | 40.658117245607 | 58.6379324818317 | 43.5369248778286 | 70.3445020693621 | 45.9882442244021 | 44.0309603546589 | 41.6431236222455 | 35.8223436340608 | Rrm2 |
| **ENSMUSG00000020650** | 0.233900242736513 | 0.140181537336716 | 0.779711473022872 | 1096.86565191482 | 1033.49355999228 | 1141.42459571003 | 1058.91923781746 | 799.319482947941 | 735.07242147639 | 1031.70838774113 | 1114.83475491456 | Bcap29 |
| **ENSMUSG00000020652** | -0.118662562834731 | 0.660847353943687 | 0.939611397743053 | 37.0440623793308 | 33.3910448854875 | 35.965285768641 | 29.075727522003 | 44.893286028583 | 30.5770558018465 | 29.1501865355718 | 42.3354970220718 | Cenpo |
| **ENSMUSG00000020653** | -0.226423411011073 | 0.194827821082265 | 0.809984565948116 | 329.782506547701 | 307.849145529616 | 345.456034356683 | 335.777756544422 | 506.965644664242 | 376.709327478749 | 351.884394607974 | 307.203734801188 | Klf11 |
| **ENSMUSG00000020654** | -1.0449045872523 | 0.00073221158039640 | 0.556246415579446 | 14.4562194651047 | 26.8757190541728 | 22.7149173275627 | 15.9447538023887 | 50.3680770076785 | 35.4693847301419 | 40.6020455316893 | 39.0789203280663 | Adcy3 |
| **ENSMUSG00000020657** | 0.265821952331065 | 0.306914337664643 | 0.851785577030345 | 381.282788392137 | 428.382673408937 | 436.315703666934 | 413.625672167849 | 286.879047304603 | 194.470074899744 | 454.951125573032 | 442.894430384751 | Dnajc27 |
| **ENSMUSG00000020658** | -0.00818490969268593 | 0.969969360683845 | 0.9951105526746 | 165.343010132135 | 180.800291818981 | 184.558703286447 | 257.929840920994 | 198.187433443257 | 200.585486060113 | 146.792010768415 | 247.49982874442 | Efr3b |
| **ENSMUSG00000020659** | -0.121119027107539 | 0.345477090007241 | 0.861266047293794 | 177.088688447533 | 215.820168162297 | 186.451613063744 | 192.274972322923 | 208.042057205628 | 207.923979452556 | 214.462086654564 | 209.506433981022 | Cbll1 |
| **ENSMUSG00000020661** | 0.264320852275599 | 0.307107486613748 | 0.851785577030345 | 545.722284807703 | 769.622863824041 | 728.770264259304 | 579.638697051544 | 492.731188118594 | 314.332133642982 | 683.988305495382 | 692.565310258508 | Dnmt3a |
| **ENSMUSG00000020664** | 0.25479508316981 | 0.20185563337809 | 0.813395633194566 | 13006.983463728 | 13912.6638970435 | 14361.5064803515 | 13020.2983696918 | 10556.4919658919 | 7487.70942475617 | 13325.7995591185 | 14138.9704798073 | Dld |
| **ENSMUSG00000020668** | 0.0530077513403251 | 0.765610877908402 | 0.958345041873658 | 258.404922938747 | 264.685111897157 | 213.898804834549 | 192.274972322923 | 225.561388338734 | 177.34692365071 | 235.283648465687 | 257.269558826436 | Kif3c |
| **ENSMUSG00000020669** | -0.0816647645170209 | 0.766075740958053 | 0.958370894524665 | 29.8159526467785 | 23.6180561385155 | 29.3401015481019 | 26.2619474392285 | 27.3739548954774 | 29.3539735697726 | 29.1501865355718 | 29.3091902460497 | Sh3yl1 |
| **ENSMUSG00000020671** | 0.0697489130752974 | 0.715568443032189 | 0.951176382858794 | 3037.61311510513 | 3817.16652142146 | 3518.91927599493 | 2765.94582136732 | 2727.54086578537 | 2365.44103683084 | 3796.81179625823 | 3628.91196268682 | Rab10 |
| **ENSMUSG00000020674** | -0.531249260330071 | 0.0194189340549696 | 0.642273034804458 | 355.984404328204 | 368.115909469277 | 373.849681016137 | 460.522006880757 | 775.230402639921 | 631.110431750112 | 416.431236222455 | 429.868123608729 | Pxdn |
| **ENSMUSG00000020676** | 0.194223492128037 | 0.702733317765126 | 0.946451777392559 | 15.3597331816738 | 22.8036404096012 | 32.1794662140472 | 21.5723139679377 | 38.3235368536684 | 19.5693157131818 | 6.24646854333682 | 16.2828834700276 | Ccl11 |
| **ENSMUSG00000020677** | -0.0379525245149766 | 0.792777150531657 | 0.961804768755116 | 331.58953398084 | 319.250965734417 | 381.421320125324 | 417.377378944882 | 414.989156215438 | 390.163232031561 | 330.021754706295 | 352.795808517265 | Ddx52 |
| **ENSMUSG00000020680** | 0.254999506015933 | 0.0344241721669919 | 0.678229434609245 | 1438.39383677792 | 1338.89945833516 | 1305.16129144621 | 1672.3232958623 | 1239.49267766722 | 1087.32010431366 | 1283.64928565572 | 1210.36100460539 | Taf15 |
| **ENSMUSG00000020681** | -1.18373803259973 | 0.0102922699076497 | 0.635862253526407 | 423.747933070882 | 529.370223794314 | 484.584902988005 | 470.839200517597 | 1525.276766776 | 1987.50862712002 | 440.376032305246 | 383.190524327983 | Ace |
| **ENSMUSG00000020682** | -0.494725986246583 | 0.249041725104691 | 0.835314899646652 | 78.6056933415069 | 102.616381843205 | 116.413951303759 | 90.0409626487835 | 180.668102310151 | 240.94719971855 | 63.5057635239243 | 61.8749571861049 | Mmp28 |
| **ENSMUSG00000020684** | 0.122427661388456 | 0.487214577509704 | 0.899236955296196 | 2955.39336689735 | 2791.0027029894 | 2784.47028240373 | 2338.2512487856 | 2043.19199339843 | 2037.65499863505 | 3106.57702221951 | 2796.31385458608 | Rasl10b |
| **ENSMUSG00000020687** | -0.0312748217177566 | 0.863051229307722 | 0.974868041734905 | 321.65088309858 | 302.96265115613 | 335.04503058155 | 298.260688774095 | 283.594172717146 | 238.501035254403 | 386.239971596327 | 375.591845375304 | Cdc27 |
| **ENSMUSG00000020689** | 0.0908412296329363 | 0.911895590281208 | 0.986226496507248 | 76.7986659083688 | 76.5550785179469 | 247.971180825893 | 41.2687745473591 | 120.445401540101 | 222.600966237442 | 24.9858741733473 | 47.7631248454143 | Itgb3 |
| **ENSMUSG00000020691** | 0.065910448399424 | 0.737475893037121 | 0.954383574136061 | 145.465708367616 | 169.39847161418 | 167.522515290775 | 203.530092654021 | 193.80760065998 | 119.862058743238 | 159.284947855089 | 181.282769299641 | Mettl2 |
| **ENSMUSG00000020692** | 0.0254021540662815 | 0.906065390211995 | 0.98504921011344 | 106.614618555147 | 100.987550385377 | 123.039135524298 | 137.87522405595 | 139.059690869025 | 128.423634367755 | 111.395355689507 | 81.4144173501381 | Nle1 |
| **ENSMUSG00000020694** | 0.123983371136632 | 0.532511880508331 | 0.909082872831377 | 891.768038253647 | 979.742121883937 | 892.506959995485 | 773.789522762983 | 708.437952694956 | 572.402484610566 | 996.311732662223 | 968.288803684309 | Tlk2 |
| **ENSMUSG00000020695** | -1.22495820601036 | 0.0055642878868656 | 0.635862253526407 | 106.614618555147 | 118.904696421492 | 119.253315969704 | 122.868396947819 | 394.184950494875 | 470.886659348436 | 109.313199508394 | 119.407812113536 | Mrc2 |
| **ENSMUSG00000020696** | 0.132668115861672 | 0.643989456416738 | 0.934847999069355 | 90.3513716569045 | 66.7820897709749 | 63.4124775394459 | 121.930470253561 | 93.0714466446232 | 69.71568722821 | 93.6970281500523 | 55.3618037980939 | Rffl |

| | | | | | | | | | | | | |
|---|---|---|---|---|---|---|---|---|---|---|---|---|
| ENSMUSG00000020697 | 0.342352843182019 | 0.200447647504495 | 0.812921911423739 | 283.70330700268 | 378.703313945163 | 384.260684791269 | 280.44008158319 | 229.94122112201 | 151.662196777159 | 359.171941241867 | 305.032683671851 | Lig3 |
| ENSMUSG00000020700 | -0.00723528732600271 | 0.969557008674999 | 0.9951105526746 | 617.099868416658 | 846.177942341987 | 947.401343537095 | 643.417712261099 | 775.230402639921 | 612.764198269004 | 884.916376972716 | 795.690238902016 | Map3k3 |
| ENSMUSG00000020705 | 0.120071842990838 | 0.375485807408287 | 0.873821423608136 | 970.373731595154 | 1086.43058237171 | 1095.04830616625 | 966.064495085906 | 895.675804180021 | 783.995710759344 | 1017.13329447335 | 1090.95319249185 | Ddx42 |
| ENSMUSG00000020706 | 0.239942039303938 | 0.0484527963811055 | 0.703396849069134 | 1019.16347228988 | 1031.86472853445 | 1025.95709929492 | 1023.27802343565 | 967.943045104082 | 716.726187995282 | 944.257828134416 | 841.282312618093 | Ftsj3 |
| ENSMUSG00000020707 | -0.356514415116648 | 0.092259048431477 | 0.754058637722175 | 70.4740698923855 | 66.7820897709749 | 50.1621090983677 | 75.9720622349111 | 97.4512794278996 | 90.5080851734656 | 75.998700610598 | 73.8157383974585 | Rnf135 |
| ENSMUSG00000020708 | 0.052222489561637 | 0.719002614601571 | 0.951674101666722 | 3694.46758705082 | 3071.16171373593 | 3333.41411781983 | 3711.37592917954 | 3313.34350054859 | 4092.43314851913 | 3054.52311769171 | 2860.35986290152 | Psmc5 |
| ENSMUSG00000020709 | -0.35732445510641 | 0.165000855140005 | 0.795216960550936 | 77.7021796249379 | 109.13170767452 | 107.895857305923 | 62.8410885152968 | 143.439523652302 | 129.646716599829 | 96.8202624217207 | 89.0130963028176 | Adap2 |
| ENSMUSG00000020715 | 0.316164790877327 | 0.280047471591043 | 0.848943557485723 | 261.115464088454 | 368.115909469277 | 419.279515671262 | 245.736793895638 | 193.80760065998 | 168.785348026193 | 303.994802442392 | 372.335268681298 | Ern1 |
| ENSMUSG00000020716 | 0.113405802115528 | 0.576860569795382 | 0.922450244597442 | 419.230364488037 | 514.710740673856 | 512.03209475881 | 425.818719193205 | 419.368988998714 | 288.647406769431 | 517.4158110064 | 503.683862006187 | Nf1 |
| ENSMUSG00000020717 | -0.632435121434746 | 0.0531591866276146 | 0.703396849069134 | 3175.85071374019 | 2777.15763559786 | 3093.96103099177 | 2962.91042716153 | 6737.2777788749 | 6662.12891810631 | 2654.74913091815 | 2564.01138374701 | Pecam1 |
| ENSMUSG00000020718 | -0.15223522075408 | 0.486941777665267 | 0.899236955296196 | 74.0881247586617 | 66.7820897709749 | 61.519567762149 | 82.5375490947182 | 82.1218646864323 | 101.51582526213 | 69.7522320672611 | 64.046008315442 | Polg2 |
| ENSMUSG00000020719 | 0.13672540723981 | 0.323148043268953 | 0.855977678491063 | 4025.1536073151 | 4286.26998127611 | 5275.53954932644 | 4455.15179772627 | 4353.55378657673 | 3412.39942748607 | 4529.73077200975 | 4113.05636452897 | Ddx5 |
| ENSMUSG00000020720 | 0.181283937539847 | 0.0342138770970622 | 0.678229434609245 | 3111.70123986379 | 3087.45002831422 | 3270.94809516903 | 3385.91536627196 | 2956.38712871156 | 2608.83440101354 | 2986.85304180556 | 2783.28754781005 | Psmd12 |
| ENSMUSG00000020721 | 0.00049879495443738: | 0.996812207389831 | 0.999262355603273 | 681.24934229306 | 759.849875077069 | 691.858523602015 | 665.927952923294 | 710.627869086594 | 578.517895770936 | 767.274552739873 | 740.328435103922 | Helz |
| ENSMUSG00000020728 | 0.248865458720888 | 0.151652666493722 | 0.783412422888014 | 142.755167217909 | 139.26508964435 | 125.878500190243 | 124.744250336335 | 98.5462376237187 | 138.208292224346 | 97.8613405122768 | 115.065709854862 | Cep112 |
| ENSMUSG00000020732 | -0.525053127215952 | 0.218356014552671 | 0.827129763070499 | 18.0702743313809 | 12.2162359337149 | 13.2503684410783 | 8.44134024832345 | 24.0890803080201 | 24.4616446414772 | 11.4518589961175 | 15.1973579053591 | Rab37 |
| ENSMUSG00000020733 | -0.421917804848436 | 0.10921503779393 | 0.763154315676794 | 75.8951521917998 | 107.502876216691 | 115.467496415111 | 105.047789756914 | 192.712642464161 | 145.546785616789 | 95.7791843311646 | 107.467030902182 | Slc9a3r1 |
| ENSMUSG00000020734 | 0.328850409956888 | 0.479153424345267 | 0.897732059433541 | 13.5527057485357 | 17.9171460361152 | 31.2330113253987 | 14.0689004138724 | 19.7092475247437 | 13.4539045528125 | 7.28754663389296 | 20.6249857287016 | Grin2c |
| ENSMUSG00000020736 | -0.141032340077286 | 0.605975125135115 | 0.926859923029135 | 116.553269437407 | 113.203786319092 | 100.324218196735 | 147.254490998531 | 131.394983498292 | 192.023910435596 | 79.1219348822664 | 124.835439936878 | Nt5c |
| ENSMUSG00000020737 | 0.077399236585283 | 0.703035271651548 | 0.946451777392559 | 1875.69447559734 | 1494.45286255779 | 1158.4607837057 | 1936.8186236431 | 1700.47007810706 | 1799.15396338065 | 1355.48367390409 | 1273.32148735616 | Hn1 |
| ENSMUSG00000020738 | 0.0433679722816376 | 0.577887130360224 | 0.922450244597442 | 1378.76193148436 | 1319.35348084121 | 1313.67938544404 | 1371.24882700543 | 1378.55236853624 | 1327.04422180014 | 1270.11527047849 | 1248.35439936878 | Sumo2 |
| ENSMUSG00000020739 | -0.0783242470635041 | 0.532422101900365 | 0.909082872831377 | 374.958192376154 | 286.674336577844 | 327.473391472363 | 313.267515882226 | 352.576539053749 | 336.347613820311 | 350.843316517418 | 334.3418739179 | Nup85 |
| ENSMUSG00000020740 | 0.0643680894685247 | 0.749816212030828 | 0.956197192463153 | 249.369785773056 | 297.26174105373 | 205.380710836713 | 181.019851991825 | 208.042057205628 | 195.693157131818 | 271.721381635152 | 217.105112933702 | Gga3 |
| ENSMUSG00000020741 | 0.193640685502966 | 0.405331698448591 | 0.881158207349486 | 5554.80232946649 | 6809.3299094527 | 6714.15098007208 | 5240.19644082035 | 4764.16311000889 | 3291.31428651076 | 6434.90367772748 | 6772.59399796682 | Cluh |
| ENSMUSG00000020743 | -0.105091070019398 | 0.261275385802113 | 0.841102613761057 | 717.389890955822 | 657.233493233863 | 642.642869392295 | 739.086235075431 | 763.185862485911 | 725.287763619799 | 732.91897575152 | 742.499486233259 | Mif4gd |
| ENSMUSG00000020744 | -0.0557750391435402 | 0.731901395419728 | 0.953219349759111 | 475.248214915318 | 459.330471107681 | 357.759947909113 | 448.328959855401 | 394.184950494875 | 446.425014706959 | 411.225845769674 | 557.960140239613 | Slc25a19 |
| ENSMUSG00000020745 | 0.0805188129127441 | 0.62413362622434 | 0.931145988862144 | 4102.85578694003 | 4614.47952002859 | 4331.92402534394 | 3749.83092364413 | 3703.14861826019 | 3008.78229090169 | 4620.30456588813 | 4553.77974378439 | Pafah1b1 |
| ENSMUSG00000020747 | 0.00543262245245982 | 0.977170538164578 | 0.996179425433073 | 1447.42897394361 | 1738.77758123209 | 1637.36695736181 | 1431.27613543795 | 1432.20532013138 | 1097.10476217025 | 1795.85970620934 | 1905.09736599323 | Tmem94 |
| ENSMUSG00000020752 | -0.164180332434427 | 0.382353864882757 | 0.875126835008975 | 139.141112351633 | 122.162359337149 | 101.270673085384 | 101.296082979881 | 144.534481848121 | 105.185071958352 | 140.545542225078 | 129.177542195552 | Recql5 |
| ENSMUSG00000020755 | -0.0436381185484022 | 0.698933376259015 | 0.945141109561958 | 399.353062723518 | 350.198763433161 | 403.189782564238 | 393.929211588428 | 436.88832013182 | 399.947889888152 | 379.99350305299 | 376.677370939972 | Sap30bp |
| ENSMUSG00000020758 | -1.38411253027058 | 0.0079330053454464€ | 0.635862253526407 | 15.3597331816738 | 26.8757190541728 | 20.8220075502658 | 15.0068271081306 | 45.9882442244021 | 110.077400886647 | 27.0680303544596 | 21.7105112933701 | Itgb4 |
| ENSMUSG00000020766 | -0.141568645911768 | 0.587288476559192 | 0.923058967588378 | 287.317361868956 | 275.272516373043 | 267.84673348751 | 405.184331919526 | 390.900075907418 | 475.778988276731 | 229.03717992235 | 268.124814473121 | Galk1 |
| ENSMUSG00000020770 | 0.0287523614415443 | 0.872738710497111 | 0.976162349063696 | 116.553269437407 | 144.151584017836 | 132.503684410783 | 117.24083678227 | 109.49581958191 | 114.969729814943 | 155.120635492864 | 120.493337678204 | Unk |
| ENSMUSG00000020773 | -0.895422355637889 | 0.0262384134529351 | 0.664353746352133 | 352.370349461928 | 313.550055632016 | 306.651383922097 | 430.508352664496 | 806.984190318674 | 1179.0512717192 | 319.610973800734 | 305.032683671851 | Trim47 |
| ENSMUSG00000020774 | -0.238733150757851 | 0.531337390163962 | 0.909082872831377 | 25.2983840639333 | 10.5874044758863 | 18.9290977729689 | 22.5102406621959 | 30.6588294829347 | 20.7923979452556 | 17.6983275394543 | 21.7105112933701 | Aspa |
| ENSMUSG00000020775 | 0.0676689927068933 | 0.549188418896775 | 0.914699131434573 | 1742.87795926169 | 1489.56636818431 | 1664.81414913262 | 1820.51571355509 | 1684.04570516977 | 1724.54594722414 | 1580.35654146422 | 1420.95296415108 | Mrpl38 |
| ENSMUSG00000020776 | -0.0258864528837785 | 0.865339854664937 | 0.974868041734905 | 309.905204783182 | 319.250965734417 | 258.382184601026 | 260.743621003769 | 281.404256325508 | 243.393364182698 | 327.939598525183 | 315.887939318536 | Fbf1 |
| ENSMUSG00000020777 | 0.307797268498041 | 0.19077841108025 | 0.808824684419004 | 10727.4183568243 | 13259.5024824542 | 15242.6559816832 | 11272.9409382888 | 8793.60927062317 | 6547.15918829137 | 12712.604563781 | 12745.1556547729 | Acox1 |
| ENSMUSG00000020778 | 0.0544096874310545 | 0.766580151492111 | 0.958624690460551 | 113.8427282877 | 97.7298874697194 | 106.949402417274 | 139.751077444466 | 118.255485148462 | 118.638976511164 | 94.7381062406084 | 109.638082031519 | Ten1 |
| ENSMUSG00000020780 | 0.218400893682238 | 0.0274882807653921 | 0.664931733862672 | 2235.29293479182 | 2240.45767024332 | 2654.80596265889 | 2494.88500672671 | 2078.23065566465 | 1928.80067998048 | 2161.27811599454 | 2102.6630187629 | Srp68 |
| ENSMUSG00000020781 | -0.0682657544572079 | 0.6919020922194 | 0.94469184542218 | 181.606257030378 | 146.594831204579 | 163.736695736181 | 175.392291826276 | 202.567266226533 | 176.123841418636 | 135.340151772298 | 185.624871558315 | Tsen54 |
| ENSMUSG00000020782 | 0.0634251303944341 | 0.724650311919799 | 0.951674101666722 | 256.597895505609 | 277.715763559786 | 308.544293699394 | 250.426427366929 | 269.359716171498 | 193.24699266767 | 322.734208072402 | 259.440609955773 | Llgl2 |
| ENSMUSG00000020783 | 0.0799788085662771 | 0.474750848594764 | 0.897549741595057 | 359.59845919448 | 416.980853204136 | 370.063861461543 | 371.418970926232 | 350.386622662111 | 335.124531588238 | 366.45948787576 | 384.276049892652 | Ncbp3 |
| ENSMUSG00000020785 | -0.661493867811014 | 0.203137951890228 | 0.815723085214011 | 13.5527057485357 | 4.88649437348597 | 11.3574586637814 | 16.8826804966469 | 16.4243729372865 | 33.0232202659942 | 11.4518589961175 | 13.0263067760221 | Camkk1 |
| ENSMUSG00000020787 | 0.744634237775858 | 0.170389694524995 | 0.799531027639052 | 11.7456783153976 | 17.1027303072009 | 26.5007368821565 | 8.44134024832345 | 7.66470737073368 | 13.4539045528125 | 4.16431236222455 | 13.0263067760221 | P2rx1 |
| ENSMUSG00000020788 | -0.534651267154698 | 0.180721364982367 | 0.802808628271642 | 338.817643713392 | 404.764617270421 | 487.42426765395 | 329.212269684615 | 713.912743674051 | 996.812019140196 | 257.146288367366 | 293.091902460497 | Atp2a3 |
| ENSMUSG00000020790 | -0.0384486673719076 | 0.834162061172755 | 0.968512902280839 | 286.413848152387 | 349.384347704247 | 383.314229902621 | 282.315934971707 | 401.849657865609 | 255.624186503437 | 334.18606706852 | 344.111603999917 | Ankfy1 |
| ENSMUSG00000020792 | 0.00666514544103359 | 0.93150853408904 | 0.987783121884152 | 867.373167906283 | 897.48613326359 | 869.792042667923 | 902.285479876351 | 878.156473046916 | 898.965440574287 | 898.450392149946 | 845.624414876767 | Exoc7 |
| ENSMUSG00000020794 | 0.209193849063707 | 0.164954966611629 | 0.795216960550936 | 2372.62701971031 | 2519.80226526093 | 2469.3008044838 | 2433.91977159993 | 1907.41717711687 | 1671.95341124497 | 2362.20618747187 | 2530.36009124229 | Ube2g1 |
| ENSMUSG00000020801 | -0.145648232154086 | 0.394888702083979 | 0.879215782741314 | 268.343573821006 | 180.800291818981 | 190.237432618338 | 229.79204009325 | 235.416012101106 | 265.408844360028 | 244.653351280692 | 216.019587369033 | Med31 |
| ENSMUSG00000020802 | 0.212402023680612 | 0.30859354249318 | 0.851785577030345 | 1057.11104838578 | 1162.17124516075 | 1742.42345000179 | 899.471699793577 | 992.032125412102 | 1155.8127093098 | 1028.58515346946 | 1019.30850522373 | Ube2o |
| ENSMUSG00000020803 | 0.142747036769782 | 0.331483234186656 | 0.856333478432716 | 925.198045766702 | 671.078560625407 | 777.039463580375 | 943.55425442371 | 731.432074807157 | 829.249753346077 | 687.11153976705 | 756.61131857395 | Txndc17 |
| ENSMUSG00000020806 | -0.438717995240389 | 0.0271473843583152 | 0.664353746352133 | 202.387072511466 | 203.603932228582 | 210.112985279955 | 160.385464718146 | 316.442918591719 | 303.324393554317 | 242.57119509958 | 191.052499381657 | Rhbdf2 |
| ENSMUSG00000020807 | -0.17910968996129 | 0.507054062090789 | 0.901698316485955 | 40.658117245607 | 39.0919549878878 | 61.519567762149 | 36.5791410760683 | 39.4184950494875 | 46.4771248188067 | 55.1771387994752 | 59.7039060567679 | 4933427D14I |
| ENSMUSG00000020810 | -0.693325171938463 | 0.042116869765637 | 0.685406501171951 | 486.090379514146 | 551.359448475 | 591.534305405279 | 746.589648629496 | 1123.42710891039 | 1604.6838884809 | 527.826591911961 | 586.183804920994 | Cygb |
| ENSMUSG00000020811 | -0.711250904872171 | 0.0174810928879468 | 0.642273034804458 | 107.518132271716 | 72.4829998733752 | 81.3951204237664 | 95.6685228143324 | 178.478185918513 | 210.370143916704 | 105.14888714617 | 91.1841474321546 | Wscd1 |
| ENSMUSG00000020812 | 0.728236100506395 | 0.0922017726571354 | 0.754058637722175 | 29.8159526467785 | 19.5459774939439 | 13.2503684410783 | 30.0136542162612 | 15.3294147414674 | 20.7923979452556 | 11.4518589961175 | 8.68420451734806 | 1810032O08I |
| ENSMUSG00000020814 | -0.579830220963958 | 0.0737174510987557 | 0.732971689437116 | 332.493047697409 | 246.767965861042 | 251.757000380487 | 292.633128608546 | 519.010184818252 | 648.233582999146 | 211.338852382896 | 301.776106977845 | Mxra7 |
| ENSMUSG00000020817 | 0.209553566833587 | 0.310317623821675 | 0.851785577030345 | 1376.95490405122 | 1689.09822176832 | 1553.1324722721 | 1450.03466932312 | 999.696832782835 | 996.812019140196 | 1572.02791673977 | 1679.30804854218 | Rabep1 |
| ENSMUSG00000020818 | -0.0723362705617258 | 0.486564604243602 | 0.899236955296196 | 544.818771091134 | 503.308920469055 | 493.102996985841 | 494.287367874051 | 504.775728272604 | 588.302553627526 | 507.005030100839 | 541.677256769585 | Mfsd11 |
| ENSMUSG00000020821 | 0.218147426977325 | 0.292152375878826 | 0.851785577030345 | 22539.956687248 | 25124.725237007 | 25015.7491618671 | 21010.4958780771 | 17147.0453465271 | 13790.2521666328 | 25414.7983466564 | 24189.851683073 | Kif1c |
| ENSMUSG00000020823 | -0.199063895015683 | 0.319142490710893 | 0.854670142081901 | 1671.50037565273 | 1773.79745757541 | 1399.80678031105 | 1133.95337335812 | 2113.26931793086 | 1895.77745971448 | 1422.11267169968 | 1433.9792709271 | Sec14l1 |
| ENSMUSG00000020826 | -0.337088238276368 | 0.229687874934782 | 0.832048466795696 | 91.2548853734735 | 77.3694942468612 | 58.6802030962037 | 117.24083678227 | 152.199189218854 | 106.408154190426 | 83.2862472444909 | 93.3551985614916 | Nos2 |
| ENSMUSG00000020827 | 0.132928370511181 | 0.582999150875809 | 0.922450244597442 | 1073.37429528403 | 1315.28140219664 | 1110.19158438463 | 889.154506156737 | 856.257309130534 | 631.110431750112 | 1281.5671294746 | 1232.07151589876 | Mink1 |
| ENSMUSG00000020828 | -0.45257843210183 | 0.0449816079475846 | 0.693563614452513 | 108.421645988285 | 127.048853710635 | 121.146225747001 | 122.868396947819 | 178.478185918513 | 226.270212933664 | 130.134761319517 | 122.664388807541 | Pld2 |
| ENSMUSG00000020829 | -0.557952851670419 | 0.0505158154637815 | 0.703396849069134 | 38.8510898124689 | 50.4937751926884 | 26.5007368821565 | 30.9515809105193 | 60.2227007700503 | 62.3771938357668 | 40.6020455316893 | 54.2762782334254 | Slc46a1 |
| ENSMUSG00000020831 | -0.171823498271748 | 0.650429202775844 | 0.936463929813731 | 26.2018977805023 | 18.7315617650296 | 17.036187995672 | 22.5102406621959 | 24.0890803080201 | 36.6924669622158 | 21.8626399016789 | 13.0263067760221 | 0610010K14I |
| ENSMUSG00000020832 | -0.0743530485047845 | 0.506734414273446 | 0.901698316485955 | 431.879556520004 | 416.980853204136 | 398.457508120996 | 387.363724728621 | 477.401773377126 | 403.617136584374 | 440.376032305246 | 399.473407798011 | Eral1 |
| ENSMUSG00000020834 | 0.0718282506931969 | 0.70060351603217 | 0.945712617851005 | 121.070838020252 | 95.2866402829764 | 108.842312194571 | 123.806323642077 | 128.110108910834 | 108.854318654574 | 102.025652874501 | 87.9275707381491 | Dhrs13 |
| ENSMUSG00000020836 | 0.322705987794388 | 0.0757160188794705 | 0.735399943172214 | 4570.8758921228 | 4804.23838486562 | 4521.21500307363 | 4869.71539658837 | 3388.89561606011 | 2656.53460806442 | 4550.55233382087 | 4407.23379255414 | Coro6 |
| ENSMUSG00000020840 | -0.0579532829262921 | 0.512083879813548 | 0.902394714365738 | 764.372604217412 | 842.92027942633 | 827.201572678743 | 801.927323590728 | 859.542183717991 | 785.218792991418 | 875.546674157711 | 847.795466006104 | Blmh |
| ENSMUSG00000020841 | -0.0709467105432876 | 0.578888503745344 | 0.922450244597442 | 1220.64703108478 | 1522.95741306979 | 1520.95300605805 | 1278.39408427387 | 1552.65072167148 | 1355.17511313784 | 1586.60301000755 | 1327.59776558958 | Cpd |
| ENSMUSG00000020843 | -0.0563292454670141 | 0.722767674442535 | 0.951674101666722 | 668.600150261093 | 541.586459728029 | 458.084166105848 | 630.286738541484 | 543.099265126272 | 661.687487551958 | 544.483841360859 | 641.545608719088 | Timm22 |
| ENSMUSG00000020844 | -0.38310783679625 | 0.181619620382198 | 0.803406537295772 | 430.072529086865 | 312.735639903102 | 346.402489245332 | 356.412143818101 | 630.6959207918 | 645.787418534998 | 291.501865355718 | 318.058990447873 | Nxn |
| ENSMUSG00000020846 | 0.0386551601854136 | 0.825173463109566 | 0.96672635991852 | 1635.35982698997 | 1448.03116600968 | 1834.22957420069 | 2420.78879788031 | 1771.6423608353 | 1713.53820713548 | 1680.3000381576 | 1977.82757882602 | Rflnb |
| ENSMUSG00000020847 | 0.123763254826357 | 0.489845450423907 | 0.899236955296196 | 115.649755720838 | 136.821842457607 | 115.467496415111 | 135.061443973175 | 137.964732673206 | 105.185071958352 | 122.847214685624 | 95.5262496908287 | Rph3al |
| ENSMUSG00000020848 | -1.22254232398035 | 0.11100931922852 | 0.764116514278315 | 6.32459601598331 | 15.4738988493722 | 3.78581955459379 | 0.937926694258161 | 18.6142893289246 | 29.3539735697726 | 8.32862472444909 | 6.51315338801105 | Doc2b |
| ENSMUSG00000020849 | 0.114843367451637 | 0.327509754879986 | 0.855977678491063 | 8998.99661702769 | 8495.17046830536 | 8752.81481022084 | 8363.49233270002 | 7304.4661243092 | 6923.86851577012 | 8722.15224267931 | 9009.86218674861 | Ywhae |
| ENSMUSG00000020850 | -0.0309470094789812 | 0.847916115203438 | 0.972467578249175 | 2045.55505431232 | 2561.33746743556 | 2689.82479353889 | 1846.77766099432 | 2430.8071947184 | 1978.9470514955 | 2571.46288367366 | 2359.93257758934 | Prpf8 |
| ENSMUSG00000020857 | 0.06256730863771 | 0.691903567505165 | 0.94469184542218 | 7131.43376487947 | 4901.96827233534 | 5662.63959878366 | 7374.91759695192 | 5930.29358855623 | 6123.97273599382 | 5456.29027260471 | 6495.78497897635 | Nme2 |
| ENSMUSG00000020859 | 0.299297423929159 | 0.280378429477674 | 0.849093714741928 | 3080.98177350044 | 4229.26088025211 | 4490.92844663688 | 3162.68881303852 | 2549.06267986686 | 1713.53820713548 | 4153.90158131899 | 3742.89214697701 | Spag9 |
| ENSMUSG00000020863 | -0.0954247292429119 | 0.551583326592017 | 0.914873920695527 | 1605.54387434319 | 1274.56061575092 | 1105.45930994139 | 1372.18675369969 | 1413.59103080245 | 1719.65361829585 | 1270.11527047849 | 1322.17013776624 | Luc7l3 |
| ENSMUSG00000020864 | 0.0858035581754478 | 0.407693700524231 | 0.881573350402261 | 7573.25197228174 | 7221.42426828335 | 6530.53873167428 | 7250.17334661559 | 6021.17511880921 | 6443.19719856509 | 7216.75332373514 | 7243.71209303295 | Ankrd40 |
| ENSMUSG00000020865 | -0.434571868922709 | 0.242672636739047 | 0.832048466795696 | 40.658117245607 | 88.7713144516618 | 75.7163910918757 | 59.0893817382642 | 70.0773245324222 | 75.8310983885793 | 56.2182168900314 | 155.230155747597 | Abcc3 |
| ENSMUSG00000020866 | -0.595482054949177 | 0.0237249233277946 | 0.654124840968651 | 201.483558794897 | 178.357044632238 | 146.700507740509 | 128.495957113368 | 171.908436743598 | 368.147751854232 | 212.379930473452 | 238.815624227072 | Cacna1g |
| ENSMUSG00000020868 | -0.190367568457397 | 0.384685929054596 | 0.875126835008975 | 135.527057485357 | 145.780415475665 | 132.503684410783 | 187.585338851632 | 216.801722772181 | 195.693157131818 | 136.381229862854 | 137.8617467129 | Xylt2 |
| ENSMUSG00000020869 | 0.0550468433212141 | 0.506152398390528 | 0.901494649536949 | 1073.37429528403 | 1097.83240257652 | 1112.08449416193 | 1082.36740517392 | 1099.33802860237 | 1064.08154190426 | 967.161546126651 | 1072.49925789249 | Lrrc59 |

| | | | | | | | | | | | | |
|---|---|---|---|---|---|---|---|---|---|---|---|---|
| **ENSMUSG00000020873** | 0.0170906351594638 | 0.922165219536656 | 0.986713918026868 | 525.844983043184 | 472.361122770311 | 459.030620994497 | 526.176875478828 | 538.719432342996 | 615.210362733151 | 383.116737324658 | 424.440495785386 | Slc35b1 |
| **ENSMUSG00000020876** | -0.0256464468661391 | 0.882546666507174 | 0.978884707960125 | 195.158962778914 | 147.409246933493 | 198.755526616174 | 239.171307035831 | 203.662224422352 | 192.023910435596 | 196.76375911511 | 200.822229463674 | Snx11 |
| **ENSMUSG00000020877** | -0.224608084782658 | 0.22572468003913 | 0.832048466795696 | 373.151164943016 | 307.849145529616 | 300.026199701558 | 377.984457786039 | 469.737066006393 | 467.217412652214 | 319.610973800734 | 332.170822788563 | Scrn2 |
| **ENSMUSG00000020882** | 0.254946713047893 | 0.192937864870813 | 0.808824684419004 | 185.220311896654 | 131.935348084121 | 112.628131749165 | 157.571684635371 | 111.685735973548 | 130.869798831903 | 123.88829277618 | 125.920965501547 | Cacnb1 |
| **ENSMUSG00000020883** | 0.0652746127714622 | 0.748320795552616 | 0.956197192463153 | 393.931980424104 | 524.483729420828 | 555.569019636638 | 472.715053906113 | 468.642107810573 | 316.77829810713 | 524.703357640293 | 549.275935722265 | Fbxl20 |
| **ENSMUSG00000020884** | -0.215394134283903 | 0.512205811683844 | 0.902402172030517 | 121.070838020252 | 204.418347957496 | 124.932045301595 | 170.702658354985 | 119.350443344282 | 171.23151249034 | 127.011527047849 | 303.947158107182 | Asgr1 |
| **ENSMUSG00000020886** | -0.601215537368997 | 0.0628529588412078 | 0.715615581523459 | 46.9827132615903 | 36.6487078011448 | 33.1259211026956 | 32.8274342990356 | 58.0327843784121 | 81.9465095489486 | 57.2592949805875 | 30.3947158107182 | Dlg4 |
| **ENSMUSG00000020888** | -0.00375500177462441 | 0.983747612340542 | 0.996940244108351 | 734.556651570634 | 705.284021239809 | 721.198625150116 | 874.147679048606 | 805.889232122855 | 993.142772443974 | 630.893322877019 | 614.407469602375 | Dvl2 |
| **ENSMUSG00000020889** | -0.471162024820237 | 0.00021931461467876 | 0.399150048388892 | 1470.01681685784 | 1527.84390744328 | 1655.34960024613 | 1607.60635395849 | 2519.49880857974 | 2282.27144504982 | 1827.09204892602 | 2050.55779165881 | Nr1d1 |
| **ENSMUSG00000020893** | 0.0236175932673566 | 0.944432564260534 | 0.990505437194639 | 1188.12053728829 | 1277.00386293767 | 874.524317111165 | 438.011766218561 | 1225.25822112157 | 1093.43551547403 | 807.876598271562 | 590.525907179668 | Per1 |
| **ENSMUSG00000020894** | 0.0219556350730444 | 0.872923868806347 | 0.976162349063696 | 1211.61189391909 | 1316.09581792556 | 1451.86179918672 | 1321.53871220975 | 1320.51958415783 | 1084.87393984951 | 1551.20635492864 | 1262.46623170947 | Vamp2 |
| **ENSMUSG00000020895** | -0.12055517168144 | 0.718010071185811 | 0.951674101666722 | 60.535419010126 | 62.7100111264033 | 68.1447519826882 | 79.7237690119437 | 94.1664048404423 | 110.077400886647 | 52.0539045278068 | 39.0789203280663 | Tmem107 |
| **ENSMUSG00000020898** | -0.0199481701264574 | 0.899522252262934 | 0.984802943363903 | 243.948703473642 | 300.519403969387 | 307.597838810745 | 244.79886720138 | 316.442918591719 | 288.647406769431 | 231.119336103462 | 276.809018990469 | Ctc1 |
| **ENSMUSG00000020899** | 0.134446367324962 | 0.570590450546537 | 0.920927417858634 | 115.649755720838 | 135.193010999779 | 156.165056626994 | 96.6064495085906 | 99.6411958195378 | 88.0619207093179 | 151.997401221196 | 118.322286548867 | Pfas |
| **ENSMUSG00000020900** | 0.220829461610451 | 0.234696490795408 | 0.832048466795696 | 822.197482077831 | 941.464582624964 | 971.062715753306 | 973.567908639971 | 682.158955995297 | 590.748718091674 | 983.818795575549 | 923.7822555329 | Myh10 |
| **ENSMUSG00000020901** | 0.274571791225494 | 0.475835013486434 | 0.897549741595057 | 21.6843291976571 | 15.4738988493722 | 21.7684624389143 | 23.448167356454 | 9.85462376237187 | 22.0154801773295 | 13.5340151772298 | 22.7960368580387 | Pik3r5 |
| **ENSMUSG00000020902** | 0.0698609466253905 | 0.776949393085227 | 0.959738279197945 | 1197.15567445398 | 1835.6930529729 | 2911.29523748262 | 1834.58461396896 | 1982.96929262838 | 1641.37635544312 | 1965.55543496999 | 1820.42637194909 | Ntn1 |
| **ENSMUSG00000020903** | -0.0218340457982902 | 0.887072470341561 | 0.980672278091991 | 579.152292320758 | 448.743066631795 | 495.942361651786 | 577.762843663027 | 589.087509350674 | 594.417964787896 | 440.376032305246 | 510.197015394199 | Stx8 |
| **ENSMUSG00000020905** | -0.374063882867057 | 0.246137100995546 | 0.832048466795696 | 30.7194663633475 | 34.2054606144018 | 50.1621090983677 | 43.1446279358754 | 66.7924499449649 | 66.0464405319884 | 31.2323427166841 | 41.2499714574033 | Usp43 |
| **ENSMUSG00000020910** | -0.110789316377662 | 0.487140551238899 | 0.899236955296196 | 137.334084918495 | 112.389370590177 | 150.486327295103 | 120.054616865045 | 134.679858085749 | 150.439114545085 | 138.463386043966 | 137.8617467129 | Adprm |
| **ENSMUSG00000020914** | -0.567603550038103 | 0.145546530746117 | 0.780607990324772 | 20.780815481088 | 26.0613033252585 | 13.2503684410783 | 15.9447538023887 | 42.7033696369448 | 17.123151249034 | 29.1501865355718 | 23.8815624227072 | Top2a |
| **ENSMUSG00000020917** | -0.123015758154234 | 0.591917782246098 | 0.923935529355527 | 519.520387027201 | 609.182965227918 | 867.899132890626 | 578.700770357285 | 884.72622222183 | 775.434135134827 | 497.635327285833 | 646.973236542431 | Acly |
| **ENSMUSG00000020918** | 0.142508099345597 | 0.167721942134849 | 0.798648534882772 | 379.475760958999 | 416.166437475222 | 397.511053232348 | 398.618845059719 | 367.905953795217 | 337.570696052385 | 383.116737324658 | 352.795808517265 | Kat2a |
| **ENSMUSG00000020919** | 0.268834084820221 | 0.0804033665374691 | 0.739855948845396 | 1545.91196904964 | 1755.88031153929 | 1857.8909464169 | 1541.95148536042 | 1390.59690869025 | 1067.75078860048 | 1525.17940266474 | 1577.26864546334 | Stat5b |
| **ENSMUSG00000020921** | -0.349381043992319 | 0.0357762173172259 | 0.679566075569698 | 237.624107457659 | 215.005752433383 | 198.755526616174 | 191.337045628665 | 262.789966996583 | 337.570696052385 | 236.324726556243 | 238.815624227072 | Tmem101 |
| **ENSMUSG00000020922** | 0.150838551803011 | 0.21527196428615 | 0.824338622350122 | 866.469654189714 | 742.747144769868 | 741.074177811734 | 832.878904501247 | 778.515277227378 | 620.102691661447 | 698.563398763168 | 768.552099785303 | Lsm12 |
| **ENSMUSG00000020923** | 0.217128755106539 | 0.0186565574212311 | 0.642273034804458 | 2556.04030417383 | 2654.1808605318 | 3042.85246700476 | 2867.2419043472 | 2357.44499559852 | 2284.71760951397 | 2475.68369934249 | 2446.77462276282 | Ubtf |
| **ENSMUSG00000020925** | 0.0332194141191723 | 0.850940037762662 | 0.972692784625411 | 409.291713605777 | 315.178887089845 | 338.830850136144 | 329.212269684615 | 303.30342024189 | 267.855008824175 | 403.938299135781 | 384.276049892652 | Ccdc43 |
| **ENSMUSG00000020926** | -0.0915179516645435 | 0.622903591135726 | 0.931059568526633 | 217.74680569314 | 241.881471487556 | 230.934992830221 | 181.957778686083 | 170.813478547779 | 233.608706326107 | 251.940897914585 | 273.552442296464 | Adam11 |
| **ENSMUSG00000020928** | -0.182650176909338 | 0.518186526914453 | 0.905473752527583 | 121.974351736821 | 90.4001459094905 | 88.966759532954 | 98.4823028971069 | 155.484063806312 | 133.315963296051 | 63.5057635239243 | 102.03940307884 | Higd1b |
| **ENSMUSG00000020929** | -0.0389133538486745 | 0.693271049812247 | 0.94469184542218 | 771.600713949964 | 774.509358197526 | 820.576388458203 | 780.35500962279 | 863.922016501267 | 865.942220308293 | 741.247600475969 | 763.124471961961 | Eftud2 |
| **ENSMUSG00000020935** | -0.114167446124427 | 0.365190712707282 | 0.86924083181804 | 778.828823682517 | 701.211942595237 | 585.855576073389 | 733.458674909882 | 746.761489548624 | 765.649477278236 | 697.522320672611 | 820.657326889392 | Dcakd |
| **ENSMUSG00000020936** | 0.0367014040647368 | 0.824257251110719 | 0.966637987288264 | 2707.83060855743 | 2402.52640029727 | 2330.17193585248 | 2898.19348525772 | 2738.49044774356 | 3101.73654053931 | 2058.21138502948 | 2181.9063849837 | Nmt1 |
| **ENSMUSG00000020937** | 0.238341544579874 | 0.285474036237193 | 0.851275833092507 | 136.430571201926 | 118.090280692578 | 174.147699511314 | 215.723139679377 | 147.819356435578 | 150.439114545085 | 122.847214685624 | 124.835439936878 | Plcd3 |
| **ENSMUSG00000020941** | 0.105184853748286 | 0.635918242592062 | 0.932595977875587 | 53.3073092775737 | 76.5550785179469 | 64.3589324280944 | 79.7237690119437 | 61.3176589658694 | 52.592535979176 | 67.6700758861489 | 72.73021283279 | Map3k14 |
| **ENSMUSG00000020946** | 0.0170631940067711 | 0.879183866925486 | 0.978393501240173 | 928.812100632978 | 932.506009606906 | 1148.99623481921 | 930.423280704096 | 1024.88087128667 | 979.688867891162 | 929.68273486663 | 959.604599166961 | Gosr2 |
| **ENSMUSG00000020948** | -0.474197441658398 | 0.123729116837169 | 0.770326056866107 | 44.2721721118832 | 76.5550785179469 | 53.001473764313 | 40.3308478531009 | 55.8428679867739 | 55.0387004433237 | 100.984574783945 | 85.7565196088121 | Klhl28 |
| **ENSMUSG00000020949** | 0.145589768668957 | 0.255321647830466 | 0.837406774112203 | 5092.20330658314 | 4296.04297002308 | 4578.00229639254 | 5202.67937305002 | 4278.00167106521 | 3625.21573586692 | 4678.60493895928 | 4744.83224316605 | Fkbp3 |
| **ENSMUSG00000020952** | 0.101309922938927 | 0.364092811085858 | 0.86924083181804 | 421.037391921175 | 392.548381336706 | 431.583429223692 | 422.067012416173 | 404.039574257247 | 340.016860516533 | 394.568596320776 | 413.585240138701 | Scfd1 |
| **ENSMUSG00000020954** | 0.208615127349043 | 0.264418654658655 | 0.842510961779757 | 3081.88528721701 | 3149.34562371171 | 2972.81480524477 | 2878.4970246783 | 2111.07940153922 | 1976.50088703136 | 3166.95955147177 | 3200.12936464276 | Strn3 |
| **ENSMUSG00000020955** | 0.00956255331363012 | 0.941898481392061 | 0.989934163630124 | 1255.88406603097 | 1109.23422278132 | 1260.67791167973 | 1539.13770527764 | 1284.3859636958 | 1383.30600447554 | 1173.29500805677 | 1289.60437082619 | Ap4s1 |
| **ENSMUSG00000020956** | 0.0376288835412898 | 0.818316996679677 | 0.966315773308182 | 171.667606148119 | 149.038078391322 | 168.468970179424 | 203.530092654021 | 145.62944004394 | 177.34692365071 | 189.476212481217 | 161.743309135608 | Dtd2 |
| **ENSMUSG00000020961** | -0.69792973352044 | 0.183114405216241 | 0.805716353538343 | 18.9737880479499 | 18.7315617650296 | 12.3039135524298 | 12.1930470253561 | 8.75966556655277 | 15.9000690169602 | 48.9306702561384 | 27.1381391167127 | Ston2 |
| **ENSMUSG00000020962** | 0.0266876423498469 | 0.821935019036545 | 0.966637987288264 | 404.774145022932 | 354.270842077733 | 424.958245003153 | 409.873965390816 | 423.748821781991 | 368.147751854232 | 364.377331694648 | 407.07208675069 | Gtf2a1 |
| **ENSMUSG00000020964** | -0.00960272989206491 | 0.950595355994203 | 0.991696503889296 | 1264.91920319666 | 1715.15952509358 | 1583.41902870885 | 1254.94591691742 | 1399.35657425681 | 1257.32853457193 | 1570.98683864921 | 1629.37387256743 | Sel1l |
| **ENSMUSG00000020973** | -0.279220565847443 | 0.224638546220491 | 0.832048466795696 | 58.7283915769879 | 65.1532583131463 | 59.6266579848522 | 62.8410885152968 | 78.836990098975 | 100.292743030056 | 64.5468416144805 | 56.4473293627624 | Dnaaf2 |
| **ENSMUSG00000020978** | 0.204247446064729 | 0.243628473913395 | 0.832048466795696 | 3380.0448136848 | 3480.81282537984 | 3728.08580638623 | 3515.34925007959 | 2785.57365016378 | 2211.33267558954 | 3685.41644056872 | 3558.35280098337 | Klhdc2 |
| **ENSMUSG00000020982** | 0.050747804417609 | 0.722852634530675 | 0.951674101666722 | 1041.75131520411 | 890.970807432275 | 882.095956220352 | 1027.96765690694 | 891.295971396745 | 1139.91264029284 | 842.232175259915 | 838.025735924088 | Nemf |
| **ENSMUSG00000020986** | 0.109411569248074 | 0.657886201360493 | 0.938763867173667 | 870.08370905599 | 1205.33527879321 | 1084.63730239112 | 768.161962597434 | 744.571573156986 | 611.54111603693 | 1180.58255469066 | 1103.97949926787 | Sec23a |
| **ENSMUSG00000020988** | 0.250723267134189 | 0.388680884791298 | 0.876405748044371 | 1042.65482892068 | 1259.08671690155 | 1328.82266366242 | 1081.42947847966 | 708.437952694956 | 551.610086665311 | 1356.52475199465 | 1342.79512349494 | L2hgdh |
| **ENSMUSG00000020990** | 0.711623376460604 | 0.00323105777658545 | 0.635862253526407 | 247.562758339918 | 331.467201668132 | 371.010316350191 | 290.75727522003 | 185.047935093427 | 116.192812047017 | 243.612273190136 | 211.677485110359 | Cdkl1 |
| **ENSMUSG00000020993** | 0.152041383942304 | 0.138888234275463 | 0.779711473022872 | 1471.82384429097 | 1397.53739081699 | 1299.48256211432 | 1455.66222948867 | 1293.14562926235 | 1110.55866672306 | 1342.99073681742 | 1313.48593324889 | Trappc6b |
| **ENSMUSG00000020994** | 0.193712547809198 | 0.19203697154568 | 0.808824684419004 | 1158.30458464152 | 1256.64346971481 | 1302.32192678026 | 1122.69825302702 | 1015.0262475243 | 826.803588881929 | 1214.93813167901 | 1173.45313540666 | Pnn |
| **ENSMUSG00000021000** | 0.126397773625087 | 0.555138487163418 | 0.916239900758826 | 1358.88462971984 | 1562.8637837866 | 1274.87473500946 | 1286.8354245222 | 1065.39432453198 | 850.042151291332 | 1619.91750890535 | 1487.17002359586 | Mia2 |
| **ENSMUSG00000021003** | 0.0890161538780718 | 0.722326429270703 | 0.951674101666722 | 194.255449062345 | 283.416673662186 | 231.881447718869 | 172.578511743502 | 169.71852035196 | 145.546785616789 | 247.776585552361 | 265.953763343784 | Galc |
| **ENSMUSG00000021007** | -0.452621128876584 | 0.0412152963608089 | 0.683365309556438 | 51.5002818444356 | 53.7514381083457 | 70.9841166486335 | 73.1582821521366 | 72.2672409240604 | 88.0619207093179 | 103.066730965058 | 77.072315091464 | Spata7 |
| **ENSMUSG00000021009** | 0.256908633932295 | 0.175012905206125 | 0.799889021902745 | 991.154547076242 | 1168.68657099206 | 1049.61847151113 | 1010.14704971604 | 713.912743674051 | 695.933790050026 | 949.463218587197 | 1171.28208427732 | Ptpn21 |
| **ENSMUSG00000021012** | 0.154663925359762 | 0.296133260140944 | 0.851785577030345 | 1638.97388185625 | 1646.74860386477 | 1649.67087091424 | 1546.64111883171 | 1262.48679977942 | 1197.39750520031 | 1733.39502077597 | 1628.28834700276 | Zc3h14 |
| **ENSMUSG00000021013** | -0.0372627384164534 | 0.90699735477457 | 0.98504921011344 | 26.2018977805023 | 21.1748089517725 | 25.5542819935081 | 33.7653609932938 | 19.7092475247437 | 23.2385624094033 | 31.2323427166841 | 34.7368180693922 | Ttc8 |
| **ENSMUSG00000021018** | 0.113464683106068 | 0.582089374499775 | 0.922450244597442 | 202.387072511466 | 139.26508964435 | 122.09268063565 | 184.771558768858 | 173.003394939417 | 147.992950080937 | 141.586620315635 | 136.776221148232 | Polr2h |
| **ENSMUSG00000021022** | 0.233049876032201 | 0.0813596950183028 | 0.739855948845396 | 627.942033015486 | 544.844122643686 | 495.942361651786 | 557.128456389348 | 419.368988998714 | 467.217412652214 | 546.565997541972 | 460.262839419447 | Ppp2r3c |
| **ENSMUSG00000021023** | 0.122596972226101 | 0.494044865399552 | 0.900141680336539 | 213.229237110295 | 236.180561385155 | 259.328639489674 | 287.005568442997 | 260.600050604945 | 178.570005882784 | 263.392756910703 | 210.59195954569 | 1110008L16R |
| **ENSMUSG00000021024** | 0.0855854850748003 | 0.451825265221759 | 0.89111066611284 | 5273.80956361352 | 4088.36695914993 | 4586.52039039037 | 4979.45281981658 | 4640.43283388133 | 4574.32754795624 | 4056.04024080671 | 4566.80605056041 | Psma6 |
| **ENSMUSG00000021025** | -0.41051734742824 | 0.352754897605761 | 0.862728204353539 | 821.293968361262 | 746.004807685525 | 468.495169880981 | 458.646153492241 | 1097.14811221074 | 1484.82182973767 | 364.377331694648 | 370.164217551961 | Nfkbia |
| **ENSMUSG00000021027** | 0.0478199665338339 | 0.841714838689378 | 0.97064500700668 | 847.495866141764 | 1035.93680717903 | 886.828230663595 | 771.913669374467 | 675.589206820383 | 572.402484610566 | 1187.87010132455 | 989.999314977679 | Ralgapa1 |
| **ENSMUSG00000021028** | -0.112598968670448 | 0.644447842639131 | 0.934857223264556 | 60.535419010126 | 50.4937751926884 | 39.7511053232348 | 54.3997482669734 | 43.7983278327639 | 50.1463715150282 | 60.3825292522559 | 67.3025850094475 | Mbip |
| **ENSMUSG00000021033** | -0.00826491257622691 | 0.948090997642746 | 0.991192530276814 | 937.847237798669 | 927.61951523342 | 960.651711978173 | 921.044013761514 | 833.263187018333 | 825.580506649855 | 1009.84574783945 | 1098.55187144453 | Gstz1 |
| **ENSMUSG00000021036** | -0.258526657004128 | 0.254425178846267 | 0.837406774112203 | 421.037391921175 | 385.218639776477 | 454.298346551254 | 318.895076047775 | 574.853052805026 | 593.194882555822 | 398.732908683 | 323.486618271215 | Sptlc2 |
| **ENSMUSG00000021037** | 0.0814357553314111 | 0.481109447886317 | 0.897732059433541 | 2682.53222449349 | 2309.68300720104 | 2237.41935676493 | 2806.27666922042 | 2178.96680968 | 2360.54870790255 | 2538.14838477586 | 2406.61017687008 | Ahsa1 |
| **ENSMUSG00000021038** | -0.0271648671056805 | 0.822307128128385 | 0.966637987288264 | 380.379274675568 | 460.144886836596 | 460.923530771794 | 387.363724728621 | 416.084114411257 | 415.847958905112 | 460.156516025812 | 428.78259804406 | Vipas39 |
| **ENSMUSG00000021039** | 0.00621392499519413 | 0.955499776622922 | 0.992371505557225 | 853.820462157748 | 834.776122137187 | 807.326020017125 | 800.98939689647 | 911.005218921489 | 727.733928083946 | 880.752064610492 | 762.038946397292 | Snw1 |
| **ENSMUSG00000021040** | 0.16511904816685 | 0.405233036044856 | 0.881158207349486 | 707.451240073562 | 497.608010366655 | 493.102996985841 | 701.569167305105 | 571.568178217569 | 653.125911927441 | 431.006329490241 | 485.229927406823 | Slirp |
| **ENSMUSG00000021044** | -0.00316712714425805 | 0.978722618676201 | 0.996664169100645 | 634.26662903147 | 619.770369703804 | 584.90912118474 | 574.949063580253 | 590.182467546493 | 514.917619703095 | 652.755962778698 | 659.999543318453 | Adck1 |
| **ENSMUSG00000021047** | 0.234685232706131 | 0.544158319561746 | 0.913837378403038 | 61.4389327266951 | 48.8649437348597 | 19.8755526616174 | 72.2203554578784 | 38.3235368536684 | 56.2617826753975 | 33.3144988977964 | 44.5065481514088 | Nova1 |
| **ENSMUSG00000021048** | 0.125254079168625 | 0.337449919125292 | 0.858570960161243 | 516.809845877494 | 611.626212414661 | 570.712297855013 | 553.376749612315 | 575.948011000845 | 431.748027922072 | 552.812466085309 | 503.683862006187 | Mthfd1 |
| **ENSMUSG00000021051** | 0.109440832120303 | 0.566993093878569 | 0.919931558693298 | 849.302893574902 | 1078.28642508257 | 1066.6546595068 | 825.375490947182 | 796.034608360484 | 659.24132308781 | 1065.02288663893 | 1019.30850522373 | Ppp2r5e |
| **ENSMUSG00000021054** | 0.100593030849776 | 0.352317715259518 | 0.862728204353539 | 589.994456919586 | 667.006481980835 | 583.016211407443 | 610.590277962063 | 561.713554455197 | 517.363784167243 | 581.96265262088 | 624.177199684392 | Sgpp1 |
| **ENSMUSG00000021057** | 0.313730081652556 | 0.559094824195968 | 0.916574868225118 | 18.9737880479499 | 21.1748089517725 | 17.036187995672 | 35.6412143818101 | 7.66470737073368 | 11.0077400886647 | 15.616171358342 | 40.1644458927348 | Akap5 |
| **ENSMUSG00000021061** | 0.432356594522538 | 0.252053290895881 | 0.83723604018117 | 3811.02085648823 | 5753.84712477973 | 5863.28803517713 | 3319.32257097963 | 2304.8870021992 | 1504.39114545085 | 5072.1324571895 | 5010.78600650983 | Sptb |
| **ENSMUSG00000021062** | 0.238524023977401 | 0.65317697690259 | 0.936991241819063 | 18.0702743313809 | 20.3603932228582 | 17.036187995672 | 16.8826804966469 | 9.85462376237187 | 17.123151249034 | 4.16431236222455 | 30.3947158107182 | Rab15 |
| **ENSMUSG00000021065** | 0.135080402586095 | 0.264835949989677 | 0.842510961779757 | 486.090379514146 | 486.206190161854 | 456.191256328551 | 404.246405225267 | 402.944616061428 | 376.709327478749 | 437.252798033577 | 451.578634902099 | Fut8 |
| **ENSMUSG00000021066** | 0.215774815510393 | 0.488181119935283 | 0.899236955296196 | 46.0791995450213 | 88.7713144516618 | 79.5022106464695 | 40.3308478531009 | 66.7924499449649 | 46.4771248188067 | 55.1771387994752 | 51.0197015394199 | Atl1 |
| **ENSMUSG00000021067** | -0.00215545639805998 | 0.989137345099737 | 0.998437672973644 | 355.080890611635 | 453.629561005281 | 534.747012086372 | 439.887619607078 | 511.345477447518 | 431.748027922072 | 426.842017128016 | 415.756291268038 | Sav1 |
| **ENSMUSG00000021068** | 0.341132681067123 | 0.120811905067466 | 0.769868040577702 | 401.160090156656 | 443.856572258309 | 429.690519446395 | 396.742991671202 | 300.018545654433 | 206.700897220482 | 390.404283958551 | 421.183919091381 | Nin |
| **ENSMUSG00000021069** | -0.0441942081170402 | 0.782891554796174 | 0.960887487507189 | 142.755167217909 | 160.439898596123 | 141.968233297267 | 145.378637610015 | 143.439523652302 | 129.646716599829 | 151.997401221196 | 183.453820428978 | Pygl |
| **ENSMUSG00000021072** | 0.112314754652355 | 0.342542226847767 | 0.860070775653557 | 930.619128066116 | 842.105863697416 | 869.792042667923 | 930.423280704096 | 916.480009900584 | 750.97249049335 | 736.042210023189 | 900.986218674861 | Tmx1 |
| **ENSMUSG00000021076** | 0.0771518681308584 | 0.471206041891419 | 0.897091462240771 | 1295.63866956001 | 1112.49188569697 | 1250.2669079046 | 1325.29041898678 | 1289.8607546749 | 1204.73599859275 | 1078.55690181616 | 1150.65709854862 | Actr10 |

| | | | | | | | | | | | | |
|---|---|---|---|---|---|---|---|---|---|---|---|---|
| **ENSMUSG00000021079** | 0.0749034186985977 | 0.593567094162648 | 0.923935529355527 | 249.369785773056 | 210.933673788811 | 195.916161950228 | 215.723139679377 | 187.237851485066 | 205.477814988408 | 208.215618111227 | 226.874843015718 | Timm9 |
| **ENSMUSG00000021087** | -1.187081721312 | 0.0194028149111946 | 0.642273034804458 | 13.5527057485357 | 11.4018202048006 | 15.1432782183751 | 24.3860940507122 | 62.4126171616885 | 45.2540425867328 | 27.0680303544596 | 11.9407812113536 | Rtn1 |
| **ENSMUSG00000021091** | -1.09134762440734 | 0.0107283623638345 | 0.635862253526407 | 188.83436676293 | 469.103459854653 | 299.079744812909 | 212.909359596603 | 614.271547854513 | 1191.28209403994 | 367.500565966316 | 321.315567141878 | Serpina3n |
| **ENSMUSG00000021094** | -0.11848840387385 | 0.572322952919889 | 0.921574199900435 | 860.145058173731 | 881.197818685304 | 645.482234058241 | 943.55425442371 | 1056.63465896543 | 1132.57414690039 | 682.947227404826 | 744.670537362596 | Dhrs7 |
| **ENSMUSG00000021096** | 0.240178312390728 | 0.084872032388107 | 0.743541556158628 | 5297.30092024431 | 5344.19601313582 | 5308.66547042914 | 5057.30073544001 | 3974.69825082332 | 3671.69286068573 | 5314.70365228908 | 4822.99008382218 | Ppm1a |
| **ENSMUSG00000021097** | -0.215511134692887 | 0.410205184335544 | 0.882192126163777 | 100.290022539164 | 78.1839099757755 | 72.8770264259304 | 45.0204813243917 | 78.836990098975 | 81.9465095489486 | 102.025652874501 | 81.4144173501381 | Clmn |
| **ENSMUSG00000021102** | -0.0567903589069419 | 0.702626328428126 | 0.946451777392559 | 3826.38058966991 | 2978.3183206397 | 3089.22875654853 | 3822.05127910201 | 3535.62001429986 | 4255.10308538496 | 3163.8363172001 | 3313.02402336829 | Glrx5 |
| **ENSMUSG00000021103** | 0.0620498958094134 | 0.651162497776313 | 0.936463929813731 | 390.317925557827 | 343.683437601847 | 427.797609669098 | 424.880792498947 | 421.558905390352 | 396.278643191931 | 375.829190690765 | 325.657669400552 | Mnat1 |
| **ENSMUSG00000021108** | -0.412365631935134 | 0.0752905137988203 | 0.735399943172214 | 481.572810931301 | 466.66021266791 | 469.44162476963 | 397.68091836546 | 843.117810780705 | 688.595296657583 | 448.704657029695 | 436.38127699674 | Prkch |
| **ENSMUSG00000021109** | 0.189779219590864 | 0.353812730270965 | 0.86286441023679 | 691.187993175319 | 898.300548992504 | 1097.8876708322 | 1148.96020046625 | 888.011096809288 | 634.779678446333 | 877.628830338823 | 961.775650296298 | Hif1a |
| **ENSMUSG00000021111** | 0.141980061233321 | 0.404495846777851 | 0.881158207349486 | 2046.45856802889 | 2224.98377139395 | 2198.61470633034 | 1951.82545075123 | 1547.17593069238 | 1552.09135250173 | 2286.20748686128 | 2245.95239329914 | Papola |
| **ENSMUSG00000021112** | 0.146014493631248 | 0.437915789122388 | 0.888759286175347 | 531.266065342598 | 574.163088884602 | 575.444572298256 | 498.039074651084 | 435.793361936001 | 353.470765069345 | 590.291277345329 | 588.354856050331 | Mpp5 |
| **ENSMUSG00000021113** | 0.0210118571918442 | 0.89181808218922 | 0.982421229269187 | 361.405486627618 | 277.715763559786 | 271.632553042104 | 294.508981997063 | 260.600050604945 | 275.193502216618 | 332.103910887408 | 319.144516012541 | Snapc1 |
| **ENSMUSG00000021114** | 0.0645884826518014 | 0.379767181232815 | 0.87432013076003 | 2449.42568561868 | 2326.78573750824 | 2465.5149849292 | 2428.29221143438 | 2263.27859075807 | 2454.72603977224 | 2215.41417670346 | 2314.34050387326 | Atp6v1d |
| **ENSMUSG00000021115** | 0.0261204855623298 | 0.840420576741108 | 0.969957774070128 | 179.79922959724 | 179.171460361152 | 176.98706417726 | 200.716312571247 | 190.522726072523 | 167.562265794119 | 182.188665847324 | 182.368294864309 | Vrk1 |
| **ENSMUSG00000021116** | 0.20445850702128 | 0.154427216338915 | 0.786853994984691 | 1254.07703859783 | 1056.29720040188 | 1297.58965233702 | 1363.74541345137 | 1139.85148184768 | 868.38838477244 | 1178.50039850955 | 1125.69001056124 | Eif2s1 |
| **ENSMUSG00000021120** | 0.0855058514803537 | 0.779066954231695 | 0.960290688328569 | 51.5002818444356 | 22.8036404096012 | 46.3762895437739 | 40.3308478531009 | 37.2285786578493 | 33.0232202659942 | 40.6020455316893 | 40.1644458927348 | Pigh |
| **ENSMUSG00000021124** | 0.0357181993323681 | 0.793173524635335 | 0.961804768755116 | 3492.98402825593 | 2707.93229864014 | 3073.13902344151 | 3747.01714356135 | 2995.80562376105 | 3581.18477551226 | 3048.27664914837 | 3077.46497583522 | Vti1b |
| **ENSMUSG00000021127** | -0.467445698407961 | 0.0118619300841394 | 0.635862253526407 | 584.573374620172 | 501.680089011226 | 623.713771619326 | 690.314046974007 | 1033.64053685323 | 916.088591823321 | 712.097413940397 | 656.742966624447 | Zfp36l1 |
| **ENSMUSG00000021130** | -0.504397136444747 | 0.20097057824599 | 0.812921911423739 | 40.658117245607 | 47.2361122770311 | 62.4660226507975 | 50.6480414899407 | 90.881530252985 | 117.415894279091 | 42.6842017128016 | 34.7368180693922 | Galnt16 |
| **ENSMUSG00000021131** | 0.0856438178947245 | 0.633761283257914 | 0.9325328237529 | 854.723975874317 | 602.667639396603 | 552.729654970693 | 865.706338800283 | 678.87408140784 | 727.733928083946 | 640.263025692024 | 663.256120012458 | Erh |
| **ENSMUSG00000021133** | 0.0794048475407622 | 0.661537208553417 | 0.939611397743053 | 1777.21148049131 | 2144.35661423143 | 2271.49173275627 | 1750.17121148573 | 1712.51461826107 | 1438.34470491886 | 2344.50785993242 | 2021.24860141276 | Susd6 |
| **ENSMUSG00000021134** | 0.0953792113217679 | 0.616630784090079 | 0.928896975098882 | 796.899098013898 | 985.443031986338 | 1131.96004682354 | 1258.69762369445 | 954.803546754252 | 804.7881087046 | 1220.14352213179 | 924.867781097568 | Srsf5 |
| **ENSMUSG00000021135** | -0.319204618098671 | 0.397552793480662 | 0.88020150974241 | 34.3335212296237 | 92.8433930962335 | 53.001473764313 | 60.0273084325223 | 44.893286028583 | 66.0464405319884 | 70.7933101578173 | 118.322286548867 | Slc10a1 |
| **ENSMUSG00000021136** | -0.42837831422479 | 0.305385189718301 | 0.851785577030345 | 63.2459601598332 | 58.6379324818317 | 43.5369248778286 | 44.0825546301336 | 118.255485148462 | 92.9542496376133 | 32.2734208072402 | 39.0789203280663 | Smoc1 |
| **ENSMUSG00000021139** | -0.0641223036465901 | 0.768457788658515 | 0.95879680569409 | 45.1756858284523 | 61.0811796685746 | 53.9479286529615 | 64.7169419038131 | 58.0327843784121 | 63.6002760678407 | 51.0128264372507 | 62.9604827507734 | Gm20498 |
| **ENSMUSG00000021140** | 0.0911853265932066 | 0.694612337628527 | 0.944771261777587 | 1651.62307388821 | 2110.15115361703 | 1763.24545755206 | 1423.77272188389 | 1423.44565456483 | 1060.41229520804 | 2048.84168221448 | 1989.76836003737 | Pcnx |
| **ENSMUSG00000021143** | 0.155778667006945 | 0.473214764772808 | 0.89728597737823 | 1181.79594127231 | 1283.51918876898 | 1440.50434052294 | 1138.64300682941 | 921.95480087968 | 782.77262852727 | 1428.35914024302 | 1393.81482503436 | Pacs2 |
| **ENSMUSG00000021144** | 0.106542970399113 | 0.175140445328172 | 0.799889021902745 | 2075.3710069591 | 1939.12385054502 | 2171.16751455954 | 2018.41824604356 | 1966.5449196911 | 1926.35451551633 | 1913.50153044218 | 1812.82769299641 | Mta1 |
| **ENSMUSG00000021147** | 0.092652612902667 | 0.56617924939158 | 0.919931558693298 | 677.635287426784 | 792.426504233642 | 744.859997366328 | 653.734905897938 | 573.758094609207 | 557.72549782568 | 757.904849924867 | 800.03234116069 | Wdr37 |
| **ENSMUSG00000021149** | 0.269156091136365 | 0.0122481731670484 | 0.635862253526407 | 1426.64815846252 | 1533.54481754568 | 1508.64909250562 | 1640.43378825752 | 1258.10696699614 | 1113.00483118721 | 1400.250031798 | 1296.1175242142 | Gtpbp4 |
| **ENSMUSG00000021156** | 0.262716827533774 | 0.156363211890171 | 0.787275540343856 | 3402.63265659902 | 3849.74315057803 | 3673.19142284462 | 3220.84026808253 | 2428.61727832676 | 2248.02514255175 | 3634.40361413147 | 3479.10943476257 | Zmynd11 |
| **ENSMUSG00000021171** | -0.0524655639341704 | 0.683871658865313 | 0.944452327903597 | 629.749060448624 | 752.52013351684 | 818.683478680906 | 639.666005484066 | 798.224524752122 | 710.610776834912 | 746.45299092875 | 690.394259129171 | Esyt2 |
| **ENSMUSG00000021177** | 0.168404832722788 | 0.26258744475427 | 0.841952818014037 | 173.474633581257 | 173.470550258752 | 163.736695736181 | 164.137171495178 | 171.908436743598 | 162.669936865823 | 130.134761319517 | 136.776221148232 | Tdp1 |
| **ENSMUSG00000021178** | 0.162916942095014 | 0.123233575064082 | 0.770251097497048 | 5186.16873310632 | 4682.89044125739 | 5243.36008311239 | 5837.6557450628 | 4730.2194059385 | 5105.14523667629 | 4465.18393039527 | 4412.66142037748 | Psmc1 |
| **ENSMUSG00000021179** | -0.174336161445817 | 0.179399209577107 | 0.801755154770274 | 192.448421629207 | 206.047179415325 | 215.791714611846 | 229.79204009325 | 260.600050604945 | 227.493295165738 | 220.708555197901 | 243.157726485746 | Nrde2 |
| **ENSMUSG00000021180** | 0.0193830417135922 | 0.946502798572683 | 0.990510945542179 | 127.395434036235 | 166.140808698523 | 177.933519065908 | 121.930470253561 | 146.724398239759 | 89.2850029413918 | 224.872867560126 | 123.74991437221 | Rps6ka5 |
| **ENSMUSG00000021182** | -0.215628365707301 | 0.338500513146237 | 0.858570960161243 | 64.1494738764022 | 61.0811796685746 | 61.519567762149 | 80.6616957062019 | 87.5966556655277 | 53.8156182112498 | 83.2862472444909 | 84.6709940441436 | Ccdc88c |
| **ENSMUSG00000021185** | 0.16324667521652 | 0.406230920329821 | 0.881192654171779 | 1112.22538509649 | 1336.45621114841 | 1253.10627257054 | 1115.19483947295 | 908.81530252985 | 777.880299598975 | 1324.25133118741 | 1289.60437082619 | Dglucy |
| **ENSMUSG00000021186** | -0.925280765214973 | 0.0430023380837871 | 0.687730553581215 | 280.992765852973 | 306.220314071788 | 344.509579468035 | 351.72251034681 | 825.598479647599 | 1136.24339359662 | 238.406882737355 | 237.730098662403 | Fbln5 |
| **ENSMUSG00000021188** | 0.161656808773075 | 0.502239324361406 | 0.900927059989498 | 885.443442237664 | 1134.48111037766 | 1056.24365573167 | 920.106087067256 | 756.616113310996 | 550.387004433237 | 1163.92530524176 | 1100.72292257387 | Trip11 |
| **ENSMUSG00000021189** | 0.12263773120185 | 0.499436532500978 | 0.900927059989498 | 145.465708367616 | 144.96599974675 | 169.415425068072 | 148.192417692789 | 119.350443344282 | 112.523565350795 | 160.326025945645 | 164.999885829613 | Atxn3 |
| **ENSMUSG00000021190** | -0.11400276010102 | 0.402414023393015 | 0.881158207349486 | 1407.67437041457 | 1490.38078391322 | 1287.17864856189 | 1353.42821981453 | 1652.29191749102 | 1722.09978275999 | 1367.97661099076 | 1253.78202719213 | Lgmn |
| **ENSMUSG00000021192** | 0.264074948134603 | 0.0585039957764678 | 0.711712402279519 | 803.223694029881 | 837.21936932393 | 798.807926019289 | 776.603302845758 | 653.690042904001 | 527.148442023833 | 772.479943192653 | 722.960026069226 | Golga5 |
| **ENSMUSG00000021193** | 0.304668927567026 | 0.22619678797904 | 0.832048466795696 | 1030.90915060528 | 1251.75697534132 | 1391.28868631322 | 1021.40217004714 | 777.420319031559 | 587.079471395453 | 1220.14352213179 | 1215.78863242873 | Pitrm1 |
| **ENSMUSG00000021196** | -0.193500825077296 | 0.0756727571833105 | 0.735399943172214 | 1370.63030803524 | 1277.00386293767 | 1187.8008852538 | 1357.17992659156 | 1547.17593069238 | 1423.66771813397 | 1637.6158364448 | 1328.68329115425 | Pfkp |
| **ENSMUSG00000021200** | -0.177129879915846 | 0.163631399532394 | 0.793968226938511 | 11627.318018527 | 10990.5402616989 | 8941.15933306188 | 11165.0793684492 | 12403.6864422387 | 13491.8201020067 | 11702.7588159415 | 10708.7096954548 | Asb2 |
| **ENSMUSG00000021203** | -0.0590932992233739 | 0.805035188955882 | 0.963461896828986 | 110.228673421423 | 113.203786319092 | 102.217127974032 | 71.2824287636203 | 134.679858085749 | 80.7234273168747 | 87.4505596067155 | 110.723607596188 | Otub2 |
| **ENSMUSG00000021209** | -1.18079292856536 | 0.0173831466875009 | 0.642273034804458 | 9.93865088225949 | 9.77298874697194 | 7.57163910918757 | 1.87585338851632 | 13.1394983498292 | 18.3462334811079 | 15.616171358342 | 19.5394601640331 | Ppp4r4 |
| **ENSMUSG00000021210** | -0.314709522324695 | 0.288738942818567 | 0.851785577030345 | 259.308436655316 | 232.922898469498 | 227.149173275627 | 292.633128608546 | 252.935343234211 | 380.37857417497 | 164.49033830787 | 461.348364984116 | Akr1c6 |
| **ENSMUSG00000021211** | -0.24941528762685 | 0.493202513875684 | 0.899931765958683 | 16.2632468982428 | 26.8757190541728 | 17.036187995672 | 25.3240207449704 | 21.8991639163819 | 23.2385624094033 | 18.7394056300105 | 37.9933947633978 | Akr1c12 |
| **ENSMUSG00000021213** | -0.139370227132394 | 0.676666215965661 | 0.94409858171788 | 45.1756858284523 | 64.338842584232 | 62.4660226507975 | 51.5859681841989 | 53.6529515951357 | 57.4848649074714 | 34.3555769883525 | 100.953877514171 | Akr1c13 |
| **ENSMUSG00000021215** | 0.280101051327618 | 0.0612138186460565 | 0.71282992338528 | 302.67709505063 | 315.178887089845 | 250.810545491838 | 276.688374806158 | 221.181555555458 | 254.401104271363 | 201.96914956789 | 267.039288908453 | Net1 |
| **ENSMUSG00000021217** | -0.533189358314013 | 0.107144243771442 | 0.762287160109383 | 25.2983840639333 | 45.6072808192024 | 27.447191770805 | 45.9584080186499 | 72.2672409240604 | 55.0387004433237 | 34.3555769883525 | 47.7631248454143 | Tshz3 |
| **ENSMUSG00000021218** | 0.121779089173515 | 0.074365617735178 | 0.73518749490507 | 5418.37175826456 | 5405.2771928044 | 5860.44867051118 | 5429.6576330605 | 5121.11948184592 | 4903.3366683841 | 5215.80123368624 | 5081.34516821328 | Gdi2 |
| **ENSMUSG00000021219** | -0.309476362841251 | 0.126348540713392 | 0.771949105062466 | 215.939778260002 | 297.26174105373 | 272.579007930753 | 215.723139679377 | 224.466430142915 | 300.878229090169 | 367.500565966316 | 348.453706258591 | Rgs6 |
| **ENSMUSG00000021221** | -0.307747599546565 | 0.21926331326605 | 0.827535829146756 | 95.7724539563188 | 104.245213301034 | 104.110037751329 | 63.779015209555 | 94.1664048404423 | 91.7311674055395 | 155.120635492864 | 113.980184290193 | Dpf3 |
| **ENSMUSG00000021222** | 0.0797682601820884 | 0.558017320304208 | 0.916502374249196 | 253.887354355902 | 254.097707421271 | 283.936466594534 | 249.488500672671 | 270.454674367317 | 204.254732756335 | 252.981976005141 | 256.184033261768 | Dcaf4 |
| **ENSMUSG00000021223** | -0.734337602691634 | 0.0248200491719884 | 0.655303666217042 | 80.412720774645 | 149.852494120236 | 88.0203046443056 | 76.9099889291692 | 244.175677667659 | 185.908499275227 | 110.35427759895 | 118.322286548867 | Papln |
| **ENSMUSG00000021224** | -0.21533787505237 | 0.229194170284243 | 0.832048466795696 | 353.273863178497 | 415.352021746308 | 446.726707442067 | 378.922384480297 | 567.188345434292 | 517.363784167243 | 360.213019332423 | 407.07208675069 | Numb |
| **ENSMUSG00000021226** | 0.394206857011193 | 0.0221933309901385 | 0.648762042745232 | 5182.55467824004 | 5213.07508078062 | 7524.31636475515 | 6479.19760393538 | 5639.03470846835 | 4052.0714348607 | 4471.43039893861 | 4401.8061647308 | Acot2 |
| **ENSMUSG00000021228** | 0.101703180816703 | 0.75598264367142 | 0.956566265296272 | 29.8159526467785 | 33.3910448854875 | 41.6440151005317 | 25.3240207449704 | 27.3739548954774 | 22.0154801773295 | 26.0269522639034 | 45.5920737160773 | Acot3 |
| **ENSMUSG00000021234** | -0.0626924382168574 | 0.824022843380892 | 0.966637987288264 | 208.711668527449 | 210.933673788811 | 278.257737262643 | 174.454365132018 | 186.142893289246 | 155.33144347338 | 361.254097422979 | 207.335382851685 | Fam161b |
| **ENSMUSG00000021235** | 0.0427648943365681 | 0.584464381096519 | 0.922450244597442 | 1416.70950758026 | 1297.36425616053 | 1347.75176143539 | 1427.52442866092 | 1344.60866446585 | 1308.69798831903 | 1300.30653510461 | 1374.27536487033 | Coq6 |
| **ENSMUSG00000021236** | 0.0152671076210977 | 0.941410640952539 | 0.989934163630124 | 3492.98402825593 | 4428.79273383612 | 3747.0149041592 | 3118.60625840839 | 2799.80810670943 | 2891.3663966226 | 4455.81422758026 | 4484.30610764561 | Entpd5 |
| **ENSMUSG00000021238** | 0.239415885410019 | 0.419129140519819 | 0.882550938324998 | 4850.9651442592 | 6354.88593271851 | 6711.31161540614 | 4867.83954319986 | 3860.82259845814 | 2497.53391789482 | 6610.84587503147 | 6330.78509314674 | Aldh6a1 |
| **ENSMUSG00000021240** | -0.204043723205606 | 0.329676773726485 | 0.855977678491063 | 103.90407740544 | 79.8127414336042 | 106.002947528626 | 111.613276616721 | 122.635317931739 | 140.654456688494 | 86.4094815161593 | 112.894658725525 | Abcd4 |
| **ENSMUSG00000021241** | 0.044355289430041 | 0.741736252789435 | 0.954383574136061 | 993.86508822595 | 819.302223287815 | 731.609628925249 | 906.037186653384 | 770.850569856644 | 785.218792991418 | 873.464517976599 | 916.18357658022 | Isca2 |
| **ENSMUSG00000021242** | -0.0912353124552213 | 0.589010989364659 | 0.92352104528132 | 1553.14007878219 | 1367.40400884716 | 1435.77206607969 | 1608.54428065275 | 1664.33645764503 | 2040.1011630992 | 1287.81359801794 | 1363.42010922365 | Npc2 |
| **ENSMUSG00000021243** | 0.151488962391453 | 0.315702419427981 | 0.853277176422981 | 518.616873310632 | 466.66021266791 | 476.066808990169 | 618.093691516128 | 523.390017601528 | 513.694537471021 | 416.431236222455 | 419.012867962044 | Fcf1 |
| **ENSMUSG00000021244** | 0.264547608145727 | 0.10426668767008 | 0.761198544536855 | 1042.65482892068 | 1191.49021140166 | 1217.1409868019 | 981.071322194037 | 866.111932892906 | 714.280023531134 | 1068.1461209106 | 1039.93349095243 | Ylpm1 |
| **ENSMUSG00000021245** | -0.00965467276273479 | 0.971271021691033 | 0.995122272737203 | 201.483558794897 | 229.665235553841 | 216.738169500494 | 179.143998603309 | 152.199189218854 | 130.869798831903 | 266.515991182371 | 282.236646813812 | Mlh3 |
| **ENSMUSG00000021248** | 0.081022054190222 | 0.282682335741529 | 0.850587941933635 | 3352.93940218773 | 3482.44165683767 | 3666.56623862408 | 3369.03268577532 | 3149.09977117572 | 3365.92230266726 | 3416.81829320524 | 3181.6754300434 | Tmed10 |
| **ENSMUSG00000021250** | -0.0916505874846733 | 0.82201074899485 | 0.966637987288264 | 39.754603529038 | 29.3189662409158 | 17.036187995672 | 25.3240207449704 | 43.7983278327639 | 33.0232202659942 | 28.1091084450157 | 14.1118323406906 | Fos |
| **ENSMUSG00000021252** | -0.0986342764727294 | 0.713145138565507 | 0.950540828584099 | 79.509207058076 | 67.5965054998893 | 68.1447519826882 | 105.047789756914 | 104.021028602814 | 108.854318654574 | 71.8343882483734 | 58.6183804920994 | Erg28 |
| **ENSMUSG00000021253** | -0.146524507207701 | 0.357027915343025 | 0.865163598066054 | 439.107666252556 | 419.424100390879 | 302.865564367503 | 391.115431505653 | 412.7992398238 | 499.017550686135 | 415.390158131898 | 392.96025441 | Tgfb3 |
| **ENSMUSG00000021254** | 0.16747137058063 | 0.517299278510744 | 0.905197321821966 | 237.624107457659 | 323.323044378988 | 359.65285768641 | 206.343872736795 | 216.801722772181 | 173.677676954488 | 331.062832796851 | 281.151121249143 | Gpatch2l |
| **ENSMUSG00000021255** | 0.195496990027597 | 0.472851191897236 | 0.89728597737823 | 470.730646332472 | 598.595560752032 | 731.609628925249 | 558.066383083606 | 442.363111110915 | 292.316653465652 | 689.193695948162 | 635.032455331077 | Esrrb |
| **ENSMUSG00000021256** | -0.15297813082476 | 0.659085457471742 | 0.938989601151841 | 185.220311896654 | 178.357044632238 | 148.593417517806 | 138.813150750208 | 127.015150715015 | 83.1695917810225 | 239.447960827911 | 273.552442296464 | Vash1 |
| **ENSMUSG00000021257** | 0.0626734839234549 | 0.721831626067734 | 0.951674101666722 | 122.87786545339 | 107.502876216691 | 130.610774633486 | 99.4202295913651 | 123.730276127558 | 92.9542496376133 | 109.313199508394 | 113.980184290193 | Angel1 |
| **ENSMUSG00000021258** | 0.219353957950516 | 0.123932200529995 | 0.771244742545532 | 763.469090500843 | 850.250020986559 | 903.864418659267 | 768.161962597434 | 642.74046094581 | 607.871869340708 | 850.560799984364 | 719.703449375221 | Ccnk |
| **ENSMUSG00000021259** | 0.65716988346151 | 0.11549291917413 | 0.769071703867842 | 14.4562194651047 | 21.9892246806869 | 16.0897331070236 | 15.0068271081306 | 7.66470737073368 | 8.56157562451702 | 14.5750932677859 | 11.9407812113536 | Cyp46a1 |
| **ENSMUSG00000021260** | -0.440309281478949 | 0.299700530408509 | 0.851785577030345 | 19.877301764519 | 35.8342920722305 | 33.1259211026956 | 12.1930470253561 | 45.9882442244021 | 44.0309603546589 | 27.0680303544596 | 20.6249857287016 | Hhipl1 |
| **ENSMUSG00000021262** | -0.1636861885372 | 0.297696560586618 | 0.851785577030345 | 216.843291976571 | 240.252640029727 | 251.757000380487 | 307.639955716677 | 309.873169416804 | 302.101311322243 | 279.008928269045 | 247.49982874442 | Evl |
| **ENSMUSG00000021264** | 0.190295650902343 | 0.117352791323076 | 0.769868040577702 | 1076.08483643373 | 1232.21099784738 | 1267.30309590027 | 1051.4158242634 | 964.658170516624 | 902.634687270508 | 1103.5427759895 | 1083.35451353917 | Yy1 |
| **ENSMUSG00000021265** | -0.372202308150327 | 0.0148844174745332 | 0.635862253526407 | 196.062476495483 | 150.666909849151 | 170.36187995672 | 157.571684635371 | 243.08071947184 | 220.154801773295 | 219.667477107345 | 189.966973816989 | Slc25a29 |
| **ENSMUSG00000021266** | -0.168838763731657 | 0.2355538031683 | 0.832048466795696 | 1092.34808333198 | 1081.54408799823 | 1219.0338965792 | 1143.3326403007 | 1432.20532013138 | 1465.25251402448 | 1161.84314906065 | 1041.0190165171 | Wars |

| | | | | | | | | | | | | |
|---|---|---|---|---|---|---|---|---|---|---|---|---|
| **ENSMUSG00000021268** | 0.516603010069796 | 0.461764042928597 | 0.895328112556867 | 15.3597331816738 | 12.2162359337149 | 5.67872933189068 | 15.0068271081306 | 2.18991639163819 | 17.123151249034 | 12.4929370866736 | 2.17105112933702 | Meg3 |
| **ENSMUSG00000021270** | 0.209933290753116 | 0.0720352233344041 | 0.728760020516246 | 9224.87504616995 | 8375.45135615496 | 8050.54528284369 | 9742.24457325952 | 7279.28208580536 | 8060.11190936673 | 6790.95238469768 | 8470.35598110837 | Hsp90aa1 |
| **ENSMUSG00000021271** | -0.224746109307268 | 0.0440654241129153 | 0.691603097094229 | 336.107102563685 | 333.910448854875 | 353.974128354519 | 332.963976461647 | 404.039574257247 | 359.586176229715 | 438.293876124134 | 382.104998763315 | Zfp839 |
| **ENSMUSG00000021273** | 0.390991689256676 | 0.0221709695756114 | 0.648762042745232 | 1277.56839522863 | 1451.28882892533 | 1994.18045038228 | 1681.70256280488 | 1358.8431210115 | 1023.71982824582 | 1235.75969349013 | 1264.63728283881 | Fdft1 |
| **ENSMUSG00000021275** | 0.140495409882642 | 0.410749999042122 | 0.882192126163777 | 195.158962778914 | 177.542628903324 | 177.933519065908 | 147.254490998531 | 155.484063806312 | 125.977469903608 | 174.901119213431 | 175.855141476298 | Tecpr2 |
| **ENSMUSG00000021276** | 0.0654806695530649 | 0.572094978684682 | 0.921574199900435 | 327.975479114563 | 316.807718547674 | 288.668741037776 | 342.343243404229 | 300.018545654433 | 283.755077841135 | 313.364505257397 | 321.315567141878 | Cinp |
| **ENSMUSG00000021277** | 0.121590924720544 | 0.643212074738109 | 0.934616627390525 | 506.871194995234 | 528.555808065399 | 376.689045682082 | 395.805064976944 | 338.342082508101 | 253.178022039289 | 524.703357640293 | 544.933833463591 | Traf3 |
| **ENSMUSG00000021279** | -0.0415790283914078 | 0.810647889482102 | 0.964527941143258 | 1091.44456961541 | 1273.74620002201 | 1336.39430277161 | 994.202295913651 | 1366.50782838223 | 906.30393396673 | 1201.40411650178 | 1357.9924814003 | Cdc42bpb |
| **ENSMUSG00000021280** | -0.0576461193325143 | 0.878484823326521 | 0.978050192049164 | 46.0791995450213 | 27.6901347830872 | 23.6613722162112 | 40.3308478531009 | 24.0890803080201 | 56.2617826753975 | 39.5609674411332 | 23.8815624227072 | Exoc3l4 |
| **ENSMUSG00000021281** | -0.296159532665073 | 0.0544597922286926 | 0.703396849069134 | 468.020105182765 | 445.485403716138 | 443.887342776122 | 447.391033161143 | 571.568178217569 | 431.748027922072 | 533.031982364742 | 678.453477917817 | Tnfaip2 |
| **ENSMUSG00000021282** | 0.176264422519335 | 0.23851502100588 | 0.832048466795696 | 3575.20377646371 | 3904.30900441529 | 4281.76191624557 | 3802.35481852259 | 2867.69551485021 | 3033.24393554317 | 3701.03261192707 | 4170.58921945641 | Eif5 |
| **ENSMUSG00000021285** | -0.0789745908873761 | 0.604686698428329 | 0.926448151609826 | 741.784761303186 | 716.685841444609 | 726.877354482007 | 649.045272426648 | 909.91026072567 | 673.918309872697 | 801.630129728225 | 607.894316214364 | Ppp1r13b |
| **ENSMUSG00000021286** | -0.129922032347475 | 0.269573835854009 | 0.843793744782845 | 761.662063067705 | 636.873100011005 | 675.768790494991 | 698.75538722233 | 826.693437843418 | 709.387694602839 | 814.123066814899 | 682.795580176491 | Zfyve21 |
| **ENSMUSG00000021287** | 0.0597791329755302 | 0.885834116814628 | 0.979948700426504 | 26.2018977805023 | 17.1027303072009 | 14.1968233297267 | 28.1378008277448 | 18.6142893289246 | 9.78465785659088 | 24.9858741733473 | 28.2236646813812 | Xrcc3 |
| **ENSMUSG00000021288** | -0.0911862575978516 | 0.331606783094849 | 0.856333478432716 | 1630.84225840713 | 1521.32858161197 | 1630.74177314127 | 1702.33695007856 | 1730.03394939417 | 1892.10821301826 | 1705.28591233095 | 1581.61074772202 | Klc1 |
| **ENSMUSG00000021290** | 0.164511245375132 | 0.322457062813028 | 0.855977678491063 | 5621.6623444926 | 3769.93040914443 | 4252.42181469747 | 5193.30010610744 | 4380.92774147221 | 4865.42111918981 | 3696.86829956484 | 3864.47101021989 | 2010107E04F |
| **ENSMUSG00000021294** | -0.0855909002189393 | 0.581910319260663 | 0.922450244597442 | 845.688838708626 | 589.636987733974 | 757.163910918757 | 671.555513088843 | 851.877476347257 | 650.679747463293 | 733.960053842076 | 801.117866725359 | Kif26a |
| **ENSMUSG00000021302** | 0.250529158828869 | 0.278481459669191 | 0.848075053184273 | 254.790868072471 | 304.591482613959 | 409.814966784777 | 325.460562907582 | 226.656346534553 | 196.916239363891 | 324.816364253515 | 338.683976176574 | Ggps1 |
| **ENSMUSG00000021306** | -0.44701243888465 | 0.140194892878563 | 0.779711473022872 | 58.7283915769879 | 61.0811796685746 | 51.1085639870161 | 53.4618215727152 | 100.736154015357 | 105.185071958352 | 51.0128264372507 | 49.9341759747513 | Gpr137b |
| **ENSMUSG00000021311** | 0.141307712641796 | 0.676434066110667 | 0.94409858171788 | 894.478579403355 | 1413.82570539527 | 1516.22073161481 | 844.134024832345 | 695.298454345127 | 599.310293716191 | 1464.79687341248 | 1473.05819125516 | Mtr |
| **ENSMUSG00000021313** | 0.330951220877147 | 0.308753151033862 | 0.851785577030345 | 29631.6358485984 | 47067.5282211458 | 34623.2127365375 | 24855.9953245355 | 20238.1123333244 | 14143.7229317021 | 37343.4711082486 | 36537.7049811773 | Ryr2 |
| **ENSMUSG00000021314** | 0.484722069013796 | 0.12310956673639 | 0.770251097497048 | 68.6670424592474 | 114.83261777692 | 141.968233297267 | 61.9031618210386 | 70.0773245324222 | 53.8156182112498 | 87.4505596067155 | 65.1315338801105 | Amph |
| **ENSMUSG00000021318** | -0.00717500142680797 | 0.988347030717393 | 0.998437672973644 | 6.32459601598331 | 10.5874044758863 | 12.3039135524298 | 15.0068271081306 | 10.949581958191 | 13.4539045528125 | 14.5750932677859 | 5.42762782334254 | Gli3 |
| **ENSMUSG00000021322** | -0.947180888515411 | 0.0997765221702487 | 0.759800953500741 | 4.51756858284523 | 8.14415728914329 | 7.57163910918757 | 12.1930470253561 | 8.75966556655277 | 31.8001380339204 | 9.36970281500523 | 13.0263067760221 | Aoah |
| **ENSMUSG00000021326** | -0.0197220525451059 | 0.869450365398916 | 0.975613767803763 | 422.844419354313 | 399.063707168021 | 375.742590793433 | 416.439452250624 | 419.368988998714 | 462.325083723919 | 363.336253604092 | 392.96025441 | Trim27 |
| **ENSMUSG00000021327** | 0.119250445091415 | 0.455071308258471 | 0.892775943845882 | 360.501972911049 | 360.786167909048 | 363.438677241004 | 343.281170098487 | 295.638712871156 | 256.847268735511 | 386.239971596327 | 374.506319810635 | Zkscan3 |
| **ENSMUSG00000021338** | 0.0190536656194475 | 0.937816434105305 | 0.988970340396118 | 61.4389327266951 | 66.7820897709749 | 76.6628459805242 | 45.9584080186499 | 75.5521155115177 | 67.2695227640623 | 54.1360607089191 | 51.0197015394199 | Carmil1 |
| **ENSMUSG00000021339** | 0.0695863704005318 | 0.843881307131252 | 0.971721411930626 | 255.69438178904 | 409.651111643907 | 352.081218577222 | 273.874594723383 | 225.561388338734 | 136.985209992272 | 458.0743598447 | 409.243137880027 | Mrs2 |
| **ENSMUSG00000021340** | 0.0769209160375234 | 0.81483439842446 | 0.966177091475091 | 299.063040184354 | 378.703313945163 | 328.419846361011 | 254.178134143962 | 212.421889988905 | 151.662196777159 | 359.171941241867 | 471.118095066132 | Gpld1 |
| **ENSMUSG00000021357** | 0.147993941211568 | 0.30547506856515 | 0.851785577030345 | 374.958192376154 | 452.000729547452 | 517.7108240907 | 407.060185308042 | 379.950493949227 | 355.916929533493 | 415.390158131898 | 428.78259804406 | Exoc2 |
| **ENSMUSG00000021360** | -0.457326259330843 | 0.0996126742042809 | 0.759800953500741 | 188.83436676293 | 191.387696294867 | 247.971180825893 | 184.771558768858 | 359.146288228664 | 382.824738639118 | 193.640524843441 | 181.282769299641 | Gcnt2 |
| **ENSMUSG00000021361** | -0.0897828851073464 | 0.324628237920058 | 0.855977678491063 | 1693.18470485039 | 1528.65832317219 | 1412.11069386348 | 1571.96513957668 | 1643.53225192446 | 1690.29964472607 | 1583.47977573588 | 1687.99225305953 | Tmem14c |
| **ENSMUSG00000021364** | -0.777955508582791 | 0.0991528346135397 | 0.759800953500741 | 29.8159526467785 | 79.8127414336042 | 31.2330113253987 | 44.0825546301336 | 35.0386622662111 | 53.8156182112498 | 85.3684034256032 | 143.289374536243 | Elovl2 |
| **ENSMUSG00000021365** | -0.0635140663183365 | 0.791475588133872 | 0.961804768755116 | 505.064167562096 | 742.747144769868 | 516.764369202052 | 378.922384480297 | 708.437952694956 | 611.54111603693 | 439.33495421469 | 481.973350712817 | Nedd9 |
| **ENSMUSG00000021366** | 0.278169055597382 | 0.275892368490976 | 0.846508725058461 | 755.337467051722 | 961.010560118908 | 913.328967545751 | 625.597105070194 | 537.624474147177 | 434.19419238622 | 865.13589325215 | 846.709940441436 | Hivep1 |
| **ENSMUSG00000021367** | -0.620648844559192 | 0.0656904032711817 | 0.718586813809351 | 66.8600150261093 | 34.2054606144018 | 66.2518422053913 | 91.9168160372998 | 95.2613630362614 | 146.769867848863 | 72.8754663389296 | 83.5854684794751 | Edn1 |
| **ENSMUSG00000021368** | 0.070539881516641 | 0.647544102188544 | 0.935161944208659 | 435.49361138628 | 333.09603312596 | 309.490748588042 | 392.053358199911 | 323.012667766634 | 321.670627035425 | 381.034581143546 | 373.420794245967 | Tbc1d7 |
| **ENSMUSG00000021371** | 0.15323936517224 | 0.300012005693753 | 0.851785577030345 | 1883.82609904646 | 2066.17270425565 | 2073.68266102875 | 1754.86084495702 | 1575.64484378368 | 1431.00621152642 | 2017.60933949779 | 1969.14337430867 | Mcur1 |
| **ENSMUSG00000021373** | 0.104570684952506 | 0.603318920200308 | 0.926448151609826 | 3251.74586593199 | 3760.15742039746 | 3586.11757308896 | 3139.24064568207 | 2523.87864136302 | 2385.01035254403 | 4092.47797397617 | 3774.3723883524 | Cap2 |
| **ENSMUSG00000021374** | 0.24494233303899 | 0.312752318118896 | 0.852613848149207 | 475.248214915318 | 636.873100011005 | 681.447519826882 | 423.942865804689 | 439.078236523458 | 305.770558018465 | 562.182168900314 | 563.387768062955 | Nup153 |
| **ENSMUSG00000021375** | 0.286903888645813 | 0.339655347086833 | 0.858570960161243 | 2865.04199524044 | 4026.47136375244 | 3622.08285885761 | 2488.3195198669 | 2025.67266226533 | 1468.92176072071 | 3722.89525182874 | 3438.94498886983 | Kif13a |
| **ENSMUSG00000021376** | 0.390088339997418 | 0.155306482618883 | 0.787275540343856 | 38.8510898124689 | 60.2667639396603 | 67.1982970940397 | 53.4618215727152 | 36.1336204620302 | 34.2463024980681 | 41.6431236222455 | 55.3618037980939 | Tpmt |
| **ENSMUSG00000021377** | -0.00113411573971193 | 0.989739018510229 | 0.998628446692557 | 1060.72510325206 | 1060.36927904646 | 1131.01359193489 | 1183.6634881538 | 1122.33215071457 | 1170.48969609468 | 1051.4888714617 | 1095.29529475052 | Dek |
| **ENSMUSG00000021379** | -1.82643530464104 | 0.00361923572977884 | 0.635862253526407 | 5.42108229941427 | 7.32974156022896 | 6.62518422053913 | 3.75170677703264 | 16.4243729372865 | 45.2540425867328 | 13.5340151772298 | 7.59867895267955 | Id4 |
| **ENSMUSG00000021384** | -0.758834441496343 | 0.0830269769918225 | 0.741377052226143 | 7.22810973255236 | 7.32974156022896 | 13.2503684410783 | 15.9447538023887 | 21.8991639163819 | 23.2385624094033 | 14.5750932677859 | 14.1118323406906 | Susd3 |
| **ENSMUSG00000021385** | -0.0461181069047331 | 0.831075218225139 | 0.967583158900775 | 142.755167217909 | 143.337168288922 | 121.146225747001 | 106.92364314543 | 130.300025302473 | 100.292743030056 | 176.983275394543 | 122.664388807541 | Ippk |
| **ENSMUSG00000021388** | -0.308116303796964 | 0.350549734942413 | 0.861774532843376 | 453.563885717661 | 295.632909595901 | 253.649910157784 | 377.984457786039 | 531.054724972262 | 643.34125407085 | 234.242570375131 | 301.776106977845 | Aspn |
| **ENSMUSG00000021390** | -0.264573634253733 | 0.100668861922646 | 0.759800953500741 | 1351.65651998729 | 1223.25242482932 | 963.491076644119 | 1212.7392156758 | 1369.79270296969 | 1722.09978275999 | 1188.91117941511 | 1428.55164310376 | Ogn |
| **ENSMUSG00000021391** | 0.526290554951378 | 0.116979613771371 | 0.769868040577702 | 84.0267756409212 | 46.4216965481167 | 46.3762895437739 | 102.23400967414 | 60.2227007700503 | 46.4771248188067 | 52.0539045278068 | 34.7368180693922 | Cenpp |
| **ENSMUSG00000021392** | 0.375075798028143 | 0.00445382405423241 | 0.635862253526407 | 704.740698923855 | 816.044560372157 | 613.302767844194 | 811.30659053331 | 534.339599559719 | 546.717757737015 | 599.660980160335 | 590.525907179668 | Nol8 |
| **ENSMUSG00000021395** | 0.168539437809261 | 0.223701435216335 | 0.832048466795696 | 1071.56726785089 | 1121.45045871503 | 1181.17570103326 | 950.119741283517 | 992.032125412102 | 803.565026472526 | 930.723812957186 | 1120.2623827379 | Spin1 |
| **ENSMUSG00000021400** | -0.00441025813528026 | 0.971477336900098 | 0.995122272737203 | 643.30176619716 | 603.482055125517 | 595.320124959873 | 689.376120279749 | 738.001823982071 | 593.194882555822 | 632.975479058131 | 574.243023709641 | Wrnip1 |
| **ENSMUSG00000021408** | -0.498360375688417 | 0.0941995968093242 | 0.757085157577544 | 57.8248778604189 | 84.6992358070902 | 83.2880302010633 | 60.9652351267805 | 160.958854785407 | 77.0541806206532 | 77.0397787011541 | 90.0986218674861 | Ripk1 |
| **ENSMUSG00000021411** | -0.444481452191575 | 0.202989665589192 | 0.815599205968724 | 170.764092431549 | 175.913797445495 | 195.916161950228 | 207.281799431054 | 365.716037403578 | 375.486245246675 | 131.175839410073 | 148.717002359586 | Pxdc1 |
| **ENSMUSG00000021413** | 0.132862399915465 | 0.578345215723572 | 0.922450244597442 | 555.660935689963 | 644.202841571234 | 654.946782944725 | 556.19052969509 | 487.256397139498 | 330.232202659942 | 646.509494235361 | 733.815281715911 | Prpf4b |
| **ENSMUSG00000021417** | 0.212488749910528 | 0.0340991188559153 | 0.67781375479544 | 6400.49116817511 | 5846.69051787597 | 7017.96299932823 | 6505.45955137461 | 5424.42290208781 | 5067.229687482 | 5810.2568233938 | 5936.73931317207 | Eci2 |
| **ENSMUSG00000021418** | 0.183986696106028 | 0.352060062618982 | 0.862728204353539 | 114.746242004269 | 92.8433930962335 | 109.78876708322 | 127.55803041911 | 112.780694169367 | 78.277262852727 | 111.395355689507 | 87.9275707381491 | Rpp40 |
| **ENSMUSG00000021420** | 0.137548737437698 | 0.120608638522304 | 0.769868040577702 | 1826.90473490261 | 1693.17030041289 | 1751.88799888827 | 1906.80496942684 | 1754.12302970219 | 1616.91471080164 | 1527.26155884585 | 1627.20282143809 | Fars2 |
| **ENSMUSG00000021423** | -0.618821103024851 | 0.057469840755514 | 0.710862211775517 | 31.6229800799166 | 26.8757190541728 | 33.1259211026956 | 42.2067012416173 | 74.4571573156986 | 57.4848649074714 | 31.2323427166841 | 42.3354970220718 | Ly86 |
| **ENSMUSG00000021427** | -0.114796055832341 | 0.334471679060807 | 0.85842786809952 | 992.058060792811 | 977.298874697194 | 1115.87031371652 | 1042.03655732082 | 1194.59939163863 | 1254.88237010778 | 1035.87270010336 | 984.571687154336 | Ssr1 |
| **ENSMUSG00000021428** | 0.0971965494060205 | 0.493619974238721 | 0.900117970535229 | 161.728955265859 | 159.625482867208 | 140.07532351997 | 178.206071909051 | 151.104231023035 | 150.439114545085 | 147.833088858971 | 148.717002359586 | Riok1 |
| **ENSMUSG00000021431** | 0.030982219755511 | 0.815725431295774 | 0.966177091475091 | 344.238726012806 | 289.931999493501 | 300.972654590206 | 349.846656958294 | 326.297542354091 | 349.801518373124 | 299.830490080167 | 282.236646813812 | Snrnp48 |
| **ENSMUSG00000021432** | -0.339873783725686 | 0.23742531701377 | 0.832048466795696 | 28.0089252136404 | 25.2468875963442 | 20.8220075502658 | 32.8274342990356 | 31.7537876787538 | 34.2463024980681 | 28.1091084450157 | 41.2499714574033 | Slc35b3 |
| **ENSMUSG00000021451** | 0.318332430434695 | 0.370400928187992 | 0.873195670830145 | 611.678786117243 | 727.273245920495 | 920.900606654939 | 634.976372012775 | 447.837902090011 | 234.831788558181 | 787.055036460439 | 851.05204270011 | Sema4d |
| **ENSMUSG00000021453** | 0.148657450988785 | 0.57006244243653 | 0.920872458148558 | 206.904641094311 | 175.913797445495 | 288.668741037776 | 353.598363735327 | 204.757182618171 | 311.885969178834 | 201.96914956789 | 206.249857287016 | Gadd45g |
| **ENSMUSG00000021456** | 0.229004018430861 | 0.107201615785307 | 0.762287160109383 | 1402.25328811516 | 1305.50841344967 | 1669.54642357586 | 1603.85464718146 | 1348.98849724913 | 1464.02943179241 | 1084.80337035949 | 1206.01890234671 | Fbp2 |
| **ENSMUSG00000021457** | 0.269533654912798 | 0.284750195540239 | 0.851125503777472 | 581.862833470465 | 773.694942468612 | 838.559031342524 | 553.376749612315 | 481.781606160403 | 385.270903103266 | 810.99983254323 | 600.295637261685 | Syk |
| **ENSMUSG00000021458** | 0.427861133697292 | 0.00483956600087225 | 0.635862253526407 | 869.180195339421 | 902.372627637076 | 787.450467355508 | 1093.62252550502 | 574.853052805026 | 658.018240855737 | 706.892023487617 | 775.065253173314 | 2010111I01R |
| **ENSMUSG00000021460** | 0.205303452458406 | 0.0390113279018814 | 0.679566075569698 | 2933.70903769969 | 2681.87099531488 | 2827.06075239291 | 3112.97869824284 | 2404.52819801874 | 2282.27144504982 | 2657.87236518982 | 2675.82051690787 | Auh |
| **ENSMUSG00000021461** | 0.0732649935127697 | 0.762894727464212 | 0.957842155335615 | 72.2810973255236 | 64.338842584232 | 61.519567762149 | 65.6548685980713 | 79.9319482947941 | 74.6080161565054 | 43.7252798033577 | 53.1907526687569 | Fancc |
| **ENSMUSG00000021464** | -1.19053637512395 | 0.114252098458993 | 0.767518180080285 | 2.71054114970714 | 4.07207864457164 | 6.62518422053913 | 15.0068271081306 | 28.4689130912965 | 25.6847268735511 | 5.20539045278068 | 5.42762782334254 | Ror2 |
| **ENSMUSG00000021466** | 0.0325691685775189 | 0.881319478881781 | 0.978695839596094 | 180.702743313809 | 225.593156909269 | 297.186835035612 | 218.536919762152 | 244.175677667659 | 157.777607937528 | 255.064132186253 | 243.157726485746 | Ptch1 |
| **ENSMUSG00000021468** | -0.0477726014048627 | 0.685675373186187 | 0.944452327903597 | 354.177376895066 | 379.517729674077 | 417.386605893965 | 415.501525556365 | 433.603445544362 | 379.155491942897 | 377.911346871878 | 427.697072479392 | Sptlc1 |
| **ENSMUSG00000021470** | 0.229644339087595 | 0.368511396642857 | 0.871495599696183 | 241.238162323935 | 280.973426475443 | 293.401015481019 | 265.43325447506 | 157.67398019795 | 160.223772401676 | 315.446661438509 | 287.664274637155 | Ercc6l2 |
| **ENSMUSG00000021474** | -0.263402617027121 | 0.084010586619687 | 0.742565114108635 | 143.658680934478 | 149.038078391322 | 153.325691961048 | 145.378637610015 | 185.047935093427 | 192.023910435596 | 142.627698406191 | 191.052499381657 | Sfxn1 |
| **ENSMUSG00000021476** | 0.129058384394925 | 0.275602925371827 | 0.846289746011912 | 2149.45913171776 | 2119.924142364 | 1811.51465687313 | 2054.99738711963 | 1713.60957645689 | 1670.73032901289 | 2085.27941538394 | 1969.14337430867 | Habp4 |
| **ENSMUSG00000021477** | 0.0468461585242704 | 0.81937012742027 | 0.966398681144067 | 4305.2428594515 | 4182.83918370399 | 6475.64434813267 | 4958.8184325429 | 5291.9329603937 | 5945.40273011103 | 4403.76032305246 | 3645.19484615685 | Ctsl |
| **ENSMUSG00000021481** | 0.29787516139129 | 0.236911183745686 | 0.832048466795696 | 401.160090156656 | 481.319695788368 | 782.718192912266 | 611.528204656321 | 405.134532453066 | 357.140011765567 | 616.318229609233 | 472.203620630801 | Zfp346 |
| **ENSMUSG00000021482** | 0.27878217746654 | 0.274934969775687 | 0.846286635233373 | 388.510898124689 | 554.617111390658 | 633.178320505811 | 406.122258613784 | 374.475702970131 | 260.516515431732 | 520.539045278068 | 477.631248454143 | Aaed1 |
| **ENSMUSG00000021483** | -0.609575953456513 | 0.012375788573108 | 0.635862253526407 | 40.658117245607 | 30.9477976987445 | 25.5542819935081 | 35.6412143818101 | 49.2731188118594 | 51.3694537471021 | 51.0128264372507 | 51.0197015394199 | Cdk20 |
| **ENSMUSG00000021484** | -0.0280249231475133 | 0.810280582969896 | 0.964412705343932 | 1718.48308891432 | 2058.84296269542 | 2090.71884902442 | 1637.62000817475 | 1904.13230252941 | 1828.50793695042 | 1971.80190351332 | 1948.51838857997 | Lman2 |
| **ENSMUSG00000021486** | -0.400852440370526 | 0.0723252709264631 | 0.72957913253658 | 628.845546732055 | 573.348673155687 | 582.069756518795 | 654.672832592197 | 849.687559955619 | 1157.03579154187 | 584.044808801993 | 630.690353072403 | Prelid1 |
| **ENSMUSG00000021488** | 0.175838192308264 | 0.372526957035781 | 0.873700364743909 | 1217.0329762185 | 1369.8472560339 | 1312.7329305554 | 1128.32581319257 | 1019.40608030758 | 761.980230582015 | 1395.04464134522 | 1273.32148735616 | Nsd1 |
| **ENSMUSG00000021492** | -0.367914181499221 | 0.42227547036372 | 0.883344319781939 | 42.4651446787451 | 116.461449234749 | 41.6440151005317 | 53.4618215727152 | 49.2731188118594 | 62.3771938357668 | 61.4236073428121 | 155.230155747597 | F12 |
| **ENSMUSG00000021493** | -0.357487013197546 | 0.0824968566133646 | 0.739855948845396 | 663.179067961679 | 540.772043999114 | 483.638448099356 | 602.14893771374 | 730.337116611338 | 1005.37359476471 | 562.182168900314 | 637.203506460414 | Pdlim7 |
| **ENSMUSG00000021494** | 0.0281383406269944 | 0.824344347941253 | 0.966637987288264 | 243.045189757073 | 253.283291692356 | 252.703455269135 | 279.502154888932 | 267.16979977986 | 272.747337752471 | 245.694429371248 | 223.618266321713 | Ddx41 |
| **ENSMUSG00000021495** | 0.244313547525541 | 0.233271637205903 | 0.832048466795696 | 89.4478579403355 | 91.2145616384048 | 130.610774633486 | 100.358156285623 | 82.1218646864323 | 85.6157562451702 | 103.066730965058 | 75.9867895267955 | Fam193b |

| | | | | | | | | | | | | |
|---|---|---|---|---|---|---|---|---|---|---|---|---|
| **ENSMUSG00000021496** | -0.0308229161808085 | 0.888002139851463 | 0.980862800996964 | 485.186865797577 | 343.683437601847 | 323.687571917769 | 479.28054076592 | 373.380744774312 | 565.063991218123 | 353.966550789086 | 375.591845375304 | Pcbd2 |
| **ENSMUSG00000021497** | 0.0122264006672855 | 0.92228429364864 | 0.986713918026868 | 978.505355044276 | 975.670043239366 | 1092.20894150031 | 1119.88447294424 | 1102.62290318983 | 1172.93586055883 | 909.902251146063 | 946.578292390939 | Txndc15 |
| **ENSMUSG00000021500** | 0.185936437208919 | 0.121532026735237 | 0.769868040577702 | 990.251033359673 | 939.021335438221 | 989.045358637627 | 930.423280704096 | 828.883354235056 | 703.272283442469 | 930.723812957186 | 918.354627709557 | Ddx46 |
| **ENSMUSG00000021501** | -0.111989372851839 | 0.54447480996581 | 0.913937916428821 | 202.387072511466 | 176.728213174409 | 187.398067952392 | 251.364354061187 | 211.326931793086 | 275.193502216618 | 186.352978209548 | 211.677485110359 | Caml |
| **ENSMUSG00000021504** | -0.0154366650597785 | 0.927469405970885 | 0.986790505471895 | 123.781379169959 | 114.018202048006 | 138.182413742673 | 120.992543559303 | 105.115986798633 | 147.992950080937 | 131.175839410073 | 118.322286548867 | B4galt7 |
| **ENSMUSG00000021508** | -0.979686063421746 | 0.0137927222219826 | 0.635862253526407 | 291.834930451802 | 133.56417954195 | 155.218601738345 | 298.260688774095 | 517.915226622433 | 714.280023531134 | 260.269522639034 | 240.986675356409 | Cxcl14 |
| **ENSMUSG00000021510** | -0.204939274462309 | 0.410525849355998 | 0.882192126163777 | 65.9565013095403 | 56.1946852950887 | 69.0912068713366 | 47.8342614071662 | 61.3176589658694 | 52.592535979176 | 67.6700758861489 | 93.3551985614916 | Zfp729a |
| **ENSMUSG00000021514** | -0.010259695454937 | 0.970543795505255 | 0.9951105526746 | 36.1405486627618 | 54.56585383726 | 66.2518422053913 | 68.4686486808458 | 49.2731188118594 | 63.6002760678407 | 44.7663578939139 | 69.4736361387845 | Zfp369 |
| **ENSMUSG00000021518** | 0.114353268737635 | 0.200115423047145 | 0.812921911423739 | 1823.29068003633 | 1803.11642381632 | 1630.74177314127 | 1813.95022669528 | 1515.42214301363 | 1602.23772401676 | 1691.75189715372 | 1722.72907112892 | Ptdss1 |
| **ENSMUSG00000021519** | 0.0656582576581156 | 0.68517516728191 | 0.944452327903597 | 308.098177350044 | 276.086932101957 | 318.008842585878 | 324.522636213324 | 306.588294829347 | 218.931719541221 | 336.268223249632 | 308.289260365856 | Mterf3 |
| **ENSMUSG00000021520** | 0.153568579605313 | 0.198903107574312 | 0.812921911423739 | 17580.5698970005 | 14025.0532676337 | 14911.3967706563 | 17315.0647026999 | 14639.5910781013 | 15808.3378495546 | 13048.8727870306 | 13890.3851254982 | Uqcrb |
| **ENSMUSG00000021532** | 0.0423336014011205 | 0.861712930094706 | 0.974868041734905 | 144.562194651047 | 131.120932355207 | 126.824955078892 | 135.999370667433 | 109.49581958191 | 81.9465095489486 | 169.69572876065 | 160.657783570939 | Fastkd3 |
| **ENSMUSG00000021537** | 0.259961907554443 | 0.0491767108030726 | 0.703396849069134 | 683.959883442767 | 607.554133770089 | 694.69788826796 | 793.485983342404 | 590.182467546493 | 659.24132308781 | 536.15521663641 | 536.249628946243 | Cetn3 |
| **ENSMUSG00000021539** | -0.0760403024735588 | 0.852285555496109 | 0.972979893633677 | 28.9124389302094 | 34.2054606144018 | 22.7149173275627 | 29.075727522003 | 27.3739548954774 | 30.5770558018465 | 12.4929370866736 | 51.0197015394199 | Lect2 |
| **ENSMUSG00000021540** | 0.0128074515718978 | 0.927448200583242 | 0.986790505471895 | 659.565013095403 | 772.066111010784 | 797.861471130641 | 634.038445318517 | 666.82954125383 | 610.318033804856 | 782.890724098215 | 777.236304302651 | Smad5 |
| **ENSMUSG00000021546** | -0.00460458249883931 | 0.953926918728802 | 0.992371505557225 | 3593.27405079509 | 3791.91963382511 | 3799.06992303487 | 3587.56960553747 | 4022.87641143936 | 3671.69286068573 | 3567.77461633588 | 3557.2672754187 | Hnrnpk |
| **ENSMUSG00000021548** | 0.108225754875011 | 0.402868359735103 | 0.881158207349486 | 575.538237454482 | 557.874774306315 | 564.087113634474 | 575.886990274511 | 593.467342133951 | 426.855698993777 | 520.539045278068 | 566.644344756961 | Ccnh |
| **ENSMUSG00000021549** | 0.0437580953495283 | 0.802565435094085 | 0.963461896828986 | 459.888481733644 | 542.400875456943 | 583.016211407443 | 469.901273823339 | 445.647985698372 | 388.940149799487 | 564.264325081426 | 593.782483873674 | Rasa1 |
| **ENSMUSG00000021550** | -0.00850891615524935 | 0.956644662230562 | 0.992371505557225 | 699.319616624441 | 581.492830444831 | 688.072704047421 | 812.244517227568 | 788.36990098975 | 777.880299598975 | 626.729010514794 | 604.637739520359 | 2210016F16F |
| **ENSMUSG00000021552** | 0.152695784919552 | 0.117665181384383 | 0.769868040577702 | 651.433389646282 | 595.337897836374 | 629.392500951217 | 669.679659700327 | 575.948011000845 | 599.310293716191 | 587.168043073661 | 527.565424428895 | Gkap1 |
| **ENSMUSG00000021555** | 0.113853071055625 | 0.263272690693919 | 0.842510961779757 | 1306.48083415884 | 1382.06349196762 | 1448.07597963212 | 1364.68334014562 | 1283.29100549998 | 1119.12024234758 | 1283.64928565572 | 1395.9858761637 | Naa35 |
| **ENSMUSG00000021556** | 0.268303354217414 | 0.175496030020714 | 0.799889021902745 | 159.018414116152 | 215.005752433383 | 184.558703286447 | 135.999370667433 | 160.958854785407 | 119.862058743238 | 135.340151772298 | 160.657783570939 | Golm1 |
| **ENSMUSG00000021557** | 0.275063628978577 | 0.240035109445933 | 0.832048466795696 | 2684.33925192663 | 3660.79870146991 | 4247.68954025423 | 2842.85581029649 | 2359.63491199015 | 1941.03150230122 | 3409.53074657135 | 3392.26738958909 | Agtpbp1 |
| **ENSMUSG00000021559** | 0.015029274003318 | 0.958521252072715 | 0.993052897090327 | 1496.21871463834 | 1724.93251384055 | 1624.11658892073 | 1405.01418799873 | 1419.06582178155 | 746.080161565054 | 2245.60544132959 | 1773.74877266834 | Dapk1 |
| **ENSMUSG00000021567** | -1.0447431667354 | 0.136239709747904 | 0.776456211338551 | 6.32459601598331 | 16.2883145782866 | 3.78581955459379 | 8.44134024832345 | 18.6142893289246 | 40.3617136584374 | 8.32862472444909 | 5.42762782334254 | Nkd2 |
| **ENSMUSG00000021573** | -0.273193247317695 | 0.265259309446199 | 0.842510961779757 | 227.685456575399 | 228.850819824926 | 318.008842585878 | 235.419600258798 | 289.068963696242 | 187.131581507301 | 399.773986773556 | 343.026078435248 | Tppp |
| **ENSMUSG00000021575** | 0.631321732969893 | 0.152057103366398 | 0.783412422888014 | 27.1054114970714 | 35.8342920722305 | 39.7511053232348 | 44.0825546301336 | 22.994122112201 | 6.1154111603693 | 27.0680303544596 | 37.9933947633978 | Ahrr |
| **ENSMUSG00000021576** | 0.0927368777056221 | 0.408037082054438 | 0.881573350402261 | 853.820462157748 | 700.397526866323 | 764.735550027945 | 880.713165908413 | 787.274942793931 | 725.287763619799 | 751.658381381531 | 734.90080728058 | Pdcd6 |
| **ENSMUSG00000021577** | 0.244945797822789 | 0.237297238705998 | 0.832048466795696 | 40277.7379709315 | 47661.2372875243 | 51028.115321481 | 41555.7801158021 | 35395.6186380481 | 25123.3321290291 | 45670.0136765166 | 46143.5207029289 | Sdha |
| **ENSMUSG00000021578** | 0.124018797699595 | 0.370885282602281 | 0.873323500696802 | 805.934235179588 | 829.889627763701 | 858.434584004141 | 768.161962597434 | 617.556422441971 | 687.372214425509 | 815.164144905455 | 872.76255399348 | Ccdc127 |
| **ENSMUSG00000021583** | 0.0496254992992499 | 0.773343863004413 | 0.959170127433816 | 477.958756065025 | 586.379324818317 | 609.5169482896 | 468.025420434822 | 512.440435643337 | 395.055560959857 | 573.634027896431 | 587.269330485663 | Erap1 |
| **ENSMUSG00000021585** | 0.19714903480537 | 0.197203554875901 | 0.812565226112804 | 3058.39393058622 | 3471.85425236178 | 3407.23759913441 | 3291.18477015189 | 2717.686242023 | 2222.3404156782 | 3418.90044938635 | 3178.41885334939 | Cast |
| **ENSMUSG00000021589** | 0.113464893177714 | 0.688111855062708 | 0.944453919646345 | 224.071401709123 | 245.953550132127 | 249.86409060319 | 200.716312571247 | 188.332809680885 | 110.077400886647 | 302.953724351836 | 248.585354309088 | Rhobtb3 |
| **ENSMUSG00000021591** | -0.207984895551519 | 0.325693269112401 | 0.855977678491063 | 224.974915425692 | 238.623808571898 | 133.450139299431 | 182.895705380341 | 189.427767876704 | 221.377884005369 | 249.858741733473 | 240.986675356409 | Glrx |
| **ENSMUSG00000021592** | -0.17261082111106 | 0.564054960646687 | 0.918598124443955 | 56.9213641438498 | 71.6685841444609 | 47.3227444324223 | 62.8410885152968 | 77.7420319031559 | 97.8465785659088 | 43.7252798033577 | 51.0197015394199 | Arsk |
| **ENSMUSG00000021594** | -0.457901495209612 | 0.390093723072886 | 0.877515569464664 | 14.4562194651047 | 14.6594831204579 | 5.67872933189068 | 14.0689004138724 | 6.56974917491458 | 25.6847268735511 | 11.4518589961175 | 23.8815624227072 | Srd5a1 |
| **ENSMUSG00000021595** | 0.255757411788137 | 0.0175514279175979 | 0.642273034804458 | 1865.75582471508 | 1696.42796332855 | 1859.7838561942 | 2113.14884216364 | 1686.23562156141 | 1451.79860947167 | 1541.83665211364 | 1629.37387256743 | Nsun2 |
| **ENSMUSG00000021596** | 0.0637071630554707 | 0.853134335174799 | 0.973120292022893 | 25.2983840639333 | 26.8757190541728 | 45.4298346551254 | 29.075727522003 | 29.5638712871156 | 45.2540425867328 | 22.903717992235 | 23.8815624227072 | Mctp1 |
| **ENSMUSG00000021597** | 0.0673326734345978 | 0.746661645120744 | 0.956085046483146 | 637.880683897746 | 631.172189908605 | 541.372196306912 | 490.535661097018 | 478.496731572945 | 384.047820871192 | 670.454290318152 | 662.17059444779 | Slf1 |
| **ENSMUSG00000021598** | -0.106544906592931 | 0.648502199120833 | 0.936253564680217 | 69.5705561758165 | 47.2361122770311 | 48.2691993210708 | 71.2824287636203 | 56.937826182593 | 59.9310293716191 | 71.8343882483734 | 65.1315338801105 | Med10 |
| **ENSMUSG00000021604** | 0.025631421406944 | 0.897090912766433 | 0.984332015045905 | 436.397125102849 | 456.072808192024 | 522.443098533943 | 610.590277962063 | 447.837902090011 | 370.593916318379 | 621.523620062014 | 548.190410157596 | Irx4 |
| **ENSMUSG00000021606** | 0.0404453977610039 | 0.822080301798519 | 0.966637987288264 | 11976.9778268393 | 8537.52008620891 | 9275.25790875478 | 11232.6100904357 | 9953.16999999559 | 12744.5168582096 | 8539.96357683199 | 8651.638750408 | Ndufs6 |
| **ENSMUSG00000021607** | -0.0394161003861505 | 0.807704406119691 | 0.964026716319739 | 769.793686516826 | 623.842448348376 | 619.927952064733 | 795.361836730921 | 803.699315731217 | 834.142082274372 | 631.934400967575 | 617.664046296381 | Mrpl36 |
| **ENSMUSG00000021608** | -0.5013884199035 | 0.0337521057102621 | 0.672620872762612 | 154.500845533307 | 114.018202048006 | 119.253315969704 | 151.944124469822 | 243.08071947184 | 229.939459629886 | 140.545542225078 | 150.888053488923 | Lpcat1 |
| **ENSMUSG00000021610** | 0.0823001062753623 | 0.379272699096211 | 0.87422797717331 | 1432.06924076194 | 1478.16454797951 | 1512.43491206022 | 1625.42696114939 | 1313.94983498292 | 1492.16032313011 | 1514.76862175918 | 1391.64377390503 | Clptm1l |
| **ENSMUSG00000021614** | -0.928062620198218 | 0.0422953705623633 | 0.685510731751723 | 78.6056933415069 | 67.5965054998893 | 55.8408384302584 | 93.7926694258161 | 179.573144114332 | 260.516515431732 | 57.2592949805875 | 66.217059444779 | Vcan |
| **ENSMUSG00000021615** | 0.023891988354319 | 0.873361123669794 | 0.976415486781506 | 200.580045078328 | 174.284965987666 | 182.66579350915 | 178.206071909051 | 205.85214081399 | 194.470074899744 | 170.736806851206 | 153.05910461826 | Xrcc4 |
| **ENSMUSG00000021619** | 0.0938807243390495 | 0.416254316794817 | 0.882192126163777 | 608.968244967536 | 601.853223667689 | 644.535779169592 | 627.47295845871 | 522.295059405709 | 528.371524255907 | 645.468416144805 | 628.519301943066 | Atg10 |
| **ENSMUSG00000021620** | -0.478288745322846 | 0.256123734795591 | 0.837517407767513 | 20.780815481088 | 42.3496179035451 | 16.0897331070236 | 23.448167356454 | 25.1840385038392 | 30.5770558018465 | 28.1091084450157 | 59.7039060567679 | Acot12 |
| **ENSMUSG00000021621** | -0.122331869309172 | 0.436283971800447 | 0.888701139843674 | 279.185738419835 | 240.252640029727 | 231.881447718869 | 304.826175633902 | 329.582416941548 | 305.770558018465 | 255.064132186253 | 259.440609955773 | Zcchc9 |
| **ENSMUSG00000021622** | -0.0353139313137345 | 0.775687165364071 | 0.9595799687189 | 49297.5154034402 | 45241.6081569199 | 44528.8096011321 | 52221.8824829059 | 52499.9606149382 | 55871.619443366 | 43879.35936076 | 43780.3315486456 | Ckmt2 |
| **ENSMUSG00000021629** | -0.00444903857942197 | 0.967475063696207 | 0.994284908685797 | 477.958756065025 | 476.433201414882 | 465.655805215036 | 401.432625142493 | 443.458069306734 | 451.317343635254 | 447.663578939139 | 485.229927406823 | Slc30a5 |
| **ENSMUSG00000021635** | 0.0755991249902234 | 0.716717826568043 | 0.951377160840035 | 362.309000344187 | 381.146561131906 | 306.651383922097 | 273.874594723383 | 278.119381738051 | 226.270212933664 | 376.870268781321 | 374.506319810635 | Rad17 |
| **ENSMUSG00000021639** | 0.151268335091869 | 0.336200953719279 | 0.858570960161243 | 313.519259649459 | 279.344595017615 | 326.526936583714 | 287.943495137256 | 289.068963696242 | 205.477814988408 | 299.830490080167 | 290.92085133116 | Gtf2h2 |
| **ENSMUSG00000021643** | -0.0313480859284217 | 0.886295363106798 | 0.980068672363141 | 76.7986659083688 | 65.1532583131463 | 68.1447519826882 | 88.1651092602672 | 66.7924499449649 | 94.1773318696872 | 83.2862472444909 | 60.7894316214364 | Serf1 |
| **ENSMUSG00000021645** | 0.019753250777409 | 0.931296174030159 | 0.987775860606096 | 156.307872966445 | 115.647033505835 | 95.5919437534931 | 162.261318106662 | 155.484063806312 | 143.100621152642 | 111.395355689507 | 112.894658725525 | Smn1 |
| **ENSMUSG00000021646** | 0.0263497682823476 | 0.855254338422458 | 0.973268281968411 | 1240.5243328493 | 1271.30295283527 | 1173.60406192407 | 1203.35994873322 | 1272.34142354179 | 905.080851734656 | 1301.34761319517 | 1319.99908663691 | Mccc2 |
| **ENSMUSG00000021650** | -0.00618398297079371 | 0.952646400587697 | 0.992000114323903 | 937.847237798669 | 902.372627637076 | 936.043884873314 | 963.250715003132 | 994.22204180374 | 807.234273168747 | 993.188498390554 | 958.519073602292 | Ptcd2 |
| **ENSMUSG00000021660** | 0.163248100321466 | 0.253258797396356 | 0.837406774112203 | 5648.76775598967 | 4257.76543076411 | 4681.16587925522 | 5774.8146565475 | 4633.86308470642 | 5139.39153917436 | 4056.04024080671 | 4355.12856545005 | Btf3 |
| **ENSMUSG00000021661** | 0.348183309229126 | 0.0376241263015503 | 0.679566075569698 | 306.291149916906 | 288.303168035672 | 317.06238769723 | 332.026049767389 | 208.042057205628 | 201.808568292187 | 305.035880532948 | 260.526135520442 | Ankra2 |
| **ENSMUSG00000021662** | 0.0161472910259962 | 0.922831993723243 | 0.986713918026868 | 219.553833126278 | 225.593156909269 | 156.165056626994 | 172.578511743502 | 195.997517051618 | 182.239252579005 | 197.804837205666 | 189.966973816989 | Arhgef28 |
| **ENSMUSG00000021665** | -0.246518235488735 | 0.174286690296424 | 0.799889021902745 | 273.764656120421 | 206.047179415325 | 230.934992830221 | 184.771558768858 | 289.068963696242 | 309.439804714686 | 225.913945650682 | 238.815624227072 | Hexb |
| **ENSMUSG00000021666** | 0.220289756451027 | 0.276316725024107 | 0.846688575024234 | 2013.02856051583 | 2100.37816487005 | 2197.66825144169 | 2026.85958629189 | 1582.21459295859 | 1196.17442296823 | 2147.74410081731 | 2229.66950982911 | Gfm2 |
| **ENSMUSG00000021668** | -0.0809421152106479 | 0.733897006372222 | 0.954182918936468 | 78.6056933415069 | 56.1946852950887 | 53.001473764313 | 45.9584080186499 | 63.5075753575076 | 68.4926049961361 | 62.4646854333682 | 53.1907526687569 | Polk |
| **ENSMUSG00000021669** | 0.0619757558832756 | 0.751657751876731 | 0.956197192463153 | 1203.48027046997 | 1394.27972790133 | 1271.08891545486 | 1034.53314376675 | 1039.11532783232 | 848.819069059259 | 1447.09854587303 | 1361.24905809431 | Col4a3bp |
| **ENSMUSG00000021670** | -0.0413740455868422 | 0.816593510304562 | 0.966315773308182 | 106.614618555147 | 103.43079757212 | 87.0738497556571 | 95.6685228143324 | 107.305903190271 | 114.969729814943 | 83.2862472444909 | 99.8683519495027 | Hmgcr |
| **ENSMUSG00000021671** | -0.050769757730585 | 0.799859699670796 | 0.963461896828986 | 191.544907912638 | 167.769640156352 | 176.040609288611 | 150.068271081306 | 177.383227722694 | 121.085140975312 | 205.092383839559 | 205.164331722348 | Poc5 |
| **ENSMUSG00000021676** | -0.512674578685699 | 0.0998762090248377 | 0.759800953500741 | 91.2548853734735 | 157.182235680465 | 73.8234813145789 | 74.0962088463947 | 141.249607260664 | 161.446854633749 | 90.5737938783839 | 173.684090346961 | Iqgap2 |
| **ENSMUSG00000021678** | -0.588590805654703 | 0.378338635708085 | 0.87422797717331 | 10.8421645988285 | 12.2162359337149 | 7.57163910918757 | 10.3171936368398 | 22.994122112201 | 30.5770558018465 | 4.16431236222455 | 4.34210225867403 | F2rl1 |
| **ENSMUSG00000021681** | 0.146604932995825 | 0.334239178539135 | 0.858148279095856 | 833.03964667666 | 951.237571371936 | 884.935320886298 | 846.009878220861 | 662.449708470554 | 680.033721033066 | 920.313032051625 | 911.841474321546 | Aggf1 |
| **ENSMUSG00000021684** | -0.268606716037494 | 0.393491365797117 | 0.878942305112115 | 37.0440623793308 | 38.2775392589734 | 24.6078271048596 | 65.6548685980713 | 53.6529515951357 | 50.1463715150282 | 49.9717483466946 | 45.5920737160773 | Pde8b |
| **ENSMUSG00000021686** | 0.285796916406122 | 0.0346806411244381 | 0.679566075569698 | 1105.90078908051 | 1216.73709899801 | 1421.57524274997 | 1185.53934154232 | 963.563212320805 | 864.719138076219 | 1107.70708835173 | 1106.15055039721 | Ap3b1 |
| **ENSMUSG00000021687** | -0.0368377808997208 | 0.765492961063443 | 0.958345041873658 | 728.23205555465 | 894.228470347933 | 886.828230663595 | 797.237690119437 | 796.034608360484 | 772.987970670679 | 950.504296677753 | 871.677028428812 | Scamp1 |
| **ENSMUSG00000021690** | 0.239836822380919 | 0.345197803484405 | 0.861266047293794 | 1484.47303632294 | 2123.99622100857 | 1902.37432618338 | 1385.3177274193 | 1184.74476787626 | 950.334894321389 | 1949.93926361164 | 1754.20931250431 | Jmy |
| **ENSMUSG00000021692** | 0.224473461836661 | 0.208179574012565 | 0.820372930586885 | 112.035700854562 | 96.1010560118908 | 111.681676860517 | 119.116690170786 | 106.210944994452 | 100.292743030056 | 77.0397787011541 | 92.2696729968231 | Dimt1 |
| **ENSMUSG00000021693** | 0.0672181633487213 | 0.552686981568953 | 0.914977712327965 | 468.020105182765 | 495.979178908826 | 486.477812765302 | 490.535661097018 | 502.585811880965 | 399.947889888152 | 453.910047482476 | 494.999657488839 | Kif2a |
| **ENSMUSG00000021694** | -0.0320433479615749 | 0.910490461730374 | 0.986166948757759 | 47.8862269781594 | 70.8541684155466 | 81.3951204237664 | 50.6480414899407 | 49.2731188118594 | 58.7079471395453 | 90.5737938783839 | 57.5328549274309 | Ercc8 |
| **ENSMUSG00000021699** | -0.0441551858770549 | 0.905104189021755 | 0.98504921011344 | 243.948703473642 | 381.146561131906 | 296.240380146964 | 195.088752405698 | 238.700886688563 | 117.415894279091 | 434.129563761909 | 360.394487469945 | Pde4d |
| **ENSMUSG00000021701** | -0.485433719040146 | 0.0655762959086035 | 0.718339708336358 | 348.756294595651 | 259.798617523671 | 401.296872786941 | 594.645524159674 | 727.05224202388 | 474.555906044658 | 553.853544175865 | 489.572029665497 | Plk2 |
| **ENSMUSG00000021702** | -0.791286406710874 | 0.0812050603269158 | 0.739855948845396 | 110.228673421423 | 83.0704043492615 | 47.3227444324223 | 113.489130005238 | 116.065568756824 | 320.447544803351 | 95.7791843311646 | 81.4144173501381 | Thbs4 |
| **ENSMUSG00000021703** | -0.0832820499694613 | 0.645036477289962 | 0.935063240511793 | 234.913566307952 | 292.375246680244 | 276.364827485346 | 233.543746870282 | 305.493336633528 | 198.139321595965 | 291.501865355718 | 302.861632542514 | Serinc5 |
| **ENSMUSG00000021704** | -0.0622127885319458 | 0.820563855472457 | 0.966398681144067 | 77.7021796249379 | 85.5136515360045 | 89.9132144216024 | 76.9099889291692 | 62.4126171616885 | 56.2617826753975 | 119.723980413956 | 105.295979772845 | Mtx3 |
| **ENSMUSG00000021706** | -0.0223378369043849 | 0.945744882508009 | 0.990505437194639 | 174.378147297826 | 241.067055758641 | 219.57753416644 | 148.192417692789 | 145.62944004394 | 102.738907494204 | 268.598147363483 | 277.894544555138 | Zfyve16 |
| **ENSMUSG00000021707** | 0.181575661670024 | 0.383840485526557 | 0.875126835008975 | 98.4829951060259 | 79.8127414336042 | 77.6093008691726 | 101.296082979881 | 85.4067392738896 | 89.2850029413918 | 59.3414511616998 | 81.4144173501381 | Dhfr |
| **ENSMUSG00000021708** | -0.201498971403638 | 0.245673351524963 | 0.832048466795696 | 147.272735800754 | 159.625482867208 | 170.36187995672 | 166.013024883695 | 231.036179317829 | 192.023910435596 | 151.997401221196 | 164.999885829613 | Rasgrf2 |
| **ENSMUSG00000021709** | 0.250800637315524 | 0.252507949470445 | 0.83723604018117 | 1484.47303632294 | 1847.90928890661 | 2011.21663837795 | 1419.0830884126 | 1230.73301210066 | 990.696607979826 | 1795.85970620934 | 1665.19621620149 | Erbin |
| **ENSMUSG00000021710** | 0.324251835267023 | 0.068183654624793 | 0.722112580225498 | 403.870631306363 | 398.249291439107 | 413.600786339371 | 355.474217123843 | 307.683253025166 | 222.600966237442 | 348.761160336306 | 374.506319810635 | Nln |

| | | | | | | | | | | | | |
|---|---|---|---|---|---|---|---|---|---|---|---|---|
| **ENSMUSG00000021711** | 0.109533298697706 | 0.494033323383633 | 0.900141680336539 | 701.126644057579 | 691.438953848265 | 689.019158936069 | 644.355638955357 | 540.909348734634 | 510.025290774799 | 703.768789215948 | 770.72315091464 | Trappc13 |
| **ENSMUSG00000021712** | 0.311556864115424 | 0.159137811462163 | 0.789393312599749 | 475.248214915318 | 513.081909216027 | 486.477812765302 | 453.018593326692 | 393.089992299056 | 231.162541861959 | 489.306702561384 | 438.552328126077 | Trim23 |
| **ENSMUSG00000021713** | 0.00661307774420152 | 0.975963629498954 | 0.99575536949131 | 91.2548853734735 | 105.874044758863 | 93.6990339761962 | 72.2203554578784 | 91.9764884488041 | 70.9387694602839 | 115.559668051731 | 82.4999429148066 | Ppwd1 |
| **ENSMUSG00000021715** | 0.122006710917766 | 0.230361027791433 | 0.832048466795696 | 513.195791011218 | 500.865673282312 | 456.191256328551 | 466.149567046306 | 475.211856985488 | 443.978850242811 | 426.842017128016 | 434.210225867403 | Cwc27 |
| **ENSMUSG00000021716** | -0.114705928534915 | 0.431605409324475 | 0.886559861988535 | 440.011179969125 | 373.002403842763 | 305.704929033448 | 411.749818779333 | 395.279908690694 | 439.086521314516 | 392.486440139664 | 430.953649173397 | Srek1ip1 |
| **ENSMUSG00000021719** | -0.148249846959717 | 0.505972175851349 | 0.901494649536949 | 95.7724539563188 | 159.625482867208 | 127.77140996754 | 101.296082979881 | 124.825234323377 | 110.077400886647 | 143.668776496747 | 158.486732441602 | Rgs7bp |
| **ENSMUSG00000021725** | 0.0961663642396084 | 0.59263854015155 | 0.923935529355527 | 124.684892886528 | 135.193010999779 | 129.664319744837 | 164.137171495178 | 121.54035973592 | 105.185071958352 | 146.792010768415 | 143.289374536243 | Parp8 |
| **ENSMUSG00000021728** | -0.940697093994532 | 0.103309130061371 | 0.759800953500741 | 14.4562194651047 | 21.9892246806869 | 16.0897331070236 | 12.1930470253561 | 42.7033696369448 | 53.8156182112498 | 22.903717992235 | 5.42762782334254 | Emb |
| **ENSMUSG00000021731** | 0.137992130982409 | 0.306743672667541 | 0.851785577030345 | 835.750187826367 | 710.170515613295 | 842.344850897118 | 937.926694258161 | 861.732100109629 | 772.987970670679 | 678.782915042601 | 708.848193728535 | Mrps30 |
| **ENSMUSG00000021733** | 0.00999777132236052 | 0.966972990355084 | 0.994284908685797 | 259.308436655316 | 340.425774686189 | 329.36630124966 | 241.047160424347 | 241.98576127602 | 195.693157131818 | 338.350379430744 | 385.36157545732 | Slc4a7 |
| **ENSMUSG00000021737** | 0.193755303581153 | 0.0465290713110173 | 0.698690595145979 | 2351.84620422922 | 2026.26633353885 | 2242.15163120817 | 2478.94025292432 | 2074.94578107719 | 1965.49314694269 | 2021.77365186002 | 1892.07105921721 | Psmd6 |
| **ENSMUSG00000021738** | 0.159952699646085 | 0.450552459138358 | 0.89108031640712 | 465.309564033058 | 503.308920469055 | 484.584902988005 | 401.432625142493 | 364.621079207759 | 280.085831144914 | 526.785513821405 | 487.40097853616 | Atxn7 |
| **ENSMUSG00000021745** | -0.386256276181454 | 0.0234561716933799 | 0.654124840968651 | 725.521514404943 | 829.889627763701 | 764.735550027945 | 710.948434247686 | 1274.53133993343 | 1037.17373279863 | 815.164144905455 | 835.854684794751 | Ptprg |
| **ENSMUSG00000021747** | 0.14921401037653 | 0.6608794715817 | 0.939611397743053 | 46.9827132615903 | 52.122606650517 | 32.1794662140472 | 53.4618215727152 | 63.5075753575076 | 22.0154801773295 | 41.6431236222455 | 39.0789203280663 | 4930452B06I |
| **ENSMUSG00000021748** | 0.105027599568107 | 0.452519204366927 | 0.891110666112841 | 20882.0090173438 | 20309.0850319366 | 19626.6350259028 | 20905.4480883202 | 18094.1841859106 | 14919.1570668369 | 21320.2382164991 | 21649.7218617487 | Pdhb |
| **ENSMUSG00000021750** | -0.335548974839351 | 0.618798945845763 | 0.929568738467614 | 461.695509166782 | 491.907100264254 | 421.172425448559 | 106.92364314543 | 704.058119911679 | 755.864819421645 | 258.187366457922 | 151.973579053591 | Fam107a |
| **ENSMUSG00000021751** | 0.101639403311489 | 0.754720959985087 | 0.956497786093635 | 63.2459601598332 | 111.574954861263 | 41.6440151005317 | 84.4134024832345 | 63.5075753575076 | 63.6002760678407 | 59.3414511616998 | 94.4407241261602 | Acox2 |
| **ENSMUSG00000021752** | 0.0974513755530365 | 0.539652954827032 | 0.913837378403038 | 137.334084918495 | 147.409246933493 | 156.165056626994 | 185.709485463116 | 134.679858085749 | 141.877538920568 | 148.874166949528 | 159.572258006271 | Kctd6 |
| **ENSMUSG00000021754** | -0.0478013088615972 | 0.697246214947674 | 0.944933386973863 | 274.66816983699 | 300.519403969387 | 307.597838810745 | 308.577882410935 | 328.487458745729 | 275.193502216618 | 291.501865355718 | 335.427399482569 | Map3k1 |
| **ENSMUSG00000021756** | 0.276216036388767 | 0.0981977675879752 | 0.759800953500741 | 4462.45424613451 | 6099.97380956832 | 6549.46782944725 | 5110.76255701272 | 4506.8479339914 | 3745.07779461016 | 5188.73320333179 | 4908.74660343099 | Il6st |
| **ENSMUSG00000021759** | -0.140728575666602 | 0.576276135369921 | 0.922450244597442 | 1278.4719089452 | 1136.9243575644 | 1356.26985543322 | 1396.5728477504 | 1779.30706820603 | 1960.6008180144 | 981.736639394437 | 976.973008201657 | Plpp1 |
| **ENSMUSG00000021760** | -0.130650684404514 | 0.478553533582782 | 0.897674307047416 | 385.800356974982 | 284.231089391101 | 320.848207251823 | 360.163850595134 | 367.905953795217 | 475.778988276731 | 298.789411989611 | 337.598450611906 | Gpx8 |
| **ENSMUSG00000021763** | 0.346742857644379 | 0.266304686326693 | 0.843513595393324 | 56.0178504272808 | 81.4415728914329 | 81.3951204237664 | 50.6480414899407 | 45.9882442244021 | 31.8001380339204 | 75.998700610598 | 57.5328549274309 | BC067074 |
| **ENSMUSG00000021764** | 0.0939552988699596 | 0.516509741512525 | 0.904619578708276 | 5390.36283305092 | 3850.55756630695 | 4475.78516841851 | 5561.9052969509 | 4615.24879537749 | 4829.95173445967 | 4152.86050322843 | 4464.76664748157 | Ndufs4 |
| **ENSMUSG00000021765** | -0.324231438709798 | 0.59538550000972 | 0.924461858177432 | 12.6491920319666 | 7.32974156022896 | 3.78581955459379 | 15.0068271081306 | 12.0445401540101 | 22.0154801773295 | 10.4107809055614 | 4.34210225867403 | Fst |
| **ENSMUSG00000021767** | 0.0683761831271437 | 0.79426734685216 | 0.961804768755116 | 338.817643713392 | 449.557482360709 | 377.63550057073 | 304.826175633902 | 244.175677667659 | 245.839528646846 | 474.731609293598 | 437.466802561409 | Kat6b |
| **ENSMUSG00000021768** | -0.0333054090182371 | 0.93674620244669 | 0.988852035581444 | 11.7456783153976 | 10.5874044758863 | 13.2503684410783 | 16.8826804966469 | 15.3294147414674 | 18.3462334811079 | 10.4107809055614 | 9.76973008201657 | Dusp13 |
| **ENSMUSG00000021770** | 0.0203296314521986 | 0.885687842173885 | 0.979948700426504 | 466.213077749627 | 504.937751926884 | 466.602260103684 | 490.535661097018 | 529.959766776443 | 369.370834086306 | 487.224546380272 | 513.453592088204 | Samd8 |
| **ENSMUSG00000021771** | 0.076805737302602 | 0.302046149625484 | 0.851785577030345 | 11346.3252526741 | 10082.4667239594 | 10165.871958973 | 10848.0601457899 | 10316.6961210075 | 10169.9287596941 | 9890.2418602833 | 9865.2563317074 | Vdac2 |
| **ENSMUSG00000021772** | 0.0710677448700471 | 0.649203495372819 | 0.936461253686283 | 959.531566996326 | 700.397526866323 | 643.589324280944 | 826.31341764144 | 777.420319031559 | 711.833859066986 | 807.876598271562 | 681.710054611823 | Nkiras1 |
| **ENSMUSG00000021773** | -0.217641001917141 | 0.452762395400287 | 0.891153418181945 | 68.6670424592474 | 54.56585383726 | 37.8581955459379 | 58.151455044006 | 67.887408140784 | 89.2850029413918 | 53.094982618363 | 45.5920737160773 | Comtd1 |
| **ENSMUSG00000021774** | 0.0220060975730302 | 0.89970389046268 | 0.984802943363903 | 435.49361138628 | 446.299819445052 | 424.011790114504 | 377.984457786039 | 390.900075907418 | 303.324393554317 | 483.060234018047 | 479.80229958348 | Ube2e1 |
| **ENSMUSG00000021775** | 0.21654791223188 | 0.31591040872182 | 0.853356198600137 | 1567.59629824729 | 1925.27878315347 | 1873.98067952392 | 1539.13770527764 | 1330.3742079202 | 974.796538962866 | 1840.62606410325 | 1796.54480952638 | Nr1d2 |
| **ENSMUSG00000021779** | 0.00145605460576308 | 0.995388004369974 | 0.999262355603273 | 345.142239729375 | 360.786167909048 | 338.830850136144 | 313.267515882226 | 240.890803080201 | 227.493295165738 | 418.513392403567 | 468.947043936795 | Thrb |
| **ENSMUSG00000021782** | 0.0895203385592837 | 0.63242057769263 | 0.931825800800958 | 172.571119864688 | 225.593156909269 | 210.112985279955 | 184.771558768858 | 160.958854785407 | 149.216032313011 | 223.831789469569 | 210.59195954569 | Dlg5 |
| **ENSMUSG00000021785** | 0.296129070923736 | 0.0241538183926184 | 0.654124840968651 | 890.864524537078 | 936.578088251478 | 965.383986421416 | 969.816201862939 | 776.32536083574 | 609.094951572782 | 843.273253350471 | 833.683633665414 | Ngly1 |
| **ENSMUSG00000021786** | 0.15440397736073 | 0.568661477563711 | 0.92059532906293 | 452.660372001092 | 526.926976607571 | 478.906173656114 | 358.287997206618 | 331.772333333186 | 237.277953022329 | 538.237372817523 | 524.308847734889 | Oxsm |
| **ENSMUSG00000021792** | 0.307773199458561 | 0.131785128695838 | 0.77303252008843 | 680.345828576491 | 573.348673155687 | 1042.04683240194 | 671.555513088843 | 567.188345434292 | 688.595296657583 | 601.743136341447 | 539.506205640248 | Fam213a |
| **ENSMUSG00000021794** | 0.12091485546915 | 0.491092358676709 | 0.899236955296196 | 4153.4525550679 | 4535.4811943239 | 4274.19027713639 | 3922.40943538763 | 3440.3586512636 | 2876.68940983772 | 4445.4034466747 | 4764.37170333008 | Glud1 |
| **ENSMUSG00000021796** | 0.196263409073905 | 0.417832829770195 | 0.882468496126562 | 1189.92756472143 | 1492.82403109996 | 1409.27132919754 | 1281.20786435665 | 840.927894389066 | 812.126602097043 | 1456.46824868804 | 1579.43969659268 | Bmpr1a |
| **ENSMUSG00000021798** | 0.0851970935742476 | 0.507119430863465 | 0.901698316485955 | 21652.7062175772 | 22110.5726242951 | 21975.7360595283 | 20528.4015572284 | 18442.380892181 | 17296.8289259885 | 24065.5611412957 | 21514.0311661652 | Ldb3 |
| **ENSMUSG00000021799** | -0.515043567010981 | 0.0841966882144167 | 0.742565114108635 | 30.7194663633475 | 21.1748089517725 | 23.6613722162112 | 24.3860940507122 | 32.8487458745729 | 45.2540425867328 | 26.0269522639034 | 39.0789203280663 | Opn4 |
| **ENSMUSG00000021806** | -1.02147019367333 | 0.0144492388525411 | 0.635862253526407 | 93.0619128066116 | 146.594831204579 | 104.110037751329 | 142.564857527241 | 361.336204620302 | 391.386314263635 | 105.14888714617 | 130.263067760221 | Nid2 |
| **ENSMUSG00000021807** | 0.111131519335945 | 0.435869961096004 | 0.888687115594624 | 2253.3632091232 | 1599.51249158774 | 1978.09071727525 | 2236.95516580571 | 1801.20623212241 | 2079.23979452556 | 1742.76472359097 | 1846.47898550113 | Rtraf |
| **ENSMUSG00000021809** | 0.195782290974089 | 0.226836437949512 | 0.832048466795696 | 690.28447945875 | 600.22439220986 | 663.464876942561 | 662.176246146262 | 528.864808580624 | 436.640356850368 | 687.11153976705 | 629.604827507734 | Nudt13 |
| **ENSMUSG00000021810** | 0.0184382933953681 | 0.865728895297208 | 0.974868041734905 | 842.978297558919 | 872.239245667246 | 856.541674226844 | 851.63743838641 | 888.011096809288 | 905.080851734656 | 873.464517976599 | 714.275821551878 | Ecd |
| **ENSMUSG00000021811** | 0.128257314504928 | 0.304888222113656 | 0.851785577030345 | 355.080890611635 | 317.622134276588 | 379.528410348027 | 332.026049767389 | 338.342082508101 | 289.870489001505 | 341.473613702413 | 295.262953589834 | Dnajc9 |
| **ENSMUSG00000021814** | 0.063674567801601 | 0.449588073689568 | 0.89108031640712 | 2054.59019147801 | 1961.92749095462 | 2038.66383014875 | 2256.65162638514 | 2068.37603190227 | 1986.28554488795 | 2003.03424623001 | 1894.24211034655 | Anxa7 |
| **ENSMUSG00000021816** | 0.137476853028267 | 0.521512665031592 | 0.90609507420585 | 1899.18583222813 | 2124.81063673748 | 1922.249878845 | 1691.08182974746 | 1502.2826446638 | 1153.36654484565 | 2064.45785357282 | 2222.07083087644 | Ppp3cb |
| **ENSMUSG00000021819** | 0.0707251895839684 | 0.561431773911507 | 0.917031459819748 | 697.512589191303 | 842.92027942633 | 816.79056890361 | 683.7485601142 | 780.705193619016 | 653.125911927441 | 729.795741479852 | 731.644230586574 | Zswim8 |
| **ENSMUSG00000021820** | 0.0765086448850576 | 0.680818969008717 | 0.944452327903597 | 842.07478384235 | 890.156391703361 | 858.434584004141 | 785.044643094081 | 672.304332232926 | 609.094951572782 | 869.300205614374 | 1049.70322103445 | Camk2g |
| **ENSMUSG00000021822** | -0.678663667860223 | 0.0407811618296528 | 0.683226273624098 | 42.4651446787451 | 50.4937751926884 | 43.5369248778286 | 90.0409626487835 | 130.300025302473 | 94.1773318696872 | 72.8754663389296 | 65.1315338801105 | Plau |
| **ENSMUSG00000021823** | 0.169131855493661 | 0.677293614895239 | 0.944214159784677 | 665.889609111386 | 1156.47033505835 | 1308.00065611215 | 542.121629281217 | 615.366506050332 | 336.347613820311 | 1229.5132249468 | 1084.44003910384 | Vcl |
| **ENSMUSG00000021824** | -0.0165539829729636 | 0.918110067796288 | 0.986713918026868 | 405.677658739501 | 495.164763179912 | 567.872933189068 | 420.191159027656 | 450.027818481649 | 421.963370065482 | 560.100012719201 | 477.631248454143 | Ap3m1 |
| **ENSMUSG00000021830** | 0.145140206838825 | 0.265484446555043 | 0.842603400964081 | 392.124952990966 | 464.216965481167 | 500.674636095028 | 423.942865804689 | 384.330326732503 | 363.255422925936 | 432.047407580797 | 429.868123608729 | Txndc16 |
| **ENSMUSG00000021831** | 0.107534415728436 | 0.377933787037539 | 0.87422797717331 | 262.922491521592 | 315.178887089845 | 298.133289924261 | 283.253861665965 | 284.689130912965 | 264.185762127954 | 279.008928269045 | 248.585354309088 | Ero1l |
| **ENSMUSG00000021832** | 0.226073273323147 | 0.0402758430522533 | 0.679680259824687 | 2333.77592989784 | 2119.924142364 | 2189.15015744386 | 2396.4027038296 | 1761.78773707293 | 1757.56916749014 | 2071.74540020671 | 2135.22878570295 | Psmc6 |
| **ENSMUSG00000021835** | -0.921311355831533 | 0.0594563746339033 | 0.712406493939829 | 88.5443442237664 | 59.452348210746 | 53.9479286529615 | 99.4202295913651 | 215.706764576362 | 240.94719971855 | 64.5468416144805 | 49.9341759747513 | Bmp4 |
| **ENSMUSG00000021838** | 0.422992807535486 | 0.121095408921931 | 0.769868040577702 | 1755.52715129365 | 2216.02519837589 | 1854.10512686231 | 1750.17121148573 | 1050.06490979051 | 837.811328970594 | 1713.6145370554 | 2048.38674052947 | Samd4 |
| **ENSMUSG00000021840** | -0.0264453036301499 | 0.877882941379223 | 0.977771955397114 | 378.57224724243 | 469.103459854653 | 412.654331450723 | 360.163850595134 | 500.395895489327 | 318.001380339203 | 429.965251399684 | 401.644458927348 | Mapk1ip1l |
| **ENSMUSG00000021843** | 0.251660711482935 | 0.101674390486821 | 0.759800953500741 | 9064.04960462066 | 10097.1262070798 | 9951.97315413842 | 9349.25328836535 | 7281.47200219699 | 6334.34287991052 | 9562.30226175812 | 9126.01342216814 | Ktn1 |
| **ENSMUSG00000021846** | 0.00582846824482727 | 0.960044429154052 | 0.993625746680416 | 517.713359594063 | 561.132437221972 | 575.444572298256 | 465.211640352048 | 549.669014301187 | 511.248373006873 | 491.388858742496 | 559.045665804281 | Peli2 |
| **ENSMUSG00000021866** | 0.0511250081732067 | 0.778370350579036 | 0.960174437792435 | 599.029594085277 | 750.076886330097 | 732.556083813898 | 631.224665235743 | 623.031213421066 | 479.448234972953 | 803.712285909337 | 711.019244857872 | Anxa11 |
| **ENSMUSG00000021868** | 0.214775375666844 | 0.129979934373322 | 0.77303252008843 | 4132.67173958681 | 3927.92706055381 | 3685.49533639705 | 4064.03636622061 | 3369.18636853536 | 2639.41145681539 | 3760.37406308877 | 3852.53022900853 | Ppif |
| **ENSMUSG00000021870** | 0.334615764731343 | 0.161557580272112 | 0.792786631432503 | 7064.57374985336 | 8322.51433377552 | 7730.64353048051 | 7794.17082928532 | 4408.30169636768 | 4258.77233208118 | 7178.23343438456 | 8666.83610831336 | Slmap |
| **ENSMUSG00000021871** | -0.688590024504712 | 0.0516234850281498 | 0.703396849069134 | 249.369785773056 | 217.448999620126 | 274.47191770805 | 190.399118934407 | 503.680770076785 | 584.633306931305 | 192.599446752885 | 221.447215192376 | Pnp |
| **ENSMUSG00000021876** | -0.52896007728106 | 0.0506358014805156 | 0.703396849069134 | 903.513716569045 | 787.540009860156 | 736.341903368492 | 945.430107812227 | 1228.54309570903 | 1861.53115721641 | 694.399086400943 | 1083.35451353917 | Rnase4 |
| **ENSMUSG00000021877** | -0.0148839979700833 | 0.902365473690722 | 0.984902747274457 | 1309.19137530855 | 1218.36593045584 | 1214.30162213596 | 1365.62126683988 | 1442.05994389375 | 1383.30600447554 | 1094.1730731745 | 1241.84124598077 | Arf4 |
| **ENSMUSG00000021879** | 0.0608307797477153 | 0.909137780918337 | 0.98549154890268 | 5.42108229941427 | 14.6594831204579 | 14.1968233297267 | 12.1930470253561 | 16.4243729372865 | 12.2308223207386 | 4.16431236222455 | 11.9407812113536 | Dnah12 |
| **ENSMUSG00000021884** | 0.143826174114399 | 0.394226822235311 | 0.878942305112115 | 238.527621174228 | 311.106808445274 | 283.936466594534 | 249.488500672671 | 244.175677667659 | 203.031650524261 | 233.201492284575 | 299.605055848508 | Hacl1 |
| **ENSMUSG00000021886** | -0.33665248990958 | 0.445665741428401 | 0.89108031640712 | 6.32459601598331 | 11.4018202048006 | 16.0897331070236 | 13.1309737196143 | 14.2344565456483 | 20.7923979452556 | 12.4929370866736 | 11.9407812113536 | Gpr65 |
| **ENSMUSG00000021890** | 0.0800744020191544 | 0.631686633944452 | 0.931825800800958 | 332.493047697409 | 432.454752053509 | 449.566072108012 | 318.895076047775 | 349.291664466292 | 314.332133642982 | 392.486440139664 | 394.045779974668 | Eaf1 |
| **ENSMUSG00000021891** | 0.113248187713768 | 0.490218146383585 | 0.899236955296196 | 165.343010132135 | 157.182235680465 | 128.717864856189 | 137.87522405595 | 127.015150715015 | 121.085140975312 | 155.120635492864 | 141.118323406906 | Mettl6 |
| **ENSMUSG00000021892** | 0.0360035117449864 | 0.784091579111361 | 0.961141387070613 | 1058.91807581892 | 1031.86472853445 | 1183.06861081056 | 1314.03529865568 | 1214.30863916338 | 956.450305481758 | 1128.52865016285 | 1173.45313540666 | Sh3bp5 |
| **ENSMUSG00000021893** | 0.274144847317734 | 0.12593335713941 | 0.771949105062466 | 975.794813894569 | 1168.68657099206 | 1165.08596792624 | 964.18864169739 | 747.856447744443 | 697.1568722821 | 1000.47604502445 | 1087.69661579784 | Capn7 |
| **ENSMUSG00000021895** | -0.299532349769889 | 0.0175537507094951 | 0.642273034804458 | 564.696072855653 | 503.308920469055 | 568.819388077717 | 560.88016316638 | 752.23628052772 | 639.672007374629 | 733.960053842076 | 577.499600403646 | Arhgef3 |
| **ENSMUSG00000021898** | 0.207403960037857 | 0.133803892432918 | 0.77303252008843 | 2097.95884987332 | 2108.5223221592 | 2252.5626349833 | 2174.11407729042 | 1723.46420021926 | 1512.95272107536 | 2063.41677548226 | 2175.39323159569 | Asb14 |
| **ENSMUSG00000021900** | -0.436733617358969 | 0.113559929436083 | 0.767518180080285 | 200.580045078328 | 210.933673788811 | 154.272146849697 | 187.585338851632 | 306.588294829347 | 368.147751854232 | 175.942197303987 | 170.427513652956 | Btd |
| **ENSMUSG00000021901** | 0.148863669589239 | 0.395148714308943 | 0.879215782741314 | 1472.72735800754 | 1492.82403109996 | 1610.86622047966 | 1309.34566518439 | 1264.67671617106 | 973.573456730792 | 1650.10877353148 | 1418.78191302174 | Bap1 |
| **ENSMUSG00000021902** | 0.118626984035109 | 0.502758722643739 | 0.900927059989498 | 160.82544154929 | 146.594831204579 | 183.612248397799 | 186.647412157374 | 188.332809680885 | 145.546785616789 | 162.408182126757 | 127.006491066215 | Phf7 |
| **ENSMUSG00000021903** | -0.284788950231027 | 0.224046614220772 | 0.832048466795696 | 852.01343472461 | 961.824975847822 | 1030.68937373816 | 859.140851940476 | 1376.3624521446 | 1515.39888553951 | 823.492769629904 | 797.861290031353 | Galnt15 |
| **ENSMUSG00000021904** | -0.694696257094952 | 0.0435422962880193 | 0.688841110980169 | 614.389327266951 | 659.676740420606 | 654.946782944725 | 540.245775892701 | 1434.39523652302 | 1472.59100741693 | 595.49666779811 | 494.999657488839 | Sema3g |
| **ENSMUSG00000021905** | 0.0998771792337474 | 0.479516161947342 | 0.897732059433541 | 1354.367061137 | 1013.13316676942 | 1084.63730239112 | 1287.77335121646 | 997.506916391197 | 1264.66702796437 | 1071.26935518226 | 1089.86766692718 | Dph3 |
| **ENSMUSG00000021906** | 0.0579686620380822 | 0.78716750508338 | 0.961141387070613 | 1303.77029300913 | 1318.5390651123 | 1157.51432881705 | 1036.40899715527 | 1013.93128932848 | 771.764888438605 | 1446.05746778247 | 1393.81482503436 | Oxnad1 |
| **ENSMUSG00000021910** | -0.0102526147992697 | 0.934670294568612 | 0.98818087308472 | 2723.1903417391 | 3415.65956706669 | 3667.51269351273 | 2878.4970246783 | 3354.95191198971 | 3124.97510294871 | 3217.97237790902 | 3077.46497583522 | Nisch |
| **ENSMUSG00000021911** | 0.237479018720045 | 0.404138014740391 | 0.881158207349486 | 377.668733525861 | 460.144886836596 | 438.208613444231 | 365.791410760683 | 270.454674367317 | 189.577745971448 | 475.772687384154 | 455.920737160773 | Parg |
| **ENSMUSG00000021913** | 0.310842032117029 | 0.132654705975672 | 0.77303252008843 | 4601.59535848615 | 4920.69983410037 | 6021.34600158142 | 4788.11577418791 | 4120.32769086726 | 2650.41919690405 | 4964.90141386222 | 4653.64809573389 | Ogdhl |

| | | | | | | | | | | | | |
|---|---|---|---|---|---|---|---|---|---|---|---|---|
| ENSMUSG00000021916 | 0.0610042333977607 | 0.610878432890023 | 0.927932112008768 | 261.115464088454 | 300.519403969387 | 266.900278598862 | 294.508981997063 | 249.650468646754 | 284.978160073209 | 279.008928269045 | 263.782712214447 | Glt8d1 |
| ENSMUSG00000021917 | 0.0163256973783733 | 0.887371322884532 | 0.980702840906172 | 1194.44513330428 | 1058.74044758863 | 1078.01211817058 | 1264.32518386 | 1193.50443344282 | 1249.99004117948 | 1054.61210573337 | 1046.44664434044 | Spcs1 |
| ENSMUSG00000021918 | 0.319101468481976 | 0.272754457106108 | 0.844738346035895 | 100.290022539164 | 111.574954861263 | 115.467496415111 | 84.4134024832345 | 61.3176589658694 | 48.9232892829544 | 107.231043327282 | 111.809133160856 | Nek4 |
| ENSMUSG00000021922 | -0.474451170413833 | 0.270021292839873 | 0.843793744782845 | 159.018414116152 | 567.647763053287 | 186.451613063744 | 197.902532488472 | 263.884925192402 | 352.247682837272 | 298.789411989611 | 629.604827507734 | Itih4 |
| ENSMUSG00000021928 | -0.285518290069075 | 0.298887453514233 | 0.851785577030345 | 81.316234491214 | 89.5857301805761 | 78.5557557578211 | 82.5375490947182 | 77.7420319031559 | 129.646716599829 | 62.4646854333682 | 135.690695583563 | Ebpl |
| ENSMUSG00000021929 | 0.102139534252766 | 0.615603251973823 | 0.928422072203769 | 1802.50986455524 | 2159.01609735189 | 1867.35549530339 | 1592.59952685036 | 1555.93559625894 | 1196.17442296823 | 2048.84168221448 | 2112.43274884492 | Kpna3 |
| ENSMUSG00000021930 | 0.0752107657676982 | 0.566160376291925 | 0.919931558693298 | 833.943160393229 | 819.302223287815 | 816.79056890361 | 772.851596068725 | 678.87408140784 | 665.35673424818 | 863.053737071037 | 869.505977299475 | Spryd7 |
| ENSMUSG00000021932 | -0.106071984616249 | 0.587319841826035 | 0.923058967588378 | 185.220311896654 | 155.553404222637 | 147.646962629158 | 237.295453647315 | 191.617684268342 | 227.493295165738 | 187.394056300105 | 174.76961591163 | Rnaseh2b |
| ENSMUSG00000021936 | 0.00384683704678059 | 0.983801599007034 | 0.996940244108351 | 365.019541493894 | 435.712414969166 | 446.726707442067 | 333.901903155905 | 352.576539053749 | 305.770558018465 | 495.553171104721 | 422.269444656049 | Mapk8 |
| ENSMUSG00000021938 | 0.114717144968156 | 0.470549480123519 | 0.896509979960267 | 249.369785773056 | 289.931999493501 | 321.794662140472 | 275.750448111899 | 233.226095709468 | 227.493295165738 | 313.364505257397 | 274.637967861132 | Pspc1 |
| ENSMUSG00000021939 | -0.0803542094981815 | 0.281255566463532 | 0.850048154552447 | 11639.0636968424 | 12090.0014957332 | 11936.6890556342 | 10941.8528152157 | 12036.8754466393 | 13099.210705511 | 11986.9731346634 | 12156.8007987226 | Ctsb |
| ENSMUSG00000021943 | -1.12895760981727 | 0.0475593627691979 | 0.703396849069134 | 18.9737880479499 | 20.3603932228582 | 8.51809399783602 | 12.1930470253561 | 42.7033696369448 | 59.9310293716191 | 20.8215618111227 | 8.68420451734806 | Gdf10 |
| ENSMUSG00000021944 | 0.00349797243382279 | 0.983647321240763 | 0.996940244108351 | 1447.42897394361 | 1608.4710646058 | 1880.60586374446 | 1973.39776471917 | 1486.95322992233 | 1439.56778715093 | 2044.67736985225 | 1920.29472389859 | Gata4 |
| ENSMUSG00000021945 | 0.0340181366950821 | 0.869801082383223 | 0.975613767803763 | 436.397125102849 | 544.844122643686 | 530.961192531779 | 449.266886549659 | 454.407651264925 | 324.116791499573 | 621.523620062014 | 514.539117652873 | Zmym2 |
| ENSMUSG00000021947 | 0.14106882485792 | 0.513375199254939 | 0.903087831516097 | 76.7986659083688 | 85.5136515360045 | 104.110037751329 | 116.302910088012 | 105.115986798633 | 85.6157562451702 | 85.3684034256032 | 70.559161703453 | Cryl1 |
| ENSMUSG00000021948 | -0.178274758595772 | 0.328645207907328 | 0.855977678491063 | 130.105975185942 | 154.738988493722 | 164.68315062483 | 167.888878272211 | 151.104231023035 | 215.262472844999 | 186.352978209548 | 146.545951230249 | Prkcd |
| ENSMUSG00000021951 | 0.13402034620238 | 0.480109579023784 | 0.897732059433541 | 688.477452025612 | 464.216965481167 | 384.260684791269 | 536.494069115668 | 453.312693069106 | 490.455975061618 | 446.622500848583 | 499.341759747513 | Eef1akmt1 |
| ENSMUSG00000021952 | 0.505890610814382 | 0.182963558070964 | 0.805716353538343 | 91.2548853734735 | 150.666909849151 | 151.432782183751 | 77.8479156234274 | 59.1277425742312 | 40.3617136584374 | 112.436433780063 | 119.407812113536 | Xpo4 |
| ENSMUSG00000021957 | -0.389178205438803 | 0.105604614381832 | 0.762094734041179 | 395.739007857242 | 378.703313945163 | 381.421320125324 | 419.253232333398 | 627.411046204342 | 705.718447906617 | 349.802238426862 | 381.019473198646 | Tkt |
| ENSMUSG00000021958 | -0.0372482237011924 | 0.856076308795631 | 0.97363563840968 | 74.9916384752307 | 73.2974156022896 | 75.7163910918757 | 72.2203554578784 | 82.1218646864323 | 95.400414101761 | 67.6700758861489 | 59.7039060567679 | Pinx1 |
| ENSMUSG00000021959 | -0.103958341342291 | 0.487892510197047 | 0.899236955296196 | 578.248778604189 | 678.408302185636 | 722.145080038765 | 494.287367874051 | 722.672409240604 | 612.764198269004 | 654.83811895981 | 667.598222271132 | Lats2 |
| ENSMUSG00000021962 | -0.0619610043252845 | 0.763035295322422 | 0.957845481430497 | 104.807591122009 | 166.140808698523 | 139.128868631322 | 153.819977858338 | 178.478185918513 | 134.539045528125 | 153.038479311752 | 122.664388807541 | Dcp1a |
| ENSMUSG00000021963 | 0.0201176006042388 | 0.866236210258221 | 0.974868041734905 | 2565.07544133952 | 2198.10805233977 | 2431.44260893786 | 2564.29158210181 | 2512.92905940483 | 2743.37344654167 | 2232.07142615236 | 2136.31431126762 | Sap18 |
| ENSMUSG00000021967 | 0.121491473594739 | 0.503780100163964 | 0.900927059989498 | 1055.30402095264 | 669.449729167578 | 758.110365807406 | 973.567908639971 | 767.565695269187 | 936.880989768576 | 708.974179668729 | 764.209997526629 | Mrpl57 |
| ENSMUSG00000021969 | -0.199451039080837 | 0.356697024987223 | 0.864965376583892 | 167.150037565273 | 127.048853710635 | 159.004421292939 | 151.944124469822 | 197.092475247437 | 225.04713070159 | 129.093683228961 | 144.374900100912 | Zdhhc20 |
| ENSMUSG00000021972 | 0.298022200260716 | 0.10641737237141 | 0.762094734041179 | 317.133314515735 | 303.777066885045 | 348.295399022628 | 257.929840920994 | 261.695008800764 | 183.462334811079 | 301.91264626128 | 249.670879873757 | Hmbox1 |
| ENSMUSG00000021973 | 0.241509787296459 | 0.134068366302786 | 0.77303252008843 | 3405.34319774873 | 3544.33725223516 | 3511.34763688574 | 3431.87377429061 | 2473.51056435534 | 2345.87172111766 | 3519.8850241703 | 3410.72132418845 | Micu2 |
| ENSMUSG00000021974 | -0.15807052238079 | 0.429083262498181 | 0.88571355566945 | 224.974915425692 | 179.171460361152 | 140.07532351997 | 239.171307035831 | 209.137015401448 | 199.362403828039 | 214.462086654564 | 250.756405438425 | Fgf9 |
| ENSMUSG00000021975 | -0.16175397919086 | 0.27523040744278 | 0.846286635233373 | 152.693818100169 | 175.913797445495 | 197.809071727525 | 179.143998603309 | 225.561388338734 | 185.908499275227 | 189.476212481217 | 187.795922687652 | Ints9 |
| ENSMUSG00000021978 | -0.143416713484161 | 0.25774744654031 | 0.838169889266919 | 983.92643734369 | 1207.77852597995 | 1148.99623481921 | 1038.28485054378 | 1098.24307040655 | 1092.21243324196 | 1342.99073681742 | 1302.63067760221 | Extl3 |
| ENSMUSG00000021981 | 0.00537036613800661 | 0.970430718890472 | 0.9951105526746 | 575.538237454482 | 500.051257553398 | 527.175372977185 | 519.611388619021 | 453.312693069106 | 477.002070508805 | 646.509494235361 | 536.249628946243 | Cab39l |
| ENSMUSG00000021982 | 0.216754355773754 | 0.264779099990324 | 0.842510961779757 | 1048.07591122009 | 1095.38915538977 | 1091.26248661166 | 1086.11911195095 | 710.627869086594 | 721.618516923577 | 1100.41954171784 | 1184.30839105334 | Cdadc1 |
| ENSMUSG00000021983 | 0.286732378532208 | 0.304660781588264 | 0.851785577030345 | 205.097613661173 | 241.881471487556 | 151.432782183751 | 236.357526953057 | 133.58489988993 | 108.854318654574 | 243.612273190136 | 197.565652769668 | Atp8a2 |
| ENSMUSG00000021987 | 0.0957460213494471 | 0.649558667917184 | 0.936461253686283 | 942.364806381514 | 1041.63771728143 | 1049.61847151113 | 902.285479876351 | 689.823663366031 | 693.487625585878 | 1124.36433780063 | 1174.53866097133 | Mtmr6 |
| ENSMUSG00000021990 | -0.340241567045927 | 0.0571056014573982 | 0.708844329082449 | 135.527057485357 | 192.202112023782 | 215.791714611846 | 178.206071909051 | 239.795844884382 | 215.262472844999 | 252.981976005141 | 205.164331722348 | Spata13 |
| ENSMUSG00000021991 | -0.277968223259906 | 0.541452644356347 | 0.913837378403038 | 43.3686583953142 | 39.9063707168021 | 16.0897331070236 | 17.8206071909051 | 20.8042057205628 | 57.4848649074714 | 38.5198893505771 | 26.0526135520442 | Cacna2d3 |
| ENSMUSG00000021993 | 0.19339830044967 | 0.159613707573564 | 0.790303302997254 | 3019.54284077375 | 2988.90572511559 | 2931.17079014424 | 3005.11712840315 | 2473.51056435534 | 2069.45513666897 | 2991.01735416778 | 2910.29403887627 | Mipep |
| ENSMUSG00000021994 | -0.25757742279067 | 0.338786341514698 | 0.858570960161243 | 124.684892886528 | 143.337168288922 | 106.002947528626 | 109.737423228205 | 130.300025302473 | 81.9465095489486 | 186.352978209548 | 179.111718170304 | Wnt5a |
| ENSMUSG00000021996 | 0.0131411939204207 | 0.910179930627345 | 0.98603671070003 | 3096.34150668212 | 2912.35064659764 | 2775.00573351725 | 3489.08730264036 | 2832.656852584 | 3389.16086507667 | 2857.75935857659 | 3082.89260365856 | Esd |
| ENSMUSG00000021998 | -0.110466261253794 | 0.571482564044138 | 0.921130164032017 | 183.413284463516 | 270.386021999557 | 291.508105703722 | 254.178134143962 | 236.510970296925 | 331.455284892016 | 237.365804646799 | 274.637967861132 | Lcp1 |
| ENSMUSG00000021999 | -0.180729076813542 | 0.570829835618637 | 0.921099723134627 | 46.0791995450213 | 82.2559886203472 | 49.2156542097192 | 72.2203554578784 | 48.1781606160403 | 68.4926049961361 | 59.3414511616998 | 107.467030902182 | Cpb2 |
| ENSMUSG00000022000 | 0.112298445064507 | 0.321062834445003 | 0.855469551206808 | 1405.86734298143 | 1351.93010999779 | 1194.42606947434 | 1268.07689063703 | 1309.57000219964 | 1055.51996627974 | 1181.62363278122 | 1282.00569187351 | Zc3h13 |
| ENSMUSG00000022003 | -0.0492519596490225 | 0.800139592180938 | 0.963461896828986 | 81.316234491214 | 104.245213301034 | 94.6454888648447 | 71.2824287636203 | 87.5966556655277 | 102.738907494204 | 82.2451691539348 | 92.2696729968231 | Slc25a30 |
| ENSMUSG00000022008 | 0.0199154305877707 | 0.918511904196762 | 0.986713918026868 | 347.852780879082 | 450.371898089624 | 512.03209475881 | 352.660437041069 | 393.089992299056 | 316.77829810713 | 482.019155927491 | 447.236532643425 | Gpalpp1 |
| ENSMUSG00000022009 | 0.05401765988679 | 0.724489718396424 | 0.951674101666722 | 124.684892886528 | 128.677685168464 | 128.717864856189 | 156.633757941113 | 130.300025302473 | 135.762127760198 | 131.175839410073 | 121.578863242873 | Nufip1 |
| ENSMUSG00000022010 | -0.188178800133531 | 0.244820748827591 | 0.832048466795696 | 7317.5575904927 | 5825.51570892419 | 6138.70640777383 | 5159.53474511415 | 6721.94836413344 | 8615.39124272827 | 6201.70218544291 | 6309.07458185337 | Tsc22d1 |
| ENSMUSG00000022012 | 0.363060253177922 | 0.507222580130703 | 0.901698316485955 | 17.1667606148119 | 15.4738988493722 | 28.3936466594534 | 12.1930470253561 | 6.56974917491458 | 30.5770558018465 | 8.32862472444909 | 11.9407812113536 | Enox1 |
| ENSMUSG00000022013 | 0.198220512662202 | 0.236062867552322 | 0.832048466795696 | 2428.64487013759 | 1533.54481754568 | 1835.17602908934 | 2153.47969001674 | 1594.25913311261 | 1635.26094428275 | 1724.02531796096 | 1975.65652769668 | Dnajc15 |
| ENSMUSG00000022014 | -0.450951195110891 | 0.236174687549781 | 0.832048466795696 | 9.03513716569045 | 21.1748089517725 | 29.3401015481019 | 26.2619474392285 | 31.7537876787538 | 24.4616446414772 | 27.0680303544596 | 33.6512925047237 | Epsti1 |
| ENSMUSG00000022016 | 0.221234790357191 | 0.318481074685244 | 0.854351922245337 | 778.828823682517 | 968.340301679137 | 941.722614205205 | 726.893188050075 | 649.310210120724 | 484.340563901248 | 914.066563508288 | 881.446758510828 | Akap11 |
| ENSMUSG00000022018 | -0.430320711710746 | 0.346931752939823 | 0.861266047293794 | 1032.71617803842 | 829.889627763701 | 567.872933189068 | 563.693943249155 | 1323.80445874529 | 1867.64656837678 | 414.349080041342 | 429.868123608729 | Rgcc |
| ENSMUSG00000022019 | 0.0296911270876235 | 0.791956897694907 | 0.961804768755116 | 515.002818444356 | 497.608010366655 | 443.887342776122 | 468.963347129081 | 439.078236523458 | 451.317343635254 | 471.60837502193 | 524.308847734889 | Tdrd3 |
| ENSMUSG00000022020 | -0.121287187949953 | 0.613169991681637 | 0.928075610917191 | 79.509207058076 | 52.9370223794314 | 52.0550188756646 | 54.3997482669734 | 64.6025335533267 | 75.8310983885793 | 68.711153976705 | 51.0197015394199 | Naa16 |
| ENSMUSG00000022022 | 0.154603872934187 | 0.184404875020595 | 0.805963874499992 | 352.370349461928 | 374.631235300591 | 363.438677241004 | 332.963976461647 | 303.30342024189 | 300.878229090169 | 359.171941241867 | 314.802413753867 | Mtrf1 |
| ENSMUSG00000022023 | -0.0339135280098436 | 0.738823658084766 | 0.954383574136061 | 763.469090500843 | 732.159740293981 | 640.749959614998 | 729.70696813285 | 700.773245324222 | 780.326464063122 | 708.974179668729 | 745.756062927265 | Wbp4 |
| ENSMUSG00000022024 | 0.12656178777595 | 0.335173106281735 | 0.858570960161243 | 1226.97162710076 | 1033.49355999228 | 1340.1801223262 | 1316.84907873846 | 1232.9229284923 | 1201.06675189653 | 1068.1461209106 | 1001.94009618903 | Sugt1 |
| ENSMUSG00000022025 | -0.884866489815749 | 0.0639541897118908 | 0.716991331547963 | 19.877301764519 | 17.9171460361152 | 22.7149173275627 | 30.0136542162612 | 53.6529515951357 | 72.1618516923577 | 14.5750932677859 | 27.1381391167127 | Cnmd |
| ENSMUSG00000022031 | 0.0682245525317437 | 0.559923576647502 | 0.916574868225118 | 842.978297558919 | 991.143942088738 | 1020.27836996303 | 876.961459131381 | 936.189257425328 | 814.572766561191 | 966.120468036095 | 841.282312618093 | Elp3 |
| ENSMUSG00000022032 | -0.920059585235648 | 0.0248614193434792 | 0.655303666217042 | 221.360860559416 | 238.623808571898 | 315.169477919933 | 300.136542162612 | 849.687559955619 | 714.280023531134 | 232.160414194018 | 238.815624227072 | Scara5 |
| ENSMUSG00000022035 | 0.0515792020442361 | 0.708520913960447 | 0.949174314815223 | 565.599586572222 | 487.020605890769 | 572.60520763231 | 472.715053906113 | 457.692525852383 | 452.540425867328 | 533.031982364742 | 579.670651532983 | Ccdc25 |
| ENSMUSG00000022037 | -0.0526027141647599 | 0.678195127292962 | 0.944452327903597 | 18158.8186756047 | 15676.6883658719 | 17885.1580307897 | 19355.9931894057 | 17647.4412420164 | 21878.4949673372 | 17438.0580168153 | 16752.9160395291 | Clu |
| ENSMUSG00000022040 | 0.204595058516294 | 0.0844423732869727 | 0.743285529122972 | 10029.0022539164 | 10099.5694542666 | 13164.2410462112 | 11467.0917640003 | 10167.7818063761 | 9913.08149095863 | 9352.00448746578 | 9408.25006898196 | Ephx2 |
| ENSMUSG00000022041 | 0.728734110495529 | 0.0069688335305625 | 0.635862253526407 | 221.360860559416 | 227.221988367098 | 422.118880337207 | 246.674720589896 | 143.439523652302 | 125.977469903608 | 200.928071477334 | 202.993280593011 | Chrna2 |
| ENSMUSG00000022043 | 0.163153965341625 | 0.363464916704117 | 0.86924083181804 | 1987.7301764519 | 2450.57692830322 | 2037.71737526011 | 1981.83910496749 | 1611.77846424571 | 1484.82182973767 | 2055.08815075781 | 2401.18254904674 | Trim35 |
| ENSMUSG00000022048 | -0.821317354042254 | 0.0661305189383728 | 0.719725956118125 | 193.351935345776 | 131.935348084121 | 134.396594188079 | 235.419600258798 | 421.558905390352 | 532.040770952129 | 115.559668051731 | 159.572258006271 | Dpysl2 |
| ENSMUSG00000022051 | 0.135465152759889 | 0.433497282251353 | 0.887868762931002 | 1677.82497166872 | 1475.72130079276 | 2262.97363875844 | 1546.64111883171 | 1431.11036193556 | 1871.31581507301 | 1484.57735713305 | 1552.30155747597 | Bnip3l |
| ENSMUSG00000022052 | -0.0503098928821079 | 0.59904193193916 | 0.924848033480823 | 1223.35757223449 | 1259.08671690155 | 1103.56640016409 | 1169.59458773993 | 1243.87251045049 | 1121.56640681173 | 1246.1704743957 | 1312.40040768423 | Ppp2r2a |
| ENSMUSG00000022053 | -0.725835092536918 | 0.0430456367406968 | 0.687730553581215 | 36.1405486627618 | 42.3496179035451 | 50.1621090983677 | 72.2203554578784 | 118.255485148462 | 107.6312364225 | 48.9306702561384 | 57.5328549274309 | Ebf2 |
| ENSMUSG00000022064 | 0.231605171129111 | 0.429004375427791 | 0.885682338687873 | 133.720030052219 | 192.202112023782 | 128.717864856189 | 135.999370667433 | 124.825234323377 | 66.0464405319884 | 178.024353485099 | 133.519644454226 | Pibf1 |
| ENSMUSG00000022066 | 0.675869199162517 | 0.0968268190362847 | 0.757809915356977 | 16.2632468982428 | 21.9892246806869 | 17.9826428843205 | 14.0689004138724 | 7.66470737073368 | 9.78465785659088 | 14.5750932677859 | 11.9407812113536 | Entpd4b |
| ENSMUSG00000022070 | 0.090851529892802 | 0.83118801061263 | 0.967583158900775 | 18.0702743313809 | 13.0306516626293 | 10.4110037751329 | 14.0689004138724 | 15.3294147414674 | 13.4539045528125 | 7.28754663389296 | 16.2828834700276 | Bora |
| ENSMUSG00000022074 | -0.618900301221324 | 0.0498846265376668 | 0.703396849069134 | 36.1405486627618 | 37.4631235300591 | 27.447191770805 | 48.7721881014244 | 82.1218646864323 | 62.3771938357668 | 48.9306702561384 | 36.9078691987293 | Tnfrsf10b |
| ENSMUSG00000022075 | -0.0267120213490572 | 0.80976254759869 | 0.964412705343932 | 524.941469326615 | 530.999055252142 | 535.693466975021 | 537.431995809926 | 535.434557755538 | 467.217412652214 | 568.428637443651 | 595.953535003011 | Rhobtb2 |
| ENSMUSG00000022089 | 0.22097105392604 | 0.241893528121629 | 0.832048466795696 | 186.123825613223 | 145.780415475665 | 187.398067952392 | 178.206071909051 | 168.623562156141 | 179.793088114857 | 133.257995591185 | 117.236760984199 | Bin3 |
| ENSMUSG00000022090 | -0.54547335687186 | 0.126691693929274 | 0.77215780108715 | 120.167324303683 | 76.5550785179469 | 92.7525790875478 | 165.075098189436 | 173.003394939417 | 273.970419984545 | 107.231043327282 | 109.638082031519 | Pdlim2 |
| ENSMUSG00000022091 | -0.55763966947212 | 0.126135028491997 | 0.771949105062466 | 851.10992100804 | 705.284021239809 | 589.641395627982 | 574.011136885995 | 1455.19944224358 | 1479.92950080937 | 610.071761065896 | 459.177313854779 | Sorbs3 |
| ENSMUSG00000022092 | 0.235319431900685 | 0.181742827818334 | 0.80356567265522 | 798.706125447036 | 631.986605637519 | 717.412805595523 | 694.065753751039 | 577.042969196664 | 447.648096939033 | 748.535147109862 | 639.374557589751 | Ppp3cc |
| ENSMUSG00000022094 | 0.0368215815716778 | 0.859604473875315 | 0.974868041734905 | 1370.63030803524 | 1724.11809811163 | 1601.40167159317 | 1264.32518386 | 1434.39523652302 | 988.250443515679 | 1886.43350008772 | 1500.19633037188 | Slc39a14 |
| ENSMUSG00000022095 | 0.0977420710929348 | 0.302076042590324 | 0.851785577030345 | 687.573938309043 | 704.469605510894 | 668.197151385803 | 727.831114744333 | 619.746338833609 | 611.54111603693 | 691.275852129275 | 681.710054611823 | Fam160b2 |
| ENSMUSG00000022096 | 0.117528305114096 | 0.449830596581254 | 0.89108031640712 | 374.054678659585 | 426.753841951108 | 335.04503058155 | 450.204813243917 | 374.475702970131 | 300.878229090169 | 411.225845769674 | 374.506319810635 | Hr |
| ENSMUSG00000022098 | -0.548198113128918 | 0.0144367633619341 | 0.635862253526407 | 245.75573090678 | 262.241864710414 | 225.25626349833 | 231.667893481766 | 401.849657865609 | 481.894399437101 | 274.84461590682 | 254.012982132431 | Bmp1 |
| ENSMUSG00000022099 | -0.705479133996349 | 0.0136908442490564 | 0.635862253526407 | 52.4037955610046 | 83.8848200781758 | 78.5557557578211 | 44.0825546301336 | 129.205067106653 | 128.423634367755 | 85.3684034256032 | 80.3288917854696 | Dmtn |
| ENSMUSG00000022100 | 0.191923758761327 | 0.36232675797287 | 0.86924083181804 | 959.531566996326 | 1231.39658211846 | 1329.76911855107 | 894.782066322286 | 823.408563255961 | 731.403174780168 | 1174.33608614732 | 1135.45974064326 | Xpo7 |
| ENSMUSG00000022102 | -1.02261514922789 | 0.0612621162068692 | 0.71282992338528 | 8.13162344912141 | 4.88649437348597 | 17.9826428843205 | 9.37926694258161 | 17.5193311331055 | 36.6924669622158 | 10.4107809055614 | 17.3684090346961 | Dok2 |
| ENSMUSG00000022103 | -1.56153783916956 | 0.066582638248872 | 0.719725956118125 | 0.903513716569045 | 8.14415728914329 | 13.2503684410783 | 6.56548685980713 | 50.3680770076785 | 23.2385624094033 | 8.32862472444909 | 3.25657669400552 | Gfra2 |
| ENSMUSG00000022105 | 0.042385764561673 | 0.824595441926673 | 0.966657594633313 | 357.791431761342 | 443.856572258309 | 435.369248778286 | 333.901903155905 | 352.576539053749 | 282.531995609062 | 448.704657029695 | 440.723379255414 | Rb1 |
| ENSMUSG00000022106 | 0.178125306926771 | 0.133388351939579 | 0.77303252008843 | 496.932544112975 | 424.310594764365 | 477.013263878817 | 503.666634816633 | 379.950493949227 | 408.509465512669 | 428.924173309128 | 462.433890548784 | Rcbtb2 |

| | | | | | | | | | | | | |
|---|---|---|---|---|---|---|---|---|---|---|---|---|
| ENSMUSG00000022108 | 0.00918704288566098 | 0.950558277222193 | 0.991696503889296 | 18400.9603516452 | 17220.8205878935 | 19509.2746197104 | 19993.7833415012 | 20793.2561386047 | 22148.7961406255 | 15767.1276814727 | 15939.8573915924 | Itm2b |
| ENSMUSG00000022109 | 0.140424527587514 | 0.170085949605823 | 0.79907358295209 | 551.143367107117 | 561.946852950887 | 577.337482075553 | 617.15576482187 | 568.283303630111 | 523.479195327612 | 519.497967187512 | 481.973350712817 | Med4 |
| ENSMUSG00000022110 | 0.23878281844714 | 0.179898576039858 | 0.802731406993634 | 17288.7349665487 | 17891.08473279 | 18224.9353358145 | 18056.0267911639 | 14112.9161859123 | 10634.7000078822 | 17480.7422185281 | 18330.1846849924 | Sucla2 |
| ENSMUSG00000022111 | 0.150512064420199 | 0.253550611223812 | 0.837406774112203 | 1020.97049972302 | 835.590537866101 | 980.527264639791 | 1170.53251443419 | 953.708588558433 | 922.20400298369 | 828.698160082685 | 905.328320933535 | Uchl3 |
| ENSMUSG00000022114 | -0.297007726231338 | 0.0551272939556773 | 0.703396849069134 | 196.965990212052 | 184.872370463553 | 226.202718386979 | 241.985087118606 | 302.208462046071 | 265.408844360028 | 241.530117009024 | 234.473521968398 | Spry2 |
| ENSMUSG00000022119 | 0.128862582060684 | 0.581565755294735 | 0.922450244597442 | 342.431698579668 | 384.404224047563 | 324.634026806417 | 269.184961252092 | 267.16979977986 | 194.470074899744 | 363.336253604092 | 382.104998763315 | Rbm26 |
| ENSMUSG00000022120 | 0.122209722951447 | 0.624155797799137 | 0.931145988862144 | 75.8951521917998 | 65.1532583131463 | 60.5731128735006 | 50.6480414899407 | 66.7924499449649 | 37.9155491942896 | 63.5057635239243 | 62.9604827507734 | Rnf219 |
| ENSMUSG00000022122 | -0.848271972258012 | 0.0177151332749689 | 0.642273034804458 | 386.703870691551 | 274.458100644129 | 251.757000380487 | 330.150196378873 | 911.005218921489 | 709.387694602839 | 322.734208072402 | 295.262953589834 | Ednrb |
| ENSMUSG00000022124 | 0.191494382530374 | 0.334789085131141 | 0.858570960161243 | 1355.27057485357 | 1557.1628736842 | 1637.36695736181 | 1269.01481733129 | 1062.10944994452 | 942.996400928946 | 1553.28851110976 | 1536.01867400594 | Fbxl3 |
| ENSMUSG00000022125 | 0.190256618959936 | 0.17233800259158 | 0.799889021902745 | 635.170142748039 | 674.336223541064 | 689.965613824718 | 649.045272426648 | 561.713554455197 | 462.325083723919 | 621.523620062014 | 674.111375659143 | Cln5 |
| ENSMUSG00000022129 | 0.148703628237928 | 0.560582466051723 | 0.916762264997884 | 303.580608767199 | 341.240190415104 | 186.451613063744 | 248.550573978413 | 211.326931793086 | 276.416584448692 | 315.446661438509 | 171.513039217624 | Dct |
| ENSMUSG00000022130 | 0.119722944139053 | 0.481205477051944 | 0.897732059433541 | 185.220311896654 | 175.913797445495 | 145.754052851861 | 154.757904552597 | 151.104231023035 | 128.423634367755 | 145.750932677859 | 183.453820428978 | Tgds |
| ENSMUSG00000022131 | -0.0688257892376067 | 0.531847215497589 | 0.909082872831377 | 521.327414460339 | 447.114235173966 | 441.994432998825 | 483.032247542953 | 502.585811880965 | 479.448234972953 | 534.073060455298 | 468.947043936795 | Gpr180 |
| ENSMUSG00000022136 | -0.0562127637703592 | 0.613739526652701 | 0.928075610917191 | 1279.37542266177 | 1588.11067138294 | 1528.52464516724 | 1403.13833461021 | 1541.70113971329 | 1624.25320419409 | 1380.46954807744 | 1484.99897246652 | Dnajc3 |
| ENSMUSG00000022139 | 0.156976600529859 | 0.554961242817956 | 0.916239900758826 | 3920.34601619309 | 5842.61843923139 | 5874.64549384091 | 3643.84520719296 | 3403.13007260575 | 2919.4972879603 | 5848.77671274438 | 5121.50961410602 | Mbnl2 |
| ENSMUSG00000022141 | 0.152086997246769 | 0.260293241232437 | 0.840316047847749 | 1127.58511827817 | 1228.95333493172 | 1253.10627257054 | 1050.47789756914 | 989.842209020464 | 874.50379593281 | 1176.41824232843 | 1151.74262411329 | Nipbl |
| ENSMUSG00000022142 | 0.0153337422561606 | 0.92216969878413 | 0.986713918026868 | 160.82544154929 | 199.531853584011 | 156.165056626994 | 163.19924480092 | 186.142893289246 | 144.323703384715 | 172.818963032319 | 169.341988088287 | Nup155 |
| ENSMUSG00000022146 | -0.598859542850725 | 0.0341102818287986 | 0.67781375479544 | 251.176813206195 | 266.313943354985 | 245.131816159948 | 201.654239265505 | 493.826146314413 | 488.00981059747 | 257.146288367366 | 222.532740757044 | Osmr |
| ENSMUSG00000022148 | 0.281452518926413 | 0.280424780177191 | 0.849093714741928 | 37.0440623793308 | 43.1640336324594 | 63.4124775394459 | 47.8342614071662 | 35.0386622662111 | 39.1386314263635 | 38.5198893505771 | 44.5065481514088 | Fyb |
| ENSMUSG00000022149 | -0.320473245296856 | 0.405109167373818 | 0.881158207349486 | 64.1494738764022 | 122.976775066064 | 46.3762895437739 | 75.0341355406529 | 72.2672409240604 | 81.9465095489486 | 66.6289977955927 | 164.999885829613 | C9 |
| ENSMUSG00000022150 | -0.313939780689041 | 0.0790219395427092 | 0.739855948845396 | 199.676531361759 | 233.737314198412 | 256.489274823729 | 284.191788360223 | 343.816873487196 | 319.224462571277 | 233.201492284575 | 314.802413753867 | Dab2 |
| ENSMUSG00000022151 | 0.132459792287571 | 0.196819028057577 | 0.812065429583558 | 732.749624137496 | 779.395852571012 | 842.344850897118 | 846.009878220861 | 719.387534653147 | 669.025980944401 | 765.19239655876 | 764.209997526629 | Ttc33 |
| ENSMUSG00000022159 | -0.297994974413557 | 0.340700148222572 | 0.859077629258968 | 18.0702743313809 | 20.3603932228582 | 26.5007368821565 | 21.5723139679377 | 21.8991639163819 | 24.4616446414772 | 26.0269522639034 | 33.6512925047237 | Rab2b |
| ENSMUSG00000022160 | 0.165992125546774 | 0.230307042502222 | 0.832048466795696 | 262.018977805023 | 237.809392842984 | 262.16800415562 | 232.605820176024 | 249.650468646754 | 190.800828203522 | 235.283648465687 | 209.506433981022 | Mettl3 |
| ENSMUSG00000022174 | -0.148239723015753 | 0.355765558968761 | 0.864662986238123 | 1223.35757223449 | 1066.07018914886 | 968.223351087361 | 1208.04958220451 | 1243.87251045049 | 1539.86053018099 | 1145.18589961175 | 1021.47955635307 | Dad1 |
| ENSMUSG00000022175 | -0.175372778386317 | 0.229222501337222 | 0.832048466795696 | 1317.32299875767 | 1494.45286255779 | 1505.80972783968 | 1308.40773849013 | 1780.40202640185 | 1819.9463613259 | 1466.8790295936 | 1287.43331969685 | Lrp10 |
| ENSMUSG00000022177 | -0.467193034559483 | 0.125840598538726 | 0.771949105062466 | 30.7194663633475 | 43.9784493613737 | 26.5007368821565 | 38.4549944645846 | 36.1336204620302 | 69.71568722821 | 39.5609674411332 | 48.8486504100828 | Haus4 |
| ENSMUSG00000022178 | -0.341382280368624 | 0.108181275136962 | 0.762287160109383 | 109.325159704854 | 87.9568987227475 | 126.824955078892 | 125.682177030594 | 185.047935093427 | 140.654456688494 | 118.6829023234 | 124.835439936878 | Ajuba |
| ENSMUSG00000022179 | -0.217517310506787 | 0.271948019179459 | 0.84423521828958 | 82.2197482077831 | 91.2145616384048 | 59.6266579848522 | 71.2824287636203 | 83.2168228822514 | 94.1773318696872 | 85.3684034256032 | 92.2696729968231 | 4931414P19 |
| ENSMUSG00000022180 | -0.279372602638989 | 0.355011540340037 | 0.864197387893991 | 90.3513716569045 | 88.7713144516618 | 109.78876708322 | 54.3997482669734 | 120.445401540101 | 88.0619207093179 | 141.586620315635 | 66.217059444779 | Slc7a8 |
| ENSMUSG00000022181 | -0.311626966844723 | 0.582167616766872 | 0.922450244597442 | 18.0702743313809 | 49.679359463774 | 14.1968233297267 | 27.1998741334867 | 15.3294147414674 | 26.9078091056249 | 20.8215618111227 | 72.73021283279 | C6 |
| ENSMUSG00000022184 | 0.0380398022350279 | 0.834227051157371 | 0.968512902280839 | 173.474633581257 | 140.893921102179 | 163.736695736181 | 145.378637610015 | 188.332809680885 | 151.662196777159 | 119.723980413956 | 147.631476794917 | Fbxo4 |
| ENSMUSG00000022185 | 0.0986753486531647 | 0.221869217862887 | 0.831327997735572 | 1673.30740308587 | 1765.65330028626 | 1579.63320915426 | 1572.90306627094 | 1525.276766776 | 1498.27573429048 | 1546.00096447586 | 1587.03837554536 | Acin1 |
| ENSMUSG00000022186 | 0.15568356680848 | 0.385665068989049 | 0.875126835008975 | 26720.5146538129 | 26305.6280439328 | 23560.1015431258 | 24764.0785084982 | 20573.169541245 | 16328.147798186 | 26609.9559946149 | 27470.3099395013 | Oxct1 |
| ENSMUSG00000022191 | 0.279616091907474 | 0.0481110912548031 | 0.703396849069134 | 1787.15013137357 | 2074.31686154479 | 2137.09513856819 | 1826.14327372064 | 1478.19356435578 | 1360.06744206613 | 1899.96751526495 | 1706.44618765889 | Drosha |
| ENSMUSG00000022193 | -0.123724371463888 | 0.526124505686388 | 0.90806673740897 | 3684.52893616857 | 2680.24216385706 | 2910.34878259397 | 3566.93521826379 | 3697.67382728109 | 4563.31980786757 | 2839.01995294658 | 2892.92562984157 | Psmb5 |
| ENSMUSG00000022194 | -0.149308001256933 | 0.345356811530931 | 0.861266047293794 | 339.721157429961 | 271.200437728471 | 269.739643264807 | 358.287997206618 | 340.531998899739 | 393.832478727783 | 341.473613702413 | 298.51953028384 | Pabpn1 |
| ENSMUSG00000022195 | 0.100848075311251 | 0.591031726060568 | 0.923935529355527 | 380.379274675568 | 372.187988113848 | 395.618143455051 | 370.481044231974 | 309.873169416804 | 254.401104271363 | 422.677704765791 | 427.697072479392 | 6030458C11 |
| ENSMUSG00000022197 | 0.0100157194569927 | 0.944973268805103 | 0.990505437194639 | 628.845546732055 | 884.455481600961 | 839.505486231172 | 720.327701190268 | 835.453103409971 | 693.487625585878 | 791.219348822664 | 731.644230586574 | Pdzd2 |
| ENSMUSG00000022199 | -1.50035664596406 | 0.0126143479642203 | 0.635862253526407 | 9.93865088225949 | 4.88649437348597 | 13.2503684410783 | 13.1309737196143 | 18.6142893289246 | 67.2695227640623 | 16.6572494488982 | 14.1118323406906 | Slc22a17 |
| ENSMUSG00000022200 | 0.0751862005038598 | 0.641969213094824 | 0.934616627390525 | 1516.99953011943 | 1704.57212061769 | 1637.36695736181 | 1357.17992659156 | 1431.11036193556 | 1121.56640681173 | 1705.28591233095 | 1641.31465377878 | Golph3 |
| ENSMUSG00000022201 | 0.292073308866794 | 0.246928257343933 | 0.833064552804398 | 1612.77198407575 | 1976.58697407508 | 2032.98510081686 | 1450.03466932312 | 1168.32039493898 | 903.857769502582 | 1898.92643717439 | 1804.14348847906 | Zfr |
| ENSMUSG00000022203 | -1.31007085653443 | 0.0091109244354247 | 0.635862253526407 | 9.03513716569045 | 20.3603932228582 | 26.5007368821565 | 37.5170677703264 | 68.9823663366031 | 95.400414101761 | 23.9447960827911 | 43.4210225867403 | Efs |
| ENSMUSG00000022204 | -0.0292417580381197 | 0.86731282757803 | 0.974868041734905 | 546.625798524272 | 403.950201541507 | 463.762895437739 | 572.135283497478 | 577.042969196664 | 573.62556684264 | 415.390158131898 | 461.348364984116 | Ngdn |
| ENSMUSG00000022205 | 0.0178160152408278 | 0.908323690762606 | 0.985057469135899 | 1627.22820354085 | 1303.87958199184 | 1600.45521670452 | 1947.13581727994 | 1679.66587238649 | 1788.14622329198 | 1546.00096447586 | 1385.13062051702 | Sub1 |
| ENSMUSG00000022206 | 0.0259709121687113 | 0.879731805150463 | 0.978393501240173 | 419.230364488037 | 452.815145276367 | 357.759947909113 | 407.060185308042 | 516.820268426614 | 380.37857417497 | 384.157815415214 | 326.743194965221 | Npr3 |
| ENSMUSG00000022210 | 0.17360541332782 | 0.0960251917526694 | 0.757809915356977 | 3474.91375392455 | 3093.96535414553 | 3417.64860290954 | 3586.63167884321 | 2672.79295599442 | 2925.61269912067 | 3247.12256444459 | 3187.10305786674 | Dhrs4 |
| ENSMUSG00000022211 | 0.623867988790225 | 0.0158621838300166 | 0.63777231095375 | 57.8248778604189 | 58.6379324818317 | 56.7872933189068 | 54.3997482669734 | 48.1781606160403 | 24.4616446414772 | 31.2323427166841 | 43.4210225867403 | Carmil3 |
| ENSMUSG00000022214 | 0.125003591745154 | 0.385920950824082 | 0.875126835008975 | 3868.84573434865 | 3838.34133037323 | 4144.52595739155 | 3690.74154190586 | 3292.53929482802 | 2860.78934082076 | 4265.29693700849 | 3831.90524327983 | Dcaf11 |
| ENSMUSG00000022215 | -0.201136428933635 | 0.228517275403908 | 0.832048466795696 | 2379.85512944286 | 1952.96891793656 | 1852.21221708501 | 2246.3344327483 | 2393.57861606055 | 3100.51345830723 | 2079.0329468406 | 2121.11695336226 | Fitm1 |
| ENSMUSG00000022216 | -0.0150607595283888 | 0.901495278968469 | 0.984802943363903 | 2691.56736165919 | 2387.0525014479 | 2525.14164291406 | 2839.10410351945 | 2707.83161826063 | 3039.35934670354 | 2319.52198575907 | 2486.93906865555 | Psme1 |
| ENSMUSG00000022217 | -0.0136431191409284 | 0.940269172253443 | 0.989269716906553 | 904.417230285614 | 730.530908836153 | 705.108892043093 | 936.988767563903 | 892.390929592564 | 1028.61215717412 | 717.302804393178 | 670.854798965138 | Emc9 |
| ENSMUSG00000022219 | -0.0288202224148531 | 0.920805418643314 | 0.986713918026868 | 29.8159526467785 | 43.9784493613737 | 44.483379766477 | 42.2067012416173 | 31.7537876787538 | 37.9155491942896 | 35.3966550789086 | 58.6183804920994 | Cideb |
| ENSMUSG00000022220 | -0.515738070248191 | 0.132673566826799 | 0.77303252008843 | 357.791431761342 | 350.198763433161 | 435.369248778286 | 384.549944645846 | 711.722827282413 | 869.611467004514 | 296.707255808499 | 307.203734801188 | Adcy4 |
| ENSMUSG00000022221 | -0.762889696756333 | 0.0374518465395236 | 0.679566075569698 | 18.9737880479499 | 27.6901347830872 | 26.5007368821565 | 35.6412143818101 | 64.6025335533267 | 59.9310293716191 | 33.3144988977964 | 27.1381391167127 | Ripk3 |
| ENSMUSG00000022223 | 0.00417911184183986 | 0.964833914622036 | 0.994284908685797 | 2387.98675289199 | 2093.04842330982 | 1997.96626993687 | 2113.14884216364 | 2083.70544664374 | 2052.33198541994 | 2279.96101831794 | 2150.42614360831 | Sdr39u1 |
| ENSMUSG00000022228 | -0.0207339054362508 | 0.923851203081763 | 0.986713918026868 | 259.308436655316 | 293.189662409158 | 366.278041906949 | 281.378008277448 | 233.226095709468 | 238.501035254403 | 382.075659234102 | 362.565538599282 | Zscan26 |
| ENSMUSG00000022231 | 0.259823476435367 | 0.537436882161387 | 0.912099403004057 | 58.7283915769879 | 99.3587189275481 | 211.059440168604 | 125.682177030594 | 132.489941694111 | 77.0541806206532 | 147.833088858971 | 55.3618037980939 | Sema5a |
| ENSMUSG00000022234 | 0.14600943352079 | 0.129008161338439 | 0.77303252008843 | 3343.90426502204 | 3032.88417447696 | 3179.14197097013 | 3705.748369014 | 3028.65436963562 | 3132.31359634115 | 2866.08798330104 | 2958.05716372168 | Cct5 |
| ENSMUSG00000022235 | 0.281705236166862 | 0.00611882026861535 | 0.635862253526407 | 1204.38378418654 | 1039.19447009468 | 1254.99918234784 | 1264.32518386 | 969.038003299901 | 1023.71982824582 | 938.011359591079 | 986.742738283673 | Cmbl |
| ENSMUSG00000022237 | 0.292183985037608 | 0.264114607162959 | 0.842510961779757 | 722.810973255236 | 1047.33862738383 | 1068.5475692841 | 848.823658303636 | 610.986673267056 | 474.555906044658 | 852.642956165476 | 1072.49925789249 | Ankrd33b |
| ENSMUSG00000022241 | 0.0937082153874844 | 0.429874013361341 | 0.886017087288711 | 601.740135234984 | 776.138189655355 | 738.234813145789 | 711.886360941944 | 672.304332232926 | 615.210362733151 | 659.002431322034 | 703.420565905193 | Tars |
| ENSMUSG00000022244 | 0.0377445866244599 | 0.760323424467366 | 0.957357521291862 | 679.442314859922 | 716.685841444609 | 663.464876942561 | 643.417712261099 | 665.734583058011 | 541.82542880872 | 708.974179668729 | 715.361347116547 | Amacr |
| ENSMUSG00000022246 | -0.0958851346389323 | 0.577011233901017 | 0.922450244597442 | 206.001127377742 | 179.171460361152 | 147.646962629158 | 152.88205116408 | 199.282391639076 | 179.793088114857 | 198.845915296222 | 155.230155747597 | Rai14 |
| ENSMUSG00000022247 | 0.088946290528219 | 0.441289052642916 | 0.89108031640712 | 546.625798524272 | 458.516055378767 | 458.084166105848 | 471.777127211855 | 440.173194719277 | 447.648096939033 | 433.088485671353 | 498.256234182845 | Brix1 |
| ENSMUSG00000022248 | -0.170539086945387 | 0.363133670262928 | 0.86924083181804 | 68.6670424592474 | 79.8127414336042 | 96.5383986421416 | 84.4134024832345 | 101.831112211176 | 95.400414101761 | 81.2040910633787 | 92.2696729968231 | Rad1 |
| ENSMUSG00000022253 | 0.224144956010827 | 0.426550002703391 | 0.885066977635295 | 1925.38773000863 | 2540.16265848379 | 3024.86982412044 | 2008.10105240672 | 1816.53564686388 | 1103.22017333062 | 2480.88908979527 | 2730.0967951413 | Nadk2 |
| ENSMUSG00000022255 | 0.0118772682133572 | 0.928328733686285 | 0.986901071486195 | 2270.52996973801 | 2023.00867062319 | 2344.3687591822 | 2487.38159317264 | 2407.81307260619 | 2606.3882365494 | 1958.26788833609 | 2078.78145634019 | Mtdh |
| ENSMUSG00000022257 | 0.00792037719521413 | 0.928428793899731 | 0.986901071486195 | 1359.78814343641 | 1292.47776178704 | 1346.80530654674 | 1317.78700543272 | 1358.8431210115 | 1286.6825081417 | 1427.31806215246 | 1213.61758129939 | Laptm4b |
| ENSMUSG00000022261 | -0.298864246864009 | 0.165290062045649 | 0.795216960550936 | 750.819898468876 | 772.880526739698 | 821.522843346852 | 840.382318055312 | 1113.57248514802 | 1339.27504412088 | 720.426038664847 | 746.841588491933 | Sdc2 |
| ENSMUSG00000022263 | 0.0746051945933902 | 0.786420105150256 | 0.961141387070613 | 478.862269781594 | 702.840774053066 | 593.427215182576 | 445.515179772627 | 516.820268426614 | 283.755077841135 | 648.591650416473 | 658.914017753784 | Trio |
| ENSMUSG00000022265 | 0.142891625921166 | 0.405502378760869 | 0.881158207349486 | 5248.51117954958 | 6107.30355112855 | 5681.56869655663 | 4808.75016146159 | 4371.07311770984 | 3825.80122192703 | 5919.57002290219 | 5668.61449869895 | Ank |
| ENSMUSG00000022270 | 0.387275165191025 | 0.0387618003889378 | 0.679566075569698 | 8043.98261861421 | 9495.27298341216 | 12119.3548491434 | 10894.0185538085 | 7960.34608360483 | 5721.57868164152 | 8410.86989360303 | 8911.07936036378 | Fam134b |
| ENSMUSG00000022272 | -0.0122496204805006 | 0.943139539334023 | 0.990505437194639 | 1162.82215322436 | 1279.44711012441 | 1023.11773462897 | 1198.67031526193 | 1387.3120341028 | 881.842289325253 | 1084.80337035949 | 1349.30827688295 | Myo10 |
| ENSMUSG00000022280 | 0.0172126883837283 | 0.863879312704005 | 0.974868041734905 | 500.546598979251 | 509.82424630037 | 571.658752743662 | 505.542488205149 | 500.395895489327 | 507.579126310652 | 505.963952010282 | 548.190410157596 | Rnf19a |
| ENSMUSG00000022283 | -0.00549647316240459 | 0.976044425597577 | 0.99575536949131 | 4624.18320140037 | 4362.82505979406 | 5893.57459161388 | 4492.66886549659 | 4870.37405500334 | 6352.68911339163 | 4064.36886553116 | 4160.81948937439 | Pabpc1 |
| ENSMUSG00000022285 | -0.3139148038622 | 0.0387769960254094 | 0.679566075569698 | 1779.01850792445 | 1851.16695182227 | 1538.93564894237 | 1662.94402891972 | 2544.68284708358 | 2323.85624094033 | 1772.9559882171 | 1852.99213888914 | Ywhaz |
| ENSMUSG00000022286 | -0.017396383564969 | 0.979501860188569 | 0.996872525500296 | 11.7456783153976 | 7.32974156022896 | 15.1432782183751 | 15.0068271081306 | 6.56974917491458 | 1.22308223207386 | 17.6983275394543 | 23.8815624227072 | Grhl2 |
| ENSMUSG00000022292 | 0.192416569401243 | 0.393909447634089 | 0.878942305112115 | 692.091506891888 | 776.95260538427 | 771.360734248484 | 666.865879617553 | 468.642107810573 | 461.102001491845 | 854.725112346588 | 758.782369703287 | Rrm2b |
| ENSMUSG00000022295 | 0.211481648306811 | 0.0565049624418742 | 0.70510438385647 | 1522.42061241884 | 1535.98806473242 | 1621.27722425479 | 1539.13770527764 | 1249.34730142959 | 1186.38976511164 | 1526.2204807553 | 1406.84113181039 | Atp6v1c1 |
| ENSMUSG00000022297 | -0.0249401778472232 | 0.887459065190641 | 0.980702840906172 | 434.590097669711 | 412.09435883065 | 551.783200082044 | 597.459304242449 | 508.060602860061 | 610.318033804856 | 468.485140750262 | 443.97995594942 | Fzd6 |
| ENSMUSG00000022299 | 0.197837571337093 | 0.364818753370499 | 0.86924083181804 | 155.404359249876 | 250.025628776699 | 205.380710836713 | 151.944124469822 | 159.863896589588 | 135.762127760198 | 198.845915296222 | 170.427513652956 | Slc25a32 |
| ENSMUSG00000022300 | -0.0598667656488307 | 0.64942111601598 | 0.936461253686283 | 335.203588847116 | 356.714089264476 | 330.312756138308 | 345.157023487003 | 365.716037403578 | 402.3940543523 | 365.418409785204 | 293.091902460497 | Dcaf13 |
| ENSMUSG00000022304 | -0.167714283722819 | 0.729203151980205 | 0.952797867741259 | 34.3335212296237 | 87.1424829938332 | 36.9117406572894 | 50.6480414899407 | 40.5134532453066 | 42.8078781225851 | 30.191264626128 | 121.578863242873 | Dpys |
| ENSMUSG00000022305 | 0.081123633525743 | 0.707053225994358 | 0.948710970699398 | 102.097049972302 | 118.904696421492 | 120.199770858353 | 100.358156285623 | 84.3117810780705 | 81.9465095489486 | 137.42230795341 | 112.894658725525 | Lrp12 |
| ENSMUSG00000022306 | 0.193918625056202 | 0.262966619251187 | 0.842390761392756 | 209.615182244018 | 242.69588721647 | 214.845259723197 | 226.040333316217 | 168.623562156141 | 163.893019097897 | 249.858741733473 | 197.565652769668 | Zfpm2 |
| ENSMUSG00000022307 | 0.0636680461348691 | 0.749906897714505 | 0.956197192463153 | 1048.97942493666 | 1332.38413250384 | 1269.19600567757 | 972.629981945713 | 921.95480087968 | 845.149822363037 | 1308.63515982906 | 1347.13722575362 | Oxr1 |

| | | | | | | | | | | | | |
|---|---|---|---|---|---|---|---|---|---|---|---|---|
| **ENSMUSG00000022309** | 0.0208171905080071 | 0.917683426889573 | 0.986713918026868 | 235.817080024521 | 283.416673662186 | 257.435729712378 | 196.026679099956 | 197.092475247437 | 204.254732756335 | 243.612273190136 | 313.716888189199 | Angpt1 |
| **ENSMUSG00000022312** | 0.11576498145446 | 0.419564324211623 | 0.882568531013176 | 8380.08972117789 | 6131.73602299598 | 6467.12625413484 | 8142.1416328551 | 6463.53822992013 | 7592.89449671452 | 6367.23360184133 | 6452.36395638961 | Eif3h |
| **ENSMUSG00000022313** | -0.125490258450839 | 0.423521368309662 | 0.884482395183227 | 95.7724539563188 | 110.760539132349 | 117.360406192407 | 119.116690170786 | 127.015150715015 | 123.53130543946 | 118.6829023234 | 113.980184290193 | Utp23 |
| **ENSMUSG00000022314** | 0.0183261580948432 | 0.849357108474118 | 0.972640876584422 | 1741.07093182855 | 1849.53812036444 | 1815.30047642772 | 1646.06134842307 | 1867.99868206738 | 1597.34539508846 | 1659.47847634648 | 1837.79478098378 | Rad21 |
| **ENSMUSG00000022323** | -0.068316309974516 | 0.70992033749308 | 0.949571160524938 | 336.107102563685 | 377.074482487334 | 334.098575692902 | 341.405316709971 | 268.264757975679 | 385.270903103266 | 335.227145159076 | 467.861518372127 | Rida |
| **ENSMUSG00000022324** | -1.32686567644564 | 0.00482539808223697 | 0.635862253526407 | 133.720030052219 | 146.594831204579 | 90.8596693102509 | 126.620103724852 | 441.268152915096 | 561.394744521902 | 137.42230795341 | 109.638082031519 | Matn2 |
| **ENSMUSG00000022325** | 0.110169276847413 | 0.652560897862465 | 0.936854870419601 | 83.1232619243521 | 54.56585383726 | 53.9479286529615 | 50.6480414899407 | 51.4630352034976 | 66.0464405319884 | 52.0539045278068 | 55.3618037980939 | Pop1 |
| **ENSMUSG00000022329** | -0.101673696137968 | 0.551822441150981 | 0.914873920695527 | 119.263810587114 | 146.594831204579 | 131.557229522134 | 154.757904552597 | 152.199189218854 | 174.900759186562 | 140.545542225078 | 125.920965501547 | Stk3 |
| **ENSMUSG00000022332** | 0.0506249101368323 | 0.672229097276552 | 0.943216235131458 | 1524.22763985198 | 1249.31372815458 | 1327.87620877377 | 1331.85590584659 | 1132.18677447695 | 1300.13641269451 | 1392.96248516411 | 1419.86743858641 | Khdrbs3 |
| **ENSMUSG00000022335** | -0.202794762313777 | 0.42378678598739 | 0.884584010063004 | 53.3073092775737 | 72.4829998733752 | 52.0550188756646 | 36.5791410760683 | 62.4126171616885 | 69.71568722821 | 59.3414511616998 | 56.4473293627624 | Zfat |
| **ENSMUSG00000022336** | 0.114030090387914 | 0.333121717862746 | 0.856918526232803 | 3784.81895870773 | 3075.2337923805 | 3373.16522314306 | 4022.76759167325 | 3279.3997964782 | 3561.61545979908 | 3108.65917840062 | 3222.9254015008 | Eif3e |
| **ENSMUSG00000022337** | 0.170837709949038 | 0.111659387194089 | 0.764116514278315 | 2686.14627935977 | 2393.56782727921 | 2511.89127447298 | 2649.64291127931 | 2263.27859075807 | 1960.6008180144 | 2429.87626335802 | 2441.34699493947 | Emc2 |
| **ENSMUSG00000022338** | 0.065048519895602 | 0.591356639041805 | 0.923935529355527 | 801.416666596743 | 639.316347197748 | 627.49959117392 | 690.314046974007 | 660.259792078915 | 706.941530138691 | 656.920275140922 | 613.321944037707 | Eny2 |
| **ENSMUSG00000022339** | 0.0259152214277645 | 0.886043910638238 | 0.980068672363141 | 458.984968017075 | 499.236841824483 | 385.207139679918 | 432.384206053012 | 420.463947194533 | 330.232202659942 | 438.293876124134 | 554.703563545607 | Ebag9 |
| **ENSMUSG00000022340** | -0.228561794233894 | 0.357742056820116 | 0.865834559821921 | 60.535419010126 | 45.6072808192024 | 53.9479286529615 | 60.9652351267805 | 77.7420319031559 | 78.277262852727 | 45.80743598447 | 57.5328549274309 | Sybu |
| **ENSMUSG00000022346** | 0.318358546013171 | 0.268074137856212 | 0.843793744782845 | 54.2108229941427 | 79.8127414336042 | 96.5383986421416 | 128.495957113368 | 64.6025335533267 | 75.8310983885793 | 86.4094815161593 | 60.7894316214364 | Myc |
| **ENSMUSG00000022350** | 0.112324361070061 | 0.497029348503859 | 0.900927059989498 | 663.179067961679 | 771.251695281869 | 660.625512276616 | 672.493439783102 | 620.841297029428 | 475.778988276731 | 756.863771834311 | 705.59161703453 | Washc5 |
| **ENSMUSG00000022351** | -0.486149876982237 | 0.179244306361313 | 0.801755154770274 | 31.6229800799166 | 23.6180561385155 | 18.9290977729689 | 27.1998741334867 | 33.943704070392 | 45.2540425867328 | 17.6983275394543 | 45.5920737160773 | Sqle |
| **ENSMUSG00000022353** | -0.217493095271048 | 0.17875867855355 | 0.801755154770274 | 177.992202164102 | 175.099381716581 | 184.558703286447 | 214.785212985119 | 232.131137513649 | 210.370143916704 | 178.024353485099 | 254.012982132431 | Mtss1 |
| **ENSMUSG00000022354** | 0.130775125752703 | 0.400380254853723 | 0.880487015130533 | 18838.2609904646 | 13526.6308415381 | 14833.7874697871 | 18544.6865988724 | 14790.6953091244 | 17475.3989318713 | 13149.8573618146 | 14630.7135606021 | Ndufb9 |
| **ENSMUSG00000022357** | -0.177827786314364 | 0.346781329959107 | 0.861266047293794 | 366.826568927032 | 397.434875710192 | 386.153594568566 | 298.260688774095 | 482.876564356222 | 332.67836712409 | 488.265624470828 | 334.3418739179 | Klhl38 |
| **ENSMUSG00000022358** | 0.240515696289442 | 0.236300820455153 | 0.832048466795696 | 7568.73440369889 | 8231.29977213712 | 8479.28934740143 | 7382.42101050599 | 6414.26511110827 | 4383.52671975271 | 8697.16636850596 | 7303.41599908972 | Fbxo32 |
| **ENSMUSG00000022359** | 0.330549538586392 | 0.064177938767177 | 0.716991331547963 | 230.395997725106 | 210.933673788811 | 234.720812384815 | 271.998741334867 | 195.997517051618 | 138.208292224346 | 198.845915296222 | 219.276164063039 | Wdyhv1 |
| **ENSMUSG00000022360** | 0.0618553397316373 | 0.786149585131128 | 0.961141387070613 | 46.9827132615903 | 52.9370223794314 | 54.8943835416099 | 39.3929211588428 | 50.3680770076785 | 48.9232892829544 | 43.7252798033577 | 43.4210225867403 | Atad2 |
| **ENSMUSG00000022361** | 0.0555299936199075 | 0.770807818100891 | 0.959021909120511 | 814.969372345279 | 871.424829938332 | 772.307189137133 | 689.376120279749 | 635.075753575076 | 569.956320146419 | 924.477344413849 | 898.815167545524 | Zhx1 |
| **ENSMUSG00000022362** | 0.121008265146989 | 0.782286089570324 | 0.960742105865155 | 14.4562194651047 | 12.2162359337149 | 10.4110037751329 | 25.3240207449704 | 15.3294147414674 | 11.0077400886647 | 18.7394056300105 | 11.9407812113536 | Gm29394 |
| **ENSMUSG00000022364** | 0.189478236405895 | 0.505993957235326 | 0.901494649536949 | 63.2459601598332 | 106.688460487777 | 68.1447519826882 | 63.779015209555 | 45.9882442244021 | 56.2617826753975 | 82.2451691539348 | 80.3288917854696 | Tbc1d31 |
| **ENSMUSG00000022365** | -0.0484590357099163 | 0.58777739547356 | 0.923241888853076 | 1496.21871463834 | 1483.86545808191 | 1558.81120160399 | 1494.11722395325 | 1653.38687568684 | 1662.16875338838 | 1493.94705994806 | 1430.72269423309 | Derl1 |
| **ENSMUSG00000022369** | -0.540204014551873 | 0.0806194442604152 | 0.739855948845396 | 18.9737880479499 | 16.2883145782866 | 21.7684624389143 | 22.5102406621959 | 25.1840385038392 | 33.0232202659942 | 33.3144988977964 | 23.8815624227072 | Mtbp |
| **ENSMUSG00000022370** | -0.00750706506687346 | 0.956763833132052 | 0.992403427270738 | 2124.16074765382 | 1654.89276115392 | 1693.20779579207 | 2143.1624963799 | 1859.23901650083 | 2163.63246853866 | 1738.60041122875 | 1894.24211034655 | Mrpl13 |
| **ENSMUSG00000022371** | -1.37132512500704 | 0.0208559758925896 | 0.648762042745232 | 37.9475760958999 | 57.009101024003 | 38.8046504345863 | 42.2067012416173 | 121.54035973592 | 267.855008824175 | 38.5198893505771 | 28.2236646813812 | Col14a1 |
| **ENSMUSG00000022372** | 0.424642793552111 | 0.403835825756788 | 0.881158207349486 | 14.4562194651047 | 34.2054606144018 | 52.0550188756646 | 14.0689004138724 | 17.5193311331055 | 34.2463024980681 | 14.5750932677859 | 19.5394601640331 | Sla |
| **ENSMUSG00000022376** | -0.243161028734687 | 0.616805268349729 | 0.929020296831244 | 11.7456783153976 | 7.32974156022896 | 19.8755526616174 | 13.1309737196143 | 13.1394983498292 | 7.33849339244316 | 17.6983275394543 | 22.7960368580387 | Adcy8 |
| **ENSMUSG00000022377** | -0.0722604757943324 | 0.628119965181786 | 0.931825800800958 | 494.222002963268 | 550.545032746086 | 550.836745193396 | 423.004939110431 | 621.936255225247 | 506.356044078578 | 536.15521663641 | 458.09178829011 | Asap1 |
| **ENSMUSG00000022378** | 0.211493986656145 | 0.378472667146556 | 0.87422797717331 | 387.60738440812 | 360.786167909048 | 429.690519446395 | 421.129085721914 | 285.784089108784 | 214.039390612925 | 442.458188486358 | 437.466802561409 | Fam49b |
| **ENSMUSG00000022383** | 0.0447592504298219 | 0.920735242939519 | 0.986713918026868 | 380.379274675568 | 849.435605257645 | 658.732602499319 | 226.978260010475 | 391.995034103237 | 223.824048469516 | 844.314331441027 | 590.525907179668 | Ppara |
| **ENSMUSG00000022386** | 0.124739400311011 | 0.40470660705967 | 0.881158207349486 | 248.466272056487 | 211.748089517725 | 254.596365046432 | 227.916186704733 | 186.142893289246 | 196.916239363891 | 244.653351280692 | 235.559047533066 | Trmu |
| **ENSMUSG00000022387** | 0.12858360702751 | 0.368386027450688 | 0.871495599696183 | 767.083145367119 | 827.446380576958 | 868.845587779274 | 704.382947387879 | 685.443830582755 | 595.64104701997 | 815.164144905455 | 800.03234116069 | Brd1 |
| **ENSMUSG00000022389** | 0.166678250228333 | 0.385408038636948 | 0.875126835008975 | 2290.40727150253 | 2555.63655733316 | 2639.66268444052 | 2233.20345902868 | 1892.0877623754 | 1539.86053018099 | 2604.77738257145 | 2620.45871310978 | Tef |
| **ENSMUSG00000022390** | 0.297162642841276 | 0.200893662712096 | 0.812921911423739 | 2496.40839888027 | 3267.43590440429 | 3110.99721898744 | 2413.28538432625 | 1852.66926732591 | 1558.2067636621 | 2947.29207436442 | 2827.79409596146 | Zc3h7b |
| **ENSMUSG00000022391** | 0.00363747645827773 | 0.966544130001655 | 0.994284908685797 | 766.17963165055 | 838.848200781758 | 789.343377132805 | 784.106716399823 | 834.358145214152 | 752.195572725424 | 784.972880279327 | 798.946815596022 | Rangap1 |
| **ENSMUSG00000022394** | 0.0560445535072848 | 0.750042133984968 | 0.956197192463153 | 271.054114970713 | 261.427448981499 | 293.401015481019 | 212.909359596603 | 224.466430142915 | 203.031650524261 | 287.337552993494 | 283.322172378481 | L3mbtl2 |
| **ENSMUSG00000022400** | -0.089302662663032 | 0.589904029181759 | 0.923810046070101 | 1730.22876722972 | 1251.75697534132 | 1160.353693483 | 1562.5858726341 | 1408.11623982336 | 1752.67683856184 | 1375.26415762466 | 1533.8476228766 | Rbx1 |
| **ENSMUSG00000022401** | 0.0466011253721873 | 0.860140144115755 | 0.974868041734905 | 177.992202164102 | 208.490426602068 | 193.076797284283 | 167.888878272211 | 159.863896589588 | 100.292743030056 | 223.831789469569 | 238.815624227072 | Xpnpep3 |
| **ENSMUSG00000022403** | 0.179786108704348 | 0.0919106175185675 | 0.754058637722175 | 3492.98402825593 | 3435.20554456064 | 3146.96250475609 | 3424.37036073655 | 2853.46105830457 | 2613.72672994184 | 3151.34338011343 | 3297.82666546293 | St13 |
| **ENSMUSG00000022404** | 0.1504569146237 | 0.372187611633201 | 0.873700364743909 | 239.431134890797 | 250.840044505613 | 302.865564367503 | 209.15765281957 | 197.092475247437 | 225.04713070159 | 264.433835001259 | 216.019587369033 | Slc25a17 |
| **ENSMUSG00000022407** | 0.208256382275502 | 0.0147817108688967 | 0.635862253526407 | 1535.97331816738 | 1509.11234567825 | 1524.73882561265 | 1508.18612436712 | 1359.93807920732 | 1212.07449198519 | 1415.86620315635 | 1271.15043622682 | Adsl |
| **ENSMUSG00000022412** | -0.0487902263203882 | 0.772053375679471 | 0.959045638244974 | 511.388763578079 | 566.018931595458 | 598.159489625818 | 540.245775892701 | 557.33372167192 | 423.186452297555 | 614.236073428121 | 695.821886952513 | Mief1 |
| **ENSMUSG00000022414** | -0.067641512918765 | 0.712926286989547 | 0.950522761209907 | 301.773581334061 | 302.96265115613 | 334.098575692902 | 340.467390015713 | 238.700886688563 | 315.555215875056 | 388.322127777439 | 397.302356668674 | Tab1 |
| **ENSMUSG00000022415** | 0.104975404579259 | 0.379994102014483 | 0.87432013076003 | 1999.4758547673 | 1624.75937918409 | 1764.1919124407 | 2007.16312571246 | 1794.6364829475 | 1888.43896632204 | 1620.9585869959 | 1572.92654320467 | Syngr1 |
| **ENSMUSG00000022419** | 0.233444331757665 | 0.0297560987254787 | 0.665956123973759 | 645.108793630298 | 794.05533569147 | 699.430162711202 | 676.245146560134 | 616.461464246152 | 601.756458180339 | 568.428637443651 | 608.979841779033 | Deptor |
| **ENSMUSG00000022420** | -0.129700137925524 | 0.428659961662711 | 0.885378691567738 | 470.730646332472 | 369.744740927105 | 393.725233677754 | 462.397860269274 | 451.122776677468 | 579.740978003009 | 427.883095218572 | 398.387882233342 | Dnal4 |
| **ENSMUSG00000022421** | -0.703199063380192 | 0.0993163093892185 | 0.759800953500741 | 51.5002818444356 | 59.452348210746 | 58.6802030962037 | 59.0893817382642 | 132.489941694111 | 151.662196777159 | 54.1360607089191 | 34.7368180693922 | Nptxr |
| **ENSMUSG00000022425** | -0.76026347987353 | 0.00676097066929877 | 0.635862253526407 | 119.263810587114 | 146.594831204579 | 125.878500190243 | 113.489130005238 | 259.505092409126 | 305.770558018465 | 129.093683228961 | 162.828834700276 | Enpp2 |
| **ENSMUSG00000022426** | -0.0346474536103357 | 0.847052368523987 | 0.972334438148346 | 668.600150261093 | 837.21936932393 | 717.412805595523 | 664.052099534778 | 682.158955995297 | 550.387004433237 | 830.780316263797 | 893.387539722182 | Josd1 |
| **ENSMUSG00000022427** | 0.068138192922758 | 0.625407551990285 | 0.931573470007391 | 1698.6057871498 | 1274.56061575092 | 1176.44342659002 | 1465.04149643125 | 1261.3918415836 | 1340.49812635295 | 1357.5658300852 | 1395.9858761637 | Tomm22 |
| **ENSMUSG00000022428** | -0.220778560560547 | 0.273706808605977 | 0.845204865395511 | 310.808718499751 | 298.890572511559 | 191.183887506986 | 238.233380341573 | 287.974005500422 | 375.486245246675 | 281.091084450157 | 268.124814473121 | Cby1 |
| **ENSMUSG00000022433** | -0.082387737554897 | 0.405761079481247 | 0.881158207349486 | 1034.52320547156 | 1061.18369477537 | 1062.86883995221 | 1193.04275509638 | 1212.11872277174 | 1220.63606760971 | 1069.18719900115 | 1106.15055039721 | Csnk1e |
| **ENSMUSG00000022434** | -0.0128148297310746 | 0.947366927643946 | 0.990957649293953 | 101.193536255733 | 106.688460487777 | 83.2880302010633 | 115.364983393754 | 86.5016974697087 | 108.854318654574 | 92.6559500594962 | 122.664388807541 | Fam118a |
| **ENSMUSG00000022436** | -0.489872529115488 | 0.0532973167387121 | 0.703396849069134 | 57.8248778604189 | 30.9477976987445 | 54.8943835416099 | 45.9584080186499 | 72.2672409240604 | 73.3849339244316 | 59.3414511616998 | 60.7894316214364 | Sh3bp1 |
| **ENSMUSG00000022437** | 0.0589768841887587 | 0.400189118299357 | 0.880487015130533 | 11137.6135841466 | 10267.3390944229 | 10809.4612832539 | 11574.0154071457 | 10522.5482618215 | 10703.1926128783 | 10514.888714617 | 10292.9534041868 | Samm50 |
| **ENSMUSG00000022438** | -0.0143207853216532 | 0.874670486566266 | 0.976778666860061 | 2209.09103701132 | 2329.22898469498 | 2090.71884902442 | 2033.42507315169 | 2034.43232783188 | 2202.77109996502 | 2336.17923520797 | 2176.47875716036 | Parvb |
| **ENSMUSG00000022440** | -0.30440105015472 | 0.292667676591194 | 0.851785577030345 | 106.614618555147 | 116.461449234749 | 83.2880302010633 | 103.171936368398 | 128.110108910834 | 200.585486060113 | 90.5737938783839 | 87.9275707381491 | C1qtnf6 |
| **ENSMUSG00000022442** | 0.0206843882499071 | 0.862652608361527 | 0.974868041734905 | 1378.76193148436 | 1291.66334605813 | 1211.46225747001 | 1406.89004138724 | 1410.306156215 | 1451.79860947167 | 1183.70578896233 | 1169.11103314798 | Ttll1 |
| **ENSMUSG00000022443** | -0.295427742116176 | 0.109692252356702 | 0.763154315676794 | 2411.47810952278 | 2554.00772587533 | 2247.83036054006 | 2289.47906068417 | 3819.21418701701 | 3176.34455669581 | 2234.15358233347 | 2433.74831598679 | Myh9 |
| **ENSMUSG00000022445** | -0.309953134150768 | 0.465738905937508 | 0.895328112556867 | 86.7373167906283 | 219.892246806869 | 92.7525790875478 | 132.247663890401 | 95.2613630362614 | 129.646716599829 | 119.723980413956 | 314.802413753867 | Cyp2d26 |
| **ENSMUSG00000022450** | 0.0608653997509449 | 0.735156584940907 | 0.954383574136061 | 6537.82525309361 | 5060.77933947364 | 5868.02030962037 | 6656.46574915017 | 6067.16336303362 | 7440.00921770529 | 4897.23133797607 | 4723.12173187268 | Ndufa6 |
| **ENSMUSG00000022451** | -0.225825669137663 | 0.168900363325169 | 0.79907358295209 | 267.440060104437 | 265.499527626071 | 321.794662140472 | 353.598363735327 | 410.609323432161 | 352.247682837272 | 292.542943446274 | 357.137910775939 | Twf1 |
| **ENSMUSG00000022452** | -0.096612658351763 | 0.664412918663529 | 0.941415427954342 | 3934.80223565819 | 2774.71438841112 | 3075.97838810745 | 4205.66329705359 | 3633.07129372776 | 5233.56887104405 | 3019.1264626128 | 3075.29392470588 | Smdt1 |
| **ENSMUSG00000022453** | -0.151141530598895 | 0.526142627981813 | 0.90806673740897 | 305.387636200337 | 330.652785939217 | 341.670214802089 | 272.936668029125 | 454.407651264925 | 434.19419238622 | 262.351678820146 | 238.815624227072 | Naga |
| **ENSMUSG00000022462** | 0.516541149403225 | 0.00124941965789062 | 0.596732488826387 | 3863.42465204924 | 5044.49102489535 | 5679.67578677933 | 5411.83702586959 | 3262.97542354091 | 3140.87517196567 | 3432.43446456358 | 4143.45108033969 | Slc38a2 |
| **ENSMUSG00000022463** | -0.0848144890198347 | 0.4885338457304 | 0.899236955296196 | 371.344137509877 | 456.072808192024 | 460.923530771794 | 413.625672167849 | 475.211856985488 | 480.671317205027 | 425.80093903746 | 424.440495785386 | Srebf2 |
| **ENSMUSG00000022464** | -0.00957211226883887 | 0.97060891055691 | 0.9951105526746 | 262.922491521592 | 450.371898089624 | 301.919109478855 | 341.405316709971 | 220.086597359638 | 352.247682837272 | 321.693129981846 | 472.203620630801 | Slc38a4 |
| **ENSMUSG00000022466** | 0.331155693113385 | 0.0376772778941914 | 0.679566075569698 | 554.757421973394 | 562.761268679801 | 643.589324280944 | 681.872706725683 | 478.496731572945 | 395.055560959857 | 598.619902069779 | 467.861518372127 | Rpap3 |
| **ENSMUSG00000022468** | -0.179511244121863 | 0.521704576672348 | 0.906152189339124 | 56.9213641438498 | 34.2054606144018 | 38.8046504345863 | 63.779015209555 | 53.6529515951357 | 59.9310293716191 | 42.6842017128016 | 62.9604827507734 | Endou |
| **ENSMUSG00000022469** | -0.237330298761719 | 0.313870275655554 | 0.853228028313558 | 421.037391921175 | 362.414999366876 | 420.22597055991 | 417.377378944882 | 638.360628162534 | 585.856389163379 | 325.857442344071 | 361.480013034613 | Rapgef3 |
| **ENSMUSG00000022471** | 0.157237218205375 | 0.116684777664141 | 0.769868040577702 | 652.336903362851 | 681.665965101293 | 624.660226507975 | 673.43136647736 | 620.841297029428 | 528.371524255907 | 620.482541971457 | 589.440381615 | Xrcc6 |
| **ENSMUSG00000022472** | 0.0584536595163661 | 0.589069161274332 | 0.92352104528132 | 554.757421973394 | 513.896324944941 | 532.854102309076 | 571.19735680322 | 515.725310230795 | 464.771248188067 | 531.990904274186 | 572.071972580304 | Desi1 |
| **ENSMUSG00000022474** | 0.00040013488935572 | 0.998690141450269 | 0.999409658843245 | 691.187993175319 | 600.22439220986 | 803.540200462531 | 860.078778634734 | 826.693437843418 | 1023.71982824582 | 625.687932424238 | 478.716774018812 | Pmm1 |
| **ENSMUSG00000022475** | -0.378505540037169 | 0.0916276915867223 | 0.753823363185192 | 527.652010476322 | 382.775392589734 | 498.781726317731 | 557.128456389348 | 805.889232122855 | 775.434135134827 | 501.799639648058 | 473.289146195469 | Hdac7 |
| **ENSMUSG00000022476** | 0.0680974666899502 | 0.650519709150948 | 0.936463929813731 | 221.360860559416 | 193.83094348161 | 247.024725937245 | 244.79886720138 | 216.801722772181 | 239.724117486476 | 223.831789469569 | 184.539345993646 | Polr3h |
| **ENSMUSG00000022477** | 0.216536753779775 | 0.0629375501650035 | 0.715615581523459 | 72948.7939620681 | 74961.2669364615 | 84877.1279591041 | 74827.7916679161 | 66685.1440417746 | 55741.9727267662 | 70899.500123054 | 71413.4703228471 | Aco2 |
| **ENSMUSG00000022488** | 0.0585748711621517 | 0.844035561499104 | 0.971721411930626 | 43.3686583953142 | 78.9983257046899 | 80.448665535118 | 74.0962088463947 | 89.7865720571659 | 69.71568722821 | 42.6842017128016 | 64.046008315442 | Nckap1l |
| **ENSMUSG00000022489** | -0.908759202480743 | 0.0137503591591018 | 0.635862253526407 | 16.2632468982428 | 15.4738988493722 | 15.1432782183751 | 24.3860940507122 | 45.9882442244021 | 39.1386314263635 | 29.1501865355718 | 19.5394601640331 | Pde1b |
| **ENSMUSG00000022490** | 0.10299096784266 | 0.745726545399351 | 0.956085046483146 | 38.8510898124689 | 35.0198763433161 | 32.1794662140472 | 61.9031618210386 | 42.7033696369448 | 51.3694537471021 | 34.3555769883525 | 28.2236646813812 | Ppp1r1a |
| **ENSMUSG00000022498** | -0.0289307733053412 | 0.810425463682008 | 0.964412705343932 | 213.229237110295 | 227.221988367098 | 212.005895057252 | 217.598993067893 | 218.991639163819 | 239.724117486476 | 224.872867560126 | 205.164331722348 | Txndc11 |
| **ENSMUSG00000022500** | -0.621413750733057 | 0.0535367153694033 | 0.703396849069134 | 315.326287082597 | 258.169786065842 | 286.775831260479 | 287.943495137256 | 646.025335533267 | 606.648787108634 | 259.228444548478 | 255.098507697099 | Litaf |

| | | | | | | | | | | | | |
|---|---|---|---|---|---|---|---|---|---|---|---|---|
| ENSMUSG00000022503 | -0.121796099060074 | 0.394002185220923 | 0.878942305112115 | 146.369222084185 | 158.811067138294 | 151.432782183751 | 166.013024883695 | 150.009272827216 | 168.785348026193 | 167.613572579538 | 191.052499381657 | Nubp1 |
| ENSMUSG00000022504 | -0.561373269763384 | 0.259594604439454 | 0.84030973098672 | 11.7456783153976 | 13.8450673915436 | 9.46454888648447 | 15.0068271081306 | 12.0445401540101 | 13.4539045528125 | 11.4518589961175 | 36.9078691987293 | Ciita |
| ENSMUSG00000022505 | -0.122258652638856 | 0.245926363799015 | 0.832048466795696 | 925.198045766702 | 872.239245667246 | 849.916490006305 | 974.50583533423 | 1107.00273597311 | 954.004141017611 | 925.518422504405 | 955.262496908287 | Emp2 |
| ENSMUSG00000022507 | 0.115183526466277 | 0.408554141070261 | 0.882113132676277 | 1127.58511827817 | 1388.57881779893 | 1221.87326124514 | 1104.87764583611 | 1043.4951606156 | 952.781058785537 | 1206.60950695456 | 1267.89385953282 | 1810013L24F |
| ENSMUSG00000022508 | 0.0972654911030594 | 0.75707582486338 | 0.956676055981837 | 270.150601254144 | 399.878122896935 | 158.057966404291 | 263.557401086543 | 168.623562156141 | 362.032340693862 | 219.667477107345 | 271.381391167127 | Bcl6 |
| ENSMUSG00000022514 | -0.327392930268334 | 0.169273392359718 | 0.79907358295209 | 80.412720774645 | 99.3587189275481 | 64.3589324280944 | 61.9031618210386 | 84.3117810780705 | 125.977469903608 | 85.3684034256032 | 90.0986218674861 | Il1rap |
| ENSMUSG00000022515 | -0.165850656316876 | 0.314191236804104 | 0.853228028313558 | 122.87786545339 | 123.791190794978 | 101.270673085384 | 122.868396947819 | 119.350443344282 | 134.539045528125 | 121.806136595068 | 153.05910461826 | Anks3 |
| ENSMUSG00000022516 | -0.378915270872808 | 0.101254798262883 | 0.759800953500741 | 230.395997725106 | 132.749763813036 | 160.897331070236 | 197.902532488472 | 234.321053905287 | 308.216722482613 | 192.599446752885 | 204.078806157679 | Nudt16l1 |
| ENSMUSG00000022517 | 0.168027487039812 | 0.348060335941284 | 0.861266047293794 | 4830.18432877811 | 6003.05833782752 | 6046.90028357493 | 4590.21324169944 | 4285.66637843595 | 3716.94690327246 | 5689.49176488929 | 5416.77256769585 | Mgrn1 |
| ENSMUSG00000022519 | 0.0530837538766557 | 0.823605028470711 | 0.966637987288264 | 21067.2293292404 | 24354.2879574541 | 20580.6615536605 | 19229.3730856808 | 15646.9526182549 | 13590.8897628047 | 26547.4913091815 | 26366.3304402334 | Srl |
| ENSMUSG00000022521 | -0.284927918342918 | 0.0787241737149162 | 0.739855948845396 | 489.704434380422 | 463.402549752253 | 417.386605893965 | 521.487242007538 | 723.767367436423 | 580.964060235083 | 485.14239019916 | 515.624643217541 | Crebbp |
| ENSMUSG00000022523 | -3.40656863046284 | NA | NA | 4.51756858284523 | 1.62883145782866 | 1.89290977729689 | 3.75170677703264 | 1.0949581958191 | 118.638976511164 | 1.04107809055614 | 4.34210225867403 | Fgf12 |
| ENSMUSG00000022525 | 0.11824051338224 | 0.552932796912737 | 0.914977712327965 | 671.3106914108 | 636.873100011005 | 540.425741418263 | 647.169419038131 | 470.832024202212 | 425.632616761703 | 701.686633034836 | 700.163989211187 | Hrasls |
| ENSMUSG00000022526 | 0.052169114854765 | 0.830916482996388 | 0.967583158900775 | 260.211950371885 | 354.270842077733 | 388.992959234512 | 307.639955716677 | 241.98576127602 | 223.824048469516 | 381.034581143546 | 416.841816832707 | Zfp251 |
| ENSMUSG00000022528 | -0.412561172368109 | 0.168466868969137 | 0.79907358295209 | 386.703870691551 | 240.252640029727 | 406.975602118832 | 529.928582255861 | 651.500126512363 | 682.479885497214 | 417.472314313011 | 329.999771659226 | Hes1 |
| ENSMUSG00000022529 | 0.0791275759713639 | 0.61659547217566 | 0.928896975098882 | 132.81651633565 | 147.409246933493 | 168.468970179424 | 135.999370667433 | 136.869774477387 | 143.100621152642 | 121.806136595068 | 151.973579053591 | Zfp263 |
| ENSMUSG00000022533 | 0.228373997968479 | 0.297156690849344 | 0.851785577030345 | 880.925873654819 | 1114.93513288372 | 1165.08596792624 | 926.671573927063 | 675.589206820383 | 645.787418534998 | 1037.95485628447 | 1128.94658725525 | Atp13a3 |
| ENSMUSG00000022536 | 0.155508203923563 | 0.360656562022939 | 0.868539734006824 | 991.154547076242 | 1215.92268326909 | 1269.19600567757 | 1005.45741624475 | 970.13296149572 | 766.87255951031 | 1131.65188443452 | 1153.91367524262 | Glyr1 |
| ENSMUSG00000022537 | -0.265445520214311 | 0.376694637223776 | 0.87422797717331 | 129.202461469373 | 83.0704043492615 | 126.824955078892 | 151.944124469822 | 178.478185918513 | 204.254732756335 | 118.6829023234 | 89.0130963028176 | Tmem44 |
| ENSMUSG00000022538 | -0.0180759209413966 | 0.856538216095012 | 0.97363563840968 | 437.300638819418 | 424.310594764365 | 385.207139679918 | 424.880792498947 | 437.983278327639 | 419.517205601334 | 408.102611498006 | 427.697072479392 | Lsg1 |
| ENSMUSG00000022540 | 0.128585095409135 | 0.424506798468095 | 0.884903615282587 | 213.229237110295 | 179.985876090067 | 202.541346170768 | 226.040333316217 | 174.098353135236 | 205.477814988408 | 215.50316474512 | 156.315681312265 | Rogdi |
| ENSMUSG00000022543 | -0.0803284055585371 | 0.851400739045261 | 0.972692784625411 | 15.3597331816738 | 7.32974156022896 | 16.0897331070236 | 17.8206071909051 | 12.0445401540101 | 12.2308223207386 | 19.7804837205666 | 15.1973579053591 | 4930451G09 |
| ENSMUSG00000022544 | 0.148257458729433 | 0.314341745210362 | 0.853228028313558 | 269.247087537575 | 230.479651282755 | 224.309808609682 | 282.315934971707 | 217.896680968 | 245.839528646846 | 245.694429371248 | 198.651178334337 | Eef2kmt |
| ENSMUSG00000022545 | 0.0693096110800003 | 0.63557288997851 | 0.932560352417921 | 286.413848152387 | 265.499527626071 | 299.079744812909 | 229.79204009325 | 266.074841584041 | 222.600966237442 | 287.337552993494 | 252.927456567762 | Ercc4 |
| ENSMUSG00000022546 | -0.030457261779301 | 0.747947752277591 | 0.956197192463153 | 784.249905981931 | 750.891302059011 | 683.340429604179 | 696.879533833814 | 765.375778877549 | 705.718447906617 | 741.247600475969 | 765.295523091298 | Gpt |
| ENSMUSG00000022548 | -1.00804736334277 | 0.0677510849365464 | 0.722112580225498 | 80.412720774645 | 96.9154717408051 | 100.324218196735 | 95.6685228143324 | 194.902558855799 | 437.863439082442 | 65.5879197050366 | 53.1907526687569 | Apod |
| ENSMUSG00000022550 | 0.0673367511240602 | 0.768219277003032 | 0.95879680569409 | 86.7373167906283 | 79.8127414336042 | 86.1273948670087 | 104.109863062656 | 118.255485148462 | 79.5003450848009 | 67.6700758861489 | 74.901263962127 | Adck5 |
| ENSMUSG00000022551 | 0.034266610048297 | 0.743086060140836 | 0.954383574136061 | 24919.8118166908 | 20786.3326490804 | 21386.0946639003 | 24298.8668681462 | 23665.3314862381 | 23496.6327603709 | 20935.0393229934 | 21149.2945764365 | Cyc1 |
| ENSMUSG00000022552 | -0.0063358729560606 | 0.964558241045039 | 0.994284908685797 | 421.940905637744 | 430.82592059568 | 424.958245003153 | 476.466760683146 | 474.116898789669 | 496.571386221987 | 443.499266576914 | 348.453706258591 | Sharpin |
| ENSMUSG00000022553 | 0.0794144679201332 | 0.508950995919798 | 0.902394714365738 | 2934.61255141626 | 2776.34321986895 | 2876.27640660263 | 3320.26049767389 | 3035.22411881054 | 3140.87517196567 | 2564.17533703976 | 2530.36009124229 | Maf1 |
| ENSMUSG00000022554 | 0.18383992546225 | 0.463443604807925 | 0.895328112556867 | 46.0791995450213 | 76.5550785179469 | 85.1809399783602 | 87.227182566009 | 68.9823663366031 | 56.2617826753975 | 62.4646854333682 | 71.6446872681215 | Hgh1 |
| ENSMUSG00000022555 | 0.0906257209265389 | 0.443544727362965 | 0.89108031640712 | 412.002254755485 | 360.786167909048 | 368.170951684246 | 380.798237868813 | 396.374866886513 | 358.363093997641 | 319.610973800734 | 354.966859646602 | Dgat1 |
| ENSMUSG00000022556 | 0.101960245082829 | 0.322651616805665 | 0.855977678491063 | 818.583427211555 | 796.498582878213 | 788.396922244156 | 796.299763425179 | 768.660653465006 | 656.795158623663 | 823.492769629904 | 730.558705021906 | Hsf1 |
| ENSMUSG00000022557 | 0.120764331290673 | 0.245552564566671 | 0.832048466795696 | 496.932544112975 | 501.680089011226 | 525.282463199888 | 580.576623745802 | 496.016062706051 | 457.432754795624 | 501.799639648058 | 478.716774018812 | Bop1 |
| ENSMUSG00000022558 | 0.0931234133651747 | 0.533413620902474 | 0.909205447826158 | 964.049135579171 | 1070.14226779343 | 961.598166866822 | 936.988767563903 | 882.536305830192 | 716.726187995282 | 1014.01006020168 | 1072.49925789249 | Mroh1 |
| ENSMUSG00000022559 | -0.169823601832509 | 0.293411582848385 | 0.851785577030345 | 177.992202164102 | 193.83094348161 | 163.736695736181 | 210.095579513828 | 213.516848184724 | 250.731857575141 | 195.722681024554 | 180.197243734972 | Fbxl6 |
| ENSMUSG00000022560 | -0.111849923227896 | 0.507327022558512 | 0.901698316485955 | 112.035700854562 | 100.987550385377 | 113.574586637814 | 104.109863062656 | 124.825234323377 | 121.085140975312 | 125.970448957293 | 93.3551985614916 | Slc52a2 |
| ENSMUSG00000022561 | -0.175471257460076 | 0.262025364638285 | 0.841137531367976 | 370.440623793308 | 339.611358957275 | 377.63550057073 | 357.350070512359 | 424.84377997781 | 507.579126310652 | 338.350379430744 | 362.565538599282 | Gpaa1 |
| ENSMUSG00000022562 | 0.112188482022097 | 0.420924601915381 | 0.882568531013176 | 2241.6175308078 | 2765.75581539306 | 2955.7786172491 | 2100.01786844402 | 2366.20466116507 | 2200.32493550087 | 2288.28964304239 | 2455.45882728016 | Oplah |
| ENSMUSG00000022564 | -0.0550535177486673 | 0.757168229579973 | 0.956693827734505 | 1280.27893637834 | 1219.18034618475 | 1370.46667876295 | 1253.0700635289 | 1498.99777007634 | 1681.73806910156 | 1120.2000254384 | 1022.56508191773 | Grina |
| ENSMUSG00000022565 | 0.193154625492385 | 0.379141167918559 | 0.87422797717331 | 7978.02611730467 | 9582.41546640599 | 6883.56640514015 | 7142.3117767759 | 6845.67864026099 | 4477.7040516224 | 7278.17693107795 | 9026.14507021864 | Plec |
| ENSMUSG00000022568 | 0.12444354776727 | 0.260588107336499 | 0.840316047847749 | 711.968808656408 | 807.0859873541 | 830.987392233336 | 775.665376151499 | 707.342994499137 | 653.125911927441 | 702.727711125392 | 803.288917854696 | Scrib |
| ENSMUSG00000022570 | 0.151077772337167 | 0.373425732492151 | 0.873700364743909 | 420.133878204606 | 298.890572511559 | 420.22597055991 | 407.9981120023 | 346.006789878835 | 409.732547744743 | 344.596847974081 | 293.091902460497 | Tsta3 |
| ENSMUSG00000022571 | -0.102181736693417 | 0.427574214940994 | 0.885066977635295 | 273.764656120421 | 244.324718674299 | 235.667267273463 | 250.426427366929 | 291.25888008788 | 273.970419984545 | 234.242570375131 | 278.980070119806 | Pycrl |
| ENSMUSG00000022574 | -0.157762481761033 | 0.413123501124173 | 0.882192126163777 | 381.282788392137 | 402.321370083678 | 377.63550057073 | 336.71568323868 | 469.737066006393 | 536.933099880424 | 314.405583347953 | 351.710282952596 | Naprt |
| ENSMUSG00000022575 | -0.477138057002102 | 0.108040812179214 | 0.762287160109383 | 55.1143367107117 | 36.6487078011448 | 43.5369248778286 | 60.9652351267805 | 79.9319482947941 | 91.7311674055395 | 57.2592949805875 | 44.5065481514088 | Gsdmd |
| ENSMUSG00000022579 | -0.315856240034514 | 0.304920166376361 | 0.851785577030345 | 2494.60137144713 | 1576.70885117814 | 1847.47994264177 | 2304.4858877923 | 3337.43258085661 | 3690.03909416683 | 1493.94705994806 | 1715.13039217624 | Gpihbp1 |
| ENSMUSG00000022587 | -0.755893201805752 | 0.0395276534142982 | 0.679566075569698 | 2010.31801936613 | 1791.71460361152 | 1803.94301776394 | 1969.64605794214 | 4431.29581847989 | 5132.05304578191 | 1481.45412286138 | 1748.78168468097 | Ly6e |
| ENSMUSG00000022594 | 0.0321860079862054 | 0.881718079195684 | 0.978854883720278 | 7156.73214894341 | 7490.99587455399 | 6027.97118580196 | 6193.12996218664 | 4977.67995819361 | 4832.39789892382 | 8264.07788283461 | 8201.14564107057 | Lynx1 |
| ENSMUSG00000022595 | -0.492639956875293 | 0.243060867590148 | 0.832048466795696 | 16.2632468982428 | 9.77298874697194 | 16.0897331070236 | 13.1309737196143 | 16.4243729372865 | 31.8001380339204 | 12.4929370866736 | 17.3684090346961 | Lypd2 |
| ENSMUSG00000022601 | 0.096467039402732 | 0.689577685911226 | 0.94469184542218 | 212.325723393726 | 299.704988240473 | 252.703455269135 | 195.088752405698 | 218.991639163819 | 141.877538920568 | 280.050006359601 | 256.184033261768 | Zbtb11 |
| ENSMUSG00000022602 | 0.0929997681016024 | 0.787132667352319 | 0.961141387070613 | 82.2197482077831 | 73.2974156022896 | 58.6802030962037 | 155.695831246855 | 67.887408140784 | 99.0696607979826 | 114.518589961175 | 65.1315338801105 | Arc |
| ENSMUSG00000022604 | 0.0222191500797376 | 0.913164792770318 | 0.986226496507248 | 90.3513716569045 | 78.9983257046899 | 90.8596693102509 | 85.3513291774927 | 64.6025335533267 | 77.0541806206532 | 93.6970281500523 | 104.210454208177 | Cep97 |
| ENSMUSG00000022607 | 0.0838894019504089 | 0.430499036884633 | 0.886017087288711 | 524.037955610046 | 583.121661902659 | 604.784673846357 | 547.749189446766 | 556.238763476101 | 502.686797382356 | 501.799639648058 | 570.986447015635 | Ptk2 |
| ENSMUSG00000022610 | -0.0754689355724947 | 0.770347985226196 | 0.958992744413128 | 376.765219809292 | 320.879797192245 | 337.884395247495 | 471.777127211855 | 404.039574257247 | 596.864129252044 | 280.050006359601 | 308.289260365856 | Mapk12 |
| ENSMUSG00000022614 | -0.354691627453516 | 0.0711583403643696 | 0.726501344493452 | 554.757421973394 | 575.79192034243 | 534.747012086372 | 597.459304242449 | 896.77076237584 | 879.396124861105 | 550.730309904196 | 567.72987032163 | Lmf2 |
| ENSMUSG00000022615 | -0.161406600534476 | 0.4103533362988 | 0.882192126163777 | 84.0267756409212 | 67.5965054998893 | 64.3589324280944 | 90.0409626487835 | 85.4067392738896 | 80.7234273168747 | 80.1630129728225 | 95.5262496908287 | Tymp |
| ENSMUSG00000022617 | 0.147878556318139 | 0.0963363384970744 | 0.757809915356977 | 880.02235993825 | 781.024684028841 | 844.237760674415 | 843.196098138087 | 764.28082068173 | 786.441875223492 | 717.302804393178 | 754.440267444613 | Chkb |
| ENSMUSG00000022620 | -0.564976664759276 | 0.0445388944272142 | 0.693563614452513 | 176.185174730964 | 167.769640156352 | 190.237432618338 | 217.598993067893 | 356.956371837026 | 387.717067567413 | 173.860041122875 | 194.309076075663 | Arsa |
| ENSMUSG00000022621 | 0.317131966277717 | 0.173672164952165 | 0.799889021902745 | 49.6932544112975 | 51.3081909216027 | 53.001473764313 | 41.2687745473591 | 42.7033696369448 | 33.0232202659942 | 41.6431236222455 | 39.0789203280663 | Rabl2 |
| ENSMUSG00000022623 | -0.132531203414974 | 0.528049279690778 | 0.908232038364594 | 758.048008201429 | 745.190391956611 | 859.38103889279 | 804.741103673502 | 1227.44813751321 | 894.073111645991 | 651.714884688142 | 699.078463646519 | Shank3 |
| ENSMUSG00000022629 | 0.211042920386134 | 0.443331080568066 | 0.89108031640712 | 983.022923627121 | 1242.79840232327 | 958.758802200877 | 837.568537972538 | 633.980795379257 | 534.486935416277 | 1117.07679116673 | 1188.65049331202 | Kif21a |
| ENSMUSG00000022634 | 0.0276915615523083 | 0.844013781020743 | 0.971721411930626 | 451.756858284523 | 368.930325198191 | 393.725233677754 | 459.584080186499 | 362.431162816121 | 421.963370065482 | 471.60837502193 | 385.36157545732 | Yaf2 |
| ENSMUSG00000022635 | -0.0460361206334085 | 0.785108317523246 | 0.961141387070613 | 2240.71401709123 | 1794.97226652718 | 1953.48289017039 | 2542.71926813388 | 2129.69369086814 | 2751.93502216618 | 1952.02141979276 | 1975.65652769668 | Zcrb1 |
| ENSMUSG00000022636 | -0.664424274588578 | 0.215390451342797 | 0.824338622350122 | 18.0702743313809 | 5.7009101024003 | 31.2330113253987 | 22.5102406621959 | 31.7537876787538 | 52.592535979176 | 15.616171358342 | 22.7960368580387 | Alcam |
| ENSMUSG00000022637 | 0.0789053931750848 | 0.691776792358095 | 0.94469184542218 | 320.747369382011 | 294.004078138073 | 384.260684791269 | 331.088123073131 | 326.297542354091 | 247.06261087892 | 420.595548584679 | 263.782712214447 | Cblb |
| ENSMUSG00000022639 | -0.28238834937668 | 0.154764156370133 | 0.787275540343856 | 71.3775836089546 | 73.2974156022896 | 92.7525790875478 | 75.9720622349111 | 85.4067392738896 | 118.638976511164 | 84.3273253350471 | 93.3551985614916 | Dubr |
| ENSMUSG00000022641 | -0.0157993308530923 | 0.920647752134232 | 0.986713918026868 | 262.018977805023 | 302.148235427216 | 222.416898832385 | 241.985087118606 | 258.410134213307 | 223.824048469516 | 255.064132186253 | 302.861632542514 | Bbx |
| ENSMUSG00000022651 | 0.569601032707376 | 0.515921155456102 | 0.904245406849874 | 24.3948703473642 | 7.32974156022896 | 47.3227444324223 | 3.75170677703264 | 13.1394983498292 | 35.4693847301419 | 4.16431236222455 | 3.25657669400552 | Retnlg |
| ENSMUSG00000022656 | -0.150650189564594 | 0.392551640980715 | 0.878537584780694 | 113.8427282877 | 122.976775066064 | 116.413951303759 | 116.302910088012 | 162.053812981226 | 130.869798831903 | 106.189965236726 | 122.664388807541 | Nectin3 |
| ENSMUSG00000022661 | -0.329726653894712 | 0.294776706280498 | 0.851785577030345 | 611.678786117243 | 633.615437095348 | 497.835271429083 | 581.51455044006 | 842.022852584886 | 1158.25887377394 | 359.171941241867 | 563.387768062955 | Cd200 |
| ENSMUSG00000022663 | -0.0130735532148292 | 0.889650367038087 | 0.981434643646972 | 733.653137854065 | 772.880526739698 | 708.894711597687 | 710.948434247686 | 715.00770186987 | 688.595296657583 | 793.301505003776 | 755.525793009281 | Atg3 |
| ENSMUSG00000022664 | -0.152462906343332 | 0.294987167799851 | 0.851785577030345 | 140.044626068202 | 127.86326943955 | 135.343049076728 | 139.751077444466 | 150.009272827216 | 155.33144347338 | 132.216917500629 | 166.085411394282 | Slc35a5 |
| ENSMUSG00000022665 | -0.904562860886913 | 0.0492712605306465 | 0.703396849069134 | 657.757985662265 | 689.810122390436 | 671.036516051749 | 852.575365080669 | 1615.06333883317 | 2703.01173288323 | 529.908748093074 | 527.565424428895 | Ccdc80 |
| ENSMUSG00000022668 | 0.168001625375033 | 0.2443717042228 | 0.832048466795696 | 238.527621174228 | 247.582381589956 | 232.827902607518 | 221.350699844926 | 214.611806380543 | 170.008430258266 | 239.447960827911 | 211.677485110359 | Gtpbp8 |
| ENSMUSG00000022671 | -0.114488163341389 | 0.486158991578767 | 0.899236955296196 | 293.64195788494 | 267.942774812814 | 241.345996605354 | 315.143369270742 | 337.247124312282 | 351.024600605198 | 255.064132186253 | 268.124814473121 | Mzt2 |
| ENSMUSG00000022672 | 0.189092776198672 | 0.345322742954565 | 0.861266047293794 | 149.079763233892 | 184.057954734638 | 161.843785958884 | 192.274972322923 | 159.863896589588 | 156.554525705454 | 181.147587756768 | 105.295979772845 | Prkdc |
| ENSMUSG00000022673 | -0.343359338075121 | 0.0903565083529675 | 0.753823363185192 | 112.035700854562 | 104.245213301034 | 108.842312194571 | 101.296082979881 | 153.294147414674 | 163.893019097897 | 130.134761319517 | 94.4407241261602 | Mcm4 |
| ENSMUSG00000022674 | 0.197365554511639 | 0.249081884183527 | 0.835314899646652 | 146.369222084185 | 194.645359210525 | 193.076797284283 | 149.130344387048 | 133.58489988993 | 144.323703384715 | 150.95632313064 | 167.17093695895 | Ube2v2 |
| ENSMUSG00000022676 | -0.505198872679813 | 0.114931745117469 | 0.767518180080285 | 44.2721721118832 | 41.5352021746308 | 52.0550188756646 | 54.3997482669734 | 100.736154015357 | 79.5003450848009 | 37.4788112600209 | 55.3618037980939 | Snai2 |
| ENSMUSG00000022677 | 0.108467249059927 | 0.529622173856529 | 0.908620485539384 | 1409.48139784771 | 1134.48111037766 | 1108.29867460733 | 1267.13896394278 | 909.91026072567 | 1002.92743030056 | 1229.5132249468 | 1419.86743858641 | Fopnl |
| ENSMUSG00000022678 | -0.00828870868309818 | 0.947635179711454 | 0.991041028853253 | 899.899661702769 | 922.733020859934 | 1012.70673085384 | 726.893188050075 | 884.72622222183 | 914.865509591247 | 915.107641598844 | 868.420451734806 | Nde1 |
| ENSMUSG00000022679 | -0.279956021981307 | 0.323285789589471 | 0.855977678491063 | 36.1405486627618 | 49.679359463774 | 53.001473764313 | 62.8410885152968 | 51.4630352034976 | 45.2540425867328 | 85.3684034256032 | 61.8749571861049 | Mpv17l |
| ENSMUSG00000022680 | 0.239403500565833 | 0.152977189990141 | 0.783412422888014 | 2624.70734663308 | 2857.78479276038 | 3106.2649445442 | 2818.46971624577 | 2119.83906710577 | 1863.97732168056 | 2918.14188782885 | 2759.40598538735 | Pdxdc1 |
| ENSMUSG00000022681 | 0.0849553725567235 | 0.490350682523558 | 0.899236955296196 | 526.748496759753 | 466.66021266791 | 512.978549647458 | 586.204183911351 | 479.591689768764 | 556.502415593606 | 472.649453112486 | 464.604941678121 | Ntan1 |
| ENSMUSG00000022682 | 0.0862978483904722 | 0.509536771676649 | 0.902394714365738 | 1172.76080410662 | 1277.00386293767 | 1237.01653946352 | 1171.47044112844 | 1131.09181628113 | 918.534756287469 | 1272.1974266596 | 1252.69650162746 | Rrn3 |
| ENSMUSG00000022684 | 0.106705901153801 | 0.258835509128646 | 0.838606660810377 | 1598.31576461064 | 1587.29625565403 | 1802.99656287529 | 1579.46855313074 | 1433.3002783272 | 1583.89149053565 | 1617.83535272424 | 1464.37398673782 | Bfar |

| | | | | | | | | | | | | |
|---|---|---|---|---|---|---|---|---|---|---|---|---|
| **ENSMUSG00000022685** | 0.183467535579484 | 0.188639253485446 | 0.808677608936066 | 286.413848152387 | 372.187988113848 | 351.134763688574 | 317.957149353517 | 319.727793179176 | 304.547475786391 | 286.296474902938 | 259.440609955773 | Parn |
| **ENSMUSG00000022687** | -0.994958779060506 | 0.00977802511684966 | 0.635862253526407 | 42.4651446787451 | 42.3496179035451 | 35.965285768641 | 51.5859681841989 | 108.400861386091 | 139.43137445642 | 55.1771387994752 | 41.2499714574033 | Boc |
| **ENSMUSG00000022698** | 0.140856017243281 | 0.390743041334739 | 0.877797322358001 | 2167.52940604914 | 2311.31183865886 | 2029.19928126227 | 1996.84593207563 | 1786.97177557677 | 1450.5755272396 | 2199.79800534512 | 2275.26158354519 | Naa50 |
| **ENSMUSG00000022701** | 0.133236705496296 | 0.534818586514978 | 0.910706162399539 | 164.439496415566 | 138.450673915436 | 109.78876708322 | 132.247663890401 | 108.400861386091 | 162.669936865823 | 127.011527047849 | 99.8683519495027 | Ccdc191 |
| **ENSMUSG00000022702** | 0.092993042919072 | 0.488886595397849 | 0.899236955296196 | 262.018977805023 | 228.036404096012 | 207.27362061401 | 260.743621003769 | 224.466430142915 | 214.039390612925 | 225.913945650682 | 233.387996403729 | Hira |
| **ENSMUSG00000022704** | -0.100873675800822 | 0.735720529342526 | 0.954383574136061 | 31.6229800799166 | 51.3081909216027 | 66.2518422053913 | 48.7721881014244 | 42.7033696369448 | 42.8078781225851 | 70.7933101578173 | 55.3618037980939 | Qtrt2 |
| **ENSMUSG00000022706** | -0.0774592209376263 | 0.479869656977304 | 0.897732059433541 | 511.388763578079 | 429.197089137851 | 501.621090983677 | 504.604561510891 | 543.099265126272 | 527.148442023833 | 488.265624470828 | 494.999657488839 | Mrpl40 |
| **ENSMUSG00000022707** | 0.0564001244221091 | 0.789635269071006 | 0.961659329450003 | 1545.91196904964 | 2011.60685041839 | 1582.4725738202 | 1671.38536916804 | 1364.31791199059 | 1161.92812047017 | 2131.08685136841 | 1892.07105921721 | Gbe1 |
| **ENSMUSG00000022708** | 0.0294846558269831 | 0.839417513799133 | 0.969820257058002 | 7382.61057808567 | 6794.67042633224 | 6047.84673846358 | 7210.78042545674 | 7275.9972112179 | 7899.88813696506 | 5507.30309904196 | 6199.43649982185 | Zbtb20 |
| **ENSMUSG00000022710** | 0.171974884625244 | 0.357955613853853 | 0.865834559821921 | 1289.31407354403 | 1454.54649184099 | 1477.41608118023 | 1209.92543559303 | 1077.43886468599 | 874.50379593281 | 1377.34631380577 | 1490.42660028986 | Usp7 |
| **ENSMUSG00000022711** | 0.0918171773116811 | 0.311828129701251 | 0.851785577030345 | 629.749060448624 | 590.451403462888 | 611.409858066897 | 626.535031764452 | 546.384139713729 | 565.063991218123 | 581.96265262088 | 612.236418473038 | Pmm2 |
| **ENSMUSG00000022718** | -0.084644057561293 | 0.616645212950739 | 0.928896975098882 | 227.685456575399 | 231.294067011669 | 198.755526616174 | 189.461192240149 | 206.947099009809 | 188.354663739374 | 222.790711379013 | 280.065595684475 | Dgcr8 |
| **ENSMUSG00000022721** | -0.195986836471015 | 0.329357676168224 | 0.855977678491063 | 234.010052591383 | 182.42912327681 | 238.506631939409 | 215.723139679377 | 293.448796479518 | 305.770558018465 | 196.76375911511 | 201.907755028342 | Trmt2a |
| **ENSMUSG00000022722** | -0.00255790854210874 | 0.991347677451861 | 0.998628446692557 | 79.509207058076 | 61.895595397489 | 43.5369248778286 | 70.3445020693621 | 63.5075753575076 | 63.6002760678407 | 59.3414511616998 | 69.4736361387845 | Arl6 |
| **ENSMUSG00000022723** | 0.297245313177659 | 0.110097000544064 | 0.763647454385034 | 1443.81491907733 | 1468.39155923253 | 1412.11069386348 | 1308.40773849013 | 1096.05315401492 | 781.549546295196 | 1378.38739189632 | 1326.51224002492 | Crybg3 |
| **ENSMUSG00000022724** | 0.191149273674036 | 0.352088320824124 | 0.862728204353539 | 373.151164943016 | 434.897999240251 | 398.457508120996 | 404.246405225267 | 402.944616061428 | 223.824048469516 | 427.883095218572 | 354.966859646602 | Riox2 |
| **ENSMUSG00000022742** | 0.0528911676967312 | 0.821865652536775 | 0.966637987288264 | 1285.70001867775 | 1414.64012112419 | 1479.30899095752 | 1117.07069286147 | 1050.06490979051 | 824.357424417781 | 1626.16397744869 | 1604.40678458005 | Cpox |
| **ENSMUSG00000022744** | 0.180827434015542 | 0.300560511306735 | 0.851785577030345 | 911.645340018166 | 987.886279173081 | 948.347798425744 | 927.609500621321 | 713.912743674051 | 631.110431750112 | 964.038311854983 | 1020.3940307884 | Cldnd1 |
| **ENSMUSG00000022747** | 0.0793008687383994 | 0.548323121814791 | 0.914699131434573 | 702.030157774148 | 693.882201035008 | 644.535779169592 | 609.652351267805 | 640.550544554172 | 508.802208542726 | 665.248899865371 | 692.565310258508 | St3gal6 |
| **ENSMUSG00000022748** | -0.0347774621774493 | 0.7407766745154 | 0.954383574136061 | 1087.83051474913 | 997.659267920053 | 906.703783325212 | 886.340726073962 | 960.278337733348 | 1018.82749931753 | 964.038311854983 | 1031.24928643508 | Cmss1 |
| **ENSMUSG00000022749** | 0.19906980147796 | 0.240430419500103 | 0.832048466795696 | 230.395997725106 | 285.859920848929 | 290.561650815073 | 298.260688774095 | 225.561388338734 | 190.800828203522 | 279.008928269045 | 265.953763343784 | Tbc1d23 |
| **ENSMUSG00000022750** | 0.107733133812567 | 0.368361014045888 | 0.871495599696183 | 543.011743657996 | 556.245942848486 | 699.430162711202 | 574.011136885995 | 569.37826182593 | 546.717757737015 | 545.524919451416 | 539.506205640248 | Klhl22 |
| **ENSMUSG00000022751** | 0.166934722039893 | 0.201739307174869 | 0.813395633194566 | 878.215332505112 | 667.820897709749 | 867.899132890626 | 891.968286239511 | 751.1413223319 | 720.395434691503 | 687.11153976705 | 784.834983255331 | Nit2 |
| **ENSMUSG00000022752** | 0.199172454305824 | 0.320667908719111 | 0.855228644171892 | 1826.90473490261 | 2048.25555821954 | 1947.8041608385 | 1699.52316999579 | 1433.3002783272 | 1130.12798243625 | 1939.52848270608 | 2048.38674052947 | Tomm70a |
| **ENSMUSG00000022754** | -0.735407385551883 | 0.0977694473122836 | 0.759517686335357 | 30.7194663633475 | 26.8757190541728 | 17.036187995672 | 22.5102406621959 | 48.1781606160403 | 70.9387694602839 | 23.9447960827911 | 19.5394601640331 | Tmem45a |
| **ENSMUSG00000022756** | -0.254461846847828 | 0.468276739114178 | 0.895328112556867 | 29.8159526467785 | 25.2468875963442 | 17.9826428843205 | 15.0068271081306 | 32.8487458745729 | 28.1308913376988 | 26.0269522639034 | 18.4539345993646 | Slc7a4 |
| **ENSMUSG00000022757** | 0.0268144356170303 | 0.783515393368708 | 0.961051227381536 | 873.697763922267 | 897.48613326359 | 831.933847121985 | 907.9130400419 | 912.100177117308 | 774.211052902753 | 911.984407327176 | 846.709940441436 | Tfg |
| **ENSMUSG00000022758** | -0.392942003924379 | 0.0679499798849504 | 0.722112580225498 | 60.535419010126 | 76.5550785179469 | 50.1621090983677 | 61.9031618210386 | 78.836990098975 | 99.0696607979826 | 79.1219348822664 | 71.6446872681215 | P2rx6 |
| **ENSMUSG00000022760** | -0.142313738907723 | 0.473216777067157 | 0.89728597737823 | 231.299511441676 | 203.603932228582 | 209.166530391307 | 237.295453647315 | 258.410134213307 | 324.116791499573 | 187.394056300105 | 204.078806157679 | Thap7 |
| **ENSMUSG00000022761** | 0.0722535051182552 | 0.552293911460357 | 0.914977712327965 | 565.599586572222 | 667.820897709749 | 742.967087589031 | 740.024161769689 | 642.74046094581 | 658.018240855737 | 648.591650416473 | 633.946929766408 | Lztr1 |
| **ENSMUSG00000022765** | 0.0620021218933676 | 0.681713855819902 | 0.944452327903597 | 487.897406947284 | 576.606336071345 | 539.479286529615 | 473.652980600371 | 573.758094609207 | 409.732547744743 | 454.951125573032 | 551.446986851602 | Snap29 |
| **ENSMUSG00000022766** | -0.575749909737492 | 0.0637156774004071 | 0.716991331547963 | 66.8600150261093 | 101.801966114291 | 57.7337482075553 | 68.4686486808458 | 79.9319482947941 | 83.1695917810225 | 103.066730965058 | 173.684090346961 | Serpind1 |
| **ENSMUSG00000022769** | -0.618685394234368 | 0.13985124355293 | 0.779711473022872 | 105.711104838578 | 101.801966114291 | 57.7337482075553 | 108.799496533947 | 144.534481848121 | 260.516515431732 | 54.1360607089191 | 116.15123541953 | Sdf2l1 |
| **ENSMUSG00000022770** | 0.219064498111813 | 0.360965255707841 | 0.868539734006824 | 2037.4234308632 | 2396.82549019487 | 2237.41935676493 | 1832.70876058045 | 1501.18768646798 | 1147.25113368528 | 2339.30246947964 | 2317.59708056726 | Dlg1 |
| **ENSMUSG00000022771** | 0.0450946517951517 | 0.716594991335202 | 0.951348124093194 | 624.32797814921 | 537.514381083457 | 524.33600831124 | 548.687116141024 | 577.042969196664 | 602.979540412413 | 493.471014923609 | 493.914131924171 | Ppil2 |
| **ENSMUSG00000022772** | 0.240235436473302 | 0.606155753068709 | 0.926859923029135 | 133.720030052219 | 211.748089517725 | 239.453086828057 | 112.551203310979 | 84.3117810780705 | 48.9232892829544 | 235.283648465687 | 221.447215192376 | Senp5 |
| **ENSMUSG00000022773** | 0.0722084020644702 | 0.866839620900982 | 0.974868041734905 | 17.1667606148119 | 13.8450673915436 | 33.1259211026956 | 17.8206071909051 | 19.7092475247437 | 9.78465785659088 | 26.0269522639034 | 21.7105112933701 | Ypel1 |
| **ENSMUSG00000022774** | 0.0697746469120804 | 0.518394375557034 | 0.905504473025266 | 665.889609111386 | 559.503605764144 | 592.480760293928 | 643.417712261099 | 560.618596259378 | 637.225842910481 | 580.921574530324 | 566.644344756961 | Ncbp2 |
| **ENSMUSG00000022779** | -0.0478964283989949 | 0.64238590977767 | 0.934616627390525 | 376.765219809292 | 374.631235300591 | 360.599312575058 | 359.225923900876 | 363.52612101194 | 365.701587390084 | 377.911346871878 | 413.585240138701 | Top3b |
| **ENSMUSG00000022781** | -0.10034226758723 | 0.501716872277324 | 0.900927059989498 | 795.09207058076 | 1022.9061555164 | 928.472245764126 | 785.982569788339 | 1054.44474257379 | 801.118862008378 | 885.957455063272 | 1045.36111877577 | Pak2 |
| **ENSMUSG00000022787** | 0.0456653032119078 | 0.796141766618525 | 0.962200030474509 | 171.667606148119 | 122.976775066064 | 123.039135524298 | 135.061443973175 | 140.154649064844 | 141.877538920568 | 137.42230795341 | 116.15123541953 | Wdr53 |
| **ENSMUSG00000022788** | 0.00426769829759492 | 0.988373054734151 | 0.998437672973644 | 417.423337054899 | 609.182965227918 | 451.458981885309 | 382.67409125733 | 365.716037403578 | 260.516515431732 | 591.332355435886 | 637.203506460414 | Fgd4 |
| **ENSMUSG00000022789** | 0.0622790252125728 | 0.753330694280432 | 0.956497786093635 | 2397.92540377425 | 2606.94474825477 | 2350.99394340274 | 2082.19726125312 | 1853.76422552173 | 1658.49950669215 | 2778.63742369433 | 2747.46520417599 | Dnm1l |
| **ENSMUSG00000022790** | -0.0352898203261543 | 0.933646866288117 | 0.98818087308472 | 23.4913566307952 | 14.6594831204579 | 11.3574586637814 | 15.9447538023887 | 9.85462376237187 | 13.4539045528125 | 22.903717992235 | 20.6249857287016 | Igsf11 |
| **ENSMUSG00000022791** | 0.112286468333648 | 0.393720155019412 | 0.878942305112115 | 380.379274675568 | 397.434875710192 | 395.618143455051 | 348.908730264036 | 330.677375137367 | 420.740287833408 | 331.062832796851 | 327.828720529889 | Tnk2 |
| **ENSMUSG00000022792** | 0.167381383782194 | 0.225867099393695 | 0.832048466795696 | 553.853908256825 | 500.865673282312 | 480.799083433411 | 554.314676306573 | 442.363111110915 | 395.055560959857 | 551.771387994752 | 470.032569501464 | Yars2 |
| **ENSMUSG00000022793** | -0.378130788324445 | 0.355825315287049 | 0.864662986238123 | 350.563322028789 | 325.766291565731 | 306.651383922097 | 211.971432902344 | 475.211856985488 | 702.049201210395 | 201.96914956789 | 174.76961591163 | B4galt4 |
| **ENSMUSG00000022797** | 0.0548944212026661 | 0.880508099806693 | 0.978433072023387 | 287.317361868956 | 339.611358957275 | 186.451613063744 | 171.640585049243 | 139.059690869025 | 143.100621152642 | 292.542943446274 | 373.420794245967 | Tfrc |
| **ENSMUSG00000022799** | -0.312553281689684 | 0.237030202335906 | 0.832048466795696 | 1026.39158202244 | 1090.50266101629 | 1049.61847151113 | 816.934150698859 | 1664.33645764503 | 1613.24546410542 | 814.123066814899 | 856.479670523452 | Arhgap31 |
| **ENSMUSG00000022800** | 0.192784221382306 | 0.300855533985858 | 0.851785577030345 | 1637.16685442311 | 1703.75770488878 | 1844.64057797582 | 1576.65477304797 | 1351.17841364077 | 1039.61989726278 | 1717.77884941763 | 1806.3145396084 | Fyttd1 |
| **ENSMUSG00000022801** | 0.023711364432878 | 0.865809648772044 | 0.974868041734905 | 326.168451681425 | 380.332145402991 | 352.081218577222 | 288.881421831514 | 371.190828382674 | 306.993640250539 | 342.514691792969 | 305.032683671851 | Lrch3 |
| **ENSMUSG00000022802** | 0.274626047349377 | 0.34322608674053 | 0.861167771001038 | 60.535419010126 | 59.452348210746 | 63.4124775394459 | 37.5170677703264 | 39.4184950494875 | 33.0232202659942 | 45.80743598447 | 64.046008315442 | Lmln |
| **ENSMUSG00000022803** | 0.0239014803300222 | 0.837102277967187 | 0.968924788429433 | 10702.1199727603 | 10350.4094987722 | 9674.66187176442 | 10316.2557101455 | 9326.85391198707 | 8742.59179486395 | 10951.10043456 | 11346.9987274799 | Popdc2 |
| **ENSMUSG00000022807** | 0.190475958632501 | 0.30837750986762 | 0.851785577030345 | 495.125516679837 | 584.750493360488 | 704.162437154444 | 502.728708122374 | 505.870686468423 | 374.263163014601 | 584.044808801993 | 538.42068007558 | Osbpl11 |
| **ENSMUSG00000022808** | 0.19869766998226 | 0.142828632498193 | 0.780607990324772 | 842.07478384235 | 987.071863444166 | 991.884723303572 | 827.251344335698 | 747.856447744443 | 680.033721033066 | 889.080689334941 | 860.821772782126 | Snx4 |
| **ENSMUSG00000022809** | -0.321656918217764 | 0.583048860176119 | 0.922450244597442 | 7.22810973255236 | 23.6180561385155 | 4.73227444324223 | 12.1930470253561 | 9.85462376237187 | 17.123151249034 | 9.36970281500523 | 23.8815624227072 | Nr1i2 |
| **ENSMUSG00000022811** | 0.00937157495303867 | 0.954324323180991 | 0.992371505557225 | 543.011743657996 | 645.017257300148 | 541.372196306912 | 507.418341593665 | 487.256397139498 | 456.20967256355 | 618.400385790345 | 659.999543318453 | Zfp148 |
| **ENSMUSG00000022812** | 0.109882888647901 | 0.597811648663455 | 0.924496535139492 | 1297.44569699315 | 1553.90521076854 | 1343.96594188079 | 1156.46361402031 | 1046.78003520306 | 884.2884537894 | 1424.19482788079 | 1603.32125901539 | Gsk3b |
| **ENSMUSG00000022814** | 0.22231175919267 | 0.0842536083394679 | 0.742565114108635 | 215.939778260002 | 231.294067011669 | 238.506631939409 | 233.543746870282 | 214.611806380543 | 204.254732756335 | 195.722681024554 | 173.684090346961 | Umps |
| **ENSMUSG00000022816** | -0.882751676274666 | 0.0561648346093371 | 0.703396849069134 | 625.231491865779 | 548.101785559343 | 374.796135904785 | 632.162591930001 | 1163.9405621557 | 2026.64725854639 | 378.952424962434 | 451.578634902099 | Fstl1 |
| **ENSMUSG00000022817** | 0.0694714961531295 | 0.715256691937746 | 0.951176382858794 | 3079.17474606731 | 3387.15501655469 | 4596.93139416551 | 4520.80666632434 | 4304.28066776487 | 4211.0721250303 | 3282.5192195235 | 3053.58341341251 | Itgb5 |
| **ENSMUSG00000022820** | 0.20244935689125 | 0.239389254384587 | 0.832048466795696 | 4123.63660242112 | 2658.25293917637 | 3188.60651985662 | 3919.59565530486 | 3071.35773927257 | 3420.96100311059 | 2572.50396176421 | 3006.90581413177 | Ndufb4 |
| **ENSMUSG00000022821** | -0.479912358113893 | 0.14200431582512 | 0.780237320732165 | 59.631905293557 | 105.874044758863 | 59.6266579848522 | 77.8479156234274 | 81.0269064906132 | 80.7234273168747 | 87.4505596067155 | 173.684090346961 | Hgd |
| **ENSMUSG00000022822** | -0.0690205633472218 | 0.64470823194033 | 0.934857223264556 | 235.817080024521 | 204.418347957496 | 201.594891282119 | 209.15765281957 | 245.270635863478 | 178.570005882784 | 237.365804646799 | 230.131419709724 | Abcc5 |
| **ENSMUSG00000022827** | -0.155895772004386 | 0.537475422144579 | 0.912099403004057 | 73.1846110420926 | 98.5443031986338 | 86.1273948670087 | 68.4686486808458 | 86.5016974697087 | 57.4848649074714 | 107.231043327282 | 111.809133160856 | Rabl3 |
| **ENSMUSG00000022828** | -0.0254137601059209 | 0.868356865120199 | 0.975106533657166 | 143.658680934478 | 127.86326943955 | 120.199770858353 | 132.247663890401 | 128.110108910834 | 119.862058743238 | 149.915245040084 | 134.605170018895 | Gtf2e1 |
| **ENSMUSG00000022831** | -0.288870371632791 | 0.173243461345918 | 0.799889021902745 | 159.018414116152 | 188.13003337921 | 206.327165725361 | 156.633757941113 | 259.505092409126 | 265.408844360028 | 162.408182126757 | 181.282769299641 | Hcls1 |
| **ENSMUSG00000022833** | -0.527321273660307 | 0.0815907944484564 | 0.739855948845396 | 20.780815481088 | 21.9892246806869 | 25.5542819935081 | 28.1378008277448 | 27.3739548954774 | 26.9078091056249 | 40.6020455316893 | 43.4210225867403 | Ccdc14 |
| **ENSMUSG00000022836** | -0.236058669578945 | 0.280816759994852 | 0.849606795105313 | 750.819898468876 | 823.374301932386 | 741.074177811734 | 599.335157630965 | 1000.79179097865 | 1126.45873574002 | 683.988305495382 | 623.091674119723 | Mylk |
| **ENSMUSG00000022837** | 0.135043169651495 | 0.342426495102455 | 0.860056790709335 | 226.78194285883 | 216.634583891211 | 261.221549266971 | 254.178134143962 | 236.510970296925 | 187.131581507301 | 212.379930473452 | 235.559047533066 | Iqcb1 |
| **ENSMUSG00000022838** | -0.164721046833684 | 0.720289237740916 | 0.951674101666722 | 16.2632468982428 | 21.1748089517725 | 15.1432782183751 | 6.56548685980713 | 10.949581958191 | 18.3462334811079 | 13.5340151772298 | 23.8815624227072 | Eaf2 |
| **ENSMUSG00000022840** | 0.146708755318865 | 0.532521134889222 | 0.909082872831377 | 2173.85400206512 | 2739.6945120678 | 2268.65236809033 | 2050.30775364834 | 1490.23810450979 | 1525.1835433961 | 2764.06233042654 | 2559.66928148834 | Adcy5 |
| **ENSMUSG00000022841** | -0.00438162593901388 | 0.961696783485145 | 0.993822507448007 | 3690.85353218455 | 3462.08126361481 | 3469.70362178521 | 3639.15557372167 | 3512.62589218766 | 3999.47889888152 | 3467.83111964249 | 3327.13585570898 | Ap2m1 |
| **ENSMUSG00000022842** | 0.103694719282531 | 0.575105536444964 | 0.922450244597442 | 146.369222084185 | 106.688460487777 | 124.932045301595 | 162.261318106662 | 139.059690869025 | 118.638976511164 | 128.052605138405 | 116.15123541953 | Ece2 |
| **ENSMUSG00000022843** | 0.235934510693744 | 0.341331496901244 | 0.859283395807012 | 36.1405486627618 | 42.3496179035451 | 44.483379766477 | 53.4618215727152 | 40.5134532453066 | 37.9155491942896 | 32.2734208072402 | 39.0789203280663 | Clcn2 |
| **ENSMUSG00000022844** | -0.549695265707654 | 0.0855033483675593 | 0.744136283725187 | 40.658117245607 | 54.56585383726 | 58.6802030962037 | 55.3376749612315 | 85.4067392738896 | 101.51582526213 | 36.4377331694648 | 83.5854684794751 | Pdia5 |
| **ENSMUSG00000022848** | 0.0401805095795055 | 0.672291286294223 | 0.943216235131458 | 1502.54331065432 | 1566.93586243117 | 1679.95742735099 | 1525.06880486377 | 1572.35996919622 | 1388.19833340383 | 1646.98553925981 | 1492.5976514192 | Dirc2 |
| **ENSMUSG00000022849** | 0.179057681277408 | 0.51571700800113 | 0.904239892322932 | 45.1756858284523 | 39.9063707168021 | 33.1259211026956 | 55.3376749612315 | 39.4184950494875 | 28.1308913376988 | 38.5198893505771 | 46.6775992807458 | Hspbap1 |
| **ENSMUSG00000022853** | 0.210184446346572 | 0.471819312689054 | 0.897171173241185 | 402.063603873225 | 509.82424630037 | 713.626986040929 | 657.486612674971 | 452.217734873287 | 272.747337752471 | 686.070461676494 | 561.216716933618 | Ehhadh |
| **ENSMUSG00000022855** | 0.17463838081645 | 0.387843819753683 | 0.875951433019341 | 524.941469326615 | 542.400875456943 | 556.515474525287 | 483.032247542953 | 401.849657865609 | 322.893709267499 | 564.264325081426 | 576.414074838978 | Senp2 |
| **ENSMUSG00000022856** | -0.192467210544543 | 0.198899214899608 | 0.812921911423739 | 257.501409222178 | 226.407572638183 | 191.183887506986 | 231.667893481766 | 294.543754675337 | 247.06261087892 | 263.392756910703 | 231.216945274392 | Tmem41a |
| **ENSMUSG00000022858** | 0.178997823333817 | 0.203733762809634 | 0.816074141171399 | 834.846674109798 | 829.889627763701 | 714.573440929577 | 793.485983342404 | 646.025335533267 | 750.97249049335 | 585.085886892549 | 821.74285245406 | Tra2b |
| **ENSMUSG00000022861** | 0.299685439163145 | 0.539801559489122 | 0.913837378403038 | 9.03513716569045 | 9.77298874697194 | 17.9826428843205 | 21.5723139679377 | 16.4243729372865 | 6.1154111603693 | 13.5340151772298 | 10.8552556466851 | Dgkg |
| **ENSMUSG00000022863** | 0.184285522377518 | 0.391051114393113 | 0.877896341705281 | 93.9654265231807 | 71.6685841444609 | 92.7525790875478 | 113.489130005238 | 93.0714466446232 | 91.7311674055395 | 71.8343882483734 | 70.559161703453 | Btg3 |
| **ENSMUSG00000022864** | -0.0738710147573467 | 0.789181240654968 | 0.961659329450003 | 26.2018977805023 | 40.7207864457164 | 31.2330113253987 | 32.8274342990356 | 42.7033696369448 | 30.5770558018465 | 33.3144988977964 | 31.4802413753867 | D16Ertd472e |
| **ENSMUSG00000022865** | 0.0141594181195868 | 0.952429490549102 | 0.992000114323903 | 516.809845877494 | 582.307246173745 | 463.762895437739 | 463.335786963532 | 455.502609460744 | 302.101311322243 | 645.468416144805 | 602.466688391022 | Cxadr |
| **ENSMUSG00000022867** | 0.145104296422875 | 0.510182973240111 | 0.902394714365738 | 1368.8232806021 | 1658.15042406957 | 1607.08040092506 | 1388.13150750208 | 1193.50443344282 | 878.173042629031 | 1693.83405333483 | 1679.30804854218 | Usp25 |

| | | | | | | | | | | | | |
|---|---|---|---|---|---|---|---|---|---|---|---|---|
| **ENSMUSG00000022868** | -0.425803342325764 | 0.151105058548625 | 0.783412422888014 | 887.250469670802 | 1453.73207611208 | 697.537252933905 | 909.788893430416 | 834.358145214152 | 1132.57414690039 | 1409.61973461301 | 1927.89340285127 | Ahsg |
| **ENSMUSG00000022871** | -0.522177652381442 | 0.197917993546083 | 0.812921911423739 | 78.6056933415069 | 209.304842330982 | 61.519567762149 | 91.9168160372998 | 105.115986798633 | 190.800828203522 | 117.641824232843 | 221.447215192376 | Fetub |
| **ENSMUSG00000022875** | -0.241512539754174 | 0.426718838853321 | 0.885066977635295 | 227.685456575399 | 457.701639649853 | 258.382184601026 | 317.019222659259 | 259.505092409126 | 288.647406769431 | 371.664878328541 | 570.986447015635 | Kng1 |
| **ENSMUSG00000022877** | -0.338967313406276 | 0.347249403279745 | 0.861266047293794 | 110.228673421423 | 221.521078264697 | 92.7525790875478 | 166.013024883695 | 129.205067106653 | 149.216032313011 | 154.079557402308 | 314.802413753867 | Hrg |
| **ENSMUSG00000022881** | 0.157062325648896 | 0.407439682557417 | 0.881573350402261 | 184.316798180085 | 129.492100897378 | 164.68315062483 | 162.261318106662 | 153.294147414674 | 161.446854633749 | 109.313199508394 | 150.888053488923 | Rfc4 |
| **ENSMUSG00000022883** | -1.12328098420693 | 0.0644381727963714 | 0.716991331547963 | 3.61405486627618 | 9.77298874697194 | 7.57163910918757 | 17.8206071909051 | 17.5193311331055 | 42.8078781225851 | 10.4107809055614 | 14.1118323406906 | Robo1 |
| **ENSMUSG00000022884** | 0.265144009175915 | 0.101714809204128 | 0.759800953500741 | 10515.9961471471 | 10355.2959931457 | 12206.428698899 | 12035.4753407207 | 8685.20840923708 | 7184.38503120185 | 10763.7063782599 | 10904.1042970952 | Eif4a2 |
| **ENSMUSG00000022885** | -0.247256846736039 | 0.1262021456021 | 0.771949105062466 | 149.079763233892 | 144.151584017836 | 162.790240847533 | 191.337045628665 | 224.466430142915 | 176.123841418636 | 182.188665847324 | 184.539345993646 | St6gal1 |
| **ENSMUSG00000022887** | -0.0599282862821518 | 0.841640892966856 | 0.970640319105748 | 409.291713605777 | 359.971752180133 | 176.98706417726 | 298.260688774095 | 262.789966996583 | 373.040080782527 | 438.293876124134 | 223.618266321713 | Masp1 |
| **ENSMUSG00000022889** | -0.0259042033093333 | 0.759973572483549 | 0.957357521291862 | 1495.31520092177 | 1566.93586243117 | 1400.7532351997 | 1447.22088924034 | 1575.64484378368 | 1404.09840242079 | 1521.01509030252 | 1516.4792138419 | Mrpl39 |
| **ENSMUSG00000022890** | 0.028188539901016 | 0.874297061908047 | 0.97671366425653 | 24370.4754770169 | 18290.148439958 | 19524.4178979288 | 24219.1430991342 | 20993.6334884395 | 27301.6415843527 | 18427.0822028436 | 18011.0401689799 | Atp5j |
| **ENSMUSG00000022892** | -0.528498021868373 | 0.0426411478078627 | 0.687730553581215 | 4020.63603873225 | 4153.52021746308 | 4139.79368294831 | 3988.0643039857 | 7515.79305610228 | 8203.21253051938 | 3958.17890029443 | 3838.41839666784 | App |
| **ENSMUSG00000022893** | 0.0921580426913103 | 0.725547015637973 | 0.951727359509341 | 573.731210021344 | 759.035459348154 | 909.543147991157 | 620.907471598903 | 870.491765676182 | 853.711397987554 | 437.252798033577 | 525.394373299558 | Adamts1 |
| **ENSMUSG00000022894** | -0.619412584643682 | 0.121014636203905 | 0.769868040577702 | 254.790868072471 | 207.676010873154 | 322.74111702912 | 427.694572581722 | 659.164833883096 | 725.287763619799 | 245.694429371248 | 233.387996403729 | Adamts5 |
| **ENSMUSG00000022895** | -0.306252956935224 | 0.1033523258681 | 0.759800953500741 | 1524.22763985198 | 1557.97728941311 | 1841.80121330988 | 1856.1569279369 | 2734.11061496029 | 2256.58671817627 | 1633.45152408258 | 1759.63694032765 | Ets2 |
| **ENSMUSG00000022897** | 0.158429426657085 | 0.471946069805372 | 0.897171173241185 | 1440.20086421106 | 1828.36331141267 | 1850.31930730771 | 1493.17929725899 | 1374.17253575297 | 947.888729857241 | 1792.73647193767 | 1808.48559073773 | Dyrk1a |
| **ENSMUSG00000022898** | 0.105499147737684 | 0.270691603482366 | 0.844129257012535 | 545.722284807703 | 554.617111390658 | 598.159489625818 | 543.997482669733 | 519.010184818252 | 495.348303989913 | 513.251498644175 | 555.789089110276 | Dscr3 |
| **ENSMUSG00000022899** | 0.142961179961202 | 0.798810155151519 | 0.963240258978423 | 36.1405486627618 | 22.8036404096012 | 20.8220075502658 | 89.1030359545253 | 71.1722827282413 | 33.0232202659942 | 32.2734208072402 | 16.2828834700276 | Slc15a2 |
| **ENSMUSG00000022901** | 0.130824838331978 | 0.668592201613699 | 0.942055056494206 | 44.2721721118832 | 39.9063707168021 | 39.7511053232348 | 27.1998741334867 | 40.5134532453066 | 26.9078091056249 | 24.9858741733473 | 45.5920737160773 | Cd86 |
| **ENSMUSG00000022905** | 0.148822155154999 | 0.514816111896614 | 0.904145806837306 | 2474.72406968261 | 2972.6174105373 | 2762.70181996482 | 2263.21711324494 | 2059.61636633572 | 1498.27573429048 | 2924.38835637219 | 2963.48479154503 | Kpna1 |
| **ENSMUSG00000022906** | -0.08550786370043 | 0.610204197526115 | 0.927932112008768 | 249.369785773056 | 303.777066885045 | 267.84673348751 | 262.619474392285 | 303.30342024189 | 353.470765069345 | 229.03717992235 | 265.953763343784 | Parp9 |
| **ENSMUSG00000022911** | -0.00038871147518804 | 0.998200717819132 | 0.999262355603273 | 140.044626068202 | 165.326392969609 | 145.754052851861 | 121.930470253561 | 158.768938393769 | 138.208292224346 | 157.202791673977 | 119.407812113536 | Arl13b |
| **ENSMUSG00000022912** | -0.489872883264111 | 0.119348941306512 | 0.769868040577702 | 337.010616280254 | 377.074482487334 | 377.63550057073 | 364.853484066425 | 709.532910890775 | 735.07242147639 | 295.666177717943 | 306.118209236519 | Pros1 |
| **ENSMUSG00000022913** | -0.00724245342678217 | 0.962180245425772 | 0.994128583764339 | 527.652010476322 | 430.011504866766 | 437.262158555582 | 560.88016316638 | 526.674892188986 | 557.72549782568 | 422.677704765791 | 459.177313854779 | Psmg1 |
| **ENSMUSG00000022914** | 0.196772719290543 | 0.360707998483644 | 0.868539734006824 | 917.96993603415 | 1001.73134656462 | 919.95415176629 | 815.058297310342 | 602.227007700503 | 587.079471395453 | 1015.05113829223 | 983.486161589668 | Brwd1 |
| **ENSMUSG00000022940** | 0.166254912313973 | 0.324826059767587 | 0.855977678491063 | 1740.16741811198 | 1147.51176204029 | 1372.35958854025 | 1670.44744247379 | 1139.85148184768 | 1444.46011607923 | 1253.45802102959 | 1447.00557770312 | Pigp |
| **ENSMUSG00000022941** | -0.441717401085321 | 0.176358745489836 | 0.799889021902745 | 118.360296870545 | 63.5244268553176 | 121.146225747001 | 127.55803041911 | 213.516848184724 | 167.562265794119 | 116.600746142287 | 86.8420451734806 | Ripply3 |
| **ENSMUSG00000022946** | -0.00473679744578229 | 0.974317101902178 | 0.995510898988495 | 251.176813206195 | 312.735639903102 | 309.490748588042 | 240.109233730089 | 295.638712871156 | 248.285693110993 | 281.091084450157 | 292.006376895829 | Dopey2 |
| **ENSMUSG00000022947** | -0.678891323574663 | 0.0409449821083528 | 0.683226273624098 | 64.1494738764022 | 47.2361122770311 | 41.6440151005317 | 56.2756016554897 | 102.926070406995 | 117.415894279091 | 41.6431236222455 | 73.8157383974585 | Cbr3 |
| **ENSMUSG00000022948** | 0.219576407509763 | 0.626863299204825 | 0.931825800800958 | 11.7456783153976 | 19.5459774939439 | 13.2503684410783 | 19.6964605794214 | 9.85462376237187 | 15.9000690169602 | 21.8626399016789 | 7.59867895267955 | Setd4 |
| **ENSMUSG00000022951** | 0.547451533312269 | 0.0412163564991829 | 0.683365309556438 | 3423.41347208011 | 3749.57001592157 | 4404.80105176987 | 6630.20380171094 | 2357.44499559852 | 2669.98851261724 | 2924.38835637219 | 4506.01661893897 | Rcan1 |
| **ENSMUSG00000022952** | 0.198729383621513 | 0.593670852166031 | 0.923935529355527 | 17.1667606148119 | 26.8757190541728 | 23.6613722162112 | 17.8206071909051 | 24.0890803080201 | 23.2385624094033 | 11.4518589961175 | 16.2828834700276 | Runx1 |
| **ENSMUSG00000022956** | 0.0388083514439791 | 0.815876890617998 | 0.966177091475091 | 34263.0471597313 | 23710.0851158828 | 26753.4403374256 | 32651.1040805151 | 28512.7114191293 | 34098.3095479871 | 24194.6548245246 | 27457.2836327252 | Atp5o |
| **ENSMUSG00000022957** | 0.117897745262444 | 0.602430080843809 | 0.926448151609826 | 1526.03466728512 | 1686.65497458157 | 1603.29458137047 | 1362.80748675711 | 1081.81869746927 | 993.142772443974 | 1831.25636128824 | 1786.77507944436 | Itsn1 |
| **ENSMUSG00000022960** | -0.00630075752774952 | 0.981356515977945 | 0.996940244108351 | 49.6932544112975 | 33.3910448854875 | 61.519567762149 | 45.0204813243917 | 56.937826182593 | 51.3694537471021 | 40.6020455316893 | 41.2499714574033 | Donson |
| **ENSMUSG00000022961** | -0.0817689774746393 | 0.413429165939031 | 0.882192126163777 | 2533.4524612596 | 2681.05657958597 | 3051.37056100259 | 2784.70435525248 | 3137.05523102171 | 2985.54372849229 | 2872.33445184438 | 2699.70207933058 | Son |
| **ENSMUSG00000022962** | 0.175159323472795 | 0.213085454486462 | 0.824215250018647 | 1020.97049972302 | 1094.57473966086 | 1128.17422726895 | 1039.22277723804 | 950.423713970976 | 737.518585940537 | 1074.39258945393 | 1029.07823530575 | Gart |
| **ENSMUSG00000022964** | -0.103487220934029 | 0.400138787372286 | 0.880487015130533 | 919.776963467288 | 833.147290679358 | 866.952678001977 | 846.94780491512 | 879.251431242735 | 796.226533080083 | 1072.31043327282 | 974.80195707232 | Tmem50b |
| **ENSMUSG00000022965** | 0.139521158258347 | 0.526988781401659 | 0.908217276872524 | 4096.53119092405 | 4013.44071208981 | 3237.82217406634 | 3491.90108272313 | 2885.21484598332 | 2246.80206031968 | 3841.57815415214 | 4497.33241442163 | Ifngr2 |
| **ENSMUSG00000022967** | -0.192782659448789 | 0.181053472781752 | 0.803056849297055 | 242.141676040504 | 259.798617523671 | 315.169477919933 | 293.571055302804 | 337.247124312282 | 275.193502216618 | 310.241270985729 | 345.197129564585 | Ifnar1 |
| **ENSMUSG00000022969** | 0.138972486889937 | 0.248509209106209 | 0.834621660998471 | 3018.63932705718 | 2453.83459121887 | 2658.59178221349 | 3157.06125287297 | 2677.17278877769 | 2323.85624094033 | 2511.0803544214 | 2737.69547409398 | Il10rb |
| **ENSMUSG00000022971** | -0.339789883501195 | 0.0939002300570833 | 0.756539556550399 | 561.082017989377 | 474.804369957054 | 510.139184981513 | 626.535031764452 | 744.571573156986 | 912.419345127099 | 524.703357640293 | 568.815395886298 | Ifnar2 |
| **ENSMUSG00000022972** | -0.0694430084138414 | 0.651277290280823 | 0.936463929813731 | 667.696636544524 | 574.977504613516 | 575.444572298256 | 695.003680445298 | 630.6959207918 | 796.226533080083 | 672.536446499264 | 538.42068007558 | 1110004E09F |
| **ENSMUSG00000022973** | 0.0808943869631853 | 0.505480270182724 | 0.901494649536949 | 333.396561413978 | 408.022280186079 | 350.188308799925 | 331.088123073131 | 321.917709570815 | 342.463024980681 | 352.92547269853 | 328.914246094558 | Synj1 |
| **ENSMUSG00000022974** | 0.110991223752557 | 0.608264571740118 | 0.927932112008768 | 110.228673421423 | 136.821842457607 | 139.128868631322 | 120.992543559303 | 110.590777777729 | 88.0619207093179 | 160.326025945645 | 109.638082031519 | Paxbp1 |
| **ENSMUSG00000022978** | -0.587194610704846 | 0.0655510508893666 | 0.718339708336358 | 45.1756858284523 | 31.7622134276588 | 26.5007368821565 | 48.7721881014244 | 56.937826182593 | 83.1695917810225 | 48.9306702561384 | 40.1644458927348 | Mis18a |
| **ENSMUSG00000022982** | 0.0189700819273245 | 0.924577841231587 | 0.986713918026868 | 9860.04518891799 | 8057.82922187837 | 9287.56182230721 | 10970.9285427377 | 9737.46323541923 | 12806.8940520454 | 7449.95481601971 | 7684.43547228837 | Sod1 |
| **ENSMUSG00000022983** | 0.24325623064913 | 0.180842972679789 | 0.802808628271642 | 489.704434380422 | 625.471279806204 | 527.175372977185 | 364.853484066425 | 430.318570956905 | 373.040080782527 | 450.786813210807 | 441.808904820083 | Scaf4 |
| **ENSMUSG00000022987** | 0.18608905812421 | 0.348513317308892 | 0.861266047293794 | 194.255449062345 | 232.922898469498 | 190.237432618338 | 200.716312571247 | 137.964732673206 | 147.992950080937 | 214.462086654564 | 218.19063849837 | Zfp641 |
| **ENSMUSG00000022992** | 0.084938436598564 | 0.559675796125399 | 0.916574868225118 | 845.688838708626 | 928.433930962335 | 984.313084194385 | 943.55425442371 | 763.185862485911 | 747.303243797128 | 1058.77641809559 | 919.440153274226 | Kansl2 |
| **ENSMUSG00000022993** | 0.404389778551791 | 0.420176118393271 | 0.882568531013176 | 8.13162344912141 | 12.2162359337149 | 17.9826428843205 | 14.0689004138724 | 5.47479097909548 | 6.1154111603693 | 13.5340151772298 | 14.1118323406906 | 4930415O20 |
| **ENSMUSG00000022994** | 0.206357831848588 | 0.512125784090596 | 0.902394714365738 | 3260.78100309768 | 4462.18377872161 | 5126.94613180864 | 3276.17794304376 | 2714.40136743554 | 1780.80772989954 | 4886.8205570705 | 4593.94418967712 | Adcy6 |
| **ENSMUSG00000022995** | 0.398589687101306 | 0.0930194169329103 | 0.754058637722175 | 4522.98966514464 | 5650.41632720761 | 4524.05436773958 | 4876.28088344818 | 2662.93833223204 | 2619.84214110221 | 4896.19025988551 | 4668.84545363925 | Enah |
| **ENSMUSG00000022999** | -0.424820787344639 | 0.0217290395868498 | 0.648762042745232 | 78.6056933415069 | 83.0704043492615 | 100.324218196735 | 106.92364314543 | 125.920192519196 | 112.523565350795 | 143.668776496747 | 111.809133160856 | Lmbr1l |
| **ENSMUSG00000023000** | -0.63071542066228 | 0.145178417861265 | 0.780607990324772 | 13.5527057485357 | 9.77298874697194 | 12.3039135524298 | 7.50341355406529 | 22.994122112201 | 19.5693157131818 | 13.5340151772298 | 10.8552556466851 | Dhh |
| **ENSMUSG00000023004** | -0.614751443001357 | 0.0212995036499179 | 0.648762042745232 | 1212.51540763566 | 925.176268046677 | 926.579335986829 | 1192.10482840212 | 1726.74907480672 | 2457.17220423638 | 993.188498390554 | 1341.70959793028 | Tuba1b |
| **ENSMUSG00000023007** | 0.280747801888042 | 0.0434327718121081 | 0.688841110980169 | 267.440060104437 | 274.458100644129 | 245.131816159948 | 259.805694309511 | 188.332809680885 | 194.470074899744 | 239.447960827911 | 238.815624227072 | Prpf40b |
| **ENSMUSG00000023008** | -0.315769320055045 | 0.0852744203715403 | 0.743541556158628 | 496.029030396406 | 420.238516119794 | 503.514000760974 | 474.59090729463 | 753.331238723539 | 634.779678446333 | 442.458188486358 | 527.565424428895 | Fmnl3 |
| **ENSMUSG00000023009** | -0.501377738656846 | 0.190053323882225 | 0.808824684419004 | 60.535419010126 | 66.7820897709749 | 49.2156542097192 | 70.3445020693621 | 124.825234323377 | 125.977469903608 | 63.5057635239243 | 35.8223436340608 | Nckap5l |
| **ENSMUSG00000023010** | -0.112063537977941 | 0.240909914024873 | 0.832048466795696 | 5506.91610248833 | 5783.16609102065 | 6168.99296421058 | 5653.8221129882 | 6729.61307150417 | 6645.00576685728 | 5798.80496439768 | 5807.56177097652 | Tmbim6 |
| **ENSMUSG00000023015** | -0.909845962915294 | 0.0246058394303193 | 0.655303666217042 | 19.877301764519 | 12.2162359337149 | 9.46454888648447 | 25.3240207449704 | 33.943704070392 | 44.0309603546589 | 22.903717992235 | 24.9670879873757 | Racgap1 |
| **ENSMUSG00000023017** | 0.06562807396538 | 0.720774957352975 | 0.951674101666722 | 126.491920319666 | 118.904696421492 | 96.5383986421416 | 115.364983393754 | 112.780694169367 | 85.6157562451702 | 124.929370866736 | 112.894658725525 | Asic1 |
| **ENSMUSG00000023018** | -0.144365307746863 | 0.298480156200713 | 0.851785577030345 | 380.379274675568 | 385.218639776477 | 461.869985660442 | 384.549944645846 | 388.710159515779 | 414.624876673038 | 510.128264372507 | 466.775992807458 | Smarcd1 |
| **ENSMUSG00000023019** | 0.33024175380494 | 0.195414423607334 | 0.811094945724481 | 2215.4156330273 | 1879.67150233427 | 3025.81627900908 | 2208.81736497797 | 1567.98013641295 | 1161.92812047017 | 2413.21901390912 | 2276.34710910986 | Gpd1 |
| **ENSMUSG00000023020** | -0.0136616854820561 | 0.930330317630676 | 0.987529878729394 | 3953.77602370614 | 2987.27689365776 | 3055.15638055719 | 3905.52675489098 | 3295.82416941548 | 4231.86452297555 | 3319.99803078352 | 3187.10305786674 | Cox14 |
| **ENSMUSG00000023021** | -0.179943768999365 | 0.363598608235818 | 0.86924083181804 | 333.396561413978 | 320.879797192245 | 344.509579468035 | 337.653609932938 | 462.072358635659 | 453.763508099402 | 338.350379430744 | 260.526135520442 | Cers5 |
| **ENSMUSG00000023022** | -0.12449554113512 | 0.495727812150916 | 0.900927059989498 | 398.449549006949 | 451.186313818538 | 482.691993210708 | 543.997482669733 | 618.65138063779 | 573.62556684264 | 395.609674411332 | 458.09178829011 | Lima1 |
| **ENSMUSG00000023025** | 0.213984012400846 | 0.319491149778767 | 0.854670142081901 | 739.977733870048 | 968.340301679137 | 816.79056890361 | 702.507093999363 | 629.600962595981 | 469.663577116362 | 810.99983254323 | 871.677028428812 | Larp4 |
| **ENSMUSG00000023026** | -0.0954378428224963 | 0.572977361671985 | 0.922101211171807 | 412.002254755485 | 472.361122770311 | 441.047978110176 | 366.729337454941 | 406.229490648885 | 359.586176229715 | 500.758561557502 | 540.591731204917 | Dip2b |
| **ENSMUSG00000023027** | 0.249743955908277 | 0.275679413125368 | 0.846289746011912 | 1227.87514081733 | 1325.86880667253 | 1370.46667876295 | 1121.76032633276 | 927.429591858775 | 670.249063176475 | 1201.40411650178 | 1443.74900100912 | Atf1 |
| **ENSMUSG00000023030** | 0.00959705341291358 | 0.961035010362507 | 0.993822507448007 | 483.379838364439 | 606.739718041175 | 468.495169880981 | 394.867138282686 | 439.078236523458 | 374.263163014601 | 581.96265262088 | 544.933833463591 | Slc11a2 |
| **ENSMUSG00000023031** | -0.161431559706568 | 0.703936553005442 | 0.946931219685582 | 16.2632468982428 | 13.0306516626293 | 17.036187995672 | 15.9447538023887 | 12.0445401540101 | 26.9078091056249 | 9.36970281500523 | 21.7105112933701 | Cela1 |
| **ENSMUSG00000023032** | -0.312584802722482 | 0.553835049921295 | 0.915567635607008 | 25.2983840639333 | 50.4937751926884 | 51.1085639870161 | 12.1930470253561 | 76.6470737073368 | 47.7002070508805 | 28.1091084450157 | 20.6249857287016 | Slc4a8 |
| **ENSMUSG00000023034** | -0.194775655264954 | 0.614377770449246 | 0.928075610917191 | 345.142239729375 | 460.95930256551 | 191.183887506986 | 192.274972322923 | 439.078236523458 | 480.671317205027 | 281.091084450157 | 161.743309135608 | Nr4a1 |
| **ENSMUSG00000023043** | 0.057471724155035 | 0.849251896499577 | 0.972640876584422 | 40.658117245607 | 69.2253369577179 | 35.965285768641 | 65.6548685980713 | 39.4184950494875 | 64.8233582999146 | 54.1360607089191 | 45.5920737160773 | Krt18 |
| **ENSMUSG00000023044** | -0.0287206358713643 | 0.858495793659137 | 0.974702188484994 | 211.422209677157 | 229.665235553841 | 177.933519065908 | 266.371181169318 | 222.276513751277 | 220.154801773295 | 213.421008564008 | 247.49982874442 | Csad |
| **ENSMUSG00000023046** | -1.11101850336785 | 0.027316900175959 | 0.664353746352133 | 199.676531361759 | 103.43079757212 | 166.576060402127 | 197.902532488472 | 508.060602860061 | 673.918309872697 | 133.257995591185 | 127.006491066215 | Igfbp6 |
| **ENSMUSG00000023048** | -0.35092246267659 | 0.150288886198746 | 0.782641594375979 | 583.669860903603 | 443.856572258309 | 589.641395627982 | 691.251973668265 | 792.749733773026 | 1040.84297949485 | 597.578823979222 | 513.453592088204 | Prr13 |
| **ENSMUSG00000023050** | -0.188895970897359 | 0.532949016088258 | 0.909082872831377 | 75.8951521917998 | 49.679359463774 | 60.5731128735006 | 69.4065753751039 | 106.210944994452 | 84.3926740130963 | 53.094982618363 | 47.7631248454143 | Map3k12 |
| **ENSMUSG00000023051** | -0.00044261572084957 | 0.997310556629769 | 0.999262355603273 | 899.899661702769 | 686.552459474779 | 698.483707822554 | 875.085605742864 | 792.749733773026 | 829.249753346077 | 773.52102128321 | 765.295523091298 | Tarbp2 |
| **ENSMUSG00000023055** | 0.234591878673822 | 0.080523146281504 | 0.739855948845396 | 4722.6661965064 | 5141.40649663616 | 5857.60930584524 | 4743.09529286352 | 4279.09662926103 | 3670.46977845365 | 5002.38022512224 | 4439.7995594942 | Calcoco1 |
| **ENSMUSG00000023057** | 0.0509217107999423 | 0.879409604691714 | 0.978393501240173 | 64.1494738764022 | 38.2775392589734 | 67.1982970940397 | 57.2135283497478 | 37.2285786578493 | 66.0464405319884 | 35.3966550789086 | 80.3288917854696 | Fabp2 |
| **ENSMUSG00000023064** | -0.383394314369664 | 0.265108491957919 | 0.842510961779757 | 104.807591122009 | 101.801966114291 | 72.8770264259304 | 108.799496533947 | 163.148771177045 | 194.470074899744 | 75.998700610598 | 73.8157383974585 | Sncg |
| **ENSMUSG00000023066** | 0.141278987921851 | 0.560981421831665 | 0.916762264997884 | 39.754603529038 | 49.679359463774 | 42.5904699891801 | 47.8342614071662 | 43.7983278327639 | 30.5770558018465 | 41.6431236222455 | 46.6775992807458 | Rttn |
| **ENSMUSG00000023067** | -0.218401131500072 | 0.315511990928999 | 0.853277176422981 | 3769.45922552606 | 3498.72997141596 | 4811.7766538887 | 2976.04140088115 | 4780.58748294618 | 5445.16209719282 | 4269.46124937072 | 3022.10317203713 | Cdkn1a |
| **ENSMUSG00000023068** | 0.173380194806726 | 0.278657789971272 | 0.848075053184273 | 2056.39721891115 | 2174.48999620126 | 2025.41346170768 | 1983.71495835601 | 1585.49946754605 | 1437.12162268679 | 2146.70302272675 | 2136.31431126762 | Nus1 |
| **ENSMUSG00000023070** | -0.383313787261802 | 0.275699143319216 | 0.846289746011912 | 152.693818100169 | 263.870696168242 | 167.522515290775 | 252.302280755445 | 222.276513751277 | 253.178022039289 | 144.709854587303 | 471.118095066132 | Rgn |

| | | | | | | | | | | | | |
|---|---|---|---|---|---|---|---|---|---|---|---|---|
| ENSMUSG00000023072 | 0.191873532942223 | 0.133150814475634 | 0.77303252008843 | 408.388199889208 | 341.240190415104 | 425.904699891801 | 453.95652002095 | 342.721915291377 | 353.470765069345 | 366.45948787576 | 362.565538599282 | Cep89 |
| ENSMUSG00000023073 | -0.156029065790974 | 0.757649024277227 | 0.95677573228595 | 12.6491920319666 | 14.6594831204579 | 5.67872933189068 | 9.37926694258161 | 10.949581958191 | 8.56157562451702 | 8.32862472444909 | 19.5394601640331 | Slc10a2 |
| ENSMUSG00000023074 | 0.314225881343908 | 0.0831195493958006 | 0.741509660313212 | 1078.79537758344 | 1153.21267214269 | 1414.95005852943 | 1083.30533186818 | 912.100177117308 | 697.1568722821 | 1098.33738553672 | 1095.29529475052 | Mospd1 |
| ENSMUSG00000023075 | 0.0474730259490641 | 0.776838949866267 | 0.959738279197945 | 671.3106914108 | 677.593886456721 | 677.661700272288 | 584.328330522834 | 601.132049504684 | 468.440494884288 | 744.370834747638 | 711.019244857872 | Akirin1 |
| ENSMUSG00000023079 | 0.056853989514248 | 0.741223196864062 | 0.954383574136061 | 159.921927832721 | 172.656134529838 | 192.130342395635 | 163.19924480092 | 155.484063806312 | 129.646716599829 | 196.76375911511 | 178.026192605635 | Gtf2ird1 |
| ENSMUSG00000023087 | -0.108410083527028 | 0.754955852368784 | 0.956497786093635 | 919.776963467288 | 1211.85060462452 | 632.231865617162 | 349.846656958294 | 1033.64053685323 | 746.080161565054 | 985.900951756661 | 591.611432744337 | Noct |
| ENSMUSG00000023088 | 0.318007525632485 | 0.017416954114009 | 0.642273034804458 | 629.749060448624 | 777.767021113184 | 603.838218957709 | 679.996853337167 | 542.004306930453 | 467.217412652214 | 591.332355435886 | 557.960140239613 | Abcc1 |
| ENSMUSG00000023089 | 0.0351867638485344 | 0.8665005260269 | 0.974868041734905 | 14303.5256470046 | 10444.0673075974 | 11910.1883187521 | 14610.0841164594 | 12716.844486243 | 17167.1822093887 | 10322.2892678641 | 9827.262936944 | Ndufa5 |
| ENSMUSG00000023092 | 0.174755488522191 | 0.356113147408431 | 0.864965376583892 | 2290.40727150253 | 2720.14853457386 | 2244.04454098547 | 2050.30775364834 | 1675.28603960322 | 1692.74580919022 | 2729.70675343819 | 2144.99851578497 | Fhl1 |
| ENSMUSG00000023094 | 0.0898983364076952 | 0.41182197022927 | 0.882192126163777 | 1555.8506199319 | 1447.21675028076 | 1365.73440431971 | 1595.41330693313 | 1356.65320461986 | 1253.65928787571 | 1439.81099923914 | 1552.30155747597 | Msrb2 |
| ENSMUSG00000023104 | -0.0386202794263066 | 0.804674896822596 | 0.963461896828986 | 364.116027777325 | 326.580707294646 | 311.383658365339 | 415.501525556365 | 402.944616061428 | 404.840218816448 | 340.432535611857 | 308.289260365856 | Rfc2 |
| ENSMUSG00000023106 | 0.105927588449848 | 0.449958787023617 | 0.89108031640712 | 2218.12617417701 | 1756.69472726821 | 1939.28606684067 | 2242.58272597126 | 1825.29531243043 | 2262.70212933664 | 1754.21658258709 | 1737.92642903428 | Denr |
| ENSMUSG00000023110 | 0.101299648061305 | 0.309834142751144 | 0.851785577030345 | 918.873449750719 | 1005.8034252092 | 1104.51285505274 | 1022.3400967414 | 921.95480087968 | 886.734618253548 | 974.449092760544 | 992.170366107016 | Prmt5 |
| ENSMUSG00000023118 | 0.17098327773348 | 0.17138305319838 | 0.799531027639052 | 3393.59751943333 | 3795.99171246969 | 4044.20173919481 | 3663.54166777238 | 3070.26278107675 | 2865.68166974905 | 3790.56532771489 | 3504.07652274994 | Sympk |
| ENSMUSG00000023143 | -0.328109110593486 | 0.168616440863151 | 0.79907358295209 | 241.238162323935 | 250.025628776699 | 226.202718386979 | 291.695201914288 | 391.995034103237 | 412.178712208891 | 251.940897914585 | 211.677485110359 | Nagpa |
| ENSMUSG00000023147 | 0.173027367986578 | 0.237827087782362 | 0.832048466795696 | 280.992765852973 | 315.99330281876 | 318.008842585878 | 318.895076047775 | 269.359716171498 | 221.377884005369 | 281.091084450157 | 321.315567141878 | Wrb |
| ENSMUSG00000023150 | 0.318980425292027 | 0.254949432982773 | 0.837406774112203 | 34736.4883472135 | 39397.3608862307 | 34187.8434877592 | 37991.6586776211 | 25126.0057194608 | 14712.4561696165 | 38761.4194675861 | 38689.2166503503 | Ivns1abp |
| ENSMUSG00000023153 | -0.223809499141732 | 0.524286782452223 | 0.907437243979788 | 52.4037955610046 | 25.2468875963442 | 45.4298346551254 | 50.6480414899407 | 76.6470737073368 | 53.8156182112498 | 38.5198893505771 | 33.6512925047237 | Tmem52 |
| ENSMUSG00000023156 | 0.123745917238878 | 0.452583217612501 | 0.891110666112841 | 927.00507319984 | 846.992358070902 | 867.899132890626 | 915.416453595965 | 684.348872386936 | 655.572076391589 | 942.175671953304 | 981.315110460331 | Rpp14 |
| ENSMUSG00000023169 | -0.0309483676498536 | 0.928520299715099 | 0.986901071486195 | 103.90407740544 | 105.874044758863 | 60.5731128735006 | 118.178763476528 | 76.6470737073368 | 162.669936865823 | 100.984574783945 | 57.5328549274309 | Slc38a1 |
| ENSMUSG00000023170 | 0.0771901726825376 | 0.560841053443566 | 0.916762264997884 | 681.24934229306 | 653.975830318206 | 636.017685171756 | 617.15576482187 | 646.025335533267 | 726.510845851873 | 542.401685179747 | 540.591731204917 | Gps2 |
| ENSMUSG00000023175 | 0.179966131876947 | 0.161862065362406 | 0.793318309062746 | 50571.4697438026 | 39530.9250657726 | 47133.4534546927 | 52515.4535382087 | 44980.8826842485 | 44942.156617554 | 37248.733002008 | 40326.1892018704 | Bsg |
| ENSMUSG00000023176 | -0.376867416156062 | 0.411521656435533 | 0.882192126163777 | 36.1405486627618 | 92.8433930962335 | 29.3401015481019 | 51.5859681841989 | 41.6084114411257 | 44.0309603546589 | 63.5057635239243 | 123.74991437221 | Cpn2 |
| ENSMUSG00000023186 | 0.0760904258954643 | 0.526724094158589 | 0.908217276872524 | 541.204716224858 | 597.781145023117 | 638.857049837702 | 631.224665235743 | 629.600962595981 | 521.033030863464 | 609.03068297534 | 524.308847734889 | Vwa5a |
| ENSMUSG00000023191 | -0.990123872924822 | 0.0424090565892505 | 0.685752963940267 | 98.4829951060259 | 78.9983257046899 | 90.8596693102509 | 112.551203310979 | 266.074841584041 | 340.016860516533 | 100.984574783945 | 49.9341759747513 | P3h3 |
| ENSMUSG00000023206 | 0.133847037525157 | 0.460721667702161 | 0.894774044236998 | 252.080326922764 | 210.933673788811 | 231.881447718869 | 172.578511743502 | 213.516848184724 | 154.108361241306 | 208.215618111227 | 213.848536239696 | Il15ra |
| ENSMUSG00000023216 | 2.25557376578358 | NA | NA | 6.32459601598331 | 4.07207864457164 | 80.448665535118 | 0 | 0 | 15.9000690169602 | 1.04107809055614 | 2.17105112933702 | Epb42 |
| ENSMUSG00000023224 | -1.05683258986252 | 0.0161576691887057 | 0.639401780715449 | 821.293968361262 | 733.78857175181 | 661.571967165264 | 763.472329126143 | 1913.98692629178 | 2997.77455081303 | 627.77008860535 | 661.085068883121 | Serping1 |
| ENSMUSG00000023232 | 0.134555382361758 | 0.690509138433302 | 0.94469184542218 | 23.4913566307952 | 32.5766291565731 | 30.2865564367503 | 19.6964605794214 | 22.994122112201 | 28.1308913376988 | 15.616171358342 | 30.3947158107182 | Serinc2 |
| ENSMUSG00000023235 | 0.20548168619529 | 0.599382273166486 | 0.924951380294376 | 17.1667606148119 | 30.9477976987445 | 17.9826428843205 | 30.0136542162612 | 18.6142893289246 | 17.123151249034 | 32.2734208072402 | 15.1973579053591 | Ccl25 |
| ENSMUSG00000023249 | 0.0244611911509642 | 0.89938291775883 | 0.984802943363903 | 929.715614349547 | 915.403279299705 | 662.518422053913 | 600.273084325223 | 669.019457645468 | 695.933790050026 | 923.436266323293 | 767.466574220635 | Parp3 |
| ENSMUSG00000023259 | -0.20467319392459 | 0.227577665193945 | 0.832048466795696 | 134.623543768788 | 169.39847161418 | 140.07532351997 | 164.137171495178 | 154.389105610493 | 159.000690169602 | 211.338852382896 | 175.855141476298 | Slc26a6 |
| ENSMUSG00000023262 | -0.0973763536845481 | 0.581724817882386 | 0.922450244597442 | 173.474633581257 | 201.160685041839 | 192.130342395635 | 186.647412157374 | 257.315176017488 | 187.131581507301 | 201.96914956789 | 159.572258006271 | Acy1 |
| ENSMUSG00000023266 | 0.138043844516489 | 0.499245925386354 | 0.900927059989498 | 74.9916384752307 | 70.8541684155466 | 70.9841166486335 | 92.854742731558 | 63.5075753575076 | 74.6080161565054 | 82.2451691539348 | 60.7894316214364 | Frs3 |
| ENSMUSG00000023267 | 0.483530107963397 | 0.10584493960891 | 0.762094734041179 | 48.7897406947284 | 57.8235167529173 | 72.8770264259304 | 44.0825546301336 | 22.994122112201 | 48.9232892829544 | 44.7663578939139 | 43.4210225867403 | Gabrr2 |
| ENSMUSG00000023272 | -0.18110240308385 | 0.525014992756144 | 0.907989757056366 | 354.177376895066 | 365.672662282534 | 376.689045682082 | 377.984457786039 | 481.781606160403 | 647.010500767072 | 247.776585552361 | 296.348479154503 | Creld2 |
| ENSMUSG00000023277 | 0.324304564119613 | 0.0356335777388417 | 0.679566075569698 | 8861.66253210919 | 8789.98896217235 | 9337.72393140558 | 11452.0849368921 | 9471.38839383519 | 7318.92407672998 | 6667.0640919215 | 7244.79761859762 | Twf2 |
| ENSMUSG00000023284 | -0.0725725973478194 | 0.800658500034781 | 0.963461896828986 | 39.754603529038 | 29.3189662409158 | 40.6975602118832 | 34.703287687552 | 31.7537876787538 | 26.9078091056249 | 49.9717483466946 | 42.3354970220718 | Zfp605 |
| ENSMUSG00000023286 | -0.0433234451473162 | 0.672818566340994 | 0.943302739919217 | 1010.12833512419 | 911.331200655134 | 1000.40281730141 | 994.202295913651 | 1080.72373927345 | 1086.09702208159 | 942.175671953304 | 927.038832226905 | Ube2j2 |
| ENSMUSG00000023307 | 0.129002572859499 | 0.11732665374485 | 0.769868040577702 | 2182.88913923081 | 2109.33673788811 | 2240.25872143087 | 2243.52065266552 | 1978.58945984511 | 1901.89287087485 | 2185.22291207733 | 1957.20259309732 | March5 |
| ENSMUSG00000023328 | -1.27593238414025 | 0.0589915652730864 | 0.712165592539891 | 16.2632468982428 | 17.1027303072009 | 16.0897331070236 | 12.1930470253561 | 21.8991639163819 | 102.738907494204 | 16.6572494488982 | 8.68420451734806 | Ache |
| ENSMUSG00000023330 | -0.0136379820138318 | 0.952635620256273 | 0.992000114323903 | 83.1232619243521 | 74.1118313312039 | 83.2880302010633 | 97.5443762028488 | 71.1722827282413 | 118.638976511164 | 68.711153976705 | 83.5854684794751 | Dtwd1 |
| ENSMUSG00000023336 | -0.015352383344785 | 0.957191894195757 | 0.992403427270738 | 311.712232216321 | 216.634583891211 | 113.574586637814 | 171.640585049243 | 176.288269526875 | 211.593226148778 | 215.50316474512 | 219.276164063039 | Wfdc1 |
| ENSMUSG00000023341 | -0.0968073407862431 | 0.869850982893355 | 0.975613767803763 | 18.9737880479499 | 13.0306516626293 | 3.78581955459379 | 4.68963347129081 | 9.85462376237187 | 14.6769867848863 | 9.36970281500523 | 9.76973008201657 | Mx2 |
| ENSMUSG00000023345 | -0.174934047257848 | 0.395801275426672 | 0.879215782741314 | 139.141112351633 | 89.5857301805761 | 104.110037751329 | 114.427056699496 | 114.970610561005 | 151.662196777159 | 105.14888714617 | 133.519644454226 | Poc1a |
| ENSMUSG00000023348 | -0.312685627541856 | 0.0531213066130254 | 0.703396849069134 | 234.913566307952 | 260.613033252585 | 218.631079277791 | 241.985087118606 | 350.386622662111 | 331.455284892016 | 246.735507461804 | 260.526135520442 | Trip6 |
| ENSMUSG00000023349 | -0.445075594526187 | 0.120674670417809 | 0.769868040577702 | 27.1054114970714 | 39.0919549878878 | 20.8220075502658 | 37.5170677703264 | 42.7033696369448 | 48.9232892829544 | 36.4377331694648 | 42.3354970220718 | Clec4n |
| ENSMUSG00000023353 | 0.0733290289997882 | 0.436814617374399 | 0.888701139843674 | 1553.14007878219 | 1585.6674241962 | 1768.92418688395 | 1698.58524330153 | 1640.24737733701 | 1657.27642446008 | 1533.50802738919 | 1448.09110326779 | Agap3 |
| ENSMUSG00000023367 | -0.744194596853308 | 0.0240544902125316 | 0.654124840968651 | 110.228673421423 | 111.574954861263 | 109.78876708322 | 129.433883807626 | 181.76306050597 | 342.463024980681 | 114.518589961175 | 134.605170018895 | Tmem176a |
| ENSMUSG00000023411 | -0.464128494560679 | 0.167851144522987 | 0.798989855183099 | 57.8248778604189 | 40.7207864457164 | 44.483379766477 | 68.4686486808458 | 84.3117810780705 | 111.300483118721 | 49.9717483466946 | 46.6775992807458 | Nfatc4 |
| ENSMUSG00000023439 | 0.618986651378524 | 0.0274063856486517 | 0.664353746352133 | 324.361424248287 | 197.903022126182 | 252.703455269135 | 374.232751009006 | 165.338687568684 | 116.192812047017 | 230.078258012906 | 235.559047533066 | Gnb3 |
| ENSMUSG00000023452 | 0.121318286388864 | 0.334463797239197 | 0.85842786809952 | 466.213077749627 | 445.485403716138 | 399.403963009645 | 456.770300103725 | 395.279908690694 | 348.57843614105 | 436.211719943021 | 443.97995594942 | Pisd |
| ENSMUSG00000023456 | 0.0262770819533915 | 0.667054058202125 | 0.942024383071524 | 21144.9315088654 | 19919.7943135156 | 19747.7812516498 | 20031.3004092715 | 20388.1216061516 | 19816.3783240607 | 19902.2898571617 | 19277.848502948 | Tpi1 |
| ENSMUSG00000023460 | 0.267759179260557 | 0.0705696221122174 | 0.72606739345142 | 5943.31322759118 | 6259.59929243553 | 7344.48993591195 | 6595.50051402339 | 5098.12535973372 | 4363.95740403953 | 6350.57635239243 | 5899.83144397334 | Rab12 |
| ENSMUSG00000023473 | 0.595009004745442 | 0.189068690385341 | 0.808677608936066 | 51.5002818444356 | 25.2468875963442 | 30.2865564367503 | 30.9515809105193 | 14.2344565456483 | 9.78465785659088 | 38.5198893505771 | 28.2236646813812 | Celsr3 |
| ENSMUSG00000023495 | 0.0469462745582509 | 0.715676341055238 | 0.951176382858794 | 1918.15962027608 | 1592.99716575643 | 1413.05714875213 | 1673.26122255656 | 1548.2708888882 | 1669.50724678082 | 1726.10747414207 | 1442.66347544445 | Pcbp4 |
| ENSMUSG00000023505 | 0.869134369721722 | 0.169556374086986 | 0.79907358295209 | 16.2632468982428 | 15.4738988493722 | 7.57163910918757 | 22.5102406621959 | 13.1394983498292 | 13.4539045528125 | 1.04107809055614 | 6.51315338801105 | Cdca3 |
| ENSMUSG00000023571 | 0.0501443045974655 | 0.809824479994319 | 0.964412705343932 | 219.553833126278 | 247.582381589956 | 183.612248397799 | 197.902532488472 | 248.555510450935 | 245.839528646846 | 151.997401221196 | 174.76961591163 | C1qtnf12 |
| ENSMUSG00000023572 | 0.0396779888106826 | 0.804525287526367 | 0.963461896828986 | 790.574501997914 | 773.694942468612 | 956.86589242358 | 819.747930781633 | 786.179984598112 | 1024.94291047789 | 770.397787011541 | 669.769273400469 | Ccndbp1 |
| ENSMUSG00000023707 | -0.00174966615642069 | 0.991324167769835 | 0.998628446692557 | 400.256576440087 | 358.342920722305 | 333.152120804253 | 407.9981120023 | 349.291664466292 | 477.002070508805 | 344.596847974081 | 332.170822788563 | Ogfod2 |
| ENSMUSG00000023723 | 0.07244334311131 | 0.500901506866728 | 0.900927059989498 | 4103.7593006566 | 3648.58246553619 | 3689.28115595165 | 3853.94078670678 | 3506.05614301275 | 3176.34455669581 | 3901.9606834044 | 3959.99725991072 | Mrps23 |
| ENSMUSG00000023755 | 0.0861794468083848 | 0.81901160919372 | 0.966315773308182 | 17.1667606148119 | 12.2162359337149 | 17.036187995672 | 18.7585338851632 | 16.4243729372865 | 19.5693157131818 | 13.5340151772298 | 11.9407812113536 | Rhebl1 |
| ENSMUSG00000023764 | 0.169185433223738 | 0.450796103390807 | 0.89108031640712 | 63.2459601598332 | 69.2253369577179 | 68.1447519826882 | 55.3376749612315 | 60.2227007700503 | 40.3617136584374 | 66.6289977955927 | 59.7039060567679 | Sfi1 |
| ENSMUSG00000023791 | 0.0767345368598241 | 0.559660384254281 | 0.916574868225118 | 296.352499034647 | 263.870696168242 | 245.131816159948 | 289.819348525772 | 278.119381738051 | 270.301173288323 | 229.03717992235 | 261.61166108511 | Pigx |
| ENSMUSG00000023800 | -0.0882745429066869 | 0.793647450574646 | 0.961804768755116 | 111.132187137993 | 159.625482867208 | 105.056492639978 | 85.3513291774927 | 91.9764884488041 | 67.2695227640623 | 187.394056300105 | 143.289374536243 | Tiam2 |
| ENSMUSG00000023805 | 0.0726635408144945 | 0.702652725006367 | 0.946451777392559 | 1017.35644485674 | 1194.74787431732 | 1131.01359193489 | 1051.4158242634 | 905.530427942393 | 753.418654957498 | 1215.97920976957 | 1302.63067760221 | Synj2 |
| ENSMUSG00000023806 | 0.101345432101767 | 0.415454208857985 | 0.882192126163777 | 897.189120553062 | 860.837425462445 | 816.79056890361 | 967.002421780164 | 722.672409240604 | 958.896469945906 | 826.616003901573 | 794.604713337348 | Rsph3b |
| ENSMUSG00000023809 | 0.221959085158449 | 0.444697796500965 | 0.89108031640712 | 1713.06200661491 | 2391.93899582138 | 1845.58703286447 | 1560.71001924558 | 1376.3624521446 | 835.365164506446 | 2092.56696201783 | 2135.22878570295 | Rps6ka2 |
| ENSMUSG00000023826 | 0.370047159142522 | 0.239347212277993 | 0.832048466795696 | 176.185174730964 | 228.036404096012 | 233.774357496166 | 197.902532488472 | 154.389105610493 | 69.71568722821 | 215.50316474512 | 206.249857287016 | Park2 |
| ENSMUSG00000023827 | -0.732500909681848 | 0.0854464403082579 | 0.744107245066898 | 56.0178504272808 | 57.009101024003 | 59.6266579848522 | 51.5859681841989 | 111.685735973548 | 172.454594722414 | 44.7663578939139 | 44.5065481514088 | Agpat4 |
| ENSMUSG00000023828 | -0.00069794209889103 | 0.997880746113055 | 0.999262355603273 | 216.843291976571 | 292.375246680244 | 205.380710836713 | 151.944124469822 | 178.478185918513 | 155.33144347338 | 279.008928269045 | 254.012982132431 | Slc22a3 |
| ENSMUSG00000023829 | -0.550317387356036 | 0.0699666331750217 | 0.724916894583216 | 140.044626068202 | 166.140808698523 | 121.146225747001 | 136.937297361692 | 133.58489988993 | 271.524255520397 | 128.052605138405 | 294.177428025166 | Slc22a1 |
| ENSMUSG00000023830 | 0.194978546926469 | 0.435402181526063 | 0.888588936437999 | 3910.40736531083 | 5304.28964241902 | 5968.34452781711 | 4004.94698448235 | 3452.40319141761 | 2751.93502216618 | 5178.32242242622 | 5378.77917293246 | Igf2r |
| ENSMUSG00000023832 | -0.168219907042366 | 0.500482656907222 | 0.900927059989498 | 65.0529875929712 | 57.8235167529173 | 38.8046504345863 | 61.9031618210386 | 68.9823663366031 | 75.8310983885793 | 57.2592949805875 | 49.9341759747513 | Acat2 |
| ENSMUSG00000023845 | 0.297807399880721 | 0.379206857286634 | 0.87422797717331 | 650.529875929712 | 850.250020986559 | 895.346324661431 | 551.500896223799 | 503.680770076785 | 260.516515431732 | 858.889424708813 | 773.979727608646 | Lnpep |
| ENSMUSG00000023852 | 0.188943282677764 | 0.16669154144166 | 0.796745160868404 | 1147.46242004269 | 1145.88293058246 | 1032.58228351546 | 969.816201862939 | 881.441347634373 | 787.664957455566 | 1042.11916864669 | 1056.21637442246 | Chd1 |
| ENSMUSG00000023861 | 0.091236258327876 | 0.416918921992143 | 0.882192126163777 | 8062.95640666216 | 6590.25207837475 | 7189.2713341736 | 8431.02305468661 | 6795.31056325332 | 6821.12960827592 | 7138.67246694343 | 7661.63943543033 | Mpc1 |
| ENSMUSG00000023868 | -0.230515211491284 | 0.510776717705349 | 0.902394714365738 | 40.658117245607 | 26.0613033252585 | 25.5542819935081 | 35.6412143818101 | 43.7983278327639 | 53.8156182112498 | 32.2734208072402 | 20.6249857287016 | Pde10a |
| ENSMUSG00000023882 | 0.491496343547148 | 0.208124684903365 | 0.820372930586885 | 23.4913566307952 | 26.8757190541728 | 29.3401015481019 | 15.0068271081306 | 25.1840385038392 | 14.6769867848863 | 16.6572494488982 | 10.8552556466851 | Zfp54 |
| ENSMUSG00000023883 | 0.256566172685002 | 0.149232083363661 | 0.782024490202776 | 1253.17352488127 | 1267.2308741907 | 1353.43049076728 | 1246.5045766691 | 916.480009900584 | 802.341944240452 | 1254.49909912014 | 1311.31488211956 | Phf10 |
| ENSMUSG00000023885 | -0.673817937545152 | 0.0880829090346768 | 0.750135871546083 | 161.728955265859 | 196.274190668353 | 405.082692341535 | 256.053987532478 | 585.802634763217 | 579.740978003009 | 226.955023741238 | 233.387996403729 | Thbs2 |
| ENSMUSG00000023886 | -0.567369395128281 | 0.046973343067407 | 0.702324795701059 | 851.10992100804 | 824.188717661301 | 714.573440929577 | 859.140851940476 | 1745.36336413564 | 1536.19128348477 | 714.17957012151 | 819.571801324723 | Smoc2 |
| ENSMUSG00000023892 | 0.125655489679283 | 0.415106880650741 | 0.882192126163777 | 206.904641094311 | 197.903022126182 | 181.719338620502 | 163.19924480092 | 163.148771177045 | 149.216032313011 | 180.106509666212 | 194.309076075663 | Zfp51 |
| ENSMUSG00000023904 | -0.00241720247451118 | 0.988860293454193 | 0.998437672973644 | 2649.10221698044 | 2251.0450747192 | 2160.7565107844 | 2911.32445897733 | 2487.74502090099 | 3181.23688562411 | 2142.53871036453 | 2178.64980828969 | Hcfc1r1 |
| ENSMUSG00000023905 | 0.628621365870182 | 0.0119751538443707 | 0.635862253526407 | 2631.03194264906 | 2172.86116474343 | 2555.42819935081 | 3602.5764326456 | 1251.53721782123 | 2268.81754049701 | 1349.23720536075 | 2220.98530531177 | Tnfrsf12a |
| ENSMUSG00000023908 | -0.0080574905806314 | 0.980307725560354 | 0.996888334226108 | 23.4913566307952 | 25.2468875963442 | 21.7684624389143 | 23.448167356454 | 17.5193311331055 | 29.3539735697726 | 17.6983275394543 | 30.3947158107182 | Pkmyt1 |
| ENSMUSG00000023909 | -0.349914896473324 | 0.0176221008006943 | 0.642273034804458 | 468.020105182765 | 431.640336324594 | 385.207139679918 | 372.35689762049 | 551.858930692825 | 615.210362733151 | 445.581422758026 | 501.51281087685 | Paqr4 |

| | | | | | | | | | | | | |
|---|---|---|---|---|---|---|---|---|---|---|---|---|
| **ENSMUSG00000023912** | 0.360017875744636 | 0.263145181489694 | 0.842510961779757 | 46.9827132615903 | 52.122606650517 | 39.7511053232348 | 46.8963347129081 | 18.6142893289246 | 35.4693847301419 | 36.4377331694648 | 54.2762782334254 | Slc25a27 |
| **ENSMUSG00000023913** | -0.302334871111834 | 0.303318011522577 | 0.851785577030345 | 75.8951521917998 | 146.594831204579 | 176.98706417726 | 154.757904552597 | 197.092475247437 | 220.154801773295 | 136.381229862854 | 130.263067760221 | Pla2g7 |
| **ENSMUSG00000023915** | -0.804583878070542 | 0.0078827410874469 | 0.635862253526407 | 137.334084918495 | 120.533527879321 | 82.3415753124149 | 102.23400967414 | 278.119381738051 | 233.608706326107 | 143.668776496747 | 118.322286548867 | Tnfrsf21 |
| **ENSMUSG00000023919** | 0.0826428496393007 | 0.669500758763858 | 0.942334932906175 | 112.035700854562 | 105.059629029948 | 147.646962629158 | 130.371810501884 | 111.685735973548 | 95.400414101761 | 139.504464134522 | 119.407812113536 | Cenpq |
| **ENSMUSG00000023921** | 0.137999344512241 | 0.582352501201867 | 0.922450244597442 | 2381.662156876 | 2876.51635452541 | 2674.68151532051 | 2193.81053786984 | 1954.50037953709 | 1352.72894867369 | 2993.09951034889 | 2901.60983435892 | Mut |
| **ENSMUSG00000023923** | 0.0197354384766342 | 0.923893113959829 | 0.986713918026868 | 425.55496050402 | 601.853223667689 | 512.03209475881 | 435.197986135787 | 451.122776677468 | 349.801518373124 | 613.194995337564 | 532.993052252237 | Tbc1d5 |
| **ENSMUSG00000023927** | 0.162643854920743 | 0.42075351162739 | 0.882568531013176 | 676.731773710215 | 767.994032366212 | 547.050925638802 | 654.672832592197 | 478.496731572945 | 456.20967256355 | 726.672507208183 | 702.335040340524 | Satb1 |
| **ENSMUSG00000023932** | 0.0512685248613345 | 0.661461575173562 | 0.939611397743053 | 929.715614349547 | 952.866402829765 | 870.738497556571 | 909.788893430416 | 943.853964796062 | 747.303243797128 | 980.695561303881 | 861.907298346795 | Cdc5l |
| **ENSMUSG00000023938** | -0.0864867873298892 | 0.546595509742802 | 0.914393789031376 | 702.933671490717 | 643.38842584232 | 622.767316730678 | 676.245146560134 | 735.811907590433 | 546.717757737015 | 806.835520181006 | 717.532398245884 | Aars2 |
| **ENSMUSG00000023939** | 0.0510362032457458 | 0.780038539941841 | 0.960643510321816 | 3108.08718499752 | 2133.76920975554 | 2360.45849228923 | 2765.00789467306 | 2564.39209460832 | 3149.43674759019 | 2065.49893166338 | 2228.58398426445 | Mrpl14 |
| **ENSMUSG00000023942** | -0.346089361069585 | 0.0587688774200598 | 0.711712402279519 | 906.224257718752 | 938.206919709307 | 863.166858447383 | 884.464872685446 | 1244.96746864631 | 1477.48333634522 | 840.150019078802 | 1005.19667288304 | Slc29a1 |
| **ENSMUSG00000023944** | 0.19565618649613 | 0.0129063837233168 | 0.635862253526407 | 62326.1831963659 | 58701.456908687 | 57205.6263796894 | 62363.6838279194 | 50663.7157205496 | 49699.9465003213 | 54013.2134942335 | 55704.8298765291 | Hsp90ab1 |
| **ENSMUSG00000023947** | -0.469315795228307 | 0.330065978833559 | 0.855977678491063 | 13.5527057485357 | 9.77298874697194 | 21.7684624389143 | 25.3240207449704 | 40.5134532453066 | 28.1308913376988 | 15.616171358342 | 13.0263067760221 | Nfkbie |
| **ENSMUSG00000023951** | 0.133482677553111 | 0.52778132678358 | 0.908232038364594 | 2219.02968789357 | 2367.50652395395 | 2651.0201431043 | 2369.20282969612 | 1859.23901650083 | 1488.49107643389 | 2501.7106516064 | 2907.03746218226 | Vegfa |
| **ENSMUSG00000023952** | 0.279976057093308 | 0.0718031830381096 | 0.728065810245271 | 1123.06754969532 | 1228.13891920281 | 1301.37547189161 | 1202.42202203896 | 900.055636963298 | 791.334204151787 | 1198.28088223011 | 1107.23607596188 | Gtpbp2 |
| **ENSMUSG00000023953** | -0.266954362072722 | 0.222120323232221 | 0.831327997735572 | 79.509207058076 | 109.13170767452 | 124.932045301595 | 85.3513291774927 | 106.210944994452 | 99.0696607979826 | 139.504464134522 | 134.605170018895 | Polh |
| **ENSMUSG00000023959** | 0.262927821090479 | 0.18780999846533 | 0.808642553838633 | 7535.30439618584 | 8980.5622427383 | 10163.9790491957 | 9274.2191528247 | 6605.88279537661 | 5339.97702523447 | 9507.12512295864 | 8509.43490143643 | Clic5 |
| **ENSMUSG00000023960** | -0.069454477352624 | 0.582248086398144 | 0.922450244597442 | 313.519259649459 | 352.642010619904 | 334.098575692902 | 316.081295965 | 399.65974147397 | 340.016860516533 | 312.323427166841 | 329.999771659226 | Enpp5 |
| **ENSMUSG00000023961** | -0.0252190310443627 | 0.876417660597913 | 0.977469019294424 | 663.179067961679 | 793.240919962556 | 871.68495244522 | 615.279911433354 | 708.437952694956 | 654.348994159515 | 886.998533153828 | 744.670537362596 | Enpp4 |
| **ENSMUSG00000023963** | 0.0038248507984106 | 0.97307542255402 | 0.995480246308729 | 224.974915425692 | 241.881471487556 | 236.613722162112 | 242.923013812864 | 240.890803080201 | 240.94719971855 | 226.955023741238 | 235.559047533066 | Cyp39a1 |
| **ENSMUSG00000023965** | 0.157128570169993 | 0.37368815791591 | 0.873700364743909 | 673.117718843939 | 860.837425462445 | 721.198625150116 | 618.093691516128 | 610.986673267056 | 500.240632918209 | 776.644255554878 | 688.223207999834 | Fbxl17 |
| **ENSMUSG00000023967** | 0.00418096720988974 | 0.973641970045918 | 0.995510898988495 | 2054.59019147801 | 1645.11977240694 | 1618.43785958884 | 1987.46666513304 | 1772.73731903112 | 2025.42417631431 | 1695.91620951595 | 1791.11718170304 | Mrps18a |
| **ENSMUSG00000023971** | -0.0106574546121917 | 0.956451623866323 | 0.992371505557225 | 230.395997725106 | 210.119258059897 | 202.541346170768 | 266.371181169318 | 222.276513751277 | 305.770558018465 | 186.352978209548 | 202.993280593011 | Rrp36 |
| **ENSMUSG00000023972** | -0.742642989132015 | 0.0777405936730905 | 0.739855948845396 | 25.2983840639333 | 21.1748089517725 | 16.0897331070236 | 30.0136542162612 | 44.893286028583 | 64.8233582999146 | 19.7804837205666 | 26.0526135520442 | Ptk7 |
| **ENSMUSG00000023973** | -0.259486633850766 | 0.32820016692131 | 0.855977678491063 | 364.116027777325 | 379.517729674077 | 348.295399022628 | 393.929211588428 | 520.105143014071 | 659.24132308781 | 298.789411989611 | 301.776106977845 | Cnpy3 |
| **ENSMUSG00000023977** | 0.313795424009999 | 0.236263885178165 | 0.832048466795696 | 3322.21993582438 | 4385.62870020366 | 4896.95759386706 | 3218.02648799975 | 2766.95936083486 | 1893.33129525033 | 4326.7205443513 | 3741.80662141235 | Ubr2 |
| **ENSMUSG00000023980** | 0.0504896855567307 | 0.699385018923693 | 0.945280238056015 | 223.167887992554 | 245.953550132127 | 282.043556817237 | 239.171307035831 | 240.890803080201 | 220.154801773295 | 247.776585552361 | 246.414303179751 | Taf8 |
| **ENSMUSG00000023984** | -0.205479913664098 | 0.676519124576623 | 0.94409858171788 | 10.8421645988285 | 5.7009101024003 | 12.3039135524298 | 14.0689004138724 | 7.66470737073368 | 19.5693157131818 | 12.4929370866736 | 9.76973008201657 | Gm20517 |
| **ENSMUSG00000023988** | 0.0112720271721002 | 0.923081559638952 | 0.986713918026868 | 711.065294939838 | 820.116639016729 | 754.324546252812 | 773.789522762983 | 823.408563255961 | 683.702967729287 | 832.862472444909 | 694.736361387845 | Bysl |
| **ENSMUSG00000023990** | -0.179970773579558 | 0.289860387472502 | 0.851785577030345 | 868.276681622852 | 825.003133390215 | 759.056820696054 | 796.299763425179 | 1028.16574587413 | 1146.02805145321 | 763.110240377648 | 744.670537362596 | Tfeb |
| **ENSMUSG00000023991** | 0.163056590531321 | 0.222707106494279 | 0.832048466795696 | 374.054678659585 | 415.352021746308 | 482.691993210708 | 376.108604397523 | 350.386622662111 | 364.47850515801 | 406.020455316893 | 350.624757387928 | Foxp4 |
| **ENSMUSG00000023992** | -0.347574073129643 | 0.355268045013165 | 0.864215962387542 | 10.8421645988285 | 23.6180561385155 | 21.7684624389143 | 35.6412143818101 | 29.5638712871156 | 34.2463024980681 | 26.0269522639034 | 27.1381391167127 | Trem2 |
| **ENSMUSG00000023993** | 0.602965111915032 | 0.459706260150235 | 0.894774044236998 | 16.2632468982428 | 15.4738988493722 | 82.3415753124149 | 3.75170677703264 | 14.2344565456483 | 35.4693847301419 | 5.20539045278068 | 22.7960368580387 | Treml1 |
| **ENSMUSG00000023994** | -0.279784885864966 | 0.0987504689379527 | 0.759800953500741 | 228.588970291968 | 253.283291692356 | 230.934992830221 | 178.206071909051 | 302.208462046071 | 302.101311322243 | 249.858741733473 | 229.045894145055 | Nfya |
| **ENSMUSG00000023999** | 0.00575085379294012 | 0.985056650648921 | 0.99735301701069 | 28.0089252136404 | 39.9063707168021 | 26.5007368821565 | 20.6343872736795 | 27.3739548954774 | 28.1308913376988 | 27.0680303544596 | 32.5657669400552 | Kif6 |
| **ENSMUSG00000024002** | -0.0459570626044085 | 0.656483616783022 | 0.938316262308724 | 908.03128515189 | 1023.72057124531 | 958.758802200877 | 878.837312519897 | 1058.82457535707 | 1001.70434806849 | 911.984407327176 | 920.525678838894 | Brd4 |
| **ENSMUSG00000024006** | -0.00392965884185032 | 0.982842108991967 | 0.996940244108351 | 410.195227322346 | 453.629561005281 | 406.975602118832 | 358.287997206618 | 355.861413641207 | 310.66288694676 | 482.019155927491 | 484.144401842154 | Stk38 |
| **ENSMUSG00000024007** | -0.298787937086054 | 0.0410571723880708 | 0.683226273624098 | 533.073092775737 | 447.114235173966 | 388.046504345863 | 445.515179772627 | 506.965644664242 | 651.902829695367 | 529.908748093074 | 543.848307898922 | Ppil1 |
| **ENSMUSG00000024008** | -2.25093694814833 | NA | NA | 7.22810973255236 | 0.814415728914329 | 11.3574586637814 | 9.37926694258161 | 5.47479097909548 | 127.200552135681 | 2.08215618111227 | 2.17105112933702 | Cpne5 |
| **ENSMUSG00000024011** | -1.32206479874571 | 0.0503063711949371 | 0.703396849069134 | 422.844419354313 | 254.912123150185 | 354.920583243168 | 394.867138282686 | 1142.04139823932 | 1877.43122623337 | 288.37863108405 | 261.61166108511 | Pi16 |
| **ENSMUSG00000024012** | -0.187499390913016 | 0.304582072757272 | 0.851785577030345 | 3493.8875419725 | 3110.25366872382 | 3028.65564367503 | 3498.46656958294 | 4044.77557535574 | 4864.19803695774 | 3183.61680092067 | 2862.53091403085 | Mtch1 |
| **ENSMUSG00000024013** | -0.737021304054248 | 0.104440632693202 | 0.761198544536855 | 6.32459601598331 | 7.32974156022896 | 13.2503684410783 | 9.37926694258161 | 17.5193311331055 | 19.5693157131818 | 12.4929370866736 | 10.8552556466851 | Fgd2 |
| **ENSMUSG00000024014** | -0.396573886638005 | 0.0889283490755899 | 0.751206603737452 | 348.756294595651 | 237.809392842984 | 388.992959234512 | 532.742362338636 | 453.312693069106 | 473.332823812584 | 530.94982618363 | 526.479898864226 | Pim1 |
| **ENSMUSG00000024018** | -0.00064724277254572 | 0.997724795965089 | 0.999262355603273 | 111.132187137993 | 121.347943608235 | 86.1273948670087 | 112.551203310979 | 82.1218646864323 | 114.969729814943 | 91.61487196894 | 143.289374536243 | Ccdc167 |
| **ENSMUSG00000024019** | 0.178812169980188 | 0.502160480990857 | 0.900927059989498 | 259.308436655316 | 315.99330281876 | 334.098575692902 | 219.47484645641 | 192.712642464161 | 161.446854633749 | 338.350379430744 | 303.947158107182 | Cmtr1 |
| **ENSMUSG00000024026** | 0.0186273101741901 | 0.856290872436893 | 0.97363563840968 | 2494.60137144713 | 2344.70288354435 | 2138.04159345684 | 2196.62431795261 | 2143.92814741379 | 2081.68595898971 | 2379.90451501133 | 2450.03119945682 | Glo1 |
| **ENSMUSG00000024030** | -0.837865303132778 | 0.0278643388292693 | 0.665956123973759 | 70.4740698923855 | 61.0811796685746 | 35.965285768641 | 45.0204813243917 | 134.679858085749 | 134.539045528125 | 54.1360607089191 | 57.5328549274309 | Abcg1 |
| **ENSMUSG00000024033** | 0.185778401945425 | 0.616532611695227 | 0.928896975098882 | 19.877301764519 | 16.2883145782866 | 17.9826428843205 | 14.0689004138724 | 13.1394983498292 | 17.123151249034 | 11.4518589961175 | 18.4539345993646 | Rsph1 |
| **ENSMUSG00000024036** | 0.0577970957327524 | 0.668630614289385 | 0.942055056494206 | 290.027903018663 | 337.982527499446 | 361.545767463707 | 302.012395551128 | 315.3479603959 | 284.978160073209 | 348.761160336306 | 290.92085133116 | Slc37a1 |
| **ENSMUSG00000024037** | 0.207005828819873 | 0.229550494575175 | 0.832048466795696 | 120.167324303683 | 138.450673915436 | 154.272146849697 | 168.826804966469 | 137.964732673206 | 108.854318654574 | 128.052605138405 | 128.092016630884 | Wdr4 |
| **ENSMUSG00000024038** | 0.0515617057904516 | 0.79114885064555 | 0.961804768755116 | 7738.59498241387 | 5668.33347324373 | 5692.92615522041 | 7004.43655271995 | 6543.47017821492 | 8268.03588881929 | 5113.77558081174 | 5263.71346307759 | Ndufv3 |
| **ENSMUSG00000024039** | -0.144581093298054 | 0.818237487522109 | 0.966315773308182 | 16.2632468982428 | 94.4722245540621 | 24.6078271048596 | 34.703287687552 | 36.1336204620302 | 20.7923979452556 | 27.0680303544596 | 104.210454208177 | Cbs |
| **ENSMUSG00000024042** | 0.0653755398770331 | 0.862130669523259 | 0.974868041734905 | 1014.64590370704 | 1490.38078391322 | 515.817914313404 | 482.094320848695 | 1227.44813751321 | 850.042151291332 | 689.193695948162 | 581.84170266232 | Sik1 |
| **ENSMUSG00000024043** | -0.513974555322119 | 0.185601058116573 | 0.805963874499992 | 22.5878429142261 | 12.2162359337149 | 23.6613722162112 | 35.6412143818101 | 43.7983278327639 | 39.1386314263635 | 22.903717992235 | 28.2236646813812 | Arhgap28 |
| **ENSMUSG00000024044** | -0.110472151557468 | 0.578981031328065 | 0.922450244597442 | 112.035700854562 | 129.492100897378 | 131.557229522134 | 92.854742731558 | 109.49581958191 | 107.6312364225 | 154.079557402308 | 131.348593324889 | Epb41l3 |
| **ENSMUSG00000024045** | 0.222080667141351 | 0.188302404909241 | 0.808677608936066 | 589.090943203017 | 632.801021366433 | 707.948256709038 | 549.625042835282 | 544.194223322091 | 402.3940543523 | 640.263025692024 | 537.335154510911 | Akap8 |
| **ENSMUSG00000024048** | 0.268398588094179 | 0.0694627887900428 | 0.724352955175446 | 26528.0662321837 | 23473.9045544977 | 19919.0895864952 | 30079.3090848592 | 22850.6825885487 | 19779.6858570985 | 20668.523331811 | 19725.0850355915 | Myl12a |
| **ENSMUSG00000024049** | 0.295857418848364 | 0.142865133209545 | 0.780607990324772 | 56372.0278041759 | 62595.1785086264 | 54017.0198598328 | 54943.7457496431 | 39480.9076666492 | 32118.1394142596 | 58398.2344116559 | 55669.0075328951 | Myom1 |
| **ENSMUSG00000024050** | -0.0681837669880273 | 0.618369256513096 | 0.929568738467614 | 352.370349461928 | 414.537606017393 | 359.65285768641 | 322.646782824807 | 421.558905390352 | 397.501725424004 | 369.582722147429 | 332.170822788563 | Wiz |
| **ENSMUSG00000024052** | -0.206968373879337 | 0.347328654607222 | 0.861266047293794 | 131.913002619081 | 179.985876090067 | 124.932045301595 | 90.0409626487835 | 144.534481848121 | 157.777607937528 | 156.16171358342 | 150.888053488923 | Lpin2 |
| **ENSMUSG00000024053** | -0.325231720689747 | 0.183611435655622 | 0.805963874499992 | 376.765219809292 | 405.579032999336 | 394.671688566402 | 208.219726125312 | 499.300937293508 | 516.140701935169 | 372.705956419097 | 348.453706258591 | Emilin2 |
| **ENSMUSG00000024054** | 0.0580583916299908 | 0.750857405487374 | 0.956197192463153 | 311.712232216321 | 382.775392589734 | 353.974128354519 | 279.502154888932 | 323.012667766634 | 237.277953022329 | 376.870268781321 | 337.598450611906 | Smchd1 |
| **ENSMUSG00000024055** | -0.143533076328221 | 0.531344402233157 | 0.909082872831377 | 54.2108229941427 | 65.9676740420606 | 52.0550188756646 | 67.5307219865876 | 65.6974917491458 | 85.6157562451702 | 62.4646854333682 | 52.1052271040884 | Cyp4f13 |
| **ENSMUSG00000024059** | 0.0547077072690048 | 0.557576601259279 | 0.916341990969984 | 5002.7554486428 | 5190.27144037102 | 4717.13116502386 | 4859.39820295153 | 4314.13529152724 | 4577.99679465246 | 4968.02464813388 | 5173.61484121011 | Clip4 |
| **ENSMUSG00000024063** | 0.3127655787791 | 0.155062817175808 | 0.787275540343856 | 8460.50244195254 | 9167.06344465968 | 9065.14492347482 | 10073.3326963327 | 5944.52804510188 | 5160.18393711961 | 8102.71077879841 | 10391.7362305716 | Lbh |
| **ENSMUSG00000024065** | -0.659192848575369 | 0.0608867833692948 | 0.71282992338528 | 60.535419010126 | 68.4109212288036 | 57.7337482075553 | 46.8963347129081 | 120.445401540101 | 138.208292224346 | 64.5468416144805 | 46.6775992807458 | Ehd3 |
| **ENSMUSG00000024066** | -0.420638460328665 | 0.18888391894093 | 0.808677608936066 | 2974.3671549453 | 2912.35064659764 | 3303.12756138308 | 2155.35554340525 | 5293.02791858951 | 5393.79264344572 | 2281.0020964085 | 2218.81425418243 | Xdh |
| **ENSMUSG00000024067** | 0.175638601425928 | 0.263176775092961 | 0.842510961779757 | 498.739571546113 | 368.115909469277 | 347.34894413398 | 407.9981120023 | 359.146288228664 | 407.286383280595 | 336.268223249632 | 334.3418739179 | Dpy30 |
| **ENSMUSG00000024068** | 0.207519694301949 | 0.0546608218968106 | 0.703396849069134 | 939.654265231807 | 925.176268046677 | 978.634354862494 | 1049.53997087488 | 816.838814081046 | 750.97249049335 | 930.723812957186 | 870.591502864143 | Spast |
| **ENSMUSG00000024069** | -0.229937168186906 | 0.0915056666181798 | 0.753823363185192 | 144.562194651047 | 144.151584017836 | 135.343049076728 | 135.999370667433 | 176.288269526875 | 166.339183562045 | 150.95632313064 | 163.914360264945 | Slc30a6 |
| **ENSMUSG00000024070** | -0.0620785528190154 | 0.610453280775721 | 0.927932112008768 | 379.475760958999 | 423.496179035451 | 486.477812765302 | 424.880792498947 | 480.686647964584 | 459.878919259771 | 426.842017128016 | 422.269444656049 | Prkd3 |
| **ENSMUSG00000024072** | -0.00788653993425677 | 0.925135125042524 | 0.986713918026868 | 973.987786461431 | 1092.94590820303 | 1020.27836996303 | 982.947175582553 | 1045.68507700724 | 1012.71208815716 | 994.22957648111 | 1041.0190165171 | Yipf4 |
| **ENSMUSG00000024073** | 0.132022259146558 | 0.393115245936186 | 0.878770935869648 | 819.486940928124 | 1067.69902060668 | 1020.27836996303 | 829.127197724215 | 858.447225522172 | 692.264543353805 | 943.21675004386 | 915.098051015552 | Birc6 |
| **ENSMUSG00000024074** | 0.0902584297839261 | 0.59451711396188 | 0.924419871591907 | 1111.32187137993 | 1505.85468276259 | 1467.95153229374 | 1081.42947847966 | 1470.52885698505 | 1150.9203803815 | 1135.81619679675 | 1096.38082031519 | Crim1 |
| **ENSMUSG00000024076** | -0.760273981598716 | 0.158292145801904 | 0.789230661864335 | 38.8510898124689 | 18.7315617650296 | 35.0188308799925 | 42.2067012416173 | 68.9823663366031 | 113.746647582869 | 17.6983275394543 | 28.2236646813812 | Vit |
| **ENSMUSG00000024077** | 0.111610206268875 | 0.607789955310442 | 0.927510709653048 | 751.723412185445 | 1028.6070656188 | 827.201572678743 | 723.141481273042 | 757.711071506815 | 503.90987961443 | 942.175671953304 | 878.190181816823 | Strn |
| **ENSMUSG00000024078** | 0.0801887398095395 | 0.594708877226774 | 0.924461858177432 | 316.229800799166 | 261.427448981499 | 284.882921483182 | 294.508981997063 | 295.638712871156 | 254.401104271363 | 316.487739529066 | 226.874843015718 | Ttc27 |
| **ENSMUSG00000024079** | 0.126303307235725 | 0.373562784103068 | 0.873700364743909 | 252.080326922764 | 230.479651282755 | 209.166530391307 | 204.468019348279 | 203.662224422352 | 179.793088114857 | 211.338852382896 | 225.78931745105 | Eif2ak2 |
| **ENSMUSG00000024081** | 0.230921070155268 | 0.00862910358324224 | 0.635862253526407 | 817.679913494986 | 920.289773673191 | 890.614050218188 | 925.733647232805 | 770.850569856644 | 763.203312814088 | 759.98700610598 | 734.90080728058 | Cebpz |
| **ENSMUSG00000024082** | 0.0703628250285818 | 0.493803074878112 | 0.900117970535229 | 717.389890955822 | 684.109212288036 | 724.037989816062 | 762.534402431885 | 655.879959295639 | 624.995020589742 | 740.206522385413 | 727.3021283279 | Ndufaf7 |
| **ENSMUSG00000024083** | 0.165772632528393 | 0.5226317786097 | 0.90648515894888 | 3341.19372387233 | 4517.56404828778 | 3925.89487811376 | 3165.50259312129 | 2763.6744862474 | 1981.39321595965 | 4364.19935561132 | 4217.26681873715 | Pja2 |
| **ENSMUSG00000024084** | -0.702581385125952 | 0.106350635729478 | 0.762094734041179 | 62.3424464432641 | 53.7514381083457 | 43.5369248778286 | 37.5170677703264 | 85.4067392738896 | 143.100621152642 | 27.0680303544596 | 66.217059444779 | Qpct |
| **ENSMUSG00000024085** | -0.433602750389566 | 0.125393918969149 | 0.771949105062466 | 491.511461813561 | 628.728942721862 | 561.247748968529 | 345.157023487003 | 1028.16574587413 | 722.841599155651 | 476.813765474711 | 510.197015394199 | Man2a1 |
| **ENSMUSG00000024087** | -0.660391283943621 | 0.176586401117905 | 0.799889021902745 | 180.702743313809 | 215.820168162297 | 159.004421292939 | 105.985716451172 | 404.039574257247 | 439.086521314516 | 121.806136595068 | 81.4144173501381 | Cyp1b1 |
| **ENSMUSG00000024091** | -0.0400251931656023 | 0.612484843011801 | 0.928055903343287 | 1737.45687696227 | 1751.80823289472 | 1642.09923180506 | 1745.48157801444 | 1721.27428382762 | 1668.28416454874 | 1864.57086018604 | 1814.99874412574 | Vapa |
| **ENSMUSG00000024095** | 0.0740953826137612 | 0.738512682370199 | 0.954383574136061 | 507.774708711803 | 620.584785432718 | 672.929425829046 | 484.908100931469 | 513.535393839156 | 360.809258461789 | 708.974179668729 | 587.269330485663 | Hnrnpll |
| **ENSMUSG00000024096** | 0.0262156624727207 | 0.847285687036368 | 0.972386862177572 | 1854.01014639968 | 1888.63007535233 | 1915.62469462446 | 1803.63303305844 | 1996.10879097821 | 2167.30171523488 | 1614.71211845257 | 1551.2160319113 | Ralbp1 |

| | | | | | | | | | | | | |
|---|---|---|---|---|---|---|---|---|---|---|---|---|
| **ENSMUSG00000024097** | -0.0762561295438674 | 0.3920489846162 | 0.878285927324016 | 565.599586572222 | 532.627886709971 | 539.479286529615 | 551.500896223799 | 563.903470846835 | 590.748718091674 | 600.702058250891 | 552.53251241627 | Srsf7 |
| **ENSMUSG00000024098** | 0.0804707364296833 | 0.680528251817355 | 0.944452327903597 | 1460.07816597558 | 1863.38318775598 | 1897.64205174014 | 1507.24819767287 | 1315.04479317874 | 1229.19764323423 | 1975.96621587555 | 1842.13688324246 | Twsg1 |
| **ENSMUSG00000024099** | 0.100031381666716 | 0.39051389651629 | 0.877797322358001 | 16805.3551281842 | 13215.5240330928 | 13600.5567498782 | 16148.2838950428 | 13515.0690109951 | 14297.8312929434 | 13133.2001123657 | 14819.5950088545 | Ndufv2 |
| **ENSMUSG00000024101** | -0.0161640996639952 | 0.899760407859486 | 0.984802943363903 | 589.090943203017 | 538.328796812371 | 580.176846741498 | 545.87333605825 | 621.936255225247 | 643.34125407085 | 485.14239019916 | 529.736475558232 | Washc1 |
| **ENSMUSG00000024104** | 0.185720533372583 | 0.348415314508856 | 0.861266047293794 | 464.406050316489 | 593.709066378546 | 647.375143835538 | 468.025420434822 | 430.318570956905 | 360.809258461789 | 540.319528998635 | 578.585125968315 | Washc2 |
| **ENSMUSG00000024109** | -0.779245561971613 | 0.105193782913996 | 0.762094734041179 | 11.7456783153976 | 9.77298874697194 | 6.62518422053913 | 8.44134024832345 | 13.1394983498292 | 25.6847268735511 | 8.32862472444909 | 16.2828834700276 | Nrxn1 |
| **ENSMUSG00000024112** | -0.705762435146063 | 0.0473725988346426 | 0.703396849069134 | 94.8689402397497 | 75.7406627890326 | 119.253315969704 | 150.068271081306 | 130.300025302473 | 322.893709267499 | 130.134761319517 | 134.605170018895 | Cacna1h |
| **ENSMUSG00000024118** | -0.3579584557704 | 0.121837018294667 | 0.769868040577702 | 56.0178504272808 | 60.2667639396603 | 44.483379766477 | 58.151455044006 | 84.3117810780705 | 78.277262852727 | 66.6289977955927 | 52.1052271040884 | Tedc2 |
| **ENSMUSG00000024120** | 0.304916354822367 | 0.0915621880866931 | 0.753823363185192 | 4686.52564784364 | 5503.82149600303 | 5765.80318164634 | 4864.08783642282 | 3994.40749834807 | 3008.78229090169 | 4954.49063295665 | 4894.6347710903 | Lrpprc |
| **ENSMUSG00000024121** | -0.0327165749305791 | 0.760628670593978 | 0.957357521291862 | 1479.05195402353 | 1378.80582905196 | 1287.17864856189 | 1465.04149643125 | 1470.52885698505 | 1587.56073723187 | 1403.37326606967 | 1278.7491151795 | Atp6v0c |
| **ENSMUSG00000024122** | 0.305074225206861 | 0.26201477869545 | 0.841137531367976 | 1008.32130769105 | 1410.56804247962 | 1639.25986713911 | 1052.35375095766 | 951.518672166795 | 595.64104701997 | 1337.78534636464 | 1250.52545049812 | Pdpk1 |
| **ENSMUSG00000024127** | 0.298130908798226 | 0.434090577860822 | 0.887868762931002 | 1367.01625316897 | 2194.85038942412 | 2120.05895057252 | 1324.35249229252 | 928.524550054594 | 605.425704876561 | 2119.63499237229 | 2044.0446382708 | Prepl |
| **ENSMUSG00000024130** | 0.0644969752146126 | 0.717084010696872 | 0.95140862628518 | 280.089252136404 | 447.928650902881 | 339.777305024792 | 306.702029022419 | 363.52612101194 | 299.655146858096 | 316.487739529066 | 335.427399482569 | Abca3 |
| **ENSMUSG00000024131** | -0.253531091072036 | 0.578766901329528 | 0.922450244597442 | 7.22810973255236 | 17.1027303072009 | 13.2503684410783 | 7.50341355406529 | 12.0445401540101 | 11.0077400886647 | 14.5750932677859 | 16.2828834700276 | Slc3a1 |
| **ENSMUSG00000024132** | 0.164144352537008 | 0.211786058359089 | 0.823370267313127 | 11804.4067069746 | 9637.79573597216 | 12542.4201843692 | 13268.8489436702 | 11180.6181375088 | 11291.4951665059 | 9581.04166738812 | 10118.1837882752 | Eci1 |
| **ENSMUSG00000024135** | 0.182290125341526 | 0.368970610458494 | 0.871495599696183 | 158.114900399583 | 148.223662662408 | 186.451613063744 | 142.564857527241 | 151.104231023035 | 123.53130543946 | 178.024353485099 | 106.381505337514 | Srbd1 |
| **ENSMUSG00000024137** | 0.0673909804576365 | 0.618652199170056 | 0.929568738467614 | 184.316798180085 | 196.274190668353 | 198.755526616174 | 209.15765281957 | 205.85214081399 | 200.585486060113 | 167.613572579538 | 179.111718170304 | E4f1 |
| **ENSMUSG00000024140** | -0.087326836367116 | 0.539267839450785 | 0.913689959742792 | 9551.04349785137 | 10096.3117913509 | 10637.2064935199 | 8922.49664247789 | 12839.4798041747 | 10451.2376730711 | 9291.62195821352 | 9071.73714393472 | Epas1 |
| **ENSMUSG00000024142** | -0.0152189437822175 | 0.933670124790622 | 0.98818087308472 | 160.82544154929 | 192.202112023782 | 149.539872406455 | 189.461192240149 | 166.433645764503 | 134.539045528125 | 194.681602933998 | 202.993280593011 | Mlst8 |
| **ENSMUSG00000024143** | 0.208599277744073 | 0.375143914710776 | 0.873700364743909 | 2693.37438909232 | 3191.69524161525 | 2858.29376371831 | 2784.70435525248 | 1870.18859845902 | 1679.29190463741 | 3089.91977277061 | 3335.82006022632 | Rhoq |
| **ENSMUSG00000024145** | -0.157380987206975 | 0.687906709964053 | 0.944452327903597 | 40.658117245607 | 15.4738988493722 | 18.9290977729689 | 32.8274342990356 | 35.0386622662111 | 35.4693847301419 | 30.191264626128 | 19.5394601640331 | Pigf |
| **ENSMUSG00000024146** | 0.0298252085082504 | 0.760469298516869 | 0.957357521291862 | 712.872322372977 | 646.646088757977 | 667.250696497155 | 726.893188050075 | 641.645502749991 | 724.064681387725 | 688.152617857606 | 643.716659848425 | Cript |
| **ENSMUSG00000024150** | -0.0121962466491919 | 0.90926338806141 | 0.985537921199608 | 1973.27395698679 | 1624.75937918409 | 1673.33224313045 | 1919.93594314646 | 1819.82052145134 | 1681.73806910156 | 1837.50282983158 | 1911.61051938124 | Mcfd2 |
| **ENSMUSG00000024151** | 0.111083214561047 | 0.422263522048879 | 0.883344319781939 | 373.151164943016 | 416.166437475222 | 444.83379766477 | 407.060185308042 | 386.520243124141 | 309.439804714686 | 428.924173309128 | 392.96025441 | Msh2 |
| **ENSMUSG00000024158** | 0.1356828916459 | 0.374336748930467 | 0.873700364743909 | 2149.45913171776 | 1545.7610534794 | 1899.53496151743 | 2257.58955307939 | 1887.70792959212 | 1986.28554488795 | 1638.65691453536 | 1634.80150039077 | Hagh |
| **ENSMUSG00000024160** | 0.0133136834162157 | 0.925070112576802 | 0.986713918026868 | 262.018977805023 | 246.767965861042 | 264.060913932917 | 197.902532488472 | 248.555510450935 | 225.04713070159 | 239.447960827911 | 248.585354309088 | Spsb3 |
| **ENSMUSG00000024163** | 0.0406305196434147 | 0.85889235650463 | 0.974833700919212 | 875.504791355405 | 1107.60539132349 | 989.991813526275 | 961.374861614615 | 762.090904290091 | 649.456665231219 | 1307.59408173851 | 1105.06502483254 | Mapk8ip3 |
| **ENSMUSG00000024164** | -1.04740649495021 | 0.0019520143922521 | 0.635862253526407 | 1289.31407354403 | 2028.70958072559 | 1697.94007023531 | 1416.26930832982 | 3487.44185368382 | 5649.41682994916 | 1620.9585869959 | 2537.95877019497 | C3 |
| **ENSMUSG00000024165** | 0.0761354593714968 | 0.652669861987224 | 0.936914467032094 | 117.456783153976 | 97.7298874697194 | 89.9132144216024 | 98.4823028971069 | 98.5462376237187 | 94.1773318696872 | 98.902418602833 | 91.1841474321546 | Jpt2 |
| **ENSMUSG00000024168** | -0.773971363825408 | 0.0381783753036505 | 0.679566075569698 | 496.932544112975 | 451.186313818538 | 428.744064557746 | 433.32213274727 | 1029.26070406995 | 1284.23634367755 | 352.92547269853 | 429.868123608729 | Tmem204 |
| **ENSMUSG00000024169** | -0.192879808762789 | 0.381992769896795 | 0.875126835008975 | 59.631905293557 | 78.9983257046899 | 74.7699362032273 | 67.5307219865876 | 97.4512794278996 | 88.0619207093179 | 59.3414511616998 | 77.072315091464 | Ift140 |
| **ENSMUSG00000024170** | -0.129994032716562 | 0.310079166693451 | 0.851785577030345 | 185.220311896654 | 209.304842330982 | 172.254789734017 | 194.150825711439 | 203.662224422352 | 206.700897220482 | 209.256696201783 | 213.848536239696 | Telo2 |
| **ENSMUSG00000024174** | -0.14504774581847 | 0.604152053060642 | 0.926448151609826 | 39.754603529038 | 33.3910448854875 | 22.7149173275627 | 28.1378008277448 | 37.2285786578493 | 35.4693847301419 | 32.2734208072402 | 32.5657669400552 | Pot1b |
| **ENSMUSG00000024177** | 0.213570826247302 | 0.271912411085987 | 0.84423521828958 | 126.491920319666 | 109.946123403434 | 125.878500190243 | 163.19924480092 | 136.869774477387 | 107.6312364225 | 109.313199508394 | 98.7828263848342 | Nme4 |
| **ENSMUSG00000024180** | 0.33692420716178 | 0.0519738315227893 | 0.703396849069134 | 228.588970291968 | 275.272516373043 | 313.276568142636 | 268.247034557834 | 211.326931793086 | 166.339183562045 | 242.57119509958 | 237.730098662403 | Tmem8 |
| **ENSMUSG00000024181** | 0.00330916649698159 | 0.981691333768892 | 0.996940244108351 | 4076.65388915953 | 3044.28599468176 | 3053.26347077989 | 3799.54103843981 | 3449.11831683016 | 3941.99403397405 | 3098.24839749506 | 3453.05682121052 | Mrpl28 |
| **ENSMUSG00000024182** | 0.0880127379884199 | 0.428526547585256 | 0.885378691567738 | 559.274990556239 | 583.936077631574 | 638.857049837702 | 503.666634816633 | 535.434557755538 | 545.494675504941 | 518.456889096956 | 551.446986851602 | Axin1 |
| **ENSMUSG00000024187** | 0.0575914544689627 | 0.575205732130404 | 0.922450244597442 | 2848.7787483422 | 3100.48067997685 | 3322.05665915605 | 2860.67641748739 | 2837.03668536728 | 2672.43467708138 | 3205.47944082234 | 2940.68875468699 | Fam234a |
| **ENSMUSG00000024188** | -0.0374944543117655 | 0.803650960099572 | 0.963461896828986 | 142.755167217909 | 165.326392969609 | 163.736695736181 | 139.751077444466 | 173.003394939417 | 167.562265794119 | 142.627698406191 | 145.46042566558 | Luc7l |
| **ENSMUSG00000024190** | -0.414768795204451 | 0.0636147918007638 | 0.716991331547963 | 894.478579403355 | 855.136515360045 | 807.326020017125 | 737.210381686915 | 1491.33306270561 | 1232.86688993045 | 952.586452858865 | 715.361347116547 | Dusp1 |
| **ENSMUSG00000024191** | 0.0181144654593032 | 0.90761945926522 | 0.98504921011344 | 439.107666252556 | 364.858246553619 | 296.240380146964 | 420.191159027656 | 389.805117711598 | 397.501725424004 | 345.637926064637 | 369.078691987293 | Bnip1 |
| **ENSMUSG00000024193** | -0.0393822147874177 | 0.779488348878663 | 0.960444627083598 | 366.826568927032 | 397.434875710192 | 429.690519446395 | 439.887619607078 | 434.698403740182 | 495.348303989913 | 381.034581143546 | 369.078691987293 | Phf1 |
| **ENSMUSG00000024194** | -0.242261845842972 | 0.236957574092362 | 0.832048466795696 | 814.06585862871 | 671.078560625407 | 600.052399403115 | 804.741103673502 | 905.530427942393 | 1135.02031136454 | 649.632728507029 | 729.473179457237 | Cuta |
| **ENSMUSG00000024197** | 0.179187152019311 | 0.135922334408385 | 0.776087404131513 | 6308.33276908507 | 5695.2091922979 | 6877.88767580826 | 7020.38130652234 | 6127.38606380367 | 4975.49852007646 | 6187.12709217512 | 5583.9435046548 | Plin3 |
| **ENSMUSG00000024201** | -0.0535976945176058 | 0.720965781539546 | 0.951674101666722 | 657.757985662265 | 650.718167402549 | 784.611102689562 | 639.666005484066 | 796.034608360484 | 567.510155682271 | 790.178270732108 | 680.624529047154 | Kdm4b |
| **ENSMUSG00000024206** | 0.281248151565837 | 0.434980016695917 | 0.888379878975956 | 38.8510898124689 | 50.4937751926884 | 43.5369248778286 | 23.448167356454 | 49.2731188118594 | 33.0232202659942 | 23.9447960827911 | 22.7960368580387 | Rfx2 |
| **ENSMUSG00000024208** | 0.111175556667961 | 0.567814948456144 | 0.920429362097163 | 2868.65605010672 | 1834.87863724398 | 1941.17897661796 | 2476.12647284155 | 2037.71720241934 | 2636.96529235124 | 1743.80580168153 | 2026.6762292361 | Uqcc2 |
| **ENSMUSG00000024210** | -0.0994136841220497 | 0.783398588690428 | 0.961051227381536 | 131.913002619081 | 71.6685841444609 | 51.1085639870161 | 76.9099889291692 | 82.1218646864323 | 51.3694537471021 | 133.257995591185 | 87.9275707381491 | Ip6k3 |
| **ENSMUSG00000024212** | 0.11990739721054 | 0.413682652665771 | 0.882192126163777 | 661.372040528541 | 781.839099757755 | 771.360734248484 | 657.486612674971 | 567.188345434292 | 588.302553627526 | 773.52102128321 | 713.190295987209 | Mllt1 |
| **ENSMUSG00000024213** | 0.0113253094808091 | 0.931925086700096 | 0.987906435243901 | 4916.92164556874 | 4368.52596989646 | 4345.17439378502 | 4287.26291945406 | 3657.16037403578 | 4196.39513824541 | 4995.09267848834 | 4928.28606359502 | Nudt3 |
| **ENSMUSG00000024217** | -0.115263536131361 | 0.47309705913093 | 0.89728597737823 | 550.239853390548 | 434.083583511337 | 513.925004536107 | 572.135283497478 | 597.847174917227 | 671.472145408549 | 479.936999746379 | 493.914131924171 | Snrpc |
| **ENSMUSG00000024218** | 0.0695578175717901 | 0.515786027240092 | 0.904239892322932 | 551.143367107117 | 488.649437348597 | 448.619617219364 | 511.170048370698 | 475.211856985488 | 473.332823812584 | 466.402984569149 | 490.657555230165 | Taf11 |
| **ENSMUSG00000024219** | -0.0449509242766073 | 0.802899037828305 | 0.963461896828986 | 1211.61189391909 | 1436.62934580488 | 1405.48550964294 | 1103.93971914186 | 1201.16914081355 | 1012.71208815716 | 1619.91750890535 | 1486.08449803119 | Anks1 |
| **ENSMUSG00000024220** | 0.216523061911639 | 0.245761817383595 | 0.832048466795696 | 234.010052591383 | 229.665235553841 | 300.972654590206 | 240.109233730089 | 160.958854785407 | 261.739597663806 | 214.462086654564 | 227.960368580387 | Zfp523 |
| **ENSMUSG00000024222** | 0.0793607756116853 | 0.720566286930727 | 0.951674101666722 | 2345.52160821324 | 2479.89589454413 | 3036.22728278422 | 1917.12216306368 | 2327.8811243114 | 1634.03786205068 | 3120.11103739674 | 2172.13665490168 | Fkbp5 |
| **ENSMUSG00000024228** | -0.0288462952026488 | 0.903544635384323 | 0.985027469232262 | 370.440623793308 | 465.031381210082 | 426.85115478045 | 415.501525556365 | 404.039574257247 | 255.624186503437 | 581.96265262088 | 468.947043936795 | Nudt12 |
| **ENSMUSG00000024231** | 0.169157856354778 | 0.361133327385636 | 0.868539734006824 | 1075.18132271716 | 965.897054492394 | 965.383986421416 | 961.374861614615 | 757.711071506815 | 647.010500767072 | 1029.62623156002 | 1093.12424362119 | Cul2 |
| **ENSMUSG00000024232** | 0.14424377817678 | 0.488833889543136 | 0.899236955296196 | 449.046317134815 | 499.236841824483 | 276.364827485346 | 475.528833988888 | 354.766455445387 | 456.20967256355 | 375.829190690765 | 352.795808517265 | Bambi |
| **ENSMUSG00000024234** | 0.020738896046029 | 0.880892007703172 | 0.97846486749867 | 352.370349461928 | 402.321370083678 | 362.492222352355 | 332.963976461647 | 373.380744774312 | 294.7628179298 | 361.254097422979 | 399.473407798011 | Mtpap |
| **ENSMUSG00000024235** | -0.417290690290617 | 0.139050163468194 | 0.779711473022872 | 24.3948703473642 | 21.1748089517725 | 20.8220075502658 | 28.1378008277448 | 33.943704070392 | 28.1308913376988 | 31.2323427166841 | 32.5657669400552 | Map3k8 |
| **ENSMUSG00000024236** | 0.477934440062218 | 0.136817599173121 | 0.776456211338551 | 7116.97754541437 | 9505.04597215913 | 9262.95399520235 | 6897.51290957452 | 4540.7916380618 | 2754.38118663033 | 8178.70947940901 | 8063.28389435767 | Svil |
| **ENSMUSG00000024238** | 0.108260050595511 | 0.584971946538927 | 0.922695927484181 | 1384.18301378378 | 1571.00794107574 | 1350.59112610133 | 1154.5877606318 | 1065.39432453198 | 939.327154232724 | 1524.13832457418 | 1536.01867400594 | Zeb1 |
| **ENSMUSG00000024240** | 0.143835071123461 | 0.362971866726451 | 0.86924083181804 | 411.098741038915 | 434.083583511337 | 427.797609669098 | 332.963976461647 | 348.196706270473 | 302.101311322243 | 370.623800237985 | 432.039174738066 | Epc1 |
| **ENSMUSG00000024241** | 0.162089713612484 | 0.421594586508403 | 0.882568531013176 | 286.413848152387 | 327.39512302356 | 335.04503058155 | 296.384835385579 | 273.739548954774 | 182.239252579005 | 332.103910887408 | 323.486618271215 | Sos1 |
| **ENSMUSG00000024242** | -0.0122201198798853 | 0.922944539753786 | 0.986713918026868 | 429.169015370296 | 492.721515993169 | 500.674636095028 | 475.528833988888 | 547.479097909548 | 441.532685778663 | 428.924173309128 | 496.085183053508 | Map4k3 |
| **ENSMUSG00000024246** | -0.312409811952113 | 0.254758152069379 | 0.837406774112203 | 48.7897406947284 | 27.6901347830872 | 29.3401015481019 | 43.1446279358754 | 41.6084114411257 | 45.2540425867328 | 52.0539045278068 | 45.5920737160773 | Thumpd2 |
| **ENSMUSG00000024247** | 0.0232495512771172 | 0.850722448266364 | 0.972692784625411 | 666.793122827955 | 688.995706661522 | 747.699362032273 | 846.94780491512 | 702.96316171586 | 659.24132308781 | 791.219348822664 | 747.927114056602 | Pkdcc |
| **ENSMUSG00000024248** | 0.0840412632709453 | 0.405522488889215 | 0.881158207349486 | 6079.7437987931 | 5116.97402476873 | 5181.84051535025 | 6149.04740755651 | 5218.57076127382 | 5519.77011334933 | 5136.67929880398 | 5377.69364736779 | Cox7a2l |
| **ENSMUSG00000024251** | -0.00636944105531518 | 0.976715369887476 | 0.996134981677394 | 107.518132271716 | 188.13003337921 | 144.807597963212 | 117.24083678227 | 142.344565456483 | 132.092881063977 | 161.367104036201 | 124.835439936878 | Thada |
| **ENSMUSG00000024253** | 0.0655975733519153 | 0.798892932506766 | 0.963240258978423 | 63.2459601598332 | 71.6685841444609 | 51.1085639870161 | 62.8410885152968 | 85.4067392738896 | 55.0387004433237 | 51.0128264372507 | 46.6775992807458 | Dync2li1 |
| **ENSMUSG00000024258** | -0.124910971173261 | 0.388735479748119 | 0.876405748044371 | 473.44118748218 | 388.476302692135 | 352.081218577222 | 378.922384480297 | 474.116898789669 | 464.771248188067 | 376.870268781321 | 422.269444656049 | Polr2d |
| **ENSMUSG00000024259** | 0.123316412903943 | 0.55977302898463 | 0.916574868225118 | 1544.1049416165 | 1839.76513161747 | 1963.89389394553 | 1510.9999044499 | 1373.07757755715 | 1083.65085761744 | 1996.78777768667 | 1842.13688324246 | Slc25a46 |
| **ENSMUSG00000024260** | 0.0271183762219043 | 0.84110111255801 | 0.970420002017028 | 309.905204783182 | 394.177212794535 | 389.93941412316 | 323.584709519066 | 360.241246424483 | 314.332133642982 | 358.130863151311 | 358.223436340608 | Sap130 |
| **ENSMUSG00000024269** | -0.0894027027642089 | 0.534639966783391 | 0.910706162399539 | 181.606257030378 | 157.99665140938 | 161.843785958884 | 182.895705380341 | 155.484063806312 | 190.800828203522 | 190.517290571773 | 191.052499381657 | Tpgs2 |
| **ENSMUSG00000024270** | -0.107526063592282 | 0.628867241457131 | 0.931825800800958 | 110.228673421423 | 144.96599974675 | 136.289503965376 | 135.061443973175 | 179.573144114332 | 168.785348026193 | 113.477511870619 | 106.381505337514 | Slc39a6 |
| **ENSMUSG00000024271** | -0.239221551778909 | 0.14019177892477 | 0.779711473022872 | 273.764656120421 | 377.888898216248 | 292.45456059237 | 328.274342990356 | 393.089992299056 | 440.309603546589 | 339.391457521301 | 331.085297223895 | Elp2 |
| **ENSMUSG00000024273** | -0.176702828674668 | 0.540906398842779 | 0.913837378403038 | 68.6670424592474 | 44.7928650902881 | 31.2330113253987 | 55.3376749612315 | 55.8428679867739 | 70.9387694602839 | 52.0539045278068 | 47.7631248454143 | 2700062C07 |
| **ENSMUSG00000024276** | 0.241886700460039 | 0.337386344600858 | 0.858570960161243 | 304.484122483768 | 327.39512302356 | 361.545767463707 | 281.378008277448 | 205.85214081399 | 173.677676954488 | 369.582722147429 | 327.828720529889 | Zfp397 |
| **ENSMUSG00000024277** | 0.0216771429358915 | 0.879695878419826 | 0.978393501240173 | 2567.78598248923 | 2920.49480388678 | 2239.31226654223 | 2577.42255582143 | 2194.29622442147 | 2287.16377397812 | 2717.21381635152 | 2952.62953589834 | Mapre2 |
| **ENSMUSG00000024283** | 0.215920045591626 | 0.162639709414797 | 0.793968226938511 | 1058.01456210235 | 1272.9317842931 | 1259.73145679108 | 1022.3400967414 | 946.0438811877 | 802.341944240452 | 1164.96638333232 | 1057.30189998713 | Wac |
| **ENSMUSG00000024286** | 0.0777491503825159 | 0.531062655763503 | 0.909082872831377 | 2888.53335187124 | 3058.1310620733 | 3022.97691434314 | 2572.73292235014 | 2693.59716171498 | 2294.50226737056 | 2953.53854290776 | 2993.87950735574 | Ccny |
| **ENSMUSG00000024287** | 0.104066561768839 | 0.365861623530972 | 0.869253398153585 | 460.791995450213 | 399.878122896935 | 380.474865236676 | 423.004939110431 | 355.861413641207 | 387.717067567413 | 402.897221045225 | 401.644458927348 | Thoc1 |
| **ENSMUSG00000024290** | -0.0194724226090343 | 0.843126998268363 | 0.971386364129692 | 1859.43122869909 | 1973.32931115942 | 1638.31341225046 | 1698.58524330153 | 1831.86506160535 | 1922.68526882011 | 1749.01119213431 | 1766.15009371566 | Rock1 |
| **ENSMUSG00000024292** | -0.316021045909545 | 0.550643892234555 | 0.914699131434573 | 17.1667606148119 | 29.3189662409158 | 8.51809399783602 | 23.448167356454 | 19.7092475247437 | 14.6769867848863 | 14.5750932677859 | 48.8486504100828 | Cyp4f14 |
| **ENSMUSG00000024293** | 0.162443098468145 | 0.336338044766387 | 0.858570960161243 | 341.528184863099 | 317.622134276588 | 301.919109478855 | 316.081295965 | 237.605928492744 | 243.393364182698 | 293.584021536831 | 365.822115293287 | Esco1 |
| **ENSMUSG00000024294** | 0.279363192770595 | 0.120131480158059 | 0.769868040577702 | 742.688275019755 | 953.680818558679 | 987.15244886033 | 905.099259959126 | 746.761489548624 | 533.263853184203 | 853.684034256032 | 821.74285245406 | Mib1 |
| **ENSMUSG00000024298** | -0.068539261598984 | 0.866811514903416 | 0.974868041734905 | 46.9827132615903 | 99.3587189275481 | 106.949402417274 | 43.1446279358754 | 66.7924499449649 | 41.5847958905112 | 122.847214685624 | 79.2433662208011 | Zfp871 |

| | | | | | | | | | | | | |
|---|---|---|---|---|---|---|---|---|---|---|---|---|
| ENSMUSG00000024299 | 0.0452153330220586 | 0.849893114638983 | 0.972640876584422 | 58.7283915769879 | 65.9676740420606 | 72.8770264259304 | 78.7858423176855 | 50.3680770076785 | 91.7311674055395 | 65.5879197050366 | 60.7894316214364 | Adamts10 |
| ENSMUSG00000024300 | -0.31172707823417 | 0.32400756278376 | 0.855977678491063 | 26.2018977805023 | 35.0198763433161 | 37.8581955459379 | 34.703287687552 | 24.0890803080201 | 53.8156182112498 | 36.4377331694648 | 52.1052271040884 | Myo1f |
| ENSMUSG00000024302 | 0.12693445339782 | 0.597637334654332 | 0.924496535139492 | 966.759676728878 | 1003.36017802245 | 996.616997746814 | 962.312788308873 | 713.912743674051 | 562.617826753975 | 1231.59538112791 | 1088.78214136251 | Dtna |
| ENSMUSG00000024304 | 0.174605213466896 | 0.341065453489888 | 0.859283395807012 | 5730.98750419745 | 6689.6107973023 | 6201.17243042462 | 5093.87987651607 | 4287.85629482758 | 4191.50280931712 | 6208.9897320768 | 6323.18641419406 | Cdh2 |
| ENSMUSG00000024308 | -0.486049932271204 | 0.0771975675408951 | 0.739855948845396 | 1120.35700854562 | 1115.74954861263 | 1227.55199057704 | 1148.02227377199 | 2162.54243674272 | 2210.10959335746 | 1122.28218161952 | 965.032226990303 | Tapbp |
| ENSMUSG00000024309 | -0.0705747112634593 | 0.740375446039029 | 0.954383574136061 | 1311.90191645825 | 1006.61784093811 | 1035.4216481814 | 1385.3177274193 | 1280.00613091252 | 1697.63813811852 | 1052.52994955225 | 947.663817955607 | Pfdn6 |
| ENSMUSG00000024312 | 0.0257722861713939 | 0.863788185927652 | 0.974868041734905 | 273.764656120421 | 287.488752306758 | 292.45456059237 | 349.846656958294 | 289.068963696242 | 340.016860516533 | 305.035880532948 | 248.585354309088 | Wdr46 |
| ENSMUSG00000024317 | -0.0318516529492744 | 0.867493613049622 | 0.974868041734905 | 97.5794813894569 | 96.9154717408051 | 108.842312194571 | 98.4823028971069 | 119.350443344282 | 121.085140975312 | 86.4094815161593 | 84.6709940441436 | Rnf138 |
| ENSMUSG00000024319 | 0.0448971392087419 | 0.542082393532667 | 0.913837378403038 | 1316.4194850411 | 1303.06516626293 | 1294.75028767108 | 1326.22834568104 | 1257.01200880032 | 1290.35175483792 | 1323.21025309685 | 1209.27547904072 | Vps52 |
| ENSMUSG00000024325 | 0.126481730851296 | 0.311050634669618 | 0.851785577030345 | 380.379274675568 | 386.033055505392 | 386.153594568566 | 409.873965390816 | 363.52612101194 | 417.071041137186 | 335.227145159076 | 316.973464883204 | Ring1 |
| ENSMUSG00000024327 | 0.0891689001026898 | 0.693832779805503 | 0.94469184542218 | 344.238726012806 | 401.506954354764 | 384.260684791269 | 310.453735799451 | 307.683253025166 | 216.485555077073 | 442.458188486358 | 386.447101021989 | Slc39a7 |
| ENSMUSG00000024331 | 0.0479813698724256 | 0.853896616719092 | 0.9731539929935 | 340.62467114653 | 429.197089137851 | 442.940887887473 | 337.653609932938 | 309.873169416804 | 225.04713070159 | 512.210420553619 | 451.578634902099 | Dsc2 |
| ENSMUSG00000024335 | 0.041287318968462 | 0.687863968269217 | 0.944452327903597 | 1295.63866956001 | 1538.43131191917 | 1341.12657721485 | 1318.72493212697 | 1361.03303740314 | 1257.32853457193 | 1439.81099923914 | 1280.92016630884 | Brd2 |
| ENSMUSG00000024338 | -0.339451804929663 | 0.309519117205027 | 0.851785577030345 | 289.124389302094 | 193.016527752696 | 198.755526616174 | 292.633128608546 | 382.140410340865 | 477.002070508805 | 180.106509666212 | 193.223550510994 | Psmb8 |
| ENSMUSG00000024339 | -0.414356739073363 | 0.125779196326668 | 0.771949105062466 | 524.037955610046 | 518.782819318427 | 535.693466975021 | 579.638697051544 | 928.524550054594 | 1006.59667699679 | 463.279750297481 | 478.716774018812 | Tap2 |
| ENSMUSG00000024346 | 0.128836487414137 | 0.422378003468231 | 0.883425759399749 | 2038.32694457977 | 1486.30870526865 | 1613.7055851456 | 2123.46603580048 | 1692.80537073632 | 1919.01602212389 | 1492.9059818575 | 1537.10419957061 | Pfdn1 |
| ENSMUSG00000024349 | -0.802269511454293 | 0.0270880716646482 | 0.664353746352133 | 37.9475760958999 | 30.1333819698302 | 45.4298346551254 | 51.5859681841989 | 99.6411958195378 | 100.292743030056 | 47.8895921655823 | 40.1644458927348 | Tmem173 |
| ENSMUSG00000024350 | 0.0650842888906375 | 0.590155883156403 | 0.923935529355527 | 542.108229941427 | 547.287369830429 | 502.567545872325 | 581.51455044006 | 486.161438943679 | 463.548165955993 | 591.332355435886 | 535.164103381574 | Dnajc18 |
| ENSMUSG00000024352 | 0.199809619642326 | 0.324743107265795 | 0.855977678491063 | 125.588406603097 | 92.0289773673191 | 106.002947528626 | 113.489130005238 | 71.1722827282413 | 92.9542496376133 | 105.14888714617 | 110.723607596188 | Spata24 |
| ENSMUSG00000024353 | 0.125974115274583 | 0.674955607535909 | 0.944023556644042 | 32.5264937964856 | 36.6487078011448 | 20.8220075502658 | 36.5791410760683 | 35.0386622662111 | 26.9078091056249 | 24.9858741733473 | 29.3091902460497 | Mzb1 |
| ENSMUSG00000024357 | -0.0332511164288533 | 0.840547594815802 | 0.97002376969022 | 646.012307346867 | 564.39010013763 | 579.230391852849 | 651.859052509422 | 620.841297029428 | 807.234273168747 | 550.730309904196 | 521.052271040884 | Sil1 |
| ENSMUSG00000024359 | 0.287012210012139 | 0.0983809168653496 | 0.759800953500741 | 22586.9394005096 | 24155.570519599 | 23701.1233215344 | 22766.2946497283 | 17915.7059999921 | 13583.5512694123 | 22788.1583241833 | 22105.6425989095 | Hspa9 |
| ENSMUSG00000024360 | 0.174802716221705 | 0.308949905212963 | 0.851785577030345 | 3565.26512558145 | 3854.62964495152 | 3726.19289660893 | 3588.50753223172 | 2741.77532233102 | 2520.77248030422 | 3924.86440139664 | 3864.47101021989 | Etf1 |
| ENSMUSG00000024366 | -1.41425158776115 | 0.0836256924594412 | 0.742148948523533 | 7.22810973255236 | 17.1027303072009 | 1.89290977729689 | 12.1930470253561 | 33.943704070392 | 57.4848649074714 | 6.24646854333682 | 5.42762782334254 | Gfra3 |
| ENSMUSG00000024369 | 0.0441658015913235 | 0.768900410484262 | 0.95879680569409 | 369.537110076739 | 347.755516246418 | 352.081218577222 | 416.439452250624 | 405.134532453066 | 410.955629976817 | 319.610973800734 | 306.118209236519 | Nelfe |
| ENSMUSG00000024370 | 0.0550765952491656 | 0.687577654246041 | 0.944452327903597 | 501.45011269582 | 536.699965354543 | 538.532831640966 | 474.59090729463 | 464.262275027297 | 408.509465512669 | 542.401685179747 | 557.960140239613 | Cdc23 |
| ENSMUSG00000024371 | -0.615337818236105 | 0.203808680799828 | 0.816074141171399 | 7.22810973255236 | 13.8450673915436 | 6.62518422053913 | 6.56548685980713 | 9.85462376237187 | 11.0077400886647 | 13.5340151772298 | 18.4539345993646 | C2 |
| ENSMUSG00000024376 | -0.179578731412905 | 0.413896879480586 | 0.882192126163777 | 115.649755720838 | 103.43079757212 | 67.1982970940397 | 91.9168160372998 | 118.255485148462 | 85.6157562451702 | 108.272121417838 | 116.15123541953 | Epb41l4a |
| ENSMUSG00000024378 | -0.152732564154039 | 0.466610263910938 | 0.895328112556867 | 117.456783153976 | 136.007426728693 | 132.503684410783 | 182.895705380341 | 174.098353135236 | 189.577745971448 | 124.929370866736 | 144.374900100912 | Stard4 |
| ENSMUSG00000024381 | -0.31039328921127 | 0.295531494360292 | 0.851785577030345 | 139.141112351633 | 89.5857301805761 | 182.66579350915 | 146.316564304273 | 209.137015401448 | 234.831788558181 | 122.847214685624 | 124.835439936878 | Bin1 |
| ENSMUSG00000024382 | 0.00506129579143551 | 0.956648989158477 | 0.992371505557225 | 414.712795905192 | 408.836695914993 | 396.564598343699 | 394.867138282686 | 399.65974147397 | 401.170972120226 | 407.061533407449 | 401.644458927348 | Ercc3 |
| ENSMUSG00000024383 | -0.147516020039603 | 0.394056673441009 | 0.878942305112115 | 146.369222084185 | 205.232763686411 | 175.094154399963 | 151.944124469822 | 171.908436743598 | 172.454594722414 | 187.394056300105 | 220.361689627707 | Map3k2 |
| ENSMUSG00000024384 | 0.0382886593889398 | 0.768999373918254 | 0.95879680569409 | 506.871194995234 | 645.017257300148 | 595.320124959873 | 535.55614242141 | 571.568178217569 | 480.671317205027 | 584.044808801993 | 586.183804920994 | Iws1 |
| ENSMUSG00000024386 | -0.134877548403807 | 0.72977460617047 | 0.952921973914044 | 61.4389327266951 | 119.719112150406 | 64.3589324280944 | 77.8479156234274 | 59.1277425742312 | 55.0387004433237 | 77.0397787011541 | 163.914360264945 | Proc |
| ENSMUSG00000024387 | -0.00497933934960068 | 0.971775142897386 | 0.995123114262614 | 1918.15962027608 | 1598.69807585883 | 1693.20779579207 | 2058.74909389666 | 2020.19787128623 | 2021.75492961809 | 1675.09464770482 | 1577.26864546334 | Csnk2b |
| ENSMUSG00000024391 | -0.158623307330905 | 0.637570863081619 | 0.932981383080877 | 37.0440623793308 | 45.6072808192024 | 31.2330113253987 | 28.1378008277448 | 21.8991639163819 | 46.4771248188067 | 35.3966550789086 | 55.3618037980939 | Apom |
| ENSMUSG00000024392 | 0.0676439907389828 | 0.451893515990383 | 0.891110666112841 | 4892.52677522138 | 5366.99965354543 | 5545.27919259125 | 4988.83208675916 | 5113.45477447518 | 4540.08124545817 | 5224.12985841069 | 4961.93735609975 | Bag6 |
| ENSMUSG00000024393 | -0.0111472065892013 | 0.955815552230848 | 0.992371505557225 | 2984.30580582756 | 3883.13419546352 | 3694.95988528354 | 2726.55290020847 | 3080.11740483912 | 2418.03357281002 | 4063.3277874406 | 3829.73419215049 | Prrc2a |
| ENSMUSG00000024395 | -0.593912921729592 | 0.100018482439904 | 0.759800953500741 | 1524.22763985198 | 1369.03284030499 | 1458.48698340726 | 1366.55919353414 | 2658.55849944877 | 3724.2853966649 | 1150.39129006453 | 1098.55187144453 | Lims2 |
| ENSMUSG00000024400 | 0.147022301532533 | 0.206728116497265 | 0.819960256425221 | 1047.17239750352 | 1082.35850372714 | 1074.22629861599 | 959.499008226099 | 964.658170516624 | 792.557286383861 | 1031.70838774113 | 969.374329248977 | Wdr33 |
| ENSMUSG00000024403 | 0.0901395803682286 | 0.666630106049298 | 0.942024383071524 | 75.8951521917998 | 67.5965054998893 | 64.3589324280944 | 60.0273084325223 | 52.5579933993167 | 56.2617826753975 | 68.711153976705 | 73.8157383974585 | Atp6v1g2 |
| ENSMUSG00000024404 | 0.262330708082134 | 0.0566611002394384 | 0.705217457281182 | 2434.06595243701 | 2271.40546794206 | 2513.78418425027 | 2340.12710217411 | 1796.82639933914 | 1647.49176660349 | 2305.98797058184 | 2217.72872861776 | Riok3 |
| ENSMUSG00000024410 | 0.061383489667193 | 0.593393931015898 | 0.923935529355527 | 487.897406947284 | 526.926976607571 | 513.925004536107 | 494.287367874051 | 560.618596259378 | 456.20967256355 | 448.704657029695 | 473.289146195469 | 3110002H16 |
| ENSMUSG00000024411 | -0.0821388100975503 | 0.867309184306614 | 0.974868041734905 | 28.9124389302094 | 48.0505280059454 | 35.965285768641 | 7.50341355406529 | 31.7537876787538 | 39.1386314263635 | 37.4788112600209 | 19.5394601640331 | Aqp4 |
| ENSMUSG00000024413 | 0.216751860227602 | 0.187211922910269 | 0.807540361382744 | 1101.38322049767 | 1549.01871639505 | 1393.18159609051 | 1035.47107046101 | 1136.56660726022 | 945.442565393093 | 1166.00746142287 | 1122.43343386724 | Npc1 |
| ENSMUSG00000024414 | -0.0119771821657092 | 0.946599047026074 | 0.990510945542179 | 915.259394884443 | 719.129088631352 | 667.250696497155 | 790.67220325963 | 742.381656765348 | 1015.1582526213 | 694.399086400943 | 667.598222271132 | Mrpl27 |
| ENSMUSG00000024420 | -0.336390293121774 | 0.0602931135255456 | 0.71282992338528 | 119.263810587114 | 119.719112150406 | 130.610774633486 | 146.316564304273 | 158.768938393769 | 199.362403828039 | 130.134761319517 | 163.914360264945 | Zfp521 |
| ENSMUSG00000024421 | -0.452455701170099 | 0.140405900642175 | 0.779711473022872 | 42.4651446787451 | 68.4109212288036 | 40.6975602118832 | 45.0204813243917 | 82.1218646864323 | 90.5080851734656 | 49.9717483466946 | 47.7631248454143 | Lama3 |
| ENSMUSG00000024422 | 0.106199631308335 | 0.286762035000793 | 0.851785577030345 | 661.372040528541 | 672.707392083235 | 705.108892043093 | 634.976372012775 | 644.930377337448 | 558.948580057754 | 657.961353231478 | 620.920622990386 | Dhx16 |
| ENSMUSG00000024423 | 0.00303652268730395 | 0.984236253956646 | 0.996940244108351 | 414.712795905192 | 467.474628396825 | 388.992959234512 | 498.039074651084 | 382.140410340865 | 391.386314263635 | 476.813765474711 | 514.539117652873 | Impact |
| ENSMUSG00000024424 | 0.0557941722166286 | 0.800398389241839 | 0.963461896828986 | 194.255449062345 | 205.232763686411 | 200.648436393471 | 200.716312571247 | 168.623562156141 | 125.977469903608 | 219.667477107345 | 255.098507697099 | Ttc39c |
| ENSMUSG00000024425 | 0.0381421504723296 | 0.582848170963418 | 0.922450244597442 | 2449.42568561868 | 2387.0525014479 | 2323.54675163194 | 2536.15378127407 | 2358.53995379433 | 2419.25665504209 | 2301.82365821962 | 2364.27467984801 | Ndfip1 |
| ENSMUSG00000024426 | -0.0346693410511136 | 0.882343372337054 | 0.978854883720278 | 73.1846110420926 | 118.904696421492 | 79.5022106464695 | 81.59962240046 | 85.4067392738896 | 101.51582526213 | 70.7933101578173 | 105.295979772845 | Atat1 |
| ENSMUSG00000024427 | -0.232861038553722 | 0.416111802653809 | 0.882192126163777 | 89.4478579403355 | 89.5857301805761 | 129.664319744837 | 146.316564304273 | 202.567266226533 | 119.862058743238 | 98.902418602833 | 112.894658725525 | Spry4 |
| ENSMUSG00000024429 | -0.00794158408078971 | 0.921815679295592 | 0.986713918026868 | 1438.39383677792 | 1391.83648071459 | 1474.57671651428 | 1545.70319213745 | 1509.94735203453 | 1526.40662562818 | 1447.09854587303 | 1399.24245285771 | Gnl1 |
| ENSMUSG00000024431 | -0.0898128118097901 | 0.278238536195323 | 0.848075053184273 | 1084.21645988285 | 1108.4198070524 | 1117.76322349382 | 1113.31898608444 | 1204.45401540101 | 1088.54318654574 | 1163.92530524176 | 1249.43992493345 | Nr3c1 |
| ENSMUSG00000024436 | 0.0858422545742268 | 0.523539679200971 | 0.906728945648564 | 658.661499378834 | 513.896324944941 | 539.479286529615 | 663.11417284052 | 605.511882287961 | 519.80994863139 | 534.073060455298 | 577.499600403646 | Mrps18b |
| ENSMUSG00000024440 | -0.742104372107844 | 0.0155266678732724 | 0.635862253526407 | 268.343573821006 | 198.717437855096 | 192.130342395635 | 274.812521417641 | 630.6959207918 | 414.624876673038 | 246.735507461804 | 270.295865602458 | Pcdh12 |
| ENSMUSG00000024442 | 0.161849643708115 | 0.309242057936045 | 0.851785577030345 | 3449.61536986061 | 3425.43255581367 | 3506.6153624425 | 3402.79804676861 | 2734.11061496029 | 2375.22569468744 | 3519.8850241703 | 3690.78691987293 | 0610009O20 |
| ENSMUSG00000024446 | -0.102891279379913 | 0.620621000936854 | 0.930228305062361 | 349.65980831222 | 266.313943354985 | 251.757000380487 | 295.446908691321 | 351.48158085793 | 398.724807656078 | 264.433835001259 | 235.559047533066 | Rpp21 |
| ENSMUSG00000024451 | -0.248777229883858 | 0.164492099878697 | 0.795216960550936 | 829.425591810383 | 736.231818938553 | 847.07712534036 | 923.857793844289 | 1225.25822112157 | 1126.45873574002 | 784.972880279327 | 828.256005842071 | Arap3 |
| ENSMUSG00000024454 | 0.210288277532882 | 0.0641452138645144 | 0.716991331547963 | 1222.45405851792 | 1032.67914426337 | 1162.24660326029 | 1129.26373988683 | 1002.98170737029 | 847.595986827185 | 1058.77641809559 | 1018.22297965906 | Hdac3 |
| ENSMUSG00000024456 | 0.0792543700134707 | 0.5973669625188 | 0.924496535139492 | 749.012871035738 | 943.907829811707 | 931.311610430072 | 780.35500962279 | 717.197618261509 | 698.379954514174 | 917.189797779956 | 889.045437463508 | Diaph1 |
| ENSMUSG00000024457 | -0.0605376620336592 | 0.579916962688037 | 0.922450244597442 | 411.098741038915 | 456.072808192024 | 396.564598343699 | 390.177504811395 | 427.033696369448 | 402.3940543523 | 458.0743598447 | 437.466802561409 | Trim26 |
| ENSMUSG00000024462 | -0.0561532320544935 | 0.745207891256876 | 0.95596456017978 | 116.553269437407 | 105.874044758863 | 140.07532351997 | 106.92364314543 | 137.964732673206 | 123.53130543946 | 117.641824232843 | 108.552556466851 | Gabbr1 |
| ENSMUSG00000024471 | 0.58869115789207 | 0.00239606102578646 | 0.635862253526407 | 2962.6214766299 | 3384.71176936795 | 3587.06402797761 | 3873.63724728621 | 1761.78773707293 | 1963.04698247854 | 3082.63222613672 | 2372.95888436536 | Myot |
| ENSMUSG00000024472 | 0.0855465756309203 | 0.708406879566034 | 0.949174314815223 | 268.343573821006 | 331.467201668132 | 344.509579468035 | 218.536919762152 | 251.840385038392 | 190.800828203522 | 327.939598525183 | 324.572143835884 | Dcp2 |
| ENSMUSG00000024474 | 0.00630082509216658 | 0.951604277396367 | 0.991915011884503 | 1994.05477246788 | 1870.71292931621 | 2107.75503702009 | 2130.96944935454 | 2132.9785654556 | 2122.04767264815 | 2048.84168221448 | 1763.97904258632 | Ik |
| ENSMUSG00000024477 | 0.127943167502709 | 0.723201684839788 | 0.951674101666722 | 224.071401709123 | 319.250965734417 | 307.597838810745 | 164.137171495178 | 169.71852035196 | 111.300483118721 | 347.72008224575 | 299.605055848508 | Pggt1b |
| ENSMUSG00000024479 | -0.0922617205428403 | 0.851400707689081 | 0.972692784625411 | 16.2632468982428 | 17.9171460361152 | 5.67872933189068 | 15.9447538023887 | 12.0445401540101 | 12.2308223207386 | 10.4107809055614 | 24.9670879873757 | Mal2 |
| ENSMUSG00000024480 | -0.143827620955393 | 0.329234881169494 | 0.855977678491063 | 309.905204783182 | 289.931999493501 | 244.185361271299 | 345.157023487003 | 331.772333333186 | 357.140011765567 | 294.625099627387 | 331.085297223895 | Ap3s1 |
| ENSMUSG00000024481 | 0.0468152634990367 | 0.898477521105041 | 0.984462340635154 | 40.658117245607 | 43.9784493613737 | 60.5731128735006 | 29.075727522003 | 65.6974917491458 | 42.8078781225851 | 37.4788112600209 | 22.7960368580387 | Lvrn |
| ENSMUSG00000024483 | 0.0724201585273788 | 0.623283466145659 | 0.931059568526633 | 721.003945822098 | 856.765346817874 | 901.97150888197 | 745.651721935238 | 729.242158415519 | 629.887349518038 | 849.519721893808 | 857.565196088121 | Ankhd1 |
| ENSMUSG00000024486 | 0.414422200864221 | 0.0366758754062762 | 0.679566075569698 | 744.495302452893 | 906.444706281648 | 1081.79793772517 | 1071.11228484282 | 643.835419141629 | 543.048511040794 | 770.397787011541 | 895.558590851519 | Hbegf |
| ENSMUSG00000024487 | -0.07226679478032 | 0.682123327649772 | 0.944452327903597 | 305.387636200337 | 258.169786065842 | 300.972654590206 | 305.764102328161 | 358.051330032845 | 365.701587390084 | 243.612273190136 | 263.782712214447 | Yipf5 |
| ENSMUSG00000024491 | 0.00755679309189795 | 0.961805606734839 | 0.9938610234747 | 346.045753445944 | 423.496179035451 | 383.314229902621 | 327.336416296098 | 323.012667766634 | 320.447544803351 | 395.609674411332 | 433.124700302735 | Rbm27 |
| ENSMUSG00000024493 | 0.239120106841388 | 0.00542114929513476 | 0.635862253526407 | 1986.82666273533 | 2059.65737842434 | 1950.64352550445 | 1964.95642447085 | 1599.7339240917 | 1596.12231285639 | 1846.87253264659 | 1702.10408540022 | Lars |
| ENSMUSG00000024498 | 0.190391130885715 | 0.325427716928055 | 0.855977678491063 | 453.563885717661 | 579.863998987002 | 537.586376752318 | 407.060185308042 | 404.039574257247 | 322.893709267499 | 492.429936833053 | 513.453592088204 | Tcerg1 |
| ENSMUSG00000024501 | -0.462503886200976 | 0.111555485027803 | 0.764116514278315 | 274.66816983699 | 301.333819698302 | 251.757000380487 | 413.625672167849 | 395.279908690694 | 692.264543353805 | 307.11803671406 | 316.973464883204 | Dpysl3 |
| ENSMUSG00000024505 | -0.0656031197773721 | 0.855270889107734 | 0.973268281968411 | 21.6843291976571 | 47.2361122770311 | 30.2865564367503 | 32.8274342990356 | 20.8042057205628 | 50.1463715150282 | 33.3144988977964 | 34.7368180693922 | Dtwd2 |
| ENSMUSG00000024507 | 0.16410511637161 | 0.284727442673346 | 0.851125503777472 | 3313.18479865869 | 3593.20219597002 | 3565.2955655387 | 3077.33748386103 | 3059.31319911856 | 2270.04062272908 | 3337.69635832297 | 3423.74763096447 | Hsd17b4 |
| ENSMUSG00000024511 | -0.143327901405386 | 0.604189968609823 | 0.926448151609826 | 32.5264937964856 | 35.0198763433161 | 23.6613722162112 | 32.8274342990356 | 26.2789966996583 | 36.6924669622158 | 38.5198893505771 | 35.8223436340608 | Rab27b |
| ENSMUSG00000024513 | 0.159590587945014 | 0.14731505698939 | 0.780607990324772 | 2092.53776757391 | 2163.08817599646 | 2138.04159345684 | 2465.80927920471 | 1893.18272057122 | 2248.02514255175 | 1958.26788833609 | 1833.45267872511 | Mbd2 |
| ENSMUSG00000024515 | 0.0790904086875202 | 0.582051623965031 | 0.922450244597442 | 1083.31294616628 | 1286.77685168464 | 1265.41018612297 | 1034.53314376675 | 1001.88674917447 | 955.227223249684 | 1274.27958284071 | 1188.65049331202 | Smad4 |
| ENSMUSG00000024516 | 0.0983791317963194 | 0.636138990910901 | 0.932595977875587 | 402.063603873225 | 263.056280439328 | 323.687571917769 | 445.515179772627 | 320.822751374995 | 419.517205601334 | 292.542943446274 | 307.203734801188 | Sec11c |
| ENSMUSG00000024524 | 0.0601669273025603 | 0.731719884468686 | 0.953164137228739 | 140.044626068202 | 126.234437981721 | 122.09268063565 | 118.178763476528 | 114.970610561005 | 125.977469903608 | 98.902418602833 | 146.545951230249 | Gnal |
| ENSMUSG00000024525 | 0.251719061055004 | 0.0555346046585194 | 0.703396849069134 | 620.713923282934 | 604.296470854432 | 646.428688946889 | 543.059555975475 | 529.959766776443 | 448.871179171106 | 461.197594116369 | 587.269330485663 | Impa2 |

| | | | | | | | | | | | | |
|---|---|---|---|---|---|---|---|---|---|---|---|---|
| **ENSMUSG00000024526** | 0.0999877316224891 | 0.595852092286309 | 0.924461858177432 | 1270.34028549608 | 1106.79097559457 | 1251.21336279325 | 1262.44933047149 | 1250.44225962541 | 1500.72189875463 | 927.600578685518 | 885.788860769502 | Cidea |
| **ENSMUSG00000024527** | 0.228512408061828 | 0.212315326465953 | 0.823370267313127 | 5130.15088267904 | 5323.02120418405 | 5261.34272599672 | 5021.6595210582 | 4142.22685478364 | 3049.14400456013 | 5339.68952646242 | 5166.01616225743 | Afg3l2 |
| **ENSMUSG00000024528** | 0.168563722316858 | 0.198426727354842 | 0.812921911423739 | 237.624107457659 | 231.294067011669 | 214.845259723197 | 235.419600258798 | 192.712642464161 | 232.385624094033 | 208.215618111227 | 185.624871558315 | Srfbp1 |
| **ENSMUSG00000024529** | -0.51154135537618 | 0.322530401667008 | 0.855977678491063 | 69.5705561758165 | 63.5244268553176 | 135.343049076728 | 92.854742731558 | 182.858018701789 | 236.054870790255 | 55.1771387994752 | 41.2499714574033 | Lox |
| **ENSMUSG00000024533** | 0.470805779214324 | 0.016222695298578 | 0.640151243458091 | 271.957628687283 | 272.014853457386 | 359.65285768641 | 287.005568442997 | 237.605928492744 | 147.992950080937 | 233.201492284575 | 238.815624227072 | Spire1 |
| **ENSMUSG00000024534** | -0.360584789057221 | 0.118629959420624 | 0.769868040577702 | 149.079763233892 | 120.533527879321 | 151.432782183751 | 122.868396947819 | 191.617684268342 | 236.054870790255 | 121.806136595068 | 149.802527924254 | Sncaip |
| **ENSMUSG00000024535** | 0.382459514245912 | 0.014633461510198 | 0.635862253526407 | 195.158962778914 | 238.623808571898 | 191.183887506986 | 226.978260010475 | 137.964732673206 | 168.785348026193 | 183.22974393788 | 163.914360264945 | Snx24 |
| **ENSMUSG00000024537** | 0.0345451470238009 | 0.831862550346017 | 0.967583158900775 | 543.011743657996 | 461.773718294424 | 398.457508120996 | 588.080037299867 | 540.909348734634 | 507.579126310652 | 475.772687384154 | 420.098393526712 | Psmg2 |
| **ENSMUSG00000024538** | -0.188997446972558 | 0.34664998537131 | 0.861266047293794 | 325.264937964856 | 256.540954608014 | 251.757000380487 | 325.460562907582 | 379.950493949227 | 398.724807656078 | 239.447960827911 | 303.947158107182 | Ppic |
| **ENSMUSG00000024539** | 0.152058202403811 | 0.302377663433401 | 0.851785577030345 | 237.624107457659 | 274.458100644129 | 270.686098153456 | 268.247034557834 | 243.08071947184 | 190.800828203522 | 238.406882737355 | 272.466916731795 | Ptpn2 |
| **ENSMUSG00000024542** | -0.102042451124706 | 0.691629950741188 | 0.94469184542218 | 42.4651446787451 | 79.8127414336042 | 60.5731128735006 | 52.523894878457 | 56.937826182593 | 69.71568722821 | 54.1360607089191 | 72.73021283279 | Cep192 |
| **ENSMUSG00000024544** | 0.136821842502253 | 0.396997409525908 | 0.879613019351548 | 175.281661014395 | 199.531853584011 | 189.290977729689 | 144.440710915757 | 146.724398239759 | 166.339183562045 | 153.038479311752 | 179.111718170304 | Ldlrad4 |
| **ENSMUSG00000024548** | 0.0164376745793776 | 0.925427872860214 | 0.986713918026868 | 160.82544154929 | 185.686786192467 | 161.843785958884 | 142.564857527241 | 144.534481848121 | 133.315963296051 | 192.599446752885 | 172.598564782293 | Setbp1 |
| **ENSMUSG00000024556** | 0.159858307235993 | 0.218600643514573 | 0.827129763070499 | 220.457346842847 | 201.160685041839 | 196.862616838877 | 226.978260010475 | 188.332809680885 | 184.685417043153 | 205.092383839559 | 178.026192605635 | Me2 |
| **ENSMUSG00000024558** | 0.499937187650787 | 0.00126094825212092 | 0.596732488826387 | 178.895715880671 | 216.634583891211 | 222.416898832385 | 213.847286290861 | 134.679858085749 | 132.092881063977 | 170.736806851206 | 149.802527924254 | Mapk4 |
| **ENSMUSG00000024560** | -0.0633276994242441 | 0.57672449833253 | 0.922450244597442 | 686.670424592474 | 705.284021239809 | 734.448993591195 | 636.852225401292 | 782.895110010654 | 735.07242147639 | 745.411912838194 | 624.177199684392 | Cxxc1 |
| **ENSMUSG00000024561** | 0.164018618954213 | 0.617705852921184 | 0.929368966320976 | 1579.34197656269 | 1853.61019900901 | 2269.59882297898 | 1263.38725716574 | 1673.09612321158 | 754.641737189571 | 2461.10860607471 | 1327.59776558958 | Mbd1 |
| **ENSMUSG00000024563** | -0.0318448917758308 | 0.837417048186437 | 0.968924788429433 | 325.264937964856 | 354.270842077733 | 337.884395247495 | 281.378008277448 | 296.733671066975 | 280.085831144914 | 377.911346871878 | 372.335268681298 | Smad2 |
| **ENSMUSG00000024566** | 0.0287836160474918 | 0.87612085962731 1 | 0.977469019294424 | 396.642521573811 | 542.400875456943 | 494.049451874489 | 455.832373409466 | 396.374866886513 | 370.593916318379 | 573.634027896431 | 510.197015394199 | Atp9b |
| **ENSMUSG00000024570** | 0.0230036885675995 | 0.856519281823864 | 0.97363563840968 | 1183.60296870545 | 1074.214346438 | 1085.58375727977 | 1231.49774956097 | 1194.59939163863 | 1262.22086350022 | 1112.91247880451 | 933.551985614916 | Rbfa |
| **ENSMUSG00000024571** | 0.0459942615901544 | 0.652213216972005 | 0.936720465895231 | 865.566140473145 | 921.91860513102 | 1023.11773462897 | 1002.64363616197 | 880.346389438554 | 876.949960396957 | 960.915077583314 | 973.716431507651 | Gm16286 |
| **ENSMUSG00000024576** | 0.0498465511522022 | 0.760623980396261 | 0.957357521291862 | 3726.99408084731 | 4251.2501049328 | 4192.79515671262 | 3463.76328189539 | 3441.45360945942 | 2931.72811028104 | 4380.85660506022 | 4348.61541206204 | Csnk1a1 |
| **ENSMUSG00000024579** | -0.429882200827597 | 0.202770079077123 | 0.815403452654488 | 22.5878429142261 | 20.3603932228582 | 21.7684624389143 | 22.5102406621959 | 35.0386622662111 | 40.3617136584374 | 17.6983275394543 | 24.9670879873757 | Pcyox1l |
| **ENSMUSG00000024580** | 0.212683696559383 | 0.247237114263345 | 0.833120697990749 | 396.642521573811 | 340.425774686189 | 407.922057007481 | 358.287997206618 | 289.068963696242 | 242.170281950624 | 404.979377226337 | 359.308961905276 | Grpel2 |
| **ENSMUSG00000024581** | 0.15319884905057 | 0.360508508084086 | 0.868539734006824 | 337.010616280254 | 432.454752053509 | 454.298346551254 | 434.260059441529 | 385.425284928322 | 291.093571233579 | 432.047407580797 | 381.019473198646 | Napg |
| **ENSMUSG00000024583** | 0.0886205163484911 | 0.394905078236813 | 0.879215782741314 | 2248.84564054035 | 1926.9076146113 | 1966.73325861147 | 2281.97564713011 | 2086.9903212312 | 1893.33129525033 | 2076.95079065949 | 1863.84739453583 | Txnl1 |
| **ENSMUSG00000024587** | 0.140224997697243 | 0.114536703900141 | 0.767518180080285 | 3278.85127742906 | 3404.25774686189 | 3130.87277164906 | 3398.10841329732 | 2796.52323212197 | 2865.68166974905 | 3283.56029761405 | 3041.64263220116 | Nars |
| **ENSMUSG00000024588** | 0.2335055815939 | 0.233165174879906 | 0.832048466795696 | 3322.21993582438 | 3600.53193753025 | 4272.29736735909 | 3460.01157511836 | 2533.73326512539 | 2360.54870790255 | 3735.38818891542 | 3834.07629440917 | Fech |
| **ENSMUSG00000024589** | -0.0415637892430529 | 0.736350202467514 | 0.954383574136061 | 568.310127721929 | 508.195414842541 | 457.1377112172 | 519.611388619021 | 579.232885588302 | 477.002070508805 | 558.017856538089 | 498.256234182845 | Nedd4l |
| **ENSMUSG00000024590** | 0.0402015988343358 | 0.858875549357999 | 0.974833700919212 | 64.1494738764022 | 67.5965054998893 | 68.1447519826882 | 61.9031618210386 | 72.2672409240604 | 45.2540425867328 | 57.2592949805875 | 79.2433662208011 | Lmnb1 |
| **ENSMUSG00000024592** | 0.16463941798556 | 0.500670339107317 | 0.900927059989498 | 577.34526488762 | 538.328796812371 | 485.531357876653 | 467.087493740564 | 360.241246424483 | 297.208982393948 | 628.811166695907 | 557.960140239613 | C330018D20 |
| **ENSMUSG00000024594** | 0.00204268016971722 | 0.983938301971971 | 0.996940244108351 | 734.556651570634 | 697.139863950665 | 616.142132510139 | 660.300392757745 | 700.773245324222 | 697.1568722821 | 671.495368408708 | 636.117980895745 | Prrc1 |
| **ENSMUSG00000024597** | 0.112400648988333 | 0.350078254576474 | 0.861774532843376 | 543.915257374565 | 616.512706788147 | 670.0900611631 | 565.569796637671 | 542.004306930453 | 496.571386221987 | 585.085886892549 | 591.611432744337 | Slc12a2 |
| **ENSMUSG00000024601** | 0.117205663827637 | 0.685108937029106 | 0.944452327903597 | 1019.16347228988 | 1259.08671690155 | 1460.37989318455 | 1026.09180351843 | 879.251431242735 | 593.194882555822 | 1436.68776496747 | 1482.82792133718 | Isoc1 |
| **ENSMUSG00000024603** | 0.0767421700670465 | 0.445525446602283 | 0.89108031640712 | 961.338594429464 | 1033.49355999228 | 1061.92238506356 | 970.754128557197 | 969.038003299901 | 854.934480219628 | 952.586452858865 | 1041.0190165171 | Dctn4 |
| **ENSMUSG00000024604** | -0.198250486224618 | 0.288695643978905 | 0.851785577030345 | 357.791431761342 | 336.353696041618 | 325.580481695066 | 388.301651422879 | 516.820268426614 | 447.648096939033 | 323.775286162958 | 327.828720529889 | Rbm22 |
| **ENSMUSG00000024608** | 0.106065673447488 | 0.54316206163611 | 0.913837378403038 | 16321.0717761032 | 10807.2967226931 | 13170.8662304318 | 15690.5756682448 | 13464.7009339874 | 15386.3744794892 | 10857.4034064099 | 12313.1164800349 | Rps14 |
| **ENSMUSG00000024610** | -0.756808697629003 | 0.0300448829291122 | 0.665956123973759 | 853.820462157748 | 1109.23422278132 | 1224.71262591109 | 1295.27676477052 | 2891.78459515823 | 2588.04200306829 | 946.339984315528 | 1149.57157298395 | Cd74 |
| **ENSMUSG00000024613** | 0.23703593432141 | 0.0266648376909457 | 0.664353746352133 | 460.791995450213 | 485.39177443294 | 469.44162476963 | 503.666634816633 | 376.665619361769 | 398.724807656078 | 451.827891301363 | 400.558933362679 | Tcof1 |
| **ENSMUSG00000024614** | 0.0161860387308486 | 0.867187781374895 | 0.974868041734905 | 524.037955610046 | 548.916201288258 | 507.299820315568 | 560.88016316638 | 522.295059405709 | 563.840908986049 | 534.073060455298 | 498.256234182845 | Tmx3 |
| **ENSMUSG00000024617** | 0.190667065949513 | 0.349727735225278 | 0.861774532843376 | 830.329105526952 | 1125.5225373596 | 961.598166866822 | 836.63061127828 | 696.393412540946 | 606.648787108634 | 987.983107937774 | 997.597993930358 | Camk2a |
| **ENSMUSG00000024620** | -0.993522775982815 | 0.00091108338382739 | 0.556246415579446 | 308.098177350044 | 359.971752180133 | 399.403963009645 | 361.101777289392 | 1065.39432453198 | 894.073111645991 | 440.376032305246 | 445.065481514088 | Pdgfrb |
| **ENSMUSG00000024621** | -0.600350160971613 | 0.0427713878893563 | 0.687730553581215 | 249.369785773056 | 346.12668478859 | 419.279515671262 | 377.984457786039 | 677.779123212021 | 728.95701031602 | 308.159114804616 | 397.302356668674 | Csf1r |
| **ENSMUSG00000024622** | 0.180894833209931 | 0.436191010731977 | 0.888701139843674 | 277.378710986697 | 348.569931975333 | 318.008842585878 | 240.109233730089 | 237.605928492744 | 170.008430258266 | 331.062832796851 | 305.032683671851 | Hmgxb3 |
| **ENSMUSG00000024639** | -0.105432877980622 | 0.464775584680037 | 0.895328112556867 | 598.126080368708 | 611.626212414661 | 573.551662520959 | 597.459304242449 | 706.248036303317 | 757.087901653719 | 529.908748093074 | 569.900921450967 | Gnaq |
| **ENSMUSG00000024640** | 0.123970519330248 | 0.649392807903483 | 0.936461253686283 | 144.562194651047 | 92.0289773673191 | 87.0738497556571 | 96.6064495085906 | 96.3563212320805 | 134.539045528125 | 65.5879197050366 | 90.0986218674861 | Psat1 |
| **ENSMUSG00000024642** | -0.130529594310362 | 0.429522055316625 | 0.885743677088682 | 131.009488902512 | 157.99665140938 | 149.539872406455 | 116.302910088012 | 156.579022002131 | 136.985209992272 | 144.709854587303 | 169.341988088287 | Tle4 |
| **ENSMUSG00000024644** | -0.0105703621086635 | 0.951239360278166 | 0.991915011884503 | 767.083145367119 | 665.377650523007 | 816.79056890361 | 782.230863011306 | 874.871598459458 | 925.873249679912 | 635.057635239243 | 618.749571861049 | Cndp2 |
| **ENSMUSG00000024645** | 0.159780890305327 | 0.347543310133303 | 0.861266047293794 | 724.618000688374 | 640.130762926662 | 631.285410728514 | 599.335157630965 | 626.316088008523 | 413.401794440965 | 632.975479058131 | 649.144287671768 | Timm21 |
| **ENSMUSG00000024646** | -0.0685001026270959 | 0.679929756976711 | 0.944452327903597 | 1285.70001867775 | 1397.53739081699 | 1168.87178748083 | 1318.72493212697 | 1306.28512761218 | 1742.89218070525 | 1062.94073045782 | 1312.40040768423 | Cyb5a |
| **ENSMUSG00000024654** | 0.240616862413429 | 0.0745213213158622 | 0.735399943172214 | 2050.07262289516 | 2044.99789530388 | 1981.87653682985 | 2153.47969001674 | 1496.80785368471 | 1509.28347437914 | 1966.59651306054 | 1991.93941116671 | Asrgl1 |
| **ENSMUSG00000024659** | -0.60089833255451 | 0.0912573292683415 | 0.753823363185192 | 330.68602026427 | 252.468875963442 | 242.292451494002 | 332.026049767389 | 598.942133113046 | 678.810638800992 | 235.283648465687 | 243.157726485746 | Anxa1 |
| **ENSMUSG00000024660** | -0.33257221148699 | 0.132256100237759 | 0.77303252008843 | 73.1846110420926 | 46.4216965481167 | 56.7872933189068 | 75.9720622349111 | 74.4571573156986 | 85.6157562451702 | 81.2040910633787 | 75.9867895267955 | Incenp |
| **ENSMUSG00000024661** | 0.00559573276902556 | 0.973539733014184 | 0.995510898988495 | 50881.3749485858 | 46901.3874124473 | 57269.9853121175 | 57822.2427743214 | 52886.4808580624 | 68857.0835012942 | 44460.2809352904 | 45848.2577493391 | Fth1 |
| **ENSMUSG00000024663** | -0.292208632349233 | 0.395225954519899 | 0.879215782741314 | 67.7635287426784 | 86.3280672649188 | 86.1273948670087 | 115.364983393754 | 170.813478547779 | 130.869798831903 | 63.5057635239243 | 70.559161703453 | Rab3il1 |
| **ENSMUSG00000024664** | 0.117265295073536 | 0.291568046802426 | 0.851785577030345 | 1627.22820354085 | 1551.4619635818 | 1267.30309590027 | 1563.52379932835 | 1324.89941694111 | 1357.62127760198 | 1425.23590597135 | 1432.89374536243 | Fads3 |
| **ENSMUSG00000024665** | -0.690189063652347 | 0.0188850388447068 | 0.642273034804458 | 40.658117245607 | 68.4109212288036 | 53.001473764313 | 59.0893817382642 | 93.0714466446232 | 83.1695917810225 | 55.1771387994752 | 125.920965501547 | Fads2 |
| **ENSMUSG00000024666** | -0.541731743313086 | 0.0790312417863871 | 0.739855948845396 | 18.0702743313809 | 23.6180561385155 | 21.7684624389143 | 25.3240207449704 | 28.4689130912965 | 41.5847958905112 | 23.9447960827911 | 35.8223436340608 | Tmem138 |
| **ENSMUSG00000024667** | -0.0944234861891829 | 0.679051650704046 | 0.944452327903597 | 67.7635287426784 | 63.5244268553176 | 87.0738497556571 | 57.2135283497478 | 56.937826182593 | 73.3849339244316 | 88.4916376972716 | 74.901263962127 | Tmem216 |
| **ENSMUSG00000024668** | 0.092408491635155 | 0.552939014923034 | 0.914977712327965 | 1472.72735800754 | 1238.72632367869 | 1191.58670480839 | 1258.69762369445 | 1047.87499339888 | 1018.82749931753 | 1319.04594073463 | 1454.6042566558 | Sdhaf2 |
| **ENSMUSG00000024672** | -0.996878450981035 | 0.0323066306827947 | 0.671606027118506 | 8.13162344912141 | 12.2162359337149 | 8.51809399783602 | 19.6964605794214 | 21.8991639163819 | 39.1386314263635 | 13.5340151772298 | 22.7960368580387 | Ms4a7 |
| **ENSMUSG00000024677** | -0.286231036258284 | 0.276458975442948 | 0.846750864146095 | 70.4740698923855 | 105.874044758863 | 72.8770264259304 | 57.2135283497478 | 97.4512794278996 | 108.854318654574 | 64.5468416144805 | 104.210454208177 | Ms4a6b |
| **ENSMUSG00000024678** | -0.613418950548905 | 0.09202616295198 4 | 0.754058637722175 | 93.9654265231807 | 96.9154717408051 | 135.343049076728 | 163.19924480092 | 235.416012101106 | 295.985900161874 | 112.436433780063 | 105.295979772845 | Ms4a4d |
| **ENSMUSG00000024679** | -0.552079524945331 | 0.0855655391066961 | 0.744211232430814 | 45.1756858284523 | 57.8235167529173 | 55.8408384302584 | 51.5859681841989 | 101.831112211176 | 108.854318654574 | 53.094982618363 | 45.5920737160773 | Ms4a6d |
| **ENSMUSG00000024683** | 0.083909891250459 | 0.443444425161433 | 0.89108031640712 | 1925.38773000863 | 1711.90186217792 | 1592.88357759534 | 1872.10168173929 | 1571.2650110004 | 1569.21450375076 | 1834.37959555991 | 1724.90012225826 | Mrpl16 |
| **ENSMUSG00000024687** | 0.190395573696356 | 0.437821480273324 | 0.888759286175347 | 2320.22322414931 | 2888.73259045912 | 2923.59915103505 | 2368.26490300186 | 1923.84155005415 | 1402.87532018872 | 2874.41660802549 | 3000.39266074375 | Osbp |
| **ENSMUSG00000024691** | -0.398010332726887 | 0.0652514246184737 | 0.718339708336358 | 93.9654265231807 | 61.0811796685746 | 70.0376617599851 | 104.109863062656 | 118.255485148462 | 106.408154190426 | 103.066730965058 | 105.295979772845 | Fam111a |
| **ENSMUSG00000024694** | -0.150546754643982 | 0.772680178497843 | 0.959045638244974 | 30.7194663633475 | 26.0613033252585 | 13.2503684410783 | 32.8274342990356 | 25.1840385038392 | 23.2385624094033 | 9.36970281500523 | 56.4473293627624 | Keg1 |
| **ENSMUSG00000024695** | 0.112506504920037 | 0.469899442571075 | 0.895897084741774 | 4819.34216417929 | 5112.90194612416 | 4474.83871352986 | 4270.38023895741 | 3577.22842574099 | 3687.59292970269 | 4982.59974140167 | 5028.15441554453 | Zfp91 |
| **ENSMUSG00000024696** | -0.465676018699119 | 0.357811853961207 | 0.865834559821921 | 4.51756858284523 | 8.95857301805762 | 10.4110037751329 | 13.1309737196143 | 18.6142893289246 | 11.0077400886647 | 7.28754663389296 | 14.1118323406906 | Lpxn |
| **ENSMUSG00000024712** | 0.154272622822409 | 0.232637499463466 | 0.832048466795696 | 2309.38105955048 | 2065.35828852674 | 2015.00245793254 | 2061.56287397944 | 1627.10787898718 | 1724.54594722414 | 2076.95079065949 | 2164.537975949 | Rfk |
| **ENSMUSG00000024713** | -0.87175590500305 | 0.0449374443620614 | 0.693563614452513 | 20.780815481088 | 21.9892246806869 | 15.1432782183751 | 22.5102406621959 | 40.5134532453066 | 64.8233582999146 | 16.6572494488982 | 26.0526135520442 | Pcsk5 |
| **ENSMUSG00000024725** | -0.0870786250253629 | 0.668304857866818 | 0.942024383071524 | 826.715050660676 | 596.966729294203 | 672.929425829046 | 844.134024832345 | 860.63714191381 | 999.258183604343 | 612.153917247008 | 652.400864365773 | Ostf1 |
| **ENSMUSG00000024726** | 0.207526404662367 | 0.163368597750394 | 0.793968226938511 | 1451.94654252646 | 1363.33193020259 | 1508.64909250562 | 1452.84844940589 | 1167.22543674316 | 976.01962119494 | 1465.83795150304 | 1391.64377390503 | Carnmt1 |
| **ENSMUSG00000024732** | 0.209696020321588 | 0.132315606923126 | 0.77303252008843 | 261.115464088454 | 225.593156909269 | 259.328639489674 | 251.364354061187 | 254.03030143003 | 201.808568292187 | 206.133461930115 | 199.736703899005 | Ccdc86 |
| **ENSMUSG00000024735** | 0.151397998639053 | 0.160314875230232 | 0.79125089991166 | 6133.05110807068 | 6951.85266201271 | 6814.47519826882 | 6586.12124708081 | 6394.55586358353 | 5163.85318381584 | 6461.97170808194 | 5824.93018001121 | Prpf19 |
| **ENSMUSG00000024736** | -0.451386968111588 | 0.0749474824778856 | 0.735399943172214 | 169.86057871498 | 133.56417954195 | 92.7525790875478 | 88.1651092602672 | 159.863896589588 | 174.900759186562 | 196.76375911511 | 131.348593324889 | Tmem132a |
| **ENSMUSG00000024737** | -0.289248753617015 | 0.384932947196655 | 0.875126835008975 | 34.3335212296237 | 32.5766291565731 | 72.8770264259304 | 59.0893817382642 | 68.9823663366031 | 75.8310983885793 | 43.7252798033577 | 54.2762782334254 | Slc15a3 |
| **ENSMUSG00000024740** | 0.174842649321782 | 0.248583859722871 | 0.834621660998471 | 4052.25901881217 | 4849.84566568483 | 5355.04175997291 | 4250.68377837799 | 4395.16219801786 | 3254.62181954854 | 4493.29303884029 | 4250.91811124188 | Ddb1 |
| **ENSMUSG00000024742** | -0.316656803147332 | 0.343752887246354 | 0.861266047293794 | 29.8159526467785 | 17.9171460361152 | 19.8755526616174 | 15.0068271081306 | 29.5638712871156 | 23.2385624094033 | 23.9447960827911 | 26.0526135520442 | Fen1 |
| **ENSMUSG00000024743** | 0.0133143685756325 | 0.945518751384695 | 0.990505437194639 | 701.126644057579 | 905.630290552733 | 801.647290685234 | 570.259430108962 | 692.013579757669 | 582.187142467157 | 900.532548331058 | 776.150778737983 | Syt7 |
| **ENSMUSG00000024747** | -0.224080634919581 | 0.626573292036091 | 0.931698409533928 | 24.3948703473642 | 64.338842584232 | 33.1259211026956 | 53.4618215727152 | 39.4184950494875 | 44.0309603546589 | 24.9858741733473 | 96.6117752554972 | Aldh1a7 |
| **ENSMUSG00000024750** | 0.00713223112580241 | 0.955684573365056 | 0.992371505557225 | 2912.9282222186 | 3349.69189302463 | 3332.46766293118 | 3101.72357791174 | 2954.19721231992 | 2671.21159484931 | 3632.32145795036 | 3374.89898055439 | Zfand5 |
| **ENSMUSG00000024754** | -0.101586647296137 | 0.662390303402053 | 0.939611397743053 | 146.369222084185 | 199.531853584011 | 158.057966404291 | 122.868396947819 | 213.516848184724 | 167.562265794119 | 174.901119213431 | 117.236760984199 | Tmem2 |
| **ENSMUSG00000024758** | -0.116971964437626 | 0.391189680734342 | 0.877948726030055 | 1824.1941937529 | 1956.22658085222 | 1920.3569690677 | 1897.42570248426 | 2301.60212761174 | 2388.67959924025 | 1816.68126802046 | 1734.66985234028 | Rtn3 |
| **ENSMUSG00000024759** | -0.385208444175431 | 0.037605561700921 | 0.679566075569698 | 206.001127377742 | 254.097707421271 | 244.185361271299 | 229.79204009325 | 387.61520131996 | 330.232202659942 | 243.612273190136 | 259.440609955773 | Atl3 |
| **ENSMUSG00000024761** | 0.120017614681597 | 0.794304913439825 | 0.961804768755116 | 9.03513716569045 | 14.6594831204579 | 27.447191770805 | 13.1309737196143 | 17.5193311331055 | 18.3462334811079 | 13.5340151772298 | 9.76973008201657 | Gm16437 |

| | | | | | | | | | | | | |
|---|---|---|---|---|---|---|---|---|---|---|---|---|
| **ENSMUSG00000024764** | 0.117428004640779 | 0.641439802387373 | 0.934544499347759 | 130.105975185942 | 160.439898596123 | 189.290977729689 | 154.757904552597 | 137.964732673206 | 86.838838477244 | 185.311900118992 | 173.684090346961 | Naa40 |
| **ENSMUSG00000024766** | 0.0376360529905197 | 0.816826146692617 | 0.966315773308182 | 191.544907912638 | 153.924572764808 | 169.415425068072 | 208.219726125312 | 168.623562156141 | 189.577745971448 | 153.038479311752 | 193.223550510994 | Lipo3 |
| **ENSMUSG00000024767** | -0.113465243555984 | 0.397560268139988 | 0.88020150974241 | 1538.68385931708 | 1295.7354247027 | 1183.06861081056 | 1414.39345494131 | 1381.8372431237 | 1723.32286499207 | 1366.93553290021 | 1405.75560624572 | Otub1 |
| **ENSMUSG00000024769** | 0.369305852425457 | 0.0587371903789799 | 0.711712402279519 | 731.846110420926 | 879.568987227475 | 841.398396008469 | 657.486612674971 | 544.194223322091 | 435.417274618294 | 690.234774038719 | 737.071858409917 | Cdc42bpg |
| **ENSMUSG00000024772** | -0.234281173054948 | 0.200382526761112 | 0.812921911423739 | 3829.09113081961 | 3570.39855556042 | 3794.33764859162 | 3655.10032752405 | 5276.60354565223 | 5320.40770952129 | 3512.5974775364 | 3358.61609708436 | Ehd1 |
| **ENSMUSG00000024773** | 0.0431149968252489 | 0.675156948286675 | 0.944023556644042 | 846.592352425195 | 898.300548992504 | 978.634354862494 | 865.706338800283 | 886.916138613469 | 798.67269754423 | 947.381062406084 | 848.880991570773 | Atg2a |
| **ENSMUSG00000024776** | -0.0886833829067903 | 0.804794583838251 | 0.963461896828986 | 28.9124389302094 | 32.5766291565731 | 18.9290977729689 | 32.8274342990356 | 16.4243729372865 | 44.0309603546589 | 29.1501865355718 | 31.4802413753867 | Stambpl1 |
| **ENSMUSG00000024777** | 0.0615319983188837 | 0.580822368247186 | 0.922450244597442 | 832.13613296009 | 797.312998607128 | 774.20009891443 | 885.402799379704 | 753.331238723539 | 738.741668172611 | 902.61470451217 | 755.525793009281 | Ppp2r5b |
| **ENSMUSG00000024778** | -0.112230118652148 | 0.656724627068357 | 0.938371059559662 | 391.221439274397 | 341.240190415104 | 335.04503058155 | 253.240207449704 | 465.357233223116 | 447.648096939033 | 255.064132186253 | 260.526135520442 | Fas |
| **ENSMUSG00000024780** | 0.136271947068501 | 0.643533003913822 | 0.934616627390525 | 2350.03917679609 | 2395.19665873704 | 2653.85950777024 | 2237.89309249997 | 1621.63308800808 | 1166.82044939846 | 3336.65528023242 | 2642.16922440315 | Cdc37l1 |
| **ENSMUSG00000024781** | -0.314922039187064 | 0.24523921879128 | 0.832048466795696 | 649.626362213143 | 772.066111010784 | 732.556083813898 | 482.094320848695 | 989.842209020464 | 1148.47421591735 | 572.592949805875 | 569.900921450967 | Lipa |
| **ENSMUSG00000024782** | 0.128726037119761 | 0.415494631536172 | 0.882192126163777 | 2430.45189757073 | 2498.62745630916 | 2580.98248134431 | 2353.25807589373 | 1996.10879097821 | 1758.79224972221 | 2642.25619383147 | 2622.62976423911 | Ak3 |
| **ENSMUSG00000024785** | -0.200889493831127 | 0.373107012534276 | 0.873700364743909 | 82.2197482077831 | 92.8433930962335 | 96.5383986421416 | 132.247663890401 | 88.6916138613469 | 138.208292224346 | 108.272121417838 | 129.177542195552 | Rcl1 |
| **ENSMUSG00000024787** | -0.109213161516493 | 0.433648750115888 | 0.887868762931002 | 245.75573090678 | 242.69588721647 | 212.005895057252 | 269.184961252092 | 283.594172717146 | 282.531995609062 | 232.160414194018 | 248.585354309088 | Snx15 |
| **ENSMUSG00000024789** | 0.164226892386524 | 0.348154339417759 | 0.861266047293794 | 608.064731250967 | 772.880526739698 | 758.110365807406 | 670.617586394585 | 564.998429042654 | 484.340563901248 | 761.028084196536 | 695.821886952513 | Jak2 |
| **ENSMUSG00000024790** | -0.341515681167286 | 0.149964886369048 | 0.782641594375979 | 211.422209677157 | 138.450673915436 | 154.272146849697 | 166.013024883695 | 237.605928492744 | 283.755077841135 | 160.326025945645 | 168.256462523619 | Sac3d1 |
| **ENSMUSG00000024792** | -0.281304349728194 | 0.245205637229562 | 0.832048466795696 | 193.351935345776 | 185.686786192467 | 179.826428843205 | 228.854113398991 | 275.929465346412 | 324.116791499573 | 206.133461930115 | 151.973579053591 | Zfpl1 |
| **ENSMUSG00000024793** | 0.223845545528515 | 0.69033617242784 | 0.94469184542218 | 13.5527057485357 | 18.7315617650296 | 16.0897331070236 | 15.9447538023887 | 24.0890803080201 | 20.7923979452556 | 7.28754663389296 | 3.25657669400552 | Tnfrsf25 |
| **ENSMUSG00000024797** | -0.235928270253057 | 0.145480955926416 | 0.780607990324772 | 265.633032671299 | 274.458100644129 | 287.722286149128 | 287.005568442997 | 395.279908690694 | 366.924669622158 | 279.008928269045 | 272.466916731795 | Vps51 |
| **ENSMUSG00000024799** | -0.330388631311267 | 0.195939922077961 | 0.811193545163524 | 44.2721721118832 | 64.338842584232 | 40.6975602118832 | 50.6480414899407 | 77.7420319031559 | 52.592535979176 | 51.0128264372507 | 70.559161703453 | Tm7sf2 |
| **ENSMUSG00000024800** | -0.0251345239838742 | 0.884277090733186 | 0.979396283413121 | 262.922491521592 | 203.603932228582 | 213.898804834549 | 243.860940507122 | 231.036179317829 | 182.239252579005 | 274.84461590682 | 250.756405438425 | Rpp30 |
| **ENSMUSG00000024803** | 0.430910780124488 | 0.148702851598381 | 0.78151662300109 | 68385.1461796779 | 51148.5654387355 | 26685.295585443 | 80193.670285767 | 33727.9973058156 | 45290.735053695 | 42586.3403722893 | 46346.5139835219 | Ankrd1 |
| **ENSMUSG00000024805** | 0.185269675453027 | 0.314468118500061 | 0.853228028313558 | 933.329669215824 | 980.556537612852 | 922.793516432236 | 899.471699793577 | 683.253914191116 | 617.656527197299 | 1031.70838774113 | 952.005920214281 | Pcgf5 |
| **ENSMUSG00000024807** | -0.0587632196757583 | 0.650998840727409 | 0.936463929813731 | 389.414411841258 | 441.413325071566 | 427.797609669098 | 364.853484066425 | 411.70428162798 | 365.701587390084 | 461.197594116369 | 451.578634902099 | Syvn1 |
| **ENSMUSG00000024810** | -0.51216808350106 | 0.180848682351474 | 0.802808628271642 | 27.1054114970714 | 30.1333819698302 | 36.9117406572894 | 22.5102406621959 | 55.8428679867739 | 59.9310293716191 | 20.8215618111227 | 30.3947158107182 | Il33 |
| **ENSMUSG00000024811** | 0.108210781305108 | 0.618541941125961 | 0.929568738467614 | 1855.81717383282 | 2159.8305130808 | 2084.09366480388 | 1606.66842726423 | 1525.276766776 | 1216.96682091349 | 2182.09967780566 | 2224.24188200577 | Tnks2 |
| **ENSMUSG00000024812** | -0.32084119925126 | 0.0111793127333436 | 0.635862253526407 | 374.054678659585 | 343.683437601847 | 384.260684791269 | 412.687745473591 | 513.535393839156 | 514.917619703095 | 445.581422758026 | 417.927342397375 | Tjp2 |
| **ENSMUSG00000024816** | -0.677999403901141 | 0.00876468860134199 | 0.635862253526407 | 102.097049972302 | 101.801966114291 | 112.628131749165 | 106.92364314543 | 223.371471947096 | 214.039390612925 | 106.189965236726 | 134.605170018895 | Frmd8 |
| **ENSMUSG00000024817** | -0.0370551838613342 | 0.774073467913388 | 0.959203117135841 | 421.037391921175 | 492.721515993169 | 491.210087208544 | 386.425798034362 | 468.642107810573 | 418.29412336926 | 486.183468289716 | 464.604941678121 | Uhrf2 |
| **ENSMUSG00000024818** | -0.308677734872663 | 0.365071060816229 | 0.86924083181804 | 25.2983840639333 | 30.9477976987445 | 30.2865564367503 | 58.151455044006 | 43.7983278327639 | 61.154111603693 | 35.3966550789086 | 39.0789203280663 | Slc25a45 |
| **ENSMUSG00000024824** | -0.335949272136415 | 0.110641991658123 | 0.764116514278315 | 55.1143367107117 | 61.0811796685746 | 66.2518422053913 | 56.2756016554897 | 73.3621991198795 | 92.9542496376133 | 75.998700610598 | 59.7039060567679 | Rad9a |
| **ENSMUSG00000024826** | 0.0387316971981808 | 0.819505760333947 | 0.966398681144067 | 804.12720774645 | 950.423155643021 | 1037.3145579587 | 721.265627884526 | 820.123688668504 | 694.710707817952 | 985.900951756661 | 918.354627709557 | Dpf2 |
| **ENSMUSG00000024827** | -0.492782736253661 | 0.242018666919894 | 0.832048466795696 | 28.0089252136404 | 42.3496179035451 | 28.3936466594534 | 15.0068271081306 | 20.8042057205628 | 50.1463715150282 | 31.2323427166841 | 58.6183804920994 | Gldc |
| **ENSMUSG00000024829** | 0.0801357881783913 | 0.556561867833522 | 0.916239900758826 | 1036.33023290469 | 868.981582751589 | 946.454888648447 | 1131.13959327534 | 1019.40608030758 | 1040.84297949485 | 885.957455063272 | 821.74285245406 | Mrpl21 |
| **ENSMUSG00000024830** | 0.188083582900512 | 0.228880350897852 | 0.832048466795696 | 280.089252136404 | 215.820168162297 | 208.220075502658 | 241.047160424347 | 177.383227722694 | 215.262472844999 | 215.50316474512 | 221.447215192376 | Rps6kb2 |
| **ENSMUSG00000024831** | -0.246005939081606 | 0.236271637903358 | 0.832048466795696 | 73.1846110420926 | 54.56585383726 | 60.5731128735006 | 53.4618215727152 | 79.9319482947941 | 69.71568722821 | 74.9576225200418 | 61.8749571861049 | Ighmbp2 |
| **ENSMUSG00000024833** | -0.229918547145146 | 0.324891840736382 | 0.855977678491063 | 93.0619128066116 | 78.9983257046899 | 111.681676860517 | 117.24083678227 | 140.154649064844 | 125.977469903608 | 79.1219348822664 | 124.835439936878 | Pola2 |
| **ENSMUSG00000024835** | -0.422968192322413 | 0.123550047591149 | 0.770251097497048 | 507.774708711803 | 456.072808192024 | 511.085639870161 | 644.355638955357 | 906.625386138212 | 987.027361283605 | 468.485140750262 | 479.80229958348 | Coro1b |
| **ENSMUSG00000024841** | -0.0774800785596029 | 0.452751570459815 | 0.891153418181945 | 475.248214915318 | 513.896324944941 | 506.353365426919 | 468.963347129081 | 467.547149614754 | 523.479195327612 | 536.15521663641 | 546.019359028259 | Eif1ad |
| **ENSMUSG00000024843** | 0.0711849409817458 | 0.710977622849064 | 0.94990235491128 | 254.790868072471 | 262.241864710414 | 283.936466594534 | 279.502154888932 | 268.264757975679 | 170.008430258266 | 300.871568170723 | 287.664274637155 | Chka |
| **ENSMUSG00000024844** | -0.0434484013997447 | 0.809483047334548 | 0.964381124236801 | 2018.44964281525 | 1474.90688506385 | 1532.31046472184 | 2062.5008006737 | 1778.21211001021 | 2294.50226737056 | 1646.98553925981 | 1585.95284998069 | Banf1 |
| **ENSMUSG00000024845** | -0.107610227770544 | 0.493245538822423 | 0.899931765958683 | 2321.12673786588 | 1926.09319888239 | 2058.53938281037 | 2285.72735390714 | 2183.34664246328 | 2972.08982393948 | 2019.69149567891 | 2083.12355859887 | Tmem134 |
| **ENSMUSG00000024846** | 0.191851626108378 | 0.467215483858012 | 0.895328112556867 | 105.711104838578 | 74.9262470601182 | 80.448665535118 | 147.254490998531 | 82.1218646864323 | 108.854318654574 | 91.61487196894 | 74.901263962127 | Cst6 |
| **ENSMUSG00000024847** | -0.127907935590017 | 0.335294509646392 | 0.858570960161243 | 1433.87626819507 | 1254.20022252807 | 1186.85443036515 | 1387.19358080782 | 1397.16665786517 | 1719.65361829585 | 1342.99073681742 | 1291.77542195552 | Aip |
| **ENSMUSG00000024851** | -0.050727606231473 | 0.708261755088734 | 0.949174314815223 | 399.353062723518 | 539.9576282702 | 539.479286529615 | 453.018593326692 | 492.731188118594 | 525.92535979176 | 490.34778065194 | 492.828606359502 | Pitpnm1 |
| **ENSMUSG00000024853** | 0.133839220222403 | 0.109832713166741 | 0.763169777831932 | 3647.48487378923 | 3621.70674648202 | 4062.18438207913 | 3870.82346720343 | 3688.91416171454 | 3314.55284892016 | 3502.18669663084 | 3347.76084143768 | Sf3b2 |
| **ENSMUSG00000024854** | -0.0718748549485855 | 0.74704521105028 | 0.956119556124081 | 164.439496415566 | 136.821842457607 | 170.36187995672 | 188.52326554589 | 153.294147414674 | 248.285693110993 | 151.997401221196 | 141.118323406906 | Pold4 |
| **ENSMUSG00000024855** | 0.0075912562113283 | 0.958849809435842 | 0.993101678702062 | 684.863397159336 | 814.415728914329 | 807.326020017125 | 594.645524159674 | 721.577451044785 | 623.771938357668 | 765.19239655876 | 775.065253173314 | Pacs1 |
| **ENSMUSG00000024856** | -0.305647762640634 | 0.170917903773553 | 0.799531027639052 | 355.984404328204 | 318.436550005503 | 305.704929033448 | 363.915557372167 | 462.072358635659 | 574.848649074714 | 324.816364253515 | 300.690581413177 | Cdk2ap2 |
| **ENSMUSG00000024858** | 0.102193485589179 | 0.339792777422087 | 0.858570960161243 | 1656.14064247106 | 1619.8728848106 | 1779.33519065908 | 1497.86893073028 | 1537.32130693001 | 1342.9442908171 | 1600.13702518478 | 1622.86071917942 | Grk2 |
| **ENSMUSG00000024862** | 0.0753164938591436 | 0.693293039820537 | 0.94469184542218 | 2178.37157064797 | 2197.29363661086 | 1905.21369084932 | 1809.26059322399 | 1595.35409130842 | 1459.13710286411 | 2468.3961527086 | 2154.76824586699 | Klc2 |
| **ENSMUSG00000024863** | 0.0270188470503876 | 0.951772990251307 | 0.991915011884503 | 40.658117245607 | 73.2974156022896 | 22.7149173275627 | 61.9031618210386 | 31.7537876787538 | 46.4771248188067 | 31.2323427166841 | 85.7565196088121 | Mbl2 |
| **ENSMUSG00000024866** | 0.0517556868747557 | 0.663538724892638 | 0.940272688100259 | 560.178504272808 | 441.413325071566 | 547.050925638802 | 522.425168701796 | 528.864808580624 | 501.463715150282 | 494.512093014165 | 472.203620630801 | Acy3 |
| **ENSMUSG00000024867** | 0.0293257518104178 | 0.858585265309575 | 0.974724097682876 | 579.152292320758 | 636.058684282091 | 597.21303473717 | 558.066383083606 | 510.250519251699 | 454.986590331476 | 699.604476853724 | 656.742966624447 | Pip5k1b |
| **ENSMUSG00000024869** | -0.300149048532738 | 0.218745592782862 | 0.827219244147549 | 41.5616309621761 | 30.9477976987445 | 42.5904699891801 | 42.2067012416173 | 42.7033696369448 | 42.8078781225851 | 57.2592949805875 | 49.9341759747513 | Nudt8 |
| **ENSMUSG00000024870** | -0.0609773611106157 | 0.377843733838189 | 0.87422797717331 | 4316.08502405033 | 4135.60307142696 | 4140.74013783696 | 3979.62296373738 | 4150.9865203502 | 4477.7040516224 | 4408.96571350524 | 4250.91811124188 | Rab1b |
| **ENSMUSG00000024871** | 0.0514897170148368 | 0.771361029014574 | 0.959045638244974 | 3779.39787640832 | 2446.50484965864 | 3022.03045945449 | 4059.34673274932 | 3365.9014939479 | 3279.08346419002 | 3360.60007631521 | 2834.30724934947 | Doc2g |
| **ENSMUSG00000024875** | 0.110158928720294 | 0.317316532075111 | 0.854351922245337 | 1658.85118362077 | 1395.90855935916 | 1363.84149454241 | 1603.85464718146 | 1515.42214301363 | 1357.62127760198 | 1321.12809691574 | 1385.13062051702 | Yif1a |
| **ENSMUSG00000024878** | 0.171758709935584 | 0.205991203521296 | 0.819960256425221 | 268.343573821006 | 259.798617523671 | 279.204192151292 | 233.543746870282 | 217.896680968 | 204.254732756335 | 242.57119509958 | 258.355084391105 | Cbwd1 |
| **ENSMUSG00000024883** | -0.384998953662565 | 0.406132098160648 | 0.881158207349486 | 18.9737880479499 | 13.8450673915436 | 7.57163910918757 | 30.0136542162612 | 20.8042057205628 | 34.2463024980681 | 20.8215618111227 | 16.2828834700276 | Rin1 |
| **ENSMUSG00000024885** | -0.880157761295965 | 0.0536507777239278 | 0.703396849069134 | 19.877301764519 | 16.2883145782866 | 17.9826428843205 | 24.3860940507122 | 53.6529515951357 | 55.0387004433237 | 16.6572494488982 | 19.5394601640331 | Aldh3b1 |
| **ENSMUSG00000024887** | 0.144229141546399 | 0.534405944296852 | 0.910449965200942 | 377.668733525861 | 495.979178908826 | 451.458981885309 | 338.591536627196 | 327.39250054991 | 251.954939807215 | 472.649453112486 | 452.664160466768 | Asah2 |
| **ENSMUSG00000024889** | -0.153338814748411 | 0.546070671171499 | 0.914211323944814 | 128.298947752804 | 117.275864963663 | 100.324218196735 | 171.640585049243 | 175.193311331055 | 177.34692365071 | 102.025652874501 | 121.578863242873 | Rce1 |
| **ENSMUSG00000024891** | -0.373852089576604 | 0.0615322590452808 | 0.71282992338528 | 77.7021796249379 | 95.2866402829764 | 103.163582862681 | 72.2203554578784 | 117.160526952643 | 134.539045528125 | 104.107809055614 | 96.6117752554972 | Slc29a2 |
| **ENSMUSG00000024892** | -0.0390872176961761 | 0.829823358889185 | 0.967535167881372 | 674.021232560508 | 848.62118952873 | 739.181268034437 | 771.913669374467 | 794.939650164664 | 543.048511040794 | 867.218049433262 | 910.755948756878 | Pcx |
| **ENSMUSG00000024896** | 0.107592139361531 | 0.345540956936795 | 0.861266047293794 | 303.580608767199 | 335.539280312703 | 344.509579468035 | 303.888248939644 | 302.208462046071 | 284.978160073209 | 318.569895710178 | 288.749800201823 | Minpp1 |
| **ENSMUSG00000024897** | -0.305001324192488 | 0.2060831494248 | 0.819960256425221 | 92.1583990900426 | 98.5443031986338 | 57.7337482075553 | 84.4134024832345 | 110.590777777729 | 129.646716599829 | 80.1630129728225 | 92.2696729968231 | Apba1 |
| **ENSMUSG00000024899** | -0.486923464234697 | 0.0668303107418287 | 0.719725956118125 | 281.896279569542 | 326.580707294646 | 335.04503058155 | 304.826175633902 | 589.087509350674 | 574.848649074714 | 297.748333899055 | 288.749800201823 | Papss2 |
| **ENSMUSG00000024900** | -0.142457190923233 | 0.399530364869832 | 0.880459952732461 | 606.257703817829 | 914.588863570791 | 967.276896198713 | 779.417082928532 | 928.524550054594 | 907.527016198804 | 801.630129728225 | 969.374329248977 | Cpt1a |
| **ENSMUSG00000024901** | -0.108463429009141 | 0.692253886822448 | 0.94469184542218 | 38.8510898124689 | 47.2361122770311 | 51.1085639870161 | 46.8963347129081 | 51.4630352034976 | 29.3539735697726 | 63.5057635239243 | 53.1907526687569 | Peli3 |
| **ENSMUSG00000024902** | 0.14597390895912 | 0.319492577793075 | 0.854670142081901 | 1434.77978191164 | 1007.43225666702 | 1194.42606947434 | 1459.4139362657 | 1172.70022772225 | 1220.63606760971 | 1129.56972825341 | 1082.2689879745 | Mrpl11 |
| **ENSMUSG00000024905** | 0.256340135075563 | 0.629079659025938 | 0.931825800800958 | 24.3948703473642 | 17.1027303072009 | 14.1968233297267 | 10.3171936368398 | 14.2344565456483 | 3.66924669622158 | 23.9447960827911 | 13.0263067760221 | Tesmin |
| **ENSMUSG00000024906** | -0.0757666350439964 | 0.535576688861009 | 0.91096005719335 | 375.861706092723 | 365.672662282534 | 362.492222352355 | 441.763472995594 | 432.508487348543 | 425.632616761703 | 400.815064864113 | 370.164217551961 | Mus81 |
| **ENSMUSG00000024908** | 0.113660249479204 | 0.571111389683328 | 0.921126010531982 | 1523.32412613541 | 1797.41551371392 | 1951.5899803931 | 1428.46235535518 | 1505.56751925126 | 1065.30462413633 | 1888.51565626883 | 1732.49880121094 | Ppp6r3 |
| **ENSMUSG00000024909** | -0.686617163310653 | 0.0912355397884395 | 0.753823363185192 | 69.5705561758165 | 87.9568987227475 | 83.2880302010633 | 118.178763476528 | 181.76306050597 | 248.285693110993 | 83.2862472444909 | 65.1315338801105 | Efemp2 |
| **ENSMUSG00000024911** | 0.12998396628241 | 0.296198079955391 | 0.851785577030345 | 486.993893230715 | 457.701639649853 | 452.405436773958 | 558.066383083606 | 452.217734873287 | 496.571386221987 | 417.472314313011 | 421.183919091381 | Fibp |
| **ENSMUSG00000024912** | -0.144890880142615 | 0.720195774012554 | 0.951674101666722 | 13.5527057485357 | 30.1333819698302 | 40.6975602118832 | 41.2687745473591 | 48.1781606160403 | 26.9078091056249 | 39.5609674411332 | 23.8815624227072 | Fosl1 |
| **ENSMUSG00000024913** | -0.0936281622385176 | 0.475260728454036 | 0.897549741595057 | 2023.87072511466 | 2351.21820937567 | 2548.80301513027 | 1965.89435116511 | 2676.07783058187 | 2270.04062272908 | 2391.35637400745 | 2148.25509247898 | Lrp5 |
| **ENSMUSG00000024914** | 0.00215320897541198 | 0.990184592579361 | 0.998628446692557 | 7620.23468554333 | 6231.90915765244 | 6530.53873167428 | 7675.99206580879 | 7186.21063916073 | 9097.28564216537 | 5998.69195778446 | 5735.91708370839 | Drap1 |
| **ENSMUSG00000024921** | 0.219033041745497 | 0.0605319644100161 | 0.71282992338528 | 3501.11565170505 | 3774.002487789 | 3852.07139679918 | 3449.69438148152 | 3262.97542354091 | 2635.74221011917 | 3503.2277747214 | 3119.80047285729 | Smarca2 |
| **ENSMUSG00000024924** | 0.412564148055975 | 0.19022644577524 | 0.808824684419004 | 10149.1695782201 | 12982.6011346233 | 15783.0817231015 | 10338.7659508077 | 7608.8645027469 | 4452.01932474885 | 12771.9460149427 | 12169.8271054986 | Vldlr |
| **ENSMUSG00000024925** | -0.160213461927516 | 0.475795510296763 | 0.897549741595057 | 432.783070236573 | 318.436550005503 | 328.419846361011 | 388.301651422879 | 441.268152915096 | 565.063991218123 | 302.953724351836 | 332.170822788563 | Rnaseh2c |
| **ENSMUSG00000024926** | 0.062877807505868 | 0.555993361647138 | 0.916239900758826 | 437.300638819418 | 434.897999240251 | 497.835271429083 | 488.659807708502 | 450.027818481649 | 437.863439082442 | 420.595548584679 | 470.032569501464 | Kat5 |
| **ENSMUSG00000024927** | -0.187522710368226 | 0.272452268999437 | 0.844738346035895 | 731.846110420926 | 822.559886203472 | 892.506959995485 | 897.59584640506 | 1140.9464400435 | 1095.88167993818 | 793.301505003776 | 779.407355431988 | Rela |
| **ENSMUSG00000024935** | 0.0253330622748072 | 0.951427564800624 | 0.991915011884503 | 36.1405486627618 | 47.2361122770311 | 22.7149173275627 | 78.7858423176855 | 68.9823663366031 | 33.0232202659942 | 45.80743598447 | 33.6512925047237 | Slc1a1 |
| **ENSMUSG00000024937** | 0.245253470309248 | 0.233945303472797 | 0.832048466795696 | 8615.90680120241 | 8925.18197317213 | 7409.79532322869 | 8633.61522064637 | 6182.13397359462 | 4784.69769187294 | 8927.24462651887 | 8438.87573973298 | Ehbp1l1 |
| **ENSMUSG00000024939** | -0.936247409296224 | 0.0215253143676967 | 0.648762042745232 | 9.03513716569045 | 8.14415728914329 | 10.4110037751329 | 16.8826804966469 | 17.5193311331055 | 26.9078091056249 | 16.6572494488982 | 23.8815624227072 | Fam89b |

| | | | | | | | | | | | | |
|---|---|---|---|---|---|---|---|---|---|---|---|---|
| **ENSMUSG00000024940** | -0.216904171557214 | 0.238530482839235 | 0.832048466795696 | 738.17070643691 | 904.815874823819 | 965.383986421416 | 822.561710864407 | 1097.14811221074 | 1258.551616804 | 850.560799984364 | 782.663932125994 | Ltbp3 |
| **ENSMUSG00000024941** | 0.130490822559466 | 0.12164112285034 | 0.769868040577702 | 1754.62363757709 | 1672.80990719003 | 1898.58850662878 | 1778.30901231347 | 1653.38687568684 | 1536.19128348477 | 1701.12159996873 | 1596.80810562737 | Scyl1 |
| **ENSMUSG00000024942** | -0.332129274079101 | 0.0380201631205317 | 0.679566075569698 | 539.39768879172 | 645.017257300148 | 444.83379766477 | 483.032247542953 | 687.633746974393 | 758.310983885793 | 642.345181873136 | 573.157498144972 | Capn1 |
| **ENSMUSG00000024943** | 0.26987851347228 | 0.112331979832823 | 0.764848627391134 | 556.564449406532 | 613.255043872489 | 676.71524538364 | 640.603932178324 | 546.384139713729 | 368.147751854232 | 579.880496439768 | 566.644344756961 | Smc5 |
| **ENSMUSG00000024944** | 0.0323112477997084 | 0.842761447024521 | 0.97128746259298 | 1844.97500923399 | 1533.54481754568 | 1572.06157004507 | 1935.88069694884 | 1696.09024532378 | 2114.7091792557 | 1502.27568467251 | 1422.03848971574 | Arl2 |
| **ENSMUSG00000024947** | -0.239672749454318 | 0.210297950179428 | 0.82236445044827 | 302.67709505063 | 295.632909595901 | 388.992959234512 | 296.384835385579 | 409.514365236342 | 480.671317205027 | 305.035880532948 | 321.315567141878 | Men1 |
| **ENSMUSG00000024948** | -0.037231279490013 | 0.825030820915865 | 0.96672635991852 | 477.055242348456 | 423.496179035451 | 628.446046062569 | 461.459933575015 | 590.182467546493 | 503.90987961443 | 511.169342463063 | 436.38127699674 | Map4k2 |
| **ENSMUSG00000024949** | 0.114190550472075 | 0.489523303872347 | 0.899236955296196 | 1704.02686944922 | 2104.45024351463 | 2464.56853004056 | 1869.28790165652 | 2181.15672607164 | 1523.96046116403 | 1911.41937426107 | 1905.09736599323 | Sf1 |
| **ENSMUSG00000024952** | -0.0578684058607416 | 0.686356506610895 | 0.944452327903597 | 544.818771091134 | 498.422426095569 | 557.461929413935 | 528.990655561603 | 631.790878987619 | 629.887349518038 | 490.34778065194 | 465.69046724279 | Rps6ka4 |
| **ENSMUSG00000024953** | -0.0584947601435529 | 0.763644886624623 | 0.95785116771606 | 14870.9322610099 | 10831.7291945606 | 11525.9276339608 | 14833.3106696928 | 14188.4683014239 | 17595.2609906145 | 11164.521443124 | 11268.8408868238 | Prdx5 |
| **ENSMUSG00000024955** | 0.0226801355404911 | 0.8125462779804 | 0.965379767964637 | 4934.99191990012 | 4234.14737462559 | 4409.53332621311 | 4228.17353771579 | 4145.5117293711 | 4179.27198699638 | 4681.72817323095 | 4521.21397684433 | Esrra |
| **ENSMUSG00000024958** | -0.0664829417133658 | 0.619796602175058 | 0.929594515086012 | 807.741262612726 | 730.530908836153 | 634.124775394459 | 646.231492343873 | 676.684165016202 | 852.48831575548 | 710.015257759285 | 714.275821551878 | Gpr137 |
| **ENSMUSG00000024959** | 0.0365508477108741 | 0.871094059302726 | 0.975613767803763 | 394.835494140673 | 261.427448981499 | 314.223023031284 | 347.03287687552 | 319.727793179176 | 448.871179171106 | 249.858741733473 | 267.039288908453 | Bad |
| **ENSMUSG00000024960** | -0.0429870423467471 | 0.714329875368618 | 0.951176382858794 | 346.045753445944 | 397.434875710192 | 368.170951684246 | 319.833002742033 | 372.285786578493 | 346.132271676902 | 381.034581143546 | 375.591845375304 | Plcb3 |
| **ENSMUSG00000024962** | 0.0350008594877702 | 0.77923910723681 | 0.960307976179513 | 7442.24248337922 | 7411.9975488493 | 7255.52317637899 | 8703.95972271574 | 8022.75870076652 | 8316.95917810224 | 7401.02414576358 | 6335.12719540541 | Vegfb |
| **ENSMUSG00000024963** | 0.0363707672068467 | 0.765008577430443 | 0.958197631858494 | 864.662626756576 | 696.325448221751 | 721.198625150116 | 804.741103673502 | 686.538788778574 | 835.365164506446 | 725.631429117627 | 763.124471961961 | Dnajc4 |
| **ENSMUSG00000024965** | 0.136374177858814 | 0.693949557158937 | 0.94469184542218 | 58.7283915769879 | 88.7713144516618 | 125.878500190243 | 56.2756016554897 | 83.2168228822514 | 106.408154190426 | 46.8485140750261 | 64.046008315442 | Fermt3 |
| **ENSMUSG00000024966** | 0.0645762649324188 | 0.549084698320819 | 0.914699131434573 | 2924.673900534 | 2592.28526513431 | 2571.51793245783 | 2699.35302607499 | 2362.91978657761 | 2383.78727031195 | 2670.36530227649 | 2897.26773210025 | Stip1 |
| **ENSMUSG00000024968** | -0.386010857370822 | 0.278612816809325 | 0.848075053184273 | 32.5264937964856 | 45.6072808192024 | 24.6078271048596 | 47.8342614071662 | 48.1781606160403 | 25.6847268735511 | 52.0539045278068 | 70.559161703453 | Rcor2 |
| **ENSMUSG00000024969** | 0.0685939783997309 | 0.47327382573932 | 0.89728597737823 | 1078.79537758344 | 1078.28642508257 | 1031.63582862681 | 916.354380290224 | 940.569090208604 | 1000.48126583642 | 964.038311854983 | 1010.62430070638 | Mark2 |
| **ENSMUSG00000024970** | -0.0691326844284755 | 0.609111805185192 | 0.927932112008768 | 202.387072511466 | 186.501201921381 | 175.094154399963 | 189.461192240149 | 215.706764576362 | 198.139321595965 | 203.010227658447 | 173.684090346961 | AI846148 |
| **ENSMUSG00000024974** | 0.0720055369244554 | 0.448062600255806 | 0.89108031640712 | 964.95264929574 | 1092.94590820303 | 957.812347312228 | 946.368034506485 | 949.328755775157 | 896.519276110139 | 929.68273486663 | 994.341417236353 | Smc3 |
| **ENSMUSG00000024975** | -0.0731943762768662 | 0.504768762860948 | 0.90122958035533 | 546.625798524272 | 494.350347450997 | 448.619617219364 | 477.404687377404 | 528.864808580624 | 544.271593272868 | 514.292576734731 | 483.058876277486 | Pdcd4 |
| **ENSMUSG00000024976** | 0.0438459565379783 | 0.661339589501733 | 0.939611397743053 | 848.399379858333 | 867.35275129376 | 881.149501331704 | 766.286109208918 | 798.224524752122 | 766.87255951031 | 805.79444209045 | 891.216488592845 | Shoc2 |
| **ENSMUSG00000024978** | 0.330786657122179 | 0.624041204496142 | 0.931145988862144 | 1601.02630576035 | 3269.87915159103 | 3511.34763688574 | 1485.67588370493 | 1175.98510230971 | 413.401794440965 | 3115.94672503452 | 3140.42545858599 | Gpam |
| **ENSMUSG00000024981** | -0.40125851565386 | 0.0448800714801781 | 0.693563614452513 | 287.317361868956 | 283.416673662186 | 246.078271048596 | 295.446908691321 | 425.938738173629 | 468.440494884288 | 290.460787265162 | 285.493223507818 | Acsl5 |
| **ENSMUSG00000024982** | 0.276729096589668 | 0.0421679776412852 | 0.685406501171951 | 668.600150261093 | 661.305571878435 | 687.126249158772 | 677.183073254392 | 513.535393839156 | 454.986590331476 | 641.30410378258 | 612.236418473038 | Zdhhc6 |
| **ENSMUSG00000024983** | 0.189865341218166 | 0.196434941666568 | 0.812065429583558 | 365.019541493894 | 408.836695914993 | 424.958245003153 | 366.729337454941 | 354.766455445387 | 269.078091056249 | 372.705956419097 | 374.506319810635 | Vti1a |
| **ENSMUSG00000024985** | -0.219583366888009 | 0.300955602969719 | 0.851785577030345 | 113.8427282877 | 108.317291945606 | 107.895857305923 | 137.87522405595 | 181.76306050597 | 136.985209992272 | 120.765058504512 | 105.295979772845 | Tcf7l2 |
| **ENSMUSG00000024986** | -0.51987106082251 | 0.0853781548706397 | 0.743979028326967 | 101.193536255733 | 68.4109212288036 | 75.7163910918757 | 107.861569839689 | 185.047935093427 | 144.323703384715 | 98.902418602833 | 78.1578406561325 | Hhex |
| **ENSMUSG00000024990** | -0.194968046266688 | 0.487846869585536 | 0.899236955296196 | 827.618564377245 | 1124.70812163069 | 749.59227180957 | 906.037186653384 | 716.102660065689 | 922.20400298369 | 795.383661184888 | 1696.67645757688 | Rbp4 |
| **ENSMUSG00000024991** | 0.316341748217105 | 0.00686495794267715 | 0.635862253526407 | 8716.19682374158 | 9770.5454997852 | 9613.14230400227 | 8888.73128148459 | 7446.81068976568 | 6196.13458768617 | 8095.42323216452 | 7965.58659353751 | Eif3a |
| **ENSMUSG00000024993** | -0.0481011088331741 | 0.706498153751405 | 0.948415903702233 | 215.036264543433 | 197.088606397268 | 212.005895057252 | 228.854113398991 | 220.086597359638 | 203.031650524261 | 215.50316474512 | 242.072200921077 | Fam45a |
| **ENSMUSG00000024997** | 0.186660385980245 | 0.184990994245013 | 0.805963874499992 | 8125.29885310542 | 7544.74731266234 | 7453.33224810652 | 7624.40609762459 | 6104.39194169147 | 5509.98545549274 | 7403.10630194469 | 7997.0668349129 | Prdx3 |
| **ENSMUSG00000024998** | 0.221804834023046 | 0.362777374400781 | 0.86924083181804 | 758.951521917998 | 846.992358070902 | 655.893237833374 | 501.790781428116 | 446.742943894192 | 448.871179171106 | 718.343882483734 | 755.525793009281 | Plce1 |
| **ENSMUSG00000024999** | 0.0365173426215168 | 0.864291479291862 | 0.974868041734905 | 160.82544154929 | 212.56250524664 | 207.27362061401 | 187.585338851632 | 168.623562156141 | 129.646716599829 | 219.667477107345 | 230.131419709724 | Noc3l |
| **ENSMUSG00000025006** | 0.332621397187411 | 0.653162066595312 | 0.936991241819063 | 5518.66178080373 | 10710.3812509523 | 11029.0388174204 | 4250.68377837799 | 2828.27701980073 | 1191.28209403994 | 11785.0039850955 | 9216.11204403563 | Sorbs1 |
| **ENSMUSG00000025007** | -0.163134598492689 | 0.274830205450301 | 0.846286635233373 | 149.983276950461 | 168.584055885266 | 159.950876181588 | 155.695831246855 | 151.104231023035 | 205.477814988408 | 184.270822028436 | 170.427513652956 | Aldh18a1 |
| **ENSMUSG00000025008** | -0.331505737453102 | 0.0721950004814276 | 0.72957913253658 | 93.0619128066116 | 82.2559886203472 | 100.324218196735 | 74.0962088463947 | 105.115986798633 | 128.423634367755 | 110.35427759895 | 96.6117752554972 | Tctn3 |
| **ENSMUSG00000025010** | 0.15416023938053 | 0.427259833554735 | 0.885066977635295 | 117.456783153976 | 96.1010560118908 | 109.78876708322 | 127.55803041911 | 86.5016974697087 | 85.6157562451702 | 109.313199508394 | 122.664388807541 | Ccnj |
| **ENSMUSG00000025016** | 0.023739281093257 | 0.851312497314471 | 0.972692784625411 | 2213.60860559416 | 2339.00197344195 | 2236.47290187628 | 2087.82482141867 | 2042.09703520262 | 1829.73101918249 | 2362.20618747187 | 2496.70879873757 | Tm9sf3 |
| **ENSMUSG00000025017** | -0.566825078166118 | 0.108154638634834 | 0.762287160109383 | 15.3597331816738 | 18.7315617650296 | 22.7149173275627 | 26.2619474392285 | 29.5638712871156 | 25.6847268735511 | 21.8626399016789 | 45.5920737160773 | Pik3ap1 |
| **ENSMUSG00000025019** | -0.148924586701304 | 0.645445050120977 | 0.935063240511793 | 58.7283915769879 | 87.1424829938332 | 60.5731128735006 | 34.703287687552 | 51.4630352034976 | 55.0387004433237 | 92.6559500594962 | 68.388110574116 | Lcor |
| **ENSMUSG00000025024** | 0.110271778995286 | 0.322754933151033 | 0.855977678491063 | 769.793686516826 | 711.799347071123 | 749.59227180957 | 807.554883756277 | 736.906865786252 | 673.918309872697 | 778.72641173599 | 624.177199684392 | Smndc1 |
| **ENSMUSG00000025025** | 0.0716230691236077 | 0.557405893432035 | 0.916341990969984 | 1438.39383677792 | 1642.6765252202 | 1847.47994264177 | 1472.54490998531 | 1618.34821342063 | 1610.79929964127 | 1475.20765431805 | 1387.30167164635 | Mxi1 |
| **ENSMUSG00000025026** | -0.327333620496573 | 0.0213696354413737 | 0.648762042745232 | 707.451240073562 | 824.188717661301 | 683.340429604179 | 754.093062183562 | 906.625386138212 | 1136.24339359662 | 805.79444209045 | 879.275707381491 | Add3 |
| **ENSMUSG00000025027** | -0.0070701500217782 | 0.934376700161182 | 0.98818087308472 | 1103.1902479308 | 1061.18369477537 | 1037.3145579587 | 1039.22277723804 | 1136.56660726022 | 1028.61215717412 | 1095.21415126506 | 1000.85457062436 | Xpnpep1 |
| **ENSMUSG00000025034** | -0.147445155585602 | 0.172709218221929 | 0.799889021902745 | 1015.54941742361 | 896.671717534676 | 870.738497556571 | 993.264369219393 | 1098.24307040655 | 1100.77400886647 | 1052.52994955225 | 931.380934485579 | Trim8 |
| **ENSMUSG00000025035** | -0.0282275802408008 | 0.867572722806332 | 0.974868041734905 | 1474.53438544068 | 1133.66669464875 | 1221.87326124514 | 1452.84844940589 | 1317.23470957037 | 1717.2074538317 | 1156.63775860787 | 1197.33469782936 | Arl3 |
| **ENSMUSG00000025036** | -0.0617223735133248 | 0.855905191954009 | 0.973591279685626 | 20.780815481088 | 27.6901347830872 | 29.3401015481019 | 32.8274342990356 | 27.3739548954774 | 42.8078781225851 | 17.6983275394543 | 28.2236646813812 | Sfxn2 |
| **ENSMUSG00000025037** | -0.185907896704223 | 0.51122683553278 | 0.902394714365738 | 74.9916384752307 | 83.0704043492615 | 69.0912068713366 | 111.613276616721 | 98.5462376237187 | 143.100621152642 | 69.7522320672611 | 74.901263962127 | Maoa |
| **ENSMUSG00000025040** | 0.0171422082433078 | 0.900762146697618 | 0.984802943363903 | 574.634723737913 | 516.339572131684 | 521.496643645294 | 516.797608536247 | 458.787484048202 | 458.655837027697 | 586.126964983105 | 599.210111697016 | Fundc1 |
| **ENSMUSG00000025041** | -0.0302022984593278 | 0.74172684541778 | 0.954383574136061 | 978.505355044276 | 948.794324185193 | 969.16980597601 | 882.58901929693 | 969.038003299901 | 1035.95065056656 | 958.832921402202 | 896.644116416187 | Nt5c2 |
| **ENSMUSG00000025044** | -0.467026256502318 | 0.133355204691625 | 0.77303252008843 | 17.1667606148119 | 33.3910448854875 | 23.6613722162112 | 30.0136542162612 | 37.2285786578493 | 39.1386314263635 | 27.0680303544596 | 41.2499714574033 | Msr1 |
| **ENSMUSG00000025047** | 0.150484182197921 | 0.224074175986591 | 0.832048466795696 | 616.196354700089 | 683.294796559122 | 620.874406953381 | 584.328330522834 | 602.227007700503 | 478.225152740879 | 620.482541971457 | 554.703563545607 | Pdcd11 |
| **ENSMUSG00000025049** | -0.0609929273486749 | 0.799409476434229 | 0.963342665478963 | 67.7635287426784 | 49.679359463774 | 56.7872933189068 | 55.3376749612315 | 79.9319482947941 | 57.4848649074714 | 54.1360607089191 | 47.7631248454143 | Taf5 |
| **ENSMUSG00000025050** | -0.290845455870952 | 0.116993844545902 | 0.769868040577702 | 68.6670424592474 | 66.7820897709749 | 68.1447519826882 | 58.151455044006 | 73.3621991198795 | 88.0619207093179 | 84.3273253350471 | 74.901263962127 | Pcgf6 |
| **ENSMUSG00000025059** | 0.138398236900605 | 0.613882869867845 | 0.928075610917191 | 499.643085262682 | 552.173864203915 | 496.888816540435 | 474.59090729463 | 366.810995599397 | 254.401104271363 | 550.730309904196 | 665.427171141795 | Gk |
| **ENSMUSG00000025060** | 0.229839843359631 | 0.38005630447432 | 0.87432013076003 | 3015.92878590747 | 4161.66437475222 | 3107.21139943285 | 2638.38779094821 | 2235.9046358626 | 1700.08430258266 | 3551.11736688698 | 3532.30018743132 | Slk |
| **ENSMUSG00000025066** | 0.0521217663692611 | 0.551561812387508 | 0.914873920695527 | 2205.47698214504 | 2164.71700745429 | 2045.28901436929 | 2057.81116720241 | 1870.18859845902 | 2095.13986354252 | 2014.48610522612 | 2193.84716619505 | Sfr1 |
| **ENSMUSG00000025068** | 0.15724540406657 | 0.166426044977702 | 0.796299608935682 | 4955.77273538121 | 4219.48789150514 | 4356.5318524488 | 5268.33424164809 | 4522.17734873287 | 4258.77233208118 | 4250.72184374071 | 3826.47761545649 | Gsto1 |
| **ENSMUSG00000025069** | 0.813142842232268 | 0.0562064598346096 | 0.703396849069134 | 21.6843291976571 | 18.7315617650296 | 26.5007368821565 | 38.4549944645846 | 12.0445401540101 | 18.3462334811079 | 21.8626399016789 | 7.59867895267955 | Gsto2 |
| **ENSMUSG00000025075** | 0.451677781389647 | 0.035313114748337 | 0.679566075569698 | 206.001127377742 | 166.140808698523 | 200.648436393471 | 280.44008158319 | 142.344565456483 | 168.785348026193 | 125.970448957293 | 186.710397122983 | Habp2 |
| **ENSMUSG00000025076** | -0.342023541659788 | 0.0910260201519296 | 0.753823363185192 | 110.228673421423 | 100.173134656462 | 87.0738497556571 | 112.551203310979 | 143.439523652302 | 160.223772401676 | 99.9434966933891 | 117.236760984199 | Casp7 |
| **ENSMUSG00000025077** | 0.0659989470775326 | 0.738740376911551 | 0.954383574136061 | 164.439496415566 | 156.367819951551 | 148.593417517806 | 136.937297361692 | 132.489941694111 | 111.300483118721 | 141.586620315635 | 193.223550510994 | Dclre1a |
| **ENSMUSG00000025078** | 0.1520280256534 | 0.586425010018566 | 0.923058967588378 | 401.160090156656 | 596.966729294203 | 526.228918088536 | 386.425798034362 | 360.241246424483 | 249.508775343067 | 589.250199254773 | 519.966745476215 | Nhlrc2 |
| **ENSMUSG00000025083** | -0.524984593339415 | 0.194215799157501 | 0.809162446871198 | 18.0702743313809 | 32.5766291565731 | 24.6078271048596 | 50.6480414899407 | 42.7033696369448 | 72.1618516923577 | 37.4788112600209 | 29.3091902460497 | Afap1l2 |
| **ENSMUSG00000025085** | 0.0192230130328994 | 0.920238496369561 | 0.986713918026868 | 9649.5264929574 | 11877.4389904866 | 12450.6140601703 | 9054.74430636829 | 9780.16660505617 | 8022.19636017245 | 13591.2744722104 | 11068.0186573601 | Ablim1 |
| **ENSMUSG00000025086** | 0.394531153114715 | 0.0989760628249976 | 0.759800953500741 | 92.1583990900426 | 114.83261777692 | 68.1447519826882 | 109.737423228205 | 66.7924499449649 | 58.7079471395453 | 77.0397787011541 | 90.0986218674861 | Trub1 |
| **ENSMUSG00000025089** | -0.487464331822343 | 0.010819386259159 | 0.635862253526407 | 183.413284463516 | 232.108482740584 | 220.523989055088 | 204.468019348279 | 356.956371837026 | 330.232202659942 | 213.421008564008 | 278.980070119806 | Gfra1 |
| **ENSMUSG00000025092** | -0.192251538225596 | 0.545682519381646 | 0.914211323944814 | 340.62467114653 | 465.845796938996 | 257.435729712378 | 228.854113398991 | 258.410134213307 | 242.170281950624 | 442.458188486358 | 534.078577816906 | Hspa12a |
| **ENSMUSG00000025094** | -0.281168591811274 | 0.372492392902253 | 0.873700364743909 | 64.1494738764022 | 43.9784493613737 | 25.5542819935081 | 47.8342614071662 | 62.4126171616885 | 66.0464405319884 | 40.6020455316893 | 52.1052271040884 | Slc18a2 |
| **ENSMUSG00000025102** | 0.244783348569791 | 0.175196480768687 | 0.799889021902745 | 361.405486627618 | 297.26174105373 | 268.793188376159 | 320.770929436291 | 246.365594059297 | 314.332133642982 | 291.501865355718 | 201.907755028342 | 3110040N11 |
| **ENSMUSG00000025103** | 0.143794964571272 | 0.281664090922785 | 0.850048154552447 | 5403.91553879946 | 5237.50755264805 | 4956.58425185192 | 4912.86002452425 | 4057.91507370557 | 3983.57882986456 | 5281.38915339128 | 5240.91742621955 | Btbd1 |
| **ENSMUSG00000025104** | 0.119892895014649 | 0.509577245871518 | 0.902394714365738 | 224.974915425692 | 196.274190668353 | 282.043556817237 | 222.288626539184 | 189.427767876704 | 266.631926592101 | 205.092383839559 | 191.052499381657 | Hdgfl3 |
| **ENSMUSG00000025127** | -0.336533998284462 | 0.316693068419596 | 0.854012936553746 | 35.2370349461928 | 42.3496179035451 | 37.8581955459379 | 48.7721881014244 | 35.0386622662111 | 61.154111603693 | 32.2734208072402 | 79.2433662208011 | Gcgr |
| **ENSMUSG00000025130** | 0.0208011468947614 | 0.818667257195231 | 0.966315773308182 | 5545.7671923008 | 5222.03365379868 | 5346.52366597508 | 5616.30504521787 | 5347.77582838047 | 5814.53293127913 | 4791.04137273934 | 5467.79226923527 | P4hb |
| **ENSMUSG00000025132** | -0.203492462120124 | 0.284480349766187 | 0.851125503777472 | 2972.56012751216 | 2976.68948918187 | 3391.14786602739 | 3462.82535520113 | 4135.65710560873 | 4783.47460964087 | 2930.63482491552 | 2894.01115540624 | Arhgdia |
| **ENSMUSG00000025133** | 0.137865923203723 | 0.28858185679693 | 0.851785577030345 | 1378.76193148436 | 1429.29960424465 | 1445.23661496618 | 1248.38043005761 | 1317.23470957037 | 1056.74304851181 | 1448.13962396359 | 1176.70971210066 | Ints4 |
| **ENSMUSG00000025134** | -0.0153528637221837 | 0.925111642136132 | 0.986713918026868 | 388.510898124689 | 379.517729674077 | 358.706402797761 | 487.721881014244 | 474.116898789669 | 439.086521314516 | 376.870268781321 | 341.94055287058 | Alyref |
| **ENSMUSG00000025135** | -0.0913221357484845 | 0.651211530083005 | 0.936463929813731 | 1550.42953763248 | 1115.74954861263 | 1236.07008457487 | 1563.52379932835 | 1423.44565456483 | 1964.27006471062 | 1188.91117941511 | 1247.26887380412 | Anapc11 |
| **ENSMUSG00000025137** | 0.0198764089419433 | 0.823051235493111 | 0.966637987288264 | 993.86508822595 | 990.329526359824 | 1036.36810307005 | 995.140222607909 | 1009.55145654521 | 916.088591823321 | 1071.26935518226 | 961.775650296298 | Pcyt2 |
| **ENSMUSG00000025138** | -0.177419655551109 | 0.529038872463779 | 0.908549695662944 | 112.939214571131 | 100.987550385377 | 131.557229522134 | 101.296082979881 | 142.344565456483 | 189.577745971448 | 89.5327157878278 | 84.6709940441436 | Sirt7 |
| **ENSMUSG00000025139** | 0.0709937433442424 | 0.667546344313638 | 0.942024383071524 | 1496.21871463834 | 1666.29458135872 | 1978.09071727525 | 1560.71001924558 | 1503.37760285962 | 1251.21312341156 | 1886.43350008772 | 1736.84090346961 | Tollip |
| **ENSMUSG00000025140** | 0.299228804592294 | 0.311819035377246 | 0.851785577030345 | 36.1405486627618 | 42.3496179035451 | 30.2865564367503 | 41.2687745473591 | 28.4689130912965 | 37.9155491942896 | 36.4377331694648 | 19.5394601640331 | Pycr1 |
| **ENSMUSG00000025141** | -0.222530035470085 | 0.0837789453768826 | 0.742148948523533 | 1197.15567445398 | 1048.96745884166 | 934.150975096017 | 1094.56045219927 | 1253.72713421287 | 1375.96751108309 | 1290.93683228961 | 1068.15715563381 | Myadml2 |
| **ENSMUSG00000025142** | 0.10816823683415 | 0.346514323968583 | 0.861266047293794 | 792.381529431053 | 706.098436968723 | 845.184215563063 | 830.065124418473 | 796.034608360484 | 716.726187995282 | 773.52102128321 | 656.742966624447 | Aspscr1 |

| | | | | | | | | | | | | |
|---|---|---|---|---|---|---|---|---|---|---|---|---|
| **ENSMUSG00000025144** | -0.101432574763667 | 0.578868708984194 | 0.922450244597442 | 468.923618899334 | 316.807718547674 | 355.867038131816 | 394.867138282686 | 341.626957095558 | 502.686797382356 | 431.006329490241 | 373.420794245967 | Cenpx |
| **ENSMUSG00000025145** | 0.0123709775473473 | 0.964159850683835 | 0.994284908685797 | 56.0178504272808 | 45.6072808192024 | 37.8581955459379 | 57.2135283497478 | 43.7983278327639 | 58.7079471395453 | 33.3144988977964 | 59.7039060567679 | Lrrc45 |
| **ENSMUSG00000025147** | -0.0277436011214519 | 0.728829825905321 | 0.952436379887563 | 1100.4797067811 | 1048.96745884166 | 1026.90355418356 | 1072.05021153708 | 1146.42123102259 | 1048.1814728873 | 1082.72121417838 | 1052.95979772845 | Mob2 |
| **ENSMUSG00000025150** | -0.776985062884537 | 0.0089588177154486 | 0.635862253526407 | 97.5794813894569 | 105.874044758863 | 83.2880302010633 | 82.5375490947182 | 178.478185918513 | 233.608706326107 | 85.3684034256032 | 136.776221148232 | Cbr2 |
| **ENSMUSG00000025151** | -0.0966709271089419 | 0.277361204924365 | 0.848075053184273 | 1709.44795174863 | 1811.26058110547 | 1683.74324690559 | 1845.83973430006 | 1938.0760065998 | 2027.87034077846 | 1823.96881465435 | 1750.9527358103 | Maged1 |
| **ENSMUSG00000025153** | -0.359109046091933 | 0.00600951380833158 | 0.635862253526407 | 283.70330700268 | 329.023954481389 | 290.561650815073 | 247.612647284155 | 387.61520131996 | 375.486245246675 | 374.788112600209 | 339.769501741243 | Fasn |
| **ENSMUSG00000025154** | 0.0177899703066966 | 0.955689769086086 | 0.992371505557225 | 26.2018977805023 | 51.3081909216027 | 44.483379766477 | 34.703287687552 | 33.943704070392 | 26.9078091056249 | 41.6431236222455 | 52.1052271040884 | Arhgap19 |
| **ENSMUSG00000025155** | 0.140912421691956 | 0.206992707043513 | 0.819960256425221 | 551.143367107117 | 459.330471107681 | 477.013263878817 | 519.611388619021 | 475.211856985488 | 429.301863457925 | 438.293876124134 | 476.545722889475 | Dus1l |
| **ENSMUSG00000025156** | 0.0151672148466409 | 0.849474365559623 | 0.972640876584422 | 1178.18188640603 | 1171.1298181788 | 1211.46225747001 | 1124.57410641554 | 1169.4153531348 | 1087.32010431366 | 1204.52735077345 | 1173.45313540666 | Gps1 |
| **ENSMUSG00000025157** | 0.10058774478871 | 0.51731870360244 | 0.905197321821966 | 229.492484008537 | 216.634583891211 | 202.541346170768 | 213.847286290861 | 232.131137513649 | 162.669936865823 | 186.352978209548 | 222.532740757044 | Zdhhc16 |
| **ENSMUSG00000025158** | 0.0559652661576096 | 0.653099214682673 | 0.936991241819063 | 254.790868072471 | 290.746415222415 | 256.489274823729 | 274.812521417641 | 243.08071947184 | 245.839528646846 | 289.419709174606 | 257.269558826436 | Rfng |
| **ENSMUSG00000025159** | 0.0516031047118383 | 0.723705334472576 | 0.951674101666722 | 236.72059374109 | 310.292392716359 | 298.133289924261 | 317.957149353517 | 295.638712871156 | 265.408844360028 | 256.10521027681 | 305.032683671851 | Mms19 |
| **ENSMUSG00000025161** | -0.645242702808426 | 0.0952441326060575 | 0.757085157577544 | 61.4389327266951 | 51.3081909216027 | 57.7337482075553 | 63.779015209555 | 139.059690869025 | 132.092881063977 | 46.8485140750261 | 48.8486504100828 | Slc16a3 |
| **ENSMUSG00000025162** | 0.234199315201315 | 0.102516673047605 | 0.759800953500741 | 1672.4038893693 | 1997.76178302685 | 2013.10954815525 | 1566.33757941113 | 1680.76083058231 | 1273.22860358889 | 1636.57475835425 | 1571.84101764 | Csnk1d |
| **ENSMUSG00000025169** | -0.129298600093396 | 0.486552669527565 | 0.899236955296196 | 358.694945477911 | 315.99330281876 | 332.205665915605 | 371.418970926232 | 356.956371837026 | 508.802208542726 | 346.679004155194 | 296.348479154503 | Ogfod3 |
| **ENSMUSG00000025170** | -0.234263996614946 | 0.280574990440651 | 0.849093714741928 | 108.421645988285 | 97.7298874697194 | 71.930571537282 | 94.7305961200743 | 93.0714466446232 | 100.292743030056 | 102.025652874501 | 143.289374536243 | Rab40b |
| **ENSMUSG00000025171** | -0.140708217268112 | 0.400623600698284 | 0.880487015130533 | 292.738444168371 | 249.211213047785 | 253.649910157784 | 333.901903155905 | 319.727793179176 | 374.263163014601 | 267.557069272927 | 284.407697943149 | Ubtd1 |
| **ENSMUSG00000025172** | 0.233809786363161 | 0.629481960312878 | 0.931825800800958 | 14.4562194651047 | 21.1748089517725 | 7.57163910918757 | 12.1930470253561 | 6.56974917491458 | 11.0077400886647 | 16.6572494488982 | 13.0263067760221 | Ankrd2 |
| **ENSMUSG00000025173** | 0.235064425501246 | 0.160684045600694 | 0.792016108372479 | 786.056933415069 | 803.828324438442 | 845.184215563063 | 756.906842266336 | 620.841297029428 | 510.025290774799 | 837.026784807134 | 742.499486233259 | Wdr45b |
| **ENSMUSG00000025175** | 0.0295993002366038 | 0.842990214328734 | 0.971386364129692 | 577.34526488762 | 465.845796938996 | 461.869985660442 | 477.404687377404 | 515.725310230795 | 569.956320146419 | 425.80093903746 | 432.039174738066 | Fn3k |
| **ENSMUSG00000025176** | -0.030015714778648 | 0.941161455996745 | 0.989833620337305 | 20.780815481088 | 26.0613033252585 | 18.9290977729689 | 36.5791410760683 | 26.2789966996583 | 25.6847268735511 | 12.4929370866736 | 40.1644458927348 | Hoga1 |
| **ENSMUSG00000025178** | 0.138606855326982 | 0.320213341504395 | 0.855228644171892 | 653.24041707942 | 660.491156149521 | 767.57491469389 | 620.907471598903 | 603.321965896322 | 525.92535979176 | 726.672507208183 | 597.039060567679 | Pi4k2a |
| **ENSMUSG00000025184** | 0.206494828732592 | 0.256021750333395 | 0.837517407767513 | 263.826005238161 | 228.850819824926 | 303.812019256151 | 258.867767615252 | 216.801722772181 | 171.23151249034 | 268.598147363483 | 256.184033261768 | R3hcc1l |
| **ENSMUSG00000025188** | -0.346239294190249 | 0.348163155016074 | 0.861266047293794 | 11.7456783153976 | 21.1748089517725 | 25.5542819935081 | 13.1309737196143 | 22.994122112201 | 26.9078091056249 | 21.8626399016789 | 19.5394601640331 | Hps1 |
| **ENSMUSG00000025190** | 0.0785206439181546 | 0.617028551284507 | 0.929187015773942 | 22564.3515575953 | 24087.1595983702 | 26637.9728410105 | 21289.998032966 | 22514.5304224323 | 17227.1132387603 | 26787.9803481 | 23038.1090589597 | Got1 |
| **ENSMUSG00000025192** | -0.512188064267544 | 0.248006102229007 | 0.834346263461314 | 13.5527057485357 | 13.8450673915436 | 16.0897331070236 | 16.8826804966469 | 33.943704070392 | 24.4616446414772 | 8.32862472444909 | 19.5394601640331 | Entpd7 |
| **ENSMUSG00000025193** | 0.173153489369071 | 0.200474767877571 | 0.812921911423739 | 511.388763578079 | 437.341246426994 | 484.584902988005 | 549.625042835282 | 512.440435643337 | 424.409534529629 | 393.52751823022 | 427.697072479392 | Cutc |
| **ENSMUSG00000025194** | -0.632172604622458 | 0.27087378217597 | 0.844129257012535 | 19.877301764519 | 61.0811796685746 | 18.9290977729689 | 41.2687745473591 | 26.2789966996583 | 29.3539735697726 | 38.5198893505771 | 124.835439936878 | Abcc2 |
| **ENSMUSG00000025195** | 0.00564078389571328 | 0.980662754580335 | 0.996888334226108 | 179.79922959724 | 228.036404096012 | 235.667267273463 | 166.950951577953 | 179.573144114332 | 135.762127760198 | 243.612273190136 | 247.49982874442 | Dnmbp |
| **ENSMUSG00000025196** | -0.278075936480628 | 0.38960000566872 | 0.877215768963936 | 36.1405486627618 | 51.3081909216027 | 32.1794662140472 | 37.5170677703264 | 37.2285786578493 | 48.9232892829544 | 32.2734208072402 | 72.73021283279 | Cpn1 |
| **ENSMUSG00000025197** | -0.325243325680678 | 0.485642887632069 | 0.899236955296196 | 18.0702743313809 | 39.9063707168021 | 15.1432782183751 | 23.448167356454 | 18.6142893289246 | 17.123151249034 | 34.3555769883525 | 51.0197015394199 | Cyp2c23 |
| **ENSMUSG00000025198** | 0.0856595156579505 | 0.602578313176425 | 0.926448151609826 | 271.957628687283 | 262.241864710414 | 240.399541716705 | 245.736793895638 | 234.321053905287 | 188.354663739374 | 239.447960827911 | 298.51953028384 | Erlin1 |
| **ENSMUSG00000025199** | 0.169473906044297 | 0.45519360938204 | 0.892775943845882 | 355.080890611635 | 470.732291312482 | 435.369248778286 | 362.977630677908 | 316.442918591719 | 238.501035254403 | 422.677704765791 | 465.69046724279 | Chuk |
| **ENSMUSG00000025200** | -0.377855868007485 | 0.0237989091360086 | 0.654124840968651 | 84.0267756409212 | 77.3694942468612 | 90.8596693102509 | 86.2892558717508 | 124.825234323377 | 103.961989726278 | 106.189965236726 | 104.210454208177 | Cwf19l1 |
| **ENSMUSG00000025203** | -1.03033296350913 | 0.00609519886197523 | 0.635862253526407 | 144.562194651047 | 158.811067138294 | 78.5557557578211 | 105.047789756914 | 323.012667766634 | 386.49398533534 | 153.038479311752 | 133.519644454226 | Scd2 |
| **ENSMUSG00000025204** | 0.0414842032054011 | 0.800748977132244 | 0.963461896828986 | 22393.5874651638 | 15508.1043099866 | 16737.1082508591 | 20843.5449264991 | 18529.9775478466 | 21560.493586998 | 15762.9633691105 | 17489.987897939 | Ndufb8 |
| **ENSMUSG00000025207** | -0.250683425610586 | 0.649975635120661 | 0.936463929813731 | 11.7456783153976 | 23.6180561385155 | 5.67872933189068 | 24.3860940507122 | 14.2344565456483 | 12.2308223207386 | 15.616171358342 | 35.8223436340608 | Sema4g |
| **ENSMUSG00000025208** | -0.0576239219787304 | 0.597780165524466 | 0.924496535139492 | 1227.87514081733 | 1032.67914426337 | 1070.44047906139 | 1149.89812716051 | 1223.06830472993 | 1243.87463001912 | 1038.99593437502 | 1158.2557775013 | Mrpl43 |
| **ENSMUSG00000025209** | 0.15283658610057 | 0.441920899689711 | 0.89108031640712 | 890.864524537078 | 1014.76199822725 | 973.902080419252 | 863.830485411767 | 693.108537953488 | 612.764198269004 | 995.270654571667 | 1064.90057893981 | Twnk |
| **ENSMUSG00000025212** | -0.655775286191145 | 0.0546162594910127 | 0.703396849069134 | 104.807591122009 | 116.461449234749 | 123.039135524298 | 151.006197775564 | 231.036179317829 | 318.001380339203 | 106.189965236726 | 125.920965501547 | Sfxn3 |
| **ENSMUSG00000025213** | -0.961064019748957 | 0.045648745048601 | 0.697906909182231 | 51.5002818444356 | 52.9370223794314 | 75.7163910918757 | 80.6616957062019 | 160.958854785407 | 237.277953022329 | 70.7933101578173 | 39.0789203280663 | Kazald1 |
| **ENSMUSG00000025217** | 0.0204956784061413 | 0.834553683566672 | 0.968516656512387 | 562.889045422515 | 526.112560878656 | 534.747012086372 | 535.55614242141 | 519.010184818252 | 495.348303989913 | 579.880496439768 | 532.993052252237 | Btrc |
| **ENSMUSG00000025218** | -0.147058619771759 | 0.203593867914608 | 0.816074141171399 | 298.159526467785 | 294.818493866987 | 261.221549266971 | 304.826175633902 | 337.247124312282 | 335.124531588238 | 311.282349076285 | 300.690581413177 | Poll |
| **ENSMUSG00000025220** | 0.327432261162116 | 0.229675752287483 | 0.832048466795696 | 2543.39111214186 | 3325.2594211572 | 3887.09022767917 | 2689.97375913241 | 1966.5449196911 | 1482.37566527352 | 3303.34078133462 | 3165.39254657337 | Mgea5 |
| **ENSMUSG00000025221** | 0.38321942378936 | 0.135467526308759 | 0.776010327117824 | 3211.08774868639 | 4525.70820557692 | 3863.42885546296 | 4015.26417811919 | 3053.83840813946 | 1605.90697071298 | 3841.57815415214 | 3470.42523024522 | Kcnip2 |
| **ENSMUSG00000025223** | -0.0154257189530375 | 0.914355415402155 | 0.986591837173614 | 773.407741383103 | 792.426504233642 | 871.68495244522 | 792.548056648146 | 921.95480087968 | 941.773318696872 | 691.275852129275 | 711.019244857872 | Ldb1 |
| **ENSMUSG00000025224** | -0.133868579354954 | 0.410265063613543 | 0.882192126163777 | 769.793686516826 | 1009.06108812485 | 910.489602879806 | 727.831114744333 | 891.295971396745 | 770.541806206532 | 1048.36563719003 | 1038.84796538776 | Gbf1 |
| **ENSMUSG00000025225** | -0.0239671438420941 | 0.851797340131511 | 0.972742337780703 | 494.222002963268 | 532.627886709971 | 597.21303473717 | 530.866508950119 | 589.087509350674 | 612.764198269004 | 505.963952010282 | 484.144401842154 | Nfkb2 |
| **ENSMUSG00000025226** | -0.563591216649056 | 0.0396985742101833 | 0.679566075569698 | 92.1583990900426 | 68.4109212288036 | 78.5557557578211 | 72.2203554578784 | 117.160526952643 | 172.454594722414 | 75.998700610598 | 95.5262496908287 | Fbxl15 |
| **ENSMUSG00000025227** | -0.0246321829272097 | 0.900267619614823 | 0.984802943363903 | 68.6670424592474 | 64.338842584232 | 75.7163910918757 | 61.9031618210386 | 63.5075753575076 | 64.8233582999146 | 81.2040910633787 | 65.1315338801105 | Mfsd13a |
| **ENSMUSG00000025228** | 0.0758548917698056 | 0.363457302764526 | 0.86924083181804 | 2418.70621925533 | 2418.81471487556 | 2651.0201431043 | 2476.12647284155 | 2499.789561055 | 2299.39459629886 | 2440.28704426358 | 2213.38662635909 | Actr1a |
| **ENSMUSG00000025229** | 0.131518903703044 | 0.743190127174231 | 0.954383574136061 | 28.9124389302094 | 14.6594831204579 | 16.0897331070236 | 15.9447538023887 | 22.994122112201 | 19.5693157131818 | 12.4929370866736 | 14.1118323406906 | Pitx3 |
| **ENSMUSG00000025231** | -0.385076517592263 | 0.00898383783501557 | 0.635862253526407 | 164.439496415566 | 132.749763813036 | 174.147699511314 | 143.502784221499 | 210.231973597267 | 204.254732756335 | 201.96914956789 | 185.624871558315 | Sufu |
| **ENSMUSG00000025232** | 0.0858092085717824 | 0.504212405410813 | 0.900927059989498 | 1710.3514654652 | 1757.50914299712 | 1899.53496151743 | 1992.15629860433 | 1816.53564686388 | 2013.19335399357 | 1620.9585869959 | 1484.99897246652 | Hexa |
| **ENSMUSG00000025234** | 0.280063946322572 | 0.277861423172445 | 0.848075053184273 | 2277.75807947056 | 3142.01588215148 | 2985.1187187972 | 2115.96262224641 | 1708.13478547779 | 1393.09066233213 | 2867.1290613916 | 2695.3599770719 | Arih1 |
| **ENSMUSG00000025235** | -0.231061088588465 | 0.437345766862168 | 0.888722970093854 | 40.658117245607 | 27.6901347830872 | 55.8408384302584 | 52.523894878457 | 39.4184950494875 | 53.8156182112498 | 65.5879197050366 | 47.7631248454143 | Bbs4 |
| **ENSMUSG00000025236** | -0.167193611053876 | 0.235705059755688 | 0.832048466795696 | 132.81651633565 | 151.481325578065 | 146.700507740509 | 125.682177030594 | 160.958854785407 | 155.33144347338 | 153.038479311752 | 156.315681312265 | Adpgk |
| **ENSMUSG00000025237** | 0.19788098510065 | 0.118516679272611 | 0.769868040577702 | 360.501972911049 | 333.09603312596 | 309.490748588042 | 375.170677703265 | 328.487458745729 | 278.86274891284 | 279.008928269045 | 314.802413753867 | Parp6 |
| **ENSMUSG00000025239** | 0.0918265480938394 | 0.619925526145351 | 0.929679074487901 | 1820.58013888663 | 2053.95646832194 | 3040.95955722746 | 2308.23759456933 | 2015.81803850296 | 2010.74718952943 | 2578.75043030755 | 2048.38674052947 | Limd1 |
| **ENSMUSG00000025240** | 0.0574017258747493 | 0.871002942358261 | 0.975613767803763 | 391.221439274397 | 574.163088884602 | 532.854102309076 | 306.702029022419 | 321.917709570815 | 204.254732756335 | 567.387559353095 | 640.46008315442 | Sacm1l |
| **ENSMUSG00000025241** | 0.415590895368767 | 0.264401444782266 | 0.842510961779757 | 8165.95697035103 | 13180.5041567495 | 12186.5531462374 | 7929.2322732585 | 5014.90853685146 | 3453.98422337658 | 11735.0322367488 | 10880.2227346725 | Fyco1 |
| **ENSMUSG00000025245** | 0.232960924534662 | 0.189835310628819 | 0.808824684419004 | 386.703870691551 | 424.310594764365 | 491.210087208544 | 391.115431505653 | 331.772333333186 | 273.970419984545 | 434.129563761909 | 399.473407798011 | Lztfl1 |
| **ENSMUSG00000025246** | -0.100672612801754 | 0.338918559315594 | 0.858570960161243 | 719.19691838896 | 855.136515360045 | 823.415753124149 | 707.196727470654 | 861.732100109629 | 821.911259953634 | 816.205222996011 | 830.427056971408 | Tbl1x |
| **ENSMUSG00000025257** | 0.431440116150946 | 0.340214151528256 | 0.858570960161243 | 15.3597331816738 | 9.77298874697194 | 19.8755526616174 | 15.0068271081306 | 17.5193311331055 | 9.78465785659088 | 8.32862472444909 | 8.68420451734806 | Ribc1 |
| **ENSMUSG00000025260** | 0.0456529334582202 | 0.724058007244489 | 0.951674101666722 | 4213.98797407803 | 3417.28839852452 | 3935.35942700024 | 4411.06924309613 | 4123.61256545472 | 4312.58795029243 | 3453.25602637471 | 3590.91856792342 | Hsd17b10 |
| **ENSMUSG00000025261** | 0.322127142078203 | 0.281620161493716 | 0.850048154552447 | 3955.58305113928 | 5821.44363027962 | 4970.78107518164 | 3495.65278950017 | 2866.6005566544 | 1994.84712051247 | 4932.62799305498 | 4798.0229958348 | Huwe1 |
| **ENSMUSG00000025262** | 0.39244171259797 | 0.392231912950616 | 0.87840402004898 | 180.702743313809 | 364.858246553619 | 256.489274823729 | 114.427056699496 | 114.970610561005 | 67.2695227640623 | 279.008928269045 | 236.644573097735 | Fam120c |
| **ENSMUSG00000025264** | 0.0768315115536138 | 0.526065855469009 | 0.90806673740897 | 292.738444168371 | 301.333819698302 | 291.508105703722 | 304.826175633902 | 277.024423542232 | 248.285693110993 | 317.528817619622 | 284.407697943149 | Tsr2 |
| **ENSMUSG00000025265** | 0.323751290302683 | 0.120233512853254 | 0.769868040577702 | 140.044626068202 | 151.481325578065 | 148.593417517806 | 147.254490998531 | 129.205067106653 | 75.8310983885793 | 145.750932677859 | 117.236760984199 | Fgd1 |
| **ENSMUSG00000025266** | 0.079441608271683 | 0.609091614289244 | 0.927932112008768 | 728.23205555465 | 823.374301932386 | 825.308662901446 | 687.500266891232 | 664.639624862192 | 580.964060235083 | 834.944628626022 | 818.486275760055 | Gnl3l |
| **ENSMUSG00000025268** | -0.227889562416362 | 0.380349270127308 | 0.87432013076003 | 246.659244623349 | 316.807718547674 | 248.917635714542 | 230.729966787508 | 326.297542354091 | 452.540425867328 | 244.653351280692 | 199.736703899005 | Maged2 |
| **ENSMUSG00000025269** | -0.472449175234836 | 0.0463394762771463 | 0.69805115405055 | 103.90407740544 | 98.5443031986338 | 112.628131749165 | 63.779015209555 | 131.394983498292 | 121.085140975312 | 167.613572579538 | 105.295979772845 | Apex2 |
| **ENSMUSG00000025270** | 0.0960591144374996 | 0.924247449087985 | 0.986713918026868 | 1057.11104838578 | 605.110886583346 | 4176.7054236056 | 412.687745473591 | 666.82954125383 | 4129.12561548135 | 401.856142954669 | 651.315338801105 | Alas2 |
| **ENSMUSG00000025271** | -0.0931029424257542 | 0.735897758232555 | 0.954383574136061 | 94.8689402397497 | 105.874044758863 | 101.270673085384 | 74.0962088463947 | 75.5521155115177 | 69.71568722821 | 148.874166949528 | 106.381505337514 | Pfkfb1 |
| **ENSMUSG00000025277** | 0.112728121525164 | 0.414584615300521 | 0.882192126163777 | 411.098741038915 | 305.405898342873 | 374.796135904785 | 361.101777289392 | 327.39250054991 | 302.101311322243 | 357.089785060755 | 354.966859646602 | Abhd6 |
| **ENSMUSG00000025278** | -0.309368862345991 | 0.199240768981746 | 0.812921911423739 | 656.854471945696 | 741.932729040953 | 786.504012466859 | 621.845398293161 | 1102.62290318983 | 1136.24339359662 | 584.044808801993 | 656.742966624447 | Flnb |
| **ENSMUSG00000025280** | 0.0261373172078466 | 0.862218567036688 | 0.974868041734905 | 225.878429142261 | 285.859920848929 | 227.149173275627 | 256.053987532478 | 246.365594059297 | 204.254732756335 | 264.433835001259 | 261.61166108511 | Polr3a |
| **ENSMUSG00000025283** | -0.168721140495985 | 0.295889641211647 | 0.851785577030345 | 321.65088309858 | 285.859920848929 | 301.919109478855 | 390.177504811395 | 396.374866886513 | 415.847958905112 | 308.159114804616 | 340.855027305911 | Sat1 |
| **ENSMUSG00000025287** | 0.0190753853729118 | 0.882999104365835 | 0.979073805351756 | 561.082017989377 | 430.011504866766 | 498.781726317731 | 570.259430108962 | 540.909348734634 | 512.471455238947 | 490.34778065194 | 488.486504100828 | Acot9 |
| **ENSMUSG00000025289** | -0.104783972952762 | 0.718158520312059 | 0.951674101666722 | 259.308436655316 | 175.913797445495 | 209.166530391307 | 320.770929436291 | 226.656346534553 | 407.286383280595 | 184.270822028436 | 220.361689627707 | Prdx4 |
| **ENSMUSG00000025290** | 0.171308380219858 | 0.276693069523029 | 0.847094277204072 | 9525.74511378744 | 6462.3888089352 | 7576.37138363082 | 9508.70082638924 | 7019.77699339623 | 8199.54328382316 | 6762.84327625266 | 7388.08699313386 | Rps24 |
| **ENSMUSG00000025314** | -0.236295096000131 | 0.364544651167804 | 0.86924083181804 | 80.412720774645 | 83.0704043492615 | 132.503684410783 | 111.613276616721 | 151.104231023035 | 143.100621152642 | 94.7381062406084 | 91.1841474321546 | Ptprj |
| **ENSMUSG00000025316** | 0.125943520134531 | 0.700144838172143 | 0.94561797945347 | 254.790868072471 | 338.796943228361 | 166.576060402127 | 198.84045918273 | 217.896680968 | 353.470765069345 | 137.42230795341 | 171.513039217624 | Banp |
| **ENSMUSG00000025317** | -0.273699967313532 | 0.559262067029525 | 0.916574868225118 | 9.93865088225949 | 13.8450673915436 | 6.62518422053913 | 15.9447538023887 | 10.949581958191 | 9.78465785659088 | 13.5340151772298 | 21.7105112933701 | Car5a |
| **ENSMUSG00000025321** | -1.09718171390419 | 0.0594608452504429 | 0.712406493939829 | 6.32459601598331 | 7.32974156022896 | 2.83936466594534 | 10.3171936368398 | 16.4243729372865 | 24.4616446414772 | 10.4107809055614 | 6.51315338801105 | Itgb8 |
| **ENSMUSG00000025323** | -0.18524133936767 | 0.213477095113823 | 0.824248286808591 | 137.334084918495 | 133.56417954195 | 136.289503965376 | 132.247663890401 | 150.009272827216 | 129.646716599829 | 171.777884941763 | 160.657783570939 | Sp4 |
| **ENSMUSG00000025324** | -1.14891400553268 | 0.00495182837357282 | 0.635862253526407 | 9.03513716569045 | 21.1748089517725 | 16.0897331070236 | 10.3171936368398 | 38.3235368536684 | 41.5847958905112 | 27.0680303544596 | 19.5394601640331 | Atp10a |

| | | | | | | | | | | | | |
|---|---|---|---|---|---|---|---|---|---|---|---|---|
| **ENSMUSG00000025326** | -0.0215363435032684 | 0.836003526200265 | 0.968924788429433 | 1889.24718134587 | 1848.72370463553 | 1653.45669046884 | 1714.52999710392 | 1597.54400770006 | 1782.03081213161 | 1920.78907707607 | 1912.69604494591 | Ube3a |
| **ENSMUSG00000025330** | 0.0570222119863683 | 0.92422392585717 | 0.986713918026868 | 31.6229800799166 | 5.7009101024003 | 7.57163910918757 | 15.9447538023887 | 10.949581958191 | 14.6769867848863 | 22.903717992235 | 9.76973008201657 | Padi4 |
| **ENSMUSG00000025332** | 0.292851498725858 | 0.174370066732751 | 0.799889021902745 | 747.2058436026 | 1050.59629029948 | 945.508433759798 | 712.824287636203 | 653.690042904001 | 491.679057293692 | 835.985706716578 | 839.111261488756 | Kdm5c |
| **ENSMUSG00000025337** | 0.139638809822052 | 0.350738965286532 | 0.861774532843376 | 1291.12110097717 | 1241.98398659435 | 1681.85033712829 | 1538.19977858338 | 1410.306156215 | 1461.58326732826 | 1211.81489740734 | 1138.71631733726 | Sbds |
| **ENSMUSG00000025340** | 0.167999459692132 | 0.37163349725153 | 0.873433041763747 | 601.740135234984 | 662.119987607349 | 667.250696497155 | 628.410885152968 | 508.060602860061 | 404.840218816448 | 648.591650416473 | 715.361347116547 | Rabgef1 |
| **ENSMUSG00000025347** | -0.219138635070053 | 0.501704688481992 | 0.900927059989498 | 63.2459601598332 | 119.719112150406 | 74.7699362032273 | 107.861569839689 | 84.3117810780705 | 77.0541806206532 | 91.61487196894 | 172.598564782293 | Mettl7b |
| **ENSMUSG00000025348** | 0.0361807133566953 | 0.716910987667481 | 0.95140862628518 | 3947.45142769016 | 4455.66845289029 | 3954.28852477321 | 3619.45911314224 | 3593.65279867828 | 3966.45567861553 | 3952.97350984165 | 4069.63534194223 | Itga7 |
| **ENSMUSG00000025350** | -0.104400773980228 | 0.743488748005021 | 0.954383574136061 | 45.1756858284523 | 50.4937751926884 | 44.483379766477 | 31.8895076047775 | 35.0386622662111 | 74.6080161565054 | 40.6020455316893 | 35.8223436340608 | Rdh5 |
| **ENSMUSG00000025351** | 0.02682140118951 | 0.880112484895815 | 0.978393501240173 | 4963.90435883033 | 3962.13252116821 | 4481.4638977504 | 5655.69796637671 | 4968.92029262706 | 5808.41752011876 | 3832.20845133714 | 4103.28663444696 | Cd63 |
| **ENSMUSG00000025352** | 0.0450509802720382 | 0.86029608989183 | 0.974868041734905 | 53.3073092775737 | 70.0397526866323 | 52.0550188756646 | 48.7721881014244 | 72.2672409240604 | 44.0309603546589 | 43.7252798033577 | 57.5328549274309 | Gdf11 |
| **ENSMUSG00000025353** | -0.149980938888636 | 0.447454776124736 | 0.89108031640712 | 78.6056933415069 | 59.452348210746 | 63.4124775394459 | 75.0341355406529 | 81.0269064906132 | 85.6157562451702 | 69.7522320672611 | 70.559161703453 | Ormdl2 |
| **ENSMUSG00000025354** | -0.018444532727377 | 0.83990417214836 | 0.969879194284716 | 932.426155499254 | 981.370953341766 | 956.86589242358 | 967.940348474422 | 903.340511550755 | 922.20400298369 | 1044.20132482781 | 1017.13745409439 | Dnajc14 |
| **ENSMUSG00000025355** | -0.364652361527467 | 0.203579768711191 | 0.816074141171399 | 31.6229800799166 | 31.7622134276588 | 33.1259211026956 | 30.0136542162612 | 45.9882442244021 | 51.3694537471021 | 24.9858741733473 | 41.2499714574033 | Mmp19 |
| **ENSMUSG00000025357** | -0.0672325074000457 | 0.696352181655262 | 0.944933386973863 | 200.580045078328 | 213.376920975554 | 262.16800415562 | 228.854113398991 | 294.543754675337 | 236.054870790255 | 218.626399016789 | 198.651178334337 | Dgka |
| **ENSMUSG00000025358** | -0.170966887866927 | 0.408697803732739 | 0.882113132676277 | 104.807591122009 | 94.4722245540621 | 105.056492639978 | 113.489130005238 | 108.400861386091 | 89.2850029413918 | 156.16171358342 | 115.065709854862 | Cdk2 |
| **ENSMUSG00000025362** | 0.0495481931692113 | 0.810187365221121 | 0.964412705343932 | 3165.00854914136 | 2378.09392842984 | 2571.51793245783 | 3261.17111593563 | 2939.96275577428 | 3666.80053175743 | 2201.88016152623 | 2184.07743611304 | Rps26 |
| **ENSMUSG00000025364** | 0.173641420589994 | 0.0392796922502836 | 0.679566075569698 | 1858.52771498253 | 1711.90186217792 | 1914.67823973581 | 1940.57033042014 | 1694.99528712796 | 1627.92245089031 | 1696.9572876065 | 1562.07128755798 | Pa2g4 |
| **ENSMUSG00000025366** | -0.302030525900092 | 0.133726716273494 | 0.77303252008843 | 502.353626412389 | 543.215291185857 | 645.482234058241 | 515.859681841989 | 770.850569856644 | 880.619207093179 | 529.908748093074 | 540.591731204917 | Esyt1 |
| **ENSMUSG00000025369** | 0.277519385700726 | 0.0582091796656292 | 0.710862211775517 | 2648.19870326387 | 3025.55443291673 | 2912.24169237127 | 2637.44986425395 | 2032.24241144024 | 1927.5775977484 | 2722.4192068043 | 2575.95216495837 | Smarcc2 |
| **ENSMUSG00000025371** | 0.00617172408284475 | 0.955937223105883 | 0.992371505557225 | 580.959319753896 | 532.627886709971 | 547.050925638802 | 625.597105070194 | 579.232885588302 | 623.771938357668 | 518.456889096956 | 555.789089110276 | Chmp6 |
| **ENSMUSG00000025372** | -0.138224124814194 | 0.435205482583615 | 0.888448582170255 | 262.018977805023 | 242.69588721647 | 280.15064703994 | 230.729966787508 | 283.594172717146 | 348.57843614105 | 275.885693997376 | 210.59195954569 | Baiap2 |
| **ENSMUSG00000025373** | 0.00125185176970796 | 0.990800314758197 | 0.998628446692557 | 378.57224724243 | 394.991628523449 | 332.205665915605 | 368.605190843457 | 363.52612101194 | 385.270903103266 | 365.418409785204 | 360.394487469945 | Rnf41 |
| **ENSMUSG00000025374** | -0.0305457581342802 | 0.757444533937705 | 0.956693827734505 | 944.171833814652 | 774.509358197526 | 839.505486231172 | 894.782066322286 | 897.865720571659 | 884.2884537894 | 880.752064610492 | 862.992823911464 | Nabp2 |
| **ENSMUSG00000025375** | -0.830122047532754 | 0.0067109185852945 | 0.635862253526407 | 34.3335212296237 | 31.7622134276588 | 20.8220075502658 | 35.6412143818101 | 61.3176589658694 | 74.6080161565054 | 41.6431236222455 | 41.2499714574033 | Aatk |
| **ENSMUSG00000025377** | -0.159452182074067 | 0.339794857533306 | 0.858570960161243 | 93.0619128066116 | 118.904696421492 | 127.77140996754 | 113.489130005238 | 130.300025302473 | 113.746647582869 | 127.011527047849 | 134.605170018895 | Tepsin |
| **ENSMUSG00000025381** | -0.0163062804218627 | 0.927410425783508 | 0.986790505471895 | 991.154547076242 | 803.013908709528 | 795.022106464695 | 957.623154837583 | 979.987585258092 | 1126.45873574002 | 733.960053842076 | 747.927114056602 | Cnpy2 |
| **ENSMUSG00000025384** | -0.464616282785956 | 0.0031632513522134 | 0.635862253526407 | 164.439496415566 | 203.603932228582 | 159.950876181588 | 167.888878272211 | 238.700886688563 | 280.085831144914 | 206.133461930115 | 237.730098662403 | Faap100 |
| **ENSMUSG00000025393** | 0.125672657396409 | 0.117410390785909 | 0.769868040577702 | 159064.493315697 | 154426.252853819 | 173476.662995262 | 172737.959281526 | 157010.435531284 | 142831.543061585 | 150302.526089771 | 154523.478604997 | Atp5b |
| **ENSMUSG00000025395** | 0.339236783649125 | 0.43208835124434 | 0.8866807012241 | 35.2370349461928 | 19.5459774939439 | 27.447191770805 | 28.1378008277448 | 18.6142893289246 | 40.3617136584374 | 10.4107809055614 | 18.4539345993646 | Prim1 |
| **ENSMUSG00000025396** | -0.58702137233789 | 0.0751061794724298 | 0.735399943172214 | 28.0089252136404 | 24.4324718674299 | 22.7149173275627 | 20.6343872736795 | 20.8042057205628 | 37.9155491942896 | 35.3966550789086 | 49.9341759747513 | Hsd17b6 |
| **ENSMUSG00000025402** | -0.848115472694708 | 0.0110259535194486 | 0.635862253526407 | 113.8427282877 | 105.874044758863 | 86.1273948670087 | 104.109863062656 | 238.700886688563 | 272.747337752471 | 139.504464134522 | 87.9275707381491 | Nab2 |
| **ENSMUSG00000025403** | 0.0862573793589152 | 0.386364868162548 | 0.875126835008975 | 920.680477183857 | 1004.17459375137 | 1026.90355418356 | 926.671573927063 | 930.714466446232 | 819.465095489486 | 970.284780398319 | 931.380934485579 | Shmt2 |
| **ENSMUSG00000025404** | 0.24538398757161 | 0.234152253551661 | 0.832048466795696 | 6086.97190852566 | 7104.9628190486 | 6390.46340815431 | 5509.38140207244 | 4181.64534983313 | 3910.19389594013 | 6626.46204638981 | 6448.02185413094 | R3hdm2 |
| **ENSMUSG00000025405** | -0.22794894606962 | 0.735574866496726 | 0.954383574136061 | 4.51756858284523 | 19.5459774939439 | 8.51809399783602 | 9.37926694258161 | 5.47479097909548 | 17.123151249034 | 4.16431236222455 | 22.7960368580387 | Inhbc |
| **ENSMUSG00000025408** | -0.245051717987557 | 0.244639041328849 | 0.832048466795696 | 281.896279569542 | 228.850819824926 | 294.347470369667 | 348.908730264036 | 296.733671066975 | 467.217412652214 | 288.37863108405 | 315.887939318536 | Ddit3 |
| **ENSMUSG00000025409** | -0.0210998825542005 | 0.901187800449087 | 0.984802943363903 | 244.852217190211 | 218.26341534904 | 274.47191770805 | 280.44008158319 | 292.353838283699 | 199.362403828039 | 252.981976005141 | 286.578749072486 | Mbd6 |
| **ENSMUSG00000025410** | 0.145108248953605 | 0.103258789293175 | 0.759800953500741 | 2747.58521208647 | 2460.34991705019 | 2810.02456439724 | 2846.60751707352 | 2474.60552255116 | 2519.54939807215 | 2538.14838477586 | 2291.54446701522 | Dctn2 |
| **ENSMUSG00000025413** | 0.0226677414314747 | 0.849366609863357 | 0.972640876584422 | 311.712232216321 | 287.488752306758 | 346.402489245332 | 313.267515882226 | 307.683253025166 | 283.755077841135 | 330.021754706295 | 315.887939318536 | Ttc4 |
| **ENSMUSG00000025417** | 0.119199160192676 | 0.348901481668819 | 0.861554058734205 | 1239.62081913273 | 1339.71387406407 | 1592.88357759534 | 1282.14579105091 | 1224.16326292575 | 1106.88942002684 | 1340.9085806363 | 1348.22275131829 | Pip4k2c |
| **ENSMUSG00000025421** | -0.0023954148515800 | 0.984665385652026 | 0.99711600049115 | 1256.78757974754 | 1112.49188569697 | 1066.6546595068 | 1085.18118525669 | 1010.64641474103 | 1021.27366378167 | 1246.1704743957 | 1249.43992493345 | Hdhd2 |
| **ENSMUSG00000025422** | -0.78078180685058 | 0.0643778050714792 | 0.716991331547963 | 53.3073092775737 | 88.7713144516618 | 80.448665535118 | 67.5307219865876 | 203.662224422352 | 172.454594722414 | 71.8343882483734 | 51.0197015394199 | Agap2 |
| **ENSMUSG00000025423** | 0.277520531781591 | 0.122411311997154 | 0.770251097497048 | 465.309564033058 | 542.400875456943 | 564.087113634474 | 490.535661097018 | 364.621079207759 | 330.232202659942 | 476.813765474711 | 528.650949993563 | Pias2 |
| **ENSMUSG00000025428** | 0.120845347880236 | 0.177293096648966 | 0.800325353413757 | 132938.490687386 | 134151.373282497 | 143524.205134205 | 137141.76538104 | 132214.012228764 | 111293.144625329 | 129670.440491129 | 130562.672816069 | Atp5a1 |
| **ENSMUSG00000025429** | 0.364240100728144 | 0.324776987308632 | 0.855977678491063 | 362.309000344187 | 457.701639649853 | 424.958245003153 | 307.639955716677 | 202.567266226533 | 124.754387671534 | 458.0743598447 | 420.098393526712 | Pstpip2 |
| **ENSMUSG00000025436** | -0.146519481205141 | 0.523486756704475 | 0.906728945648564 | 104.807591122009 | 70.8541684155466 | 63.4124775394459 | 105.985716451172 | 91.9764884488041 | 100.292743030056 | 84.3273253350471 | 105.295979772845 | Atp23 |
| **ENSMUSG00000025437** | 0.160465002413483 | 0.576369950075523 | 0.922450244597442 | 263.826005238161 | 349.384347704247 | 347.34894413398 | 236.357526953057 | 231.036179317829 | 145.546785616789 | 332.103910887408 | 361.480013034613 | Usp33 |
| **ENSMUSG00000025439** | 0.113320728799329 | 0.213611720812684 | 0.824248286808591 | 646.915821063436 | 574.977504613516 | 602.891764069061 | 611.528204656321 | 551.858930692825 | 554.056251129458 | 583.003730711437 | 562.302242498287 | Clns1a |
| **ENSMUSG00000025451** | 0.257480390290657 | 0.0419977764606373 | 0.685406501171951 | 2590.37382540345 | 2343.07405208652 | 2339.63648473896 | 2750.93899425919 | 1962.16508690782 | 1843.18492373531 | 2451.7389032597 | 2126.54458118561 | Paip1 |
| **ENSMUSG00000025453** | 0.408265581686241 | 0.139168751098149 | 0.779711473022872 | 11960.714579941 | 14180.6066718563 | 16174.914047002 | 12652.6311055426 | 9274.29591858775 | 5338.7539430024 | 13464.2629451625 | 13342.1947153406 | Nnt |
| **ENSMUSG00000025464** | 0.136592732929531 | 0.557871033190068 | 0.916370464878199 | 59.631905293557 | 83.8848200781758 | 87.0738497556571 | 74.0962088463947 | 62.4126171616885 | 69.71568722821 | 55.1771387994752 | 90.0986218674861 | Paox |
| **ENSMUSG00000025465** | 0.127845755452136 | 0.35030477553145 | 0.861774532843376 | 7908.45556112885 | 7415.25521176496 | 7939.81006087182 | 7946.11495375514 | 6774.50635753275 | 5679.993885751 | 8026.71207818781 | 8079.5667778277 | Echs1 |
| **ENSMUSG00000025466** | -0.0330657853055203 | 0.8718898423166 | 0.975948900779883 | 651.433389646282 | 495.979178908826 | 388.046504345863 | 544.935409363992 | 531.054724972262 | 678.810638800992 | 444.54034466747 | 475.460197324806 | Fuom |
| **ENSMUSG00000025470** | -0.0234085541552795 | 0.912938958386561 | 0.986226496507248 | 112.035700854562 | 97.7298874697194 | 100.324218196735 | 141.626930832982 | 104.021028602814 | 150.439114545085 | 97.8613405122768 | 107.467030902182 | Zfp511 |
| **ENSMUSG00000025474** | 0.0944134998002171 | 0.502657239228629 | 0.900927059989498 | 537.590661358582 | 614.069459601404 | 670.0900611631 | 625.597105070194 | 637.265669966714 | 470.886659348436 | 626.729010514794 | 555.789089110276 | Tubgcp2 |
| **ENSMUSG00000025477** | 0.103263789319909 | 0.51530795149307 | 0.90419750457791 | 9877.2119495328 | 10097.9406228088 | 9582.85574756552 | 8997.53077801854 | 7417.24681847856 | 7488.93250698824 | 11044.7974627101 | 9940.15759566952 | Inpp5a |
| **ENSMUSG00000025478** | -0.242938084659432 | 0.383691068968615 | 0.875126835008975 | 54.2108229941427 | 45.6072808192024 | 43.5369248778286 | 31.8895076047775 | 58.0327843784121 | 33.0232202659942 | 58.3003730711437 | 57.5328549274309 | Dpysl4 |
| **ENSMUSG00000025479** | -0.387615344332569 | 0.280035102154695 | 0.848943557485723 | 1374.24436290152 | 2580.88344492951 | 1299.48256211432 | 1713.59207040966 | 1548.2708888882 | 2159.96322184244 | 1339.86750254575 | 4068.54981637757 | Cyp2e1 |
| **ENSMUSG00000025481** | -0.0463882710260995 | 0.867604187495634 | 0.974868041734905 | 47.8862269781594 | 61.895595397489 | 52.0550188756646 | 58.151455044006 | 48.1781606160403 | 55.0387004433237 | 39.5609674411332 | 84.6709940441436 | Urah |
| **ENSMUSG00000025484** | -0.43595009310633 | 0.0301443569763976 | 0.665956123973759 | 189.737880479499 | 167.769640156352 | 191.183887506986 | 239.171307035831 | 324.107625962453 | 306.993640250539 | 213.421008564008 | 221.447215192376 | Bet1l |
| **ENSMUSG00000025485** | 0.0210445248492694 | 0.863947720904322 | 0.974868041734905 | 480.669297214732 | 487.020605890769 | 556.515474525287 | 498.977001345342 | 525.579933993166 | 558.948580057754 | 441.417110395802 | 468.947043936795 | Ric8a |
| **ENSMUSG00000025486** | 0.0549199730313225 | 0.595151416392627 | 0.924461858177432 | 894.478579403355 | 795.684167149299 | 797.861471130641 | 794.423910036663 | 790.559817381388 | 710.610776834912 | 853.684034256032 | 803.288917854696 | Sirt3 |
| **ENSMUSG00000025487** | 0.0609112481809868 | 0.603764123311617 | 0.926448151609826 | 2302.15294981793 | 1901.66072701496 | 2115.32667612928 | 2353.25807589373 | 2092.46511221029 | 2329.9716521007 | 1896.84428099328 | 1995.19598786072 | Psmd13 |
| **ENSMUSG00000025488** | 0.167083524654359 | 0.319632462671076 | 0.854773759434202 | 15804.2619302257 | 10671.2892959644 | 12644.6373123432 | 14569.7532686063 | 12403.6864422387 | 14274.592730534 | 10471.1634348136 | 10669.6307751268 | Cox8b |
| **ENSMUSG00000025491** | -0.711179172982811 | 0.212499070661202 | 0.823370267313127 | 15.3597331816738 | 13.8450673915436 | 23.6613722162112 | 11.2551203310979 | 35.0386622662111 | 48.9232892829544 | 7.28754663389296 | 14.1118323406906 | Ifitm1 |
| **ENSMUSG00000025492** | -0.558793389929473 | 0.126256280623292 | 0.771949105062466 | 807.741262612726 | 732.159740293981 | 708.894711597687 | 835.692684584022 | 1444.24986028539 | 1925.13143328426 | 546.565997541972 | 628.519301943066 | Ifitm3 |
| **ENSMUSG00000025494** | -0.50425313324644 | 0.168113604459325 | 0.79907358295209 | 85.8338030740593 | 78.1839099757755 | 64.3589324280944 | 85.3513291774927 | 151.104231023035 | 168.785348026193 | 51.0128264372507 | 74.901263962127 | Sigirr |
| **ENSMUSG00000025495** | 0.201837697199255 | 0.0547517112794761 | 0.703396849069134 | 760.758549351136 | 651.532583131463 | 790.289832021453 | 790.67220325963 | 650.405168316544 | 669.025980944401 | 631.934400967575 | 650.229813236436 | Ptdss2 |
| **ENSMUSG00000025498** | -0.324296766915564 | 0.328196572716987 | 0.855977678491063 | 164.439496415566 | 148.223662662408 | 199.701981504822 | 181.019851991825 | 229.94122112201 | 370.593916318379 | 118.6829023234 | 149.802527924254 | Irf7 |
| **ENSMUSG00000025499** | -0.030711169396025 | 0.845943300091981 | 0.972180580805179 | 1591.08765487809 | 1368.21842457607 | 1341.12657721485 | 1587.90989337907 | 1623.82300439972 | 1826.06177248627 | 1342.99073681742 | 1223.38731138141 | Hras |
| **ENSMUSG00000025503** | -0.106778267444217 | 0.467262867937383 | 0.895328112556867 | 2260.59131885575 | 2093.04842330982 | 2287.5814658633 | 2243.52065266552 | 2477.89039713862 | 2976.98215286777 | 2011.36287095446 | 2102.6630187629 | Taldo1 |
| **ENSMUSG00000025504** | -0.621506366903805 | 0.097732564997513 | 0.759517686335357 | 32.5264937964856 | 60.2667639396603 | 54.8943835416099 | 49.7101147956825 | 68.9823663366031 | 135.762127760198 | 46.8485140750261 | 53.1907526687569 | Eps8l2 |
| **ENSMUSG00000025505** | -0.0423590838905545 | 0.804322934139597 | 0.963461896828986 | 248.466272056487 | 202.789516499668 | 197.809071727525 | 211.971432902344 | 167.528603960322 | 239.724117486476 | 233.201492284575 | 246.414303179751 | Tmem80 |
| **ENSMUSG00000025508** | 0.0731631884830118 | 0.679429537714399 | 0.944452327903597 | 5810.49671125553 | 4374.22687999886 | 5102.33830470378 | 6072.13741862734 | 5306.16741693934 | 6383.26616919347 | 4227.81812574847 | 4386.60880682544 | Rplp2 |
| **ENSMUSG00000025509** | 0.291552931725389 | 0.107464942126432 | 0.762287160109383 | 15687.7086607883 | 16634.4412630752 | 24475.3234204488 | 21900.5883109281 | 18829.996093501 | 15962.4462107959 | 15704.6629960393 | 13800.2865036307 | Pnpla2 |
| **ENSMUSG00000025510** | -0.0858135458095354 | 0.478565748285538 | 0.897674307047416 | 6269.4816792726 | 6029.11964115277 | 5403.31095929398 | 6062.75815168475 | 6430.68948404556 | 7345.8318858356 | 5952.88452179999 | 5493.84488278732 | Cd151 |
| **ENSMUSG00000025511** | -0.101769412828018 | 0.703067717474341 | 0.946451777392559 | 463.50253659992 | 395.806044252364 | 457.1377112172 | 563.693943249155 | 706.248036303317 | 621.325773893521 | 330.021754706295 | 360.394487469945 | Tspan4 |
| **ENSMUSG00000025512** | 0.0818495068300975 | 0.50042724326884 | 0.900927059989498 | 743.591788736324 | 605.92530231226 | 583.016211407443 | 619.031618210386 | 594.56230032977 | 556.502415593606 | 644.427338054249 | 614.407469602375 | Chid1 |
| **ENSMUSG00000025521** | 0.16491407469077 | 0.249241036676322 | 0.835314899646652 | 475.248214915318 | 498.422426095569 | 512.03209475881 | 564.631869943413 | 480.686647964584 | 363.255422925936 | 463.279750297481 | 519.966745476215 | Tmem192 |
| **ENSMUSG00000025525** | 0.00083958776274966 | 0.99505372027514 | 0.999262355603273 | 1076.08483643373 | 867.35275129376 | 897.239234438728 | 989.51266244236 | 906.625386138212 | 813.349684329117 | 1062.94073045782 | 1043.19006764644 | Apool |
| **ENSMUSG00000025531** | 0.272101680308838 | 0.0993487745328599 | 0.759800953500741 | 318.036828232304 | 390.919549878878 | 306.651383922097 | 346.094950181261 | 249.650468646754 | 239.724117486476 | 345.637926064637 | 292.006376895829 | Chm |
| **ENSMUSG00000025532** | 0.0654686894301611 | 0.641075085927717 | 0.934544499347759 | 266.536546387868 | 241.067055758641 | 241.345996605354 | 288.881421831514 | 240.890803080201 | 278.86274891284 | 214.462086654564 | 258.355084391105 | Crcp |
| **ENSMUSG00000025533** | -0.0495608749903177 | 0.790075970405519 | 0.961659329450003 | 200.580045078328 | 254.097707421271 | 173.201244622666 | 263.557401086543 | 192.712642464161 | 228.716377397812 | 256.10521027681 | 245.328777615083 | Asl |
| **ENSMUSG00000025534** | -0.409697702820891 | 0.0122687911656021 | 0.635862253526407 | 303.580608767199 | 305.405898342873 | 284.882921483182 | 356.412143818101 | 439.078236523458 | 506.356044078578 | 376.870268781321 | 339.769501741243 | Gusb |
| **ENSMUSG00000025537** | 0.110301952042935 | 0.691125451784557 | 0.94469184542218 | 248.466272056487 | 457.701639649853 | 520.550188756646 | 260.743621003769 | 355.861413641207 | 392.609396495709 | 374.788112600209 | 255.098507697099 | Phkg1 |
| **ENSMUSG00000025538** | -0.34481565932288 | 0.130086320831987 | 0.77303252008843 | 96.6759676728878 | 90.4001459094905 | 104.110037751329 | 120.992543559303 | 152.199189218854 | 166.339183562045 | 93.6970281500523 | 111.809133160856 | Sumf2 |
| **ENSMUSG00000025544** | 0.13008451598603 | 0.256001912752697 | 0.837517407767513 | 3681.81839501886 | 4089.18137487884 | 4350.85312311691 | 3593.19716570302 | 3542.18976347478 | 3172.67530999959 | 3927.9876356683 | 3715.7540078603 | Tm9sf2 |
| **ENSMUSG00000025545** | 0.0977415598305975 | 0.255705573935612 | 0.837479467570303 | 2939.1301199991 | 2660.69618636311 | 2544.07074068703 | 2829.72483657687 | 2460.37106600551 | 2492.64158896653 | 2637.05080337869 | 2663.87973569652 | Clybl |

| | | | | | | | | | | | | |
|---|---|---|---|---|---|---|---|---|---|---|---|---|
| **ENSMUSG00000025551** | 0.564009622177656 | 0.269618894106316 | 0.843793744782845 | 9.03513716569045 | 25.2468875963442 | 30.2865564367503 | 12.1930470253561 | 18.6142893289246 | 9.78465785659088 | 8.32862472444909 | 15.1973579053591 | Fgf14 |
| **ENSMUSG00000025555** | -0.103412440924703 | 0.636999312713869 | 0.932931968211785 | 115.649755720838 | 163.69756151178 | 161.843785958884 | 113.489130005238 | 171.908436743598 | 174.900759186562 | 114.518589961175 | 135.690695583563 | Farp1 |
| **ENSMUSG00000025558** | -0.2556258563107 | 0.301678223030858 | 0.851785577030345 | 596.31905293557 | 696.325448221751 | 679.554610049585 | 528.990655561603 | 1098.24307040655 | 802.341944240452 | 570.510793624763 | 515.624643217541 | Dock9 |
| **ENSMUSG00000025571** | -0.062770979121854 | 0.700768570016165 | 0.945843497961766 | 918.873449750719 | 1021.27732405857 | 933.204520207369 | 750.341355406529 | 1077.43886468599 | 736.295503708463 | 1036.91377819391 | 933.551985614916 | Tnrc6c |
| **ENSMUSG00000025572** | -0.661992492731989 | 0.0393197607054077 | 0.679566075569698 | 165.343010132135 | 182.42912327681 | 154.272146849697 | 163.19924480092 | 323.012667766634 | 413.401794440965 | 142.627698406191 | 174.76961591163 | Tmc6 |
| **ENSMUSG00000025574** | -0.0819863224494533 | 0.747098965936681 | 0.956119556124081 | 46.0791995450213 | 39.9063707168021 | 38.8046504345863 | 56.2756016554897 | 39.4184950494875 | 51.3694537471021 | 59.3414511616998 | 41.2499714574033 | Tk1 |
| **ENSMUSG00000025575** | -0.0836217124573136 | 0.545324825570246 | 0.914211323944814 | 424.651446787451 | 474.804369957054 | 501.621090983677 | 437.073839524303 | 573.758094609207 | 505.132961846504 | 428.924173309128 | 440.723379255414 | Cant1 |
| **ENSMUSG00000025577** | -0.59517276691637 | 0.0967628423709528 | 0.757809915356977 | 22.5878429142261 | 44.7928650902881 | 59.6266579848522 | 42.2067012416173 | 95.2613630362614 | 57.4848649074714 | 61.4236073428121 | 41.2499714574033 | Cbx2 |
| **ENSMUSG00000025578** | -0.698640293329602 | 0.00221212696093417 | 0.635862253526407 | 65.0529875929712 | 74.9262470601182 | 42.5904699891801 | 60.0273084325223 | 100.736154015357 | 108.854318654574 | 106.189965236726 | 79.2433662208011 | Cbx8 |
| **ENSMUSG00000025579** | -0.0627637018944549 | 0.569201536106531 | 0.920872458148558 | 4176.9439116987 | 4778.17708154037 | 5068.26592871243 | 4413.88302317891 | 4951.40096149396 | 5012.19098703868 | 5070.05030100839 | 4223.77997212516 | Gaa |
| **ENSMUSG00000025580** | -0.0861778339217151 | 0.614011167081064 | 0.928075610917191 | 1084.21645988285 | 1062.8125262332 | 953.080072868986 | 1089.87081872798 | 1275.62629812925 | 1353.95203090576 | 955.709687130533 | 864.078349476132 | Eif4a3 |
| **ENSMUSG00000025583** | 0.0463796575896652 | 0.829218423598053 | 0.967483872587428 | 818.583427211555 | 1090.50266101629 | 958.758802200877 | 725.017334661559 | 775.230402639921 | 615.210362733151 | 1052.52994955225 | 1035.59138869376 | Rptor |
| **ENSMUSG00000025584** | 0.22182408134921 | 0.108203311640635 | 0.762287160109383 | 682.152856009629 | 728.902077378324 | 569.765842966365 | 740.024161769689 | 572.663136413388 | 500.240632918209 | 630.893322877019 | 628.519301943066 | Pde8a |
| **ENSMUSG00000025586** | 0.279843294107076 | 0.0654321032859087 | 0.718339708336358 | 197.869503928621 | 228.036404096012 | 229.042083052924 | 235.419600258798 | 176.288269526875 | 151.662196777159 | 192.599446752885 | 211.677485110359 | Cpeb1 |
| **ENSMUSG00000025591** | 0.0489546681896173 | 0.730708899233769 | 0.952921973914044 | 224.071401709123 | 180.800291818981 | 184.558703286447 | 224.164479927701 | 201.472308030714 | 196.916239363891 | 204.051305749003 | 183.453820428978 | Tma16 |
| **ENSMUSG00000025597** | -0.350777999915876 | 0.196548790394396 | 0.812065429583558 | 57.8248778604189 | 52.9370223794314 | 75.7163910918757 | 84.4134024832345 | 108.400861386091 | 106.408154190426 | 66.6289977955927 | 64.046008315442 | Klhl4 |
| **ENSMUSG00000025602** | 0.373615238650781 | 0.249103265886657 | 0.835314899646652 | 57.8248778604189 | 61.895595397489 | 57.7337482075553 | 57.2135283497478 | 35.0386622662111 | 22.0154801773295 | 61.4236073428121 | 61.8749571861049 | Zfp202 |
| **ENSMUSG00000025607** | 0.0865429366444782 | 0.585418029171632 | 0.922964550512322 | 894.478579403355 | 899.929380450333 | 932.25806531872 | 877.899385825639 | 781.800151814835 | 644.564336302924 | 1000.47604502445 | 966.117752554972 | Copg2 |
| **ENSMUSG00000025608** | -0.554148454584179 | 0.0631044793619447 | 0.715615581523459 | 2227.1613113427 | 2222.5405242072 | 2408.7276916103 | 2486.44366647839 | 4755.40344444234 | 4806.71317205027 | 2008.23963668279 | 2151.51166917298 | Podxl |
| **ENSMUSG00000025609** | 0.158803524230104 | 0.307358536042426 | 0.851785577030345 | 1194.44513330428 | 1314.46698646773 | 1271.08891545486 | 1099.25008567056 | 1032.54557865741 | 845.149822363037 | 1235.75969349013 | 1255.95307832146 | Mkln1 |
| **ENSMUSG00000025610** | -0.136250834167315 | 0.707162117130536 | 0.948710970699398 | 44.2721721118832 | 39.0919549878878 | 20.8220075502658 | 45.0204813243917 | 51.4630352034976 | 53.8156182112498 | 36.4377331694648 | 22.7960368580387 | Map3k7cl |
| **ENSMUSG00000025612** | -0.0772375801671019 | 0.71142423608707 | 0.950020479830613 | 504.160653845527 | 759.849875077069 | 738.234813145789 | 564.631869943413 | 659.164833883096 | 489.232892829544 | 812.040910633787 | 746.841588491933 | Bach1 |
| **ENSMUSG00000025613** | 0.121130903746999 | 0.127604606559508 | 0.772837268577874 | 3854.38951488355 | 3665.68519584339 | 3453.61388867818 | 3891.45785447711 | 3400.94015621412 | 3438.08415435962 | 3564.65138206421 | 3264.1753729582 | Cct8 |
| **ENSMUSG00000025616** | 0.257468486296587 | 0.152311665235944 | 0.783412422888014 | 1067.04969926804 | 1148.3261777692 | 1178.33633636732 | 1148.02227377199 | 806.984190318674 | 710.610776834912 | 1152.47344624564 | 1127.86106169058 | Usp16 |
| **ENSMUSG00000025626** | -0.148961576886678 | 0.366024800329629 | 0.869253398153585 | 103.90407740544 | 96.9154717408051 | 96.5383986421416 | 88.1651092602672 | 93.0714466446232 | 111.300483118721 | 107.231043327282 | 116.15123541953 | Phf6 |
| **ENSMUSG00000025630** | 0.0495029790660158 | 0.710562421967256 | 0.949886733585872 | 1535.97331816738 | 1158.09916651618 | 1202.94416347218 | 1447.22088924034 | 1152.99098019751 | 1314.8133994794 | 1342.99073681742 | 1352.56485357696 | Hprt |
| **ENSMUSG00000025645** | -0.0418992386299832 | 0.840052944437377 | 0.969879194284716 | 161.728955265859 | 143.337168288922 | 134.396594188079 | 119.116690170786 | 193.80760065998 | 124.754387671534 | 118.6829023234 | 137.8617467129 | Ccdc51 |
| **ENSMUSG00000025647** | -0.86812171532871 | 0.0275259717041783 | 0.664931733862672 | 518.616873310632 | 430.011504866766 | 424.958245003153 | 559.004309777864 | 1204.45401540101 | 1478.7064185773 | 394.568596320776 | 450.493109337431 | Shisa5 |
| **ENSMUSG00000025648** | -0.12411413833092 | 0.651234866179764 | 0.936463929813731 | 48.7897406947284 | 51.3081909216027 | 28.3936466594534 | 45.9584080186499 | 45.9882442244021 | 61.154111603693 | 40.6020455316893 | 43.4210225867403 | Pfkfb4 |
| **ENSMUSG00000025651** | -0.0321127419669296 | 0.777903253887155 | 0.960109845076647 | 38223.1477794535 | 33538.454132421 | 34741.5195976185 | 38848.923676173 | 39466.6732101035 | 41526.0879433717 | 33549.7825462621 | 34082.2461538971 | Uqcrc1 |
| **ENSMUSG00000025656** | -0.119174764428131 | 0.541675390437648 | 0.913837378403038 | 280.992765852973 | 313.550055632016 | 212.005895057252 | 228.854113398991 | 232.131137513649 | 245.839528646846 | 307.11803671406 | 339.769501741243 | Arhgef9 |
| **ENSMUSG00000025658** | 0.546152202101985 | 0.0908604370342938 | 0.753823363185192 | 55.1143367107117 | 52.9370223794314 | 62.4660226507975 | 72.2203554578784 | 19.7092475247437 | 55.0387004433237 | 37.4788112600209 | 54.2762782334254 | Ksr2 |
| **ENSMUSG00000025666** | -0.0992939011803287 | 0.376226538178615 | 0.8741697248747 | 295.448985318078 | 316.807718547674 | 325.580481695066 | 299.198615468353 | 323.012667766634 | 305.770558018465 | 336.268223249632 | 359.308961905276 | Tmem47 |
| **ENSMUSG00000025701** | -0.0159899438139134 | 0.944406034347916 | 0.990505437194639 | 229.492484008537 | 174.284965987666 | 284.882921483182 | 197.902532488472 | 254.03030143003 | 174.900759186562 | 183.22974393788 | 283.322172378481 | Alox5 |
| **ENSMUSG00000025702** | 0.12537767169328 | 0.543351316225343 | 0.913837378403038 | 1265.82271691323 | 1531.10157035894 | 1409.27132919754 | 1198.67031526193 | 973.417836083177 | 928.319414144059 | 1553.28851110976 | 1499.11080480721 | March8 |
| **ENSMUSG00000025722** | 0.178207386663776 | 0.25199396709012 | 0.83723604018117 | 174.378147297826 | 171.841718800923 | 158.057966404291 | 151.944124469822 | 159.863896589588 | 159.000690169602 | 142.627698406191 | 119.407812113536 | Wdr73 |
| **ENSMUSG00000025723** | -0.896178330579347 | 0.0711468064395478 | 0.726501344493452 | 49.6932544112975 | 38.2775392589734 | 24.6078271048596 | 44.0825546301336 | 121.54035973592 | 110.077400886647 | 33.3144988977964 | 27.1381391167127 | Nmb |
| **ENSMUSG00000025724** | -0.139796327467188 | 0.518835387401897 | 0.905696057686609 | 626.135005582348 | 448.743066631795 | 493.102996985841 | 602.14893771374 | 666.82954125383 | 781.549546295196 | 429.965251399684 | 513.453592088204 | Sec11a |
| **ENSMUSG00000025728** | -0.0395978229585161 | 0.792760284014671 | 0.961804768755116 | 543.011743657996 | 637.687515739919 | 597.21303473717 | 476.466760683146 | 505.870686468423 | 522.256113095538 | 655.879197050366 | 632.86140420174 | Pigq |
| **ENSMUSG00000025730** | 0.00726919876080238 | 0.96073002451701 | 0.993784632152314 | 303.580608767199 | 265.499527626071 | 271.632553042104 | 316.081295965 | 349.291664466292 | 266.631926592101 | 271.721381635152 | 262.697186649779 | Rab40c |
| **ENSMUSG00000025731** | -0.00281447353599428 | 0.987552142766697 | 0.998437672973644 | 297.256012751216 | 254.097707421271 | 283.936466594534 | 365.791410760683 | 280.309298129689 | 380.37857417497 | 264.433835001259 | 278.980070119806 | Mettl26 |
| **ENSMUSG00000025732** | -0.0607346213141941 | 0.73564562096916 | 0.954383574136061 | 4336.86583953142 | 2900.13441066392 | 3388.30850136144 | 4020.89173828474 | 4056.82011550975 | 4635.48165955993 | 3268.98520434627 | 3315.19507449762 | Mcrip2 |
| **ENSMUSG00000025733** | 0.0928957671708424 | 0.556375329938376 | 0.916239900758826 | 2925.57741425057 | 2816.24959058575 | 2829.90011705886 | 2610.24999012046 | 2206.34076457548 | 2140.39390612925 | 3022.24969688446 | 3114.37284503395 | Rhot2 |
| **ENSMUSG00000025736** | -0.174286785168267 | 0.228049074540336 | 0.832048466795696 | 562.889045422515 | 546.472954101514 | 494.049451874489 | 483.970174237211 | 603.321965896322 | 704.495365674543 | 563.22324699087 | 486.315452971491 | Jmjd8 |
| **ENSMUSG00000025737** | 0.0219603553445295 | 0.865421367849314 | 0.974868041734905 | 402.063603873225 | 372.187988113848 | 364.385132129652 | 346.094950181261 | 378.855535753408 | 306.993640250539 | 407.061533407449 | 367.993166422624 | Wdr24 |
| **ENSMUSG00000025738** | 0.265799762170894 | 0.318064544676536 | 0.854351922245337 | 58.7283915769879 | 54.56585383726 | 61.519567762149 | 51.5859681841989 | 52.5579933993167 | 39.1386314263635 | 64.5468416144805 | 31.4802413753867 | Fbxl16 |
| **ENSMUSG00000025742** | 0.110376293850459 | 0.564834797345291 | 0.919517890403047 | 301.773581334061 | 485.39177443294 | 465.655805215036 | 333.901903155905 | 339.43704070392 | 392.609396495709 | 400.815064864113 | 337.598450611906 | Prps2 |
| **ENSMUSG00000025743** | -0.674289054777496 | 0.0352237589902046 | 0.679566075569698 | 881.829387371388 | 934.949256793649 | 710.787621374984 | 869.458045577315 | 1870.18859845902 | 1974.05472256721 | 799.547973547113 | 778.32182986732 | Sdc3 |
| **ENSMUSG00000025745** | 0.266702091116285 | 0.0885741020660759 | 0.751206603737452 | 61303.4056692097 | 65612.588784254 | 73814.963220581 | 65640.7996976574 | 54443.5114125171 | 41817.1815146053 | 59848.4561918006 | 65301.9613937634 | Hadha |
| **ENSMUSG00000025747** | -0.585412853509699 | 0.305462131807481 | 0.851785577030345 | 18.0702743313809 | 21.1748089517725 | 4.73227444324223 | 7.50341355406529 | 25.1840385038392 | 29.3539735697726 | 9.36970281500523 | 14.1118323406906 | Tyms |
| **ENSMUSG00000025757** | 0.288052531228829 | 0.231039866451248 | 0.832048466795696 | 1052.59347980294 | 1448.84558173859 | 1278.66055456405 | 1185.53934154232 | 815.743855885227 | 660.464405319884 | 1251.37586484848 | 1338.45302123627 | Hspa4l |
| **ENSMUSG00000025758** | 0.302989125989433 | 0.335051505335828 | 0.858570960161243 | 40.658117245607 | 35.8342920722305 | 39.7511053232348 | 22.5102406621959 | 22.994122112201 | 23.2385624094033 | 29.1501865355718 | 36.9078691987293 | Plk4 |
| **ENSMUSG00000025759** | 0.013590522391134 | 0.949525784322538 | 0.991267322147698 | 281.896279569542 | 325.766291565731 | 301.919109478855 | 326.39848960184 | 260.600050604945 | 206.700897220482 | 371.664878328541 | 384.276049892652 | Mfsd8 |
| **ENSMUSG00000025762** | 0.0398632422192771 | 0.636762028122574 | 0.932878870437987 | 1769.07985704219 | 1636.9756151178 | 1545.56083316291 | 1676.07500263933 | 1586.59442574187 | 1561.87601035832 | 1704.2448342404 | 1593.55152893337 | Larp1b |
| **ENSMUSG00000025764** | -0.137594134995884 | 0.147573401552015 | 0.780607990324772 | 663.179067961679 | 638.501931468834 | 718.359260484171 | 710.010507553428 | 715.00770186987 | 726.510845851873 | 763.110240377648 | 796.775764466685 | Jade1 |
| **ENSMUSG00000025766** | 0.250959635934428 | 0.230322235467059 | 0.832048466795696 | 517.713359594063 | 478.062032872711 | 438.208613444231 | 444.577253078368 | 355.861413641207 | 260.516515431732 | 474.731609293598 | 486.315452971491 | D3Ertd751e |
| **ENSMUSG00000025776** | -0.263627876449254 | 0.350338903565454 | 0.861774532843376 | 45.1756858284523 | 66.7820897709749 | 42.5904699891801 | 33.7653609932938 | 48.1781606160403 | 70.9387694602839 | 60.3825292522559 | 47.7631248454143 | Crispld1 |
| **ENSMUSG00000025779** | -0.253867176395042 | 0.43955027172665 | 0.889733791064291 | 51.5002818444356 | 26.0613033252585 | 34.0723759913441 | 33.7653609932938 | 51.4630352034976 | 57.4848649074714 | 35.3966550789086 | 29.3091902460497 | Ly96 |
| **ENSMUSG00000025780** | -1.04486266737737 | 0.0183304745349883 | 0.642273034804458 | 69.5705561758165 | 78.1839099757755 | 67.1982970940397 | 53.4618215727152 | 179.573144114332 | 247.06261087892 | 49.9717483466946 | 78.1578406561325 | Itih5 |
| **ENSMUSG00000025781** | 0.100407526907313 | 0.32808164300175 | 0.855977678491063 | 33833.878144361 | 27966.2217151891 | 31884.1722887889 | 35284.802237992 | 30292.0184873353 | 31335.3667857323 | 28323.5705316703 | 30346.9526858728 | Atp5c1 |
| **ENSMUSG00000025782** | 0.0528632393183586 | 0.668848602676408 | 0.942055056494206 | 507.774708711803 | 435.712414969166 | 439.155068332879 | 392.053358199911 | 441.268152915096 | 457.432754795624 | 404.979377226337 | 408.157612315359 | Taf3 |
| **ENSMUSG00000025784** | -1.05553402648658 | 0.0312706324259605 | 0.669064667366769 | 269.247087537575 | 244.324718674299 | 297.186835035612 | 370.481044231974 | 773.040486248282 | 1229.19764323423 | 184.270822028436 | 269.21034003779 | Clec3b |
| **ENSMUSG00000025785** | 0.0111114424434166 | 0.934705367017768 | 0.98818087308472 | 438.204152535987 | 390.105134149963 | 341.670214802089 | 450.204813243917 | 407.324448844704 | 436.640356850368 | 401.856142954669 | 362.565538599282 | Exosc7 |
| **ENSMUSG00000025786** | 0.0393174038607517 | 0.840038590236642 | 0.969879194284716 | 973.084272744862 | 1147.51176204029 | 1152.78205437381 | 900.409626487835 | 847.497643563981 | 771.764888438605 | 1272.1974266596 | 1169.11103314798 | Zdhhc3 |
| **ENSMUSG00000025787** | -0.245727974265131 | 0.613085895523605 | 0.928075610917191 | 11.7456783153976 | 8.95857301805762 | 14.1968233297267 | 8.44134024832345 | 19.7092475247437 | 15.9000690169602 | 9.36970281500523 | 6.51315338801105 | Tgm4 |
| **ENSMUSG00000025790** | -0.0328177320859728 | 0.801183101170252 | 0.963461896828986 | 3090.01691066613 | 3154.23211808519 | 3674.13787773327 | 3768.58945752929 | 3062.59807370601 | 3612.98491354618 | 4014.39711718446 | 3310.85297223895 | Slco3a1 |
| **ENSMUSG00000025791** | 0.0728008050873706 | 0.442339324074801 | 0.89108031640712 | 7210.94297193755 | 7603.38524514417 | 7672.90978227296 | 7449.95173249257 | 7330.74512100885 | 6223.0423967918 | 7695.64924539096 | 7212.23185165756 | Pgm2 |
| **ENSMUSG00000025792** | -0.203931775150717 | 0.185509024933874 | 0.805963874499992 | 303.580608767199 | 298.076156782644 | 227.149173275627 | 263.557401086543 | 268.264757975679 | 311.885969178834 | 323.775286162958 | 354.966859646602 | Slc25a10 |
| **ENSMUSG00000025793** | 0.192347380876899 | 0.142214888337759 | 0.780607990324772 | 782.442878548793 | 826.631964848044 | 1035.4216481814 | 773.789522762983 | 787.274942793931 | 697.1568722821 | 749.576225200418 | 756.61131857395 | Hgs |
| **ENSMUSG00000025794** | 0.00680746743243721 | 0.968316396356599 | 0.994697802893254 | 6142.08624523637 | 4538.73885723955 | 5471.45571127667 | 6317.87421252297 | 5471.50610450803 | 7081.64612370765 | 4857.67037053493 | 4954.33867714707 | Rpl14 |
| **ENSMUSG00000025795** | 0.0629091441141605 | 0.766062878452525 | 0.958370894524665 | 4553.70913150799 | 4878.35021619683 | 3875.73276901539 | 4063.09843952635 | 3199.4678481834 | 3077.27489589783 | 5197.06182805623 | 5155.16090661074 | Rassf3 |
| **ENSMUSG00000025804** | 0.762295315620994 | 0.229089130429656 | 0.832048466795696 | 13.5527057485357 | 7.32974156022896 | 38.8046504345863 | 15.0068271081306 | 14.2344565456483 | 15.9000690169602 | 3.12323427166841 | 10.8552556466851 | Ccr1 |
| **ENSMUSG00000025809** | 0.0735281616226978 | 0.611225862588622 | 0.927932112008768 | 8944.78579403355 | 10218.4741506881 | 10107.1917558768 | 8109.31419855606 | 7902.31329922642 | 7638.14853930125 | 10098.4574783945 | 9882.62474074209 | Itgb1 |
| **ENSMUSG00000025810** | -0.449756996290725 | 0.0344792538273708 | 0.678352458445015 | 917.96993603415 | 912.960032112962 | 947.401343537095 | 1073.92606492559 | 1661.05158305757 | 1625.47628642616 | 938.011359591079 | 1037.76243982309 | Nrp1 |
| **ENSMUSG00000025812** | 0.277406397612257 | 0.207961517519763 | 0.820372930586885 | 964.95264929574 | 1215.92268326909 | 1208.62289280407 | 951.995594672034 | 810.269064906132 | 584.633306931305 | 1071.26935518226 | 1114.83475491456 | Pard3 |
| **ENSMUSG00000025813** | 0.198383137785059 | 0.338274837815582 | 0.858570960161243 | 2436.77649358671 | 2208.69545681566 | 1768.92418688395 | 2217.25870522629 | 1449.72465126448 | 1477.48333634522 | 2141.49763227397 | 2453.28777615083 | Homer2 |
| **ENSMUSG00000025815** | 0.538179312705909 | 0.150472919862587 | 0.782641594375979 | 33.4300075130547 | 48.8649437348597 | 56.7872933189068 | 52.523894878457 | 41.6084114411257 | 12.2308223207386 | 43.7252798033577 | 33.6512925047237 | Dhtkd1 |
| **ENSMUSG00000025816** | 0.174475727014836 | 0.539349639788858 | 0.913717099239814 | 110.228673421423 | 144.96599974675 | 141.968233297267 | 100.358156285623 | 88.6916138613469 | 68.4926049961361 | 162.408182126757 | 120.493337678204 | Sec61a2 |
| **ENSMUSG00000025817** | 0.00461597394640071 | 0.98233627260203 | 0.996940244108351 | 197.869503928621 | 151.481325578065 | 108.842312194571 | 175.392291826276 | 168.623562156141 | 146.769867848863 | 145.750932677859 | 170.427513652956 | Nudt5 |
| **ENSMUSG00000025821** | -0.187340285389001 | 0.281117365216826 | 0.850048154552447 | 141.85165350134 | 131.935348084121 | 142.914688185915 | 156.633757941113 | 194.902558855799 | 176.123841418636 | 129.093683228961 | 153.05910461826 | Zfp282 |
| **ENSMUSG00000025823** | -0.457027437416281 | 0.0420046075548868 | 0.685406501171951 | 538.494175075151 | 515.52515640277 | 482.691993210708 | 502.728708122374 | 832.168228822514 | 918.534756287469 | 445.581422758026 | 604.637739520359 | Pdia4 |
| **ENSMUSG00000025825** | -0.014310412028643 | 0.923522307631594 | 0.986713918026868 | 4007.98684670028 | 3428.69021872932 | 3906.01932545214 | 4218.79427077321 | 3800.59989768809 | 4927.79831302558 | 3592.76049050923 | 3396.60949184776 | Iscu |
| **ENSMUSG00000025854** | -0.443750601634171 | 0.00052708421784431 | 0.522942841846969 | 268.343573821006 | 320.879797192245 | 281.097101928589 | 325.460562907582 | 424.84377997781 | 445.201932474885 | 374.788112600209 | 383.190524327983 | Fam20c |
| **ENSMUSG00000025855** | -0.575614375213186 | 0.191257198568615 | 0.808824684419004 | 36.1405486627618 | 23.6180561385155 | 29.3401015481019 | 16.8826804966469 | 38.3235368536684 | 68.4926049961361 | 35.3966550789086 | 16.2828834700276 | Prkar1b |
| **ENSMUSG00000025856** | -0.204950757781541 | 0.414190096779795 | 0.882192126163777 | 378.57224724243 | 280.973426475443 | 365.3315870183 | 565.569796637671 | 479.591689768764 | 599.310293716191 | 355.007628879643 | 399.473407798011 | Pdgfa |
| **ENSMUSG00000025857** | 0.0998890902757028 | 0.480370706773223 | 0.897732059433541 | 357.791431761342 | 390.919549878878 | 424.958245003153 | 330.150196378873 | 308.778211220985 | 330.232202659942 | 396.650752501888 | 366.907640857956 | Dnaaf5 |
| **ENSMUSG00000025858** | 0.0959751183491951 | 0.234291713110676 | 0.832048466795696 | 1100.4797067811 | 1052.22512175731 | 1160.353693483 | 1106.75349922463 | 1017.21616391594 | 1081.20469315329 | 1001.517123115 | 1035.59138869376 | Get4 |

| | | | | | | | | | | | | |
|---|---|---|---|---|---|---|---|---|---|---|---|---|
| **ENSMUSG00000025860** | 0.0206327182330112 | 0.884012388769163 | 0.979384300778265 | 807.741262612726 | 825.003133390215 | 1056.24365573167 | 923.857793844289 | 950.423713970976 | 779.103381831049 | 831.821394354353 | 998.683519495027 | Xiap |
| **ENSMUSG00000025862** | 0.114366351996278 | 0.33382504825018 | 0.85787787607678 | 1020.06698600645 | 1053.85395321514 | 1070.44047906139 | 984.823028971069 | 866.111932892906 | 839.034411202668 | 1047.32455909947 | 1060.55847668113 | Stag2 |
| **ENSMUSG00000025867** | -0.273181190648567 | 0.144190030658618 | 0.780607990324772 | 127.395434036235 | 134.378595270864 | 88.0203046443056 | 115.364983393754 | 139.059690869025 | 140.654456688494 | 123.88829277618 | 159.572258006271 | Cplx2 |
| **ENSMUSG00000025868** | 0.0206124562770042 | 0.866416439169821 | 0.974868041734905 | 1851.29960524997 | 1586.48183992511 | 1469.84444207104 | 1794.25376611586 | 1612.87342244153 | 1867.64656837678 | 1600.13702518478 | 1527.33446948859 | Higd2a |
| **ENSMUSG00000025869** | 0.093914375224572 | 0.476601771189837 | 0.897553735056661 | 458.984968017075 | 395.806044252364 | 402.24332767559 | 479.28054076592 | 466.452191418935 | 387.717067567413 | 402.897221045225 | 369.078691987293 | Nop16 |
| **ENSMUSG00000025870** | -0.268282421036383 | 0.343559611785063 | 0.861266047293794 | 36.1405486627618 | 31.7622134276588 | 36.9117406572894 | 23.448167356454 | 37.2285786578493 | 30.5770558018465 | 36.4377331694648 | 49.9341759747513 | Arl10 |
| **ENSMUSG00000025871** | 0.130959396463677 | 0.427677044082574 | 0.885066977635295 | 2539.77705727559 | 2584.95552357408 | 2632.09104533133 | 2473.31269275877 | 1932.60121562071 | 1871.31581507301 | 2749.48723715876 | 2787.62965006873 | 4833439L19R |
| **ENSMUSG00000025872** | 0.135543998200716 | 0.214755547153369 | 0.824248286808591 | 478.862269781594 | 409.651111643907 | 422.118880337207 | 451.142739938176 | 420.463947194533 | 415.847958905112 | 374.788112600209 | 392.96025441 | Thoc3 |
| **ENSMUSG00000025873** | -0.0338020198839813 | 0.784997740795108 | 0.961141387070613 | 320.747369382011 | 351.013179162076 | 338.830850136144 | 279.502154888932 | 336.152166116463 | 348.57843614105 | 331.062832796851 | 306.118209236519 | Faf2 |
| **ENSMUSG00000025875** | -0.147714901516315 | 0.463051651808983 | 0.895328112556867 | 352.370349461928 | 298.076156782644 | 231.881447718869 | 332.026049767389 | 298.923587458613 | 445.201932474885 | 303.994802442392 | 298.51953028384 | Tspan17 |
| **ENSMUSG00000025876** | -0.690334176262033 | 0.0652891886007829 | 0.718339708336358 | 34.3335212296237 | 19.5459774939439 | 16.0897331070236 | 40.3308478531009 | 53.6529515951357 | 57.4848649074714 | 33.3144988977964 | 33.6512925047237 | Unc5a |
| **ENSMUSG00000025877** | -1.22003703301432 | 0.01745179395027 | 0.642273034804458 | 2.71054114970714 | 4.88649437348597 | 12.3039135524298 | 8.44134024832345 | 12.0445401540101 | 18.3462334811079 | 15.616171358342 | 19.5394601640331 | Hk3 |
| **ENSMUSG00000025878** | 0.170155131287853 | 0.18430808630225 | 0.805963874499992 | 334.300075130547 | 300.519403969387 | 325.580481695066 | 364.853484066425 | 275.929465346412 | 272.747337752471 | 295.666177717943 | 332.170822788563 | Uimc1 |
| **ENSMUSG00000025880** | -0.552839237832626 | 0.080574728322984 | 0.739855948845396 | 455.370913150799 | 426.753841951108 | 457.1377112172 | 424.880792498947 | 1055.53970076961 | 757.087901653719 | 416.431236222455 | 359.308961905276 | Smad7 |
| **ENSMUSG00000025885** | 0.242935991850472 | 0.443758066796244 | 0.89108031640712 | 1455.56059739273 | 1747.73615425015 | 1428.20042697051 | 1192.10482840212 | 919.764884488041 | 598.087211484117 | 1717.77884941763 | 1684.73567636552 | Myo5b |
| **ENSMUSG00000025887** | -0.0679828584992012 | 0.799204546766395 | 0.96328975655894 | 141.85165350134 | 131.120932355207 | 98.4313084194385 | 191.337045628665 | 136.869774477387 | 206.700897220482 | 137.42230795341 | 109.638082031519 | Casp12 |
| **ENSMUSG00000025888** | -0.0287870242670537 | 0.944753467706235 | 0.990505437194639 | 29.8159526467785 | 23.6180561385155 | 31.2330113253987 | 50.6480414899407 | 47.0832024202212 | 42.8078781225851 | 13.5340151772298 | 34.7368180693922 | Casp1 |
| **ENSMUSG00000025889** | 0.317896267944812 | 0.764809736220549 | 0.958167875539229 | 218.650319409709 | 135.193010999779 | 1009.86736618789 | 70.3445020693621 | 168.623562156141 | 806.011190936674 | 72.8754663389296 | 103.124928643508 | Snca |
| **ENSMUSG00000025893** | 0.0687906228114119 | 0.675881793672981 | 0.94409858171788 | 156.307872966445 | 144.96599974675 | 134.396594188079 | 152.88205116408 | 120.445401540101 | 147.992950080937 | 125.970448957293 | 167.17093695895 | Kbtbd3 |
| **ENSMUSG00000025894** | 0.0405637911492462 | 0.740822284081568 | 0.954383574136061 | 686.670424592474 | 570.09101024003 | 580.176846741498 | 639.666005484066 | 585.802634763217 | 532.040770952129 | 666.289977955927 | 622.006148555055 | Aasdhppt |
| **ENSMUSG00000025898** | 0.03295569678796 | 0.743022173265593 | 0.954383574136061 | 482.47632464787 | 499.236841824483 | 493.102996985841 | 486.783954319986 | 442.363111110915 | 467.217412652214 | 480.978077836935 | 526.479898864226 | Cwf19l2 |
| **ENSMUSG00000025899** | 0.0295435345898093 | 0.907721292352539 | 0.98504921011344 | 200.580045078328 | 258.169786065842 | 232.827902607518 | 202.592165959763 | 173.003394939417 | 138.208292224346 | 290.460787265162 | 273.552442296464 | Alkbh8 |
| **ENSMUSG00000025900** | -0.433309782066971 | 0.441757821897061 | 0.89108031640712 | 7.22810973255236 | 24.4324718674299 | 16.0897331070236 | 12.1930470253561 | 32.8487458745729 | 23.2385624094033 | 19.7804837205666 | 5.42762782334254 | Rp1 |
| **ENSMUSG00000025902** | -0.979165483665876 | 0.0501061563959967 | 0.703396849069134 | 224.071401709123 | 263.056280439328 | 222.416898832385 | 161.323391412404 | 513.535393839156 | 886.734618253548 | 192.599446752885 | 124.835439936878 | Sox17 |
| **ENSMUSG00000025903** | -0.00665085876858182 | 0.96665752229483 | 0.994284908685797 | 1808.83446057123 | 1571.00794107574 | 1577.74029937696 | 1688.26804966469 | 1462.86414961431 | 1335.60579742465 | 1852.07792309937 | 2024.50517810677 | Lypla1 |
| **ENSMUSG00000025907** | 0.150486726205964 | 0.330629905729931 | 0.856303993370193 | 1509.77142038687 | 1531.10157035894 | 1698.88652512396 | 1458.47600957144 | 1277.81621452089 | 1095.88167993818 | 1653.23200780315 | 1555.55813416997 | Rb1cc1 |
| **ENSMUSG00000025911** | 0.328666824276399 | 0.126886728192366 | 0.772551378490123 | 1337.20030052219 | 1531.91598608785 | 1641.15277691641 | 1321.53871220975 | 986.557334433006 | 783.995710759344 | 1406.49650034134 | 1465.45951230249 | Adhfe1 |
| **ENSMUSG00000025915** | -0.0126232804469609 | 0.931170602438992 | 0.987775860606096 | 199.676531361759 | 236.99497711407 | 246.078271048596 | 216.661066373635 | 239.795844884382 | 188.354663739374 | 244.653351280692 | 233.387996403729 | Sgk3 |
| **ENSMUSG00000025917** | 0.185622930478842 | 0.0639094859507821 | 0.716991331547963 | 2012.12504679926 | 1745.29290706341 | 1881.55231863311 | 1852.40522115987 | 1478.19356435578 | 1612.02238187335 | 1752.13442640598 | 1743.35405685762 | Cops5 |
| **ENSMUSG00000025920** | 0.191636733499287 | 0.326554414417408 | 0.855977678491063 | 1758.23769244336 | 1783.57044632238 | 1764.1919124407 | 1780.18486570199 | 1289.8607546749 | 1113.00483118721 | 2008.23963668279 | 1792.20270726771 | Stau2 |
| **ENSMUSG00000025921** | 0.116727679575905 | 0.560085967274263 | 0.916574868225118 | 188.83436676293 | 196.274190668353 | 192.130342395635 | 172.578511743502 | 170.813478547779 | 116.192812047017 | 186.352978209548 | 217.105112933702 | Rdh10 |
| **ENSMUSG00000025925** | -0.280867321608746 | 0.244958620448571 | 0.832048466795696 | 81.316234491214 | 71.6685841444609 | 47.3227444324223 | 66.5927952923295 | 95.2613630362614 | 92.9542496376133 | 72.8754663389296 | 64.046008315442 | Terf1 |
| **ENSMUSG00000025930** | -0.878778192985037 | 0.0174076415047339 | 0.642273034804458 | 21.6843291976571 | 22.8036404096012 | 33.1259211026956 | 29.075727522003 | 68.9823663366031 | 64.8233582999146 | 36.4377331694648 | 26.0526135520442 | Msc |
| **ENSMUSG00000025931** | -0.390590806234022 | 0.0842165630699664 | 0.742565114108635 | 43.3686583953142 | 50.4937751926884 | 49.2156542097192 | 51.5859681841989 | 79.9319482947941 | 67.2695227640623 | 56.2182168900314 | 52.1052271040884 | Paqr8 |
| **ENSMUSG00000025932** | -0.30964933576 | 0.464292009172455 | 0.895328112556867 | 7.22810973255236 | 16.2883145782866 | 17.9826428843205 | 13.1309737196143 | 10.949581958191 | 20.7923979452556 | 17.6983275394543 | 18.4539345993646 | Eya1 |
| **ENSMUSG00000025933** | -0.170406996879271 | 0.489377347838718 | 0.899236955296196 | 48.7897406947284 | 58.6379324818317 | 35.0188308799925 | 54.3997482669734 | 62.4126171616885 | 58.7079471395453 | 44.7663578939139 | 56.4473293627624 | Tmem14a |
| **ENSMUSG00000025934** | -0.317428513334894 | 0.215991349036218 | 0.825342458903182 | 451.756858284523 | 594.52348210746 | 427.797609669098 | 559.004309777864 | 651.500126512363 | 776.657217366901 | 348.761160336306 | 757.696844138618 | Gsta3 |
| **ENSMUSG00000025935** | -0.0627764734085554 | 0.714831517588403 | 0.951176382858794 | 795.09207058076 | 706.912852697637 | 763.789095139297 | 747.527575323754 | 930.714466446232 | 913.642427359173 | 577.798340258656 | 726.216602763232 | Tram1 |
| **ENSMUSG00000025936** | -0.336971878323278 | 0.193943764145971 | 0.809162446871198 | 29.8159526467785 | 28.5045505120015 | 35.965285768641 | 31.8895076047775 | 39.4184950494875 | 33.0232202659942 | 37.4788112600209 | 48.8486504100828 | Gm4956 |
| **ENSMUSG00000025937** | 0.115778278132897 | 0.406924577176615 | 0.881296485258502 | 535.783633925444 | 582.307246173745 | 507.299820315568 | 491.473587791276 | 462.072358635659 | 409.732547744743 | 508.046108191395 | 573.157498144972 | Lactb2 |
| **ENSMUSG00000025938** | 0.276490720828182 | 0.281430391742648 | 0.850048154552447 | 308.098177350044 | 394.177212794535 | 299.079744812909 | 256.991914226736 | 239.795844884382 | 154.108361241306 | 342.514691792969 | 301.776106977845 | Slco5a1 |
| **ENSMUSG00000025939** | -0.143197437933943 | 0.206496634340435 | 0.819960256425221 | 350.563322028789 | 338.796943228361 | 312.330113253987 | 354.536290429585 | 371.190828382674 | 344.909189444828 | 369.582722147429 | 411.414189009364 | Ube2w |
| **ENSMUSG00000025940** | 0.058631160566886 | 0.671544859436715 | 0.943173644832837 | 2354.55674537893 | 2167.16025464103 | 2122.89831523847 | 2117.83847563493 | 1912.89196809596 | 1741.66909847318 | 2271.63239359349 | 2485.85354309088 | Tmem70 |
| **ENSMUSG00000025949** | 0.0868145366011972 | 0.631976266871491 | 0.931825800800958 | 138.237598635064 | 175.913797445495 | 177.933519065908 | 151.944124469822 | 127.015150715015 | 132.092881063977 | 166.572494488982 | 180.197243734972 | Pikfyve |
| **ENSMUSG00000025950** | -0.101025699883483 | 0.447113710303479 | 0.89108031640712 | 1196.25216073742 | 1281.07594158224 | 986.205993971682 | 1161.1532474916 | 1123.42710891039 | 1124.01257127588 | 1322.16917500629 | 1390.55824834036 | Idh1 |
| **ENSMUSG00000025956** | -0.349296963803979 | 0.0519005210357126 | 0.703396849069134 | 85.8338030740593 | 83.8848200781758 | 94.6454888648447 | 100.358156285623 | 127.015150715015 | 95.400414101761 | 132.216917500629 | 108.552556466851 | Mettl21a |
| **ENSMUSG00000025958** | -0.0813748516658716 | 0.468750956148543 | 0.895328112556867 | 520.42390074377 | 526.926976607571 | 522.443098533943 | 444.577253078368 | 547.479097909548 | 576.071731306788 | 513.251498644175 | 496.085183053508 | Creb1 |
| **ENSMUSG00000025959** | -0.435160734202717 | 0.0622243329914349 | 0.715615581523459 | 237.624107457659 | 197.903022126182 | 219.57753416644 | 326.39848960184 | 441.268152915096 | 352.247682837272 | 272.762459725708 | 260.526135520442 | Klf7 |
| **ENSMUSG00000025962** | 0.0916597576742443 | 0.65983232673075 | 0.939611397743053 | 671.3106914108 | 629.543358450776 | 635.071230283108 | 532.742362338636 | 547.479097909548 | 377.932409710823 | 698.563398763168 | 691.479784693839 | Fastkd2 |
| **ENSMUSG00000025964** | -0.583534861218506 | 0.111229243338229 | 0.764116514278315 | 51.5002818444356 | 57.8235167529173 | 60.5731128735006 | 36.5791410760683 | 117.160526952643 | 101.51582526213 | 45.80743598447 | 45.5920737160773 | Adam23 |
| **ENSMUSG00000025967** | 0.0715957426730694 | 0.640567047822882 | 0.934417517272849 | 4794.94729383192 | 3677.08701604819 | 4194.68806648992 | 5412.77495256385 | 4158.65122772093 | 4887.43659936714 | 3800.97610862045 | 4357.29961657939 | Eef1b2 |
| **ENSMUSG00000025968** | 0.261790333926128 | 0.222087957553676 | 0.831327997735572 | 17905.8348349653 | 20101.4090210635 | 20272.1172599611 | 17693.9870871802 | 13747.2001485088 | 10333.821778792 | 19414.0242326908 | 19869.4599356924 | Ndufs1 |
| **ENSMUSG00000025969** | -0.311299497588642 | 0.154380234482354 | 0.786853994984691 | 403.870631306363 | 362.414999366876 | 415.493696116668 | 422.067012416173 | 725.957283828061 | 472.10974158051 | 368.541644056872 | 423.354970220718 | Nrp2 |
| **ENSMUSG00000025971** | 0.0679964367845137 | 0.672875705336184 | 0.943302739919217 | 581.862833470465 | 484.577358704026 | 437.262158555582 | 567.445650026188 | 430.318570956905 | 437.863439082442 | 519.497967187512 | 587.269330485663 | Maip1 |
| **ENSMUSG00000025978** | -0.27546638200652 | 0.169738158654672 | 0.79907358295209 | 70.4740698923855 | 90.4001459094905 | 88.0203046443056 | 93.7926694258161 | 121.54035973592 | 117.415894279091 | 85.3684034256032 | 91.1841474321546 | Rftn2 |
| **ENSMUSG00000025979** | 0.00954335642534578 | 0.924338158919073 | 0.986713918026868 | 915.259394884443 | 872.239245667246 | 892.506959995485 | 1036.40899715527 | 880.346389438554 | 977.242703427014 | 892.203923606609 | 942.236190132265 | Mob4 |
| **ENSMUSG00000025980** | 0.231533305588063 | 0.217115520364395 | 0.825996545635915 | 11158.3943996277 | 11222.6487444394 | 10826.4974712496 | 11034.7075579473 | 8551.62350934715 | 6520.25137918575 | 10759.5420658977 | 11849.5970639214 | Hspd1 |
| **ENSMUSG00000025981** | 0.0835660830174718 | 0.653028022215183 | 0.936991241819063 | 730.039082987788 | 858.394178275702 | 611.409858066897 | 604.024791102256 | 802.604357535398 | 549.163922201163 | 726.672507208183 | 567.72987032163 | Coq10b |
| **ENSMUSG00000025982** | 0.0262868325100532 | 0.868073822975328 | 0.974898560893217 | 1735.64984952914 | 2044.99789530388 | 2260.13427409249 | 1920.87386984071 | 1901.94238613777 | 1564.32217482247 | 2383.027749283 | 1966.97232317934 | Sf3b1 |
| **ENSMUSG00000025986** | -0.139249160166613 | 0.399661627212251 | 0.880459952732461 | 194.255449062345 | 157.99665140938 | 201.594891282119 | 167.888878272211 | 194.902558855799 | 238.501035254403 | 192.599446752885 | 169.341988088287 | Slc39a10 |
| **ENSMUSG00000025991** | -0.51758695105958 | 0.125190519103921 | 0.771949105062466 | 234.010052591383 | 505.752167655798 | 236.613722162112 | 317.957149353517 | 278.119381738051 | 357.140011765567 | 530.94982618363 | 687.137682435165 | Cps1 |
| **ENSMUSG00000025993** | 0.107107368795309 | 0.590790590509438 | 0.923935529355527 | 1122.16403597875 | 1400.79505373265 | 1835.17602908934 | 1160.21532079735 | 1255.9170506045 | 1020.0505815496 | 1496.02921612917 | 1350.39380244762 | Slc40a1 |
| **ENSMUSG00000025995** | 0.139653490036048 | 0.37792948816439 | 0.87422797717331 | 317.133314515735 | 365.672662282534 | 304.7584741448 | 323.584709519066 | 266.074841584041 | 243.393364182698 | 347.72008224575 | 332.170822788563 | Wdr75 |
| **ENSMUSG00000025997** | -0.383212587104946 | 0.0494761405028092 | 0.703396849069134 | 55.1143367107117 | 51.3081909216027 | 52.0550188756646 | 52.523894878457 | 68.9823663366031 | 62.3771938357668 | 75.998700610598 | 67.3025850094475 | Ikzf2 |
| **ENSMUSG00000026000** | -0.0316572245731521 | 0.82419308426726 | 0.966637987288264 | 599.029594085277 | 509.009830571455 | 604.784673846357 | 744.71379524098 | 655.879959295639 | 621.325773893521 | 651.714884688142 | 581.84170266232 | Lancl1 |
| **ENSMUSG00000026003** | 0.22182284246736 | 0.0292795063910875 | 0.665956123973759 | 39339.8907331328 | 37742.4681250767 | 41288.1480623998 | 44850.716592731 | 33850.6326237474 | 31759.7763202619 | 36669.8935836588 | 37676.4212985146 | Acadl |
| **ENSMUSG00000026004** | 0.0635392445706616 | 0.807970817362104 | 0.964026716319739 | 371.344137509877 | 421.052931848708 | 376.689045682082 | 317.019222659259 | 304.398378437709 | 201.808568292187 | 460.156516025812 | 454.835211596105 | Kansl1l |
| **ENSMUSG00000026005** | -0.197529527700392 | 0.257176958738012 | 0.837517407767513 | 149.983276950461 | 169.39847161418 | 194.023252172932 | 148.192417692789 | 163.148771177045 | 221.377884005369 | 166.572494488982 | 208.420908416353 | Rpe |
| **ENSMUSG00000026011** | 0.157382293981832 | 0.691775024526423 | 0.94469184542218 | 34.3335212296237 | 26.8757190541728 | 14.1968233297267 | 31.8895076047775 | 26.2789966996583 | 12.2308223207386 | 31.2323427166841 | 26.0526135520442 | Ctla4 |
| **ENSMUSG00000026012** | 0.0836076389483215 | 0.781375886223252 | 0.960742105865155 | 68.6670424592474 | 62.7100111264033 | 93.6990339761962 | 81.59962240046 | 43.7983278327639 | 56.2617826753975 | 83.2862472444909 | 105.295979772845 | Cd28 |
| **ENSMUSG00000026014** | 0.0934780486305244 | 0.701211493304433 | 0.945920819861844 | 1765.46580217591 | 2006.72035604491 | 1860.73031108285 | 1420.95894180111 | 1470.52885698505 | 1015.1582526213 | 2270.59131550293 | 1854.07766445381 | Raph1 |
| **ENSMUSG00000026017** | 0.194002597070861 | 0.527516045734235 | 0.908217276872524 | 65.0529875929712 | 77.3694942468612 | 50.1621090983677 | 70.3445020693621 | 38.3235368536684 | 44.0309603546589 | 57.2592949805875 | 90.0986218674861 | Carf |
| **ENSMUSG00000026019** | -0.011555961962142 | 0.926648618554216 | 0.986790505471895 | 543.915257374565 | 473.989954228139 | 449.566072108012 | 583.390403828576 | 492.731188118594 | 536.933099880424 | 541.360607089191 | 496.085183053508 | Wdr12 |
| **ENSMUSG00000026020** | 0.143122057695362 | 0.359507994246831 | 0.867338970126053 | 456.274426867368 | 554.617111390658 | 394.671688566402 | 507.418341593665 | 442.363111110915 | 500.240632918209 | 374.788112600209 | 416.841816832707 | Nop58 |
| **ENSMUSG00000026021** | 0.0301726039636827 | 0.821835291205496 | 0.966637987288264 | 1606.44738805976 | 1241.16957086544 | 1281.49991923 | 1539.13770527764 | 1296.43050384981 | 1578.99916160735 | 1314.8816283724 | 1361.24905809431 | Sumo1 |
| **ENSMUSG00000026024** | 0.126268352695002 | 0.607538304662468 | 0.927510709653048 | 430.072529086865 | 549.730617017172 | 475.12035410152 | 407.9981120023 | 395.279908690694 | 249.508775343067 | 519.497967187512 | 541.677256769585 | Als2 |
| **ENSMUSG00000026027** | 0.211379934842294 | 0.185000550478275 | 0.805963874499992 | 1794.37824110612 | 1865.82643494273 | 1838.96184864393 | 1975.27361810769 | 1485.85827172651 | 1223.08223207386 | 1907.25506189884 | 1837.79478098378 | Stradb |
| **ENSMUSG00000026028** | 0.223449689093087 | 0.437519969824262 | 0.888722970093854 | 6080.64731250967 | 7363.13260511445 | 6231.45898686137 | 5079.8109761022 | 3876.1520131996 | 2997.77455081303 | 7080.37209387229 | 7248.05419529162 | Trak2 |
| **ENSMUSG00000026029** | -0.331959518966771 | 0.235421225226782 | 0.832048466795696 | 115.649755720838 | 128.677685168464 | 134.396594188079 | 151.006197775564 | 217.896680968 | 226.270212933664 | 107.231043327282 | 116.15123541953 | Casp8 |
| **ENSMUSG00000026031** | 0.17088363477123 | 0.327733289265527 | 0.855977678491063 | 922.487504616995 | 926.805099504506 | 927.525790875478 | 756.906842266336 | 677.779123212021 | 629.887349518038 | 831.821394354353 | 998.683519495027 | Cflar |
| **ENSMUSG00000026032** | 0.0371796667077466 | 0.849582350665724 | 0.972640876584422 | 6461.02658718524 | 4655.2003064743 | 4529.73309707147 | 6336.63274640814 | 5287.55312761042 | 6949.55324264367 | 4695.26218840818 | 4491.90478659828 | Ndufb3 |
| **ENSMUSG00000026034** | 0.224047832387002 | 0.0393382285101888 | 0.679566075569698 | 387.60738440812 | 343.683437601847 | 388.046504345863 | 392.053358199911 | 318.632834983357 | 338.793778284459 | 328.980676615739 | 307.203734801188 | Clk1 |
| **ENSMUSG00000026035** | -0.120142392383124 | 0.340724165557066 | 0.859077629258968 | 217.74680569314 | 210.933673788811 | 195.916161950228 | 235.419600258798 | 245.270635863478 | 242.170281950624 | 217.585320926233 | 230.131419709724 | Ppil3 |
| **ENSMUSG00000026036** | -0.204910858563707 | 0.173832376980384 | 0.799889021902745 | 269.247087537575 | 282.602257933272 | 210.112985279955 | 234.48167356454 | 267.16979977986 | 294.7628179298 | 323.775286162958 | 263.782712214447 | Nif3l1 |
| **ENSMUSG00000026037** | 0.142210435786656 | 0.451107007123078 | 0.891094723422636 | 300.870067617492 | 280.973426475443 | 271.632553042104 | 240.109233730089 | 194.902558855799 | 204.254732756335 | 294.625099627387 | 296.348479154503 | Orc2 |
| **ENSMUSG00000026039** | 0.0440596551708083 | 0.850107211544968 | 0.972692784625411 | 126.491920319666 | 142.522752560008 | 126.824955078892 | 149.130344387048 | 135.774816281568 | 78.277262852727 | 160.326025945645 | 153.05910461826 | Sgol2a |
| **ENSMUSG00000026042** | -0.96532632216361 | 0.0303946139986353 | 0.665956123973759 | 130.105975185942 | 124.605606523892 | 112.628131749165 | 135.061443973175 | 325.202584158272 | 451.317343635254 | 100.984574783945 | 104.210454208177 | Col5a2 |

| | | | | | | | | | | | | |
|---|---|---|---|---|---|---|---|---|---|---|---|---|
| **ENSMUSG00000026043** | -1.54867554815203 | 0.0204251917205811 | 0.646254927104491 | 533.976606492306 | 672.707392083235 | 440.101523221528 | 634.976372012775 | 2205.24580637966 | 3438.08415435962 | 478.895921655823 | 553.618037980939 | Col3a1 |
| **ENSMUSG00000026047** | -0.0570374739458547 | 0.760704041618553 | 0.957357521291862 | 149.983276950461 | 122.976775066064 | 106.002947528626 | 108.799496533947 | 118.255485148462 | 118.638976511164 | 151.997401221196 | 118.322286548867 | Kdelc1 |
| **ENSMUSG00000026048** | 0.180125897053225 | 0.225630323877711 | 0.832048466795696 | 438.204152535987 | 412.09435883065 | 466.602260103684 | 385.487871340104 | 378.855535753408 | 348.57843614105 | 454.951125573032 | 319.144516012541 | Ercc5 |
| **ENSMUSG00000026049** | 0.0081648006242731 | 0.971851779532435 | 0.995123114262614 | 97.5794813894569 | 69.2253369577179 | 86.1273948670087 | 95.6685228143324 | 100.736154015357 | 63.6002760678407 | 74.9576225200418 | 106.381505337514 | Tex30 |
| **ENSMUSG00000026062** | 0.0713030168917091 | 0.835543919508401 | 0.968865818776815 | 98.4829951060259 | 195.459774939439 | 122.09268063565 | 66.5927952923295 | 127.015150715015 | 79.5003450848009 | 151.997401221196 | 100.953877514171 | Slc9a2 |
| **ENSMUSG00000026064** | 0.549568723023924 | 0.0158567279268405 | 0.63777231095375 | 2531.64543382646 | 2737.25126488106 | 3937.25233677754 | 3915.84394852782 | 2454.89627502642 | 1498.27573429048 | 2391.35637400745 | 2619.37318754511 | Ptp4a1 |
| **ENSMUSG00000026072** | -0.368855405384872 | 0.110969566514194 | 0.764116514278315 | 149.079763233892 | 152.295741306979 | 162.790240847533 | 140.689004138724 | 241.98576127602 | 251.954939807215 | 140.545542225078 | 147.631476794917 | Il1r1 |
| **ENSMUSG00000026074** | 0.347098543652127 | 0.0970876635336925 | 0.757809915356977 | 3656.52001095493 | 4136.41748715588 | 4340.44211934178 | 3537.85949074178 | 2647.60891749058 | 2103.70143916704 | 3958.17890029443 | 3609.37250252279 | Map4k4 |
| **ENSMUSG00000026077** | -0.084187495991491 | 0.786649889388005 | 0.961141387070613 | 27.1054114970714 | 46.4216965481167 | 58.6802030962037 | 30.9515809105193 | 35.0386622662111 | 46.4771248188067 | 41.6431236222455 | 49.9341759747513 | Npas2 |
| **ENSMUSG00000026078** | 0.153207124949875 | 0.271612794560399 | 0.844129257012535 | 2058.20424634428 | 1769.72537893084 | 1636.42050247316 | 2215.38285183778 | 1713.60957645689 | 1959.37773578232 | 1547.04204256642 | 1686.90672749486 | Pdcl3 |
| **ENSMUSG00000026082** | 0.348224483954802 | 0.269206233289083 | 0.843793744782845 | 279.185738419835 | 287.488752306758 | 266.900278598862 | 142.564857527241 | 148.914314631397 | 116.192812047017 | 262.351678820146 | 238.815624227072 | Rev1 |
| **ENSMUSG00000026083** | 0.0915837448455763 | 0.562306197940982 | 0.917686886311859 | 7952.72773324073 | 6674.95131418184 | 6527.69936700834 | 8788.37312519897 | 7002.25766226312 | 8544.45247326798 | 6151.73043709621 | 6404.60083154419 | Eif5b |
| **ENSMUSG00000026087** | 0.111307688843691 | 0.418680157629521 | 0.882468496126562 | 3879.68789894748 | 2869.18661296518 | 3079.76420766205 | 3941.16796927279 | 3114.06110890951 | 3405.06093409363 | 3040.98910251448 | 3187.10305786674 | Mrpl30 |
| **ENSMUSG00000026088** | -0.602052594461218 | 0.101940598039095 | 0.759800953500741 | 41.5616309621761 | 25.2468875963442 | 34.0723759913441 | 25.3240207449704 | 51.4630352034976 | 77.0541806206532 | 35.3966550789086 | 28.2236646813812 | Mitd1 |
| **ENSMUSG00000026090** | -0.120070727177702 | 0.746917112074091 | 0.95610346389357 | 47.8862269781594 | 52.122606650517 | 39.7511053232348 | 55.3376749612315 | 32.8487458745729 | 28.1308913376988 | 87.4505596067155 | 62.9604827507734 | 2010300C02I |
| **ENSMUSG00000026094** | -0.321966449381788 | 0.11000379877462 | 0.763594585197136 | 126.491920319666 | 184.872370463553 | 159.950876181588 | 161.323391412404 | 212.421889988905 | 249.508775343067 | 167.613572579538 | 162.828834700276 | Stk17b |
| **ENSMUSG00000026095** | 0.0730135007683004 | 0.685769569396253 | 0.944452327903597 | 636.073656464608 | 485.39177443294 | 530.961192531779 | 740.024161769689 | 535.434557755538 | 697.1568722821 | 530.94982618363 | 511.282540958867 | Asnsd1 |
| **ENSMUSG00000026096** | 0.172543352654173 | 0.252424457624942 | 0.83723604018117 | 1260.40163461382 | 1075.02876216691 | 1079.90502794788 | 1175.22214790548 | 912.100177117308 | 830.472835578151 | 1132.69296252508 | 1196.2491722647 | Osgepl1 |
| **ENSMUSG00000026097** | 0.0850684626471576 | 0.681561609227194 | 0.944452327903597 | 251.176813206195 | 203.603932228582 | 193.076797284283 | 267.309107863576 | 195.997517051618 | 170.008430258266 | 208.215618111227 | 287.664274637155 | Ormdl1 |
| **ENSMUSG00000026098** | -0.0760839535463595 | 0.75248597173247 | 0.956497786093635 | 64.1494738764022 | 52.122606650517 | 37.8581955459379 | 45.9584080186499 | 51.4630352034976 | 48.9232892829544 | 51.0128264372507 | 59.7039060567679 | Pms1 |
| **ENSMUSG00000026100** | 0.167631749524074 | 0.769827544187262 | 0.958815818668915 | 15.3597331816738 | 20.3603932228582 | 5.67872933189068 | 5.62756016554897 | 6.56974917491458 | 13.4539045528125 | 13.5340151772298 | 8.68420451734806 | Mstn |
| **ENSMUSG00000026102** | -0.106814411331859 | 0.440783277337745 | 0.890970307297912 | 387.60738440812 | 314.364471360931 | 367.224496795597 | 355.474217123843 | 338.342082508101 | 408.509465512669 | 350.843316517418 | 436.38127699674 | Inpp1 |
| **ENSMUSG00000026103** | 0.150678702852447 | 0.455888062237818 | 0.892805623137304 | 1178.18188640603 | 1404.86713237722 | 1206.72998302677 | 1128.32581319257 | 1007.36154015357 | 749.749408261276 | 1329.45672164019 | 1342.79512349494 | Gls |
| **ENSMUSG00000026104** | -0.173044806840821 | 0.241460191016028 | 0.832048466795696 | 365.019541493894 | 452.000729547452 | 407.922057007481 | 406.122258613784 | 534.339599559719 | 479.448234972953 | 373.747034509653 | 452.664160466768 | Stat1 |
| **ENSMUSG00000026107** | 0.272157884848231 | 0.335174077278397 | 0.858570960161243 | 721.907459538667 | 793.240919962556 | 591.534305405279 | 673.43136647736 | 428.128654565267 | 329.009120427868 | 721.467116755403 | 822.828378018729 | Nabp1 |
| **ENSMUSG00000026109** | -1.29206831039488 | 0.0110599039775063 | 0.635862253526407 | 6.32459601598331 | 6.51532583131463 | 4.73227444324223 | 8.44134024832345 | 22.994122112201 | 20.7923979452556 | 9.36970281500523 | 10.8552556466851 | Tmeff2 |
| **ENSMUSG00000026110** | -0.134047277018262 | 0.483553187952578 | 0.89863290032536 | 239.431134890797 | 256.540954608014 | 265.953823710214 | 223.226553233442 | 332.867291529005 | 308.216722482613 | 195.722681024554 | 245.328777615083 | Mgat4a |
| **ENSMUSG00000026111** | 0.0865065259415554 | 0.482151271661211 | 0.897732059433541 | 411.098741038915 | 342.054606144018 | 414.54724122802 | 429.570425970238 | 406.229490648885 | 368.147751854232 | 374.788112600209 | 353.881334081933 | Unc50 |
| **ENSMUSG00000026112** | 0.144327135247667 | 0.496262661440742 | 0.900927059989498 | 4182.36499399811 | 4728.49772207659 | 4097.20321295913 | 3889.58200108859 | 2842.51147634638 | 2873.0201631415 | 4742.1107024832 | 4830.58876277486 | Coa5 |
| **ENSMUSG00000026113** | 0.237661466566331 | 0.386578634977303 | 0.875379400038268 | 411.098741038915 | 557.874774306315 | 581.123301630146 | 451.142739938176 | 308.778211220985 | 266.631926592101 | 593.414511616998 | 527.565424428895 | Inpp4a |
| **ENSMUSG00000026114** | -0.0256500129256838 | 0.933904371168798 | 0.98818087308472 | 131.009488902512 | 177.542628903324 | 105.056492639978 | 74.0962088463947 | 104.021028602814 | 88.0619207093179 | 166.572494488982 | 137.8617467129 | Cnga3 |
| **ENSMUSG00000026116** | 0.0880247506329235 | 0.5873887017743 | 0.923058967588378 | 1056.20753466921 | 1288.40568314247 | 1101.67349038679 | 865.706338800283 | 1089.48340484 | 853.711397987554 | 1147.26805579286 | 966.117752554972 | Tmem131 |
| **ENSMUSG00000026121** | -0.74678066517404 | 0.0113201860190259 | 0.635862253526407 | 297.256012751216 | 249.211213047785 | 265.953823710214 | 208.219726125312 | 699.678287128403 | 434.19419238622 | 292.542943446274 | 286.578749072486 | Sema4c |
| **ENSMUSG00000026123** | 0.065108898383774 | 0.810748676972434 | 0.964565235387333 | 2794.56792534806 | 3313.8576009524 | 2934.95660969883 | 2437.67147837696 | 2177.87185148418 | 1545.97594134136 | 3595.8837247809 | 3653.8790506742 | Plekhb2 |
| **ENSMUSG00000026126** | -0.109322546779095 | 0.753322788491668 | 0.956497786093635 | 25.2983840639333 | 17.1027303072009 | 40.6975602118832 | 30.9515809105193 | 27.3739548954774 | 36.6924669622158 | 22.903717992235 | 35.8223436340608 | Ptpn18 |
| **ENSMUSG00000026127** | 0.0132396844221741 | 0.920523585817033 | 0.986713918026868 | 442.721721118832 | 479.69086433054 | 520.550188756646 | 440.825546301336 | 456.597567656563 | 393.832478727783 | 488.265624470828 | 526.479898864226 | Imp4 |
| **ENSMUSG00000026131** | 0.53275616113368 | 0.0504131672935482 | 0.703396849069134 | 10087.7306454934 | 14135.813806766 | 11984.9582549553 | 10001.1123408748 | 6861.00805500246 | 4394.53445984138 | 10666.8861158382 | 10018.3154363257 | Dst |
| **ENSMUSG00000026134** | -0.991251823527533 | 0.0151825722563982 | 0.635862253526407 | 18.0702743313809 | 17.1027303072009 | 17.9826428843205 | 18.7585338851632 | 44.893286028583 | 56.2617826753975 | 24.9858741733473 | 17.3684090346961 | Prim2 |
| **ENSMUSG00000026135** | 0.0914015263696635 | 0.734729210924695 | 0.954383574136061 | 118.360296870545 | 167.769640156352 | 163.736695736181 | 107.861569839689 | 119.350443344282 | 81.9465095489486 | 144.709854587303 | 176.940667040967 | Zfp142 |
| **ENSMUSG00000026142** | -0.146167654356814 | 0.338394908128344 | 0.858570960161243 | 191.544907912638 | 241.881471487556 | 203.487801059416 | 211.971432902344 | 259.505092409126 | 248.285693110993 | 239.447960827911 | 193.223550510994 | Rhbdd1 |
| **ENSMUSG00000026150** | 0.167624112250197 | 0.295246832821696 | 0.851785577030345 | 2048.26559546202 | 2120.73855809291 | 2290.42083052924 | 1998.72178546414 | 1509.94735203453 | 1614.46854633749 | 2220.61956715624 | 2184.07743611304 | Mff |
| **ENSMUSG00000026153** | 0.308607856537179 | 0.244727016184796 | 0.832048466795696 | 318.036828232304 | 415.352021746308 | 394.671688566402 | 309.515809105193 | 235.416012101106 | 172.454594722414 | 388.322127777439 | 363.65106416395 | Fam135a |
| **ENSMUSG00000026154** | -0.0312055975974202 | 0.865043345751393 | 0.974868041734905 | 860.145058173731 | 640.130762926662 | 502.567545872325 | 725.017334661559 | 650.405168316544 | 796.226533080083 | 636.098713329799 | 705.59161703453 | Sdhaf4 |
| **ENSMUSG00000026155** | -0.0127471094927079 | 0.949661213521916 | 0.991267322147698 | 137.334084918495 | 117.275864963663 | 175.094154399963 | 125.682177030594 | 156.579022002131 | 163.893019097897 | 124.929370866736 | 115.065709854862 | Smap1 |
| **ENSMUSG00000026156** | 0.241880595585555 | 0.577374497865395 | 0.922450244597442 | 22.5878429142261 | 16.2883145782866 | 10.4110037751329 | 32.8274342990356 | 19.7092475247437 | 13.4539045528125 | 21.8626399016789 | 14.1118323406906 | B3gat2 |
| **ENSMUSG00000026158** | -0.014990564364016 | 0.926247485417099 | 0.986713918026868 | 894.478579403355 | 697.139863950665 | 736.341903368492 | 1008.27119632752 | 817.933772276865 | 984.581196819457 | 753.740537562643 | 815.229699066049 | Ogfrl1 |
| **ENSMUSG00000026159** | 0.100290201207506 | 0.484787235799476 | 0.899236955296196 | 539.39768879172 | 631.986605637519 | 663.464876942561 | 605.900644490772 | 570.47322002175 | 454.986590331476 | 627.77008860535 | 622.006148555055 | Agfg1 |
| **ENSMUSG00000026162** | 0.0648689206441139 | 0.807230750446748 | 0.964026716319739 | 67.7635287426784 | 61.0811796685746 | 54.8943835416099 | 48.7721881014244 | 63.5075753575076 | 68.4926049961361 | 57.2592949805875 | 33.6512925047237 | Nhej1 |
| **ENSMUSG00000026170** | -0.0273285637860347 | 0.848791076713243 | 0.972640876584422 | 326.168451681425 | 356.714089264476 | 399.403963009645 | 385.487871340104 | 418.274030802895 | 413.401794440965 | 318.569895710178 | 346.282655129254 | Cyp27a1 |
| **ENSMUSG00000026171** | -0.00392156265801166 | 0.978223010503089 | 0.996580229842593 | 242.141676040504 | 186.501201921381 | 222.416898832385 | 235.419600258798 | 229.94122112201 | 236.054870790255 | 224.872867560126 | 197.565652769668 | Rnf25 |
| **ENSMUSG00000026172** | -0.0389395136851779 | 0.771360162315448 | 0.959045638244974 | 742.688275019755 | 650.718167402549 | 564.087113634474 | 713.762214330461 | 649.310210120724 | 733.849339244316 | 616.318229609233 | 745.756062927265 | Bcs1l |
| **ENSMUSG00000026173** | 0.0858585991432084 | 0.84806757301766 | 0.972560979955028 | 13.5527057485357 | 13.0306516626293 | 7.57163910918757 | 10.3171936368398 | 7.66470737073368 | 11.0077400886647 | 11.4518589961175 | 11.9407812113536 | Plcd4 |
| **ENSMUSG00000026174** | 0.0777059700546534 | 0.525786739601273 | 0.90806673740897 | 254.790868072471 | 249.211213047785 | 238.506631939409 | 241.985087118606 | 229.94122112201 | 234.831788558181 | 210.29777429234 | 258.355084391105 | Cnot9 |
| **ENSMUSG00000026176** | 0.0566101591486832 | 0.483560838199056 | 0.89863290032536 | 2930.99849654998 | 3112.69691591056 | 3299.34174182849 | 2906.63482550604 | 2913.68375907462 | 2843.66618957172 | 3060.76958623504 | 2959.14268928635 | Ctdsp1 |
| **ENSMUSG00000026177** | -0.710001883030094 | 0.0234449227062063 | 0.654124840968651 | 29.8159526467785 | 44.7928650902881 | 63.4124775394459 | 60.9652351267805 | 98.5462376237187 | 97.8465785659088 | 52.0539045278068 | 77.072315091464 | Slc11a1 |
| **ENSMUSG00000026179** | -0.119014586300101 | 0.271101445881076 | 0.844129257012535 | 1769.98337075876 | 1421.96986268442 | 1502.02390828509 | 1631.05452131494 | 1677.47595599486 | 1868.86965060886 | 1646.98553925981 | 1676.05147184818 | Pnkd |
| **ENSMUSG00000026180** | -0.205804553842632 | 0.790508201931701 | 0.961804768755116 | 15.3597331816738 | 8.95857301805762 | 21.7684624389143 | 1.87585338851632 | 13.1394983498292 | 31.8001380339204 | 4.16431236222455 | 6.51315338801105 | Cxcr2 |
| **ENSMUSG00000026181** | -0.277054577670284 | 0.0187783403148486 | 0.642273034804458 | 739.074220153479 | 688.995706661522 | 731.609628925249 | 603.086864407998 | 935.094299229509 | 837.811328970594 | 784.972880279327 | 790.262611078674 | Ppm1f |
| **ENSMUSG00000026185** | -0.477180294655146 | 0.0516144122966552 | 0.703396849069134 | 2680.72519706036 | 2163.90259172537 | 1577.74029937696 | 1405.01418799873 | 2658.55849944877 | 3619.10032470655 | 2472.56046507082 | 2147.16956691431 | Igfbp5 |
| **ENSMUSG00000026187** | 0.13671351146661 | 0.263814184405169 | 0.842510961779757 | 627.038519298917 | 628.728942721862 | 702.269527377147 | 735.334528298398 | 545.28918151791 | 577.294813538862 | 681.906149314269 | 643.716659848425 | Xrcc5 |
| **ENSMUSG00000026189** | 0.0739780794283043 | 0.566251255296367 | 0.919931558693298 | 298.159526467785 | 289.117583764587 | 291.508105703722 | 256.053987532478 | 266.074841584041 | 249.508775343067 | 254.023054095697 | 308.289260365856 | Pecr |
| **ENSMUSG00000026192** | 0.111643858635343 | 0.336490965438737 | 0.858570960161243 | 837.557215259505 | 968.340301679137 | 910.489602879806 | 969.816201862939 | 937.284215621147 | 755.864819421645 | 911.984407327176 | 805.459968984033 | Atic |
| **ENSMUSG00000026193** | -0.919929548469461 | 0.00175156004887313 | 0.635862253526407 | 406.58117245607 | 601.038807938775 | 462.81644054909 | 397.68091836546 | 1131.09181628113 | 1224.30531430593 | 490.34778065194 | 690.394259129171 | Fn1 |
| **ENSMUSG00000026196** | 0.104246984728356 | 0.8160261477087 | 0.966177091475091 | 9.93865088225949 | 12.2162359337149 | 12.3039135524298 | 9.37926694258161 | 9.85462376237187 | 7.33849339244316 | 9.36970281500523 | 14.1118323406906 | Bard1 |
| **ENSMUSG00000026197** | -0.167617015925344 | 0.338829797414212 | 0.858570960161243 | 196.965990212052 | 185.686786192467 | 148.593417517806 | 206.343872736795 | 234.321053905287 | 233.608706326107 | 175.942197303987 | 185.624871558315 | Zfand2b |
| **ENSMUSG00000026198** | -0.231667845735327 | 0.311560458178136 | 0.851785577030345 | 300.870067617492 | 245.953550132127 | 189.290977729689 | 250.426427366929 | 303.30342024189 | 395.055560959857 | 226.955023741238 | 234.473521968398 | Abcb6 |
| **ENSMUSG00000026199** | -0.0732365778239542 | 0.591388990604707 | 0.923935529355527 | 179.79922959724 | 195.459774939439 | 196.862616838877 | 174.454365132018 | 221.181555555458 | 199.362403828039 | 186.352978209548 | 179.111718170304 | Ankzf1 |
| **ENSMUSG00000026200** | -0.281083412191021 | 0.463990647218451 | 0.895328112556867 | 10.8421645988285 | 13.8450673915436 | 12.3039135524298 | 15.0068271081306 | 19.7092475247437 | 15.9000690169602 | 12.4929370866736 | 15.1973579053591 | Glb1l |
| **ENSMUSG00000026201** | -0.120328172763087 | 0.415623496285749 | 0.882192126163777 | 458.081454300506 | 400.69253862585 | 357.759947909113 | 439.887619607078 | 390.900075907418 | 535.71001764835 | 425.80093903746 | 449.407583772762 | Stk16 |
| **ENSMUSG00000026202** | -0.287646118614302 | 0.0868070168503439 | 0.747269672026173 | 5870.12861654909 | 4144.56164444502 | 4344.22793889637 | 4944.74953212903 | 4947.02112871068 | 6711.05220738927 | 5229.33524886347 | 6675.98222271132 | Tuba4a |
| **ENSMUSG00000026203** | 0.184812344365827 | 0.0964110285963233 | 0.757809915356977 | 3138.80665136086 | 3219.38537639834 | 3640.06550174193 | 3452.50816156429 | 3002.37537293596 | 2644.30378574368 | 3350.18929540965 | 2834.30724934947 | Dnajb2 |
| **ENSMUSG00000026204** | -0.0804227454941164 | 0.721221024644707 | 0.951674101666722 | 272.861142403852 | 215.820168162297 | 171.308334845369 | 266.371181169318 | 248.555510450935 | 284.978160073209 | 276.926772087932 | 169.341988088287 | Ptprn |
| **ENSMUSG00000026207** | 0.200867704954918 | 0.42750270548305 | 0.885066977635295 | 4229.3477072597 | 5024.94504740141 | 4611.12821749523 | 3867.0717604264 | 2740.6803641352 | 2561.13419396266 | 5126.26851789842 | 4998.84522529848 | Speg |
| **ENSMUSG00000026208** | 0.153969336823058 | 0.107890901935068 | 0.762287160109383 | 81873.7024543372 | 74222.5918703363 | 77245.8621919316 | 88533.7144511106 | 71306.962586327 | 77865.0841405181 | 72765.1120613306 | 67356.8612876809 | Des |
| **ENSMUSG00000026209** | 0.0709942541371353 | 0.40568713305979 | 0.881158207349486 | 1694.99173228353 | 1695.61354759963 | 1750.94154399963 | 1844.9018076058 | 1704.84991089033 | 1737.99985177695 | 1523.09724648363 | 1685.82120193019 | Dnpep |
| **ENSMUSG00000026211** | -0.0288290915515607 | 0.819114021412593 | 0.966315773308182 | 2474.72406968261 | 2284.43611960469 | 2091.66530391307 | 2431.10599151715 | 2250.13909240824 | 2096.3629457746 | 2802.58221977712 | 2318.68260613193 | Obsl1 |
| **ENSMUSG00000026213** | 0.214478968675484 | 0.258054370961294 | 0.838310965110319 | 178.895715880671 | 241.881471487556 | 288.668741037776 | 256.053987532478 | 234.321053905287 | 172.454594722414 | 201.96914956789 | 222.532740757044 | Stk11ip |
| **ENSMUSG00000026219** | 0.190649016339098 | 0.333180165582918 | 0.856918526232803 | 3446.00131499434 | 3957.24602679472 | 4046.09464897211 | 3090.46845758064 | 2933.39300659936 | 2227.2327446065 | 3888.42666822717 | 3689.70139430826 | Trip12 |
| **ENSMUSG00000026222** | -0.341839944693102 | 0.13188317796244 | 0.77303252008843 | 320.747369382011 | 317.622134276588 | 276.364827485346 | 323.584709519066 | 439.078236523458 | 545.494675504941 | 275.885693997376 | 310.460311495193 | Sp100 |
| **ENSMUSG00000026223** | -0.616510054574766 | 0.0152044325502242 | 0.635862253526407 | 937.847237798669 | 1056.29720040188 | 910.489602879806 | 945.430107812227 | 1600.82888228752 | 2217.44808674991 | 1151.43236815509 | 934.637511179585 | Itm2c |
| **ENSMUSG00000026227** | -0.161578288347265 | 0.814830480461398 | 0.966177091475091 | 9.93865088225949 | 16.2883145782866 | 8.51809399783602 | 13.1309737196143 | 3.28487458745729 | 6.1154111603693 | 9.36970281500523 | 34.7368180693922 | 2810459M11 |
| **ENSMUSG00000026229** | 0.058504733315132 | 0.358007313385719 | 0.865834559821921 | 4317.89205148347 | 4356.30973396274 | 4419.94432998825 | 4439.20704392388 | 4045.87053355156 | 4401.87295323382 | 4222.61273529569 | 4167.3326427624 | Psmd1 |
| **ENSMUSG00000026234** | 0.23976643626558 | 0.0123809588254379 | 0.635862253526407 | 9264.62964969899 | 9284.33930962335 | 9491.99607825527 | 9928.89198541689 | 7824.57126732327 | 7111.00009727742 | 8532.6760301981 | 8685.29004291273 | Ncl |
| **ENSMUSG00000026235** | -0.299179385413269 | 0.101032386874483 | 0.759800953500741 | 364.116027777325 | 491.09268453534 | 378.581955459379 | 317.019222659259 | 387.61520131996 | 489.232892829544 | 559.058934628645 | 473.289146195469 | Epha4 |
| **ENSMUSG00000026238** | -0.26216212375959 | 0.248330953221387 | 0.834621660998471 | 3512.86133002045 | 3213.68446629594 | 3215.10725673877 | 3537.85949074178 | 4505.75297579558 | 5724.02484610566 | 2759.89801806432 | 3177.33332778472 | Ptma |
| **ENSMUSG00000026239** | -0.104292269466392 | 0.437021889621294 | 0.888701139843674 | 899.899661702769 | 701.211942595237 | 697.537252933905 | 906.975113347642 | 810.269064906132 | 856.157562451702 | 901.573626421614 | 877.104656252154 | Pde6d |
| **ENSMUSG00000026240** | 0.288995012103362 | 0.105260480255789 | 0.762094734041179 | 267.440060104437 | 268.757190541728 | 273.525462819401 | 288.881421831514 | 212.421889988905 | 171.23151249034 | 293.584021536831 | 220.361689627707 | Cops7b |

| | | | | | | | | | | | | |
|---|---|---|---|---|---|---|---|---|---|---|---|---|
| **ENSMUSG00000026245** | 0.132975534611109 | 0.330717089908415 | 0.856303993370193 | 967.663190445447 | 1012.31875104051 | 1064.7617497295 | 998.891929384942 | 961.373295929167 | 726.510845851873 | 1060.8585742767 | 936.808562308922 | Farsb |
| **ENSMUSG00000026248** | 0.00363191125236902 | 0.971376311706875 | 0.995122272737203 | 1192.63810587114 | 1009.87550385377 | 1102.61994527544 | 1118.00861955573 | 1182.55485148462 | 1011.48900592508 | 1078.55690181616 | 1137.6307917726 | Mrpl44 |
| **ENSMUSG00000026249** | -0.765881117587776 | 0.0477160240923379 | 0.703396849069134 | 312.61574593289 | 332.281617397046 | 277.311282373995 | 307.639955716677 | 656.974917491458 | 920.980920751616 | 259.228444548478 | 255.098507697099 | Serpine2 |
| **ENSMUSG00000026254** | 0.171109329520984 | 0.282772778945154 | 0.850587941933635 | 1203.48027046997 | 1439.07259299162 | 1364.78794943106 | 1329.98005245807 | 1085.10357205672 | 923.427085215764 | 1309.67623791962 | 1420.95296415108 | Eif4e2 |
| **ENSMUSG00000026255** | -0.575583503755556 | 0.245202816746676 | 0.832048466795696 | 32.5264937964856 | 57.009101024003 | 58.6802030962037 | 45.9584080186499 | 88.6916138613469 | 140.654456688494 | 26.0269522639034 | 34.7368180693922 | Efhd1 |
| **ENSMUSG00000026260** | 0.12202644314569 | 0.102788703739368 | 0.759800953500741 | 17912.1594309813 | 16102.6277920941 | 16123.8054830149 | 17788.7176833003 | 15268.0970825015 | 15560.0521564436 | 15679.677121866 | 15909.4626757816 | Ndufa10 |
| **ENSMUSG00000026269** | -0.0580311929330945 | 0.546420619970472 | 0.914211323944814 | 1506.1573655206 | 1685.02614312375 | 1717.81562289693 | 1566.33757941113 | 1670.90620681994 | 1538.63744794892 | 1805.22940902434 | 1724.90012225826 | Rnpepl1 |
| **ENSMUSG00000026270** | -0.259622749805656 | 0.18355238949389 | 0.805963874499992 | 152.693818100169 | 95.2866402829764 | 158.057966404291 | 126.620103724852 | 169.71852035196 | 147.992950080937 | 164.49033830787 | 154.144630182928 | Capn10 |
| **ENSMUSG00000026271** | -0.959630348430389 | 0.0299104104807956 | 0.665956123973759 | 7.22810973255236 | 7.32974156022896 | 9.46454888648447 | 5.62756016554897 | 16.4243729372865 | 14.6769867848863 | 13.5340151772298 | 13.0263067760221 | Gpr35 |
| **ENSMUSG00000026272** | -0.150021691885902 | 0.714182214522544 | 0.951176382858794 | 39.754603529038 | 48.8649437348597 | 30.2865564367503 | 52.523894878457 | 33.943704070392 | 30.5770558018465 | 33.3144988977964 | 92.2696729968231 | Agxt |
| **ENSMUSG00000026273** | -0.105974859795575 | 0.463812639425236 | 0.895328112556867 | 211.422209677157 | 222.335493993612 | 180.772883731853 | 208.219726125312 | 249.650468646754 | 223.824048469516 | 222.790711379013 | 189.966973816989 | Mterf4 |
| **ENSMUSG00000026275** | 0.116779406000496 | 0.146690810788595 | 0.780607990324772 | 936.9437240821 | 919.475357944277 | 979.580809751142 | 1007.33326963327 | 865.016974697087 | 897.742358342213 | 900.532548331058 | 880.36123294616 | Ppp1r7 |
| **ENSMUSG00000026276** | 0.0198518911697457 | 0.821038315220632 | 0.96650690096489 | 2506.34704976253 | 2414.74263623098 | 2356.67267273463 | 2344.8167356454 | 2368.39457755671 | 2145.28623505755 | 2421.54763863357 | 2554.241653665 | Sept2 |
| **ENSMUSG00000026277** | 0.165349396417908 | 0.091663618460748 | 0.753823363185192 | 1456.4641111093 | 1332.38413250384 | 1380.87768253808 | 1484.73795701067 | 1283.29100549998 | 1132.57414690039 | 1370.05876717188 | 1253.78202719213 | Stk25 |
| **ENSMUSG00000026278** | -0.315160238752964 | 0.397869818125294 | 0.880282219458479 | 67.7635287426784 | 48.8649437348597 | 55.8408384302584 | 83.4754757889764 | 105.115986798633 | 121.085140975312 | 40.6020455316893 | 52.1052271040884 | Bok |
| **ENSMUSG00000026279** | 0.0386727016202522 | 0.785464678675974 | 0.961141387070613 | 917.066422317581 | 823.374301932386 | 815.844114014961 | 991.388515830876 | 958.08842134171 | 979.688867891162 | 777.685333645434 | 739.242909539254 | Thap4 |
| **ENSMUSG00000026280** | -0.151032182749837 | 0.163263316827303 | 0.793968226938511 | 447.239289701677 | 447.928650902881 | 493.102996985841 | 439.887619607078 | 563.903470846835 | 501.463715150282 | 472.649453112486 | 491.743080794834 | Atg4b |
| **ENSMUSG00000026281** | 0.00183070509831314 | 0.991268053712846 | 0.998628446692557 | 296.352499034647 | 269.571606270643 | 312.330113253987 | 343.281170098487 | 306.588294829347 | 384.047820871192 | 267.557069272927 | 262.697186649779 | Dtymk |
| **ENSMUSG00000026283** | 0.0762763735650786 | 0.695375453288958 | 0.944933386973863 | 218.650319409709 | 316.807718547674 | 324.634026806417 | 332.963976461647 | 313.158044004262 | 215.262472844999 | 312.323427166841 | 289.835325766492 | Ing5 |
| **ENSMUSG00000026288** | -0.422581477955959 | 0.0875138281471941 | 0.74812051555524 | 119.263810587114 | 140.893921102179 | 147.646962629158 | 102.23400967414 | 227.751304730372 | 200.585486060113 | 128.052605138405 | 128.092016630884 | Inpp5d |
| **ENSMUSG00000026289** | 0.161725533466449 | 0.403075445767763 | 0.881158207349486 | 786.960447131638 | 829.889627763701 | 826.255117790094 | 754.093062183562 | 670.114415841287 | 480.671317205027 | 876.587752248267 | 829.34153140674 | Atg16l1 |
| **ENSMUSG00000026295** | -0.254351107036799 | 0.54106329964716 | 0.913837378403038 | 17.1667606148119 | 26.8757190541728 | 23.6613722162112 | 23.448167356454 | 15.3294147414674 | 28.1308913376988 | 17.6983275394543 | 47.7631248454143 | Spp2 |
| **ENSMUSG00000026303** | -0.921671286657189 | 0.0558020998930015 | 0.703396849069134 | 14.4562194651047 | 19.5459774939439 | 9.46454888648447 | 19.6964605794214 | 37.2285786578493 | 51.3694537471021 | 16.6572494488982 | 15.1973579053591 | Mlph |
| **ENSMUSG00000026304** | 0.0178275473860689 | 0.949601271714542 | 0.991267322147698 | 27.1054114970714 | 28.5045505120015 | 29.3401015481019 | 31.8895076047775 | 21.8991639163819 | 31.8001380339204 | 27.0680303544596 | 34.7368180693922 | Rab17 |
| **ENSMUSG00000026305** | 0.132353308105745 | 0.205219582900465 | 0.819390360194292 | 1154.69052977524 | 1120.63604298612 | 1032.58228351546 | 1254.00799022316 | 997.506916391197 | 972.350374498718 | 1082.72121417838 | 1108.32160152655 | Lrrfip1 |
| **ENSMUSG00000026307** | -0.143454728571042 | 0.212807722077331 | 0.823370267313127 | 221.360860559416 | 226.407572638183 | 229.988537941573 | 244.79886720138 | 248.555510450935 | 247.06261087892 | 270.680303544596 | 251.841931003094 | Scly |
| **ENSMUSG00000026308** | -0.118603065941209 | 0.405903336394278 | 0.881158207349486 | 1037.23374662126 | 950.423155643021 | 742.020632700382 | 815.9962240046 | 895.675804180021 | 900.188522806361 | 987.983107937774 | 1065.98610450447 | Klhl30 |
| **ENSMUSG00000026309** | 0.0856001705181411 | 0.544180643065549 | 0.913837378403038 | 945.97886124779 | 759.849875077069 | 883.988865997649 | 902.285479876351 | 845.307727172343 | 963.788798874201 | 796.424739275444 | 686.052156870497 | Ilkap |
| **ENSMUSG00000026311** | 0.119423449940045 | 0.543580752779976 | 0.913837378403038 | 395.739007857242 | 477.247617143797 | 469.44162476963 | 331.088123073131 | 379.950493949227 | 280.085831144914 | 447.663578939139 | 432.039174738066 | Asb1 |
| **ENSMUSG00000026313** | -0.0349661641218183 | 0.828404905140794 | 0.967424258651894 | 453.563885717661 | 469.103459854653 | 410.761421673426 | 363.915557372167 | 519.010184818252 | 415.847958905112 | 453.910047482476 | 350.624757387928 | Hdac4 |
| **ENSMUSG00000026315** | 0.0857565690124983 | 0.84698008432778 | 0.972334438148346 | 9.93865088225949 | 17.1027303072009 | 30.2865564367503 | 29.075727522003 | 14.2344565456483 | 26.9078091056249 | 24.9858741733473 | 15.1973579053591 | Serpinb8 |
| **ENSMUSG00000026317** | -0.0556333006863132 | 0.61968064023754 | 0.929594515086012 | 522.230928176908 | 485.39177443294 | 469.44162476963 | 428.63249927598 | 521.20010120989 | 456.20967256355 | 515.333654825288 | 487.40097853616 | Cln8 |
| **ENSMUSG00000026319** | -0.0384172696660863 | 0.836431536488118 | 0.968924788429433 | 225.878429142261 | 284.231089391101 | 247.024725937245 | 221.350699844926 | 237.605928492744 | 185.908499275227 | 281.091084450157 | 299.605055848508 | 2310035C23 |
| **ENSMUSG00000026321** | -0.409141805293153 | 0.344705797649569 | 0.861266047293794 | 10.8421645988285 | 27.6901347830872 | 17.036187995672 | 13.1309737196143 | 14.2344565456483 | 33.0232202659942 | 18.7394056300105 | 26.0526135520442 | Tnfrsf11a |
| **ENSMUSG00000026333** | -0.186345802305777 | 0.452507093487148 | 0.891110666112841 | 115.649755720838 | 74.1118313312039 | 111.681676860517 | 88.1651092602672 | 85.4067392738896 | 91.7311674055395 | 148.874166949528 | 116.15123541953 | Gin1 |
| **ENSMUSG00000026335** | 0.00679915037258743 | 0.966010618906251 | 0.994284908685797 | 15783.4811147446 | 17869.9099238382 | 15994.1411632701 | 15626.7966530352 | 11932.8544180365 | 17624.6149641843 | 19054.852291449 | 16355.6136828604 | Pam |
| **ENSMUSG00000026339** | 0.0366074200659965 | 0.871069662748873 | 0.975613767803763 | 252.080326922764 | 354.270842077733 | 262.16800415562 | 272.936668029125 | 250.745426842573 | 187.131581507301 | 322.734208072402 | 351.710282952596 | Ccdc93 |
| **ENSMUSG00000026341** | 0.0269723052859589 | 0.754840163581113 | 0.956497786093635 | 1997.66882733416 | 2044.18347957496 | 1950.64352550445 | 2067.19043414499 | 1954.50037953709 | 1894.55437748241 | 1878.10487536327 | 2182.99191054837 | Actr3 |
| **ENSMUSG00000026342** | 0.256444322318187 | 0.125044736831891 | 0.771949105062466 | 3539.96674151752 | 3928.74147628272 | 3693.06697550624 | 3453.44608825855 | 2505.26435203409 | 2481.63384887786 | 3606.29450568646 | 3640.85274389817 | Slc35f5 |
| **ENSMUSG00000026347** | -5.89022843344038 | NA | NA | 0.903513716569045 | 0 | 0 | 0.937926694258161 | 0 | 108.854318654574 | 0 | 0 | Tmem163 |
| **ENSMUSG00000026349** | 0.00080560597507754 | 0.994705420019856 | 0.999262355603273 | 251.176813206195 | 274.458100644129 | 254.596365046432 | 293.571055302804 | 252.935343234211 | 260.516515431732 | 268.598147363483 | 290.92085133116 | Ccnt2 |
| **ENSMUSG00000026353** | 0.208910444782636 | 0.0432740463520505 | 0.687730553581215 | 2322.93376529901 | 2337.37314198412 | 2572.46438734648 | 2464.87135251045 | 1926.03146644579 | 1931.24684444462 | 2315.35767339685 | 2215.55767748842 | Ubxn4 |
| **ENSMUSG00000026355** | -0.422318770818105 | 0.131665292964771 | 0.77303252008843 | 121.974351736821 | 104.245213301034 | 103.163582862681 | 110.675349922463 | 153.294147414674 | 229.939459629886 | 93.6970281500523 | 113.980184290193 | Mcm6 |
| **ENSMUSG00000026356** | 0.232437234901816 | 0.0699865770796323 | 0.724916894583216 | 2098.86236358989 | 1989.6176257377 | 2127.63058968171 | 2070.00421422776 | 1566.88517821713 | 1511.72963884329 | 1993.664543415 | 1978.91310439069 | Dars |
| **ENSMUSG00000026357** | 1.02081530068784 | 0.0919160830975696 | 0.754058637722175 | 46.0791995450213 | 35.8342920722305 | 124.932045301595 | 15.9447538023887 | 18.6142893289246 | 45.2540425867328 | 12.4929370866736 | 33.6512925047237 | Rgs18 |
| **ENSMUSG00000026360** | 0.0494116284842252 | 0.844121062697268 | 0.971721411930626 | 1030.90915060528 | 1237.91190794978 | 1049.61847151113 | 622.783324987419 | 1288.76579647908 | 957.673387713832 | 848.478643803251 | 714.275821551878 | Rgs2 |
| **ENSMUSG00000026361** | 0.154656775303755 | 0.399781678316845 | 0.880487015130533 | 1192.63810587114 | 1268.85970564852 | 1322.19747944188 | 1045.78826409785 | 1052.25482618215 | 765.649477278236 | 1302.38869128573 | 1216.8741579934 | Cdc73 |
| **ENSMUSG00000026365** | -0.469206358196613 | 0.175613738493653 | 0.799889021902745 | 1727.51822608001 | 1248.49931242567 | 1303.26838166891 | 1335.60761262362 | 2443.94669306822 | 3161.66756991093 | 1210.77381931679 | 957.433548037624 | Cfh |
| **ENSMUSG00000026368** | -0.475823260813327 | 0.0887163297449406 | 0.751206603737452 | 38.8510898124689 | 46.4216965481167 | 43.5369248778286 | 44.0825546301336 | 43.7983278327639 | 59.9310293716191 | 47.8895921655823 | 89.0130963028176 | F13b |
| **ENSMUSG00000026374** | -0.060378217677112 | 0.59594188605595 | 0.924461858177432 | 1169.14674924034 | 1068.5134363356 | 1047.72556173383 | 1138.64300682941 | 1149.70610561005 | 1336.82887965673 | 1075.43366754449 | 1052.95979772845 | Tsn |
| **ENSMUSG00000026377** | 0.105517875673182 | 0.365703859702995 | 0.869253398153585 | 371.344137509877 | 408.022280186079 | 328.419846361011 | 389.239578117137 | 355.861413641207 | 330.232202659942 | 353.966550789086 | 351.710282952596 | Nifk |
| **ENSMUSG00000026380** | 0.655254882078373 | 0.217470796788376 | 0.825996545635915 | 18.0702743313809 | 9.77298874697194 | 13.2503684410783 | 14.0689004138724 | 4.37983278327639 | 4.89232892829544 | 16.6572494488982 | 8.68420451734806 | Tfcp2l1 |
| **ENSMUSG00000026383** | 0.187248308409419 | 0.462458658810371 | 0.895328112556867 | 110.228673421423 | 145.780415475665 | 163.736695736181 | 136.937297361692 | 107.305903190271 | 77.0541806206532 | 163.449260217313 | 140.032797842237 | Epb41l5 |
| **ENSMUSG00000026384** | 0.125706321257889 | 0.675953150910334 | 0.94409858171788 | 573.731210021344 | 683.294796559122 | 704.162437154444 | 592.769670771158 | 432.508487348543 | 309.439804714686 | 707.933101578173 | 890.130963028176 | Ptpn4 |
| **ENSMUSG00000026385** | 0.0740491042122738 | 0.66323705592905 | 0.940031968637737 | 3193.92098807157 | 2400.08315311053 | 2593.28639489674 | 3046.38590295051 | 2402.3382816271 | 3387.93778284459 | 2242.48220705792 | 2639.99817327381 | Dbi |
| **ENSMUSG00000026388** | -0.277776095175921 | 0.585436085107213 | 0.922964550512322 | 23.4913566307952 | 18.7315617650296 | 5.67872933189068 | 19.6964605794214 | 12.0445401540101 | 35.4693847301419 | 19.7804837205666 | 15.1973579053591 | 3110009E18 |
| **ENSMUSG00000026389** | -0.166397425439663 | 0.194657918374195 | 0.809762948852223 | 730.942596704357 | 781.839099757755 | 682.39397471553 | 604.962717796514 | 771.945528052463 | 894.073111645991 | 725.631429117627 | 753.354741879944 | Steap3 |
| **ENSMUSG00000026393** | 0.276035077347034 | 0.192361894047133 | 0.808824684419004 | 3496.5980831222 | 3951.54511669232 | 4447.39152175905 | 3435.62548106764 | 2781.19381738051 | 2136.72465943303 | 3886.34451204606 | 3855.78680570254 | Nek7 |
| **ENSMUSG00000026395** | -0.0959085506680347 | 0.743377248873241 | 0.954383574136061 | 47.8862269781594 | 37.4631235300591 | 79.5022106464695 | 50.6480414899407 | 55.8428679867739 | 70.9387694602839 | 43.7252798033577 | 59.7039060567679 | Ptprc |
| **ENSMUSG00000026398** | -0.610146139501897 | 0.145469091620481 | 0.780607990324772 | 10.8421645988285 | 13.0306516626293 | 6.62518422053913 | 13.1309737196143 | 16.4243729372865 | 20.7923979452556 | 11.4518589961175 | 18.4539345993646 | Nr5a2 |
| **ENSMUSG00000026399** | -0.605131280444619 | 0.0947674476419581 | 0.757085157577544 | 78.6056933415069 | 48.0505280059454 | 55.8408384302584 | 98.4823028971069 | 134.679858085749 | 157.777607937528 | 61.4236073428121 | 73.8157383974585 | Cd55 |
| **ENSMUSG00000026404** | 0.129583214486997 | 0.354303537343791 | 0.863443308527191 | 263.826005238161 | 245.139134403213 | 279.204192151292 | 244.79886720138 | 234.321053905287 | 220.154801773295 | 277.967850178489 | 210.59195954569 | Ddx59 |
| **ENSMUSG00000026405** | -0.329679530364752 | 0.246111008921328 | 0.832048466795696 | 141.85165350134 | 208.490426602068 | 159.950876181588 | 167.888878272211 | 192.712642464161 | 185.908499275227 | 135.340151772298 | 338.683976176574 | C4bp |
| **ENSMUSG00000026407** | 0.113311532917327 | 0.7116299644881 | 0.950020479830613 | 361.405486627618 | 430.82592059568 | 424.958245003153 | 367.667264149199 | 290.163921892061 | 177.34692365071 | 529.908748093074 | 466.775992807458 | Cacna1s |
| **ENSMUSG00000026409** | 0.0937808136298926 | 0.815153768448133 | 0.966177091475091 | 259.308436655316 | 446.299819445052 | 473.227444324223 | 291.695201914288 | 193.80760065998 | 156.554525705454 | 504.922873919726 | 522.137796605552 | Pfkfb2 |
| **ENSMUSG00000026411** | -0.0316763000013801 | 0.840398569451816 | 0.969957774070128 | 324.361424248287 | 262.241864710414 | 283.936466594534 | 294.508981997063 | 304.398378437709 | 362.032340693862 | 277.967850178489 | 247.49982874442 | Tmem9 |
| **ENSMUSG00000026414** | 0.075838023190555 | 0.491760498700196 | 0.899236955296196 | 257243.907812956 | 221474.656568149 | 212005.895057252 | 244909.542551303 | 204072.833745784 | 246820.440596969 | 225318.448982884 | 211510.3141734 | Tnnt2 |
| **ENSMUSG00000026417** | -0.324632052325731 | 0.45340156295749 | 0.891651261195907 | 121.070838020252 | 349.384347704247 | 151.432782183751 | 181.957778686083 | 163.148771177045 | 156.554525705454 | 199.886993386778 | 487.40097853616 | Pigr |
| **ENSMUSG00000026421** | -0.406778141382533 | 0.12517820628114 | 0.771949105062466 | 765.276117933981 | 886.08431305879 | 700.376617599851 | 717.513921107493 | 1213.21368096756 | 1479.92950080937 | 664.207821774815 | 713.190295987209 | Csrp1 |
| **ENSMUSG00000026425** | -0.105817747078495 | 0.500670618974374 | 0.900927059989498 | 353.273863178497 | 428.382673408937 | 328.419846361011 | 352.660437041069 | 479.591689768764 | 346.132271676902 | 388.322127777439 | 360.394487469945 | Srgap2 |
| **ENSMUSG00000026426** | -0.0363234033448964 | 0.766673724788311 | 0.958624690460551 | 464.406050316489 | 469.103459854653 | 441.994432998825 | 528.990655561603 | 523.390017601528 | 464.771248188067 | 530.94982618363 | 433.124700302735 | Arl8a |
| **ENSMUSG00000026427** | 0.0624271798888688 | 0.557839567286948 | 0.916370464878199 | 848.399379858333 | 693.067785306094 | 766.628459805242 | 846.009878220861 | 743.476614961167 | 781.549546295196 | 732.91897575152 | 762.038946397292 | Eif2d |
| **ENSMUSG00000026429** | 0.269664180441378 | 0.260517016652841 | 0.840316047847749 | 46.0791995450213 | 50.4937751926884 | 44.483379766477 | 45.0204813243917 | 33.943704070392 | 31.8001380339204 | 41.6431236222455 | 46.6775992807458 | Ube2t |
| **ENSMUSG00000026430** | 0.0843997424618089 | 0.837244877169453 | 0.968924788429433 | 14.4562194651047 | 26.8757190541728 | 41.6440151005317 | 30.9515809105193 | 40.5134532453066 | 20.7923979452556 | 17.6983275394543 | 28.2236646813812 | Rassf5 |
| **ENSMUSG00000026433** | -0.107724748790059 | 0.549691328154252 | 0.914699131434573 | 168.957064998411 | 147.409246933493 | 125.878500190243 | 181.957778686083 | 187.237851485066 | 187.131581507301 | 144.709854587303 | 154.144630182928 | Rab29 |
| **ENSMUSG00000026434** | -0.0837307304420421 | 0.409439250157913 | 0.882192126163777 | 1120.35700854562 | 1064.44135769103 | 1007.02800152195 | 1171.47044112844 | 1227.44813751321 | 1066.52770636841 | 1103.5427759895 | 1225.55836251075 | Nucks1 |
| **ENSMUSG00000026435** | -0.34013762351072 | 0.0787138765380037 | 0.739855948845396 | 323.457910531718 | 270.386021999557 | 281.097101928589 | 192.274972322923 | 408.419407040523 | 321.670627035425 | 298.789411989611 | 322.401092706547 | Slc45a3 |
| **ENSMUSG00000026436** | 0.0189548311515712 | 0.895219183234337 | 0.983733637008786 | 739.977733870048 | 672.707392083235 | 638.857049837702 | 817.872077393117 | 819.028730472685 | 631.110431750112 | 627.77008860535 | 753.354741879944 | Elk4 |
| **ENSMUSG00000026437** | 0.431394401841091 | 0.0421922669677083 | 0.685406501171951 | 647.819334780005 | 696.325448221751 | 664.41133183121 | 661.238319452004 | 519.010184818252 | 297.208982393948 | 594.455589707554 | 567.72987032163 | Cdk18 |
| **ENSMUSG00000026439** | -0.20500695288481 | 0.258358163303055 | 0.838495963179748 | 290.027903018663 | 298.076156782644 | 315.169477919933 | 245.736793895638 | 269.359716171498 | 282.531995609062 | 352.92547269853 | 419.012867962044 | Rbbp5 |
| **ENSMUSG00000026443** | -0.65851784030832 | 0.214194219254012 | 0.824248286808591 | 36.1405486627618 | 42.3496179035451 | 39.7511053232348 | 45.0204813243917 | 55.8428679867739 | 149.216032313011 | 19.7804837205666 | 33.6512925047237 | Lrrn2 |
| **ENSMUSG00000026447** | 0.146072653769889 | 0.499372955353071 | 0.900927059989498 | 517.713359594063 | 479.69086433054 | 491.210087208544 | 505.542488205149 | 512.440435643337 | 265.408844360028 | 500.758561557502 | 522.137796605552 | Pik3c2b |
| **ENSMUSG00000026455** | 0.241266308487029 | 0.270410868783233 | 0.844129257012535 | 702.030157774148 | 778.581436842098 | 775.146553803078 | 642.47978556684 | 558.428679867739 | 388.940149799487 | 765.19239655876 | 738.157383974585 | Klhl12 |
| **ENSMUSG00000026456** | -0.0424781633509298 | 0.742003419521264 | 0.954383574136061 | 748.109357319169 | 633.615437095348 | 585.855576073389 | 761.596475737627 | 704.058119911679 | 760.757148349941 | 649.632728507029 | 696.907412517182 | Cyb5r1 |
| **ENSMUSG00000026457** | 0.0832377865365466 | 0.382783446804912 | 0.875126835008975 | 5263.87091273126 | 5100.68571019044 | 6207.79761464516 | 5527.20200926334 | 4983.15474917271 | 5128.38379908569 | 5351.14138545854 | 5396.14758196715 | Adipor1 |
| **ENSMUSG00000026458** | -0.376112395790065 | 0.122744431227469 | 0.770251097497048 | 35.2370349461928 | 43.1640336324594 | 31.2330113253987 | 50.6480414899407 | 50.3680770076785 | 51.3694537471021 | 48.9306702561384 | 57.5328549274309 | Ppfia4 |
| **ENSMUSG00000026463** | -0.533752843458391 | 0.0565504883812893 | 0.70510438385647 | 262.922491521592 | 240.252640029727 | 309.490748588042 | 235.419600258798 | 542.004306930453 | 464.771248188067 | 288.37863108405 | 222.532740757044 | Atp2b4 |

| | | | | | | | | | | | | |
|---|---|---|---|---|---|---|---|---|---|---|---|---|
| **ENSMUSG00000026466** | 0.190732510205834 | 0.350448986419787 | 0.861774532843376 | 2242.52104452437 | 3068.71846654919 | 3073.13902344151 | 2300.73418101527 | 1989.5390418033 | 1779.58464766747 | 2916.05973164774 | 2675.82051690787 | Tor1aip1 |
| **ENSMUSG00000026469** | 0.223460586319675 | 0.339391431233989 | 0.858570960161243 | 1159.20809835808 | 1521.32858161197 | 1478.36253606887 | 1214.61506906432 | 831.073270626695 | 852.48831575548 | 1547.04204256642 | 1371.01878817632 | Xpr1 |
| **ENSMUSG00000026470** | -0.0473681637342471 | 0.735224166612632 | 0.954383574136061 | 347.852780879082 | 355.899673535562 | 347.34894413398 | 438.949692912819 | 399.65974147397 | 434.19419238622 | 357.089785060755 | 349.539231823259 | Stx6 |
| **ENSMUSG00000026471** | 0.0649897510510198 | 0.691980724787039 | 0.94469184542218 | 227.685456575399 | 222.335493993612 | 202.541346170768 | 260.743621003769 | 170.813478547779 | 245.839528646846 | 234.242570375131 | 222.532740757044 | Mr1 |
| **ENSMUSG00000026473** | 0.0555436224057567 | 0.644247271925513 | 0.934857223264556 | 17358.3055227245 | 18668.8517539032 | 20030.7712633557 | 15609.9139725386 | 17511.6664257348 | 15343.5666013666 | 19093.3721807995 | 17011.2711239202 | Glul |
| **ENSMUSG00000026475** | -0.0947640527971935 | 0.818857331516191 | 0.966315773308182 | 10.8421645988285 | 12.2162359337149 | 18.9290977729689 | 17.8206071909051 | 13.1394983498292 | 22.0154801773295 | 10.4107809055614 | 18.4539345993646 | Rgs16 |
| **ENSMUSG00000026478** | -0.310027967979739 | 0.14548321640559 | 0.780607990324772 | 1695.8952460001 | 2305.61092855646 | 2322.60029674329 | 1912.43252959239 | 3137.05523102171 | 3138.42900750152 | 1940.56956079664 | 1996.28151342539 | Lamc1 |
| **ENSMUSG00000026479** | 0.350650217793337 | 0.0309815739110385 | 0.667184591665619 | 393.931980424104 | 394.991628523449 | 407.922057007481 | 396.742991671202 | 241.98576127602 | 278.86274891284 | 367.500565966316 | 360.394487469945 | Lamc2 |
| **ENSMUSG00000026480** | -0.198075492569232 | 0.485921649884441 | 0.899236955296196 | 60.535419010126 | 52.122606650517 | 66.2518422053913 | 73.1582821521366 | 68.9823663366031 | 110.077400886647 | 47.8895921655823 | 62.9604827507734 | Ncf2 |
| **ENSMUSG00000026482** | -0.135077067582898 | 0.386404094968326 | 0.875126835008975 | 421.940905637744 | 458.516055378767 | 454.298346551254 | 577.762843663027 | 595.657258525589 | 545.494675504941 | 523.662279549737 | 435.295751432072 | Rgl1 |
| **ENSMUSG00000026483** | -0.354067764231217 | 0.0578737033720294 | 0.710862211775517 | 228.588970291968 | 283.416673662186 | 268.793188376159 | 235.419600258798 | 383.235368536684 | 387.717067567413 | 261.31060072959 | 268.124814473121 | Fam129a |
| **ENSMUSG00000026484** | 0.0209104546596211 | 0.870347806679357 | 0.975613767803763 | 271.054114970713 | 296.447325324816 | 307.597838810745 | 269.184961252092 | 277.024423542232 | 324.116791499573 | 264.433835001259 | 263.782712214447 | Rnf2 |
| **ENSMUSG00000026489** | 0.168425726479666 | 0.111507562735707 | 0.764116514278315 | 13979.1642227563 | 15337.0770069146 | 16148.4133101198 | 16283.3453390159 | 14839.9684279362 | 12093.8371107463 | 13554.8367390409 | 14453.7728935612 | Coq8a |
| **ENSMUSG00000026490** | 0.0310074888268483 | 0.854859377391782 | 0.973268281968411 | 1077.89186386687 | 1268.85970564852 | 1042.04683240194 | 867.582192188799 | 1073.05903190271 | 813.349684329117 | 1182.66471087177 | 1096.38082031519 | Cdc42bpa |
| **ENSMUSG00000026491** | 0.224854669651506 | 0.226790082815271 | 0.832048466795696 | 456.274426867368 | 560.318021493058 | 514.871459424755 | 437.073839524303 | 404.039574257247 | 302.101311322243 | 501.799639648058 | 475.460197324806 | Ahctf1 |
| **ENSMUSG00000026492** | 0.376799184483359 | 0.100540403211887 | 0.759800953500741 | 1563.98224338102 | 1825.10564849701 | 1702.67234467856 | 1722.97133735224 | 1015.0262475243 | 876.949960396957 | 1706.32699042151 | 1648.91333273146 | Tfb2m |
| **ENSMUSG00000026494** | 0.0716661889137799 | 0.827809986371043 | 0.967319591893432 | 91.2548853734735 | 90.4001459094905 | 45.4298346551254 | 45.0204813243917 | 50.3680770076785 | 51.3694537471021 | 78.0808567917102 | 79.2433662208011 | Kif26b |
| **ENSMUSG00000026495** | 0.101818159246658 | 0.395377779427702 | 0.879215782741314 | 4673.87645581167 | 4919.07100264254 | 4368.83576600123 | 4372.61424863155 | 3771.03602640097 | 3885.73225129865 | 4887.86163516106 | 4539.6679114437 | Efcab2 |
| **ENSMUSG00000026496** | 0.168516410984016 | 0.345878382249749 | 0.861266047293794 | 1301.05975185942 | 1424.41310987116 | 1359.10922009917 | 1220.24262922987 | 1081.81869746927 | 850.042151291332 | 1364.8533767191 | 1422.03848971574 | Parp1 |
| **ENSMUSG00000026499** | 0.0150108073194107 | 0.918409548085758 | 0.986713918026868 | 526.748496759753 | 688.995706661522 | 650.214508501483 | 538.369922504185 | 643.835419141629 | 502.686797382356 | 598.619902069779 | 633.946929766408 | Acbd3 |
| **ENSMUSG00000026500** | -0.00862782018351197 | 0.963363225216063 | 0.994284908685797 | 513.195791011218 | 368.115909469277 | 441.994432998825 | 453.95652002095 | 408.419407040523 | 594.417964787896 | 376.870268781321 | 409.243137880027 | Cox20 |
| **ENSMUSG00000026502** | 0.0888063920413737 | 0.563116412415523 | 0.918253929144354 | 364.116027777325 | 416.980853204136 | 305.704929033448 | 347.03287687552 | 292.353838283699 | 311.885969178834 | 365.418409785204 | 378.848422069309 | Desi2 |
| **ENSMUSG00000026504** | 0.128268910823603 | 0.255303526997773 | 0.837406774112203 | 256.597895505609 | 261.427448981499 | 259.328639489674 | 249.488500672671 | 222.276513751277 | 248.285693110993 | 239.447960827911 | 230.131419709724 | Sdccag8 |
| **ENSMUSG00000026509** | -0.138019067554051 | 0.395220334814935 | 0.879215782741314 | 2431.3554112873 | 2630.56280439328 | 2739.0404477486 | 2471.43683937025 | 3174.28380967956 | 3431.96874319925 | 2500.66957351584 | 2198.18926845373 | Capn2 |
| **ENSMUSG00000026510** | -0.104163337515904 | 0.542952145529104 | 0.913837378403038 | 308.098177350044 | 267.942774812814 | 289.615195926425 | 237.295453647315 | 274.834507150593 | 232.385624094033 | 338.350379430744 | 338.683976176574 | Trp53bp2 |
| **ENSMUSG00000026511** | 0.0256131825283169 | 0.86038539058393 | 0.974868041734905 | 1917.25610655951 | 1571.82235680465 | 1683.74324690559 | 1898.36362917852 | 1517.61205940527 | 2126.94000157644 | 1574.11007292088 | 1729.24222451693 | Srp9 |
| **ENSMUSG00000026516** | 0.183347766786274 | 0.264755614220915 | 0.842510961779757 | 351.466835745359 | 400.69253862585 | 358.706402797761 | 322.646782824807 | 274.834507150593 | 262.96267989588 | 386.239971596327 | 337.598450611906 | Nvl |
| **ENSMUSG00000026519** | -0.15753810123798 | 0.385871524484618 | 0.875126835008975 | 327.975479114563 | 388.476302692135 | 358.706402797761 | 383.612017951588 | 505.870686468423 | 462.325083723919 | 331.062832796851 | 328.914246094558 | Tmem63a |
| **ENSMUSG00000026520** | -0.166135080948461 | 0.292941023592635 | 0.851785577030345 | 266.536546387868 | 272.014853457386 | 224.309808609682 | 259.805694309511 | 330.677375137367 | 319.224462571277 | 264.433835001259 | 234.473521968398 | Pycr2 |
| **ENSMUSG00000026525** | -0.323004456825366 | 0.342370181130915 | 0.860056790709335 | 18.9737880479499 | 21.1748089517725 | 42.5904699891801 | 36.5791410760683 | 44.893286028583 | 39.1386314263635 | 34.3555769883525 | 30.3947158107182 | Opn3 |
| **ENSMUSG00000026526** | 0.118108017673295 | 0.180089992981164 | 0.802731406993634 | 14186.0688638506 | 12822.1612360272 | 13733.060434289 | 15258.1914621918 | 13411.0479823923 | 12197.7991004726 | 12756.3298435843 | 13230.3855821798 | Fh1 |
| **ENSMUSG00000026527** | -0.331904729032434 | 0.447679957139134 | 0.89108031640712 | 10.8421645988285 | 12.2162359337149 | 9.46454888648447 | 23.448167356454 | 15.3294147414674 | 12.2308223207386 | 22.903717992235 | 19.5394601640331 | Rgs7 |
| **ENSMUSG00000026532** | -0.0528670439840395 | 0.824247438072314 | 0.966637987288264 | 202.387072511466 | 276.086932101957 | 318.008842585878 | 166.013024883695 | 226.656346534553 | 201.808568292187 | 288.37863108405 | 281.151121249143 | Spta1 |
| **ENSMUSG00000026536** | -0.336187598549252 | 0.371345867710097 | 0.873374202289323 | 56.0178504272808 | 81.4415728914329 | 80.448665535118 | 59.0893817382642 | 118.255485148462 | 129.646716599829 | 37.4788112600209 | 65.1315338801105 | Mnda |
| **ENSMUSG00000026542** | -0.158571546032684 | 0.716367038615291 | 0.951348124093194 | 9.03513716569045 | 34.2054606144018 | 21.7684624389143 | 21.5723139679377 | 16.4243729372865 | 20.7923979452556 | 28.1091084450157 | 31.4802413753867 | Apcs |
| **ENSMUSG00000026544** | -0.11407870075821 | 0.564703029016873 | 0.919437940809327 | 355.080890611635 | 308.663561258531 | 405.082692341535 | 403.308478531009 | 483.971522552041 | 473.332823812584 | 326.898520434627 | 309.374785930525 | Dusp23 |
| **ENSMUSG00000026547** | -0.725989516271336 | 0.0381981107960754 | 0.679566075569698 | 786.056933415069 | 699.583111137408 | 837.612576453875 | 822.561710864407 | 1747.55328052728 | 2067.00897220482 | 662.125665593703 | 727.3021283279 | Tagln2 |
| **ENSMUSG00000026548** | 0.185573680506929 | 0.688362389001352 | 0.944584424505182 | 23.4913566307952 | 32.5766291565731 | 11.3574586637814 | 27.1998741334867 | 19.7092475247437 | 36.6924669622158 | 15.616171358342 | 11.9407812113536 | Slamf9 |
| **ENSMUSG00000026553** | 0.121544948222348 | 0.566311203968198 | 0.919931558693298 | 2686.14627935977 | 3777.26015070466 | 4012.02227298077 | 2654.3325447506 | 2826.08710340909 | 2216.22500451783 | 3502.18669663084 | 3523.61598291398 | Copa |
| **ENSMUSG00000026554** | 0.227431913771099 | 0.157753210187209 | 0.789230661864335 | 4401.91882712439 | 4946.76113742563 | 5122.21385736539 | 4425.13814351 | 3684.53432893126 | 3107.85195169968 | 4732.7409996682 | 4613.48364984116 | Dcaf8 |
| **ENSMUSG00000026556** | -0.359082104978262 | 0.448620505422451 | 0.89108031640712 | 13.5527057485357 | 15.4738988493722 | 10.4110037751329 | 24.3860940507122 | 37.2285786578493 | 14.6769867848863 | 12.4929370866736 | 17.3684090346961 | Vangl2 |
| **ENSMUSG00000026558** | 0.556537641625092 | 0.00849019174948175 | 0.635862253526407 | 1829.61527605232 | 2049.88438967737 | 2538.39201135513 | 2270.72052679901 | 1288.76579647908 | 1040.84297949485 | 1628.2461336298 | 1948.51838857997 | Uck2 |
| **ENSMUSG00000026563** | -0.18085459349242 | 0.432972294963447 | 0.887804130062338 | 77.7021796249379 | 65.9676740420606 | 75.7163910918757 | 82.5375490947182 | 59.1277425742312 | 77.0541806206532 | 98.902418602833 | 106.381505337514 | Tada1 |
| **ENSMUSG00000026564** | 0.38897461860479 | 0.0764512881367535 | 0.737436383485768 | 1067.04969926804 | 1181.71722265469 | 1472.68380673698 | 1798.0054728929 | 904.435469746574 | 839.034411202668 | 1123.32325971007 | 1347.13722575362 | Dusp27 |
| **ENSMUSG00000026565** | 0.0226943663232805 | 0.872765256763503 | 0.976162349063696 | 139.141112351633 | 144.96599974675 | 141.968233297267 | 132.247663890401 | 129.205067106653 | 136.985209992272 | 150.95632313064 | 132.434118889558 | Pou2f1 |
| **ENSMUSG00000026566** | -0.565254424274961 | 0.0913636375784498 | 0.753823363185192 | 135.527057485357 | 108.317291945606 | 95.5919437534931 | 166.950951577953 | 231.036179317829 | 284.978160073209 | 114.518589961175 | 119.407812113536 | Mpzl1 |
| **ENSMUSG00000026568** | 0.0750640245614547 | 0.550149181290406 | 0.914699131434573 | 9672.11433587163 | 7264.58830191581 | 7786.48436891077 | 9157.91624273669 | 8051.22761385782 | 8746.26104156017 | 7771.64794600156 | 7594.33685042088 | Mpc2 |
| **ENSMUSG00000026571** | 0.303171146039421 | 0.1021257609449 | 0.759800953500741 | 2925.57741425057 | 3356.20721885595 | 3169.67742208365 | 2916.01409244862 | 2109.9844433434 | 1871.31581507301 | 3153.42553629454 | 2887.49800201823 | Dcaf6 |
| **ENSMUSG00000026574** | -0.386688356778946 | 0.120668366688849 | 0.769868040577702 | 1274.85785407892 | 1097.0179868476 | 1040.15392262464 | 1250.25628344613 | 1696.09024532378 | 2244.35589585553 | 1058.77641809559 | 1097.46634587986 | Dpt |
| **ENSMUSG00000026575** | -0.453221935330528 | 0.0653532603410511 | 0.718339708336358 | 52.4037955610046 | 43.9784493613737 | 38.8046504345863 | 35.6412143818101 | 64.6025335533267 | 66.0464405319884 | 60.3825292522559 | 43.4210225867403 | Nme7 |
| **ENSMUSG00000026576** | 0.0816008704283374 | 0.357385970261978 | 0.865729181537996 | 4280.84798910414 | 4370.9692170832 | 4268.5115478045 | 3933.66455571873 | 3787.46039933826 | 3742.63163014601 | 4334.0080909852 | 4062.03666298956 | Atp1b1 |
| **ENSMUSG00000026577** | 0.222780145350783 | 0.0852260018574854 | 0.743541556158628 | 739.074220153479 | 772.066111010784 | 739.181268034437 | 720.327701190268 | 674.494248624564 | 505.132961846504 | 691.275852129275 | 673.025850094475 | Blzf1 |
| **ENSMUSG00000026578** | 0.14766161437124 | 0.228144697008997 | 0.832048466795696 | 339.721157429961 | 314.364471360931 | 343.563124579386 | 381.736164563072 | 280.309298129689 | 331.455284892016 | 325.857442344071 | 307.203734801188 | Ccdc181 |
| **ENSMUSG00000026579** | -0.202394511076501 | 0.594391368716027 | 0.924327822600271 | 54.2108229941427 | 99.3587189275481 | 97.48485353079 | 71.2824287636203 | 66.7924499449649 | 72.1618516923577 | 59.3414511616998 | 172.598564782293 | F5 |
| **ENSMUSG00000026581** | 0.0212320868191015 | 0.964478840050322 | 0.994284908685797 | 16.2632468982428 | 10.5874044758863 | 23.6613722162112 | 7.50341355406529 | 15.3294147414674 | 19.5693157131818 | 9.36970281500523 | 13.0263067760221 | Sell |
| **ENSMUSG00000026584** | 0.102175357327913 | 0.543663089715729 | 0.913837378403038 | 138.237598635064 | 106.688460487777 | 131.557229522134 | 143.502784221499 | 133.58489988993 | 118.638976511164 | 122.847214685624 | 108.552556466851 | Scyl3 |
| **ENSMUSG00000026585** | 0.18160168010657 | 0.164636329851752 | 0.795216960550936 | 786.960447131638 | 883.641065872047 | 841.398396008469 | 815.058297310342 | 720.482492848966 | 595.64104701997 | 797.465817366001 | 818.486275760055 | Kifap3 |
| **ENSMUSG00000026586** | 0.045664223248812 | 0.805738438547736 | 0.963475137341661 | 800.513152880174 | 858.394178275702 | 562.194203857177 | 756.906842266336 | 597.847174917227 | 634.779678446333 | 856.8072685277 | 795.690238902016 | Prrx1 |
| **ENSMUSG00000026587** | -1.30864142258491 | 0.0434455687141994 | 0.688841110980169 | 9.03513716569045 | 11.4018202048006 | 2.83936466594534 | 12.1930470253561 | 47.0832024202212 | 22.0154801773295 | 10.4107809055614 | 8.68420451734806 | Astn1 |
| **ENSMUSG00000026589** | -0.452684383696976 | 0.306071427428832 | 0.851785577030345 | 10.8421645988285 | 24.4324718674299 | 6.62518422053913 | 21.5723139679377 | 25.1840385038392 | 19.5693157131818 | 21.8626399016789 | 20.6249857287016 | Sec16b |
| **ENSMUSG00000026594** | 0.236935538441026 | 0.44249626865709 | 0.89108031640712 | 140.044626068202 | 167.769640156352 | 119.253315969704 | 107.861569839689 | 87.5966556655277 | 64.8233582999146 | 172.818963032319 | 128.092016630884 | Ralgps2 |
| **ENSMUSG00000026596** | 0.244600568816802 | 0.405439847029835 | 0.881158207349486 | 504.160653845527 | 796.498582878213 | 710.787621374984 | 481.156394154437 | 397.469825082332 | 318.001380339203 | 673.57752458982 | 714.275821551878 | Abl2 |
| **ENSMUSG00000026600** | -0.575207695614644 | 0.149840792659205 | 0.782641594375979 | 39.754603529038 | 24.4324718674299 | 41.6440151005317 | 45.9584080186499 | 73.3621991198795 | 88.0619207093179 | 29.1501865355718 | 35.8223436340608 | Soat1 |
| **ENSMUSG00000026603** | 0.214729216461045 | 0.117711484567956 | 0.769868040577702 | 2199.15238612906 | 1882.11474952101 | 1984.71590149579 | 2251.96199291385 | 1916.17684268342 | 1431.00621152642 | 1971.80190351332 | 1846.47898550113 | Smyd2 |
| **ENSMUSG00000026604** | 0.237040944508669 | 0.148550589589752 | 0.781423388079902 | 835.750187826367 | 1140.18202048006 | 1053.40429106572 | 963.250715003132 | 848.5926017598 | 658.018240855737 | 969.243702307763 | 910.755948756878 | Ptpn14 |
| **ENSMUSG00000026605** | -0.451967569888633 | 0.0612796657277972 | 0.71282992338528 | 140.044626068202 | 139.26508964435 | 184.558703286447 | 99.4202295913651 | 212.421889988905 | 141.877538920568 | 239.447960827911 | 175.855141476298 | Cenpf |
| **ENSMUSG00000026608** | 0.264493357143175 | 0.0711856248996104 | 0.726501344493452 | 225.878429142261 | 266.313943354985 | 278.257737262643 | 245.736793895638 | 209.137015401448 | 177.34692365071 | 220.708555197901 | 237.730098662403 | Kctd3 |
| **ENSMUSG00000026610** | 0.201982223855087 | 0.418226620382473 | 0.882468496126562 | 950.496429830635 | 1344.60036843756 | 1340.1801223262 | 832.878904501247 | 790.559817381388 | 699.603036746248 | 1322.16917500629 | 1071.41373232782 | Esrrg |
| **ENSMUSG00000026611** | 0.34473587916243 | 0.442047648327729 | 0.89108031640712 | 13.5527057485357 | 13.8450673915436 | 15.1432782183751 | 21.5723139679377 | 17.5193311331055 | 4.89232892829544 | 15.616171358342 | 11.9407812113536 | Spata17 |
| **ENSMUSG00000026615** | 0.227708243931956 | 0.00101755802357017 | 0.556246415579446 | 4232.96176212598 | 4280.56907117371 | 4301.63746890719 | 4188.78061655695 | 3861.91755665395 | 3457.6534700728 | 3577.14431915089 | 3623.48433486348 | Eprs |
| **ENSMUSG00000026617** | 0.0980119907344261 | 0.562276082470484 | 0.917686886311859 | 364.116027777325 | 393.362797065621 | 268.793188376159 | 295.446908691321 | 283.594172717146 | 305.770558018465 | 289.419709174606 | 357.137910775939 | Bpnt1 |
| **ENSMUSG00000026618** | 0.220057886741771 | 0.171338682119248 | 0.799531027639052 | 2951.77931203107 | 3171.3348483924 | 3110.99721898744 | 3073.58577708399 | 2475.70048074698 | 1961.82390024647 | 3086.79653849895 | 3040.55710663649 | Iars2 |
| **ENSMUSG00000026620** | 0.305163755640595 | 0.0463598759133836 | 0.69805115405055 | 247.562758339918 | 268.757190541728 | 246.078271048596 | 193.212899017181 | 195.997517051618 | 172.454594722414 | 204.051305749003 | 200.822229463674 | Mark1 |
| **ENSMUSG00000026621** | -0.114742338807446 | 0.665215749754162 | 0.941592455323073 | 151.7903043836 | 208.490426602068 | 176.98706417726 | 193.212899017181 | 150.009272827216 | 195.693157131818 | 140.545542225078 | 305.032683671851 | Marc1 |
| **ENSMUSG00000026623** | 0.159704685203016 | 0.566477223534169 | 0.919931558693298 | 2479.24163826546 | 3009.26611833844 | 2474.03307892704 | 2249.14821283107 | 1636.96250274955 | 1358.84435983406 | 2874.41660802549 | 3270.68852634621 | Lpgat1 |
| **ENSMUSG00000026626** | 0.00643965907627413 | 0.943652574201569 | 0.990505437194639 | 2644.58464839759 | 2734.80801769432 | 2721.05780486428 | 2546.47097491091 | 2816.23247964672 | 2745.81961100581 | 2682.85823936316 | 2355.59047533066 | Ppp2r5a |
| **ENSMUSG00000026627** | -0.267220960740575 | 0.186918738879496 | 0.807540361382744 | 74.9916384752307 | 66.7820897709749 | 62.4660226507975 | 61.9031618210386 | 85.4067392738896 | 95.400414101761 | 64.5468416144805 | 75.9867895267955 | Tmem206 |
| **ENSMUSG00000026628** | 0.526067126701052 | 0.0110746354875663 | 0.635862253526407 | 185.220311896654 | 285.859920848929 | 210.112985279955 | 186.647412157374 | 129.205067106653 | 146.769867848863 | 140.545542225078 | 186.710397122983 | Atf3 |
| **ENSMUSG00000026632** | 0.0115403493136367 | 0.948747932807881 | 0.991233613135933 | 80.412720774645 | 95.2866402829764 | 75.7163910918757 | 91.9168160372998 | 95.2613630362614 | 83.1695917810225 | 82.2451691539348 | 80.3288917854696 | Tatdn3 |
| **ENSMUSG00000026634** | 0.0446519440764191 | 0.640950086469335 | 0.934544499347759 | 623.424464432641 | 616.512706788147 | 673.875880717694 | 588.080037299867 | 569.37826182593 | 628.664267285964 | 609.03068297534 | 618.749571861049 | Angel2 |
| **ENSMUSG00000026637** | -0.511497222823608 | 0.190912901406897 | 0.808824684419004 | 50.5967681278665 | 35.8342920722305 | 31.2330113253987 | 37.5170677703264 | 71.1722827282413 | 85.6157562451702 | 24.9858741733473 | 40.1644458927348 | Traf5 |
| **ENSMUSG00000026638** | -0.537745800706984 | 0.162497307979792 | 0.793968226938511 | 17.1667606148119 | 15.4738988493722 | 11.3574586637814 | 9.37926694258161 | 14.2344565456483 | 24.4616446414772 | 19.7804837205666 | 19.5394601640331 | Irf6 |
| **ENSMUSG00000026639** | -0.0384922534076409 | 0.86485024605646 | 0.974868041734905 | 180.702743313809 | 219.892246806869 | 280.15064703994 | 151.006197775564 | 168.623562156141 | 214.039390612925 | 247.776585552361 | 223.618266321713 | Lamb3 |
| **ENSMUSG00000026640** | 0.199283361894286 | 0.324839706991112 | 0.855977678491063 | 819.486940928124 | 1132.03786319092 | 1052.45783617707 | 701.569167305105 | 976.702710670634 | 638.448925142555 | 757.904849924867 | 854.308619394115 | Plxna2 |
| **ENSMUSG00000026641** | -0.171347960168759 | 0.319339226087527 | 0.854670142081901 | 618.003382133227 | 448.743066631795 | 479.852628544763 | 608.714424573547 | 617.556422441971 | 730.180092548094 | 510.128264372507 | 569.900921450967 | Usf1 |
| **ENSMUSG00000026643** | 0.00636153714237186 | 0.96710044979176 | 0.994284908685797 | 221.360860559416 | 172.656134529838 | 167.522515290775 | 192.274972322923 | 171.908436743598 | 203.031650524261 | 200.928071477334 | 174.76961591163 | Nmt2 |
| **ENSMUSG00000026644** | -0.402699527899776 | 0.399652582972309 | 0.880459952732461 | 9.93865088225949 | 4.88649437348597 | 8.51809399783602 | 12.1930470253561 | 10.949581958191 | 14.6769867848863 | 12.4929370866736 | 8.68420451734806 | Acbd7 |

| | | | | | | | | | | | | |
|---|---|---|---|---|---|---|---|---|---|---|---|---|
| **ENSMUSG00000026646** | 0.0575116695114342 | 0.855521774899487 | 0.973387788552416 | 32.5264937964856 | 24.4324718674299 | 22.7149173275627 | 23.448167356454 | 20.8042057205628 | 19.5693157131818 | 32.2734208072402 | 26.0526135520442 | Suv39h2 |
| **ENSMUSG00000026648** | -0.358769629049919 | 0.202882241351614 | 0.815403452654488 | 23.4913566307952 | 23.6180561385155 | 27.447191770805 | 34.703287687552 | 29.5638712871156 | 37.9155491942896 | 32.2734208072402 | 40.1644458927348 | Dclre1c |
| **ENSMUSG00000026649** | 0.360370810241513 | 0.256333360152813 | 0.837517407767513 | 33.4300075130547 | 45.6072808192024 | 27.447191770805 | 36.5791410760683 | 27.3739548954774 | 39.1386314263635 | 18.7394056300105 | 27.1381391167127 | Cfap126 |
| **ENSMUSG00000026650** | 0.557149108735826 | 0.0955587125850827 | 0.757085157577544 | 55.1143367107117 | 29.3189662409158 | 42.5904699891801 | 53.4618215727152 | 28.4689130912965 | 34.2463024980681 | 18.7394056300105 | 41.2499714574033 | Meig1 |
| **ENSMUSG00000026655** | -0.33703471352612 | 0.219862086912043 | 0.828432177683914 | 64.1494738764022 | 81.4415728914329 | 88.0203046443056 | 44.0825546301336 | 105.115986798633 | 102.738907494204 | 66.6289977955927 | 77.072315091464 | Fam107b |
| **ENSMUSG00000026656** | -0.266352056832197 | 0.305474120004445 | 0.851785577030345 | 124.684892886528 | 134.378595270864 | 138.182413742673 | 160.385464718146 | 220.086597359638 | 216.485555077073 | 118.6829023234 | 116.15123541953 | Fcgr2b |
| **ENSMUSG00000026657** | -0.25316600075825 | 0.192173814706843 | 0.808824684419004 | 148.176249517323 | 221.521078264697 | 193.076797284283 | 241.047160424347 | 232.131137513649 | 196.916239363891 | 262.351678820146 | 265.953763343784 | Frmd4a |
| **ENSMUSG00000026659** | 0.10585585440032 | 0.546405629515817 | 0.914211323944814 | 125.588406603097 | 97.7298874697194 | 106.949402417274 | 105.047789756914 | 108.400861386091 | 85.6157562451702 | 97.8613405122768 | 111.809133160856 | Dusp12 |
| **ENSMUSG00000026662** | 0.0333633565011243 | 0.794718022390386 | 0.961804768755116 | 328.878992831132 | 297.26174105373 | 365.3315870183 | 334.839829850164 | 293.448796479518 | 304.547475786391 | 342.514691792969 | 353.881334081933 | Sephs1 |
| **ENSMUSG00000026663** | 0.194957976991721 | 0.450280633385785 | 0.89108031640712 | 2709.63763599057 | 3931.99913919838 | 4581.78811594713 | 2906.63482550604 | 2420.95257095602 | 2156.29397514621 | 4092.47797397617 | 3673.41851083823 | Atf6 |
| **ENSMUSG00000026664** | 0.225478025968033 | 0.0541398936669292 | 0.703396849069134 | 14669.448702215 | 14314.1708513982 | 15253.0669854584 | 15451.4043612089 | 12110.2376457592 | 10776.5775468028 | 14012.9110988856 | 14149.825735454 | Phyh |
| **ENSMUSG00000026667** | 0.155698405741008 | 0.499030566834514 | 0.900927059989498 | 969.470217878585 | 1247.68489669675 | 1149.94268970786 | 770.975742680209 | 777.420319031559 | 669.025980944401 | 1169.13069569454 | 1098.55187144453 | Uhmk1 |
| **ENSMUSG00000026669** | -0.73339393905488 | 0.0368251103049933 | 0.679566075569698 | 26.2018977805023 | 15.4738988493722 | 16.0897331070236 | 19.6964605794214 | 30.6588294829347 | 30.5770558018465 | 21.8626399016789 | 45.5920737160773 | Mcm10 |
| **ENSMUSG00000026670** | -0.0952592865439181 | 0.307335397555675 | 0.851785577030345 | 824.004509510969 | 839.662616510673 | 747.699362032273 | 816.934150698859 | 817.933772276865 | 830.472835578151 | 877.628830338823 | 922.696729968231 | Uap1 |
| **ENSMUSG00000026672** | 0.324706800931475 | 0.078231372668303 | 0.739855948845396 | 3715.24840253191 | 3795.17729674077 | 4531.62600684876 | 3855.8166400953 | 3304.58383498203 | 2157.51705737829 | 3704.15584619873 | 3525.78703404331 | Optn |
| **ENSMUSG00000026674** | -0.107886327123038 | 0.532121363142167 | 0.909082872831377 | 438.204152535987 | 650.718167402549 | 467.548714992333 | 414.563598862107 | 493.826146314413 | 500.240632918209 | 562.182168900314 | 568.815395886298 | Ddr2 |
| **ENSMUSG00000026675** | -0.114550527016195 | 0.715978148301512 | 0.951348124093194 | 135.527057485357 | 175.913797445495 | 171.308334845369 | 105.985716451172 | 137.964732673206 | 83.1695917810225 | 233.201492284575 | 182.368294864309 | Hsd17b7 |
| **ENSMUSG00000026676** | -0.952054725805116 | 0.0516917964475454 | 0.703396849069134 | 19.877301764519 | 28.5045505120015 | 23.6613722162112 | 23.448167356454 | 51.4630352034976 | 91.7311674055395 | 23.9447960827911 | 18.4539345993646 | Ccdc3 |
| **ENSMUSG00000026678** | -0.129292036671216 | 0.254974816394894 | 0.837406774112203 | 12830.798288997 | 11884.7687320468 | 11058.3789189685 | 9877.3060172327 | 13666.1732420181 | 11964.1903941465 | 12104.6149588962 | 12196.9652446154 | Rgs5 |
| **ENSMUSG00000026687** | 0.100235442314554 | 0.505567492953263 | 0.901494649536949 | 618.003382133227 | 670.264144896492 | 874.524317111165 | 771.913669374467 | 707.342994499137 | 591.971800323748 | 751.658381381531 | 684.966631305828 | Aldh9a1 |
| **ENSMUSG00000026688** | 0.0904542533957681 | 0.556859963629722 | 0.916239900758826 | 9453.46401646192 | 7167.67283017501 | 7652.08777472269 | 10093.9670836063 | 8238.46546534289 | 9190.23989180298 | 7009.57878371447 | 7840.75115360063 | Mgst3 |
| **ENSMUSG00000026694** | -0.0043112677735279 | 0.978933323258058 | 0.996668984706452 | 126.491920319666 | 142.522752560008 | 111.681676860517 | 143.502784221499 | 119.350443344282 | 123.53130543946 | 136.381229862854 | 146.545951230249 | Mettl13 |
| **ENSMUSG00000026696** | -0.00563744323114103 | 0.964255421447312 | 0.994284908685797 | 518.616873310632 | 416.166437475222 | 462.81644054909 | 538.369922504185 | 502.585811880965 | 516.140701935169 | 465.361906478593 | 459.177313854779 | Vamp4 |
| **ENSMUSG00000026697** | -1.23312217264313 | 0.0371000350950428 | 0.679566075569698 | 77.7021796249379 | 54.56585383726 | 58.6802030962037 | 81.59962240046 | 130.300025302473 | 412.178712208891 | 49.9717483466946 | 48.8486504100828 | Myoc |
| **ENSMUSG00000026698** | -0.367575028331283 | 0.00669289300343051 | 0.635862253526407 | 196.062476495483 | 158.811067138294 | 159.004421292939 | 174.454365132018 | 212.421889988905 | 234.831788558181 | 224.872867560126 | 216.019587369033 | Pigc |
| **ENSMUSG00000026701** | 0.0810990859695756 | 0.399618330445796 | 0.880459952732461 | 6220.69193857787 | 5583.63423743664 | 5308.66547042914 | 6441.68053616505 | 5436.46744224182 | 5398.68497237402 | 5634.31462608981 | 5796.70651532983 | Prdx6 |
| **ENSMUSG00000026705** | 0.190417655650332 | 0.270769763586401 | 0.844129257012535 | 232.203025158245 | 235.366145656241 | 185.505158175096 | 212.909359596603 | 173.003394939417 | 155.33144347338 | 197.804837205666 | 232.302470839061 | Klhl20 |
| **ENSMUSG00000026707** | 0.281313557834726 | 0.171517244013667 | 0.799531027639052 | 61.4389327266951 | 58.6379324818317 | 76.6628459805242 | 64.7169419038131 | 49.2731188118594 | 55.0387004433237 | 57.2592949805875 | 53.1907526687569 | Nsun6 |
| **ENSMUSG00000026708** | -0.620768504995066 | 0.0997525417468387 | 0.759800953500741 | 19.877301764519 | 10.5874044758863 | 17.036187995672 | 12.1930470253561 | 22.994122112201 | 30.5770558018465 | 16.6572494488982 | 21.7105112933701 | Cenpl |
| **ENSMUSG00000026709** | 0.187968029273337 | 0.330185984621838 | 0.85612905103553 | 515.002818444356 | 563.575684408715 | 483.638448099356 | 471.777127211855 | 440.173194719277 | 299.655146858096 | 526.785513821405 | 517.795694346878 | Dars2 |
| **ENSMUSG00000026712** | -0.322179804703589 | 0.188971556958592 | 0.808677608936066 | 184.316798180085 | 272.014853457386 | 353.027673465871 | 279.502154888932 | 409.514365236342 | 412.178712208891 | 267.557069272927 | 272.466916731795 | Mrc1 |
| **ENSMUSG00000026715** | -0.336202933164627 | 0.246200086287279 | 0.832048466795696 | 289.124389302094 | 484.577358704026 | 280.15064703994 | 374.232751009006 | 314.253002200081 | 393.832478727783 | 383.116737324658 | 712.104770422541 | Serpinc1 |
| **ENSMUSG00000026718** | 0.0625143152851989 | 0.663301175278507 | 0.940031968637737 | 337.914129996823 | 346.12668478859 | 308.544293699394 | 284.191788360223 | 318.632834983357 | 260.516515431732 | 290.460787265162 | 352.795808517265 | Stam |
| **ENSMUSG00000026721** | 0.0899972597727306 | 0.512017460666699 | 0.902394714365738 | 1075.18132271716 | 1185.78930129926 | 1270.14246056622 | 1123.63617972128 | 940.569090208604 | 973.573456730792 | 1268.03311429737 | 1189.73601887668 | Rabgap1l |
| **ENSMUSG00000026723** | 0.115990417231727 | 0.657264981615604 | 0.938389079274206 | 127.395434036235 | 136.007426728693 | 102.217127974032 | 103.171936368398 | 107.305903190271 | 63.6002760678407 | 112.436433780063 | 148.717002359586 | Trdmt1 |
| **ENSMUSG00000026726** | 0.0376498274593145 | 0.915651604947208 | 0.986713918026868 | 52.4037955610046 | 65.1532583131463 | 35.0188308799925 | 58.151455044006 | 27.3739548954774 | 40.3617136584374 | 59.3414511616998 | 78.1578406561325 | Cubn |
| **ENSMUSG00000026727** | -0.156698078237097 | 0.434405893548929 | 0.888121557461676 | 412.002254755485 | 383.589808318649 | 388.992959234512 | 438.949692912819 | 539.814390538815 | 567.510155682271 | 326.898520434627 | 376.677370939972 | Rsu1 |
| **ENSMUSG00000026728** | -0.888681755318399 | 0.02624727730072 | 0.664353746352133 | 2793.66441163149 | 2685.12865823054 | 2423.87096982867 | 2752.8148476477 | 5922.6288811855 | 9187.79372733883 | 2362.20618747187 | 2256.80764894583 | Vim |
| **ENSMUSG00000026730** | 0.193766956376053 | 0.302202895596626 | 0.851785577030345 | 285.510334435818 | 317.622134276588 | 245.131816159948 | 368.605190843457 | 261.695008800764 | 206.700897220482 | 312.323427166841 | 282.236646813812 | Pter |
| **ENSMUSG00000026737** | -0.0799577337212536 | 0.662470887484883 | 0.939611397743053 | 232.203025158245 | 244.324718674299 | 260.275094378323 | 211.033506208086 | 304.398378437709 | 287.424324537357 | 213.421008564008 | 197.565652769668 | Pip4k2a |
| **ENSMUSG00000026739** | 0.195723899919633 | 0.310960744567566 | 0.851785577030345 | 1398.63923324888 | 1414.64012112419 | 1688.47552134883 | 1330.91797915233 | 1084.00861386091 | 940.550236464798 | 1471.04334195582 | 1595.72258006271 | Bmi1 |
| **ENSMUSG00000026740** | 0.0403386655958231 | 0.650281346710014 | 0.936463929813731 | 795.09207058076 | 812.7868974565 | 901.97150888197 | 874.147679048606 | 802.604357535398 | 834.142082274372 | 844.314331441027 | 808.716545678038 | Dnajc1 |
| **ENSMUSG00000026743** | 0.106795171753554 | 0.520700485128149 | 0.905962179970576 | 253.887354355902 | 274.458100644129 | 251.757000380487 | 262.619474392285 | 220.086597359638 | 195.693157131818 | 308.159114804616 | 243.157726485746 | Mllt10 |
| **ENSMUSG00000026748** | -0.582359449040311 | 0.0982035797051838 | 0.759800953500741 | 190.641394196068 | 188.13003337921 | 189.290977729689 | 181.957778686083 | 410.609323432161 | 409.732547744743 | 165.531416398426 | 137.8617467129 | Plxdc2 |
| **ENSMUSG00000026749** | -0.0607784001852738 | 0.763798382550258 | 0.95785116771606 | 106.614618555147 | 103.43079757212 | 95.5919437534931 | 142.564857527241 | 121.54035973592 | 139.43137445642 | 110.35427759895 | 96.6117752554972 | Nek6 |
| **ENSMUSG00000026750** | 0.114660332024017 | 0.175842718823242 | 0.799889021902745 | 5049.73816190439 | 4395.40168895063 | 4566.64483772876 | 4996.33550031322 | 4284.57142024013 | 4529.0735053695 | 4499.53950738362 | 4242.23390672453 | Psmb7 |
| **ENSMUSG00000026753** | 0.0225263911218322 | 0.792505734165563 | 0.961804768755116 | 902.610202852476 | 961.824975847822 | 904.810873547915 | 953.87144806055 | 916.480009900584 | 872.057631468662 | 896.368235968834 | 980.229584895662 | Ppp6c |
| **ENSMUSG00000026754** | 0.283606506415756 | 0.31084869072527 | 0.851785577030345 | 269.247087537575 | 324.137460107903 | 326.526936583714 | 260.743621003769 | 154.389105610493 | 163.893019097897 | 340.432535611857 | 310.460311495193 | Golga1 |
| **ENSMUSG00000026755** | 0.228615836594998 | 0.118009492539238 | 0.769868040577702 | 3170.42963144078 | 2358.5479509359 | 2272.43818764492 | 2974.16554749263 | 2057.42644994408 | 2298.17151406678 | 2389.27421782633 | 2451.11672502149 | Arpc5l |
| **ENSMUSG00000026761** | -0.0292319230818631 | 0.829276349975198 | 0.967483872587428 | 412.002254755485 | 386.033055505392 | 348.295399022628 | 374.232751009006 | 341.626957095558 | 351.024600605198 | 412.26692386023 | 446.151007078757 | Orc4 |
| **ENSMUSG00000026764** | -0.648494495097401 | 0.17847178525618 | 0.801696999069105 | 12.6491920319666 | 16.2883145782866 | 4.73227444324223 | 21.5723139679377 | 17.5193311331055 | 18.3462334811079 | 33.3144988977964 | 17.3684090346961 | Kif5c |
| **ENSMUSG00000026766** | 0.0749676727349954 | 0.397225656363122 | 0.8799282544021 | 3217.41234470237 | 2817.06400631466 | 2835.57884639075 | 3127.98552535097 | 2798.71314851361 | 2704.2348151153 | 2893.1560136555 | 2992.79398179108 | Mmadhc |
| **ENSMUSG00000026767** | -0.060737818289948 | 0.641608220321168 | 0.934557275614621 | 226.78194285883 | 206.047179415325 | 197.809071727525 | 217.598993067893 | 204.757182618171 | 218.931719541221 | 212.379930473452 | 248.585354309088 | Mindy3 |
| **ENSMUSG00000026768** | -0.465182508625208 | 0.195620451272643 | 0.811094945724481 | 30.7194663633475 | 69.2253369577179 | 31.2330113253987 | 46.8963347129081 | 82.1218646864323 | 74.6080161565054 | 51.0128264372507 | 39.0789203280663 | Itga8 |
| **ENSMUSG00000026770** | -0.427729143681066 | 0.345491156094488 | 0.861266047293794 | 20.780815481088 | 16.2883145782866 | 17.9826428843205 | 13.1309737196143 | 43.7983278327639 | 17.123151249034 | 17.6983275394543 | 13.0263067760221 | Il2ra |
| **ENSMUSG00000026771** | 0.225900197901797 | 0.299157721152523 | 0.851785577030345 | 295.448985318078 | 347.755516246418 | 309.490748588042 | 299.198615468353 | 221.181555555458 | 183.462334811079 | 345.637926064637 | 319.144516012541 | Spopl |
| **ENSMUSG00000026773** | -0.340423151278848 | 0.299194124732636 | 0.851785577030345 | 123.781379169959 | 148.223662662408 | 109.78876708322 | 93.7926694258161 | 169.71852035196 | 242.170281950624 | 87.4505596067155 | 104.210454208177 | Pfkfb3 |
| **ENSMUSG00000026775** | 0.0804336176186999 | 0.59051178776496 | 0.923935529355527 | 1601.02630576035 | 1604.39898596123 | 1698.88652512396 | 1480.04832353938 | 1507.7574356429 | 1144.80496922113 | 1671.97141343316 | 1711.87381548224 | Yme1l1 |
| **ENSMUSG00000026778** | -0.0141439901632508 | 0.956052137881078 | 0.992371505557225 | 114.746242004269 | 105.874044758863 | 148.593417517806 | 83.4754757889764 | 113.875652365186 | 103.961989726278 | 155.120635492864 | 83.5854684794751 | Prkcq |
| **ENSMUSG00000026781** | 0.026511910382407 | 0.887758761591548 | 0.980702840906172 | 685.766910875905 | 763.92195372164 | 736.341903368492 | 602.14893771374 | 559.523638063559 | 536.933099880424 | 843.273253350471 | 796.775764466685 | Acbd5 |
| **ENSMUSG00000026782** | -0.170833971748333 | 0.139472048320472 | 0.779711473022872 | 262.922491521592 | 268.757190541728 | 287.722286149128 | 255.11606083822 | 316.442918591719 | 286.201242305283 | 321.693129981846 | 284.407697943149 | Abi2 |
| **ENSMUSG00000026784** | 0.10482544236158 | 0.680981170668291 | 0.944452327903597 | 262.922491521592 | 200.346269312925 | 193.076797284283 | 238.233380341573 | 170.813478547779 | 129.646716599829 | 266.515991182371 | 263.782712214447 | Pdss1 |
| **ENSMUSG00000026785** | -0.543392011888094 | 0.0975167876497904 | 0.758828112300049 | 122.87786545339 | 87.1424829938332 | 110.735221971868 | 136.937297361692 | 209.137015401448 | 251.954939807215 | 96.8202624217207 | 109.638082031519 | Pkn3 |
| **ENSMUSG00000026786** | -0.61446373628091 | 0.0921449583856123 | 0.754058637722175 | 50.5967681278665 | 36.6487078011448 | 66.2518422053913 | 73.1582821521366 | 116.065568756824 | 123.53130543946 | 51.0128264372507 | 56.4473293627624 | Apbb1ip |
| **ENSMUSG00000026788** | 0.218946156309926 | 0.285070971457909 | 0.851275833092507 | 245.75573090678 | 341.240190415104 | 321.794662140472 | 261.681547698027 | 229.94122112201 | 184.685417043153 | 313.364505257397 | 276.809018990469 | Zbtb43 |
| **ENSMUSG00000026790** | 0.238180904176235 | 0.128428882356593 | 0.77303252008843 | 489.704434380422 | 454.443976734195 | 535.693466975021 | 505.542488205149 | 439.078236523458 | 346.132271676902 | 519.497967187512 | 376.677370939972 | Odf2 |
| **ENSMUSG00000026791** | 0.198229326810831 | 0.255524744059478 | 0.837479467570303 | 355.080890611635 | 375.445651029505 | 530.01473764313 | 412.687745473591 | 381.045452145046 | 423.186452297555 | 333.144988977964 | 321.315567141878 | Slc2a8 |
| **ENSMUSG00000026792** | 0.0542556827866389 | 0.788329006912788 | 0.961444367900485 | 322.554396815149 | 292.375246680244 | 290.561650815073 | 220.412773150668 | 254.03030143003 | 196.916239363891 | 313.364505257397 | 319.144516012541 | Lrsam1 |
| **ENSMUSG00000026796** | -0.134759690484451 | 0.145739767303356 | 0.780607990324772 | 974.891300178 | 967.525885950222 | 959.705257089525 | 1000.76778277346 | 1054.44474257379 | 1194.95134073616 | 1014.01006020168 | 1023.6506074824 | Fam129b |
| **ENSMUSG00000026797** | -0.27249262232899 | 0.130669157814673 | 0.77303252008843 | 204.194099944604 | 256.540954608014 | 201.594891282119 | 164.137171495178 | 250.745426842573 | 289.870489001505 | 245.694429371248 | 213.848536239696 | Stxbp1 |
| **ENSMUSG00000026798** | -0.0924356701891339 | 0.472052282901759 | 0.897171173241185 | 405.677658739501 | 316.807718547674 | 367.224496795597 | 377.046531091781 | 356.956371837026 | 406.063301048521 | 375.829190690765 | 424.440495785386 | Coq4 |
| **ENSMUSG00000026799** | 0.00043103104926646 | 0.998351238971074 | 0.999262355603273 | 201.483558794897 | 178.357044632238 | 194.96970706158 | 174.454365132018 | 218.991639163819 | 240.94719971855 | 143.668776496747 | 146.545951230249 | Med27 |
| **ENSMUSG00000026803** | 0.0895056778008299 | 0.570382177784812 | 0.920872458148558 | 249.369785773056 | 210.119258059897 | 229.042083052924 | 172.578511743502 | 191.617684268342 | 192.023910435596 | 206.133461930115 | 219.276164063039 | Ttf1 |
| **ENSMUSG00000026806** | 0.229564165944259 | 0.216633073591778 | 0.825523564386777 | 75.8951521917998 | 79.8127414336042 | 80.448665535118 | 86.2892558717508 | 70.0773245324222 | 59.9310293716191 | 79.1219348822664 | 65.1315338801105 | Ddx31 |
| **ENSMUSG00000026810** | 0.0945704716619977 | 0.604395495152042 | 0.926448151609826 | 170.764092431549 | 112.389370590177 | 133.450139299431 | 139.751077444466 | 141.249607260664 | 140.654456688494 | 119.723980413956 | 119.407812113536 | Dpm2 |
| **ENSMUSG00000026811** | 0.0194629641023152 | 0.905857089642254 | 0.98504921011344 | 1760.0447198765 | 1927.72203034022 | 1932.66088262013 | 1656.37854205991 | 1517.61205940527 | 1461.58326732826 | 2210.20878625068 | 1988.68283447271 | St6galnac6 |
| **ENSMUSG00000026812** | 0.439161698514298 | 0.199315063255019 | 0.812921911423739 | 795.09207058076 | 1170.31540244989 | 1184.01506569921 | 759.720622349111 | 522.295059405709 | 341.239942748607 | 1019.21545065446 | 999.769045059695 | Tsc1 |
| **ENSMUSG00000026814** | -0.746125755149251 | 0.045783092868592 | 0.697906909182231 | 2364.49539626119 | 2062.91504133999 | 2538.39201135513 | 2946.96567335914 | 5679.54816171366 | 6757.52933220807 | 2164.40135026621 | 2025.59070367144 | Eng |
| **ENSMUSG00000026816** | -0.172170897868529 | 0.196719093650872 | 0.812065429583558 | 252.983840639333 | 240.252640029727 | 192.130342395635 | 236.357526953057 | 257.315176017488 | 264.185762127954 | 250.899819824029 | 267.039288908453 | Gtf3c5 |
| **ENSMUSG00000026817** | -0.0574436525187659 | 0.592330225647178 | 0.923935529355527 | 14407.42972441 | 13050.1976401232 | 11223.0620695933 | 13205.0699284607 | 12427.7755225467 | 13721.7595616366 | 13969.1858190822 | 13875.1877675929 | Ak1 |
| **ENSMUSG00000026819** | 0.102455246242211 | 0.739141706629039 | 0.954383574136061 | 238.527621174228 | 298.076156782644 | 130.610774633486 | 139.751077444466 | 241.98576127602 | 209.14706168463 | 159.284947855089 | 142.203848971575 | Slc25a25 |
| **ENSMUSG00000026820** | -0.0860294817082779 | 0.325057030078529 | 0.855977678491063 | 1299.25272442629 | 1184.97488557035 | 1118.70967838246 | 1133.95337335812 | 1280.00613091252 | 1265.89011019644 | 1239.92400585236 | 1242.92677154544 | Ptges2 |
| **ENSMUSG00000026821** | -0.503986818512496 | 0.00306568046027161 | 0.635862253526407 | 164.439496415566 | 161.254314325037 | 160.897331070236 | 181.957778686083 | 282.499214521327 | 266.631926592101 | 198.845915296222 | 200.822229463674 | Ralgds |
| **ENSMUSG00000026822** | -0.404421464200717 | 0.153655992230755 | 0.785239783695802 | 181.606257030378 | 152.295741306979 | 117.360406192407 | 124.744250336335 | 210.231973597267 | 245.839528646846 | 206.133461930115 | 100.953877514171 | Lcn2 |
| **ENSMUSG00000026824** | -0.0050914985284990 | 0.978797403218279 | 0.996664169100645 | 1005.61076654135 | 1185.78930129926 | 873.577862222516 | 741.900015158206 | 732.527033002976 | 958.896469945906 | 1071.26935518226 | 1058.38742555179 | Kcnj3 |
| **ENSMUSG00000026825** | -0.926072840078981 | 0.0332547451980362 | 0.671606027118506 | 47.8862269781594 | 49.679359463774 | 49.2156542097192 | 76.9099889291692 | 125.920192519196 | 189.577745971448 | 39.5609674411332 | 70.559161703453 | Dnm1 |
| **ENSMUSG00000026826** | -0.449402057451357 | 0.113758202847515 | 0.767518180080285 | 23.4913566307952 | 33.3910448854875 | 33.1259211026956 | 26.2619474392285 | 49.2731188118594 | 44.0309603546589 | 30.191264626128 | 35.8223436340608 | Nr4a2 |

| | | | | | | | | | | | | |
|---|---|---|---|---|---|---|---|---|---|---|---|---|
| **ENSMUSG00000026827** | -0.16852778790614 | 0.228914758772791 | 0.832048466795696 | 196.062476495483 | 214.191336704468 | 193.076797284283 | 185.709485463116 | 252.935343234211 | 194.470074899744 | 209.256696201783 | 230.131419709724 | Gpd2 |
| **ENSMUSG00000026832** | -0.0960195370784475 | 0.861869880462064 | 0.974868041734905 | 27.1054114970714 | 29.3189662409158 | 11.3574586637814 | 15.9447538023887 | 35.0386622662111 | 31.8001380339204 | 17.6983275394543 | 5.42762782334254 | Cytip |
| **ENSMUSG00000026833** | -0.690152164222653 | 0.0945552031653238 | 0.757085157577544 | 74.9916384752307 | 56.1946852950887 | 74.7699362032273 | 79.7237690119437 | 137.964732673206 | 207.923979452556 | 63.5057635239243 | 52.1052271040884 | Olfm1 |
| **ENSMUSG00000026836** | -0.27417492805023 | 0.0416307462516152 | 0.684320787496965 | 213.229237110295 | 188.944449108124 | 178.879973954556 | 189.461192240149 | 249.650468646754 | 218.931719541221 | 212.379930473452 | 250.756405438425 | Acvr1 |
| **ENSMUSG00000026837** | -1.39765268418761 | 0.00096528983341887 | 0.556246415579446 | 157.211386683014 | 181.614707547895 | 149.539872406455 | 158.509611329629 | 597.847174917227 | 732.626257012242 | 195.722681024554 | 179.111718170304 | Col5a1 |
| **ENSMUSG00000026839** | -0.21583612066037 | 0.412478788294891 | 0.882192126163777 | 142.755167217909 | 129.492100897378 | 83.2880302010633 | 102.23400967414 | 98.5462376237187 | 176.123841418636 | 105.14888714617 | 153.05910461826 | Upp2 |
| **ENSMUSG00000026842** | -0.100749116467875 | 0.570192002181787 | 0.920872458148558 | 526.748496759753 | 708.541684155466 | 700.376617599851 | 519.611388619021 | 598.942133113046 | 545.494675504941 | 772.479943192653 | 715.361347116547 | Abl1 |
| **ENSMUSG00000026843** | 0.00947377576605879 | 0.957221325845185 | 0.992403427270738 | 151.7903043836 | 179.985876090067 | 194.96970706158 | 175.392291826276 | 203.662224422352 | 139.43137445642 | 157.202791673977 | 196.480127205 | Fubp3 |
| **ENSMUSG00000026848** | 0.0370251097066734 | 0.796506564619663 | 0.962278883153809 | 327.975479114563 | 444.670987987223 | 385.207139679918 | 372.35689762049 | 395.279908690694 | 348.57843614105 | 412.26692386023 | 335.427399482569 | Tor1b |
| **ENSMUSG00000026849** | -0.0647248760773294 | 0.754879833697597 | 0.956497786093635 | 190.641394196068 | 134.378595270864 | 154.272146849697 | 175.392291826276 | 187.237851485066 | 210.370143916704 | 124.929370866736 | 162.828834700276 | Tor1a |
| **ENSMUSG00000026851** | 0.0093773351554054 | 0.933546105451685 | 0.98818087308472 | 766.17963165055 | 743.561560498782 | 769.467824471187 | 877.899385825639 | 759.900987898453 | 739.964750404685 | 884.916376972716 | 750.098165185939 | BC005624 |
| **ENSMUSG00000026853** | 0.171972068714094 | 0.289113521082965 | 0.851785577030345 | 17320.3579466286 | 18435.9288554337 | 19952.2155075979 | 17149.9896045105 | 15100.5684785412 | 12189.2375248481 | 18133.4981813068 | 19246.3682615726 | Crat |
| **ENSMUSG00000026854** | -0.0818110415007706 | 0.682882450830803 | 0.944452327903597 | 805.934235179588 | 1008.24667239594 | 912.382512657103 | 849.761584997894 | 782.895110010654 | 693.487625585878 | 1199.32196032067 | 1108.32160152655 | Usp20 |
| **ENSMUSG00000026856** | -0.127863409290678 | 0.506405270074396 | 0.901689186353298 | 98.4829951060259 | 100.173134656462 | 100.324218196735 | 117.24083678227 | 136.869774477387 | 108.854318654574 | 86.4094815161593 | 122.664388807541 | Dolpp1 |
| **ENSMUSG00000026857** | -0.0529908070252082 | 0.737033889885277 | 0.954383574136061 | 334.300075130547 | 312.735639903102 | 296.240380146964 | 358.287997206618 | 293.448796479518 | 425.632616761703 | 311.282349076285 | 321.315567141878 | Ntmt1 |
| **ENSMUSG00000026858** | -0.0162981400933746 | 0.924473831218803 | 0.986713918026868 | 1115.83943996277 | 1125.5225373596 | 1214.30162213596 | 1039.22277723804 | 986.557334433006 | 869.611467004514 | 1334.66211209297 | 1353.65037914163 | Miga2 |
| **ENSMUSG00000026860** | -0.0170150061865106 | 0.870707219741228 | 0.975613767803763 | 1299.25272442629 | 1161.35682943183 | 1082.74439261382 | 1216.49092245284 | 1143.13635643514 | 1325.82113956806 | 1148.30913388342 | 1200.59127452337 | Sh3glb2 |
| **ENSMUSG00000026864** | -0.0246198622313924 | 0.792688010895139 | 0.961804768755116 | 11104.1835766336 | 10567.8584983923 | 11133.1488551717 | 9608.1210559806 | 9991.49353684926 | 10563.7612384219 | 10983.3738553672 | 11604.2682863063 | Hspa5 |
| **ENSMUSG00000026866** | -0.247732179219302 | 0.528068313062349 | 0.908232038364594 | 19.877301764519 | 54.56585383726 | 25.5542819935081 | 23.448167356454 | 33.943704070392 | 25.6847268735511 | 39.5609674411332 | 47.7631248454143 | Kynu |
| **ENSMUSG00000026867** | 0.0442169218706463 | 0.763780142001459 | 0.95785116771606 | 765.276117933981 | 993.587189275481 | 972.009170641955 | 752.217208795045 | 878.156473046916 | 724.064681387725 | 878.669908429379 | 896.644116416187 | Gapvd1 |
| **ENSMUSG00000026869** | -0.171041253602938 | 0.122194419513128 | 0.770251097497048 | 349.65980831222 | 337.982527499446 | 295.293925258315 | 320.770929436291 | 381.045452145046 | 353.470765069345 | 357.089785060755 | 376.677370939972 | Psmd5 |
| **ENSMUSG00000026870** | -0.249231069267242 | 0.570901809282893 | 0.921108854796072 | 17.1667606148119 | 15.4738988493722 | 14.1968233297267 | 15.0068271081306 | 14.2344565456483 | 20.7923979452556 | 8.32862472444909 | 30.3947158107182 | Cutal |
| **ENSMUSG00000026872** | 0.0866794699643027 | 0.604049956886433 | 0.926448151609826 | 445.432262268539 | 514.710740673856 | 536.639921863669 | 545.87333605825 | 391.995034103237 | 410.955629976817 | 553.853544175865 | 565.558819192293 | Zeb2 |
| **ENSMUSG00000026873** | 0.16128714811908 | 0.646319754266542 | 0.935063240511793 | 20.780815481088 | 25.2468875963442 | 26.5007368821565 | 14.0689004138724 | 24.0890803080201 | 19.5693157131818 | 17.6983275394543 | 16.2828834700276 | Phf19 |
| **ENSMUSG00000026874** | -0.393468129193859 | 0.362839064089025 | 0.86924083181804 | 60.535419010126 | 176.728213174409 | 65.3053873167428 | 85.3513291774927 | 79.9319482947941 | 91.7311674055395 | 109.313199508394 | 229.045894145055 | Hc |
| **ENSMUSG00000026875** | -1.13668151796062 | 0.0313845101292996 | 0.669064667366769 | 9.93865088225949 | 7.32974156022896 | 8.51809399783602 | 10.3171936368398 | 21.8991639163819 | 34.2463024980681 | 6.24646854333682 | 17.3684090346961 | Traf1 |
| **ENSMUSG00000026878** | 0.0299407791964254 | 0.785760732703514 | 0.961141387070613 | 2613.86518203425 | 2700.60255707991 | 2770.273459074 | 2608.37413673195 | 2676.07783058187 | 2189.31719541221 | 2816.11623495435 | 2789.80070119806 | Rab14 |
| **ENSMUSG00000026879** | -1.51224305710569 | 0.0170795424588615 | 0.642273034804458 | 9507.67483945606 | 8653.98153544366 | 7833.80711334319 | 11175.396562086 | 38784.5142541082 | 52725.851942472 | 7239.65704172737 | 7281.70548779635 | Gsn |
| **ENSMUSG00000026880** | -0.0493459240678248 | 0.600113422518517 | 0.925516139867954 | 4383.84855279301 | 4124.20125122216 | 4500.39299552336 | 3843.62359306994 | 4268.14704730284 | 4038.61753030788 | 4506.82705401752 | 4623.25337992317 | Stom |
| **ENSMUSG00000026883** | 0.0158622878088027 | 0.912559520342684 | 0.986226496507248 | 1232.39270940018 | 1656.52159261174 | 1613.7055851456 | 1258.69762369445 | 1570.17005280458 | 1443.23703384715 | 1289.89575419905 | 1395.9858761637 | Dab2ip |
| **ENSMUSG00000026885** | 0.460678984006961 | 0.0664287483462256 | 0.719725956118125 | 89.4478579403355 | 75.7406627890326 | 141.021778408619 | 88.1651092602672 | 75.5521155115177 | 61.154111603693 | 65.5879197050366 | 83.5854684794751 | Ttll11 |
| **ENSMUSG00000026887** | 0.0888559997444943 | 0.54775595390375 | 0.914699131434573 | 1115.83943996277 | 1152.39825641378 | 1059.08302039761 | 1167.71873435141 | 1036.92541144068 | 807.234273168747 | 1190.99333559622 | 1189.73601887668 | Mrrf |
| **ENSMUSG00000026888** | 0.041540567636377 | 0.697635720509091 | 0.944933386973863 | 3942.03034539074 | 4112.79943101736 | 3535.00900910195 | 3309.94330403705 | 3375.75611771028 | 3554.27696640664 | 3708.32015856096 | 3839.50392223251 | Grb14 |
| **ENSMUSG00000026889** | 0.124698891987349 | 0.372046176502886 | 0.873663802472542 | 787.863960848207 | 851.064436715473 | 720.252170261468 | 779.417082928532 | 659.164833883096 | 599.310293716191 | 795.383661184888 | 823.913903583397 | Rbm18 |
| **ENSMUSG00000026890** | -0.234154586880062 | 0.521412859984513 | 0.90609507420585 | 56.0178504272808 | 54.56585383726 | 151.432782183751 | 121.930470253561 | 134.679858085749 | 134.539045528125 | 84.3273253350471 | 97.6973008201657 | Lhx6 |
| **ENSMUSG00000026893** | 0.172231071918722 | 0.550667317468438 | 0.914699131434573 | 237.624107457659 | 287.488752306758 | 249.86409060319 | 219.47484645641 | 168.623562156141 | 121.085140975312 | 294.625099627387 | 297.434004719171 | Gca |
| **ENSMUSG00000026895** | -0.045853867254414 | 0.794716024130377 | 0.961804768755116 | 15862.9903218027 | 11887.2119792335 | 11732.2547996861 | 15051.847589455 | 14319.8632849222 | 17647.8535265937 | 11669.4443170437 | 12659.3991351641 | Ndufa8 |
| **ENSMUSG00000026896** | -0.021185396619592 | 0.938797283399562 | 0.989269716906553 | 157.211386683014 | 159.625482867208 | 159.004421292939 | 188.52326554589 | 224.466430142915 | 226.270212933664 | 112.436433780063 | 111.809133160856 | Ifih1 |
| **ENSMUSG00000026914** | 0.103497387386936 | 0.447826472962629 | 0.89108031640712 | 1314.61245760796 | 1060.36927904646 | 946.454888648447 | 1176.16007459973 | 1004.07666556611 | 974.796538962866 | 1043.16024673725 | 1163.68340532464 | Psmd14 |
| **ENSMUSG00000026915** | 0.204810268539906 | 0.354308099077424 | 0.863443308527191 | 213.229237110295 | 197.088606397268 | 263.114459044268 | 223.226553233442 | 167.528603960322 | 139.43137445642 | 266.515991182371 | 202.993280593011 | Strbp |
| **ENSMUSG00000026917** | 0.186150356738032 | 0.127879002966801 | 0.772950109316303 | 601.740135234984 | 708.541684155466 | 760.949730473351 | 713.762214330461 | 633.980795379257 | 536.933099880424 | 635.057635239243 | 640.46008315442 | Wdr5 |
| **ENSMUSG00000026918** | 0.0935453485884453 | 0.352789947946346 | 0.862728204353539 | 891.768038253647 | 794.05533569147 | 803.540200462531 | 855.389145163443 | 851.877476347257 | 763.203312814088 | 719.38496057429 | 800.03234116069 | Brd3 |
| **ENSMUSG00000026921** | -0.388830730281306 | 0.145171643603631 | 0.780607990324772 | 829.425591810383 | 518.782819318427 | 696.590798045257 | 843.196098138087 | 1161.75064576406 | 1273.22860358889 | 722.508194845959 | 624.177199684392 | Egfl7 |
| **ENSMUSG00000026922** | 0.11345975310962 | 0.365378878897918 | 0.86924083181804 | 1132.10268686101 | 965.897054492394 | 1190.64024991975 | 1350.61443973175 | 1055.53970076961 | 1053.07380181559 | 1104.58385408006 | 1073.58478345715 | Agpat2 |
| **ENSMUSG00000026923** | -0.405700814867995 | 0.113489245916065 | 0.767518180080285 | 645.108793630298 | 646.646088757977 | 573.551662520959 | 480.218467460179 | 1212.11872277174 | 704.495365674543 | 659.002431322034 | 531.907526687569 | Notch1 |
| **ENSMUSG00000026924** | 0.0639098950201851 | 0.774939412819326 | 0.959435639901991 | 958.628053279757 | 1269.67412137744 | 1316.51875010999 | 925.733647232805 | 1077.43886468599 | 703.272283442469 | 1359.64798626631 | 1135.45974064326 | Sec16a |
| **ENSMUSG00000026925** | 0.094704315263394 | 0.673710074492864 | 0.943734776094628 | 395.739007857242 | 491.09268453534 | 366.278041906949 | 336.71568323868 | 292.353838283699 | 286.201242305283 | 504.922873919726 | 404.901035621353 | Inpp5e |
| **ENSMUSG00000026926** | 0.182426866691856 | 0.348105840401402 | 0.861266047293794 | 3235.48261903375 | 3598.0886903435 | 3729.03226127488 | 3411.23938701693 | 2516.21393399228 | 2256.58671817627 | 3766.6205316321 | 3773.28686278773 | Pmpca |
| **ENSMUSG00000026927** | 0.121375844011603 | 0.307897231499997 | 0.851785577030345 | 679.442314859922 | 565.204515866544 | 686.179794270124 | 719.38977449601 | 587.992551154855 | 611.54111603693 | 660.043509412591 | 575.328549274309 | Sdccag3 |
| **ENSMUSG00000026928** | 0.223930896290796 | 0.587321737205325 | 0.923058967588378 | 17.1667606148119 | 21.1748089517725 | 11.3574586637814 | 18.7585338851632 | 7.66470737073368 | 18.3462334811079 | 17.6983275394543 | 15.1973579053591 | Card9 |
| **ENSMUSG00000026930** | 0.0202035020607435 | 0.884195761825734 | 0.979384300778265 | 2793.66441163149 | 2404.96964748401 | 2333.01130051842 | 2509.89183383484 | 2397.95844884382 | 1996.07020274454 | 2764.06233042654 | 2742.03757635265 | Gpsm1 |
| **ENSMUSG00000026932** | -0.0515716382469485 | 0.793402274819928 | 0.961804768755116 | 1114.9359262462 | 1162.17124516075 | 1004.188636856 | 850.699511692152 | 832.168228822514 | 873.280713700736 | 1284.69036374627 | 1291.77542195552 | Nacc2 |
| **ENSMUSG00000026933** | 0.189168214198925 | 0.412782499132875 | 0.882192126163777 | 1001.99671167507 | 1209.40735743778 | 1054.35074595437 | 857.264998551959 | 760.995946094272 | 594.417964787896 | 1117.07679116673 | 1143.05841959594 | Camsap1 |
| **ENSMUSG00000026938** | -0.631466164242309 | 0.173488003638146 | 0.799889021902745 | 15.3597331816738 | 13.8450673915436 | 34.0723759913441 | 27.1998741334867 | 29.5638712871156 | 52.592535979176 | 14.5750932677859 | 43.4210225867403 | Fcna |
| **ENSMUSG00000026939** | -0.030959490817484 | 0.882646406819293 | 0.978917166298306 | 674.924746277077 | 469.103459854653 | 379.528410348027 | 558.066383083606 | 478.496731572945 | 660.464405319884 | 462.238672206925 | 526.479898864226 | Tmem141 |
| **ENSMUSG00000026942** | -0.128782516501418 | 0.383687436514609 | 0.875126835008975 | 297.256012751216 | 241.067055758641 | 260.275094378323 | 247.612647284155 | 305.493336633528 | 326.562955963721 | 243.612273190136 | 269.21034003779 | Traf2 |
| **ENSMUSG00000026944** | 0.145377537084041 | 0.546811323862484 | 0.914451747770672 | 1569.40332568043 | 1978.2158055329 | 1694.15425068072 | 1494.11722395325 | 1257.01200880032 | 957.673387713832 | 2024.89688613169 | 1849.73556219514 | Abca2 |
| **ENSMUSG00000026946** | 0.0282176315560681 | 0.905627777616245 | 0.98504921011344 | 190.641394196068 | 165.326392969609 | 160.897331070236 | 244.79886720138 | 206.947099009809 | 249.508775343067 | 145.750932677859 | 145.46042566558 | Nmi |
| **ENSMUSG00000026950** | 1.37221518548646 | 0.00145393048975175 | 0.631096703207871 | 44.2721721118832 | 232.108482740584 | 103.163582862681 | 150.068271081306 | 59.1277425742312 | 57.4848649074714 | 42.6842017128016 | 45.5920737160773 | Neb |
| **ENSMUSG00000026956** | 0.037604639657273 | 0.867434614666944 | 0.974868041734905 | 165.343010132135 | 155.553404222637 | 143.861143074564 | 189.461192240149 | 200.377349834895 | 189.577745971448 | 140.545542225078 | 107.467030902182 | Uap1l1 |
| **ENSMUSG00000026958** | -0.881055634280629 | 0.0432128226665196 | 0.687730553581215 | 41.5616309621761 | 26.8757190541728 | 29.3401015481019 | 42.2067012416173 | 82.1218646864323 | 110.077400886647 | 28.1091084450157 | 37.9933947633978 | Dpp7 |
| **ENSMUSG00000026960** | -0.11206177965755 | 0.546230370333878 | 0.914211323944814 | 227.685456575399 | 267.1283590839 | 169.415425068072 | 220.412773150668 | 258.410134213307 | 190.800828203522 | 254.023054095697 | 252.927456567762 | Arl6ip6 |
| **ENSMUSG00000026965** | -0.0319527367633675 | 0.793334833748669 | 0.961804768755116 | 1086.92700103256 | 1184.16046984143 | 1131.01359193489 | 1146.14642038347 | 1297.52546204563 | 1291.57483707 | 1036.91377819391 | 1025.82165861174 | Anapc2 |
| **ENSMUSG00000026966** | -0.210782869728051 | 0.18122840670642 | 0.803056849297055 | 481.572810931301 | 365.672662282534 | 345.456034356683 | 412.687745473591 | 474.116898789669 | 541.82542880872 | 417.472314313011 | 425.526021350055 | Ssna1 |
| **ENSMUSG00000026970** | -0.183995211766215 | 0.229650062957372 | 0.832048466795696 | 735.460165287203 | 697.95427967958 | 586.802030962037 | 783.168789705565 | 814.648897689408 | 955.227223249684 | 690.234774038719 | 726.216602763232 | Rbms1 |
| **ENSMUSG00000026971** | -0.0417101371587033 | 0.895236318776344 | 0.983733637008786 | 130.105975185942 | 118.090280692578 | 82.3415753124149 | 117.24083678227 | 84.3117810780705 | 75.8310983885793 | 107.231043327282 | 193.223550510994 | Itgb6 |
| **ENSMUSG00000026972** | -0.163604522227948 | 0.697277104958296 | 0.944933386973863 | 17.1667606148119 | 30.9477976987445 | 31.2330113253987 | 27.1998741334867 | 39.4184950494875 | 46.4771248188067 | 15.616171358342 | 18.4539345993646 | Arrdc1 |
| **ENSMUSG00000026974** | 0.235347920630619 | 0.0216597425050298 | 0.648762042745232 | 422.844419354313 | 468.289044125739 | 466.602260103684 | 463.335786963532 | 413.894198019619 | 369.370834086306 | 374.788112600209 | 388.618152151326 | Zmynd19 |
| **ENSMUSG00000026975** | -0.447047734387423 | 0.167006418396132 | 0.797702596809586 | 37.0440623793308 | 36.6487078011448 | 45.4298346551254 | 51.5859681841989 | 71.1722827282413 | 84.3926740130963 | 37.4788112600209 | 40.1644458927348 | Dph7 |
| **ENSMUSG00000026977** | 0.0831367959722341 | 0.618584965550469 | 0.929568738467614 | 398.449549006949 | 456.887223920938 | 558.408384302584 | 504.604561510891 | 440.173194719277 | 371.816998550453 | 542.401685179747 | 454.835211596105 | March7 |
| **ENSMUSG00000026979** | -0.34078599455744 | 0.446108533798443 | 0.89108031640712 | 8.13162344912141 | 12.2162359337149 | 16.0897331070236 | 10.3171936368398 | 17.5193311331055 | 20.7923979452556 | 12.4929370866736 | 8.68420451734806 | Psd4 |
| **ENSMUSG00000026986** | 0.139681065088626 | 0.404838818737195 | 0.881158207349486 | 215.036264543433 | 258.169786065842 | 331.259211026956 | 309.515809105193 | 244.175677667659 | 240.94719971855 | 260.269522639034 | 264.868237779116 | Hnmt |
| **ENSMUSG00000026987** | 0.157302282604673 | 0.262461893864788 | 0.841952818014037 | 470.730646332472 | 508.195414842541 | 456.191256328551 | 463.335786963532 | 450.027818481649 | 335.124531588238 | 434.129563761909 | 481.973350712817 | Baz2b |
| **ENSMUSG00000026988** | -0.132447027242185 | 0.237079657293244 | 0.832048466795696 | 268.343573821006 | 284.231089391101 | 306.651383922097 | 264.495327780801 | 307.683253025166 | 315.555215875056 | 312.323427166841 | 296.348479154503 | Wdsub1 |
| **ENSMUSG00000026991** | 0.154538503949657 | 0.155625426647573 | 0.787275540343856 | 7257.02217148257 | 6850.05069589842 | 7477.94007521138 | 6211.8884960718 | 5950.00283608097 | 5678.77080351893 | 6899.22450611552 | 6443.67975187226 | Pkp4 |
| **ENSMUSG00000026999** | -0.256230087085163 | 0.216434903131373 | 0.825447777181429 | 76.7986659083688 | 54.56585383726 | 63.4124775394459 | 75.9720622349111 | 95.2613630362614 | 72.1618516923577 | 79.1219348822664 | 75.9867895267955 | Nup35 |
| **ENSMUSG00000027001** | 0.111228022693603 | 0.500445923612044 | 0.900927059989498 | 257.501409222178 | 200.346269312925 | 192.130342395635 | 196.026679099956 | 171.908436743598 | 190.800828203522 | 227.996101831794 | 192.138024946326 | Dusp19 |
| **ENSMUSG00000027002** | 0.061962121879403 | 0.742658051882745 | 0.954383574136061 | 1933.51935345776 | 2307.23976001429 | 2170.22105967089 | 1790.50205933883 | 1734.41378217745 | 1443.23703384715 | 2403.84931109412 | 2274.17605798052 | Nckap1 |
| **ENSMUSG00000027004** | -0.947862194390545 | 0.101075277027572 | 0.759800953500741 | 37.9475760958999 | 31.7622134276588 | 60.5731128735006 | 60.9652351267805 | 101.831112211176 | 203.031650524261 | 43.7252798033577 | 20.6249857287016 | Frzb |
| **ENSMUSG00000027006** | 0.0168738330147804 | 0.854049472612726 | 0.9731539929935 | 764.372604217412 | 752.52013351684 | 860.327493781438 | 762.534402431885 | 798.224524752122 | 783.995710759344 | 772.479943192653 | 747.927114056602 | Dnajc10 |
| **ENSMUSG00000027007** | -0.00980416282334418 | 0.938684441848603 | 0.989269716906553 | 662.27555424511 | 791.612088504727 | 745.806452254976 | 594.645524159674 | 745.666531352805 | 647.010500767072 | 712.097413940397 | 708.848193728535 | Ssfa2 |
| **ENSMUSG00000027009** | -0.361437859074501 | 0.336483715972032 | 0.858570960161243 | 15.3597331816738 | 21.1748089517725 | 12.3039135524298 | 10.3171936368398 | 17.5193311331055 | 20.7923979452556 | 17.6983275394543 | 20.6249857287016 | Itga4 |
| **ENSMUSG00000027010** | 0.107247716865653 | 0.503218001077724 | 0.900927059989498 | 6031.85757181495 | 6503.92401110983 | 6464.28688946889 | 5646.31869943413 | 5448.51198239583 | 4280.78781225851 | 6615.01018739369 | 6534.86389930442 | Slc25a12 |
| **ENSMUSG00000027011** | 0.12244339222008l | 0.14263345882893 | 0.780607990324772 | 1150.17296119239 | 1063.62694196211 | 1059.08302039761 | 1051.4158242634 | 990.937167216283 | 995.588936908122 | 945.298906224972 | 1041.0190165171 | Ube2e3 |
| **ENSMUSG00000027012** | -0.032648774776299 | 0.676714627413242 | 0.94409858171788 | 1407.67437041457 | 1427.67077278682 | 1510.54200228292 | 1356.2419998973 | 1465.05406600595 | 1514.17580330744 | 1410.66081270357 | 1443.74900100912 | Dync1i2 |
| **ENSMUSG00000027014** | 0.267639293889861 | 0.0398252760195885 | 0.679566075569698 | 297.256012751216 | 292.375246680244 | 247.024725937245 | 287.005568442997 | 235.416012101106 | 210.370143916704 | 231.119336103462 | 256.184033261768 | Cwc22 |
| **ENSMUSG00000027016** | 0.222691646236872 | 0.386173027293557 | 0.875126835008975 | 112.035700854562 | 144.151584017836 | 86.1273948670087 | 148.192417692789 | 71.1722827282413 | 101.51582526213 | 121.806136595068 | 125.920965501547 | Zfp385b |

| | | | | | | | | | | | | |
|---|---|---|---|---|---|---|---|---|---|---|---|---|
| **ENSMUSG00000027018** | 0.0849382074639629 | 0.450512884196007 | 0.89108031640712 | 551.143367107117 | 557.874774306315 | 519.603733867997 | 491.473587791276 | 486.161438943679 | 448.871179171106 | 553.853544175865 | 509.11148982953 | Hat1 |
| **ENSMUSG00000027022** | 0.39515272456089 | 0.202259506367883 | 0.81455046200345 | 25497.1570815785 | 28081.868748695 | 16706.8216944224 | 24431.1145320366 | 12123.377144109 | 10404.7605482523 | 24959.8472210834 | 24535.0488126376 | Xirp2 |
| **ENSMUSG00000027030** | -0.0386563151161181 | 0.844125172764099 | 0.971721411930626 | 1845.87852295056 | 2194.85038942412 | 1688.47552134883 | 1278.39408427387 | 1674.1910814074 | 1575.32991491113 | 2251.85190987292 | 1696.67645757688 | Stk39 |
| **ENSMUSG00000027035** | -0.0543244802127986 | 0.852989967740781 | 0.973120292022893 | 69.5705561758165 | 81.4415728914329 | 77.6093008691726 | 40.3308478531009 | 63.5075753575076 | 48.9232892829544 | 91.61487196894 | 74.901263962127 | Cers6 |
| **ENSMUSG00000027048** | -0.0743468442033716 | 0.856694642539394 | 0.973691889769428 | 48.7897406947284 | 114.018202048006 | 46.3762895437739 | 63.779015209555 | 60.2227007700503 | 52.592535979176 | 46.8485140750261 | 128.092016630884 | Abcb11 |
| **ENSMUSG00000027067** | -0.0488793573583978 | 0.704210020277218 | 0.946931219685582 | 866.469654189714 | 899.114964721419 | 943.615523982501 | 953.87144806055 | 1055.53970076961 | 1067.75078860048 | 843.273253350471 | 823.913903583397 | Ssrp1 |
| **ENSMUSG00000027071** | 0.159195798084117 | 0.794558850392414 | 0.961804768755116 | 13.5527057485357 | 15.4738988493722 | 11.3574586637814 | 6.56548685980713 | 8.75966556655277 | 2.44616446414772 | 21.8626399016789 | 8.68420451734806 | P2rx3 |
| **ENSMUSG00000027074** | -0.728163267780928 | 0.0502527961203244 | 0.703396849069134 | 1302.86677929256 | 1176.8307282812 | 1363.84149454241 | 1220.24262922987 | 3047.26865896455 | 3247.2833261561 | 1159.76099287954 | 934.637511179585 | Slc43a3 |
| **ENSMUSG00000027075** | -0.773100041866478 | 0.0507875819339576 | 0.703396849069134 | 25.2983840639333 | 23.6180561385155 | 29.3401015481019 | 30.0136542162612 | 74.4571573156986 | 57.4848649074714 | 22.903717992235 | 30.3947158107182 | Slc43a1 |
| **ENSMUSG00000027076** | 0.210132368110792 | 0.0698631431676925 | 0.724916894583216 | 1489.89411862236 | 1162.98566088966 | 1235.12362968622 | 1451.91052271163 | 1195.69434983445 | 1138.68955806076 | 1138.93943106841 | 1141.97289403127 | Timm10 |
| **ENSMUSG00000027078** | -0.018233882743145 | 0.979682210248477 | 0.996872525500296 | 434.590097669711 | 241.881471487556 | 1020.27836996303 | 249.488500672671 | 369.000911991036 | 1113.00483118721 | 211.338852382896 | 277.894544555138 | Ube2l6 |
| **ENSMUSG00000027079** | 0.00076900761740484 | 0.996633613455079 | 0.999262355603273 | 122.87786545339 | 127.86326943955 | 171.308334845369 | 159.447538023887 | 152.199189218854 | 165.116101329971 | 122.847214685624 | 141.118323406906 | Clp1 |
| **ENSMUSG00000027080** | -0.0644116396773365 | 0.600219538367929 | 0.925516139867954 | 288.220875585525 | 283.416673662186 | 259.328639489674 | 297.322762079837 | 272.644590758955 | 332.67836712409 | 295.666177717943 | 280.065595684475 | Med19 |
| **ENSMUSG00000027082** | 0.319178893128634 | 0.132172029073731 | 0.77303252008843 | 838.460728976074 | 849.435605257645 | 1407.37841942024 | 1085.18118525669 | 733.621991198795 | 694.710707817952 | 947.381062406084 | 973.716431507651 | Tfpi |
| **ENSMUSG00000027086** | 0.264908233172616 | 0.267438449296505 | 0.843793744782845 | 452.660372001092 | 496.79359463774 | 468.495169880981 | 360.163850595134 | 393.089992299056 | 212.816308380852 | 426.842017128016 | 446.151007078757 | Fastkd1 |
| **ENSMUSG00000027087** | -0.244606487290336 | 0.0661798530802035 | 0.719725956118125 | 375.861706092723 | 427.568257680022 | 338.830850136144 | 347.970803569778 | 502.585811880965 | 436.640356850368 | 416.431236222455 | 411.414189009364 | Itgav |
| **ENSMUSG00000027088** | 0.142642225448769 | 0.404874751153036 | 0.881158207349486 | 849.302893574902 | 720.757920089181 | 810.16538468307 | 814.120370616084 | 678.87408140784 | 530.817688720055 | 851.60187807492 | 830.427056971408 | Phospho2 |
| **ENSMUSG00000027091** | 0.171914574491204 | 0.0220389351600589 | 0.648762042745232 | 1957.91422380512 | 1866.64085067164 | 1953.48289017039 | 1946.19789058568 | 1691.7104125405 | 1692.74580919022 | 1827.09204892602 | 1643.48570490812 | Zc3h15 |
| **ENSMUSG00000027099** | 0.110551457801122 | 0.447423091655622 | 0.89108031640712 | 1923.5807025755 | 1636.16119938889 | 1502.02390828509 | 1761.42633181683 | 1509.94735203453 | 1290.35175483792 | 1730.2717865043 | 1787.86060500903 | Mtx2 |
| **ENSMUSG00000027102** | -1.3123092004729 | 0.0184527157516351 | 0.642273034804458 | 4.51756858284523 | 7.32974156022896 | 6.62518422053913 | 11.2551203310979 | 17.5193311331055 | 35.4693847301419 | 10.4107809055614 | 10.8552556466851 | Hoxd8 |
| **ENSMUSG00000027104** | 0.162744399710097 | 0.240589487233235 | 0.832048466795696 | 692.995020608457 | 808.714818811928 | 782.718192912266 | 710.010507553428 | 584.707676567398 | 585.856389163379 | 762.069162287092 | 741.413960668591 | Atf2 |
| **ENSMUSG00000027108** | 0.123405410573821 | 0.312037891857754 | 0.851810228496481 | 1030.90915060528 | 976.48445896828 | 857.488129115493 | 956.685228143324 | 826.693437843418 | 770.541806206532 | 925.518422504405 | 984.571687154336 | Ola1 |
| **ENSMUSG00000027109** | 0.0507156669453277 | 0.676532839730867 | 0.94409858171788 | 657.757985662265 | 639.316347197748 | 736.341903368492 | 615.279911433354 | 675.589206820383 | 549.163922201163 | 637.139791420356 | 693.650835823176 | Sp3 |
| **ENSMUSG00000027111** | -0.47493610212771 | 0.037911585795723 | 0.679566075569698 | 672.214205127369 | 611.626212414661 | 551.783200082044 | 757.844768960594 | 1260.29688338778 | 982.135032355309 | 655.879197050366 | 706.677142599198 | Itga6 |
| **ENSMUSG00000027115** | -0.168690268628169 | 0.724067203703929 | 0.951674101666722 | 6.32459601598331 | 16.2883145782866 | 12.3039135524298 | 9.37926694258161 | 16.4243729372865 | 14.6769867848863 | 11.4518589961175 | 7.59867895267955 | Kif18a |
| **ENSMUSG00000027122** | -0.0610512654096277 | 0.72145362435664 | 0.951674101666722 | 121.070838020252 | 118.904696421492 | 145.754052851861 | 125.682177030594 | 112.780694169367 | 127.200552135681 | 133.257995591185 | 159.572258006271 | Arl14ep |
| **ENSMUSG00000027130** | 0.0449188373442901 | 0.753259209700634 | 0.956497786093635 | 428.265501653727 | 426.753841951108 | 411.707876562074 | 455.832373409466 | 416.084114411257 | 327.786038195794 | 476.813765474711 | 447.236532643425 | Slc12a6 |
| **ENSMUSG00000027131** | -0.0749259046331521 | 0.635604669282971 | 0.932560352417921 | 1181.79594127231 | 934.949256793649 | 1065.70820461815 | 1308.40773849013 | 1169.4153531348 | 1432.22929375849 | 1056.69426191448 | 1072.49925789249 | Emc4 |
| **ENSMUSG00000027132** | -0.60016362939617 | 0.00397750812185222 | 0.635862253526407 | 72.2810973255236 | 55.3802695661743 | 70.0376617599851 | 59.0893817382642 | 112.780694169367 | 102.738907494204 | 74.9576225200418 | 98.7828263848342 | Katnbl1 |
| **ENSMUSG00000027133** | 0.0632661499679763 | 0.755550289568648 | 0.956566265296272 | 1187.21702357173 | 860.023009733531 | 1028.79646396086 | 1305.59395840736 | 1076.34390649017 | 1364.95977099443 | 945.298906224972 | 807.63102011337 | Nop10 |
| **ENSMUSG00000027134** | -0.528050367342881 | 0.135941011825196 | 0.776087404131513 | 52.4037955610046 | 44.7928650902881 | 41.6440151005317 | 25.3240207449704 | 73.3621991198795 | 85.6157562451702 | 39.5609674411332 | 39.0789203280663 | Lpcat4 |
| **ENSMUSG00000027160** | -0.174828109639945 | 0.513533672013571 | 0.903136226008053 | 168.957064998411 | 138.450673915436 | 143.861143074564 | 151.006197775564 | 160.958854785407 | 270.301173288323 | 122.847214685624 | 127.006491066215 | Ccdc34 |
| **ENSMUSG00000027162** | 0.155146104172846 | 0.268965895832503 | 0.843793744782845 | 1013.74238999047 | 1133.66669464875 | 1113.03094905057 | 1042.97448401508 | 855.162350934715 | 820.68817772156 | 1033.79054392224 | 1153.91367524262 | Lin7c |
| **ENSMUSG00000027163** | -0.213777774540851 | 0.198534235068247 | 0.812921911423739 | 394.835494140673 | 294.818493866987 | 335.04503058155 | 389.239578117137 | 379.950493949227 | 508.802208542726 | 369.582722147429 | 382.104998763315 | Commd9 |
| **ENSMUSG00000027164** | -0.0282312333132341 | 0.907031387832003 | 0.98504921011344 | 206.001127377742 | 294.004078138073 | 265.953823710214 | 193.212899017181 | 217.896680968 | 163.893019097897 | 283.173240631269 | 312.63136262453 | Traf6 |
| **ENSMUSG00000027165** | 0.070885446045931 | 0.512050015056547 | 0.902394714365738 | 1816.06257030378 | 1571.82235680465 | 1641.15277691641 | 1943.38411050291 | 1593.16417491679 | 1802.82321007687 | 1603.26025945645 | 1639.14360264945 | B230118H07 |
| **ENSMUSG00000027166** | 0.0347972420847242 | 0.843537964579736 | 0.971654415184575 | 86.7373167906283 | 110.760539132349 | 97.48485353079 | 105.047789756914 | 107.305903190271 | 83.1695917810225 | 104.107809055614 | 95.5262496908287 | Dnajc24 |
| **ENSMUSG00000027167** | -0.223288095064894 | 0.264544923269299 | 0.842510961779757 | 159.921927832721 | 120.533527879321 | 128.717864856189 | 132.247663890401 | 170.813478547779 | 110.077400886647 | 171.777884941763 | 178.026192605635 | Elp4 |
| **ENSMUSG00000027170** | 0.217062955304581 | 0.025310648174902 | 0.658361241852789 | 3712.53786138221 | 3089.89327550096 | 3401.55886980252 | 3650.41069405276 | 2947.62746314501 | 2832.65844948306 | 2959.7850114511 | 3177.33332778472 | Eif3m |
| **ENSMUSG00000027171** | 0.209306115976736 | 0.619684188931816 | 0.929594515086012 | 20.780815481088 | 27.6901347830872 | 30.2865564367503 | 24.3860940507122 | 19.7092475247437 | 7.33849339244316 | 33.3144988977964 | 28.2236646813812 | Prrg4 |
| **ENSMUSG00000027173** | -0.649340671204462 | 0.0754150836637437 | 0.735399943172214 | 30.7194663633475 | 23.6180561385155 | 11.3574586637814 | 37.5170677703264 | 35.0386622662111 | 41.5847958905112 | 35.3966550789086 | 49.9341759747513 | Depdc7 |
| **ENSMUSG00000027175** | 0.0703638029251166 | 0.78142292980886 1 | 0.960742105865155 | 80.412720774645 | 68.4109212288036 | 61.519567762149 | 84.4134024832345 | 52.5579933993167 | 99.0696607979826 | 57.2592949805875 | 72.73021283279 | Tcp11l1 |
| **ENSMUSG00000027176** | -0.146423501747296 | 0.261292856614756 | 0.841102613761057 | 182.509770746947 | 178.357044632238 | 169.415425068072 | 179.143998603309 | 202.567266226533 | 177.34692365071 | 214.462086654564 | 189.966973816989 | Cstf3 |
| **ENSMUSG00000027177** | 0.266784971823677 | 0.339468587345102 | 0.858570960161243 | 6154.73543726833 | 8133.5698846674 | 7854.62912089346 | 5582.53968422458 | 4400.63698899695 | 3384.26853614837 | 7967.37062702611 | 7291.47521787837 | Hipk3 |
| **ENSMUSG00000027180** | 0.191043453094418 | 0.375699175074715 | 0.873821423608136 | 1854.91366011625 | 2182.6341534904 | 1991.34108571633 | 1687.33012297043 | 1559.22047084639 | 1084.87393984951 | 2087.36157156505 | 2026.6762292361 | Fbxo3 |
| **ENSMUSG00000027184** | 0.123466912475731 | 0.419155844538177 | 0.882550938324998 | 3156.87692569224 | 3541.89400504842 | 3114.78303854204 | 2850.35922385055 | 2626.80471177001 | 2345.87172111766 | 3367.8876229491 | 3283.71483312224 | Caprin1 |
| **ENSMUSG00000027185** | 0.0759748166822734 | 0.721076529787315 | 0.951674101666722 | 230.395997725106 | 320.065381463331 | 335.991485470199 | 280.44008158319 | 274.834507150593 | 188.354663739374 | 327.939598525183 | 314.802413753867 | Nat10 |
| **ENSMUSG00000027187** | 0.199705018029162 | 0.114602679431547 | 0.767518180080285 | 9748.91300178 | 11634.7431032701 | 11684.9320552537 | 11333.9061734156 | 9980.54395489107 | 8193.42787266279 | 11040.6331503478 | 9446.24346374535 | Cat |
| **ENSMUSG00000027188** | -1.32864622935598 | 0.141765770006718 | 0.779852097185473 | 19.877301764519 | 4.88649437348597 | 15.1432782183751 | 14.0689004138724 | 25.1840385038392 | 99.0696607979826 | 10.4107809055614 | 1.08552556466851 | Pamr1 |
| **ENSMUSG00000027189** | 0.319137292763687 | 0.064570659792583 | 0.716991331547963 | 1301.05975185942 | 1440.70142444945 | 1585.31193848615 | 1214.61506906432 | 1015.0262475243 | 852.48831575548 | 1313.84055028184 | 1259.20965501547 | Trim44 |
| **ENSMUSG00000027193** | -0.0741485603032072 | 0.602269693106429 | 0.926448151609826 | 748.109357319169 | 943.907829811707 | 956.86589242358 | 725.955261355817 | 856.257309130534 | 797.449615312156 | 935.929203409967 | 962.861175860966 | Api5 |
| **ENSMUSG00000027194** | 0.081057423770564 | 0.644254496122304 | 0.934857223264556 | 594.512025502432 | 743.561560498782 | 694.69788826796 | 550.562969529541 | 613.176589658694 | 457.432754795624 | 693.358008310387 | 677.367952353149 | Ttc17 |
| **ENSMUSG00000027195** | -0.190952343950395 | 0.147053958601602 | 0.780607990324772 | 388.510898124689 | 345.312269059675 | 300.026199701558 | 348.908730264036 | 395.279908690694 | 443.978850242811 | 372.705956419097 | 367.993166422624 | Hsd17b12 |
| **ENSMUSG00000027198** | -0.105381347860934 | 0.298695835660647 | 0.851785577030345 | 1148.36593375926 | 1151.58384068486 | 1162.24660326029 | 1103.0017924476 | 1327.08933333275 | 1297.69024823037 | 1215.97920976957 | 1071.41373232782 | Ext2 |
| **ENSMUSG00000027199** | 0.189568361685373 | 0.628783673073155 | 0.931825800800958 | 18.9737880479499 | 17.9171460361152 | 17.036187995672 | 36.5791410760683 | 19.7092475247437 | 26.9078091056249 | 17.6983275394543 | 15.1973579053591 | Gatm |
| **ENSMUSG00000027200** | -0.460364347004002 | 0.0027724051446851 | 0.635862253526407 | 244.852217190211 | 237.809392842984 | 222.416898832385 | 259.805694309511 | 390.900075907418 | 264.185762127954 | 347.72008224575 | 323.486618271215 | Sema6d |
| **ENSMUSG00000027201** | -0.0777993192888448 | 0.665920769109236 | 0.942024383071524 | 75.8951521917998 | 74.1118313312039 | 84.2344850897118 | 95.6685228143324 | 95.2613630362614 | 84.3926740130963 | 83.2862472444909 | 84.6709940441436 | Myef2 |
| **ENSMUSG00000027203** | -0.0281842002169306 | 0.849697829982878 | 0.972640876584422 | 439.107666252556 | 378.703313945163 | 435.369248778286 | 508.356268287923 | 373.380744774312 | 491.679057293692 | 442.458188486358 | 488.486504100828 | Dut |
| **ENSMUSG00000027204** | -0.798025311709331 | 0.0285106691409379 | 0.665956123973759 | 300.870067617492 | 271.200437728471 | 414.54724122802 | 405.184331919526 | 869.396807480363 | 898.965440574287 | 343.555769883525 | 308.289260365856 | Fbn1 |
| **ENSMUSG00000027206** | 0.177723013855726 | 0.10060467027679 | 0.759800953500741 | 1966.04584725424 | 2141.91336704468 | 2324.49320652059 | 2120.6522557177 | 1757.40790428965 | 1731.88444061659 | 2084.23833729339 | 1986.51178334337 | Cops2 |
| **ENSMUSG00000027207** | 0.0971531820469861 | 0.552104357144585 | 0.914957699363693 | 169.86057871498 | 175.099381716581 | 182.66579350915 | 202.592165959763 | 193.80760065998 | 193.24699266767 | 156.16171358342 | 140.032797842237 | Galk2 |
| **ENSMUSG00000027208** | -0.287178710779744 | 0.54547134078198 | 0.914211323944814 | 26.2018977805023 | 11.4018202048006 | 34.0723759913441 | 9.37926694258161 | 35.0386622662111 | 22.0154801773295 | 17.6983275394543 | 23.8815624227072 | Fgf7 |
| **ENSMUSG00000027210** | 0.151164960203402 | 0.455723106138128 | 0.892805623137304 | 499.643085262682 | 523.669313691913 | 500.674636095028 | 478.342614071662 | 435.793361936001 | 289.870489001505 | 523.662279549737 | 552.53251241627 | Meis2 |
| **ENSMUSG00000027215** | -0.325541903182932 | 0.146524860919307 | 0.780607990324772 | 219.553833126278 | 236.99497711407 | 276.364827485346 | 262.619474392285 | 391.995034103237 | 388.940149799487 | 242.57119509958 | 224.703791886381 | Cd82 |
| **ENSMUSG00000027217** | -0.566547693241339 | 0.101149547588091 | 0.759800953500741 | 122.87786545339 | 152.295741306979 | 160.897331070236 | 190.399118934407 | 285.784089108784 | 373.040080782527 | 140.545542225078 | 129.177542195552 | Tspan18 |
| **ENSMUSG00000027219** | -0.16704001951033 | 0.474893158985802 | 0.897549741595057 | 273.764656120421 | 201.160685041839 | 182.66579350915 | 299.198615468353 | 304.398378437709 | 336.347613820311 | 206.133461930115 | 227.960368580387 | Slc28a2 |
| **ENSMUSG00000027221** | -1.06048938131706 | 0.0314060592601248 | 0.669064667366769 | 33.4300075130547 | 14.6594831204579 | 18.9290977729689 | 35.6412143818101 | 82.1218646864323 | 77.0541806206532 | 35.3966550789086 | 19.5394601640331 | Chst1 |
| **ENSMUSG00000027222** | 0.104648185008809 | 0.491348892306708 | 0.899236955296196 | 601.740135234984 | 618.95595397489 | 662.518422053913 | 707.196727470654 | 704.058119911679 | 677.587556568918 | 527.826591911961 | 500.427285312182 | Pex16 |
| **ENSMUSG00000027223** | -0.551940189011619 | 0.123303222980563 | 0.770251097497048 | 77.7021796249379 | 61.0811796685746 | 54.8943835416099 | 72.2203554578784 | 135.774816281568 | 143.100621152642 | 55.1771387994752 | 56.4473293627624 | Mapk8ip1 |
| **ENSMUSG00000027227** | 0.176852090097265 | 0.195888734730728 | 0.811193545163524 | 7521.7516904373 | 7582.2104361924 | 8310.82037722201 | 7613.1509772935 | 6580.69875687277 | 5552.79333361532 | 7131.38492030954 | 8181.60618090654 | Sord |
| **ENSMUSG00000027230** | -0.61152066645131 | 0.0391940099275528 | 0.679566075569698 | 56.9213641438498 | 55.3802695661743 | 65.3053873167428 | 71.2824287636203 | 134.679858085749 | 117.415894279091 | 67.6700758861489 | 60.7894316214364 | Creb3l1 |
| **ENSMUSG00000027236** | 0.197176269892848 | 0.054196378744518 | 0.703396849069134 | 3013.21824475777 | 2891.17583764587 | 3102.47912498961 | 3175.81978675813 | 2404.52819801874 | 2466.95686209297 | 2828.60917204102 | 2924.40587121696 | Eif3j1 |
| **ENSMUSG00000027238** | 0.126577485439664 | 0.605152690144497 | 0.926448151609826 | 2704.21655369115 | 3265.80707294646 | 3694.01343039489 | 3350.27415189015 | 2023.48274587369 | 2212.55575782161 | 3565.69246015477 | 4118.48399235232 | Frmd5 |
| **ENSMUSG00000027239** | -1.01584812590698 | 0.0215395099504806 | 0.648762042745232 | 17.1667606148119 | 11.4018202048006 | 8.51809399783602 | 11.2551203310979 | 27.3739548954774 | 39.1386314263635 | 15.616171358342 | 16.2828834700276 | Mdk |
| **ENSMUSG00000027242** | -0.505049810187017 | 0.125180753248297 | 0.771949105062466 | 24.3948703473642 | 21.1748089517725 | 20.8220075502658 | 16.8826804966469 | 41.6084114411257 | 23.2385624094033 | 22.903717992235 | 30.3947158107182 | Wdr76 |
| **ENSMUSG00000027243** | 0.040736175564068 | 0.789824427451951 | 0.961659329450003 | 252.983840639333 | 270.386021999557 | 273.525462819401 | 262.619474392285 | 281.404256325508 | 217.708637309147 | 306.076958623504 | 223.618266321713 | Harbi1 |
| **ENSMUSG00000027244** | 0.226910231496856 | 0.340070267476544 | 0.858570960161243 | 2094.34479500705 | 2984.0192307421 | 3707.26379883597 | 2491.13329994968 | 2273.13321452045 | 1750.23067409769 | 3278.35490716127 | 2332.79443847262 | Atg13 |
| **ENSMUSG00000027245** | 0.054198725750103 | 0.772279788063668 | 0.959045638244974 | 474.344701198749 | 385.218639776477 | 369.117406572894 | 487.721881014244 | 452.217734873287 | 513.694537471021 | 330.021754706295 | 358.223436340608 | Hypk |
| **ENSMUSG00000027247** | -0.325673866572085 | 0.0609848731420093 | 0.71282992338528 | 257.501409222178 | 303.777066885045 | 263.114459044268 | 254.178134143962 | 419.368988998714 | 368.147751854232 | 279.008928269045 | 286.578749072486 | Arhgap1 |
| **ENSMUSG00000027248** | -0.236344659672893 | 0.21224441546362 | 0.823370267313127 | 3457.74699330974 | 2812.17751194118 | 3044.74537678205 | 3350.27415189015 | 4256.10250714883 | 4692.9665244674 | 2723.46028489485 | 3247.89248948817 | Pdia3 |
| **ENSMUSG00000027249** | -0.292482259388731 | 0.331840479879681 | 0.856422197236858 | 293.64195788494 | 568.462178782201 | 326.526936583714 | 390.177504811395 | 331.772333333186 | 413.401794440965 | 425.80093903746 | 763.124471961961 | F2 |
| **ENSMUSG00000027253** | -0.0659128788131242 | 0.753744968751334 | 0.956497786093635 | 152.693818100169 | 159.625482867208 | 135.343049076728 | 114.427056699496 | 186.142893289246 | 106.408154190426 | 146.792010768415 | 148.717002359586 | Lrp4 |
| **ENSMUSG00000027254** | -0.53678515129261 | 0.0144949938286639 | 0.635862253526407 | 116.553269437407 | 130.306516626293 | 113.574586637814 | 123.806323642077 | 200.377349834895 | 232.385624094033 | 130.134761319517 | 141.118323406906 | Map1a |
| **ENSMUSG00000027255** | 0.198417728644886 | 0.370894815599225 | 0.873323500696802 | 622.520950716072 | 790.797672775813 | 904.810873547915 | 663.11417284052 | 593.467342133951 | 436.640356850368 | 802.671207818781 | 764.209997526629 | Arfgap2 |
| **ENSMUSG00000027257** | 0.0713658431832729 | 0.522769642080338 | 0.90648515894888 | 3979.97792148664 | 3570.39855556042 | 3110.99721898744 | 4053.71917258377 | 3462.25781517998 | 3573.84628211982 | 3535.50119552864 | 3433.51736104649 | Pacsin3 |
| **ENSMUSG00000027259** | -0.213371588394512 | 0.428729811337985 | 0.885378691567738 | 146.369222084185 | 165.326392969609 | 172.254789734017 | 163.19924480092 | 133.58489988993 | 119.862058743238 | 232.160414194018 | 263.782712214447 | Adal |
| **ENSMUSG00000027261** | -0.260123551195124 | 0.294495033212341 | 0.851785577030345 | 65.0529875929712 | 83.8848200781758 | 70.9841166486335 | 81.59962240046 | 83.2168228822514 | 95.400414101761 | 60.3825292522559 | 122.664388807541 | Hao1 |

| | | | | | | | | | | | | |
|---|---|---|---|---|---|---|---|---|---|---|---|---|
| **ENSMUSG00000027263** | 0.165660021478824 | 0.628877715276083 | 0.931825800800958 | 217.74680569314 | 267.1283590839 | 243.238906382651 | 151.944124469822 | 113.875652365186 | 117.415894279091 | 272.762459725708 | 280.065595684475 | Tubgcp4 |
| **ENSMUSG00000027272** | 0.343057895011697 | 0.164126907174144 | 0.794328481062321 | 448.142803418246 | 678.408302185636 | 769.467824471187 | 484.908100931469 | 453.312693069106 | 332.67836712409 | 627.77008860535 | 462.433890548784 | Ubr1 |
| **ENSMUSG00000027274** | -0.0188544974405102 | 0.879595099575753 | 0.978393501240173 | 1800.70283712211 | 1470.02039069036 | 1573.00802493372 | 1724.84719074076 | 1408.11623982336 | 1763.68457865051 | 1689.66974097261 | 1793.28823283237 | Mkks |
| **ENSMUSG00000027276** | -0.669481119765405 | 0.018209432226594 | 0.642273034804458 | 141.85165350134 | 140.079505373265 | 183.612248397799 | 125.682177030594 | 340.531998899739 | 280.085831144914 | 155.120635492864 | 164.999885829613 | Jag1 |
| **ENSMUSG00000027281** | -0.40112035537435 | 0.201120452196961 | 0.813027672006923 | 22.5878429142261 | 26.0613033252585 | 17.9826428843205 | 17.8206071909051 | 20.8042057205628 | 31.8001380339204 | 30.191264626128 | 29.3091902460497 | Slx4ip |
| **ENSMUSG00000027282** | 0.125334904843684 | 0.244641228985355 | 0.832048466795696 | 3187.59639205559 | 2685.94307395946 | 2889.52677504371 | 3199.26795411459 | 2678.26774697351 | 2534.22638485704 | 2707.84411353651 | 3044.89920889516 | Mtch2 |
| **ENSMUSG00000027284** | -0.136857384883408 | 0.44467720676015 | 0.89108031640712 | 128.298947752804 | 117.275864963663 | 134.396594188079 | 118.178763476528 | 164.243729372865 | 108.854318654574 | 123.88829277618 | 149.802527924254 | Cdan1 |
| **ENSMUSG00000027285** | 0.179433422592237 | 0.266783941223496 | 0.843513595393324 | 339.721157429961 | 386.033055505392 | 353.974128354519 | 357.350070512359 | 273.739548954774 | 258.070350967584 | 378.952424962434 | 357.137910775939 | Haus2 |
| **ENSMUSG00000027286** | 0.175497620312574 | 0.124385807037922 | 0.771949105062466 | 361.405486627618 | 347.755516246418 | 355.867038131816 | 345.157023487003 | 297.828629262794 | 280.085831144914 | 342.514691792969 | 326.743194965221 | Lrrc57 |
| **ENSMUSG00000027287** | 0.19980448684812 | 0.247463038523751 | 0.833398011296137 | 976.698327611138 | 972.412380323708 | 1041.10037751329 | 987.636809053844 | 740.19174037371 | 660.464405319884 | 1049.40671528059 | 1011.70982627105 | Snap23 |
| **ENSMUSG00000027288** | 0.332717331579995 | 0.238949628335906 | 0.832048466795696 | 18972.8845342334 | 22685.5501289086 | 20316.6006397276 | 17108.7208299631 | 11914.2401287076 | 8632.5143939773 | 21067.256240494 | 21180.7748178119 | Zfp106 |
| **ENSMUSG00000027291** | 0.162309380374945 | 0.407682940413032 | 0.881573350402261 | 593.608511785863 | 705.284021239809 | 645.482234058241 | 528.052728867345 | 485.06648074786 | 408.509465512669 | 705.850945397061 | 608.979841779033 | Vps39 |
| **ENSMUSG00000027293** | 0.225622391357029 | 0.20629467776894 | 0.819960256425221 | 8289.73834952099 | 9854.43031986338 | 10000.2423534595 | 9879.18187062121 | 7684.41661825842 | 5951.5181412714 | 10332.7000487697 | 8548.5138217645 | Ehd4 |
| **ENSMUSG00000027298** | -0.106552960741897 | 0.707400973616535 | 0.948710970699398 | 27.1054114970714 | 49.679359463774 | 48.2691993210708 | 33.7653609932938 | 44.893286028583 | 50.1463715150282 | 36.4377331694648 | 40.1644458927348 | Tyro3 |
| **ENSMUSG00000027300** | 0.0564620582933874 | 0.733630683353161 | 0.953953397470081 | 113.8427282877 | 128.677685168464 | 145.754052851861 | 132.247663890401 | 129.205067106653 | 106.408154190426 | 121.806136595068 | 142.203848971575 | Ubox5 |
| **ENSMUSG00000027303** | -0.111149047728998 | 0.281471981865266 | 0.850048154552447 | 817.679913494986 | 759.035459348154 | 713.626986040929 | 771.913669374467 | 861.732100109629 | 900.188522806361 | 779.767489826546 | 767.466574220635 | Ptpra |
| **ENSMUSG00000027304** | -0.00053354621455939 | 0.994727240239697 | 0.999262355603273 | 1386.89355493348 | 1380.43466050979 | 1283.39282900729 | 1406.89004138724 | 1311.75991859128 | 1411.43689581323 | 1321.12809691574 | 1416.6108618924 | Rtf1 |
| **ENSMUSG00000027305** | 0.0148406300164005 | 0.918113233146759 | 0.986713918026868 | 1223.35757223449 | 1097.83240257652 | 952.133617980337 | 1048.60204418062 | 1024.88087128667 | 879.396124861105 | 1157.67883669842 | 1214.70310686406 | Ndufaf1 |
| **ENSMUSG00000027306** | 0.112543414928752 | 0.851679481141525 | 0.972732497553372 | 38.8510898124689 | 15.4738988493722 | 55.8408384302584 | 10.3171936368398 | 14.2344565456483 | 51.3694537471021 | 14.5750932677859 | 31.4802413753867 | Nusap1 |
| **ENSMUSG00000027309** | 0.113750413335097 | 0.402108313873217 | 0.881158207349486 | 1192.63810587114 | 1527.02949171437 | 1600.45521670452 | 1223.05640931264 | 1381.8372431237 | 1216.96682091349 | 1237.84184967125 | 1286.34779413218 | 4930402H24I |
| **ENSMUSG00000027312** | 0.117559247548656 | 0.614702677492073 | 0.928075610917191 | 540.301202508289 | 803.013908709528 | 756.217456030109 | 499.9149280396 | 551.858930692825 | 420.740287833408 | 744.370834747638 | 678.453477917817 | Atrn |
| **ENSMUSG00000027313** | -0.210215181604952 | 0.469624963941956 | 0.895614823680169 | 223.167887992554 | 231.294067011669 | 159.950876181588 | 126.620103724852 | 223.371471947096 | 306.993640250539 | 193.640524843441 | 134.605170018895 | Chac1 |
| **ENSMUSG00000027314** | -0.791477956871905 | 0.0198055659712405 | 0.642273034804458 | 393.028466707535 | 219.077831077954 | 250.810545491838 | 331.088123073131 | 844.212768976524 | 584.633306931305 | 315.446661438509 | 322.401092706547 | Dll4 |
| **ENSMUSG00000027316** | -0.0414515170512561 | 0.828658960561401 | 0.967476923778924 | 133.720030052219 | 145.780415475665 | 191.183887506986 | 139.751077444466 | 122.635317931739 | 168.785348026193 | 178.024353485099 | 158.486732441602 | Gfra4 |
| **ENSMUSG00000027322** | -0.00130322922371109 | 0.997666959062876 | 0.999262355603273 | 16.2632468982428 | 23.6180561385155 | 36.9117406572894 | 26.2619474392285 | 22.994122112201 | 46.4771248188067 | 12.4929370866736 | 21.7105112933701 | Siglec1 |
| **ENSMUSG00000027324** | -0.163706129321199 | 0.408136871729504 | 0.881651811558757 | 64.1494738764022 | 64.338842584232 | 72.8770264259304 | 66.5927952923295 | 68.9823663366031 | 61.154111603693 | 84.3273253350471 | 84.6709940441436 | Rpusd2 |
| **ENSMUSG00000027327** | -0.116290533410307 | 0.433365288012651 | 0.887868762931002 | 197.869503928621 | 203.603932228582 | 228.095628164276 | 196.964605794214 | 233.226095709468 | 264.185762127954 | 198.845915296222 | 200.822229463674 | 1700037H04I |
| **ENSMUSG00000027329** | -0.0345468617076929 | 0.837858523805124 | 0.969013729862879 | 297.256012751216 | 219.077831077954 | 291.508105703722 | 313.267515882226 | 324.107625962453 | 289.870489001505 | 237.365804646799 | 296.348479154503 | Spef1 |
| **ENSMUSG00000027330** | -0.0874641531972392 | 0.803824160649443 | 0.963461896828986 | 64.1494738764022 | 56.1946852950887 | 119.253315969704 | 62.8410885152968 | 86.5016974697087 | 123.53130543946 | 54.1360607089191 | 57.5328549274309 | Cdc25b |
| **ENSMUSG00000027331** | -0.0463285962220746 | 0.919765306374861 | 0.986713918026868 | 13.5527057485357 | 13.8450673915436 | 8.51809399783602 | 15.0068271081306 | 7.66470737073368 | 22.0154801773295 | 10.4107809055614 | 13.0263067760221 | Knstrn |
| **ENSMUSG00000027332** | 0.2999394612907 | 0.315584329413936 | 0.853277176422981 | 8574.34517024024 | 9138.55889414768 | 9459.81661204123 | 8049.28689012354 | 5385.00440703832 | 3610.53874908203 | 10386.8361094786 | 9226.96729968231 | Ivd |
| **ENSMUSG00000027333** | -0.140518198675629 | 0.508799834534076 | 0.902394714365738 | 433.686583953142 | 390.105134149963 | 606.677583623654 | 344.219096792745 | 532.149683168081 | 580.964060235083 | 434.129563761909 | 409.243137880027 | Smox |
| **ENSMUSG00000027335** | -0.795930446084332 | 0.00305638528341399 | 0.635862253526407 | 44.2721721118832 | 57.8235167529173 | 42.5904699891801 | 46.8963347129081 | 83.2168228822514 | 50.1463715150282 | 94.7381062406084 | 104.210454208177 | Adra1d |
| **ENSMUSG00000027339** | -0.385212722188212 | 0.131268137481661 | 0.77303252008843 | 103.90407740544 | 134.378595270864 | 97.48485353079 | 83.4754757889764 | 185.047935093427 | 150.439114545085 | 117.641824232843 | 95.5262496908287 | Rassf2 |
| **ENSMUSG00000027340** | -0.0628317247278894 | 0.707265998407389 | 0.948710970699398 | 210.518695960587 | 267.1283590839 | 278.257737262643 | 249.488500672671 | 307.683253025166 | 207.923979452556 | 265.474913091815 | 268.124814473121 | Slc23a2 |
| **ENSMUSG00000027341** | 0.218348052601385 | 0.141729697098073 | 0.779852097185473 | 477.055242348456 | 498.422426095569 | 499.72818120638 | 391.115431505653 | 383.235368536684 | 338.793778284459 | 463.279750297481 | 417.927342397375 | Tmem230 |
| **ENSMUSG00000027342** | -0.0252942522663648 | 0.838547370724893 | 0.969245483845283 | 822.197482077831 | 771.251695281869 | 842.344850897118 | 884.464872685446 | 969.038003299901 | 892.850029413917 | 740.206522385413 | 777.236304302651 | Pcna |
| **ENSMUSG00000027346** | 0.330157034510415 | 0.165483008526117 | 0.795216960550936 | 3681.81839501886 | 5042.86219343752 | 4809.88374411141 | 3819.23749901923 | 2750.53498789757 | 2312.84850085167 | 4367.32258988299 | 4372.49697448475 | Gpcpd1 |
| **ENSMUSG00000027349** | -0.0383376165636118 | 0.71269279589437 | 0.950456121590218 | 637.880683897746 | 664.563234794092 | 636.964140060405 | 601.211011019481 | 648.215251924905 | 580.964060235083 | 711.056335849841 | 667.598222271132 | Fam98b |
| **ENSMUSG00000027351** | -0.223699125053222 | 0.134484751020957 | 0.773512701181749 | 319.843855665442 | 316.807718547674 | 321.794662140472 | 416.439452250624 | 387.61520131996 | 456.20967256355 | 353.966550789086 | 408.157612315359 | Spred1 |
| **ENSMUSG00000027353** | 0.462148833316486 | 0.053672406634632 | 0.703396849069134 | 93.0619128066116 | 63.5244268553176 | 117.360406192407 | 92.854742731558 | 66.7924499449649 | 56.2617826753975 | 77.0397787011541 | 65.1315338801105 | Mcm8 |
| **ENSMUSG00000027357** | 0.259491246807162 | 0.0501843945406947 | 0.703396849069134 | 576.441751171051 | 553.802695661743 | 529.068282754482 | 673.43136647736 | 503.680770076785 | 415.847958905112 | 517.4158110064 | 510.197015394199 | Crls1 |
| **ENSMUSG00000027358** | -0.769668440416246 | 0.0905824703084034 | 0.753823363185192 | 33.4300075130547 | 27.6901347830872 | 36.9117406572894 | 30.9515809105193 | 44.893286028583 | 116.192812047017 | 29.1501865355718 | 30.3947158107182 | Bmp2 |
| **ENSMUSG00000027359** | -0.38237038793948 | 0.25033152881499 | 0.835923538390114 | 211.422209677157 | 403.135785812593 | 258.382184601026 | 244.79886720138 | 249.650468646754 | 287.424324537357 | 286.296474902938 | 633.946929766408 | Slc27a2 |
| **ENSMUSG00000027360** | 0.164153423839063 | 0.712882501728349 | 0.950522761209907 | 43.3686583953142 | 21.9892246806869 | 60.5731128735006 | 31.8895076047775 | 50.3680770076785 | 50.1463715150282 | 17.6983275394543 | 22.7960368580387 | Hdc |
| **ENSMUSG00000027361** | -0.132497257263936 | 0.334804583117498 | 0.858570960161243 | 199.676531361759 | 162.883145782866 | 191.183887506986 | 191.337045628665 | 191.617684268342 | 192.023910435596 | 210.29777429234 | 221.447215192376 | Gabpb1 |
| **ENSMUSG00000027363** | 0.20296816497952 | 0.373030908505516 | 0.873700364743909 | 773.407741383103 | 996.844852191138 | 976.741445085197 | 716.575994413235 | 626.316088008523 | 525.92535979176 | 1000.47604502445 | 855.394144958784 | Usp8 |
| **ENSMUSG00000027365** | 0.0868669942870499 | 0.65501277701895 | 0.937504201434612 | 412.905768472054 | 525.298145149742 | 540.425741418263 | 406.122258613784 | 414.989156215438 | 329.009120427868 | 547.607075632528 | 481.973350712817 | Trpm7 |
| **ENSMUSG00000027366** | -0.0414711134821895 | 0.670473579181688 | 0.942789837500876 | 1100.4797067811 | 1122.26487444394 | 1163.19305814894 | 1084.24325856243 | 1240.58763586304 | 1015.1582526213 | 1157.67883669842 | 1185.39391661801 | Sppl2a |
| **ENSMUSG00000027367** | 0.17718666840418 | 0.457513969918639 | 0.892966448175628 | 3160.49098055852 | 3726.76637551197 | 3536.90191887925 | 2908.51067889456 | 2457.08619141805 | 1837.06951257494 | 3661.47164448593 | 3835.16181997384 | Stard7 |
| **ENSMUSG00000027371** | 0.0145774879062905 | 0.912390782149898 | 0.986226496507248 | 868.276681622852 | 668.635313438664 | 777.039463580375 | 920.106087067256 | 765.375778877549 | 868.38838477244 | 791.219348822664 | 776.150778737983 | Fahd2a |
| **ENSMUSG00000027374** | 0.12417708355924 | 0.2105838039569 | 0.822896573073231 | 1082.40943244972 | 886.898728787704 | 972.009170641955 | 1009.20912302178 | 901.150595159117 | 897.742358342213 | 945.298906224972 | 879.275707381491 | Mrps5 |
| **ENSMUSG00000027375** | 0.0848774019087067 | 0.581054888841176 | 0.922450244597442 | 715.582863522684 | 695.511032492837 | 643.589324280944 | 684.686486808458 | 620.841297029428 | 496.571386221987 | 775.603177464322 | 688.223207999834 | Mal |
| **ENSMUSG00000027377** | -0.706440905032669 | 0.0459608210627031 | 0.697906909182231 | 93.0619128066116 | 65.1532583131463 | 82.3415753124149 | 78.7858423176855 | 181.76306050597 | 190.800828203522 | 68.711153976705 | 80.3288917854696 | Mall |
| **ENSMUSG00000027378** | 0.303193849587307 | 0.0896854760959137 | 0.753162795025539 | 344.238726012806 | 320.065381463331 | 381.421320125324 | 329.212269684615 | 256.220217821669 | 209.14706168463 | 349.802238426862 | 297.434004719171 | Nphp1 |
| **ENSMUSG00000027381** | -0.203275887230388 | 0.366015337146163 | 0.869253398153585 | 178.895715880671 | 168.584055885266 | 238.506631939409 | 192.274972322923 | 218.991639163819 | 147.992950080937 | 285.255396812381 | 242.072200921077 | Bcl2l11 |
| **ENSMUSG00000027384** | 0.0330755028480197 | 0.842206858000926 | 0.971051154447817 | 602.643648951553 | 460.95930256551 | 455.244801439903 | 577.762843663027 | 445.647985698372 | 497.794468454061 | 478.895921655823 | 626.348250813729 | Ndufaf5 |
| **ENSMUSG00000027387** | 0.0860339227437282 | 0.660548813381609 | 0.939611397743053 | 199.676531361759 | 181.614707547895 | 168.468970179424 | 159.447538023887 | 164.243729372865 | 118.638976511164 | 175.942197303987 | 208.420908416353 | Zc3h8 |
| **ENSMUSG00000027394** | 0.17114371130014 | 0.332131656683597 | 0.856655692540538 | 787.863960848207 | 948.794324185193 | 839.505486231172 | 828.189271029956 | 623.031213421066 | 604.202622644487 | 864.094815161593 | 931.380934485579 | Ttl |
| **ENSMUSG00000027395** | 0.155106774236451 | 0.283276819205055 | 0.850587941933635 | 297.256012751216 | 360.786167909048 | 310.437203476691 | 335.777756544422 | 326.297542354091 | 242.170281950624 | 283.173240631269 | 319.144516012541 | Polr1b |
| **ENSMUSG00000027397** | -0.0292260259223404 | 0.860279001405671 | 0.974868041734905 | 154.500845533307 | 211.748089517725 | 195.916161950228 | 167.888878272211 | 197.092475247437 | 194.470074899744 | 197.804837205666 | 156.315681312265 | Slc20a1 |
| **ENSMUSG00000027404** | -0.163244146287224 | 0.150243462576124 | 0.782641594375979 | 1877.50150303048 | 1757.50914299712 | 1866.40904041474 | 1914.30838298091 | 2223.86009570859 | 2327.52548763655 | 1956.18573215498 | 1797.63033509105 | Snrpb |
| **ENSMUSG00000027405** | -0.102478279469125 | 0.425074611027854 | 0.885066977635295 | 354.177376895066 | 310.292392716359 | 295.293925258315 | 367.667264149199 | 379.950493949227 | 363.255422925936 | 321.693129981846 | 360.394487469945 | Nop56 |
| **ENSMUSG00000027406** | 0.209417296218138 | 0.0592174215859111 | 0.712406493939829 | 22198.4285023849 | 19234.8706854986 | 21657.7272169424 | 22550.571510049 | 17455.823557748 | 16462.6868437141 | 20162.5593798007 | 19986.6966966766 | Idh3b |
| **ENSMUSG00000027408** | -1.17546408404799 | 0.0247764527255914 | 0.655303666217042 | 43.3686583953142 | 60.2667639396603 | 25.5542819935081 | 60.9652351267805 | 144.534481848121 | 201.808568292187 | 40.6020455316893 | 43.4210225867403 | Cpxm1 |
| **ENSMUSG00000027411** | -0.00761662819410692 | 0.963777541617938 | 0.994284908685797 | 312.61574593289 | 323.323044378988 | 359.65285768641 | 422.067012416173 | 441.268152915096 | 341.239942748607 | 316.487739529066 | 325.657669400552 | Vps16 |
| **ENSMUSG00000027412** | -0.211684620899488 | 0.36520118074068 | 0.86924083181804 | 81.316234491214 | 97.7298874697194 | 76.6628459805242 | 94.7305961200743 | 134.679858085749 | 106.408154190426 | 95.7791843311646 | 69.4736361387845 | Lpin3 |
| **ENSMUSG00000027422** | 0.0417641175566712 | 0.60512999446044 | 0.926448151609826 | 3218.31585841894 | 3199.02498317548 | 3003.10136168152 | 2838.1661768252 | 2875.36022222095 | 3074.82873143368 | 2925.42943446274 | 3035.12947881315 | Rrbp1 |
| **ENSMUSG00000027423** | 0.156102695243149 | 0.214213785014682 | 0.824248286808591 | 1647.10550530537 | 1654.89276115392 | 1656.29605513478 | 1689.20597635895 | 1299.71537843727 | 1312.36723501525 | 1704.2448342404 | 1647.82780716679 | Snx5 |
| **ENSMUSG00000027424** | -0.0407562191278117 | 0.807631720543238 | 0.964026716319739 | 392.124952990966 | 447.928650902881 | 460.923530771794 | 453.018593326692 | 356.956371837026 | 393.832478727783 | 520.539045278068 | 531.907526687569 | Mgme1 |
| **ENSMUSG00000027425** | 0.13382085133958 | 0.393455306675794 | 0.878942305112115 | 712.872322372977 | 750.076886330097 | 649.268053612834 | 581.51455044006 | 548.574056105368 | 513.694537471021 | 670.454290318152 | 721.874500504558 | Kat14 |
| **ENSMUSG00000027427** | 0.106061028349633 | 0.421489703030719 | 0.882568531013176 | 250.273299489625 | 258.984201794756 | 238.506631939409 | 234.48167356454 | 209.137015401448 | 204.254732756335 | 245.694429371248 | 252.927456567762 | Polr3f |
| **ENSMUSG00000027428** | -0.148816123008298 | 0.242869141893077 | 0.832048466795696 | 242.141676040504 | 224.778741180355 | 203.487801059416 | 214.785212985119 | 258.410134213307 | 233.608706326107 | 227.996101831794 | 261.61166108511 | Rbbp9 |
| **ENSMUSG00000027429** | -0.38450064937392 | 0.0147081215709175 | 0.635862253526407 | 205.097613661173 | 272.8292691863 | 239.453086828057 | 185.709485463116 | 316.442918591719 | 299.655146858096 | 286.296474902938 | 277.894544555138 | Sec23b |
| **ENSMUSG00000027430** | 0.0993074384799357 | 0.476300720372551 | 0.897553735056661 | 294.545471601509 | 286.674336577844 | 305.704929033448 | 362.03970398365 | 335.057207920644 | 276.416584448692 | 280.050006359601 | 273.552442296464 | Dtd1 |
| **ENSMUSG00000027433** | 0.0282948895477143 | 0.876868930983994 | 0.977515813184586 | 484.283352081008 | 629.543358450776 | 556.515474525287 | 507.418341593665 | 545.28918151791 | 379.155491942897 | 617.359307699789 | 592.696958309005 | Xrn2 |
| **ENSMUSG00000027435** | -0.609291108777306 | 0.0281161678727111 | 0.665956123973759 | 805.934235179588 | 753.334549245754 | 882.095956220352 | 1042.97448401508 | 1931.50625742489 | 1624.25320419409 | 857.848346618257 | 902.07174423953 | Cd93 |
| **ENSMUSG00000027438** | -0.186965091636767 | 0.629770635528962 | 0.931825800800958 | 30.7194663633475 | 30.9477976987445 | 52.0550188756646 | 49.7101147956825 | 25.1840385038392 | 33.0232202659942 | 75.998700610598 | 51.0197015394199 | Napb |
| **ENSMUSG00000027439** | 0.171694094653198 | 0.533305500950397 | 0.909132720692319 | 604.450676384691 | 757.406627890326 | 750.538726698218 | 637.79015209555 | 436.88832013182 | 366.924669622158 | 815.164144905455 | 821.74285245406 | Gzf1 |
| **ENSMUSG00000027447** | -0.157529495457241 | 0.521719238054617 | 0.906152189339124 | 14749.8614229897 | 12449.1588321844 | 14459.937788771 | 16528.1442062173 | 16634.6049108837 | 24597.4067692374 | 11804.784468816 | 11864.7944218268 | Cst3 |
| **ENSMUSG00000027452** | 0.407024051962723 | 0.174310740078898 | 0.799889021902745 | 8570.73111537396 | 11212.0613399636 | 15192.4938725849 | 9481.50095225575 | 8080.79148514493 | 4217.18753619067 | 10940.6896536544 | 10288.6113019281 | Acss1 |
| **ENSMUSG00000027454** | -0.00622489788337585 | 0.979794785305168 | 0.996872525500296 | 67.7635287426784 | 43.1640336324594 | 65.3053873167428 | 45.0204813243917 | 54.7479097909548 | 48.9232892829544 | 63.5057635239243 | 54.2762782334254 | Gins1 |
| **ENSMUSG00000027455** | 0.00765499040660749 | 0.939682607635796 | 0.989269716906553 | 1981.40558043592 | 1836.50746870181 | 1913.73178484716 | 2070.00421422776 | 1972.01971067019 | 2139.17082389718 | 1921.83015516663 | 1728.15669895226 | Nsfl1c |
| **ENSMUSG00000027456** | -0.522362865007028 | 0.10382482859067 | 0.759800953500741 | 16.2632468982428 | 20.3603932228582 | 23.6613722162112 | 17.8206071909051 | 33.943704070392 | 31.8001380339204 | 26.0269522639034 | 20.6249857287016 | Sdcbp2 |
| **ENSMUSG00000027459** | -0.552358884005621 | 0.115499177413126 | 0.769071703867842 | 64.1494738764022 | 36.6487078011448 | 44.483379766477 | 61.9031618210386 | 97.4512794278996 | 110.077400886647 | 49.9717483466946 | 46.6775992807458 | Fam110a |
| **ENSMUSG00000027463** | -0.604529489956594 | 0.130817158081352 | 0.77303252008843 | 35.2370349461928 | 25.2468875963442 | 39.7511053232348 | 52.523894878457 | 81.0269064906132 | 84.3926740130963 | 33.3144988977964 | 33.6512925047237 | Slc52a3 |
| **ENSMUSG00000027465** | 0.125888648724672 | 0.178446085463956 | 0.801696999069105 | 1063.43564440177 | 1079.10084081149 | 1105.45930994139 | 1108.62935261315 | 948.233797579338 | 979.688867891162 | 948.422140496641 | 1115.92028047923 | Tbc1d20 |

| | | | | | | | | | | | | |
|---|---|---|---|---|---|---|---|---|---|---|---|---|
| **ENSMUSG00000027466** | -0.253224915013714 | 0.0786511648575332 | 0.739855948845396 | 852.01343472461 | 843.734695155244 | 787.450467355508 | 847.885731609378 | 1002.98170737029 | 1235.3130543946 | 852.642956165476 | 881.446758510828 | Rbck1 |
| **ENSMUSG00000027469** | -0.757125035831937 | 0.121839489145617 | 0.769868040577702 | 13.5527057485357 | 6.51532583131463 | 6.62518422053913 | 15.0068271081306 | 22.994122112201 | 19.5693157131818 | 8.32862472444909 | 19.5394601640331 | Tpx2 |
| **ENSMUSG00000027472** | 0.0733205211995595 | 0.452531188743197 | 0.891110666112841 | 3176.75422745676 | 2763.31256820632 | 2842.20403061129 | 3167.37844650981 | 2589.57613311216 | 2965.97441277911 | 2828.60917204102 | 2973.25452162704 | Pdrg1 |
| **ENSMUSG00000027474** | -0.570387540976947 | 0.125629435061169 | 0.771949105062466 | 171.667606148119 | 139.26508964435 | 122.09268063565 | 116.302910088012 | 290.163921892061 | 309.439804714686 | 105.14888714617 | 111.809133160856 | Ccm2l |
| **ENSMUSG00000027475** | 0.0949772027606613 | 0.694514037655241 | 0.944771261777587 | 245.75573090678 | 285.045505120015 | 272.579007930753 | 199.778385876988 | 192.712642464161 | 165.116101329971 | 334.18606706852 | 246.414303179751 | Kif3b |
| **ENSMUSG00000027478** | -0.204236791427063 | 0.690803823272502 | 0.94469184542218 | 18.0702743313809 | 9.77298874697194 | 10.4110037751329 | 6.56548685980713 | 10.949581958191 | 7.33849339244316 | 11.4518589961175 | 21.7105112933701 | Dnmt3b |
| **ENSMUSG00000027479** | -0.0590972165709686 | 0.406957210687403 | 0.881296485258502 | 1983.21260786905 | 1857.68227765358 | 1815.30047642772 | 1882.41887537613 | 1942.45583938308 | 1939.80842006914 | 1945.77495124942 | 2025.59070367144 | Mapre1 |
| **ENSMUSG00000027487** | 0.149157457842956 | 0.307994915565018 | 0.851785577030345 | 272.861142403852 | 289.117583764587 | 268.793188376159 | 305.764102328161 | 298.923587458613 | 220.154801773295 | 276.926772087932 | 227.960368580387 | Cdk5rap1 |
| **ENSMUSG00000027488** | 0.0468567648873459 | 0.716611301890736 | 0.951348124093194 | 3389.07995085049 | 3085.82119685639 | 4173.86605893965 | 3509.72168991404 | 3702.05366006437 | 3209.36777696181 | 3645.85547312759 | 3146.938611974 | Snta1 |
| **ENSMUSG00000027490** | -0.0324259948396342 | 0.937848619416679 | 0.988970340396118 | 20.780815481088 | 14.6594831204579 | 19.8755526616174 | 26.2619474392285 | 20.8042057205628 | 35.4693847301419 | 15.616171358342 | 11.9407812113536 | E2f1 |
| **ENSMUSG00000027495** | -0.176953129536727 | 0.18404964098276 | 0.805963874499992 | 238.527621174228 | 226.407572638183 | 276.364827485346 | 227.916186704733 | 306.588294829347 | 255.624186503437 | 267.557069272927 | 264.868237779116 | Fam210b |
| **ENSMUSG00000027496** | 0.335267902327114 | 0.384472296078409 | 0.875126835008975 | 23.4913566307952 | 20.3603932228582 | 17.9826428843205 | 17.8206071909051 | 25.1840385038392 | 14.6769867848863 | 11.4518589961175 | 11.9407812113536 | Aurka |
| **ENSMUSG00000027498** | -0.00649624825857125 | 0.951167155331796 | 0.991915011884503 | 461.695509166782 | 403.135785812593 | 415.493696116668 | 415.501525556365 | 402.944616061428 | 450.09426140318 | 441.417110395802 | 409.243137880027 | Cstf1 |
| **ENSMUSG00000027499** | 0.0686947548811862 | 0.817394291151917 | 0.966315773308182 | 4540.15642575945 | 4805.86721632345 | 4660.34387170495 | 3869.88554050917 | 2868.79047304603 | 2413.14124388173 | 5324.07335510408 | 6438.25212404892 | Pkia |
| **ENSMUSG00000027500** | -0.487777501762644 | 0.11707212656499 | 0.769868040577702 | 213.229237110295 | 149.852494120236 | 188.344522841041 | 214.785212985119 | 293.448796479518 | 435.417274618294 | 176.983275394543 | 169.341988088287 | Stmn2 |
| **ENSMUSG00000027502** | -0.0246095590346192 | 0.825509284505015 | 0.966806404871388 | 870.987222772559 | 833.961706408273 | 972.955625530603 | 846.94780491512 | 933.99934103369 | 991.9196902119 | 823.492769629904 | 836.940210359419 | Rtf2 |
| **ENSMUSG00000027506** | 0.149315280338384 | 0.222226123512756 | 0.831327997735572 | 506.871194995234 | 515.52515640277 | 520.550188756646 | 571.19735680322 | 411.70428162798 | 453.763508099402 | 512.210420553619 | 527.565424428895 | Tpd52 |
| **ENSMUSG00000027508** | -0.0813258330887262 | 0.818999758958678 | 0.966315773308182 | 35.2370349461928 | 52.9370223794314 | 26.5007368821565 | 20.6343872736795 | 31.7537876787538 | 47.7002070508805 | 35.3966550789086 | 29.3091902460497 | Pag1 |
| **ENSMUSG00000027509** | -0.0202335628540808 | 0.865441445553716 | 0.974868041734905 | 447.239289701677 | 380.332145402991 | 415.493696116668 | 469.901273823339 | 477.401773377126 | 414.624876673038 | 420.595548584679 | 423.354970220718 | Rae1 |
| **ENSMUSG00000027510** | 0.0650273593264884 | 0.626582830166882 | 0.931698409533928 | 3934.80223565819 | 4310.70245314354 | 5219.69871089618 | 4216.91841738469 | 3840.01839273757 | 3871.05526451377 | 4750.43932720765 | 4439.7995594942 | Rbm38 |
| **ENSMUSG00000027513** | -0.172732661433356 | 0.68303935522955 | 0.944452327903597 | 149.079763233892 | 463.402549752253 | 162.790240847533 | 226.040333316217 | 144.534481848121 | 217.708637309147 | 320.65205189129 | 446.151007078757 | Pck1 |
| **ENSMUSG00000027514** | -0.153243490525784 | 0.746123742922816 | 0.956085046483146 | 7.22810973255236 | 10.5874044758863 | 14.1968233297267 | 11.2551203310979 | 12.0445401540101 | 18.3462334811079 | 11.4518589961175 | 6.51315338801105 | Zbp1 |
| **ENSMUSG00000027519** | 0.109884399770813 | 0.449595619703094 | 0.89108031640712 | 1975.08098441993 | 2107.70790643028 | 2170.22105967089 | 1860.84656140819 | 1646.81712651192 | 1574.10683267906 | 2186.26399016789 | 2110.26169771558 | Rab22a |
| **ENSMUSG00000027522** | 0.0980602059758297 | 0.544678046954484 | 0.913937916428821 | 386.703870691551 | 411.279943101736 | 409.814966784777 | 370.481044231974 | 348.196706270473 | 281.308913376988 | 398.732908683 | 445.065481514088 | Stx16 |
| **ENSMUSG00000027523** | -0.0489573431673827 | 0.732948104946262 | 0.953733898884637 | 14784.1949442193 | 13803.532189369 | 13397.0689488188 | 16528.1442062173 | 15248.3878349767 | 18367.0258790532 | 14093.0741118584 | 12825.4845465584 | Gnas |
| **ENSMUSG00000027524** | -0.0840608548827192 | 0.846702379009935 | 0.972334438148346 | 483.379838364439 | 358.342920722305 | 122.09268063565 | 291.695201914288 | 155.484063806312 | 295.985900161874 | 558.017856538089 | 321.315567141878 | Edn3 |
| **ENSMUSG00000027531** | -0.104125084228554 | 0.382575715200095 | 0.875126835008975 | 317.133314515735 | 339.611358957275 | 293.401015481019 | 351.72251034681 | 349.291664466292 | 326.562955963721 | 382.075659234102 | 340.855027305911 | Impa1 |
| **ENSMUSG00000027533** | -0.456324449707905 | 0.214686462812273 | 0.824248286808591 | 417.423337054899 | 272.014853457386 | 300.972654590206 | 505.542488205149 | 641.645502749991 | 819.465095489486 | 246.735507461804 | 345.197129564585 | Fabp5 |
| **ENSMUSG00000027534** | 0.174553469255035 | 0.282089493500396 | 0.850308824809137 | 209.615182244018 | 217.448999620126 | 194.023252172932 | 212.909359596603 | 202.567266226533 | 155.33144347338 | 159.284947855089 | 221.447215192376 | Snx16 |
| **ENSMUSG00000027536** | 0.333561978562475 | 0.203220142228549 | 0.815817276171832 | 957.724539563188 | 1162.98566088966 | 1005.13509174465 | 934.174987481129 | 632.885837183438 | 472.10974158051 | 1085.84444845005 | 1030.16376087041 | Chmp4c |
| **ENSMUSG00000027540** | -0.230501867498918 | 0.153082529601729 | 0.783412422888014 | 412.002254755485 | 454.443976734195 | 484.584902988005 | 410.811892085075 | 620.841297029428 | 573.62556684264 | 438.293876124134 | 435.295751432072 | Ptpn1 |
| **ENSMUSG00000027544** | 0.0970374586223106 | 0.674135093476433 | 0.943983909911292 | 190.641394196068 | 202.789516499668 | 262.16800415562 | 352.660437041069 | 194.902558855799 | 221.377884005369 | 250.899819824029 | 274.637967861132 | Nfatc2 |
| **ENSMUSG00000027546** | -0.137307004235046 | 0.258606118556349 | 0.838606660810377 | 1949.782600356 | 1899.21747982821 | 2009.32372860065 | 1696.70938991301 | 2141.73823102215 | 2344.64863888559 | 2057.17030693893 | 1767.23561928033 | Atp9a |
| **ENSMUSG00000027547** | -0.013131002785797 | 0.973136461698127 | 0.995480246308729 | 27.1054114970714 | 37.4631235300591 | 29.3401015481019 | 22.5102406621959 | 31.7537876787538 | 11.0077400886647 | 39.5609674411332 | 34.7368180693922 | Sall4 |
| **ENSMUSG00000027550** | 0.217433626098976 | 0.472290680590529 | 0.89728597737823 | 396.642521573811 | 386.847471234306 | 176.98706417726 | 416.439452250624 | 217.896680968 | 304.547475786391 | 413.308001950786 | 248.585354309088 | Lrrcc1 |
| **ENSMUSG00000027551** | -0.321688992871496 | 0.147040497710368 | 0.780607990324772 | 124.684892886528 | 109.946123403434 | 129.664319744837 | 137.87522405595 | 165.338687568684 | 212.816308380852 | 111.395355689507 | 138.947272277569 | Zfp64 |
| **ENSMUSG00000027552** | -0.122249065674686 | 0.605101505426155 | 0.926448151609826 | 39.754603529038 | 47.2361122770311 | 39.7511053232348 | 39.3929211588428 | 40.5134532453066 | 42.8078781225851 | 55.1771387994752 | 42.3354970220718 | E2f5 |
| **ENSMUSG00000027559** | -0.379410653840276 | 0.452338504343723 | 0.891110666112841 | 132.81651633565 | 329.023954481389 | 141.968233297267 | 232.605820176024 | 239.795844884382 | 200.585486060113 | 79.1219348822664 | 568.815395886298 | Car3 |
| **ENSMUSG00000027562** | 0.71953746311089 | NA | NA | 115.649755720838 | 91.2145616384048 | 550.836745193396 | 83.4754757889764 | 78.836990098975 | 258.070350967584 | 66.6289977955927 | 107.467030902182 | Car2 |
| **ENSMUSG00000027566** | 0.139043203974016 | 0.44670013612086 | 0.89108031640712 | 3529.12457691869 | 2522.24551244768 | 3340.98575692902 | 4054.65709927803 | 3292.53929482802 | 3599.53100899337 | 2554.80563422476 | 2764.83361321069 | Psma7 |
| **ENSMUSG00000027569** | -0.0099298061807934 | 0.950536516171651 | 0.991696503889296 | 160.82544154929 | 156.367819951551 | 150.486327295103 | 147.254490998531 | 143.439523652302 | 125.977469903608 | 171.777884941763 | 176.940667040967 | Mrgbp |
| **ENSMUSG00000027573** | -0.0800758502805839 | 0.549325225942775 | 0.914699131434573 | 887.250469670802 | 930.877178149078 | 850.862944894954 | 796.299763425179 | 918.669926292222 | 743.633997100907 | 1042.11916864669 | 957.433548037624 | Gid8 |
| **ENSMUSG00000027575** | 0.0202931830929579 | 0.845284835771138 | 0.972136081650911 | 533.976606492306 | 574.163088884602 | 650.214508501483 | 557.128456389348 | 566.093387238473 | 557.72549782568 | 578.839418349212 | 579.670651532983 | Arfgap1 |
| **ENSMUSG00000027579** | 0.399805559333207 | 0.475060983920813 | 0.897549741595057 | 9.03513716569045 | 8.95857301805762 | 18.9290977729689 | 15.0068271081306 | 16.4243729372865 | 13.4539045528125 | 4.16431236222455 | 5.42762782334254 | Srms |
| **ENSMUSG00000027580** | -0.248145604871298 | 0.335426307865168 | 0.858570960161243 | 180.702743313809 | 265.499527626071 | 234.720812384815 | 200.716312571247 | 385.425284928322 | 272.747337752471 | 206.133461930115 | 183.453820428978 | Helz2 |
| **ENSMUSG00000027582** | -0.157363772916453 | 0.200695381563798 | 0.812921911423739 | 306.291149916906 | 342.869021872932 | 338.830850136144 | 349.846656958294 | 372.285786578493 | 382.824738639118 | 412.26692386023 | 324.572143835884 | Zgpat |
| **ENSMUSG00000027583** | -0.130604864646327 | 0.435141259548625 | 0.888448051614053 | 187.930853046361 | 219.892246806869 | 270.686098153456 | 193.212899017181 | 260.600050604945 | 233.608706326107 | 209.256696201783 | 250.756405438425 | Zbtb46 |
| **ENSMUSG00000027589** | 0.210642083155918 | 0.287282781605338 | 0.851785577030345 | 1011.03184884076 | 1228.13891920281 | 1401.69969008835 | 966.064495085906 | 950.423713970976 | 722.841599155651 | 1145.18589961175 | 1161.5123541953 | Pcmtd2 |
| **ENSMUSG00000027593** | -0.0680224235227514 | 0.603233809477192 | 0.926448151609826 | 2064.52884236027 | 1921.2067045089 | 1896.69559685149 | 2132.84530274306 | 2275.32313091208 | 2438.82597075528 | 1859.36546973326 | 1830.1961020311 | Raly |
| **ENSMUSG00000027597** | -0.028601282631487 | 0.864611207436651 | 0.974868041734905 | 1132.10268686101 | 1509.92676140717 | 1257.83854701379 | 1264.32518386 | 1081.81869746927 | 1219.41298537764 | 1300.30653510461 | 1666.28174176616 | Ahcy |
| **ENSMUSG00000027598** | 0.0202979375041066 | 0.920895522842507 | 0.986713918026868 | 587.283915769879 | 709.35609988438 | 728.770264259304 | 513.983828453472 | 606.60684048378 | 436.640356850368 | 745.411912838194 | 714.275821551878 | Itch |
| **ENSMUSG00000027599** | 0.0120655273491726 | 0.945895180811261 | 0.990505437194639 | 1003.80373910821 | 947.979908456279 | 918.061241988993 | 865.706338800283 | 781.800151814835 | 710.610776834912 | 1076.47474563505 | 1134.37421507859 | Armc1 |
| **ENSMUSG00000027601** | 0.140784613536478 | 0.487504484216052 | 0.899236955296196 | 990.251033359673 | 975.670043239366 | 1001.34927219006 | 960.436934920357 | 720.482492848966 | 634.779678446333 | 1062.94073045782 | 1143.05841959594 | Mtfr1 |
| **ENSMUSG00000027602** | -0.0882006544519043 | 0.580084666433217 | 0.922450244597442 | 16200.000938083 | 14014.4658631578 | 13012.8082640275 | 15336.0393778152 | 14857.4877590693 | 20025.5253857453 | 13847.3796824872 | 13526.7340613343 | Map1lc3a |
| **ENSMUSG00000027603** | -1.03863450338097 | 0.152447040009683 | 0.783412422888014 | 9.03513716569045 | 7.32974156022896 | 4.73227444324223 | 7.50341355406529 | 29.5638712871156 | 22.0154801773295 | 3.12323427166841 | 4.34210225867403 | Ggt7 |
| **ENSMUSG00000027605** | 0.0206632675741151 | 0.845574583227823 | 0.972180580805179 | 748.109357319169 | 810.343650269757 | 851.809399783602 | 809.430737144793 | 753.331238723539 | 717.949270227356 | 840.150019078802 | 860.821772782126 | Acss2 |
| **ENSMUSG00000027610** | 0.16535919093168 | 0.290382807828902 | 0.851785577030345 | 504.160653845527 | 392.548381336706 | 493.102996985841 | 547.749189446766 | 376.665619361769 | 508.802208542726 | 432.047407580797 | 410.328663444696 | Gss |
| **ENSMUSG00000027611** | -0.146695388082475 | 0.818514364423759 | 0.966315773308182 | 12.6491920319666 | 6.51532583131463 | 21.7684624389143 | 23.448167356454 | 30.6588294829347 | 26.9078091056249 | 9.36970281500523 | 4.34210225867403 | Procr |
| **ENSMUSG00000027613** | -0.142070888369764 | 0.468618126837098 | 0.895328112556867 | 829.425591810383 | 776.138189655355 | 877.36368177711 | 934.174987481129 | 1136.56660726022 | 1169.26661386261 | 730.836819570408 | 734.90080728058 | Eif6 |
| **ENSMUSG00000027615** | 0.0754107959987065 | 0.661560879340534 | 0.939611397743053 | 114.746242004269 | 120.533527879321 | 159.004421292939 | 129.433883807626 | 132.489941694111 | 112.523565350795 | 117.641824232843 | 133.519644454226 | Hps3 |
| **ENSMUSG00000027618** | 0.198272701277243 | 0.290411206559841 | 0.851785577030345 | 2399.73243120738 | 2440.80393955624 | 2527.03455269135 | 2551.1606083822 | 1888.80288778794 | 1534.96820125269 | 2708.88519162707 | 2510.82063107826 | Nfs1 |
| **ENSMUSG00000027620** | -0.143220651407608 | 0.107087617540594 | 0.762287160109383 | 2031.09883484721 | 2000.20503021359 | 1879.65940885582 | 1863.66034149097 | 2298.31725302428 | 2109.81685032741 | 2209.16770816012 | 1969.14337430867 | Rbm39 |
| **ENSMUSG00000027624** | -0.0797326011722352 | 0.638538575465545 | 0.933504827868072 | 126.491920319666 | 171.841718800923 | 172.254789734017 | 145.378637610015 | 173.003394939417 | 177.34692365071 | 162.408182126757 | 138.947272277569 | Epb41l1 |
| **ENSMUSG00000027628** | -0.081879118409892 | 0.430429363697147 | 0.886017087288711 | 651.433389646282 | 658.862324691692 | 592.480760293928 | 547.749189446766 | 664.639624862192 | 661.687487551958 | 638.180869510912 | 630.690353072403 | Aar2 |
| **ENSMUSG00000027630** | 0.35009333492881 | 0.0295632803641403 | 0.665956123973759 | 922.487504616995 | 843.734695155244 | 994.724087969518 | 941.678401035194 | 700.773245324222 | 546.717757737015 | 868.259127523818 | 787.006034384668 | Tbl1xr1 |
| **ENSMUSG00000027634** | -0.214405744821853 | 0.179100844321672 | 0.801755154770274 | 293.64195788494 | 296.447325324816 | 316.115932808581 | 327.336416296098 | 386.520243124141 | 429.301863457925 | 337.309301340188 | 278.980070119806 | Ndrg3 |
| **ENSMUSG00000027635** | 0.63942100629954 | 0.10030638689719 | 0.759800953500741 | 21.6843291976571 | 17.9171460361152 | 17.036187995672 | 22.5102406621959 | 10.949581958191 | 12.2308223207386 | 18.7394056300105 | 8.68420451734806 | Dsn1 |
| **ENSMUSG00000027637** | 0.0498108894004832 | 0.660502920592675 | 0.939611397743053 | 1104.09376164737 | 1081.54408799823 | 1068.5475692841 | 1137.70508013515 | 984.367418041368 | 935.657907536503 | 1110.8303226234 | 1210.36100460539 | 1110008F13 |
| **ENSMUSG00000027639** | -0.0072123719437798 | 0.959995202606797 | 0.993625746680416 | 570.117155155067 | 606.739718041175 | 707.948256709038 | 659.362466063487 | 730.337116611338 | 703.272283442469 | 551.771387994752 | 572.071972580304 | Samhd1 |
| **ENSMUSG00000027641** | -0.223963627098074 | 0.446521684675861 | 0.89108031640712 | 32.5264937964856 | 42.3496179035451 | 28.3936466594534 | 30.9515809105193 | 47.0832024202212 | 50.1463715150282 | 30.191264626128 | 30.3947158107182 | Rbl1 |
| **ENSMUSG00000027642** | -0.0899671893426845 | 0.55937889613752 | 0.916574868225118 | 1491.70114605549 | 1699.6856262442 | 1791.63910421151 | 1693.89560983024 | 1906.32221892105 | 2173.41712639525 | 1479.37196668027 | 1549.04498078196 | Rpn2 |
| **ENSMUSG00000027646** | -0.560257434896609 | 0.0942478516549691 | 0.757085157577544 | 50.5967681278665 | 39.9063707168021 | 42.5904699891801 | 52.523894878457 | 78.836990098975 | 100.292743030056 | 62.4646854333682 | 32.5657669400552 | Src |
| **ENSMUSG00000027649** | 0.0222242269405128 | 0.885810168025682 | 0.979948700426504 | 378.57224724243 | 359.971752180133 | 429.690519446395 | 434.260059441529 | 444.553027502553 | 450.09426140318 | 359.171941241867 | 324.572143835884 | Ctnnbl1 |
| **ENSMUSG00000027650** | 0.0539145187577663 | 0.778193236010515 | 0.960126492111037 | 596.31905293557 | 756.592212161411 | 657.786147610671 | 516.797608536247 | 515.725310230795 | 496.571386221987 | 763.110240377648 | 658.914017753784 | Tti1 |
| **ENSMUSG00000027651** | -0.0133751368437174 | 0.940215578730862 | 0.989269716906553 | 524.941469326615 | 691.438953848265 | 639.80350472635 | 494.287367874051 | 504.775728272604 | 505.132961846504 | 692.316930219831 | 669.769273400469 | Rprd1b |
| **ENSMUSG00000027652** | 0.0991874597666916 | 0.625676666788982 | 0.931573470007391 | 840.267756409212 | 1148.3261777692 | 1088.42312194571 | 815.9962240046 | 885.821180417649 | 640.895089606702 | 1046.28348100892 | 1060.55847668113 | Ralgapb |
| **ENSMUSG00000027655** | 0.0591416053003107 | 0.749202490660651 | 0.956197192463153 | 252.983840639333 | 301.333819698302 | 309.490748588042 | 256.053987532478 | 227.751304730372 | 211.593226148778 | 316.487739529066 | 318.058990447873 | Dhx35 |
| **ENSMUSG00000027656** | -0.957372415916649 | 0.152622069132936 | 0.783412422888014 | 18.0702743313809 | 18.7315617650296 | 35.0188308799925 | 53.4618215727152 | 50.3680770076785 | 154.108361241306 | 26.0269522639034 | 13.0263067760221 | Wisp2 |
| **ENSMUSG00000027660** | -0.244180255793257 | 0.219706243324594 | 0.82837125944045 | 500.546598979251 | 464.216965481167 | 354.920583243168 | 317.957149353517 | 598.942133113046 | 432.971110154146 | 492.429936833053 | 415.756291268038 | Skil |
| **ENSMUSG00000027661** | -1.65565961309029 | 0.0078587851981017 | 0.635862253526407 | 11.7456783153976 | 8.95857301805762 | 8.51809399783602 | 9.37926694258161 | 32.8487458745729 | 62.3771938357668 | 5.20539045278068 | 21.7105112933701 | Slc2a10 |
| **ENSMUSG00000027663** | 0.034855914917238 | 0.816183839242528 | 0.966231443542036 | 210.518695960587 | 249.211213047785 | 215.791714611846 | 208.219726125312 | 208.042057205628 | 179.793088114857 | 230.078258012906 | 244.243252050414 | Zmat3 |
| **ENSMUSG00000027665** | 0.136733987283391 | 0.266626783294182 | 0.843513595393324 | 1452.85005624302 | 1638.60444657563 | 1446.18306985483 | 1391.88321427911 | 1330.3742079202 | 1141.13572252491 | 1400.250031798 | 1520.82131610058 | Pik3ca |
| **ENSMUSG00000027667** | 0.206301391800302 | 0.140978161712236 | 0.779711473022872 | 467.116591466196 | 468.289044125739 | 456.191256328551 | 486.783954319986 | 445.647985698372 | 318.001380339203 | 408.102611498006 | 454.835211596105 | Zfp639 |
| **ENSMUSG00000027668** | 0.212537531623797 | 0.255239986640061 | 0.837406774112203 | 10644.2950948999 | 11282.9155083791 | 11395.3168593273 | 10172.752925924 | 8748.71598459458 | 6441.97411633302 | 11198.8770201124 | 11146.1764980162 | Mfn1 |
| **ENSMUSG00000027669** | -0.251034593087024 | 0.336908554167103 | 0.858570960161243 | 230.395997725106 | 191.387696294867 | 182.66579350915 | 246.674720589896 | 309.873169416804 | 329.009120427868 | 145.750932677859 | 229.045894145055 | Gnb4 |
| **ENSMUSG00000027671** | 0.0217083811583421 | 0.861218161044033 | 0.974868041734905 | 500.546598979251 | 412.09435883065 | 419.279515671262 | 424.880792498947 | 465.357233223116 | 381.601656407044 | 435.170641852465 | 447.236532643425 | Actl6a |
| **ENSMUSG00000027673** | 0.0586107014317677 | 0.674476252045904 | 0.944023556644042 | 11242.4211752686 | 8558.69489516068 | 9119.09285212779 | 11168.8310752262 | 9339.9934103369 | 11087.2404337495 | 8771.08291293545 | 9295.35541025643 | Ndufb5 |

| | | | | | | | | | | | | |
|---|---|---|---|---|---|---|---|---|---|---|---|---|
| **ENSMUSG00000027677** | 0.321535696715157 | 0.159083576936721 | 0.789393312599749 | 208.711668527449 | 250.840044505613 | 222.416898832385 | 236.357526953057 | 165.338687568684 | 114.969729814943 | 236.324726556243 | 217.105112933702 | Ttc14 |
| **ENSMUSG00000027678** | -0.0608387519637285 | 0.683439714096744 | 0.944452327903597 | 621.617436999503 | 664.563234794092 | 633.178320505811 | 468.025420434822 | 710.627869086594 | 600.533375948265 | 590.291277345329 | 589.440381615 | Ncoa3 |
| **ENSMUSG00000027679** | 0.0786352609659898 | 0.545776783535967 | 0.914211323944814 | 1336.29678680562 | 1066.07018914886 | 1066.6546595068 | 1168.65666104567 | 1077.43886468599 | 1270.78243912474 | 1027.54407537891 | 1017.13745409439 | Dnajc19 |
| **ENSMUSG00000027680** | 0.223043215643357 | 0.144904918595586 | 0.780607990324772 | 5510.53015735461 | 5930.57533795414 | 6078.13329490033 | 5693.21503414704 | 4315.23024972306 | 4006.81739227396 | 5855.02318128771 | 5708.77894459168 | Fxr1 |
| **ENSMUSG00000027684** | -0.294353941875368 | 0.0871545621801584 | 0.747269672026173 | 188.83436676293 | 208.490426602068 | 279.204192151292 | 243.860940507122 | 327.39250054991 | 284.978160073209 | 250.899819824029 | 264.868237779116 | Mecom |
| **ENSMUSG00000027690** | -0.0653679367754845 | 0.88908116952423 | 0.98126431530921 | 29.8159526467785 | 55.3802695661743 | 32.1794662140472 | 30.9515809105193 | 29.5638712871156 | 25.6847268735511 | 19.7804837205666 | 80.3288917854696 | Slc2a2 |
| **ENSMUSG00000027692** | 0.09962199270164 | 0.769254154829898 | 0.95879680569409 | 535.783633925444 | 505.752167655798 | 734.448993591195 | 471.777127211855 | 359.146288228664 | 259.293433199658 | 817.246301086567 | 661.085068883121 | Tnik |
| **ENSMUSG00000027694** | -0.100486412681752 | 0.779738259254048 | 0.960582210291683 | 76.7986659083688 | 98.5443031986338 | 99.3777633080869 | 69.4065753751039 | 73.3621991198795 | 39.1386314263635 | 137.42230795341 | 118.322286548867 | Gm8325 |
| **ENSMUSG00000027695** | -0.205308226727319 | 0.319200063249202 | 0.854670142081901 | 193.351935345776 | 174.284965987666 | 163.736695736181 | 123.806323642077 | 244.175677667659 | 174.900759186562 | 175.942197303987 | 160.657783570939 | Pld1 |
| **ENSMUSG00000027698** | 0.127322470985822 | 0.388943772837935 | 0.876536136761342 | 4811.21054073017 | 5948.49248399026 | 6335.5690246127 | 5409.96117248107 | 5773.7145665541 | 4131.5717799455 | 5472.94752205361 | 5224.63454274953 | Nceh1 |
| **ENSMUSG00000027706** | -0.0683444218192211 | 0.692339150516689 | 0.94469184542218 | 3236.38613275032 | 2783.67296142918 | 2957.6715270264 | 3422.49450734803 | 3453.49814961343 | 4159.7026712832 | 2639.13295955981 | 2750.72178087 | Sec62 |
| **ENSMUSG00000027708** | 0.0808968907815512 | 0.626796921153014 | 0.931825800800958 | 992.058060792811 | 1052.22512175731 | 1000.40281730141 | 857.264998551959 | 781.800151814835 | 748.526326029202 | 1070.22827709171 | 1087.69661579784 | Dcun1d1 |
| **ENSMUSG00000027709** | 0.251677225745504 | 0.153185611271695 | 0.783412422888014 | 3113.50826729693 | 3325.2594211572 | 3319.2172944901 | 3138.30271898781 | 2598.33579867872 | 1892.10821301826 | 3170.08278574344 | 3169.73464883204 | Mccc1 |
| **ENSMUSG00000027710** | 0.114595516575577 | 0.494683902944391 | 0.900200915538898 | 920.680477183857 | 961.824975847822 | 1027.85000907221 | 941.678401035194 | 881.441347634373 | 639.672007374629 | 971.325858488875 | 1063.81505337514 | Acad9 |
| **ENSMUSG00000027712** | -0.274614282128525 | 0.192559604843311 | 0.808824684419004 | 2060.91478749399 | 1777.86953621998 | 1910.89242018121 | 2192.87261117558 | 2896.16442794151 | 3085.83647152235 | 1753.17550449653 | 1873.61712461784 | Anxa5 |
| **ENSMUSG00000027714** | 0.307741551205332 | 0.0145084040235156 | 0.635862253526407 | 432.783070236573 | 359.971752180133 | 441.047978110176 | 471.777127211855 | 332.867291529005 | 360.809258461789 | 335.227145159076 | 348.453706258591 | Exosc9 |
| **ENSMUSG00000027715** | -0.239274338961556 | 0.55785446746622 | 0.916370464878199 | 11.7456783153976 | 8.14415728914329 | 17.036187995672 | 17.8206071909051 | 19.7092475247437 | 14.6769867848863 | 14.5750932677859 | 15.1973579053591 | Ccna2 |
| **ENSMUSG00000027716** | -0.289218469514944 | 0.25516097134007 | 0.837406774112203 | 66.8600150261093 | 38.2775392589734 | 59.6266579848522 | 45.9584080186499 | 70.0773245324222 | 67.2695227640623 | 69.7522320672611 | 49.9341759747513 | Trpc3 |
| **ENSMUSG00000027722** | 0.0679381908080786 | 0.630682569136319 | 0.931825800800958 | 180.702743313809 | 176.728213174409 | 192.130342395635 | 191.337045628665 | 189.427767876704 | 181.016170346931 | 188.435134390661 | 147.631476794917 | Spata5 |
| **ENSMUSG00000027739** | -0.0838970810798324 | 0.511713786153995 | 0.902394714365738 | 250.273299489625 | 254.912123150185 | 229.988537941573 | 291.695201914288 | 273.739548954774 | 276.416584448692 | 252.981976005141 | 285.493223507818 | Rab33b |
| **ENSMUSG00000027742** | 0.155461306593506 | 0.403409695163054 | 0.881158207349486 | 1657.04415618763 | 1887.81565962341 | 1840.85475842123 | 1693.89560983024 | 1489.14314631397 | 1109.33558449099 | 1995.74669959611 | 1760.72246589232 | Cog6 |
| **ENSMUSG00000027746** | 0.0244019583989894 | 0.80471856677386 | 0.963461896828986 | 1019.16347228988 | 974.041211781537 | 989.045358637627 | 1029.84351029546 | 863.922016501267 | 1005.37359476471 | 1025.46191919779 | 1049.70322103445 | Ufm1 |
| **ENSMUSG00000027750** | -1.56218961230657 | 0.0322270688087568 | 0.671606027118506 | 224.071401709123 | 187.315617650296 | 153.325691961048 | 199.778385876988 | 521.20010120989 | 1412.65997804531 | 161.367104036201 | 162.828834700276 | Postn |
| **ENSMUSG00000027751** | 0.192464721984418 | 0.328824847634417 | 0.855977678491063 | 176.185174730964 | 188.13003337921 | 155.218601738345 | 137.87522405595 | 141.249607260664 | 110.077400886647 | 181.147587756768 | 142.203848971575 | Supt20 |
| **ENSMUSG00000027752** | -0.167159875968951 | 0.230817814383769 | 0.832048466795696 | 190.641394196068 | 169.39847161418 | 167.522515290775 | 173.51643843776 | 215.706764576362 | 168.785348026193 | 204.051305749003 | 197.565652769668 | Exosc8 |
| **ENSMUSG00000027761** | -0.524460681572879 | 0.0549828278685881 | 0.703396849069134 | 97.5794813894569 | 124.605606523892 | 77.6093008691726 | 109.737423228205 | 117.160526952643 | 140.654456688494 | 109.313199508394 | 222.532740757044 | Aadac |
| **ENSMUSG00000027763** | -0.0478546718794939 | 0.592005380841773 | 0.923935529355527 | 2699.69898510831 | 2669.65475938117 | 2301.77828919302 | 2561.47780201904 | 2718.78120021882 | 2712.79639073982 | 2505.87496396862 | 2642.16922440315 | Mbnl1 |
| **ENSMUSG00000027765** | -0.13084340064054 | 0.492137950522289 | 0.899236955296196 | 661.372040528541 | 700.397526866323 | 606.677583623654 | 473.652980600371 | 590.182467546493 | 528.371524255907 | 802.671207818781 | 752.269216315276 | P2ry1 |
| **ENSMUSG00000027770** | 0.151334212636371 | 0.381615163411628 | 0.875126835008975 | 387.60738440812 | 535.885549625628 | 526.228918088536 | 527.114802173087 | 471.926982398031 | 357.140011765567 | 424.759860946904 | 525.394373299558 | Dhx36 |
| **ENSMUSG00000027774** | 0.246995246796155 | 0.232526482756125 | 0.832048466795696 | 4546.48102177543 | 5081.1397326965 | 4986.87080828867 | 4587.39946161667 | 3659.35029042742 | 2632.07296342295 | 4965.94249195277 | 4921.77291020701 | Gfm1 |
| **ENSMUSG00000027775** | -0.0674351128498303 | 0.543337816093107 | 0.913837378403038 | 468.020105182765 | 540.772043999114 | 508.246275204216 | 500.852854733858 | 542.004306930453 | 583.410224699231 | 482.019155927491 | 509.11148982953 | Mfsd1 |
| **ENSMUSG00000027777** | 0.310584285912373 | 0.198717704251381 | 0.812921911423739 | 788.767474564776 | 703.65518978198 | 787.450467355508 | 696.879533833814 | 508.060602860061 | 359.586176229715 | 746.45299092875 | 784.834983255331 | Schip1 |
| **ENSMUSG00000027778** | -0.105163655549942 | 0.630762311381368 | 0.931825800800958 | 74.9916384752307 | 83.0704043492615 | 74.7699362032273 | 73.1582821521366 | 84.3117810780705 | 94.1773318696872 | 95.7791843311646 | 55.3618037980939 | Ift80 |
| **ENSMUSG00000027782** | 0.154273060707816 | 0.438857575903314 | 0.889583654898613 | 3225.54396815149 | 3475.11191527744 | 3196.1781589658 | 3106.41321138303 | 2425.3324037393 | 2064.56280774067 | 3566.73353824532 | 3626.74091155748 | Kpna4 |
| **ENSMUSG00000027784** | 0.0971829122825652 | 0.681823718460034 | 0.944452327903597 | 1069.76024041775 | 1374.73375040739 | 1349.64467121269 | 925.733647232805 | 1054.44474257379 | 708.164612370765 | 1470.00226386526 | 1178.88076323 | Ppm1l |
| **ENSMUSG00000027787** | 0.122702430722675 | 0.300756311984549 | 0.851785577030345 | 489.704434380422 | 463.402549752253 | 441.994432998825 | 453.95652002095 | 398.564783278151 | 375.486245246675 | 448.704657029695 | 474.374671760138 | Nmd3 |
| **ENSMUSG00000027792** | 0.122071710316631 | 0.548064781986243 | 0.914699131434573 | 291.834930451802 | 281.787842204358 | 221.470443943737 | 280.44008158319 | 208.042057205628 | 188.354663739374 | 320.65205189129 | 270.295865602458 | Bche |
| **ENSMUSG00000027796** | -0.243714551687337 | 0.295574016197414 | 0.851785577030345 | 35.2370349461928 | 46.4216965481167 | 41.6440151005317 | 40.3308478531009 | 52.5579933993167 | 52.592535979176 | 48.9306702561384 | 40.1644458927348 | Smad9 |
| **ENSMUSG00000027797** | -0.286758531248815 | 0.443380724281777 | 0.89108031640712 | 23.4913566307952 | 39.0919549878878 | 16.0897331070236 | 19.6964605794214 | 26.2789966996583 | 42.8078781225851 | 27.0680303544596 | 24.9670879873757 | Dclk1 |
| **ENSMUSG00000027799** | 0.174649786159738 | 0.431854861498519 | 0.886559861988535 | 254.790868072471 | 374.631235300591 | 336.937940358847 | 248.550573978413 | 231.036179317829 | 201.808568292187 | 320.65205189129 | 322.401092706547 | Nbea |
| **ENSMUSG00000027800** | -0.401197544897811 | 0.234891219718661 | 0.832048466795696 | 1155.59404349181 | 730.530908836153 | 905.757328436564 | 1571.02721288242 | 1571.2650110004 | 2306.7330896913 | 846.396487622139 | 1037.76243982309 | Tm4sf1 |
| **ENSMUSG00000027801** | -0.326187865978911 | 0.305734700795497 | 0.851785577030345 | 32.5264937964856 | 43.1640336324594 | 40.6975602118832 | 35.6412143818101 | 26.2789966996583 | 51.3694537471021 | 43.7252798033577 | 69.4736361387845 | Tm4sf4 |
| **ENSMUSG00000027803** | -0.02167944647232 | 0.820202144459377 | 0.966398681144067 | 1815.15905658721 | 1875.5994236897 | 1980.9300819412 | 2024.98373290337 | 1952.31046314545 | 1774.69231873917 | 2137.33331991175 | 1946.34733745063 | Wwtr1 |
| **ENSMUSG00000027804** | 0.0794837925112552 | 0.413933667270458 | 0.882192126163777 | 1182.69945498888 | 1116.56396434154 | 1129.1206821576 | 1303.71810501884 | 1080.72373927345 | 1059.18921297596 | 1150.39129006453 | 1186.47944218268 | Ppid |
| **ENSMUSG00000027805** | 0.0500981111426713 | 0.743512214235156 | 0.954383574136061 | 1060.72510325206 | 947.979908456279 | 904.810873547915 | 1024.21595012991 | 806.984190318674 | 803.565026472526 | 1072.31043327282 | 1119.17685717323 | Pfn2 |
| **ENSMUSG00000027806** | 0.146618652705005 | 0.411893921512978 | 0.882192126163777 | 566.503100288791 | 693.067785306094 | 712.68053115228 | 592.769670771158 | 535.434557755538 | 432.971110154146 | 677.741836952045 | 669.769273400469 | Tsc22d2 |
| **ENSMUSG00000027808** | -0.139720906722687 | 0.366956265009913 | 0.870096026116028 | 677.635287426784 | 655.604661776035 | 689.019158936069 | 633.100518624259 | 771.945528052463 | 854.934480219628 | 545.524919451416 | 754.440267444613 | Serp1 |
| **ENSMUSG00000027809** | 0.275407926125599 | 0.163191820324866 | 0.793968226938511 | 20934.4128129048 | 22489.2759382403 | 24473.4305106715 | 20795.710665092 | 16267.7939152843 | 12576.9545924155 | 22514.354786367 | 21918.9322017865 | Etfdh |
| **ENSMUSG00000027810** | 0.0872855677789388 | 0.341542509666595 | 0.859283395807012 | 1171.85729039005 | 1151.58384068486 | 1207.67643791542 | 1316.84907873846 | 1078.53382288181 | 1113.00483118721 | 1209.73274122623 | 1160.42682863063 | Eif2a |
| **ENSMUSG00000027811** | -0.358609764617762 | 0.25192593651266 | 0.83723604018117 | 22.5878429142261 | 32.5766291565731 | 30.2865564367503 | 28.1378008277448 | 28.4689130912965 | 30.5770558018465 | 32.2734208072402 | 54.2762782334254 | 4930579G24 |
| **ENSMUSG00000027820** | 0.111964150499577 | 0.582325882581352 | 0.922450244597442 | 242.141676040504 | 289.117583764587 | 282.990011705886 | 241.047160424347 | 252.935343234211 | 172.454594722414 | 227.996101831794 | 322.401092706547 | Mme |
| **ENSMUSG00000027822** | -0.109773302013946 | 0.431093493507296 | 0.886306782832496 | 196.062476495483 | 188.944449108124 | 186.451613063744 | 181.019851991825 | 177.383227722694 | 196.916239363891 | 201.96914956789 | 235.559047533066 | Slc33a1 |
| **ENSMUSG00000027823** | 0.145389430842252 | 0.216785255458295 | 0.825692108906787 | 1278.4719089452 | 1285.96243595572 | 1248.3739981273 | 1214.61506906432 | 1094.9581958191 | 954.004141017611 | 1239.92400585236 | 1254.86755275679 | Gmps |
| **ENSMUSG00000027827** | -0.728356310898307 | 0.163033596314823 | 0.793968226938511 | 15.3597331816738 | 26.0613033252585 | 12.3039135524298 | 9.37926694258161 | 33.943704070392 | 44.0309603546589 | 16.6572494488982 | 10.8552556466851 | Kcnab1 |
| **ENSMUSG00000027828** | 0.0744392270904686 | 0.526393004272522 | 0.908217276872524 | 3507.44024772103 | 3874.17562244546 | 3851.12494191053 | 3322.13635106241 | 3229.03171947052 | 3065.04407357709 | 3900.91960531384 | 3626.74091155748 | Ssr3 |
| **ENSMUSG00000027829** | 0.0465483666201419 | 0.772947724715477 | 0.959121750920843 | 112.035700854562 | 115.647033505835 | 112.628131749165 | 123.806323642077 | 118.255485148462 | 99.0696607979826 | 128.052605138405 | 103.124928643508 | Ccnl1 |
| **ENSMUSG00000027833** | -0.966434498680647 | 0.0482289820714137 | 0.703396849069134 | 23.4913566307952 | 10.5874044758863 | 26.5007368821565 | 16.8826804966469 | 17.5193311331055 | 72.1618516923577 | 26.0269522639034 | 35.8223436340608 | Shox2 |
| **ENSMUSG00000027834** | -0.633205658673971 | 0.0820599269673095 | 0.739855948845396 | 99.3865088225949 | 86.3280672649188 | 58.6802030962037 | 72.2203554578784 | 192.712642464161 | 156.554525705454 | 67.6700758861489 | 74.901263962127 | Serpini1 |
| **ENSMUSG00000027835** | -0.0172325793612199 | 0.904123307202315 | 0.985039825636533 | 385.800356974982 | 324.951875836817 | 300.026199701558 | 306.702029022419 | 318.632834983357 | 302.101311322243 | 327.939598525183 | 384.276049892652 | Pdcd10 |
| **ENSMUSG00000027845** | 0.104945004880467 | 0.632035756334202 | 0.931825800800958 | 100.290022539164 | 148.223662662408 | 107.895857305923 | 91.9168160372998 | 128.110108910834 | 89.2850029413918 | 96.8202624217207 | 103.124928643508 | Dclre1b |
| **ENSMUSG00000027848** | -1.60542080119795 | 0.00443857634819806 | 0.635862253526407 | 101.193536255733 | 87.1424829938332 | 75.7163910918757 | 117.24083678227 | 397.469825082332 | 600.533375948265 | 78.0808567917102 | 84.6709940441436 | Olfml3 |
| **ENSMUSG00000027852** | -0.105636108019038 | 0.332214876603736 | 0.856655692540538 | 533.073092775737 | 567.647763053287 | 598.159489625818 | 634.976372012775 | 682.158955995297 | 632.333513982185 | 578.839418349212 | 617.664046296381 | Nras |
| **ENSMUSG00000027854** | 0.132696432518724 | 0.391090519980342 | 0.877896341705281 | 580.055806037327 | 659.676740420606 | 697.537252933905 | 584.328330522834 | 531.054724972262 | 461.102001491845 | 646.509494235361 | 659.999543318453 | Sike1 |
| **ENSMUSG00000027858** | -0.294967899897849 | 0.223720974955188 | 0.832048466795696 | 130.105975185942 | 161.254314325037 | 139.128868631322 | 188.52326554589 | 187.237851485066 | 273.970419984545 | 156.16171358342 | 143.289374536243 | Tspan2 |
| **ENSMUSG00000027859** | -0.862770961452861 | 0.0597958582885969 | 0.71282992338528 | 40.658117245607 | 34.2054606144018 | 28.3936466594534 | 72.2203554578784 | 90.881530252985 | 128.423634367755 | 72.8754663389296 | 27.1381391167127 | Ngf |
| **ENSMUSG00000027860** | -0.474728660288103 | 0.0179067120302221 | 0.642273034804458 | 140.948139784771 | 149.038078391322 | 151.432782183751 | 111.613276616721 | 245.270635863478 | 149.216032313011 | 198.845915296222 | 174.76961591163 | Vangl1 |
| **ENSMUSG00000027861** | 0.00471668982382265 | 0.977043376755552 | 0.996179425433073 | 21271.423429185 | 19989.8340662022 | 15546.4680009394 | 18388.9907676255 | 15715.9349845915 | 16276.7783444389 | 21853.2701988639 | 21104.7880282851 | Casq2 |
| **ENSMUSG00000027864** | -0.564005458946386 | 0.0354900539251188 | 0.679566075569698 | 501.45011269582 | 612.440628143575 | 576.391027186904 | 483.032247542953 | 1041.30524422396 | 1099.5509266344 | 539.278450908079 | 534.078577816906 | Ptgfrn |
| **ENSMUSG00000027865** | -0.149355903628289 | 0.298132096169314 | 0.851785577030345 | 148.176249517323 | 179.985876090067 | 178.879973954556 | 178.206071909051 | 208.042057205628 | 194.470074899744 | 188.435134390661 | 169.341988088287 | Gdap2 |
| **ENSMUSG00000027868** | 0.304568071632203 | 0.471833602985366 | 0.897171173241185 | 64.1494738764022 | 66.7820897709749 | 14.1968233297267 | 49.7101147956825 | 33.943704070392 | 39.1386314263635 | 44.7663578939139 | 40.1644458927348 | Tbx15 |
| **ENSMUSG00000027875** | 0.613711263075349 | 0.0630070895246659 | 0.715615581523459 | 1589.28062744495 | 2975.87507345296 | 5241.4671733351 | 2311.05137465211 | 1620.53812981226 | 2338.53322772522 | 1675.09464770482 | 2285.03131362721 | Hmgcs2 |
| **ENSMUSG00000027878** | -0.349998959220573 | 0.221247231056207 | 0.830349645871581 | 113.8427282877 | 150.666909849151 | 141.021778408619 | 82.5375490947182 | 191.617684268342 | 205.477814988408 | 119.723980413956 | 106.381505337514 | Notch2 |
| **ENSMUSG00000027879** | -0.06996171084707 | 0.509769603598623 | 0.902394714365738 | 884.539928521095 | 803.013908709528 | 768.521369582539 | 860.078778634734 | 903.340511550755 | 950.334894321389 | 833.903550535465 | 794.604713337348 | Sec22b |
| **ENSMUSG00000027881** | -0.0865218015342706 | 0.354587657473149 | 0.863771056173629 | 746.302329886031 | 745.190391956611 | 691.858523602015 | 676.245146560134 | 747.856447744443 | 724.064681387725 | 815.164144905455 | 749.01263962127 | Prpf38b |
| **ENSMUSG00000027882** | 0.158933121355288 | 0.266438533054605 | 0.843513595393324 | 351.466835745359 | 381.96097686082 | 389.93941412316 | 344.219096792745 | 309.873169416804 | 269.078091056249 | 373.747034509653 | 360.394487469945 | Stxbp3 |
| **ENSMUSG00000027883** | -0.302317149716948 | 0.588886834295244 | 0.92352104528132 | 9.03513716569045 | 12.2162359337149 | 14.1968233297267 | 27.1998741334867 | 36.1336204620302 | 22.0154801773295 | 10.4107809055614 | 8.68420451734806 | Gpsm2 |
| **ENSMUSG00000027884** | 0.186720753388924 | 0.114824287982302 | 0.767518180080285 | 798.706125447036 | 807.900403083014 | 871.68495244522 | 843.196098138087 | 692.013579757669 | 624.995020589742 | 810.99983254323 | 788.091559949337 | Clcc1 |
| **ENSMUSG00000027889** | -0.751187753853401 | 0.0153699567382271 | 0.635862253526407 | 36.1405486627618 | 52.9370223794314 | 28.3936466594534 | 35.6412143818101 | 74.4571573156986 | 85.6157562451702 | 41.6431236222455 | 57.5328549274309 | Ampd2 |
| **ENSMUSG00000027890** | 0.136081733114224 | 0.252061408908096 | 0.83723604018117 | 335.203588847116 | 320.879797192245 | 324.634026806417 | 300.136542162612 | 270.454674367317 | 322.893709267499 | 270.680303544596 | 302.861632542514 | Gstm4 |
| **ENSMUSG00000027893** | 0.157310203162607 | 0.45104968399245 | 0.891094723422636 | 1971.46692955366 | 2381.3515913455 | 2160.7565107844 | 1790.50205933883 | 1521.99189218854 | 1344.16737304917 | 2302.86473631017 | 2276.34710910986 | Ahcyl1 |
| **ENSMUSG00000027894** | 0.034865511401931 | 0.913402638836895 | 0.986234232499696 | 72.2810973255236 | 125.420022252807 | 165.629605513478 | 62.8410885152968 | 99.6411958195378 | 86.838838477244 | 121.806136595068 | 107.467030902182 | Slc6a17 |
| **ENSMUSG00000027895** | -0.252712925155251 | 0.752470273037808 | 0.956497786093635 | 0.903513716569045 | 8.95857301805762 | 26.5007368821565 | 6.56548685980713 | 14.2344565456483 | 23.2385624094033 | 8.32862472444909 | 5.42762782334254 | Kcnc4 |
| **ENSMUSG00000027900** | 0.184700127592027 | 0.0976965288015836 | 0.759517686335357 | 1833.22933091859 | 1652.44951396717 | 2010.2701834893 | 1943.38411050291 | 1492.42802090143 | 1578.99916160735 | 1659.47847634648 | 1812.82769299641 | Dram2 |
| **ENSMUSG00000027905** | 0.00664913281413751 | 0.96633787262099 | 0.994284908685797 | 259.308436655316 | 246.767965861042 | 301.919109478855 | 259.805694309511 | 295.638712871156 | 211.593226148778 | 299.830490080167 | 254.012982132431 | Ddx20 |
| **ENSMUSG00000027907** | -0.594667936895292 | 0.0970622448426685 | 0.757809915356977 | 346.949267162513 | 240.252640029727 | 216.738169500494 | 342.343243404229 | 559.523638063559 | 681.25680326514 | 221.749633288457 | 269.21034003779 | S100a11 |
| **ENSMUSG00000027931** | -0.686574147500567 | 0.0388760211232625 | 0.679566075569698 | 184.316798180085 | 206.861595144239 | 249.86409060319 | 272.936668029125 | 475.211856985488 | 562.617826753975 | 213.421008564008 | 220.361689627707 | Npr1 |
| **ENSMUSG00000027932** | -1.02361032192923 | 0.116435663888736 | 0.769868040577702 | 9.93865088225949 | 4.88649437348597 | 5.67872933189068 | 18.7585338851632 | 26.2789966996583 | 36.6924669622158 | 10.4107809055614 | 6.51315338801105 | Slc27a3 |

| | | | | | | | | | | | | |
|---|---|---|---|---|---|---|---|---|---|---|---|---|
| **ENSMUSG00000027933** | -0.0211594218062175 | 0.898247282942152 | 0.984462340635154 | 689.380965742181 | 722.386751547009 | 807.326020017125 | 642.47978556684 | 631.790878987619 | 605.425704876561 | 914.066563508288 | 751.183690750607 | Ints3 |
| **ENSMUSG00000027935** | -0.434432498756771 | 0.0665649830897299 | 0.719725956118125 | 100.290022539164 | 102.616381843205 | 123.039135524298 | 142.564857527241 | 173.003394939417 | 212.816308380852 | 117.641824232843 | 130.263067760221 | Rab13 |
| **ENSMUSG00000027936** | -0.127379418947239 | 0.323419252524844 | 0.855977678491063 | 307.194663633475 | 333.910448854875 | 374.796135904785 | 345.157023487003 | 406.229490648885 | 399.947889888152 | 348.761160336306 | 332.170822788563 | Crtc2 |
| **ENSMUSG00000027937** | -0.0115285579562249 | 0.926171675083324 | 0.986713918026868 | 920.680477183857 | 750.891302059011 | 771.360734248484 | 946.368034506485 | 792.749733773026 | 931.988660840281 | 850.560799984364 | 841.282312618093 | Jtb |
| **ENSMUSG00000027940** | -0.247025182429763 | 0.147494145139249 | 0.780607990324772 | 1131.19917314444 | 1148.3261777692 | 1312.7329305554 | 1328.10419906956 | 1756.31294609383 | 1656.05334222801 | 1111.87140071395 | 1315.65698437823 | Tpm3 |
| **ENSMUSG00000027942** | 0.134699684661618 | 0.200844262515285 | 0.812921911423739 | 1761.85174730964 | 1513.18442432282 | 1497.29163384184 | 1760.48840512257 | 1417.97086358573 | 1412.65997804531 | 1556.41174538142 | 1562.07128755798 | 4933434E20F |
| **ENSMUSG00000027944** | 0.0104749827516961 | 0.934483761226232 | 0.98818087308472 | 577.34526488762 | 448.743066631795 | 597.21303473717 | 575.886990274511 | 556.238763476101 | 568.733237914345 | 533.031982364742 | 524.308847734889 | Hax1 |
| **ENSMUSG00000027947** | -0.493302074205392 | 0.0492664024519776 | 0.703396849069134 | 72.2810973255236 | 109.13170767452 | 88.966759532954 | 76.9099889291692 | 169.71852035196 | 117.415894279091 | 87.4505596067155 | 115.065709854862 | Il6ra |
| **ENSMUSG00000027951** | -0.277052584540645 | 0.215742511060004 | 0.825072543673859 | 137.334084918495 | 193.016527752696 | 166.576060402127 | 167.888878272211 | 226.656346534553 | 258.070350967584 | 185.311900118992 | 136.776221148232 | Adar |
| **ENSMUSG00000027952** | -0.0112575920834195 | 0.94188647028609 | 0.989934163630124 | 174.378147297826 | 166.140808698523 | 142.914688185915 | 154.757904552597 | 158.768938393769 | 189.577745971448 | 151.997401221196 | 144.374900100912 | Pmvk |
| **ENSMUSG00000027953** | -0.128945747646863 | 0.601047053510506 | 0.925969784079517 | 244.852217190211 | 194.645359210525 | 230.934992830221 | 282.315934971707 | 336.152166116463 | 319.224462571277 | 167.613572579538 | 219.276164063039 | Slc50a1 |
| **ENSMUSG00000027954** | -0.378563903971927 | 0.132253488487743 | 0.77303252008843 | 274.66816983699 | 215.820168162297 | 282.990011705886 | 411.749818779333 | 462.072358635659 | 454.986590331476 | 265.474913091815 | 358.223436340608 | Efna1 |
| **ENSMUSG00000027955** | -0.401202650139404 | 0.255700967550774 | 0.837479467570303 | 218.650319409709 | 223.149909722526 | 187.398067952392 | 489.59773440276 | 460.97740043984 | 495.348303989913 | 246.735507461804 | 274.637967861132 | Fam198b |
| **ENSMUSG00000027956** | 0.32512649429856 | 0.108033540331523 | 0.762287160109383 | 1640.78090928939 | 1922.02112023782 | 2299.88537941573 | 2106.58335530383 | 1208.83384818428 | 1298.91333046244 | 1884.35134390661 | 1968.057848744 | Tmem144 |
| **ENSMUSG00000027957** | -0.12965309382559 | 0.407913712016311 | 0.881573350402261 | 231.299511441676 | 192.202112023782 | 221.470443943737 | 246.674720589896 | 261.695008800764 | 280.085831144914 | 205.092383839559 | 229.045894145055 | Slc35a3 |
| **ENSMUSG00000027959** | -0.042647813846679 | 0.91346088751335 | 0.986234232499696 | 13.5527057485357 | 19.5459774939439 | 10.4110037751329 | 14.0689004138724 | 13.1394983498292 | 13.4539045528125 | 17.6983275394543 | 15.1973579053591 | Sass6 |
| **ENSMUSG00000027961** | 0.172385702854575 | 0.13365039013067 | 0.77303252008843 | 5038.89599730556 | 4693.47784573328 | 4783.38300722925 | 4972.88733295677 | 3723.95282398075 | 4038.61753030788 | 4652.57798669537 | 4877.2663620556 | Lrrc39 |
| **ENSMUSG00000027962** | -0.586862151210175 | 0.144817330096157 | 0.780607990324772 | 184.316798180085 | 133.56417954195 | 212.005895057252 | 162.261318106662 | 351.48158085793 | 434.19419238622 | 117.641824232843 | 136.776221148232 | Vcam1 |
| **ENSMUSG00000027963** | -0.227607057944572 | 0.114748564261357 | 0.767518180080285 | 218.650319409709 | 276.086932101957 | 232.827902607518 | 215.723139679377 | 250.745426842573 | 260.516515431732 | 308.159114804616 | 285.493223507818 | Extl2 |
| **ENSMUSG00000027968** | -0.00530498508470627 | 0.975802910216252 | 0.99575536949131 | 1123.06754969532 | 851.878852444388 | 756.217456030109 | 1047.66411748637 | 907.720344334031 | 1146.02805145321 | 815.164144905455 | 924.867781097568 | Larp7 |
| **ENSMUSG00000027981** | -0.180152018350233 | 0.31757760154596 | 0.854351922245337 | 120.167324303683 | 112.389370590177 | 138.182413742673 | 107.861569839689 | 124.825234323377 | 113.746647582869 | 164.49033830787 | 137.8617467129 | Rnpc3 |
| **ENSMUSG00000027983** | -0.0625004342636072 | 0.64624768794678 | 0.935063240511793 | 371.344137509877 | 309.477976987445 | 336.937940358847 | 335.777756544422 | 407.324448844704 | 306.993640250539 | 331.062832796851 | 366.907640857956 | Cyp2u1 |
| **ENSMUSG00000027984** | 0.225425652654582 | 0.086037666851305 | 0.746460787597018 | 23106.4597875368 | 21612.9646139285 | 23829.8411863906 | 22987.6453495733 | 18890.2187942711 | 15832.7994941961 | 20862.1638566544 | 22708.1092873005 | Hadh |
| **ENSMUSG00000027993** | -0.268516700530621 | 0.0923600454446484 | 0.754058637722175 | 151.7903043836 | 179.171460361152 | 149.539872406455 | 148.192417692789 | 228.846262926191 | 176.123841418636 | 172.818963032319 | 180.197243734972 | Trim2 |
| **ENSMUSG00000027994** | -0.471523561240004 | 0.234523004292098 | 0.832048466795696 | 24.3948703473642 | 21.1748089517725 | 17.036187995672 | 15.9447538023887 | 30.6588294829347 | 44.0309603546589 | 17.6983275394543 | 17.3684090346961 | Mcub |
| **ENSMUSG00000027995** | -0.332015208094666 | 0.279059307596918 | 0.848075053184273 | 52.4037955610046 | 43.1640336324594 | 34.0723759913441 | 50.6480414899407 | 73.3621991198795 | 74.6080161565054 | 41.6431236222455 | 37.9933947633978 | Tlr2 |
| **ENSMUSG00000027996** | -0.696352974989314 | 0.264705196005791 | 0.842510961779757 | 16.2632468982428 | 19.5459774939439 | 11.3574586637814 | 17.8206071909051 | 28.4689130912965 | 57.4848649074714 | 15.616171358342 | 4.34210225867403 | Sfrp2 |
| **ENSMUSG00000027997** | -0.0989934036786022 | 0.510031281315162 | 0.902394714365738 | 197.869503928621 | 155.553404222637 | 175.094154399963 | 155.695831246855 | 203.662224422352 | 184.685417043153 | 171.777884941763 | 172.598564782293 | Casp6 |
| **ENSMUSG00000027998** | -0.148743498034503 | 0.261549807340266 | 0.841137531367976 | 471.634160049041 | 452.000729547452 | 477.959718767466 | 496.163221262567 | 598.942133113046 | 574.848649074714 | 443.499266576914 | 487.40097853616 | Plrg1 |
| **ENSMUSG00000027999** | 0.165193065366205 | 0.299071058091665 | 0.851785577030345 | 1544.1049416165 | 1386.13557061219 | 2044.34255948065 | 1700.46109669005 | 1528.56164136346 | 1492.16032313011 | 1676.13572579538 | 1254.86755275679 | Pla2g12a |
| **ENSMUSG00000028001** | -0.286454616742703 | 0.417758042139985 | 0.882468496126562 | 431.879556520004 | 1104.34772840783 | 441.994432998825 | 582.452477134318 | 560.618596259378 | 599.310293716191 | 737.083288113745 | 1226.64388807541 | Fga |
| **ENSMUSG00000028005** | -0.452636410426845 | 0.07181066666926 | 0.728065810245271 | 156.307872966445 | 145.780415475665 | 157.111511515642 | 176.330218520534 | 295.638712871156 | 266.631926592101 | 159.284947855089 | 148.717002359586 | Gucy1b1 |
| **ENSMUSG00000028007** | -0.339393317377384 | 0.0508821444779292 | 0.703396849069134 | 131.009488902512 | 137.636258186522 | 134.396594188079 | 109.737423228205 | 173.003394939417 | 189.577745971448 | 157.202791673977 | 130.263067760221 | Snx7 |
| **ENSMUSG00000028010** | 0.338040175428653 | 0.0317470603820545 | 0.671606027118506 | 321.65088309858 | 333.910448854875 | 368.170951684246 | 399.556771753977 | 343.816873487196 | 277.639666680766 | 236.324726556243 | 268.124814473121 | Gar1 |
| **ENSMUSG00000028011** | -0.26895110995724 | 0.294116292281829 | 0.851785577030345 | 565.599586572222 | 777.767021113184 | 547.997380527451 | 630.286738541484 | 528.864808580624 | 760.757148349941 | 595.49666779811 | 1153.91367524262 | Tdo2 |
| **ENSMUSG00000028013** | 0.212114118956732 | 0.102873925662573 | 0.759800953500741 | 2234.38942107525 | 1868.26968212947 | 1891.0168675196 | 2535.21585457981 | 1842.81464356354 | 1812.60786793346 | 1764.62736349265 | 1942.00523519196 | Ppa2 |
| **ENSMUSG00000028015** | -0.125100593302672 | 0.540623576453679 | 0.913837378403038 | 210.518695960587 | 210.119258059897 | 207.27362061401 | 211.971432902344 | 292.353838283699 | 269.078091056249 | 165.531416398426 | 189.966973816989 | Ctso |
| **ENSMUSG00000028016** | 0.018845514967104 | 0.887588267713681 | 0.980702840906172 | 209.615182244018 | 215.820168162297 | 226.202718386979 | 206.343872736795 | 206.947099009809 | 183.462334811079 | 217.585320926233 | 237.730098662403 | Ints12 |
| **ENSMUSG00000028017** | 0.169877813686353 | 0.658740955905389 | 0.938989601151841 | 225.878429142261 | 346.12668478859 | 173.201244622666 | 128.495957113368 | 162.053812981226 | 103.961989726278 | 324.816364253515 | 185.624871558315 | Egf |
| **ENSMUSG00000028018** | -0.0584228337639629 | 0.867108832097824 | 0.974868041734905 | 36.1405486627618 | 23.6180561385155 | 17.036187995672 | 20.6343872736795 | 32.8487458745729 | 20.7923979452556 | 24.9858741733473 | 22.7960368580387 | Gstcd |
| **ENSMUSG00000028019** | -0.201412549090315 | 0.579071442127842 | 0.922450244597442 | 23.4913566307952 | 13.8450673915436 | 22.7149173275627 | 35.6412143818101 | 19.7092475247437 | 28.1308913376988 | 30.191264626128 | 31.4802413753867 | Pdgfc |
| **ENSMUSG00000028024** | -0.344280975598199 | 0.136310981475903 | 0.776456211338551 | 118.360296870545 | 117.275864963663 | 107.895857305923 | 90.9788893430416 | 182.858018701789 | 113.746647582869 | 99.9434966933891 | 155.230155747597 | Enpep |
| **ENSMUSG00000028028** | 0.021466476144662 | 0.959103271588499 | 0.993211395091646 | 16.2632468982428 | 9.77298874697194 | 16.0897331070236 | 12.1930470253561 | 16.4243729372865 | 15.9000690169602 | 12.4929370866736 | 8.68420451734806 | Alpk1 |
| **ENSMUSG00000028029** | 0.0905332599461547 | 0.462381644862675 | 0.895328112556867 | 2022.96721139809 | 1727.37576102729 | 1920.3569690677 | 2242.58272597126 | 1854.85918371755 | 2096.3629457746 | 1757.33981685876 | 1723.81459669359 | Aimp1 |
| **ENSMUSG00000028030** | 0.0899585370459336 | 0.728717883615687 | 0.952401994188708 | 196.965990212052 | 283.416673662186 | 282.043556817237 | 261.681547698027 | 228.846262926191 | 140.654456688494 | 330.021754706295 | 261.61166108511 | Tbck |
| **ENSMUSG00000028032** | -0.554429094152078 | 0.0457299406987537 | 0.697906909182231 | 144.562194651047 | 196.274190668353 | 165.629605513478 | 142.564857527241 | 338.342082508101 | 289.870489001505 | 169.69572876065 | 156.315681312265 | Papss1 |
| **ENSMUSG00000028034** | 0.2116111679091 | 0.21108355095566 | 0.823123035902316 | 1194.44513330428 | 1410.56804247962 | 1333.55493810566 | 1111.44313269592 | 1039.11532783232 | 824.357424417781 | 1303.42976937628 | 1192.99259557069 | Fubp1 |
| **ENSMUSG00000028035** | 0.298078652026701 | 0.0378732977980976 | 0.679566075569698 | 4064.00469712756 | 5306.73288960577 | 4319.62011179151 | 3873.63724728621 | 3116.25102530115 | 3392.83011177289 | 3787.44209344323 | 3989.30645015677 | Dnajb4 |
| **ENSMUSG00000028036** | -0.550285668224673 | 0.126014077612083 | 0.771949105062466 | 41.5616309621761 | 65.1532583131463 | 38.8046504345863 | 52.523894878457 | 54.7479097909548 | 128.423634367755 | 60.3825292522559 | 47.7631248454143 | Ptgfr |
| **ENSMUSG00000028037** | -0.0426079841393061 | 0.850512899000696 | 0.972692784625411 | 76.7986659083688 | 70.8541684155466 | 76.6628459805242 | 62.8410885152968 | 95.2613630362614 | 80.7234273168747 | 55.1771387994752 | 65.1315338801105 | Ifi44 |
| **ENSMUSG00000028041** | -0.693542901398949 | 0.0547921133477364 | 0.703396849069134 | 810.451803762433 | 789.983257046899 | 911.436057768454 | 931.361207398354 | 1932.60121562071 | 2199.1018532688 | 755.822693743755 | 681.710054611823 | Adam15 |
| **ENSMUSG00000028042** | 0.0421205133233936 | 0.769578747393323 | 0.95879680569409 | 567.40661400536 | 572.534257426773 | 759.056820696054 | 639.666005484066 | 620.841297029428 | 533.263853184203 | 677.741836952045 | 631.775878637071 | Zbtb7b |
| **ENSMUSG00000028044** | -0.0142379382082187 | 0.954529994604986 | 0.992371505557225 | 57.8248778604189 | 65.9676740420606 | 42.5904699891801 | 64.7169419038131 | 67.887408140784 | 67.2695227640623 | 42.6842017128016 | 56.4473293627624 | Cks1b |
| **ENSMUSG00000028047** | -1.21339578953563 | 0.0211929587124316 | 0.648762042745232 | 25.2983840639333 | 25.2468875963442 | 19.8755526616174 | 39.3929211588428 | 72.2672409240604 | 130.869798831903 | 26.0269522639034 | 26.0526135520442 | Thbs3 |
| **ENSMUSG00000028048** | -0.203209012317881 | 0.321493902581902 | 0.855704427283138 | 129.202461469373 | 148.223662662408 | 161.843785958884 | 159.447538023887 | 208.042057205628 | 207.923979452556 | 140.545542225078 | 133.519644454226 | Gba |
| **ENSMUSG00000028049** | -0.19518668194538 | 0.0373953085109943 | 0.679566075569698 | 1116.74295367934 | 1125.5225373596 | 1074.22629861599 | 1080.4915517854 | 1280.00613091252 | 1369.85209992272 | 1129.56972825341 | 1257.03860388613 | Scamp3 |
| **ENSMUSG00000028053** | 0.308821255990033 | 0.28328436424658 | 0.850587941933635 | 1442.91140536077 | 2085.7186817496 | 1804.88947265259 | 1214.61506906432 | 1103.71786138565 | 747.303243797128 | 1779.20245676044 | 1655.42648611947 | Ash1l |
| **ENSMUSG00000028057** | -0.082860176960026 | 0.395704031992825 | 0.879215782741314 | 1239.62081913273 | 1145.06851485355 | 1041.10037751329 | 1160.21532079735 | 1305.19016941636 | 1152.14346261358 | 1180.58255469066 | 1219.04520912273 | Rit1 |
| **ENSMUSG00000028059** | -0.278223047702463 | 0.160275533969868 | 0.79125089991166 | 642.398252480591 | 610.811796685746 | 657.786147610671 | 694.065753751039 | 1035.83045324487 | 897.742358342213 | 595.49666779811 | 630.690353072403 | Arhgef2 |
| **ENSMUSG00000028060** | 0.224812256625334 | 0.297921713821123 | 0.851785577030345 | 430.072529086865 | 620.584785432718 | 530.01473764313 | 434.260059441529 | 341.626957095558 | 341.239942748607 | 484.101312108604 | 556.874614674944 | 2810403A07F |
| **ENSMUSG00000028062** | -0.0726934027974887 | 0.72674886757228 | 0.952045814446758 | 1402.25328811516 | 1102.71889695 | 1217.1409868019 | 1494.11722395325 | 1384.02715951534 | 1898.22362417863 | 1069.18719900115 | 1135.45974064326 | Lamtor2 |
| **ENSMUSG00000028063** | -0.199284740514325 | 0.291872565175452 | 0.851785577030345 | 2987.91986069383 | 2678.61333239923 | 2716.32553042104 | 3037.94456270218 | 3661.54020681906 | 4277.11856556229 | 2657.87236518982 | 2517.33378446627 | Lmna |
| **ENSMUSG00000028064** | -0.459813374921662 | 0.149998420148863 | 0.782641594375979 | 26.2018977805023 | 45.6072808192024 | 46.3762895437739 | 38.4549944645846 | 79.9319482947941 | 55.0387004433237 | 40.6020455316893 | 40.1644458927348 | Sema4a |
| **ENSMUSG00000028066** | -0.346380322754122 | 0.07766753276738 | 0.739855948845396 | 131.009488902512 | 103.43079757212 | 113.574586637814 | 105.985716451172 | 147.819356435578 | 181.016170346931 | 139.504464134522 | 109.638082031519 | Pmf1 |
| **ENSMUSG00000028068** | -0.454032450738708 | 0.480280457600253 | 0.897732059433541 | 9.93865088225949 | 12.2162359337149 | 9.46454888648447 | 2.81378008277448 | 18.6142893289246 | 18.3462334811079 | 4.16431236222455 | 6.51315338801105 | Iqgap3 |
| **ENSMUSG00000028069** | 0.242667955110026 | 0.152686926052543 | 0.783412422888014 | 704.740698923855 | 619.770369703804 | 566.92647830042 | 910.726820124675 | 597.847174917227 | 636.002760678407 | 581.96265262088 | 552.53251241627 | Gpatch4 |
| **ENSMUSG00000028070** | -0.0614692786282784 | 0.722998050695902 | 0.951674101666722 | 2149.45913171776 | 1462.69064913013 | 1600.45521670452 | 2094.39030827847 | 1943.5507975789 | 2241.90973139138 | 1653.23200780315 | 1786.77507944436 | Naxe |
| **ENSMUSG00000028073** | -0.620839417854111 | 0.0744629724235783 | 0.735399943172214 | 147.272735800754 | 148.223662662408 | 175.094154399963 | 177.268145214792 | 352.576539053749 | 364.47850515801 | 160.326025945645 | 119.407812113536 | Pear1 |
| **ENSMUSG00000028076** | -0.097743246153647 | 0.592823489906294 | 0.923935529355527 | 85.8338030740593 | 92.8433930962335 | 101.270673085384 | 101.296082979881 | 99.6411958195378 | 99.0696607979826 | 85.3684034256032 | 123.74991437221 | Cd1d1 |
| **ENSMUSG00000028080** | 0.239104666280835 | 0.490217787510965 | 0.899236955296196 | 806.837748896157 | 1206.14969452212 | 1092.20894150031 | 669.679659700327 | 616.461464246152 | 366.924669622158 | 1070.22827709171 | 1144.14394516061 | Lrba |
| **ENSMUSG00000028081** | 0.0492146038393716 | 0.72640039147565 | 0.952038256048006 | 12877.7810022586 | 10025.4576229354 | 11880.848217204 | 13616.81974724 | 11517.8652618211 | 13653.2669566405 | 10458.6704977269 | 11148.3475491456 | Rps3a1 |
| **ENSMUSG00000028082** | -0.101844629283517 | 0.32643302687326 | 0.855977678491063 | 418.326850771468 | 457.701639649853 | 437.262158555582 | 412.687745473591 | 486.161438943679 | 420.740287833408 | 474.731609293598 | 470.032569501464 | Sh3d19 |
| **ENSMUSG00000028085** | 0.159637465760443 | 0.123630824824154 | 0.770326056866107 | 1551.33305134905 | 1390.20764925676 | 1542.72146849697 | 1715.46792379818 | 1412.49607260663 | 1274.45168582096 | 1405.45542225078 | 1455.68978222047 | Gatb |
| **ENSMUSG00000028086** | -0.15773783628585 | 0.256797593062111 | 0.837517407767513 | 802.320180313312 | 699.583111137408 | 684.286884492827 | 847.885731609378 | 721.577451044785 | 798.67269754423 | 956.75076522109 | 906.413846498204 | Fbxw7 |
| **ENSMUSG00000028088** | -0.065025236339331 | 0.743138909562524 | 0.954383574136061 | 451.756858284523 | 690.624538119351 | 733.502538702546 | 628.410885152968 | 571.568178217569 | 604.202622644487 | 609.03068297534 | 834.769159230082 | Fmo5 |
| **ENSMUSG00000028089** | 0.19017951367209 | 0.131598862782989 | 0.77303252008843 | 470.730646332472 | 497.608010366655 | 516.764369202052 | 540.245775892701 | 489.446313531136 | 370.593916318379 | 443.499266576914 | 470.032569501464 | Chd1l |
| **ENSMUSG00000028093** | -0.0390665827745527 | 0.732803360953108 | 0.953733898884637 | 405.677658739501 | 378.703313945163 | 449.566072108012 | 423.004939110431 | 411.70428162798 | 390.163232031561 | 438.293876124134 | 460.262839419447 | Acp6 |
| **ENSMUSG00000028096** | 0.199338045556996 | 0.127531921039857 | 0.772837268577874 | 467.116591466196 | 484.577358704026 | 534.747012086372 | 467.087493740564 | 377.760577557588 | 380.37857417497 | 464.320828388037 | 477.631248454143 | Gpr89 |
| **ENSMUSG00000028098** | 0.223461696146438 | 0.147648689797739 | 0.780607990324772 | 899.899661702769 | 969.154717408051 | 765.682004916593 | 956.685228143324 | 651.500126512363 | 673.918309872697 | 859.930502799369 | 890.130963028176 | Rnf115 |
| **ENSMUSG00000028099** | 0.0942708393616891 | 0.281447092105576 | 0.850048154552447 | 717.389890955822 | 678.408302185636 | 685.233339381476 | 754.093062183562 | 679.969039603659 | 648.233582999146 | 657.961353231478 | 668.683747835801 | Polr3c |
| **ENSMUSG00000028100** | -0.125756213172533 | 0.628117467025759 | 0.931825800800958 | 42.4651446787451 | 31.7622134276588 | 45.4298346551254 | 44.0825546301336 | 37.2285786578493 | 58.7079471395453 | 41.6431236222455 | 41.2499714574033 | Nudt17 |
| **ENSMUSG00000028101** | -0.149181580124319 | 0.363957620479673 | 0.86924083181804 | 177.992202164102 | 166.955224427437 | 173.201244622666 | 215.723139679377 | 204.757182618171 | 236.054870790255 | 206.133461930115 | 167.17093695895 | Pias3 |
| **ENSMUSG00000028102** | 0.156557933438125 | 0.191327804626347 | 0.808824684419004 | 448.142803418246 | 372.187988113848 | 466.602260103684 | 432.384206053012 | 402.944616061428 | 397.501725424004 | 383.116737324658 | 358.223436340608 | Pex11b |
| **ENSMUSG00000028104** | 0.225946485138726 | 0.0785136264527741 | 0.739855948845396 | 1051.68996608637 | 934.134841064735 | 1198.21188902893 | 1212.7392156758 | 933.99934103369 | 837.811328970594 | 993.188498390554 | 992.170366107016 | Polr3gl |
| **ENSMUSG00000028106** | 0.211060341681299 | 0.207119283074181 | 0.819960256425221 | 641.494738764022 | 732.974156022896 | 742.020632700382 | 680.934780031425 | 562.808512651016 | 451.317343635254 | 693.358008310387 | 707.762668163867 | Rprd2 |
| **ENSMUSG00000028107** | 0.0493553391532153 | 0.62142930271411 | 0.93078489223231 | 1240.5243328493 | 1088.87382955846 | 1025.95709929492 | 1201.4840953447 | 1116.85735973548 | 1117.89716011551 | 1120.2000254384 | 1048.61769546978 | Tars2 |
| **ENSMUSG00000028108** | -1.50272564302656 | 0.0286961901215826 | 0.665956123973759 | 302.67709505063 | 276.086932101957 | 292.45456059237 | 278.564228194674 | 979.987585258092 | 1801.6001278448 | 216.544242835676 | 260.526135520442 | Ecm1 |

| | | | | | | | | | | | | |
|---|---|---|---|---|---|---|---|---|---|---|---|---|
| **ENSMUSG00000028111** | -0.827205075080929 | 0.0562273603843011 | 0.703396849069134 | 50.5967681278665 | 48.0505280059454 | 47.3227444324223 | 30.9515809105193 | 94.1664048404423 | 140.654456688494 | 33.3144988977964 | 46.6775992807458 | Ctsk |
| **ENSMUSG00000028114** | 0.0854808937099947 | 0.520497995973288 | 0.905962179970576 | 427.361987937158 | 500.051257553398 | 417.386605893965 | 426.756645887463 | 412.7992398238 | 353.470765069345 | 438.293876124134 | 464.604941678121 | Mettl14 |
| **ENSMUSG00000028116** | 0.205060709412574 | 0.18485799149168 | 0.805963874499992 | 28577.2353413623 | 20740.7253682612 | 22153.6695785942 | 31509.6472936029 | 22792.6498041703 | 22223.404156782 | 22046.9107237073 | 22272.8135358684 | Myoz2 |
| **ENSMUSG00000028121** | -0.309542522853405 | 0.194280544299168 | 0.809162446871198 | 92.1583990900426 | 78.9983257046899 | 65.3053873167428 | 80.6616957062019 | 125.920192519196 | 110.077400886647 | 67.6700758861489 | 90.0986218674861 | Bcar3 |
| **ENSMUSG00000028124** | 0.152778057345557 | 0.302422228003194 | 0.851785577030345 | 1562.17521594788 | 1869.08409785838 | 2254.4555447606 | 1667.63366239101 | 1701.56503630288 | 1758.79224972221 | 1710.49130278373 | 1443.74900100912 | Gclm |
| **ENSMUSG00000028125** | 0.226337939095412 | 0.591948764297912 | 0.923935529355527 | 141.85165350134 | 67.5965054998893 | 198.755526616174 | 201.654239265505 | 100.736154015357 | 90.5080851734656 | 87.4505596067155 | 242.072200921077 | Abca4 |
| **ENSMUSG00000028126** | -0.0257407411092071 | 0.847846266470674 | 0.972467578249175 | 337.914129996823 | 333.910448854875 | 290.561650815073 | 297.322762079837 | 350.386622662111 | 269.078091056249 | 326.898520434627 | 335.427399482569 | Pip5k1a |
| **ENSMUSG00000028127** | 0.204753855675672 | 0.338468193070934 | 0.858570960161243 | 3052.9728482868 | 3456.38035351241 | 3491.47208422412 | 2853.17300393333 | 2558.91730362923 | 1795.48471668443 | 3277.31382907072 | 3520.35940621997 | Abcd3 |
| **ENSMUSG00000028128** | -0.290288899677501 | 0.0876337604065859 | 0.74814562510601 | 176.185174730964 | 181.614707547895 | 235.667267273463 | 243.860940507122 | 285.784089108784 | 262.96267989588 | 254.023054095697 | 220.361689627707 | F3 |
| **ENSMUSG00000028132** | -0.353809413015288 | 0.305399196325519 | 0.851785577030345 | 52.4037955610046 | 102.616381843205 | 50.1621090983677 | 65.6548685980713 | 56.937826182593 | 68.4926049961361 | 85.3684034256032 | 135.690695583563 | Tmem56 |
| **ENSMUSG00000028133** | 0.543791713652752 | 0.109465382568915 | 0.763154315676794 | 39.754603529038 | 24.4324718674299 | 24.6078271048596 | 25.3240207449704 | 21.8991639163819 | 13.4539045528125 | 19.7804837205666 | 22.7960368580387 | Rwdd3 |
| **ENSMUSG00000028134** | 0.205226391385216 | 0.360110171305468 | 0.868239937412421 | 187.930853046361 | 237.809392842984 | 298.133289924261 | 251.364354061187 | 183.952976897608 | 152.885279009232 | 258.187366457922 | 249.670879873757 | Ptbp2 |
| **ENSMUSG00000028136** | 0.0833773155039701 | 0.710825380210831 | 0.949886733585872 | 627.038519298917 | 789.983257046899 | 693.751433379311 | 531.804435644377 | 578.137927392483 | 408.509465512669 | 761.028084196536 | 745.756062927265 | Snx27 |
| **ENSMUSG00000028138** | 0.0295368636083394 | 0.782355890898374 | 0.960742105865155 | 2166.62589233257 | 1989.6176257377 | 1969.57262327742 | 2264.1550399392 | 2242.47438503751 | 2140.39390612925 | 1821.88665847324 | 2015.82097358942 | Adh5 |
| **ENSMUSG00000028140** | 0.214994358067571 | 0.127538974539264 | 0.772837268577874 | 2327.45133388186 | 2196.47922088194 | 2322.60029674329 | 2400.15441060663 | 1881.13818041721 | 1572.88375044698 | 2263.30376886904 | 2247.03791886381 | Mrpl9 |
| **ENSMUSG00000028145** | 0.0370560967243659 | 0.846056073489777 | 0.972180580805179 | 237.624107457659 | 160.439898596123 | 159.004421292939 | 218.536919762152 | 180.668102310151 | 163.893019097897 | 200.928071477334 | 209.506433981022 | Them4 |
| **ENSMUSG00000028149** | -0.0319466799649077 | 0.837183559876412 | 0.968924788429433 | 826.715050660676 | 923.547436588849 | 1013.65318574249 | 813.182443921826 | 847.497643563981 | 737.518585940537 | 999.434966933891 | 1071.41373232782 | Rap1gds1 |
| **ENSMUSG00000028150** | 0.411276086475564 | 0.00339368704493976 | 0.635862253526407 | 1664.27226592018 | 1809.63174964764 | 2144.66677767738 | 2142.22456968564 | 1531.84651595092 | 1226.75147877008 | 1470.00226386526 | 1605.49231014472 | Rorc |
| **ENSMUSG00000028152** | -0.489549826396754 | 0.159005585430902 | 0.789393312599749 | 91.2548853734735 | 83.8848200781758 | 88.0203046443056 | 61.9031618210386 | 151.104231023035 | 165.116101329971 | 53.094982618363 | 87.9275707381491 | Tspan5 |
| **ENSMUSG00000028156** | 0.0502836913931183 | 0.800822679959167 | 0.963461896828986 | 4036.89928563049 | 3612.74817346396 | 2798.66710573346 | 3382.16365949493 | 2948.72242134083 | 2498.7570001269 | 3572.98000678866 | 4335.58910528602 | Eif4e |
| **ENSMUSG00000028158** | 0.453459745960407 | 0.0073033233872450 | 0.635862253526407 | 365.019541493894 | 364.858246553619 | 476.066808990169 | 392.053358199911 | 262.789966996583 | 240.94719971855 | 312.323427166841 | 349.539231823259 | Mttp |
| **ENSMUSG00000028159** | -0.00829280416081535 | 0.961404516503524 | 0.993822507448007 | 176.185174730964 | 194.645359210525 | 154.272146849697 | 172.578511743502 | 135.774816281568 | 183.462334811079 | 179.065431575656 | 204.078806157679 | Dapp1 |
| **ENSMUSG00000028161** | -0.020169552672475 | 0.880077531516725 | 0.978393501240173 | 2137.71345340236 | 2019.75100770753 | 1944.01834128391 | 2348.56844242244 | 2258.8987579748 | 2503.64932905519 | 1858.3243916427 | 1949.60391414464 | Ppp3ca |
| **ENSMUSG00000028163** | -0.0854217146496728 | 0.378302627503745 | 0.87422797717331 | 637.880683897746 | 707.727268426552 | 682.39397471553 | 737.210381686915 | 787.274942793931 | 713.05694129906 | 727.71358529874 | 705.59161703453 | Nfkb1 |
| **ENSMUSG00000028164** | -0.213534232555979 | 0.0372322166034959 | 0.679566075569698 | 318.940341948873 | 357.52850499339 | 330.312756138308 | 338.591536627196 | 391.995034103237 | 371.816998550453 | 393.52751823022 | 402.729984492016 | Manba |
| **ENSMUSG00000028165** | 0.180005738217569 | 0.106957215678166 | 0.762287160109383 | 1796.18526853926 | 1808.81733391872 | 1716.86916800828 | 1754.86084495702 | 1420.16077997737 | 1397.98299126042 | 1645.94446116925 | 1781.34745162102 | Cisd2 |
| **ENSMUSG00000028173** | -0.0910571344841446 | 0.426439463988669 | 0.885066977635295 | 702.030157774148 | 788.35442558907 | 716.466350706874 | 644.355638955357 | 709.532910890775 | 814.572766561191 | 810.99983254323 | 703.420565905193 | Wls |
| **ENSMUSG00000028179** | -0.120644774746441 | 0.622447556625174 | 0.93078489223231 | 113.8427282877 | 153.924572764808 | 82.3415753124149 | 113.489130005238 | 104.021028602814 | 113.746647582869 | 120.765058504512 | 166.085411394282 | Cth |
| **ENSMUSG00000028180** | 0.206155076715828 | 0.259871361338906 | 0.840316047847749 | 1137.52376916043 | 1167.05773953423 | 1121.54904304841 | 1023.27802343565 | 821.218646864323 | 717.949270227356 | 1195.15764795844 | 1121.34790830257 | Zranb2 |
| **ENSMUSG00000028184** | -0.107346994316539 | 0.521160536520362 | 0.90609507420585 | 715.582863522684 | 756.592212161411 | 855.595219338196 | 757.844768960594 | 763.185862485911 | 629.887349518038 | 989.02418602833 | 940.065139002928 | Adgrl2 |
| **ENSMUSG00000028186** | -0.333369990003354 | 0.252557715286931 | 0.83723604018117 | 171.667606148119 | 307.034729800702 | 201.594891282119 | 189.461192240149 | 177.383227722694 | 233.608706326107 | 272.762459725708 | 412.499714574033 | Uox |
| **ENSMUSG00000028187** | 0.0105071139232042 | 0.931382321552054 | 0.987775860606096 | 434.590097669711 | 359.157336451219 | 392.778778789105 | 413.625672167849 | 445.647985698372 | 366.924669622158 | 389.363205867995 | 385.36157545732 | Rpf1 |
| **ENSMUSG00000028188** | 0.218303425676311 | 0.488776688364542 | 0.899236955296196 | 26.2018977805023 | 23.6180561385155 | 24.6078271048596 | 30.9515809105193 | 19.7092475247437 | 17.123151249034 | 22.903717992235 | 30.3947158107182 | Spata1 |
| **ENSMUSG00000028189** | -0.301994793375594 | 0.169474446494054 | 0.79907358295209 | 88.5443442237664 | 77.3694942468612 | 68.1447519826882 | 64.7169419038131 | 106.210944994452 | 103.961989726278 | 92.6559500594962 | 66.217059444779 | Ctbs |
| **ENSMUSG00000028191** | -0.0329725246116689 | 0.806703680189134 | 0.964026716319739 | 407.484686172639 | 380.332145402991 | 378.581955459379 | 418.31530563914 | 473.02194059385 | 429.301863457925 | 357.089785060755 | 362.565538599282 | Bcl10 |
| **ENSMUSG00000028194** | -0.185741213283297 | 0.543405241074451 | 0.913837378403038 | 136.430571201926 | 133.56417954195 | 82.3415753124149 | 119.116690170786 | 101.831112211176 | 79.5003450848009 | 164.49033830787 | 189.966973816989 | Ddah1 |
| **ENSMUSG00000028195** | 0.534623817090251 | 0.00166350207346837 | 0.635862253526407 | 1184.50648242202 | 1517.25650296739 | 1683.74324690559 | 2009.97690579524 | 1107.00273597311 | 1043.289143959 | 1100.41954171784 | 1163.68340532464 | Cyr61 |
| **ENSMUSG00000028199** | 0.201713743995805 | 0.335285415437748 | 0.858570960161243 | 246.659244623349 | 265.499527626071 | 327.473391472363 | 201.654239265505 | 205.85214081399 | 173.677676954488 | 265.474913091815 | 259.440609955773 | Cryz |
| **ENSMUSG00000028207** | 0.0583925408683008 | 0.764414063117607 | 0.958110466593105 | 5290.07281051176 | 5454.95655226817 | 4312.04847268232 | 4777.79858055107 | 3940.75454675293 | 3549.38463747834 | 5885.21444591384 | 5671.87107539295 | Asph |
| **ENSMUSG00000028211** | -0.0687274340441779 | 0.823294160890453 | 0.966637987288264 | 327.975479114563 | 456.887223920938 | 237.56017705076 | 178.206071909051 | 412.7992398238 | 353.470765069345 | 250.899819824029 | 243.157726485746 | Trp53inp1 |
| **ENSMUSG00000028212** | 0.038253472887607 | 0.889767330243336 | 0.981485722449169 | 51.5002818444356 | 57.009101024003 | 40.6975602118832 | 39.3929211588428 | 36.1336204620302 | 36.6924669622158 | 47.8895921655823 | 62.9604827507734 | Ccne2 |
| **ENSMUSG00000028214** | -0.726298351044921 | 0.100804940462726 | 0.759800953500741 | 62.3424464432641 | 53.7514381083457 | 22.7149173275627 | 39.3929211588428 | 96.3563212320805 | 118.638976511164 | 40.6020455316893 | 40.1644458927348 | Gem |
| **ENSMUSG00000028218** | 0.133208600737124 | 0.413918933906583 | 0.882192126163777 | 367.730082643601 | 317.622134276588 | 317.06238769723 | 417.377378944882 | 291.25888008788 | 282.531995609062 | 332.103910887408 | 387.532626586657 | Fam92a |
| **ENSMUSG00000028221** | 0.0126321461009546 | 0.919992542098815 | 0.986713918026868 | 543.915257374565 | 522.040482234085 | 498.781726317731 | 526.176875478828 | 501.490853685146 | 436.640356850368 | 537.196294726966 | 595.953535003011 | Tmem55a |
| **ENSMUSG00000028223** | 0.160365439743488 | 0.0735844374548856 | 0.732679452507786 | 20397.7256652628 | 19489.7828086488 | 22666.6481282417 | 21356.5908282583 | 18263.9027062625 | 18418.3953328003 | 17997.1169514439 | 20402.4529879446 | Decr1 |
| **ENSMUSG00000028224** | 0.256803555063682 | 0.111438276014849 | 0.764116514278315 | 580.055806037327 | 679.22271791455 | 649.268053612834 | 590.893817382642 | 506.965644664242 | 393.832478727783 | 583.003730711437 | 606.808790649696 | Nbn |
| **ENSMUSG00000028228** | -0.280641376041298 | 0.0359673564736131 | 0.679566075569698 | 246.659244623349 | 299.704988240473 | 286.775831260479 | 289.819348525772 | 385.425284928322 | 309.439804714686 | 353.966550789086 | 314.802413753867 | Cpne3 |
| **ENSMUSG00000028229** | 0.145316137787809 | 0.372264551160086 | 0.873700364743909 | 775.214768816241 | 585.564909089402 | 680.501064938233 | 736.272454992657 | 547.479097909548 | 569.956320146419 | 604.866370613115 | 788.091559949337 | Rmdn1 |
| **ENSMUSG00000028232** | -0.158967253626291 | 0.38481330795808 | 0.875126835008975 | 99.3865088225949 | 87.1424829938332 | 111.681676860517 | 94.7305961200743 | 90.881530252985 | 125.977469903608 | 119.723980413956 | 102.03940307884 | Tmem68 |
| **ENSMUSG00000028233** | -0.11007207566011 | 0.60506838858979 | 0.926448151609826 | 201.483558794897 | 266.313943354985 | 207.27362061401 | 220.412773150668 | 241.98576127602 | 173.677676954488 | 224.872867560126 | 325.657669400552 | Tgs1 |
| **ENSMUSG00000028234** | -0.00412155746515747 | 0.985666342760709 | 0.997588567541988 | 3489.36997338965 | 2568.66720899579 | 2970.92189546747 | 3734.824096536 | 3339.62249724825 | 4535.18891652987 | 2510.03927633085 | 2416.3799069521 | Rps20 |
| **ENSMUSG00000028240** | -0.318793051226503 | 0.395717662729322 | 0.879215782741314 | 25.2983840639333 | 34.2054606144018 | 30.2865564367503 | 31.8895076047775 | 40.5134532453066 | 22.0154801773295 | 24.9858741733473 | 64.046008315442 | Cyp7a1 |
| **ENSMUSG00000028243** | 0.0634868397949949 | 0.718872277337321 | 0.951674101666722 | 161.728955265859 | 166.140808698523 | 124.932045301595 | 137.87522405595 | 121.54035973592 | 152.885279009232 | 129.093683228961 | 162.828834700276 | Ubxn2b |
| **ENSMUSG00000028245** | 0.373047860053882 | 0.0540978325607279 | 0.703396849069134 | 1376.05139033466 | 1549.01871639505 | 1880.60586374446 | 1533.51014511209 | 1205.54897359683 | 856.157562451702 | 1515.80969984973 | 1315.65698437823 | Nsmaf |
| **ENSMUSG00000028246** | 0.125741570392552 | 0.780666589115374 | 0.960742105865155 | 8.13162344912141 | 13.8450673915436 | 13.2503684410783 | 9.37926694258161 | 9.85462376237187 | 9.78465785659088 | 8.32862472444909 | 13.0263067760221 | Faxc |
| **ENSMUSG00000028247** | 0.205964978412751 | 0.12922294951011 | 0.77303252008843 | 1596.5087371775 | 1479.79337943734 | 1527.57819027859 | 1523.19295147525 | 1242.77755225467 | 1077.53544645707 | 1550.16527683809 | 1439.40689875044 | Coq3 |
| **ENSMUSG00000028248** | 0.0848380801659487 | 0.609677973917041 | 0.927932112008768 | 309.001691066613 | 229.665235553841 | 307.597838810745 | 348.908730264036 | 250.745426842573 | 269.078091056249 | 306.076958623504 | 299.605055848508 | Pnisr |
| **ENSMUSG00000028249** | -0.110028437588537 | 0.26423740507346 | 0.842510961779757 | 1690.47416370068 | 1889.44449108124 | 1810.56820198448 | 1636.68208148049 | 1979.68441804093 | 2033.98575193883 | 1746.9290359532 | 1825.85399977243 | Sdcbp |
| **ENSMUSG00000028251** | -0.0109823381734708 | 0.931311875989951 | 0.987775860606096 | 476.151728631887 | 480.505280059454 | 436.315703666934 | 497.101147956825 | 424.84377997781 | 429.301863457925 | 544.483841360859 | 504.769387570856 | Tstd3 |
| **ENSMUSG00000028252** | 0.0750737116235675 | 0.593954689403804 | 0.923935529355527 | 160.82544154929 | 149.038078391322 | 174.147699511314 | 180.081925297567 | 168.623562156141 | 149.216032313011 | 150.95632313064 | 160.657783570939 | Ccnc |
| **ENSMUSG00000028256** | 0.127676787379238 | 0.526079516256128 | 0.90806673740897 | 91.2548853734735 | 95.2866402829764 | 101.270673085384 | 75.0341355406529 | 76.6470737073368 | 78.277262852727 | 104.107809055614 | 72.73021283279 | Odf2l |
| **ENSMUSG00000028261** | 0.0966963630395488 | 0.489916941665027 | 0.899236955296196 | 1261.30514833039 | 1127.15136881743 | 1257.83854701379 | 1199.60824195619 | 1006.26658195775 | 952.781058785537 | 1236.80077158069 | 1334.1109189776 | Ndufaf4 |
| **ENSMUSG00000028266** | -0.341984486209115 | 0.0222693326104362 | 0.648762042745232 | 726.425028121512 | 627.100111264033 | 570.712297855013 | 653.734905897938 | 810.269064906132 | 996.812019140196 | 764.151318468204 | 697.99293808185 | Lmo4 |
| **ENSMUSG00000028268** | 0.00448497093524981 | 0.981943508703152 | 0.996940244108351 | 242.141676040504 | 201.975100770753 | 194.023252172932 | 207.281799431054 | 275.929465346412 | 221.377884005369 | 187.394056300105 | 158.486732441602 | Gbp3 |
| **ENSMUSG00000028270** | -0.190813263992226 | 0.511098655494187 | 0.902394714365738 | 268.343573821006 | 243.510302945384 | 260.275094378323 | 321.708856130549 | 434.698403740182 | 414.624876673038 | 195.722681024554 | 204.078806157679 | Gbp2 |
| **ENSMUSG00000028271** | 0.0131485350781854 | 0.913021046706057 | 0.986226496507248 | 443.625234835401 | 403.135785812593 | 431.583429223692 | 431.446279358754 | 411.70428162798 | 495.348303989913 | 394.568596320776 | 394.045779974668 | Gtf2b |
| **ENSMUSG00000028273** | 0.133433715241714 | 0.497272486145644 | 0.900927059989498 | 15118.4950193498 | 15465.7546920831 | 14709.8018793742 | 14257.4236794183 | 11622.9812486197 | 9614.64942633261 | 18062.704871149 | 14988.9369969428 | Pdlim5 |
| **ENSMUSG00000028274** | -0.0249642958094411 | 0.907445812348944 | 0.98504921011344 | 220.457346842847 | 254.097707421271 | 266.900278598862 | 219.47484645641 | 201.472308030714 | 173.677676954488 | 302.953724351836 | 298.51953028384 | Rngtt |
| **ENSMUSG00000028277** | 0.139189082606753 | 0.267659911992819 | 0.843793744782845 | 767.986659083688 | 835.590537866101 | 936.043884873314 | 790.67220325963 | 811.364023101951 | 839.034411202668 | 675.659680770933 | 699.078463646519 | Ube2j1 |
| **ENSMUSG00000028278** | 0.24575387512621 | 0.477192896188701 | 0.897674307047416 | 2479.24163826546 | 3357.02163458486 | 3677.92369728786 | 2181.61749084448 | 1745.36336413564 | 1165.59736716639 | 3597.96588096201 | 3354.27399482569 | Rragd |
| **ENSMUSG00000028282** | 0.0499160703916992 | 0.76317467341079 | 0.95785116771606 | 124.684892886528 | 137.636258186522 | 137.235958854025 | 131.309737196143 | 110.590777777729 | 113.746647582869 | 153.038479311752 | 134.605170018895 | Casp8ap2 |
| **ENSMUSG00000028284** | 0.261097729573917 | 0.209492850986387 | 0.821566820583092 | 1094.15511076511 | 1211.85060462452 | 1146.15687015327 | 980.133395499779 | 778.515277227378 | 643.34125407085 | 1187.87010132455 | 1087.69661579784 | Map3k7 |
| **ENSMUSG00000028289** | -0.277346772634289 | 0.228739098702541 | 0.832048466795696 | 55.1143367107117 | 57.009101024003 | 40.6975602118832 | 71.2824287636203 | 59.1277425742312 | 74.6080161565054 | 68.711153976705 | 69.4736361387845 | Epha7 |
| **ENSMUSG00000028291** | -0.0715755617471532 | 0.720087016312909 | 0.951674101666722 | 858.338030740593 | 860.837425462445 | 811.111839571719 | 911.664746818933 | 1089.48340484 | 1053.07380181559 | 892.203923606609 | 582.927228226989 | Akirin2 |
| **ENSMUSG00000028292** | 0.205934952949126 | 0.211000403641069 | 0.823123035902316 | 429.169015370296 | 443.856572258309 | 372.903226127488 | 363.915557372167 | 359.146288228664 | 261.739597663806 | 383.116737324658 | 390.789203280663 | Rars2 |
| **ENSMUSG00000028293** | -0.120054596505425 | 0.579595101521238 | 0.922450244597442 | 323.457910531718 | 200.346269312925 | 178.879973954556 | 222.288626539184 | 257.315176017488 | 254.401104271363 | 208.215618111227 | 285.493223507818 | Slc35a1 |
| **ENSMUSG00000028295** | 0.240019863853132 | 0.0246696763834038 | 0.655303666217042 | 838.460728976074 | 701.21942595237 | 691.858523602015 | 740.962088463947 | 609.891715071237 | 634.779678446333 | 609.03068297534 | 663.256120012458 | Smim8 |
| **ENSMUSG00000028300** | 0.126411913416693 | 0.519899433544722 | 0.905962179970576 | 480.669297214732 | 527.741392336485 | 460.923530771794 | 446.453106466885 | 377.760577557588 | 313.109051410908 | 522.621201459181 | 540.591731204917 | 3110043O21 |
| **ENSMUSG00000028307** | -0.273035576726362 | 0.241751075291371 | 0.832048466795696 | 954.110484696911 | 1440.70142444945 | 1190.64024991975 | 983.885102276811 | 1042.40020241978 | 1185.16668287957 | 1309.67623791962 | 1984.34073221403 | Aldob |
| **ENSMUSG00000028309** | 0.246982196757286 | 0.160032465015649 | 0.790986484775773 | 965.856163012309 | 1157.28475078726 | 1119.65613327111 | 1006.39534293901 | 833.263187018333 | 655.572076391589 | 1074.39258945393 | 1016.05192852972 | Rnf20 |
| **ENSMUSG00000028312** | -0.116664890383153 | 0.634333922395554 | 0.932543559775713 | 57.8248778604189 | 61.0811796685746 | 50.1621090983677 | 57.2135283497478 | 66.7924499449649 | 78.277262852727 | 40.6020455316893 | 60.7894316214364 | Smc2 |
| **ENSMUSG00000028318** | 0.071615500633494 | 0.605118245299606 | 0.926448151609826 | 335.203588847116 | 279.344595017615 | 258.382184601026 | 319.833002742033 | 257.315176017488 | 282.531995609062 | 308.159114804616 | 286.578749072486 | Polr1e |
| **ENSMUSG00000028322** | -0.0534807330754599 | 0.734851552393168 | 0.954383574136061 | 450.853344567953 | 364.858246553619 | 363.438677241004 | 440.825546301336 | 452.217734873287 | 495.348303989913 | 364.377331694648 | 370.164217551961 | Exosc3 |
| **ENSMUSG00000028327** | -0.403178898734877 | 0.485421673512788 | 0.899236955296196 | 11.7456783153976 | 17.9171460361152 | 2.83936466594534 | 11.2551203310979 | 12.0445401540101 | 8.56157562451702 | 12.4929370866736 | 24.9670879873757 | Stra6l |
| **ENSMUSG00000028328** | 0.263984220311861 | 0.286842063596686 | 0.851785577030345 | 3879.68789894748 | 4500.46131798058 | 4244.85017558828 | 3724.50690289916 | 2670.60303960278 | 2096.3629457746 | 4383.97983933189 | 4463.6811219169 | Tmod1 |
| **ENSMUSG00000028329** | -0.0993192496439471 | 0.641832284318627 | 0.934589624613243 | 187.930853046361 | 204.418347957496 | 190.237432618338 | 225.102406621959 | 260.600050604945 | 273.970419984545 | 161.367104036201 | 170.427513652956 | Xpa |
| **ENSMUSG00000028330** | 0.0865310763437269 | 0.459145860841626 | 0.894716050377411 | 491.511461813561 | 577.420751800259 | 597.21303473717 | 493.349441179793 | 514.630352034976 | 497.794468454061 | 538.237372817523 | 483.058876277486 | Ncbp1 |

| | | | | | | | | | | | | |
|---|---|---|---|---|---|---|---|---|---|---|---|---|
| ENSMUSG00000028331 | -0.261826876507508 | 0.466701335992573 | 0.895328112556867 | 22.5878429142261 | 26.8757190541728 | 29.3401015481019 | 30.9515809105193 | 47.0832024202212 | 40.3617136584374 | 28.1091084450157 | 16.2828834700276 | Trmo |
| ENSMUSG00000028333 | 0.103270182595448 | 0.285474645827566 | 0.851275833092507 | 3009.60418989149 | 2823.57933214598 | 2722.95071464158 | 3194.5783206433 | 2849.08122552129 | 2915.82804126408 | 2612.06492920535 | 2562.92585818235 | Anp32b |
| ENSMUSG00000028334 | -0.205735849972902 | 0.354761496048946 | 0.863949229409685 | 299.966553900923 | 230.479651282755 | 257.435729712378 | 296.384835385579 | 388.710159515779 | 387.717067567413 | 248.817663642917 | 225.78931745105 | Nans |
| ENSMUSG00000028337 | -0.928513900870312 | 0.0693505136583531 | 0.724352955175446 | 17.1667606148119 | 15.4738988493722 | 13.2503684410783 | 2.81378008277448 | 27.3739548954774 | 33.0232202659942 | 16.6572494488982 | 16.2828834700276 | Coro2a |
| ENSMUSG00000028339 | -1.38596821016972 | 0.0320832616336474 | 0.671606027118506 | 307.194663633475 | 482.134111517283 | 177.933519065908 | 299.198615468353 | 1244.96746864631 | 1471.36792518485 | 292.542943446274 | 301.776106977845 | Col15a1 |
| ENSMUSG00000028341 | 0.0490190702872322 | 0.862392038805233 | 0.974868041734905 | 135.527057485357 | 245.953550132127 | 215.791714611846 | 127.55803041911 | 211.326931793086 | 117.415894279091 | 165.531416398426 | 206.249857287016 | Nr4a3 |
| ENSMUSG00000028343 | -0.0595735725896296 | 0.566927822197914 | 0.919931558693298 | 845.688838708626 | 802.199492980614 | 767.57491469389 | 830.065124418473 | 942.759006600242 | 863.496055844145 | 771.438865102097 | 805.459968984033 | Erp44 |
| ENSMUSG00000028344 | 0.0980861289933226 | 0.677094693826872 | 0.94409858171788 | 44.2721721118832 | 51.3081909216027 | 56.7872933189068 | 47.8342614071662 | 45.9882442244021 | 58.7079471395453 | 37.4788112600209 | 45.5920737160773 | Invs |
| ENSMUSG00000028345 | 0.0913148022613852 | 0.591342393981971 | 0.923935529355527 | 151.7903043836 | 179.985876090067 | 172.254789734017 | 160.385464718146 | 165.338687568684 | 118.638976511164 | 180.106509666212 | 158.486732441602 | Tex10 |
| ENSMUSG00000028347 | 0.435082974008384 | 0.0331458411288295 | 0.671606027118506 | 325.264937964856 | 355.085257806647 | 361.545767463707 | 311.39166249371 | 228.846262926191 | 167.562265794119 | 312.323427166841 | 290.92085133116 | Tmeff1 |
| ENSMUSG00000028348 | 0.205946503256916 | 0.285450930365592 | 0.851275833092507 | 3274.33370884622 | 3488.95698266898 | 3146.96250475609 | 3318.38464428537 | 2724.25599119791 | 1912.90061096352 | 3274.19059479905 | 3556.18174985403 | Cavin4 |
| ENSMUSG00000028356 | -0.235457628145337 | 0.38580044548846 | 0.875126835008975 | 552.046880823686 | 972.412380323708 | 553.676109859341 | 660.300392757745 | 590.182467546493 | 706.941530138691 | 724.590351027071 | 1202.76232565271 | Ambp |
| ENSMUSG00000028367 | -0.00895753963985424 | 0.946725940462057 | 0.990510945542179 | 2069.04641094311 | 1864.1976034849 | 1720.65498756288 | 2279.16186704733 | 2032.24241144024 | 2276.15603388945 | 1815.6401899299 | 1859.50529227715 | Txn1 |
| ENSMUSG00000028369 | -0.867074957515648 | 0.0863628570413195 | 0.746460787597018 | 86.7373167906283 | 73.2974156022896 | 95.5919437534931 | 113.489130005238 | 235.416012101106 | 319.224462571277 | 59.3414511616998 | 59.7039060567679 | Svep1 |
| ENSMUSG00000028373 | -0.133956799643233 | 0.67006344378375 | 0.94249140110542 | 47.8862269781594 | 30.1333819698302 | 34.0723759913441 | 33.7653609932938 | 33.943704070392 | 59.9310293716191 | 38.5198893505771 | 28.2236646813812 | Astn2 |
| ENSMUSG00000028378 | -0.589289216597344 | 0.0890012115125952 | 0.751206603737452 | 37.9475760958999 | 42.3496179035451 | 37.8581955459379 | 68.4686486808458 | 67.887408140784 | 108.854318654574 | 40.6020455316893 | 64.046008315442 | Ptgr1 |
| ENSMUSG00000028381 | -0.0240965146939006 | 0.869841860353093 | 0.975613767803763 | 272.861142403852 | 342.054606144018 | 360.599312575058 | 287.005568442997 | 352.576539053749 | 284.978160073209 | 335.227145159076 | 310.460311495193 | Ugcg |
| ENSMUSG00000028382 | -0.194417049285945 | 0.441876122148933 | 0.89108031640712 | 444.52874855197 | 460.144886836596 | 605.731128735006 | 533.680289032894 | 804.794273927036 | 706.941530138691 | 395.609674411332 | 432.039174738066 | Ptbp3 |
| ENSMUSG00000028383 | 0.330455220087308 | 0.251363650442932 | 0.83723604018117 | 6695.03663977662 | 8200.35197443838 | 9366.11757806503 | 6422.92200227989 | 4641.52779207715 | 3416.06867418229 | 8273.44758564962 | 8070.88257331035 | Hsdl2 |
| ENSMUSG00000028385 | -0.0628050392660776 | 0.849727718597546 | 0.972640876584422 | 74.9916384752307 | 112.389370590177 | 137.235958854025 | 107.861569839689 | 84.3117810780705 | 61.154111603693 | 142.627698406191 | 162.828834700276 | Snx30 |
| ENSMUSG00000028389 | -0.556857241193195 | 0.149464568758229 | 0.782641594375979 | 20.780815481088 | 20.3603932228582 | 17.036187995672 | 31.8895076047775 | 41.6084114411257 | 46.4771248188067 | 17.6983275394543 | 27.1381391167127 | Zfp37 |
| ENSMUSG00000028393 | 0.0117129411891834 | 0.926182882067339 | 0.986713918026868 | 1415.80599386369 | 1163.80007661858 | 1158.4607837057 | 1289.64920460497 | 1415.78094719409 | 1295.24408376622 | 1142.06266534008 | 1134.37421507859 | Alad |
| ENSMUSG00000028394 | 0.0172011405508095 | 0.899219658803211 | 0.984802943363903 | 608.968244967536 | 554.617111390658 | 561.247748968529 | 578.700770357285 | 533.2446413639 | 467.217412652214 | 647.550572325917 | 626.348250813729 | Pole3 |
| ENSMUSG00000028396 | 0.29335780864913 | 0.239059828981705 | 0.832048466795696 | 1834.13284463516 | 1375.5481661363 | 1669.54642357586 | 1673.26122255656 | 1195.69434983445 | 770.541806206532 | 1756.2987387682 | 1622.86071917942 | 2310002L09F |
| ENSMUSG00000028397 | -0.245298453990625 | 0.148725717069273 | 0.78151662300109 | 170.764092431549 | 215.820168162297 | 262.16800415562 | 202.592165959763 | 249.650468646754 | 221.377884005369 | 258.187366457922 | 278.980070119806 | Kdm4c |
| ENSMUSG00000028398 | -0.0225717588647568 | 0.859547845140938 | 0.974868041734905 | 796.899098013898 | 654.79024604712 | 651.160963390131 | 784.106716399823 | 781.800151814835 | 795.003450848009 | 677.741836952045 | 678.453477917817 | Dmac1 |
| ENSMUSG00000028399 | -0.069559260475202 | 0.7090070204065 | 0.949280056132338 | 117.456783153976 | 121.347943608235 | 112.628131749165 | 110.675349922463 | 90.881530252985 | 136.985209992272 | 114.518589961175 | 143.289374536243 | Ptprd |
| ENSMUSG00000028402 | 0.183629730548262 | 0.414798765246418 | 0.882192126163777 | 1254.07703859783 | 1580.78092982271 | 1524.73882561265 | 1218.36677584135 | 1047.87499339888 | 813.349684329117 | 1580.35654146422 | 1468.71608899649 | Mpdz |
| ENSMUSG00000028403 | 0.0666660549623655 | 0.78210542111505 | 0.960742105865155 | 198.77301764519 | 241.881471487556 | 154.272146849697 | 139.751077444466 | 146.724398239759 | 136.985209992272 | 205.092383839559 | 212.763010675027 | Zdhhc21 |
| ENSMUSG00000028405 | 0.134554881943711 | 0.389977936981439 | 0.877515569464664 | 781.539364832224 | 933.320425335821 | 933.204520207369 | 729.70696813285 | 791.654775577207 | 605.425704876561 | 822.451691539348 | 856.479670523452 | Aco1 |
| ENSMUSG00000028407 | -0.0178882178641001 | 0.956202158727861 | 0.992371505557225 | 68.6670424592474 | 39.9063707168021 | 54.8943835416099 | 60.0273084325223 | 62.4126171616885 | 86.838838477244 | 38.5198893505771 | 39.0789203280663 | Smim27 |
| ENSMUSG00000028409 | 0.014628477852289 | 0.892804392240771 | 0.983118202649778 | 977.601841327707 | 881.197818685304 | 962.54462175547 | 1078.61569839689 | 1013.93128932848 | 1029.83523940619 | 923.436266323293 | 893.387539722182 | Smu1 |
| ENSMUSG00000028410 | 0.0425405623728298 | 0.696361640951118 | 0.944933386973863 | 3620.37946229216 | 3495.4723085003 | 3157.37350853122 | 3191.76454056052 | 3433.78890208869 | 3365.92230266726 | 2817.15731304491 | 3458.48444903387 | Dnaja1 |
| ENSMUSG00000028411 | 0.215037945977338 | 0.0993699860014 | 0.759800953500741 | 416.51982333833 | 411.279943101736 | 440.101523221528 | 401.432625142493 | 319.727793179176 | 316.77829810713 | 402.897221045225 | 397.302356668674 | Aptx |
| ENSMUSG00000028412 | -0.112944701823248 | 0.420689566485107 | 0.882568531013176 | 807.741262612726 | 882.012234414218 | 791.236286910102 | 814.120370616084 | 1012.83633113266 | 1015.1582526213 | 756.863771834311 | 780.492880996657 | Slc44a1 |
| ENSMUSG00000028413 | 0.113870478274748 | 0.486182648559567 | 0.899236955296196 | 1385.08652750035 | 1312.02373928098 | 1763.24545755206 | 1620.7373276781 | 1403.73640704008 | 1101.99709109855 | 1483.53627904249 | 1628.28834700276 | B4galt1 |
| ENSMUSG00000028414 | -0.167215159345809 | 0.392699634143709 | 0.878640128585071 | 746.302329886031 | 824.188717661301 | 905.757328436564 | 628.410885152968 | 664.639624862192 | 757.087901653719 | 1062.94073045782 | 1000.85457062436 | Fktn |
| ENSMUSG00000028416 | 0.0347913290325982 | 0.802070737194314 | 0.963461896828986 | 7742.20903728015 | 6736.84690957933 | 6736.86589739964 | 8044.59725665225 | 6843.48872386936 | 8716.9070679904 | 6642.07821774815 | 6362.26533452212 | Bag1 |
| ENSMUSG00000028417 | -0.0578470361201818 | 0.926975362434778 | 0.986790505471895 | 11.7456783153976 | 13.0306516626293 | 12.3039135524298 | 2.81378008277448 | 3.28487458745729 | 8.56157562451702 | 17.6983275394543 | 11.9407812113536 | Tal2 |
| ENSMUSG00000028419 | 0.0698823784768178 | 0.456364562162031 | 0.892805623137304 | 891.768038253647 | 808.714818811928 | 782.718192912266 | 877.899385825639 | 825.598479647599 | 828.026671114003 | 790.178270732108 | 758.782369703287 | Chmp5 |
| ENSMUSG00000028420 | 0.141786192156389 | 0.403230853764826 | 0.881158207349486 | 1460.98167969215 | 1422.78427841333 | 1708.35107401045 | 1292.46298468775 | 1127.80694169367 | 1130.12798243625 | 1673.01249152371 | 1401.41350398704 | Tmem38b |
| ENSMUSG00000028423 | 0.162223705350621 | 0.200701373747555 | 0.812921911423739 | 659.565013095403 | 717.500257173524 | 651.160963390131 | 623.721251681677 | 542.004306930453 | 516.140701935169 | 644.427338054249 | 666.512696706464 | Nfx1 |
| ENSMUSG00000028426 | 0.224686979595168 | 0.158238766966051 | 0.789230661864335 | 3153.26287082597 | 3673.01493740362 | 3880.46504345863 | 3151.43369270742 | 2895.06946974569 | 2276.15603388945 | 3406.40751229968 | 3280.45825642823 | Rad23b |
| ENSMUSG00000028427 | -0.129606621639624 | 0.643331710974598 | 0.934616627390525 | 488.800920663853 | 426.753841951108 | 588.694940739334 | 863.830485411767 | 870.491765676182 | 731.403174780168 | 438.293876124134 | 550.361461286933 | Aqp7 |
| ENSMUSG00000028430 | -0.13134527484036 | 0.239833131821364 | 0.832048466795696 | 435.49361138628 | 478.062032872711 | 472.280989435575 | 444.577253078368 | 529.959766776443 | 452.540425867328 | 473.690531203042 | 548.190410157596 | Nol6 |
| ENSMUSG00000028431 | 0.254148191449003 | 0.100263377069734 | 0.759800953500741 | 489.704434380422 | 563.575684408715 | 512.978549647458 | 485.846027625728 | 493.826146314413 | 325.339873731647 | 444.54034466747 | 455.920737160773 | Ikbkap |
| ENSMUSG00000028433 | 0.271079572723789 | 0.175118365291232 | 0.799889021902745 | 2305.7670046842 | 2738.88009633889 | 2646.28786866106 | 2182.55541753874 | 1792.44656655586 | 1437.12162268679 | 2430.91734144858 | 2520.59036116027 | Ubap2 |
| ENSMUSG00000028434 | -0.237149895143575 | 0.497709786933402 | 0.900927059989498 | 24.3948703473642 | 55.3802695661743 | 31.2330113253987 | 44.0825546301336 | 42.7033696369448 | 57.4848649074714 | 28.1091084450157 | 55.3618037980939 | Epb41l4b |
| ENSMUSG00000028436 | 0.0156225744894882 | 0.906509446521223 | 0.98504921011344 | 758.048008201429 | 759.849875077069 | 937.936794650611 | 653.734905897938 | 783.990068206473 | 744.857079332981 | 767.274552739873 | 779.407355431988 | Dcaf12 |
| ENSMUSG00000028437 | 0.115412011210567 | 0.371723701732476 | 0.873497245316205 | 925.198045766702 | 965.082638763479 | 1025.01064440627 | 900.409626487835 | 845.307727172343 | 730.180092548094 | 997.352810752779 | 947.663817955607 | Ubap1 |
| ENSMUSG00000028439 | 0.0717043886204867 | 0.602040175231578 | 0.926448151609826 | 334.300075130547 | 415.352021746308 | 468.495169880981 | 421.129085721914 | 393.089992299056 | 362.032340693862 | 394.568596320776 | 409.243137880027 | Fam219a |
| ENSMUSG00000028443 | 0.141959790349188 | 0.385567897665496 | 0.875126835008975 | 438.204152535987 | 296.447325324816 | 322.74111702912 | 388.301651422879 | 330.677375137367 | 333.901449356164 | 290.460787265162 | 354.966859646602 | Nudt2 |
| ENSMUSG00000028444 | -1.40868852572876 | 0.00408208082141152 | 0.635862253526407 | 48.7897406947284 | 29.3189662409158 | 35.965285768641 | 36.5791410760683 | 141.249607260664 | 172.454594722414 | 29.1501865355718 | 57.5328549274309 | Cntfr |
| ENSMUSG00000028445 | -0.16088916837644 | 0.455600768107837 | 0.892805623137304 | 253.887354355902 | 182.42912327681 | 160.897331070236 | 287.943495137256 | 236.510970296925 | 287.424324537357 | 224.872867560126 | 240.986675356409 | Enho |
| ENSMUSG00000028447 | 0.128847127546653 | 0.410514098564377 | 0.882192126163777 | 1172.76080410662 | 921.91860513102 | 1024.06418951762 | 1320.60078551549 | 1115.76240153966 | 1122.7894890438 | 885.957455063272 | 935.723036744253 | Dctn3 |
| ENSMUSG00000028452 | 0.0725362595757987 | 0.329255969662154 | 0.855977678491063 | 13388.2662521201 | 14354.891637844 | 14256.4499877116 | 14417.8091441365 | 14388.8456512588 | 13340.1579052296 | 13045.7495527589 | 12876.5042480978 | Vcp |
| ENSMUSG00000028453 | -0.116952240783826 | 0.739236909872725 | 0.954383574136061 | 16.2632468982428 | 23.6180561385155 | 20.8220075502658 | 24.3860940507122 | 29.5638712871156 | 28.1308913376988 | 20.8215618111227 | 14.1118323406906 | Fancg |
| ENSMUSG00000028454 | -0.139079921293596 | 0.46293178011024 | 0.895328112556867 | 347.852780879082 | 389.290718421049 | 364.385132129652 | 284.191788360223 | 358.051330032845 | 281.308913376988 | 458.0743598447 | 427.697072479392 | Pigo |
| ENSMUSG00000028455 | 0.126325561851929 | 0.339812596060881 | 0.858570960161243 | 2051.8796503283 | 1590.55391856968 | 1686.58261157153 | 2052.18360703686 | 1848.28943454264 | 1772.24615427502 | 1569.94576055865 | 1571.84101764 | Stoml2 |
| ENSMUSG00000028456 | -0.132539534861767 | 0.487715041496738 | 0.899236955296196 | 197.869503928621 | 184.057954734638 | 224.309808609682 | 143.502784221499 | 251.840385038392 | 190.800828203522 | 199.886993386778 | 179.111718170304 | Unc13b |
| ENSMUSG00000028458 | 0.0696096781931543 | 0.68183762079971 | 0.944452327903597 | 551.143367107117 | 662.119987607349 | 675.768790494991 | 517.735535230505 | 534.339599559719 | 453.763508099402 | 673.57752458982 | 630.690353072403 | Tesk1 |
| ENSMUSG00000028461 | 0.222012781723722 | 0.226319408919276 | 0.832048466795696 | 681.24934229306 | 430.82592059568 | 494.049451874489 | 636.852225401292 | 513.535393839156 | 535.71001764835 | 458.0743598447 | 415.756291268038 | Ccdc107 |
| ENSMUSG00000028464 | -0.34723504022831 | 0.31894323844167 | 0.854670142081901 | 562.889045422515 | 535.071133896714 | 351.134763688574 | 575.886990274511 | 607.701798679599 | 1165.59736716639 | 442.458188486358 | 361.480013034613 | Tpm2 |
| ENSMUSG00000028465 | 0.00205373722948759 | 0.984098775036708 | 0.996940244108351 | 2810.8311722463 | 3335.03240990418 | 3184.82070030202 | 3011.68261526296 | 3386.70569966847 | 3150.65982982226 | 2854.63612430493 | 2934.17560129898 | Tln1 |
| ENSMUSG00000028466 | -0.208038509033625 | 0.273813619987867 | 0.845204865395511 | 627.942033015486 | 488.649437348597 | 546.104470750154 | 599.335157630965 | 731.432074807157 | 818.242013257412 | 580.921574530324 | 483.058876277486 | Creb3 |
| ENSMUSG00000028467 | -0.0109674971997011 | 0.934735105738025 | 0.98818087308472 | 293.64195788494 | 285.859920848929 | 372.903226127488 | 331.088123073131 | 341.626957095558 | 299.655146858096 | 326.898520434627 | 323.486618271215 | Gba2 |
| ENSMUSG00000028468 | 0.152551147283322 | 0.429608002343157 | 0.885743677088682 | 880.925873654819 | 1069.32785206451 | 1058.13656550896 | 896.657919710802 | 753.331238723539 | 650.679747463293 | 1120.2000254384 | 987.828263848342 | Rgp1 |
| ENSMUSG00000028469 | 0.0451585538156579 | 0.7456929710994 | 0.956085046483146 | 1221.55054480135 | 1413.01128966636 | 1297.58965233702 | 1146.14642038347 | 1221.97334653411 | 1007.81975922886 | 1425.23590597135 | 1265.72280840348 | Npr2 |
| ENSMUSG00000028470 | -0.0459848121687079 | 0.799202074151111 | 0.96328975655894 | 2057.30073262772 | 1598.69807585883 | 1824.76502531421 | 2194.7484645641 | 2047.57182618171 | 2533.00330262496 | 1603.26025945645 | 1741.18300572829 | Hint2 |
| ENSMUSG00000028475 | -0.347391699002001 | 0.294845797493918 | 0.851785577030345 | 52.4037955610046 | 56.1946852950887 | 100.324218196735 | 105.047789756914 | 123.730276127558 | 130.869798831903 | 65.5879197050366 | 79.2433662208011 | Spaar |
| ENSMUSG00000028476 | -0.698819390908408 | 0.127693414786625 | 0.772837268577874 | 46.9827132615903 | 72.4829998733752 | 49.2156542097192 | 36.5791410760683 | 118.255485148462 | 139.43137445642 | 39.5609674411332 | 36.9078691987293 | Reck |
| ENSMUSG00000028478 | -0.0153578292876191 | 0.922348306483421 | 0.986713918026868 | 1800.70283712211 | 1635.34678365997 | 1649.67087091424 | 2009.97690579524 | 1854.85918371755 | 2229.67890907065 | 1575.15115101143 | 1513.2226371479 | Clta |
| ENSMUSG00000028479 | -0.257103327617533 | 0.0495948736379618 | 0.703396849069134 | 338.817643713392 | 404.764617270421 | 435.369248778286 | 354.536290429585 | 477.401773377126 | 488.00981059747 | 412.26692386023 | 455.920737160773 | Gne |
| ENSMUSG00000028480 | -0.434398581549158 | 0.22830579527189 | 0.832048466795696 | 35.2370349461928 | 43.1640336324594 | 34.0723759913441 | 32.8274342990356 | 40.5134532453066 | 88.0619207093179 | 37.4788112600209 | 31.4802413753867 | Glipr2 |
| ENSMUSG00000028483 | 0.111077990546238 | 0.557801523774671 | 0.916370464878199 | 204.194099944604 | 176.728213174409 | 148.593417517806 | 151.006197775564 | 198.187433443257 | 139.43137445642 | 138.463386043966 | 154.144630182928 | Snapc3 |
| ENSMUSG00000028484 | -0.0236718455288122 | 0.84764796599873 | 0.972455319662159 | 386.703870691551 | 401.506954354764 | 461.869985660442 | 448.328959855401 | 440.173194719277 | 465.99433042014 | 441.417110395802 | 378.848422069309 | Psip1 |
| ENSMUSG00000028488 | 0.00832406669015999 | 0.982212335858211 | 0.996940244108351 | 29.8159526467785 | 27.6901347830872 | 17.036187995672 | 33.7653609932938 | 15.3294147414674 | 22.0154801773295 | 36.4377331694648 | 33.6512925047237 | Sh3gl2 |
| ENSMUSG00000028494 | 0.294642496638638 | 0.162218978701965 | 0.793874624603454 | 2228.06482505926 | 2203.80896244217 | 3685.49533639705 | 3655.10032752405 | 2707.83161826063 | 2233.34815576687 | 2240.40005087681 | 2415.29438138743 | Plin2 |
| ENSMUSG00000028495 | 0.0450084245641116 | 0.762045170303214 | 0.957666649379949 | 6527.88660221135 | 5006.21348563638 | 6014.72081736088 | 6640.52099534778 | 5863.50113861126 | 7049.84598567373 | 5294.92316856851 | 5238.74637509022 | Rps6 |
| ENSMUSG00000028496 | 0.197320801420228 | 0.140239059325541 | 0.779711473022872 | 317.133314515735 | 378.703313945163 | 411.707876562074 | 347.03287687552 | 304.398378437709 | 352.247682837272 | 317.528817619622 | 295.262953589834 | Mllt3 |
| ENSMUSG00000028497 | -0.0798505691699677 | 0.751711267739427 | 0.956197192463153 | 86.7373167906283 | 62.7100111264033 | 93.6990339761962 | 82.5375490947182 | 87.5966556655277 | 121.085140975312 | 66.6289977955927 | 69.4736361387845 | Hacd4 |
| ENSMUSG00000028514 | 0.255902604096924 | 0.39592089190592 | 0.87923508914506 | 2141.32750826864 | 2840.68206245318 | 2395.47732316922 | 1827.0812004149 | 1482.57339713906 | 1013.93517038923 | 2626.64002247313 | 2584.63636947572 | Usp24 |
| ENSMUSG00000028517 | -0.417465454162777 | 0.0563895501169627 | 0.704362276191198 | 1044.46185635382 | 989.515110630909 | 1042.04683240194 | 1151.77398054902 | 1852.66926732591 | 1709.86896043926 | 1081.68013608783 | 1003.0256217537 | Plpp3 |
| ENSMUSG00000028518 | 0.159577979157984 | 0.628808405583772 | 0.931825800800958 | 1949.782600356 | 3000.30754532039 | 2533.65973691189 | 1737.04023776611 | 1542.79609790911 | 1022.49674601375 | 2793.21251696211 | 2896.18220653558 | Prkaa2 |
| ENSMUSG00000028521 | 0.127843039588909 | 0.591374286181024 | 0.923935529355527 | 156.307872966445 | 197.088606397268 | 166.576060402127 | 159.447538023887 | 154.389105610493 | 92.9542496376133 | 171.777884941763 | 201.907755028342 | Slc35d1 |
| ENSMUSG00000028522 | 0.0464294990395312 | 0.671976045337987 | 0.943216235131458 | 674.924746277077 | 659.676740420606 | 669.143606274452 | 556.19052969509 | 605.511882287961 | 613.987280501078 | 595.49666779811 | 664.341645577127 | Mier1 |
| ENSMUSG00000028525 | -0.37799738909479 | 0.109942373403875 | 0.763549783289909 | 344.238726012806 | 342.054606144018 | 217.684624389143 | 296.384835385579 | 513.535393839156 | 371.816998550453 | 410.184767679118 | 264.868237779116 | Pde4b |

| | | | | | | | | | | | | |
|---|---|---|---|---|---|---|---|---|---|---|---|---|
| **ENSMUSG00000028527** | -0.0862167895761715 | 0.781459143976139 | 0.960742105865155 | 917.066422317581 | 1059.55486331754 | 732.556083813898 | 654.672832592197 | 666.82954125383 | 477.002070508805 | 1181.62363278122 | 1245.09782267478 | Ak4 |
| **ENSMUSG00000028530** | 0.307088241391098 | 0.136955919206584 | 0.776456211338551 | 3483.04537737367 | 4981.78101376895 | 5539.60046325936 | 4355.7315681349 | 4226.53863586171 | 2581.92659190792 | 3997.73986773556 | 4032.72747274351 | Jak1 |
| **ENSMUSG00000028532** | -0.033264121535465 | 0.908164816207824 | 0.98504921011344 | 43.3686583953142 | 61.895595397489 | 36.9117406572894 | 47.8342614071662 | 63.5075753575076 | 56.2617826753975 | 34.3555769883525 | 41.2499714574033 | Cachd1 |
| **ENSMUSG00000028538** | 0.105901597348052 | 0.512026738952222 | 0.902394714365738 | 1487.18357747265 | 1500.96818838911 | 1508.64909250562 | 1363.74541345137 | 1223.06830472993 | 1077.53544645707 | 1730.2717865043 | 1413.3542851984 | St3gal3 |
| **ENSMUSG00000028540** | 0.00570883099232222 | 0.980676059043848 | 0.996888334226108 | 56.9213641438498 | 88.7713144516618 | 60.5731128735006 | 66.5927952923295 | 54.7479097909548 | 81.9465095489486 | 63.5057635239243 | 72.73021283279 | Dph2 |
| **ENSMUSG00000028541** | -0.0355434403240923 | 0.809030274474167 | 0.964172502140384 | 147.272735800754 | 141.708336831093 | 134.396594188079 | 135.061443973175 | 152.199189218854 | 157.777607937528 | 137.42230795341 | 125.920965501547 | B4galt2 |
| **ENSMUSG00000028542** | -0.516684507065782 | 0.243694752321569 | 0.832048466795696 | 14.4562194651047 | 29.3189662409158 | 27.447191770805 | 9.37926694258161 | 42.7033696369448 | 29.3539735697726 | 18.7394056300105 | 24.9670879873757 | Slc6a9 |
| **ENSMUSG00000028545** | 0.0212907740356833 | 0.875376068930557 | 0.976986380159756 | 266.536546387868 | 258.169786065842 | 263.114459044268 | 271.060814640609 | 282.499214521327 | 269.078091056249 | 280.050006359601 | 211.677485110359 | Bend5 |
| **ENSMUSG00000028549** | -0.0480898901685841 | 0.812253905677115 | 0.965224347191244 | 76.7986659083688 | 65.9676740420606 | 81.3951204237664 | 78.7858423176855 | 60.2227007700503 | 75.8310983885793 | 88.4916376972716 | 87.9275707381491 | Itgb3bp |
| **ENSMUSG00000028550** | 0.0604057855954361 | 0.738401421073003 | 0.954383574136061 | 100.290022539164 | 100.173134656462 | 83.2880302010633 | 96.6064495085906 | 91.9764884488041 | 73.3849339244316 | 97.8613405122768 | 100.953877514171 | Atg4c |
| **ENSMUSG00000028551** | -0.129615454758743 | 0.433931705001374 | 0.887868762931002 | 211.422209677157 | 203.603932228582 | 227.149173275627 | 240.109233730089 | 292.353838283699 | 261.739597663806 | 203.010227658447 | 208.420908416353 | Cdkn2c |
| **ENSMUSG00000028552** | 0.0157383822578377 | 0.869216778742374 | 0.975613767803763 | 797.802611730467 | 857.579762546788 | 907.65023821386 | 771.913669374467 | 797.129566556303 | 812.126602097043 | 832.862472444909 | 856.479670523452 | Eps15 |
| **ENSMUSG00000028553** | -0.0402553313818886 | 0.867373846713795 | 0.974868041734905 | 148.176249517323 | 198.717437855096 | 171.308334845369 | 197.902532488472 | 150.009272827216 | 172.454594722414 | 141.586620315635 | 272.466916731795 | Angptl3 |
| **ENSMUSG00000028555** | 0.22417404219024 | 0.63991627199788 | 0.933921420069286 | 50.5967681278665 | 66.7820897709749 | 17.036187995672 | 50.6480414899407 | 37.2285786578493 | 14.6769867848863 | 52.0539045278068 | 54.2762782334254 | Ttc39a |
| **ENSMUSG00000028556** | 0.0657573209574549 | 0.776866844569269 | 0.959738279197945 | 94.8689402397497 | 150.666909849151 | 141.021778408619 | 87.227182566009 | 107.305903190271 | 97.8465785659088 | 125.970448957293 | 121.578863242873 | Dock7 |
| **ENSMUSG00000028557** | 0.0545416808720676 | 0.723174410498107 | 0.951674101666722 | 3196.63152922128 | 3576.91388139173 | 3283.25200872146 | 3136.42686559929 | 2644.32404290312 | 2673.65775931346 | 3686.45751865928 | 3698.38559882561 | Rnf11 |
| **ENSMUSG00000028559** | 0.173533032793844 | 0.0601938566102654 | 0.71282992338528 | 2428.64487013759 | 2582.51227638734 | 2775.00573351725 | 2592.42938292956 | 2229.33488668768 | 2120.82459041607 | 2466.31399652749 | 2383.81414001204 | Osbpl9 |
| **ENSMUSG00000028560** | 0.0515362016771961 | 0.802940706010746 | 0.963461896828986 | 412.905768472054 | 479.69086433054 | 497.835271429083 | 387.363724728621 | 425.938738173629 | 282.531995609062 | 521.580123368624 | 484.144401842154 | Usp1 |
| **ENSMUSG00000028563** | 0.271419333554106 | 0.109740401745359 | 0.763154315676794 | 891.768038253647 | 801.385077251699 | 835.719666676579 | 857.264998551959 | 646.025335533267 | 516.140701935169 | 807.876598271562 | 833.683633665414 | Tm2d1 |
| **ENSMUSG00000028565** | -0.129434473642079 | 0.478323590582833 | 0.897674307047416 | 1270.34028549608 | 1049.78187457057 | 914.2754224344 | 1288.71127791071 | 1322.70950054947 | 1483.59874750559 | 949.463218587197 | 1192.99259557069 | Nfia |
| **ENSMUSG00000028567** | -0.111590157675039 | 0.581730873351539 | 0.922450244597442 | 386.703870691551 | 319.250965734417 | 321.794662140472 | 346.094950181261 | 406.229490648885 | 475.778988276731 | 254.023054095697 | 349.539231823259 | Txndc12 |
| **ENSMUSG00000028568** | 0.151390577189479 | 0.148328274540634 | 0.781357001591751 | 1059.82158953549 | 902.372627637076 | 892.506959995485 | 1004.51948955049 | 814.648897689408 | 847.595986827185 | 919.271953961069 | 892.302014157513 | Btf3l4 |
| **ENSMUSG00000028572** | 0.25670358247456 | 0.444852277438797 | 0.89108031640712 | 275.571683553559 | 458.516055378767 | 437.262158555582 | 271.998741334867 | 270.454674367317 | 139.43137445642 | 414.349080041342 | 383.190524327983 | Hook1 |
| **ENSMUSG00000028573** | -0.111527938891342 | 0.499114493878856 | 0.900927059989498 | 143.658680934478 | 138.450673915436 | 131.557229522134 | 143.502784221499 | 143.439523652302 | 176.123841418636 | 120.765058504512 | 162.828834700276 | Fggy |
| **ENSMUSG00000028576** | 0.276868137153606 | 0.157200513988258 | 0.788998730890588 | 110.228673421423 | 132.749763813036 | 147.646962629158 | 99.4202295913651 | 112.780694169367 | 95.400414101761 | 109.313199508394 | 86.8420451734806 | Ift74 |
| **ENSMUSG00000028577** | 0.273227487998777 | 0.147941848279006 | 0.780607990324772 | 1866.65933843165 | 1927.72203034022 | 1910.89242018121 | 1767.99181867663 | 1377.45741034042 | 1078.75852868914 | 1864.57086018604 | 1861.67634340649 | Plaa |
| **ENSMUSG00000028578** | -0.217988252663101 | 0.288123478525183 | 0.851785577030345 | 63.2459601598332 | 53.7514381083457 | 66.2518422053913 | 64.7169419038131 | 81.0269064906132 | 56.2617826753975 | 72.8754663389296 | 77.072315091464 | Caap1 |
| **ENSMUSG00000028580** | 0.122175908246663 | 0.579597026474513 | 0.922450244597442 | 1404.96382926487 | 1767.28213174409 | 1484.98772028941 | 1290.58713129923 | 1085.10357205672 | 973.573456730792 | 1673.01249152371 | 1732.49880121094 | Pum1 |
| **ENSMUSG00000028581** | -0.431069558818008 | 0.132539000799042 | 0.77303252008843 | 178.895715880671 | 277.715763559786 | 307.597838810745 | 243.860940507122 | 394.184950494875 | 494.125221757839 | 238.406882737355 | 233.387996403729 | Laptm5 |
| **ENSMUSG00000028582** | 0.311181110670908 | 0.0333567137165209 | 0.671606027118506 | 402.063603873225 | 437.341246426994 | 375.742590793433 | 443.63932638411 | 338.342082508101 | 272.747337752471 | 394.568596320776 | 329.999771659226 | Cc2d1b |
| **ENSMUSG00000028583** | -0.930887459960827 | 0.0758817185292958 | 0.735399943172214 | 25.2983840639333 | 35.8342920722305 | 31.2330113253987 | 43.1446279358754 | 85.4067392738896 | 124.754387671534 | 27.0680303544596 | 21.7105112933701 | Pdpn |
| **ENSMUSG00000028597** | -0.470166295785934 | 0.156775644427622 | 0.787563725533336 | 149.983276950461 | 109.946123403434 | 109.78876708322 | 149.130344387048 | 226.656346534553 | 275.193502216618 | 115.559668051731 | 102.03940307884 | Gpx7 |
| **ENSMUSG00000028599** | -0.584856824243799 | 0.0422395943126847 | 0.685406501171951 | 96.6759676728878 | 112.389370590177 | 112.628131749165 | 115.364983393754 | 204.757182618171 | 236.054870790255 | 109.313199508394 | 106.381505337514 | Tnfrsf1b |
| **ENSMUSG00000028600** | -0.875151788964979 | 0.00726521171850836 | 0.635862253526407 | 102.097049972302 | 86.3280672649188 | 106.949402417274 | 85.3513291774927 | 167.528603960322 | 299.655146858096 | 133.257995591185 | 98.7828263848342 | Podn |
| **ENSMUSG00000028601** | 0.0665748057278385 | 0.542517984684427 | 0.913837378403038 | 673.117718843939 | 599.409976480946 | 656.839692722022 | 749.403428712271 | 648.215251924905 | 620.102691661447 | 613.194995337564 | 675.196901223812 | Echdc2 |
| **ENSMUSG00000028603** | 0.0676949860716715 | 0.499436550463904 | 0.900927059989498 | 9306.19128066116 | 8875.50261370835 | 9298.91928097099 | 9395.211696384 | 8046.84778107454 | 8279.04362890796 | 8785.65800620324 | 10072.5917145591 | Scp2 |
| **ENSMUSG00000028607** | 0.0498613637298219 | 0.406899320143276 | 0.881296485258502 | 8136.14101770425 | 8073.30312072774 | 8445.21697141009 | 8122.44517227568 | 8136.63435313171 | 7704.19497983324 | 7757.07285273377 | 8064.36941992234 | Cpt2 |
| **ENSMUSG00000028608** | 0.0570541976022426 | 0.673891924294115 | 0.943870003876702 | 564.696072855653 | 440.598909342652 | 496.888816540435 | 543.059555975475 | 459.882442244021 | 568.733237914345 | 457.033281754144 | 480.887825148149 | 0610037L13 |
| **ENSMUSG00000028609** | 0.0427370671407429 | 0.751142792421051 | 0.956197192463153 | 462.599022883351 | 352.642010619904 | 374.796135904785 | 367.667264149199 | 411.70428162798 | 382.824738639118 | 350.843316517418 | 366.907640857956 | Magoh |
| **ENSMUSG00000028613** | -0.184016010683839 | 0.676596300388154 | 0.94409858171788 | 20.780815481088 | 28.5045505120015 | 9.46454888648447 | 24.3860940507122 | 14.2344565456483 | 34.2463024980681 | 29.1501865355718 | 17.3684090346961 | Lrp8 |
| **ENSMUSG00000028614** | 0.127438930503694 | 0.473539515570475 | 0.897334770978705 | 316.229800799166 | 369.744740927105 | 376.689045682082 | 292.633128608546 | 258.410134213307 | 260.516515431732 | 348.761160336306 | 372.335268681298 | Ndc1 |
| **ENSMUSG00000028617** | -0.0845726127699783 | 0.603790376981859 | 0.926448151609826 | 128.298947752804 | 159.625482867208 | 116.413951303759 | 144.440710915757 | 133.58489988993 | 140.654456688494 | 155.120635492864 | 153.05910461826 | Lrrc42 |
| **ENSMUSG00000028618** | -0.109813845604852 | 0.417457675371778 | 0.882436023575407 | 2307.57403211734 | 2230.68468149635 | 2292.31374030654 | 2405.78197077218 | 2557.82234543341 | 3045.47475786391 | 2175.85320926233 | 2189.50506393638 | Tmem59 |
| **ENSMUSG00000028619** | -0.0410913185919297 | 0.786550735520515 | 0.961141387070613 | 240.334648607366 | 282.602257933272 | 294.347470369667 | 260.743621003769 | 234.321053905287 | 253.178022039289 | 303.994802442392 | 316.973464883204 | Tceanc2 |
| **ENSMUSG00000028621** | 0.00653755553446966 | 0.95813280445447 | 0.992835150787715 | 339.721157429961 | 351.013179162076 | 341.670214802089 | 363.915557372167 | 325.202584158272 | 302.101311322243 | 373.747034509653 | 387.532626586657 | Cyb5rl |
| **ENSMUSG00000028622** | 0.0696271752223122 | 0.402040968650656 | 0.881158207349486 | 1971.46692955366 | 1963.55632241245 | 1912.78532995851 | 1873.03960843355 | 1817.6306050597 | 1707.42279597511 | 1992.62346532445 | 1837.79478098378 | Mrpl37 |
| **ENSMUSG00000028629** | -0.142520195862091 | 0.617183542658775 | 0.929187015773942 | 42.4651446787451 | 28.5045505120015 | 31.2330113253987 | 38.4549944645846 | 32.8487458745729 | 53.8156182112498 | 35.3966550789086 | 33.6512925047237 | Exo5 |
| **ENSMUSG00000028630** | -0.111586956273729 | 0.713027280805373 | 0.950526426881904 | 125.588406603097 | 219.892246806869 | 201.594891282119 | 137.87522405595 | 141.249607260664 | 114.969729814943 | 247.776585552361 | 235.559047533066 | Dyrk2 |
| **ENSMUSG00000028631** | 0.300340876831603 | 0.216267401150373 | 0.825447777181429 | 98.4829951060259 | 105.874044758863 | 106.002947528626 | 82.5375490947182 | 64.6025335533267 | 55.0387004433237 | 103.066730965058 | 95.5262496908287 | Kcnq4 |
| **ENSMUSG00000028633** | 0.238704729460993 | 0.125218369449741 | 0.771949105062466 | 931.522641782685 | 957.75289720325 | 883.988865997649 | 1028.9055836012 | 783.990068206473 | 618.879609429373 | 843.273253350471 | 974.80195707232 | Ctps |
| **ENSMUSG00000028634** | 0.0335603331883537 | 0.831198302461338 | 0.967583158900775 | 226.78194285883 | 181.614707547895 | 163.736695736181 | 201.654239265505 | 175.193311331055 | 207.923979452556 | 175.942197303987 | 197.565652769668 | Hivep3 |
| **ENSMUSG00000028636** | 0.0671145169737185 | 0.6844149539041 | 0.944452327903597 | 331.58953398084 | 213.376920975554 | 259.328639489674 | 266.371181169318 | 249.650468646754 | 238.501035254403 | 257.146288367366 | 275.723493425801 | Ppcs |
| **ENSMUSG00000028639** | -0.0463908799483492 | 0.747961261569114 | 0.956197192463153 | 28239.3212113655 | 23751.6203180575 | 27350.6533721628 | 30219.0601623037 | 28554.3198305704 | 34417.5340105584 | 25266.9652577974 | 24903.0419790602 | Ybx1 |
| **ENSMUSG00000028641** | -0.276737206855386 | 0.0329414156327651 | 0.671606027118506 | 215.939778260002 | 261.427448981499 | 212.952349945901 | 242.923013812864 | 302.208462046071 | 286.201242305283 | 262.351678820146 | 281.151121249143 | P3h1 |
| **ENSMUSG00000028643** | -0.0468054967252588 | 0.822974696733536 | 0.966637987288264 | 344.238726012806 | 275.272516373043 | 281.097101928589 | 365.791410760683 | 366.810995599397 | 421.963370065482 | 273.803537816264 | 246.414303179751 | Svbp |
| **ENSMUSG00000028645** | -0.396623430207411 | 0.0698293546864939 | 0.724916894583216 | 369.537110076739 | 434.083583511337 | 368.170951684246 | 348.908730264036 | 683.253914191116 | 490.455975061618 | 498.676405376389 | 329.999771659226 | Slc2a1 |
| **ENSMUSG00000028646** | 0.0971549833289558 | 0.482624059015352 | 0.898371506261489 | 1063.43564440177 | 1108.4198070524 | 1148.04977993057 | 1045.78826409785 | 993.127083607921 | 808.457355400821 | 1103.5427759895 | 1174.53866097133 | Rragc |
| **ENSMUSG00000028647** | 0.0803737409675837 | 0.668714555214059 | 0.942055056494206 | 332.493047697409 | 329.838370210303 | 369.117406572894 | 362.03970398365 | 240.890803080201 | 289.870489001505 | 373.747034509653 | 412.499714574033 | Mycbp |
| **ENSMUSG00000028648** | 0.0193992904364546 | 0.918770056348386 | 0.986713918026868 | 5942.40971387461 | 4545.25418307087 | 4831.65220655032 | 6120.90960672876 | 5571.14730032757 | 6857.82207523813 | 4286.11849881961 | 4439.7995594942 | Ndufs5 |
| **ENSMUSG00000028649** | 0.339781287573602 | 0.122543899531072 | 0.770251097497048 | 3218.31585841894 | 4301.74388012548 | 4119.91813028669 | 3315.5708642026 | 2725.35094939373 | 1922.68526882011 | 3650.01978548982 | 3518.18835509063 | Macf1 |
| **ENSMUSG00000028651** | -0.0988724973557849 | 0.480776323667609 | 0.897732059433541 | 409.291713605777 | 325.766291565731 | 343.563124579386 | 401.432625142493 | 382.140410340865 | 447.648096939033 | 348.761160336306 | 407.07208675069 | Ppie |
| **ENSMUSG00000028653** | 0.0204230191427297 | 0.923375386060696 | 0.986713918026868 | 112.035700854562 | 123.791190794978 | 85.1809399783602 | 121.930470253561 | 90.881530252985 | 92.9542496376133 | 136.381229862854 | 116.15123541953 | Trit1 |
| **ENSMUSG00000028655** | -0.529750750293417 | 0.373164039809305 | 0.873700364743909 | 5.42108229941427 | 18.7315617650296 | 4.73227444324223 | 6.56548685980713 | 9.85462376237187 | 20.7923979452556 | 8.32862472444909 | 13.0263067760221 | Mfsd2a |
| **ENSMUSG00000028656** | -0.436875826935125 | 0.0474757748602614 | 0.703396849069134 | 805.934235179588 | 756.592212161411 | 863.166858447383 | 863.830485411767 | 1364.31791199059 | 1451.79860947167 | 796.424739275444 | 841.282312618093 | Cap1 |
| **ENSMUSG00000028657** | -0.00198059303953468 | 0.983326492291571 | 0.996940244108351 | 1169.14674924034 | 1140.18202048006 | 1116.81676860517 | 1027.96765690694 | 1065.39432453198 | 1044.51222619108 | 1184.74686705288 | 1164.76893088931 | Ppt1 |
| **ENSMUSG00000028668** | 0.178219272058275 | 0.203916399341173 | 0.816074141171399 | 1021.87401343959 | 1176.8307282812 | 1215.24807702461 | 1008.27119632752 | 923.049759075499 | 813.349684329117 | 1103.5427759895 | 1067.07163006914 | Eloa |
| **ENSMUSG00000028669** | 0.0272524343658581 | 0.810209770790668 | 0.964412705343932 | 475.248214915318 | 429.197089137851 | 491.210087208544 | 464.27371365779 | 416.084114411257 | 481.894399437101 | 429.965251399684 | 497.170708618176 | Pithd1 |
| **ENSMUSG00000028670** | -0.0447839868903899 | 0.780390729203317 | 0.960742105865155 | 806.837748896157 | 694.696616763922 | 765.682004916593 | 781.292936317048 | 790.559817381388 | 1006.59667699679 | 697.522320672611 | 651.315338801105 | Lypla2 |
| **ENSMUSG00000028672** | 0.0672501352294085 | 0.531222306800738 | 0.909082872831377 | 1669.6933482196 | 1431.74285143139 | 1409.27132919754 | 1509.12405106138 | 1396.07169966935 | 1301.35949492659 | 1481.45412286138 | 1565.32786425199 | Hmgcl |
| **ENSMUSG00000028673** | -0.0527637954388645 | 0.745543618817922 | 0.956085046483146 | 753.530439618584 | 645.017257300148 | 813.951204237664 | 672.493439783102 | 800.41444114376 | 912.419345127099 | 617.359307699789 | 663.256120012458 | Fuca1 |
| **ENSMUSG00000028675** | -0.122171854226115 | 0.13518581849597 | 0.775642208276055 | 931.522641782685 | 923.547436588849 | 949.294253314392 | 935.112914175387 | 1046.78003520306 | 1050.62763735145 | 1031.70838774113 | 941.150664567596 | Pnrc2 |
| **ENSMUSG00000028676** | 0.160694131945243 | 0.233731946264797 | 0.832048466795696 | 1058.91807581892 | 1134.48111037766 | 1013.65318574249 | 933.23706078687 | 778.515277227378 | 875.726878164883 | 987.983107937774 | 1061.6440022458 | Srsf10 |
| **ENSMUSG00000028677** | -0.0200124450872782 | 0.850774531718758 | 0.972692784625411 | 718.293404672391 | 733.78857175181 | 740.127722923085 | 797.237690119437 | 721.577451044785 | 793.780368615935 | 832.862472444909 | 682.795580176491 | Rnf220 |
| **ENSMUSG00000028680** | 0.0774550399258452 | 0.82625986926789 | 0.967261756842012 | 36.1405486627618 | 38.2775392589734 | 21.7684624389143 | 37.5170677703264 | 43.7983278327639 | 37.9155491942896 | 28.1091084450157 | 17.3684090346961 | Plk3 |
| **ENSMUSG00000028683** | 0.0929589982754724 | 0.572379312546335 | 0.921574199900435 | 468.923618899334 | 345.312269059675 | 381.421320125324 | 446.453106466885 | 453.312693069106 | 403.617136584374 | 360.213019332423 | 322.401092706547 | Eif2b3 |
| **ENSMUSG00000028684** | 0.00493755943738847 | 0.975024139222782 | 0.99575536949131 | 1508.86790667031 | 1320.98231229904 | 1556.91829182669 | 1636.68208148049 | 1680.76083058231 | 1784.47697659576 | 1286.77251992738 | 1251.61097606279 | Urod |
| **ENSMUSG00000028688** | -0.0256918407348807 | 0.924303176030568 | 0.986713918026868 | 67.7635287426784 | 68.4109212288036 | 50.1621090983677 | 105.047789756914 | 63.5075753575076 | 80.7234273168747 | 63.5057635239243 | 89.0130963028176 | Toe1 |
| **ENSMUSG00000028689** | -0.456986527710713 | 0.0891305198980741 | 0.751206603737452 | 30.7194663633475 | 35.8342920722305 | 28.3936466594534 | 21.5723139679377 | 38.3235368536684 | 39.1386314263635 | 37.4788112600209 | 45.5920737160773 | Ccdc163 |
| **ENSMUSG00000028690** | 0.0576597869661674 | 0.722859175517748 | 0.951674101666722 | 1030.90915060528 | 967.525885950222 | 842.344850897118 | 867.582192188799 | 834.358145214152 | 686.149132193435 | 1027.54407537891 | 1013.88087740039 | Mmachc |
| **ENSMUSG00000028691** | -0.0416034106902412 | 0.699665451699233 | 0.945280238056015 | 6386.93846242658 | 5770.13543935802 | 5194.14442890268 | 6665.84501609275 | 5991.6112475221 | 6205.91924554276 | 6036.17076904448 | 6486.01524889433 | Prdx1 |
| **ENSMUSG00000028692** | -0.14411456925234 | 0.310085236051653 | 0.851785577030345 | 5135.57196497845 | 4704.87966593808 | 4952.79843229732 | 5265.52046156532 | 5175.86739163687 | 7016.82276540773 | 4929.50475878331 | 5045.52282457922 | Akr1a1 |
| **ENSMUSG00000028693** | 0.173635265746076 | 0.211929570385759 | 0.823370267313127 | 356.887918044773 | 333.09603312596 | 361.545767463707 | 383.612017951588 | 338.342082508101 | 254.401104271363 | 350.843316517418 | 326.743194965221 | Nasp |
| **ENSMUSG00000028696** | 0.295374179110144 | 0.244351504528901 | 0.832048466795696 | 233.106538874814 | 281.787842204358 | 271.632553042104 | 184.771558768858 | 174.098353135236 | 123.53130543946 | 248.817663642917 | 244.243252050414 | Ipp |
| **ENSMUSG00000028698** | -0.0929410796579366 | 0.655595511157842 | 0.937625787683528 | 258.404922938747 | 214.191336704468 | 178.879973954556 | 220.412773150668 | 210.231973597267 | 170.008430258266 | 294.625099627387 | 254.012982132431 | Pik3r3 |
| **ENSMUSG00000028700** | 0.144370917447949 | 0.278772295973299 | 0.848075053184273 | 1438.39383677792 | 1569.37910961791 | 1666.70705890991 | 1566.33757941113 | 1211.02376457592 | 1257.32853457193 | 1589.72624427922 | 1587.03837554536 | Pomgnt1 |
| **ENSMUSG00000028701** | 0.163180887926162 | 0.3595809124105 | 0.867338970126053 | 84.9302893574902 | 83.0704043492615 | 77.6093008691726 | 77.8479156234274 | 71.1722827282413 | 70.9387694602839 | 66.6289977955927 | 80.3288917854696 | Lurap1 |
| **ENSMUSG00000028702** | -0.32174302422588 | 0.49484019624624 | 0.900200915538898 | 16.2632468982428 | 10.5874044758863 | 19.8755526616174 | 19.6964605794214 | 32.8487458745729 | 26.9078091056249 | 14.5750932677859 | 8.68420451734806 | Rad54l |

| | | | | | | | | | | | | |
|---|---|---|---|---|---|---|---|---|---|---|---|---|
| **ENSMUSG00000028703** | 0.0223760880351677 | 0.898741743277844 | 0.984540500469898 | 255.69438178904 | 340.425774686189 | 367.224496795597 | 262.619474392285 | 301.113503850252 | 256.847268735511 | 346.679004155194 | 301.776106977845 | Lrrc41 |
| **ENSMUSG00000028706** | 0.115521759672414 | 0.427881426501699 | 0.885066977635295 | 927.908586916409 | 943.093414082792 | 862.220403558735 | 790.67220325963 | 698.583328932584 | 705.718447906617 | 911.984407327176 | 935.723036744253 | Nsun4 |
| **ENSMUSG00000028708** | 0.325527074393872 | 0.0500785662715157 | 0.703396849069134 | 404.774145022932 | 398.249291439107 | 455.244801439903 | 352.660437041069 | 323.012667766634 | 261.739597663806 | 399.773986773556 | 299.605055848508 | Mknk1 |
| **ENSMUSG00000028709** | -0.204813546457955 | 0.394132479544123 | 0.878942305112115 | 61.4389327266951 | 76.5550785179469 | 74.7699362032273 | 106.92364314543 | 110.590777777729 | 102.738907494204 | 73.9165444294857 | 81.4144173501381 | Mob3c |
| **ENSMUSG00000028710** | 0.175600342710022 | 0.452514605262614 | 0.891110666112841 | 1930.80881230805 | 2014.05009760513 | 2075.57557080604 | 1854.28107454838 | 1549.36584708402 | 1034.72756833449 | 2231.0303480618 | 2155.85377143166 | Atpaf1 |
| **ENSMUSG00000028713** | -0.11663354648266 | 0.71409483935163 | 0.951176382858794 | 477.958756065025 | 421.052931848708 | 488.370722542599 | 528.990655561603 | 705.153078107498 | 742.410914868833 | 249.858741733473 | 381.019473198646 | Cyp4b1 |
| **ENSMUSG00000028715** | -0.291030281468334 | 0.491134053151169 | 0.899236955296196 | 172.571119864688 | 438.155662155909 | 188.344522841041 | 177.268145214792 | 121.54035973592 | 249.508775343067 | 433.088485671353 | 390.789203280663 | Cyp4a14 |
| **ENSMUSG00000028717** | -0.237889311782591 | 0.463567999905458 | 0.895328112556867 | 71.3775836089546 | 54.56585383726 | 66.2518422053913 | 68.4686486808458 | 109.49581958191 | 102.738907494204 | 48.9306702561384 | 46.6775992807458 | Tal1 |
| **ENSMUSG00000028719** | 0.0384855290641423 | 0.731022110486421 | 0.952984317124008 | 844.785324992057 | 777.767021113184 | 872.631407333868 | 838.506464666796 | 790.559817381388 | 743.633997100907 | 774.562099373766 | 935.723036744253 | Cmpk1 |
| **ENSMUSG00000028729** | 0.0181707798010737 | 0.910291486055182 | 0.98603671070003 | 749.012871035738 | 739.48948185421 | 788.396922244156 | 878.837312519897 | 886.916138613469 | 924.650167447838 | 667.331056046484 | 638.289032025082 | Ebna1bp2 |
| **ENSMUSG00000028737** | 0.142448831603256 | 0.481988814673302 | 0.897732059433541 | 1962.43179238797 | 2071.87361435805 | 2053.80710836713 | 2066.25250745073 | 1732.22386578581 | 1191.28209403994 | 2286.20748686128 | 2176.47875716036 | Aldh4a1 |
| **ENSMUSG00000028741** | 0.094655232645634 | 0.572360412239845 | 0.921574199900435 | 642.398252480591 | 536.699965354543 | 437.262158555582 | 673.43136647736 | 551.858930692825 | 594.417964787896 | 480.978077836935 | 517.795694346878 | Mrto4 |
| **ENSMUSG00000028743** | -0.0747160124324831 | 0.680514356102207 | 0.944452327903597 | 1380.5689589175 | 1184.16046984143 | 1235.12362968622 | 1682.64048949914 | 1541.70113971329 | 1779.58464766747 | 1167.04853951343 | 1286.34779413218 | Akr7a5 |
| **ENSMUSG00000028744** | -0.437451014307363 | 0.0265764832383704 | 0.664353746352133 | 88.5443442237664 | 72.4829998733752 | 87.0738497556571 | 104.109863062656 | 141.249607260664 | 127.200552135681 | 108.272121417838 | 99.8683519495027 | Pqlc2 |
| **ENSMUSG00000028745** | -0.00858227793715063 | 0.889108557377936 | 0.98126431530921 | 5610.82017989377 | 5779.90842810499 | 5531.08236926152 | 5608.8016316638 | 5563.48259295683 | 5891.58711189978 | 5698.86146770429 | 5513.38434295135 | Capzb |
| **ENSMUSG00000028755** | -0.113353891015268 | 0.841170397716704 | 0.970420002017028 | 10.8421645988285 | 13.8450673915436 | 11.3574586637814 | 20.6343872736795 | 6.56974917491458 | 33.0232202659942 | 14.5750932677859 | 7.59867895267955 | Cda |
| **ENSMUSG00000028756** | 0.124219714088854 | 0.17002725720107 | 0.79907358295209 | 25975.1158376435 | 25878.8742019817 | 29132.8279274878 | 27127.6537780288 | 23229.5381243021 | 23455.0479644804 | 25899.9407368556 | 26608.4026411545 | Pink1 |
| **ENSMUSG00000028757** | -0.119378280532069 | 0.505428412428638 | 0.901494649536949 | 1697.70227343324 | 1337.27062687733 | 1492.5593593986 | 1640.43378825752 | 1849.38439273845 | 2098.80911023874 | 1452.30393632581 | 1300.45962647287 | Ddost |
| **ENSMUSG00000028759** | 0.119021002332077 | 0.466598222922666 | 0.895328112556867 | 2093.44128129048 | 2540.16265848379 | 2623.57295133349 | 2026.85958629189 | 1991.72895819494 | 1704.97663151096 | 2470.47830888971 | 2380.55756331804 | Hp1bp3 |
| **ENSMUSG00000028760** | 0.253947524826507 | 0.00468137016275147 | 0.635862253526407 | 1188.12053728829 | 1364.1463459315 | 1298.53610722567 | 1213.67714237006 | 1117.9523179313 | 1032.28140387034 | 1040.03701246558 | 1057.30189998713 | Eif4g3 |
| **ENSMUSG00000028763** | -0.492903360322991 | 0.0838627114808085 | 0.742295439672922 | 3849.8719463007 | 4864.50514880528 | 4083.0063896294 | 3926.16114216466 | 8487.02097579382 | 7573.32518100134 | 3651.06086358037 | 3824.30656432715 | Hspg2 |
| **ENSMUSG00000028766** | -0.298351154054019 | 0.0387729614910503 | 0.679566075569698 | 770.697200233395 | 725.644414462667 | 608.570493400951 | 683.7485601142 | 879.251431242735 | 983.358114587383 | 871.382361795486 | 696.907412517182 | Alpl |
| **ENSMUSG00000028771** | -0.0997502620315122 | 0.300293310371045 | 0.851785577030345 | 601.740135234984 | 644.202841571234 | 600.052399403115 | 585.266257217093 | 705.153078107498 | 607.871869340708 | 649.632728507029 | 642.631134283757 | Ptpn12 |
| **ENSMUSG00000028772** | -0.0981480043079545 | 0.386083911144534 | 0.875126835008975 | 703.837185207286 | 558.689190035229 | 578.283936964201 | 649.045272426648 | 653.690042904001 | 693.487625585878 | 663.166743684259 | 654.57191549511 | Zcchc17 |
| **ENSMUSG00000028773** | 0.118715595557684 | 0.285352081829557 | 0.851275833092507 | 124384.926332627 | 99822.9358930293 | 113139.217389035 | 117564.421491789 | 104363.750518106 | 117217.754957495 | 98303.7987007632 | 99092.2011707647 | Fabp3 |
| **ENSMUSG00000028776** | -0.636146201253588 | 0.090245608750655 | 0.753823363185192 | 1470.92033057441 | 1403.23830091939 | 1511.48845717157 | 1287.77335121646 | 3131.58044004262 | 3509.0229238199 | 1193.07549177733 | 984.571687154336 | Tinagl1 |
| **ENSMUSG00000028777** | 0.363136915793986 | 0.37870428419808 | 0.87422797717331 | 16.2632468982428 | 11.4018202048006 | 22.7149173275627 | 22.5102406621959 | 18.6142893289246 | 11.0077400886647 | 10.4107809055614 | 16.2828834700276 | Gnat3 |
| **ENSMUSG00000028779** | -0.0367997652756838 | 0.797694330005565 | 0.962555316113048 | 674.021232560508 | 595.337897836374 | 660.625512276616 | 642.47978556684 | 736.906865786252 | 764.426395046162 | 588.209121164217 | 550.361461286933 | Pef1 |
| **ENSMUSG00000028780** | -0.153314634007108 | 0.448365874383841 | 0.89108031640712 | 227.685456575399 | 283.416673662186 | 159.950876181588 | 207.281799431054 | 213.516848184724 | 229.939459629886 | 250.899819824029 | 283.322172378481 | Sema3c |
| **ENSMUSG00000028782** | -0.0601495758354628 | 0.856103816446186 | 0.97363563840968 | 30.7194663633475 | 37.4631235300591 | 35.0188308799925 | 18.7585338851632 | 20.8042057205628 | 34.2463024980681 | 32.2734208072402 | 40.1644458927348 | Adgrb2 |
| **ENSMUSG00000028788** | -0.00607360369317377 | 0.941553412829082 | 0.989934163630124 | 6629.98365218365 | 6371.17424729679 | 6265.53136285272 | 6550.480032699 | 5827.36751814923 | 6653.5673424818 | 6687.88565373262 | 6757.39664006146 | Ptp4a2 |
| **ENSMUSG00000028789** | 0.0685948649744434 | 0.799550227833255 | 0.963355472513329 | 26.2018977805023 | 34.2054606144018 | 35.0188308799925 | 38.4549944645846 | 35.0386622662111 | 35.4693847301419 | 30.191264626128 | 27.1381391167127 | Azin2 |
| **ENSMUSG00000028790** | -0.0272082014868137 | 0.751432355233188 | 0.956197192463153 | 930.619128066116 | 939.835751167135 | 1013.65318574249 | 935.112914175387 | 1035.83045324487 | 941.773318696872 | 977.572327032212 | 935.723036744253 | Khdrbs1 |
| **ENSMUSG00000028792** | 0.0342758198378256 | 0.820638764873598 | 0.966398681144067 | 1322.74408105708 | 1177.64514401012 | 1225.65908079974 | 1401.26248122169 | 1389.50195049443 | 1482.37566527352 | 1045.24240291836 | 1090.95319249185 | Ak2 |
| **ENSMUSG00000028793** | 0.206283030287384 | 0.0556604619226277 | 0.703396849069134 | 824.908023227538 | 912.145616384048 | 919.95415176629 | 877.899385825639 | 842.022852584886 | 673.918309872697 | 794.342583094332 | 752.269216315276 | Rnf19b |
| **ENSMUSG00000028795** | -0.152553011873289 | 0.611068504270915 | 0.927932112008768 | 98.4829951060259 | 74.1118313312039 | 60.5731128735006 | 94.7305961200743 | 95.2613630362614 | 135.762127760198 | 56.2182168900314 | 78.1578406561325 | Ccdc28b |
| **ENSMUSG00000028796** | -0.204710421457273 | 0.163764724874833 | 0.793968226938511 | 684.863397159336 | 658.862324691692 | 769.467824471187 | 693.127827056781 | 872.68168206782 | 963.788798874201 | 670.454290318152 | 728.387653892569 | Phc2 |
| **ENSMUSG00000028797** | -0.0708420573798257 | 0.6577690702291 | 0.938704652777376 | 1226.97162710076 | 1202.07761587755 | 1312.7329305554 | 1407.8279680815 | 1296.43050384981 | 1742.89218070525 | 1114.99463498562 | 1255.95307832146 | Tmem234 |
| **ENSMUSG00000028798** | 0.0677521038827884 | 0.623336279194838 | 0.931059568526633 | 2362.68836882805 | 1776.24070476215 | 1977.14426238661 | 2227.57589886313 | 2010.34324752386 | 2262.70212933664 | 1732.35394268541 | 1956.11706753265 | Eif3i |
| **ENSMUSG00000028799** | 0.00627874503924071 | 0.965075924934872 | 0.994284908685797 | 197.869503928621 | 227.221988367098 | 250.810545491838 | 201.654239265505 | 238.700886688563 | 210.370143916704 | 197.804837205666 | 226.874843015718 | Zfp362 |
| **ENSMUSG00000028800** | -0.217531117041042 | 0.179333666773675 | 0.801755154770274 | 389.414411841258 | 352.642010619904 | 353.974128354519 | 348.908730264036 | 491.636229922775 | 486.786728365396 | 342.514691792969 | 360.394487469945 | Hdac1 |
| **ENSMUSG00000028803** | 0.18300699361824 | 0.240342754191299 | 0.832048466795696 | 922.487504616995 | 1134.48111037766 | 1212.40871235866 | 976.381688722746 | 909.91026072567 | 755.864819421645 | 1079.59797990671 | 993.255891671684 | Nipal3 |
| **ENSMUSG00000028807** | -0.56651301417691 | 0.131949479421789 | 0.77303252008843 | 25.2983840639333 | 18.7315617650296 | 32.1794662140472 | 39.3929211588428 | 63.5075753575076 | 50.1463715150282 | 30.191264626128 | 27.1381391167127 | Zbtb8a |
| **ENSMUSG00000028809** | 0.176642686101981 | 0.282260791831594 | 0.850587941933635 | 914.355881167874 | 1022.09173978748 | 1074.22629861599 | 967.002421780164 | 877.061514851097 | 642.118171838776 | 991.106342209442 | 1007.36772401237 | Srrm1 |
| **ENSMUSG00000028811** | 0.00266877143114501 | 0.977460719166836 | 0.996179425433073 | 706.547726356993 | 780.210268299927 | 772.307189137133 | 736.272454992657 | 798.224524752122 | 703.272283442469 | 717.302804393178 | 770.72315091464 | Yars |
| **ENSMUSG00000028820** | 0.0885980116843072 | 0.523098639227463 | 0.90648515894888 | 1650.71956017165 | 1911.43371576193 | 1860.73031108285 | 1465.97942312551 | 1824.20035423462 | 1736.77676954488 | 1475.20765431805 | 1443.74900100912 | Sfpq |
| **ENSMUSG00000028821** | 0.0822052569672612 | 0.514846582656177 | 0.904145806837306 | 1945.26503177315 | 1570.19352534683 | 1808.67529220718 | 1940.57033042014 | 1815.44068866806 | 1923.90835105218 | 1566.82252628699 | 1556.64365973464 | Syf2 |
| **ENSMUSG00000028822** | -0.176389935952178 | 0.470019489813818 | 0.895920229657462 | 1532.3592633011 | 1317.72464938338 | 1478.36253606887 | 1622.61318106662 | 2046.47686798589 | 2323.85624094033 | 1169.13069569454 | 1186.47944218268 | Tmem50a |
| **ENSMUSG00000028825** | 0.0877591859404194 | 0.606032595121912 | 0.926859923029135 | 436.397125102849 | 347.755516246418 | 291.508105703722 | 423.004939110431 | 344.911831683016 | 407.286383280595 | 333.144988977964 | 325.657669400552 | Rhd |
| **ENSMUSG00000028826** | 0.165550029648339 | 0.11340698333913 | 0.767518180080285 | 1097.76916563139 | 1171.1298181788 | 1107.35221971868 | 1039.22277723804 | 1015.0262475243 | 881.842289325253 | 1088.96768272172 | 949.834869084944 | Tmem57 |
| **ENSMUSG00000028830** | 0.175566566838242 | 0.395495782525017 | 0.879215782741314 | 1133.90971429415 | 1347.0436156243 | 1220.9268063565 | 1005.45741624475 | 935.094299229509 | 713.05694129906 | 1260.74556766348 | 1258.1241294508 | AU040320 |
| **ENSMUSG00000028832** | 0.103179248833761 | 0.626926202131302 | 0.931825800800958 | 322.554396815149 | 285.859920848929 | 429.690519446395 | 396.742991671202 | 404.039574257247 | 390.163232031561 | 251.940897914585 | 289.835325766492 | Stmn1 |
| **ENSMUSG00000028833** | -0.213631636121271 | 0.0393409195187672 | 0.679566075569698 | 425.55496050402 | 492.721515993169 | 416.440151005317 | 431.446279358754 | 515.725310230795 | 524.702277559686 | 509.087186281951 | 500.427285312182 | Ncdn |
| **ENSMUSG00000028834** | 0.0550583130081019 | 0.758283831817246 | 0.95677573228595 | 4264.58474220589 | 4543.62535161304 | 5645.60341078799 | 5694.1529608413 | 4054.63019911812 | 4082.64849066254 | 6003.89734823724 | 5250.68715630157 | Trim63 |
| **ENSMUSG00000028836** | -0.0266887414998598 | 0.940847071206643 | 0.989652845063254 | 26.2018977805023 | 17.9171460361152 | 19.8755526616174 | 20.6343872736795 | 20.8042057205628 | 15.9000690169602 | 16.6572494488982 | 32.5657669400552 | Slc30a2 |
| **ENSMUSG00000028837** | 0.0361702606927208 | 0.798411856957719 | 0.963086469226463 | 1661.56172477047 | 1358.4454358291 | 1232.28426502028 | 1651.68890858862 | 1538.41626512583 | 1560.65292812624 | 1353.40151772298 | 1305.88725429621 | Psmb2 |
| **ENSMUSG00000028838** | -0.0397993861367465 | 0.809785276692687 | 0.964412705343932 | 739.074220153479 | 802.199492980614 | 685.233339381476 | 624.659178375935 | 642.74046094581 | 596.864129252044 | 878.669908429379 | 811.973122372044 | Extl1 |
| **ENSMUSG00000028840** | 0.251014986817534 | 0.565638201658061 | 0.919917953383268 | 19.877301764519 | 17.9171460361152 | 15.1432782183751 | 31.8895076047775 | 24.0890803080201 | 9.78465785659088 | 24.9858741733473 | 11.9407812113536 | Zfp593 |
| **ENSMUSG00000028841** | 0.387278910860177 | 0.0545374195594943 | 0.703396849069134 | 608.968244967536 | 476.433201414882 | 522.443098533943 | 733.458674909882 | 384.330326732503 | 368.147751854232 | 569.469715534207 | 466.775992807458 | Cnksr1 |
| **ENSMUSG00000028842** | -0.121521373064266 | 0.582214223665274 | 0.922450244597442 | 154.500845533307 | 229.665235553841 | 193.076797284283 | 135.061443973175 | 176.288269526875 | 154.108361241306 | 216.544242835676 | 227.960368580387 | Ago3 |
| **ENSMUSG00000028843** | -0.518355751735495 | 0.1693217252106 | 0.79907358295209 | 208.711668527449 | 197.903022126182 | 279.204192151292 | 209.15765281957 | 341.626957095558 | 591.971800323748 | 162.408182126757 | 186.710397122983 | Sh3bgrl3 |
| **ENSMUSG00000028847** | 0.0348959970614939 | 0.75947683075562 | 0.957357521291862 | 1165.53269437407 | 924.361852317763 | 971.062715753306 | 1113.31898608444 | 1024.88087128667 | 1081.20469315329 | 981.736639394437 | 986.742738283673 | Trappc3 |
| **ENSMUSG00000028848** | -0.391015561514667 | 0.0813603154282913 | 0.739855948845396 | 142.755167217909 | 101.801966114291 | 127.77140996754 | 119.116690170786 | 175.193311331055 | 204.254732756335 | 109.313199508394 | 156.315681312265 | Gpn2 |
| **ENSMUSG00000028849** | -0.130773316896954 | 0.311453443937021 | 0.851785577030345 | 7341.04894712349 | 7600.12758222851 | 6002.41690380845 | 7034.45020693621 | 7740.2594862452 | 6430.96637624435 | 8293.22806937019 | 8167.49434856585 | Map7d1 |
| **ENSMUSG00000028850** | -0.399659679733826 | 0.0588008640150439 | 0.711712402279519 | 55.1143367107117 | 49.679359463774 | 69.0912068713366 | 67.5307219865876 | 90.881530252985 | 78.277262852727 | 82.2451691539348 | 66.217059444779 | Gpatch3 |
| **ENSMUSG00000028851** | 0.0656448349765191 | 0.686600857274591 | 0.944452327903597 | 2556.04030417383 | 2410.67055758641 | 2823.27493283832 | 3173.00600667536 | 2812.94760505926 | 3184.90613232033 | 2251.85190987292 | 2226.41293313511 | Nudc |
| **ENSMUSG00000028854** | -0.231895792305042 | 0.18437968269241 | 0.805963874499992 | 266.536546387868 | 361.600583637962 | 339.777305024792 | 254.178134143962 | 353.671497249568 | 430.524945689999 | 321.693129981846 | 331.085297223895 | Slc9a1 |
| **ENSMUSG00000028857** | 0.0326399914110607 | 0.806162393221116 | 0.963604452772887 | 603.547162668122 | 465.031381210082 | 578.283936964201 | 640.603932178324 | 560.618596259378 | 589.5256358596 | 564.264325081426 | 521.052271040884 | Tmem222 |
| **ENSMUSG00000028859** | 0.308211470501404 | 0.624249924282526 | 0.931145988862144 | 39.754603529038 | 16.2883145782866 | 27.447191770805 | 12.1930470253561 | 12.0445401540101 | 47.7002070508805 | 9.36970281500523 | 8.68420451734806 | Csf3r |
| **ENSMUSG00000028861** | 0.0313376870027671 | 0.827094698256603 | 0.967319591893432 | 1342.6213828216 | 1078.28642508257 | 1207.67643791542 | 1436.9036956035 | 1282.19604730416 | 1438.34470491886 | 1133.73404061563 | 1102.8939737032 | Mrps15 |
| **ENSMUSG00000028862** | -0.607873219480241 | 0.173531786200792 | 0.799889021902745 | 537.590661358582 | 564.39010013763 | 485.531357876653 | 189.461192240149 | 1013.93128932848 | 1024.94291047789 | 348.761160336306 | 321.315567141878 | Map3k6 |
| **ENSMUSG00000028863** | -0.0510464418652022 | 0.769451916442506 | 0.95879680569409 | 244.852217190211 | 214.191336704468 | 245.131816159948 | 267.309107863576 | 287.974005500422 | 298.432064626022 | 215.50316474512 | 205.164331722348 | Meaf6 |
| **ENSMUSG00000028868** | -0.327646428204451 | 0.118105374593664 | 0.769868040577702 | 909.838312585028 | 1046.52421165491 | 912.382512657103 | 753.155135489303 | 1530.7515577551 | 1230.4207254663 | 872.423439886043 | 912.926999886215 | Wasf2 |
| **ENSMUSG00000028869** | 0.20592785630334 | 0.153138989928945 | 0.783412422888014 | 948.689402397497 | 954.495234287593 | 1120.60258815976 | 1103.0017924476 | 1076.34390649017 | 919.757838519542 | 812.040910633787 | 769.637625349972 | Gnl2 |
| **ENSMUSG00000028871** | -0.0177170867220894 | 0.977828209424522 | 0.996261558637615 | 5.42108229941427 | 7.32974156022896 | 18.9290977729689 | 18.7585338851632 | 17.5193311331055 | 22.0154801773295 | 7.28754663389296 | 4.34210225867403 | Rspo1 |
| **ENSMUSG00000028878** | 0.0258901165724521 | 0.795576403311834 | 0.962043367573119 | 1024.5845545893 | 1004.17459375137 | 972.955625530603 | 949.181814589259 | 1035.83045324487 | 854.934480219628 | 1024.42084110724 | 963.946701425635 | Fam76a |
| **ENSMUSG00000028879** | 0.0931880036271745 | 0.326562575703106 | 0.855977678491063 | 1124.87457712846 | 1214.29385181126 | 1313.67938544404 | 1262.44933047149 | 1213.21368096756 | 1073.86619976085 | 1185.78794514344 | 1133.28868951392 | Stx12 |
| **ENSMUSG00000028882** | -0.0449639551079174 | 0.740497623199347 | 0.954383574136061 | 353.273863178497 | 277.715763559786 | 303.812019256151 | 300.136542162612 | 354.766455445387 | 281.308913376988 | 316.487739529066 | 320.23004157721 | Ppp1r8 |
| **ENSMUSG00000028883** | -0.305328700756874 | 0.51359777172337 | 0.903136226008053 | 16.2632468982428 | 35.0198763433161 | 12.3039135524298 | 13.1309737196143 | 20.8042057205628 | 29.3539735697726 | 32.2734208072402 | 13.0263067760221 | Sema3a |
| **ENSMUSG00000028884** | 0.0335876613694708 | 0.803458369629567 | 0.963461896828986 | 349.65980831222 | 296.447325324816 | 348.295399022628 | 315.143369270742 | 351.48158085793 | 303.324393554317 | 346.679004155194 | 276.809018990469 | Rpa2 |
| **ENSMUSG00000028886** | 0.0924058847431353 | 0.583386077721873 | 0.922450244597442 | 469.827132615903 | 633.615437095348 | 604.784673846357 | 456.770300103725 | 485.06648074786 | 426.855698993777 | 580.921574530324 | 537.335154510911 | Eya3 |
| **ENSMUSG00000028889** | 0.130731206426212 | 0.289658314762789 | 0.851785577030345 | 328.878992831132 | 309.477976987445 | 296.240380146964 | 353.598363735327 | 318.632834983357 | 306.993640250539 | 283.173240631269 | 268.124814473121 | Yrdc |
| **ENSMUSG00000028890** | 0.101708441784851 | 0.550353975003555 | 0.914699131434573 | 551.143367107117 | 537.514381083457 | 555.569019636638 | 454.894446715208 | 436.88832013182 | 380.37857417497 | 555.935700356977 | 581.84170266232 | Mtf1 |
| **ENSMUSG00000028893** | -0.0879902890860298 | 0.643373159066315 | 0.934616627390525 | 104.807591122009 | 86.3280672649188 | 83.2880302010633 | 85.3513291774927 | 116.065568756824 | 94.1773318696872 | 81.2040910633787 | 91.1841474321546 | Sesn2 |
| **ENSMUSG00000028894** | -0.294726115518705 | 0.0634132145675377 | 0.716991331547963 | 113.8427282877 | 110.760539132349 | 127.77140996754 | 136.937297361692 | 162.053812981226 | 163.893019097897 | 136.381229862854 | 137.8617467129 | Inpp5b |
| **ENSMUSG00000028896** | -0.058262618729849 | 0.738957262547317 | 0.954383574136061 | 128.298947752804 | 122.162359337149 | 117.360406192407 | 98.4823028971069 | 99.6411958195378 | 136.985209992272 | 127.011527047849 | 122.664388807541 | Rcc1 |
| **ENSMUSG00000028898** | 0.147881260153674 | 0.443712092575475 | 0.89108031640712 | 487.897406947284 | 408.022280186079 | 523.389553422591 | 591.8317440769 | 486.161438943679 | 577.294813538862 | 378.952424962434 | 373.420794245967 | Trnau1ap |

| | | | | | | | | | | | | |
|---|---|---|---|---|---|---|---|---|---|---|---|---|
| **ENSMUSG00000028899** | 0.0204713531817288 | 0.878810934224552 | 0.978116690177517 | 544.818771091134 | 393.362797065621 | 462.81644054909 | 496.163221262567 | 498.205979097689 | 451.317343635254 | 447.663578939139 | 472.203620630801 | Taf12 |
| **ENSMUSG00000028901** | 0.25103431547501 | 0.137704204428731 | 0.779425998172399 | 227.685456575399 | 270.386021999557 | 243.238906382651 | 216.661066373635 | 162.053812981226 | 182.239252579005 | 224.872867560126 | 235.559047533066 | Gmeb1 |
| **ENSMUSG00000028902** | -0.144654436301592 | 0.384343124145447 | 0.875126835008975 | 287.317361868956 | 303.777066885045 | 306.651383922097 | 345.157023487003 | 395.279908690694 | 403.617136584374 | 287.337552993494 | 288.749800201823 | Sf3a3 |
| **ENSMUSG00000028906** | 0.266386316436004 | 0.242069522079781 | 0.832048466795696 | 3077.36771863417 | 3988.19382449347 | 4939.54806385624 | 2259.46540646791 | 2776.81398459723 | 2940.28968590556 | 3164.87739529066 | 2977.59662388572 | Epb41 |
| **ENSMUSG00000028907** | 0.120881101324509 | 0.439368320797054 | 0.889583654898613 | 498.739571546113 | 422.681763306537 | 444.83379766477 | 528.052728867345 | 441.268152915096 | 535.71001764835 | 386.239971596327 | 379.933947633978 | Utp11 |
| **ENSMUSG00000028909** | -0.419087700896937 | 0.273104954990388 | 0.844738346035895 | 44.2721721118832 | 63.5244268553176 | 111.681676860517 | 35.6412143818101 | 110.590777777729 | 92.9542496376133 | 84.3273253350471 | 53.1907526687569 | Ptpru |
| **ENSMUSG00000028910** | 0.095399625166614 | 0.380774160922792 | 0.874640829372842 | 505.064167562096 | 510.638662029284 | 451.458981885309 | 490.535661097018 | 503.680770076785 | 458.655837027697 | 451.827891301363 | 419.012867962044 | Mecr |
| **ENSMUSG00000028911** | 0.0136451714063739 | 0.920887657959138 | 0.986713918026868 | 346.045753445944 | 395.806044252364 | 410.761421673426 | 330.150196378873 | 377.760577557588 | 324.116791499573 | 414.349080041342 | 351.710282952596 | Srsf4 |
| **ENSMUSG00000028914** | -0.112376638038537 | 0.468035556853794 | 0.895328112556867 | 174.378147297826 | 149.038078391322 | 189.290977729689 | 183.8336320746 | 199.282391639076 | 156.554525705454 | 203.010227658447 | 192.138024946326 | Casp9 |
| **ENSMUSG00000028917** | 0.135011170028631 | 0.416503559022248 | 0.882192126163777 | 2197.34535869592 | 2442.43277101407 | 2494.85508647731 | 2329.80990853727 | 1999.39366556567 | 1599.79155955261 | 2445.49243471637 | 2572.69558826436 | Plekhm2 |
| **ENSMUSG00000028919** | -0.293743568386686 | 0.0771926880655704 | 0.739855948845396 | 564.696072855653 | 544.844122643686 | 433.476339000989 | 456.770300103725 | 535.434557755538 | 516.140701935169 | 679.823993133157 | 719.703449375221 | Arhgef19 |
| **ENSMUSG00000028920** | -0.0555433134785958 | 0.751494399240514 | 0.956197192463153 | 273.764656120421 | 393.362797065621 | 332.205665915605 | 248.550573978413 | 326.297542354091 | 293.539735697726 | 325.857442344071 | 351.710282952596 | Fbxo42 |
| **ENSMUSG00000028923** | -0.342715587255709 | 0.227973281634896 | 0.832048466795696 | 188.83436676293 | 171.027303072009 | 227.149173275627 | 256.991914226736 | 325.202584158272 | 381.601656407044 | 168.654650670094 | 195.394601640331 | Necap2 |
| **ENSMUSG00000028926** | -0.0352946857194464 | 0.846239263280486 | 0.972230220592717 | 168.053551281842 | 184.057954734638 | 201.594891282119 | 152.88205116408 | 210.231973597267 | 207.923979452556 | 161.367104036201 | 145.46042566558 | Cdk14 |
| **ENSMUSG00000028927** | -0.407546626720732 | 0.325592438518616 | 0.855977678491063 | 14.4562194651047 | 16.2883145782866 | 12.3039135524298 | 18.7585338851632 | 21.8991639163819 | 33.0232202659942 | 11.4518589961175 | 16.2828834700276 | Padi2 |
| **ENSMUSG00000028931** | -0.208940905866088 | 0.640590300407834 | 0.934417517272849 | 18.9737880479499 | 19.5459774939439 | 26.5007368821565 | 17.8206071909051 | 27.3739548954774 | 40.3617136584374 | 19.7804837205666 | 8.68420451734806 | Kcnab2 |
| **ENSMUSG00000028932** | -0.0185739765742398 | 0.895008344799451 | 0.983733637008786 | 3041.22716997141 | 2647.66553470048 | 2926.438515701 | 3181.44734692368 | 3353.85695379389 | 3413.62250971814 | 2655.7902090087 | 2527.10351454829 | Psmc2 |
| **ENSMUSG00000028933** | -0.042544052910271 | 0.808324431410788 | 0.964026716319739 | 216.843291976571 | 205.232763686411 | 198.755526616174 | 164.137171495178 | 195.997517051618 | 156.554525705454 | 223.831789469569 | 231.216945274392 | Xrcc2 |
| **ENSMUSG00000028936** | 0.219379398057099 | 0.0200219023808733 | 0.642273034804458 | 4267.2952833556 | 3681.97351042168 | 3979.84280676672 | 4375.42802871432 | 3397.65528162666 | 3762.20094585919 | 3389.75026285078 | 3455.22787233986 | Rpl22 |
| **ENSMUSG00000028937** | -0.107293659625948 | 0.380003660016361 | 0.87432013076003 | 1984.11612158562 | 1752.62264862364 | 1861.67676597149 | 2156.29347009951 | 2187.72647524656 | 2347.09480334974 | 1844.79037646547 | 1974.57100213202 | Acot7 |
| **ENSMUSG00000028943** | -0.0371908636370885 | 0.854281137835436 | 0.973258284492635 | 70.4740698923855 | 78.1839099757755 | 89.9132144216024 | 67.5307219865876 | 71.1722827282413 | 91.7311674055395 | 85.3684034256032 | 66.217059444779 | Espn |
| **ENSMUSG00000028944** | 0.00791706368100088 | 0.96474268710568 | 0.994284908685797 | 736.363679003772 | 769.622863824041 | 634.124775394459 | 617.15576482187 | 548.574056105368 | 577.294813538862 | 772.479943192653 | 843.45336374743 | Prkag2 |
| **ENSMUSG00000028945** | 0.0916171245748342 | 0.235316349405811 | 0.832048466795696 | 3102.6661026981 | 2809.73426475443 | 3105.31848965555 | 3021.06188220554 | 2765.86440263904 | 2762.94276225485 | 2983.72980753389 | 2783.28754781005 | Rheb |
| **ENSMUSG00000028948** | -0.00824107939342586 | 0.955598499705176 | 0.992371505557225 | 207.80815481088 | 265.499527626071 | 235.667267273463 | 211.033506208086 | 249.650468646754 | 198.139321595965 | 235.283648465687 | 242.072200921077 | Nol9 |
| **ENSMUSG00000028949** | -0.053388870952418 | 0.693367355287832 | 0.94469184542218 | 1241.42784656587 | 1086.43058237171 | 959.705257089525 | 1130.20166658108 | 1032.54557865741 | 1352.72894867369 | 1111.87140071395 | 1088.78214136251 | Smarcd3 |
| **ENSMUSG00000028952** | 0.144062986905774 | 0.528461390106196 | 0.908232038364594 | 62.3424464432641 | 65.9676740420606 | 53.001473764313 | 51.5859681841989 | 62.4126171616885 | 59.9310293716191 | 43.7252798033577 | 45.5920737160773 | Zbtb48 |
| **ENSMUSG00000028953** | 0.182572265533684 | 0.097229183002899 | 0.757809915356977 | 1562.17521594788 | 1616.61522189494 | 1667.65351379856 | 1639.49586156327 | 1355.55824642404 | 1236.53613662667 | 1553.28851110976 | 1567.49891538132 | Abcf2 |
| **ENSMUSG00000028954** | 0.191551238078392 | 0.13101614911996 | 0.77303252008843 | 1332.68273193934 | 1551.4619635818 | 1784.06746510232 | 1448.1588159346 | 1365.41287018641 | 1197.39750520031 | 1429.40021833358 | 1362.33458365898 | Nub1 |
| **ENSMUSG00000028955** | 0.0064173479980729 | 0.946118258060885 | 0.990505437194639 | 1327.26164963993 | 1252.57139107024 | 1232.28426502028 | 1426.58650196666 | 1303.00025302473 | 1210.85140975312 | 1316.96378455351 | 1382.95956938768 | Vamp3 |
| **ENSMUSG00000028957** | 0.0935097954609392 | 0.726709904056631 | 0.952045814446758 | 103.90407740544 | 118.090280692578 | 96.5383986421416 | 99.4202295913651 | 89.7865720571659 | 57.4848649074714 | 142.627698406191 | 100.953877514171 | Per3 |
| **ENSMUSG00000028958** | -0.0349295105413043 | 0.796904329547956 | 0.962278883153809 | 212.325723393726 | 188.13003337921 | 217.684624389143 | 212.909359596603 | 225.561388338734 | 229.939459629886 | 211.338852382896 | 184.539345993646 | Tmub1 |
| **ENSMUSG00000028959** | 0.00519226827588892 | 0.949492394683609 | 0.991267322147698 | 6167.3846293003 | 5450.07005789469 | 5539.60046325936 | 5679.14613373317 | 5518.58930692825 | 5522.21627781348 | 6074.69065839506 | 5637.13425732356 | Fastk |
| **ENSMUSG00000028960** | 0.0889850570437855 | 0.507561962825331 | 0.90183410140733 | 872.794250205697 | 946.35107699845 | 980.527264639791 | 990.450589136618 | 856.257309130534 | 724.064681387725 | 990.065264118886 | 991.084840542347 | Ube4b |
| **ENSMUSG00000028961** | -0.247097053451352 | 0.118106742083919 | 0.769868040577702 | 688.477452025612 | 781.839099757755 | 708.894711597687 | 740.962088463947 | 942.759006600242 | 1046.95839065522 | 806.835520181006 | 670.854798965138 | Pgd |
| **ENSMUSG00000028962** | -0.080736819174786 | 0.500858693395149 | 0.900927059989498 | 543.011743657996 | 589.636987733974 | 550.836745193396 | 486.783954319986 | 614.271547854513 | 623.771938357668 | 544.483841360859 | 514.539117652873 | Slc4a2 |
| **ENSMUSG00000028964** | -0.109511058457289 | 0.573634942600508 | 0.922207298536842 | 8372.86161144534 | 6928.23460587419 | 6290.13918995758 | 8915.93115561808 | 8204.52176127249 | 10923.3474146516 | 6833.63658641048 | 6952.79124170179 | Park7 |
| **ENSMUSG00000028967** | -0.506992107077454 | 0.160898535948362 | 0.792229941270026 | 385.800356974982 | 571.719841697859 | 263.114459044268 | 233.543746870282 | 696.393412540946 | 704.495365674543 | 292.542943446274 | 374.506319810635 | Errfi1 |
| **ENSMUSG00000028969** | -0.038821222464957 | 0.800556587348997 | 0.963461896828986 | 426.458474220589 | 382.775392589734 | 340.723759913441 | 355.474217123843 | 384.330326732503 | 469.663577116362 | 368.541644056872 | 325.657669400552 | Cdk5 |
| **ENSMUSG00000028970** | 0.0969465873377573 | 0.540897117934467 | 0.913837378403038 | 151.7903043836 | 176.728213174409 | 200.648436393471 | 199.778385876988 | 179.573144114332 | 165.116101329971 | 186.352978209548 | 149.802527924254 | Abcb1b |
| **ENSMUSG00000028973** | 0.0784150260139539 | 0.467913734686979 | 0.895328112556867 | 2127.7748025201 | 2118.29531090617 | 1934.55379239743 | 1891.79814231871 | 1726.74907480672 | 1784.47697659576 | 2033.22551085613 | 2100.49196763356 | Abcb8 |
| **ENSMUSG00000028974** | 0.0253143706583269 | 0.907473489481541 | 0.98504921011344 | 703.837185207286 | 816.858976101072 | 840.451941119821 | 678.120999948651 | 669.019457645468 | 488.00981059747 | 962.997233764426 | 865.163875040801 | Dffa |
| **ENSMUSG00000028975** | -0.00281698977412118 | 0.97451128103287 | 0.995510898988495 | 626.135005582348 | 601.853223667689 | 638.857049837702 | 664.990026229036 | 629.600962595981 | 654.348994159515 | 642.345181873136 | 610.065367343701 | Pex14 |
| **ENSMUSG00000028977** | -0.030667075039589 | 0.843920796016193 | 0.971721411930626 | 598.126080368708 | 735.417403209639 | 840.451941119821 | 936.050840869645 | 793.844691968845 | 787.664957455566 | 761.028084196536 | 833.683633665414 | Casz1 |
| **ENSMUSG00000028978** | -0.562133765676061 | 0.0531644559247116 | 0.703396849069134 | 639.687711330884 | 488.649437348597 | 494.995906763138 | 553.376749612315 | 1198.97922442191 | 976.01962119494 | 574.675105986987 | 464.604941678121 | Nos3 |
| **ENSMUSG00000028979** | -0.0299725937081452 | 0.944535637463631 | 0.990505437194639 | 17.1667606148119 | 30.9477976987445 | 28.3936466594534 | 27.1998741334867 | 15.3294147414674 | 20.7923979452556 | 19.7804837205666 | 49.9341759747513 | Masp2 |
| **ENSMUSG00000028980** | -0.192410389578899 | 0.198264733610724 | 0.812921911423739 | 621.617436999503 | 739.48948185421 | 849.916490006305 | 648.107345732389 | 919.764884488041 | 830.472835578151 | 695.440164491499 | 821.74285245406 | H6pd |
| **ENSMUSG00000028982** | 0.260262453140815 | 0.129045259582726 | 0.77303252008843 | 656.854471945696 | 689.810122390436 | 724.037989816062 | 783.168789705565 | 780.705193619016 | 566.287073450197 | 560.100012719201 | 475.460197324806 | Slc25a33 |
| **ENSMUSG00000028986** | 0.220155465349532 | 0.234322279358697 | 0.832048466795696 | 1121.26052226218 | 1245.24164951001 | 1664.81414913262 | 1129.26373988683 | 1023.78591309086 | 892.850029413917 | 1288.8546761085 | 1223.38731138141 | Klhl7 |
| **ENSMUSG00000028988** | -0.17773147267308 | 0.2474994094022 | 0.833398011296137 | 588.187429486448 | 541.586459728029 | 529.068282754482 | 661.238319452004 | 803.699315731217 | 579.740978003009 | 574.675105986987 | 665.427171141795 | Ctnnbip1 |
| **ENSMUSG00000028989** | -1.52225480165444 | 0.0448048813526453 | 0.693563614452513 | 10.8421645988285 | 15.4738988493722 | 12.3039135524298 | 17.8206071909051 | 29.5638712871156 | 116.192812047017 | 10.4107809055614 | 6.51315338801105 | Angptl7 |
| **ENSMUSG00000028990** | 0.21811481414119 | 0.13635721979655 | 0.776456211338551 | 533.073092775737 | 574.977504613516 | 561.247748968529 | 574.011136885995 | 443.458069306734 | 385.270903103266 | 540.319528998635 | 557.960140239613 | Lzic |
| **ENSMUSG00000028991** | 0.3840940007913 | 0.0959968785904501 | 0.757809915356977 | 2200.95941356219 | 2576.81136628494 | 2876.27640660263 | 2234.14138572294 | 1984.0642508242 | 1106.88942002684 | 2227.90711379013 | 2256.80764894583 | Mtor |
| **ENSMUSG00000028992** | 0.081425024798158 | 0.590188324765246 | 0.923935529355527 | 515.002818444356 | 447.114235173966 | 381.421320125324 | 434.260059441529 | 395.279908690694 | 355.916929533493 | 454.951125573032 | 473.289146195469 | Nmnat1 |
| **ENSMUSG00000028995** | 0.128829544475726 | 0.466499942032781 | 0.895328112556867 | 573.731210021344 | 737.860650396382 | 667.250696497155 | 543.997482669733 | 487.256397139498 | 481.894399437101 | 679.823993133157 | 657.828492189116 | Fam126a |
| **ENSMUSG00000028996** | -0.0762417336544508 | 0.7314132137253 | 0.953029200153788 | 683.056369726198 | 460.144886836596 | 563.140658745826 | 743.775868546722 | 758.806029702634 | 786.441875223492 | 473.690531203042 | 564.473293627624 | Rbp7 |
| **ENSMUSG00000028998** | 0.103761823844312 | 0.589515452916199 | 0.923594634030771 | 1237.81379169959 | 841.291447968501 | 1087.47666705707 | 1216.49092245284 | 997.506916391197 | 1329.49038626429 | 876.587752248267 | 876.019130687486 | Tomm7 |
| **ENSMUSG00000028999** | -0.00358313808426359 | 0.9852193886163 | 0.99735301701069 | 215.036264543433 | 175.913797445495 | 215.791714611846 | 168.826804966469 | 151.104231023035 | 171.23151249034 | 216.544242835676 | 237.730098662403 | Rint1 |
| **ENSMUSG00000029001** | 0.115693754375622 | 0.433500921708276 | 0.887868762931002 | 286.413848152387 | 324.137460107903 | 335.04503058155 | 316.081295965 | 257.315176017488 | 267.855008824175 | 357.089785060755 | 281.151121249143 | Fbxo44 |
| **ENSMUSG00000029003** | -0.035087453056157 | 0.844662257866945 | 0.971865371253468 | 156.307872966445 | 133.56417954195 | 160.897331070236 | 170.702658354985 | 170.813478547779 | 194.470074899744 | 136.381229862854 | 135.690695583563 | Mad2l2 |
| **ENSMUSG00000029004** | -0.0303693785187789 | 0.815887765358327 | 0.966177091475091 | 908.03128515189 | 1108.4198070524 | 940.776159316556 | 879.775239214155 | 971.227919691539 | 854.934480219628 | 1110.8303226234 | 981.315110460331 | Kmt2e |
| **ENSMUSG00000029007** | 0.103489885814935 | 0.284619814866917 | 0.851125503777472 | 1276.66488151206 | 1407.31037956396 | 1263.51727634568 | 1242.75286989206 | 1128.90189988949 | 1143.58188698906 | 1275.32066093127 | 1283.09121743818 | Agtrap |
| **ENSMUSG00000029009** | 0.106007502588944 | 0.565266784503296 | 0.919692942798738 | 153.597331816738 | 140.079505373265 | 209.166530391307 | 165.075098189436 | 154.389105610493 | 129.646716599829 | 159.284947855089 | 175.855141476298 | Mthfr |
| **ENSMUSG00000029012** | 0.0279679308816988 | 0.865473257575928 | 0.974868041734905 | 121.974351736821 | 124.605606523892 | 151.432782183751 | 132.247663890401 | 139.059690869025 | 106.408154190426 | 134.299073681742 | 138.947272277569 | Orc5 |
| **ENSMUSG00000029014** | 0.286310793703416 | 0.0214855202293024 | 0.648762042745232 | 1105.90078908051 | 1126.33695308852 | 1071.38693395004 | 1344.9868795662 | 999.696832782835 | 1013.93517038923 | 971.325858488875 | 827.170480277403 | Dnajc2 |
| **ENSMUSG00000029016** | 0.0876475872595898 | 0.561792333216973 | 0.917279359161133 | 197.869503928621 | 184.057954734638 | 234.720812384815 | 213.847286290861 | 205.85214081399 | 163.893019097897 | 212.379930473452 | 197.565652769668 | Clcn6 |
| **ENSMUSG00000029017** | 0.118801082142873 | 0.230036255079724 | 0.832048466795696 | 4643.15698944832 | 4319.6610261616 | 4077.32766029751 | 4783.42614071662 | 3916.66546644491 | 3798.89341282141 | 4340.25455952853 | 4357.29961657939 | Pmpcb |
| **ENSMUSG00000029019** | 1.17073217470051 | 0.0849737377414648 | 0.743541556158628 | 3876.0738440812 | 3351.32072448246 | 3685.49533639705 | 14456.264138601 | 1094.9581958191 | 2958.63591938667 | 2039.47197939947 | 5175.78589233944 | Nppb |
| **ENSMUSG00000029020** | 0.217333994714948 | 0.369792805098352 | 0.87265070724025 | 23001.6521964147 | 27738.999726822 | 25760.6091592334 | 20292.0440302753 | 16824.0326787604 | 13238.6420799675 | 26809.8429880016 | 26383.6988492681 | Mfn2 |
| **ENSMUSG00000029022** | 0.041536903974795 | 0.855108598983602 | 0.973268281968411 | 88.5443442237664 | 80.6271571625185 | 88.0203046443056 | 81.59962240046 | 100.736154015357 | 102.738907494204 | 62.4646854333682 | 64.046008315442 | Miip |
| **ENSMUSG00000029028** | 0.199626684887534 | 0.134270680755902 | 0.77303252008843 | 955.91751213005 | 1171.94423390772 | 1279.6070094527 | 1065.48472467727 | 1056.63465896543 | 850.042151291332 | 1000.47604502445 | 986.742738283673 | Lrrc47 |
| **ENSMUSG00000029029** | -0.28863061943046 | 0.0681847582531899 | 0.722112580225498 | 199.676531361759 | 191.387696294867 | 160.897331070236 | 173.51643843776 | 239.795844884382 | 254.401104271363 | 186.352978209548 | 207.335382851685 | Wrap73 |
| **ENSMUSG00000029030** | 0.00931296011395836 | 0.919932977006503 | 0.986713918026868 | 2697.89195767517 | 2769.82789403763 | 3141.2837754242 | 2943.21396658211 | 2724.25599119791 | 3101.73654053931 | 2823.40378158824 | 2828.87962152613 | Tprgl |
| **ENSMUSG00000029033** | -0.168224000259122 | 0.209734066903145 | 0.821566820583092 | 159.921927832721 | 180.800291818981 | 168.468970179424 | 170.702658354985 | 188.332809680885 | 178.570005882784 | 183.22974393788 | 213.848536239696 | Acap3 |
| **ENSMUSG00000029034** | 0.0533263239019579 | 0.756219981325002 | 0.956566265296272 | 355.080890611635 | 291.56083095133 | 419.279515671262 | 426.756645887463 | 416.084114411257 | 373.040080782527 | 313.364505257397 | 335.427399482569 | Ints11 |
| **ENSMUSG00000029036** | 0.226249211904547 | 0.114550107543547 | 0.767518180080285 | 852.916948441179 | 991.958357817652 | 971.062715753306 | 939.802547646678 | 855.162350934715 | 618.879609429373 | 855.766190437144 | 879.275707381491 | Atad3a |
| **ENSMUSG00000029038** | 0.0663982013150054 | 0.63590642735717 | 0.932595977875587 | 1047.17239750352 | 787.540009860156 | 858.434584004141 | 1044.85033740359 | 959.183379537529 | 957.673387713832 | 819.328457267679 | 833.683633665414 | Ssu72 |
| **ENSMUSG00000029047** | 0.113828823677752 | 0.43696398287725 | 0.888701139843674 | 299.063040184354 | 285.859920848929 | 330.312756138308 | 350.784583652552 | 314.253002200081 | 271.524255520397 | 333.144988977964 | 249.670879873757 | Pex10 |
| **ENSMUSG00000029048** | 0.0672345604715283 | 0.462427184548576 | 0.895328112556867 | 2013.02856051583 | 1799.04434517175 | 1957.26870972499 | 2131.9073760488 | 1911.79700990014 | 1953.26232462195 | 1766.70951967376 | 1909.4394682519 | Rer1 |
| **ENSMUSG00000029049** | 0.0206792058350095 | 0.966537313000641 | 0.994284908685797 | 4.51756858284523 | 14.6594831204579 | 18.9290977729689 | 11.2551203310979 | 10.949581958191 | 12.2308223207386 | 14.5750932677859 | 10.8552556466851 | Morn1 |
| **ENSMUSG00000029050** | -0.173510403581966 | 0.191395867656454 | 0.808824684419004 | 1187.21702357173 | 1534.3592332746 | 1615.5984949229 | 1288.71127791071 | 1730.03394939417 | 1533.74511902062 | 1564.74037010587 | 1516.4792138419 | Ski |
| **ENSMUSG00000029053** | -0.251248801302998 | 0.310807741764604 | 0.851785577030345 | 74.0881247586617 | 54.56585383726 | 74.7699362032273 | 66.5927952923295 | 90.881530252985 | 73.3849339244316 | 102.025652874501 | 54.2762782334254 | Prkcz |
| **ENSMUSG00000029056** | -0.0646698360160554 | 0.555872318686706 | 0.916239900758826 | 674.924746277077 | 667.820897709749 | 711.734076263632 | 725.017334661559 | 704.058119911679 | 634.779678446333 | 777.685333645434 | 788.091559949337 | Pank4 |
| **ENSMUSG00000029059** | 0.0316086296554536 | 0.871459777810332 | 0.975613767803763 | 5899.0410554793 | 4241.47711618582 | 3858.69658101972 | 6319.75006591149 | 5149.58839493721 | 5841.44074038475 | 4403.76032305246 | 4484.30610764561 | Fam213b |
| **ENSMUSG00000029060** | -0.0733316943916634 | 0.43403566820573 | 0.887868762931002 | 723.714486971805 | 695.511032492837 | 725.930899593359 | 751.279282100787 | 747.856447744443 | 799.895779776304 | 802.671207818781 | 696.907412517182 | Mib2 |
| **ENSMUSG00000029061** | -0.842188175998326 | 0.132469415202581 | 0.77303252008843 | 62.3424464432641 | 54.56585383726 | 39.7511053232348 | 88.1651092602672 | 124.825234323377 | 238.501035254403 | 46.8485140750261 | 29.3091902460497 | Mmp23 |
| **ENSMUSG00000029062** | 0.127000889420965 | 0.176793448629188 | 0.799889021902745 | 2201.86292727876 | 1935.86618762936 | 1997.96626993687 | 2155.35554340525 | 1797.92135753496 | 1819.9463613259 | 2066.54000975393 | 1906.1828915579 | Cdk11b |
| **ENSMUSG00000029063** | -0.0559908863959534 | 0.630551089183509 | 0.931825800800958 | 631.556087881762 | 720.757920089181 | 661.571967165264 | 634.038445318517 | 785.085026402292 | 689.818378889657 | 659.002431322034 | 619.835097425718 | Nadk |

| | | | | | | | | | | | | |
|---|---|---|---|---|---|---|---|---|---|---|---|---|
| ENSMUSG00000029064 | -0.130077954158682 | 0.328590526054173 | 0.855977678491063 | 2630.12842893249 | 2666.39709646551 | 2488.22990225677 | 2478.94025292432 | 3270.64013091164 | 3082.16722482613 | 2379.90451501133 | 2501.05090099624 | Gnb1 |
| ENSMUSG00000029066 | 0.107363852797656 | 0.62423703854301 | 0.931145988862144 | 765.276117933981 | 456.072808192024 | 562.194203857177 | 641.541858872582 | 581.42280197994 | 739.964750404685 | 479.936999746379 | 450.493109337431 | Mrpl20 |
| ENSMUSG00000029068 | -0.0193077337051051 | 0.894068431612228 | 0.983730237253949 | 412.002254755485 | 362.414999366876 | 431.583429223692 | 527.114802173087 | 420.463947194533 | 418.29412336926 | 458.0743598447 | 458.09178829011 | Ccnl2 |
| ENSMUSG00000029070 | -1.27068331767799 | 0.00212112054248658 | 0.635862253526407 | 152.693818100169 | 163.69756151178 | 156.165056626994 | 176.330218520534 | 563.903470846835 | 649.456665231219 | 173.860041122875 | 179.111718170304 | Mxra8 |
| ENSMUSG00000029071 | 0.150240006965581 | 0.115279529634835 | 0.768714674329266 | 891.768038253647 | 886.08431305879 | 948.347798425744 | 931.361207398354 | 745.666531352805 | 806.011190936674 | 875.546674157711 | 867.334926170138 | Dvl1 |
| ENSMUSG00000029073 | -0.28296754825871 | 0.113557155378001 | 0.767518180080285 | 82.2197482077831 | 81.4415728914329 | 69.0912068713366 | 67.5307219865876 | 84.3117810780705 | 100.292743030056 | 91.61487196894 | 90.0986218674861 | Cptp |
| ENSMUSG00000029076 | -0.0440621764541617 | 0.774420598627516 | 0.959391911785248 | 2216.31914674387 | 1988.80321000879 | 2224.16898832385 | 2298.85832762675 | 2382.62903410236 | 2793.5198180567 | 1940.56956079664 | 1883.38685469986 | Sdf4 |
| ENSMUSG00000029082 | -0.0666123788179789 | 0.904967498027701 | 0.98504921011344 | 9.93865088225949 | 13.0306516626293 | 19.8755526616174 | 15.9447538023887 | 15.3294147414674 | 30.5770558018465 | 4.16431236222455 | 11.9407812113536 | Bst1 |
| ENSMUSG00000029084 | -0.375109891166253 | 0.20948585009881 | 0.821566820583092 | 134.623543768788 | 110.760539132349 | 114.521041526462 | 187.585338851632 | 233.226095709468 | 229.939459629886 | 107.231043327282 | 140.032797842237 | Cd38 |
| ENSMUSG00000029086 | -0.221188969254616 | 0.578241719246343 | 0.922450244597442 | 37.0440623793308 | 27.6901347830872 | 33.1259211026956 | 17.8206071909051 | 56.937826182593 | 28.1308913376988 | 30.191264626128 | 19.5394601640331 | Prom1 |
| ENSMUSG00000029089 | -0.538324990066562 | 0.0322072198096921 | 0.671606027118506 | 37.9475760958999 | 29.3189662409158 | 34.0723759913441 | 32.8274342990356 | 39.4184950494875 | 61.154111603693 | 47.8895921655823 | 46.6775992807458 | 5730480H06l |
| ENSMUSG00000029090 | 0.0882824797214072 | 0.596889507126747 | 0.924496535139492 | 575.538237454482 | 551.359448475 | 464.709350326387 | 449.266886549659 | 450.027818481649 | 387.717067567413 | 585.085886892549 | 496.085183053508 | Adgra3 |
| ENSMUSG00000029093 | 0.147374890203451 | 0.550762235524941 | 0.914699131434573 | 199.676531361759 | 252.468875963442 | 355.867038131816 | 234.48167356454 | 270.454674367317 | 152.885279009232 | 250.899819824029 | 265.953763343784 | Sorcs2 |
| ENSMUSG00000029094 | -0.0502771948993569 | 0.750857096954862 | 0.956197192463153 | 174.378147297826 | 188.944449108124 | 238.506631939409 | 187.585338851632 | 204.757182618171 | 216.485555077073 | 176.983275394543 | 219.276164063039 | Afap1 |
| ENSMUSG00000029095 | 0.169862388192939 | 0.393755950610175 | 0.878942305112115 | 478.862269781594 | 712.613762800038 | 737.28835825714 | 494.287367874051 | 446.742943894192 | 500.240632918209 | 591.332355435886 | 615.492995167044 | Ablim2 |
| ENSMUSG00000029096 | -1.3696597443164 | 0.00359167008025774 | 0.635862253526407 | 795.995584297329 | 793.240919962556 | 866.006223113329 | 1051.4158242634 | 2895.06946974569 | 4469.14247599788 | 885.957455063272 | 811.973122372044 | Htra3 |
| ENSMUSG00000029097 | 0.0561779413422086 | 0.852886565807188 | 0.973120292022893 | 30.7194663633475 | 53.7514381083457 | 61.519567762149 | 31.8895076047775 | 41.6084114411257 | 37.9155491942896 | 42.6842017128016 | 48.8486504100828 | Trmt44 |
| ENSMUSG00000029098 | 0.0477851402913287 | 0.76131155587104 | 0.957479700684068 | 481.572810931301 | 573.348673155687 | 563.140658745826 | 464.27371365779 | 458.787484048202 | 414.624876673038 | 548.648153723084 | 591.611432744337 | Acox3 |
| ENSMUSG00000029101 | -0.178892821066328 | 0.341690248009526 | 0.859283395807012 | 379.475760958999 | 411.279943101736 | 439.155068332879 | 372.35689762049 | 480.686647964584 | 599.310293716191 | 370.623800237985 | 364.736589728619 | Rgs12 |
| ENSMUSG00000029102 | -0.360853034135649 | 0.471736567764631 | 0.897171173241185 | 11.7456783153976 | 29.3189662409158 | 17.036187995672 | 15.9447538023887 | 17.5193311331055 | 12.2308223207386 | 18.7394056300105 | 46.6775992807458 | Hgfac |
| ENSMUSG00000029103 | -0.032888624275351 | 0.812049882232233 | 0.965224347191244 | 1569.40332568043 | 1275.37503147984 | 1437.66497585699 | 1495.05515064751 | 1595.35409130842 | 1712.3151249034 | 1276.36173902182 | 1327.59776558958 | Lrpap1 |
| ENSMUSG00000029104 | 0.144393654094745 | 0.481444180163586 | 0.897732059433541 | 363.212514060756 | 423.496179035451 | 421.172425448559 | 337.653609932938 | 321.917709570815 | 239.724117486476 | 439.33495421469 | 396.216831104005 | Htt |
| ENSMUSG00000029106 | 0.0185526264767393 | 0.866271772464148 | 0.974868041734905 | 2905.70011248605 | 3256.03408419949 | 3151.69477919933 | 2761.25618789603 | 2862.22072387112 | 2650.41919690405 | 3228.38315881458 | 3178.41885334939 | Add1 |
| ENSMUSG00000029108 | -0.0100687828877004 | 0.965620579914415 | 0.994284908685797 | 1316.4194850411 | 1541.68897483482 | 1477.41608118023 | 1229.62189617245 | 1058.82457535707 | 971.127292266645 | 1914.54260853274 | 1658.68306281348 | Pcdh7 |
| ENSMUSG00000029110 | 0.122357613524342 | 0.480859628757177 | 0.897732059433541 | 510.48524986151 | 668.635313438664 | 708.894711597687 | 570.259430108962 | 597.847174917227 | 465.99433042014 | 510.128264372507 | 683.88110574116 | Rnf4 |
| ENSMUSG00000029111 | -0.153431625172021 | 0.303030915750824 | 0.851785577030345 | 200.580045078328 | 181.614707547895 | 199.701981504822 | 201.654239265505 | 212.421889988905 | 227.493295165738 | 251.940897914585 | 179.111718170304 | Nelfa |
| ENSMUSG00000029119 | 0.0105441152506355 | 0.944876738608551 | 0.990505437194639 | 392.124952990966 | 340.425774686189 | 364.385132129652 | 384.549944645846 | 411.70428162798 | 428.078781225851 | 330.021754706295 | 301.776106977845 | Man2b2 |
| ENSMUSG00000029122 | -0.800683923804669 | 0.0329759381543134 | 0.671606027118506 | 82.2197482077831 | 77.3694942468612 | 62.4660226507975 | 69.4065753751039 | 180.668102310151 | 192.023910435596 | 71.8343882483734 | 64.046008315442 | Evc |
| ENSMUSG00000029123 | -1.04724103793261 | 0.221510982636183 | 0.831114951057963 | 11.7456783153976 | 14.6594831204579 | 4.73227444324223 | 2.81378008277448 | 17.5193311331055 | 42.8078781225851 | 9.36970281500523 | 1.08552556466851 | Stk32b |
| ENSMUSG00000029125 | -0.0439707998622308 | 0.752830649421594 | 0.956497786093635 | 299.966553900923 | 274.458100644129 | 271.632553042104 | 285.129715054481 | 349.291664466292 | 292.316653465652 | 267.557069272927 | 257.269558826436 | Stx18 |
| ENSMUSG00000029126 | -0.92374745404557 | 0.0199883886400747 | 0.642273034804458 | 37.0440623793308 | 43.1640336324594 | 61.519567762149 | 61.9031618210386 | 123.730276127558 | 156.554525705454 | 51.0128264372507 | 55.3618037980939 | Nsg1 |
| ENSMUSG00000029127 | -0.235163695748243 | 0.370633129431444 | 0.873323500696802 | 26.2018977805023 | 30.9477976987445 | 30.2865564367503 | 30.9515809105193 | 41.6084114411257 | 31.8001380339204 | 34.3555769883525 | 31.4802413753867 | Zbtb49 |
| ENSMUSG00000029128 | 0.066532263126261 | 0.601316152484053 | 0.926043912368388 | 2157.59075516688 | 2045.81231103279 | 2152.23841678657 | 2100.95579513828 | 1862.52389108828 | 1684.1842335657 | 2235.19466042403 | 2291.54446701522 | Rab28 |
| ENSMUSG00000029131 | 0.232024197101227 | 0.195205450908201 | 0.81082646923293 | 1838.65041321801 | 2150.87194006274 | 2298.93892452708 | 1887.10850884742 | 1604.11375687498 | 1289.12867260585 | 2137.33331991175 | 1928.97892841594 | Dnajb6 |
| ENSMUSG00000029134 | -0.108307648202301 | 0.721874648459344 | 0.951674101666722 | 27.1054114970714 | 32.5766291565731 | 23.6613722162112 | 21.5723139679377 | 25.1840385038392 | 23.2385624094033 | 36.4377331694648 | 28.2236646813812 | Plb1 |
| ENSMUSG00000029135 | -0.361219168009205 | 0.0889510672456053 | 0.751206603737452 | 421.940905637744 | 529.370223794314 | 426.85115478045 | 359.225923900876 | 675.589206820383 | 616.433444965225 | 567.387559353095 | 373.420794245967 | Fosl2 |
| ENSMUSG00000029136 | 0.14789400305418 | 0.593211863815971 | 0.923935529355527 | 78.6056933415069 | 64.338842584232 | 77.6093008691726 | 61.9031618210386 | 64.6025335533267 | 96.6234963338349 | 44.7663578939139 | 49.9341759747513 | Rbks |
| ENSMUSG00000029141 | 0.0575869265317933 | 0.629254767803772 | 0.931825800800958 | 480.669297214732 | 401.506954354764 | 459.030620994497 | 449.266886549659 | 470.832024202212 | 439.086521314516 | 426.842017128016 | 383.190524327983 | Slc4a1ap |
| ENSMUSG00000029145 | -0.0666639366558017 | 0.54116129257697 | 0.913837378403038 | 1260.40163461382 | 1173.57306536555 | 1146.15687015327 | 1403.13833461021 | 1356.65320461986 | 1395.53682679627 | 1250.33478675792 | 1216.8741579934 | Eif2b4 |
| ENSMUSG00000029146 | -0.0635868060442406 | 0.667461143613842 | 0.942024383071524 | 1776.30796677474 | 1618.24405335277 | 1672.38578824181 | 1880.54302198761 | 1881.13818041721 | 2233.34815576687 | 1641.78014880703 | 1505.62395819522 | Snx17 |
| ENSMUSG00000029147 | 0.0511523858355555 | 0.56599036202826 | 0.919931558693298 | 1324.55110849022 | 1447.21675028076 | 1466.05862251644 | 1463.16564304273 | 1419.06582178155 | 1443.23703384715 | 1377.34631380577 | 1263.55175727414 | Ppm1g |
| ENSMUSG00000029148 | 0.212673027020541 | 0.0363949551721896 | 0.679566075569698 | 1859.43122869909 | 1899.21747982821 | 2099.23694302226 | 1812.07437330677 | 1650.10200109938 | 1505.61422768292 | 1821.88665847324 | 1639.14360264945 | Nrbp1 |
| ENSMUSG00000029149 | -0.506447071667657 | 0.256262648424779 | 0.837517407767513 | 11.7456783153976 | 13.8450673915436 | 13.2503684410783 | 7.50341355406529 | 15.3294147414674 | 9.78465785659088 | 14.5750932677859 | 26.0526135520442 | Krtcap3 |
| ENSMUSG00000029152 | 0.0692768789577618 | 0.51439799773484 | 0.90408556099417 | 3112.60475358036 | 2898.5055792061 | 2798.66710573346 | 2868.17983104146 | 2784.47869196796 | 2424.14898397039 | 3115.94672503452 | 2803.91253353876 | Ociad1 |
| ENSMUSG00000029153 | -0.0607913933324541 | 0.656875450436329 | 0.938389079274206 | 158.114900399583 | 184.057954734638 | 175.094154399963 | 155.695831246855 | 179.573144114332 | 165.116101329971 | 170.736806851206 | 186.710397122983 | Ociad2 |
| ENSMUSG00000029156 | 0.121658418345889 | 0.460493829835315 | 0.894774044236998 | 3343.90426502204 | 3601.34635325916 | 3717.6748026111 | 3315.5708642026 | 2784.47869196796 | 2509.76474021556 | 3908.20715194774 | 3644.10932059218 | Sgcb |
| ENSMUSG00000029158 | 0.106186251598388 | 0.269744065106454 | 0.843793744782845 | 2150.36264543433 | 1922.02112023782 | 1989.44817593904 | 2190.05883109281 | 2114.36427612668 | 1841.96184150323 | 1835.42067365047 | 1873.61712461784 | Yipf7 |
| ENSMUSG00000029161 | 0.103084832100028 | 0.766888867756814 | 0.958655025595455 | 46.9827132615903 | 36.6487078011448 | 46.3762895437739 | 59.0893817382642 | 45.9882442244021 | 70.9387694602839 | 23.9447960827911 | 35.8223436340608 | Cgref1 |
| ENSMUSG00000029162 | -0.168375119197716 | 0.301450292159893 | 0.851785577030345 | 549.336339673979 | 527.741392336485 | 574.498117409607 | 585.266257217093 | 618.65138063779 | 743.633997100907 | 464.320828388037 | 688.223207999834 | Khk |
| ENSMUSG00000029163 | -1.0416026363502 | 0.00837696895256961 | 0.635862253526407 | 201.483558794897 | 212.56250524664 | 182.66579350915 | 241.985087118606 | 648.215251924905 | 666.579816480253 | 206.133461930115 | 206.249857287016 | Emilin1 |
| ENSMUSG00000029165 | 0.103405232185686 | 0.704727408278536 | 0.947228924036081 | 93.9654265231807 | 128.677685168464 | 94.6454888648447 | 76.9099889291692 | 70.0773245324222 | 64.8233582999146 | 113.477511870619 | 118.322286548867 | Agbl5 |
| ENSMUSG00000029166 | 0.049537854005788 | 0.607274902408304 | 0.927437976300313 | 1143.84836517641 | 1044.89538019708 | 1131.96004682354 | 1176.16007459973 | 1021.59599669922 | 1196.17442296823 | 1087.92660463116 | 1039.93349095243 | Mapre3 |
| ENSMUSG00000029167 | 0.160531497872291 | 0.555169103473262 | 0.916239900758826 | 1254.07703859783 | 1881.3003337921 | 2586.6612106762 | 1534.44807180635 | 1370.88766116551 | 1113.00483118721 | 2013.44502713557 | 1994.11046229605 | Ppargc1a |
| ENSMUSG00000029169 | 0.232184836928359 | 0.250225322003818 | 0.835923538390114 | 1523.32412613541 | 1758.32355872604 | 1768.92418688395 | 1517.5653913097 | 1277.81621452089 | 938.10407200065 | 1634.49260217313 | 1740.09748016362 | Dhx15 |
| ENSMUSG00000029171 | -0.426784968435239 | 0.215050358418281 | 0.824338622350122 | 72.2810973255236 | 57.8235167529173 | 55.8408384302584 | 84.4134024832345 | 135.774816281568 | 116.192812047017 | 49.9717483466946 | 61.8749571861049 | Pgm1 |
| ENSMUSG00000029173 | -0.012779591509597 | 0.956524309328649 | 0.992371505557225 | 147.272735800754 | 180.800291818981 | 174.147699511314 | 174.454365132018 | 147.819356435578 | 108.854318654574 | 217.585320926233 | 207.335382851685 | Sepsecs |
| ENSMUSG00000029174 | -0.552960480568389 | 0.0219939066166211 | 0.648762042745232 | 147.272735800754 | 138.450673915436 | 167.522515290775 | 155.695831246855 | 209.137015401448 | 337.570696052385 | 168.654650670094 | 179.111718170304 | Tbc1d1 |
| ENSMUSG00000029175 | -0.168630111201703 | 0.145248389461965 | 0.780607990324772 | 683.056369726198 | 722.386751547009 | 769.467824471187 | 775.665376151499 | 868.301849284544 | 931.988660840281 | 742.288678566525 | 775.065253173314 | Slc35f6 |
| ENSMUSG00000029176 | 0.145234567681799 | 0.067603216871499 | 0.722038584319139 | 1046.26888378695 | 1133.66669464875 | 1089.36957683436 | 1137.70508013515 | 999.696832782835 | 989.473525747752 | 1030.66730965058 | 965.032226990303 | Anapc4 |
| ENSMUSG00000029177 | 0.0868587042312459 | 0.476476987556008 | 0.897553735056661 | 556.564449406532 | 469.103459854653 | 459.030620994497 | 571.19735680322 | 469.737066006393 | 510.025290774799 | 462.238672206925 | 493.914131924171 | Cenpa |
| ENSMUSG00000029178 | -0.284487875364696 | 0.104485385862569 | 0.761198544536855 | 740.881247586617 | 688.995706661522 | 656.839692722022 | 617.15576482187 | 1100.43298679819 | 760.757148349941 | 753.740537562643 | 678.453477917817 | Klf3 |
| ENSMUSG00000029179 | 0.111272044238375 | 0.42329644552763 | 0.884281490205862 | 156.307872966445 | 157.99665140938 | 152.3792370724 | 138.813150750208 | 134.679858085749 | 134.539045528125 | 142.627698406191 | 148.717002359586 | Zcchc4 |
| ENSMUSG00000029185 | -0.377014884531132 | 0.0582226997533007 | 0.710862211775517 | 132.81651633565 | 171.841718800923 | 153.325691961048 | 167.888878272211 | 221.181555555458 | 260.516515431732 | 168.654650670094 | 163.914360264945 | Fam114a1 |
| ENSMUSG00000029186 | -0.366157485316419 | 0.24640215801841 | 0.832123990973914 | 27.1054114970714 | 21.9892246806869 | 33.1259211026956 | 31.8895076047775 | 33.943704070392 | 53.8156182112498 | 26.0269522639034 | 33.6512925047237 | Pi4k2b |
| ENSMUSG00000029189 | 0.0671427182114443 | 0.86896996517971 | 0.975431777626165 | 14.4562194651047 | 17.9171460361152 | 9.46454888648447 | 15.0068271081306 | 10.949581958191 | 17.123151249034 | 11.4518589961175 | 15.1973579053591 | Sel1l3 |
| ENSMUSG00000029190 | 0.184113904040366 | 0.49107885663536 | 0.899236955296196 | 2204.57346842847 | 2787.74504007375 | 2434.2819736038 | 1925.563503312 | 1450.8196094603 | 1334.38271519258 | 2701.59764499317 | 2744.20862748199 | D5Ertd579e |
| ENSMUSG00000029191 | 0.289404701859095 | 0.0326923852314765 | 0.671606027118506 | 1029.10212317214 | 1156.47033505835 | 1339.23366743755 | 1219.30470253561 | 992.032125412102 | 810.903519864969 | 1058.77641809559 | 1018.22297965906 | Rfc1 |
| ENSMUSG00000029192 | -0.118531594255177 | 0.483986976538332 | 0.89898089116307 | 188.83436676293 | 219.892246806869 | 250.810545491838 | 184.771558768858 | 216.801722772181 | 190.800828203522 | 255.064132186253 | 252.927456567762 | Tbc1d14 |
| ENSMUSG00000029196 | 0.0876714670955425 | 0.623344188357539 | 0.931059568526633 | 348.756294595651 | 465.845796938996 | 464.709350326387 | 345.157023487003 | 327.39250054991 | 338.793778284459 | 453.910047482476 | 408.157612315359 | Tada2b |
| ENSMUSG00000029198 | 0.171014832985999 | 0.22291069203221 | 0.832048466795696 | 2736.74304748764 | 2422.88679352013 | 2581.92893623296 | 2391.71307035831 | 2022.38778767787 | 1870.09273284093 | 2489.21771451972 | 2617.20213641577 | Grpel1 |
| ENSMUSG00000029199 | 0.120516951244064 | 0.290434648725399 | 0.851785577030345 | 1094.15511076511 | 1067.69902060668 | 1231.33781013163 | 1133.01544666386 | 1085.10357205672 | 908.750098430878 | 1157.67883669842 | 1009.53877514171 | Lias |
| ENSMUSG00000029201 | -0.240996983585857 | 0.0628073827013992 | 0.715615581523459 | 236.72059374109 | 254.097707421271 | 216.738169500494 | 247.612647284155 | 295.638712871156 | 293.539735697726 | 244.653351280692 | 296.348479154503 | Ugdh |
| ENSMUSG00000029202 | 0.147604784405683 | 0.502007275182761 | 0.900927059989498 | 1016.45293114018 | 1131.223447462 | 1148.04977993057 | 1011.0849764103 | 849.687559955619 | 620.102691661447 | 1207.65058504512 | 1209.27547904072 | Pds5a |
| ENSMUSG00000029203 | 0.0606675897795272 | 0.597289078455907 | 0.924496535139492 | 1753.72012386052 | 1620.68730053951 | 1686.58261157153 | 1796.12961950438 | 1560.31542904221 | 1413.88306027738 | 1812.51695565823 | 1785.6895538797 | Ube2k |
| ENSMUSG00000029207 | -0.100451373220244 | 0.550677770522456 | 0.914699131434573 | 430.072529086865 | 473.989954228139 | 449.566072108012 | 652.79697920368 | 524.484975797347 | 534.486935416277 | 488.265624470828 | 603.55221395569 | Apbb2 |
| ENSMUSG00000029208 | 0.257841389740739 | 0.177171386268867 | 0.800237299431217 | 655.950958229127 | 735.417403209639 | 659.679057387967 | 746.589648629496 | 498.205979097689 | 428.078781225851 | 739.165444294857 | 673.025850094475 | Guf1 |
| ENSMUSG00000029209 | -0.0413872792393169 | 0.82902559831774 | 0.967483872587428 | 118.360296870545 | 139.26508964435 | 141.968233297267 | 124.744250336335 | 166.433645764503 | 99.0696607979826 | 133.257995591185 | 140.032797842237 | Gnpda2 |
| ENSMUSG00000029213 | 0.0586046150426089 | 0.628131566446993 | 0.931825800800958 | 609.871758684105 | 552.988279932829 | 652.10741827878 | 681.872706725683 | 628.506004400162 | 585.856389163379 | 648.591650416473 | 532.993052252237 | Commd8 |
| ENSMUSG00000029217 | 0.374407083267801 | 0.206792368526112 | 0.819960256425221 | 318.036828232304 | 473.175538499225 | 398.457508120996 | 427.694572581722 | 234.321053905287 | 176.123841418636 | 365.418409785204 | 471.118095066132 | Tec |
| ENSMUSG00000029221 | 0.228570896697566 | 0.286362798061147 | 0.851785577030345 | 3988.10954493576 | 4053.34708280661 | 3679.81660706516 | 3528.4802237992 | 2509.64418481737 | 2318.96391201204 | 4021.68466381836 | 4164.0760660684 | Slc30a9 |
| ENSMUSG00000029223 | -0.317518330932862 | 0.231270146123429 | 0.832048466795696 | 351.466835745359 | 257.355370336928 | 198.755526616174 | 316.081295965 | 286.879047304603 | 527.148442023833 | 302.953724351836 | 284.407697943149 | Uchl1 |
| ENSMUSG00000029227 | 0.152803325723972 | 0.23905065001617 | 0.832048466795696 | 805.030721463019 | 866.538335564846 | 820.576388458203 | 754.093062183562 | 779.610235423197 | 587.079471395453 | 769.356708920985 | 782.663932125994 | Fip1l1 |
| ENSMUSG00000029228 | -0.251374123704141 | 0.486656785605026 | 0.899236955296196 | 306.291149916906 | 240.252640029727 | 150.486327295103 | 241.047160424347 | 320.822751374995 | 448.871179171106 | 217.585320926233 | 130.263067760221 | Lnx1 |
| ENSMUSG00000029229 | 0.030957476544394 | 0.80849152040379 | 0.964026716319739 | 340.62467114653 | 340.425774686189 | 345.456034356683 | 384.549944645846 | 382.140410340865 | 359.586176229715 | 293.584021536831 | 346.282655129254 | Chic2 |
| ENSMUSG00000029231 | -0.764850793518839 | 0.0368471961044159 | 0.679566075569698 | 234.010052591383 | 285.045505120015 | 293.401015481019 | 371.418970926232 | 704.058119911679 | 772.987970670679 | 279.008928269045 | 256.184033261768 | Pdgfra |
| ENSMUSG00000029233 | -0.394352130187345 | 0.189914461990737 | 0.808824684419004 | 68.6670424592474 | 90.4001459094905 | 64.3589324280944 | 65.6548685980713 | 121.54035973592 | 132.092881063977 | 62.4646854333682 | 65.1315338801105 | Srd5a3 |
| ENSMUSG00000029234 | -0.340735605915221 | 0.0560520590726151 | 0.703396849069134 | 274.66816983699 | 223.149909722526 | 248.917635714542 | 269.184961252092 | 351.48158085793 | 396.278643191931 | 249.858741733473 | 289.835325766492 | Tmem165 |

| | | | | | | | | | | | | |
|---|---|---|---|---|---|---|---|---|---|---|---|---|
| **ENSMUSG00000029238** | 0.11993564842337 | 0.647185132297718 | 0.935063240511793 | 788.767474564776 | 1091.3170767452 | 1087.47666705707 | 697.817460528072 | 667.924499449649 | 555.279333361532 | 1099.37846362728 | 1049.70322103445 | Clock |
| **ENSMUSG00000029246** | 0.0768359769516171 | 0.584158768321601 | 0.922450244597442 | 309.905204783182 | 323.323044378988 | 335.04503058155 | 320.770929436291 | 326.297542354091 | 245.839528646846 | 349.802238426862 | 298.51953028384 | Ppat |
| **ENSMUSG00000029247** | 0.155129007252042 | 0.322022196721541 | 0.855977678491063 | 3267.10559911367 | 3713.73572384934 | 3871.9469494608 | 3357.77756544422 | 3121.72581628025 | 2419.25665504209 | 3727.05956419097 | 3492.13574153859 | Paics |
| **ENSMUSG00000029249** | -0.0284561171809499 | 0.857313686400335 | 0.974029039597056 | 280.089252136404 | 232.922898469498 | 212.952349945901 | 241.985087118606 | 211.326931793086 | 227.493295165738 | 261.31060072959 | 286.578749072486 | Rest |
| **ENSMUSG00000029250** | 0.0356087234761052 | 0.81018505078185 | 0.964412705343932 | 1068.85672670118 | 1155.65591932943 | 1134.79941148949 | 903.223406570609 | 1135.4716490644 | 826.803588881929 | 1134.77511870619 | 1060.55847668113 | Polr2b |
| **ENSMUSG00000029253** | -0.290523162690915 | 0.293387176902764 | 0.851785577030345 | 53.3073092775737 | 76.5550785179469 | 47.3227444324223 | 62.8410885152968 | 61.3176589658694 | 50.1463715150282 | 86.4094815161593 | 95.5262496908287 | Cenpc1 |
| **ENSMUSG00000029260** | -0.236010847541147 | 0.586834549373694 | 0.923058967588378 | 84.9302893574902 | 113.203786319092 | 56.7872933189068 | 104.109863062656 | 72.2672409240604 | 96.6234963338349 | 45.80743598447 | 208.420908416353 | Ugt2b34 |
| **ENSMUSG00000029263** | 0.219458534862911 | 0.266653595249878 | 0.843513595393324 | 390.317925557827 | 397.434875710192 | 478.906173656114 | 365.791410760683 | 298.923587458613 | 275.193502216618 | 470.567296931374 | 356.05238521127 | Pigg |
| **ENSMUSG00000029265** | 0.137555164085126 | 0.297606489863034 | 0.851785577030345 | 601.740135234984 | 715.05700998678 | 724.98444470471 | 641.541858872582 | 580.327843784121 | 530.817688720055 | 702.727711125392 | 624.177199684392 | Dr1 |
| **ENSMUSG00000029267** | -0.217624187795869 | 0.317050489996125 | 0.854283473529811 | 203.290586228035 | 150.666909849151 | 129.664319744837 | 173.51643843776 | 248.555510450935 | 156.554525705454 | 196.76375911511 | 161.743309135608 | Mtf2 |
| **ENSMUSG00000029270** | -0.272797052013942 | 0.114932986491102 | 0.767518180080285 | 107.518132271716 | 112.389370590177 | 138.182413742673 | 100.358156285623 | 157.67398019795 | 122.308223207386 | 136.381229862854 | 136.776221148232 | Fam69a |
| **ENSMUSG00000029273** | -0.470247564570338 | 0.121095689434412 | 0.769868040577702 | 99.3865088225949 | 81.4415728914329 | 99.3777633080869 | 90.9788893430416 | 87.5966556655277 | 147.992950080937 | 80.1630129728225 | 198.651178334337 | Sult1d1 |
| **ENSMUSG00000029276** | -0.0614616922210282 | 0.8242365746538 | 0.966637987288264 | 57.8248778604189 | 44.7928650902881 | 44.483379766477 | 52.523894878457 | 45.9882442244021 | 33.0232202659942 | 70.7933101578173 | 57.5328549274309 | Glmn |
| **ENSMUSG00000029279** | 0.020178774813378 | 0.967301125469659 | 0.994284908685797 | 13.5527057485357 | 5.7009101024003 | 14.1968233297267 | 8.44134024832345 | 9.85462376237187 | 11.0077400886647 | 7.28754663389296 | 13.0263067760221 | Brdt |
| **ENSMUSG00000029287** | 0.00817897553088694 | 0.968720790179288 | 0.994790165280964 | 664.986095394817 | 916.21769502862 | 829.094482456039 | 559.942236472122 | 843.117810780705 | 544.271593272868 | 830.780316263797 | 734.90080728058 | Tgfbr3 |
| **ENSMUSG00000029290** | 0.0482543039859201 | 0.759200674421035 | 0.957357521291862 | 275.571683553559 | 278.5301792887 | 269.739643264807 | 261.681547698027 | 226.656346534553 | 217.708637309147 | 314.405583347953 | 289.835325766492 | Zfp326 |
| **ENSMUSG00000029291** | 0.197543823388479 | 0.241512419430212 | 0.832048466795696 | 633.363115314901 | 726.458830191581 | 605.731128735006 | 639.666005484066 | 503.680770076785 | 442.755768010737 | 630.893322877019 | 693.650835823176 | Rufy3 |
| **ENSMUSG00000029298** | 0.0769254807880211 | 0.678674637588671 | 0.944452327903597 | 223.167887992554 | 224.778741180355 | 280.15064703994 | 299.198615468353 | 312.063085808443 | 217.708637309147 | 217.585320926233 | 225.78931745105 | Gbp9 |
| **ENSMUSG00000029309** | -0.758967659864065 | 0.0132516429961693 | 0.635862253526407 | 4330.54124351543 | 4230.88971170994 | 3520.81218577222 | 3913.96809513931 | 9497.66739053485 | 9503.34894321389 | 4035.21867899559 | 4034.89852387284 | Sparcl1 |
| **ENSMUSG00000029310** | 0.11598488588215 | 0.405190582805422 | 0.881158207349486 | 1957.01071008855 | 1548.20430066614 | 1821.92566064826 | 2164.73481034784 | 1769.45244444366 | 1945.92383122951 | 1603.26025945645 | 1594.63705449804 | Nudt9 |
| **ENSMUSG00000029311** | 0.181762408067247 | 0.296336920416791 | 0.851785577030345 | 1610.06144292604 | 1590.55391856968 | 2310.29638319086 | 1971.52191133066 | 2007.0583729364 | 1643.82251990727 | 1468.96118577471 | 1476.31476794917 | Hsd17b11 |
| **ENSMUSG00000029312** | 0.101494254453395 | 0.529672581699679 | 0.908620485539384 | 216.843291976571 | 241.881471487556 | 279.204192151292 | 256.991914226736 | 231.036179317829 | 187.131581507301 | 234.242570375131 | 273.552442296464 | Klhl8 |
| **ENSMUSG00000029313** | 0.184400986031339 | 0.27908138148193 | 0.848075053184273 | 1139.33079659357 | 1515.62767150957 | 1285.28573878459 | 996.078149302167 | 1019.40608030758 | 913.642427359173 | 1224.30783449402 | 1186.47944218268 | Aff1 |
| **ENSMUSG00000029314** | 0.163256894335987 | 0.412675327852686 | 0.882192126163777 | 365.923055210463 | 304.591482613959 | 375.742590793433 | 432.384206053012 | 352.576539053749 | 232.385624094033 | 327.939598525183 | 405.986561186022 | Gpat3 |
| **ENSMUSG00000029319** | 0.13544502313779 | 0.258192534826634 | 0.838310965110319 | 2669.88303246153 | 2413.92822050207 | 2611.26903778106 | 2868.17983104146 | 2207.4357227713 | 2130.60924827266 | 2703.67980117429 | 2572.69558826436 | Coq2 |
| **ENSMUSG00000029321** | -0.376233458478572 | 0.475503262301871 | 0.897549741595057 | 68.6670424592474 | 77.3694942468612 | 51.1085639870161 | 16.8826804966469 | 76.6470737073368 | 130.869798831903 | 37.4788112600209 | 33.6512925047237 | Slc10a6 |
| **ENSMUSG00000029326** | -0.0857970451535095 | 0.58314723301644 | 0.922450244597442 | 168.053551281842 | 171.841718800923 | 156.165056626994 | 170.702658354985 | 204.757182618171 | 189.577745971448 | 170.736806851206 | 143.289374536243 | Enoph1 |
| **ENSMUSG00000029328** | 0.185371486789699 | 0.178283654186375 | 0.801696999069105 | 1219.74351736821 | 1113.30630142589 | 1310.8400207781 | 1347.80065964898 | 954.803546754252 | 988.250443515679 | 1291.97791038017 | 1152.82814967796 | Hnrnpdl |
| **ENSMUSG00000029330** | 0.474535134338104 | 0.0312837608753327 | 0.669064667366769 | 269.247087537575 | 315.178887089845 | 316.115932808581 | 242.923013812864 | 195.997517051618 | 133.315963296051 | 261.31060072959 | 231.216945274392 | Cds1 |
| **ENSMUSG00000029338** | -0.0522124897369444 | 0.787109181522051 | 0.961141387070613 | 1980.50206671935 | 2692.45839979077 | 2098.29048813361 | 1704.21280346708 | 1942.45583938308 | 1751.45375632977 | 2555.84671231532 | 2537.95877019497 | Antxr2 |
| **ENSMUSG00000029343** | -0.527620752677821 | 0.059978146343773 | 0.71282992338528 | 68.6670424592474 | 49.679359463774 | 51.1085639870161 | 49.7101147956825 | 63.5075753575076 | 110.077400886647 | 53.094982618363 | 90.0986218674861 | Crybb1 |
| **ENSMUSG00000029344** | -0.32676962761387 | 0.171861367260144 | 0.799889021902745 | 719.19691838896 | 537.514381083457 | 494.049451874489 | 506.480414899407 | 837.643019801609 | 941.773318696872 | 538.237372817523 | 514.539117652873 | Tpst2 |
| **ENSMUSG00000029345** | -0.0914652024783743 | 0.432375546535533 | 0.887104334029329 | 642.398252480591 | 635.244268553176 | 715.519895818226 | 623.721251681677 | 719.387534653147 | 787.664957455566 | 654.83811895981 | 627.433776378397 | Tfip11 |
| **ENSMUSG00000029346** | -0.168357560194609 | 0.512023452162982 | 0.902394714365738 | 118.360296870545 | 93.6578088251478 | 94.6454888648447 | 135.061443973175 | 168.623562156141 | 141.877538920568 | 88.4916376972716 | 97.6973008201657 | Srrd |
| **ENSMUSG00000029352** | 0.0845901206517675 | 0.882941605411237 | 0.979073805351756 | 17.1667606148119 | 17.1027303072009 | 4.73227444324223 | 13.1309737196143 | 17.5193311331055 | 18.3462334811079 | 9.36970281500523 | 4.34210225867403 | Crybb3 |
| **ENSMUSG00000029359** | -0.0764759027447397 | 0.665019116082849 | 0.941544702132496 | 5643.34667369026 | 3939.32888075861 | 4029.05846097644 | 5274.8997285079 | 5094.84048514626 | 5983.31827930532 | 4141.40864423231 | 4695.98359275596 | Tesc |
| **ENSMUSG00000029363** | -0.0471480844666573 | 0.836694241567544 | 0.968924788429433 | 59.631905293557 | 65.1532583131463 | 71.930571537282 | 51.5859681841989 | 49.2731188118594 | 74.6080161565054 | 58.3003730711437 | 74.901263962127 | Rfc5 |
| **ENSMUSG00000029364** | 0.0554872477720114 | 0.593760103475266 | 0.923935529355527 | 2482.85569313174 | 2475.00940017064 | 2303.67119897032 | 2460.18171903916 | 2267.65842354135 | 2063.3397255086 | 2632.88649101647 | 2389.24176783538 | Wsb2 |
| **ENSMUSG00000029366** | -0.114104799492533 | 0.677468561296911 | 0.944214159784677 | 50.5967681278665 | 69.2253369577179 | 53.9479286529615 | 38.4549944645846 | 74.4571573156986 | 44.0309603546589 | 62.4646854333682 | 48.8486504100828 | Dck |
| **ENSMUSG00000029368** | -0.416703766047966 | 0.175260902429224 | 0.799889021902745 | 6760.0896273696 | 12378.3046637689 | 6753.90208539532 | 7828.87411697287 | 7035.1064081377 | 8349.98239836824 | 12627.2361603554 | 17001.5013938382 | Alb |
| **ENSMUSG00000029369** | 0.149393783949799 | 0.726997787737206 | 0.95219229341535 | 23.4913566307952 | 39.9063707168021 | 18.9290977729689 | 18.7585338851632 | 12.0445401540101 | 19.5693157131818 | 23.9447960827911 | 35.8223436340608 | Afm |
| **ENSMUSG00000029372** | 0.473566542695359 | 0.59803326523024 | 0.924505515197332 | 89.4478579403355 | 144.96599974675 | 425.904699891801 | 33.7653609932938 | 90.881530252985 | 282.531995609062 | 33.3144988977964 | 93.3551985614916 | Ppbp |
| **ENSMUSG00000029373** | 0.0968280708623147 | 0.836300469669876 | 0.968924788429433 | 123.781379169959 | 101.801966114291 | 316.115932808581 | 99.4202295913651 | 136.869774477387 | 275.193502216618 | 75.998700610598 | 111.809133160856 | Pf4 |
| **ENSMUSG00000029376** | 0.0356924584256865 | 0.831886493163214 | 0.967583158900775 | 117.456783153976 | 117.275864963663 | 129.664319744837 | 131.309737196143 | 99.6411958195378 | 116.192812047017 | 124.929370866736 | 142.203848971575 | Mthfd2l |
| **ENSMUSG00000029381** | 0.129830058428107 | 0.371396509222193 | 0.873374202289323 | 914.355881167874 | 1162.17124516075 | 1194.42606947434 | 1095.49837889353 | 903.340511550755 | 873.280713700736 | 1155.59668051731 | 1057.30189998713 | Shroom3 |
| **ENSMUSG00000029385** | 0.293902698603532 | 0.140374882182842 | 0.779711473022872 | 541.204716224858 | 586.379324818317 | 871.68495244522 | 523.363095396054 | 574.853052805026 | 428.078781225851 | 540.319528998635 | 513.453592088204 | Ccng2 |
| **ENSMUSG00000029386** | -0.253701782800801 | 0.190176073285565 | 0.808824684419004 | 83.1232619243521 | 70.8541684155466 | 62.4660226507975 | 68.4686486808458 | 75.5521155115177 | 100.292743030056 | 83.2862472444909 | 81.4144173501381 | Tctn2 |
| **ENSMUSG00000029387** | 0.0388867540318079 | 0.839815412581607 | 0.969879194284716 | 252.983840639333 | 242.69588721647 | 234.720812384815 | 223.226553233442 | 238.700886688563 | 155.33144347338 | 254.023054095697 | 278.980070119806 | Gtf2h3 |
| **ENSMUSG00000029388** | 0.120616611489512 | 0.229105679798444 | 0.832048466795696 | 700.22313034101 | 583.936077631574 | 651.160963390131 | 661.238319452004 | 593.467342133951 | 595.64104701997 | 619.441463880901 | 578.585125968315 | Eif2b1 |
| **ENSMUSG00000029389** | -0.382782300678519 | 0.0750772933969105 | 0.735399943172214 | 55.1143367107117 | 74.9262470601182 | 54.8943835416099 | 79.7237690119437 | 94.1664048404423 | 94.1773318696872 | 72.8754663389296 | 84.6709940441436 | Ddx55 |
| **ENSMUSG00000029390** | 0.0886229694567643 | 0.381446017544865 | 0.875126835008975 | 2472.01352853291 | 2182.6341534904 | 2312.18929296816 | 2644.01535111376 | 2224.9550539044 | 2087.80137015008 | 2388.23313973578 | 2334.96548960196 | Tmed2 |
| **ENSMUSG00000029392** | 0.161724052532883 | 0.263823551588285 | 0.842510961779757 | 4684.7186204105 | 4560.72808192024 | 4975.51334962488 | 5115.45219048401 | 4069.95961385958 | 3431.96874319925 | 5159.58301679621 | 4622.1678543585 | Rilpl1 |
| **ENSMUSG00000029394** | 0.194558360780788 | 0.254185652922668 | 0.837406774112203 | 734.556651570634 | 657.233493233863 | 762.842640250648 | 911.664746818933 | 701.868203520041 | 793.780368615935 | 665.248899865371 | 518.881219911547 | Cdk2ap1 |
| **ENSMUSG00000029397** | 0.00449294904352219 | 0.963544378527912 | 0.994284908685797 | 733.653137854065 | 683.294796559122 | 733.502538702546 | 754.093062183562 | 748.951405940262 | 755.864819421645 | 648.591650416473 | 742.499486233259 | Rchy1 |
| **ENSMUSG00000029401** | -0.511078156490197 | 0.0794190067449796 | 0.739855948845396 | 191.544907912638 | 95.2866402829764 | 114.521041526462 | 135.061443973175 | 211.326931793086 | 267.855008824175 | 146.792010768415 | 138.947272277569 | Rilpl2 |
| **ENSMUSG00000029402** | -0.150916801640213 | 0.285350722728904 | 0.851275833092507 | 160.82544154929 | 158.811067138294 | 153.325691961048 | 164.137171495178 | 163.148771177045 | 205.477814988408 | 176.983275394543 | 162.828834700276 | Snrnp35 |
| **ENSMUSG00000029403** | 0.0291239994623723 | 0.906847043697567 | 0.98504921011344 | 155.404359249876 | 176.728213174409 | 183.612248397799 | 139.751077444466 | 121.54035973592 | 110.077400886647 | 213.421008564008 | 196.480127205 | Cdkl2 |
| **ENSMUSG00000029404** | -0.0499365426724686 | 0.7961529841788 | 0.962200030474509 | 730.039082987788 | 564.39010013763 | 626.553136285272 | 763.472329126143 | 678.87408140784 | 930.765578608207 | 606.948526794228 | 563.387768062955 | Arl6ip4 |
| **ENSMUSG00000029405** | -0.0907666124692975 | 0.399029208166938 | 0.880459952732461 | 584.573374620172 | 661.305571878435 | 637.910594949053 | 574.949063580253 | 616.461464246152 | 610.318033804856 | 696.481242582055 | 694.736361387845 | G3bp2 |
| **ENSMUSG00000029406** | 0.0409472120753904 | 0.806094037731358 | 0.963604452772887 | 1104.09376164737 | 1265.60204273287 | 1234.17717479757 | 1038.28485054378 | 1124.52206710621 | 829.249753346077 | 1277.40281711238 | 1279.83464074417 | Pitpnm2 |
| **ENSMUSG00000029407** | 0.25988275565018 | 0.101661518667505 | 0.759800953500741 | 1419.42004872997 | 1534.3592332746 | 1601.40167159317 | 1373.12468039395 | 1148.61114741423 | 958.896469945906 | 1499.15245040084 | 1342.79512349494 | Uso1 |
| **ENSMUSG00000029408** | -0.0839659739352283 | 0.558321718964497 | 0.916574868225118 | 281.896279569542 | 338.796943228361 | 276.364827485346 | 246.674720589896 | 327.39250054991 | 275.193502216618 | 297.748333899055 | 312.63136262453 | Abcb9 |
| **ENSMUSG00000029413** | -0.115151319395901 | 0.468737701375995 | 0.895328112556867 | 261.115464088454 | 315.99330281876 | 252.703455269135 | 237.295453647315 | 266.074841584041 | 346.132271676902 | 288.37863108405 | 257.269558826436 | Naaa |
| **ENSMUSG00000029415** | 0.373232385964617 | 0.0492483184193325 | 0.703396849069134 | 324.361424248287 | 376.26006675842 | 393.725233677754 | 327.336416296098 | 281.404256325508 | 188.354663739374 | 298.789411989611 | 327.828720529889 | Sdad1 |
| **ENSMUSG00000029416** | -0.0945466714801504 | 0.503493742807627 | 0.900927059989498 | 346.045753445944 | 376.26006675842 | 413.600786339371 | 294.508981997063 | 402.944616061428 | 375.486245246675 | 351.884394607974 | 397.302356668674 | Slc15a4 |
| **ENSMUSG00000029417** | -0.269211561044318 | 0.478767058708243 | 0.897732059433541 | 30.7194663633475 | 29.3189662409158 | 44.483379766477 | 31.8895076047775 | 66.7924499449649 | 47.7002070508805 | 22.903717992235 | 27.1381391167127 | Cxcl9 |
| **ENSMUSG00000029422** | 0.00340687593494631 | 0.97436601344513 | 0.995510898988495 | 788.767474564776 | 784.282346944498 | 757.163910918757 | 795.361836730921 | 718.292576457328 | 711.833859066986 | 815.164144905455 | 871.677028428812 | Rsrc2 |
| **ENSMUSG00000029426** | 0.120889904171871 | 0.449432427378274 | 0.89108031640712 | 4127.2506572874 | 4882.4222948414 | 5944.68315560089 | 4420.44851003871 | 4271.4319218903 | 3796.44724835726 | 5233.4995612257 | 4514.70082345632 | Scarb2 |
| **ENSMUSG00000029427** | -0.254433673295467 | 0.0229725559454983 | 0.650514839131011 | 262.922491521592 | 278.5301792887 | 253.649910157784 | 241.985087118606 | 312.063085808443 | 303.324393554317 | 322.734208072402 | 299.605055848508 | Zcchc8 |
| **ENSMUSG00000029428** | -0.287362605817853 | 0.110996373318158 | 0.764116514278315 | 351.466835745359 | 320.065381463331 | 329.36630124966 | 354.536290429585 | 476.306815181307 | 511.248373006873 | 325.857442344071 | 341.94055287058 | Stx2 |
| **ENSMUSG00000029430** | 0.0626037930218739 | 0.491085539825572 | 0.899236955296196 | 3201.14909780413 | 2786.93062434483 | 2781.63091773779 | 3074.52370377825 | 2826.08710340909 | 2655.31152583235 | 2908.77218501385 | 2949.37295920434 | Ran |
| **ENSMUSG00000029432** | 0.196449579660057 | 0.352054253377157 | 0.862728204353539 | 24610.8101256242 | 25951.3572018551 | 27525.7475265628 | 23861.7930286219 | 19048.9877326648 | 14754.040965507 | 27631.2536014504 | 27535.4414733814 | Nipsnap2 |
| **ENSMUSG00000029433** | -0.013975459890809 | 0.891070068549672 | 0.982142775127356 | 1528.74520843482 | 1279.44711012441 | 1263.51727634568 | 1371.24882700543 | 1287.67083828326 | 1389.4214156359 | 1438.76992114858 | 1379.70299269367 | Diablo |
| **ENSMUSG00000029434** | 0.144443949281451 | 0.506776847122996 | 0.901698316485955 | 555.660935689963 | 724.829998733753 | 796.915016241992 | 575.886990274511 | 504.775728272604 | 440.309603546589 | 688.152617857606 | 766.381048655966 | Vps33a |
| **ENSMUSG00000029436** | -0.442124755292295 | 0.284053163486853 | 0.851125503777472 | 60.535419010126 | 61.895595397489 | 53.001473764313 | 53.4618215727152 | 97.4512794278996 | 136.985209992272 | 42.6842017128016 | 34.7368180693922 | Mmp17 |
| **ENSMUSG00000029438** | -0.0114866533840608 | 0.960497465953105 | 0.993694458032251 | 215.036264543433 | 233.737314198412 | 243.238906382651 | 212.909359596603 | 190.522726072523 | 146.769867848863 | 292.542943446274 | 281.151121249143 | Bcl7a |
| **ENSMUSG00000029439** | 0.0785701693069739 | 0.517678998965974 | 0.905270212216717 | 496.029030396406 | 454.443976734195 | 506.353365426919 | 521.487242007538 | 480.686647964584 | 414.624876673038 | 531.990904274186 | 443.97995594942 | Sfswap |
| **ENSMUSG00000029440** | -0.0870688321135956 | 0.440328569823359 | 0.890530552540252 | 420.133878204606 | 359.971752180133 | 399.403963009645 | 422.067012416173 | 464.262275027297 | 421.963370065482 | 406.020455316893 | 408.157612315359 | Psmd9 |
| **ENSMUSG00000029442** | -0.0316841419624687 | 0.931932182937425 | 0.987906435243901 | 25.2983840639333 | 16.2883145782866 | 27.447191770805 | 41.2687745473591 | 18.6142893289246 | 34.2463024980681 | 26.0269522639034 | 33.6512925047237 | Wdr66 |
| **ENSMUSG00000029445** | -0.250449054836952 | 0.416515007893583 | 0.882192126163777 | 356.887918044773 | 680.037133643464 | 345.456034356683 | 400.494698448235 | 355.861413641207 | 502.686797382356 | 443.499266576914 | 819.571801324723 | Hpd |
| **ENSMUSG00000029446** | -0.189112844161598 | 0.423221321839216 | 0.884281490205862 | 90.3513716569045 | 87.1424829938332 | 61.519567762149 | 86.2892558717508 | 87.5966556655277 | 125.977469903608 | 71.8343882483734 | 86.8420451734806 | Psph |
| **ENSMUSG00000029447** | 0.0804466549382611 | 0.288050769390008 | 0.851785577030345 | 2265.1088874386 | 2286.06495106252 | 2168.32814989359 | 2284.78942721288 | 2268.75338173717 | 2014.41643622565 | 2120.67607046285 | 2111.34722328025 | Cct6a |
| **ENSMUSG00000029452** | 0.297855250242477 | 0.153932151453187 | 0.785495071155314 | 181.606257030378 | 156.367819951551 | 209.166530391307 | 211.971432902344 | 162.053812981226 | 103.961989726278 | 169.69572876065 | 180.197243734972 | Tmem116 |
| **ENSMUSG00000029454** | -0.0411817411904014 | 0.750866917079203 | 0.956197192463153 | 598.126080368708 | 475.618785685968 | 576.391027186904 | 558.066383083606 | 502.585811880965 | 642.118171838776 | 567.387559353095 | 560.13119136895 | Mapkapk5 |
| **ENSMUSG00000029455** | 0.0213627136345533 | 0.799456508637582 | 0.963342665478963 | 8178.606162383 | 8687.37258032914 | 9023.50090837429 | 9416.78401035194 | 9043.25973926992 | 8164.07389909301 | 9001.16117094836 | 8576.73748644588 | Aldh2 |
| **ENSMUSG00000029456** | 0.0894881642396958 | 0.675994148974195 | 0.94409858171788 | 64.1494738764022 | 60.2667639396603 | 55.8408384302584 | 45.9584080186499 | 53.6529515951357 | 57.4848649074714 | 51.0128264372507 | 51.0197015394199 | Acad10 |
| **ENSMUSG00000029458** | 0.211938839443206 | 0.0571980001183097 | 0.709357340752966 | 838.460728976074 | 874.682492853989 | 904.810873547915 | 900.409626487835 | 697.488370736765 | 680.033721033066 | 819.328457267679 | 839.111261488756 | Brap |

| | | | | | | | | | | | | |
|---|---|---|---|---|---|---|---|---|---|---|---|---|
| **ENSMUSG00000029461** | 0.0840240727089795 | 0.565323135033522 | 0.919692942798738 | 884.539928521095 | 1090.50266101629 | 1016.49255040843 | 853.513291774927 | 911.005218921489 | 783.995710759344 | 1077.5158237256 | 854.308619394115 | Fam168a |
| **ENSMUSG00000029462** | 0.102183118051816 | 0.451954099926382 | 0.891110666112841 | 1087.83051474913 | 843.734695155244 | 808.272474905774 | 1000.76778277346 | 831.073270626695 | 815.795848793264 | 879.710986519935 | 957.433548037624 | Vps29 |
| **ENSMUSG00000029463** | 0.0532131592673308 | 0.792973970330686 | 0.961804768755116 | 114.746242004269 | 82.2559886203472 | 82.3415753124149 | 111.613276616721 | 100.736154015357 | 101.51582526213 | 78.0808567917102 | 96.6117752554972 | Fam216a |
| **ENSMUSG00000029464** | 0.129483772769928 | 0.338125486920832 | 0.858570960161243 | 446.335775985108 | 427.568257680022 | 460.923530771794 | 481.156394154437 | 396.374866886513 | 369.370834086306 | 502.840717738614 | 389.703677715994 | Gpn3 |
| **ENSMUSG00000029465** | -0.0592940582326245 | 0.747217393461596 | 0.956119556124081 | 1742.87795926169 | 1521.32858161197 | 1551.2395624948 | 1725.78511743502 | 1830.77010340953 | 2243.13281362346 | 1371.09984526243 | 1372.10431374099 | Arpc3 |
| **ENSMUSG00000029466** | -0.0611317300874243 | 0.614245475156213 | 0.928075610917191 | 912.548853734736 | 950.423155643021 | 959.705257089525 | 1077.67777170263 | 1136.56660726022 | 1090.98935100988 | 938.011359591079 | 904.242795368867 | Anapc7 |
| **ENSMUSG00000029467** | 0.245490144568782 | 0.392537175748866 | 0.878537584780694 | 292194.528910996 | 395743.334241237 | 401114.206993432 | 256284.717499266 | 208947.587633571 | 171350.151466852 | 382600.362591742 | 371932.538696806 | Atp2a2 |
| **ENSMUSG00000029468** | -0.413247056016697 | 0.230256806604401 | 0.832048466795696 | 25.2983840639333 | 37.4631235300591 | 35.0188308799925 | 30.9515809105193 | 50.3680770076785 | 58.7079471395453 | 20.8215618111227 | 42.3354970220718 | P2rx7 |
| **ENSMUSG00000029469** | 0.21115240153339 | 0.127673862875346 | 0.772837268577874 | 882.732901087957 | 921.91860513102 | 1165.08596792624 | 1005.45741624475 | 788.36990098975 | 779.103381831049 | 888.039611244385 | 976.973008201657 | Ift81 |
| **ENSMUSG00000029470** | 0.212358885900214 | 0.131838599611971 | 0.77303252008843 | 333.396561413978 | 392.548381336706 | 411.707876562074 | 440.825546301336 | 330.677375137367 | 390.163232031561 | 308.159114804616 | 334.3418739179 | P2rx4 |
| **ENSMUSG00000029471** | 0.042932496497006 | 0.801268887674057 | 0.963461896828986 | 140.948139784771 | 175.913797445495 | 190.237432618338 | 152.88205116408 | 151.104231023035 | 166.339183562045 | 185.311900118992 | 137.8617467129 | Camkk2 |
| **ENSMUSG00000029472** | 0.150319962259939 | 0.11913938725322 | 0.769868040577702 | 4813.0175681633 | 4711.39499176939 | 5063.53365426919 | 5133.27279767492 | 4161.93610230839 | 4058.18684602107 | 4830.60234018047 | 4716.60857848467 | Anapc5 |
| **ENSMUSG00000029474** | 0.103233328495548 | 0.593567849995523 | 0.923935529355527 | 666.793122827955 | 697.95427967958 | 772.307189137133 | 604.024791102256 | 527.769850384805 | 481.894399437101 | 775.603177464322 | 765.295523091298 | Rnf34 |
| **ENSMUSG00000029475** | 0.11503286068084 | 0.542328311810084 | 0.913837378403038 | 185.220311896654 | 232.922898469498 | 253.649910157784 | 189.461192240149 | 170.813478547779 | 166.339183562045 | 219.667477107345 | 237.730098662403 | Kdm2b |
| **ENSMUSG00000029478** | 0.0863686068598093 | 0.578082922639278 | 0.922450244597442 | 1660.6582110539 | 2153.31518724949 | 1864.51613063744 | 1575.71684635371 | 1742.07848954818 | 1385.75216893968 | 1946.81602933998 | 1757.46588919831 | Ncor2 |
| **ENSMUSG00000029480** | 0.376408135393689 | 0.0773476798000639 | 0.739855948845396 | 492.41497553013 | 609.182965227918 | 668.197151385803 | 658.424539369229 | 479.591689768764 | 302.101311322243 | 511.169342463063 | 576.414074838978 | Dhx37 |
| **ENSMUSG00000029482** | -0.0526179927341031 | 0.76461538957143 | 0.958110466593105 | 86.7373167906283 | 84.6992358070902 | 96.5383986421416 | 77.8479156234274 | 91.9764884488041 | 99.0696607979826 | 79.1219348822664 | 89.0130963028176 | Aacs |
| **ENSMUSG00000029484** | -0.472080190105697 | 0.0773085430155104 | 0.739855948845396 | 344.238726012806 | 258.984201794756 | 303.812019256151 | 312.329589187968 | 580.327843784121 | 536.933099880424 | 269.639225454039 | 305.032683671851 | Anxa3 |
| **ENSMUSG00000029486** | 0.0864333842178007 | 0.605292798837954 | 0.926448151609826 | 916.162908601012 | 702.026358324151 | 658.732602499319 | 816.934150698859 | 730.337116611338 | 584.633306931305 | 739.165444294857 | 858.650721652789 | Mrpl1 |
| **ENSMUSG00000029490** | -0.0204792793418665 | 0.945469509370168 | 0.990505437194639 | 22.5878429142261 | 30.1333819698302 | 21.7684624389143 | 24.3860940507122 | 29.5638712871156 | 22.0154801773295 | 24.9858741733473 | 23.8815624227072 | Mfsd7a |
| **ENSMUSG00000029499** | 0.0242635790753891 | 0.856575054094046 | 0.97363563840968 | 1757.33417872679 | 1346.22919989539 | 1520.95300605805 | 1808.32266652973 | 1727.84403300253 | 1657.27642446008 | 1462.71471723137 | 1477.40029351384 | Pxmp2 |
| **ENSMUSG00000029500** | 0.110608917188597 | 0.624821396684632 | 0.931529302402539 | 329.782506547701 | 359.157336451219 | 394.671688566402 | 338.591536627196 | 291.25888008788 | 207.923979452556 | 422.677704765791 | 394.045779974668 | Pgam5 |
| **ENSMUSG00000029501** | 0.113017325705411 | 0.464227336730774 | 0.895328112556867 | 680.345828576491 | 733.78857175181 | 791.236286910102 | 667.803806311811 | 638.360628162534 | 517.363784167243 | 783.931802188771 | 715.361347116547 | Ankle2 |
| **ENSMUSG00000029502** | 0.286376326634721 | 0.0906430604773676 | 0.753823363185192 | 2709.63763599057 | 2951.44260158553 | 2855.45439905236 | 2819.40764294003 | 2038.81216061516 | 1757.56916749014 | 2847.34857767103 | 2649.76790335583 | Golga3 |
| **ENSMUSG00000029504** | 0.297827948861018 | 0.0947512385804341 | 0.757085157577544 | 172.571119864688 | 140.079505373265 | 195.916161950228 | 198.84045918273 | 156.579022002131 | 118.638976511164 | 147.833088858971 | 150.888053488923 | Ddx51 |
| **ENSMUSG00000029505** | 0.12931082770452 | 0.506009012526035 | 0.901494649536949 | 515.002818444356 | 724.015583004838 | 662.518422053913 | 488.659807708502 | 560.618596259378 | 408.509465512669 | 588.209121164217 | 627.433776378397 | Ep400 |
| **ENSMUSG00000029507** | -0.0216440754936038 | 0.87321873178935 | 0.976334877610406 | 549.336339673979 | 460.144886836596 | 484.584902988005 | 561.818089860639 | 594.56230032977 | 512.471455238947 | 531.990904274186 | 447.236532643425 | Pus1 |
| **ENSMUSG00000029512** | 0.248025937310775 | 0.25717272717184 | 0.837517407767513 | 1586.57008629524 | 2106.89349070137 | 2364.24431184382 | 1649.81305520011 | 1472.71877337669 | 1131.35106466832 | 1986.37699678111 | 1898.58421260522 | Ulk1 |
| **ENSMUSG00000029513** | -0.0196152630203596 | 0.876625743221789 | 0.977469019294424 | 391.221439274397 | 449.557482360709 | 533.800557197724 | 477.404687377404 | 473.02194059385 | 469.663577116362 | 473.690531203042 | 460.262839419447 | Prkab1 |
| **ENSMUSG00000029516** | 0.0274484775325 | 0.938835486200623 | 0.989269716906553 | 16.2632468982428 | 13.0306516626293 | 18.9290977729689 | 18.7585338851632 | 15.3294147414674 | 18.3462334811079 | 14.5750932677859 | 17.3684090346961 | Cit |
| **ENSMUSG00000029518** | 0.0308470879161114 | 0.774641050210513 | 0.959414024438738 | 843.881811275488 | 990.329526359824 | 1036.36810307005 | 936.050840869645 | 997.506916391197 | 905.080851734656 | 952.586452858865 | 870.591502864143 | Rab35 |
| **ENSMUSG00000029521** | -0.250545661633997 | 0.492359623347703 | 0.899236955296196 | 36.1405486627618 | 42.3496179035451 | 20.8220075502658 | 50.6480414899407 | 60.2227007700503 | 51.3694537471021 | 24.9858741733473 | 42.3354970220718 | Chek2 |
| **ENSMUSG00000029524** | 0.442902991883599 | 0.00939417187068007 | 0.635862253526407 | 257.501409222178 | 175.913797445495 | 220.523989055088 | 214.785212985119 | 152.199189218854 | 165.116101329971 | 180.106509666212 | 141.118323406906 | Sirt4 |
| **ENSMUSG00000029528** | -0.338638646168514 | 0.115825194425705 | 0.769868040577702 | 619.810409566365 | 653.161414589292 | 742.967087589031 | 643.417712261099 | 1100.43298679819 | 1001.70434806849 | 605.907448703672 | 655.657441059779 | Pxn |
| **ENSMUSG00000029534** | 0.0876823508789467 | 0.472991890850555 | 0.89728597737823 | 286.413848152387 | 287.488752306758 | 316.115932808581 | 283.253861665965 | 268.264757975679 | 244.616446414772 | 300.871568170723 | 288.749800201823 | St7 |
| **ENSMUSG00000029535** | 0.0356390117649888 | 0.793092748130704 | 0.961804768755116 | 409.291713605777 | 377.888898216248 | 361.545767463707 | 404.246405225267 | 344.911831683016 | 331.455284892016 | 391.445362049107 | 446.151007078757 | Triap1 |
| **ENSMUSG00000029536** | 0.0532167598831889 | 0.749300917821615 | 0.956197192463153 | 520.42390074377 | 589.636987733974 | 531.907647420427 | 508.356268287923 | 451.122776677468 | 412.178712208891 | 638.180869510912 | 569.900921450967 | Gatc |
| **ENSMUSG00000029538** | -0.280291089003298 | 0.270003485133121 | 0.843793744782845 | 255.69438178904 | 211.748089517725 | 225.25626349833 | 321.708856130549 | 372.285786578493 | 410.955629976817 | 232.160414194018 | 217.105112933702 | Srsf9 |
| **ENSMUSG00000029544** | -0.0839971719689469 | 0.737328141829518 | 0.954383574136061 | 46.9827132615903 | 31.7622134276588 | 51.1085639870161 | 41.2687745473591 | 50.3680770076785 | 47.7002070508805 | 39.5609674411332 | 43.4210225867403 | Cabp1 |
| **ENSMUSG00000029545** | 0.136281705346884 | 0.214144813360236 | 0.824248286808591 | 7186.54810159018 | 6644.00351648309 | 6856.11921336935 | 7365.53833000934 | 6359.51720131732 | 5418.2542880872 | 6954.40164491499 | 6788.87688143685 | Acads |
| **ENSMUSG00000029547** | -0.0458707399436043 | 0.638352412116799 | 0.933338421505509 | 1010.12833512419 | 1128.78020027526 | 978.634354862494 | 947.305961200743 | 1101.52794499401 | 1068.97387083255 | 1018.1743725639 | 1009.53877514171 | Ints1 |
| **ENSMUSG00000029550** | -0.114067568764067 | 0.333204964736024 | 0.856918526232803 | 473.44118748218 | 474.804369957054 | 410.761421673426 | 467.087493740564 | 529.959766776443 | 429.301863457925 | 501.799639648058 | 514.539117652873 | Sppl3 |
| **ENSMUSG00000029551** | -0.171408286311757 | 0.515905817530005 | 0.904245406849874 | 124.684892886528 | 71.6685841444609 | 84.2344850897118 | 108.799496533947 | 128.110108910834 | 139.43137445642 | 80.1630129728225 | 91.1841474321546 | Psmg3 |
| **ENSMUSG00000029552** | -0.461505263309991 | 0.198670652559779 | 0.812921911423739 | 79.509207058076 | 96.1010560118908 | 70.0376617599851 | 73.1582821521366 | 178.478185918513 | 135.762127760198 | 60.3825292522559 | 65.1315338801105 | Tes |
| **ENSMUSG00000029554** | -0.0826149558715132 | 0.659103676896269 | 0.938989601151841 | 311.712232216321 | 262.241864710414 | 208.220075502658 | 259.805694309511 | 271.549632563136 | 332.67836712409 | 284.214318721825 | 216.019587369033 | Mad1l1 |
| **ENSMUSG00000029557** | -0.265579345297886 | 0.155816881215539 | 0.787275540343856 | 318.940341948873 | 201.975100770753 | 211.059440168604 | 267.309107863576 | 321.917709570815 | 258.070350967584 | 294.625099627387 | 325.657669400552 | Mrm2 |
| **ENSMUSG00000029559** | -0.0287764944705502 | 0.86967429174987 | 0.975613767803763 | 475.248214915318 | 384.404224047563 | 329.36630124966 | 456.770300103725 | 448.93286028583 | 496.571386221987 | 366.45948787576 | 367.993166422624 | 2210016L21[illegible] |
| **ENSMUSG00000029560** | 0.112364859392078 | 0.323871516453612 | 0.855977678491063 | 373.151164943016 | 352.642010619904 | 367.224496795597 | 351.72251034681 | 341.626957095558 | 300.878229090169 | 321.693129981846 | 371.24974311663 | Snx8 |
| **ENSMUSG00000029561** | -0.509935228064166 | 0.15935180992525 | 0.789652743439786 | 57.8248778604189 | 79.8127414336042 | 98.4313084194385 | 90.9788893430416 | 165.338687568684 | 165.116101329971 | 75.998700610598 | 59.7039060567679 | Oasl2 |
| **ENSMUSG00000029563** | -0.9563571166637 | 0.0496461008493766 | 0.703396849069134 | 28.0089252136404 | 44.7928650902881 | 18.9290977729689 | 28.1378008277448 | 59.1277425742312 | 116.192812047017 | 28.1091084450157 | 30.3947158107182 | Foxp2 |
| **ENSMUSG00000029569** | -0.0863554809002988 | 0.395633377366484 | 0.879215782741314 | 356.887918044773 | 355.085257806647 | 349.241853911277 | 329.212269684615 | 374.475702970131 | 348.57843614105 | 368.541644056872 | 384.276049892652 | Tmem168 |
| **ENSMUSG00000029570** | -0.468683181951541 | 0.331192581264903 | 0.856333478432716 | 85.8338030740593 | 82.2559886203472 | 98.4313084194385 | 63.779015209555 | 167.528603960322 | 200.585486060113 | 37.4788112600209 | 52.1052271040884 | Lfng |
| **ENSMUSG00000029571** | 0.148379686882683 | 0.447692352115268 | 0.89108031640712 | 3074.65717748446 | 3441.72087039195 | 3676.03078751057 | 2946.96567335914 | 2589.57613311216 | 2131.83233050474 | 3802.01718671101 | 3330.39243240298 | Tmem106b |
| **ENSMUSG00000029575** | 0.13542046217136 | 0.588468808148357 | 0.92352104528132 | 886.346955954233 | 966.711470221308 | 815.844114014961 | 762.534402431885 | 559.523638063559 | 530.817688720055 | 1101.46061980839 | 931.380934485579 | Mmab |
| **ENSMUSG00000029576** | -0.954858686054921 | 0.0805632744042012 | 0.739855948845396 | 8.13162344912141 | 13.8450673915436 | 16.0897331070236 | 5.62756016554897 | 20.8042057205628 | 39.1386314263635 | 16.6572494488982 | 8.68420451734806 | Radil |
| **ENSMUSG00000029577** | 0.149169458168466 | 0.501406571021981 | 0.900927059989498 | 3515.57187117015 | 4310.70245314354 | 4100.98903251372 | 3250.85392229879 | 2703.45178547735 | 2419.25665504209 | 4197.62686112234 | 4365.98382109674 | Ube3b |
| **ENSMUSG00000029578** | 0.0752601709250701 | 0.484944804620124 | 0.899236955296196 | 1103.1902479308 | 1113.30630142589 | 1276.76764478675 | 1262.44933047149 | 1183.64980968044 | 1006.59667699679 | 1183.70578896233 | 1137.6307917726 | Wipi2 |
| **ENSMUSG00000029580** | -0.639212560868536 | 0.0348632748541109 | 0.679566075569698 | 3247.22829734915 | 2940.85519710964 | 3200.91043340905 | 4048.09161241822 | 7376.73336523326 | 7120.78475513401 | 2941.04560582109 | 3489.96469040925 | Actb |
| **ENSMUSG00000029581** | -0.839382603191235 | 0.00637123322510974 | 0.635862253526407 | 341.528184863099 | 258.984201794756 | 280.15064703994 | 425.818719193205 | 829.978312430875 | 759.534066117867 | 370.623800237985 | 377.762896504641 | Fscn1 |
| **ENSMUSG00000029587** | -0.182143135100223 | 0.325791676539784 | 0.855977678491063 | 121.070838020252 | 113.203786319092 | 175.094154399963 | 143.502784221499 | 146.724398239759 | 145.546785616789 | 170.736806851206 | 162.828834700276 | Zfp12 |
| **ENSMUSG00000029591** | -0.0146398016244497 | 0.937846750416587 | 0.988970340396118 | 261.115464088454 | 264.685111897157 | 229.042083052924 | 239.171307035831 | 215.706764576362 | 189.577745971448 | 311.282349076285 | 286.578749072486 | Ung |
| **ENSMUSG00000029592** | -0.0364223496718155 | 0.753533824970403 | 0.956497786093635 | 278.282224703266 | 299.704988240473 | 272.579007930753 | 256.991914226736 | 290.163921892061 | 302.101311322243 | 271.721381635152 | 273.552442296464 | Usp30 |
| **ENSMUSG00000029594** | 0.060550830783399 | 0.611020693824526 | 0.927932112008768 | 282.799793286111 | 277.715763559786 | 268.793188376159 | 271.060814640609 | 259.505092409126 | 237.277953022329 | 263.392756910703 | 294.177428025166 | Rbm19 |
| **ENSMUSG00000029596** | -0.173080365206514 | 0.424692695703306 | 0.885066977635295 | 224.974915425692 | 195.459774939439 | 190.237432618338 | 251.364354061187 | 266.074841584041 | 321.670627035425 | 210.29777429234 | 174.76961591163 | Sdsl |
| **ENSMUSG00000029597** | -0.0927338093003914 | 0.766174847915855 | 0.958408558857279 | 31.6229800799166 | 65.1532583131463 | 34.0723759913441 | 36.5791410760683 | 38.3235368536684 | 56.2617826753975 | 41.6431236222455 | 43.4210225867403 | Sds |
| **ENSMUSG00000029598** | 0.0343623291897022 | 0.782304808892712 | 0.960742105865155 | 1219.74351736821 | 1496.89610974454 | 1662.92123935532 | 1331.85590584659 | 1373.07757755715 | 1454.24477393582 | 1415.86620315635 | 1334.1109189776 | Plbd2 |
| **ENSMUSG00000029599** | 0.0455304238445658 | 0.692752725469111 | 0.94469184542218 | 860.145058173731 | 986.257447715252 | 967.276896198713 | 921.981940455772 | 1047.87499339888 | 835.365164506446 | 882.834220791604 | 853.223093829447 | Ddx54 |
| **ENSMUSG00000029600** | -0.184464168881173 | 0.173943074896441 | 0.799889021902745 | 211.422209677157 | 179.985876090067 | 213.898804834549 | 177.268145214792 | 229.94122112201 | 209.14706168463 | 235.283648465687 | 213.848536239696 | Rita1 |
| **ENSMUSG00000029603** | -1.46033168845681 | 0.00301661098138811 | 0.635862253526407 | 9.03513716569045 | 7.32974156022896 | 6.62518422053913 | 6.56548685980713 | 29.5638712871156 | 25.6847268735511 | 17.6983275394543 | 8.68420451734806 | Dtx1 |
| **ENSMUSG00000029610** | 0.021084458390622 | 0.902054461006758 | 0.984871597499125 | 599.933107801846 | 466.66021266791 | 438.208613444231 | 627.47295845871 | 614.271547854513 | 543.048511040794 | 465.361906478593 | 478.716774018812 | Aimp2 |
| **ENSMUSG00000029613** | 0.0222515459729169 | 0.907534040225389 | 0.98504921011344 | 882.732901087957 | 1089.68824528737 | 1082.74439261382 | 822.561710864407 | 806.984190318674 | 742.410914868833 | 1158.71991478898 | 1109.40712709121 | Eif2ak1 |
| **ENSMUSG00000029614** | 0.0370073411544975 | 0.824460780086685 | 0.966637987288264 | 8925.8120059856 | 6662.73507824812 | 7700.35697404376 | 9461.80449167633 | 8356.72095049135 | 9634.21874204579 | 6906.51205274941 | 7024.43592896991 | Rpl6 |
| **ENSMUSG00000029616** | -0.0575562335463871 | 0.73124518496316 | 0.952992645818943 | 910.741826301597 | 697.95427967958 | 836.666121565227 | 907.9130400419 | 989.842209020464 | 1023.71982824582 | 754.781615653199 | 721.874500504558 | Erp29 |
| **ENSMUSG00000029617** | 0.146172526693675 | 0.212065363597013 | 0.823370267313127 | 801.416666596743 | 693.882201035008 | 729.716719147953 | 847.885731609378 | 722.672409240604 | 753.418654957498 | 673.57752458982 | 627.433776378397 | Ccz1 |
| **ENSMUSG00000029621** | -0.0184995178048102 | 0.888815699553107 | 0.981215233412228 | 1245.94541514871 | 1051.4107060284 | 1122.49549793706 | 1288.71127791071 | 1258.10696699614 | 1369.85209992272 | 1082.72121417838 | 1059.47295111646 | Arpc1a |
| **ENSMUSG00000029622** | -0.454071923766866 | 0.118445059017872 | 0.769868040577702 | 894.478579403355 | 864.909504107017 | 897.239234438728 | 1077.67777170263 | 1554.84063806312 | 1942.25458453329 | 791.219348822664 | 828.256005842071 | Arpc1b |
| **ENSMUSG00000029623** | 0.0655373584255361 | 0.642412125959338 | 0.934616627390525 | 1621.80712124144 | 1404.0527166483 | 1516.22073161481 | 1703.27487677282 | 1403.73640704008 | 1843.18492373531 | 1376.30523571521 | 1346.05170018895 | Pdap1 |
| **ENSMUSG00000029624** | -0.0319829118347522 | 0.830297619302656 | 0.967583158900775 | 430.976042803434 | 397.434875710192 | 433.476339000989 | 405.184331919526 | 407.324448844704 | 333.901449356164 | 466.402984569149 | 494.999657488839 | Ptcd1 |
| **ENSMUSG00000029625** | -0.0552423965293649 | 0.697954143437625 | 0.944933386973863 | 181.606257030378 | 159.625482867208 | 151.432782183751 | 177.268145214792 | 166.433645764503 | 192.023910435596 | 160.326025945645 | 178.026192605635 | Cpsf4 |
| **ENSMUSG00000029627** | -0.191189251235513 | 0.382568110643013 | 0.875126835008975 | 147.272735800754 | 112.389370590177 | 100.324218196735 | 140.689004138724 | 166.433645764503 | 171.23151249034 | 124.929370866736 | 109.638082031519 | Zkscan14 |
| **ENSMUSG00000029629** | 0.111373996526682 | 0.277879835287492 | 0.848075053184273 | 934.233182932393 | 887.713144516618 | 866.006223113329 | 967.002421780164 | 790.559817381388 | 791.334204151787 | 938.011359591079 | 861.907298346795 | Phf14 |
| **ENSMUSG00000029630** | -0.152371109241935 | 0.682997381134683 | 0.944452327903597 | 65.9565013095403 | 98.5443031986338 | 99.3777633080869 | 91.9168160372998 | 39.4184950494875 | 94.1773318696872 | 97.8613405122768 | 163.914360264945 | Cyp3a25 |
| **ENSMUSG00000029632** | 0.125740196799813 | 0.463790000559904 | 0.895328112556867 | 26562.3997534134 | 18048.2669684704 | 19637.9924845666 | 25482.5303563 | 20807.4905951503 | 24177.8895636361 | 17582.7678714026 | 19674.065334052 | Ndufa4 |
| **ENSMUSG00000029634** | 0.0722014592455051 | 0.704043940287375 | 0.946931219685582 | 801.416666596743 | 979.742121883937 | 819.629933569555 | 710.948434247686 | 730.337116611338 | 572.402484610566 | 938.011359591079 | 908.584897627541 | Rnf6 |
| **ENSMUSG00000029635** | 0.259147849172564 | 0.0561009993853294 | 0.703396849069134 | 244.852217190211 | 320.065381463331 | 300.026199701558 | 294.508981997063 | 226.656346534553 | 242.170281950624 | 261.31060072959 | 238.815624227072 | Cdk8 |
| **ENSMUSG00000029636** | -0.112428429515673 | 0.682569371167177 | 0.944452327903597 | 43.3686583953142 | 49.679359463774 | 45.4298346551254 | 28.1378008277448 | 42.7033696369448 | 34.2463024980681 | 54.1360607089191 | 48.8486504100828 | Wasf3 |
| **ENSMUSG00000029640** | 0.0825931728751642 | 0.692903974464861 | 0.94469184542218 | 604.450676384691 | 673.52180781215 | 580.176846741498 | 542.121629281217 | 488.351355335317 | 387.717067567413 | 706.892023487617 | 682.795580176491 | Usp12 |

| | | | | | | | | | | | | |
|---|---|---|---|---|---|---|---|---|---|---|---|---|
| **ENSMUSG00000029641** | -0.369083554688027 | 0.515443529302603 | 0.904239892322932 | 26.2018977805023 | 7.32974156022896 | 13.2503684410783 | 20.6343872736795 | 33.943704070392 | 33.0232202659942 | 9.36970281500523 | 10.8552556466851 | Rasl11a |
| **ENSMUSG00000029642** | 0.120628638832596 | 0.327664870887191 | 0.855977678491063 | 1303.77029300913 | 1128.78020027526 | 1139.53168593273 | 1385.3177274193 | 1090.57836303582 | 1260.99778126815 | 1020.25652874501 | 1188.65049331202 | Polr1d |
| **ENSMUSG00000029647** | -0.0626743443873377 | 0.658789225804438 | 0.938989601151841 | 168.957064998411 | 184.057954734638 | 201.594891282119 | 180.081925297567 | 214.611806380543 | 179.793088114857 | 171.777884941763 | 200.822229463674 | Pan3 |
| **ENSMUSG00000029648** | -0.318169330808146 | 0.151613784842891 | 0.783412422888014 | 4338.67286696455 | 4129.90216132456 | 4622.48567615901 | 3557.55595132121 | 7072.33498679555 | 5945.40273011103 | 3945.68596320776 | 3793.91184851643 | Flt1 |
| **ENSMUSG00000029649** | -0.0535515880781232 | 0.777287786392188 | 0.95974290923544 | 3099.05204783182 | 2150.05752433383 | 2251.61618009465 | 3127.04759865671 | 2641.03916831566 | 3480.8920324822 | 2422.58871672413 | 2485.85354309088 | Pomp |
| **ENSMUSG00000029650** | 0.0969249655179842 | 0.460150645664252 | 0.894774044236998 | 290.027903018663 | 271.200437728471 | 267.84673348751 | 270.12288794635 | 264.979883388221 | 221.377884005369 | 293.584021536831 | 246.414303179751 | Slc46a3 |
| **ENSMUSG00000029651** | -0.0245462324703872 | 0.867054695624842 | 0.974868041734905 | 2604.83004486856 | 3164.81952256108 | 3150.74832431068 | 2496.76086011523 | 2351.97020461942 | 2972.08982393948 | 3182.57572283011 | 3106.77416608127 | Mtus2 |
| **ENSMUSG00000029655** | 0.0451559115331222 | 0.716518329631484 | 0.951348124093194 | 477.055242348456 | 526.926976607571 | 481.745538322059 | 529.928582255861 | 490.541271726955 | 408.509465512669 | 531.990904274186 | 521.052271040884 | N4bp2l2 |
| **ENSMUSG00000029656** | -0.388330201822004 | 0.248413720580375 | 0.834621660998471 | 68.6670424592474 | 64.338842584232 | 28.3936466594534 | 59.0893817382642 | 45.9882442244021 | 73.3849339244316 | 69.7522320672611 | 99.8683519495027 | C8b |
| **ENSMUSG00000029657** | 0.31081917055416 | 0.075864247053178 | 0.735399943172214 | 2680.72519706036 | 2670.46917511008 | 2415.35287583084 | 2752.8148476477 | 1903.03734433359 | 1592.45306616017 | 2258.09837841626 | 2725.75469288262 | Hsph1 |
| **ENSMUSG00000029659** | -0.41916815820862 | 0.135344815598434 | 0.776010327117824 | 191.544907912638 | 212.56250524664 | 207.27362061401 | 256.053987532478 | 309.873169416804 | 453.763508099402 | 201.96914956789 | 195.394601640331 | Medag |
| **ENSMUSG00000029661** | -1.52804672771726 | 0.0228587098624649 | 0.650119841310877 | 570.117155155067 | 627.914526992947 | 505.406910538271 | 735.334528298398 | 2414.38282178111 | 3562.83854203115 | 526.785513821405 | 529.736475558232 | Col1a2 |
| **ENSMUSG00000029664** | -0.570082253444326 | 0.121714926522433 | 0.769868040577702 | 22.5878429142261 | 22.8036404096012 | 19.8755526616174 | 32.8274342990356 | 35.0386622662111 | 47.7002070508805 | 17.6983275394543 | 45.5920737160773 | Tfpi2 |
| **ENSMUSG00000029669** | -0.0309301516931612 | 0.81510734826301 | 0.966177091475091 | 867.373167906283 | 784.282346944498 | 621.82086184203 | 770.03781598595 | 769.755611660825 | 724.064681387725 | 777.685333645434 | 838.025735924088 | Tspan12 |
| **ENSMUSG00000029670** | -0.299365811216971 | 0.0540065061727679 | 0.703396849069134 | 192.448421629207 | 189.758864837039 | 159.950876181588 | 192.274972322923 | 217.896680968 | 273.970419984545 | 199.886993386778 | 213.848536239696 | Ing3 |
| **ENSMUSG00000029672** | -0.0457554811354823 | 0.6094102611162 | 0.927932112008768 | 636.977170181177 | 641.759594384491 | 633.178320505811 | 652.79697920368 | 711.722827282413 | 670.249063176475 | 629.852244786463 | 636.117980895745 | Fam3c |
| **ENSMUSG00000029673** | 0.00127082261016618 | 0.995110237009014 | 0.999262355603273 | 385.800356974982 | 373.002403842763 | 408.868511896129 | 392.99128489417 | 308.778211220985 | 284.978160073209 | 449.745735120251 | 514.539117652873 | Auts2 |
| **ENSMUSG00000029674** | -0.205999947165946 | 0.52225933009999 | 0.90648515894888 | 86.7373167906283 | 103.43079757212 | 106.949402417274 | 64.7169419038131 | 119.350443344282 | 146.769867848863 | 100.984574783945 | 51.0197015394199 | Limk1 |
| **ENSMUSG00000029675** | -1.1417252477957 | 0.0812447704962029 | 0.739855948845396 | 403.870631306363 | 437.341246426994 | 618.981497176084 | 440.825546301336 | 1194.59939163863 | 2305.51000745923 | 355.007628879643 | 339.769501741243 | Eln |
| **ENSMUSG00000029676** | -0.279890272954274 | 0.136742280232848 | 0.776456211338551 | 144.562194651047 | 132.749763813036 | 136.289503965376 | 124.744250336335 | 139.059690869025 | 140.654456688494 | 215.50316474512 | 157.401206876934 | Pot1a |
| **ENSMUSG00000029681** | 0.0660911741701525 | 0.478078894271654 | 0.897674307047416 | 1289.31407354403 | 1219.18034618475 | 1342.0730321035 | 1258.69762369445 | 1129.99685808531 | 1202.2898341286 | 1205.56842886401 | 1341.70959793028 | Bcl7b |
| **ENSMUSG00000029683** | 0.33942376482902 | 0.0622170185589936 | 0.715615581523459 | 26277.7929326941 | 28603.909230929 | 23495.7426106977 | 25320.2690381933 | 17325.5235324456 | 15467.097906806 | 24688.1258394482 | 24476.4304321455 | Lmod2 |
| **ENSMUSG00000029684** | -0.0371495970308869 | 0.734868243271443 | 0.954383574136061 | 489.704434380422 | 602.667639396603 | 529.068282754482 | 521.487242007538 | 532.149683168081 | 545.494675504941 | 537.196294726966 | 585.098279356326 | Wasl |
| **ENSMUSG00000029685** | 0.0902683087766094 | 0.793814398543121 | 0.961804768755116 | 836.653701542936 | 1238.72632367869 | 1093.15539638896 | 696.879533833814 | 569.37826182593 | 488.00981059747 | 1210.77381931679 | 1362.33458365898 | Asb15 |
| **ENSMUSG00000029686** | 0.211344478079927 | 0.183933249048915 | 0.805963874499992 | 1913.64205169324 | 2078.38894018937 | 2227.95480787844 | 1958.39093761104 | 1667.62133223248 | 1338.0519618888 | 2061.33461930115 | 1995.19598786072 | Cul1 |
| **ENSMUSG00000029687** | -0.0677703974910001 | 0.808985035894565 | 0.964172502140384 | 80.412720774645 | 39.9063707168021 | 59.6266579848522 | 85.3513291774927 | 73.3621991198795 | 70.9387694602839 | 55.1771387994752 | 78.1578406561325 | Ezh2 |
| **ENSMUSG00000029695** | -0.199031243106178 | 0.475421093665276 | 0.897549741595057 | 64.1494738764022 | 103.43079757212 | 58.6802030962037 | 79.7237690119437 | 68.9823663366031 | 81.9465095489486 | 74.9576225200418 | 125.920965501547 | Aass |
| **ENSMUSG00000029699** | -0.328678265451008 | 0.547389079099579 | 0.914699131434573 | 9.93865088225949 | 13.8450673915436 | 6.62518422053913 | 5.62756016554897 | 16.4243729372865 | 13.4539045528125 | 11.4518589961175 | 4.34210225867403 | Ssc4d |
| **ENSMUSG00000029701** | 0.189387481914972 | 0.138780090462404 | 0.779711473022872 | 351.466835745359 | 407.207864457164 | 431.583429223692 | 397.68091836546 | 316.442918591719 | 322.893709267499 | 382.075659234102 | 370.164217551961 | Rbm28 |
| **ENSMUSG00000029703** | -0.198934357221823 | 0.365420189166622 | 0.86924083181804 | 120.167324303683 | 115.647033505835 | 125.878500190243 | 136.937297361692 | 180.668102310151 | 172.454594722414 | 114.518589961175 | 105.295979772845 | Lrwd1 |
| **ENSMUSG00000029705** | 0.0541477932948747 | 0.748286421823042 | 0.956197192463153 | 2286.79321663625 | 2456.27783840562 | 2403.04896227841 | 1970.5839846364 | 1869.0936402632 | 1763.68457865051 | 2578.75043030755 | 2568.35348600569 | Cux1 |
| **ENSMUSG00000029708** | -0.0874744074401951 | 0.568021864719038 | 0.920432752255281 | 245.75573090678 | 280.973426475443 | 325.580481695066 | 244.79886720138 | 295.638712871156 | 251.954939807215 | 290.460787265162 | 326.743194965221 | Gcc1 |
| **ENSMUSG00000029710** | -0.642944325290034 | 0.0284332665316991 | 0.665956123973759 | 1002.90022539164 | 925.990683775592 | 1138.58523104408 | 997.954002690684 | 2391.38869966891 | 1969.16239363891 | 1056.69426191448 | 931.380934485579 | Ephb4 |
| **ENSMUSG00000029713** | -0.248535355759441 | 0.264026414457952 | 0.842510961779757 | 2847.87523462563 | 2460.34991705019 | 2548.80301513027 | 3051.0755364218 | 3688.91416171454 | 4452.01932474885 | 2484.01232406694 | 2334.96548960196 | Gnb2 |
| **ENSMUSG00000029714** | -0.0100876939514678 | 0.954873301523407 | 0.992371505557225 | 630.652574165193 | 606.739718041175 | 636.964140060405 | 495.225294568309 | 560.618596259378 | 443.978850242811 | 696.481242582055 | 683.88110574116 | Gigyf1 |
| **ENSMUSG00000029715** | 0.00124586402552794 | 0.995824499148259 | 0.999262355603273 | 186.123825613223 | 128.677685168464 | 169.415425068072 | 151.944124469822 | 177.383227722694 | 206.700897220482 | 148.874166949528 | 103.124928643508 | Pop7 |
| **ENSMUSG00000029716** | -0.391146008433989 | 0.4181248019827 | 0.882468496126562 | 37.9475760958999 | 109.13170767452 | 46.3762895437739 | 47.8342614071662 | 39.4184950494875 | 57.4848649074714 | 61.4236073428121 | 158.486732441602 | Tfr2 |
| **ENSMUSG00000029718** | -1.18438245421346 | 0.0134835407545755 | 0.635862253526407 | 461.695509166782 | 422.681763306537 | 322.74111702912 | 454.894446715208 | 1208.83384818428 | 1871.31581507301 | 333.144988977964 | 364.736589728619 | Pcolce |
| **ENSMUSG00000029722** | -0.191430846442832 | 0.414770523799718 | 0.882192126163777 | 283.70330700268 | 399.063707168021 | 453.351891662606 | 263.557401086543 | 476.306815181307 | 469.663577116362 | 339.391457521301 | 313.716888189199 | Agfg2 |
| **ENSMUSG00000029723** | -0.0298453411314567 | 0.690411826114868 | 0.94469184542218 | 3130.67502791174 | 3002.75079250713 | 2831.79302683615 | 2931.02091955675 | 2921.34846644535 | 2979.42831733192 | 3180.493566649 | 3063.35314349453 | Tsc22d4 |
| **ENSMUSG00000029725** | -0.334140278642758 | 0.261621844328414 | 0.841137531367976 | 175.281661014395 | 109.13170767452 | 132.503684410783 | 190.399118934407 | 249.650468646754 | 251.954939807215 | 141.586620315635 | 122.664388807541 | Ppp1r35 |
| **ENSMUSG00000029726** | -0.173994942960706 | 0.294246956901897 | 0.851785577030345 | 691.187993175319 | 693.882201035008 | 708.894711597687 | 679.996853337167 | 877.061514851097 | 968.681127802497 | 660.043509412591 | 625.26272524906 | Mepce |
| **ENSMUSG00000029727** | -0.303678234304403 | 0.27513589871626 | 0.846286635233373 | 56.0178504272808 | 83.0704043492615 | 55.8408384302584 | 54.3997482669734 | 50.3680770076785 | 67.2695227640623 | 92.6559500594962 | 97.6973008201657 | Cyp3a13 |
| **ENSMUSG00000029729** | 0.03535253758309 | 0.758149053235972 | 0.95677573228595 | 302.67709505063 | 330.652785939217 | 296.240380146964 | 315.143369270742 | 317.537876787538 | 287.424324537357 | 283.173240631269 | 326.743194965221 | Zkscan1 |
| **ENSMUSG00000029730** | -0.103695378636821 | 0.592293938713212 | 0.923935529355527 | 188.83436676293 | 200.346269312925 | 240.399541716705 | 244.79886720138 | 257.315176017488 | 291.093571233579 | 178.024353485099 | 213.848536239696 | Mcm7 |
| **ENSMUSG00000029735** | -0.175167651415246 | 0.548610696949874 | 0.914699131434573 | 32.5264937964856 | 30.1333819698302 | 21.7684624389143 | 33.7653609932938 | 28.4689130912965 | 31.8001380339204 | 43.7252798033577 | 29.3091902460497 | Tpk1 |
| **ENSMUSG00000029752** | 0.0656467672475291 | 0.815444300148524 | 0.966177091475091 | 74.0881247586617 | 78.1839099757755 | 92.7525790875478 | 55.3376749612315 | 75.5521155115177 | 101.51582526213 | 46.8485140750261 | 64.046008315442 | Asns |
| **ENSMUSG00000029759** | -0.00499367617527355 | 0.980807210567286 | 0.996888334226108 | 272.861142403852 | 400.69253862585 | 427.797609669098 | 301.07446885687 | 278.119381738051 | 319.224462571277 | 411.225845769674 | 398.387882233342 | Pon3 |
| **ENSMUSG00000029761** | -0.167717961934221 | 0.297013785462993 | 0.851785577030345 | 1014.64590370704 | 1206.96411025103 | 1093.15539638896 | 951.995594672034 | 1319.42462596201 | 1440.79086938301 | 1034.8316220128 | 999.769045059695 | Cald1 |
| **ENSMUSG00000029762** | -0.460790629677054 | 0.31734222538926 | 0.854351922245337 | 53.3073092775737 | 54.56585383726 | 31.2330113253987 | 44.0825546301336 | 68.9823663366031 | 123.53130543946 | 30.191264626128 | 30.3947158107182 | Akr1b8 |
| **ENSMUSG00000029763** | 0.123223053829043 | 0.436492234293795 | 0.888701139843674 | 865.566140473145 | 984.628616257423 | 1008.92091129924 | 876.023532437123 | 744.571573156986 | 706.941530138691 | 1047.32455909947 | 929.209883356242 | Exoc4 |
| **ENSMUSG00000029765** | 0.286790384748151 | 0.154314467769265 | 0.786853994984691 | 195.158962778914 | 289.117583764587 | 183.612248397799 | 168.826804966469 | 170.813478547779 | 163.893019097897 | 170.736806851206 | 181.282769299641 | Plxna4 |
| **ENSMUSG00000029767** | 0.150405989629207 | 0.304140428744056 | 0.851785577030345 | 6635.40473448307 | 7701.11513261389 | 7372.8835825714 | 6270.97787781007 | 5542.67838723627 | 5342.42318969862 | 7230.28733891237 | 7093.9095651087 | Calu |
| **ENSMUSG00000029769** | -0.171125786135748 | 0.48165036360847 | 0.897732059433541 | 46.0791995450213 | 43.1640336324594 | 43.5369248778286 | 47.8342614071662 | 64.6025335533267 | 56.2617826753975 | 43.7252798033577 | 39.0789203280663 | Ccdc136 |
| **ENSMUSG00000029771** | -0.487896251075791 | 0.140980295414435 | 0.779711473022872 | 29.8159526467785 | 28.5045505120015 | 34.0723759913441 | 37.5170677703264 | 52.5579933993167 | 67.2695227640623 | 26.0269522639034 | 36.9078691987293 | Irf5 |
| **ENSMUSG00000029772** | 0.262756924222421 | 0.317481592495959 | 0.854351922245337 | 605.35419010126 | 668.635313438664 | 603.838218957709 | 510.23212167644 | 365.716037403578 | 310.66288694676 | 641.30410378258 | 671.940324529806 | Ahcyl2 |
| **ENSMUSG00000029775** | 0.0602179802279357 | 0.806691388771702 | 0.964026716319739 | 944.171833814652 | 1093.76032393194 | 1098.83412572085 | 813.182443921826 | 734.716949394614 | 620.102691661447 | 1218.06136595068 | 1214.70310686406 | Klhdc10 |
| **ENSMUSG00000029776** | 0.190207574926376 | 0.168424179496267 | 0.79907358295209 | 7160.34620380968 | 6910.31745983808 | 7208.20043194657 | 7256.73883347539 | 6023.36503520085 | 4900.89050391996 | 7022.07172080114 | 7062.42932373331 | Hibadh |
| **ENSMUSG00000029777** | 0.124989030002259 | 0.225009108882453 | 0.832048466795696 | 2116.0291242047 | 1926.09319888239 | 2207.13280032818 | 2346.69258903392 | 2105.60461056012 | 2041.32424533127 | 1831.25636128824 | 1904.01184042856 | Gars |
| **ENSMUSG00000029778** | -0.449537511020246 | 0.0123983708840803 | 0.635862253526407 | 56.9213641438498 | 65.9676740420606 | 67.1982970940397 | 69.4065753751039 | 95.2613630362614 | 85.6157562451702 | 84.3273253350471 | 89.0130963028176 | Adcyap1r1 |
| **ENSMUSG00000029780** | 0.189695121538779 | 0.0751436151184456 | 0.735399943172214 | 1623.61414867457 | 1487.93753672648 | 1520.95300605805 | 1611.35806073552 | 1253.72713421287 | 1259.77469903608 | 1414.82512506579 | 1544.70287852329 | Nt5c3 |
| **ENSMUSG00000029781** | -0.028410075155325 | 0.635192102825454 | 0.932543559775713 | 3455.9399658766 | 3582.61479149413 | 3470.65007667385 | 3445.00474801023 | 3570.65867656607 | 3479.66895025013 | 3541.74766407198 | 3639.76721833351 | Fkbp9 |
| **ENSMUSG00000029782** | 0.0682989461319427 | 0.642954965359657 | 0.934616627390525 | 187.027339329792 | 201.975100770753 | 173.201244622666 | 196.964605794214 | 185.047935093427 | 154.108361241306 | 208.215618111227 | 175.855141476298 | Tmem209 |
| **ENSMUSG00000029787** | -0.116829306193502 | 0.503369560348397 | 0.900927059989498 | 110.228673421423 | 119.719112150406 | 123.985590412947 | 126.620103724852 | 118.255485148462 | 107.6312364225 | 158.243869764533 | 135.690695583563 | Avl9 |
| **ENSMUSG00000029790** | -0.139583131761882 | 0.68474625570256 | 0.944452327903597 | 18.0702743313809 | 17.9171460361152 | 16.0897331070236 | 21.5723139679377 | 22.994122112201 | 22.0154801773295 | 13.5340151772298 | 22.7960368580387 | Cep41 |
| **ENSMUSG00000029798** | -0.0183502646266578 | 0.939852093165013 | 0.989269716906553 | 40.658117245607 | 47.2361122770311 | 47.3227444324223 | 41.2687745473591 | 35.0386622662111 | 52.592535979176 | 39.5609674411332 | 52.1052271040884 | Herc6 |
| **ENSMUSG00000029802** | -0.211360451084492 | 0.419251357933207 | 0.882568531013176 | 552.950394540256 | 549.730617017172 | 469.44162476963 | 583.390403828576 | 787.274942793931 | 858.60372691585 | 385.198893505771 | 465.69046724279 | Abcg2 |
| **ENSMUSG00000029804** | 0.0724651459403596 | 0.814476460971059 | 0.966177091475091 | 477.958756065025 | 632.801021366433 | 639.80350472635 | 558.066383083606 | 416.084114411257 | 273.970419984545 | 736.042210023189 | 768.552099785303 | Herc3 |
| **ENSMUSG00000029810** | -0.726847538383737 | 0.0291444141314232 | 0.665956123973759 | 256.597895505609 | 295.632909595901 | 229.042083052924 | 268.247034557834 | 399.65974147397 | 790.111121919713 | 294.625099627387 | 254.012982132431 | Tmem176b |
| **ENSMUSG00000029815** | 0.0377628656844867 | 0.852614277972775 | 0.973120292022893 | 503.257140128958 | 320.065381463331 | 395.618143455051 | 510.23212167644 | 457.692525852383 | 503.90987961443 | 378.952424962434 | 344.111603999917 | Malsu1 |
| **ENSMUSG00000029816** | -1.28835424989813 | 0.0185718465964991 | 0.642273034804458 | 25.2983840639333 | 16.2883145782866 | 23.6613722162112 | 23.448167356454 | 54.7479097909548 | 119.862058743238 | 21.8626399016789 | 20.6249857287016 | Gpnmb |
| **ENSMUSG00000029817** | -0.0106925957986405 | 0.921231119335735 | 0.986713918026868 | 427.361987937158 | 377.074482487334 | 362.492222352355 | 377.046531091781 | 399.65974147397 | 379.155491942897 | 401.856142954669 | 374.506319810635 | Tra2a |
| **ENSMUSG00000029821** | 0.00565456771368121 | 0.980629502450828 | 0.996888334226108 | 168.053551281842 | 105.059629029948 | 87.0738497556571 | 144.440710915757 | 121.54035973592 | 136.985209992272 | 128.052605138405 | 116.15123541953 | Gsdme |
| **ENSMUSG00000029822** | 0.112998985824363 | 0.634998377209831 | 0.932543559775713 | 321.65088309858 | 468.289044125739 | 412.654331450723 | 295.446908691321 | 293.448796479518 | 244.616446414772 | 435.170641852465 | 411.414189009364 | Osbpl3 |
| **ENSMUSG00000029823** | 0.069138872321569 | 0.727253307941995 | 0.952237115597464 | 1974.17747070336 | 2083.27543456285 | 1935.50024728607 | 1630.11659462068 | 1492.42802090143 | 1345.39045528125 | 2243.52328514847 | 2184.07743611304 | Luc7l2 |
| **ENSMUSG00000029826** | -0.399085002419603 | 0.0657397452365035 | 0.718586813809351 | 254.790868072471 | 334.724864583789 | 333.152120804253 | 257.929840920994 | 500.395895489327 | 448.871179171106 | 314.405583347953 | 294.177428025166 | Zc3hav1 |
| **ENSMUSG00000029828** | -0.811388676693009 | 0.117993811155987 | 0.769868040577702 | 9.93865088225949 | 4.88649437348597 | 10.4110037751329 | 10.3171936368398 | 14.2344565456483 | 26.9078091056249 | 7.28754663389296 | 14.1118323406906 | 4921507P07R |
| **ENSMUSG00000029833** | 0.159137699007611 | 0.23332746008395 | 0.832048466795696 | 572.827696304775 | 614.069459601404 | 688.072704047421 | 625.597105070194 | 589.087509350674 | 453.763508099402 | 597.578823979222 | 597.039060567679 | Trim24 |
| **ENSMUSG00000029836** | 0.139786640865493 | 0.202795664975981 | 0.815403452654488 | 889.05749710394 | 811.972481727586 | 860.327493781438 | 906.037186653384 | 803.699315731217 | 676.364474336844 | 851.60187807492 | 813.058647936712 | Cbx3 |
| **ENSMUSG00000029838** | -0.939186724853701 | 0.0738460229335022 | 0.733705544272914 | 49.6932544112975 | 73.2974156022896 | 27.447191770805 | 63.779015209555 | 154.389105610493 | 177.34692365071 | 36.4377331694648 | 43.4210225867403 | Ptn |
| **ENSMUSG00000029840** | 0.00551386006571373 | 0.958813565288712 | 0.993101678702062 | 1143.84836517641 | 1227.32450347389 | 1221.87326124514 | 1145.20849368921 | 1098.24307040655 | 1086.09702208159 | 1186.829023234 | 1348.22275131829 | Mtpn |
| **ENSMUSG00000029851** | -0.724484220365349 | 0.0683869735722962 | 0.72237439886039 | 24.3948703473642 | 26.8757190541728 | 21.7684624389143 | 15.0068271081306 | 51.4630352034976 | 50.1463715150282 | 20.8215618111227 | 23.8815624227072 | Tcaf2 |
| **ENSMUSG00000029859** | -0.13746373015667 | 0.787595262410673 | 0.961224689823763 | 22.5878429142261 | 24.4324718674299 | 16.0897331070236 | 4.68963347129081 | 21.8991639163819 | 23.2385624094033 | 10.4107809055614 | 19.5394601640331 | Epha1 |
| **ENSMUSG00000029860** | -0.25793125163057 | 0.26984130995162 | 0.843793744782845 | 1294.73515584344 | 1439.88700872053 | 1427.25397208186 | 1290.58713129923 | 1677.47595599486 | 2430.26439513076 | 1257.62233339181 | 1156.08472637196 | Zyx |
| **ENSMUSG00000029862** | 0.215042813475015 | 0.562442271457322 | 0.917693310295102 | 37.0440623793308 | 53.7514381083457 | 46.3762895437739 | 41.2687745473591 | 32.8487458745729 | 15.9000690169602 | 44.7663578939139 | 59.7039060567679 | Clcn1 |
| **ENSMUSG00000029863** | -0.284568837592455 | 0.131765669057885 | 0.77303252008843 | 78.6056933415069 | 101.801966114291 | 88.966759532954 | 82.5375490947182 | 100.736154015357 | 130.869798831903 | 106.189965236726 | 92.2696729968231 | Casp2 |
| **ENSMUSG00000029864** | -0.0483050651918749 | 0.726549809917709 | 0.952045814446758 | 1835.9398720683 | 1473.27805360602 | 1581.52611893155 | 1843.02595421729 | 1812.15581408061 | 2003.40869613698 | 1651.14985162203 | 1496.93975367787 | Gstk1 |

| | | | | | | | | | | | | |
|---|---|---|---|---|---|---|---|---|---|---|---|---|
| **ENSMUSG00000029869** | -0.963829364789977 | 0.084517609283909 | 0.743476626316337 | 8.13162344912141 | 4.88649437348597 | 13.2503684410783 | 15.0068271081306 | 32.8487458745729 | 22.0154801773295 | 18.7394056300105 | 6.51315338801105 | Ephb6 |
| **ENSMUSG00000029875** | -1.78039295139619 | 0.00076978793205157 | 0.556246415579446 | 5.42108229941427 | 4.07207864457164 | 8.51809399783602 | 4.68963347129081 | 22.994122112201 | 30.5770558018465 | 13.5340151772298 | 10.8552556466851 | Ccdc184 |
| **ENSMUSG00000029910** | 0.081472654707989 | 0.741475779647396 | 0.954383574136061 | 84.9302893574902 | 83.0704043492615 | 69.0912068713366 | 112.551203310979 | 54.7479097909548 | 92.9542496376133 | 91.61487196894 | 91.1841474321546 | Mad2l1 |
| **ENSMUSG00000029911** | 0.147501603431822 | 0.245269135900728 | 0.832048466795696 | 791.478015714483 | 624.65686407729 | 686.179794270124 | 753.155135489303 | 562.808512651016 | 692.264543353805 | 675.659680770933 | 646.973236542431 | Ssbp1 |
| **ENSMUSG00000029913** | 0.377798412190511 | 0.0319133330463339 | 0.671606027118506 | 137.334084918495 | 112.389370590177 | 151.432782183751 | 133.185590584659 | 110.590777777729 | 95.400414101761 | 89.5327157878278 | 115.065709854862 | Prdm5 |
| **ENSMUSG00000029915** | 0.357290494247782 | 0.468553093792652 | 0.895328112556867 | 9.93865088225949 | 8.14415728914329 | 24.6078271048596 | 13.1309737196143 | 8.75966556655277 | 13.4539045528125 | 11.4518589961175 | 9.76973008201657 | Clec5a |
| **ENSMUSG00000029916** | 0.0275401118572813 | 0.901978154207571 | 0.984871597499125 | 225.878429142261 | 269.571606270643 | 238.506631939409 | 172.578511743502 | 204.757182618171 | 157.777607937528 | 285.255396812381 | 240.986675356409 | Agk |
| **ENSMUSG00000029918** | 0.154581926506499 | 0.289365118176518 | 0.851785577030345 | 920.680477183857 | 636.873100011005 | 684.286884492827 | 825.375490947182 | 679.969039603659 | 728.95701031602 | 676.700758861489 | 669.769273400469 | Mrps33 |
| **ENSMUSG00000029920** | 0.0624694964167708 | 0.633199439993019 | 0.932281134354785 | 194.255449062345 | 202.789516499668 | 204.434255948065 | 181.957778686083 | 181.76306050597 | 171.23151249034 | 191.558368662329 | 205.164331722348 | Smarcad1 |
| **ENSMUSG00000029922** | 0.0224166703262005 | 0.954387373068102 | 0.992371505557225 | 1116.74295367934 | 1103.53331267892 | 2439.9607029357 | 959.499008226099 | 1209.9288063801 | 2502.42624682312 | 894.286079787721 | 927.038832226905 | Mkrn1 |
| **ENSMUSG00000029924** | -0.0799817824333349 | 0.570415985608759 | 0.920872458148558 | 346.045753445944 | 356.714089264476 | 323.687571917769 | 268.247034557834 | 317.537876787538 | 320.447544803351 | 379.99350305299 | 350.624757387928 | Slc37a3 |
| **ENSMUSG00000029925** | -0.334312866859533 | 0.542735556220635 | 0.913837378403038 | 18.0702743313809 | 8.14415728914329 | 50.1621090983677 | 21.5723139679377 | 28.4689130912965 | 48.9232892829544 | 15.616171358342 | 30.3947158107182 | Tbxas1 |
| **ENSMUSG00000029992** | 0.156897835752659 | 0.520048050463031 | 0.905962179970576 | 431.879556520004 | 578.235167529173 | 509.192730092864 | 413.625672167849 | 356.956371837026 | 281.308913376988 | 573.634027896431 | 521.052271040884 | Gfpt1 |
| **ENSMUSG00000029993** | 0.11331400478347 | 0.300025470345969 | 0.851785577030345 | 1651.62307388821 | 1512.37000859391 | 1504.86327295103 | 1773.61937884218 | 1460.67423322268 | 1647.49176660349 | 1501.23460658195 | 1347.13722575362 | Nfu1 |
| **ENSMUSG00000029994** | -0.0367176842790788 | 0.777329570115097 | 0.95974290923544 | 566.503100288791 | 626.285695535119 | 507.299820315568 | 609.652351267805 | 594.56230032977 | 678.810638800992 | 574.675105986987 | 523.223322170221 | Anxa4 |
| **ENSMUSG00000029998** | 0.136392397179592 | 0.300569316006321 | 0.851785577030345 | 2337.38998476412 | 2456.27783840562 | 2353.83330806869 | 2358.88563605928 | 1892.0877623754 | 1852.9695815919 | 2437.16380999192 | 2465.22855736218 | Pcyox1 |
| **ENSMUSG00000029999** | -0.424222636490824 | 0.320045023149731 | 0.855228644171892 | 20.780815481088 | 8.95857301805762 | 16.0897331070236 | 11.2551203310979 | 28.4689130912965 | 15.9000690169602 | 14.5750932677859 | 17.3684090346961 | Tgfa |
| **ENSMUSG00000030002** | 0.0140518563159291 | 0.920960849992804 | 0.986713918026868 | 403.870631306363 | 383.589808318649 | 442.940887887473 | 510.23212167644 | 492.731188118594 | 403.617136584374 | 401.856142954669 | 424.440495785386 | Dusp11 |
| **ENSMUSG00000030007** | 0.0762397055535916 | 0.462029872491898 | 0.895328112556867 | 4761.51728631887 | 4316.40336324594 | 4507.0181797439 | 5069.49378246536 | 4424.72606930497 | 4910.67516177655 | 4010.23280482224 | 4349.70093762671 | Cct7 |
| **ENSMUSG00000030008** | 0.0214691270140748 | 0.887786718614414 | 0.980702840906172 | 186.123825613223 | 153.110157035894 | 156.165056626994 | 175.392291826276 | 158.768938393769 | 179.793088114857 | 179.065431575656 | 143.289374536243 | Pradc1 |
| **ENSMUSG00000030016** | 0.316115135709688 | 0.0830397952797582 | 0.741377052226143 | 887.250469670802 | 1020.46290832965 | 979.580809751142 | 788.796349871114 | 738.001823982071 | 523.479195327612 | 866.176971342706 | 823.913903583397 | Zfp638 |
| **ENSMUSG00000030019** | -0.0505214677821373 | 0.647310201313418 | 0.935063240511793 | 309.001691066613 | 337.982527499446 | 321.794662140472 | 337.653609932938 | 314.253002200081 | 325.339873731647 | 363.336253604092 | 349.539231823259 | Fbxl14 |
| **ENSMUSG00000030020** | -0.150257245296145 | 0.569772672846843 | 0.920872458148558 | 103.000563688871 | 93.6578088251478 | 169.415425068072 | 159.447538023887 | 158.768938393769 | 177.34692365071 | 143.668776496747 | 103.124928643508 | Prickle2 |
| **ENSMUSG00000030022** | -0.0206712205415975 | 0.925056363703412 | 0.986713918026868 | 326.168451681425 | 312.735639903102 | 371.95677123884 | 333.901903155905 | 492.731188118594 | 253.178022039289 | 337.309301340188 | 280.065595684475 | Adamts9 |
| **ENSMUSG00000030029** | -0.0476920912986894 | 0.812251378032316 | 0.965224347191244 | 867.373167906283 | 1044.08096446817 | 996.616997746814 | 860.078778634734 | 906.625386138212 | 683.702967729287 | 1301.34761319517 | 1001.94009618903 | Lrig1 |
| **ENSMUSG00000030030** | 0.0975116444950577 | 0.789648345558308 | 0.961659329450003 | 33.4300075130547 | 33.3910448854875 | 26.5007368821565 | 23.448167356454 | 19.7092475247437 | 15.9000690169602 | 42.6842017128016 | 30.3947158107182 | 1700003E16R |
| **ENSMUSG00000030031** | 0.0824253438677318 | 0.816337680726326 | 0.966315773308182 | 16.2632468982428 | 14.6594831204579 | 24.6078271048596 | 19.6964605794214 | 17.5193311331055 | 17.123151249034 | 20.8215618111227 | 15.1973579053591 | Kbtbd8 |
| **ENSMUSG00000030032** | -0.425074471263705 | 0.430737391089746 | 0.886017087288711 | 16.2632468982428 | 25.2468875963442 | 11.3574586637814 | 23.448167356454 | 28.4689130912965 | 50.1463715150282 | 8.32862472444909 | 16.2828834700276 | Wdr54 |
| **ENSMUSG00000030034** | -0.170635215482942 | 0.46662856202328 | 0.895328112556867 | 346.045753445944 | 348.569931975333 | 320.848207251823 | 329.212269684615 | 451.122776677468 | 511.248373006873 | 309.200192895173 | 243.157726485746 | Ino80b |
| **ENSMUSG00000030035** | -0.105418045226562 | 0.369045080012934 | 0.871495599696183 | 1337.20030052219 | 1175.20189682338 | 1256.89209212514 | 1326.22834568104 | 1228.54309570903 | 1591.22998392809 | 1287.81359801794 | 1375.360890435 | Wbp1 |
| **ENSMUSG00000030036** | -0.055818938611593 | 0.766498208277707 | 0.958624690460551 | 158.114900399583 | 213.376920975554 | 188.344522841041 | 175.392291826276 | 222.276513751277 | 216.485555077073 | 182.188665847324 | 144.374900100912 | Mogs |
| **ENSMUSG00000030037** | -0.196800210733396 | 0.498742505499173 | 0.900927059989498 | 51.5002818444356 | 38.2775392589734 | 25.5542819935081 | 57.2135283497478 | 54.7479097909548 | 44.0309603546589 | 52.0539045278068 | 46.6775992807458 | Mrpl53 |
| **ENSMUSG00000030042** | 0.0178060638439557 | 0.904708632758525 | 0.98504921011344 | 598.126080368708 | 456.072808192024 | 415.493696116668 | 557.128456389348 | 502.585811880965 | 478.225152740879 | 530.94982618363 | 489.572029665497 | Pole4 |
| **ENSMUSG00000030045** | -0.0529053373505529 | 0.702826807054477 | 0.946451777392559 | 655.047444512558 | 509.82424630037 | 456.191256328551 | 562.756016554897 | 540.909348734634 | 578.517895770936 | 564.264325081426 | 581.84170266232 | Mrpl19 |
| **ENSMUSG00000030047** | -0.950769295950134 | 0.00570236250153579 | 0.635862253526407 | 34.3335212296237 | 46.4216965481167 | 31.2330113253987 | 39.3929211588428 | 108.400861386091 | 91.7311674055395 | 49.9717483466946 | 43.4210225867403 | Arhgap25 |
| **ENSMUSG00000030051** | -0.0256398056253346 | 0.91663105805057 | 0.986713918026868 | 69.5705561758165 | 72.4829998733752 | 72.8770264259304 | 94.7305961200743 | 108.400861386091 | 79.5003450848009 | 69.7522320672611 | 57.5328549274309 | Aplf |
| **ENSMUSG00000030054** | 0.433516674685435 | 0.600364116134603 | 0.925533581921158 | 24.3948703473642 | 32.5766291565731 | 136.289503965376 | 16.8826804966469 | 12.0445401540101 | 103.961989726278 | 6.24646854333682 | 33.6512925047237 | Gp9 |
| **ENSMUSG00000030055** | 0.0376610712349852 | 0.922780341273274 | 0.986713918026868 | 24.3948703473642 | 39.9063707168021 | 35.965285768641 | 17.8206071909051 | 21.8991639163819 | 24.4616446414772 | 45.80743598447 | 22.7960368580387 | Rab43 |
| **ENSMUSG00000030056** | 0.0503386092131496 | 0.662157043831087 | 0.939611397743053 | 470.730646332472 | 434.897999240251 | 414.54724122802 | 468.963347129081 | 386.520243124141 | 418.29412336926 | 469.526218840818 | 452.664160466768 | Isy1 |
| **ENSMUSG00000030057** | 0.115344158621982 | 0.20020664402194 | 0.812921911423739 | 9886.24708669849 | 9356.00789376781 | 9518.49681513743 | 10546.985676933 | 9010.41099339535 | 8269.25897105137 | 9744.49092760544 | 9260.61859218704 | Cnbp |
| **ENSMUSG00000030058** | -0.164766892636583 | 0.131310631362683 | 0.77303252008843 | 965.856163012309 | 976.48445896828 | 954.026527757634 | 965.126568391648 | 1183.64980968044 | 1187.61284734372 | 970.284780398319 | 988.91378941301 | Copg1 |
| **ENSMUSG00000030059** | 0.21182933019358 | 0.219305931395911 | 0.827535829146756 | 909.838312585028 | 876.311324311818 | 867.899132890626 | 903.223406570609 | 655.879959295639 | 587.079471395453 | 885.957455063272 | 941.150664567596 | Tmf1 |
| **ENSMUSG00000030060** | -0.149587891534775 | 0.531077665089122 | 0.909082872831377 | 114.746242004269 | 83.0704043492615 | 73.8234813145789 | 123.806323642077 | 130.300025302473 | 122.308223207386 | 86.4094815161593 | 99.8683519495027 | Hmces |
| **ENSMUSG00000030061** | 0.0175191109666242 | 0.932149025378607 | 0.987982116552499 | 1259.49812089725 | 1196.37670577515 | 1039.20746773599 | 931.361207398354 | 909.91026072567 | 803.565026472526 | 1366.93553290021 | 1291.77542195552 | Uba3 |
| **ENSMUSG00000030062** | -0.193124122753113 | 0.230995209737536 | 0.832048466795696 | 1345.33192397131 | 1206.96411025103 | 1252.1598176819 | 1256.82177030594 | 1489.14314631397 | 1859.08499275227 | 1207.65058504512 | 1232.07151589876 | Rpn1 |
| **ENSMUSG00000030064** | -0.470470813876207 | 0.0678585587040732 | 0.722112580225498 | 128.298947752804 | 161.254314325037 | 157.111511515642 | 153.819977858338 | 273.739548954774 | 266.631926592101 | 138.463386043966 | 154.144630182928 | Frmd4b |
| **ENSMUSG00000030067** | -0.214906559687726 | 0.319532184869065 | 0.854670142081901 | 354.177376895066 | 415.352021746308 | 499.72818120638 | 391.115431505653 | 609.891715071237 | 572.402484610566 | 362.295175513536 | 383.190524327983 | Foxp1 |
| **ENSMUSG00000030068** | 0.137787639565531 | 0.753050386288344 | 0.956497786093635 | 28.9124389302094 | 20.3603932228582 | 17.9826428843205 | 17.8206071909051 | 14.2344565456483 | 9.78465785659088 | 34.3555769883525 | 18.4539345993646 | Gm20696 |
| **ENSMUSG00000030074** | -0.939010352669704 | 0.0640073845009361 | 0.716991331547963 | 18.9737880479499 | 21.1748089517725 | 32.1794662140472 | 36.5791410760683 | 78.836990098975 | 85.6157562451702 | 24.9858741733473 | 19.5394601640331 | Gxylt2 |
| **ENSMUSG00000030077** | -1.81441836627242 | 0.0228324199275144 | 0.650119841310877 | 5.42108229941427 | 12.2162359337149 | 1.89290977729689 | 2.81378008277448 | 36.1336204620302 | 31.8001380339204 | 6.24646854333682 | 5.42762782334254 | Chl1 |
| **ENSMUSG00000030079** | -0.0824909246313996 | 0.543550427176907 | 0.913837378403038 | 570.117155155067 | 491.09268453534 | 500.674636095028 | 580.576623745802 | 614.271547854513 | 632.333513982185 | 477.854843565267 | 544.933833463591 | Ruvbl1 |
| **ENSMUSG00000030082** | -0.216733444972967 | 0.245898567386632 | 0.832048466795696 | 519.520387027201 | 563.575684408715 | 655.893237833374 | 524.301022090312 | 722.672409240604 | 839.034411202668 | 522.621201459181 | 547.104884592928 | Sec61a1 |
| **ENSMUSG00000030083** | -0.0334123962141938 | 0.851718692242197 | 0.972732497553372 | 573.731210021344 | 624.65686407729 | 646.428688946889 | 612.466131350579 | 803.699315731217 | 688.595296657583 | 543.442763270303 | 479.80229958348 | Abtb1 |
| **ENSMUSG00000030084** | -0.0358594655651389 | 0.888514343732412 | 0.98103849240407 | 318.036828232304 | 426.753841951108 | 363.438677241004 | 256.991914226736 | 295.638712871156 | 228.716377397812 | 410.184767679118 | 464.604941678121 | Plxna1 |
| **ENSMUSG00000030086** | 0.163815463089806 | 0.264960062731296 | 0.842510961779757 | 321.65088309858 | 242.69588721647 | 261.221549266971 | 312.329589187968 | 250.745426842573 | 277.639666680766 | 255.064132186253 | 232.302470839061 | Chchd6 |
| **ENSMUSG00000030087** | 0.00816950275520022 | 0.966010990863102 | 0.994284908685797 | 1317.32299875767 | 1628.01704209974 | 1409.27132919754 | 951.057667977776 | 1385.12211771116 | 1154.58962707772 | 1563.69929201532 | 1172.36760984199 | Klf15 |
| **ENSMUSG00000030088** | -0.488019582111873 | 0.123129873240208 | 0.770251097497048 | 128.298947752804 | 263.056280439328 | 148.593417517806 | 129.433883807626 | 229.94122112201 | 220.154801773295 | 148.874166949528 | 340.855027305911 | Aldh1l1 |
| **ENSMUSG00000030089** | -0.332850395903242 | 0.0155022192990051 | 0.635862253526407 | 636.073656464608 | 724.829998733753 | 706.055346931741 | 600.273084325223 | 685.443830582755 | 936.880989768576 | 853.684034256032 | 884.703335204834 | Slc41a3 |
| **ENSMUSG00000030091** | 0.214223259811203 | 0.522320964865713 | 0.90648515894888 | 40.658117245607 | 61.895595397489 | 58.6802030962037 | 36.5791410760683 | 49.2731188118594 | 20.7923979452556 | 47.8895921655823 | 52.1052271040884 | Nup210 |
| **ENSMUSG00000030094** | 0.248867538641588 | 0.051635245557299 | 0.703396849069134 | 373.151164943016 | 432.454752053509 | 494.995906763138 | 452.080666632434 | 382.140410340865 | 341.239942748607 | 370.623800237985 | 379.933947633978 | Xpc |
| **ENSMUSG00000030095** | -0.362867451799511 | 0.107212363364838 | 0.762287160109383 | 468.923618899334 | 507.380999113627 | 565.980023411771 | 579.638697051544 | 827.788396039237 | 880.619207093179 | 568.428637443651 | 452.664160466768 | Tmem43 |
| **ENSMUSG00000030096** | -0.128959250754477 | 0.551354758101146 | 0.914873920695527 | 1499.83276950461 | 1652.44951396717 | 2533.65973691189 | 2432.98184490567 | 2411.09794719365 | 2658.98077252857 | 1883.31026581605 | 1924.63682615726 | Slc6a6 |
| **ENSMUSG00000030098** | 0.237588788059569 | 0.496239566655141 | 0.900927059989498 | 56.0178504272808 | 61.0811796685746 | 111.681676860517 | 126.620103724852 | 62.4126171616885 | 52.592535979176 | 74.9576225200418 | 110.723607596188 | Grip2 |
| **ENSMUSG00000030101** | -0.106683250614367 | 0.436570643755714 | 0.888701139843674 | 446.335775985108 | 377.074482487334 | 420.22597055991 | 489.59773440276 | 475.211856985488 | 532.040770952129 | 442.458188486358 | 416.841816832707 | Sumf1 |
| **ENSMUSG00000030102** | -0.125369479964813 | 0.46750663158901 | 0.895328112556867 | 373.151164943016 | 464.216965481167 | 377.63550057073 | 305.764102328161 | 491.636229922775 | 359.586176229715 | 422.677704765791 | 385.36157545732 | Itpr1 |
| **ENSMUSG00000030103** | -0.523254690705028 | 0.00924467949947137 | 0.635862253526407 | 698.416102907872 | 758.22104361924 | 787.450467355508 | 600.273084325223 | 1393.88178327771 | 966.234963338349 | 970.284780398319 | 757.696844138618 | Bhlhe40 |
| **ENSMUSG00000030104** | -0.291210063571441 | 0.109218985600512 | 0.763154315676794 | 164.439496415566 | 214.191336704468 | 206.327165725361 | 202.592165959763 | 295.638712871156 | 262.96267989588 | 204.051305749003 | 201.907755028342 | Edem1 |
| **ENSMUSG00000030105** | 0.141208667813303 | 0.22869122426984 | 0.832048466795696 | 2705.12006740772 | 2729.10710759192 | 2895.2055043756 | 2572.73292235014 | 2241.37942684169 | 2192.98644210843 | 2633.92756910703 | 2815.85331475011 | Arl8b |
| **ENSMUSG00000030107** | -0.420249541327956 | 0.258502672748227 | 0.838547517499604 | 52.4037955610046 | 39.0919549878878 | 40.6975602118832 | 31.8895076047775 | 70.0773245324222 | 80.7234273168747 | 24.9858741733473 | 44.5065481514088 | Usp18 |
| **ENSMUSG00000030108** | -0.416062288435019 | 0.370020950632113 | 0.872916429836026 | 17.1667606148119 | 26.0613033252585 | 16.0897331070236 | 14.0689004138724 | 20.8042057205628 | 17.123151249034 | 13.5340151772298 | 46.6775992807458 | Slc6a13 |
| **ENSMUSG00000030110** | -1.16522600987569 | 0.007489136861009 | 0.635862253526407 | 19.877301764519 | 30.1333819698302 | 33.1259211026956 | 29.075727522003 | 79.9319482947941 | 107.6312364225 | 34.3555769883525 | 30.3947158107182 | Ret |
| **ENSMUSG00000030116** | -1.06408040027142 | 0.0344117904474612 | 0.678229434609245 | 276.475197270128 | 164.511977240694 | 126.824955078892 | 203.530092654021 | 482.876564356222 | 824.357424417781 | 149.915245040084 | 156.315681312265 | Mfap5 |
| **ENSMUSG00000030120** | -0.101679071543843 | 0.226392414815554 | 0.832048466795696 | 4380.23449792673 | 4208.90048702925 | 4375.46095022177 | 4597.71665525351 | 4505.75297579558 | 5151.6223614951 | 4718.16590640041 | 4470.19427530491 | Mlf2 |
| **ENSMUSG00000030122** | -0.36735809748825 | 0.152743527536209 | 0.783412422888014 | 3873.3633029315 | 3870.9179595298 | 3711.99607327921 | 4238.49073135263 | 5143.0186457623 | 7968.3807419612 | 3588.596178147 | 3547.49754533668 | Ptms |
| **ENSMUSG00000030123** | -0.244433385510004 | 0.112135399633602 | 0.764116514278315 | 1308.28786159198 | 1351.11569426887 | 1098.83412572085 | 1211.80128898154 | 1835.14993619281 | 1248.76695894741 | 1324.25133118741 | 1479.57134464318 | Plxnd1 |
| **ENSMUSG00000030126** | 0.0684511494272731 | 0.680621110369258 | 0.944452327903597 | 674.924746277077 | 751.705717787925 | 617.088587398787 | 532.742362338636 | 558.428679867739 | 527.148442023833 | 623.605776243126 | 747.927114056602 | Tmcc1 |
| **ENSMUSG00000030127** | 0.12144422282336 | 0.146854485544328 | 0.780607990324772 | 2034.71288971349 | 1934.23735617153 | 1924.14278862229 | 1951.82545075123 | 1711.41966006525 | 1724.54594722414 | 1965.55543496999 | 1808.48559073773 | Cops7a |
| **ENSMUSG00000030131** | -0.694575152895264 | 0.149254638122934 | 0.782024490202776 | 19.877301764519 | 61.0811796685746 | 29.3401015481019 | 40.3308478531009 | 38.3235368536684 | 57.4848649074714 | 31.2323427166841 | 117.236760984199 | Mug2 |
| **ENSMUSG00000030134** | -0.582645421756389 | 0.252239572836582 | 0.83723604018117 | 22.5878429142261 | 13.0306516626293 | 11.3574586637814 | 7.50341355406529 | 9.85462376237187 | 33.0232202659942 | 24.9858741733473 | 14.1118323406906 | Rasgef1a |
| **ENSMUSG00000030137** | -0.0289332812282949 | 0.833834295515145 | 0.96835936700622 | 4911.50056326933 | 3641.25272397596 | 4592.19911972226 | 4936.3081918807 | 4140.03693839201 | 4658.72022196933 | 4459.97853994249 | 5187.7266735508 | Tuba8 |
| **ENSMUSG00000030138** | 0.220165453650307 | 0.162469866294163 | 0.793968226938511 | 908.03128515189 | 932.506009606906 | 803.540200462531 | 821.623784170149 | 692.013579757669 | 578.517895770936 | 883.87529888216 | 819.571801324723 | Bms1 |
| **ENSMUSG00000030144** | -0.648373455398419 | 0.130906649203687 | 0.77303252008843 | 19.877301764519 | 26.8757190541728 | 27.447191770805 | 25.3240207449704 | 49.2731188118594 | 66.0464405319884 | 22.903717992235 | 18.4539345993646 | Clec4d |
| **ENSMUSG00000030145** | 0.0877660381210111 | 0.760716699963879 | 0.957357521291862 | 33.4300075130547 | 34.2054606144018 | 36.9117406572894 | 51.5859681841989 | 48.1781606160403 | 28.1308913376988 | 38.5198893505771 | 31.4802413753867 | Zfp248 |
| **ENSMUSG00000030148** | -0.450631049224013 | 0.346880558790449 | 0.861266047293794 | 8.13162344912141 | 19.5459774939439 | 23.6613722162112 | 15.9447538023887 | 31.7537876787538 | 24.4616446414772 | 27.0680303544596 | 8.68420451734806 | Clec4a2 |
| **ENSMUSG00000030157** | 0.0961611654310839 | 0.566192017136491 | 0.919931558693298 | 155.404359249876 | 140.079505373265 | 187.398067952392 | 163.19924480092 | 166.433645764503 | 165.116101329971 | 146.792010768415 | 125.920965501547 | Clec2d |
| **ENSMUSG00000030159** | 0.497436784562127 | 0.439339006893973 | 0.889583654898613 | 39.754603529038 | 53.7514381083457 | 156.165056626994 | 16.8826804966469 | 36.1336204620302 | 92.9542496376133 | 17.6983275394543 | 42.3354970220718 | Clec1b |
| **ENSMUSG00000030161** | 0.327455752734496 | 0.0529573955154175 | 0.703396849069134 | 5090.39627915 | 5854.83467516511 | 7999.43671885667 | 7194.83567165435 | 5886.49526072347 | 4678.28953768251 | 5367.79863490744 | 4897.89134778431 | Gabarapl1 |

| | | | | | | | | | | | | |
|---|---|---|---|---|---|---|---|---|---|---|---|---|
| ENSMUSG00000030166 | -0.276617461290067 | 0.241239010645912 | 0.832048466795696 | 59.631905293557 | 66.7820897709749 | 51.1085639870161 | 80.6616957062019 | 75.5521155115177 | 99.0696607979826 | 78.0808567917102 | 60.7894316214364 | Rad52 |
| ENSMUSG00000030168 | 0.207150158928177 | 0.158009645612914 | 0.789230661864335 | 2605.73355858513 | 2947.37052294096 | 3416.70214802089 | 2393.58892374683 | 2270.94329812881 | 2237.01740246309 | 2632.88649101647 | 2701.87313045992 | Adipor2 |
| ENSMUSG00000030170 | 0.243662352966438 | 0.416684047893513 | 0.882192126163777 | 56.9213641438498 | 62.7100111264033 | 65.3053873167428 | 45.0204813243917 | 54.7479097909548 | 58.7079471395453 | 24.9858741733473 | 56.4473293627624 | Wnt5b |
| ENSMUSG00000030172 | 0.151638128328707 | 0.533490770764943 | 0.909225374960129 | 793.285043147622 | 1040.82330155251 | 846.130670451711 | 608.714424573547 | 635.075753575076 | 505.132961846504 | 844.314331441027 | 975.887482636988 | Erc1 |
| ENSMUSG00000030177 | 0.0620342154334398 | 0.74662700969757 | 0.956085046483146 | 76.7986659083688 | 73.2974156022896 | 79.5022106464695 | 72.2203554578784 | 64.6025335533267 | 78.277262852727 | 84.3273253350471 | 61.8749571861049 | Ccdc77 |
| ENSMUSG00000030180 | -0.0087322614892282 | 0.937024513486174 | 0.988852035581444 | 459.888481733644 | 576.606336071345 | 514.871459424755 | 524.301022090312 | 510.250519251699 | 503.90987961443 | 539.278450908079 | 535.164103381574 | Kdm5a |
| ENSMUSG00000030188 | -0.0091764633342490 | 0.967825190574927 | 0.994406701590602 | 93.0619128066116 | 86.3280672649188 | 86.1273948670087 | 109.737423228205 | 118.255485148462 | 111.300483118721 | 80.1630129728225 | 68.388110574116 | Magohb |
| ENSMUSG00000030189 | 0.0905752884698241 | 0.589218530963202 | 0.92352104528132 | 5121.11574551335 | 4545.25418307087 | 5714.69461765932 | 6038.37205763404 | 5413.47332012961 | 6137.42664054663 | 4529.73077200975 | 4035.98404943751 | Ybx3 |
| ENSMUSG00000030199 | -0.0395888671948594 | 0.792241777678585 | 0.961804768755116 | 243.948703473642 | 277.715763559786 | 248.917635714542 | 196.964605794214 | 247.460552255116 | 225.04713070159 | 251.940897914585 | 270.295865602458 | Etv6 |
| ENSMUSG00000030201 | 0.182873212089836 | 0.431747346744966 | 0.886559861988535 | 1161.91863950779 | 1461.06181767231 | 1335.44784788296 | 1010.14704971604 | 906.625386138212 | 735.07242147639 | 1370.05876717188 | 1364.50563478831 | Lrp6 |
| ENSMUSG00000030203 | 0.23174771723212 | 0.151272051416542 | 0.783412422888014 | 829.425591810383 | 890.970807432275 | 887.774685552243 | 799.113543507953 | 631.790878987619 | 589.5256358596 | 762.069162287092 | 917.269102144889 | Dusp16 |
| ENSMUSG00000030204 | -0.0530517745891718 | 0.705891635376003 | 0.947784902404319 | 810.451803762433 | 742.747144769868 | 880.203046443056 | 875.085605742864 | 971.227919691539 | 962.565716642128 | 764.151318468204 | 734.90080728058 | Ddx47 |
| ENSMUSG00000030207 | -0.153874051878035 | 0.342162658255419 | 0.859912121266038 | 171.667606148119 | 219.077831077954 | 167.522515290775 | 181.019851991825 | 195.997517051618 | 193.24699266767 | 244.653351280692 | 188.88144825232 | Fam234b |
| ENSMUSG00000030208 | -0.807877594854407 | 0.0237105560912003 | 0.654124840968651 | 518.616873310632 | 554.617111390658 | 469.44162476963 | 596.521377548191 | 1127.80694169367 | 1610.79929964127 | 482.019155927491 | 525.394373299558 | Emp1 |
| ENSMUSG00000030213 | -0.0497990548984198 | 0.827257269736086 | 0.967319591893432 | 952.303457263773 | 1098.64681830543 | 1048.67201662248 | 961.374861614615 | 855.162350934715 | 682.479885497214 | 1349.23720536075 | 1315.65698437823 | Atf7ip |
| ENSMUSG00000030214 | 0.285931059091514 | 0.0063281368451377 | 0.635862253526407 | 1935.32638089089 | 1663.03691844306 | 1894.80268707419 | 1965.89435116511 | 1416.87590538991 | 1511.72963884329 | 1497.07029421972 | 1691.24882975353 | Plbd1 |
| ENSMUSG00000030216 | 0.0873977242587587 | 0.524471288313156 | 0.907437243979788 | 1449.23600137675 | 1521.32858161197 | 1341.12657721485 | 1303.71810501884 | 1301.90529482891 | 1062.85845967218 | 1410.66081270357 | 1508.88053488923 | Wbp11 |
| ENSMUSG00000030217 | -0.274998195563006 | 0.116508413905571 | 0.769868040577702 | 270.150601254144 | 314.364471360931 | 310.437203476691 | 284.191788360223 | 331.772333333186 | 435.417274618294 | 392.486440139664 | 268.124814473121 | Art4 |
| ENSMUSG00000030218 | -0.675026509371113 | 0.146932185863233 | 0.780607990324772 | 1263.11217576352 | 777.767021113184 | 654.000328056077 | 977.319615417004 | 1551.55576347566 | 3049.14400456013 | 719.38496057429 | 543.848307898922 | Mgp |
| ENSMUSG00000030220 | -0.234392804626515 | 0.312566447577844 | 0.852613848149207 | 256.597895505609 | 229.665235553841 | 300.972654590206 | 301.07446885687 | 362.431162816121 | 431.748027922072 | 216.544242835676 | 270.295865602458 | Arhgdib |
| ENSMUSG00000030222 | -0.365227226715502 | 0.335201182144835 | 0.858570960161243 | 50.5967681278665 | 26.0613033252585 | 60.5731128735006 | 46.8963347129081 | 91.9764884488041 | 68.4926049961361 | 38.5198893505771 | 37.9933947633978 | Rerg |
| ENSMUSG00000030223 | 0.0164079672733087 | 0.956425021860736 | 0.992371505557225 | 34.3335212296237 | 85.5136515360045 | 79.5022106464695 | 72.2203554578784 | 68.9823663366031 | 75.8310983885793 | 69.7522320672611 | 54.2762782334254 | Ptpro |
| ENSMUSG00000030224 | 0.160704401190882 | 0.0541013735397292 | 0.703396849069134 | 4082.97848517551 | 3950.73070096341 | 4021.48682186725 | 4230.98731779857 | 3537.8099306915 | 3346.35298695408 | 3854.07109123882 | 3828.64866658583 | Strap |
| ENSMUSG00000030225 | 0.0436924385056702 | 0.81771866871109 | 0.966315773308182 | 106.614618555147 | 135.193010999779 | 95.5919437534931 | 89.1030359545253 | 99.6411958195378 | 100.292743030056 | 108.272121417838 | 106.381505337514 | Dera |
| ENSMUSG00000030231 | 0.127219406000637 | 0.576218071940298 | 0.922450244597442 | 306.291149916906 | 350.198763433161 | 367.224496795597 | 250.426427366929 | 274.834507150593 | 190.800828203522 | 360.213019332423 | 339.769501741243 | Plekha5 |
| ENSMUSG00000030232 | 0.300829046855421 | 0.0981813281354244 | 0.759800953500741 | 1432.97275447851 | 1478.97896370842 | 1442.39725030023 | 1310.28359187865 | 976.702710670634 | 856.157562451702 | 1358.60690817576 | 1405.75560624572 | Aebp2 |
| ENSMUSG00000030236 | -0.417310787764377 | 0.138429858490581 | 0.779711473022872 | 333.396561413978 | 345.312269059675 | 318.955297474527 | 392.053358199911 | 367.905953795217 | 470.886659348436 | 275.885693997376 | 741.413960668591 | Slco1b2 |
| ENSMUSG00000030237 | -0.392791752647891 | 0.436846520259386 | 0.888701139843674 | 4.51756858284523 | 8.95857301805762 | 15.1432782183751 | 9.37926694258161 | 18.6142893289246 | 9.78465785659088 | 10.4107809055614 | 10.8552556466851 | Slco1a4 |
| ENSMUSG00000030243 | 0.152179506200489 | 0.29023787136985 | 0.851785577030345 | 180.702743313809 | 190.573280565953 | 199.701981504822 | 190.399118934407 | 175.193311331055 | 140.654456688494 | 178.024353485099 | 189.966973816989 | Recql |
| ENSMUSG00000030244 | -0.570350279316055 | 0.270026147988462 | 0.843793744782845 | 11.7456783153976 | 23.6180561385155 | 9.46454888648447 | 17.8206071909051 | 15.3294147414674 | 13.4539045528125 | 17.6983275394543 | 46.6775992807458 | Gys2 |
| ENSMUSG00000030245 | 0.0493176370480432 | 0.722271144424458 | 0.951674101666722 | 285.510334435818 | 269.571606270643 | 240.399541716705 | 242.923013812864 | 256.220217821669 | 288.647406769431 | 224.872867560126 | 235.559047533066 | Golt1b |
| ENSMUSG00000030246 | -0.0464697965732368 | 0.704003233941368 | 0.946931219685582 | 41699.8685608111 | 35005.2168601957 | 37940.5371212503 | 41612.0557174576 | 41477.0164576274 | 46568.8559862122 | 36240.9694103497 | 37086.9809168996 | Ldhb |
| ENSMUSG00000030247 | -0.605942564698554 | 0.0722454512887189 | 0.72957913253658 | 473.44118748218 | 297.26174105373 | 356.813493020464 | 492.411514485535 | 952.613630362614 | 770.541806206532 | 330.021754706295 | 412.499714574033 | Kcnj8 |
| ENSMUSG00000030249 | 0.184127259890518 | 0.421344028299336 | 0.882568531013176 | 4498.59479479728 | 5614.58203513538 | 6734.97298762235 | 4211.29085721914 | 4224.34871947008 | 3173.89839223167 | 5430.26332034081 | 5706.60789346234 | Abcc9 |
| ENSMUSG00000030254 | -0.259485352860092 | 0.483557880193548 | 0.89863290032536 | 17.1667606148119 | 15.4738988493722 | 19.8755526616174 | 13.1309737196143 | 22.994122112201 | 19.5693157131818 | 23.9447960827911 | 11.9407812113536 | Rad18 |
| ENSMUSG00000030255 | 0.158913800015174 | 0.486320937061572 | 0.899236955296196 | 5827.66347187034 | 6345.92735970045 | 5474.29507594262 | 4793.74333435346 | 3935.27975577383 | 3444.19956551999 | 6141.31965619065 | 6579.37044745582 | Sspn |
| ENSMUSG00000030256 | -0.0531643476229035 | 0.791201497683833 | 0.961804768755116 | 564.696072855653 | 456.887223920938 | 487.42426765395 | 648.107345732389 | 476.306815181307 | 426.855698993777 | 651.714884688142 | 681.710054611823 | Bhlhe41 |
| ENSMUSG00000030257 | 0.19722908242155 | 0.684727027414418 | 0.944452327903597 | 18.9737880479499 | 11.4018202048006 | 9.46454888648447 | 8.44134024832345 | 8.75966556655277 | 13.4539045528125 | 13.5340151772298 | 6.51315338801105 | Srgap3 |
| ENSMUSG00000030259 | 0.234020836923452 | 0.31360804402464 | 0.85295001595893 | 718.293404672391 | 987.886279173081 | 877.36368177711 | 668.741733006069 | 547.479097909548 | 501.463715150282 | 797.465817366001 | 918.354627709557 | Rassf8 |
| ENSMUSG00000030263 | -0.165895171993971 | 0.680474987553798 | 0.944452327903597 | 10.8421645988285 | 14.6594831204579 | 14.1968233297267 | 16.8826804966469 | 20.8042057205628 | 18.3462334811079 | 9.36970281500523 | 15.1973579053591 | Lrmp |
| ENSMUSG00000030264 | 0.342008117041003 | 0.0048299749682186 | 0.635862253526407 | 680.345828576491 | 649.08933594472 | 620.874406953381 | 672.493439783102 | 548.574056105368 | 426.855698993777 | 530.94982618363 | 561.216716933618 | Thumpd3 |
| ENSMUSG00000030265 | -0.0009802714659909 | 0.9934929702436 | 0.999133450854041 | 973.084272744862 | 1010.68991958268 | 979.580809751142 | 914.478526901707 | 897.865720571659 | 839.034411202668 | 1054.61210573337 | 1087.69661579784 | Kras |
| ENSMUSG00000030268 | -1.68957552613591 | 0.0285635930760169 | 0.665956123973759 | 7.22810973255236 | 4.88649437348597 | 3.78581955459379 | 4.68963347129081 | 20.8042057205628 | 33.0232202659942 | 1.04107809055614 | 11.9407812113536 | Bcat1 |
| ENSMUSG00000030269 | 0.179750806636725 | 0.201466209629198 | 0.813395633194566 | 482.47632464787 | 548.101785559343 | 555.569019636638 | 471.777127211855 | 515.725310230795 | 373.040080782527 | 484.101312108604 | 442.894430384751 | Mtmr14 |
| ENSMUSG00000030271 | 0.0201603264957304 | 0.931353859321404 | 0.987775860606096 | 116.553269437407 | 78.9983257046899 | 71.930571537282 | 108.799496533947 | 101.831112211176 | 69.71568722821 | 102.025652874501 | 96.6117752554972 | Ogg1 |
| ENSMUSG00000030272 | -0.0964176982950624 | 0.481932608205721 | 0.897732059433541 | 1488.99060490579 | 1105.97655986566 | 1082.74439261382 | 1278.39408427387 | 1321.61454235365 | 1451.79860947167 | 1251.37586484848 | 1274.40701292083 | Camk1 |
| ENSMUSG00000030275 | 0.116783941002449 | 0.56153104522266 | 0.917031459819748 | 889.961010820509 | 982.999784799595 | 1040.15392262464 | 759.720622349111 | 840.927894389066 | 572.402484610566 | 1009.84574783945 | 962.861175860966 | Etnk1 |
| ENSMUSG00000030278 | -1.67441893664781 | 0.0127124420708495 | 0.635862253526407 | 4.51756858284523 | 11.4018202048006 | 4.73227444324223 | 10.3171936368398 | 12.0445401540101 | 58.7079471395453 | 8.32862472444909 | 20.6249857287016 | Cidec |
| ENSMUSG00000030279 | 0.271465691409843 | 0.268970745708489 | 0.843793744782845 | 262.922491521592 | 302.148235427216 | 291.508105703722 | 257.929840920994 | 165.338687568684 | 160.223772401676 | 281.091084450157 | 315.887939318536 | C2cd5 |
| ENSMUSG00000030281 | -0.287058118171812 | 0.25094241069652 | 0.836703579022786 | 104.807591122009 | 105.059629029948 | 84.2344850897118 | 146.316564304273 | 158.768938393769 | 167.562265794119 | 100.984574783945 | 110.723607596188 | Il17rc |
| ENSMUSG00000030282 | 0.14786276417632 | 0.159662366150352 | 0.790303302997254 | 1030.90915060528 | 953.680818558679 | 1128.17422726895 | 1181.78763476528 | 946.0438811877 | 982.135032355309 | 1009.84574783945 | 936.808562308922 | Cmas |
| ENSMUSG00000030283 | 0.291732043597146 | 0.353998588355751 | 0.86309292474309 | 36.1405486627618 | 37.4631235300591 | 40.6975602118832 | 40.3308478531009 | 25.1840385038392 | 37.9155491942896 | 44.7663578939139 | 18.4539345993646 | St8sia1 |
| ENSMUSG00000030284 | -0.106585269376061 | 0.473123528608107 | 0.89728597737823 | 1308.28786159198 | 1165.4289080764 | 1087.47666705707 | 1418.14516171834 | 1411.40111441082 | 1577.77607937528 | 1218.06136595068 | 1154.99920080729 | Creld1 |
| ENSMUSG00000030286 | 0.0107280910987563 | 0.883275023072189 | 0.97910597465636 | 1448.33248766018 | 1402.42388519047 | 1454.70116385266 | 1483.80003031641 | 1507.7574356429 | 1429.78312929434 | 1398.16787561689 | 1410.09770850439 | Emc3 |
| ENSMUSG00000030287 | -0.20159268315797 | 0.393082048210851 | 0.878770935869648 | 359.59845919448 | 373.002403842763 | 492.156542097192 | 341.405316709971 | 597.847174917227 | 529.594606487981 | 331.062832796851 | 343.026078435248 | Itpr2 |
| ENSMUSG00000030291 | 0.138909536710563 | 0.395241161451297 | 0.879215782741314 | 499.643085262682 | 356.714089264476 | 367.224496795597 | 484.908100931469 | 337.247124312282 | 398.724807656078 | 407.061533407449 | 408.157612315359 | Med21 |
| ENSMUSG00000030298 | -0.0364010847652884 | 0.761965764518353 | 0.957666649379949 | 963.145621862602 | 830.704043492615 | 803.540200462531 | 923.857793844289 | 854.067392738896 | 1038.39681503071 | 848.478643803251 | 871.677028428812 | Sec13 |
| ENSMUSG00000030301 | 0.168125167431132 | 0.0840631125846955 | 0.742565114108635 | 1092.34808333198 | 1046.52421165491 | 1161.30014837164 | 1084.24325856243 | 1005.17162376193 | 864.719138076219 | 1021.29760683557 | 1008.45324957704 | Ccdc91 |
| ENSMUSG00000030304 | 0.0427463428391986 | 0.755766151654851 | 0.956566265296272 | 756.240980768291 | 727.273245920495 | 708.894711597687 | 733.458674909882 | 651.500126512363 | 598.087211484117 | 727.71358529874 | 861.907298346795 | Ergic2 |
| ENSMUSG00000030306 | 0.602331380483674 | 0.0354301807430253 | 0.679566075569698 | 2571.4000373555 | 4386.44311593257 | 6312.85410728514 | 3307.12952395428 | 2518.40385038392 | 2007.0779428332 | 3541.74766407198 | 2851.67565838417 | Tmtc1 |
| ENSMUSG00000030309 | 0.431783074727314 | 0.165616741691438 | 0.795216960550936 | 76.7986659083688 | 79.8127414336042 | 109.78876708322 | 71.2824287636203 | 45.9882442244021 | 42.8078781225851 | 100.984574783945 | 59.7039060567679 | Caprin2 |
| ENSMUSG00000030313 | -0.0548309104741372 | 0.638156951804112 | 0.933198322011444 | 387.60738440812 | 412.09435883065 | 385.207139679918 | 358.287997206618 | 448.93286028583 | 399.947889888152 | 364.377331694648 | 390.789203280663 | Dennd5b |
| ENSMUSG00000030314 | 0.0484446659322742 | 0.622156983723898 | 0.93078489223231 | 505.064167562096 | 517.968403589513 | 528.121827865833 | 532.742362338636 | 504.775728272604 | 549.163922201163 | 496.594249195277 | 465.69046724279 | Atg7 |
| ENSMUSG00000030315 | -0.234446959047574 | 0.213838867011312 | 0.824248286808591 | 196.965990212052 | 198.717437855096 | 225.25626349833 | 209.15765281957 | 308.778211220985 | 273.970419984545 | 195.722681024554 | 198.651178334337 | Vgll4 |
| ENSMUSG00000030316 | -0.0255276668137305 | 0.86278466548621 | 0.974868041734905 | 183.413284463516 | 169.39847161418 | 205.380710836713 | 214.785212985119 | 223.371471947096 | 189.577745971448 | 191.558368662329 | 181.282769299641 | Tamm41 |
| ENSMUSG00000030317 | -0.328794955460237 | 0.255489458106049 | 0.837479467570303 | 1491.70114605549 | 1255.01463825698 | 1459.43343829591 | 1296.21469146478 | 2527.16351595048 | 2196.65568880465 | 985.900951756661 | 1201.67680008804 | Timp4 |
| ENSMUSG00000030319 | 0.108875923122132 | 0.475624290785343 | 0.897549741595057 | 2655.42681299642 | 3208.79797192245 | 2747.55854174644 | 2356.0718559765 | 2396.863490648 | 2100.03219247082 | 2738.03537816264 | 2935.26112686364 | Cand2 |
| ENSMUSG00000030322 | -0.760146587667293 | 0.0869404238082608 | 0.747269672026173 | 7.22810973255236 | 12.2162359337149 | 15.1432782183751 | 10.3171936368398 | 25.1840385038392 | 25.6847268735511 | 12.4929370866736 | 13.0263067760221 | Mbd4 |
| ENSMUSG00000030323 | 0.151755593699797 | 0.419807849081807 | 0.882568531013176 | 171.667606148119 | 162.883145782866 | 204.434255948065 | 131.309737196143 | 180.668102310151 | 143.100621152642 | 142.627698406191 | 136.776221148232 | Ift122 |
| ENSMUSG00000030327 | 0.00969531884217815 | 0.943883277549784 | 0.990505437194639 | 332.493047697409 | 344.497853330761 | 301.919109478855 | 302.950322245386 | 315.3479603959 | 266.631926592101 | 323.775286162958 | 366.907640857956 | Necap1 |
| ENSMUSG00000030329 | -0.876503762669117 | 0.230861870953258 | 0.832048466795696 | 10.8421645988285 | 7.32974156022896 | 2.83936466594534 | 13.1309737196143 | 22.994122112201 | 28.1308913376988 | 2.08215618111227 | 9.76973008201657 | Pianp |
| ENSMUSG00000030330 | -0.133574810842228 | 0.806814594213691 | 0.964026716319739 | 18.0702743313809 | 19.5459774939439 | 8.51809399783602 | 15.9447538023887 | 14.2344565456483 | 3.66924669622158 | 26.0269522639034 | 23.8815624227072 | Ing4 |
| ENSMUSG00000030335 | -0.0166575480036605 | 0.912887673754248 | 0.986226496507248 | 4620.5691465341 | 3727.58079124088 | 3561.50974598411 | 4397.93826937652 | 4114.85289988817 | 5023.19872712734 | 3703.11476810818 | 3657.1356273682 | Mrpl51 |
| ENSMUSG00000030337 | 0.0578204441666183 | 0.786790589415898 | 0.961141387070613 | 112.035700854562 | 151.481325578065 | 128.717864856189 | 131.309737196143 | 93.0714466446232 | 113.746647582869 | 164.49033830787 | 131.348593324889 | Vamp1 |
| ENSMUSG00000030339 | -0.200301476745556 | 0.108553203937823 | 0.762287160109383 | 650.529875929712 | 607.554133770089 | 599.105944514467 | 628.410885152968 | 781.800151814835 | 803.565026472526 | 616.318229609233 | 655.657441059779 | Ltbr |
| ENSMUSG00000030340 | -0.0004444565943271 | 0.999158090886774 | 0.999709467616704 | 12.6491920319666 | 18.7315617650296 | 15.1432782183751 | 8.44134024832345 | 13.1394983498292 | 13.4539045528125 | 17.6983275394543 | 10.8552556466851 | Scnn1a |
| ENSMUSG00000030341 | -0.587302161525892 | 0.0356190606875927 | 0.679566075569698 | 592.704998069294 | 712.613762800038 | 682.39397471553 | 849.761584997894 | 1354.46328822822 | 1504.39114545085 | 674.618602680377 | 730.558705021906 | Tnfrsf1a |
| ENSMUSG00000030342 | -0.362646815178727 | 0.354548362981544 | 0.863771056173629 | 681.24934229306 | 523.669313691913 | 756.217456030109 | 506.480414899407 | 1017.21616391594 | 1367.40593545857 | 362.295175513536 | 426.611546914723 | Cd9 |
| ENSMUSG00000030347 | 0.194036279939288 | 0.218549104731112 | 0.827129763070499 | 271.957628687283 | 227.221988367098 | 219.57753416644 | 270.12288794635 | 223.371471947096 | 174.900759186562 | 222.790711379013 | 242.072200921077 | D6Wsu163e |
| ENSMUSG00000030351 | -0.535051701242975 | 0.487761327341514 | 0.899236955296196 | 8.13162344912141 | 9.77298874697194 | 8.51809399783602 | 9.37926694258161 | 29.5638712871156 | 17.123151249034 | 1.04107809055614 | 4.34210225867403 | Tspan11 |
| ENSMUSG00000030352 | -0.249659342375016 | 0.190916951970256 | 0.808824684419004 | 1212.51540763566 | 1238.72632367869 | 748.645816920921 | 919.168160372998 | 1204.45401540101 | 1276.89785028511 | 1044.20132482781 | 1373.18983930566 | Tspan9 |
| ENSMUSG00000030353 | -0.101806870777353 | 0.654065189488197 | 0.937077409179068 | 51.5002818444356 | 52.9370223794314 | 63.4124775394459 | 48.7721881014244 | 68.9823663366031 | 56.2617826753975 | 62.4646854333682 | 44.5065481514088 | Tead4 |
| ENSMUSG00000030357 | -0.0537918933442437 | 0.633832409939869 | 0.9325328237529 | 11305.6671354285 | 9474.09817446039 | 8468.8783436263 | 9892.31284434083 | 9820.68005830148 | 10410.8759594127 | 9816.32531585381 | 10580.6176788239 | Fkbp4 |
| ENSMUSG00000030359 | -0.544454304024133 | 0.207776642319784 | 0.820372930586885 | 248.466272056487 | 623.842448348376 | 252.703455269135 | 332.963976461647 | 261.695008800764 | 365.701587390084 | 475.772687384154 | 1023.6506074824 | Pzp |
| ENSMUSG00000030374 | 0.119039768974306 | 0.435822898764101 | 0.888687115594624 | 1320.03353990737 | 1447.21675028076 | 1467.00507740509 | 1284.02164443942 | 1163.9405621557 | 1001.70434806849 | 1434.60560878636 | 1479.57134464318 | Strn4 |
| ENSMUSG00000030376 | 0.0692529211332295 | 0.805196524600752 | 0.963475137341661 | 40.658117245607 | 48.0505280059454 | 27.447191770805 | 37.5170677703264 | 36.1336204620302 | 45.2540425867328 | 29.1501865355718 | 36.9078691987293 | Slc8a2 |
| ENSMUSG00000030378 | -0.344147590867746 | 0.300708544900577 | 0.851785577030345 | 282.799793286111 | 236.180561385155 | 224.309808609682 | 303.888248939644 | 208.042057205628 | 363.255422925936 | 192.599446752885 | 565.558819192293 | Sult2a8 |

| | | | | | | | | | | | | |
|---|---|---|---|---|---|---|---|---|---|---|---|---|
| **ENSMUSG00000030382** | -0.388127322007148 | 0.282654608681175 | 0.850587941933635 | 108.421645988285 | 211.748089517725 | 99.3777633080869 | 134.123517278917 | 117.160526952643 | 143.100621152642 | 148.874166949528 | 315.887939318536 | Slc27a5 |
| **ENSMUSG00000030386** | 0.136407912710278 | 0.513467005082454 | 0.903134950056386 | 154.500845533307 | 159.625482867208 | 153.325691961048 | 208.219726125312 | 113.875652365186 | 139.43137445642 | 169.69572876065 | 191.052499381657 | Zfp606 |
| **ENSMUSG00000030393** | -0.131139320701505 | 0.69002663460976 | 0.94469184542218 | 15.3597331816738 | 20.3603932228582 | 23.6613722162112 | 25.3240207449704 | 24.0890803080201 | 17.123151249034 | 24.9858741733473 | 26.0526135520442 | Zik1 |
| **ENSMUSG00000030397** | 0.195085440359992 | 0.166710976424274 | 0.796745160868404 | 992.961574509381 | 1093.76032393194 | 1265.41018612297 | 1134.89130005238 | 1067.58424092362 | 787.664957455566 | 1018.1743725639 | 1044.2755932111 | Mark4 |
| **ENSMUSG00000030399** | 0.0396694445512383 | 0.724920710263331 | 0.951674101666722 | 67170.8237446091 | 55912.8974528843 | 58751.1872128523 | 66381.7617861214 | 67563.3005148216 | 60987.7724201309 | 55007.4430707146 | 57925.8151818409 | Ckm |
| **ENSMUSG00000030400** | -0.0750437185432012 | 0.657945734120637 | 0.938763867173667 | 166.246523848704 | 179.171460361152 | 194.023252172932 | 182.895705380341 | 163.148771177045 | 167.562265794119 | 188.435134390661 | 240.986675356409 | Ercc2 |
| **ENSMUSG00000030401** | 0.0130905133547828 | 0.939361950040651 | 0.989269716906553 | 2615.67220946739 | 2301.53884991189 | 2878.16931637993 | 3176.75771345239 | 3119.53589988861 | 3182.45996785618 | 2334.09707902686 | 2237.26818878179 | Rtn2 |
| **ENSMUSG00000030403** | -0.411071492133564 | 0.0579610566102763 | 0.710862211775517 | 341.528184863099 | 333.910448854875 | 362.492222352355 | 398.618845059719 | 539.814390538815 | 647.010500767072 | 374.788112600209 | 349.539231823259 | Vasp |
| **ENSMUSG00000030407** | -0.0805872695900544 | 0.709736770117065 | 0.949571160524938 | 77.7021796249379 | 79.8127414336042 | 76.6628459805242 | 73.1582821521366 | 100.736154015357 | 57.4848649074714 | 73.9165444294857 | 92.2696729968231 | Qpctl |
| **ENSMUSG00000030409** | 0.0767685658479693 | 0.686663284556358 | 0.944452327903597 | 13143.4140349299 | 13833.6655713388 | 11365.9767577792 | 11250.4306976266 | 8975.37233112914 | 9752.85771855696 | 15067.523204619 | 13227.1290054858 | Dmpk |
| **ENSMUSG00000030410** | 0.0923394057402922 | 0.376897156899502 | 0.87422797717331 | 1821.48365260319 | 1675.25315437677 | 1966.73325861147 | 1824.26742033212 | 1824.20035423462 | 1818.72327909383 | 1666.76602298037 | 1526.24894392392 | Dmwd |
| **ENSMUSG00000030411** | -0.566681106855809 | 0.0664123792725665 | 0.719725956118125 | 89.4478579403355 | 103.43079757212 | 91.8061241988993 | 85.3513291774927 | 201.472308030714 | 172.454594722414 | 97.8613405122768 | 77.072315091464 | Nova2 |
| **ENSMUSG00000030413** | -0.994565615398707 | 0.221027732493937 | 0.829974372625247 | 7.22810973255236 | 0.81441572891432§ | 15.1432782183751 | 10.3171936368398 | 22.994122112201 | 33.0232202659942 | 5.20539045278068 | 5.42762782334254 | Pglyrp1 |
| **ENSMUSG00000030417** | 0.112533518308646 | 0.609536111488814 | 0.927932112008768 | 2195.53833126278 | 1501.78260411802 | 2012.1630932666 | 2440.48525845974 | 1851.57430913009 | 2585.59583860414 | 1614.71211845257 | 1487.17002359586 | Pdcd5 |
| **ENSMUSG00000030421** | 0.273466786175055 | 0.173204813277422 | 0.799889021902745 | 369.537110076739 | 430.011504866766 | 441.047978110176 | 413.625672167849 | 289.068963696242 | 244.616446414772 | 407.061533407449 | 426.611546914723 | Uri1 |
| **ENSMUSG00000030423** | 0.00906067054356227 | 0.947964449652977 | 0.991192530276814 | 245.75573090678 | 224.778741180355 | 282.990011705886 | 256.053987532478 | 273.739548954774 | 269.078091056249 | 225.913945650682 | 234.473521968398 | Pop4 |
| **ENSMUSG00000030424** | 0.488247275767188 | 0.196896695211534 | 0.812065429583558 | 30.7194663633475 | 23.6180561385155 | 32.1794662140472 | 21.5723139679377 | 22.994122112201 | 9.78465785659088 | 15.616171358342 | 28.2236646813812 | Zfp939 |
| **ENSMUSG00000030428** | -0.787145747397084 | 0.172635857658773 | 0.799889021902745 | 13.5527057485357 | 6.51532583131463 | 12.3039135524298 | 22.5102406621959 | 39.4184950494875 | 33.0232202659942 | 14.5750932677859 | 7.59867895267955 | Ttyh1 |
| **ENSMUSG00000030431** | 0.023446621307822 | 0.935605833665782 | 0.98818087308472 | 122.87786545339 | 100.987550385377 | 120.199770858353 | 151.944124469822 | 129.205067106653 | 189.577745971448 | 85.3684034256032 | 84.6709940441436 | Tmem238 |
| **ENSMUSG00000030432** | -0.0639372422765158 | 0.775746830003206 | 0.9595799687189 | 7949.11367837446 | 5808.41297861699 | 7037.83855198985 | 8249.06527600053 | 7939.54187788427 | 10690.9617905576 | 6033.04753477281 | 5697.923688945 | Rpl28 |
| **ENSMUSG00000030433** | -0.343692623498192 | 0.248526494091515 | 0.834621660998471 | 56.0178504272808 | 96.9154717408051 | 123.985590412947 | 69.4065753751039 | 112.780694169367 | 147.992950080937 | 96.8202624217207 | 82.4999429148066 | Sbk2 |
| **ENSMUSG00000030435** | -0.0101089714622749 | 0.911337656134481 | 0.986226496507248 | 1221.55054480135 | 1232.21099784738 | 1356.26985543322 | 1206.173728816 | 1356.65320461986 | 1204.73599859275 | 1277.40281711238 | 1211.44653017005 | U2af2 |
| **ENSMUSG00000030443** | -0.441155021563251 | 0.365474403717806 | 0.86924083181804 | 17.1667606148119 | 6.51532583131463 | 9.46454888648447 | 11.2551203310979 | 16.4243729372865 | 15.9000690169602 | 7.28754663389296 | 20.6249857287016 | Zfp583 |
| **ENSMUSG00000030446** | -0.248365263990602 | 0.474966216036081 | 0.897549741595057 | 18.9737880479499 | 17.1027303072009 | 31.2330113253987 | 33.7653609932938 | 25.1840385038392 | 28.1308913376988 | 26.0269522639034 | 40.1644458927348 | Zfp273 |
| **ENSMUSG00000030447** | -0.185630367935054 | 0.135797972795572 | 0.776087404131513 | 669.503663977662 | 716.685841444609 | 779.87882824632 | 668.741733006069 | 928.524550054594 | 820.68817772156 | 764.151318468204 | 711.019244857872 | Cyfip1 |
| **ENSMUSG00000030451** | 0.25965551854403 | 0.226659453565208 | 0.832048466795696 | 1171.85729039005 | 1687.46939031049 | 1555.0253820494 | 1157.40154071457 | 1150.80106380587 | 795.003450848009 | 1370.05876717188 | 1337.3674956716 | Herc2 |
| **ENSMUSG00000030452** | 0.0939535526218145 | 0.716138642399349 | 0.951348124093194 | 383.993329541844 | 520.411650776256 | 471.334534546927 | 399.556771753977 | 336.152166116463 | 255.624186503437 | 575.716184077544 | 494.999657488839 | Nipa2 |
| **ENSMUSG00000030465** | -0.115145908693037 | 0.695758624972606 | 0.944933386973863 | 352.370349461928 | 554.617111390658 | 391.832323900457 | 289.819348525772 | 294.543754675337 | 302.101311322243 | 602.784214432003 | 521.052271040884 | Psd3 |
| **ENSMUSG00000030469** | -0.0467992471040637 | 0.823041051217463 | 0.966637987288264 | 69.5705561758165 | 57.8235167529173 | 50.1621090983677 | 61.9031618210386 | 60.2227007700503 | 53.8156182112498 | 64.5468416144805 | 68.388110574116 | Zfp719 |
| **ENSMUSG00000030470** | 0.106782590518235 | 0.501492416611796 | 0.900927059989498 | 36946.4828979414 | 26132.971909403 | 25993.4370618409 | 37562.0882516508 | 28487.5273806254 | 30678.5716271086 | 29489.5779930931 | 28944.4536563211 | Csrp3 |
| **ENSMUSG00000030471** | 0.129571694045973 | 0.635333899653005 | 0.932543559775713 | 138.237598635064 | 175.913797445495 | 154.272146849697 | 113.489130005238 | 114.970610561005 | 78.277262852727 | 167.613572579538 | 170.427513652956 | Zdhhc13 |
| **ENSMUSG00000030474** | -0.00580633496174485 | 0.989937663190545 | 0.998628446692557 | 16.2632468982428 | 21.9892246806869 | 21.7684624389143 | 23.448167356454 | 20.8042057205628 | 37.9155491942896 | 7.28754663389296 | 18.4539345993646 | Siglece |
| **ENSMUSG00000030483** | 0.062467537018863 | 0.827438125380075 | 0.967319591893432 | 182.509770746947 | 93.6578088251478 | 97.48485353079 | 79.7237690119437 | 101.831112211176 | 107.6312364225 | 130.134761319517 | 94.4407241261602 | Cyp2b10 |
| **ENSMUSG00000030486** | -0.387866337508947 | 0.269174443395677 | 0.843793744782845 | 17.1667606148119 | 12.2162359337149 | 21.7684624389143 | 19.6964605794214 | 19.7092475247437 | 29.3539735697726 | 24.9858741733473 | 18.4539345993646 | Zfp108 |
| **ENSMUSG00000030493** | -0.75115128905079 | 0.0226963103527901 | 0.650119841310877 | 21.6843291976571 | 19.5459774939439 | 19.8755526616174 | 20.6343872736795 | 40.5134532453066 | 47.7002070508805 | 23.9447960827911 | 26.0526135520442 | Faap24 |
| **ENSMUSG00000030494** | -0.126695167424022 | 0.630644203628282 | 0.931825800800958 | 137.334084918495 | 140.079505373265 | 126.824955078892 | 120.054616865045 | 171.908436743598 | 203.031650524261 | 99.9434966933891 | 98.7828263848342 | Rhpn2 |
| **ENSMUSG00000030498** | -0.148208979940789 | 0.56079342176797 | 0.916762264997884 | 64.1494738764022 | 53.7514381083457 | 55.8408384302584 | 67.5307219865876 | 74.4571573156986 | 73.3849339244316 | 40.6020455316893 | 79.2433662208011 | Gas2 |
| **ENSMUSG00000030499** | -0.641631913732279 | 0.0464362322780927 | 0.69805115405055 | 39.754603529038 | 35.8342920722305 | 22.7149173275627 | 34.703287687552 | 68.9823663366031 | 64.8233582999146 | 34.3555769883525 | 40.1644458927348 | Kctd15 |
| **ENSMUSG00000030505** | 0.114712449280422 | 0.46065838636793 | 0.894774044236998 | 491.511461813561 | 487.020605890769 | 510.139184981513 | 480.218467460179 | 487.256397139498 | 332.67836712409 | 504.922873919726 | 491.743080794834 | Prmt3 |
| **ENSMUSG00000030509** | -0.0699321566486366 | 0.749320763441759 | 0.956197192463153 | 156.307872966445 | 159.625482867208 | 164.68315062483 | 142.564857527241 | 134.679858085749 | 116.192812047017 | 217.585320926233 | 184.539345993646 | Asb7 |
| **ENSMUSG00000030512** | -0.0657782580170972 | 0.697655796926204 | 0.944933386973863 | 439.107666252556 | 418.609684661965 | 471.334534546927 | 491.473587791276 | 503.680770076785 | 601.756458180339 | 378.952424962434 | 422.269444656049 | Snrpa1 |
| **ENSMUSG00000030513** | -0.928091323074283 | 0.0102682690919672 | 0.635862253526407 | 352.370349461928 | 420.238516119794 | 477.013263878817 | 451.142739938176 | 1224.16326292575 | 1163.15120270224 | 394.568596320776 | 454.835211596105 | Pcsk6 |
| **ENSMUSG00000030515** | 0.326300624224746 | 0.155519781289606 | 0.787275540343856 | 1204.38378418654 | 1530.28715463002 | 1535.14982938778 | 1182.72556145954 | 970.13296149572 | 692.264543353805 | 1352.36043963242 | 1333.02539341293 | Tarsl2 |
| **ENSMUSG00000030516** | -0.0606000587062896 | 0.622466584877317 | 0.93078489223231 | 1398.63923324888 | 1690.72705322615 | 1620.33076936614 | 1343.11102617769 | 1728.93899119835 | 1479.92950080937 | 1646.98553925981 | 1456.77530778514 | Tjp1 |
| **ENSMUSG00000030521** | 0.270817678806638 | 0.0609901645630375 | 0.71282992338528 | 481.572810931301 | 370.55915665602 | 417.386605893965 | 440.825546301336 | 328.487458745729 | 319.224462571277 | 351.884394607974 | 416.841816832707 | Mphosph10 |
| **ENSMUSG00000030522** | -0.175213753412808 | 0.4423072322187 | 0.89108031640712 | 330.68602026427 | 348.569931975333 | 301.919109478855 | 198.84045918273 | 419.368988998714 | 363.255422925936 | 268.598147363483 | 282.236646813812 | Mtmr10 |
| **ENSMUSG00000030527** | -0.0701625206716956 | 0.660913707688667 | 0.939611397743053 | 217.74680569314 | 252.468875963442 | 305.704929033448 | 268.247034557834 | 323.012667766634 | 269.078091056249 | 252.981976005141 | 250.756405438425 | Crtc3 |
| **ENSMUSG00000030528** | 0.209593486011397 | 0.589143449254712 | 0.92352104528132 | 18.9737880479499 | 44.7928650902881 | 36.9117406572894 | 18.7585338851632 | 24.0890803080201 | 23.2385624094033 | 34.3555769883525 | 21.7105112933701 | Blm |
| **ENSMUSG00000030530** | 0.154292067961771 | 0.498330826598374 | 0.900927059989498 | 4014.31144271627 | 4780.62032872711 | 4786.2223718952 | 3158.93710626149 | 3473.20739713818 | 2605.16515431732 | 5116.89881508341 | 3846.01707562052 | Furin |
| **ENSMUSG00000030532** | 0.0135086097337143 | 0.957360181989396 | 0.992403427270738 | 34.3335212296237 | 39.9063707168021 | 43.5369248778286 | 47.8342614071662 | 30.6588294829347 | 44.0309603546589 | 45.80743598447 | 43.4210225867403 | Hddc3 |
| **ENSMUSG00000030533** | -0.309451466539807 | 0.160076530504298 | 0.790986484775773 | 274.66816983699 | 290.746415222415 | 234.720812384815 | 253.240207449704 | 406.229490648885 | 406.063301048521 | 267.557069272927 | 226.874843015718 | Unc45a |
| **ENSMUSG00000030534** | 0.0188358150671622 | 0.903908396102967 | 0.985039825636533 | 111.132187137993 | 116.461449234749 | 127.77140996754 | 118.178763476528 | 118.255485148462 | 105.185071958352 | 131.175839410073 | 111.809133160856 | Vps33b |
| **ENSMUSG00000030536** | -0.468512905005814 | 0.023068406238995 | 0.65126049321065 | 496.932544112975 | 567.647763053287 | 538.532831640966 | 513.045901759214 | 959.183379537529 | 852.48831575548 | 546.565997541972 | 570.986447015635 | Iqgap1 |
| **ENSMUSG00000030538** | -0.158322250371894 | 0.4349940734127 | 0.888379878975956 | 310.808718499751 | 237.809392842984 | 245.131816159948 | 327.336416296098 | 301.113503850252 | 418.29412336926 | 262.351678820146 | 270.295865602458 | Cib1 |
| **ENSMUSG00000030539** | -0.0894270326394151 | 0.721348950713895 | 0.951674101666722 | 177.088688447533 | 223.149909722526 | 231.881447718869 | 151.944124469822 | 199.282391639076 | 143.100621152642 | 296.707255808499 | 194.309076075663 | Sema4b |
| **ENSMUSG00000030541** | 0.193072619214072 | 0.095771855624579 | 0.757475844708466 | 74202.871000666 | 63476.3763273117 | 69192.477544422 | 80580.0960838014 | 64175.4998569573 | 55607.433681238 | 67954.2902048707 | 63706.2388137007 | Idh2 |
| **ENSMUSG00000030545** | 0.244871091364646 | 0.110768282549535 | 0.764116514278315 | 1058.91807581892 | 1014.76199822725 | 1065.70820461815 | 1013.89875649307 | 768.660653465006 | 730.180092548094 | 891.162845516053 | 1113.74922934989 | Pex11a |
| **ENSMUSG00000030551** | -0.76845722470185 | 0.0146908890812183 | 0.635862253526407 | 147.272735800754 | 144.151584017836 | 154.272146849697 | 162.261318106662 | 346.006789878835 | 379.155491942897 | 160.326025945645 | 150.888053488923 | Nr2f2 |
| **ENSMUSG00000030554** | -0.0474404660766204 | 0.790692913974313 | 0.961804768755116 | 255.69438178904 | 374.631235300591 | 257.435729712378 | 328.274342990356 | 296.733671066975 | 302.101311322243 | 286.296474902938 | 372.335268681298 | Synm |
| **ENSMUSG00000030555** | -0.226590489119954 | 0.0793497556123432 | 0.739855948845396 | 150.886790667031 | 173.470550258752 | 171.308334845369 | 167.888878272211 | 203.662224422352 | 196.916239363891 | 188.435134390661 | 187.795922687652 | Ttc23 |
| **ENSMUSG00000030556** | 0.0102915923356439 | 0.955003590385536 | 0.992371505557225 | 177.992202164102 | 177.542628903324 | 124.932045301595 | 194.150825711439 | 179.573144114332 | 151.662196777159 | 184.270822028436 | 154.144630182928 | Lrrc28 |
| **ENSMUSG00000030557** | 0.0969029864490256 | 0.63430316412151 | 0.932543559775713 | 2916.54227708488 | 3734.91053280111 | 3434.68479090521 | 2408.59575085496 | 2480.08031353025 | 2277.37911612153 | 3419.94152747691 | 3505.16204831461 | Mef2a |
| **ENSMUSG00000030560** | -0.156966689188814 | 0.344572964917114 | 0.861266047293794 | 325.264937964856 | 393.362797065621 | 483.638448099356 | 461.459933575015 | 532.149683168081 | 428.078781225851 | 420.595548584679 | 473.289146195469 | Ctsc |
| **ENSMUSG00000030562** | -0.554118843894055 | 0.199906746560976 | 0.812921911423739 | 26.2018977805023 | 21.1748089517725 | 18.9290977729689 | 23.448167356454 | 24.0890803080201 | 64.8233582999146 | 18.7394056300105 | 24.9670879873757 | Nox4 |
| **ENSMUSG00000030579** | -0.394517191842702 | 0.230062971742195 | 0.832048466795696 | 74.9916384752307 | 52.122606650517 | 79.5022106464695 | 97.5443762028488 | 102.926070406995 | 160.223772401676 | 68.711153976705 | 68.388110574116 | Tyrobp |
| **ENSMUSG00000030583** | -0.207442253172114 | 0.338148417655821 | 0.858570960161243 | 92.1583990900426 | 105.874044758863 | 82.3415753124149 | 71.2824287636203 | 117.160526952643 | 107.6312364225 | 74.9576225200418 | 107.467030902182 | Sipa1l3 |
| **ENSMUSG00000030587** | -0.779267717131518 | 0.103250427896873 | 0.759800953500741 | 20.780815481088 | 17.9171460361152 | 13.2503684410783 | 21.5723139679377 | 32.8487458745729 | 61.154111603693 | 17.6983275394543 | 15.1973579053591 | 2200002D01I |
| **ENSMUSG00000030588** | -0.213331164763236 | 0.292548133273499 | 0.851785577030345 | 1325.45462220679 | 961.824975847822 | 1027.85000907221 | 1202.42202203896 | 1495.71289548889 | 1654.83025999593 | 1112.91247880451 | 974.80195707232 | Yif1b |
| **ENSMUSG00000030589** | -0.00519617888068355 | 0.990749515231116 | 0.998628446692557 | 11.7456783153976 | 11.4018202048006 | 9.46454888648447 | 10.3171936368398 | 13.1394983498292 | 8.56157562451702 | 7.28754663389296 | 14.1118323406906 | Rasgrp4 |
| **ENSMUSG00000030590** | -0.0714782019657869 | 0.629699726938951 | 0.931825800800958 | 245.75573090678 | 187.315617650296 | 212.005895057252 | 227.916186704733 | 231.036179317829 | 254.401104271363 | 232.160414194018 | 199.736703899005 | Fam98c |
| **ENSMUSG00000030591** | 0.11797954686851 | 0.273275364131081 | 0.844738346035895 | 2479.24163826546 | 2170.41791755669 | 2021.62764215308 | 2505.20220036355 | 2200.86597359638 | 2162.40938630658 | 2102.9777429234 | 1989.76836003737 | Psmd8 |
| **ENSMUSG00000030592** | -0.658813256665493 | 0.244521210559591 | 0.832048466795696 | 16.2632468982428 | 30.1333819698302 | 11.3574586637814 | 20.6343872736795 | 7.66470737073368 | 19.5693157131818 | 52.0539045278068 | 44.5065481514088 | Ryr1 |
| **ENSMUSG00000030595** | -0.135610477244857 | 0.404375688669891 | 0.881158207349486 | 553.853908256825 | 465.845796938996 | 490.263632319895 | 602.14893771374 | 594.56230032977 | 710.610776834912 | 537.196294726966 | 478.716774018812 | Nfkbib |
| **ENSMUSG00000030598** | -0.366813990702023 | 0.405786414084605 | 0.881158207349486 | 14.4562194651047 | 15.4738988493722 | 11.3574586637814 | 14.0689004138724 | 27.3739548954774 | 23.2385624094033 | 11.4518589961175 | 9.76973008201657 | Fbxo17 |
| **ENSMUSG00000030602** | 0.0341388742798286 | 0.79118969189225 | 0.961804768755116 | 187.930853046361 | 197.088606397268 | 198.755526616174 | 210.095579513828 | 174.098353135236 | 201.808568292187 | 207.174540020671 | 192.138024946326 | Pak4 |
| **ENSMUSG00000030603** | -0.0217861152556896 | 0.866254634114531 | 0.974868041734905 | 1759.14120615993 | 1457.80415475665 | 1562.59702115859 | 1863.66034149097 | 1705.94486908615 | 1905.56211757107 | 1644.9033830787 | 1488.25554916052 | Psmc4 |
| **ENSMUSG00000030604** | 0.0209734714911525 | 0.924920304205223 | 0.986713918026868 | 112.035700854562 | 71.6685841444609 | 80.448665535118 | 107.861569839689 | 108.400861386091 | 81.9465095489486 | 79.1219348822664 | 96.6117752554972 | Zfp626 |
| **ENSMUSG00000030605** | -0.391078986427013 | 0.12147516481861 | 0.769868040577702 | 2997.85851157609 | 3216.12771348268 | 2941.58179391937 | 3438.43926115042 | 5329.16153905154 | 5567.47032040021 | 2742.19969052486 | 2877.72827193621 | Mfge8 |
| **ENSMUSG00000030609** | 0.196973143156212 | 0.188795326399048 | 0.808677608936066 | 775.214768816241 | 772.066111010784 | 648.321598724186 | 875.085605742864 | 591.277425742312 | 591.971800323748 | 767.274552739873 | 727.3021283279 | Aen |
| **ENSMUSG00000030610** | 0.0811900214325308 | 0.482292178122435 | 0.897874058989495 | 502.353626412389 | 446.299819445052 | 493.102996985841 | 468.963347129081 | 458.787484048202 | 390.163232031561 | 488.265624470828 | 466.775992807458 | Det1 |
| **ENSMUSG00000030611** | -0.0705583107238459 | 0.677679812162245 | 0.944224354592595 | 468.020105182765 | 348.569931975333 | 405.082692341535 | 431.446279358754 | 428.128654565267 | 533.263853184203 | 346.679004155194 | 428.78259804406 | Mrps11 |
| **ENSMUSG00000030612** | -0.0230585916011436 | 0.853127509645638 | 0.973120292022893 | 860.145058173731 | 715.05700998678 | 641.696414503647 | 741.900015158206 | 696.393412540946 | 804.7881087046 | 752.699459472087 | 753.354741879944 | Mrpl46 |
| **ENSMUSG00000030613** | 0.27445448101469 | 0.117899919927756 | 0.769868040577702 | 874.601277638836 | 818.4878075589 | 926.579335986829 | 846.94780491512 | 609.891715071237 | 547.940839969089 | 833.903550535465 | 872.76255399348 | Ccdc90b |
| **ENSMUSG00000030614** | -0.0216288204401333 | 0.898632836293947 | 0.984540500469898 | 885.443442237664 | 808.714818811928 | 640.749959614998 | 799.113543507953 | 653.690042904001 | 699.603036746248 | 900.532548331058 | 927.038832226905 | Tmem126b |
| **ENSMUSG00000030615** | 0.0402891750000862 | 0.768253233818761 | 0.95879680569409 | 1573.92089426328 | 1207.77852597995 | 1147.10332504192 | 1458.47600957144 | 1203.35905720519 | 1296.46716599829 | 1352.36043963242 | 1386.21614608168 | Tmem126a |
| **ENSMUSG00000030616** | -0.247844679593183 | 0.669539485446439 | 0.942334932906175 | 8.13162344912141 | 12.2162359337149 | 12.3039135524298 | 8.44134024832345 | 12.0445401540101 | 25.6847268735511 | 7.28754663389296 | 4.34210225867403 | Sytl2 |
| **ENSMUSG00000030619** | 0.0792387118057548 | 0.4758499853928 | 0.897549741595057 | 481.572810931301 | 437.341246426994 | 432.52988411234 | 425.818719193205 | 395.279908690694 | 388.940149799487 | 448.704657029695 | 448.322058208094 | Eed |

| | | | | | | | | | | | | |
|---|---|---|---|---|---|---|---|---|---|---|---|---|
| ENSMUSG00000030621 | 0.0659339772320038 | 0.456283054382447 | 0.892805623137304 | 2487.37326171458 | 2394.38224300813 | 2324.49320652059 | 2644.95327780801 | 2335.54583168213 | 2189.31719541221 | 2370.53481219632 | 2514.07720777226 | Me3 |
| ENSMUSG00000030629 | 0.0771425161930703 | 0.500340439810537 | 0.900927059989498 | 1299.25272442629 | 1339.71387406407 | 1361.00212987647 | 1272.76652410832 | 1078.53382288181 | 1177.82818948713 | 1402.33218797912 | 1338.45302123627 | Zfand6 |
| ENSMUSG00000030630 | 0.216632819754772 | 0.193796495420155 | 0.809162446871198 | 511.388763578079 | 526.112560878656 | 636.017685171756 | 619.031618210386 | 526.674892188986 | 464.771248188067 | 387.281049686883 | 593.782483873674 | Fah |
| ENSMUSG00000030643 | -0.0187544633642019 | 0.935365993056466 | 0.98818087308472 | 182.509770746947 | 132.749763813036 | 158.057966404291 | 104.109863062656 | 152.199189218854 | 106.408154190426 | 154.079557402308 | 171.513039217624 | Rab30 |
| ENSMUSG00000030647 | -0.104614757954989 | 0.520712065736305 | 0.905962179970576 | 4482.33154789903 | 3458.82360069915 | 3298.39528693984 | 4031.20893192158 | 4144.41677117528 | 5053.77578292919 | 3603.17127141479 | 3619.1422326048 | Ndufc2 |
| ENSMUSG00000030649 | -0.150322597306952 | 0.367978439081273 | 0.871495599696183 | 161.728955265859 | 153.110157035894 | 130.610774633486 | 166.013024883695 | 153.294147414674 | 207.923979452556 | 163.449260217313 | 155.230155747597 | Anapc15 |
| ENSMUSG00000030652 | -0.0810810910277597 | 0.659117970570947 | 0.938989601151841 | 3681.81839501886 | 2928.63896117593 | 2614.10840244701 | 3552.86631784992 | 3463.3527733758 | 4293.01863457925 | 2999.34597889223 | 2761.57703651668 | Coq7 |
| ENSMUSG00000030654 | -0.353517429976318 | 0.110490449053113 | 0.764116514278315 | 588.187429486448 | 570.09101024003 | 590.587850516631 | 649.983199120906 | 904.435469746574 | 999.258183604343 | 489.306702561384 | 673.025850094475 | Arl6ip1 |
| ENSMUSG00000030655 | 0.491305033399603 | 0.16555112448287 | 0.795216960550936 | 585.476888336741 | 982.999784799595 | 1058.13656550896 | 632.162591930001 | 446.742943894192 | 266.631926592101 | 845.355409531583 | 758.782369703287 | Smg1 |
| ENSMUSG00000030657 | -0.562164069036606 | 0.171118894556022 | 0.799531027639052 | 29.8159526467785 | 28.5045505120015 | 30.2865564367503 | 21.5723139679377 | 62.4126171616885 | 56.2617826753975 | 24.9858741733473 | 19.5394601640331 | Xylt1 |
| ENSMUSG00000030659 | -0.619642454873816 | 0.0875062650527833 | 0.74812051555524 | 118.360296870545 | 109.13170767452 | 96.5383986421416 | 147.254490998531 | 192.712642464161 | 320.447544803351 | 96.8202624217207 | 115.065709854862 | Nucb2 |
| ENSMUSG00000030660 | -0.264362336198701 | 0.235216145245051 | 0.832048466795696 | 138.237598635064 | 149.852494120236 | 159.004421292939 | 105.047789756914 | 226.656346534553 | 141.877538920568 | 140.545542225078 | 154.144630182928 | Pik3c2a |
| ENSMUSG00000030662 | 0.24886402004783 | 0.17465754719201 | 0.799889021902745 | 6695.94015349319 | 7509.72743631902 | 6764.31308917045 | 6920.96107693097 | 5109.07494169191 | 4267.3339077057 | 7330.23083560576 | 6763.90979344947 | Ipo5 |
| ENSMUSG00000030663 | 0.023136839360078 | 0.795265405570904 | 0.962032824235414 | 1301.96326557599 | 1255.01463825698 | 1355.32340054458 | 1469.73112990254 | 1325.99437513693 | 1375.96751108309 | 1278.44389520294 | 1315.65698437823 | 1110004F10R |
| ENSMUSG00000030664 | -0.208007234264149 | 0.649589933712551 | 0.936461253686283 | 19.877301764519 | 14.6594831204579 | 22.7149173275627 | 6.56548685980713 | 13.1394983498292 | 15.9000690169602 | 27.0680303544596 | 17.3684090346961 | Sox6os |
| ENSMUSG00000030671 | -0.129715277419206 | 0.595446314091297 | 0.924461858177432 | 137.334084918495 | 200.346269312925 | 117.360406192407 | 121.930470253561 | 129.205067106653 | 144.323703384715 | 146.792010768415 | 211.677485110359 | Pde3b |
| ENSMUSG00000030672 | -0.17749648192799 | 0.718590688863013 | 0.951674101666722 | 16.2632468982428 | 8.95857301805762 | 8.51809399783602 | 27.1998741334867 | 17.5193311331055 | 19.5693157131818 | 21.8626399016789 | 9.76973008201657 | Mylpf |
| ENSMUSG00000030674 | 0.217282571027604 | 0.640822848211037 | 0.934495223272154 | 14.4562194651047 | 27.6901347830872 | 25.5542819935081 | 21.5723139679377 | 6.56974917491458 | 24.4616446414772 | 15.616171358342 | 30.3947158107182 | Qprt |
| ENSMUSG00000030678 | 0.00674309172464288 | 0.944377936066985 | 0.990505437194639 | 1270.34028549608 | 1282.70477304007 | 1301.37547189161 | 1311.22151857291 | 1373.07757755715 | 1395.53682679627 | 1183.70578896233 | 1190.82154444135 | Maz |
| ENSMUSG00000030681 | 0.0283741522178706 | 0.865663664651096 | 0.974868041734905 | 2484.66272056487 | 2400.08315311053 | 2376.54822539625 | 2829.72483657687 | 3025.36949504816 | 2772.72742011144 | 2153.99056936065 | 1943.09076075663 | Mvp |
| ENSMUSG00000030682 | -0.289013673488319 | 0.101514465382411 | 0.759800953500741 | 650.529875929712 | 649.903751673634 | 609.5169482896 | 725.955261355817 | 902.245553354936 | 989.473525747752 | 731.877897660964 | 598.124586132348 | Cdipt |
| ENSMUSG00000030683 | -0.953835505540026 | 0.0284364321914829 | 0.665956123973759 | 58.7283915769879 | 66.7820897709749 | 33.1259211026956 | 62.8410885152968 | 171.908436743598 | 146.769867848863 | 41.6431236222455 | 69.4736361387845 | Sez6l2 |
| ENSMUSG00000030685 | 0.0308745078623221 | 0.81483306568757 | 0.966177091475091 | 176.185174730964 | 189.758864837039 | 162.790240847533 | 179.143998603309 | 165.338687568684 | 166.339183562045 | 181.147587756768 | 180.197243734972 | Kctd13 |
| ENSMUSG00000030688 | -0.432559864298114 | 0.117914570524999 | 0.769868040577702 | 1182.69945498888 | 1195.56229004623 | 1152.78205437381 | 1394.69699436189 | 1328.18429152856 | 2793.5198180567 | 1107.70708835173 | 1419.86743858641 | Stard10 |
| ENSMUSG00000030689 | -0.30129401268849 | 0.249714993754524 | 0.835314899646652 | 269.247087537575 | 234.551729927327 | 217.684624389143 | 317.019222659259 | 346.006789878835 | 465.99433042014 | 262.351678820146 | 206.249857287016 | Ino80e |
| ENSMUSG00000030691 | 0.00164723089236851 | 0.993593023425217 | 0.999133450854041 | 396.642521573811 | 358.342920722305 | 294.347470369667 | 292.633128608546 | 306.588294829347 | 240.94719971855 | 415.390158131898 | 376.677370939972 | Fchsd2 |
| ENSMUSG00000030695 | -0.0892647338384566 | 0.531084570944831 | 0.909082872831377 | 93232.6768990432 | 80628.7859939764 | 82787.3555649683 | 90471.471001448 | 100837.985127568 | 109021.880920368 | 82822.9674941934 | 76594.6838430099 | Aldoa |
| ENSMUSG00000030697 | -0.0709984876227675 | 0.627352382462355 | 0.931825800800958 | 587.283915769879 | 527.741392336485 | 547.997380527451 | 571.19735680322 | 641.645502749991 | 698.379954514174 | 514.292576734731 | 493.914131924171 | Ppp4c |
| ENSMUSG00000030701 | -0.480582133545524 | 0.167495843237316 | 0.798119129525325 | 49.6932544112975 | 68.4109212288036 | 30.2865564367503 | 48.7721881014244 | 90.881530252985 | 86.838838477244 | 60.3825292522559 | 37.9933947633978 | Plekhb1 |
| ENSMUSG00000030704 | -0.0035309426468842 | 0.974858626814535 | 0.995719268067791 | 2397.92540377425 | 2466.8652428815 | 2176.84624389143 | 2252.8999196081 | 2037.71720241934 | 2210.10959335746 | 2546.47700950031 | 2522.76141228961 | Rab6a |
| ENSMUSG00000030706 | -0.0040819403309261 | 0.971577160531729 | 0.995123114262614 | 1516.99953011943 | 1352.7445257267 | 1169.81824236948 | 1395.63492105614 | 1265.77167436688 | 1455.46785616789 | 1392.96248516411 | 1337.3674956716 | Mrpl48 |
| ENSMUSG00000030707 | -0.388690680452786 | 0.207705738260842 | 0.820372930586885 | 74.9916384752307 | 85.5136515360045 | 129.664319744837 | 82.5375490947182 | 128.110108910834 | 183.462334811079 | 73.9165444294857 | 103.124928643508 | Coro1a |
| ENSMUSG00000030711 | -0.48633052900768 | 0.0536041729192788 | 0.703396849069134 | 805.030721463019 | 613.255043872489 | 669.143606274452 | 560.88016316638 | 1225.25822112157 | 1174.15894279091 | 660.043509412591 | 651.315338801105 | Sult1a1 |
| ENSMUSG00000030714 | -0.19577172554694 | 0.250433483137263 | 0.835923538390114 | 197.869503928621 | 176.728213174409 | 176.040609288611 | 201.654239265505 | 220.086597359638 | 269.078091056249 | 199.886993386778 | 173.684090346961 | Sgf29 |
| ENSMUSG00000030717 | -0.884475491866909 | 0.0741101355503516 | 0.734229516971743 | 41.5616309621761 | 47.2361122770311 | 38.8046504345863 | 60.9652351267805 | 99.6411958195378 | 176.123841418636 | 43.7252798033577 | 29.3091902460497 | Nupr1 |
| ENSMUSG00000030718 | 0.0672914706556146 | 0.559564554027365 | 0.916574868225118 | 824.908023227538 | 880.383402956389 | 915.221877323048 | 778.479156234274 | 731.432074807157 | 759.534066117867 | 855.766190437144 | 896.644116416187 | Ppme1 |
| ENSMUSG00000030720 | 0.0178977160980912 | 0.933346396824443 | 0.98818087308472 | 75.8951521917998 | 77.3694942468612 | 81.3951204237664 | 90.0409626487835 | 76.6470737073368 | 102.738907494204 | 83.2862472444909 | 58.6183804920994 | Cln3 |
| ENSMUSG00000030722 | -0.181402842881844 | 0.503744488277132 | 0.900927059989498 | 37.9475760958999 | 48.0505280059454 | 57.7337482075553 | 40.3308478531009 | 58.0327843784121 | 35.4693847301419 | 65.5879197050366 | 48.8486504100828 | Nfatc2ip |
| ENSMUSG00000030725 | 0.0958644485893727 | 0.574062295291746 | 0.922280709351328 | 140.044626068202 | 109.946123403434 | 127.77140996754 | 144.440710915757 | 110.590777777729 | 112.523565350795 | 140.545542225078 | 123.74991437221 | Lipt2 |
| ENSMUSG00000030726 | 0.0165772473764746 | 0.90392797515291 | 0.985039825636533 | 146.369222084185 | 140.079505373265 | 146.700507740509 | 151.006197775564 | 143.439523652302 | 151.662196777159 | 149.915245040084 | 132.434118889558 | Pold3 |
| ENSMUSG00000030727 | 0.040742168618557 | 0.808281640074325 | 0.964026716319739 | 143.658680934478 | 126.234437981721 | 112.628131749165 | 127.55803041911 | 136.869774477387 | 135.762127760198 | 119.723980413956 | 104.210454208177 | Rabep2 |
| ENSMUSG00000030729 | -0.0026959013458880 | 0.986812557094221 | 0.998220569123292 | 204.194099944604 | 260.613033252585 | 293.401015481019 | 226.040333316217 | 259.505092409126 | 272.747337752471 | 237.365804646799 | 217.105112933702 | Pgm2l1 |
| ENSMUSG00000030730 | -0.214653129396887 | 0.582753274837924 | 0.922450244597442 | 14.4562194651047 | 19.5459774939439 | 22.7149173275627 | 20.6343872736795 | 24.0890803080201 | 35.4693847301419 | 17.6983275394543 | 13.0263067760221 | Atp2a1 |
| ENSMUSG00000030731 | 0.115445895196107 | 0.610154257743625 | 0.927932112008768 | 58.7283915769879 | 68.4109212288036 | 53.9479286529615 | 76.9099889291692 | 48.1781606160403 | 67.2695227640623 | 68.711153976705 | 54.2762782334254 | Syt3 |
| ENSMUSG00000030733 | 0.135997423294326 | 0.316109942480358 | 0.853356198600137 | 1290.2175872606 | 1310.39490782315 | 1332.60848321701 | 1216.49092245284 | 998.601874587016 | 1038.39681503071 | 1385.67493853022 | 1262.46623170947 | Sh2b1 |
| ENSMUSG00000030735 | 0.323690786551423 | 0.195203978096291 | 0.81082646923293 | 61.4389327266951 | 72.4829998733752 | 64.3589324280944 | 68.4686486808458 | 39.4184950494875 | 40.3617136584374 | 69.7522320672611 | 62.9604827507734 | Gm9755 |
| ENSMUSG00000030737 | -0.696087231510238 | 0.0250435671734869 | 0.655303666217042 | 336.107102563685 | 329.023954481389 | 356.813493020464 | 402.370551836751 | 739.09678217789 | 869.611467004514 | 317.528817619622 | 382.104998763315 | Slco2b1 |
| ENSMUSG00000030738 | 0.144242965013905 | 0.0830353674511581 | 0.741377052226143 | 3872.45978921493 | 4107.91293664387 | 4454.96316086824 | 4206.60122374785 | 3947.32429592784 | 3833.13971531948 | 3564.65138206421 | 3713.58295673096 | Eif3c |
| ENSMUSG00000030739 | 0.35667018381296 | 0.0038337365802242 | 0.635862253526407 | 6742.01935303821 | 7780.9278740475 | 8286.21255011715 | 8032.40420962689 | 6655.15591418847 | 5154.06852595924 | 6403.6713350108 | 5871.60777929196 | Myh14 |
| ENSMUSG00000030741 | -0.0161981226510272 | 0.911653127740169 | 0.986226496507248 | 738.17070643691 | 685.738043745865 | 784.611102689562 | 653.734905897938 | 785.085026402292 | 828.026671114003 | 705.850945397061 | 576.414074838978 | Spns1 |
| ENSMUSG00000030744 | 0.0708488254450869 | 0.636311175282988 | 0.932595977875587 | 14051.4453200818 | 9723.30938750817 | 11449.2647879803 | 13469.5652562415 | 11314.2030373987 | 13029.4950182828 | 10350.3983763091 | 11666.1432434925 | Rps3 |
| ENSMUSG00000030747 | 0.198978318391361 | 0.0473659791706763 | 0.703396849069134 | 12283.2689767562 | 10734.8137228198 | 11611.1085739391 | 11361.1060475491 | 10492.9843905344 | 8806.19207093179 | 10764.7474563505 | 9998.77597616162 | Dgat2 |
| ENSMUSG00000030748 | -0.68878427026631 | 0.0460246745004264 | 0.697906909182231 | 290.027903018663 | 315.178887089845 | 278.257737262643 | 253.240207449704 | 619.746338833609 | 709.387694602839 | 267.557069272927 | 236.644573097735 | Il4ra |
| ENSMUSG00000030750 | 0.222420525811442 | 0.236024786463066 | 0.832048466795696 | 1372.43733546838 | 1119.00721152829 | 1231.33781013163 | 1668.57158908527 | 1442.05994389375 | 1224.30531430593 | 973.408014669988 | 981.315110460331 | Nsmce1 |
| ENSMUSG00000030751 | 0.0684010158824179 | 0.48340890112635 | 0.89863290032536 | 2984.30580582756 | 2607.75916398368 | 2721.05780486428 | 3215.21270791698 | 2796.52323212197 | 2709.1271440436 | 2792.17143887156 | 2695.3599770719 | Psma1 |
| ENSMUSG00000030752 | 0.517965490143631 | 0.020084452816095 | 0.642795045197142 | 85.8338030740593 | 99.3587189275481 | 82.3415753124149 | 62.8410885152968 | 48.1781606160403 | 56.2617826753975 | 61.4236073428121 | 65.1315338801105 | Kdm8 |
| ENSMUSG00000030753 | 0.152770843829637 | 0.473447441708319 | 0.89728597737823 | 2630.12842893249 | 2595.54292804997 | 2624.51940622214 | 2408.59575085496 | 1931.50625742489 | 1541.08361241306 | 2750.52831524931 | 3003.64923743776 | Thap12 |
| ENSMUSG00000030754 | -0.0766886565929906 | 0.481156577296591 | 0.897732059433541 | 699.319616624441 | 697.95427967958 | 702.269527377147 | 691.251973668265 | 808.079148514494 | 793.780368615935 | 692.316930219831 | 650.229813236436 | Copb1 |
| ENSMUSG00000030759 | 0.0656263123531083 | 0.681408931024744 | 0.944452327903597 | 241.238162323935 | 324.951875836817 | 297.186835035612 | 225.102406621959 | 269.359716171498 | 238.501035254403 | 265.474913091815 | 267.039288908453 | Far1 |
| ENSMUSG00000030760 | 0.152268440563098 | 0.489200365121628 | 0.899236955296196 | 115.649755720838 | 136.007426728693 | 121.146225747001 | 120.054616865045 | 110.590777777729 | 72.1618516923577 | 115.559668051731 | 144.374900100912 | Acer3 |
| ENSMUSG00000030761 | 0.0322948302621539 | 0.872780742519558 | 0.976162349063696 | 118.360296870545 | 131.120932355207 | 140.07532351997 | 166.950951577953 | 174.098353135236 | 107.6312364225 | 128.052605138405 | 133.519644454226 | Myo7a |
| ENSMUSG00000030762 | 0.25574308757301 | 0.578288598996971 | 0.922450244597442 | 228.588970291968 | 102.616381843205 | 72.8770264259304 | 280.44008158319 | 229.94122112201 | 116.192812047017 | 71.8343882483734 | 155.230155747597 | Aqp8 |
| ENSMUSG00000030763 | -0.0698466777633117 | 0.60683717785645 | 0.927177252274348 | 436.397125102849 | 391.733965607792 | 360.599312575058 | 473.652980600371 | 445.647985698372 | 477.002070508805 | 436.211719943021 | 386.447101021989 | Lcmt1 |
| ENSMUSG00000030766 | -0.15744469781522 | 0.419992706872162 | 0.882568531013176 | 160.82544154929 | 209.304842330982 | 228.095628164276 | 202.592165959763 | 258.410134213307 | 237.277953022329 | 161.367104036201 | 236.644573097735 | Arhgap17 |
| ENSMUSG00000030768 | -0.184528162662032 | 0.327834707491344 | 0.855977678491063 | 139.141112351633 | 176.728213174409 | 147.646962629158 | 106.92364314543 | 156.579022002131 | 150.439114545085 | 182.188665847324 | 159.572258006271 | Disp1 |
| ENSMUSG00000030770 | -0.120313911631226 | 0.220208870481574 | 0.828720352121776 | 674.021232560508 | 778.581436842098 | 726.877354482007 | 789.734276565372 | 785.085026402292 | 843.926740130963 | 772.479943192653 | 827.170480277403 | Parva |
| ENSMUSG00000030771 | 0.510229423963038 | 0.4445129735169 | 0.89108031640712 | 20.780815481088 | 6.51532583131463 | 3.78581955459379 | 15.9447538023887 | 12.0445401540101 | 3.66924669622158 | 5.20539045278068 | 11.9407812113536 | Micalcl |
| ENSMUSG00000030772 | -3.56571391212248 | NA | NA | 44.2721721118832 | 25.2468875963442 | 20.8220075502658 | 45.9584080186499 | 58.0327843784121 | 1501.9449809867 | 30.191264626128 | 23.8815624227072 | Dkk3 |
| ENSMUSG00000030774 | -0.630768973081939 | 0.320834617579608 | 0.855228644171892 | 4.51756858284523 | 19.5459774939439 | 13.2503684410783 | 15.0068271081306 | 36.1336204620302 | 28.1308913376988 | 5.20539045278068 | 11.9407812113536 | Pak1 |
| ENSMUSG00000030779 | 0.250697250385787 | 0.0487589140793014 | 0.703396849069134 | 1011.93536255733 | 1232.21099784738 | 1057.19011062032 | 917.292306984482 | 885.821180417649 | 969.904210034571 | 874.505596067155 | 817.400750195386 | Rbbp6 |
| ENSMUSG00000030780 | -0.299177650653974 | 0.284493903866677 | 0.851125503777472 | 76.7986659083688 | 100.987550385377 | 76.6628459805242 | 74.0962088463947 | 131.394983498292 | 130.869798831903 | 80.1630129728225 | 62.9604827507734 | BC017158 |
| ENSMUSG00000030782 | -0.780527541159699 | 0.0409782106934997 | 0.683226273624098 | 183.413284463516 | 156.367819951551 | 156.165056626994 | 166.950951577953 | 378.855535753408 | 474.555906044658 | 138.463386043966 | 147.631476794917 | Tgfb1i1 |
| ENSMUSG00000030785 | 0.024291460744254 | 0.893951988223015 | 0.983730237253949 | 37392.8186739265 | 27842.4305243942 | 28437.1835843312 | 35916.0269032278 | 31833.7196270486 | 41126.1400534835 | 27973.7682932434 | 26492.2514057349 | Cox6a2 |
| ENSMUSG00000030786 | -0.250437621788453 | 0.52121295186652 | 0.90609507420585 | 33.4300075130547 | 21.1748089517725 | 35.965285768641 | 39.3929211588428 | 48.1781606160403 | 59.9310293716191 | 22.903717992235 | 23.8815624227072 | Itgam |
| ENSMUSG00000030787 | -0.676536862267577 | 0.105821358561166 | 0.762094734041179 | 364.116027777325 | 269.571606270643 | 424.011790114504 | 269.184961252092 | 776.32536083574 | 857.380644683776 | 225.913945650682 | 261.61166108511 | Lyve1 |
| ENSMUSG00000030788 | 0.0701015072097761 | 0.737081408227783 | 0.954383574136061 | 681.24934229306 | 816.044560372157 | 712.68053115228 | 629.348811847226 | 571.568178217569 | 478.225152740879 | 873.464517976599 | 780.492880996657 | Rnf141 |
| ENSMUSG00000030790 | -0.248543884464263 | 0.500175234546794 | 0.900927059989498 | 158.114900399583 | 136.007426728693 | 136.289503965376 | 85.3513291774927 | 195.997517051618 | 243.393364182698 | 88.4916376972716 | 85.7565196088121 | Adm |
| ENSMUSG00000030793 | -0.218269603715425 | 0.33135144064018 | 0.856333478432716 | 76.7986659083688 | 47.2361122770311 | 56.7872933189068 | 64.7169419038131 | 76.6470737073368 | 78.277262852727 | 69.7522320672611 | 60.7894316214364 | Pycard |
| ENSMUSG00000030795 | 0.235559016007513 | 0.0834802961833995 | 0.742148948523533 | 2211.80157816102 | 2596.35734377888 | 3118.56885809663 | 2624.31889053433 | 2287.36767106609 | 2007.0779428332 | 2464.23184034638 | 2201.44584514773 | Fus |
| ENSMUSG00000030796 | -0.512605928826259 | 0.14047033002589 | 0.779711473022872 | 98.4829951060259 | 66.7820897709749 | 75.7163910918757 | 73.1582821521366 | 136.869774477387 | 177.34692365071 | 74.9576225200418 | 59.7039060567679 | Tead2 |
| ENSMUSG00000030798 | -0.546556727097832 | 0.244575260795511 | 0.832048466795696 | 17.1667606148119 | 17.9171460361152 | 7.57163910918757 | 20.6343872736795 | 22.994122112201 | 40.3617136584374 | 13.5340151772298 | 16.2828834700276 | Cd37 |
| ENSMUSG00000030801 | -0.0395710590628725 | 0.757325136258234 | 0.956693827734505 | 305.387636200337 | 292.375246680244 | 295.293925258315 | 331.088123073131 | 340.531998899739 | 309.439804714686 | 338.350379430744 | 269.21034003779 | Kat8 |
| ENSMUSG00000030802 | 0.0483276413712526 | 0.511554755230679 | 0.902394714365738 | 3110.79772614722 | 2936.78311846507 | 2994.58326768369 | 3206.77136766865 | 2860.03080747948 | 2886.47406769431 | 3003.51029125445 | 3093.74785930525 | Bckdk |
| ENSMUSG00000030805 | 0.142343992711139 | 0.329995862677624 | 0.855977678491063 | 1601.92981947692 | 1315.28140219664 | 1367.62731409701 | 1634.80622809198 | 1368.69774477387 | 1592.45306616017 | 1274.27958284071 | 1128.94658725525 | Stx4a |
| ENSMUSG00000030806 | -0.152904915391781 | 0.790794914980452 | 0.961804768755116 | 6.32459601598331 | 13.8450673915436 | 17.9826428843205 | 8.44134024832345 | 16.4243729372865 | 22.0154801773295 | 9.36970281500523 | 4.34210225867403 | Stx1b |
| ENSMUSG00000030811 | -0.193992914666404 | 0.414115288289366 | 0.882192126163777 | 47.8862269781594 | 68.4109212288036 | 46.3762895437739 | 52.523894878457 | 64.6025335533267 | 56.2617826753975 | 51.0128264372507 | 74.901263962127 | Fbxl19 |
| ENSMUSG00000030814 | -0.170219867979127 | 0.45242693483035 | 0.891110666112841 | 668.600150261093 | 478.876448601625 | 510.139184981513 | 696.879533833814 | 704.058119911679 | 895.296193878065 | 482.019155927491 | 568.815395886298 | Bcl7c |

| | | | | | | | | | | | | |
|---|---|---|---|---|---|---|---|---|---|---|---|---|
| **ENSMUSG00000030815** | -0.0307334559422103 | 0.862143106835266 | 0.974868041734905 | 129.202461469373 | 127.048853710635 | 137.235958854025 | 153.819977858338 | 146.724398239759 | 124.754387671534 | 116.600746142287 | 170.427513652956 | Phkg2 |
| **ENSMUSG00000030816** | 0.108043550524259 | 0.466432403123947 | 0.895328112556867 | 1187.21702357173 | 1242.79840232327 | 1284.33928389594 | 1008.27119632752 | 1000.79179097865 | 913.642427359173 | 1246.1704743957 | 1220.1307346874 | Rnf40 |
| **ENSMUSG00000030822** | -0.190507003289549 | 0.08917567627787 | 0.751206603737452 | 398.449549006949 | 329.838370210303 | 366.278041906949 | 367.667264149199 | 445.647985698372 | 406.063301048521 | 418.513392403567 | 397.302356668674 | Prr14 |
| **ENSMUSG00000030823** | -0.161537866407338 | 0.595625284078241 | 0.924461858177432 | 36.1405486627618 | 28.5045505120015 | 65.3053873167428 | 51.5859681841989 | 43.7983278327639 | 59.9310293716191 | 48.9306702561384 | 49.9341759747513 | 9130019O22R |
| **ENSMUSG00000030824** | -0.0585582759363991 | 0.62892720905464 | 0.931825800800958 | 2433.16243872044 | 2627.30514147762 | 2871.54413215939 | 2470.498912676 | 2827.18206160491 | 3073.60564920161 | 2587.079055032 | 2346.90627081331 | Nucb1 |
| **ENSMUSG00000030826** | 0.075560191249651 | 0.507286061588599 | 0.901698316485955 | 2985.20931954412 | 2642.779040327 | 2556.37465423945 | 2768.75960145009 | 2564.39209460832 | 2261.47904710457 | 2927.51159064386 | 2638.91264770914 | Bcat2 |
| **ENSMUSG00000030830** | 0.247232196370678 | 0.649533312608404 | 0.936461253686283 | 12.6491920319666 | 30.1333819698302 | 30.2865564367503 | 10.3171936368398 | 13.1394983498292 | 34.2463024980681 | 9.36970281500523 | 14.1118323406906 | Itgal |
| **ENSMUSG00000030834** | -0.112052006820917 | 0.86381482332417 | 0.974868041734905 | 8.13162344912141 | 18.7315617650296 | 7.57163910918757 | 17.8206071909051 | 7.66470737073368 | 3.66924669622158 | 13.5340151772298 | 31.4802413753867 | Abcc6 |
| **ENSMUSG00000030835** | 0.195622945654129 | 0.246980549204357 | 0.833064552804398 | 4740.73647083778 | 5462.2862938284 | 5230.10971467132 | 4641.79920988364 | 3923.23521561982 | 3326.7836712409 | 5051.31089537837 | 5226.80559387886 | Nomo1 |
| **ENSMUSG00000030838** | -0.483308881657866 | 0.28127065571964 | 0.850048154552447 | 19.877301764519 | 5.7009101024003 | 19.8755526616174 | 22.5102406621959 | 21.8991639163819 | 15.9000690169602 | 27.0680303544596 | 29.3091902460497 | Ush1c |
| **ENSMUSG00000030839** | -0.263229099971593 | 0.217698350058183 | 0.826183082597859 | 110.228673421423 | 100.173134656462 | 64.3589324280944 | 99.4202295913651 | 95.2613630362614 | 127.200552135681 | 113.477511870619 | 113.980184290193 | Sergef |
| **ENSMUSG00000030842** | 0.0460341333941538 | 0.736058534409503 | 0.954383574136061 | 2359.07431396178 | 1789.27135642478 | 1619.38431447749 | 2009.97690579524 | 1834.05497799699 | 1932.4699266767 | 1901.00859335551 | 1866.01844566516 | Lamtor1 |
| **ENSMUSG00000030844** | 0.119665736555798 | 0.639305810917182 | 0.933801135937988 | 77.7021796249379 | 76.5550785179469 | 113.574586637814 | 75.0341355406529 | 72.2672409240604 | 91.7311674055395 | 54.1360607089191 | 97.6973008201657 | Rgs10 |
| **ENSMUSG00000030846** | 0.0285752550602557 | 0.83456989645765 | 0.968516656512387 | 638.784197614315 | 586.379324818317 | 637.910594949053 | 620.907471598903 | 571.568178217569 | 507.579126310652 | 728.754663389296 | 625.26272524906 | Tial1 |
| **ENSMUSG00000030847** | 0.12400103342352 | 0.24526404294107 | 0.832048466795696 | 8166.8604840676 | 9829.99784799595 | 8921.28378040026 | 8852.15214040853 | 9013.69586798281 | 7392.30901065441 | 8512.89554647753 | 7904.79716191607 | Bag3 |
| **ENSMUSG00000030850** | 0.0355004395343846 | 0.831411516988211 | 0.967583158900775 | 681.24934229306 | 826.631964848044 | 824.362208012797 | 640.603932178324 | 677.779123212021 | 574.848649074714 | 832.862472444909 | 814.144173501381 | Ate1 |
| **ENSMUSG00000030852** | 0.414875455717051 | 0.119577789923323 | 0.769868040577702 | 29480.7490579314 | 34910.7446356416 | 45615.3398133005 | 34575.7296571329 | 25002.2754433333 | 15133.1964574499 | 36988.463479369 | 31323.9256940745 | Tacc2 |
| **ENSMUSG00000030861** | 0.255779563109112 | 0.266637084466953 | 0.843513595393324 | 2854.19983064161 | 3414.03073560887 | 3772.56918615271 | 3142.05442576484 | 2206.34076457548 | 1846.85417043153 | 3579.226475332 | 3407.46474749445 | Acadsb |
| **ENSMUSG00000030862** | 0.334163109970393 | 0.102385618340022 | 0.759800953500741 | 2365.39890997776 | 1398.3518065459 | 1428.20042697051 | 2165.67273704209 | 1534.03643234255 | 1254.88237010778 | 1536.63126166086 | 1509.96606045389 | Cpxm2 |
| **ENSMUSG00000030865** | -0.539020226489302 | 0.324867365901568 | 0.855977678491063 | 3.61405486627618 | 13.8450673915436 | 22.7149173275627 | 37.5170677703264 | 25.1840385038392 | 26.9078091056249 | 34.3555769883525 | 26.0526135520442 | Chp2 |
| **ENSMUSG00000030868** | -0.147865563029841 | 0.249449436114169 | 0.835314899646652 | 290.027903018663 | 286.674336577844 | 299.079744812909 | 314.205442576484 | 346.006789878835 | 360.809258461789 | 336.268223249632 | 275.723493425801 | Dctn5 |
| **ENSMUSG00000030869** | 0.129491340608035 | 0.216828664174906 | 0.825692108906787 | 12943.7375035681 | 10712.0100824102 | 10578.5262904237 | 12581.348676779 | 11087.5466908642 | 10975.9399506308 | 10228.592239714 | 10504.6308892971 | Ndufab1 |
| **ENSMUSG00000030870** | 0.0647179409483533 | 0.545636732826044 | 0.914211323944814 | 829.425591810383 | 753.334549245754 | 752.431636475515 | 755.968915572078 | 799.319482947941 | 644.564336302924 | 761.028084196536 | 749.01263962127 | Ubfd1 |
| **ENSMUSG00000030871** | 0.282423745530654 | 0.0302239123563271 | 0.665956123973759 | 268.343573821006 | 252.468875963442 | 272.579007930753 | 264.495327780801 | 250.745426842573 | 205.477814988408 | 218.626399016789 | 194.309076075663 | Ears2 |
| **ENSMUSG00000030872** | 0.0233766482688258 | 0.897489319939718 | 0.984332015045905 | 1030.00563688871 | 1135.29552610657 | 1039.20746773599 | 930.423280704096 | 943.853964796062 | 728.95701031602 | 1224.30783449402 | 1170.19655871265 | Gga2 |
| **ENSMUSG00000030876** | 0.188602239367054 | 0.0357626439829744 | 0.679566075569698 | 2343.7145807801 | 2121.55297382183 | 2077.46848058334 | 2364.51319622482 | 1858.14405830501 | 1933.69300890877 | 1950.9803417022 | 2072.26830295218 | Mettl9 |
| **ENSMUSG00000030878** | -0.232180935762826 | 0.466273471823574 | 0.895328112556867 | 174.378147297826 | 141.708336831093 | 241.345996605354 | 126.620103724852 | 183.952976897608 | 329.009120427868 | 144.709854587303 | 146.545951230249 | Cdr2 |
| **ENSMUSG00000030879** | 0.0577973533414066 | 0.675542743336103 | 0.94409858171788 | 1339.91084167189 | 1123.89370590177 | 1253.10627257054 | 1526.00673155803 | 1298.62042024145 | 1429.78312929434 | 1142.06266534008 | 1166.93998201865 | Mrpl17 |
| **ENSMUSG00000030880** | 0.198282775988519 | 0.326420164549748 | 0.855977678491063 | 242.141676040504 | 271.200437728471 | 212.952349945901 | 247.612647284155 | 212.421889988905 | 141.877538920568 | 235.283648465687 | 258.355084391105 | Polr3e |
| **ENSMUSG00000030881** | -0.0450131284010631 | 0.741940718462661 | 0.954383574136061 | 271.957628687283 | 244.324718674299 | 226.202718386979 | 252.302280755445 | 239.795844884382 | 297.208982393948 | 257.146288367366 | 233.387996403729 | Arfip2 |
| **ENSMUSG00000030884** | 0.153793744316248 | 0.192536360427265 | 0.808824684419004 | 26510.8994715689 | 24553.8198110381 | 25772.9130727859 | 25799.5495789592 | 22491.5363003201 | 19113.1060406182 | 25193.0487133679 | 25458.8310681705 | Uqcrc2 |
| **ENSMUSG00000030887** | -0.717619375783471 | 0.0534473969997966 | 0.703396849069134 | 19.877301764519 | 17.1027303072009 | 8.51809399783602 | 20.6343872736795 | 33.943704070392 | 29.3539735697726 | 19.7804837205666 | 26.0526135520442 | Pdzd9 |
| **ENSMUSG00000030888** | 0.0859707864132038 | 0.632018716781179 | 0.931825800800958 | 176.185174730964 | 206.861595144239 | 216.738169500494 | 195.088752405698 | 173.003394939417 | 141.877538920568 | 211.338852382896 | 221.447215192376 | Rrp8 |
| **ENSMUSG00000030889** | 0.0361816363915113 | 0.823573390790399 | 0.966637987288264 | 131.913002619081 | 116.461449234749 | 142.914688185915 | 131.309737196143 | 129.205067106653 | 149.216032313011 | 115.559668051731 | 116.15123541953 | Vwa3a |
| **ENSMUSG00000030890** | -0.151747184023664 | 0.21355144185424 | 0.824248286808591 | 256.597895505609 | 299.704988240473 | 247.971180825893 | 258.867767615252 | 280.309298129689 | 286.201242305283 | 310.241270985729 | 305.032683671851 | Ilk |
| **ENSMUSG00000030894** | -0.0369649253103411 | 0.761219604891838 | 0.957479700684068 | 828.522078093814 | 958.567312932165 | 1047.72556173383 | 859.140851940476 | 951.518672166795 | 1018.82749931753 | 973.408014669988 | 846.709940441436 | Tpp1 |
| **ENSMUSG00000030895** | -0.37932738646893 | 0.231547402929853 | 0.832048466795696 | 568.310127721929 | 1262.34437981721 | 709.841166486335 | 758.782695654852 | 623.031213421066 | 923.427085215764 | 1215.97920976957 | 1529.50552061793 | Hpx |
| **ENSMUSG00000030911** | 0.0566314275470106 | 0.90425041504148 | 0.98504921011344 | 9.03513716569045 | 15.4738988493722 | 11.3574586637814 | 7.50341355406529 | 13.1394983498292 | 8.56157562451702 | 12.4929370866736 | 7.59867895267955 | Zp2 |
| **ENSMUSG00000030917** | -0.185847041913656 | 0.090676546392892 | 0.753823363185192 | 652.336903362851 | 598.595560752032 | 583.962666296092 | 652.79697920368 | 731.432074807157 | 785.218792991418 | 666.289977955927 | 648.058762107099 | Tmem159 |
| **ENSMUSG00000030921** | -0.168499144450328 | 0.46770311726529 | 0.895328112556867 | 115.649755720838 | 97.7298874697194 | 123.985590412947 | 118.178763476528 | 169.71852035196 | 145.546785616789 | 89.5327157878278 | 107.467030902182 | Trim30a |
| **ENSMUSG00000030922** | 0.163292605855162 | 0.438492493329386 | 0.889346190988072 | 65.9565013095403 | 84.6992358070902 | 85.1809399783602 | 90.9788893430416 | 76.6470737073368 | 55.0387004433237 | 84.3273253350471 | 74.901263962127 | Lyrm1 |
| **ENSMUSG00000030924** | 0.554434267810515 | 0.138130105902687 | 0.779711473022872 | 25.2983840639333 | 15.4738988493722 | 23.6613722162112 | 16.8826804966469 | 12.0445401540101 | 12.2308223207386 | 15.616171358342 | 15.1973579053591 | Rexo5 |
| **ENSMUSG00000030929** | -0.171440253508425 | 0.638185011908408 | 0.933198322011444 | 42.4651446787451 | 25.2468875963442 | 22.7149173275627 | 18.7585338851632 | 30.6588294829347 | 19.5693157131818 | 32.2734208072402 | 40.1644458927348 | Eri2 |
| **ENSMUSG00000030930** | -0.449574799213396 | 0.0799081567070126 | 0.739855948845396 | 123.781379169959 | 126.234437981721 | 106.949402417274 | 153.819977858338 | 220.086597359638 | 222.600966237442 | 112.436433780063 | 143.289374536243 | Chst15 |
| **ENSMUSG00000030934** | 0.0839457360422007 | 0.345134138138852 | 0.861266047293794 | 4402.82234084096 | 4708.95174458265 | 4759.72163501304 | 4602.4062887248 | 4524.36726512451 | 3906.52464924391 | 4343.3777938002 | 4653.64809573389 | Oat |
| **ENSMUSG00000030935** | 0.0265243902697852 | 0.939565610609432 | 0.989269716906553 | 15.3597331816738 | 26.0613033252585 | 21.7684624389143 | 18.7585338851632 | 15.3294147414674 | 18.3462334811079 | 20.8215618111227 | 26.0526135520442 | Acsm3 |
| **ENSMUSG00000030942** | 0.107730715466854 | 0.338107598502245 | 0.858570960161243 | 614.389327266951 | 675.150639269978 | 593.427215182576 | 643.417712261099 | 602.227007700503 | 518.586866399316 | 649.632728507029 | 573.157498144972 | Thumpd1 |
| **ENSMUSG00000030946** | -0.0814168869039012 | 0.579312982176764 | 0.922450244597442 | 431.879556520004 | 363.229415095791 | 311.383658365339 | 361.101777289392 | 337.247124312282 | 391.386314263635 | 394.568596320776 | 429.868123608729 | Lhpp |
| **ENSMUSG00000030956** | 0.0055994708731892 | 0.967454579596069 | 0.994284908685797 | 315.326287082597 | 339.611358957275 | 306.651383922097 | 317.019222659259 | 362.431162816121 | 260.516515431732 | 312.323427166841 | 337.598450611906 | Fam53b |
| **ENSMUSG00000030960** | 0.332917465912337 | 0.0847090577695828 | 0.743541556158628 | 90.3513716569045 | 77.3694942468612 | 101.270673085384 | 79.7237690119437 | 60.2227007700503 | 68.4926049961361 | 77.0397787011541 | 70.559161703453 | Eef1akmt2 |
| **ENSMUSG00000030965** | 0.0900177804463378 | 0.601359711655066 | 0.926043912368388 | 423.747933070882 | 548.101785559343 | 459.030620994497 | 484.908100931469 | 396.374866886513 | 357.140011765567 | 528.867670002517 | 516.71016878221 | Abraxas2 |
| **ENSMUSG00000030966** | -0.10075058358015 | 0.598679536741716 | 0.924848033480823 | 76.7986659083688 | 69.2253369577179 | 66.2518422053913 | 78.7858423176855 | 75.5521155115177 | 91.7311674055395 | 79.1219348822664 | 66.217059444779 | Trim21 |
| **ENSMUSG00000030967** | 0.0790124414409413 | 0.506068414706092 | 0.901494649536949 | 1523.32412613541 | 1407.31037956396 | 1470.79089695969 | 1745.48157801444 | 1342.41874807421 | 1553.3144347338 | 1337.78534636464 | 1585.95284998069 | Zranb1 |
| **ENSMUSG00000030970** | -0.243868517527236 | 0.134577619312275 | 0.773512701181749 | 268.343573821006 | 246.767965861042 | 299.079744812909 | 294.508981997063 | 390.900075907418 | 355.916929533493 | 264.433835001259 | 301.776106977845 | Ctbp2 |
| **ENSMUSG00000030972** | 0.060612518057694 | 0.771377815122269 | 0.959045638244974 | 332.493047697409 | 369.744740927105 | 368.170951684246 | 238.233380341573 | 282.499214521327 | 276.416584448692 | 414.349080041342 | 281.151121249143 | Acsm5 |
| **ENSMUSG00000030978** | -0.291333740994032 | 0.0399763912644133 | 0.679566075569698 | 289.124389302094 | 233.737314198412 | 294.347470369667 | 285.129715054481 | 369.000911991036 | 359.586176229715 | 330.021754706295 | 289.835325766492 | Rrm1 |
| **ENSMUSG00000030979** | 0.162451944754625 | 0.363825154652692 | 0.86924083181804 | 693.898534325027 | 553.802695661743 | 581.123301630146 | 602.14893771374 | 424.84377997781 | 463.548165955993 | 621.523620062014 | 661.085068883121 | Uros |
| **ENSMUSG00000030980** | -0.0043801829842351 | 0.971317525356676 | 0.995122272737203 | 384.896843258413 | 352.642010619904 | 346.402489245332 | 406.122258613784 | 379.950493949227 | 414.624876673038 | 336.268223249632 | 364.736589728619 | Knop1 |
| **ENSMUSG00000030982** | -0.0173287638211208 | 0.902947606989764 | 0.985027469232262 | 477.055242348456 | 580.678414715916 | 602.891764069061 | 512.107975064956 | 653.690042904001 | 514.917619703095 | 530.94982618363 | 499.341759747513 | 9030624J02R |
| **ENSMUSG00000030983** | 0.13259425722836 | 0.201831159762277 | 0.813395633194566 | 1925.38773000863 | 1717.60277228032 | 1629.79531825263 | 1967.77020455362 | 1524.18180858018 | 1720.87670052792 | 1675.09464770482 | 1684.73567636552 | Bccip |
| **ENSMUSG00000030986** | 0.308548214840863 | 0.0692913012242727 | 0.724352955175446 | 1697.70227343324 | 1939.12385054502 | 2140.88095812279 | 2010.9148324895 | 1469.43389878923 | 1177.82818948713 | 1798.982940481 | 1841.05135767779 | Dhx32 |
| **ENSMUSG00000030987** | -0.345041713608784 | 0.0261277145193546 | 0.664353746352133 | 362.309000344187 | 381.146561131906 | 356.813493020464 | 296.384835385579 | 485.06648074786 | 512.471455238947 | 408.102611498006 | 370.164217551961 | Stim1 |
| **ENSMUSG00000030990** | 0.154880255215856 | 0.279272519313525 | 0.848075053184273 | 1139.33079659357 | 1040.0088858236 | 965.383986421416 | 1184.60141484806 | 956.993463145891 | 780.326464063122 | 1060.8585742767 | 1088.78214136251 | Pgap2 |
| **ENSMUSG00000030994** | 0.562616334304772 | 0.100409833888023 | 0.759800953500741 | 39.754603529038 | 52.9370223794314 | 58.6802030962037 | 38.4549944645846 | 41.6084114411257 | 26.9078091056249 | 16.6572494488982 | 43.4210225867403 | D7Ertd443e |
| **ENSMUSG00000030996** | 0.13931988013133 | 0.171397520581758 | 0.799531027639052 | 2993.34094299325 | 2993.79221948907 | 3505.66890755385 | 3438.43926115042 | 2880.83501320004 | 3133.53667857323 | 2999.34597889223 | 2726.84021844729 | Art1 |
| **ENSMUSG00000031004** | -0.0824251009426351 | 0.852019523483484 | 0.97283610202923 | 20.780815481088 | 30.1333819698302 | 55.8408384302584 | 48.7721881014244 | 66.7924499449649 | 48.9232892829544 | 22.903717992235 | 26.0526135520442 | Mki67 |
| **ENSMUSG00000031007** | -0.303297707629828 | 0.0811048722711702 | 0.739855948845396 | 303.580608767199 | 268.757190541728 | 323.687571917769 | 269.184961252092 | 420.463947194533 | 420.740287833408 | 297.748333899055 | 299.605055848508 | Atp6ap2 |
| **ENSMUSG00000031010** | 0.301223720588094 | 0.457729295650666 | 0.892966448175628 | 1794.37824110612 | 3126.54198330211 | 2710.64680108915 | 1458.47600957144 | 1332.56412431184 | 697.1568722821 | 2856.71828048604 | 2490.19564534956 | Usp9x |
| **ENSMUSG00000031012** | -0.378913714103344 | 0.0054358718891339 | 0.635862253526407 | 159.018414116152 | 167.769640156352 | 175.094154399963 | 179.143998603309 | 249.650468646754 | 223.824048469516 | 214.462086654564 | 197.565652769668 | Cask |
| **ENSMUSG00000031015** | -0.20708945686837 | 0.243244689706704 | 0.832048466795696 | 219.553833126278 | 235.366145656241 | 254.596365046432 | 304.826175633902 | 342.721915291377 | 326.562955963721 | 244.653351280692 | 257.269558826436 | Swap70 |
| **ENSMUSG00000031016** | 0.152887686273834 | 0.577527790994958 | 0.922450244597442 | 296.352499034647 | 483.762942975111 | 330.312756138308 | 239.171307035831 | 297.828629262794 | 195.693157131818 | 396.650752501888 | 323.486618271215 | Wee1 |
| **ENSMUSG00000031021** | -0.0530000626368244 | 0.615940201249657 | 0.928422072203769 | 857.434517024024 | 768.808448095126 | 763.789095139297 | 862.892558717508 | 872.68168206782 | 914.865509591247 | 819.328457267679 | 768.552099785303 | Tmem9b |
| **ENSMUSG00000031023** | 0.0583733932301671 | 0.75074634780233 | 0.956197192463153 | 293.64195788494 | 192.202112023782 | 200.648436393471 | 241.047160424347 | 252.935343234211 | 222.600966237442 | 222.790711379013 | 192.138024946326 | Akip1 |
| **ENSMUSG00000031024** | -0.159531225561193 | 0.278936021579756 | 0.848075053184273 | 206.904641094311 | 244.324718674299 | 253.649910157784 | 290.75727522003 | 280.309298129689 | 269.078091056249 | 257.146288367366 | 305.032683671851 | St5 |
| **ENSMUSG00000031029** | -0.0391446618051413 | 0.756874743744095 | 0.956596013703865 | 3459.55402074287 | 2999.49312959147 | 3446.04224956899 | 3717.00348934509 | 3710.81332563092 | 3964.00951415138 | 3273.14951670849 | 3049.24131115384 | Eif3f |
| **ENSMUSG00000031059** | -0.00091754403092375 | 0.996350136944348 | 0.999262355603273 | 15664.2173041575 | 11204.7315984033 | 12733.6040718762 | 15394.1908328592 | 13979.3312860224 | 18448.9723886021 | 10945.8950441072 | 11658.5445645398 | Ndufb11 |
| **ENSMUSG00000031060** | -0.0814043328569254 | 0.427901555513817 | 0.885066977635295 | 485.186865797577 | 513.081909216027 | 541.372196306912 | 485.846027625728 | 543.099265126272 | 582.187142467157 | 514.292576734731 | 504.769387570856 | Rbm10 |
| **ENSMUSG00000031065** | 0.185860235874951 | 0.344382269240238 | 0.861266047293794 | 3333.96561413978 | 3616.00583637962 | 3345.71803137226 | 2983.54481443521 | 2322.4063333323 | 2192.98644210843 | 3641.69116076537 | 3516.0173039613 | Cdk16 |
| **ENSMUSG00000031066** | 0.0753800769168491 | 0.758325704588626 | 0.95677573228595 | 77.7021796249379 | 92.0289773673191 | 109.78876708322 | 72.2203554578784 | 81.0269064906132 | 105.185071958352 | 57.2592949805875 | 91.1841474321546 | Usp11 |
| **ENSMUSG00000031068** | 0.101652084494027 | 0.313519045831377 | 0.85295001595893 | 3429.7380680961 | 2890.36142191695 | 3287.03782827606 | 3576.31448520637 | 3010.0400803067 | 3102.95962277138 | 2918.14188782885 | 3254.40564287619 | Glrx3 |
| **ENSMUSG00000031070** | -1.31870920397585 | 0.0160545231322229 | 0.63777231095375 | 14.4562194651047 | 17.1027303072009 | 14.1968233297267 | 28.1378008277448 | 68.9823663366031 | 79.5003450848009 | 16.6572494488982 | 19.5394601640331 | Mrgprf |
| **ENSMUSG00000031072** | 0.223416709535916 | 0.128812642539826 | 0.77303252008843 | 391.221439274397 | 353.456426348819 | 374.796135904785 | 411.749818779333 | 267.16979977986 | 310.66288694676 | 385.198893505771 | 347.368180693922 | Oraov1 |
| **ENSMUSG00000031075** | -0.194776255450952 | 0.372890959336718 | 0.873700364743909 | 73.1846110420926 | 70.0397526866323 | 79.5022106464695 | 86.2892558717508 | 109.49581958191 | 99.0696607979826 | 81.2040910633787 | 64.046008315442 | Ano1 |
| **ENSMUSG00000031077** | 0.077587967159535 | 0.490608611038337 | 0.899236955296196 | 325.264937964856 | 359.157336451219 | 371.010316350191 | 389.239578117137 | 346.006789878835 | 348.57843614105 | 320.65205189129 | 353.881334081933 | Fadd |
| **ENSMUSG00000031078** | 0.0453676260876033 | 0.592098058341066 | 0.923935529355527 | 944.171833814652 | 1053.03953748623 | 1064.7617497295 | 1045.78826409785 | 1024.88087128667 | 993.142772443974 | 992.147420299998 | 970.459854813646 | Cttn |
| **ENSMUSG00000031090** | -0.1853363762687 | 0.435043911936246 | 0.888379878975956 | 56.0178504272808 | 73.2974156022896 | 42.5904699891801 | 52.523894878457 | 61.3176589658694 | 64.8233582999146 | 56.2182168900314 | 73.8157383974585 | Nadsyn1 |
| **ENSMUSG00000031093** | -0.633903612530559 | 0.0070966576803812 | 0.635862253526407 | 28.9124389302094 | 37.4631235300591 | 42.5904699891801 | 46.8963347129081 | 65.6974917491458 | 61.154111603693 | 60.3825292522559 | 54.2762782334254 | Dock11 |

| | | | | | | | | | | | | |
|---|---|---|---|---|---|---|---|---|---|---|---|---|
| ENSMUSG00000031095 | 0.0382622752482314 | 0.823914257299209 | 0.966637987288264 | 252.983840639333 | 267.1283590839 | 262.16800415562 | 226.978260010475 | 258.410134213307 | 178.570005882784 | 258.187366457922 | 286.578749072486 | Cul4b |
| ENSMUSG00000031097 | -0.257537804850914 | 0.459979072302251 | 0.894774044236998 | 29.8159526467785 | 20.3603932228582 | 28.3936466594534 | 21.5723139679377 | 31.7537876787538 | 37.9155491942896 | 15.616171358342 | 34.7368180693922 | Tnni2 |
| ENSMUSG00000031099 | -0.240638975722786 | 0.28341767933515 | 0.850804315099468 | 56.9213641438498 | 45.6072808192024 | 43.5369248778286 | 42.2067012416173 | 52.5579933993167 | 53.8156182112498 | 51.0128264372507 | 65.1315338801105 | Smarca1 |
| ENSMUSG00000031101 | 0.141714621081564 | 0.776921023774333 | 0.959738279197945 | 13.5527057485357 | 9.77298874697194 | 22.7149173275627 | 6.56548685980713 | 12.0445401540101 | 9.78465785659088 | 9.36970281500523 | 16.2828834700276 | Sash3 |
| ENSMUSG00000031103 | -0.110570360942821 | 0.602279072582293 | 0.926448151609826 | 99.3865088225949 | 112.389370590177 | 141.968233297267 | 105.047789756914 | 163.148771177045 | 114.969729814943 | 104.107809055614 | 112.894658725525 | Elf4 |
| ENSMUSG00000031105 | -0.588101620388523 | 0.0578667097363415 | 0.710862211775517 | 32.5264937964856 | 23.6180561385155 | 40.6975602118832 | 38.4549944645846 | 40.5134532453066 | 74.6080161565054 | 39.5609674411332 | 48.8486504100828 | Slc25a14 |
| ENSMUSG00000031107 | -0.219989859121997 | 0.456571400104227 | 0.892805623137304 | 61.4389327266951 | 57.8235167529173 | 42.5904699891801 | 61.9031618210386 | 79.9319482947941 | 89.2850029413918 | 51.0128264372507 | 41.2499714574033 | Rbmx2 |
| ENSMUSG00000031109 | 0.150953314919947 | 0.350630660024865 | 0.861774532843376 | 230.395997725106 | 206.861595144239 | 212.952349945901 | 210.095579513828 | 175.193311331055 | 156.554525705454 | 233.201492284575 | 208.420908416353 | Enox2 |
| ENSMUSG00000031111 | -0.119716137182189 | 0.595675524621327 | 0.924461858177432 | 680.345828576491 | 505.752167655798 | 345.456034356683 | 405.184331919526 | 441.268152915096 | 533.263853184203 | 607.989604884784 | 522.137796605552 | Igsf1 |
| ENSMUSG00000031112 | -0.856330383253952 | 0.0585893353005126 | 0.711712402279519 | 6.32459601598331 | 11.4018202048006 | 10.4110037751329 | 4.68963347129081 | 16.4243729372865 | 14.6769867848863 | 15.616171358342 | 13.0263067760221 | Stk26 |
| ENSMUSG00000031119 | -0.00227975299288972 | 0.985140838279518 | 0.99735301701069 | 305.387636200337 | 333.910448854875 | 368.170951684246 | 294.508981997063 | 317.537876787538 | 340.016860516533 | 335.227145159076 | 311.545837059862 | Gpc4 |
| ENSMUSG00000031125 | 0.231160377145343 | 0.585969371878953 | 0.923058967588378 | 36.1405486627618 | 41.5352021746308 | 47.3227444324223 | 13.1309737196143 | 20.8042057205628 | 23.2385624094033 | 31.2323427166841 | 42.3354970220718 | 3830403N18 |
| ENSMUSG00000031129 | -0.314827566306013 | 0.368287481587819 | 0.871495599696183 | 27.1054114970714 | 21.9892246806869 | 35.0188308799925 | 21.5723139679377 | 35.0386622662111 | 48.9232892829544 | 20.8215618111227 | 27.1381391167127 | Slc9a9 |
| ENSMUSG00000031133 | 0.215610937415556 | 0.358994886480316 | 0.866754558180356 | 1168.24323552378 | 1517.25650296739 | 1409.27132919754 | 1245.56664997484 | 938.379173816966 | 747.303243797128 | 1529.34371502696 | 1382.95956938768 | Arhgef6 |
| ENSMUSG00000031134 | 0.154706547986391 | 0.324827310705401 | 0.855977678491063 | 327.071965397994 | 403.950201541507 | 336.937940358847 | 262.619474392285 | 281.404256325508 | 306.993640250539 | 302.953724351836 | 305.032683671851 | Rbmx |
| ENSMUSG00000031137 | 0.179742660928431 | 0.402063515612975 | 0.881158207349486 | 980.312382477414 | 1170.31540244989 | 1209.56934769271 | 1008.27119632752 | 908.81530252985 | 615.210362733151 | 1149.35021197397 | 1182.137339924 | Fgf13 |
| ENSMUSG00000031138 | -0.367080584022944 | 0.299033617124892 | 0.851785577030345 | 51.5002818444356 | 68.4109212288036 | 31.2330113253987 | 50.6480414899407 | 43.7983278327639 | 48.9232892829544 | 62.4646854333682 | 105.295979772845 | F9 |
| ENSMUSG00000031144 | -0.357447580878235 | 0.234493116396215 | 0.832048466795696 | 59.631905293557 | 64.338842584232 | 63.4124775394459 | 81.59962240046 | 93.0714466446232 | 130.869798831903 | 51.0128264372507 | 70.559161703453 | Syp |
| ENSMUSG00000031145 | -0.0347229738988128 | 0.778460346176337 | 0.960200178349109 | 440.914693685694 | 494.350347450997 | 419.279515671262 | 422.067012416173 | 503.680770076785 | 472.10974158051 | 400.815064864113 | 445.065481514088 | Prickle3 |
| ENSMUSG00000031146 | -0.0171058844531443 | 0.885461321871756 | 0.979948700426504 | 864.662626756576 | 695.511032492837 | 820.576388458203 | 822.561710864407 | 831.073270626695 | 886.734618253548 | 797.465817366001 | 726.216602763232 | Plp2 |
| ENSMUSG00000031147 | 0.162128135751715 | 0.16140508660179 | 0.792786631432503 | 855.627489590886 | 851.878852444388 | 918.061241988993 | 1041.09863062656 | 826.693437843418 | 793.780368615935 | 899.491470240502 | 755.525793009281 | Magix |
| ENSMUSG00000031148 | 0.0635405119723564 | 0.726700886610971 | 0.952045814446758 | 463.50253659992 | 478.062032872711 | 459.977075883145 | 393.929211588428 | 404.039574257247 | 313.109051410908 | 535.114138545854 | 464.604941678121 | Gpkow |
| ENSMUSG00000031149 | -0.608633573751548 | 0.204978984023142 | 0.819090359056803 | 56.9213641438498 | 38.2775392589734 | 25.5542819935081 | 49.7101147956825 | 73.3621991198795 | 124.754387671534 | 36.4377331694648 | 26.0526135520442 | Praf2 |
| ENSMUSG00000031153 | 0.200674634736219 | 0.250476153115886 | 0.835923538390114 | 460.791995450213 | 572.534257426773 | 543.265106084208 | 500.852854733858 | 441.268152915096 | 329.009120427868 | 520.539045278068 | 515.624643217541 | Gripap1 |
| ENSMUSG00000031154 | 0.00880897581228072 | 0.934406302226762 | 0.98818087308472 | 509.581736144941 | 523.669313691913 | 590.587850516631 | 518.673461924763 | 579.232885588302 | 514.917619703095 | 534.073060455298 | 500.427285312182 | Otud5 |
| ENSMUSG00000031155 | 0.226350180825004 | 0.456942958443063 | 0.892805623137304 | 41.5616309621761 | 23.6180561385155 | 35.0188308799925 | 34.703287687552 | 29.5638712871156 | 24.4616446414772 | 23.9447960827911 | 36.9078691987293 | Pim2 |
| ENSMUSG00000031156 | -0.192563098735164 | 0.138925401277137 | 0.779711473022872 | 199.676531361759 | 206.861595144239 | 229.042083052924 | 223.226553233442 | 251.840385038392 | 250.731857575141 | 263.392756910703 | 214.934061804364 | Slc35a2 |
| ENSMUSG00000031157 | -0.273935564808337 | 0.206001273562586 | 0.819960256425221 | 425.55496050402 | 373.002403842763 | 420.22597055991 | 408.936038696558 | 529.959766776443 | 688.595296657583 | 392.486440139664 | 358.223436340608 | Pqbp1 |
| ENSMUSG00000031158 | -0.122196839720169 | 0.338985904321691 | 0.858570960161243 | 764.372604217412 | 688.181290932608 | 698.483707822554 | 843.196098138087 | 735.811907590433 | 941.773318696872 | 803.712285909337 | 778.32182986732 | Timm17b |
| ENSMUSG00000031161 | 0.278256180376406 | 0.114379525138893 | 0.767518180080285 | 398.449549006949 | 460.144886836596 | 443.887342776122 | 439.887619607078 | 372.285786578493 | 253.178022039289 | 433.088485671353 | 376.677370939972 | Hdac6 |
| ENSMUSG00000031165 | -0.582962401033098 | 0.125517995700177 | 0.771949105062466 | 17.1667606148119 | 26.0613033252585 | 30.2865564367503 | 27.1998741334867 | 45.9882442244021 | 31.8001380339204 | 18.7394056300105 | 54.2762782334254 | Was |
| ENSMUSG00000031166 | 0.240237501306392 | 0.314893363714623 | 0.853277176422981 | 985.733464776828 | 1171.1298181788 | 1091.26248661166 | 970.754128557197 | 660.259792078915 | 609.094951572782 | 1218.06136595068 | 1083.35451353917 | Wdr13 |
| ENSMUSG00000031167 | 0.268949838433542 | 0.0802793023785908 | 0.739855948845396 | 1084.21645988285 | 808.714818811928 | 1085.58375727977 | 1060.79509120598 | 884.72622222183 | 973.573456730792 | 746.45299092875 | 747.927114056602 | Rbm3 |
| ENSMUSG00000031168 | -0.158468137479518 | 0.467426849740989 | 0.895328112556867 | 212.325723393726 | 237.809392842984 | 216.738169500494 | 267.309107863576 | 297.828629262794 | 344.909189444828 | 189.476212481217 | 211.677485110359 | Ebp |
| ENSMUSG00000031169 | -0.0122525320055335 | 0.966031316659359 | 0.994284908685797 | 49.6932544112975 | 52.9370223794314 | 47.3227444324223 | 53.4618215727152 | 55.8428679867739 | 77.0541806206532 | 37.4788112600209 | 35.8223436340608 | Porcn |
| ENSMUSG00000031171 | -0.0125031450335783 | 0.945338625097867 | 0.990505437194639 | 280.089252136404 | 193.83094348161 | 205.380710836713 | 218.536919762152 | 248.555510450935 | 237.277953022329 | 178.024353485099 | 242.072200921077 | Ftsj1 |
| ENSMUSG00000031173 | -0.465523426349629 | 0.0911890283916516 | 0.753823363185192 | 98.4829951060259 | 94.4722245540621 | 70.0376617599851 | 93.7926694258161 | 95.2613630362614 | 114.969729814943 | 91.61487196894 | 191.052499381657 | Otc |
| ENSMUSG00000031174 | 0.325850336805859 | 0.235649856414975 | 0.832048466795696 | 84.0267756409212 | 54.56585383726 | 106.002947528626 | 69.4065753751039 | 48.1781606160403 | 53.8156182112498 | 74.9576225200418 | 72.73021283279 | Rpgr |
| ENSMUSG00000031176 | 0.12864584273379 | 0.242018367518627 | 0.832048466795696 | 1388.70058236662 | 1232.21099784738 | 1247.42754323865 | 1360.93163336859 | 1051.15986798633 | 1157.03579154187 | 1296.14222274239 | 1277.66358961483 | Dynlt3 |
| ENSMUSG00000031189 | 0.701670286751925 | 0.414227227505651 | 0.882192126163777 | 8.13162344912141 | 14.6594831204579 | 29.3401015481019 | 0.937926694258161 | 4.37983278327639 | 2.44616446414772 | 9.36970281500523 | 16.2828834700276 | Aff2 |
| ENSMUSG00000031196 | 0.0229127954866938 | 0.934213611771532 | 0.98818087308472 | 28.0089252136404 | 35.0198763433161 | 41.6440151005317 | 30.9515809105193 | 28.4689130912965 | 28.1308913376988 | 40.6020455316893 | 35.8223436340608 | F8 |
| ENSMUSG00000031197 | 0.15298025009654 | 0.193932433629878 | 0.809162446871198 | 904.417230285614 | 811.158065998671 | 872.631407333868 | 987.636809053844 | 732.527033002976 | 748.526326029202 | 844.314331441027 | 889.045437463508 | Vbp1 |
| ENSMUSG00000031198 | 0.175106190380836 | 0.18801498435281 | 0.808677608936066 | 4192.30364488037 | 3856.25847640935 | 4157.77632583263 | 4330.40754738993 | 3169.90397689629 | 3200.80620133729 | 4010.23280482224 | 4263.9444180179 | Fundc2 |
| ENSMUSG00000031200 | 0.262364884209012 | 0.546351675533129 | 0.914211323944814 | 16.2632468982428 | 11.4018202048006 | 14.1968233297267 | 15.0068271081306 | 7.66470737073368 | 9.78465785659088 | 18.7394056300105 | 10.8552556466851 | Mtcp1 |
| ENSMUSG00000031201 | 0.0955000820008596 | 0.433990150566484 | 0.887868762931002 | 725.521514404943 | 587.193740547231 | 657.786147610671 | 746.589648629496 | 591.277425742312 | 660.464405319884 | 606.948526794228 | 683.88110574116 | Brcc3 |
| ENSMUSG00000031204 | 0.133446198090305 | 0.50508383560754 | 0.901404296286819 | 245.75573090678 | 217.448999620126 | 192.130342395635 | 257.929840920994 | 203.662224422352 | 149.216032313011 | 262.351678820146 | 216.019587369033 | Asb12 |
| ENSMUSG00000031207 | -0.139446423526712 | 0.225884522577364 | 0.832048466795696 | 2741.26061607048 | 3318.74409532589 | 3269.05518539174 | 2876.62117128978 | 3668.10995599397 | 3495.56901926709 | 3282.5192195235 | 2999.30713517909 | Msn |
| ENSMUSG00000031209 | -0.457846822734501 | 0.224169395133041 | 0.832048466795696 | 52.4037955610046 | 36.6487078011448 | 41.6440151005317 | 44.0825546301336 | 100.736154015357 | 68.4926049961361 | 40.6020455316893 | 30.3947158107182 | Heph |
| ENSMUSG00000031214 | -0.263223676063493 | 0.231727935516664 | 0.832048466795696 | 58.7283915769879 | 81.4415728914329 | 61.519567762149 | 80.6616957062019 | 97.4512794278996 | 97.8465785659088 | 71.8343882483734 | 72.73021283279 | Ophn1 |
| ENSMUSG00000031216 | -0.180683579218378 | 0.176489868241424 | 0.799889021902745 | 543.011743657996 | 587.193740547231 | 639.80350472635 | 537.431995809926 | 751.1413223319 | 639.672007374629 | 552.812466085309 | 671.940324529806 | Stard8 |
| ENSMUSG00000031217 | -0.500940603228269 | 0.10840820367472 | 0.762287160109383 | 516.809845877494 | 530.184639523228 | 869.792042667923 | 712.824287636203 | 1132.18677447695 | 1391.86758010005 | 603.825292522559 | 593.782483873674 | Efnb1 |
| ENSMUSG00000031221 | 0.0789214108118506 | 0.511899986170293 | 0.902394714365738 | 1206.19081161968 | 1054.66836894406 | 1084.63730239112 | 1319.66285882123 | 1092.76827942746 | 1237.75921885875 | 1055.65318382392 | 1031.24928643508 | Igbp1 |
| ENSMUSG00000031226 | -0.055973732551772 | 0.71945053465033 | 0.951674101666722 | 137.334084918495 | 118.904696421492 | 114.521041526462 | 119.116690170786 | 128.110108910834 | 117.415894279091 | 142.627698406191 | 120.493337678204 | Pbdc1 |
| ENSMUSG00000031227 | -0.935756305068996 | 0.0041739287675072 | 0.635862253526407 | 33.4300075130547 | 50.4937751926884 | 45.4298346551254 | 36.5791410760683 | 97.4512794278996 | 114.969729814943 | 57.2592949805875 | 48.8486504100828 | Magee1 |
| ENSMUSG00000031229 | 0.154098077275759 | 0.377855690093057 | 0.87422797717331 | 1038.13726033783 | 1355.18777291344 | 1349.64467121269 | 1062.6709445945 | 974.512794278996 | 863.496055844145 | 1284.69036374627 | 1195.16364670003 | Atrx |
| ENSMUSG00000031230 | -0.179986400383655 | 0.422040986983741 | 0.883119811570378 | 278.282224703266 | 237.809392842984 | 154.272146849697 | 174.454365132018 | 279.21433993387 | 194.470074899744 | 242.57119509958 | 240.986675356409 | Fgf16 |
| ENSMUSG00000031231 | 0.151517054143502 | 0.2831222868874 | 0.850587941933635 | 14907.0728096727 | 10679.4334532536 | 12283.0915448795 | 14385.9196365317 | 11374.4257381688 | 13326.7040006768 | 10971.9219963711 | 11373.051341032 | Cox7b |
| ENSMUSG00000031232 | -0.133826792964664 | 0.227701198060941 | 0.832048466795696 | 836.653701542936 | 812.7868974565 | 779.87882824632 | 693.127827056781 | 813.553939493589 | 947.888729857241 | 805.79444209045 | 860.821772782126 | Magt1 |
| ENSMUSG00000031239 | -0.342161913392313 | 0.267078563111901 | 0.843793744782845 | 153.597331816738 | 89.5857301805761 | 105.056492639978 | 105.047789756914 | 218.991639163819 | 133.315963296051 | 82.2451691539348 | 140.032797842237 | Itm2a |
| ENSMUSG00000031242 | 0.142124983167538 | 0.464600235942268 | 0.895328112556867 | 147.272735800754 | 213.376920975554 | 237.56017705076 | 191.337045628665 | 201.472308030714 | 150.439114545085 | 195.722681024554 | 167.17093695895 | 2610002M06 |
| ENSMUSG00000031245 | 0.0429037927532759 | 0.741845108519361 | 0.954383574136061 | 1188.12053728829 | 991.958357817652 | 957.812347312228 | 1282.14579105091 | 1051.15986798633 | 1139.91264029284 | 1012.96898211112 | 1086.61109023318 | Hmgn5 |
| ENSMUSG00000031246 | -0.0876724944381485 | 0.593417717160527 | 0.923935529355527 | 856.531003307455 | 835.590537866101 | 785.557557578211 | 1015.77460988159 | 913.195135313127 | 1147.25113368528 | 737.083288113745 | 916.18357658022 | Sh3bgrl |
| ENSMUSG00000031253 | -0.659130305246848 | 0.0925007250599859 | 0.754058637722175 | 47.8862269781594 | 43.9784493613737 | 56.7872933189068 | 62.8410885152968 | 111.685735973548 | 133.315963296051 | 40.6020455316893 | 48.8486504100828 | Srpx2 |
| ENSMUSG00000031255 | 0.173194210093811 | 0.620857310679589 | 0.930381707340543 | 39.754603529038 | 48.8649437348597 | 34.0723759913441 | 19.6964605794214 | 22.994122112201 | 26.9078091056249 | 38.5198893505771 | 37.9933947633978 | Sytl4 |
| ENSMUSG00000031256 | -0.309533598057035 | 0.0369033246057838 | 0.679566075569698 | 159.921927832721 | 157.182235680465 | 193.076797284283 | 144.440710915757 | 197.092475247437 | 187.131581507301 | 218.626399016789 | 207.335382851685 | Cstf2 |
| ENSMUSG00000031258 | 0.0342173291433576 | 0.928662755392325 | 0.986901071486195 | 25.2983840639333 | 19.5459774939439 | 15.1432782183751 | 15.0068271081306 | 15.3294147414674 | 20.7923979452556 | 12.4929370866736 | 24.9670879873757 | Xkrx |
| ENSMUSG00000031266 | -0.171591829491492 | 0.460248814708595 | 0.894774044236998 | 44.2721721118832 | 50.4937751926884 | 48.2691993210708 | 51.5859681841989 | 58.0327843784121 | 64.8233582999146 | 39.5609674411332 | 57.5328549274309 | Gla |
| ENSMUSG00000031271 | 0.260235334803475 | 0.519249194416381 | 0.905962179970576 | 26.2018977805023 | 26.0613033252585 | 38.8046504345863 | 37.5170677703264 | 21.8991639163819 | 31.8001380339204 | 11.4518589961175 | 42.3354970220718 | Serpina7 |
| ENSMUSG00000031274 | 0.105513640240105 | 0.611169790782854 | 0.927932112008768 | 490.607948096991 | 567.647763053287 | 424.958245003153 | 363.915557372167 | 413.894198019619 | 310.66288694676 | 515.333654825288 | 476.545722889475 | Col4a5 |
| ENSMUSG00000031278 | -0.0415399536424377 | 0.797440204051041 | 0.962499516359833 | 177.992202164102 | 223.96432545144 | 234.720812384815 | 220.412773150668 | 220.086597359638 | 188.354663739374 | 211.338852382896 | 261.61166108511 | Acsl4 |
| ENSMUSG00000031284 | 0.608967748934321 | 0.283102718006261 | 0.850587941933635 | 10.8421645988285 | 20.3603932228582 | 13.2503684410783 | 8.44134024832345 | 6.56974917491458 | 2.44616446414772 | 14.5750932677859 | 10.8552556466851 | Pak3 |
| ENSMUSG00000031286 | 0.111997040185116 | 0.722678120493399 | 0.951674101666722 | 137.334084918495 | 193.016527752696 | 113.574586637814 | 62.8410885152968 | 131.394983498292 | 127.200552135681 | 114.518589961175 | 96.6117752554972 | Glt28d2 |
| ENSMUSG00000031290 | -0.00455167490181225 | 0.989190720209051 | 0.998437672973644 | 22.5878429142261 | 17.1027303072009 | 20.8220075502658 | 28.1378008277448 | 28.4689130912965 | 20.7923979452556 | 16.6572494488982 | 22.7960368580387 | Lrch2 |
| ENSMUSG00000031292 | -0.375970547007108 | 0.0620968344238648 | 0.715615581523459 | 83.1232619243521 | 103.43079757212 | 94.6454888648447 | 95.6685228143324 | 112.780694169367 | 90.5080851734656 | 139.504464134522 | 145.46042566558 | Cdkl5 |
| ENSMUSG00000031295 | 0.131048990937637 | 0.334230483395361 | 0.858148279095856 | 297.256012751216 | 285.045505120015 | 318.008842585878 | 271.998741334867 | 266.074841584041 | 223.824048469516 | 296.707255808499 | 282.236646813812 | Phka2 |
| ENSMUSG00000031299 | 0.320707457667727 | 0.133333363933974 | 0.77303252008843 | 30411.3681859975 | 34636.2865349975 | 36347.653543655 | 30310.976978341 | 22153.194217812 | 17804.4080522992 | 33100.0368111418 | 32395.3394264023 | Pdha1 |
| ENSMUSG00000031303 | 0.33455222021098 | 0.506173993645692 | 0.901494649536949 | 13.5527057485357 | 21.1748089517725 | 32.1794662140472 | 29.075727522003 | 13.1394983498292 | 9.78465785659088 | 38.5198893505771 | 14.1118323406906 | Map3k15 |
| ENSMUSG00000031304 | -0.778943617495477 | 0.0329171108051939 | 0.671606027118506 | 68.6670424592474 | 52.122606650517 | 62.4660226507975 | 47.8342614071662 | 127.015150715015 | 156.554525705454 | 58.3003730711437 | 55.3618037980939 | Il2rg |
| ENSMUSG00000031309 | -0.0160555627061678 | 0.908245157229865 | 0.98504921011344 | 683.056369726198 | 785.911178402327 | 760.949730473351 | 715.638067718977 | 712.817785478232 | 602.979540412413 | 854.725112346588 | 806.545494548701 | Rps6ka3 |
| ENSMUSG00000031310 | -0.0377280689306018 | 0.848672122702817 | 0.972640876584422 | 205.097613661173 | 217.448999620126 | 229.988537941573 | 226.040333316217 | 222.276513751277 | 152.885279009232 | 286.296474902938 | 238.815624227072 | Zmym3 |
| ENSMUSG00000031311 | -0.00908646510613753 | 0.922847959727302 | 0.986713918026868 | 1674.21091680244 | 1826.73447995484 | 2020.68118726443 | 1815.8260800838 | 1924.93650824997 | 1871.31581507301 | 1838.54390792214 | 1748.78168468097 | Nono |
| ENSMUSG00000031312 | 0.0289529635486272 | 0.815694097647081 | 0.966177091475091 | 3359.26399820371 | 2788.55945580266 | 2490.12281203406 | 2802.52496244339 | 3000.18545654433 | 2945.18201483385 | 2601.65414829979 | 2667.13631239052 | Itgb1bp2 |
| ENSMUSG00000031314 | 0.0714532678600754 | 0.723113780148881 | 0.951674101666722 | 449.949830851384 | 562.761268679801 | 489.317177431247 | 401.432625142493 | 434.698403740182 | 314.332133642982 | 533.031982364742 | 528.650949993563 | Taf1 |
| ENSMUSG00000031320 | -0.0160582339260134 | 0.912937091733839 | 0.986226496507248 | 8282.51023978844 | 6978.72838106688 | 8156.54823037231 | 9464.61827175911 | 8881.2059262887 | 9631.77257758165 | 7124.09737367564 | 7613.87631058491 | Rps4x |
| ENSMUSG00000031327 | -0.642711152626896 | 0.123463473697529 | 0.770251097497048 | 17.1667606148119 | 11.4018202048006 | 8.51809399783602 | 15.9447538023887 | 22.994122112201 | 19.5693157131818 | 28.1091084450157 | 11.9407812113536 | Chic1 |
| ENSMUSG00000031328 | -0.546786352672444 | 0.102252333166751 | 0.759800953500741 | 1030.00563688871 | 1413.01128966636 | 1198.21188902893 | 973.567908639971 | 2345.40045544451 | 2479.18768441371 | 995.270654571667 | 922.696729968231 | Flna |
| ENSMUSG00000031333 | 0.0861330113206481 | 0.736204629141714 | 0.954383574136061 | 1014.64590370704 | 1224.88125628715 | 1028.79646396086 | 840.382318055312 | 786.179984598112 | 591.971800323748 | 1235.75969349013 | 1255.95307832146 | Abcb7 |

| | | | | | | | | | | | | |
|---|---|---|---|---|---|---|---|---|---|---|---|---|
| **ENSMUSG00000031337** | 0.346020965628865 | 0.228273898720541 | 0.832048466795696 | 344.238726012806 | 399.878122896935 | 483.638448099356 | 288.881421831514 | 216.801722772181 | 187.131581507301 | 398.732908683 | 389.703677715994 | Mtm1 |
| **ENSMUSG00000031342** | -0.194444760106253 | 0.370083400974246 | 0.872916429836026 | 254.790868072471 | 313.550055632016 | 240.399541716705 | 310.453735799451 | 384.330326732503 | 403.617136584374 | 249.858741733473 | 244.243252050414 | Gpm6b |
| **ENSMUSG00000031343** | -0.635961320656558 | 0.0450391922416247 | 0.693563614452513 | 31.6229800799166 | 36.6487078011448 | 37.8581955459379 | 24.3860940507122 | 47.0832024202212 | 78.277262852727 | 42.6842017128016 | 35.8223436340608 | Gabra3 |
| **ENSMUSG00000031347** | -0.0338888048786095 | 0.731618124579113 | 0.953120967023437 | 579.152292320758 | 526.926976607571 | 540.425741418263 | 579.638697051544 | 537.624474147177 | 611.54111603693 | 548.648153723084 | 581.84170266232 | Cetn2 |
| **ENSMUSG00000031349** | -0.300730786439183 | 0.101321902115483 | 0.759800953500741 | 70.4740698923855 | 84.6992358070902 | 63.4124775394459 | 84.4134024832345 | 93.0714466446232 | 100.292743030056 | 88.4916376972716 | 92.2696729968231 | Nsdhl |
| **ENSMUSG00000031352** | 0.0126508712772172 | 0.955145116389843 | 0.992371505557225 | 864.662626756576 | 1022.9061555164 | 1047.72556173383 | 814.120370616084 | 791.654775577207 | 613.987280501078 | 1196.198726049 | 1113.74922934989 | Hccs |
| **ENSMUSG00000031353** | 0.096697820145175 | 0.511350232847738 | 0.902394714365738 | 3298.72857919358 | 3414.03073560887 | 3445.09579468035 | 3016.37224873425 | 2957.48208690738 | 2389.90268147232 | 3510.51532135529 | 3460.6555001632 | Rbbp7 |
| **ENSMUSG00000031355** | -0.500537466427963 | 0.097983418755896 | 0.759800953500741 | 40.658117245607 | 49.679359463774 | 40.6975602118832 | 30.9515809105193 | 51.4630352034976 | 79.5003450848009 | 35.3966550789086 | 64.046008315442 | Arhgap6 |
| **ENSMUSG00000031357** | 0.161332744233281 | 0.179230155840711 | 0.801755154770274 | 778.828823682517 | 723.201167275924 | 716.466350706874 | 728.769041438591 | 625.221129812704 | 563.840908986049 | 744.370834747638 | 700.163989211187 | Syap1 |
| **ENSMUSG00000031358** | 0.0878784203482779 | 0.581545542429625 | 0.922450244597442 | 1081.50591873315 | 1174.38748109446 | 1288.12510345054 | 1127.38788649831 | 966.848086908263 | 876.949960396957 | 1269.07419238793 | 1280.92016630884 | Msl3 |
| **ENSMUSG00000031360** | 0.0918653764217918 | 0.630758558961033 | 0.931825800800958 | 116.553269437407 | 140.079505373265 | 184.558703286447 | 159.447538023887 | 153.294147414674 | 155.33144347338 | 130.134761319517 | 124.835439936878 | Ctps2 |
| **ENSMUSG00000031365** | -0.17380776262706 | 0.31603942807243 | 0.853356198600137 | 96.6759676728878 | 117.275864963663 | 130.610774633486 | 102.23400967414 | 136.869774477387 | 111.300483118721 | 119.723980413956 | 135.690695583563 | Zfp275 |
| **ENSMUSG00000031367** | -0.0750078937038453 | 0.541305796515726 | 0.913837378403038 | 683.056369726198 | 719.129088631352 | 600.998854291764 | 703.445020693621 | 694.203496149307 | 817.018931025338 | 707.933101578173 | 633.946929766408 | Ap1s2 |
| **ENSMUSG00000031370** | 0.174969771856686 | 0.0941516691621263 | 0.757085157577544 | 549.336339673979 | 513.896324944941 | 528.121827865833 | 579.638697051544 | 523.390017601528 | 479.448234972953 | 443.499266576914 | 476.545722889475 | Zrsr2 |
| **ENSMUSG00000031371** | -0.135996586420319 | 0.518328055275595 | 0.905504473025266 | 102.097049972302 | 88.7713144516618 | 88.0203046443056 | 123.806323642077 | 136.869774477387 | 118.638976511164 | 92.6559500594962 | 94.4407241261602 | Haus7 |
| **ENSMUSG00000031375** | -1.04028220704794 | 0.0110379307018747 | 0.635862253526407 | 2199.15238612906 | 2383.79483853224 | 1856.94449152825 | 2224.76211878036 | 5216.38084488218 | 8489.41377282466 | 2159.19595981343 | 1956.11706753265 | Bgn |
| **ENSMUSG00000031376** | 0.484782659880885 | 0.382676390338851 | 0.875126835008975 | 21.6843291976571 | 13.0306516626293 | 6.62518422053913 | 9.37926694258161 | 4.37983278327639 | 7.33849339244316 | 11.4518589961175 | 13.0263067760221 | Atp2b3 |
| **ENSMUSG00000031377** | -0.489654886823942 | 0.313600267937786 | 0.85295001595893 | 20.780815481088 | 17.9171460361152 | 8.51809399783602 | 10.3171936368398 | 24.0890803080201 | 33.0232202659942 | 12.4929370866736 | 11.9407812113536 | Bmx |
| **ENSMUSG00000031378** | 0.028526530368346 | 0.852690611101091 | 0.973120292022893 | 706.547726356993 | 889.341975974447 | 745.806452254976 | 698.75538722233 | 742.381656765348 | 594.417964787896 | 784.972880279327 | 858.650721652789 | Abcd1 |
| **ENSMUSG00000031379** | 0.0374098010302028 | 0.847100002594178 | 0.972334438148346 | 211.422209677157 | 280.159010746529 | 300.026199701558 | 362.03970398365 | 302.208462046071 | 316.77829810713 | 266.515991182371 | 238.815624227072 | Pir |
| **ENSMUSG00000031380** | -0.139916285879874 | 0.682246699986359 | 0.944452327903597 | 112.035700854562 | 87.1424829938332 | 38.8046504345863 | 87.227182566009 | 72.2672409240604 | 124.754387671534 | 95.7791843311646 | 66.217059444779 | Vegfd |
| **ENSMUSG00000031381** | 0.0416378596955924 | 0.890952604794427 | 0.982142775127356 | 22.5878429142261 | 37.4631235300591 | 29.3401015481019 | 25.3240207449704 | 28.4689130912965 | 29.3539735697726 | 32.2734208072402 | 21.7105112933701 | Piga |
| **ENSMUSG00000031382** | 0.052427242090817 | 0.709327623464161 | 0.949394580728696 | 2933.70903769969 | 2918.05155670004 | 4004.45063387158 | 3440.31511453894 | 3371.376284927 | 3423.40716757473 | 3181.53464473955 | 2845.16250499616 | Asb11 |
| **ENSMUSG00000031385** | 0.240186319618317 | 0.348288348249969 | 0.861266047293794 | 35.2370349461928 | 46.4216965481167 | 38.8046504345863 | 48.7721881014244 | 36.1336204620302 | 29.3539735697726 | 42.6842017128016 | 34.7368180693922 | Plxnb3 |
| **ENSMUSG00000031386** | 0.0140712623980265 | 0.957395235131983 | 0.992403427270738 | 638.784197614315 | 963.453807305651 | 995.670542858166 | 577.762843663027 | 722.672409240604 | 512.471455238947 | 1026.50299728835 | 882.532284075497 | Hcfc1 |
| **ENSMUSG00000031387** | -0.286604784125206 | 0.470651603590372 | 0.896509979960267 | 31.6229800799166 | 19.5459774939439 | 45.4298346551254 | 58.151455044006 | 60.2227007700503 | 62.3771938357668 | 30.191264626128 | 35.8223436340608 | Renbp |
| **ENSMUSG00000031388** | -0.0493887044035963 | 0.801651523543466 | 0.963461896828986 | 548.43282595741 | 440.598909342652 | 482.691993210708 | 622.783324987419 | 583.612718371579 | 681.25680326514 | 401.856142954669 | 501.51281087685 | Naa10 |
| **ENSMUSG00000031391** | -1.25088132339526 | 0.0102377018050516 | 0.635862253526407 | 26.2018977805023 | 30.9477976987445 | 25.5542819935081 | 19.6964605794214 | 111.685735973548 | 79.5003450848009 | 27.0680303544596 | 26.0526135520442 | L1cam |
| **ENSMUSG00000031392** | 0.0874603300387236 | 0.277850618368439 | 0.848075053184273 | 1852.20311896654 | 1839.76513161747 | 1791.63910421151 | 1887.10850884742 | 1623.82300439972 | 1674.39957570911 | 1798.982940481 | 1838.88030654845 | Irak1 |
| **ENSMUSG00000031393** | 0.12977133789983 | 0.314423130521549 | 0.853228028313558 | 727.328541838081 | 769.622863824041 | 892.506959995485 | 778.479156234274 | 789.464859185569 | 607.871869340708 | 725.631429117627 | 770.72315091464 | Mecp2 |
| **ENSMUSG00000031399** | -0.0499092558968564 | 0.735003367347574 | 0.954383574136061 | 293.64195788494 | 351.013179162076 | 356.813493020464 | 333.901903155905 | 375.57066116595 | 391.386314263635 | 333.144988977964 | 283.322172378481 | Fam3a |
| **ENSMUSG00000031400** | -0.369556922596789 | 0.159093131432109 | 0.789393312599749 | 101.193536255733 | 115.647033505835 | 123.985590412947 | 144.440710915757 | 147.819356435578 | 239.724117486476 | 114.518589961175 | 125.920965501547 | G6pdx |
| **ENSMUSG00000031402** | -0.119649097943446 | 0.726166799144769 | 0.952001589440381 | 149.983276950461 | 114.83261777692 | 274.47191770805 | 146.316564304273 | 188.332809680885 | 289.870489001505 | 128.052605138405 | 138.947272277569 | Mpp1 |
| **ENSMUSG00000031403** | 0.0496322507023535 | 0.70573700112526 | 0.947760509100828 | 534.880120208875 | 467.474628396825 | 406.975602118832 | 500.852854733858 | 488.351355335317 | 462.325083723919 | 410.184767679118 | 485.229927406823 | Dkc1 |
| **ENSMUSG00000031422** | 0.0587921183884099 | 0.502780906774057 | 0.900927059989498 | 1150.17296119239 | 1099.46123403434 | 1010.81382107654 | 1072.05021153708 | 1054.44474257379 | 1088.54318654574 | 984.859873666105 | 1033.42033756442 | Morf4l2 |
| **ENSMUSG00000031425** | -0.726898388078428 | 0.0943885303902575 | 0.757085157577544 | 40.658117245607 | 74.1118313312039 | 38.8046504345863 | 27.1998741334867 | 105.115986798633 | 108.854318654574 | 35.3966550789086 | 51.0197015394199 | Plp1 |
| **ENSMUSG00000031429** | 0.0973167261702731 | 0.472154671543687 | 0.897171173241185 | 415.616309621761 | 346.12668478859 | 334.098575692902 | 357.350070512359 | 333.962249724825 | 368.147751854232 | 293.584021536831 | 363.65106416395 | Psmd10 |
| **ENSMUSG00000031431** | -0.180228986053909 | 0.584839182690853 | 0.922651176944186 | 2571.4000373555 | 2313.75508584561 | 1414.95005852943 | 1194.9186084849 | 2671.6979977986 | 2987.98989295644 | 1460.63256105026 | 1373.18983930566 | Tsc22d3 |
| **ENSMUSG00000031432** | 0.240614235417261 | 0.126923127841214 | 0.772551378490123 | 659.565013095403 | 688.995706661522 | 795.022106464695 | 695.003680445298 | 551.858930692825 | 474.555906044658 | 711.056335849841 | 663.256120012458 | Prps1 |
| **ENSMUSG00000031433** | 0.337273205395314 | 0.450589354448765 | 0.89108031640712 | 43.3686583953142 | 59.452348210746 | 58.6802030962037 | 44.0825546301336 | 16.4243729372865 | 20.7923979452556 | 71.8343882483734 | 53.1907526687569 | Rbm41 |
| **ENSMUSG00000031434** | 0.158779077918383 | 0.49086390480578 | 0.899236955296196 | 58.7283915769879 | 74.1118313312039 | 88.0203046443056 | 77.8479156234274 | 61.3176589658694 | 50.1463715150282 | 78.0808567917102 | 77.072315091464 | Morc4 |
| **ENSMUSG00000031438** | -0.0415818633807032 | 0.711660409837115 | 0.950020479830613 | 1438.39383677792 | 1411.38245820853 | 1329.76911855107 | 1265.26311055426 | 1320.51958415783 | 1428.56004706227 | 1595.97271282256 | 1259.20965501547 | Rnf128 |
| **ENSMUSG00000031441** | 0.135722457786968 | 0.651691207672844 | 0.936720465895231 | 1598.31576461064 | 2084.08985029177 | 1867.35549530339 | 1344.04895287195 | 1119.04727612712 | 884.2884537894 | 2049.88276030503 | 2220.98530531177 | Atp11a |
| **ENSMUSG00000031442** | -0.233021791456701 | 0.427941671835755 | 0.885066977635295 | 1675.11443051901 | 1677.69640156352 | 1374.25249831754 | 936.050840869645 | 2348.68533003196 | 2048.66273872371 | 1157.67883669842 | 1101.80844813854 | Mcf2l |
| **ENSMUSG00000031443** | -0.533296013379507 | 0.227158980201732 | 0.832048466795696 | 9.93865088225949 | 22.8036404096012 | 10.4110037751329 | 13.1309737196143 | 15.3294147414674 | 19.5693157131818 | 15.616171358342 | 31.4802413753867 | F7 |
| **ENSMUSG00000031444** | -0.375749123769832 | 0.231587761442124 | 0.832048466795696 | 55.1143367107117 | 118.090280692578 | 81.3951204237664 | 71.2824287636203 | 68.9823663366031 | 108.854318654574 | 93.6970281500523 | 151.973579053591 | F10 |
| **ENSMUSG00000031445** | -0.436220158247738 | 0.237406776772015 | 0.832048466795696 | 37.9475760958999 | 81.4415728914329 | 40.6975602118832 | 53.4618215727152 | 58.0327843784121 | 51.3694537471021 | 58.3003730711437 | 121.578863242873 | Proz |
| **ENSMUSG00000031446** | 0.24803468761782 | 0.172182146469017 | 0.799889021902745 | 3126.1574593289 | 3376.56761207881 | 3128.03340698312 | 2962.91042716153 | 2350.8752464236 | 1912.90061096352 | 3232.5474711768 | 3106.77416608127 | Cul4a |
| **ENSMUSG00000031447** | 0.0335968632200355 | 0.558330714619365 | 0.916574868225118 | 9172.47125060895 | 8930.06846754561 | 8980.91043838511 | 8950.63444330563 | 8725.72186248238 | 8998.21598136739 | 8977.21637486557 | 8504.00727361309 | Lamp1 |
| **ENSMUSG00000031448** | 0.148615656480472 | 0.312932649789713 | 0.852615634676169 | 25086.0583405395 | 21145.4899855316 | 19281.1789915462 | 27990.5463367463 | 20893.99229262 | 23643.4026282198 | 20287.4887506674 | 19526.4338572571 | Adprhl1 |
| **ENSMUSG00000031451** | -0.257398477135376 | 0.11721321182275 | 0.769868040577702 | 2876.78767355584 | 2590.65643367648 | 2372.76240584166 | 2810.96630269171 | 3588.17800769918 | 3852.70903103266 | 2639.13295955981 | 2653.02448004983 | Gas6 |
| **ENSMUSG00000031453** | -0.360879475656873 | 0.0500759468248501 | 0.703396849069134 | 254.790868072471 | 233.737314198412 | 300.026199701558 | 289.819348525772 | 439.078236523458 | 365.701587390084 | 301.91264626128 | 277.894544555138 | Rasa3 |
| **ENSMUSG00000031458** | 0.00079387336571881 | 0.99572776248168 | 0.999262355603273 | 257.501409222178 | 250.025628776699 | 212.952349945901 | 287.943495137256 | 249.650468646754 | 288.647406769431 | 240.489038918468 | 230.131419709724 | Coprs |
| **ENSMUSG00000031461** | 0.52333357910293 | 0.0356782469707001 | 0.679566075569698 | 35068.0778811943 | 35899.4453305436 | 47166.5793757953 | 46943.231047621 | 27890.775163904 | 16467.5791726424 | 37231.0346744686 | 33263.7598781371 | Myom2 |
| **ENSMUSG00000031465** | -1.15543213006734 | 0.0146808780065828 | 0.635862253526407 | 36.1405486627618 | 17.9171460361152 | 13.2503684410783 | 39.3929211588428 | 87.5966556655277 | 81.9465095489486 | 28.1091084450157 | 40.1644458927348 | Angpt2 |
| **ENSMUSG00000031467** | 0.0824130106078606 | 0.475340002777035 | 0.897549741595057 | 556.564449406532 | 494.350347450997 | 598.159489625818 | 584.328330522834 | 481.781606160403 | 508.802208542726 | 558.017856538089 | 559.045665804281 | Agpat5 |
| **ENSMUSG00000031478** | 0.139433227551305 | 0.469257642866378 | 0.895328112556867 | 78.6056933415069 | 94.4722245540621 | 100.324218196735 | 94.7305961200743 | 97.4512794278996 | 91.7311674055395 | 75.998700610598 | 69.4736361387845 | Nek3 |
| **ENSMUSG00000031479** | 0.157024420730966 | 0.305134337543638 | 0.851785577030345 | 881.829387371388 | 1055.48278467297 | 962.54462175547 | 1037.34692384953 | 856.257309130534 | 683.702967729287 | 1017.13329447335 | 972.630905942983 | Vps36 |
| **ENSMUSG00000031480** | -0.461588698458118 | 0.145873506122531 | 0.780607990324772 | 140.948139784771 | 92.0289773673191 | 126.824955078892 | 118.178763476528 | 222.276513751277 | 228.716377397812 | 93.6970281500523 | 113.980184290193 | Thsd1 |
| **ENSMUSG00000031482** | 0.0996725382180897 | 0.637405005996094 | 0.932931968211785 | 412.905768472054 | 520.411650776256 | 428.744064557746 | 364.853484066425 | 356.956371837026 | 297.208982393948 | 414.349080041342 | 542.762782334254 | Slc25a15 |
| **ENSMUSG00000031483** | -0.00495052970785582 | 0.977183996966499 | 0.996179425433073 | 464.406050316489 | 547.287369830429 | 520.550188756646 | 425.818719193205 | 475.211856985488 | 374.263163014601 | 515.333654825288 | 599.210111697016 | Erlin2 |
| **ENSMUSG00000031485** | 0.149757846193696 | 0.435888402319977 | 0.888687115594624 | 1602.83333319349 | 1790.08577215369 | 1948.75061572715 | 1687.33012297043 | 1516.51710120945 | 1081.20469315329 | 1977.0072939661 | 1759.63694032765 | Plpbp |
| **ENSMUSG00000031486** | -0.782597116879645 | 0.0401754697110024 | 0.679566075569698 | 135.527057485357 | 149.852494120236 | 147.646962629158 | 105.985716451172 | 302.208462046071 | 374.263163014601 | 157.202791673977 | 94.4407241261602 | Adgra2 |
| **ENSMUSG00000031487** | -0.417969646268696 | 0.0907035361407978 | 0.753823363185192 | 94.8689402397497 | 72.4829998733752 | 89.9132144216024 | 125.682177030594 | 128.110108910834 | 172.454594722414 | 110.35427759895 | 100.953877514171 | Brf2 |
| **ENSMUSG00000031490** | 0.3511013893402 | 0.0344201023518229 | 0.678229434609245 | 2875.88415983927 | 2653.36644480288 | 3251.07254250741 | 3939.29211588428 | 2803.09298129689 | 2759.27351555863 | 2205.0033957979 | 2204.70242184174 | Eif4ebp1 |
| **ENSMUSG00000031497** | -0.456424213446742 | 0.353273621377858 | 0.862728204353539 | 16.2632468982428 | 13.0306516626293 | 11.3574586637814 | 15.0068271081306 | 29.5638712871156 | 28.1308913376988 | 11.4518589961175 | 7.59867895267955 | Tnfsf13b |
| **ENSMUSG00000031502** | -0.658751226570518 | 0.0180956111904832 | 0.642273034804458 | 4605.20941335242 | 5374.32939510565 | 5395.7393201848 | 5269.27216834235 | 11963.5132475195 | 10165.0364307658 | 5374.04510345078 | 5090.02937273063 | Col4a1 |
| **ENSMUSG00000031503** | -0.433672649932809 | 0.0410722777880577 | 0.683226273624098 | 4146.22444533535 | 4984.22426095569 | 4734.16735301953 | 4336.03510755548 | 8566.95292408861 | 6636.44419123276 | 4744.19285866431 | 4636.2796866992 | Col4a2 |
| **ENSMUSG00000031504** | 0.0983501528398218 | 0.629126409979877 | 0.931825800800958 | 369.537110076739 | 286.674336577844 | 417.386605893965 | 554.314676306573 | 372.285786578493 | 381.601656407044 | 368.541644056872 | 397.302356668674 | Rab20 |
| **ENSMUSG00000031505** | 0.0109944589905306 | 0.945412150484013 | 0.990505437194639 | 939.654265231807 | 763.107537992726 | 831.933847121985 | 1062.6709445945 | 983.272459845549 | 1017.60441708545 | 818.287379177123 | 751.183690750607 | Naxd |
| **ENSMUSG00000031508** | 0.0143011585691548 | 0.921116840483325 | 0.986713918026868 | 300.870067617492 | 289.931999493501 | 263.114459044268 | 302.012395551128 | 263.884925192402 | 256.847268735511 | 274.84461590682 | 348.453706258591 | Ankrd10 |
| **ENSMUSG00000031511** | 0.0654913078854463 | 0.684930777553006 | 0.944452327903597 | 2521.7067829442 | 2560.52305170665 | 2214.70443943737 | 2099.07994174977 | 1915.0818844876 | 1852.9695815919 | 2600.61307020923 | 2609.60345746309 | Arhgef7 |
| **ENSMUSG00000031513** | 0.0263126284356223 | 0.787969663512944 | 0.961343752849784 | 677.635287426784 | 645.017257300148 | 709.841166486335 | 630.286738541484 | 706.248036303317 | 638.448925142555 | 653.797040869254 | 615.492995167044 | Leprotl1 |
| **ENSMUSG00000031516** | 0.0629300744926811 | 0.606593139204411 | 0.927110332696882 | 1366.1127394524 | 1083.98733518497 | 1249.32045301595 | 1267.13896394278 | 1165.03552035152 | 1351.50586644161 | 1164.96638333232 | 1073.58478345715 | Dctn6 |
| **ENSMUSG00000031517** | -0.773081742231549 | 0.181421968737043 | 0.803056849297055 | 32.5264937964856 | 10.5874044758863 | 13.2503684410783 | 15.9447538023887 | 50.3680770076785 | 42.8078781225851 | 22.903717992235 | 7.59867895267955 | Gpm6a |
| **ENSMUSG00000031519** | 0.368669897366553 | 0.125774400250897 | 0.771949105062466 | 608.064731250967 | 448.743066631795 | 351.134763688574 | 539.307849198443 | 348.196706270473 | 264.185762127954 | 394.568596320776 | 500.427285312182 | Asb5 |
| **ENSMUSG00000031520** | -0.632979573541749 | 0.119441472665797 | 0.769868040577702 | 55.1143367107117 | 55.3802695661743 | 59.6266579848522 | 79.7237690119437 | 119.350443344282 | 165.116101329971 | 39.5609674411332 | 64.046008315442 | Vegfc |
| **ENSMUSG00000031521** | 0.030508057713853 | 0.755887565053724 | 0.956566265296272 | 565.599586572222 | 496.79359463774 | 508.246275204216 | 489.59773440276 | 505.870686468423 | 508.802208542726 | 493.471014923609 | 509.11148982953 | Aga |
| **ENSMUSG00000031523** | -0.12330598651301 | 0.429931932146605 | 0.886017087288711 | 1627.22820354085 | 1643.49094094912 | 2238.36581165358 | 1615.10976751255 | 2230.4298448835 | 1691.52272695815 | 1977.0072939661 | 1859.50529227715 | Dlc1 |
| **ENSMUSG00000031527** | -0.0272344632625104 | 0.846498340431038 | 0.972262781013841 | 278.282224703266 | 247.582381589956 | 298.133289924261 | 249.488500672671 | 241.98576127602 | 261.739597663806 | 315.446661438509 | 273.552442296464 | Eri1 |
| **ENSMUSG00000031529** | 0.335788013819758 | 0.217299930354976 | 0.825996545635915 | 1030.90915060528 | 1391.02206498567 | 1264.46373123432 | 982.009248888295 | 733.621991198795 | 541.82542880872 | 1275.32066093127 | 1147.40052185461 | Tnks |
| **ENSMUSG00000031530** | -0.505136869516895 | 0.169130658978358 | 0.79907358295209 | 26.2018977805023 | 48.0505280059454 | 52.0550188756646 | 45.9584080186499 | 48.1781606160403 | 42.8078781225851 | 105.14888714617 | 47.7631248454143 | Dusp4 |
| **ENSMUSG00000031532** | 0.121392849175188 | 0.327752721704295 | 0.855977678491063 | 11858.6175299687 | 11477.5608675896 | 14194.9304199494 | 14581.9463156316 | 13298.2672882229 | 11999.6597788766 | 11534.1041652714 | 11074.5318107481 | Saraf |
| **ENSMUSG00000031533** | 0.107084678070555 | 0.253450191307284 | 0.837406774112203 | 897.189120553062 | 908.073537739476 | 908.596693102509 | 910.726820124675 | 848.5926017598 | 775.434135134827 | 929.68273486663 | 809.802071242707 | Mrps31 |
| **ENSMUSG00000031534** | 0.178800157568577 | 0.0319961362438696 | 0.671606027118506 | 1111.32187137993 | 1162.17124516075 | 1211.46225747001 | 1140.51886021792 | 1064.29936633616 | 1038.39681503071 | 1029.62623156002 | 954.176971343618 | Smim19 |
| **ENSMUSG00000031536** | 0.213901049611274 | 0.0523138485884202 | 0.703396849069134 | 649.626362213143 | 569.276594511116 | 660.625512276616 | 698.75538722233 | 558.428679867739 | 519.80994863139 | 593.414511616998 | 549.275935722265 | Polb |
| **ENSMUSG00000031537** | 0.0488963912972011 | 0.727531654160619 | 0.952237115597464 | 346.045753445944 | 408.836695914993 | 413.600786339371 | 332.963976461647 | 350.386622662111 | 315.555215875056 | 382.075659234102 | 402.729984492016 | Ikbkb |

| | | | | | | | | | | | | |
|---|---|---|---|---|---|---|---|---|---|---|---|---|
| **ENSMUSG00000031538** | -0.631366303286889 | 0.0866632768752998 | 0.747269672026173 | 244.852217190211 | 177.542628903324 | 158.057966404291 | 224.164479927701 | 429.223612761086 | 483.117481669175 | 168.654650670094 | 166.085411394282 | Plat |
| **ENSMUSG00000031539** | -0.41313405257503 | 0.125557728135413 | 0.771949105062466 | 82.2197482077831 | 51.3081909216027 | 62.4660226507975 | 62.8410885152968 | 104.021028602814 | 111.300483118721 | 61.4236073428121 | 68.388110574116 | Ap3m2 |
| **ENSMUSG00000031540** | 0.128133192927565 | 0.549848184265022 | 0.914699131434573 | 600.836621518415 | 828.260796305872 | 835.719666676579 | 518.673461924763 | 594.56230032977 | 486.786728365396 | 747.494069019306 | 717.532398245884 | Kat6a |
| **ENSMUSG00000031543** | -0.00096479073504813 | 0.994810658599338 | 0.999262355603273 | 9593.50864253012 | 8210.93937891426 | 7947.38169998101 | 8635.49107403489 | 7053.72069746662 | 7709.08730876154 | 10030.7874025084 | 9615.58545183364 | Ank1 |
| **ENSMUSG00000031545** | 0.0500606124482649 | 0.768896165467952 | 0.95879680569409 | 1598.31576461064 | 1814.51824402112 | 1627.90240847533 | 1437.84162229776 | 1503.37760285962 | 1174.15894279091 | 1932.24093607219 | 1646.74228160213 | Gpat4 |
| **ENSMUSG00000031546** | 0.184606511798604 | 0.147946714740298 | 0.780607990324772 | 463.50253659992 | 360.786167909048 | 422.118880337207 | 419.253232333398 | 348.196706270473 | 404.840218816448 | 368.541644056872 | 344.111603999917 | Gins4 |
| **ENSMUSG00000031548** | -0.986100882527006 | 0.0368869515415836 | 0.679566075569698 | 104.807591122009 | 119.719112150406 | 120.199770858353 | 116.302910088012 | 286.879047304603 | 448.871179171106 | 96.8202624217207 | 81.4144173501381 | Sfrp1 |
| **ENSMUSG00000031549** | -0.0375106139305235 | 0.949389478705332 | 0.991267322147698 | 12.6491920319666 | 21.1748089517725 | 8.51809399783602 | 6.56548685980713 | 6.56974917491458 | 7.33849339244316 | 11.4518589961175 | 24.9670879873757 | Ido2 |
| **ENSMUSG00000031555** | 0.185500109869293 | 0.405944307965551 | 0.881158207349486 | 1347.13895140445 | 1601.14132304557 | 1518.11364139211 | 1206.173728816 | 962.468254124986 | 897.742358342213 | 1635.53368026369 | 1491.51212585453 | Adam9 |
| **ENSMUSG00000031556** | 0.0307716723469932 | 0.832079323366008 | 0.967583158900775 | 990.251033359673 | 745.190391956611 | 833.826756899282 | 860.078778634734 | 914.290093508946 | 951.557976553463 | 756.863771834311 | 734.90080728058 | Tm2d2 |
| **ENSMUSG00000031557** | -0.187896961517971 | 0.189320448010247 | 0.808677608936066 | 523.134441893477 | 538.328796812371 | 460.923530771794 | 446.453106466885 | 644.930377337448 | 463.548165955993 | 542.401685179747 | 591.611432744337 | Plekha2 |
| **ENSMUSG00000031558** | -0.619248457759935 | 0.054506326835907 | 0.703396849069134 | 17.1667606148119 | 17.1027303072009 | 30.2865564367503 | 34.703287687552 | 42.7033696369448 | 36.6924669622158 | 35.3966550789086 | 36.9078691987293 | Slit2 |
| **ENSMUSG00000031561** | -0.837159759909617 | 0.0791988875222782 | 0.739855948845396 | 19.877301764519 | 45.6072808192024 | 22.7149173275627 | 14.0689004138724 | 49.2731188118594 | 79.5003450848009 | 29.1501865355718 | 26.0526135520442 | Tenm3 |
| **ENSMUSG00000031562** | -0.32071109781553 | 0.445779691758027 | 0.89108031640712 | 9.93865088225949 | 10.5874044758863 | 10.4110037751329 | 15.0068271081306 | 16.4243729372865 | 11.0077400886647 | 18.7394056300105 | 10.8552556466851 | Dctd |
| **ENSMUSG00000031563** | 0.141802259130827 | 0.391859801998982 | 0.878285927324016 | 1269.43677177951 | 1698.87121051529 | 1504.86327295103 | 1227.74604278393 | 1283.29100549998 | 1026.16599270997 | 1395.04464134522 | 1462.20293560848 | Wwc2 |
| **ENSMUSG00000031565** | -0.691832869555779 | 0.017695362690432 | 0.642273034804458 | 243.948703473642 | 297.26174105373 | 318.955297474527 | 285.129715054481 | 546.384139713729 | 706.941530138691 | 313.364505257397 | 284.407697943149 | Fgfr1 |
| **ENSMUSG00000031568** | 0.035096623232996 | 0.870194038195455 | 0.975613767803763 | 840.267756409212 | 800.570661522785 | 719.30571537282 | 761.596475737627 | 712.817785478232 | 479.448234972953 | 961.95615567387 | 891.216488592845 | Rwdd4a |
| **ENSMUSG00000031570** | -0.228466668745295 | 0.255285925346636 | 0.837406774112203 | 128.298947752804 | 109.13170767452 | 105.056492639978 | 137.87522405595 | 165.338687568684 | 165.116101329971 | 111.395355689507 | 121.578863242873 | Plpp5 |
| **ENSMUSG00000031574** | -0.0499409889378409 | 0.907362290844584 | 0.98504921011344 | 64.1494738764022 | 74.9262470601182 | 30.2865564367503 | 62.8410885152968 | 26.2789966996583 | 107.6312364225 | 54.1360607089191 | 53.1907526687569 | Star |
| **ENSMUSG00000031575** | 0.164991436418828 | 0.145277522661686 | 0.780607990324772 | 852.01343472461 | 906.444706281648 | 930.365155541423 | 935.112914175387 | 794.939650164664 | 689.818378889657 | 864.094815161593 | 881.446758510828 | Ash2l |
| **ENSMUSG00000031577** | -0.205926424119098 | 0.140552394054217 | 0.779711473022872 | 204.194099944604 | 185.686786192467 | 203.487801059416 | 182.895705380341 | 238.700886688563 | 250.731857575141 | 212.379930473452 | 194.309076075663 | Tti2 |
| **ENSMUSG00000031578** | 0.191245046722289 | 0.177846615334166 | 0.801696999069105 | 315.326287082597 | 267.1283590839 | 264.060913932917 | 273.874594723383 | 240.890803080201 | 210.370143916704 | 245.694429371248 | 283.322172378481 | Mak16 |
| **ENSMUSG00000031583** | -0.299159837000249 | 0.231663374134095 | 0.832048466795696 | 66.8600150261093 | 102.616381843205 | 71.930571537282 | 81.59962240046 | 114.970610561005 | 124.754387671534 | 90.5737938783839 | 68.388110574116 | Wrn |
| **ENSMUSG00000031584** | 0.0863327714581481 | 0.563725675195966 | 0.918590066040684 | 738.17070643691 | 934.134841064735 | 777.985918469023 | 825.375490947182 | 732.527033002976 | 626.218102821816 | 841.191097169358 | 884.703335204834 | Gsr |
| **ENSMUSG00000031585** | -0.0499910139656406 | 0.693725038866204 | 0.94469184542218 | 296.352499034647 | 268.757190541728 | 238.506631939409 | 256.991914226736 | 277.024423542232 | 298.432064626022 | 265.474913091815 | 258.355084391105 | Gtf2e2 |
| **ENSMUSG00000031586** | 0.145616872120755 | 0.127161472520504 | 0.772837268577874 | 2655.42681299642 | 2726.66386040517 | 3096.80039565772 | 3166.44051981555 | 2619.14000439928 | 2542.78796048155 | 2696.39225454039 | 2667.13631239052 | Rbpms |
| **ENSMUSG00000031590** | -0.119023896758981 | 0.447003615043154 | 0.89108031640712 | 352.370349461928 | 294.004078138073 | 317.06238769723 | 331.088123073131 | 379.950493949227 | 421.963370065482 | 295.666177717943 | 309.374785930525 | Frg1 |
| **ENSMUSG00000031591** | -0.145983113279039 | 0.235372598690257 | 0.832048466795696 | 466.213077749627 | 467.474628396825 | 473.227444324223 | 526.176875478828 | 601.132049504684 | 569.956320146419 | 478.895921655823 | 489.572029665497 | Asah1 |
| **ENSMUSG00000031592** | 0.288408035626988 | 0.311153012276444 | 0.851785577030345 | 985.733464776828 | 1307.1372449075 | 1344.91239676944 | 930.423280704096 | 718.292576457328 | 539.379264344572 | 1286.77251992738 | 1195.16364670003 | Pcm1 |
| **ENSMUSG00000031594** | -0.0945629205543052 | 0.765750904432285 | 0.958345041873658 | 125.588406603097 | 302.96265115613 | 159.004421292939 | 143.502784221499 | 131.394983498292 | 214.039390612925 | 185.311900118992 | 250.756405438425 | Fgl1 |
| **ENSMUSG00000031595** | -1.09912917168426 | 0.0415191499209145 | 0.683543358336265 | 28.0089252136404 | 24.4324718674299 | 14.1968233297267 | 28.1378008277448 | 67.887408140784 | 92.9542496376133 | 13.5340151772298 | 29.3091902460497 | Pdgfrl |
| **ENSMUSG00000031596** | -0.349544908870352 | 0.211102742393033 | 0.823123035902316 | 67.7635287426784 | 140.079505373265 | 65.3053873167428 | 96.6064495085906 | 97.4512794278996 | 113.746647582869 | 120.765058504512 | 140.032797842237 | Slc7a2 |
| **ENSMUSG00000031600** | 0.0931412411535106 | 0.642617156127967 | 0.934616627390525 | 503.257140128958 | 531.813470981057 | 590.587850516631 | 515.859681841989 | 462.072358635659 | 337.570696052385 | 625.687932424238 | 580.756177097652 | Vps37a |
| **ENSMUSG00000031601** | 0.26070891919953 | 0.184274630186801 | 0.805963874499992 | 1024.5845545893 | 1026.97823416097 | 1101.67349038679 | 992.326442525135 | 738.001823982071 | 612.764198269004 | 991.106342209442 | 1117.00580604389 | Cnot7 |
| **ENSMUSG00000031604** | -0.399058917280257 | 0.0404494680924788 | 0.681021953702461 | 103.90407740544 | 99.3587189275481 | 76.6628459805242 | 83.4754757889764 | 101.831112211176 | 112.523565350795 | 118.6829023234 | 146.545951230249 | Msmo1 |
| **ENSMUSG00000031605** | 0.0904738147695714 | 0.669849182835824 | 0.942388853397102 | 511.388763578079 | 533.442302438885 | 536.639921863669 | 296.384835385579 | 513.535393839156 | 441.532685778663 | 395.609674411332 | 413.585240138701 | Klhl2 |
| **ENSMUSG00000031608** | -0.140340443016395 | 0.421264705378969 | 0.882568531013176 | 140.948139784771 | 198.717437855096 | 191.183887506986 | 170.702658354985 | 164.243729372865 | 212.816308380852 | 216.544242835676 | 180.197243734972 | Galnt7 |
| **ENSMUSG00000031609** | -0.0544792290749445 | 0.733171934707712 | 0.953733898884637 | 176.185174730964 | 161.254314325037 | 180.772883731853 | 217.598993067893 | 208.042057205628 | 196.916239363891 | 161.367104036201 | 197.565652769668 | Sap30 |
| **ENSMUSG00000031613** | -0.107069040599836 | 0.709394346586099 | 0.949394580728696 | 65.0529875929712 | 76.5550785179469 | 100.324218196735 | 128.495957113368 | 82.1218646864323 | 112.523565350795 | 72.8754663389296 | 131.348593324889 | Hpgd |
| **ENSMUSG00000031616** | -0.136735340727094 | 0.381841809869377 | 0.875126835008975 | 766.17963165055 | 842.92027942633 | 856.541674226844 | 676.245146560134 | 809.174106710313 | 732.626257012242 | 1072.31043327282 | 839.111261488756 | Ednra |
| **ENSMUSG00000031617** | 0.253329007464967 | 0.1727201316842 | 0.799889021902745 | 285.510334435818 | 254.097707421271 | 224.309808609682 | 253.240207449704 | 151.104231023035 | 218.931719541221 | 249.858741733473 | 233.387996403729 | Tmem184c |
| **ENSMUSG00000031618** | -0.111042387735795 | 0.510384856200762 | 0.902394714365738 | 214.132750826864 | 215.820168162297 | 171.308334845369 | 158.509611329629 | 186.142893289246 | 185.908499275227 | 217.585320926233 | 231.216945274392 | Nr3c2 |
| **ENSMUSG00000031622** | -0.0904339940961437 | 0.575990865031484 | 0.922450244597442 | 1054.40050723608 | 889.341975974447 | 949.294253314392 | 1118.00861955573 | 1169.4153531348 | 1287.90559037377 | 936.970281500523 | 877.104656252154 | Sin3b |
| **ENSMUSG00000031626** | 0.128284524099557 | 0.562545007153741 | 0.917753130064067 | 9077.6023103692 | 10905.0266101629 | 7439.13542477679 | 8157.14845996323 | 6453.68360615776 | 6020.01074626754 | 10761.6242220788 | 9315.98039598513 | Sorbs2 |
| **ENSMUSG00000031627** | -0.130408066781035 | 0.353158650737203 | 0.862728204353539 | 388.510898124689 | 333.09603312596 | 419.279515671262 | 375.170677703265 | 435.793361936001 | 478.225152740879 | 377.911346871878 | 367.993166422624 | Irf2 |
| **ENSMUSG00000031628** | -0.180022040055275 | 0.513087848722538 | 0.903078589555164 | 38.8510898124689 | 43.9784493613737 | 48.2691993210708 | 61.9031618210386 | 74.4571573156986 | 50.1463715150282 | 39.5609674411332 | 54.2762782334254 | Casp3 |
| **ENSMUSG00000031631** | -0.0666794048710546 | 0.722813795400418 | 0.951674101666722 | 254.790868072471 | 309.477976987445 | 300.972654590206 | 246.674720589896 | 223.371471947096 | 256.847268735511 | 358.130863151311 | 325.657669400552 | Cfap97 |
| **ENSMUSG00000031633** | 0.0545181612931228 | 0.712061735798056 | 0.950190941515516 | 144798.915244788 | 114014.94438509 | 130801.012066104 | 155892.795852649 | 137632.960339873 | 153080.972166364 | 118663.121839679 | 115901.564539657 | Slc25a4 |
| **ENSMUSG00000031634** | 0.0983535775018086 | 0.389027007362073 | 0.876536136761342 | 445.432262268539 | 456.887223920938 | 462.81644054909 | 518.673461924763 | 419.368988998714 | 494.125221757839 | 421.636626675235 | 425.526021350055 | Ufsp2 |
| **ENSMUSG00000031636** | -0.265248704760004 | 0.449196916447864 | 0.89108031640712 | 225.878429142261 | 298.076156782644 | 268.793188376159 | 231.667893481766 | 280.309298129689 | 557.72549782568 | 247.776585552361 | 146.545951230249 | Pdlim3 |
| **ENSMUSG00000031637** | 0.0243840832606705 | 0.943581858311715 | 0.990505437194639 | 34.3335212296237 | 52.122606650517 | 53.9479286529615 | 46.8963347129081 | 20.8042057205628 | 45.2540425867328 | 59.3414511616998 | 58.6183804920994 | Lrp2bp |
| **ENSMUSG00000031639** | -0.259333232104572 | 0.250119611048351 | 0.835923538390114 | 89.4478579403355 | 111.574954861263 | 61.519567762149 | 82.5375490947182 | 120.445401540101 | 101.51582526213 | 104.107809055614 | 87.9275707381491 | Tlr3 |
| **ENSMUSG00000031641** | 0.0325290005176912 | 0.772013169286467 | 0.959045638244974 | 1079.69889130001 | 1023.72057124531 | 925.632881098181 | 1209.92543559303 | 1000.79179097865 | 1018.82749931753 | 1069.18719900115 | 1055.13084885779 | Cbr4 |
| **ENSMUSG00000031642** | -0.222639879910532 | 0.327280358568382 | 0.855977678491063 | 110.228673421423 | 107.502876216691 | 166.576060402127 | 190.399118934407 | 185.047935093427 | 157.777607937528 | 151.997401221196 | 174.76961591163 | Sh3rf1 |
| **ENSMUSG00000031644** | 0.0546048045158357 | 0.833137716509295 | 0.967868401766821 | 271.054114970713 | 315.99330281876 | 273.525462819401 | 211.033506208086 | 187.237851485066 | 181.016170346931 | 363.336253604092 | 299.605055848508 | Nek1 |
| **ENSMUSG00000031647** | -0.0240130562229333 | 0.929103648470329 | 0.987017183728709 | 247.562758339918 | 325.766291565731 | 176.040609288611 | 209.15765281957 | 162.053812981226 | 206.700897220482 | 296.707255808499 | 309.374785930525 | Mfap3l |
| **ENSMUSG00000031652** | -0.0217425765686538 | 0.906908108053337 | 0.98504921011344 | 465.309564033058 | 681.665965101293 | 721.198625150116 | 513.983828453472 | 592.372383938131 | 501.463715150282 | 644.427338054249 | 679.539003482486 | N4bp1 |
| **ENSMUSG00000031657** | 0.278473933559552 | 0.233490381958592 | 0.832048466795696 | 639.687711330884 | 720.757920089181 | 686.179794270124 | 597.459304242449 | 488.351355335317 | 327.786038195794 | 677.741836952045 | 684.966631305828 | Heatr3 |
| **ENSMUSG00000031659** | -0.569959580256583 | 0.106568340201775 | 0.76222154758118 | 41.5616309621761 | 64.338842584232 | 39.7511053232348 | 60.9652351267805 | 85.4067392738896 | 122.308223207386 | 43.7252798033577 | 56.4473293627624 | Adcy7 |
| **ENSMUSG00000031660** | 0.190332911550497 | 0.173262989394001 | 0.799889021902745 | 2310.28457326705 | 2404.96964748401 | 2236.47290187628 | 2359.82356275353 | 1867.99868206738 | 1667.06108231667 | 2449.65674707859 | 2174.30770603102 | Brd7 |
| **ENSMUSG00000031661** | -0.921502795914829 | 0.0317707236188124 | 0.671606027118506 | 21.6843291976571 | 26.0613033252585 | 27.447191770805 | 13.1309737196143 | 41.6084114411257 | 74.6080161565054 | 28.1091084450157 | 23.8815624227072 | Nkd1 |
| **ENSMUSG00000031662** | -0.650888573169813 | 0.178982996829287 | 0.801755154770274 | 14.4562194651047 | 6.51532583131463 | 11.3574586637814 | 21.5723139679377 | 22.994122112201 | 33.0232202659942 | 16.6572494488982 | 11.9407812113536 | Snx20 |
| **ENSMUSG00000031665** | 0.0808775038272397 | 0.89345948179028 | 0.983527674914169 | 17.1667606148119 | 14.6594831204579 | 12.3039135524298 | 5.62756016554897 | 19.7092475247437 | 17.123151249034 | 7.28754663389296 | 3.25657669400552 | Sall1 |
| **ENSMUSG00000031666** | 0.386182125171061 | 0.163995309077508 | 0.793968226938511 | 1042.65482892068 | 1453.73207611208 | 1745.26281466774 | 1049.53997087488 | 848.5926017598 | 617.656527197299 | 1347.15504917964 | 1234.24256702809 | Rbl2 |
| **ENSMUSG00000031667** | 0.10555871859993 | 0.386842352102802 | 0.875691048029321 | 2013.02856051583 | 1755.88031153929 | 1663.86769424397 | 2053.12153373112 | 1544.98601430075 | 1675.62265794119 | 1886.43350008772 | 1849.73556219514 | Aktip |
| **ENSMUSG00000031668** | -0.10822988664164 | 0.520367289544482 | 0.905962179970576 | 103.90407740544 | 100.173134656462 | 128.717864856189 | 113.489130005238 | 136.869774477387 | 116.192812047017 | 118.6829023234 | 108.552556466851 | Eif2ak3 |
| **ENSMUSG00000031669** | -0.521975524853531 | 0.222858595341246 | 0.832048466795696 | 25.2983840639333 | 13.8450673915436 | 13.2503684410783 | 17.8206071909051 | 28.4689130912965 | 11.0077400886647 | 23.9447960827911 | 36.9078691987293 | Gins3 |
| **ENSMUSG00000031671** | -0.0602748761306846 | 0.705728126765224 | 0.947760509100828 | 241.238162323935 | 211.748089517725 | 214.845259723197 | 212.909359596603 | 180.668102310151 | 269.078091056249 | 247.776585552361 | 221.447215192376 | Setd6 |
| **ENSMUSG00000031672** | 0.114908685879942 | 0.386236153376642 | 0.875126835008975 | 22876.9673035282 | 25326.7003377778 | 26581.1855476916 | 23907.7514366405 | 21873.9798778781 | 18607.9730787717 | 24604.8395922037 | 26048.2714497855 | Got2 |
| **ENSMUSG00000031673** | -0.384782473361634 | 0.39909706165798 | 0.880459952732461 | 61.4389327266951 | 26.0613033252585 | 27.447191770805 | 28.1378008277448 | 54.7479097909548 | 66.0464405319884 | 49.9717483466946 | 16.2828834700276 | Cdh11 |
| **ENSMUSG00000031681** | -0.282977955995405 | 0.149091394562191 | 0.782024490202776 | 207.80815481088 | 208.490426602068 | 247.024725937245 | 228.854113398991 | 321.917709570815 | 326.562955963721 | 195.722681024554 | 242.072200921077 | Smad1 |
| **ENSMUSG00000031683** | 0.0462050857063136 | 0.808315380053953 | 0.964026716319739 | 506.871194995234 | 455.25839246311 | 459.030620994497 | 647.169419038131 | 454.407651264925 | 655.572076391589 | 441.417110395802 | 452.664160466768 | Lsm6 |
| **ENSMUSG00000031684** | -0.205852269489689 | 0.384656278779266 | 0.875126835008975 | 86.7373167906283 | 90.4001459094905 | 61.519567762149 | 73.1582821521366 | 117.160526952643 | 96.6234963338349 | 69.7522320672611 | 77.072315091464 | Slc10a7 |
| **ENSMUSG00000031691** | 0.0973335279241875 | 0.614829653314058 | 0.928075610917191 | 571.020668871636 | 805.457155896271 | 714.573440929577 | 608.714424573547 | 631.790878987619 | 450.09426140318 | 726.672507208183 | 714.275821551878 | Tnpo2 |
| **ENSMUSG00000031696** | 0.191822403840966 | 0.332664916293653 | 0.856918526232803 | 2236.19644850839 | 2386.23808571898 | 2345.31521407085 | 1984.65288505027 | 1675.28603960322 | 1388.19833340383 | 2310.15228294407 | 2463.05750623284 | Vps35 |
| **ENSMUSG00000031697** | 0.0385464598901887 | 0.880578916345065 | 0.978433072023387 | 64.1494738764022 | 44.7928650902881 | 46.3762895437739 | 63.779015209555 | 47.0832024202212 | 70.9387694602839 | 43.7252798033577 | 52.1052271040884 | Orc6 |
| **ENSMUSG00000031698** | 0.155863265390065 | 0.399401504451286 | 0.880459952732461 | 16358.1158384826 | 18037.6795639945 | 17616.3648424135 | 15759.9822436199 | 13933.343041798 | 10768.0159711783 | 19281.8073151902 | 16847.3567636552 | Mylk3 |
| **ENSMUSG00000031700** | -0.00062654480647439 | 0.997600910033357 | 0.999262355603273 | 1000.18968424193 | 1155.65591932943 | 954.972982646283 | 793.485983342404 | 770.850569856644 | 760.757148349941 | 1223.26675640346 | 1150.65709854862 | Gpt2 |
| **ENSMUSG00000031701** | 0.220200670104861 | 0.150629000460815 | 0.783020695097675 | 9443.52536557966 | 10040.9315217848 | 10192.3726958551 | 9083.82003389029 | 8066.55702859929 | 6290.31191955586 | 9762.18925514489 | 9153.15156128486 | Dnaja2 |
| **ENSMUSG00000031703** | -0.00995640465902965 | 0.925962749651408 | 0.986713918026868 | 1098.67267934796 | 1202.07761587755 | 1218.08744169055 | 1057.04338442895 | 1065.39432453198 | 1086.09702208159 | 1275.32066093127 | 1179.96628879467 | Itfg1 |
| **ENSMUSG00000031706** | -0.132670763347267 | 0.414424525426996 | 0.882192126163777 | 205.097613661173 | 184.057954734638 | 240.399541716705 | 169.764731660727 | 243.08071947184 | 218.931719541221 | 222.790711379013 | 191.052499381657 | Rfx1 |
| **ENSMUSG00000031708** | -0.0285525454714094 | 0.782687901325262 | 0.960786612650028 | 11338.1936292249 | 10190.784015905 | 9289.4547320845 | 10032.0639217853 | 9329.04382837871 | 10775.3544645707 | 10951.10043456 | 10612.0979201993 | Tecr |
| **ENSMUSG00000031709** | -0.095467472566391 | 0.720860021297512 | 0.951674101666722 | 93.9654265231807 | 89.5857301805761 | 91.8061241988993 | 58.151455044006 | 119.350443344282 | 101.51582526213 | 65.5879197050366 | 70.559161703453 | Tbc1d9 |
| **ENSMUSG00000031711** | 0.164119182541078 | 0.230458301138703 | 0.832048466795696 | 530.362551626029 | 465.845796938996 | 427.797609669098 | 512.107975064956 | 459.882442244021 | 386.49398533534 | 391.445362049107 | 489.572029665497 | Zfp330 |
| **ENSMUSG00000031712** | 0.0736414231932326 | 0.554247335559939 | 0.915833391735373 | 435.49361138628 | 486.206190161854 | 409.814966784777 | 469.901273823339 | 480.686647964584 | 377.932409710823 | 414.349080041342 | 438.552328126077 | Il15 |
| **ENSMUSG00000031714** | -0.17203030118275 | 0.416805556044872 | 0.882192126163777 | 756.240980768291 | 1004.17459375137 | 828.148027567391 | 668.741733006069 | 819.028730472685 | 656.795158623663 | 1152.47344624564 | 1041.0190165171 | Gab1 |
| **ENSMUSG00000031715** | 0.0629636917295787 | 0.676244296082729 | 0.94409858171788 | 1214.3224350688 | 1288.40568314247 | 1272.98182523216 | 1057.04338442895 | 1101.52794499401 | 934.434825304429 | 1395.04464134522 | 1194.07812113536 | Smarca5 |

| | | | | | | | | | | | | |
|---|---|---|---|---|---|---|---|---|---|---|---|---|
| **ENSMUSG00000031722** | -0.288922944715821 | 0.331302122651409 | 0.856333478432716 | 893.575065686785 | 1618.24405335277 | 822.4692982355 | 838.506464666796 | 859.542183717991 | 1246.32079448326 | 1111.87140071395 | 1881.21580357052 | Hp |
| **ENSMUSG00000031723** | -0.136966584029385 | 0.49092932542516 | 0.899236955296196 | 121.070838020252 | 101.801966114291 | 88.0203046443056 | 115.364983393754 | 110.590777777729 | 138.208292224346 | 91.61487196894 | 129.177542195552 | Txnl4b |
| **ENSMUSG00000031725** | -0.198254687717224 | 0.416888405309411 | 0.882192126163777 | 67.7635287426784 | 87.9568987227475 | 61.519567762149 | 81.59962240046 | 70.0773245324222 | 81.9465095489486 | 71.8343882483734 | 119.407812113536 | Ces1f |
| **ENSMUSG00000031728** | -0.178302640198275 | 0.349104803364207 | 0.861774532843376 | 116.553269437407 | 115.647033505835 | 114.521041526462 | 151.944124469822 | 163.148771177045 | 157.777607937528 | 113.477511870619 | 130.263067760221 | Zfp821 |
| **ENSMUSG00000031729** | 0.0893236545270195 | 0.350015205713586 | 0.861774532843376 | 759.855035634567 | 829.075212034787 | 787.450467355508 | 710.948434247686 | 741.286698569529 | 709.387694602839 | 693.358008310387 | 758.782369703287 | Ist1 |
| **ENSMUSG00000031730** | -0.0167561803182639 | 0.896270180189601 | 0.984095879090375 | 376.765219809292 | 290.746415222415 | 330.312756138308 | 315.143369270742 | 359.146288228664 | 325.339873731647 | 316.487739529066 | 326.743194965221 | Dhodh |
| **ENSMUSG00000031731** | 0.11503893820233 | 0.574276444537566 | 0.922280709351328 | 711.968808656408 | 1022.9061555164 | 946.454888648447 | 672.493439783102 | 728.147200219699 | 582.187142467157 | 910.94332923662 | 874.933605122817 | Ap1g1 |
| **ENSMUSG00000031732** | 0.0324517501890555 | 0.874556456577877 | 0.9767298530085 | 67.7635287426784 | 91.2145616384048 | 88.966759532954 | 93.7926694258161 | 93.0714466446232 | 64.8233582999146 | 82.2451691539348 | 93.3551985614916 | Phlpp2 |
| **ENSMUSG00000031734** | 0.0199476946891941 | 0.876921285892249 | 0.977515813184586 | 817.679913494986 | 688.181290932608 | 943.615523982501 | 916.354380290224 | 860.63714191381 | 826.803588881929 | 807.876598271562 | 822.828378018729 | Irx3 |
| **ENSMUSG00000031736** | -0.177499120627094 | 0.592934598202711 | 0.923935529355527 | 53.3073092775737 | 32.5766291565731 | 57.7337482075553 | 94.7305961200743 | 70.0773245324222 | 84.3926740130963 | 56.2182168900314 | 58.6183804920994 | Crnde |
| **ENSMUSG00000031737** | -0.0138572314937089 | 0.964012671364118 | 0.994284908685797 | 227.685456575399 | 107.502876216691 | 220.523989055088 | 344.219096792745 | 222.276513751277 | 236.054870790255 | 196.76375911511 | 252.927456567762 | Irx5 |
| **ENSMUSG00000031740** | -1.4159867000409 | 0.00367073038544936 | 0.635862253526407 | 266.536546387868 | 329.838370210303 | 299.079744812909 | 295.446908691321 | 1216.49855555502 | 1410.21381358116 | 244.653351280692 | 307.203734801188 | Mmp2 |
| **ENSMUSG00000031748** | -0.123621068114942 | 0.581795930532707 | 0.922450244597442 | 690.28447945875 | 710.170515613295 | 485.531357876653 | 695.003680445298 | 531.054724972262 | 956.450305481758 | 756.863771834311 | 568.815395886298 | Gnao1 |
| **ENSMUSG00000031749** | 0.100419077362055 | 0.364807057211107 | 0.86924083181804 | 665.889609111386 | 846.992358070902 | 795.022106464695 | 777.541229540016 | 728.147200219699 | 689.818378889657 | 707.933101578173 | 752.269216315276 | St3gal2 |
| **ENSMUSG00000031750** | -1.06502413602688 | 0.0501004710500524 | 0.703396849069134 | 47.8862269781594 | 27.6901347830872 | 31.2330113253987 | 30.9515809105193 | 99.6411958195378 | 138.208292224346 | 29.1501865355718 | 21.7105112933701 | Il34 |
| **ENSMUSG00000031751** | 0.104615455623772 | 0.326489020445178 | 0.855977678491063 | 4753.38566286975 | 4939.4313958654 | 4818.40183810924 | 4486.10337863679 | 4144.41677117528 | 3902.85540254769 | 4886.8205570705 | 4732.89146195469 | Amfr |
| **ENSMUSG00000031753** | 0.142383640778877 | 0.0742085727197676 | 0.734338471687791 | 2025.6777525478 | 1983.9167156353 | 1997.96626993687 | 2081.25933455886 | 1912.89196809596 | 1719.65361829585 | 1931.19985798163 | 1762.89351702166 | Cog4 |
| **ENSMUSG00000031754** | -0.0967177523704275 | 0.3530195586987 | 0.862728204353539 | 455.370913150799 | 401.506954354764 | 417.386605893965 | 404.246405225267 | 479.591689768764 | 426.855698993777 | 437.252798033577 | 450.493109337431 | Nudt21 |
| **ENSMUSG00000031755** | 0.141064672495938 | 0.273160709231874 | 0.844738346035895 | 241.238162323935 | 225.593156909269 | 256.489274823729 | 244.79886720138 | 191.617684268342 | 220.154801773295 | 236.324726556243 | 229.045894145055 | Bbs2 |
| **ENSMUSG00000031756** | 0.353175577191424 | 0.478120450950305 | 0.897674307047416 | 24.3948703473642 | 9.77298874697194 | 8.51809399783602 | 17.8206071909051 | 10.949581958191 | 9.78465785659088 | 17.6983275394543 | 8.68420451734806 | Cenpn |
| **ENSMUSG00000031758** | -0.137926059861745 | 0.650323184446151 | 0.936463929813731 | 40.658117245607 | 42.3496179035451 | 28.3936466594534 | 30.9515809105193 | 36.1336204620302 | 24.4616446414772 | 44.7663578939139 | 51.0197015394199 | Cdyl2 |
| **ENSMUSG00000031760** | -0.550698814136211 | 0.30664842606653 | 0.851785577030345 | 22.5878429142261 | 7.32974156022896 | 16.0897331070236 | 21.5723139679377 | 30.6588294829347 | 44.0309603546589 | 11.4518589961175 | 13.0263067760221 | Mt3 |
| **ENSMUSG00000031762** | -0.206971986141066 | 0.457555121963906 | 0.892966448175628 | 1030.90915060528 | 1268.04528991961 | 1272.98182523216 | 984.823028971069 | 1897.56255335449 | 1330.71346849636 | 1358.60690817576 | 673.025850094475 | Mt2 |
| **ENSMUSG00000031765** | -0.143837412372766 | 0.536830579759525 | 0.91199738788281 | 6169.19165673344 | 5528.25396787046 | 6578.80793099535 | 5804.82831076376 | 8492.49576677292 | 7742.11052902753 | 6552.54550196032 | 3818.87893650381 | Mt1 |
| **ENSMUSG00000031767** | 0.180875333814743 | 0.0476036056045914 | 0.703396849069134 | 1617.28955265859 | 1562.8637837866 | 1632.63468291857 | 1420.02101510686 | 1331.46916611602 | 1369.85209992272 | 1324.25133118741 | 1473.05819125516 | Nudt7 |
| **ENSMUSG00000031770** | -0.2278153976961 | 0.200998494719874 | 0.812921911423739 | 8504.77461406442 | 7844.45230090281 | 10655.1891364042 | 8038.03176979244 | 12127.7569768923 | 10439.0068507504 | 10899.0465300322 | 7570.45528799817 | Herpud1 |
| **ENSMUSG00000031774** | -0.0349374201960289 | 0.758246177427825 | 0.95677573228595 | 479.765783498163 | 448.743066631795 | 395.618143455051 | 469.901273823339 | 483.971522552041 | 479.448234972953 | 444.54034466747 | 430.953649173397 | Fam192a |
| **ENSMUSG00000031776** | 0.0573229169541108 | 0.688966872141741 | 0.94469184542218 | 1669.6933482196 | 1522.14299734088 | 1466.05862251644 | 1496.93100403603 | 1284.3859636958 | 1251.21312341156 | 1652.19092971259 | 1725.98564782293 | Arl2bp |
| **ENSMUSG00000031778** | -0.48423039177345 | 0.103852370984768 | 0.759800953500741 | 142.755167217909 | 155.553404222637 | 123.985590412947 | 237.295453647315 | 296.733671066975 | 299.655146858096 | 150.95632313064 | 175.855141476298 | Cx3cl1 |
| **ENSMUSG00000031781** | -0.0291494933456989 | 0.819500001048048 | 0.966398681144067 | 1143.84836517641 | 1247.68489669675 | 1468.89798718239 | 1084.24325856243 | 1189.12460065954 | 1262.22086350022 | 1361.73014244743 | 1232.07151589876 | Ciapin1 |
| **ENSMUSG00000031782** | 0.134068975648801 | 0.336074128455165 | 0.858570960161243 | 14238.4726594116 | 13122.6806399966 | 13111.2395724469 | 13484.5720833496 | 11693.0585731521 | 9705.15751150608 | 13786.9971532349 | 13981.5692729304 | Coq9 |
| **ENSMUSG00000031783** | 0.0576117374351075 | 0.650819461208158 | 0.936463929813731 | 1662.46523848704 | 1406.49596383505 | 1611.81267536831 | 1738.91609115463 | 1600.82888228752 | 1772.24615427502 | 1405.45542225078 | 1390.55824834036 | Polr2c |
| **ENSMUSG00000031785** | -0.853246829282481 | 0.0316093959541814 | 0.671337170953485 | 1488.08709118922 | 1391.83648071459 | 1361.00212987647 | 1080.4915517854 | 3443.64352585106 | 3928.54012942124 | 1105.62493217062 | 1136.54526620793 | Adgrg1 |
| **ENSMUSG00000031787** | 0.0490125802267641 | 0.719427570545685 | 0.951674101666722 | 234.010052591383 | 197.903022126182 | 205.380710836713 | 196.964605794214 | 224.466430142915 | 200.585486060113 | 196.76375911511 | 184.539345993646 | Katnb1 |
| **ENSMUSG00000031788** | 0.00326946445458785 | 0.981350486583162 | 0.996940244108351 | 2209.99455072788 | 2177.74765911691 | 2134.25577390225 | 2038.11470662298 | 1990.63399999912 | 1706.19971374303 | 2395.52068636967 | 2446.77462276282 | Kifc3 |
| **ENSMUSG00000031789** | 0.283751711384021 | 0.483939127719082 | 0.89898089116307 | 34.3335212296237 | 24.4324718674299 | 23.6613722162112 | 21.5723139679377 | 10.949581958191 | 15.9000690169602 | 35.3966550789086 | 22.7960368580387 | Cngb1 |
| **ENSMUSG00000031790** | -0.289518658312208 | 0.0327296019882044 | 0.671606027118506 | 1620.90360752487 | 1825.10564849701 | 1338.2872125489 | 1394.69699436189 | 1854.85918371755 | 1955.7084890861 | 1706.32699042151 | 2037.53148488279 | Mmp15 |
| **ENSMUSG00000031791** | 0.15483717915699 | 0.243755327643731 | 0.832048466795696 | 12155.8735427199 | 11515.0239911197 | 11919.6528676385 | 11098.4865731568 | 9873.2380517008 | 8499.19843068125 | 11878.7010132455 | 11684.5971780918 | Tmem38a |
| **ENSMUSG00000031792** | -0.149671305368986 | 0.392277907585588 | 0.87840402004898 | 149.079763233892 | 167.769640156352 | 166.576060402127 | 169.764731660727 | 136.869774477387 | 218.931719541221 | 191.558368662329 | 178.026192605635 | Usb1 |
| **ENSMUSG00000031796** | -0.115144061517504 | 0.502121189580824 | 0.900927059989498 | 196.965990212052 | 207.676010873154 | 162.790240847533 | 184.771558768858 | 224.466430142915 | 239.724117486476 | 164.49033830787 | 187.795922687652 | Cfap20 |
| **ENSMUSG00000031799** | -0.133380698394333 | 0.285128620177988 | 0.851275833092507 | 2254.26672283977 | 2083.27543456285 | 1994.18045038228 | 2002.47349224117 | 2639.94421011984 | 2431.48747736283 | 1994.70562150556 | 2076.61040521085 | Tpm4 |
| **ENSMUSG00000031803** | -0.940473348696081 | 0.0139690818094056 | 0.635862253526407 | 203.290586228035 | 148.223662662408 | 131.557229522134 | 176.330218520534 | 470.832024202212 | 469.663577116362 | 153.038479311752 | 172.598564782293 | B3gnt3 |
| **ENSMUSG00000031805** | -0.308017401060109 | 0.0824672970562331 | 0.739855948845396 | 254.790868072471 | 217.448999620126 | 282.043556817237 | 281.378008277448 | 349.291664466292 | 391.386314263635 | 275.885693997376 | 265.953763343784 | Jak3 |
| **ENSMUSG00000031807** | -0.188367174458228 | 0.233102539996044 | 0.832048466795696 | 883.636414804526 | 710.984931342209 | 797.861471130641 | 892.90621293377 | 953.708588558433 | 1158.25887377394 | 841.191097169358 | 791.348136643342 | Pgls |
| **ENSMUSG00000031808** | 0.298790540464705 | 0.0500034485558255 | 0.703396849069134 | 4656.70969519686 | 5704.98218104487 | 6901.54904802447 | 4844.3913758434 | 4767.44798459635 | 4143.80260226624 | 4773.34304519989 | 4286.74045487594 | Slc27a1 |
| **ENSMUSG00000031812** | 0.17322883794467 | 0.149607126077635 | 0.782641594375979 | 6473.67577921721 | 5911.0293604602 | 8076.0995648372 | 6895.637056186 | 6114.24656545384 | 6380.82000472933 | 5860.22857174049 | 5905.25907179668 | Map1lc3b |
| **ENSMUSG00000031813** | -0.267153032032385 | 0.201641107248672 | 0.813395633194566 | 436.397125102849 | 341.240190415104 | 333.152120804253 | 448.328959855401 | 501.490853685146 | 621.325773893521 | 381.034581143546 | 373.420794245967 | Mvb12a |
| **ENSMUSG00000031816** | 0.129658385131551 | 0.475580380671065 | 0.897549741595057 | 151.7903043836 | 144.96599974675 | 139.128868631322 | 126.620103724852 | 114.970610561005 | 111.300483118721 | 165.531416398426 | 121.578863242873 | Mthfsd |
| **ENSMUSG00000031818** | -0.00393100246230222 | 0.982261510221375 | 0.996940244108351 | 84317.7070576564 | 59254.4451886198 | 65698.1660955319 | 83631.1716202232 | 76145.5828536516 | 89426.8804803123 | 61123.7768527319 | 67005.1510047283 | Cox4i1 |
| **ENSMUSG00000031819** | -0.0595232180819686 | 0.647041650336457 | 0.935063240511793 | 586.38040205331 | 505.752167655798 | 501.621090983677 | 607.776497879289 | 571.568178217569 | 658.018240855737 | 543.442763270303 | 522.137796605552 | Emc8 |
| **ENSMUSG00000031820** | -0.0664963028901922 | 0.539816720264673 | 0.913837378403038 | 1198.96270188712 | 1002.54576229354 | 1031.63582862681 | 1208.98750889877 | 1144.23131463096 | 1248.76695894741 | 1121.24110352896 | 1137.6307917726 | Babam1 |
| **ENSMUSG00000031821** | 0.0667054548723149 | 0.906755698644978 | 0.98504921011344 | 13.5527057485357 | 3.25766291565731 | 9.46454888648447 | 15.9447538023887 | 14.2344565456483 | 11.0077400886647 | 6.24646854333682 | 8.68420451734806 | Gins2 |
| **ENSMUSG00000031822** | 0.0860280537022674 | 0.78755676104209 | 0.961224689823763 | 165.343010132135 | 241.881471487556 | 254.596365046432 | 144.440710915757 | 182.858018701789 | 94.1773318696872 | 257.146288367366 | 224.703791886381 | Gse1 |
| **ENSMUSG00000031823** | 0.0250379330039635 | 0.830308411831506 | 0.967583158900775 | 940.557778948376 | 1057.92603185971 | 991.884723303572 | 921.044013761514 | 889.106055005107 | 854.934480219628 | 1046.28348100892 | 1052.95979772845 | Zdhhc7 |
| **ENSMUSG00000031824** | 0.0838572592465086 | 0.634781665120269 | 0.932543559775713 | 740.881247586617 | 959.381728661079 | 826.255117790094 | 778.479156234274 | 643.835419141629 | 684.926049961361 | 1014.01006020168 | 775.065253173314 | 6430548M08 |
| **ENSMUSG00000031825** | -0.82149332602509 | 0.0555809065364459 | 0.703396849069134 | 644.205279913729 | 738.675066125296 | 727.823809370656 | 838.506464666796 | 1851.57430913009 | 2265.14829380079 | 544.483841360859 | 551.446986851602 | Crispld2 |
| **ENSMUSG00000031826** | 0.0447321183409525 | 0.666861105382809 | 0.942024383071524 | 1103.1902479308 | 1225.69567201606 | 1195.37252436299 | 1088.93289203373 | 1116.85735973548 | 988.250443515679 | 1178.50039850955 | 1187.56496774735 | Usp10 |
| **ENSMUSG00000031827** | -0.700157436476504 | 0.0185111243508426 | 0.642273034804458 | 99.3865088225949 | 113.203786319092 | 132.503684410783 | 110.675349922463 | 222.276513751277 | 275.193502216618 | 109.313199508394 | 134.605170018895 | Cotl1 |
| **ENSMUSG00000031828** | -0.761031903930344 | 0.0135139774002241 | 0.635862253526407 | 40.658117245607 | 60.2667639396603 | 42.5904699891801 | 48.7721881014244 | 95.2613630362614 | 117.415894279091 | 64.5468416144805 | 49.9341759747513 | Klhl36 |
| **ENSMUSG00000031832** | -0.0921312476902522 | 0.727699946770091 | 0.952263232836493 | 77.7021796249379 | 129.492100897378 | 138.182413742673 | 82.5375490947182 | 141.249607260664 | 128.423634367755 | 92.6559500594962 | 94.4407241261602 | Taf1c |
| **ENSMUSG00000031833** | -0.252801958398689 | 0.308659864038946 | 0.851785577030345 | 121.070838020252 | 122.976775066064 | 126.824955078892 | 91.9168160372998 | 181.76306050597 | 155.33144347338 | 128.052605138405 | 86.8420451734806 | Mast3 |
| **ENSMUSG00000031834** | -0.132815648874311 | 0.244873331326517 | 0.832048466795696 | 583.669860903603 | 588.008156276145 | 611.409858066897 | 589.017963994125 | 702.96316171586 | 719.172352459429 | 610.071761065896 | 569.900921450967 | Pik3r2 |
| **ENSMUSG00000031835** | -0.0958484153921257 | 0.323714065093067 | 0.855977678491063 | 781.539364832224 | 875.496908582903 | 844.237760674415 | 808.492810450535 | 974.512794278996 | 862.272973612071 | 841.191097169358 | 859.736247217458 | Mbtps1 |
| **ENSMUSG00000031838** | -0.782199683946463 | 0.0492787617095729 | 0.703396849069134 | 67.7635287426784 | 70.0397526866323 | 62.4660226507975 | 53.4618215727152 | 150.009272827216 | 174.900759186562 | 64.5468416144805 | 47.7631248454143 | Ifi30 |
| **ENSMUSG00000031839** | 0.0765320629851657 | 0.511353688470648 | 0.902394714365738 | 3606.82675654363 | 2956.32909595901 | 3171.57033186095 | 3641.96935380444 | 3005.66024752342 | 3545.71539078212 | 2962.90824572277 | 3171.90569996138 | Hsbp1 |
| **ENSMUSG00000031840** | -0.165806099962428 | 0.281889160154358 | 0.850258509130083 | 687.573938309043 | 609.997380956832 | 540.425741418263 | 598.397230936707 | 656.974917491458 | 847.595986827185 | 650.673806597585 | 579.670651532983 | Rab3a |
| **ENSMUSG00000031841** | -0.244414571894635 | 0.0397639129722652 | 0.679566075569698 | 2512.67164577851 | 2546.67798431511 | 2477.81889848163 | 2539.9054880511 | 2990.33083278195 | 3550.60771971041 | 2786.96604841878 | 2611.77450859243 | Cdh13 |
| **ENSMUSG00000031842** | -0.60096086861444 | 0.0331886126573031 | 0.671606027118506 | 22.5878429142261 | 18.7315617650296 | 33.1259211026956 | 28.1378008277448 | 37.2285786578493 | 36.6924669622158 | 40.6020455316893 | 40.1644458927348 | Pde4c |
| **ENSMUSG00000031843** | 0.0344587805794105 | 0.871310285371712 | 0.975613767803763 | 104.807591122009 | 83.8848200781758 | 86.1273948670087 | 137.87522405595 | 109.49581958191 | 102.738907494204 | 91.61487196894 | 98.7828263848342 | Mphosph6 |
| **ENSMUSG00000031844** | -0.145126921916118 | 0.767760318015637 | 0.95879680569409 | 26.2018977805023 | 34.2054606144018 | 17.9826428843205 | 31.8895076047775 | 12.0445401540101 | 30.5770558018465 | 18.7394056300105 | 60.7894316214364 | Hsd17b2 |
| **ENSMUSG00000031848** | -0.0696818959692984 | 0.716627460215053 | 0.951348124093194 | 1837.74689950144 | 1501.78260411802 | 1565.43638582453 | 2163.79688365358 | 1858.14405830501 | 2414.3643261138 | 1628.2461336298 | 1518.65026497124 | Lsm4 |
| **ENSMUSG00000031849** | -2.97442995513421 | 0.0145703196037072 | 0.635862253526407 | 27.1054114970714 | 6.51532583131463 | 10.4110037751329 | 36.5791410760683 | 65.6974917491458 | 532.040770952129 | 28.1091084450157 | 7.59867895267955 | Comp |
| **ENSMUSG00000031851** | 0.0825666041902166 | 0.589121366006075 | 0.92352104528132 | 319.843855665442 | 278.5301792887 | 286.775831260479 | 331.088123073131 | 262.789966996583 | 349.801518373124 | 290.460787265162 | 246.414303179751 | Ntpcr |
| **ENSMUSG00000031853** | 0.176194210734149 | 0.712299600377604 | 0.950233477782287 | 29.8159526467785 | 24.4324718674299 | 56.7872933189068 | 26.2619474392285 | 27.3739548954774 | 9.78465785659088 | 49.9717483466946 | 33.6512925047237 | Map3k21 |
| **ENSMUSG00000031858** | 0.212368512275707 | 0.365503034836122 | 0.86924083181804 | 502.353626412389 | 669.449729167578 | 628.446046062569 | 548.687116141024 | 417.179072607076 | 332.67836712409 | 661.084587503147 | 615.492995167044 | Mau2 |
| **ENSMUSG00000031862** | -0.0554419900450104 | 0.714797787524684 | 0.951176382858794 | 313.519259649459 | 371.373572384934 | 380.474865236676 | 278.564228194674 | 393.089992299056 | 351.024600605198 | 340.432535611857 | 312.63136262453 | Atp13a1 |
| **ENSMUSG00000031864** | -0.0149829682701001 | 0.935029822090253 | 0.98818087308472 | 284.606820719249 | 286.674336577844 | 260.275094378323 | 241.985087118606 | 236.510970296925 | 205.477814988408 | 324.816364253515 | 316.973464883204 | Ints10 |
| **ENSMUSG00000031865** | 0.241583548500253 | 0.0952491267430348 | 0.757085157577544 | 5109.37006719795 | 5720.45607989424 | 6016.61372713818 | 5319.92020983229 | 4428.01094389243 | 3735.29313675357 | 5446.92056978971 | 5136.70697201138 | Dctn1 |
| **ENSMUSG00000031871** | -0.659230165767877 | 0.0410253236804847 | 0.683226273624098 | 2318.41619671617 | 2316.19833303235 | 2546.91010535297 | 2751.87692095345 | 5749.62548624608 | 5463.50833067393 | 2208.12663006957 | 2266.57737902784 | Cdh5 |
| **ENSMUSG00000031872** | -0.843919597017846 | 0.0971919200723582 | 0.757809915356977 | 17.1667606148119 | 21.1748089517725 | 12.3039135524298 | 31.8895076047775 | 53.6529515951357 | 59.9310293716191 | 18.7394056300105 | 16.2828834700276 | Bean1 |
| **ENSMUSG00000031875** | -0.576379986293688 | 0.0868914744450102 | 0.747269672026173 | 130.105975185942 | 109.13170767452 | 123.039135524298 | 123.806323642077 | 216.801722772181 | 293.539735697726 | 92.6559500594962 | 122.664388807541 | Cmtm3 |
| **ENSMUSG00000031877** | -1.00805645618986 | 0.00732778781031745 | 0.635862253526407 | 37.9475760958999 | 48.0505280059454 | 30.2865564367503 | 45.9584080186499 | 118.255485148462 | 112.523565350795 | 56.2182168900314 | 40.1644458927348 | Ces2g |
| **ENSMUSG00000031878** | 0.116140500474805 | 0.394205948138985 | 0.878942305112115 | 521.327414460339 | 535.885549625628 | 596.266579848521 | 576.824916968769 | 514.630352034976 | 413.401794440965 | 564.264325081426 | 563.387768062955 | Nae1 |
| **ENSMUSG00000031879** | 0.00198877656997137 | 0.991259759562283 | 0.998628446692557 | 1671.50037565273 | 1312.02373928098 | 1295.69674255972 | 1647.93720181159 | 1498.99777007634 | 1906.78519980315 | 1315.92270646296 | 1198.42022339403 | Fam96b |
| **ENSMUSG00000031880** | -0.087356874215628 | 0.786636382038545 | 0.961141387070613 | 3329.44804555693 | 2750.28191654369 | 5093.82021070594 | 8197.47930781633 | 3921.04529922819 | 4931.4675597218 | 6256.87932424238 | 5469.96332036461 | Rrad |
| **ENSMUSG00000031881** | 0.273683196113891 | 0.38494102805695 | 0.875126835008975 | 21.6843291976571 | 25.2468875963442 | 35.0188308799925 | 28.1378008277448 | 24.0890803080201 | 22.0154801773295 | 17.6983275394543 | 27.1381391167127 | Cdh16 |
| **ENSMUSG00000031883** | -0.207333519594124 | 0.61259720350871 | 0.928055903343287 | 60.535419010126 | 27.6901347830872 | 35.965285768641 | 68.4686486808458 | 55.8428679867739 | 91.7311674055395 | 44.7663578939139 | 30.3947158107182 | Car7 |

| | | | | | | | | | | | | |
|---|---|---|---|---|---|---|---|---|---|---|---|---|
| ENSMUSG00000031885 | -0.105093949050829 | 0.443209216599468 | 0.89108031640712 | 361.405486627618 | 411.279943101736 | 442.940887887473 | 372.35689762049 | 451.122776677468 | 355.916929533493 | 466.402984569149 | 433.124700302735 | Cbfb |
| ENSMUSG00000031886 | -0.578199572920595 | 0.102719562269959 | 0.759800953500741 | 44.2721721118832 | 59.452348210746 | 29.3401015481019 | 45.9584080186499 | 59.1277425742312 | 66.0464405319884 | 37.4788112600209 | 105.295979772845 | Ces2e |
| ENSMUSG00000031887 | -0.315896814256858 | 0.222124449343511 | 0.831327997735572 | 49.6932544112975 | 67.5965054998893 | 60.5731128735006 | 59.0893817382642 | 85.4067392738896 | 96.6234963338349 | 48.9306702561384 | 65.1315338801105 | Tradd |
| ENSMUSG00000031889 | 0.0982161882076217 | 0.62732260629615 | 0.931825800800958 | 282.799793286111 | 410.465527372822 | 402.24332767559 | 345.157023487003 | 343.816873487196 | 237.277953022329 | 391.445362049107 | 372.335268681298 | D230025D16 |
| ENSMUSG00000031897 | -0.091217697221557 | 0.69475677609973 | 0.944771261777587 | 1131.19917314444 | 793.240919962556 | 636.964140060405 | 1082.36740517392 | 1108.09769416893 | 1185.16668287957 | 767.274552739873 | 821.74285245406 | Psmb10 |
| ENSMUSG00000031901 | 0.317319532285777 | 0.0206142759404417 | 0.646348290773668 | 314.422773366028 | 307.034729800702 | 363.438677241004 | 356.412143818101 | 295.638712871156 | 260.516515431732 | 231.119336103462 | 288.749800201823 | Dus2 |
| ENSMUSG00000031902 | 0.0490268842572542 | 0.735886280724843 | 0.954383574136061 | 449.046317134815 | 574.977504613516 | 504.460455649622 | 423.942865804689 | 455.502609460744 | 413.401794440965 | 509.087186281951 | 509.11148982953 | Nfatc3 |
| ENSMUSG00000031903 | -0.378044144625443 | 0.0962346381714308 | 0.757809915356977 | 159.018414116152 | 147.409246933493 | 227.149173275627 | 283.253861665965 | 261.695008800764 | 286.201242305283 | 271.721381635152 | 240.986675356409 | Pla2g15 |
| ENSMUSG00000031904 | 0.161520673706703 | 0.455139603919383 | 0.892775943845882 | 76.7986659083688 | 116.461449234749 | 107.895857305923 | 84.4134024832345 | 84.3117810780705 | 69.71568722821 | 89.5327157878278 | 100.953877514171 | Slc7a6 |
| ENSMUSG00000031906 | -1.1496013277097 | 0.0307847485323752 | 0.667184591665619 | 12.6491920319666 | 18.7315617650296 | 15.1432782183751 | 17.8206071909051 | 55.8428679867739 | 58.7079471395453 | 12.4929370866736 | 16.2828834700276 | Smpd3 |
| ENSMUSG00000031907 | -0.31161783847461 | 0.469156842452025 | 0.895328112556867 | 20.780815481088 | 10.5874044758863 | 15.1432782183751 | 8.44134024832345 | 16.4243729372865 | 11.0077400886647 | 20.8215618111227 | 19.5394601640331 | Zfp90 |
| ENSMUSG00000031913 | 0.127871689586795 | 0.229714111767907 | 0.832048466795696 | 1715.77254776462 | 1794.15785079827 | 1878.71295396717 | 1854.28107454838 | 1676.38099779904 | 1416.32922474153 | 1750.05227022487 | 1783.51850275036 | Vps4a |
| ENSMUSG00000031916 | -0.0280982257548741 | 0.820585918101966 | 0.966398681144067 | 713.775836089546 | 621.399201161633 | 661.571967165264 | 665.927952923294 | 782.895110010654 | 686.149132193435 | 584.044808801993 | 662.17059444779 | Cog8 |
| ENSMUSG00000031917 | 0.0205007553038322 | 0.908078140835403 | 0.98504921011344 | 379.475760958999 | 275.272516373043 | 242.292451494002 | 350.784583652552 | 326.297542354091 | 331.455284892016 | 289.419709174606 | 283.322172378481 | Nip7 |
| ENSMUSG00000031918 | 0.0511406975133251 | 0.61932470579622 | 0.929568738467614 | 383.089815825275 | 381.146561131906 | 387.100049457215 | 396.742991671202 | 393.089992299056 | 338.793778284459 | 385.198893505771 | 375.591845375304 | Mtmr2 |
| ENSMUSG00000031921 | 0.0171253461677717 | 0.907958158105886 | 0.98504921011344 | 256.597895505609 | 304.591482613959 | 316.115932808581 | 256.053987532478 | 239.795844884382 | 267.855008824175 | 310.241270985729 | 301.776106977845 | Terf2 |
| ENSMUSG00000031922 | 0.196064269865419 | 0.171991318069063 | 0.799889021902745 | 201.483558794897 | 184.057954734638 | 220.523989055088 | 212.909359596603 | 165.338687568684 | 159.000690169602 | 194.681602933998 | 194.309076075663 | Cep57 |
| ENSMUSG00000031924 | 0.0226276862922199 | 0.832908853540669 | 0.967854847250989 | 1446.52546022704 | 1465.94831204579 | 1474.57671651428 | 1498.80685742454 | 1409.21119801918 | 1248.76695894741 | 1516.85077794029 | 1617.43309135608 | Cyb5b |
| ENSMUSG00000031925 | -0.348774095317605 | 0.219901327510482 | 0.828432177683914 | 93.9654265231807 | 88.7713144516618 | 104.110037751329 | 74.0962088463947 | 159.863896589588 | 139.43137445642 | 68.711153976705 | 92.2696729968231 | Maml2 |
| ENSMUSG00000031928 | 0.145763762634408 | 0.477545682993321 | 0.897674307047416 | 186.123825613223 | 210.119258059897 | 194.023252172932 | 157.571684635371 | 142.344565456483 | 128.423634367755 | 214.462086654564 | 189.966973816989 | Mre11a |
| ENSMUSG00000031930 | 0.162593261898926 | 0.191158040642937 | 0.808824684419004 | 563.792559139084 | 683.294796559122 | 767.57491469389 | 695.941607139556 | 605.511882287961 | 627.44118505389 | 609.03068297534 | 579.670651532983 | Wwp2 |
| ENSMUSG00000031931 | -0.0486565795771468 | 0.769513241669171 | 0.95879680569409 | 149.079763233892 | 133.56417954195 | 113.574586637814 | 133.185590584659 | 121.54035973592 | 124.754387671534 | 146.792010768415 | 154.144630182928 | Ankrd49 |
| ENSMUSG00000031934 | -0.516980064395377 | 0.356147177698717 | 0.864965376583892 | 30.7194663633475 | 18.7315617650296 | 31.2330113253987 | 30.9515809105193 | 42.7033696369448 | 86.838838477244 | 19.7804837205666 | 10.8552556466851 | Panx1 |
| ENSMUSG00000031935 | -0.0616108126497135 | 0.60124653110184 | 0.926043912368388 | 240.334648607366 | 225.593156909269 | 222.416898832385 | 243.860940507122 | 244.175677667659 | 223.824048469516 | 251.940897914585 | 251.841931003094 | Med17 |
| ENSMUSG00000031938 | 0.0351227331169998 | 0.850757102222245 | 0.972692784625411 | 161.728955265859 | 172.656134529838 | 134.396594188079 | 183.8336320746 | 144.534481848121 | 138.208292224346 | 150.95632313064 | 202.993280593011 | 4931406C07 |
| ENSMUSG00000031939 | -0.0381411108362016 | 0.874301479446921 | 0.97671366425653 | 109.325159704854 | 104.245213301034 | 134.396594188079 | 166.013024883695 | 118.255485148462 | 182.239252579005 | 106.189965236726 | 121.578863242873 | Taf1d |
| ENSMUSG00000031948 | 0.0421768071733822 | 0.709956551623577 | 0.949571160524938 | 3497.50159683877 | 3130.61406194668 | 3478.22171578304 | 3860.50627356659 | 3652.78054125251 | 3660.68512059706 | 3083.67330422728 | 3167.56359770271 | Kars |
| ENSMUSG00000031949 | 0.0838615445204423 | 0.742606741185454 | 0.954383574136061 | 34.3335212296237 | 40.7207864457164 | 48.2691993210708 | 33.7653609932938 | 42.7033696369448 | 33.0232202659942 | 35.3966550789086 | 36.9078691987293 | Adat1 |
| ENSMUSG00000031950 | 0.169470126238877 | 0.133030139201021 | 0.77303252008843 | 1899.18583222813 | 1582.40976128054 | 1814.35402153907 | 1756.73669834554 | 1382.93220131952 | 1631.59169758653 | 1534.54910547975 | 1721.64354556425 | Gabarapl2 |
| ENSMUSG00000031951 | -0.303032650424724 | 0.303175813136525 | 0.851785577030345 | 48.7897406947284 | 57.009101024003 | 35.0188308799925 | 42.2067012416173 | 83.2168228822514 | 53.8156182112498 | 41.6431236222455 | 47.7631248454143 | Tmem231 |
| ENSMUSG00000031953 | 0.0238731796278198 | 0.927840109188086 | 0.986901071486195 | 61.4389327266951 | 48.0505280059454 | 59.6266579848522 | 80.6616957062019 | 43.7983278327639 | 67.2695227640623 | 56.2182168900314 | 78.1578406561325 | Tmem170 |
| ENSMUSG00000031954 | -0.0460028398787205 | 0.762967751103238 | 0.957845481430497 | 798.706125447036 | 705.284021239809 | 670.0900611631 | 877.899385825639 | 775.230402639921 | 956.450305481758 | 726.672507208183 | 693.650835823176 | Cfdp1 |
| ENSMUSG00000031955 | -0.245150967795779 | 0.0561435957060048 | 0.703396849069134 | 463.50253659992 | 473.175538499225 | 443.887342776122 | 384.549944645846 | 566.093387238473 | 571.179402378492 | 475.772687384154 | 480.887825148149 | Bcar1 |
| ENSMUSG00000031958 | 0.31076367963524 | 0.147122246504159 | 0.780607990324772 | 1238.71730541616 | 1222.43800910041 | 1367.62731409701 | 1450.97259601738 | 893.485887788383 | 717.949270227356 | 1363.81229862854 | 1279.83464074417 | Ldhd |
| ENSMUSG00000031959 | -0.0414264496740228 | 0.823838219283818 | 0.966637987288264 | 152.693818100169 | 172.656134529838 | 152.3792370724 | 165.075098189436 | 166.433645764503 | 117.415894279091 | 178.024353485099 | 198.651178334337 | Wdr59 |
| ENSMUSG00000031960 | 0.0431222799686904 | 0.58870548641494 | 0.92352104528132 | 1912.73853797667 | 1902.47514274387 | 2073.68266102875 | 1904.92911603833 | 1930.41129922907 | 1768.5769075788 | 1911.41937426107 | 1951.77496527398 | Aars |
| ENSMUSG00000031963 | -0.953905609464282 | 0.0467721709987787 | 0.701439656975469 | 45.1756858284523 | 47.2361122770311 | 35.0188308799925 | 39.3929211588428 | 153.294147414674 | 101.51582526213 | 40.6020455316893 | 28.2236646813812 | Bmper |
| ENSMUSG00000031965 | -0.101687426723847 | 0.411826446404929 | 0.882192126163777 | 1001.99671167507 | 1192.30462713058 | 1117.76322349382 | 1136.76715344089 | 1007.36154015357 | 1210.85140975312 | 1341.94965872686 | 1213.61758129939 | Tbx20 |
| ENSMUSG00000031967 | 0.160733825511011 | 0.163432289144675 | 0.793968226938511 | 2635.5495112319 | 2733.17918623649 | 2588.5541204535 | 2815.655936163 | 2337.73574807377 | 2033.98575193883 | 2570.4218055831 | 2693.18892594257 | Afg3l1 |
| ENSMUSG00000031969 | 0.192008013390518 | 0.166087855862012 | 0.796071270505327 | 1300.15623814286 | 1274.56061575092 | 1364.78794943106 | 1423.77272188389 | 1102.62290318983 | 945.442565393093 | 1311.75839410073 | 1333.02539341293 | Acad8 |
| ENSMUSG00000031970 | -0.573331492006824 | 0.381097739389725 | 0.875094660294807 | 15.3597331816738 | 17.9171460361152 | 6.62518422053913 | 12.1930470253561 | 5.47479097909548 | 6.1154111603693 | 35.3966550789086 | 30.3947158107182 | Dbndd1 |
| ENSMUSG00000031971 | -0.578196681456088 | 0.171053753437942 | 0.799531027639052 | 14.4562194651047 | 15.4738988493722 | 12.3039135524298 | 18.7585338851632 | 35.0386622662111 | 13.4539045528125 | 28.1091084450157 | 14.1118323406906 | Ccsap |
| ENSMUSG00000031972 | 0.611191994575789 | 0.178794975450635 | 0.801755154770274 | 22230.9549961814 | 15744.2848713718 | 3527.43736999276 | 25699.1914226736 | 11207.9920924043 | 12713.9398024078 | 12309.7073427358 | 7762.5933129445 | Acta1 |
| ENSMUSG00000031974 | 0.17855469794888 | 0.390098961865143 | 0.877515569464664 | 1083.31294616628 | 1150.76942495595 | 1047.72556173383 | 1053.29167765192 | 866.111932892906 | 623.771938357668 | 1117.07679116673 | 1222.30178581674 | Abcb10 |
| ENSMUSG00000031976 | 0.141408581215695 | 0.486700501960655 | 0.899236955296196 | 104.807591122009 | 170.212887343095 | 131.557229522134 | 113.489130005238 | 122.635317931739 | 102.738907494204 | 128.052605138405 | 118.322286548867 | Urb2 |
| ENSMUSG00000031979 | 0.139464584229402 | 0.314960063011598 | 0.853277176422981 | 507.774708711803 | 480.505280059454 | 475.12035410152 | 446.453106466885 | 402.944616061428 | 359.586176229715 | 467.444062659705 | 502.598336441519 | Cog2 |
| ENSMUSG00000031980 | -0.409603063635223 | 0.0337424001813377 | 0.672620872762612 | 248.466272056487 | 349.384347704247 | 244.185361271299 | 213.847286290861 | 339.43704070392 | 420.740287833408 | 307.11803671406 | 337.598450611906 | Agt |
| ENSMUSG00000031982 | 0.287743613145522 | 0.503387586724179 | 0.900927059989498 | 28.0089252136404 | 30.9477976987445 | 39.7511053232348 | 16.8826804966469 | 10.949581958191 | 24.4616446414772 | 38.5198893505771 | 20.6249857287016 | Arv1 |
| ENSMUSG00000031983 | 0.141652819834574 | 0.0657540561808269 | 0.718586813809351 | 1885.6331264796 | 1905.73280565953 | 1917.51760440175 | 1899.30155587278 | 1658.86166666593 | 1713.53820713548 | 1661.56063252759 | 1862.76186897116 | 2310022B05 |
| ENSMUSG00000031984 | 0.261973691099056 | 0.0506965653573535 | 0.703396849069134 | 2163.91535118286 | 2343.88846781544 | 2478.76535337028 | 2468.62305928748 | 1711.41966006525 | 1753.89992079391 | 2337.22031329853 | 2080.95250746953 | 2810004N23 |
| ENSMUSG00000031985 | 0.187380537411466 | 0.329938392638853 | 0.855977678491063 | 5931.56754927578 | 6016.90340521906 | 5691.97970033176 | 5385.57507843036 | 4225.44367766589 | 3647.23121604425 | 6135.07318764731 | 6212.46280659787 | Gnpat |
| ENSMUSG00000031986 | 0.0605203502372619 | 0.66219929977848 | 0.939611397743053 | 306.291149916906 | 343.683437601847 | 391.832323900457 | 322.646782824807 | 337.247124312282 | 287.424324537357 | 365.418409785204 | 316.973464883204 | Sprtn |
| ENSMUSG00000031987 | 0.0771110192723621 | 0.564550096493766 | 0.919296698745417 | 15601.8748577143 | 15410.3744225169 | 15593.7907453718 | 13602.7508468261 | 12652.2419526897 | 12233.2684852027 | 16188.7643081479 | 15999.5612976491 | Egln1 |
| ENSMUSG00000031988 | 0.0178481205953626 | 0.907588228412577 | 0.98504921011344 | 706.547726356993 | 896.671717534676 | 999.45636241276 | 748.465502018013 | 792.749733773026 | 730.180092548094 | 930.723812957186 | 855.394144958784 | Vps26b |
| ENSMUSG00000031990 | -0.416472607477908 | 0.088555119886788 | 0.751206603737452 | 249.369785773056 | 210.933673788811 | 148.593417517806 | 197.902532488472 | 337.247124312282 | 329.009120427868 | 196.76375911511 | 214.934061804364 | Jam3 |
| ENSMUSG00000031993 | 0.0852578434912358 | 0.623702603373543 | 0.931145988862144 | 1026.39158202244 | 1192.30462713058 | 1118.70967838246 | 976.381688722746 | 906.625386138212 | 772.987970670679 | 1231.59538112791 | 1153.91367524262 | Snx19 |
| ENSMUSG00000031994 | -0.941500897530989 | 0.134541337151108 | 0.773512701181749 | 9.03513716569045 | 14.6594831204579 | 9.46454888648447 | 17.8206071909051 | 22.994122112201 | 57.4848649074714 | 9.36970281500523 | 8.68420451734806 | Adamts8 |
| ENSMUSG00000031996 | 0.022688307188263 | 0.896112573608595 | 0.984073355116573 | 9514.90294918861 | 11831.0172939385 | 10068.3871054422 | 7993.94921516231 | 9004.93620241625 | 7723.76429554642 | 11114.5496947773 | 10949.6963708112 | Aplp2 |
| ENSMUSG00000032000 | -0.214696277879687 | 0.33224816727523 | 0.856655692540538 | 163.535982698997 | 142.522752560008 | 131.557229522134 | 95.6685228143324 | 178.478185918513 | 177.34692365071 | 119.723980413956 | 144.374900100912 | Birc3 |
| ENSMUSG00000032002 | 0.13812925129756 | 0.221607032359703 | 0.831250791108906 | 1031.81266432185 | 897.48613326359 | 1075.17275350464 | 1144.27056699496 | 951.518672166795 | 1012.71208815716 | 911.984407327176 | 893.387539722182 | Dcun1d5 |
| ENSMUSG00000032006 | -0.200490761801919 | 0.399922139364031 | 0.880487015130533 | 509.581736144941 | 408.022280186079 | 444.83379766477 | 519.611388619021 | 734.716949394614 | 636.002760678407 | 356.048706970199 | 436.38127699674 | Pdgfd |
| ENSMUSG00000032009 | 0.128150707352478 | 0.671564820059439 | 0.943173644832837 | 413.809282188623 | 591.265819191803 | 425.904699891801 | 334.839829850164 | 302.208462046071 | 233.608706326107 | 548.648153723084 | 530.8220011229 | Sesn3 |
| ENSMUSG00000032010 | -0.0415719684578467 | 0.824404983475218 | 0.966637987288264 | 3437.86969154522 | 3575.2850499339 | 2870.59767727074 | 2204.12773150668 | 3268.45021452 | 2647.97303243991 | 3666.67703493871 | 2858.18881177218 | Usp2 |
| ENSMUSG00000032011 | -1.08036990963725 | 0.0158940565632265 | 0.63777231095375 | 68.6670424592474 | 60.2667639396603 | 46.3762895437739 | 45.0204813243917 | 151.104231023035 | 206.700897220482 | 65.5879197050366 | 43.4210225867403 | Thy1 |
| ENSMUSG00000032012 | -0.778306527154042 | 0.0561187530828568 | 0.703396849069134 | 30.7194663633475 | 36.6487078011448 | 29.3401015481019 | 43.1446279358754 | 88.6916138613469 | 88.0619207093179 | 33.3144988977964 | 30.3947158107182 | Nectin1 |
| ENSMUSG00000032014 | -0.658687150092129 | 0.0728648162680777 | 0.73022532320606 | 160.82544154929 | 185.686786192467 | 129.664319744837 | 257.929840920994 | 351.48158085793 | 472.10974158051 | 161.367104036201 | 174.76961591163 | Oaf |
| ENSMUSG00000032018 | 0.0857552557056251 | 0.662405696974958 | 0.939611397743053 | 460.791995450213 | 475.618785685968 | 497.835271429083 | 476.466760683146 | 381.045452145046 | 320.447544803351 | 530.94982618363 | 566.644344756961 | Sc5d |
| ENSMUSG00000032020 | -0.425017813363688 | 0.13082181694551 | 0.77303252008843 | 32.5264937964856 | 26.0613033252585 | 40.6975602118832 | 24.3860940507122 | 45.9882442244021 | 47.7002070508805 | 33.3144988977964 | 39.0789203280663 | Ubash3b |
| ENSMUSG00000032024 | -0.473361259156019 | 0.179232310037093 | 0.801755154770274 | 65.0529875929712 | 61.0811796685746 | 82.3415753124149 | 103.171936368398 | 139.059690869025 | 161.446854633749 | 74.9576225200418 | 57.5328549274309 | Clmp |
| ENSMUSG00000032026 | 0.0471969804626511 | 0.610242689050951 | 0.927932112008768 | 1501.63979693775 | 1311.20932355207 | 1257.83854701379 | 1384.37980072505 | 1300.81033663309 | 1361.29052429821 | 1334.66211209297 | 1283.09121743818 | Rexo2 |
| ENSMUSG00000032028 | 0.375059210649551 | 0.269936660707794 | 0.843793744782845 | 37.0440623793308 | 36.6487078011448 | 38.8046504345863 | 40.3308478531009 | 37.2285786578493 | 44.0309603546589 | 16.6572494488982 | 20.6249857287016 | Nxpe2 |
| ENSMUSG00000032030 | 0.0710988280679277 | 0.751565579125293 | 0.956197192463153 | 1131.19917314444 | 1363.33193020259 | 1136.69232126678 | 1014.83668318733 | 970.13296149572 | 715.503105763208 | 1331.5388778213 | 1404.67008068105 | Cul5 |
| ENSMUSG00000032034 | -0.0439073980195379 | 0.793237204797063 | 0.961804768755116 | 1104.99727536394 | 1266.41645846178 | 1233.23071990893 | 959.499008226099 | 928.524550054594 | 1086.09702208159 | 1390.880328983 | 1299.3741009082 | Kcnj5 |
| ENSMUSG00000032035 | -0.191501387916323 | 0.299056154417434 | 0.851785577030345 | 1011.03184884076 | 1082.35850372714 | 1352.48403587863 | 1527.88258494654 | 1666.52637403667 | 1510.50655661122 | 1134.77511870619 | 1367.76221148232 | Ets1 |
| ENSMUSG00000032038 | 0.0431077132769953 | 0.789929552613937 | 0.961659329450003 | 489.704434380422 | 351.82759489099 | 409.814966784777 | 487.721881014244 | 436.88832013182 | 348.57843614105 | 439.33495421469 | 461.348364984116 | St3gal4 |
| ENSMUSG00000032040 | -0.134848769467032 | 0.412795796241722 | 0.882192126163777 | 240.334648607366 | 202.789516499668 | 160.897331070236 | 218.536919762152 | 235.416012101106 | 245.839528646846 | 219.667477107345 | 202.993280593011 | Dcps |
| ENSMUSG00000032041 | -0.0954474447915635 | 0.571789470005764 | 0.921363774754067 | 95.7724539563188 | 87.9568987227475 | 88.966759532954 | 97.5443762028488 | 91.9764884488041 | 113.746647582869 | 89.5327157878278 | 100.953877514171 | Tirap |
| ENSMUSG00000032042 | 0.12974190163237 | 0.426182059045543 | 0.885066977635295 | 1872.08042073106 | 2326.78573750824 | 2730.52235375077 | 1935.88069694884 | 2026.76762046115 | 1693.9688914223 | 2216.45525479401 | 2164.537975949 | Srpr |
| ENSMUSG00000032044 | 0.1792566416189 | 0.189299551539067 | 0.808677608936066 | 321.65088309858 | 300.519403969387 | 256.489274823729 | 301.07446885687 | 244.175677667659 | 233.608706326107 | 276.926772087932 | 286.578749072486 | Rpusd4 |
| ENSMUSG00000032046 | 0.0934857148487599 | 0.208878696221628 | 0.821566820583092 | 1123.97106341189 | 1134.48111037766 | 1149.94268970786 | 1148.96020046625 | 1039.11532783232 | 1061.63537744011 | 1121.24110352896 | 1048.61769546978 | Abhd12 |
| ENSMUSG00000032047 | 0.289220730299318 | 0.131422766678385 | 0.77303252008843 | 21568.6794419362 | 20311.5282791234 | 22234.1182441293 | 21142.7435419675 | 15236.3432948227 | 12165.9989624387 | 21337.9365440386 | 21027.7157131937 | Acat1 |
| ENSMUSG00000032050 | 0.096199798632646 | 0.501811094619308 | 0.900927059989498 | 3255.35992079827 | 3829.38275735517 | 4230.65335225856 | 3149.55783931891 | 3119.53589988861 | 3002.66687974133 | 3685.41644056872 | 3723.35268681298 | Rdx |
| ENSMUSG00000032051 | 0.106635180843221 | 0.301721025399444 | 0.851785577030345 | 1418.5165350134 | 1204.52086306429 | 1296.64319744837 | 1394.69699436189 | 1344.60866446585 | 1256.10545233985 | 1175.37716423788 | 1159.34130306597 | Fdx1 |
| ENSMUSG00000032058 | -0.0142700977703573 | 0.927213190664602 | 0.986790505471895 | 121.070838020252 | 109.13170767452 | 116.413951303759 | 133.185590584659 | 128.110108910834 | 107.6312364225 | 120.765058504512 | 127.006491066215 | Ppp2r1b |
| ENSMUSG00000032059 | -0.202447242728834 | 0.208427810062133 | 0.820595884853464 | 194.255449062345 | 193.83094348161 | 187.398067952392 | 235.419600258798 | 256.220217821669 | 233.608706326107 | 256.10521027681 | 186.710397122983 | Alg9 |
| ENSMUSG00000032060 | -0.0206606249949068 | 0.903725633969045 | 0.985039825636533 | 50529.0045991238 | 38433.907078925 | 35666.2060238281 | 49468.1297085639 | 44317.3380175821 | 53660.2867677764 | 39700.4719052677 | 38931.2888512714 | Cryab |
| ENSMUSG00000032062 | -0.22255297669121 | 0.264863172467454 | 0.842510961779757 | 59.631905293557 | 80.6271571625185 | 84.2344850897118 | 69.4065753751039 | 72.2672409240604 | 88.0619207093179 | 94.7381062406084 | 87.9275707381491 | 2310030G06 |

| | | | | | | | | | | | | |
|---|---|---|---|---|---|---|---|---|---|---|---|---|
| **ENSMUSG00000032064** | -0.537698167510424 | 0.0152221809222893 | 0.635862253526407 | 113.8427282877 | 150.666909849151 | 145.754052851861 | 119.116690170786 | 191.617684268342 | 145.546785616789 | 262.351678820146 | 168.256462523619 | Dixdc1 |
| **ENSMUSG00000032067** | -0.014544976400807 | 0.94136823318227 | 0.989934163630124 | 680.345828576491 | 434.083583511337 | 409.814966784777 | 615.279911433354 | 486.161438943679 | 558.948580057754 | 508.046108191395 | 607.894316214364 | Pts |
| **ENSMUSG00000032076** | -0.468781833756273 | 0.115798628460711 | 0.769868040577702 | 38.8510898124689 | 31.7622134276588 | 27.447191770805 | 33.7653609932938 | 41.6084114411257 | 68.4926049961361 | 38.5198893505771 | 34.7368180693922 | Cadm1 |
| **ENSMUSG00000032077** | 0.085088015686751 | 0.59844194362284 | 0.924848033480823 | 183.413284463516 | 247.582381589956 | 218.631079277791 | 236.357526953057 | 233.226095709468 | 185.908499275227 | 229.03717992235 | 186.710397122983 | Bud13 |
| **ENSMUSG00000032078** | 0.376103621592198 | 0.025939692894498 | 0.664353746352133 | 675.828259993646 | 696.325448221751 | 864.113313336032 | 969.816201862939 | 711.722827282413 | 639.672007374629 | 598.619902069779 | 519.966745476215 | Zpr1 |
| **ENSMUSG00000032079** | -0.134833516128946 | 0.693065078647441 | 0.94469184542218 | 209.615182244018 | 351.013179162076 | 228.095628164276 | 255.11606083822 | 175.193311331055 | 226.270212933664 | 225.913945650682 | 518.881219911547 | Apoa5 |
| **ENSMUSG00000032080** | -0.654366525969065 | 0.160541911264287 | 0.791877538160847 | 63.2459601598332 | 157.182235680465 | 237.56017705076 | 133.185590584659 | 164.243729372865 | 88.0619207093179 | 352.92547269853 | 324.572143835884 | Apoa4 |
| **ENSMUSG00000032081** | -0.109073372535791 | 0.671474774320327 | 0.943173644832837 | 433.686583953142 | 610.811796685746 | 459.977075883145 | 528.052728867345 | 405.134532453066 | 571.179402378492 | 391.445362049107 | 824.999429148066 | Apoc3 |
| **ENSMUSG00000032083** | -0.339320933485934 | 0.192915671149117 | 0.808824684419004 | 2068.14289722654 | 2742.13775925454 | 1798.26428843205 | 1988.4045918273 | 1666.52637403667 | 2413.14124388173 | 2830.69132822214 | 3966.51041329873 | Apoa1 |
| **ENSMUSG00000032085** | -0.603950656856122 | 0.133862382220578 | 0.77303252008843 | 297.256012751216 | 412.09435883065 | 212.005895057252 | 186.647412157374 | 463.167316831478 | 755.864819421645 | 236.324726556243 | 230.131419709724 | Tagln |
| **ENSMUSG00000032086** | 0.121371906995995 | 0.598845831202973 | 0.924848033480823 | 552.046880823686 | 751.705717787925 | 708.894711597687 | 530.866508950119 | 523.390017601528 | 385.270903103266 | 731.877897660964 | 696.907412517182 | Bace1 |
| **ENSMUSG00000032089** | -0.523160651692007 | 0.0688040255319377 | 0.723457921755197 | 18.9737880479499 | 30.1333819698302 | 31.2330113253987 | 39.3929211588428 | 48.1781606160403 | 45.2540425867328 | 41.6431236222455 | 36.9078691987293 | Il10ra |
| **ENSMUSG00000032096** | 0.0556949427349998 | 0.724455295348295 | 0.951674101666722 | 2033.80937599692 | 2385.42366999007 | 2411.56705627624 | 1865.53619487948 | 1880.04322222139 | 1731.88444061659 | 2374.69912455855 | 2379.47203775337 | Arcn1 |
| **ENSMUSG00000032097** | 0.0423447576142495 | 0.725300362868533 | 0.951674101666722 | 3011.41121732463 | 3349.69189302463 | 3171.57033186095 | 2913.20031236585 | 3025.36949504816 | 2509.76474021556 | 3270.02628243683 | 3279.37273086356 | Ddx6 |
| **ENSMUSG00000032105** | 0.274062192476006 | 0.510602596430757 | 0.902394714365738 | 248.466272056487 | 179.985876090067 | 45.4298346551254 | 187.585338851632 | 109.49581958191 | 118.638976511164 | 158.243869764533 | 160.657783570939 | Pdzd3 |
| **ENSMUSG00000032109** | -0.160165500281849 | 0.232663623806264 | 0.832048466795696 | 398.449549006949 | 441.413325071566 | 361.545767463707 | 370.481044231974 | 494.921104510232 | 463.548165955993 | 385.198893505771 | 414.67076570337 | Nlrx1 |
| **ENSMUSG00000032112** | -0.0934021186274865 | 0.319276758258529 | 0.854670142081901 | 584.573374620172 | 557.874774306315 | 577.337482075553 | 641.541858872582 | 651.500126512363 | 627.44118505389 | 613.194995337564 | 626.348250813729 | Trappc4 |
| **ENSMUSG00000032114** | -0.0587267485309664 | 0.626055729595202 | 0.931698409533928 | 402.063603873225 | 359.971752180133 | 365.3315870183 | 352.660437041069 | 335.057207920644 | 393.832478727783 | 421.636626675235 | 390.789203280663 | Slc37a4 |
| **ENSMUSG00000032115** | -0.11406640398359 | 0.411052539534271 | 0.882192126163777 | 880.02235993825 | 1057.92603185971 | 1076.11920839328 | 800.051470202212 | 958.08842134171 | 1121.56640681173 | 962.997233764426 | 1086.61109023318 | Hyou1 |
| **ENSMUSG00000032116** | -0.194312606658009 | 0.275384994037736 | 0.846286635233373 | 688.477452025612 | 695.511032492837 | 599.105944514467 | 623.721251681677 | 839.832936193247 | 929.542496376133 | 573.634027896431 | 641.545608719088 | Stt3a |
| **ENSMUSG00000032118** | -0.119851903272895 | 0.759887313854635 | 0.957357521291862 | 35.2370349461928 | 25.2468875963442 | 38.8046504345863 | 49.7101147956825 | 41.6084114411257 | 69.71568722821 | 24.9858741733473 | 26.0526135520442 | Fez1 |
| **ENSMUSG00000032119** | 0.156877886789704 | 0.58024546573476 | 0.922450244597442 | 67.7635287426784 | 65.1532583131463 | 93.6990339761962 | 49.7101147956825 | 62.4126171616885 | 52.592535979176 | 46.8485140750261 | 85.7565196088121 | Hinfp |
| **ENSMUSG00000032120** | -0.15496960858243 | 0.180515437353329 | 0.802808628271642 | 326.168451681425 | 332.281617397046 | 367.224496795597 | 343.281170098487 | 429.223612761086 | 370.593916318379 | 360.213019332423 | 363.65106416395 | C2cd2l |
| **ENSMUSG00000032121** | 0.0550882707370591 | 0.616656516010568 | 0.928896975098882 | 441.818207402263 | 381.146561131906 | 401.296872786941 | 420.191159027656 | 363.52612101194 | 408.509465512669 | 419.554470494123 | 390.789203280663 | Tmem218 |
| **ENSMUSG00000032122** | -0.088593279952131 | 0.629632268982131 | 0.931825800800958 | 93.0619128066116 | 82.2559886203472 | 82.3415753124149 | 80.6616957062019 | 94.1664048404423 | 72.1618516923577 | 89.5327157878278 | 103.124928643508 | Slc37a2 |
| **ENSMUSG00000032123** | -0.0491264277422261 | 0.704233225295325 | 0.946931219685582 | 459.888481733644 | 453.629561005281 | 459.977075883145 | 394.867138282686 | 516.820268426614 | 488.00981059747 | 409.143689588562 | 416.841816832707 | Dpagt1 |
| **ENSMUSG00000032125** | -0.686689641762948 | 0.0371972478281313 | 0.679566075569698 | 610.775272400674 | 495.164763179912 | 506.353365426919 | 603.086864407998 | 1413.59103080245 | 1128.90490020417 | 504.922873919726 | 518.881219911547 | Robo4 |
| **ENSMUSG00000032126** | -0.103350821546765 | 0.351264313201461 | 0.862329676629249 | 507.774708711803 | 526.112560878656 | 495.942361651786 | 494.287367874051 | 494.921104510232 | 500.240632918209 | 595.49666779811 | 582.927228226989 | Hmbs |
| **ENSMUSG00000032127** | -0.0378062933250979 | 0.801630876931492 | 0.963461896828986 | 399.353062723518 | 460.95930256551 | 507.299820315568 | 387.363724728621 | 502.585811880965 | 488.00981059747 | 382.075659234102 | 429.868123608729 | Vps11 |
| **ENSMUSG00000032135** | -0.893660720848846 | 0.00089486791235119 | 0.556246415579446 | 327.071965397994 | 302.148235427216 | 254.596365046432 | 313.267515882226 | 769.755611660825 | 708.164612370765 | 375.829190690765 | 371.24974311663 | Mcam |
| **ENSMUSG00000032171** | -0.0807876943080119 | 0.541089091445851 | 0.913837378403038 | 1550.42953763248 | 1311.20932355207 | 1254.05272745919 | 1589.78574676758 | 1521.99189218854 | 1717.2074538317 | 1436.68776496747 | 1359.07800696497 | Pin1 |
| **ENSMUSG00000032175** | 0.120060643438315 | 0.491789385511272 | 0.899236955296196 | 206.904641094311 | 292.375246680244 | 264.060913932917 | 210.095579513828 | 197.092475247437 | 205.477814988408 | 245.694429371248 | 247.49982874442 | Tyk2 |
| **ENSMUSG00000032177** | -0.197706110869875 | 0.421692362156557 | 0.882568531013176 | 983.92643734369 | 1250.94255961241 | 904.810873547915 | 793.485983342404 | 905.530427942393 | 759.534066117867 | 1416.9072812469 | 1428.55164310376 | Pde4a |
| **ENSMUSG00000032178** | 0.244849427793827 | 0.118705750237404 | 0.769868040577702 | 674.021232560508 | 783.467931215584 | 713.626986040929 | 619.969544904645 | 551.858930692825 | 467.217412652214 | 655.879197050366 | 679.539003482486 | Ilf3 |
| **ENSMUSG00000032180** | -0.138372060166956 | 0.427135140368205 | 0.885066977635295 | 945.075347531221 | 694.696616763922 | 580.176846741498 | 787.858423176855 | 809.174106710313 | 951.557976553463 | 748.535147109862 | 802.203392290027 | Tmed1 |
| **ENSMUSG00000032182** | -0.226834649763042 | 0.269107117363785 | 0.843793744782845 | 55.1143367107117 | 83.0704043492615 | 71.930571537282 | 60.0273084325223 | 84.3117810780705 | 78.277262852727 | 73.9165444294857 | 80.3288917854696 | Yipf2 |
| **ENSMUSG00000032184** | 0.024213126797666 | 0.854795296686634 | 0.973268281968411 | 185.220311896654 | 186.501201921381 | 191.183887506986 | 198.84045918273 | 191.617684268342 | 206.700897220482 | 186.352978209548 | 164.999885829613 | Lysmd2 |
| **ENSMUSG00000032185** | 0.335914255584495 | 0.176499820508601 | 0.799889021902745 | 1045.36537007039 | 1286.77685168464 | 1299.48256211432 | 968.878275168681 | 786.179984598112 | 551.610086665311 | 1210.77381931679 | 1095.29529475052 | Carm1 |
| **ENSMUSG00000032186** | -0.528375899124483 | 0.0933967065196822 | 0.754786063592529 | 77.7021796249379 | 49.679359463774 | 55.8408384302584 | 45.0204813243917 | 100.736154015357 | 112.523565350795 | 67.6700758861489 | 48.8486504100828 | Tmod2 |
| **ENSMUSG00000032187** | 0.0418595498968989 | 0.628541737396401 | 0.931825800800958 | 2335.58295733098 | 2308.05417574321 | 1994.18045038228 | 2212.569071755 | 2188.82143344237 | 2076.79363006141 | 2166.48350644732 | 2165.62350151367 | Smarca4 |
| **ENSMUSG00000032192** | -0.100553026646299 | 0.361666237532143 | 0.868824634957016 | 527.652010476322 | 525.298145149742 | 441.994432998825 | 506.480414899407 | 504.775728272604 | 529.594606487981 | 577.798340258656 | 534.078577816906 | Gnb5 |
| **ENSMUSG00000032193** | -0.336009118591585 | 0.1522800173877 | 0.783412422888014 | 48.7897406947284 | 82.2559886203472 | 70.9841166486335 | 81.59962240046 | 102.926070406995 | 69.71568722821 | 89.5327157878278 | 95.5262496908287 | Ldlr |
| **ENSMUSG00000032194** | 0.0578315676229802 | 0.75093786471264 | 0.956197192463153 | 2656.33032671299 | 2536.90499556813 | 2280.00982675411 | 2100.01786844402 | 1834.05497799699 | 1873.76197953715 | 2692.22794217817 | 2796.31385458608 | Kank2 |
| **ENSMUSG00000032198** | -0.135922604066909 | 0.458820881491018 | 0.894333713711794 | 531.266065342598 | 483.762942975111 | 600.052399403115 | 498.039074651084 | 700.773245324222 | 683.702967729287 | 478.895921655823 | 459.177313854779 | Dock6 |
| **ENSMUSG00000032199** | 0.0838244440652487 | 0.582183484182887 | 0.922450244597442 | 7531.69034131956 | 7534.97432391537 | 7217.66498083306 | 6913.45766337691 | 5656.55403960145 | 5859.78697386586 | 7878.87898932884 | 8153.38251622516 | Polr2m |
| **ENSMUSG00000032202** | -0.466540381326273 | 0.279270032889677 | 0.848075053184273 | 89.4478579403355 | 52.9370223794314 | 58.6802030962037 | 48.7721881014244 | 145.62944004394 | 112.523565350795 | 36.4377331694648 | 51.0197015394199 | Rab27a |
| **ENSMUSG00000032204** | -0.515768474600762 | 0.193130924073423 | 0.808863491502674 | 48.7897406947284 | 98.5443031986338 | 48.2691993210708 | 65.6548685980713 | 78.836990098975 | 85.6157562451702 | 46.8485140750261 | 162.828834700276 | Aqp9 |
| **ENSMUSG00000032207** | -0.477039538619954 | 0.187197315076607 | 0.807540361382744 | 20.780815481088 | 27.6901347830872 | 17.036187995672 | 19.6964605794214 | 26.2789966996583 | 20.7923979452556 | 24.9858741733473 | 46.6775992807458 | Lipc |
| **ENSMUSG00000032212** | 0.0548865926823421 | 0.590643699157688 | 0.923935529355527 | 1001.99671167507 | 862.466256920274 | 860.327493781438 | 1012.96082979881 | 893.485887788383 | 924.650167447838 | 871.382361795486 | 908.584897627541 | Sltm |
| **ENSMUSG00000032215** | 0.0655417556797831 | 0.653600298199057 | 0.936991241819063 | 856.531003307455 | 727.273245920495 | 691.858523602015 | 879.775239214155 | 833.263187018333 | 624.995020589742 | 732.91897575152 | 822.828378018729 | Rsl24d1 |
| **ENSMUSG00000032216** | 0.247830664247234 | 0.240299716868862 | 0.832048466795696 | 7955.43827439044 | 11313.8633060779 | 13283.494362181 | 8261.25832302588 | 7801.57714521107 | 6893.29145996827 | 10192.1545065446 | 9484.23685850875 | Nedd4 |
| **ENSMUSG00000032217** | 0.0676614621071298 | 0.684291105564373 | 0.944452327903597 | 248.466272056487 | 332.281617397046 | 304.7584741448 | 238.233380341573 | 268.264757975679 | 226.270212933664 | 279.008928269045 | 298.51953028384 | Rnf111 |
| **ENSMUSG00000032220** | -0.217480461558606 | 0.383510570544368 | 0.875126835008975 | 185.220311896654 | 205.232763686411 | 162.790240847533 | 237.295453647315 | 273.739548954774 | 310.66288694676 | 167.613572579538 | 168.256462523619 | Myo1e |
| **ENSMUSG00000032224** | -0.0238861723833687 | 0.908866545110535 | 0.985338457039129 | 271.957628687283 | 214.191336704468 | 198.755526616174 | 257.929840920994 | 170.813478547779 | 226.270212933664 | 258.187366457922 | 302.861632542514 | Fam81a |
| **ENSMUSG00000032228** | -0.0304934327882652 | 0.771540549317354 | 0.959045638244974 | 416.51982333833 | 470.732291312482 | 450.512526996661 | 410.811892085075 | 477.401773377126 | 434.19419238622 | 432.047407580797 | 442.894430384751 | Tcf12 |
| **ENSMUSG00000032231** | -0.663392068097104 | 0.0652487370605287 | 0.718339708336358 | 983.022923627121 | 779.395852571012 | 888.721140440892 | 1027.02973021269 | 1927.12642464161 | 2371.55644799121 | 758.945928015424 | 768.552099785303 | Anxa2 |
| **ENSMUSG00000032232** | 0.13399698329431 | 0.382754947940904 | 0.875126835008975 | 1293.83164212687 | 1250.12814388349 | 1171.71115214678 | 1015.77460988159 | 1148.61114741423 | 829.249753346077 | 1169.13069569454 | 1163.68340532464 | Cgnl1 |
| **ENSMUSG00000032235** | 0.0914079808909657 | 0.719546062252115 | 0.951674101666722 | 61.4389327266951 | 73.2974156022896 | 83.2880302010633 | 51.5859681841989 | 52.5579933993167 | 48.9232892829544 | 71.8343882483734 | 79.2433662208011 | Ice2 |
| **ENSMUSG00000032238** | 0.118850071023144 | 0.292460239847006 | 0.851785577030345 | 770.697200233395 | 829.889627763701 | 926.579335986829 | 767.224035903176 | 810.269064906132 | 691.041461121731 | 740.206522385413 | 791.348136643342 | Rora |
| **ENSMUSG00000032239** | -0.116880590679958 | 0.51210939227802 | 0.902394714365738 | 1289.31407354403 | 1053.85395321514 | 1085.58375727977 | 1289.64920460497 | 1287.67083828326 | 1676.84574017326 | 1049.40671528059 | 1103.97949926787 | Rp9 |
| **ENSMUSG00000032243** | -0.690177760414077 | 0.279375355237323 | 0.848201898195939 | 19.877301764519 | 32.5766291565731 | 20.8220075502658 | 31.8895076047775 | 52.5579933993167 | 95.400414101761 | 10.4107809055614 | 11.9407812113536 | Itga11 |
| **ENSMUSG00000032244** | -0.1025337255046 | 0.450725005002555 | 0.89108031640712 | 349.65980831222 | 442.22774080048 | 406.975602118832 | 344.219096792745 | 448.93286028583 | 424.409534529629 | 414.349080041342 | 370.164217551961 | Fem1b |
| **ENSMUSG00000032245** | 0.329710463883205 | 0.0907250545005769 | 0.753823363185192 | 533.976606492306 | 594.52348210746 | 804.48665535118 | 792.548056648146 | 536.529515951357 | 418.29412336926 | 615.277151518677 | 597.039060567679 | Cln6 |
| **ENSMUSG00000032249** | 0.00942800435960572 | 0.913561551042704 | 0.986234232499696 | 4597.0777899033 | 4103.8408579993 | 4071.64893096562 | 4578.02019467409 | 4178.36047524567 | 4544.97357438646 | 4143.49080041342 | 4371.41144892008 | Anp32a |
| **ENSMUSG00000032251** | 0.118343306056081 | 0.536325815749584 | 0.911291453479697 | 103.000563688871 | 106.688460487777 | 90.8596693102509 | 129.433883807626 | 96.3563212320805 | 118.638976511164 | 85.3684034256032 | 96.6117752554972 | Irak1bp1 |
| **ENSMUSG00000032252** | -0.0546272542889619 | 0.68637526649996 | 0.944452327903597 | 206.904641094311 | 240.252640029727 | 242.292451494002 | 208.219726125312 | 240.890803080201 | 207.923979452556 | 227.996101831794 | 255.098507697099 | Glce |
| **ENSMUSG00000032253** | -0.153219718418966 | 0.151082852882252 | 0.783412422888014 | 383.089815825275 | 394.177212794535 | 359.65285768641 | 398.618845059719 | 464.262275027297 | 435.417274618294 | 403.938299135781 | 404.901035621353 | Phip |
| **ENSMUSG00000032254** | -0.612880398600796 | 0.256545923483097 | 0.837517407767513 | 4.51756858284523 | 17.9171460361152 | 12.3039135524298 | 15.0068271081306 | 25.1840385038392 | 28.1308913376988 | 15.616171358342 | 7.59867895267955 | Kif23 |
| **ENSMUSG00000032258** | -0.862949754760557 | 0.0381994575155217 | 0.679566075569698 | 15.3597331816738 | 21.9892246806869 | 8.51809399783602 | 15.9447538023887 | 37.2285786578493 | 14.6769867848863 | 27.0680303544596 | 33.6512925047237 | Lca5 |
| **ENSMUSG00000032259** | 0.348689368885011 | 0.62607950948213 | 0.931698409533928 | 27.1054114970714 | 0.814415728914329 | 23.6613722162112 | 30.0136542162612 | 25.1840385038392 | 11.0077400886647 | 13.5340151772298 | 14.1118323406906 | Drd2 |
| **ENSMUSG00000032261** | -0.321015189937234 | 0.220644704487945 | 0.829144114546069 | 110.228673421423 | 124.605606523892 | 140.07532351997 | 90.9788893430416 | 165.338687568684 | 198.139321595965 | 93.6970281500523 | 125.920965501547 | Sh3bgrl2 |
| **ENSMUSG00000032263** | 0.0811226795840896 | 0.48472452446046 | 0.899236955296196 | 1480.85898145666 | 1373.10491894956 | 1552.18601738345 | 1485.67588370493 | 1503.37760285962 | 1185.16668287957 | 1529.34371502696 | 1349.30827688295 | Bckdhb |
| **ENSMUSG00000032264** | 0.143536572139058 | 0.418064391392667 | 0.882468496126562 | 369.537110076739 | 386.847471234306 | 433.476339000989 | 379.860311174555 | 327.39250054991 | 265.408844360028 | 445.581422758026 | 381.019473198646 | Zw10 |
| **ENSMUSG00000032265** | -0.0867590299775589 | 0.672508725287721 | 0.943216235131458 | 296.352499034647 | 352.642010619904 | 407.922057007481 | 334.839829850164 | 363.52612101194 | 507.579126310652 | 296.707255808499 | 311.545837059862 | Fam46a |
| **ENSMUSG00000032267** | 0.375525925563809 | 0.0931580001678367 | 0.754058637722175 | 2624.70734663308 | 3110.25366872382 | 3050.42410611394 | 2895.37970517494 | 2059.61636633572 | 1385.75216893968 | 2633.92756910703 | 2923.32034565229 | Usp28 |
| **ENSMUSG00000032271** | -0.418418662292365 | 0.0418741644045674 | 0.685406501171951 | 158.114900399583 | 177.542628903324 | 146.700507740509 | 148.192417692789 | 193.80760065998 | 261.739597663806 | 147.833088858971 | 240.986675356409 | Nnmt |
| **ENSMUSG00000032279** | 0.142869524241619 | 0.225632037734858 | 0.832048466795696 | 17338.42822096 | 17111.688880219 | 17295.5166351617 | 15910.0505147012 | 14577.1784609396 | 12926.7561107886 | 16618.7295595476 | 17152.3894473271 | Idh3a |
| **ENSMUSG00000032280** | -0.476540280280141 | 0.0676293879193089 | 0.722038584319139 | 179.79922959724 | 183.243539005724 | 176.98706417726 | 171.640585049243 | 308.778211220985 | 340.016860516533 | 190.517290571773 | 151.973579053591 | Tle3 |
| **ENSMUSG00000032285** | 0.309561932737124 | 0.11495543474855 | 0.767518180080285 | 3701.69569678338 | 3777.26015070466 | 3588.01048286626 | 4003.07113109383 | 2784.47869196796 | 2033.98575193883 | 3697.9093776554 | 3641.93826946284 | Dnaja4 |
| **ENSMUSG00000032288** | -0.0893405307163334 | 0.596324469389814 | 0.92448662081833 | 719.19691838896 | 597.781145023117 | 686.179794270124 | 742.837941852464 | 826.693437843418 | 874.50379593281 | 643.386259963692 | 577.499600403646 | Imp3 |
| **ENSMUSG00000032289** | -0.22357817309365 | 0.294278168275739 | 0.851785577030345 | 58.7283915769879 | 65.1532583131463 | 50.1621090983677 | 49.7101147956825 | 55.8428679867739 | 62.3771938357668 | 74.9576225200418 | 68.388110574116 | Thsd4 |
| **ENSMUSG00000032290** | -0.106840255639344 | 0.573464981030718 | 0.922207298536842 | 202.387072511466 | 231.294067011669 | 231.881447718869 | 163.19924480092 | 268.264757975679 | 242.170281950624 | 181.147587756768 | 201.907755028342 | Ptpn9 |
| **ENSMUSG00000032293** | 0.183954801796848 | 0.4913174708421 | 0.899236955296196 | 644.205279913729 | 737.860650396382 | 841.398396008469 | 614.341984739096 | 527.769850384805 | 353.470765069345 | 853.684034256032 | 762.038946397292 | Ireb2 |
| **ENSMUSG00000032294** | -0.137274589753048 | 0.0913139565297166 | 0.753823363185192 | 41241.7871065106 | 38824.8266288039 | 36739.4858675554 | 36830.5054301295 | 43428.231962577 | 44514.0778363281 | 40759.2483233633 | 40274.0839747663 | Pkm |
| **ENSMUSG00000032295** | 0.0978342884437758 | 0.421442572669081 | 0.882568531013176 | 594.512025502432 | 598.595560752032 | 704.162437154444 | 531.804435644377 | 566.093387238473 | 545.494675504941 | 604.866370613115 | 552.53251241627 | Man2c1 |
| **ENSMUSG00000032298** | -0.204826696737814 | 0.318944918004771 | 0.854670142081901 | 95.7724539563188 | 76.5550785179469 | 60.5731128735006 | 93.7926694258161 | 93.0714466446232 | 84.3926740130963 | 102.025652874501 | 96.6117752554972 | Neil1 |
| **ENSMUSG00000032299** | 0.062100024422592 | 0.739989133666309 | 0.954383574136061 | 341.528184863099 | 250.025628776699 | 302.865564367503 | 352.660437041069 | 298.923587458613 | 379.155491942897 | 271.721381635152 | 245.328777615083 | Commd4 |

| | | | | | | | | | | | | |
|---|---|---|---|---|---|---|---|---|---|---|---|---|
| ENSMUSG00000032300 | 0.121789136098936 | 0.570700945383486 | 0.920998737234417 | 256.597895505609 | 368.930325198191 | 419.279515671262 | 270.12288794635 | 367.905953795217 | 309.439804714686 | 291.501865355718 | 239.90114979174 | 1700017B05I |
| ENSMUSG00000032301 | 0.0746286925183105 | 0.517552369289972 | 0.905207722478398 | 2425.93432898789 | 2102.8214120568 | 2138.04159345684 | 2503.32634697503 | 2178.96680968 | 2454.72603977224 | 1979.08945014722 | 2096.14986537489 | Psma4 |
| ENSMUSG00000032305 | 0.0915712059772018 | 0.491176305630664 | 0.899236955296196 | 388.510898124689 | 352.642010619904 | 331.259211026956 | 432.384206053012 | 319.727793179176 | 358.363093997641 | 353.966550789086 | 379.933947633978 | Fam219b |
| ENSMUSG00000032306 | -0.0368227551410217 | 0.681040628942299 | 0.944452327903597 | 2730.41845147165 | 2369.13535541178 | 2338.69002985031 | 2393.58892374683 | 2428.61727832676 | 2562.35727619474 | 2629.7632567448 | 2465.22855736218 | Mpi |
| ENSMUSG00000032307 | 0.106708042672069 | 0.540888260107523 | 0.913837378403038 | 861.0485718903 | 961.010560118908 | 963.491076644119 | 916.354380290224 | 766.470737073368 | 639.672007374629 | 1058.77641809559 | 971.545380378314 | Ube2q2 |
| ENSMUSG00000032308 | 0.44228356771073 | 0.0123150377536459 | 0.635862253526407 | 148.176249517323 | 162.068730053951 | 144.807597963212 | 147.254490998531 | 104.021028602814 | 85.6157562451702 | 134.299073681742 | 118.322286548867 | Ulk3 |
| ENSMUSG00000032309 | 0.00038354572688115 | 0.99723723566185 | 0.999262355603273 | 790.574501997914 | 781.024684028841 | 778.932373357672 | 763.472329126143 | 791.654775577207 | 656.795158623663 | 846.396487622139 | 816.315224630718 | Fbxo22 |
| ENSMUSG00000032310 | -0.342596533164228 | 0.451899543499253 | 0.891110666112841 | 38.8510898124689 | 83.8848200781758 | 30.2865564367503 | 62.8410885152968 | 45.9882442244021 | 55.0387004433237 | 40.6020455316893 | 132.434118889558 | Cyp1a2 |
| ENSMUSG00000032311 | 0.158407624243439 | 0.635409307770544 | 0.932543559775713 | 37.0440623793308 | 38.2775392589734 | 44.483379766477 | 34.703287687552 | 18.6142893289246 | 29.3539735697726 | 53.094982618363 | 36.9078691987293 | Nrg4 |
| ENSMUSG00000032312 | -0.0367524318432997 | 0.767784220137846 | 0.95879680569409 | 538.494175075151 | 489.463853077511 | 477.959718767466 | 455.832373409466 | 572.663136413388 | 521.033030863464 | 453.910047482476 | 465.69046724279 | Csk |
| ENSMUSG00000032313 | 0.0738938360161524 | 0.760471746202122 | 0.957357521291862 | 101.193536255733 | 103.43079757212 | 67.1982970940397 | 78.7858423176855 | 74.4571573156986 | 63.6002760678407 | 86.4094815161593 | 108.552556466851 | Tmem266 |
| ENSMUSG00000032314 | 0.237055559442199 | 0.0862626289465592 | 0.746460787597018 | 24547.5641654644 | 24219.0949464543 | 26106.0651935901 | 25248.9866094297 | 20219.4980439954 | 16781.9113062854 | 23884.4135535389 | 24062.8451920068 | Etfa |
| ENSMUSG00000032316 | 0.0855856626268641 | 0.449190721912913 | 0.89108031640712 | 340.62467114653 | 331.467201668132 | 365.3315870183 | 380.798237868813 | 306.588294829347 | 354.693847301419 | 333.144988977964 | 341.94055287058 | Clk3 |
| ENSMUSG00000032320 | -0.088464743279481 | 0.25837024639376 | 0.838495963179748 | 868.276681622852 | 917.032110757534 | 879.256591554407 | 924.795720538547 | 969.038003299901 | 979.688867891162 | 926.559500594962 | 942.236190132265 | Rcn2 |
| ENSMUSG00000032322 | -0.597410353302046 | 0.136897331535367 | 0.776456211338551 | 35.2370349461928 | 22.8036404096012 | 13.2503684410783 | 16.8826804966469 | 45.9882442244021 | 39.1386314263635 | 27.0680303544596 | 21.7105112933701 | Pstpip1 |
| ENSMUSG00000032324 | -0.0881486527203942 | 0.374878262884354 | 0.873700364743909 | 4210.37391921175 | 4010.18304917415 | 3854.91076146512 | 3709.50007579103 | 3686.7242453229 | 4250.21075645666 | 4419.3764944108 | 4423.51667602417 | Tspan3 |
| ENSMUSG00000032328 | 0.128011899179146 | 0.253001630305255 | 0.837406774112203 | 2340.10052591383 | 2313.75508584561 | 2420.08515027408 | 2290.41698737843 | 1993.91887458658 | 1860.30807498434 | 2347.63109420409 | 2365.36020541268 | Tmem30a |
| ENSMUSG00000032329 | -0.13484216985195 | 0.270963570098307 | 0.844129257012535 | 265.633032671299 | 256.540954608014 | 256.489274823729 | 264.495327780801 | 281.404256325508 | 250.731857575141 | 319.610973800734 | 292.006376895829 | Hmg20a |
| ENSMUSG00000032330 | 0.132485000756396 | 0.41319580561578 | 0.882192126163777 | 8549.95029989287 | 6299.50566315233 | 7417.36696233788 | 9251.7089121625 | 7156.64676787362 | 8515.09849969821 | 6226.68805961625 | 6855.09394088163 | Cox7a2 |
| ENSMUSG00000032332 | -2.00399038240642 | 0.013747045674164 | 0.635862253526407 | 3.61405486627618 | 6.51532583131463 | 5.67872933189068 | 6.56548685980713 | 20.8042057205628 | 59.9310293716191 | 6.24646854333682 | 3.25657669400552 | Col12a1 |
| ENSMUSG00000032333 | -0.0219583652243628 | 0.902771785936963 | 0.985027469232262 | 187.930853046361 | 166.955224427437 | 156.165056626994 | 172.578511743502 | 140.154649064844 | 209.14706168463 | 196.76375911511 | 148.717002359586 | Stoml1 |
| ENSMUSG00000032334 | -1.09068180253542 | 0.0142652923161423 | 0.635862253526407 | 297.256012751216 | 311.106808445274 | 273.525462819401 | 378.922384480297 | 844.212768976524 | 1283.01326144548 | 286.296474902938 | 272.466916731795 | Loxl1 |
| ENSMUSG00000032336 | -0.0609850818276738 | 0.551296353510458 | 0.914873920695527 | 1342.6213828216 | 1360.88868301584 | 1559.75765649264 | 1294.33883807626 | 1455.19944224358 | 1567.99142151869 | 1384.63386043966 | 1390.55824834036 | Nptn |
| ENSMUSG00000032338 | -0.248582199440538 | 0.192667431584638 | 0.808824684419004 | 164.439496415566 | 163.69756151178 | 155.218601738345 | 141.626930832982 | 135.774816281568 | 187.131581507301 | 184.270822028436 | 235.559047533066 | Hcn4 |
| ENSMUSG00000032340 | 0.220738004851222 | 0.328183916476571 | 0.855977678491063 | 1282.08596381147 | 1835.6930529729 | 1701.72588978991 | 1226.80811608967 | 1119.04727612712 | 919.757838519542 | 1630.32828981091 | 1518.65026497124 | Neo1 |
| ENSMUSG00000032342 | 0.197705901139862 | 0.185298385067635 | 0.805963874499992 | 701.126644057579 | 729.716493107238 | 765.682004916593 | 712.824287636203 | 584.707676567398 | 506.356044078578 | 675.659680770933 | 768.552099785303 | Mto1 |
| ENSMUSG00000032344 | -0.459781316071205 | 0.427454690475679 | 0.885066977635295 | 16.2632468982428 | 6.51532583131463 | 5.67872933189068 | 12.1930470253561 | 25.1840385038392 | 15.9000690169602 | 7.28754663389296 | 7.59867895267955 | Mb21d1 |
| ENSMUSG00000032348 | -0.15028851657097 | 0.227468243064068 | 0.832048466795696 | 3872.45978921493 | 3945.84420658992 | 3090.17521143718 | 3667.29337454941 | 3750.23182068041 | 4491.15795617521 | 4240.31106283514 | 3696.21454769627 | Gsta4 |
| ENSMUSG00000032349 | -0.448228189198947 | 0.110446464941921 | 0.764116514278315 | 366.826568927032 | 409.651111643907 | 325.580481695066 | 416.439452250624 | 671.209374037106 | 711.833859066986 | 291.501865355718 | 398.387882233342 | Elovl5 |
| ENSMUSG00000032350 | -0.0281693564875164 | 0.890845119565399 | 0.982142775127356 | 245.75573090678 | 418.609684661965 | 283.936466594534 | 324.522636213324 | 289.068963696242 | 276.416584448692 | 386.239971596327 | 346.282655129254 | Gclc |
| ENSMUSG00000032352 | 0.0507521068909282 | 0.782616211581751 | 0.960786612650028 | 357.791431761342 | 406.39344872825 | 291.508105703722 | 378.922384480297 | 333.962249724825 | 259.293433199658 | 387.281049686883 | 403.815510056685 | Lrrc1 |
| ENSMUSG00000032353 | -0.162661504204028 | 0.446593982477777 | 0.89108031640712 | 466.213077749627 | 373.816819571677 | 405.082692341535 | 471.777127211855 | 558.428679867739 | 628.664267285964 | 364.377331694648 | 371.24974311663 | Tmed3 |
| ENSMUSG00000032355 | 0.217401429071629 | 0.106238548006495 | 0.762094734041179 | 2053.68667776144 | 2005.09152458708 | 1634.52759269587 | 1888.98436223594 | 1501.18768646798 | 1413.88306027738 | 1787.53108148489 | 1818.25532081975 | Mlip |
| ENSMUSG00000032359 | -0.101734375272179 | 0.403354348396635 | 0.881158207349486 | 801.416666596743 | 743.561560498782 | 910.489602879806 | 757.844768960594 | 917.574968096403 | 945.442565393093 | 799.547973547113 | 785.920508819999 | Ctsh |
| ENSMUSG00000032363 | -0.15361607888891 | 0.497598650614221 | 0.900927059989498 | 257.501409222178 | 252.468875963442 | 170.36187995672 | 171.640585049243 | 195.997517051618 | 195.693157131818 | 300.871568170723 | 255.098507697099 | Adamts7 |
| ENSMUSG00000032366 | -0.0225168308049434 | 0.853566355339271 | 0.9731539929935 | 250971.715592533 | 218569.635663112 | 207555.664170827 | 256102.759720579 | 236408.044226518 | 270905.375910967 | 220828.279178315 | 219738.597953587 | Tpm1 |
| ENSMUSG00000032369 | 0.454251383303992 | 0.0273532937328345 | 0.664353746352133 | 370.440623793308 | 385.218639776477 | 564.087113634474 | 643.417712261099 | 382.140410340865 | 344.909189444828 | 341.473613702413 | 363.65106416395 | Plscr1 |
| ENSMUSG00000032370 | 0.0688335456594719 | 0.683077906274622 | 0.944452327903597 | 465.309564033058 | 479.69086433054 | 445.780252553418 | 479.28054076592 | 420.463947194533 | 329.009120427868 | 488.265624470828 | 543.848307898922 | Lactb |
| ENSMUSG00000032372 | 0.082459922189052 | 0.599847294676212 | 0.925250296840931 | 288.220875585525 | 290.746415222415 | 334.098575692902 | 388.301651422879 | 323.012667766634 | 325.339873731647 | 256.10521027681 | 324.572143835884 | Plscr2 |
| ENSMUSG00000032374 | 0.0512731848592482 | 0.778788709230326 | 0.960254744040417 | 444.52874855197 | 568.462178782201 | 506.353365426919 | 326.39848960184 | 471.926982398031 | 423.186452297555 | 433.088485671353 | 453.749686031436 | Plod2 |
| ENSMUSG00000032375 | -0.192164257159247 | 0.193718576036338 | 0.809162446871198 | 121.974351736821 | 146.594831204579 | 134.396594188079 | 118.178763476528 | 155.484063806312 | 151.662196777159 | 141.586620315635 | 147.631476794917 | Aph1b |
| ENSMUSG00000032376 | -0.00693603452001507 | 0.960504690018772 | 0.993694458032251 | 309.001691066613 | 356.714089264476 | 369.117406572894 | 347.970803569778 | 393.089992299056 | 308.216722482613 | 379.99350305299 | 307.203734801188 | Usp3 |
| ENSMUSG00000032377 | -0.397962235248379 | 0.140337024464225 | 0.779711473022872 | 139.141112351633 | 152.295741306979 | 119.253315969704 | 163.19924480092 | 255.12525962585 | 234.831788558181 | 153.038479311752 | 113.980184290193 | Plscr4 |
| ENSMUSG00000032380 | -0.00345028642131333 | 0.988846717029703 | 0.998437672973644 | 157.211386683014 | 93.6578088251478 | 116.413951303759 | 127.55803041911 | 150.009272827216 | 152.885279009232 | 94.7381062406084 | 98.7828263848342 | Dapk2 |
| ENSMUSG00000032381 | 0.185964316591669 | 0.255855052938969 | 0.837517407767513 | 1378.76193148436 | 1097.0179868476 | 1019.33191507438 | 1311.22151857291 | 880.346389438554 | 980.911950123235 | 1094.1730731745 | 1268.97938509749 | Fam96a |
| ENSMUSG00000032382 | -0.137133170232426 | 0.384274857850095 | 0.875126835008975 | 401.160090156656 | 403.950201541507 | 377.63550057073 | 409.873965390816 | 543.099265126272 | 465.99433042014 | 364.377331694648 | 378.848422069309 | Snx1 |
| ENSMUSG00000032383 | -0.337267345680396 | 0.177215977901398 | 0.800237299431217 | 651.433389646282 | 505.752167655798 | 511.085639870161 | 612.466131350579 | 820.123688668504 | 1031.05832163826 | 482.019155927491 | 549.275935722265 | Ppib |
| ENSMUSG00000032384 | -0.0264291735145853 | 0.890629694128403 | 0.982124996145087 | 116.553269437407 | 124.605606523892 | 130.610774633486 | 110.675349922463 | 124.825234323377 | 86.838838477244 | 137.42230795341 | 141.118323406906 | Csnk1g1 |
| ENSMUSG00000032386 | 0.0302334129810454 | 0.81510972568792 | 0.966177091475091 | 336.107102563685 | 417.795268933051 | 335.04503058155 | 346.094950181261 | 339.43704070392 | 332.67836712409 | 386.239971596327 | 347.368180693922 | Trip4 |
| ENSMUSG00000032387 | -0.0855598760200309 | 0.593266008573196 | 0.923935529355527 | 564.696072855653 | 688.995706661522 | 643.589324280944 | 501.790781428116 | 581.42280197994 | 540.602346576646 | 681.906149314269 | 741.413960668591 | Rbpms2 |
| ENSMUSG00000032388 | -0.0115239864918651 | 0.953027494923096 | 0.992049433957868 | 317.133314515735 | 264.685111897157 | 319.901752363175 | 317.957149353517 | 339.43704070392 | 399.947889888152 | 245.694429371248 | 245.328777615083 | Spg21 |
| ENSMUSG00000032392 | -0.148895560140661 | 0.501647241065691 | 0.900927059989498 | 60.535419010126 | 63.5244268553176 | 64.3589324280944 | 90.9788893430416 | 75.5521155115177 | 94.1773318696872 | 67.6700758861489 | 72.73021283279 | Parp16 |
| ENSMUSG00000032393 | 0.19731503902148 | 0.467612754176902 | 0.895328112556867 | 1485.37655003951 | 1945.63917637633 | 1992.28754060498 | 1421.89686849537 | 1105.90777777729 | 928.319414144059 | 2032.18443276558 | 1902.92631486389 | Dpp8 |
| ENSMUSG00000032396 | 0.186177035990209 | 0.0420300352051543 | 0.685406501171951 | 908.03128515189 | 902.372627637076 | 928.472245764126 | 900.409626487835 | 787.274942793931 | 725.287763619799 | 824.53384772046 | 859.736247217458 | Dis3l |
| ENSMUSG00000032397 | 0.185588828392143 | 0.271653973875544 | 0.844129257012535 | 114.746242004269 | 127.048853710635 | 116.413951303759 | 135.999370667433 | 113.875652365186 | 118.638976511164 | 114.518589961175 | 87.9275707381491 | Tipin |
| ENSMUSG00000032398 | 0.0060021907031801 | 0.972521631591585 | 0.995322826578593 | 482.47632464787 | 382.775392589734 | 401.296872786941 | 507.418341593665 | 379.950493949227 | 560.171662289828 | 429.965251399684 | 397.302356668674 | Snapc5 |
| ENSMUSG00000032399 | 0.0561706404118783 | 0.684807198901008 | 0.944452327903597 | 11840.5472556373 | 9969.2629376403 | 11743.6122583499 | 13122.5323793659 | 11309.8232046155 | 13270.4422180014 | 10310.837408868 | 10003.1180784203 | Rpl4 |
| ENSMUSG00000032402 | -0.296703688049851 | 0.0799619900210717 | 0.739855948845396 | 154.500845533307 | 193.83094348161 | 211.059440168604 | 151.944124469822 | 201.472308030714 | 199.362403828039 | 247.776585552361 | 224.703791886381 | Smad3 |
| ENSMUSG00000032403 | -0.043686818867485 | 0.83199819688466 | 0.967583158900775 | 162.632468982428 | 102.616381843205 | 112.628131749165 | 121.930470253561 | 143.439523652302 | 145.546785616789 | 110.35427759895 | 116.15123541953 | 2300009A05I |
| ENSMUSG00000032405 | -0.229222484650266 | 0.0573235005494936 | 0.710279592000416 | 458.984968017075 | 513.896324944941 | 551.783200082044 | 467.087493740564 | 595.657258525589 | 558.948580057754 | 648.591650416473 | 530.8220011229 | Pias1 |
| ENSMUSG00000032407 | 0.0244806174950428 | 0.875348258331047 | 0.976986380159756 | 677.635287426784 | 767.179616637298 | 691.858523602015 | 638.728078789808 | 612.081631462875 | 552.833168897385 | 822.451691539348 | 740.328435103922 | U2surp |
| ENSMUSG00000032409 | 0.343039623678408 | 0.0903013217174486 | 0.753823363185192 | 185.220311896654 | 197.903022126182 | 166.576060402127 | 170.702658354985 | 136.869774477387 | 102.738907494204 | 189.476212481217 | 137.8617467129 | Atr |
| ENSMUSG00000032410 | 0.11845431052588 | 0.559961829502685 | 0.916574868225118 | 355.080890611635 | 493.535931722083 | 481.745538322059 | 325.460562907582 | 325.202584158272 | 320.447544803351 | 458.0743598447 | 421.183919091381 | Xrn1 |
| ENSMUSG00000032411 | 0.20530105713084 | 0.398399363588562 | 0.880459952732461 | 2406.05702722337 | 2726.66386040517 | 3119.51531298528 | 2091.5765281957 | 1748.6482387231 | 1525.1835433961 | 2998.30490080167 | 2698.61655376591 | Tfdp2 |
| ENSMUSG00000032412 | -0.126701315684387 | 0.467758705706638 | 0.895328112556867 | 1143.84836517641 | 998.473683648967 | 937.936794650611 | 1128.32581319257 | 1310.66496039546 | 1401.65223795664 | 955.709687130533 | 928.124357791574 | Atp1b3 |
| ENSMUSG00000032413 | -0.232338190126162 | 0.361227294095578 | 0.868539734006824 | 37.9475760958999 | 39.9063707168021 | 37.8581955459379 | 36.5791410760683 | 55.8428679867739 | 51.3694537471021 | 35.3966550789086 | 36.9078691987293 | Rasa2 |
| ENSMUSG00000032415 | 0.0585281256420013 | 0.897733474917316 | 0.984332015045905 | 31.6229800799166 | 13.0306516626293 | 9.46454888648447 | 26.2619474392285 | 24.0890803080201 | 14.6769867848863 | 21.8626399016789 | 16.2828834700276 | Ube2cbp |
| ENSMUSG00000032418 | 0.171544094751811 | 0.344640563009253 | 0.861266047293794 | 5498.78447903921 | 6080.42783207438 | 5714.69461765932 | 5420.27836611791 | 4232.01342684081 | 3779.32409710823 | 6012.22597296169 | 6142.98917045908 | Me1 |
| ENSMUSG00000032419 | -0.441047833042852 | 0.0850334686010238 | 0.743541556158628 | 42.4651446787451 | 35.8342920722305 | 46.3762895437739 | 62.8410885152968 | 64.6025335533267 | 75.8310983885793 | 49.9717483466946 | 64.046008315442 | Tbx18 |
| ENSMUSG00000032420 | 0.45468972767658 | 0.0685380125832476 | 0.72237439886039 | 322.554396815149 | 502.494504740141 | 341.670214802089 | 227.916186704733 | 285.784089108784 | 210.370143916704 | 291.501865355718 | 230.131419709724 | Nt5e |
| ENSMUSG00000032422 | 0.0257216893902171 | 0.894270322402502 | 0.983733637008786 | 436.397125102849 | 495.164763179912 | 496.888816540435 | 419.253232333398 | 395.279908690694 | 327.786038195794 | 536.15521663641 | 554.703563545607 | Snx14 |
| ENSMUSG00000032423 | 0.192502564279503 | 0.0945239471742816 | 0.757085157577544 | 2170.23994719885 | 2171.2323332856 | 2126.68413479306 | 2151.60383662822 | 1713.60957645689 | 1648.71484883556 | 2064.45785357282 | 2114.60379997425 | Syncrip |
| ENSMUSG00000032425 | -0.129431783917362 | 0.549072206292077 | 0.914699131434573 | 193.351935345776 | 109.13170767452 | 131.557229522134 | 141.626930832982 | 176.288269526875 | 127.200552135681 | 163.449260217313 | 161.743309135608 | Zfp949 |
| ENSMUSG00000032431 | -0.502721412582946 | 0.0733021638118035 | 0.730864975478592 | 355.984404328204 | 365.672662282534 | 330.312756138308 | 383.612017951588 | 610.986673267056 | 757.087901653719 | 367.500565966316 | 299.605055848508 | Crtap |
| ENSMUSG00000032434 | -0.296427572282322 | 0.0469283067842726 | 0.702324795701059 | 520.42390074377 | 526.112560878656 | 650.214508501483 | 484.908100931469 | 757.711071506815 | 699.603036746248 | 561.141090809758 | 661.085068883121 | Cmtm6 |
| ENSMUSG00000032435 | 0.280428632602006 | 0.10343889322606 | 0.759800953500741 | 2160.30129631659 | 2368.32093968287 | 2211.86507477142 | 2274.47223357604 | 1716.89445104434 | 1345.39045528125 | 2101.93666483284 | 2256.80764894583 | Dync1li1 |
| ENSMUSG00000032436 | -0.214639072160866 | 0.211203059752041 | 0.823123035902316 | 150.886790667031 | 114.83261777692 | 140.07532351997 | 161.323391412404 | 166.433645764503 | 190.800828203522 | 147.833088858971 | 153.05910461826 | Cmtm7 |
| ENSMUSG00000032437 | -0.010754398982608 | 0.919615391795909 | 0.986713918026868 | 1404.96382926487 | 1540.060143377 | 1386.55641186997 | 1257.75969700019 | 1333.65908250766 | 1325.82113956806 | 1535.5901835703 | 1436.15032205644 | Stt3b |
| ENSMUSG00000032440 | -0.400805105490914 | 0.083179637886532 | 0.74156942891138 | 795.995584297329 | 923.547436588849 | 1124.38840771435 | 792.548056648146 | 1348.98849724913 | 1637.7071087469 | 930.723812957186 | 884.703335204834 | Tgfbr2 |
| ENSMUSG00000032449 | 0.205544478728846 | 0.326165997121041 | 0.855977678491063 | 727.328541838081 | 886.898728787704 | 964.437531532767 | 822.561710864407 | 631.790878987619 | 514.917619703095 | 924.477344413849 | 877.104656252154 | Slc25a36 |
| ENSMUSG00000032456 | 0.126213007729788 | 0.281667466683571 | 0.850048154552447 | 417.423337054899 | 429.197089137851 | 418.333060782613 | 390.177504811395 | 384.330326732503 | 324.116791499573 | 401.856142954669 | 404.901035621353 | Nmnat3 |
| ENSMUSG00000032458 | 0.135154461031017 | 0.194033853606952 | 0.809162446871198 | 2941.84066114881 | 3089.89327550096 | 3132.76568142636 | 2770.63545483861 | 2807.47281408016 | 2349.54096781388 | 2865.04690521049 | 2844.07697943149 | Copb2 |
| ENSMUSG00000032459 | -0.123337774450744 | 0.403280110057195 | 0.881158207349486 | 365.923055210463 | 329.838370210303 | 305.704929033448 | 329.212269684615 | 316.442918591719 | 314.332133642982 | 399.773986773556 | 417.927342397375 | Mrps22 |
| ENSMUSG00000032462 | 0.155520724189094 | 0.518399681029295 | 0.905504473025266 | 398.449549006949 | 574.977504613516 | 567.872933189068 | 441.763472995594 | 429.223612761086 | 273.970419984545 | 520.539045278068 | 555.789089110276 | Pik3cb |
| ENSMUSG00000032463 | 0.30864873756468 | 0.14125297787363 | 0.779711473022872 | 109.325159704854 | 99.3587189275481 | 85.1809399783602 | 90.9788893430416 | 97.4512794278996 | 84.3926740130963 | 68.711153976705 | 60.7894316214364 | Faim |
| ENSMUSG00000032468 | 0.148598709093137 | 0.438775466396362 | 0.889583654898613 | 586.38040205331 | 682.480380830207 | 660.625512276616 | 538.369922504185 | 504.775728272604 | 398.724807656078 | 671.495368408708 | 650.229813236436 | Armc8 |
| ENSMUSG00000032469 | -0.181935475637995 | 0.468760373804419 | 0.895328112556867 | 110.228673421423 | 131.935348084121 | 100.324218196735 | 115.364983393754 | 179.573144114332 | 150.439114545085 | 92.6559500594962 | 97.6973008201657 | Dbr1 |

| | | | | | | | | | | | | |
|---|---|---|---|---|---|---|---|---|---|---|---|---|
| **ENSMUSG00000032470** | 0.196372515767923 | 0.178573716310309 | 0.801696999069105 | 149.079763233892 | 161.254314325037 | 135.343049076728 | 137.87522405595 | 124.825234323377 | 135.762127760198 | 123.88829277618 | 125.920965501547 | Mras |
| **ENSMUSG00000032475** | -0.0542046612098934 | 0.723928875598219 | 0.951674101666722 | 744.495302452893 | 561.132437221972 | 635.071230283108 | 715.638067718977 | 698.583328932584 | 821.911259953634 | 591.332355435886 | 646.973236542431 | Nck1 |
| **ENSMUSG00000032477** | -0.13580864252349 | 0.445439140067125 | 0.89108031640712 | 130.105975185942 | 111.574954861263 | 131.557229522134 | 136.937297361692 | 121.54035973592 | 117.415894279091 | 155.120635492864 | 164.999885829613 | Cdc25a |
| **ENSMUSG00000032478** | 0.131405952954474 | 0.391103136247104 | 0.877896341705281 | 289.124389302094 | 246.767965861042 | 266.900278598862 | 300.136542162612 | 251.840385038392 | 293.539735697726 | 204.051305749003 | 258.355084391105 | Nme6 |
| **ENSMUSG00000032479** | 0.304260279803682 | 0.164913203662074 | 0.795216960550936 | 14161.6739935032 | 17636.9870253687 | 15005.0958046325 | 12554.1488026455 | 9832.72459845549 | 8439.26740130963 | 15360.0661480652 | 14438.5755356558 | Map4 |
| **ENSMUSG00000032480** | 0.0364630476223703 | 0.805809263534779 | 0.963475137341661 | 801.416666596743 | 1020.46290832965 | 1006.0815466333 | 847.885731609378 | 1021.59599669922 | 733.849339244316 | 918.230875870512 | 909.670423192209 | Dhx30 |
| **ENSMUSG00000032481** | 0.174994171706865 | 0.170491282332472 | 0.799531027639052 | 824.004509510969 | 846.992358070902 | 795.022106464695 | 752.217208795045 | 658.069875687277 | 606.648787108634 | 792.26042691322 | 792.433662208011 | Smarcc1 |
| **ENSMUSG00000032485** | 0.0610670440979607 | 0.596102128893793 | 0.924461858177432 | 758.951521917998 | 924.361852317763 | 962.54462175547 | 911.664746818933 | 917.574968096403 | 796.226533080083 | 838.06786289769 | 857.565196088121 | Scap |
| **ENSMUSG00000032492** | -0.495925499150925 | 0.169676827484107 | 0.79907358295209 | 346.949267162513 | 320.879797192245 | 477.959718767466 | 313.267515882226 | 585.802634763217 | 896.519276110139 | 288.37863108405 | 287.664274637155 | Pth1r |
| **ENSMUSG00000032495** | 0.209911387260785 | 0.0906385928374175 | 0.753823363185192 | 16458.4058610217 | 15700.3064220104 | 16355.6869307338 | 16801.0808742464 | 13204.1008833825 | 11795.4050461203 | 16248.1057593096 | 15221.2394677818 | Lrrc2 |
| **ENSMUSG00000032497** | 0.237991494877332 | 0.11316325988994 | 0.767518180080285 | 2794.56792534806 | 2891.99025337478 | 2526.0880978027 | 2532.40207449704 | 2140.64327282633 | 1785.70005882783 | 2670.36530227649 | 2512.9916822076 | Lrrfip2 |
| **ENSMUSG00000032498** | -0.158812201650034 | 0.348365886555403 | 0.861266047293794 | 168.957064998411 | 145.780415475665 | 115.467496415111 | 141.626930832982 | 158.768938393769 | 144.323703384715 | 156.16171358342 | 179.111718170304 | Mlh1 |
| **ENSMUSG00000032501** | 0.132650156750039 | 0.569031796045358 | 0.920872458148558 | 529.45903790946 | 653.975830318206 | 397.511053232348 | 334.839829850164 | 458.787484048202 | 527.148442023833 | 382.075659234102 | 381.019473198646 | Trib1 |
| **ENSMUSG00000032503** | -0.283407298691621 | 0.532307882241634 | 0.909082872831377 | 12.6491920319666 | 23.6180561385155 | 13.2503684410783 | 8.44134024832345 | 22.994122112201 | 19.5693157131818 | 18.7394056300105 | 9.76973008201657 | Arpp21 |
| **ENSMUSG00000032504** | 0.0797002248001691 | 0.597377448554623 | 0.924496535139492 | 1575.72792169641 | 1996.13295156902 | 1974.30489772066 | 1593.53745354462 | 1698.28016171542 | 1371.0751821548 | 1901.00859335551 | 1784.60402831503 | Pdcd6ip |
| **ENSMUSG00000032507** | -0.224819073218679 | 0.615751044963065 | 0.928422072203769 | 9.03513716569045 | 10.5874044758863 | 15.1432782183751 | 10.3171936368398 | 12.0445401540101 | 20.7923979452556 | 10.4107809055614 | 9.76973008201657 | Fbxl2 |
| **ENSMUSG00000032508** | 0.0350767767119021 | 0.846000614531112 | 0.972180580805179 | 267.440060104437 | 235.366145656241 | 351.134763688574 | 312.329589187968 | 328.487458745729 | 308.216722482613 | 262.351678820146 | 238.815624227072 | Myd88 |
| **ENSMUSG00000032511** | 0.0450593208211397 | 0.879637682348712 | 0.978393501240173 | 4694.65727129276 | 5580.37657452098 | 6078.13329490033 | 4145.63598862107 | 3730.52257315566 | 2668.76543038516 | 6631.66743684259 | 6836.64000628226 | Scn5a |
| **ENSMUSG00000032512** | 0.136520400541217 | 0.415752158650071 | 0.882192126163777 | 461.695509166782 | 532.627886709971 | 523.389553422591 | 413.625672167849 | 388.710159515779 | 359.586176229715 | 524.703357640293 | 483.058876277486 | Wdr48 |
| **ENSMUSG00000032513** | 0.159941321897214 | 0.319131716122719 | 0.854670142081901 | 659.565013095403 | 728.08766164941 | 830.987392233336 | 668.741733006069 | 591.277425742312 | 523.479195327612 | 777.685333645434 | 690.394259129171 | Gorasp1 |
| **ENSMUSG00000032515** | 0.135730293804601 | 0.601006506251477 | 0.925969784079517 | 263.826005238161 | 446.299819445052 | 285.829376371831 | 196.026679099956 | 295.638712871156 | 319.224462571277 | 248.817663642917 | 222.532740757044 | Csrnp1 |
| **ENSMUSG00000032518** | -0.0160314502040509 | 0.93132417247756 | 0.987775860606096 | 14517.6583978314 | 9866.64655579709 | 11878.9553074267 | 14125.1760155279 | 12738.7436501594 | 16203.3934105145 | 10527.3816517037 | 11482.6894230635 | Rpsa |
| **ENSMUSG00000032519** | 0.107550267065402 | 0.504795117701739 | 0.90122958035533 | 543.915257374565 | 394.177212794535 | 498.781726317731 | 510.23212167644 | 417.179072607076 | 382.824738639118 | 530.94982618363 | 474.374671760138 | Slc25a38 |
| **ENSMUSG00000032523** | -0.151727536525032 | 0.206801203914263 | 0.819960256425221 | 3379.14129996823 | 3474.29749954853 | 3317.32438471281 | 2921.64165261417 | 3729.42761495984 | 4169.48732913979 | 3458.46141682749 | 3189.27410899608 | Hhatl |
| **ENSMUSG00000032525** | 0.207338343474755 | 0.556040284860413 | 0.916239900758826 | 253.887354355902 | 359.157336451219 | 273.525462819401 | 166.950951577953 | 155.484063806312 | 125.977469903608 | 358.130863151311 | 272.466916731795 | Nktr |
| **ENSMUSG00000032526** | 0.0709076432379023 | 0.63746992874716 | 0.932931968211785 | 1165.53269437407 | 796.498582878213 | 813.951204237664 | 932.299134092612 | 854.067392738896 | 945.442565393093 | 846.396487622139 | 884.703335204834 | Ss18l2 |
| **ENSMUSG00000032527** | 0.0586539599737935 | 0.423010266958224 | 0.884281490205862 | 4640.44644829861 | 4243.10594764365 | 4491.87490152553 | 4797.49504113049 | 4423.63111110915 | 4247.76459199251 | 4398.55493259968 | 4376.83907674342 | Pccb |
| **ENSMUSG00000032528** | -0.104321591649215 | 0.77309895968913 | 0.959121750920843 | 34.3335212296237 | 55.3802695661743 | 41.6440151005317 | 25.3240207449704 | 41.6084114411257 | 23.2385624094033 | 41.6431236222455 | 61.8749571861049 | Vipr1 |
| **ENSMUSG00000032531** | -0.057144631827488 | 0.790093923179327 | 0.961659329450003 | 1056.20753466921 | 1270.48853710635 | 1939.28606684067 | 1349.67651303749 | 1761.78773707293 | 1199.84366966446 | 1553.28851110976 | 1326.51224002492 | Amotl2 |
| **ENSMUSG00000032534** | 0.144347197850419 | 0.172402155728488 | 0.799889021902745 | 1395.02517838261 | 1311.20932355207 | 1221.87326124514 | 1384.37980072505 | 1171.60526952643 | 1066.52770636841 | 1279.48497329349 | 1287.43331969685 | Cep63 |
| **ENSMUSG00000032536** | 0.191760567937572 | 0.300667437996853 | 0.851785577030345 | 10401.2499051428 | 11116.7746996806 | 9969.00934213409 | 9386.77035613568 | 7854.13513861038 | 6452.98185642168 | 10703.3238490076 | 10774.9267548996 | Trak1 |
| **ENSMUSG00000032537** | -0.170070960810245 | 0.594591473376392 | 0.924432008640932 | 69.5705561758165 | 37.4631235300591 | 31.2330113253987 | 52.523894878457 | 56.937826182593 | 70.9387694602839 | 41.6431236222455 | 45.5920737160773 | Ephb1 |
| **ENSMUSG00000032540** | 0.07326973590241 | 0.655008570886941 | 0.937504201434612 | 997.479143092226 | 1061.99811050428 | 958.758802200877 | 918.23023367874 | 835.453103409971 | 722.841599155651 | 1061.89965236726 | 1120.2623827379 | Abhd5 |
| **ENSMUSG00000032547** | -0.0341269265096777 | 0.739412918860995 | 0.954383574136061 | 1095.96213819825 | 1126.33695308852 | 1095.9947610549 | 1048.60204418062 | 1103.71786138565 | 1268.33627466059 | 1091.04983890283 | 1010.62430070638 | Ryk |
| **ENSMUSG00000032548** | -0.837445984479 | 0.0328559368757965 | 0.671606027118506 | 42.4651446787451 | 36.6487078011448 | 54.8943835416099 | 38.4549944645846 | 120.445401540101 | 103.961989726278 | 40.6020455316893 | 43.4210225867403 | Slco2a1 |
| **ENSMUSG00000032549** | 0.106579078056996 | 0.527478099019111 | 0.908217276872524 | 515.002818444356 | 517.153987860599 | 460.923530771794 | 426.756645887463 | 581.42280197994 | 413.401794440965 | 393.52751823022 | 395.131305539337 | Rab6b |
| **ENSMUSG00000032551** | 0.139851900432977 | 0.301758919499965 | 0.851785577030345 | 215.939778260002 | 226.407572638183 | 214.845259723197 | 221.350699844926 | 190.522726072523 | 170.008430258266 | 219.667477107345 | 216.019587369033 | 1110059G10 |
| **ENSMUSG00000032553** | -0.174883066494303 | 0.35369206156475 | 0.86286441023679 | 659.565013095403 | 583.936077631574 | 563.140658745826 | 664.990026229036 | 792.749733773026 | 886.734618253548 | 550.730309904196 | 561.216716933618 | Srprb |
| **ENSMUSG00000032554** | -0.315400350609247 | 0.260419777635912 | 0.840316047847749 | 568.310127721929 | 1250.94255961241 | 668.197151385803 | 767.224035903176 | 785.085026402292 | 889.180782717696 | 960.915077583314 | 1415.52533632773 | Trf |
| **ENSMUSG00000032555** | -0.0412512674897873 | 0.806160718774725 | 0.963604452772887 | 225.878429142261 | 259.798617523671 | 239.453086828057 | 202.592165959763 | 215.706764576362 | 195.693157131818 | 285.255396812381 | 257.269558826436 | Topbp1 |
| **ENSMUSG00000032556** | -0.309720889659565 | 0.57110262835609 | 0.921126010531982 | 28.0089252136404 | 9.77298874697194 | 8.51809399783602 | 12.1930470253561 | 7.66470737073368 | 26.9078091056249 | 23.9447960827911 | 14.1118323406906 | Bfsp2 |
| **ENSMUSG00000032557** | -0.00506741197241407 | 0.975938774981564 | 0.99575536949131 | 400.256576440087 | 393.362797065621 | 416.440151005317 | 459.584080186499 | 418.274030802895 | 541.82542880872 | 346.679004155194 | 370.164217551961 | Uba5 |
| **ENSMUSG00000032558** | -0.0588189226001091 | 0.836854953124454 | 0.968924788429433 | 35.2370349461928 | 43.1640336324594 | 31.2330113253987 | 23.448167356454 | 36.1336204620302 | 29.3539735697726 | 34.3555769883525 | 39.0789203280663 | Nphp3 |
| **ENSMUSG00000032560** | 0.20594812252196 | 0.466721593783958 | 0.895328112556867 | 778.828823682517 | 985.443031986338 | 866.952678001977 | 619.969544904645 | 584.707676567398 | 391.386314263635 | 947.381062406084 | 894.47306528685 | Dnajc13 |
| **ENSMUSG00000032562** | -0.265699071818969 | 0.0424835878185944 | 0.686159342791019 | 5278.32713219636 | 4826.22760954631 | 4918.72605630598 | 5075.12134263091 | 6424.11973487064 | 7172.15420888111 | 5276.1837629385 | 5291.93712775897 | Gnai2 |
| **ENSMUSG00000032563** | 0.220541018543536 | 0.208115468696657 | 0.820372930586885 | 2387.08323917542 | 2311.31183865886 | 2160.7565107844 | 2198.50017134113 | 1963.26004510364 | 1341.72120858502 | 2206.04447388845 | 2261.1497512045 | Mrpl3 |
| **ENSMUSG00000032565** | 0.3678207534072 | 0.0918646291518516 | 0.754058637722175 | 168.053551281842 | 146.594831204579 | 165.629605513478 | 195.088752405698 | 118.255485148462 | 167.562265794119 | 147.833088858971 | 90.0986218674861 | Nudt16 |
| **ENSMUSG00000032567** | -0.212906544217554 | 0.354915196888041 | 0.864114300574038 | 45.1756858284523 | 71.6685841444609 | 72.8770264259304 | 51.5859681841989 | 64.6025335533267 | 68.4926049961361 | 69.7522320672611 | 77.072315091464 | Aste1 |
| **ENSMUSG00000032570** | 0.123409103524963 | 0.457708779117586 | 0.892966448175628 | 1003.80373910821 | 1241.16957086544 | 1268.24955078892 | 1081.42947847966 | 1013.93128932848 | 802.341944240452 | 1187.87010132455 | 1212.53205573472 | Atp2c1 |
| **ENSMUSG00000032571** | -0.213858965326663 | 0.101283199763975 | 0.759800953500741 | 337.010616280254 | 359.157336451219 | 423.065335225856 | 345.157023487003 | 479.591689768764 | 396.278643191931 | 408.102611498006 | 413.585240138701 | Pik3r4 |
| **ENSMUSG00000032575** | -0.222532645565821 | 0.335774206504272 | 0.858570960161243 | 835.750187826367 | 637.687515739919 | 739.181268034437 | 831.003051112731 | 885.821180417649 | 1298.91333046244 | 639.221947601468 | 728.387653892569 | Manf |
| **ENSMUSG00000032577** | 0.125955561076177 | 0.475831273671598 | 0.897549741595057 | 541.204716224858 | 499.236841824483 | 554.62256474799 | 476.466760683146 | 434.698403740182 | 348.57843614105 | 566.346481262538 | 547.104884592928 | Mapkapk3 |
| **ENSMUSG00000032578** | -0.318576486200813 | 0.168020531401568 | 0.79907358295209 | 1133.00620057758 | 868.981582751589 | 1237.01653946352 | 914.478526901707 | 1897.56255335449 | 1138.68955806076 | 985.900951756661 | 1157.17025193663 | Cish |
| **ENSMUSG00000032579** | 0.188212270356765 | 0.121467142933523 | 0.769868040577702 | 348.756294595651 | 313.550055632016 | 276.364827485346 | 329.212269684615 | 282.499214521327 | 278.86274891284 | 272.762459725708 | 278.980070119806 | Hemk1 |
| **ENSMUSG00000032580** | 0.179847112594073 | 0.348660833976458 | 0.861266047293794 | 749.012871035738 | 922.733020859934 | 542.31865119556 | 569.321503414704 | 620.841297029428 | 672.695227640623 | 574.675105986987 | 590.525907179668 | Rbm5 |
| **ENSMUSG00000032582** | 0.336712685354184 | 0.0499001040117191 | 0.703396849069134 | 648.722848496574 | 738.675066125296 | 776.093008691726 | 646.231492343873 | 548.574056105368 | 404.840218816448 | 637.139791420356 | 632.86140420174 | Rbm6 |
| **ENSMUSG00000032583** | -0.149015836691079 | 0.310148457191409 | 0.851785577030345 | 313.519259649459 | 284.231089391101 | 252.703455269135 | 273.874594723383 | 297.828629262794 | 358.363093997641 | 330.021754706295 | 261.61166108511 | Mon1a |
| **ENSMUSG00000032584** | -0.571100953474283 | 0.289638329857265 | 0.851785577030345 | 7.22810973255236 | 8.14415728914329 | 10.4110037751329 | 8.44134024832345 | 12.0445401540101 | 22.0154801773295 | 4.16431236222455 | 13.0263067760221 | Mst1r |
| **ENSMUSG00000032590** | 0.171741983950591 | 0.171098130605686 | 0.799531027639052 | 838.460728976074 | 914.588863570791 | 1036.36810307005 | 1034.53314376675 | 952.613630362614 | 780.326464063122 | 872.423439886043 | 788.091559949337 | Apeh |
| **ENSMUSG00000032591** | -0.579253398788714 | 0.360968933980476 | 0.868539734006824 | 2.71054114970714 | 16.2883145782866 | 7.57163910918757 | 15.0068271081306 | 7.66470737073368 | 8.56157562451702 | 18.7394056300105 | 27.1381391167127 | Mst1 |
| **ENSMUSG00000032594** | -0.277424706926184 | 0.0760173048058161 | 0.735399943172214 | 577.34526488762 | 517.968403589513 | 667.250696497155 | 668.741733006069 | 854.067392738896 | 783.995710759344 | 708.974179668729 | 599.210111697016 | Ip6k1 |
| **ENSMUSG00000032596** | -0.297657695231889 | 0.150769986261785 | 0.783020695097675 | 109.325159704854 | 119.719112150406 | 125.878500190243 | 114.427056699496 | 173.003394939417 | 176.123841418636 | 107.231043327282 | 121.578863242873 | Uba7 |
| **ENSMUSG00000032598** | -0.1338808418613 | 0.312085115393707 | 0.851810228496481 | 208.711668527449 | 216.634583891211 | 189.290977729689 | 176.330218520534 | 209.137015401448 | 218.931719541221 | 231.119336103462 | 209.506433981022 | Nckipsd |
| **ENSMUSG00000032599** | -0.103763809982872 | 0.594006067469032 | 0.923935529355527 | 256.597895505609 | 233.737314198412 | 167.522515290775 | 236.357526953057 | 255.12525962585 | 293.539735697726 | 201.96914956789 | 211.677485110359 | Ip6k2 |
| **ENSMUSG00000032601** | 0.0905192391157203 | 0.60513328033592 | 0.926448151609826 | 2683.43573821006 | 3212.87005056703 | 2746.61208685779 | 2462.99549912193 | 2211.81555555458 | 2079.23979452556 | 2996.22274462056 | 3142.59650971533 | Prkar2a |
| **ENSMUSG00000032602** | 0.243589420281059 | 0.0468131434136976 | 0.701439656975469 | 10164.5293114018 | 9540.88026423136 | 11103.8087536236 | 9811.65114863462 | 8209.99655225159 | 7251.65455396591 | 9194.8016957918 | 9651.4077954677 | Slc25a20 |
| **ENSMUSG00000032604** | 0.0912177182818685 | 0.445797545666052 | 0.89108031640712 | 1260.40163461382 | 1359.25985155801 | 1606.13394603641 | 1395.63492105614 | 1481.47843894324 | 1284.23634367755 | 1222.2256783129 | 1288.51884526152 | Qars |
| **ENSMUSG00000032606** | 0.0551870475271084 | 0.721616240453008 | 0.951674101666722 | 449.046317134815 | 493.535931722083 | 472.280989435575 | 453.018593326692 | 377.760577557588 | 387.717067567413 | 475.772687384154 | 555.789089110276 | Nicn1 |
| **ENSMUSG00000032607** | -0.246478456939119 | 0.29644979375012 | 0.851785577030345 | 54.2108229941427 | 54.56585383726 | 34.0723759913441 | 58.151455044006 | 55.8428679867739 | 56.2617826753975 | 62.4646854333682 | 64.046008315442 | Amt |
| **ENSMUSG00000032609** | -0.313204974058552 | 0.250446898865757 | 0.835923538390114 | 74.9916384752307 | 58.6379324818317 | 55.8408384302584 | 86.2892558717508 | 99.6411958195378 | 114.969729814943 | 64.5468416144805 | 64.046008315442 | Klhdc8b |
| **ENSMUSG00000032611** | 0.423486284946551 | 0.353757889455607 | 0.86286441023679 | 9.93865088225949 | 12.2162359337149 | 12.3039135524298 | 13.1309737196143 | 6.56974917491458 | 11.0077400886647 | 11.4518589961175 | 6.51315338801105 | 1700102P08 |
| **ENSMUSG00000032612** | 0.213103959958722 | 0.0254825637767381 | 0.661594038988582 | 2472.91704224948 | 2567.85279326688 | 2786.36319218103 | 2580.2363359042 | 2204.15084818384 | 2018.08568292187 | 2359.08295320021 | 2394.66939565873 | Usp4 |
| **ENSMUSG00000032615** | 0.127447327428385 | 0.323837178247644 | 0.855977678491063 | 375.861706092723 | 346.12668478859 | 305.704929033448 | 362.977630677908 | 286.879047304603 | 302.101311322243 | 335.227145159076 | 348.453706258591 | Nt5m |
| **ENSMUSG00000032621** | 0.0393776720679132 | 0.766568918655497 | 0.958624690460551 | 386.703870691551 | 424.310594764365 | 443.887342776122 | 453.95652002095 | 343.816873487196 | 424.409534529629 | 436.211719943021 | 458.09178829011 | Srek1 |
| **ENSMUSG00000032624** | -0.355676385775402 | 0.0713581723778791 | 0.727489215531785 | 75.8951521917998 | 88.7713144516618 | 66.2518422053913 | 89.1030359545253 | 112.780694169367 | 81.9465095489486 | 114.518589961175 | 99.8683519495027 | Eml4 |
| **ENSMUSG00000032625** | -0.278185666395107 | 0.384210336326106 | 0.875126835008975 | 109.325159704854 | 138.450673915436 | 116.413951303759 | 99.4202295913651 | 195.997517051618 | 193.24699266767 | 95.7791843311646 | 78.1578406561325 | Thsd7a |
| **ENSMUSG00000032633** | -0.0158597045779202 | 0.94692045328125 | 0.990639835510776 | 599.029594085277 | 783.467931215584 | 779.87882824632 | 721.265627884526 | 627.411046204342 | 452.540425867328 | 917.189797779956 | 917.269102144889 | Flcn |
| **ENSMUSG00000032637** | 0.0810143675361219 | 0.5516810499264 | 0.914873920695527 | 667.696636544524 | 811.972481727586 | 864.113313336032 | 635.914298707033 | 738.001823982071 | 724.064681387725 | 693.358008310387 | 662.17059444779 | Atxn2l |
| **ENSMUSG00000032640** | 0.0952454271708304 | 0.537209232708626 | 0.912099403004057 | 245.75573090678 | 263.870696168242 | 206.327165725361 | 217.598993067893 | 256.220217821669 | 221.377884005369 | 200.928071477334 | 196.480127205 | Chsy1 |
| **ENSMUSG00000032641** | -0.0312986782844852 | 0.903090471953992 | 0.985027469232262 | 62.3424464432641 | 41.5352021746308 | 36.9117406572894 | 48.7721881014244 | 58.0327843784121 | 44.0309603546589 | 42.6842017128016 | 48.8486504100828 | Gpr19 |
| **ENSMUSG00000032643** | -0.248178152750303 | 0.309696323115592 | 0.851785577030345 | 88.5443442237664 | 74.9262470601182 | 62.4660226507975 | 129.433883807626 | 109.49581958191 | 102.738907494204 | 103.066730965058 | 106.381505337514 | Fhl3 |
| **ENSMUSG00000032648** | 0.108935507180763 | 0.351057545694757 | 0.86217195868522 | 40949.9521760588 | 44524.1078997463 | 46080.9956185156 | 40016.6424105245 | 40326.2153938215 | 33349.7832219579 | 42761.2414915028 | 42654.6415380843 | Pygm |
| **ENSMUSG00000032649** | -0.550512810728302 | 0.103501675470247 | 0.759800953500741 | 82.2197482077831 | 95.2866402829764 | 61.519567762149 | 82.5375490947182 | 148.914314631397 | 179.793088114857 | 72.8754663389296 | 70.559161703453 | Colgalt2 |
| **ENSMUSG00000032652** | -0.0112883677891526 | 0.951946901602698 | 0.991915011884503 | 447.239289701677 | 462.588134023339 | 506.353365426919 | 452.080666632434 | 398.564783278151 | 352.247682837272 | 596.537745888666 | 534.078577816906 | Crebl2 |
| **ENSMUSG00000032656** | 0.0341224680695913 | 0.861063158182319 | 0.974868041734905 | 79.509207058076 | 105.874044758863 | 82.3415753124149 | 80.6616957062019 | 93.0714466446232 | 69.71568722821 | 89.5327157878278 | 87.9275707381491 | March3 |
| **ENSMUSG00000032657** | -0.226752354332165 | 0.459808838524304 | 0.894774044236998 | 53.3073092775737 | 52.9370223794314 | 31.2330113253987 | 44.0825546301336 | 68.9823663366031 | 64.8233582999146 | 45.80743598447 | 33.6512925047237 | Fam189b |
| **ENSMUSG00000032661** | 0.204517321055017 | 0.819084504074661 | 0.966315773308182 | 2.71054114970714 | 6.51532583131463 | 34.0723759913441 | 4.68963347129081 | 7.66470737073368 | 25.6847268735511 | 5.20539045278068 | 3.25657669400552 | Oas3 |

| | | | | | | | | | | | | |
|---|---|---|---|---|---|---|---|---|---|---|---|---|
| ENSMUSG00000032666 | 0.0104745417323111 | 0.948356811470766 | 0.991217716191233 | 823.1009957944 | 833.961706408273 | 883.042411109001 | 923.857793844289 | 741.286698569529 | 752.195572725424 | 1125.40541589118 | 818.486275760055 | 1700025G04 |
| ENSMUSG00000032667 | -0.0613055314767575 | 0.526776934050992 | 0.908217276872524 | 872.794250205697 | 828.260796305872 | 865.05976822468 | 917.292306984482 | 859.542183717991 | 1009.04284146093 | 895.327157878277 | 871.677028428812 | Pon2 |
| ENSMUSG00000032671 | 0.854322854004648 | 0.0582102726479427 | 0.710862211775517 | 9.93865088225949 | 17.1027303072009 | 16.0897331070236 | 15.0068271081306 | 6.56974917491458 | 9.78465785659088 | 7.28754663389296 | 8.68420451734806 | A930018P22 |
| ENSMUSG00000032673 | 0.109536231734351 | 0.414063798923128 | 0.882192126163777 | 473.44118748218 | 355.085257806647 | 398.457508120996 | 461.459933575015 | 416.084114411257 | 380.37857417497 | 377.911346871878 | 389.703677715994 | Prorsd1 |
| ENSMUSG00000032679 | 0.0159613027884702 | 0.908509687443324 | 0.985182259238642 | 2971.65661379559 | 2421.2579620623 | 2437.12133826975 | 2614.0016968975 | 2699.07195269407 | 3047.92092232806 | 2328.89168857408 | 2254.63659781649 | Cd59a |
| ENSMUSG00000032680 | 0.0428523791888419 | 0.845167286559701 | 0.972136081650911 | 111.132187137993 | 112.389370590177 | 72.8770264259304 | 91.9168160372998 | 79.9319482947941 | 118.638976511164 | 87.4505596067155 | 92.2696729968231 | 6820408C15 |
| ENSMUSG00000032688 | 0.00756994892203271 | 0.975977924444655 | 0.99575536949131 | 49.6932544112975 | 61.895595397489 | 56.7872933189068 | 83.4754757889764 | 73.3621991198795 | 72.1618516923577 | 48.9306702561384 | 56.4473293627624 | Malt1 |
| ENSMUSG00000032690 | -0.203559732749297 | 0.517933806457054 | 0.905473752527583 | 94.8689402397497 | 103.43079757212 | 93.6990339761962 | 79.7237690119437 | 132.489941694111 | 161.446854633749 | 61.4236073428121 | 73.8157383974585 | Oas2 |
| ENSMUSG00000032698 | -0.0450932630331258 | 0.714884024204072 | 0.951176382858794 | 510.48524986151 | 416.980853204136 | 422.118880337207 | 492.411514485535 | 516.820268426614 | 484.340563901248 | 448.704657029695 | 450.493109337431 | Lmo2 |
| ENSMUSG00000032702 | 0.14290778846476 | 0.53330032331422 | 0.909132720692319 | 1121.26052226218 | 1491.19519964214 | 1352.48403587863 | 1127.38788649831 | 914.290093508946 | 806.011190936674 | 1547.04204256642 | 1343.88064905961 | Kank1 |
| ENSMUSG00000032705 | 0.147019034010881 | 0.407670395722065 | 0.881573350402261 | 290.931416735233 | 331.467201668132 | 279.204192151292 | 307.639955716677 | 268.264757975679 | 195.693157131818 | 315.446661438509 | 311.545837059862 | Exd2 |
| ENSMUSG00000032712 | 0.249619296096917 | 0.453114785759295 | 0.891468041670908 | 1028.19860945557 | 1178.45955973903 | 540.425741418263 | 397.68091836546 | 752.23628052772 | 792.557286383861 | 572.592949805875 | 528.650949993563 | 2810474O19 |
| ENSMUSG00000032714 | -0.312161032437003 | 0.124350149760065 | 0.771949105062466 | 140.948139784771 | 154.738988493722 | 186.451613063744 | 226.040333316217 | 263.884925192402 | 231.162541861959 | 187.394056300105 | 196.480127205 | Syde1 |
| ENSMUSG00000032715 | -0.711538419013455 | 0.072481271107566 | 0.729721356194038 | 90.3513716569045 | 51.3081909216027 | 80.448665535118 | 95.6685228143324 | 131.394983498292 | 239.724117486476 | 78.0808567917102 | 71.6446872681215 | Trib3 |
| ENSMUSG00000032718 | -0.546874070577118 | 0.133905813282517 | 0.77303252008843 | 18.0702743313809 | 29.3189662409158 | 19.8755526616174 | 29.075727522003 | 47.0832024202212 | 46.4771248188067 | 21.8626399016789 | 26.0526135520442 | Mansc1 |
| ENSMUSG00000032724 | 0.260270113692021 | 0.32508707056454 | 0.855977678491063 | 552.950394540256 | 788.35442558907 | 794.075651576047 | 712.824287636203 | 475.211856985488 | 359.586176229715 | 742.288678566525 | 800.03234116069 | Abtb2 |
| ENSMUSG00000032725 | -0.629024133938083 | 0.0540204047058359 | 0.703396849069134 | 39.754603529038 | 35.0198763433161 | 52.0550188756646 | 70.3445020693621 | 97.4512794278996 | 95.400414101761 | 46.8485140750261 | 65.1315338801105 | Folr2 |
| ENSMUSG00000032727 | -0.0672104044615372 | 0.672599864507678 | 0.943216235131458 | 134.623543768788 | 171.027303072009 | 175.094154399963 | 158.509611329629 | 190.522726072523 | 173.677676954488 | 154.079557402308 | 151.973579053591 | Mier3 |
| ENSMUSG00000032733 | -0.387735716896816 | 0.163002836156741 | 0.793968226938511 | 102.097049972302 | 74.9262470601182 | 132.503684410783 | 133.185590584659 | 171.908436743598 | 181.016170346931 | 91.61487196894 | 134.605170018895 | Snx33 |
| ENSMUSG00000032735 | -0.0312855101200665 | 0.755319753701818 | 0.956566265296272 | 2494.60137144713 | 2356.91911947807 | 2305.56410874762 | 2015.60446596079 | 2482.27022992189 | 2375.22569468744 | 2250.81083178237 | 2266.57737902784 | Ablim3 |
| ENSMUSG00000032737 | 0.0513094635991402 | 0.783303838377268 | 0.961051227381536 | 4345.90097669711 | 4687.77693563088 | 3780.1408252619 | 3604.45228603411 | 3457.87798239671 | 2969.64365947533 | 4717.12482830985 | 4699.24016944997 | Inppl1 |
| ENSMUSG00000032740 | -0.322635631898721 | 0.129784121943466 | 0.77303252008843 | 108.421645988285 | 110.760539132349 | 105.056492639978 | 123.806323642077 | 134.679858085749 | 194.470074899744 | 123.88829277618 | 108.552556466851 | Ccdc88a |
| ENSMUSG00000032741 | -0.111215544446726 | 0.444765278332604 | 0.89108031640712 | 831.232619243521 | 981.370953341766 | 908.596693102509 | 698.75538722233 | 870.491765676182 | 819.465095489486 | 1015.05113829223 | 988.91378941301 | Tpcn1 |
| ENSMUSG00000032743 | 0.145094729968309 | 0.536343242058785 | 0.911291453479697 | 183.413284463516 | 180.800291818981 | 264.060913932917 | 158.509611329629 | 151.104231023035 | 139.43137445642 | 234.242570375131 | 185.624871558315 | D430042O09 |
| ENSMUSG00000032744 | -0.968030976171624 | 0.00186579203912117 | 0.635862253526407 | 174.378147297826 | 150.666909849151 | 174.147699511314 | 205.405946042537 | 498.205979097689 | 450.09426140318 | 244.653351280692 | 185.624871558315 | Heyl |
| ENSMUSG00000032745 | 0.148595257657897 | 0.350245181626892 | 0.861774532843376 | 1841.36095436771 | 1876.41383941861 | 1943.07188639526 | 1678.88878272211 | 1486.95322992233 | 1287.90559037377 | 1930.15877989108 | 1914.86709607525 | Gpbp1 |
| ENSMUSG00000032750 | -1.18827853718972 | 0.0104582446076267 | 0.635862253526407 | 10.8421645988285 | 15.4738988493722 | 5.67872933189068 | 11.2551203310979 | 19.7092475247437 | 42.8078781225851 | 18.7394056300105 | 18.4539345993646 | Gab3 |
| ENSMUSG00000032754 | -0.605911920370726 | 0.0298703991506652 | 0.665956123973759 | 98.4829951060259 | 113.203786319092 | 103.163582862681 | 133.185590584659 | 240.890803080201 | 207.923979452556 | 120.765058504512 | 112.894658725525 | Slc8b1 |
| ENSMUSG00000032757 | -0.057566777760415 | 0.648883611447282 | 0.936461253686283 | 365.019541493894 | 298.890572511559 | 320.848207251823 | 377.984457786039 | 349.291664466292 | 349.801518373124 | 338.350379430744 | 379.933947633978 | Bet1 |
| ENSMUSG00000032763 | -0.0640028220098139 | 0.592441581545309 | 0.923935529355527 | 692.995020608457 | 610.811796685746 | 698.483707822554 | 657.486612674971 | 715.00770186987 | 600.533375948265 | 781.849646007659 | 680.624529047154 | Ilvbl |
| ENSMUSG00000032766 | -0.100084144350954 | 0.725678760773174 | 0.951727359509341 | 514.099304727787 | 402.321370083678 | 707.00180182039 | 572.135283497478 | 605.511882287961 | 883.065371557327 | 352.92547269853 | 512.368066523536 | Gng11 |
| ENSMUSG00000032777 | 0.0121009570083376 | 0.94011690560105 | 0.989269716906553 | 1254.07703859783 | 1442.33025590728 | 1237.96299435217 | 1151.77398054902 | 1172.70022772225 | 978.465785659088 | 1454.38609250692 | 1437.2358476211 | Gtf3c1 |
| ENSMUSG00000032782 | -0.0806040160617011 | 0.789440310559205 | 0.961659329450003 | 37.9475760958999 | 26.8757190541728 | 32.1794662140472 | 31.8895076047775 | 38.3235368536684 | 19.5693157131818 | 40.6020455316893 | 36.9078691987293 | Cntrob |
| ENSMUSG00000032786 | -0.0489403446791332 | 0.743219386496444 | 0.954383574136061 | 2146.74859056805 | 2152.50077152057 | 2069.89684147415 | 2146.91420315693 | 2204.15084818384 | 1659.72258892423 | 2602.69522639034 | 2341.47864298997 | Alas1 |
| ENSMUSG00000032788 | 0.0463437211347899 | 0.812726306679354 | 0.965402199310489 | 411.098741038915 | 557.874774306315 | 374.796135904785 | 333.901903155905 | 477.401773377126 | 342.463024980681 | 410.184767679118 | 395.131305539337 | Pdxk |
| ENSMUSG00000032802 | 0.0584053014168769 | 0.793824646409741 | 0.961804768755116 | 1286.60353239432 | 2312.94067011669 | 1379.93122764944 | 1398.44870113892 | 1667.62133223248 | 1295.24408376622 | 1823.96881465435 | 1338.45302123627 | Srxn1 |
| ENSMUSG00000032803 | 0.302166289894362 | 0.0522970469479524 | 0.703396849069134 | 5717.43479844892 | 6030.7484726106 | 5309.61192531779 | 5901.43476027235 | 4201.35459735787 | 3630.10806479522 | 5469.82428778194 | 5317.98974131102 | Cdv3 |
| ENSMUSG00000032806 | -0.0613145211033485 | 0.763782083151425 | 0.95785116771606 | 82.2197482077831 | 83.0704043492615 | 88.0203046443056 | 76.9099889291692 | 102.926070406995 | 61.154111603693 | 92.6559500594962 | 86.8420451734806 | Slc10a3 |
| ENSMUSG00000032812 | -0.0475237364666418 | 0.741647472809201 | 0.954383574136061 | 525.844983043184 | 588.008156276145 | 518.657278979349 | 419.253232333398 | 578.137927392483 | 562.617826753975 | 510.128264372507 | 471.118095066132 | Arap1 |
| ENSMUSG00000032815 | 0.0977097068629975 | 0.710767065540067 | 0.949886733585872 | 53.3073092775737 | 58.6379324818317 | 50.1621090983677 | 37.5170677703264 | 44.893286028583 | 34.2463024980681 | 60.3825292522559 | 46.6775992807458 | Fanca |
| ENSMUSG00000032816 | 0.0343236765677106 | 0.907878527973643 | 0.98504921011344 | 245.75573090678 | 383.589808318649 | 300.972654590206 | 216.661066373635 | 231.036179317829 | 161.446854633749 | 387.281049686883 | 339.769501741243 | Igdcc4 |
| ENSMUSG00000032826 | 0.196849195783901 | 0.529345511101477 | 0.908560090446144 | 1722.0971437806 | 2501.88511922482 | 2154.13132656387 | 1523.19295147525 | 1347.89353905331 | 879.396124861105 | 2390.31529591689 | 2275.26158354519 | Ank2 |
| ENSMUSG00000032827 | 0.435173684400157 | 0.190708847031235 | 0.808824684419004 | 239.431134890797 | 394.991628523449 | 300.026199701558 | 227.916186704733 | 154.389105610493 | 114.969729814943 | 322.734208072402 | 267.039288908453 | Ppp1r9a |
| ENSMUSG00000032834 | 0.220254555598441 | 0.158189641688 | 0.789230661864335 | 533.976606492306 | 588.008156276145 | 459.977075883145 | 548.687116141024 | 524.484975797347 | 355.916929533493 | 479.936999746379 | 467.861518372127 | Pwp2 |
| ENSMUSG00000032839 | -1.63437341438056E-0 | 0.999994597936908 | 0.999994597936908 | 115.649755720838 | 103.43079757212 | 159.004421292939 | 110.675349922463 | 99.6411958195378 | 92.9542496376133 | 161.367104036201 | 133.519644454226 | Trpc1 |
| ENSMUSG00000032840 | 0.0583094045333022 | 0.79518148640602 | 0.962032824235414 | 502.353626412389 | 628.728942721862 | 587.748485850685 | 507.418341593665 | 471.926982398031 | 344.909189444828 | 628.811166695907 | 691.479784693839 | 2410131K14 |
| ENSMUSG00000032842 | 0.0163127577565703 | 0.946353024196286 | 0.990510945542179 | 56.0178504272808 | 71.6685841444609 | 59.6266579848522 | 41.2687745473591 | 65.6974917491458 | 48.9232892829544 | 53.094982618363 | 58.6183804920994 | Abcc10 |
| ENSMUSG00000032845 | 0.23124966222795 | 0.294490712789859 | 0.851785577030345 | 3712.53786138221 | 5117.78844049764 | 5177.108240907 | 4500.17227905066 | 3298.01408580712 | 2756.82735109448 | 5173.11703197344 | 4537.49686031436 | Alpk2 |
| ENSMUSG00000032846 | -0.270636024843119 | 0.0786688511832765 | 0.739855948845396 | 109.325159704854 | 104.245213301034 | 97.48485353079 | 91.9168160372998 | 121.54035973592 | 125.977469903608 | 115.559668051731 | 123.74991437221 | Zswim6 |
| ENSMUSG00000032849 | -0.0539860710379944 | 0.75124814537717 | 0.956197192463153 | 428.265501653727 | 535.071133896714 | 427.797609669098 | 362.977630677908 | 557.33372167192 | 421.963370065482 | 428.924173309128 | 413.585240138701 | Abcc4 |
| ENSMUSG00000032850 | -0.35253465204465 | 0.367130035096984 | 0.870359479006162 | 18.0702743313809 | 17.1027303072009 | 17.036187995672 | 14.0689004138724 | 19.7092475247437 | 12.2308223207386 | 18.7394056300105 | 33.6512925047237 | Rnft2 |
| ENSMUSG00000032855 | -0.338681127883111 | 0.0604604311807044 | 0.71282992338528 | 430.072529086865 | 592.080234920717 | 554.62256474799 | 410.811892085075 | 716.102660065689 | 708.164612370765 | 538.237372817523 | 552.53251241627 | Pkd1 |
| ENSMUSG00000032860 | 0.0900234454036356 | 0.495311106593897 | 0.900742507277983 | 986.636978493397 | 1040.0088858236 | 1226.60553568839 | 1052.35375095766 | 973.417836083177 | 929.542496376133 | 1197.23980413956 | 943.321715696933 | P2ry2 |
| ENSMUSG00000032867 | 0.0166071303891569 | 0.887647146197489 | 0.980702840906172 | 1589.28062744495 | 1713.53069363575 | 1707.4046191218 | 1459.4139362657 | 1828.58018701789 | 1631.59169758653 | 1472.08442004638 | 1464.37398673782 | Fbxw8 |
| ENSMUSG00000032869 | -0.0466398437859877 | 0.739952189248554 | 0.954383574136061 | 421.940905637744 | 448.743066631795 | 520.550188756646 | 453.018593326692 | 535.434557755538 | 511.248373006873 | 463.279750297481 | 395.131305539337 | Psmf1 |
| ENSMUSG00000032870 | 0.0808302936928181 | 0.5676475386131 | 0.920429362097163 | 1792.57121367299 | 1813.70382829221 | 1683.74324690559 | 1724.84719074076 | 1634.77258635791 | 1292.79791930207 | 1942.65171697775 | 1760.72246589232 | Smap2 |
| ENSMUSG00000032872 | 0.158812140776645 | 0.161920344855403 | 0.793324017651341 | 907.127771435321 | 838.033785052844 | 888.721140440892 | 970.754128557197 | 744.571573156986 | 737.518585940537 | 907.820094964951 | 836.940210359419 | Cyb5r4 |
| ENSMUSG00000032875 | -0.209148957196239 | 0.185195452243892 | 0.805963874499992 | 904.417230285614 | 1015.57641395617 | 1089.36957683436 | 893.844139628028 | 1048.96995159469 | 912.419345127099 | 1387.75709471133 | 1161.5123541953 | Arhgef17 |
| ENSMUSG00000032878 | -0.105184599520729 | 0.600641158011048 | 0.925755180289997 | 548.43282595741 | 531.813470981057 | 506.353365426919 | 379.860311174555 | 653.690042904001 | 595.64104701997 | 468.485140750262 | 398.387882233342 | Ccdc85a |
| ENSMUSG00000032883 | 0.116835566131695 | 0.621636925674378 | 0.93078489223231 | 123.781379169959 | 138.450673915436 | 170.36187995672 | 130.371810501884 | 116.065568756824 | 89.2850029413918 | 179.065431575656 | 133.519644454226 | Acsl3 |
| ENSMUSG00000032897 | -0.0118491810566624 | 0.909993176777874 | 0.985944245354498 | 467.116591466196 | 489.463853077511 | 495.942361651786 | 476.466760683146 | 476.306815181307 | 437.863439082442 | 533.031982364742 | 496.085183053508 | Nfyc |
| ENSMUSG00000032898 | 0.178159080639161 | 0.383309331753617 | 0.875126835008975 | 1053.49699351951 | 1379.62024478087 | 1419.68233297267 | 1023.27802343565 | 984.367418041368 | 779.103381831049 | 1305.5119255574 | 1239.67019485144 | Fbxo21 |
| ENSMUSG00000032902 | 0.15406951725405 | 0.195994639133822 | 0.811193545163524 | 3969.13575688781 | 4433.6792282096 | 4335.70984489854 | 4261.93889870909 | 3693.29399449781 | 3221.59859928255 | 4011.27388291279 | 4350.78646319138 | Slc16a1 |
| ENSMUSG00000032905 | 0.156424866589731 | 0.309917337158033 | 0.851785577030345 | 862.855599323438 | 822.559886203472 | 847.07712534036 | 749.403428712271 | 696.393412540946 | 568.733237914345 | 818.287379177123 | 859.736247217458 | Atg12 |
| ENSMUSG00000032911 | -0.69831939411905 | 0.0290830798855019 | 0.665956123973759 | 146.369222084185 | 169.39847161418 | 146.700507740509 | 235.419600258798 | 387.61520131996 | 387.717067567413 | 191.558368662329 | 166.085411394282 | Cspg4 |
| ENSMUSG00000032913 | 0.0596755640165627 | 0.767586428015428 | 0.95879680569409 | 138.237598635064 | 140.893921102179 | 160.897331070236 | 138.813150750208 | 152.199189218854 | 94.1773318696872 | 169.69572876065 | 137.8617467129 | Lrig2 |
| ENSMUSG00000032925 | -0.669579977604429 | 0.077365251137882 | 0.739855948845396 | 166.246523848704 | 155.553404222637 | 124.932045301595 | 167.888878272211 | 290.163921892061 | 430.524945689999 | 137.42230795341 | 120.493337678204 | Itgbl1 |
| ENSMUSG00000032932 | -0.0695479688231095 | 0.583681300079205 | 0.922450244597442 | 319.843855665442 | 392.548381336706 | 331.259211026956 | 317.019222659259 | 378.855535753408 | 325.339873731647 | 352.92547269853 | 371.24974311663 | Hspa13 |
| ENSMUSG00000032939 | -0.0869739516841614 | 0.625010604537745 | 0.931529302402539 | 271.054114970713 | 279.344595017615 | 189.290977729689 | 226.978260010475 | 255.12525962585 | 211.593226148778 | 283.173240631269 | 276.809018990469 | Nup93 |
| ENSMUSG00000032942 | 0.508417012159234 | 0.127291692967668 | 0.772837268577874 | 1644.39496415566 | 1790.08577215369 | 3154.53414386527 | 2958.22079369024 | 2477.89039713862 | 837.811328970594 | 1842.70822028436 | 1552.30155747597 | Ucp3 |
| ENSMUSG00000032946 | -0.21143951828132 | 0.359458736754511 | 0.867338970126053 | 390.317925557827 | 296.447325324816 | 413.600786339371 | 332.963976461647 | 493.826146314413 | 540.602346576646 | 292.542943446274 | 333.256348353232 | Rasgrp2 |
| ENSMUSG00000032952 | -0.106324974519643 | 0.573935057404957 | 0.922280709351328 | 103.90407740544 | 100.173134656462 | 91.8061241988993 | 90.0409626487835 | 125.920192519196 | 83.1695917810225 | 112.436433780063 | 93.3551985614916 | Ap4b1 |
| ENSMUSG00000032959 | -0.0189028663161615 | 0.889253154208215 | 0.98126431530921 | 10147.3625507869 | 8089.59143530603 | 8302.30228322417 | 9952.34015277335 | 9029.02528272427 | 10868.3087142083 | 8462.92379813083 | 8613.64535564461 | Pebp1 |
| ENSMUSG00000032965 | 0.112420957080824 | 0.416751142883961 | 0.882192126163777 | 337.914129996823 | 270.386021999557 | 293.401015481019 | 314.205442576484 | 282.499214521327 | 313.109051410908 | 287.337552993494 | 242.072200921077 | Ift57 |
| ENSMUSG00000032966 | -0.101416700990702 | 0.327620873323418 | 0.855977678491063 | 3470.3961853417 | 3303.27019647652 | 3247.28672295282 | 3690.74154190586 | 3785.27048294662 | 4102.21780637573 | 3338.73743641353 | 3485.62258815058 | Fkbp1a |
| ENSMUSG00000032968 | -0.404097240270311 | 0.205273488909674 | 0.819390360194292 | 34.3335212296237 | 17.9171460361152 | 29.3401015481019 | 23.448167356454 | 36.1336204620302 | 25.6847268735511 | 43.7252798033577 | 32.5657669400552 | Inha |
| ENSMUSG00000032977 | -0.168409086559089 | 0.304157754127239 | 0.851785577030345 | 271.054114970713 | 223.96432545144 | 260.275094378323 | 245.736793895638 | 284.689130912965 | 349.801518373124 | 232.160414194018 | 259.440609955773 | Fam207a |
| ENSMUSG00000032988 | 0.154769451320794 | 0.583455477460557 | 0.922450244597442 | 33.4300075130547 | 38.2775392589734 | 36.9117406572894 | 24.3860940507122 | 24.0890803080201 | 33.0232202659942 | 31.2323427166841 | 31.4802413753867 | Slc16a8 |
| ENSMUSG00000032997 | -0.294787819453422 | 0.147918008655034 | 0.780607990324772 | 521.327414460339 | 537.514381083457 | 520.550188756646 | 543.997482669733 | 776.32536083574 | 834.142082274372 | 530.94982618363 | 464.604941678121 | Chpf |
| ENSMUSG00000032998 | 0.0261128830630979 | 0.800842869736017 | 0.963461896828986 | 749.916384752307 | 740.303897583125 | 784.611102689562 | 686.562340196974 | 773.040486248282 | 649.456665231219 | 724.590351027071 | 759.867895267955 | Foxj3 |
| ENSMUSG00000033004 | 0.322574268814921 | 0.10712614213744 | 0.762287160109383 | 1040.84780148754 | 1361.70309874476 | 1151.83559948516 | 915.416453595965 | 802.604357535398 | 660.464405319884 | 1061.89965236726 | 1048.61769546978 | Mycbp2 |
| ENSMUSG00000033006 | -2.31139383629467 | 0.00040637653867561 | 0.522942841846969 | 5.42108229941427 | 4.07207864457164 | 5.67872933189068 | 9.37926694258161 | 48.1781606160403 | 51.3694537471021 | 14.5750932677859 | 7.59867895267955 | Sox10 |
| ENSMUSG00000033009 | 0.178346736603781 | 0.515006585667875 | 0.904145806837306 | 550.239853390548 | 712.613762800038 | 762.842640250648 | 551.500896223799 | 452.217734873287 | 332.67836712409 | 732.91897575152 | 758.782369703287 | Ogfod1 |
| ENSMUSG00000033014 | 0.187710655246205 | 0.300160574900514 | 0.851785577030345 | 506.871194995234 | 542.400875456943 | 433.476339000989 | 435.197986135787 | 395.279908690694 | 311.885969178834 | 508.046108191395 | 467.861518372127 | Trim33 |
| ENSMUSG00000033016 | -0.203310399375144 | 0.0839023643114821 | 0.742295439672922 | 355.984404328204 | 393.362797065621 | 412.654331450723 | 347.970803569778 | 471.926982398031 | 448.871179171106 | 406.020455316893 | 412.499714574033 | Nfatc1 |
| ENSMUSG00000033020 | -0.142855789446311 | 0.535096915984263 | 0.910813988293389 | 1939.84394947374 | 1325.86880667253 | 1386.55641186997 | 1842.08802752303 | 1915.0818844876 | 2464.51069762883 | 1457.50932677859 | 1334.1109189776 | Polr2f |
| ENSMUSG00000033021 | 0.0939618771962477 | 0.490513909714748 | 0.899236955296196 | 486.993893230715 | 403.950201541507 | 510.139184981513 | 480.218467460179 | 419.368988998714 | 511.248373006873 | 438.293876124134 | 394.045779974668 | Gmppa |

| | | | | | | | | | | | | |
|---|---|---|---|---|---|---|---|---|---|---|---|---|
| **ENSMUSG00000033022** | -0.134266154603462 | 0.615293890855618 | 0.928422072203769 | 284.606820719249 | 569.276594511116 | 465.655805215036 | 356.412143818101 | 327.39250054991 | 522.256113095538 | 392.486440139664 | 598.124586132348 | Cdo1 |
| **ENSMUSG00000033024** | -0.217846412247363 | 0.761694209431205 | 0.957666649379949 | 21.6843291976571 | 5.7009101024003 | 0.946454888648447 | 10.3171936368398 | 8.75966556655277 | 8.56157562451702 | 16.6572494488982 | 10.8552556466851 | Klra9 |
| **ENSMUSG00000033029** | 0.2798540430568 | 0.479650662256926 | 0.897732059433541 | 15.3597331816738 | 23.6180561385155 | 21.7684624389143 | 16.8826804966469 | 9.85462376237187 | 17.123151249034 | 23.9447960827911 | 13.0263067760221 | 1700088E04R |
| **ENSMUSG00000033031** | -0.223695906531301 | 0.548710815822166 | 0.914699131434573 | 12.6491920319666 | 26.0613033252585 | 23.6613722162112 | 28.1378008277448 | 17.5193311331055 | 22.0154801773295 | 32.2734208072402 | 33.6512925047237 | C330027C09 |
| **ENSMUSG00000033032** | -0.083932829088031 | 0.460507072659911 | 0.894774044236998 | 1561.27170223131 | 1365.77517738933 | 1259.73145679108 | 1341.23517278917 | 1636.96250274955 | 1364.95977099443 | 1453.34501441637 | 1403.58455511638 | Afap1l1 |
| **ENSMUSG00000033033** | -0.913417749519451 | 0.0301447867979954 | 0.665956123973759 | 40.658117245607 | 52.9370223794314 | 50.1621090983677 | 44.0825546301336 | 119.350443344282 | 149.216032313011 | 43.7252798033577 | 42.3354970220718 | Calhm2 |
| **ENSMUSG00000033039** | 0.250200508533714 | 0.371418356543635 | 0.873374202289323 | 1174.56783153976 | 1480.60779516625 | 1384.66350209268 | 1131.13959327534 | 748.951405940262 | 664.133652016106 | 1458.55040486915 | 1475.2292423845 | Micall1 |
| **ENSMUSG00000033044** | 0.0116924672246833 | 0.935666707820735 | 0.98818087308472 | 2846.06820719249 | 2323.52807459258 | 1977.14426238661 | 2742.49765401086 | 2355.25507920688 | 2685.8885816342 | 2281.0020964085 | 2488.02459422022 | Dhrs7c |
| **ENSMUSG00000033047** | 0.0477884323396732 | 0.615433290293952 | 0.928422072203769 | 1842.26446808428 | 1678.51081729243 | 1831.39020953474 | 1812.07437330677 | 1801.20623212241 | 1890.88513078619 | 1641.78014880703 | 1597.89363119204 | Eif3l |
| **ENSMUSG00000033054** | 0.0141466457953 | 0.945656411473975 | 0.990505437194639 | 116.553269437407 | 120.533527879321 | 157.111511515642 | 94.7305961200743 | 107.305903190271 | 106.408154190426 | 140.545542225078 | 129.177542195552 | Npat |
| **ENSMUSG00000033055** | -0.539259273884077 | 0.0240678237855582 | 0.654124840968651 | 113.8427282877 | 98.5443031986338 | 78.5557557578211 | 91.9168160372998 | 166.433645764503 | 173.677676954488 | 121.806136595068 | 95.5262496908287 | Ankrd54 |
| **ENSMUSG00000033059** | 0.204685226661364 | 0.200525731804914 | 0.812921911423739 | 25715.8074009882 | 31685.6583491409 | 30448.4002227092 | 28285.0553187434 | 24202.9559603853 | 19113.1060406182 | 29080.4343035046 | 28375.6382604348 | Pygb |
| **ENSMUSG00000033060** | 0.179074523673316 | 0.506853259579319 | 0.901698316485955 | 7062.76672242023 | 7879.47217724613 | 7842.32520734103 | 6694.92074361476 | 5633.55991748925 | 3456.43038784073 | 9204.1713986068 | 7743.05385278046 | Lmo7 |
| **ENSMUSG00000033065** | 0.0559280928914605 | 0.589284882961004 | 0.92352104528132 | 29362.3887610608 | 30282.4200482215 | 29595.6443680369 | 28215.6487433683 | 27723.2465599437 | 24290.4131289869 | 30272.4687171913 | 30701.9195455194 | Pfkm |
| **ENSMUSG00000033066** | -0.512177371159297 | 0.0550411518521743 | 0.703396849069134 | 67.7635287426784 | 67.5965054998893 | 92.7525790875478 | 95.6685228143324 | 119.350443344282 | 149.216032313011 | 69.7522320672611 | 123.74991437221 | Gas7 |
| **ENSMUSG00000033068** | 0.0741949474862658 | 0.597320117014246 | 0.924496535139492 | 390.317925557827 | 332.281617397046 | 366.278041906949 | 398.618845059719 | 383.235368536684 | 305.770558018465 | 400.815064864113 | 321.315567141878 | Entpd6 |
| **ENSMUSG00000033075** | 0.027710751678098 | 0.868596722648295 | 0.975147618296101 | 210.518695960587 | 206.047179415325 | 261.221549266971 | 186.647412157374 | 234.321053905287 | 177.34692365071 | 203.010227658447 | 232.302470839061 | Senp1 |
| **ENSMUSG00000033082** | 0.00602898900337977 | 0.974131392721101 | 0.995510898988495 | 241.238162323935 | 230.479651282755 | 253.649910157784 | 213.847286290861 | 252.935343234211 | 300.878229090169 | 185.311900118992 | 197.565652769668 | Clec1a |
| **ENSMUSG00000033083** | 0.445671382297492 | 0.0864331161723338 | 0.746460787597018 | 1536.87683188395 | 2514.91577088745 | 2886.68741037776 | 1642.30964164604 | 1365.41287018641 | 1138.68955806076 | 1840.62606410325 | 1955.03154196798 | Tbc1d4 |
| **ENSMUSG00000033088** | -0.428721058715597 | 0.080203666744797 | 0.739855948845396 | 620.713923282934 | 549.730617017172 | 535.693466975021 | 538.369922504185 | 959.183379537529 | 1006.59667699679 | 552.812466085309 | 503.683862006187 | Triobp |
| **ENSMUSG00000033096** | -0.126181151122482 | 0.240425714611638 | 0.832048466795696 | 851.10992100804 | 806.271571625185 | 845.184215563063 | 711.886360941944 | 913.195135313127 | 930.765578608207 | 860.971580889925 | 804.374443419364 | Apmap |
| **ENSMUSG00000033099** | -0.0965682198641231 | 0.553398199458983 | 0.915192402724762 | 362.309000344187 | 276.086932101957 | 284.882921483182 | 377.046531091781 | 352.576539053749 | 399.947889888152 | 306.076958623504 | 332.170822788563 | Nol12 |
| **ENSMUSG00000033102** | -0.0173624223246718 | 0.952508974775522 | 0.992000114323903 | 159.921927832721 | 193.016527752696 | 116.413951303759 | 141.626930832982 | 120.445401540101 | 102.738907494204 | 154.079557402308 | 240.986675356409 | Cdc14b |
| **ENSMUSG00000033105** | 0.0842260120242277 | 0.819936495448354 | 0.966398681144067 | 18.9737880479499 | 24.4324718674299 | 36.9117406572894 | 22.5102406621959 | 18.6142893289246 | 17.123151249034 | 24.9858741733473 | 35.8223436340608 | Lss |
| **ENSMUSG00000033106** | -0.158267071860678 | 0.268336632625493 | 0.843793744782845 | 147.272735800754 | 145.780415475665 | 160.897331070236 | 168.826804966469 | 190.522726072523 | 182.239252579005 | 156.16171358342 | 166.085411394282 | Slc7a6os |
| **ENSMUSG00000033107** | -0.282175063530506 | 0.278730956496459 | 0.848075053184273 | 43.3686583953142 | 61.895595397489 | 49.2156542097192 | 51.5859681841989 | 76.6470737073368 | 78.277262852727 | 47.8895921655823 | 48.8486504100828 | Rnf125 |
| **ENSMUSG00000033111** | 0.188027149583814 | 0.236500629682764 | 0.832048466795696 | 421.037391921175 | 368.930325198191 | 371.95677123884 | 417.377378944882 | 269.359716171498 | 327.786038195794 | 401.856142954669 | 386.447101021989 | 3830406C13I |
| **ENSMUSG00000033114** | 0.0822185549235883 | 0.822462913842787 | 0.966637987288264 | 25.2983840639333 | 16.2883145782866 | 17.9826428843205 | 24.3860940507122 | 22.994122112201 | 24.4616446414772 | 11.4518589961175 | 20.6249857287016 | Slc35d2 |
| **ENSMUSG00000033124** | 0.0525012085416433 | 0.869778214367435 | 0.975613767803763 | 703.837185207286 | 1046.52421165491 | 1065.70820461815 | 619.031618210386 | 707.342994499137 | 413.401794440965 | 1136.8572748873 | 1054.04532329312 | Atg9a |
| **ENSMUSG00000033126** | -0.155965963977127 | 0.444245811580302 | 0.89108031640712 | 94.8689402397497 | 80.6271571625185 | 90.8596693102509 | 81.59962240046 | 107.305903190271 | 77.0541806206532 | 119.723980413956 | 82.4999429148066 | Ybey |
| **ENSMUSG00000033128** | 0.0671753453309926 | 0.420400536744671 | 0.882568531013176 | 1773.59742562504 | 1753.43706435255 | 1836.12248397799 | 1792.37791272735 | 1595.35409130842 | 1656.05334222801 | 1854.16007928048 | 1722.72907112892 | Gga1 |
| **ENSMUSG00000033147** | -0.0849457442146948 | 0.767388257475552 | 0.95879680569409 | 33.4300075130547 | 36.6487078011448 | 57.7337482075553 | 64.7169419038131 | 61.3176589658694 | 47.7002070508805 | 47.8895921655823 | 46.6775992807458 | Slc22a15 |
| **ENSMUSG00000033149** | -0.11640654816718 | 0.490522631780113 | 0.899236955296196 | 257.501409222178 | 210.933673788811 | 204.434255948065 | 292.633128608546 | 296.733671066975 | 270.301173288323 | 236.324726556243 | 243.157726485746 | Phldb2 |
| **ENSMUSG00000033152** | -0.0221763999282677 | 0.88416605958641 | 0.979384300778265 | 310.808718499751 | 286.674336577844 | 270.686098153456 | 345.157023487003 | 313.158044004262 | 359.586176229715 | 303.994802442392 | 256.184033261768 | Podxl2 |
| **ENSMUSG00000033157** | -0.0776296698324217 | 0.660636261703489 | 0.939611397743053 | 193.351935345776 | 188.13003337921 | 204.434255948065 | 215.723139679377 | 162.053812981226 | 196.916239363891 | 263.392756910703 | 222.532740757044 | Abhd10 |
| **ENSMUSG00000033159** | -0.0930781219894268 | 0.550404208801017 | 0.914699131434573 | 645.108793630298 | 714.242594257866 | 757.163910918757 | 686.562340196974 | 883.631264026011 | 836.58824673852 | 674.618602680377 | 595.953535003011 | Cnppd1 |
| **ENSMUSG00000033161** | 0.0713565726916221 | 0.701302563206427 | 0.945920819861844 | 29829.505352527 | 32667.0293024826 | 36679.8592095706 | 27203.6258402637 | 28921.1308261698 | 22065.6265488445 | 38249.2090470325 | 31043.86009839 | Atp1a1 |
| **ENSMUSG00000033166** | 0.0713097949242927 | 0.696902245956625 | 0.944933386973863 | 252.983840639333 | 243.510302945384 | 247.024725937245 | 233.543746870282 | 212.421889988905 | 167.562265794119 | 270.680303544596 | 277.894544555138 | Dis3 |
| **ENSMUSG00000033170** | -0.38886689069392 | 0.030397133376484 | 0.665956123973759 | 551.143367107117 | 468.289044125739 | 502.567545872325 | 507.418341593665 | 780.705193619016 | 807.234273168747 | 534.073060455298 | 536.249628946243 | Card10 |
| **ENSMUSG00000033174** | -0.339862584874192 | 0.216576612181751 | 0.825523564386777 | 3782.10841755802 | 3537.82192640384 | 4432.24824354068 | 4698.07481153913 | 6389.08107260443 | 7519.50956279009 | 3162.79523910954 | 3749.40530036503 | Mgll |
| **ENSMUSG00000033182** | 0.192579380605649 | 0.443653911288879 | 0.89108031640712 | 344.238726012806 | 325.766291565731 | 322.74111702912 | 301.07446885687 | 223.371471947096 | 172.454594722414 | 373.747034509653 | 361.480013034613 | Kbtbd12 |
| **ENSMUSG00000033184** | -0.070678580474984 | 0.364904635602508 | 0.86924083181804 | 1220.64703108478 | 1284.3336044979 | 1284.33928389594 | 1285.89749782794 | 1415.78094719409 | 1291.57483707 | 1334.66211209297 | 1287.43331969685 | Tmed7 |
| **ENSMUSG00000033186** | 0.254968630653513 | 0.226725112502044 | 0.832048466795696 | 1312.80543017482 | 1151.58384068486 | 1228.49844546568 | 1311.22151857291 | 908.81530252985 | 693.487625585878 | 1280.52605138405 | 1309.14383099022 | Mzt1 |
| **ENSMUSG00000033191** | -0.772405966655285 | 0.019565169689446 | 0.642273034804458 | 1338.10381423876 | 1316.09581792556 | 1322.19747944188 | 1507.24819767287 | 3356.04687018553 | 3394.05319400496 | 1217.02028786012 | 1400.32797842237 | Tie1 |
| **ENSMUSG00000033192** | -0.16558816670696 | 0.723778136419456 | 0.951674101666722 | 9.03513716569045 | 15.4738988493722 | 13.2503684410783 | 14.0689004138724 | 20.8042057205628 | 19.5693157131818 | 6.24646854333682 | 11.9407812113536 | Lpcat2 |
| **ENSMUSG00000033207** | -1.15077335008376 | 0.00649457008388675 | 0.635862253526407 | 23.4913566307952 | 32.5766291565731 | 21.7684624389143 | 23.448167356454 | 94.1664048404423 | 70.9387694602839 | 30.191264626128 | 30.3947158107182 | Mamdc2 |
| **ENSMUSG00000033208** | -0.814438081917781 | 0.175465733808137 | 0.799889021902745 | 15.3597331816738 | 26.0613033252585 | 9.46454888648447 | 10.3171936368398 | 30.6588294829347 | 52.592535979176 | 18.7394056300105 | 6.51315338801105 | S100b |
| **ENSMUSG00000033209** | -0.181590392653097 | 0.300763636078342 | 0.851785577030345 | 383.993329541844 | 454.443976734195 | 390.885869011809 | 372.35689762049 | 563.903470846835 | 499.017550686135 | 395.609674411332 | 359.308961905276 | Ttc28 |
| **ENSMUSG00000033216** | 0.0530008355777947 | 0.780848457698676 | 0.960742105865155 | 116.553269437407 | 115.647033505835 | 92.7525790875478 | 96.6064495085906 | 116.065568756824 | 116.192812047017 | 89.5327157878278 | 85.7565196088121 | Eefsec |
| **ENSMUSG00000033220** | -0.374057858921915 | 0.175968193635739 | 0.799889021902745 | 32.5264937964856 | 33.3910448854875 | 30.2865564367503 | 32.8274342990356 | 26.2789966996583 | 47.7002070508805 | 48.9306702561384 | 44.5065481514088 | Rac2 |
| **ENSMUSG00000033222** | 0.211736228465099 | 0.271334035708767 | 0.844129257012535 | 107.518132271716 | 104.245213301034 | 100.324218196735 | 103.171936368398 | 90.881530252985 | 63.6002760678407 | 103.066730965058 | 99.8683519495027 | Ttf2 |
| **ENSMUSG00000033228** | 0.197359918586535 | 0.141405999889395 | 0.779725819159865 | 1733.842822096 | 1966.8139853281 | 1855.9980366396 | 1733.28853098908 | 1539.51122332165 | 1284.23634367755 | 1769.83275394543 | 1762.89351702166 | Scaf11 |
| **ENSMUSG00000033233** | 0.0092272311688928 | 0.960075676472128 | 0.993625746680416 | 132.81651633565 | 151.481325578065 | 103.163582862681 | 112.551203310979 | 120.445401540101 | 140.654456688494 | 111.395355689507 | 125.920965501547 | Trim45 |
| **ENSMUSG00000033237** | -0.0845619786530729 | 0.645603770221395 | 0.935063240511793 | 352.370349461928 | 513.896324944941 | 430.636974335043 | 333.901903155905 | 433.603445544362 | 386.49398533534 | 516.374732915844 | 392.96025441 | Arid2 |
| **ENSMUSG00000033249** | -0.166175558267944 | 0.380403858385379 | 0.87432013076003 | 67.7635287426784 | 66.7820897709749 | 83.2880302010633 | 62.8410885152968 | 77.7420319031559 | 83.1695917810225 | 82.2451691539348 | 71.6446872681215 | Hsf4 |
| **ENSMUSG00000033253** | -0.065995942255122 | 0.702140462826078 | 0.94640766896208 | 201.483558794897 | 219.892246806869 | 209.166530391307 | 203.530092654021 | 208.042057205628 | 163.893019097897 | 267.557069272927 | 232.302470839061 | Szt2 |
| **ENSMUSG00000033256** | -0.099142274031853 | 0.715571803783966 | 0.951176382858794 | 93.9654265231807 | 62.7100111264033 | 73.8234813145789 | 69.4065753751039 | 98.5462376237187 | 103.961989726278 | 52.0539045278068 | 67.3025850094475 | Shf |
| **ENSMUSG00000033257** | 0.15818015286721 | 0.647951152961441 | 0.9356525121774 | 94.8689402397497 | 147.409246933493 | 121.146225747001 | 136.937297361692 | 77.7420319031559 | 57.4848649074714 | 182.188665847324 | 130.263067760221 | Ttll4 |
| **ENSMUSG00000033272** | -0.00360315643955375 | 0.979887149109283 | 0.996872525500296 | 896.285606836493 | 1077.47200935366 | 1055.29720084302 | 771.913669374467 | 879.251431242735 | 911.196262895025 | 1042.11916864669 | 978.058533766325 | Slc35a4 |
| **ENSMUSG00000033278** | 0.0134782433152089 | 0.937318836405186 | 0.98894578968943 | 2088.92371270763 | 2523.05992817659 | 2271.49173275627 | 1823.32949363787 | 2153.78277117616 | 1629.14553312238 | 2501.7106516064 | 2340.3931174253 | Ptprm |
| **ENSMUSG00000033282** | -0.271712437200286 | 0.254607552722235 | 0.837406774112203 | 43.3686583953142 | 40.7207864457164 | 53.9479286529615 | 42.2067012416173 | 48.1781606160403 | 44.0309603546589 | 58.3003730711437 | 66.217059444779 | Rpgrip1l |
| **ENSMUSG00000033285** | 0.0426786451125555 | 0.793877476172363 | 0.961804768755116 | 209.615182244018 | 267.942774812814 | 240.399541716705 | 226.040333316217 | 251.840385038392 | 179.793088114857 | 256.10521027681 | 227.960368580387 | Wdr3 |
| **ENSMUSG00000033287** | -0.684918070450365 | 0.015564662793303 | 0.635862253526407 | 140.044626068202 | 140.893921102179 | 129.664319744837 | 216.661066373635 | 318.632834983357 | 332.67836712409 | 167.613572579538 | 189.966973816989 | Kctd17 |
| **ENSMUSG00000033294** | -0.14960271058255 | 0.462246063076855 | 0.895328112556867 | 134.623543768788 | 134.378595270864 | 148.593417517806 | 171.640585049243 | 198.187433443257 | 135.762127760198 | 191.558368662329 | 127.006491066215 | Noc4l |
| **ENSMUSG00000033295** | 0.25871279234776 | 0.411998288915368 | 0.882192126163777 | 59.631905293557 | 80.6271571625185 | 143.861143074564 | 90.0409626487835 | 58.0327843784121 | 63.6002760678407 | 90.5737938783839 | 99.8683519495027 | Ptprf |
| **ENSMUSG00000033306** | -0.149775505374533 | 0.446801926159896 | 0.89108031640712 | 546.625798524272 | 556.245942848486 | 418.333060782613 | 520.54931531328 | 654.78500109982 | 702.049201210395 | 425.80093903746 | 484.144401842154 | Lpp |
| **ENSMUSG00000033307** | -0.126660088018516 | 0.529502802891399 | 0.908560090446144 | 1717.57957519775 | 1420.34103122659 | 1345.85885165809 | 1827.0812004149 | 1744.26840593982 | 2327.52548763655 | 1468.96118577471 | 1350.39380244762 | Mif |
| **ENSMUSG00000033308** | 0.299031778445637 | 0.226191037081871 | 0.832048466795696 | 322.554396815149 | 447.928650902881 | 515.817914313404 | 321.708856130549 | 245.270635863478 | 260.516515431732 | 366.45948787576 | 434.210225867403 | Dpyd |
| **ENSMUSG00000033313** | -0.454796747364921 | 0.162260770053004 | 0.793874624603454 | 36.1405486627618 | 33.3910448854875 | 23.6613722162112 | 32.8274342990356 | 39.4184950494875 | 68.4926049961361 | 33.3144988977964 | 32.5657669400552 | Fbxl8 |
| **ENSMUSG00000033318** | -0.211177336997635 | 0.476019628265815 | 0.897553735056661 | 94.8689402397497 | 67.5965054998893 | 59.6266579848522 | 119.116690170786 | 123.730276127558 | 111.300483118721 | 63.5057635239243 | 96.6117752554972 | Gstt2 |
| **ENSMUSG00000033319** | -0.0905931362483821 | 0.427160297407195 | 0.885066977635295 | 468.923618899334 | 517.153987860599 | 462.81644054909 | 404.246405225267 | 490.541271726955 | 483.117481669175 | 487.224546380272 | 513.453592088204 | Fem1c |
| **ENSMUSG00000033323** | -0.109426485035741 | 0.403676901381185 | 0.881158207349486 | 225.878429142261 | 216.634583891211 | 242.292451494002 | 199.778385876988 | 266.074841584041 | 229.939459629886 | 232.160414194018 | 225.78931745105 | Ctdp1 |
| **ENSMUSG00000033326** | 0.114108910798016 | 0.549171120889078 | 0.914699131434573 | 1074.27780900059 | 1224.88125628715 | 1220.9268063565 | 880.713165908413 | 999.696832782835 | 738.741668172611 | 1236.80077158069 | 1089.86766692718 | Kdm4a |
| **ENSMUSG00000033327** | -1.479329788341 | 0.0336814350301653 | 0.672620872762612 | 325.264937964856 | 327.39512302356 | 387.100049457215 | 498.977001345342 | 1623.82300439972 | 2062.11664327653 | 332.103910887408 | 272.466916731795 | Tnxb |
| **ENSMUSG00000033335** | -0.0299270000990199 | 0.730835032790742 | 0.952921973914044 | 1481.76249517323 | 1531.10157035894 | 1604.24103625912 | 1494.11722395325 | 1645.7221683161 | 1646.26868437142 | 1466.8790295936 | 1481.74239577251 | Dnm2 |
| **ENSMUSG00000033350** | -1.0316751960481 | 0.0174929534868884 | 0.642273034804458 | 19.877301764519 | 13.8450673915436 | 15.1432782183751 | 27.1998741334867 | 54.7479097909548 | 56.2617826753975 | 21.8626399016789 | 22.7960368580387 | Chst2 |
| **ENSMUSG00000033352** | 0.124269974185147 | 0.531861399179936 | 0.909082872831377 | 883.636414804526 | 987.886279173081 | 1041.10037751329 | 889.154506156737 | 820.123688668504 | 588.302553627526 | 1108.74816644229 | 969.374329248977 | Map2k4 |
| **ENSMUSG00000033355** | -0.339058875198727 | 0.238024899971465 | 0.832048466795696 | 127.395434036235 | 109.13170767452 | 86.1273948670087 | 121.930470253561 | 176.288269526875 | 195.693157131818 | 94.7381062406084 | 96.6117752554972 | Rtp4 |
| **ENSMUSG00000033356** | -0.121184977207777 | 0.650088885132444 | 0.936463929813731 | 26.2018977805023 | 39.0919549878878 | 36.9117406572894 | 35.6412143818101 | 32.8487458745729 | 31.8001380339204 | 42.6842017128016 | 42.3354970220718 | Pus7l |
| **ENSMUSG00000033361** | -0.403218505085373 | 0.206488296399809 | 0.819960256425221 | 175.281661014395 | 234.551729927327 | 230.934992830221 | 151.006197775564 | 353.671497249568 | 368.147751854232 | 163.449260217313 | 162.828834700276 | Prrg3 |
| **ENSMUSG00000033364** | -0.114964823445872 | 0.556993333529477 | 0.916239900758826 | 114.746242004269 | 127.048853710635 | 106.949402417274 | 116.302910088012 | 153.294147414674 | 89.2850029413918 | 131.175839410073 | 129.177542195552 | Usp37 |
| **ENSMUSG00000033365** | 0.151551297523517 | 0.275879748829111 | 0.846508725058461 | 1638.97388185625 | 1750.99381716581 | 1523.792370724 | 1746.4195047087 | 1486.95322992233 | 1185.16668287957 | 1634.49260217313 | 1687.99225305953 | Ipo13 |
| **ENSMUSG00000033373** | 0.0671413407184029 | 0.699672543845468 | 0.945280238056015 | 179.79922959724 | 205.232763686411 | 258.382184601026 | 213.847286290861 | 212.421889988905 | 218.931719541221 | 222.790711379013 | 163.914360264945 | Fntb |
| **ENSMUSG00000033377** | -0.0643814345639197 | 0.4915812820209 | 0.899236955296196 | 1521.51709870227 | 1553.09079503962 | 1601.40167159317 | 1788.62620595031 | 1732.22386578581 | 1724.54594722414 | 1726.10747414207 | 1576.18311989867 | Palmd |
| **ENSMUSG00000033379** | 0.143208934697945 | 0.299523961163049 | 0.851785577030345 | 2425.03081527132 | 1735.51991831643 | 2030.14573615092 | 2300.73418101527 | 1946.83567216635 | 2079.23979452556 | 1719.86100559874 | 1943.09076075663 | Atp6v0b |
| **ENSMUSG00000033382** | 0.179825946859807 | 0.493860824082072 | 0.900117970535229 | 826.715050660676 | 1017.205245414 | 948.347798425744 | 763.472329126143 | 660.259792078915 | 446.425014706959 | 1032.74946583169 | 998.683519495027 | Trappc8 |
| **ENSMUSG00000033386** | 0.140866734385269 | 0.596024410578918 | 0.924461858177432 | 56.0178504272808 | 54.56585383726 | 46.3762895437739 | 57.2135283497478 | 51.4630352034976 | 28.1308913376988 | 53.094982618363 | 60.7894316214364 | Frrs1 |

| | | | | | | | | | | | | |
|---|---|---|---|---|---|---|---|---|---|---|---|---|
| **ENSMUSG00000033389** | 0.196105578237002 | 0.356591665992351 | 0.864965376583892 | 1081.50591873315 | 1317.72464938338 | 1226.60553568839 | 1000.76778277346 | 774.135444444101 | 764.426395046162 | 1210.77381931679 | 1288.51884526152 | Arhgap44 |
| **ENSMUSG00000033392** | 0.0653468932628383 | 0.718247444326276 | 0.951674101666722 | 905.320744002183 | 1237.91190794978 | 1039.20746773599 | 839.444391361054 | 999.696832782835 | 744.857079332981 | 1133.73404061563 | 965.032226990303 | Clasp2 |
| **ENSMUSG00000033396** | 0.32242685043476 | 0.264486574484524 | 0.842510961779757 | 345.142239729375 | 442.22774080048 | 446.726707442067 | 308.577882410935 | 266.074841584041 | 162.669936865823 | 413.308001950786 | 390.789203280663 | Spg11 |
| **ENSMUSG00000033400** | 0.268256164399849 | 0.292787352058867 | 0.851785577030345 | 5871.03213026565 | 7719.03227865001 | 7377.61585701464 | 5652.88418629394 | 4476.18910450847 | 3385.49161838044 | 7142.83677930565 | 7098.25166736737 | Agl |
| **ENSMUSG00000033405** | -0.0988504427504244 | 0.776487135083513 | 0.959733396167384 | 63.2459601598332 | 87.1424829938332 | 91.8061241988993 | 65.6548685980713 | 50.3680770076785 | 47.7002070508805 | 105.14888714617 | 125.920965501547 | Nudt15 |
| **ENSMUSG00000033411** | 0.0210164947637107 | 0.906200689832526 | 0.98504921011344 | 130.105975185942 | 178.357044632238 | 152.3792370724 | 132.247663890401 | 168.623562156141 | 123.53130543946 | 141.586620315635 | 150.888053488923 | Ctdspl2 |
| **ENSMUSG00000033416** | 0.238561864119977 | 0.186981967035688 | 0.807540361382744 | 677.635287426784 | 756.592212161411 | 977.687899973846 | 781.292936317048 | 614.271547854513 | 533.263853184203 | 768.315630830429 | 789.177085514005 | Gucd1 |
| **ENSMUSG00000033417** | 0.07688324411823 | 0.710607741828886 | 0.949886733585872 | 842.978297558919 | 1026.16381843205 | 944.56197887115 | 750.341355406529 | 731.432074807157 | 600.533375948265 | 989.02418602833 | 1057.30189998713 | Cacul1 |
| **ENSMUSG00000033419** | 0.117996780590882 | 0.685791575908749 | 0.944452327903597 | 131.009488902512 | 141.708336831093 | 146.700507740509 | 115.364983393754 | 100.736154015357 | 64.8233582999146 | 162.408182126757 | 163.914360264945 | Snap91 |
| **ENSMUSG00000033420** | -0.760065570179433 | 0.0312874010474255 | 0.669064667366769 | 60.535419010126 | 82.2559886203472 | 63.4124775394459 | 69.4065753751039 | 137.964732673206 | 192.023910435596 | 73.9165444294857 | 64.046008315442 | Antxr1 |
| **ENSMUSG00000033423** | 0.00631931003920921 | 0.948772168965375 | 0.991233613135933 | 1587.47360001181 | 1456.17532329882 | 1512.43491206022 | 1540.0756319719 | 1567.98013641295 | 1682.96115133363 | 1427.31806215246 | 1392.7292994697 | Eri3 |
| **ENSMUSG00000033427** | 0.00413674582222777 | 0.991320465200267 | 0.998628446692557 | 55.1143367107117 | 114.83261777692 | 48.2691993210708 | 64.7169419038131 | 55.8428679867739 | 56.2617826753975 | 51.0128264372507 | 119.407812113536 | Upb1 |
| **ENSMUSG00000033429** | 0.054050905933869 | 0.738581669045039 | 0.954383574136061 | 1748.2990415611 | 1312.8381550099 | 1363.84149454241 | 1817.70193347232 | 1538.41626512583 | 1760.01533195428 | 1312.79947219129 | 1402.49902955171 | Mcee |
| **ENSMUSG00000033430** | -0.0157482356822182 | 0.914960986841417 | 0.986713918026868 | 575.538237454482 | 577.420751800259 | 624.660226507975 | 559.004309777864 | 695.298454345127 | 465.99433042014 | 638.180869510912 | 561.216716933618 | Terf2ip |
| **ENSMUSG00000033434** | -0.0759841548024888 | 0.722484260967486 | 0.951674101666722 | 188.83436676293 | 144.151584017836 | 151.432782183751 | 259.805694309511 | 189.427767876704 | 215.262472844999 | 188.435134390661 | 191.052499381657 | Gtpbp6 |
| **ENSMUSG00000033436** | -0.757168382799139 | 0.0603788872738535 | 0.71282992338528 | 59.631905293557 | 43.9784493613737 | 50.1621090983677 | 63.779015209555 | 98.5462376237187 | 170.008430258266 | 52.0539045278068 | 47.7631248454143 | Armcx2 |
| **ENSMUSG00000033439** | -0.00679550173099025 | 0.972262423980869 | 0.995254967835651 | 189.737880479499 | 164.511977240694 | 160.897331070236 | 224.164479927701 | 191.617684268342 | 136.985209992272 | 194.681602933998 | 218.19063849837 | Trmt13 |
| **ENSMUSG00000033444** | 0.281319900527034 | 0.258038361382532 | 0.838310965110319 | 1490.79763233892 | 1944.82476064742 | 2006.48436393471 | 1324.35249229252 | 1101.52794499401 | 942.996400928946 | 1809.39372138657 | 1712.9593410469 | Specc1l |
| **ENSMUSG00000033446** | -0.156126576116183 | 0.649750161953692 | 0.936463929813731 | 75.8951521917998 | 76.5550785179469 | 109.78876708322 | 209.15765281957 | 145.62944004394 | 136.985209992272 | 96.8202624217207 | 145.46042566558 | Lpar6 |
| **ENSMUSG00000033453** | 0.112705405052928 | 0.772669692446667 | 0.959045638244974 | 82.2197482077831 | 78.1839099757755 | 265.953823710214 | 201.654239265505 | 162.053812981226 | 134.539045528125 | 118.6829023234 | 164.999885829613 | Adamts15 |
| **ENSMUSG00000033454** | -0.130997736018755 | 0.468046823801227 | 0.895328112556867 | 226.78194285883 | 235.366145656241 | 265.953823710214 | 249.488500672671 | 264.979883388221 | 192.023910435596 | 287.337552993494 | 324.572143835884 | Zbtb1 |
| **ENSMUSG00000033458** | -0.118899451261528 | 0.751597800596546 | 0.956197192463153 | 108.421645988285 | 115.647033505835 | 102.217127974032 | 61.9031618210386 | 65.6974917491458 | 55.0387004433237 | 133.257995591185 | 167.17093695895 | Fan1 |
| **ENSMUSG00000033460** | -0.371817582143382 | 0.35036861199436 | 0.861774532843376 | 28.0089252136404 | 21.1748089517725 | 27.447191770805 | 35.6412143818101 | 60.2227007700503 | 39.1386314263635 | 17.6983275394543 | 28.2236646813812 | Armcx1 |
| **ENSMUSG00000033467** | -0.636290796479468 | 0.0997185188863247 | 0.759800953500741 | 116.553269437407 | 76.5550785179469 | 86.1273948670087 | 121.930470253561 | 192.712642464161 | 261.739597663806 | 94.7381062406084 | 74.901263962127 | Crlf2 |
| **ENSMUSG00000033475** | 0.0780736491495494 | 0.812314283543886 | 0.965224347191244 | 46.0791995450213 | 46.4216965481167 | 42.5904699891801 | 64.7169419038131 | 39.4184950494875 | 79.5003450848009 | 35.3966550789086 | 35.8223436340608 | Tomm6 |
| **ENSMUSG00000033478** | -0.0735930121774223 | 0.63619186184156 | 0.932595977875587 | 280.992765852973 | 330.652785939217 | 322.74111702912 | 273.874594723383 | 328.487458745729 | 261.739597663806 | 301.91264626128 | 378.848422069309 | Fam160b1 |
| **ENSMUSG00000033486** | 0.41407882659365 | 0.0614576916287412 | 0.71282992338528 | 108.421645988285 | 109.13170767452 | 141.021778408619 | 144.440710915757 | 78.836990098975 | 72.1618516923577 | 113.477511870619 | 111.809133160856 | Catsper2 |
| **ENSMUSG00000033487** | -0.240117649254057 | 0.0956031221303825 | 0.757085157577544 | 172.571119864688 | 210.933673788811 | 228.095628164276 | 185.709485463116 | 247.460552255116 | 247.06261087892 | 215.50316474512 | 232.302470839061 | Fndc3a |
| **ENSMUSG00000033488** | -0.243672404651943 | 0.0952873617150102 | 0.757085157577544 | 176.185174730964 | 171.027303072009 | 167.522515290775 | 201.654239265505 | 223.371471947096 | 236.054870790255 | 182.188665847324 | 207.335382851685 | Cryzl2 |
| **ENSMUSG00000033499** | -0.0049387656653166 | 0.971480058914073 | 0.995122272737203 | 1827.80824861918 | 1853.61019900901 | 1500.13099850779 | 1526.00673155803 | 1467.24398239759 | 1577.77607937528 | 1742.76472359097 | 1943.09076075663 | Larp4b |
| **ENSMUSG00000033502** | -0.582412941215518 | 0.140436977878244 | 0.779711473022872 | 28.0089252136404 | 11.4018202048006 | 25.5542819935081 | 16.8826804966469 | 43.7983278327639 | 19.5693157131818 | 29.1501865355718 | 29.3091902460497 | Cdc14a |
| **ENSMUSG00000033526** | 0.311946076654143 | 0.236972146951922 | 0.832048466795696 | 2234.38942107525 | 2521.43109671876 | 2583.82184601026 | 2008.10105240672 | 1354.46328822822 | 1181.49743618335 | 2547.51808759087 | 2445.68909719815 | Ppip5k1 |
| **ENSMUSG00000033530** | 0.184065329220028 | 0.191602556425723 | 0.808824684419004 | 2209.99455072788 | 2302.35326564081 | 2250.66972520601 | 2221.94833869758 | 1797.92135753496 | 1598.56847732053 | 2335.13815711741 | 2175.39323159569 | Ttc7b |
| **ENSMUSG00000033533** | -0.269351650619282 | 0.480137294877204 | 0.897732059433541 | 38.8510898124689 | 70.8541684155466 | 34.0723759913441 | 73.1582821521366 | 58.0327843784121 | 59.9310293716191 | 38.5198893505771 | 105.295979772845 | Acsm1 |
| **ENSMUSG00000033538** | -0.0882964281171833 | 0.856254729189642 | 0.97363563840968 | 26.2018977805023 | 15.4738988493722 | 13.2503684410783 | 47.8342614071662 | 18.6142893289246 | 20.7923979452556 | 24.9858741733473 | 44.5065481514088 | Casp4 |
| **ENSMUSG00000033540** | -0.128392603971601 | 0.346460506230082 | 0.861266047293794 | 296.352499034647 | 351.013179162076 | 367.224496795597 | 346.094950181261 | 418.274030802895 | 354.693847301419 | 390.404283958551 | 323.486618271215 | Idua |
| **ENSMUSG00000033542** | 0.254228384664049 | 0.18932719183946 | 0.808677608936066 | 630.652574165193 | 680.851549372379 | 784.611102689562 | 593.707597465416 | 538.719432342996 | 395.055560959857 | 686.070461676494 | 633.946929766408 | Arhgef5 |
| **ENSMUSG00000033543** | 0.132582563381325 | 0.285448286248241 | 0.851275833092507 | 562.889045422515 | 492.721515993169 | 562.194203857177 | 649.045272426648 | 516.820268426614 | 492.902139525765 | 495.553171104721 | 561.216716933618 | Gtf2a2 |
| **ENSMUSG00000033544** | -0.227131283781357 | 0.505238314445624 | 0.901494649536949 | 44.2721721118832 | 32.5766291565731 | 40.6975602118832 | 20.6343872736795 | 51.4630352034976 | 47.7002070508805 | 24.9858741733473 | 37.9933947633978 | Angptl1 |
| **ENSMUSG00000033545** | -0.074851876199225 | 0.531355635835336 | 0.909082872831377 | 795.995584297329 | 759.035459348154 | 629.392500951217 | 677.183073254392 | 686.538788778574 | 757.087901653719 | 805.79444209045 | 765.295523091298 | Znrf1 |
| **ENSMUSG00000033554** | 0.157422032643443 | 0.45182653469238 | 0.891110666112841 | 175.281661014395 | 154.738988493722 | 124.932045301595 | 210.095579513828 | 174.098353135236 | 132.092881063977 | 127.011527047849 | 162.828834700276 | Dph5 |
| **ENSMUSG00000033557** | 0.17357815111642 | 0.548933775600033 | 0.914699131434573 | 2660.84789529584 | 3304.89902793435 | 3144.12314009014 | 2324.18234837172 | 1804.49110670987 | 1464.02943179241 | 3420.98260556746 | 3447.62919338718 | Fam20b |
| **ENSMUSG00000033565** | 0.00899267822619434 | 0.96714460697169 | 0.994284908685797 | 1053.49699351951 | 1333.19854823276 | 1216.19453191325 | 1153.64983393754 | 1009.55145654521 | 786.441875223492 | 1394.00356325467 | 1536.01867400594 | Rbfox2 |
| **ENSMUSG00000033577** | 0.0322535099923364 | 0.796890799001481 | 0.962278883153809 | 273.764656120421 | 298.890572511559 | 282.990011705886 | 324.522636213324 | 318.632834983357 | 292.316653465652 | 261.31060072959 | 282.236646813812 | Myo6 |
| **ENSMUSG00000033578** | -0.156299922464218 | 0.7400437491804 | 0.954383574136061 | 16.2632468982428 | 19.5459774939439 | 21.7684624389143 | 27.1998741334867 | 16.4243729372865 | 47.7002070508805 | 11.4518589961175 | 19.5394601640331 | Tmem35a |
| **ENSMUSG00000033581** | 0.282882523304543 | 0.384406269166204 | 0.875126835008975 | 171.667606148119 | 158.811067138294 | 236.613722162112 | 210.095579513828 | 113.875652365186 | 90.5080851734656 | 167.613572579538 | 265.953763343784 | Igf2bp2 |
| **ENSMUSG00000033585** | -0.894636245637982 | 0.0654383521190826 | 0.718339708336358 | 65.0529875929712 | 50.4937751926884 | 46.3762895437739 | 58.151455044006 | 148.914314631397 | 181.016170346931 | 37.4788112600209 | 42.3354970220718 | Ndn |
| **ENSMUSG00000033589** | -0.506738937861642 | 0.148422734250073 | 0.781357001591751 | 33.4300075130547 | 32.5766291565731 | 32.1794662140472 | 53.4618215727152 | 70.0773245324222 | 70.9387694602839 | 28.1091084450157 | 46.6775992807458 | Reep4 |
| **ENSMUSG00000033590** | 0.115886581675148 | 0.763992473116448 | 0.958008075434456 | 44.2721721118832 | 26.0613033252585 | 43.5369248778286 | 28.1378008277448 | 15.3294147414674 | 30.5770558018465 | 49.9717483466946 | 34.7368180693922 | Myo5c |
| **ENSMUSG00000033594** | -0.431365771637458 | 0.156749480782937 | 0.787563725533336 | 74.0881247586617 | 83.0704043492615 | 42.5904699891801 | 43.1446279358754 | 100.736154015357 | 79.5003450848009 | 91.61487196894 | 56.4473293627624 | Spata2l |
| **ENSMUSG00000033595** | -0.0878096614574447 | 0.811939551151282 | 0.965224347191244 | 27.1054114970714 | 20.3603932228582 | 20.8220075502658 | 15.9447538023887 | 17.5193311331055 | 35.4693847301419 | 18.7394056300105 | 18.4539345993646 | Lgi3 |
| **ENSMUSG00000033596** | 0.0321933216708177 | 0.861015015314399 | 0.974868041734905 | 327.071965397994 | 377.888898216248 | 395.618143455051 | 395.805064976944 | 302.208462046071 | 293.539735697726 | 400.815064864113 | 465.69046724279 | Rfwd3 |
| **ENSMUSG00000033610** | 0.124958824306405 | 0.567830036271461 | 0.920429362097163 | 1486.28006375608 | 1896.77423264147 | 2024.46700681903 | 1487.55173709344 | 1401.54649064844 | 1075.08928199292 | 1923.91231134774 | 1921.38024946326 | Pank1 |
| **ENSMUSG00000033618** | 0.0394106951565348 | 0.915120068929174 | 0.986713918026868 | 11.7456783153976 | 18.7315617650296 | 25.5542819935081 | 17.8206071909051 | 18.6142893289246 | 14.6769867848863 | 18.7394056300105 | 19.5394601640331 | Map3k13 |
| **ENSMUSG00000033623** | 0.128216432886684 | 0.41459359942674 | 0.882192126163777 | 711.065294939838 | 859.208594004617 | 693.751433379311 | 651.859052509422 | 620.841297029428 | 547.940839969089 | 719.38496057429 | 779.407355431988 | Pcgf3 |
| **ENSMUSG00000033624** | 0.230570476374932 | 0.758213754133534 | 0.95677573228595 | 793.285043147622 | 1928.53644606913 | 1758.51318310881 | 589.955890688383 | 466.452191418935 | 238.501035254403 | 1770.87383203599 | 1845.39345993646 | Pdpr |
| **ENSMUSG00000033628** | -0.0195319893411685 | 0.853632746883998 | 0.9731539929935 | 527.652010476322 | 561.946852950887 | 545.158015861505 | 505.542488205149 | 580.327843784121 | 525.92535979176 | 573.634027896431 | 489.572029665497 | Pik3c3 |
| **ENSMUSG00000033629** | -0.0974279347409149 | 0.474931279177081 | 0.897549741595057 | 458.081454300506 | 420.238516119794 | 365.3315870183 | 336.71568323868 | 434.698403740182 | 390.163232031561 | 435.170641852465 | 430.953649173397 | Hacd3 |
| **ENSMUSG00000033632** | -0.0844876087932322 | 0.680733811971982 | 0.944452327903597 | 108.421645988285 | 88.7713144516618 | 124.932045301595 | 96.6064495085906 | 107.305903190271 | 90.5080851734656 | 140.545542225078 | 104.210454208177 | AW554918 |
| **ENSMUSG00000033633** | 0.470975335411605 | 0.181351016411627 | 0.803056849297055 | 27.1054114970714 | 36.6487078011448 | 66.2518422053913 | 65.6548685980713 | 31.7537876787538 | 40.3617136584374 | 40.6020455316893 | 28.2236646813812 | Clec18a |
| **ENSMUSG00000033634** | -0.432755067113826 | 0.227356637116492 | 0.832048466795696 | 156.307872966445 | 248.39679731887 | 130.610774633486 | 192.274972322923 | 154.389105610493 | 193.24699266767 | 184.270822028436 | 450.493109337431 | Nat8f2 |
| **ENSMUSG00000033653** | 0.168585625952907 | 0.146141353914191 | 0.780607990324772 | 788.767474564776 | 821.745470474558 | 810.16538468307 | 873.209752354348 | 685.443830582755 | 639.672007374629 | 804.753363999894 | 798.946815596022 | Vps8 |
| **ENSMUSG00000033658** | 0.376902720578853 | 0.145412648821665 | 0.780607990324772 | 326.168451681425 | 335.539280312703 | 329.36630124966 | 326.39848960184 | 200.377349834895 | 147.992950080937 | 338.350379430744 | 326.743194965221 | Ddx19b |
| **ENSMUSG00000033669** | -0.509204764715544 | 0.296474138798312 | 0.851785577030345 | 16.2632468982428 | 6.51532583131463 | 6.62518422053913 | 15.9447538023887 | 22.994122112201 | 14.6769867848863 | 10.4107809055614 | 16.2828834700276 | Zfp7 |
| **ENSMUSG00000033671** | 0.351706486061219 | 0.257284865294743 | 0.837517407767513 | 637.880683897746 | 918.660942215363 | 795.968561353344 | 553.376749612315 | 427.033696369448 | 306.993640250539 | 788.096114550995 | 754.440267444613 | Cep350 |
| **ENSMUSG00000033684** | -0.212681081959202 | 0.372030012735101 | 0.873663802472542 | 1020.06698600645 | 1067.69902060668 | 1119.65613327111 | 1065.48472467727 | 1693.90032893214 | 1497.0526520584 | 795.383661184888 | 966.117752554972 | Qsox1 |
| **ENSMUSG00000033685** | -0.409977893830473 | 0.291072587518347 | 0.851785577030345 | 1576.63143541298 | 2102.8214120568 | 2700.23579731402 | 1450.97259601738 | 2907.1140098997 | 4699.08193562777 | 1512.68646557807 | 1286.34779413218 | Ucp2 |
| **ENSMUSG00000033688** | -0.506999231737235 | 0.108648919950531 | 0.762573773680082 | 104.807591122009 | 172.656134529838 | 105.056492639978 | 120.054616865045 | 124.825234323377 | 152.885279009232 | 138.463386043966 | 298.51953028384 | 1300017J02F |
| **ENSMUSG00000033697** | 0.230979712289179 | 0.360054873654013 | 0.868239937412421 | 112.035700854562 | 131.935348084121 | 107.895857305923 | 122.868396947819 | 114.970610561005 | 57.4848649074714 | 98.902418602833 | 132.434118889558 | Arhgap39 |
| **ENSMUSG00000033701** | -0.0522724208576797 | 0.7850372396268 | 0.961141387070613 | 1032.71617803842 | 941.464582624964 | 1042.04683240194 | 1223.9943360069 | 1165.03552035152 | 1451.79860947167 | 926.559500594962 | 854.308619394115 | Acbd6 |
| **ENSMUSG00000033703** | -0.220404458292606 | 0.322716222325988 | 0.855977678491063 | 69.5705561758165 | 68.4109212288036 | 58.6802030962037 | 93.7926694258161 | 81.0269064906132 | 103.961989726278 | 82.2451691539348 | 71.6446872681215 | Fuk |
| **ENSMUSG00000033705** | 0.281290803799299 | 0.190259386174588 | 0.808824684419004 | 91.2548853734735 | 92.8433930962335 | 113.574586637814 | 132.247663890401 | 87.5966556655277 | 70.9387694602839 | 109.313199508394 | 84.6709940441436 | Stard9 |
| **ENSMUSG00000033706** | 0.0661239588159901 | 0.635197804872323 | 0.932543559775713 | 260.211950371885 | 294.004078138073 | 227.149173275627 | 256.991914226736 | 245.270635863478 | 220.154801773295 | 254.023054095697 | 272.466916731795 | Smyd5 |
| **ENSMUSG00000033707** | 0.0753031956991657 | 0.308202020392887 | 0.851785577030345 | 1916.35259284294 | 1958.66982803896 | 1985.66235638444 | 1898.36362917852 | 1772.73731903112 | 1783.25389436369 | 1952.02141979276 | 1855.16319001848 | Lrrc24 |
| **ENSMUSG00000033712** | 0.0073868483203781 | 0.948465032443419 | 0.991217716191233 | 437.300638819418 | 479.69086433054 | 436.315703666934 | 505.542488205149 | 433.603445544362 | 430.524945689999 | 485.14239019916 | 499.341759747513 | Ccar2 |
| **ENSMUSG00000033713** | -0.0494218945180612 | 0.768923622914902 | 0.95879680569409 | 268.343573821006 | 381.96097686082 | 362.492222352355 | 278.564228194674 | 286.879047304603 | 342.463024980681 | 360.213019332423 | 347.368180693922 | Foxn3 |
| **ENSMUSG00000033715** | -0.21313011203219 | 0.365110369808244 | 0.86924083181804 | 288.220875585525 | 275.272516373043 | 216.738169500494 | 314.205442576484 | 323.012667766634 | 332.67836712409 | 198.845915296222 | 414.67076570337 | Akr1c14 |
| **ENSMUSG00000033720** | -0.495457323456257 | 0.0992217109352117 | 0.759800953500741 | 44.2721721118832 | 27.6901347830872 | 19.8755526616174 | 41.2687745473591 | 45.9882442244021 | 50.1463715150282 | 41.6431236222455 | 49.9341759747513 | Sfxn5 |
| **ENSMUSG00000033721** | -0.431767577224342 | 0.108287321971497 | 0.762287160109383 | 59.631905293557 | 53.7514381083457 | 42.5904699891801 | 45.0204813243917 | 91.9764884488041 | 75.8310983885793 | 55.1771387994752 | 48.8486504100828 | Vav3 |
| **ENSMUSG00000033722** | 0.269326092125063 | 0.145028275911323 | 0.780607990324772 | 650.529875929712 | 781.839099757755 | 735.395448479843 | 682.810633419941 | 529.959766776443 | 435.417274618294 | 754.781615653199 | 643.716659848425 | BC034090 |
| **ENSMUSG00000033728** | -0.133486806476697 | 0.2847191489687 | 0.851125503777472 | 285.510334435818 | 289.117583764587 | 300.026199701558 | 275.750448111899 | 327.39250054991 | 341.239942748607 | 326.898520434627 | 267.039288908453 | Lrrc14 |
| **ENSMUSG00000033732** | -0.0409113997254516 | 0.789132897274069 | 0.961659329450003 | 964.049135579171 | 1149.14059349812 | 1148.99623481921 | 851.63743838641 | 1058.82457535707 | 878.173042629031 | 1178.50039850955 | 1115.92028047923 | Sf3b3 |
| **ENSMUSG00000033735** | -0.205107869883794 | 0.198005241751241 | 0.812921911423739 | 2875.88415983927 | 2544.23473712836 | 2443.74652249029 | 3026.68944237109 | 3537.8099306915 | 3710.83149211209 | 2703.67980117429 | 2603.09030407508 | Spr |
| **ENSMUSG00000033739** | -0.143223858086441 | 0.520735438349594 | 0.905962179970576 | 128.298947752804 | 93.6578088251478 | 101.270673085384 | 153.819977858338 | 135.774816281568 | 162.669936865823 | 110.35427759895 | 118.322286548867 | Fkbpl |
| **ENSMUSG00000033751** | -0.00624066054658048 | 0.970534109610781 | 0.9951105526746 | 1451.94654252646 | 1126.33695308852 | 1116.81676860517 | 1455.66222948867 | 1351.17841364077 | 1575.32991491113 | 1106.66601026117 | 1140.8873684666 | Gadd45gip1 |
| **ENSMUSG00000033752** | 0.552476223414632 | 0.197560959921191 | 0.812921911423739 | 49.6932544112975 | 27.6901347830872 | 19.8755526616174 | 60.9652351267805 | 27.3739548954774 | 40.3617136584374 | 17.6983275394543 | 22.7960368580387 | Mnd1 |
| **ENSMUSG00000033760** | -0.0132641594821427 | 0.932737366092177 | 0.988114361264825 | 141.85165350134 | 133.56417954195 | 175.094154399963 | 157.571684635371 | 152.199189218854 | 139.43137445642 | 170.736806851206 | 149.802527924254 | Rbm4b |

| | | | | | | | | | | | | |
|---|---|---|---|---|---|---|---|---|---|---|---|---|
| **ENSMUSG00000033762** | -0.24094894560342 | 0.555551297704108 | 0.916239900758826 | 11.7456783153976 | 10.5874044758863 | 13.2503684410783 | 15.9447538023887 | 9.85462376237187 | 14.6769867848863 | 20.8215618111227 | 15.1973579053591 | Recql4 |
| **ENSMUSG00000033763** | 0.259800162087999 | 0.13651669996872 | 0.776456211338551 | 1397.73571953231 | 1443.9590873651 | 1447.12952474348 | 1238.06323642077 | 1071.9640737069 | 841.480575666815 | 1388.79817280189 | 1312.40040768423 | Mtss1l |
| **ENSMUSG00000033767** | -0.0540008939877712 | 0.761897349613816 | 0.957666649379949 | 421.037391921175 | 361.600583637962 | 358.706402797761 | 313.267515882226 | 377.760577557588 | 277.639666680766 | 425.80093903746 | 427.697072479392 | Tmem131l |
| **ENSMUSG00000033768** | -0.500156149020492 | 0.0685449573576669 | 0.72237439886039 | 121.974351736821 | 211.748089517725 | 170.36187995672 | 159.447538023887 | 249.650468646754 | 303.324393554317 | 252.981976005141 | 133.519644454226 | Nrxn2 |
| **ENSMUSG00000033769** | 0.127261866659396 | 0.356201152554518 | 0.864965376583892 | 471.634160049041 | 534.2567181678 | 600.052399403115 | 448.328959855401 | 437.983278327639 | 436.640356850368 | 500.758561557502 | 504.769387570856 | Exoc6b |
| **ENSMUSG00000033773** | 0.0755936967808234 | 0.614665636966854 | 0.928075610917191 | 220.457346842847 | 188.13003337921 | 206.327165725361 | 166.950951577953 | 179.573144114332 | 166.339183562045 | 201.96914956789 | 193.223550510994 | Rpap2 |
| **ENSMUSG00000033777** | -0.528610261904138 | 0.138030502021622 | 0.779711473022872 | 13.5527057485357 | 15.4738988493722 | 25.5542819935081 | 15.0068271081306 | 19.7092475247437 | 30.5770558018465 | 21.8626399016789 | 28.2236646813812 | Tlr13 |
| **ENSMUSG00000033781** | 0.0514491213043711 | 0.693227897585132 | 0.94469184542218 | 299.966553900923 | 317.622134276588 | 280.15064703994 | 316.081295965 | 304.398378437709 | 245.839528646846 | 298.789411989611 | 321.315567141878 | Asb13 |
| **ENSMUSG00000033788** | -0.147229685614321 | 0.22717775310173 | 0.832048466795696 | 1758.23769244336 | 1813.70382829221 | 1400.7532351997 | 1622.61318106662 | 1672.00116501576 | 2015.63951845772 | 1794.81862811878 | 1823.68294864309 | Dysf |
| **ENSMUSG00000033790** | -0.121347769021893 | 0.430081317589803 | 0.886017087288711 | 132.81651633565 | 131.120932355207 | 138.182413742673 | 114.427056699496 | 133.58489988993 | 124.754387671534 | 154.079557402308 | 148.717002359586 | Tubgcp5 |
| **ENSMUSG00000033792** | 0.165045409526137 | 0.758242173970626 | 0.95677573228595 | 33.4300075130547 | 100.173134656462 | 121.146225747001 | 33.7653609932938 | 42.7033696369448 | 25.6847268735511 | 108.272121417838 | 80.3288917854696 | Atp7a |
| **ENSMUSG00000033793** | 0.0603339004557133 | 0.687651319116203 | 0.944452327903597 | 730.039082987788 | 639.316347197748 | 729.716719147953 | 747.527575323754 | 764.28082068173 | 799.895779776304 | 590.291277345329 | 576.414074838978 | Atp6v1h |
| **ENSMUSG00000033799** | 0.119755582359112 | 0.66918036881175 | 0.942250071683273 | 162.632468982428 | 207.676010873154 | 234.720812384815 | 159.447538023887 | 192.712642464161 | 90.5080851734656 | 219.667477107345 | 199.736703899005 | Fam208b |
| **ENSMUSG00000033808** | -0.0126167925135828 | 0.931346535273834 | 0.987775860606096 | 270.150601254144 | 250.840044505613 | 349.241853911277 | 285.129715054481 | 290.163921892061 | 318.001380339203 | 275.885693997376 | 281.151121249143 | Tmem87a |
| **ENSMUSG00000033809** | -0.317403721610551 | 0.0323127354970242 | 0.671606027118506 | 143.658680934478 | 177.542628903324 | 139.128868631322 | 167.888878272211 | 191.617684268342 | 216.485555077073 | 186.352978209548 | 189.966973816989 | Alg3 |
| **ENSMUSG00000033813** | 0.27760975639526 | 0.0876294543711177 | 0.74814562510601 | 960.435080712895 | 925.176268046677 | 1002.29572707871 | 880.713165908413 | 735.811907590433 | 579.740978003009 | 875.546674157711 | 916.18357658022 | Tcea1 |
| **ENSMUSG00000033819** | 0.0397301050195585 | 0.801373330063255 | 0.963461896828986 | 543.915257374565 | 475.618785685968 | 507.299820315568 | 479.28054076592 | 419.368988998714 | 402.3940543523 | 545.524919451416 | 582.927228226989 | Ppp1r16a |
| **ENSMUSG00000033831** | -0.367095903552342 | 0.297308902762347 | 0.851785577030345 | 494.222002963268 | 1097.0179868476 | 554.62256474799 | 600.273084325223 | 562.808512651016 | 648.233582999146 | 825.574925811016 | 1505.62395819522 | Fgb |
| **ENSMUSG00000033845** | 0.0749850107437515 | 0.518183681747008 | 0.905473752527583 | 1712.15849289834 | 1396.72297508807 | 1512.43491206022 | 1777.37108561922 | 1632.58266996627 | 1537.41436571684 | 1416.9072812469 | 1487.17002359586 | Mrpl15 |
| **ENSMUSG00000033849** | 0.227682066157015 | 0.583527360743113 | 0.922450244597442 | 116.553269437407 | 161.254314325037 | 40.6975602118832 | 182.895705380341 | 77.7420319031559 | 111.300483118721 | 87.4505596067155 | 151.973579053591 | B3galt2 |
| **ENSMUSG00000033855** | -0.00074400948620353 | 0.997487511477451 | 0.999262355603273 | 217.74680569314 | 284.231089391101 | 305.704929033448 | 168.826804966469 | 320.822751374995 | 218.931719541221 | 231.119336103462 | 206.249857287016 | Ston1 |
| **ENSMUSG00000033857** | -0.0327623186714008 | 0.879741094157643 | 0.978393501240173 | 103.000563688871 | 83.0704043492615 | 110.735221971868 | 146.316564304273 | 130.300025302473 | 114.969729814943 | 98.902418602833 | 108.552556466851 | Engase |
| **ENSMUSG00000033860** | -0.314846206398252 | 0.245608211452838 | 0.832048466795696 | 684.863397159336 | 1239.54073940761 | 665.357786719858 | 775.665376151499 | 770.850569856644 | 889.180782717696 | 1024.42084110724 | 1502.36738150121 | Fgg |
| **ENSMUSG00000033862** | 0.00148731096842664 | 0.992158220289183 | 0.998990770555763 | 236.72059374109 | 181.614707547895 | 212.005895057252 | 230.729966787508 | 202.567266226533 | 222.600966237442 | 192.599446752885 | 242.072200921077 | Cdk10 |
| **ENSMUSG00000033863** | -0.307853414816257 | 0.0500533202171092 | 0.703396849069134 | 2765.65548641785 | 2816.24959058575 | 2735.25462819401 | 2432.98184490567 | 3485.25193729219 | 4143.80260226624 | 3035.78371206169 | 2644.34027553248 | Klf9 |
| **ENSMUSG00000033871** | 0.186250711181405 | 0.571463949537876 | 0.921130164032017 | 176.185174730964 | 145.780415475665 | 318.008842585878 | 371.418970926232 | 191.617684268342 | 161.446854633749 | 235.283648465687 | 299.605055848508 | Ppargc1b |
| **ENSMUSG00000033880** | -0.642202712341362 | 0.0452539570750235 | 0.695534138838186 | 314.422773366028 | 385.218639776477 | 401.296872786941 | 354.536290429585 | 743.476614961167 | 861.049891379997 | 320.65205189129 | 347.368180693922 | Lgals3bp |
| **ENSMUSG00000033883** | -0.195064849435369 | 0.242503932388012 | 0.832048466795696 | 90.3513716569045 | 88.7713144516618 | 100.324218196735 | 103.171936368398 | 124.825234323377 | 108.854318654574 | 105.14888714617 | 98.7828263848342 | D3Ertd254e |
| **ENSMUSG00000033885** | 0.0506376132022421 | 0.604454243944029 | 0.926448151609826 | 554.757421973394 | 610.811796685746 | 606.677583623654 | 545.87333605825 | 537.624474147177 | 540.602346576646 | 573.634027896431 | 586.183804920994 | Pxk |
| **ENSMUSG00000033898** | -0.390750381334746 | 0.173673875157193 | 0.799889021902745 | 197.869503928621 | 241.067055758641 | 174.147699511314 | 224.164479927701 | 186.142893289246 | 277.639666680766 | 193.640524843441 | 440.723379255414 | Cfhr2 |
| **ENSMUSG00000033900** | -0.514918802935979 | 0.255116957843469 | 0.837406774112203 | 35.2370349461928 | 27.6901347830872 | 7.57163910918757 | 30.9515809105193 | 21.8991639163819 | 50.1463715150282 | 35.3966550789086 | 37.9933947633978 | Map9 |
| **ENSMUSG00000033902** | 0.286111552428352 | 0.255181134944023 | 0.837406774112203 | 182.509770746947 | 171.841718800923 | 224.309808609682 | 143.502784221499 | 130.300025302473 | 94.1773318696872 | 183.22974393788 | 183.453820428978 | Mapkbp1 |
| **ENSMUSG00000033904** | -0.246407822606017 | 0.457087183616081 | 0.892805623137304 | 23.4913566307952 | 25.2468875963442 | 35.965285768641 | 41.2687745473591 | 40.5134532453066 | 22.0154801773295 | 47.8895921655823 | 37.9933947633978 | Ccp110 |
| **ENSMUSG00000033909** | 0.0386945651521801 | 0.83513783701882 | 0.968771050291583 | 335.203588847116 | 528.555808065399 | 452.405436773958 | 408.936038696558 | 419.368988998714 | 331.455284892016 | 454.951125573032 | 473.289146195469 | Usp36 |
| **ENSMUSG00000033910** | -0.775397073047748 | 0.0150763764696457 | 0.635862253526407 | 181.606257030378 | 166.140808698523 | 169.415425068072 | 166.950951577953 | 407.324448844704 | 418.29412336926 | 181.147587756768 | 164.999885829613 | Gucy1a1 |
| **ENSMUSG00000033916** | -0.0644098966750114 | 0.690272547598698 | 0.94469184542218 | 2436.77649358671 | 1863.38318775598 | 2153.18487167522 | 2383.27173010999 | 2482.27022992189 | 2797.18906475292 | 1949.93926361164 | 2011.47887133074 | Chmp2a |
| **ENSMUSG00000033917** | -0.0707437813480922 | 0.556174464423964 | 0.916239900758826 | 702.030157774148 | 688.181290932608 | 705.108892043093 | 795.361836730921 | 747.856447744443 | 883.065371557327 | 697.522320672611 | 708.848193728535 | Gde1 |
| **ENSMUSG00000033918** | 0.0664941844364981 | 0.563286808409094 | 0.918316170047219 | 1545.91196904964 | 1301.4363348051 | 1310.8400207781 | 1640.43378825752 | 1398.26161606099 | 1445.6831983113 | 1316.96378455351 | 1376.44641599967 | Parl |
| **ENSMUSG00000033931** | 0.133556471014645 | 0.485096241291392 | 0.899236955296196 | 88.5443442237664 | 101.801966114291 | 91.8061241988993 | 101.296082979881 | 104.021028602814 | 85.6157562451702 | 92.6559500594962 | 67.3025850094475 | Rbm34 |
| **ENSMUSG00000033933** | -0.131959773392876 | 0.4305280358371 | 0.886017087288711 | 243.045189757073 | 184.872370463553 | 171.308334845369 | 208.219726125312 | 246.365594059297 | 187.131581507301 | 231.119336103462 | 219.276164063039 | Vhl |
| **ENSMUSG00000033938** | 0.0407639031543782 | 0.84776540430447 | 0.972455319662159 | 9296.2526297789 | 6346.74177542936 | 6694.27542741046 | 8804.31787900136 | 7709.60065676226 | 10184.605746479 | 5992.44548924112 | 6388.31794807417 | Ndufb7 |
| **ENSMUSG00000033940** | 0.063305673129703 | 0.597963811221772 | 0.92450103938896 | 1619.09658009173 | 1277.00386293767 | 1322.19747944188 | 1573.84099296519 | 1315.04479317874 | 1439.56778715093 | 1323.21025309685 | 1465.45951230249 | Brk1 |
| **ENSMUSG00000033943** | 0.202729611198149 | 0.295938188332714 | 0.851785577030345 | 523.134441893477 | 642.574010113405 | 600.052399403115 | 532.742362338636 | 434.698403740182 | 362.032340693862 | 613.194995337564 | 586.183804920994 | Mga |
| **ENSMUSG00000033948** | -0.567709690781173 | 0.135931700799909 | 0.776087404131513 | 16.2632468982428 | 30.9477976987445 | 34.0723759913441 | 15.9447538023887 | 41.6084114411257 | 24.4616446414772 | 29.1501865355718 | 48.8486504100828 | Zswim5 |
| **ENSMUSG00000033949** | 0.0712424406797521 | 0.86083580730919 | 0.974868041734905 | 12.6491920319666 | 13.0306516626293 | 16.0897331070236 | 13.1309737196143 | 9.85462376237187 | 17.123151249034 | 15.616171358342 | 9.76973008201657 | Trim36 |
| **ENSMUSG00000033953** | 0.0517394935816953 | 0.730405523920548 | 0.952921973914044 | 1726.61471236345 | 1871.52734504513 | 1943.07188639526 | 1607.60635395849 | 1552.65072167148 | 1415.10614250946 | 2049.88276030503 | 1877.95922687652 | Ppp3r1 |
| **ENSMUSG00000033955** | -0.23155714283709 | 0.182426105609251 | 0.804505478598291 | 929.715614349547 | 822.559886203472 | 1041.10037751329 | 837.568537972538 | 1320.51958415783 | 1170.48969609468 | 896.368235968834 | 876.019130687486 | Tnks1bp1 |
| **ENSMUSG00000033960** | 0.0818360911655445 | 0.630493873422597 | 0.931825800800958 | 3499.30862427191 | 3676.27260031928 | 3197.12461385445 | 2199.43809803539 | 3031.93924422308 | 2827.76612055476 | 3039.94802442392 | 2979.76767501505 | Jcad |
| **ENSMUSG00000033961** | -0.179282949024119 | 0.644631307262874 | 0.934857223264556 | 25.2983840639333 | 50.4937751926884 | 51.1085639870161 | 32.8274342990356 | 31.7537876787538 | 28.1308913376988 | 74.9576225200418 | 45.5920737160773 | Zfp446 |
| **ENSMUSG00000033964** | 0.17845735373543 | 0.447402599316924 | 0.89108031640712 | 479.765783498163 | 619.770369703804 | 477.959718767466 | 429.570425970238 | 350.386622662111 | 314.332133642982 | 507.005030100839 | 601.381162826353 | Zbtb41 |
| **ENSMUSG00000033965** | -0.682816360943219 | 0.029351553047886 | 0.665956123973759 | 77.7021796249379 | 127.048853710635 | 108.842312194571 | 142.564857527241 | 191.617684268342 | 283.755077841135 | 111.395355689507 | 146.545951230249 | Slc16a2 |
| **ENSMUSG00000033970** | -0.175903958060988 | 0.578288418622595 | 0.922450244597442 | 58.7283915769879 | 54.56585383726 | 28.3936466594534 | 85.3513291774927 | 67.887408140784 | 69.71568722821 | 59.3414511616998 | 59.7039060567679 | Rfc3 |
| **ENSMUSG00000033972** | 0.224701767410447 | 0.308249452486165 | 0.851785577030345 | 112.035700854562 | 86.3280672649188 | 93.6990339761962 | 97.5443762028488 | 66.7924499449649 | 64.8233582999146 | 103.066730965058 | 97.6973008201657 | Zfp944 |
| **ENSMUSG00000033983** | 0.0872489078880708 | 0.609996394926022 | 0.927932112008768 | 111.132187137993 | 107.502876216691 | 118.306861081056 | 118.178763476528 | 124.825234323377 | 88.0619207093179 | 114.518589961175 | 99.8683519495027 | Coil |
| **ENSMUSG00000033985** | -0.315753282145352 | 0.310440400290338 | 0.851785577030345 | 35.2370349461928 | 48.8649437348597 | 23.6613722162112 | 25.3240207449704 | 39.4184950494875 | 36.6924669622158 | 51.0128264372507 | 39.0789203280663 | Tesk2 |
| **ENSMUSG00000033991** | 0.212337352640395 | 0.485800314982133 | 0.899236955296196 | 192.448421629207 | 254.912123150185 | 276.364827485346 | 183.8336320746 | 154.389105610493 | 105.185071958352 | 260.269522639034 | 262.697186649779 | Ttc37 |
| **ENSMUSG00000033998** | 0.0130897199688739 | 0.972087049661336 | 0.995230273442615 | 101.193536255733 | 30.9477976987445 | 58.6802030962037 | 50.6480414899407 | 68.9823663366031 | 37.9155491942896 | 67.6700758861489 | 64.046008315442 | Kcnk1 |
| **ENSMUSG00000034006** | 0.086191358695799 | 0.44684151265434 | 0.89108031640712 | 769.793686516826 | 756.592212161411 | 789.343377132805 | 902.285479876351 | 727.05224202388 | 687.372214425509 | 792.26042691322 | 822.828378018729 | Pqlc1 |
| **ENSMUSG00000034007** | 0.357894772179018 | 0.0549551987903178 | 0.703396849069134 | 712.872322372977 | 794.869751420385 | 750.538726698218 | 667.803806311811 | 487.256397139498 | 423.186452297555 | 697.522320672611 | 674.111375659143 | Scaper |
| **ENSMUSG00000034021** | 0.182055882137901 | 0.300620682861102 | 0.851785577030345 | 355.080890611635 | 410.465527372822 | 461.869985660442 | 333.901903155905 | 325.202584158272 | 271.524255520397 | 378.952424962434 | 399.473407798011 | Pds5b |
| **ENSMUSG00000034022** | 0.133774795900491 | 0.347103976362548 | 0.861266047293794 | 942.364806381514 | 1083.98733518497 | 1060.97593017491 | 973.567908639971 | 955.898504950072 | 737.518585940537 | 1086.88552654061 | 919.440153274226 | Cpsf1 |
| **ENSMUSG00000034024** | 0.171248793384589 | 0.0623097069878722 | 0.715615581523459 | 3696.27461448396 | 3221.82862358508 | 3482.95399022628 | 3864.25798034362 | 3312.24854235277 | 3206.92161249766 | 3042.03018060503 | 3106.77416608127 | Cct2 |
| **ENSMUSG00000034028** | -0.139859402688213 | 0.844626117872026 | 0.971865371253468 | 6.32459601598331 | 8.14415728914329 | 30.2865564367503 | 5.62756016554897 | 8.75966556655277 | 22.0154801773295 | 4.16431236222455 | 20.6249857287016 | Cd226 |
| **ENSMUSG00000034032** | 0.0992154784343884 | 0.590576090353371 | 0.923935529355527 | 311.712232216321 | 370.55915665602 | 323.687571917769 | 271.998741334867 | 317.537876787538 | 214.039390612925 | 340.432535611857 | 320.23004157721 | Rpap1 |
| **ENSMUSG00000034035** | 0.335031541717168 | 0.43697447393096 | 0.888701139843674 | 14.4562194651047 | 25.2468875963442 | 23.6613722162112 | 18.7585338851632 | 28.4689130912965 | 11.0077400886647 | 10.4107809055614 | 15.1973579053591 | Ccdc17 |
| **ENSMUSG00000034037** | -0.285960270629099 | 0.0552710398019 | 0.703396849069134 | 599.029594085277 | 644.202841571234 | 653.053873167428 | 579.638697051544 | 901.150595159117 | 826.803588881929 | 657.961353231478 | 633.946929766408 | Fgd5 |
| **ENSMUSG00000034040** | 0.0283809900404883 | 0.910871037972692 | 0.986166948757759 | 99.3865088225949 | 156.367819951551 | 179.826428843205 | 99.4202295913651 | 125.920192519196 | 102.738907494204 | 146.792010768415 | 148.717002359586 | Galnt17 |
| **ENSMUSG00000034041** | -0.715178236685789 | 0.0622819767851191 | 0.715615581523459 | 59.631905293557 | 41.5352021746308 | 63.4124775394459 | 62.8410885152968 | 124.825234323377 | 145.546785616789 | 45.80743598447 | 57.5328549274309 | Lyl1 |
| **ENSMUSG00000034042** | 0.14899945800331 | 0.425429087994188 | 0.885066977635295 | 921.583990900426 | 908.887953468391 | 859.38103889279 | 920.106087067256 | 721.577451044785 | 576.071731306788 | 952.586452858865 | 1004.11114731837 | Gpbp1l1 |
| **ENSMUSG00000034055** | 0.202422823280803 | 0.484893380405096 | 0.899236955296196 | 1092.34808333198 | 1379.62024478087 | 1394.12805097916 | 936.050840869645 | 750.046364136081 | 658.018240855737 | 1632.41044599202 | 1132.20316394925 | Phka1 |
| **ENSMUSG00000034057** | 0.480464660059088 | 0.131164279053674 | 0.77303252008843 | 39.754603529038 | 48.0505280059454 | 41.6440151005317 | 43.1446279358754 | 26.2789966996583 | 15.9000690169602 | 37.4788112600209 | 43.4210225867403 | Myrfl |
| **ENSMUSG00000034063** | 0.0863385581130279 | 0.880436476274721 | 0.978433072023387 | 12.6491920319666 | 13.8450673915436 | 14.1968233297267 | 2.81378008277448 | 12.0445401540101 | 9.78465785659088 | 5.20539045278068 | 14.1118323406906 | 4930590J08F |
| **ENSMUSG00000034064** | 0.105411280145931 | 0.476857882672965 | 0.897622440754504 | 247.562758339918 | 232.922898469498 | 277.311282373995 | 309.515809105193 | 258.410134213307 | 272.747337752471 | 241.530117009024 | 219.276164063039 | Poglut1 |
| **ENSMUSG00000034066** | 0.345630430381153 | 0.192930632923219 | 0.808824684419004 | 383.993329541844 | 544.844122643686 | 562.194203857177 | 422.067012416173 | 329.582416941548 | 226.270212933664 | 535.114138545854 | 413.585240138701 | Farp2 |
| **ENSMUSG00000034071** | -0.318059365688439 | 0.633999152794715 | 0.932543559775713 | 4.51756858284523 | 17.1027303072009 | 12.3039135524298 | 7.50341355406529 | 8.75966556655277 | 2.44616446414772 | 22.903717992235 | 17.3684090346961 | Zfp551 |
| **ENSMUSG00000034075** | 0.245354572453682 | 0.214179996742184 | 0.824248286808591 | 2315.70565556646 | 2799.96127600746 | 2783.52382751508 | 2281.03772043585 | 1841.71968536772 | 1552.09135250173 | 2669.32422418593 | 2523.84693785428 | Zdhhc5 |
| **ENSMUSG00000034083** | 0.015141686133663 | 0.907074368128756 | 0.98504921011344 | 269.247087537575 | 237.809392842984 | 251.757000380487 | 289.819348525772 | 245.270635863478 | 273.970419984545 | 239.447960827911 | 278.980070119806 | Ccdc174 |
| **ENSMUSG00000034088** | 0.226002766051078 | 0.287833192778181 | 0.851785577030345 | 15603.6818851474 | 18568.6786192467 | 19373.9315706337 | 14566.0015618292 | 12157.3208481794 | 10266.552256028 | 17755.5868344349 | 18055.5467171313 | Hdlbp |
| **ENSMUSG00000034101** | -0.24476207316303 | 0.112169443672677 | 0.764116514278315 | 274.66816983699 | 256.540954608014 | 324.634026806417 | 268.247034557834 | 360.241246424483 | 386.49398533534 | 289.419709174606 | 296.348479154503 | Ctnnd1 |
| **ENSMUSG00000034105** | 0.215148390828144 | 0.237262348390006 | 0.832048466795696 | 281.896279569542 | 308.663561258531 | 319.901752363175 | 285.129715054481 | 255.12525962585 | 184.685417043153 | 323.775286162958 | 264.868237779116 | Tldc1 |
| **ENSMUSG00000034108** | -0.0726735110253981 | 0.685470082329084 | 0.944452327903597 | 571.020668871636 | 444.670987987223 | 513.925004536107 | 598.397230936707 | 615.366506050332 | 687.372214425509 | 452.868969391919 | 483.058876277486 | Ccs |
| **ENSMUSG00000034109** | 0.0202518045836171 | 0.8706664997007 | 0.975613767803763 | 623.424464432641 | 686.552459474779 | 684.286884492827 | 521.487242007538 | 604.416924092141 | 600.533375948265 | 649.632728507029 | 626.348250813729 | Golim4 |
| **ENSMUSG00000034110** | -0.483838458299984 | 0.114395743484375 | 0.767518180080285 | 18.9737880479499 | 28.5045505120015 | 35.965285768641 | 46.8963347129081 | 48.1781606160403 | 47.7002070508805 | 44.7663578939139 | 41.2499714574033 | Kctd7 |
| **ENSMUSG00000034111** | -0.0616333767781548 | 0.749191512510906 | 0.956197192463153 | 159.921927832721 | 198.717437855096 | 199.701981504822 | 138.813150750208 | 195.997517051618 | 140.654456688494 | 185.311900118992 | 205.164331722348 | Tmed8 |
| **ENSMUSG00000034116** | -0.447389788133817 | 0.12111548920597 | 0.769868040577702 | 24.3948703473642 | 29.3189662409158 | 31.2330113253987 | 36.5791410760683 | 35.0386622662111 | 45.2540425867328 | 31.2323427166841 | 54.2762782334254 | Vav1 |
| **ENSMUSG00000034118** | -0.39444473577396 | 0.234671541592169 | 0.832048466795696 | 189.737880479499 | 158.811067138294 | 185.505158175096 | 348.908730264036 | 394.184950494875 | 365.701587390084 | 172.818963032319 | 227.960368580387 | Tpst1 |
| **ENSMUSG00000034120** | -0.221624844701455 | 0.071417132652929 | 0.727489215531785 | 711.968808656408 | 754.963380703583 | 670.0900611631 | 826.31341764144 | 744.571573156986 | 900.188522806361 | 936.970281500523 | 873.848079558149 | Srsf2 |

| | | | | | | | | | | | | |
|---|---|---|---|---|---|---|---|---|---|---|---|---|
| **ENSMUSG00000034121** | 0.170802022231424 | 0.60494783373019 | 0.926448151609826 | 43.3686583953142 | 52.122606650517 | 32.1794662140472 | 57.2135283497478 | 45.9882442244021 | 28.1308913376988 | 60.3825292522559 | 29.3091902460497 | Mks1 |
| **ENSMUSG00000034126** | -0.0393765500583883 | 0.787391823575397 | 0.961224689823763 | 234.010052591383 | 281.787842204358 | 273.525462819401 | 204.468019348279 | 263.884925192402 | 244.616446414772 | 260.269522639034 | 252.927456567762 | Pomt2 |
| **ENSMUSG00000034135** | 0.350910080340962 | 0.149137567764316 | 0.782024490202776 | 2299.44240866822 | 2792.63153444723 | 4054.61274296995 | 2738.74594723383 | 2367.29961936089 | 1538.63744794892 | 3035.78371206169 | 2376.21546105936 | Sik3 |
| **ENSMUSG00000034152** | 0.146169469676833 | 0.308161209078621 | 0.851785577030345 | 1803.41337827181 | 1852.7957832801 | 1968.62616838877 | 1583.22025990778 | 1519.80197579691 | 1339.27504412088 | 1877.06379727271 | 1775.91982379768 | Exoc3 |
| **ENSMUSG00000034154** | 0.214183792729385 | 0.320770406082163 | 0.855228644171892 | 306.291149916906 | 362.414999366876 | 324.634026806417 | 257.929840920994 | 223.371471947096 | 195.693157131818 | 332.103910887408 | 326.743194965221 | Ino80 |
| **ENSMUSG00000034157** | 0.329794044102868 | 0.130228108272653 | 0.77303252008843 | 1091.44456961541 | 1337.27062687733 | 1396.02096075646 | 1218.36677584135 | 871.586723872001 | 661.687487551958 | 1251.37586484848 | 1226.64388807541 | Cipc |
| **ENSMUSG00000034158** | 0.180192562915736 | 0.243741536100695 | 0.832048466795696 | 1952.49314150571 | 1993.68970438228 | 2006.48436393471 | 2031.54921976318 | 2039.90711881098 | 1287.90559037377 | 1819.80450229213 | 1897.49868704055 | Lrrc58 |
| **ENSMUSG00000034160** | 0.110880402017194 | 0.552475838414382 | 0.914977712327965 | 456.274426867368 | 564.39010013763 | 486.477812765302 | 513.045901759214 | 452.217734873287 | 326.562955963721 | 536.15521663641 | 554.703563545607 | Ogt |
| **ENSMUSG00000034161** | 0.0623453787827545 | 0.787031681884678 | 0.961141387070613 | 248.466272056487 | 179.171460361152 | 205.380710836713 | 249.488500672671 | 198.187433443257 | 304.547475786391 | 168.654650670094 | 174.76961591163 | Scx |
| **ENSMUSG00000034163** | 0.134483632076315 | 0.612305621746389 | 0.928055903343287 | 175.281661014395 | 201.975100770753 | 184.558703286447 | 151.006197775564 | 127.015150715015 | 107.6312364225 | 246.735507461804 | 167.17093695895 | Zfc3h1 |
| **ENSMUSG00000034164** | -1.07083525756446 | 0.050313694521858 | 0.703396849069134 | 4.51756858284523 | 4.88649437348597 | 13.2503684410783 | 11.2551203310979 | 24.0890803080201 | 25.6847268735511 | 10.4107809055614 | 10.8552556466851 | Emid1 |
| **ENSMUSG00000034165** | -0.141475747162058 | 0.282911857603375 | 0.850587941933635 | 1039.94428777097 | 1021.27732405857 | 1104.51285505274 | 951.995594672034 | 1237.30276127558 | 1306.25182385488 | 1007.76359165834 | 992.170366107016 | Ccnd3 |
| **ENSMUSG00000034168** | -0.704809372981334 | 0.00026560770640537 | 0.409921226885629 | 765.276117933981 | 724.829998733753 | 605.731128735006 | 711.886360941944 | 1345.70362266167 | 1405.32148465286 | 994.22957648111 | 832.598108100745 | Irf2bpl |
| **ENSMUSG00000034171** | -0.535417260750632 | 0.287381388692283 | 0.851785577030345 | 12.6491920319666 | 26.8757190541728 | 12.3039135524298 | 14.0689004138724 | 19.7092475247437 | 15.9000690169602 | 13.5340151772298 | 46.6775992807458 | Faah |
| **ENSMUSG00000034173** | -0.480754848477908 | 0.252474884271097 | 0.83723604018117 | 22.5878429142261 | 21.1748089517725 | 9.46454888648447 | 15.0068271081306 | 27.3739548954774 | 29.3539735697726 | 27.0680303544596 | 11.9407812113536 | Zbed5 |
| **ENSMUSG00000034175** | 0.0749113579276282 | 0.527316151109646 | 0.908217276872524 | 482.47632464787 | 481.319695788368 | 430.636974335043 | 467.087493740564 | 481.781606160403 | 424.409534529629 | 387.281049686883 | 474.374671760138 | Rhbdd3 |
| **ENSMUSG00000034187** | -0.0850059509076275 | 0.638600638905862 | 0.933504827868072 | 297.256012751216 | 390.105134149963 | 335.991485470199 | 272.936668029125 | 333.962249724825 | 266.631926592101 | 381.034581143546 | 392.96025441 | Nsf |
| **ENSMUSG00000034189** | 0.0120178072271873 | 0.922313286262906 | 0.986713918026868 | 578.248778604189 | 507.380999113627 | 495.942361651786 | 447.391033161143 | 529.959766776443 | 473.332823812584 | 473.690531203042 | 535.164103381574 | Hsdl1 |
| **ENSMUSG00000034190** | -0.0251907493082749 | 0.715028824504395 | 0.951176382858794 | 1451.04302880989 | 1465.94831204579 | 1409.27132919754 | 1401.26248122169 | 1462.86414961431 | 1416.32922474153 | 1482.49520095194 | 1466.54503786715 | Chmp7 |
| **ENSMUSG00000034192** | 0.0376926106318039 | 0.825063814673568 | 0.96672635991852 | 607.161217534398 | 386.847471234306 | 432.52988411234 | 556.19052969509 | 502.585811880965 | 500.240632918209 | 450.786813210807 | 477.631248454143 | Lsm3 |
| **ENSMUSG00000034194** | 0.0424486744319244 | 0.673334852482713 | 0.943631400280823 | 635.170142748039 | 587.193740547231 | 643.589324280944 | 696.879533833814 | 593.467342133951 | 636.002760678407 | 604.866370613115 | 653.486389930442 | R3hcc1 |
| **ENSMUSG00000034201** | -0.116883494513448 | 0.347667717191954 | 0.861266047293794 | 524.941469326615 | 544.844122643686 | 575.444572298256 | 445.515179772627 | 590.182467546493 | 583.410224699231 | 587.168043073661 | 506.940438700193 | Gas2l1 |
| **ENSMUSG00000034203** | 0.179023506392757 | 0.236733980398287 | 0.832048466795696 | 939.654265231807 | 743.561560498782 | 791.236286910102 | 975.443762028488 | 747.856447744443 | 650.679747463293 | 748.535147109862 | 898.815167545524 | Chchd4 |
| **ENSMUSG00000034205** | -1.05080245818002 | 0.0116871731719881 | 0.635862253526407 | 64.1494738764022 | 64.338842584232 | 58.6802030962037 | 80.6616957062019 | 194.902558855799 | 226.270212933664 | 57.2592949805875 | 77.072315091464 | Loxl2 |
| **ENSMUSG00000034206** | 0.168150689732845 | 0.655157886134211 | 0.937579130270368 | 255.69438178904 | 287.488752306758 | 119.253315969704 | 177.268145214792 | 136.869774477387 | 94.1773318696872 | 244.653351280692 | 271.381391167127 | Polq |
| **ENSMUSG00000034209** | -0.471620617593033 | 0.106413610643439 | 0.762094734041179 | 410.195227322346 | 228.850819824926 | 171.308334845369 | 253.240207449704 | 348.196706270473 | 335.124531588238 | 511.169342463063 | 280.065595684475 | Rasl10a |
| **ENSMUSG00000034210** | 0.0675206528223678 | 0.733139113861702 | 0.953733898884637 | 850.206407291471 | 979.742121883937 | 975.794990196549 | 722.203554578784 | 720.482492848966 | 631.110431750112 | 978.613405122768 | 1035.59138869376 | Efcab14 |
| **ENSMUSG00000034211** | 0.0347683009745003 | 0.775050928667777 | 0.959488181746473 | 880.925873654819 | 777.767021113184 | 667.250696497155 | 766.286109208918 | 802.604357535398 | 691.041461121731 | 748.535147109862 | 776.150778737983 | Mrps17 |
| **ENSMUSG00000034216** | -0.421655666157447 | 0.0152558328213868 | 0.635862253526407 | 324.361424248287 | 332.281617397046 | 264.060913932917 | 367.667264149199 | 487.256397139498 | 502.686797382356 | 382.075659234102 | 354.966859646602 | Vps18 |
| **ENSMUSG00000034218** | 0.289602215288321 | 0.505331013282286 | 0.901494649536949 | 108.421645988285 | 162.068730053951 | 144.807597963212 | 78.7858423176855 | 70.0773245324222 | 34.2463024980681 | 157.202791673977 | 142.203848971575 | Atm |
| **ENSMUSG00000034220** | 0.0351845870308499 | 0.827385396623078 | 0.967319591893432 | 13528.3108781883 | 15969.0636125522 | 13999.0142579992 | 12805.5131567067 | 12437.6301463091 | 10788.8083691235 | 16373.0351301764 | 15344.989382154 | Gpc1 |
| **ENSMUSG00000034235** | 0.33672386910505 | 0.136118599601924 | 0.776456211338551 | 232.203025158245 | 296.447325324816 | 229.042083052924 | 216.661066373635 | 167.528603960322 | 133.315963296051 | 256.10521027681 | 213.848536239696 | Usp54 |
| **ENSMUSG00000034243** | 0.391599424462306 | 0.0860803936431783 | 0.746460787597018 | 1792.57121367299 | 2365.87769249612 | 2418.19224049678 | 1837.39839405174 | 1384.02715951534 | 1062.85845967218 | 1995.74669959611 | 1970.22889987334 | Golgb1 |
| **ENSMUSG00000034245** | 0.122376920992094 | 0.500707393344998 | 0.900927059989498 | 465.309564033058 | 508.195414842541 | 572.60520763231 | 542.121629281217 | 362.431162816121 | 418.29412336926 | 558.017856538089 | 578.585125968315 | Hdac11 |
| **ENSMUSG00000034247** | -0.0759556423462756 | 0.455812519536366 | 0.892805623137304 | 345.142239729375 | 375.445651029505 | 357.759947909113 | 337.653609932938 | 369.000911991036 | 360.809258461789 | 388.322127777439 | 374.506319810635 | Plekhm1 |
| **ENSMUSG00000034248** | 0.254726962469713 | 0.4813493328644 | 0.897732059433541 | 247.562758339918 | 215.820168162297 | 489.317177431247 | 183.8336320746 | 152.199189218854 | 371.816998550453 | 199.886993386778 | 229.045894145055 | Slc25a37 |
| **ENSMUSG00000034252** | 0.271125889332922 | 0.142688332719911 | 0.780607990324772 | 1891.95772249558 | 2145.98544568926 | 2013.10954815525 | 1709.84036363263 | 1439.87002750211 | 1170.48969609468 | 1963.47327878887 | 1856.24871558315 | Senp6 |
| **ENSMUSG00000034254** | 0.0226015658548479 | 0.789981088523385 | 0.961659329450003 | 1732.93930837943 | 1582.40976128054 | 1681.85033712829 | 1750.17121148573 | 1595.35409130842 | 1791.8154699882 | 1642.82122689758 | 1613.0909890974 | Agpat1 |
| **ENSMUSG00000034255** | -0.412003870490164 | 0.242392704566883 | 0.832048466795696 | 316.229800799166 | 330.652785939217 | 372.903226127488 | 226.978260010475 | 621.936255225247 | 569.956320146419 | 259.228444548478 | 208.420908416353 | Arhgap27 |
| **ENSMUSG00000034258** | -0.186857750823722 | 0.618630573610548 | 0.929568738467614 | 103.90407740544 | 112.389370590177 | 100.324218196735 | 77.8479156234274 | 152.199189218854 | 40.3617136584374 | 160.326025945645 | 95.5262496908287 | Flvcr2 |
| **ENSMUSG00000034259** | -0.336429123793144 | 0.109677293041105 | 0.763154315676794 | 603.547162668122 | 493.535931722083 | 573.551662520959 | 640.603932178324 | 825.598479647599 | 967.458045570423 | 577.798340258656 | 548.190410157596 | Exosc4 |
| **ENSMUSG00000034263** | -0.0505183214480966 | 0.676943884705787 | 0.94409858171788 | 373.151164943016 | 364.043830824705 | 331.259211026956 | 334.839829850164 | 344.911831683016 | 324.116791499573 | 385.198893505771 | 398.387882233342 | Ints14 |
| **ENSMUSG00000034265** | 0.116880891445267 | 0.592392180453299 | 0.923935529355527 | 171.667606148119 | 258.169786065842 | 242.292451494002 | 211.033506208086 | 235.416012101106 | 240.94719971855 | 197.804837205666 | 141.118323406906 | Zdhhc14 |
| **ENSMUSG00000034269** | 0.0976715927294479 | 0.526537947704159 | 0.908217276872524 | 590.897970636155 | 567.647763053287 | 571.658752743662 | 529.928582255861 | 516.820268426614 | 401.170972120226 | 621.523620062014 | 570.986447015635 | Setd5 |
| **ENSMUSG00000034271** | -0.316768057114845 | 0.287033004973075 | 0.851785577030345 | 262.922491521592 | 223.96432545144 | 186.451613063744 | 175.392291826276 | 339.43704070392 | 374.263163014601 | 167.613572579538 | 176.940667040967 | Jdp2 |
| **ENSMUSG00000034274** | -0.0475594780838253 | 0.639999456348669 | 0.933921420069286 | 397.54603529038 | 395.806044252364 | 417.386605893965 | 441.763472995594 | 441.268152915096 | 417.071041137186 | 443.499266576914 | 404.901035621353 | Thoc5 |
| **ENSMUSG00000034275** | 0.0573218055469725 | 0.871034393265954 | 0.975613767803763 | 53.3073092775737 | 28.5045505120015 | 49.2156542097192 | 54.3997482669734 | 45.9882442244021 | 22.0154801773295 | 62.4646854333682 | 46.6775992807458 | Igsf9b |
| **ENSMUSG00000034278** | -0.207675190021398 | 0.383538535825137 | 0.875126835008975 | 125.588406603097 | 96.9154717408051 | 97.48485353079 | 158.509611329629 | 157.67398019795 | 168.785348026193 | 112.436433780063 | 113.980184290193 | Dnajc17 |
| **ENSMUSG00000034282** | -0.30441783598224 | 0.633043222004342 | 0.93224876522904 | 18.9737880479499 | 4.88649437348597 | 13.2503684410783 | 14.0689004138724 | 27.3739548954774 | 20.7923979452556 | 3.12323427166841 | 11.9407812113536 | Evpl |
| **ENSMUSG00000034285** | -0.169283550629945 | 0.619444797930615 | 0.929568738467614 | 58.7283915769879 | 100.173134656462 | 66.2518422053913 | 75.9720622349111 | 67.887408140784 | 84.3926740130963 | 47.8895921655823 | 138.947272277569 | Nipsnap1 |
| **ENSMUSG00000034290** | 0.26588125746918 | 0.307604457508211 | 0.851785577030345 | 5796.94400550699 | 7280.06220076518 | 7179.80678528712 | 5163.28645189118 | 4062.29490648885 | 3310.88360222394 | 6968.97673818278 | 6798.64661151886 | Nek9 |
| **ENSMUSG00000034292** | 0.298473709984862 | 0.12222461677227 | 0.770251097497048 | 215.939778260002 | 210.933673788811 | 185.505158175096 | 205.405946042537 | 171.908436743598 | 111.300483118721 | 189.476212481217 | 191.052499381657 | Traf3ip1 |
| **ENSMUSG00000034295** | 0.137719384877371 | 0.547155022789228 | 0.914699131434573 | 5967.70809793854 | 6806.88666226596 | 7273.50581926331 | 5272.08594842512 | 4973.30012541034 | 3768.31635701956 | 7661.29366840261 | 6610.85068883121 | Fhod3 |
| **ENSMUSG00000034297** | 0.107039876699683 | 0.487285949451122 | 0.899236955296196 | 721.907459538667 | 823.374301932386 | 768.521369582539 | 600.273084325223 | 646.025335533267 | 563.840908986049 | 724.590351027071 | 770.72315091464 | Med13 |
| **ENSMUSG00000034300** | 0.0676411978017117 | 0.64308039555057 | 0.934616627390525 | 376.765219809292 | 482.948527246197 | 447.673162330715 | 419.253232333398 | 365.716037403578 | 365.701587390084 | 444.54034466747 | 471.118095066132 | Fam53c |
| **ENSMUSG00000034303** | 0.556287119558793 | 0.0547233359670534 | 0.703396849069134 | 30.7194663633475 | 39.9063707168021 | 44.483379766477 | 34.703287687552 | 19.7092475247437 | 25.6847268735511 | 23.9447960827911 | 32.5657669400552 | Ccdc15 |
| **ENSMUSG00000034308** | 0.0345815714011433 | 0.890518781927528 | 0.982080657481013 | 72.2810973255236 | 70.8541684155466 | 87.0738497556571 | 109.737423228205 | 78.836990098975 | 61.154111603693 | 79.1219348822664 | 111.809133160856 | Sdr42e1 |
| **ENSMUSG00000034312** | 0.0168610256200941 | 0.940119792054188 | 0.989269716906553 | 2063.6253286437 | 2606.94474825477 | 2457.94334582002 | 1888.98436223594 | 2042.09703520262 | 1433.45237599056 | 2670.36530227649 | 2765.91913877536 | Iqsec1 |
| **ENSMUSG00000034317** | -0.165874111096108 | 0.778813894700519 | 0.960254744040417 | 7.22810973255236 | 7.32974156022896 | 26.5007368821565 | 14.0689004138724 | 22.994122112201 | 18.3462334811079 | 5.20539045278068 | 15.1973579053591 | Trim59 |
| **ENSMUSG00000034320** | -0.212100542350576 | 0.19996879830497 | 0.812921911423739 | 117.456783153976 | 110.760539132349 | 117.360406192407 | 149.130344387048 | 132.489941694111 | 141.877538920568 | 138.463386043966 | 159.572258006271 | Slc26a2 |
| **ENSMUSG00000034321** | 0.165460343203377 | 0.351633544007029 | 0.862685236001802 | 205.097613661173 | 198.717437855096 | 258.382184601026 | 273.874594723383 | 229.94122112201 | 205.477814988408 | 169.69572876065 | 229.045894145055 | Exosc1 |
| **ENSMUSG00000034327** | 0.308120031181445 | 0.168920049583624 | 0.79907358295209 | 2212.70509187759 | 2392.7534115503 | 2381.28049983949 | 2239.76894588849 | 1525.276766776 | 1207.1821630569 | 2345.54893802298 | 2372.95888436536 | Kctd9 |
| **ENSMUSG00000034330** | -0.0371553705053898 | 0.854674495570523 | 0.973268281968411 | 313.519259649459 | 251.654460234528 | 204.434255948065 | 225.102406621959 | 308.778211220985 | 194.470074899744 | 275.885693997376 | 240.986675356409 | Plcg2 |
| **ENSMUSG00000034333** | 0.114044803623865 | 0.728336376073712 | 0.952401994188708 | 73.1846110420926 | 92.0289773673191 | 81.3951204237664 | 69.4065753751039 | 36.1336204620302 | 57.4848649074714 | 88.4916376972716 | 109.638082031519 | Zbed4 |
| **ENSMUSG00000034334** | -0.102341092566477 | 0.798555379027169 | 0.963175947784593 | 19.877301764519 | 17.1027303072009 | 25.5542819935081 | 13.1309737196143 | 28.4689130912965 | 23.2385624094033 | 18.7394056300105 | 10.8552556466851 | Fam151b |
| **ENSMUSG00000034341** | -0.00886511125937046 | 0.930258578924105 | 0.987529878729394 | 2588.56679797031 | 2582.51227638734 | 2949.15343302856 | 2967.60006063282 | 2954.19721231992 | 2903.59721894334 | 2758.85693997376 | 2539.04429575964 | Wbp2 |
| **ENSMUSG00000034342** | 0.0360249002241942 | 0.827805554872152 | 0.967319591893432 | 302.67709505063 | 315.99330281876 | 311.383658365339 | 249.488500672671 | 339.43704070392 | 226.270212933664 | 306.076958623504 | 277.894544555138 | Cbl |
| **ENSMUSG00000034343** | 0.0953518903478544 | 0.426322831941276 | 0.885066977635295 | 869.180195339421 | 754.148964974668 | 760.949730473351 | 815.9962240046 | 709.532910890775 | 653.125911927441 | 824.53384772046 | 806.545494548701 | Ube2f |
| **ENSMUSG00000034345** | 0.0609576309870455 | 0.715581439196366 | 0.951176382858794 | 866.469654189714 | 677.593886456721 | 742.967087589031 | 968.878275168681 | 676.684165016202 | 950.334894321389 | 739.165444294857 | 755.525793009281 | Gtf2h5 |
| **ENSMUSG00000034349** | -0.179159523824916 | 0.311892112654884 | 0.851785577030345 | 430.976042803434 | 321.69421292116 | 343.563124579386 | 420.191159027656 | 427.033696369448 | 536.933099880424 | 398.732908683 | 354.966859646602 | Smc4 |
| **ENSMUSG00000034353** | 0.374756806156548 | 0.132016623139399 | 0.77303252008843 | 125.588406603097 | 65.9676740420606 | 120.199770858353 | 120.054616865045 | 84.3117810780705 | 92.9542496376133 | 68.711153976705 | 86.8420451734806 | Ramp1 |
| **ENSMUSG00000034354** | 0.138250741443502 | 0.525519439884156 | 0.908006595346551 | 1534.16629073424 | 1837.32188443073 | 1724.44080711747 | 1376.87638717098 | 1271.24646534597 | 991.9196902119 | 1721.94316177985 | 1895.32763591121 | Mtmr3 |
| **ENSMUSG00000034361** | -0.517620812676163 | 0.189467754654228 | 0.808824684419004 | 46.0791995450213 | 30.9477976987445 | 46.3762895437739 | 42.2067012416173 | 67.887408140784 | 102.738907494204 | 32.2734208072402 | 34.7368180693922 | Cpne2 |
| **ENSMUSG00000034371** | 0.0668873455172766 | 0.762065477004793 | 0.957666649379949 | 161.728955265859 | 201.160685041839 | 158.057966404291 | 207.281799431054 | 155.484063806312 | 129.646716599829 | 169.69572876065 | 239.90114979174 | Tkfc |
| **ENSMUSG00000034377** | 0.288478964889394 | 0.206152821235997 | 0.819960256425221 | 1166.43620809064 | 1435.81493007596 | 1442.39725030023 | 1208.04958220451 | 884.72622222183 | 708.164612370765 | 1334.66211209297 | 1372.10431374099 | Tulp4 |
| **ENSMUSG00000034379** | -0.196498188686997 | 0.487083979253338 | 0.899236955296196 | 37.9475760958999 | 43.9784493613737 | 35.965285768641 | 66.5927952923295 | 51.4630352034976 | 41.5847958905112 | 65.5879197050366 | 52.1052271040884 | Wdr5b |
| **ENSMUSG00000034382** | -0.0701943596572094 | 0.901573268196028 | 0.984810686948948 | 11.7456783153976 | 19.5459774939439 | 12.3039135524298 | 15.0068271081306 | 21.8991639163819 | 18.3462334811079 | 2.08215618111227 | 19.5394601640331 | AI661453 |
| **ENSMUSG00000034387** | -0.966508540223434 | 0.0198943742133911 | 0.642273034804458 | 56.0178504272808 | 39.9063707168021 | 26.5007368821565 | 32.8274342990356 | 56.937826182593 | 145.546785616789 | 52.0539045278068 | 49.9341759747513 | Ssu2 |
| **ENSMUSG00000034390** | -0.516711570477215 | 0.10366187104038 | 0.759800953500741 | 274.66816983699 | 249.211213047785 | 278.257737262643 | 214.785212985119 | 466.452191418935 | 552.833168897385 | 224.872867560126 | 211.677485110359 | Cmip |
| **ENSMUSG00000034401** | -0.418420416103126 | 0.0857376155733782 | 0.744775159671184 | 128.298947752804 | 107.502876216691 | 143.861143074564 | 120.054616865045 | 193.80760065998 | 226.270212933664 | 131.175839410073 | 117.236760984199 | Spata6 |
| **ENSMUSG00000034403** | -0.190944891708966 | 0.0856215076170698 | 0.744232003004443 | 431.879556520004 | 483.762942975111 | 438.208613444231 | 488.659807708502 | 580.327843784121 | 529.594606487981 | 486.183468289716 | 508.025964264862 | Pja1 |
| **ENSMUSG00000034412** | -0.491159263031915 | 0.120357433305455 | 0.769868040577702 | 106.614618555147 | 96.1010560118908 | 92.7525790875478 | 100.358156285623 | 183.952976897608 | 201.808568292187 | 77.0397787011541 | 94.4407241261602 | Tbc1d10a |
| **ENSMUSG00000034413** | -0.352264197235613 | 0.297924030660844 | 0.851785577030345 | 75.8951521917998 | 64.338842584232 | 66.2518422053913 | 85.3513291774927 | 163.148771177045 | 83.1695917810225 | 71.8343882483734 | 54.2762782334254 | Neurl1b |
| **ENSMUSG00000034422** | -0.133688677608489 | 0.390112213206933 | 0.877515569464664 | 382.186302108706 | 430.82592059568 | 436.315703666934 | 403.308478531009 | 558.428679867739 | 461.102001491845 | 364.377331694648 | 429.868123608729 | Parp14 |
| **ENSMUSG00000034424** | 0.0671602338067719 | 0.467523787563563 | 0.895328112556867 | 1645.29847787223 | 1509.11234567825 | 1483.09481051212 | 1754.86084495702 | 1489.14314631397 | 1508.06039214707 | 1524.13832457418 | 1579.43969659268 | Gcsh |
| **ENSMUSG00000034429** | -0.00192209447432106 | 0.99462036712328 | 0.999262355603273 | 74.0881247586617 | 40.7207864457164 | 81.3951204237664 | 84.4134024832345 | 64.6025335533267 | 92.9542496376133 | 59.3414511616998 | 64.046008315442 | Zfp707 |
| **ENSMUSG00000034430** | 0.0465519395154081 | 0.771225733930356 | 0.959045638244974 | 318.036828232304 | 452.000729547452 | 428.744064557746 | 384.549944645846 | 398.564783278151 | 319.224462571277 | 421.636626675235 | 392.96025441 | Zxdc |

| | | | | | | | | | | | | |
|---|---|---|---|---|---|---|---|---|---|---|---|---|
| **ENSMUSG00000034432** | 0.0614360429463094 | 0.416251216336398 | 0.882192126163777 | 1893.76474992872 | 1873.97059223187 | 1774.60291621584 | 1916.18423636942 | 1738.79361496073 | 1713.53820713548 | 1853.11900118992 | 1841.05135767779 | Cops8 |
| **ENSMUSG00000034435** | 0.0497829553246697 | 0.918964861898095 | 0.986713918026868 | 13.5527057485357 | 16.2883145782866 | 14.1968233297267 | 15.9447538023887 | 8.75966556655277 | 6.1154111603693 | 15.616171358342 | 27.1381391167127 | Tmem30b |
| **ENSMUSG00000034442** | -0.11915858295864 | 0.482136403202126 | 0.897732059433541 | 636.073656464608 | 616.512706788147 | 483.638448099356 | 476.466760683146 | 568.283303630111 | 584.633306931305 | 736.042210023189 | 514.539117652873 | Trmt5 |
| **ENSMUSG00000034445** | -0.0851300132916412 | 0.635458948400081 | 0.932543559775713 | 130.105975185942 | 150.666909849151 | 147.646962629158 | 115.364983393754 | 133.58489988993 | 117.415894279091 | 162.408182126757 | 162.828834700276 | Cyb561a3 |
| **ENSMUSG00000034449** | 0.0540304144234141 | 0.637964627350092 | 0.933164350662676 | 1979.59855300278 | 1635.34678365997 | 1997.96626993687 | 1607.60635395849 | 1663.24149944921 | 1701.30738481474 | 1814.59911183935 | 1774.83429823301 | Dhrs11 |
| **ENSMUSG00000034450** | -0.408414043814595 | 0.444938776662279 | 0.89108031640712 | 31.6229800799166 | 87.1424829938332 | 25.5542819935081 | 46.8963347129081 | 32.8487458745729 | 42.8078781225851 | 40.6020455316893 | 137.8617467129 | Gulo |
| **ENSMUSG00000034453** | 0.162659586934372 | 0.482058404644408 | 0.897732059433541 | 260.211950371885 | 329.838370210303 | 312.330113253987 | 253.240207449704 | 204.757182618171 | 181.016170346931 | 305.035880532948 | 340.855027305911 | Polr3b |
| **ENSMUSG00000034456** | -0.0749649831007138 | 0.890368232235335 | 0.981992595342925 | 19.877301764519 | 63.5244268553176 | 26.5007368821565 | 28.1378008277448 | 19.7092475247437 | 19.5693157131818 | 28.1091084450157 | 78.1578406561325 | Uroc1 |
| **ENSMUSG00000034457** | -0.237797874305095 | 0.403841909657626 | 0.881158207349486 | 24.3948703473642 | 32.5766291565731 | 26.5007368821565 | 31.8895076047775 | 41.6084114411257 | 30.5770558018465 | 26.0269522639034 | 37.9933947633978 | Eda2r |
| **ENSMUSG00000034459** | -0.249020404988242 | 0.461517887034611 | 0.895328112556867 | 65.9565013095403 | 86.3280672649188 | 55.8408384302584 | 66.5927952923295 | 82.1218646864323 | 138.208292224346 | 49.9717483466946 | 57.5328549274309 | Ifit1 |
| **ENSMUSG00000034460** | 0.296202870102676 | 0.1581489499555 | 0.789230661864335 | 172.571119864688 | 200.346269312925 | 212.952349945901 | 216.661066373635 | 146.724398239759 | 112.523565350795 | 203.010227658447 | 189.966973816989 | Six4 |
| **ENSMUSG00000034462** | -0.145527009250087 | 0.350057865596525 | 0.861774532843376 | 581.862833470465 | 658.047908962778 | 599.105944514467 | 546.811262752508 | 748.951405940262 | 759.534066117867 | 610.071761065896 | 522.137796605552 | Pkd2 |
| **ENSMUSG00000034463** | -0.98801003842764 | 0.0680699548701769 | 0.722112580225498 | 24.3948703473642 | 20.3603932228582 | 10.4110037751329 | 26.2619474392285 | 39.4184950494875 | 85.6157562451702 | 18.7394056300105 | 18.4539345993646 | Scara3 |
| **ENSMUSG00000034471** | -0.288205564418732 | 0.170033757522033 | 0.79907358295209 | 1113.12889881306 | 1225.69567201606 | 1317.46520499864 | 978.257542111262 | 1843.90960175936 | 1623.03012196201 | 1157.67883669842 | 1035.59138869376 | Caskin2 |
| **ENSMUSG00000034472** | 0.207249973290067 | 0.438805240007573 | 0.889583654898613 | 65.9565013095403 | 108.317291945606 | 144.807597963212 | 105.047789756914 | 94.1664048404423 | 68.4926049961361 | 104.107809055614 | 99.8683519495027 | Rasd2 |
| **ENSMUSG00000034473** | -0.117961873289589 | 0.411688671798714 | 0.882192126163777 | 247.562758339918 | 186.501201921381 | 196.862616838877 | 209.15765281957 | 208.042057205628 | 233.608706326107 | 238.406882737355 | 231.216945274392 | Sec22a |
| **ENSMUSG00000034480** | 0.310985490266824 | 0.144365645300943 | 0.780607990324772 | 282.799793286111 | 416.980853204136 | 410.761421673426 | 320.770929436291 | 309.873169416804 | 211.593226148778 | 363.336253604092 | 268.124814473121 | Diaph2 |
| **ENSMUSG00000034484** | 0.122030624588251 | 0.291107478068289 | 0.851785577030345 | 573.731210021344 | 549.730617017172 | 662.518422053913 | 685.624413502716 | 561.713554455197 | 556.502415593606 | 594.455589707554 | 556.874614674944 | Snx2 |
| **ENSMUSG00000034485** | 0.191397975486275 | 0.0772055997225041 | 0.739855948845396 | 2931.90201026655 | 2791.0027029894 | 2960.51089169234 | 3263.04696932414 | 2596.14588228708 | 2278.6021983536 | 2825.48593776935 | 2759.40598538735 | Uaca |
| **ENSMUSG00000034487** | -0.321038074350129 | 0.114942321972723 | 0.767518180080285 | 174.378147297826 | 225.593156909269 | 161.843785958884 | 159.447538023887 | 281.404256325508 | 242.170281950624 | 174.901119213431 | 204.078806157679 | Kdelc2 |
| **ENSMUSG00000034501** | -0.0173041772268765 | 0.925165007798737 | 0.986713918026868 | 113.8427282877 | 128.677685168464 | 157.111511515642 | 109.737423228205 | 125.920192519196 | 117.415894279091 | 150.95632313064 | 120.493337678204 | Pcnx4 |
| **ENSMUSG00000034509** | -0.255741554287311 | 0.234667789297121 | 0.832048466795696 | 330.68602026427 | 306.220314071788 | 251.757000380487 | 304.826175633902 | 427.033696369448 | 447.648096939033 | 249.858741733473 | 301.776106977845 | Mad2l1bp |
| **ENSMUSG00000034518** | 0.15302773362443 | 0.326083165288149 | 0.855977678491063 | 208.711668527449 | 230.479651282755 | 208.220075502658 | 180.081925297567 | 179.573144114332 | 157.777607937528 | 190.517290571773 | 216.019587369033 | Hmgxb4 |
| **ENSMUSG00000034520** | -0.261181854975761 | 0.0904147673266614 | 0.753823363185192 | 234.010052591383 | 197.088606397268 | 225.25626349833 | 253.240207449704 | 302.208462046071 | 299.655146858096 | 262.351678820146 | 225.78931745105 | Gjc1 |
| **ENSMUSG00000034522** | -0.207054226347391 | 0.220630897204519 | 0.829144114546069 | 176.185174730964 | 177.542628903324 | 209.166530391307 | 247.612647284155 | 243.08071947184 | 199.362403828039 | 232.160414194018 | 259.440609955773 | Zfp395 |
| **ENSMUSG00000034525** | 0.181431467799106 | 0.57701740378355 | 0.922450244597442 | 563.792559139084 | 697.95427967958 | 832.880302010633 | 482.094320848695 | 468.642107810573 | 271.524255520397 | 799.547973547113 | 731.644230586574 | Ice1 |
| **ENSMUSG00000034528** | -0.280748440835352 | 0.363876887204687 | 0.86924083181804 | 84.9302893574902 | 132.749763813036 | 105.056492639978 | 104.109863062656 | 71.1722827282413 | 105.185071958352 | 148.874166949528 | 193.223550510994 | Hsd17b13 |
| **ENSMUSG00000034533** | 0.393172159359556 | 0.393094580647863 | 0.878770935869648 | 21.6843291976571 | 6.51532583131463 | 18.9290977729689 | 21.5723139679377 | 13.1394983498292 | 13.4539045528125 | 14.5750932677859 | 10.8552556466851 | Scn10a |
| **ENSMUSG00000034538** | 0.0805086180813909 | 0.820911522392019 | 0.96650690096489 | 15.3597331816738 | 19.5459774939439 | 18.9290977729689 | 20.6343872736795 | 19.7092475247437 | 11.0077400886647 | 20.8215618111227 | 18.4539345993646 | Zfp418 |
| **ENSMUSG00000034543** | 0.220884920757927 | 0.138822136224118 | 0.779711473022872 | 446.335775985108 | 482.948527246197 | 435.369248778286 | 462.397860269274 | 402.944616061428 | 297.208982393948 | 436.211719943021 | 429.868123608729 | Morc2a |
| **ENSMUSG00000034544** | -0.0365463645362342 | 0.815312338601681 | 0.966177091475091 | 316.229800799166 | 322.508628650074 | 321.794662140472 | 292.633128608546 | 307.683253025166 | 407.286383280595 | 303.994802442392 | 268.124814473121 | Rsrc1 |
| **ENSMUSG00000034557** | 0.144322162535884 | 0.352985210649836 | 0.862728204353539 | 979.408868760845 | 1097.83240257652 | 1083.69084750247 | 1023.27802343565 | 940.569090208604 | 705.718447906617 | 1079.59797990671 | 1058.38742555179 | Zfyve9 |
| **ENSMUSG00000034560** | 0.00626663417547864 | 0.972836195091836 | 0.995410294646453 | 280.992765852973 | 320.879797192245 | 281.097101928589 | 261.681547698027 | 279.21433993387 | 201.808568292187 | 334.18606706852 | 323.486618271215 | Washc4 |
| **ENSMUSG00000034563** | 0.136604796584237 | 0.649167277472831 | 0.936461253686283 | 250.273299489625 | 368.115909469277 | 377.63550057073 | 219.47484645641 | 217.896680968 | 165.116101329971 | 377.911346871878 | 344.111603999917 | Ccpg1 |
| **ENSMUSG00000034566** | 0.0541294276730228 | 0.752406317889769 | 0.956497786093635 | 32595.1608389449 | 23157.9112516789 | 25877.9695654258 | 31707.5498260914 | 27422.1330560935 | 33598.0689150689 | 22900.5947579633 | 25244.9825319308 | Atp5h |
| **ENSMUSG00000034570** | -0.069381975899123 | 0.707248444580929 | 0.948710970699398 | 590.897970636155 | 535.071133896714 | 373.849681016137 | 481.156394154437 | 422.653863586171 | 517.363784167243 | 525.744435730849 | 613.321944037707 | Inpp5j |
| **ENSMUSG00000034573** | -0.0556785351696928 | 0.880589764821049 | 0.978433072023387 | 18.0702743313809 | 44.7928650902881 | 46.3762895437739 | 29.075727522003 | 49.2731188118594 | 34.2463024980681 | 35.3966550789086 | 24.9670879873757 | Ptpn13 |
| **ENSMUSG00000034574** | 0.203765499721561 | 0.508893131435835 | 0.902394714365738 | 1674.21091680244 | 2076.76010873154 | 1517.16718650346 | 1386.25565411356 | 905.530427942393 | 878.173042629031 | 1803.14725284323 | 2190.59058950105 | Daam1 |
| **ENSMUSG00000034575** | 0.177134009206788 | 0.190336671996856 | 0.808824684419004 | 365.923055210463 | 390.919549878878 | 330.312756138308 | 375.170677703265 | 331.772333333186 | 265.408844360028 | 345.637926064637 | 349.539231823259 | Papd7 |
| **ENSMUSG00000034586** | -0.595064969451619 | 0.13269787152754 | 0.77303252008843 | 30.7194663633475 | 12.2162359337149 | 26.5007368821565 | 41.2687745473591 | 50.3680770076785 | 53.8156182112498 | 30.191264626128 | 32.5657669400552 | Hid1 |
| **ENSMUSG00000034587** | 0.172638428429105 | 0.610629174292231 | 0.927932112008768 | 24.3948703473642 | 17.9171460361152 | 31.2330113253987 | 17.8206071909051 | 19.7092475247437 | 20.7923979452556 | 21.8626399016789 | 18.4539345993646 | 8430429K09I |
| **ENSMUSG00000034591** | -0.407564161190178 | 0.188153612552625 | 0.808677608936066 | 26.2018977805023 | 33.3910448854875 | 18.9290977729689 | 17.8206071909051 | 31.7537876787538 | 34.2463024980681 | 33.3144988977964 | 29.3091902460497 | Slc41a2 |
| **ENSMUSG00000034593** | 0.0298835507152011 | 0.89386990030675 | 0.983730237253949 | 73.1846110420926 | 122.162359337149 | 112.628131749165 | 86.2892558717508 | 112.780694169367 | 91.7311674055395 | 82.2451691539348 | 99.8683519495027 | Myo5a |
| **ENSMUSG00000034595** | -0.364672890137138 | 0.10645013570499 | 0.762094734041179 | 272.861142403852 | 261.427448981499 | 281.097101928589 | 230.729966787508 | 413.894198019619 | 434.19419238622 | 238.406882737355 | 261.61166108511 | Ppp1r18 |
| **ENSMUSG00000034601** | -0.158210233094041 | 0.455896347064354 | 0.892805623137304 | 58.7283915769879 | 68.4109212288036 | 87.0738497556571 | 63.779015209555 | 77.7420319031559 | 80.7234273168747 | 87.4505596067155 | 64.046008315442 | 2700049A03I |
| **ENSMUSG00000034602** | 0.161878876686832 | 0.399374461660643 | 0.880459952732461 | 712.872322372977 | 974.041211781537 | 983.366629305736 | 760.658549043369 | 743.476614961167 | 567.510155682271 | 927.600578685518 | 827.170480277403 | Mon2 |
| **ENSMUSG00000034610** | 0.182542310802833 | 0.511335079462219 | 0.902394714365738 | 355.984404328204 | 460.95930256551 | 409.814966784777 | 306.702029022419 | 246.365594059297 | 210.370143916704 | 447.663578939139 | 446.151007078757 | Zcchc11 |
| **ENSMUSG00000034612** | -0.174503322435438 | 0.54793344742327 | 0.914699131434573 | 83.1232619243521 | 65.9676740420606 | 108.842312194571 | 75.9720622349111 | 117.160526952643 | 127.200552135681 | 66.6289977955927 | 66.217059444779 | Chst11 |
| **ENSMUSG00000034613** | 0.00694105582314863 | 0.970201784660756 | 0.9951105526746 | 116.553269437407 | 133.56417954195 | 104.110037751329 | 164.137171495178 | 122.635317931739 | 132.092881063977 | 136.381229862854 | 124.835439936878 | Ppm1h |
| **ENSMUSG00000034614** | -0.265246033376297 | 0.0725654188630396 | 0.729721356194038 | 1076.08483643373 | 881.197818685304 | 963.491076644119 | 1254.00799022316 | 1189.12460065954 | 1391.86758010005 | 1320.08701882518 | 1115.92028047923 | Pik3ip1 |
| **ENSMUSG00000034616** | 0.266460561873992 | 0.247813342626192 | 0.834050721850692 | 100.290022539164 | 122.976775066064 | 162.790240847533 | 115.364983393754 | 105.115986798633 | 83.1695917810225 | 135.340151772298 | 92.2696729968231 | Ssh3 |
| **ENSMUSG00000034617** | 0.139776709312151 | 0.569540287148573 | 0.920872458148558 | 312.61574593289 | 379.517729674077 | 373.849681016137 | 320.770929436291 | 312.063085808443 | 178.570005882784 | 419.554470494123 | 347.368180693922 | Mtrr |
| **ENSMUSG00000034620** | 0.241859458778739 | 0.075271206278379 | 0.735399943172214 | 688.477452025612 | 623.842448348376 | 752.431636475515 | 809.430737144793 | 678.87408140784 | 532.040770952129 | 640.263025692024 | 577.499600403646 | Tmem5 |
| **ENSMUSG00000034621** | 0.0974490055668912 | 0.685726056505952 | 0.944452327903597 | 255.69438178904 | 378.703313945163 | 385.207139679918 | 284.191788360223 | 275.929465346412 | 201.808568292187 | 388.322127777439 | 351.710282952596 | Gpatch8 |
| **ENSMUSG00000034636** | 0.0846949514189843 | 0.776492721823546 | 0.959733396167384 | 1030.90915060528 | 1429.29960424465 | 1297.58965233702 | 957.623154837583 | 836.54806160579 | 607.871869340708 | 1475.20765431805 | 1526.24894392392 | Zyg11b |
| **ENSMUSG00000034639** | -0.138711874325428 | 0.52259103352072 | 0.90648515894888 | 49.6932544112975 | 53.7514381083457 | 44.483379766477 | 53.4618215727152 | 56.937826182593 | 63.6002760678407 | 56.2182168900314 | 45.5920737160773 | Setmar |
| **ENSMUSG00000034640** | 0.431912068541608 | 0.0885157211946122 | 0.751206603737452 | 406.58117245607 | 453.629561005281 | 269.739643264807 | 215.723139679377 | 297.828629262794 | 250.731857575141 | 221.749633288457 | 227.960368580387 | Tiparp |
| **ENSMUSG00000034641** | -0.14627758168689 | 0.576242635648131 | 0.922450244597442 | 62.3424464432641 | 53.7514381083457 | 53.9479286529615 | 45.0204813243917 | 79.9319482947941 | 67.2695227640623 | 46.8485140750261 | 44.5065481514088 | Cd300ld |
| **ENSMUSG00000034647** | -0.0374406747169203 | 0.820571197045709 | 0.966398681144067 | 584.573374620172 | 548.916201288258 | 464.709350326387 | 618.093691516128 | 594.56230032977 | 698.379954514174 | 491.388858742496 | 491.743080794834 | Ankrd12 |
| **ENSMUSG00000034652** | -0.00445998062336032 | 0.989860029230026 | 0.998628446692557 | 28.0089252136404 | 28.5045505120015 | 50.1621090983677 | 60.9652351267805 | 45.9882442244021 | 55.0387004433237 | 29.1501865355718 | 37.9933947633978 | Cd300a |
| **ENSMUSG00000034653** | -0.00366517807913223 | 0.990802229485256 | 0.998628446692557 | 101.193536255733 | 125.420022252807 | 176.040609288611 | 101.296082979881 | 94.1664048404423 | 85.6157562451702 | 203.010227658447 | 121.578863242873 | Ythdc2 |
| **ENSMUSG00000034656** | -0.0433826224176467 | 0.897390200722637 | 0.984332015045905 | 27.1054114970714 | 41.5352021746308 | 35.0188308799925 | 30.9515809105193 | 41.6084114411257 | 31.8001380339204 | 46.8485140750261 | 18.4539345993646 | Cacna1a |
| **ENSMUSG00000034659** | -0.237628800521248 | 0.047159502560151 | 0.702838509185083 | 2949.97228459793 | 2969.35974762164 | 2769.32700418536 | 3097.03394444045 | 3744.75702970131 | 3938.32478727783 | 3229.42423690514 | 2985.1953028384 | Tmem109 |
| **ENSMUSG00000034663** | -0.0591013232137917 | 0.77009412587482 | 0.958992744413128 | 74.0881247586617 | 97.7298874697194 | 76.6628459805242 | 104.109863062656 | 75.5521155115177 | 94.1773318696872 | 94.7381062406084 | 103.124928643508 | Bmp2k |
| **ENSMUSG00000034664** | 0.465603597865992 | 0.583156792416267 | 0.922450244597442 | 70.4740698923855 | 93.6578088251478 | 282.043556817237 | 25.3240207449704 | 84.3117810780705 | 162.669936865823 | 20.8215618111227 | 73.8157383974585 | Itga2b |
| **ENSMUSG00000034667** | 0.055293395420151 | 0.723162062556422 | 0.951674101666722 | 809.548290045864 | 1036.75122290794 | 903.864418659267 | 827.251344335698 | 759.900987898453 | 725.287763619799 | 993.188498390554 | 963.946701425635 | Xpot |
| **ENSMUSG00000034673** | -0.136765214955772 | 0.277641336498175 | 0.848075053184273 | 362.309000344187 | 336.353696041618 | 409.814966784777 | 392.99128489417 | 454.407651264925 | 432.971110154146 | 384.157815415214 | 378.848422069309 | Pbx2 |
| **ENSMUSG00000034674** | -0.0275244009924635 | 0.897809851112532 | 0.984338056038603 | 173.474633581257 | 215.005752433383 | 237.56017705076 | 222.288626539184 | 204.757182618171 | 146.769867848863 | 239.447960827911 | 272.466916731795 | Tdg |
| **ENSMUSG00000034675** | -0.868468478733944 | 0.0267257129315252 | 0.664353746352133 | 100.290022539164 | 97.7298874697194 | 76.6628459805242 | 101.296082979881 | 233.226095709468 | 280.085831144914 | 94.7381062406084 | 79.2433662208011 | Dbn1 |
| **ENSMUSG00000034681** | 0.135053941769467 | 0.241520685998134 | 0.832048466795696 | 1154.69052977524 | 1268.04528991961 | 1181.17570103326 | 1150.83605385476 | 1031.45062046159 | 933.211743072355 | 1205.56842886401 | 1158.2557775013 | Rnps1 |
| **ENSMUSG00000034684** | -0.706311724841238 | 0.0177541713873819 | 0.642273034804458 | 227.685456575399 | 227.221988367098 | 212.952349945901 | 218.536919762152 | 505.870686468423 | 490.455975061618 | 246.735507461804 | 204.078806157679 | Sema3f |
| **ENSMUSG00000034685** | -0.489598482932731 | 0.0758653065873478 | 0.735399943172214 | 70.4740698923855 | 50.4937751926884 | 69.0912068713366 | 75.0341355406529 | 119.350443344282 | 116.192812047017 | 77.0397787011541 | 59.7039060567679 | Fam171a2 |
| **ENSMUSG00000034686** | -0.77582925978526 | 0.113434867094732 | 0.767518180080285 | 8.13162344912141 | 10.5874044758863 | 10.4110037751329 | 3.75170677703264 | 20.8042057205628 | 13.4539045528125 | 11.4518589961175 | 10.8552556466851 | Prr7 |
| **ENSMUSG00000034707** | 0.0925974787862104 | 0.538360022703499 | 0.912708496930502 | 2709.63763599057 | 3317.92967959697 | 3664.67332884679 | 2570.85706896162 | 2932.29804840354 | 2526.88789146459 | 3156.54877056621 | 2884.24142532422 | Gns |
| **ENSMUSG00000034708** | -0.244466286872083 | 0.211332840737827 | 0.823363318105013 | 1659.75469733734 | 1618.24405335277 | 2025.41346170768 | 1822.39156694361 | 2508.54922662155 | 2628.40371672672 | 1627.20505553924 | 1678.22252297751 | Grn |
| **ENSMUSG00000034709** | -0.148643390262638 | 0.336239971836379 | 0.858570960161243 | 194.255449062345 | 216.634583891211 | 199.701981504822 | 195.088752405698 | 261.695008800764 | 244.616446414772 | 189.476212481217 | 198.651178334337 | Ppp1r21 |
| **ENSMUSG00000034714** | -0.707562670958765 | 0.0220910246524459 | 0.648762042745232 | 18.0702743313809 | 28.5045505120015 | 38.8046504345863 | 40.3308478531009 | 64.6025335533267 | 48.9232892829544 | 48.9306702561384 | 42.3354970220718 | Ttyh2 |
| **ENSMUSG00000034723** | 0.0531800396784027 | 0.693925951533554 | 0.94469184542218 | 1423.93761731282 | 1563.67819951551 | 1328.82266366242 | 1290.58713129923 | 1211.02376457592 | 1180.27435395127 | 1553.28851110976 | 1458.94635891447 | Tmx4 |
| **ENSMUSG00000034724** | -0.0580797009017771 | 0.677113756016812 | 0.94409858171788 | 618.906895849796 | 710.170515613295 | 722.145080038765 | 606.83857118503 | 724.862325632242 | 556.502415593606 | 746.45299092875 | 738.157383974585 | Cnot6l |
| **ENSMUSG00000034729** | 0.169706328355245 | 0.215125014393754 | 0.824338622350122 | 801.416666596743 | 679.22271791455 | 611.409858066897 | 815.058297310342 | 698.583328932584 | 629.887349518038 | 585.085886892549 | 670.854798965138 | Mrps10 |
| **ENSMUSG00000034731** | 0.576170009787385 | 0.073920349250835 | 0.733705544272914 | 33.4300075130547 | 48.0505280059454 | 53.9479286529615 | 35.6412143818101 | 36.1336204620302 | 25.6847268735511 | 35.3966550789086 | 17.3684090346961 | Dgkh |
| **ENSMUSG00000034738** | -0.290265461857817 | 0.391913269542151 | 0.878285927324016 | 153.597331816738 | 95.2866402829764 | 97.48485353079 | 87.227182566009 | 181.76306050597 | 183.462334811079 | 86.4094815161593 | 79.2433662208011 | Nostrin |
| **ENSMUSG00000034744** | -0.0705104245012315 | 0.626500310855709 | 0.931698409533928 | 491.511461813561 | 408.022280186079 | 474.173899212872 | 538.369922504185 | 528.864808580624 | 577.294813538862 | 459.115437935256 | 442.894430384751 | Nagk |
| **ENSMUSG00000034748** | -0.0406514458645039 | 0.837173848757286 | 0.968924788429433 | 75.8951521917998 | 104.245213301034 | 89.9132144216024 | 96.6064495085906 | 88.6916138613469 | 100.292743030056 | 111.395355689507 | 77.072315091464 | Sirt6 |
| **ENSMUSG00000034751** | -0.228224117176138 | 0.12307110956023 | 0.770251097497048 | 278.282224703266 | 364.043830824705 | 330.312756138308 | 298.260688774095 | 432.508487348543 | 380.37857417497 | 339.391457521301 | 337.598450611906 | Mast4 |
| **ENSMUSG00000034757** | -0.103494120888498 | 0.450928980851379 | 0.89108031640712 | 462.599022883351 | 460.144886836596 | 502.567545872325 | 472.715053906113 | 512.440435643337 | 609.094951572782 | 496.594249195277 | 422.269444656049 | Tmub2 |
| **ENSMUSG00000034758** | -0.0823944016080592 | 0.726477835462252 | 0.952045814446758 | 88.5443442237664 | 82.2559886203472 | 91.8061241988993 | 95.6685228143324 | 129.205067106653 | 103.961989726278 | 80.1630129728225 | 66.217059444779 | Tle6 |

| | | | | | | | | | | | | |
|---|---|---|---|---|---|---|---|---|---|---|---|---|
| ENSMUSG00000034761 | 0.15697609539155 | 0.455431505496457 | 0.892805623137304 | 458.984968017075 | 650.718167402549 | 682.39397471553 | 511.170048370698 | 536.529515951357 | 355.916929533493 | 584.044808801993 | 588.354856050331 | Map4k5 |
| ENSMUSG00000034765 | -0.368646001050665 | 0.124253979246416 | 0.771949105062466 | 53.3073092775737 | 64.338842584232 | 71.930571537282 | 86.2892558717508 | 104.021028602814 | 73.3849339244316 | 106.189965236726 | 71.6446872681215 | Dusp5 |
| ENSMUSG00000034768 | 0.103345757733905 | 0.634535838950823 | 0.932543559775713 | 52.4037955610046 | 57.009101024003 | 52.0550188756646 | 50.6480414899407 | 41.6084114411257 | 58.7079471395453 | 46.8485140750261 | 51.0197015394199 | Asb16 |
| ENSMUSG00000034771 | 0.110321396649209 | 0.655416276580321 | 0.937625787683528 | 47.8862269781594 | 62.7100111264033 | 43.5369248778286 | 39.3929211588428 | 43.7983278327639 | 51.3694537471021 | 40.6020455316893 | 44.5065481514088 | Tle2 |
| ENSMUSG00000034780 | 0.112178813485157 | 0.813005654253703 | 0.965596283675411 | 9.93865088225949 | 19.5459774939439 | 10.4110037751329 | 17.8206071909051 | 9.85462376237187 | 17.123151249034 | 7.28754663389296 | 19.5394601640331 | B3galt1 |
| ENSMUSG00000034781 | -0.0328109994300073 | 0.739395915019242 | 0.954383574136061 | 1867.56285214822 | 1793.34343506935 | 1913.73178484716 | 1813.95022669528 | 2108.88948514758 | 1939.80842006914 | 1707.36806851206 | 1803.05796291439 | Gna11 |
| ENSMUSG00000034785 | -0.408009619726829 | 0.470299885578719 | 0.89620872694312 | 11.7456783153976 | 39.0919549878878 | 23.6613722162112 | 30.0136542162612 | 15.3294147414674 | 24.4616446414772 | 20.8215618111227 | 78.1578406561325 | Dio1 |
| ENSMUSG00000034786 | -0.555443493841754 | 0.120889359227414 | 0.769868040577702 | 28.0089252136404 | 44.7928650902881 | 40.6975602118832 | 62.8410885152968 | 82.1218646864323 | 90.5080851734656 | 47.8895921655823 | 39.0789203280663 | Gpsm3 |
| ENSMUSG00000034789 | 0.123184573046283 | 0.439915059295766 | 0.890212729256729 | 636.977170181177 | 500.865673282312 | 605.731128735006 | 692.189900362523 | 577.042969196664 | 669.025980944401 | 487.224546380272 | 503.683862006187 | Rab24 |
| ENSMUSG00000034793 | 0.0606803104460213 | 0.560702646066909 | 0.916762264997884 | 1091.44456961541 | 1086.43058237171 | 941.722614205205 | 1176.16007459973 | 1055.53970076961 | 982.135032355309 | 1074.39258945393 | 1006.28219844771 | G6pc3 |
| ENSMUSG00000034795 | 0.466047765190492 | 0.0694216880838066 | 0.724352955175446 | 72.2810973255236 | 49.679359463774 | 43.5369248778286 | 60.0273084325223 | 42.7033696369448 | 33.0232202659942 | 45.80743598447 | 41.2499714574033 | Ccdc122 |
| ENSMUSG00000034799 | -0.0596974505622363 | 0.75535638147346 | 0.956566265296272 | 62.3424464432641 | 68.4109212288036 | 57.7337482075553 | 69.4065753751039 | 61.3176589658694 | 67.2695227640623 | 67.6700758861489 | 72.73021283279 | Unc13a |
| ENSMUSG00000034800 | 0.448533600230609 | 0.205847677644273 | 0.819960256425221 | 21.6843291976571 | 30.1333819698302 | 42.5904699891801 | 44.0825546301336 | 31.7537876787538 | 15.9000690169602 | 22.903717992235 | 30.3947158107182 | Zfp661 |
| ENSMUSG00000034801 | 0.270534657274181 | 0.246471174798971 | 0.832154744277519 | 987.540492209966 | 1135.29552610657 | 1014.59964063114 | 852.575365080669 | 724.862325632242 | 513.694537471021 | 1034.8316220128 | 1033.42033756442 | Sos2 |
| ENSMUSG00000034807 | 0.0442631481912301 | 0.74522564849724 | 0.95596456017978 | 765.276117933981 | 957.75289720325 | 1080.85148283653 | 875.085605742864 | 948.233797579338 | 813.349684329117 | 910.94332923662 | 894.47306528685 | Colgalt1 |
| ENSMUSG00000034810 | -0.654601332550284 | 0.0723276073076221 | 0.72957913253658 | 140.948139784771 | 171.027303072009 | 148.593417517806 | 138.813150750208 | 327.39250054991 | 365.701587390084 | 137.42230795341 | 113.980184290193 | Scn7a |
| ENSMUSG00000034820 | 0.0636274564409935 | 0.676963098964158 | 0.94409858171788 | 440.011179969125 | 363.229415095791 | 413.600786339371 | 328.274342990356 | 418.274030802895 | 315.555215875056 | 351.884394607974 | 391.874728845331 | Cpsf7 |
| ENSMUSG00000034826 | 0.065825553899778 | 0.643179973312411 | 0.934616627390525 | 203.290586228035 | 219.892246806869 | 231.881447718869 | 199.778385876988 | 186.142893289246 | 181.016170346931 | 221.749633288457 | 226.874843015718 | Nup54 |
| ENSMUSG00000034832 | -0.121353525799803 | 0.512724830869717 | 0.902972980953514 | 201.483558794897 | 178.357044632238 | 179.826428843205 | 228.854113398991 | 227.751304730372 | 167.562265794119 | 265.474913091815 | 195.394601640331 | Tet3 |
| ENSMUSG00000034837 | -0.0618709987346836 | 0.874404582862381 | 0.976717302449415 | 32.5264937964856 | 27.6901347830872 | 17.9826428843205 | 12.1930470253561 | 24.0890803080201 | 19.5693157131818 | 31.2323427166841 | 19.5394601640331 | Gnat1 |
| ENSMUSG00000034839 | -0.91704686780074 | 0.0265308975245337 | 0.664353746352133 | 13.5527057485357 | 21.1748089517725 | 21.7684624389143 | 31.8895076047775 | 63.5075753575076 | 51.3694537471021 | 23.9447960827911 | 28.2236646813812 | Larp6 |
| ENSMUSG00000034842 | 0.0712219471736822 | 0.606068218110314 | 0.926859923029135 | 5181.65116452347 | 4620.9948458599 | 6105.58048667113 | 5918.317440769 | 4939.35642133995 | 4772.4668695522 | 6084.06036121006 | 4977.13471400511 | Art3 |
| ENSMUSG00000034845 | -0.812784055629434 | 0.0825244341293142 | 0.739855948845396 | 93.9654265231807 | 77.3694942468612 | 72.8770264259304 | 110.675349922463 | 190.522726072523 | 302.101311322243 | 74.9576225200418 | 56.4473293627624 | Plvap |
| ENSMUSG00000034848 | 0.0882258773088422 | 0.644096137587741 | 0.934847999069355 | 148.176249517323 | 194.645359210525 | 177.933519065908 | 177.268145214792 | 174.098353135236 | 117.415894279091 | 193.640524843441 | 170.427513652956 | Ttc21b |
| ENSMUSG00000034850 | 0.0746271030654932 | 0.556921064677353 | 0.916239900758826 | 759.855035634567 | 832.332874950444 | 849.916490006305 | 728.769041438591 | 730.337116611338 | 634.779678446333 | 814.123066814899 | 830.427056971408 | Tmem127 |
| ENSMUSG00000034853 | -0.191449133496763 | 0.449271060600825 | 0.89108031640712 | 442.721721118832 | 493.535931722083 | 494.995906763138 | 337.653609932938 | 469.737066006393 | 298.432064626022 | 652.755962778698 | 598.124586132348 | Acot11 |
| ENSMUSG00000034854 | -0.198363995750813 | 0.269482896759253 | 0.843793744782845 | 125.588406603097 | 162.883145782866 | 108.842312194571 | 141.626930832982 | 152.199189218854 | 173.677676954488 | 158.243869764533 | 135.690695583563 | Mfsd12 |
| ENSMUSG00000034858 | -0.0166687230770784 | 0.946696724873727 | 0.990510945542179 | 550.239853390548 | 668.635313438664 | 530.01473764313 | 495.225294568309 | 520.105143014071 | 381.601656407044 | 865.13589325215 | 502.598336441519 | Fam214a |
| ENSMUSG00000034863 | 0.0321850699991115 | 0.754629895760607 | 0.956497786093635 | 702.933671490717 | 709.35609988438 | 600.052399403115 | 665.927952923294 | 652.595084708182 | 616.433444965225 | 693.358008310387 | 656.742966624447 | Ano8 |
| ENSMUSG00000034867 | -0.149526546737122 | 0.299497296581993 | 0.851785577030345 | 226.78194285883 | 203.603932228582 | 256.489274823729 | 226.040333316217 | 262.789966996583 | 288.647406769431 | 223.831789469569 | 237.730098662403 | Ankrd27 |
| ENSMUSG00000034868 | -0.438380244868479 | 0.128031305653197 | 0.77303252008843 | 641.494738764022 | 622.213616890547 | 636.017685171756 | 797.237690119437 | 1168.32039493898 | 1313.59031724733 | 531.990904274186 | 641.545608719088 | Myl12b |
| ENSMUSG00000034872 | -0.0717790378974074 | 0.742121808213144 | 0.954383574136061 | 70.4740698923855 | 57.8235167529173 | 72.8770264259304 | 90.0409626487835 | 87.5966556655277 | 84.3926740130963 | 67.6700758861489 | 66.217059444779 | Gipc3 |
| ENSMUSG00000034875 | 0.0877325591557064 | 0.460672019431504 | 0.894774044236998 | 1252.2700111647 | 1065.25577341994 | 1055.29720084302 | 1165.84288096289 | 1024.88087128667 | 938.10407200065 | 1130.61080634396 | 1175.62418653599 | Nudt19 |
| ENSMUSG00000034880 | -0.0696685755546476 | 0.623883948412429 | 0.931145988862144 | 1305.57732044227 | 1105.97655986566 | 916.168332211697 | 1136.76715344089 | 1129.99685808531 | 1328.26730403221 | 1068.1461209106 | 1160.42682863063 | Mrpl34 |
| ENSMUSG00000034881 | -0.37276788122412 | 0.330290230052292 | 0.85614138351386 | 23.4913566307952 | 25.2468875963442 | 25.5542819935081 | 36.5791410760683 | 30.6588294829347 | 62.3771938357668 | 20.8215618111227 | 30.3947158107182 | Tbxa2r |
| ENSMUSG00000034889 | -0.0727664143628883 | 0.649588332185626 | 0.936461253686283 | 477.958756065025 | 478.876448601625 | 524.33600831124 | 498.977001345342 | 633.980795379257 | 576.071731306788 | 419.554470494123 | 453.749686031436 | Cactin |
| ENSMUSG00000034892 | 0.180604772791751 | 0.405594898001863 | 0.881158207349486 | 2631.93545636563 | 1890.25890681016 | 2278.11691697681 | 2598.99486978936 | 2217.29034653367 | 2837.55077841135 | 1699.03944378762 | 1540.36077626461 | Rps29 |
| ENSMUSG00000034893 | 0.155692667583417 | 0.462040836912443 | 0.895328112556867 | 268.343573821006 | 356.714089264476 | 387.100049457215 | 299.198615468353 | 256.220217821669 | 215.262472844999 | 350.843316517418 | 353.881334081933 | Cog3 |
| ENSMUSG00000034898 | 0.306456730577964 | 0.273781709987273 | 0.845204865395511 | 2530.7419201099 | 3376.56761207881 | 3234.03635451174 | 2629.00852400563 | 1761.78773707293 | 1385.75216893968 | 3367.8876229491 | 3001.47818630842 | Filip1 |
| ENSMUSG00000034902 | -0.0055824804598991 | 0.957249099301281 | 0.992403427270738 | 545.722284807703 | 634.429852824262 | 632.231865617162 | 617.15576482187 | 616.461464246152 | 650.679747463293 | 566.346481262538 | 606.808790649696 | Pip5k1c |
| ENSMUSG00000034903 | 0.147086121058195 | 0.585772320282397 | 0.923015034455189 | 1363.40219830269 | 1869.08409785838 | 1903.32078107203 | 1181.78763476528 | 1327.08933333275 | 821.911259953634 | 1796.90078429989 | 1758.55141476298 | Cobll1 |
| ENSMUSG00000034906 | -0.974379291260059 | 0.0309104312498322 | 0.667184591665619 | 15.3597331816738 | 9.77298874697194 | 8.51809399783602 | 12.1930470253561 | 37.2285786578493 | 23.2385624094033 | 16.6572494488982 | 13.0263067760221 | Ncaph |
| ENSMUSG00000034908 | 0.245016932739398 | 0.316310099611693 | 0.85344741328796 | 1204.38378418654 | 1505.04026703368 | 2100.1833979109 | 1284.02164443942 | 1140.9464400435 | 898.965440574287 | 1665.72494488982 | 1435.06479649177 | Sidt2 |
| ENSMUSG00000034910 | -0.0033712826305390 | 0.981779191764393 | 0.996940244108351 | 1228.7786545339 | 1318.5390651123 | 1337.34075766026 | 999.8298560792 | 1097.14811221074 | 1084.87393984951 | 1385.67493853022 | 1327.59776558958 | Pygo1 |
| ENSMUSG00000034911 | -0.202904922143487 | 0.298683077763266 | 0.851785577030345 | 438.204152535987 | 421.052931848708 | 408.868511896129 | 487.721881014244 | 586.897592959036 | 640.895089606702 | 386.239971596327 | 408.157612315359 | Ushbp1 |
| ENSMUSG00000034926 | -0.461204211905948 | 0.190908475260275 | 0.808824684419004 | 87.6408305071974 | 162.883145782866 | 89.9132144216024 | 120.992543559303 | 134.679858085749 | 136.985209992272 | 91.61487196894 | 272.466916731795 | Dhcr24 |
| ENSMUSG00000034928 | 0.0127980498988815 | 0.937100289025335 | 0.988856872640121 | 932.426155499254 | 1214.29385181126 | 1077.06566328193 | 903.223406570609 | 965.753128712444 | 825.580506649855 | 1155.59668051731 | 1143.05841959594 | Rnf44 |
| ENSMUSG00000034930 | -0.613120735391749 | 0.065170859335028 | 0.718339708336358 | 20.780815481088 | 26.8757190541728 | 16.0897331070236 | 34.703287687552 | 33.943704070392 | 50.1463715150282 | 29.1501865355718 | 37.9933947633978 | Rtkn |
| ENSMUSG00000034931 | 0.135936137747688 | 0.343596735956539 | 0.861266047293794 | 367.730082643601 | 419.424100390879 | 445.780252553418 | 317.019222659259 | 352.576539053749 | 327.786038195794 | 369.582722147429 | 360.394487469945 | Dhx8 |
| ENSMUSG00000034932 | -0.0170480762763047 | 0.935819381177567 | 0.98818087308472 | 1121.26052226218 | 771.251695281869 | 800.700835796586 | 1072.98813823134 | 954.803546754252 | 1278.12093251718 | 799.547973547113 | 779.407355431988 | Mrpl54 |
| ENSMUSG00000034936 | -0.768807981986346 | 0.028195919595714 | 0.665956123973759 | 214.132750826864 | 123.791190794978 | 166.576060402127 | 204.468019348279 | 409.514365236342 | 446.425014706959 | 188.435134390661 | 163.914360264945 | Arl4d |
| ENSMUSG00000034940 | 0.170886651060961 | 0.298756220220542 | 0.851785577030345 | 792.381529431053 | 1022.9061555164 | 978.634354862494 | 785.982569788339 | 753.331238723539 | 644.564336302924 | 943.21675004386 | 838.025735924088 | Synrg |
| ENSMUSG00000034947 | -0.401386198949848 | 0.241609396543891 | 0.832048466795696 | 47.8862269781594 | 70.0397526866323 | 53.9479286529615 | 60.9652351267805 | 95.2613630362614 | 118.638976511164 | 44.7663578939139 | 49.9341759747513 | Tmem106a |
| ENSMUSG00000034949 | 0.259872851793961 | 0.433310474656086 | 0.887868762931002 | 29.8159526467785 | 21.1748089517725 | 23.6613722162112 | 26.2619474392285 | 15.3294147414674 | 20.7923979452556 | 18.7394056300105 | 29.3091902460497 | Zfr2 |
| ENSMUSG00000034951 | -0.0264536847704489 | 0.819803413438822 | 0.966398681144067 | 601.740135234984 | 721.572335818095 | 714.573440929577 | 664.052099534778 | 728.147200219699 | 610.318033804856 | 732.91897575152 | 679.539003482486 | Cog7 |
| ENSMUSG00000034957 | -0.425493098161658 | 0.193055002458825 | 0.808863491502674 | 107.518132271716 | 239.438224300813 | 154.272146849697 | 144.440710915757 | 191.617684268342 | 302.101311322243 | 120.765058504512 | 254.012982132431 | Cebpa |
| ENSMUSG00000034959 | -0.28445095505064 | 0.5598759059477 | 0.916574868225118 | 5.42108229941427 | 8.14415728914329 | 15.1432782183751 | 17.8206071909051 | 10.949581958191 | 14.6769867848863 | 18.7394056300105 | 11.9407812113536 | Rubcnl |
| ENSMUSG00000034968 | -0.0475714281728889 | 0.894845024439869 | 0.983733637008786 | 28.0089252136404 | 17.1027303072009 | 34.0723759913441 | 27.1998741334867 | 33.943704070392 | 28.1308913376988 | 32.2734208072402 | 15.1973579053591 | Lbx2 |
| ENSMUSG00000034973 | 0.166634116172291 | 0.420543787192262 | 0.882568531013176 | 248.466272056487 | 329.023954481389 | 349.241853911277 | 287.943495137256 | 248.555510450935 | 203.031650524261 | 359.171941241867 | 270.295865602458 | Dopey1 |
| ENSMUSG00000034974 | -0.139224510863117 | 0.35336155185998 | 0.862751266538077 | 678.538801143353 | 673.52180781215 | 638.857049837702 | 764.410255820401 | 833.263187018333 | 895.296193878065 | 681.906149314269 | 625.26272524906 | Dapk3 |
| ENSMUSG00000034981 | 0.028825924487133 | 0.821259137364397 | 0.96663752376845 | 1853.10663268311 | 1963.55632241245 | 2095.45112346766 | 1879.60509529336 | 1754.12302970219 | 1621.80703972994 | 2102.9777429234 | 2156.93929699632 | Parm1 |
| ENSMUSG00000034987 | 0.0897946865419981 | 0.8362374859188 | 0.968924788429433 | 13.5527057485357 | 21.9892246806869 | 15.1432782183751 | 45.0204813243917 | 24.0890803080201 | 23.2385624094033 | 20.8215618111227 | 21.7105112933701 | Hrh2 |
| ENSMUSG00000034993 | -0.730084106735021 | 0.0528631008204724 | 0.703396849069134 | 374.054678659585 | 434.083583511337 | 500.674636095028 | 473.652980600371 | 919.764884488041 | 1283.01326144548 | 401.856142954669 | 352.795808517265 | Vat1 |
| ENSMUSG00000034994 | -0.0107682925645402 | 0.928451082707268 | 0.986901071486195 | 60298.6984163849 | 55807.0234081255 | 63538.3560396362 | 66214.8108345434 | 64205.0637282444 | 71262.8862517834 | 57254.0895901347 | 54979.6987993306 | Eef2 |
| ENSMUSG00000034998 | 0.0226817101675267 | 0.90123919969231 | 0.984802943363903 | 154.500845533307 | 175.099381716581 | 214.845259723197 | 182.895705380341 | 222.276513751277 | 157.777607937528 | 154.079557402308 | 181.282769299641 | Foxn2 |
| ENSMUSG00000035000 | -0.195615546936029 | 0.559318548085437 | 0.916574868225118 | 22.5878429142261 | 39.9063707168021 | 37.8581955459379 | 37.5170677703264 | 39.4184950494875 | 30.5770558018465 | 28.1091084450157 | 59.7039060567679 | Dpp4 |
| ENSMUSG00000035004 | 0.426879121335843 | 0.184376810667122 | 0.805963874499992 | 27.1054114970714 | 32.5766291565731 | 49.2156542097192 | 33.7653609932938 | 20.8042057205628 | 36.6924669622158 | 22.903717992235 | 26.0526135520442 | Igsf6 |
| ENSMUSG00000035007 | -0.0449632956433295 | 0.713953723530418 | 0.951176382858794 | 388.510898124689 | 395.806044252364 | 472.280989435575 | 407.9981120023 | 454.407651264925 | 380.37857417497 | 450.786813210807 | 429.868123608729 | Rundc1 |
| ENSMUSG00000035011 | -0.0973398111979835 | 0.317000576627621 | 0.854283473529811 | 1273.05082664578 | 1511.55559286499 | 1469.84444207104 | 1343.11102617769 | 1492.42802090143 | 1596.12231285639 | 1449.18070205414 | 1452.43320552646 | Zbtb7a |
| ENSMUSG00000035021 | -0.0157799539885279 | 0.913016436656081 | 0.986226496507248 | 172.571119864688 | 203.603932228582 | 217.684624389143 | 182.895705380341 | 213.516848184724 | 174.900759186562 | 204.051305749003 | 192.138024946326 | Baz1a |
| ENSMUSG00000035024 | 0.0550428032243822 | 0.793416410463166 | 0.961804768755116 | 216.843291976571 | 252.468875963442 | 195.916161950228 | 184.771558768858 | 170.813478547779 | 154.108361241306 | 256.10521027681 | 236.644573097735 | Ncapd3 |
| ENSMUSG00000035027 | -0.133475248187667 | 0.385629806975472 | 0.875126835008975 | 2748.48872580303 | 2329.22898469498 | 2223.2225334352 | 2780.95264847545 | 2777.90894279305 | 3403.83785186155 | 2378.86343692077 | 2499.96537543157 | Map2k2 |
| ENSMUSG00000035031 | -0.16146788797164 | 0.715818858986605 | 0.951274775289317 | 56.0178504272808 | 145.780415475665 | 40.6975602118832 | 93.7926694258161 | 65.6974917491458 | 69.71568722821 | 74.9576225200418 | 166.085411394282 | C8a |
| ENSMUSG00000035041 | -0.173144043658826 | 0.692275265624942 | 0.94469184542218 | 33.4300075130547 | 82.2559886203472 | 51.1085639870161 | 64.7169419038131 | 53.6529515951357 | 44.0309603546589 | 36.4377331694648 | 127.006491066215 | Creb3l3 |
| ENSMUSG00000035047 | 0.43598409307531 | 0.0153026097085361 | 0.635862253526407 | 145.465708367616 | 165.326392969609 | 164.68315062483 | 148.192417692789 | 121.54035973592 | 83.1695917810225 | 124.929370866736 | 130.263067760221 | Kri1 |
| ENSMUSG00000035048 | 0.0098225440336615 | 0.963512355514769 | 0.994284908685797 | 2829.80496029425 | 2085.7186817496 | 2441.85361271299 | 2784.70435525248 | 2469.13073157206 | 3577.51552881604 | 2049.88276030503 | 1977.82757882602 | Anapc13 |
| ENSMUSG00000035049 | 0.320110088429869 | 0.0462467587799732 | 0.69805115405055 | 898.9961479862 | 973.226796052623 | 910.489602879806 | 1027.96765690694 | 832.168228822514 | 552.833168897385 | 851.60187807492 | 814.144173501381 | Rrp12 |
| ENSMUSG00000035051 | 0.116866128438206 | 0.505633918824819 | 0.901494649536949 | 158.114900399583 | 166.955224427437 | 134.396594188079 | 136.937297361692 | 143.439523652302 | 124.754387671534 | 165.531416398426 | 116.15123541953 | Dhx57 |
| ENSMUSG00000035062 | -0.624261474139506 | 0.123408006338333 | 0.770251097497048 | 17.1667606148119 | 17.9171460361152 | 7.57163910918757 | 19.6964605794214 | 33.943704070392 | 23.2385624094033 | 22.903717992235 | 16.2828834700276 | Zc4h2 |
| ENSMUSG00000035064 | 0.132390515749398 | 0.315505735022463 | 0.853277176422981 | 769.793686516826 | 906.444706281648 | 923.739971320884 | 763.472329126143 | 824.50352145178 | 686.149132193435 | 853.684034256032 | 703.420565905193 | Eef2k |
| ENSMUSG00000035067 | 0.325601061838863 | 0.368849787711225 | 0.871495599696183 | 23.4913566307952 | 23.6180561385155 | 15.1432782183751 | 17.8206071909051 | 20.8042057205628 | 13.4539045528125 | 15.616171358342 | 14.1118323406906 | Xkr6 |
| ENSMUSG00000035069 | -0.0822331685424865 | 0.472932510409137 | 0.89728597737823 | 829.425591810383 | 784.282346944498 | 637.910594949053 | 763.472329126143 | 820.123688668504 | 759.534066117867 | 799.547973547113 | 813.058647936712 | Oma1 |
| ENSMUSG00000035078 | 0.126869179246668 | 0.641783545345993 | 0.934589624613243 | 182.509770746947 | 276.901347830872 | 312.330113253987 | 166.013024883695 | 197.092475247437 | 145.546785616789 | 255.064132186253 | 260.526135520442 | Mtmr9 |
| ENSMUSG00000035085 | -0.924231080340984 | 0.0414619852758463 | 0.683543358336265 | 37.9475760958999 | 40.7207864457164 | 32.1794662140472 | 38.4549944645846 | 87.5966556655277 | 130.869798831903 | 37.4788112600209 | 28.2236646813812 | 1700020L24R |
| ENSMUSG00000035086 | 0.108688241289219 | 0.147424998697134 | 0.780607990324772 | 1558.5611610816 | 1572.63677253357 | 1556.91829182669 | 1632.93037470346 | 1396.07169966935 | 1433.45237599056 | 1501.23460658195 | 1530.5910461826 | Becn1 |
| ENSMUSG00000035093 | 0.0108082236436701 | 0.963751554522777 | 0.994284908685797 | 482.47632464787 | 688.181290932608 | 650.214508501483 | 433.32213274727 | 539.814390538815 | 368.147751854232 | 685.029383585938 | 643.716659848425 | Secisbp2l |
| ENSMUSG00000035104 | -0.323959734514239 | 0.196795100102728 | 0.812065429583558 | 58.7283915769879 | 58.6379324818317 | 67.1982970940397 | 75.0341355406529 | 86.5016974697087 | 112.523565350795 | 58.3003730711437 | 68.388110574116 | Eva1a |

| | | | | | | | | | | | | |
|---|---|---|---|---|---|---|---|---|---|---|---|---|
| **ENSMUSG00000035105** | -0.154485053991649 | 0.393925995037695 | 0.878942305112115 | 2191.9242763965 | 2227.42701858069 | 2001.75208949147 | 1380.62809394801 | 2090.27519581866 | 1893.33129525033 | 2527.7376038703 | 2172.13665490168 | Egln3 |
| **ENSMUSG00000035107** | -0.132503238668089 | 0.521452445342822 | 0.90609507420585 | 164.439496415566 | 184.872370463553 | 205.380710836713 | 157.571684635371 | 180.668102310151 | 143.100621152642 | 257.146288367366 | 198.651178334337 | Dcbld2 |
| **ENSMUSG00000035109** | 0.399549290936355 | 0.237909756938484 | 0.832048466795696 | 31.6229800799166 | 41.5352021746308 | 33.1259211026956 | 28.1378008277448 | 16.4243729372865 | 18.3462334811079 | 36.4377331694648 | 30.3947158107182 | Shc4 |
| **ENSMUSG00000035112** | 0.220895199166589 | 0.126435148597344 | 0.771949105062466 | 219.553833126278 | 249.211213047785 | 247.971180825893 | 231.667893481766 | 244.175677667659 | 185.908499275227 | 197.804837205666 | 185.624871558315 | Wnk4 |
| **ENSMUSG00000035121** | 0.831448820820613 | 0.0139614490241707 | 0.635862253526407 | 28.0089252136404 | 54.56585383726 | 68.1447519826882 | 47.8342614071662 | 35.0386622662111 | 24.4616446414772 | 30.191264626128 | 21.7105112933701 | Neil2 |
| **ENSMUSG00000035125** | -0.0115609115917377 | 0.971362369151848 | 0.995122272737203 | 27.1054114970714 | 27.6901347830872 | 45.4298346551254 | 23.448167356454 | 35.0386622662111 | 22.0154801773295 | 32.2734208072402 | 34.7368180693922 | Gcfc2 |
| **ENSMUSG00000035126** | -0.477146652179985 | 0.294706203125977 | 0.851785577030345 | 13.5527057485357 | 8.95857301805762 | 7.57163910918757 | 6.56548685980713 | 9.85462376237187 | 15.9000690169602 | 12.4929370866736 | 13.0263067760221 | Wdr78 |
| **ENSMUSG00000035133** | 0.0989313961757998 | 0.714871260642728 | 0.951176382858794 | 887.250469670802 | 1246.87048096784 | 1180.22924614461 | 783.168789705565 | 689.823663366031 | 648.233582999146 | 1236.80077158069 | 1250.52545049812 | Arhgap5 |
| **ENSMUSG00000035139** | 0.105371170848063 | 0.480865200601057 | 0.897732059433541 | 180.702743313809 | 206.861595144239 | 205.380710836713 | 174.454365132018 | 168.623562156141 | 155.33144347338 | 203.010227658447 | 185.624871558315 | Secisbp2 |
| **ENSMUSG00000035142** | 0.0575680054730587 | 0.678193693860297 | 0.944452327903597 | 316.229800799166 | 344.497853330761 | 371.010316350191 | 344.219096792745 | 359.146288228664 | 267.855008824175 | 361.254097422979 | 332.170822788563 | Nubpl |
| **ENSMUSG00000035150** | 0.16441745845427 | 0.588588058221975 | 0.92352104528132 | 909.838312585028 | 1317.72464938338 | 1183.06861081056 | 874.147679048606 | 747.856447744443 | 500.240632918209 | 1306.55300364795 | 1267.89385953282 | Eif2s3x |
| **ENSMUSG00000035151** | -0.0442322976657222 | 0.825893128017846 | 0.967092863612197 | 177.088688447533 | 229.665235553841 | 224.309808609682 | 167.888878272211 | 211.326931793086 | 147.992950080937 | 221.749633288457 | 242.072200921077 | Elmod2 |
| **ENSMUSG00000035152** | -0.113242218480021 | 0.453910335584758 | 0.891964629399053 | 772.504227666534 | 1097.0179868476 | 945.508433759798 | 904.161333264867 | 1124.52206710621 | 1093.43551547403 | 883.87529888216 | 922.696729968231 | Ap2b1 |
| **ENSMUSG00000035158** | -0.0152189036347364 | 0.927239779298308 | 0.986790505471895 | 971.277245311723 | 797.312998607128 | 836.666121565227 | 912.602673513191 | 763.185862485911 | 714.280023531134 | 1055.65318382392 | 1020.3940307884 | Mitf |
| **ENSMUSG00000035161** | 0.127709458433092 | 0.61648729505839 | 0.928896975098882 | 211.422209677157 | 232.922898469498 | 227.149173275627 | 192.274972322923 | 203.662224422352 | 105.185071958352 | 232.160414194018 | 248.585354309088 | Ints6 |
| **ENSMUSG00000035164** | -0.104416007978122 | 0.605479904887029 | 0.92658295801349 | 112.939214571131 | 118.090280692578 | 128.717864856189 | 105.985716451172 | 128.110108910834 | 85.6157562451702 | 147.833088858971 | 137.8617467129 | Zc3h12c |
| **ENSMUSG00000035168** | -0.0372562813836106 | 0.74651828711119 | 0.956085046483146 | 832.13613296009 | 888.527560245533 | 892.506959995485 | 697.817460528072 | 882.536305830192 | 854.934480219628 | 807.876598271562 | 853.223093829447 | Tanc1 |
| **ENSMUSG00000035171** | -0.0597731554236873 | 0.697078206651926 | 0.944933386973863 | 253.887354355902 | 188.944449108124 | 233.774357496166 | 230.729966787508 | 224.466430142915 | 223.824048469516 | 220.708555197901 | 275.723493425801 | 1110059E24F |
| **ENSMUSG00000035172** | -0.267994743197481 | 0.0825055979928109 | 0.739855948845396 | 397.54603529038 | 357.52850499339 | 524.33600831124 | 488.659807708502 | 545.28918151791 | 565.063991218123 | 524.703357640293 | 492.828606359502 | Plekhh3 |
| **ENSMUSG00000035173** | -0.0581018900738306 | 0.72988655376565 | 0.952921973914044 | 550.239853390548 | 626.285695535119 | 572.60520763231 | 541.183702586959 | 527.769850384805 | 459.878919259771 | 692.316930219831 | 703.420565905193 | Ccdc186 |
| **ENSMUSG00000035181** | 0.0500648508691579 | 0.795949137658992 | 0.962200030474509 | 483.379838364439 | 665.377650523007 | 592.480760293928 | 478.342614071662 | 492.731188118594 | 402.3940543523 | 615.277151518677 | 632.86140420174 | Heatr5a |
| **ENSMUSG00000035184** | 0.106034820922543 | 0.344981622532482 | 0.861266047293794 | 267.440060104437 | 280.159010746529 | 296.240380146964 | 276.688374806158 | 263.884925192402 | 270.301173288323 | 242.57119509958 | 264.868237779116 | Fam124a |
| **ENSMUSG00000035189** | 0.110090617993842 | 0.778136322294025 | 0.960126492111037 | 106.614618555147 | 106.688460487777 | 68.1447519826882 | 74.0962088463947 | 62.4126171616885 | 31.8001380339204 | 121.806136595068 | 112.894658725525 | Ano4 |
| **ENSMUSG00000035198** | -0.0484772676928876 | 0.720372845270086 | 0.951674101666722 | 508.678222428372 | 484.577358704026 | 409.814966784777 | 454.894446715208 | 487.256397139498 | 552.833168897385 | 468.485140750262 | 414.67076570337 | Tubg1 |
| **ENSMUSG00000035199** | -0.0500059932635207 | 0.584738245273015 | 0.922641625223466 | 2341.0040396304 | 2122.36738955074 | 2267.70591320168 | 2465.80927920471 | 2424.23744554348 | 2544.01104271363 | 2327.85061048352 | 2225.32740757044 | Arl6ip5 |
| **ENSMUSG00000035202** | 0.399637940611896 | 0.201614503330286 | 0.813395633194566 | 9613.38594429464 | 8188.13573850466 | 11035.6640016409 | 16912.6941508632 | 13980.4262442182 | 7674.84100626347 | 5570.80886256589 | 7454.30405257864 | Lars2 |
| **ENSMUSG00000035203** | -0.0028877624666181 | 0.977235627915096 | 0.996179425433073 | 5561.12692548247 | 5491.60526006932 | 5196.98379356862 | 5111.70048370698 | 4721.45974037195 | 5736.2556684264 | 5796.72280821657 | 5150.81880435207 | Epn1 |
| **ENSMUSG00000035206** | -0.154858406102242 | 0.302356197138582 | 0.851785577030345 | 203.290586228035 | 255.726538879099 | 269.739643264807 | 205.405946042537 | 243.08071947184 | 253.178022039289 | 283.173240631269 | 260.526135520442 | Sppl2b |
| **ENSMUSG00000035208** | 0.0118343028399498 | 0.974071557469177 | 0.995510898988495 | 28.9124389302094 | 30.9477976987445 | 37.8581955459379 | 36.5791410760683 | 32.8487458745729 | 57.4848649074714 | 20.8215618111227 | 22.7960368580387 | Slfn8 |
| **ENSMUSG00000035212** | -0.185525700710724 | 0.373193801095409 | 0.873700364743909 | 618.906895849796 | 575.79192034243 | 642.642869392295 | 710.010507553428 | 836.54806160579 | 947.888729857241 | 593.414511616998 | 519.966745476215 | Leprot |
| **ENSMUSG00000035215** | 0.166674565874009 | 0.361143777554667 | 0.868539734006824 | 428.265501653727 | 302.96265115613 | 318.955297474527 | 377.984457786039 | 319.727793179176 | 395.055560959857 | 295.666177717943 | 262.697186649779 | Lsm7 |
| **ENSMUSG00000035226** | 0.437174384671627 | 0.111991763728242 | 0.764116514278315 | 118.360296870545 | 134.378595270864 | 101.270673085384 | 106.92364314543 | 86.5016974697087 | 44.0309603546589 | 111.395355689507 | 97.6973008201657 | Rims4 |
| **ENSMUSG00000035227** | -0.0702627829669442 | 0.682273747054077 | 0.944452327903597 | 1436.58680934478 | 1262.34437981721 | 1220.9268063565 | 1420.95894180111 | 1486.95322992233 | 1769.79998981087 | 1072.31043327282 | 1279.83464074417 | Spcs2 |
| **ENSMUSG00000035228** | 0.154902836758028 | 0.748383025644981 | 0.956197192463153 | 14.4562194651047 | 16.2883145782866 | 10.4110037751329 | 9.37926694258161 | 4.37983278327639 | 14.6769867848863 | 11.4518589961175 | 15.1973579053591 | Ccdc106 |
| **ENSMUSG00000035232** | -0.461805346142994 | 0.23475013667351 | 0.832048466795696 | 18.9737880479499 | 16.2883145782866 | 25.5542819935081 | 23.448167356454 | 33.943704070392 | 45.2540425867328 | 18.7394056300105 | 18.4539345993646 | Pdk3 |
| **ENSMUSG00000035234** | -0.0015766690602553 | 0.99406860472904 | 0.999262355603273 | 192.448421629207 | 187.315617650296 | 178.879973954556 | 159.447538023887 | 154.389105610493 | 125.977469903608 | 229.03717992235 | 208.420908416353 | Abraxas1 |
| **ENSMUSG00000035235** | -0.0784261723374936 | 0.67585655361324 | 0.94409858171788 | 102.097049972302 | 103.43079757212 | 72.8770264259304 | 90.9788893430416 | 88.6916138613469 | 110.077400886647 | 95.7791843311646 | 96.6117752554972 | Trim13 |
| **ENSMUSG00000035236** | 0.166785326740668 | 0.500823217085303 | 0.900927059989498 | 44.2721721118832 | 46.4216965481167 | 36.9117406572894 | 50.6480414899407 | 37.2285786578493 | 31.8001380339204 | 41.6431236222455 | 47.7631248454143 | Scai |
| **ENSMUSG00000035237** | -0.122113763492755 | 0.702309133588754 | 0.946434671388391 | 66.8600150261093 | 129.492100897378 | 76.6628459805242 | 85.3513291774927 | 76.6470737073368 | 105.185071958352 | 63.5057635239243 | 145.46042566558 | Lcat |
| **ENSMUSG00000035239** | -0.182549707511154 | 0.425793396906108 | 0.885066977635295 | 89.4478579403355 | 74.1118313312039 | 79.5022106464695 | 65.6548685980713 | 93.0714466446232 | 63.6002760678407 | 113.477511870619 | 79.2433662208011 | Neu3 |
| **ENSMUSG00000035242** | -0.0551908155082899 | 0.53175721663941 | 0.909082872831377 | 12446.8049594552 | 10716.8965767837 | 10552.0255535415 | 11215.7274099391 | 11269.3097513701 | 12053.4753970879 | 11319.6420786169 | 12041.7350888678 | Oaz1 |
| **ENSMUSG00000035245** | -0.29782485029228 | 0.0336848443884843 | 0.672620872762612 | 224.974915425692 | 215.005752433383 | 231.881447718869 | 229.79204009325 | 332.867291529005 | 271.524255520397 | 251.940897914585 | 251.841931003094 | Eogt |
| **ENSMUSG00000035247** | 0.35984894217918 | 0.227117222531655 | 0.832048466795696 | 2827.09441914454 | 4199.12749828228 | 4675.48714992333 | 2785.64228194674 | 2478.98535533444 | 1530.0758723244 | 3844.70138842381 | 3434.60288661116 | Hectd1 |
| **ENSMUSG00000035248** | 0.215476836494413 | 0.274803145284646 | 0.846286635233373 | 898.9961479862 | 1154.84150360052 | 1043.93974217924 | 902.285479876351 | 794.939650164664 | 601.756458180339 | 1014.01006020168 | 1033.42033756442 | Zcchc6 |
| **ENSMUSG00000035258** | -0.862710314461574 | 0.0812057384776548 | 0.739855948845396 | 39.754603529038 | 37.4631235300591 | 22.7149173275627 | 44.0825546301336 | 88.6916138613469 | 118.638976511164 | 28.1091084450157 | 27.1381391167127 | Abi3bp |
| **ENSMUSG00000035266** | 0.334363030846878 | 0.168793118473864 | 0.79907358295209 | 53.3073092775737 | 49.679359463774 | 53.001473764313 | 61.9031618210386 | 29.5638712871156 | 45.2540425867328 | 48.9306702561384 | 48.8486504100828 | Helq |
| **ENSMUSG00000035268** | -0.0864806515079826 | 0.578100531778577 | 0.922450244597442 | 4512.14750054581 | 3871.73237525872 | 3772.56918615271 | 4697.13688484487 | 4206.82938833697 | 5654.30915887745 | 4043.54730372003 | 3991.4775012861 | Pkig |
| **ENSMUSG00000035273** | 0.148069265894512 | 0.804528916488182 | 0.963461896828986 | 19.877301764519 | 13.8450673915436 | 53.001473764313 | 11.2551203310979 | 24.0890803080201 | 35.4693847301419 | 7.28754663389296 | 21.7105112933701 | Hpse |
| **ENSMUSG00000035275** | 0.133361864729289 | 0.565660132458351 | 0.919917953383268 | 132.81651633565 | 123.791190794978 | 145.754052851861 | 166.013024883695 | 165.338687568684 | 81.9465095489486 | 143.668776496747 | 125.920965501547 | Raver2 |
| **ENSMUSG00000035278** | 0.0203999069779335 | 0.897037342396254 | 0.984332015045905 | 468.020105182765 | 387.66188696322 | 430.636974335043 | 500.852854733858 | 490.541271726955 | 510.025290774799 | 379.99350305299 | 382.104998763315 | Plekhj1 |
| **ENSMUSG00000035279** | -1.38368980908846 | 0.0194409564094641 | 0.642273034804458 | 35.2370349461928 | 45.6072808192024 | 34.0723759913441 | 52.523894878457 | 116.065568756824 | 256.847268735511 | 35.3966550789086 | 29.3091902460497 | Ssc5d |
| **ENSMUSG00000035283** | -0.128366720462216 | 0.447743715434124 | 0.89108031640712 | 360.501972911049 | 378.703313945163 | 354.920583243168 | 413.625672167849 | 406.229490648885 | 376.709327478749 | 530.94982618363 | 333.256348353232 | Adrb1 |
| **ENSMUSG00000035284** | 0.468047699711212 | 0.200089618167433 | 0.812921911423739 | 777.925309965948 | 1452.91766038316 | 1500.13099850779 | 811.30659053331 | 641.645502749991 | 379.155491942897 | 1166.00746142287 | 1096.38082031519 | Vps13c |
| **ENSMUSG00000035285** | -0.52762867865135 | 0.144414137645461 | 0.780607990324772 | 47.8862269781594 | 47.2361122770311 | 41.6440151005317 | 34.703287687552 | 79.9319482947941 | 92.9542496376133 | 44.7663578939139 | 30.3947158107182 | Nat14 |
| **ENSMUSG00000035293** | -0.539631138218417 | 0.0082694401364517 | 0.635862253526407 | 49.6932544112975 | 54.56585383726 | 42.5904699891801 | 47.8342614071662 | 79.9319482947941 | 69.71568722821 | 64.5468416144805 | 69.4736361387845 | G2e3 |
| **ENSMUSG00000035296** | 0.075779561264143 | 0.348321642894785 | 0.861266047293794 | 5905.36565149528 | 5803.52648424351 | 6383.83822393377 | 6178.12313507851 | 5496.69014301187 | 5985.76444376947 | 6072.60850221395 | 5473.21989705862 | Sgcg |
| **ENSMUSG00000035297** | 0.178638933084916 | 0.0785045596036082 | 0.739855948845396 | 2038.32694457977 | 1995.31853584011 | 2234.57999209898 | 2160.9831035708 | 1786.97177557677 | 1671.95341124497 | 2035.30766703725 | 1950.68943970931 | Cops4 |
| **ENSMUSG00000035299** | -0.463875112280052 | 0.138259228504717 | 0.779711473022872 | 19.877301764519 | 31.7622134276588 | 19.8755526616174 | 18.7585338851632 | 25.1840385038392 | 34.2463024980681 | 34.3555769883525 | 31.4802413753867 | Mid1 |
| **ENSMUSG00000035305** | -0.352030011865645 | 0.154944265792462 | 0.787275540343856 | 54.2108229941427 | 54.56585383726 | 35.965285768641 | 37.5170677703264 | 62.4126171616885 | 67.2695227640623 | 54.1360607089191 | 49.9341759747513 | Ror1 |
| **ENSMUSG00000035310** | 0.0643962768805961 | 0.752326544099229 | 0.956497786093635 | 162.632468982428 | 179.985876090067 | 175.094154399963 | 135.999370667433 | 163.148771177045 | 107.6312364225 | 188.435134390661 | 164.999885829613 | Lin54 |
| **ENSMUSG00000035311** | 0.325889266229999 | 0.114383845683245 | 0.767518180080285 | 735.460165287203 | 956.124065745422 | 849.916490006305 | 805.679030367761 | 610.986673267056 | 454.986590331476 | 865.13589325215 | 738.157383974585 | Gnptab |
| **ENSMUSG00000035314** | -0.287916920795638 | 0.0624195185285398 | 0.715615581523459 | 316.229800799166 | 337.982527499446 | 252.703455269135 | 240.109233730089 | 344.911831683016 | 376.709327478749 | 357.089785060755 | 323.486618271215 | Gdpd5 |
| **ENSMUSG00000035325** | 0.163008299111567 | 0.269394599950104 | 0.843793744782845 | 1838.65041321801 | 2323.52807459258 | 2275.27755231087 | 1836.46046735748 | 1811.06085588479 | 1539.86053018099 | 2112.3474457384 | 1925.72235172193 | Sec31a |
| **ENSMUSG00000035329** | -0.0852550878638986 | 0.642653085324058 | 0.934616627390525 | 92.1583990900426 | 84.6992358070902 | 80.448665535118 | 84.4134024832345 | 83.2168228822514 | 112.523565350795 | 85.3684034256032 | 82.4999429148066 | Fbxo33 |
| **ENSMUSG00000035342** | -0.0647373595772125 | 0.467625581943021 | 0.895328112556867 | 1936.22989460746 | 1947.26800783416 | 2127.63058968171 | 2069.0662875335 | 2157.06764576362 | 2233.34815576687 | 2141.49763227397 | 1919.20919833392 | Lzts2 |
| **ENSMUSG00000035349** | -0.0341694370570228 | 0.950643552864361 | 0.991696503889296 | 9.03513716569045 | 21.9892246806869 | 7.57163910918757 | 7.50341355406529 | 7.66470737073368 | 11.0077400886647 | 9.36970281500523 | 19.5394601640331 | NA |
| **ENSMUSG00000035351** | -0.0514162424967264 | 0.764091682419741 | 0.958041307101859 | 88.5443442237664 | 91.2145616384048 | 90.8596693102509 | 96.6064495085906 | 100.736154015357 | 92.9542496376133 | 106.189965236726 | 80.3288917854696 | Nup37 |
| **ENSMUSG00000035354** | 0.06735221635847 | 0.645815552660399 | 0.935063240511793 | 543.011743657996 | 625.471279806204 | 738.234813145789 | 614.341984739096 | 618.65138063779 | 645.787418534998 | 643.386259963692 | 498.256234182845 | Uvrag |
| **ENSMUSG00000035356** | -0.028587091237745 | 0.926566120173629 | 0.986790505471895 | 22.5878429142261 | 17.9171460361152 | 27.447191770805 | 23.448167356454 | 20.8042057205628 | 22.0154801773295 | 24.9858741733473 | 24.9670879873757 | Nfkbiz |
| **ENSMUSG00000035357** | 0.193708080283597 | 0.338995260253099 | 0.858570960161243 | 1278.4719089452 | 1318.5390651123 | 1688.47552134883 | 1273.70445080258 | 1193.50443344282 | 843.926740130963 | 1567.86360437754 | 1253.78202719213 | Pdzrn3 |
| **ENSMUSG00000035367** | 0.159192257871554 | 0.284066780073443 | 0.851125503777472 | 145.465708367616 | 124.605606523892 | 141.021778408619 | 155.695831246855 | 125.920192519196 | 123.53130543946 | 124.929370866736 | 132.434118889558 | Rmi1 |
| **ENSMUSG00000035372** | -0.24598429898448 | 0.357995942796669 | 0.865834559821921 | 59.631905293557 | 72.4829998733752 | 59.6266579848522 | 77.8479156234274 | 97.4512794278996 | 106.408154190426 | 61.4236073428121 | 55.3618037980939 | 1810055G02 |
| **ENSMUSG00000035376** | -0.202735899653912 | 0.34635652587792 | 0.861266047293794 | 175.281661014395 | 168.584055885266 | 131.557229522134 | 155.695831246855 | 178.478185918513 | 231.162541861959 | 123.8882927618 | 194.309076075663 | Hacd2 |
| **ENSMUSG00000035378** | 0.0204412414524591 | 0.933752154962543 | 0.98818087308472 | 81.316234491214 | 98.5443031986338 | 74.7699362032273 | 88.1651092602672 | 101.831112211176 | 51.3694537471021 | 83.2862472444909 | 100.953877514171 | Shq1 |
| **ENSMUSG00000035382** | -0.109032878981253 | 0.469471978773089 | 0.895491044377672 | 239.431134890797 | 325.766291565731 | 279.204192151292 | 241.985087118606 | 283.594172717146 | 284.978160073209 | 326.898520434627 | 276.809018990469 | Pcsk7 |
| **ENSMUSG00000035385** | -0.832189729943054 | 0.189331863517645 | 0.808677608936066 | 4.51756858284523 | 6.51532583131463 | 6.62518422053913 | 17.8206071909051 | 26.2789966996583 | 22.0154801773295 | 7.28754663389296 | 7.59867895267955 | Ccl2 |
| **ENSMUSG00000035390** | -0.772091463294611 | 0.130963494928524 | 0.77303252008843 | 12.6491920319666 | 12.2162359337149 | 12.3039135524298 | 9.37926694258161 | 14.2344565456483 | 41.5847958905112 | 13.5340151772298 | 10.8552556466851 | Brsk1 |
| **ENSMUSG00000035392** | 0.0998418962885156 | 0.630371933693673 | 0.931825800800958 | 397.54603529038 | 483.762942975111 | 480.799083433411 | 317.019222659259 | 372.285786578493 | 287.424324537357 | 457.033281754144 | 449.407583772762 | Dennd1a |
| **ENSMUSG00000035397** | -0.528745856456877 | 0.0587042305174503 | 0.711712402279519 | 110.228673421423 | 78.1839099757755 | 93.6990339761962 | 112.551203310979 | 169.71852035196 | 200.585486060113 | 105.14888714617 | 94.4407241261602 | Klf16 |
| **ENSMUSG00000035399** | -0.0532436801775542 | 0.680453021020088 | 0.944452327903597 | 234.913566307952 | 227.221988367098 | 238.506631939409 | 215.723139679377 | 250.745426842573 | 264.185762127954 | 212.379930473452 | 224.703791886381 | Oser1 |
| **ENSMUSG00000035401** | -0.062987939170107 | 0.625423415174527 | 0.931573470007391 | 443.625234835401 | 393.362797065621 | 381.421320125324 | 430.508352664496 | 497.11102090187 | 436.640356850368 | 383.116737324658 | 405.986561186022 | Emsy |
| **ENSMUSG00000035407** | -0.748844819099513 | 0.147323724318328 | 0.780607990324772 | 9.93865088225949 | 13.8450673915436 | 16.0897331070236 | 20.6343872736795 | 39.4184950494875 | 39.1386314263635 | 11.4518589961175 | 11.9407812113536 | Kank4 |
| **ENSMUSG00000035413** | 0.0145164726962622 | 0.925938775319028 | 0.986713918026868 | 271.054114970713 | 337.168111770532 | 279.204192151292 | 306.702029022419 | 316.442918591719 | 347.355353908976 | 246.735507461804 | 273.552442296464 | Tmem98 |
| **ENSMUSG00000035437** | 0.229115373121935 | 0.158574984583313 | 0.789230661864335 | 824.908023227538 | 898.300548992504 | 844.237760674415 | 769.099889291692 | 612.081631462875 | 572.402484610566 | 862.012658980481 | 798.946815596022 | Rabgap1 |
| **ENSMUSG00000035439** | 0.0559611509740936 | 0.74913385755069 | 0.956197192463153 | 567.40661400536 | 513.081909216027 | 356.813493020464 | 554.314676306573 | 527.769850384805 | 431.748027922072 | 458.0743598447 | 498.256234182845 | Haus8 |
| **ENSMUSG00000035441** | -0.40467571993804 | 0.204739195675554 | 0.818367605160705 | 133.720030052219 | 165.326392969609 | 148.593417517806 | 144.440710915757 | 267.16979977986 | 280.085831144914 | 133.257995591185 | 104.210454208177 | Myo1d |
| **ENSMUSG00000035443** | 0.0780404249326543 | 0.559980990053627 | 0.916574868225118 | 432.783070236573 | 372.187988113848 | 312.330113253987 | 395.805064976944 | 358.051330032845 | 346.132271676902 | 362.295175513536 | 366.907640857956 | Thyn1 |

| | | | | | | | | | | | | |
|---|---|---|---|---|---|---|---|---|---|---|---|---|
| **ENSMUSG00000035458** | -0.129006486449984 | 0.485631431161692 | 0.899236955296196 | 94098.2430395163 | 73292.5291079161 | 81152.8279722724 | 92793.7774964312 | 98945.8973651927 | 121711.359078134 | 76522.3628901477 | 76087.7434043097 | Tnni3 |
| **ENSMUSG00000035459** | 0.727055475977574 | 0.0911950552316861 | 0.753823363185192 | 37.9475760958999 | 38.2775392589734 | 61.519567762149 | 59.0893817382642 | 47.0832024202212 | 8.56157562451702 | 33.3144988977964 | 29.3091902460497 | Stab2 |
| **ENSMUSG00000035469** | 0.0365131462634402 | 0.741594252277201 | 0.954383574136061 | 274.66816983699 | 260.613033252585 | 268.793188376159 | 267.309107863576 | 272.644590758955 | 245.839528646846 | 254.023054095697 | 271.381391167127 | Rcbtb1 |
| **ENSMUSG00000035473** | 0.0763141577834144 | 0.666986414985321 | 0.942024383071524 | 459.888481733644 | 524.483729420828 | 448.619617219364 | 366.729337454941 | 388.710159515779 | 340.016860516533 | 489.306702561384 | 488.486504100828 | Galm |
| **ENSMUSG00000035476** | 0.163880016005647 | 0.374690141014639 | 0.873700364743909 | 233.106538874814 | 255.726538879099 | 247.971180825893 | 208.219726125312 | 199.282391639076 | 157.777607937528 | 218.626399016789 | 267.039288908453 | Tab3 |
| **ENSMUSG00000035478** | -0.0818258997314833 | 0.575622031620364 | 0.922450244597442 | 3161.39449427509 | 2679.42774812814 | 2673.73506043186 | 3313.69501081408 | 2972.81150164885 | 3838.03204424777 | 2870.25229566327 | 2838.64935160815 | Mbd3 |
| **ENSMUSG00000035486** | 0.0979520701272753 | 0.808878744318205 | 0.964157363647118 | 40.658117245607 | 30.1333819698302 | 27.447191770805 | 28.1378008277448 | 41.6084114411257 | 17.123151249034 | 43.7252798033577 | 15.1973579053591 | Plk5 |
| **ENSMUSG00000035493** | -0.735138800891546 | 0.0647833409873406 | 0.718339708336358 | 217.74680569314 | 256.540954608014 | 255.542819935081 | 271.060814640609 | 499.300937293508 | 760.757148349941 | 182.188665847324 | 224.703791886381 | Tgfbi |
| **ENSMUSG00000035495** | 0.35416269566302 | 0.325179738567508 | 0.855977678491063 | 191.544907912638 | 269.571606270643 | 258.382184601026 | 151.944124469822 | 135.774816281568 | 73.3849339244316 | 234.242570375131 | 237.730098662403 | Tstd2 |
| **ENSMUSG00000035504** | -0.0499475068956231 | 0.861739106832606 | 0.974868041734905 | 290.027903018663 | 380.332145402991 | 262.16800415562 | 282.315934971707 | 232.131137513649 | 227.493295165738 | 276.926772087932 | 521.052271040884 | Reep6 |
| **ENSMUSG00000035505** | -0.260710773440528 | 0.12460963283617 | 0.771949105062466 | 145.465708367616 | 157.182235680465 | 123.039135524298 | 152.88205116408 | 193.80760065998 | 190.800828203522 | 139.504464134522 | 170.427513652956 | Cox18 |
| **ENSMUSG00000035506** | -0.219003781445096 | 0.741584658487524 | 0.954383574136061 | 9.03513716569045 | 9.77298874697194 | 14.1968233297267 | 7.50341355406529 | 10.949581958191 | 28.1308913376988 | 3.12323427166841 | 5.42762782334254 | Slc12a8 |
| **ENSMUSG00000035513** | -0.133746342194222 | 0.523011530358234 | 0.90648515894888 | 307.194663633475 | 325.766291565731 | 175.094154399963 | 315.143369270742 | 317.537876787538 | 330.232202659942 | 295.666177717943 | 289.835325766492 | Ntng2 |
| **ENSMUSG00000035517** | -0.141538556445439 | 0.290915341090377 | 0.851785577030345 | 269.247087537575 | 266.313943354985 | 312.330113253987 | 250.426427366929 | 330.677375137367 | 309.439804714686 | 265.474913091815 | 306.118209236519 | Tdrd7 |
| **ENSMUSG00000035521** | -0.0476427325040384 | 0.818451124848389 | 0.966315773308182 | 241.238162323935 | 171.841718800923 | 232.827902607518 | 243.860940507122 | 270.454674367317 | 277.639666680766 | 190.517290571773 | 181.282769299641 | Gnptg |
| **ENSMUSG00000035529** | 0.152507567914107 | 0.416616251335553 | 0.882192126163777 | 332.493047697409 | 394.177212794535 | 414.54724122802 | 310.453735799451 | 315.3479603959 | 239.724117486476 | 387.281049686883 | 362.565538599282 | Prdm4 |
| **ENSMUSG00000035530** | 0.119514032135813 | 0.300651541994482 | 0.851785577030345 | 18420.8376534097 | 15684.8325231611 | 18141.6473056134 | 19800.570442484 | 17643.0614092331 | 18003.7704561272 | 15547.4602043653 | 15125.713218091 | Eif1 |
| **ENSMUSG00000035540** | -0.2679389764737 | 0.400334892792398 | 0.880487015130533 | 704.740698923855 | 1398.3518065459 | 763.789095139297 | 950.119741283517 | 762.090904290091 | 954.004141017611 | 981.736639394437 | 1898.58421260522 | Gc |
| **ENSMUSG00000035545** | -0.0142125518686253 | 0.953419499727805 | 0.992207750896771 | 347.852780879082 | 262.241864710414 | 311.383658365339 | 330.150196378873 | 314.253002200081 | 185.908499275227 | 425.80093903746 | 336.512925047237 | Leng8 |
| **ENSMUSG00000035547** | -0.594174186632199 | 0.143151710835478 | 0.780607990324772 | 30.7194663633475 | 39.0919549878878 | 24.6078271048596 | 24.3860940507122 | 48.1781606160403 | 73.3849339244316 | 18.7394056300105 | 40.1644458927348 | Capn5 |
| **ENSMUSG00000035559** | -0.0515278955260462 | 0.817755784551126 | 0.966315773308182 | 473.44118748218 | 362.414999366876 | 412.654331450723 | 483.032247542953 | 513.535393839156 | 601.756458180339 | 343.555769883525 | 336.512925047237 | Mpv17l2 |
| **ENSMUSG00000035561** | -0.0305503547515599 | 0.829784594488901 | 0.967535167881372 | 296.352499034647 | 325.766291565731 | 320.848207251823 | 285.129715054481 | 279.21433993387 | 371.816998550453 | 326.898520434627 | 277.894544555138 | Aldh1b1 |
| **ENSMUSG00000035566** | -0.107588755788093 | 0.708763337270671 | 0.949267137065756 | 58.7283915769879 | 56.1946852950887 | 96.5383986421416 | 105.047789756914 | 97.4512794278996 | 106.408154190426 | 69.7522320672611 | 67.3025850094475 | Pcdh17 |
| **ENSMUSG00000035569** | 0.227589373741701 | 0.147705804492821 | 0.780607990324772 | 3511.05430258731 | 4140.48956580045 | 3644.79777618517 | 3051.0755364218 | 2704.54674367317 | 2579.48042744377 | 3485.52944718195 | 3483.45153702124 | Ankrd11 |
| **ENSMUSG00000035572** | 0.0349942488543989 | 0.804893674605474 | 0.963461896828986 | 275.571683553559 | 359.971752180133 | 312.330113253987 | 297.322762079837 | 341.626957095558 | 272.747337752471 | 312.323427166841 | 288.749800201823 | Dcaf10 |
| **ENSMUSG00000035575** | 0.0688872885592166 | 0.618249082911148 | 0.929568738467614 | 969.470217878585 | 1019.64849260074 | 998.509907524111 | 895.719993016544 | 929.619508250413 | 731.403174780168 | 987.983107937774 | 1051.87427216378 | Utp6 |
| **ENSMUSG00000035578** | 0.389414328582677 | 0.466576207197919 | 0.895328112556867 | 6.32459601598331 | 10.5874044758863 | 13.2503684410783 | 15.9447538023887 | 8.75966556655277 | 3.66924669622158 | 8.32862472444909 | 14.1118323406906 | Iqcg |
| **ENSMUSG00000035585** | -0.139694554741583 | 0.326667888219272 | 0.855977678491063 | 1552.23656506562 | 1373.91933467847 | 1433.8791563024 | 1497.86893073028 | 1768.35748624784 | 1894.55437748241 | 1480.41304477083 | 1311.31488211956 | Tsen34 |
| **ENSMUSG00000035595** | -0.0354554536958821 | 0.880010138743506 | 0.978393501240173 | 88.5443442237664 | 72.4829998733752 | 88.966759532954 | 115.364983393754 | 101.831112211176 | 119.862058743238 | 74.9576225200418 | 78.1578406561325 | 1600002K03[illegible] |
| **ENSMUSG00000035596** | 0.0598373833415315 | 0.639675227990362 | 0.933896249399425 | 404.774145022932 | 523.669313691913 | 481.745538322059 | 493.349441179793 | 501.490853685146 | 417.071041137186 | 460.156516025812 | 447.236532643425 | Mboat7 |
| **ENSMUSG00000035597** | 0.366190497904789 | 0.0689848189575027 | 0.724157781586334 | 288.220875585525 | 315.99330281876 | 256.489274823729 | 225.102406621959 | 189.427767876704 | 161.446854633749 | 217.585320926233 | 273.552442296464 | Prpf39 |
| **ENSMUSG00000035601** | 0.113330504500899 | 0.62444714171546 | 0.931259048662274 | 120.167324303683 | 83.8848200781758 | 74.7699362032273 | 118.178763476528 | 111.685735973548 | 81.9465095489486 | 93.6970281500523 | 79.2433662208011 | Trmt10b |
| **ENSMUSG00000035606** | -0.492701034376653 | 0.127439648852172 | 0.772837268577874 | 107.518132271716 | 100.173134656462 | 84.2344850897118 | 91.9168160372998 | 90.881530252985 | 73.3849339244316 | 188.435134390661 | 186.710397122983 | Ky |
| **ENSMUSG00000035614** | -0.141929867830557 | 0.495254620834417 | 0.900742507277983 | 145.465708367616 | 239.438224300813 | 244.185361271299 | 159.447538023887 | 202.567266226533 | 215.262472844999 | 232.160414194018 | 220.361689627707 | Togaram1 |
| **ENSMUSG00000035620** | 0.20193205316305 | 0.0914802841487472 | 0.753823363185192 | 1225.16459966762 | 1221.62359337149 | 1232.28426502028 | 1197.73238856767 | 932.904382837871 | 952.781058785537 | 1131.65188443452 | 1221.21626025207 | Ric8b |
| **ENSMUSG00000035621** | 0.00437618921745397 | 0.988112553451954 | 0.998437672973644 | 1787.15013137357 | 2567.03837753796 | 1764.1919124407 | 868.520118883057 | 2097.93990318939 | 1878.65430846545 | 1589.72624427922 | 1400.32797842237 | Midn |
| **ENSMUSG00000035623** | 0.186825624831288 | 0.105535921183097 | 0.762094734041179 | 468.020105182765 | 504.937751926884 | 468.495169880981 | 509.294194982182 | 427.033696369448 | 375.486245246675 | 478.895921655823 | 430.953649173397 | Rsf1 |
| **ENSMUSG00000035629** | 0.140658648419615 | 0.423887053346246 | 0.884584010063004 | 553.853908256825 | 710.984931342209 | 704.162437154444 | 568.383576720446 | 605.511882287961 | 424.409534529629 | 619.441463880901 | 651.315338801105 | Rubcn |
| **ENSMUSG00000035632** | -0.0808755677458484 | 0.449522433852572 | 0.89108031640712 | 318.036828232304 | 357.52850499339 | 341.670214802089 | 345.157023487003 | 377.760577557588 | 375.486245246675 | 341.473613702413 | 347.368180693922 | Cnot3 |
| **ENSMUSG00000035637** | -0.0155467496533797 | 0.886796951527812 | 0.980467217760193 | 1401.34977439859 | 1390.20764925676 | 1467.00507740509 | 1380.62809394801 | 1659.95662486175 | 1377.19059331517 | 1290.93683228961 | 1372.10431374099 | Grhpr |
| **ENSMUSG00000035640** | 0.175550957951258 | 0.497685767717438 | 0.900927059989498 | 37.9475760958999 | 36.6487078011448 | 43.5369248778286 | 40.3308478531009 | 42.7033696369448 | 28.1308913376988 | 39.5609674411332 | 29.3091902460497 | Cbarp |
| **ENSMUSG00000035642** | 0.0287844577150639 | 0.821782246053676 | 0.966637987288264 | 1194.44513330428 | 978.113290426109 | 961.598166866822 | 1170.53251443419 | 1008.45649834939 | 1188.83592957579 | 970.284780398319 | 1052.95979772845 | Aamdc |
| **ENSMUSG00000035649** | 0.141590962453669 | 0.435996207606637 | 0.888687115594624 | 111.132187137993 | 91.2145616384048 | 96.5383986421416 | 116.302910088012 | 78.836990098975 | 106.408154190426 | 98.902418602833 | 92.2696729968231 | Zcchc7 |
| **ENSMUSG00000035666** | 0.14471096432051 | 0.522822650083401 | 0.90648515894888 | 356.887918044773 | 481.319695788368 | 505.406910538271 | 355.474217123843 | 347.101748074654 | 262.96267989588 | 457.033281754144 | 468.947043936795 | Gtf3c4 |
| **ENSMUSG00000035671** | 0.0223361046552855 | 0.903079452275908 | 0.985027469232262 | 149.079763233892 | 178.357044632238 | 230.934992830221 | 226.040333316217 | 202.567266226533 | 170.008430258266 | 197.804837205666 | 200.822229463674 | Zswim4 |
| **ENSMUSG00000035673** | -0.0397853946266035 | 0.827568227213419 | 0.967319591893432 | 234.010052591383 | 288.303168035672 | 345.456034356683 | 375.170677703265 | 369.000911991036 | 284.978160073209 | 313.364505257397 | 309.374785930525 | Sbno2 |
| **ENSMUSG00000035674** | 0.214361904590593 | 0.214748643899143 | 0.824248286808591 | 10185.3101268828 | 7008.86176303671 | 8188.72769658636 | 8940.31724966879 | 7031.82153355024 | 9381.0407200065 | 6472.3824889875 | 6699.86378513403 | Ndufa3 |
| **ENSMUSG00000035686** | 0.00248868246707643 | 0.991582688033387 | 0.998628446692557 | 928.812100632978 | 623.842448348376 | 636.964140060405 | 617.15576482187 | 738.001823982071 | 949.111812089315 | 482.019155927491 | 633.946929766408 | Thrsp |
| **ENSMUSG00000035692** | 0.148189293292978 | 0.764665026141352 | 0.958110466593105 | 74.0881247586617 | 52.9370223794314 | 167.522515290775 | 62.8410885152968 | 89.7865720571659 | 143.100621152642 | 29.1501865355718 | 60.7894316214364 | Isg15 |
| **ENSMUSG00000035696** | 0.0268219024717852 | 0.864525062275429 | 0.974868041734905 | 329.782506547701 | 330.652785939217 | 358.706402797761 | 283.253861665965 | 308.778211220985 | 255.624186503437 | 339.391457521301 | 373.420794245967 | Rnf38 |
| **ENSMUSG00000035697** | -0.182554234407409 | 0.486301319022007 | 0.899236955296196 | 56.0178504272808 | 67.5965054998893 | 98.4313084194385 | 66.5927952923295 | 90.881530252985 | 105.185071958352 | 62.4646854333682 | 69.4736361387845 | Arhgap45 |
| **ENSMUSG00000035704** | 0.1023385201159 | 0.631662283759318 | 0.931825800800958 | 55.1143367107117 | 65.1532583131463 | 60.5731128735006 | 56.2756016554897 | 58.0327843784121 | 66.0464405319884 | 52.0539045278068 | 45.5920737160773 | Alg8 |
| **ENSMUSG00000035711** | -0.00477280893660115 | 0.991845339411814 | 0.998820544075263 | 18.0702743313809 | 21.1748089517725 | 19.8755526616174 | 25.3240207449704 | 25.1840385038392 | 37.9155491942896 | 14.5750932677859 | 7.59867895267955 | Dok3 |
| **ENSMUSG00000035713** | -0.538775854204585 | 0.295117151818162 | 0.851785577030345 | 13.5527057485357 | 4.88649437348597 | 7.57163910918757 | 8.44134024832345 | 12.0445401540101 | 14.6769867848863 | 16.6572494488982 | 6.51315338801105 | Usp35 |
| **ENSMUSG00000035722** | -0.130336351993007 | 0.465171672239356 | 0.895328112556867 | 206.904641094311 | 284.231089391101 | 308.544293699394 | 226.978260010475 | 317.537876787538 | 302.101311322243 | 254.023054095697 | 250.756405438425 | Abca7 |
| **ENSMUSG00000035725** | -0.0660973166940185 | 0.501136884499085 | 0.900927059989498 | 558.37147683967 | 518.782819318427 | 554.62256474799 | 486.783954319986 | 545.28918151791 | 540.602346576646 | 549.68923181364 | 581.84170266232 | Prkx |
| **ENSMUSG00000035726** | 0.144451362453333 | 0.444753437264344 | 0.89108031640712 | 481.572810931301 | 526.926976607571 | 569.765842966365 | 425.818719193205 | 460.97740043984 | 316.77829810713 | 517.4158110064 | 516.71016878221 | Supt16 |
| **ENSMUSG00000035735** | -0.0436778439215123 | 0.875714799810584 | 0.977159268104838 | 65.9565013095403 | 80.6271571625185 | 48.2691993210708 | 39.3929211588428 | 49.2731188118594 | 58.7079471395453 | 62.4646854333682 | 71.6446872681215 | Dagla |
| **ENSMUSG00000035745** | 0.164806226041446 | 0.484531030823766 | 0.899236955296196 | 58.7283915769879 | 67.5965054998893 | 70.0376617599851 | 72.2203554578784 | 66.7924499449649 | 37.9155491942896 | 68.711153976705 | 65.1315338801105 | Grin3b |
| **ENSMUSG00000035754** | -0.0617339365438579 | 0.660332173719376 | 0.939611397743053 | 1214.3224350688 | 1073.39993070909 | 1025.01064440627 | 1271.82859741407 | 1246.06242684213 | 1384.52908670761 | 1155.59668051731 | 999.769045059695 | Wdr18 |
| **ENSMUSG00000035757** | 0.0857750216330636 | 0.511754716073498 | 0.902394714365738 | 294.545471601509 | 245.139134403213 | 300.026199701558 | 287.943495137256 | 296.733671066975 | 256.847268735511 | 255.064132186253 | 252.927456567762 | Selenoo |
| **ENSMUSG00000035759** | -0.27047960483993 | 0.445415116525402 | 0.89108031640712 | 23.4913566307952 | 26.0613033252585 | 25.5542819935081 | 37.5170677703264 | 33.943704070392 | 22.0154801773295 | 53.094982618363 | 26.0526135520442 | Bbs10 |
| **ENSMUSG00000035762** | 0.0492282528367715 | 0.822554475790923 | 0.966637987288264 | 118.360296870545 | 145.780415475665 | 186.451613063744 | 123.806323642077 | 133.58489988993 | 103.961989726278 | 162.408182126757 | 154.144630182928 | Tmem161b |
| **ENSMUSG00000035764** | 0.0839631678918167 | 0.578190260908584 | 0.922450244597442 | 489.704434380422 | 477.247617143797 | 491.210087208544 | 434.260059441529 | 422.653863586171 | 354.693847301419 | 471.60837502193 | 535.164103381574 | Fbxo45 |
| **ENSMUSG00000035765** | 0.0878603843421408 | 0.233991342741787 | 0.832048466795696 | 2531.64543382646 | 2624.04747856197 | 2449.42525182218 | 2669.33937185873 | 2419.8576127602 | 2294.50226737056 | 2469.43723079916 | 2482.59696639688 | Dym |
| **ENSMUSG00000035769** | -0.241229017268465 | 0.673727432667698 | 0.943734776094628 | 16.2632468982428 | 28.5045505120015 | 14.1968233297267 | 24.3860940507122 | 15.3294147414674 | 11.0077400886647 | 13.5340151772298 | 58.6183804920994 | Xylb |
| **ENSMUSG00000035770** | 0.181373240893986 | 0.439170513188978 | 0.889583654898613 | 2347.32863564638 | 2659.06735490528 | 2471.19371426109 | 2217.25870522629 | 1647.91208470774 | 1408.99073134909 | 2772.39095515099 | 2719.24153949461 | Dync1li2 |
| **ENSMUSG00000035772** | 0.0139233511745878 | 0.926706493792529 | 0.986790505471895 | 672.214205127369 | 640.945178655577 | 576.391027186904 | 649.045272426648 | 594.56230032977 | 494.125221757839 | 687.11153976705 | 737.071858409917 | Mrps2 |
| **ENSMUSG00000035776** | 0.148445390723354 | 0.380508653019754 | 0.87432013076003 | 2142.23102198521 | 2511.65810797179 | 2551.64237979621 | 2278.22394035307 | 1867.99868206738 | 1658.49950669215 | 2531.90191623252 | 2496.70879873757 | Cd99l2 |
| **ENSMUSG00000035778** | -0.597492800680293 | 0.0475184263530348 | 0.703396849069134 | 427.361987937158 | 314.364471360931 | 387.100049457215 | 407.060185308042 | 808.079148514494 | 792.557286383861 | 336.268223249632 | 387.532626586657 | Ggta1 |
| **ENSMUSG00000035780** | -0.355109729879366 | 0.141917306757758 | 0.78006782384854 | 65.0529875929712 | 76.5550785179469 | 50.1621090983677 | 56.2756016554897 | 71.1722827282413 | 77.0541806206532 | 64.5468416144805 | 105.295979772845 | Ugt2a3 |
| **ENSMUSG00000035781** | 0.02249859115318 | 0.815899111316657 | 0.966177091475091 | 1657.9476699042 | 1517.25650296739 | 1678.0645175737 | 1737.04023776611 | 1518.70701760109 | 1791.8154699882 | 1577.23330719255 | 1601.15020788605 | R3hdm4 |
| **ENSMUSG00000035783** | -0.812344217987007 | 0.0375195357410462 | 0.679566075569698 | 1175.47134525633 | 1786.82810923804 | 773.253644025781 | 908.850966736158 | 2321.31137513649 | 3528.59223953308 | 1225.34891258457 | 1082.2689879745 | Acta2 |
| **ENSMUSG00000035790** | 0.00934271278923385 | 0.951877071512143 | 0.991915011884503 | 287.317361868956 | 269.571606270643 | 239.453086828057 | 279.502154888932 | 254.03030143003 | 214.039390612925 | 295.666177717943 | 303.947158107182 | Cep19 |
| **ENSMUSG00000035798** | -0.147227684686593 | 0.467120437599194 | 0.895328112556867 | 271.957628687283 | 339.611358957275 | 324.634026806417 | 245.736793895638 | 287.974005500422 | 243.393364182698 | 390.404283958551 | 386.447101021989 | Zdhhc17 |
| **ENSMUSG00000035799** | -0.953744980030781 | 0.0862378597507058 | 0.746460787597018 | 15.3597331816738 | 17.9171460361152 | 23.6613722162112 | 25.3240207449704 | 68.9823663366031 | 59.9310293716191 | 20.8215618111227 | 9.76973008201657 | Twist1 |
| **ENSMUSG00000035811** | -0.277239281204669 | 0.416380450705116 | 0.882192126163777 | 46.0791995450213 | 59.452348210746 | 59.6266579848522 | 60.9652351267805 | 48.1781606160403 | 56.2617826753975 | 47.8895921655823 | 121.578863242873 | Ugt2b35 |
| **ENSMUSG00000035824** | 0.147737553563286 | 0.337901785349375 | 0.858570960161243 | 202.387072511466 | 193.016527752696 | 244.185361271299 | 247.612647284155 | 204.757182618171 | 171.23151249034 | 208.215618111227 | 214.934061804364 | Tk2 |
| **ENSMUSG00000035828** | -0.15117011543238 | 0.530434616677049 | 0.908923862033581 | 2834.32252887709 | 2738.06568060997 | 2644.39495888376 | 2054.05946042537 | 3987.83774917315 | 3228.93709267499 | 2303.90581440073 | 1885.5579058292 | Pim3 |
| **ENSMUSG00000035829** | 0.330672110775837 | 0.361468582740268 | 0.868564573435346 | 66.8600150261093 | 65.9676740420606 | 47.3227444324223 | 73.1582821521366 | 36.1336204620302 | 23.2385624094033 | 69.7522320672611 | 71.6446872681215 | Ppp1r26 |
| **ENSMUSG00000035834** | 0.198745072414855 | 0.381696823975348 | 0.875126835008975 | 182.509770746947 | 228.850819824926 | 164.68315062483 | 220.412773150668 | 127.015150715015 | 141.877538920568 | 225.913945650682 | 198.651178334337 | Polr3g |
| **ENSMUSG00000035836** | -0.357371768594635 | 0.311273792655497 | 0.851785577030345 | 84.0267756409212 | 113.203786319092 | 63.4124775394459 | 112.551203310979 | 73.3621991198795 | 102.738907494204 | 92.6559500594962 | 209.506433981022 | Ugt2b1 |
| **ENSMUSG00000035840** | 0.0963936321456249 | 0.674857892455099 | 0.944023556644042 | 196.062476495483 | 289.931999493501 | 269.739643264807 | 210.095579513828 | 173.003394939417 | 178.570005882784 | 282.132162540713 | 269.21034003779 | Lysmd3 |
| **ENSMUSG00000035842** | 0.321097736245088 | 0.365528170981191 | 0.86924083181804 | 15.3597331816738 | 23.6180561385155 | 19.8755526616174 | 22.5102406621959 | 13.1394983498292 | 14.6769867848863 | 16.6572494488982 | 20.6249857287016 | Ddx11 |
| **ENSMUSG00000035845** | 0.0893636135554356 | 0.643916933256301 | 0.934847999069355 | 96.6759676728878 | 92.8433930962335 | 131.557229522134 | 96.6064495085906 | 96.3563212320805 | 103.961989726278 | 108.272121417838 | 83.5854684794751 | Alg12 |
| **ENSMUSG00000035847** | -0.143520201179083 | 0.312319719305971 | 0.852283084707256 | 470.730646332472 | 609.182965227918 | 616.142132510139 | 478.342614071662 | 607.701798679599 | 549.163922201163 | 588.209121164217 | 656.742966624447 | Ids |
| **ENSMUSG00000035849** | 0.16425700391894 | 0.563921667219922 | 0.918590066040684 | 507.774708711803 | 456.072808192024 | 399.403963009645 | 536.494069115668 | 297.828629262794 | 254.401104271363 | 528.867670002517 | 613.321944037707 | Krt222 |

| | | | | | | | | | | | | |
|---|---|---|---|---|---|---|---|---|---|---|---|---|
| **ENSMUSG00000035851** | 0.0832264283307927 | 0.521865061165909 | 0.906201487635264 | 980.312382477414 | 1101.09006549217 | 1104.51285505274 | 887.278652768221 | 930.714466446232 | 850.042151291332 | 966.120468036095 | 1097.46634587986 | Ythdc1 |
| **ENSMUSG00000035863** | -0.677934089944132 | 0.0197591003763684 | 0.642273034804458 | 798.706125447036 | 699.583111137408 | 905.757328436564 | 763.472329126143 | 1653.38687568684 | 1821.16944355798 | 810.99983254323 | 782.663932125994 | Palm |
| **ENSMUSG00000035873** | -0.247445347294751 | 0.440802588319025 | 0.890970307297912 | 36.1405486627618 | 46.4216965481167 | 39.7511053232348 | 43.1446279358754 | 68.9823663366031 | 62.3771938357668 | 37.4788112600209 | 28.2236646813812 | Pawr |
| **ENSMUSG00000035875** | -0.427084038911337 | 0.206054740768755 | 0.819960256425221 | 64.1494738764022 | 79.8127414336042 | 42.5904699891801 | 64.7169419038131 | 55.8428679867739 | 74.6080161565054 | 66.6289977955927 | 141.118323406906 | AI182371 |
| **ENSMUSG00000035877** | 0.00205573109436287 | 0.99037149602783 | 0.998628446692557 | 709.2582675067 | 739.48948185421 | 574.498117409607 | 532.742362338636 | 779.610235423197 | 638.448925142555 | 560.100012719201 | 575.328549274309 | Zhx3 |
| **ENSMUSG00000035878** | -0.520879798497021 | 0.155999486856779 | 0.787275540343856 | 15.3597331816738 | 23.6180561385155 | 12.3039135524298 | 19.6964605794214 | 32.8487458745729 | 15.9000690169602 | 27.0680303544596 | 26.0526135520442 | Hykk |
| **ENSMUSG00000035885** | 0.071251428246506 | 0.697379660997619 | 0.944933386973863 | 2493.69785773056 | 2087.34751320742 | 2386.95922917138 | 2758.44240781325 | 2362.91978657761 | 3054.03633348843 | 1927.03554561941 | 1914.86709607525 | Cox8a |
| **ENSMUSG00000035890** | 0.129892537751886 | 0.33970770075673 | 0.858570960161243 | 724.618000688374 | 614.883875330318 | 750.538726698218 | 702.507093999363 | 706.248036303317 | 706.941530138691 | 598.619902069779 | 540.591731204917 | Rnf126 |
| **ENSMUSG00000035891** | 0.126023150834787 | 0.364278359820187 | 0.86924083181804 | 814.969372345279 | 943.907829811707 | 1018.38546018573 | 847.885731609378 | 829.978312430875 | 682.479885497214 | 901.573626421614 | 906.413846498204 | Cerk |
| **ENSMUSG00000035896** | 1.83171863896354 | 0.00452172863795305 | 0.635862253526407 | 28.0089252136404 | 45.6072808192024 | 168.468970179424 | 81.59962240046 | 48.1781606160403 | 19.5693157131818 | 17.6983275394543 | 5.42762782334254 | Rnase1 |
| **ENSMUSG00000035898** | -0.28249870172106 | 0.0810234824728103 | 0.739855948845396 | 177.088688447533 | 162.068730053951 | 166.576060402127 | 140.689004138724 | 220.086597359638 | 218.931719541221 | 185.311900118992 | 162.828834700276 | Uba6 |
| **ENSMUSG00000035900** | 0.00732118959898943 | 0.971057159728498 | 0.995122272737203 | 530.362551626029 | 574.163088884602 | 602.891764069061 | 483.032247542953 | 525.579933993166 | 362.032340693862 | 667.331056046484 | 623.091674119723 | Gramd4 |
| **ENSMUSG00000035901** | 0.131466474549348 | 0.380030954596314 | 0.87432013076003 | 3455.9399658766 | 3873.36120671655 | 3752.69363349109 | 3521.9147369394 | 2981.5711672154 | 2667.54234815309 | 3807.22257716379 | 3874.2407403019 | Dennd5a |
| **ENSMUSG00000035914** | -0.140765138448177 | 0.622213576090645 | 0.93078489223231 | 36.1405486627618 | 43.1640336324594 | 31.2330113253987 | 26.2619474392285 | 39.4184950494875 | 44.0309603546589 | 27.0680303544596 | 41.2499714574033 | Cd276 |
| **ENSMUSG00000035919** | 0.120198226364368 | 0.4415240844474 | 0.89108031640712 | 202.387072511466 | 196.274190668353 | 226.202718386979 | 168.826804966469 | 193.80760065998 | 154.108361241306 | 197.804837205666 | 183.453820428978 | Bbs9 |
| **ENSMUSG00000035929** | -0.132617667514297 | 0.675392803656636 | 0.94409858171788 | 582.766347187034 | 528.555808065399 | 695.644343156608 | 552.438822918057 | 849.687559955619 | 962.565716642128 | 433.088485671353 | 341.94055287058 | H2-Q4 |
| **ENSMUSG00000035933** | 0.338752746297174 | 0.290543694145152 | 0.851785577030345 | 420.133878204606 | 619.770369703804 | 636.017685171756 | 362.03970398365 | 279.21433993387 | 237.277953022329 | 555.935700356977 | 538.42068007558 | Cog5 |
| **ENSMUSG00000035934** | -0.239264543612857 | 0.227746891754291 | 0.832048466795696 | 142.755167217909 | 162.068730053951 | 134.396594188079 | 90.9788893430416 | 154.389105610493 | 160.223772401676 | 162.408182126757 | 149.802527924254 | Pknox2 |
| **ENSMUSG00000035936** | 0.177801506158157 | 0.409823641383452 | 0.882192126163777 | 916.162908601012 | 931.691593877992 | 1026.90355418356 | 926.671573927063 | 753.331238723539 | 534.486935416277 | 1063.98180854837 | 1007.36772401237 | Aldh5a1 |
| **ENSMUSG00000035941** | 0.11219644966386 | 0.665117133673767 | 0.941548816421589 | 687.573938309043 | 766.365200908383 | 659.679057387967 | 621.845398293161 | 451.122776677468 | 413.401794440965 | 753.740537562643 | 911.841474321546 | Ibtk |
| **ENSMUSG00000035944** | 0.023751920122855 | 0.844412441901646 | 0.971806333564415 | 633.363115314901 | 705.284021239809 | 713.626986040929 | 601.211011019481 | 722.672409240604 | 577.294813538862 | 679.823993133157 | 629.604827507734 | Ttc38 |
| **ENSMUSG00000035948** | -0.13488265294494 | 0.812096175220535 | 0.965224347191244 | 7.22810973255236 | 10.5874044758863 | 33.1259211026956 | 15.9447538023887 | 18.6142893289246 | 7.33849339244316 | 18.7394056300105 | 28.2236646813812 | Acss3 |
| **ENSMUSG00000035949** | -0.0158444150671596 | 0.8679062781925 | 0.974868041734905 | 919.776963467288 | 914.588863570791 | 981.473719528439 | 883.526945991188 | 975.607752474815 | 919.757838519542 | 851.60187807492 | 993.255891671684 | Fbxw2 |
| **ENSMUSG00000035953** | -0.169658130735721 | 0.278968418943374 | 0.848075053184273 | 247.562758339918 | 273.643684915214 | 297.186835035612 | 275.750448111899 | 366.810995599397 | 324.116791499573 | 292.542943446274 | 247.49982874442 | Tmem55b |
| **ENSMUSG00000035954** | 0.0405526562157072 | 0.859277589807627 | 0.974868041734905 | 118.360296870545 | 157.99665140938 | 149.539872406455 | 94.7305961200743 | 133.58489988993 | 95.400414101761 | 151.997401221196 | 124.835439936878 | Dock4 |
| **ENSMUSG00000035958** | -0.0990436734921388 | 0.431588552942058 | 0.886559861988535 | 221.360860559416 | 248.39679731887 | 247.971180825893 | 261.681547698027 | 235.416012101106 | 259.293433199658 | 272.762459725708 | 281.151121249143 | Tdp2 |
| **ENSMUSG00000035960** | -0.112912154915184 | 0.392590553544548 | 0.878537584780694 | 252.080326922764 | 263.056280439328 | 218.631079277791 | 251.364354061187 | 297.828629262794 | 242.170281950624 | 251.940897914585 | 273.552442296464 | Apex1 |
| **ENSMUSG00000035967** | -0.118041315851225 | 0.708322302067699 | 0.949174314815223 | 37.0440623793308 | 23.6180561385155 | 31.2330113253987 | 42.2067012416173 | 27.3739548954774 | 29.3539735697726 | 49.9717483466946 | 37.9933947633978 | Ints6l |
| **ENSMUSG00000035969** | 0.204628023962349 | 0.0637875170536241 | 0.716991331547963 | 2077.17803439223 | 2640.33579314025 | 2485.39053759082 | 2667.46351847021 | 2159.25756215526 | 2146.50931728962 | 2055.08815075781 | 2204.70242184174 | Rusc2 |
| **ENSMUSG00000035984** | 0.240475095474471 | 0.440677360658192 | 0.890970307297912 | 32.5264937964856 | 31.7622134276588 | 59.6266579848522 | 33.7653609932938 | 37.2285786578493 | 36.6924669622158 | 33.3144988977964 | 26.0526135520442 | Nme5 |
| **ENSMUSG00000035992** | 0.0493273888031708 | 0.844023419544934 | 0.971721411930626 | 829.425591810383 | 1056.29720040188 | 1194.42606947434 | 1054.22960434617 | 791.654775577207 | 634.779678446333 | 1309.67623791962 | 1258.1241294508 | Fnip1 |
| **ENSMUSG00000036002** | 0.327058889618092 | 0.0796295377549445 | 0.739855948845396 | 655.950958229127 | 709.35609988438 | 751.485181586867 | 634.038445318517 | 452.217734873287 | 426.855698993777 | 710.015257759285 | 602.466688391022 | Fam214b |
| **ENSMUSG00000036006** | 0.103057621352995 | 0.767897173600934 | 0.95879680569409 | 46.9827132615903 | 43.1640336324594 | 48.2691993210708 | 75.0341355406529 | 42.7033696369448 | 24.4616446414772 | 67.6700758861489 | 62.9604827507734 | Ripor2 |
| **ENSMUSG00000036009** | 0.0349192302132992 | 0.865074581892565 | 0.974868041734905 | 66.8600150261093 | 77.3694942468612 | 90.8596693102509 | 70.3445020693621 | 79.9319482947941 | 86.838838477244 | 61.4236073428121 | 70.559161703453 | Mettl25 |
| **ENSMUSG00000036019** | 0.018463751854625 | 0.926026014091456 | 0.986713918026868 | 70.4740698923855 | 94.4722245540621 | 81.3951204237664 | 89.1030359545253 | 74.4571573156986 | 102.738907494204 | 79.1219348822664 | 75.9867895267955 | Tmtc2 |
| **ENSMUSG00000036022** | -0.180561390933039 | 0.36893561294219 | 0.871495599696183 | 53.3073092775737 | 68.4109212288036 | 54.8943835416099 | 54.3997482669734 | 64.6025335533267 | 64.8233582999146 | 62.4646854333682 | 70.559161703453 | Fam122b |
| **ENSMUSG00000036023** | -0.00186645704457769 | 0.989508631160667 | 0.998596768760793 | 154.500845533307 | 162.068730053951 | 156.165056626994 | 171.640585049243 | 145.62944004394 | 150.439114545085 | 180.106509666212 | 168.256462523619 | Parp2 |
| **ENSMUSG00000036026** | 0.19660065281524 | 0.270568150335816 | 0.844129257012535 | 2812.63819967944 | 3078.49145529616 | 3217.94662140472 | 2566.16743549033 | 2281.892880087 | 1899.4467064107 | 3049.31772723892 | 2955.88611259235 | Tmem63b |
| **ENSMUSG00000036040** | -1.57478009828828 | 0.0381047274279105 | 0.679566075569698 | 161.728955265859 | 100.173134656462 | 72.8770264259304 | 144.440710915757 | 526.674892188986 | 717.949270227356 | 107.231043327282 | 75.9867895267955 | Adamtsl2 |
| **ENSMUSG00000036046** | 0.0976215195458844 | 0.60995007378433 | 0.927932112008768 | 1266.7262306298 | 1657.33600834066 | 1476.46962629158 | 1308.40773849013 | 1062.10944994452 | 1068.97387083255 | 1568.9046824681 | 1634.80150039077 | 5031439G07 |
| **ENSMUSG00000036052** | 0.166330104173496 | 0.389987203344591 | 0.877515569464664 | 1981.40558043592 | 2297.46677126732 | 2593.28639489674 | 2312.92722804063 | 1599.7339240917 | 1647.49176660349 | 2297.65934585739 | 2638.91264770914 | Dnajb5 |
| **ENSMUSG00000036053** | -0.0669824466530692 | 0.746125816558934 | 0.956085046483146 | 130.105975185942 | 148.223662662408 | 171.308334845369 | 107.861569839689 | 179.573144114332 | 146.769867848863 | 138.463386043966 | 119.407812113536 | Fmnl2 |
| **ENSMUSG00000036054** | 0.0864800364710716 | 0.661240737107529 | 0.939611397743053 | 553.853908256825 | 649.903751673634 | 570.712297855013 | 439.887619607078 | 482.876564356222 | 385.270903103266 | 640.263025692024 | 576.414074838978 | Sugp2 |
| **ENSMUSG00000036057** | -0.135134946213172 | 0.294506087219091 | 0.851785577030345 | 346.949267162513 | 431.640336324594 | 432.52988411234 | 373.294824314748 | 473.02194059385 | 437.863439082442 | 393.52751823022 | 436.38127699674 | Ptpn23 |
| **ENSMUSG00000036061** | 0.0934534511965793 | 0.597528815025004 | 0.924496535139492 | 195.158962778914 | 164.511977240694 | 178.879973954556 | 146.316564304273 | 158.768938393769 | 124.754387671534 | 174.901119213431 | 182.368294864309 | Smug1 |
| **ENSMUSG00000036062** | 0.0190024910616308 | 0.940234636461004 | 0.989269716906553 | 77.7021796249379 | 61.0811796685746 | 67.1982970940397 | 64.7169419038131 | 70.0773245324222 | 41.5847958905112 | 65.5879197050366 | 89.0130963028176 | Phf24 |
| **ENSMUSG00000036073** | 0.0754483386864375 | 0.482785512159599 | 0.898414997395275 | 971.277245311723 | 935.763672522564 | 900.078599104673 | 1047.66411748637 | 998.601874587016 | 961.342634410054 | 857.848346618257 | 841.282312618093 | Galt |
| **ENSMUSG00000036078** | -0.183630838740549 | 0.312809227513921 | 0.852613848149207 | 346.949267162513 | 357.52850499339 | 300.972654590206 | 330.150196378873 | 444.553027502553 | 459.878919259771 | 313.364505257397 | 300.690581413177 | Sigmar1 |
| **ENSMUSG00000036083** | 0.0829237709070563 | 0.853078462122893 | 0.973120292022893 | 11.7456783153976 | 13.8450673915436 | 16.0897331070236 | 12.1930470253561 | 10.949581958191 | 9.78465785659088 | 8.32862472444909 | 21.7105112933701 | Slc17a3 |
| **ENSMUSG00000036086** | 0.106984546580965 | 0.709414424289642 | 0.949394580728696 | 91.2548853734735 | 115.647033505835 | 95.5919437534931 | 80.6616957062019 | 72.2672409240604 | 58.7079471395453 | 139.504464134522 | 84.6709940441436 | Zranb3 |
| **ENSMUSG00000036087** | 0.292238698919227 | 0.220162094178395 | 0.828720352121776 | 5683.10127721929 | 6600.83948285063 | 5738.35598987553 | 5455.91958049972 | 3481.96706270473 | 3280.30654642209 | 6070.52634603283 | 6339.46929766408 | Slain2 |
| **ENSMUSG00000036093** | 0.0872222856477997 | 0.593936530083504 | 0.923935529355527 | 869.180195339421 | 918.660942215363 | 879.256591554407 | 831.003051112731 | 734.716949394614 | 645.787418534998 | 871.382361795486 | 1039.93349095243 | Arl5a |
| **ENSMUSG00000036095** | -0.998154173880658 | 0.0663809183597098 | 0.719725956118125 | 9.93865088225949 | 3.25766291565731 | 7.57163910918757 | 8.44134024832345 | 22.994122112201 | 15.9000690169602 | 7.28754663389296 | 11.9407812113536 | Dgkb |
| **ENSMUSG00000036097** | -0.0879371852061216 | 0.680635604068685 | 0.944452327903597 | 293.64195788494 | 350.198763433161 | 334.098575692902 | 244.79886720138 | 270.454674367317 | 243.393364182698 | 373.747034509653 | 411.414189009364 | Slf2 |
| **ENSMUSG00000036098** | 0.249078519430071 | 0.510037103072395 | 0.902394714365738 | 153.597331816738 | 184.872370463553 | 149.539872406455 | 97.5443762028488 | 67.887408140784 | 68.4926049961361 | 154.079557402308 | 201.907755028342 | Myrf |
| **ENSMUSG00000036099** | 0.271704534536922 | 0.00371278654995519 | 0.635862253526407 | 1083.31294616628 | 1049.78187457057 | 1113.03094905057 | 1208.98750889877 | 888.011096809288 | 875.726878164883 | 964.038311854983 | 960.690124731629 | Vezt |
| **ENSMUSG00000036103** | -0.804454627123817 | 0.01729067291516 | 0.642273034804458 | 120.167324303683 | 132.749763813036 | 147.646962629158 | 119.116690170786 | 306.588294829347 | 341.239942748607 | 124.929370866736 | 135.690695583563 | Colec12 |
| **ENSMUSG00000036104** | 0.147257705392908 | 0.398988707984395 | 0.880459952732461 | 1014.64590370704 | 1274.56061575092 | 1343.01948699215 | 968.878275168681 | 1032.54557865741 | 806.011190936674 | 1151.43236815509 | 1163.68340532464 | Rab3gap1 |
| **ENSMUSG00000036106** | -0.151205513740282 | 0.403807138426431 | 0.881158207349486 | 86.7373167906283 | 88.7713144516618 | 85.1809399783602 | 86.2892558717508 | 119.350443344282 | 91.7311674055395 | 85.3684034256032 | 89.0130963028176 | Prr5 |
| **ENSMUSG00000036110** | -0.234771302229728 | 0.402768977014757 | 0.881158207349486 | 40.658117245607 | 45.6072808192024 | 30.2865564367503 | 38.4549944645846 | 38.3235368536684 | 44.0309603546589 | 36.4377331694648 | 64.046008315442 | Slc17a2 |
| **ENSMUSG00000036111** | -0.12415734539667 | 0.639442153087147 | 0.933801135937988 | 100.290022539164 | 55.3802695661743 | 91.8061241988993 | 87.227182566009 | 84.3117810780705 | 124.754387671534 | 86.4094815161593 | 69.4736361387845 | Lmo1 |
| **ENSMUSG00000036112** | 0.213215153432384 | 0.0083154361271415 | 0.635862253526407 | 3662.84460697091 | 3585.05803868087 | 3671.29851306733 | 3646.65898727573 | 3084.4972376224 | 2870.57399867735 | 3343.94282686631 | 3263.08984739353 | Metap2 |
| **ENSMUSG00000036114** | -0.191187795821766 | 0.331610110424144 | 0.856333478432716 | 301.773581334061 | 277.715763559786 | 238.506631939409 | 346.094950181261 | 351.48158085793 | 424.409534529629 | 290.460787265162 | 263.782712214447 | Rpp25l |
| **ENSMUSG00000036120** | 0.167677062662759 | 0.305277059231807 | 0.851785577030345 | 318.940341948873 | 333.910448854875 | 321.794662140472 | 379.860311174555 | 325.202584158272 | 354.693847301419 | 296.707255808499 | 230.131419709724 | Rfxank |
| **ENSMUSG00000036138** | -0.00236714643875782 | 0.983587735826308 | 0.996940244108351 | 1874.79096188077 | 1918.76345732216 | 1825.71148020285 | 2031.54921976318 | 2019.10291309041 | 2163.63246853866 | 1648.02661735036 | 1834.53820428978 | Acaa1a |
| **ENSMUSG00000036144** | -0.473575642906953 | 0.105125359923213 | 0.762094734041179 | 220.457346842847 | 157.99665140938 | 193.076797284283 | 341.405316709971 | 462.072358635659 | 293.539735697726 | 276.926772087932 | 234.473521968398 | Meox2 |
| **ENSMUSG00000036151** | -0.403996948179287 | 0.413153874054015 | 0.882192126163777 | 9.93865088225949 | 12.2162359337149 | 8.51809399783602 | 9.37926694258161 | 4.37983278327639 | 15.9000690169602 | 17.6983275394543 | 15.1973579053591 | Tm6sf2 |
| **ENSMUSG00000036155** | -0.282365422587493 | 0.148577333903457 | 0.781423388079902 | 159.921927832721 | 218.26341534904 | 128.717864856189 | 167.888878272211 | 194.902558855799 | 193.24699266767 | 192.599446752885 | 240.986675356409 | Mgat5 |
| **ENSMUSG00000036158** | -0.0608817209652233 | 0.782071343732554 | 0.960742105865155 | 330.68602026427 | 298.076156782644 | 397.511053232348 | 566.507723331929 | 440.173194719277 | 469.663577116362 | 379.99350305299 | 371.24974311663 | Prickle1 |
| **ENSMUSG00000036160** | 0.173444916891231 | 0.199474352063258 | 0.812921911423739 | 182.509770746947 | 170.212887343095 | 180.772883731853 | 184.771558768858 | 166.433645764503 | 140.654456688494 | 157.202791673977 | 171.513039217624 | Surf6 |
| **ENSMUSG00000036167** | 0.172112315685184 | 0.28302095970583 | 0.850587941933635 | 262.922491521592 | 351.82759489099 | 310.437203476691 | 260.743621003769 | 289.068963696242 | 220.154801773295 | 270.680303544596 | 272.466916731795 | Pphln1 |
| **ENSMUSG00000036180** | -0.0807268599141091 | 0.538788163862713 | 0.913211421116911 | 550.239853390548 | 678.408302185636 | 682.39397471553 | 634.038445318517 | 775.230402639921 | 680.033721033066 | 625.687932424238 | 611.15089290837 | Gatad2a |
| **ENSMUSG00000036181** | 0.475689271891835 | 0.111765652816196 | 0.764116514278315 | 2231.67887992554 | 3231.60161233206 | 6378.15949460188 | 3185.19905370072 | 3150.19472937154 | 2343.42555665351 | 2992.05843225834 | 2319.7681316966 | Hist1h1c |
| **ENSMUSG00000036186** | 0.0133986762382138 | 0.952791470496413 | 0.992000114323903 | 259.308436655316 | 190.573280565953 | 152.3792370724 | 280.44008158319 | 195.997517051618 | 261.739597663806 | 193.640524843441 | 223.618266321713 | Fam69b |
| **ENSMUSG00000036188** | -0.0650072694168979 | 0.584676932482425 | 0.922641625223466 | 311.712232216321 | 293.189662409158 | 274.47191770805 | 310.453735799451 | 301.113503850252 | 287.424324537357 | 344.596847974081 | 310.460311495193 | Ankmy2 |
| **ENSMUSG00000036192** | 0.48188218289797 | 0.259793499382219 | 0.840316047847749 | 37.0440623793308 | 19.5459774939439 | 11.3574586637814 | 26.2619474392285 | 20.8042057205628 | 15.9000690169602 | 17.6983275394543 | 13.0263067760221 | Rorb |
| **ENSMUSG00000036197** | 0.315497727380515 | 0.127064537974662 | 0.77273486535379 | 372.247651226447 | 437.341246426994 | 503.514000760974 | 361.101777289392 | 293.448796479518 | 250.731857575141 | 433.088485671353 | 366.907640857956 | Gxylt1 |
| **ENSMUSG00000036199** | -0.161589921947451 | 0.525218678934746 | 0.908006595346551 | 11149.359262462 | 7900.6469861979 | 8307.98101255607 | 11441.7677432553 | 10946.2970836035 | 16286.5630022955 | 8118.32695015675 | 8048.08653645232 | Ndufa13 |
| **ENSMUSG00000036202** | 0.198380800374859 | 0.491088393266819 | 0.899236955296196 | 379.475760958999 | 493.535931722083 | 423.065335225856 | 326.39848960184 | 271.549632563136 | 200.585486060113 | 482.019155927491 | 459.177313854779 | Rif1 |
| **ENSMUSG00000036206** | -0.485452172693855 | 0.0742268692764333 | 0.734338471687791 | 243.948703473642 | 238.623808571898 | 229.042083052924 | 269.184961252092 | 468.642107810573 | 450.09426140318 | 219.667477107345 | 235.559047533066 | Sh3bp4 |
| **ENSMUSG00000036208** | 0.42704580682666 | 0.134181637669887 | 0.77303252008843 | 87.6408305071974 | 93.6578088251478 | 115.467496415111 | 61.9031618210386 | 59.1277425742312 | 44.0309603546589 | 71.8343882483734 | 91.1841474321546 | Nepro |
| **ENSMUSG00000036214** | 0.0165637331880195 | 0.90976958227891 | 0.985888646122408 | 376.765219809292 | 342.054606144018 | 298.133289924261 | 389.239578117137 | 308.778211220985 | 373.040080782527 | 389.363205867995 | 319.144516012541 | Znrd1as |
| **ENSMUSG00000036216** | -0.497767497964023 | 0.139178812717435 | 0.779711473022872 | 30.7194663633475 | 28.5045505120015 | 29.3401015481019 | 23.448167356454 | 28.4689130912965 | 35.4693847301419 | 30.191264626128 | 64.046008315442 | Leap2 |
| **ENSMUSG00000036218** | -0.265753922414467 | 0.557551010634225 | 0.916341990969984 | 11.7456783153976 | 14.6594831204579 | 6.62518422053913 | 11.2551203310979 | 15.3294147414674 | 18.3462334811079 | 11.4518589961175 | 8.68420451734806 | Pdzrn4 |
| **ENSMUSG00000036225** | -0.0561138080114588 | 0.797871650705369 | 0.962602034942898 | 266.536546387868 | 206.861595144239 | 234.720812384815 | 170.702658354985 | 192.712642464161 | 173.677676954488 | 271.721381635152 | 274.637967861132 | Kctd1 |
| **ENSMUSG00000036241** | 0.0299129309909187 | 0.77203635659445 | 0.959045638244974 | 4666.64834607912 | 4190.16892526422 | 4397.22941266068 | 4740.28151278075 | 4216.68401209934 | 5029.31413828771 | 4138.28540996064 | 4242.23390672453 | Ube2r2 |
| **ENSMUSG00000036242** | -0.0127889523304426 | 0.9451162891322 | 0.990505437194639 | 431.879556520004 | 630.35777417969 | 422.118880337207 | 499.9149280396 | 408.419407040523 | 556.502415593606 | 575.716184077544 | 462.433890548784 | 3632451O06 |

| | | | | | | | | | | | | |
|---|---|---|---|---|---|---|---|---|---|---|---|---|
| **ENSMUSG00000036246** | -0.327836827047862 | 0.321209711448194 | 0.855590711009608 | 13.5527057485357 | 31.7622134276588 | 28.3936466594534 | 25.3240207449704 | 29.5638712871156 | 36.6924669622158 | 27.0680303544596 | 31.4802413753867 | Gmip |
| **ENSMUSG00000036249** | -0.230012433121186 | 0.406706918960625 | 0.881296485258502 | 69.5705561758165 | 65.1532583131463 | 56.7872933189068 | 61.9031618210386 | 99.6411958195378 | 94.1773318696872 | 49.9717483466946 | 54.2762782334254 | Rbm43 |
| **ENSMUSG00000036256** | -1.08597181843259 | 0.0175118152714017 | 0.642273034804458 | 1445.62194651047 | 1237.09749222087 | 875.470771999813 | 1429.40028204944 | 3489.63177007546 | 4975.49852007646 | 1091.04983890283 | 1032.33481199975 | Igfbp7 |
| **ENSMUSG00000036257** | 0.274407583086098 | 0.272865930535382 | 0.844738346035895 | 3642.06379148982 | 4460.55494726378 | 4613.02112727253 | 3429.9979209021 | 2785.57365016378 | 2027.87034077846 | 4190.33931448845 | 4344.27330980337 | Pnpla8 |
| **ENSMUSG00000036264** | 0.12194505829444 | 0.567760929420283 | 0.920429362097163 | 132.81651633565 | 136.007426728693 | 200.648436393471 | 151.006197775564 | 133.58489988993 | 176.123841418636 | 149.915245040084 | 110.723607596188 | Fstl4 |
| **ENSMUSG00000036270** | -0.060308350912489 | 0.575496764267225 | 0.922450244597442 | 437.300638819418 | 502.494504740141 | 514.871459424755 | 508.356268287923 | 547.479097909548 | 503.90987961443 | 514.292576734731 | 480.887825148149 | Edc4 |
| **ENSMUSG00000036273** | 0.160683053630053 | 0.37646503534349 | 0.87422797717331 | 253.887354355902 | 320.879797192245 | 245.131816159948 | 244.79886720138 | 247.460552255116 | 176.123841418636 | 260.269522639034 | 268.124814473121 | Lrrk2 |
| **ENSMUSG00000036275** | 0.0482419174490895 | 0.537632902515679 | 0.912143766452032 | 2193.73130382964 | 2252.67390617703 | 2365.19076673247 | 2296.04454754398 | 2127.50377447651 | 2183.20178425184 | 2367.41157792465 | 2128.71563231494 | 9530068E07R |
| **ENSMUSG00000036278** | -0.0317405050167366 | 0.812184458580537 | 0.965224347191244 | 10171.7574211343 | 8710.99063646766 | 9363.27821339908 | 10696.1160213201 | 10121.7935621517 | 11825.9821019221 | 8963.68235968834 | 8898.05305358776 | Macrod1 |
| **ENSMUSG00000036281** | -0.0248070358614033 | 0.912761531790706 | 0.986226496507248 | 51.5002818444356 | 70.8541684155466 | 88.0203046443056 | 66.5927952923295 | 59.1277425742312 | 77.0541806206532 | 72.8754663389296 | 72.73021283279 | Snapc4 |
| **ENSMUSG00000036282** | 0.102348397487272 | 0.497457119441807 | 0.900927059989498 | 552.950394540256 | 622.213616890547 | 595.320124959873 | 515.859681841989 | 508.060602860061 | 423.186452297555 | 581.96265262088 | 615.492995167044 | Naa30 |
| **ENSMUSG00000036285** | 0.0553361260324926 | 0.636919310063799 | 0.932931968211785 | 482.47632464787 | 473.175538499225 | 474.173899212872 | 498.977001345342 | 482.876564356222 | 388.940149799487 | 479.936999746379 | 502.598336441519 | Noa1 |
| **ENSMUSG00000036291** | -0.0220583149240628 | 0.925032740861692 | 0.986713918026868 | 119.263810587114 | 171.027303072009 | 159.950876181588 | 148.192417692789 | 143.439523652302 | 97.8465785659088 | 186.352978209548 | 179.111718170304 | Ap5m1 |
| **ENSMUSG00000036295** | 0.506710087017092 | 0.531358845018919 | 0.909082872831377 | 8.13162344912141 | 15.4738988493722 | 20.8220075502658 | 2.81378008277448 | 9.85462376237187 | 18.3462334811079 | 1.04107809055614 | 4.34210225867403 | Lrrn3 |
| **ENSMUSG00000036299** | -0.131905443883583 | 0.136701200429603 | 0.776456211338551 | 797.802611730467 | 702.026358324151 | 757.163910918757 | 765.34818251466 | 847.497643563981 | 848.819069059259 | 822.451691539348 | 792.433662208011 | BC031181 |
| **ENSMUSG00000036305** | -0.308247977429054 | 0.504417236483579 | 0.901138960097352 | 13.5527057485357 | 9.77298874697194 | 10.4110037751329 | 13.1309737196143 | 12.0445401540101 | 6.1154111603693 | 19.7804837205666 | 19.5394601640331 | Rpl39-ps |
| **ENSMUSG00000036306** | -0.722813203545325 | 0.0471454878787305 | 0.702838509185083 | 14.4562194651047 | 17.9171460361152 | 15.1432782183751 | 31.8895076047775 | 37.2285786578493 | 22.0154801773295 | 36.4377331694648 | 34.7368180693922 | Lzts1 |
| **ENSMUSG00000036309** | 0.0937523576402571 | 0.378094035114661 | 0.87422797717331 | 6414.04387392365 | 5703.35334958704 | 5415.61487284641 | 6112.46826648044 | 4903.22280087792 | 5612.72436298694 | 5605.16443955424 | 6036.60766512157 | Skp1a |
| **ENSMUSG00000036315** | 0.0116701591876168 | 0.951995610038323 | 0.991915011884503 | 215.036264543433 | 206.861595144239 | 195.916161950228 | 258.867767615252 | 220.086597359638 | 286.201242305283 | 183.22974393788 | 181.282769299641 | Znrd1 |
| **ENSMUSG00000036323** | 0.162819302691515 | 0.163771446010039 | 0.793968226938511 | 972.180759028292 | 1123.89370590177 | 1069.49402417274 | 1005.45741624475 | 914.290093508946 | 817.018931025338 | 1060.8585742767 | 932.466460050248 | Srp72 |
| **ENSMUSG00000036327** | -0.0705497845860633 | 0.662344089373677 | 0.939611397743053 | 278.282224703266 | 302.148235427216 | 406.975602118832 | 375.170677703265 | 344.911831683016 | 325.339873731647 | 373.747034509653 | 385.36157545732 | Qsox2 |
| **ENSMUSG00000036333** | 0.136837914907638 | 0.576703347593833 | 0.922450244597442 | 1172.76080410662 | 1544.94663775048 | 1402.646144977 | 1126.44995980405 | 1062.10944994452 | 717.949270227356 | 1540.79557402308 | 1450.26215439713 | Kidins220 |
| **ENSMUSG00000036334** | -1.00785749409369 | 0.0668519307153266 | 0.719725956118125 | 9.93865088225949 | 19.5459774939439 | 19.8755526616174 | 15.9447538023887 | 38.3235368536684 | 66.0464405319884 | 14.5750932677859 | 13.0263067760221 | Igsf10 |
| **ENSMUSG00000036339** | 0.128712889875218 | 0.455897327447536 | 0.892805623137304 | 226.78194285883 | 247.582381589956 | 228.095628164276 | 191.337045628665 | 211.326931793086 | 155.33144347338 | 211.338852382896 | 238.815624227072 | Tmem260 |
| **ENSMUSG00000036352** | 0.0267303504339135 | 0.751470490958525 | 0.956197192463153 | 1717.57957519775 | 1570.19352534683 | 1677.11806268505 | 1589.78574676758 | 1715.79949284852 | 1533.74511902062 | 1639.69799262592 | 1543.61735295862 | Ubac1 |
| **ENSMUSG00000036353** | 0.0716327834889622 | 0.879880874233493 | 0.978393501240173 | 27.1054114970714 | 38.2775392589734 | 52.0550188756646 | 15.9447538023887 | 24.0890803080201 | 57.4848649074714 | 16.6572494488982 | 29.3091902460497 | P2ry12 |
| **ENSMUSG00000036356** | -0.393962482013667 | 0.188365618134026 | 0.808677608936066 | 42.4651446787451 | 44.7928650902881 | 51.1085639870161 | 42.2067012416173 | 78.836990098975 | 77.0541806206532 | 36.4377331694648 | 45.5920737160773 | Csgalnact1 |
| **ENSMUSG00000036362** | -0.462359369542525 | 0.30702817206812 | 0.851785577030345 | 9.03513716569045 | 6.51532583131463 | 8.51809399783602 | 13.1309737196143 | 14.2344565456483 | 12.2308223207386 | 9.36970281500523 | 15.1973579053591 | P2ry13 |
| **ENSMUSG00000036368** | 0.063180215895674 | 0.762420429639685 | 0.957766099999568 | 204.194099944604 | 243.510302945384 | 229.988537941573 | 204.468019348279 | 195.997517051618 | 150.439114545085 | 287.337552993494 | 209.506433981022 | Rmdn2 |
| **ENSMUSG00000036371** | 0.0959514172426904 | 0.145521204986177 | 0.780607990324772 | 8911.35578652049 | 8431.64604145004 | 8574.88129115493 | 9060.37186653384 | 8524.24955445167 | 8001.40396222719 | 8210.98290021625 | 7989.46815596022 | Serbp1 |
| **ENSMUSG00000036372** | 0.00533572514348241 | 0.977536125496423 | 0.996183182916017 | 1199.86621560369 | 897.48613326359 | 1018.38546018573 | 1239.93908980929 | 1094.9581958191 | 1415.10614250946 | 961.95615567387 | 868.420451734806 | Tmem258 |
| **ENSMUSG00000036376** | -0.0702150488305486 | 0.75865800476518 | 0.957021132157692 | 374.054678659585 | 319.250965734417 | 300.972654590206 | 347.03287687552 | 431.413529152724 | 458.655837027697 | 272.762459725708 | 246.414303179751 | Abt1 |
| **ENSMUSG00000036377** | -0.112458948504204 | 0.641019160876557 | 0.934544499347759 | 50.5967681278665 | 73.2974156022896 | 50.1621090983677 | 47.8342614071662 | 53.6529515951357 | 56.2617826753975 | 73.9165444294857 | 56.4473293627624 | C530008M17 |
| **ENSMUSG00000036381** | -0.554433300216464 | 0.151581523103269 | 0.783412422888014 | 52.4037955610046 | 39.0919549878878 | 28.3936466594534 | 37.5170677703264 | 95.2613630362614 | 67.2695227640623 | 34.3555769883525 | 34.7368180693922 | P2ry14 |
| **ENSMUSG00000036390** | 0.202705289442788 | 0.326962583958295 | 0.855977678491063 | 494.222002963268 | 399.878122896935 | 371.95677123884 | 556.19052969509 | 353.671497249568 | 443.978850242811 | 299.830490080167 | 486.315452971491 | Gadd45a |
| **ENSMUSG00000036391** | 0.113570236245086 | 0.607073362380434 | 0.92742133338909 | 231.299511441676 | 291.56083095133 | 295.293925258315 | 222.288626539184 | 211.326931793086 | 166.339183562045 | 301.91264626128 | 281.151121249143 | Sec24a |
| **ENSMUSG00000036398** | 0.000846475522433323 | 0.996432577768422 | 0.999262355603273 | 1733.842822096 | 1273.74620002201 | 1423.46815252726 | 1859.90863471393 | 1659.95662486175 | 1976.50088703136 | 1306.55300364795 | 1344.96617462428 | Ppp1r11 |
| **ENSMUSG00000036402** | 0.0674197795117029 | 0.519738657240142 | 0.905962179970576 | 648.722848496574 | 793.240919962556 | 775.146553803078 | 754.093062183562 | 700.773245324222 | 730.180092548094 | 724.590351027071 | 680.624529047154 | Gng12 |
| **ENSMUSG00000036403** | 0.344987088186837 | 0.207456153559849 | 0.820327948642971 | 112.939214571131 | 144.96599974675 | 113.574586637814 | 98.4823028971069 | 63.5075753575076 | 66.0464405319884 | 124.929370866736 | 115.065709854862 | Cep135 |
| **ENSMUSG00000036427** | -0.0654463581159635 | 0.575294631260717 | 0.922450244597442 | 18199.4767928503 | 16873.0650716471 | 17895.5690345648 | 18290.5084647284 | 19025.9936105526 | 21872.3795561768 | 17310.0054116769 | 16358.8702595544 | Gpi1 |
| **ENSMUSG00000036430** | -0.382424772510292 | 0.114764423369512 | 0.767518180080285 | 311.712232216321 | 258.169786065842 | 287.722286149128 | 348.908730264036 | 462.072358635659 | 540.602346576646 | 283.173240631269 | 287.664274637155 | Tbcc |
| **ENSMUSG00000036432** | -0.0801991491166445 | 0.6821418358189 | 0.944452327903597 | 176.185174730964 | 207.676010873154 | 176.040609288611 | 159.447538023887 | 235.416012101106 | 215.262472844999 | 164.49033830787 | 146.545951230249 | Siah2 |
| **ENSMUSG00000036435** | -0.0568653969324439 | 0.717056028110065 | 0.95140862628518 | 195.158962778914 | 205.232763686411 | 223.363353721033 | 192.274972322923 | 260.600050604945 | 209.14706168463 | 195.722681024554 | 183.453820428978 | Exoc1 |
| **ENSMUSG00000036437** | -0.553606145361778 | 0.400229300107125 | 0.880487015130533 | 6.32459601598331 | 38.2775392589734 | 29.3401015481019 | 25.3240207449704 | 54.7479097909548 | 66.0464405319884 | 15.616171358342 | 9.76973008201657 | Npy1r |
| **ENSMUSG00000036438** | -0.242901986702532 | 0.102656095667235 | 0.759800953500741 | 1769.98337075876 | 1711.90186217792 | 1467.00507740509 | 1724.84719074076 | 2042.09703520262 | 2418.03357281002 | 1664.68386679926 | 1774.83429823301 | Calm2 |
| **ENSMUSG00000036442** | -0.0872627924656305 | 0.507091344912307 | 0.901698316485955 | 543.011743657996 | 504.123336197969 | 476.066808990169 | 588.080037299867 | 527.769850384805 | 654.348994159515 | 552.812466085309 | 509.11148982953 | Thap11 |
| **ENSMUSG00000036446** | -0.994332085150621 | 0.0258656780541891 | 0.664353746352133 | 539.39768879172 | 416.166437475222 | 209.166530391307 | 423.942865804689 | 1031.45062046159 | 1385.75216893968 | 314.405583347953 | 434.210225867403 | Lum |
| **ENSMUSG00000036450** | 0.190760961721292 | 0.586552980604616 | 0.923058967588378 | 1504.35033808746 | 2419.62913060447 | 2186.31079277791 | 1364.68334014562 | 1051.15986798633 | 847.595986827185 | 2351.79540656631 | 2298.05762040323 | Hif1an |
| **ENSMUSG00000036452** | -0.0105231948722145 | 0.96548839101818 | 0.994284908685797 | 1158.30458464152 | 1154.0270878716 | 1361.00212987647 | 1129.26373988683 | 1069.77415731526 | 739.964750404685 | 1751.09334831542 | 1275.4925384855 | Arhgap26 |
| **ENSMUSG00000036459** | -0.516746164978304 | 0.0817734619176943 | 0.739855948845396 | 84.9302893574902 | 78.9983257046899 | 79.5022106464695 | 82.5375490947182 | 133.58489988993 | 179.793088114857 | 79.1219348822664 | 74.901263962127 | Wtip |
| **ENSMUSG00000036461** | -0.1312638397643 | 0.388416583307751 | 0.876358159057779 | 252.080326922764 | 258.984201794756 | 319.901752363175 | 272.936668029125 | 325.202584158272 | 344.909189444828 | 283.173240631269 | 256.184033261768 | Elf1 |
| **ENSMUSG00000036466** | 0.374587069777367 | 0.369052348265871 | 0.871495599696183 | 31.6229800799166 | 18.7315617650296 | 46.3762895437739 | 17.8206071909051 | 28.4689130912965 | 14.6769867848863 | 18.7394056300105 | 26.0526135520442 | Megf11 |
| **ENSMUSG00000036469** | -0.303824744476147 | 0.366877187500499 | 0.870096026116028 | 15.3597331816738 | 22.8036404096012 | 16.0897331070236 | 24.3860940507122 | 21.8991639163819 | 20.7923979452556 | 22.903717992235 | 31.4802413753867 | March1 |
| **ENSMUSG00000036473** | 0.0124968902051642 | 0.946585594344675 | 0.990510945542179 | 253.887354355902 | 293.189662409158 | 297.186835035612 | 222.288626539184 | 271.549632563136 | 192.023910435596 | 302.953724351836 | 289.835325766492 | Tbc1d24 |
| **ENSMUSG00000036478** | -0.297229843667196 | 0.174735351278688 | 0.799889021902745 | 412.002254755485 | 404.764617270421 | 465.655805215036 | 574.949063580253 | 619.746338833609 | 765.649477278236 | 434.129563761909 | 463.519416113453 | Btg1 |
| **ENSMUSG00000036492** | 0.421053897615708 | 0.41548474532818 | 0.882192126163777 | 25.2983840639333 | 27.6901347830872 | 83.2880302010633 | 129.433883807626 | 43.7983278327639 | 47.7002070508805 | 71.8343882483734 | 34.7368180693922 | Rnf39 |
| **ENSMUSG00000036499** | 0.361543426828668 | 0.146490992694175 | 0.780607990324772 | 810.451803762433 | 1018.83407687183 | 943.615523982501 | 772.851596068725 | 529.959766776443 | 448.871179171106 | 946.339984315528 | 833.683633665414 | Eea1 |
| **ENSMUSG00000036501** | 0.238252029178313 | 0.323349298719552 | 0.855977678491063 | 169.86057871498 | 249.211213047785 | 197.809071727525 | 155.695831246855 | 151.104231023035 | 110.077400886647 | 194.681602933998 | 198.651178334337 | Fam13b |
| **ENSMUSG00000036503** | 0.241084200152231 | 0.215480306392877 | 0.824338622350122 | 2436.77649358671 | 2584.14110784516 | 2609.37612800377 | 2292.29284076695 | 1845.00455995518 | 1456.69093839997 | 2560.01102467754 | 2532.53114237163 | Rnf13 |
| **ENSMUSG00000036504** | 0.0820379307367331 | 0.626448059150417 | 0.931698409533928 | 1101.38322049767 | 759.849875077069 | 842.344850897118 | 1070.17435814856 | 917.574968096403 | 1024.94291047789 | 758.945928015424 | 864.078349476132 | Phpt1 |
| **ENSMUSG00000036513** | -0.00417247120029836 | 0.981733698846706 | 0.996940244108351 | 258.404922938747 | 172.656134529838 | 226.202718386979 | 220.412773150668 | 212.421889988905 | 187.131581507301 | 212.379930473452 | 267.039288908453 | Commd2 |
| **ENSMUSG00000036523** | 0.535146510727183 | 0.315657175652133 | 0.853277176422981 | 8.13162344912141 | 13.0306516626293 | 35.0188308799925 | 23.448167356454 | 9.85462376237187 | 8.56157562451702 | 18.7394056300105 | 17.3684090346961 | Greb1 |
| **ENSMUSG00000036528** | 0.103816148253298 | 0.666154032874607 | 0.942024383071524 | 416.51982333833 | 541.586459728029 | 562.194203857177 | 362.977630677908 | 348.196706270473 | 322.893709267499 | 490.34778065194 | 590.525907179668 | Ppfibp2 |
| **ENSMUSG00000036529** | 0.0266517610969327 | 0.897972295310807 | 0.984360759361306 | 1170.05026295691 | 1343.78595270864 | 1172.65760703543 | 953.87144806055 | 996.411958195378 | 801.118862008378 | 1375.26415762466 | 1381.87404382301 | Sbf1 |
| **ENSMUSG00000036533** | -0.158027637886942 | 0.446126993245975 | 0.89108031640712 | 440.914693685694 | 328.209538752474 | 474.173899212872 | 640.603932178324 | 545.28918151791 | 580.964060235083 | 475.772687384154 | 499.341759747513 | Cdc42ep3 |
| **ENSMUSG00000036534** | -0.0718208506726224 | 0.625078437138156 | 0.931529302402539 | 188.83436676293 | 224.778741180355 | 226.202718386979 | 194.150825711439 | 223.371471947096 | 189.577745971448 | 213.421008564008 | 249.670879873757 | Slc38a7 |
| **ENSMUSG00000036537** | -0.199259894760193 | 0.490008667859372 | 0.899236955296196 | 49.6932544112975 | 27.6901347830872 | 49.2156542097192 | 52.523894878457 | 59.1277425742312 | 62.3771938357668 | 41.6431236222455 | 42.3354970220718 | Rnf113a1 |
| **ENSMUSG00000036545** | -1.04940171391661 | 0.00744877422096307 | 0.635862253526407 | 182.509770746947 | 199.531853584011 | 191.183887506986 | 216.661066373635 | 450.027818481649 | 771.764888438605 | 219.667477107345 | 194.309076075663 | Adamts2 |
| **ENSMUSG00000036550** | 0.221272122203404 | 0.41044439856508 | 0.882192126163777 | 1947.07205920629 | 2749.46750081477 | 2722.95071464158 | 1849.59144107709 | 1681.85578877813 | 1179.0512717192 | 2608.94169493368 | 2480.42591526754 | Cnot1 |
| **ENSMUSG00000036552** | -0.19311220351944 | 0.450066162193191 | 0.89108031640712 | 56.0178504272808 | 37.4631235300591 | 53.001473764313 | 69.4065753751039 | 75.5521155115177 | 61.154111603693 | 53.094982618363 | 56.4473293627624 | Ermard |
| **ENSMUSG00000036553** | -0.591463226319205 | 0.0398490572913534 | 0.679566075569698 | 106.614618555147 | 116.461449234749 | 147.646962629158 | 139.751077444466 | 264.979883388221 | 247.06261087892 | 120.765058504512 | 136.776221148232 | Sh3tc1 |
| **ENSMUSG00000036555** | -0.350122286671442 | 0.233789971260739 | 0.832048466795696 | 28.0089252136404 | 49.679359463774 | 36.9117406572894 | 57.2135283497478 | 71.1722827282413 | 51.3694537471021 | 52.0539045278068 | 44.5065481514088 | Iqce |
| **ENSMUSG00000036560** | -0.719816266122185 | 0.226710348006035 | 0.832048466795696 | 4.51756858284523 | 22.8036404096012 | 5.67872933189068 | 16.8826804966469 | 14.2344565456483 | 36.6924669622158 | 13.5340151772298 | 18.4539345993646 | Lgi4 |
| **ENSMUSG00000036561** | 0.272547022284534 | 0.173149113708029 | 0.799889021902745 | 1219.74351736821 | 1209.40735743778 | 1087.47666705707 | 1095.49837889353 | 909.91026072567 | 623.771938357668 | 1161.84314906065 | 1121.34790830257 | Ppp6r2 |
| **ENSMUSG00000036564** | -0.0374095172371018 | 0.830587246449606 | 0.967583158900775 | 2461.17136393408 | 2544.23473712836 | 2206.18634543953 | 2565.22950879607 | 2004.86845654477 | 2049.88582095579 | 3105.53594412896 | 2872.30064411287 | Ndrg4 |
| **ENSMUSG00000036565** | -0.521208860175966 | 0.0163659952308926 | 0.641523953292557 | 140.948139784771 | 122.162359337149 | 158.057966404291 | 144.440710915757 | 244.175677667659 | 251.954939807215 | 147.833088858971 | 168.256462523619 | Ttyh3 |
| **ENSMUSG00000036568** | -0.141333877568921 | 0.321648799634917 | 0.855704427283138 | 224.071401709123 | 240.252640029727 | 243.238906382651 | 232.605820176024 | 263.884925192402 | 207.923979452556 | 271.721381635152 | 292.006376895829 | Bicral |
| **ENSMUSG00000036570** | -0.083662011044528 | 0.675025285449272 | 0.944023556644042 | 6769.12476453529 | 5375.14381083457 | 5249.98526733293 | 6904.07839643433 | 6348.56761935912 | 8769.49960396957 | 5504.17986477029 | 5128.02276749403 | Fxyd1 |
| **ENSMUSG00000036572** | -0.0454806477937833 | 0.756659017312294 | 0.956566265296272 | 283.70330700268 | 249.211213047785 | 265.007368821565 | 322.646782824807 | 252.935343234211 | 297.208982393948 | 275.885693997376 | 329.999771659226 | Upf3b |
| **ENSMUSG00000036580** | 0.145731231906217 | 0.367937042608566 | 0.871495599696183 | 808.644776329295 | 1104.34772840783 | 1059.08302039761 | 954.809374754808 | 821.218646864323 | 726.510845851873 | 1009.84574783945 | 991.084840542347 | Spg20 |
| **ENSMUSG00000036585** | 0.209043949492711 | 0.396439708778208 | 0.879359239049715 | 4691.94673014305 | 5121.86051914221 | 5219.69871089618 | 4157.82903564643 | 3504.96118481693 | 2534.22638485704 | 5921.6521790833 | 4640.62178895787 | Fgf1 |
| **ENSMUSG00000036591** | 0.222792841048219 | 0.335453863668252 | 0.858570960161243 | 971.277245311723 | 1234.65424503412 | 1125.334862603 | 899.471699793577 | 697.488370736765 | 639.672007374629 | 1164.96638333232 | 1122.43343386724 | Arhgap21 |
| **ENSMUSG00000036594** | -0.527908493319513 | 0.109061131393306 | 0.763154315676794 | 364.116027777325 | 443.856572258309 | 522.443098533943 | 531.804435644377 | 997.506916391197 | 898.965440574287 | 359.171941241867 | 429.868123608729 | H2-Aa |
| **ENSMUSG00000036599** | -1.17465096507327 | 0.0065964629247461 | 0.635862253526407 | 52.4037955610046 | 43.9784493613737 | 45.4298346551254 | 54.3997482669734 | 137.964732673206 | 200.585486060113 | 57.2592949805875 | 47.7631248454143 | Chst12 |
| **ENSMUSG00000036606** | 0.170181328646596 | 0.404065076711241 | 0.881158207349486 | 2609.3476134514 | 3158.30419672977 | 3086.38939188259 | 2420.78879788031 | 2208.53068096712 | 1769.79998981087 | 3205.47944082234 | 2835.39277491414 | Plxnb2 |
| **ENSMUSG00000036611** | -0.428129103461179 | 0.0115867988302954 | 0.635862253526407 | 260.211950371885 | 204.418347957496 | 193.076797284283 | 211.033506208086 | 331.772333333186 | 320.447544803351 | 235.283648465687 | 282.236646813812 | Eepd1 |
| **ENSMUSG00000036613** | 0.0753067353456821 | 0.660217365322864 | 0.939611397743053 | 190.641394196068 | 126.234437981721 | 171.308334845369 | 148.192417692789 | 150.009272827216 | 149.216032313011 | 149.915245040084 | 154.144630182928 | Eipr1 |

| | | | | | | | | | | | | |
|---|---|---|---|---|---|---|---|---|---|---|---|---|
| ENSMUSG00000036615 | -0.13145331235245 | 0.280586043964395 | 0.849093714741928 | 318.940341948873 | 295.632909595901 | 305.704929033448 | 355.474217123843 | 359.146288228664 | 377.932409710823 | 343.555769883525 | 316.973464883204 | Rfxap |
| ENSMUSG00000036617 | 0.0929197222535528 | 0.662648501114446 | 0.939682254260302 | 627.038519298917 | 911.331200655134 | 710.787621374984 | 650.921125815164 | 602.227007700503 | 489.232892829544 | 866.176971342706 | 760.953420832624 | Etl4 |
| ENSMUSG00000036620 | 0.00604936008207167 | 0.939327194851457 | 0.989269716906553 | 2032.90586228035 | 1844.65162599095 | 1845.58703286447 | 1920.87386984071 | 1917.27180087924 | 1953.26232462195 | 1795.85970620934 | 1946.34733745063 | Mgat4b |
| ENSMUSG00000036622 | -0.107513238828973 | 0.499122863916665 | 0.900927059989498 | 436.397125102849 | 448.743066631795 | 463.762895437739 | 430.508352664496 | 519.010184818252 | 572.402484610566 | 465.361906478593 | 361.480013034613 | Atp13a2 |
| ENSMUSG00000036632 | 0.0663014495359987 | 0.74283673487757 | 0.954383574136061 | 345.142239729375 | 249.211213047785 | 334.098575692902 | 316.081295965 | 333.962249724825 | 369.370834086306 | 267.557069272927 | 218.19063849837 | Alg5 |
| ENSMUSG00000036636 | -0.111074348330459 | 0.455932828693898 | 0.892805623137304 | 480.669297214732 | 639.316347197748 | 596.266579848521 | 471.777127211855 | 534.339599559719 | 556.502415593606 | 627.77008860535 | 644.802185413094 | Clcn7 |
| ENSMUSG00000036639 | -0.0419768144242333 | 0.823812233422756 | 0.966637987288264 | 183.413284463516 | 131.120932355207 | 134.396594188079 | 145.378637610015 | 146.724398239759 | 184.685417043153 | 155.120635492864 | 125.920965501547 | Nudt1 |
| ENSMUSG00000036644 | 0.00130717204402649 | 0.993371088000834 | 0.999133450854041 | 1456.4641111093 | 1613.35755897928 | 1646.8315062483 | 1269.01481733129 | 1318.32966776619 | 1253.65928787571 | 1732.35394268541 | 1674.96594628351 | Tbc1d9b |
| ENSMUSG00000036646 | -0.163367043613143 | 0.0826406781215735 | 0.739855948845396 | 618.003382133227 | 631.986605637519 | 662.518422053913 | 595.583450853932 | 711.722827282413 | 650.679747463293 | 701.686633034836 | 743.585011797928 | Man1b1 |
| ENSMUSG00000036655 | -0.583031976333313 | 0.0727229684923251 | 0.729806017191742 | 220.457346842847 | 198.717437855096 | 193.076797284283 | 260.743621003769 | 433.603445544362 | 488.00981059747 | 193.640524843441 | 193.223550510994 | Colec11 |
| ENSMUSG00000036661 | -0.368169859253055 | 0.119997902234508 | 0.769868040577702 | 198.77301764519 | 188.13003337921 | 189.290977729689 | 195.088752405698 | 346.006789878835 | 288.647406769431 | 176.983275394543 | 184.539345993646 | Dennd3 |
| ENSMUSG00000036667 | -0.529796944058566 | 0.0601265530411644 | 0.71282992338528 | 112.035700854562 | 104.245213301034 | 124.932045301595 | 126.620103724852 | 199.282391639076 | 249.508775343067 | 114.518589961175 | 112.894658725525 | Tcaf1 |
| ENSMUSG00000036672 | -0.340675518783243 | 0.424258099130429 | 0.884586455610141 | 27.1054114970714 | 19.5459774939439 | 18.9290977729689 | 28.1378008277448 | 33.943704070392 | 51.3694537471021 | 18.7394056300105 | 15.1973579053591 | Cenpt |
| ENSMUSG00000036676 | -0.00404513592804516 | 0.982556364767127 | 0.996940244108351 | 206.904641094311 | 313.550055632016 | 291.508105703722 | 216.661066373635 | 235.416012101106 | 234.831788558181 | 275.885693997376 | 285.493223507818 | Tmtc3 |
| ENSMUSG00000036678 | 0.126594418211126 | 0.456197465220967 | 0.892805623137304 | 150.886790667031 | 132.749763813036 | 172.254789734017 | 183.8336320746 | 135.774816281568 | 135.762127760198 | 168.654650670094 | 144.374900100912 | Aaas |
| ENSMUSG00000036686 | -0.112895337921269 | 0.478248663226442 | 0.897674307047416 | 259.308436655316 | 325.766291565731 | 308.544293699394 | 317.019222659259 | 375.57066116595 | 369.370834086306 | 282.132162540713 | 283.322172378481 | Cc2d1a |
| ENSMUSG00000036693 | 0.31814151773894 | 0.0481273098209229 | 0.703396849069134 | 646.915821063436 | 701.211942595237 | 693.751433379311 | 770.975742680209 | 568.283303630111 | 425.632616761703 | 681.906149314269 | 578.585125968315 | Nop14 |
| ENSMUSG00000036698 | 0.182518805238581 | 0.449384290580634 | 0.89108031640712 | 1057.11104838578 | 1310.39490782315 | 1196.31897925164 | 870.395972271574 | 902.245553354936 | 605.425704876561 | 1122.28218161952 | 1276.57806405016 | Ago2 |
| ENSMUSG00000036707 | 0.0893800842313254 | 0.615766361696343 | 0.928422072203769 | 2930.99849654998 | 3271.50798304886 | 3328.68184337659 | 2960.09664707876 | 2460.37106600551 | 2261.47904710457 | 3666.67703493871 | 3351.01741813168 | Cab39 |
| ENSMUSG00000036712 | 0.331377570211905 | 0.155252690655945 | 0.787275540343856 | 981.215896193983 | 1162.17124516075 | 1182.12215592191 | 1040.1607039323 | 733.621991198795 | 541.82542880872 | 1110.8303226234 | 1082.2689879745 | Cyld |
| ENSMUSG00000036718 | -0.278718459979109 | 0.510224605462231 | 0.902394714365738 | 38.8510898124689 | 17.1027303072009 | 26.5007368821565 | 33.7653609932938 | 45.9882442244021 | 53.8156182112498 | 20.8215618111227 | 20.6249857287016 | Micall2 |
| ENSMUSG00000036721 | -0.0391137122290854 | 0.827941050091948 | 0.967319591893432 | 140.044626068202 | 132.749763813036 | 98.4313084194385 | 135.061443973175 | 119.350443344282 | 119.862058743238 | 130.134761319517 | 150.888053488923 | Zscan12 |
| ENSMUSG00000036737 | 0.263598346050542 | 0.302267872079781 | 0.851785577030345 | 1942.55449062345 | 2498.62745630916 | 2339.63648473896 | 1870.22582835077 | 1389.50195049443 | 1128.90490020417 | 2343.46678184186 | 2343.64969411931 | Oxsr1 |
| ENSMUSG00000036743 | 0.608011188391998 | 0.14467492160787 | 0.780607990324772 | 28.9124389302094 | 14.6594831204579 | 18.9290977729689 | 21.5723139679377 | 20.8042057205628 | 8.56157562451702 | 14.5750932677859 | 10.8552556466851 | Psma8 |
| ENSMUSG00000036745 | 0.194977403153894 | 0.208060504788297 | 0.820372930586885 | 606.257703817829 | 721.572335818095 | 561.247748968529 | 629.348811847226 | 592.372383938131 | 436.640356850368 | 634.016557148687 | 536.249628946243 | Ttll7 |
| ENSMUSG00000036748 | -0.0129244947996229 | 0.944441359284178 | 0.990505437194639 | 1803.41337827181 | 1413.82570539527 | 1479.30899095752 | 1860.84656140819 | 1805.58606490569 | 2086.578287918 | 1334.66211209297 | 1390.55824834036 | Cuedc2 |
| ENSMUSG00000036751 | 0.154298635571711 | 0.274095679083148 | 0.845290626657401 | 15638.9189200936 | 11527.2402270534 | 13225.7606139734 | 15125.0058716071 | 12566.8352134158 | 14390.785542581 | 11359.203046058 | 11569.531468237 | Cox6b1 |
| ENSMUSG00000036752 | -0.276762560296242 | 0.136189809232641 | 0.776456211338551 | 4017.92549758254 | 3007.63728688062 | 3809.48092681 | 4102.4913606852 | 5053.23207370513 | 5629.84751423598 | 3616.70528659202 | 3797.16842521044 | Tubb4b |
| ENSMUSG00000036764 | 0.109001942985287 | 0.481992704898087 | 0.897732059433541 | 237.624107457659 | 184.872370463553 | 225.25626349833 | 243.860940507122 | 205.85214081399 | 207.923979452556 | 179.065431575656 | 233.387996403729 | Dnajc12 |
| ENSMUSG00000036769 | -0.0331000674503611 | 0.886254993779528 | 0.980068672363141 | 175.281661014395 | 199.531853584011 | 173.201244622666 | 156.633757941113 | 121.54035973592 | 149.216032313011 | 217.585320926233 | 232.302470839061 | Wdr44 |
| ENSMUSG00000036775 | 0.202623816536188 | 0.266647242552079 | 0.843513595393324 | 383.089815825275 | 385.218639776477 | 349.241853911277 | 421.129085721914 | 316.442918591719 | 259.293433199658 | 316.487739529066 | 443.97995594942 | Decr2 |
| ENSMUSG00000036777 | -0.561306770395193 | 0.173950116462748 | 0.799889021902745 | 15.3597331816738 | 29.3189662409158 | 11.3574586637814 | 17.8206071909051 | 32.8487458745729 | 35.4693847301419 | 15.616171358342 | 26.0526135520442 | Anln |
| ENSMUSG00000036779 | 0.226208974289115 | 0.25815522429971 | 0.838310965110319 | 619.810409566365 | 790.797672775813 | 680.501064938233 | 605.900644490772 | 483.971522552041 | 426.855698993777 | 707.933101578173 | 686.052156870497 | Papd5 |
| ENSMUSG00000036781 | 0.26102473579182 | 0.128976425381495 | 0.77303252008843 | 1367.91976688553 | 911.331200655134 | 999.45636241276 | 1260.57347708297 | 903.340511550755 | 1126.45873574002 | 885.957455063272 | 872.76255399348 | Rps27l |
| ENSMUSG00000036782 | 0.057105047801274 | 0.699595075367664 | 0.945280238056015 | 257.501409222178 | 325.766291565731 | 324.634026806417 | 268.247034557834 | 301.113503850252 | 314.332133642982 | 263.392756910703 | 252.927456567762 | Klhl13 |
| ENSMUSG00000036792 | 0.192892284852481 | 0.455158690630204 | 0.892775943845882 | 280.089252136404 | 371.373572384934 | 302.865564367503 | 271.998741334867 | 211.326931793086 | 170.008430258266 | 372.705956419097 | 318.058990447873 | Mbd5 |
| ENSMUSG00000036810 | 0.0411123202014288 | 0.769774624144201 | 0.958815818668915 | 798.706125447036 | 776.138189655355 | 711.734076263632 | 664.052099534778 | 625.221129812704 | 634.779678446333 | 774.562099373766 | 832.598108100745 | Cnep1r1 |
| ENSMUSG00000036817 | -0.0137290278291866 | 0.913412134074782 | 0.986234232499696 | 903.513716569045 | 991.958357817652 | 1109.24512949598 | 847.885731609378 | 1050.06490979051 | 873.280713700736 | 1008.8046697489 | 956.348022472955 | Sun1 |
| ENSMUSG00000036819 | 0.124444764129891 | 0.649107499595358 | 0.936461253686283 | 205.097613661173 | 252.468875963442 | 271.632553042104 | 214.785212985119 | 147.819356435578 | 143.100621152642 | 280.050006359601 | 294.177428025166 | Jmjd4 |
| ENSMUSG00000036820 | 0.143340199159953 | 0.447975280800152 | 0.89108031640712 | 168.957064998411 | 146.594831204579 | 162.790240847533 | 192.274972322923 | 179.573144114332 | 176.123841418636 | 123.88829277618 | 128.092016630884 | Amdhd2 |
| ENSMUSG00000036822 | -0.257475157543936 | 0.0963869969808795 | 0.757809915356977 | 243.045189757073 | 249.211213047785 | 310.437203476691 | 263.557401086543 | 285.784089108784 | 384.047820871192 | 315.446661438509 | 289.835325766492 | Topors |
| ENSMUSG00000036825 | 0.139558548049607 | 0.629727788319603 | 0.931825800800958 | 458.984968017075 | 452.815145276367 | 405.082692341535 | 317.019222659259 | 293.448796479518 | 203.031650524261 | 500.758561557502 | 485.229927406823 | Ssx2ip |
| ENSMUSG00000036833 | 0.54835817746927 | 0.0457406266517777 | 0.697906909182231 | 680.345828576491 | 1023.72057124531 | 1724.44080711747 | 1314.03529865568 | 1008.45649834939 | 692.264543353805 | 834.944628626022 | 706.677142599198 | Pnpla7 |
| ENSMUSG00000036835 | -0.0440950658899015 | 0.827955129105728 | 0.967319591893432 | 975.794813894569 | 835.590537866101 | 824.362208012797 | 1052.35375095766 | 998.601874587016 | 1290.35175483792 | 749.576225200418 | 765.295523091298 | Psenen |
| ENSMUSG00000036840 | 0.0192494175083594 | 0.913729093499469 | 0.986299122529346 | 337.914129996823 | 356.714089264476 | 466.602260103684 | 534.618215727152 | 457.692525852383 | 413.401794440965 | 399.773986773556 | 401.644458927348 | Siah1a |
| ENSMUSG00000036845 | -0.165400915354087 | 0.454633854911289 | 0.892775943845882 | 221.360860559416 | 157.99665140938 | 141.021778408619 | 194.150825711439 | 201.472308030714 | 264.185762127954 | 175.942197303987 | 160.657783570939 | Lin37 |
| ENSMUSG00000036850 | 0.14413636807985 | 0.273997534952319 | 0.845204865395511 | 1731.13228094629 | 1253.38580679915 | 1342.0730321035 | 1485.67588370493 | 1207.73888998846 | 1369.85209992272 | 1308.63515982906 | 1373.18983930566 | Mrpl41 |
| ENSMUSG00000036854 | 0.0820993589828168 | 0.609623478390976 | 0.927932112008768 | 31633.8222445154 | 27665.7023112197 | 20160.4355831006 | 31697.2326324546 | 24919.058620451 | 26708.4467017969 | 27658.3216318049 | 25722.613780385 | Hspb6 |
| ENSMUSG00000036860 | -0.00373237207751144 | 0.981240198411738 | 0.996940244108351 | 2304.86349096763 | 1639.41886230454 | 1564.48993093588 | 1935.88069694884 | 1943.5507975789 | 2100.03219247082 | 1690.71081906317 | 1730.3277500816 | Mrpl55 |
| ENSMUSG00000036862 | -0.899980140560372 | 0.00651991988889032 | 0.635862253526407 | 112.035700854562 | 89.5857301805761 | 97.48485353079 | 139.751077444466 | 296.733671066975 | 272.747337752471 | 125.970448957293 | 123.74991437221 | Dchs1 |
| ENSMUSG00000036863 | -0.100044547992535 | 0.606043477347906 | 0.926859923029135 | 343.335212296237 | 389.290718421049 | 364.385132129652 | 289.819348525772 | 279.21433993387 | 326.562955963721 | 460.156516025812 | 420.098393526712 | Syde2 |
| ENSMUSG00000036864 | -0.116920375225174 | 0.701627100123698 | 0.945991110533699 | 31.6229800799166 | 43.1640336324594 | 38.8046504345863 | 40.3308478531009 | 63.5075753575076 | 36.6924669622158 | 38.5198893505771 | 28.2236646813812 | Proser3 |
| ENSMUSG00000036867 | -0.656342389502845 | 0.0219090869910167 | 0.648762042745232 | 318.940341948873 | 257.355370336928 | 318.008842585878 | 297.322762079837 | 646.025335533267 | 631.110431750112 | 301.91264626128 | 299.605055848508 | Smad6 |
| ENSMUSG00000036873 | 0.0718053603264997 | 0.520692764842694 | 0.905962179970576 | 533.976606492306 | 424.310594764365 | 478.906173656114 | 500.852854733858 | 454.407651264925 | 457.432754795624 | 466.402984569149 | 464.604941678121 | 2410004B18I |
| ENSMUSG00000036879 | 0.200575794334796 | 0.377218628313885 | 0.87422797717331 | 1773.59742562504 | 2362.62002958047 | 2075.57557080604 | 1864.59826818522 | 1356.65320461986 | 1236.53613662667 | 2192.51045871122 | 2241.61029104047 | Phkb |
| ENSMUSG00000036880 | 0.0555721955497006 | 0.612387074663031 | 0.928055903343287 | 36332.997084391 | 32933.3432458376 | 38838.7228105777 | 38048.8722059708 | 36917.6105302367 | 38930.707446911 | 31441.5994128859 | 33341.9177187932 | Acaa2 |
| ENSMUSG00000036885 | -0.984360221982945 | 0.0119671550966544 | 0.635862253526407 | 12.6491920319666 | 15.4738988493722 | 34.0723759913441 | 20.6343872736795 | 49.2731188118594 | 39.1386314263635 | 24.9858741733473 | 49.9341759747513 | Arhgef26 |
| ENSMUSG00000036887 | -0.658970071445648 | 0.0586260042921078 | 0.711712402279519 | 522.230928176908 | 513.081909216027 | 621.82086184203 | 619.031618210386 | 1075.24894829435 | 1517.84505000366 | 438.293876124134 | 563.387768062955 | C1qa |
| ENSMUSG00000036890 | 0.321188991772708 | 0.0444989632886408 | 0.693563614452513 | 271.054114970713 | 315.99330281876 | 323.687571917769 | 306.702029022419 | 235.416012101106 | 190.800828203522 | 257.146288367366 | 289.835325766492 | Gtdc1 |
| ENSMUSG00000036892 | -0.443089106878453 | 0.308144299162568 | 0.851785577030345 | 25.2983840639333 | 41.5352021746308 | 25.5542819935081 | 29.075727522003 | 19.7092475247437 | 52.592535979176 | 23.9447960827911 | 69.4736361387845 | Prodh2 |
| ENSMUSG00000036893 | -0.040193467639329 | 0.83826346388492 | 0.969135716755098 | 340.62467114653 | 393.362797065621 | 383.314229902621 | 303.888248939644 | 351.48158085793 | 253.178022039289 | 425.80093903746 | 429.868123608729 | Ehmt1 |
| ENSMUSG00000036894 | -0.468292751333409 | 0.0940656416776592 | 0.757085157577544 | 38.8510898124689 | 43.9784493613737 | 51.1085639870161 | 60.9652351267805 | 78.836990098975 | 89.2850029413918 | 54.1360607089191 | 47.7631248454143 | Rap2b |
| ENSMUSG00000036896 | -0.729020348338793 | 0.0270689435297138 | 0.664353746352133 | 336.107102563685 | 342.054606144018 | 395.618143455051 | 409.873965390816 | 713.912743674051 | 1026.16599270997 | 349.802238426862 | 370.164217551961 | C1qc |
| ENSMUSG00000036898 | -0.14142934766163 | 0.51011906502437 | 0.902394714365738 | 112.035700854562 | 92.0289773673191 | 91.8061241988993 | 123.806323642077 | 89.7865720571659 | 99.0696607979826 | 138.463386043966 | 134.605170018895 | Zfp157 |
| ENSMUSG00000036902 | -0.107653753724276 | 0.839670950050084 | 0.969879194284716 | 19.877301764519 | 20.3603932228582 | 8.51809399783602 | 10.3171936368398 | 5.47479097909548 | 26.9078091056249 | 17.6983275394543 | 14.1118323406906 | Neto2 |
| ENSMUSG00000036904 | -0.0590459146437802 | 0.803101362282756 | 0.963461896828986 | 113.8427282877 | 137.636258186522 | 193.076797284283 | 163.19924480092 | 193.80760065998 | 184.685417043153 | 134.299073681742 | 120.493337678204 | Fzd8 |
| ENSMUSG00000036905 | -0.678726304750804 | 0.0301498897847195 | 0.665956123973759 | 314.422773366028 | 360.786167909048 | 387.100049457215 | 436.135912830045 | 718.292576457328 | 924.650167447838 | 307.11803671406 | 449.407583772762 | C1qb |
| ENSMUSG00000036908 | -0.679291377955699 | 0.0173380199404211 | 0.642273034804458 | 116.553269437407 | 109.946123403434 | 154.272146849697 | 140.689004138724 | 277.024423542232 | 271.524255520397 | 121.806136595068 | 164.999885829613 | Unc93b1 |
| ENSMUSG00000036913 | -0.0340330050866327 | 0.902344646740268 | 0.984902747274457 | 100.290022539164 | 65.9676740420606 | 61.519567762149 | 93.7926694258161 | 53.6529515951357 | 85.6157562451702 | 109.313199508394 | 80.3288917854696 | Trim67 |
| ENSMUSG00000036916 | -0.0714960850102225 | 0.85763738428174 | 0.974046056228403 | 30.7194663633475 | 21.9892246806869 | 17.9826428843205 | 41.2687745473591 | 43.7983278327639 | 15.9000690169602 | 27.0680303544596 | 30.3947158107182 | Zfp280c |
| ENSMUSG00000036918 | -0.0292430881650384 | 0.886377323484533 | 0.980081278713594 | 416.51982333833 | 495.979178908826 | 551.783200082044 | 366.729337454941 | 453.312693069106 | 336.347613820311 | 562.182168900314 | 515.624643217541 | Ttc7 |
| ENSMUSG00000036923 | -0.586331224971928 | 0.243032848661102 | 0.832048466795696 | 18.9737880479499 | 15.4738988493722 | 12.3039135524298 | 15.9447538023887 | 29.5638712871156 | 42.8078781225851 | 9.36970281500523 | 13.0263067760221 | Stox1 |
| ENSMUSG00000036932 | 0.184002921522418 | 0.163682892252933 | 0.793968226938511 | 2342.81106706353 | 2472.5661529839 | 2455.10398115407 | 2490.19537325542 | 2041.0020770068 | 1733.10752284866 | 2429.87626335802 | 2385.98519114138 | Aifm1 |
| ENSMUSG00000036934 | 0.171758503427634 | 0.21450579351665 | 0.824248286808591 | 364.116027777325 | 317.622134276588 | 391.832323900457 | 384.549944645846 | 301.113503850252 | 309.439804714686 | 302.953724351836 | 379.933947633978 | 4921524J17F |
| ENSMUSG00000036940 | 0.134488235586589 | 0.341656308349714 | 0.859283395807012 | 590.897970636155 | 613.255043872489 | 565.033568523123 | 542.121629281217 | 532.149683168081 | 418.29412336926 | 614.236073428121 | 539.506205640248 | Kdm1a |
| ENSMUSG00000036941 | 0.0836221430163887 | 0.647356886348151 | 0.935063240511793 | 187.930853046361 | 247.582381589956 | 266.900278598862 | 229.79204009325 | 217.896680968 | 171.23151249034 | 227.996101831794 | 261.61166108511 | Elac1 |
| ENSMUSG00000036943 | -0.207064688043472 | 0.209362385081807 | 0.821566820583092 | 187.027339329792 | 201.160685041839 | 224.309808609682 | 225.102406621959 | 290.163921892061 | 254.401104271363 | 226.955023741238 | 195.394601640331 | Rab8b |
| ENSMUSG00000036944 | 0.581754383938684 | 0.00436878655195811 | 0.635862253526407 | 259.308436655316 | 276.901347830872 | 336.937940358847 | 365.791410760683 | 274.834507150593 | 171.23151249034 | 176.983275394543 | 204.078806157679 | Tmem71 |
| ENSMUSG00000036948 | 0.428554914383852 | 0.230181877565537 | 0.832048466795696 | 36.1405486627618 | 21.1748089517725 | 19.8755526616174 | 28.1378008277448 | 15.3294147414674 | 15.9000690169602 | 26.0269522639034 | 20.6249857287016 | BC037034 |
| ENSMUSG00000036955 | 0.103400976595515 | 0.379243912006813 | 0.87422797717331 | 924.294532050133 | 1057.92603185971 | 1069.49402417274 | 909.788893430416 | 942.759006600242 | 815.795848793264 | 1018.1743725639 | 909.670423192209 | Kif1bp |
| ENSMUSG00000036957 | -0.598031982712809 | 0.140944419427566 | 0.779711473022872 | 18.9737880479499 | 25.2468875963442 | 11.3574586637814 | 20.6343872736795 | 30.6588294829347 | 44.0309603546589 | 15.616171358342 | 26.0526135520442 | Lrfn3 |
| ENSMUSG00000036959 | -0.327943845406868 | 0.0624439940487485 | 0.715615581523459 | 136.430571201926 | 144.96599974675 | 125.878500190243 | 114.427056699496 | 195.997517051618 | 140.654456688494 | 142.627698406191 | 175.855141476298 | Bcorl1 |
| ENSMUSG00000036966 | -0.0962406796914336 | 0.410497049139959 | 0.882192126163777 | 412.002254755485 | 420.238516119794 | 440.101523221528 | 396.742991671202 | 479.591689768764 | 488.00981059747 | 422.677704765791 | 395.131305539337 | Spryd3 |
| ENSMUSG00000036968 | -0.616331718732287 | 0.111617175512608 | 0.764116514278315 | 93.0619128066116 | 65.9676740420606 | 71.930571537282 | 83.4754757889764 | 166.433645764503 | 190.800828203522 | 63.5057635239243 | 61.8749571861049 | Cnpy4 |
| ENSMUSG00000036975 | -0.158201997284722 | 0.270584406773677 | 0.844129257012535 | 325.264937964856 | 302.148235427216 | 283.936466594534 | 286.067641748739 | 344.911831683016 | 271.524255520397 | 386.239971596327 | 332.170822788563 | Tmem177 |
| ENSMUSG00000036977 | 0.0723094486049109 | 0.739940334871642 | 0.954383574136061 | 186.123825613223 | 116.461449234749 | 110.735221971868 | 167.888878272211 | 146.724398239759 | 141.877538920568 | 118.6829023234 | 145.46042566558 | Anapc10 |
| ENSMUSG00000036980 | 0.102867506246607 | 0.631700246630026 | 0.931825800800958 | 643.30176619716 | 646.646088757977 | 615.19567762149 | 534.618215727152 | 545.28918151791 | 364.47850515801 | 746.45299092875 | 614.407469602375 | Taf6 |

| | | | | | | | | | | | | |
|---|---|---|---|---|---|---|---|---|---|---|---|---|
| ENSMUSG00000036983 | -0.0250909022232288 | 0.885809974517415 | 0.979948700426504 | 160.82544154929 | 179.171460361152 | 137.235958854025 | 138.813150750208 | 139.059690869025 | 143.100621152642 | 155.120635492864 | 189.966973816989 | Tfb1m |
| ENSMUSG00000036985 | 0.0962628150609492 | 0.625369103367301 | 0.931573470007391 | 701.126644057579 | 778.581436842098 | 803.540200462531 | 583.390403828576 | 556.238763476101 | 538.156182112498 | 881.793142701048 | 704.506091469861 | Zdhhc9 |
| ENSMUSG00000036986 | -0.442196247010974 | 0.115165260242137 | 0.768321548877656 | 73.1846110420926 | 118.090280692578 | 114.521041526462 | 122.868396947819 | 192.712642464161 | 176.123841418636 | 92.6559500594962 | 121.578863242873 | Pml |
| ENSMUSG00000036989 | -0.0116407684862641 | 0.901148896687972 | 0.984802943363903 | 489.704434380422 | 536.699965354543 | 530.961192531779 | 499.9149280396 | 543.099265126272 | 516.140701935169 | 500.758561557502 | 514.539117652873 | Trim3 |
| ENSMUSG00000036990 | 0.23630935194957 | 0.377450266023381 | 0.87422797717331 | 932.426155499254 | 1292.47776178704 | 1153.72850926246 | 906.037186653384 | 816.838814081046 | 506.356044078578 | 1091.04983890283 | 1222.30178581674 | Otud4 |
| ENSMUSG00000036992 | -0.269073846122469 | 0.249812677691303 | 0.835314899646652 | 85.8338030740593 | 66.7820897709749 | 85.1809399783602 | 118.178763476528 | 105.115986798633 | 135.762127760198 | 93.6970281500523 | 94.4407241261602 | Nxt1 |
| ENSMUSG00000036995 | -0.541363893112441 | 0.050806912323112 | 0.703396849069134 | 103.90407740544 | 86.3280672649188 | 54.8943835416099 | 108.799496533947 | 142.344565456483 | 102.738907494204 | 169.69572876065 | 99.8683519495027 | Asap3 |
| ENSMUSG00000037001 | -0.199782144789758 | 0.341440388464761 | 0.859283395807012 | 63.2459601598332 | 61.0811796685746 | 65.3053873167428 | 87.227182566009 | 67.887408140784 | 77.0541806206532 | 80.1630129728225 | 92.2696729968231 | Zfp39 |
| ENSMUSG00000037003 | -0.0631663698454125 | 0.659046473544919 | 0.938989601151841 | 2390.69729404169 | 2755.98282664609 | 3378.84395247496 | 2760.31826120177 | 3404.22503080157 | 2810.64296930573 | 2896.27924792717 | 2679.07709360188 | Tns2 |
| ENSMUSG00000037007 | -0.207894337805121 | 0.280245246246366 | 0.849093714741928 | 95.7724539563188 | 115.647033505835 | 99.3777633080869 | 92.854742731558 | 124.825234323377 | 85.6157562451702 | 122.847214685624 | 132.434118889558 | Zfp113 |
| ENSMUSG00000037010 | -0.555377962701504 | 0.133369251818567 | 0.77303252008843 | 94.8689402397497 | 74.9262470601182 | 29.3401015481019 | 91.9168160372998 | 136.869774477387 | 66.0464405319884 | 103.066730965058 | 121.578863242873 | Apln |
| ENSMUSG00000037012 | -0.0953081303864572 | 0.340295487873479 | 0.858620222808832 | 3530.02809063526 | 3680.34467896385 | 3252.01899739606 | 3037.94456270218 | 3445.8334422427 | 3568.95395319152 | 3896.75529295162 | 3511.67520170262 | Hk1 |
| ENSMUSG00000037013 | -0.0224844446315249 | 0.846008071530768 | 0.972180580805179 | 549.336339673979 | 561.132437221972 | 477.959718767466 | 469.901273823339 | 540.909348734634 | 491.679057293692 | 499.717483466946 | 559.045665804281 | Ss18 |
| ENSMUSG00000037014 | 0.265148775885951 | 0.618909690926972 | 0.929568738467614 | 12.6491920319666 | 15.4738988493722 | 44.483379766477 | 10.3171936368398 | 10.949581958191 | 19.5693157131818 | 16.6572494488982 | 21.7105112933701 | Sstr4 |
| ENSMUSG00000037016 | 0.560736468724977 | 0.22068724200697 | 0.829144114546069 | 22.5878429142261 | 26.0613033252585 | 53.001473764313 | 81.59962240046 | 22.994122112201 | 22.0154801773295 | 37.4788112600209 | 41.2499714574033 | Frem2 |
| ENSMUSG00000037017 | -0.0260764787385222 | 0.846840654051358 | 0.972334438148346 | 261.115464088454 | 239.438224300813 | 217.684624389143 | 251.364354061187 | 266.074841584041 | 225.04713070159 | 269.639225454039 | 225.78931745105 | Zscan21 |
| ENSMUSG00000037020 | -0.169303345898262 | 0.672841685787549 | 0.943302739919217 | 18.0702743313809 | 13.8450673915436 | 20.8220075502658 | 11.2551203310979 | 24.0890803080201 | 13.4539045528125 | 12.4929370866736 | 21.7105112933701 | Wdr62 |
| ENSMUSG00000037022 | 0.0291147841575149 | 0.79193142742498 | 0.961804768755116 | 1084.21645988285 | 931.691593877992 | 906.703783325212 | 1127.38788649831 | 959.183379537529 | 1005.37359476471 | 985.900951756661 | 1018.22297965906 | Mmaa |
| ENSMUSG00000037029 | 0.0857760851155189 | 0.720914050737547 | 0.951674101666722 | 548.43282595741 | 604.296470854432 | 456.191256328551 | 430.508352664496 | 347.101748074654 | 355.916929533493 | 629.852244786463 | 588.354856050331 | Zfp146 |
| ENSMUSG00000037031 | -0.595854111045604 | 0.0222280447860066 | 0.648762042745232 | 169.86057871498 | 110.760539132349 | 122.09268063565 | 172.578511743502 | 283.594172717146 | 259.293433199658 | 151.997401221196 | 174.76961591163 | Tspan15 |
| ENSMUSG00000037032 | -0.0721360194806842 | 0.427091174346314 | 0.885066977635295 | 4275.42690680472 | 3642.06713970488 | 4025.27264142184 | 3707.62422240251 | 4089.66886138433 | 4357.84199287916 | 3939.43949466442 | 4066.37876524823 | Apbb1 |
| ENSMUSG00000037035 | -0.682066436498959 | 0.0532335281811037 | 0.703396849069134 | 93.9654265231807 | 56.1946852950887 | 90.8596693102509 | 103.171936368398 | 188.332809680885 | 194.470074899744 | 97.8613405122768 | 71.6446872681215 | Inhbb |
| ENSMUSG00000037049 | -0.0985143885800833 | 0.328551538070915 | 0.855977678491063 | 4291.69015370296 | 4054.16149853553 | 3829.35647947162 | 3513.47339669107 | 4426.91598569661 | 3855.15519549681 | 4294.44712354406 | 4220.52339543116 | Smpd1 |
| ENSMUSG00000037053 | -0.389917637108967 | 0.108511145899698 | 0.762287160109383 | 318.036828232304 | 466.66021266791 | 323.687571917769 | 389.239578117137 | 388.710159515779 | 458.655837027697 | 389.363205867995 | 726.216602763232 | Azgp1 |
| ENSMUSG00000037058 | 0.132985980846823 | 0.155758771645576 | 0.787275540343856 | 4768.74539605142 | 4277.31140825805 | 4745.52481168331 | 5186.73461924763 | 4051.34532453066 | 4389.64213091308 | 4517.23783492308 | 4347.52988649737 | Paip2 |
| ENSMUSG00000037060 | -0.57226170971388 | 0.100560415342159 | 0.759800953500741 | 337.010616280254 | 320.065381463331 | 263.114459044268 | 448.328959855401 | 589.087509350674 | 852.48831575548 | 297.748333899055 | 296.348479154503 | Cavin3 |
| ENSMUSG00000037062 | 0.1297797691295 | 0.631336325217983 | 0.931825800800958 | 8901.41713563823 | 8241.07276088409 | 9325.42001785315 | 4853.77064278598 | 8462.9318954858 | 4224.52602958311 | 8081.88921698729 | 7857.03403707066 | Sh3glb1 |
| ENSMUSG00000037070 | -0.279670118316581 | 0.229147030075896 | 0.832048466795696 | 215.939778260002 | 182.42912327681 | 177.933519065908 | 188.52326554589 | 269.359716171498 | 314.332133642982 | 160.326025945645 | 185.624871558315 | Rbmxl1 |
| ENSMUSG00000037071 | -0.533073100936964 | 0.180423554126622 | 0.802808628271642 | 459.888481733644 | 956.124065745422 | 780.825283134969 | 719.38977449601 | 687.633746974393 | 1029.83523940619 | 495.553171104721 | 2007.13676907207 | Scd1 |
| ENSMUSG00000037072 | -0.159414268961511 | 0.157476688107242 | 0.789197677076549 | 2056.39721891115 | 1766.46771601518 | 1796.37137865475 | 1940.57033042014 | 2096.84494499357 | 2410.69507941758 | 1911.41937426107 | 2025.59070367144 | Selenof |
| ENSMUSG00000037075 | 0.0486685119334195 | 0.690299924144111 | 0.94469184542218 | 1083.31294616628 | 1065.25577341994 | 1025.01064440627 | 1029.84351029546 | 1051.15986798633 | 823.134342185708 | 1106.66601026117 | 1081.18346240983 | Rnf139 |
| ENSMUSG00000037085 | -0.0890065677505537 | 0.602985257987768 | 0.926448151609826 | 167.150037565273 | 142.522752560008 | 152.3792370724 | 211.971432902344 | 195.997517051618 | 174.900759186562 | 176.983275394543 | 168.256462523619 | Trmt12 |
| ENSMUSG00000037089 | -0.104197676151932 | 0.494883671062277 | 0.900200915538898 | 550.239853390548 | 471.546707041396 | 501.621090983677 | 556.19052969509 | 594.56230032977 | 653.125911927441 | 437.252798033577 | 551.446986851602 | Slc35b2 |
| ENSMUSG00000037095 | -0.121370495413185 | 0.237380434374167 | 0.832048466795696 | 2044.65154059575 | 2032.78165937016 | 2099.23694302226 | 1764.2401118996 | 2079.32561386047 | 2188.09411318014 | 2359.08295320021 | 2011.47887133074 | Lrg1 |
| ENSMUSG00000037096 | -0.470892258306946 | 0.0448942243399094 | 0.693563614452513 | 37.0440623793308 | 36.6487078011448 | 32.1794662140472 | 42.2067012416173 | 52.5579933993167 | 41.5847958905112 | 55.1771387994752 | 55.3618037980939 | Gm9762 |
| ENSMUSG00000037098 | 0.296365635076633 | 0.275320797268284 | 0.846286635233373 | 1151.07647490896 | 1421.1554469555 | 1491.61290450995 | 1224.93226270116 | 809.174106710313 | 632.333513982185 | 1436.68776496747 | 1427.46611753909 | Rab11fip3 |
| ENSMUSG00000037103 | -0.297770522552512 | 0.109772351149661 | 0.763154315676794 | 74.0881247586617 | 88.7713144516618 | 101.270673085384 | 105.985716451172 | 116.065568756824 | 129.646716599829 | 102.025652874501 | 107.467030902182 | Dcaf15 |
| ENSMUSG00000037104 | -0.0297459974400754 | 0.858831602739094 | 0.974833700919212 | 511.388763578079 | 569.276594511116 | 494.995906763138 | 369.543117537716 | 488.351355335317 | 429.301863457925 | 507.005030100839 | 561.216716933618 | Socs5 |
| ENSMUSG00000037106 | 0.87062740904947 | 0.0401735919820048 | 0.679566075569698 | 12.6491920319666 | 21.9892246806869 | 20.8220075502658 | 16.8826804966469 | 10.949581958191 | 6.1154111603693 | 12.4929370866736 | 9.76973008201657 | Fer1l6 |
| ENSMUSG00000037110 | 0.272523654598294 | 0.455370808453303 | 0.892805623137304 | 992.961574509381 | 1557.1628736842 | 1169.81824236948 | 725.955261355817 | 664.639624862192 | 428.078781225851 | 1178.50039850955 | 1409.01218293972 | Ralgapa2 |
| ENSMUSG00000037111 | 0.383058182116607 | 0.136198447320224 | 0.776456211338551 | 1592.89468231123 | 1982.28788417748 | 1878.71295396717 | 1603.85464718146 | 983.272459845549 | 865.942220308293 | 1800.02401857156 | 1761.80799145699 | Setd7 |
| ENSMUSG00000037112 | 0.00298350683470365 | 0.993358255830016 | 0.999133450854041 | 91.2548853734735 | 140.079505373265 | 141.021778408619 | 62.8410885152968 | 85.4067392738896 | 63.6002760678407 | 167.613572579538 | 117.236760984199 | Sik2 |
| ENSMUSG00000037119 | -0.105484676527516 | 0.525586009515627 | 0.908008665693042 | 203.290586228035 | 223.96432545144 | 261.221549266971 | 244.79886720138 | 312.063085808443 | 227.493295165738 | 219.667477107345 | 244.243252050414 | Fam91a1 |
| ENSMUSG00000037126 | -0.138318590464826 | 0.519068767403606 | 0.905875760677986 | 84.0267756409212 | 89.5857301805761 | 87.0738497556571 | 106.92364314543 | 93.0714466446232 | 136.985209992272 | 81.2040910633787 | 94.4407241261602 | Psd |
| ENSMUSG00000037129 | -0.804471528052346 | 0.150044409192417 | 0.782641594375979 | 9.93865088225949 | 22.8036404096012 | 9.46454888648447 | 2.81378008277448 | 14.2344565456483 | 28.1308913376988 | 19.7804837205666 | 17.3684090346961 | Tmprss13 |
| ENSMUSG00000037134 | 0.0856875310889832 | 0.481045109750789 | 0.897732059433541 | 332.493047697409 | 385.218639776477 | 394.671688566402 | 319.833002742033 | 339.43704070392 | 343.686107212755 | 343.555769883525 | 323.486618271215 | Prmt9 |
| ENSMUSG00000037138 | -0.706701325505994 | 0.110630331097858 | 0.764116514278315 | 18.9737880479499 | 17.9171460361152 | 24.6078271048596 | 19.6964605794214 | 40.5134532453066 | 57.4848649074714 | 18.7394056300105 | 16.2828834700276 | Aff3 |
| ENSMUSG00000037139 | 0.208424508876682 | 0.553888757523808 | 0.915567635607008 | 73.1846110420926 | 41.5352021746308 | 67.1982970940397 | 77.8479156234274 | 27.3739548954774 | 55.0387004433237 | 84.3273253350471 | 57.5328549274309 | Myom3 |
| ENSMUSG00000037143 | -0.196233824946812 | 0.726396037113286 | 0.952038256048006 | 9.93865088225949 | 17.9171460361152 | 11.3574586637814 | 10.3171936368398 | 12.0445401540101 | 12.2308223207386 | 28.1091084450157 | 4.34210225867403 | Cfap61 |
| ENSMUSG00000037145 | -0.440797045475548 | 0.170535274079378 | 0.799531027639052 | 74.0881247586617 | 122.162359337149 | 106.949402417274 | 72.2203554578784 | 78.836990098975 | 201.808568292187 | 128.052605138405 | 102.03940307884 | 2210407C18I |
| ENSMUSG00000037148 | 0.185409858225523 | 0.0835456705754534 | 0.742148948523533 | 1636.26334070654 | 1594.62599721426 | 1690.36843112613 | 1620.7373276781 | 1362.12799559896 | 1256.10545233985 | 1556.41174538142 | 1576.18311989867 | Arhgap10 |
| ENSMUSG00000037149 | 0.318942336076958 | 0.0102498272640457 | 0.635862253526407 | 3193.92098807157 | 3140.38705069365 | 3798.12346814622 | 3632.59008686186 | 2730.82574037283 | 2364.21795459877 | 3061.8106643256 | 2875.55722080688 | Ddx1 |
| ENSMUSG00000037151 | 0.0619719569789998 | 0.697077112313394 | 0.944933386973863 | 883.636414804526 | 1104.34772840783 | 916.168332211697 | 866.644265494541 | 770.850569856644 | 785.218792991418 | 965.079389945539 | 1090.95319249185 | Lrrc20 |
| ENSMUSG00000037152 | 0.0286006868544705 | 0.878223358348111 | 0.977934951033036 | 6463.73712833495 | 4877.53580046791 | 4956.58425185192 | 6164.99216135889 | 5795.61373047048 | 7086.53845263594 | 4481.84117984417 | 4659.07572355723 | Ndufc1 |
| ENSMUSG00000037166 | -0.741909683991857 | 0.0540575147905574 | 0.703396849069134 | 84.0267756409212 | 70.0397526866323 | 44.483379766477 | 78.7858423176855 | 146.724398239759 | 185.908499275227 | 59.3414511616998 | 72.73021283279 | Ppp1r14a |
| ENSMUSG00000037169 | -1.1084531932296 | 0.00866789792161137 | 0.635862253526407 | 26.2018977805023 | 13.8450673915436 | 32.1794662140472 | 42.2067012416173 | 89.7865720571659 | 69.71568722821 | 29.1501865355718 | 57.5328549274309 | Mycn |
| ENSMUSG00000037172 | 0.0860769235617648 | 0.687224733110296 | 0.944452327903597 | 96.6759676728878 | 109.13170767452 | 89.9132144216024 | 84.4134024832345 | 76.6470737073368 | 72.1618516923577 | 117.641824232843 | 91.1841474321546 | E330009J07F |
| ENSMUSG00000037174 | 0.0291240180302913 | 0.876578829235092 | 0.977469019294424 | 930.619128066116 | 1091.3170767452 | 989.045358637627 | 765.34818251466 | 932.904382837871 | 667.802898712327 | 1095.21415126506 | 1004.11114731837 | Elf2 |
| ENSMUSG00000037185 | 0.19756267367964 | 0.358052823010591 | 0.865834559821921 | 78.6056933415069 | 109.946123403434 | 123.039135524298 | 93.7926694258161 | 88.6916138613469 | 72.1618516923577 | 106.189965236726 | 85.7565196088121 | Krt80 |
| ENSMUSG00000037190 | -0.00337177284264078 | 0.988585213750106 | 0.998437672973644 | 100.290022539164 | 74.9262470601182 | 63.4124775394459 | 69.4065753751039 | 93.0714466446232 | 81.9465095489486 | 58.3003730711437 | 75.9867895267955 | Cyb561d2 |
| ENSMUSG00000037196 | 0.12088478391002 | 0.766929699145277 | 0.958655025595455 | 23.4913566307952 | 30.9477976987445 | 24.6078271048596 | 25.3240207449704 | 20.8042057205628 | 45.2540425867328 | 14.5750932677859 | 16.2828834700276 | Pacrg |
| ENSMUSG00000037197 | 0.208632878004851 | 0.0163960405629127 | 0.641523953292557 | 1076.9883501503 | 1009.87550385377 | 1072.33338883869 | 1135.82922674663 | 936.189257425328 | 983.358114587383 | 890.121767425497 | 907.499372062872 | Rbm17 |
| ENSMUSG00000037204 | -0.113979209659468 | 0.604990492623301 | 0.926448151609826 | 624.32797814921 | 465.845796938996 | 575.444572298256 | 637.79015209555 | 728.147200219699 | 807.234273168747 | 521.580123368624 | 436.38127699674 | Atg101 |
| ENSMUSG00000037206 | -1.26195775960992 | 0.0135875504062078 | 0.635862253526407 | 282.799793286111 | 201.975100770753 | 195.916161950228 | 276.688374806158 | 718.292576457328 | 1188.83592957579 | 183.22974393788 | 206.249857287016 | Islr |
| ENSMUSG00000037210 | 0.00974517433606957 | 0.953952767009469 | 0.992371505557225 | 338.817643713392 | 469.917875583568 | 409.814966784777 | 306.702029022419 | 418.274030802895 | 341.239942748607 | 371.664878328541 | 384.276049892652 | Fam193a |
| ENSMUSG00000037211 | -0.374971294582648 | 0.309475895149264 | 0.851785577030345 | 270.150601254144 | 294.004078138073 | 251.757000380487 | 236.357526953057 | 481.781606160403 | 533.263853184203 | 181.147587756768 | 169.341988088287 | Spry1 |
| ENSMUSG00000037214 | -0.19193231888077 | 0.19380240911897 | 0.809162446871198 | 224.974915425692 | 209.304842330982 | 184.558703286447 | 232.605820176024 | 235.416012101106 | 273.970419984545 | 209.256696201783 | 255.098507697099 | Thap1 |
| ENSMUSG00000037216 | -0.0093287105544083 | 0.955659842472818 | 0.992371505557225 | 153.597331816738 | 147.409246933493 | 134.396594188079 | 177.268145214792 | 183.952976897608 | 134.539045528125 | 150.95632313064 | 146.545951230249 | Lipt1 |
| ENSMUSG00000037221 | -0.343612904222684 | 0.00865345780414651 | 0.635862253526407 | 195.158962778914 | 216.634583891211 | 189.290977729689 | 218.536919762152 | 267.16979977986 | 286.201242305283 | 234.242570375131 | 254.012982132431 | Mospd3 |
| ENSMUSG00000037224 | -0.173635973311145 | 0.509049371379494 | 0.902394714365738 | 32.5264937964856 | 33.3910448854875 | 40.6975602118832 | 44.0825546301336 | 30.6588294829347 | 48.9232892829544 | 47.8895921655823 | 42.3354970220718 | Zfyve28 |
| ENSMUSG00000037225 | -0.0459848067785646 | 0.857366537261366 | 0.974029039597056 | 299.966553900923 | 330.652785939217 | 429.690519446395 | 270.12288794635 | 282.499214521327 | 228.716377397812 | 474.731609293598 | 386.447101021989 | Fgf2 |
| ENSMUSG00000037234 | 0.178756534272423 | 0.205375342733912 | 0.819495406657294 | 740.881247586617 | 758.22104361924 | 783.664647800914 | 738.148308381173 | 717.197618261509 | 510.025290774799 | 700.64555494428 | 739.242909539254 | Hook3 |
| ENSMUSG00000037235 | -0.160127797267914 | 0.435987824289226 | 0.888687115594624 | 2326.54782016529 | 2341.44522062869 | 2272.43818764492 | 1764.2401118996 | 2893.97451154987 | 3044.25167563184 | 1801.06509666212 | 1988.68283447271 | Mxd4 |
| ENSMUSG00000037236 | 0.256367203829383 | 0.0943746188722305 | 0.757085157577544 | 1952.49314150571 | 2150.87194006274 | 2391.69150361463 | 2246.3344327483 | 1701.56503630288 | 1442.01395161508 | 2083.19725920283 | 2089.63671198688 | Matr3 |
| ENSMUSG00000037242 | -0.0296792764170802 | 0.848358624705255 | 0.972640876584422 | 8119.87777080601 | 10418.820420001 | 10069.3335603308 | 8004.26640879915 | 8808.93868536464 | 7735.99511786716 | 10764.7474563505 | 10062.8219844771 | Clic4 |
| ENSMUSG00000037243 | 0.0894569703414983 | 0.768015344004701 | 0.95879680569409 | 25.2983840639333 | 27.6901347830872 | 26.5007368821565 | 30.9515809105193 | 21.8991639163819 | 20.7923979452556 | 24.9858741733473 | 35.8223436340608 | Zfp692 |
| ENSMUSG00000037251 | -0.184822245800477 | 0.269428897607033 | 0.843793744782845 | 175.281661014395 | 161.254314325037 | 178.879973954556 | 123.806323642077 | 170.813478547779 | 167.562265794119 | 196.76375911511 | 191.052499381657 | Pomk |
| ENSMUSG00000037253 | 0.0430576234008412 | 0.756364767288968 | 0.956566265296272 | 265.633032671299 | 268.757190541728 | 330.312756138308 | 276.688374806158 | 259.505092409126 | 255.624186503437 | 314.405583347953 | 276.809018990469 | Mex3c |
| ENSMUSG00000037254 | -0.483281111823969 | 0.187006933553217 | 0.807540361382744 | 86.7373167906283 | 180.800291818981 | 67.1982970940397 | 112.551203310979 | 100.736154015357 | 123.53130543946 | 155.120635492864 | 246.414303179751 | Itih2 |
| ENSMUSG00000037257 | 0.202120770043679 | 0.179025912358093 | 0.801755154770274 | 524.037955610046 | 496.79359463774 | 497.835271429083 | 455.832373409466 | 352.576539053749 | 385.270903103266 | 464.320828388037 | 513.453592088204 | Aagab |
| ENSMUSG00000037259 | -1.45790682709811 | 0.00475839594142045 | 0.635862253526407 | 12.6491920319666 | 1.62883145782866 | 11.3574586637814 | 5.62756016554897 | 20.8042057205628 | 19.5693157131818 | 21.8626399016789 | 22.7960368580387 | Dzank1 |
| ENSMUSG00000037260 | -0.0569841602656973 | 0.522431680361739 | 0.90648515894888 | 481.572810931301 | 509.82424630037 | 487.42426765395 | 498.039074651084 | 522.295059405709 | 516.140701935169 | 494.512093014165 | 524.308847734889 | Hgsnat |
| ENSMUSG00000037262 | -0.157987563800523 | 0.366581938300658 | 0.869802378373101 | 173.474633581257 | 171.841718800923 | 134.396594188079 | 194.150825711439 | 165.338687568684 | 225.04713070159 | 176.983275394543 | 185.624871558315 | Kin |
| ENSMUSG00000037266 | -0.582027369839602 | 0.0462687498951448 | 0.69805115405055 | 1602.83333319349 | 867.35275129376 | 936.043884873314 | 762.534402431885 | 2150.49789658871 | 1242.65154778704 | 1764.62736349265 | 1082.2689879745 | Rsrp1 |
| ENSMUSG00000037270 | 0.423974681241916 | 0.131920445738906 | 0.77303252008843 | 714.679349806115 | 1052.22512175731 | 1083.69084750247 | 766.286109208918 | 582.51776017576 | 381.601656407044 | 953.627530949421 | 777.236304302651 | 4932438A13F |
| ENSMUSG00000037275 | 0.249689805226287 | 0.119317179223992 | 0.769868040577702 | 326.168451681425 | 402.321370083678 | 424.958245003153 | 367.667264149199 | 360.241246424483 | 258.070350967584 | 358.130863151311 | 301.776106977845 | Gemin5 |
| ENSMUSG00000037278 | 0.0222867924786525 | 0.917313338925776 | 0.986713918026868 | 93.0619128066116 | 76.5550785179469 | 99.3777633080869 | 90.0409626487835 | 104.021028602814 | 103.961989726278 | 64.5468416144805 | 81.4144173501381 | Tmem97 |

| | | | | | | | | | | | | |
|---|---|---|---|---|---|---|---|---|---|---|---|---|
| ENSMUSG00000037286 | -0.0151168574759665 | 0.903013384109697 | 0.985027469232262 | 394.835494140673 | 509.009830571455 | 487.42426765395 | 419.253232333398 | 459.882442244021 | 439.086521314516 | 468.485140750262 | 462.433890548784 | Stag1 |
| ENSMUSG00000037287 | 0.00808625576919883 | 0.950002977037981 | 0.991548537174153 | 753.530439618584 | 950.423155643021 | 842.344850897118 | 811.30659053331 | 705.153078107498 | 883.065371557327 | 872.423439886043 | 879.275707381491 | Tbcel |
| ENSMUSG00000037295 | -0.287404082034755 | 0.0189474956829636 | 0.642273034804458 | 303.580608767199 | 327.39512302356 | 363.438677241004 | 311.39166249371 | 433.603445544362 | 368.147751854232 | 418.513392403567 | 372.335268681298 | Ldlrap1 |
| ENSMUSG00000037296 | 0.101535537111836 | 0.504209043400925 | 0.900927059989498 | 208.711668527449 | 168.584055885266 | 177.933519065908 | 229.79204009325 | 186.142893289246 | 173.677676954488 | 186.352978209548 | 184.539345993646 | Lsm1 |
| ENSMUSG00000037300 | 0.247319583768605 | 0.120421288137579 | 0.769868040577702 | 393.028466707535 | 403.135785812593 | 388.992959234512 | 402.370551836751 | 298.923587458613 | 265.408844360028 | 412.26692386023 | 359.308961905276 | Ttc13 |
| ENSMUSG00000037306 | -0.147774335164367 | 0.147571687346242 | 0.780607990324772 | 704.740698923855 | 770.437279552955 | 650.214508501483 | 715.638067718977 | 741.286698569529 | 791.334204151787 | 766.233474649317 | 849.966517135441 | Man1c1 |
| ENSMUSG00000037313 | -0.418802548499936 | 0.165104038900246 | 0.795216960550936 | 37.0440623793308 | 32.5766291565731 | 37.8581955459379 | 25.3240207449704 | 59.1277425742312 | 41.5847958905112 | 28.1091084450157 | 48.8486504100828 | Tacc3 |
| ENSMUSG00000037315 | -0.240454297594703 | 0.449449913757549 | 0.89108031640712 | 39.754603529038 | 23.6180561385155 | 27.447191770805 | 25.3240207449704 | 41.6084114411257 | 42.8078781225851 | 27.0680303544596 | 26.0526135520442 | Jade3 |
| ENSMUSG00000037316 | -0.154113083847926 | 0.35827788583403 | 0.866227995515174 | 90.3513716569045 | 87.9568987227475 | 81.3951204237664 | 103.171936368398 | 99.6411958195378 | 94.1773318696872 | 102.025652874501 | 107.467030902182 | Bag4 |
| ENSMUSG00000037321 | -0.455772564450184 | 0.279597670876192 | 0.848522228930484 | 138.237598635064 | 101.801966114291 | 139.128868631322 | 145.378637610015 | 286.879047304603 | 269.078091056249 | 78.0808567917102 | 85.7565196088121 | Tap1 |
| ENSMUSG00000037325 | 0.0592825229566726 | 0.795224820779861 | 0.962032824235414 | 70.4740698923855 | 88.7713144516618 | 82.3415753124149 | 75.0341355406529 | 75.5521155115177 | 52.592535979176 | 98.902418602833 | 75.9867895267955 | Bbs7 |
| ENSMUSG00000037326 | 0.0800227663029988 | 0.619274044867457 | 0.929568738467614 | 108.421645988285 | 127.86326943955 | 107.895857305923 | 107.861569839689 | 110.590777777729 | 106.408154190426 | 114.518589961175 | 96.6117752554972 | Capn15 |
| ENSMUSG00000037331 | 0.223024314218858 | 0.103172057787895 | 0.759800953500741 | 3788.43301357401 | 3936.88563357186 | 3479.16817067169 | 3835.18225282162 | 3298.01408580712 | 2515.88015137593 | 3456.37926064637 | 3613.71460478146 | Larp1 |
| ENSMUSG00000037336 | 0.488296238239767 | 0.381860022692933 | 0.875126835008975 | 11.7456783153976 | 16.2883145782866 | 39.7511053232348 | 12.1930470253561 | 13.1394983498292 | 20.7923979452556 | 17.6983275394543 | 5.42762782334254 | Mfsd2b |
| ENSMUSG00000037337 | 0.42683666090759 | 0.409649704864075 | 0.882192126163777 | 12.6491920319666 | 14.6594831204579 | 17.036187995672 | 37.5170677703264 | 17.5193311331055 | 23.2385624094033 | 6.24646854333682 | 14.1118323406906 | Map4k1 |
| ENSMUSG00000037339 | 0.142021775832088 | 0.243814880758073 | 0.832048466795696 | 778.828823682517 | 742.747144769868 | 770.414279359836 | 876.961459131381 | 760.995946094272 | 796.226533080083 | 695.440164491499 | 619.835097425718 | Fam53a |
| ENSMUSG00000037341 | -0.164185592554344 | 0.711547073762998 | 0.950020479830613 | 12.6491920319666 | 10.5874044758863 | 23.6613722162112 | 17.8206071909051 | 19.7092475247437 | 9.78465785659088 | 27.0680303544596 | 15.1973579053591 | Slc9a7 |
| ENSMUSG00000037343 | -0.0728456520888997 | 0.747375447301219 | 0.956119556124081 | 290.931416735233 | 344.497853330761 | 299.079744812909 | 214.785212985119 | 257.315176017488 | 220.154801773295 | 373.747034509653 | 357.137910775939 | Taf2 |
| ENSMUSG00000037344 | -0.560794177294805 | 0.153061529644292 | 0.783412422888014 | 17.1667606148119 | 18.7315617650296 | 15.1432782183751 | 29.075727522003 | 28.4689130912965 | 47.7002070508805 | 19.7804837205666 | 22.7960368580387 | Slc12a9 |
| ENSMUSG00000037347 | -0.7451273393845 | 0.0809984100544386 | 0.739855948845396 | 51.5002818444356 | 39.9063707168021 | 26.5007368821565 | 56.2756016554897 | 109.49581958191 | 105.185071958352 | 34.3555769883525 | 43.4210225867403 | Chst7 |
| ENSMUSG00000037348 | -0.200235093728657 | 0.277984626128824 | 0.848075053184273 | 474.344701198749 | 398.249291439107 | 281.097101928589 | 367.667264149199 | 394.184950494875 | 458.655837027697 | 398.732908683 | 497.170708618176 | Paqr7 |
| ENSMUSG00000037349 | -0.232494990116539 | 0.243001137546031 | 0.832048466795696 | 309.905204783182 | 209.304842330982 | 236.613722162112 | 256.053987532478 | 354.766455445387 | 338.793778284459 | 267.557069272927 | 227.960368580387 | Nudt22 |
| ENSMUSG00000037351 | 0.111201890730091 | 0.420209333423397 | 0.882568531013176 | 2758.42737668529 | 2848.01180401341 | 2592.3399400081 | 2678.71863880131 | 2123.12394169323 | 2208.88651112539 | 2807.7876102299 | 2929.8334990403 | Actr1b |
| ENSMUSG00000037353 | -0.0687235113710731 | 0.676855570256046 | 0.94409858171788 | 107.518132271716 | 123.791190794978 | 134.396594188079 | 117.24083678227 | 140.154649064844 | 132.092881063977 | 128.052605138405 | 106.381505337514 | Letmd1 |
| ENSMUSG00000037355 | 0.193132445442629 | 0.344285077587143 | 0.861266047293794 | 77.7021796249379 | 87.1424829938332 | 110.735221971868 | 93.7926694258161 | 75.5521155115177 | 80.7234273168747 | 97.8613405122768 | 68.388110574116 | Uvssa |
| ENSMUSG00000037358 | 0.217930578579977 | 0.420654736787459 | 0.882568531013176 | 68.6670424592474 | 83.8848200781758 | 118.306861081056 | 94.7305961200743 | 87.5966556655277 | 59.9310293716191 | 106.189965236726 | 59.7039060567679 | 4930578C19l |
| ENSMUSG00000037361 | 0.0431907647473601 | 0.717449825955538 | 0.951530419413962 | 401.160090156656 | 334.724864583789 | 347.34894413398 | 395.805064976944 | 342.721915291377 | 376.709327478749 | 341.473613702413 | 374.506319810635 | Sf3b6 |
| ENSMUSG00000037362 | -1.30614888345522 | 0.0248955975694444 | 0.655303666217042 | 93.9654265231807 | 70.0397526866323 | 30.2865564367503 | 83.4754757889764 | 215.706764576362 | 359.586176229715 | 65.5879197050366 | 46.6775992807458 | Nov |
| ENSMUSG00000037363 | 0.0956136571692692 | 0.603728066465816 | 0.926448151609826 | 151.7903043836 | 101.801966114291 | 116.413951303759 | 114.427056699496 | 101.831112211176 | 116.192812047017 | 112.436433780063 | 122.664388807541 | Letm2 |
| ENSMUSG00000037364 | 0.0581194409598427 | 0.528268709773997 | 0.908232038364594 | 973.987786461431 | 937.392503980392 | 1042.04683240194 | 1100.18801236482 | 975.607752474815 | 955.227223249684 | 1005.68143547723 | 955.262496908287 | Srrt |
| ENSMUSG00000037366 | -0.137292234735803 | 0.520814111379773 | 0.905962179970576 | 227.685456575399 | 237.809392842984 | 201.594891282119 | 155.695831246855 | 185.047935093427 | 183.462334811079 | 254.023054095697 | 282.236646813812 | Pafah2 |
| ENSMUSG00000037369 | -0.0206308868660847 | 0.937326948974899 | 0.98894578968943 | 137.334084918495 | 182.42912327681 | 221.470443943737 | 158.509611329629 | 129.205067106653 | 129.646716599829 | 238.406882737355 | 211.677485110359 | Kdm6a |
| ENSMUSG00000037370 | -0.179994381160939 | 0.374668251535607 | 0.873700364743909 | 75.8951521917998 | 61.895595397489 | 87.0738497556571 | 78.7858423176855 | 73.3621991198795 | 85.6157562451702 | 81.2040910633787 | 103.124928643508 | Enpp1 |
| ENSMUSG00000037373 | 0.0452445938543981 | 0.550118105480798 | 0.914699131434573 | 4476.91046559962 | 4386.44311593257 | 4417.1049653223 | 4505.79983921621 | 4269.24200549866 | 3970.12492531175 | 4461.01961803305 | 4534.24028362036 | Ctbp1 |
| ENSMUSG00000037375 | -0.0960177802606642 | 0.684964799451756 | 0.944452327903597 | 99.3865088225949 | 69.2253369577179 | 67.1982970940397 | 64.7169419038131 | 73.3621991198795 | 64.8233582999146 | 102.025652874501 | 80.3288917854696 | Hhat |
| ENSMUSG00000037376 | 0.141744140071088 | 0.329758367016988 | 0.855977678491063 | 211.422209677157 | 234.551729927327 | 201.594891282119 | 216.661066373635 | 185.047935093427 | 166.339183562045 | 223.831789469569 | 207.335382851685 | Trmt6 |
| ENSMUSG00000037379 | -0.612928655502187 | 0.310656404723762 | 0.851785577030345 | 7.22810973255236 | 8.95857301805762 | 10.4110037751329 | 31.8895076047775 | 30.6588294829347 | 35.4693847301419 | 12.4929370866736 | 10.8552556466851 | Spon2 |
| ENSMUSG00000037395 | 0.0937688105728891 | 0.560461685281509 | 0.916762264997884 | 425.55496050402 | 408.022280186079 | 534.747012086372 | 468.025420434822 | 409.514365236342 | 344.909189444828 | 471.60837502193 | 492.828606359502 | Rcor3 |
| ENSMUSG00000037400 | 0.15412357154308 | 0.54887270864413 | 0.914699131434573 | 568.310127721929 | 747.633639143354 | 696.590798045257 | 523.363095396054 | 445.647985698372 | 363.255422925936 | 723.549272936515 | 745.756062927265 | Atp11b |
| ENSMUSG00000037405 | -0.882718919567088 | 0.0298756346185447 | 0.665956123973759 | 162.632468982428 | 121.347943608235 | 159.004421292939 | 149.130344387048 | 387.61520131996 | 447.648096939033 | 133.257995591185 | 123.74991437221 | Icam1 |
| ENSMUSG00000037406 | -1.28506177033546 | 0.0695668039122604 | 0.724893403106749 | 20.780815481088 | 7.32974156022896 | 5.67872933189068 | 22.5102406621959 | 27.3739548954774 | 85.6157562451702 | 10.4107809055614 | 14.1118323406906 | Htra4 |
| ENSMUSG00000037408 | -0.0672456989066155 | 0.719021632739374 | 0.951674101666722 | 126.491920319666 | 140.893921102179 | 126.824955078892 | 112.551203310979 | 117.160526952643 | 122.308223207386 | 171.777884941763 | 119.407812113536 | Cnnm4 |
| ENSMUSG00000037410 | -0.406469974713982 | 0.100811846424348 | 0.759800953500741 | 135.527057485357 | 201.160685041839 | 182.66579350915 | 150.068271081306 | 257.315176017488 | 298.432064626022 | 159.284947855089 | 173.684090346961 | Tbc1d2b |
| ENSMUSG00000037411 | 0.621331393186273 | 0.00140234377972495 | 0.628340487109018 | 2057.30073262772 | 2933.52545554941 | 3513.24054666303 | 3282.74342990356 | 2088.08527942702 | 1541.08361241306 | 2126.92253900619 | 1905.09736599323 | Serpine1 |
| ENSMUSG00000037415 | 0.198061083726282 | 0.1822421249804 | 0.804505478598291 | 498.739571546113 | 606.739718041175 | 642.642869392295 | 630.286738541484 | 476.306815181307 | 447.648096939033 | 540.319528998635 | 607.894316214364 | Ranbp10 |
| ENSMUSG00000037416 | 0.064853973316892 | 0.798761514076062 | 0.963240258978423 | 635.170142748039 | 762.293122263812 | 773.253644025781 | 543.997482669733 | 493.826146314413 | 428.078781225851 | 868.259127523818 | 804.374443419364 | Dmxl1 |
| ENSMUSG00000037419 | -0.433215450280621 | 0.152543079092069 | 0.783412422888014 | 129.202461469373 | 136.007426728693 | 122.09268063565 | 169.764731660727 | 241.98576127602 | 269.078091056249 | 131.175839410073 | 110.723607596188 | Endod1 |
| ENSMUSG00000037426 | 0.0846386773030324 | 0.681885822917166 | 0.944452327903597 | 427.361987937158 | 412.09435883065 | 405.082692341535 | 365.791410760683 | 327.39250054991 | 260.516515431732 | 477.854843565267 | 451.578634902099 | Depdc5 |
| ENSMUSG00000037432 | 0.29531438018933 | 0.207297913583942 | 0.820099122666178 | 157.211386683014 | 131.120932355207 | 166.576060402127 | 113.489130005238 | 91.9764884488041 | 89.2850029413918 | 128.052605138405 | 153.05910461826 | Fer1l5 |
| ENSMUSG00000037434 | -0.405986279669206 | 0.0141496196396149 | 0.635862253526407 | 414.712795905192 | 390.919549878878 | 394.671688566402 | 397.68091836546 | 697.488370736765 | 500.240632918209 | 453.910047482476 | 465.69046724279 | Slc30a1 |
| ENSMUSG00000037440 | -0.302429063170633 | 0.452257337526743 | 0.891110666112841 | 22.5878429142261 | 35.0198763433161 | 22.7149173275627 | 31.8895076047775 | 25.1840385038392 | 42.8078781225851 | 16.6572494488982 | 54.2762782334254 | Vnn1 |
| ENSMUSG00000037443 | 0.21918359279818 | 0.282042209725603 | 0.850308824809137 | 1011.03184884076 | 1374.73375040739 | 1201.05125369488 | 1228.68396947819 | 839.832936193247 | 780.326464063122 | 1179.5414766001 | 1336.28197010693 | Cep85 |
| ENSMUSG00000037447 | 0.0884855185252139 | 0.683750345046092 | 0.944452327903597 | 165.343010132135 | 204.418347957496 | 262.16800415562 | 321.708856130549 | 216.801722772181 | 233.608706326107 | 233.201492284575 | 212.763010675027 | Arid5a |
| ENSMUSG00000037455 | -0.305887545955345 | 0.142713158442606 | 0.780607990324772 | 60.535419010126 | 51.3081909216027 | 76.6628459805242 | 58.151455044006 | 66.7924499449649 | 74.6080161565054 | 80.1630129728225 | 82.4999429148066 | Slc18b1 |
| ENSMUSG00000037458 | 0.12399444897459 | 0.425142424168405 | 0.885066977635295 | 1296.54218327658 | 1395.90855935916 | 1663.86769424397 | 1551.330752303 | 1246.06242684213 | 1111.78174895514 | 1439.81099923914 | 1621.77519361475 | Azin1 |
| ENSMUSG00000037461 | -0.0545059386618656 | 0.823131944331457 | 0.966637987288264 | 148.176249517323 | 122.162359337149 | 161.843785958884 | 130.371810501884 | 158.768938393769 | 88.0619207093179 | 191.558368662329 | 144.374900100912 | Ints7 |
| ENSMUSG00000037465 | -0.300928734760369 | 0.317654479895885 | 0.854351922245337 | 471.634160049041 | 280.973426475443 | 629.392500951217 | 653.734905897938 | 845.307727172343 | 692.264543353805 | 550.730309904196 | 419.012867962044 | Klf10 |
| ENSMUSG00000037470 | 0.144116974328349 | 0.3362513909857 | 0.858570960161243 | 1011.03184884076 | 1184.97488557035 | 1299.48256211432 | 885.402799379704 | 959.183379537529 | 900.188522806361 | 1048.36563719003 | 1056.21637442246 | Uggt1 |
| ENSMUSG00000037475 | -0.190050821865104 | 0.156319078150055 | 0.787275540343856 | 577.34526488762 | 510.638662029284 | 611.409858066897 | 600.273084325223 | 670.114415841287 | 770.541806206532 | 576.7572621681 | 606.808790649696 | Thoc2 |
| ENSMUSG00000037486 | -0.0721324935032876 | 0.727180467147284 | 0.952237115597464 | 166.246523848704 | 248.39679731887 | 250.810545491838 | 236.357526953057 | 217.896680968 | 179.793088114857 | 275.885693997376 | 273.552442296464 | Asxl2 |
| ENSMUSG00000037487 | 0.347173003049238 | 0.244234522668613 | 0.832048466795696 | 4710.01700447443 | 6110.56121404421 | 5106.12412425837 | 4220.67012416173 | 3083.40227942658 | 2057.22431434823 | 5692.61499916096 | 5004.27285312182 | Ubr5 |
| ENSMUSG00000037490 | 0.12271283488182 | 0.724631535683851 | 0.951674101666722 | 104.807591122009 | 188.944449108124 | 134.396594188079 | 89.1030359545253 | 130.300025302473 | 59.9310293716191 | 175.942197303987 | 108.552556466851 | Slc2a12 |
| ENSMUSG00000037493 | -0.853633424675538 | 0.0392975146285059 | 0.679566075569698 | 22.5878429142261 | 8.95857301805762 | 15.1432782183751 | 18.7585338851632 | 28.4689130912965 | 42.8078781225851 | 16.6572494488982 | 30.3947158107182 | Cib2 |
| ENSMUSG00000037499 | -0.120600832901412 | 0.584441537195882 | 0.922450244597442 | 1617.28955265859 | 1170.31540244989 | 1235.12362968622 | 1653.56476197714 | 1508.85239383872 | 2162.40938630658 | 1197.23980413956 | 1303.71620316688 | Nenf |
| ENSMUSG00000037503 | 0.0893201027998198 | 0.532556527181448 | 0.909082872831377 | 883.636414804526 | 1075.84317789583 | 1117.76322349382 | 876.961459131381 | 958.08842134171 | 774.211052902753 | 984.859873666105 | 998.683519495027 | Fam168b |
| ENSMUSG00000037514 | 0.0256790755718401 | 0.832954416417446 | 0.967854847250989 | 370.440623793308 | 447.114235173966 | 403.189782564238 | 450.204813243917 | 398.564783278151 | 431.748027922072 | 374.788112600209 | 437.466802561409 | Pank2 |
| ENSMUSG00000037519 | 0.35194323578256 | 0.15543284029653 | 0.787275540343856 | 641.494738764022 | 916.21769502862 | 986.205993971682 | 813.182443921826 | 481.781606160403 | 472.10974158051 | 823.492769629904 | 852.137568264778 | Ppfia1 |
| ENSMUSG00000037523 | -0.00084549379125879 | 0.996885264307002 | 0.999262355603273 | 1130.29565942788 | 1307.1372449075 | 1451.86179918672 | 1025.15387682417 | 1097.14811221074 | 842.703657898889 | 1581.39761955477 | 1394.90035059903 | Mavs |
| ENSMUSG00000037525 | -0.154981065879238 | 0.437029177572371 | 0.888701139843674 | 73.1846110420926 | 64.338842584232 | 52.0550188756646 | 61.9031618210386 | 74.4571573156986 | 72.1618516923577 | 69.7522320672611 | 64.046008315442 | Bcdin3d |
| ENSMUSG00000037526 | 0.0757161509277964 | 0.617141141006262 | 0.929187015773942 | 236.72059374109 | 300.519403969387 | 265.007368821565 | 264.495327780801 | 240.890803080201 | 215.262472844999 | 258.187366457922 | 297.434004719171 | Atg14 |
| ENSMUSG00000037531 | 0.0361840216629082 | 0.737489539184656 | 0.954383574136061 | 581.862833470465 | 514.710740673856 | 477.959718767466 | 506.480414899407 | 506.965644664242 | 484.340563901248 | 535.114138545854 | 502.598336441519 | Mrpl47 |
| ENSMUSG00000037533 | -0.0451122548876859 | 0.801567305620502 | 0.963461896828986 | 221.360860559416 | 283.416673662186 | 301.919109478855 | 235.419600258798 | 261.695008800764 | 209.14706168463 | 310.241270985729 | 293.091902460497 | Rapgef6 |
| ENSMUSG00000037536 | -0.148128359819714 | 0.246129944545371 | 0.832048466795696 | 215.939778260002 | 199.531853584011 | 194.023252172932 | 229.79204009325 | 247.460552255116 | 214.039390612925 | 236.324726556243 | 231.216945274392 | Fbxo34 |
| ENSMUSG00000037542 | -0.473169319737443 | 0.223100136365184 | 0.832048466795696 | 33.4300075130547 | 74.1118313312039 | 34.0723759913441 | 61.9031618210386 | 55.8428679867739 | 58.7079471395453 | 48.9306702561384 | 119.407812113536 | Aldh8a1 |
| ENSMUSG00000037552 | -0.483307269349213 | 0.128134748940584 | 0.77303252008843 | 162.632468982428 | 201.975100770753 | 187.398067952392 | 137.87522405595 | 363.52612101194 | 303.324393554317 | 155.120635492864 | 143.289374536243 | Plekhg2 |
| ENSMUSG00000037553 | -0.109410935687455 | 0.450722066436747 | 0.89108031640712 | 645.108793630298 | 612.440628143575 | 650.214508501483 | 764.410255820401 | 840.927894389066 | 759.534066117867 | 681.906149314269 | 600.295637261685 | Zdhhc18 |
| ENSMUSG00000037563 | 0.0545485508468838 | 0.802047294223093 | 0.963461896828986 | 5477.10014984155 | 3576.91388139173 | 4254.31472447477 | 5290.84448231029 | 4527.65213971197 | 6101.95725581649 | 3592.76049050923 | 3687.53034317892 | Rps16 |
| ENSMUSG00000037570 | -0.0747013647696075 | 0.534757503301055 | 0.910706162399539 | 901.706689135907 | 771.251695281869 | 892.506959995485 | 915.416453595965 | 938.379173816966 | 1042.06606172693 | 860.971580889925 | 824.999429148066 | Mcrs1 |
| ENSMUSG00000037572 | -0.212541028734838 | 0.716616541868505 | 0.951348124093194 | 10.8421645988285 | 5.7009101024003 | 19.8755526616174 | 4.68963347129081 | 9.85462376237187 | 8.56157562451702 | 9.36970281500523 | 19.5394601640331 | Wdhd1 |
| ENSMUSG00000037573 | -0.0358086052489359 | 0.795920576222476 | 0.962200030474509 | 1197.15567445398 | 1213.47943608235 | 1346.80530654674 | 1031.71936368398 | 1219.78343014247 | 1050.62763735145 | 1427.31806215246 | 1210.36100460539 | Tob1 |
| ENSMUSG00000037577 | -0.281869944156217 | 0.582000548629688 | 0.922450244597442 | 11.7456783153976 | 8.95857301805762 | 9.46454888648447 | 17.8206071909051 | 20.8042057205628 | 20.7923979452556 | 11.4518589961175 | 5.42762782334254 | Ephx3 |
| ENSMUSG00000037580 | -0.272137407984637 | 0.168858278863771 | 0.79907358295209 | 150.886790667031 | 171.841718800923 | 124.932045301595 | 162.261318106662 | 210.231973597267 | 220.154801773295 | 140.545542225078 | 167.17093695895 | Gch1 |
| ENSMUSG00000037583 | -0.371703587640464 | 0.179367336248815 | 0.801755154770274 | 177.088688447533 | 119.719112150406 | 84.2344850897118 | 100.358156285623 | 164.243729372865 | 205.477814988408 | 122.847214685624 | 131.348593324889 | Nr0b2 |
| ENSMUSG00000037594 | -0.0586864840146024 | 0.86098548193723 | 0.974868041734905 | 36.1405486627618 | 26.8757190541728 | 29.3401015481019 | 60.9652351267805 | 41.6084114411257 | 50.1463715150282 | 36.4377331694648 | 31.4802413753867 | BC022687 |
| ENSMUSG00000037601 | -0.0568062204942802 | 0.755832581058482 | 0.956566265296272 | 276.475197270128 | 230.479651282755 | 179.826428843205 | 261.681547698027 | 278.119381738051 | 264.185762127954 | 206.133461930115 | 238.815624227072 | Nme1 |
| ENSMUSG00000037605 | -0.611535715717805 | 0.185784314729828 | 0.805963874499992 | 6.32459601598331 | 9.77298874697194 | 9.46454888648447 | 9.37926694258161 | 16.4243729372865 | 17.123151249034 | 12.4929370866736 | 7.59867895267955 | Adgrl3 |
| ENSMUSG00000037606 | -0.0928681938876514 | 0.628335455919279 | 0.931825800800958 | 192.448421629207 | 228.850819824926 | 198.755526616174 | 237.295453647315 | 289.068963696242 | 253.178022039289 | 188.435134390661 | 184.539345993646 | Osbpl5 |

| | | | | | | | | | | | | |
|---|---|---|---|---|---|---|---|---|---|---|---|---|
| **ENSMUSG00000037608** | 0.353732971252977 | 0.0794061277670184 | 0.739855948845396 | 1385.08652750035 | 1779.49836767781 | 1892.90977729689 | 1364.68334014562 | 1173.79518591807 | 923.427085215764 | 1608.46564990923 | 1318.91356107224 | Bclaf1 |
| **ENSMUSG00000037613** | -0.301122516043853 | 0.121109642234072 | 0.769868040577702 | 70.4740698923855 | 66.7820897709749 | 57.7337482075553 | 55.3376749612315 | 86.5016974697087 | 73.3849339244316 | 70.7933101578173 | 78.1578406561325 | Tnfrsf23 |
| **ENSMUSG00000037621** | -0.258732958464305 | 0.411133642417158 | 0.882192126163777 | 293.64195788494 | 286.674336577844 | 277.311282373995 | 411.749818779333 | 528.864808580624 | 489.232892829544 | 317.528817619622 | 183.453820428978 | Atoh8 |
| **ENSMUSG00000037622** | 0.298105344382338 | 0.211786041234521 | 0.823370267313127 | 2603.92653115199 | 3313.04318522349 | 4053.6662880813 | 2460.18171903916 | 2277.51304730372 | 1706.19971374303 | 3053.48203960115 | 3072.03734801188 | Wdtc1 |
| **ENSMUSG00000037624** | -0.735166974123189 | 0.0460205026966855 | 0.697906909182231 | 25.2983840639333 | 35.0198763433161 | 24.6078271048596 | 41.2687745473591 | 78.836990098975 | 64.8233582999146 | 34.3555769883525 | 32.5657669400552 | Kcnk2 |
| **ENSMUSG00000037628** | 0.267167525140256 | 0.666886809360287 | 0.942024383071524 | 16.2632468982428 | 9.77298874697194 | 2.83936466594534 | 15.9447538023887 | 12.0445401540101 | 3.66924669622158 | 8.32862472444909 | 13.0263067760221 | Cdkn3 |
| **ENSMUSG00000037638** | -0.614143415063536 | 0.0011941953306471 | 0.596732488826387 | 60.535419010126 | 73.2974156022896 | 69.0912068713366 | 63.779015209555 | 112.780694169367 | 108.854318654574 | 82.2451691539348 | 105.295979772845 | Zbtb42 |
| **ENSMUSG00000037640** | 0.0522894475765721 | 0.807860965596725 | 0.964026716319739 | 184.316798180085 | 207.676010873154 | 173.201244622666 | 159.447538023887 | 171.908436743598 | 113.746647582869 | 206.133461930115 | 206.249857287016 | Zfp60 |
| **ENSMUSG00000037643** | -0.0136328768494094 | 0.94345023369789 | 0.990505437194639 | 231.299511441676 | 255.726538879099 | 253.649910157784 | 189.461192240149 | 205.85214081399 | 188.354663739374 | 244.653351280692 | 299.605055848508 | Prkci |
| **ENSMUSG00000037646** | 0.289751305813073 | 0.173219115959411 | 0.799889021902745 | 362.309000344187 | 432.454752053509 | 446.726707442067 | 314.205442576484 | 305.493336633528 | 216.485555077073 | 365.418409785204 | 384.276049892652 | Vps13b |
| **ENSMUSG00000037649** | -0.560093582919025 | 0.154094542415837 | 0.786034959293418 | 48.7897406947284 | 38.2775392589734 | 53.001473764313 | 41.2687745473591 | 95.2613630362614 | 96.6234963338349 | 27.0680303544596 | 48.8486504100828 | H2-DMa |
| **ENSMUSG00000037652** | 0.277348959903648 | 0.173808487048592 | 0.799889021902745 | 415.616309621761 | 475.618785685968 | 569.765842966365 | 398.618845059719 | 325.202584158272 | 293.539735697726 | 471.60837502193 | 442.894430384751 | Phc3 |
| **ENSMUSG00000037653** | -0.0602747507642196 | 0.901154479822975 | 0.984802943363903 | 9.03513716569045 | 15.4738988493722 | 9.46454888648447 | 14.0689004138724 | 12.0445401540101 | 4.89232892829544 | 18.7394056300105 | 14.1118323406906 | Kctd8 |
| **ENSMUSG00000037656** | 0.176935639965215 | 0.387616590669545 | 0.875951433019341 | 4957.57976281435 | 6147.20992184535 | 6378.15949460188 | 4794.68126104772 | 4726.93453135104 | 3370.81463159556 | 6206.90757589569 | 5400.48968422582 | Slc20a2 |
| **ENSMUSG00000037661** | -0.499140976191816 | 0.294409630563982 | 0.851785577030345 | 58.7283915769879 | 76.5550785179469 | 97.48485353079 | 66.5927952923295 | 145.62944004394 | 192.023910435596 | 43.7252798033577 | 42.3354970220718 | Gpr160 |
| **ENSMUSG00000037664** | -0.566488902640273 | 0.112476171505861 | 0.765080324297948 | 380.379274675568 | 274.458100644129 | 395.618143455051 | 513.983828453472 | 686.538788778574 | 951.557976553463 | 288.37863108405 | 390.789203280663 | Cdkn1c |
| **ENSMUSG00000037669** | 0.0213739782592824 | 0.895373856020254 | 0.983733637008786 | 432.783070236573 | 485.39177443294 | 445.780252553418 | 390.177504811395 | 411.70428162798 | 332.67836712409 | 495.553171104721 | 487.40097853616 | Ldah |
| **ENSMUSG00000037674** | 0.0291670557657792 | 0.905621393834987 | 0.98504921011344 | 150.886790667031 | 175.099381716581 | 141.021778408619 | 123.806323642077 | 101.831112211176 | 118.638976511164 | 205.092383839559 | 153.05910461826 | Rfx7 |
| **ENSMUSG00000037679** | -0.449210005786408 | 0.109595124784913 | 0.763154315676794 | 362.309000344187 | 369.744740927105 | 457.1377112172 | 343.281170098487 | 696.393412540946 | 713.05694129906 | 337.309301340188 | 346.282655129254 | Inf2 |
| **ENSMUSG00000037685** | 0.211542080938306 | 0.322543222932362 | 0.855977678491063 | 719.19691838896 | 966.711470221308 | 862.220403558735 | 805.679030367761 | 686.538788778574 | 469.663577116362 | 886.998533153828 | 852.137568264778 | Atp8a1 |
| **ENSMUSG00000037692** | 0.0473562231604349 | 0.815700358149723 | 0.966177091475091 | 617.099868416658 | 817.673391829986 | 771.360734248484 | 548.687116141024 | 658.069875687277 | 508.802208542726 | 857.848346618257 | 640.46008315442 | Ahdc1 |
| **ENSMUSG00000037697** | 0.290604528006868 | 0.125172309721862 | 0.771949105062466 | 183.413284463516 | 218.26341534904 | 220.523989055088 | 221.350699844926 | 142.344565456483 | 146.769867848863 | 227.996101831794 | 171.513039217624 | Ddhd1 |
| **ENSMUSG00000037703** | -0.0197774729775667 | 0.926117385019607 | 0.986713918026868 | 144.562194651047 | 176.728213174409 | 165.629605513478 | 115.364983393754 | 121.54035973592 | 132.092881063977 | 187.394056300105 | 169.341988088287 | Lzts3 |
| **ENSMUSG00000037706** | -0.268661136892246 | 0.185655086407621 | 0.805963874499992 | 5843.02320505201 | 5126.7470135157 | 5080.56984226486 | 6022.42730383165 | 7853.04018041456 | 8556.68329558872 | 5008.62669366557 | 5173.61484121011 | Cd81 |
| **ENSMUSG00000037709** | 0.276658787455275 | 0.314025806041844 | 0.853228028313558 | 513.195791011218 | 758.22104361924 | 609.5169482896 | 747.527575323754 | 365.716037403578 | 371.816998550453 | 665.248899865371 | 766.381048655966 | Fam13a |
| **ENSMUSG00000037710** | 0.0654702225301999 | 0.515134859407931 | 0.904145806837306 | 5743.63669622942 | 4818.08345225717 | 4761.61454479034 | 5522.51237579205 | 4902.1278426821 | 5184.64558176109 | 5101.28264372507 | 4732.89146195469 | Cisd1 |
| **ENSMUSG00000037712** | 0.173323397420733 | 0.303862567445967 | 0.851785577030345 | 4167.908774533 | 4581.90289087201 | 5374.91731263453 | 4661.49567046306 | 3832.35368536684 | 3192.24462571277 | 5003.42130321279 | 4629.76653331118 | Fermt2 |
| **ENSMUSG00000037720** | -0.0148389362214942 | 0.901741078931958 | 0.984871597499125 | 875.504791355405 | 829.889627763701 | 836.666121565227 | 876.023532437123 | 764.28082068173 | 812.126602097043 | 854.725112346588 | 1021.47955635307 | Tmem33 |
| **ENSMUSG00000037722** | -0.0358482804787853 | 0.836248695392471 | 0.968924788429433 | 265.633032671299 | 239.438224300813 | 194.96970706158 | 225.102406621959 | 221.181555555458 | 200.585486060113 | 232.160414194018 | 294.177428025166 | Gnpnat1 |
| **ENSMUSG00000037730** | 0.0889192706306213 | 0.598574058141615 | 0.924848033480823 | 214.132750826864 | 254.912123150185 | 251.757000380487 | 183.8336320746 | 194.902558855799 | 188.354663739374 | 220.708555197901 | 246.414303179751 | Mynn |
| **ENSMUSG00000037731** | 0.0562281568754444 | 0.88092297144559 | 0.97846486749867 | 25.2983840639333 | 26.8757190541728 | 51.1085639870161 | 53.4618215727152 | 39.4184950494875 | 55.0387004433237 | 24.9858741733473 | 31.4802413753867 | Themis2 |
| **ENSMUSG00000037736** | 0.273197613347455 | 0.32005778685346 | 0.855228644171892 | 4902.46542610364 | 6349.18502261611 | 6728.34780340181 | 5208.30693321557 | 3727.23769856821 | 2793.5198180567 | 6884.64941284773 | 5781.50915742447 | Limch1 |
| **ENSMUSG00000037740** | 0.00173824405203929 | 0.98686572031685 | 0.998220569123292 | 793.285043147622 | 688.995706661522 | 768.521369582539 | 826.31341764144 | 788.36990098975 | 825.580506649855 | 723.549272936515 | 735.986332845248 | Mrps26 |
| **ENSMUSG00000037742** | -0.21269357658217 | 0.300537063180042 | 0.851785577030345 | 29138.3173593517 | 21294.5280639229 | 22747.0967937768 | 26452.3465581629 | 31244.6321176979 | 38084.3345423158 | 20939.2036353556 | 25191.7917792621 | Eef1a1 |
| **ENSMUSG00000037750** | -0.114683515088629 | 0.492191947763414 | 0.899236955296196 | 228.588970291968 | 313.550055632016 | 248.917635714542 | 237.295453647315 | 284.689130912965 | 225.04713070159 | 305.035880532948 | 298.51953028384 | Fam222b |
| **ENSMUSG00000037752** | -0.592837057289375 | 0.0378310964078832 | 0.679566075569698 | 34.3335212296237 | 42.3496179035451 | 38.8046504345863 | 30.0136542162612 | 63.5075753575076 | 72.1618516923577 | 48.9306702561384 | 35.8223436340608 | Xkr8 |
| **ENSMUSG00000037754** | -0.416314034112799 | 0.0151588958154535 | 0.635862253526407 | 221.360860559416 | 192.202112023782 | 165.629605513478 | 197.902532488472 | 318.632834983357 | 247.06261087892 | 215.50316474512 | 256.184033261768 | Ppp1r16b |
| **ENSMUSG00000037761** | -0.249918436413567 | 0.191443680540036 | 0.808824684419004 | 77.7021796249379 | 72.4829998733752 | 92.7525790875478 | 105.047789756914 | 106.210944994452 | 114.969729814943 | 98.902418602833 | 93.3551985614916 | Actr5 |
| **ENSMUSG00000037762** | -0.665198204049286 | 0.024227282673576 | 0.654702249680876 | 73.1846110420926 | 52.122606650517 | 42.5904699891801 | 81.59962240046 | 87.5966556655277 | 64.8233582999146 | 121.806136595068 | 120.493337678204 | Slc16a9 |
| **ENSMUSG00000037772** | -0.078657100418506 | 0.714682392777754 | 0.951176382858794 | 1670.59686193616 | 1260.71554835938 | 1459.43343829591 | 2014.66653926653 | 1740.98353135236 | 2259.03288264042 | 1421.07159360913 | 1343.88064905961 | Mrpl23 |
| **ENSMUSG00000037773** | 0.0136049423868216 | 0.949255044076716 | 0.991267322147698 | 101.193536255733 | 157.182235680465 | 140.07532351997 | 113.489130005238 | 131.394983498292 | 118.638976511164 | 155.120635492864 | 102.03940307884 | Pced1a |
| **ENSMUSG00000037780** | -0.475892000687107 | 0.178717476238239 | 0.801755154770274 | 43.3686583953142 | 62.7100111264033 | 27.447191770805 | 54.3997482669734 | 68.9823663366031 | 59.9310293716191 | 37.4788112600209 | 95.5262496908287 | Mbl1 |
| **ENSMUSG00000037784** | -0.332733335093253 | 0.426087688768946 | 0.885066977635295 | 40.658117245607 | 28.5045505120015 | 17.9826428843205 | 52.523894878457 | 30.6588294829347 | 30.5770558018465 | 72.8754663389296 | 41.2499714574033 | Dzip1l |
| **ENSMUSG00000037787** | 0.0515213209143922 | 0.676757416059603 | 0.94409858171788 | 772.504227666534 | 583.936077631574 | 693.751433379311 | 744.71379524098 | 669.019457645468 | 720.395434691503 | 660.043509412591 | 646.973236542431 | Apopt1 |
| **ENSMUSG00000037788** | -0.0075247964876214 | 0.952881690670867 | 0.992019688458877 | 377.668733525861 | 357.52850499339 | 407.922057007481 | 311.39166249371 | 338.342082508101 | 385.270903103266 | 366.45948787576 | 372.335268681298 | Vopp1 |
| **ENSMUSG00000037791** | 0.0236623956442454 | 0.905798686477586 | 0.98504921011344 | 404.774145022932 | 502.494504740141 | 563.140658745826 | 430.508352664496 | 445.647985698372 | 332.67836712409 | 572.592949805875 | 517.795694346878 | Phf12 |
| **ENSMUSG00000037795** | 0.422136479003596 | 0.229017126235125 | 0.832048466795696 | 112.939214571131 | 238.623808571898 | 216.738169500494 | 126.620103724852 | 112.780694169367 | 69.71568722821 | 144.709854587303 | 191.052499381657 | N4bp2 |
| **ENSMUSG00000037797** | -0.471475940797062 | 0.365528387202908 | 0.86924083181804 | 20.780815481088 | 28.5045505120015 | 17.9826428843205 | 17.8206071909051 | 16.4243729372865 | 31.8001380339204 | 10.4107809055614 | 59.7039060567679 | Adh4 |
| **ENSMUSG00000037798** | -0.144553288517701 | 0.701252088408375 | 0.945920819861844 | 471.634160049041 | 1099.46123403434 | 475.12035410152 | 547.749189446766 | 402.944616061428 | 534.486935416277 | 732.91897575152 | 1197.33469782936 | Mat1a |
| **ENSMUSG00000037805** | 0.0632857229928696 | 0.691786608860381 | 0.94469184542218 | 3507.44024772103 | 2600.42942242345 | 3117.62240320798 | 3917.71980191634 | 3125.0106908677 | 3665.57744952536 | 2830.69132822214 | 2958.05716372168 | Rpl10a |
| **ENSMUSG00000037808** | -0.177398184757465 | 0.263454987768314 | 0.842510961779757 | 139.141112351633 | 110.760539132349 | 124.932045301595 | 134.123517278917 | 158.768938393769 | 128.423634367755 | 135.340151772298 | 151.973579053591 | Fam76b |
| **ENSMUSG00000037813** | -1.31031108161843 | 0.0006055657126763 | 0.525706734317154 | 43.3686583953142 | 43.9784493613737 | 34.0723759913441 | 38.4549944645846 | 130.300025302473 | 157.777607937528 | 55.1771387994752 | 54.2762782334254 | D630003M21 |
| **ENSMUSG00000037815** | 0.0444849022038701 | 0.725456181638535 | 0.951698749807259 | 8352.08079596425 | 10501.0764086214 | 10148.8357709773 | 8513.56060378133 | 8742.14623541967 | 8143.28150114776 | 10099.4985564851 | 9390.88165994726 | Ctnna1 |
| **ENSMUSG00000037816** | -0.340870097115943 | 0.297161792186205 | 0.851785577030345 | 33.4300075130547 | 31.7622134276588 | 43.5369248778286 | 38.4549944645846 | 58.0327843784121 | 64.8233582999146 | 26.0269522639034 | 37.9933947633978 | Fbxw17 |
| **ENSMUSG00000037818** | 0.561941115287962 | 0.0798811211324276 | 0.739855948845396 | 654.143930795989 | 903.18704336599 | 1628.84886336398 | 812.244517227568 | 598.942133113046 | 419.517205601334 | 888.039611244385 | 801.117866725359 | Abhd18 |
| **ENSMUSG00000037820** | 0.0162432955214634 | 0.875343843697366 | 0.976986380159756 | 8223.78184821145 | 8250.03133390215 | 9491.04962336663 | 7606.58549043369 | 8753.09581737786 | 8504.09075960955 | 8297.39238173241 | 7641.01444970162 | Tgm2 |
| **ENSMUSG00000037822** | -0.0972156699691213 | 0.28563743989926 | 0.851362869828931 | 1351.65651998729 | 1318.5390651123 | 1176.44342659002 | 1352.49029312027 | 1303.00025302473 | 1440.79086938301 | 1406.49650034134 | 1412.26875963373 | Smim14 |
| **ENSMUSG00000037824** | -0.635945078485031 | 0.0509868683071874 | 0.703396849069134 | 194.255449062345 | 219.077831077954 | 239.453086828057 | 176.330218520534 | 489.446313531136 | 418.29412336926 | 195.722681024554 | 185.624871558315 | Tspan14 |
| **ENSMUSG00000037826** | 0.101363195886731 | 0.705693219814022 | 0.947760509100828 | 1609.15792920947 | 1917.94904159324 | 1767.03127710665 | 1474.42076337383 | 1403.73640704008 | 839.034411202668 | 2129.0046951873 | 1936.57760736862 | Ppm1k |
| **ENSMUSG00000037827** | -0.0115477088256556 | 0.972231622526362 | 0.995254967835651 | 19.877301764519 | 19.5459774939439 | 16.0897331070236 | 28.1378008277448 | 22.994122112201 | 22.0154801773295 | 19.7804837205666 | 19.5394601640331 | Gm5884 |
| **ENSMUSG00000037845** | -0.0852952844137655 | 0.629681731023143 | 0.931825800800958 | 93.9654265231807 | 92.8433930962335 | 82.3415753124149 | 85.3513291774927 | 85.4067392738896 | 84.3926740130963 | 110.35427759895 | 95.5262496908287 | Fdxacb1 |
| **ENSMUSG00000037847** | 0.100601416146457 | 0.631146852318173 | 0.931825800800958 | 63.2459601598332 | 52.122606650517 | 51.1085639870161 | 58.151455044006 | 47.0832024202212 | 55.0387004433237 | 52.0539045278068 | 55.3618037980939 | Nmrk1 |
| **ENSMUSG00000037851** | 0.352943742982123 | 0.0737193268641938 | 0.732971689437116 | 1536.87683188395 | 1954.59774939439 | 1942.12543150661 | 1689.20597635895 | 1385.12211771116 | 933.211743072355 | 1632.41044599202 | 1625.03177030876 | Iars |
| **ENSMUSG00000037852** | 0.0620357399404327 | 0.568813153373187 | 0.920733562563053 | 7788.28823682517 | 9125.52824248505 | 7919.9345082102 | 8604.53949312437 | 7690.98636743334 | 7119.56167290194 | 8675.30372860429 | 8544.17171950582 | Cpe |
| **ENSMUSG00000037857** | 0.151468213907047 | 0.490296055460464 | 0.899236955296196 | 425.55496050402 | 675.150639269978 | 522.443098533943 | 460.522006880757 | 389.805117711598 | 360.809258461789 | 547.607075632528 | 577.499600403646 | Nufip2 |
| **ENSMUSG00000037860** | 1.22649762786429 | 0.0111958819465235 | 0.635862253526407 | 20.780815481088 | 12.2162359337149 | 17.9826428843205 | 11.2551203310979 | 5.47479097909548 | 6.1154111603693 | 6.24646854333682 | 8.68420451734806 | Aim2 |
| **ENSMUSG00000037876** | 0.30869248113468 | 0.187379595733375 | 0.807540361382744 | 641.494738764022 | 825.817549119129 | 785.557557578211 | 687.500266891232 | 460.97740043984 | 402.3940543523 | 770.397787011541 | 739.242909539254 | Jmjd1c |
| **ENSMUSG00000037885** | -0.0067929309175366 | 0.975490941502897 | 0.99575536949131 | 238.527621174228 | 366.487078011448 | 312.330113253987 | 256.991914226736 | 307.683253025166 | 199.362403828039 | 348.761160336306 | 323.486618271215 | Stk35 |
| **ENSMUSG00000037887** | 0.289678424020563 | 0.141215042912239 | 0.779711473022872 | 1390.50760979976 | 1619.8728848106 | 1618.43785958884 | 1526.94465825229 | 913.195135313127 | 1165.59736716639 | 1222.2256783129 | 1734.66985234028 | Dusp8 |
| **ENSMUSG00000037890** | -0.29230856875869 | 0.406660359656727 | 0.881296485258502 | 19.877301764519 | 30.1333819698302 | 21.7684624389143 | 16.8826804966469 | 38.3235368536684 | 18.3462334811079 | 28.1091084450157 | 23.8815624227072 | Wdr19 |
| **ENSMUSG00000037892** | -0.436026926073979 | 0.256802212988756 | 0.837517407767513 | 28.9124389302094 | 27.6901347830872 | 39.7511053232348 | 48.7721881014244 | 61.3176589658694 | 75.8310983885793 | 30.191264626128 | 29.3091902460497 | Pcdh18 |
| **ENSMUSG00000037894** | -0.168250188727234 | 0.178993722027974 | 0.801755154770274 | 1070.66375413432 | 951.237571371936 | 850.862944894954 | 1008.27119632752 | 1103.71786138565 | 1224.30531430593 | 955.709687130533 | 1079.0124112805 | H2afz |
| **ENSMUSG00000037896** | 0.131038344012564 | 0.613310065862633 | 0.928075610917191 | 259.308436655316 | 407.207864457164 | 378.581955459379 | 297.322762079837 | 287.974005500422 | 184.685417043153 | 403.938299135781 | 348.453706258591 | Rcor1 |
| **ENSMUSG00000037902** | -0.078944503329557 | 0.464774357859321 | 0.895328112556867 | 624.32797814921 | 711.799347071123 | 626.553136285272 | 760.658549043369 | 747.856447744443 | 706.941530138691 | 715.220648212066 | 706.677142599198 | Sirpa |
| **ENSMUSG00000037904** | -0.309720519386738 | 0.0897462771912517 | 0.75321800011268 | 1451.04302880989 | 1181.71722265469 | 1392.23514120187 | 994.202295913651 | 1642.43729372865 | 1845.63108819945 | 1540.79557402308 | 1192.99259557069 | Ankrd9 |
| **ENSMUSG00000037905** | 0.120789475149423 | 0.362819285905157 | 0.86924083181804 | 1777.21148049131 | 1483.05104235299 | 1853.15867197366 | 1941.50825711439 | 1804.49110670987 | 1731.88444061659 | 1471.04334195582 | 1480.65687020784 | Bri3bp |
| **ENSMUSG00000037907** | 0.152542681265975 | 0.311763265823881 | 0.851785577030345 | 327.975479114563 | 329.023954481389 | 287.722286149128 | 390.177504811395 | 275.929465346412 | 266.631926592101 | 341.473613702413 | 315.887939318536 | Ankrd13b |
| **ENSMUSG00000037916** | 0.0495353868546664 | 0.660890980204006 | 0.939611397743053 | 25613.7103510159 | 22472.987623662 | 25201.2543200422 | 27536.5898167254 | 25949.4142827168 | 26717.0082774214 | 22239.5101704602 | 22515.9712623542 | Ndufv1 |
| **ENSMUSG00000037921** | 0.23536900919189 | 0.604418113050389 | 0.926448151609826 | 24.3948703473642 | 15.4738988493722 | 14.1968233297267 | 10.3171936368398 | 12.0445401540101 | 7.33849339244316 | 19.7804837205666 | 15.1973579053591 | Ddx60 |
| **ENSMUSG00000037926** | 0.039817580704193 | 0.688107139430195 | 0.944453919646345 | 1565.78927081416 | 1820.21915412352 | 1827.60438998015 | 1872.10168173929 | 1795.73144114332 | 1704.97663151096 | 1602.21918136589 | 1790.03165613837 | Ssh2 |
| **ENSMUSG00000037933** | 0.222282840243745 | 0.308110673680473 | 0.851785577030345 | 1813.35202915407 | 2325.97132177932 | 2387.90568406003 | 2068.12836083925 | 1699.37511991124 | 1191.28209403994 | 2353.87756274742 | 2122.20247892693 | Bicd2 |
| **ENSMUSG00000037935** | 0.107466908522015 | 0.185426099740466 | 0.805963874499992 | 1527.84169471826 | 1390.20764925676 | 1536.09628427643 | 1541.01355866616 | 1438.77506930629 | 1340.49812635295 | 1363.81229862854 | 1419.86743858641 | Smarce1 |
| **ENSMUSG00000037936** | -0.48458456260909 | 0.0512754713203704 | 0.703396849069134 | 381.282788392137 | 388.476302692135 | 377.63550057073 | 425.818719193205 | 699.678287128403 | 731.403174780168 | 345.637926064637 | 425.526021350055 | Scarb1 |
| **ENSMUSG00000037938** | -0.0520765474234188 | 0.794186803840114 | 0.961804768755116 | 190.641394196068 | 166.955224427437 | 161.843785958884 | 206.343872736795 | 181.76306050597 | 243.393364182698 | 138.463386043966 | 189.966973816989 | Chchd5 |
| **ENSMUSG00000037940** | 0.330025629077188 | 0.0561354947528792 | 0.703396849069134 | 812.258831195571 | 1131.223447462 | 1041.10037751329 | 957.623154837583 | 734.716949394614 | 612.764198269004 | 867.218049433262 | 920.525678838894 | Inpp4b |
| **ENSMUSG00000037942** | -0.105213705954848 | 0.768551211367291 | 0.95879680569409 | 61.4389327266951 | 135.193010999779 | 65.3053873167428 | 79.7237690119437 | 65.6974917491458 | 72.1618516923577 | 79.1219348822664 | 150.888053488923 | Crp |
| **ENSMUSG00000037946** | -0.0373801993378619 | 0.870933496836834 | 0.975613767803763 | 60.535419010126 | 66.7820897709749 | 51.1085639870161 | 45.9584080186499 | 68.9823663366031 | 47.7002070508805 | 55.1771387994752 | 58.6183804920994 | Fgd3 |

| | | | | | | | | | | | | |
|---|---|---|---|---|---|---|---|---|---|---|---|---|
| **ENSMUSG00000037949** | -0.0517186853472872 | 0.676366758963158 | 0.94409858171788 | 1330.8757045062 | 1237.09749222087 | 1686.58261157153 | 1404.07626130447 | 1496.80785368471 | 1527.62970786025 | 1394.00356325467 | 1445.92005213845 | Ano10 |
| **ENSMUSG00000037957** | -0.0349464093873847 | 0.809355051725433 | 0.964311345725361 | 293.64195788494 | 325.766291565731 | 288.668741037776 | 281.378008277448 | 312.063085808443 | 242.170281950624 | 324.816364253515 | 338.683976176574 | Wdr20 |
| **ENSMUSG00000037958** | -0.00974491462992242 | 0.953641725916294 | 0.992364666839775 | 565.599586572222 | 513.896324944941 | 425.904699891801 | 515.859681841989 | 519.010184818252 | 633.556596214259 | 418.513392403567 | 465.69046724279 | Nsrp1 |
| **ENSMUSG00000037960** | -0.0499726028938731 | 0.756764396820539 | 0.956566265296272 | 2147.65210428462 | 1905.73280565953 | 2055.70001814443 | 2220.07248530907 | 2178.96680968 | 2661.42693699272 | 2157.11380363232 | 1626.11729587342 | Card19 |
| **ENSMUSG00000037965** | 0.170149626441004 | 0.383560770620667 | 0.875126835008975 | 252.983840639333 | 244.324718674299 | 335.04503058155 | 313.267515882226 | 249.650468646754 | 188.354663739374 | 310.241270985729 | 268.124814473121 | Zc3h7a |
| **ENSMUSG00000037966** | 0.00184562366299728 | 0.98875352828813 | 0.998437672973644 | 2964.42850406304 | 2441.61835528516 | 2954.83216236045 | 3088.59260419213 | 3001.28041474014 | 3161.66756991093 | 2862.96474902938 | 2408.78122799942 | Ninj1 |
| **ENSMUSG00000037971** | -0.00820389583828948 | 0.963466866704491 | 0.994284908685797 | 274.66816983699 | 232.922898469498 | 270.686098153456 | 233.543746870282 | 202.567266226533 | 330.232202659942 | 239.447960827911 | 246.414303179751 | 1110032A03 |
| **ENSMUSG00000037972** | -0.0259817378738113 | 0.855161435913236 | 0.973268281968411 | 301.773581334061 | 319.250965734417 | 308.544293699394 | 283.253861665965 | 280.309298129689 | 264.185762127954 | 324.816364253515 | 364.736589728619 | Snn |
| **ENSMUSG00000037977** | -0.0588391812702182 | 0.779068651399958 | 0.960290688328569 | 84.0267756409212 | 71.6685841444609 | 64.3589324280944 | 81.59962240046 | 65.6974917491458 | 68.4926049961361 | 96.8202624217207 | 82.4999429148066 | 6430571L13 |
| **ENSMUSG00000037979** | -0.569134793953864 | 0.218739574966725 | 0.827219244147549 | 18.0702743313809 | 31.7622134276588 | 33.1259211026956 | 13.1309737196143 | 24.0890803080201 | 67.2695227640623 | 27.0680303544596 | 24.9670879873757 | Ccdc92 |
| **ENSMUSG00000037989** | 0.104111890624937 | 0.686137626417839 | 0.944452327903597 | 1902.79988709441 | 2102.00699632788 | 2375.6017705076 | 1611.35806073552 | 1740.98353135236 | 1062.85845967218 | 2553.7645561342 | 2076.61040521085 | Wnk2 |
| **ENSMUSG00000037990** | 0.652496559564711 | 0.18737065795445 | 0.807540361382744 | 24.3948703473642 | 21.1748089517725 | 6.62518422053913 | 30.0136542162612 | 10.949581958191 | 15.9000690169602 | 9.36970281500523 | 16.2828834700276 | Sh3rf3 |
| **ENSMUSG00000037992** | -0.300988238375903 | 0.261950105523005 | 0.841137531367976 | 160.82544154929 | 184.872370463553 | 233.774357496166 | 211.971432902344 | 314.253002200081 | 321.670627035425 | 180.106509666212 | 159.572258006271 | Rara |
| **ENSMUSG00000037993** | -0.118627087043514 | 0.31013544556033 | 0.851785577030345 | 477.958756065025 | 569.276594511116 | 565.033568523123 | 488.659807708502 | 619.746338833609 | 567.510155682271 | 525.744435730849 | 568.815395886298 | Dhx38 |
| **ENSMUSG00000037996** | -0.102533217471346 | 0.845437754469641 | 0.972136081650911 | 16.2632468982428 | 21.9892246806869 | 16.0897331070236 | 45.0204813243917 | 17.5193311331055 | 52.592535979176 | 23.9447960827911 | 13.0263067760221 | Slc24a2 |
| **ENSMUSG00000037997** | 0.0349218253411677 | 0.849575575594928 | 0.972640876584422 | 105.711104838578 | 109.946123403434 | 93.6990339761962 | 101.296082979881 | 88.6916138613469 | 85.6157562451702 | 123.88829277618 | 102.03940307884 | Parp11 |
| **ENSMUSG00000037999** | 0.0459162967806579 | 0.862723089622074 | 0.974868041734905 | 225.878429142261 | 242.69588721647 | 212.005895057252 | 182.895705380341 | 173.003394939417 | 122.308223207386 | 287.337552993494 | 252.927456567762 | Arap2 |
| **ENSMUSG00000038000** | -0.0269267122439601 | 0.870358811269033 | 0.975613767803763 | 150.886790667031 | 169.39847161418 | 131.557229522134 | 155.695831246855 | 140.154649064844 | 181.016170346931 | 163.449260217313 | 135.690695583563 | Acd |
| **ENSMUSG00000038002** | 0.076479579737472 | 0.55532481728701 | 0.916239900758826 | 196.062476495483 | 209.304842330982 | 216.738169500494 | 185.709485463116 | 204.757182618171 | 177.34692365071 | 193.640524843441 | 189.966973816989 | Cramp1l |
| **ENSMUSG00000038005** | 0.152406377797394 | 0.31058625840229 | 0.851785577030345 | 602.643648951553 | 491.907100264254 | 561.247748968529 | 649.983199120906 | 533.2446413639 | 604.202622644487 | 433.088485671353 | 504.769387570856 | Hpf1 |
| **ENSMUSG00000038007** | -0.400000168985729 | 0.265914838892555 | 0.843277879501733 | 694.802048041596 | 725.644414462667 | 795.022106464695 | 453.95652002095 | 1325.99437513693 | 1223.08223207386 | 527.826591911961 | 446.151007078757 | Acer2 |
| **ENSMUSG00000038009** | 0.0221504898376653 | 0.967063024335232 | 0.994284908685797 | 14.4562194651047 | 20.3603932228582 | 11.3574586637814 | 20.6343872736795 | 8.75966556655277 | 19.5693157131818 | 6.24646854333682 | 31.4802413753867 | Dnajc22 |
| **ENSMUSG00000038013** | -0.130316734504411 | 0.38057202497173 | 0.874321109304885 | 214.132750826864 | 265.499527626071 | 185.505158175096 | 219.47484645641 | 234.321053905287 | 243.393364182698 | 254.023054095697 | 237.730098662403 | Wipf2 |
| **ENSMUSG00000038014** | 0.133512186857889 | 0.580140843075494 | 0.922450244597442 | 2695.18141652546 | 3155.86094954302 | 2953.8857074718 | 2462.05757242767 | 2107.79452695176 | 1593.67614839224 | 3302.29970324407 | 3266.34642408754 | Fam120a |
| **ENSMUSG00000038020** | 0.0202953797411642 | 0.962263208642172 | 0.994128583764339 | 16.2632468982428 | 30.1333819698302 | 24.6078271048596 | 14.0689004138724 | 9.85462376237187 | 22.0154801773295 | 21.8626399016789 | 30.3947158107182 | Rapgefl1 |
| **ENSMUSG00000038022** | -0.331996396213931 | 0.175879411318438 | 0.799889021902745 | 70.4740698923855 | 47.2361122770311 | 41.6440151005317 | 51.5859681841989 | 75.5521155115177 | 74.6080161565054 | 60.3825292522559 | 55.3618037980939 | Mindy4 |
| **ENSMUSG00000038023** | 0.0600267183797948 | 0.783817076935745 | 0.961141387070613 | 1427.55167217909 | 1747.73615425015 | 1647.77796113695 | 1273.70445080258 | 1185.83972607208 | 1042.06606172693 | 1742.76472359097 | 1876.87370131185 | Atp6v0a2 |
| **ENSMUSG00000038024** | 0.298786277469459 | 0.351436404391336 | 0.862447289221848 | 584.573374620172 | 925.176268046677 | 844.237760674415 | 522.425168701796 | 459.882442244021 | 308.216722482613 | 763.110240377648 | 806.545494548701 | Dennd4c |
| **ENSMUSG00000038025** | 0.199420001368293 | 0.245124271769535 | 0.832048466795696 | 1143.84836517641 | 1403.23830091939 | 1493.50581428725 | 1077.67777170263 | 1056.63465896543 | 886.734618253548 | 1268.03311429737 | 1245.09782267478 | Phf2 |
| **ENSMUSG00000038028** | 0.245983379527027 | 0.207252505173889 | 0.820099122666178 | 1499.83276950461 | 1676.8819858346 | 1852.21221708501 | 1603.85464718146 | 1294.24058745817 | 960.11955217798 | 1593.89055664145 | 1743.35405685762 | Tigar |
| **ENSMUSG00000038034** | -0.654104728368263 | 0.109468596123296 | 0.763154315676794 | 106.614618555147 | 99.3587189275481 | 128.717864856189 | 123.806323642077 | 214.611806380543 | 326.562955963721 | 109.313199508394 | 71.6446872681215 | Igsf8 |
| **ENSMUSG00000038039** | 0.191150506464625 | 0.482841465482774 | 0.898414997395275 | 481.572810931301 | 596.966729294203 | 735.395448479843 | 528.990655561603 | 431.413529152724 | 297.208982393948 | 705.850945397061 | 616.578520731712 | Gcc2 |
| **ENSMUSG00000038042** | 0.21096227015429 | 0.212662151038143 | 0.823370267313127 | 104.807591122009 | 133.56417954195 | 117.360406192407 | 132.247663890401 | 99.6411958195378 | 94.1773318696872 | 115.559668051731 | 111.809133160856 | Ptpdc1 |
| **ENSMUSG00000038046** | 0.00649005440117591 | 0.953869865638884 | 0.992371505557225 | 467.116591466196 | 406.39344872825 | 389.93941412316 | 468.025420434822 | 427.033696369448 | 439.086521314516 | 436.211719943021 | 421.183919091381 | Mrm3 |
| **ENSMUSG00000038047** | -0.0187543336139659 | 0.931828599804207 | 0.987906435243901 | 97.5794813894569 | 100.173134656462 | 78.5557557578211 | 76.9099889291692 | 86.5016974697087 | 63.6002760678407 | 104.107809055614 | 103.124928643508 | Haus6 |
| **ENSMUSG00000038055** | 0.092256173237589 | 0.647324222736302 | 0.935063240511793 | 769.793686516826 | 658.862324691692 | 622.767316730678 | 819.747930781633 | 638.360628162534 | 874.50379593281 | 706.892023487617 | 474.374671760138 | Dexi |
| **ENSMUSG00000038056** | 0.158484409972335 | 0.409270765869094 | 0.882192126163777 | 636.073656464608 | 903.18704336599 | 780.825283134969 | 649.045272426648 | 595.657258525589 | 512.471455238947 | 780.808567917102 | 770.72315091464 | Kmt2c |
| **ENSMUSG00000038058** | -0.268525824157144 | 0.034994473971313 | 0.679566075569698 | 270.150601254144 | 280.159010746529 | 280.15064703994 | 256.991914226736 | 367.905953795217 | 322.893709267499 | 282.132162540713 | 337.598450611906 | Nod1 |
| **ENSMUSG00000038059** | 0.00786308947286319 | 0.976771666698783 | 0.996134981677394 | 83.1232619243521 | 103.43079757212 | 173.201244622666 | 129.433883807626 | 91.9764884488041 | 154.108361241306 | 113.477511870619 | 127.006491066215 | Smim3 |
| **ENSMUSG00000038065** | 0.0597516008246092 | 0.669263910313937 | 0.942250071683273 | 555.660935689963 | 636.873100011005 | 748.645816920921 | 629.348811847226 | 593.467342133951 | 544.271593272868 | 701.686633034836 | 625.26272524906 | Mturn |
| **ENSMUSG00000038068** | 0.290020407586294 | 0.245436122416534 | 0.832048466795696 | 1012.8388762739 | 1288.40568314247 | 1148.04977993057 | 621.845398293161 | 862.827058305448 | 587.079471395453 | 903.655782602727 | 975.887482636988 | Rnf144b |
| **ENSMUSG00000038069** | -0.240896196706208 | 0.121742413251156 | 0.769868040577702 | 105.711104838578 | 131.935348084121 | 131.557229522134 | 133.185590584659 | 160.958854785407 | 134.539045528125 | 144.709854587303 | 153.05910461826 | Cdkn2aip |
| **ENSMUSG00000038070** | -0.0366561527471732 | 0.872548149042544 | 0.976162349063696 | 63.2459601598332 | 68.4109212288036 | 69.0912068713366 | 77.8479156234274 | 79.9319482947941 | 48.9232892829544 | 88.4916376972716 | 67.3025850094475 | Cntln |
| **ENSMUSG00000038072** | 0.145988947753821 | 0.39580534841479 | 0.879215782741314 | 394.835494140673 | 418.609684661965 | 446.726707442067 | 378.922384480297 | 381.045452145046 | 264.185762127954 | 410.184767679118 | 424.440495785386 | Galnt11 |
| **ENSMUSG00000038074** | -0.381591994008587 | 0.131953383242872 | 0.77303252008843 | 217.74680569314 | 171.027303072009 | 141.968233297267 | 131.309737196143 | 277.024423542232 | 258.070350967584 | 164.49033830787 | 163.914360264945 | Fkbp14 |
| **ENSMUSG00000038079** | -0.489006866481043 | 0.233191608939956 | 0.832048466795696 | 39.754603529038 | 25.2468875963442 | 35.965285768641 | 46.8963347129081 | 58.0327843784121 | 90.5080851734656 | 32.2734208072402 | 27.1381391167127 | Tmem237 |
| **ENSMUSG00000038080** | 0.111817659605728 | 0.620976616230194 | 0.93041735818988 | 432.783070236573 | 489.463853077511 | 435.369248778286 | 354.536290429585 | 323.012667766634 | 273.970419984545 | 510.128264372507 | 476.545722889475 | Kdm1b |
| **ENSMUSG00000038084** | 0.262436131252118 | 0.37592376130984 | 0.874078513608115 | 4601.59535848615 | 5950.12131544808 | 5436.43688039668 | 4253.49755846076 | 3204.9426391625 | 2227.2327446065 | 5706.14901433819 | 5735.91708370839 | Opa1 |
| **ENSMUSG00000038085** | -0.0467488880892463 | 0.84972316476088 | 0.972640876584422 | 121.070838020252 | 99.3587189275481 | 116.413951303759 | 130.371810501884 | 116.065568756824 | 178.570005882784 | 102.025652874501 | 86.8420451734806 | Cnbd2 |
| **ENSMUSG00000038086** | -0.162952557956414 | 0.426909047539283 | 0.885066977635295 | 2009.41450564956 | 1500.96818838911 | 1465.1121676278 | 1760.48840512257 | 1800.1112739266 | 2617.39597663806 | 1601.17810327534 | 1524.07789279458 | Hspb2 |
| **ENSMUSG00000038092** | -0.0953664569255171 | 0.897663148429712 | 0.984332015045905 | 15.3597331816738 | 12.2162359337149 | 2.83936466594534 | 13.1309737196143 | 5.47479097909548 | 9.78465785659088 | 3.12323427166841 | 28.2236646813812 | Hsd3b5 |
| **ENSMUSG00000038095** | 0.123147302060264 | 0.391030137064476 | 0.877896341705281 | 1365.20922573583 | 1460.24740194339 | 1327.87620877377 | 1263.38725716574 | 1143.13635643514 | 1005.37359476471 | 1414.82512506579 | 1409.01218293972 | Sbno1 |
| **ENSMUSG00000038102** | 0.108241212660238 | 0.418266607672698 | 0.882468496126562 | 512.292277294649 | 579.049583258088 | 617.088587398787 | 527.114802173087 | 523.390017601528 | 434.19419238622 | 585.085886892549 | 529.736475558232 | Trappc11 |
| **ENSMUSG00000038116** | -0.0253249266516419 | 0.812614848599175 | 0.965379767964637 | 533.976606492306 | 530.999055252142 | 575.444572298256 | 483.032247542953 | 513.535393839156 | 544.271593272868 | 584.044808801993 | 518.881219911547 | Phf20 |
| **ENSMUSG00000038119** | -0.498758631861791 | 0.0140414022453072 | 0.635862253526407 | 47.8862269781594 | 68.4109212288036 | 67.1982970940397 | 62.8410885152968 | 89.7865720571659 | 84.3926740130963 | 97.8613405122768 | 75.9867895267955 | Cdon |
| **ENSMUSG00000038121** | 0.232772076989286 | 0.428320780937831 | 0.885378691567738 | 3840.83680913501 | 5120.23168768438 | 4723.7563492444 | 3399.04633999158 | 2677.17278877769 | 2048.66273872371 | 4811.86293455046 | 4999.93075086315 | Fam210a |
| **ENSMUSG00000038122** | 0.00500210029348442 | 0.981028364124373 | 0.996940244108351 | 110.228673421423 | 96.1010560118908 | 97.48485353079 | 97.5443762028488 | 95.2613630362614 | 68.4926049961361 | 113.477511870619 | 121.578863242873 | Tbc1d32 |
| **ENSMUSG00000038126** | 0.148649550969212 | 0.567458608619129 | 0.9203643243484 | 69.5705561758165 | 84.6992358070902 | 94.6454888648447 | 76.9099889291692 | 56.937826182593 | 50.1463715150282 | 93.6970281500523 | 92.2696729968231 | Mphosph9 |
| **ENSMUSG00000038127** | -0.0709283612372861 | 0.503675077790704 | 0.900927059989498 | 497.836057829544 | 547.287369830429 | 482.691993210708 | 470.839200517597 | 561.713554455197 | 505.132961846504 | 539.278450908079 | 493.914131924171 | Ccdc50 |
| **ENSMUSG00000038132** | 0.327131415584453 | 0.0750244107113094 | 0.735399943172214 | 3494.79105568907 | 4204.82840838468 | 4427.51596909743 | 3881.14066084027 | 2736.30053135192 | 2441.27213521942 | 4062.28670935004 | 3519.2738806553 | Rbm24 |
| **ENSMUSG00000038141** | 0.0391239035846921 | 0.737109257149845 | 0.954383574136061 | 318.036828232304 | 329.023954481389 | 318.955297474527 | 315.143369270742 | 327.39250054991 | 271.524255520397 | 337.309301340188 | 309.374785930525 | Tmem181a |
| **ENSMUSG00000038143** | 0.0479299469516395 | 0.811410048695307 | 0.965065430498408 | 215.036264543433 | 280.973426475443 | 270.686098153456 | 251.364354061187 | 205.85214081399 | 185.908499275227 | 293.584021536831 | 298.51953028384 | Stox2 |
| **ENSMUSG00000038145** | -0.405497026112688 | 0.138986097949687 | 0.779711473022872 | 2753.00629438588 | 2639.52137741134 | 2465.5149849292 | 2046.55604687131 | 4430.20086028407 | 4377.41130859234 | 2242.48220705792 | 2070.09725182284 | Snrk |
| **ENSMUSG00000038146** | -0.508949985288265 | 0.0691653402598614 | 0.724352955175446 | 303.580608767199 | 403.950201541507 | 395.618143455051 | 318.895076047775 | 725.957283828061 | 626.218102821816 | 347.72008224575 | 324.572143835884 | Notch3 |
| **ENSMUSG00000038147** | -0.109705944881845 | 0.827280894512258 | 0.967319591893432 | 27.1054114970714 | 20.3603932228582 | 74.7699362032273 | 15.9447538023887 | 39.4184950494875 | 46.4771248188067 | 21.8626399016789 | 41.2499714574033 | Cd84 |
| **ENSMUSG00000038150** | 0.0132506582508923 | 0.897547249598659 | 0.984332015045905 | 1440.20086421106 | 1306.32282917858 | 1389.39577653592 | 1417.20723502408 | 1221.97334653411 | 1521.51429669988 | 1423.15374979024 | 1336.28197010693 | Ormdl3 |
| **ENSMUSG00000038151** | -0.00924763775219095 | 0.98165849643098 | 0.996940244108351 | 26.2018977805023 | 19.5459774939439 | 20.8220075502658 | 31.8895076047775 | 36.1336204620302 | 12.2308223207386 | 17.6983275394543 | 32.5657669400552 | Prdm1 |
| **ENSMUSG00000038152** | 0.145846982108952 | 0.563466314366307 | 0.918501010039666 | 52.4037955610046 | 50.4937751926884 | 42.5904699891801 | 52.523894878457 | 37.2285786578493 | 39.1386314263635 | 40.6020455316893 | 61.8749571861049 | 5033430I15 |
| **ENSMUSG00000038155** | 0.357221933222349 | 0.130003164501994 | 0.77303252008843 | 153.597331816738 | 126.234437981721 | 121.146225747001 | 183.8336320746 | 122.635317931739 | 132.092881063977 | 127.011527047849 | 74.901263962127 | Gstp2 |
| **ENSMUSG00000038156** | -0.987754123653234 | 0.0186168918682481 | 0.642273034804458 | 224.974915425692 | 229.665235553841 | 247.971180825893 | 283.253861665965 | 715.00770186987 | 809.680437632895 | 229.03717992235 | 201.907755028342 | Spon1 |
| **ENSMUSG00000038160** | 0.0956836795918765 | 0.504574218334613 | 0.901187590673496 | 364.116027777325 | 337.168111770532 | 312.330113253987 | 353.598363735327 | 279.21433993387 | 287.424324537357 | 331.062832796851 | 381.019473198646 | Atg5 |
| **ENSMUSG00000038167** | -0.488665762675157 | 0.273369248771155 | 0.844738346035895 | 20.780815481088 | 20.3603932228582 | 26.5007368821565 | 19.6964605794214 | 49.2731188118594 | 41.5847958905112 | 12.4929370866736 | 19.5394601640331 | Plekhg6 |
| **ENSMUSG00000038168** | -0.332736574231294 | 0.207493415247557 | 0.820327948642971 | 104.807591122009 | 87.9568987227475 | 59.6266579848522 | 105.047789756914 | 125.920192519196 | 140.654456688494 | 108.272121417838 | 75.9867895267955 | P3h2 |
| **ENSMUSG00000038170** | 0.372427924088297 | 0.333981037332275 | 0.857961274005048 | 40088.000090452 | 60651.1681637079 | 51045.1515094767 | 35167.5614012098 | 21671.4126116516 | 15440.1900977004 | 56635.6892043444 | 50669.0767820319 | Pde4dip |
| **ENSMUSG00000038172** | 0.0996794991361821 | 0.662940622416846 | 0.939808659458052 | 211.422209677157 | 338.796943228361 | 335.04503058155 | 257.929840920994 | 245.270635863478 | 196.916239363891 | 280.050006359601 | 344.111603999917 | Ttc39b |
| **ENSMUSG00000038173** | -0.820434202203312 | 0.0796399523563353 | 0.739855948845396 | 40.658117245607 | 42.3496179035451 | 19.8755526616174 | 40.3308478531009 | 68.9823663366031 | 116.192812047017 | 45.80743598447 | 22.7960368580387 | Enpp6 |
| **ENSMUSG00000038174** | 0.190573402397126 | 0.321705436952667 | 0.855704427283138 | 297.256012751216 | 377.888898216248 | 322.74111702912 | 360.163850595134 | 255.12525962585 | 223.824048469516 | 351.884394607974 | 358.223436340608 | Fam126b |
| **ENSMUSG00000038175** | -0.0643140717419038 | 0.78774469645429 | 0.961258876168556 | 674.021232560508 | 782.65351548667 | 766.628459805242 | 458.646153492241 | 874.871598459458 | 828.026671114003 | 598.619902069779 | 503.683862006187 | Mylip |
| **ENSMUSG00000038178** | 0.241889042333852 | 0.352212246884101 | 0.862728204353539 | 368.63359636017 | 579.863998987002 | 749.59227180957 | 455.832373409466 | 431.413529152724 | 332.67836712409 | 603.825292522559 | 452.664160466768 | Slc43a2 |
| **ENSMUSG00000038180** | -0.315201664543826 | 0.415205210303997 | 0.882192126163777 | 24.3948703473642 | 25.2468875963442 | 10.4110037751329 | 14.0689004138724 | 21.8991639163819 | 22.0154801773295 | 29.1501865355718 | 19.5394601640331 | Spag4 |
| **ENSMUSG00000038181** | -0.30355229005837 | 0.35348227172391 | 0.86286441023679 | 71.3775836089546 | 92.8433930962335 | 88.0203046443056 | 103.171936368398 | 120.445401540101 | 170.008430258266 | 95.7791843311646 | 53.1907526687569 | Chpf2 |
| **ENSMUSG00000038187** | 0.0547026431522414 | 0.763659993597959 | 0.95785116771606 | 244.852217190211 | 237.809392842984 | 203.487801059416 | 210.095579513828 | 179.573144114332 | 189.577745971448 | 211.338852382896 | 282.236646813812 | Btbd10 |
| **ENSMUSG00000038188** | -0.499686225331676 | 0.126245764495839 | 0.771949105062466 | 87.6408305071974 | 55.3802695661743 | 77.6093008691726 | 123.806323642077 | 148.914314631397 | 166.339183562045 | 95.7791843311646 | 75.9867895267955 | Scarf1 |
| **ENSMUSG00000038193** | 0.103399941802404 | 0.817629404756284 | 0.966315773308182 | 4183.26850771468 | 4984.22426095569 | 1540.82855871967 | 1125.51203310979 | 3363.71157755627 | 3512.69217051612 | 2630.80433483536 | 1508.88053488923 | Hand2 |
| **ENSMUSG00000038195** | -0.141573206359012 | 0.361495449894149 | 0.868564573435346 | 894.478579403355 | 596.966729294203 | 648.321598724186 | 791.610129953888 | 869.396807480363 | 799.895779776304 | 802.671207818781 | 760.953420832624 | Rilp |
| **ENSMUSG00000038201** | 0.165172039129787 | 0.617554391183696 | 0.929314954729851 | 158.114900399583 | 207.676010873154 | 177.933519065908 | 124.744250336335 | 120.445401540101 | 70.9387694602839 | 194.681602933998 | 209.506433981022 | Kcna7 |
| **ENSMUSG00000038204** | -0.0282795132825006 | 0.880970599923569 | 0.97846486749867 | 2188.31022153023 | 2239.6432545144 | 1836.12248397799 | 1771.74352545367 | 1811.06085588479 | 1515.39888553951 | 2334.09707902686 | 2533.6166679363 | Asb10 |

| | | | | | | | | | | | | |
|---|---|---|---|---|---|---|---|---|---|---|---|---|
| **ENSMUSG00000038205** | 0.255380944594513 | 0.381806550698577 | 0.875126835008975 | 1361.59517086955 | 1595.44041294317 | 1642.09923180506 | 1323.41456559827 | 874.871598459458 | 692.264543353805 | 1644.9033830787 | 1748.78168468097 | Prkab2 |
| **ENSMUSG00000038206** | 0.137044731377141 | 0.345997889048091 | 0.861266047293794 | 1108.61133023022 | 1130.40903173309 | 1139.53168593273 | 1140.51886021792 | 943.853964796062 | 812.126602097043 | 1182.66471087177 | 1169.11103314798 | Fbxo8 |
| **ENSMUSG00000038208** | -0.0629151495451529 | 0.723677970514212 | 0.951674101666722 | 162.632468982428 | 188.13003337921 | 229.042083052924 | 192.274972322923 | 187.237851485066 | 226.270212933664 | 229.03717992235 | 163.914360264945 | Pgap3 |
| **ENSMUSG00000038212** | -0.226171186580016 | 0.192808926965056 | 0.808824684419004 | 345.142239729375 | 277.715763559786 | 279.204192151292 | 312.329589187968 | 451.122776677468 | 360.809258461789 | 311.282349076285 | 297.434004719171 | Mfsd14b |
| **ENSMUSG00000038213** | -0.361278392441803 | 0.203594257779641 | 0.816074141171399 | 136.430571201926 | 111.574954861263 | 94.6454888648447 | 133.185590584659 | 169.71852035196 | 226.270212933664 | 94.7381062406084 | 121.578863242873 | Tapbpl |
| **ENSMUSG00000038214** | 0.0548926791177807 | 0.824616691059045 | 0.966657594633313 | 142.755167217909 | 154.738988493722 | 174.147699511314 | 182.895705380341 | 127.015150715015 | 102.738907494204 | 184.270822028436 | 214.934061804364 | Bend3 |
| **ENSMUSG00000038215** | 0.0348439456006258 | 0.853059450564769 | 0.973120292022893 | 103.000563688871 | 83.0704043492615 | 87.0738497556571 | 106.92364314543 | 76.6470737073368 | 106.408154190426 | 95.7791843311646 | 92.2696729968231 | Cep44 |
| **ENSMUSG00000038216** | -0.0438091100323917 | 0.919669406161554 | 0.986713918026868 | 22.5878429142261 | 12.2162359337149 | 9.46454888648447 | 19.6964605794214 | 17.5193311331055 | 23.2385624094033 | 14.5750932677859 | 10.8552556466851 | Pnmt |
| **ENSMUSG00000038217** | -0.0466906105473057 | 0.827657522264502 | 0.967319591893432 | 65.9565013095403 | 57.009101024003 | 49.2156542097192 | 48.7721881014244 | 56.937826182593 | 61.154111603693 | 51.0128264372507 | 59.7039060567679 | Tlcd2 |
| **ENSMUSG00000038224** | -0.352835838429759 | 0.384062140185853 | 0.875126835008975 | 65.0529875929712 | 162.068730053951 | 78.5557557578211 | 71.2824287636203 | 73.3621991198795 | 84.3926740130963 | 114.518589961175 | 209.506433981022 | Serpinf2 |
| **ENSMUSG00000038225** | 0.0819419056721324 | 0.808393388544951 | 0.964026716319739 | 18.0702743313809 | 26.0613033252585 | 21.7684624389143 | 22.5102406621959 | 16.4243729372865 | 15.9000690169602 | 27.0680303544596 | 23.8815624227072 | Primpol |
| **ENSMUSG00000038235** | -0.321267306288956 | 0.100754111407502 | 0.759800953500741 | 937.847237798669 | 801.385077251699 | 890.614050218188 | 981.071322194037 | 1351.17841364077 | 1401.65223795664 | 866.176971342706 | 893.387539722182 | F11r |
| **ENSMUSG00000038239** | 0.114011950922289 | 0.280536416048877 | 0.849093714741928 | 37099.1767160416 | 35524.814095243 | 32226.7889584796 | 32582.6354318343 | 29656.9427337602 | 28858.6252657827 | 34346.2072855375 | 34127.8382276132 | Hrc |
| **ENSMUSG00000038240** | 0.025726560352075 | 0.846941319729795 | 0.972334438148346 | 681.24934229306 | 706.098436968723 | 633.178320505811 | 678.120999948651 | 606.60684048378 | 557.72549782568 | 726.672507208183 | 758.782369703287 | Pdss2 |
| **ENSMUSG00000038241** | -0.130947682717422 | 0.478305002600262 | 0.897674307047416 | 161.728955265859 | 180.800291818981 | 169.415425068072 | 172.578511743502 | 249.650468646754 | 157.777607937528 | 181.147587756768 | 160.657783570939 | Cep250 |
| **ENSMUSG00000038244** | 0.275354755215293 | 0.210202031567254 | 0.822220844401339 | 1647.10550530537 | 1939.12385054502 | 1706.45816423315 | 1762.36425851108 | 1323.80445874529 | 908.750098430878 | 1773.99706630766 | 1821.51189751376 | Mical2 |
| **ENSMUSG00000038248** | -0.024832837562678 | 0.882022106551544 | 0.978854883720278 | 592.704998069294 | 508.195414842541 | 441.994432998825 | 484.908100931469 | 437.983278327639 | 451.317343635254 | 539.278450908079 | 633.946929766408 | Sobp |
| **ENSMUSG00000038250** | 0.00090470291963799 | 0.996095878331159 | 0.999262355603273 | 376.765219809292 | 345.312269059675 | 398.457508120996 | 308.577882410935 | 341.626957095558 | 258.070350967584 | 409.143689588562 | 417.927342397375 | Usp38 |
| **ENSMUSG00000038252** | -0.0616252523203717 | 0.749912299843413 | 0.956197192463153 | 208.711668527449 | 196.274190668353 | 145.754052851861 | 150.068271081306 | 195.997517051618 | 141.877538920568 | 195.722681024554 | 197.565652769668 | Ncapd2 |
| **ENSMUSG00000038256** | 0.109997031692441 | 0.683792933158392 | 0.944452327903597 | 388.510898124689 | 526.926976607571 | 416.440151005317 | 295.446908691321 | 305.493336633528 | 262.96267989588 | 560.100012719201 | 378.848422069309 | Bcl9 |
| **ENSMUSG00000038260** | 0.14440155270352 | 0.297293412350505 | 0.851785577030345 | 343.335212296237 | 405.579032999336 | 429.690519446395 | 393.929211588428 | 330.677375137367 | 309.439804714686 | 385.198893505771 | 396.216831104005 | Trpm4 |
| **ENSMUSG00000038264** | -0.533895438501721 | 0.176174080421072 | 0.799889021902745 | 738.17070643691 | 762.293122263812 | 581.123301630146 | 361.101777289392 | 1175.98510230971 | 1431.00621152642 | 447.663578939139 | 483.058876277486 | Sema7a |
| **ENSMUSG00000038267** | 0.201410011624562 | 0.490839305953639 | 0.899236955296196 | 219.553833126278 | 352.642010619904 | 275.418372596698 | 183.8336320746 | 195.997517051618 | 134.539045528125 | 265.474913091815 | 300.690581413177 | Slc22a23 |
| **ENSMUSG00000038268** | -0.092602607227197 | 0.693391576335759 | 0.94469184542218 | 103.90407740544 | 65.9676740420606 | 81.3951204237664 | 80.6616957062019 | 98.5462376237187 | 111.300483118721 | 71.8343882483734 | 72.73021283279 | Ovca2 |
| **ENSMUSG00000038271** | -0.447644342420597 | 0.0265229252145821 | 0.664353746352133 | 77.7021796249379 | 65.9676740420606 | 82.3415753124149 | 75.0341355406529 | 100.736154015357 | 113.746647582869 | 118.6829023234 | 77.072315091464 | Iffo1 |
| **ENSMUSG00000038274** | 0.0256426853987648 | 0.902001696520584 | 0.984871597499125 | 5597.26747414523 | 3562.25439827127 | 4339.49566445313 | 5121.07975064956 | 4454.28994059209 | 6136.20355831455 | 3723.9363299193 | 3978.45119451008 | Fau |
| **ENSMUSG00000038279** | 0.246166484242298 | 0.0594068666742194 | 0.712406493939829 | 515.906332160925 | 653.975830318206 | 547.997380527451 | 507.418341593665 | 476.306815181307 | 458.655837027697 | 429.965251399684 | 512.368066523536 | Nop2 |
| **ENSMUSG00000038280** | -0.0356377832744677 | 0.778693225472254 | 0.960254744040417 | 619.810409566365 | 530.184639523228 | 584.90912118474 | 506.480414899407 | 591.277425742312 | 532.040770952129 | 648.591650416473 | 524.308847734889 | Ostm1 |
| **ENSMUSG00000038286** | 0.0772993567716217 | 0.613970359351658 | 0.928075610917191 | 636.073656464608 | 530.184639523228 | 431.583429223692 | 622.783324987419 | 521.20010120989 | 524.702277559686 | 493.471014923609 | 565.558819192293 | Bphl |
| **ENSMUSG00000038290** | 0.198138575375437 | 0.246842642323246 | 0.833003960609788 | 547.529312240841 | 630.35777417969 | 715.519895818226 | 581.51455044006 | 520.105143014071 | 407.286383280595 | 619.441463880901 | 608.979841779033 | Smg6 |
| **ENSMUSG00000038291** | 0.0552666210990873 | 0.697197863447342 | 0.944933386973863 | 201.483558794897 | 197.088606397268 | 235.667267273463 | 235.419600258798 | 232.131137513649 | 184.685417043153 | 212.379930473452 | 206.249857287016 | Snx25 |
| **ENSMUSG00000038295** | -0.64276786419071 | 0.121776482836878 | 0.769868040577702 | 12.6491920319666 | 17.1027303072009 | 6.62518422053913 | 10.3171936368398 | 18.6142893289246 | 23.2385624094033 | 15.616171358342 | 16.2828834700276 | Atg9b |
| **ENSMUSG00000038296** | -0.664506361557736 | 0.0413797956766283 | 0.683543358336265 | 166.246523848704 | 145.780415475665 | 154.272146849697 | 218.536919762152 | 331.772333333186 | 421.963370065482 | 172.818963032319 | 159.572258006271 | Galnt18 |
| **ENSMUSG00000038299** | 0.10195628987564 | 0.448471544442522 | 0.89108031640712 | 513.195791011218 | 564.39010013763 | 578.283936964201 | 534.618215727152 | 547.479097909548 | 406.063301048521 | 552.812466085309 | 532.993052252237 | Wdr36 |
| **ENSMUSG00000038301** | 0.114776578875755 | 0.321231501649756 | 0.855590711009608 | 320.747369382011 | 366.487078011448 | 315.169477919933 | 324.522636213324 | 283.594172717146 | 305.770558018465 | 313.364505257397 | 323.486618271215 | Snx10 |
| **ENSMUSG00000038302** | 0.179028790625598 | 0.339816173222589 | 0.858570960161243 | 922.487504616995 | 969.969133136965 | 877.36368177711 | 771.913669374467 | 648.215251924905 | 604.202622644487 | 983.818795575549 | 891.216488592845 | Afg1l |
| **ENSMUSG00000038305** | 0.185031451500728 | 0.26875611169795 | 0.843793744782845 | 125.588406603097 | 169.39847161418 | 161.843785958884 | 145.378637610015 | 145.62944004394 | 122.308223207386 | 119.723980413956 | 142.203848971575 | Spats2l |
| **ENSMUSG00000038312** | -0.110993586539974 | 0.666905782420317 | 0.942024383071524 | 141.85165350134 | 140.079505373265 | 183.612248397799 | 196.964605794214 | 239.795844884382 | 214.039390612925 | 119.723980413956 | 142.203848971575 | Edem2 |
| **ENSMUSG00000038319** | -0.227493329055266 | 0.141261456106539 | 0.779711473022872 | 1445.62194651047 | 1544.94663775048 | 1390.34223142457 | 1148.02227377199 | 1766.1675698562 | 1486.04491196974 | 1872.89948491049 | 1348.22275131829 | Kcnh2 |
| **ENSMUSG00000038323** | -0.00085890961757599 | 0.997387951837121 | 0.999262355603273 | 44.2721721118832 | 74.1118313312039 | 51.1085639870161 | 65.6548685980713 | 53.6529515951357 | 53.8156182112498 | 49.9717483466946 | 78.1578406561325 | 1700066M21 |
| **ENSMUSG00000038324** | 0.188870605552246 | 0.0892523633918327 | 0.751342622734882 | 3100.85907526496 | 3286.16746616932 | 3811.3738365873 | 3318.38464428537 | 3143.62498019663 | 2622.28830556636 | 3116.98780312507 | 2973.25452162704 | Trpc4ap |
| **ENSMUSG00000038332** | 0.174340929719729 | 0.31468009446193 | 0.853277176422981 | 3609.53729769333 | 3505.24529724727 | 4023.37973164455 | 3334.32939808776 | 2870.98038943767 | 2408.24891495343 | 3920.70008903441 | 3623.48433486348 | Sesn1 |
| **ENSMUSG00000038335** | 0.388784797911675 | 0.0312075768513422 | 0.669064667366769 | 623.424464432641 | 776.95260538427 | 652.10741827878 | 687.500266891232 | 442.363111110915 | 410.955629976817 | 584.044808801993 | 654.57191549511 | Tsr1 |
| **ENSMUSG00000038342** | 0.155713710346399 | 0.552657799575914 | 0.914977712327965 | 780.635851115655 | 1126.33695308852 | 1062.86883995221 | 800.051470202212 | 794.939650164664 | 484.340563901248 | 1137.89835297786 | 966.117752554972 | Mlxip |
| **ENSMUSG00000038344** | 0.256056910526558 | 0.169737909112244 | 0.79907358295209 | 149.079763233892 | 213.376920975554 | 229.988537941573 | 208.219726125312 | 152.199189218854 | 147.992950080937 | 192.599446752885 | 176.940667040967 | Txlng |
| **ENSMUSG00000038346** | 0.0439432768422099 | 0.730233328351137 | 0.952921973914044 | 444.52874855197 | 551.359448475 | 502.567545872325 | 460.522006880757 | 505.870686468423 | 413.401794440965 | 502.840717738614 | 477.631248454143 | Zfp384 |
| **ENSMUSG00000038349** | -0.374576573285809 | 0.261628826495328 | 0.841137531367976 | 46.9827132615903 | 47.2361122770311 | 79.5022106464695 | 53.4618215727152 | 94.1664048404423 | 102.738907494204 | 44.7663578939139 | 53.1907526687569 | Plcl1 |
| **ENSMUSG00000038351** | 0.0201276030439337 | 0.925652442931988 | 0.986713918026868 | 150.886790667031 | 223.96432545144 | 169.415425068072 | 156.633757941113 | 125.920192519196 | 167.562265794119 | 212.379930473452 | 185.624871558315 | Sgsm2 |
| **ENSMUSG00000038354** | 0.186737342541805 | 0.272547165834487 | 0.844738346035895 | 120.167324303683 | 97.7298874697194 | 96.5383986421416 | 108.799496533947 | 101.831112211176 | 88.0619207093179 | 94.7381062406084 | 86.8420451734806 | Ankrd35 |
| **ENSMUSG00000038365** | -0.0721938564631463 | 0.617100323890001 | 0.929187015773942 | 288.220875585525 | 332.281617397046 | 291.508105703722 | 316.081295965 | 332.867291529005 | 266.631926592101 | 377.911346871878 | 312.63136262453 | Fbxo25 |
| **ENSMUSG00000038366** | -0.588587580786137 | 0.0263342424484629 | 0.664353746352133 | 523.134441893477 | 592.080234920717 | 692.804978490663 | 683.7485601142 | 1258.10696699614 | 1212.07449198519 | 692.316930219831 | 585.098279356326 | Lasp1 |
| **ENSMUSG00000038368** | 0.0203713081071113 | 0.932296624757617 | 0.987982116552499 | 345.142239729375 | 418.609684661965 | 313.276568142636 | 293.571055302804 | 291.25888008788 | 218.931719541221 | 395.609674411332 | 445.065481514088 | Focad |
| **ENSMUSG00000038369** | 0.204766389418725 | 0.305423142053952 | 0.851785577030345 | 366.826568927032 | 423.496179035451 | 447.673162330715 | 297.322762079837 | 349.291664466292 | 244.616446414772 | 400.815064864113 | 336.512925047237 | Ncoa6 |
| **ENSMUSG00000038370** | 0.13281162900059 | 0.459966792998973 | 0.894774044236998 | 4080.26794402581 | 4309.07362168571 | 6514.44899856726 | 4724.33675897836 | 4068.86465566376 | 5023.19872712734 | 4845.17743344826 | 3964.33936216939 | Pcp4l1 |
| **ENSMUSG00000038371** | 0.14549274986326 | 0.551500288921213 | 0.914873920695527 | 1053.49699351951 | 1246.87048096784 | 1255.94563723649 | 760.658549043369 | 900.055636963298 | 633.556596214259 | 1198.28088223011 | 1170.19655871265 | Sbf2 |
| **ENSMUSG00000038372** | 0.0670329648358856 | 0.738536203906684 | 0.954383574136061 | 99.3865088225949 | 113.203786319092 | 81.3951204237664 | 105.047789756914 | 109.49581958191 | 101.51582526213 | 97.8613405122768 | 72.73021283279 | Gmds |
| **ENSMUSG00000038374** | -0.115123992111355 | 0.483106340228273 | 0.89863290032536 | 944.171833814652 | 884.455481600961 | 927.525790875478 | 956.685228143324 | 1065.39432453198 | 1284.23634367755 | 829.739238173241 | 843.45336374743 | Rbm8a |
| **ENSMUSG00000038375** | 0.120680829143268 | 0.637210220198319 | 0.932931968211785 | 14634.2116672688 | 16728.0990719003 | 14723.0522478152 | 12027.0340004724 | 9576.50438063382 | 8851.44611351852 | 18250.0989274491 | 16770.2844485638 | Trp53inp2 |
| **ENSMUSG00000038383** | -0.103565436361611 | 0.427738121246325 | 0.885066977635295 | 415.616309621761 | 431.640336324594 | 476.066808990169 | 395.805064976944 | 405.134532453066 | 507.579126310652 | 505.963952010282 | 428.78259804406 | Pigu |
| **ENSMUSG00000038384** | -0.0336407605810871 | 0.888898732300565 | 0.981228911360951 | 341.528184863099 | 425.939426222194 | 427.797609669098 | 364.853484066425 | 378.855535753408 | 233.608706326107 | 512.210420553619 | 471.118095066132 | Setd1b |
| **ENSMUSG00000038387** | -0.3088443959777 | 0.249662308185518 | 0.835314899646652 | 886.346955954233 | 834.776122137187 | 928.472245764126 | 1104.87764583611 | 1267.96159075851 | 1780.80772989954 | 848.478643803251 | 754.440267444613 | Rras |
| **ENSMUSG00000038388** | 0.132021216850067 | 0.0891821230786939 | 0.751206603737452 | 1761.85174730964 | 1729.00459248512 | 1659.13541980073 | 1791.43998603309 | 1559.22047084639 | 1559.42984589417 | 1686.54650670094 | 1528.41999505326 | Mpp6 |
| **ENSMUSG00000038393** | -0.0460175506475115 | 0.802898452547988 | 0.963461896828986 | 12249.8389692431 | 11818.8010580047 | 18748.3248892371 | 16404.3378825752 | 17951.8396204541 | 14290.492799551 | 15219.5206058402 | 13677.6221148232 | Txnip |
| **ENSMUSG00000038398** | -0.0829711042021734 | 0.600316606788565 | 0.925533581921158 | 444.52874855197 | 387.66188696322 | 365.3315870183 | 333.901903155905 | 417.179072607076 | 473.332823812584 | 403.938299135781 | 328.914246094558 | Upf3a |
| **ENSMUSG00000038400** | -0.300994769085123 | 0.10275704816671 | 0.759800953500741 | 756.240980768291 | 658.047908962778 | 681.447519826882 | 815.9962240046 | 918.669926292222 | 1188.83592957579 | 729.795741479852 | 751.183690750607 | Pmepa1 |
| **ENSMUSG00000038403** | -0.00378669757948167 | 0.974068235054764 | 0.995510898988495 | 6408.62279162424 | 6102.41705675506 | 5597.33421146691 | 5339.61667041171 | 5019.28836963474 | 6050.58780206938 | 6119.45701628897 | 6321.01536306472 | Hfe2 |
| **ENSMUSG00000038406** | -0.00048288723578821 | 0.994802458062233 | 0.999262355603273 | 1671.50037565273 | 1661.40808698523 | 1689.42197623748 | 1779.24693900773 | 1743.173447744 | 1687.85348026193 | 1751.09334831542 | 1620.68966805008 | Scaf1 |
| **ENSMUSG00000038412** | 0.0655801672332318 | 0.58101247664829 | 0.922450244597442 | 3296.92155176045 | 2536.90499556813 | 2481.60471803623 | 2858.80056409888 | 2625.70975357419 | 2750.71193993411 | 2628.72217865424 | 2672.56394021387 | Higd1a |
| **ENSMUSG00000038416** | 0.112407946838047 | 0.2985006899611 | 0.851785577030345 | 750.819898468876 | 708.541684155466 | 750.538726698218 | 759.720622349111 | 693.108537953488 | 590.748718091674 | 746.45299092875 | 714.275821551878 | Cdc16 |
| **ENSMUSG00000038417** | -0.00909138002849111 | 0.934272171611651 | 0.98818087308472 | 959.531566996326 | 1043.26654873925 | 944.56197887115 | 823.499637558666 | 994.22204180374 | 963.788798874201 | 959.873999492758 | 878.190181816823 | Fig4 |
| **ENSMUSG00000038418** | -0.406027394171403 | 0.249793187176749 | 0.835314899646652 | 162.632468982428 | 134.378595270864 | 112.628131749165 | 164.137171495178 | 274.834507150593 | 267.855008824175 | 127.011527047849 | 91.1841474321546 | Egr1 |
| **ENSMUSG00000038422** | -0.641932970291459 | 0.0690791260332536 | 0.724157781586334 | 32.5264937964856 | 35.0198763433161 | 13.2503684410783 | 35.6412143818101 | 47.0832024202212 | 58.7079471395453 | 32.2734208072402 | 44.5065481514088 | Hdhd3 |
| **ENSMUSG00000038425** | 0.00102249469325802 | 0.997622840070126 | 0.999262355603273 | 25.2983840639333 | 38.2775392589734 | 24.6078271048596 | 34.703287687552 | 38.3235368536684 | 41.5847958905112 | 18.7394056300105 | 24.9670879873757 | Poli |
| **ENSMUSG00000038429** | 0.0744319694512416 | 0.514918362088212 | 0.904145806837306 | 1530.55223586796 | 1733.07667112969 | 1530.41755494454 | 1616.04769420681 | 1456.2944004394 | 1328.26730403221 | 1706.32699042151 | 1595.72258006271 | Usp5 |
| **ENSMUSG00000038437** | 0.223301046738891 | 0.180018437374447 | 0.802731406993634 | 1986.82666273533 | 2283.62170387578 | 2517.57000380487 | 2079.38348117034 | 1663.24149944921 | 1515.39888553951 | 2240.40005087681 | 2175.39323159569 | Mllt6 |
| **ENSMUSG00000038446** | 0.100078263206584 | 0.621800289844221 | 0.93078489223231 | 356.887918044773 | 424.310594764365 | 332.205665915605 | 353.598363735327 | 309.873169416804 | 242.170281950624 | 445.581422758026 | 370.164217551961 | Cdc40 |
| **ENSMUSG00000038451** | -0.446232220776458 | 0.210620259813698 | 0.822896573073231 | 28.0089252136404 | 37.4631235300591 | 60.5731128735006 | 84.4134024832345 | 66.7924499449649 | 94.1773318696872 | 58.3003730711437 | 67.3025850094475 | Spsb2 |
| **ENSMUSG00000038456** | -0.684805575586397 | 0.0044036509289894 | 0.635862253526407 | 67.7635287426784 | 94.4722245540621 | 106.002947528626 | 94.7305961200743 | 182.858018701789 | 172.454594722414 | 109.313199508394 | 119.407812113536 | Dennd2a |
| **ENSMUSG00000038457** | -0.604989861297386 | 0.183941938076265 | 0.805963874499992 | 101.193536255733 | 62.7100111264033 | 49.2156542097192 | 62.8410885152968 | 140.154649064844 | 182.239252579005 | 42.6842017128016 | 55.3618037980939 | Tmem255b |
| **ENSMUSG00000038459** | 0.0371055635588242 | 0.723303810609721 | 0.951674101666722 | 592.704998069294 | 702.840774053066 | 667.250696497155 | 617.15576482187 | 666.82954125383 | 587.079471395453 | 630.893322877019 | 629.604827507734 | Abhd17c |
| **ENSMUSG00000038462** | 0.0470573800093572 | 0.608823087419927 | 0.927932112008768 | 21388.880212339 | 18819.5186637523 | 18425.583772208 | 21371.5976553665 | 19873.4912541166 | 18266.7331360231 | 19237.0409572963 | 20059.4269095094 | Uqcrfs1 |
| **ENSMUSG00000038463** | -1.23869779401182 | 0.0200122302957893 | 0.642273034804458 | 68.6670424592474 | 90.4001459094905 | 35.965285768641 | 32.8274342990356 | 137.964732673206 | 264.185762127954 | 100.984574783945 | 35.8223436340608 | Olfml2b |
| **ENSMUSG00000038467** | 0.138485227485136 | 0.078684570070533 | 0.739855948845396 | 3123.44691817919 | 3144.45912933822 | 3305.02047116038 | 3458.13572172984 | 2985.95099999868 | 2844.8892718038 | 3122.19319357785 | 2883.15589975956 | Chmp4b |
| **ENSMUSG00000038473** | 0.0911466912897551 | 0.877153261281883 | 0.977569405470617 | 78.6056933415069 | 125.420022252807 | 115.467496415111 | 47.8342614071662 | 36.1336204620302 | 17.123151249034 | 144.709854587303 | 146.545951230249 | Nos1ap |
| **ENSMUSG00000038481** | -0.0665080590070182 | 0.637157363349387 | 0.932931968211785 | 593.608511785863 | 754.963380703583 | 761.896185362 | 641.541858872582 | 676.684165016202 | 632.333513982185 | 809.958754452674 | 762.038946397292 | Cdk19 |
| **ENSMUSG00000038482** | 0.129614294855807 | 0.386092391066904 | 0.875126835008975 | 496.932544112975 | 458.516055378767 | 476.066808990169 | 467.087493740564 | 395.279908690694 | 347.355353908976 | 479.936999746379 | 511.282540958867 | Tfdp1 |
| **ENSMUSG00000038485** | 0.198290635710796 | 0.438992608666708 | 0.889583654898613 | 936.9437240821 | 1080.72967226931 | 1015.54609551978 | 877.899385825639 | 593.467342133951 | 565.063991218123 | 1137.89835297786 | 1111.57817822055 | Socs7 |

| | | | | | | | | | | | | |
|---|---|---|---|---|---|---|---|---|---|---|---|---|
| **ENSMUSG00000038486** | -0.363462711329114 | 0.390406680433477 | 0.877797322358001 | 14.4562194651047 | 11.4018202048006 | 9.46454888648447 | 14.0689004138724 | 13.1394983498292 | 23.2385624094033 | 17.6983275394543 | 9.76973008201657 | Sv2a |
| **ENSMUSG00000038489** | 0.0396174307223776 | 0.802992339825246 | 0.963461896828986 | 1172.76080410662 | 978.927706155023 | 1017.43900529708 | 1280.26993766239 | 1110.28761056056 | 1322.15189287184 | 942.175671953304 | 955.262496908287 | Polr2l |
| **ENSMUSG00000038495** | 0.262750843023725 | 0.196965329896833 | 0.812065429583558 | 1739.26390439541 | 2258.37481627943 | 2368.97658628706 | 1880.54302198761 | 1553.7456798673 | 1208.40524528897 | 2064.45785357282 | 2046.21568940014 | Otud7b |
| **ENSMUSG00000038497** | -0.0898441830721965 | 0.543869471973877 | 0.913837378403038 | 651.433389646282 | 623.028032619461 | 549.890290304748 | 563.693943249155 | 534.339599559719 | 568.733237914345 | 750.617303290975 | 687.137682435165 | Tmco3 |
| **ENSMUSG00000038500** | -0.219541757029366 | 0.424234765594596 | 0.884586455610141 | 37.9475760958999 | 34.2054606144018 | 52.0550188756646 | 37.5170677703264 | 32.8487458745729 | 59.9310293716191 | 49.9717483466946 | 45.5920737160773 | Prr3 |
| **ENSMUSG00000038502** | -0.0409788086252803 | 0.805689349297289 | 0.963475137341661 | 2400.63594492395 | 2214.39636691806 | 2482.55117292488 | 2572.73292235014 | 2646.51395929476 | 3153.10599428641 | 2199.79800534512 | 1950.68943970931 | Ptov1 |
| **ENSMUSG00000038503** | -0.189243552082578 | 0.131418704117767 | 0.77303252008843 | 933.329669215824 | 828.260796305872 | 789.343377132805 | 973.567908639971 | 1074.15399009853 | 1103.22017333062 | 946.339984315528 | 895.558590851519 | Mesd |
| **ENSMUSG00000038506** | 0.0466658258139764 | 0.773596950853843 | 0.959203117135841 | 1604.64036062662 | 1711.087446449 | 1513.38136694887 | 1464.10356973699 | 1401.54649064844 | 1164.37428493431 | 1700.08052187817 | 1825.85399977243 | Dcun1d2 |
| **ENSMUSG00000038507** | -0.0160042485669332 | 0.905013664460672 | 0.98504921011344 | 215.036264543433 | 241.881471487556 | 226.202718386979 | 232.605820176024 | 268.264757975679 | 206.700897220482 | 225.913945650682 | 224.703791886381 | Parp12 |
| **ENSMUSG00000038508** | 0.269114825104605 | 0.578809784455137 | 0.922450244597442 | 117.456783153976 | 70.0397526866323 | 83.2880302010633 | 308.577882410935 | 91.9764884488041 | 209.14706168463 | 83.2862472444909 | 96.6117752554972 | Gdf15 |
| **ENSMUSG00000038510** | 0.246299816097334 | 0.0704378352557471 | 0.72606739345142 | 314.422773366028 | 292.375246680244 | 274.47191770805 | 356.412143818101 | 256.220217821669 | 248.285693110993 | 288.37863108405 | 249.670879873757 | Rpf2 |
| **ENSMUSG00000038515** | 0.212012412834578 | 0.486519061931651 | 0.899236955296196 | 35.2370349461928 | 43.1640336324594 | 47.3227444324223 | 52.523894878457 | 35.0386622662111 | 42.8078781225851 | 53.094982618363 | 22.7960368580387 | Grtp1 |
| **ENSMUSG00000038517** | -0.444110397821078 | 0.0837170665865676 | 0.742148948523533 | 402.967117589794 | 352.642010619904 | 385.207139679918 | 387.363724728621 | 721.577451044785 | 649.456665231219 | 360.213019332423 | 348.453706258591 | Tbkbp1 |
| **ENSMUSG00000038518** | 0.119392181625749 | 0.57178177717134 | 0.921363774754067 | 256.597895505609 | 258.169786065842 | 234.720812384815 | 262.619474392285 | 241.98576127602 | 150.439114545085 | 232.160414194018 | 306.118209236519 | Jarid2 |
| **ENSMUSG00000038520** | -0.193271371690035 | 0.245410979160142 | 0.832048466795696 | 715.582863522684 | 732.974156022896 | 723.091534927413 | 718.451847801752 | 932.904382837871 | 1015.1582526213 | 715.220648212066 | 642.631134283757 | Tbc1d17 |
| **ENSMUSG00000038521** | -1.00341614893376 | 0.0058067021511708 | 0.635862253526407 | 279.185738419835 | 359.971752180133 | 293.401015481019 | 265.43325447506 | 780.705193619016 | 978.465785659088 | 279.008928269045 | 364.736589728619 | C1s1 |
| **ENSMUSG00000038522** | -0.177214692192168 | 0.490634801260497 | 0.899236955296196 | 104.807591122009 | 127.86326943955 | 115.467496415111 | 113.489130005238 | 107.305903190271 | 80.7234273168747 | 155.120635492864 | 178.026192605635 | Mfsd4b1 |
| **ENSMUSG00000038524** | -0.592001241089284 | 0.0871394836198465 | 0.747269672026173 | 12.6491920319666 | 17.9171460361152 | 20.8220075502658 | 22.5102406621959 | 36.1336204620302 | 30.5770558018465 | 19.7804837205666 | 24.9670879873757 | Fchsd1 |
| **ENSMUSG00000038525** | 0.101533821244141 | 0.49183058821012 | 0.899236955296196 | 255.69438178904 | 214.191336704468 | 221.470443943737 | 226.040333316217 | 220.086597359638 | 249.508775343067 | 204.051305749003 | 182.368294864309 | Armc10 |
| **ENSMUSG00000038526** | 0.00187194495158371 | 0.989045300105379 | 0.998437672973644 | 1952.49314150571 | 2023.00867062319 | 1930.76797284283 | 1821.45364024935 | 1789.1616919684 | 1610.79929964127 | 2334.09707902686 | 1982.16968108469 | Car14 |
| **ENSMUSG00000038527** | -0.128216067598538 | 0.703524784717168 | 0.946803532576442 | 42.4651446787451 | 71.6685841444609 | 30.2865564367503 | 41.2687745473591 | 63.5075753575076 | 53.8156182112498 | 32.2734208072402 | 54.2762782334254 | C1rl |
| **ENSMUSG00000038528** | 0.550096563086643 | 0.269367256327661 | 0.843793744782845 | 32.5264937964856 | 57.009101024003 | 47.3227444324223 | 28.1378008277448 | 19.7092475247437 | 7.33849339244316 | 36.4377331694648 | 48.8486504100828 | Mfsd4b5 |
| **ENSMUSG00000038530** | -0.796404984784879 | 0.0142922615425271 | 0.635862253526407 | 186.123825613223 | 182.42912327681 | 178.879973954556 | 224.164479927701 | 523.390017601528 | 420.740287833408 | 198.845915296222 | 197.565652769668 | Rgs4 |
| **ENSMUSG00000038533** | 0.116083217031145 | 0.444288399221752 | 0.89108031640712 | 291.834930451802 | 359.971752180133 | 318.955297474527 | 297.322762079837 | 267.16979977986 | 260.516515431732 | 287.337552993494 | 354.966859646602 | Cbfa2t2 |
| **ENSMUSG00000038534** | 0.0216150593059804 | 0.919452464352957 | 0.986713918026868 | 65.0529875929712 | 57.8235167529173 | 64.3589324280944 | 59.0893817382642 | 56.937826182593 | 51.3694537471021 | 77.0397787011541 | 56.4473293627624 | Osbpl7 |
| **ENSMUSG00000038535** | -0.0838970846811727 | 0.664710249826649 | 0.941513723457524 | 189.737880479499 | 196.274190668353 | 162.790240847533 | 141.626930832982 | 152.199189218854 | 159.000690169602 | 191.558368662329 | 229.045894145055 | Zfp280d |
| **ENSMUSG00000038538** | 0.0461958428843501 | 0.659649887338094 | 0.939458313865081 | 352.370349461928 | 332.281617397046 | 366.278041906949 | 334.839829850164 | 351.48158085793 | 331.455284892016 | 339.391457521301 | 319.144516012541 | Ubn2 |
| **ENSMUSG00000038539** | -0.107247328889877 | 0.547448321762049 | 0.914699131434573 | 2767.46251385099 | 2650.10878188723 | 2482.55117292488 | 2381.39587672147 | 2503.07443564246 | 3797.67033058933 | 2416.34224818079 | 2359.93257758934 | Atf5 |
| **ENSMUSG00000038542** | -0.107001472904912 | 0.49925046757778 | 0.900927059989498 | 192.448421629207 | 207.676010873154 | 282.990011705886 | 225.102406621959 | 241.98576127602 | 234.831788558181 | 260.269522639034 | 239.90114979174 | Pcid2 |
| **ENSMUSG00000038543** | -0.0373269757807163 | 0.896015460402601 | 0.984073355116573 | 94.8689402397497 | 71.6685841444609 | 53.001473764313 | 71.2824287636203 | 85.4067392738896 | 81.9465095489486 | 43.7252798033577 | 87.9275707381491 | BC028528 |
| **ENSMUSG00000038544** | 0.00036988289074643 | 0.998042262151091 | 0.999262355603273 | 138.237598635064 | 156.367819951551 | 162.790240847533 | 160.385464718146 | 145.62944004394 | 172.454594722414 | 135.340151772298 | 164.999885829613 | Inip |
| **ENSMUSG00000038545** | 0.189117461022768 | 0.0628160375849764 | 0.715615581523459 | 692.995020608457 | 746.004807685525 | 687.126249158772 | 685.624413502716 | 661.354750274735 | 636.002760678407 | 550.730309904196 | 619.835097425718 | Cul7 |
| **ENSMUSG00000038546** | 0.27943313607013 | 0.121937342640097 | 0.769868040577702 | 1132.10268686101 | 1283.51918876898 | 1356.26985543322 | 1164.90495426864 | 1076.34390649017 | 722.841599155651 | 1266.99203620682 | 999.769045059695 | Ranbp9 |
| **ENSMUSG00000038550** | -0.221557396755989 | 0.516257286767332 | 0.904492143440747 | 150.886790667031 | 122.976775066064 | 84.2344850897118 | 60.9652351267805 | 162.053812981226 | 121.085140975312 | 136.381229862854 | 69.4736361387845 | Ciart |
| **ENSMUSG00000038552** | -0.376898040192464 | 0.149145317990971 | 0.782024490202776 | 37.0440623793308 | 43.1640336324594 | 53.9479286529615 | 41.2687745473591 | 60.2227007700503 | 75.8310983885793 | 48.9306702561384 | 43.4210225867403 | Fndc4 |
| **ENSMUSG00000038563** | -0.0235093461088533 | 0.861812574560238 | 0.974868041734905 | 218.650319409709 | 227.221988367098 | 250.810545491838 | 195.088752405698 | 232.131137513649 | 207.923979452556 | 225.913945650682 | 239.90114979174 | Efl1 |
| **ENSMUSG00000038564** | 0.158574278032456 | 0.158441119225738 | 0.789230661864335 | 997.479143092226 | 1092.94590820303 | 1201.99770858353 | 1132.0775199696 | 1075.24894829435 | 883.065371557327 | 1042.11916864669 | 961.775650296298 | Ift172 |
| **ENSMUSG00000038569** | 0.323779084464589 | 0.217595428962791 | 0.826183082597859 | 39.754603529038 | 44.7928650902881 | 65.3053873167428 | 66.5927952923295 | 38.3235368536684 | 41.5847958905112 | 45.80743598447 | 46.6775992807458 | Rad9b |
| **ENSMUSG00000038578** | 0.306172949278465 | 0.409977623590103 | 0.882192126163777 | 27.1054114970714 | 25.2468875963442 | 35.965285768641 | 32.8274342990356 | 37.2285786578493 | 23.2385624094033 | 11.4518589961175 | 26.0526135520442 | Susd1 |
| **ENSMUSG00000038582** | 0.129408208269136 | 0.701728358120223 | 0.946035804551092 | 3056.58690315308 | 3882.3197797346 | 3650.47650551706 | 2592.42938292956 | 2078.23065566465 | 1419.99847143775 | 4241.3521409257 | 4310.62201729864 | Pptc7 |
| **ENSMUSG00000038583** | 0.10995207423733 | 0.535161128671837 | 0.910813988293389 | 107450.368742974 | 109563.348010845 | 110639.630028115 | 101479.916611956 | 81636.7982056844 | 76638.332661748 | 120962.863341718 | 118403.700966217 | Pln |
| **ENSMUSG00000038587** | -0.212193832713093 | 0.176186773263239 | 0.799889021902745 | 811.355317479002 | 803.013908709528 | 745.806452254976 | 837.568537972538 | 1112.4775269522 | 1033.50448610241 | 823.492769629904 | 735.986332845248 | Akap12 |
| **ENSMUSG00000038593** | -0.0451494167919907 | 0.853389364804237 | 0.973120292022893 | 41.5616309621761 | 74.1118313312039 | 54.8943835416099 | 56.2756016554897 | 65.6974917491458 | 57.4848649074714 | 61.4236073428121 | 49.9341759747513 | Tctn1 |
| **ENSMUSG00000038594** | -0.0572440647456221 | 0.796685207958179 | 0.962278883153809 | 74.0881247586617 | 94.4722245540621 | 101.270673085384 | 77.8479156234274 | 85.4067392738896 | 73.3849339244316 | 119.723980413956 | 82.4999429148066 | Cep85l |
| **ENSMUSG00000038604** | -0.156790936079913 | 0.245521819000528 | 0.832048466795696 | 1386.89355493348 | 1531.10157035894 | 1684.68970179424 | 1417.20723502408 | 2013.62812211132 | 1488.49107643389 | 1565.78144819643 | 1642.40017934345 | Ripor1 |
| **ENSMUSG00000038605** | -0.042063121456702 | 0.88833718079468 | 0.980942551992548 | 42.4651446787451 | 38.2775392589734 | 44.483379766477 | 27.1998741334867 | 25.1840385038392 | 51.3694537471021 | 38.5198893505771 | 42.3354970220718 | Samd10 |
| **ENSMUSG00000038607** | -0.110793828735411 | 0.631792227289512 | 0.931825800800958 | 67.7635287426784 | 57.009101024003 | 76.6628459805242 | 88.1651092602672 | 78.836990098975 | 81.9465095489486 | 58.3003730711437 | 93.3551985614916 | Gng10 |
| **ENSMUSG00000038608** | -0.662307346190907 | 0.0190588342886047 | 0.642273034804458 | 22.5878429142261 | 31.7622134276588 | 18.9290977729689 | 33.7653609932938 | 44.893286028583 | 46.4771248188067 | 38.5198893505771 | 40.1644458927348 | Dock10 |
| **ENSMUSG00000038611** | 0.0379586725563558 | 0.832087345110581 | 0.967583158900775 | 407.484686172639 | 549.730617017172 | 513.925004536107 | 345.157023487003 | 504.775728272604 | 412.178712208891 | 419.554470494123 | 433.124700302735 | Phrf1 |
| **ENSMUSG00000038612** | 0.0463443554118074 | 0.532588695642046 | 0.909082872831377 | 4662.13077749627 | 4383.18545301692 | 4557.18028884227 | 4440.14497061814 | 4594.44458965693 | 4071.64075057388 | 4362.11719943021 | 4441.97061062353 | Mcl1 |
| **ENSMUSG00000038615** | 0.259928571307684 | 0.23049755197271 | 0.832048466795696 | 23859.0867134388 | 27176.2384581422 | 27418.7981241455 | 21863.0712431577 | 16783.5192255151 | 14637.84815346 | 26252.8662095541 | 26102.5477280189 | Nfe2l1 |
| **ENSMUSG00000038619** | 0.151323659200207 | 0.338553904525617 | 0.858570960161243 | 3226.44748186806 | 3146.90237652497 | 2866.81185771615 | 3189.88868717201 | 2299.4122112201 | 2348.31788558181 | 3360.60007631521 | 3182.76095560806 | Ensa |
| **ENSMUSG00000038622** | -0.0567178848064716 | 0.728072658325508 | 0.952401994188708 | 314.422773366028 | 254.097707421271 | 244.185361271299 | 284.191788360223 | 331.772333333186 | 313.109051410908 | 239.447960827911 | 257.269558826436 | Med30 |
| **ENSMUSG00000038623** | -0.455899791609881 | 0.125332729257153 | 0.771949105062466 | 74.0881247586617 | 85.5136515360045 | 97.48485353079 | 74.0962088463947 | 152.199189218854 | 152.885279009232 | 72.8754663389296 | 77.072315091464 | Tm6sf1 |
| **ENSMUSG00000038624** | -0.326669116251041 | 0.550639856557545 | 0.914699131434573 | 34.3335212296237 | 15.4738988493722 | 16.0897331070236 | 30.0136542162612 | 56.937826182593 | 37.9155491942896 | 11.4518589961175 | 14.1118323406906 | Nepn |
| **ENSMUSG00000038628** | -0.0919569067361929 | 0.630672136026057 | 0.931825800800958 | 295.448985318078 | 251.654460234528 | 293.401015481019 | 222.288626539184 | 203.662224422352 | 287.424324537357 | 308.159114804616 | 333.256348353232 | Polr3k |
| **ENSMUSG00000038633** | -0.181189465013852 | 0.207532930995781 | 0.820327948642971 | 908.03128515189 | 985.443031986338 | 695.644343156608 | 819.747930781633 | 1096.05315401492 | 918.534756287469 | 949.463218587197 | 902.07174423953 | Degs1 |
| **ENSMUSG00000038637** | 0.392809429532588 | 0.199919658172278 | 0.812921911423739 | 46.9827132615903 | 52.122606650517 | 41.6440151005317 | 31.8895076047775 | 26.2789966996583 | 23.2385624094033 | 45.80743598447 | 35.8223436340608 | Lrrc56 |
| **ENSMUSG00000038641** | -0.510177323507248 | 0.23181508512004 | 0.832048466795696 | 19.877301764519 | 38.2775392589734 | 16.0897331070236 | 20.6343872736795 | 32.8487458745729 | 34.2463024980681 | 16.6572494488982 | 52.1052271040884 | Akr1d1 |
| **ENSMUSG00000038642** | -0.442244508572331 | 0.155587997546849 | 0.787275540343856 | 128.298947752804 | 177.542628903324 | 228.095628164276 | 225.102406621959 | 281.404256325508 | 387.717067567413 | 145.750932677859 | 217.105112933702 | Ctss |
| **ENSMUSG00000038644** | 0.0973853715406807 | 0.608567497446393 | 0.927932112008768 | 262.922491521592 | 221.521078264697 | 257.435729712378 | 228.854113398991 | 252.935343234211 | 287.424324537357 | 174.901119213431 | 193.223550510994 | Pold1 |
| **ENSMUSG00000038646** | 0.0888131034365328 | 0.46533718488805 | 0.895328112556867 | 634.26662903147 | 640.945178655577 | 766.628459805242 | 684.686486808458 | 636.170711770895 | 567.510155682271 | 714.17957012151 | 643.716659848425 | Fam103a1 |
| **ENSMUSG00000038648** | -0.444932043256234 | 0.0295768565330668 | 0.665956123973759 | 402.967117589794 | 416.980853204136 | 372.903226127488 | 328.274342990356 | 564.998429042654 | 675.14139210477 | 468.485140750262 | 363.65106416395 | Creb3l2 |
| **ENSMUSG00000038650** | -0.101215580340742 | 0.546140102795109 | 0.914211323944814 | 1282.98947752804 | 1154.84150360052 | 1139.53168593273 | 1340.29724609491 | 1436.58515291466 | 1647.49176660349 | 1129.56972825341 | 1062.72952781047 | Rnh1 |
| **ENSMUSG00000038658** | -0.0073600841032746 | 0.972769187461041 | 0.995410294646453 | 323.457910531718 | 374.631235300591 | 381.421320125324 | 395.805064976944 | 352.576539053749 | 234.831788558181 | 441.417110395802 | 452.664160466768 | Ric1 |
| **ENSMUSG00000038663** | 0.111857566201986 | 0.673550295833251 | 0.943676982965892 | 7903.03447882944 | 9805.56537612852 | 7787.43082379942 | 6305.68116549762 | 5364.20020131776 | 4768.79762285598 | 9728.8747562471 | 9566.73680142356 | Fsd2 |
| **ENSMUSG00000038664** | 0.32209258698037 | 0.123522888587876 | 0.770251097497048 | 1161.01512579122 | 1448.84558173859 | 1524.73882561265 | 1107.69142591889 | 1119.04727612712 | 695.933790050026 | 1273.23850475016 | 1103.97949926787 | Herc1 |
| **ENSMUSG00000038668** | -0.348222848471129 | 0.347229573693678 | 0.861266047293794 | 129.202461469373 | 127.048853710635 | 80.448665535118 | 114.427056699496 | 154.389105610493 | 254.401104271363 | 88.4916376972716 | 78.1578406561325 | Lpar1 |
| **ENSMUSG00000038670** | 0.295831353456011 | 0.369688743502158 | 0.872553381010192 | 1282.08596381147 | 1362.51751447367 | 424.958245003153 | 1277.45615757962 | 762.090904290091 | 940.550236464798 | 1096.25522935561 | 742.499486233259 | Mybpc2 |
| **ENSMUSG00000038671** | 0.0108411420998883 | 0.908021889072191 | 0.98504921011344 | 1076.9883501503 | 939.021335438221 | 955.919437534931 | 1093.62252550502 | 1002.98170737029 | 1015.1582526213 | 1006.72251356778 | 1009.53877514171 | Arfrp1 |
| **ENSMUSG00000038676** | -0.0646630274654364 | 0.849721762842425 | 0.972640876584422 | 28.0089252136404 | 27.6901347830872 | 24.6078271048596 | 20.6343872736795 | 15.3294147414674 | 23.2385624094033 | 36.4377331694648 | 30.3947158107182 | Ucn |
| **ENSMUSG00000038677** | -1.09496201782824 | 0.174186342321357 | 0.799889021902745 | 5.42108229941427 | 13.0306516626293 | 2.83936466594534 | 5.62756016554897 | 22.994122112201 | 25.6847268735511 | 8.32862472444909 | 1.08552556466851 | Scube3 |
| **ENSMUSG00000038679** | -0.350912332802001 | 0.296800620740182 | 0.851785577030345 | 43.3686583953142 | 41.5352021746308 | 51.1085639870161 | 75.0341355406529 | 90.881530252985 | 88.0619207093179 | 43.7252798033577 | 46.6775992807458 | Trps1 |
| **ENSMUSG00000038683** | 0.075817188127905 | 0.492287998752 | 0.899236955296196 | 743.591788736324 | 661.305571878435 | 682.39397471553 | 741.900015158206 | 743.476614961167 | 658.018240855737 | 682.947227404826 | 599.210111697016 | Pak1ip1 |
| **ENSMUSG00000038685** | 0.315078092266748 | 0.377434306463238 | 0.87422797717331 | 50.5967681278665 | 52.122606650517 | 35.965285768641 | 38.4549944645846 | 15.3294147414674 | 40.3617136584374 | 51.0128264372507 | 35.8223436340608 | Rtel1 |
| **ENSMUSG00000038690** | 0.153541017286326 | 0.38313155905684 | 0.875126835008975 | 15516.0410546402 | 11058.9511829277 | 12395.7196766287 | 14801.421162088 | 12559.170506045 | 15062.2576879896 | 10192.1545065446 | 10530.6835028492 | Atp5j2 |
| **ENSMUSG00000038692** | -1.06013031645323 | 0.105097231650878 | 0.762094734041179 | 1.80702743313809 | 10.5874044758863 | 7.57163910918757 | 16.8826804966469 | 14.2344565456483 | 35.4693847301419 | 19.7804837205666 | 7.59867895267955 | Hoxb4 |
| **ENSMUSG00000038695** | -0.0269288370440273 | 0.803866106366778 | 0.963461896828986 | 1203.48027046997 | 995.21602073331 | 1076.11920839328 | 1208.04958220451 | 1111.38256875638 | 1241.42846555497 | 1114.99463498562 | 1099.6373970092 | Josd2 |
| **ENSMUSG00000038696** | 0.0308034906835094 | 0.775562606468549 | 0.9595799687189 | 1151.97998862553 | 1221.62359337149 | 1053.40429106572 | 1059.85716451172 | 1018.31112211176 | 1021.27366378167 | 1167.04853951343 | 1185.39391661801 | Mapkap1 |
| **ENSMUSG00000038697** | -0.0852435570027786 | 0.535545283782949 | 0.91096005719335 | 281.896279569542 | 260.613033252585 | 239.453086828057 | 226.978260010475 | 250.745426842573 | 286.201242305283 | 241.530117009024 | 293.091902460497 | Taf5l |
| **ENSMUSG00000038702** | -0.343406441486165 | 0.123334994031333 | 0.770251097497048 | 49.6932544112975 | 49.679359463774 | 53.001473764313 | 48.7721881014244 | 65.6974917491458 | 80.7234273168747 | 49.9717483466946 | 59.7039060567679 | Dsel |
| **ENSMUSG00000038705** | 0.0475818000396847 | 0.837430887501781 | 0.968924788429433 | 139.141112351633 | 133.56417954195 | 169.415425068072 | 103.171936368398 | 155.484063806312 | 161.446854633749 | 109.313199508394 | 102.03940307884 | Gmeb2 |
| **ENSMUSG00000038708** | 0.464502545566596 | 0.0730319919823137 | 0.730383296285387 | 20704.9203288962 | 23191.3022965644 | 21808.2135442375 | 20436.4847411911 | 13348.6353652306 | 8548.12171996421 | 19958.5080740517 | 20571.7949760329 | Golga4 |
| **ENSMUSG00000038712** | 0.0694322675067241 | 0.560059310310451 | 0.916574868225118 | 513.195791011218 | 593.709066378546 | 560.301294079881 | 567.445650026188 | 571.568178217569 | 589.5256358596 | 487.224546380272 | 483.058876277486 | Mindy1 |
| **ENSMUSG00000038717** | 0.186307939836445 | 0.143093538052755 | 0.780607990324772 | 9998.28278755305 | 7592.79784066829 | 8533.2372760544 | 9378.32901588736 | 8121.30493839024 | 8670.42994317159 | 7144.91893548677 | 7265.42260432632 | Atp5l |
| **ENSMUSG00000038718** | -0.177432796929269 | 0.282464304879699 | 0.850587941933635 | 135.527057485357 | 155.553404222637 | 164.68315062483 | 127.55803041911 | 155.484063806312 | 141.877538920568 | 172.818963032319 | 188.88144825232 | Pbx3 |
| **ENSMUSG00000038722** | 0.0993174011059421 | 0.434530841111895 | 0.888246266820341 | 930.619128066116 | 734.602987480724 | 773.253644025781 | 933.23706078687 | 815.743855885227 | 845.149822363037 | 750.617303290975 | 735.986332845248 | Bud31 |

| | | | | | | | | | | | | |
|---|---|---|---|---|---|---|---|---|---|---|---|---|
| **ENSMUSG00000038729** | 0.277203189118135 | 0.371945895363904 | 0.873663802472542 | 46.9827132615903 | 92.0289773673191 | 103.163582862681 | 52.523894878457 | 70.0773245324222 | 47.7002070508805 | 67.6700758861489 | 57.5328549274309 | Akap2 |
| **ENSMUSG00000038732** | -0.102777970763881 | 0.777973670620123 | 0.960111442462328 | 21.6843291976571 | 25.2468875963442 | 14.1968233297267 | 38.4549944645846 | 27.3739548954774 | 20.7923979452556 | 27.0680303544596 | 31.4802413753867 | Mboat1 |
| **ENSMUSG00000038733** | 0.155002530858912 | 0.293338146174279 | 0.851785577030345 | 2721.38331430596 | 3104.55275862142 | 3092.06812121448 | 2679.65656549557 | 2496.50468646754 | 2052.33198541994 | 3006.63352552612 | 2859.27433733685 | Wdr26 |
| **ENSMUSG00000038736** | -0.10881525876075 | 0.546005611901718 | 0.914211323944814 | 74.0881247586617 | 92.8433930962335 | 77.6093008691726 | 82.5375490947182 | 77.7420319031559 | 89.2850029413918 | 88.4916376972716 | 97.6973008201657 | Nudcd1 |
| **ENSMUSG00000038738** | -0.031022227196003 | 0.91592885537377 | 0.986713918026868 | 44.2721721118832 | 52.122606650517 | 29.3401015481019 | 33.7653609932938 | 30.6588294829347 | 37.9155491942896 | 42.6842017128016 | 52.1052271040884 | Shank1 |
| **ENSMUSG00000038740** | 0.00443283032786769 | 0.983260075441977 | 0.996940244108351 | 535.783633925444 | 667.820897709749 | 616.142132510139 | 495.225294568309 | 500.395895489327 | 403.617136584374 | 699.604476853724 | 703.420565905193 | Mvb12b |
| **ENSMUSG00000038742** | 0.0977089854598518 | 0.738965139443495 | 0.954383574136061 | 88.5443442237664 | 57.009101024003 | 76.6628459805242 | 103.171936368398 | 75.5521155115177 | 111.300483118721 | 66.6289977955927 | 51.0197015394199 | Angptl6 |
| **ENSMUSG00000038754** | -0.205832587824702 | 0.551695588413455 | 0.914873920695527 | 41.5616309621761 | 93.6578088251478 | 50.1621090983677 | 69.4065753751039 | 61.3176589658694 | 64.8233582999146 | 53.094982618363 | 115.065709854862 | Elovl3 |
| **ENSMUSG00000038759** | -0.018789998432492 | 0.901455182008818 | 0.984802943363903 | 210.518695960587 | 238.623808571898 | 205.380710836713 | 173.51643843776 | 227.751304730372 | 187.131581507301 | 205.092383839559 | 219.276164063039 | Nup205 |
| **ENSMUSG00000038762** | 0.199074791404733 | 0.0122049446140017 | 0.635862253526407 | 3775.78382154204 | 3558.99673535562 | 3825.57065991702 | 4019.01588489622 | 3326.48299889842 | 3107.85195169968 | 3428.27015220136 | 3357.53057151969 | Abcf1 |
| **ENSMUSG00000038763** | 0.334679067854454 | 0.260667742244142 | 0.840316047847749 | 4474.19992444991 | 5625.16943961127 | 4545.82283017849 | 3895.20956125414 | 2665.12824862368 | 2005.85486060113 | 4782.71274801489 | 5247.43057960757 | Alpk3 |
| **ENSMUSG00000038764** | 0.245463726295517 | 0.340620213163767 | 0.859077629258968 | 1938.0369220406 | 2361.80561385155 | 1761.35254777476 | 1616.98562090107 | 1473.8137315725 | 922.20400298369 | 2016.56826140724 | 2063.58409843483 | Ptpn3 |
| **ENSMUSG00000038766** | 0.122562367518768 | 0.507598200489741 | 0.90183410140733 | 260.211950371885 | 245.953550132127 | 301.919109478855 | 226.978260010475 | 216.801722772181 | 183.462334811079 | 255.064132186253 | 294.177428025166 | Gabpb2 |
| **ENSMUSG00000038773** | 0.103110054150672 | 0.627369110746832 | 0.931825800800958 | 730.942596704357 | 912.960032112962 | 859.38103889279 | 645.293565649615 | 712.817785478232 | 488.00981059747 | 853.684034256032 | 876.019130687486 | Kdm3b |
| **ENSMUSG00000038774** | 0.291112446972836 | 0.20952487236485 | 0.821566820583092 | 265.633032671299 | 288.303168035672 | 264.060913932917 | 218.536919762152 | 194.902558855799 | 132.092881063977 | 273.803537816264 | 245.328777615083 | Ascc3 |
| **ENSMUSG00000038776** | -0.50242482829322 | 0.159296701054768 | 0.789652743439786 | 985.733464776828 | 1517.25650296739 | 1319.35811477593 | 1267.13896394278 | 2155.9726875678 | 3080.94414259405 | 1005.68143547723 | 968.288803684309 | Ephx1 |
| **ENSMUSG00000038777** | -0.563810927304212 | 0.0391749162731072 | 0.679566075569698 | 42.4651446787451 | 61.895595397489 | 70.9841166486335 | 89.1030359545253 | 106.210944994452 | 108.854318654574 | 108.272121417838 | 67.3025850094475 | Sema6c |
| **ENSMUSG00000038780** | 0.0701322650243893 | 0.720350613157422 | 0.951674101666722 | 710.161781223269 | 955.309650016507 | 900.078599104673 | 656.548685980713 | 705.153078107498 | 585.856389163379 | 849.519721893808 | 928.124357791574 | Smurf1 |
| **ENSMUSG00000038781** | -0.150881050740165 | 0.428871023213841 | 0.88553865206485 | 198.77301764519 | 171.841718800923 | 142.914688185915 | 220.412773150668 | 223.371471947096 | 236.054870790255 | 175.942197303987 | 180.197243734972 | Stap2 |
| **ENSMUSG00000038784** | 0.102694086577162 | 0.443110805017345 | 0.89108031640712 | 531.266065342598 | 540.772043999114 | 550.836745193396 | 529.928582255861 | 496.016062706051 | 399.947889888152 | 564.264325081426 | 542.762782334254 | Cnot4 |
| **ENSMUSG00000038793** | -0.542667802368457 | 0.244056875565699 | 0.832048466795696 | 10.8421645988285 | 10.5874044758863 | 4.73227444324223 | 15.0068271081306 | 19.7092475247437 | 17.123151249034 | 12.4929370866736 | 10.8552556466851 | Lefty1 |
| **ENSMUSG00000038797** | -0.131831932868309 | 0.45177966152518 | 0.891110666112841 | 106.614618555147 | 78.1839099757755 | 88.966759532954 | 95.6685228143324 | 100.736154015357 | 100.292743030056 | 94.7381062406084 | 108.552556466851 | Zscan2 |
| **ENSMUSG00000038801** | 0.471479439377003 | 0.0887109518545341 | 0.751206603737452 | 88.5443442237664 | 51.3081909216027 | 59.6266579848522 | 99.4202295913651 | 47.0832024202212 | 47.7002070508805 | 57.2592949805875 | 62.9604827507734 | Scgb1c1 |
| **ENSMUSG00000038803** | -0.0387942602231073 | 0.803059981368042 | 0.963461896828986 | 1104.99727536394 | 891.78522316119 | 941.722614205205 | 1128.32581319257 | 1002.98170737029 | 1294.02100153414 | 901.573626421614 | 980.229584895662 | Ost4 |
| **ENSMUSG00000038806** | -0.0428010172758498 | 0.761011602225497 | 0.957468401713057 | 222.264374275985 | 250.840044505613 | 263.114459044268 | 233.543746870282 | 269.359716171498 | 275.193502216618 | 244.653351280692 | 210.59195954569 | Sde2 |
| **ENSMUSG00000038807** | 0.116057140825062 | 0.767829921478568 | 0.95879680569409 | 1551.33305134905 | 2302.35326564081 | 1717.81562289693 | 1099.25008567056 | 1028.16574587413 | 620.102691661447 | 2201.88016152623 | 2304.57077379124 | Rap1gap2 |
| **ENSMUSG00000038811** | 0.03609208109378 | 0.887409681682072 | 0.980702840906172 | 210.518695960587 | 127.048853710635 | 191.183887506986 | 241.047160424347 | 197.092475247437 | 251.954939807215 | 146.792010768415 | 155.230155747597 | Gngt2 |
| **ENSMUSG00000038812** | -0.0275926890178614 | 0.893983224499052 | 0.983730237253949 | 677.635287426784 | 569.276594511116 | 592.480760293928 | 765.34818251466 | 706.248036303317 | 869.611467004514 | 613.194995337564 | 466.775992807458 | Trmt112 |
| **ENSMUSG00000038816** | 0.306876369063103 | 0.147006862960068 | 0.780607990324772 | 695.705561758165 | 792.426504233642 | 1190.64024991975 | 743.775868546722 | 602.227007700503 | 617.656527197299 | 862.012658980481 | 683.88110574116 | Ctnnal1 |
| **ENSMUSG00000038822** | 0.100113402878619 | 0.573219582087559 | 0.922207298536842 | 105.711104838578 | 137.636258186522 | 138.182413742673 | 109.737423228205 | 127.015150715015 | 100.292743030056 | 109.313199508394 | 121.578863242873 | Hace1 |
| **ENSMUSG00000038827** | -0.174958385130401 | 0.586359574153078 | 0.923058967588378 | 32.5264937964856 | 55.3802695661743 | 37.8581955459379 | 36.5791410760683 | 31.7537876787538 | 68.4926049961361 | 36.4377331694648 | 47.7631248454143 | Fam206a |
| **ENSMUSG00000038828** | -0.00259822239058536 | 0.980110303950033 | 0.996872525500296 | 770.697200233395 | 802.199492980614 | 877.36368177711 | 818.810004087375 | 894.580845984202 | 793.780368615935 | 845.355409531583 | 740.328435103922 | Tmem214 |
| **ENSMUSG00000038831** | 0.468250214907373 | 0.179657028023216 | 0.802391035126198 | 116.553269437407 | 144.96599974675 | 142.914688185915 | 79.7237690119437 | 54.7479097909548 | 53.8156182112498 | 100.984574783945 | 140.032797842237 | Ralgps1 |
| **ENSMUSG00000038836** | 0.287629591480003 | 0.438018469518798 | 0.888835141214916 | 26.2018977805023 | 30.9477976987445 | 24.6078271048596 | 38.4549944645846 | 19.7092475247437 | 12.2308223207386 | 31.2323427166841 | 34.7368180693922 | Agbl3 |
| **ENSMUSG00000038838** | 0.137893243045499 | 0.417363270616168 | 0.882436023575407 | 622.520950716072 | 713.428178528952 | 596.266579848521 | 584.328330522834 | 582.51776017576 | 414.624876673038 | 649.632728507029 | 639.374557589751 | Vars2 |
| **ENSMUSG00000038843** | 0.196574697925931 | 0.522212069046549 | 0.90648515894888 | 103.90407740544 | 71.6685841444609 | 65.3053873167428 | 45.0204813243917 | 56.937826182593 | 73.3849339244316 | 43.7252798033577 | 75.9867895267955 | Gcnt1 |
| **ENSMUSG00000038844** | 0.321808274610218 | 0.279205092449371 | 0.848075053184273 | 1625.42117610771 | 2261.63247919509 | 1860.73031108285 | 1494.11722395325 | 1127.80694169367 | 769.318723974458 | 1927.03554561941 | 1969.14337430867 | Kif16b |
| **ENSMUSG00000038845** | 0.0269779204953825 | 0.815846382655505 | 0.966177091475091 | 10475.3380299015 | 8364.86395167907 | 8651.54413713545 | 10203.7045068345 | 9495.47747414321 | 9965.67402693781 | 8627.41413643871 | 8908.90830923444 | Phb |
| **ENSMUSG00000038848** | 0.0457184319212361 | 0.712246525779493 | 0.950233477782287 | 619.810409566365 | 751.705717787925 | 731.609628925249 | 614.341984739096 | 727.05224202388 | 627.44118505389 | 668.37213413704 | 610.065367343701 | Ythdf1 |
| **ENSMUSG00000038855** | -0.39160345647562 | 0.0542566357189845 | 0.703396849069134 | 1451.04302880989 | 1497.71052547345 | 1665.76060402127 | 1658.25439544843 | 2814.04256325508 | 2227.2327446065 | 1606.38349372812 | 1581.61074772202 | Itpkb |
| **ENSMUSG00000038859** | 0.428013335940901 | 0.0015808503671826 | 0.635862253526407 | 480.669297214732 | 614.069459601404 | 620.874406953381 | 561.818089860639 | 395.279908690694 | 402.3940543523 | 482.019155927491 | 412.499714574033 | Baiap2l1 |
| **ENSMUSG00000038860** | 0.400820347579131 | 0.290596146079969 | 0.851785577030345 | 27.1054114970714 | 26.0613033252585 | 56.7872933189068 | 26.2619474392285 | 36.1336204620302 | 23.2385624094033 | 22.903717992235 | 20.6249857287016 | Garnl3 |
| **ENSMUSG00000038861** | 0.209293201086948 | 0.0951360577269672 | 0.757085157577544 | 590.897970636155 | 602.667639396603 | 585.855576073389 | 574.011136885995 | 499.300937293508 | 418.29412336926 | 547.607075632528 | 568.815395886298 | Pi4kb |
| **ENSMUSG00000038866** | 0.0910909970254476 | 0.661299639019092 | 0.939611397743053 | 253.887354355902 | 250.840044505613 | 274.47191770805 | 258.867767615252 | 194.902558855799 | 177.34692365071 | 315.446661438509 | 285.493223507818 | Zcchc2 |
| **ENSMUSG00000038871** | 0.126347690770986 | 0.841043073579651 | 0.970420002017028 | 908.934798868459 | 732.974156022896 | 2476.87244359299 | 586.204183911351 | 725.957283828061 | 2188.09411318014 | 670.454290318152 | 726.216602763232 | Bpgm |
| **ENSMUSG00000038872** | -0.268930561637996 | 0.185866065837599 | 0.805963874499992 | 165.343010132135 | 155.553404222637 | 142.914688185915 | 188.52326554589 | 228.846262926191 | 240.94719971855 | 147.833088858971 | 169.341988088287 | Zfhx3 |
| **ENSMUSG00000038876** | 0.118419821213284 | 0.57008387345674 | 0.920872458148558 | 454.46739943423 | 562.761268679801 | 586.802030962037 | 423.004939110431 | 408.419407040523 | 343.686107212755 | 599.660980160335 | 514.539117652873 | Rnf146 |
| **ENSMUSG00000038879** | 0.180870466694936 | 0.720024479270355 | 0.951674101666722 | 13.5527057485357 | 4.88649437348597 | 14.1968233297267 | 10.3171936368398 | 9.85462376237187 | 8.56157562451702 | 7.28754663389296 | 11.9407812113536 | Nipal2 |
| **ENSMUSG00000038880** | -0.0674121172796542 | 0.693377522069075 | 0.94469184542218 | 1839.55392693458 | 1356.00218864236 | 1310.8400207781 | 1769.86767206515 | 1693.90032893214 | 1965.49314694269 | 1475.20765431805 | 1442.66347544445 | Mrps34 |
| **ENSMUSG00000038884** | -0.394749210200394 | 0.301078469656146 | 0.851785577030345 | 34.3335212296237 | 18.7315617650296 | 24.6078271048596 | 44.0825546301336 | 37.2285786578493 | 63.6002760678407 | 30.191264626128 | 29.3091902460497 | A230050P20 |
| **ENSMUSG00000038886** | 0.0781213252462796 | 0.743634807466338 | 0.954452732924361 | 1568.49981196386 | 2276.29196231555 | 1980.9300819412 | 1559.77209255132 | 1389.50195049443 | 1213.29757421727 | 2245.60544132959 | 2147.16956691431 | Man2a2 |
| **ENSMUSG00000038888** | -0.344834180573215 | 0.236390486951861 | 0.832048466795696 | 39.754603529038 | 67.5965054998893 | 51.1085639870161 | 82.5375490947182 | 84.3117810780705 | 99.0696607979826 | 59.3414511616998 | 64.046008315442 | Ctu1 |
| **ENSMUSG00000038893** | -0.386934231358737 | 0.405671443380371 | 0.881158207349486 | 28.9124389302094 | 26.8757190541728 | 81.3951204237664 | 32.8274342990356 | 67.887408140784 | 85.6157562451702 | 40.6020455316893 | 28.2236646813812 | Fam117a |
| **ENSMUSG00000038894** | 0.0815337675954132 | 0.696523777889201 | 0.944933386973863 | 126.491920319666 | 194.645359210525 | 182.66579350915 | 229.79204009325 | 174.098353135236 | 145.546785616789 | 188.435134390661 | 184.539345993646 | Irs2 |
| **ENSMUSG00000038895** | 0.0144009397003911 | 0.944926487973027 | 0.990505437194639 | 84.9302893574902 | 81.4415728914329 | 65.3053873167428 | 99.4202295913651 | 70.0773245324222 | 74.6080161565054 | 94.7381062406084 | 87.9275707381491 | Zfp653 |
| **ENSMUSG00000038900** | 0.0625981609627655 | 0.752513344425905 | 0.956497786093635 | 6804.36179948148 | 4341.65025084229 | 5257.55690644212 | 6537.34905897938 | 5301.78758415607 | 7071.86146585106 | 4552.63449000199 | 5041.18072232055 | Rpl12 |
| **ENSMUSG00000038902** | -0.0195382080508905 | 0.922842038968515 | 0.986713918026868 | 246.659244623349 | 293.189662409158 | 247.971180825893 | 177.268145214792 | 269.359716171498 | 193.24699266767 | 283.173240631269 | 232.302470839061 | Pogz |
| **ENSMUSG00000038903** | 0.0190201193345322 | 0.957284233023197 | 0.992403427270738 | 68.6670424592474 | 53.7514381083457 | 21.7684624389143 | 59.0893817382642 | 53.6529515951357 | 56.2617826753975 | 55.1771387994752 | 35.8223436340608 | Ccdc68 |
| **ENSMUSG00000038909** | 0.00010607993884828 | 0.999329448034391 | 0.999709467616704 | 537.590661358582 | 662.119987607349 | 664.41133183121 | 541.183702586959 | 644.930377337448 | 552.833168897385 | 607.989604884784 | 599.210111697016 | Kat7 |
| **ENSMUSG00000038910** | 0.0235977673088536 | 0.897277035017139 | 0.984332015045905 | 297.256012751216 | 278.5301792887 | 222.416898832385 | 287.943495137256 | 277.024423542232 | 309.439804714686 | 196.76375911511 | 286.578749072486 | Plcl2 |
| **ENSMUSG00000038914** | 0.251873959092028 | 0.278241293336955 | 0.848075053184273 | 608.064731250967 | 912.960032112962 | 822.4692982355 | 527.114802173087 | 578.137927392483 | 429.301863457925 | 703.768789215948 | 699.078463646519 | Dido1 |
| **ENSMUSG00000038930** | -0.558622569988391 | 0.0800940068616514 | 0.739855948845396 | 22.5878429142261 | 22.8036404096012 | 25.5542819935081 | 15.9447538023887 | 21.8991639163819 | 30.5770558018465 | 39.5609674411332 | 35.8223436340608 | Rccd1 |
| **ENSMUSG00000038936** | 0.0887215824687641 | 0.563202180639577 | 0.91828598298905 | 1150.17296119239 | 1096.20357111869 | 1020.27836996303 | 1148.02227377199 | 930.714466446232 | 825.580506649855 | 1169.13069569454 | 1224.47283694608 | Sccpdh |
| **ENSMUSG00000038943** | -0.622353186789773 | 0.192629557738767 | 0.808824684419004 | 8.13162344912141 | 9.77298874697194 | 7.57163910918757 | 15.0068271081306 | 24.0890803080201 | 17.123151249034 | 12.4929370866736 | 8.68420451734806 | Prc1 |
| **ENSMUSG00000038949** | -0.0736019428809364 | 0.741358460988889 | 0.954383574136061 | 1094.15511076511 | 1302.25075053401 | 1339.23366743755 | 867.582192188799 | 967.943045104082 | 938.10407200065 | 1602.21918136589 | 1335.19644454226 | Cnst |
| **ENSMUSG00000038954** | 0.224892919026097 | 0.0614818969215749 | 0.71282992338528 | 483.379838364439 | 422.681763306537 | 445.780252553418 | 512.107975064956 | 414.989156215438 | 429.301863457925 | 359.171941241867 | 391.874728845331 | Supt3 |
| **ENSMUSG00000038957** | -0.180418453791558 | 0.329728840624827 | 0.855977678491063 | 163.535982698997 | 166.955224427437 | 154.272146849697 | 165.075098189436 | 218.991639163819 | 214.039390612925 | 168.654650670094 | 135.690695583563 | Edc3 |
| **ENSMUSG00000038965** | 0.00281997167121162 | 0.980811805294024 | 0.996888334226108 | 2815.34874082914 | 2741.32334352563 | 2688.87833865024 | 2624.31889053433 | 2310.36179317829 | 2606.3882365494 | 2793.21251696211 | 3138.25440745666 | Ube2l3 |
| **ENSMUSG00000038967** | 0.162751946856567 | 0.153070959844512 | 0.783412422888014 | 11916.4424078291 | 13200.0501342434 | 14550.7974580812 | 12943.3883807626 | 11498.1560142963 | 10276.3369138846 | 12878.1359801794 | 12343.5111958456 | Pdk2 |
| **ENSMUSG00000038970** | 0.162334219185222 | 0.537563147538918 | 0.912136833534763 | 755.337467051722 | 1002.54576229354 | 879.256591554407 | 733.458674909882 | 619.746338833609 | 439.086521314516 | 980.695561303881 | 971.545380378314 | Lmtk2 |
| **ENSMUSG00000038975** | 0.144814377751059 | 0.180312886738942 | 0.802808628271642 | 2420.51324668847 | 2068.61595144239 | 2301.77828919302 | 2542.71926813388 | 2012.5331639155 | 1985.06246265587 | 2115.47068001007 | 2327.36681064928 | Rabggtb |
| **ENSMUSG00000038976** | -0.0216964308652372 | 0.786577908144566 | 0.961141387070613 | 1701.31632829951 | 1747.73615425015 | 1629.79531825263 | 1812.07437330677 | 1791.35160836004 | 1827.28485471835 | 1674.05356961427 | 1704.27513652956 | Ppp1r9b |
| **ENSMUSG00000038982** | -0.0313865399309531 | 0.821660799347251 | 0.966637987288264 | 475.248214915318 | 451.186313818538 | 386.153594568566 | 474.59090729463 | 411.70428162798 | 404.840218816448 | 492.429936833053 | 516.71016878221 | Bloc1s5 |
| **ENSMUSG00000038984** | -0.0458693304828343 | 0.832170768460916 | 0.967591827718242 | 135.527057485357 | 199.531853584011 | 147.646962629158 | 147.254490998531 | 150.009272827216 | 124.754387671534 | 212.379930473452 | 162.828834700276 | Tspyl5 |
| **ENSMUSG00000038987** | -0.277543346392843 | 0.552636050160573 | 0.914977712327965 | 11.7456783153976 | 9.77298874697194 | 9.46454888648447 | 5.62756016554897 | 9.85462376237187 | 14.6769867848863 | 11.4518589961175 | 8.68420451734806 | Cfap157 |
| **ENSMUSG00000038990** | -0.174301993971317 | 0.245742077063995 | 0.832048466795696 | 199.676531361759 | 210.119258059897 | 236.613722162112 | 228.854113398991 | 203.662224422352 | 286.201242305283 | 256.10521027681 | 242.072200921077 | Cables2 |
| **ENSMUSG00000038991** | -0.250791793395806 | 0.165501466111763 | 0.795216960550936 | 865.566140473145 | 820.931054745643 | 815.844114014961 | 880.713165908413 | 1167.22543674316 | 1257.32853457193 | 790.178270732108 | 811.973122372044 | Txndc5 |
| **ENSMUSG00000039000** | 0.129200212338294 | 0.643709451043745 | 0.934775146366713 | 906.224257718752 | 1338.89945833516 | 1237.96299435217 | 882.58901929693 | 885.821180417649 | 552.833168897385 | 1213.89705358846 | 1338.45302123627 | Ube3c |
| **ENSMUSG00000039001** | 0.218477962582617 | 0.171533654597867 | 0.799531027639052 | 3442.38726012806 | 2423.70120924904 | 3227.4111702912 | 3602.5764326456 | 2822.80222882163 | 3124.97510294871 | 2412.17793581857 | 2552.07060253566 | Rps21 |
| **ENSMUSG00000039004** | -0.6096300122423 | 0.0822792773545223 | 0.739855948845396 | 714.679349806115 | 715.871425715695 | 889.66759532954 | 877.899385825639 | 1910.70205170432 | 1646.26868437142 | 736.042210023189 | 587.269330485663 | Bmp6 |
| **ENSMUSG00000039005** | 0.141714777797005 | 0.589276319048682 | 0.92352104528132 | 327.975479114563 | 366.487078011448 | 421.172425448559 | 345.157023487003 | 231.036179317829 | 218.931719541221 | 422.677704765791 | 450.493109337431 | Tlr4 |
| **ENSMUSG00000039007** | -0.216720938898033 | 0.22602665139881 | 0.832048466795696 | 861.952085606869 | 803.828324438442 | 671.036516051749 | 891.968286239511 | 1063.20440814034 | 1137.46647582869 | 749.576225200418 | 803.288917854696 | Cpq |
| **ENSMUSG00000039016** | 0.177602249740937 | 0.236707259607308 | 0.832048466795696 | 2301.24943610136 | 1563.67819951551 | 1849.37285241907 | 2160.04517687655 | 1616.15829702899 | 1868.86965060886 | 1625.12289935813 | 1851.90661332447 | Timm8b |
| **ENSMUSG00000039018** | -0.0136065942463257 | 0.897055632617284 | 0.984332015045905 | 539.39768879172 | 509.82424630037 | 514.871459424755 | 585.266257217093 | 498.205979097689 | 533.263853184203 | 569.469715534207 | 567.72987032163 | Mtg1 |
| **ENSMUSG00000039031** | 0.15541412436605 | 0.585269065482972 | 0.922964550512322 | 151.7903043836 | 151.481325578065 | 209.166530391307 | 347.970803569778 | 223.371471947096 | 174.900759186562 | 172.818963032319 | 200.822229463674 | Arhgap18 |
| **ENSMUSG00000039033** | 0.253616057486527 | 0.0799676358017322 | 0.739855948845396 | 165.343010132135 | 185.686786192467 | 180.772883731853 | 176.330218520534 | 134.679858085749 | 133.315963296051 | 161.367104036201 | 163.914360264945 | Tasp1 |
| **ENSMUSG00000039041** | -0.067067008998019 | 0.684097941116094 | 0.944452327903597 | 1955.20368265541 | 1802.30200808741 | 1838.01539375528 | 2025.92165959763 | 2327.8811243114 | 2377.67185915158 | 1696.9572876065 | 1582.69627328668 | Adrm1 |

| | | | | | | | | | | | | |
|---|---|---|---|---|---|---|---|---|---|---|---|---|
| **ENSMUSG00000039043** | 0.0825813167650326 | 0.510049719246993 | 0.902394714365738 | 274.66816983699 | 266.313943354985 | 272.579007930753 | 231.667893481766 | 260.600050604945 | 255.624186503437 | 246.735507461804 | 224.703791886381 | Arpin |
| **ENSMUSG00000039046** | -0.132086972625129 | 0.450381708386896 | 0.89108031640712 | 128.298947752804 | 167.769640156352 | 168.468970179424 | 146.316564304273 | 197.092475247437 | 178.570005882784 | 143.668776496747 | 150.888053488923 | Usp6nl |
| **ENSMUSG00000039047** | -0.269704772701835 | 0.0470852604154916 | 0.702838509185083 | 426.458474220589 | 399.878122896935 | 353.027673465871 | 380.798237868813 | 509.15556105588 | 506.356044078578 | 478.895921655823 | 387.532626586657 | Pigk |
| **ENSMUSG00000039048** | 0.212849561315523 | 0.244546320050596 | 0.832048466795696 | 1510.67493410344 | 1492.00961537105 | 1599.50876181588 | 1442.53125576905 | 1270.15150715015 | 889.180782717696 | 1501.23460658195 | 1553.38708304063 | Foxred1 |
| **ENSMUSG00000039050** | 0.106282349420138 | 0.584326463843547 | 0.922450244597442 | 1133.00620057758 | 1225.69567201606 | 1258.78500190243 | 914.478526901707 | 959.183379537529 | 764.426395046162 | 1229.5132249468 | 1255.95307832146 | Osbpl2 |
| **ENSMUSG00000039062** | -0.837423117909396 | 0.0245058537035798 | 0.655303666217042 | 65.9565013095403 | 78.1839099757755 | 82.3415753124149 | 82.5375490947182 | 146.724398239759 | 237.277953022329 | 57.2592949805875 | 111.809133160856 | Anpep |
| **ENSMUSG00000039063** | 0.174006612152313 | 0.0905918905850538 | 0.753823363185192 | 730.942596704357 | 714.242594257866 | 702.269527377147 | 779.417082928532 | 624.126171616885 | 616.433444965225 | 625.687932424238 | 727.3021283279 | Echdc3 |
| **ENSMUSG00000039065** | 0.107365581210608 | 0.356431083081557 | 0.864965376583892 | 240.334648607366 | 226.407572638183 | 244.185361271299 | 244.79886720138 | 228.846262926191 | 228.716377397812 | 208.215618111227 | 221.447215192376 | Fam173b |
| **ENSMUSG00000039067** | 0.177480070731082 | 0.0624639851038948 | 0.715615581523459 | 4366.68179217819 | 4225.18880160754 | 4486.19617219364 | 4938.18404526922 | 4252.81763256137 | 4141.35643780209 | 3843.66031033326 | 3692.95797100226 | Psmd7 |
| **ENSMUSG00000039068** | 0.164445823569152 | 0.374253207920627 | 0.873700364743909 | 546.625798524272 | 700.397526866323 | 584.90912118474 | 548.687116141024 | 505.870686468423 | 384.047820871192 | 630.893322877019 | 602.466688391022 | Zzz3 |
| **ENSMUSG00000039069** | 0.0726245220261272 | 0.470596898989104 | 0.896509979960267 | 561.082017989377 | 535.885549625628 | 518.657278979349 | 600.273084325223 | 560.618596259378 | 508.802208542726 | 526.785513821405 | 510.197015394199 | Mtg2 |
| **ENSMUSG00000039081** | -0.910091187020203 | 0.0196305763696841 | 0.642273034804458 | 36.1405486627618 | 17.1027303072009 | 25.5542819935081 | 47.8342614071662 | 77.7420319031559 | 78.277262852727 | 43.7252798033577 | 37.9933947633978 | Zfp503 |
| **ENSMUSG00000039086** | 0.191424699115959 | 0.513073362969977 | 0.903078589555164 | 47.8862269781594 | 38.2775392589734 | 41.6440151005317 | 65.6548685980713 | 41.6084114411257 | 30.5770558018465 | 57.2592949805875 | 39.0789203280663 | Ss18l1 |
| **ENSMUSG00000039087** | 0.0151047586924297 | 0.962442991265901 | 0.994128583764339 | 571.924182588206 | 686.552459474779 | 947.401343537095 | 736.272454992657 | 577.042969196664 | 357.140011765567 | 1080.63905799727 | 895.558590851519 | Rreb1 |
| **ENSMUSG00000039089** | -0.169214203177707 | 0.300786388205734 | 0.851785577030345 | 155.404359249876 | 171.841718800923 | 139.128868631322 | 150.068271081306 | 150.009272827216 | 157.777607937528 | 207.174540020671 | 178.026192605635 | L3mbtl3 |
| **ENSMUSG00000039096** | 0.288845988883932 | 0.197893207536439 | 0.812921911423739 | 150.886790667031 | 145.780415475665 | 170.36187995672 | 141.626930832982 | 111.685735973548 | 81.9465095489486 | 139.504464134522 | 163.914360264945 | Rsad1 |
| **ENSMUSG00000039100** | 0.260486772396335 | 0.320425596175019 | 0.855228644171892 | 4440.76991693686 | 5970.48170867094 | 5954.14770448738 | 4169.08415597753 | 3529.05026512495 | 2594.15741422866 | 5564.56239402255 | 5453.68043689458 | March6 |
| **ENSMUSG00000039103** | -0.0563129909441469 | 0.683972210250746 | 0.944452327903597 | 28107.4082087464 | 23762.2077225334 | 22509.536616726 | 28149.0559480759 | 23043.3952310129 | 31164.1352732419 | 25421.0448151997 | 26981.8234354004 | Nexn |
| **ENSMUSG00000039105** | -0.0565146706811236 | 0.727737524590307 | 0.952263232836493 | 2767.46251385099 | 2101.19258059897 | 2267.70591320168 | 2729.36668029125 | 2565.48705280414 | 3176.34455669581 | 2264.3448469596 | 2254.63659781649 | Atp6v1g1 |
| **ENSMUSG00000039108** | 0.189500616897002 | 0.300341181814705 | 0.851785577030345 | 1155.59404349181 | 1281.89035731115 | 1303.26838166891 | 1074.86399161985 | 976.702710670634 | 769.318723974458 | 1336.74426827408 | 1138.71631733726 | Lsm14b |
| **ENSMUSG00000039109** | -0.257764405553989 | 0.236268443319079 | 0.832048466795696 | 355.080890611635 | 431.640336324594 | 602.891764069061 | 413.625672167849 | 577.042969196664 | 639.672007374629 | 389.363205867995 | 550.361461286933 | F13a1 |
| **ENSMUSG00000039110** | 0.450829068607743 | 0.317962993121718 | 0.854351922245337 | 17.1667606148119 | 17.1027303072009 | 10.4110037751329 | 9.37926694258161 | 10.949581958191 | 7.33849339244316 | 9.36970281500523 | 11.9407812113536 | Mycbpap |
| **ENSMUSG00000039114** | -0.0150160239375849 | 0.959495010672928 | 0.99332083910315 | 258.404922938747 | 439.784493613737 | 196.862616838877 | 374.232751009006 | 274.834507150593 | 385.270903103266 | 404.979377226337 | 218.19063849837 | Nrn1 |
| **ENSMUSG00000039115** | -0.0633053314523413 | 0.76005517684595 | 0.957357521291862 | 857.434517024024 | 1094.57473966086 | 804.48665535118 | 702.507093999363 | 763.185862485911 | 699.603036746248 | 1048.36563719003 | 1102.8939737032 | Itga9 |
| **ENSMUSG00000039116** | -0.425145784727652 | 0.404751028897964 | 0.881158207349486 | 9.03513716569045 | 17.9171460361152 | 8.51809399783602 | 20.6343872736795 | 24.0890803080201 | 29.3539735697726 | 8.32862472444909 | 14.1118323406906 | Adgrg6 |
| **ENSMUSG00000039117** | 0.194542680681038 | 0.339015959810853 | 0.858570960161243 | 318.036828232304 | 389.290718421049 | 412.654331450723 | 370.481044231974 | 281.404256325508 | 232.385624094033 | 397.691830592444 | 389.703677715994 | Taf4 |
| **ENSMUSG00000039126** | 0.210349765329101 | 0.480572621701853 | 0.897732059433541 | 2564.17192762295 | 3422.17489289801 | 2763.64827485346 | 2492.07122664393 | 1593.16417491679 | 1456.69093839997 | 3685.41644056872 | 2980.85320057972 | Prune2 |
| **ENSMUSG00000039128** | 0.177034772488847 | 0.148834695993465 | 0.78151662300109 | 1348.04246512102 | 1247.68489669675 | 1513.38136694887 | 1533.51014511209 | 1139.85148184768 | 1169.26661386261 | 1271.15634856904 | 1409.01218293972 | Cdc123 |
| **ENSMUSG00000039130** | -0.149565330088164 | 0.331904535926664 | 0.856428386405603 | 320.747369382011 | 300.519403969387 | 303.812019256151 | 361.101777289392 | 358.051330032845 | 437.863439082442 | 327.939598525183 | 303.947158107182 | Zc3hc1 |
| **ENSMUSG00000039131** | 0.11073935695655 | 0.633645530522207 | 0.9325328237529 | 53.3073092775737 | 45.6072808192024 | 50.1621090983677 | 53.4618215727152 | 39.4184950494875 | 48.9232892829544 | 40.6020455316893 | 58.6183804920994 | Gipc2 |
| **ENSMUSG00000039137** | 0.147360586438555 | 0.495401586327336 | 0.900789112984252 | 524.941469326615 | 634.429852824262 | 669.143606274452 | 569.321503414704 | 459.882442244021 | 370.593916318379 | 705.850945397061 | 627.433776378397 | Whrn |
| **ENSMUSG00000039145** | -0.129256113775194 | 0.707370676355811 | 0.948710970699398 | 34.3335212296237 | 43.9784493613737 | 42.5904699891801 | 44.0825546301336 | 41.6084114411257 | 74.6080161565054 | 23.9447960827911 | 41.2499714574033 | Camk1d |
| **ENSMUSG00000039148** | 0.169765641862393 | 0.244860971466452 | 0.832048466795696 | 1905.51042824412 | 1753.43706435255 | 1989.44817593904 | 2157.23139679377 | 1939.17096479562 | 1998.51636720869 | 1561.6171358342 | 1440.49242431511 | Sart1 |
| **ENSMUSG00000039156** | -0.295174452345298 | 0.0795119665776306 | 0.739855948845396 | 244.852217190211 | 195.459774939439 | 253.649910157784 | 252.302280755445 | 360.241246424483 | 281.308913376988 | 258.187366457922 | 260.526135520442 | Stim2 |
| **ENSMUSG00000039157** | -0.552506014533887 | 0.00513425370734778 | 0.635862253526407 | 245.75573090678 | 240.252640029727 | 286.775831260479 | 211.033506208086 | 486.161438943679 | 347.355353908976 | 295.666177717943 | 313.716888189199 | Fam102a |
| **ENSMUSG00000039158** | -0.363258334614795 | 0.177429348826277 | 0.800680199869068 | 45.1756858284523 | 54.56585383726 | 62.4660226507975 | 53.4618215727152 | 100.736154015357 | 64.8233582999146 | 48.9306702561384 | 62.9604827507734 | Akna |
| **ENSMUSG00000039159** | 0.094379712538381 | 0.445123911025582 | 0.89108031640712 | 1878.40501674704 | 2263.26131065292 | 2192.93597699845 | 2060.62494728518 | 1769.45244444366 | 1793.03855222028 | 2253.93406605404 | 2046.21568940014 | Ube2h |
| **ENSMUSG00000039163** | 0.267864256004758 | 0.0358628644313917 | 0.679566075569698 | 469.827132615903 | 364.858246553619 | 412.654331450723 | 443.63932638411 | 331.772333333186 | 366.924669622158 | 332.103910887408 | 373.420794245967 | Cmc1 |
| **ENSMUSG00000039164** | -0.104110557511756 | 0.608628710983217 | 0.927932112008768 | 69.5705561758165 | 104.245213301034 | 109.78876708322 | 88.1651092602672 | 98.5462376237187 | 96.6234963338349 | 90.5737938783839 | 113.980184290193 | Naif1 |
| **ENSMUSG00000039166** | 0.377829635220947 | 0.0393544093801948 | 0.679566075569698 | 606.257703817829 | 763.107537992726 | 830.987392233336 | 641.541858872582 | 499.300937293508 | 424.409534529629 | 667.331056046484 | 594.868009438342 | Akap7 |
| **ENSMUSG00000039167** | -0.409454164377865 | 0.197795218130672 | 0.812921911423739 | 1607.35090177633 | 1366.58959311824 | 1765.13836732935 | 1847.71558768858 | 3196.18297359594 | 2975.7590706357 | 1196.198726049 | 1380.78851825834 | Adgrl4 |
| **ENSMUSG00000039168** | -0.262390992245183 | 0.305366949779839 | 0.851785577030345 | 341.528184863099 | 355.899673535562 | 329.36630124966 | 332.963976461647 | 429.223612761086 | 607.871869340708 | 242.57119509958 | 352.795808517265 | Dap |
| **ENSMUSG00000039176** | 0.15500437292559 | 0.498776141603061 | 0.900927059989498 | 1591.99116859466 | 1887.81565962341 | 1855.05158175096 | 1498.80685742454 | 1459.57927502686 | 927.096331911986 | 1950.9803417022 | 1798.71586065572 | Polg |
| **ENSMUSG00000039178** | 0.141668034590581 | 0.443498780950202 | 0.89108031640712 | 299.966553900923 | 294.004078138073 | 276.364827485346 | 244.79886720138 | 208.042057205628 | 203.031650524261 | 313.364505257397 | 285.493223507818 | Tbc1d19 |
| **ENSMUSG00000039182** | 0.0184171729810525 | 0.895421754821937 | 0.983733637008786 | 325.264937964856 | 244.324718674299 | 257.435729712378 | 287.943495137256 | 256.220217821669 | 269.078091056249 | 294.625099627387 | 280.065595684475 | AW209491 |
| **ENSMUSG00000039183** | -0.0536721526311265 | 0.725683546764826 | 0.951727359509341 | 395.739007857242 | 348.569931975333 | 362.492222352355 | 377.046531091781 | 379.950493949227 | 479.448234972953 | 317.528817619622 | 364.736589728619 | Nubp2 |
| **ENSMUSG00000039191** | 0.0845279839761104 | 0.64730748339816 | 0.935063240511793 | 402.967117589794 | 466.66021266791 | 537.586376752318 | 425.818719193205 | 378.855535753408 | 333.901449356164 | 516.374732915844 | 498.256234182845 | Rbpj |
| **ENSMUSG00000039195** | -0.107907274682881 | 0.586296756161076 | 0.923058967588378 | 507.774708711803 | 397.434875710192 | 354.920583243168 | 492.411514485535 | 446.742943894192 | 626.218102821816 | 419.554470494123 | 397.302356668674 | 1110008P14R |
| **ENSMUSG00000039196** | -0.0830440564015007 | 0.782408859246069 | 0.960742105865155 | 133.720030052219 | 265.499527626071 | 192.130342395635 | 171.640585049243 | 175.193311331055 | 238.501035254403 | 119.723980413956 | 275.723493425801 | Orm1 |
| **ENSMUSG00000039197** | 0.0716014114331615 | 0.641504044364172 | 0.934544499347759 | 3365.58859421969 | 3282.90980325366 | 3239.71508384363 | 3137.36479229355 | 2888.49972057078 | 2362.9948723667 | 3506.35100899307 | 3635.42511607483 | Adk |
| **ENSMUSG00000039199** | 0.0673805919937576 | 0.591882190556618 | 0.923935529355527 | 364.116027777325 | 412.908774559565 | 403.189782564238 | 364.853484066425 | 384.330326732503 | 396.278643191931 | 313.364505257397 | 382.104998763315 | Zdhhc1 |
| **ENSMUSG00000039201** | 0.124317613816246 | 0.486163024082551 | 0.899236955296196 | 81.316234491214 | 100.987550385377 | 81.3951204237664 | 94.7305961200743 | 76.6470737073368 | 88.0619207093179 | 79.1219348822664 | 85.7565196088121 | Tbc1d25 |
| **ENSMUSG00000039202** | -0.384247379786112 | 0.0201841000051518 | 0.64449919326795 | 327.975479114563 | 436.52683069808 | 359.65285768641 | 307.639955716677 | 556.238763476101 | 468.440494884288 | 408.102611498006 | 437.466802561409 | Abhd2 |
| **ENSMUSG00000039205** | 0.00709822706819347 | 0.956504058366248 | 0.992371505557225 | 577.34526488762 | 562.761268679801 | 472.280989435575 | 447.391033161143 | 499.300937293508 | 551.610086665311 | 520.539045278068 | 479.80229958348 | Ciz1 |
| **ENSMUSG00000039206** | -0.0705795114004739 | 0.621735474489412 | 0.93078489223231 | 179.79922959724 | 215.820168162297 | 178.879973954556 | 205.405946042537 | 194.902558855799 | 193.24699266767 | 233.201492284575 | 197.565652769668 | Daglb |
| **ENSMUSG00000039208** | -0.42077372495 | 0.0588596139797508 | 0.711712402279519 | 209.615182244018 | 166.140808698523 | 179.826428843205 | 191.337045628665 | 290.163921892061 | 319.224462571277 | 164.49033830787 | 226.874843015718 | Metrnl |
| **ENSMUSG00000039210** | 0.198823602663671 | 0.337275656612392 | 0.858570960161243 | 162.632468982428 | 181.614707547895 | 207.27362061401 | 190.399118934407 | 136.869774477387 | 119.862058743238 | 210.29777429234 | 178.026192605635 | Gpatch2 |
| **ENSMUSG00000039218** | 0.179203810630733 | 0.303925506185991 | 0.851785577030345 | 1775.40445305817 | 2360.99119812264 | 2653.85950777024 | 1994.97007868711 | 2119.83906710577 | 1523.96046116403 | 2110.26528955729 | 2003.88019237806 | Srrm2 |
| **ENSMUSG00000039219** | -0.0772900474338685 | 0.510161919374434 | 0.902394714365738 | 692.995020608457 | 541.586459728029 | 571.658752743662 | 655.610759286455 | 633.980795379257 | 669.025980944401 | 637.139791420356 | 656.742966624447 | Arid4b |
| **ENSMUSG00000039220** | -0.0249512719763174 | 0.914809057294925 | 0.986713918026868 | 463.50253659992 | 457.701639649853 | 404.136237452887 | 402.370551836751 | 639.455586358353 | 464.771248188067 | 311.282349076285 | 343.026078435248 | Ppp1r10 |
| **ENSMUSG00000039221** | 0.0644686539972748 | 0.764673975041144 | 0.958110466593105 | 691.187993175319 | 504.123336197969 | 641.696414503647 | 846.009878220861 | 646.025335533267 | 830.472835578151 | 485.14239019916 | 604.637739520359 | Rpl22l1 |
| **ENSMUSG00000039230** | -0.015603197906657 | 0.913597974771316 | 0.986234232499696 | 264.72951895473 | 332.281617397046 | 297.186835035612 | 267.309107863576 | 290.163921892061 | 273.970419984545 | 269.639225454039 | 340.855027305911 | Tbcd |
| **ENSMUSG00000039231** | -0.0308048405582753 | 0.853207739421748 | 0.973120292022893 | 91.2548853734735 | 92.8433930962335 | 102.217127974032 | 85.3513291774927 | 104.021028602814 | 91.7311674055395 | 94.7381062406084 | 89.0130963028176 | Suv39h1 |
| **ENSMUSG00000039232** | 0.0975788691146163 | 0.788614597385632 | 0.961623804555037 | 50.5967681278665 | 39.0919549878878 | 59.6266579848522 | 31.8895076047775 | 52.5579933993167 | 61.154111603693 | 34.3555769883525 | 21.7105112933701 | Stx11 |
| **ENSMUSG00000039233** | 0.0413659844996109 | 0.750896147092407 | 0.956197192463153 | 451.756858284523 | 482.134111517283 | 504.460455649622 | 456.770300103725 | 459.882442244021 | 376.709327478749 | 503.88179582917 | 499.341759747513 | Tbce |
| **ENSMUSG00000039234** | 0.0848508088534265 | 0.556792057930668 | 0.916239900758826 | 617.099868416658 | 786.725594131241 | 670.0900611631 | 574.011136885995 | 697.488370736765 | 552.833168897385 | 652.755962778698 | 593.782483873674 | Sec24d |
| **ENSMUSG00000039236** | -0.103457169920367 | 0.905284883152275 | 0.98504921011344 | 197.869503928621 | 92.0289773673191 | 536.639921863669 | 118.178763476528 | 166.433645764503 | 664.133652016106 | 74.9576225200418 | 109.638082031519 | Isg20 |
| **ENSMUSG00000039239** | 0.0221464330412413 | 0.948772562033278 | 0.991233613135933 | 198.77301764519 | 249.211213047785 | 144.807597963212 | 322.646782824807 | 185.047935093427 | 386.49398533534 | 181.147587756768 | 149.802527924254 | Tgfb2 |
| **ENSMUSG00000039242** | 0.0762975303756356 | 0.746370480989462 | 0.956085046483146 | 693.898534325027 | 866.538335564846 | 692.804978490663 | 642.47978556684 | 510.250519251699 | 486.786728365396 | 886.998533153828 | 861.907298346795 | B3galnt2 |
| **ENSMUSG00000039244** | -0.0859801116482876 | 0.430638424002748 | 0.886017087288711 | 397.54603529038 | 364.043830824705 | 380.474865236676 | 396.742991671202 | 454.407651264925 | 384.047820871192 | 399.773986773556 | 394.045779974668 | E130309D02R |
| **ENSMUSG00000039246** | 0.63852504802431 | 0.0781540104841146 | 0.739855948845396 | 28.0089252136404 | 34.2054606144018 | 26.5007368821565 | 26.2619474392285 | 20.8042057205628 | 18.3462334811079 | 26.0269522639034 | 8.68420451734806 | Lyplal1 |
| **ENSMUSG00000039252** | -0.29417199284802 | 0.346921760123225 | 0.861266047293794 | 42.4651446787451 | 34.2054606144018 | 31.2330113253987 | 19.6964605794214 | 42.7033696369448 | 30.5770558018465 | 34.3555769883525 | 48.8486504100828 | Lgi2 |
| **ENSMUSG00000039253** | -0.221908068760752 | 0.264629196111405 | 0.842510961779757 | 65.9565013095403 | 67.5965054998893 | 57.7337482075553 | 62.8410885152968 | 71.1722827282413 | 63.6002760678407 | 89.5327157878278 | 71.6446872681215 | Fn3krp |
| **ENSMUSG00000039254** | 0.0100975956532721 | 0.959432103313482 | 0.99332083910315 | 178.895715880671 | 229.665235553841 | 236.613722162112 | 151.006197775564 | 201.472308030714 | 182.239252579005 | 235.283648465687 | 171.513039217624 | Pomt1 |
| **ENSMUSG00000039262** | 0.130618278870742 | 0.646915878362264 | 0.935063240511793 | 2106.99398703901 | 3241.37460107903 | 3022.03045945449 | 1772.68145214792 | 1946.83567216635 | 1410.21381358116 | 2973.31902662833 | 2934.17560129898 | Prrc2b |
| **ENSMUSG00000039263** | 0.105507841079171 | 0.222532304318968 | 0.832048466795696 | 2086.21317155792 | 2156.57285016514 | 2385.06631939409 | 2423.60257796309 | 2104.5096523643 | 2086.578287918 | 2133.16900754952 | 2087.46566085754 | Npepl1 |
| **ENSMUSG00000039270** | -0.205913032906903 | 0.146270168788693 | 0.780607990324772 | 338.817643713392 | 422.681763306537 | 352.081218577222 | 296.384835385579 | 396.374866886513 | 393.832478727783 | 398.732908683 | 438.552328126077 | Megf9 |
| **ENSMUSG00000039275** | 0.205359840409995 | 0.155847086976009 | 0.787275540343856 | 658.661499378834 | 653.161414589292 | 734.448993591195 | 586.204183911351 | 523.390017601528 | 480.671317205027 | 662.125665593703 | 615.492995167044 | Foxk2 |
| **ENSMUSG00000039285** | 0.217177458235898 | 0.244261221562462 | 0.832048466795696 | 1443.81491907733 | 1537.61689619025 | 1515.27427672616 | 1460.35186295996 | 1053.34978437797 | 955.227223249684 | 1543.91880829475 | 1570.75549207533 | Azi2 |
| **ENSMUSG00000039286** | -0.0322432402637562 | 0.866772207414619 | 0.974868041734905 | 544.818771091134 | 904.001459094905 | 753.378091364164 | 537.431995809926 | 704.058119911679 | 680.033721033066 | 751.658381381531 | 666.512696706464 | Fndc3b |
| **ENSMUSG00000039294** | -0.520328086087182 | 0.0159912947518901 | 0.63777231095375 | 126.491920319666 | 120.533527879321 | 108.842312194571 | 128.495957113368 | 208.042057205628 | 218.931719541221 | 135.340151772298 | 133.519644454226 | BC017643 |
| **ENSMUSG00000039298** | 0.173298602293411 | 0.231803744093908 | 0.832048466795696 | 263.826005238161 | 258.169786065842 | 261.221549266971 | 230.729966787508 | 206.947099009809 | 190.800828203522 | 258.187366457922 | 242.072200921077 | Cdk5rap2 |
| **ENSMUSG00000039304** | -0.168926806354118 | 0.460146303292909 | 0.894774044236998 | 187.027339329792 | 157.182235680465 | 147.646962629158 | 241.985087118606 | 258.410134213307 | 223.824048469516 | 155.120635492864 | 187.795922687652 | Tnfsf10 |
| **ENSMUSG00000039307** | -0.022342883105678 | 0.895850041406374 | 0.983975729490316 | 112.939214571131 | 99.3587189275481 | 108.842312194571 | 106.92364314543 | 97.4512794278996 | 108.854318654574 | 130.134761319517 | 97.6973008201657 | Hexdc |
| **ENSMUSG00000039308** | -0.240858324227826 | 0.183786256389784 | 0.805963874499992 | 110.228673421423 | 127.048853710635 | 138.182413742673 | 153.819977858338 | 173.003394939417 | 179.793088114857 | 138.463386043966 | 134.605170018895 | Ndst2 |
| **ENSMUSG00000039316** | -0.158007323210367 | 0.360305191044636 | 0.868408659311123 | 901.706689135907 | 989.515110630909 | 775.146553803078 | 876.023532437123 | 1136.56660726022 | 1179.0512717192 | 772.479943192653 | 866.249400605469 | Rftn1 |
| **ENSMUSG00000039318** | 0.174600128390764 | 0.353615387773333 | 0.86286441023679 | 527.652010476322 | 618.95595397489 | 586.802030962037 | 440.825546301336 | 464.262275027297 | 355.916929533493 | 578.839418349212 | 526.479898864226 | Rab3gap2 |

| | | | | | | | | | | | | |
|---|---|---|---|---|---|---|---|---|---|---|---|---|
| **ENSMUSG00000039321** | -0.237429574678363 | 0.739125801025656 | 0.954383574136061 | 5.42108229941427 | 3.25766291565731 | 3.78581955459379 | 25.3240207449704 | 8.75966556655277 | 11.0077400886647 | 9.36970281500523 | 15.1973579053591 | Uts2r |
| **ENSMUSG00000039323** | -0.49363576345326 | 0.132138717298837 | 0.77303252008843 | 127.395434036235 | 147.409246933493 | 71.930571537282 | 98.4823028971069 | 84.3117810780705 | 136.985209992272 | 206.133461930115 | 199.736703899005 | Igfbp2 |
| **ENSMUSG00000039328** | 0.0702943696802135 | 0.767534785437801 | 0.95879680569409 | 84.9302893574902 | 92.0289773673191 | 61.519567762149 | 105.985716451172 | 96.3563212320805 | 92.9542496376133 | 63.5057635239243 | 75.9867895267955 | Rnf122 |
| **ENSMUSG00000039345** | -0.00494767623122291 | 0.971303100736378 | 0.995122272737203 | 765.276117933981 | 567.647763053287 | 626.553136285272 | 715.638067718977 | 653.690042904001 | 746.080161565054 | 676.700758861489 | 607.894316214364 | Mettl22 |
| **ENSMUSG00000039347** | -0.136754733858757 | 0.527713891165389 | 0.908232038364594 | 185.220311896654 | 201.975100770753 | 189.290977729689 | 208.219726125312 | 214.611806380543 | 308.216722482613 | 163.449260217313 | 178.026192605635 | Atp6v0e2 |
| **ENSMUSG00000039349** | 0.0283011178130442 | 0.792183070611845 | 0.961804768755116 | 826.715050660676 | 798.127414336042 | 758.110365807406 | 906.975113347642 | 756.616113310996 | 771.764888438605 | 810.99983254323 | 885.788860769502 | C130074G19 |
| **ENSMUSG00000039354** | 0.0296159757982055 | 0.808396635539114 | 0.964026716319739 | 558.37147683967 | 637.687515739919 | 664.41133183121 | 644.355638955357 | 605.511882287961 | 530.817688720055 | 681.906149314269 | 633.946929766408 | Smarcal1 |
| **ENSMUSG00000039356** | -0.0929327462196196 | 0.719332306659021 | 0.951674101666722 | 143.658680934478 | 142.522752560008 | 131.557229522134 | 106.92364314543 | 143.439523652302 | 209.14706168463 | 100.984574783945 | 107.467030902182 | Exosc2 |
| **ENSMUSG00000039357** | -0.675560215305145 | 0.0238811295608437 | 0.654124840968651 | 85.8338030740593 | 100.173134656462 | 103.163582862681 | 117.24083678227 | 228.846262926191 | 210.370143916704 | 104.107809055614 | 106.381505337514 | Fut11 |
| **ENSMUSG00000039361** | 0.14031943103586 | 0.398355533111677 | 0.880459952732461 | 2900.27903018663 | 3996.33798178261 | 4521.21500307363 | 3233.03331510788 | 3508.24605940439 | 3646.00813381218 | 3108.65917840062 | 3030.78737655447 | Picalm |
| **ENSMUSG00000039367** | 0.122644393602411 | 0.34699577566084 | 0.861266047293794 | 1629.93874469056 | 1877.22825514753 | 1953.48289017039 | 1621.67525437236 | 1585.49946754605 | 1374.74442885102 | 1743.80580168153 | 1799.80138622039 | Sec24c |
| **ENSMUSG00000039376** | 0.522592691594701 | 0.0384876987233328 | 0.679566075569698 | 5249.41469326615 | 4972.82244075089 | 3967.53889321429 | 6449.18394971912 | 2471.3206479637 | 2865.68166974905 | 3834.29060751825 | 5195.32535250348 | Synpo2l |
| **ENSMUSG00000039377** | -0.647958787957457 | 0.0630488417045248 | 0.715615581523459 | 149.079763233892 | 83.8848200781758 | 71.930571537282 | 128.495957113368 | 227.751304730372 | 231.162541861959 | 121.806136595068 | 98.7828263848342 | Hlx |
| **ENSMUSG00000039382** | 0.32650653512702 | 0.0213762353640323 | 0.648762042745232 | 867.373167906283 | 846.992358070902 | 1111.13803927328 | 1069.2364314543 | 888.011096809288 | 748.526326029202 | 744.370834747638 | 724.045551633895 | Wdr45 |
| **ENSMUSG00000039384** | 0.166227030797511 | 0.682422727771545 | 0.944452327903597 | 46.0791995450213 | 46.4216965481167 | 117.360406192407 | 145.378637610015 | 56.937826182593 | 73.3849339244316 | 100.984574783945 | 84.6709940441436 | Dusp10 |
| **ENSMUSG00000039395** | 0.372270025747411 | 0.0630509658952573 | 0.715615581523459 | 1361.59517086955 | 1314.46698646773 | 1167.92533259218 | 1358.11785328582 | 932.904382837871 | 684.926049961361 | 1070.22827709171 | 1329.76881671892 | Mreg |
| **ENSMUSG00000039405** | -0.772740242595773 | 0.0550159237343958 | 0.703396849069134 | 128.298947752804 | 108.317291945606 | 106.949402417274 | 128.495957113368 | 249.650468646754 | 359.586176229715 | 89.5327157878278 | 108.552556466851 | Prss23 |
| **ENSMUSG00000039410** | -0.167227583987896 | 0.304441191139579 | 0.851785577030345 | 206.904641094311 | 250.025628776699 | 240.399541716705 | 238.233380341573 | 309.873169416804 | 284.978160073209 | 209.256696201783 | 247.49982874442 | Prdm16 |
| **ENSMUSG00000039414** | -0.0584865103538149 | 0.786744549177307 | 0.961141387070613 | 735.460165287203 | 898.300548992504 | 848.970035117657 | 615.279911433354 | 769.755611660825 | 557.72549782568 | 1059.81749618615 | 838.025735924088 | Heatr5b |
| **ENSMUSG00000039427** | -0.223093112273886 | 0.252948691371899 | 0.837406774112203 | 397.54603529038 | 365.672662282534 | 348.295399022628 | 463.335786963532 | 535.434557755538 | 562.617826753975 | 391.445362049107 | 349.539231823259 | Alg1 |
| **ENSMUSG00000039428** | 0.139425453653696 | 0.385858185994582 | 0.875126835008975 | 1064.33915811834 | 1177.64514401012 | 1281.49991923 | 962.312788308873 | 1081.81869746927 | 779.103381831049 | 1113.95355689507 | 1096.38082031519 | Tmem135 |
| **ENSMUSG00000039438** | -0.117270833988826 | 0.77647150117993 | 0.959733396167384 | 44.2721721118832 | 95.2866402829764 | 52.0550188756646 | 63.779015209555 | 41.6084114411257 | 57.4848649074714 | 46.8485140750261 | 131.348593324889 | Ttc36 |
| **ENSMUSG00000039449** | 0.151898973249118 | 0.416888321571822 | 0.882192126163777 | 981.215896193983 | 1110.04863851023 | 1016.49255040843 | 964.18864169739 | 846.402685368162 | 638.448925142555 | 1112.91247880451 | 1065.98610450447 | Prpf18 |
| **ENSMUSG00000039450** | -0.0374175077593936 | 0.803401484043108 | 0.963461896828986 | 271.054114970713 | 253.283291692356 | 296.240380146964 | 292.633128608546 | 328.487458745729 | 315.555215875056 | 240.489038918468 | 258.355084391105 | Dcxr |
| **ENSMUSG00000039452** | -0.442858339230826 | 0.43764971928886 | 0.888722970093854 | 10.8421645988285 | 4.88649437348597 | 6.62518422053913 | 19.6964605794214 | 21.8991639163819 | 17.123151249034 | 10.4107809055614 | 7.59867895267955 | Snx22 |
| **ENSMUSG00000039456** | -0.0171805983468106 | 0.932976611591696 | 0.98813739334255 | 214.132750826864 | 244.324718674299 | 245.131816159948 | 170.702658354985 | 251.840385038392 | 157.777607937528 | 254.023054095697 | 220.361689627707 | Morc3 |
| **ENSMUSG00000039457** | -0.340141769813074 | 0.289025901592141 | 0.851785577030345 | 111.132187137993 | 128.677685168464 | 166.576060402127 | 96.6064495085906 | 216.801722772181 | 193.24699266767 | 149.915245040084 | 77.072315091464 | Ppl |
| **ENSMUSG00000039458** | -0.112976376814402 | 0.586887326108128 | 0.923058967588378 | 140.044626068202 | 189.758864837039 | 185.505158175096 | 114.427056699496 | 192.712642464161 | 174.900759186562 | 169.69572876065 | 144.374900100912 | Mtmr12 |
| **ENSMUSG00000039461** | 0.218299314403449 | 0.133008067570123 | 0.77303252008843 | 1623.61414867457 | 1426.04194132899 | 1615.5984949229 | 1786.7503525618 | 1174.89014411389 | 1243.87463001912 | 1488.74166949528 | 1636.97255152011 | Tcta |
| **ENSMUSG00000039463** | 0.140515468081095 | 0.340568917788544 | 0.859077629258968 | 503.257140128958 | 644.202841571234 | 642.642869392295 | 516.797608536247 | 497.11102090187 | 463.548165955993 | 601.743136341447 | 529.736475558232 | Slc9a8 |
| **ENSMUSG00000039470** | -0.259947092054399 | 0.185840779471635 | 0.805963874499992 | 79.509207058076 | 70.0397526866323 | 70.0376617599851 | 87.227182566009 | 110.590777777729 | 95.400414101761 | 84.3273253350471 | 77.072315091464 | Zdhhc2 |
| **ENSMUSG00000039473** | -0.0192334918305634 | 0.900947677868021 | 0.984802943363903 | 655.047444512558 | 767.179616637298 | 772.307189137133 | 649.045272426648 | 759.900987898453 | 556.502415593606 | 737.083288113745 | 827.170480277403 | Ubn1 |
| **ENSMUSG00000039474** | 0.0640083185699221 | 0.65380416754438 | 0.937077409179068 | 2702.40952625801 | 3585.05803868087 | 3513.24054666303 | 2846.60751707352 | 2683.74253795261 | 2868.1278342132 | 3390.79134094134 | 3155.62281649135 | Wfs1 |
| **ENSMUSG00000039477** | -0.154876588779649 | 0.4041656308785 | 0.881158207349486 | 840.267756409212 | 946.35107699845 | 953.080072868986 | 759.720622349111 | 948.233797579338 | 686.149132193435 | 1161.84314906065 | 1098.55187144453 | Tnrc18 |
| **ENSMUSG00000039478** | 0.314168063338189 | 0.0989554973305702 | 0.759800953500741 | 128.298947752804 | 135.193010999779 | 150.486327295103 | 123.806323642077 | 82.1218646864323 | 100.292743030056 | 131.175839410073 | 118.322286548867 | Micu3 |
| **ENSMUSG00000039480** | 0.0823095139601524 | 0.622248706763335 | 0.93078489223231 | 172.571119864688 | 212.56250524664 | 248.917635714542 | 189.461192240149 | 190.522726072523 | 172.454594722414 | 219.667477107345 | 194.309076075663 | Nt5dc1 |
| **ENSMUSG00000039481** | -0.407750368230938 | 0.206981789487063 | 0.819960256425221 | 674.924746277077 | 531.813470981057 | 520.550188756646 | 713.762214330461 | 731.432074807157 | 1442.01395161508 | 586.126964983105 | 479.80229958348 | Nrtn |
| **ENSMUSG00000039483** | -0.183691818417367 | 0.327459653863733 | 0.855977678491063 | 137.334084918495 | 144.151584017836 | 173.201244622666 | 180.081925297567 | 217.896680968 | 194.470074899744 | 143.668776496747 | 164.999885829613 | Asb6 |
| **ENSMUSG00000039485** | -0.313736279602352 | 0.215371762629814 | 0.824338622350122 | 155.404359249876 | 171.841718800923 | 218.631079277791 | 187.585338851632 | 170.813478547779 | 170.008430258266 | 338.350379430744 | 231.216945274392 | Tspyl4 |
| **ENSMUSG00000039488** | -0.0190399876614488 | 0.939395358321934 | 0.989269716906553 | 65.0529875929712 | 83.8848200781758 | 70.9841166486335 | 60.0273084325223 | 49.2731188118594 | 61.154111603693 | 90.5737938783839 | 82.4999429148066 | Cntn5 |
| **ENSMUSG00000039496** | 0.302751875682397 | 0.198027901048508 | 0.812921911423739 | 760.758549351136 | 925.176268046677 | 963.491076644119 | 785.982569788339 | 502.585811880965 | 501.463715150282 | 893.245001697165 | 886.874386334171 | Cdnf |
| **ENSMUSG00000039497** | -0.487327683193588 | 0.170928094238306 | 0.799531027639052 | 39.754603529038 | 64.338842584232 | 42.5904699891801 | 61.9031618210386 | 84.3117810780705 | 105.185071958352 | 33.3144988977964 | 70.559161703453 | Dse |
| **ENSMUSG00000039501** | -0.0247450838702733 | 0.854533037881754 | 0.973268281968411 | 1049.88293865323 | 1295.7354247027 | 1185.9079754765 | 986.698882359586 | 1298.62042024145 | 1020.0505815496 | 1107.70708835173 | 1170.19655871265 | Znfx1 |
| **ENSMUSG00000039509** | 0.0746918669000437 | 0.604219801012017 | 0.926448151609826 | 371.344137509877 | 376.26006675842 | 439.155068332879 | 362.03970398365 | 416.084114411257 | 298.432064626022 | 377.911346871878 | 376.677370939972 | Nup133 |
| **ENSMUSG00000039512** | 0.0767865030270106 | 0.788891675922976 | 0.961659329450003 | 368.63359636017 | 443.856572258309 | 374.796135904785 | 317.019222659259 | 278.119381738051 | 195.693157131818 | 462.238672206925 | 489.572029665497 | Uhrf1bp1 |
| **ENSMUSG00000039515** | -0.0732212113756948 | 0.450849749310777 | 0.89108031640712 | 1520.6135849857 | 1466.76272777471 | 1308.9471110008 | 1509.12405106138 | 1391.69186688607 | 1576.5529971432 | 1556.41174538142 | 1583.78179885135 | Ptpa |
| **ENSMUSG00000039519** | -0.506379706215786 | 0.258506873285335 | 0.838547517499604 | 38.8510898124689 | 71.6685841444609 | 28.3936466594534 | 43.1446279358754 | 32.8487458745729 | 68.4926049961361 | 39.5609674411332 | 118.322286548867 | Cyp7b1 |
| **ENSMUSG00000039523** | 0.231375563284354 | 0.372740648540071 | 0.873700364743909 | 1383.27950006721 | 1666.29458135872 | 1516.22073161481 | 1195.85653517916 | 879.251431242735 | 804.7881087046 | 1623.04074317702 | 1600.06468232138 | Cep104 |
| **ENSMUSG00000039529** | -0.249860894759352 | 0.16144283732259 | 0.792786631432503 | 70.4740698923855 | 79.8127414336042 | 72.8770264259304 | 85.3513291774927 | 95.2613630362614 | 85.6157562451702 | 102.025652874501 | 83.5854684794751 | Atp8b1 |
| **ENSMUSG00000039530** | 0.109686189057634 | 0.47321154962204 | 0.89728597737823 | 225.878429142261 | 246.767965861042 | 272.579007930753 | 277.626301500416 | 275.929465346412 | 228.716377397812 | 197.804837205666 | 245.328777615083 | Tusc3 |
| **ENSMUSG00000039531** | -0.262469398555206 | 0.310228918836977 | 0.851785577030345 | 43.3686583953142 | 37.4631235300591 | 60.5731128735006 | 53.4618215727152 | 75.5521155115177 | 47.7002070508805 | 52.0539045278068 | 57.5328549274309 | Zufsp |
| **ENSMUSG00000039536** | 0.0762168945336932 | 0.349261333513504 | 0.861774532843376 | 1385.08652750035 | 1536.80248046134 | 1410.21778408619 | 1420.95894180111 | 1422.35069636901 | 1331.93655072843 | 1390.880328983 | 1312.40040768423 | Stau1 |
| **ENSMUSG00000039542** | -1.49967813749731 | 0.00581643530346587 | 0.635862253526407 | 17.1667606148119 | 32.5766291565731 | 19.8755526616174 | 22.5102406621959 | 49.2731188118594 | 150.439114545085 | 22.903717992235 | 39.0789203280663 | Ncam1 |
| **ENSMUSG00000039556** | 0.641052029197159 | 0.157877951852792 | 0.789230661864335 | 28.9124389302094 | 36.6487078011448 | 60.5731128735006 | 20.6343872736795 | 21.8991639163819 | 17.123151249034 | 40.6020455316893 | 14.1118323406906 | Ppp1r3f |
| **ENSMUSG00000039568** | -0.130067013625544 | 0.449977402914579 | 0.89108031640712 | 533.073092775737 | 570.905425968944 | 636.017685171756 | 670.617586394585 | 652.595084708182 | 852.48831575548 | 591.332355435886 | 542.762782334254 | Ubald1 |
| **ENSMUSG00000039578** | -0.446271025162844 | 0.278896819862706 | 0.848075053184273 | 20.780815481088 | 15.4738988493722 | 20.8220075502658 | 15.9447538023887 | 33.943704070392 | 9.78465785659088 | 32.2734208072402 | 22.7960368580387 | Ccser1 |
| **ENSMUSG00000039585** | 0.159679058932691 | 0.537268836667055 | 0.912099403004057 | 280.992765852973 | 434.897999240251 | 448.619617219364 | 304.826175633902 | 281.404256325508 | 216.485555077073 | 395.609674411332 | 421.183919091381 | Myo9a |
| **ENSMUSG00000039599** | 0.0913228691141843 | 0.467642928437522 | 0.895328112556867 | 608.968244967536 | 645.017257300148 | 678.608155160936 | 656.548685980713 | 533.2446413639 | 563.840908986049 | 714.17957012151 | 617.664046296381 | Fam149b |
| **ENSMUSG00000039601** | 0.16980683199857 | 0.376509720206088 | 0.87422797717331 | 16477.3796490697 | 18004.2885191091 | 17161.1200409736 | 15525.5005700553 | 11805.8392673215 | 11389.3417450718 | 18867.4582351489 | 17646.3035792513 | Rcan2 |
| **ENSMUSG00000039607** | -0.487670760061881 | 0.253232691446136 | 0.837406774112203 | 72.2810973255236 | 88.7713144516618 | 59.6266579848522 | 95.6685228143324 | 125.920192519196 | 209.14706168463 | 55.1771387994752 | 54.2762782334254 | Rbms3 |
| **ENSMUSG00000039611** | -0.206900904612382 | 0.0351377130581161 | 0.679566075569698 | 1088.7340284657 | 917.846526486448 | 1023.11773462897 | 951.995594672034 | 1216.49855555502 | 1158.25887377394 | 1133.73404061563 | 1086.61109023318 | Tmem246 |
| **ENSMUSG00000039615** | -0.127494308927554 | 0.306460470682169 | 0.851785577030345 | 2012.12504679926 | 1776.24070476215 | 1725.38726200612 | 2089.70067480718 | 2158.16260395944 | 2359.32562567048 | 1932.24093607219 | 1857.33424114782 | Stub1 |
| **ENSMUSG00000039616** | 0.0955438368122807 | 0.576439207020456 | 0.922450244597442 | 144.562194651047 | 114.83261777692 | 162.790240847533 | 138.813150750208 | 133.58489988993 | 147.992950080937 | 121.806136595068 | 121.578863242873 | Mocos |
| **ENSMUSG00000039620** | -0.19254087733782 | 0.627927693747601 | 0.931825800800958 | 10.8421645988285 | 29.3189662409158 | 17.036187995672 | 20.6343872736795 | 25.1840385038392 | 19.5693157131818 | 28.1091084450157 | 16.2828834700276 | 6430573F11R |
| **ENSMUSG00000039621** | -0.336486639025776 | 0.105478165613513 | 0.762094734041179 | 98.4829951060259 | 114.83261777692 | 141.021778408619 | 103.171936368398 | 174.098353135236 | 160.223772401676 | 111.395355689507 | 132.434118889558 | Prex1 |
| **ENSMUSG00000039623** | 0.0643381602966144 | 0.715268836422727 | 0.951176382858794 | 98.4829951060259 | 109.13170767452 | 128.717864856189 | 130.371810501884 | 113.875652365186 | 94.1773318696872 | 116.600746142287 | 120.493337678204 | Ap5z1 |
| **ENSMUSG00000039629** | -0.0623070450289351 | 0.823735391262619 | 0.966637987288264 | 1106.80430279708 | 1216.73709899801 | 752.431636475515 | 698.75538722233 | 822.313605060142 | 618.879609429373 | 1325.29240927796 | 1174.53866097133 | Strip2 |
| **ENSMUSG00000039630** | 0.218065756382266 | 0.0447154515892087 | 0.693563614452513 | 4457.0331638351 | 4960.60620481718 | 5424.13296684425 | 5038.54220155484 | 4522.17734873287 | 3747.52395907431 | 4451.64991521804 | 4368.15487222607 | Hnrnpu |
| **ENSMUSG00000039633** | 0.31463383571334 | 0.161382996303912 | 0.792786631432503 | 319.843855665442 | 335.539280312703 | 261.221549266971 | 249.488500672671 | 220.086597359638 | 156.554525705454 | 248.817663642917 | 311.545837059862 | Lonrf1 |
| **ENSMUSG00000039634** | 0.039652729848995 | 0.862700615013824 | 0.974868041734905 | 93.0619128066116 | 153.110157035894 | 118.306861081056 | 88.1651092602672 | 117.160526952643 | 118.638976511164 | 89.5327157878278 | 116.15123541953 | Zfp189 |
| **ENSMUSG00000039637** | -0.0548350390270383 | 0.720853179446744 | 0.951674101666722 | 172.571119864688 | 171.027303072009 | 187.398067952392 | 148.192417692789 | 180.668102310151 | 203.031650524261 | 168.654650670094 | 154.144630182928 | Coro7 |
| **ENSMUSG00000039639** | -0.94049332654288 | 0.0207354931078342 | 0.648684683035624 | 42.4651446787451 | 13.8450673915436 | 28.3936466594534 | 29.075727522003 | 36.1336204620302 | 86.838838477244 | 55.1771387994752 | 40.1644458927348 | Kcne1 |
| **ENSMUSG00000039640** | 0.0300283317885773 | 0.818844372170935 | 0.966315773308182 | 4580.81454300506 | 3531.30660057253 | 3194.28524918851 | 3955.23686968667 | 3680.15449614798 | 3796.44724835726 | 3682.29320629706 | 3788.48422069309 | Mrpl12 |
| **ENSMUSG00000039646** | -0.633095583937693 | 0.205770164405671 | 0.819960256425221 | 133.720030052219 | 181.614707547895 | 119.253315969704 | 61.9031618210386 | 293.448796479518 | 315.555215875056 | 98.902418602833 | 62.9604827507734 | Vasn |
| **ENSMUSG00000039648** | 0.104287464419301 | 0.298242406731136 | 0.851785577030345 | 587.283915769879 | 548.916201288258 | 536.639921863669 | 528.052728867345 | 549.669014301187 | 489.232892829544 | 483.060234018047 | 525.394373299558 | Kyat1 |
| **ENSMUSG00000039652** | 0.041903523321448 | 0.845980748396437 | 0.972180580805179 | 870.987222772559 | 989.515110630909 | 883.042411109001 | 824.437564252924 | 699.678287128403 | 598.087211484117 | 1083.76229226894 | 1083.35451353917 | Cpeb3 |
| **ENSMUSG00000039653** | -0.578589725280642 | 0.0833596719925856 | 0.742148948523533 | 46.0791995450213 | 63.5244268553176 | 38.8046504345863 | 60.9652351267805 | 56.937826182593 | 59.9310293716191 | 64.5468416144805 | 131.348593324889 | Baat |
| **ENSMUSG00000039656** | -0.0393442806190947 | 0.698758842172469 | 0.945141109561958 | 1295.63866956001 | 1439.07259299162 | 1498.23808873049 | 1354.36614650878 | 1597.54400770006 | 1342.9442908171 | 1397.12679752634 | 1403.58455511638 | Rxrb |
| **ENSMUSG00000039660** | -0.139640190502216 | 0.438504437033302 | 0.889346190988072 | 134.623543768788 | 131.935348084121 | 120.199770858353 | 156.633757941113 | 165.338687568684 | 176.123841418636 | 136.381229862854 | 121.578863242873 | Spout1 |
| **ENSMUSG00000039661** | 0.0337780608974283 | 0.955767597270475 | 0.992371505557225 | 7.22810973255236 | 4.88649437348597 | 11.3574586637814 | 20.6343872736795 | 9.85462376237187 | 19.5693157131818 | 8.32862472444909 | 5.42762782334254 | Dusp26 |
| **ENSMUSG00000039662** | 0.0475825403819196 | 0.728263023051614 | 0.952401994188708 | 999.286170525364 | 1027.79264988988 | 930.365155541423 | 925.733647232805 | 827.788396039237 | 802.341944240452 | 1053.57102764281 | 1072.49925789249 | Icmt |
| **ENSMUSG00000039670** | -0.0290524337382022 | 0.852978765111816 | 0.973120292022893 | 483.379838364439 | 373.002403842763 | 407.922057007481 | 557.128456389348 | 485.06648074786 | 491.679057293692 | 434.129563761909 | 447.236532643425 | Oxld1 |
| **ENSMUSG00000039671** | 0.169147632399296 | 0.225246150668122 | 0.832048466795696 | 727.328541838081 | 702.840774053066 | 743.913542477679 | 680.934780031425 | 676.684165016202 | 494.125221757839 | 718.343882483734 | 648.058762107099 | Zmynd8 |
| **ENSMUSG00000039678** | -0.0252618649654873 | 0.830666563638219 | 0.967583158900775 | 393.931980424104 | 345.312269059675 | 418.333060782613 | 341.405316709971 | 393.089992299056 | 366.924669622158 | 375.829190690765 | 388.618152151326 | Tbc1d13 |
| **ENSMUSG00000039680** | 0.0678232465386769 | 0.612053440433977 | 0.928055903343287 | 766.17963165055 | 614.883875330318 | 579.230391852849 | 751.279282100787 | 656.974917491458 | 692.264543353805 | 620.482541971457 | 617.664046296381 | Mrps6 |
| **ENSMUSG00000039682** | 0.18492729329751 | 0.0664123119063996 | 0.719725956118125 | 2029.29180741408 | 1929.35086179804 | 2018.78827748714 | 2125.34188918899 | 1739.88857315654 | 1571.66066821491 | 1868.73517254827 | 1945.26181188597 | Lap3 |
| **ENSMUSG00000039683** | -1.39266261667316 | 0.0333034139652615 | 0.671606027118506 | 6.32459601598331 | 9.77298874697194 | 1.89290977729689 | 7.50341355406529 | 10.949581958191 | 35.4693847301419 | 7.28754663389296 | 14.1118323406906 | Sdk1 |

| | | | | | | | | | | | | |
|---|---|---|---|---|---|---|---|---|---|---|---|---|
| **ENSMUSG00000039684** | 0.419860546994715 | 0.324183442358427 | 0.855977678491063 | 11.7456783153976 | 17.9171460361152 | 17.036187995672 | 12.1930470253561 | 8.75966556655277 | 9.78465785659088 | 15.616171358342 | 9.76973008201657 | Gm5422 |
| **ENSMUSG00000039686** | 0.102750656020068 | 0.569277289312702 | 0.920872458148558 | 572.827696304775 | 706.098436968723 | 766.628459805242 | 630.286738541484 | 665.734583058011 | 458.655837027697 | 750.617303290975 | 615.492995167044 | Zer1 |
| **ENSMUSG00000039693** | 0.0738216775953654 | 0.572384494063431 | 0.921574199900435 | 375.861706092723 | 395.806044252364 | 353.027673465871 | 349.846656958294 | 307.683253025166 | 321.670627035425 | 396.650752501888 | 374.506319810635 | Msantd3 |
| **ENSMUSG00000039697** | -0.392949606674049 | 0.241707405909893 | 0.832048466795696 | 28.0089252136404 | 33.3910448854875 | 48.2691993210708 | 26.2619474392285 | 31.7537876787538 | 62.3771938357668 | 52.0539045278068 | 32.5657669400552 | Ncoa7 |
| **ENSMUSG00000039699** | 0.101812748061905 | 0.809903276840166 | 0.964412705343932 | 22.5878429142261 | 13.0306516626293 | 14.1968233297267 | 22.5102406621959 | 27.3739548954774 | 11.0077400886647 | 14.5750932677859 | 14.1118323406906 | Batf2 |
| **ENSMUSG00000039701** | 0.366715273804826 | 0.254394590143535 | 0.837406774112203 | 112.035700854562 | 156.367819951551 | 229.988537941573 | 167.888878272211 | 101.831112211176 | 75.8310983885793 | 191.558368662329 | 146.545951230249 | Usp53 |
| **ENSMUSG00000039703** | 0.235882234987233 | 0.181959962154926 | 0.803888000741704 | 1811.54500172094 | 2265.70455783966 | 2434.2819736038 | 1979.02532488472 | 1762.88269526875 | 1340.49812635295 | 1962.43220069832 | 2142.82746465563 | Nploc4 |
| **ENSMUSG00000039704** | 0.0524776930661166 | 0.92320448628806 | 0.986713918026868 | 46.9827132615903 | 128.677685168464 | 127.77140996754 | 50.6480414899407 | 40.5134532453066 | 29.3539735697726 | 156.16171358342 | 115.065709854862 | Lmbrd2 |
| **ENSMUSG00000039706** | -0.216555600650118 | 0.561013625088698 | 0.916762264997884 | 81.316234491214 | 83.8848200781758 | 72.8770264259304 | 87.227182566009 | 136.869774477387 | 140.654456688494 | 44.7663578939139 | 56.4473293627624 | Ldb2 |
| **ENSMUSG00000039713** | -0.31878905606273 | 0.122503009015117 | 0.770251097497048 | 321.65088309858 | 352.642010619904 | 313.276568142636 | 338.591536627196 | 546.384139713729 | 469.663577116362 | 339.391457521301 | 299.605055848508 | Plekhg5 |
| **ENSMUSG00000039715** | 0.214367285241262 | 0.107514325152198 | 0.762287160109383 | 193.351935345776 | 197.903022126182 | 212.005895057252 | 190.399118934407 | 171.908436743598 | 174.900759186562 | 151.997401221196 | 185.624871558315 | Wdr34 |
| **ENSMUSG00000039725** | -0.0943621699889184 | 0.464781327009821 | 0.895328112556867 | 261.115464088454 | 240.252640029727 | 238.506631939409 | 245.736793895638 | 301.113503850252 | 267.855008824175 | 239.447960827911 | 244.243252050414 | Trp53rka |
| **ENSMUSG00000039735** | 0.129902216978531 | 0.346137653992776 | 0.861266047293794 | 498.739571546113 | 544.029706914772 | 660.625512276616 | 487.721881014244 | 511.345477447518 | 452.540425867328 | 514.292576734731 | 523.223322170221 | Fnbp1l |
| **ENSMUSG00000039737** | -0.148028666791267 | 0.301467386838722 | 0.851785577030345 | 238.527621174228 | 210.119258059897 | 194.96970706158 | 220.412773150668 | 227.751304730372 | 282.531995609062 | 224.872867560126 | 223.618266321713 | Prkrip1 |
| **ENSMUSG00000039738** | 0.0884776957147616 | 0.680743159876507 | 0.944452327903597 | 71.3775836089546 | 86.3280672649188 | 77.6093008691726 | 56.2756016554897 | 66.7924499449649 | 57.4848649074714 | 73.9165444294857 | 75.9867895267955 | Slx4 |
| **ENSMUSG00000039740** | -0.303994423426417 | 0.206214078985501 | 0.819960256425221 | 105.711104838578 | 85.5136515360045 | 117.360406192407 | 116.302910088012 | 143.439523652302 | 179.793088114857 | 103.066730965058 | 98.7828263848342 | Alg2 |
| **ENSMUSG00000039741** | -0.27098207456384 | 0.163914180176279 | 0.793968226938511 | 175.281661014395 | 162.068730053951 | 214.845259723197 | 190.399118934407 | 193.80760065998 | 181.016170346931 | 291.501865355718 | 227.960368580387 | Bahcc1 |
| **ENSMUSG00000039745** | 0.00661983492788159 | 0.96152548366064 | 0.993822507448007 | 1116.74295367934 | 1108.4198070524 | 1071.38693395004 | 1342.17309948343 | 1219.78343014247 | 1331.93655072843 | 1042.11916864669 | 1024.73613304707 | Htatip2 |
| **ENSMUSG00000039747** | -0.776766875092635 | 0.111616640232636 | 0.764116514278315 | 11.7456783153976 | 19.5459774939439 | 12.3039135524298 | 11.2551203310979 | 44.893286028583 | 19.5693157131818 | 11.4518589961175 | 18.4539345993646 | Orai2 |
| **ENSMUSG00000039753** | 0.289809867981212 | 0.135536557254949 | 0.776011038858713 | 1073.37429528403 | 1086.43058237171 | 1330.71557343972 | 1082.36740517392 | 871.586723872001 | 653.125911927441 | 1043.16024673725 | 1171.28208427732 | Fbxl5 |
| **ENSMUSG00000039754** | -0.40317357733942 | 0.0931042753041212 | 0.754058637722175 | 56.9213641438498 | 53.7514381083457 | 31.2330113253987 | 55.3376749612315 | 66.7924499449649 | 64.8233582999146 | 64.5468416144805 | 65.1315338801105 | Alkbh4 |
| **ENSMUSG00000039756** | 0.169665949770547 | 0.0808932614299195 | 0.739855948845396 | 1002.90022539164 | 1036.75122290794 | 958.758802200877 | 1057.98131112321 | 924.144717271318 | 835.365164506446 | 986.942029847218 | 858.650721652789 | Dnttip2 |
| **ENSMUSG00000039759** | -0.119385511298221 | 0.700306653005265 | 0.94561797945347 | 161.728955265859 | 151.481325578065 | 163.736695736181 | 195.088752405698 | 213.516848184724 | 287.424324537357 | 105.14888714617 | 124.835439936878 | Thap3 |
| **ENSMUSG00000039763** | 0.244592374245117 | 0.432597286434288 | 0.887428195033564 | 899.899661702769 | 1211.03618889561 | 1248.3739981273 | 917.292306984482 | 729.242158415519 | 435.417274618294 | 1256.58125530126 | 1187.56496774735 | Dnajc28 |
| **ENSMUSG00000039765** | -0.296524103767186 | 0.291526697925176 | 0.851785577030345 | 66.8600150261093 | 132.749763813036 | 63.4124775394459 | 70.3445020693621 | 88.6916138613469 | 100.292743030056 | 99.9434966933891 | 121.578863242873 | Cc2d2a |
| **ENSMUSG00000039768** | 0.131992779716364 | 0.325273339901081 | 0.855977678491063 | 1656.14064247106 | 1711.087446449 | 1522.84591583535 | 1428.46235535518 | 1428.92044554392 | 1174.15894279091 | 1580.35654146422 | 1581.61074772202 | Dnajc11 |
| **ENSMUSG00000039770** | 0.129108205314948 | 0.332910292462419 | 0.856918526232803 | 1287.50704611089 | 977.298874697194 | 1142.37105059868 | 1261.51140377723 | 997.506916391197 | 1190.05901180787 | 974.449092760544 | 1107.23607596188 | Ypel5 |
| **ENSMUSG00000039771** | -0.0146991464501395 | 0.941798961569544 | 0.989934163630124 | 677.635287426784 | 487.835021619683 | 558.408384302584 | 744.71379524098 | 647.120293729086 | 796.226533080083 | 552.812466085309 | 498.256234182845 | Polr2j |
| **ENSMUSG00000039781** | 0.160257338527623 | 0.252064136550845 | 0.83723604018117 | 298.159526467785 | 288.303168035672 | 318.955297474527 | 304.826175633902 | 306.588294829347 | 226.270212933664 | 297.748333899055 | 250.756405438425 | Cep131 |
| **ENSMUSG00000039782** | 0.197660162318204 | 0.478038631957812 | 0.897674307047416 | 475.248214915318 | 597.781145023117 | 594.373670071225 | 461.459933575015 | 384.330326732503 | 256.847268735511 | 543.442763270303 | 670.854798965138 | Cpeb2 |
| **ENSMUSG00000039783** | -0.403219929040116 | 0.250245689653491 | 0.835923538390114 | 39.754603529038 | 95.2866402829764 | 53.9479286529615 | 48.7721881014244 | 62.4126171616885 | 78.277262852727 | 56.2182168900314 | 118.322286548867 | Kmo |
| **ENSMUSG00000039787** | -0.891587986671732 | 0.128725369649304 | 0.77303252008843 | 20.780815481088 | 28.5045505120015 | 18.9290977729689 | 25.3240207449704 | 53.6529515951357 | 92.9542496376133 | 13.5340151772298 | 14.1118323406906 | Cercam |
| **ENSMUSG00000039789** | 0.040033809420368 | 0.911567110962029 | 0.986226496507248 | 90.3513716569045 | 112.389370590177 | 119.253315969704 | 88.1651092602672 | 67.887408140784 | 45.2540425867328 | 136.381229862854 | 148.717002359586 | Zfp597 |
| **ENSMUSG00000039795** | 0.256572649321738 | 0.120898931332709 | 0.769868040577702 | 527.652010476322 | 518.782819318427 | 548.943835416099 | 561.818089860639 | 370.095870186855 | 371.816998550453 | 515.333654825288 | 547.104884592928 | Zfand1 |
| **ENSMUSG00000039801** | 0.215946203333011 | 0.327081487683502 | 0.855977678491063 | 103.000563688871 | 153.924572764808 | 172.254789734017 | 107.861569839689 | 99.6411958195378 | 125.977469903608 | 115.559668051731 | 121.578863242873 | 2410089E03 |
| **ENSMUSG00000039804** | 0.171610544157288 | 0.365563743049523 | 0.86924083181804 | 543.915257374565 | 690.624538119351 | 787.450467355508 | 466.149567046306 | 512.440435643337 | 495.348303989913 | 601.743136341447 | 599.210111697016 | Ncoa5 |
| **ENSMUSG00000039810** | -0.0465663744730102 | 0.761102650975067 | 0.957479700684068 | 192.448421629207 | 188.13003337921 | 198.755526616174 | 224.164479927701 | 232.131137513649 | 228.716377397812 | 197.804837205666 | 171.513039217624 | Zc3h10 |
| **ENSMUSG00000039813** | -0.445161070433194 | 0.290903568283729 | 0.851785577030345 | 18.9737880479499 | 13.8450673915436 | 29.3401015481019 | 32.8274342990356 | 39.4184950494875 | 46.4771248188067 | 27.0680303544596 | 16.2828834700276 | Tbc1d2 |
| **ENSMUSG00000039824** | 0.0577241622382153 | 0.854122481216673 | 0.97315735084075 | 87.6408305071974 | 74.1118313312039 | 58.6802030962037 | 92.854742731558 | 56.937826182593 | 121.085140975312 | 74.9576225200418 | 48.8486504100828 | Myl6b |
| **ENSMUSG00000039826** | 0.109974541140785 | 0.532742647180434 | 0.909082872831377 | 422.844419354313 | 395.806044252364 | 426.85115478045 | 467.087493740564 | 326.297542354091 | 321.670627035425 | 440.376032305246 | 497.170708618176 | Trub2 |
| **ENSMUSG00000039828** | 0.0314662564219754 | 0.808809883373439 | 0.964157363647118 | 224.071401709123 | 206.861595144239 | 183.612248397799 | 219.47484645641 | 192.712642464161 | 207.923979452556 | 206.133461930115 | 209.506433981022 | Wdr70 |
| **ENSMUSG00000039831** | -0.223654943618354 | 0.121626223303335 | 0.769868040577702 | 787.863960848207 | 671.892976354321 | 859.38103889279 | 912.602673513191 | 1089.48340484 | 951.557976553463 | 831.821394354353 | 899.900693110193 | Arhgap29 |
| **ENSMUSG00000039834** | 0.0722011960715809 | 0.72012424040277 | 0.951674101666722 | 141.85165350134 | 143.337168288922 | 147.646962629158 | 120.054616865045 | 122.635317931739 | 92.9542496376133 | 156.16171358342 | 153.05910461826 | Zfp335 |
| **ENSMUSG00000039835** | -0.141650944203323 | 0.529865077960332 | 0.908620485539384 | 193.351935345776 | 182.42912327681 | 145.754052851861 | 161.323391412404 | 158.768938393769 | 133.315963296051 | 250.899819824029 | 209.506433981022 | Nhsl1 |
| **ENSMUSG00000039840** | 0.0809260724102832 | 0.770175301966326 | 0.958992744413128 | 171.667606148119 | 249.211213047785 | 216.738169500494 | 142.564857527241 | 143.439523652302 | 127.200552135681 | 260.269522639034 | 206.249857287016 | Epg5 |
| **ENSMUSG00000039841** | -0.0906747488502821 | 0.644707651083271 | 0.934857223264556 | 72.2810973255236 | 76.5550785179469 | 63.4124775394459 | 76.9099889291692 | 82.1218646864323 | 63.6002760678407 | 71.8343882483734 | 90.0986218674861 | Zfp800 |
| **ENSMUSG00000039842** | 0.153413595388972 | 0.477874235017452 | 0.897674307047416 | 130.105975185942 | 174.284965987666 | 175.094154399963 | 147.254490998531 | 141.249607260664 | 95.400414101761 | 173.860041122875 | 151.973579053591 | Mcph1 |
| **ENSMUSG00000039844** | 0.229147289855011 | 0.186154250325649 | 0.806376426354616 | 5353.31877067159 | 5977.81145023117 | 5972.1303473717 | 5229.87924718351 | 4733.50428052596 | 3420.96100311059 | 5799.84604248824 | 5268.05556533627 | Rapgef1 |
| **ENSMUSG00000039849** | -0.155068865749737 | 0.408982215955844 | 0.882192126163777 | 529.45903790946 | 614.069459601404 | 630.338955839866 | 672.493439783102 | 707.342994499137 | 898.965440574287 | 589.250199254773 | 529.736475558232 | Pcif1 |
| **ENSMUSG00000039850** | -0.106536078741993 | 0.494120202736066 | 0.900141680336539 | 101.193536255733 | 118.090280692578 | 101.270673085384 | 113.489130005238 | 116.065568756824 | 121.085140975312 | 108.272121417838 | 122.664388807541 | Endov |
| **ENSMUSG00000039852** | 0.018551771654722 | 0.932486917317653 | 0.98804205366864 | 831.232619243521 | 1247.68489669675 | 1244.58817857271 | 853.513291774927 | 1131.09181628113 | 731.403174780168 | 1236.80077158069 | 1023.6506074824 | Rere |
| **ENSMUSG00000039853** | -0.408133719356046 | 0.323961188555592 | 0.855977678491063 | 16.2632468982428 | 34.2054606144018 | 28.3936466594534 | 16.8826804966469 | 28.4689130912965 | 53.8156182112498 | 21.8626399016789 | 23.8815624227072 | Trim14 |
| **ENSMUSG00000039873** | -0.109077352075986 | 0.458820592595281 | 0.894333713711794 | 791.478015714483 | 712.613762800038 | 684.286884492827 | 885.402799379704 | 872.68168206782 | 969.904210034571 | 750.617303290975 | 722.960026069226 | Neurl2 |
| **ENSMUSG00000039878** | 0.0362559053587873 | 0.936876087731302 | 0.988852035581444 | 18.0702743313809 | 13.0306516626293 | 6.62518422053913 | 15.9447538023887 | 13.1394983498292 | 18.3462334811079 | 12.4929370866736 | 8.68420451734806 | Slc39a5 |
| **ENSMUSG00000039879** | 0.308929376382578 | 0.254638488302214 | 0.837406774112203 | 806.837748896157 | 1050.59629029948 | 1241.74881390676 | 758.782695654852 | 640.550544554172 | 480.671317205027 | 959.873999492758 | 1032.33481199975 | Heca |
| **ENSMUSG00000039883** | -0.603818016595485 | 0.309613192091845 | 0.851785577030345 | 12.6491920319666 | 8.95857301805762 | 7.57163910918757 | 11.2551203310979 | 19.7092475247437 | 29.3539735697726 | 4.16431236222455 | 8.68420451734806 | Lrrc17 |
| **ENSMUSG00000039886** | 0.048828960146202 | 0.703692143295079 | 0.946931219685582 | 631.556087881762 | 485.39177443294 | 553.676109859341 | 586.204183911351 | 564.998429042654 | 479.448234972953 | 561.141090809758 | 574.243023709641 | Tmem120a |
| **ENSMUSG00000039887** | 0.00504450891211174 | 0.977117711274355 | 0.996179425433073 | 268.343573821006 | 227.221988367098 | 231.881447718869 | 263.557401086543 | 192.712642464161 | 238.501035254403 | 312.323427166841 | 243.157726485746 | Alg14 |
| **ENSMUSG00000039891** | 0.392844259849187 | 0.106033131238909 | 0.762094734041179 | 22932.9851539555 | 28617.7542983206 | 29980.8515077168 | 23586.04258051 | 16641.1746600586 | 12452.200204744 | 26278.893161818 | 24687.0223916912 | Txlnb |
| **ENSMUSG00000039899** | -0.0492366758603655 | 0.842618993740136 | 0.97128746259298 | 348.756294595651 | 422.681763306537 | 415.493696116668 | 445.515179772627 | 502.585811880965 | 582.187142467157 | 338.350379430744 | 267.039288908453 | Fgl2 |
| **ENSMUSG00000039901** | 0.125882690020859 | 0.548499195285367 | 0.914699131434573 | 379.475760958999 | 464.216965481167 | 438.208613444231 | 335.777756544422 | 329.582416941548 | 265.408844360028 | 408.102611498006 | 478.716774018812 | 9130011E15 |
| **ENSMUSG00000039903** | 0.0382819159194856 | 0.901449801153843 | 0.984802943363903 | 62.3424464432641 | 54.56585383726 | 49.2156542097192 | 45.0204813243917 | 70.0773245324222 | 67.2695227640623 | 36.4377331694648 | 32.5657669400552 | Eva1c |
| **ENSMUSG00000039908** | -0.439160610272586 | 0.126649253671637 | 0.77215780108715 | 29.8159526467785 | 46.4216965481167 | 26.5007368821565 | 29.075727522003 | 45.9882442244021 | 36.6924669622158 | 55.1771387994752 | 41.2499714574033 | Slc26a11 |
| **ENSMUSG00000039910** | -0.0084233366479283 | 0.972326127683929 | 0.995254967835651 | 979.408868760845 | 1242.79840232327 | 783.664647800914 | 694.065753751039 | 961.373295929167 | 1251.21312341156 | 820.369535358236 | 690.394259129171 | Cited2 |
| **ENSMUSG00000039911** | -0.126961310239506 | 0.394593350215774 | 0.879215782741314 | 438.204152535987 | 490.278268806426 | 548.943835416099 | 395.805064976944 | 574.853052805026 | 513.694537471021 | 514.292576734731 | 442.894430384751 | Spsb1 |
| **ENSMUSG00000039914** | 0.180537706124263 | 0.120594323343876 | 0.769868040577702 | 6426.69306595562 | 5606.43787784624 | 5617.20976412853 | 6940.65753751039 | 5760.57506820427 | 5138.16845694228 | 5807.13358912213 | 4991.2465463458 | Coq10a |
| **ENSMUSG00000039917** | 0.111251275709844 | 0.425788412296935 | 0.885066977635295 | 372.247651226447 | 488.649437348597 | 479.852628544763 | 422.067012416173 | 389.805117711598 | 388.940149799487 | 464.320828388037 | 388.618152151326 | Rhbdd2 |
| **ENSMUSG00000039929** | 0.17505475388854 | 0.466791936301591 | 0.895328112556867 | 126.491920319666 | 184.057954734638 | 164.68315062483 | 127.55803041911 | 136.869774477387 | 97.8465785659088 | 115.559668051731 | 183.453820428978 | Urb1 |
| **ENSMUSG00000039934** | -0.51042298105799 | 0.109411809015596 | 0.763154315676794 | 32.5264937964856 | 26.8757190541728 | 18.9290977729689 | 16.8826804966469 | 26.2789966996583 | 34.2463024980681 | 37.4788112600209 | 37.9933947633978 | Gsap |
| **ENSMUSG00000039936** | -0.41292009658863 | 0.158563467435489 | 0.789230661864335 | 56.9213641438498 | 38.2775392589734 | 71.930571537282 | 51.5859681841989 | 105.115986798633 | 69.71568722821 | 58.3003730711437 | 57.5328549274309 | Pik3cd |
| **ENSMUSG00000039942** | -0.383001750615803 | 0.255231460770614 | 0.837406774112203 | 65.0529875929712 | 44.7928650902881 | 70.0376617599851 | 91.9168160372998 | 143.439523652302 | 75.8310983885793 | 77.0397787011541 | 57.5328549274309 | Ptger4 |
| **ENSMUSG00000039943** | -0.0096753659941705 | 0.936335596164937 | 0.988592177836394 | 1145.65539260955 | 1276.18944720875 | 1242.69526879541 | 1101.12593905908 | 1228.54309570903 | 1024.94291047789 | 1357.5658300852 | 1185.39391661801 | Plcb4 |
| **ENSMUSG00000039952** | -0.0066151125814583 | 0.972608551508056 | 0.995324009463412 | 7466.63735372659 | 9799.0500502972 | 9801.48682684331 | 6548.60417931048 | 7800.48218701525 | 6813.79111488347 | 10471.1634348136 | 8684.20451734806 | Dag1 |
| **ENSMUSG00000039953** | -0.148622685299981 | 0.429545709015863 | 0.885743677088682 | 1714.86903404805 | 1887.0012438945 | 1927.92860817689 | 1568.21343279965 | 2488.83997909681 | 2251.69438924798 | 1568.9046824681 | 1559.90023642865 | Clstn1 |
| **ENSMUSG00000039954** | 0.372829384892505 | 0.523028512253003 | 0.90648515894888 | 17.1667606148119 | 12.2162359337149 | 3.78581955459379 | 15.0068271081306 | 14.2344565456483 | 3.66924669622158 | 9.36970281500523 | 9.76973008201657 | Stk32a |
| **ENSMUSG00000039956** | -0.0777535696323165 | 0.683422351388874 | 0.944452327903597 | 239.431134890797 | 183.243539005724 | 141.968233297267 | 158.509611329629 | 200.377349834895 | 195.693157131818 | 184.270822028436 | 183.453820428978 | Mrap |
| **ENSMUSG00000039958** | 0.207475893365311 | 0.425377269050178 | 0.885066977635295 | 325.264937964856 | 373.002403842763 | 566.92647830042 | 431.446279358754 | 395.279908690694 | 243.393364182698 | 507.005030100839 | 322.401092706547 | Etfbkmt |
| **ENSMUSG00000039959** | -0.112520252792432 | 0.444636793867653 | 0.89108031640712 | 829.425591810383 | 995.21602073331 | 1051.51138128842 | 801.927323590728 | 1121.23719251876 | 1073.86619976085 | 890.121767425497 | 892.302014157513 | Hip1 |
| **ENSMUSG00000039960** | 0.341070046656537 | 0.0270093745511303 | 0.664353746352133 | 674.924746277077 | 646.646088757977 | 505.406910538271 | 781.292936317048 | 519.010184818252 | 485.563646133322 | 501.799639648058 | 552.53251241627 | Rhou |
| **ENSMUSG00000039967** | 0.0373403131365921 | 0.851987908725924 | 0.97283610202923 | 258.404922938747 | 359.971752180133 | 321.794662140472 | 302.950322245386 | 259.505092409126 | 228.716377397812 | 370.623800237985 | 351.710282952596 | Zfp292 |
| **ENSMUSG00000039968** | -0.0569217166389873 | 0.593891802103564 | 0.923935529355527 | 464.406050316489 | 459.330471107681 | 443.887342776122 | 493.349441179793 | 474.116898789669 | 483.117481669175 | 534.073060455298 | 443.97995594942 | Rsbn1l |
| **ENSMUSG00000039976** | 0.15806605241011 | 0.694804783229521 | 0.944771261777587 | 1748.2990415611 | 2578.44019774276 | 2846.93630505453 | 1507.24819767287 | 1211.02376457592 | 777.880299598975 | 2890.03277938384 | 2900.52430879425 | Tbc1d16 |
| **ENSMUSG00000039982** | -0.478219606965293 | 0.0516914158326422 | 0.703396849069134 | 97.5794813894569 | 102.616381843205 | 152.3792370724 | 146.316564304273 | 221.181555555458 | 195.693157131818 | 127.011527047849 | 150.888053488923 | Dtx4 |
| **ENSMUSG00000039983** | -0.152441107050244 | 0.269628120340141 | 0.843793744782845 | 318.940341948873 | 263.056280439328 | 284.882921483182 | 260.743621003769 | 358.051330032845 | 314.332133642982 | 302.953724351836 | 277.894544555138 | Ccdc32 |
| **ENSMUSG00000039985** | -0.224642753019537 | 0.540363920355802 | 0.913837378403038 | 29.8159526467785 | 21.1748089517725 | 13.2503684410783 | 38.4549944645846 | 30.6588294829347 | 34.2463024980681 | 31.2323427166841 | 23.8815624227072 | Fam60a |
| **ENSMUSG00000039987** | 0.155773277581362 | 0.581425985509945 | 0.922450244597442 | 1557.65764736503 | 2116.66647944834 | 1977.14426238661 | 1482.86210362215 | 1228.54309570903 | 917.311674055395 | 2185.22291207733 | 2072.26830295218 | Phtf2 |
| **ENSMUSG00000039988** | 0.104448128915947 | 0.464226815630355 | 0.895328112556867 | 575.538237454482 | 697.95427967958 | 660.625512276616 | 602.14893771374 | 542.004306930453 | 495.348303989913 | 668.37213413704 | 652.400864365773 | Ankrd13c |

| | | | | | | | | | | | | |
|---|---|---|---|---|---|---|---|---|---|---|---|---|
| **ENSMUSG00000039989** | -0.361685916396155 | 0.119956048937531 | 0.769868040577702 | 122.87786545339 | 107.502876216691 | 141.021778408619 | 174.454365132018 | 195.997517051618 | 223.824048469516 | 138.463386043966 | 143.289374536243 | Cbx4 |
| **ENSMUSG00000039990** | 0.0658340515766094 | 0.780514398377444 | 0.960742105865155 | 242.141676040504 | 255.726538879099 | 233.774357496166 | 185.709485463116 | 185.047935093427 | 150.439114545085 | 234.242570375131 | 306.118209236519 | Edrf1 |
| **ENSMUSG00000039994** | 0.0114226105068777 | 0.955673075003211 | 0.992371505557225 | 256.597895505609 | 209.304842330982 | 220.523989055088 | 332.026049767389 | 283.594172717146 | 260.516515431732 | 191.558368662329 | 274.637967861132 | Timeless |
| **ENSMUSG00000039997** | -0.268374901979282 | 0.218379064771769 | 0.827129763070499 | 222.264374275985 | 257.355370336928 | 279.204192151292 | 250.426427366929 | 307.683253025166 | 397.501725424004 | 195.722681024554 | 315.887939318536 | Ifi203 |
| **ENSMUSG00000040003** | 0.108134810669238 | 0.634291728697189 | 0.932543559775713 | 163.535982698997 | 220.706662535783 | 178.879973954556 | 141.626930832982 | 159.863896589588 | 112.523565350795 | 175.942197303987 | 205.164331722348 | Magi2 |
| **ENSMUSG00000040006** | 0.202377256060154 | 0.0544200534201636 | 0.703396849069134 | 2068.14289722654 | 1747.73615425015 | 2076.52202569469 | 2151.60383662822 | 1769.45244444366 | 1866.42348614471 | 1631.36936790147 | 1723.81459669359 | Ginm1 |
| **ENSMUSG00000040007** | -0.00829574983701132 | 0.949220684035836 | 0.991267322147698 | 534.880120208875 | 562.761268679801 | 598.159489625818 | 476.466760683146 | 614.271547854513 | 500.240632918209 | 573.634027896431 | 496.085183053508 | Bahd1 |
| **ENSMUSG00000040010** | 0.0801962364067586 | 0.696184912503448 | 0.944933386973863 | 185.220311896654 | 258.169786065842 | 262.16800415562 | 170.702658354985 | 211.326931793086 | 251.954939807215 | 180.106509666212 | 186.710397122983 | Slc7a5 |
| **ENSMUSG00000040017** | -0.271797822575686 | 0.448104065218508 | 0.89108031640712 | 138.237598635064 | 288.303168035672 | 126.824955078892 | 117.24083678227 | 160.958854785407 | 183.462334811079 | 144.709854587303 | 321.315567141878 | Saa4 |
| **ENSMUSG00000040018** | 0.107816141715445 | 0.687829280565253 | 0.944452327903597 | 719.19691838896 | 913.774447841877 | 888.721140440892 | 693.127827056781 | 607.701798679599 | 431.748027922072 | 1046.28348100892 | 896.644116416187 | Cox15 |
| **ENSMUSG00000040021** | -0.00883089383007214 | 0.954218804830229 | 0.992371505557225 | 198.77301764519 | 271.200437728471 | 225.25626349833 | 247.612647284155 | 217.896680968 | 216.485555077073 | 264.433835001259 | 249.670879873757 | Lats1 |
| **ENSMUSG00000040022** | 0.398767556990132 | 0.176494833299039 | 0.799889021902745 | 168.053551281842 | 173.470550258752 | 194.023252172932 | 103.171936368398 | 114.970610561005 | 69.71568722821 | 166.572494488982 | 132.434118889558 | Rab11fip2 |
| **ENSMUSG00000040025** | -0.107644707195748 | 0.498294582761737 | 0.900927059989498 | 559.274990556239 | 478.876448601625 | 465.655805215036 | 537.431995809926 | 638.360628162534 | 618.879609429373 | 437.252798033577 | 505.854913135525 | Ythdf2 |
| **ENSMUSG00000040028** | -0.0682453281855844 | 0.563912444346497 | 0.918590066040684 | 608.968244967536 | 715.871425715695 | 620.874406953381 | 592.769670771158 | 736.906865786252 | 617.656527197299 | 620.482541971457 | 687.137682435165 | Elavl1 |
| **ENSMUSG00000040029** | 0.231270754121476 | 0.145814321357408 | 0.780607990324772 | 597.222566652139 | 708.541684155466 | 654.000328056077 | 592.769670771158 | 520.105143014071 | 419.517205601334 | 595.49666779811 | 638.289032025082 | Ipo8 |
| **ENSMUSG00000040033** | -0.182857663335414 | 0.261396212175075 | 0.841137531367976 | 121.974351736821 | 141.708336831093 | 119.253315969704 | 136.937297361692 | 147.819356435578 | 176.123841418636 | 135.340151772298 | 132.434118889558 | Stat2 |
| **ENSMUSG00000040034** | -0.161889020108519 | 0.524857753794447 | 0.907879726052909 | 72.2810973255236 | 53.7514381083457 | 40.6975602118832 | 58.151455044006 | 66.7924499449649 | 78.277262852727 | 56.2182168900314 | 51.0197015394199 | Nup43 |
| **ENSMUSG00000040037** | -0.681952608335152 | 0.0535713062017549 | 0.703396849069134 | 26.2018977805023 | 35.8342920722305 | 29.3401015481019 | 33.7653609932938 | 64.6025335533267 | 73.3849339244316 | 35.3966550789086 | 28.2236646813812 | Negr1 |
| **ENSMUSG00000040040** | 0.253747018368551 | 0.117454326979827 | 0.769868040577702 | 146.369222084185 | 152.295741306979 | 192.130342395635 | 154.757904552597 | 128.110108910834 | 121.085140975312 | 147.833088858971 | 143.289374536243 | Ift88 |
| **ENSMUSG00000040043** | 0.0331745299074252 | 0.713621078692616 | 0.951025296130537 | 2313.89862813332 | 2308.86859147212 | 2008.377273712 | 2100.95579513828 | 2039.90711881098 | 2079.23979452556 | 2181.05859971511 | 2234.01161208779 | Rbms2 |
| **ENSMUSG00000040044** | 0.193838284732812 | 0.122029794220791 | 0.770101700012172 | 647.819334780005 | 662.934403336263 | 744.859997366328 | 647.169419038131 | 559.523638063559 | 512.471455238947 | 615.277151518677 | 674.111375659143 | Orc3 |
| **ENSMUSG00000040048** | 0.0754088113146847 | 0.604094328534851 | 0.926448151609826 | 19115.6397014513 | 14387.4682670005 | 15392.1958540897 | 18214.5364024935 | 16384.954442237 | 18589.6268452906 | 14245.0715130796 | 14473.3123537252 | Ndufb10 |
| **ENSMUSG00000040054** | 0.147183447664053 | 0.465417813891848 | 0.895328112556867 | 621.617436999503 | 890.970807432275 | 767.57491469389 | 629.348811847226 | 602.227007700503 | 484.340563901248 | 741.247600475969 | 798.946815596022 | Baz2a |
| **ENSMUSG00000040061** | -0.342732927688649 | 0.191902831910915 | 0.808824684419004 | 34.3335212296237 | 34.2054606144018 | 36.9117406572894 | 24.3860940507122 | 35.0386622662111 | 39.1386314263635 | 41.6431236222455 | 48.8486504100828 | Plcb2 |
| **ENSMUSG00000040078** | 0.140368287421826 | 0.382098280387347 | 0.875126835008975 | 206.001127377742 | 212.56250524664 | 250.810545491838 | 226.040333316217 | 180.668102310151 | 170.008430258266 | 230.078258012906 | 230.131419709724 | Ptges3-ps |
| **ENSMUSG00000040084** | -0.513068211714946 | 0.237162801517597 | 0.832048466795696 | 7.22810973255236 | 16.2883145782866 | 7.57163910918757 | 17.8206071909051 | 16.4243729372865 | 20.7923979452556 | 15.616171358342 | 17.3684090346961 | Bub1b |
| **ENSMUSG00000040086** | -0.0154048026932572 | 0.911198754339792 | 0.986226496507248 | 2673.4970873278 | 2835.79556807969 | 2329.22548096383 | 2344.8167356454 | 2430.8071947184 | 2156.29397514621 | 2938.96344963997 | 2765.91913877536 | Tnni3k |
| **ENSMUSG00000040093** | -0.323598953658909 | 0.28527369668835 | 0.851275833092507 | 59.631905293557 | 67.5965054998893 | 70.9841166486335 | 93.7926694258161 | 104.021028602814 | 135.762127760198 | 66.6289977955927 | 59.7039060567679 | Bmf |
| **ENSMUSG00000040097** | 0.100572251801771 | 0.58656262962879 | 0.923058967588378 | 459.888481733644 | 658.047908962778 | 547.050925638802 | 484.908100931469 | 430.318570956905 | 425.632616761703 | 627.77008860535 | 521.052271040884 | Flywch1 |
| **ENSMUSG00000040102** | -0.343448232532843 | 0.119298305120976 | 0.769868040577702 | 101.193536255733 | 73.2974156022896 | 113.574586637814 | 101.296082979881 | 157.67398019795 | 113.746647582869 | 123.88829277618 | 97.6973008201657 | Klhl42 |
| **ENSMUSG00000040105** | -0.156742064394204 | 0.406842399336523 | 0.881296485258502 | 228.588970291968 | 193.016527752696 | 173.201244622666 | 191.337045628665 | 234.321053905287 | 277.639666680766 | 176.983275394543 | 188.88144825232 | Plpp6 |
| **ENSMUSG00000040111** | 0.168281711538903 | 0.532734674160378 | 0.909082872831377 | 435.49361138628 | 436.52683069808 | 618.981497176084 | 474.59090729463 | 355.861413641207 | 256.847268735511 | 565.305403171982 | 569.900921450967 | Gramd1b |
| **ENSMUSG00000040112** | 0.0576105011607095 | 0.606380118023388 | 0.927090791342307 | 1968.75638840395 | 1555.53404222637 | 1775.54937110449 | 2008.10105240672 | 1743.173447744 | 1815.05403239761 | 1702.16267805928 | 1760.72246589232 | Mrps35 |
| **ENSMUSG00000040113** | 0.236568105176674 | 0.419484937394733 | 0.882568531013176 | 44.2721721118832 | 48.8649437348597 | 25.5542819935081 | 43.1446279358754 | 36.1336204620302 | 26.9078091056249 | 40.6020455316893 | 33.6512925047237 | Mettl11b |
| **ENSMUSG00000040118** | 0.212029959321255 | 0.436380795570651 | 0.888701139843674 | 3350.22886103802 | 4765.14642987774 | 4118.02522050939 | 3242.41258205046 | 2397.95844884382 | 2129.38616604059 | 4354.82965279632 | 4477.79295425759 | Cacna2d1 |
| **ENSMUSG00000040121** | -0.132508263770643 | 0.738254074977227 | 0.954383574136061 | 16.2632468982428 | 14.6594831204579 | 10.4110037751329 | 22.5102406621959 | 21.8991639163819 | 19.5693157131818 | 16.6572494488982 | 11.9407812113536 | Rep15 |
| **ENSMUSG00000040123** | 0.254001949584609 | 0.217265673295181 | 0.825996545635915 | 613.485813550382 | 624.65686407729 | 725.930899593359 | 542.121629281217 | 482.876564356222 | 362.032340693862 | 676.700758861489 | 578.585125968315 | Zmym5 |
| **ENSMUSG00000040124** | 0.218838961298269 | 0.464341089573106 | 0.895328112556867 | 61.4389327266951 | 37.4631235300591 | 35.0188308799925 | 71.2824287636203 | 50.3680770076785 | 42.8078781225851 | 41.6431236222455 | 41.2499714574033 | Gorab |
| **ENSMUSG00000040127** | -0.23135521490595 | 0.625110484355224 | 0.931529302402539 | 18.9737880479499 | 20.3603932228582 | 6.62518422053913 | 15.9447538023887 | 15.3294147414674 | 14.6769867848863 | 12.4929370866736 | 30.3947158107182 | Sdr9c7 |
| **ENSMUSG00000040128** | -0.101396937581963 | 0.528298136532066 | 0.908232038364594 | 3274.33370884622 | 3008.45170260953 | 2869.65122238209 | 3352.15000527867 | 3749.13686248459 | 4015.37896789848 | 3064.93389859727 | 2586.80742060505 | Pnrc1 |
| **ENSMUSG00000040134** | -0.311277782238973 | 0.252323233833208 | 0.83723604018117 | 382.186302108706 | 439.784493613737 | 367.224496795597 | 416.439452250624 | 366.810995599397 | 490.455975061618 | 333.144988977964 | 802.203392290027 | Rdh7 |
| **ENSMUSG00000040136** | 0.0424215647597186 | 0.759770181044313 | 0.957357521291862 | 618.003382133227 | 659.676740420606 | 609.5169482896 | 508.356268287923 | 566.093387238473 | 496.571386221987 | 652.755962778698 | 610.065367343701 | Abcc8 |
| **ENSMUSG00000040139** | 0.145266583759309 | 0.485614329705035 | 0.899236955296196 | 180.702743313809 | 152.295741306979 | 129.664319744837 | 181.957778686083 | 182.858018701789 | 159.000690169602 | 113.477511870619 | 128.092016630884 | 9430038I01R |
| **ENSMUSG00000040146** | 0.124862176833219 | 0.589003830485036 | 0.92352104528132 | 207.80815481088 | 184.872370463553 | 223.363353721033 | 221.350699844926 | 168.623562156141 | 122.308223207386 | 252.981976005141 | 222.532740757044 | Rgl3 |
| **ENSMUSG00000040147** | 0.201068506048131 | 0.353842741591703 | 0.86286441023679 | 1269.43677177951 | 1783.57044632238 | 1760.40609288611 | 1072.98813823134 | 1071.9640737069 | 1066.52770636841 | 1529.34371502696 | 1452.43320552646 | Maob |
| **ENSMUSG00000040151** | 0.0434319545008012 | 0.788448378889689 | 0.961505529655644 | 241.238162323935 | 236.99497711407 | 194.96970706158 | 208.219726125312 | 246.365594059297 | 232.385624094033 | 174.901119213431 | 202.993280593011 | Hs2st1 |
| **ENSMUSG00000040152** | -0.548365547578444 | 0.297445615824757 | 0.851785577030345 | 91.2548853734735 | 88.7713144516618 | 168.468970179424 | 115.364983393754 | 278.119381738051 | 284.978160073209 | 65.5879197050366 | 49.9341759747513 | Thbs1 |
| **ENSMUSG00000040158** | -0.108379392709869 | 0.462793156987023 | 0.895328112556867 | 467.116591466196 | 442.22774080048 | 441.994432998825 | 468.025420434822 | 529.959766776443 | 590.748718091674 | 411.225845769674 | 430.953649173397 | Tax1bp3 |
| **ENSMUSG00000040167** | -0.274313490832138 | 0.073951602734491 | 0.733705544272914 | 157.211386683014 | 154.738988493722 | 188.344522841041 | 187.585338851632 | 190.522726072523 | 190.800828203522 | 240.489038918468 | 208.420908416353 | Ikzf5 |
| **ENSMUSG00000040170** | -0.436852344530999 | 0.187043142394319 | 0.807540361382744 | 907.127771435321 | 886.08431305879 | 1224.71262591109 | 854.451218469185 | 1733.31882398163 | 1994.84712051247 | 787.055036460439 | 727.3021283279 | Fmo2 |
| **ENSMUSG00000040174** | 0.245406882076537 | 0.0401905801033681 | 0.679566075569698 | 681.24934229306 | 522.854897962999 | 633.178320505811 | 643.417712261099 | 547.479097909548 | 521.033030863464 | 520.539045278068 | 502.598336441519 | Alkbh3 |
| **ENSMUSG00000040177** | 0.383427162716436 | 0.0129689121668368 | 0.635862253526407 | 255.69438178904 | 273.643684915214 | 341.670214802089 | 263.557401086543 | 187.237851485066 | 216.485555077073 | 233.201492284575 | 232.302470839061 | 2310057M21 |
| **ENSMUSG00000040181** | -0.571608550064718 | 0.0823035961858994 | 0.739855948845396 | 836.653701542936 | 759.035459348154 | 926.579335986829 | 882.58901929693 | 1719.08436743598 | 1905.56211757107 | 657.961353231478 | 778.32182986732 | Fmo1 |
| **ENSMUSG00000040183** | 0.0018334585242298 | 0.995198908387047 | 0.999262355603273 | 32.5264937964856 | 49.679359463774 | 50.1621090983677 | 43.1446279358754 | 39.4184950494875 | 25.6847268735511 | 53.094982618363 | 56.4473293627624 | Ankrd6 |
| **ENSMUSG00000040187** | -0.200037437793897 | 0.684509833602647 | 0.944452327903597 | 22.5878429142261 | 13.8450673915436 | 17.9826428843205 | 15.9447538023887 | 7.66470737073368 | 18.3462334811079 | 15.616171358342 | 39.0789203280663 | Arntl2 |
| **ENSMUSG00000040188** | -0.14080507678608 | 0.304371016772862 | 0.851785577030345 | 366.826568927032 | 435.712414969166 | 428.744064557746 | 386.425798034362 | 467.547149614754 | 508.802208542726 | 432.047407580797 | 376.677370939972 | Scamp2 |
| **ENSMUSG00000040189** | 0.67896601997573 | 0.121357560605751 | 0.769868040577702 | 26.2018977805023 | 13.0306516626293 | 16.0897331070236 | 12.1930470253561 | 12.0445401540101 | 11.0077400886647 | 9.36970281500523 | 9.76973008201657 | Ccdc114 |
| **ENSMUSG00000040195** | 0.125188897419944 | 0.599383636915251 | 0.924951380294376 | 91.2548853734735 | 68.4109212288036 | 86.1273948670087 | 86.2892558717508 | 87.5966556655277 | 52.592535979176 | 96.8202624217207 | 66.217059444779 | Nemp1 |
| **ENSMUSG00000040205** | 0.379583158343689 | 0.416387881547108 | 0.882192126163777 | 27.1054114970714 | 30.9477976987445 | 9.46454888648447 | 19.6964605794214 | 8.75966556655277 | 17.123151249034 | 23.9447960827911 | 17.3684090346961 | Cuzd1 |
| **ENSMUSG00000040209** | 0.328130995272419 | 0.0863225741532988 | 0.746460787597018 | 615.29284098352 | 754.148964974668 | 679.554610049585 | 685.624413502716 | 451.122776677468 | 408.509465512669 | 666.289977955927 | 651.315338801105 | Zfp704 |
| **ENSMUSG00000040212** | -0.544166087242089 | 0.123520285836652 | 0.770251097497048 | 106.614618555147 | 109.13170767452 | 91.8061241988993 | 137.87522405595 | 189.427767876704 | 270.301173288323 | 84.3273253350471 | 106.381505337514 | Emp3 |
| **ENSMUSG00000040213** | 0.0236666998395851 | 0.835555143193514 | 0.968865818776815 | 616.196354700089 | 675.150639269978 | 697.537252933905 | 640.603932178324 | 673.399290428745 | 555.279333361532 | 700.64555494428 | 655.657441059779 | Kyat3 |
| **ENSMUSG00000040219** | 0.15473665669466 | 0.223902701809827 | 0.832048466795696 | 590.897970636155 | 680.037133643464 | 715.519895818226 | 752.217208795045 | 539.814390538815 | 628.664267285964 | 675.659680770933 | 615.492995167044 | Ttc12 |
| **ENSMUSG00000040220** | 0.133187179630708 | 0.425956028117226 | 0.885066977635295 | 165.343010132135 | 174.284965987666 | 147.646962629158 | 172.578511743502 | 166.433645764503 | 177.34692365071 | 131.175839410073 | 128.092016630884 | Gas8 |
| **ENSMUSG00000040225** | 0.256689191455307 | 0.219785351915451 | 0.828432177683914 | 805.030721463019 | 1032.67914426337 | 1034.47519329275 | 722.203554578784 | 812.45898129777 | 507.579126310652 | 848.478643803251 | 839.111261488756 | Prrc2c |
| **ENSMUSG00000040229** | -0.17724501969481 | 0.738814234091006 | 0.954383574136061 | 10.8421645988285 | 9.77298874697194 | 22.7149173275627 | 38.4549944645846 | 9.85462376237187 | 28.1308913376988 | 27.0680303544596 | 27.1381391167127 | Gpr34 |
| **ENSMUSG00000040234** | -0.149435050775133 | 0.343309185304738 | 0.861167771001038 | 317.133314515735 | 392.548381336706 | 365.3315870183 | 348.908730264036 | 478.496731572945 | 418.29412336926 | 336.268223249632 | 347.368180693922 | Tm7sf3 |
| **ENSMUSG00000040236** | -0.0373573505586114 | 0.742004427938227 | 0.954383574136061 | 923.391018333564 | 781.024684028841 | 776.093008691726 | 956.685228143324 | 863.922016501267 | 928.319414144059 | 853.684034256032 | 881.446758510828 | Trappc5 |
| **ENSMUSG00000040242** | 0.122387324445293 | 0.415318390130682 | 0.882192126163777 | 1972.37044327023 | 1957.85541231005 | 1844.64057797582 | 1809.26059322399 | 1466.14902420177 | 1468.92176072071 | 1943.69279506831 | 2087.46566085754 | Fgfr1op2 |
| **ENSMUSG00000040247** | 0.0486001632627432 | 0.818856392015747 | 0.966315773308182 | 149.983276950461 | 210.119258059897 | 229.988537941573 | 202.592165959763 | 244.175677667659 | 193.24699266767 | 143.668776496747 | 185.624871558315 | Tbc1d10c |
| **ENSMUSG00000040249** | -1.21950557045739 | 0.00435716814400479 | 0.635862253526407 | 763.469090500843 | 1057.92603185971 | 911.436057768454 | 939.802547646678 | 2826.08710340909 | 3894.29382692317 | 901.573626421614 | 931.380934485579 | Lrp1 |
| **ENSMUSG00000040250** | 0.061050236937157 | 0.681187521833706 | 0.944452327903597 | 570.117155155067 | 531.813470981057 | 539.479286529615 | 524.301022090312 | 458.787484048202 | 424.409534529629 | 600.702058250891 | 590.525907179668 | Ints13 |
| **ENSMUSG00000040253** | -0.0592854707777967 | 0.770669859050874 | 0.958995611819782 | 232.203025158245 | 241.067055758641 | 274.47191770805 | 227.916186704733 | 346.006789878835 | 262.96267989588 | 209.256696201783 | 198.651178334337 | Gbp7 |
| **ENSMUSG00000040254** | -0.696563884875019 | 0.178299995175229 | 0.801696999069105 | 22.5878429142261 | 13.0306516626293 | 19.8755526616174 | 20.6343872736795 | 36.1336204620302 | 59.9310293716191 | 11.4518589961175 | 16.2828834700276 | Sema3d |
| **ENSMUSG00000040260** | -0.254502928093851 | 0.0963580963405774 | 0.757809915356977 | 166.246523848704 | 201.160685041839 | 164.68315062483 | 153.819977858338 | 211.326931793086 | 188.354663739374 | 230.078258012906 | 188.88144825232 | Daam2 |
| **ENSMUSG00000040263** | -0.231371046758063 | 0.311778751802313 | 0.851785577030345 | 319.843855665442 | 332.281617397046 | 301.919109478855 | 363.915557372167 | 391.995034103237 | 563.840908986049 | 274.84461590682 | 318.058990447873 | Klhdc4 |
| **ENSMUSG00000040265** | -0.504792126406477 | 0.152215191920699 | 0.783412422888014 | 58.7283915769879 | 57.009101024003 | 90.8596693102509 | 54.3997482669734 | 122.635317931739 | 134.539045528125 | 60.3825292522559 | 53.1907526687569 | Dnm3 |
| **ENSMUSG00000040268** | 0.216688146415844 | 0.13056402127185 | 0.77303252008843 | 1139.33079659357 | 1312.02373928098 | 1198.21188902893 | 1162.09117418586 | 866.111932892906 | 935.657907536503 | 1088.96768272172 | 1249.43992493345 | Plekha1 |
| **ENSMUSG00000040269** | -0.0204770045574508 | 0.904055381436169 | 0.985039825636533 | 1277.56839522863 | 999.288099377881 | 994.724087969518 | 1349.67651303749 | 1189.12460065954 | 1417.5523069736 | 978.613405122768 | 1102.8939737032 | Mrps28 |
| **ENSMUSG00000040270** | -0.150031614911529 | 0.574311800579895 | 0.922280709351328 | 32.5264937964856 | 47.2361122770311 | 38.8046504345863 | 57.2135283497478 | 60.2227007700503 | 48.9232892829544 | 44.7663578939139 | 41.2499714574033 | Bach2 |
| **ENSMUSG00000040272** | 0.225888649035937 | 0.309022948483719 | 0.851785577030345 | 71.3775836089546 | 83.8848200781758 | 94.6454888648447 | 84.4134024832345 | 76.6470737073368 | 50.1463715150282 | 88.4916376972716 | 69.4736361387845 | Accs |
| **ENSMUSG00000040274** | 0.00998020077816893 | 0.970246523959006 | 0.9951105526746 | 91.2548853734735 | 175.913797445495 | 176.040609288611 | 108.799496533947 | 159.863896589588 | 156.554525705454 | 123.88829277618 | 108.552556466851 | Cdk6 |
| **ENSMUSG00000040276** | -0.548660012901226 | 0.389316835985199 | 0.876720306717642 | 9.93865088225949 | 17.1027303072009 | 12.3039135524298 | 0.937926694258161 | 9.85462376237187 | 12.2308223207386 | 22.903717992235 | 14.1118323406906 | Pacsin1 |
| **ENSMUSG00000040280** | -0.909778697866631 | 0.00800564461793294 | 0.635862253526407 | 183.413284463516 | 177.542628903324 | 164.68315062483 | 178.206071909051 | 453.312693069106 | 502.686797382356 | 195.722681024554 | 171.513039217624 | Ndufa4l2 |
| **ENSMUSG00000040282** | 0.312177899108332 | 0.0710774764669843 | 0.726501344493452 | 154.500845533307 | 132.749763813036 | 130.610774633486 | 116.302910088012 | 109.49581958191 | 89.2850029413918 | 111.395355689507 | 119.407812113536 | BC052040 |
| **ENSMUSG00000040283** | -0.209177124478423 | 0.361811917722828 | 0.868897239438884 | 1534.16629073424 | 1481.42221089516 | 1876.82004418987 | 1714.52999710392 | 2207.4357227713 | 2621.06522333428 | 1326.33348736852 | 1483.91344690185 | Btnl9 |
| **ENSMUSG00000040289** | -1.06501069885171 | 0.0077031491119115 | 0.635862253526407 | 402.967117589794 | 268.757190541728 | 266.900278598862 | 185.709485463116 | 979.987585258092 | 760.757148349941 | 307.11803671406 | 305.032683671851 | Hey1 |

| | | | | | | | | | | | | |
|---|---|---|---|---|---|---|---|---|---|---|---|---|
| **ENSMUSG00000040296** | -0.280949702793385 | 0.280330146542964 | 0.849093714741928 | 106.614618555147 | 147.409246933493 | 110.735221971868 | 116.302910088012 | 181.76306050597 | 178.570005882784 | 85.3684034256032 | 140.032797842237 | Ddx58 |
| **ENSMUSG00000040297** | 0.0852996768302621 | 0.558485731714438 | 0.916574868225118 | 268.343573821006 | 280.159010746529 | 259.328639489674 | 241.047160424347 | 237.605928492744 | 206.700897220482 | 291.501865355718 | 251.841931003094 | Suco |
| **ENSMUSG00000040302** | -0.119485221030565 | 0.559646923874991 | 0.916574868225118 | 98.4829951060259 | 71.6685841444609 | 95.5919437534931 | 74.0962088463947 | 106.210944994452 | 100.292743030056 | 78.0808567917102 | 84.6709940441436 | Rbm48 |
| **ENSMUSG00000040310** | -0.59543068008383 | 0.299988992549164 | 0.851785577030345 | 18.0702743313809 | 7.32974156022896 | 5.67872933189068 | 9.37926694258161 | 4.37983278327639 | 19.5693157131818 | 17.6983275394543 | 19.5394601640331 | Alx4 |
| **ENSMUSG00000040312** | -0.290124103178754 | 0.551810242337085 | 0.914873920695527 | 18.0702743313809 | 39.0919549878878 | 35.965285768641 | 22.5102406621959 | 45.9882442244021 | 61.154111603693 | 20.8215618111227 | 14.1118323406906 | Cchcr1 |
| **ENSMUSG00000040321** | 0.234395199036237 | 0.470844521067651 | 0.896754476570639 | 440.914693685694 | 511.453077758198 | 535.693466975021 | 475.528833988888 | 321.917709570815 | 187.131581507301 | 537.196294726966 | 622.006148555055 | Zfp770 |
| **ENSMUSG00000040322** | -0.49644494325409 | 0.102811352828996 | 0.759800953500741 | 82.2197482077831 | 101.801966114291 | 86.1273948670087 | 100.358156285623 | 181.76306050597 | 174.900759186562 | 82.2451691539348 | 84.6709940441436 | Slc25a24 |
| **ENSMUSG00000040325** | 0.249736777598912 | 0.224316947767524 | 0.832048466795696 | 740.881247586617 | 804.642740167357 | 800.700835796586 | 739.086235075431 | 595.657258525589 | 421.963370065482 | 794.342583094332 | 781.578406561325 | Dcaf1 |
| **ENSMUSG00000040327** | 0.169515924231693 | 0.264744409615818 | 0.842510961779757 | 900.803175419338 | 949.608739914107 | 883.042411109001 | 932.299134092612 | 843.117810780705 | 602.979540412413 | 918.230875870512 | 893.387539722182 | Cul9 |
| **ENSMUSG00000040331** | 0.0610282850294864 | 0.638954621855482 | 0.933801135937988 | 660.468526811972 | 595.337897836374 | 697.537252933905 | 711.886360941944 | 699.678287128403 | 709.387694602839 | 589.250199254773 | 556.874614674944 | Nsmce4a |
| **ENSMUSG00000040339** | -0.206652008270714 | 0.357867887551514 | 0.865834559821921 | 285.510334435818 | 274.458100644129 | 312.330113253987 | 217.598993067893 | 423.748821781991 | 331.455284892016 | 217.585320926233 | 285.493223507818 | Fam102b |
| **ENSMUSG00000040345** | 0.474631007446344 | 0.0458065161072615 | 0.697906909182231 | 67.7635287426784 | 67.5965054998893 | 55.8408384302584 | 88.1651092602672 | 59.1277425742312 | 44.0309603546589 | 55.1771387994752 | 42.3354970220718 | Arhgap9 |
| **ENSMUSG00000040350** | 0.0342845510565585 | 0.882921143060053 | 0.979073805351756 | 433.686583953142 | 365.672662282534 | 350.188308799925 | 407.060185308042 | 337.247124312282 | 243.393364182698 | 529.908748093074 | 408.157612315359 | Trim7 |
| **ENSMUSG00000040351** | 0.234419981138231 | 0.250900984075119 | 0.836703579022786 | 316.229800799166 | 394.991628523449 | 347.34894413398 | 291.695201914288 | 270.454674367317 | 198.139321595965 | 333.144988977964 | 345.197129564585 | Ankib1 |
| **ENSMUSG00000040354** | -0.048932163273785 | 0.731479071478728 | 0.953029200153788 | 1260.40163461382 | 1146.69734631137 | 1409.27132919754 | 1431.27613543795 | 1529.65659955928 | 1512.95272107536 | 1187.87010132455 | 1198.42022339403 | Mars |
| **ENSMUSG00000040356** | 0.0172834767973836 | 0.891828589414929 | 0.982421229269187 | 604.450676384691 | 634.429852824262 | 681.447519826882 | 708.134654164912 | 708.437952694956 | 724.064681387725 | 593.414511616998 | 572.071972580304 | Skiv2l |
| **ENSMUSG00000040359** | 0.176222877103057 | 0.345496336252344 | 0.861266047293794 | 479.765783498163 | 581.492830444831 | 560.301294079881 | 515.859681841989 | 378.855535753408 | 382.824738639118 | 610.071761065896 | 518.881219911547 | Ufl1 |
| **ENSMUSG00000040363** | -0.0907604848336231 | 0.520718725902507 | 0.905962179970576 | 475.248214915318 | 522.854897962999 | 566.92647830042 | 512.107975064956 | 527.769850384805 | 459.878919259771 | 654.83811895981 | 567.72987032163 | Bcor |
| **ENSMUSG00000040364** | 0.197070527032896 | 0.731968148634318 | 0.953219349759111 | 4.51756858284523 | 13.8450673915436 | 24.6078271048596 | 12.1930470253561 | 13.1394983498292 | 4.89232892829544 | 15.616171358342 | 14.1118323406906 | Sec1 |
| **ENSMUSG00000040365** | 0.107256357945192 | 0.533021429543946 | 0.909094751518347 | 708.354753790131 | 706.098436968723 | 729.716719147953 | 605.900644490772 | 522.295059405709 | 529.594606487981 | 792.26042691322 | 707.762668163867 | Trim41 |
| **ENSMUSG00000040370** | 0.223055609336089 | 0.108442287096173 | 0.762287160109383 | 1545.00845533307 | 1485.49428953974 | 1367.62731409701 | 1402.20040791595 | 1091.67332123164 | 1071.4200352967 | 1312.79947219129 | 1492.5976514192 | Etfrf1 |
| **ENSMUSG00000040374** | 0.0331987686858957 | 0.803150202648052 | 0.963461896828986 | 874.601277638836 | 775.323773926441 | 754.324546252812 | 715.638067718977 | 696.393412540946 | 677.587556568918 | 781.849646007659 | 892.302014157513 | Pex2 |
| **ENSMUSG00000040383** | 0.0175054772056816 | 0.87762124554459 | 0.977664000726269 | 475.248214915318 | 592.080234920717 | 570.712297855013 | 538.369922504185 | 556.238763476101 | 499.017550686135 | 550.730309904196 | 543.848307898922 | Aqr |
| **ENSMUSG00000040385** | 0.0655335307689968 | 0.564884532671649 | 0.919517890403047 | 7674.44550853747 | 6353.25710126068 | 7416.42050744923 | 7777.28814878867 | 7075.619861383 | 7797.14922947086 | 6351.61743048299 | 6699.86378513403 | Ppp1ca |
| **ENSMUSG00000040387** | 0.25316408818105 | 0.457103315072319 | 0.892805623137304 | 45.1756858284523 | 56.1946852950887 | 51.1085639870161 | 61.9031618210386 | 24.0890803080201 | 73.3849339244316 | 34.3555769883525 | 48.8486504100828 | Klhl32 |
| **ENSMUSG00000040389** | -0.0449458689652125 | 0.820298407263899 | 0.966398681144067 | 166.246523848704 | 179.985876090067 | 177.933519065908 | 139.751077444466 | 166.433645764503 | 121.085140975312 | 197.804837205666 | 198.651178334337 | Wdr47 |
| **ENSMUSG00000040390** | -0.003260997789522 | 0.973970151046361 | 0.995510898988495 | 662.27555424511 | 684.92362801695 | 655.893237833374 | 634.976372012775 | 626.316088008523 | 611.54111603693 | 725.631429117627 | 679.539003482486 | Map3k10 |
| **ENSMUSG00000040396** | -0.0059360288451192 | 0.964943258082932 | 0.994284908685797 | 259.308436655316 | 287.488752306758 | 317.06238769723 | 274.812521417641 | 294.543754675337 | 316.77829810713 | 286.296474902938 | 246.414303179751 | Abhd13 |
| **ENSMUSG00000040407** | 0.311622252549364 | 0.0535692441651605 | 0.703396849069134 | 1264.01568948009 | 1594.62599721426 | 1481.20190073482 | 1260.57347708297 | 1097.14811221074 | 874.50379593281 | 1253.45802102959 | 1286.34779413218 | Akap9 |
| **ENSMUSG00000040410** | 0.120036333530346 | 0.383627667130453 | 0.875126835008975 | 205.097613661173 | 175.913797445495 | 203.487801059416 | 198.84045918273 | 160.958854785407 | 181.016170346931 | 198.845915296222 | 179.111718170304 | Fbxl4 |
| **ENSMUSG00000040413** | -0.22970831105541 | 0.635074306325683 | 0.932543559775713 | 12.6491920319666 | 13.8450673915436 | 13.2503684410783 | 9.37926694258161 | 5.47479097909548 | 15.9000690169602 | 12.4929370866736 | 23.8815624227072 | Timd2 |
| **ENSMUSG00000040414** | 0.185235553907351 | 0.374177953616378 | 0.873700364743909 | 736.363679003772 | 653.975830318206 | 935.097429984665 | 918.23023367874 | 832.168228822514 | 884.2884537894 | 552.812466085309 | 584.012753791657 | Slc25a28 |
| **ENSMUSG00000040415** | 0.100136765377653 | 0.565527108554724 | 0.919917032184696 | 272.861142403852 | 209.304842330982 | 291.508105703722 | 246.674720589896 | 222.276513751277 | 294.7628179298 | 227.996101831794 | 207.335382851685 | Dtx3 |
| **ENSMUSG00000040423** | 0.160676540238685 | 0.384874825588844 | 0.875126835008975 | 945.075347531221 | 1093.76032393194 | 927.525790875478 | 741.900015158206 | 889.106055005107 | 604.202622644487 | 893.245001697165 | 930.295408920911 | Rc3h1 |
| **ENSMUSG00000040424** | -0.210498540326545 | 0.604185119442545 | 0.926448151609826 | 13.5527057485357 | 21.9892246806869 | 22.7149173275627 | 16.8826804966469 | 10.949581958191 | 18.3462334811079 | 26.0269522639034 | 31.4802413753867 | Hipk4 |
| **ENSMUSG00000040428** | -0.40266217550836 | 0.121245155754547 | 0.769868040577702 | 65.9565013095403 | 39.9063707168021 | 36.9117406572894 | 45.0204813243917 | 59.1277425742312 | 75.8310983885793 | 56.2182168900314 | 57.5328549274309 | Plekha4 |
| **ENSMUSG00000040429** | -0.128324502200658 | 0.636430147152485 | 0.932595977875587 | 32.5264937964856 | 30.1333819698302 | 31.2330113253987 | 29.075727522003 | 30.6588294829347 | 25.6847268735511 | 41.6431236222455 | 35.8223436340608 | Mterf1a |
| **ENSMUSG00000040430** | -0.0010769384185415 | 0.992961659591873 | 0.999133450854041 | 1440.20086421106 | 1579.15209836488 | 1341.12657721485 | 1205.23580212174 | 1361.03303740314 | 1260.99778126815 | 1454.38609250692 | 1493.68317698387 | Pitpnc1 |
| **ENSMUSG00000040433** | 0.0797057550114775 | 0.436090092144921 | 0.888687115594624 | 857.434517024024 | 1016.39082968508 | 907.65023821386 | 856.327071857701 | 806.984190318674 | 858.60372691585 | 901.573626421614 | 876.019130687486 | Zbtb38 |
| **ENSMUSG00000040435** | 0.168108401660474 | 0.350126006811151 | 0.861774532843376 | 420.133878204606 | 495.979178908826 | 668.197151385803 | 597.459304242449 | 493.826146314413 | 453.763508099402 | 563.22324699087 | 429.868123608729 | Ppp1r15a |
| **ENSMUSG00000040441** | -0.722874836099491 | 0.0862061442025371 | 0.746460787597018 | 153.597331816738 | 102.616381843205 | 139.128868631322 | 166.013024883695 | 270.454674367317 | 431.748027922072 | 97.8613405122768 | 127.006491066215 | Slc26a10 |
| **ENSMUSG00000040446** | 0.155483495072276 | 0.668058394414084 | 0.942024383071524 | 655.950958229127 | 838.848200781758 | 820.576388458203 | 602.14893771374 | 413.894198019619 | 291.093571233579 | 1018.1743725639 | 895.558590851519 | Rprd1a |
| **ENSMUSG00000040447** | -0.343764564318507 | 0.224169290723938 | 0.832048466795696 | 185.220311896654 | 193.016527752696 | 238.506631939409 | 236.357526953057 | 306.588294829347 | 412.178712208891 | 178.024353485099 | 186.710397122983 | Spns2 |
| **ENSMUSG00000040451** | 0.125352759656413 | 0.261556066907352 | 0.841137531367976 | 1397.73571953231 | 1422.78427841333 | 1441.45079541158 | 1261.51140377723 | 1351.17841364077 | 1339.27504412088 | 1290.93683228961 | 1083.35451353917 | Sgms1 |
| **ENSMUSG00000040455** | 0.00357401639544544 | 0.987487353657128 | 0.998437672973644 | 180.702743313809 | 176.728213174409 | 159.950876181588 | 173.51643843776 | 142.344565456483 | 114.969729814943 | 205.092383839559 | 225.78931745105 | Usp45 |
| **ENSMUSG00000040459** | 0.113639922421876 | 0.260359584346192 | 0.840316047847749 | 977.601841327707 | 926.805099504506 | 830.040937344688 | 983.885102276811 | 822.313605060142 | 823.134342185708 | 901.573626421614 | 889.045437463508 | Arglu1 |
| **ENSMUSG00000040462** | -0.275394587049427 | 0.180653681457729 | 0.802808628271642 | 929.715614349547 | 1040.0088858236 | 1241.74881390676 | 1040.1607039323 | 1476.00364796414 | 1671.95341124497 | 1005.68143547723 | 993.255891671684 | Os9 |
| **ENSMUSG00000040463** | 0.218148359719491 | 0.0378913545901686 | 0.679566075569698 | 1891.95772249558 | 2027.89516499668 | 1981.87653682985 | 2015.60446596079 | 1947.93063036217 | 1547.19902357343 | 1718.81992750818 | 1591.38047780403 | Mybbp1a |
| **ENSMUSG00000040464** | -0.0218274970339475 | 0.91602418414515 | 0.986713918026868 | 254.790868072471 | 307.849145529616 | 285.829376371831 | 271.060814640609 | 231.036179317829 | 207.923979452556 | 334.18606706852 | 362.565538599282 | Gtpbp10 |
| **ENSMUSG00000040466** | -0.0533478316185121 | 0.832045754192191 | 0.967583158900775 | 306.291149916906 | 290.746415222415 | 421.172425448559 | 358.287997206618 | 319.727793179176 | 524.702277559686 | 236.324726556243 | 348.453706258591 | Blvrb |
| **ENSMUSG00000040472** | -0.0984085454633051 | 0.402820795744031 | 0.881158207349486 | 330.68602026427 | 321.69421292116 | 307.597838810745 | 307.639955716677 | 367.905953795217 | 365.701587390084 | 314.405583347953 | 310.460311495193 | Rabggta |
| **ENSMUSG00000040479** | -0.0718288263939218 | 0.632338675886246 | 0.931825800800958 | 591.801484352725 | 579.049583258088 | 646.428688946889 | 732.520748215624 | 731.432074807157 | 770.541806206532 | 620.482541971457 | 557.960140239613 | Dgkz |
| **ENSMUSG00000040481** | 0.160606819116754 | 0.271650770511963 | 0.844129257012535 | 1148.36593375926 | 1228.95333493172 | 1132.90650171219 | 934.174987481129 | 901.150595159117 | 854.934480219628 | 1102.50169789895 | 1117.00580604389 | Bptf |
| **ENSMUSG00000040482** | -0.134962339195604 | 0.530338216913347 | 0.908923862033581 | 133.720030052219 | 141.708336831093 | 145.754052851861 | 139.751077444466 | 181.76306050597 | 199.362403828039 | 117.641824232843 | 118.322286548867 | Dxo |
| **ENSMUSG00000040483** | -0.127033106543405 | 0.738723192476998 | 0.954383574136061 | 84.9302893574902 | 49.679359463774 | 97.48485353079 | 88.1651092602672 | 107.305903190271 | 138.208292224346 | 41.6431236222455 | 62.9604827507734 | Xaf1 |
| **ENSMUSG00000040485** | -0.419189806617447 | 0.193373344543457 | 0.809022817984525 | 65.0529875929712 | 76.5550785179469 | 102.217127974032 | 53.4618215727152 | 129.205067106653 | 102.738907494204 | 114.518589961175 | 51.0197015394199 | Lrrc52 |
| **ENSMUSG00000040488** | -1.1343984857687 | 0.0051114084842381 | 0.635862253526407 | 2967.13904521274 | 3693.37533062648 | 3082.60357232799 | 3026.68944237109 | 11066.7424851436 | 10675.0617215406 | 3342.90174877575 | 2949.37295920434 | Ltbp4 |
| **ENSMUSG00000040489** | 0.149426880454716 | 0.742497289740162 | 0.954383574136061 | 21.6843291976571 | 8.95857301805762 | 20.8220075502658 | 24.3860940507122 | 12.0445401540101 | 22.0154801773295 | 10.4107809055614 | 23.8815624227072 | Sox30 |
| **ENSMUSG00000040498** | -0.278770851168743 | 0.463235165499843 | 0.895328112556867 | 32.5264937964856 | 50.4937751926884 | 55.8408384302584 | 29.075727522003 | 30.6588294829347 | 86.838838477244 | 45.80743598447 | 41.2499714574033 | Igsf23 |
| **ENSMUSG00000040502** | 0.0741398467946009 | 0.65730248620115 | 0.938389079274206 | 168.957064998411 | 163.69756151178 | 160.897331070236 | 151.944124469822 | 159.863896589588 | 179.793088114857 | 157.202791673977 | 117.236760984199 | March9 |
| **ENSMUSG00000040506** | 0.075734768155844 | 0.64020735452195 | 0.934084049822467 | 423.747933070882 | 491.09268453534 | 583.016211407443 | 482.094320848695 | 509.15556105588 | 395.055560959857 | 548.648153723084 | 424.440495785386 | Ambra1 |
| **ENSMUSG00000040511** | 0.166322659067302 | 0.202837848496647 | 0.815403452654488 | 576.441751171051 | 700.397526866323 | 744.859997366328 | 652.79697920368 | 627.411046204342 | 518.586866399316 | 644.427338054249 | 591.611432744337 | Pvr |
| **ENSMUSG00000040520** | -0.292774142620265 | 0.114243586412134 | 0.767518180080285 | 188.83436676293 | 248.39679731887 | 227.149173275627 | 235.419600258798 | 278.119381738051 | 211.593226148778 | 347.72008224575 | 263.782712214447 | Manea |
| **ENSMUSG00000040521** | 0.00246735168412627 | 0.980750027450255 | 0.996888334226108 | 853.820462157748 | 798.127414336042 | 708.894711597687 | 795.361836730921 | 825.598479647599 | 768.095641742384 | 740.206522385413 | 817.400750195386 | Tsfm |
| **ENSMUSG00000040522** | -0.191250092092183 | 0.701075727714957 | 0.945920819861844 | 18.0702743313809 | 23.6180561385155 | 19.8755526616174 | 6.56548685980713 | 30.6588294829347 | 9.78465785659088 | 14.5750932677859 | 22.7960368580387 | Tlr8 |
| **ENSMUSG00000040524** | 0.285321108951116 | 0.390198490695579 | 0.877567525220465 | 515.002818444356 | 713.428178528952 | 603.838218957709 | 461.459933575015 | 367.905953795217 | 214.039390612925 | 631.934400967575 | 667.598222271132 | Zfp609 |
| **ENSMUSG00000040525** | 0.490052255077476 | 0.396302964141374 | 0.879359239049715 | 11.7456783153976 | 17.1027303072009 | 10.4110037751329 | 13.1309737196143 | 6.56974917491458 | 18.3462334811079 | 2.08215618111227 | 10.8552556466851 | Cblc |
| **ENSMUSG00000040532** | -0.0098098568939660 | 0.946111030202334 | 0.990505437194639 | 1198.96270188712 | 923.547436588849 | 896.292779550079 | 1264.32518386 | 1062.10944994452 | 1072.64311752877 | 1081.68013608783 | 1095.29529475052 | Abhd11 |
| **ENSMUSG00000040536** | 0.0552168674839716 | 0.928472886692413 | 0.986901071486195 | 7.22810973255236 | 11.4018202048006 | 8.51809399783602 | 14.0689004138724 | 5.47479097909548 | 6.1154111603693 | 5.20539045278068 | 22.7960368580387 | Necab1 |
| **ENSMUSG00000040537** | 0.136920882029994 | 0.343314806589296 | 0.861167771001038 | 194.255449062345 | 216.634583891211 | 224.309808609682 | 189.461192240149 | 181.76306050597 | 163.893019097897 | 213.421008564008 | 189.966973816989 | Adam22 |
| **ENSMUSG00000040543** | 0.303426558571217 | 0.433844003412476 | 0.887868762931002 | 31.6229800799166 | 44.7928650902881 | 46.3762895437739 | 23.448167356454 | 31.7537876787538 | 15.9000690169602 | 23.9447960827911 | 46.6775992807458 | Pitpnm3 |
| **ENSMUSG00000040548** | 0.181332818965569 | 0.476718188111496 | 0.897602769807331 | 1646.2019915888 | 1876.41383941861 | 1666.70705890991 | 1536.32392519487 | 1162.84560395988 | 889.180782717696 | 1887.47457817828 | 1990.85388560204 | Tex2 |
| **ENSMUSG00000040549** | 0.179564488889204 | 0.420220020283449 | 0.882568531013176 | 941.461292664945 | 1135.29552610657 | 1075.17275350464 | 813.182443921826 | 786.179984598112 | 574.848649074714 | 1052.52994955225 | 1086.61109023318 | Ckap5 |
| **ENSMUSG00000040550** | 0.264701052097542 | 0.311855817185152 | 0.851785577030345 | 2000.37936848387 | 2495.3697933935 | 2349.10103362544 | 1949.94959736272 | 1332.56412431184 | 1152.14346261358 | 2363.24726556243 | 2471.74171075019 | Otud6b |
| **ENSMUSG00000040552** | -0.364739048425274 | 0.0788671412839544 | 0.739855948845396 | 65.9565013095403 | 58.6379324818317 | 79.5022106464695 | 86.2892558717508 | 106.210944994452 | 99.0696607979826 | 78.0808567917102 | 90.0986218674861 | C3ar1 |
| **ENSMUSG00000040557** | 0.139015630899429 | 0.59354453147025 | 0.923935529355527 | 39.754603529038 | 47.2361122770311 | 40.6975602118832 | 47.8342614071662 | 39.4184950494875 | 29.3539735697726 | 53.094982618363 | 36.9078691987293 | Mettl27 |
| **ENSMUSG00000040560** | -0.1210074350876 | 0.518766419725382 | 0.905689488434585 | 329.782506547701 | 462.588134023339 | 382.367775013973 | 352.660437041069 | 340.531998899739 | 348.57843614105 | 478.895921655823 | 492.828606359502 | Wdr7 |
| **ENSMUSG00000040562** | -0.179129075530886 | 0.226703758874399 | 0.832048466795696 | 2341.0040396304 | 2592.28526513431 | 1985.66235638444 | 2467.68513259322 | 2477.89039713862 | 3219.1524348184 | 2644.33835001259 | 2288.28789032121 | Gstm2 |
| **ENSMUSG00000040563** | 0.111207144019582 | 0.727308415400737 | 0.952237115597464 | 37.0440623793308 | 39.9063707168021 | 36.9117406572894 | 38.4549944645846 | 37.2285786578493 | 53.8156182112498 | 31.2323427166841 | 19.5394601640331 | Plppr2 |
| **ENSMUSG00000040564** | -0.125500132783497 | 0.674880085855374 | 0.944023556644042 | 739.977733870048 | 913.774447841877 | 822.4692982355 | 858.202925246218 | 598.942133113046 | 1015.1582526213 | 549.68923181364 | 1474.14371681983 | Apoc1 |
| **ENSMUSG00000040565** | 0.146004624068757 | 0.630717520797931 | 0.931825800800958 | 199.676531361759 | 323.323044378988 | 337.884395247495 | 189.461192240149 | 236.510970296925 | 135.762127760198 | 333.144988977964 | 243.157726485746 | Btaf1 |
| **ENSMUSG00000040584** | -0.3177618444327 | 0.220495593219461 | 0.829144114546069 | 364.116027777325 | 425.939426222194 | 490.263632319895 | 391.115431505653 | 753.331238723539 | 588.302553627526 | 417.472314313011 | 324.572143835884 | Abcb1a |
| **ENSMUSG00000040586** | 0.1078815775417 | 0.490168576322261 | 0.899236955296196 | 228.588970291968 | 253.283291692356 | 223.363353721033 | 241.985087118606 | 238.700886688563 | 168.785348026193 | 247.776585552361 | 222.532740757044 | Ofd1 |
| **ENSMUSG00000040591** | -0.0529216688951592 | 0.686789185732306 | 0.944452327903597 | 226.78194285883 | 216.634583891211 | 211.059440168604 | 197.902532488472 | 231.036179317829 | 199.362403828039 | 209.256696201783 | 244.243252050414 | 1110051M20 |
| **ENSMUSG00000040596** | 0.169884267905879 | 0.478361061809095 | 0.897674307047416 | 254.790868072471 | 378.703313945163 | 364.385132129652 | 268.247034557834 | 229.94122112201 | 200.585486060113 | 340.432535611857 | 353.881334081933 | Pogk |
| **ENSMUSG00000040599** | -0.22787747975903 | 0.0749211233076999 | 0.735399943172214 | 223.167887992554 | 223.149909722526 | 209.166530391307 | 195.088752405698 | 249.650468646754 | 243.393364182698 | 227.996101831794 | 275.723493425801 | Mis12 |
| **ENSMUSG00000040605** | -0.653245155707395 | 0.10474205902379 | 0.762094734041179 | 74.9916384752307 | 31.7622134276588 | 41.6440151005317 | 56.2756016554897 | 84.3117810780705 | 138.208292224346 | 52.0539045278068 | 47.7631248454143 | Bace2 |

| | | | | | | | | | | | | |
|---|---|---|---|---|---|---|---|---|---|---|---|---|
| **ENSMUSG00000040606** | -0.0972063412116396 | 0.717183132419006 | 0.951449255902579 | 43.3686583953142 | 39.0919549878878 | 45.4298346551254 | 49.7101147956825 | 65.6974917491458 | 48.9232892829544 | 41.6431236222455 | 33.6512925047237 | Kazn |
| **ENSMUSG00000040612** | -0.392840522086015 | 0.368767220747167 | 0.871495599696183 | 14.4562194651047 | 12.2162359337149 | 17.036187995672 | 10.3171936368398 | 14.2344565456483 | 30.5770558018465 | 11.4518589961175 | 15.1973579053591 | Ildr2 |
| **ENSMUSG00000040613** | -0.172887885962604 | 0.491489121090419 | 0.899236955296196 | 49.6932544112975 | 55.3802695661743 | 63.4124775394459 | 63.779015209555 | 66.7924499449649 | 91.7311674055395 | 53.094982618363 | 51.0197015394199 | Apobec1 |
| **ENSMUSG00000040616** | -0.937871004394996 | 0.0886086876962251 | 0.751206603737452 | 33.4300075130547 | 18.7315617650296 | 23.6613722162112 | 38.4549944645846 | 49.2731188118594 | 123.53130543946 | 28.1091084450157 | 18.4539345993646 | Tmem51 |
| **ENSMUSG00000040618** | -0.718742118014485 | 0.0482852170673663 | 0.703396849069134 | 133.720030052219 | 141.708336831093 | 159.004421292939 | 180.081925297567 | 339.43704070392 | 398.724807656078 | 120.765058504512 | 153.05910461826 | Pck2 |
| **ENSMUSG00000040620** | 0.152615976282523 | 0.53888778632352 | 0.91326883260538 | 138.237598635064 | 225.593156909269 | 217.684624389143 | 175.392291826276 | 134.679858085749 | 128.423634367755 | 191.558368662329 | 225.78931745105 | Dhx33 |
| **ENSMUSG00000040621** | -0.238541200420971 | 0.440287087324551 | 0.890530552540252 | 25.2983840639333 | 17.9171460361152 | 17.9826428843205 | 24.3860940507122 | 22.994122112201 | 26.9078091056249 | 27.0680303544596 | 23.8815624227072 | Gemin8 |
| **ENSMUSG00000040624** | -0.320898361568479 | 0.0894420569596203 | 0.751593644682828 | 197.869503928621 | 198.717437855096 | 202.541346170768 | 155.695831246855 | 305.493336633528 | 236.054870790255 | 197.804837205666 | 204.078806157679 | Plekhg1 |
| **ENSMUSG00000040631** | -0.404506977082007 | 0.0553309303942614 | 0.703396849069134 | 362.309000344187 | 264.685111897157 | 272.579007930753 | 362.03970398365 | 557.33372167192 | 397.501725424004 | 317.528817619622 | 397.302356668674 | Dok4 |
| **ENSMUSG00000040640** | 0.237885826346152 | 0.690493115468391 | 0.94469184542218 | 10.8421645988285 | 9.77298874697194 | 9.46454888648447 | 18.7585338851632 | 3.28487458745729 | 4.89232892829544 | 18.7394056300105 | 14.1118323406906 | Erc2 |
| **ENSMUSG00000040648** | 0.201417170826199 | 0.583855966142454 | 0.922450244597442 | 1755.52715129365 | 2269.77663648423 | 2626.41231599944 | 1429.40028204944 | 1191.31451705118 | 807.234273168747 | 2908.77218501385 | 2120.0314277976 | Ppip5k2 |
| **ENSMUSG00000040651** | 0.0602187564139754 | 0.719852096365113 | 0.951674101666722 | 252.983840639333 | 237.809392842984 | 239.453086828057 | 194.150825711439 | 213.516848184724 | 178.570005882784 | 264.433835001259 | 229.045894145055 | Fam208a |
| **ENSMUSG00000040652** | -0.0696587757155183 | 0.464349188152889 | 0.895328112556867 | 4681.10456554422 | 4398.65935186629 | 4026.21909631049 | 4379.17973549136 | 4195.87980637878 | 4461.80398260544 | 4922.21721214941 | 4769.79933115342 | Oaz2 |
| **ENSMUSG00000040653** | 0.0205484381228643 | 0.92215981649122 | 0.986713918026868 | 2273.24051088772 | 2138.65570412903 | 1657.24251002343 | 2025.92165959763 | 1621.63308800808 | 1523.96046116403 | 2096.73127438006 | 2737.69547409398 | Ppp1r14c |
| **ENSMUSG00000040658** | 0.372011141078975 | 0.373624278013098 | 0.873700364743909 | 29.8159526467785 | 17.9171460361152 | 16.0897331070236 | 15.0068271081306 | 22.994122112201 | 13.4539045528125 | 10.4107809055614 | 14.1118323406906 | Dnph1 |
| **ENSMUSG00000040659** | -0.0202112132882393 | 0.932057301707743 | 0.987963669163657 | 1019.16347228988 | 890.970807432275 | 1321.25102455323 | 1297.15261815904 | 1082.91365566509 | 1658.49950669215 | 1023.37976301668 | 828.256005842071 | Efhd2 |
| **ENSMUSG00000040661** | 0.130693449463376 | 0.686334052584277 | 0.944452327903597 | 196.062476495483 | 363.229415095791 | 301.919109478855 | 242.923013812864 | 206.947099009809 | 128.423634367755 | 341.473613702413 | 331.085297223895 | Rad54l2 |
| **ENSMUSG00000040666** | 0.0155361266486097 | 0.923547749046938 | 0.986713918026868 | 7000.42427597696 | 6259.59929243553 | 4734.16735301953 | 7056.02252090415 | 5483.55064466204 | 5736.2556684264 | 6859.66353867438 | 6702.03483626337 | Sh3bgr |
| **ENSMUSG00000040667** | 0.0460055008600058 | 0.685651618413153 | 0.944452327903597 | 854.723975874317 | 847.806773799816 | 819.629933569555 | 804.741103673502 | 826.693437843418 | 675.14139210477 | 878.669908429379 | 840.196787053425 | Nup88 |
| **ENSMUSG00000040669** | 0.0972080889545616 | 0.427619425150375 | 0.885066977635295 | 310.808718499751 | 308.663561258531 | 313.276568142636 | 275.750448111899 | 250.745426842573 | 281.308913376988 | 288.37863108405 | 309.374785930525 | Phc1 |
| **ENSMUSG00000040675** | -0.372934338983504 | 0.440257669676415 | 0.890530552540252 | 18.9737880479499 | 18.7315617650296 | 10.4110037751329 | 10.3171936368398 | 22.994122112201 | 26.9078091056249 | 19.7804837205666 | 6.51315338801105 | Mthfd1l |
| **ENSMUSG00000040681** | -0.00245006901174201 | 0.978632693279108 | 0.996664169100645 | 814.969372345279 | 827.446380576958 | 895.346324661431 | 924.795720538547 | 873.776640263639 | 906.30393396673 | 859.930502799369 | 828.256005842071 | Hmgn1 |
| **ENSMUSG00000040687** | 0.105409000759033 | 0.585615452815893 | 0.922964550512322 | 181.606257030378 | 229.665235553841 | 218.631079277791 | 166.950951577953 | 212.421889988905 | 138.208292224346 | 180.106509666212 | 209.506433981022 | Madd |
| **ENSMUSG00000040688** | -0.178978973598982 | 0.15045003797993 | 0.782641594375979 | 394.835494140673 | 374.631235300591 | 355.867038131816 | 378.922384480297 | 489.446313531136 | 440.309603546589 | 379.99350305299 | 394.045779974668 | Tbl3 |
| **ENSMUSG00000040690** | -1.65275728586521 | 0.00793150809982732 | 0.635862253526407 | 18.9737880479499 | 48.0505280059454 | 19.8755526616174 | 34.703287687552 | 142.344565456483 | 185.908499275227 | 32.2734208072402 | 22.7960368580387 | Col16a1 |
| **ENSMUSG00000040694** | -0.181994011008688 | 0.224144280713264 | 0.832048466795696 | 3808.31031533853 | 3208.79797192245 | 3364.64712914523 | 3837.05810621014 | 3905.71588448672 | 5050.10653623297 | 3721.85417373819 | 3454.14234677519 | Apobec2 |
| **ENSMUSG00000040697** | 0.13858000858142 | 0.622227962890464 | 0.93078489223231 | 814.06585862871 | 1037.56563863685 | 1037.3145579587 | 703.445020693621 | 626.316088008523 | 479.448234972953 | 1096.25522935561 | 1060.55847668113 | Dnajc16 |
| **ENSMUSG00000040699** | -0.201949504143393 | 0.480087186203371 | 0.897732059433541 | 110.228673421423 | 117.275864963663 | 112.628131749165 | 147.254490998531 | 158.768938393769 | 211.593226148778 | 97.8613405122768 | 93.3551985614916 | Limd2 |
| **ENSMUSG00000040701** | 0.293917981538087 | 0.0799449316386708 | 0.739855948845396 | 101.193536255733 | 104.245213301034 | 101.270673085384 | 102.23400967414 | 86.5016974697087 | 90.5080851734656 | 86.4094815161593 | 70.559161703453 | Ap1g2 |
| **ENSMUSG00000040705** | -0.0316157243290914 | 0.933003737486505 | 0.98813739334255 | 31.6229800799166 | 18.7315617650296 | 14.1968233297267 | 21.5723139679377 | 16.4243729372865 | 30.5770558018465 | 21.8626399016789 | 19.5394601640331 | A930016O22 |
| **ENSMUSG00000040706** | -0.0993746583339215 | 0.740265955540083 | 0.954383574136061 | 28.9124389302094 | 43.1640336324594 | 24.6078271048596 | 44.0825546301336 | 32.8487458745729 | 42.8078781225851 | 31.2323427166841 | 44.5065481514088 | Agmat |
| **ENSMUSG00000040710** | -0.130157175931777 | 0.640825558075397 | 0.934495223272154 | 127.395434036235 | 118.090280692578 | 134.396594188079 | 249.488500672671 | 210.231973597267 | 188.354663739374 | 149.915245040084 | 140.032797842237 | St8sia4 |
| **ENSMUSG00000040711** | -0.347104376867081 | 0.311042804196599 | 0.851785577030345 | 61.4389327266951 | 103.43079757212 | 67.1982970940397 | 71.2824287636203 | 128.110108910834 | 136.985209992272 | 69.7522320672611 | 52.1052271040884 | Sh3pxd2b |
| **ENSMUSG00000040712** | 0.0330758156300093 | 0.810026433624102 | 0.964412705343932 | 1250.46298373156 | 1247.68489669675 | 1256.89209212514 | 1068.29850476005 | 1158.4657711766 | 951.557976553463 | 1282.60820756516 | 1319.99908663691 | Camta2 |
| **ENSMUSG00000040713** | 0.180042079182417 | 0.0888737177349471 | 0.751206603737452 | 5123.82628666305 | 5618.65411377995 | 6487.9482616851 | 5738.23551547143 | 4834.24043454131 | 4820.16707660308 | 5224.12985841069 | 5393.97653083781 | Creg1 |
| **ENSMUSG00000040717** | -0.273679425056062 | 0.359611600356799 | 0.867338970126053 | 109.325159704854 | 127.048853710635 | 123.039135524298 | 90.9788893430416 | 191.617684268342 | 174.900759186562 | 82.2451691539348 | 96.6117752554972 | Il17rd |
| **ENSMUSG00000040720** | 0.0751438481930349 | 0.540958699060038 | 0.913837378403038 | 462.599022883351 | 507.380999113627 | 551.783200082044 | 461.459933575015 | 517.915226622433 | 414.624876673038 | 478.895921655823 | 470.032569501464 | Virma |
| **ENSMUSG00000040721** | 0.261021996091423 | 0.313467131066842 | 0.85295001595893 | 212.325723393726 | 320.879797192245 | 269.739643264807 | 255.11606083822 | 249.650468646754 | 125.977469903608 | 290.460787265162 | 216.019587369033 | Zfhx2 |
| **ENSMUSG00000040722** | 0.00713879975110694 | 0.969006111324739 | 0.994908125383341 | 161.728955265859 | 226.407572638183 | 188.344522841041 | 141.626930832982 | 186.142893289246 | 173.677676954488 | 192.599446752885 | 162.828834700276 | Scamp5 |
| **ENSMUSG00000040723** | -0.0272045880905403 | 0.660085540853373 | 0.939611397743053 | 1960.62476495483 | 1948.08242356307 | 1959.16161950228 | 1921.81179653497 | 2003.77349834895 | 2005.85486060113 | 1974.92513778499 | 1953.94601640331 | Rcsd1 |
| **ENSMUSG00000040724** | -0.577935719482345 | 0.178501342077896 | 0.801696999069105 | 15.3597331816738 | 33.3910448854875 | 22.7149173275627 | 34.703287687552 | 47.0832024202212 | 62.3771938357668 | 31.2323427166841 | 18.4539345993646 | Kcna2 |
| **ENSMUSG00000040725** | 0.0471975831426555 | 0.683624258355418 | 0.944452327903597 | 1628.13171725742 | 1782.75603059347 | 1831.39020953474 | 1530.69636502932 | 1653.38687568684 | 1417.5523069736 | 1718.81992750818 | 1763.97904258632 | Hnrnpul1 |
| **ENSMUSG00000040729** | 0.310501998936127 | 0.352260671212469 | 0.862728204353539 | 93.9654265231807 | 88.7713144516618 | 105.056492639978 | 71.2824287636203 | 51.4630352034976 | 37.9155491942896 | 111.395355689507 | 87.9275707381491 | Cep126 |
| **ENSMUSG00000040731** | 0.0246529295142397 | 0.793381963465247 | 0.961804768755116 | 3044.84122483768 | 3460.45243215698 | 3711.04961839056 | 3451.57023487003 | 3552.04438723715 | 3297.42969767113 | 3346.02498304742 | 3240.2938105355 | Eif4h |
| **ENSMUSG00000040732** | -0.0333435637664813 | 0.793576595691973 | 0.961804768755116 | 1059.82158953549 | 871.424829938332 | 936.043884873314 | 1171.47044112844 | 973.417836083177 | 995.588936908122 | 1057.73534000503 | 1105.06502483254 | Erg |
| **ENSMUSG00000040734** | 0.07262916786135 | 0.593806549175002 | 0.923935529355527 | 969.470217878585 | 974.041211781537 | 1294.75028767108 | 1214.61506906432 | 1009.55145654521 | 1148.47421591735 | 1036.91377819391 | 1038.84796538776 | Ppp1r13l |
| **ENSMUSG00000040738** | 0.0564622878488091 | 0.753981684986747 | 0.956497786093635 | 309.905204783182 | 263.870696168242 | 265.953823710214 | 239.171307035831 | 255.12525962585 | 190.800828203522 | 283.173240631269 | 307.203734801188 | Ints8 |
| **ENSMUSG00000040740** | 0.36176005449891 | 0.0989731779279392 | 0.759800953500741 | 5773.4526488762 | 6118.70537133335 | 8236.05044101878 | 6375.08774087272 | 4406.11177997605 | 3598.3079267613 | 6709.7482936343 | 5909.60117405535 | Slc25a34 |
| **ENSMUSG00000040746** | -0.100358161585366 | 0.443659481947565 | 0.89108031640712 | 960.435080712895 | 934.134841064735 | 933.204520207369 | 987.636809053844 | 1131.09181628113 | 1172.93586055883 | 920.313032051625 | 867.334926170138 | Rnf167 |
| **ENSMUSG00000040747** | -0.487756222008905 | 0.0845838846336719 | 0.743541556158628 | 55.1143367107117 | 52.9370223794314 | 70.0376617599851 | 82.5375490947182 | 108.400861386091 | 123.53130543946 | 69.7522320672611 | 64.046008315442 | Cd53 |
| **ENSMUSG00000040749** | 0.102392523479523 | 0.641532167118854 | 0.934544499347759 | 67.7635287426784 | 69.2253369577179 | 70.9841166486335 | 47.8342614071662 | 65.6974917491458 | 51.3694537471021 | 63.5057635239243 | 57.5328549274309 | Siah1b |
| **ENSMUSG00000040752** | 0.224533168131433 | 0.0259589193897275 | 0.664353746352133 | 587456.486889744 | 615337.504891323 | 597789.425764357 | 627211.276911012 | 556980.050174671 | 445751.096397086 | 522440.053871424 | 552708.367557747 | Myh6 |
| **ENSMUSG00000040760** | 0.138082071739033 | 0.34888110599374 | 0.861554058734205 | 582.766347187034 | 629.543358450776 | 641.696414503647 | 613.404058044837 | 536.529515951357 | 436.640356850368 | 622.56469815257 | 644.802185413094 | Appl1 |
| **ENSMUSG00000040761** | 0.0603772315945573 | 0.786843145814753 | 0.961141387070613 | 637.880683897746 | 892.599638890104 | 833.826756899282 | 543.997482669733 | 762.090904290091 | 483.117481669175 | 820.369535358236 | 722.960026069226 | Spen |
| **ENSMUSG00000040767** | -0.19981440241069 | 0.376662239899526 | 0.87422797717331 | 206.904641094311 | 134.378595270864 | 151.432782183751 | 225.102406621959 | 199.282391639076 | 243.393364182698 | 157.202791673977 | 224.703791886381 | Snrnp25 |
| **ENSMUSG00000040771** | 0.0065118783674766 | 0.965496359017616 | 0.994284908685797 | 477.958756065025 | 432.454752053509 | 444.83379766477 | 588.080037299867 | 499.300937293508 | 541.82542880872 | 477.854843565267 | 415.756291268038 | Oard1 |
| **ENSMUSG00000040774** | -0.16874965439432 | 0.216362518764263 | 0.825447777181429 | 291.834930451802 | 326.580707294646 | 300.972654590206 | 286.067641748739 | 374.475702970131 | 377.932409710823 | 312.323427166841 | 292.006376895829 | Cept1 |
| **ENSMUSG00000040782** | 0.181141965352798 | 0.0958160589742819 | 0.757475844708466 | 2344.61809449667 | 2430.21653508036 | 2269.59882297898 | 2362.63734283631 | 1840.6247271719 | 1908.00828203522 | 2260.18053459737 | 2287.20236475655 | Cop1 |
| **ENSMUSG00000040785** | 0.225942972429656 | 0.309091336992322 | 0.851785577030345 | 2158.49426888345 | 2467.67965861042 | 2262.97363875844 | 1869.28790165652 | 1500.09272827216 | 1291.57483707 | 2308.07012676295 | 2388.15624227072 | Ttc3 |
| **ENSMUSG00000040794** | -0.168642097964338 | 0.560883156919842 | 0.916762264997884 | 108.421645988285 | 96.9154717408051 | 123.985590412947 | 135.061443973175 | 154.389105610493 | 192.023910435596 | 93.6970281500523 | 82.4999429148066 | C1qtnf4 |
| **ENSMUSG00000040795** | -0.00339866787177112 | 0.984203909090539 | 0.996940244108351 | 126.491920319666 | 112.389370590177 | 99.3777633080869 | 138.813150750208 | 114.970610561005 | 121.085140975312 | 131.175839410073 | 110.723607596188 | Iqcc |
| **ENSMUSG00000040797** | -0.395424417204084 | 0.336190508164575 | 0.858570960161243 | 20.780815481088 | 13.0306516626293 | 18.9290977729689 | 17.8206071909051 | 19.7092475247437 | 34.2463024980681 | 28.1091084450157 | 10.8552556466851 | Iqsec3 |
| **ENSMUSG00000040811** | 0.135096418316458 | 0.217391056116746 | 0.825996545635915 | 679.442314859922 | 736.231818938553 | 732.556083813898 | 679.996853337167 | 732.527033002976 | 615.210362733151 | 639.221947601468 | 588.354856050331 | Eml2 |
| **ENSMUSG00000040812** | 0.971804297579397 | 0.0245605973149896 | 0.655303666217042 | 12.6491920319666 | 18.7315617650296 | 20.8220075502658 | 15.0068271081306 | 10.949581958191 | 7.33849339244316 | 8.32862472444909 | 7.59867895267955 | Agbl2 |
| **ENSMUSG00000040813** | -0.00779356965169477 | 0.966113171363294 | 0.994284908685797 | 992.058060792811 | 885.269897329875 | 1091.26248661166 | 1199.60824195619 | 1181.45989328881 | 1291.57483707 | 865.13589325215 | 853.223093829447 | Tex264 |
| **ENSMUSG00000040818** | -0.143782267740615 | 0.346601915406222 | 0.861266047293794 | 227.685456575399 | 257.355370336928 | 304.7584741448 | 219.47484645641 | 278.119381738051 | 253.178022039289 | 275.885693997376 | 307.203734801188 | Dennd6a |
| **ENSMUSG00000040820** | 0.212764779115568 | 0.26530906744598 | 0.842510961779757 | 204.194099944604 | 232.108482740584 | 224.309808609682 | 176.330218520534 | 141.249607260664 | 156.554525705454 | 216.544242835676 | 207.335382851685 | Hlcs |
| **ENSMUSG00000040822** | -0.287382999281237 | 0.106059495174602 | 0.762094734041179 | 229.492484008537 | 193.016527752696 | 160.897331070236 | 228.854113398991 | 245.270635863478 | 299.655146858096 | 223.831789469569 | 223.618266321713 | 1700123O20 |
| **ENSMUSG00000040824** | -0.0910158058723691 | 0.682880517712233 | 0.944452327903597 | 1858.52771498253 | 1222.43800910041 | 1446.18306985483 | 1828.95705380341 | 1750.83815511474 | 2306.7330896913 | 1310.71731601018 | 1402.49902955171 | Snrpd2 |
| **ENSMUSG00000040829** | 0.0251003646938287 | 0.954440591204805 | 0.992371505557225 | 11.7456783153976 | 7.32974156022896 | 17.9826428843205 | 15.0068271081306 | 15.3294147414674 | 11.0077400886647 | 13.5340151772298 | 10.8552556466851 | Zmynd15 |
| **ENSMUSG00000040836** | -0.393140030501319 | 0.218526513742294 | 0.827129763070499 | 18.9737880479499 | 23.6180561385155 | 17.9826428843205 | 23.448167356454 | 36.1336204620302 | 24.4616446414772 | 29.1501865355718 | 20.6249857287016 | Gpr161 |
| **ENSMUSG00000040841** | -0.490395001324708 | 0.00780008914399616 | 0.635862253526407 | 84.9302893574902 | 110.760539132349 | 89.9132144216024 | 120.054616865045 | 119.350443344282 | 147.992950080937 | 153.038479311752 | 149.802527924254 | Six5 |
| **ENSMUSG00000040842** | 0.066260286110799 | 0.629943486682132 | 0.931825800800958 | 1947.97557292286 | 2027.08074926776 | 1966.73325861147 | 1749.23328479147 | 1606.30367326662 | 1582.66840830357 | 2129.0046951873 | 2026.6762292361 | Szrd1 |
| **ENSMUSG00000040843** | -0.0254180943990778 | 0.834964348688552 | 0.9687316073575 | 553.853908256825 | 522.854897962999 | 500.674636095028 | 547.749189446766 | 540.909348734634 | 461.102001491845 | 616.318229609233 | 542.762782334254 | Tiprl |
| **ENSMUSG00000040848** | -0.175590478235503 | 0.170715523740054 | 0.799531027639052 | 542.108229941427 | 732.159740293981 | 657.786147610671 | 616.217838127612 | 745.666531352805 | 737.518585940537 | 652.755962778698 | 743.585011797928 | Sft2d2 |
| **ENSMUSG00000040850** | 0.219069909969348 | 0.412443383751859 | 0.882192126163777 | 7544.33953335153 | 8737.86635552183 | 7400.33077434221 | 6651.77611567888 | 4948.1160869065 | 3829.47046862325 | 8600.34610608425 | 8682.03346621872 | Psme4 |
| **ENSMUSG00000040852** | -0.193151016985096 | 0.449361201118256 | 0.89108031640712 | 30.7194663633475 | 44.7928650902881 | 33.1259211026956 | 41.2687745473591 | 50.3680770076785 | 41.5847958905112 | 36.4377331694648 | 43.4210225867403 | Plekhh2 |
| **ENSMUSG00000040855** | -0.573714680413817 | 0.119635090144829 | 0.769868040577702 | 18.9737880479499 | 27.6901347830872 | 12.3039135524298 | 30.0136542162612 | 30.6588294829347 | 22.0154801773295 | 41.6431236222455 | 37.9933947633978 | Reps2 |
| **ENSMUSG00000040857** | -0.219966057636602 | 0.188317502069812 | 0.808677608936066 | 386.703870691551 | 474.804369957054 | 459.030620994497 | 332.026049767389 | 508.060602860061 | 558.948580057754 | 440.376032305246 | 419.012867962044 | Erf |
| **ENSMUSG00000040859** | 0.247321667986795 | 0.185766514970643 | 0.805963874499992 | 1674.21091680244 | 1917.13462586433 | 2062.32520236497 | 1642.30964164604 | 1448.62969306867 | 1103.22017333062 | 1928.07662370997 | 1665.19621620149 | Bsdc1 |
| **ENSMUSG00000040860** | -0.0973373639094935 | 0.623669791575001 | 0.931145988862144 | 78.6056933415069 | 83.8848200781758 | 62.4660226507975 | 65.6548685980713 | 72.2672409240604 | 86.838838477244 | 83.2862472444909 | 69.4736361387845 | Crocc |
| **ENSMUSG00000040865** | -0.0635398560736746 | 0.651921091725815 | 0.936720465895231 | 248.466272056487 | 236.99497711407 | 209.166530391307 | 219.47484645641 | 262.789966996583 | 209.14706168463 | 261.31060072959 | 221.447215192376 | Ino80d |
| **ENSMUSG00000040875** | -0.192825589153566 | 0.408779371442138 | 0.882113132676277 | 45.1756858284523 | 52.122606650517 | 50.1621090983677 | 45.9584080186499 | 50.3680770076785 | 41.5847958905112 | 67.6700758861489 | 60.7894316214364 | Osbpl10 |
| **ENSMUSG00000040877** | -0.0795067053292972 | 0.622431735640251 | 0.93078489223231 | 103.90407740544 | 100.987550385377 | 96.5383986421416 | 101.296082979881 | 94.1664048404423 | 102.738907494204 | 119.723980413956 | 108.552556466851 | Wdr25 |
| **ENSMUSG00000040883** | -0.125236015111875 | 0.539885592973541 | 0.913837378403038 | 1707.6409243155 | 1184.16046984143 | 1318.41165988729 | 1863.66034149097 | 1673.09612321158 | 2128.16308380852 | 1385.67493853022 | 1438.32137318577 | Tmem205 |
| **ENSMUSG00000040888** | 0.108553989723029 | 0.451419035384821 | 0.891110666112841 | 473.44118748218 | 396.620459981278 | 459.030620994497 | 509.294194982182 | 440.173194719277 | 443.978850242811 | 349.802238426862 | 471.118095066132 | Gfer |
| **ENSMUSG00000040891** | -0.694559295283526 | 0.125685550712664 | 0.771949105062466 | 9.93865088225949 | 14.6594831204579 | 8.51809399783602 | 13.1309737196143 | 10.949581958191 | 15.9000690169602 | 16.6572494488982 | 31.4802413753867 | Foxa3 |

| | | | | | | | | | | | | |
|---|---|---|---|---|---|---|---|---|---|---|---|---|
| **ENSMUSG00000040896** | 0.233394008519343 | 0.468673459925868 | 0.895328112556867 | 209.615182244018 | 320.065381463331 | 253.649910157784 | 247.612647284155 | 151.104231023035 | 118.638976511164 | 337.309301340188 | 269.21034003779 | Kcnd3 |
| **ENSMUSG00000040913** | 0.115351951512022 | 0.305844656369686 | 0.851785577030345 | 851.10992100804 | 953.680818558679 | 989.045358637627 | 985.760955665328 | 872.68168206782 | 764.426395046162 | 956.75076522109 | 893.387539722182 | Fbxw4 |
| **ENSMUSG00000040918** | -0.0170275494028203 | 0.930121597917451 | 0.987529878729394 | 187.027339329792 | 228.036404096012 | 226.202718386979 | 180.081925297567 | 168.623562156141 | 171.23151249034 | 257.146288367366 | 233.387996403729 | Slc19a2 |
| **ENSMUSG00000040928** | 0.216041853772257 | 0.253637627924069 | 0.837406774112203 | 104.807591122009 | 157.99665140938 | 137.235958854025 | 151.006197775564 | 122.635317931739 | 107.6312364225 | 135.340151772298 | 108.552556466851 | S100pbp |
| **ENSMUSG00000040929** | -0.391852616562208 | 0.368180593003086 | 0.871495599696183 | 10.8421645988285 | 19.5459774939439 | 9.46454888648447 | 27.1998741334867 | 28.4689130912965 | 14.6769867848863 | 20.8215618111227 | 23.8815624227072 | Rfx3 |
| **ENSMUSG00000040940** | -0.200927220685995 | 0.153784915431701 | 0.785320762994975 | 575.538237454482 | 605.110886583346 | 598.159489625818 | 597.459304242449 | 813.553939493589 | 738.741668172611 | 579.880496439768 | 600.295637261685 | Arhgef1 |
| **ENSMUSG00000040943** | 0.300146074161475 | 0.277600336570776 | 0.848075053184273 | 103.90407740544 | 155.553404222637 | 133.450139299431 | 130.371810501884 | 90.881530252985 | 61.154111603693 | 133.257995591185 | 138.947272277569 | Tet2 |
| **ENSMUSG00000040945** | -0.186355866390302 | 0.213719989877458 | 0.824248286808591 | 234.913566307952 | 299.704988240473 | 229.988537941573 | 281.378008277448 | 329.582416941548 | 315.555215875056 | 267.557069272927 | 278.980070119806 | Rcc2 |
| **ENSMUSG00000040950** | -0.581784938626464 | 0.166707719950484 | 0.796745160868404 | 37.9475760958999 | 43.9784493613737 | 79.5022106464695 | 54.3997482669734 | 117.160526952643 | 118.638976511164 | 36.4377331694648 | 51.0197015394199 | Mgl2 |
| **ENSMUSG00000040952** | 0.0373291836437507 | 0.862063571453439 | 0.974868041734905 | 4566.35832353995 | 2948.18493866987 | 3527.43736999276 | 4472.03447822291 | 3676.86962156053 | 5130.82996354984 | 3111.78241267229 | 3199.04383907809 | Rps19 |
| **ENSMUSG00000040957** | 0.0427526840062119 | 0.841367740126234 | 0.970567055091221 | 159.921927832721 | 189.758864837039 | 193.076797284283 | 157.571684635371 | 224.466430142915 | 183.462334811079 | 121.806136595068 | 150.888053488923 | Cables1 |
| **ENSMUSG00000040963** | -0.238270130678432 | 0.62560293174625 | 0.931573470007391 | 24.3948703473642 | 59.452348210746 | 20.8220075502658 | 21.5723139679377 | 19.7092475247437 | 26.9078091056249 | 36.4377331694648 | 66.217059444779 | Asgr2 |
| **ENSMUSG00000040964** | 0.0490414997117879 | 0.735639877538231 | 0.954383574136061 | 1888.3436676293 | 1859.31110911141 | 1710.24398378774 | 1934.00484356033 | 1511.04231023035 | 1534.96820125269 | 1987.41807487166 | 2110.26169771558 | Arhgef10l |
| **ENSMUSG00000040972** | -0.223739275783049 | 0.687722861861534 | 0.944452327903597 | 18.0702743313809 | 20.3603932228582 | 3.78581955459379 | 10.3171936368398 | 19.7092475247437 | 22.0154801773295 | 9.36970281500523 | 10.8552556466851 | Igsf21 |
| **ENSMUSG00000040985** | 0.278081001667553 | 0.544803529628176 | 0.91403805127858 | 9.03513716569045 | 21.9892246806869 | 11.3574586637814 | 21.5723139679377 | 10.949581958191 | 9.78465785659088 | 13.5340151772298 | 18.4539345993646 | Sun3 |
| **ENSMUSG00000040987** | -0.0510464194626534 | 0.809297307581273 | 0.964311345725361 | 81.316234491214 | 100.173134656462 | 85.1809399783602 | 69.4065753751039 | 108.400861386091 | 84.3926740130963 | 85.3684034256032 | 70.559161703453 | Mill2 |
| **ENSMUSG00000040990** | -0.16697124776959 | 0.378873857329942 | 0.87422797717331 | 1739.26390439541 | 1952.15450220765 | 1414.95005852943 | 1662.00610222546 | 1569.07509460877 | 1519.06813223573 | 2083.19725920283 | 2427.23516259878 | Sh3kbp1 |
| **ENSMUSG00000040997** | 0.0407426756612934 | 0.734322323801247 | 0.954227437328031 | 802.320180313312 | 1009.87550385377 | 1021.22482485167 | 957.623154837583 | 905.530427942393 | 962.565716642128 | 977.572327032212 | 840.196787053425 | Abhd4 |
| **ENSMUSG00000040998** | -1.19245381544337 | 0.0284657126052175 | 0.665956123973759 | 23.4913566307952 | 31.7622134276588 | 16.0897331070236 | 14.0689004138724 | 76.6470737073368 | 78.277262852727 | 27.0680303544596 | 14.1118323406906 | Npnt |
| **ENSMUSG00000041000** | -0.19304421282471 | 0.684768964590526 | 0.944452327903597 | 22.5878429142261 | 17.1027303072009 | 21.7684624389143 | 44.0825546301336 | 42.7033696369448 | 45.2540425867328 | 18.7394056300105 | 14.1118323406906 | Trim62 |
| **ENSMUSG00000041012** | -0.136514882842106 | 0.421633388403766 | 0.882568531013176 | 449.046317134815 | 373.816819571677 | 358.706402797761 | 312.329589187968 | 490.541271726955 | 428.078781225851 | 348.761160336306 | 375.591845375304 | Cmtm8 |
| **ENSMUSG00000041020** | -0.355342160587427 | 0.492245482912177 | 0.899236955296196 | 20.780815481088 | 20.3603932228582 | 13.2503684410783 | 11.2551203310979 | 21.8991639163819 | 41.5847958905112 | 9.36970281500523 | 11.9407812113536 | Map7d2 |
| **ENSMUSG00000041025** | -0.0735912167769882 | 0.650115534034165 | 0.936463929813731 | 156.307872966445 | 175.099381716581 | 179.826428843205 | 168.826804966469 | 208.042057205628 | 147.992950080937 | 160.326025945645 | 198.651178334337 | Iffo2 |
| **ENSMUSG00000041028** | 0.363503507691731 | 0.0655157185059294 | 0.718339708336358 | 15684.9981196386 | 15929.9716575643 | 18143.5402153907 | 16350.8760610025 | 10477.6549757929 | 9333.34051295562 | 16039.8901411984 | 15532.7853048417 | Ghitm |
| **ENSMUSG00000041035** | -0.257098975512937 | 0.188385376870077 | 0.808677608936066 | 409.291713605777 | 426.753841951108 | 431.583429223692 | 423.004939110431 | 514.630352034976 | 698.379954514174 | 397.691830592444 | 411.414189009364 | Gm17018 |
| **ENSMUSG00000041037** | -0.00857730226459401 | 0.973355282019559 | 0.995501426054906 | 531.266065342598 | 607.554133770089 | 566.92647830042 | 403.308478531009 | 442.363111110915 | 327.786038195794 | 704.809867306505 | 645.887710977762 | Irgq |
| **ENSMUSG00000041040** | -0.471923220629488 | 0.338076837414856 | 0.858570960161243 | 835.750187826367 | 991.143942088738 | 724.037989816062 | 304.826175633902 | 1486.95322992233 | 1651.16101329971 | 415.390158131898 | 408.157612315359 | Fam117b |
| **ENSMUSG00000041046** | -0.464484305375132 | 0.169773774666901 | 0.79907358295209 | 217.74680569314 | 131.935348084121 | 138.182413742673 | 173.51643843776 | 312.063085808443 | 325.339873731647 | 125.970448957293 | 149.802527924254 | Ramp3 |
| **ENSMUSG00000041057** | 0.310464670842798 | 0.00646857091139063 | 0.635862253526407 | 556.564449406532 | 646.646088757977 | 562.194203857177 | 581.51455044006 | 460.97740043984 | 423.186452297555 | 498.676405376389 | 509.11148982953 | Wdr43 |
| **ENSMUSG00000041058** | 0.264641947473238 | 0.268807769706467 | 0.843793744782845 | 2321.12673786588 | 2393.56782727921 | 2304.61765385897 | 2054.05946042537 | 1420.16077997737 | 1242.65154778704 | 2345.54893802298 | 2543.38639801831 | Wwp1 |
| **ENSMUSG00000041073** | -0.240950239588985 | 0.502443978531197 | 0.900927059989498 | 20.780815481088 | 21.9892246806869 | 20.8220075502658 | 16.8826804966469 | 21.8991639163819 | 37.9155491942896 | 18.7394056300105 | 17.3684090346961 | Nacad |
| **ENSMUSG00000041075** | -0.256479057446016 | 0.113143655522732 | 0.767518180080285 | 210.518695960587 | 239.438224300813 | 182.66579350915 | 250.426427366929 | 257.315176017488 | 295.985900161874 | 280.050006359601 | 222.532740757044 | Fzd7 |
| **ENSMUSG00000041079** | 0.0250337524250584 | 0.89190889824604 | 0.982421229269187 | 158.114900399583 | 132.749763813036 | 106.002947528626 | 125.682177030594 | 150.009272827216 | 114.969729814943 | 128.052605138405 | 120.493337678204 | Rwdd2b |
| **ENSMUSG00000041084** | -0.0290339053495592 | 0.860128191646202 | 0.974868041734905 | 881.829387371388 | 697.139863950665 | 653.053873167428 | 873.209752354348 | 743.476614961167 | 934.434825304429 | 664.207821774815 | 827.170480277403 | Ostc |
| **ENSMUSG00000041096** | 0.111825383470726 | 0.529463051375743 | 0.908560090446144 | 284.606820719249 | 293.189662409158 | 334.098575692902 | 294.508981997063 | 282.499214521327 | 207.923979452556 | 352.92547269853 | 271.381391167127 | Tspyl2 |
| **ENSMUSG00000041112** | -0.575608414072857 | 0.061454563885359 | 0.71282992338528 | 97.5794813894569 | 100.987550385377 | 94.6454888648447 | 135.999370667433 | 197.092475247437 | 236.054870790255 | 103.066730965058 | 104.210454208177 | Elmo1 |
| **ENSMUSG00000041115** | 0.00111749630230487 | 0.991393052124668 | 0.998628446692557 | 1387.79706865005 | 1326.68322240144 | 1293.80383278243 | 1190.22897501361 | 1183.64980968044 | 1243.87463001912 | 1411.70189079412 | 1354.7359047063 | Iqsec2 |
| **ENSMUSG00000041117** | -0.442065723133564 | 0.311096792972454 | 0.851785577030345 | 56.0178504272808 | 66.7820897709749 | 35.965285768641 | 47.8342614071662 | 106.210944994452 | 100.292743030056 | 51.0128264372507 | 23.8815624227072 | Ccdc8 |
| **ENSMUSG00000041119** | -0.154287122870562 | 0.671896171607069 | 0.943216235131458 | 33.4300075130547 | 22.8036404096012 | 43.5369248778286 | 51.5859681841989 | 55.8428679867739 | 48.9232892829544 | 40.6020455316893 | 22.7960368580387 | Pde9a |
| **ENSMUSG00000041120** | -1.03343734831556 | 0.0359490058803259 | 0.679566075569698 | 214.132750826864 | 183.243539005724 | 179.826428843205 | 170.702658354985 | 469.737066006393 | 785.218792991418 | 155.120635492864 | 121.578863242873 | Nbl1 |
| **ENSMUSG00000041124** | 0.0943065049760325 | 0.632156151122548 | 0.931825800800958 | 1230.58568196704 | 1225.69567201606 | 1120.60258815976 | 1022.3400967414 | 901.150595159117 | 782.77262852727 | 1281.5671294746 | 1341.70959793028 | Msantd4 |
| **ENSMUSG00000041126** | -0.136246284553907 | 0.434677996447673 | 0.888379878975956 | 1653.43010132135 | 1403.23830091939 | 1474.57671651428 | 1792.37791272735 | 1781.49698459767 | 2240.68664915931 | 1497.07029421972 | 1431.80821979776 | H2afv |
| **ENSMUSG00000041130** | 0.0563571624852635 | 0.670668784797346 | 0.942873423161451 | 460.791995450213 | 443.042156529395 | 447.673162330715 | 349.846656958294 | 427.033696369448 | 373.040080782527 | 406.020455316893 | 429.868123608729 | Zfp598 |
| **ENSMUSG00000041132** | 0.188807201996266 | 0.375987500034822 | 0.874078513608115 | 54.2108229941427 | 70.0397526866323 | 77.6093008691726 | 75.9720622349111 | 70.0773245324222 | 57.4848649074714 | 54.1360607089191 | 61.8749571861049 | N4bp2l1 |
| **ENSMUSG00000041133** | 0.0801527855013136 | 0.425308704024957 | 0.885066977635295 | 1102.28673421423 | 1245.24164951001 | 1231.33781013163 | 1219.30470253561 | 1267.96159075851 | 1070.19695306463 | 1093.13199508394 | 1107.23607596188 | Smc1a |
| **ENSMUSG00000041134** | 0.0691770525400825 | 0.723345156471785 | 0.951674101666722 | 646.012307346867 | 874.682492853989 | 1050.56492639978 | 952.933521366292 | 1025.97582948249 | 881.842289325253 | 681.906149314269 | 769.637625349972 | Cyyr1 |
| **ENSMUSG00000041135** | -0.438845957274517 | 0.0453173790963685 | 0.695534138838186 | 70.4740698923855 | 69.2253369577179 | 58.6802030962037 | 65.6548685980713 | 97.4512794278996 | 114.969729814943 | 69.7522320672611 | 77.072315091464 | Ripk2 |
| **ENSMUSG00000041143** | 0.161706149882928 | 0.303419715319361 | 0.851785577030345 | 393.931980424104 | 474.804369957054 | 541.372196306912 | 404.246405225267 | 385.425284928322 | 366.924669622158 | 475.772687384154 | 392.96025441 | Tmco4 |
| **ENSMUSG00000041147** | 0.3825646787711 | 0.411274126002323 | 0.882192126163777 | 28.9124389302094 | 28.5045505120015 | 34.0723759913441 | 11.2551203310979 | 12.0445401540101 | 15.9000690169602 | 34.3555769883525 | 16.2828834700276 | Brca2 |
| **ENSMUSG00000041153** | 0.0418964409695032 | 0.845191327972451 | 0.972136081650911 | 172.571119864688 | 152.295741306979 | 116.413951303759 | 135.061443973175 | 125.920192519196 | 103.96198972627 8 | 168.654650670094 | 160.657783570939 | Osgin2 |
| **ENSMUSG00000041161** | 0.211028604453706 | 0.228589674081385 | 0.832048466795696 | 231.299511441676 | 276.901347830872 | 262.16800415562 | 239.171307035831 | 165.338687568684 | 205.477814988408 | 237.365804646799 | 263.782712214447 | Otud3 |
| **ENSMUSG00000041164** | 0.0512878603383335 | 0.753790462813652 | 0.956497786093635 | 1151.07647490896 | 1253.38580679915 | 1136.69232126678 | 995.140222607909 | 932.904382837871 | 898.965440574287 | 1279.48497329349 | 1265.72280840348 | Zmiz2 |
| **ENSMUSG00000041168** | 0.188151444718017 | 0.0671994023328132 | 0.721898114207542 | 6742.92286675478 | 6521.84115714594 | 6424.53578414566 | 6588.93502716358 | 5272.22371286895 | 5202.9918152422 | 6295.39921359296 | 6292.79169838334 | Lonp1 |
| **ENSMUSG00000041180** | 0.184665024406534 | 0.685642916444281 | 0.944452327903597 | 23.4913566307952 | 17.9171460361152 | 17.036187995672 | 24.3860940507122 | 14.2344565456483 | 6.1154111603693 | 30.191264626128 | 21.7105112933701 | Hectd2 |
| **ENSMUSG00000041187** | -0.303316875897936 | 0.298072555807692 | 0.851785577030345 | 309.905204783182 | 302.96265115613 | 301.919109478855 | 227.916186704733 | 512.440435643337 | 442.755768010737 | 234.242570375131 | 221.447215192376 | Prkd2 |
| **ENSMUSG00000041189** | -0.452550898446488 | 0.27395497360575 | 0.845204865395511 | 32.5264937964856 | 31.7622134276588 | 38.8046504345863 | 67.5307219865876 | 96.3563212320805 | 66.0464405319884 | 40.6020455316893 | 30.3947158107182 | Chrnb1 |
| **ENSMUSG00000041193** | -0.195005221970324 | 0.38288914289318 | 0.875126835008975 | 625.231491865779 | 698.768695408494 | 510.139184981513 | 363.915557372167 | 665.734583058011 | 497.794468454061 | 730.836819570408 | 622.006148555055 | Pla2g5 |
| **ENSMUSG00000041199** | -0.386289275823714 | 0.0846722545756979 | 0.743541556158628 | 74.9916384752307 | 62.7100111264033 | 59.6266579848522 | 86.2892558717508 | 86.5016974697087 | 119.862058743238 | 75.998700610598 | 89.0130963028176 | Rpusd1 |
| **ENSMUSG00000041202** | -0.466809405699267 | 0.314275774493041 | 0.853228028313558 | 8.13162344912141 | 16.2883145782866 | 17.9826428843205 | 13.1309737196143 | 31.7537876787538 | 20.7923979452556 | 13.5340151772298 | 10.8552556466851 | Pla2g2d |
| **ENSMUSG00000041203** | -0.126080441660524 | 0.57704979182417 | 0.922450244597442 | 1017.35644485674 | 949.608739914107 | 1030.68937373816 | 1154.5877606318 | 1208.83384818428 | 1591.22998392809 | 994.22957648111 | 738.157383974585 | Trir |
| **ENSMUSG00000041215** | -0.157211580916393 | 0.459251130007879 | 0.894774044236998 | 60.535419010126 | 67.5965054998893 | 63.4124775394459 | 90.0409626487835 | 77.7420319031559 | 68.4926049961361 | 75.998700610598 | 91.1841474321546 | Yeats2 |
| **ENSMUSG00000041219** | -0.238756279872058 | 0.283611740467381 | 0.8510190268075 | 58.7283915769879 | 55.3802695661743 | 51.1085639870161 | 45.0204813243917 | 52.5579933993167 | 64.8233582999146 | 55.1771387994752 | 75.9867895267955 | Arhgap11a |
| **ENSMUSG00000041220** | -0.161392529078238 | 0.745918759587362 | 0.956085046483146 | 7.22810973255236 | 22.8036404096012 | 7.57163910918757 | 10.3171936368398 | 15.3294147414674 | 11.0077400886647 | 13.5340151772298 | 14.1118323406906 | Elovl6 |
| **ENSMUSG00000041225** | 0.108174645289894 | 0.652981772725048 | 0.936991241819063 | 561.985531705946 | 692.253369577179 | 559.354839191232 | 490.535661097018 | 419.368988998714 | 364.47850515801 | 618.400385790345 | 734.90080728058 | Arhgap12 |
| **ENSMUSG00000041229** | -0.162768285013542 | 0.30277166278591 | 0.851785577030345 | 199.676531361759 | 225.593156909269 | 248.917635714542 | 178.206071909051 | 254.03030143003 | 204.254732756335 | 248.817663642917 | 246.414303179751 | Phf8 |
| **ENSMUSG00000041231** | 0.189483328610583 | 0.428691550815895 | 0.885378691567738 | 451.756858284523 | 510.638662029284 | 547.050925638802 | 422.067012416173 | 328.487458745729 | 282.531995609062 | 546.565997541972 | 535.164103381574 | Ublcp1 |
| **ENSMUSG00000041235** | 0.272596228092805 | 0.0995590816459608 | 0.759800953500741 | 580.055806037327 | 650.718167402549 | 613.302767844194 | 661.238319452004 | 436.88832013182 | 421.963370065482 | 575.716184077544 | 638.289032025082 | Chd7 |
| **ENSMUSG00000041236** | 0.0906987763763983 | 0.583724384700279 | 0.922450244597442 | 1118.54998111248 | 1296.54984043161 | 1157.51432881705 | 979.19546880552 | 913.195135313127 | 889.180782717696 | 1252.41694293903 | 1219.04520912273 | Vps41 |
| **ENSMUSG00000041237** | -0.839757724478145 | 0.274273212639637 | 0.845462699415127 | 9.03513716569045 | 18.7315617650296 | 11.3574586637814 | 6.56548685980713 | 8.75966556655277 | 6.1154111603693 | 7.28754663389296 | 59.7039060567679 | Pklr |
| **ENSMUSG00000041238** | -0.234486324442539 | 0.22021647369917 | 0.828720352121776 | 70.4740698923855 | 78.1839099757755 | 61.519567762149 | 64.7169419038131 | 83.2168228822514 | 86.838838477244 | 68.711153976705 | 85.7565196088121 | Rbbp8 |
| **ENSMUSG00000041241** | 0.0452221771042683 | 0.65685727929178 | 0.938389079274206 | 1620.90360752487 | 1646.74860386477 | 1785.96037487962 | 1502.55856420157 | 1585.49946754605 | 1429.78312929434 | 1678.21788197649 | 1658.68306281348 | Mul1 |
| **ENSMUSG00000041258** | 0.288859498295718 | 0.143572472481897 | 0.780607990324772 | 299.063040184354 | 406.39344872825 | 387.100049457215 | 290.75727522003 | 262.789966996583 | 214.039390612925 | 325.857442344071 | 328.914246094558 | Zfp236 |
| **ENSMUSG00000041261** | -0.140997908006585 | 0.544289174336093 | 0.913837378403038 | 420.133878204606 | 335.539280312703 | 424.011790114504 | 657.486612674971 | 595.657258525589 | 535.71001764835 | 391.445362049107 | 502.598336441519 | Car8 |
| **ENSMUSG00000041263** | -0.248505849206646 | 0.0848218762987425 | 0.743541556158628 | 203.290586228035 | 226.407572638183 | 248.917635714542 | 247.612647284155 | 251.840385038392 | 321.670627035425 | 251.940897914585 | 275.723493425801 | Rusc1 |
| **ENSMUSG00000041264** | 0.209402843580996 | 0.235535153635601 | 0.832048466795696 | 518.616873310632 | 506.566583384712 | 511.085639870161 | 513.045901759214 | 371.190828382674 | 346.132271676902 | 486.183468289716 | 567.72987032163 | Uspl1 |
| **ENSMUSG00000041268** | 0.44995917206253 | 0.174152732562448 | 0.799889021902745 | 604.450676384691 | 832.332874950444 | 900.078599104673 | 551.500896223799 | 379.950493949227 | 267.855008824175 | 726.672507208183 | 739.242909539254 | Dmxl2 |
| **ENSMUSG00000041272** | -0.24970110631118 | 0.672492544024854 | 0.943216235131458 | 5.42108229941427 | 4.07207864457164 | 15.1432782183751 | 14.0689004138724 | 16.4243729372865 | 12.2308223207386 | 5.20539045278068 | 11.9407812113536 | Tox |
| **ENSMUSG00000041278** | 0.115771808588476 | 0.270140914484815 | 0.843865788665109 | 1331.77921822277 | 1149.95500922703 | 1166.97887770353 | 1183.6634881538 | 1017.21616391594 | 1066.52770636841 | 1201.40411650178 | 1173.45313540666 | Ttc1 |
| **ENSMUSG00000041297** | 0.132563004762716 | 0.523025380917132 | 0.90648515894888 | 415.616309621761 | 586.379324818317 | 565.033568523123 | 400.494698448235 | 407.324448844704 | 338.793778284459 | 521.580123368624 | 526.479898864226 | Cdk13 |
| **ENSMUSG00000041298** | 0.134422265509771 | 0.385198900538453 | 0.875126835008975 | 435.49361138628 | 483.762942975111 | 473.227444324223 | 450.204813243917 | 379.950493949227 | 332.67836712409 | 490.34778065194 | 474.374671760138 | Katnal1 |
| **ENSMUSG00000041303** | 0.0910562575346802 | 0.671746140576287 | 0.943216235131458 | 213.229237110295 | 294.818493866987 | 221.470443943737 | 236.357526953057 | 266.074841584041 | 147.992950080937 | 244.653351280692 | 247.49982874442 | Gtf3c3 |
| **ENSMUSG00000041308** | -0.234289673015446 | 0.145857270088458 | 0.780607990324772 | 158.114900399583 | 180.800291818981 | 174.147699511314 | 178.206071909051 | 244.175677667659 | 205.477814988408 | 199.886993386778 | 163.914360264945 | Sntb2 |
| **ENSMUSG00000041313** | -0.0137050666802711 | 0.957206147527118 | 0.992403427270738 | 460.791995450213 | 615.698291059232 | 979.580809751142 | 649.983199120906 | 773.040486248282 | 481.894399437101 | 800.589051637669 | 675.196901223812 | Slc7a1 |
| **ENSMUSG00000041319** | -0.0363817832235339 | 0.792265876157716 | 0.961804768755116 | 177.992202164102 | 171.027303072009 | 146.700507740509 | 167.888878272211 | 181.76306050597 | 170.008430258266 | 170.736806851206 | 158.486732441602 | Thoc6 |
| **ENSMUSG00000041324** | -0.702976105230769 | 0.150741734971131 | 0.783020695097675 | 16.2632468982428 | 7.32974156022896 | 12.3039135524298 | 23.448167356454 | 13.1394983498292 | 37.9155491942896 | 29.1501865355718 | 16.2828834700276 | Inhba |
| **ENSMUSG00000041328** | 0.238122298353457 | 0.201798411845955 | 0.813395633194566 | 421.037391921175 | 523.669313691913 | 495.942361651786 | 335.777756544422 | 410.609323432161 | 297.208982393948 | 406.020455316893 | 391.874728845331 | Pcf11 |
| **ENSMUSG00000041329** | -0.747632194799696 | 0.0234268585002049 | 0.654124840968651 | 174.378147297826 | 189.758864837039 | 156.165056626994 | 150.068271081306 | 414.989156215438 | 384.047820871192 | 159.284947855089 | 168.256462523619 | Atp1b2 |

| | | | | | | | | | | | | |
|---|---|---|---|---|---|---|---|---|---|---|---|---|
| **ENSMUSG00000041341** | 0.154336659790396 | 0.461206267555148 | 0.895328112556867 | 460.791995450213 | 663.748819065178 | 611.409858066897 | 469.901273823339 | 521.20010120989 | 340.016860516533 | 563.22324699087 | 556.874614674944 | Atg2b |
| **ENSMUSG00000041346** | -0.16918771454797 | 0.249258209633558 | 0.835314899646652 | 184.316798180085 | 166.955224427437 | 183.612248397799 | 147.254490998531 | 182.858018701789 | 184.685417043153 | 182.188665847324 | 217.105112933702 | Wrap53 |
| **ENSMUSG00000041351** | 0.125953289824882 | 0.409451382691961 | 0.882192126163777 | 251.176813206195 | 221.521078264697 | 236.613722162112 | 208.219726125312 | 218.991639163819 | 181.016170346931 | 191.558368662329 | 248.585354309088 | Rap1gap |
| **ENSMUSG00000041353** | -0.0801472514900752 | 0.795110366965049 | 0.962032824235414 | 25.2983840639333 | 21.1748089517725 | 20.8220075502658 | 26.2619474392285 | 19.7092475247437 | 30.5770558018465 | 27.0680303544596 | 21.7105112933701 | Tmem29 |
| **ENSMUSG00000041354** | -0.28813595457991 | 0.00761896290548483 | 0.635862253526407 | 488.800920663853 | 469.917875583568 | 462.81644054909 | 468.963347129081 | 626.316088008523 | 605.425704876561 | 570.510793624763 | 506.940438700193 | Rgl2 |
| **ENSMUSG00000041355** | -0.217080900490189 | 0.195397392712282 | 0.811094945724481 | 726.425028121512 | 680.037133643464 | 602.891764069061 | 742.837941852464 | 875.966556655277 | 958.896469945906 | 606.948526794228 | 758.782369703287 | Ssr2 |
| **ENSMUSG00000041360** | 0.282157531555598 | 0.00162635887337783 | 0.635862253526407 | 716.486377239253 | 784.282346944498 | 794.075651576047 | 741.900015158206 | 618.65138063779 | 655.572076391589 | 617.359307699789 | 606.808790649696 | Pum3 |
| **ENSMUSG00000041361** | 0.248104161485876 | 0.159013015613457 | 0.789393312599749 | 5887.2953771639 | 5653.67399012327 | 7373.83003746005 | 6482.01138401815 | 4925.1219647943 | 4012.93280343433 | 6261.04363660461 | 6183.15361635182 | Myzap |
| **ENSMUSG00000041362** | -0.40458747976914 | 0.13297867099144 | 0.77303252008843 | 74.9916384752307 | 87.1424829938332 | 102.217127974032 | 109.737423228205 | 93.0714466446232 | 183.462334811079 | 128.052605138405 | 91.1841474321546 | Shtn1 |
| **ENSMUSG00000041375** | 0.0562526133863322 | 0.646087922029344 | 0.935063240511793 | 601.740135234984 | 652.346998860377 | 711.734076263632 | 576.824916968769 | 618.65138063779 | 541.82542880872 | 674.618602680377 | 608.979841779033 | Ccdc9 |
| **ENSMUSG00000041378** | -1.04102762575657 | 0.0212268664156541 | 0.648762042745232 | 783.346392265362 | 317.622134276588 | 477.959718767466 | 1008.27119632752 | 2551.2525962585 | 1318.48264617562 | 814.123066814899 | 639.374557589751 | Cldn5 |
| **ENSMUSG00000041390** | -0.213890433915841 | 0.134169484917363 | 0.77303252008843 | 318.940341948873 | 307.034729800702 | 338.830850136144 | 347.970803569778 | 451.122776677468 | 401.170972120226 | 340.432535611857 | 329.999771659226 | Mdfic |
| **ENSMUSG00000041396** | -0.00145712362359551 | 0.99427445654777 | 0.999262355603273 | 107.518132271716 | 77.3694942468612 | 81.3951204237664 | 72.2203554578784 | 83.2168228822514 | 74.6080161565054 | 95.7791843311646 | 84.6709940441436 | Mettl18 |
| **ENSMUSG00000041408** | 0.23823025230988 | 0.262030388641413 | 0.841137531367976 | 1220.64703108478 | 1469.20597496145 | 1620.33076936614 | 1239.93908980929 | 1038.0203696365 | 804.7881087046 | 1453.34501441637 | 1407.92665737505 | Wapl |
| **ENSMUSG00000041415** | 0.202019281078075 | 0.399210040714017 | 0.880459952732461 | 522.230928176908 | 554.617111390658 | 583.016211407443 | 525.23894878457 | 430.318570956905 | 275.193502216618 | 615.277151518677 | 577.499600403646 | Dicer1 |
| **ENSMUSG00000041417** | 0.0510173820342622 | 0.853346544252384 | 0.973120292022893 | 5827.66347187034 | 5947.67806826134 | 5849.0912118474 | 3131.737232128 | 4431.29581847989 | 3227.71401044292 | 6830.51335213881 | 5544.86458432674 | Pik3r1 |
| **ENSMUSG00000041420** | -0.0962077179773785 | 0.750394240946436 | 0.956197192463153 | 48.7897406947284 | 49.679359463774 | 58.6802030962037 | 72.2203554578784 | 82.1218646864323 | 78.277262852727 | 39.5609674411332 | 45.5920737160773 | Meis3 |
| **ENSMUSG00000041426** | 0.155300072501987 | 0.421590585058006 | 0.882568531013176 | 2428.64487013759 | 2374.02184978527 | 2169.27460478224 | 2239.76894588849 | 1755.21798789801 | 1475.03717188107 | 2568.33964940199 | 2471.74171075019 | Hibch |
| **ENSMUSG00000041429** | -0.209627915630686 | 0.408024324609043 | 0.881573350402261 | 43.3686583953142 | 49.679359463774 | 50.1621090983677 | 37.5170677703264 | 54.7479097909548 | 66.0464405319884 | 52.0539045278068 | 36.9078691987293 | Nthl1 |
| **ENSMUSG00000041438** | 0.0975568743868743 | 0.511393369303575 | 0.902394714365738 | 502.353626412389 | 572.534257426773 | 577.337482075553 | 576.824916968769 | 474.116898789669 | 426.855698993777 | 617.359307699789 | 563.387768062955 | Utp4 |
| **ENSMUSG00000041439** | -0.0345253774291854 | 0.770640775089891 | 0.958995611819782 | 638.784197614315 | 746.004807685525 | 694.69788826796 | 657.486612674971 | 678.87408140784 | 649.456665231219 | 808.917676362118 | 665.427171141795 | Mfsd6 |
| **ENSMUSG00000041440** | 0.543320834589622 | 0.141648199054083 | 0.779852097185473 | 28.9124389302094 | 32.5766291565731 | 33.1259211026956 | 23.448167356454 | 28.4689130912965 | 8.56157562451702 | 23.9447960827911 | 19.5394601640331 | Gk5 |
| **ENSMUSG00000041444** | -0.177673616059023 | 0.2425384744445 | 0.832048466795696 | 168.053551281842 | 175.913797445495 | 161.843785958884 | 190.399118934407 | 187.237851485066 | 165.116101329971 | 207.174540020671 | 226.874843015718 | Arhgap32 |
| **ENSMUSG00000041445** | -0.668702800570943 | 0.0631122453107442 | 0.715615581523459 | 624.32797814921 | 494.350347450997 | 491.210087208544 | 716.575994413235 | 1358.8431210115 | 1351.50586644161 | 494.512093014165 | 493.914131924171 | Mmrn2 |
| **ENSMUSG00000041453** | -0.0136096060919576 | 0.943887914118024 | 0.990505437194639 | 6912.78344546976 | 5473.6881140332 | 6460.5010699143 | 7489.34465365142 | 6672.67524532158 | 8954.18502101273 | 5324.07335510408 | 5636.04873175889 | Rpl21 |
| **ENSMUSG00000041459** | 0.21872935536503 | 0.174920560701395 | 0.799889021902745 | 1571.21035311357 | 1634.53236793106 | 1676.1716077964 | 1600.10294040442 | 1277.81621452089 | 1053.07380181559 | 1692.79297524428 | 1544.70287852329 | Tardbp |
| **ENSMUSG00000041471** | 0.587153869715212 | 0.00434518190016938 | 0.635862253526407 | 108.421645988285 | 142.522752560008 | 123.039135524298 | 109.737423228205 | 76.6470737073368 | 61.154111603693 | 99.9434966933891 | 83.5854684794751 | Fam35a |
| **ENSMUSG00000041476** | 0.110956322859506 | 0.416331839080769 | 0.882192126163777 | 5254.83577556557 | 3927.11264482489 | 3992.14672031915 | 5031.97671469504 | 3832.35368536684 | 4173.15657583601 | 4433.95158767859 | 4418.08904820083 | Smpx |
| **ENSMUSG00000041477** | -0.259626079057357 | 0.447220761103281 | 0.89108031640712 | 42.4651446787451 | 41.5352021746308 | 56.7872933189068 | 40.3308478531009 | 52.5579933993167 | 91.7311674055395 | 36.4377331694648 | 36.9078691987293 | Dcp1b |
| **ENSMUSG00000041481** | 0.427599686972219 | 0.330667686559717 | 0.856303993370193 | 27.1054114970714 | 13.8450673915436 | 24.6078271048596 | 24.3860940507122 | 28.4689130912965 | 15.9000690169602 | 8.32862472444909 | 14.1118323406906 | Serpina3g |
| **ENSMUSG00000041483** | -0.240069784548367 | 0.0607755693065771 | 0.71282992338528 | 209.615182244018 | 223.149909722526 | 219.57753416644 | 199.778385876988 | 248.555510450935 | 271.524255520397 | 221.749633288457 | 265.953763343784 | Zfp281 |
| **ENSMUSG00000041488** | 0.154115685649168 | 0.309387259022844 | 0.851785577030345 | 311.712232216321 | 235.366145656241 | 234.720812384815 | 266.371181169318 | 244.175677667659 | 214.039390612925 | 260.269522639034 | 222.532740757044 | Stx3 |
| **ENSMUSG00000041491** | 0.0553638964227292 | 0.858045590282745 | 0.974350331019239 | 59.631905293557 | 49.679359463774 | 63.4124775394459 | 44.0825546301336 | 33.943704070392 | 36.6924669622158 | 62.4646854333682 | 74.901263962127 | Cep78 |
| **ENSMUSG00000041506** | 0.132758173525319 | 0.286569470814681 | 0.851785577030345 | 520.42390074377 | 424.310594764365 | 520.550188756646 | 571.19735680322 | 476.306815181307 | 456.20967256355 | 463.279750297481 | 460.262839419447 | Rrp9 |
| **ENSMUSG00000041515** | -0.490812666064044 | 0.0236679979957571 | 0.654124840968651 | 70.4740698923855 | 71.6685841444609 | 76.6628459805242 | 92.854742731558 | 141.249607260664 | 112.523565350795 | 85.3684034256032 | 98.7828263848342 | Irf8 |
| **ENSMUSG00000041528** | 0.110713584716792 | 0.665046441513457 | 0.941544702132496 | 1680.53551281842 | 2133.76920975554 | 1851.26576219636 | 1585.09611329629 | 1438.77506930629 | 971.127292266645 | 2208.12663006957 | 2096.14986537489 | Rnf123 |
| **ENSMUSG00000041530** | -0.0858793294285376 | 0.634899776714138 | 0.932543559775713 | 326.168451681425 | 359.157336451219 | 337.884395247495 | 269.184961252092 | 336.152166116463 | 253.178022039289 | 383.116737324658 | 398.387882233342 | Ago1 |
| **ENSMUSG00000041540** | -0.786707825032949 | 0.061319608674407 | 0.71282992338528 | 13.5527057485357 | 13.8450673915436 | 9.46454888648447 | 17.8206071909051 | 21.8991639163819 | 39.1386314263635 | 18.7394056300105 | 15.1973579053591 | Sox5 |
| **ENSMUSG00000041548** | 0.0302062566569662 | 0.719993729638708 | 0.951674101666722 | 15126.626642799 | 13718.8329535619 | 13128.2757604426 | 15362.3013252544 | 13920.2035434482 | 14275.8158127661 | 14145.1280163862 | 13806.7996570187 | Hspb8 |
| **ENSMUSG00000041559** | -1.5900714413972 | NA | NA | 4.51756858284523 | 6.51532583131463 | 10.4110037751329 | 26.2619474392285 | 13.1394983498292 | 113.746647582869 | 11.4518589961175 | 5.42762782334254 | Fmod |
| **ENSMUSG00000041560** | -0.121898197024119 | 0.517045334885026 | 0.905197321821966 | 3289.69344202789 | 2831.72348943512 | 3333.41411781983 | 3469.39084206094 | 3619.93179537793 | 4735.77440258998 | 2941.04560582109 | 2768.09018990469 | Nop53 |
| **ENSMUSG00000041566** | 0.64568006761442 | 0.102300268469306 | 0.759800953500741 | 19.877301764519 | 20.3603932228582 | 15.1432782183751 | 16.8826804966469 | 15.3294147414674 | 7.33849339244316 | 12.4929370866736 | 10.8552556466851 | Tssk1 |
| **ENSMUSG00000041567** | -0.313914077253183 | 0.585567873536397 | 0.922964550512322 | 25.2983840639333 | 22.8036404096012 | 19.8755526616174 | 30.9515809105193 | 21.8991639163819 | 33.0232202659942 | 5.20539045278068 | 62.9604827507734 | Serpina12 |
| **ENSMUSG00000041570** | 0.120550773961182 | 0.607197235841641 | 0.92742133338909 | 668.600150261093 | 823.374301932386 | 748.645816920921 | 514.921755147731 | 607.701798679599 | 408.509465512669 | 784.972880279327 | 732.729756151243 | Camsap2 |
| **ENSMUSG00000041571** | -0.271026045695595 | 0.254430435963791 | 0.837406774112203 | 3502.01916542162 | 2526.31759109225 | 2816.64974861778 | 3600.70057925708 | 3578.32338393681 | 5571.13956709643 | 3054.52311769171 | 2814.76778918544 | Selenow |
| **ENSMUSG00000041577** | -1.2897566527546 | 0.00467216999589625 | 0.635862253526407 | 267.440060104437 | 337.168111770532 | 396.564598343699 | 375.170677703265 | 1107.00273597311 | 1583.89149053565 | 316.487739529066 | 358.223436340608 | Prelp |
| **ENSMUSG00000041592** | -0.100544655831505 | 0.798955437270728 | 0.963240258978423 | 37.9475760958999 | 44.7928650902881 | 55.8408384302584 | 36.5791410760683 | 54.7479097909548 | 80.7234273168747 | 28.1091084450157 | 24.9670879873757 | Sdk2 |
| **ENSMUSG00000041594** | 0.228472669343022 | 0.334185696134651 | 0.858148279095856 | 123.781379169959 | 151.481325578065 | 170.36187995672 | 103.171936368398 | 93.0714466446232 | 95.400414101761 | 142.627698406191 | 136.776221148232 | Tmtc4 |
| **ENSMUSG00000041598** | -0.78161739894728 | 0.0697488191848751 | 0.724916894583216 | 367.730082643601 | 356.714089264476 | 319.901752363175 | 263.557401086543 | 797.129566556303 | 963.788798874201 | 207.174540020671 | 281.151121249143 | Cdc42ep4 |
| **ENSMUSG00000041607** | -0.312547084854509 | 0.264181684160858 | 0.842510961779757 | 51.5002818444356 | 66.7820897709749 | 46.3762895437739 | 48.7721881014244 | 71.1722827282413 | 95.400414101761 | 48.9306702561384 | 51.0197015394199 | Mbp |
| **ENSMUSG00000041609** | -0.340863717335776 | 0.159994548534976 | 0.790986484775773 | 63.2459601598332 | 78.9983257046899 | 54.8943835416099 | 41.2687745473591 | 76.6470737073368 | 68.4926049961361 | 71.8343882483734 | 85.7565196088121 | Bicdl1 |
| **ENSMUSG00000041616** | -2.24271369920512 | 0.0195027221962084 | 0.642273034804458 | 1913.64205169324 | 728.902077378324 | 532.854102309076 | 1902.11533595555 | 1560.31542904221 | 18077.1553900516 | 3284.60137570461 | 1109.40712709121 | Nppa |
| **ENSMUSG00000041623** | -0.168535561524657 | 0.226612837798724 | 0.832048466795696 | 151.7903043836 | 146.594831204579 | 135.343049076728 | 135.061443973175 | 168.623562156141 | 168.785348026193 | 155.120635492864 | 147.631476794917 | D11Wsu47e |
| **ENSMUSG00000041624** | -0.214198769351347 | 0.689507666603307 | 0.94469184542218 | 6.32459601598331 | 16.2883145782866 | 16.0897331070236 | 5.62756016554897 | 8.75966556655277 | 8.56157562451702 | 15.616171358342 | 18.4539345993646 | Gucy1a2 |
| **ENSMUSG00000041625** | -0.0442940970890375 | 0.753034550616101 | 0.956497786093635 | 388.510898124689 | 327.39512302356 | 352.081218577222 | 389.239578117137 | 305.493336633528 | 395.055560959857 | 392.486440139664 | 409.243137880027 | Ggact |
| **ENSMUSG00000041629** | 0.0294207612664652 | 0.81005737660634 | 0.964412705343932 | 726.425028121512 | 570.905425968944 | 609.5169482896 | 628.410885152968 | 604.416924092141 | 688.595296657583 | 617.359307699789 | 574.243023709641 | Fam104a |
| **ENSMUSG00000041632** | 0.0234204103257849 | 0.855025482118134 | 0.973268281968411 | 908.934798868459 | 804.642740167357 | 819.629933569555 | 886.340726073962 | 847.497643563981 | 689.818378889657 | 879.710986519935 | 945.49276682627 | Mrps27 |
| **ENSMUSG00000041633** | -0.338366054382539 | 0.147886306272238 | 0.780607990324772 | 107.518132271716 | 116.461449234749 | 154.272146849697 | 199.778385876988 | 201.472308030714 | 205.477814988408 | 150.95632313064 | 172.598564782293 | Kctd12b |
| **ENSMUSG00000041638** | 0.14665970368053 | 0.399842196463142 | 0.880487015130533 | 1144.75187889298 | 1393.46531217242 | 1687.52906646018 | 1178.03592798825 | 1378.55236853624 | 977.242703427014 | 1279.48497329349 | 1245.09782267478 | Gcn1l1 |
| **ENSMUSG00000041645** | 0.0911698413893373 | 0.358748110389174 | 0.866648170843829 | 2497.31191259684 | 2518.98784953202 | 2978.49353457666 | 2546.47097491091 | 2505.26435203409 | 2577.03426297962 | 2519.40897914585 | 2293.71551814456 | Ddx24 |
| **ENSMUSG00000041650** | 0.22520674181938 | 0.144404176128667 | 0.780607990324772 | 2358.17080024521 | 2340.63080489978 | 2297.99246963843 | 2405.78197077218 | 1919.46171727088 | 1515.39888553951 | 2209.16770816012 | 2397.92597235273 | Pcca |
| **ENSMUSG00000041654** | -0.188507234592758 | 0.446386914985471 | 0.89108031640712 | 65.9565013095403 | 68.4109212288036 | 51.1085639870161 | 55.3376749612315 | 73.3621991198795 | 92.9542496376133 | 54.1360607089191 | 55.3618037980939 | Slc39a11 |
| **ENSMUSG00000041658** | 0.392273548665423 | 0.148103573523658 | 0.78100176015323 | 39.754603529038 | 50.4937751926884 | 61.519567762149 | 47.8342614071662 | 48.1781606160403 | 26.9078091056249 | 41.6431236222455 | 34.7368180693922 | Rragb |
| **ENSMUSG00000041660** | -0.395195027939925 | 0.276026898879062 | 0.846578349250793 | 32.5264937964856 | 39.9063707168021 | 41.6440151005317 | 47.8342614071662 | 38.3235368536684 | 48.9232892829544 | 32.2734208072402 | 93.3551985614916 | Bbox1 |
| **ENSMUSG00000041671** | 0.162824730578785 | 0.215071851157368 | 0.824338622350122 | 301.773581334061 | 236.99497711407 | 273.525462819401 | 261.681547698027 | 247.460552255116 | 222.600966237442 | 235.283648465687 | 252.927456567762 | Pyroxd1 |
| **ENSMUSG00000041684** | 0.22015884220668 | 0.23970875460816 | 0.832048466795696 | 174.378147297826 | 166.955224427437 | 173.201244622666 | 164.137171495178 | 159.863896589588 | 99.0696607979826 | 153.038479311752 | 169.341988088287 | Bivm |
| **ENSMUSG00000041685** | 0.204242586325206 | 0.202340518231556 | 0.814640521227916 | 664.082581678248 | 834.776122137187 | 863.166858447383 | 728.769041438591 | 690.91862156185 | 517.363784167243 | 732.91897575152 | 740.328435103922 | Fcho2 |
| **ENSMUSG00000041688** | 0.329524360886603 | 0.202588190767211 | 0.815165112907463 | 539.39768879172 | 530.184639523228 | 426.85115478045 | 467.087493740564 | 296.733671066975 | 242.170281950624 | 502.840717738614 | 519.966745476215 | Amot |
| **ENSMUSG00000041695** | -0.277326127227803 | 0.225155416023403 | 0.832048466795696 | 664.986095394817 | 582.307246173745 | 754.324546252812 | 591.8317440769 | 812.45898129777 | 472.10974158051 | 1001.517123115 | 855.394144958784 | Kcnj2 |
| **ENSMUSG00000041696** | -0.821178012283171 | 0.00887326116924681 | 0.635862253526407 | 74.9916384752307 | 74.1118313312039 | 59.6266579848522 | 81.59962240046 | 163.148771177045 | 187.131581507301 | 82.2451691539348 | 81.4144173501381 | Rasl12 |
| **ENSMUSG00000041697** | -0.211664706039964 | 0.344435761396037 | 0.861266047293794 | 2680.72519706036 | 2167.97467036994 | 2226.06189810115 | 2637.44986425395 | 3130.4854818468 | 3915.08622486842 | 2098.81343056117 | 2103.74854432757 | Cox6a1 |
| **ENSMUSG00000041698** | -0.485022379601929 | 0.0952231568405703 | 0.757085157577544 | 74.0881247586617 | 61.895595397489 | 40.6975602118832 | 64.7169419038131 | 67.887408140784 | 64.8233582999146 | 80.1630129728225 | 124.835439936878 | Slco1a1 |
| **ENSMUSG00000041702** | -0.140347488194073 | 0.693685421037233 | 0.94469184542218 | 159.018414116152 | 263.870696168242 | 255.542819935081 | 150.068271081306 | 197.092475247437 | 102.738907494204 | 344.596847974081 | 268.124814473121 | Btbd7 |
| **ENSMUSG00000041712** | 0.19322128275637 | 0.128196001788552 | 0.77303252008843 | 734.556651570634 | 818.4878075589 | 866.006223113329 | 719.38977449601 | 669.019457645468 | 594.417964787896 | 773.52102128321 | 706.677142599198 | Ubr7 |
| **ENSMUSG00000041716** | -0.366341046216129 | 0.0547012944198091 | 0.703396849069134 | 56.0178504272808 | 75.7406627890326 | 72.8770264259304 | 75.9720622349111 | 91.9764884488041 | 90.5080851734656 | 98.902418602833 | 80.3288917854696 | Gm20604 |
| **ENSMUSG00000041718** | 0.174367883690739 | 0.377365579472731 | 0.87422797717331 | 77.7021796249379 | 77.3694942468612 | 99.3777633080869 | 73.1582821521366 | 74.4571573156986 | 68.4926049961361 | 64.5468416144805 | 82.4999429148066 | Alg13 |
| **ENSMUSG00000041720** | 0.0380462159846363 | 0.854038713014586 | 0.9731539929935 | 980.312382477414 | 1285.96243595572 | 1334.50139299431 | 1002.64363616197 | 1054.44474257379 | 779.103381831049 | 1365.89445480965 | 1283.09121743818 | Pi4ka |
| **ENSMUSG00000041729** | -0.565269318001063 | 0.0522589279701409 | 0.703396849069134 | 169.86057871498 | 184.057954734638 | 268.793188376159 | 176.330218520534 | 385.425284928322 | 390.163232031561 | 174.901119213431 | 232.302470839061 | Coro2b |
| **ENSMUSG00000041731** | 0.0611864266953973 | 0.772022731717492 | 0.959045638244974 | 2869.55956382329 | 3413.21631987995 | 2640.60913932917 | 2315.7410081234 | 2264.37354895389 | 1980.17013372758 | 3456.37926064637 | 3070.95182244721 | Pgm5 |
| **ENSMUSG00000041733** | 0.109507759488103 | 0.475716369213861 | 0.897549741595057 | 3307.76371635927 | 3174.59251130805 | 2956.72507213775 | 3052.95138981032 | 2651.98875027385 | 2248.02514255175 | 3400.16104375634 | 3277.20167973422 | Coq5 |
| **ENSMUSG00000041734** | -0.797325455943978 | 0.00794288147355256 | 0.635862253526407 | 68.6670424592474 | 83.0704043492615 | 91.8061241988993 | 75.0341355406529 | 171.908436743598 | 198.139321595965 | 104.107809055614 | 80.3288917854696 | Kirrel |
| **ENSMUSG00000041736** | -0.279266836293967 | 0.297049824896035 | 0.851785577030345 | 931.522641782685 | 793.240919962556 | 776.093008691726 | 973.567908639971 | 1066.4892827278 | 1669.50724678082 | 712.097413940397 | 769.637625349972 | Tspo |
| **ENSMUSG00000041740** | 0.207013979141339 | 0.163832508274136 | 0.793968226938511 | 12846.1580221787 | 13424.8288754238 | 15097.84838372 | 12033.5994873322 | 9993.6834532409 | 9897.18142194167 | 13449.6878518947 | 12922.0963218139 | Rnf10 |
| **ENSMUSG00000041741** | 0.164623372593212 | 0.525240791902421 | 0.908006595346551 | 1602.83333319349 | 2346.33171500218 | 2052.86065347848 | 1435.02784221499 | 1415.78094719409 | 1060.41229520804 | 2273.7145497746 | 1884.47238026453 | Pde3a |
| **ENSMUSG00000041747** | 0.179527111810028 | 0.256139481427603 | 0.837517407767513 | 140.948139784771 | 140.079505373265 | 132.503684410783 | 166.013024883695 | 145.62944004394 | 122.308223207386 | 116.600746142287 | 127.006491066215 | Utp15 |
| **ENSMUSG00000041757** | -0.109968834748749 | 0.628491669417316 | 0.931825800800958 | 589.090943203017 | 746.819223414439 | 686.179794270124 | 646.231492343873 | 548.574056105368 | 514.917619703095 | 985.900951756661 | 829.34153140674 | Plekha6 |
| **ENSMUSG00000041762** | -0.00357067869960877 | 0.985997244716968 | 0.997705378387025 | 258.404922938747 | 422.681763306537 | 311.383658365339 | 265.43325447506 | 309.873169416804 | 259.293433199658 | 369.582722147429 | 322.401092706547 | Gpr155 |

| | | | | | | | | | | | | |
|---|---|---|---|---|---|---|---|---|---|---|---|---|
| **ENSMUSG00000041763** | 0.182172708287165 | 0.135915898867531 | 0.776087404131513 | 1049.88293865323 | 1182.53163838361 | 1163.19305814894 | 1075.80191831411 | 931.809424642051 | 853.711397987554 | 1136.8572748873 | 1017.13745409439 | Tpp2 |
| **ENSMUSG00000041765** | 0.029232253315139 | 0.790448497882371 | 0.961804768755116 | 821.293968361262 | 776.138189655355 | 830.987392233336 | 832.878904501247 | 902.245553354936 | 765.649477278236 | 821.410613448792 | 705.59161703453 | Ubac2 |
| **ENSMUSG00000041769** | 0.172256146777682 | 0.0666493361449258 | 0.719725956118125 | 1245.04190143214 | 1111.67746996806 | 1276.76764478675 | 1259.63555038871 | 1057.72961716125 | 1024.94291047789 | 1100.41954171784 | 1157.17025193663 | Ppp2r2d |
| **ENSMUSG00000041773** | -0.413176020688475 | 0.298092088502977 | 0.851785577030345 | 163.535982698997 | 196.274190668353 | 154.272146849697 | 105.985716451172 | 283.594172717146 | 329.009120427868 | 121.806136595068 | 92.2696729968231 | Enc1 |
| **ENSMUSG00000041774** | 0.199371168580342 | 0.460472975003398 | 0.894774044236998 | 61.4389327266951 | 36.6487078011448 | 40.6975602118832 | 45.9584080186499 | 49.2731188118594 | 39.1386314263635 | 33.3144988977964 | 39.0789203280663 | Ydjc |
| **ENSMUSG00000041775** | 0.00259634394173739 | 0.988804265297156 | 0.998437672973644 | 240.334648607366 | 241.881471487556 | 248.917635714542 | 235.419600258798 | 270.454674367317 | 210.370143916704 | 302.953724351836 | 180.197243734972 | Mapk1ip1 |
| **ENSMUSG00000041777** | -0.105342500647107 | 0.65165576629036 | 0.936720465895231 | 493.318489246699 | 445.485403716138 | 446.726707442067 | 549.625042835282 | 544.194223322091 | 749.749408261276 | 438.293876124134 | 350.624757387928 | Cir1 |
| **ENSMUSG00000041779** | -0.107429606469993 | 0.631969828924111 | 0.931825800800958 | 93.9654265231807 | 132.749763813036 | 126.824955078892 | 143.502784221499 | 168.623562156141 | 150.439114545085 | 104.107809055614 | 112.894658725525 | Tram2 |
| **ENSMUSG00000041781** | 0.195041052456982 | 0.303430082751383 | 0.851785577030345 | 599.933107801846 | 626.285695535119 | 606.677583623654 | 447.391033161143 | 467.547149614754 | 370.593916318379 | 583.003730711437 | 569.900921450967 | Cpsf2 |
| **ENSMUSG00000041782** | 0.461896596351702 | 0.243666823918808 | 0.832048466795696 | 25.2983840639333 | 26.0613033252585 | 61.519567762149 | 30.0136542162612 | 16.4243729372865 | 33.0232202659942 | 26.0269522639034 | 28.2236646813812 | Lad1 |
| **ENSMUSG00000041794** | -0.439606099671834 | 0.159870405471749 | 0.790811941596364 | 190.641394196068 | 197.088606397268 | 194.023252172932 | 107.861569839689 | 262.789966996583 | 352.247682837272 | 175.942197303987 | 145.46042566558 | Myrip |
| **ENSMUSG00000041797** | -0.393116705940712 | 0.21417726844986 | 0.824248286808591 | 131.009488902512 | 193.016527752696 | 212.952349945901 | 170.702658354985 | 294.543754675337 | 318.001380339203 | 112.436433780063 | 205.164331722348 | Abca9 |
| **ENSMUSG00000041798** | 0.085299811703386 | 0.683127774200134 | 0.944452327903597 | 1138.427282877 | 939.835751167135 | 719.30571537282 | 1123.63617972128 | 738.001823982071 | 949.111812089315 | 816.205222996011 | 1192.99259557069 | Gck |
| **ENSMUSG00000041801** | -0.766470710128414 | 0.0777309588563836 | 0.739855948845396 | 197.869503928621 | 147.409246933493 | 90.8596693102509 | 217.598993067893 | 362.431162816121 | 472.10974158051 | 119.723980413956 | 158.486732441602 | Phlda3 |
| **ENSMUSG00000041815** | -0.171366417436639 | 0.211053023712797 | 0.823123035902316 | 416.51982333833 | 427.568257680022 | 424.958245003153 | 360.163850595134 | 524.484975797347 | 491.679057293692 | 399.773986773556 | 420.098393526712 | Poldip3 |
| **ENSMUSG00000041817** | 0.471925279205148 | 0.245729768701649 | 0.832048466795696 | 25.2983840639333 | 27.6901347830872 | 41.6440151005317 | 43.1446279358754 | 27.3739548954774 | 8.56157562451702 | 33.3144988977964 | 29.3091902460497 | Fam169a |
| **ENSMUSG00000041827** | -0.286296763292658 | 0.515148302038189 | 0.904145806837306 | 18.9737880479499 | 36.6487078011448 | 17.9826428843205 | 24.3860940507122 | 29.5638712871156 | 53.8156182112498 | 19.7804837205666 | 17.3684090346961 | Oasl1 |
| **ENSMUSG00000041828** | -0.700307460718412 | 0.0346112616172533 | 0.679566075569698 | 294.545471601509 | 311.921224174188 | 274.47191770805 | 369.543117537716 | 697.488370736765 | 750.97249049335 | 274.84461590682 | 309.374785930525 | Abca8a |
| **ENSMUSG00000041836** | 0.289001185007756 | 0.338932441674472 | 0.858570960161243 | 73.1846110420926 | 45.6072808192024 | 70.0376617599851 | 105.047789756914 | 61.3176589658694 | 52.592535979176 | 46.8485140750261 | 79.2433662208011 | Ptpre |
| **ENSMUSG00000041837** | 0.0288880092536293 | 0.829937212001617 | 0.967566911862526 | 387.60738440812 | 418.609684661965 | 393.725233677754 | 519.611388619021 | 405.134532453066 | 419.517205601334 | 415.390158131898 | 445.065481514088 | Pdcd7 |
| **ENSMUSG00000041840** | 0.0164866105415601 | 0.905347923920164 | 0.98504921011344 | 446.335775985108 | 350.198763433161 | 365.3315870183 | 471.777127211855 | 416.084114411257 | 404.840218816448 | 409.143689588562 | 384.276049892652 | Haus1 |
| **ENSMUSG00000041841** | 0.148197238807336 | 0.397853981362823 | 0.880282219458479 | 2496.40839888027 | 1835.6930529729 | 2145.61323256603 | 2567.10536218459 | 2073.85082288137 | 2563.58035842681 | 1767.75059776432 | 1757.46588919831 | Rpl37 |
| **ENSMUSG00000041845** | -0.336853874048892 | 0.175079175190846 | 0.799889021902745 | 55.1143367107117 | 51.3081909216027 | 48.2691993210708 | 81.59962240046 | 81.0269064906132 | 88.0619207093179 | 69.7522320672611 | 59.7039060567679 | Rhod |
| **ENSMUSG00000041846** | -0.0219022782333246 | 0.863506517120196 | 0.974868041734905 | 488.800920663853 | 539.9576282702 | 555.569019636638 | 444.577253078368 | 501.490853685146 | 456.20967256355 | 541.360607089191 | 560.13119136895 | Ppp4r3a |
| **ENSMUSG00000041849** | -0.254370050924206 | 0.141761698363385 | 0.779852097185473 | 95.7724539563188 | 98.5443031986338 | 132.503684410783 | 97.5443762028488 | 124.825234323377 | 121.085140975312 | 125.970448957293 | 133.519644454226 | Card6 |
| **ENSMUSG00000041852** | 0.117584098253161 | 0.560617301243826 | 0.916762264997884 | 343.335212296237 | 370.55915665602 | 385.207139679918 | 377.046531091781 | 291.25888008788 | 237.277953022329 | 426.842017128016 | 403.815510056685 | Tcf20 |
| **ENSMUSG00000041859** | -0.566576247325206 | 0.0714395457152883 | 0.727489215531785 | 86.7373167906283 | 61.0811796685746 | 74.7699362032273 | 60.9652351267805 | 140.154649064844 | 144.323703384715 | 67.6700758861489 | 68.388110574116 | Mcm3 |
| **ENSMUSG00000041870** | -0.111440137702185 | 0.392096377297995 | 0.878285927324016 | 720.100432105529 | 754.148964974668 | 852.755854672251 | 710.948434247686 | 874.871598459458 | 930.765578608207 | 763.110240377648 | 714.275821551878 | Ankrd13a |
| **ENSMUSG00000041879** | 0.0877739236671518 | 0.520741516818788 | 0.905962179970576 | 699.31961662441 | 802.199492980614 | 733.502538702546 | 662.176246146262 | 673.399290428745 | 556.502415593606 | 750.617303290975 | 744.670537362596 | Ipo9 |
| **ENSMUSG00000041881** | 0.146644719304845 | 0.349814007790546 | 0.861774532843376 | 5162.67737647552 | 3537.82192640384 | 4220.24234848342 | 4933.49441179793 | 3844.39822552085 | 4722.32049803717 | 3675.00565966316 | 3887.26704707793 | Ndufa7 |
| **ENSMUSG00000041889** | 0.189312888278713 | 0.606732082257172 | 0.927177252274348 | 118.360296870545 | 64.338842584232 | 106.002947528626 | 209.15765281957 | 91.9764884488041 | 165.116101329971 | 87.4505596067155 | 92.2696729968231 | Shisa4 |
| **ENSMUSG00000041890** | -0.181426964986072 | 0.247596917300769 | 0.83352428049144 | 404.774145022932 | 473.989954228139 | 440.101523221528 | 383.612017951588 | 569.37826182593 | 532.040770952129 | 426.842017128016 | 403.815510056685 | Git2 |
| **ENSMUSG00000041891** | -0.0136144342462974 | 0.91928500687456 | 0.986713918026868 | 1461.88519340871 | 1589.73950284077 | 1579.63320915426 | 1183.6634881538 | 1450.8196094603 | 1305.02874162281 | 1590.76732236978 | 1522.99236722992 | Lman1 |
| **ENSMUSG00000041895** | 0.178235615692324 | 0.216976441132867 | 0.825699388311104 | 3136.09611021116 | 3532.93543203036 | 3481.06108044899 | 3484.39766916907 | 2642.13412651148 | 2502.42624682312 | 3536.5422736192 | 3367.30030160171 | Wipi1 |
| **ENSMUSG00000041912** | 0.341066703186111 | 0.109648522838999 | 0.763154315676794 | 142.755167217909 | 122.976775066064 | 193.076797284283 | 126.620103724852 | 128.110108910834 | 96.6234963338349 | 133.257995591185 | 103.124928643508 | Tdrkh |
| **ENSMUSG00000041915** | -0.241007544321055 | 0.215210912000393 | 0.824338622350122 | 168.957064998411 | 228.036404096012 | 239.453086828057 | 155.695831246855 | 232.131137513649 | 205.477814988408 | 221.749633288457 | 276.809018990469 | Ammecr1l |
| **ENSMUSG00000041920** | 0.170931159913969 | 0.494397471510976 | 0.900141680336539 | 271.957628687283 | 283.416673662186 | 331.259211026956 | 339.529463321454 | 257.315176017488 | 154.108361241306 | 355.007628879643 | 321.315567141878 | Slc16a6 |
| **ENSMUSG00000041921** | 0.153192398831574 | 0.233970704825236 | 0.832048466795696 | 456.274426867368 | 368.115909469277 | 402.24332767559 | 436.135912830045 | 351.48158085793 | 337.570696052385 | 402.897221045225 | 401.644458927348 | Metap1d |
| **ENSMUSG00000041926** | -0.239413348736004 | 0.063991558987318 | 0.716991331547963 | 516.809845877494 | 514.710740673856 | 488.370722542599 | 490.535661097018 | 671.209374037106 | 620.102691661447 | 598.619902069779 | 484.144401842154 | Rnpep |
| **ENSMUSG00000041930** | -0.242950621508491 | 0.551489837444025 | 0.914873920695527 | 28.0089252136404 | 48.0505280059454 | 24.6078271048596 | 12.1930470253561 | 37.2285786578493 | 36.6924669622158 | 35.3966550789086 | 24.9670879873757 | Fam222a |
| **ENSMUSG00000041935** | -0.205336355018617 | 0.196237864733812 | 0.811906300001476 | 310.808718499751 | 329.838370210303 | 329.36630124966 | 315.143369270742 | 342.721915291377 | 286.201242305283 | 425.80093903746 | 425.526021350055 | AW549877 |
| **ENSMUSG00000041936** | -0.81718107425865 | 0.00264003004374854 | 0.635862253526407 | 262.018977805023 | 317.622134276588 | 295.293925258315 | 245.736793895638 | 688.728705170212 | 624.995020589742 | 335.227145159076 | 326.743194965221 | Agrn |
| **ENSMUSG00000041939** | -0.213634408814881 | 0.180830370480891 | 0.802808628271642 | 119.263810587114 | 118.904696421492 | 134.396594188079 | 128.495957113368 | 140.154649064844 | 174.900759186562 | 132.216917500629 | 134.605170018895 | Mvk |
| **ENSMUSG00000041949** | 0.167133324868044 | 0.436751859136306 | 0.888701139843674 | 215.036264543433 | 308.663561258531 | 321.794662140472 | 263.557401086543 | 254.03030143003 | 168.785348026193 | 293.584021536831 | 270.295865602458 | Tango6 |
| **ENSMUSG00000041957** | 0.322234675344162 | 0.00951628664596463 | 0.635862253526407 | 4885.29866548883 | 5500.56383308738 | 5507.42099704531 | 5565.65700372793 | 4292.23612761086 | 3620.32340693862 | 5001.33914703168 | 4247.66153454787 | Pkp2 |
| **ENSMUSG00000041958** | -0.0881260653149019 | 0.412087906367727 | 0.882192126163777 | 527.652010476322 | 574.977504613516 | 578.283936964201 | 498.039074651084 | 571.568178217569 | 545.494675504941 | 631.934400967575 | 566.644344756961 | Pigs |
| **ENSMUSG00000041959** | -0.286530033456429 | 0.243116209132185 | 0.832048466795696 | 709.2582675067 | 525.298145149742 | 594.373670071225 | 667.803806311811 | 857.352267326353 | 1064.08154190426 | 478.895921655823 | 645.887710977762 | S100a10 |
| **ENSMUSG00000041961** | 0.43119649454975 | 0.0555048715252805 | 0.703396849069134 | 71.3775836089546 | 67.5965054998893 | 104.110037751329 | 90.9788893430416 | 63.5075753575076 | 51.3694537471021 | 62.4646854333682 | 69.4736361387845 | Znrf3 |
| **ENSMUSG00000041966** | 0.11135566608035 | 0.58317210817756 | 0.922450244597442 | 104.807591122009 | 74.1118313312039 | 92.7525790875478 | 102.23400967414 | 73.3621991198795 | 81.9465095489486 | 104.107809055614 | 85.7565196088121 | Dcaf17 |
| **ENSMUSG00000041974** | -0.145015347126376 | 0.613941081340247 | 0.928075610917191 | 30.7194663633475 | 61.0811796685746 | 47.3227444324223 | 34.703287687552 | 55.8428679867739 | 48.9232892829544 | 47.8895921655823 | 40.1644458927348 | Spidr |
| **ENSMUSG00000041975** | 0.15523680443193 | 0.419882906483307 | 0.882568531013176 | 82.2197482077831 | 123.791190794978 | 98.4313084194385 | 105.047789756914 | 87.5966556655277 | 85.6157562451702 | 90.5737938783839 | 104.210454208177 | Mettl8 |
| **ENSMUSG00000041977** | 0.202986661592663 | 0.44447807711777 | 0.89108031640712 | 550.239853390548 | 698.768695408494 | 624.660226507975 | 463.335786963532 | 423.748821781991 | 299.655146858096 | 655.879197050366 | 650.229813236436 | Arhgef11 |
| **ENSMUSG00000041992** | -0.152504200866797 | 0.509900892064136 | 0.902394714365738 | 261.115464088454 | 284.231089391101 | 340.723759913441 | 187.585338851632 | 366.810995599397 | 220.154801773295 | 330.021754706295 | 275.723493425801 | Rapgef5 |
| **ENSMUSG00000041995** | -0.0829140278463067 | 0.602881398816191 | 0.926448151609826 | 266.536546387868 | 302.96265115613 | 363.438677241004 | 283.253861665965 | 308.778211220985 | 278.86274891284 | 379.99350305299 | 319.144516012541 | Zbed3 |
| **ENSMUSG00000041997** | 0.163793114969171 | 0.460122030805086 | 0.894774044236998 | 787.863960848207 | 1050.59629029948 | 968.223351087361 | 759.720622349111 | 679.969039603659 | 552.833168897385 | 1033.79054392224 | 916.18357658022 | Tlk1 |
| **ENSMUSG00000042010** | 0.367848504496391 | 0.184705609930795 | 0.805963874499992 | 9682.95650047046 | 12871.840595491 | 15706.418877121 | 9923.26442525135 | 8581.18738063426 | 5180.97633506487 | 11893.2761065133 | 11683.5116525271 | Acacb |
| **ENSMUSG00000042015** | 0.275968446665245 | 0.181829151127096 | 0.80356567265522 | 700.22313034101 | 761.478706534897 | 738.234813145789 | 710.948434247686 | 537.624474147177 | 404.840218816448 | 808.917676362118 | 651.315338801105 | Wdr41 |
| **ENSMUSG00000042029** | -0.110495493462977 | 0.676185275360319 | 0.94409858171788 | 29.8159526467785 | 39.0919549878878 | 27.447191770805 | 33.7653609932938 | 30.6588294829347 | 40.3617136584374 | 36.4377331694648 | 33.6512925047237 | Ncapg2 |
| **ENSMUSG00000042032** | 0.059576965478026 | 0.432105231575661 | 0.8866807012241 | 1944.36151805658 | 1968.44281678593 | 1956.32225483634 | 1880.54302198761 | 1975.30458525765 | 1750.23067409769 | 1844.79037646547 | 1864.9329201005 | Mat2b |
| **ENSMUSG00000042035** | -0.238669141154415 | 0.107700636255503 | 0.762287160109383 | 200.580045078328 | 195.459774939439 | 231.881447718869 | 177.268145214792 | 267.16979977986 | 206.700897220482 | 236.324726556243 | 238.815624227072 | Igsf3 |
| **ENSMUSG00000042042** | 0.10952205862824 | 0.288943163064995 | 0.851785577030345 | 592.704998069294 | 650.718167402549 | 577.337482075553 | 559.004309777864 | 529.959766776443 | 523.479195327612 | 563.22324699087 | 589.440381615 | Csgalnact2 |
| **ENSMUSG00000042043** | 0.123558371677935 | 0.47161395068827 | 0.897171173241185 | 1306.48083415884 | 924.361852317763 | 919.007696877642 | 1178.03592798825 | 995.316999999559 | 1187.61284734372 | 851.60187807492 | 938.979613438259 | Tbca |
| **ENSMUSG00000042045** | -5.86150753739434 | NA | NA | 18.0702743313809 | 4.07207864457164 | 0 | 18.7585338851632 | 3.28487458745729 | 2369.11028352707 | 1.04107809055614 | 4.34210225867403 | Sln |
| **ENSMUSG00000042046** | 0.0987326904199579 | 0.590776451389803 | 0.923935529355527 | 379.475760958999 | 466.66021266791 | 530.01473764313 | 376.108604397523 | 342.721915291377 | 353.470765069345 | 473.690531203042 | 465.69046724279 | Dstyk |
| **ENSMUSG00000042050** | 0.0407099316330877 | 0.803318565330473 | 0.963461896828986 | 97.5794813894569 | 112.389370590177 | 115.467496415111 | 126.620103724852 | 110.590777777729 | 117.415894279091 | 99.9434966933891 | 111.809133160856 | Wdr60 |
| **ENSMUSG00000042055** | 0.262046685697693 | 0.117499039942286 | 0.769868040577702 | 593.608511785863 | 716.685841444609 | 776.093008691726 | 676.245146560134 | 529.959766776443 | 456.20967256355 | 700.64555494428 | 615.492995167044 | Wdr11 |
| **ENSMUSG00000042063** | 0.0353681430663259 | 0.84104884459021 | 0.970420002017028 | 207.80815481088 | 215.820168162297 | 210.112985279955 | 259.805694309511 | 233.226095709468 | 160.223772401676 | 245.694429371248 | 231.216945274392 | Zfp386 |
| **ENSMUSG00000042066** | 0.0257717350820734 | 0.818086847417732 | 0.966315773308182 | 1313.70894389139 | 1339.71387406407 | 1243.64172368406 | 1180.84970807103 | 1162.84560395988 | 1113.00483118721 | 1329.45672164019 | 1381.87404382301 | Tmcc2 |
| **ENSMUSG00000042073** | -0.255465521725857 | 0.120848969187694 | 0.769868040577702 | 129.202461469373 | 137.636258186522 | 125.878500190243 | 123.806323642077 | 150.009272827216 | 189.577745971448 | 131.175839410073 | 147.631476794917 | Abhd14b |
| **ENSMUSG00000042079** | -0.0551985025920462 | 0.612437219874624 | 0.928055903343287 | 1402.25328811516 | 1372.29050322064 | 1542.72146849697 | 1645.12342172881 | 1721.27428382762 | 1519.06813223573 | 1452.30393632581 | 1501.28185593655 | Hnrnpf |
| **ENSMUSG00000042082** | -0.363414913360696 | 0.303917314709126 | 0.851785577030345 | 35.2370349461928 | 85.5136515360045 | 110.735221971868 | 44.0825546301336 | 96.3563212320805 | 79.5003450848009 | 87.4505596067155 | 91.1841474321546 | Arsb |
| **ENSMUSG00000042087** | 0.261818019930499 | 0.393344540785377 | 0.878942305112115 | 31.6229800799166 | 60.2667639396603 | 35.965285768641 | 46.8963347129081 | 31.7537876787538 | 40.3617136584374 | 45.80743598447 | 28.2236646813812 | 4933440N22 |
| **ENSMUSG00000042099** | -0.380884090243455 | 0.194230545921117 | 0.809162446871198 | 912.548853734736 | 826.631964848044 | 1004.188636856 | 991.388515830876 | 1641.34233553283 | 1714.76128936755 | 738.124366204301 | 769.637625349972 | Kank3 |
| **ENSMUSG00000042102** | -0.302443747963133 | 0.457476596914685 | 0.892966448175628 | 55.1143367107117 | 117.275864963663 | 49.2156542097192 | 70.3445020693621 | 60.2227007700503 | 77.0541806206532 | 59.3414511616998 | 163.914360264945 | Dmgdh |
| **ENSMUSG00000042104** | 0.0719190490230671 | 0.740055786986048 | 0.954383574136061 | 149.079763233892 | 151.481325578065 | 140.07532351997 | 146.316564304273 | 134.679858085749 | 92.9542496376133 | 188.435134390661 | 141.118323406906 | Uggt2 |
| **ENSMUSG00000042105** | -0.201022378506002 | 0.329444904087171 | 0.855977678491063 | 93.9654265231807 | 107.502876216691 | 90.8596693102509 | 111.613276616721 | 135.774816281568 | 133.315963296051 | 113.477511870619 | 82.4999429148066 | Inpp5f |
| **ENSMUSG00000042106** | -0.46312660257697 | 0.203724946809611 | 0.816074141171399 | 62.3424464432641 | 36.6487078011448 | 77.6093008691726 | 83.4754757889764 | 110.590777777729 | 132.092881063977 | 57.2592949805875 | 58.6183804920994 | Fam212a |
| **ENSMUSG00000042109** | 0.128385326873255 | 0.444714109328741 | 0.89108031640712 | 2977.077696095 | 3187.62316297068 | 3617.35058441436 | 2994.79993476631 | 2390.29374147309 | 2652.8653613682 | 3810.34581143546 | 2834.30724934947 | Csdc2 |
| **ENSMUSG00000042111** | 0.144844474921536 | 0.173675425352088 | 0.799889021902745 | 337.010616280254 | 312.735639903102 | 316.115932808581 | 350.784583652552 | 295.638712871156 | 300.878229090169 | 299.830490080167 | 294.177428025166 | Ccdc115 |
| **ENSMUSG00000042115** | -0.145818984663649 | 0.62812991575455 | 0.931825800800958 | 376.765219809292 | 351.013179162076 | 149.539872406455 | 250.426427366929 | 226.656346534553 | 294.7628179298 | 318.569895710178 | 408.157612315359 | Klhdc8a |
| **ENSMUSG00000042116** | -1.06856142516704 | 0.0273206030667625 | 0.664353746352133 | 235.817080024521 | 211.748089517725 | 151.432782183751 | 243.860940507122 | 618.65138063779 | 821.911259953634 | 149.915245040084 | 178.026192605635 | Vwa1 |
| **ENSMUSG00000042118** | -0.285956326449101 | 0.464994151793652 | 0.895328112556867 | 30.7194663633475 | 59.452348210746 | 31.2330113253987 | 31.8895076047775 | 37.2285786578493 | 34.2463024980681 | 34.3555769883525 | 81.4144173501381 | Bhmt2 |
| **ENSMUSG00000042121** | -0.183119743287171 | 0.169411565479599 | 0.79907358295209 | 196.965990212052 | 208.490426602068 | 194.96970706158 | 235.419600258798 | 258.410134213307 | 244.616446414772 | 226.955023741238 | 219.276164063039 | Ssh1 |
| **ENSMUSG00000042129** | -0.142262650944056 | 0.632718706685802 | 0.93206732801631 | 207.80815481088 | 212.56250524664 | 306.651383922097 | 150.068271081306 | 343.816873487196 | 280.085831144914 | 170.736806851206 | 173.684090346961 | Rassf4 |
| **ENSMUSG00000042133** | 0.0330196298492252 | 0.783963164052658 | 0.961141387070613 | 1535.06980445081 | 1414.64012112419 | 1319.35811477593 | 1642.30964164604 | 1469.43389878923 | 1620.58395749786 | 1295.10114465183 | 1393.81482503436 | Ppig |
| **ENSMUSG00000042138** | -0.211159316081955 | 0.47734613914248 | 0.897674307047416 | 43.3686583953142 | 26.8757190541728 | 24.6078271048596 | 32.8274342990356 | 31.7537876787538 | 45.2540425867328 | 40.6020455316893 | 30.3947158107182 | Msantd2 |
| **ENSMUSG00000042148** | -0.0866332837154755 | 0.329948011837565 | 0.855977678491063 | 1740.16741811198 | 1513.18442432282 | 1607.08040092506 | 1649.81305520011 | 1618.34821342063 | 1829.73101918249 | 1752.13442640598 | 1712.9593410469 | Cox10 |

| | | | | | | | | | | | | |
|---|---|---|---|---|---|---|---|---|---|---|---|---|
| **ENSMUSG00000042155** | -0.0768896832673058 | 0.774125968429168 | 0.959203117135841 | 585.476888336741 | 682.480380830207 | 684.286884492827 | 466.149567046306 | 468.642107810573 | 410.955629976817 | 843.273253350471 | 827.170480277403 | Klhl23 |
| **ENSMUSG00000042156** | -0.185458724154678 | 0.573518067058269 | 0.922207298536842 | 43.3686583953142 | 30.1333819698302 | 27.447191770805 | 21.5723139679377 | 38.3235368536684 | 46.4771248188067 | 30.191264626128 | 24.9670879873757 | Dzip1 |
| **ENSMUSG00000042165** | 0.0419207361941933 | 0.91261273016741 | 0.986226496507248 | 27.1054114970714 | 16.2883145782866 | 25.5542819935081 | 15.9447538023887 | 28.4689130912965 | 22.0154801773295 | 12.4929370866736 | 19.5394601640331 | Gm9774 |
| **ENSMUSG00000042167** | 0.152462253421594 | 0.426774819616917 | 0.885066977635295 | 160.82544154929 | 183.243539005724 | 159.950876181588 | 185.709485463116 | 147.819356435578 | 114.969729814943 | 157.202791673977 | 199.736703899005 | Papd4 |
| **ENSMUSG00000042178** | -0.259174319615893 | 0.0450389519158883 | 0.693563614452513 | 256.597895505609 | 240.252640029727 | 247.024725937245 | 229.79204009325 | 333.962249724825 | 295.985900161874 | 264.433835001259 | 271.381391167127 | Armc5 |
| **ENSMUSG00000042182** | -0.0122397543769037 | 0.974235263279397 | 0.995510898988495 | 25.2983840639333 | 28.5045505120015 | 22.7149173275627 | 24.3860940507122 | 15.3294147414674 | 19.5693157131818 | 43.7252798033577 | 22.7960368580387 | Bend6 |
| **ENSMUSG00000042185** | 0.119136573183708 | 0.479584112566026 | 0.897732059433541 | 150.886790667031 | 154.738988493722 | 173.201244622666 | 155.695831246855 | 131.394983498292 | 118.638976511164 | 160.326025945645 | 172.598564782293 | Nfrkb |
| **ENSMUSG00000042190** | -0.731709091636412 | 0.0238904929035436 | 0.654124840968651 | 164.439496415566 | 201.975100770753 | 241.345996605354 | 159.447538023887 | 404.039574257247 | 480.671317205027 | 191.558368662329 | 198.651178334337 | Cmklr1 |
| **ENSMUSG00000042197** | -0.0223404805287247 | 0.900815605613159 | 0.984802943363903 | 223.167887992554 | 254.912123150185 | 295.293925258315 | 217.598993067893 | 228.846262926191 | 209.14706168463 | 257.146288367366 | 310.460311495193 | Zfp451 |
| **ENSMUSG00000042198** | 0.0480138483118231 | 0.716434943066922 | 0.951348124093194 | 1436.58680934478 | 1159.727997974 | 1402.646144977 | 1411.57967485853 | 1306.28512761218 | 1534.96820125269 | 1211.81489740734 | 1181.05181435934 | Chchd7 |
| **ENSMUSG00000042202** | -0.0974462132592956 | 0.694608750136134 | 0.944771261777587 | 74.9916384752307 | 131.120932355207 | 71.930571537282 | 73.1582821521366 | 88.6916138613469 | 88.0619207093179 | 98.902418602833 | 100.953877514171 | Slc35e2 |
| **ENSMUSG00000042203** | 0.137485692916276 | 0.401664206276246 | 0.881158207349486 | 337.010616280254 | 356.714089264476 | 367.224496795597 | 320.770929436291 | 255.12525962585 | 271.524255520397 | 379.99350305299 | 348.453706258591 | Tbc1d22b |
| **ENSMUSG00000042207** | -0.153453533210845 | 0.292385163617836 | 0.851785577030345 | 229.492484008537 | 249.211213047785 | 258.382184601026 | 195.088752405698 | 289.068963696242 | 262.96267989588 | 250.899819824029 | 234.473521968398 | Kdm5b |
| **ENSMUSG00000042208** | 0.13132815145154 | 0.587414471092228 | 0.923058967588378 | 206.001127377742 | 255.726538879099 | 273.525462819401 | 218.536919762152 | 204.757182618171 | 130.869798831903 | 252.981976005141 | 281.151121249143 | 0610010F05R |
| **ENSMUSG00000042210** | 0.0039491477997672 | 0.982732090261792 | 0.996940244108351 | 247.562758339918 | 228.036404096012 | 227.149173275627 | 296.384835385579 | 248.555510450935 | 318.001380339203 | 204.051305749003 | 226.874843015718 | Abhd14a |
| **ENSMUSG00000042211** | 0.254351064487312 | 0.289150494351929 | 0.851785577030345 | 434.590097669711 | 690.624538119351 | 720.252170261468 | 533.680289032894 | 489.446313531136 | 327.786038195794 | 563.22324699087 | 613.321944037707 | Fbxo38 |
| **ENSMUSG00000042213** | 0.740706865363108 | 0.108736443018351 | 0.76280262299237 | 16.2632468982428 | 9.77298874697194 | 20.8220075502658 | 15.9447538023887 | 12.0445401540101 | 4.89232892829544 | 9.36970281500523 | 10.8552556466851 | Zfand4 |
| **ENSMUSG00000042215** | 0.0706322752622286 | 0.535304924492565 | 0.910861252626585 | 1282.08596381147 | 1333.19854823276 | 1109.24512949598 | 1259.63555038871 | 1182.55485148462 | 1038.39681503071 | 1287.81359801794 | 1236.41361815743 | Bag2 |
| **ENSMUSG00000042216** | -0.529373424327606 | 0.271607889224199 | 0.844129257012535 | 10.8421645988285 | 8.95857301805762 | 9.46454888648447 | 12.1930470253561 | 14.2344565456483 | 26.9078091056249 | 8.32862472444909 | 10.8552556466851 | Sgsm1 |
| **ENSMUSG00000042225** | 0.35311689333184 | 0.118665912975922 | 0.769868040577702 | 360.501972911049 | 370.55915665602 | 425.904699891801 | 308.577882410935 | 209.137015401448 | 220.154801773295 | 365.418409785204 | 351.710282952596 | Ammecr1 |
| **ENSMUSG00000042228** | -0.442832072679303 | 0.0724398955204437 | 0.729721356194038 | 200.580045078328 | 203.603932228582 | 200.648436393471 | 254.178134143962 | 408.419407040523 | 335.124531588238 | 200.928071477334 | 223.618266321713 | Lyn |
| **ENSMUSG00000042229** | -0.129996948864105 | 0.412688481205665 | 0.882192126163777 | 243.948703473642 | 186.501201921381 | 224.309808609682 | 234.48167356454 | 238.700886688563 | 277.639666680766 | 201.96914956789 | 255.098507697099 | Rabif |
| **ENSMUSG00000042246** | -0.0225090864552687 | 0.94464571158201 | 0.990505437194639 | 178.895715880671 | 202.789516499668 | 163.736695736181 | 109.737423228205 | 136.869774477387 | 86.838838477244 | 242.57119509958 | 198.651178334337 | Tmc7 |
| **ENSMUSG00000042248** | -0.145917483856035 | 0.595137870415638 | 0.924461858177432 | 75.8951521917998 | 126.234437981721 | 74.7699362032273 | 95.6685228143324 | 72.2672409240604 | 110.077400886647 | 89.5327157878278 | 141.118323406906 | Cyp2c37 |
| **ENSMUSG00000042249** | -0.0680625330178688 | 0.768007174977196 | 0.95879680569409 | 124.684892886528 | 170.212887343095 | 166.576060402127 | 133.185590584659 | 105.115986798633 | 141.877538920568 | 190.517290571773 | 185.624871558315 | Grk3 |
| **ENSMUSG00000042251** | 0.263626108073307 | 0.356636436216663 | 0.864965376583892 | 214.132750826864 | 308.663561258531 | 239.453086828057 | 184.771558768858 | 213.516848184724 | 101.51582526213 | 229.03717992235 | 244.243252050414 | Pm20d1 |
| **ENSMUSG00000042254** | -2.13840118260014 | 0.0127755622806171 | 0.635862253526407 | 110.228673421423 | 79.8127414336042 | 53.9479286529615 | 74.0962088463947 | 281.404256325508 | 969.904210034571 | 80.1630129728225 | 69.4736361387845 | Cilp |
| **ENSMUSG00000042271** | -0.0541860700757389 | 0.724718884009064 | 0.951674101666722 | 160.82544154929 | 171.841718800923 | 151.432782183751 | 155.695831246855 | 143.439523652302 | 150.439114545085 | 179.065431575656 | 191.052499381657 | Nxt2 |
| **ENSMUSG00000042272** | 0.286267481241861 | 0.14646627063927 | 0.780607990324772 | 421.940905637744 | 527.741392336485 | 538.532831640966 | 443.63932638411 | 356.956371837026 | 283.755077841135 | 482.019155927491 | 460.262839419447 | Sestd1 |
| **ENSMUSG00000042275** | -0.195801338875716 | 0.398981741945231 | 0.880459952732461 | 389.414411841258 | 331.467201668132 | 312.330113253987 | 446.453106466885 | 512.440435643337 | 543.048511040794 | 341.473613702413 | 298.51953028384 | Pelo |
| **ENSMUSG00000042284** | -0.206215593489343 | 0.236890413291462 | 0.832048466795696 | 335.203588847116 | 452.000729547452 | 358.706402797761 | 333.901903155905 | 531.054724972262 | 423.186452297555 | 389.363205867995 | 364.736589728619 | Itga1 |
| **ENSMUSG00000042286** | -0.648322888339191 | 0.0125182416391196 | 0.635862253526407 | 301.773581334061 | 405.579032999336 | 375.742590793433 | 414.563598862107 | 776.32536083574 | 752.195572725424 | 375.829190690765 | 443.97995594942 | Stab1 |
| **ENSMUSG00000042292** | -0.403680747970346 | 0.0835243814922932 | 0.742148948523533 | 101.193536255733 | 137.636258186522 | 145.754052851861 | 93.7926694258161 | 191.617684268342 | 184.685417043153 | 121.806136595068 | 135.690695583563 | Mkl1 |
| **ENSMUSG00000042293** | 0.246991676803096 | 0.0685222823498054 | 0.72237439886039 | 440.914693685694 | 338.796943228361 | 394.671688566402 | 407.060185308042 | 292.353838283699 | 363.255422925936 | 342.514691792969 | 334.3418739179 | Gm5617 |
| **ENSMUSG00000042298** | 0.1767838764045 | 0.337945325688166 | 0.858570960161243 | 1012.8388762739 | 985.443031986338 | 1045.83265195653 | 877.899385825639 | 837.643019801609 | 611.54111603693 | 1096.25522935561 | 922.696729968231 | Ttc19 |
| **ENSMUSG00000042302** | 0.2886708515518 | 0.293816710305901 | 0.851785577030345 | 1168.24323552378 | 1331.56971677493 | 1136.69232126678 | 943.55425442371 | 649.310210120724 | 590.748718091674 | 1218.06136595068 | 1290.68989639086 | Ehbp1 |
| **ENSMUSG00000042303** | -0.0498870556459427 | 0.793039137678115 | 0.961804768755116 | 282.799793286111 | 204.418347957496 | 176.040609288611 | 266.371181169318 | 247.460552255116 | 228.716377397812 | 269.639225454039 | 216.019587369033 | Sgsm3 |
| **ENSMUSG00000042305** | -0.00826395693568295 | 0.953777453342131 | 0.992371505557225 | 365.019541493894 | 325.766291565731 | 318.008842585878 | 311.39166249371 | 364.621079207759 | 284.978160073209 | 299.830490080167 | 377.762896504641 | Tmem183a |
| **ENSMUSG00000042308** | 0.0924076568822564 | 0.661073094587754 | 0.939611397743053 | 167.150037565273 | 192.202112023782 | 259.328639489674 | 177.268145214792 | 168.623562156141 | 145.546785616789 | 222.790711379013 | 208.420908416353 | Setd1a |
| **ENSMUSG00000042312** | -0.00129827833801899 | 0.995104482683045 | 0.999262355603273 | 552.950394540256 | 371.373572384934 | 401.296872786941 | 528.990655561603 | 477.401773377126 | 609.094951572782 | 385.198893505771 | 385.36157545732 | S100a13 |
| **ENSMUSG00000042320** | -0.446768916330238 | 0.322683042761646 | 0.855977678491063 | 6.32459601598331 | 15.4738988493722 | 15.1432782183751 | 11.2551203310979 | 18.6142893289246 | 8.56157562451702 | 18.7394056300105 | 19.5394601640331 | Prox2 |
| **ENSMUSG00000042323** | -0.0598581673538816 | 0.527252266589067 | 0.908217276872524 | 920.680477183857 | 1055.48278467297 | 980.527264639791 | 890.092432850995 | 1012.83633113266 | 963.788798874201 | 1002.55820120556 | 1031.24928643508 | Pbrm1 |
| **ENSMUSG00000042328** | 0.0245490013336552 | 0.875081985465576 | 0.976986380159756 | 165.343010132135 | 201.160685041839 | 188.344522841041 | 179.143998603309 | 177.383227722694 | 146.769867848863 | 193.640524843441 | 202.993280593011 | Hps4 |
| **ENSMUSG00000042331** | 0.123302089507939 | 0.812153423603453 | 0.965224347191244 | 60.535419010126 | 50.4937751926884 | 176.040609288611 | 36.5791410760683 | 63.5075753575076 | 123.53130543946 | 35.3966550789086 | 74.901263962127 | Specc1 |
| **ENSMUSG00000042340** | 0.0500294084429306 | 0.773133378001065 | 0.959121750920843 | 152.693818100169 | 168.584055885266 | 134.396594188079 | 124.744250336335 | 145.62944004394 | 122.308223207386 | 162.408182126757 | 130.263067760221 | Ctf1 |
| **ENSMUSG00000042348** | -0.355320106149904 | 0.0894445136544791 | 0.751593644682828 | 97.5794813894569 | 118.904696421492 | 88.966759532954 | 104.109863062656 | 145.62944004394 | 163.893019097897 | 99.9434966933891 | 116.15123541953 | Arl15 |
| **ENSMUSG00000042350** | 0.315858576920079 | 0.264032977425757 | 0.842510961779757 | 1259.49812089725 | 1701.31445770203 | 1792.58555910016 | 1215.55299575858 | 1007.36154015357 | 661.687487551958 | 1522.05616839307 | 1603.32125901539 | Arel1 |
| **ENSMUSG00000042351** | -0.0132272680633704 | 0.986850896602306 | 0.998220569123292 | 9.03513716569045 | 7.32974156022896 | 27.447191770805 | 2.81378008277448 | 7.66470737073368 | 24.4616446414772 | 2.08215618111227 | 13.0263067760221 | Grap2 |
| **ENSMUSG00000042354** | 0.278691740113573 | 0.0754297260319362 | 0.735399943172214 | 588.187429486448 | 627.914526992947 | 548.943835416099 | 601.211011019481 | 525.579933993166 | 365.701587390084 | 487.224546380272 | 570.986447015635 | Gnl3 |
| **ENSMUSG00000042359** | 0.141950475079678 | 0.576224831912438 | 0.922450244597442 | 363.212514060756 | 465.031381210082 | 384.260684791269 | 297.322762079837 | 247.460552255116 | 244.616446414772 | 437.252798033577 | 438.552328126077 | Osbpl6 |
| **ENSMUSG00000042363** | 0.102555314483055 | 0.42748932129401 | 0.885066977635295 | 653.24041707942 | 677.593886456721 | 761.896185362 | 582.452477134318 | 646.025335533267 | 541.82542880872 | 664.207821774815 | 638.289032025082 | Lgalsl |
| **ENSMUSG00000042364** | -0.251018921845338 | 0.0730383296285387 | 0.730383296285387 | 355.080890611635 | 394.991628523449 | 430.636974335043 | 460.522006880757 | 536.529515951357 | 532.040770952129 | 454.951125573032 | 429.868123608729 | Snx18 |
| **ENSMUSG00000042369** | 0.123777235944678 | 0.473931850600595 | 0.897549741595057 | 326.168451681425 | 370.55915665602 | 403.189782564238 | 438.011766218561 | 333.962249724825 | 282.531995609062 | 436.211719943021 | 357.137910775939 | Rbm45 |
| **ENSMUSG00000042377** | -0.569339291515169 | 0.222983408996047 | 0.832048466795696 | 8.13162344912141 | 8.14415728914329 | 11.3574586637814 | 9.37926694258161 | 17.5193311331055 | 19.5693157131818 | 9.36970281500523 | 8.68420451734806 | Fam83g |
| **ENSMUSG00000042379** | -0.567564873397333 | 0.29380904056431 | 0.851785577030345 | 19.877301764519 | 11.4018202048006 | 5.67872933189068 | 25.3240207449704 | 10.949581958191 | 36.6924669622158 | 16.6572494488982 | 28.2236646813812 | Esm1 |
| **ENSMUSG00000042380** | -0.0512162304238567 | 0.720597678568996 | 0.951674101666722 | 854.723975874317 | 657.233493233863 | 706.055346931741 | 815.9962240046 | 821.218646864323 | 903.857769502582 | 700.64555494428 | 718.617923810552 | Smim12 |
| **ENSMUSG00000042389** | 0.261473693408588 | 0.141537613144027 | 0.779832386580936 | 218.650319409709 | 227.221988367098 | 160.897331070236 | 226.978260010475 | 200.377349834895 | 185.908499275227 | 160.326025945645 | 149.802527924254 | Tsen2 |
| **ENSMUSG00000042390** | 0.0489474454662544 | 0.59685107619891 | 0.924496535139492 | 833.03964667666 | 933.320425335821 | 875.470771999813 | 924.795720538547 | 879.251431242735 | 796.226533080083 | 899.491470240502 | 871.677028428812 | Gatad2b |
| **ENSMUSG00000042396** | 0.0315272915730751 | 0.834100740944985 | 0.968512902280839 | 510.48524986151 | 379.517729674077 | 455.244801439903 | 464.27371365779 | 478.496731572945 | 491.679057293692 | 368.541644056872 | 432.039174738066 | Rbm7 |
| **ENSMUSG00000042404** | 0.110227260730966 | 0.693579421480385 | 0.94469184542218 | 1269.43677177951 | 1649.19185105152 | 1500.13099850779 | 1139.58093352367 | 1099.33802860237 | 689.818378889657 | 1682.38219433872 | 1677.13699741284 | Dennd4b |
| **ENSMUSG00000042406** | 0.22452576787111 | 0.018173133353517 | 0.642273034804458 | 1877.50150303048 | 1726.56134529838 | 1948.75061572715 | 1777.37108561922 | 1476.00364796414 | 1722.09978275999 | 1493.94705994806 | 1582.69627328668 | Atf4 |
| **ENSMUSG00000042408** | 0.187273076900619 | 0.457101055141392 | 0.892805623137304 | 135.527057485357 | 141.708336831093 | 170.36187995672 | 126.620103724852 | 107.305903190271 | 78.277262852727 | 163.449260217313 | 154.144630182928 | Zmym6 |
| **ENSMUSG00000042410** | 0.201113612072843 | 0.377704775690743 | 0.87422797717331 | 606.257703817829 | 648.274920215806 | 745.806452254976 | 557.128456389348 | 476.306815181307 | 363.255422925936 | 663.166743684259 | 720.788974939889 | Agps |
| **ENSMUSG00000042419** | -0.183079249960772 | 0.457890376243321 | 0.893021247686005 | 196.965990212052 | 146.594831204579 | 164.68315062483 | 199.778385876988 | 223.371471947096 | 276.416584448692 | 131.175839410073 | 173.684090346961 | Nfkbil1 |
| **ENSMUSG00000042423** | -0.0591204837270163 | 0.577121339052995 | 0.922450244597442 | 548.43282595741 | 589.636987733974 | 630.338955839866 | 572.135283497478 | 659.164833883096 | 549.163922201163 | 611.112839156452 | 617.664046296381 | Fbrs |
| **ENSMUSG00000042425** | -0.62792295790494 | 0.0169711393590424 | 0.642273034804458 | 48.7897406947284 | 62.7100111264033 | 49.2156542097192 | 56.2756016554897 | 65.6974917491458 | 63.6002760678407 | 85.3684034256032 | 120.493337678204 | Frmpd3 |
| **ENSMUSG00000042426** | 0.299632044515341 | 0.0487460356319312 | 0.703396849069134 | 779.732337399086 | 767.994032366212 | 869.792042667923 | 757.844768960594 | 604.416924092141 | 503.90987961443 | 740.206522385413 | 729.473179457237 | Dhx29 |
| **ENSMUSG00000042428** | -0.313676706258045 | 0.329795935363576 | 0.855977678491063 | 77.7021796249379 | 78.1839099757755 | 63.4124775394459 | 47.8342614071662 | 91.9764884488041 | 40.3617136584374 | 86.4094815161593 | 112.894658725525 | Mgat3 |
| **ENSMUSG00000042429** | -0.214806522580173 | 0.0826013896087429 | 0.739855948845396 | 615.29284098352 | 579.049583258088 | 553.676109859341 | 537.431995809926 | 731.432074807157 | 660.464405319884 | 702.727711125392 | 557.960140239613 | Adora1 |
| **ENSMUSG00000042436** | -1.94353485851302 | 0.0166540358113658 | 0.642273034804458 | 51.5002818444356 | 50.4937751926884 | 38.8046504345863 | 77.8479156234274 | 180.668102310151 | 557.72549782568 | 54.1360607089191 | 48.8486504100828 | Mfap4 |
| **ENSMUSG00000042439** | 0.0430841839012875 | 0.872441530686979 | 0.976162349063696 | 306.291149916906 | 390.919549878878 | 297.186835035612 | 287.943495137256 | 263.884925192402 | 177.34692365071 | 421.636626675235 | 381.019473198646 | Zfp532 |
| **ENSMUSG00000042444** | 0.00547846868227908 | 0.974387373396967 | 0.995510898988495 | 990.251033359673 | 1176.01631255229 | 1004.188636856 | 837.568537972538 | 947.138839383519 | 781.549546295196 | 1084.80337035949 | 1178.88076323 | Mindy2 |
| **ENSMUSG00000042446** | 0.211366351975075 | 0.360234802875454 | 0.868389693151693 | 497.836057829544 | 622.213616890547 | 600.998854291764 | 438.011766218561 | 399.65974147397 | 311.885969178834 | 568.428637443651 | 584.012753791657 | Zmym4 |
| **ENSMUSG00000042447** | 0.233519643084143 | 0.138826597845644 | 0.779711473022872 | 744.495302452893 | 741.118313312039 | 813.951204237664 | 649.045272426648 | 623.031213421066 | 478.225152740879 | 700.64555494428 | 704.506091469861 | Mios |
| **ENSMUSG00000042453** | -0.557576148693358 | 0.0566326949044524 | 0.705217457281182 | 70.4740698923855 | 88.7713144516618 | 70.0376617599851 | 39.3929211588428 | 120.445401540101 | 107.6312364225 | 102.025652874501 | 66.217059444779 | Reln |
| **ENSMUSG00000042460** | -0.0323638132026392 | 0.860841875063637 | 0.974868041734905 | 400.256576440087 | 483.762942975111 | 442.940887887473 | 375.170677703265 | 413.894198019619 | 319.224462571277 | 469.526218840818 | 537.335154510911 | C1galt1 |
| **ENSMUSG00000042462** | 0.000938870122748602 | 0.995299292239206 | 0.999262355603273 | 501.45011269582 | 352.642010619904 | 406.029147230184 | 409.873965390816 | 411.70428162798 | 489.232892829544 | 409.143689588562 | 359.308961905276 | Dctpp1 |
| **ENSMUSG00000042472** | 0.0449769759407232 | 0.760339166965913 | 0.957357521291862 | 454.46739943423 | 397.434875710192 | 427.797609669098 | 371.418970926232 | 377.760577557588 | 330.232202659942 | 465.361906478593 | 425.526021350055 | Zfp410 |
| **ENSMUSG00000042473** | -0.104365798356092 | 0.550995969290152 | 0.914814010690916 | 126.491920319666 | 142.522752560008 | 159.004421292939 | 109.737423228205 | 137.964732673206 | 128.423634367755 | 161.367104036201 | 149.802527924254 | Tbc1d8b |
| **ENSMUSG00000042476** | 0.303520228443742 | 0.164379997253062 | 0.794999359973894 | 1577.53494912955 | 1512.37000859391 | 1239.85590412947 | 1523.19295147525 | 1001.88674917447 | 799.895779776304 | 1552.2474330192 | 1387.30167164635 | Abcb4 |
| **ENSMUSG00000042485** | -0.31375817208884 | 0.395940639496209 | 0.87923508914506 | 156.307872966445 | 155.553404222637 | 156.165056626994 | 126.620103724852 | 261.695008800764 | 284.978160073209 | 100.984574783945 | 92.2696729968231 | Mustn1 |
| **ENSMUSG00000042487** | 0.0497438024100219 | 0.729380391378321 | 0.952818017249783 | 462.599022883351 | 435.712414969166 | 410.761421673426 | 398.618845059719 | 381.045452145046 | 348.57843614105 | 494.512093014165 | 424.440495785386 | Leo1 |
| **ENSMUSG00000042492** | 0.0117944136710702 | 0.912448508672438 | 0.986226496507248 | 719.19691838896 | 735.417403209639 | 720.252170261468 | 652.79697920368 | 787.274942793931 | 703.272283442469 | 650.673806597585 | 664.341645577127 | Tbc1d10b |
| **ENSMUSG00000042496** | 0.1168508357101 | 0.602358990422391 | 0.926448151609826 | 87.6408305071974 | 105.059629029948 | 79.5022106464695 | 94.7305961200743 | 66.7924499449649 | 68.4926049961361 | 110.35427759895 | 92.2696729968231 | Prdm10 |
| **ENSMUSG00000042500** | 0.152324683831532 | 0.640028653537763 | 0.933921420069286 | 463.50253659992 | 576.606336071345 | 470.388079658278 | 378.922384480297 | 342.721915291377 | 193.24699266767 | 564.264325081426 | 599.210111697016 | Ago4 |
| **ENSMUSG00000042502** | 0.10903075409822 | 0.199436430143045 | 0.812921911423739 | 862.855599323438 | 825.003133390215 | 883.988865997649 | 859.140851940476 | 783.990068206473 | 763.203312814088 | 849.519721893808 | 782.663932125994 | Cd2bp2 |
| **ENSMUSG00000042505** | 0.112090495535882 | 0.588180883064726 | 0.92352104528132 | 269.247087537575 | 230.479651282755 | 224.309808609682 | 243.860940507122 | 225.561388338734 | 152.885279009232 | 218.626399016789 | 297.434004719171 | Sdhaf3 |

| | | | | | | | | | | | | |
|---|---|---|---|---|---|---|---|---|---|---|---|---|
| **ENSMUSG00000042506** | -0.00196798631099666 | 0.988990635756965 | 0.998437672973644 | 488.800920663853 | 570.09101024003 | 633.178320505811 | 517.735535230505 | 570.47322002175 | 464.771248188067 | 623.605776243126 | 552.53251241627 | Usp22 |
| **ENSMUSG00000042507** | 0.0373853339751783 | 0.829989903156371 | 0.967566911862526 | 383.089815825275 | 500.051257553398 | 525.282463199888 | 478.342614071662 | 496.016062706051 | 343.686107212755 | 511.169342463063 | 486.315452971491 | Elmsan1 |
| **ENSMUSG00000042508** | 0.305445182110156 | 0.0819366575413578 | 0.739855948845396 | 224.071401709123 | 267.1283590839 | 254.596365046432 | 223.226553233442 | 183.952976897608 | 149.216032313011 | 214.462086654564 | 235.559047533066 | Dmtf1 |
| **ENSMUSG00000042510** | -0.749585216908864 | 0.0489847481834376 | 0.703396849069134 | 10.8421645988285 | 10.5874044758863 | 17.036187995672 | 11.2551203310979 | 24.0890803080201 | 22.0154801773295 | 14.5750932677859 | 22.7960368580387 | AA986860 |
| **ENSMUSG00000042515** | 0.240729498860356 | 0.596141458869349 | 0.924461858177432 | 14.4562194651047 | 26.0613033252585 | 13.2503684410783 | 21.5723139679377 | 6.56974917491458 | 14.6769867848863 | 19.7804837205666 | 22.7960368580387 | Mum1l1 |
| **ENSMUSG00000042520** | 0.126085710287004 | 0.512136611471931 | 0.902394714365738 | 1677.82497166872 | 1898.4030640993 | 2214.70443943737 | 1641.37171495178 | 1509.94735203453 | 1262.22086350022 | 2091.52588392728 | 1945.26181188597 | Ubap2l |
| **ENSMUSG00000042523** | -0.0458453026997899 | 0.859673657712358 | 0.974868041734905 | 103.000563688871 | 140.893921102179 | 118.306861081056 | 83.4754757889764 | 90.881530252985 | 85.6157562451702 | 154.079557402308 | 129.177542195552 | Dnal1 |
| **ENSMUSG00000042524** | 0.109836661839873 | 0.510498133117238 | 0.902394714365738 | 2320.22322414931 | 2801.59010746529 | 3490.52562933547 | 2561.47780201904 | 2806.37785588435 | 2159.96322184244 | 2907.73110692329 | 2479.34038970287 | Sun2 |
| **ENSMUSG00000042529** | -0.0932415737944808 | 0.722692830139114 | 0.951674101666722 | 523.134441893477 | 491.907100264254 | 485.531357876653 | 358.287997206618 | 460.97740043984 | 286.201242305283 | 698.563398763168 | 536.249628946243 | Kcnj12 |
| **ENSMUSG00000042532** | 0.390580825917291 | 0.303016961020206 | 0.851785577030345 | 17.1667606148119 | 13.0306516626293 | 19.8755526616174 | 16.8826804966469 | 13.1394983498292 | 12.2308223207386 | 13.5340151772298 | 11.9407812113536 | Golga7b |
| **ENSMUSG00000042535** | 0.057912466138737 | 0.753481724685787 | 0.956497786093635 | 670.407177694231 | 776.95260538427 | 841.398396008469 | 621.845398293161 | 662.449708470554 | 525.92535979176 | 867.218049433262 | 739.242909539254 | Gtpbp1 |
| **ENSMUSG00000042541** | 0.0240453608284582 | 0.912056525355824 | 0.986226496507248 | 3096.34150668212 | 2284.43611960469 | 2446.58588715624 | 3211.46100113994 | 2775.71902640141 | 3763.42402809127 | 2186.26399016789 | 2131.97220900895 | Sem1 |
| **ENSMUSG00000042548** | -0.0707052338486155 | 0.723121892568291 | 0.951674101666722 | 366.826568927032 | 505.752167655798 | 543.265106084208 | 445.515179772627 | 505.870686468423 | 348.57843614105 | 514.292576734731 | 585.098279356326 | Asxl1 |
| **ENSMUSG00000042549** | 0.47024549791178 | 0.0101038527666264 | 0.635862253526407 | 192.448421629207 | 180.800291818981 | 168.468970179424 | 173.51643843776 | 98.5462376237187 | 133.315963296051 | 120.765058504512 | 163.914360264945 | Map2k3os |
| **ENSMUSG00000042557** | -0.118270433356539 | 0.391903604556564 | 0.878285927324016 | 502.353626412389 | 496.79359463774 | 563.140658745826 | 475.528833988888 | 540.909348734634 | 450.09426140318 | 611.112839156452 | 607.894316214364 | Sin3a |
| **ENSMUSG00000042558** | -0.296208663212336 | 0.122782786854025 | 0.770251097497048 | 231.299511441676 | 165.326392969609 | 204.434255948065 | 196.964605794214 | 257.315176017488 | 306.993640250539 | 190.517290571773 | 225.78931745105 | Adprhl2 |
| **ENSMUSG00000042569** | -0.0152852577421764 | 0.920029283880861 | 0.986713918026868 | 624.32797814921 | 471.546707041396 | 431.583429223692 | 521.487242007538 | 506.965644664242 | 587.079471395453 | 494.512093014165 | 483.058876277486 | Dhrs7b |
| **ENSMUSG00000042570** | 0.0559302151851672 | 0.752179893089854 | 0.956497786093635 | 251.176813206195 | 254.912123150185 | 229.042083052924 | 233.543746870282 | 247.460552255116 | 163.893019097897 | 271.721381635152 | 247.49982874442 | Mier2 |
| **ENSMUSG00000042572** | 0.00459560587016919 | 0.971032161826215 | 0.995122272737203 | 635.170142748039 | 739.48948185421 | 693.751433379311 | 680.934780031425 | 789.464859185569 | 721.618516923577 | 614.236073428121 | 616.578520731712 | Ube2q1 |
| **ENSMUSG00000042579** | 0.349497064208543 | 0.11104566687692 | 0.764116514278315 | 182.509770746947 | 197.088606397268 | 214.845259723197 | 167.888878272211 | 113.875652365186 | 113.746647582869 | 187.394056300105 | 182.368294864309 | 4632404H12 |
| **ENSMUSG00000042589** | -0.0218783437315853 | 0.930317978470138 | 0.987529878729394 | 183.413284463516 | 200.346269312925 | 159.004421292939 | 155.695831246855 | 171.908436743598 | 101.51582526213 | 211.338852382896 | 223.618266321713 | Cux2 |
| **ENSMUSG00000042590** | 0.202595154944234 | 0.138308833304265 | 0.779711473022872 | 622.520950716072 | 711.799347071123 | 827.201572678743 | 668.741733006069 | 566.093387238473 | 563.840908986049 | 639.221947601468 | 689.308733564502 | Ipo11 |
| **ENSMUSG00000042594** | -0.244602634505028 | 0.154975962995791 | 0.787275540343856 | 628.845546732055 | 648.274920215806 | 618.981497176084 | 702.507093999363 | 1011.74137293685 | 765.649477278236 | 686.070461676494 | 615.492995167044 | Sh2b3 |
| **ENSMUSG00000042595** | 0.0261418213570196 | 0.906629628724571 | 0.98504921011344 | 117.456783153976 | 146.594831204579 | 121.146225747001 | 125.682177030594 | 109.49581958191 | 88.0619207093179 | 136.381229862854 | 167.17093695895 | Fam199x |
| **ENSMUSG00000042599** | 0.081148379994708 | 0.53617458895169 | 0.911291453479697 | 548.43282595741 | 562.761268679801 | 619.927952064733 | 498.039074651084 | 515.725310230795 | 458.655837027697 | 603.825292522559 | 527.565424428895 | Kdm7a |
| **ENSMUSG00000042605** | 0.172743343238473 | 0.353265496039709 | 0.862728204353539 | 961.338594429464 | 1249.31372815458 | 1125.334862603 | 961.374861614615 | 867.206891088725 | 706.941530138691 | 1142.06266534008 | 1095.29529475052 | Atxn2 |
| **ENSMUSG00000042606** | 0.236718916275896 | 0.227276868171773 | 0.832048466795696 | 163.535982698997 | 124.605606523892 | 116.413951303759 | 148.192417692789 | 114.970610561005 | 130.869798831903 | 132.216917500629 | 91.1841474321546 | Hirip3 |
| **ENSMUSG00000042607** | -0.153035188197136 | 0.577514251398257 | 0.922450244597442 | 47.8862269781594 | 61.895595397489 | 88.966759532954 | 85.3513291774927 | 66.7924499449649 | 59.9310293716191 | 93.6970281500523 | 94.4407241261602 | Asb4 |
| **ENSMUSG00000042608** | 0.0976653137382481 | 0.588255404125942 | 0.92352104528132 | 2023.87072511466 | 2380.53717561658 | 2250.66972520601 | 1962.14264438807 | 1737.69865676491 | 1536.19128348477 | 2240.40005087681 | 2537.95877019497 | Stk40 |
| **ENSMUSG00000042613** | -0.0253191468667193 | 0.795607787135523 | 0.962043367573119 | 3234.57910531718 | 3212.87005056703 | 2969.97544057883 | 3395.29463321454 | 3020.98966226489 | 3631.33114702729 | 3263.77981389349 | 3125.22810068063 | Pbxip1 |
| **ENSMUSG00000042616** | 0.104577094873916 | 0.644684232788306 | 0.934857223264556 | 74.9916384752307 | 61.0811796685746 | 45.4298346551254 | 60.9652351267805 | 62.4126171616885 | 53.8156182112498 | 56.2182168900314 | 53.1907526687569 | Oscp1 |
| **ENSMUSG00000042622** | -0.201033053706666 | 0.501128952121328 | 0.900927059989498 | 568.310127721929 | 490.278268806426 | 544.211560972857 | 331.088123073131 | 730.337116611338 | 719.172352459429 | 477.854843565267 | 296.348479154503 | Maff |
| **ENSMUSG00000042625** | 0.103210791436033 | 0.292888325327293 | 0.851785577030345 | 691.187993175319 | 632.801021366433 | 646.428688946889 | 666.865879617553 | 676.684165016202 | 602.979540412413 | 594.455589707554 | 580.756177097652 | Safb2 |
| **ENSMUSG00000042626** | -0.0286815417221573 | 0.838283490628342 | 0.969135716755098 | 522.230928176908 | 576.606336071345 | 691.858523602015 | 496.163221262567 | 636.170711770895 | 573.62556684264 | 577.798340258656 | 544.933833463591 | Shc1 |
| **ENSMUSG00000042628** | 0.0967741273777206 | 0.530639092344552 | 0.909082872831377 | 328.878992831132 | 436.52683069808 | 443.887342776122 | 391.115431505653 | 346.006789878835 | 329.009120427868 | 431.006329490241 | 389.703677715994 | Zfyve1 |
| **ENSMUSG00000042632** | -0.0509564242459073 | 0.712635371449611 | 0.950456121590218 | 192.448421629207 | 193.016527752696 | 176.98706417726 | 185.709485463116 | 178.478185918513 | 193.24699266767 | 179.065431575656 | 224.703791886381 | Pla2g6 |
| **ENSMUSG00000042642** | -0.0580918711387493 | 0.504857261680693 | 0.90122958035533 | 1286.60353239432 | 1186.60371702818 | 1100.72703549814 | 1211.80128898154 | 1234.01788668812 | 1205.95908082483 | 1279.48497329349 | 1262.46623170947 | Flad1 |
| **ENSMUSG00000042644** | -0.405973557110809 | 0.109775545192385 | 0.763154315676794 | 155.404359249876 | 140.079505373265 | 196.862616838877 | 146.316564304273 | 308.778211220985 | 218.931719541221 | 169.69572876065 | 148.717002359586 | Itpr3 |
| **ENSMUSG00000042647** | 0.355808754791027 | 0.0924281521669406 | 0.754058637722175 | 2479.24163826546 | 3071.16171373593 | 3487.68626466953 | 3120.4821117969 | 2239.18951045005 | 1571.66066821491 | 2831.73240631269 | 2857.10328620751 | Acad12 |
| **ENSMUSG00000042650** | 0.0885887987002384 | 0.680577666594199 | 0.944452327903597 | 2572.30355107207 | 3381.45410645229 | 3462.13198267602 | 2669.33937185873 | 2586.29125852471 | 1914.12369319559 | 3376.21624767355 | 3487.79363927991 | Alkbh5 |
| **ENSMUSG00000042659** | 0.0615754038459326 | 0.699115488797774 | 0.945173655771957 | 324.361424248287 | 255.726538879099 | 261.221549266971 | 318.895076047775 | 266.074841584041 | 236.054870790255 | 326.898520434627 | 281.151121249143 | Arrdc4 |
| **ENSMUSG00000042660** | 0.326027128443524 | 0.0284332318716662 | 0.665956123973759 | 383.089815825275 | 377.074482487334 | 415.493696116668 | 460.522006880757 | 371.190828382674 | 362.032340693862 | 295.666177717943 | 276.809018990469 | Wdr55 |
| **ENSMUSG00000042670** | 0.144248048806058 | 0.362768253746387 | 0.86924083181804 | 2405.1535135068 | 1604.39898596123 | 1968.62616838877 | 2370.14075639037 | 1739.88857315654 | 1866.42348614471 | 1835.42067365047 | 2111.34722328025 | Immp1l |
| **ENSMUSG00000042675** | 0.183860362323225 | 0.413703724246783 | 0.882192126163777 | 1021.87401343959 | 1146.69734631137 | 1573.00802493372 | 1567.27550610539 | 1289.8607546749 | 1465.25251402448 | 1102.50169789895 | 816.315224630718 | Ypel3 |
| **ENSMUSG00000042677** | -0.23853432199613 | 0.342169495625442 | 0.859912121266038 | 121.070838020252 | 104.245213301034 | 70.0376617599851 | 72.2203554578784 | 116.065568756824 | 96.6234963338349 | 134.299073681742 | 86.8420451734806 | Zc3h12a |
| **ENSMUSG00000042680** | 0.159163936224379 | 0.681533039411916 | 0.944452327903597 | 329.782506547701 | 523.669313691913 | 428.744064557746 | 279.502154888932 | 241.98576127602 | 141.877538920568 | 474.731609293598 | 539.506205640248 | Garem1 |
| **ENSMUSG00000042682** | -0.020675393273564 | 0.907638724713655 | 0.98504921011344 | 2507.2505634791 | 2001.83386167142 | 2256.3484545379 | 2609.3120634262 | 2242.47438503751 | 3153.10599428641 | 1972.84298160388 | 2142.82746465563 | Selenok |
| **ENSMUSG00000042684** | -0.0860599666934097 | 0.870217829375018 | 0.975613767803763 | 12.6491920319666 | 4.88649437348597 | 11.3574586637814 | 24.3860940507122 | 12.0445401540101 | 13.4539045528125 | 19.7804837205666 | 10.8552556466851 | Npl |
| **ENSMUSG00000042686** | 0.389069278352436 | 0.0946876478619062 | 0.757085157577544 | 849.302893574902 | 914.588863570791 | 997.563452635463 | 921.981940455772 | 635.075753575076 | 418.29412336926 | 876.587752248267 | 881.446758510828 | Jph1 |
| **ENSMUSG00000042688** | 0.127956168829014 | 0.576975067145861 | 0.922450244597442 | 453.563885717661 | 562.761268679801 | 473.227444324223 | 415.501525556365 | 331.772333333186 | 316.77829810713 | 507.005030100839 | 587.269330485663 | Mapk6 |
| **ENSMUSG00000042694** | -0.124353519558219 | 0.41696121729771 | 0.882192126163777 | 314.422773366028 | 313.550055632016 | 264.060913932917 | 366.729337454941 | 377.760577557588 | 357.140011765567 | 349.802238426862 | 287.664274637155 | Stn1 |
| **ENSMUSG00000042699** | 0.261070063932617 | 0.167350372284293 | 0.797973453837565 | 1123.06754969532 | 1269.67412137744 | 1398.8603254224 | 1064.54679798301 | 925.239675467137 | 738.741668172611 | 1248.25263057681 | 1138.71631733726 | Dhx9 |
| **ENSMUSG00000042700** | 0.288990327457117 | 0.139260596693567 | 0.779711473022872 | 371.344137509877 | 403.950201541507 | 377.63550057073 | 312.329589187968 | 269.359716171498 | 216.485555077073 | 374.788112600209 | 337.598450611906 | Sipa1l1 |
| **ENSMUSG00000042705** | 0.103294689914398 | 0.481790481162671 | 0.897732059433541 | 207.80815481088 | 159.625482867208 | 173.201244622666 | 187.585338851632 | 177.383227722694 | 154.108361241306 | 174.901119213431 | 170.427513652956 | Commd10 |
| **ENSMUSG00000042709** | 0.0358782667894685 | 0.835261930700288 | 0.968814461162998 | 726.425028121512 | 837.21936932393 | 659.679057387967 | 740.962088463947 | 708.437952694956 | 529.594606487981 | 804.753363999894 | 847.795466006104 | Atpaf2 |
| **ENSMUSG00000042712** | -0.256768509620515 | 0.40934140176701 | 0.882192126163777 | 349.65980831222 | 218.26341534904 | 217.684624389143 | 280.44008158319 | 381.045452145046 | 478.225152740879 | 189.476212481217 | 225.78931745105 | Tceal9 |
| **ENSMUSG00000042717** | 0.0853805130737345 | 0.667487471562494 | 0.942024383071524 | 3674.59028528631 | 4018.3272064633 | 4187.11642738073 | 2991.04822798928 | 3290.34937843639 | 2506.09549351934 | 4465.18393039527 | 3753.7474026237 | Ppp1r3a |
| **ENSMUSG00000042719** | 0.31836972148691 | 0.233634933850442 | 0.832048466795696 | 513.195791011218 | 734.602987480724 | 743.913542477679 | 596.521377548191 | 462.072358635659 | 292.316653465652 | 678.782915042601 | 641.545608719088 | Naa25 |
| **ENSMUSG00000042726** | 0.3120672925205 | 0.0360511352189213 | 0.679566075569698 | 1493.50817348863 | 1635.34678365997 | 1994.18045038228 | 1602.9167204872 | 1511.04231023035 | 1146.02805145321 | 1507.48107512529 | 1251.61097606279 | Trafd1 |
| **ENSMUSG00000042729** | -0.474248449676998 | 0.317811657769811 | 0.854351922245337 | 32.5264937964856 | 13.0306516626293 | 26.5007368821565 | 54.3997482669734 | 47.0832024202212 | 67.2695227640623 | 40.6020455316893 | 20.6249857287016 | Wdr74 |
| **ENSMUSG00000042734** | -1.14088866502387 | 0.00010011390109083 | 0.399150048388892 | 18.9737880479499 | 28.5045505120015 | 26.5007368821565 | 24.3860940507122 | 68.9823663366031 | 63.6002760678407 | 40.6020455316893 | 44.5065481514088 | Ttc9 |
| **ENSMUSG00000042737** | 0.106571593133358 | 0.55322702418077 | 0.915115571755578 | 198.77301764519 | 158.811067138294 | 165.629605513478 | 174.454365132018 | 167.528603960322 | 204.254732756335 | 140.545542225078 | 136.776221148232 | Dpm3 |
| **ENSMUSG00000042742** | -0.0443138765101093 | 0.725117165026622 | 0.951674101666722 | 271.957628687283 | 317.622134276588 | 272.579007930753 | 320.770929436291 | 297.828629262794 | 282.531995609062 | 307.11803671406 | 332.170822788563 | Bmt2 |
| **ENSMUSG00000042743** | -0.0458229087540228 | 0.905786277929989 | 0.98504921011344 | 28.0089252136404 | 14.6594831204579 | 25.5542819935081 | 15.0068271081306 | 28.4689130912965 | 24.4616446414772 | 14.5750932677859 | 18.4539345993646 | Sgtb |
| **ENSMUSG00000042744** | 0.0536411293580382 | 0.798289034544753 | 0.963021946311154 | 540.301202508289 | 837.21936932393 | 734.448993591195 | 572.135283497478 | 647.120293729086 | 463.548165955993 | 744.370834747638 | 730.558705021906 | Hectd4 |
| **ENSMUSG00000042745** | -0.714778409878236 | 0.0514108697353248 | 0.703396849069134 | 718.293404672391 | 509.009830571455 | 555.569019636638 | 581.51455044006 | 1442.05994389375 | 1438.34470491886 | 512.210420553619 | 488.486504100828 | Id1 |
| **ENSMUSG00000042747** | -0.0590957750820475 | 0.773735349457893 | 0.959203117135841 | 605.35419010126 | 421.867347577622 | 460.923530771794 | 605.900644490772 | 593.467342133951 | 688.595296657583 | 477.854843565267 | 422.269444656049 | Krtcap2 |
| **ENSMUSG00000042757** | -0.519856006126894 | 0.183086687961813 | 0.805716353538343 | 17.1667606148119 | 14.6594831204579 | 17.036187995672 | 23.448167356454 | 32.8487458745729 | 31.8001380339204 | 27.0680303544596 | 11.9407812113536 | Tmem108 |
| **ENSMUSG00000042759** | 0.0388824758840571 | 0.928781195570017 | 0.986901071486195 | 12.6491920319666 | 16.2883145782866 | 13.2503684410783 | 21.5723139679377 | 19.7092475247437 | 17.123151249034 | 18.7394056300105 | 6.51315338801105 | Apobr |
| **ENSMUSG00000042763** | -0.580787460149412 | 0.287514290455852 | 0.851785577030345 | 5.42108229941427 | 12.2162359337149 | 7.57163910918757 | 8.44134024832345 | 7.66470737073368 | 12.2308223207386 | 22.903717992235 | 7.59867895267955 | Maneal |
| **ENSMUSG00000042770** | 0.0334797914674909 | 0.773647615857151 | 0.959203117135841 | 411.098741038915 | 418.609684661965 | 406.029147230184 | 453.018593326692 | 388.710159515779 | 462.325083723919 | 373.747034509653 | 426.611546914723 | Hebp1 |
| **ENSMUSG00000042772** | 0.0959340384809274 | 0.693040419829967 | 0.94469184542218 | 799.609639163605 | 945.536661269536 | 1010.81382107654 | 676.245146560134 | 616.461464246152 | 563.840908986049 | 956.75076522109 | 1073.58478345715 | Smg7 |
| **ENSMUSG00000042787** | -0.277720987086165 | 0.325479728376927 | 0.855977678491063 | 70.4740698923855 | 43.9784493613737 | 53.001473764313 | 40.3308478531009 | 50.3680770076785 | 47.7002070508805 | 73.9165444294857 | 79.2433662208011 | Exog |
| **ENSMUSG00000042790** | 0.187511513296381 | 0.475508502631152 | 0.897549741595057 | 567.40661400536 | 647.460504486891 | 627.49959117392 | 492.411514485535 | 384.330326732503 | 320.447544803351 | 722.508194845959 | 622.006148555055 | Rnf214 |
| **ENSMUSG00000042793** | 0.322735306751617 | 0.172350886783221 | 0.799889021902745 | 554.757421973394 | 857.579762546788 | 980.527264639791 | 606.83857118503 | 566.093387238473 | 436.640356850368 | 685.029383585938 | 709.933719293204 | Lgr6 |
| **ENSMUSG00000042797** | -0.0824473599182946 | 0.80870189623553 | 0.964157363647118 | 42.4651446787451 | 61.895595397489 | 29.3401015481019 | 37.5170677703264 | 51.4630352034976 | 51.3694537471021 | 54.1360607089191 | 24.9670879873757 | Aqp11 |
| **ENSMUSG00000042804** | -0.979173064811727 | 0.0103635025616192 | 0.635862253526407 | 95.7724539563188 | 111.574954861263 | 109.78876708322 | 130.371810501884 | 304.398378437709 | 352.247682837272 | 116.600746142287 | 109.638082031519 | Gpr153 |
| **ENSMUSG00000042807** | -0.0576094563496857 | 0.772663449737995 | 0.959045638244974 | 430.976042803434 | 563.575684408715 | 425.904699891801 | 339.529463321454 | 512.440435643337 | 534.486935416277 | 420.595548584679 | 365.822115293287 | Hecw2 |
| **ENSMUSG00000042810** | 0.150655727737823 | 0.488037544686656 | 0.899236955296196 | 192.448421629207 | 219.892246806869 | 264.060913932917 | 220.412773150668 | 171.908436743598 | 147.992950080937 | 271.721381635152 | 214.934061804364 | Krba1 |
| **ENSMUSG00000042814** | 0.0544225356373158 | 0.817263854645975 | 0.966315773308182 | 183.413284463516 | 101.801966114291 | 128.717864856189 | 165.075098189436 | 158.768938393769 | 151.662196777159 | 141.586620315635 | 105.295979772845 | Mcts2 |
| **ENSMUSG00000042821** | -0.922460475319773 | 0.0550866899240191 | 0.703396849069134 | 24.3948703473642 | 29.3189662409158 | 17.036187995672 | 27.1998741334867 | 61.3176589658694 | 80.7234273168747 | 28.1091084450157 | 16.2828834700276 | Snai1 |
| **ENSMUSG00000042826** | -0.315440457555026 | 0.152605688226353 | 0.783412422888014 | 47.8862269781594 | 53.7514381083457 | 53.9479286529615 | 49.7101147956825 | 82.1218646864323 | 56.2617826753975 | 56.2182168900314 | 60.7894316214364 | Fgf11 |
| **ENSMUSG00000042828** | -0.0862473687718729 | 0.58075328098477 | 0.922450244597442 | 14729.9841212251 | 13544.5479875742 | 10389.235312694 | 12377.818584125 | 12112.4275621508 | 11778.2818948713 | 15559.953141452 | 14734.9240148103 | Trim72 |
| **ENSMUSG00000042831** | 0.121996856156258 | 0.486087680149683 | 0.899236955296196 | 421.940905637744 | 307.849145529616 | 364.385132129652 | 458.646153492241 | 355.861413641207 | 428.078781225851 | 313.364505257397 | 329.999771659226 | Alkbh6 |
| **ENSMUSG00000042834** | -0.0487506362727995 | 0.859852735173627 | 0.974868041734905 | 199.676531361759 | 222.335493993612 | 160.897331070236 | 297.322762079837 | 195.997517051618 | 209.14706168463 | 164.49033830787 | 340.855027305911 | Nrep |
| **ENSMUSG00000042842** | 0.362219789510474 | 0.0277906793962625 | 0.665956123973759 | 1284.79650496118 | 1276.18944720875 | 924.686426209533 | 1322.47663890401 | 1000.79179097865 | 770.541806206532 | 1057.73534000503 | 910.755948756878 | Serpinb6b |
| **ENSMUSG00000042851** | -0.102388765619595 | 0.654604374030398 | 0.937269843859626 | 110.228673421423 | 92.0289773673191 | 110.735221971868 | 119.116690170786 | 98.5462376237187 | 154.108361241306 | 129.093683228961 | 82.4999429148066 | Zc3h6 |

| | | | | | | | | | | | | |
|---|---|---|---|---|---|---|---|---|---|---|---|---|
| **ENSMUSG00000042854** | -0.174115843462537 | 0.5765684892806 | 0.922450244597442 | 48.7897406947284 | 44.7928650902881 | 34.0723759913441 | 57.2135283497478 | 59.1277425742312 | 75.8310983885793 | 36.4377331694648 | 37.9933947633978 | Trp53rkb |
| **ENSMUSG00000042857** | 0.209215008543847 | 0.722601805896311 | 0.951674101666722 | 12.6491920319666 | 3.25766291565731 | 18.9290977729689 | 22.5102406621959 | 15.3294147414674 | 6.1154111603693 | 12.4929370866736 | 15.1973579053591 | Gm9776 |
| **ENSMUSG00000042870** | 0.1389538998191 | 0.28864795370232 | 0.851785577030345 | 845.688838708626 | 961.824975847822 | 1065.70820461815 | 859.140851940476 | 881.441347634373 | 720.395434691503 | 898.450392149946 | 887.959911898839 | Tom1 |
| **ENSMUSG00000042895** | 0.554590033035754 | 0.0188309609965288 | 0.642273034804458 | 2847.87523462563 | 3616.00583637962 | 3470.65007667385 | 5320.85813652655 | 1919.46171727088 | 2353.21021451011 | 2802.58221977712 | 3310.85297223895 | Abra |
| **ENSMUSG00000042901** | -0.00950769346110648 | 0.927575408337439 | 0.9868276977487 | 1011.03184884076 | 1003.36017802245 | 1013.65318574249 | 1106.75349922463 | 1025.97582948249 | 1157.03579154187 | 926.559500594962 | 1054.04532329312 | Aida |
| **ENSMUSG00000042903** | 0.243399383373016 | 0.0888831922843323 | 0.751206603737452 | 2167.52940604914 | 2410.67055758641 | 2526.0880978027 | 2235.0793124172 | 1770.54740263948 | 1632.8147798186 | 2283.08425258961 | 2201.44584514773 | Foxo4 |
| **ENSMUSG00000042918** | -0.379599193036076 | 0.21830960019401 | 0.827129763070499 | 23.4913566307952 | 27.6901347830872 | 44.483379766477 | 39.3929211588428 | 42.7033696369448 | 39.1386314263635 | 58.3003730711437 | 34.7368180693922 | Mamstr |
| **ENSMUSG00000042938** | 0.250124352098066 | 0.540112838866929 | 0.913837378403038 | 12.6491920319666 | 16.2883145782866 | 13.2503684410783 | 14.0689004138724 | 9.85462376237187 | 8.56157562451702 | 13.5340151772298 | 15.1973579053591 | Gm14117 |
| **ENSMUSG00000042961** | -0.217134237812695 | 0.293214828665073 | 0.851785577030345 | 210.518695960587 | 199.531853584011 | 290.561650815073 | 257.929840920994 | 356.956371837026 | 267.855008824175 | 276.926772087932 | 211.677485110359 | Egflam |
| **ENSMUSG00000042962** | -0.0743830297734819 | 0.784375359685945 | 0.961141387070613 | 51.5002818444356 | 40.7207864457164 | 42.5904699891801 | 40.3308478531009 | 40.5134532453066 | 63.6002760678407 | 32.2734208072402 | 48.8486504100828 | Gm5436 |
| **ENSMUSG00000042978** | 0.193523734023782 | 0.338214161015393 | 0.858570960161243 | 2060.01127377742 | 2771.45672549546 | 2552.58883468486 | 2113.14884216364 | 1736.60369856909 | 1549.64518803758 | 2553.7645561342 | 2464.14303179751 | Sbk1 |
| **ENSMUSG00000042992** | -0.0327439744954926 | 0.82513410229718 | 0.96672635991852 | 230.395997725106 | 177.542628903324 | 183.612248397799 | 195.088752405698 | 199.282391639076 | 200.585486060113 | 182.188665847324 | 222.532740757044 | Borcs5 |
| **ENSMUSG00000042997** | -0.131093582659432 | 0.41325155140414 | 0.882192126163777 | 112.035700854562 | 127.86326943955 | 130.610774633486 | 120.054616865045 | 142.344565456483 | 129.646716599829 | 150.95632313064 | 113.980184290193 | Nhlrc3 |
| **ENSMUSG00000043003** | 0.516744501384806 | 0.109158021588508 | 0.763154315676794 | 25.2983840639333 | 33.3910448854875 | 35.965285768641 | 28.1378008277448 | 16.4243729372865 | 15.9000690169602 | 28.1091084450157 | 24.9670879873757 | Rasef |
| **ENSMUSG00000043004** | -0.534461726090539 | 0.0836556893020876 | 0.742148948523533 | 58.7283915769879 | 83.0704043492615 | 77.6093008691726 | 55.3376749612315 | 89.7865720571659 | 159.000690169602 | 87.4505596067155 | 62.9604827507734 | Gng2 |
| **ENSMUSG00000043008** | -0.281648737290521 | 0.332362672937434 | 0.856655692540538 | 84.9302893574902 | 73.2974156022896 | 61.519567762149 | 139.751077444466 | 102.926070406995 | 138.208292224346 | 85.3684034256032 | 110.723607596188 | Klhl6 |
| **ENSMUSG00000043015** | -0.237475775759928 | 0.643110218698643 | 0.934616627390525 | 6.32459601598331 | 7.32974156022896 | 18.9290977729689 | 9.37926694258161 | 10.949581958191 | 18.3462334811079 | 10.4107809055614 | 9.76973008201657 | Nemp2 |
| **ENSMUSG00000043017** | -0.503306862387666 | 0.220541296529759 | 0.829144114546069 | 37.0440623793308 | 49.679359463774 | 50.1621090983677 | 83.4754757889764 | 122.635317931739 | 97.8465785659088 | 57.2592949805875 | 34.7368180693922 | Ptgir |
| **ENSMUSG00000043019** | 0.207764136415903 | 0.381773343503842 | 0.875126835008975 | 535.783633925444 | 741.118313312039 | 681.447519826882 | 477.404687377404 | 460.97740043984 | 355.916929533493 | 668.37213413704 | 623.091674119723 | Edem3 |
| **ENSMUSG00000043051** | 0.372770351486921 | 0.252617440201532 | 0.83723604018117 | 21.6843291976571 | 21.1748089517725 | 30.2865564367503 | 27.1998741334867 | 20.8042057205628 | 14.6769867848863 | 18.7394056300105 | 22.7960368580387 | Disc1 |
| **ENSMUSG00000043059** | -0.0941922965497005 | 0.619315605468177 | 0.929568738467614 | 108.421645988285 | 84.6992358070902 | 128.717864856189 | 100.358156285623 | 107.305903190271 | 129.646716599829 | 109.313199508394 | 104.210454208177 | Zfp513 |
| **ENSMUSG00000043061** | -0.0637761259979093 | 0.69849528228308 | 0.945012697918882 | 147.272735800754 | 157.182235680465 | 148.593417517806 | 154.757904552597 | 177.383227722694 | 124.754387671534 | 182.188665847324 | 149.802527924254 | Tmem18 |
| **ENSMUSG00000043065** | -0.16603347294463 | 0.445153526646214 | 0.89108031640712 | 114.746242004269 | 106.688460487777 | 98.4313084194385 | 84.4134024832345 | 108.400861386091 | 154.108361241306 | 90.5737938783839 | 102.03940307884 | Spice1 |
| **ENSMUSG00000043067** | -0.380724928152424 | 0.0202796447977306 | 0.644586421602923 | 159.921927832721 | 152.295741306979 | 129.664319744837 | 184.771558768858 | 199.282391639076 | 236.054870790255 | 194.681602933998 | 186.710397122983 | Dpy19l1 |
| **ENSMUSG00000043068** | -0.769482432874715 | 0.0690273970833657 | 0.724157781586334 | 38.8510898124689 | 26.8757190541728 | 31.2330113253987 | 35.6412143818101 | 93.0714466446232 | 74.6080161565054 | 24.9858741733473 | 33.6512925047237 | Fam89a |
| **ENSMUSG00000043079** | 0.176345762098811 | 0.295294511161747 | 0.851785577030345 | 5441.86311489536 | 6505.55284256766 | 6339.3548441673 | 5666.95308670781 | 4752.11856985488 | 4074.08691503803 | 6496.32728507029 | 5873.77883042129 | Synpo |
| **ENSMUSG00000043085** | 0.288260021441134 | 0.278979645804586 | 0.848075053184273 | 606.257703817829 | 518.782819318427 | 560.301294079881 | 513.983828453472 | 406.229490648885 | 237.277953022329 | 605.907448703672 | 550.361461286933 | Tmem82 |
| **ENSMUSG00000043088** | 0.600816315118028 | 0.20973870921958 | 0.821566820583092 | 11.7456783153976 | 18.7315617650296 | 10.4110037751329 | 12.1930470253561 | 6.56974917491458 | 6.1154111603693 | 13.5340151772298 | 8.68420451734806 | Il17re |
| **ENSMUSG00000043090** | 0.0745505643085407 | 0.714126397569218 | 0.951176382858794 | 211.422209677157 | 254.097707421271 | 253.649910157784 | 193.212899017181 | 201.472308030714 | 154.108361241306 | 261.31060072959 | 248.585354309088 | Zfp866 |
| **ENSMUSG00000043091** | -0.0533610601212866 | 0.788294678926645 | 0.961444367900485 | 500.546598979251 | 371.373572384934 | 604.784673846357 | 606.83857118503 | 533.2446413639 | 631.110431750112 | 435.170641852465 | 562.302242498287 | Tuba1c |
| **ENSMUSG00000043099** | -1.02185262906617 | 0.00572561953767438 | 0.635862253526407 | 168.957064998411 | 109.946123403434 | 222.416898832385 | 172.578511743502 | 567.188345434292 | 406.063301048521 | 211.338852382896 | 183.453820428978 | Hic1 |
| **ENSMUSG00000043122** | -0.173615980788008 | 0.443158865439864 | 0.89108031640712 | 818.583427211555 | 877.940155769646 | 1111.13803927328 | 664.990026229036 | 842.022852584886 | 754.641737189571 | 1339.86750254575 | 979.144059330994 | A530016L24 |
| **ENSMUSG00000043126** | -0.161630106846083 | 0.302613520802329 | 0.851785577030345 | 177.088688447533 | 149.038078391322 | 131.557229522134 | 164.137171495178 | 173.003394939417 | 171.23151249034 | 193.640524843441 | 157.401206876934 | D830039M14 |
| **ENSMUSG00000043131** | -0.213608927232438 | 0.144781488631244 | 0.780607990324772 | 532.169579059168 | 596.966729294203 | 570.712297855013 | 548.687116141024 | 767.565695269187 | 722.841599155651 | 551.771387994752 | 566.644344756961 | Mob1a |
| **ENSMUSG00000043140** | -0.00684541768132712 | 0.97478115628054 | 0.995713359371724 | 604.450676384691 | 618.141538245975 | 510.139184981513 | 475.528833988888 | 493.826146314413 | 371.816998550453 | 693.358008310387 | 658.914017753784 | Tmem186 |
| **ENSMUSG00000043153** | 0.123160280417898 | 0.55586473327432 | 0.916239900758826 | 89.4478579403355 | 87.1424829938332 | 90.8596693102509 | 81.59962240046 | 70.0773245324222 | 69.71568722821 | 108.272121417838 | 71.6446872681215 | Ispd |
| **ENSMUSG00000043154** | 0.124201815344408 | 0.525461449819143 | 0.908006595346551 | 7866.89393016668 | 8846.99806319635 | 7452.38579321787 | 6977.23667858646 | 6216.07767766501 | 5084.35283873103 | 8688.83774378152 | 8584.33616539856 | Ppp2r3a |
| **ENSMUSG00000043155** | -0.170019108651479 | 0.626039802287751 | 0.931698409533928 | 33.4300075130547 | 30.1333819698302 | 17.9826428843205 | 50.6480414899407 | 27.3739548954774 | 42.8078781225851 | 40.6020455316893 | 37.9933947633978 | Hpdl |
| **ENSMUSG00000043157** | -0.67366681849696 | 0.125019420984263 | 0.771949105062466 | 11.7456783153976 | 11.4018202048006 | 17.9826428843205 | 4.68963347129081 | 15.3294147414674 | 18.3462334811079 | 18.7394056300105 | 20.6249857287016 | Arl11 |
| **ENSMUSG00000043162** | 0.0655274475436106 | 0.477879918195423 | 0.897674307047416 | 1234.19973683332 | 1053.85395321514 | 1081.79793772517 | 1144.27056699496 | 1025.97582948249 | 1092.21243324196 | 1091.04983890283 | 1103.97949926787 | Pyurf |
| **ENSMUSG00000043165** | -0.446542717871686 | 0.311322881563769 | 0.851785577030345 | 17.1667606148119 | 13.0306516626293 | 18.9290977729689 | 13.1309737196143 | 35.0386622662111 | 24.4616446414772 | 14.5750932677859 | 10.8552556466851 | Lor |
| **ENSMUSG00000043183** | 0.263688691820078 | 0.42007943327306 | 0.882568531013176 | 28.9124389302094 | 28.5045505120015 | 31.2330113253987 | 18.7585338851632 | 17.5193311331055 | 23.2385624094033 | 29.1501865355718 | 19.5394601640331 | Simc1 |
| **ENSMUSG00000043190** | 0.00961362589631651 | 0.957465682422978 | 0.992403427270738 | 204.194099944604 | 160.439898596123 | 169.415425068072 | 213.847286290861 | 156.579022002131 | 229.939459629886 | 182.188665847324 | 174.76961591163 | Rfesd |
| **ENSMUSG00000043192** | -0.115968820633664 | 0.568097191150794 | 0.920432752255281 | 53.3073092775737 | 52.9370223794314 | 55.8408384302584 | 48.7721881014244 | 54.7479097909548 | 63.6002760678407 | 54.1360607089191 | 56.4473293627624 | Gm1840 |
| **ENSMUSG00000043207** | -0.065045737769449 | 0.739261893507437 | 0.954383574136061 | 627.038519298917 | 750.891302059011 | 672.929425829046 | 443.63932638411 | 546.384139713729 | 593.194882555822 | 755.822693743755 | 714.275821551878 | Zmpste24 |
| **ENSMUSG00000043230** | 0.00499664498241372 | 0.990937237401474 | 0.998628446692557 | 19.877301764519 | 10.5874044758863 | 31.2330113253987 | 17.8206071909051 | 18.6142893289246 | 13.4539045528125 | 30.191264626128 | 16.2828834700276 | Fam124b |
| **ENSMUSG00000043241** | 0.120540381502148 | 0.295032403592735 | 0.851785577030345 | 1092.34808333198 | 1070.14226779343 | 986.205993971682 | 946.368034506485 | 829.978312430875 | 927.096331911986 | 948.422140496641 | 1061.6440022458 | Upf2 |
| **ENSMUSG00000043251** | -0.593418827227645 | 0.0955937329578855 | 0.757085157577544 | 35.2370349461928 | 21.1748089517725 | 25.5542819935081 | 51.5859681841989 | 62.4126171616885 | 63.6002760678407 | 39.5609674411332 | 35.8223436340608 | Exoc3l |
| **ENSMUSG00000043252** | 0.11385000191105 | 0.553285858286778 | 0.915115571755578 | 733.653137854065 | 995.21602073331 | 1002.29572707871 | 783.168789705565 | 811.364023101951 | 591.971800323748 | 858.889424708813 | 984.571687154336 | Tmem64 |
| **ENSMUSG00000043257** | 0.145143440460696 | 0.444880991982174 | 0.89108031640712 | 255.69438178904 | 328.209538752474 | 303.812019256151 | 213.847286290861 | 229.94122112201 | 218.931719541221 | 307.11803671406 | 239.90114979174 | Pigv |
| **ENSMUSG00000043259** | -0.602824143547279 | 0.00664511661160182 | 0.635862253526407 | 60.535419010126 | 73.2974156022896 | 77.6093008691726 | 60.0273084325223 | 109.49581958191 | 123.53130543946 | 107.231043327282 | 72.73021283279 | Fam13c |
| **ENSMUSG00000043262** | 0.0737013716416337 | 0.790539879306793 | 0.961804768755116 | 192.448421629207 | 184.057954734638 | 196.862616838877 | 134.123517278917 | 150.009272827216 | 92.9542496376133 | 212.379930473452 | 216.019587369033 | Uevld |
| **ENSMUSG00000043263** | 0.24368922052003 | 0.595228770524361 | 0.924461858177432 | 9.93865088225949 | 26.0613033252585 | 19.8755526616174 | 10.3171936368398 | 14.2344565456483 | 9.78465785659088 | 14.5750932677859 | 17.3684090346961 | Ifi209 |
| **ENSMUSG00000043279** | 0.0347534444011469 | 0.825232141796537 | 0.96672635991852 | 758.048008201429 | 800.570661522785 | 761.896185362 | 728.769041438591 | 810.269064906132 | 540.602346576646 | 829.739238173241 | 794.604713337348 | Trim56 |
| **ENSMUSG00000043284** | 0.0551301562825002 | 0.728651519119268 | 0.952401994188708 | 1319.13002619081 | 1059.55486331754 | 1160.353693483 | 1425.64857527241 | 1419.06582178155 | 1276.89785028511 | 1053.57102764281 | 1029.07823530575 | Tmem11 |
| **ENSMUSG00000043290** | -0.0736033803720113 | 0.84307554799906 | 0.971386364129692 | 11.7456783153976 | 25.2468875963442 | 17.9826428843205 | 20.6343872736795 | 16.4243729372865 | 23.2385624094033 | 23.9447960827911 | 16.2828834700276 | Zfp784 |
| **ENSMUSG00000043300** | -0.556522539890556 | 0.0850650030601702 | 0.743541556158628 | 23.4913566307952 | 25.2468875963442 | 21.7684624389143 | 30.9515809105193 | 45.9882442244021 | 46.4771248188067 | 35.3966550789086 | 21.7105112933701 | B3galnt1 |
| **ENSMUSG00000043311** | -0.160396142955889 | 0.229537664241127 | 0.832048466795696 | 243.045189757073 | 212.56250524664 | 243.238906382651 | 261.681547698027 | 298.923587458613 | 271.524255520397 | 260.269522639034 | 242.072200921077 | D17H6S53E |
| **ENSMUSG00000043313** | -0.343929459110528 | 0.257275560038026 | 0.837517407767513 | 26.2018977805023 | 31.7622134276588 | 30.2865564367503 | 21.5723139679377 | 26.2789966996583 | 35.4693847301419 | 30.191264626128 | 47.7631248454143 | Pcdhb19 |
| **ENSMUSG00000043323** | 0.0171486341904777 | 0.912175363842113 | 0.986226496507248 | 635.170142748039 | 719.943504360267 | 724.98444470471 | 649.983199120906 | 612.081631462875 | 536.933099880424 | 801.630129728225 | 745.756062927265 | Fbrsl1 |
| **ENSMUSG00000043336** | -0.159308280507268 | 0.227481955486633 | 0.832048466795696 | 384.896843258413 | 453.629561005281 | 414.54724122802 | 484.908100931469 | 466.452191418935 | 499.017550686135 | 544.483841360859 | 430.953649173397 | Filip1l |
| **ENSMUSG00000043372** | 0.0173050689905488 | 0.966249466120734 | 0.994284908685797 | 16.2632468982428 | 25.2468875963442 | 10.4110037751329 | 30.0136542162612 | 25.1840385038392 | 22.0154801773295 | 15.616171358342 | 18.4539345993646 | Hexim2 |
| **ENSMUSG00000043384** | 0.234426375586404 | 0.219076576606535 | 0.827535829146756 | 484.283352081008 | 566.018931595458 | 574.498117409607 | 585.266257217093 | 419.368988998714 | 337.570696052385 | 531.990904274186 | 588.354856050331 | Gprasp1 |
| **ENSMUSG00000043391** | -0.0020252564247591 | 0.991240216359139 | 0.998628446692557 | 316.229800799166 | 364.858246553619 | 347.34894413398 | 300.136542162612 | 298.923587458613 | 245.839528646846 | 385.198893505771 | 399.473407798011 | 2510009E07 |
| **ENSMUSG00000043411** | 0.176058294690667 | 0.153031826509764 | 0.783412422888014 | 457.177940583937 | 519.597235047342 | 538.532831640966 | 528.990655561603 | 464.262275027297 | 432.971110154146 | 401.856142954669 | 510.197015394199 | Usp48 |
| **ENSMUSG00000043415** | 0.103071276948304 | 0.639140270167269 | 0.933801135937988 | 540.301202508289 | 728.902077378324 | 375.742590793433 | 465.211640352048 | 505.870686468423 | 582.187142467157 | 429.965251399684 | 448.322058208094 | Otud1 |
| **ENSMUSG00000043421** | -0.0226589218405394 | 0.941058369275081 | 0.989800147601914 | 104.807591122009 | 46.4216965481167 | 72.8770264259304 | 93.7926694258161 | 105.115986798633 | 83.1695917810225 | 55.1771387994752 | 79.2433662208011 | Hilpda |
| **ENSMUSG00000043424** | -0.0285939719124745 | 0.888186762976965 | 0.980942551992548 | 56.9213641438498 | 54.56585383726 | 70.0376617599851 | 56.2756016554897 | 58.0327843784121 | 58.7079471395453 | 66.6289977955927 | 58.6183804920994 | Eif3j2 |
| **ENSMUSG00000043432** | -0.116974747020131 | 0.610013330235402 | 0.927932112008768 | 101.193536255733 | 71.6685841444609 | 87.0738497556571 | 94.7305961200743 | 114.970610561005 | 107.6312364225 | 96.8202624217207 | 65.1315338801105 | Leng9 |
| **ENSMUSG00000043439** | -0.730327266338756 | 0.0868474840833855 | 0.747269672026173 | 8.13162344912141 | 15.4738988493722 | 27.447191770805 | 11.2551203310979 | 25.1840385038392 | 25.6847268735511 | 19.7804837205666 | 32.5657669400552 | Epop |
| **ENSMUSG00000043445** | -0.121388240321684 | 0.301647178719872 | 0.851785577030345 | 1034.52320547156 | 1009.06108812485 | 1122.49549793706 | 1099.25008567056 | 1169.4153531348 | 1342.9442908171 | 1016.09221638279 | 1112.66370378522 | Pgp |
| **ENSMUSG00000043448** | -1.12159681537686 | 0.0803392658680613 | 0.739855948845396 | 6.32459601598331 | 5.7009101024003 | 13.2503684410783 | 10.3171936368398 | 39.4184950494875 | 19.5693157131818 | 4.16431236222455 | 14.1118323406906 | Gjc2 |
| **ENSMUSG00000043456** | -0.367593185584471 | 0.110878701249642 | 0.764116514278315 | 54.2108229941427 | 56.1946852950887 | 44.483379766477 | 53.4618215727152 | 74.4571573156986 | 61.154111603693 | 51.0128264372507 | 82.4999429148066 | Zfp536 |
| **ENSMUSG00000043460** | -0.480347398116445 | 0.378003078373584 | 0.87422797717331 | 22.5878429142261 | 8.95857301805762 | 11.3574586637814 | 4.68963347129081 | 9.85462376237187 | 15.9000690169602 | 14.5750932677859 | 26.0526135520442 | Elfn2 |
| **ENSMUSG00000043467** | 0.434754157642224 | 0.168126534644658 | 0.79907358295209 | 84.9302893574902 | 119.719112150406 | 135.343049076728 | 109.737423228205 | 64.6025335533267 | 44.0309603546589 | 110.35427759895 | 112.894658725525 | Zbtb37 |
| **ENSMUSG00000043483** | 0.125902138522105 | 0.66877413106378 | 0.942055056494206 | 45.1756858284523 | 36.6487078011448 | 43.5369248778286 | 25.3240207449704 | 35.0386622662111 | 42.8078781225851 | 26.0269522639034 | 34.7368180693922 | Gm6863 |
| **ENSMUSG00000043487** | 0.274840860044019 | 0.158984592479418 | 0.789393312599749 | 77.7021796249379 | 83.0704043492615 | 101.270673085384 | 75.9720622349111 | 74.4571573156986 | 73.3849339244316 | 71.8343882483734 | 59.7039060567679 | Acot6 |
| **ENSMUSG00000043496** | -0.909634634896978 | 0.053161583628269 | 0.703396849069134 | 93.0619128066116 | 104.245213301034 | 88.0203046443056 | 148.192417692789 | 275.929465346412 | 370.593916318379 | 70.7933101578173 | 97.6973008201657 | Tril |
| **ENSMUSG00000043505** | -0.273986521102676 | 0.31931927109387 | 0.854670142081901 | 73.1846110420926 | 47.2361122770311 | 43.5369248778286 | 37.5170677703264 | 75.5521155115177 | 59.9310293716191 | 56.2182168900314 | 52.1052271040884 | Gimap5 |
| **ENSMUSG00000043510** | 0.29639415232943 | 0.0307484038458806 | 0.667184591665619 | 437.300638819418 | 357.52850499339 | 424.011790114504 | 476.466760683146 | 358.051330032845 | 379.155491942897 | 313.364505257397 | 329.999771659226 | Hscb |
| **ENSMUSG00000043518** | 0.0732406783419892 | 0.502302961706561 | 0.900927059989498 | 1299.25272442629 | 1265.60204273287 | 1261.62436656838 | 1223.05640931264 | 1174.89014411389 | 1040.84297949485 | 1361.73014244743 | 1220.1307346874 | Rai2 |
| **ENSMUSG00000043535** | 0.225387165401671 | 0.384451204951725 | 0.875126835008975 | 894.478579403355 | 1185.78930129926 | 1205.78352813812 | 822.561710864407 | 777.420319031559 | 522.256113095538 | 1134.77511870619 | 1079.0124112805 | Setx |
| **ENSMUSG00000043541** | -0.0392303866931403 | 0.842304934554295 | 0.971083627237645 | 114.746242004269 | 81.4415728914329 | 98.4313084194385 | 97.5443762028488 | 85.4067392738896 | 89.2850029413918 | 119.723980413956 | 107.467030902182 | Casc1 |
| **ENSMUSG00000043542** | 0.152600482603763 | 0.310143034759609 | 0.851785577030345 | 198.77301764519 | 236.99497711407 | 204.434255948065 | 194.150825711439 | 189.427767876704 | 173.677676954488 | 168.654650670094 | 219.276164063039 | Zc2hc1a |
| **ENSMUSG00000043556** | -0.0758393959421371 | 0.850147440196324 | 0.972692784625411 | 10.8421645988285 | 16.2883145782866 | 12.3039135524298 | 21.5723139679377 | 12.0445401540101 | 17.123151249034 | 14.5750932677859 | 20.6249857287016 | Fbxl7 |
| **ENSMUSG00000043557** | -0.142410617949494 | 0.599499055624956 | 0.924958767906592 | 160.82544154929 | 137.636258186522 | 70.9841166486335 | 150.068271081306 | 163.148771177045 | 116.192812047017 | 129.093683228961 | 164.999885829613 | Mdga1 |
| **ENSMUSG00000043572** | -0.144193032899691 | 0.471921401555006 | 0.897171173241185 | 80.412720774645 | 70.0397526866323 | 85.1809399783602 | 92.854742731558 | 94.1664048404423 | 68.4926049961361 | 95.7791843311646 | 103.124928643508 | Pars2 |

| | | | | | | | | | | | | |
|---|---|---|---|---|---|---|---|---|---|---|---|---|
| **ENSMUSG00000043587** | -0.348899975872106 | 0.0330936629975317 | 0.671606027118506 | 96.6759676728878 | 111.574954861263 | 83.2880302010633 | 103.171936368398 | 118.255485148462 | 124.754387671534 | 130.134761319517 | 130.263067760221 | Pxylp1 |
| **ENSMUSG00000043602** | -0.00097227159073627 | 0.995333720442663 | 0.999262355603273 | 134.623543768788 | 147.409246933493 | 121.146225747001 | 153.819977858338 | 160.958854785407 | 134.539045528125 | 143.668776496747 | 118.322286548867 | Zfp3 |
| **ENSMUSG00000043613** | -0.83410979273612 | 0.157281544417618 | 0.788998730890588 | 83.1232619243521 | 34.2054606144018 | 113.574586637814 | 65.6548685980713 | 119.350443344282 | 314.332133642982 | 56.2182168900314 | 39.0789203280663 | Mmp3 |
| **ENSMUSG00000043621** | -0.141423227872658 | 0.777778459779961 | 0.960041127374359 | 49.6932544112975 | 97.7298874697194 | 127.77140996754 | 23.448167356454 | 120.445401540101 | 42.8078781225851 | 105.14888714617 | 60.7894316214364 | Ubxn10 |
| **ENSMUSG00000043629** | -0.178857751095424 | 0.66978256378176 | 0.942388853397102 | 40.658117245607 | 21.9892246806869 | 24.6078271048596 | 33.7653609932938 | 37.2285786578493 | 58.7079471395453 | 16.6572494488982 | 24.9670879873757 | 1700019D03I |
| **ENSMUSG00000043631** | -0.508413243180495 | 0.169439830534675 | 0.79907358295209 | 255.69438178904 | 259.798617523671 | 182.66579350915 | 210.095579513828 | 429.223612761086 | 517.363784167243 | 147.833088858971 | 198.651178334337 | Ecm2 |
| **ENSMUSG00000043633** | -0.234074309009773 | 0.229575843156575 | 0.832048466795696 | 121.070838020252 | 112.389370590177 | 101.270673085384 | 128.495957113368 | 129.205067106653 | 178.570005882784 | 112.436433780063 | 125.920965501547 | Fam221b |
| **ENSMUSG00000043635** | -0.419680492308962 | 0.253294295034102 | 0.837406774112203 | 20.780815481088 | 17.1027303072009 | 35.0188308799925 | 22.5102406621959 | 39.4184950494875 | 17.123151249034 | 35.3966550789086 | 34.7368180693922 | Adamts3 |
| **ENSMUSG00000043639** | 0.278792702249927 | 0.414566501777021 | 0.882192126163777 | 2036.51991714663 | 2703.04580426666 | 2769.32700418536 | 2199.43809803539 | 1246.06242684213 | 987.027361283605 | 2744.28184670598 | 3024.27422316646 | Rbm20 |
| **ENSMUSG00000043644** | -0.00226537371803387 | 0.990939788275065 | 0.998628446692557 | 298.159526467785 | 193.83094348161 | 228.095628164276 | 270.12288794635 | 243.08071947184 | 309.439804714686 | 198.845915296222 | 240.986675356409 | 0610009L18F |
| **ENSMUSG00000043668** | 0.236990578824261 | 0.30169834242843 | 0.851785577030345 | 59.631905293557 | 78.9983257046899 | 87.0738497556571 | 105.985716451172 | 67.887408140784 | 67.2695227640623 | 64.5468416144805 | 81.4144173501381 | Tox3 |
| **ENSMUSG00000043671** | -0.187978588294978 | 0.652027200677322 | 0.936720465895231 | 15.3597331816738 | 8.95857301805762 | 25.5542819935081 | 20.6343872736795 | 27.3739548954774 | 19.5693157131818 | 16.6572494488982 | 16.2828834700276 | Dpy19l3 |
| **ENSMUSG00000043683** | 0.0398455124170536 | 0.823615827242457 | 0.966637987288264 | 7804.55148372341 | 8620.59049055817 | 8201.03161013879 | 6989.42972561182 | 7035.1064081377 | 5635.96292539634 | 9523.78237240754 | 8558.28355184651 | Fem1a |
| **ENSMUSG00000043687** | 0.15003959319658 | 0.586656321155459 | 0.923058967588378 | 56.0178504272808 | 41.5352021746308 | 33.1259211026956 | 63.779015209555 | 49.2731188118594 | 45.2540425867328 | 42.6842017128016 | 37.9933947633978 | 1190005I06R |
| **ENSMUSG00000043702** | -0.0391171657462391 | 0.750288409462943 | 0.956197192463153 | 242.141676040504 | 280.973426475443 | 251.757000380487 | 227.916186704733 | 263.884925192402 | 260.516515431732 | 248.817663642917 | 258.355084391105 | Pde12 |
| **ENSMUSG00000043716** | 0.0207322454397988 | 0.903194178320685 | 0.985027469232262 | 8361.11593312994 | 6441.21399998343 | 7376.66940212599 | 9109.14405463526 | 8055.6074466411 | 9449.53332500264 | 6394.30163219579 | 6943.02151161977 | Rpl7 |
| **ENSMUSG00000043719** | -0.838359818819446 | 0.0866758362859178 | 0.747269672026173 | 34.3335212296237 | 39.9063707168021 | 47.3227444324223 | 45.0204813243917 | 135.774816281568 | 102.738907494204 | 27.0680303544596 | 32.5657669400552 | Col6a6 |
| **ENSMUSG00000043733** | 0.217766305731121 | 0.461699726061827 | 0.895328112556867 | 10019.9671167507 | 12282.203607757 | 12642.744402566 | 8922.49664247789 | 6400.03065456262 | 5574.80881379265 | 13900.4746651055 | 11845.2549616628 | Ptpn11 |
| **ENSMUSG00000043795** | -0.0449883193873156 | 0.835968882795031 | 0.968924788429433 | 214.132750826864 | 198.717437855096 | 168.468970179424 | 131.309737196143 | 154.389105610493 | 152.885279009232 | 200.928071477334 | 226.874843015718 | Prr33 |
| **ENSMUSG00000043801** | -0.097175152563889 | 0.68577420717853 | 0.944452327903597 | 140.948139784771 | 97.7298874697194 | 97.48485353079 | 110.675349922463 | 90.881530252985 | 103.961989726278 | 116.600746142287 | 166.085411394282 | Oaz1-ps |
| **ENSMUSG00000043811** | -0.424182828969171 | 0.34770924904397 | 0.861266047293794 | 27.1054114970714 | 32.5766291565731 | 52.0550188756646 | 38.4549944645846 | 17.5193311331055 | 57.4848649074714 | 38.5198893505771 | 87.9275707381491 | Rtn4r |
| **ENSMUSG00000043822** | 0.0856437490682984 | 0.621845161559649 | 0.93078489223231 | 2485.56623428144 | 2303.16768136972 | 1646.8315062483 | 2110.33506208086 | 1765.07261166038 | 1736.77676954488 | 2350.75432847576 | 2200.36031958306 | Adamtsl5 |
| **ENSMUSG00000043831** | 0.0545917083132394 | 0.698138023856498 | 0.944933386973863 | 340.62467114653 | 352.642010619904 | 317.06238769723 | 300.136542162612 | 305.493336633528 | 260.516515431732 | 352.92547269853 | 341.94055287058 | Lysmd4 |
| **ENSMUSG00000043832** | -0.370507821681498 | 0.31357254352573 | 0.85295001595893 | 13.5527057485357 | 12.2162359337149 | 13.2503684410783 | 16.8826804966469 | 21.8991639163819 | 19.5693157131818 | 16.6572494488982 | 14.1118323406906 | Clec4a3 |
| **ENSMUSG00000043857** | 0.210404697337861 | 0.624526856608286 | 0.931259048662274 | 41.5616309621761 | 35.0198763433161 | 9.46454888648447 | 32.8274342990356 | 19.7092475247437 | 26.9078091056249 | 28.1091084450157 | 28.2236646813812 | Mgat5b |
| **ENSMUSG00000043866** | -0.0368926895357516 | 0.814246756520553 | 0.966177091475091 | 1033.61969175499 | 829.075212034787 | 966.330441310064 | 1036.40899715527 | 923.049759075499 | 1254.88237010778 | 870.34128370493 | 918.354627709557 | Taf10 |
| **ENSMUSG00000043872** | -0.151295033850841 | 0.70045198908958 | 0.94561797945347 | 51.5002818444356 | 48.0505280059454 | 40.6975602118832 | 28.1378008277448 | 30.6588294829347 | 23.2385624094033 | 70.7933101578173 | 61.8749571861049 | Zmym1 |
| **ENSMUSG00000043881** | -0.0862321785975012 | 0.579601113781586 | 0.922450244597442 | 147.272735800754 | 185.686786192467 | 185.505158175096 | 180.081925297567 | 187.237851485066 | 198.139321595965 | 156.16171358342 | 200.822229463674 | Kbtbd7 |
| **ENSMUSG00000043885** | -0.0739126361668102 | 0.67141024937709 | 0.943173644832837 | 122.87786545339 | 131.120932355207 | 159.004421292939 | 138.813150750208 | 169.71852035196 | 117.415894279091 | 140.545542225078 | 151.973579053591 | Slc36a4 |
| **ENSMUSG00000043895** | -0.987231284830888 | 0.0137995751973818 | 0.635862253526407 | 49.6932544112975 | 29.3189662409158 | 39.7511053232348 | 58.151455044006 | 125.920192519196 | 127.200552135681 | 51.0128264372507 | 46.6775992807458 | S1pr2 |
| **ENSMUSG00000043903** | 0.699757980318186 | 0.0890074239690972 | 0.751206603737452 | 19.877301764519 | 35.8342920722305 | 26.5007368821565 | 33.7653609932938 | 29.5638712871156 | 11.0077400886647 | 19.7804837205666 | 10.8552556466851 | Zfp469 |
| **ENSMUSG00000043909** | 0.205220649050087 | 0.37833249692797 | 0.87422797717331 | 487.897406947284 | 548.101785559343 | 552.729654970693 | 390.177504811395 | 394.184950494875 | 278.86274891284 | 576.7572621681 | 465.69046724279 | Trp53bp1 |
| **ENSMUSG00000043923** | -0.332374166249024 | 0.247156020347456 | 0.833120697990749 | 38.8510898124689 | 38.2775392589734 | 19.8755526616174 | 31.8895076047775 | 43.7983278327639 | 39.1386314263635 | 35.3966550789086 | 44.5065481514088 | Ccdc84 |
| **ENSMUSG00000043929** | -0.178623451416791 | 0.373541168228449 | 0.873700364743909 | 73.1846110420926 | 65.9676740420606 | 61.519567762149 | 58.151455044006 | 77.7420319031559 | 84.3926740130963 | 70.7933101578173 | 60.7894316214364 | Klhl15 |
| **ENSMUSG00000043940** | 0.205380561836022 | 0.409064245888921 | 0.882192126163777 | 531.266065342598 | 774.509358197526 | 827.201572678743 | 485.846027625728 | 524.484975797347 | 386.49398533534 | 672.536446499264 | 687.137682435165 | Wdfy3 |
| **ENSMUSG00000043943** | -0.596074340183004 | 0.138387111433509 | 0.779711473022872 | 103.000563688871 | 114.018202048006 | 135.343049076728 | 101.296082979881 | 257.315176017488 | 261.739597663806 | 72.8754663389296 | 94.4407241261602 | Naalad2 |
| **ENSMUSG00000043953** | -0.24386349218549 | 0.350790007825518 | 0.861774532843376 | 355.080890611635 | 316.807718547674 | 229.042083052924 | 220.412773150668 | 422.653863586171 | 365.701587390084 | 339.391457521301 | 200.822229463674 | Ccrl2 |
| **ENSMUSG00000043962** | 0.199619641403715 | 0.294260011985253 | 0.851785577030345 | 2449.42568561868 | 3146.08796079605 | 3069.35320388691 | 2246.3344327483 | 2259.99371617062 | 1757.56916749014 | 2801.54114168656 | 2681.24814473121 | Thrap3 |
| **ENSMUSG00000043964** | 0.0733085424426108 | 0.570490100962444 | 0.920872458148558 | 364.116027777325 | 370.55915665602 | 397.511053232348 | 434.260059441529 | 407.324448844704 | 402.3940543523 | 328.980676615739 | 350.624757387928 | Orai3 |
| **ENSMUSG00000043987** | 0.345458234366445 | 0.143947869705829 | 0.780607990324772 | 144.562194651047 | 218.26341534904 | 184.558703286447 | 119.116690170786 | 107.305903190271 | 117.415894279091 | 143.668776496747 | 156.315681312265 | Cep164 |
| **ENSMUSG00000043991** | -0.158275296090708 | 0.501873480771011 | 0.900927059989498 | 579.152292320758 | 509.009830571455 | 476.066808990169 | 639.666005484066 | 752.23628052772 | 809.680437632895 | 466.402984569149 | 432.039174738066 | Pura |
| **ENSMUSG00000043993** | 0.0334591196663639 | 0.912359796430727 | 0.986226496507248 | 42.4651446787451 | 44.7928650902881 | 53.9479286529615 | 45.9584080186499 | 37.2285786578493 | 30.5770558018465 | 71.8343882483734 | 42.3354970220718 | 2900052L18F |
| **ENSMUSG00000043998** | -0.197587346010127 | 0.154481702255094 | 0.786853994984691 | 358.694945477911 | 403.950201541507 | 405.082692341535 | 351.72251034681 | 502.585811880965 | 465.99433042014 | 379.99350305299 | 395.131305539337 | Mgat2 |
| **ENSMUSG00000044005** | 0.029658162358489 | 0.928150202286096 | 0.986901071486195 | 40.658117245607 | 70.8541684155466 | 44.483379766477 | 41.2687745473591 | 29.5638712871156 | 40.3617136584374 | 57.2592949805875 | 66.217059444779 | Gls2 |
| **ENSMUSG00000044017** | -1.13980703967466 | 0.0288579001060732 | 0.665956123973759 | 18.9737880479499 | 25.2468875963442 | 44.483379766477 | 21.5723139679377 | 78.836990098975 | 108.854318654574 | 18.7394056300105 | 36.9078691987293 | Adgrd1 |
| **ENSMUSG00000044018** | 0.0613167039413685 | 0.66724289064661 | 0.942024383071524 | 1718.48308891432 | 1654.89276115392 | 1513.38136694887 | 1585.09611329629 | 1333.65908250766 | 1323.37497510392 | 1803.14725284323 | 1741.18300572829 | Mrpl50 |
| **ENSMUSG00000044026** | -0.148089151755713 | 0.612256408595469 | 0.928055903343287 | 60.535419010126 | 61.895595397489 | 39.7511053232348 | 34.703287687552 | 45.9882442244021 | 42.8078781225851 | 70.7933101578173 | 58.6183804920994 | Slc35g1 |
| **ENSMUSG00000044030** | -0.336231684030392 | 0.108103669154785 | 0.762287160109383 | 556.564449406532 | 586.379324818317 | 539.479286529615 | 568.383576720446 | 850.782518151438 | 916.088591823321 | 578.839418349212 | 497.170708618176 | Irf2bp1 |
| **ENSMUSG00000044033** | 0.245589161515828 | 0.350569503203331 | 0.861774532843376 | 1700.41281458294 | 2162.27376026754 | 2305.56410874762 | 1667.63366239101 | 1228.54309570903 | 1056.74304851181 | 2249.76975369181 | 2073.35382851685 | Ccdc141 |
| **ENSMUSG00000044037** | -0.122256892164342 | 0.455140960111878 | 0.892775943845882 | 138.237598635064 | 147.409246933493 | 183.612248397799 | 189.461192240149 | 185.047935093427 | 189.577745971448 | 181.147587756768 | 160.657783570939 | Als2cl |
| **ENSMUSG00000044042** | 0.471338219334269 | 0.25894772989314 | 0.838606660810377 | 35.2370349461928 | 55.3802695661743 | 28.3936466594534 | 24.3860940507122 | 10.949581958191 | 26.9078091056249 | 35.3966550789086 | 30.3947158107182 | Fmn1 |
| **ENSMUSG00000044043** | -0.274078019112293 | 0.473626846850197 | 0.897377834231242 | 28.9124389302094 | 35.8342920722305 | 35.0188308799925 | 29.075727522003 | 55.8428679867739 | 55.0387004433237 | 27.0680303544596 | 18.4539345993646 | Pcdhb14 |
| **ENSMUSG00000044060** | 0.222659289726507 | 0.542573895448495 | 0.913837378403038 | 21.6843291976571 | 25.2468875963442 | 12.3039135524298 | 24.3860940507122 | 15.3294147414674 | 18.3462334811079 | 21.8626399016789 | 16.2828834700276 | NA |
| **ENSMUSG00000044066** | 0.233523762785395 | 0.346855114136481 | 0.861266047293794 | 1064.33915811834 | 1303.06516626293 | 1257.83854701379 | 1015.77460988159 | 792.749733773026 | 622.548856125595 | 1380.46954807744 | 1150.65709854862 | Cep68 |
| **ENSMUSG00000044067** | -0.398527373215829 | 0.137168137377723 | 0.777073872865278 | 658.661499378834 | 402.321370083678 | 343.563124579386 | 337.653609932938 | 432.508487348543 | 480.671317205027 | 737.083288113745 | 645.887710977762 | Gpr22 |
| **ENSMUSG00000044068** | 0.215923572458101 | 0.030113111187748 | 0.665956123973759 | 731.846110420926 | 753.334549245754 | 788.396922244156 | 825.375490947182 | 661.354750274735 | 705.718447906617 | 694.399086400943 | 606.808790649696 | Zrsr1 |
| **ENSMUSG00000044072** | 0.274210758600133 | 0.490017460515149 | 0.899236955296196 | 103.90407740544 | 171.841718800923 | 129.664319744837 | 110.675349922463 | 77.7420319031559 | 39.1386314263635 | 153.038479311752 | 156.315681312265 | Eml6 |
| **ENSMUSG00000044080** | 0.0856204492240894 | 0.626242237451737 | 0.931698409533928 | 18285.3105959243 | 12244.7404842269 | 13632.7362160922 | 18276.4395643145 | 14445.7834774413 | 17475.3989318713 | 13307.0601534885 | 13613.5761065078 | S100a1 |
| **ENSMUSG00000044086** | 0.0320933900060648 | 0.889910488521799 | 0.981565686140537 | 2024.77423883123 | 1886.18682816558 | 1116.81676860517 | 1764.2401118996 | 1515.42214301363 | 1234.08997216252 | 1778.16137866988 | 2114.60379997425 | Lmod3 |
| **ENSMUSG00000044098** | -0.2927036616633 | 0.070841023160492 | 0.726501344493452 | 108.421645988285 | 121.347943608235 | 139.128868631322 | 127.55803041911 | 170.813478547779 | 150.439114545085 | 156.16171358342 | 130.263067760221 | Rsbn1 |
| **ENSMUSG00000044122** | 0.131702083260645 | 0.709068841868182 | 0.949280056132338 | 25.2983840639333 | 13.0306516626293 | 25.5542819935081 | 22.5102406621959 | 19.7092475247437 | 17.123151249034 | 18.7394056300105 | 22.7960368580387 | Proca1 |
| **ENSMUSG00000044134** | -0.144948383130694 | 0.472161358991581 | 0.897171173241185 | 171.667606148119 | 155.553404222637 | 154.272146849697 | 242.923013812864 | 189.427767876704 | 228.716377397812 | 214.462086654564 | 168.256462523619 | Fam109a |
| **ENSMUSG00000044147** | -0.209280100778243 | 0.250950835868173 | 0.836703579022786 | 679.442314859922 | 725.644414462667 | 850.862944894954 | 903.223406570609 | 1065.39432453198 | 1084.87393984951 | 756.863771834311 | 745.756062927265 | Arf6 |
| **ENSMUSG00000044148** | 0.125374913264287 | 0.443607141536265 | 0.89108031640712 | 327.071965397994 | 341.240190415104 | 381.421320125324 | 348.908730264036 | 286.879047304603 | 254.401104271363 | 350.843316517418 | 388.618152151326 | 1810030O07I |
| **ENSMUSG00000044149** | 0.0659770114607248 | 0.815251455198723 | 0.966177091475091 | 57.8248778604189 | 61.895595397489 | 46.3762895437739 | 75.9720622349111 | 66.7924499449649 | 33.0232202659942 | 66.6289977955927 | 64.046008315442 | Nkrf |
| **ENSMUSG00000044150** | 0.356493876396561 | 0.168738218438748 | 0.79907358295209 | 46.9827132615903 | 39.9063707168021 | 54.8943835416099 | 37.5170677703264 | 40.5134532453066 | 37.9155491942896 | 29.1501865355718 | 32.5657669400552 | Bclaf3 |
| **ENSMUSG00000044155** | 0.099070633957557 | 0.566442255562562 | 0.919931558693298 | 277.378710986697 | 271.200437728471 | 316.115932808581 | 311.39166249371 | 318.632834983357 | 325.339873731647 | 229.03717992235 | 225.78931745105 | Lsm8 |
| **ENSMUSG00000044162** | -0.166209036877956 | 0.626564343719866 | 0.931698409533928 | 28.0089252136404 | 47.2361122770311 | 30.2865564367503 | 33.7653609932938 | 28.4689130912965 | 29.3539735697726 | 61.4236073428121 | 36.9078691987293 | Tnip3 |
| **ENSMUSG00000044167** | -0.24524571846562 | 0.165627261833971 | 0.795216960550936 | 540.301202508289 | 594.52348210746 | 702.269527377147 | 625.597105070194 | 857.352267326353 | 846.372904595111 | 660.043509412591 | 555.789089110276 | Foxo1 |
| **ENSMUSG00000044177** | 0.256841026475024 | 0.152143398732022 | 0.783412422888014 | 194.255449062345 | 293.189662409158 | 260.275094378323 | 254.178134143962 | 192.712642464161 | 178.570005882784 | 227.996101831794 | 238.815624227072 | Wfikkn2 |
| **ENSMUSG00000044197** | -0.132651787525597 | 0.298796531798173 | 0.851785577030345 | 1194.44513330428 | 1295.7354247027 | 1403.59259986565 | 1322.47663890401 | 1655.57679207847 | 1231.64380769838 | 1503.31676276306 | 1326.51224002492 | Gpr146 |
| **ENSMUSG00000044199** | -0.511472306584413 | 0.329140048375354 | 0.855977678491063 | 23.4913566307952 | 15.4738988493722 | 32.1794662140472 | 41.2687745473591 | 59.1277425742312 | 66.0464405319884 | 14.5750932677859 | 20.6249857287016 | S1pr4 |
| **ENSMUSG00000044206** | -0.417217792713506 | 0.371102574015995 | 0.873374202289323 | 15.3597331816738 | 11.4018202048006 | 12.3039135524298 | 15.9447538023887 | 9.85462376237187 | 34.2463024980681 | 12.4929370866736 | 17.3684090346961 | Vsig4 |
| **ENSMUSG00000044216** | 0.170646531426929 | 0.553998964771617 | 0.915640840156802 | 33.4300075130547 | 34.2054606144018 | 34.0723759913441 | 43.1446279358754 | 28.4689130912965 | 22.0154801773295 | 42.6842017128016 | 34.7368180693922 | Kcnj4 |
| **ENSMUSG00000044221** | 0.275473380769615 | 0.180195574298499 | 0.802731406993634 | 5102.1419574654 | 5501.37824881629 | 5473.34862105397 | 4820.94320848695 | 3746.94694609295 | 2923.16653465652 | 5613.49306427869 | 4980.39129069911 | Grsf1 |
| **ENSMUSG00000044224** | 0.237329827645412 | 0.0272267455484791 | 0.664353746352133 | 1709.44795174863 | 1603.58457023231 | 1646.8315062483 | 1775.4952322307 | 1421.25573817319 | 1230.4207254663 | 1563.69929201532 | 1495.8542281132 | Dnajc21 |
| **ENSMUSG00000044229** | -0.581152746237205 | 0.0545831549351692 | 0.703396849069134 | 112.939214571131 | 84.6992358070902 | 116.413951303759 | 149.130344387048 | 247.460552255116 | 207.923979452556 | 106.189965236726 | 131.348593324889 | Nxpe4 |
| **ENSMUSG00000044231** | 0.181079983290416 | 0.4540159648007 | 0.891964629399053 | 149.079763233892 | 195.459774939439 | 151.432782183751 | 164.137171495178 | 131.394983498292 | 90.5080851734656 | 170.736806851206 | 188.88144825232 | Nhlrc1 |
| **ENSMUSG00000044244** | -0.158348961712772 | 0.661506583592904 | 0.939611397743053 | 19.877301764519 | 14.6594831204579 | 25.5542819935081 | 24.3860940507122 | 24.0890803080201 | 34.2463024980681 | 18.7394056300105 | 17.3684090346961 | Il20rb |
| **ENSMUSG00000044250** | 0.31071329781371 | 0.331049041441535 | 0.856333478432716 | 27.1054114970714 | 19.5459774939439 | 24.6078271048596 | 25.3240207449704 | 21.8991639163819 | 22.0154801773295 | 17.6983275394543 | 16.2828834700276 | Pced1b |
| **ENSMUSG00000044252** | 0.127270764920113 | 0.509583974953957 | 0.902394714365738 | 1529.64872215139 | 1555.53404222637 | 1556.91829182669 | 1409.70382147002 | 1123.42710891039 | 1024.94291047789 | 1703.20375614984 | 1687.99225305953 | Osbpl1a |
| **ENSMUSG00000044258** | 0.326150433392229 | 0.456393386557283 | 0.892805623137304 | 718.293404672391 | 642.574010113405 | 1827.60438998015 | 468.963347129081 | 720.482492848966 | 1192.50517627201 | 391.445362049107 | 613.321944037707 | Ctla2a |
| **ENSMUSG00000044285** | -0.4249309213055 | 0.133985756403343 | 0.77303252008843 | 370.440623793308 | 281.787842204358 | 325.580481695066 | 401.432625142493 | 467.547149614754 | 736.295503708463 | 289.419709174606 | 359.308961905276 | Gm1821 |
| **ENSMUSG00000044296** | 0.296974487030337 | 0.453824065448831 | 0.891964629399053 | 24.3948703473642 | 21.9892246806869 | 26.5007368821565 | 12.1930470253561 | 17.5193311331055 | 13.4539045528125 | 24.9858741733473 | 13.0263067760221 | Zfp879 |
| **ENSMUSG00000044308** | 0.283773200619822 | 0.344223129201666 | 0.861266047293794 | 4154.35606878447 | 6240.05331494159 | 5663.58605367231 | 3774.21701769484 | 3278.30483828238 | 2215.00192228576 | 5560.39808166033 | 5236.57532396088 | Ubr3 |
| **ENSMUSG00000044317** | -0.645459590108408 | 0.0264879223145842 | 0.664353746352133 | 179.79922959724 | 145.780415475665 | 175.094154399963 | 180.081925297567 | 361.336204620302 | 362.032340693862 | 175.942197303987 | 166.085411394282 | Gpr4 |
| **ENSMUSG00000044328** | -0.359863816026054 | 0.151756209891815 | 0.783412422888014 | 252.983840639333 | 219.892246806869 | 254.596365046432 | 216.661066373635 | 402.944616061428 | 385.270903103266 | 210.29777429234 | 213.848536239696 | Trp53i13 |

| | | | | | | | | | | | | |
|---|---|---|---|---|---|---|---|---|---|---|---|---|
| **ENSMUSG00000044337** | 0.145687884659316 | 0.534968731815794 | 0.910813988293389 | 1328.1651633565 | 1140.99643620897 | 1282.44637411865 | 2207.87943828371 | 1134.37669086858 | 1660.9456711563 | 1203.48627268289 | 1388.38719721102 | Ackr3 |
| **ENSMUSG00000044338** | -0.365263580831823 | 0.304627861519565 | 0.851785577030345 | 114.746242004269 | 97.7298874697194 | 147.646962629158 | 305.764102328161 | 208.042057205628 | 233.608706326107 | 155.120635492864 | 260.526135520442 | Aplnr |
| **ENSMUSG00000044339** | -0.511953779485492 | 0.262804390371446 | 0.842065278491208 | 15.3597331816738 | 9.77298874697194 | 7.57163910918757 | 16.8826804966469 | 16.4243729372865 | 25.6847268735511 | 9.36970281500523 | 19.5394601640331 | Alkbh2 |
| **ENSMUSG00000044340** | -0.0105294072403606 | 0.931585733263884 | 0.987783121884152 | 430.072529086865 | 479.69086433054 | 427.797609669098 | 373.294824314748 | 418.274030802895 | 403.617136584374 | 435.170641852465 | 466.775992807458 | Phlpp1 |
| **ENSMUSG00000044345** | -0.229236601869654 | 0.156548239195805 | 0.787275540343856 | 604.450676384691 | 762.293122263812 | 653.053873167428 | 628.410885152968 | 889.106055005107 | 898.965440574287 | 680.865071223713 | 637.203506460414 | Marveld1 |
| **ENSMUSG00000044348** | 0.165594763918814 | 0.664873058540354 | 0.941513723457524 | 27.1054114970714 | 23.6180561385155 | 23.6613722162112 | 18.7585338851632 | 25.1840385038392 | 23.2385624094033 | 26.0269522639034 | 8.68420451734806 | Slc25a53 |
| **ENSMUSG00000044350** | -8.66196716719554E-0 | 0.999691619493903 | 0.999835584300858 | 63.2459601598332 | 49.679359463774 | 45.4298346551254 | 57.2135283497478 | 54.7479097909548 | 51.3694537471021 | 62.4646854333682 | 46.6775992807458 | Lacc1 |
| **ENSMUSG00000044364** | 0.0284822918886373 | 0.895487984617645 | 0.983733637008786 | 93.0619128066116 | 88.7713144516618 | 72.8770264259304 | 69.4065753751039 | 84.3117810780705 | 57.4848649074714 | 83.2862472444909 | 92.2696729968231 | Tmem74b |
| **ENSMUSG00000044365** | -0.0450970758479498 | 0.877811677601186 | 0.977770986518081 | 34.3335212296237 | 38.2775392589734 | 58.6802030962037 | 64.7169419038131 | 42.7033696369448 | 66.0464405319884 | 47.8895921655823 | 45.5920737160773 | Cxxc4 |
| **ENSMUSG00000044367** | -0.518725425446892 | 0.0708541629455949 | 0.726501344493452 | 38.8510898124689 | 32.5766291565731 | 47.3227444324223 | 47.8342614071662 | 78.836990098975 | 59.9310293716191 | 62.4646854333682 | 36.9078691987293 | Slc16a13 |
| **ENSMUSG00000044393** | 0.362113966353938 | 0.160795633080019 | 0.792229941270026 | 1478.14844030696 | 2096.30608622548 | 2188.20370255521 | 1392.82114097337 | 1157.37081298079 | 903.857769502582 | 1781.28461294155 | 1723.81459669359 | Dsg2 |
| **ENSMUSG00000044405** | 0.509884998337208 | 0.135480564669598 | 0.776010327117824 | 69.5705561758165 | 41.5352021746308 | 39.7511053232348 | 49.7101147956825 | 47.0832024202212 | 46.4771248188067 | 26.0269522639034 | 21.7105112933701 | Adig |
| **ENSMUSG00000044408** | 0.0802729027818805 | 0.503124105702697 | 0.900927059989498 | 741.784761303186 | 704.469605510894 | 736.341903368492 | 740.024161769689 | 626.316088008523 | 620.102691661447 | 708.974179668729 | 807.63102011337 | Sptssa |
| **ENSMUSG00000044424** | -0.443708944872304 | 0.284716214286192 | 0.851125503777472 | 10.8421645988285 | 8.14415728914329 | 10.4110037751329 | 14.0689004138724 | 12.0445401540101 | 17.123151249034 | 12.4929370866736 | 17.3684090346961 | Gm9493 |
| **ENSMUSG00000044433** | -0.750247791168854 | 0.0693879033226467 | 0.724352955175446 | 11.7456783153976 | 13.0306516626293 | 15.1432782183751 | 8.44134024832345 | 12.0445401540101 | 28.1308913376988 | 17.6983275394543 | 23.8815624227072 | Camsap3 |
| **ENSMUSG00000044442** | 0.190013329711267 | 0.13020509696129 | 0.77303252008843 | 728.23205555465 | 565.204515866544 | 692.804978490663 | 767.224035903176 | 607.701798679599 | 595.64104701997 | 617.359307699789 | 591.611432744337 | N6amt1 |
| **ENSMUSG00000044447** | 0.20484228131913 | 0.56693035242414 | 0.919931558693298 | 262.922491521592 | 344.497853330761 | 239.453086828057 | 214.785212985119 | 178.478185918513 | 95.400414101761 | 337.309301340188 | 309.374785930525 | Dock5 |
| **ENSMUSG00000044452** | 0.113197500605291 | 0.765090064987696 | 0.958197631858494 | 101.193536255733 | 143.337168288922 | 166.576060402127 | 94.7305961200743 | 78.836990098975 | 53.8156182112498 | 178.024353485099 | 156.315681312265 | Zfp507 |
| **ENSMUSG00000044456** | -0.396297501030639 | 0.191440981185756 | 0.808824684419004 | 503.257140128958 | 536.699965354543 | 513.925004536107 | 315.143369270742 | 897.865720571659 | 757.087901653719 | 398.732908683 | 407.07208675069 | Rin3 |
| **ENSMUSG00000044461** | -1.01044503089491 | 0.0130494486542273 | 0.635862253526407 | 21.6843291976571 | 19.5459774939439 | 12.3039135524298 | 9.37926694258161 | 29.5638712871156 | 45.2540425867328 | 34.3555769883525 | 18.4539345993646 | Shisa2 |
| **ENSMUSG00000044465** | 0.192219705090193 | 0.162469629550814 | 0.793968226938511 | 421.037391921175 | 509.009830571455 | 486.477812765302 | 399.556771753977 | 404.039574257247 | 344.909189444828 | 437.252798033577 | 402.729984492016 | Fam160a2 |
| **ENSMUSG00000044468** | 0.114421054046638 | 0.908844235642085 | 0.985338457039129 | 194.255449062345 | 106.688460487777 | 848.970035117657 | 80.6616957062019 | 186.142893289246 | 752.195572725424 | 78.0808567917102 | 120.493337678204 | Fam46c |
| **ENSMUSG00000044469** | -0.508052800836346 | 0.173936331557832 | 0.799889021902745 | 25.2983840639333 | 18.7315617650296 | 15.1432782183751 | 41.2687745473591 | 39.4184950494875 | 44.0309603546589 | 33.3144988977964 | 26.0526135520442 | Tnfaip8l1 |
| **ENSMUSG00000044475** | -0.15101593224018 | 0.261776342724703 | 0.841137531367976 | 573.731210021344 | 563.575684408715 | 657.786147610671 | 692.189900362523 | 670.114415841287 | 801.118862008378 | 682.947227404826 | 607.894316214364 | Ascc1 |
| **ENSMUSG00000044477** | 0.059162297212694 | 0.52466695725519 | 0.907662727148411 | 2799.98900764747 | 3010.08053406736 | 3158.31996341987 | 2750.93899425919 | 2690.31228712752 | 2682.21933493797 | 3052.44096151059 | 2822.36646813812 | Zfand3 |
| **ENSMUSG00000044496** | -0.0597679283716011 | 0.59244386601441 | 0.923935529355527 | 1004.70725282478 | 1124.70812163069 | 1148.99623481921 | 1074.86399161985 | 1227.44813751321 | 1179.0512717192 | 1156.63775860787 | 974.80195707232 | 2510039O18 |
| **ENSMUSG00000044499** | 0.596121823402869 | 0.212661407631642 | 0.823370267313127 | 11.7456783153976 | 22.8036404096012 | 27.447191770805 | 37.5170677703264 | 16.4243729372865 | 7.33849339244316 | 15.616171358342 | 26.0526135520442 | Hs3st5 |
| **ENSMUSG00000044501** | -0.0756983968435575 | 0.723985608214872 | 0.951674101666722 | 173.474633581257 | 182.42912327681 | 160.897331070236 | 110.675349922463 | 139.059690869025 | 143.100621152642 | 198.845915296222 | 180.197243734972 | Zfp758 |
| **ENSMUSG00000044502** | 0.0853666696539937 | 0.455603572945225 | 0.892805623137304 | 3267.10559911367 | 2623.23306283305 | 2851.66857949777 | 3360.59134552699 | 2651.98875027385 | 2937.84352144141 | 2862.96474902938 | 2953.71506146301 | Bod1 |
| **ENSMUSG00000044505** | -0.629439797263642 | 0.192153556516168 | 0.808824684419004 | 5.42108229941427 | 5.7009101024003 | 9.46454888648447 | 14.0689004138724 | 13.1394983498292 | 13.4539045528125 | 10.4107809055614 | 16.2828834700276 | Lingo4 |
| **ENSMUSG00000044522** | -0.220239703891953 | 0.518938847877785 | 0.905762829482588 | 20.780815481088 | 13.0306516626293 | 17.036187995672 | 16.8826804966469 | 19.7092475247437 | 20.7923979452556 | 19.7804837205666 | 18.4539345993646 | A730020M07 |
| **ENSMUSG00000044533** | -0.0550246435331212 | 0.730820841031124 | 0.952921973914044 | 24243.0800429806 | 18705.5004617043 | 20853.2405615912 | 24042.8128806137 | 22847.3977139613 | 28429.3234023248 | 19454.6262782225 | 20529.4594790108 | Rps2 |
| **ENSMUSG00000044534** | -1.28111606354648 | 0.0461295940519097 | 0.69805115405055 | 12.6491920319666 | 16.2883145782866 | 13.2503684410783 | 15.0068271081306 | 48.1781606160403 | 72.1618516923577 | 7.28754663389296 | 11.9407812113536 | Ackr2 |
| **ENSMUSG00000044548** | -0.268705124540939 | 0.437413885576777 | 0.888722970093854 | 40.658117245607 | 34.2054606144018 | 79.5022106464695 | 71.2824287636203 | 63.5075753575076 | 91.7311674055395 | 71.8343882483734 | 44.5065481514088 | Dact1 |
| **ENSMUSG00000044556** | 0.521088598338349 | 0.181175626798914 | 0.803056849297055 | 24.3948703473642 | 26.0613033252585 | 18.9290977729689 | 28.1378008277448 | 25.1840385038392 | 7.33849339244316 | 18.7394056300105 | 16.2828834700276 | Tex38 |
| **ENSMUSG00000044562** | -0.624987978212889 | 0.0761117653565274 | 0.735694099375202 | 591.801484352725 | 487.020605890769 | 543.265106084208 | 606.83857118503 | 1213.21368096756 | 1311.14415278318 | 464.320828388037 | 449.407583772762 | Rasip1 |
| **ENSMUSG00000044566** | 0.983241835118467 | 0.0150254985506038 | 0.635862253526407 | 34.3335212296237 | 31.7622134276588 | 21.7684624389143 | 18.7585338851632 | 7.66470737073368 | 13.4539045528125 | 19.7804837205666 | 13.0263067760221 | Cage1 |
| **ENSMUSG00000044573** | -0.085004659453801 | 0.498761197185208 | 0.900927059989498 | 795.995584297329 | 719.943504360267 | 757.163910918757 | 870.395972271574 | 821.218646864323 | 973.573456730792 | 757.904849924867 | 782.663932125994 | Acp1 |
| **ENSMUSG00000044583** | -0.0382202561424751 | 0.897925753639419 | 0.984360759361306 | 21.6843291976571 | 28.5045505120015 | 27.447191770805 | 26.2619474392285 | 29.5638712871156 | 33.0232202659942 | 22.903717992235 | 21.7105112933701 | Tlr7 |
| **ENSMUSG00000044595** | -0.817424029846742 | 0.0240817366852632 | 0.654124840968651 | 19.877301764519 | 17.9171460361152 | 17.036187995672 | 15.0068271081306 | 44.893286028583 | 20.7923979452556 | 35.3966550789086 | 21.7105112933701 | Dnd1 |
| **ENSMUSG00000044600** | 0.0819991866925266 | 0.445228579366366 | 0.89108031640712 | 2693.37438909232 | 2269.77663648423 | 2533.65973691189 | 2574.60877573865 | 2281.892880087 | 2158.74013961036 | 2425.7119509958 | 2646.51132666182 | Smim7 |
| **ENSMUSG00000044617** | -0.0408969694040162 | 0.851473835007946 | 0.972692784625411 | 113.8427282877 | 179.985876090067 | 164.68315062483 | 120.054616865045 | 173.003394939417 | 114.969729814943 | 156.16171358342 | 150.888053488923 | Zbtb39 |
| **ENSMUSG00000044627** | -0.0526347090256815 | 0.783545748252172 | 0.961051227381536 | 3572.493235314 | 2660.69618636311 | 2924.5456059237 | 3736.69994992451 | 3258.59559075763 | 4416.54994001871 | 2968.11363617555 | 2731.18232070597 | Swi5 |
| **ENSMUSG00000044628** | 0.574610169380624 | 0.3443101748101 | 0.861266047293794 | 3.61405486627618 | 17.1027303072009 | 21.7684624389143 | 10.3171936368398 | 6.56974917491458 | 11.0077400886647 | 12.4929370866736 | 5.42762782334254 | Rnf208 |
| **ENSMUSG00000044629** | -0.568777210666052 | 0.0440172210693077 | 0.691603097094229 | 28.0089252136404 | 28.5045505120015 | 28.3936466594534 | 30.0136542162612 | 53.6529515951357 | 51.3694537471021 | 31.2323427166841 | 34.7368180693922 | Cnrip1 |
| **ENSMUSG00000044636** | -0.33472055479919 | 0.131802697897749 | 0.77303252008843 | 38.8510898124689 | 60.2667639396603 | 59.6266579848522 | 60.9652351267805 | 68.9823663366031 | 70.9387694602839 | 62.4646854333682 | 74.901263962127 | Csrnp2 |
| **ENSMUSG00000044641** | 0.257858151254327 | 0.28088499416555 | 0.849628172682816 | 46.0791995450213 | 61.0811796685746 | 61.519567762149 | 43.1446279358754 | 38.3235368536684 | 48.9232892829544 | 45.80743598447 | 44.5065481514088 | Pard6b |
| **ENSMUSG00000044646** | -0.72140426053986 | 0.0288856183815659 | 0.665956123973759 | 23.4913566307952 | 17.9171460361152 | 18.9290977729689 | 22.5102406621959 | 28.4689130912965 | 51.3694537471021 | 31.2323427166841 | 26.0526135520442 | Zbtb7c |
| **ENSMUSG00000044674** | -0.576226359435624 | 0.0227694880214035 | 0.650119841310877 | 111.132187137993 | 79.8127414336042 | 123.985590412947 | 84.4134024832345 | 166.433645764503 | 196.916239363891 | 125.970448957293 | 106.381505337514 | Fzd1 |
| **ENSMUSG00000044676** | -0.156977530259284 | 0.51528699910966 | 0.90419750457791 | 179.79922959724 | 170.212887343095 | 188.344522841041 | 181.957778686083 | 153.294147414674 | 147.992950080937 | 300.871568170723 | 199.736703899005 | Zfp612 |
| **ENSMUSG00000044700** | 0.07696584650266 | 0.57037384088053 | 0.920872458148558 | 2823.48036427827 | 3272.32239877777 | 3222.67889584796 | 2684.34619896686 | 2789.95348294706 | 2376.44877691951 | 3273.14951670849 | 2938.51770355765 | Tmem201 |
| **ENSMUSG00000044707** | -0.647598534322772 | 0.275178278149301 | 0.846286635233373 | 9.93865088225949 | 8.14415728914329 | 17.9826428843205 | 2.81378008277448 | 18.6142893289246 | 23.2385624094033 | 7.28754663389296 | 11.9407812113536 | Ccnjl |
| **ENSMUSG00000044709** | -0.0531144646534941 | 0.687701848441861 | 0.944452327903597 | 668.600150261093 | 557.060358577401 | 532.854102309076 | 603.086864407998 | 614.271547854513 | 705.718447906617 | 548.648153723084 | 582.927228226989 | Gemin7 |
| **ENSMUSG00000044712** | 0.0691463035666386 | 0.847744802975928 | 0.972455319662159 | 22.5878429142261 | 17.9171460361152 | 29.3401015481019 | 20.6343872736795 | 14.2344565456483 | 20.7923979452556 | 31.2323427166841 | 19.5394601640331 | Slc38a6 |
| **ENSMUSG00000044715** | -0.0678270448718204 | 0.649197139706943 | 0.936461253686283 | 250.273299489625 | 236.99497711407 | 237.56017705076 | 236.357526953057 | 277.024423542232 | 292.316653465652 | 206.133461930115 | 233.387996403729 | Gskip |
| **ENSMUSG00000044716** | 0.369345823498372 | 0.00073724931475952 | 0.556246415579446 | 994.768601942519 | 1071.77109925126 | 1193.47961458569 | 1095.49837889353 | 793.844691968845 | 806.011190936674 | 826.616003901573 | 944.407241261602 | Dok7 |
| **ENSMUSG00000044730** | 0.141063045636143 | 0.479666931789212 | 0.897732059433541 | 126.491920319666 | 117.275864963663 | 133.450139299431 | 114.427056699496 | 90.881530252985 | 88.0619207093179 | 130.134761319517 | 135.690695583563 | 9930104L06 |
| **ENSMUSG00000044734** | -0.404314447739873 | 0.126657244708888 | 0.77215780108715 | 43.3686583953142 | 49.679359463774 | 36.9117406572894 | 46.8963347129081 | 60.2227007700503 | 73.3849339244316 | 37.4788112600209 | 64.046008315442 | Serpinb1a |
| **ENSMUSG00000044737** | -1.72493071001017 | 0.0180646676861759 | 0.642273034804458 | 5.42108229941427 | 4.88649437348597 | 9.46454888648447 | 4.68963347129081 | 24.0890803080201 | 45.2540425867328 | 4.16431236222455 | 7.59867895267955 | Klk14 |
| **ENSMUSG00000044749** | -0.190013681123896 | 0.432064603810805 | 0.8866807012241 | 50.5967681278665 | 73.2974156022896 | 82.3415753124149 | 81.59962240046 | 98.5462376237187 | 59.9310293716191 | 82.2451691539348 | 86.8420451734806 | Abca6 |
| **ENSMUSG00000044751** | 0.0915629339798041 | 0.741685875324203 | 0.954383574136061 | 55.1143367107117 | 36.6487078011448 | 41.6440151005317 | 45.9584080186499 | 59.1277425742312 | 35.4693847301419 | 34.3555769883525 | 39.0789203280663 | Gm12231 |
| **ENSMUSG00000044763** | 0.190717142095733 | 0.305927782285981 | 0.851785577030345 | 414.712795905192 | 366.487078011448 | 334.098575692902 | 302.950322245386 | 272.644590758955 | 242.170281950624 | 364.377331694648 | 362.565538599282 | Trmt10c |
| **ENSMUSG00000044768** | -0.336360973508588 | 0.163977097297864 | 0.793968226938511 | 52.4037955610046 | 63.5244268553176 | 73.8234813145789 | 53.4618215727152 | 63.5075753575076 | 105.185071958352 | 67.6700758861489 | 71.6446872681215 | D1Ertd622e |
| **ENSMUSG00000044770** | -0.386654149368934 | 0.539981976412808 | 0.913837378403038 | 11.7456783153976 | 16.2883145782866 | 4.73227444324223 | 15.9447538023887 | 19.7092475247437 | 31.8001380339204 | 6.24646854333682 | 6.51315338801105 | Scml4 |
| **ENSMUSG00000044783** | -0.123959973977788 | 0.328480887246225 | 0.855977678491063 | 189.737880479499 | 197.088606397268 | 194.023252172932 | 179.143998603309 | 227.751304730372 | 194.470074899744 | 207.174540020671 | 198.651178334337 | Hjurp |
| **ENSMUSG00000044786** | -0.103201023875537 | 0.707238810483873 | 0.948710970699398 | 408.388199889208 | 504.937751926884 | 368.170951684246 | 286.067641748739 | 538.719432342996 | 532.040770952129 | 360.213019332423 | 254.012982132431 | Zfp36 |
| **ENSMUSG00000044788** | 0.135081818311575 | 0.602635099452721 | 0.926448151609826 | 46.9827132615903 | 65.1532583131463 | 50.1621090983677 | 64.7169419038131 | 39.4184950494875 | 42.8078781225851 | 67.6700758861489 | 56.4473293627624 | Fads6 |
| **ENSMUSG00000044791** | 0.248489356298856 | 0.338472455111399 | 0.858570960161243 | 773.407741383103 | 963.453807305651 | 898.185689327376 | 677.183073254392 | 596.752216721408 | 410.955629976817 | 957.791843311646 | 821.74285245406 | Setd2 |
| **ENSMUSG00000044792** | 0.165761297499535 | 0.358763641638014 | 0.866648170843829 | 7408.81247586617 | 7240.9702457773 | 7944.54233531506 | 6710.86549741714 | 5571.14730032757 | 4905.78283284825 | 7543.65184416977 | 8102.36281468574 | Isca1 |
| **ENSMUSG00000044795** | 0.258941768716921 | 0.423293087456668 | 0.884281490205862 | 260.211950371885 | 395.806044252364 | 432.52988411234 | 273.874594723383 | 188.332809680885 | 168.785348026193 | 373.747034509653 | 407.07208675069 | Cyb5d1 |
| **ENSMUSG00000044807** | -0.39428097842644 | 0.211779674801817 | 0.823370267313127 | 28.9124389302094 | 27.6901347830872 | 37.8581955459379 | 24.3860940507122 | 51.4630352034976 | 47.7002070508805 | 30.191264626128 | 27.1381391167127 | Zfp354c |
| **ENSMUSG00000044811** | -0.470348204335146 | 0.172109987913405 | 0.799889021902745 | 14.4562194651047 | 20.3603932228582 | 15.1432782183751 | 13.1309737196143 | 20.8042057205628 | 26.9078091056249 | 19.7804837205666 | 20.6249857287016 | Cd300c2 |
| **ENSMUSG00000044813** | 0.218149026639457 | 0.220652530006972 | 0.829144114546069 | 589.994456919586 | 684.109212288036 | 573.551662520959 | 619.969544904645 | 544.194223322091 | 662.910569784032 | 383.116737324658 | 532.993052252237 | Shb |
| **ENSMUSG00000044835** | 0.193890289370429 | 0.552136460479948 | 0.914957699363693 | 110.228673421423 | 141.708336831093 | 79.5022106464695 | 88.1651092602672 | 52.5579933993167 | 78.277262852727 | 143.668776496747 | 92.2696729968231 | Ankrd45 |
| **ENSMUSG00000044847** | 0.156710555751577 | 0.340180743909984 | 0.858570960161243 | 166.246523848704 | 168.584055885266 | 177.933519065908 | 165.075098189436 | 152.199189218854 | 116.192812047017 | 160.326025945645 | 178.026192605635 | Lsm11 |
| **ENSMUSG00000044854** | 0.38289034186079 | 0.209307385712203 | 0.821566820583092 | 28.0089252136404 | 43.1640336324594 | 35.0188308799925 | 34.703287687552 | 27.3739548954774 | 37.9155491942896 | 20.8215618111227 | 22.7960368580387 | 1700056E22 |
| **ENSMUSG00000044857** | 0.0883045181380237 | 0.604451598476758 | 0.926448151609826 | 814.06585862871 | 908.073537739476 | 1127.2277723803 | 1005.45741624475 | 1056.63465896543 | 968.681127802497 | 903.655782602727 | 696.907412517182 | Lemd2 |
| **ENSMUSG00000044864** | 0.0343304312684632 | 0.874300114409945 | 0.97671366425653 | 803.223694029881 | 1129.59461600417 | 941.722614205205 | 720.327701190268 | 813.553939493589 | 618.879609429373 | 1075.43366754449 | 1001.94009618903 | Ankrd50 |
| **ENSMUSG00000044876** | -0.302441159601259 | 0.121764298358062 | 0.769868040577702 | 183.413284463516 | 149.852494120236 | 159.950876181588 | 199.778385876988 | 227.751304730372 | 267.855008824175 | 196.76375911511 | 162.828834700276 | Zfp444 |
| **ENSMUSG00000044881** | -0.0665110789876496 | 0.757986586588021 | 0.95677573228595 | 207.80815481088 | 131.935348084121 | 144.807597963212 | 232.605820176024 | 176.288269526875 | 192.023910435596 | 175.942197303987 | 206.249857287016 | Coa4 |
| **ENSMUSG00000044894** | -0.0114515120098476 | 0.958168120684616 | 0.992835150787715 | 24669.5385172012 | 16386.0444657563 | 18572.2842799485 | 23583.2288004272 | 21618.8546182522 | 28554.0777899963 | 16784.260975946 | 16917.9159253587 | Uqcrq |
| **ENSMUSG00000044906** | 0.173232300139819 | 0.636177222264706 | 0.932595977875587 | 37.0440623793308 | 44.7928650902881 | 51.1085639870161 | 55.3376749612315 | 63.5075753575076 | 52.592535979176 | 19.7804837205666 | 31.4802413753867 | 4930503L19 |
| **ENSMUSG00000044921** | -0.0254250136404034 | 0.939011387980703 | 0.989269716906553 | 137.334084918495 | 95.2866402829764 | 251.757000380487 | 292.633128608546 | 218.991639163819 | 215.262472844999 | 179.065431575656 | 176.940667040967 | Rassf9 |
| **ENSMUSG00000044927** | -0.646928755897489 | 0.197787030576739 | 0.812921911423739 | 16.2632468982428 | 15.4738988493722 | 21.7684624389143 | 24.3860940507122 | 29.5638712871156 | 61.154111603693 | 20.8215618111227 | 10.8552556466851 | H1fx |
| **ENSMUSG00000044933** | -0.0735231950852937 | 0.849198309707923 | 0.972640876584422 | 34.3335212296237 | 37.4631235300591 | 14.1968233297267 | 26.2619474392285 | 30.6588294829347 | 22.0154801773295 | 42.6842017128016 | 22.7960368580387 | Sstr3 |
| **ENSMUSG00000044934** | 0.104334693375994 | 0.642413758813767 | 0.934616627390525 | 577.34526488762 | 535.071133896714 | 467.548714992333 | 519.611388619021 | 347.101748074654 | 373.040080782527 | 581.96265262088 | 650.229813236436 | Zfp367 |
| **ENSMUSG00000044937** | 0.347013419416192 | 0.341083186637126 | 0.859283395807012 | 26.2018977805023 | 30.1333819698302 | 53.001473764313 | 31.8895076047775 | 32.8487458745729 | 14.6769867848863 | 28.1091084450157 | 34.7368180693922 | Ttc41 |

| | | | | | | | | | | | | |
|---|---|---|---|---|---|---|---|---|---|---|---|---|
| **ENSMUSG00000044938** | 0.127907219332628 | 0.406089720176247 | 0.881158207349486 | 2627.41788778278 | 2344.70288354435 | 1984.71590149579 | 3082.96504402658 | 2503.07443564246 | 2241.90973139138 | 2140.45655418342 | 2302.3997226619 | Klhl31 |
| **ENSMUSG00000044948** | -0.0988665803267993 | 0.568354721983395 | 0.92059532906293 | 225.878429142261 | 290.746415222415 | 362.492222352355 | 345.157023487003 | 313.158044004262 | 332.67836712409 | 349.802238426862 | 314.802413753867 | Cfap43 |
| **ENSMUSG00000044949** | 0.00064886125594281 | 0.998034532025492 | 0.999262355603273 | 28.9124389302094 | 37.4631235300591 | 35.965285768641 | 34.703287687552 | 33.943704070392 | 29.3539735697726 | 31.2323427166841 | 42.3354970220718 | Ubtd2 |
| **ENSMUSG00000044950** | -0.105819376688349 | 0.602301543475352 | 0.926448151609826 | 108.421645988285 | 89.5857301805761 | 127.77140996754 | 131.309737196143 | 113.875652365186 | 102.738907494204 | 150.95632313064 | 122.664388807541 | Pwwp2a |
| **ENSMUSG00000044951** | 0.34020219657015 | 0.295327856897003 | 0.851785577030345 | 1765.46580217591 | 1917.94904159324 | 1640.20632202776 | 1263.38725716574 | 1011.74137293685 | 613.987280501078 | 1574.11007292088 | 2002.7946668134 | Mylk4 |
| **ENSMUSG00000044952** | 0.104064736168164 | 0.563641609491398 | 0.918571155207735 | 106.614618555147 | 92.0289773673191 | 86.1273948670087 | 106.92364314543 | 84.3117810780705 | 79.5003450848009 | 98.902418602833 | 100.953877514171 | Kctd21 |
| **ENSMUSG00000044968** | 0.0641940256997747 | 0.781563957298279 | 0.960742105865155 | 165.343010132135 | 193.016527752696 | 217.684624389143 | 165.075098189436 | 123.730276127558 | 144.323703384715 | 218.626399016789 | 221.447215192376 | Napepld |
| **ENSMUSG00000044982** | -0.18183641009126 | 0.411325534536632 | 0.882192126163777 | 95.7724539563188 | 100.173134656462 | 116.413951303759 | 103.171936368398 | 153.294147414674 | 134.539045528125 | 88.4916376972716 | 95.5262496908287 | Sft2d3 |
| **ENSMUSG00000044986** | -0.0513888826322914 | 0.721061123429576 | 0.951674101666722 | 345.142239729375 | 327.39512302356 | 294.347470369667 | 321.708856130549 | 375.57066116595 | 369.370834086306 | 274.84461590682 | 316.973464883204 | Tst |
| **ENSMUSG00000044991** | 0.145122183537742 | 0.254833527695915 | 0.837406774112203 | 658.661499378834 | 539.143212541285 | 542.31865119556 | 618.093691516128 | 583.612718371579 | 523.479195327612 | 555.935700356977 | 468.947043936795 | 1110034G24 |
| **ENSMUSG00000045005** | -0.13238897857857 | 0.612982625670917 | 0.928075610917191 | 65.9565013095403 | 57.8235167529173 | 88.966759532954 | 77.8479156234274 | 105.115986798633 | 52.592535979176 | 72.8754663389296 | 86.8420451734806 | Fzd5 |
| **ENSMUSG00000045007** | -0.278623984865195 | 0.398691994745167 | 0.880459952732461 | 24.3948703473642 | 26.8757190541728 | 25.5542819935081 | 52.523894878457 | 40.5134532453066 | 41.5847958905112 | 31.2323427166841 | 43.4210225867403 | Tubg2 |
| **ENSMUSG00000045008** | -0.551597508424776 | 0.0213957312870449 | 0.648762042745232 | 64.1494738764022 | 55.3802695661743 | 39.7511053232348 | 47.8342614071662 | 58.0327843784121 | 74.6080161565054 | 82.2451691539348 | 89.0130963028176 | 9030612E09 |
| **ENSMUSG00000045031** | 0.19355524007739 | 0.711905783964641 | 0.950074110229522 | 8.13162344912141 | 17.9171460361152 | 6.62518422053913 | 14.0689004138724 | 8.75966556655277 | 13.4539045528125 | 13.5340151772298 | 5.42762782334254 | Cetn4 |
| **ENSMUSG00000045038** | -0.00684364447731261 | 0.968518256349498 | 0.994790165280964 | 1176.3748589729 | 1190.67579567275 | 1003.24218196735 | 908.850966736158 | 913.195135313127 | 897.742358342213 | 1164.96638333232 | 1323.25566333091 | Prkce |
| **ENSMUSG00000045039** | -0.00737427049242111 | 0.956195444719459 | 0.992371505557225 | 1356.17408857014 | 1781.12719913564 | 1603.29458137047 | 1358.11785328582 | 1443.15490208957 | 1400.42915572457 | 1665.72494488982 | 1620.68966805008 | Megf8 |
| **ENSMUSG00000045045** | -0.110410595212779 | 0.527564253276453 | 0.908217276872524 | 140.948139784771 | 159.625482867208 | 149.539872406455 | 183.8336320746 | 210.231973597267 | 172.454594722414 | 149.915245040084 | 151.973579053591 | Lrfn4 |
| **ENSMUSG00000045064** | -0.124190807661785 | 0.501534977881411 | 0.900927059989498 | 93.0619128066116 | 96.9154717408051 | 92.7525790875478 | 96.6064495085906 | 119.350443344282 | 78.277262852727 | 99.9434966933891 | 115.065709854862 | Zc2hc1c |
| **ENSMUSG00000045071** | -0.188651200351937 | 0.377181342534432 | 0.87422797717331 | 70.4740698923855 | 72.4829998733752 | 64.3589324280944 | 49.7101147956825 | 70.0773245324222 | 61.154111603693 | 79.1219348822664 | 82.4999429148066 | E130308A19 |
| **ENSMUSG00000045078** | 0.12472217885629 | 0.342261782002371 | 0.859988450074698 | 618.906895849796 | 778.581436842098 | 808.272474905774 | 698.75538722233 | 697.488370736765 | 583.410224699231 | 706.892023487617 | 675.196901223812 | Rnf216 |
| **ENSMUSG00000045087** | -0.290920836336915 | 0.60527820129699 | 0.926448151609826 | 20.780815481088 | 28.5045505120015 | 19.8755526616174 | 10.3171936368398 | 4.37983278327639 | 39.1386314263635 | 28.1091084450157 | 26.0526135520442 | S1pr5 |
| **ENSMUSG00000045092** | -0.723719185616064 | 0.103646201168581 | 0.759800953500741 | 1749.20255527767 | 1740.40641268992 | 1433.8791563024 | 1146.14642038347 | 3517.00572497094 | 4452.01932474885 | 1104.58385408006 | 950.920394649613 | S1pr1 |
| **ENSMUSG00000045094** | 0.121108874350332 | 0.730661986975895 | 0.952921973914044 | 50.5967681278665 | 77.3694942468612 | 76.6628459805242 | 53.4618215727152 | 50.3680770076785 | 26.9078091056249 | 80.1630129728225 | 79.2433662208011 | Arhgef37 |
| **ENSMUSG00000045095** | 0.0805423758440183 | 0.480231839303714 | 0.897732059433541 | 312.61574593289 | 321.69421292116 | 345.456034356683 | 300.136542162612 | 308.778211220985 | 319.224462571277 | 306.076958623504 | 276.809018990469 | Magi1 |
| **ENSMUSG00000045098** | 0.164596471876318 | 0.361193901812398 | 0.868539734006824 | 586.38040205331 | 584.750493360488 | 570.712297855013 | 473.652980600371 | 463.167316831478 | 362.032340693862 | 595.49666779811 | 554.703563545607 | Kmt5b |
| **ENSMUSG00000045100** | -0.0544905825983376 | 0.739989054706504 | 0.954383574136061 | 908.03128515189 | 843.734695155244 | 636.017685171756 | 839.444391361054 | 677.779123212021 | 824.357424417781 | 948.422140496641 | 900.986218674861 | Slc25a26 |
| **ENSMUSG00000045102** | 0.221010841587312 | 0.326585699840924 | 0.855977678491063 | 121.974351736821 | 78.1839099757755 | 137.235958854025 | 106.92364314543 | 100.736154015357 | 96.6234963338349 | 77.0397787011541 | 106.381505337514 | Poln |
| **ENSMUSG00000045103** | 0.430744170078467 | 0.19851121476725 | 0.812921911423739 | 2116.0291242047 | 3420.54606144018 | 2704.96807175726 | 2065.31458075647 | 1325.99437513693 | 974.796538962866 | 2875.45768611605 | 2469.57065962085 | Dmd |
| **ENSMUSG00000045104** | 0.0891551392199248 | 0.711024822463109 | 0.94990235491128 | 75.8951521917998 | 48.0505280059454 | 72.8770264259304 | 83.4754757889764 | 60.2227007700503 | 74.6080161565054 | 72.8754663389296 | 55.3618037980939 | Gm5514 |
| **ENSMUSG00000045106** | 0.218764722032572 | 0.371001569377249 | 0.873374202289323 | 53.3073092775737 | 62.7100111264033 | 40.6975602118832 | 67.5307219865876 | 53.6529515951357 | 51.3694537471021 | 46.8485140750261 | 41.2499714574033 | Ccdc73 |
| **ENSMUSG00000045107** | -0.088937147368555 | 0.645783074661913 | 0.935063240511793 | 177.088688447533 | 132.749763813036 | 133.450139299431 | 176.330218520534 | 192.712642464161 | 181.016170346931 | 154.079557402308 | 131.348593324889 | Saysd1 |
| **ENSMUSG00000045128** | -0.0764205664548834 | 0.728365561773332 | 0.952401994188708 | 9826.61518140493 | 7382.67858260839 | 9176.82660033534 | 11014.0731706736 | 10212.6750924047 | 13725.4288083329 | 7743.53883755654 | 7753.90910842715 | Rpl18a |
| **ENSMUSG00000045136** | -0.239866369177397 | 0.552050619219857 | 0.914957699363693 | 51.5002818444356 | 51.3081909216027 | 44.483379766477 | 31.8895076047775 | 83.2168228822514 | 69.71568722821 | 36.4377331694648 | 22.7960368580387 | Tubb2b |
| **ENSMUSG00000045140** | 0.180697347117487 | 0.52707418993884 | 0.908217276872524 | 74.9916384752307 | 80.6271571625185 | 106.949402417274 | 60.0273084325223 | 73.3621991198795 | 41.5847958905112 | 89.5327157878278 | 79.2433662208011 | Pigw |
| **ENSMUSG00000045160** | 0.0345196420042892 | 0.82026763038184 | 0.966398681144067 | 4803.98243099761 | 3510.13179162076 | 3511.34763688574 | 4196.28403011101 | 3667.01499779816 | 4634.25857732785 | 3633.36253604092 | 3709.24085447229 | Bola3 |
| **ENSMUSG00000045165** | -0.0586570679714921 | 0.877202629034497 | 0.977569405470617 | 21.6843291976571 | 15.4738988493722 | 23.6613722162112 | 15.0068271081306 | 28.4689130912965 | 20.7923979452556 | 16.6572494488982 | 13.0263067760221 | AI467606 |
| **ENSMUSG00000045176** | -0.195954540159231 | 0.230523624793231 | 0.832048466795696 | 408.388199889208 | 350.198763433161 | 336.937940358847 | 395.805064976944 | 431.413529152724 | 532.040770952129 | 398.732908683 | 347.368180693922 | Borcs6 |
| **ENSMUSG00000045180** | -0.350841106430142 | 0.278509212378995 | 0.848075053184273 | 73.1846110420926 | 39.9063707168021 | 53.001473764313 | 38.4549944645846 | 93.0714466446232 | 41.5847958905112 | 57.2592949805875 | 68.388110574116 | Shroom2 |
| **ENSMUSG00000045193** | 0.254654822090192 | 0.013975671157282 | 0.635862253526407 | 702.030157774148 | 685.738043745865 | 598.159489625818 | 714.700141024719 | 597.847174917227 | 554.056251129458 | 549.68923181364 | 562.302242498287 | Cirbp |
| **ENSMUSG00000045201** | -0.380679359861312 | 0.100099020331463 | 0.759800953500741 | 163.535982698997 | 196.274190668353 | 113.574586637814 | 105.985716451172 | 189.427767876704 | 215.262472844999 | 188.435134390661 | 162.828834700276 | Lrrc3b |
| **ENSMUSG00000045205** | 0.299909118160895 | 0.314849029885734 | 0.853277176422981 | 471.634160049041 | 647.460504486891 | 622.767316730678 | 434.260059441529 | 297.828629262794 | 270.301173288323 | 599.660980160335 | 599.210111697016 | Dpy19l4 |
| **ENSMUSG00000045210** | -0.107185092480287 | 0.499562784961957 | 0.900927059989498 | 318.940341948873 | 406.39344872825 | 395.618143455051 | 311.39166249371 | 360.241246424483 | 333.901449356164 | 449.745735120251 | 398.387882233342 | Vcpip1 |
| **ENSMUSG00000045211** | 0.237927428849527 | 0.23502527037788 | 0.832048466795696 | 684.863397159336 | 561.132437221972 | 500.674636095028 | 770.975742680209 | 580.327843784121 | 659.24132308781 | 470.567296931374 | 425.526021350055 | Nudt18 |
| **ENSMUSG00000045216** | 0.241736442858724 | 0.0966055771767365 | 0.757809915356977 | 811.355317479002 | 880.383402956389 | 1056.24365573167 | 942.616327729452 | 790.559817381388 | 631.110431750112 | 841.191097169358 | 856.479670523452 | Hs6st1 |
| **ENSMUSG00000045237** | -0.0914543672472501 | 0.641678260920156 | 0.934561292249236 | 278.282224703266 | 200.346269312925 | 197.809071727525 | 262.619474392285 | 295.638712871156 | 282.531995609062 | 218.626399016789 | 204.078806157679 | 1110012L19 |
| **ENSMUSG00000045248** | -0.0169937504741653 | 0.934907031648367 | 0.98818087308472 | 187.027339329792 | 168.584055885266 | 152.3792370724 | 136.937297361692 | 187.237851485066 | 204.254732756335 | 134.299073681742 | 128.092016630884 | Med26 |
| **ENSMUSG00000045251** | -0.371649686529342 | 0.380365570493684 | 0.87432013076003 | 37.0440623793308 | 26.0613033252585 | 33.1259211026956 | 48.7721881014244 | 49.2731188118594 | 85.6157562451702 | 23.9447960827911 | 29.3091902460497 | Zfp688 |
| **ENSMUSG00000045252** | -0.162921032402149 | 0.348630055444383 | 0.861266047293794 | 227.685456575399 | 249.211213047785 | 288.668741037776 | 294.508981997063 | 342.721915291377 | 327.786038195794 | 287.337552993494 | 229.045894145055 | Zfp574 |
| **ENSMUSG00000045257** | 0.287822095232153 | 0.057918957272632 | 0.710862211775517 | 138.237598635064 | 132.749763813036 | 134.396594188079 | 130.371810501884 | 102.926070406995 | 112.523565350795 | 99.9434966933891 | 123.74991437221 | Morn2 |
| **ENSMUSG00000045268** | -0.0539573881735492 | 0.613990958474396 | 0.928075610917191 | 489.704434380422 | 430.011504866766 | 424.011790114504 | 405.184331919526 | 446.742943894192 | 443.978850242811 | 461.197594116369 | 463.519416113453 | Zfp691 |
| **ENSMUSG00000045281** | -0.965806264165453 | 0.0415342400601734 | 0.683543358336265 | 5.42108229941427 | 13.8450673915436 | 9.46454888648447 | 7.50341355406529 | 26.2789966996583 | 19.5693157131818 | 12.4929370866736 | 13.0263067760221 | Gpr20 |
| **ENSMUSG00000045282** | 0.233515423705688 | 0.29741421806681 | 0.851785577030345 | 59.631905293557 | 58.6379324818317 | 62.4660226507975 | 66.5927952923295 | 53.6529515951357 | 36.6924669622158 | 61.4236073428121 | 57.5328549274309 | Tmem86b |
| **ENSMUSG00000045284** | 0.188457666152995 | 0.432891697964628 | 0.887769922446285 | 105.711104838578 | 156.367819951551 | 113.574586637814 | 90.0409626487835 | 114.970610561005 | 73.3849339244316 | 103.066730965058 | 117.236760984199 | Dcaf12l1 |
| **ENSMUSG00000045287** | -0.519090880481253 | 0.104868839322878 | 0.762094734041179 | 56.9213641438498 | 57.009101024003 | 44.483379766477 | 50.6480414899407 | 73.3621991198795 | 123.53130543946 | 57.2592949805875 | 46.6775992807458 | Rtn4rl1 |
| **ENSMUSG00000045294** | 0.0472009802542 | 0.800795189504989 | 0.963461896828986 | 853.820462157748 | 666.192066251921 | 772.307189137133 | 946.368034506485 | 912.100177117308 | 563.840908986049 | 806.835520181006 | 849.966517135441 | Insig1 |
| **ENSMUSG00000045302** | 0.0830595234630733 | 0.445187996943306 | 0.89108031640712 | 1097.76916563139 | 1120.63604298612 | 1196.31897925164 | 1090.80874542224 | 1073.05903190271 | 911.196262895025 | 1159.76099287954 | 1107.23607596188 | Preb |
| **ENSMUSG00000045312** | -1.03784029646353 | 0.0199299488498929 | 0.642273034804458 | 26.2018977805023 | 21.9892246806869 | 17.9826428843205 | 30.9515809105193 | 64.6025335533267 | 81.9465095489486 | 19.7804837205666 | 33.6512925047237 | Lhfpl2 |
| **ENSMUSG00000045316** | -0.0813011778998718 | 0.410597291662136 | 0.882192126163777 | 1803.41337827181 | 1765.65330028626 | 1507.70263761698 | 1649.81305520011 | 1831.86506160535 | 1714.76128936755 | 1883.31026581605 | 1686.90672749486 | Fahd1 |
| **ENSMUSG00000045318** | -0.863447605989605 | 0.125454643991395 | 0.771949105062466 | 17.1667606148119 | 16.2883145782866 | 7.57163910918757 | 7.50341355406529 | 26.2789966996583 | 41.5847958905112 | 10.4107809055614 | 10.8552556466851 | Adra2c |
| **ENSMUSG00000045319** | -0.242465039993596 | 0.103393856822041 | 0.759800953500741 | 323.457910531718 | 309.477976987445 | 330.312756138308 | 247.612647284155 | 379.950493949227 | 402.3940543523 | 330.021754706295 | 321.315567141878 | Proser2 |
| **ENSMUSG00000045333** | -0.510094736831425 | 0.106652624306404 | 0.762287160109383 | 48.7897406947284 | 35.0198763433161 | 35.0188308799925 | 45.9584080186499 | 70.0773245324222 | 84.3926740130963 | 38.5198893505771 | 42.3354970220718 | Zfp423 |
| **ENSMUSG00000045348** | -0.237137088901466 | 0.617259062527898 | 0.929199997671237 | 7.22810973255236 | 18.7315617650296 | 8.51809399783602 | 11.2551203310979 | 10.949581958191 | 15.9000690169602 | 17.6983275394543 | 9.76973008201657 | Nyap1 |
| **ENSMUSG00000045374** | -0.175485620639808 | 0.19596223266376 | 0.811193545163524 | 217.74680569314 | 228.850819824926 | 265.007368821565 | 226.040333316217 | 297.828629262794 | 264.185762127954 | 237.365804646799 | 259.440609955773 | Wdr81 |
| **ENSMUSG00000045377** | -0.771283005652055 | 0.0449767662942997 | 0.693563614452513 | 259.308436655316 | 201.975100770753 | 269.739643264807 | 346.094950181261 | 650.405168316544 | 713.05694129906 | 207.174540020671 | 268.124814473121 | Tmem88 |
| **ENSMUSG00000045382** | -0.500064384717319 | 0.161599746669968 | 0.792786631432503 | 54.2108229941427 | 39.9063707168021 | 33.1259211026956 | 97.5443762028488 | 61.3176589658694 | 85.6157562451702 | 69.7522320672611 | 100.953877514171 | Cxcr4 |
| **ENSMUSG00000045392** | -0.357026460745916 | 0.0271031851437763 | 0.664353746352133 | 571.020668871636 | 516.339572131684 | 610.463403178248 | 527.114802173087 | 828.883354235056 | 815.795848793264 | 646.509494235361 | 559.045665804281 | Olfr1033 |
| **ENSMUSG00000045409** | 0.172411435240407 | 0.268231897693016 | 0.843793744782845 | 200.580045078328 | 193.016527752696 | 234.720812384815 | 219.47484645641 | 187.237851485066 | 212.816308380852 | 153.038479311752 | 199.736703899005 | Trim39 |
| **ENSMUSG00000045410** | 0.0552690875440441 | 0.64320157910929 | 0.934616627390525 | 510.48524986151 | 437.341246426994 | 441.994432998825 | 424.880792498947 | 407.324448844704 | 406.063301048521 | 464.320828388037 | 467.861518372127 | Akr1e1 |
| **ENSMUSG00000045411** | -0.0342708453398979 | 0.839463567358413 | 0.969820257058002 | 98.4829951060259 | 85.5136515360045 | 97.48485353079 | 110.675349922463 | 97.4512794278996 | 105.185071958352 | 106.189965236726 | 92.2696729968231 | 2410002F23 |
| **ENSMUSG00000045414** | 0.154127798312838 | 0.314486459031958 | 0.853228028313558 | 1205.28729790311 | 1318.5390651123 | 1015.54609551978 | 1263.38725716574 | 898.960678767479 | 1028.61215717412 | 1079.59797990671 | 1309.14383099022 | 1190002N15 |
| **ENSMUSG00000045427** | 0.0949677796110926 | 0.460547817919705 | 0.894774044236998 | 1128.48863199474 | 1347.85803135321 | 1385.60995698133 | 1238.06323642077 | 1206.64393179264 | 1010.26592369301 | 1340.9085806363 | 1215.78863242873 | Hnrnph2 |
| **ENSMUSG00000045435** | -0.0556370137187318 | 0.728608594050627 | 0.952401994188708 | 458.081454300506 | 322.508628650074 | 325.580481695066 | 439.887619607078 | 376.665619361769 | 413.401794440965 | 406.020455316893 | 410.328663444696 | Tmem60 |
| **ENSMUSG00000045438** | 0.0596656486607551 | 0.738442988870357 | 0.954383574136061 | 1542.29791418336 | 1238.72632367869 | 959.705257089525 | 1526.94465825229 | 1121.23719251876 | 1423.66771813397 | 1214.93813167901 | 1295.03199864953 | Cox19 |
| **ENSMUSG00000045464** | -0.163128453961639 | 0.524343958447233 | 0.907437243979788 | 79.509207058076 | 62.7100111264033 | 71.930571537282 | 48.7721881014244 | 64.6025335533267 | 57.4848649074714 | 99.9434966933891 | 71.6446872681215 | 2810002D19 |
| **ENSMUSG00000045466** | -0.8235679021746 | 0.0112644793871886 | 0.635862253526407 | 27.1054114970714 | 19.5459774939439 | 14.1968233297267 | 27.1998741334867 | 48.1781606160403 | 44.0309603546589 | 29.1501865355718 | 34.7368180693922 | Zfp956 |
| **ENSMUSG00000045482** | 0.235905734228188 | 0.231547183757301 | 0.832048466795696 | 535.783633925444 | 729.716493107238 | 662.518422053913 | 527.114802173087 | 519.010184818252 | 370.593916318379 | 576.7572621681 | 617.664046296381 | Trrap |
| **ENSMUSG00000045503** | -0.127397426922701 | 0.479295245546021 | 0.897732059433541 | 658.661499378834 | 443.856572258309 | 529.068282754482 | 626.535031764452 | 608.796756875418 | 761.980230582015 | 562.182168900314 | 534.078577816906 | Sys1 |
| **ENSMUSG00000045519** | -0.0474928436293101 | 0.828554656901203 | 0.967436464724085 | 129.202461469373 | 74.9262470601182 | 94.6454888648447 | 93.7926694258161 | 116.065568756824 | 108.854318654574 | 89.5327157878278 | 91.1841474321546 | Zfp560 |
| **ENSMUSG00000045534** | -0.768383469208078 | 0.0994598176629838 | 0.759800953500741 | 660.468526811972 | 566.018931595458 | 420.22597055991 | 329.212269684615 | 1155.18089658915 | 1527.62970786025 | 357.089785060755 | 326.743194965221 | Kcna5 |
| **ENSMUSG00000045538** | -0.0163605846190252 | 0.925485608167736 | 0.986713918026868 | 326.168451681425 | 324.951875836817 | 336.937940358847 | 348.908730264036 | 440.173194719277 | 348.57843614105 | 268.598147363483 | 295.262953589834 | Ddx28 |
| **ENSMUSG00000045555** | -0.654460012348727 | 0.138998881675441 | 0.779711473022872 | 12.6491920319666 | 13.0306516626293 | 4.73227444324223 | 13.1309737196143 | 14.2344565456483 | 18.3462334811079 | 13.5340151772298 | 22.7960368580387 | Mettl24 |
| **ENSMUSG00000045573** | -1.30717494848996 | 0.0228159318714895 | 0.650119841310877 | 69.5705561758165 | 60.2667639396603 | 62.4660226507975 | 93.7926694258161 | 266.074841584041 | 338.793778284459 | 68.711153976705 | 34.7368180693922 | Penk |
| **ENSMUSG00000045576** | 0.0472514372468788 | 0.65331395670406 | 0.936991241819063 | 348.756294595651 | 345.312269059675 | 342.616669690738 | 322.646782824807 | 332.867291529005 | 305.770558018465 | 330.021754706295 | 346.282655129254 | St7l |
| **ENSMUSG00000045594** | -0.373741489756456 | 0.018125711224706 | 0.642273034804458 | 234.010052591383 | 248.39679731887 | 210.112985279955 | 214.785212985119 | 300.018545654433 | 359.586176229715 | 271.721381635152 | 246.414303179751 | Glb1 |
| **ENSMUSG00000045598** | -0.147910130967463 | 0.271200261567206 | 0.844129257012535 | 187.930853046361 | 209.304842330982 | 219.57753416644 | 195.088752405698 | 256.220217821669 | 211.593226148778 | 215.50316474512 | 216.019587369033 | Zfp553 |
| **ENSMUSG00000045613** | -0.0957397249698203 | 0.721902876979727 | 0.951674101666722 | 1211.61189391909 | 1679.32523302135 | 1378.03831787214 | 993.264369219393 | 924.144717271318 | 1082.42777538537 | 1995.74669959611 | 1620.68966805008 | Chrm2 |
| **ENSMUSG00000045624** | -0.0935938003672603 | 0.490559582790621 | 0.899236955296196 | 685.766910875905 | 623.842448348376 | 536.639921863669 | 683.7485601142 | 755.521155115177 | 715.503105763208 | 621.523620062014 | 607.894316214364 | Esf1 |

| | | | | | | | | | | | | |
|---|---|---|---|---|---|---|---|---|---|---|---|---|
| **ENSMUSG00000045625** | -0.691281048581763 | 0.0654515128983005 | 0.718339708336358 | 15.3597331816738 | 10.5874044758863 | 10.4110037751329 | 12.1930470253561 | 22.994122112201 | 14.6769867848863 | 19.7804837205666 | 20.6249857287016 | Pigz |
| **ENSMUSG00000045629** | -0.604277711555477 | 0.0607163339917532 | 0.71282992338528 | 56.0178504272808 | 38.2775392589734 | 29.3401015481019 | 61.9031618210386 | 95.2613630362614 | 80.7234273168747 | 54.1360607089191 | 52.1052271040884 | Sh3tc2 |
| **ENSMUSG00000045636** | 0.370977136456212 | 0.0476108251472173 | 0.703396849069134 | 5295.49389281117 | 6014.46015803232 | 6153.8496859922 | 5840.46952514557 | 4272.52688008612 | 3080.94414259405 | 5417.77038325413 | 5247.43057960757 | Mtus1 |
| **ENSMUSG00000045639** | 0.0244952429631092 | 0.88712362665413 | 0.980672278091991 | 276.475197270128 | 349.384347704247 | 353.027673465871 | 304.826175633902 | 308.778211220985 | 239.724117486476 | 346.679004155194 | 365.822115293287 | Zfp629 |
| **ENSMUSG00000045658** | -0.388951333325419 | 0.127278147122357 | 0.772837268577874 | 106.614618555147 | 139.26508964435 | 144.807597963212 | 118.178763476528 | 227.751304730372 | 193.24699266767 | 109.313199508394 | 136.776221148232 | Pid1 |
| **ENSMUSG00000045659** | -0.764281853805669 | 0.282695949600842 | 0.850587941933635 | 6.32459601598331 | 9.77298874697194 | 19.8755526616174 | 5.62756016554897 | 12.0445401540101 | 44.0309603546589 | 5.20539045278068 | 9.76973008201657 | Plekha7 |
| **ENSMUSG00000045664** | -0.591174156654745 | 0.0385150373385513 | 0.679566075569698 | 276.475197270128 | 184.872370463553 | 176.040609288611 | 261.681547698027 | 400.754699669789 | 480.671317205027 | 218.626399016789 | 255.098507697099 | Cdc42ep2 |
| **ENSMUSG00000045665** | -0.364124519567237 | 0.0331328807268981 | 0.671606027118506 | 449.046317134815 | 414.537606017393 | 424.011790114504 | 439.887619607078 | 682.158955995297 | 633.556596214259 | 457.033281754144 | 451.578634902099 | Mfsd5 |
| **ENSMUSG00000045667** | -0.763942255520764 | 0.0229951609800811 | 0.650514839131011 | 172.571119864688 | 105.059629029948 | 124.932045301595 | 135.061443973175 | 305.493336633528 | 330.232202659942 | 157.202791673977 | 120.493337678204 | Smtnl2 |
| **ENSMUSG00000045671** | -0.406846511493399 | 0.0227201558840068 | 0.650119841310877 | 370.440623793308 | 274.458100644129 | 324.634026806417 | 446.453106466885 | 539.814390538815 | 425.632616761703 | 466.402984569149 | 443.97995594942 | Spred2 |
| **ENSMUSG00000045672** | 0.00570206431777542 | 0.987997998334801 | 0.998437672973644 | 14.4562194651047 | 17.9171460361152 | 14.1968233297267 | 27.1998741334867 | 14.2344565456483 | 23.2385624094033 | 16.6572494488982 | 19.5394601640331 | Col27a1 |
| **ENSMUSG00000045679** | -0.688459964795852 | 0.156368777481879 | 0.787275540343856 | 60.535419010126 | 33.3910448854875 | 25.5542819935081 | 52.523894878457 | 105.115986798633 | 108.854318654574 | 35.3966550789086 | 28.2236646813812 | Pqlc3 |
| **ENSMUSG00000045680** | -0.829701735026946 | 0.0132657800331026 | 0.635862253526407 | 85.8338030740593 | 74.1118313312039 | 70.9841166486335 | 89.1030359545253 | 156.579022002131 | 236.054870790255 | 98.902418602833 | 78.1578406561325 | Tcf21 |
| **ENSMUSG00000045690** | 0.147101860057882 | 0.612579714649044 | 0.928055903343287 | 48.7897406947284 | 43.9784493613737 | 49.2156542097192 | 63.779015209555 | 56.937826182593 | 62.3771938357668 | 32.2734208072402 | 34.7368180693922 | Wdr89 |
| **ENSMUSG00000045691** | 0.145190482805891 | 0.423201030861887 | 0.884281490205862 | 311.712232216321 | 395.806044252364 | 328.419846361011 | 285.129715054481 | 255.12525962585 | 250.731857575141 | 322.734208072402 | 365.822115293287 | Thtpa |
| **ENSMUSG00000045725** | -0.456335484963862 | 0.388164832061271 | 0.876114643700205 | 9.93865088225949 | 10.5874044758863 | 15.1432782183751 | 10.3171936368398 | 18.6142893289246 | 26.9078091056249 | 13.5340151772298 | 4.34210225867403 | Prr15 |
| **ENSMUSG00000045730** | -0.584481081892086 | 0.143165630040965 | 0.780607990324772 | 28.9124389302094 | 19.5459774939439 | 26.5007368821565 | 22.5102406621959 | 61.3176589658694 | 40.3617136584374 | 24.9858741733473 | 19.5394601640331 | Adrb2 |
| **ENSMUSG00000045752** | -0.300936762265469 | 0.181482753775627 | 0.803056849297055 | 216.843291976571 | 177.542628903324 | 185.505158175096 | 244.79886720138 | 284.689130912965 | 338.793778284459 | 195.722681024554 | 197.565652769668 | Tssc4 |
| **ENSMUSG00000045757** | -0.509288363436104 | 0.0955681213220207 | 0.757085157577544 | 46.0791995450213 | 31.7622134276588 | 51.1085639870161 | 42.2067012416173 | 81.0269064906132 | 74.6080161565054 | 49.9717483466946 | 37.9933947633978 | Zfp764 |
| **ENSMUSG00000045761** | -0.057810168192859 | 0.803764806830696 | 0.963461896828986 | 125.588406603097 | 100.987550385377 | 67.1982970940397 | 77.8479156234274 | 100.736154015357 | 102.738907494204 | 86.4094815161593 | 97.6973008201657 | Togaram2 |
| **ENSMUSG00000045763** | -0.332177290740925 | 0.404176980951858 | 0.881158207349486 | 14.4562194651047 | 14.6594831204579 | 25.5542819935081 | 29.075727522003 | 19.7092475247437 | 39.1386314263635 | 28.1091084450157 | 18.4539345993646 | Basp1 |
| **ENSMUSG00000045767** | 0.0511379084160041 | 0.773580350677392 | 0.959203117135841 | 1114.9359262462 | 1162.98566088966 | 1282.44637411865 | 1067.36057806579 | 997.506916391197 | 826.803588881929 | 1330.49779973074 | 1310.22935655489 | B230219D22 |
| **ENSMUSG00000045776** | 0.118719219234499 | 0.622257884813802 | 0.93078489223231 | 13961.9974621415 | 18067.8129459644 | 11474.8190699738 | 9379.26694258161 | 10686.7919911944 | 9390.82537786309 | 16040.931219289 | 12587.754447896 | Lrtm1 |
| **ENSMUSG00000045777** | -0.646822904490132 | 0.190707185150911 | 0.808824684419004 | 37.9475760958999 | 26.8757190541728 | 37.8581955459379 | 25.3240207449704 | 53.6529515951357 | 102.738907494204 | 21.8626399016789 | 22.7960368580387 | Ifitm10 |
| **ENSMUSG00000045790** | 0.178136856627078 | 0.268813707758593 | 0.843793744782845 | 265.633032671299 | 219.892246806869 | 230.934992830221 | 223.226553233442 | 185.047935093427 | 173.677676954488 | 243.612273190136 | 226.874843015718 | Ccdc149 |
| **ENSMUSG00000045795** | 0.0764432349741663 | 0.651353678051845 | 0.936476823117702 | 253.887354355902 | 238.623808571898 | 226.202718386979 | 296.384835385579 | 274.834507150593 | 270.301173288323 | 224.872867560126 | 193.223550510994 | Whamm |
| **ENSMUSG00000045799** | -0.070242268181639 | 0.771001985409536 | 0.959045638244974 | 51.5002818444356 | 54.56585383726 | 50.1621090983677 | 63.779015209555 | 47.0832024202212 | 79.5003450848009 | 51.0128264372507 | 54.2762782334254 | Gm9800 |
| **ENSMUSG00000045817** | -0.569868402797382 | 0.00431256175307284 | 0.635862253526407 | 361.405486627618 | 334.724864583789 | 420.22597055991 | 453.95652002095 | 708.437952694956 | 689.818378889657 | 448.704657029695 | 484.144401842154 | Zfp36l2 |
| **ENSMUSG00000045822** | -0.0704513040579402 | 0.800889277026727 | 0.963461896828986 | 32.5264937964856 | 50.4937751926884 | 43.5369248778286 | 44.0825546301336 | 48.1781606160403 | 58.7079471395453 | 42.6842017128016 | 30.3947158107182 | Zswim3 |
| **ENSMUSG00000045827** | 0.00883685569044097 | 0.959093677568128 | 0.993211395091646 | 292.738444168371 | 277.715763559786 | 317.06238769723 | 405.184331919526 | 314.253002200081 | 382.824738639118 | 286.296474902938 | 301.776106977845 | Serpinb9 |
| **ENSMUSG00000045838** | 0.0443793370838561 | 0.828903836581738 | 0.967483872587428 | 982.119409910552 | 820.116639016729 | 668.197151385803 | 790.67220325963 | 642.74046094581 | 616.433444965225 | 1006.72251356778 | 895.558590851519 | A430105I19F |
| **ENSMUSG00000045854** | 0.166770224703531 | 0.176908500740477 | 0.799889021902745 | 439.107666252556 | 390.105134149963 | 401.296872786941 | 392.053358199911 | 323.012667766634 | 332.67836712409 | 395.609674411332 | 392.96025441 | Lyrm2 |
| **ENSMUSG00000045867** | -0.064379759272129 | 0.735553546666282 | 0.954383574136061 | 300.870067617492 | 241.067055758641 | 274.47191770805 | 282.315934971707 | 262.789966996583 | 391.386314263635 | 260.269522639034 | 235.559047533066 | Cradd |
| **ENSMUSG00000045868** | -0.466331315697759 | 0.217434637339083 | 0.825996545635915 | 28.0089252136404 | 35.0198763433161 | 25.5542819935081 | 42.2067012416173 | 45.9882442244021 | 77.0541806206532 | 27.0680303544596 | 31.4802413753867 | Gvin1 |
| **ENSMUSG00000045875** | 0.164825654493625 | 0.650734906879833 | 0.936463929813731 | 283.70330700268 | 483.762942975111 | 527.175372977185 | 263.557401086543 | 241.98576127602 | 179.793088114857 | 472.649453112486 | 494.999657488839 | Adra1a |
| **ENSMUSG00000045896** | 0.306166173677528 | 0.25689741460793 | 0.837517407767513 | 1442.91140536077 | 1579.15209836488 | 1693.20779579207 | 1381.56602064227 | 933.99934103369 | 711.833859066986 | 1667.80710107093 | 1616.34756579141 | Paip2b |
| **ENSMUSG00000045917** | 0.0173321450313937 | 0.930801149869773 | 0.987762852142344 | 372.247651226447 | 346.12668478859 | 318.008842585878 | 278.564228194674 | 293.448796479518 | 233.608706326107 | 367.500565966316 | 403.815510056685 | Tmem268 |
| **ENSMUSG00000045930** | -0.557687450363412 | 0.102374192391661 | 0.759800953500741 | 311.712232216321 | 261.427448981499 | 417.386605893965 | 477.404687377404 | 774.135444444101 | 741.187832636759 | 315.446661438509 | 329.999771659226 | Clec14a |
| **ENSMUSG00000045932** | 0.0885348273690976 | 0.687325785953729 | 0.944452327903597 | 277.378710986697 | 310.292392716359 | 210.112985279955 | 322.646782824807 | 313.158044004262 | 322.893709267499 | 206.133461930115 | 212.763010675027 | Ifit2 |
| **ENSMUSG00000045934** | -0.820095623983174 | 0.0421463538225387 | 0.685406501171951 | 21.6843291976571 | 24.4324718674299 | 13.2503684410783 | 37.5170677703264 | 50.3680770076785 | 58.7079471395453 | 39.5609674411332 | 22.7960368580387 | Mtmr11 |
| **ENSMUSG00000045948** | 0.0238544151733094 | 0.891998286837884 | 0.982421229269187 | 960.435080712895 | 753.334549245754 | 846.130670451711 | 1108.62935261315 | 866.111932892906 | 1128.90490020417 | 818.287379177123 | 795.690238902016 | Mrps12 |
| **ENSMUSG00000045954** | -0.226157071811779 | 0.237543684445588 | 0.832048466795696 | 2142.23102198521 | 1975.77255834616 | 2482.55117292488 | 2801.58703574913 | 3512.62589218766 | 2921.94345242445 | 2146.70302272675 | 2416.3799069521 | Cavin2 |
| **ENSMUSG00000045962** | 0.183732305340489 | 0.562162786127 | 0.917668480350692 | 9487.79753769154 | 12375.8614165821 | 9225.09579965641 | 7796.98460936809 | 6128.48102199949 | 4406.76528216212 | 11904.7279655094 | 11795.320785688 | Wnk1 |
| **ENSMUSG00000045968** | -0.238824658431774 | 0.373483937792798 | 0.873700364743909 | 30.7194663633475 | 38.2775392589734 | 42.5904699891801 | 40.3308478531009 | 48.1781606160403 | 29.3539735697726 | 53.094982618363 | 47.7631248454143 | Teddm2 |
| **ENSMUSG00000045969** | -0.0239015962351596 | 0.857482152353693 | 0.974029039597056 | 556.564449406532 | 469.103459854653 | 600.998854291764 | 624.659178375935 | 586.897592959036 | 609.094951572782 | 583.003730711437 | 509.11148982953 | Ing1 |
| **ENSMUSG00000045973** | 0.106964827652727 | 0.488987740296164 | 0.899236955296196 | 1721.19363006403 | 1819.40473839461 | 1762.29900266341 | 1630.11659462068 | 1430.01540373974 | 1272.00552135681 | 1856.24223546159 | 1877.95922687652 | Slc25a51 |
| **ENSMUSG00000045975** | -0.0418600431927404 | 0.708566484544208 | 0.949174314815223 | 675.828259993646 | 768.808448095126 | 758.110365807406 | 634.038445318517 | 777.420319031559 | 739.964750404685 | 731.877897660964 | 671.940324529806 | C2cd2 |
| **ENSMUSG00000045980** | -0.128833258085219 | 0.284386469934354 | 0.851125503777472 | 238.527621174228 | 237.809392842984 | 227.149173275627 | 225.102406621959 | 281.404256325508 | 242.170281950624 | 255.064132186253 | 236.644573097735 | Tmem104 |
| **ENSMUSG00000045983** | 0.165188427285327 | 0.395743554819014 | 0.879215782741314 | 13107.2734862671 | 15214.1002318486 | 13642.2007649787 | 12967.7744748133 | 10983.5256622614 | 8449.05205916622 | 15175.7953260368 | 14378.8716295991 | Eif4g1 |
| **ENSMUSG00000045996** | 0.238853187501569 | 0.157563256198566 | 0.789230661864335 | 548.43282595741 | 361.600583637962 | 366.278041906949 | 479.28054076592 | 377.760577557588 | 384.047820871192 | 367.500565966316 | 358.223436340608 | Polr2k |
| **ENSMUSG00000046010** | 0.0682726227935379 | 0.54558242998528 | 0.914211323944814 | 299.063040184354 | 309.477976987445 | 276.364827485346 | 286.067641748739 | 289.068963696242 | 259.293433199658 | 292.542943446274 | 275.723493425801 | Zfp830 |
| **ENSMUSG00000046020** | -0.16902381146018 | 0.185308058976434 | 0.805963874499992 | 258.404922938747 | 299.704988240473 | 287.722286149128 | 233.543746870282 | 312.063085808443 | 302.101311322243 | 308.159114804616 | 292.006376895829 | Pofut1 |
| **ENSMUSG00000046027** | -0.218550646285555 | 0.176732679750604 | 0.799889021902745 | 99.3865088225949 | 117.275864963663 | 106.949402417274 | 120.054616865045 | 131.394983498292 | 143.100621152642 | 111.395355689507 | 131.348593324889 | Stard5 |
| **ENSMUSG00000046032** | 0.122884104344552 | 0.306217902395319 | 0.851785577030345 | 639.687711330884 | 680.851549372379 | 652.10741827878 | 612.466131350579 | 556.238763476101 | 513.694537471021 | 655.879197050366 | 646.973236542431 | Snx12 |
| **ENSMUSG00000046034** | 0.454312799398287 | 0.0264300442598559 | 0.664353746352133 | 921.583990900426 | 934.134841064735 | 1198.21188902893 | 1478.17247015086 | 919.764884488041 | 909.973180662952 | 853.684034256032 | 624.177199684392 | Otulin |
| **ENSMUSG00000046058** | -0.00833846233646082 | 0.970607089026805 | 0.9951105526746 | 208.711668527449 | 162.883145782866 | 173.201244622666 | 259.805694309511 | 206.947099009809 | 259.293433199658 | 194.681602933998 | 148.717002359586 | Eid2 |
| **ENSMUSG00000046062** | -0.209343920951917 | 0.295516520969394 | 0.851785577030345 | 453.563885717661 | 484.577358704026 | 545.158015861505 | 481.156394154437 | 724.862325632242 | 631.110431750112 | 387.281049686883 | 528.650949993563 | Ppp1r15b |
| **ENSMUSG00000046070** | -0.00930333543015546 | 0.962934522203144 | 0.994284908685797 | 68.6670424592474 | 74.1118313312039 | 82.3415753124149 | 93.7926694258161 | 71.1722827282413 | 85.6157562451702 | 70.7933101578173 | 93.3551985614916 | Igfals |
| **ENSMUSG00000046079** | -0.10543416164077 | 0.527057758314834 | 0.908217276872524 | 128.298947752804 | 152.295741306979 | 162.790240847533 | 175.392291826276 | 194.902558855799 | 147.992950080937 | 161.367104036201 | 160.657783570939 | Lrrc8d |
| **ENSMUSG00000046080** | -0.11609112241473 | 0.769536310059394 | 0.95879680569409 | 50.5967681278665 | 56.1946852950887 | 70.0376617599851 | 28.1378008277448 | 77.7420319031559 | 77.0541806206532 | 37.4788112600209 | 30.3947158107182 | Clec9a |
| **ENSMUSG00000046096** | 0.0244418404065366 | 0.849188089116534 | 0.972640876584422 | 417.423337054899 | 467.474628396825 | 477.959718767466 | 475.528833988888 | 389.805117711598 | 425.632616761703 | 509.087186281951 | 481.973350712817 | BC030336 |
| **ENSMUSG00000046111** | 0.187219624197623 | 0.432743489604243 | 0.88759702755507 | 98.4829951060259 | 89.5857301805761 | 81.3951204237664 | 56.2756016554897 | 66.7924499449649 | 57.4848649074714 | 80.1630129728225 | 81.4144173501381 | Cep295 |
| **ENSMUSG00000046138** | 0.128322852166349 | 0.510419687110363 | 0.902394714365738 | 618.003382133227 | 674.336223541064 | 530.01473764313 | 449.266886549659 | 402.944616061428 | 468.440494884288 | 587.168043073661 | 619.835097425718 | 9930021J03F |
| **ENSMUSG00000046139** | 0.20060130815688 | 0.476005974137658 | 0.897553735056661 | 599.933107801846 | 778.581436842098 | 755.271001141461 | 551.500896223799 | 453.312693069106 | 335.124531588238 | 748.535147109862 | 798.946815596022 | Patl1 |
| **ENSMUSG00000046152** | -0.879579161786623 | 0.00100411705393582 | 0.556246415579446 | 36.1405486627618 | 32.5766291565731 | 38.8046504345863 | 34.703287687552 | 86.5016974697087 | 69.71568722821 | 43.7252798033577 | 61.8749571861049 | Fut10 |
| **ENSMUSG00000046157** | -0.334539007476029 | 0.374510472376317 | 0.873700364743909 | 39.754603529038 | 63.5244268553176 | 65.3053873167428 | 78.7858423176855 | 91.9764884488041 | 122.308223207386 | 36.4377331694648 | 61.8749571861049 | Tmem229b |
| **ENSMUSG00000046169** | -0.550502452333428 | 0.239479033716036 | 0.832048466795696 | 6.32459601598331 | 10.5874044758863 | 9.46454888648447 | 10.3171936368398 | 9.85462376237187 | 9.78465785659088 | 20.8215618111227 | 13.0263067760221 | Adamts6 |
| **ENSMUSG00000046179** | -0.262085580388791 | 0.531634630352734 | 0.909082872831377 | 9.03513716569045 | 9.77298874697194 | 12.3039135524298 | 15.0068271081306 | 14.2344565456483 | 11.0077400886647 | 17.6983275394543 | 11.9407812113536 | E2f8 |
| **ENSMUSG00000046182** | -0.396383900241903 | 0.0635665109974973 | 0.716991331547963 | 287.317361868956 | 242.69588721647 | 223.363353721033 | 179.143998603309 | 214.611806380543 | 329.009120427868 | 348.761160336306 | 335.427399482569 | Gsg1l |
| **ENSMUSG00000046185** | -0.118076450080012 | 0.396187086656083 | 0.879359239049715 | 160.82544154929 | 151.481325578065 | 161.843785958884 | 140.689004138724 | 176.288269526875 | 163.893019097897 | 175.942197303987 | 150.888053488923 | Zfp84 |
| **ENSMUSG00000046186** | -0.728536433550738 | 0.162835754337379 | 0.793968226938511 | 7.22810973255236 | 11.4018202048006 | 6.62518422053913 | 12.1930470253561 | 14.2344565456483 | 29.3539735697726 | 9.36970281500523 | 9.76973008201657 | Cd109 |
| **ENSMUSG00000046191** | 0.0410783487311159 | 0.883194219583017 | 0.97910597465636 | 42.4651446787451 | 67.5965054998893 | 69.0912068713366 | 45.9584080186499 | 51.4630352034976 | 75.8310983885793 | 46.8485140750261 | 45.5920737160773 | Pcdhb20 |
| **ENSMUSG00000046201** | 0.145254040094496 | 0.467867470504668 | 0.895328112556867 | 604.450676384691 | 706.098436968723 | 786.504012466859 | 576.824916968769 | 586.897592959036 | 420.740287833408 | 741.247600475969 | 667.598222271132 | Scaf8 |
| **ENSMUSG00000046207** | -0.275856280444335 | 0.356554385056989 | 0.864965376583892 | 89.4478579403355 | 68.4109212288036 | 86.1273948670087 | 91.9168160372998 | 142.344565456483 | 125.977469903608 | 56.2182168900314 | 82.4999429148066 | Pik3r6 |
| **ENSMUSG00000046223** | -0.724347031905373 | 0.111702539643546 | 0.764116514278315 | 37.0440623793308 | 17.1027303072009 | 27.447191770805 | 30.0136542162612 | 85.4067392738896 | 48.9232892829544 | 24.9858741733473 | 24.9670879873757 | Plaur |
| **ENSMUSG00000046229** | -0.367908394663915 | 0.200767069850404 | 0.812921911423739 | 990.251033359673 | 824.188717661301 | 797.861471130641 | 1120.8223996385 | 1278.91117271671 | 1921.46218658803 | 821.410613448792 | 796.775764466685 | Scand1 |
| **ENSMUSG00000046230** | 0.2052549775536 | 0.515140146759968 | 0.904145806837306 | 558.37147683967 | 820.116639016729 | 601.945309180412 | 481.156394154437 | 416.084114411257 | 275.193502216618 | 747.494069019306 | 695.821886952513 | Vps13a |
| **ENSMUSG00000046269** | 0.0717350002031353 | 0.903463743980678 | 0.985027469232262 | 12.6491920319666 | 3.25766291565731 | 18.9290977729689 | 19.6964605794214 | 15.3294147414674 | 19.5693157131818 | 10.4107809055614 | 6.51315338801105 | Usp27x |
| **ENSMUSG00000046275** | -0.646593499456538 | 0.121526705083559 | 0.769868040577702 | 32.5264937964856 | 45.6072808192024 | 71.930571537282 | 60.0273084325223 | 125.920192519196 | 112.523565350795 | 38.5198893505771 | 52.1052271040884 | Tusc5 |
| **ENSMUSG00000046280** | -0.39308631109168 | 0.125016863440488 | 0.771949105062466 | 496.029030396406 | 408.836695914993 | 453.351891662606 | 426.756645887463 | 875.966556655277 | 644.564336302924 | 434.129563761909 | 389.703677715994 | She |
| **ENSMUSG00000046311** | 0.137785092699948 | 0.373467372720822 | 0.873700364743909 | 501.45011269582 | 538.328796812371 | 504.460455649622 | 487.721881014244 | 411.70428162798 | 368.147751854232 | 539.278450908079 | 526.479898864226 | Zfp62 |
| **ENSMUSG00000046312** | -0.269742693403509 | 0.110819695082919 | 0.764116514278315 | 224.071401709123 | 247.582381589956 | 215.791714611846 | 203.530092654021 | 340.531998899739 | 222.600966237442 | 262.351678820146 | 248.585354309088 | AI464131 |
| **ENSMUSG00000046314** | -0.209090343132626 | 0.521771433429396 | 0.906152189339124 | 60.535419010126 | 56.1946852950887 | 55.8408384302584 | 47.8342614071662 | 84.3117810780705 | 86.838838477244 | 31.2323427166841 | 53.1907526687569 | Stxbp6 |
| **ENSMUSG00000046318** | -0.339460372025907 | 0.067200450804201 | 0.721898114207542 | 119.263810587114 | 118.904696421492 | 131.557229522134 | 94.7305961200743 | 150.009272827216 | 114.969729814943 | 168.654650670094 | 153.05910461826 | Ccbe1 |
| **ENSMUSG00000046324** | -0.0200217760606409 | 0.889410210196736 | 0.981325587388408 | 273.764656120421 | 321.69421292116 | 297.186835035612 | 256.053987532478 | 295.638712871156 | 267.855008824175 | 338.350379430744 | 262.697186649779 | Ermp1 |
| **ENSMUSG00000046329** | 0.0232856060967052 | 0.837461912621749 | 0.968924788429433 | 1040.84780148754 | 1110.86305423914 | 918.061241988993 | 1034.53314376675 | 903.340511550755 | 988.250443515679 | 1117.07679116673 | 1030.16376087041 | Slc25a23 |
| **ENSMUSG00000046330** | 0.15070213073715 | 0.426314260497761 | 0.885066977635295 | 3724.2835396976 | 2713.63320874254 | 3325.84247871064 | 3777.03079777762 | 3165.52414411301 | 3957.89410299101 | 2457.98537180304 | 2617.20213641577 | Rpl37a |

| | | | | | | | | | | | | |
|---|---|---|---|---|---|---|---|---|---|---|---|---|
| **ENSMUSG00000046345** | 0.018393827881172 | 0.914646145168347 | 0.986675594624755 | 340.62467114653 | 288.303168035672 | 248.917635714542 | 278.564228194674 | 301.113503850252 | 217.708637309147 | 306.076958623504 | 315.887939318536 | Smco1 |
| **ENSMUSG00000046351** | 0.0320794153607844 | 0.90347764228829 | 0.985027469232262 | 240.334648607366 | 271.200437728471 | 221.470443943737 | 228.854113398991 | 209.137015401448 | 128.423634367755 | 310.241270985729 | 292.006376895829 | Zfp322a |
| **ENSMUSG00000046352** | -0.164058124241205 | 0.686000090718756 | 0.944452327903597 | 32.5264937964856 | 82.2559886203472 | 35.0188308799925 | 39.3929211588428 | 44.893286028583 | 37.9155491942896 | 41.6431236222455 | 87.9275707381491 | Gjb2 |
| **ENSMUSG00000046364** | 0.0961479868930601 | 0.599525110688484 | 0.924958767906592 | 5163.58089019209 | 3684.41675760842 | 4330.03111556664 | 5344.306303883 | 4432.3907766757 | 5464.73141290601 | 3655.2251759426 | 3776.54343948174 | Rpl27a |
| **ENSMUSG00000046380** | -0.189247793131684 | 0.426966490810958 | 0.885066977635295 | 42.4651446787451 | 57.009101024003 | 66.2518422053913 | 62.8410885152968 | 59.1277425742312 | 53.8156182112498 | 65.5879197050366 | 81.4144173501381 | Jrk |
| **ENSMUSG00000046387** | -0.29347052092207 | 0.402335238816872 | 0.881158207349486 | 24.3948703473642 | 57.8235167529173 | 47.3227444324223 | 30.9515809105193 | 43.7983278327639 | 72.1618516923577 | 43.7252798033577 | 37.9933947633978 | Pcdhb17 |
| **ENSMUSG00000046402** | -0.365327378490855 | 0.323650910222711 | 0.855977678491063 | 120.167324303683 | 92.8433930962335 | 90.8596693102509 | 130.371810501884 | 150.009272827216 | 249.508775343067 | 74.9576225200418 | 85.7565196088121 | Rbp1 |
| **ENSMUSG00000046404** | -0.396378803062301 | 0.141089391541725 | 0.779711473022872 | 80.412720774645 | 63.5244268553176 | 53.9479286529615 | 89.1030359545253 | 122.635317931739 | 110.077400886647 | 79.1219348822664 | 66.217059444779 | Yod1 |
| **ENSMUSG00000046408** | -0.33069973620497 | 0.426679735942503 | 0.885066977635295 | 9.93865088225949 | 11.4018202048006 | 11.3574586637814 | 10.3171936368398 | 12.0445401540101 | 11.0077400886647 | 12.4929370866736 | 18.4539345993646 | 1700067K01I |
| **ENSMUSG00000046410** | -0.249412080574028 | 0.494334317434438 | 0.900141680336539 | 56.9213641438498 | 53.7514381083457 | 76.6628459805242 | 47.8342614071662 | 114.970610561005 | 75.8310983885793 | 55.1771387994752 | 33.6512925047237 | Kcnk6 |
| **ENSMUSG00000046415** | 0.275666643726158 | 0.371892085493141 | 0.873663802472542 | 53.3073092775737 | 32.5766291565731 | 43.5369248778286 | 64.7169419038131 | 37.2285786578493 | 42.8078781225851 | 28.1091084450157 | 52.1052271040884 | B430212C06 |
| **ENSMUSG00000046417** | -0.312405319457763 | 0.238467062769729 | 0.832048466795696 | 56.9213641438498 | 61.895595397489 | 62.4660226507975 | 48.7721881014244 | 70.0773245324222 | 102.738907494204 | 47.8895921655823 | 66.217059444779 | Lrrc75a |
| **ENSMUSG00000046432** | -0.569260008896692 | 0.147972702557604 | 0.780607990324772 | 50.5967681278665 | 31.7622134276588 | 39.7511053232348 | 33.7653609932938 | 58.0327843784121 | 103.961989726278 | 38.5198893505771 | 31.4802413753867 | Bex3 |
| **ENSMUSG00000046434** | 0.110056120592024 | 0.350492586601575 | 0.861774532843376 | 3044.84122483768 | 2888.73259045912 | 3198.0710687431 | 3552.86631784992 | 2801.99802310107 | 2630.84988119087 | 3308.5461717874 | 3009.0768652611 | Hnrnpa1 |
| **ENSMUSG00000046441** | -0.0544690039091066 | 0.852634674268341 | 0.973120292022893 | 37.9475760958999 | 60.2667639396603 | 50.1621090983677 | 30.0136542162612 | 44.893286028583 | 52.592535979176 | 35.3966550789086 | 53.1907526687569 | Cmtr2 |
| **ENSMUSG00000046442** | 0.471105771153385 | 0.0756462885341672 | 0.735399943172214 | 169.86057871498 | 305.405898342873 | 278.257737262643 | 277.626301500416 | 136.869774477387 | 140.654456688494 | 225.913945650682 | 239.90114979174 | Ppm1e |
| **ENSMUSG00000046447** | -0.050221991803156 | 0.829290689628335 | 0.967483872587428 | 1197.15567445398 | 1124.70812163069 | 1645.88505135965 | 1326.22834568104 | 1549.36584708402 | 1870.09273284093 | 1061.89965236726 | 1000.85457062436 | Camk2n1 |
| **ENSMUSG00000046470** | -0.931989944863023 | 0.0221141194664586 | 0.648762042745232 | 820.390454644693 | 623.028032619461 | 818.683478680906 | 912.602673513191 | 2348.68533003196 | 2331.19473433278 | 687.11153976705 | 690.394259129171 | Sox18 |
| **ENSMUSG00000046480** | 0.299714151294038 | 0.266791267132159 | 0.843513595393324 | 2122.35372022069 | 2366.69210822504 | 2657.64532732484 | 1813.01230000103 | 1435.49019471884 | 1084.87393984951 | 2164.40135026621 | 2593.32057399306 | Scn4b |
| **ENSMUSG00000046491** | -0.88617651547764 | 0.145359331407234 | 0.780607990324772 | 17.1667606148119 | 17.1027303072009 | 16.0897331070236 | 28.1378008277448 | 56.937826182593 | 66.0464405319884 | 8.32862472444909 | 14.1118323406906 | C1qtnf2 |
| **ENSMUSG00000046516** | 0.0603904881007913 | 0.785261027649151 | 0.961141387070613 | 1999.4758547673 | 1458.61857048556 | 1576.79384448831 | 2194.7484645641 | 1646.81712651192 | 2430.26439513076 | 1372.14092335299 | 1484.99897246652 | Cox17 |
| **ENSMUSG00000046519** | -0.199463373175931 | 0.0818050101282338 | 0.739855948845396 | 236.72059374109 | 265.499527626071 | 244.185361271299 | 260.743621003769 | 295.638712871156 | 302.101311322243 | 290.460787265162 | 269.21034003779 | Golph3l |
| **ENSMUSG00000046532** | -0.698568156515935 | 0.104101969784236 | 0.760854717370327 | 31.6229800799166 | 52.9370223794314 | 36.9117406572894 | 29.075727522003 | 88.6916138613469 | 92.9542496376133 | 35.3966550789086 | 28.2236646813812 | Ar |
| **ENSMUSG00000046541** | -0.0448876450982613 | 0.891916085296346 | 0.982421229269187 | 91.2548853734735 | 131.120932355207 | 135.343049076728 | 76.9099889291692 | 95.2613630362614 | 58.7079471395453 | 162.408182126757 | 131.348593324889 | Zfp526 |
| **ENSMUSG00000046546** | -0.718666621409461 | 0.0112151561083211 | 0.635862253526407 | 451.756858284523 | 390.919549878878 | 451.458981885309 | 599.335157630965 | 1005.17162376193 | 1061.63537744011 | 459.115437935256 | 590.525907179668 | Fam43a |
| **ENSMUSG00000046556** | -0.143416400739868 | 0.348021111356608 | 0.861266047293794 | 219.553833126278 | 286.674336577844 | 293.401015481019 | 233.543746870282 | 304.398378437709 | 306.993640250539 | 254.023054095697 | 276.809018990469 | Zfp319 |
| **ENSMUSG00000046562** | -0.535903440161475 | 0.0276392684179915 | 0.665956123973759 | 485.186865797577 | 517.968403589513 | 617.088587398787 | 479.28054076592 | 923.049759075499 | 1033.50448610241 | 568.428637443651 | 519.966745476215 | Unc119b |
| **ENSMUSG00000046567** | -0.0619056437975679 | 0.787001370351751 | 0.961141387070613 | 61.4389327266951 | 75.7406627890326 | 57.7337482075553 | 53.4618215727152 | 64.6025335533267 | 66.0464405319884 | 79.1219348822664 | 49.9341759747513 | 4930430F08I |
| **ENSMUSG00000046572** | -0.161174061555808 | 0.588936588100308 | 0.92352104528132 | 75.8951521917998 | 127.048853710635 | 95.5919437534931 | 85.3513291774927 | 79.9319482947941 | 78.277262852727 | 167.613572579538 | 103.124928643508 | Zfp518b |
| **ENSMUSG00000046573** | 0.122295781842953 | 0.356877325562576 | 0.865163598066054 | 295.448985318078 | 289.117583764587 | 226.202718386979 | 276.688374806158 | 256.220217821669 | 248.285693110993 | 248.817663642917 | 246.414303179751 | Lyrm4 |
| **ENSMUSG00000046574** | -0.0739966390469215 | 0.581989089752154 | 0.922450244597442 | 557.467963123101 | 590.451403462888 | 631.285410728514 | 494.287367874051 | 684.348872386936 | 552.833168897385 | 538.237372817523 | 617.664046296381 | Prr12 |
| **ENSMUSG00000046589** | 0.181298001522237 | 0.74342825810401 | 0.954383574136061 | 16.2632468982428 | 6.51532583131463 | 19.8755526616174 | 45.0204813243917 | 13.1394983498292 | 15.9000690169602 | 22.903717992235 | 24.9670879873757 | Lrrc8e |
| **ENSMUSG00000046598** | 0.0838593851466045 | 0.607732595267263 | 0.927510709653048 | 3631.22162689099 | 3973.53434137301 | 3202.80334318634 | 3437.50133445616 | 3164.42918591719 | 2558.68802949851 | 4003.9863362789 | 3712.4974311663 | Bdh1 |
| **ENSMUSG00000046603** | 0.220756991831908 | 0.320243312564461 | 0.855228644171892 | 1191.73459215457 | 1258.27230117264 | 1354.37694565593 | 963.250715003132 | 896.77076237584 | 680.033721033066 | 1238.8829277618 | 1274.40701292083 | Tcaim |
| **ENSMUSG00000046605** | 0.18026632813134 | 0.502675721106331 | 0.900927059989498 | 30.7194663633475 | 48.0505280059454 | 38.8046504345863 | 55.3376749612315 | 39.4184950494875 | 35.4693847301419 | 36.4377331694648 | 41.2499714574033 | B3gntl1 |
| **ENSMUSG00000046618** | -0.951731873148255 | 0.0971378585530605 | 0.757809915356977 | 53.3073092775737 | 87.9568987227475 | 29.3401015481019 | 45.0204813243917 | 122.635317931739 | 221.377884005369 | 43.7252798033577 | 30.3947158107182 | Olfml2a |
| **ENSMUSG00000046658** | -0.0655468560561694 | 0.725245753102302 | 0.951674101666722 | 278.282224703266 | 298.076156782644 | 208.220075502658 | 246.674720589896 | 238.700886688563 | 238.501035254403 | 344.596847974081 | 257.269558826436 | Zfp316 |
| **ENSMUSG00000046667** | -0.102161353182446 | 0.671571911318716 | 0.943173644832837 | 103.000563688871 | 85.5136515360045 | 73.8234813145789 | 55.3376749612315 | 95.2613630362614 | 97.8465785659088 | 75.998700610598 | 72.73021283279 | Rbm12b1 |
| **ENSMUSG00000046668** | -0.53667249523258 | 0.00226072880343293 | 0.635862253526407 | 321.65088309858 | 265.499527626071 | 375.742590793433 | 347.03287687552 | 540.909348734634 | 540.602346576646 | 437.252798033577 | 381.019473198646 | Cxxc5 |
| **ENSMUSG00000046671** | 0.156590984496132 | 0.318346937713351 | 0.854351922245337 | 4007.98684670028 | 3962.94693689712 | 3725.24644172029 | 3487.21144925184 | 2919.15855005371 | 2766.61200895107 | 3925.90547948719 | 4008.8459103208 | Mtfr1l |
| **ENSMUSG00000046675** | -0.0142073847715082 | 0.922727712626462 | 0.986713918026868 | 159.921927832721 | 184.872370463553 | 200.648436393471 | 181.019851991825 | 177.383227722694 | 172.454594722414 | 172.818963032319 | 210.59195954569 | Tmem251 |
| **ENSMUSG00000046679** | -0.180094159517135 | 0.271560081740423 | 0.844129257012535 | 131.009488902512 | 184.872370463553 | 160.897331070236 | 161.323391412404 | 180.668102310151 | 171.23151249034 | 206.133461930115 | 164.999885829613 | C87436 |
| **ENSMUSG00000046683** | -0.163097644313166 | 0.707248190260273 | 0.948710970699398 | 9.03513716569045 | 33.3910448854875 | 16.0897331070236 | 20.6343872736795 | 25.1840385038392 | 26.9078091056249 | 19.7804837205666 | 17.3684090346961 | 0610025J13I |
| **ENSMUSG00000046688** | -0.0235772159323815 | 0.924432977008317 | 0.986713918026868 | 44.2721721118832 | 46.4216965481167 | 54.8943835416099 | 62.8410885152968 | 40.5134532453066 | 68.4926049961361 | 49.9717483466946 | 53.1907526687569 | Tifa |
| **ENSMUSG00000046691** | 0.0284305541172419 | 0.819719506303267 | 0.966398681144067 | 731.846110420926 | 889.341975974447 | 891.560505106837 | 732.520748215624 | 863.922016501267 | 716.726187995282 | 792.26042691322 | 808.716545678038 | Chtf8 |
| **ENSMUSG00000046694** | -0.961183125057648 | 0.0929166798597573 | 0.754058637722175 | 10.8421645988285 | 9.77298874697194 | 21.7684624389143 | 4.68963347129081 | 29.5638712871156 | 35.4693847301419 | 8.32862472444909 | 18.4539345993646 | Fam46b |
| **ENSMUSG00000046699** | -0.073804321992898 | 0.863735106858812 | 0.974868041734905 | 16.2632468982428 | 14.6594831204579 | 9.46454888648447 | 30.9515809105193 | 18.6142893289246 | 15.9000690169602 | 22.903717992235 | 17.3684090346961 | Slitrk4 |
| **ENSMUSG00000046707** | 0.0766225678064863 | 0.460816465192472 | 0.89483303530315 | 452.660372001092 | 448.743066631795 | 493.102996985841 | 475.528833988888 | 473.02194059385 | 453.763508099402 | 442.458188486358 | 403.815510056685 | Csnk2a2 |
| **ENSMUSG00000046709** | -0.0583872947054298 | 0.750786824808384 | 0.956197192463153 | 378.57224724243 | 452.000729547452 | 300.972654590206 | 378.922384480297 | 296.733671066975 | 425.632616761703 | 421.636626675235 | 429.868123608729 | Mapk10 |
| **ENSMUSG00000046711** | 0.0825409303371079 | 0.450329702125955 | 0.89108031640712 | 1392.3146372329 | 1304.69399772075 | 1416.84296830672 | 1476.29661676235 | 1152.99098019751 | 1404.09840242079 | 1276.36173902182 | 1445.92005213845 | Hmga1 |
| **ENSMUSG00000046714** | -1.41885972535416 | 0.0271540066138421 | 0.664353746352133 | 9.03513716569045 | 17.1027303072009 | 17.9826428843205 | 16.8826804966469 | 55.8428679867739 | 78.277262852727 | 23.9447960827911 | 5.42762782334254 | Foxc2 |
| **ENSMUSG00000046718** | -0.49438435271932 | 0.163753779516648 | 0.793968226938511 | 424.651446787451 | 291.56083095133 | 282.990011705886 | 379.860311174555 | 673.399290428745 | 722.841599155651 | 227.996101831794 | 319.144516012541 | Bst2 |
| **ENSMUSG00000046721** | 0.165694739009187 | 0.58428014273275 | 0.922450244597442 | 41.5616309621761 | 38.2775392589734 | 24.6078271048596 | 46.8963347129081 | 28.4689130912965 | 36.6924669622158 | 42.6842017128016 | 27.1381391167127 | Rpl14-ps1 |
| **ENSMUSG00000046722** | 0.0333129529294404 | 0.830312274477137 | 0.967583158900775 | 331.58953398084 | 420.238516119794 | 424.958245003153 | 442.701399689852 | 388.710159515779 | 475.778988276731 | 366.45948787576 | 352.795808517265 | Cdc42se1 |
| **ENSMUSG00000046727** | 0.0616505155653092 | 0.719582366030544 | 0.951674101666722 | 1207.99783905281 | 1030.23589707663 | 1008.92091129924 | 1257.75969700019 | 1065.39432453198 | 1389.4214156359 | 987.983107937774 | 874.933605122817 | Cystm1 |
| **ENSMUSG00000046731** | -0.64081045125941 | 0.0600900695786642 | 0.71282992338528 | 57.8248778604189 | 51.3081909216027 | 69.0912068713366 | 88.1651092602672 | 146.724398239759 | 138.208292224346 | 69.7522320672611 | 60.7894316214364 | Kctd11 |
| **ENSMUSG00000046733** | -0.518278708165841 | 0.433472788881052 | 0.887868762931002 | 16.2632468982428 | 14.6594831204579 | 11.3574586637814 | 20.6343872736795 | 24.0890803080201 | 53.8156182112498 | 7.28754663389296 | 5.42762782334254 | Gprc5a |
| **ENSMUSG00000046743** | -0.306547790004658 | 0.215202452376518 | 0.824338622350122 | 143.658680934478 | 120.533527879321 | 134.396594188079 | 164.137171495178 | 224.466430142915 | 217.708637309147 | 120.765058504512 | 133.519644454226 | Fat4 |
| **ENSMUSG00000046747** | -0.406342520066561 | 0.253844245061772 | 0.837406774112203 | 14.4562194651047 | 26.0613033252585 | 15.1432782183751 | 13.1309737196143 | 24.0890803080201 | 22.0154801773295 | 23.9447960827911 | 21.7105112933701 | Gm9812 |
| **ENSMUSG00000046753** | 0.105592792142024 | 0.614265901239545 | 0.928075610917191 | 287.317361868956 | 273.643684915214 | 237.56017705076 | 248.550573978413 | 220.086597359638 | 168.785348026193 | 324.816364253515 | 258.355084391105 | Ccdc66 |
| **ENSMUSG00000046756** | 0.132656396949211 | 0.374536964994807 | 0.873700364743909 | 2112.41506933843 | 1658.96483979849 | 1681.85033712829 | 2263.21711324494 | 1786.97177557677 | 2003.40869613698 | 1608.46564990923 | 1640.22912821412 | Mrps7 |
| **ENSMUSG00000046761** | -0.232944203455468 | 0.632920697020652 | 0.932167159539483 | 16.2632468982428 | 17.9171460361152 | 9.46454888648447 | 7.50341355406529 | 21.8991639163819 | 7.33849339244316 | 12.4929370866736 | 18.4539345993646 | Fam83h |
| **ENSMUSG00000046768** | -0.705913080864654 | 0.0334910256077242 | 0.671782526713461 | 555.660935689963 | 564.39010013763 | 726.877354482007 | 563.693943249155 | 1208.83384818428 | 1591.22998392809 | 568.428637443651 | 564.473293627624 | Rhoj |
| **ENSMUSG00000046785** | 0.211983552272805 | 0.418745271444238 | 0.882471828305336 | 310.808718499751 | 443.042156529395 | 489.317177431247 | 367.667264149199 | 293.448796479518 | 214.039390612925 | 438.293876124134 | 443.97995594942 | Epm2aip1 |
| **ENSMUSG00000046791** | -0.316627408775925 | 0.166225341521491 | 0.79616206680466 | 87.6408305071974 | 73.2974156022896 | 76.6628459805242 | 76.9099889291692 | 124.825234323377 | 112.523565350795 | 86.4094815161593 | 68.388110574116 | Riox1 |
| **ENSMUSG00000046792** | 0.00673588474466031 | 0.955548631823971 | 0.992371505557225 | 602.643648951553 | 634.429852824262 | 614.249222732842 | 581.51455044006 | 628.506004400162 | 700.826118978322 | 556.976778447533 | 537.335154510911 | Zfp787 |
| **ENSMUSG00000046794** | -0.113947466298397 | 0.610646486801272 | 0.927932112008768 | 382.186302108706 | 434.083583511337 | 404.136237452887 | 348.908730264036 | 329.582416941548 | 300.878229090169 | 509.087186281951 | 557.960140239613 | Ppp1r3b |
| **ENSMUSG00000046798** | 0.0292908689976816 | 0.878826227027747 | 0.978116690177517 | 339.721157429961 | 375.445651029505 | 408.868511896129 | 327.336416296098 | 339.43704070392 | 258.070350967584 | 362.295175513536 | 461.348364984116 | Cldn12 |
| **ENSMUSG00000046805** | -0.381514148142426 | 0.150499330351205 | 0.782641594375979 | 125.588406603097 | 181.614707547895 | 233.774357496166 | 230.729966787508 | 260.600050604945 | 344.909189444828 | 183.22974393788 | 217.105112933702 | Mpeg1 |
| **ENSMUSG00000046806** | -0.120208942279378 | 0.698516449321263 | 0.945012697918882 | 28.0089252136404 | 33.3910448854875 | 21.7684624389143 | 39.3929211588428 | 22.994122112201 | 41.5847958905112 | 35.3966550789086 | 33.6512925047237 | 3110062M04 |
| **ENSMUSG00000046808** | -0.0592010033662888 | 0.828212529615702 | 0.967362263400782 | 74.9916384752307 | 64.338842584232 | 50.1621090983677 | 40.3308478531009 | 58.0327843784121 | 48.9232892829544 | 78.0808567917102 | 54.2762782334254 | Atp10d |
| **ENSMUSG00000046811** | 0.427552159547639 | 0.421127887844176 | 0.882568531013176 | 16.2632468982428 | 24.4324718674299 | 11.3574586637814 | 22.5102406621959 | 5.47479097909548 | 14.6769867848863 | 8.32862472444909 | 27.1381391167127 | Gltpd2 |
| **ENSMUSG00000046814** | 0.0546219226650492 | 0.755802456313943 | 0.956566265296272 | 120.167324303683 | 111.574954861263 | 134.396594188079 | 117.24083678227 | 112.780694169367 | 122.308223207386 | 93.6970281500523 | 136.776221148232 | Gchfr |
| **ENSMUSG00000046818** | -0.00387677582905747 | 0.990957831598103 | 0.998628446692557 | 20.780815481088 | 32.5766291565731 | 47.3227444324223 | 41.2687745473591 | 30.6588294829347 | 51.3694537471021 | 29.1501865355718 | 31.4802413753867 | Ddit4l |
| **ENSMUSG00000046822** | -0.218653404549535 | 0.126558240430833 | 0.77215780108715 | 408.388199889208 | 373.816819571677 | 424.011790114504 | 383.612017951588 | 480.686647964584 | 534.486935416277 | 466.402984569149 | 369.078691987293 | Slc39a3 |
| **ENSMUSG00000046836** | 0.170006439438214 | 0.311743851144797 | 0.851785577030345 | 588.187429486448 | 630.35777417969 | 690.912068713366 | 551.500896223799 | 534.339599559719 | 407.286383280595 | 629.852244786463 | 614.407469602375 | Brox |
| **ENSMUSG00000046841** | 0.0743026534947777 | 0.613958471491459 | 0.928075610917191 | 724.618000688374 | 679.22271791455 | 699.430162711202 | 883.526945991188 | 716.102660065689 | 804.7881087046 | 586.126964983105 | 730.558705021906 | Ckap4 |
| **ENSMUSG00000046861** | 0.158626511013367 | 0.262185923949355 | 0.841442348349478 | 564.696072855653 | 706.098436968723 | 683.340429604179 | 637.79015209555 | 531.054724972262 | 497.794468454061 | 651.714884688142 | 640.46008315442 | Hectd3 |
| **ENSMUSG00000046862** | -0.23315872092331 | 0.0811507164120774 | 0.739855948845396 | 150.886790667031 | 147.409246933493 | 149.539872406455 | 143.502784221499 | 190.522726072523 | 168.785348026193 | 173.860041122875 | 161.743309135608 | Pramef8 |
| **ENSMUSG00000046865** | 0.0398918800910996 | 0.797047472080813 | 0.962278883153809 | 892.671551970216 | 726.458830191581 | 794.075651576047 | 878.837312519897 | 890.201013200926 | 942.996400928946 | 686.070461676494 | 683.88110574116 | Fbl |
| **ENSMUSG00000046873** | 0.179738003797317 | 0.245213377007134 | 0.832048466795696 | 261.115464088454 | 280.973426475443 | 322.74111702912 | 252.302280755445 | 234.321053905287 | 212.816308380852 | 292.542943446274 | 245.328777615083 | Mbtps2 |
| **ENSMUSG00000046876** | 0.178943353930897 | 0.412616355469018 | 0.882192126163777 | 390.317925557827 | 487.835021619683 | 302.865564367503 | 403.308478531009 | 411.70428162798 | 247.06261087892 | 324.816364253515 | 415.756291268038 | Atxn1 |
| **ENSMUSG00000046879** | -0.270343954559953 | 0.103106413284497 | 0.759800953500741 | 253.887354355902 | 301.333819698302 | 299.079744812909 | 301.07446885687 | 413.894198019619 | 390.163232031561 | 309.200192895173 | 281.151121249143 | Irgm1 |
| **ENSMUSG00000046897** | 0.0258877557195295 | 0.832428219760093 | 0.967671708900743 | 780.635851115655 | 806.271571625185 | 866.952678001977 | 764.410255820401 | 729.242158415519 | 688.595296657583 | 882.834220791604 | 858.650721652789 | Zfp740 |
| **ENSMUSG00000046909** | -0.0227561121158311 | 0.900751910410538 | 0.984802943363903 | 432.783070236573 | 390.919549878878 | 307.597838810745 | 390.177504811395 | 382.140410340865 | 291.093571233579 | 408.102611498006 | 463.519416113453 | Tefm |
| **ENSMUSG00000046916** | -0.485590219120658 | 0.113783000503 | 0.767518180080285 | 377.668733525861 | 235.366145656241 | 280.15064703994 | 477.404687377404 | 578.137927392483 | 669.025980944401 | 299.830490080167 | 372.335268681298 | Myct1 |
| **ENSMUSG00000046947** | 0.0630314643755161 | 0.656362700811137 | 0.938239982943985 | 239.431134890797 | 215.820168162297 | 237.56017705076 | 220.412773150668 | 250.745426842573 | 218.931719541221 | 183.22974393788 | 221.447215192376 | Adck2 |

| | | | | | | | | | | | | |
|---|---|---|---|---|---|---|---|---|---|---|---|---|
| **ENSMUSG00000046949** | -0.0483339186245393 | 0.76895468639019 | 0.95879680569409 | 734.556651570634 | 719.943504360267 | 614.249222732842 | 627.47295845871 | 551.858930692825 | 623.771938357668 | 786.013958369883 | 826.084954712734 | Nqo2 |
| **ENSMUSG00000046959** | -0.266565404019644 | 0.59718205749784 | 0.924496535139492 | 12.6491920319666 | 36.6487078011448 | 22.7149173275627 | 22.5102406621959 | 22.994122112201 | 13.4539045528125 | 20.8215618111227 | 56.4473293627624 | Slc26a1 |
| **ENSMUSG00000046962** | 0.0192999236056397 | 0.91101323902162 | 0.986203249162988 | 180.702743313809 | 175.099381716581 | 137.235958854025 | 136.937297361692 | 154.389105610493 | 172.454594722414 | 163.449260217313 | 132.434118889558 | Zbtb21 |
| **ENSMUSG00000046982** | 0.0335863349504424 | 0.76513351847219 | 0.958197631858494 | 799.609639163605 | 826.631964848044 | 906.703783325212 | 921.981940455772 | 858.447225522172 | 949.111812089315 | 781.849646007659 | 787.006034384668 | Tshz1 |
| **ENSMUSG00000046985** | 0.238484648925793 | 0.246538122787849 | 0.832178499519616 | 526.748496759753 | 661.305571878435 | 713.626986040929 | 578.700770357285 | 526.674892188986 | 348.57843614105 | 600.702058250891 | 625.26272524906 | Tapt1 |
| **ENSMUSG00000046994** | 0.128395416383722 | 0.521147793573101 | 0.90609507420585 | 291.834930451802 | 315.178887089845 | 304.7584741448 | 243.860940507122 | 263.884925192402 | 181.016170346931 | 328.980676615739 | 282.236646813812 | Mars2 |
| **ENSMUSG00000046997** | 0.521953697895399 | 0.276196571674176 | 0.846688575024234 | 94.8689402397497 | 59.452348210746 | 105.056492639978 | 286.067641748739 | 60.2227007700503 | 67.2695227640623 | 99.9434966933891 | 151.973579053591 | Spsb4 |
| **ENSMUSG00000047003** | -0.516075637094631 | 0.228570523104712 | 0.832048466795696 | 18.9737880479499 | 22.8036404096012 | 11.3574586637814 | 15.9447538023887 | 41.6084114411257 | 14.6769867848863 | 17.6983275394543 | 24.9670879873757 | Zfp41 |
| **ENSMUSG00000047022** | 0.228634467248753 | 0.255227405857428 | 0.837406774112203 | 74.0881247586617 | 87.9568987227475 | 108.842312194571 | 82.5375490947182 | 74.4571573156986 | 64.8233582999146 | 78.0808567917102 | 83.5854684794751 | Mipol1 |
| **ENSMUSG00000047030** | 0.00779428694034097 | 0.961389172867522 | 0.993822507448007 | 277.378710986697 | 325.766291565731 | 276.364827485346 | 216.661066373635 | 298.923587458613 | 243.393364182698 | 275.885693997376 | 272.466916731795 | Spata2 |
| **ENSMUSG00000047036** | 0.320661148978186 | 0.214269811545747 | 0.824248286808591 | 768.890172800257 | 939.021335438221 | 1001.34927219006 | 672.493439783102 | 573.758094609207 | 421.963370065482 | 959.873999492758 | 751.183690750607 | Zfp445 |
| **ENSMUSG00000047037** | 0.206912411747885 | 0.240054344488581 | 0.832048466795696 | 180.702743313809 | 171.027303072009 | 172.254789734017 | 146.316564304273 | 153.294147414674 | 107.6312364225 | 163.449260217313 | 155.230155747597 | Nipa1 |
| **ENSMUSG00000047044** | -0.0980396228522459 | 0.59316193067952 | 0.923935529355527 | 228.588970291968 | 145.780415475665 | 192.130342395635 | 156.633757941113 | 193.80760065998 | 211.593226148778 | 192.599446752885 | 175.855141476298 | D030056L22 |
| **ENSMUSG00000047045** | -0.113747676261596 | 0.659945505172665 | 0.939611397743053 | 433.686583953142 | 706.912852697637 | 583.016211407443 | 428.63249927598 | 557.33372167192 | 360.809258461789 | 735.001131932632 | 675.196901223812 | Tmem164 |
| **ENSMUSG00000047061** | -0.56406021996553 | 0.161153892598056 | 0.792786631432503 | 11.7456783153976 | 11.4018202048006 | 12.3039135524298 | 12.1930470253561 | 18.6142893289246 | 24.4616446414772 | 10.4107809055614 | 17.3684090346961 | Gm9817 |
| **ENSMUSG00000047067** | -0.054367812193732 | 0.676473123584167 | 0.94409858171788 | 383.993329541844 | 375.445651029505 | 388.992959234512 | 334.839829850164 | 325.202584158272 | 426.855698993777 | 382.075659234102 | 407.07208675069 | Dusp28 |
| **ENSMUSG00000047084** | 0.152438236447528 | 0.25167275402364 | 0.83723604018117 | 496.932544112975 | 430.82592059568 | 409.814966784777 | 444.577253078368 | 434.698403740182 | 391.386314263635 | 437.252798033577 | 339.769501741243 | Ngrn |
| **ENSMUSG00000047085** | 0.0746236329203881 | 0.749447091909473 | 0.956197192463153 | 82.2197482077831 | 73.2974156022896 | 86.1273948670087 | 98.4823028971069 | 73.3621991198795 | 55.0387004433237 | 102.025652874501 | 91.1841474321546 | Lrrc4b |
| **ENSMUSG00000047090** | -0.130535776685893 | 0.683611183501528 | 0.944452327903597 | 64.1494738764022 | 73.2974156022896 | 50.1621090983677 | 47.8342614071662 | 56.937826182593 | 101.51582526213 | 62.4646854333682 | 37.9933947633978 | Tmem198b |
| **ENSMUSG00000047098** | -0.0931219729671528 | 0.524410593607574 | 0.907437243979788 | 517.713359594063 | 539.9576282702 | 605.731128735006 | 638.728078789808 | 731.432074807157 | 604.202622644487 | 604.866370613115 | 514.539117652873 | Rnf31 |
| **ENSMUSG00000047117** | 0.0738620364309524 | 0.816537827132424 | 0.966315773308182 | 28.0089252136404 | 30.1333819698302 | 41.6440151005317 | 24.3860940507122 | 39.4184950494875 | 20.7923979452556 | 31.2323427166841 | 26.0526135520442 | Ankdd1b |
| **ENSMUSG00000047123** | -0.0319171736506089 | 0.880041906001076 | 0.978393501240173 | 256.597895505609 | 234.551729927327 | 136.289503965376 | 198.84045918273 | 222.276513751277 | 226.270212933664 | 207.174540020671 | 189.966973816989 | Ticam1 |
| **ENSMUSG00000047126** | -0.0665700155357184 | 0.632100890389358 | 0.931825800800958 | 1981.40558043592 | 2563.78071462231 | 2346.2616689595 | 2003.41141893543 | 2391.38869966891 | 1977.72396926343 | 2342.42570375131 | 2603.09030407508 | Cltc |
| **ENSMUSG00000047129** | 0.253972060638064 | 0.647336691925668 | 0.935063240511793 | 14.4562194651047 | 17.1027303072009 | 4.73227444324223 | 15.0068271081306 | 4.37983278327639 | 11.0077400886647 | 10.4107809055614 | 17.3684090346961 | 1700113H08 |
| **ENSMUSG00000047139** | 0.113987166196786 | 0.882116214889136 | 0.978854883720278 | 129.202461469373 | 68.4109212288036 | 371.95677123884 | 60.9652351267805 | 112.780694169367 | 318.001380339203 | 74.9576225200418 | 77.072315091464 | Cd24a |
| **ENSMUSG00000047141** | 0.110409506455511 | 0.60368718524909 | 0.926448151609826 | 327.071965397994 | 350.198763433161 | 335.04503058155 | 329.212269684615 | 269.359716171498 | 205.477814988408 | 374.788112600209 | 391.874728845331 | Zfp654 |
| **ENSMUSG00000047146** | -0.366146001316094 | 0.420090430673927 | 0.882568531013176 | 9.03513716569045 | 11.4018202048006 | 16.0897331070236 | 21.5723139679377 | 12.0445401540101 | 12.2308223207386 | 22.903717992235 | 27.1381391167127 | Tet1 |
| **ENSMUSG00000047153** | -0.248985907520986 | 0.200871373723028 | 0.812921911423739 | 166.246523848704 | 144.96599974675 | 195.916161950228 | 166.013024883695 | 195.997517051618 | 150.439114545085 | 252.981976005141 | 198.651178334337 | Khnyn |
| **ENSMUSG00000047171** | 0.138091791458752 | 0.784069786224137 | 0.961141387070613 | 15.3597331816738 | 13.0306516626293 | 14.1968233297267 | 11.2551203310979 | 10.949581958191 | 3.66924669622158 | 21.8626399016789 | 11.9407812113536 | Helt |
| **ENSMUSG00000047180** | -0.240004776959178 | 0.41113574799714 | 0.882192126163777 | 64.1494738764022 | 38.2775392589734 | 47.3227444324223 | 72.2203554578784 | 90.881530252985 | 63.6002760678407 | 55.1771387994752 | 52.1052271040884 | Neurl3 |
| **ENSMUSG00000047181** | 0.0650168762407672 | 0.804192917887353 | 0.963461896828986 | 71.3775836089546 | 119.719112150406 | 141.968233297267 | 88.1651092602672 | 102.926070406995 | 127.200552135681 | 87.4505596067155 | 85.7565196088121 | Samd14 |
| **ENSMUSG00000047187** | 0.0671602737687205 | 0.394301277688667 | 0.878965776679486 | 3774.88030782547 | 3760.97183612637 | 4048.93401363806 | 3743.26543678432 | 3630.88137733613 | 3395.27627623703 | 3807.22257716379 | 3794.9973740811 | Rab2a |
| **ENSMUSG00000047193** | -0.00666288135945352 | 0.980089405143151 | 0.996872525500296 | 36.1405486627618 | 63.5244268553176 | 53.9479286529615 | 40.3308478531009 | 45.9882442244021 | 44.0309603546589 | 59.3414511616998 | 45.5920737160773 | Dync2h1 |
| **ENSMUSG00000047205** | 0.0332310344525302 | 0.916139138560614 | 0.986713918026868 | 1129.39214571131 | 1638.60444657563 | 2068.9503865855 | 1133.01544666386 | 1367.60278657805 | 806.011190936674 | 2258.09837841626 | 1401.41350398704 | Dusp18 |
| **ENSMUSG00000047213** | 0.0638455005780545 | 0.754659468618344 | 0.956497786093635 | 1114.03241252963 | 1358.4454358291 | 1292.85737789378 | 962.312788308873 | 1033.64053685323 | 798.67269754423 | 1376.30523571521 | 1313.48593324889 | Ythdf3 |
| **ENSMUSG00000047215** | 0.157403914474744 | 0.312416740151508 | 0.852380381202996 | 8427.97594815605 | 5524.99630495481 | 6995.24808200067 | 8179.65870062542 | 6187.60876457372 | 6740.40618095904 | 6117.37486010786 | 7071.11352825066 | Rpl9 |
| **ENSMUSG00000047216** | -0.46299771580949 | 0.166064790887445 | 0.796071270505327 | 57.8248778604189 | 51.3081909216027 | 46.3762895437739 | 48.7721881014244 | 109.49581958191 | 81.9465095489486 | 37.4788112600209 | 53.1907526687569 | Cdh19 |
| **ENSMUSG00000047221** | 0.166870299555408 | 0.510643386165202 | 0.902394714365738 | 522.230928176908 | 660.491156149521 | 574.498117409607 | 577.762843663027 | 386.520243124141 | 333.901449356164 | 644.427338054249 | 714.275821551878 | Fam185a |
| **ENSMUSG00000047227** | 0.0992423749056732 | 0.733528387925805 | 0.953909681517594 | 28.0089252136404 | 21.1748089517725 | 28.3936466594534 | 27.1998741334867 | 24.0890803080201 | 25.6847268735511 | 22.903717992235 | 24.9670879873757 | Gm527 |
| **ENSMUSG00000047230** | -0.305513390236734 | 0.524418731071496 | 0.907437243979788 | 9.93865088225949 | 17.9171460361152 | 14.1968233297267 | 10.3171936368398 | 18.6142893289246 | 8.56157562451702 | 10.4107809055614 | 27.1381391167127 | Cldn2 |
| **ENSMUSG00000047238** | -0.115025778974151 | 0.572795092071437 | 0.921914696277202 | 97.5794813894569 | 98.5443031986338 | 99.3777633080869 | 88.1651092602672 | 129.205067106653 | 116.192812047017 | 93.6970281500523 | 77.072315091464 | Mageh1 |
| **ENSMUSG00000047242** | 0.0752837824280393 | 0.55501744742121 | 0.916239900758826 | 172.571119864688 | 197.903022126182 | 188.344522841041 | 181.957778686083 | 178.478185918513 | 181.016170346931 | 173.860041122875 | 170.427513652956 | Taf9b |
| **ENSMUSG00000047246** | 0.048227119020143 | 0.77413008949552 | 0.959203117135841 | 149.079763233892 | 160.439898596123 | 200.648436393471 | 159.447538023887 | 163.148771177045 | 133.315963296051 | 171.777884941763 | 178.026192605635 | Hist1h2be |
| **ENSMUSG00000047248** | 0.204957464402599 | 0.236911457923636 | 0.832048466795696 | 193.351935345776 | 244.324718674299 | 219.57753416644 | 200.716312571247 | 203.662224422352 | 139.43137445642 | 207.174540020671 | 193.223550510994 | C2cd3 |
| **ENSMUSG00000047250** | -0.541631096652988 | 0.193957249682889 | 0.809162446871198 | 154.500845533307 | 100.987550385377 | 153.325691961048 | 89.1030359545253 | 318.632834983357 | 218.931719541221 | 111.395355689507 | 75.9867895267955 | Ptgs1 |
| **ENSMUSG00000047260** | 0.10143618497832 | 0.389171947833492 | 0.876536136761342 | 1619.09658009173 | 1257.45788544372 | 1369.5202238743 | 1569.1513594939 | 1327.08933333275 | 1415.10614250946 | 1406.49650034134 | 1271.15043622682 | Emc6 |
| **ENSMUSG00000047264** | 0.0149494838520401 | 0.88215399389998 | 0.978854883720278 | 882.732901087957 | 803.013908709528 | 920.900606654939 | 865.706338800283 | 903.340511550755 | 916.088591823321 | 786.013958369883 | 831.512582536077 | Zfp358 |
| **ENSMUSG00000047281** | -0.442715754707445 | 0.173566235935411 | 0.799889021902745 | 32.5264937964856 | 19.5459774939439 | 28.3936466594534 | 40.3308478531009 | 48.1781606160403 | 50.1463715150282 | 38.5198893505771 | 27.1381391167127 | Sfn |
| **ENSMUSG00000047284** | 0.219701994969141 | 0.252810620122688 | 0.837406774112203 | 1065.2426718349 | 1202.89203160646 | 1219.0338965792 | 1075.80191831411 | 987.652292628825 | 647.010500767072 | 1130.61080634396 | 1151.74262411329 | Neurl4 |
| **ENSMUSG00000047298** | 0.00867031199495725 | 0.965128907396813 | 0.994284908685797 | 225.878429142261 | 284.231089391101 | 242.292451494002 | 207.281799431054 | 178.478185918513 | 217.708637309147 | 257.146288367366 | 300.690581413177 | Kcnv2 |
| **ENSMUSG00000047330** | -0.685256606894489 | 0.216240316071603 | 0.825447777181429 | 30.7194663633475 | 20.3603932228582 | 16.0897331070236 | 12.1930470253561 | 40.5134532453066 | 61.154111603693 | 12.4929370866736 | 14.1118323406906 | Kcne4 |
| **ENSMUSG00000047342** | -0.777155025225799 | 0.165596467661603 | 0.795216960550936 | 16.2632468982428 | 5.7009101024003 | 4.73227444324223 | 3.75170677703264 | 10.949581958191 | 15.9000690169602 | 13.5340151772298 | 11.9407812113536 | Zfp286 |
| **ENSMUSG00000047344** | 0.0578328081023575 | 0.849540776386241 | 0.972640876584422 | 60.535419010126 | 83.8848200781758 | 74.7699362032273 | 32.8274342990356 | 64.6025335533267 | 48.9232892829544 | 61.4236073428121 | 67.3025850094475 | Lancl3 |
| **ENSMUSG00000047368** | -0.0989224836661285 | 0.417440314895764 | 0.882436023575407 | 176.185174730964 | 175.913797445495 | 177.933519065908 | 181.957778686083 | 189.427767876704 | 196.916239363891 | 183.22974393788 | 193.223550510994 | Abhd17b |
| **ENSMUSG00000047371** | -0.0173281316045268 | 0.854425839504373 | 0.973268281968411 | 876.408305071974 | 923.547436588849 | 859.38103889279 | 853.513291774927 | 874.871598459458 | 984.581196819457 | 846.396487622139 | 852.137568264778 | Zfp768 |
| **ENSMUSG00000047379** | -0.0455635839147893 | 0.737023828126485 | 0.954383574136061 | 1193.54161958771 | 1243.61281805218 | 1228.49844546568 | 1216.49092245284 | 1407.02128162754 | 1457.91402063204 | 1116.03571307618 | 1059.47295111646 | B4gat1 |
| **ENSMUSG00000047388** | 0.21143371194116 | 0.146283784588096 | 0.780607990324772 | 533.073092775737 | 636.058684282091 | 632.231865617162 | 583.390403828576 | 458.787484048202 | 436.640356850368 | 572.592949805875 | 590.525907179668 | Atmin |
| **ENSMUSG00000047407** | -0.564658394332517 | 0.0930789609836982 | 0.754058637722175 | 31.6229800799166 | 33.3910448854875 | 32.1794662140472 | 33.7653609932938 | 61.3176589658694 | 68.4926049961361 | 39.5609674411332 | 24.9670879873757 | Tgif1 |
| **ENSMUSG00000047409** | -0.535141154700372 | 0.0489329027644804 | 0.703396849069134 | 100.290022539164 | 93.6578088251478 | 70.0376617599851 | 106.92364314543 | 168.623562156141 | 174.900759186562 | 87.4505596067155 | 107.467030902182 | Ctdspl |
| **ENSMUSG00000047412** | 0.206927858006583 | 0.137176755670674 | 0.777073872865278 | 727.328541838081 | 793.240919962556 | 653.053873167428 | 701.569167305105 | 685.443830582755 | 507.579126310652 | 604.866370613115 | 692.565310258508 | Zbtb44 |
| **ENSMUSG00000047414** | -0.735712946007423 | 0.169138216615367 | 0.79907358295209 | 18.0702743313809 | 29.3189662409158 | 18.9290977729689 | 45.9584080186499 | 50.3680770076785 | 91.7311674055395 | 30.191264626128 | 15.1973579053591 | Flrt2 |
| **ENSMUSG00000047417** | 0.0639602292736546 | 0.704143981440802 | 0.946931219685582 | 306.291149916906 | 395.806044252364 | 383.314229902621 | 330.150196378873 | 329.582416941548 | 262.96267989588 | 389.363205867995 | 371.24974311663 | Rexo1 |
| **ENSMUSG00000047419** | 0.409923572117595 | 0.220768209950555 | 0.829224022772635 | 20485.36649577 | 28805.8843316998 | 29455.569044517 | 18756.6580317747 | 12638.007496144 | 9202.47071412372 | 26368.4258776058 | 25177.6799469214 | Cmya5 |
| **ENSMUSG00000047423** | -0.209199712857607 | 0.268074414597973 | 0.843793744782845 | 775.214768816241 | 743.561560498782 | 747.699362032273 | 801.927323590728 | 1034.73549504905 | 1113.00483118721 | 755.822693743755 | 644.802185413094 | AI837181 |
| **ENSMUSG00000047434** | -0.0307551011149765 | 0.808823217121815 | 0.964157363647118 | 207.80815481088 | 210.119258059897 | 194.96970706158 | 188.52326554589 | 185.047935093427 | 209.14706168463 | 216.544242835676 | 208.420908416353 | Xxylt1 |
| **ENSMUSG00000047446** | 0.00972609046274331 | 0.939972418529389 | 0.989269716906553 | 582.766347187034 | 453.629561005281 | 474.173899212872 | 454.894446715208 | 494.921104510232 | 517.363784167243 | 446.622500848583 | 493.914131924171 | Arl4a |
| **ENSMUSG00000047454** | 0.0832567657926631 | 0.558923897080431 | 0.916574868225118 | 604.450676384691 | 749.262470601182 | 684.286884492827 | 607.776497879289 | 571.568178217569 | 536.933099880424 | 688.152617857606 | 700.163989211187 | Gphn |
| **ENSMUSG00000047459** | -0.012259451699828 | 0.939984576531366 | 0.989269716906553 | 3634.83568175727 | 3207.98355619354 | 3178.19551608148 | 3837.05810621014 | 3347.28720461898 | 4527.85042313743 | 3130.5218183023 | 2972.16899606237 | Dynlrb1 |
| **ENSMUSG00000047466** | 0.138874564349331 | 0.734081977808142 | 0.954182918936468 | 284.606820719249 | 472.361122770311 | 412.654331450723 | 237.295453647315 | 204.757182618171 | 127.200552135681 | 486.183468289716 | 459.177313854779 | 8030462N17 |
| **ENSMUSG00000047473** | -0.0797668670401775 | 0.584324352205065 | 0.922450244597442 | 214.132750826864 | 195.459774939439 | 189.290977729689 | 156.633757941113 | 198.187433443257 | 203.031650524261 | 188.435134390661 | 209.506433981022 | Zfp30 |
| **ENSMUSG00000047485** | 0.465657894044152 | 0.0851748047404539 | 0.743541556158628 | 323.457910531718 | 324.951875836817 | 419.279515671262 | 409.873965390816 | 202.567266226533 | 166.339183562045 | 309.200192895173 | 390.789203280663 | Klhl34 |
| **ENSMUSG00000047492** | -0.5227017138591 | 0.387705723641914 | 0.875951433019341 | 9.03513716569045 | 22.8036404096012 | 7.57163910918757 | 13.1309737196143 | 9.85462376237187 | 11.0077400886647 | 12.4929370866736 | 42.3354970220718 | Inhbe |
| **ENSMUSG00000047496** | -0.146954621678589 | 0.586849606942248 | 0.923058967588378 | 36.1405486627618 | 57.009101024003 | 42.5904699891801 | 51.5859681841989 | 44.893286028583 | 37.9155491942896 | 64.5468416144805 | 59.7039060567679 | Rnf152 |
| **ENSMUSG00000047511** | -0.128571481993961 | 0.644370984781182 | 0.934857223264556 | 57.8248778604189 | 45.6072808192024 | 58.6802030962037 | 46.8963347129081 | 43.7983278327639 | 80.7234273168747 | 40.6020455316893 | 64.046008315442 | Olfr1396 |
| **ENSMUSG00000047514** | 0.0959529687198219 | 0.506607046727462 | 0.901698316485955 | 2392.50432147483 | 2764.12698393523 | 3146.96250475609 | 2727.49082690273 | 2597.2408404829 | 2089.02445238215 | 2915.01865355718 | 2718.15601392994 | Tspyl1 |
| **ENSMUSG00000047539** | 0.0554291511091616 | 0.657437154713647 | 0.938389079274206 | 423.747933070882 | 493.535931722083 | 499.72818120638 | 460.522006880757 | 427.033696369448 | 408.509465512669 | 517.4158110064 | 452.664160466768 | Fbxo28 |
| **ENSMUSG00000047547** | -0.0297372568708369 | 0.864452344942923 | 0.974868041734905 | 2732.22547890479 | 2403.34081602618 | 2405.88832694435 | 3087.65467749787 | 2832.656852584 | 3453.98422337658 | 2304.94689249129 | 2260.06422563983 | Cltb |
| **ENSMUSG00000047554** | 0.0683541192249952 | 0.548828958666601 | 0.914699131434573 | 576.441751171051 | 533.442302438885 | 548.943835416099 | 563.693943249155 | 558.428679867739 | 447.648096939033 | 561.141090809758 | 550.361461286933 | Tmem41b |
| **ENSMUSG00000047557** | -0.51222415922196 | 0.176717928053964 | 0.799889021902745 | 31.6229800799166 | 21.9892246806869 | 20.8220075502658 | 36.5791410760683 | 52.5579933993167 | 53.8156182112498 | 20.8215618111227 | 31.4802413753867 | Lxn |
| **ENSMUSG00000047583** | -0.298947557023459 | 0.407301747208459 | 0.881573350402261 | 32.5264937964856 | 49.679359463774 | 24.6078271048596 | 19.6964605794214 | 30.6588294829347 | 46.4771248188067 | 48.9306702561384 | 30.3947158107182 | Tyw3 |
| **ENSMUSG00000047603** | 0.204739673746502 | 0.379452293773803 | 0.87432013076003 | 52.4037955610046 | 54.56585383726 | 62.4660226507975 | 74.0962088463947 | 63.5075753575076 | 42.8078781225851 | 47.8895921655823 | 56.4473293627624 | Zfp235 |
| **ENSMUSG00000047604** | -0.336509079271225 | 0.212279273505091 | 0.823370267313127 | 75.8951521917998 | 37.4631235300591 | 60.5731128735006 | 65.6548685980713 | 86.5016974697087 | 62.3771938357668 | 88.4916376972716 | 64.046008315442 | Frat2 |
| **ENSMUSG00000047613** | -0.15453868618544 | 0.38805116185086 | 0.876000428751576 | 381.282788392137 | 322.508628650074 | 282.990011705886 | 344.219096792745 | 364.621079207759 | 457.432754795624 | 284.214318721825 | 376.677370939972 | A430005L14 |
| **ENSMUSG00000047617** | -0.0532688790803134 | 0.755545115497248 | 0.956566265296272 | 262.018977805023 | 219.892246806869 | 280.15064703994 | 321.708856130549 | 244.175677667659 | 336.347613820311 | 275.885693997376 | 268.124814473121 | Paxx |
| **ENSMUSG00000047631** | -0.312251256643323 | 0.315093302870117 | 0.853277176422981 | 110.228673421423 | 177.542628903324 | 106.002947528626 | 121.930470253561 | 105.115986798633 | 141.877538920568 | 133.257995591185 | 260.526135520442 | Apof |
| **ENSMUSG00000047632** | -0.215588422198444 | 0.380003657120537 | 0.87432013076003 | 39.754603529038 | 38.2775392589734 | 41.6440151005317 | 38.4549944645846 | 49.2731188118594 | 45.2540425867328 | 55.1771387994752 | 33.6512925047237 | Fgfbp3 |
| **ENSMUSG00000047635** | -0.0576128453501989 | 0.714267589281818 | 0.951176382858794 | 287.317361868956 | 236.99497711407 | 228.095628164276 | 209.15765281957 | 224.466430142915 | 229.939459629886 | 284.214318721825 | 261.61166108511 | 2810006K23 |

| | | | | | | | | | | | | |
|---|---|---|---|---|---|---|---|---|---|---|---|---|
| ENSMUSG00000047638 | -0.36643879330383 | 0.383476668831934 | 0.875126835008975 | 22.5878429142261 | 35.8342920722305 | 22.7149173275627 | 21.5723139679377 | 25.1840385038392 | 18.3462334811079 | 28.1091084450157 | 60.7894316214364 | Nr1h4 |
| ENSMUSG00000047648 | 0.140902135166099 | 0.375310811291655 | 0.873821423608136 | 539.39768879172 | 636.873100011005 | 671.982970940397 | 635.914298707033 | 462.072358635659 | 497.794468454061 | 626.729010514794 | 665.427171141795 | Fbxo30 |
| ENSMUSG00000047649 | 0.120205252842421 | 0.516525405236525 | 0.904619578708276 | 432.783070236573 | 379.517729674077 | 346.402489245332 | 532.742362338636 | 410.609323432161 | 464.771248188067 | 334.18606706852 | 347.368180693922 | Cd3eap |
| ENSMUSG00000047656 | -0.0164542429376969 | 0.911292046213333 | 0.986226496507248 | 340.62467114653 | 300.519403969387 | 273.525462819401 | 314.205442576484 | 333.962249724825 | 359.586176229715 | 268.598147363483 | 282.236646813812 | Trpt1 |
| ENSMUSG00000047658 | -0.368593815536462 | 0.132574608493884 | 0.77303252008843 | 175.281661014395 | 130.306516626293 | 135.343049076728 | 136.937297361692 | 144.534481848121 | 132.092881063977 | 213.421008564008 | 255.098507697099 | Gal3st3 |
| ENSMUSG00000047669 | -0.942978645383641 | 0.0592701361272307 | 0.712406493939829 | 11.7456783153976 | 7.32974156022896 | 19.8755526616174 | 17.8206071909051 | 31.7537876787538 | 46.4771248188067 | 13.5340151772298 | 17.3684090346961 | Msl3l2 |
| ENSMUSG00000047675 | 0.0489214705207092 | 0.777302385463017 | 0.95974290923544 | 7395.25977011763 | 5395.50420405743 | 6196.44015598138 | 7827.93619027861 | 6658.44078877593 | 7925.57286383861 | 5608.28767382591 | 5729.40393032038 | Rps8 |
| ENSMUSG00000047676 | -0.041014458448047 | 0.877571083978632 | 0.977664000726269 | 61.4389327266951 | 35.8342920722305 | 50.1621090983677 | 58.151455044006 | 64.6025335533267 | 55.0387004433237 | 39.5609674411332 | 52.1052271040884 | Rpsa-ps10 |
| ENSMUSG00000047694 | -0.0581664449105378 | 0.638830830252704 | 0.933743052952758 | 336.107102563685 | 377.888898216248 | 374.796135904785 | 320.770929436291 | 325.202584158272 | 369.370834086306 | 399.773986773556 | 373.420794245967 | Yipf6 |
| ENSMUSG00000047710 | -0.0548994484134629 | 0.748790710017124 | 0.956197192463153 | 126.491920319666 | 176.728213174409 | 157.111511515642 | 138.813150750208 | 173.003394939417 | 162.669936865823 | 154.079557402308 | 133.519644454226 | Champ1 |
| ENSMUSG00000047712 | -1.02487491229261 | 0.0112978441509379 | 0.635862253526407 | 19.877301764519 | 16.2883145782866 | 10.4110037751329 | 25.3240207449704 | 36.1336204620302 | 56.2617826753975 | 20.8215618111227 | 33.6512925047237 | Ust |
| ENSMUSG00000047714 | -0.0257169700669464 | 0.6853801980152 | 0.944452327903597 | 2955.39336689735 | 2768.1990625798 | 2868.70476749344 | 2903.82104542327 | 2883.02492959168 | 2890.14331439053 | 2973.31902662833 | 2954.80058702768 | Ppp1r2 |
| ENSMUSG00000047719 | 0.0207860869250954 | 0.907756800739426 | 0.98504921011344 | 337.914129996823 | 478.062032872711 | 394.671688566402 | 285.129715054481 | 384.330326732503 | 362.032340693862 | 340.432535611857 | 388.618152151326 | Ubiad1 |
| ENSMUSG00000047721 | 0.0506180842545621 | 0.831575431763712 | 0.967583158900775 | 764.372604217412 | 519.597235047342 | 568.819388077717 | 822.561710864407 | 684.348872386936 | 885.511536021474 | 505.963952010282 | 508.025964264862 | Bola2 |
| ENSMUSG00000047728 | 0.0233473765524524 | 0.948471898192777 | 0.991217716191233 | 59.631905293557 | 71.6685841444609 | 68.1447519826882 | 57.2135283497478 | 37.2285786578493 | 58.7079471395453 | 41.6431236222455 | 115.065709854862 | BC025446 |
| ENSMUSG00000047731 | -0.173593341537163 | 0.336814209090024 | 0.858570960161243 | 1848.58906410027 | 1869.08409785838 | 1449.02243452077 | 1232.43567625522 | 2166.92226952599 | 1583.89149053565 | 1834.37959555991 | 1632.63044926144 | Wbp1l |
| ENSMUSG00000047735 | -0.242755797467861 | 0.243387532763015 | 0.832048466795696 | 233.106538874814 | 159.625482867208 | 220.523989055088 | 202.592165959763 | 286.879047304603 | 282.531995609062 | 179.065431575656 | 217.105112933702 | Samd9l |
| ENSMUSG00000047746 | 0.0974645867365619 | 0.619986844275121 | 0.929679074487901 | 6554.99201370842 | 8440.6046144681 | 7410.74177811734 | 7074.78105478931 | 5243.75479977766 | 5544.23175799081 | 8018.38345346336 | 8748.2505256635 | Fbxo40 |
| ENSMUSG00000047747 | 0.139671810119752 | 0.514071351721098 | 0.903740169017346 | 418.326850771468 | 561.132437221972 | 418.333060782613 | 410.811892085075 | 326.297542354091 | 313.109051410908 | 510.128264372507 | 491.743080794834 | Rnf150 |
| ENSMUSG00000047749 | -0.181486491257131 | 0.693643160531613 | 0.94469184542218 | 11.7456783153976 | 8.95857301805762 | 14.1968233297267 | 22.5102406621959 | 12.0445401540101 | 13.4539045528125 | 26.0269522639034 | 13.0263067760221 | Zc3hav1l |
| ENSMUSG00000047757 | -0.127182236636602 | 0.756980933158529 | 0.95664317728796 | 14.4562194651047 | 18.7315617650296 | 12.3039135524298 | 11.2551203310979 | 21.8991639163819 | 17.123151249034 | 10.4107809055614 | 13.0263067760221 | Fancb |
| ENSMUSG00000047759 | -1.01167357754426 | 0.115469996218813 | 0.769071703867842 | 2.71054114970714 | 11.4018202048006 | 9.46454888648447 | 10.3171936368398 | 17.5193311331055 | 35.4693847301419 | 9.36970281500523 | 6.51315338801105 | Hs3st3a1 |
| ENSMUSG00000047766 | 0.0479525307347315 | 0.78082180528563 | 0.960742105865155 | 88.5443442237664 | 96.1010560118908 | 80.448665535118 | 90.0409626487835 | 89.7865720571659 | 94.1773318696872 | 80.1630129728225 | 80.3288917854696 | Lrrc49 |
| ENSMUSG00000047767 | -0.17153853733096 | 0.55415953108119 | 0.915797250055649 | 37.9475760958999 | 43.9784493613737 | 46.3762895437739 | 59.0893817382642 | 65.6974917491458 | 67.2695227640623 | 43.7252798033577 | 34.7368180693922 | Atg16l2 |
| ENSMUSG00000047777 | -0.0517473664878749 | 0.817890005394202 | 0.966315773308182 | 124.684892886528 | 129.492100897378 | 146.700507740509 | 150.068271081306 | 201.472308030714 | 144.323703384715 | 112.436433780063 | 112.894658725525 | Phf13 |
| ENSMUSG00000047786 | 0.227058914993205 | 0.426576471536799 | 0.885066977635295 | 61.4389327266951 | 65.1532583131463 | 52.0550188756646 | 50.6480414899407 | 43.7983278327639 | 34.2463024980681 | 74.9576225200418 | 42.3354970220718 | Lix1 |
| ENSMUSG00000047789 | 0.12477306867403 | 0.358695185441891 | 0.866648170843829 | 263.826005238161 | 228.850819824926 | 221.470443943737 | 243.860940507122 | 250.745426842573 | 204.254732756335 | 205.092383839559 | 218.19063849837 | Slc38a9 |
| ENSMUSG00000047793 | -0.0524476699284025 | 0.883733654722215 | 0.979250426961432 | 28.0089252136404 | 38.2775392589734 | 45.4298346551254 | 18.7585338851632 | 22.994122112201 | 29.3539735697726 | 43.7252798033577 | 39.0789203280663 | Sned1 |
| ENSMUSG00000047797 | -0.132980408458246 | 0.707810782946354 | 0.949077302357839 | 103.000563688871 | 166.140808698523 | 92.7525790875478 | 129.433883807626 | 81.0269064906132 | 100.292743030056 | 117.641824232843 | 239.90114979174 | Gjb1 |
| ENSMUSG00000047804 | -0.0518692880845572 | 0.794678194151502 | 0.961804768755116 | 268.343573821006 | 312.735639903102 | 262.16800415562 | 226.040333316217 | 252.935343234211 | 201.808568292187 | 341.473613702413 | 311.545837059862 | Akap10 |
| ENSMUSG00000047810 | -0.217882532429902 | 0.474178739479121 | 0.897549741595057 | 31.6229800799166 | 31.7622134276588 | 26.5007368821565 | 31.8895076047775 | 28.4689130912965 | 23.2385624094033 | 41.6431236222455 | 47.7631248454143 | Ccdc88b |
| ENSMUSG00000047821 | -0.129174658359303 | 0.520770242216749 | 0.905962179970576 | 181.606257030378 | 262.241864710414 | 174.147699511314 | 280.44008158319 | 221.181555555458 | 229.939459629886 | 261.31060072959 | 270.295865602458 | Trim16 |
| ENSMUSG00000047824 | -0.332069305066068 | 0.0580760122366508 | 0.710862211775517 | 318.036828232304 | 254.912123150185 | 260.275094378323 | 249.488500672671 | 366.810995599397 | 419.517205601334 | 296.707255808499 | 281.151121249143 | Pygo2 |
| ENSMUSG00000047832 | -0.270445862487153 | 0.211093773678599 | 0.823123035902316 | 73.1846110420926 | 78.1839099757755 | 58.6802030962037 | 67.5307219865876 | 105.115986798633 | 86.838838477244 | 65.5879197050366 | 78.1578406561325 | Cdca4 |
| ENSMUSG00000047843 | 0.0521249665245834 | 0.768382434431625 | 0.95879680569409 | 3978.17089405351 | 2861.85687140495 | 3115.72949343069 | 4528.3100798784 | 3899.1461353118 | 3713.27765657624 | 3317.91587460241 | 3039.47158107182 | Bri3 |
| ENSMUSG00000047866 | 0.127100859652063 | 0.240530039322271 | 0.832048466795696 | 2446.71514446897 | 2748.65308508586 | 3040.95955722746 | 2747.18728748215 | 2682.64757975679 | 2262.70212933664 | 2602.69522639034 | 2507.56405438425 | Lonp2 |
| ENSMUSG00000047867 | -0.450771096610411 | 0.180743751525478 | 0.802808628271642 | 508.678222428372 | 335.539280312703 | 290.561650815073 | 515.859681841989 | 693.108537953488 | 863.496055844145 | 331.062832796851 | 369.078691987293 | Gimap6 |
| ENSMUSG00000047875 | -0.0659093002374158 | 0.7591517223038 | 0.957357521291862 | 841.171270125781 | 1018.83407687183 | 1054.35074595437 | 855.389145163443 | 885.821180417649 | 651.902829695367 | 1235.75969349013 | 1171.28208427732 | Gpr157 |
| ENSMUSG00000047878 | -0.656937275742747 | 0.122589669997843 | 0.770251097497048 | 66.8600150261093 | 57.8235167529173 | 58.6802030962037 | 54.3997482669734 | 143.439523652302 | 143.100621152642 | 53.094982618363 | 35.8223436340608 | A4galt |
| ENSMUSG00000047879 | 0.150245860447485 | 0.299471926864118 | 0.851785577030345 | 1331.77921822277 | 1451.28882892533 | 1502.02390828509 | 1383.44187403079 | 1195.69434983445 | 1015.1582526213 | 1496.02921612917 | 1399.24245285771 | Usp14 |
| ENSMUSG00000047880 | -0.266456180529221 | 0.522396644504891 | 0.90648515894888 | 39.754603529038 | 29.3189662409158 | 21.7684624389143 | 43.1446279358754 | 54.7479097909548 | 61.154111603693 | 20.8215618111227 | 24.9670879873757 | Cxcr5 |
| ENSMUSG00000047881 | -0.296513586465048 | 0.108324715754734 | 0.762287160109383 | 225.878429142261 | 217.448999620126 | 210.112985279955 | 230.729966787508 | 328.487458745729 | 321.670627035425 | 218.626399016789 | 218.19063849837 | Rell1 |
| ENSMUSG00000047888 | -0.0507666355772744 | 0.710425358794454 | 0.949886733585872 | 923.391018333564 | 890.156391703361 | 760.003275584703 | 826.31341764144 | 1034.73549504905 | 748.526326029202 | 837.026784807134 | 900.986218674861 | Tnrc6b |
| ENSMUSG00000047905 | 0.140785227357487 | 0.681014796911067 | 0.944452327903597 | 23.4913566307952 | 17.1027303072009 | 33.1259211026956 | 30.9515809105193 | 28.4689130912965 | 26.9078091056249 | 17.6983275394543 | 21.7105112933701 | Gm8566 |
| ENSMUSG00000047907 | 0.0247282757067381 | 0.867260006457475 | 0.974868041734905 | 200.580045078328 | 159.625482867208 | 184.558703286447 | 207.281799431054 | 195.997517051618 | 185.908499275227 | 191.558368662329 | 164.999885829613 | Tshz2 |
| ENSMUSG00000047909 | 0.0907504530801055 | 0.607146744613572 | 0.92742133338909 | 137.334084918495 | 119.719112150406 | 130.610774633486 | 134.123517278917 | 100.736154015357 | 111.300483118721 | 150.95632313064 | 125.920965501547 | Ankrd16 |
| ENSMUSG00000047921 | 0.0912152760436292 | 0.404630144935231 | 0.881158207349486 | 1000.18968424193 | 1114.93513288372 | 1170.76469725813 | 1072.05021153708 | 998.601874587016 | 931.988660840281 | 1152.47344624564 | 1006.28219844771 | Trappc9 |
| ENSMUSG00000047945 | -0.616349447120288 | 0.0690626844796728 | 0.724157781586334 | 40.658117245607 | 35.0198763433161 | 53.9479286529615 | 56.2756016554897 | 81.0269064906132 | 101.51582526213 | 35.3966550789086 | 67.3025850094475 | Marcksl1 |
| ENSMUSG00000047953 | 0.269284236219887 | 0.75616243591549 | 0.956566265296272 | 5.42108229941427 | 10.5874044758863 | 27.447191770805 | 1.87585338851632 | 7.66470737073368 | 17.123151249034 | 1.04107809055614 | 11.9407812113536 | Gp5 |
| ENSMUSG00000047963 | -0.311290103471407 | 0.158584793179508 | 0.789230661864335 | 268.343573821006 | 272.014853457386 | 208.220075502658 | 209.15765281957 | 260.600050604945 | 423.186452297555 | 237.365804646799 | 269.21034003779 | Stbd1 |
| ENSMUSG00000047976 | -1.49943636045457 | 0.104507244379538 | 0.761198544536855 | 11.7456783153976 | 12.2162359337149 | 0.946454888648447 | 0.937926694258161 | 20.8042057205628 | 40.3617136584374 | 9.36970281500523 | 3.25657669400552 | Kcna1 |
| ENSMUSG00000047989 | 0.158879141066098 | 0.141291128696801 | 0.779711473022872 | 658.661499378834 | 643.38842584232 | 593.427215182576 | 623.721251681677 | 522.295059405709 | 518.586866399316 | 611.112839156452 | 603.55221395569 | Ino80c |
| ENSMUSG00000047996 | 0.00865352483828798 | 0.956291184799753 | 0.992371505557225 | 118.360296870545 | 151.481325578065 | 138.182413742673 | 124.744250336335 | 134.679858085749 | 125.977469903608 | 123.88829277618 | 145.46042566558 | Prrg1 |
| ENSMUSG00000048000 | 0.226769454625712 | 0.280359076427356 | 0.849093714741928 | 1262.20866204696 | 1463.50506485905 | 1325.03684410783 | 1072.98813823134 | 983.272459845549 | 742.410914868833 | 1298.2243789235 | 1353.65037914163 | Gigyf2 |
| ENSMUSG00000048004 | 0.620516958391648 | 0.287294732728045 | 0.851785577030345 | 14.4562194651047 | 23.6180561385155 | 12.3039135524298 | 9.37926694258161 | 7.66470737073368 | 2.44616446414772 | 18.7394056300105 | 9.76973008201657 | Tmem196 |
| ENSMUSG00000048007 | -0.0670049252118034 | 0.65550688537563 | 0.937625787683528 | 164.439496415566 | 155.553404222637 | 143.861143074564 | 192.274972322923 | 186.142893289246 | 162.669936865823 | 168.654650670094 | 169.341988088287 | Timm8a1 |
| ENSMUSG00000048027 | -0.240238500023903 | 0.222204060028033 | 0.831327997735572 | 133.720030052219 | 187.315617650296 | 191.183887506986 | 168.826804966469 | 254.03030143003 | 172.454594722414 | 207.174540020671 | 170.427513652956 | Rgmb |
| ENSMUSG00000048039 | 0.0173718179897339 | 0.900706391931049 | 0.984802943363903 | 161.728955265859 | 166.955224427437 | 164.68315062483 | 159.447538023887 | 160.958854785407 | 184.685417043153 | 155.120635492864 | 145.46042566558 | Isg20l2 |
| ENSMUSG00000048047 | 0.130505960917956 | 0.425416639274612 | 0.885066977635295 | 176.185174730964 | 199.531853584011 | 227.149173275627 | 165.075098189436 | 183.952976897608 | 150.439114545085 | 192.599446752885 | 173.684090346961 | Zbtb33 |
| ENSMUSG00000048058 | 0.117258869831555 | 0.719088758153045 | 0.951674101666722 | 91.2548853734735 | 53.7514381083457 | 80.448665535118 | 104.109863062656 | 89.7865720571659 | 111.300483118721 | 49.9717483466946 | 53.1907526687569 | Ldlrad3 |
| ENSMUSG00000048065 | 0.250055578097053 | 0.374806914977532 | 0.873700364743909 | 145.465708367616 | 171.027303072009 | 169.415425068072 | 181.019851991825 | 97.4512794278996 | 86.838838477244 | 176.983275394543 | 198.651178334337 | Cyb5r2 |
| ENSMUSG00000048070 | -0.582793159587688 | 0.117867491478972 | 0.769868040577702 | 35.2370349461928 | 25.2468875963442 | 32.1794662140472 | 35.6412143818101 | 30.6588294829347 | 79.5003450848009 | 54.1360607089191 | 28.2236646813812 | Pirt |
| ENSMUSG00000048076 | -0.00014916657568499 | 0.998327749384948 | 0.999262355603273 | 6300.20114563595 | 6312.53631481496 | 6080.97265956627 | 5907.0623204379 | 5905.10955005239 | 5939.28731895066 | 6376.60330465634 | 6381.80479468616 | Arf1 |
| ENSMUSG00000048078 | 0.30560226894204 | 0.441826380791364 | 0.89108031640712 | 18.0702743313809 | 19.5459774939439 | 21.7684624389143 | 20.6343872736795 | 24.0890803080201 | 20.7923979452556 | 10.4107809055614 | 9.76973008201657 | Tenm4 |
| ENSMUSG00000048096 | -0.787293621215974 | 0.0655205461503357 | 0.718339708336358 | 122.87786545339 | 192.202112023782 | 111.681676860517 | 81.59962240046 | 312.063085808443 | 340.016860516533 | 137.42230795341 | 89.0130963028176 | Lmod1 |
| ENSMUSG00000048100 | 0.229669306734665 | 0.185383045843592 | 0.805963874499992 | 593.608511785863 | 483.762942975111 | 520.550188756646 | 632.162591930001 | 410.609323432161 | 415.847958905112 | 463.279750297481 | 611.15089290837 | Taf13 |
| ENSMUSG00000048106 | -0.0190727514307539 | 0.925121872879354 | 0.986713918026868 | 103.90407740544 | 121.347943608235 | 146.700507740509 | 115.364983393754 | 134.679858085749 | 110.077400886647 | 97.8613405122768 | 150.888053488923 | 4632415L05R |
| ENSMUSG00000048109 | 0.227939689343412 | 0.0936291198945225 | 0.755231402633518 | 308.098177350044 | 408.022280186079 | 355.867038131816 | 317.019222659259 | 306.588294829347 | 281.308913376988 | 292.542943446274 | 306.118209236519 | Rbm15 |
| ENSMUSG00000048118 | -0.104058196983432 | 0.288621395046297 | 0.851785577030345 | 435.49361138628 | 457.701639649853 | 441.994432998825 | 429.570425970238 | 477.401773377126 | 440.309603546589 | 471.60837502193 | 506.940438700193 | Arid4a |
| ENSMUSG00000048120 | -0.432640436722286 | 0.195563222077867 | 0.811094945724481 | 282.799793286111 | 270.386021999557 | 374.796135904785 | 291.695201914288 | 602.227007700503 | 572.402484610566 | 232.160414194018 | 239.90114979174 | Entpd1 |
| ENSMUSG00000048126 | -1.24311258084258 | 0.0047035559633227€ | 0.635862253526407 | 163.535982698997 | 197.088606397268 | 141.021778408619 | 145.378637610015 | 580.327843784121 | 623.771938357668 | 137.42230795341 | 191.052499381657 | Col6a3 |
| ENSMUSG00000048142 | 0.210260523816274 | 0.385979465790618 | 0.875126835008975 | 56.0178504272808 | 62.7100111264033 | 38.8046504345863 | 59.0893817382642 | 47.0832024202212 | 40.3617136584374 | 53.094982618363 | 46.6775992807458 | Nat8l |
| ENSMUSG00000048154 | 0.283926063420952 | 0.145613419964115 | 0.780607990324772 | 978.505355044276 | 1114.1207171548 | 978.634354862494 | 889.154506156737 | 944.948922991881 | 534.486935416277 | 842.232175259915 | 930.295408920911 | Kmt2d |
| ENSMUSG00000048163 | -0.435334865805021 | 0.241910686223149 | 0.832048466795696 | 59.631905293557 | 40.7207864457164 | 51.1085639870161 | 30.0136542162612 | 76.6470737073368 | 92.9542496376133 | 40.6020455316893 | 35.8223436340608 | Selplg |
| ENSMUSG00000048170 | 0.0163889814154511 | 0.897411160374478 | 0.984332015045905 | 430.072529086865 | 466.66021266791 | 425.904699891801 | 449.266886549659 | 513.535393839156 | 428.078781225851 | 379.99350305299 | 430.953649173397 | Mcmbp |
| ENSMUSG00000048174 | -0.19900002869064 | 0.593050445114555 | 0.923935529355527 | 23.4913566307952 | 17.1027303072009 | 18.9290977729689 | 34.703287687552 | 38.3235368536684 | 29.3539735697726 | 19.7804837205666 | 20.6249857287016 | Tmem81 |
| ENSMUSG00000048175 | 0.0910435737599497 | 0.620845419805568 | 0.930381707340543 | 1648.00901902194 | 1764.02446882844 | 1894.80268707419 | 1622.61318106662 | 1434.39523652302 | 1191.28209403994 | 2062.37569739171 | 1816.08426969041 | Asb8 |
| ENSMUSG00000048186 | 0.0700180214104529 | 0.735304333082152 | 0.954383574136061 | 117.456783153976 | 143.337168288922 | 198.755526616174 | 130.371810501884 | 130.300025302473 | 124.754387671534 | 151.997401221196 | 154.144630182928 | Bend7 |
| ENSMUSG00000048200 | -0.478002542736218 | 0.166390766647304 | 0.796299608935682 | 173.474633581257 | 155.553404222637 | 173.201244622666 | 269.184961252092 | 344.911831683016 | 408.509465512669 | 147.833088858971 | 173.684090346961 | Cracr2b |
| ENSMUSG00000048217 | -0.0827824755598145 | 0.862342928728555 | 0.974868041734905 | 11.7456783153976 | 18.7315617650296 | 6.62518422053913 | 15.9447538023887 | 8.75966556655277 | 13.4539045528125 | 12.4929370866736 | 21.7105112933701 | Nags |
| ENSMUSG00000048218 | -0.750472982136646 | 0.0091259376981301¢ | 0.635862253526407 | 102.097049972302 | 70.8541684155466 | 92.7525790875478 | 72.2203554578784 | 194.902558855799 | 166.339183562045 | 125.970448957293 | 81.4144173501381 | Amigo2 |
| ENSMUSG00000048222 | 0.0641073412582752 | 0.517806943832994 | 0.905379965991979 | 634.26662903147 | 530.184639523228 | 583.016211407443 | 583.390403828576 | 551.858930692825 | 568.733237914345 | 558.017856538089 | 550.361461286933 | Mfap1b |
| ENSMUSG00000048232 | -0.130971123274563 | 0.595246860179171 | 0.924461858177432 | 151.7903043836 | 175.913797445495 | 133.450139299431 | 115.364983393754 | 118.255485148462 | 118.638976511164 | 200.928071477334 | 193.223550510994 | Fbxo10 |
| ENSMUSG00000048234 | -0.0115463813037095 | 0.932839713410054 | 0.988114361264825 | 159.921927832721 | 178.357044632238 | 189.290977729689 | 171.640585049243 | 166.433645764503 | 168.785348026193 | 173.860041122875 | 195.394601640331 | Rnf149 |
| ENSMUSG00000048240 | 0.0660650011225609 | 0.867764980782005 | 0.974868041734905 | 22.5878429142261 | 24.4324718674299 | 13.2503684410783 | 12.1930470253561 | 15.3294147414674 | 17.123151249034 | 22.903717992235 | 14.1118323406906 | Gng7 |
| ENSMUSG00000048249 | 0.0889551888766418 | 0.513214877593931 | 0.903078589555164 | 777.021796249379 | 942.278998353878 | 963.491076644119 | 793.485983342404 | 777.420319031559 | 708.164612370765 | 876.587752248267 | 905.328320933535 | Crebrf |
| ENSMUSG00000048251 | 0.250969026121555 | 0.546158576719167 | 0.914211323944814 | 39.754603529038 | 13.8450673915436 | 32.1794662140472 | 34.703287687552 | 22.994122112201 | 14.6769867848863 | 26.0269522639034 | 36.9078691987293 | Rit1 |

| | | | | | | | | | | | | |
|---|---|---|---|---|---|---|---|---|---|---|---|---|
| ENSMUSG00000048264 | 0.203916218721219 | 0.433446556042022 | 0.887868762931002 | 876.408305071974 | 1114.1207171548 | 871.68495244522 | 697.817460528072 | 606.60684048378 | 494.125221757839 | 998.393888843335 | 991.084840542347 | Dip2c |
| ENSMUSG00000048271 | 0.299398462243264 | 0.212792366052898 | 0.823370267313127 | 897.189120553062 | 1014.76199822725 | 1049.61847151113 | 860.078778634734 | 650.405168316544 | 479.448234972953 | 1046.28348100892 | 928.124357791574 | Rbm33 |
| ENSMUSG00000048277 | -0.133965773252741 | 0.277086479301428 | 0.847924916831206 | 2596.69842141944 | 2426.14445643578 | 2557.3211091281 | 2500.51256689226 | 2885.21484598332 | 3184.90613232033 | 2712.00842589874 | 2280.68921136853 | Syngr2 |
| ENSMUSG00000048279 | 0.188996822781843 | 0.520503589675698 | 0.905962179970576 | 338.817643713392 | 487.020605890769 | 382.367775013973 | 386.425798034362 | 280.309298129689 | 184.685417043153 | 494.512093014165 | 438.552328126077 | Sacs |
| ENSMUSG00000048280 | -0.268625155833749 | 0.333956734049318 | 0.857961274005048 | 33.4300075130547 | 23.6180561385155 | 45.4298346551254 | 41.2687745473591 | 47.0832024202212 | 37.9155491942896 | 42.6842017128016 | 44.5065481514088 | Zfp738 |
| ENSMUSG00000048285 | 0.226432815806958 | 0.191265884579502 | 0.808824684419004 | 632.459601598332 | 725.644414462667 | 730.663174036601 | 726.893188050075 | 489.446313531136 | 500.240632918209 | 649.632728507029 | 766.381048655966 | Frmd6 |
| ENSMUSG00000048307 | 0.0338618443726665 | 0.748701062009073 | 0.956197192463153 | 672.214205127369 | 768.808448095126 | 809.218929794422 | 728.769041438591 | 729.242158415519 | 705.718447906617 | 786.013958369883 | 688.223207999834 | Ankrd46 |
| ENSMUSG00000048310 | 0.279632031236873 | 0.227670746071502 | 0.832048466795696 | 511.388763578079 | 663.748819065178 | 665.357786719858 | 461.459933575015 | 407.324448844704 | 324.116791499573 | 601.743136341447 | 562.302242498287 | Pskh1 |
| ENSMUSG00000048330 | -0.105232801175553 | 0.612653225267529 | 0.928055903343287 | 57.8248778604189 | 57.8235167529173 | 67.1982970940397 | 46.8963347129081 | 58.0327843784121 | 66.0464405319884 | 63.5057635239243 | 59.7039060567679 | Ric3 |
| ENSMUSG00000048332 | -0.518041313121639 | 0.0359996949254842 | 0.679566075569698 | 501.45011269582 | 460.144886836596 | 425.904699891801 | 507.418341593665 | 834.358145214152 | 930.765578608207 | 464.320828388037 | 485.229927406823 | Lhfp |
| ENSMUSG00000048347 | -0.434393540846408 | 0.413741727865376 | 0.882192126163777 | 8.13162344912141 | 19.5459774939439 | 7.57163910918757 | 6.56548685980713 | 19.7092475247437 | 18.3462334811079 | 8.32862472444909 | 10.8552556466851 | Pcdhb18 |
| ENSMUSG00000048351 | -0.0473352631377181 | 0.753892229017008 | 0.956497786093635 | 406.58117245607 | 368.930325198191 | 312.330113253987 | 347.970803569778 | 325.202584158272 | 329.009120427868 | 416.431236222455 | 412.499714574033 | Coa7 |
| ENSMUSG00000048368 | -0.263164307394141 | 0.53736412726477 | 0.912099403004057 | 9.93865088225949 | 12.2162359337149 | 19.8755526616174 | 13.1309737196143 | 12.0445401540101 | 25.6847268735511 | 13.5340151772298 | 15.1973579053591 | Omd |
| ENSMUSG00000048371 | -0.292743408665929 | 0.227172497489314 | 0.832048466795696 | 412.905768472054 | 445.485403716138 | 565.980023411771 | 577.762843663027 | 650.405168316544 | 376.709327478749 | 823.492769629904 | 600.295637261685 | Pdp2 |
| ENSMUSG00000048376 | -0.503315850027028 | 0.0355106045692693 | 0.679566075569698 | 396.642521573811 | 360.786167909048 | 358.706402797761 | 407.9981120023 | 689.823663366031 | 697.1568722821 | 425.80093903746 | 348.453706258591 | F2r |
| ENSMUSG00000048379 | 0.068374978464528 | 0.606534455119625 | 0.927110332696882 | 383.993329541844 | 375.445651029505 | 392.778778789105 | 357.350070512359 | 330.677375137367 | 306.993640250539 | 394.568596320776 | 405.986561186022 | Socs4 |
| ENSMUSG00000048387 | -0.611208689597 | 0.348537620100789 | 0.861266047293794 | 19.877301764519 | 17.9171460361152 | 17.9826428843205 | 25.3240207449704 | 48.1781606160403 | 61.154111603693 | 6.24646854333682 | 8.68420451734806 | Osr1 |
| ENSMUSG00000048388 | -0.798963767497429 | 0.110991606694414 | 0.764116514278315 | 13.5527057485357 | 6.51532583131463 | 7.57163910918757 | 11.2551203310979 | 20.8042057205628 | 26.9078091056249 | 9.36970281500523 | 10.8552556466851 | Fam171b |
| ENSMUSG00000048406 | -0.0928223698923576 | 0.658376102640584 | 0.938989601151841 | 124.684892886528 | 145.780415475665 | 159.004421292939 | 169.764731660727 | 127.015150715015 | 211.593226148778 | 167.613572579538 | 133.519644454226 | B330016D10 |
| ENSMUSG00000048410 | 0.257532952815046 | 0.379669358433861 | 0.87432013076003 | 147.272735800754 | 221.521078264697 | 255.542819935081 | 180.081925297567 | 157.67398019795 | 91.7311674055395 | 193.640524843441 | 229.045894145055 | Zfp407 |
| ENSMUSG00000048416 | 0.136136251290373 | 0.300545812133979 | 0.851785577030345 | 916.162908601012 | 789.983257046899 | 919.95415176629 | 1022.3400967414 | 896.77076237584 | 890.40386494977 | 710.015257759285 | 822.828378018729 | Mlf1 |
| ENSMUSG00000048429 | 0.170988693563659 | 0.438334732065335 | 0.889346190988072 | 1666.07929335332 | 1843.02279453313 | 1898.58850662878 | 1454.72430279441 | 1270.15150715015 | 1031.05832163826 | 1847.91361073714 | 1945.26181188597 | Timm29 |
| ENSMUSG00000048439 | 0.0577102458089728 | 0.78221074698439 | 0.960742105865155 | 75.8951521917998 | 122.162359337149 | 119.253315969704 | 120.992543559303 | 120.445401540101 | 96.6234963338349 | 106.189965236726 | 97.6973008201657 | Nupl2 |
| ENSMUSG00000048440 | 0.016171695712945 | 0.95481513244805 | 0.992371505557225 | 47.8862269781594 | 52.122606650517 | 57.7337482075553 | 72.2203554578784 | 78.836990098975 | 62.3771938357668 | 36.4377331694648 | 49.9341759747513 | Cyp4f16 |
| ENSMUSG00000048442 | -0.142945796465026 | 0.517398129870059 | 0.905197321821966 | 139.141112351633 | 96.9154717408051 | 154.272146849697 | 196.026679099956 | 166.433645764503 | 159.000690169602 | 162.408182126757 | 158.486732441602 | Smim5 |
| ENSMUSG00000048445 | 0.116644894044634 | 0.744866174356834 | 0.955679952135269 | 26.2018977805023 | 17.1027303072009 | 25.5542819935081 | 15.0068271081306 | 17.5193311331055 | 17.123151249034 | 24.9858741733473 | 17.3684090346961 | Ccdc57 |
| ENSMUSG00000048450 | -1.08617502142098 | 0.0302636271868083 | 0.665956123973759 | 49.6932544112975 | 26.0613033252585 | 35.0188308799925 | 66.5927952923295 | 117.160526952643 | 174.900759186562 | 49.9717483466946 | 34.7368180693922 | Msx1 |
| ENSMUSG00000048458 | -0.057474395135995 | 0.701083753073117 | 0.945920819861844 | 384.896843258413 | 331.467201668132 | 308.544293699394 | 401.432625142493 | 385.425284928322 | 429.301863457925 | 338.350379430744 | 332.170822788563 | Fam212b |
| ENSMUSG00000048478 | 0.0896965731970138 | 0.675115224254526 | 0.944023556644042 | 156.307872966445 | 105.874044758863 | 112.628131749165 | 163.19924480092 | 148.914314631397 | 130.869798831903 | 104.107809055614 | 121.578863242873 | Spata33 |
| ENSMUSG00000048481 | -0.309981381946275 | 0.265284670501984 | 0.842510961779757 | 55.1143367107117 | 48.8649437348597 | 37.8581955459379 | 47.8342614071662 | 84.3117810780705 | 59.9310293716191 | 48.9306702561384 | 42.3354970220718 | Mypop |
| ENSMUSG00000048482 | 0.0833139012507759 | 0.815044747101066 | 0.966177091475091 | 176.185174730964 | 140.893921102179 | 313.276568142636 | 452.080666632434 | 223.371471947096 | 190.800828203522 | 310.241270985729 | 296.348479154503 | Bdnf |
| ENSMUSG00000048486 | 0.0421294857281882 | 0.814761448261301 | 0.966177091475091 | 4933.18489246699 | 5937.90507951437 | 7072.85738286984 | 4970.073552874 | 5786.85406490393 | 4395.75754207345 | 6721.20015263042 | 5349.46998268641 | Fitm2 |
| ENSMUSG00000048489 | -0.814685819156423 | 0.209798237048828 | 0.821566820583092 | 848.399379858333 | 1102.71889695 | 880.203046443056 | 488.659807708502 | 1719.08436743598 | 3105.40578723553 | 465.361906478593 | 550.361461286933 | 8430408G22 |
| ENSMUSG00000048490 | 0.128614724441947 | 0.490162641446952 | 0.899236955296196 | 205.097613661173 | 268.757190541728 | 292.45456059237 | 332.963976461647 | 245.270635863478 | 222.600966237442 | 296.707255808499 | 239.90114979174 | Nrip1 |
| ENSMUSG00000048495 | 0.107536936116992 | 0.746630066719235 | 0.956085046483146 | 40.658117245607 | 26.8757190541728 | 22.7149173275627 | 52.523894878457 | 31.7537876787538 | 28.1308913376988 | 38.5198893505771 | 33.6512925047237 | Tyw5 |
| ENSMUSG00000048497 | -0.359710750921793 | 0.118513635995862 | 0.769868040577702 | 60.535419010126 | 49.679359463774 | 67.1982970940397 | 54.3997482669734 | 88.6916138613469 | 86.838838477244 | 64.5468416144805 | 57.5328549274309 | Mmgt2 |
| ENSMUSG00000048503 | 0.420534121124776 | 0.0248660284602302 | 0.655303666217042 | 104.807591122009 | 142.522752560008 | 137.235958854025 | 157.571684635371 | 106.210944994452 | 94.1773318696872 | 91.61487196894 | 112.894658725525 | Tmem136 |
| ENSMUSG00000048534 | -0.675712413340417 | 0.268185566728666 | 0.843793744782845 | 16.2632468982428 | 12.2162359337149 | 16.0897331070236 | 6.56548685980713 | 31.7537876787538 | 35.4693847301419 | 8.32862472444909 | 6.51315338801105 | Jaml |
| ENSMUSG00000048537 | 0.080766846675268 | 0.641288759147644 | 0.934544499347759 | 2338.29349848069 | 2724.22061321843 | 2262.02718386979 | 2032.48714645744 | 2054.14157535663 | 1682.96115133363 | 2486.09448024805 | 2623.71528980378 | Phldb1 |
| ENSMUSG00000048546 | -0.028486485524674 | 0.909149654977606 | 0.98549154890268 | 1511.57844782001 | 1800.67317662958 | 1341.12657721485 | 833.816831195505 | 1587.68938393769 | 1587.56073723187 | 1409.61973461301 | 1012.79535183572 | Tob2 |
| ENSMUSG00000048550 | 0.0232614975922394 | 0.883451056901955 | 0.97910597465636 | 146.369222084185 | 155.553404222637 | 169.415425068072 | 145.378637610015 | 134.679858085749 | 132.092881063977 | 168.654650670094 | 170.427513652956 | Thnsl1 |
| ENSMUSG00000048572 | 0.111059118848991 | 0.550835049732779 | 0.914699131434573 | 820.390454644693 | 785.911178402327 | 500.674636095028 | 707.196727470654 | 628.506004400162 | 561.394744521902 | 761.028084196536 | 654.57191549511 | Tmem252 |
| ENSMUSG00000048578 | 0.095159113920111 | 0.580339535866451 | 0.922450244597442 | 2557.84733160697 | 3094.77976987445 | 2836.5253012794 | 2250.08613952533 | 2125.31385808487 | 2096.3629457746 | 2922.30620019108 | 2909.2085133116 | Mlec |
| ENSMUSG00000048581 | 0.0344835951669921 | 0.877645242984582 | 0.977664000726269 | 204.194099944604 | 201.975100770753 | 142.914688185915 | 191.337045628665 | 169.71852035196 | 122.308223207386 | 222.790711379013 | 207.335382851685 | E130311K13 |
| ENSMUSG00000048582 | 0.0779702638854596 | 0.791646846758098 | 0.961804768755116 | 154.500845533307 | 207.676010873154 | 205.380710836713 | 160.385464718146 | 131.394983498292 | 96.6234963338349 | 241.530117009024 | 219.276164063039 | Gja3 |
| ENSMUSG00000048583 | -0.616148217819973 | 0.102933614419217 | 0.759800953500741 | 456.274426867368 | 373.002403842763 | 338.830850136144 | 348.908730264036 | 971.227919691539 | 739.964750404685 | 377.911346871878 | 236.644573097735 | Igf2 |
| ENSMUSG00000048612 | -0.602363673229807 | 0.0913169518444299 | 0.753823363185192 | 64.1494738764022 | 95.2866402829764 | 72.8770264259304 | 79.7237690119437 | 156.579022002131 | 182.239252579005 | 69.7522320672611 | 66.217059444779 | Myof |
| ENSMUSG00000048620 | -0.106092856379217 | 0.849090552212665 | 0.972640876584422 | 18.0702743313809 | 12.2162359337149 | 10.4110037751329 | 9.37926694258161 | 3.28487458745729 | 14.6769867848863 | 23.9447960827911 | 11.9407812113536 | Olfr1336 |
| ENSMUSG00000048644 | -0.624180577992787 | 0.172448581714117 | 0.799889021902745 | 8.13162344912141 | 7.32974156022896 | 8.51809399783602 | 9.37926694258161 | 13.1394983498292 | 18.3462334811079 | 11.4518589961175 | 8.68420451734806 | Ctxn1 |
| ENSMUSG00000048647 | -0.0401199385727494 | 0.908123775106168 | 0.98504921011344 | 23.4913566307952 | 22.8036404096012 | 16.0897331070236 | 15.0068271081306 | 22.994122112201 | 22.0154801773295 | 19.7804837205666 | 15.1973579053591 | Exd1 |
| ENSMUSG00000048652 | 0.0589383161205229 | 0.850507277044662 | 0.972692784625411 | 21.6843291976571 | 26.0613033252585 | 27.447191770805 | 22.5102406621959 | 19.7092475247437 | 22.0154801773295 | 31.2323427166841 | 20.6249857287016 | Samd13 |
| ENSMUSG00000048661 | 0.229606007337 | 0.329219486107642 | 0.855977678491063 | 332.493047697409 | 452.000729547452 | 396.564598343699 | 277.626301500416 | 284.689130912965 | 210.370143916704 | 369.582722147429 | 378.848422069309 | Lemd3 |
| ENSMUSG00000048668 | -0.064054951909308 | 0.749035358993275 | 0.956197192463153 | 97.5794813894569 | 98.5443031986338 | 88.966759532954 | 112.551203310979 | 124.825234323377 | 108.854318654574 | 106.189965236726 | 75.9867895267955 | Rhno1 |
| ENSMUSG00000048677 | -0.599634844903244 | 0.189123515634591 | 0.808677608936066 | 6.32459601598331 | 14.6594831204579 | 17.9826428843205 | 14.0689004138724 | 10.949581958191 | 29.3539735697726 | 22.903717992235 | 17.3684090346961 | Tpcn2 |
| ENSMUSG00000048696 | -0.112168195347119 | 0.45967923994416 | 0.894774044236998 | 169.86057871498 | 177.542628903324 | 231.881447718869 | 187.585338851632 | 221.181555555458 | 214.039390612925 | 196.76375911511 | 196.480127205 | Mex3d |
| ENSMUSG00000048701 | 0.0726749820744281 | 0.631930690854871 | 0.931825800800958 | 739.977733870048 | 833.147290679358 | 678.608155160936 | 639.666005484066 | 697.488370736765 | 560.171662289828 | 688.152617857606 | 803.288917854696 | Ccdc6 |
| ENSMUSG00000048706 | -0.323314091460132 | 0.248021290268666 | 0.834346263461314 | 49.6932544112975 | 56.1946852950887 | 30.2865564367503 | 48.7721881014244 | 45.9882442244021 | 74.6080161565054 | 47.8895921655823 | 64.046008315442 | Lurap1l |
| ENSMUSG00000048707 | -0.187383014852094 | 0.398166220614369 | 0.880459952732461 | 144.562194651047 | 141.708336831093 | 133.450139299431 | 114.427056699496 | 187.237851485066 | 188.354663739374 | 116.600746142287 | 117.236760984199 | Tprn |
| ENSMUSG00000048720 | -0.0716976931455968 | 0.75405817844787 | 0.956497786093635 | 171.667606148119 | 217.448999620126 | 214.845259723197 | 155.695831246855 | 182.858018701789 | 135.762127760198 | 220.708555197901 | 258.355084391105 | Tbc1d12 |
| ENSMUSG00000048728 | -0.211589880838031 | 0.682286108258576 | 0.944452327903597 | 4.51756858284523 | 11.4018202048006 | 8.51809399783602 | 15.9447538023887 | 7.66470737073368 | 14.6769867848863 | 9.36970281500523 | 15.1973579053591 | Zfp454 |
| ENSMUSG00000048731 | -0.0402828076336074 | 0.862871491757894 | 0.974868041734905 | 211.422209677157 | 158.811067138294 | 178.879973954556 | 217.598993067893 | 215.706764576362 | 269.078091056249 | 139.504464134522 | 164.999885829613 | Ggnbp1 |
| ENSMUSG00000048732 | -0.147910010879147 | 0.60897195040964 | 0.927932112008768 | 28.9124389302094 | 28.5045505120015 | 29.3401015481019 | 21.5723139679377 | 33.943704070392 | 22.0154801773295 | 33.3144988977964 | 30.3947158107182 | Klhl11 |
| ENSMUSG00000048755 | -0.0316306595645368 | 0.801953438691176 | 0.963461896828986 | 968.566704162016 | 780.210268299927 | 767.57491469389 | 855.389145163443 | 868.301849284544 | 757.087901653719 | 880.752064610492 | 938.979613438259 | Mcat |
| ENSMUSG00000048756 | -0.0199730790921255 | 0.940552187068305 | 0.989492530362702 | 2172.04697463198 | 3145.27354506714 | 2707.80743642321 | 1474.42076337383 | 2325.69120791976 | 1690.29964472607 | 3248.16364253515 | 2367.53125654201 | Foxo3 |
| ENSMUSG00000048758 | 0.0464193745523763 | 0.762155413949026 | 0.957693025126829 | 4085.68902632522 | 3396.11358957275 | 3866.26822012891 | 4481.41374516549 | 4026.16128602682 | 4615.91234384675 | 3249.2047206257 | 3437.85946330516 | Rpl29 |
| ENSMUSG00000048772 | -0.085634860046315 | 0.637685874270829 | 0.933051384559339 | 564.696072855653 | 412.09435883065 | 368.170951684246 | 524.301022090312 | 520.105143014071 | 577.294813538862 | 432.047407580797 | 454.835211596105 | Tmem53 |
| ENSMUSG00000048776 | -0.743249467853396 | 0.252916565518011 | 0.837406774112203 | 29.8159526467785 | 22.8036404096012 | 8.51809399783602 | 9.37926694258161 | 39.4184950494875 | 47.7002070508805 | 27.0680303544596 | 4.34210225867403 | Pthlh |
| ENSMUSG00000048779 | -0.52191744091041 | 0.172843107235627 | 0.799889021902745 | 37.0440623793308 | 51.3081909216027 | 50.1621090983677 | 52.523894878457 | 76.6470737073368 | 119.862058743238 | 38.5198893505771 | 40.1644458927348 | P2ry6 |
| ENSMUSG00000048787 | 0.0352755073351564 | 0.831214035743975 | 0.967583158900775 | 226.78194285883 | 216.634583891211 | 225.25626349833 | 226.040333316217 | 266.074841584041 | 173.677676954488 | 193.640524843441 | 238.815624227072 | Dcun1d3 |
| ENSMUSG00000048796 | -0.0930270997421177 | 0.65085740250294 | 0.936463929813731 | 108.421645988285 | 115.647033505835 | 135.343049076728 | 135.061443973175 | 135.774816281568 | 96.6234963338349 | 168.654650670094 | 124.835439936878 | Cyb561d1 |
| ENSMUSG00000048799 | 0.2119043213381 | 0.367209883218863 | 0.870400218073379 | 643.30176619716 | 908.073537739476 | 854.648764449547 | 701.569167305105 | 605.511882287961 | 426.855698993777 | 870.34128370493 | 779.407355431988 | Cep120 |
| ENSMUSG00000048807 | -0.330286656770903 | 0.184652859280937 | 0.805963874499992 | 158.114900399583 | 148.223662662408 | 144.807597963212 | 226.978260010475 | 246.365594059297 | 266.631926592101 | 200.928071477334 | 138.947272277569 | Slc35e4 |
| ENSMUSG00000048814 | 0.139429155007517 | 0.579094024474156 | 0.922450244597442 | 181.606257030378 | 193.016527752696 | 206.327165725361 | 147.254490998531 | 137.964732673206 | 108.854318654574 | 235.283648465687 | 178.026192605635 | Lonrf2 |
| ENSMUSG00000048827 | -0.197864264138787 | 0.686416594194005 | 0.944452327903597 | 5.42108229941427 | 11.4018202048006 | 17.036187995672 | 10.3171936368398 | 9.85462376237187 | 8.56157562451702 | 14.5750932677859 | 17.3684090346961 | Pkd1l3 |
| ENSMUSG00000048832 | 0.207984022485971 | 0.0373002190757447 | 0.679566075569698 | 710.161781223269 | 697.95427967958 | 746.752907143624 | 770.03781598595 | 652.595084708182 | 568.733237914345 | 673.57752458982 | 635.032455331077 | Vps37c |
| ENSMUSG00000048833 | -0.0155255014278804 | 0.91819324489057 | 0.986713918026868 | 292.738444168371 | 404.764617270421 | 396.564598343699 | 304.826175633902 | 356.956371837026 | 346.132271676902 | 345.637926064637 | 365.822115293287 | Slc39a9 |
| ENSMUSG00000048856 | 0.051267855255511 | 0.834287752036735 | 0.968512902280839 | 211.422209677157 | 239.438224300813 | 233.774357496166 | 261.681547698027 | 155.484063806312 | 218.931719541221 | 194.681602933998 | 344.111603999917 | Slc25a47 |
| ENSMUSG00000048865 | -0.516518111972343 | 0.0875230984720854 | 0.74812051555524 | 28.9124389302094 | 43.9784493613737 | 56.7872933189068 | 49.7101147956825 | 50.3680770076785 | 66.0464405319884 | 49.9717483466946 | 90.0986218674861 | Arhgap30 |
| ENSMUSG00000048874 | 0.217934867570969 | 0.231796392224937 | 0.832048466795696 | 931.522641782685 | 1149.14059349812 | 1003.24218196735 | 873.209752354348 | 835.453103409971 | 610.318033804856 | 1004.64035738667 | 950.920394649613 | Phf3 |
| ENSMUSG00000048878 | -0.344062226793019 | 0.0609607545611595 | 0.71282992338528 | 612.582299833812 | 547.287369830429 | 497.835271429083 | 622.783324987419 | 810.269064906132 | 907.527016198804 | 574.675105986987 | 603.55221395569 | Hexim1 |
| ENSMUSG00000048895 | -0.822886549775716 | 0.126380009666385 | 0.771949105062466 | 7.22810973255236 | 3.25766291565731 | 10.4110037751329 | 10.3171936368398 | 18.6142893289246 | 18.3462334811079 | 10.4107809055614 | 7.59867895267955 | Cdk5r1 |
| ENSMUSG00000048897 | 0.110590837806523 | 0.465590640880047 | 0.895328112556867 | 587.283915769879 | 530.184639523228 | 550.836745193396 | 519.611388619021 | 529.959766776443 | 381.601656407044 | 574.675105986987 | 538.42068007558 | Zfp710 |
| ENSMUSG00000048911 | -0.207022355461481 | 0.48882315743293 | 0.899236955296196 | 41.5616309621761 | 49.679359463774 | 38.8046504345863 | 34.703287687552 | 49.2731188118594 | 25.6847268735511 | 58.3003730711437 | 56.4473293627624 | Rnf24 |
| ENSMUSG00000048915 | 0.320050829532093 | 0.0532650996527777 | 0.703396849069134 | 318.940341948873 | 329.838370210303 | 232.827902607518 | 257.929840920994 | 199.282391639076 | 226.270212933664 | 243.612273190136 | 244.243252050414 | Efna5 |
| ENSMUSG00000048916 | 0.286602010386868 | 0.546790636362013 | 0.914451747770672 | 14.4562194651047 | 17.1027303072009 | 10.4110037751329 | 14.0689004138724 | 9.85462376237187 | 4.89232892829544 | 11.4518589961175 | 19.5394601640331 | Gm5540 |
| ENSMUSG00000048920 | 0.0677567370513238 | 0.650709532575985 | 0.936463929813731 | 1383.27950006721 | 1526.21507598545 | 1525.6852805013 | 1264.32518386 | 1273.43638173761 | 1089.76626877781 | 1566.82252628699 | 1506.70948375989 | Fkrp |
| ENSMUSG00000048921 | -0.357108751495476 | 0.27132041537963 | 0.844129257012535 | 19.877301764519 | 26.0613033252585 | 38.8046504345863 | 25.3240207449704 | 33.943704070392 | 48.9232892829544 | 29.1501865355718 | 29.3091902460497 | Zfp689 |

| | | | | | | | | | | | | |
|---|---|---|---|---|---|---|---|---|---|---|---|---|
| **ENSMUSG00000048930** | -0.143571044089632 | 0.271434915192964 | 0.844129257012535 | 377.668733525861 | 370.55915665602 | 294.347470369667 | 316.081295965 | 371.190828382674 | 403.617136584374 | 370.623800237985 | 357.137910775939 | Tada3 |
| **ENSMUSG00000048949** | 0.140195613629255 | 0.66774261095135 | 0.942024383071524 | 28.0089252136404 | 20.3603932228582 | 19.8755526616174 | 31.8895076047775 | 17.5193311331055 | 23.2385624094033 | 27.0680303544596 | 22.7960368580387 | Gm6206 |
| **ENSMUSG00000048960** | -0.0815700865880832 | 0.675733900268303 | 0.94409858171788 | 387.60738440812 | 337.168111770532 | 423.065335225856 | 338.591536627196 | 542.004306930453 | 336.347613820311 | 349.802238426862 | 344.111603999917 | Prex2 |
| **ENSMUSG00000048965** | -0.280562209549504 | 0.408794977324492 | 0.882113132676277 | 29.8159526467785 | 31.7622134276588 | 28.3936466594534 | 41.2687745473591 | 47.0832024202212 | 57.4848649074714 | 21.8626399016789 | 33.6512925047237 | Mrgpre |
| **ENSMUSG00000048967** | -0.303814554612484 | 0.174235612493249 | 0.799889021902745 | 240.334648607366 | 189.758864837039 | 210.112985279955 | 316.081295965 | 304.398378437709 | 381.601656407044 | 232.160414194018 | 262.697186649779 | Yjefn3 |
| **ENSMUSG00000048970** | -0.166256140852288 | 0.374164061270138 | 0.873700364743909 | 160.82544154929 | 130.306516626293 | 141.021778408619 | 147.254490998531 | 192.712642464161 | 188.354663739374 | 141.586620315635 | 128.092016630884 | C1galt1c1 |
| **ENSMUSG00000048997** | -0.378249614270893 | 0.272909891501174 | 0.844738346035895 | 21.6843291976571 | 16.2883145782866 | 19.8755526616174 | 27.1998741334867 | 18.6142893289246 | 25.6847268735511 | 38.5198893505771 | 27.1381391167127 | Atxn7l2 |
| **ENSMUSG00000049001** | -0.47558161317513 | 0.0949009396616084 | 0.757085157577544 | 52.4037955610046 | 70.0397526866323 | 70.9841166486335 | 52.523894878457 | 53.6529515951357 | 124.754387671534 | 87.4505596067155 | 77.072315091464 | Ndnf |
| **ENSMUSG00000049037** | -0.194275087774403 | 0.480051912002579 | 0.897732059433541 | 34.3335212296237 | 38.2775392589734 | 36.9117406572894 | 42.2067012416173 | 53.6529515951357 | 47.7002070508805 | 27.0680303544596 | 45.5920737160773 | Clec4a1 |
| **ENSMUSG00000049038** | 0.0973231121004891 | 0.644094656238961 | 0.934847999069355 | 361.405486627618 | 294.818493866987 | 282.990011705886 | 294.508981997063 | 254.03030143003 | 200.585486060113 | 370.623800237985 | 326.743194965221 | Mterf2 |
| **ENSMUSG00000049044** | 0.242660254034755 | 0.19333096906797 | 0.809022817984525 | 896.285606836493 | 1127.96578454635 | 830.987392233336 | 871.333898965832 | 687.633746974393 | 606.648787108634 | 923.436266323293 | 931.380934485579 | Rapgef4 |
| **ENSMUSG00000049047** | -0.222091333337239 | 0.405731140665122 | 0.881158207349486 | 89.4478579403355 | 95.2866402829764 | 159.004421292939 | 150.068271081306 | 142.344565456483 | 194.470074899744 | 115.559668051731 | 123.74991437221 | Armcx3 |
| **ENSMUSG00000049076** | -0.327898790292316 | 0.0931479639112567 | 0.754058637722175 | 263.826005238161 | 303.777066885045 | 356.813493020464 | 258.867767615252 | 491.636229922775 | 357.140011765567 | 321.693129981846 | 314.802413753867 | Acap2 |
| **ENSMUSG00000049086** | -0.111306446275239 | 0.614573607809142 | 0.928075610917191 | 155.404359249876 | 111.574954861263 | 130.610774633486 | 173.51643843776 | 200.377349834895 | 145.546785616789 | 150.95632313064 | 119.407812113536 | Bmyc |
| **ENSMUSG00000049090** | -0.0443255629272175 | 0.835948183632255 | 0.968924788429433 | 1697.70227343324 | 1767.28213174409 | 1555.0253820494 | 1295.27676477052 | 1293.14562926235 | 1204.73599859275 | 1891.6388905405 | 2122.20247892693 | Zadh2 |
| **ENSMUSG00000049091** | -0.0329045440310773 | 0.837323118508611 | 0.968924788429433 | 743.591788736324 | 860.837425462445 | 731.609628925249 | 914.478526901707 | 700.773245324222 | 783.995710759344 | 797.465817366001 | 1043.19006764644 | Sephs2 |
| **ENSMUSG00000049092** | 0.475070052235818 | 0.245842013266307 | 0.832048466795696 | 59.631905293557 | 97.7298874697194 | 75.7163910918757 | 67.5307219865876 | 52.5579933993167 | 20.7923979452556 | 98.902418602833 | 43.4210225867403 | Gpr137c |
| **ENSMUSG00000049097** | -0.562977906946052 | 0.107602207654736 | 0.762287160109383 | 22.5878429142261 | 21.9892246806869 | 42.5904699891801 | 55.3376749612315 | 56.937826182593 | 48.9232892829544 | 61.4236073428121 | 42.3354970220718 | Ankrd34a |
| **ENSMUSG00000049103** | 0.135575589643842 | 0.704190356976311 | 0.946931219685582 | 21.6843291976571 | 30.9477976987445 | 57.7337482075553 | 36.5791410760683 | 41.6084114411257 | 36.6924669622158 | 24.9858741733473 | 30.3947158107182 | Ccr2 |
| **ENSMUSG00000049106** | 0.135770085824725 | 0.536930438495915 | 0.91199738788281 | 1349.84949255415 | 1528.65832317219 | 1248.3739981273 | 1221.18055592413 | 1024.88087128667 | 813.349684329117 | 1448.13962396359 | 1580.52522215735 | Dcaf5 |
| **ENSMUSG00000049107** | -0.574653835734075 | 0.117137522626483 | 0.769868040577702 | 27.1054114970714 | 21.1748089517725 | 22.7149173275627 | 30.9515809105193 | 47.0832024202212 | 55.0387004433237 | 19.7804837205666 | 30.3947158107182 | Ntf3 |
| **ENSMUSG00000049115** | 0.314998664768624 | 0.111984254881659 | 0.764116514278315 | 470.730646332472 | 535.885549625628 | 593.427215182576 | 458.646153492241 | 358.051330032845 | 302.101311322243 | 502.840717738614 | 490.657555230165 | Agtr1a |
| **ENSMUSG00000049119** | 0.258180494454188 | 0.358877800334159 | 0.866754558180356 | 153.597331816738 | 118.904696421492 | 168.468970179424 | 216.661066373635 | 94.1664048404423 | 144.323703384715 | 111.395355689507 | 199.736703899005 | Fam110b |
| **ENSMUSG00000049124** | 0.359637435859647 | 0.192369509842879 | 0.808824684419004 | 62.3424464432641 | 41.5352021746308 | 60.5731128735006 | 78.7858423176855 | 56.937826182593 | 53.8156182112498 | 36.4377331694648 | 42.3354970220718 | Gm8186 |
| **ENSMUSG00000049130** | -0.594249370452092 | 0.104455876807372 | 0.761198544536855 | 44.2721721118832 | 53.7514381083457 | 76.6628459805242 | 53.4618215727152 | 99.6411958195378 | 139.43137445642 | 61.4236073428121 | 44.5065481514088 | C5ar1 |
| **ENSMUSG00000049134** | 0.343666113972564 | 0.0794838510656045 | 0.739855948845396 | 34293.7666260947 | 37430.5469009025 | 29048.5934423981 | 38383.712035821 | 23482.4734675364 | 20081.7871684207 | 33417.5656287614 | 32677.5760732161 | Nrap |
| **ENSMUSG00000049152** | -0.583583598676425 | 0.101201672133523 | 0.759800953500741 | 65.9565013095403 | 116.461449234749 | 71.930571537282 | 85.3513291774927 | 78.836990098975 | 127.200552135681 | 84.3273253350471 | 219.276164063039 | Ugt3a2 |
| **ENSMUSG00000049164** | -0.129814561265295 | 0.573565614286004 | 0.922207298536842 | 118.360296870545 | 149.852494120236 | 133.450139299431 | 135.999370667433 | 131.394983498292 | 95.400414101761 | 172.818963032319 | 187.795922687652 | Zfp518a |
| **ENSMUSG00000049184** | 0.016623187576788 | 0.930054145350857 | 0.987529878729394 | 64.1494738764022 | 75.7406627890326 | 63.4124775394459 | 63.779015209555 | 68.9823663366031 | 63.6002760678407 | 65.5879197050366 | 66.217059444779 | Purg |
| **ENSMUSG00000049191** | -0.103410422357898 | 0.734077669706876 | 0.954182918936468 | 36.1405486627618 | 38.2775392589734 | 32.1794662140472 | 28.1378008277448 | 45.9882442244021 | 44.0309603546589 | 21.8626399016789 | 33.6512925047237 | Rtl5 |
| **ENSMUSG00000049225** | -0.0302399850873504 | 0.860261882575439 | 0.974868041734905 | 461.695509166782 | 539.9576282702 | 450.512526996661 | 477.404687377404 | 482.876564356222 | 365.701587390084 | 515.333654825288 | 605.723265085027 | Pdp1 |
| **ENSMUSG00000049232** | -0.0606435522271404 | 0.652063049006999 | 0.936720465895231 | 216.843291976571 | 233.737314198412 | 243.238906382651 | 255.11606083822 | 249.650468646754 | 211.593226148778 | 272.762459725708 | 254.012982132431 | Tigd2 |
| **ENSMUSG00000049233** | -0.19366970719065 | 0.497964562089591 | 0.900927059989498 | 39.754603529038 | 28.5045505120015 | 34.0723759913441 | 48.7721881014244 | 30.6588294829347 | 41.5847958905112 | 47.8895921655823 | 52.1052271040884 | Apoo-ps |
| **ENSMUSG00000049265** | 0.0202777641136084 | 0.876815168076626 | 0.977515813184586 | 7512.71655327161 | 7947.88309847493 | 9168.3085063375 | 7676.92999250305 | 6871.95763696065 | 7326.26257012242 | 8839.79406691216 | 8815.55311067295 | Kcnk3 |
| **ENSMUSG00000049281** | -0.707778942950487 | 0.163747770327983 | 0.793968226938511 | 9.93865088225949 | 8.14415728914329 | 7.57163910918757 | 9.37926694258161 | 16.4243729372865 | 23.2385624094033 | 12.4929370866736 | 5.42762782334254 | Scn3b |
| **ENSMUSG00000049285** | -0.151012434928124 | 0.624517638730011 | 0.931259048662274 | 24.3948703473642 | 24.4324718674299 | 18.9290977729689 | 31.8895076047775 | 35.0386622662111 | 24.4616446414772 | 24.9858741733473 | 26.0526135520442 | Mblac1 |
| **ENSMUSG00000049287** | 0.141149423094597 | 0.365130704296532 | 0.86924083181804 | 858.338030740593 | 895.857301805761 | 940.776159316556 | 834.754757889763 | 878.156473046916 | 612.764198269004 | 951.545374768309 | 756.61131857395 | Iba57 |
| **ENSMUSG00000049288** | 0.180372621501417 | 0.152037916285369 | 0.783412422888014 | 950.496429830635 | 1168.68657099206 | 1214.30162213596 | 1158.33946740883 | 1096.05315401492 | 979.688867891162 | 1009.84574783945 | 878.190181816823 | Lix1l |
| **ENSMUSG00000049295** | -0.187956606425621 | 0.0901874791187502 | 0.753823363185192 | 405.677658739501 | 460.95930256551 | 435.369248778286 | 436.135912830045 | 457.692525852383 | 540.602346576646 | 519.497967187512 | 463.519416113453 | Zfp219 |
| **ENSMUSG00000049299** | -0.127688540394275 | 0.380270177840981 | 0.87432013076003 | 599.933107801846 | 512.267493487113 | 488.370722542599 | 604.962717796514 | 616.461464246152 | 695.933790050026 | 596.537745888666 | 501.51281087685 | Trappc1 |
| **ENSMUSG00000049300** | -0.111164439062968 | 0.692728505680437 | 0.94469184542218 | 41.5616309621761 | 48.0505280059454 | 63.4124775394459 | 78.7858423176855 | 62.4126171616885 | 53.8156182112498 | 82.2451691539348 | 51.0197015394199 | Prmt6 |
| **ENSMUSG00000049303** | 0.258410956103266 | 0.40531761916056 | 0.881158207349486 | 163.535982698997 | 136.821842457607 | 236.613722162112 | 188.52326554589 | 195.997517051618 | 102.738907494204 | 212.379930473452 | 94.4407241261602 | Syt12 |
| **ENSMUSG00000049305** | -0.136306312997584 | 0.342157570212245 | 0.859912121266038 | 416.51982333833 | 410.465527372822 | 500.674636095028 | 438.011766218561 | 548.574056105368 | 527.148442023833 | 461.197594116369 | 403.815510056685 | Ccdc71 |
| **ENSMUSG00000049307** | 0.231286267982229 | 0.62162087361841 | 0.93078489223231 | 23.4913566307952 | 31.7622134276588 | 7.57163910918757 | 27.1998741334867 | 13.1394983498292 | 15.9000690169602 | 27.0680303544596 | 20.6249857287016 | Fut4 |
| **ENSMUSG00000049313** | 0.231461654680733 | 0.216136694252683 | 0.825444325292919 | 252.983840639333 | 286.674336577844 | 210.112985279955 | 282.315934971707 | 194.902558855799 | 225.04713070159 | 277.967850178489 | 181.282769299641 | Sorl1 |
| **ENSMUSG00000049321** | -0.112143327641385 | 0.698269967379727 | 0.944950296853508 | 78.6056933415069 | 61.0811796685746 | 64.3589324280944 | 40.3308478531009 | 50.3680770076785 | 52.592535979176 | 70.7933101578173 | 90.0986218674861 | Zfp2 |
| **ENSMUSG00000049323** | 0.0564514819849519 | 0.789655711000067 | 0.961659329450003 | 308.098177350044 | 433.269167782423 | 420.22597055991 | 350.784583652552 | 384.330326732503 | 244.616446414772 | 446.622500848583 | 377.762896504641 | Smcr8 |
| **ENSMUSG00000049327** | 0.139005372039739 | 0.447109801838263 | 0.89108031640712 | 4519.37561027836 | 4614.47952002859 | 4143.5795025029 | 4168.14622928327 | 3243.26617601616 | 3022.23619545451 | 4728.57668730597 | 4847.95717180955 | Kmt5a |
| **ENSMUSG00000049339** | -0.131815751530589 | 0.416072597200717 | 0.882192126163777 | 541.204716224858 | 642.574010113405 | 652.10741827878 | 560.88016316638 | 769.755611660825 | 747.303243797128 | 537.196294726966 | 573.157498144972 | Retreg2 |
| **ENSMUSG00000049354** | 0.00481950677050443 | 0.9808943529113 | 0.996899287476254 | 1010.12833512419 | 1448.84558173859 | 1308.00065611215 | 923.857793844289 | 1158.4657711766 | 875.726878164883 | 1308.63515982906 | 1331.93986784826 | Dcaf7 |
| **ENSMUSG00000049382** | -0.370215881427911 | 0.306135994752481 | 0.851785577030345 | 28.9124389302094 | 50.4937751926884 | 27.447191770805 | 38.4549944645846 | 27.3739548954774 | 46.4771248188067 | 40.6020455316893 | 73.8157383974585 | Krt8 |
| **ENSMUSG00000049401** | 0.0949695653651206 | 0.308504754400656 | 0.851785577030345 | 905.320744002183 | 904.001459094905 | 1033.5287384041 | 911.664746818933 | 924.144717271318 | 856.157562451702 | 882.834220791604 | 851.05204270011 | Ogfr |
| **ENSMUSG00000049404** | -0.430072369185544 | 0.303831635715251 | 0.851785577030345 | 28.0089252136404 | 38.2775392589734 | 29.3401015481019 | 31.8895076047775 | 50.3680770076785 | 35.4693847301419 | 15.616171358342 | 70.559161703453 | Rarres1 |
| **ENSMUSG00000049411** | -0.462135847859414 | 0.0555981337784974 | 0.703396849069134 | 36.1405486627618 | 51.3081909216027 | 31.2330113253987 | 45.0204813243917 | 60.2227007700503 | 59.9310293716191 | 53.094982618363 | 53.1907526687569 | Tmem241 |
| **ENSMUSG00000049421** | 0.112701209714621 | 0.277157168316507 | 0.847954420245878 | 465.309564033058 | 542.400875456943 | 501.621090983677 | 509.294194982182 | 452.217734873287 | 452.540425867328 | 501.799639648058 | 460.262839419447 | Zfp260 |
| **ENSMUSG00000049422** | -0.0372782510327465 | 0.807970725438102 | 0.964026716319739 | 23893.4202346684 | 19045.1118206616 | 19147.7288522467 | 22820.6943979953 | 22674.3943190219 | 26407.5684727067 | 18891.4030312317 | 19157.3551652698 | Chchd10 |
| **ENSMUSG00000049439** | -0.266548380808453 | 0.0992811218326449 | 0.759800953500741 | 154.500845533307 | 190.573280565953 | 162.790240847533 | 176.330218520534 | 228.846262926191 | 226.270212933664 | 204.051305749003 | 164.999885829613 | Cyp20a1 |
| **ENSMUSG00000049470** | 0.23418071822128 | 0.403346919829215 | 0.881158207349486 | 1191.73459215457 | 1909.8048843041 | 1835.17602908934 | 1229.62189617245 | 1110.28761056056 | 775.434135134827 | 1688.62866288205 | 1667.36726733083 | Aff4 |
| **ENSMUSG00000049482** | -0.115794942689834 | 0.507177951702461 | 0.901698316485955 | 122.87786545339 | 114.83261777692 | 139.128868631322 | 162.261318106662 | 164.243729372865 | 143.100621152642 | 147.833088858971 | 128.092016630884 | Ctu2 |
| **ENSMUSG00000049488** | -0.336981742243587 | 0.176054060864363 | 0.799889021902745 | 29.8159526467785 | 46.4216965481167 | 36.9117406572894 | 49.7101147956825 | 53.6529515951357 | 48.9232892829544 | 46.8485140750261 | 56.4473293627624 | Tmem67 |
| **ENSMUSG00000049489** | 0.0116914518512681 | 0.935525889763768 | 0.98818087308472 | 355.984404328204 | 303.777066885045 | 296.240380146964 | 388.301651422879 | 343.816873487196 | 371.816998550453 | 316.487739529066 | 301.776106977845 | Fam58b |
| **ENSMUSG00000049502** | -0.0375463248973865 | 0.789118701451967 | 0.961659329450003 | 343.335212296237 | 392.548381336706 | 284.882921483182 | 343.281170098487 | 370.095870186855 | 341.239942748607 | 317.528817619622 | 372.335268681298 | Dtx3l |
| **ENSMUSG00000049504** | 0.0956632419228611 | 0.647239285685956 | 0.935063240511793 | 122.87786545339 | 135.193010999779 | 144.807597963212 | 115.364983393754 | 154.389105610493 | 84.3926740130963 | 127.011527047849 | 118.322286548867 | Proser1 |
| **ENSMUSG00000049516** | -0.0620052975501283 | 0.77520436160366 | 0.959523366491736 | 177.088688447533 | 173.470550258752 | 215.791714611846 | 239.171307035831 | 260.600050604945 | 220.154801773295 | 217.585320926233 | 142.203848971575 | Spty2d1 |
| **ENSMUSG00000049517** | 0.0692390782808548 | 0.73263174674831 | 0.953733898884637 | 5178.94062337377 | 3629.85090377116 | 4281.76191624557 | 5369.63032462797 | 4640.43283388133 | 5733.80950396225 | 3720.81309564763 | 3500.81994605594 | Rps23 |
| **ENSMUSG00000049521** | -0.521415421063207 | 0.0811839436755946 | 0.739855948845396 | 721.003945822098 | 643.38842584232 | 761.896185362 | 880.713165908413 | 1526.37172497182 | 1445.6831983113 | 619.441463880901 | 725.131077198563 | Cdc42ep1 |
| **ENSMUSG00000049526** | -0.986618751356295 | 0.113419063346151 | 0.767518180080285 | 3.61405486627618 | 5.7009101024003 | 7.57163910918757 | 11.2551203310979 | 8.75966556655277 | 9.78465785659088 | 29.1501865355718 | 7.59867895267955 | Tmem202 |
| **ENSMUSG00000049532** | -0.510417690523498 | 0.158400468878999 | 0.789230661864335 | 19.877301764519 | 14.6594831204579 | 33.1259211026956 | 30.9515809105193 | 31.7537876787538 | 50.1463715150282 | 30.191264626128 | 28.2236646813812 | Sall2 |
| **ENSMUSG00000049536** | -0.00143928082359156 | 0.991352406684869 | 0.998628446692557 | 163.535982698997 | 171.841718800923 | 185.505158175096 | 184.771558768858 | 189.427767876704 | 179.793088114857 | 163.449260217313 | 173.684090346961 | Tceal1 |
| **ENSMUSG00000049537** | 0.249931414787981 | 0.224678421361633 | 0.832048466795696 | 1182.69945498888 | 1174.38748109446 | 1724.44080711747 | 1280.26993766239 | 977.797668866454 | 879.396124861105 | 1464.79687341248 | 1185.39391661801 | Tecrl |
| **ENSMUSG00000049550** | 0.375695760896211 | 0.14149605528986 | 0.779832386580936 | 7416.04058559872 | 10651.7433184705 | 9527.96136402391 | 8421.64378774403 | 5912.77425742312 | 4132.79486217757 | 9491.5089516003 | 8221.77062679928 | Clip1 |
| **ENSMUSG00000049551** | -0.668726704595605 | 0.0217324977382794 | 0.648762042745232 | 23.4913566307952 | 17.9171460361152 | 23.6613722162112 | 32.8274342990356 | 45.9882442244021 | 36.6924669622158 | 36.4377331694648 | 35.8223436340608 | Fzd9 |
| **ENSMUSG00000049553** | 0.219118168292872 | 0.274999081308436 | 0.846286635233373 | 1292.02461469373 | 1417.89778403985 | 1338.2872125489 | 1069.2364314543 | 1078.53382288181 | 733.849339244316 | 1286.77251992738 | 1296.1175242142 | Polr1a |
| **ENSMUSG00000049555** | -0.288319590569353 | 0.283599337051986 | 0.8510190268075 | 40.658117245607 | 68.4109212288036 | 45.4298346551254 | 49.7101147956825 | 61.3176589658694 | 55.0387004433237 | 83.2862472444909 | 49.9341759747513 | Tmie |
| **ENSMUSG00000049562** | -0.178127725162626 | 0.587926143259206 | 0.923266719035655 | 18.0702743313809 | 39.0919549878878 | 28.3936466594534 | 28.1378008277448 | 41.6084114411257 | 24.4616446414772 | 32.2734208072402 | 30.3947158107182 | Ap5b1 |
| **ENSMUSG00000049577** | -0.243411056785295 | 0.0702486080129098 | 0.726006819419135 | 557.467963123101 | 483.762942975111 | 533.800557197724 | 528.990655561603 | 723.767367436423 | 655.572076391589 | 587.168043073661 | 524.308847734889 | Zfpm1 |
| **ENSMUSG00000049580** | -0.85415979252313 | 0.0205770824906416 | 0.646348290773668 | 32.5264937964856 | 46.4216965481167 | 31.2330113253987 | 56.2756016554897 | 74.4571573156986 | 113.746647582869 | 35.3966550789086 | 78.1578406561325 | Tsku |
| **ENSMUSG00000049588** | -0.389844369780254 | 0.0979547464380329 | 0.759800953500741 | 62.3424464432641 | 39.9063707168021 | 43.5369248778286 | 61.9031618210386 | 67.887408140784 | 58.7079471395453 | 69.7522320672611 | 74.901263962127 | Ccdc69 |
| **ENSMUSG00000049600** | -0.25739320037151 | 0.305619312365917 | 0.851785577030345 | 126.491920319666 | 107.502876216691 | 126.824955078892 | 126.620103724852 | 157.67398019795 | 205.477814988408 | 130.134761319517 | 90.0986218674861 | Zbtb45 |
| **ENSMUSG00000049606** | 0.223850442632036 | 0.258921401396253 | 0.838606660810377 | 428.265501653727 | 537.514381083457 | 430.636974335043 | 348.908730264036 | 346.006789878835 | 283.755077841135 | 453.910047482476 | 410.328663444696 | Zfp644 |
| **ENSMUSG00000049624** | 0.0600047827974013 | 0.727167368029558 | 0.952237115597464 | 114.746242004269 | 142.522752560008 | 136.289503965376 | 113.489130005238 | 125.920192519196 | 122.308223207386 | 136.381229862854 | 102.03940307884 | Slc17a5 |
| **ENSMUSG00000049625** | 0.141911116254758 | 0.727941741560171 | 0.952401994188708 | 13.5527057485357 | 21.9892246806869 | 31.2330113253987 | 32.8274342990356 | 12.0445401540101 | 28.1308913376988 | 21.8626399016789 | 28.2236646813812 | Tifab |
| **ENSMUSG00000049641** | -0.867905108746727 | 0.238517586522526 | 0.832048466795696 | 27.1054114970714 | 14.6594831204579 | 4.73227444324223 | 26.2619474392285 | 7.66470737073368 | 84.3926740130963 | 28.1091084450157 | 13.0263067760221 | Vgll2 |
| **ENSMUSG00000049643** | 0.126790495842618 | 0.287717687650163 | 0.851785577030345 | 1537.78034560051 | 1455.36090756991 | 1289.07155833918 | 1439.71747568628 | 1275.62629812925 | 1124.01257127588 | 1428.35914024302 | 1411.18323406906 | 2310022A10 |
| **ENSMUSG00000049657** | -0.376080772691921 | 0.0491086055487171 | 0.703396849069134 | 61.4389327266951 | 86.3280672649188 | 88.966759532954 | 81.59962240046 | 104.021028602814 | 117.415894279091 | 95.7791843311646 | 96.6117752554972 | Zbtb5 |
| **ENSMUSG00000049658** | 0.257748534900886 | 0.27325229838446 | 0.844738346035895 | 422.844419354313 | 551.359448475 | 420.22597055991 | 379.860311174555 | 316.442918591719 | 245.839528646846 | 443.499266576914 | 477.631248454143 | Bdp1 |
| **ENSMUSG00000049659** | 0.162173984310544 | 0.30062134578932 | 0.851785577030345 | 481.572810931301 | 539.9576282702 | 563.140658745826 | 439.887619607078 | 440.173194719277 | 360.809258461789 | 486.183468289716 | 521.052271040884 | Aftph |

| | | | | | | | | | | | | |
|---|---|---|---|---|---|---|---|---|---|---|---|---|
| ENSMUSG00000049670 | 0.0480587911110048 | 0.896866429982884 | 0.984332015045905 | 47.8862269781594 | 44.7928650902881 | 34.0723759913441 | 63.779015209555 | 18.6142893289246 | 55.0387004433237 | 51.0128264372507 | 59.7039060567679 | Morn4 |
| ENSMUSG00000049672 | -0.0167126978702378 | 0.926331856808521 | 0.986713918026868 | 301.773581334061 | 271.200437728471 | 292.45456059237 | 269.184961252092 | 278.119381738051 | 203.031650524261 | 339.391457521301 | 325.657669400552 | Zbtb14 |
| ENSMUSG00000049680 | 0.0802118191095304 | 0.477644634360547 | 0.897674307047416 | 1450.13951509332 | 1714.34510936466 | 1868.30195019203 | 1543.82733874893 | 1582.21459295859 | 1475.03717188107 | 1633.45152408258 | 1529.50552061793 | Urgcp |
| ENSMUSG00000049686 | -0.359709952915417 | 0.0290046727983884 | 0.665956123973759 | 214.132750826864 | 210.119258059897 | 169.415425068072 | 228.854113398991 | 304.398378437709 | 286.201242305283 | 224.872867560126 | 240.986675356409 | Orai1 |
| ENSMUSG00000049690 | 0.558748932515942 | 0.0205567196575133 | 0.646348290773668 | 109.325159704854 | 171.027303072009 | 138.182413742673 | 135.061443973175 | 75.5521155115177 | 70.9387694602839 | 104.107809055614 | 124.835439936878 | Nckap5 |
| ENSMUSG00000049709 | 0.228939063567263 | 0.422849894738182 | 0.88414647567565 | 610.775272400674 | 792.426504233642 | 676.71524538364 | 544.935409363992 | 398.564783278151 | 336.347613820311 | 799.547973547113 | 704.506091469861 | Nlrp10 |
| ENSMUSG00000049717 | -0.170569613464851 | 0.269156790785431 | 0.843793744782845 | 176.185174730964 | 176.728213174409 | 160.897331070236 | 182.895705380341 | 227.751304730372 | 184.685417043153 | 162.408182126757 | 209.506433981022 | Lig4 |
| ENSMUSG00000049728 | -0.192379649887845 | 0.296804091084681 | 0.851785577030345 | 78.6056933415069 | 79.8127414336042 | 82.3415753124149 | 63.779015209555 | 83.2168228822514 | 91.7311674055395 | 78.0808567917102 | 95.5262496908287 | Zfp668 |
| ENSMUSG00000049739 | -0.029139426797219 | 0.863104002836353 | 0.974868041734905 | 284.606820719249 | 313.550055632016 | 410.761421673426 | 279.502154888932 | 336.152166116463 | 281.308913376988 | 349.802238426862 | 346.282655129254 | Zfp646 |
| ENSMUSG00000049744 | 0.339760280809817 | 0.457189617514409 | 0.892805623137304 | 9.93865088225949 | 12.2162359337149 | 13.2503684410783 | 9.37926694258161 | 10.949581958191 | 8.56157562451702 | 9.36970281500523 | 6.51315338801105 | Arhgap15 |
| ENSMUSG00000049751 | 0.0917412081581361 | 0.635271613767766 | 0.932543559775713 | 2524.41732409391 | 1854.42461473793 | 2032.98510081686 | 2476.12647284155 | 2012.5331639155 | 2787.40440689633 | 1664.68386679926 | 1876.87370131185 | Rpl36al |
| ENSMUSG00000049755 | 0.171422416403013 | 0.238870226536647 | 0.832048466795696 | 629.749060448624 | 613.255043872489 | 671.036516051749 | 630.286738541484 | 497.11102090187 | 468.440494884288 | 628.811166695907 | 663.256120012458 | Zfp672 |
| ENSMUSG00000049760 | 0.146584293399509 | 0.301899396409433 | 0.851785577030345 | 3531.8351180684 | 2618.34656845957 | 2921.70624125776 | 3469.39084206094 | 2845.79635093383 | 3247.2833261561 | 2607.90061684312 | 2629.14291762713 | 2410015M20 |
| ENSMUSG00000049764 | -0.137803738476051 | 0.615877371968022 | 0.928422072203769 | 122.87786545339 | 149.038078391322 | 150.486327295103 | 124.744250336335 | 107.305903190271 | 97.8465785659088 | 180.106509666212 | 216.019587369033 | Zfp280b |
| ENSMUSG00000049775 | -0.0837719093213869 | 0.738325637166514 | 0.954383574136061 | 4475.10343816648 | 3359.46488177161 | 4883.70722542599 | 4672.75079079416 | 5107.97998349609 | 6374.70459356896 | 2764.06233042654 | 4184.7010517971 | Tmsb4x |
| ENSMUSG00000049791 | -0.304594228501465 | 0.277385641652076 | 0.848075053184273 | 1031.81266432185 | 1126.33695308852 | 979.580809751142 | 692.189900362523 | 1562.50534543385 | 1574.10683267906 | 808.917676362118 | 785.920508819999 | Fzd4 |
| ENSMUSG00000049792 | -0.150381654356319 | 0.461284563754635 | 0.895328112556867 | 561.985531705946 | 535.885549625628 | 559.354839191232 | 547.749189446766 | 744.571573156986 | 764.426395046162 | 422.677704765791 | 516.71016878221 | Bag5 |
| ENSMUSG00000049800 | 0.0195010631402182 | 0.826978123887781 | 0.967319591893432 | 494.222002963268 | 501.680089011226 | 502.567545872325 | 497.101147956825 | 506.965644664242 | 470.886659348436 | 505.963952010282 | 484.144401842154 | Sertad2 |
| ENSMUSG00000049804 | 0.471446919216201 | 0.163811924798599 | 0.793968226938511 | 52.4037955610046 | 57.8235167529173 | 124.932045301595 | 131.309737196143 | 64.6025335533267 | 67.2695227640623 | 59.3414511616998 | 72.73021283279 | Armcx4 |
| ENSMUSG00000049807 | -0.265708523587157 | 0.112150611430936 | 0.764116514278315 | 424.651446787451 | 460.95930256551 | 477.959718767466 | 377.984457786039 | 667.924499449649 | 507.579126310652 | 477.854843565267 | 440.723379255414 | Arhgap23 |
| ENSMUSG00000049811 | -0.377858372874735 | 0.373387870354819 | 0.873700364743909 | 8.13162344912141 | 9.77298874697194 | 17.9826428843205 | 18.7585338851632 | 21.8991639163819 | 14.6769867848863 | 19.7804837205666 | 14.1118323406906 | Fam161a |
| ENSMUSG00000049823 | 0.111523179214887 | 0.55161695079512 | 0.914873920695527 | 235.817080024521 | 254.097707421271 | 175.094154399963 | 205.405946042537 | 215.706764576362 | 152.885279009232 | 220.708555197901 | 216.019587369033 | Zbtb12 |
| ENSMUSG00000049858 | 0.13383450926674 | 0.322429280536246 | 0.855977678491063 | 581.862833470465 | 557.060358577401 | 610.463403178248 | 562.756016554897 | 535.434557755538 | 426.855698993777 | 527.826591911961 | 615.492995167044 | Suox |
| ENSMUSG00000049866 | 0.0908635872593399 | 0.731329286781844 | 0.953012833605386 | 186.123825613223 | 188.944449108124 | 210.112985279955 | 330.150196378873 | 160.958854785407 | 170.008430258266 | 255.064132186253 | 272.466916731795 | Arl4c |
| ENSMUSG00000049871 | 0.356550741212381 | 0.35013660032205 | 0.861774532843376 | 74.0881247586617 | 68.4109212288036 | 46.3762895437739 | 103.171936368398 | 24.0890803080201 | 58.7079471395453 | 84.3273253350471 | 60.7894316214364 | Nlrc3 |
| ENSMUSG00000049872 | 0.11493623909026 | 0.771884384822039 | 0.959045638244974 | 65.9565013095403 | 62.7100111264033 | 76.6628459805242 | 163.19924480092 | 45.9882442244021 | 129.646716599829 | 80.1630129728225 | 84.6709940441436 | Fam26e |
| ENSMUSG00000049878 | 0.118950288170355 | 0.601625772025463 | 0.926347630355136 | 205.097613661173 | 293.189662409158 | 258.382184601026 | 181.957778686083 | 247.460552255116 | 146.769867848863 | 237.365804646799 | 232.302470839061 | Rlf |
| ENSMUSG00000049881 | -0.238389283243797 | 0.362532059488508 | 0.86924083181804 | 920.680477183857 | 701.211942595237 | 717.412805595523 | 723.141481273042 | 1048.96995159469 | 1316.03648171147 | 650.673806597585 | 598.124586132348 | 2810025M15 |
| ENSMUSG00000049882 | 0.115423495176765 | 0.674994772709463 | 0.944023556644042 | 53.3073092775737 | 50.4937751926884 | 49.2156542097192 | 75.0341355406529 | 48.1781606160403 | 75.8310983885793 | 42.6842017128016 | 44.5065481514088 | Vcpkmt |
| ENSMUSG00000049892 | -0.538493657559015 | 0.302921146166883 | 0.851785577030345 | 71.3775836089546 | 98.5443031986338 | 19.8755526616174 | 23.448167356454 | 76.6470737073368 | 91.7311674055395 | 105.14888714617 | 36.9078691987293 | Rasd1 |
| ENSMUSG00000049904 | 0.00876459550970045 | 0.974437556302271 | 0.995510898988495 | 48.7897406947284 | 59.452348210746 | 34.0723759913441 | 60.9652351267805 | 51.4630352034976 | 37.9155491942896 | 62.4646854333682 | 49.9341759747513 | Tmem17 |
| ENSMUSG00000049907 | 0.339061962349095 | 0.336021400342043 | 0.858570960161243 | 510.48524986151 | 424.310594764365 | 1068.5475692841 | 1075.80191831411 | 448.93286028583 | 470.886659348436 | 599.660980160335 | 914.012525450883 | Rasl11b |
| ENSMUSG00000049922 | -0.188232594200964 | 0.49763230802827 | 0.900927059989498 | 80.412720774645 | 66.7820897709749 | 137.235958854025 | 99.4202295913651 | 147.819356435578 | 103.961989726278 | 93.6970281500523 | 91.1841474321546 | Slc35c1 |
| ENSMUSG00000049929 | -0.145040468856916 | 0.743109197322336 | 0.954383574136061 | 23.4913566307952 | 12.2162359337149 | 39.7511053232348 | 49.7101147956825 | 29.5638712871156 | 50.1463715150282 | 27.0680303544596 | 31.4802413753867 | Lpar4 |
| ENSMUSG00000049932 | -0.180729822737827 | 0.492476279261207 | 0.899236955296196 | 187.027339329792 | 180.800291818981 | 181.719338620502 | 222.288626539184 | 304.398378437709 | 262.96267989588 | 132.216917500629 | 175.855141476298 | H2afx |
| ENSMUSG00000049939 | -0.305267933571863 | 0.44909729219654 | 0.89108031640712 | 14.4562194651047 | 15.4738988493722 | 11.3574586637814 | 8.44134024832345 | 16.4243729372865 | 12.2308223207386 | 18.7394056300105 | 14.1118323406906 | Lrrc4 |
| ENSMUSG00000049940 | 0.0952231341586583 | 0.348328950568185 | 0.861266047293794 | 1573.92089426328 | 1483.05104235299 | 1811.51465687313 | 1776.43315892496 | 1512.13726842617 | 1590.00690169602 | 1590.76732236978 | 1526.24894392392 | Pgrmc2 |
| ENSMUSG00000049950 | 0.0831190033753058 | 0.750188423873697 | 0.956197192463153 | 66.8600150261093 | 58.6379324818317 | 38.8046504345863 | 59.0893817382642 | 64.6025335533267 | 56.2617826753975 | 39.5609674411332 | 51.0197015394199 | Rpp38 |
| ENSMUSG00000049957 | 0.177217311783161 | 0.316095461703698 | 0.853356198600137 | 99.3865088225949 | 100.987550385377 | 119.253315969704 | 130.371810501884 | 106.210944994452 | 94.1773318696872 | 89.5327157878278 | 107.467030902182 | Ccdc137 |
| ENSMUSG00000049960 | 0.0157160589400741 | 0.927201363818345 | 0.986790505471895 | 1080.60240501658 | 761.478706534897 | 795.022106464695 | 988.574735748102 | 831.073270626695 | 1098.32784440233 | 782.890724098215 | 874.933605122817 | Mrps16 |
| ENSMUSG00000049969 | -0.337802913499574 | 0.0792757583861372 | 0.739855948845396 | 179.79922959724 | 156.367819951551 | 152.3792370724 | 168.826804966469 | 247.460552255116 | 249.508775343067 | 166.572494488982 | 168.256462523619 | Plekhf2 |
| ENSMUSG00000049985 | 1.24221581864112 | 0.0012888390687395 | 0.596732488826387 | 25.2983840639333 | 28.5045505120015 | 35.965285768641 | 29.075727522003 | 9.85462376237187 | 17.123151249034 | 6.24646854333682 | 17.3684090346961 | Ankrd55 |
| ENSMUSG00000049988 | -0.248570196751417 | 0.61540918121925 | 0.928422072203769 | 20.780815481088 | 26.0613033252585 | 29.3401015481019 | 23.448167356454 | 35.0386622662111 | 57.4848649074714 | 15.616171358342 | 10.8552556466851 | Lrrc25 |
| ENSMUSG00000049999 | 0.0565698541163813 | 0.743435538022293 | 0.954383574136061 | 383.993329541844 | 442.22774080048 | 579.230391852849 | 479.28054076592 | 460.97740043984 | 364.47850515801 | 493.471014923609 | 491.743080794834 | Ppp1r3d |
| ENSMUSG00000050002 | -0.0438737093654324 | 0.802906595838763 | 0.963461896828986 | 284.606820719249 | 201.975100770753 | 238.506631939409 | 290.75727522003 | 290.163921892061 | 293.539735697726 | 227.996101831794 | 235.559047533066 | Idnk |
| ENSMUSG00000050010 | 0.272450889368299 | 0.475725944320358 | 0.897549741595057 | 65.9565013095403 | 121.347943608235 | 64.3589324280944 | 45.9584080186499 | 33.943704070392 | 51.3694537471021 | 88.4916376972716 | 72.73021283279 | Shisa3 |
| ENSMUSG00000050014 | -0.147701118871813 | 0.688358715282871 | 0.944584424505182 | 50.5967681278665 | 35.8342920722305 | 30.2865564367503 | 84.4134024832345 | 58.0327843784121 | 70.9387694602839 | 36.4377331694648 | 57.5328549274309 | Apol10b |
| ENSMUSG00000050017 | 0.23416403112552 | 0.0294568733068188 | 0.665956123973759 | 1538.68385931708 | 1598.69807585883 | 1756.62027333152 | 1518.50331800396 | 1274.53133993343 | 1242.65154778704 | 1451.26285823525 | 1481.74239577251 | Pitpnb |
| ENSMUSG00000050022 | -0.086587038772936 | 0.856560346548224 | 0.97363563840968 | 17.1667606148119 | 7.32974156022896 | 14.1968233297267 | 7.50341355406529 | 16.4243729372865 | 13.4539045528125 | 9.36970281500523 | 9.76973008201657 | Amz1 |
| ENSMUSG00000050029 | 0.190038051293745 | 0.365145035136934 | 0.86924083181804 | 697.512589191303 | 729.716493107238 | 721.198625150116 | 658.424539369229 | 485.06648074786 | 440.309603546589 | 733.960053842076 | 800.03234116069 | Rap2c |
| ENSMUSG00000050043 | -0.119070053550004 | 0.479702246126718 | 0.897732059433541 | 1770.88688447533 | 1495.26727828671 | 1232.28426502028 | 1410.64174816427 | 1295.33554565399 | 1523.96046116403 | 1686.54650670094 | 1911.61051938124 | Tmx2 |
| ENSMUSG00000050052 | -0.207104869389162 | 0.286929022339942 | 0.851785577030345 | 188.83436676293 | 146.594831204579 | 145.754052851861 | 206.343872736795 | 215.706764576362 | 232.385624094033 | 158.243869764533 | 187.795922687652 | Tdrp |
| ENSMUSG00000050064 | 0.188053649558441 | 0.259206207407504 | 0.839248070137582 | 198.77301764519 | 174.284965987666 | 250.810545491838 | 229.79204009325 | 176.288269526875 | 184.685417043153 | 212.379930473452 | 174.76961591163 | Zfp697 |
| ENSMUSG00000050071 | 0.384454687925095 | 0.440240360753338 | 0.890530552540252 | 27.1054114970714 | 28.5045505120015 | 6.62518422053913 | 45.0204813243917 | 22.994122112201 | 23.2385624094033 | 19.7804837205666 | 16.2828834700276 | Bex1 |
| ENSMUSG00000050079 | 0.0240424751345133 | 0.838542070920371 | 0.969245483845283 | 237.624107457659 | 258.984201794756 | 270.686098153456 | 267.309107863576 | 256.220217821669 | 236.054870790255 | 254.023054095697 | 270.295865602458 | Rspry1 |
| ENSMUSG00000050088 | 0.0895738662080303 | 0.595100142051867 | 0.924461858177432 | 276.475197270128 | 203.603932228582 | 183.612248397799 | 233.543746870282 | 216.801722772181 | 227.493295165738 | 208.215618111227 | 191.052499381657 | 1600012H06I |
| ENSMUSG00000050103 | -0.55089257231661 | 0.160635600191089 | 0.792016108372479 | 50.5967681278665 | 49.679359463774 | 54.8943835416099 | 45.9584080186499 | 79.9319482947941 | 121.085140975312 | 26.0269522639034 | 68.388110574116 | Agmo |
| ENSMUSG00000050105 | -0.730246284939824 | 0.0639138841601679 | 0.716991331547963 | 131.009488902512 | 85.5136515360045 | 159.004421292939 | 124.744250336335 | 267.16979977986 | 344.909189444828 | 102.025652874501 | 116.15123541953 | Grrp1 |
| ENSMUSG00000050106 | -0.654561035573639 | 0.0197040498446778 | 0.642273034804458 | 53.3073092775737 | 45.6072808192024 | 35.0188308799925 | 54.3997482669734 | 108.400861386091 | 62.3771938357668 | 57.2592949805875 | 68.388110574116 | Tmc8 |
| ENSMUSG00000050144 | -0.0360782054165523 | 0.854783336802088 | 0.973268281968411 | 440.011179969125 | 522.854897962999 | 558.408384302584 | 380.798237868813 | 431.413529152724 | 375.486245246675 | 572.592949805875 | 569.900921450967 | Slc25a44 |
| ENSMUSG00000050148 | -0.143626751089694 | 0.331132827482142 | 0.856333478432716 | 432.783070236573 | 554.617111390658 | 580.176846741498 | 508.356268287923 | 651.500126512363 | 612.764198269004 | 517.4158110064 | 512.368066523536 | Ubqln2 |
| ENSMUSG00000050174 | -0.0550356116375662 | 0.711903432697833 | 0.950074110229522 | 424.651446787451 | 315.178887089845 | 326.526936583714 | 425.818719193205 | 366.810995599397 | 401.170972120226 | 375.829190690765 | 405.986561186022 | Nudt6 |
| ENSMUSG00000050188 | -0.113004301608176 | 0.479901084420276 | 0.897732059433541 | 304.484122483768 | 329.023954481389 | 286.775831260479 | 364.853484066425 | 366.810995599397 | 418.29412336926 | 292.542943446274 | 313.716888189199 | Lsm10 |
| ENSMUSG00000050192 | -0.369798165833966 | 0.290429047210301 | 0.851785577030345 | 41.5616309621761 | 23.6180561385155 | 20.8220075502658 | 38.4549944645846 | 52.5579933993167 | 48.9232892829544 | 29.1501865355718 | 30.3947158107182 | Eif5a2 |
| ENSMUSG00000050195 | 0.431082580226701 | 0.00612995353960957 | 0.635862253526407 | 385.800356974982 | 409.651111643907 | 514.871459424755 | 416.439452250624 | 285.784089108784 | 387.717067567413 | 299.830490080167 | 308.289260365856 | Scd4 |
| ENSMUSG00000050199 | 0.0993198463361638 | 0.666252760192594 | 0.942024383071524 | 503.257140128958 | 584.750493360488 | 605.731128735006 | 520.54931531328 | 405.134532453066 | 348.57843614105 | 640.263025692024 | 671.940324529806 | Lgr4 |
| ENSMUSG00000050211 | -0.104460459176984 | 0.75039117697307 | 0.956197192463153 | 62.3424464432641 | 57.8235167529173 | 70.9841166486335 | 38.4549944645846 | 39.4184950494875 | 44.0309603546589 | 73.9165444294857 | 89.0130963028176 | Pla2g4e |
| ENSMUSG00000050212 | -0.640997762953281 | 0.117851427760926 | 0.769868040577702 | 112.035700854562 | 87.1424829938332 | 83.2880302010633 | 143.502784221499 | 231.036179317829 | 267.855008824175 | 81.2040910633787 | 84.6709940441436 | Eva1b |
| ENSMUSG00000050213 | -0.0141127103578147 | 0.952076655377068 | 0.991925048243885 | 140.948139784771 | 118.904696421492 | 138.182413742673 | 142.564857527241 | 193.80760065998 | 143.100621152642 | 111.395355689507 | 97.6973008201657 | Snip1 |
| ENSMUSG00000050222 | -0.346367003895839 | 0.229058906293959 | 0.832048466795696 | 131.913002619081 | 102.616381843205 | 87.0738497556571 | 135.061443973175 | 158.768938393769 | 212.816308380852 | 119.723980413956 | 90.0986218674861 | Il17d |
| ENSMUSG00000050229 | -0.112683759742191 | 0.543877673790101 | 0.913837378403038 | 133.720030052219 | 92.8433930962335 | 127.77140996754 | 149.130344387048 | 145.62944004394 | 130.869798831903 | 134.299073681742 | 132.434118889558 | Pigm |
| ENSMUSG00000050234 | -0.907572025463897 | 0.0368137059637315 | 0.679566075569698 | 158.114900399583 | 90.4001459094905 | 114.521041526462 | 138.813150750208 | 309.873169416804 | 414.624876673038 | 94.7381062406084 | 122.664388807541 | Gja4 |
| ENSMUSG00000050240 | 0.146767485632836 | 0.618411551818057 | 0.929568738467614 | 22.5878429142261 | 29.3189662409158 | 34.0723759913441 | 39.3929211588428 | 32.8487458745729 | 28.1308913376988 | 24.9858741733473 | 27.1381391167127 | Hic2 |
| ENSMUSG00000050244 | 0.106414163054044 | 0.525482738610989 | 0.908006595346551 | 273.764656120421 | 347.755516246418 | 318.955297474527 | 256.053987532478 | 337.247124312282 | 239.724117486476 | 269.639225454039 | 264.868237779116 | Heatr1 |
| ENSMUSG00000050248 | 0.0754745533034569 | 0.554566029990806 | 0.915945640531442 | 203.290586228035 | 221.521078264697 | 205.380710836713 | 213.847286290861 | 218.991639163819 | 196.916239363891 | 205.092383839559 | 180.197243734972 | Evc2 |
| ENSMUSG00000050270 | 0.101015923765319 | 0.632403133182549 | 0.931825800800958 | 63.2459601598332 | 60.2667639396603 | 73.8234813145789 | 50.6480414899407 | 61.3176589658694 | 52.592535979176 | 57.2592949805875 | 59.7039060567679 | Tmem220 |
| ENSMUSG00000050271 | -0.396019380294083 | 0.244071555746106 | 0.832048466795696 | 228.588970291968 | 247.582381589956 | 305.704929033448 | 271.060814640609 | 402.944616061428 | 580.964060235083 | 183.22974393788 | 219.276164063039 | Prag1 |
| ENSMUSG00000050288 | -0.888529968871194 | 0.00314606523898526 | 0.635862253526407 | 47.8862269781594 | 36.6487078011448 | 31.2330113253987 | 45.0204813243917 | 71.1722827282413 | 113.746647582869 | 58.3003730711437 | 55.3618037980939 | Fzd2 |
| ENSMUSG00000050295 | -0.759280382816169 | 0.0785437258814323 | 0.739855948845396 | 71.3775836089546 | 55.3802695661743 | 58.6802030962037 | 62.8410885152968 | 142.344565456483 | 181.016170346931 | 55.1771387994752 | 42.3354970220718 | Foxc1 |
| ENSMUSG00000050296 | -0.196627924855429 | 0.409399242880064 | 0.882192126163777 | 141.85165350134 | 161.254314325037 | 145.754052851861 | 90.0409626487835 | 142.344565456483 | 116.192812047017 | 190.517290571773 | 168.256462523619 | Abca12 |
| ENSMUSG00000050299 | 0.00532453277607054 | 0.988737884136601 | 0.998437672973644 | 46.0791995450213 | 30.9477976987445 | 43.5369248778286 | 65.6548685980713 | 35.0386622662111 | 79.5003450848009 | 29.1501865355718 | 42.3354970220718 | Gm9843 |
| ENSMUSG00000050310 | -0.0491621271499125 | 0.793522341837983 | 0.961804768755116 | 140.948139784771 | 191.387696294867 | 193.076797284283 | 143.502784221499 | 168.623562156141 | 143.100621152642 | 175.942197303987 | 204.078806157679 | Rictor |
| ENSMUSG00000050312 | 0.00012854363229048 | 0.999556196937678 | 0.999772130443173 | 96.6759676728878 | 142.522752560008 | 126.824955078892 | 119.116690170786 | 96.3563212320805 | 92.9542496376133 | 154.079557402308 | 141.118323406906 | Nsun3 |
| ENSMUSG00000050315 | 0.211856437894923 | 0.406028067586688 | 0.881158207349486 | 5021.72923669075 | 6466.46088757977 | 6045.95382868628 | 4650.24055013196 | 3911.19067546581 | 2940.28968590556 | 6612.92803121258 | 5689.23948442765 | Synpo2 |
| ENSMUSG00000050323 | 0.152093880378012 | 0.331037151369213 | 0.856333478432716 | 478.862269781594 | 346.941100517504 | 357.759947909113 | 421.129085721914 | 328.487458745729 | 332.67836712409 | 369.582722147429 | 412.499714574033 | Ndufaf6 |
| ENSMUSG00000050332 | 0.0209558379928901 | 0.903072832371086 | 0.985027469232262 | 100.290022539164 | 100.987550385377 | 97.48485353079 | 120.054616865045 | 87.5966556655277 | 102.738907494204 | 113.477511870619 | 108.552556466851 | Amer1 |
| ENSMUSG00000050335 | -0.846483616201916 | 0.102579720058204 | 0.759800953500741 | 38.8510898124689 | 33.3910448854875 | 61.519567762149 | 48.7721881014244 | 83.2168228822514 | 178.570005882784 | 35.3966550789086 | 31.4802413753867 | Lgals3 |

| | | | | | | | | | | | | |
|---|---|---|---|---|---|---|---|---|---|---|---|---|
| **ENSMUSG00000050345** | -0.135542431172999 | 0.583548661888005 | 0.922450244597442 | 57.8248778604189 | 41.5352021746308 | 50.1621090983677 | 60.9652351267805 | 61.3176589658694 | 73.3849339244316 | 44.7663578939139 | 52.1052271040884 | 4930486L24R |
| **ENSMUSG00000050373** | -0.00447124927797476 | 0.970230888855544 | 0.9951105526746 | 681.24934229306 | 589.636987733974 | 708.894711597687 | 641.541858872582 | 625.221129812704 | 752.195572725424 | 607.989604884784 | 644.802185413094 | Snx21 |
| **ENSMUSG00000050379** | -0.506004181242363 | 0.0725491957058408 | 0.729721356194038 | 51.5002818444356 | 42.3496179035451 | 45.4298346551254 | 38.4549944645846 | 89.7865720571659 | 68.4926049961361 | 51.0128264372507 | 43.4210225867403 | Sept6 |
| **ENSMUSG00000050382** | 0.207581717169513 | 0.509260546081449 | 0.902394714365738 | 23.4913566307952 | 26.8757190541728 | 34.0723759913441 | 37.5170677703264 | 33.943704070392 | 25.6847268735511 | 18.7394056300105 | 27.1381391167127 | Kif7 |
| **ENSMUSG00000050390** | -0.930351339020085 | 0.00022989203650908 | 0.399150048388892 | 64.1494738764022 | 83.8848200781758 | 67.1982970940397 | 65.6548685980713 | 171.908436743598 | 165.116101329971 | 103.066730965058 | 96.6117752554972 | C77080 |
| **ENSMUSG00000050410** | -0.186739834866767 | 0.628713047133385 | 0.931825800800958 | 22.5878429142261 | 23.6180561385155 | 30.2865564367503 | 21.5723139679377 | 44.893286028583 | 31.8001380339204 | 15.616171358342 | 19.5394601640331 | Tcf19 |
| **ENSMUSG00000050423** | 0.00221195846406342 | 0.995605434961114 | 0.999262355603273 | 15.3597331816738 | 16.2883145782866 | 17.9826428843205 | 26.2619474392285 | 14.2344565456483 | 11.0077400886647 | 26.0269522639034 | 23.8815624227072 | Ppp1r3g |
| **ENSMUSG00000050428** | -0.266351661913993 | 0.123170796374563 | 0.770251097497048 | 101.193536255733 | 109.13170767452 | 106.949402417274 | 128.495957113368 | 150.009272827216 | 145.546785616789 | 130.134761319517 | 110.723607596188 | Fbxo46 |
| **ENSMUSG00000050440** | -0.269822402162548 | 0.516392072567864 | 0.904614186904734 | 172.571119864688 | 215.005752433383 | 140.07532351997 | 135.061443973175 | 113.875652365186 | 340.016860516533 | 80.1630129728225 | 265.953763343784 | Hamp |
| **ENSMUSG00000050445** | -0.174952025577339 | 0.623385053546095 | 0.931059568526633 | 56.0178504272808 | 104.245213301034 | 70.9841166486335 | 66.5927952923295 | 56.937826182593 | 59.9310293716191 | 71.8343882483734 | 147.631476794917 | Cyp8b1 |
| **ENSMUSG00000050447** | 0.178271777636378 | 0.753026957802367 | 0.956497786093635 | 35.2370349461928 | 21.9892246806869 | 12.3039135524298 | 9.37926694258161 | 12.0445401540101 | 13.4539045528125 | 34.3555769883525 | 9.76973008201657 | Lypd6 |
| **ENSMUSG00000050471** | 0.0284367696383349 | 0.808489705217925 | 0.964026716319739 | 235.817080024521 | 215.820168162297 | 237.56017705076 | 236.357526953057 | 233.226095709468 | 214.039390612925 | 233.201492284575 | 225.78931745105 | Fam118b |
| **ENSMUSG00000050490** | 0.068599869033128 | 0.880816886800588 | 0.97846486749867 | 12.6491920319666 | 13.0306516626293 | 11.3574586637814 | 11.2551203310979 | 12.0445401540101 | 4.89232892829544 | 11.4518589961175 | 17.3684090346961 | Gm8394 |
| **ENSMUSG00000050503** | -0.29830893361734 | 0.178578150836559 | 0.801696999069105 | 1378.76193148436 | 1075.84317789583 | 678.608155160936 | 1191.16690170786 | 1244.96746864631 | 1592.45306616017 | 1175.37716423788 | 1305.88725429621 | Fbxl22 |
| **ENSMUSG00000050505** | -0.664445683932652 | 0.347515421282504 | 0.861266047293794 | 11.7456783153976 | 11.4018202048006 | 7.57163910918757 | 3.75170677703264 | 19.7092475247437 | 26.9078091056249 | 3.12323427166841 | 5.42762782334254 | Pcdh20 |
| **ENSMUSG00000050530** | -0.250682068115825 | 0.217689425423108 | 0.826183082597859 | 322.554396815149 | 332.281617397046 | 382.367775013973 | 397.68091836546 | 508.060602860061 | 534.486935416277 | 331.062832796851 | 334.3418739179 | Fam171a1 |
| **ENSMUSG00000050541** | -0.0425355123621309 | 0.771982865164633 | 0.959045638244974 | 816.776399778417 | 899.114964721419 | 823.415753124149 | 647.169419038131 | 868.301849284544 | 741.187832636759 | 927.600578685518 | 744.670537362596 | Adra1b |
| **ENSMUSG00000050545** | 0.148308713623494 | 0.690070918517777 | 0.94469184542218 | 37.0440623793308 | 22.8036404096012 | 18.9290977729689 | 27.1998741334867 | 29.5638712871156 | 17.123151249034 | 32.2734208072402 | 16.2828834700276 | Fam228b |
| **ENSMUSG00000050549** | 0.0474781731161254 | 0.790879631139249 | 0.961804768755116 | 89.4478579403355 | 74.1118313312039 | 92.7525790875478 | 91.9168160372998 | 76.6470737073368 | 84.3926740130963 | 92.6559500594962 | 82.4999429148066 | Fam241a |
| **ENSMUSG00000050552** | -0.0735568695568677 | 0.666110274493354 | 0.942024383071524 | 1032.71617803842 | 804.642740167357 | 794.075651576047 | 971.692055251455 | 972.322877887358 | 1186.38976511164 | 804.753363999894 | 829.34153140674 | Lamtor4 |
| **ENSMUSG00000050555** | -0.558288369915581 | 0.0676092251248779 | 0.722038584319139 | 34.3335212296237 | 19.5459774939439 | 21.7684624389143 | 27.1998741334867 | 37.2285786578493 | 46.4771248188067 | 39.5609674411332 | 28.2236646813812 | Hyls1 |
| **ENSMUSG00000050556** | 0.0406867699677812 | 0.843569517113808 | 0.971654415184575 | 911.645340018166 | 1077.47200935366 | 988.098903748979 | 821.623784170149 | 892.390929592564 | 606.648787108634 | 1109.78924453284 | 1083.35451353917 | Kcnb1 |
| **ENSMUSG00000050565** | -0.0997581938691019 | 0.418171197255149 | 0.882468496126562 | 893.575065686785 | 1097.83240257652 | 916.168332211697 | 887.278652768221 | 1073.05903190271 | 889.180782717696 | 1080.63905799727 | 1023.6506074824 | Tor1aip2 |
| **ENSMUSG00000050567** | -0.162306197373565 | 0.351396244118671 | 0.862447289221848 | 186.123825613223 | 180.800291818981 | 226.202718386979 | 163.19924480092 | 256.220217821669 | 209.14706168463 | 179.065431575656 | 201.907755028342 | Maml1 |
| **ENSMUSG00000050592** | -0.440800864677986 | 0.0309769860347654 | 0.667184591665619 | 187.930853046361 | 183.243539005724 | 179.826428843205 | 152.88205116408 | 158.768938393769 | 261.739597663806 | 249.858741733473 | 285.493223507818 | Fam78a |
| **ENSMUSG00000050605** | -0.0844292531041486 | 0.801617393354291 | 0.963461896828986 | 36.1405486627618 | 17.1027303072009 | 36.9117406572894 | 45.0204813243917 | 35.0386622662111 | 36.6924669622158 | 41.6431236222455 | 29.3091902460497 | Zfp61 |
| **ENSMUSG00000050608** | 0.0413950769800076 | 0.811447023959028 | 0.965065430498408 | 7132.33727859604 | 5127.56142924461 | 5687.24742588852 | 7048.51910735008 | 5921.53392298968 | 7585.55600332208 | 5112.73450272119 | 5669.70002426361 | Minos1 |
| **ENSMUSG00000050619** | 0.0443830311275432 | 0.763612406383996 | 0.95785116771606 | 191.544907912638 | 182.42912327681 | 163.736695736181 | 166.013024883695 | 170.813478547779 | 146.769867848863 | 193.640524843441 | 170.427513652956 | Zscan29 |
| **ENSMUSG00000050621** | -0.0946110685883684 | 0.752794009439126 | 0.956497786093635 | 31.6229800799166 | 18.7315617650296 | 30.2865564367503 | 31.8895076047775 | 25.1840385038392 | 33.0232202659942 | 33.3144988977964 | 28.2236646813812 | Rps27rt |
| **ENSMUSG00000050627** | 0.17087143579685 | 0.487684636992349 | 0.899236955296196 | 1004.70725282478 | 1297.36425616053 | 1472.68380673698 | 1012.02290310456 | 846.402685368162 | 715.503105763208 | 1337.78534636464 | 1351.47932801229 | Gpd1l |
| **ENSMUSG00000050628** | -0.289179711118226 | 0.129978992773263 | 0.77303252008843 | 424.651446787451 | 374.631235300591 | 337.884395247495 | 399.556771753977 | 560.618596259378 | 567.510155682271 | 390.404283958551 | 360.394487469945 | Ubald2 |
| **ENSMUSG00000050640** | 0.112746731708289 | 0.687826362250817 | 0.944452327903597 | 430.072529086865 | 368.930325198191 | 352.081218577222 | 268.247034557834 | 282.499214521327 | 184.685417043153 | 457.033281754144 | 387.532626586657 | Tmem150c |
| **ENSMUSG00000050666** | -0.238016543350585 | 0.17712565745092 | 0.800237299431217 | 146.369222084185 | 102.616381843205 | 113.574586637814 | 113.489130005238 | 155.484063806312 | 124.754387671534 | 146.792010768415 | 133.519644454226 | Vstm4 |
| **ENSMUSG00000050668** | 0.0140911284895653 | 0.924367781264283 | 0.986713918026868 | 459.888481733644 | 364.858246553619 | 336.937940358847 | 443.63932638411 | 387.61520131996 | 434.19419238622 | 358.130863151311 | 410.328663444696 | Gpatch11 |
| **ENSMUSG00000050675** | 0.345162594720988 | 0.0813042085203226 | 0.739855948845396 | 81.316234491214 | 71.6685841444609 | 79.5022106464695 | 72.2203554578784 | 51.4630352034976 | 53.8156182112498 | 64.5468416144805 | 69.4736361387845 | Gp1ba |
| **ENSMUSG00000050677** | 0.103022401374178 | 0.84438187329218 | 0.971806333564415 | 15.3597331816738 | 8.14415728914329 | 6.62518422053913 | 14.0689004138724 | 9.85462376237187 | 13.4539045528125 | 13.5340151772298 | 4.34210225867403 | Ccdc96 |
| **ENSMUSG00000050697** | -0.0128892955235811 | 0.927689706060665 | 0.98687370890577 | 301.773581334061 | 307.849145529616 | 277.311282373995 | 302.950322245386 | 356.956371837026 | 291.093571233579 | 301.91264626128 | 250.756405438425 | Prkaa1 |
| **ENSMUSG00000050705** | 0.100558305918258 | 0.261762946832813 | 0.841137531367976 | 2518.9962417945 | 2276.29196231555 | 2333.95775540707 | 2504.26427366929 | 2343.21053905287 | 2036.43191640298 | 2307.0290486724 | 2295.88656927389 | 2310061I04R |
| **ENSMUSG00000050708** | -0.192144561098798 | 0.456187685920486 | 0.892805623137304 | 9778.72895442678 | 12687.7826407563 | 13080.9530160102 | 14927.1033391186 | 16087.1258129742 | 20964.852539978 | 9555.01471512422 | 11059.3344528428 | Ftl1 |
| **ENSMUSG00000050714** | -0.197642304226812 | 0.662162763594858 | 0.939611397743053 | 72.2810973255236 | 71.6685841444609 | 45.4298346551254 | 22.5102406621959 | 44.893286028583 | 28.1308913376988 | 90.5737938783839 | 79.2433662208011 | Zbtb26 |
| **ENSMUSG00000050721** | -0.606375112341379 | 0.0244107567614557 | 0.655303666217042 | 162.632468982428 | 153.110157035894 | 165.629605513478 | 162.261318106662 | 339.43704070392 | 311.885969178834 | 170.736806851206 | 158.486732441602 | Plekho2 |
| **ENSMUSG00000050730** | -0.077935274955763 | 0.668433136868803 | 0.942024383071524 | 117.456783153976 | 141.708336831093 | 127.77140996754 | 100.358156285623 | 116.065568756824 | 117.415894279091 | 128.052605138405 | 153.05910461826 | Arhgap42 |
| **ENSMUSG00000050732** | -0.0208448640792787 | 0.9238757693642 | 0.986713918026868 | 1080.60240501658 | 791.612088504727 | 937.936794650611 | 1118.00861955573 | 1039.11532783232 | 1375.96751108309 | 752.699459472087 | 818.486275760055 | Vamp8 |
| **ENSMUSG00000050737** | -0.404889947544738 | 0.433783535372858 | 0.887868762931002 | 12.6491920319666 | 8.95857301805762 | 8.51809399783602 | 14.0689004138724 | 21.8991639163819 | 22.0154801773295 | 7.28754663389296 | 7.59867895267955 | Ptges |
| **ENSMUSG00000050761** | 0.0883718281612893 | 0.908648313995057 | 0.985255665994641 | 18.9737880479499 | 14.6594831204579 | 88.0203046443056 | 7.50341355406529 | 19.7092475247437 | 72.1618516923577 | 13.5340151772298 | 16.2828834700276 | Gp1bb |
| **ENSMUSG00000050777** | 0.120932469118703 | 0.59737087109751 | 0.924496535139492 | 77.7021796249379 | 52.122606650517 | 61.519567762149 | 65.6548685980713 | 64.6025335533267 | 56.2617826753975 | 46.8485140750261 | 68.388110574116 | Tmem37 |
| **ENSMUSG00000050786** | -0.237807810876437 | 0.239372025063378 | 0.832048466795696 | 88.5443442237664 | 80.6271571625185 | 99.3777633080869 | 80.6616957062019 | 100.736154015357 | 81.9465095489486 | 131.175839410073 | 96.6117752554972 | Ccdc126 |
| **ENSMUSG00000050796** | 0.266313977442226 | 0.183718693631691 | 0.805963874499992 | 74.9916384752307 | 82.2559886203472 | 67.1982970940397 | 60.9652351267805 | 62.4126171616885 | 55.0387004433237 | 59.3414511616998 | 60.7894316214364 | B3galt6 |
| **ENSMUSG00000050812** | 0.253198086407753 | 0.140376068908575 | 0.779711473022872 | 5619.85531705946 | 6351.62826980285 | 6200.22597553597 | 5525.32615587483 | 4493.70843564157 | 3690.03909416683 | 5875.84474309884 | 5821.67360331721 | AI314180 |
| **ENSMUSG00000050821** | -0.02634271789698 | 0.87493014467591 | 0.976986380159756 | 689.380965742181 | 543.215291185857 | 515.817914313404 | 527.114802173087 | 532.149683168081 | 474.555906044658 | 620.482541971457 | 689.308733564502 | Fam131a |
| **ENSMUSG00000050846** | 0.0216594640115462 | 0.915214318836595 | 0.986713918026868 | 132.81651633565 | 144.96599974675 | 161.843785958884 | 120.992543559303 | 177.383227722694 | 147.992950080937 | 114.518589961175 | 112.894658725525 | Zfp623 |
| **ENSMUSG00000050855** | 0.069498362292267 | 0.858183419290877 | 0.974427180082587 | 22.5878429142261 | 13.8450673915436 | 13.2503684410783 | 13.1309737196143 | 13.1394983498292 | 15.9000690169602 | 13.5340151772298 | 17.3684090346961 | Zfp940 |
| **ENSMUSG00000050856** | 0.203838991711267 | 0.242482848975355 | 0.832048466795696 | 7573.25197228174 | 5448.44122643686 | 5927.64696760522 | 7296.13175463424 | 6059.49865566288 | 6958.11481826819 | 4939.91553968887 | 4830.58876277486 | Atp5k |
| **ENSMUSG00000050860** | 0.11359160166727 | 0.673458533572311 | 0.943643602473459 | 383.089815825275 | 468.289044125739 | 512.978549647458 | 434.260059441529 | 320.822751374995 | 245.839528646846 | 573.634027896431 | 521.052271040884 | Phospho1 |
| **ENSMUSG00000050890** | -0.120355882334335 | 0.519956505994643 | 0.905962179970576 | 72.2810973255236 | 68.4109212288036 | 81.3951204237664 | 64.7169419038131 | 77.7420319031559 | 70.9387694602839 | 87.4505596067155 | 74.901263962127 | Pdik1l |
| **ENSMUSG00000050891** | 0.210441739854613 | 0.23044575064676 | 0.832048466795696 | 282.799793286111 | 219.892246806869 | 192.130342395635 | 284.191788360223 | 189.427767876704 | 209.14706168463 | 230.078258012906 | 217.105112933702 | Tatdn1 |
| **ENSMUSG00000050896** | -1.31618618322033 | 0.00955624722130705 | 0.635862253526407 | 41.5616309621761 | 71.6685841444609 | 50.1621090983677 | 33.7653609932938 | 201.472308030714 | 192.023910435596 | 60.3825292522559 | 37.9933947633978 | Rtn4rl2 |
| **ENSMUSG00000050910** | -0.388509773611501 | 0.154837531673238 | 0.787275540343856 | 52.4037955610046 | 43.9784493613737 | 66.2518422053913 | 51.5859681841989 | 86.5016974697087 | 88.0619207093179 | 59.3414511616998 | 46.6775992807458 | Cdr2l |
| **ENSMUSG00000050912** | -0.122275409713418 | 0.373705909335996 | 0.873700364743909 | 1150.17296119239 | 1311.20932355207 | 1345.85885165809 | 1175.22214790548 | 1524.18180858018 | 1512.95272107536 | 1132.69296252508 | 1254.86755275679 | Tmem123 |
| **ENSMUSG00000050914** | -0.719174393937359 | 0.0612833491353663 | 0.71282992338528 | 111.132187137993 | 52.122606650517 | 75.7163910918757 | 74.0962088463947 | 169.71852035196 | 198.139321595965 | 69.7522320672611 | 78.1578406561325 | Ankrd37 |
| **ENSMUSG00000050919** | 0.142797418852407 | 0.478339153696972 | 0.897674307047416 | 102.097049972302 | 123.791190794978 | 110.735221971868 | 99.4202295913651 | 118.255485148462 | 89.2850029413918 | 112.436433780063 | 74.901263962127 | Zfp366 |
| **ENSMUSG00000050930** | -0.0134253413458568 | 0.941705461666426 | 0.989934163630124 | 114.746242004269 | 109.13170767452 | 142.914688185915 | 136.937297361692 | 137.964732673206 | 103.961989726278 | 118.6829023234 | 146.545951230249 | Map10 |
| **ENSMUSG00000050931** | -0.489720056798819 | 0.211153096617542 | 0.823123035902316 | 17.1667606148119 | 29.3189662409158 | 24.6078271048596 | 24.3860940507122 | 30.6588294829347 | 56.2617826753975 | 17.6983275394543 | 30.3947158107182 | Sgms2 |
| **ENSMUSG00000050945** | 0.0397641906983062 | 0.838965171375422 | 0.96956148850045 | 129.202461469373 | 113.203786319092 | 108.842312194571 | 94.7305961200743 | 91.9764884488041 | 107.6312364225 | 136.381229862854 | 97.6973008201657 | Zfp438 |
| **ENSMUSG00000050947** | 0.0550359771343511 | 0.822730091247893 | 0.966637987288264 | 480.669297214732 | 604.296470854432 | 737.28835825714 | 428.63249927598 | 547.479097909548 | 351.024600605198 | 678.782915042601 | 588.354856050331 | Amigo1 |
| **ENSMUSG00000050953** | -0.135205223647327 | 0.453464687479329 | 0.891651261195907 | 2995.14797042638 | 3955.61719533689 | 3104.37203476691 | 3714.18970926232 | 2884.1198877875 | 3419.73792087851 | 4518.27891301363 | 4299.76676165196 | Gja1 |
| **ENSMUSG00000050954** | 0.438273551230669 | 0.355724669391411 | 0.864662986238123 | 10.8421645988285 | 18.7315617650296 | 15.1432782183751 | 23.448167356454 | 12.0445401540101 | 7.33849339244316 | 21.8626399016789 | 8.68420451734806 | Zfp169 |
| **ENSMUSG00000050965** | -0.011274480971575 | 0.959313897640915 | 0.993281404266293 | 518.616873310632 | 591.265819191803 | 367.224496795597 | 438.949692912819 | 460.97740043984 | 336.347613820311 | 581.96265262088 | 551.446986851602 | Prkca |
| **ENSMUSG00000050973** | -0.778666380097495 | 0.00590521320440344 | 0.635862253526407 | 30.7194663633475 | 29.3189662409158 | 49.2156542097192 | 25.3240207449704 | 54.7479097909548 | 46.4771248188067 | 60.3825292522559 | 68.388110574116 | Gdpgp1 |
| **ENSMUSG00000050989** | -0.417335240426964 | 0.0782395139532545 | 0.739855948845396 | 244.852217190211 | 267.1283590839 | 254.596365046432 | 279.502154888932 | 367.905953795217 | 512.471455238947 | 247.776585552361 | 270.295865602458 | Selenon |
| **ENSMUSG00000051000** | -0.0646697943303902 | 0.796792527607747 | 0.962278883153809 | 1382.37598635064 | 1467.57714350362 | 1394.12805097916 | 1042.03655732082 | 1304.09521122054 | 797.449615312156 | 1855.20115737104 | 1570.75549207533 | Fam160a1 |
| **ENSMUSG00000051007** | -0.0208538311609952 | 0.904852027122091 | 0.98504921011344 | 130.105975185942 | 118.904696421492 | 123.039135524298 | 159.447538023887 | 140.154649064844 | 146.769867848863 | 111.395355689507 | 141.118323406906 | Gatd1 |
| **ENSMUSG00000051022** | -0.620857139812025 | 0.2571259422068 | 0.837517407767513 | 21.6843291976571 | 18.7315617650296 | 29.3401015481019 | 45.0204813243917 | 51.4630352034976 | 89.2850029413918 | 19.7804837205666 | 16.2828834700276 | Hs3st1 |
| **ENSMUSG00000051034** | -0.184889239621145 | 0.434154401571445 | 0.887868762931002 | 37.0440623793308 | 46.4216965481167 | 42.5904699891801 | 49.7101147956825 | 51.4630352034976 | 40.3617136584374 | 47.8895921655823 | 59.7039060567679 | Zfp11 |
| **ENSMUSG00000051041** | -0.936400212426002 | 0.0273923410758477 | 0.664353746352133 | 25.2983840639333 | 11.4018202048006 | 26.5007368821565 | 30.0136542162612 | 58.0327843784121 | 64.8233582999146 | 24.9858741733473 | 30.3947158107182 | Olfml1 |
| **ENSMUSG00000051043** | -0.807253774346716 | 0.011959293261318 | 0.635862253526407 | 92.1583990900426 | 61.0811796685746 | 64.3589324280944 | 80.6616957062019 | 182.858018701789 | 171.23151249034 | 75.998700610598 | 92.2696729968231 | Gprc5c |
| **ENSMUSG00000051065** | -0.25260420232418 | 0.175553504580545 | 0.799889021902745 | 183.413284463516 | 256.540954608014 | 204.434255948065 | 229.79204009325 | 238.700886688563 | 204.254732756335 | 309.200192895173 | 288.749800201823 | Mb21d2 |
| **ENSMUSG00000051067** | -0.382732383007849 | 0.127517594600624 | 0.772837268577874 | 131.009488902512 | 157.99665140938 | 91.8061241988993 | 96.6064495085906 | 141.249607260664 | 112.523565350795 | 182.188665847324 | 186.710397122983 | Lingo3 |
| **ENSMUSG00000051098** | -0.389524557306421 | 0.0609406950824731 | 0.71282992338528 | 175.281661014395 | 185.686786192467 | 131.557229522134 | 139.751077444466 | 199.282391639076 | 265.408844360028 | 158.243869764533 | 207.335382851685 | Mblac2 |
| **ENSMUSG00000051107** | 0.100944039773088 | 0.498491365510846 | 0.900927059989498 | 461.695509166782 | 401.506954354764 | 478.906173656114 | 593.707597465416 | 478.496731572945 | 423.186452297555 | 433.088485671353 | 468.947043936795 | NA |
| **ENSMUSG00000051111** | -0.103251439942263 | 0.846328711443701 | 0.97225256818733 | 30.7194663633475 | 8.95857301805762 | 8.51809399783602 | 24.3860940507122 | 16.4243729372865 | 11.0077400886647 | 19.7804837205666 | 30.3947158107182 | Sv2c |
| **ENSMUSG00000051113** | 0.351442021997269 | 0.37145677006441 | 0.873374202289323 | 16.2632468982428 | 17.9171460361152 | 36.9117406572894 | 20.6343872736795 | 19.7092475247437 | 14.6769867848863 | 21.8626399016789 | 15.1973579053591 | Fam71e1 |
| **ENSMUSG00000051124** | -0.114755586898517 | 0.729396254673646 | 0.952818017249783 | 48.7897406947284 | 33.3910448854875 | 37.8581955459379 | 45.0204813243917 | 53.6529515951357 | 66.0464405319884 | 27.0680303544596 | 32.5657669400552 | Gimap9 |
| **ENSMUSG00000051146** | -0.597482567226213 | 0.0386540113392258 | 0.679566075569698 | 28.0089252136404 | 24.4324718674299 | 24.6078271048596 | 25.3240207449704 | 36.1336204620302 | 52.592535979176 | 38.5198893505771 | 28.2236646813812 | Camk2n2 |
| **ENSMUSG00000051147** | -0.160337903940246 | 0.612595105978648 | 0.928055903343287 | 42.4651446787451 | 24.4324718674299 | 20.8220075502658 | 38.4549944645846 | 29.5638712871156 | 37.9155491942896 | 33.3144988977964 | 40.1644458927348 | Nat2 |
| **ENSMUSG00000051149** | -0.0676226400044252 | 0.88611123877586 | 0.980068672363141 | 10.8421645988285 | 36.6487078011448 | 50.1621090983677 | 33.7653609932938 | 40.5134532453066 | 15.9000690169602 | 37.4788112600209 | 43.4210225867403 | Adnp |
| **ENSMUSG00000051154** | 0.112929277915755 | 0.350607682740382 | 0.861774532843376 | 1391.41112351633 | 1331.56971677493 | 1322.19747944188 | 1374.06260708821 | 1190.21955885536 | 1049.40455511937 | 1361.73014244743 | 1407.92665737505 | Commd3 |

| | | | | | | | | | | | | |
|---|---|---|---|---|---|---|---|---|---|---|---|---|
| **ENSMUSG00000051166** | -0.0357648444745128 | 0.93413495300802 | 0.98818087308472 | 44.2721721118832 | 54.56585383726 | 70.0376617599851 | 40.3308478531009 | 30.6588294829347 | 23.2385624094033 | 93.6970281500523 | 66.217059444779 | Eml5 |
| **ENSMUSG00000051169** | 0.185828108372459 | 0.501900770659853 | 0.900927059989498 | 55.1143367107117 | 39.0919549878878 | 32.1794662140472 | 36.5791410760683 | 38.3235368536684 | 41.5847958905112 | 30.191264626128 | 33.6512925047237 | Rpusd3 |
| **ENSMUSG00000051177** | -0.228065444211282 | 0.293730836250648 | 0.851785577030345 | 102.097049972302 | 124.605606523892 | 123.985590412947 | 135.999370667433 | 150.009272827216 | 190.800828203522 | 122.847214685624 | 107.467030902182 | Plcb1 |
| **ENSMUSG00000051184** | -0.108377588074666 | 0.614841740220302 | 0.928075610917191 | 161.728955265859 | 126.234437981721 | 111.681676860517 | 157.571684635371 | 168.623562156141 | 183.462334811079 | 136.381229862854 | 112.894658725525 | Zfp524 |
| **ENSMUSG00000051185** | 0.0812418277169073 | 0.513130762716343 | 0.903078589555164 | 1110.41835766336 | 940.65016689605 | 946.454888648447 | 1075.80191831411 | 927.429591858775 | 831.695917810225 | 1051.4888714617 | 1037.76243982309 | Fam174a |
| **ENSMUSG00000051212** | 0.110086526774564 | 0.826080312639945 | 0.967195718403264 | 4.51756858284523 | 10.5874044758863 | 14.1968233297267 | 18.7585338851632 | 9.85462376237187 | 12.2308223207386 | 10.4107809055614 | 11.9407812113536 | Gpr183 |
| **ENSMUSG00000051223** | 0.0733508629454254 | 0.661476350805248 | 0.939611397743053 | 1697.70227343324 | 1883.74358097884 | 1973.35844283201 | 1745.48157801444 | 1694.99528712796 | 1263.4439457323 | 1907.25506189884 | 2071.18277738751 | Bzw1 |
| **ENSMUSG00000051224** | -0.0979473445000512 | 0.654736243245171 | 0.937362030372647 | 116.553269437407 | 68.4109212288036 | 79.5022106464695 | 86.2892558717508 | 104.021028602814 | 100.292743030056 | 88.4916376972716 | 82.4999429148066 | Tceanc |
| **ENSMUSG00000051232** | 0.00765401820441986 | 0.970879515684596 | 0.995122272737203 | 196.062476495483 | 130.306516626293 | 130.610774633486 | 136.937297361692 | 154.389105610493 | 140.654456688494 | 176.983275394543 | 118.322286548867 | Tmem199 |
| **ENSMUSG00000051234** | 0.0371495241897918 | 0.775613556749645 | 0.9595799687189 | 1873.8874481642 | 1420.34103122659 | 1414.00360364078 | 1682.64048949914 | 1462.86414961431 | 1526.40662562818 | 1540.79557402308 | 1697.76198314155 | Rnf7 |
| **ENSMUSG00000051235** | -0.0399401854526414 | 0.945616159082403 | 0.990505437194639 | 10.8421645988285 | 8.14415728914329 | 15.1432782183751 | 10.3171936368398 | 13.1394983498292 | 1.22308223207386 | 15.616171358342 | 15.1973579053591 | Gen1 |
| **ENSMUSG00000051236** | 0.183202454310331 | 0.405011475554159 | 0.881158207349486 | 2389.79378032512 | 2933.52545554941 | 2704.02161686861 | 2362.63734283631 | 1720.1793256318 | 1671.95341124497 | 2968.11363617555 | 2789.80070119806 | Msrb3 |
| **ENSMUSG00000051238** | -0.238014766928783 | 0.147029204631926 | 0.780607990324772 | 162.632468982428 | 135.193010999779 | 152.3792370724 | 145.378637610015 | 145.62944004394 | 165.116101329971 | 208.215618111227 | 182.368294864309 | Swsap1 |
| **ENSMUSG00000051243** | -1.16964299505606 | 0.0732554188453453 | 0.730864975478592 | 9.03513716569045 | 6.51532583131463 | 9.46454888648447 | 19.6964605794214 | 19.7092475247437 | 57.4848649074714 | 6.24646854333682 | 17.3684090346961 | Islr2 |
| **ENSMUSG00000051255** | 0.0230687697232084 | 0.903538507805441 | 0.985027469232262 | 187.930853046361 | 136.821842457607 | 134.396594188079 | 156.633757941113 | 176.288269526875 | 140.654456688494 | 122.847214685624 | 166.085411394282 | Gm6563 |
| **ENSMUSG00000051256** | 0.0461643187017555 | 0.752477412133636 | 0.956497786093635 | 318.940341948873 | 285.859920848929 | 367.224496795597 | 353.598363735327 | 373.380744774312 | 322.893709267499 | 285.255396812381 | 301.776106977845 | Jagn1 |
| **ENSMUSG00000051279** | -0.749394634023573 | 0.293093250983845 | 0.851785577030345 | 7.22810973255236 | 4.07207864457164 | 3.78581955459379 | 17.8206071909051 | 24.0890803080201 | 19.5693157131818 | 7.28754663389296 | 4.34210225867403 | Gdf6 |
| **ENSMUSG00000051285** | 0.177190127970038 | 0.402903514549541 | 0.881158207349486 | 1930.80881230805 | 2433.47419799601 | 2509.05190980703 | 1948.0737439742 | 1490.23810450979 | 1504.39114545085 | 2423.62979481469 | 2382.72861444737 | Pcmtd1 |
| **ENSMUSG00000051306** | 0.0368670979920655 | 0.797324234526289 | 0.962443175247254 | 141.85165350134 | 157.99665140938 | 154.272146849697 | 143.502784221499 | 131.394983498292 | 139.43137445642 | 151.997401221196 | 159.572258006271 | Usp42 |
| **ENSMUSG00000051316** | -0.190573083605728 | 0.168681503197986 | 0.79907358295209 | 164.439496415566 | 175.913797445495 | 166.576060402127 | 199.778385876988 | 214.611806380543 | 214.039390612925 | 190.517290571773 | 187.795922687652 | Taf7 |
| **ENSMUSG00000051319** | 0.0317346058415123 | 0.815157414813686 | 0.966177091475091 | 1074.27780900059 | 802.199492980614 | 829.094482456039 | 1012.02290310456 | 895.675804180021 | 984.581196819457 | 913.025485417732 | 843.45336374743 | 1500011K16I |
| **ENSMUSG00000051323** | -0.810224053241607 | 0.0325764221363233 | 0.671606027118506 | 103.000563688871 | 91.2145616384048 | 87.0738497556571 | 130.371810501884 | 280.309298129689 | 250.731857575141 | 97.8613405122768 | 93.3551985614916 | Pcdh19 |
| **ENSMUSG00000051329** | -0.0990388856952261 | 0.664497614102841 | 0.941439398193438 | 114.746242004269 | 136.007426728693 | 176.040609288611 | 108.799496533947 | 113.875652365186 | 124.754387671534 | 179.065431575656 | 155.230155747597 | Nup160 |
| **ENSMUSG00000051331** | 0.26843140793107 | 0.314455876541184 | 0.853228028313558 | 4796.75432126506 | 6940.45084180791 | 6272.15654707326 | 4549.88239384634 | 3873.96209680796 | 2813.08913376988 | 6136.11426573787 | 5905.25907179668 | Cacna1c |
| **ENSMUSG00000051335** | 0.157871348793111 | 0.587918782442862 | 0.923266719035655 | 924.294532050133 | 1307.1372449075 | 1435.77206607969 | 858.202925246218 | 911.005218921489 | 566.287073450197 | 1448.13962396359 | 1130.03211281992 | Gfod1 |
| **ENSMUSG00000051339** | -0.265105515760346 | 0.0726480921160969 | 0.729721356194038 | 639.687711330884 | 736.231818938553 | 689.019158936069 | 699.693313916588 | 1030.35566226577 | 843.926740130963 | 719.38496057429 | 729.473179457237 | 2900026A02I |
| **ENSMUSG00000051341** | 0.114456513205544 | 0.575317157495254 | 0.922450244597442 | 137.334084918495 | 133.56417954195 | 186.451613063744 | 135.999370667433 | 109.49581958191 | 123.53130543946 | 170.736806851206 | 143.289374536243 | Zfp52 |
| **ENSMUSG00000051343** | 0.284981953122475 | 0.147383474906108 | 0.780607990324772 | 2740.35710235391 | 3596.45985888568 | 4066.91665652238 | 3346.52244511312 | 2729.73078217701 | 2010.74718952943 | 3352.27145159076 | 3191.44516012541 | Rab11fip5 |
| **ENSMUSG00000051344** | -0.0665694177930302 | 0.676482932064712 | 0.94409858171788 | 108.421645988285 | 127.048853710635 | 121.146225747001 | 114.427056699496 | 121.54035973592 | 112.523565350795 | 142.627698406191 | 116.15123541953 | Plekhm3 |
| **ENSMUSG00000051346** | 0.037363838066149 | 0.817758383989738 | 0.966315773308182 | 510.48524986151 | 396.620459981278 | 423.065335225856 | 417.377378944882 | 410.609323432161 | 333.901449356164 | 472.649453112486 | 484.144401842154 | Spryd4 |
| **ENSMUSG00000051351** | -0.295201426438618 | 0.0325642010410196 | 0.671606027118506 | 449.046317134815 | 473.175538499225 | 388.046504345863 | 485.846027625728 | 563.903470846835 | 507.579126310652 | 640.263025692024 | 491.743080794834 | Zfp46 |
| **ENSMUSG00000051355** | -0.0156776508726451 | 0.888321237276095 | 0.980942551992548 | 1627.22820354085 | 1272.11736856418 | 1431.03979163645 | 1547.57904552597 | 1470.52885698505 | 1591.22998392809 | 1413.78404697523 | 1466.54503786715 | Commd1 |
| **ENSMUSG00000051359** | -0.46565916484791 | 0.0590138658756958 | 0.712165592539891 | 93.0619128066116 | 104.245213301034 | 95.5919437534931 | 75.0341355406529 | 162.053812981226 | 154.108361241306 | 107.231043327282 | 85.7565196088121 | Ncald |
| **ENSMUSG00000051367** | 0.0119992036299338 | 0.965659712079704 | 0.994284908685797 | 52.4037955610046 | 54.56585383726 | 37.8581955459379 | 66.5927952923295 | 35.0386622662111 | 51.3694537471021 | 65.5879197050366 | 57.5328549274309 | Six1 |
| **ENSMUSG00000051373** | -0.115984456098128 | 0.398961348721046 | 0.880459952732461 | 666.793122827955 | 588.008156276145 | 580.176846741498 | 623.721251681677 | 619.746338833609 | 818.242013257412 | 623.605776243126 | 604.637739520359 | Plpp7 |
| **ENSMUSG00000051375** | -0.753932831972209 | 0.0121466307854848 | 0.635862253526407 | 358.694945477911 | 347.755516246418 | 384.260684791269 | 500.852854733858 | 1007.36154015357 | 818.242013257412 | 459.115437935256 | 399.473407798011 | Pcdh1 |
| **ENSMUSG00000051390** | -0.180059860534293 | 0.135192929174682 | 0.775642208276055 | 394.835494140673 | 362.414999366876 | 466.602260103684 | 419.253232333398 | 474.116898789669 | 491.679057293692 | 444.54034466747 | 450.493109337431 | Zbtb22 |
| **ENSMUSG00000051391** | 0.138843419622174 | 0.452283445016855 | 0.891110666112841 | 5789.71589577444 | 6758.83613426001 | 6500.25217523753 | 5757.93197605085 | 5121.11948184592 | 4008.04047450604 | 6777.41836952045 | 6622.79147004256 | Ywhag |
| **ENSMUSG00000051396** | 0.0424295574107382 | 0.917466434048706 | 0.986713918026868 | 10.8421645988285 | 17.1027303072009 | 21.7684624389143 | 18.7585338851632 | 8.75966556655277 | 18.3462334811079 | 20.8215618111227 | 18.4539345993646 | Gm45902 |
| **ENSMUSG00000051403** | 0.125415071230234 | 0.266839361998435 | 0.843513595393324 | 2233.48590735868 | 2268.14780502641 | 2449.42525182218 | 2264.1550399392 | 1969.82979427856 | 1841.96184150323 | 2314.31659530629 | 2319.7681316966 | Ppp1r37 |
| **ENSMUSG00000051412** | 0.119591801737293 | 0.543122659926559 | 0.913837378403038 | 320.747369382011 | 253.283291692356 | 280.15064703994 | 297.322762079837 | 214.611806380543 | 205.477814988408 | 327.939598525183 | 310.460311495193 | Vamp7 |
| **ENSMUSG00000051413** | -0.208630665017913 | 0.347107720165629 | 0.861266047293794 | 61.4389327266951 | 48.8649437348597 | 70.0376617599851 | 74.0962088463947 | 83.2168228822514 | 81.9465095489486 | 65.5879197050366 | 62.9604827507734 | Plagl2 |
| **ENSMUSG00000051427** | -0.573491188775746 | 0.00287489262989938 | 0.635862253526407 | 56.0178504272808 | 57.8235167529173 | 58.6802030962037 | 54.3997482669734 | 97.4512794278996 | 79.5003450848009 | 72.8754663389296 | 87.9275707381491 | Ccdc157 |
| **ENSMUSG00000051435** | 0.717867985969182 | 0.0627350152356872 | 0.715615581523459 | 26.2018977805023 | 28.5045505120015 | 39.7511053232348 | 50.6480414899407 | 10.949581958191 | 31.8001380339204 | 24.9858741733473 | 20.6249857287016 | Fhad1 |
| **ENSMUSG00000051439** | -0.351529160942945 | 0.257101829602257 | 0.837517407767513 | 131.009488902512 | 148.223662662408 | 130.610774633486 | 118.178763476528 | 205.85214081399 | 251.954939807215 | 130.134761319517 | 86.8420451734806 | Cd14 |
| **ENSMUSG00000051444** | 0.305638155054216 | 0.580580991894425 | 0.922450244597442 | 11.7456783153976 | 6.51532583131463 | 23.6613722162112 | 8.44134024832345 | 8.75966556655277 | 6.1154111603693 | 14.5750932677859 | 10.8552556466851 | Bbs12 |
| **ENSMUSG00000051451** | 0.0177126089260065 | 0.920697924035808 | 0.986713918026868 | 246.659244623349 | 280.973426475443 | 218.631079277791 | 227.916186704733 | 201.472308030714 | 203.031650524261 | 292.542943446274 | 264.868237779116 | Crebzf |
| **ENSMUSG00000051456** | -0.391538691590632 | 0.200146855098923 | 0.812921911423739 | 182.509770746947 | 114.83261777692 | 65.3053873167428 | 124.744250336335 | 123.730276127558 | 182.239252579005 | 193.640524843441 | 140.032797842237 | Hspb3 |
| **ENSMUSG00000051469** | 0.0552091899751343 | 0.72340622674186 | 0.951674101666722 | 702.933671490717 | 775.323773926441 | 658.732602499319 | 532.742362338636 | 584.707676567398 | 576.071731306788 | 690.234774038719 | 718.617923810552 | Zfp24 |
| **ENSMUSG00000051483** | -0.00354228209769401 | 0.977412270013603 | 0.996179425433073 | 666.793122827955 | 570.905425968944 | 577.337482075553 | 679.996853337167 | 631.790878987619 | 702.049201210395 | 553.853544175865 | 614.407469602375 | Cbr1 |
| **ENSMUSG00000051495** | -0.382350926563571 | 0.232809979813234 | 0.832048466795696 | 723.714486971805 | 775.323773926441 | 500.674636095028 | 548.687116141024 | 1064.29936633616 | 1243.87463001912 | 484.101312108604 | 530.8220011229 | Irf2bp2 |
| **ENSMUSG00000051498** | 0.179589464836261 | 0.79009764346173 | 0.961659329450003 | 11.7456783153976 | 8.14415728914329 | 20.8220075502658 | 5.62756016554897 | 15.3294147414674 | 18.3462334811079 | 3.12323427166841 | 4.34210225867403 | Tlr6 |
| **ENSMUSG00000051499** | 0.0724571483473509 | 0.88244930553962 | 0.978854883720278 | 12.6491920319666 | 13.0306516626293 | 22.7149173275627 | 32.8274342990356 | 18.6142893289246 | 11.0077400886647 | 13.5340151772298 | 33.6512925047237 | Zfp786 |
| **ENSMUSG00000051502** | -0.307742961775993 | 0.178193365007183 | 0.801696999069105 | 103.90407740544 | 63.5244268553176 | 65.3053873167428 | 75.9720622349111 | 97.4512794278996 | 114.969729814943 | 83.2862472444909 | 86.8420451734806 | Ufsp1 |
| **ENSMUSG00000051506** | -0.256433132552553 | 0.375662058033656 | 0.873821423608136 | 36.1405486627618 | 53.7514381083457 | 46.3762895437739 | 60.9652351267805 | 41.6084114411257 | 47.7002070508805 | 65.5879197050366 | 80.3288917854696 | Wdfy4 |
| **ENSMUSG00000051510** | 0.110289726557035 | 0.478386651793822 | 0.897674307047416 | 1215.22594878537 | 1621.50171626843 | 1484.04126540076 | 1314.03529865568 | 1244.96746864631 | 1064.08154190426 | 1397.12679752634 | 1513.2226371479 | Mafg |
| **ENSMUSG00000051515** | -0.810121000389471 | 0.0750449100346244 | 0.735399943172214 | 28.0089252136404 | 41.5352021746308 | 70.0376617599851 | 50.6480414899407 | 99.6411958195378 | 149.216032313011 | 41.6431236222455 | 43.4210225867403 | Fam181b |
| **ENSMUSG00000051518** | -0.0101796870096349 | 0.955091875027336 | 0.992371505557225 | 249.369785773056 | 211.748089517725 | 169.415425068072 | 211.971432902344 | 212.421889988905 | 249.508775343067 | 170.736806851206 | 217.105112933702 | Rps19bp1 |
| **ENSMUSG00000051550** | -0.348464793779016 | 0.128123571105462 | 0.77303252008843 | 82.2197482077831 | 71.6685841444609 | 86.1273948670087 | 93.7926694258161 | 111.685735973548 | 143.100621152642 | 92.6559500594962 | 78.1578406561325 | Zfp579 |
| **ENSMUSG00000051557** | 0.327085120961244 | 0.121879626137357 | 0.769868040577702 | 97.5794813894569 | 84.6992358070902 | 90.8596693102509 | 114.427056699496 | 75.5521155115177 | 95.400414101761 | 58.3003730711437 | 80.3288917854696 | Pusl1 |
| **ENSMUSG00000051579** | -0.171537060872181 | 0.517092106135425 | 0.905197321821966 | 116.553269437407 | 75.7406627890326 | 106.002947528626 | 143.502784221499 | 160.958854785407 | 141.877538920568 | 94.7381062406084 | 99.8683519495027 | Tceal8 |
| **ENSMUSG00000051586** | 0.429811284422263 | 0.082224080188106 | 0.739855948845396 | 1551.33305134905 | 1946.45359210525 | 1961.05452927958 | 1605.73050056997 | 1100.43298679819 | 798.67269754423 | 1554.32958920031 | 1790.03165613837 | Mical3 |
| **ENSMUSG00000051615** | -0.193501549585331 | 0.158354334681984 | 0.789230661864335 | 271.957628687283 | 269.571606270643 | 243.238906382651 | 280.44008158319 | 326.297542354091 | 254.401104271363 | 296.707255808499 | 339.769501741243 | Rap2a |
| **ENSMUSG00000051627** | -0.0595316980767938 | 0.927371601975547 | 0.986790505471895 | 13.5527057485357 | 30.9477976987445 | 44.483379766477 | 10.3171936368398 | 16.4243729372865 | 4.89232892829544 | 40.6020455316893 | 41.2499714574033 | Hist1h1e |
| **ENSMUSG00000051650** | -0.0124586738813554 | 0.962313018289737 | 0.994128583764339 | 31.6229800799166 | 46.4216965481167 | 30.2865564367503 | 34.703287687552 | 37.2285786578493 | 33.0232202659942 | 37.4788112600209 | 36.9078691987293 | B3gnt2 |
| **ENSMUSG00000051652** | -0.548354967353496 | 0.288202449669595 | 0.851785577030345 | 17.1667606148119 | 36.6487078011448 | 10.4110037751329 | 13.1309737196143 | 20.8042057205628 | 14.6769867848863 | 29.1501865355718 | 48.8486504100828 | Lrrc3 |
| **ENSMUSG00000051669** | -0.807700141392031 | 0.0648738619254871 | 0.718339708336358 | 90.3513716569045 | 53.7514381083457 | 76.6628459805242 | 126.620103724852 | 173.003394939417 | 277.639666680766 | 67.6700758861489 | 90.0986218674861 | AU021092 |
| **ENSMUSG00000051671** | -0.0428342060596316 | 0.758244261425986 | 0.95677573228595 | 411.098741038915 | 337.982527499446 | 321.794662140472 | 417.377378944882 | 354.766455445387 | 421.963370065482 | 378.952424962434 | 377.762896504641 | Coa6 |
| **ENSMUSG00000051674** | 0.25706109994959 | 0.303550863855083 | 0.851785577030345 | 293.64195788494 | 336.353696041618 | 371.010316350191 | 287.005568442997 | 231.036179317829 | 161.446854633749 | 348.761160336306 | 335.427399482569 | Dcun1d4 |
| **ENSMUSG00000051675** | -0.146277746511571 | 0.265498030560617 | 0.842603400964081 | 247.562758339918 | 263.056280439328 | 262.16800415562 | 242.923013812864 | 254.03030143003 | 269.078091056249 | 328.980676615739 | 271.381391167127 | Trim32 |
| **ENSMUSG00000051695** | -0.0360420945804438 | 0.831752637711332 | 0.967583158900775 | 3762.2311157935 | 3670.57169021688 | 3791.49828392568 | 4170.96000936604 | 4657.95216501444 | 4713.75892241266 | 3312.71048414963 | 3101.34653825793 | Pcbp1 |
| **ENSMUSG00000051703** | -0.242720468728784 | 0.610105328133421 | 0.927932112008768 | 16.2632468982428 | 17.1027303072009 | 6.62518422053913 | 6.56548685980713 | 16.4243729372865 | 14.6769867848863 | 11.4518589961175 | 13.0263067760221 | Tmem198 |
| **ENSMUSG00000051705** | -0.0544212663277758 | 0.814783686274076 | 0.966177091475091 | 113.8427282877 | 114.83261777692 | 82.3415753124149 | 88.1651092602672 | 99.6411958195378 | 70.9387694602839 | 118.6829023234 | 124.835439936878 | Senp8 |
| **ENSMUSG00000051716** | -0.343215178077613 | 0.377674782393203 | 0.87422797717331 | 59.631905293557 | 122.162359337149 | 55.8408384302584 | 84.4134024832345 | 63.5075753575076 | 80.7234273168747 | 81.2040910633787 | 183.453820428978 | Apon |
| **ENSMUSG00000051721** | 0.253057183401752 | 0.493820313948035 | 0.900117970535229 | 103.90407740544 | 118.904696421492 | 110.735221971868 | 51.5859681841989 | 49.2731188118594 | 51.3694537471021 | 109.313199508394 | 112.894658725525 | Wdcp |
| **ENSMUSG00000051730** | 0.207967890948085 | 0.0555779183679937 | 0.703396849069134 | 393.028466707535 | 382.775392589734 | 352.081218577222 | 403.308478531009 | 331.772333333186 | 341.239942748607 | 308.159114804616 | 345.197129564585 | Mettl5 |
| **ENSMUSG00000051735** | -0.109183498672931 | 0.730847932909021 | 0.952921973914044 | 25.2983840639333 | 35.0198763433161 | 39.7511053232348 | 30.0136542162612 | 45.9882442244021 | 39.1386314263635 | 20.8215618111227 | 34.7368180693922 | Rinl |
| **ENSMUSG00000051736** | 0.139974059174569 | 0.571465334047647 | 0.921130164032017 | 64.1494738764022 | 44.7928650902881 | 58.6802030962037 | 64.7169419038131 | 42.7033696369448 | 66.0464405319884 | 43.7252798033577 | 58.6183804920994 | Fam229b |
| **ENSMUSG00000051747** | 0.528835365592098 | 0.00256267525230934 | 0.635862253526407 | 69656.3899788905 | 86376.1177929248 | 68302.8099490925 | 78350.6443315498 | 58305.4289691711 | 37208.607664151 | 55022.0181639824 | 59258.8405752538 | Ttn |
| **ENSMUSG00000051748** | -0.358291052270958 | 0.352399819543757 | 0.862728204353539 | 71.3775836089546 | 120.533527879321 | 93.6990339761962 | 86.2892558717508 | 60.2227007700503 | 125.977469903608 | 75.998700610598 | 214.934061804364 | Wfdc21 |
| **ENSMUSG00000051768** | -0.0037913954845247 | 0.98363576843828 | 0.996940244108351 | 521.327414460339 | 437.341246426994 | 482.691993210708 | 513.045901759214 | 604.416924092141 | 569.956320146419 | 382.075659234102 | 403.815510056685 | Xrcc1 |
| **ENSMUSG00000051786** | 0.102074054379417 | 0.707682880677384 | 0.948997413845227 | 151.7903043836 | 251.654460234528 | 251.757000380487 | 193.212899017181 | 156.579022002131 | 132.092881063977 | 264.433835001259 | 236.644573097735 | Tubgcp6 |
| **ENSMUSG00000051790** | -0.328273369767118 | 0.231046140911704 | 0.832048466795696 | 66.8600150261093 | 92.8433930962335 | 74.7699362032273 | 61.9031618210386 | 127.015150715015 | 110.077400886647 | 67.6700758861489 | 68.388110574116 | Nlgn2 |
| **ENSMUSG00000051805** | -0.44147009348687 | 0.293273531885908 | 0.851785577030345 | 15.3597331816738 | 13.0306516626293 | 10.4110037751329 | 25.3240207449704 | 15.3294147414674 | 30.5770558018465 | 24.9858741733473 | 16.2828834700276 | A530095I07F |
| **ENSMUSG00000051811** | 0.28748525067694 | 0.487325233696752 | 0.899236955296196 | 34.3335212296237 | 25.2468875963442 | 58.6802030962037 | 34.703287687552 | 17.5193311331055 | 53.8156182112498 | 23.9447960827911 | 30.3947158107182 | Cox6b2 |
| **ENSMUSG00000051817** | 0.0633402345755486 | 0.754369191537542 | 0.956497786093635 | 497.836057829544 | 570.905425968944 | 419.279515671262 | 458.646153492241 | 395.279908690694 | 343.686107212755 | 560.100012719201 | 563.387768062955 | Sox12 |
| **ENSMUSG00000051839** | 1.14445687199388 | NA | NA | 31.6229800799166 | 22.8036404096012 | 298.133289924261 | 9.37926694258161 | 26.2789966996583 | 100.292743030056 | 16.6572494488982 | 20.6249857287016 | Gypa |

| | | | | | | | | | | | | |
|---|---|---|---|---|---|---|---|---|---|---|---|---|
| **ENSMUSG00000051851** | -0.41146740726178 | 0.155262749894892 | 0.787275540343856 | 121.070838020252 | 126.234437981721 | 104.110037751329 | 111.613276616721 | 160.958854785407 | 242.170281950624 | 122.847214685624 | 91.1841474321546 | Rtl8c |
| **ENSMUSG00000051853** | -0.200896089666759 | 0.171906306977683 | 0.799889021902745 | 614.389327266951 | 750.891302059011 | 771.360734248484 | 671.555513088843 | 929.619508250413 | 881.842289325253 | 691.275852129275 | 726.216602763232 | Arf3 |
| **ENSMUSG00000051855** | -0.537434148463167 | 0.0668390657206732 | 0.719725956118125 | 46.9827132615903 | 43.1640336324594 | 28.3936466594534 | 64.7169419038131 | 81.0269064906132 | 74.6080161565054 | 53.094982618363 | 57.5328549274309 | Mest |
| **ENSMUSG00000051864** | -0.0518398930856185 | 0.587761320646492 | 0.923241888853076 | 775.214768816241 | 797.312998607128 | 879.256591554407 | 768.161962597434 | 879.251431242735 | 781.549546295196 | 850.560799984364 | 824.999429148066 | Tbc1d22a |
| **ENSMUSG00000051890** | 0.358533023548839 | 0.155099789558666 | 0.787275540343856 | 861.952085606869 | 1031.05031280554 | 1149.94268970786 | 846.94780491512 | 699.678287128403 | 435.417274618294 | 986.942029847218 | 910.755948756878 | Klhdc1 |
| **ENSMUSG00000051910** | -0.157956645647288 | 0.541970208775786 | 0.913837378403038 | 186.123825613223 | 193.83094348161 | 216.738169500494 | 164.137171495178 | 186.142893289246 | 123.53130543946 | 288.37863108405 | 249.670879873757 | Sox6 |
| **ENSMUSG00000051934** | -0.156101820496615 | 0.292394978778698 | 0.851785577030345 | 152.693818100169 | 149.852494120236 | 127.77140996754 | 137.87522405595 | 176.288269526875 | 160.223772401676 | 155.120635492864 | 142.203848971575 | Spats2 |
| **ENSMUSG00000051950** | -0.251529204421754 | 0.123550715998807 | 0.770251097497048 | 103.90407740544 | 136.007426728693 | 126.824955078892 | 139.751077444466 | 144.534481848121 | 135.762127760198 | 157.202791673977 | 164.999885829613 | B3glct |
| **ENSMUSG00000051977** | 0.0300932560180031 | 0.953203247430745 | 0.992057028833586 | 28.9124389302094 | 27.6901347830872 | 8.51809399783602 | 13.1309737196143 | 10.949581958191 | 18.3462334811079 | 32.2734208072402 | 15.1973579053591 | Prdm9 |
| **ENSMUSG00000051978** | 0.0776070809513434 | 0.658138758618395 | 0.938942826336227 | 119.263810587114 | 89.5857301805761 | 121.146225747001 | 115.364983393754 | 105.115986798633 | 113.746647582869 | 108.272121417838 | 94.4407241261602 | Erich1 |
| **ENSMUSG00000051984** | 0.260565012655684 | 0.45967221903457 | 0.894774044236998 | 150.886790667031 | 195.459774939439 | 194.96970706158 | 133.185590584659 | 85.4067392738896 | 73.3849339244316 | 208.215618111227 | 195.394601640331 | Sec31b |
| **ENSMUSG00000051989** | -0.0676726243685956 | 0.73011845502246 | 0.952921973914044 | 1217.93648993507 | 816.044560372157 | 853.702309560899 | 1166.78080765715 | 1036.92541144068 | 1349.05970197747 | 940.093515772191 | 923.7822555329 | Smim11 |
| **ENSMUSG00000052031** | 0.0638778672310379 | 0.760609243451442 | 0.957357521291862 | 262.922491521592 | 178.357044632238 | 176.040609288611 | 202.592165959763 | 148.914314631397 | 199.362403828039 | 243.612273190136 | 192.138024946326 | Tagap1 |
| **ENSMUSG00000052033** | 0.324345001184674 | 0.176906581534658 | 0.799889021902745 | 188.83436676293 | 134.378595270864 | 122.09268063565 | 172.578511743502 | 97.4512794278996 | 156.554525705454 | 95.7791843311646 | 144.374900100912 | Pfdn4 |
| **ENSMUSG00000052040** | -0.330603824534734 | 0.207113337802767 | 0.819960256425221 | 1067.04969926804 | 1149.95500922703 | 868.845587779274 | 787.858423176855 | 1517.61205940527 | 1659.72258892423 | 866.176971342706 | 829.34153140674 | Klf13 |
| **ENSMUSG00000052056** | -0.229322907369149 | 0.25151517521137 | 0.83723604018117 | 129.202461469373 | 130.306516626293 | 149.539872406455 | 145.378637610015 | 210.231973597267 | 124.754387671534 | 175.942197303987 | 137.8617467129 | Zfp217 |
| **ENSMUSG00000052062** | 0.179889681832497 | 0.31349282009613 | 0.85295001595893 | 357.791431761342 | 399.878122896935 | 371.95677123884 | 294.508981997063 | 277.024423542232 | 253.178022039289 | 381.034581143546 | 345.197129564585 | Pard3b |
| **ENSMUSG00000052085** | 0.0786658084470111 | 0.762768805480659 | 0.957842155335615 | 359.59845919448 | 418.609684661965 | 315.169477919933 | 285.129715054481 | 310.968127612624 | 183.462334811079 | 395.609674411332 | 414.67076570337 | Dock8 |
| **ENSMUSG00000052102** | 0.217575207223143 | 0.162712092659979 | 0.793968226938511 | 220.457346842847 | 274.458100644129 | 214.845259723197 | 243.860940507122 | 183.952976897608 | 185.908499275227 | 239.447960827911 | 210.59195954569 | Gnpda1 |
| **ENSMUSG00000052105** | 0.10527031897149 | 0.683845386094442 | 0.944452327903597 | 63.2459601598332 | 106.688460487777 | 65.3053873167428 | 80.6616957062019 | 55.8428679867739 | 64.8233582999146 | 88.4916376972716 | 84.6709940441436 | Mtcl1 |
| **ENSMUSG00000052117** | 0.190144804821142 | 0.67417713364435 | 0.943983909911292 | 9.03513716569045 | 8.95857301805762 | 13.2503684410783 | 16.8826804966469 | 13.1394983498292 | 8.56157562451702 | 10.4107809055614 | 9.76973008201657 | D630039A03 |
| **ENSMUSG00000052133** | -0.709859317832088 | 0.111309225252715 | 0.764116514278315 | 11.7456783153976 | 16.2883145782866 | 15.1432782183751 | 11.2551203310979 | 28.4689130912965 | 12.2308223207386 | 13.5340151772298 | 34.7368180693922 | Sema5b |
| **ENSMUSG00000052135** | -0.386411549203735 | 0.276451910225242 | 0.846750864146095 | 13.5527057485357 | 21.1748089517725 | 17.036187995672 | 13.1309737196143 | 22.994122112201 | 18.3462334811079 | 17.6983275394543 | 26.0526135520442 | Foxo6 |
| **ENSMUSG00000052137** | -0.0957601561626957 | 0.741117294486744 | 0.954383574136061 | 60.535419010126 | 49.679359463774 | 39.7511053232348 | 33.7653609932938 | 65.6974917491458 | 51.3694537471021 | 37.4788112600209 | 42.3354970220718 | Rbm12b2 |
| **ENSMUSG00000052139** | -0.0251264832519931 | 0.823834705626988 | 0.966637987288264 | 777.021796249379 | 684.109212288036 | 688.072704047421 | 768.161962597434 | 809.174106710313 | 786.441875223492 | 704.809867306505 | 668.683747835801 | Babam2 |
| **ENSMUSG00000052143** | 0.144890386046587 | 0.769055761311285 | 0.95879680569409 | 19.877301764519 | 15.4738988493722 | 19.8755526616174 | 30.0136542162612 | 14.2344565456483 | 6.1154111603693 | 34.3555769883525 | 21.7105112933701 | Gm9869 |
| **ENSMUSG00000052144** | 0.228366330926263 | 0.135862610625223 | 0.776087404131513 | 1060.72510325206 | 1181.71722265469 | 1148.04977993057 | 997.954002690684 | 883.631264026011 | 731.403174780168 | 1045.24240291836 | 1084.44003910384 | Ppp4r2 |
| **ENSMUSG00000052146** | 0.14159100700073 | 0.337170540120504 | 0.858570960161243 | 5592.74990556239 | 4101.39761081256 | 5159.12559802268 | 5890.17963994125 | 4813.43622882075 | 5295.94606487981 | 4170.55883076788 | 4524.47055353834 | Rps10 |
| **ENSMUSG00000052151** | -0.322253386568021 | 0.260104010426578 | 0.840316047847749 | 61.4389327266951 | 50.4937751926884 | 32.1794662140472 | 34.703287687552 | 50.3680770076785 | 72.1618516923577 | 49.9717483466946 | 52.1052271040884 | Plpp2 |
| **ENSMUSG00000052154** | 0.412982072634875 | 0.383475297313462 | 0.875126835008975 | 15.3597331816738 | 34.2054606144018 | 12.3039135524298 | 17.8206071909051 | 14.2344565456483 | 7.33849339244316 | 19.7804837205666 | 18.4539345993646 | Gm9870 |
| **ENSMUSG00000052155** | 0.127781180030932 | 0.238252868956745 | 0.832048466795696 | 407.484686172639 | 449.557482360709 | 431.583429223692 | 446.453106466885 | 371.190828382674 | 379.155491942897 | 398.732908683 | 438.552328126077 | Acvr2a |
| **ENSMUSG00000052160** | -0.305876386252037 | 0.344762351111533 | 0.861266047293794 | 42.4651446787451 | 52.9370223794314 | 48.2691993210708 | 51.5859681841989 | 78.836990098975 | 85.6157562451702 | 34.3555769883525 | 43.4210225867403 | Pld4 |
| **ENSMUSG00000052188** | -0.238313002799994 | 0.262775780307168 | 0.842065278491208 | 56.9213641438498 | 59.452348210746 | 54.8943835416099 | 57.2135283497478 | 78.836990098975 | 74.6080161565054 | 65.5879197050366 | 51.0197015394199 | Gm14964 |
| **ENSMUSG00000052214** | -0.0386908725094533 | 0.768712976020179 | 0.95879680569409 | 1996.76531361759 | 1940.75268200285 | 1790.69264932286 | 2001.53556554692 | 1622.7280462039 | 1890.88513078619 | 2188.346146349 | 2237.26818878179 | Opa3 |
| **ENSMUSG00000052241** | -0.213082704527322 | 0.642100476633204 | 0.934616627390525 | 8.13162344912141 | 15.4738988493722 | 11.3574586637814 | 10.3171936368398 | 17.5193311331055 | 7.33849339244316 | 16.6572494488982 | 10.8552556466851 | A930035D04 |
| **ENSMUSG00000052248** | 0.277302263210138 | 0.244322076654189 | 0.832048466795696 | 50.5967681278665 | 57.8235167529173 | 54.8943835416099 | 68.4686486808458 | 55.8428679867739 | 56.2617826753975 | 42.6842017128016 | 36.9078691987293 | Zeb2os |
| **ENSMUSG00000052253** | -0.00612261174317287 | 0.967407588282259 | 0.994284908685797 | 1209.80486648595 | 1172.75864963663 | 1226.60553568839 | 1285.89749782794 | 1375.26749394879 | 1465.25251402448 | 1067.10504282004 | 1009.53877514171 | Zfp622 |
| **ENSMUSG00000052273** | 0.355712797542595 | 0.478182916563457 | 0.897674307047416 | 10.8421645988285 | 13.8450673915436 | 18.9290977729689 | 14.0689004138724 | 6.56974917491458 | 22.0154801773295 | 10.4107809055614 | 6.51315338801105 | Dnah3 |
| **ENSMUSG00000052291** | -0.146334521643945 | 0.740541263495933 | 0.954383574136061 | 7.22810973255236 | 10.5874044758863 | 12.3039135524298 | 12.1930470253561 | 9.85462376237187 | 15.9000690169602 | 10.4107809055614 | 10.8552556466851 | 5330438D12I |
| **ENSMUSG00000052293** | 0.0903680923470035 | 0.784651182684663 | 0.961141387070613 | 16.2632468982428 | 26.8757190541728 | 18.9290977729689 | 22.5102406621959 | 19.7092475247437 | 19.5693157131818 | 19.7804837205666 | 20.6249857287016 | Taf9 |
| **ENSMUSG00000052296** | 0.108647115382397 | 0.410672313013655 | 0.882192126163777 | 2039.23045829633 | 2152.50077152057 | 2157.91714611846 | 1957.45301091678 | 1800.1112739266 | 1646.26868437142 | 2335.13815711741 | 1921.38024946326 | Ppp6r1 |
| **ENSMUSG00000052298** | 0.0522813974026855 | 0.612407860435483 | 0.928055903343287 | 602.643648951553 | 667.820897709749 | 738.234813145789 | 662.176246146262 | 672.304332232926 | 618.879609429373 | 645.468416144805 | 638.289032025082 | Cdc42se2 |
| **ENSMUSG00000052299** | 0.124178913271546 | 0.513315223069231 | 0.903087831516097 | 447.239289701677 | 548.101785559343 | 594.373670071225 | 455.832373409466 | 485.06648074786 | 331.455284892016 | 543.442763270303 | 515.624643217541 | Ltn1 |
| **ENSMUSG00000052302** | -0.33488723951978 | 0.334843332391729 | 0.858570960161243 | 18.9737880479499 | 24.4324718674299 | 22.7149173275627 | 26.2619474392285 | 28.4689130912965 | 42.8078781225851 | 16.6572494488982 | 29.3091902460497 | Tbc1d30 |
| **ENSMUSG00000052305** | 0.307415114036691 | 0.731894507020757 | 0.953219349759111 | 30557.7374080817 | 12694.2979665876 | 81068.5934871827 | 12018.5926602241 | 14315.4834521389 | 64741.4117903656 | 10751.2134411732 | 20366.6306443105 | Hbb-bs |
| **ENSMUSG00000052310** | -0.0845991994737488 | 0.240986875695284 | 0.832048466795696 | 1465.49924827499 | 1425.22752560008 | 1458.48698340726 | 1424.71064857815 | 1608.49358965825 | 1516.62196777159 | 1525.17940266474 | 1471.9726656905 | Slc39a1 |
| **ENSMUSG00000052316** | -0.0184641699094093 | 0.962648599079098 | 0.994140449160496 | 207.80815481088 | 193.016527752696 | 101.270673085384 | 128.495957113368 | 121.54035973592 | 73.3849339244316 | 162.408182126757 | 281.151121249143 | Lrrc15 |
| **ENSMUSG00000052331** | 0.0419434606130688 | 0.822305792598716 | 0.966637987288264 | 96.6759676728878 | 114.018202048006 | 135.343049076728 | 97.5443762028488 | 117.160526952643 | 118.638976511164 | 98.902418602833 | 96.6117752554972 | Ankrd44 |
| **ENSMUSG00000052336** | 0.283074229067713 | 0.631904983152836 | 0.931825800800958 | 15.3597331816738 | 28.5045505120015 | 48.2691993210708 | 61.9031618210386 | 13.1394983498292 | 11.0077400886647 | 53.094982618363 | 48.8486504100828 | Cx3cr1 |
| **ENSMUSG00000052337** | 0.189337444513604 | 0.293597538393939 | 0.851785577030345 | 13085.5891570695 | 14012.8370316999 | 14041.6047279884 | 13281.0419906956 | 10810.5222673219 | 8464.95212818318 | 14233.6196540835 | 14217.1283204634 | Immt |
| **ENSMUSG00000052364** | -0.118566942673006 | 0.770370125425607 | 0.958992744413128 | 18.0702743313809 | 14.6594831204579 | 23.6613722162112 | 18.7585338851632 | 31.7537876787538 | 23.2385624094033 | 11.4518589961175 | 15.1973579053591 | B630019K06 |
| **ENSMUSG00000052369** | 0.104870776843897 | 0.337667859092863 | 0.858570960161243 | 333.396561413978 | 351.82759489099 | 316.115932808581 | 326.39848960184 | 298.923587458613 | 333.901449356164 | 309.200192895173 | 294.177428025166 | Tmem106c |
| **ENSMUSG00000052373** | 0.35317200839711 | 0.406526815553362 | 0.881296485258502 | 14.4562194651047 | 9.77298874697194 | 15.1432782183751 | 13.1309737196143 | 12.0445401540101 | 9.78465785659088 | 10.4107809055614 | 8.68420451734806 | Mpp3 |
| **ENSMUSG00000052374** | 0.261714482734569 | 0.0309083414018987 | 0.667184591665619 | 69177.5277091089 | 71674.2850545633 | 69627.8467932003 | 72830.9457358405 | 59902.9729768712 | 47722.2225310579 | 65203.7618896214 | 63477.1929195557 | Actn2 |
| **ENSMUSG00000052384** | -0.560079196196762 | 0.101028256123935 | 0.759800953500741 | 48.7897406947284 | 37.4631235300591 | 58.6802030962037 | 59.0893817382642 | 108.400861386091 | 96.6234963338349 | 42.6842017128016 | 53.1907526687569 | Nrros |
| **ENSMUSG00000052387** | -0.295262091684128 | 0.401297222946222 | 0.880989161802279 | 13.5527057485357 | 24.4324718674299 | 21.7684624389143 | 16.8826804966469 | 16.4243729372865 | 28.1308913376988 | 27.0680303544596 | 22.7960368580387 | Trpm3 |
| **ENSMUSG00000052392** | 0.20318731916989 | 0.556120814496868 | 0.916239900758826 | 46.9827132615903 | 56.1946852950887 | 44.483379766477 | 50.6480414899407 | 31.7537876787538 | 20.7923979452556 | 58.3003730711437 | 60.7894316214364 | Acot4 |
| **ENSMUSG00000052395** | 0.111765619094826 | 0.415216150653974 | 0.882192126163777 | 191.544907912638 | 183.243539005724 | 193.076797284283 | 188.52326554589 | 167.528603960322 | 159.000690169602 | 169.69572876065 | 202.993280593011 | Rft1 |
| **ENSMUSG00000052397** | 0.143542774436702 | 0.228192107655952 | 0.832048466795696 | 1021.87401343959 | 973.226796052623 | 1094.1018512776 | 1134.89130005238 | 1008.45649834939 | 818.242013257412 | 939.052437681635 | 1056.21637442246 | Ezr |
| **ENSMUSG00000052406** | -0.232741693686143 | 0.180080786994641 | 0.802731406993634 | 177.992202164102 | 164.511977240694 | 137.235958854025 | 185.709485463116 | 180.668102310151 | 239.724117486476 | 170.736806851206 | 192.138024946326 | Rexo4 |
| **ENSMUSG00000052407** | 0.548650469948779 | 0.270019235708175 | 0.843793744782845 | 14.4562194651047 | 18.7315617650296 | 10.4110037751329 | 9.37926694258161 | 9.85462376237187 | 8.56157562451702 | 13.5340151772298 | 4.34210225867403 | Ccdc171 |
| **ENSMUSG00000052419** | -0.0194170854447454 | 0.892597365483892 | 0.982968160356082 | 695.705561758165 | 514.710740673856 | 512.978549647458 | 580.576623745802 | 590.182467546493 | 649.456665231219 | 565.305403171982 | 530.8220011229 | 2610001J05F |
| **ENSMUSG00000052423** | -0.203333153953406 | 0.180473211546902 | 0.802808628271642 | 194.255449062345 | 187.315617650296 | 195.916161950228 | 173.51643843776 | 238.700886688563 | 181.016170346931 | 197.804837205666 | 246.414303179751 | B4galt3 |
| **ENSMUSG00000052428** | 0.0394733197210051 | 0.652180634129653 | 0.936720465895231 | 1467.30627570813 | 1388.57881779893 | 1277.7140996754 | 1435.96576890924 | 1351.17841364077 | 1419.99847143775 | 1360.68906435687 | 1288.51884526152 | Tmco1 |
| **ENSMUSG00000052446** | 0.109287387894513 | 0.644093101335022 | 0.934847999069355 | 82.2197482077831 | 72.4829998733752 | 65.3053873167428 | 69.4065753751039 | 54.7479097909548 | 57.4848649074714 | 61.4236073428121 | 94.4407241261602 | Zfp961 |
| **ENSMUSG00000052456** | 0.141374894217817 | 0.268514844442034 | 0.843793744782845 | 1596.5087371775 | 1332.38413250384 | 1431.03979163645 | 1590.72367346184 | 1419.06582178155 | 1542.30669464514 | 1195.15764795844 | 1239.67019485144 | Asna1 |
| **ENSMUSG00000052459** | -0.290482135751293 | 0.0149841843641059 | 0.635862253526407 | 365.019541493894 | 377.888898216248 | 327.473391472363 | 318.895076047775 | 468.642107810573 | 424.409534529629 | 411.225845769674 | 396.216831104005 | Atp6v1a |
| **ENSMUSG00000052485** | 0.140835723601074 | 0.623351875565777 | 0.931059568526633 | 93.9654265231807 | 74.9262470601182 | 125.878500190243 | 169.764731660727 | 85.4067392738896 | 89.2850029413918 | 118.6829023234 | 127.006491066215 | Tmem171 |
| **ENSMUSG00000052488** | -0.0129017873992056 | 0.901295398114422 | 0.984802943363903 | 546.625798524272 | 626.285695535119 | 633.178320505811 | 617.15576482187 | 664.639624862192 | 579.740978003009 | 593.414511616998 | 606.808790649696 | Cherp |
| **ENSMUSG00000052496** | -0.0121467774591868 | 0.962493966484005 | 0.994128583764339 | 112.035700854562 | 113.203786319092 | 111.681676860517 | 126.620103724852 | 128.110108910834 | 63.6002760678407 | 155.120635492864 | 119.407812113536 | Pkdrej |
| **ENSMUSG00000052512** | 0.166353438270222 | 0.442143731589412 | 0.89108031640712 | 580.959319753896 | 814.415728914329 | 674.822335606343 | 563.693943249155 | 697.488370736765 | 382.824738639118 | 625.687932424238 | 640.46008315442 | Nav2 |
| **ENSMUSG00000052516** | 0.213247041852056 | 0.730525941911301 | 0.952921973914044 | 21.6843291976571 | 34.2054606144018 | 6.62518422053913 | 9.37926694258161 | 18.6142893289246 | 14.6769867848863 | 5.20539045278068 | 23.8815624227072 | Robo2 |
| **ENSMUSG00000052520** | -0.294666939172867 | 0.396074769068721 | 0.879251804756997 | 112.939214571131 | 200.346269312925 | 111.681676860517 | 124.744250336335 | 134.679858085749 | 133.315963296051 | 110.35427759895 | 296.348479154503 | Cyp2j5 |
| **ENSMUSG00000052533** | -0.0175916686376855 | 0.918044393956509 | 0.986713918026868 | 361.405486627618 | 360.786167909048 | 355.867038131816 | 311.39166249371 | 329.582416941548 | 262.96267989588 | 416.431236222455 | 396.216831104005 | Nup188 |
| **ENSMUSG00000052534** | 0.094853101310048 | 0.252370035551819 | 0.83723604018117 | 1341.71786910503 | 1412.19687393745 | 1292.85737789378 | 1454.72430279441 | 1292.05067106653 | 1253.65928787571 | 1259.70448957293 | 1346.05170018895 | Pbx1 |
| **ENSMUSG00000052539** | 0.153566659656218 | 0.438526426355457 | 0.889346190988072 | 426.458474220589 | 519.597235047342 | 427.797609669098 | 380.798237868813 | 323.012667766634 | 311.885969178834 | 433.088485671353 | 509.11148982953 | Magi3 |
| **ENSMUSG00000052544** | -0.667001036490407 | 0.0694139091245217 | 0.724352955175446 | 48.7897406947284 | 32.5766291565731 | 34.0723759913441 | 28.1378008277448 | 62.4126171616885 | 92.9542496376133 | 34.3555769883525 | 39.0789203280663 | St6galnac3 |
| **ENSMUSG00000052557** | 0.0108898143073114 | 0.961574442131956 | 0.993822507448007 | 89.4478579403355 | 133.56417954195 | 120.199770858353 | 112.551203310979 | 141.249607260664 | 101.51582526213 | 127.011527047849 | 82.4999429148066 | Gan |
| **ENSMUSG00000052560** | -0.434401691502263 | 0.0502526447213322 | 0.703396849069134 | 78.6056933415069 | 76.5550785179469 | 92.7525790875478 | 75.9720622349111 | 129.205067106653 | 135.762127760198 | 82.2451691539348 | 91.1841474321546 | Cpne8 |
| **ENSMUSG00000052562** | -0.47182965889768 | 0.183127899792696 | 0.805716353538343 | 22.5878429142261 | 28.5045505120015 | 24.6078271048596 | 21.5723139679377 | 30.6588294829347 | 29.3539735697726 | 20.8215618111227 | 54.2762782334254 | Slc22a30 |
| **ENSMUSG00000052566** | 0.38601098431796 | 0.101405657406055 | 0.759800953500741 | 116.553269437407 | 118.090280692578 | 182.66579350915 | 127.55803041911 | 134.679858085749 | 95.400414101761 | 80.1630129728225 | 106.381505337514 | Hook2 |
| **ENSMUSG00000052572** | -0.134302689979699 | 0.677429813165927 | 0.944214159784677 | 18.9737880479499 | 16.2883145782866 | 20.8220075502658 | 20.6343872736795 | 22.994122112201 | 18.3462334811079 | 22.903717992235 | 19.5394601640331 | Dlg2 |
| **ENSMUSG00000052593** | -0.143810594022739 | 0.370093942066548 | 0.872916429836026 | 500.546598979251 | 591.265819191803 | 490.263632319895 | 383.612017951588 | 520.105143014071 | 489.232892829544 | 605.907448703672 | 556.874614674944 | Adam17 |
| **ENSMUSG00000052595** | -0.583804014553617 | 0.292816362797801 | 0.851785577030345 | 8.13162344912141 | 22.8036404096012 | 12.3039135524298 | 12.1930470253561 | 18.6142893289246 | 9.78465785659088 | 12.4929370866736 | 42.3354970220718 | A1cf |
| **ENSMUSG00000052605** | -0.0954505047796542 | 0.609441155982144 | 0.927932112008768 | 141.85165350134 | 123.791190794978 | 113.574586637814 | 140.689004138724 | 137.964732673206 | 177.34692365071 | 117.641824232843 | 123.74991437221 | Isoc2b |
| **ENSMUSG00000052609** | -0.383513433885869 | 0.0209985581550669 | 0.648762042745232 | 132.81651633565 | 162.883145782866 | 159.950876181588 | 158.509611329629 | 245.270635863478 | 194.470074899744 | 171.777884941763 | 189.966973816989 | Plekhg3 |
| **ENSMUSG00000052632** | 0.114081939736978 | 0.59652838216859 | 0.924496535139492 | 383.089815825275 | 489.463853077511 | 373.849681016137 | 394.867138282686 | 295.638712871156 | 283.755077841135 | 464.320828388037 | 472.203620630801 | Asap2 |

| | | | | | | | | | | | | |
|---|---|---|---|---|---|---|---|---|---|---|---|---|
| **ENSMUSG00000052656** | -0.142101128241514 | 0.210676754953107 | 0.822896573073231 | 529.45903790946 | 619.770369703804 | 545.158015861505 | 569.321503414704 | 560.618596259378 | 601.756458180339 | 672.536446499264 | 663.256120012458 | Rnf103 |
| **ENSMUSG00000052658** | 0.259831608747725 | 0.559726304642041 | 0.916574868225118 | 18.9737880479499 | 18.7315617650296 | 24.6078271048596 | 30.9515809105193 | 16.4243729372865 | 31.8001380339204 | 7.28754663389296 | 22.7960368580387 | 5830454E08 |
| **ENSMUSG00000052673** | -0.270688193690394 | 0.544908390158728 | 0.914103567548881 | 9.93865088225949 | 11.4018202048006 | 7.57163910918757 | 9.37926694258161 | 12.0445401540101 | 8.56157562451702 | 10.4107809055614 | 15.1973579053591 | Gm9887 |
| **ENSMUSG00000052675** | 0.228505241540796 | 0.246397997874725 | 0.832123990973914 | 73.1846110420926 | 65.1532583131463 | 70.9841166486335 | 85.3513291774927 | 67.887408140784 | 57.4848649074714 | 58.3003730711437 | 67.3025850094475 | Zfp112 |
| **ENSMUSG00000052676** | 0.0421106727498939 | 0.782181420153151 | 0.960742105865155 | 153.597331816738 | 191.387696294867 | 190.237432618338 | 154.757904552597 | 152.199189218854 | 179.793088114857 | 173.860041122875 | 164.999885829613 | Zmat1 |
| **ENSMUSG00000052681** | -0.108595268362593 | 0.426909505467272 | 0.885066977635295 | 1398.63923324888 | 1135.29552610657 | 1280.55346434135 | 1180.84970807103 | 1466.14902420177 | 1453.02169170375 | 1094.1730731745 | 1373.18983930566 | Rap1b |
| **ENSMUSG00000052684** | -0.365750760971251 | 0.0196982445913549 | 0.642273034804458 | 1234.19973683332 | 1227.32450347389 | 1258.78500190243 | 1294.33883807626 | 1864.71380747992 | 1857.86191052019 | 1524.13832457418 | 1215.78863242873 | Jun |
| **ENSMUSG00000052688** | -0.110942145315578 | 0.743851358821787 | 0.954466085361166 | 41.5616309621761 | 55.3802695661743 | 53.001473764313 | 57.2135283497478 | 82.1218646864323 | 72.1618516923577 | 39.5609674411332 | 30.3947158107182 | Rab7b |
| **ENSMUSG00000052698** | 0.324837232099088 | 0.139247667799216 | 0.779711473022872 | 2372.62701971031 | 3024.74001718782 | 2745.66563196914 | 2254.77577299662 | 1823.1053960388 | 1375.96751108309 | 2605.81846066201 | 2495.6232731729 | Tln2 |
| **ENSMUSG00000052707** | 0.0895151768697723 | 0.459867931492885 | 0.894774044236998 | 1133.90971429415 | 1143.43968339572 | 1048.67201662248 | 1068.29850476005 | 996.411958195378 | 867.165302540366 | 1158.71991478898 | 1106.15055039721 | Tnrc6a |
| **ENSMUSG00000052712** | -0.0227473287864202 | 0.815017100761474 | 0.966177091475091 | 5786.10184090816 | 4878.35021619683 | 4972.67398495894 | 4906.29453766444 | 4894.46313531136 | 5439.04668603245 | 5091.91294091006 | 5444.99623237723 | BC004004 |
| **ENSMUSG00000052713** | 0.211308495575655 | 0.469942848051013 | 0.895897084741774 | 111.132187137993 | 116.461449234749 | 219.57753416644 | 190.399118934407 | 105.115986798633 | 107.6312364225 | 167.613572579538 | 169.341988088287 | Zfp608 |
| **ENSMUSG00000052726** | -0.45749924542255 | 0.3251679337343 | 0.855977678491063 | 32.5264937964856 | 16.2883145782866 | 12.3039135524298 | 30.0136542162612 | 39.4184950494875 | 48.9232892829544 | 19.7804837205666 | 17.3684090346961 | Kcnt2 |
| **ENSMUSG00000052727** | -0.164755866034005 | 0.496455200535589 | 0.900927059989498 | 188.83436676293 | 219.077831077954 | 135.343049076728 | 212.909359596603 | 232.131137513649 | 277.639666680766 | 192.599446752885 | 146.545951230249 | Map1b |
| **ENSMUSG00000052738** | 0.0509938582886528 | 0.608107899707078 | 0.927893960994322 | 10706.6375413432 | 9269.67982650289 | 9633.01785666389 | 10964.3630558779 | 10183.1112211176 | 10440.2299329825 | 9156.28180644122 | 9385.45403212392 | Suclg1 |
| **ENSMUSG00000052748** | -0.097307710116239 | 0.656984427994896 | 0.938389079274206 | 128.298947752804 | 93.6578088251478 | 85.1809399783602 | 137.87522405595 | 141.249607260664 | 107.6312364225 | 104.107809055614 | 122.664388807541 | Swt1 |
| **ENSMUSG00000052751** | -0.0539478705812364 | 0.81672894194472 | 0.966315773308182 | 380.379274675568 | 461.773718294424 | 523.389553422591 | 408.936038696558 | 443.458069306734 | 282.531995609062 | 585.085886892549 | 529.736475558232 | Repin1 |
| **ENSMUSG00000052752** | 0.0384207204584187 | 0.714833525564973 | 0.951176382858794 | 634.26662903147 | 593.709066378546 | 674.822335606343 | 660.300392757745 | 562.808512651016 | 611.54111603693 | 664.207821774815 | 655.657441059779 | Traf7 |
| **ENSMUSG00000052763** | -0.373077366533063 | 0.0447484190309367 | 0.693563614452513 | 143.658680934478 | 166.955224427437 | 127.77140996754 | 171.640585049243 | 209.137015401448 | 243.393364182698 | 167.613572579538 | 171.513039217624 | Zfp212 |
| **ENSMUSG00000052776** | -0.459697125869261 | 0.16724943804357 | 0.797837911261499 | 32.5264937964856 | 36.6487078011448 | 32.1794662140472 | 45.0204813243917 | 55.8428679867739 | 78.277262852727 | 34.3555769883525 | 33.6512925047237 | Oas1a |
| **ENSMUSG00000052783** | 0.397059335147587 | 0.226398248930589 | 0.832048466795696 | 43.3686583953142 | 35.0198763433161 | 21.7684624389143 | 43.1446279358754 | 28.4689130912965 | 19.5693157131818 | 34.3555769883525 | 26.0526135520442 | Grk4 |
| **ENSMUSG00000052794** | -0.161816225653912 | 0.376121788663152 | 0.8741697248747 | 77.7021796249379 | 73.2974156022896 | 91.8061241988993 | 86.2892558717508 | 102.926070406995 | 91.7311674055395 | 94.7381062406084 | 78.1578406561325 | 1700030K09 |
| **ENSMUSG00000052798** | 0.00415101587109573 | 0.982181407782384 | 0.996940244108351 | 156.307872966445 | 155.553404222637 | 176.040609288611 | 127.55803041911 | 130.300025302473 | 152.885279009232 | 138.463386043966 | 192.138024946326 | Nup107 |
| **ENSMUSG00000052812** | 0.317542063087997 | 0.455077760160003 | 0.892775943845882 | 56.9213641438498 | 105.059629029948 | 119.253315969704 | 74.0962088463947 | 44.893286028583 | 29.3539735697726 | 108.272121417838 | 102.03940307884 | Atad2b |
| **ENSMUSG00000052821** | 0.0842345171551496 | 0.836368367852301 | 0.968924788429433 | 36.1405486627618 | 31.7622134276588 | 43.5369248778286 | 31.8895076047775 | 41.6084114411257 | 55.0387004433237 | 26.0269522639034 | 13.0263067760221 | Cysltr1 |
| **ENSMUSG00000052825** | -0.0377180233467059 | 0.90491322891746 | 0.98504921011344 | 34.3335212296237 | 20.3603932228582 | 29.3401015481019 | 42.2067012416173 | 35.0386622662111 | 39.1386314263635 | 26.0269522639034 | 29.3091902460497 | Gm9892 |
| **ENSMUSG00000052833** | 0.0220901075551487 | 0.797606786312895 | 0.9625446841889 | 776.11828253281 | 744.375976227696 | 705.108892043093 | 771.913669374467 | 752.23628052772 | 728.95701031602 | 711.056335849841 | 759.867895267955 | Sae1 |
| **ENSMUSG00000052837** | -0.643855852182176 | 0.0604902531805823 | 0.71282992338528 | 318.036828232304 | 241.067055758641 | 249.86409060319 | 326.39848960184 | 635.075753575076 | 624.995020589742 | 308.159114804616 | 206.249857287016 | Junb |
| **ENSMUSG00000052852** | -0.0788964669837982 | 0.762868528194005 | 0.957842155335615 | 254.790868072471 | 276.086932101957 | 226.202718386979 | 211.971432902344 | 182.858018701789 | 170.008430258266 | 323.775286162958 | 346.282655129254 | Reep1 |
| **ENSMUSG00000052889** | -0.240871525919531 | 0.590893778717975 | 0.923935529355527 | 9.93865088225949 | 11.4018202048006 | 15.1432782183751 | 17.8206071909051 | 15.3294147414674 | 26.9078091056249 | 12.4929370866736 | 9.76973008201657 | Prkcb |
| **ENSMUSG00000052906** | 0.00312598456095348 | 0.975388676478509 | 0.99575536949131 | 476.151728631887 | 447.928650902881 | 434.422793889637 | 473.652980600371 | 459.882442244021 | 492.902139525765 | 447.663578939139 | 428.78259804406 | Ubxn8 |
| **ENSMUSG00000052911** | -0.0411232183308586 | 0.649302843274681 | 0.936461253686283 | 6177.32328018256 | 6977.09954960905 | 6997.14099177797 | 6458.5632166617 | 7209.20476127293 | 7208.84667584333 | 6539.01148678309 | 6424.14029170823 | Lamb2 |
| **ENSMUSG00000052914** | -0.207235309622264 | 0.355500826272195 | 0.864479425231231 | 131.009488902512 | 150.666909849151 | 130.610774633486 | 154.757904552597 | 187.237851485066 | 214.039390612925 | 111.395355689507 | 143.289374536243 | Cyp2j6 |
| **ENSMUSG00000052915** | 0.000256256391232575 | 0.997311453612108 | 0.999262355603273 | 2934.61255141626 | 2761.68373674849 | 2758.91600041022 | 2985.42066782373 | 2804.18793949271 | 2739.70419984545 | 2879.62199847827 | 3013.41896751978 | Msl1 |
| **ENSMUSG00000052917** | 0.109644305291845 | 0.595587336082541 | 0.924461858177432 | 194.255449062345 | 241.067055758641 | 247.024725937245 | 184.771558768858 | 157.67398019795 | 167.562265794119 | 221.749633288457 | 256.184033261768 | Senp7 |
| **ENSMUSG00000052920** | 0.174477096219465 | 0.653376012729636 | 0.936991241819063 | 113.8427282877 | 190.573280565953 | 150.486327295103 | 99.4202295913651 | 93.0714466446232 | 55.0387004433237 | 214.462086654564 | 128.092016630884 | Prkg1 |
| **ENSMUSG00000052921** | -0.286367623083969 | 0.0849974847677682 | 0.743541556158628 | 1049.88293865323 | 1066.07018914886 | 1208.62289280407 | 1092.68459881076 | 1540.60618151747 | 1631.59169758653 | 1106.66601026117 | 1109.40712709121 | Arhgef15 |
| **ENSMUSG00000052926** | 0.106026380488975 | 0.51731230582907 | 0.905197321821966 | 303.580608767199 | 320.879797192245 | 352.081218577222 | 399.556771753977 | 343.816873487196 | 358.363093997641 | 326.898520434627 | 249.670879873757 | Rnaseh2a |
| **ENSMUSG00000052928** | -0.0960734011045589 | 0.541557484454998 | 0.913837378403038 | 158.114900399583 | 189.758864837039 | 188.344522841041 | 212.909359596603 | 228.846262926191 | 200.585486060113 | 173.860041122875 | 197.565652769668 | Ctif |
| **ENSMUSG00000052934** | 0.23778932761747 | 0.191785395057361 | 0.808824684419004 | 4063.101183411 | 4768.40409279339 | 6740.65171695424 | 4526.43422648989 | 4182.74030802895 | 3789.10875496482 | 4937.83338350776 | 4133.68135025768 | Fbxo31 |
| **ENSMUSG00000052949** | 0.0810613867614675 | 0.731208050189219 | 0.952992645818943 | 173.474633581257 | 138.450673915436 | 174.147699511314 | 158.509611329629 | 144.534481848121 | 96.6234963338349 | 208.215618111227 | 158.486732441602 | Rnf157 |
| **ENSMUSG00000052957** | -1.37317968399383 | 0.0041924230472559 | 0.635862253526407 | 212.325723393726 | 221.521078264697 | 174.147699511314 | 218.536919762152 | 669.019457645468 | 1070.19695306463 | 229.03717992235 | 173.684090346961 | Gas1 |
| **ENSMUSG00000052962** | 0.00689074915595102 | 0.949483417814887 | 0.991267322147698 | 662.27555424511 | 698.768695408494 | 601.945309180412 | 693.127827056781 | 684.348872386936 | 601.756458180339 | 718.343882483734 | 638.289032025082 | Mrpl35 |
| **ENSMUSG00000052974** | -0.354616466999971 | 0.374778096022436 | 0.873700364743909 | 69.5705561758165 | 187.315617650296 | 93.6990339761962 | 123.806323642077 | 94.1664048404423 | 110.077400886647 | 130.134761319517 | 272.466916731795 | Cyp2f2 |
| **ENSMUSG00000052981** | 0.179598530726384 | 0.218230692134161 | 0.827129763070499 | 335.203588847116 | 359.157336451219 | 360.599312575058 | 347.03287687552 | 294.543754675337 | 245.839528646846 | 353.966550789086 | 341.94055287058 | Ube2ql1 |
| **ENSMUSG00000052997** | -0.0364348120162377 | 0.75302687772435 | 0.956497786093635 | 590.897970636155 | 509.009830571455 | 536.639921863669 | 585.266257217093 | 637.265669966714 | 571.179402378492 | 516.374732915844 | 553.618037980939 | Uba2 |
| **ENSMUSG00000053007** | 0.211799144561287 | 0.476434882246197 | 0.897553735056661 | 192.448421629207 | 175.913797445495 | 101.270673085384 | 221.350699844926 | 150.009272827216 | 201.808568292187 | 93.6970281500523 | 151.973579053591 | Creb5 |
| **ENSMUSG00000053012** | 0.173918480204713 | 0.11167815746808 | 0.764116514278315 | 1648.00901902194 | 1518.07091869631 | 1549.34665271751 | 1504.43441759009 | 1238.3977194714 | 1279.34401474926 | 1432.52345260524 | 1562.07128755798 | Krcc1 |
| **ENSMUSG00000053024** | -0.958326833977369 | 0.38512092188548 | 0.875126835008975 | 45.1756858284523 | 2.44324718674299 | 47.3227444324223 | 127.55803041911 | 42.7033696369448 | 304.547475786391 | 40.6020455316893 | 44.5065481514088 | Cntn2 |
| **ENSMUSG00000053025** | 0.550462130948607 | 0.287450271331328 | 0.851785577030345 | 31.6229800799166 | 28.5045505120015 | 9.46454888648447 | 20.6343872736795 | 7.66470737073368 | 12.2308223207386 | 13.5340151772298 | 28.2236646813812 | Sv2b |
| **ENSMUSG00000053040** | 0.273145952703169 | 0.428334221175332 | 0.885378691567738 | 24.3948703473642 | 19.5459774939439 | 30.2865564367503 | 26.2619474392285 | 29.5638712871156 | 15.9000690169602 | 15.616171358342 | 21.7105112933701 | Aph1c |
| **ENSMUSG00000053062** | -0.0889692396383673 | 0.487371278195433 | 0.899236955296196 | 2774.69062358354 | 2647.66553470048 | 2803.3993801767 | 2344.8167356454 | 3288.15946204475 | 2874.24324537357 | 2623.51678820146 | 2457.6298784095 | Jam2 |
| **ENSMUSG00000053063** | -0.345360484156124 | 0.544555808116622 | 0.913937916428821 | 9.93865088225949 | 4.88649437348597 | 9.46454888648447 | 15.0068271081306 | 12.0445401540101 | 23.2385624094033 | 9.36970281500523 | 5.42762782334254 | Clec12a |
| **ENSMUSG00000053070** | -0.023855579612281 | 0.958298356391528 | 0.992896029410587 | 12.6491920319666 | 15.4738988493722 | 16.0897331070236 | 6.56548685980713 | 12.0445401540101 | 11.0077400886647 | 18.7394056300105 | 9.76973008201657 | 9230110C19 |
| **ENSMUSG00000053080** | -0.163387034245561 | 0.318765013550685 | 0.854670142081901 | 234.010052591383 | 310.292392716359 | 304.7584741448 | 259.805694309511 | 367.905953795217 | 324.116791499573 | 271.721381635152 | 278.980070119806 | 2700081O15 |
| **ENSMUSG00000053091** | 0.140461728554626 | 0.520139548338109 | 0.905962179970576 | 80.412720774645 | 82.2559886203472 | 88.966759532954 | 70.3445020693621 | 60.2227007700503 | 67.2695227640623 | 97.8613405122768 | 66.217059444779 | Lins1 |
| **ENSMUSG00000053093** | 0.456071168289405 | 0.139578825067888 | 0.779711473022872 | 20293.8215878573 | 12768.4097979188 | 14708.8554244855 | 28612.3917350395 | 11202.5173014252 | 13105.3261166714 | 21359.7991839403 | 10012.8878085023 | Myh7 |
| **ENSMUSG00000053094** | 0.0988035916185713 | 0.332591779131681 | 0.856918526232803 | 1008.32130769105 | 1040.0088858236 | 1173.60406192407 | 994.202295913651 | 1018.31112211176 | 1043.289143959 | 960.915077583314 | 915.098051015552 | Tmem248 |
| **ENSMUSG00000053110** | 0.221252349067514 | 0.213481812206622 | 0.824248286808591 | 2769.26954128412 | 3270.69356731994 | 3035.28082789557 | 2561.47780201904 | 2180.06176787582 | 1926.35451551633 | 3104.4948660384 | 2770.26124103403 | Yap1 |
| **ENSMUSG00000053113** | -0.351109390699698 | 0.386342799317473 | 0.875126835008975 | 93.9654265231807 | 61.0811796685746 | 62.4660226507975 | 76.9099889291692 | 119.350443344282 | 155.33144347338 | 64.5468416144805 | 36.9078691987293 | Socs3 |
| **ENSMUSG00000053119** | 0.0910241371456923 | 0.35243258587121 | 0.862728204353539 | 1853.10663268311 | 1793.34343506935 | 1761.35254777476 | 2040.92848670576 | 1739.88857315654 | 1608.35313517713 | 1744.84687977209 | 1898.58421260522 | Chmp3 |
| **ENSMUSG00000053128** | -0.776954543221239 | 0.0077064120050074 | 0.635862253526407 | 23.4913566307952 | 18.7315617650296 | 21.7684624389143 | 15.9447538023887 | 36.1336204620302 | 29.3539735697726 | 37.4788112600209 | 33.6512925047237 | Rnf26 |
| **ENSMUSG00000053134** | -0.0122742463459654 | 0.930230249049325 | 0.987529878729394 | 233.106538874814 | 258.984201794756 | 201.594891282119 | 210.095579513828 | 245.270635863478 | 237.277953022329 | 213.421008564008 | 217.105112933702 | Supt7l |
| **ENSMUSG00000053137** | -0.445990144975149 | 0.254475418516421 | 0.837406774112203 | 39.754603529038 | 27.6901347830872 | 32.1794662140472 | 41.2687745473591 | 66.7924499449649 | 59.9310293716191 | 17.6983275394543 | 47.7631248454143 | Mapk11 |
| **ENSMUSG00000053158** | -0.354935490049241 | 0.157908962142918 | 0.789230661864335 | 144.562194651047 | 143.337168288922 | 117.360406192407 | 140.689004138724 | 197.092475247437 | 245.839528646846 | 143.668776496747 | 112.894658725525 | Fes |
| **ENSMUSG00000053166** | -0.588264071591448 | 0.0026915142934412 | 0.635862253526407 | 134.623543768788 | 91.2145616384048 | 140.07532351997 | 104.109863062656 | 155.484063806312 | 199.362403828039 | 190.517290571773 | 160.657783570939 | Cdh22 |
| **ENSMUSG00000053173** | 0.257967787134863 | 0.6456696260692 | 0.935063240511793 | 8.13162344912141 | 7.32974156022896 | 9.46454888648447 | 20.6343872736795 | 6.56974917491458 | 12.2308223207386 | 5.20539045278068 | 14.1118323406906 | Rpl18-ps2 |
| **ENSMUSG00000053175** | -0.264829321517267 | 0.368509197782234 | 0.871495599696183 | 69.5705561758165 | 53.7514381083457 | 77.6093008691726 | 80.6616957062019 | 113.875652365186 | 106.408154190426 | 52.0539045278068 | 66.217059444779 | Bcl3 |
| **ENSMUSG00000053192** | 0.217302058414833 | 0.317997893867398 | 0.854351922245337 | 441.818207402263 | 381.96097686082 | 257.435729712378 | 427.694572581722 | 261.695008800764 | 287.424324537357 | 395.609674411332 | 352.795808517265 | Mllt11 |
| **ENSMUSG00000053198** | -0.452256041822313 | 0.169280393812577 | 0.79907358295209 | 26.2018977805023 | 24.4324718674299 | 44.483379766477 | 28.1378008277448 | 52.5579933993167 | 51.3694537471021 | 37.4788112600209 | 27.1381391167127 | Prx |
| **ENSMUSG00000053199** | 0.176709459320467 | 0.528231771017382 | 0.908232038364594 | 1121.26052226218 | 1512.37000859391 | 1450.91534429807 | 1021.40217004714 | 1002.98170737029 | 609.094951572782 | 1536.63126166086 | 1367.76221148232 | Arhgap20 |
| **ENSMUSG00000053205** | 0.180024521940706 | 0.500237318871407 | 0.900927059989498 | 101.193536255733 | 124.605606523892 | 118.306861081056 | 90.9788893430416 | 79.9319482947941 | 58.7079471395453 | 116.600746142287 | 128.092016630884 | Styx |
| **ENSMUSG00000053214** | 0.0996412334444036 | 0.694777376346604 | 0.944771261777587 | 367.730082643601 | 437.341246426994 | 459.030620994497 | 391.115431505653 | 292.353838283699 | 245.839528646846 | 526.785513821405 | 478.716774018812 | Gm9899 |
| **ENSMUSG00000053226** | -0.230116765846286 | 0.110730440194708 | 0.764116514278315 | 311.712232216321 | 254.912123150185 | 239.453086828057 | 286.067641748739 | 319.727793179176 | 340.016860516533 | 345.637926064637 | 275.723493425801 | Dand5 |
| **ENSMUSG00000053253** | -0.034439077981115 | 0.770676737412624 | 0.958995611819782 | 346.949267162513 | 316.807718547674 | 282.043556817237 | 330.150196378873 | 324.107625962453 | 308.216722482613 | 331.062832796851 | 343.026078435248 | Ndfip2 |
| **ENSMUSG00000053263** | 0.269225965423977 | 0.580338702750966 | 0.922450244597442 | 8.13162344912141 | 15.4738988493722 | 15.1432782183751 | 6.56548685980713 | 8.75966556655277 | 11.0077400886647 | 9.36970281500523 | 8.68420451734806 | Gm12592 |
| **ENSMUSG00000053279** | -0.549981978192526 | 0.0342760783881415 | 0.678229434609245 | 680.345828576491 | 891.78522316119 | 760.003275584703 | 841.320244749571 | 1261.3918415836 | 1559.42984589417 | 617.359307699789 | 1209.27547904072 | Aldh1a1 |
| **ENSMUSG00000053286** | 0.0103517382958623 | 0.937664484902854 | 0.988970340396118 | 488.800920663853 | 570.09101024003 | 530.961192531779 | 530.866508950119 | 594.56230032977 | 512.471455238947 | 561.141090809758 | 437.466802561409 | Trmt1l |
| **ENSMUSG00000053289** | 0.112686282616402 | 0.351657581442923 | 0.862685236001802 | 332.493047697409 | 329.838370210303 | 276.364827485346 | 295.446908691321 | 290.163921892061 | 284.978160073209 | 297.748333899055 | 269.21034003779 | Ddx10 |
| **ENSMUSG00000053291** | -0.104387750930237 | 0.681255631755349 | 0.944452327903597 | 62.3424464432641 | 50.4937751926884 | 44.483379766477 | 51.5859681841989 | 72.2672409240604 | 61.154111603693 | 39.5609674411332 | 52.1052271040884 | Rab4b |
| **ENSMUSG00000053293** | -0.0427460593473455 | 0.870286808200541 | 0.975613767803763 | 281.896279569542 | 444.670987987223 | 409.814966784777 | 247.612647284155 | 349.291664466292 | 231.162541861959 | 427.883095218572 | 416.841816832707 | Pom121 |
| **ENSMUSG00000053297** | 0.095759486132371 | 0.815259040365658 | 0.966177091475091 | 15.3597331816738 | 13.8450673915436 | 10.4110037751329 | 12.1930470253561 | 10.949581958191 | 9.78465785659088 | 12.4929370866736 | 15.1973579053591 | AI854703 |
| **ENSMUSG00000053310** | 0.26257572253618 | 0.759488531594441 | 0.957357521291862 | 159.018414116152 | 196.274190668353 | 654.000328056077 | 72.2203554578784 | 162.053812981226 | 519.80994863139 | 61.4236073428121 | 158.486732441602 | Nrgn |
| **ENSMUSG00000053317** | -0.0593425556091804 | 0.77560798531592 | 0.9595799687189 | 180.702743313809 | 162.068730053951 | 164.68315062483 | 172.578511743502 | 193.80760065998 | 236.054870790255 | 128.052605138405 | 151.973579053591 | Sec61b |
| **ENSMUSG00000053329** | 0.128067014294462 | 0.134252486054284 | 0.77303252008843 | 10780.7256661018 | 9962.74761180898 | 10418.5754142421 | 10536.6684832962 | 9266.63121121702 | 8701.00699897344 | 10121.3611963868 | 10064.9930356064 | D10Jhu81e |
| **ENSMUSG00000053332** | 0.153342898373756 | 0.536955697487805 | 0.91199738788281 | 692.091506891888 | 493.535931722083 | 658.732602499319 | 788.796349871114 | 578.137927392483 | 876.949960396957 | 446.622500848583 | 466.775992807458 | Gas5 |
| **ENSMUSG00000053333** | -0.0449968892329648 | 0.761376184552911 | 0.957479700684068 | 242.141676040504 | 283.416673662186 | 259.328639489674 | 207.281799431054 | 283.594172717146 | 229.939459629886 | 250.899819824029 | 259.440609955773 | Dis3l2 |

| | | | | | | | | | | | | |
|---|---|---|---|---|---|---|---|---|---|---|---|---|
| **ENSMUSG00000053334** | -0.242460660128575 | 0.121164085577457 | 0.769868040577702 | 195.158962778914 | 227.221988367098 | 211.059440168604 | 193.212899017181 | 269.359716171498 | 281.308913376988 | 227.996101831794 | 200.822229463674 | Ficd |
| **ENSMUSG00000053347** | 0.304303344857623 | 0.0762706596649299 | 0.736318922445191 | 382.186302108706 | 361.600583637962 | 414.54724122802 | 392.053358199911 | 310.968127612624 | 225.04713070159 | 350.843316517418 | 366.907640857956 | Zfp943 |
| **ENSMUSG00000053353** | 0.286546351976499 | 0.303601747451543 | 0.851785577030345 | 37.0440623793308 | 46.4216965481167 | 62.4660226507975 | 36.5791410760683 | 29.5638712871156 | 40.3617136584374 | 40.6020455316893 | 39.0789203280663 | 2310001K24I |
| **ENSMUSG00000053358** | 0.115014719896029 | 0.393233134599048 | 0.878842838226995 | 195.158962778914 | 205.232763686411 | 192.130342395635 | 214.785212985119 | 166.433645764503 | 187.131581507301 | 209.256696201783 | 182.368294864309 | Gm9905 |
| **ENSMUSG00000053390** | 0.0949451537847 | 0.611248729932082 | 0.927932112008768 | 145.465708367616 | 144.151584017836 | 118.306861081056 | 146.316564304273 | 143.439523652302 | 95.400414101761 | 148.874166949528 | 130.263067760221 | Zfp952 |
| **ENSMUSG00000053398** | -0.673032168409538 | 0.219265497210859 | 0.827535829146756 | 45.1756858284523 | 35.8342920722305 | 27.447191770805 | 23.448167356454 | 30.6588294829347 | 128.423634367755 | 24.9858741733473 | 27.1381391167127 | Phgdh |
| **ENSMUSG00000053411** | 0.124491157381279 | 0.40868211639213 | 0.882113132676277 | 475.248214915318 | 521.22606650517 | 644.535779169592 | 494.287367874051 | 504.775728272604 | 425.632616761703 | 559.058934628645 | 467.861518372127 | Cbx7 |
| **ENSMUSG00000053436** | -0.036328366627375 | 0.823998255397764 | 0.966637987288264 | 1299.25272442629 | 1495.26727828671 | 1507.70263761698 | 1210.86336228729 | 1328.18429152856 | 1093.43551547403 | 1618.87643081479 | 1612.00546353273 | Mapk14 |
| **ENSMUSG00000053453** | 0.151899572512194 | 0.203996209504623 | 0.816102347355766 | 1047.17239750352 | 806.271571625185 | 920.900606654939 | 1050.47789756914 | 836.54806160579 | 883.065371557327 | 851.60187807492 | 870.591502864143 | Thoc7 |
| **ENSMUSG00000053460** | -0.221397293648537 | 0.0882475804596512 | 0.751077752809164 | 304.484122483768 | 290.746415222415 | 282.043556817237 | 291.695201914288 | 346.006789878835 | 399.947889888152 | 306.076958623504 | 312.63136262453 | Ggcx |
| **ENSMUSG00000053469** | 0.294445593299867 | 0.429191526523278 | 0.885743677088682 | 18.9737880479499 | 41.5352021746308 | 54.8943835416099 | 42.2067012416173 | 33.943704070392 | 28.1308913376988 | 22.903717992235 | 43.4210225867403 | Tg |
| **ENSMUSG00000053470** | 0.0173445960885134 | 0.923334025725624 | 0.986713918026868 | 361.405486627618 | 384.404224047563 | 373.849681016137 | 302.012395551128 | 393.089992299056 | 267.855008824175 | 425.80093903746 | 316.973464883204 | Kdm3a |
| **ENSMUSG00000053475** | -1.32340205843125 | 0.0160086599861398 | 0.63777231095375 | 10.8421645988285 | 11.4018202048006 | 5.67872933189068 | 11.2551203310979 | 37.2285786578493 | 39.1386314263635 | 11.4518589961175 | 10.8552556466851 | Tnfaip6 |
| **ENSMUSG00000053477** | -0.384099497539382 | 0.139308361319334 | 0.779711473022872 | 1862.1417698488 | 1557.97728941311 | 1361.00212987647 | 2085.01104133589 | 2850.17618371711 | 2940.28968590556 | 1446.05746778247 | 1724.90012225826 | Tcf4 |
| **ENSMUSG00000053483** | -0.134396187809685 | 0.242424293529438 | 0.832048466795696 | 333.396561413978 | 312.735639903102 | 332.205665915605 | 331.088123073131 | 386.520243124141 | 337.570696052385 | 386.239971596327 | 325.657669400552 | Usp21 |
| **ENSMUSG00000053510** | 0.223951946372032 | 0.170866503643044 | 0.799531027639052 | 6564.02715087411 | 7538.23198683103 | 7282.9703681498 | 6824.35462742238 | 4998.48416391418 | 4932.69064195388 | 7228.20518273126 | 6992.95568759453 | Nrd1 |
| **ENSMUSG00000053536** | -0.108211092419591 | 0.435393995060694 | 0.888588936437999 | 456.274426867368 | 534.2567181678 | 499.72818120638 | 401.432625142493 | 582.51776017576 | 505.132961846504 | 473.690531203042 | 478.716774018812 | Cstf2t |
| **ENSMUSG00000053553** | 0.0436705136419175 | 0.805777276941397 | 0.963475137341661 | 137.334084918495 | 105.059629029948 | 105.056492639978 | 142.564857527241 | 113.875652365186 | 122.308223207386 | 127.011527047849 | 111.809133160856 | 3110082I17R |
| **ENSMUSG00000053559** | -0.276057124789019 | 0.28786539481385 | 0.851785577030345 | 197.869503928621 | 157.182235680465 | 184.558703286447 | 211.971432902344 | 297.828629262794 | 293.539735697726 | 149.915245040084 | 169.341988088287 | Smagp |
| **ENSMUSG00000053560** | -0.922487336790159 | 0.0179966515580752 | 0.642273034804458 | 229.492484008537 | 134.378595270864 | 143.861143074564 | 197.902532488472 | 387.61520131996 | 576.071731306788 | 233.201492284575 | 141.118323406906 | Ier2 |
| **ENSMUSG00000053565** | 0.0276555892048357 | 0.88240617451068 | 0.978854883720278 | 2114.22209677157 | 1544.94663775048 | 1683.74324690559 | 2172.2382239019 | 1924.93650824997 | 2325.07932317241 | 1624.08182126757 | 1499.11080480721 | Eif3k |
| **ENSMUSG00000053580** | -0.0694317849576677 | 0.754579102548804 | 0.956497786093635 | 103.000563688871 | 140.893921102179 | 141.021778408619 | 159.447538023887 | 123.730276127558 | 107.6312364225 | 174.901119213431 | 163.914360264945 | Tanc2 |
| **ENSMUSG00000053581** | 0.130703845925407 | 0.535623526788919 | 0.91096005719335 | 380.379274675568 | 318.436550005503 | 364.385132129652 | 319.833002742033 | 255.12525962585 | 227.493295165738 | 383.116737324658 | 396.216831104005 | Zfand2a |
| **ENSMUSG00000053600** | -0.190277200489606 | 0.466455581432988 | 0.895328112556867 | 56.0178504272808 | 38.2775392589734 | 37.8581955459379 | 54.3997482669734 | 67.887408140784 | 51.3694537471021 | 51.0128264372507 | 42.3354970220718 | Zfp472 |
| **ENSMUSG00000053604** | -0.0164849793906932 | 0.918361558492945 | 0.986713918026868 | 299.966553900923 | 306.220314071788 | 232.827902607518 | 301.07446885687 | 325.202584158272 | 232.385624094033 | 300.871568170723 | 294.177428025166 | Rpia |
| **ENSMUSG00000053617** | 0.0903529387443863 | 0.728368100794795 | 0.952401994188708 | 1633.55279955683 | 1960.29865949679 | 1644.938596471 | 1168.65666104567 | 1311.75991859128 | 913.642427359173 | 1864.57086018604 | 1927.89340285127 | Sh3pxd2a |
| **ENSMUSG00000053641** | -0.0363247135449643 | 0.901124649942098 | 0.984802943363903 | 93.0619128066116 | 138.450673915436 | 165.629605513478 | 83.4754757889764 | 121.54035973592 | 78.277262852727 | 141.586620315635 | 150.888053488923 | Dennd4a |
| **ENSMUSG00000053644** | 0.0323689847421037 | 0.773241886614687 | 0.959129291398286 | 1152.8835023421 | 1147.51176204029 | 1184.96152058786 | 1176.16007459973 | 1109.19265236475 | 965.011881106275 | 1243.04724012403 | 1238.58466928677 | Aldh7a1 |
| **ENSMUSG00000053646** | -0.183300239575927 | 0.501958592958308 | 0.900927059989498 | 488.800920663853 | 522.854897962999 | 448.619617219364 | 354.536290429585 | 434.698403740182 | 291.093571233579 | 685.029383585938 | 649.144287671768 | Plxnb1 |
| **ENSMUSG00000053647** | -1.2829583293341 | 0.0118942624526568 | 0.635862253526407 | 8.13162344912141 | 17.9171460361152 | 11.3574586637814 | 10.3171936368398 | 35.0386622662111 | 51.3694537471021 | 17.6983275394543 | 13.0263067760221 | Gper1 |
| **ENSMUSG00000053684** | -0.0758882610560106 | 0.76342185868593 | 0.95785116771606 | 36.1405486627618 | 39.0919549878878 | 36.9117406572894 | 45.9584080186499 | 40.5134532453066 | 36.6924669622158 | 53.094982618363 | 35.8223436340608 | BC048403 |
| **ENSMUSG00000053687** | 0.215688940689849 | 0.728486482234004 | 0.952401994188708 | 7.22810973255236 | 14.6594831204579 | 26.5007368821565 | 10.3171936368398 | 21.8991639163819 | 15.9000690169602 | 3.12323427166841 | 9.76973008201657 | Dpep2 |
| **ENSMUSG00000053702** | 0.215384257559939 | 0.4791494307499 | 0.897732059433541 | 4080.26794402581 | 6117.89095560444 | 5444.00851950587 | 3183.3232003122 | 2924.63334103281 | 2506.09549351934 | 5392.78450908079 | 5390.71995414381 | Nebl |
| **ENSMUSG00000053716** | -0.0135229166758261 | 0.916159556336872 | 0.986713918026868 | 1161.01512579122 | 986.257447715252 | 1058.13656550896 | 979.19546880552 | 1188.02964246372 | 1122.7894890438 | 1019.21545065446 | 894.47306528685 | Dusp7 |
| **ENSMUSG00000053719** | -0.174431100425602 | 0.654412130039732 | 0.937269843859626 | 28.9124389302094 | 23.6180561385155 | 21.7684624389143 | 15.0068271081306 | 19.7092475247437 | 31.8001380339204 | 35.3966550789086 | 14.1118323406906 | Klk1b26 |
| **ENSMUSG00000053730** | -0.176510488041779 | 0.430430349950692 | 0.886017087288711 | 62.3424464432641 | 74.9262470601182 | 76.6628459805242 | 81.59962240046 | 79.9319482947941 | 113.746647582869 | 70.7933101578173 | 70.559161703453 | Tmem39b |
| **ENSMUSG00000053746** | -0.308942967545572 | 0.300603690028608 | 0.851785577030345 | 58.7283915769879 | 57.009101024003 | 44.483379766477 | 60.9652351267805 | 77.7420319031559 | 96.6234963338349 | 62.4646854333682 | 37.9933947633978 | Ptrh1 |
| **ENSMUSG00000053754** | 0.00952385459229365 | 0.913099464598523 | 0.986226496507248 | 681.24934229306 | 748.448054872268 | 725.930899593359 | 723.141481273042 | 679.969039603659 | 720.395434691503 | 713.138492030954 | 746.841588491933 | Chd8 |
| **ENSMUSG00000053768** | 0.167457536634586 | 0.0810537063467382 | 0.739855948845396 | 10766.2694466367 | 9025.35510782859 | 10540.6680948778 | 11281.3822785372 | 9449.48922991881 | 9159.66283600113 | 9270.8003964024 | 9171.60549588422 | Chchd3 |
| **ENSMUSG00000053769** | 0.154265337197116 | 0.595143250055539 | 0.924461858177432 | 53.3073092775737 | 52.9370223794314 | 53.001473764313 | 85.3513291774927 | 72.2672409240604 | 50.1463715150282 | 36.4377331694648 | 60.7894316214364 | Lysmd1 |
| **ENSMUSG00000053774** | -0.0625872080248436 | 0.751505979933054 | 0.956197192463153 | 485.186865797577 | 545.6585383726 | 441.047978110176 | 324.522636213324 | 501.490853685146 | 370.593916318379 | 539.278450908079 | 464.604941678121 | Ubxn7 |
| **ENSMUSG00000053799** | 0.154702385633901 | 0.444044036923946 | 0.89108031640712 | 911.645340018166 | 952.05198710085 | 859.38103889279 | 821.623784170149 | 694.203496149307 | 544.271593272868 | 980.695561303881 | 963.946701425635 | Exoc6 |
| **ENSMUSG00000053801** | 0.00877553304588851 | 0.950447374169501 | 0.991696503889296 | 351.466835745359 | 390.919549878878 | 337.884395247495 | 399.556771753977 | 431.413529152724 | 315.555215875056 | 344.596847974081 | 378.848422069309 | Grwd1 |
| **ENSMUSG00000053819** | -0.00174470481500332 | 0.993348971139065 | 0.999133450854041 | 2059.10776006085 | 2317.82716449018 | 1982.8229917185 | 1935.88069694884 | 1688.42553795305 | 1462.80634956034 | 2477.76585552361 | 2675.82051690787 | Camk2d |
| **ENSMUSG00000053821** | -0.0163813156679213 | 0.972535738108327 | 0.995322826578593 | 18.9737880479499 | 8.95857301805762 | 9.46454888648447 | 7.50341355406529 | 8.75966556655277 | 12.2308223207386 | 12.4929370866736 | 11.9407812113536 | Gm9922 |
| **ENSMUSG00000053835** | 0.0668244189745562 | 0.724817896668279 | 0.951674101666722 | 178.895715880671 | 180.800291818981 | 240.399541716705 | 240.109233730089 | 217.896680968 | 226.270212933664 | 154.079557402308 | 204.078806157679 | H2-T24 |
| **ENSMUSG00000053838** | 0.0924697507392086 | 0.512025351192525 | 0.902394714365738 | 1800.70283712211 | 2027.89516499668 | 2141.82741301144 | 1800.81925297567 | 1624.91796259554 | 1559.42984589417 | 2115.47068001007 | 1987.59730890804 | Nudcd3 |
| **ENSMUSG00000053841** | 0.0116452787923219 | 0.939348451500424 | 0.989269716906553 | 385.800356974982 | 532.627886709971 | 561.247748968529 | 446.453106466885 | 474.116898789669 | 434.19419238622 | 510.128264372507 | 491.743080794834 | Txlna |
| **ENSMUSG00000053870** | 0.135055978552152 | 0.350703003229527 | 0.861774532843376 | 193.351935345776 | 236.99497711407 | 206.327165725361 | 195.088752405698 | 173.003394939417 | 176.123841418636 | 196.76375911511 | 211.677485110359 | Fpgt |
| **ENSMUSG00000053877** | 0.0992000713168155 | 0.807479943199724 | 0.964026716319739 | 62.3424464432641 | 127.86326943955 | 133.450139299431 | 50.6480414899407 | 65.6974917491458 | 47.7002070508805 | 123.888292776180 | 111.809133160856 | Srcap |
| **ENSMUSG00000053886** | 0.255087784227193 | 0.237829559801049 | 0.832048466795696 | 221.360860559416 | 219.077831077954 | 189.290977729689 | 210.095579513828 | 177.383227722694 | 128.423634367755 | 149.915245040084 | 247.49982874442 | Sh2d4a |
| **ENSMUSG00000053897** | 0.196253587923187 | 0.636351307505431 | 0.932595977875587 | 18.0702743313809 | 27.6901347830872 | 25.5542819935081 | 19.6964605794214 | 8.75966556655277 | 31.8001380339204 | 16.6572494488982 | 22.7960368580387 | Slc39a8 |
| **ENSMUSG00000053898** | 0.0265152361600598 | 0.861714238266622 | 0.974868041734905 | 64240.7287617757 | 50018.1564070024 | 61181.6833669015 | 67657.3420903125 | 60616.8857205452 | 71577.2183854264 | 50741.1050556155 | 55736.3101179045 | Ech1 |
| **ENSMUSG00000053907** | 0.256390678017947 | 0.159710681894406 | 0.790303302997254 | 1767.27282960905 | 2214.39636691806 | 1724.44080711747 | 1406.89004138724 | 1615.06333883317 | 1201.06675189653 | 1749.01119213431 | 1389.47272277569 | Mat2a |
| **ENSMUSG00000053929** | 0.120690324184631 | 0.254471002949548 | 0.837406774112203 | 2673.4970873278 | 2678.61333239923 | 2757.02309063293 | 2629.00852400563 | 2168.01722772181 | 2320.18699424411 | 2735.95322198153 | 2650.8534289205 | Cyhr1 |
| **ENSMUSG00000053930** | -0.249880290841157 | 0.585901253988989 | 0.923058967588378 | 9.93865088225949 | 12.2162359337149 | 11.3574586637814 | 18.7585338851632 | 10.949581958191 | 14.6769867848863 | 10.4107809055614 | 26.0526135520442 | Shisa6 |
| **ENSMUSG00000053931** | -0.457365167170671 | 0.127837367842273 | 0.772950109316303 | 703.837185207286 | 705.284021239809 | 635.071230283108 | 684.686486808458 | 1433.3002783272 | 1169.26661386261 | 564.264325081426 | 580.756177097652 | Cnn3 |
| **ENSMUSG00000053950** | -0.188541123304828 | 0.208980713732054 | 0.821566820583092 | 146.369222084185 | 161.254314325037 | 194.023252172932 | 179.143998603309 | 208.042057205628 | 184.685417043153 | 204.051305749003 | 178.026192605635 | Adnp2 |
| **ENSMUSG00000053963** | 0.202574466464646 | 0.309955517145989 | 0.851785577030345 | 163.535982698997 | 175.913797445495 | 159.950876181588 | 126.620103724852 | 164.243729372865 | 118.638976511164 | 107.231043327282 | 154.144630182928 | Stum |
| **ENSMUSG00000053964** | 0.187847049727873 | 0.488048037854542 | 0.899236955296196 | 68.6670424592474 | 43.1640336324594 | 72.8770264259304 | 66.5927952923295 | 54.7479097909548 | 74.6080161565054 | 41.6431236222455 | 49.9341759747513 | Lgals4 |
| **ENSMUSG00000053965** | -0.409847667082115 | 0.143123018071428 | 0.780607990324772 | 46.0791995450213 | 67.5965054998893 | 47.3227444324223 | 45.9584080186499 | 88.6916138613469 | 84.3926740130963 | 52.0539045278068 | 51.0197015394199 | Pde5a |
| **ENSMUSG00000053985** | -0.154848541852522 | 0.50340542602463 | 0.900927059989498 | 40.658117245607 | 54.56585383726 | 53.001473764313 | 45.9584080186499 | 64.6025335533267 | 45.2540425867328 | 57.2592949805875 | 48.8486504100828 | Zfp14 |
| **ENSMUSG00000054000** | -0.191178447430225 | 0.550519462719841 | 0.914699131434573 | 54.2108229941427 | 47.2361122770311 | 42.5904699891801 | 43.1446279358754 | 84.3117810780705 | 57.4848649074714 | 35.3966550789086 | 36.9078691987293 | Tusc1 |
| **ENSMUSG00000054008** | 0.19129935545801 | 0.614198271265539 | 0.928075610917191 | 674.021232560508 | 1161.35682943183 | 1051.51138128842 | 580.576623745802 | 521.20010120989 | 342.463024980681 | 1105.62493217062 | 1067.07163006914 | Ndst1 |
| **ENSMUSG00000054013** | 0.0775395760662893 | 0.826586882611397 | 0.967319591893432 | 91.2548853734735 | 114.018202048006 | 151.432782183751 | 78.7858423176855 | 91.9764884488041 | 46.4771248188067 | 128.052605138405 | 145.46042566558 | Tmem179 |
| **ENSMUSG00000054021** | 0.0285383838031049 | 0.795842335701023 | 0.962200030474509 | 762.565576784274 | 695.511032492837 | 632.231865617162 | 732.520748215624 | 756.616113310996 | 639.672007374629 | 696.481242582055 | 674.111375659143 | Sirt5 |
| **ENSMUSG00000054027** | 0.168542080805287 | 0.543755379307318 | 0.913837378403038 | 1522.42061241884 | 1858.4966933825 | 1859.7838561942 | 1503.49649089583 | 1167.22543674316 | 846.372904595111 | 1803.14725284323 | 2182.99191054837 | Nt5dc3 |
| **ENSMUSG00000054051** | 0.143224576538408 | 0.604221707389154 | 0.926448151609826 | 37.9475760958999 | 54.56585383726 | 49.2156542097192 | 51.5859681841989 | 38.3235368536684 | 31.8001380339204 | 42.6842017128016 | 61.8749571861049 | Ercc6 |
| **ENSMUSG00000054057** | -0.279498750497922 | 0.0878457153498987 | 0.749494463274013 | 123.781379169959 | 105.874044758863 | 110.735221971868 | 101.296082979881 | 127.015150715015 | 130.869798831903 | 156.16171358342 | 121.578863242873 | A930004D18 |
| **ENSMUSG00000054072** | 0.0294004530301793 | 0.894577518670117 | 0.983733637008786 | 330.68602026427 | 416.166437475222 | 620.874406953381 | 448.328959855401 | 563.903470846835 | 425.632616761703 | 400.815064864113 | 388.618152151326 | Iigp1 |
| **ENSMUSG00000054074** | -0.402674327029314 | 0.0820856443651204 | 0.739855948845396 | 50.5967681278665 | 44.7928650902881 | 39.7511053232348 | 64.7169419038131 | 73.3621991198795 | 57.4848649074714 | 69.7522320672611 | 62.9604827507734 | Skida1 |
| **ENSMUSG00000054079** | 0.0469888830736857 | 0.735671035890147 | 0.954383574136061 | 200.580045078328 | 207.676010873154 | 199.701981504822 | 196.026679099956 | 203.662224422352 | 160.223772401676 | 214.462086654564 | 198.651178334337 | Utp18 |
| **ENSMUSG00000054083** | -0.37232512283071 | 0.434937461173372 | 0.888379878975956 | 12.6491920319666 | 8.14415728914329 | 23.6613722162112 | 12.1930470253561 | 19.7092475247437 | 26.9078091056249 | 9.36970281500523 | 17.3684090346961 | Capn12 |
| **ENSMUSG00000054091** | 0.115284429177583 | 0.481752446433114 | 0.897732059433541 | 1099.57619306453 | 869.795998480503 | 864.113313336032 | 1143.3326403007 | 869.396807480363 | 1120.34332457966 | 892.203923606609 | 790.262611078674 | 1810037I17R |
| **ENSMUSG00000054099** | 0.401675628184678 | 0.0775049184127782 | 0.739855948845396 | 150.886790667031 | 161.254314325037 | 179.826428843205 | 136.937297361692 | 93.0714466446232 | 88.0619207093179 | 158.243869764533 | 135.690695583563 | Slc25a40 |
| **ENSMUSG00000054150** | 0.152563623105263 | 0.578652663741484 | 0.922450244597442 | 65.9565013095403 | 87.9568987227475 | 101.270673085384 | 61.9031618210386 | 50.3680770076785 | 59.9310293716191 | 79.1219348822664 | 95.5262496908287 | Syne3 |
| **ENSMUSG00000054178** | 0.714696270923295 | 0.0668946155038517 | 0.719725956118125 | 28.0089252136404 | 48.8649437348597 | 44.483379766477 | 29.075727522003 | 16.4243729372865 | 12.2308223207386 | 34.3555769883525 | 28.2236646813812 | Gm9938 |
| **ENSMUSG00000054181** | 0.1539970431002 | 0.755938086848605 | 0.956566265296272 | 14.4562194651047 | 5.7009101024003 | 14.1968233297267 | 13.1309737196143 | 13.1394983498292 | 14.6769867848863 | 7.28754663389296 | 7.59867895267955 | A930012O16 |
| **ENSMUSG00000054199** | 0.330363563874833 | 0.132403889715244 | 0.77303252008843 | 280.992765852973 | 300.519403969387 | 356.813493020464 | 299.198615468353 | 171.908436743598 | 201.808568292187 | 317.528817619622 | 292.006376895829 | Gon4l |
| **ENSMUSG00000054203** | -0.220327904479683 | 0.541890226011371 | 0.913837378403038 | 214.132750826864 | 228.850819824926 | 247.971180825893 | 151.944124469822 | 312.063085808443 | 391.386314263635 | 115.559668051731 | 163.914360264945 | Ifi205 |
| **ENSMUSG00000054204** | -1.05522806571153 | 0.0606666007698271 | 0.71282992338528 | 19.877301764519 | 10.5874044758863 | 6.62518422053913 | 3.75170677703264 | 30.6588294829347 | 28.1308913376988 | 13.5340151772298 | 13.0263067760221 | Alkal2 |
| **ENSMUSG00000054206** | 0.138751157652029 | 0.400624761384087 | 0.880487015130533 | 148.176249517323 | 127.048853710635 | 140.07532351997 | 137.87522405595 | 143.439523652302 | 139.43137445642 | 107.231043327282 | 112.894658725525 | Gzmm |
| **ENSMUSG00000054226** | 0.0527748505845117 | 0.790609117629488 | 0.961804768755116 | 287.317361868956 | 254.912123150185 | 239.453086828057 | 234.48167356454 | 176.288269526875 | 229.939459629886 | 251.940897914585 | 321.315567141878 | Tprkb |
| **ENSMUSG00000054237** | 0.136747931969142 | 0.284005878289236 | 0.851125503777472 | 459.888481733644 | 380.332145402991 | 430.636974335043 | 441.763472995594 | 370.095870186855 | 357.140011765567 | 447.663578939139 | 381.019473198646 | Fra10ac1 |
| **ENSMUSG00000054247** | -0.0933676699661119 | 0.840073188368569 | 0.969879194284716 | 9.03513716569045 | 26.8757190541728 | 11.3574586637814 | 12.1930470253561 | 19.7092475247437 | 12.2308223207386 | 17.6983275394543 | 14.1118323406906 | Gm9939 |
| **ENSMUSG00000054252** | -0.4211259388132 | 0.233534652942912 | 0.832048466795696 | 179.79922959724 | 233.737314198412 | 183.612248397799 | 103.171936368398 | 323.012667766634 | 324.116791499573 | 161.367104036201 | 130.263067760221 | Fgfr3 |
| **ENSMUSG00000054263** | -0.050828324228431 | 0.76910773770569 | 0.95879680569409 | 713.775836089546 | 833.147290679358 | 813.004749349016 | 715.638067718977 | 1024.88087128667 | 611.54111603693 | 796.424739275444 | 752.269216315276 | Lifr |

| | | | | | | | | | | | | |
|---|---|---|---|---|---|---|---|---|---|---|---|---|
| **ENSMUSG00000054277** | -0.0994293805764095 | 0.631261861078064 | 0.931825800800958 | 264.72951895473 | 258.984201794756 | 295.293925258315 | 335.777756544422 | 344.911831683016 | 403.617136584374 | 257.146288367366 | 232.302470839061 | Arfgap3 |
| **ENSMUSG00000054280** | 0.276038308375003 | 0.350481172898008 | 0.861774532843376 | 74.0881247586617 | 106.688460487777 | 69.0912068713366 | 65.6548685980713 | 72.2672409240604 | 36.6924669622158 | 64.5468416144805 | 86.8420451734806 | Prr14l |
| **ENSMUSG00000054302** | 0.189967665997656 | 0.11360063779319 | 0.767518180080285 | 662.27555424511 | 586.379324818317 | 640.749959614998 | 698.75538722233 | 597.847174917227 | 621.325773893521 | 493.471014923609 | 556.874614674944 | Eapp |
| **ENSMUSG00000054309** | -0.013239411496586 | 0.907482279677576 | 0.98504921011344 | 884.539928521095 | 846.177942341987 | 742.967087589031 | 861.95463202325 | 879.251431242735 | 929.542496376133 | 773.52102128321 | 785.920508819999 | Cpsf3 |
| **ENSMUSG00000054312** | 0.0788189274169398 | 0.69205295994 8012 | 0.94469184542218 | 2905.70011248605 | 2099.56374914114 | 2282.84919142005 | 2922.57957930843 | 2413.28786358529 | 3222.82168151462 | 2009.28071477334 | 2023.4196525421 | Mrps21 |
| **ENSMUSG00000054321** | -0.0466203845622884 | 0.883397861339897 | 0.97910597465636 | 74.9916384752307 | 84.6992358070902 | 62.4660226507975 | 41.2687745473591 | 61.3176589658694 | 40.3617136584374 | 82.2451691539348 | 87.9275707381491 | Taf4b |
| **ENSMUSG00000054364** | -0.333495690491383 | 0.194269697247556 | 0.809162446871198 | 4730.79781995552 | 4594.93354253464 | 3709.15670861326 | 4171.8979360603 | 6569.74917491458 | 7665.05634840688 | 3982.12369637722 | 3466.08312798654 | Rhob |
| **ENSMUSG00000054381** | -0.261502315572673 | 0.345435404481049 | 0.861266047293794 | 42.4651446787451 | 35.0198763433161 | 56.7872933189068 | 66.5927952923295 | 77.7420319031559 | 53.8156182112498 | 54.1360607089191 | 54.2762782334254 | Zfp747 |
| **ENSMUSG00000054387** | 0.308741168282193 | 0.120167361280393 | 0.769868040577702 | 214.132750826864 | 237.809392842984 | 229.988537941573 | 225.102406621959 | 146.724398239759 | 138.208292224346 | 225.913945650682 | 220.361689627707 | Mdm4 |
| **ENSMUSG00000054404** | -0.149945725833178 | 0.339675032920019 | 0.858570960161243 | 2154.88021401717 | 2320.27041167692 | 2540.28492113243 | 2331.68576192579 | 2909.30392629134 | 3120.08277402042 | 2143.57978845509 | 2199.2747940184 | Slfn5 |
| **ENSMUSG00000054405** | 0.113792391952983 | 0.192095963180392 | 0.808824684419004 | 1687.76362255098 | 1720.86043519598 | 1764.1919124407 | 1945.25996389143 | 1679.66587238649 | 1618.13779303372 | 1708.40914660262 | 1570.75549207533 | Dnajc8 |
| **ENSMUSG00000054408** | 0.067760701039072 | 0.574879221612184 | 0.922450244597442 | 1240.5243328493 | 1316.91023365447 | 1259.73145679108 | 1325.29041898678 | 1142.04139823932 | 1044.51222619108 | 1371.09984526243 | 1347.13722575362 | Spcs3 |
| **ENSMUSG00000054414** | -0.505926899039244 | 0.0261433094816566 | 0.664353746352133 | 76.7986659083688 | 101.801966114291 | 96.5383986421416 | 130.371810501884 | 147.819356435578 | 183.462334811079 | 130.134761319517 | 115.065709854862 | Slc30a7 |
| **ENSMUSG00000054422** | -0.30692803776515 | 0.260475287122544 | 0.840316047847749 | 505.064167562096 | 601.038807938775 | 484.584902988005 | 602.14893771374 | 504.775728272604 | 709.387694602839 | 435.170641852465 | 1063.81505337514 | Fabp1 |
| **ENSMUSG00000054426** | -0.11534839587239 | 0.541348900643497 | 0.913837378403038 | 94.8689402397497 | 98.5443031986338 | 70.9841166486335 | 82.5375490947182 | 95.2613630362614 | 107.6312364225 | 88.4916376972716 | 85.7565196088121 | A930005H10 |
| **ENSMUSG00000054428** | 0.0527843901879173 | 0.801205424953131 | 0.963461896828986 | 3120.73637702948 | 2168.78908609886 | 2624.51940622214 | 3449.69438148152 | 2823.89718701745 | 3573.84628211982 | 2437.16380999192 | 2121.11695336226 | Atpif1 |
| **ENSMUSG00000054434** | -0.159935659789867 | 0.498933346940907 | 0.900927059989498 | 94.8689402397497 | 98.5443031986338 | 103.163582862681 | 86.2892558717508 | 137.964732673206 | 111.300483118721 | 112.436433780063 | 66.217059444779 | Tmem120b |
| **ENSMUSG00000054435** | -0.364162207881997 | 0.0211367875562208 | 0.648762042745232 | 474.344701198749 | 421.052931848708 | 378.581955459379 | 438.949692912819 | 605.511882287961 | 648.233582999146 | 444.54034466747 | 508.025964264862 | Gimap4 |
| **ENSMUSG00000054452** | -0.0608159650025794 | 0.581125049275504 | 0.922450244597442 | 20422.1205356101 | 19019.8649330652 | 19234.8027020024 | 20111.0241782835 | 18854.085173809 | 24121.6277809607 | 20259.3796422224 | 18946.7632057241 | Aes |
| **ENSMUSG00000054455** | 0.113340781299095 | 0.206760782591695 | 0.819960256425221 | 4992.81679776054 | 4903.59710379317 | 4896.95759386706 | 4842.51552245489 | 4181.64534983313 | 4235.53376967178 | 4839.97204299548 | 4893.54924552563 | Vapb |
| **ENSMUSG00000054469** | 0.161467507955909 | 0.418355310623184 | 0.882468496126562 | 2217.22266046044 | 2798.33244454963 | 2695.50352287078 | 2142.22456968564 | 2001.58358195731 | 1580.22224383943 | 2841.1021091277 | 2385.98519114138 | Lclat1 |
| **ENSMUSG00000054474** | 0.269072618140218 | 0.240078385140067 | 0.832048466795696 | 128.298947752804 | 161.254314325037 | 207.27362061401 | 194.150825711439 | 159.863896589588 | 171.23151249034 | 141.586620315635 | 100.953877514171 | Thnsl2 |
| **ENSMUSG00000054477** | 0.0599417784417952 | 0.861208993784778 | 0.974868041734905 | 253.887354355902 | 254.097707421271 | 159.950876181588 | 152.88205116408 | 118.255485148462 | 123.53130543946 | 243.612273190136 | 301.776106977845 | Kcnn2 |
| **ENSMUSG00000054484** | 0.183642765549463 | 0.435951261704643 | 0.888687115594624 | 93.0619128066116 | 74.9262470601182 | 92.7525790875478 | 90.0409626487835 | 52.5579933993167 | 67.2695227640623 | 91.61487196894 | 96.6117752554972 | Tmem62 |
| **ENSMUSG00000054499** | -0.269083707218239 | 0.0439906667918033 | 0.691603097094229 | 376.765219809292 | 388.476302692135 | 360.599312575058 | 328.274342990356 | 493.826146314413 | 392.609396495709 | 473.690531203042 | 391.874728845331 | Dedd2 |
| **ENSMUSG00000054509** | 0.183369651961224 | 0.335753426199268 | 0.858570960161243 | 590.897970636155 | 749.262470601182 | 940.776159316556 | 622.783324987419 | 629.600962595981 | 518.586866399316 | 738.124366204301 | 669.769273400469 | Parp4 |
| **ENSMUSG00000054517** | -0.0429087022792522 | 0.777294925758306 | 0.95974290923544 | 187.930853046361 | 207.676010873154 | 158.057966404291 | 179.143998603309 | 203.662224422352 | 166.339183562045 | 181.147587756768 | 204.078806157679 | Trim65 |
| **ENSMUSG00000054519** | 0.14861467680789 | 0.406665237722015 | 0.881296485258502 | 200.580045078328 | 163.69756151178 | 186.451613063744 | 211.033506208086 | 159.863896589588 | 133.315963296051 | 190.517290571773 | 201.907755028342 | Zfp867 |
| **ENSMUSG00000054520** | -0.18514986923448 | 0.54279584895158 | 0.913837378403038 | 16.2632468982428 | 25.2468875963442 | 23.6613722162112 | 26.2619474392285 | 27.3739548954774 | 24.4616446414772 | 26.0269522639034 | 26.0526135520442 | Sh3bp2 |
| **ENSMUSG00000054580** | -0.0596611948893407 | 0.805768930560216 | 0.963475137341661 | 43.3686583953142 | 53.7514381083457 | 44.483379766477 | 46.8963347129081 | 60.2227007700503 | 56.2617826753975 | 40.6020455316893 | 40.1644458927348 | Pla2r1 |
| **ENSMUSG00000054594** | 0.460080238558084 | 0.260924876339343 | 0.840697409499762 | 21.6843291976571 | 13.0306516626293 | 24.6078271048596 | 30.0136542162612 | 9.85462376237187 | 23.2385624094033 | 15.616171358342 | 16.2828834700276 | Oscar |
| **ENSMUSG00000054604** | -0.0448078754293805 | 0.702933058130072 | 0.946451777392559 | 718.293404672391 | 823.374301932386 | 799.754380907938 | 694.065753751039 | 831.073270626695 | 823.134342185708 | 681.906149314269 | 796.775764466685 | Cggbp1 |
| **ENSMUSG00000054611** | 0.0651861299970754 | 0.742636332695645 | 0.954383574136061 | 760.758549351136 | 1013.13316676942 | 989.991813526275 | 685.624413502716 | 850.782518151438 | 600.533375948265 | 948.422140496641 | 896.644116416187 | Kdm2a |
| **ENSMUSG00000054612** | 0.0207651006197894 | 0.926215597752635 | 0.986713918026868 | 289.124389302094 | 176.728213174409 | 181.719338620502 | 266.371181169318 | 222.276513751277 | 283.755077841135 | 198.845915296222 | 196.480127205 | Mgmt |
| **ENSMUSG00000054619** | 0.0312317310559327 | 0.746696226367831 | 0.956085046483146 | 1312.80543017482 | 1436.62934580488 | 1209.56934769271 | 1223.05640931264 | 1288.76579647908 | 1309.9210705511 | 1200.36303841123 | 1274.40701292083 | Mettl7a1 |
| **ENSMUSG00000054626** | -0.0639697282929842 | 0.880131518934194 | 0.978393501240173 | 14.4562194651047 | 16.2883145782866 | 17.036187995672 | 25.3240207449704 | 28.4689130912965 | 13.4539045528125 | 10.4107809055614 | 23.8815624227072 | Xlr |
| **ENSMUSG00000054630** | -0.330342956668229 | 0.329314052725612 | 0.855977678491063 | 248.466272056487 | 314.364471360931 | 182.66579350915 | 278.564228194674 | 187.237851485066 | 333.901449356164 | 216.544242835676 | 550.361461286933 | Ugt2b5 |
| **ENSMUSG00000054640** | 0.225819529284922 | 0.722662115885141 | 0.951674101666722 | 1308.28786159198 | 3070.34729800702 | 2559.2140189054 | 1041.09863062656 | 821.218646864323 | 585.856389163379 | 2722.4192068043 | 2693.18892594257 | Slc8a1 |
| **ENSMUSG00000054641** | -0.935306627790462 | 0.0279336072295712 | 0.665956123973759 | 65.0529875929712 | 36.6487078011448 | 112.628131749165 | 54.3997482669734 | 168.623562156141 | 194.470074899744 | 82.2451691539348 | 68.388110574116 | Mmrn1 |
| **ENSMUSG00000054648** | 0.155725937243076 | 0.365307853075352 | 0.86924083181804 | 251.176813206195 | 280.159010746529 | 307.597838810745 | 286.067641748739 | 237.605928492744 | 192.023910435596 | 273.803537816264 | 305.032683671851 | Zfp869 |
| **ENSMUSG00000054659** | 0.118220993736406 | 0.872783994651583 | 0.976162349063696 | 302.67709505063 | 638.501931468834 | 517.7108240907 | 225.102406621959 | 168.623562156141 | 75.8310983885793 | 638.180869510912 | 668.683747835801 | Pm20d2 |
| **ENSMUSG00000054675** | -1.98150269959017 | 0.00554238960966401 | 0.635862253526407 | 13.5527057485357 | 10.5874044758863 | 11.3574586637814 | 11.2551203310979 | 51.4630352034976 | 112.523565350795 | 14.5750932677859 | 6.51315338801105 | Tmem119 |
| **ENSMUSG00000054676** | 0.0986106878721749 | 0.507305672794954 | 0.901698316485955 | 1063.43564440177 | 1004.98900948028 | 1013.65318574249 | 932.299134092612 | 809.174106710313 | 783.995710759344 | 1043.16024673725 | 1111.57817822055 | 1600014C10l |
| **ENSMUSG00000054690** | -0.504805295452591 | 0.183833461200191 | 0.805963874499992 | 386.703870691551 | 269.571606270643 | 362.492222352355 | 492.411514485535 | 763.185862485911 | 812.126602097043 | 242.57119509958 | 326.743194965221 | Emcn |
| **ENSMUSG00000054693** | -0.0989156524245118 | 0.352819383969893 | 0.862728204353539 | 738.17070643691 | 868.167167022674 | 888.721140440892 | 815.9962240046 | 954.803546754252 | 857.380644683776 | 830.780316263797 | 903.157269804198 | Adam10 |
| **ENSMUSG00000054708** | 0.0594233122994002 | 0.734194027147284 | 0.954227437328031 | 118.360296870545 | 101.801966114291 | 104.110037751329 | 113.489130005238 | 108.400861386091 | 114.969729814943 | 83.2862472444909 | 113.980184290193 | Ankrd24 |
| **ENSMUSG00000054715** | 0.344531571653694 | 0.132212424905803 | 0.77303252008843 | 66.8600150261093 | 68.4109212288036 | 79.5022106464695 | 74.0962088463947 | 44.893286028583 | 47.7002070508805 | 58.3003730711437 | 75.9867895267955 | Zscan22 |
| **ENSMUSG00000054716** | 0.0162207354152064 | 0.919264287001099 | 0.986713918026868 | 690.28447945875 | 598.595560752032 | 644.535779169592 | 746.589648629496 | 746.761489548624 | 752.195572725424 | 642.345181873136 | 509.11148982953 | Zfp771 |
| **ENSMUSG00000054717** | -0.381091291441617 | 0.0375357275429634 | 0.679566075569698 | 236.72059374109 | 171.027303072009 | 246.078271048596 | 288.881421831514 | 331.772333333186 | 273.970419984545 | 324.816364253515 | 295.262953589834 | Hmgb2 |
| **ENSMUSG00000054720** | -0.640117519035221 | 0.0687078678420753 | 0.723457921755197 | 478.862269781594 | 476.433201414882 | 447.673162330715 | 423.004939110431 | 1070.86911551108 | 1018.82749931753 | 350.843316517418 | 405.986561186022 | Lrrc8c |
| **ENSMUSG00000054723** | -0.346741985860359 | 0.13236809956401 | 0.77303252008843 | 87.6408305071974 | 51.3081909216027 | 59.6266579848522 | 70.3445020693621 | 88.6916138613469 | 99.0696607979826 | 71.8343882483734 | 82.4999429148066 | Vmac |
| **ENSMUSG00000054728** | -0.0181099035466794 | 0.938848431011736 | 0.989269716906553 | 87.6408305071974 | 111.574954861263 | 121.146225747001 | 115.364983393754 | 132.489941694111 | 134.539045528125 | 102.025652874501 | 72.73021283279 | Phactr1 |
| **ENSMUSG00000054733** | 0.00660323838584589 | 0.948315043284983 | 0.991217716191233 | 1345.33192397131 | 1224.06684055824 | 1088.42312194571 | 1286.8354245222 | 1161.75064576406 | 1249.99004117948 | 1248.25263057681 | 1262.46623170947 | Msra |
| **ENSMUSG00000054737** | 0.14757356519645 | 0.585526587435856 | 0.922964550512322 | 28.0089252136404 | 28.5045505120015 | 34.0723759913441 | 34.703287687552 | 27.3739548954774 | 26.9078091056249 | 27.0680303544596 | 31.4802413753867 | Zfp182 |
| **ENSMUSG00000054745** | -0.0384983688776013 | 0.934490132848883 | 0.98818087308472 | 8.13162344912141 | 13.0306516626293 | 8.51809399783602 | 14.0689004138724 | 9.85462376237187 | 7.33849339244316 | 16.6572494488982 | 10.8552556466851 | Gm9955 |
| **ENSMUSG00000054752** | 0.183631999798258 | 0.454806585015693 | 0.892775943845882 | 1098.67267934796 | 1330.75530104601 | 1272.98182523216 | 996.078149302167 | 948.233797579338 | 623.771938357668 | 1452.30393632581 | 1111.57817822055 | Fsd1l |
| **ENSMUSG00000054757** | -0.587793210406698 | 0.172828376548984 | 0.799889021902745 | 19.877301764519 | 23.6180561385155 | 19.8755526616174 | 27.1998741334867 | 35.0386622662111 | 36.6924669622158 | 11.4518589961175 | 53.1907526687569 | Akr1c20 |
| **ENSMUSG00000054766** | 0.143876427808926 | 0.0546214524581262 | 0.703396849069134 | 2883.11226957182 | 2826.83699506163 | 2706.86098153456 | 3059.51687667012 | 2612.57025522437 | 2523.21864476837 | 2601.65414829979 | 2648.68237779116 | Set |
| **ENSMUSG00000054770** | -0.0153893747363186 | 0.934006630340377 | 0.98818087308472 | 121.070838020252 | 121.347943608235 | 106.002947528626 | 117.24083678227 | 104.021028602814 | 140.654456688494 | 93.6970281500523 | 133.519644454226 | Kctd18 |
| **ENSMUSG00000054792** | 0.131689394915918 | 0.468137245306302 | 0.895328112556867 | 233.106538874814 | 331.467201668132 | 313.276568142636 | 248.550573978413 | 218.991639163819 | 232.385624094033 | 302.953724351836 | 273.552442296464 | Klhl18 |
| **ENSMUSG00000054793** | 0.0971432203072673 | 0.388364075452272 | 0.876358159057779 | 739.977733870048 | 684.109212288036 | 738.234813145789 | 628.410885152968 | 587.992551154855 | 636.002760678407 | 676.700758861489 | 707.762668163867 | Cadm4 |
| **ENSMUSG00000054808** | -0.42958575896582 | 0.0560111555039279 | 0.703396849069134 | 1856.72068754939 | 1870.71292931621 | 1816.24693131637 | 1837.39839405174 | 2984.85604180286 | 3365.92230266726 | 1785.44892530377 | 1806.3145396084 | Actn4 |
| **ENSMUSG00000054814** | 0.0328123665386179 | 0.837406005978356 | 0.968924788429433 | 718.293404672391 | 732.159740293981 | 741.074177811734 | 597.459304242449 | 595.657258525589 | 568.733237914345 | 736.042210023189 | 824.999429148066 | Usp46 |
| **ENSMUSG00000054823** | 0.0373771540007696 | 0.828098579711091 | 0.967319591893432 | 505.967681278665 | 618.95595397489 | 649.268053612834 | 431.446279358754 | 538.719432342996 | 451.317343635254 | 601.743136341447 | 556.874614674944 | Nsd3 |
| **ENSMUSG00000054827** | -0.407863253790773 | 0.277521634971851 | 0.848075053184273 | 94.8689402397497 | 158.811067138294 | 55.8408384302584 | 109.737423228205 | 102.926070406995 | 124.754387671534 | 95.7791843311646 | 233.387996403729 | Cyp2c50 |
| **ENSMUSG00000054836** | -0.484539350572765 | 0.0928305799610515 | 0.754058637722175 | 27.1054114970714 | 26.0613033252585 | 21.7684624389143 | 23.448167356454 | 41.6084114411257 | 39.1386314263635 | 26.0269522639034 | 31.4802413753867 | Elp6 |
| **ENSMUSG00000054843** | -0.0890739599659948 | 0.620335377541551 | 0.929900538965265 | 437.300638819418 | 627.100111264033 | 518.657278979349 | 383.612017951588 | 519.010184818252 | 481.894399437101 | 597.578823979222 | 493.914131924171 | Atrnl1 |
| **ENSMUSG00000054855** | -0.100277130039976 | 0.827735216790152 | 0.967319591893432 | 38.8510898124689 | 21.9892246806869 | 17.036187995672 | 61.9031618210386 | 38.3235368536684 | 58.7079471395453 | 29.1501865355718 | 23.8815624227072 | Rnd1 |
| **ENSMUSG00000054863** | 0.118262429201556 | 0.425926559899823 | 0.885066977635295 | 177.088688447533 | 212.56250524664 | 178.879973954556 | 167.888878272211 | 158.768938393769 | 156.554525705454 | 187.394056300105 | 175.855141476298 | Fam19a5 |
| **ENSMUSG00000054871** | -1.38600797832743 | 0.0171580787497269 | 0.642273034804458 | 14.4562194651047 | 13.8450673915436 | 11.3574586637814 | 25.3240207449704 | 51.4630352034976 | 86.838838477244 | 14.5750932677859 | 17.3684090346961 | Tmem158 |
| **ENSMUSG00000054874** | 0.0634704013534719 | 0.699806816985602 | 0.945280238056015 | 275.571683553559 | 333.910448854875 | 330.312756138308 | 235.419600258798 | 304.398378437709 | 243.393364182698 | 271.721381635152 | 305.032683671851 | Pcnx3 |
| **ENSMUSG00000054889** | 0.40001649764374 | 0.193967034255183 | 0.809162446871198 | 37357.5816389803 | 49996.9815980506 | 40842.3678098464 | 31000.3530986207 | 24186.531587448 | 15068.37309915 | 41459.8938783076 | 39932.1434218957 | Dsp |
| **ENSMUSG00000054893** | -0.0592742495211848 | 0.775962387220623 | 0.959602406551077 | 115.649755720838 | 122.976775066064 | 143.861143074564 | 135.061443973175 | 118.255485148462 | 101.51582526213 | 178.024353485099 | 140.032797842237 | Zfp667 |
| **ENSMUSG00000054894** | 0.22690254930308 | 0.143172679840106 | 0.780607990324772 | 608.064731250967 | 494.350347450997 | 603.838218957709 | 629.348811847226 | 456.597567656563 | 410.955629976817 | 574.675105986987 | 551.446986851602 | Atp5s |
| **ENSMUSG00000054920** | -0.106355433085076 | 0.547222075213193 | 0.914699131434573 | 322.554396815149 | 325.766291565731 | 342.616669690738 | 387.363724728621 | 460.97740043984 | 410.955629976817 | 302.953724351836 | 309.374785930525 | Klhl5 |
| **ENSMUSG00000054931** | -0.148531788596192 | 0.692604912454951 | 0.94469184542218 | 19.877301764519 | 12.2162359337149 | 19.8755526616174 | 11.2551203310979 | 16.4243729372865 | 19.5693157131818 | 16.6572494488982 | 17.3684090346961 | Zkscan4 |
| **ENSMUSG00000054939** | 0.164512110492393 | 0.584877149963539 | 0.922651176944186 | 44.2721721118832 | 48.8649437348597 | 42.5904699891801 | 34.703287687552 | 33.943704070392 | 22.0154801773295 | 51.0128264372507 | 44.5065481514088 | Zfp174 |
| **ENSMUSG00000054942** | 0.310669350461542 | 0.100272865373693 | 0.759800953500741 | 376.765219809292 | 474.804369957054 | 412.654331450723 | 343.281170098487 | 313.158044004262 | 236.054870790255 | 373.747034509653 | 372.335268681298 | Miga1 |
| **ENSMUSG00000054958** | -0.115726383117021 | 0.909799903959469 | 0.985888646122408 | 102.097049972302 | 234.551729927327 | 183.612248397799 | 44.0825546301336 | 37.2285786578493 | 9.78465785659088 | 282.132162540713 | 282.236646813812 | Nt5c1a |
| **ENSMUSG00000054966** | -3.08202649540422 | NA | NA | 11.7456783153976 | 0 | 0.946454888648447 | 16.8826804966469 | 5.47479097909548 | 231.162541861959 | 10.4107809055614 | 3.25657669400552 | Lmntd1 |
| **ENSMUSG00000054967** | -0.841164571324734 | 0.114753006694048 | 0.767518180080285 | 9.93865088225949 | 17.1027303072009 | 10.4110037751329 | 16.8826804966469 | 30.6588294829347 | 36.6924669622158 | 24.9858741733473 | 5.42762782334254 | Zfp647 |
| **ENSMUSG00000054978** | -0.247909713552356 | 0.476627526981852 | 0.897553735056661 | 41.5616309621761 | 32.5766291565731 | 56.7872933189068 | 69.4065753751039 | 47.0832024202212 | 35.4693847301419 | 73.9165444294857 | 80.3288917854696 | Kbtbd13 |
| **ENSMUSG00000055013** | 0.159187750644659 | 0.324526629686641 | 0.855977678491063 | 734.556651570634 | 819.302223287815 | 909.543147991157 | 802.865250284986 | 875.966556655277 | 556.502415593606 | 718.343882483734 | 772.894202043977 | Agap1 |
| **ENSMUSG00000055024** | 0.139494574593463 | 0.513005508423617 | 0.903078589555164 | 837.557215259505 | 1076.65759362474 | 972.955625530603 | 631.224665235743 | 711.722827282413 | 620.102691661447 | 960.915077583314 | 900.986218674861 | Ep300 |
| **ENSMUSG00000055027** | 0.121160535582008 | 0.584484132663937 | 0.922450244597442 | 10325.354752951 | 11800.0694962397 | 10285.1252749427 | 8852.15214040853 | 7635.14349944656 | 6521.47446141782 | 11777.7164384616 | 12003.7416941044 | Smyd1 |
| **ENSMUSG00000055041** | -0.0397380743801037 | 0.864861623538613 | 0.974868041734905 | 211.422209677157 | 124.605606523892 | 139.128868631322 | 219.47484645641 | 182.858018701789 | 205.477814988408 | 180.106509666212 | 145.46042566558 | Commd5 |
| **ENSMUSG00000055044** | -0.0199660094141716 | 0.904759091866207 | 0.98504921011344 | 1114.03241252963 | 932.506009606906 | 1032.58228351546 | 1273.70445080258 | 1309.57000219964 | 1223.08223207386 | 944.257828134416 | 936.808562308922 | Pdlim1 |

| | | | | | | | | | | | | |
|---|---|---|---|---|---|---|---|---|---|---|---|---|
| **ENSMUSG00000055053** | 0.158517498215538 | 0.119796355928012 | 0.769868040577702 | 3845.35437771786 | 4252.06452066171 | 3952.39561499591 | 3942.10589596705 | 3259.69054895345 | 3351.24531588238 | 3699.99153383651 | 4016.44458927348 | Nfic |
| **ENSMUSG00000055065** | 0.117184692614849 | 0.487654424833886 | 0.899236955296196 | 1299.25272442629 | 1483.86545808191 | 2017.84182259849 | 1428.46235535518 | 1497.90281188052 | 1281.7901792134 | 1624.08182126757 | 1338.45302123627 | Ddx17 |
| **ENSMUSG00000055067** | -0.0368083958515236 | 0.815024129263847 | 0.966177091475091 | 143.658680934478 | 137.636258186522 | 133.450139299431 | 143.502784221499 | 156.579022002131 | 159.000690169602 | 141.586620315635 | 116.15123541953 | Smyd3 |
| **ENSMUSG00000055093** | -0.397447202260746 | 0.471215212995584 | 0.897091462240771 | 11.7456783153976 | 9.77298874697194 | 5.67872933189068 | 27.1998741334867 | 24.0890803080201 | 20.7923979452556 | 8.32862472444909 | 18.4539345993646 | Gm8430 |
| **ENSMUSG00000055114** | 0.129372312913216 | 0.714375199762731 | 0.951176382858794 | 33.4300075130547 | 21.9892246806869 | 20.8220075502658 | 30.9515809105193 | 16.4243729372865 | 18.3462334811079 | 30.191264626128 | 32.5657669400552 | Anxa13 |
| **ENSMUSG00000055116** | -0.214575126393317 | 0.26067395903724 | 0.840316047847749 | 330.68602026427 | 346.12668478859 | 406.975602118832 | 283.253861665965 | 340.531998899739 | 314.332133642982 | 457.033281754144 | 473.289146195469 | Arntl |
| **ENSMUSG00000055128** | -0.0757527997342501 | 0.614539435175906 | 0.928075610917191 | 520.42390074377 | 381.96097686082 | 373.849681016137 | 445.515179772627 | 457.692525852383 | 501.463715150282 | 447.663578939139 | 408.157612315359 | Cgrrf1 |
| **ENSMUSG00000055137** | 0.056657848198366 | 0.876257738057365 | 0.977469019294424 | 18.9737880479499 | 35.8342920722305 | 23.6613722162112 | 31.8895076047775 | 40.5134532453066 | 24.4616446414772 | 22.903717992235 | 18.4539345993646 | Sugct |
| **ENSMUSG00000055148** | -0.932448292222634 | 0.0432441658317589 | 0.687730553581215 | 983.022923627121 | 842.105863697416 | 693.751433379311 | 654.672832592197 | 2111.07940153922 | 2756.82735109448 | 687.11153976705 | 502.598336441519 | Klf2 |
| **ENSMUSG00000055150** | -0.171440481133377 | 0.602812942591063 | 0.926448151609826 | 17.1667606148119 | 17.9171460361152 | 18.9290977729689 | 17.8206071909051 | 20.8042057205628 | 19.5693157131818 | 16.6572494488982 | 23.8815624227072 | Zfp78 |
| **ENSMUSG00000055172** | -1.17086605222037 | 0.0116480644209554 | 0.635862253526407 | 96.6759676728878 | 149.852494120236 | 128.717864856189 | 146.316564304273 | 387.61520131996 | 543.048511040794 | 96.8202624217207 | 147.631476794917 | C1ra |
| **ENSMUSG00000055184** | 0.138834137214849 | 0.671320015950002 | 0.943173644832837 | 25.2983840639333 | 21.9892246806869 | 22.7149173275627 | 20.6343872736795 | 22.994122112201 | 24.4616446414772 | 13.5340151772298 | 21.7105112933701 | Fam72a |
| **ENSMUSG00000055200** | -0.448281636521469 | 0.0373461619308901 | 0.679566075569698 | 92.1583990900426 | 81.4415728914329 | 67.1982970940397 | 112.551203310979 | 129.205067106653 | 141.877538920568 | 105.14888714617 | 106.381505337514 | Sertad3 |
| **ENSMUSG00000055204** | 0.171637859746757 | 0.410616703456268 | 0.882192126163777 | 2008.51099193299 | 2622.41864710414 | 2366.13722162112 | 1757.67462503979 | 1738.79361496073 | 1395.53682679627 | 2330.97384475519 | 2306.74182492058 | Ankrd17 |
| **ENSMUSG00000055228** | 0.175978480968379 | 0.467231689850967 | 0.895328112556867 | 47.8862269781594 | 44.7928650902881 | 48.2691993210708 | 60.0273084325223 | 38.3235368536684 | 36.6924669622158 | 47.8895921655823 | 54.2762782334254 | Zfp935 |
| **ENSMUSG00000055239** | 0.257467122600489 | 0.0109854261254012 | 0.635862253526407 | 4491.36668506472 | 4219.48789150514 | 4713.34534546927 | 4506.73776591046 | 3454.59310780925 | 3438.08415435962 | 3997.73986773556 | 4107.62873670563 | Kcmf1 |
| **ENSMUSG00000055240** | -0.256728842620324 | 0.433834128327308 | 0.887868762931002 | 45.1756858284523 | 30.9477976987445 | 21.7684624389143 | 56.2756016554897 | 42.7033696369448 | 37.9155491942896 | 51.0128264372507 | 52.1052271040884 | Zfp101 |
| **ENSMUSG00000055296** | 0.262620590003705 | 0.271761164074083 | 0.844129257012535 | 1233.29622311675 | 1672.80990719003 | 1957.26870972499 | 1442.53125576905 | 1239.49267766722 | 831.695917810225 | 1673.01249152371 | 1511.05158601856 | Tmem245 |
| **ENSMUSG00000055302** | -0.0382086101298117 | 0.645760009361311 | 0.935063240511793 | 6422.17549737277 | 6117.07653987552 | 5419.40069240101 | 6104.02692623211 | 6102.20202529983 | 6291.53500178793 | 6059.07448703672 | 6256.96935474928 | Mrfap1 |
| **ENSMUSG00000055305** | 0.0909418580412731 | 0.642340847114598 | 0.934616627390525 | 77.7021796249379 | 72.4829998733752 | 78.5557557578211 | 75.0341355406529 | 54.7479097909548 | 80.7234273168747 | 71.8343882483734 | 78.1578406561325 | Zfp93 |
| **ENSMUSG00000055312** | -0.701272682188754 | 0.178127435522968 | 0.801696999069105 | 4.51756858284523 | 10.5874044758863 | 13.2503684410783 | 4.68963347129081 | 12.0445401540101 | 8.56157562451702 | 15.616171358342 | 17.3684090346961 | Them7 |
| **ENSMUSG00000055319** | 0.0463222349028288 | 0.72957048016636 | 0.952921973914044 | 717.389890955822 | 789.983257046899 | 802.593745573883 | 702.507093999363 | 716.102660065689 | 595.64104701997 | 823.492769629904 | 780.492880996657 | Sec23ip |
| **ENSMUSG00000055320** | 0.255341857950084 | 0.401162417220439 | 0.880924293718604 | 3015.92878590747 | 3803.32145402991 | 3597.47503175275 | 2891.62799839791 | 1899.75246974613 | 1527.62970786025 | 3684.37536247817 | 4037.06957500218 | Tead1 |
| **ENSMUSG00000055322** | 0.268804097185652 | 0.243927036760731 | 0.832048466795696 | 31172.1267353486 | 37634.96524886 | 41505.832686789 | 28642.4053892557 | 23651.0970296925 | 19736.8779789759 | 36445.0207160987 | 35499.9425413542 | Tns1 |
| **ENSMUSG00000055334** | 0.0286675211999137 | 0.838084558361982 | 0.969115750107235 | 228.588970291968 | 210.119258059897 | 195.916161950228 | 217.598993067893 | 190.522726072523 | 190.800828203522 | 241.530117009024 | 211.677485110359 | Snupn |
| **ENSMUSG00000055371** | 0.103302617130559 | 0.472005202905945 | 0.897171173241185 | 289.124389302094 | 352.642010619904 | 377.63550057073 | 301.07446885687 | 295.638712871156 | 273.970419984545 | 332.103910887408 | 326.743194965221 | Stam2 |
| **ENSMUSG00000055401** | -0.0764863752822618 | 0.541019625900859 | 0.913837378403038 | 928.812100632978 | 826.631964848044 | 838.559031342524 | 938.864620952419 | 1007.36154015357 | 1050.62763735145 | 842.232175259915 | 826.084954712734 | Fbxo6 |
| **ENSMUSG00000055407** | -0.451257229268423 | 0.239296181129095 | 0.832048466795696 | 59.631905293557 | 62.7100111264033 | 37.8581955459379 | 89.1030359545253 | 117.160526952643 | 118.638976511164 | 42.6842017128016 | 62.9604827507734 | Map6 |
| **ENSMUSG00000055413** | 0.111677648219197 | 0.802127243940257 | 0.963461896828986 | 18.0702743313809 | 8.95857301805762 | 17.9826428843205 | 20.6343872736795 | 17.5193311331055 | 22.0154801773295 | 8.32862472444909 | 13.0263067760221 | H2-Q5 |
| **ENSMUSG00000055421** | -0.805048701487126 | 0.374995318013848 | 0.873700364743909 | 9.03513716569045 | 9.77298874697194 | 4.73227444324223 | 11.2551203310979 | 27.3739548954774 | 29.3539735697726 | 0 | 4.34210225867403 | Pcdh9 |
| **ENSMUSG00000055435** | -0.246562186384186 | 0.295074942648819 | 0.851785577030345 | 206.001127377742 | 212.56250524664 | 265.007368821565 | 279.502154888932 | 290.163921892061 | 406.063301048521 | 204.051305749003 | 243.157726485746 | Maf |
| **ENSMUSG00000055436** | -0.125040217981444 | 0.185892144941573 | 0.805963874499992 | 638.784197614315 | 590.451403462888 | 588.694940739334 | 546.811262752508 | 667.924499449649 | 632.333513982185 | 636.098713329799 | 642.631134283757 | Srsf11 |
| **ENSMUSG00000055447** | -0.0191226330239349 | 0.910590620580034 | 0.986166948757759 | 1546.81548276621 | 1442.33025590728 | 1627.90240847533 | 1740.79194454315 | 1995.01383278239 | 1800.37704561272 | 1344.03181490797 | 1303.71620316688 | Cd47 |
| **ENSMUSG00000055485** | -0.0144733691354882 | 0.957367093760281 | 0.992403427270738 | 126.491920319666 | 209.304842330982 | 277.311282373995 | 170.702658354985 | 247.460552255116 | 141.877538920568 | 200.928071477334 | 200.822229463674 | Soga1 |
| **ENSMUSG00000055489** | 0.15008626752935 | 0.612534696727642 | 0.928055903343287 | 123.781379169959 | 118.904696421492 | 106.949402417274 | 102.23400967414 | 65.6974917491458 | 64.8233582999146 | 146.792010768415 | 129.177542195552 | Ano5 |
| **ENSMUSG00000055491** | 0.361894957336774 | 0.0300846914251967 | 0.665956123973759 | 195.158962778914 | 233.737314198412 | 276.364827485346 | 202.592165959763 | 168.623562156141 | 159.000690169602 | 196.76375911511 | 181.282769299641 | Pprc1 |
| **ENSMUSG00000055493** | 0.184609521490952 | 0.348022332575684 | 0.861266047293794 | 1067.95321298461 | 1105.97655986566 | 1002.29572707871 | 1001.70570946772 | 770.850569856644 | 654.348994159515 | 1105.62493217062 | 1144.14394516061 | Epm2a |
| **ENSMUSG00000055531** | 0.205917099238511 | 0.10371511446184 | 0.759800953500741 | 566.503100288791 | 543.215291185857 | 630.338955839866 | 507.418341593665 | 485.06648074786 | 426.855698993777 | 491.388858742496 | 543.848307898922 | Cpsf6 |
| **ENSMUSG00000055538** | 0.0109853575278956 | 0.947660687201505 | 0.991041028853253 | 1081.50591873315 | 1292.47776178704 | 1103.56640016409 | 821.623784170149 | 971.227919691539 | 950.334894321389 | 1158.71991478898 | 1186.47944218268 | Zcchc24 |
| **ENSMUSG00000055541** | 0.510541929070357 | 0.292875424273933 | 0.851785577030345 | 15.3597331816738 | 11.4018202048006 | 9.46454888648447 | 18.7585338851632 | 13.1394983498292 | 4.89232892829544 | 7.28754663389296 | 13.0263067760221 | Lair1 |
| **ENSMUSG00000055553** | 0.0850581435136132 | 0.517441809594414 | 0.905197321821966 | 2322.93376529901 | 1952.15450220765 | 2008.377273712 | 2286.6652806014 | 2015.81803850296 | 2393.57192816854 | 1879.14595345383 | 1792.20270726771 | Kxd1 |
| **ENSMUSG00000055612** | -0.497155568482887 | 0.246130988056228 | 0.832048466795696 | 11.7456783153976 | 10.5874044758863 | 11.3574586637814 | 13.1309737196143 | 15.3294147414674 | 23.2385624094033 | 8.32862472444909 | 19.5394601640331 | Cdca7 |
| **ENSMUSG00000055632** | -1.35013278876802 | 0.0128031983585833 | 0.635862253526407 | 92.1583990900426 | 43.1640336324594 | 96.5383986421416 | 89.1030359545253 | 364.621079207759 | 311.885969178834 | 89.5327157878278 | 52.1052271040884 | Hmcn2 |
| **ENSMUSG00000055633** | -0.144042855721235 | 0.621038487859484 | 0.93041735818988 | 41.5616309621761 | 32.5766291565731 | 44.483379766477 | 47.8342614071662 | 47.0832024202212 | 58.7079471395453 | 51.0128264372507 | 27.1381391167127 | Zfp580 |
| **ENSMUSG00000055639** | -0.302124707822842 | 0.164975534301111 | 0.795216960550936 | 109.325159704854 | 101.801966114291 | 114.521041526462 | 114.427056699496 | 168.623562156141 | 163.893019097897 | 97.8613405122768 | 112.894658725525 | Dach1 |
| **ENSMUSG00000055652** | -0.645725558715587 | 0.0122388949718632 | 0.635862253526407 | 64.1494738764022 | 77.3694942468612 | 93.6990339761962 | 80.6616957062019 | 118.255485148462 | 182.239252579005 | 93.6970281500523 | 100.953877514171 | Klhl25 |
| **ENSMUSG00000055653** | -1.38122055426011 | 0.010785371791499 | 0.635862253526407 | 18.0702743313809 | 28.5045505120015 | 21.7684624389143 | 21.5723139679377 | 100.736154015357 | 91.7311674055395 | 23.9447960827911 | 18.4539345993646 | Gpc3 |
| **ENSMUSG00000055660** | -0.124479895918107 | 0.712739450142816 | 0.950456121590218 | 76.7986659083688 | 82.2559886203472 | 40.6975602118832 | 39.3929211588428 | 55.8428679867739 | 45.2540425867328 | 71.8343882483734 | 87.9275707381491 | Mettl4 |
| **ENSMUSG00000055670** | 0.0827495616963891 | 0.672361848868961 | 0.943216235131458 | 346.045753445944 | 437.341246426994 | 451.458981885309 | 351.72251034681 | 382.140410340865 | 260.516515431732 | 431.006329490241 | 423.354970220718 | Zzef1 |
| **ENSMUSG00000055675** | -0.869186321423965 | 0.0422366362101772 | 0.685406501171951 | 26.2018977805023 | 30.1333819698302 | 35.965285768641 | 15.9447538023887 | 68.9823663366031 | 72.1618516923577 | 34.3555769883525 | 22.7960368580387 | Kbtbd11 |
| **ENSMUSG00000055676** | 0.0652352054468135 | 0.800920365724024 | 0.963461896828986 | 54.2108229941427 | 58.6379324818317 | 42.5904699891801 | 65.6548685980713 | 68.9823663366031 | 51.3694537471021 | 53.094982618363 | 37.9933947633978 | Gm5069 |
| **ENSMUSG00000055681** | -0.091182437691089 | 0.6295371397368 | 0.931825800800958 | 2817.15576826228 | 2276.29196231555 | 2461.72916537461 | 2838.1661768252 | 2779.00390098887 | 3735.29313675357 | 2359.08295320021 | 2199.2747940184 | Cope |
| **ENSMUSG00000055692** | 0.182638595371952 | 0.668865872364712 | 0.942055056494206 | 18.9737880479499 | 12.2162359337149 | 10.4110037751329 | 15.0068271081306 | 16.4243729372865 | 11.0077400886647 | 8.32862472444909 | 14.1118323406906 | Tmem191c |
| **ENSMUSG00000055707** | 0.167097070160887 | 0.234169411684365 | 0.832048466795696 | 252.983840639333 | 241.881471487556 | 313.276568142636 | 299.198615468353 | 267.16979977986 | 238.501035254403 | 235.283648465687 | 244.243252050414 | Klhl26 |
| **ENSMUSG00000055720** | -0.00373703560202576 | 0.967512946421687 | 0.994284908685797 | 1476.34141287382 | 1405.68154810613 | 1471.73735184833 | 1414.39345494131 | 1414.68598899827 | 1613.24546410542 | 1405.45542225078 | 1351.47932801229 | Ubl7 |
| **ENSMUSG00000055723** | 0.159353794972148 | 0.242676896169005 | 0.832048466795696 | 2085.30965784136 | 1991.24645719553 | 1639.25986713911 | 2086.88689472441 | 1586.59442574187 | 1564.32217482247 | 1830.21528319769 | 2004.96571794273 | Rras2 |
| **ENSMUSG00000055725** | -0.0821436506659822 | 0.818342002773976 | 0.966315773308182 | 18.9737880479499 | 27.6901347830872 | 33.1259211026956 | 20.6343872736795 | 22.994122112201 | 15.9000690169602 | 35.3966550789086 | 31.4802413753867 | Paqr3 |
| **ENSMUSG00000055730** | 0.292229540032045 | 0.378532985784842 | 0.87422797717331 | 49.6932544112975 | 108.317291945606 | 54.8943835416099 | 78.7858423176855 | 37.2285786578493 | 57.4848649074714 | 64.5468416144805 | 79.2433662208011 | Ces2a |
| **ENSMUSG00000055733** | -0.349928270657691 | 0.418614649963582 | 0.882468496126562 | 18.9737880479499 | 7.32974156022896 | 9.46454888648447 | 15.0068271081306 | 16.4243729372865 | 19.5693157131818 | 15.616171358342 | 13.0263067760221 | Nap1l3 |
| **ENSMUSG00000055737** | 0.0992527816526187 | 0.569968457421286 | 0.920872458148558 | 2851.48928949191 | 3117.58341028405 | 2827.06075239291 | 2371.07868308463 | 2129.69369086814 | 2164.85555077073 | 2956.66177717943 | 3172.99122552605 | Ghr |
| **ENSMUSG00000055745** | 0.0350531305167853 | 0.879606528923661 | 0.978393501240173 | 143.658680934478 | 178.357044632238 | 143.861143074564 | 158.509611329629 | 131.394983498292 | 105.185071958352 | 212.379930473452 | 159.572258006271 | Rtl6 |
| **ENSMUSG00000055760** | 0.124649063517153 | 0.570231407168464 | 0.920872458148558 | 141.85165350134 | 133.56417954195 | 113.574586637814 | 195.088752405698 | 158.768938393769 | 141.877538920568 | 108.272121417838 | 127.006491066215 | Gemin6 |
| **ENSMUSG00000055762** | -0.0482700397126747 | 0.799122958514679 | 0.96328975655894 | 5292.78335166147 | 4032.98668958376 | 4417.1049653223 | 5525.32615587483 | 5109.07494169191 | 6520.25137918575 | 4045.62945990115 | 4249.83258567721 | Eef1d |
| **ENSMUSG00000055782** | -0.0374942538860952 | 0.883288669155177 | 0.97910597465636 | 72.2810973255236 | 131.935348084121 | 95.5919437534931 | 107.861569839689 | 94.1664048404423 | 83.1695917810225 | 98.902418602833 | 142.203848971575 | Abcd2 |
| **ENSMUSG00000055799** | -0.198271671334942 | 0.364950920501284 | 0.86924083181804 | 332.493047697409 | 314.364471360931 | 292.45456059237 | 281.378008277448 | 446.742943894192 | 428.078781225851 | 244.653351280692 | 282.236646813812 | Tcf7l1 |
| **ENSMUSG00000055805** | -0.291736930481302 | 0.140971897757571 | 0.779711473022872 | 293.64195788494 | 378.703313945163 | 288.668741037776 | 345.157023487003 | 467.547149614754 | 490.455975061618 | 333.144988977964 | 309.374785930525 | Fmnl1 |
| **ENSMUSG00000055809** | -0.0853174354135171 | 0.716803283479225 | 0.951399675826701 | 89.4478579403355 | 69.2253369577179 | 79.5022106464695 | 60.0273084325223 | 59.1277425742312 | 100.292743030056 | 71.8343882483734 | 85.7565196088121 | Dnaaf3 |
| **ENSMUSG00000055817** | 0.0743768395136132 | 0.601945997451973 | 0.926448151609826 | 315.326287082597 | 346.941100517504 | 368.170951684246 | 317.019222659259 | 325.202584158272 | 267.855008824175 | 374.788112600209 | 310.460311495193 | Mta3 |
| **ENSMUSG00000055835** | -0.229762427472335 | 0.189556883967848 | 0.808824684419004 | 125.588406603097 | 105.059629029948 | 143.861143074564 | 151.944124469822 | 153.294147414674 | 173.677676954488 | 151.997401221196 | 137.8617467129 | Zfp1 |
| **ENSMUSG00000055839** | -0.107075707279768 | 0.662380679438994 | 0.939611397743053 | 2954.48985318078 | 2114.2232322616 | 2202.40052588494 | 3071.70992369548 | 2751.62994609339 | 4110.77938200024 | 2182.09967780566 | 2096.14986537489 | Elob |
| **ENSMUSG00000055850** | 0.113650112855061 | 0.326864106621036 | 0.855977678491063 | 910.741826301597 | 789.168841317984 | 993.777633080869 | 907.9130400419 | 758.806029702634 | 909.973180662952 | 840.150019078802 | 819.571801324723 | Rnf181 |
| **ENSMUSG00000055865** | 0.418634582145888 | 0.30250278152066 | 0.851785577030345 | 24.3948703473642 | 35.8342920722305 | 21.7684624389143 | 30.9515809105193 | 8.75966556655277 | 18.3462334811079 | 31.2323427166841 | 26.0526135520442 | Fam19a3 |
| **ENSMUSG00000055866** | 0.209749817904274 | 0.497115432214302 | 0.900927059989498 | 230.395997725106 | 263.870696168242 | 209.166530391307 | 112.551203310979 | 262.789966996583 | 157.777607937528 | 156.16171358342 | 129.177542195552 | Per2 |
| **ENSMUSG00000055884** | 0.258561024900882 | 0.555453641447433 | 0.916239900758826 | 48.7897406947284 | 41.5352021746308 | 66.2518422053913 | 34.703287687552 | 20.8042057205628 | 18.3462334811079 | 59.3414511616998 | 60.7894316214364 | Fancm |
| **ENSMUSG00000055897** | 0.00175962322891492 | 0.99263079491605 | 0.999071668397934 | 138.237598635064 | 98.5443031986338 | 100.324218196735 | 110.675349922463 | 101.831112211176 | 134.539045528125 | 106.189965236726 | 105.295979772845 | Ppp4r1l-ps |
| **ENSMUSG00000055900** | 0.212309149529815 | 0.436660510363148 | 0.888701139843674 | 353.273863178497 | 388.476302692135 | 424.958245003153 | 323.584709519066 | 258.410134213307 | 181.016170346931 | 425.80093903746 | 420.098393526712 | Tmem69 |
| **ENSMUSG00000055912** | 0.0958738255297721 | 0.562959359507325 | 0.918213422211925 | 322.554396815149 | 366.487078011448 | 265.007368821565 | 387.363724728621 | 264.979883388221 | 319.224462571277 | 346.679004155194 | 324.572143835884 | Tmem150a |
| **ENSMUSG00000055917** | 0.0598424236961023 | 0.625649137894852 | 0.931573470007391 | 371.344137509877 | 303.777066885045 | 361.545767463707 | 345.157023487003 | 328.487458745729 | 309.439804714686 | 323.775286162958 | 362.565538599282 | Zfp277 |
| **ENSMUSG00000055923** | 0.391602467664615 | 0.129074840313169 | 0.77303252008843 | 157.211386683014 | 170.212887343095 | 195.916161950228 | 173.51643843776 | 100.736154015357 | 85.6157562451702 | 187.394056300105 | 156.315681312265 | Aasdh |
| **ENSMUSG00000055932** | 0.171309538075024 | 0.233243703144985 | 0.832048466795696 | 1649.81604645508 | 1926.9076146113 | 2120.05895057252 | 1602.9167204872 | 1551.55576347566 | 1385.75216893968 | 1760.46305113043 | 1783.51850275036 | Fto |
| **ENSMUSG00000055943** | 0.0432011647687548 | 0.676035053164168 | 0.94409858171788 | 1869.36987958135 | 1686.65497458157 | 1891.96332240825 | 2003.41141893543 | 1910.70205170432 | 1920.23910435596 | 1630.32828981091 | 1770.49219597434 | Emc7 |
| **ENSMUSG00000055963** | 0.358290334680395 | 0.138093152083655 | 0.779711473022872 | 109.325159704854 | 100.173134656462 | 70.9841166486335 | 105.047789756914 | 54.7479097909548 | 73.3849339244316 | 73.9165444294857 | 98.7828263848342 | Triqk |
| **ENSMUSG00000055978** | 0.452776165745863 | 0.0754138555575108 | 0.735399943172214 | 64.1494738764022 | 79.8127414336042 | 67.1982970940397 | 81.59962240046 | 45.9882442244021 | 52.592535979176 | 75.998700610598 | 39.0789203280663 | Fut2 |
| **ENSMUSG00000055980** | 0.0329985608101939 | 0.943503403030663 | 0.990505437194639 | 103.90407740544 | 147.409246933493 | 235.667267273463 | 101.296082979881 | 91.9764884488041 | 48.9232892829544 | 247.776585552361 | 185.624871558315 | Irs1 |
| **ENSMUSG00000055991** | 0.0552711413537902 | 0.69975158012037 | 0.945280238056015 | 306.291149916906 | 368.115909469277 | 360.599312575058 | 327.336416296098 | 302.208462046071 | 287.424324537357 | 387.281049686883 | 333.256348353232 | Zkscan5 |
| **ENSMUSG00000055994** | -0.19744745222544 | 0.618959026062663 | 0.929568738467614 | 43.3686583953142 | 44.7928650902881 | 29.3401015481019 | 42.2067012416173 | 63.5075753575076 | 61.154111603693 | 42.6842017128016 | 16.2828834700276 | Nod2 |
| **ENSMUSG00000056014** | 0.0447918564602016 | 0.868399411183524 | 0.975106533657166 | 74.0881247586617 | 80.6271571625185 | 88.966759532954 | 78.7858423176855 | 77.7420319031559 | 41.5847958905112 | 100.984574783945 | 91.1841474321546 | A430033K04 |

| | | | | | | | | | | | | |
|---|---|---|---|---|---|---|---|---|---|---|---|---|
| **ENSMUSG00000056019** | 0.156784685716952 | 0.549812696430868 | 0.914699131434573 | 56.9213641438498 | 74.9262470601182 | 89.9132144216024 | 99.4202295913651 | 55.8428679867739 | 61.154111603693 | 74.9576225200418 | 95.5262496908287 | Zfp709 |
| **ENSMUSG00000056035** | -0.0898228943128463 | 0.634970314859753 | 0.932543559775713 | 1297.44569699315 | 1478.16454797951 | 1313.67938544404 | 1521.31709808674 | 1087.29348844836 | 1319.70572840769 | 1633.45152408258 | 1930.06445398061 | Cyp3a11 |
| **ENSMUSG00000056050** | 0.220625431400383 | 0.133996236957365 | 0.77303252008843 | 1715.77254776462 | 1737.96316550318 | 1840.85475842123 | 1568.21343279965 | 1374.17253575297 | 1175.38202502298 | 1745.88795786264 | 1592.4660033687 | Mia3 |
| **ENSMUSG00000056054** | 0.699522647150549 | 0.26768679321051 | 0.843793744782845 | 58.7283915769879 | 21.1748089517725 | 84.2344850897118 | 15.9447538023887 | 21.8991639163819 | 58.7079471395453 | 20.8215618111227 | 9.76973008201657 | S100a8 |
| **ENSMUSG00000056069** | -0.554943447863781 | 0.106324057279869 | 0.762094734041179 | 15.3597331816738 | 16.2883145782866 | 25.5542819935081 | 14.0689004138724 | 30.6588294829347 | 29.3539735697726 | 20.8215618111227 | 23.8815624227072 | Fam105a |
| **ENSMUSG00000056071** | 0.467168188765368 | 0.469127903555854 | 0.895328112556867 | 100.290022539164 | 31.7622134276588 | 83.2880302010633 | 16.8826804966469 | 32.8487458745729 | 95.400414101761 | 26.0269522639034 | 14.1118323406906 | S100a9 |
| **ENSMUSG00000056076** | 0.0916072156551949 | 0.328479463857581 | 0.855977678491063 | 4785.00864294966 | 4761.07435123317 | 5187.51924468214 | 5180.16913238782 | 5222.95059405709 | 4553.53515001098 | 4522.44322537586 | 4388.77985795478 | Eif3b |
| **ENSMUSG00000056091** | 0.121745864054103 | 0.332949357006811 | 0.856918526232803 | 2003.08990963357 | 1968.44281678593 | 2038.66383014875 | 1728.59889751779 | 1539.51122332165 | 1641.37635544312 | 1956.18573215498 | 1974.57100213202 | St3gal5 |
| **ENSMUSG00000056116** | -0.280753222356553 | 0.294236940488429 | 0.851785577030345 | 420.133878204606 | 336.353696041618 | 431.583429223692 | 451.142739938176 | 528.864808580624 | 765.649477278236 | 362.295175513536 | 335.427399482569 | H2-T22 |
| **ENSMUSG00000056121** | -0.120338750404336 | 0.270998007925595 | 0.844129257012535 | 2758.42737668529 | 2641.15020886917 | 2335.85066518437 | 2327.93405514876 | 2523.87864136302 | 2924.3896168886 | 2917.10080973829 | 2574.8666393937 | Fez2 |
| **ENSMUSG00000056124** | -0.0793393053295034 | 0.557310712945332 | 0.916341990969984 | 475.248214915318 | 528.555808065399 | 469.44162476963 | 468.963347129081 | 458.787484048202 | 448.871179171106 | 598.619902069779 | 544.933833463591 | B4galt6 |
| **ENSMUSG00000056131** | 0.0362842243101811 | 0.85052672835329 | 0.972692784625411 | 75.8951521917998 | 88.7713144516618 | 79.5022106464695 | 70.3445020693621 | 75.5521155115177 | 78.277262852727 | 64.5468416144805 | 89.0130963028176 | Pgm3 |
| **ENSMUSG00000056144** | -0.0601651643049114 | 0.825508054810173 | 0.966806404871388 | 87.6408305071974 | 64.338842584232 | 62.4660226507975 | 66.5927952923295 | 84.3117810780705 | 92.9542496376133 | 42.6842017128016 | 73.8157383974585 | Trim34a |
| **ENSMUSG00000056153** | 0.0635377078606279 | 0.730649216156587 | 0.952921973914044 | 487.897406947284 | 499.236841824483 | 472.280989435575 | 557.128456389348 | 427.033696369448 | 352.247682837272 | 574.675105986987 | 574.243023709641 | Socs6 |
| **ENSMUSG00000056167** | 0.0802544187197924 | 0.46382915027897 | 0.895328112556867 | 768.890172800257 | 682.480380830207 | 833.826756899282 | 760.658549043369 | 767.565695269187 | 673.918309872697 | 754.781615653199 | 682.795580176491 | Cnot10 |
| **ENSMUSG00000056185** | 0.0897381035845558 | 0.558584066106111 | 0.916574868225118 | 134.623543768788 | 130.306516626293 | 125.878500190243 | 153.819977858338 | 133.58489988993 | 116.192812047017 | 138.463386043966 | 122.664388807541 | Snx32 |
| **ENSMUSG00000056201** | -0.411673203672873 | 0.0710469960563445 | 0.726501344493452 | 1689.57064998411 | 1496.89610974454 | 1608.02685581371 | 1808.32266652973 | 2343.21053905287 | 3169.00606330337 | 1539.75449593253 | 1732.49880121094 | Cfl1 |
| **ENSMUSG00000056204** | -0.13512225698994 | 0.252467861500181 | 0.83723604018117 | 546.625798524272 | 613.255043872489 | 546.104470750154 | 558.066383083606 | 640.550544554172 | 528.371524255907 | 638.180869510912 | 678.453477917817 | Pgpep1 |
| **ENSMUSG00000056209** | -0.0812518083950417 | 0.650571158045536 | 0.936463929813731 | 342.431698579668 | 282.602257933272 | 260.275094378323 | 356.412143818101 | 328.487458745729 | 410.955629976817 | 280.050006359601 | 295.262953589834 | Npm3 |
| **ENSMUSG00000056211** | 0.274946895973753 | 0.226582467412975 | 0.832048466795696 | 1120.35700854562 | 1294.10659324487 | 1416.84296830672 | 1085.18118525669 | 928.524550054594 | 634.779678446333 | 1272.1974266596 | 1226.64388807541 | R3hdm1 |
| **ENSMUSG00000056214** | -0.7662064432165 | 0.0333049845085896 | 0.671606027118506 | 44.2721721118832 | 27.6901347830872 | 42.5904699891801 | 37.5170677703264 | 97.4512794278996 | 74.6080161565054 | 31.2323427166841 | 55.3618037980939 | Pard6g |
| **ENSMUSG00000056216** | 0.0212549374102024 | 0.867775822633113 | 0.974868041734905 | 945.075347531221 | 864.909504107017 | 774.20009891443 | 751.279282100787 | 744.571573156986 | 901.411605038435 | 765.19239655876 | 877.104656252154 | Cebpg |
| **ENSMUSG00000056220** | -0.596892103427705 | 0.194250532538119 | 0.809162446871198 | 35.2370349461928 | 21.1748089517725 | 13.2503684410783 | 19.6964605794214 | 44.893286028583 | 52.592535979176 | 18.7394056300105 | 19.5394601640331 | Pla2g4a |
| **ENSMUSG00000056228** | 0.0518172500624606 | 0.680159594198757 | 0.944452327903597 | 1033.61969175499 | 939.021335438221 | 996.616997746814 | 1088.93289203373 | 914.290093508946 | 839.034411202668 | 1110.8303226234 | 1048.61769546978 | Cars2 |
| **ENSMUSG00000056234** | 0.125141846293231 | 0.673381889098885 | 0.943631400280823 | 516.809845877494 | 764.736369450555 | 858.434584004141 | 432.384206053012 | 503.680770076785 | 360.809258461789 | 715.220648212066 | 778.32182986732 | Ncoa4 |
| **ENSMUSG00000056260** | -0.0775629626664991 | 0.550520933928295 | 0.914699131434573 | 256.597895505609 | 211.748089517725 | 237.56017705076 | 255.11606083822 | 273.739548954774 | 261.739597663806 | 239.447960827911 | 238.815624227072 | Lrif1 |
| **ENSMUSG00000056267** | 0.177331922275067 | 0.424270336016901 | 0.884586455610141 | 135.527057485357 | 154.738988493722 | 134.396594188079 | 147.254490998531 | 112.780694169367 | 84.3926740130963 | 140.545542225078 | 167.17093695895 | Cep70 |
| **ENSMUSG00000056268** | 0.193695419306595 | 0.382814521661325 | 0.875126835008975 | 440.011179969125 | 429.197089137851 | 385.207139679918 | 327.336416296098 | 295.638712871156 | 240.94719971855 | 378.952424962434 | 466.775992807458 | Dennd1b |
| **ENSMUSG00000056271** | 0.0596561928477776 | 0.898405674381183 | 0.984462340635154 | 20.780815481088 | 17.9171460361152 | 7.57163910918757 | 24.3860940507122 | 13.1394983498292 | 23.2385624094033 | 21.8626399016789 | 9.76973008201657 | Lman1l |
| **ENSMUSG00000056305** | 0.0557650467797752 | 0.574721908741478 | 0.922450244597442 | 410.195227322346 | 423.496179035451 | 404.136237452887 | 439.887619607078 | 413.894198019619 | 375.486245246675 | 409.143689588562 | 414.67076570337 | Usp39 |
| **ENSMUSG00000056310** | 0.0578970032542068 | 0.635060850710527 | 0.932543559775713 | 294.545471601509 | 264.685111897157 | 262.16800415562 | 255.11606083822 | 268.264757975679 | 280.085831144914 | 240.489038918468 | 246.414303179751 | Tyw1 |
| **ENSMUSG00000056313** | -0.193728333513367 | 0.560717684603972 | 0.916762264997884 | 734.556651570634 | 732.159740293981 | 713.626986040929 | 554.314676306573 | 1050.06490979051 | 1181.49743618335 | 395.609674411332 | 501.51281087685 | Tcim |
| **ENSMUSG00000056342** | 0.356004937458319 | 0.258130455217088 | 0.838310965110319 | 900.803175419338 | 1147.51176204029 | 1180.22924614461 | 767.224035903176 | 629.600962595981 | 381.601656407044 | 1100.41954171784 | 1009.53877514171 | Usp34 |
| **ENSMUSG00000056366** | -0.00440218847225453 | 0.97849945578167 | 0.996653035184233 | 449.949830851384 | 361.600583637962 | 453.351891662606 | 477.404687377404 | 413.894198019619 | 547.940839969089 | 393.52751823022 | 392.96025441 | Fabp3-ps1 |
| **ENSMUSG00000056367** | 0.185218880324689 | 0.218906947710604 | 0.827378912571506 | 1428.45518589566 | 1193.93345858841 | 1130.06713704625 | 1326.22834568104 | 1064.29936633616 | 901.411605038435 | 1203.48627268289 | 1296.1175242142 | Actr3b |
| **ENSMUSG00000056383** | 0.264415431627568 | 0.0412282310589125 | 0.683365309556438 | 245.75573090678 | 214.191336704468 | 216.738169500494 | 205.405946042537 | 181.76306050597 | 173.677676954488 | 186.352978209548 | 192.138024946326 | AI987944 |
| **ENSMUSG00000056394** | 0.00180422188570397 | 0.993501586550123 | 0.999133450854041 | 73.1846110420926 | 59.452348210746 | 91.8061241988993 | 86.2892558717508 | 90.881530252985 | 83.1695917810225 | 71.8343882483734 | 64.046008315442 | Lig1 |
| **ENSMUSG00000056413** | 0.246127283858196 | 0.289714444565496 | 0.851785577030345 | 82.2197482077831 | 74.1118313312039 | 94.6454888648447 | 105.047789756914 | 94.1664048404423 | 81.9465095489486 | 70.7933101578173 | 53.1907526687569 | Adap1 |
| **ENSMUSG00000056418** | 0.198762496935209 | 0.687843603270957 | 0.944452327903597 | 14.4562194651047 | 16.2883145782866 | 5.67872933189068 | 15.9447538023887 | 6.56974917491458 | 17.123151249034 | 10.4107809055614 | 11.9407812113536 | BC043934 |
| **ENSMUSG00000056427** | -0.586557361630792 | 0.12940971236773 | 0.77303252008843 | 88.5443442237664 | 112.389370590177 | 131.557229522134 | 79.7237690119437 | 182.858018701789 | 265.408844360028 | 99.9434966933891 | 71.6446872681215 | Slit3 |
| **ENSMUSG00000056429** | 0.14390173130613 | 0.387894866250919 | 0.875951433019341 | 2703.31303997458 | 3385.52618509686 | 4065.02374674508 | 2933.83469963953 | 2812.94760505926 | 2504.87241128726 | 3302.29970324407 | 3224.01092706547 | Tgoln1 |
| **ENSMUSG00000056458** | 0.313446309689557 | 0.42150113361196 | 0.882568531013176 | 17.1667606148119 | 13.8450673915436 | 21.7684624389143 | 30.0136542162612 | 12.0445401540101 | 15.9000690169602 | 17.6983275394543 | 20.6249857287016 | Mok |
| **ENSMUSG00000056459** | 0.0277190931913919 | 0.874542481695848 | 0.9767298530085 | 92.1583990900426 | 93.6578088251478 | 84.2344850897118 | 90.0409626487835 | 95.2613630362614 | 79.5003450848009 | 99.9434966933891 | 78.1578406561325 | Zbtb25 |
| **ENSMUSG00000056476** | 0.0655910872272135 | 0.866451430478267 | 0.974868041734905 | 52.4037955610046 | 41.5352021746308 | 43.5369248778286 | 35.6412143818101 | 29.5638712871156 | 17.123151249034 | 56.2182168900314 | 61.8749571861049 | Med12l |
| **ENSMUSG00000056481** | -1.39706711923316 | 0.00990754093208805 | 0.635862253526407 | 73.1846110420926 | 52.9370223794314 | 66.2518422053913 | 69.4065753751039 | 244.175677667659 | 337.570696052385 | 61.4236073428121 | 46.6775992807458 | Cd248 |
| **ENSMUSG00000056486** | 0.164356828870687 | 0.23326156897575 | 0.832048466795696 | 559.274990556239 | 594.52348210746 | 508.246275204216 | 566.507723331929 | 411.70428162798 | 486.786728365396 | 584.044808801993 | 505.854913135525 | Chn1 |
| **ENSMUSG00000056492** | -0.193894757208136 | 0.349686190310501 | 0.861774532843376 | 6114.9808337393 | 6380.13282031485 | 7343.5434810233 | 5675.39442695613 | 9730.89348624431 | 8205.65869498352 | 5795.68173012601 | 5452.59491132991 | Adgrf5 |
| **ENSMUSG00000056493** | -0.0166361149974834 | 0.921788468075416 | 0.986713918026868 | 271.957628687283 | 403.950201541507 | 409.814966784777 | 335.777756544422 | 376.665619361769 | 314.332133642982 | 388.322127777439 | 358.223436340608 | Foxk1 |
| **ENSMUSG00000056494** | 0.0781044688854766 | 0.79543859376177 | 0.962043367573119 | 38.8510898124689 | 50.4937751926884 | 47.3227444324223 | 45.0204813243917 | 30.6588294829347 | 33.0232202659942 | 66.6289977955927 | 41.2499714574033 | Cngb3 |
| **ENSMUSG00000056498** | -1.07081464035743 | 0.0116947669442867 | 0.635862253526407 | 9.93865088225949 | 12.2162359337149 | 10.4110037751329 | 24.3860940507122 | 36.1336204620302 | 34.2463024980681 | 16.6572494488982 | 32.5657669400552 | Tmem154 |
| **ENSMUSG00000056501** | -0.0868913078551461 | 0.83467680144494 | 0.968559797165432 | 1599.21927832721 | 1972.5148954305 | 1274.87473500946 | 1099.25008567056 | 2186.63151705074 | 2584.37275637207 | 986.942029847218 | 557.960140239613 | Cebpb |
| **ENSMUSG00000056508** | 0.309404715233701 | 0.325282716713151 | 0.855977678491063 | 37.0440623793308 | 30.9477976987445 | 23.6613722162112 | 24.3860940507122 | 18.6142893289246 | 20.7923979452556 | 28.1091084450157 | 26.0526135520442 | 1700001K19I |
| **ENSMUSG00000056515** | -0.445999133905012 | 0.0577380830798477 | 0.710862211775517 | 228.588970291968 | 233.737314198412 | 151.432782183751 | 179.143998603309 | 323.012667766634 | 335.124531588238 | 196.76375911511 | 226.874843015718 | Rab31 |
| **ENSMUSG00000056529** | -0.24809968212484 | 0.585673833564838 | 0.922964550512322 | 19.877301764519 | 28.5045505120015 | 54.8943835416099 | 51.5859681841989 | 74.4571573156986 | 56.2617826753975 | 26.0269522639034 | 27.1381391167127 | Ptafr |
| **ENSMUSG00000056536** | 0.297311664456142 | 0.211742474127048 | 0.823370267313127 | 76.7986659083688 | 101.801966114291 | 121.146225747001 | 104.109863062656 | 73.3621991198795 | 58.7079471395453 | 96.8202624217207 | 98.7828263848342 | Pign |
| **ENSMUSG00000056537** | -0.11559835243405 | 0.40786062506014 | 0.881573350402261 | 423.747933070882 | 552.173864203915 | 533.800557197724 | 444.577253078368 | 514.630352034976 | 474.555906044658 | 587.168043073661 | 540.591731204917 | Rlim |
| **ENSMUSG00000056579** | 0.0126655831757426 | 0.935687677596426 | 0.98818087308472 | 521.327414460339 | 552.173864203915 | 642.642869392295 | 486.783954319986 | 571.568178217569 | 426.855698993777 | 598.619902069779 | 585.098279356326 | Tug1 |
| **ENSMUSG00000056592** | -0.411793427388142 | 0.346298807809904 | 0.861266047293794 | 8.13162344912141 | 12.2162359337149 | 11.3574586637814 | 7.50341355406529 | 12.0445401540101 | 17.123151249034 | 10.4107809055614 | 13.0263067760221 | Zfp658 |
| **ENSMUSG00000056596** | -1.21148742544262 | 0.00979129404544353 | 0.635862253526407 | 18.0702743313809 | 26.0613033252585 | 20.8220075502658 | 21.5723139679377 | 43.7983278327639 | 101.51582526213 | 35.3966550789086 | 20.6249857287016 | Trnp1 |
| **ENSMUSG00000056598** | -0.0141250394370771 | 0.94276527692257 | 0.990470440696959 | 109.325159704854 | 96.1010560118908 | 82.3415753124149 | 105.985716451172 | 112.780694169367 | 74.6080161565054 | 104.107809055614 | 105.295979772845 | Drc3 |
| **ENSMUSG00000056602** | 0.0170994927495835 | 0.942270228966317 | 0.99017502499184 | 1778.11499420788 | 2340.63080489978 | 2255.40199964925 | 1264.32518386 | 2031.14745324442 | 1477.48333634522 | 2409.0547015469 | 1630.4593981321 | Fry |
| **ENSMUSG00000056608** | -0.120927155098337 | 0.607409034333846 | 0.927459968605428 | 259.308436655316 | 315.99330281876 | 318.008842585878 | 181.957778686083 | 271.549632563136 | 212.816308380852 | 342.514691792969 | 341.94055287058 | Chd9 |
| **ENSMUSG00000056612** | -0.00194415792090246 | 0.991572598216939 | 0.998628446692557 | 908.03128515189 | 750.891302059011 | 742.020632700382 | 957.623154837583 | 885.821180417649 | 1073.86619976085 | 742.288678566525 | 662.17059444779 | Ppp1r14b |
| **ENSMUSG00000056629** | -0.150995448480004 | 0.491192690448885 | 0.899236955296196 | 430.976042803434 | 347.755516246418 | 333.152120804253 | 458.646153492241 | 457.692525852383 | 591.971800323748 | 318.569895710178 | 376.677370939972 | Fkbp2 |
| **ENSMUSG00000056665** | -0.357293733641233 | 0.0568880826975022 | 0.707207375413293 | 257.501409222178 | 228.036404096012 | 189.290977729689 | 249.488500672671 | 324.107625962453 | 365.701587390084 | 239.447960827911 | 256.184033261768 | Them6 |
| **ENSMUSG00000056666** | 0.134699648115748 | 0.196648374322927 | 0.812065429583558 | 2776.49765101668 | 3177.85017422371 | 3451.72097890089 | 3011.68261526296 | 2997.99554015269 | 2928.05886358482 | 2641.21511574092 | 2744.20862748199 | Retsat |
| **ENSMUSG00000056671** | 0.169785116579867 | 0.475241180459626 | 0.897549741595057 | 171.667606148119 | 96.9154717408051 | 98.4313084194385 | 135.061443973175 | 131.394983498292 | 114.969729814943 | 96.8202624217207 | 103.124928643508 | Prelid2 |
| **ENSMUSG00000056673** | 0.264646353758802 | 0.333300047340863 | 0.857004379408475 | 210.518695960587 | 233.737314198412 | 256.489274823729 | 230.729966787508 | 167.528603960322 | 106.408154190426 | 284.214318721825 | 216.019587369033 | Kdm5d |
| **ENSMUSG00000056687** | -0.029166295362313 | 0.922772491872828 | 0.986713918026868 | 28.0089252136404 | 28.5045505120015 | 20.8220075502658 | 24.3860940507122 | 32.8487458745729 | 24.4616446414772 | 24.9858741733473 | 21.7105112933701 | Gm11696 |
| **ENSMUSG00000056692** | 0.122602667015408 | 0.473449166897412 | 0.89728597737823 | 5955.96241962315 | 6626.90078617589 | 5804.60783208092 | 5331.17533016339 | 4633.86308470642 | 4300.35712797169 | 6518.18992497197 | 6332.95614427607 | D17Wsu92e |
| **ENSMUSG00000056698** | 0.207820723644114 | 0.104131376365802 | 0.760854717370327 | 509.581736144941 | 581.492830444831 | 668.197151385803 | 585.266257217093 | 523.390017601528 | 469.663577116362 | 556.976778447533 | 478.716774018812 | Elmod3 |
| **ENSMUSG00000056708** | -0.171440728271274 | 0.44857655569579 | 0.89108031640712 | 745.398816169462 | 528.555808065399 | 702.269527377147 | 958.561081531841 | 859.542183717991 | 1079.98161092122 | 680.865071223713 | 684.966631305828 | Ier5 |
| **ENSMUSG00000056724** | -0.150053894646218 | 0.28797596878625 | 0.851785577030345 | 228.588970291968 | 224.778741180355 | 218.631079277791 | 192.274972322923 | 281.404256325508 | 226.270212933664 | 230.078258012906 | 221.447215192376 | Nbeal2 |
| **ENSMUSG00000056737** | -0.75876778400738 | 0.0594199129777707 | 0.712406493939829 | 156.307872966445 | 145.780415475665 | 123.985590412947 | 185.709485463116 | 306.588294829347 | 472.10974158051 | 125.970448957293 | 131.348593324889 | Capg |
| **ENSMUSG00000056749** | 0.14788909845353 | 0.458650806557459 | 0.894253186844906 | 632.459601598332 | 740.303897583125 | 548.943835416099 | 489.59773440276 | 500.395895489327 | 717.949270227356 | 446.622500848583 | 513.453592088204 | Nfil3 |
| **ENSMUSG00000056753** | 0.280903315392336 | 0.504108894521524 | 0.900927059989498 | 52.4037955610046 | 80.6271571625185 | 69.0912068713366 | 40.3308478531009 | 20.8042057205628 | 33.0232202659942 | 75.998700610598 | 69.4736361387845 | C330011M18 |
| **ENSMUSG00000056763** | -0.0667032013430328 | 0.772246802246149 | 0.959045638244974 | 50.5967681278665 | 51.3081909216027 | 55.8408384302584 | 56.2756016554897 | 51.4630352034976 | 56.2617826753975 | 43.7252798033577 | 72.73021283279 | Cspp1 |
| **ENSMUSG00000056770** | 0.130838794425024 | 0.281204692538818 | 0.850048154552447 | 3159.58746684195 | 3254.40525274166 | 3198.0710687431 | 3060.45480336438 | 2565.48705280414 | 2547.68028940985 | 3324.16234314574 | 3134.99783076265 | Setd3 |
| **ENSMUSG00000056820** | 0.129973500912387 | 0.320849277521939 | 0.855228644171892 | 2013.02856051583 | 1906.54722138844 | 2131.4164092363 | 2241.64479927701 | 1679.66587238649 | 1657.27642446008 | 2104.01882101395 | 2135.22878570295 | Tsnax |
| **ENSMUSG00000056832** | -0.635632688213072 | 0.0704774690162139 | 0.72606739345142 | 18.9737880479499 | 11.4018202048006 | 17.9826428843205 | 13.1309737196143 | 18.6142893289246 | 25.6847268735511 | 28.1091084450157 | 22.7960368580387 | Ttc26 |
| **ENSMUSG00000056851** | 0.0377785918483605 | 0.613797710005228 | 0.928075610917191 | 4722.6661965064 | 4457.29728434812 | 4386.81840888555 | 4763.7296801372 | 4553.93113641162 | 4680.73570214666 | 4273.62556173294 | 4349.70093762671 | Pcbp2 |
| **ENSMUSG00000056870** | -0.465931632909621 | 0.124210165325726 | 0.771949105062466 | 59.631905293557 | 52.122606650517 | 38.8046504345863 | 68.4686486808458 | 91.9764884488041 | 101.51582526213 | 47.8895921655823 | 61.8749571861049 | Gulp1 |
| **ENSMUSG00000056888** | -1.14718950202488 | 0.0698075114571174 | 0.724916894583216 | 2.71054114970714 | 16.2883145782866 | 5.67872933189068 | 3.75170677703264 | 22.994122112201 | 19.5693157131818 | 9.36970281500523 | 11.9407812113536 | Glipr1 |
| **ENSMUSG00000056899** | 0.0801920405804192 | 0.660617218136686 | 0.939611397743053 | 779.732337399086 | 554.617111390658 | 621.82086184203 | 853.513291774927 | 689.823663366031 | 768.095641742384 | 544.483841360859 | 655.657441059779 | Immp2l |
| **ENSMUSG00000056900** | 0.205729040263123 | 0.0733495878918039 | 0.730864975478592 | 7966.28043898927 | 9144.25980425008 | 9041.48355125861 | 8864.34518743388 | 7311.03587348411 | 6525.14370811404 | 8138.10743387732 | 8386.77051262889 | Usp13 |
| **ENSMUSG00000056904** | -0.36249763436525 | 0.410500953445743 | 0.882192126163777 | 15.3597331816738 | 27.6901347830872 | 35.965285768641 | 15.0068271081306 | 16.4243729372865 | 23.2385624094033 | 34.3555769883525 | 46.6775992807458 | Gm5620 |
| **ENSMUSG00000056917** | -0.522734379904848 | 0.0250768869540971 | 0.655303666217042 | 255.69438178904 | 280.159010746529 | 290.561650815073 | 386.425798034362 | 539.814390538815 | 540.602346576646 | 321.693129981846 | 340.855027305911 | Sipa1 |

| | | | | | | | | | | | | |
|---|---|---|---|---|---|---|---|---|---|---|---|---|
| ENSMUSG00000056919 | 0.281831031370332 | 0.195804321829253 | 0.811193545163524 | 87.6408305071974 | 111.574954861263 | 95.5919437534931 | 64.7169419038131 | 77.7420319031559 | 67.2695227640623 | 72.8754663389296 | 78.1578406561325 | Cep162 |
| ENSMUSG00000056938 | -0.0195438290070907 | 0.918310007342904 | 0.986713918026868 | 180.702743313809 | 156.367819951551 | 232.827902607518 | 233.543746870282 | 171.908436743598 | 214.039390612925 | 234.242570375131 | 193.223550510994 | Acbd4 |
| ENSMUSG00000056941 | -0.0091009724120270 | 0.943725854024616 | 0.990505437194639 | 330.68602026427 | 285.859920848929 | 299.079744812909 | 331.088123073131 | 331.772333333186 | 327.786038195794 | 268.598147363483 | 326.743194965221 | Commd7 |
| ENSMUSG00000056952 | 0.126407461875518 | 0.340036575187499 | 0.858570960161243 | 1177.27837268947 | 1286.77685168464 | 1261.62436656838 | 1187.41519493083 | 966.848086908263 | 1017.60441708545 | 1342.99073681742 | 1172.36760984199 | Tatdn2 |
| ENSMUSG00000056962 | -0.0253281191257372 | 0.835082502505498 | 0.968771050291583 | 816.776399778417 | 743.561560498782 | 790.289832021453 | 885.402799379704 | 843.117810780705 | 881.842289325253 | 873.464517976599 | 694.736361387845 | Jmjd6 |
| ENSMUSG00000056973 | -0.0832080774160773 | 0.498653152602108 | 0.900927059989498 | 2802.69954879718 | 3296.7548706452 | 3037.17373767287 | 3279.92964982079 | 3877.24697139542 | 3030.79777107902 | 3010.79783788835 | 3234.86618271215 | Ces1d |
| ENSMUSG00000056999 | 0.214934543772436 | 0.128234741039153 | 0.77303252008843 | 1576.63143541298 | 1699.6856262442 | 1732.95890111531 | 1571.96513957668 | 1303.00025302473 | 1147.25113368528 | 1631.36936790147 | 1587.03837554536 | Ide |
| ENSMUSG00000057036 | 0.199484347819777 | 0.348300864628421 | 0.861266047293794 | 62.3424464432641 | 61.895595397489 | 63.4124775394459 | 76.9099889291692 | 60.2227007700503 | 69.71568722821 | 49.9717483466946 | 51.0197015394199 | Gm7536 |
| ENSMUSG00000057037 | -0.471209372322558 | 0.117856728085327 | 0.769868040577702 | 54.2108229941427 | 60.2667639396603 | 47.3227444324223 | 73.1582821521366 | 52.5579933993167 | 80.7234273168747 | 67.6700758861489 | 124.835439936878 | Cfhr1 |
| ENSMUSG00000057054 | 0.239894283704123 | 0.212382649722196 | 0.823370267313127 | 84.0267756409212 | 72.4829998733752 | 72.8770264259304 | 83.4754757889764 | 61.3176589658694 | 72.1618516923577 | 73.9165444294857 | 57.5328549274309 | Inca1 |
| ENSMUSG00000057069 | -0.338643223834201 | 0.0904582475407149 | 0.753823363185192 | 56.9213641438498 | 71.6685841444609 | 72.8770264259304 | 66.5927952923295 | 75.5521155115177 | 74.6080161565054 | 86.4094815161593 | 102.03940307884 | Ero1lb |
| ENSMUSG00000057074 | -0.433626815586317 | 0.357905771729713 | 0.865834559821921 | 28.9124389302094 | 46.4216965481167 | 42.5904699891801 | 36.5791410760683 | 37.2285786578493 | 46.4771248188067 | 18.7394056300105 | 106.381505337514 | Ces1g |
| ENSMUSG00000057093 | -0.29778646020666 | 0.513239912595282 | 0.903078589555164 | 11.7456783153976 | 23.6180561385155 | 16.0897331070236 | 6.56548685980713 | 18.6142893289246 | 17.123151249034 | 22.903717992235 | 13.0263067760221 | Zfp607b |
| ENSMUSG00000057098 | -0.429625900172378 | 0.139764000549504 | 0.779711473022872 | 175.281661014395 | 185.686786192467 | 185.505158175096 | 240.109233730089 | 318.632834983357 | 392.609396495709 | 163.449260217313 | 185.624871558315 | Ebf1 |
| ENSMUSG00000057101 | 0.150326045511843 | 0.391572562958163 | 0.878285927324016 | 556.564449406532 | 607.554133770089 | 559.354839191232 | 494.287367874051 | 462.072358635659 | 369.370834086306 | 587.168043073661 | 578.585125968315 | Zfp180 |
| ENSMUSG00000057103 | -0.0554326444227757 | 0.851428489067168 | 0.972692784625411 | 68.6670424592474 | 60.2667639396603 | 51.1085639870161 | 66.5927952923295 | 42.7033696369448 | 66.0464405319884 | 48.9306702561384 | 98.7828263848342 | Nat8f1 |
| ENSMUSG00000057110 | -0.0666058795724449 | 0.77435534340235 | 0.959391911785248 | 93.0619128066116 | 165.326392969609 | 102.217127974032 | 134.123517278917 | 106.210944994452 | 122.308223207386 | 148.874166949528 | 141.118323406906 | Cntrl |
| ENSMUSG00000057113 | 0.212967388629483 | 0.0553540957200659 | 0.703396849069134 | 4817.53513674615 | 4291.97089137851 | 4640.46831904333 | 5277.71350859067 | 3975.79320901914 | 4645.26631741652 | 3918.6179328533 | 3877.49731699591 | Npm1 |
| ENSMUSG00000057123 | -0.713300565132637 | 0.0901407830929898 | 0.753823363185192 | 100.290022539164 | 63.5244268553176 | 97.48485353079 | 126.620103724852 | 221.181555555458 | 258.070350967584 | 85.3684034256032 | 71.6446872681215 | Gja5 |
| ENSMUSG00000057130 | -0.0666185534576148 | 0.682182483673634 | 0.944452327903597 | 834.846674109798 | 579.049583258088 | 610.463403178248 | 785.044643094081 | 756.616113310996 | 820.68817772156 | 646.509494235361 | 718.617923810552 | Txnl4a |
| ENSMUSG00000057133 | 0.122998700828633 | 0.641390619110584 | 0.934544499347759 | 646.012307346867 | 719.943504360267 | 1148.99623481921 | 682.810633419941 | 604.416924092141 | 511.248373006873 | 985.900951756661 | 833.683633665414 | Chd6 |
| ENSMUSG00000057134 | -0.172198369756879 | 0.0973940916722955 | 0.758298168905933 | 469.827132615903 | 548.916201288258 | 477.959718767466 | 481.156394154437 | 549.669014301187 | 527.148442023833 | 571.551871715319 | 580.756177097652 | Ado |
| ENSMUSG00000057137 | 0.118297534595684 | 0.528892270241027 | 0.908410242815367 | 166.246523848704 | 181.614707547895 | 182.66579350915 | 190.399118934407 | 216.801722772181 | 173.677676954488 | 139.504464134522 | 134.605170018895 | Tmem140 |
| ENSMUSG00000057143 | -0.289634389933766 | 0.0384292286023618 | 0.679566075569698 | 237.624107457659 | 235.366145656241 | 200.648436393471 | 214.785212985119 | 269.359716171498 | 316.77829810713 | 244.653351280692 | 257.269558826436 | Trim12c |
| ENSMUSG00000057147 | 0.131045530015726 | 0.528476690915907 | 0.908232038364594 | 136.430571201926 | 140.893921102179 | 120.199770858353 | 93.7926694258161 | 128.110108910834 | 85.6157562451702 | 123.88829277618 | 110.723607596188 | Dph6 |
| ENSMUSG00000057156 | -0.303945206809497 | 0.25149824323155 | 0.83723604018117 | 65.0529875929712 | 81.4415728914329 | 61.519567762149 | 45.9584080186499 | 63.5075753575076 | 61.154111603693 | 102.025652874501 | 86.8420451734806 | Homez |
| ENSMUSG00000057176 | 0.0794817813994576 | 0.692043952272886 | 0.94469184542218 | 222.264374275985 | 201.160685041839 | 159.950876181588 | 200.716312571247 | 212.421889988905 | 226.270212933664 | 164.49033830787 | 140.032797842237 | Ccdc189 |
| ENSMUSG00000057177 | 0.132045269526055 | 0.424912692729852 | 0.885066977635295 | 1679.63199910185 | 1939.93826627393 | 2014.05600304389 | 1660.13024883695 | 1303.00025302473 | 1494.60648759426 | 1969.71974733221 | 1887.72895695853 | Gsk3a |
| ENSMUSG00000057181 | 0.161100480798605 | 0.453799435518217 | 0.891964629399053 | 755.337467051722 | 874.682492853989 | 853.702309560899 | 719.38977449601 | 569.37826182593 | 507.579126310652 | 891.162845516053 | 895.558590851519 | 5730455P16R |
| ENSMUSG00000057182 | -1.13318686967336 | 0.0126467434925729 | 0.635862253526407 | 10.8421645988285 | 19.5459774939439 | 14.1968233297267 | 12.1930470253561 | 45.9882442244021 | 39.1386314263635 | 28.1091084450157 | 11.9407812113536 | Scn3a |
| ENSMUSG00000057193 | -0.139467943303104 | 0.314794056081259 | 0.853277176422981 | 1140.23431031013 | 1307.1372449075 | 1343.01948699215 | 1162.09117418586 | 1468.33894059341 | 1581.4453260715 | 1262.82772384459 | 1144.14394516061 | Slc44a2 |
| ENSMUSG00000057219 | -0.188478973760709 | 0.554559530117611 | 0.915945640531442 | 22.5878429142261 | 24.4324718674299 | 28.3936466594534 | 29.075727522003 | 38.3235368536684 | 30.5770558018465 | 17.6983275394543 | 32.5657669400552 | Armc7 |
| ENSMUSG00000057228 | 0.00126039152788776 | 0.998373681477538 | 0.999262355603273 | 12.6491920319666 | 39.0919549878878 | 11.3574586637814 | 18.7585338851632 | 5.47479097909548 | 20.7923979452556 | 14.5750932677859 | 41.2499714574033 | Aadat |
| ENSMUSG00000057229 | -0.00121400446862273 | 0.991475564844228 | 0.998628446692557 | 943.268320098083 | 752.52013351684 | 752.431636475515 | 871.333898965832 | 836.54806160579 | 802.341944240452 | 839.108940988246 | 843.45336374743 | Dmac2 |
| ENSMUSG00000057230 | 0.0770786697751674 | 0.748462353333802 | 0.956197192463153 | 852.916948441179 | 1215.92268326909 | 996.616997746814 | 701.569167305105 | 806.984190318674 | 604.202622644487 | 1111.87140071395 | 1047.53216990511 | Aak1 |
| ENSMUSG00000057234 | 0.228349369157962 | 0.152569780126064 | 0.783412422888014 | 160.82544154929 | 143.337168288922 | 148.593417517806 | 149.130344387048 | 137.964732673206 | 101.51582526213 | 141.586620315635 | 131.348593324889 | Mettl15 |
| ENSMUSG00000057236 | -0.012007918861305 | 0.877547846014702 | 0.977664000726269 | 1732.03579466286 | 1750.99381716581 | 1694.15425068072 | 1801.75717966993 | 1865.80876567574 | 1658.49950669215 | 1721.94316177985 | 1790.03165613837 | Rbbp4 |
| ENSMUSG00000057265 | 0.318548310517876 | 0.140988087258107 | 0.779711473022872 | 58.7283915769879 | 57.009101024003 | 54.8943835416099 | 49.7101147956825 | 47.0832024202212 | 37.9155491942896 | 44.7663578939139 | 46.6775992807458 | Bbof1 |
| ENSMUSG00000057278 | 0.145386397229542 | 0.321686483583081 | 0.855704427283138 | 251.176813206195 | 216.634583891211 | 205.380710836713 | 232.605820176024 | 229.94122112201 | 211.593226148778 | 172.818963032319 | 205.164331722348 | Snrpg |
| ENSMUSG00000057286 | -0.521793942418877 | 0.0996313353864484 | 0.759800953500741 | 131.009488902512 | 126.234437981721 | 129.664319744837 | 162.261318106662 | 239.795844884382 | 304.547475786391 | 110.35427759895 | 134.605170018895 | St6galnac2 |
| ENSMUSG00000057315 | 0.0532523502206951 | 0.732316508208766 | 0.953583603545492 | 665.889609111386 | 609.182965227918 | 754.324546252812 | 760.658549043369 | 650.405168316544 | 550.387004433237 | 817.246301086567 | 668.683747835801 | Arhgap24 |
| ENSMUSG00000057322 | 0.258430933635114 | 0.178185461034665 | 0.801696999069105 | 1764.56228845934 | 1256.64346971481 | 1445.23661496618 | 1615.10976751255 | 1278.91117271671 | 1685.40731579778 | 1072.31043327282 | 1048.61769546978 | Rpl38 |
| ENSMUSG00000057329 | -0.0954675285643435 | 0.667840040652755 | 0.942024383071524 | 175.281661014395 | 114.018202048006 | 104.110037751329 | 163.19924480092 | 179.573144114332 | 139.43137445642 | 135.340151772298 | 140.032797842237 | Bcl2 |
| ENSMUSG00000057335 | -0.0171972213342147 | 0.899919614802194 | 0.984802943363903 | 201.483558794897 | 211.748089517725 | 258.382184601026 | 213.847286290861 | 233.226095709468 | 210.370143916704 | 233.201492284575 | 218.19063849837 | Cep170 |
| ENSMUSG00000057337 | -0.0444952674029993 | 0.859545292000712 | 0.974868041734905 | 47.8862269781594 | 41.5352021746308 | 51.1085639870161 | 33.7653609932938 | 48.1781606160403 | 50.1463715150282 | 46.8485140750261 | 34.7368180693922 | Chst3 |
| ENSMUSG00000057342 | -0.0727283422389522 | 0.689888886257624 | 0.94469184542218 | 332.493047697409 | 430.011504866766 | 559.354839191232 | 462.397860269274 | 477.401773377126 | 435.417274618294 | 538.237372817523 | 424.440495785386 | Sphk2 |
| ENSMUSG00000057346 | -0.603013461908895 | 0.208552499230571 | 0.820854127036734 | 8.13162344912141 | 6.51532583131463 | 6.62518422053913 | 12.1930470253561 | 8.75966556655277 | 9.78465785659088 | 15.616171358342 | 16.2828834700276 | Apol9a |
| ENSMUSG00000057363 | 0.0480723054503892 | 0.728367279514044 | 0.952401994188708 | 201.483558794897 | 171.027303072009 | 185.505158175096 | 188.52326554589 | 160.958854785407 | 185.908499275227 | 198.845915296222 | 175.855141476298 | Uxs1 |
| ENSMUSG00000057367 | 0.114244268843779 | 0.483342558317872 | 0.89863290032536 | 597.222566652139 | 579.049583258088 | 676.71524538364 | 545.87333605825 | 496.016062706051 | 440.309603546589 | 646.509494235361 | 631.775878637071 | Birc2 |
| ENSMUSG00000057375 | -0.138836445460827 | 0.346044988635246 | 0.861266047293794 | 324.361424248287 | 294.004078138073 | 302.865564367503 | 313.267515882226 | 369.000911991036 | 401.170972120226 | 283.173240631269 | 307.203734801188 | Yipf1 |
| ENSMUSG00000057378 | -0.077127834615841 | 0.799605848999725 | 0.963355472513329 | 50.5967681278665 | 39.0919549878878 | 56.7872933189068 | 48.7721881014244 | 30.6588294829347 | 41.5847958905112 | 63.5057635239243 | 69.4736361387845 | Ryr3 |
| ENSMUSG00000057388 | -0.0225688956994879 | 0.873975731719636 | 0.97671366425653 | 2990.63040184354 | 2290.13702970709 | 2072.7362061401 | 2718.11155996015 | 2461.46602420133 | 2781.28899573596 | 2436.12273190136 | 2552.07060253566 | Mrpl18 |
| ENSMUSG00000057396 | -0.00757257610291263 | 0.975269587287992 | 0.99575536949131 | 66.8600150261093 | 47.2361122770311 | 45.4298346551254 | 54.3997482669734 | 44.893286028583 | 45.2540425867328 | 63.5057635239243 | 60.7894316214364 | Zfp759 |
| ENSMUSG00000057400 | -0.422095656926227 | 0.146341797487101 | 0.780607990324772 | 422.844419354313 | 640.945178655577 | 348.295399022628 | 502.728708122374 | 389.805117711598 | 589.5256358596 | 616.318229609233 | 970.459854813646 | Ces1c |
| ENSMUSG00000057406 | 0.160851917405241 | 0.531710409901635 | 0.909082872831377 | 373.151164943016 | 473.989954228139 | 511.085639870161 | 359.225923900876 | 302.208462046071 | 247.06261087892 | 498.676405376389 | 487.40097853616 | Nsd2 |
| ENSMUSG00000057409 | -0.0741376671404438 | 0.79442002414081 | 0.961804768755116 | 60.535419010126 | 36.6487078011448 | 45.4298346551254 | 42.2067012416173 | 41.6084114411257 | 35.4693847301419 | 65.5879197050366 | 51.0197015394199 | Zfp53 |
| ENSMUSG00000057411 | -0.111747949537189 | 0.529823984898382 | 0.908620485539384 | 710.161781223269 | 683.294796559122 | 602.891764069061 | 859.140851940476 | 789.464859185569 | 968.681127802497 | 638.180869510912 | 690.394259129171 | Fam173a |
| ENSMUSG00000057421 | 0.241322968210427 | 0.0358771329786078 | 0.679566075569698 | 1666.07929335332 | 1623.13054772626 | 1677.11806268505 | 1636.68208148049 | 1348.98849724913 | 1182.72051841542 | 1588.68516618866 | 1463.28846117315 | Las1l |
| ENSMUSG00000057440 | 0.256578031420041 | 0.343999239905105 | 0.861266047293794 | 82.2197482077831 | 88.7713144516618 | 138.182413742673 | 119.116690170786 | 64.6025335533267 | 73.3849339244316 | 120.765058504512 | 98.7828263848342 | Mpp7 |
| ENSMUSG00000057465 | 1.00220889370584 | 0.14635754293273 | 0.780607990324772 | 15.3597331816738 | 127.86326943955 | 27.447191770805 | 10.3171936368398 | 21.8991639163819 | 20.7923979452556 | 22.903717992235 | 24.9670879873757 | Saa2 |
| ENSMUSG00000057469 | 0.128350360924971 | 0.545875471125679 | 0.914211323944814 | 2439.48703473642 | 2559.70863597773 | 2144.66677767738 | 2249.14821283107 | 1682.95074697395 | 1533.74511902062 | 2688.06362981594 | 2687.76129811922 | E2f6 |
| ENSMUSG00000057497 | 0.159596758773651 | 0.269879295804428 | 0.843793744782845 | 1122.16403597875 | 924.361852317763 | 876.417226888462 | 1163.02910088012 | 859.542183717991 | 809.680437632895 | 966.120468036095 | 1021.47955635307 | Fam136a |
| ENSMUSG00000057506 | 0.0396880382813893 | 0.836620180691132 | 0.968924788429433 | 288.220875585525 | 215.005752433383 | 269.739643264807 | 315.143369270742 | 315.3479603959 | 284.978160073209 | 199.886993386778 | 258.355084391105 | Bloc1s2 |
| ENSMUSG00000057522 | 0.160531006456024 | 0.242413797549839 | 0.832048466795696 | 5810.49671125553 | 6327.19579793542 | 6192.65433642679 | 5799.20075059821 | 4936.07154675249 | 4414.10377555456 | 5978.91147406389 | 6258.05488031395 | Spop |
| ENSMUSG00000057530 | -0.548270258767748 | 0.0530721099229513 | 0.703396849069134 | 1341.71786910503 | 1382.06349196762 | 1625.06304380938 | 1347.80065964898 | 2850.17618371711 | 2837.55077841135 | 1318.00486264407 | 1325.42671446025 | Ece1 |
| ENSMUSG00000057531 | 0.0144648757412995 | 0.923803998623918 | 0.986713918026868 | 3089.11339694956 | 2520.61668098985 | 2921.70624125776 | 3307.12952395428 | 2953.1022541241 | 3582.40785774433 | 2587.079055032 | 2598.74820181641 | Dtnbp1 |
| ENSMUSG00000057541 | 0.0915145451956162 | 0.666444505704978 | 0.942024383071524 | 205.097613661173 | 257.355370336928 | 255.542819935081 | 199.778385876988 | 174.098353135236 | 163.893019097897 | 269.639225454039 | 252.927456567762 | Pus7 |
| ENSMUSG00000057551 | 0.117982599900492 | 0.486204829900674 | 0.899236955296196 | 266.536546387868 | 295.632909595901 | 296.240380146964 | 228.854113398991 | 237.605928492744 | 200.585486060113 | 264.433835001259 | 298.51953028384 | Zfp317 |
| ENSMUSG00000057554 | 0.306925538994556 | 0.114857335191371 | 0.767518180080285 | 643.30176619716 | 821.745470474558 | 812.058294460367 | 682.810633419941 | 524.484975797347 | 450.09426140318 | 779.767489826546 | 637.203506460414 | Lgals8 |
| ENSMUSG00000057561 | 0.195778957167149 | 0.207959426764477 | 0.820372930586885 | 402.063603873225 | 516.339572131684 | 530.961192531779 | 465.211640352048 | 479.591689768764 | 340.016860516533 | 432.047407580797 | 419.012867962044 | Eif1a |
| ENSMUSG00000057572 | -0.0151572087555824 | 0.93281835206966 | 0.988114361264825 | 267.440060104437 | 243.510302945384 | 253.649910157784 | 322.646782824807 | 274.834507150593 | 346.132271676902 | 258.187366457922 | 220.361689627707 | Zbtb8os |
| ENSMUSG00000057594 | 0.0558500936223122 | 0.746575646432059 | 0.956085046483146 | 122.87786545339 | 102.616381843205 | 97.48485353079 | 118.178763476528 | 104.021028602814 | 118.638976511164 | 91.61487196894 | 110.723607596188 | Arl16 |
| ENSMUSG00000057596 | -0.314847150158514 | 0.378746041258046 | 0.87422797717331 | 16.2632468982428 | 22.8036404096012 | 28.3936466594534 | 23.448167356454 | 32.8487458745729 | 40.3617136584374 | 21.8626399016789 | 18.4539345993646 | Trim30d |
| ENSMUSG00000057604 | 0.00117898329559517 | 0.995055953000881 | 0.999262355603273 | 780.635851115655 | 664.563234794092 | 876.417226888462 | 1159.27739410309 | 918.669926292222 | 971.127292266645 | 767.274552739873 | 820.657326889392 | Lmcd1 |
| ENSMUSG00000057606 | -0.382319543241084 | 0.236121084073203 | 0.832048466795696 | 164.439496415566 | 100.987550385377 | 80.448665535118 | 99.4202295913651 | 109.49581958191 | 91.7311674055395 | 210.29777429234 | 168.256462523619 | Colq |
| ENSMUSG00000057614 | -0.269517769866875 | 0.168528680536254 | 0.79907358295209 | 85.8338030740593 | 71.6685841444609 | 88.0203046443056 | 85.3513291774927 | 94.1664048404423 | 116.192812047017 | 78.0808567917102 | 110.723607596188 | Gnai1 |
| ENSMUSG00000057637 | 0.0731389684559085 | 0.652341481282626 | 0.936720465895231 | 934.233182932393 | 1245.24164951001 | 1148.99623481921 | 898.533773099319 | 935.094299229509 | 847.595986827185 | 1122.28218161952 | 1112.66370378522 | Prdm2 |
| ENSMUSG00000057649 | -0.00044904809144926 | 0.997063308950008 | 0.999262355603273 | 400.256576440087 | 453.629561005281 | 419.279515671262 | 481.156394154437 | 412.7992398238 | 428.078781225851 | 496.594249195277 | 416.841816832707 | Brd9 |
| ENSMUSG00000057666 | -0.132358019765616 | 0.413002044718203 | 0.882192126163777 | 90768.7949939594 | 78784.9487837143 | 81824.8109432128 | 88997.9881647684 | 96332.2321517725 | 119332.46413675 | 81044.8061155236 | 76373.2366278175 | Gapdh |
| ENSMUSG00000057667 | -0.254733065481537 | 0.196624023434154 | 0.812065429583558 | 115.649755720838 | 93.6578088251478 | 79.5022106464695 | 108.799496533947 | 140.154649064844 | 103.961989726278 | 104.107809055614 | 125.920965501547 | Bloc1s3 |
| ENSMUSG00000057672 | 0.215508968287737 | 0.226805550929379 | 0.832048466795696 | 1221.55054480135 | 1279.44711012441 | 1332.60848321701 | 1106.75349922463 | 967.943045104082 | 776.657217366901 | 1266.99203620682 | 1241.84124598077 | Pkn1 |
| ENSMUSG00000057691 | 0.031134452416949 | 0.847191472631833 | 0.972359077413333 | 223.167887992554 | 290.746415222415 | 302.865564367503 | 237.295453647315 | 273.739548954774 | 287.424324537357 | 248.817663642917 | 222.532740757044 | Zfp746 |
| ENSMUSG00000057706 | -0.446199143331332 | 0.219671016190743 | 0.82837125944045 | 35.2370349461928 | 29.3189662409158 | 20.8220075502658 | 24.3860940507122 | 51.4630352034976 | 45.2540425867328 | 18.7394056300105 | 34.7368180693922 | Mex3b |
| ENSMUSG00000057719 | 0.28997932994154 | 0.241373067792154 | 0.832048466795696 | 657.757985662265 | 690.624538119351 | 825.308662901446 | 828.189271029956 | 407.324448844704 | 462.325083723919 | 847.437565712695 | 737.071858409917 | Sh3rf2 |
| ENSMUSG00000057722 | -0.062369281533994 | 0.894623099891216 | 0.983733637008786 | 12.6491920319666 | 24.4324718674299 | 24.6078271048596 | 20.6343872736795 | 22.994122112201 | 37.9155491942896 | 8.32862472444909 | 17.3684090346961 | Lepr |
| ENSMUSG00000057738 | 0.0438839056869983 | 0.762318815101826 | 0.957746336519503 | 6317.36790625076 | 6971.39863950665 | 6753.90208539532 | 5208.30693321557 | 7324.17537183394 | 5964.97204582421 | 5667.62912498761 | 5538.35143093873 | Sptan1 |
| ENSMUSG00000057751 | -0.729432424346595 | 0.230817107598646 | 0.832048466795696 | 33.4300075130547 | 18.7315617650296 | 29.3401015481019 | 11.2551203310979 | 79.9319482947941 | 46.4771248188067 | 16.6572494488982 | 10.8552556466851 | Megf6 |

| | | | | | | | | | | | | |
|---|---|---|---|---|---|---|---|---|---|---|---|---|
| ENSMUSG00000057766 | -0.689390250200151 | 0.0102895764260961 | 0.635862253526407 | 75.8951521917998 | 61.895595397489 | 61.519567762149 | 57.2135283497478 | 111.685735973548 | 149.216032313011 | 83.2862472444909 | 70.559161703453 | Ankrd29 |
| ENSMUSG00000057778 | 0.092497571448373 | 0.510083520663753 | 0.902394714365738 | 439.107666252556 | 456.887223920938 | 467.548714992333 | 449.266886549659 | 435.793361936001 | 330.232202659942 | 458.0743598447 | 474.374671760138 | Cyb5d2 |
| ENSMUSG00000057788 | -0.00497585423046174 | 0.97017600013664 | 0.9951105526746 | 454.46739943423 | 408.836695914993 | 392.778778789105 | 482.094320848695 | 504.775728272604 | 431.748027922072 | 409.143689588562 | 398.387882233342 | Ddx49 |
| ENSMUSG00000057789 | -0.266757580766922 | 0.0292946303522676 | 0.665956123973759 | 353.273863178497 | 324.137460107903 | 270.686098153456 | 317.957149353517 | 375.57066116595 | 392.609396495709 | 368.541644056872 | 387.532626586657 | Bak1 |
| ENSMUSG00000057835 | -0.0656405148071929 | 0.796304693109583 | 0.962278883153809 | 45.1756858284523 | 65.9676740420606 | 41.6440151005317 | 53.4618215727152 | 63.5075753575076 | 42.8078781225851 | 47.8895921655823 | 61.8749571861049 | Zfp119a |
| ENSMUSG00000057841 | 0.218683219552249 | 0.203397869278206 | 0.81605904225138 | 6397.78062702541 | 4170.62294777028 | 4831.65220655032 | 6150.92326094502 | 4535.3168470827 | 5374.22332773254 | 4088.31366161395 | 4522.299502409 | Rpl32 |
| ENSMUSG00000057842 | -0.40988185783323 | 0.257628981380386 | 0.838048372687018 | 29.8159526467785 | 21.1748089517725 | 14.1968233297267 | 28.1378008277448 | 45.9882442244021 | 26.9078091056249 | 23.9447960827911 | 27.1381391167127 | Zfp595 |
| ENSMUSG00000057858 | 0.20658644200843 | 0.185287983615138 | 0.805963874499992 | 362.309000344187 | 288.303168035672 | 288.668741037776 | 331.088123073131 | 273.739548954774 | 238.501035254403 | 330.021754706295 | 257.269558826436 | Fam204a |
| ENSMUSG00000057863 | 0.247142094467058 | 0.156252245227438 | 0.787275540343856 | 5142.800074711 | 3510.94620734967 | 4283.65482602287 | 5055.42488205149 | 3657.16037403578 | 4700.30501785984 | 3323.12126505519 | 3480.19496032724 | Rpl36 |
| ENSMUSG00000057880 | -0.199339147533235 | 0.481293056735233 | 0.897732059433541 | 232.203025158245 | 300.519403969387 | 132.503684410783 | 230.729966787508 | 183.952976897608 | 215.262472844999 | 327.939598525183 | 301.776106977845 | Abat |
| ENSMUSG00000057894 | -0.112170281050007 | 0.498909097976146 | 0.900927059989498 | 99.3865088225949 | 109.13170767452 | 93.6990339761962 | 86.2892558717508 | 96.3563212320805 | 105.185071958352 | 108.272121417838 | 110.723607596188 | Zfp329 |
| ENSMUSG00000057895 | 0.0451472672489688 | 0.876519010052419 | 0.977469019294424 | 27.1054114970714 | 28.5045505120015 | 33.1259211026956 | 25.3240207449704 | 33.943704070392 | 24.4616446414772 | 30.191264626128 | 21.7105112933701 | Zfp105 |
| ENSMUSG00000057897 | 0.2939490522888 | 0.23409754379548 | 0.832048466795696 | 430.976042803434 | 346.12668478859 | 367.224496795597 | 356.412143818101 | 237.605928492744 | 195.693157131818 | 395.609674411332 | 394.045779974668 | Camk2b |
| ENSMUSG00000057913 | 0.645562626438049 | 0.245549471208216 | 0.832048466795696 | 16.2632468982428 | 20.3603932228582 | 80.448665535118 | 23.448167356454 | 20.8042057205628 | 12.2308223207386 | 27.0680303544596 | 29.3091902460497 | Gm10032 |
| ENSMUSG00000057914 | 0.0916248070856467 | 0.72873494558946 | 0.952401994188708 | 626.135005582348 | 829.889627763701 | 510.139184981513 | 391.115431505653 | 466.452191418935 | 425.632616761703 | 732.91897575152 | 587.269330485663 | Cacnb2 |
| ENSMUSG00000057933 | -0.413620126145176 | 0.183693850876834 | 0.805963874499992 | 37.0440623793308 | 85.5136515360045 | 48.2691993210708 | 41.2687745473591 | 55.8428679867739 | 77.0541806206532 | 83.2862472444909 | 67.3025850094475 | Gsta2 |
| ENSMUSG00000057963 | -0.297466444451221 | 0.149909108666245 | 0.782641594375979 | 191.544907912638 | 207.676010873154 | 212.952349945901 | 192.274972322923 | 280.309298129689 | 306.993640250539 | 239.447960827911 | 162.828834700276 | Itpk1 |
| ENSMUSG00000057967 | 0.14213747185431 | 0.577906019493065 | 0.922450244597442 | 79.509207058076 | 39.9063707168021 | 61.519567762149 | 58.151455044006 | 52.5579933993167 | 57.4848649074714 | 53.094982618363 | 53.1907526687569 | Fgf18 |
| ENSMUSG00000057969 | 0.164875490566549 | 0.256195721196223 | 0.837517407767513 | 758.951521917998 | 710.984931342209 | 797.861471130641 | 845.071951526603 | 716.102660065689 | 538.156182112498 | 750.617303290975 | 769.637625349972 | Sema3b |
| ENSMUSG00000057982 | 0.116477284212722 | 0.666818575471833 | 0.942024383071524 | 271.957628687283 | 280.159010746529 | 317.06238769723 | 182.895705380341 | 216.801722772181 | 147.992950080937 | 326.898520434627 | 277.894544555138 | Zfp809 |
| ENSMUSG00000058006 | 0.373260938390689 | 0.240205559831918 | 0.832048466795696 | 256.597895505609 | 318.436550005503 | 226.202718386979 | 208.219726125312 | 182.858018701789 | 86.838838477244 | 257.146288367366 | 251.841931003094 | Mdn1 |
| ENSMUSG00000058013 | -0.0635869936015371 | 0.630030346445829 | 0.931825800800958 | 396.642521573811 | 412.908774559565 | 477.013263878817 | 411.749818779333 | 520.105143014071 | 432.971110154146 | 414.349080041342 | 407.07208675069 | Sept11 |
| ENSMUSG00000058022 | -0.062244921253741 | 0.863239833952487 | 0.974868041734905 | 44.2721721118832 | 57.8235167529173 | 44.483379766477 | 70.3445020693621 | 47.0832024202212 | 53.8156182112498 | 30.191264626128 | 95.5262496908287 | Adtrp |
| ENSMUSG00000058056 | 0.152659369958823 | 0.465582349179752 | 0.895328112556867 | 4696.4642987259 | 5555.94410265355 | 4803.25855989087 | 4026.51929845029 | 3715.1931584142 | 2986.76681072437 | 5084.62539427617 | 5378.77917293246 | Palld |
| ENSMUSG00000058064 | -0.82629675960644 | 0.0550456421949898 | 0.703396849069134 | 9.93865088225949 | 8.14415728914329 | 9.46454888648447 | 13.1309737196143 | 17.5193311331055 | 14.6769867848863 | 12.4929370866736 | 27.1381391167127 | Gm10036 |
| ENSMUSG00000058070 | 0.0202637858817199 | 0.895261401366242 | 0.983733637008786 | 2370.81999227717 | 2475.00940017064 | 1941.17897661796 | 1896.48777579 | 1986.25416721584 | 1799.15396338065 | 2459.02644989359 | 2317.59708056726 | Eml1 |
| ENSMUSG00000058076 | 0.0384021716317324 | 0.648038637114784 | 0.935681566478623 | 14232.1480633956 | 12673.9375733648 | 13559.8591896663 | 13719.0537569141 | 12319.3746611607 | 13456.3507172766 | 12828.1642318327 | 14157.4244144067 | Sdhc |
| ENSMUSG00000058093 | -0.115449596795258 | 0.542527235272514 | 0.913837378403038 | 66.8600150261093 | 90.4001459094905 | 84.2344850897118 | 71.2824287636203 | 91.9764884488041 | 75.8310983885793 | 86.4094815161593 | 84.6709940441436 | Zfp729b |
| ENSMUSG00000058099 | -0.404809663471909 | 0.153143241975703 | 0.783412422888014 | 28.0089252136404 | 29.3189662409158 | 47.3227444324223 | 28.1378008277448 | 51.4630352034976 | 46.4771248188067 | 36.4377331694648 | 41.2499714574033 | Nfam1 |
| ENSMUSG00000058126 | -0.751198700170847 | 0.057643952508794 | 0.710862211775517 | 10.8421645988285 | 17.9171460361152 | 17.036187995672 | 11.2551203310979 | 18.6142893289246 | 35.4693847301419 | 16.6572494488982 | 26.0526135520442 | Tpm3-rs7 |
| ENSMUSG00000058135 | -0.237121821272701 | 0.0779602427581395 | 0.739855948845396 | 4775.97350578397 | 5963.96638283963 | 5189.41215445943 | 5853.60049886518 | 5995.99108030537 | 7596.56374341074 | 6366.19252375078 | 5717.46314910903 | Gstm1 |
| ENSMUSG00000058173 | -0.105195912553395 | 0.484247911378 | 0.899225065379735 | 290.027903018663 | 236.99497711407 | 228.095628164276 | 282.315934971707 | 274.834507150593 | 320.447544803351 | 243.612273190136 | 277.894544555138 | Smco4 |
| ENSMUSG00000058174 | 0.495513264941089 | 0.284545029428716 | 0.851125503777472 | 11.7456783153976 | 10.5874044758863 | 9.46454888648447 | 15.0068271081306 | 9.85462376237187 | 7.33849339244316 | 9.36970281500523 | 6.51315338801105 | Gm5148 |
| ENSMUSG00000058192 | -0.052278414798781 | 0.829057972615082 | 0.967483872587428 | 65.0529875929712 | 61.895595397489 | 46.3762895437739 | 55.3376749612315 | 77.7420319031559 | 58.7079471395453 | 45.80743598447 | 55.3618037980939 | Zfp846 |
| ENSMUSG00000058207 | -0.321933926262114 | 0.266109794835691 | 0.843513595393324 | 1622.710634958 | 2648.4799504294 | 1380.87768253808 | 2002.47349224117 | 1723.46420021926 | 2064.56280774067 | 2001.99316813945 | 3778.71449061107 | Serpina3k |
| ENSMUSG00000058230 | 0.136036376433234 | 0.552600553762629 | 0.914977712327965 | 1491.70114605549 | 1851.98136755118 | 1511.48845717157 | 1255.88384361168 | 1094.9581958191 | 987.027361283605 | 1689.66974097261 | 1788.9461305737 | Arhgap35 |
| ENSMUSG00000058239 | -0.00070672577802705 | 0.992352148429626 | 0.999041193135284 | 1636.26334070654 | 1672.80990719003 | 1639.25986713911 | 1689.20597635895 | 1564.69526182549 | 1643.82251990727 | 1719.86100559874 | 1711.87381548224 | Usf2 |
| ENSMUSG00000058240 | 0.172505459222587 | 0.174768499341569 | 0.799889021902745 | 700.22313034101 | 541.586459728029 | 552.729654970693 | 622.783324987419 | 512.440435643337 | 582.187142467157 | 510.128264372507 | 540.591731204917 | Cryzl1 |
| ENSMUSG00000058254 | -0.405686460012586 | 0.0674309307872035 | 0.722038584319139 | 2221.74022904328 | 2110.15115361703 | 1939.28606684067 | 2475.18854614729 | 3568.46876017444 | 3738.96238344979 | 2215.41417670346 | 2064.6696239995 | Tspan7 |
| ENSMUSG00000058258 | -0.235794168738402 | 0.401865934634356 | 0.881158207349486 | 27.1054114970714 | 41.5352021746308 | 40.6975602118832 | 31.8895076047775 | 36.1336204620302 | 33.0232202659942 | 42.6842017128016 | 54.2762782334254 | Idi1 |
| ENSMUSG00000058267 | -0.0170456220899628 | 0.901476695421749 | 0.984802943363903 | 1351.65651998729 | 1005.8034252092 | 977.687899973846 | 1168.65666104567 | 1134.37669086858 | 1243.87463001912 | 1097.29630744617 | 1082.2689879745 | Mrps14 |
| ENSMUSG00000058290 | -0.703957256389705 | 0.241837947031696 | 0.832048466795696 | 4.51756858284523 | 4.88649437348597 | 13.2503684410783 | 9.37926694258161 | 17.5193311331055 | 20.7923979452556 | 5.20539045278068 | 8.68420451734806 | Espl1 |
| ENSMUSG00000058291 | 0.0726635527478381 | 0.566674697953004 | 0.919931558693298 | 580.959319753896 | 504.937751926884 | 571.658752743662 | 545.87333605825 | 478.496731572945 | 461.102001491845 | 563.22324699087 | 590.525907179668 | Zfp68 |
| ENSMUSG00000058297 | -0.585481714214409 | 0.102033186542426 | 0.759800953500741 | 684.863397159336 | 584.750493360488 | 870.738497556571 | 614.341984739096 | 1674.1910814074 | 1308.69798831903 | 681.906149314269 | 468.947043936795 | Spock2 |
| ENSMUSG00000058298 | 0.0477936173416674 | 0.857472587169438 | 0.974029039597056 | 35.2370349461928 | 57.8235167529173 | 51.1085639870161 | 51.5859681841989 | 45.9882442244021 | 35.4693847301419 | 46.8485140750261 | 60.7894316214364 | Mcm9 |
| ENSMUSG00000058301 | 0.0377651237828226 | 0.735236031038777 | 0.954383574136061 | 1013.74238999047 | 1125.5225373596 | 1151.83559948516 | 1022.3400967414 | 1062.10944994452 | 907.527016198804 | 1117.07679116673 | 1113.74922934989 | Upf1 |
| ENSMUSG00000058317 | 0.233059352201043 | 0.119513338415033 | 0.769868040577702 | 206.904641094311 | 192.202112023782 | 199.701981504822 | 157.571684635371 | 147.819356435578 | 166.339183562045 | 157.202791673977 | 172.598564782293 | Ube2e2 |
| ENSMUSG00000058318 | 0.0253611449187974 | 0.851298890303213 | 0.972692784625411 | 369.537110076739 | 285.045505120015 | 335.04503058155 | 388.301651422879 | 335.057207920644 | 330.232202659942 | 352.92547269853 | 334.3418739179 | Phf21a |
| ENSMUSG00000058325 | -0.0112756654652744 | 0.93552973932761 | 0.98818087308472 | 738.17070643691 | 903.18704336599 | 850.862944894954 | 728.769041438591 | 831.073270626695 | 667.802898712327 | 892.203923606609 | 854.308619394115 | Dock1 |
| ENSMUSG00000058331 | -0.0975101141401765 | 0.767969879285611 | 0.95879680569409 | 25.2983840639333 | 24.4324718674299 | 21.7684624389143 | 21.5723139679377 | 20.8042057205628 | 23.2385624094033 | 18.7394056300105 | 36.9078691987293 | Zfp85 |
| ENSMUSG00000058351 | 0.393212488338427 | 0.0273818440449346 | 0.664353746352133 | 303.580608767199 | 219.892246806869 | 282.043556817237 | 294.508981997063 | 206.947099009809 | 251.954939807215 | 167.613572579538 | 211.677485110359 | Smim4 |
| ENSMUSG00000058355 | 0.182375140273838 | 0.337062661283665 | 0.858570960161243 | 1348.04246512102 | 1542.50339056374 | 1440.50434052294 | 1364.68334014562 | 1078.53382288181 | 911.196262895025 | 1607.42457181867 | 1420.95296415108 | Abce1 |
| ENSMUSG00000058388 | -0.0620617632961256 | 0.723479876927931 | 0.951674101666722 | 151.7903043836 | 131.120932355207 | 136.289503965376 | 131.309737196143 | 116.065568756824 | 130.869798831903 | 173.860041122875 | 153.05910461826 | Phtf1 |
| ENSMUSG00000058392 | 0.031768973459179 | 0.832738401784398 | 0.967854847250989 | 126.491920319666 | 157.99665140938 | 149.539872406455 | 164.137171495178 | 155.484063806312 | 139.43137445642 | 147.833088858971 | 142.203848971575 | Rrp1b |
| ENSMUSG00000058396 | -0.236702985196086 | 0.540607280265003 | 0.913837378403038 | 126.491920319666 | 96.9154717408051 | 144.807597963212 | 90.0409626487835 | 245.270635863478 | 143.100621152642 | 83.2862472444909 | 68.388110574116 | Gpr182 |
| ENSMUSG00000058402 | 0.301832091377843 | 0.361374264405175 | 0.868564573435346 | 60.535419010126 | 47.2361122770311 | 24.6078271048596 | 44.0825546301336 | 38.3235368536684 | 42.8078781225851 | 37.4788112600209 | 24.9670879873757 | Zfp420 |
| ENSMUSG00000058407 | 0.0113930041928903 | 0.891529024885969 | 0.982421229269187 | 903.513716569045 | 823.374301932386 | 848.023580229008 | 904.161333264867 | 886.916138613469 | 867.165302540366 | 855.766190437144 | 841.282312618093 | Txndc9 |
| ENSMUSG00000058420 | -0.620414309449589 | 0.191183995467075 | 0.808824684419004 | 8.13162344912141 | 5.7009101024003 | 19.8755526616174 | 15.9447538023887 | 19.7092475247437 | 24.4616446414772 | 18.7394056300105 | 13.0263067760221 | Syt17 |
| ENSMUSG00000058440 | 0.2472286472488 | 0.0820701239247994 | 0.739855948845396 | 244.852217190211 | 302.96265115613 | 343.563124579386 | 296.384835385579 | 255.12525962585 | 248.285693110993 | 259.228444548478 | 237.730098662403 | Nrf1 |
| ENSMUSG00000058441 | 0.515624273889102 | 0.0774052764828615 | 0.739855948845396 | 41.5616309621761 | 43.9784493613737 | 54.8943835416099 | 66.5927952923295 | 28.4689130912965 | 47.7002070508805 | 41.6431236222455 | 27.1381391167127 | Panx2 |
| ENSMUSG00000058444 | -0.0273031034383037 | 0.779469014365594 | 0.960444627083598 | 455.370913150799 | 446.299819445052 | 425.904699891801 | 434.260059441529 | 483.971522552041 | 440.309603546589 | 436.211719943021 | 435.295751432072 | Map2k5 |
| ENSMUSG00000058446 | 0.203752510820049 | 0.273321270439766 | 0.844738346035895 | 750.819898468876 | 856.765346817874 | 951.187163091689 | 746.589648629496 | 613.176589658694 | 550.387004433237 | 842.232175259915 | 862.992823911464 | Znrf2 |
| ENSMUSG00000058454 | -0.412623155468744 | 0.0191785999814634 | 0.642273034804458 | 89.4478579403355 | 78.9983257046899 | 81.3951204237664 | 80.6616957062019 | 96.3563212320805 | 111.300483118721 | 102.025652874501 | 130.263067760221 | Dhcr7 |
| ENSMUSG00000058486 | -0.0356680418349187 | 0.833120995662561 | 0.967868401766821 | 165.343010132135 | 166.955224427437 | 139.128868631322 | 163.19924480092 | 132.489941694111 | 183.462334811079 | 148.874166949528 | 186.710397122983 | Wdr91 |
| ENSMUSG00000058492 | 0.452683795749203 | 0.062246197165936 | 0.715615581523459 | 69.5705561758165 | 55.3802695661743 | 60.5731128735006 | 66.5927952923295 | 49.2731188118594 | 44.0309603546589 | 32.2734208072402 | 58.6183804920994 | Scp2-ps2 |
| ENSMUSG00000058498 | 0.0721115583361566 | 0.687214858612751 | 0.944452327903597 | 275.571683553559 | 250.840044505613 | 194.023252172932 | 289.819348525772 | 289.068963696242 | 226.270212933664 | 212.379930473452 | 233.387996403729 | Rnf207 |
| ENSMUSG00000058503 | -0.215776301470244 | 0.216502754517436 | 0.825479895758216 | 274.66816983699 | 226.407572638183 | 250.810545491838 | 253.240207449704 | 287.974005500422 | 379.155491942897 | 258.187366457922 | 243.157726485746 | Fam133b |
| ENSMUSG00000058546 | 0.0973787172244495 | 0.460237752338721 | 0.894774044236998 | 5626.17991307544 | 4600.63445263704 | 5296.36155687671 | 5979.28267589578 | 5132.06906380411 | 5783.95587547728 | 4473.51255511972 | 4710.09542509665 | Rpl23a |
| ENSMUSG00000058558 | 0.0876095195343483 | 0.544012846453634 | 0.913837378403038 | 9659.46514383966 | 6790.59834768767 | 8116.79712504908 | 9840.72687615663 | 7951.58641803828 | 8758.49186388091 | 7593.62359251646 | 8076.3102011337 | Rpl5 |
| ENSMUSG00000058569 | -0.155741773463053 | 0.286129273248549 | 0.851765024736894 | 1764.56228845934 | 1450.47441319642 | 1531.36400983319 | 1608.54428065275 | 1772.73731903112 | 2137.94774166511 | 1728.18963032319 | 1441.57794987978 | Tmed9 |
| ENSMUSG00000058571 | -0.958944669489914 | 0.028582056589293 | 0.665956123973759 | 69.5705561758165 | 84.6992358070902 | 74.7699362032273 | 70.3445020693621 | 226.656346534553 | 228.716377397812 | 68.711153976705 | 58.6183804920994 | Gpc6 |
| ENSMUSG00000058586 | 0.00247885118992603 | 0.978788792006504 | 0.996664169100645 | 500.546598979251 | 469.917875583568 | 478.906173656114 | 491.473587791276 | 476.306815181307 | 464.771248188067 | 482.019155927491 | 513.453592088204 | Serhl |
| ENSMUSG00000058587 | -0.394503830486639 | 0.101298886936382 | 0.759800953500741 | 1000.18968424193 | 878.754571498561 | 995.670542858166 | 1250.25628344613 | 1761.78773707293 | 1703.75354927889 | 970.284780398319 | 986.742738283673 | Tmod3 |
| ENSMUSG00000058594 | 0.142290739187621 | 0.39139496959126 | 0.878267548888951 | 1434.77978191164 | 1599.51249158774 | 1641.15277691641 | 1454.72430279441 | 1311.75991859128 | 1035.95065056656 | 1688.62866288205 | 1516.4792138419 | Fbxo18 |
| ENSMUSG00000058600 | -0.0283896151198904 | 0.894185716121788 | 0.983733637008786 | 4476.91046559962 | 3416.47398279561 | 4110.4535814002 | 4902.54283088741 | 4384.21261605966 | 5985.76444376947 | 3565.69246015477 | 3307.59639554494 | Rpl30 |
| ENSMUSG00000058603 | 0.0917475890933482 | 0.756567214374659 | 0.956566265296272 | 30.7194663633475 | 27.6901347830872 | 24.6078271048596 | 30.9515809105193 | 27.3739548954774 | 35.4693847301419 | 20.8215618111227 | 23.8815624227072 | Rpl28-ps1 |
| ENSMUSG00000058620 | -0.700367805876663 | 0.256844331429907 | 0.837517407767513 | 11.7456783153976 | 13.0306516626293 | 6.62518422053913 | 3.75170677703264 | 21.8991639163819 | 20.7923979452556 | 4.16431236222455 | 10.8552556466851 | Adra2b |
| ENSMUSG00000058624 | -0.0106174583272442 | 0.962984486003103 | 0.994284908685797 | 413.809282188623 | 481.319695788368 | 719.30571537282 | 506.480414899407 | 572.663136413388 | 692.264543353805 | 419.554470494123 | 452.664160466768 | Gda |
| ENSMUSG00000058626 | -1.72303352456434 | 0.314508819651938 | 0.853228028313558 | 17.1667606148119 | 0 | 16.0897331070236 | 0 | 59.1277425742312 | 6.1154111603693 | 2.08215618111227 | 42.3354970220718 | Capn11 |
| ENSMUSG00000058638 | 0.0245729615389461 | 0.838086277198568 | 0.969115750107235 | 496.932544112975 | 564.39010013763 | 625.606681396623 | 611.528204656321 | 612.081631462875 | 535.71001764835 | 577.798340258656 | 532.993052252237 | Zfp110 |
| ENSMUSG00000058655 | 0.185589246523617 | 0.143405486686907 | 0.780607990324772 | 8232.81698537714 | 8896.67742266013 | 9341.50975096017 | 8533.25706436075 | 6716.47357315434 | 6832.13734836458 | 8813.76711464825 | 8414.99417731027 | Eif4b |
| ENSMUSG00000058656 | -0.00295444017485254 | 0.995605307202392 | 0.999262355603273 | 9.03513716569045 | 15.4738988493722 | 17.036187995672 | 11.2551203310979 | 4.37983278327639 | 11.0077400886647 | 12.4929370866736 | 24.9670879873757 | Samd12 |
| ENSMUSG00000058672 | -0.703307252321684 | 0.0455093972127823 | 0.697710295017159 | 403.870631306363 | 323.323044378988 | 332.205665915605 | 464.27371365779 | 794.939650164664 | 989.473525747752 | 298.789411989611 | 398.387882233342 | Tubb2a |
| ENSMUSG00000058690 | 0.191125103475445 | 0.424026007351343 | 0.884586455610141 | 5078.6506008346 | 5744.88855176167 | 4749.31063123791 | 4529.24800657266 | 3129.39052365098 | 3101.73654053931 | 5599.95904910146 | 5776.08152960113 | Ccser2 |
| ENSMUSG00000058704 | 0.174990828392473 | 0.24742491770023 | 0.833398011296137 | 1059.82158953549 | 846.992358070902 | 800.700835796586 | 833.816831195505 | 720.482492848966 | 670.249063176475 | 849.519721893808 | 895.558590851519 | Memo1 |
| ENSMUSG00000058706 | 0.0879606509619365 | 0.640637726205136 | 0.934417517272849 | 137.334084918495 | 153.110157035894 | 180.772883731853 | 151.006197775564 | 133.58489988993 | 112.523565350795 | 168.654650670094 | 169.341988088287 | 0610030E20F |
| ENSMUSG00000058709 | 0.00069521691599822 | 0.993636997213758 | 0.999133450854041 | 1573.92089426328 | 1544.13222202157 | 1533.25691961048 | 1635.74415478623 | 1685.14066336559 | 1565.54525705454 | 1601.17810327534 | 1431.80821979776 | Egln2 |

| | | | | | | | | | | | | |
|---|---|---|---|---|---|---|---|---|---|---|---|---|
| **ENSMUSG00000058715** | -0.293420408248873 | 0.389806964555696 | 0.877515569464664 | 45.1756858284523 | 58.6379324818317 | 78.5557557578211 | 72.2203554578784 | 77.7420319031559 | 128.423634367755 | 46.8485140750261 | 59.7039060567679 | Fcer1g |
| **ENSMUSG00000058729** | -0.0766199418921743 | 0.784469256375132 | 0.961141387070613 | 44.2721721118832 | 34.2054606144018 | 66.2518422053913 | 36.5791410760683 | 42.7033696369448 | 52.592535979176 | 51.0128264372507 | 44.5065481514088 | Lin9 |
| **ENSMUSG00000058743** | -0.218139136615274 | 0.645538682595749 | 0.935063240511793 | 71.3775836089546 | 57.009101024003 | 13.2503684410783 | 30.9515809105193 | 31.7537876787538 | 55.0387004433237 | 68.711153976705 | 45.5920737160773 | Kcnj14 |
| **ENSMUSG00000058748** | 0.105212541488366 | 0.6392966339886 | 0.933801135937988 | 163.535982698997 | 96.9154717408051 | 131.557229522134 | 120.054616865045 | 105.115986798633 | 95.400414101761 | 131.175839410073 | 143.289374536243 | Zfp958 |
| **ENSMUSG00000058756** | 0.00707510525874056 | 0.935514226994693 | 0.98818087308472 | 1984.11612158562 | 2219.28286129155 | 2244.04454098547 | 2034.36299984595 | 1972.01971067019 | 2161.18630407451 | 2178.97644353399 | 2128.71563231494 | Thra |
| **ENSMUSG00000058761** | -0.0639122291353317 | 0.698055584306184 | 0.944933386973863 | 213.229237110295 | 225.593156909269 | 212.952349945901 | 156.633757941113 | 212.421889988905 | 239.724117486476 | 196.76375911511 | 197.565652769668 | Rnf169 |
| **ENSMUSG00000058793** | 0.0377492498675984 | 0.836913340170633 | 0.968924788429433 | 2118.73966535441 | 3283.72421898257 | 2955.7786172491 | 2133.78322943732 | 2800.90306490525 | 2162.40938630658 | 2707.84411353651 | 2549.89955140632 | Cds2 |
| **ENSMUSG00000058794** | 0.484010060690632 | 0.521286595760589 | 0.90609507420585 | 17.1667606148119 | 11.4018202048006 | 61.519567762149 | 7.50341355406529 | 16.4243729372865 | 33.0232202659942 | 3.12323427166841 | 17.3684090346961 | Nfe2 |
| **ENSMUSG00000058799** | 0.192200286038859 | 0.0357350188883669 | 0.679566075569698 | 3210.18423496982 | 3143.64471360931 | 3316.37792982416 | 3537.85949074178 | 2796.52323212197 | 2662.65001922479 | 2984.77088562444 | 3114.37284503395 | Nap1l1 |
| **ENSMUSG00000058806** | 0.154607124279638 | 0.717431496503176 | 0.951530419413962 | 17.1667606148119 | 13.8450673915436 | 34.0723759913441 | 16.8826804966469 | 26.2789966996583 | 19.5693157131818 | 13.5340151772298 | 14.1118323406906 | Col13a1 |
| **ENSMUSG00000058809** | 0.18546440434205 | 0.47017727466528 | 0.896098016616457 | 97.5794813894569 | 75.7406627890326 | 67.1982970940397 | 77.8479156234274 | 64.6025335533267 | 45.2540425867328 | 75.998700610598 | 93.3551985614916 | Hspd1-ps3 |
| **ENSMUSG00000058818** | -0.319974191600362 | 0.46286642406441 | 0.895328112556867 | 19.877301764519 | 22.8036404096012 | 38.8046504345863 | 34.703287687552 | 28.4689130912965 | 61.154111603693 | 15.616171358342 | 40.1644458927348 | Pirb |
| **ENSMUSG00000058833** | -0.187483268111509 | 0.362546216997548 | 0.86924083181804 | 1175.47134525633 | 980.556537612852 | 1039.20746773599 | 1255.88384361168 | 1236.20780307976 | 1784.47697659576 | 993.188498390554 | 1056.21637442246 | Rex1bd |
| **ENSMUSG00000058835** | -0.188266510370489 | 0.254316491778288 | 0.837406774112203 | 448.142803418246 | 466.66021266791 | 451.458981885309 | 437.073839524303 | 572.663136413388 | 617.656527197299 | 385.198893505771 | 480.887825148149 | Abi1 |
| **ENSMUSG00000058881** | 0.116871431817526 | 0.60995799496395 | 0.927932112008768 | 170.764092431549 | 298.890572511559 | 257.435729712378 | 188.52326554589 | 186.142893289246 | 172.454594722414 | 244.653351280692 | 240.986675356409 | Zfp516 |
| **ENSMUSG00000058886** | 0.205819314415411 | 0.150497591057623 | 0.782641594375979 | 209.615182244018 | 221.521078264697 | 202.541346170768 | 178.206071909051 | 191.617684268342 | 185.908499275227 | 157.202791673977 | 170.427513652956 | Deaf1 |
| **ENSMUSG00000058900** | -0.169657404575222 | 0.559065868963423 | 0.916574868225118 | 33.4300075130547 | 39.9063707168021 | 48.2691993210708 | 30.0136542162612 | 32.8487458745729 | 33.0232202659942 | 52.0539045278068 | 52.1052271040884 | Rsl1 |
| **ENSMUSG00000058922** | -0.29535284307475 | 0.308763308303468 | 0.851785577030345 | 25.2983840639333 | 21.1748089517725 | 30.2865564367503 | 34.703287687552 | 27.3739548954774 | 35.4693847301419 | 33.3144988977964 | 40.1644458927348 | Gm10052 |
| **ENSMUSG00000058927** | 0.0872636476165294 | 0.611339579337094 | 0.927932112008768 | 149.983276950461 | 122.162359337149 | 110.735221971868 | 134.123517278917 | 124.825234323377 | 106.408154190426 | 137.42230795341 | 117.236760984199 | Gm10053 |
| **ENSMUSG00000058952** | -0.528172039840001 | 0.0954003175832729 | 0.757085157577544 | 101.193536255733 | 179.171460361152 | 88.966759532954 | 123.806323642077 | 120.445401540101 | 146.769867848863 | 166.572494488982 | 277.894544555138 | Cfi |
| **ENSMUSG00000058966** | -0.379072557777542 | 0.136578531156529 | 0.776456211338551 | 107.518132271716 | 73.2974156022896 | 76.6628459805242 | 131.309737196143 | 145.62944004394 | 150.439114545085 | 98.902418602833 | 110.723607596188 | Fam57b |
| **ENSMUSG00000058975** | 0.739849432908282 | 0.0046249327909428 | 0.635862253526407 | 89.4478579403355 | 109.13170767452 | 69.0912068713366 | 78.7858423176855 | 43.7983278327639 | 45.2540425867328 | 44.7663578939139 | 73.8157383974585 | Kcnc1 |
| **ENSMUSG00000058979** | 0.258158245612122 | 0.241273638954926 | 0.832048466795696 | 372.247651226447 | 314.364471360931 | 319.901752363175 | 402.370551836751 | 250.745426842573 | 200.585486060113 | 349.802238426862 | 375.591845375304 | Hdhd5 |
| **ENSMUSG00000058997** | 0.392154081531405 | 0.0760283886533693 | 0.735399943172214 | 10558.4612918259 | 13120.2373928098 | 14696.5515109331 | 11155.7001015066 | 8524.24955445167 | 6196.13458768617 | 11471.6394798381 | 11548.9064825083 | Vwa8 |
| **ENSMUSG00000059005** | 0.0552658338536318 | 0.568648974629171 | 0.92059532906293 | 4397.40125854154 | 4151.89138600525 | 4742.68544701737 | 4686.81969120803 | 4526.55718151615 | 3987.24807656078 | 4669.23523614427 | 4117.39846678765 | Hnrnpa3 |
| **ENSMUSG00000059013** | -0.590789900997091 | 0.110187518586275 | 0.763724866847985 | 492.41497553013 | 422.681763306537 | 381.421320125324 | 354.536290429585 | 926.334633662956 | 923.427085215764 | 319.610973800734 | 318.058990447873 | Sh2d3c |
| **ENSMUSG00000059022** | -1.28341770297135 | 0.00774940552463216 | 0.635862253526407 | 18.9737880479499 | 14.6594831204579 | 14.1968233297267 | 15.0068271081306 | 52.5579933993167 | 63.6002760678407 | 14.5750932677859 | 22.7960368580387 | Kcp |
| **ENSMUSG00000059027** | -0.097325426038615 | 0.831770807160605 | 0.967583158900775 | 16.2632468982428 | 10.5874044758863 | 9.46454888648447 | 14.0689004138724 | 16.4243729372865 | 9.78465785659088 | 19.7804837205666 | 7.59867895267955 | 9630013D21[illegible] |
| **ENSMUSG00000059058** | 0.422731287700414 | 0.315020397177725 | 0.853277176422981 | 10.8421645988285 | 16.2883145782866 | 16.0897331070236 | 17.8206071909051 | 10.949581958191 | 12.2308223207386 | 7.28754663389296 | 15.1973579053591 | Tma7-ps |
| **ENSMUSG00000059070** | -0.00292763191813116 | 0.987086856096995 | 0.998371545269589 | 7252.50460289972 | 5183.7561145397 | 5840.57311784957 | 7301.75931479978 | 6589.45842243933 | 8041.76567588563 | 5430.26332034081 | 5569.83167231411 | Rpl18 |
| **ENSMUSG00000059089** | 0.0376241833495846 | 0.924059487410089 | 0.986713918026868 | 18.9737880479499 | 19.5459774939439 | 28.3936466594534 | 12.1930470253561 | 17.5193311331055 | 25.6847268735511 | 12.4929370866736 | 21.7105112933701 | Fcgr4 |
| **ENSMUSG00000059119** | 0.106724193253082 | 0.446095815802708 | 0.89108031640712 | 4886.2021792054 | 4950.01880034129 | 4967.9417105157 | 4639.92335649512 | 4020.68649504772 | 3707.16224541587 | 5201.22614041846 | 5126.93724192936 | Nap1l4 |
| **ENSMUSG00000059142** | 0.116045463939423 | 0.431668833038399 | 0.886559861988535 | 358.694945477911 | 273.643684915214 | 330.312756138308 | 370.481044231974 | 274.834507150593 | 309.439804714686 | 300.871568170723 | 344.111603999917 | Zfp945 |
| **ENSMUSG00000059149** | 0.116312288935323 | 0.722429791564368 | 0.951674101666722 | 247.562758339918 | 368.930325198191 | 344.509579468035 | 235.419600258798 | 246.365594059297 | 129.646716599829 | 308.159114804616 | 419.012867962044 | Mfsd4a |
| **ENSMUSG00000059173** | -0.440689400079524 | 0.173818024946818 | 0.799889021902745 | 57.8248778604189 | 76.5550785179469 | 45.4298346551254 | 74.0962088463947 | 107.305903190271 | 123.53130543946 | 55.1771387994752 | 59.7039060567679 | Pde1a |
| **ENSMUSG00000059182** | 0.0182993540512889 | 0.871091831653433 | 0.975613767803763 | 308.098177350044 | 290.746415222415 | 316.115932808581 | 286.067641748739 | 292.353838283699 | 321.670627035425 | 295.666177717943 | 276.809018990469 | Skap2 |
| **ENSMUSG00000059183** | 0.0516673345364565 | 0.78242252059014 | 0.960742105865155 | 141.85165350134 | 113.203786319092 | 102.217127974032 | 137.87522405595 | 121.54035973592 | 97.8465785659088 | 121.806136595068 | 135.690695583563 | Mtfmt |
| **ENSMUSG00000059195** | -0.67847695668322 | 0.148876384770402 | 0.78151662300109 | 5.42108229941427 | 12.2162359337149 | 7.57163910918757 | 13.1309737196143 | 16.4243729372865 | 22.0154801773295 | 10.4107809055614 | 13.0263067760221 | Gm12715 |
| **ENSMUSG00000059208** | 0.00341118945414477 | 0.961430112115374 | 0.993822507448007 | 2987.01634697726 | 3036.95625312153 | 2886.68741037776 | 2921.64165261417 | 3133.77035643426 | 2898.70489001505 | 2852.55396812381 | 2920.06376895829 | Hnrnpm |
| **ENSMUSG00000059248** | -0.414982917790879 | 0.0440202169531446 | 0.691603097094229 | 661.372040528541 | 617.327122517061 | 631.285410728514 | 574.949063580253 | 1074.15399009853 | 976.01962119494 | 600.702058250891 | 663.256120012458 | Sept9 |
| **ENSMUSG00000059263** | 0.252813511736146 | 0.174795357203338 | 0.799889021902745 | 5024.43977784046 | 5630.05593398475 | 5283.11118843563 | 4931.61855840941 | 3709.7183674351 | 3213.03702365803 | 5383.41480626578 | 5207.26613371483 | Usp47 |
| **ENSMUSG00000059273** | 0.317795846578706 | 0.131953436008096 | 0.77303252008843 | 662.27555424511 | 864.095088378103 | 780.825283134969 | 571.19735680322 | 486.161438943679 | 437.863439082442 | 670.454290318152 | 714.275821551878 | Zc3h4 |
| **ENSMUSG00000059277** | -0.0484690692389318 | 0.830596170765832 | 0.967583158900775 | 37.9475760958999 | 46.4216965481167 | 40.6975602118832 | 45.9584080186499 | 47.0832024202212 | 41.5847958905112 | 43.7252798033577 | 44.5065481514088 | R74862 |
| **ENSMUSG00000059278** | -0.048580582168886 | 0.783455423355275 | 0.961051227381536 | 852.916948441179 | 670.264144896492 | 618.035042287436 | 803.803176979244 | 783.990068206473 | 945.442565393093 | 693.358008310387 | 624.177199684392 | Naa38 |
| **ENSMUSG00000059288** | -0.105565421365743 | 0.574578998152771 | 0.922450244597442 | 133.720030052219 | 167.769640156352 | 192.130342395635 | 155.695831246855 | 185.047935093427 | 135.762127760198 | 205.092383839559 | 171.513039217624 | Cdyl |
| **ENSMUSG00000059291** | 0.0611366616148989 | 0.753504554986885 | 0.956497786093635 | 6432.11414825503 | 4499.64690225167 | 5182.78697023889 | 6612.38319452004 | 5613.85066996451 | 7010.70735424736 | 4565.12742708866 | 4595.02971524179 | Rpl11 |
| **ENSMUSG00000059316** | -0.0284673562989627 | 0.769882852816012 | 0.958815818668915 | 635.170142748039 | 649.08933594472 | 628.446046062569 | 609.652351267805 | 708.437952694956 | 601.756458180339 | 615.277151518677 | 646.973236542431 | Slc27a4 |
| **ENSMUSG00000059323** | -0.10380070843787 | 0.488535642663862 | 0.899236955296196 | 1279.37542266177 | 1014.76199822725 | 1061.92238506356 | 1258.69762369445 | 1265.77167436688 | 1490.93724089803 | 1100.41954171784 | 1102.8939737032 | Tonsl |
| **ENSMUSG00000059325** | 0.0969218329231261 | 0.498659909362847 | 0.900927059989498 | 5265.67794016439 | 4239.84828472799 | 3927.78778789105 | 4860.33612964579 | 3953.89404510276 | 3798.89341282141 | 4386.061995513 | 4965.19393279375 | Hopx |
| **ENSMUSG00000059326** | -0.126848897285043 | 0.687840061553028 | 0.944452327903597 | 70.4740698923855 | 84.6992358070902 | 92.7525790875478 | 67.5307219865876 | 79.9319482947941 | 140.654456688494 | 75.998700610598 | 48.8486504100828 | Csf2ra |
| **ENSMUSG00000059327** | -0.510430947320983 | 0.190709096639559 | 0.808824684419004 | 27.1054114970714 | 30.1333819698302 | 18.9290977729689 | 19.6964605794214 | 44.893286028583 | 50.1463715150282 | 22.903717992235 | 19.5394601640331 | Eda |
| **ENSMUSG00000059343** | -0.257699321722887 | 0.521246118908414 | 0.90609507420585 | 21.6843291976571 | 31.7622134276588 | 23.6613722162112 | 12.1930470253561 | 16.4243729372865 | 29.3539735697726 | 38.5198893505771 | 22.7960368580387 | Aldoart1 |
| **ENSMUSG00000059355** | -0.179593077224414 | 0.392915475114794 | 0.878770935869648 | 786.960447131638 | 614.069459601404 | 492.156542097192 | 785.044643094081 | 724.862325632242 | 989.473525747752 | 635.057635239243 | 684.966631305828 | Wdr83os |
| **ENSMUSG00000059363** | 0.171406285612434 | 0.287383843905561 | 0.851785577030345 | 917.96993603415 | 718.314672902438 | 788.396922244156 | 1012.02290310456 | 871.586723872001 | 831.695917810225 | 678.782915042601 | 669.769273400469 | Fxn |
| **ENSMUSG00000059395** | -0.0108401775753801 | 0.982759432203005 | 0.996940244108351 | 10.8421645988285 | 4.88649437348597 | 14.1968233297267 | 11.2551203310979 | 10.949581958191 | 12.2308223207386 | 6.24646854333682 | 11.9407812113536 | Nkapl |
| **ENSMUSG00000059401** | 0.0586701058771175 | 0.793010160828997 | 0.961804768755116 | 125.588406603097 | 147.409246933493 | 123.039135524298 | 90.9788893430416 | 105.115986798633 | 90.5080851734656 | 145.750932677859 | 125.920965501547 | Mamld1 |
| **ENSMUSG00000059408** | -0.084481124571531 | 0.798927457761022 | 0.963240258978423 | 51.5002818444356 | 61.0811796685746 | 61.519567762149 | 37.5170677703264 | 50.3680770076785 | 30.5770558018465 | 57.2592949805875 | 85.7565196088121 | Mrgprh |
| **ENSMUSG00000059409** | 0.0881700311485399 | 0.38009318576629 | 0.87432013076003 | 892.671551970216 | 976.48445896828 | 1069.49402417274 | 932.299134092612 | 946.0438811877 | 920.980920751616 | 927.600578685518 | 846.709940441436 | Ppp2r5d |
| **ENSMUSG00000059412** | -0.421058990724751 | 0.439178496454603 | 0.889583654898613 | 18.0702743313809 | 12.2162359337149 | 7.57163910918757 | 18.7585338851632 | 26.2789966996583 | 31.8001380339204 | 7.28754663389296 | 10.8552556466851 | Fxyd2 |
| **ENSMUSG00000059422** | 0.201355398917432 | 0.611103358263969 | 0.927932112008768 | 18.0702743313809 | 17.9171460361152 | 26.5007368821565 | 15.9447538023887 | 14.2344565456483 | 9.78465785659088 | 18.7394056300105 | 24.9670879873757 | Gm8116 |
| **ENSMUSG00000059423** | -0.125225554109875 | 0.550539383024871 | 0.914699131434573 | 193.351935345776 | 145.780415475665 | 159.004421292939 | 163.19924480092 | 150.009272827216 | 138.208292224346 | 235.283648465687 | 196.480127205 | Zfp933 |
| **ENSMUSG00000059430** | -0.853694597795247 | 0.147100866216101 | 0.780607990324772 | 10.8421645988285 | 25.2468875963442 | 13.2503684410783 | 14.0689004138724 | 19.7092475247437 | 67.2695227640623 | 17.6983275394543 | 10.8552556466851 | Actg2 |
| **ENSMUSG00000059434** | -0.759724069632563 | 0.0715952853209102 | 0.728064275108613 | 9.03513716569045 | 22.8036404096012 | 12.3039135524298 | 18.7585338851632 | 19.7092475247437 | 20.7923979452556 | 24.9858741733473 | 41.2499714574033 | Gckr |
| **ENSMUSG00000059436** | -0.00232943700564381 | 0.989043398083492 | 0.998437672973644 | 1369.72679431867 | 1414.64012112419 | 1733.90535600395 | 1148.02227377199 | 1509.94735203453 | 1632.8147798186 | 1378.38739189632 | 1154.99920080729 | Max |
| **ENSMUSG00000059439** | 0.201230748162629 | 0.127061136176744 | 0.77273486535379 | 641.494738764022 | 746.819223414439 | 868.845587779274 | 689.376120279749 | 655.879959295639 | 680.033721033066 | 646.509494235361 | 580.756177097652 | Bcas3 |
| **ENSMUSG00000059447** | 0.27495932607457 | 0.170112353116582 | 0.79907358295209 | 45210.9228633984 | 48134.4128260236 | 57393.9709025305 | 48000.2744320499 | 37361.0685995434 | 28017.1446901159 | 47867.7295256806 | 51005.5897070792 | Hadhb |
| **ENSMUSG00000059456** | 0.077561817463685 | 0.739211519914535 | 0.954383574136061 | 205.097613661173 | 302.148235427216 | 310.437203476691 | 223.226553233442 | 175.193311331055 | 229.939459629886 | 324.816364253515 | 256.184033261768 | Ptk2b |
| **ENSMUSG00000059474** | -0.0266596129326822 | 0.894693648458283 | 0.983733637008786 | 150.886790667031 | 146.594831204579 | 199.701981504822 | 151.006197775564 | 131.394983498292 | 140.654456688494 | 188.435134390661 | 198.651178334337 | Mbtd1 |
| **ENSMUSG00000059475** | -0.127568801088766 | 0.456425167576168 | 0.892805623137304 | 149.079763233892 | 153.924572764808 | 152.3792370724 | 206.343872736795 | 192.712642464161 | 152.885279009232 | 192.599446752885 | 183.453820428978 | Zfp426 |
| **ENSMUSG00000059479** | -0.382703236745416 | 0.131495715087597 | 0.77303252008843 | 219.553833126278 | 172.656134529838 | 202.541346170768 | 208.219726125312 | 296.733671066975 | 377.932409710823 | 184.270822028436 | 188.88144825232 | B3gnt8 |
| **ENSMUSG00000059481** | -0.35086083933811 | 0.347695368340265 | 0.861266047293794 | 222.264374275985 | 547.287369830429 | 233.774357496166 | 290.75727522003 | 248.555510450935 | 362.032340693862 | 344.596847974081 | 695.821886952513 | Plg |
| **ENSMUSG00000059482** | -0.0191500767766597 | 0.917304455085447 | 0.986713918026868 | 129.202461469373 | 119.719112150406 | 134.396594188079 | 114.427056699496 | 102.926070406995 | 110.077400886647 | 134.299073681742 | 156.315681312265 | 2610301B20[illegible] |
| **ENSMUSG00000059486** | -0.0376002347526401 | 0.786727470062425 | 0.961141387070613 | 428.265501653727 | 425.939426222194 | 420.22597055991 | 336.71568323868 | 443.458069306734 | 353.470765069345 | 441.417110395802 | 414.67076570337 | Kbtbd2 |
| **ENSMUSG00000059495** | 0.366098873555948 | 0.170842337028763 | 0.799531027639052 | 4973.84300971259 | 6866.3390104767 | 6590.16538965914 | 4970.073552874 | 3481.96706270473 | 2782.51207796803 | 5982.03470833556 | 5908.51564849069 | Arhgef12 |
| **ENSMUSG00000059498** | -0.303701317709992 | 0.207144747285536 | 0.819960256425221 | 175.281661014395 | 168.584055885266 | 189.290977729689 | 201.654239265505 | 217.896680968 | 341.239942748607 | 164.49033830787 | 184.539345993646 | Fcgr3 |
| **ENSMUSG00000059518** | -0.0617447813715095 | 0.694559646893464 | 0.944771261777587 | 889.961010820509 | 683.294796559122 | 805.433110239828 | 881.651092602672 | 838.737977997428 | 1050.62763735145 | 762.069162287092 | 752.269216315276 | Znhit1 |
| **ENSMUSG00000059534** | 0.149092032152405 | 0.366749871937213 | 0.870052215406984 | 6797.13368974893 | 4966.30711491958 | 5770.53545608958 | 6568.3006398899 | 5354.34557755538 | 6795.44488140236 | 4762.93226429433 | 4824.07560938685 | Uqcr10 |
| **ENSMUSG00000059540** | 0.426953161878281 | 0.0538233934601192 | 0.703396849069134 | 98.4829951060259 | 65.9676740420606 | 90.8596693102509 | 96.6064495085906 | 55.8428679867739 | 80.7234273168747 | 64.5468416144805 | 60.7894316214364 | Tcea2 |
| **ENSMUSG00000059552** | -0.455636847570194 | 0.0751665034370629 | 0.735399943172214 | 109.325159704854 | 122.162359337149 | 144.807597963212 | 152.88205116408 | 228.846262926191 | 231.162541861959 | 148.874166949528 | 117.236760984199 | Trp53 |
| **ENSMUSG00000059554** | 0.126891137889079 | 0.399632557209196 | 0.880459952732461 | 281.896279569542 | 232.108482740584 | 295.293925258315 | 277.626301500416 | 229.94122112201 | 277.639666680766 | 214.462086654564 | 273.552442296464 | Ccdc28a |
| **ENSMUSG00000059555** | -0.757426558759692 | 0.0324210903753989 | 0.671606027118506 | 83.1232619243521 | 74.9262470601182 | 92.7525790875478 | 105.047789756914 | 197.092475247437 | 231.162541861959 | 99.9434966933891 | 73.8157383974585 | Tor4a |
| **ENSMUSG00000059586** | 0.0788212368637669 | 0.567408874088372 | 0.9203643243484 | 442.721721118832 | 395.806044252364 | 420.22597055991 | 396.742991671202 | 343.816873487196 | 343.686107212755 | 450.786813210807 | 427.697072479392 | Nsmce2 |
| **ENSMUSG00000059588** | -0.309488259421767 | 0.305962937483303 | 0.851785577030345 | 414.712795905192 | 314.364471360931 | 385.207139679918 | 385.487871340104 | 629.600962595981 | 639.672007374629 | 241.530117009024 | 348.453706258591 | Calcrl |
| **ENSMUSG00000059602** | 0.164386438067231 | 0.806194913295444 | 0.963604452772887 | 9.93865088225949 | 13.8450673915436 | 18.9290977729689 | 2.81378008277448 | 12.0445401540101 | 18.3462334811079 | 7.28754663389296 | 3.25657669400552 | Syn3 |
| **ENSMUSG00000059659** | 0.102984579984228 | 0.781799172147928 | 0.960742105865155 | 34.3335212296237 | 21.9892246806869 | 17.036187995672 | 25.3240207449704 | 14.2344565456483 | 23.2385624094033 | 21.8626399016789 | 32.5657669400552 | Gm10069 |
| **ENSMUSG00000059669** | 0.13904761655525 | 0.308219779675585 | 0.851785577030345 | 183.413284463516 | 206.861595144239 | 194.96970706158 | 196.026679099956 | 182.858018701789 | 155.33144347338 | 196.76375911511 | 173.684090346961 | Taf1b |
| **ENSMUSG00000059689** | 0.0286155873802459 | 0.832944395651794 | 0.967854847250989 | 388.510898124689 | 303.777066885045 | 358.706402797761 | 372.35689762049 | 376.665619361769 | 358.363093997641 | 302.953724351836 | 357.137910775939 | Zfp637 |

| | | | | | | | | | | | | |
|---|---|---|---|---|---|---|---|---|---|---|---|---|
| **ENSMUSG00000059708** | -0.467941722143032 | 0.0270771949731344 | 0.664353746352133 | 48.7897406947284 | 52.122606650517 | 49.2156542097192 | 43.1446279358754 | 59.1277425742312 | 62.3771938357668 | 65.5879197050366 | 80.3288917854696 | Akap17b |
| **ENSMUSG00000059713** | -0.195885306596408 | 0.363578192481057 | 0.86924083181804 | 160.82544154929 | 225.593156909269 | 161.843785958884 | 150.068271081306 | 173.003394939417 | 156.554525705454 | 219.667477107345 | 250.756405438425 | Rcan3 |
| **ENSMUSG00000059714** | 0.0391924887887177 | 0.69006582911312 | 0.94469184542218 | 6461.93010090181 | 5630.87034971367 | 5415.61487284641 | 6242.84007698232 | 5920.43896479386 | 6134.98047608248 | 5653.05403171982 | 5407.00283761384 | Flot1 |
| **ENSMUSG00000059734** | -0.0255335687383161 | 0.86779319174406 | 0.974868041734905 | 9104.70772186627 | 6690.42521303121 | 7031.21336776931 | 8831.51775313485 | 8145.39401869826 | 9376.14839107821 | 7044.97543879338 | 7657.29733317165 | Ndufs8 |
| **ENSMUSG00000059741** | 0.113456835463373 | 0.376501263986018 | 0.87422797717331 | 160764.002616563 | 121850.438112975 | 126894.992740652 | 158771.292877327 | 139225.029556594 | 129552.539267959 | 123747.747233955 | 132776.059442428 | Myl3 |
| **ENSMUSG00000059742** | 0.663846841538155 | 0.134292834483325 | 0.77303252008843 | 13.5527057485357 | 16.2883145782866 | 19.8755526616174 | 13.1309737196143 | 8.75966556655277 | 8.56157562451702 | 15.616171358342 | 6.51315338801105 | Kcnh7 |
| **ENSMUSG00000059743** | -0.159001021097314 | 0.300010985161558 | 0.851785577030345 | 271.957628687283 | 244.324718674299 | 251.757000380487 | 257.929840920994 | 309.873169416804 | 338.793778284459 | 231.119336103462 | 267.039288908453 | Fdps |
| **ENSMUSG00000059751** | -0.420940195268812 | 0.418848887817532 | 0.882550938324998 | 8.13162344912141 | 7.32974156022896 | 11.3574586637814 | 8.44134024832345 | 15.3294147414674 | 3.66924669622158 | 11.4518589961175 | 16.2828834700276 | Rps3a3 |
| **ENSMUSG00000059772** | -0.284938134313512 | 0.254643801672189 | 0.837406774112203 | 51.5002818444356 | 48.0505280059454 | 62.4660226507975 | 65.6548685980713 | 66.7924499449649 | 95.400414101761 | 63.5057635239243 | 52.1052271040884 | Slx1b |
| **ENSMUSG00000059775** | 0.22922107395573 | 0.596090331248996 | 0.924461858177432 | 10.8421645988285 | 13.0306516626293 | 12.3039135524298 | 18.7585338851632 | 8.75966556655277 | 17.123151249034 | 11.4518589961175 | 9.76973008201657 | Rps26-ps1 |
| **ENSMUSG00000059791** | -0.487098487036559 | 0.0780974528805607 | 0.739855948845396 | 46.9827132615903 | 27.6901347830872 | 49.2156542097192 | 47.8342614071662 | 58.0327843784121 | 79.5003450848009 | 53.094982618363 | 49.9341759747513 | Nrm |
| **ENSMUSG00000059796** | 0.150112916698045 | 0.1229348429679 | 0.770251097497048 | 4333.25178466514 | 4029.7290266681 | 4463.48125486607 | 4900.66697749889 | 4258.29242354047 | 4078.97924396632 | 3820.75659234102 | 3816.70788537447 | Eif4a1 |
| **ENSMUSG00000059810** | -0.39087023310265 | 0.0228875883658041 | 0.650119841310877 | 474.344701198749 | 456.072808192024 | 448.619617219364 | 526.176875478828 | 752.23628052772 | 716.726187995282 | 540.319528998635 | 489.572029665497 | Rgs3 |
| **ENSMUSG00000059811** | -0.0024907951759213[illegible] | 0.987368178313279 | 0.998437672973644 | 961.338594429464 | 1093.76032393194 | 1061.92238506356 | 891.968286239511 | 1027.07078767831 | 771.764888438605 | 1188.91117941511 | 1026.90718417641 | Atl2 |
| **ENSMUSG00000059820** | 0.0422240449619157 | 0.787844309521268 | 0.961275251163951 | 231.299511441676 | 228.036404096012 | 239.453086828057 | 257.929840920994 | 214.611806380543 | 188.354663739374 | 245.694429371248 | 278.980070119806 | AU019823 |
| **ENSMUSG00000059824** | -0.624213757100467 | 0.0149013485872486 | 0.635862253526407 | 513.195791011218 | 360.786167909048 | 288.668741037776 | 516.797608536247 | 900.055636963298 | 600.533375948265 | 606.948526794228 | 480.887825148149 | Dbp |
| **ENSMUSG00000059834** | 0.259689257613612 | 0.185202103175385 | 0.805963874499992 | 202.387072511466 | 138.450673915436 | 185.505158175096 | 146.316564304273 | 146.724398239759 | 110.077400886647 | 156.16171358342 | 147.631476794917 | Sclt1 |
| **ENSMUSG00000059839** | 0.00092367878163620[illegible] | 0.995628230415462 | 0.999262355603273 | 183.413284463516 | 172.656134529838 | 159.950876181588 | 177.268145214792 | 169.71852035196 | 132.092881063977 | 204.051305749003 | 185.624871558315 | Zfp874b |
| **ENSMUSG00000059842** | 0.169597998142044 | 0.407835003842008 | 0.881573350402261 | 94.8689402397497 | 79.8127414336042 | 106.949402417274 | 118.178763476528 | 79.9319482947941 | 89.2850029413918 | 107.231043327282 | 78.1578406561325 | Zfp341 |
| **ENSMUSG00000059851** | -0.145586986941021 | 0.481726580679139 | 0.897732059433541 | 81.316234491214 | 82.2559886203472 | 85.1809399783602 | 60.9652351267805 | 95.2613630362614 | 66.0464405319884 | 87.4505596067155 | 93.3551985614916 | Kmt5c |
| **ENSMUSG00000059852** | -0.107850635216802 | 0.412329104036358 | 0.882192126163777 | 3324.02696325752 | 3394.48475811492 | 3075.0319332188 | 3268.67452948969 | 3293.63425302384 | 4198.84130270956 | 3644.81439503703 | 2940.68875468699 | Kcng2 |
| **ENSMUSG00000059866** | 0.0203327291712253 | 0.900635440723025 | 0.984802943363903 | 181.606257030378 | 160.439898596123 | 141.968233297267 | 178.206071909051 | 183.952976897608 | 171.23151249034 | 162.408182126757 | 135.690695583563 | Tnip2 |
| **ENSMUSG00000059878** | -0.0171854993514771 | 0.923455166916426 | 0.986713918026868 | 270.150601254144 | 314.364471360931 | 326.526936583714 | 238.233380341573 | 343.816873487196 | 324.116791499573 | 252.981976005141 | 243.157726485746 | Zfp422 |
| **ENSMUSG00000059883** | -0.310588839183012 | 0.21430386759042 | 0.824248286808591 | 89.4478579403355 | 103.43079757212 | 101.270673085384 | 73.1582821521366 | 152.199189218854 | 132.092881063977 | 87.4505596067155 | 84.6709940441436 | Irak4 |
| **ENSMUSG00000059890** | 0.0593624611788534 | 0.762064487414521 | 0.957666649379949 | 520.42390074377 | 624.65686407729 | 666.304241608507 | 507.418341593665 | 494.921104510232 | 414.624876673038 | 711.056335849841 | 603.55221395569 | Ube4a |
| **ENSMUSG00000059895** | 0.127964050575094 | 0.412583473562768 | 0.882192126163777 | 12862.4212690769 | 11454.75722718 | 16411.5277691641 | 14587.5738757972 | 13171.2521375079 | 14758.9332944353 | 11687.1426445832 | 11003.9726490447 | Ptp4a3 |
| **ENSMUSG00000059897** | 0.286400449885851 | 0.151972038165216 | 0.783412422888014 | 140.044626068202 | 108.317291945606 | 131.557229522134 | 123.806323642077 | 79.9319482947941 | 100.292743030056 | 99.9434966933891 | 132.434118889558 | Zfp930 |
| **ENSMUSG00000059900** | 0.614276530481683 | 0.501504074024225 | 0.900927059989498 | 3.61405486627618 | 5.7009101024003 | 36.9117406572894 | 2.81378008277448 | 5.47479097909548 | 15.9000690169602 | 2.08215618111227 | 8.68420451734806 | Tmem40 |
| **ENSMUSG00000059901** | -1.02366620626765 | 0.169487867127771 | 0.79907358295209 | 3.61405486627618 | 8.14415728914329 | 5.67872933189068 | 13.1309737196143 | 20.8042057205628 | 26.9078091056249 | 13.5340151772298 | 1.08552556466851 | Adamts14 |
| **ENSMUSG00000059908** | -0.495816381931276 | 0.227142937850025 | 0.832048466795696 | 287.317361868956 | 590.451403462888 | 255.542819935081 | 369.543117537716 | 258.410134213307 | 377.932409710823 | 474.731609293598 | 1008.45324957704 | Mug1 |
| **ENSMUSG00000059920** | 0.0750053821458618 | 0.476063597510671 | 0.897553735056661 | 787.863960848207 | 668.635313438664 | 784.611102689562 | 772.851596068725 | 664.639624862192 | 704.495365674543 | 762.069162287092 | 728.387653892569 | 4930453N24[illegible] |
| **ENSMUSG00000059923** | -0.0229122464310618 | 0.827927921999844 | 0.967319591893432 | 2435.87297987015 | 2277.92079377338 | 1912.78532995851 | 2096.26616166699 | 2192.10630802983 | 2173.41712639525 | 2175.85320926233 | 2321.93918282594 | Grb2 |
| **ENSMUSG00000059939** | -0.0878794192651573 | 0.814898840492746 | 0.966177091475091 | 18.0702743313809 | 17.9171460361152 | 13.2503684410783 | 19.6964605794214 | 25.1840385038392 | 20.7923979452556 | 15.616171358342 | 11.9407812113536 | 9430015G10[illegible] |
| **ENSMUSG00000059970** | -0.639177247767513 | 0.0186157596112313 | 0.642273034804458 | 133.720030052219 | 83.8848200781758 | 95.5919437534931 | 82.5375490947182 | 220.086597359638 | 161.446854633749 | 115.559668051731 | 119.407812113536 | Hspa2 |
| **ENSMUSG00000059974** | -0.230869636778866 | 0.578908769873366 | 0.922450244597442 | 17.1667606148119 | 33.3910448854875 | 17.036187995672 | 19.6964605794214 | 30.6588294829347 | 31.8001380339204 | 11.4518589961175 | 29.3091902460497 | Ntm |
| **ENSMUSG00000059975** | -0.0613228953634178 | 0.71060810471286 | 0.949886733585872 | 110.228673421423 | 131.935348084121 | 123.985590412947 | 103.171936368398 | 119.350443344282 | 107.6312364225 | 131.175839410073 | 131.348593324889 | Zfp74 |
| **ENSMUSG00000059981** | -0.0365344559078993 | 0.709570504707666 | 0.949511975952744 | 892.671551970216 | 899.929380450333 | 954.026527757634 | 890.092432850995 | 1040.21028602814 | 862.272973612071 | 930.723812957186 | 895.558590851519 | Taok2 |
| **ENSMUSG00000059995** | 0.078720145326745 | 0.573954235664132 | 0.922280709351328 | 494.222002963268 | 609.997380956832 | 654.946782944725 | 544.935409363992 | 509.15556105588 | 485.563646133322 | 594.455589707554 | 591.611432744337 | Atxn7l3 |
| **ENSMUSG00000060002** | 0.163545279987285 | 0.209921561189236 | 0.82165491136391 | 11486.3698787423 | 11661.6188223243 | 11294.9926411306 | 11858.2071955059 | 9081.58327612359 | 8829.43063334119 | 11575.7472888937 | 11850.6825894861 | Chpt1 |
| **ENSMUSG00000060012** | 0.00984481042741775 | 0.969384593412798 | 0.995103983630461 | 128.298947752804 | 198.717437855096 | 195.916161950228 | 142.564857527241 | 153.294147414674 | 105.185071958352 | 199.886993386778 | 201.907755028342 | Kif13b |
| **ENSMUSG00000060019** | 0.0876146369879616 | 0.805403330683144 | 0.963475137341661 | 23.4913566307952 | 18.7315617650296 | 18.9290977729689 | 29.075727522003 | 22.994122112201 | 26.9078091056249 | 12.4929370866736 | 22.7960368580387 | Gm10073 |
| **ENSMUSG00000060032** | -0.188085995063541 | 0.490216092757233 | 0.899236955296196 | 1129.39214571131 | 852.693268173302 | 883.042411109001 | 1150.83605385476 | 1201.16914081355 | 1779.58464766747 | 819.328457267679 | 776.150778737983 | H2afj |
| **ENSMUSG00000060036** | -0.0866160111973423 | 0.667136210944021 | 0.942024383071524 | 4600.69184476958 | 3352.13514021138 | 3272.84100494633 | 4193.47025002824 | 4095.14365236342 | 5492.8623042437 | 3337.69635832297 | 3448.71471895185 | Rpl3 |
| **ENSMUSG00000060038** | 0.182865800521035 | 0.234083147203525 | 0.832048466795696 | 466.213077749627 | 386.033055505392 | 388.046504345863 | 506.480414899407 | 442.363111110915 | 398.724807656078 | 363.336253604092 | 334.3418739179 | Dhps |
| **ENSMUSG00000060063** | -0.14302419088864 | 0.626103933668051 | 0.931698409533928 | 174.378147297826 | 151.481325578065 | 227.149173275627 | 141.626930832982 | 194.902558855799 | 292.316653465652 | 113.477511870619 | 167.17093695895 | Alox5ap |
| **ENSMUSG00000060073** | 0.107223918628682 | 0.345276447462019 | 0.861266047293794 | 2443.1010896027 | 1991.24645719553 | 1982.8229917185 | 2221.01041200333 | 1864.71380747992 | 2040.1011630992 | 1921.83015516663 | 2192.76164063039 | Psma3 |
| **ENSMUSG00000060090** | 0.136608759249092 | 0.386660653763559 | 0.875422409254416 | 181.606257030378 | 201.160685041839 | 217.684624389143 | 211.033506208086 | 180.668102310151 | 146.769867848863 | 204.051305749003 | 205.164331722348 | Rp2 |
| **ENSMUSG00000060098** | 0.127673518650407 | 0.252308259594004 | 0.83723604018117 | 1260.40163461382 | 1081.54408799823 | 1188.74734014245 | 1399.38662783318 | 1190.21955885536 | 1130.12798243625 | 1103.5427759895 | 1087.69661579784 | Prmt7 |
| **ENSMUSG00000060121** | -0.0605780858573737 | 0.746454552269335 | 0.956085046483146 | 95.7724539563188 | 78.9983257046899 | 89.9132144216024 | 97.5443762028488 | 81.0269064906132 | 90.5080851734656 | 90.5737938783839 | 115.065709854862 | Gemin2 |
| **ENSMUSG00000060126** | 0.0743582597435267 | 0.65714058018985 | 0.938389079274206 | 35109.6395121565 | 25817.7930223131 | 30627.2801966637 | 36245.2391729124 | 30409.179014288 | 37492.3627419921 | 25842.681441875 | 27636.3953508955 | Tpt1 |
| **ENSMUSG00000060143** | 0.155618515358661 | 0.320760226182137 | 0.855228644171892 | 618.003382133227 | 425.12501049328 | 629.392500951217 | 594.645524159674 | 477.401773377126 | 525.92535979176 | 485.14239019916 | 546.019359028259 | Gm10076 |
| **ENSMUSG00000060147** | 0.0745463083848484 | 0.565237636140554 | 0.919692942798738 | 2788.24332933207 | 2255.93156909269 | 2400.20959761246 | 2946.96567335914 | 2499.789561055 | 2766.61200895107 | 2300.78258012906 | 2301.31419709724 | Serpinb6a |
| **ENSMUSG00000060149** | -0.0863337957768116 | 0.789724795716352 | 0.961659329450003 | 51.5002818444356 | 36.6487078011448 | 24.6078271048596 | 32.8274342990356 | 39.4184950494875 | 24.4616446414772 | 45.80743598447 | 44.5065481514088 | BC002059 |
| **ENSMUSG00000060152** | -0.146381145452051 | 0.441499517662761 | 0.89108031640712 | 575.538237454482 | 368.115909469277 | 405.082692341535 | 478.342614071662 | 564.998429042654 | 585.856389163379 | 431.006329490241 | 440.723379255414 | Pop5 |
| **ENSMUSG00000060166** | -0.180722202839909 | 0.103269230876909 | 0.759800953500741 | 546.625798524272 | 691.438953848265 | 610.463403178248 | 568.383576720446 | 687.633746974393 | 695.933790050026 | 671.495368408708 | 686.052156870497 | Zdhhc8 |
| **ENSMUSG00000060181** | 0.184686964717021 | 0.272679338460533 | 0.844738346035895 | 195.158962778914 | 221.521078264697 | 201.594891282119 | 202.592165959763 | 168.623562156141 | 140.654456688494 | 219.667477107345 | 192.138024946326 | Slc35e3 |
| **ENSMUSG00000060187** | -0.0569929380903898 | 0.656608273463639 | 0.938361205344872 | 3059.29744430279 | 2334.92989479738 | 2833.68593661345 | 3012.62054195721 | 2782.28877557633 | 3305.99127329564 | 2884.82738893105 | 2720.32706505928 | Lrrc10 |
| **ENSMUSG00000060206** | 0.019522877606876 | 0.933316300183257 | 0.98818087308472 | 64.1494738764022 | 65.9676740420606 | 77.6093008691726 | 81.59962240046 | 60.2227007700503 | 53.8156182112498 | 77.0397787011541 | 93.3551985614916 | Zfp462 |
| **ENSMUSG00000060212** | 0.193028517991004 | 0.573831333224414 | 0.922280709351328 | 24.3948703473642 | 28.5045505120015 | 21.7684624389143 | 15.9447538023887 | 15.3294147414674 | 18.3462334811079 | 21.8626399016789 | 23.8815624227072 | Pcnx2 |
| **ENSMUSG00000060216** | -0.314156594800747 | 0.194722483481863 | 0.809789010647629 | 99.3865088225949 | 99.3587189275481 | 120.199770858353 | 100.358156285623 | 142.344565456483 | 183.462334811079 | 102.025652874501 | 94.4407241261602 | Arrb2 |
| **ENSMUSG00000060227** | 0.452982776889441 | 0.0050736638196445[illegible] | 0.635862253526407 | 175.281661014395 | 196.274190668353 | 213.898804834549 | 217.598993067893 | 147.819356435578 | 117.415894279091 | 156.16171358342 | 163.914360264945 | Casc4 |
| **ENSMUSG00000060240** | -0.384929188319409 | 0.439334216871539 | 0.889583654898613 | 19.877301764519 | 13.8450673915436 | 17.036187995672 | 27.1998741334867 | 37.2285786578493 | 39.1386314263635 | 8.32862472444909 | 17.3684090346961 | Cend1 |
| **ENSMUSG00000060244** | 0.0483259689900309 | 0.893670100744032 | 0.983681567424884 | 30.7194663633475 | 26.0613033252585 | 19.8755526616174 | 40.3308478531009 | 32.8487458745729 | 40.3617136584374 | 19.7804837205666 | 20.6249857287016 | Alyref2 |
| **ENSMUSG00000060260** | -0.130522093276047 | 0.485232365802227 | 0.899236955296196 | 275.571683553559 | 249.211213047785 | 274.47191770805 | 285.129715054481 | 361.336204620302 | 349.801518373124 | 224.872867560126 | 251.841931003094 | Pwwp2b |
| **ENSMUSG00000060261** | 0.178817293993246 | 0.214745176055658 | 0.824248286808591 | 4793.14026639878 | 5773.39310227368 | 5705.23006877284 | 4850.95686270321 | 4519.98743234123 | 3827.02430415911 | 5502.09770858918 | 4809.96377704616 | Gtf2i |
| **ENSMUSG00000060275** | -0.804770734397271 | 0.0094087329901006[illegible] | 0.635862253526407 | 33.4300075130547 | 29.3189662409158 | 24.6078271048596 | 29.075727522003 | 74.4571573156986 | 31.8001380339204 | 46.8485140750261 | 49.9341759747513 | Nrg2 |
| **ENSMUSG00000060279** | 0.0410719403692793 | 0.5869994933507 | 0.923058967588378 | 1450.13951509332 | 1488.75195245539 | 1537.98919405373 | 1490.36551717622 | 1504.47256105544 | 1486.04491196974 | 1440.85207732969 | 1368.84773704699 | Ap2a1 |
| **ENSMUSG00000060288** | -0.0336706461746737 | 0.882179452795579 | 0.978854883720278 | 96.6759676728878 | 73.2974156022896 | 64.3589324280944 | 94.7305961200743 | 73.3621991198795 | 102.738907494204 | 91.61487196894 | 69.4736361387845 | Ppih |
| **ENSMUSG00000060301** | -0.0195304560689744 | 0.923505720302967 | 0.986713918026868 | 259.308436655316 | 234.551729927327 | 231.881447718869 | 253.240207449704 | 208.042057205628 | 177.34692365071 | 308.159114804616 | 297.434004719171 | 2610008E11[illegible] |
| **ENSMUSG00000060317** | 0.114730789243739 | 0.803338285951608 | 0.963461896828986 | 15.3597331816738 | 35.0198763433161 | 28.3936466594534 | 15.0068271081306 | 16.4243729372865 | 13.4539045528125 | 18.7394056300105 | 37.9933947633978 | Acnat2 |
| **ENSMUSG00000060336** | -0.24246141008194 | 0.577113416163334 | 0.922450244597442 | 14.4562194651047 | 13.0306516626293 | 10.4110037751329 | 11.2551203310979 | 15.3294147414674 | 23.2385624094033 | 10.4107809055614 | 9.76973008201657 | Zfp937 |
| **ENSMUSG00000060373** | 0.00017442486576483[illegible] | 0.998412951503125 | 0.999262355603273 | 3183.07882347275 | 3001.93637677822 | 3431.84542623927 | 3317.44671759112 | 3358.23678657717 | 2994.10530411681 | 3375.17516958299 | 3202.3004157721 | Hnrnpc |
| **ENSMUSG00000060376** | 0.0845845074820473 | 0.474917241650657 | 0.897549741595057 | 4351.32205899652 | 3986.56499303564 | 4318.67365690286 | 4508.61361929898 | 4198.06972277042 | 3329.22983570505 | 4355.87073088688 | 4301.9378127813 | Bckdha |
| **ENSMUSG00000060380** | 0.0734595830568214 | 0.839683880555704 | 0.969879194284716 | 22.5878429142261 | 23.6180561385155 | 14.1968233297267 | 22.5102406621959 | 13.1394983498292 | 26.9078091056249 | 21.8626399016789 | 17.3684090346961 | C030014I23[illegible] |
| **ENSMUSG00000060397** | 0.0920168879780027 | 0.684004650860532 | 0.944452327903597 | 53.3073092775737 | 46.4216965481167 | 57.7337482075553 | 62.8410885152968 | 56.937826182593 | 41.5847958905112 | 58.3003730711437 | 48.8486504100828 | Zfp128 |
| **ENSMUSG00000060407** | -0.607655771813095 | 0.122880027388215 | 0.770251097497048 | 50.5967681278665 | 123.791190794978 | 66.2518422053913 | 72.2203554578784 | 74.4571573156986 | 101.51582526213 | 87.4505596067155 | 213.848536239696 | Cyp2a12 |
| **ENSMUSG00000060419** | -0.0839864859187833 | 0.804804464126404 | 0.963461896828986 | 85.8338030740593 | 37.4631235300591 | 42.5904699891801 | 64.7169419038131 | 51.4630352034976 | 84.3926740130963 | 42.6842017128016 | 66.217059444779 | Rps16-ps2 |
| **ENSMUSG00000060427** | -0.191643598001375 | 0.291009187505932 | 0.851785577030345 | 116.553269437407 | 89.5857301805761 | 101.270673085384 | 130.371810501884 | 112.780694169367 | 139.43137445642 | 128.052605138405 | 119.407812113536 | Zfp868 |
| **ENSMUSG00000060429** | -0.225152158211847 | 0.512121704892381 | 0.902394714365738 | 18.9737880479499 | 21.9892246806869 | 14.1968233297267 | 29.075727522003 | 27.3739548954774 | 22.0154801773295 | 19.7804837205666 | 29.3091902460497 | Sntb1 |
| **ENSMUSG00000060441** | -0.136724457894885 | 0.746387345115015 | 0.956085046483146 | 21.6843291976571 | 22.8036404096012 | 9.46454888648447 | 37.5170677703264 | 31.7537876787538 | 20.7923979452556 | 20.8215618111227 | 27.1381391167127 | Trim5 |
| **ENSMUSG00000060450** | 0.239597949160582 | 0.313847798338705 | 0.853228028313558 | 2002.186395917 | 2233.12792868309 | 2210.91861988277 | 1795.19169281012 | 1325.99437513693 | 1163.15120270224 | 2251.85190987292 | 2238.35371434646 | Rnf14 |
| **ENSMUSG00000060459** | -0.330022183624385 | 0.37089429117859 | 0.873323500696802 | 37.9475760958999 | 67.5965054998893 | 32.1794662140472 | 48.7721881014244 | 36.1336204620302 | 48.9232892829544 | 53.094982618363 | 96.6117752554972 | Kng2 |
| **ENSMUSG00000060470** | -0.344716798361786 | 0.636434648976436 | 0.932595977875587 | 6.32459601598331 | 4.88649437348597 | 20.8220075502658 | 7.50341355406529 | 19.7092475247437 | 22.0154801773295 | 3.12323427166841 | 5.42762782334254 | Adgrg3 |
| **ENSMUSG00000060475** | -0.0150545598880891 | 0.903779613181943 | 0.985039825636533 | 700.22313034101 | 653.161414589292 | 724.037989816062 | 692.189900362523 | 661.354750274735 | 590.748718091674 | 771.438865102097 | 772.894202043977 | Wtap |
| **ENSMUSG00000060477** | -0.399618018723469 | 0.132905327377346 | 0.77303252008843 | 159.018414116152 | 166.955224427437 | 159.950876181588 | 166.013024883695 | 261.695008800764 | 304.547475786391 | 162.408182126757 | 132.434118889558 | Irak2 |
| **ENSMUSG00000060509** | 0.0371837475367298 | 0.932359221293748 | 0.987982116552499 | 18.0702743313809 | 9.77298874697194 | 12.3039135524298 | 17.8206071909051 | 12.0445401540101 | 14.6769867848863 | 20.8215618111227 | 8.68420451734806 | Xcr1 |
| **ENSMUSG00000060510** | -0.148887939335053 | 0.216915083182208 | 0.825692108906787 | 271.957628687283 | 257.355370336928 | 298.133289924261 | 255.11606083822 | 291.25888008788 | 292.316653465652 | 328.980676615739 | 286.578749072486 | Zfp266 |

| | | | | | | | | | | | | |
|---|---|---|---|---|---|---|---|---|---|---|---|---|
| **ENSMUSG00000060512** | 0.056643981292219 | 0.809330734795162 | 0.964311345725361 | 149.983276950461 | 139.26508964435 | 120.199770858353 | 90.0409626487835 | 147.819356435578 | 85.6157562451702 | 124.929370866736 | 121.578863242873 | 0610040J01F |
| **ENSMUSG00000060519** | -0.251906202734661 | 0.31771107964004 | 0.854351922245337 | 235.817080024521 | 273.643684915214 | 296.240380146964 | 318.895076047775 | 408.419407040523 | 457.432754795624 | 233.201492284575 | 240.986675356409 | Tor3a |
| **ENSMUSG00000060538** | 0.172353112659994 | 0.478513705751764 | 0.897674307047416 | 166.246523848704 | 125.420022252807 | 134.396594188079 | 189.461192240149 | 134.679858085749 | 193.24699266767 | 105.14888714617 | 113.980184290193 | Tmem219 |
| **ENSMUSG00000060548** | -0.0911996160297969 | 0.82502944375994 | 0.96672635991852 | 28.9124389302094 | 48.0505280059454 | 34.0723759913441 | 21.5723139679377 | 35.0386622662111 | 19.5693157131818 | 60.3825292522559 | 26.0526135520442 | Tnfrsf19 |
| **ENSMUSG00000060550** | -0.478855492344305 | 0.298530934405274 | 0.851785577030345 | 151.7903043836 | 76.5550785179469 | 129.664319744837 | 117.24083678227 | 244.175677667659 | 276.416584448692 | 79.1219348822664 | 62.9604827507734 | H2-Q7 |
| **ENSMUSG00000060568** | -0.245931388179371 | 0.287876678918276 | 0.851785577030345 | 93.0619128066116 | 79.8127414336042 | 96.5383986421416 | 75.0341355406529 | 114.970610561005 | 114.969729814943 | 113.477511870619 | 65.1315338801105 | Fam78b |
| **ENSMUSG00000060572** | -2.10145199599699 | 0.0010242483969922 | 0.556246415579446 | 5.42108229941427 | 8.95857301805762 | 14.1968233297267 | 9.37926694258161 | 56.937826182593 | 79.5003450848009 | 11.4518589961175 | 15.1973579053591 | Mfap2 |
| **ENSMUSG00000060579** | 0.127432935049722 | 0.668437172034049 | 0.942024383071524 | 33.4300075130547 | 25.2468875963442 | 38.8046504345863 | 28.1378008277448 | 28.4689130912965 | 36.6924669622158 | 21.8626399016789 | 28.2236646813812 | Fhit |
| **ENSMUSG00000060586** | -0.574826739614125 | 0.0554476798742666 | 0.703396849069134 | 461.695509166782 | 504.123336197969 | 623.713771619326 | 638.728078789808 | 1202.26409900937 | 1075.08928199292 | 510.128264372507 | 531.907526687569 | H2-Eb1 |
| **ENSMUSG00000060591** | -0.738599071976985 | 0.0442802582448243 | 0.691978859031708 | 957.724539563188 | 711.799347071123 | 757.163910918757 | 959.499008226099 | 1735.50874037327 | 2429.04131289869 | 758.945928015424 | 727.3021283279 | Ifitm2 |
| **ENSMUSG00000060594** | -0.627724237435113 | 0.120758364286017 | 0.769868040577702 | 49.6932544112975 | 39.9063707168021 | 34.0723759913441 | 51.5859681841989 | 97.4512794278996 | 102.738907494204 | 36.4377331694648 | 34.7368180693922 | Layn |
| **ENSMUSG00000060600** | 0.0552478266037959 | 0.653916557444092 | 0.937077409179068 | 33720.0354160733 | 26695.7331780828 | 29312.654356331 | 31965.4796670124 | 33348.0468118664 | 29179.0728105861 | 26062.3489189823 | 28531.9539417471 | Eno3 |
| **ENSMUSG00000060601** | -0.125853507439274 | 0.253833083895296 | 0.837406774112203 | 936.9437240821 | 921.91860513102 | 1064.7617497295 | 973.567908639971 | 1081.81869746927 | 1175.38202502298 | 1056.69426191448 | 938.979613438259 | Nr1h2 |
| **ENSMUSG00000060613** | 0.0305135038459217 | 0.9271845014826 | 0.986790505471895 | 44.2721721118832 | 54.56585383726 | 34.0723759913441 | 47.8342614071662 | 44.893286028583 | 48.9232892829544 | 21.8626399016789 | 61.8749571861049 | Cyp2c70 |
| **ENSMUSG00000060636** | 0.19525378196912 | 0.290736619347914 | 0.851785577030345 | 5161.77386275895 | 3453.93710632567 | 3981.73571654402 | 5022.59744775245 | 3855.34780747904 | 4792.03618526538 | 3286.68353188572 | 3456.31339790453 | Rpl35a |
| **ENSMUSG00000060639** | 0.238167206781591 | 0.341575674017918 | 0.859283395807012 | 54.2108229941427 | 51.3081909216027 | 83.2880302010633 | 66.5927952923295 | 50.3680770076785 | 67.2695227640623 | 44.7663578939139 | 54.2762782334254 | Hist1h4i |
| **ENSMUSG00000060657** | 0.208168824834764 | 0.310125252583104 | 0.851785577030345 | 738.17070643691 | 930.062762420163 | 928.472245764126 | 751.279282100787 | 628.506004400162 | 514.917619703095 | 903.655782602727 | 849.966517135441 | Marf1 |
| **ENSMUSG00000060671** | -0.321928762498355 | 0.0131450966966137 | 0.635862253526407 | 195.158962778914 | 201.160685041839 | 239.453086828057 | 235.419600258798 | 275.929465346412 | 270.301173288323 | 283.173240631269 | 258.355084391105 | Atp8b2 |
| **ENSMUSG00000060675** | -0.00646471646584711 | 0.954120823302087 | 0.992371505557225 | 4161.58417851702 | 3524.79127474121 | 3979.84280676672 | 4665.24737724009 | 3973.6032926275 | 3965.23259638345 | 4343.3777938002 | 4120.65504348165 | Pla2g16 |
| **ENSMUSG00000060679** | 0.121643985296999 | 0.29342872483908 | 0.851785577030345 | 2369.01296484404 | 1997.76178302685 | 1954.42934505904 | 2485.50573978413 | 2071.66090648973 | 1920.23910435596 | 2101.93666483284 | 1999.53809011939 | Mrps9 |
| **ENSMUSG00000060681** | 0.153444238497451 | 0.528331252353345 | 0.908232038364594 | 644.205279913729 | 900.743796179247 | 887.774685552243 | 656.548685980713 | 535.434557755538 | 499.017550686135 | 965.079389945539 | 777.236304302651 | Slc9a6 |
| **ENSMUSG00000060703** | -0.307359855008698 | 0.236044988272014 | 0.832048466795696 | 235.817080024521 | 172.656134529838 | 189.290977729689 | 251.364354061187 | 246.365594059297 | 379.155491942897 | 165.531416398426 | 260.526135520442 | Cd302 |
| **ENSMUSG00000060708** | -0.0696908280347138 | 0.689304720880373 | 0.94469184542218 | 272.861142403852 | 221.521078264697 | 295.293925258315 | 306.702029022419 | 315.3479603959 | 336.347613820311 | 262.351678820146 | 236.644573097735 | Bloc1s4 |
| **ENSMUSG00000060716** | -0.694833892311716 | 0.198066436963189 | 0.812921911423739 | 15.3597331816738 | 5.7009101024003 | 4.73227444324223 | 5.62756016554897 | 12.0445401540101 | 13.4539045528125 | 16.6572494488982 | 8.68420451734806 | Plekhh1 |
| **ENSMUSG00000060733** | 0.0669772792254694 | 0.694056236829656 | 0.94469184542218 | 489.704434380422 | 524.483729420828 | 519.603733867997 | 454.894446715208 | 388.710159515779 | 390.163232031561 | 546.565997541972 | 572.071972580304 | Ipmk |
| **ENSMUSG00000060739** | 0.0206537028357527 | 0.905172962685589 | 0.98504921011344 | 1925.38773000863 | 1386.9499863411 | 1574.90093471102 | 1876.79131521058 | 1781.49698459767 | 2027.87034077846 | 1451.26285823525 | 1407.92665737505 | Nsa2 |
| **ENSMUSG00000060743** | -0.0266043854924623 | 0.804327964825003 | 0.963461896828986 | 2743.97115722019 | 2187.52064786389 | 2372.76240584166 | 2598.99486978936 | 2527.16351595048 | 2608.83440101354 | 2333.0560009363 | 2618.28766198044 | H3f3a |
| **ENSMUSG00000060771** | 0.318618181613485 | 0.198482296257603 | 0.812921911423739 | 109.325159704854 | 140.893921102179 | 91.8061241988993 | 132.247663890401 | 78.836990098975 | 70.9387694602839 | 99.9434966933891 | 130.263067760221 | Tsga10 |
| **ENSMUSG00000060791** | -0.0861551813627406 | 0.741303948551343 | 0.954383574136061 | 37.9475760958999 | 29.3189662409158 | 40.6975602118832 | 43.1446279358754 | 40.5134532453066 | 47.7002070508805 | 40.6020455316893 | 31.4802413753867 | Gmfg |
| **ENSMUSG00000060798** | 0.369924252030759 | 0.151167283096127 | 0.783412422888014 | 507.774708711803 | 732.974156022896 | 556.515474525287 | 596.521377548191 | 437.983278327639 | 259.293433199658 | 576.7572621681 | 577.499600403646 | Intu |
| **ENSMUSG00000060802** | -0.399610459773489 | 0.232327068090324 | 0.832048466795696 | 3305.05317520957 | 2873.25869160975 | 3572.86720464789 | 3314.63293750834 | 5861.31122221963 | 6580.18240855736 | 2206.04447388845 | 2588.97847173439 | B2m |
| **ENSMUSG00000060803** | -0.184793799088325 | 0.331159612861904 | 0.856333478432716 | 2768.36602756755 | 2515.73018661636 | 2423.87096982867 | 3208.64722105717 | 3151.28968756736 | 4142.57952003416 | 2599.57199211867 | 2516.2482589016 | Gstp1 |
| **ENSMUSG00000060807** | -0.432733778433559 | 0.474777432153028 | 0.897549741595057 | 10.8421645988285 | 31.7622134276588 | 5.67872933189068 | 14.0689004138724 | 8.75966556655277 | 15.9000690169602 | 21.8626399016789 | 37.9933947633978 | Serpina6 |
| **ENSMUSG00000060843** | 0.190141951840229 | 0.566561474097538 | 0.919931558693298 | 375.861706092723 | 702.026358324151 | 591.534305405279 | 372.35689762049 | 302.208462046071 | 272.747337752471 | 645.468416144805 | 568.815395886298 | Ctnna3 |
| **ENSMUSG00000060860** | -0.16345388861083 | 0.366428976394131 | 0.869754677013001 | 842.978297558919 | 636.873100011005 | 708.894711597687 | 860.078778634734 | 848.5926017598 | 1100.77400886647 | 748.535147109862 | 717.532398245884 | Ube2s |
| **ENSMUSG00000060862** | 0.270403589584836 | 0.101994948318873 | 0.759800953500741 | 268.343573821006 | 336.353696041618 | 264.060913932917 | 239.171307035831 | 244.175677667659 | 188.354663739374 | 237.365804646799 | 248.585354309088 | Zbtb40 |
| **ENSMUSG00000060882** | 0.107086626331215 | 0.756782339045409 | 0.956566265296272 | 315.326287082597 | 633.615437095348 | 399.403963009645 | 214.785212985119 | 309.873169416804 | 226.270212933664 | 482.019155927491 | 433.124700302735 | Kcnd2 |
| **ENSMUSG00000060904** | 0.127616271105828 | 0.194629615135367 | 0.809762948852223 | 2857.81388550789 | 2718.51970311603 | 2825.16784261561 | 2938.52433311082 | 2344.30549724869 | 2449.83371084394 | 2713.04950398929 | 2871.2151185482 | Arl1 |
| **ENSMUSG00000060913** | 0.149963893930492 | 0.594077964816547 | 0.923943895566212 | 853.820462157748 | 1174.38748109446 | 897.239234438728 | 799.113543507953 | 615.366506050332 | 502.686797382356 | 1148.30913388342 | 1089.86766692718 | Trim55 |
| **ENSMUSG00000060923** | 0.104454927661456 | 0.418164784597362 | 0.882468496126562 | 2451.23271305182 | 1908.99046857519 | 1802.99656287529 | 2303.54796109804 | 1931.50625742489 | 2038.87808086712 | 1895.80320290272 | 2009.30782020141 | Acyp2 |
| **ENSMUSG00000060935** | -0.0780277054791507 | 0.544275423570315 | 0.913837378403038 | 353.273863178497 | 298.890572511559 | 331.259211026956 | 309.515809105193 | 390.900075907418 | 309.439804714686 | 325.857442344071 | 337.598450611906 | Tmem263 |
| **ENSMUSG00000060938** | 0.0651776695923146 | 0.739179341064084 | 0.954383574136061 | 5541.24962371795 | 3887.20627410809 | 4493.76781130283 | 5731.67002861162 | 4621.81854455241 | 6116.63424260137 | 3762.45621926988 | 4285.65492931127 | Rpl26 |
| **ENSMUSG00000060950** | 0.281146780721356 | 0.143102883009144 | 0.780607990324772 | 125.588406603097 | 145.780415475665 | 99.3777633080869 | 105.047789756914 | 101.831112211176 | 86.838838477244 | 94.7381062406084 | 108.552556466851 | Trmt61a |
| **ENSMUSG00000060961** | 0.361037478128727 | 0.139149375836086 | 0.779711473022872 | 524.941469326615 | 797.312998607128 | 664.41133183121 | 452.080666632434 | 512.440435643337 | 308.216722482613 | 513.251498644175 | 564.473293627624 | Slc4a4 |
| **ENSMUSG00000060969** | 0.093276443665613 | 0.767281680348786 | 0.95879680569409 | 362.309000344187 | 140.893921102179 | 335.991485470199 | 278.564228194674 | 382.140410340865 | 209.14706168463 | 232.160414194018 | 223.618266321713 | Irx1 |
| **ENSMUSG00000060981** | 0.0150995591953535 | 0.928688917222403 | 0.986901071486195 | 463.50253659992 | 375.445651029505 | 385.207139679918 | 374.232751009006 | 381.045452145046 | 502.686797382356 | 374.788112600209 | 324.572143835884 | Hist1h4h |
| **ENSMUSG00000060992** | -0.014335008591159 | 0.90007111230273 | 0.984802943363903 | 727.328541838081 | 723.201167275924 | 866.952678001977 | 749.403428712271 | 739.09678217789 | 717.949270227356 | 854.725112346588 | 783.749457690662 | Copz1 |
| **ENSMUSG00000061024** | 0.0154935269509686 | 0.911763594049972 | 0.986226496507248 | 439.107666252556 | 425.12501049328 | 508.246275204216 | 483.970174237211 | 411.70428162798 | 402.3940543523 | 530.94982618363 | 489.572029665497 | Rrs1 |
| **ENSMUSG00000061028** | 0.283642815890277 | 0.230151216776857 | 0.832048466795696 | 112.939214571131 | 133.56417954195 | 210.112985279955 | 140.689004138724 | 107.305903190271 | 108.854318654574 | 149.915245040084 | 123.74991437221 | Clasrp |
| **ENSMUSG00000061032** | 0.0606292529528915 | 0.576024361063711 | 0.922450244597442 | 3303.24614777643 | 2990.53455657341 | 3229.3040800685 | 3450.63230817578 | 3180.85355885448 | 3464.99196346524 | 3074.30360141227 | 2720.32706505928 | Rrp1 |
| **ENSMUSG00000061046** | -0.156088600578547 | 0.407974399661829 | 0.881573350402261 | 193.351935345776 | 160.439898596123 | 136.289503965376 | 165.075098189436 | 217.896680968 | 193.24699266767 | 141.586620315635 | 178.026192605635 | Haghl |
| **ENSMUSG00000061079** | -0.00458101013135216 | 0.97345392371756 | 0.995510898988495 | 221.360860559416 | 232.922898469498 | 232.827902607518 | 233.543746870282 | 263.884925192402 | 195.693157131818 | 244.653351280692 | 218.19063849837 | Zfp143 |
| **ENSMUSG00000061080** | -0.706130035818279 | 0.128195769065783 | 0.77303252008843 | 24.3948703473642 | 29.3189662409158 | 15.1432782183751 | 22.5102406621959 | 45.9882442244021 | 64.8233582999146 | 23.9447960827911 | 15.1973579053591 | Lsamp |
| **ENSMUSG00000061086** | -5.4437171248298 | NA | NA | 99.3865088225949 | 18.7315617650296 | 21.7684624389143 | 71.2824287636203 | 39.4184950494875 | 9063.0393396673 | 49.9717483466946 | 37.9933947633978 | Myl4 |
| **ENSMUSG00000061100** | -0.33933096356698 | 0.601361849500566 | 0.926043912368388 | 13.5527057485357 | 16.2883145782866 | 50.1621090983677 | 30.9515809105193 | 67.887408140784 | 50.1463715150282 | 10.4107809055614 | 11.9407812113536 | Retnla |
| **ENSMUSG00000061104** | -0.390466295350209 | 0.260746016286811 | 0.840316047847749 | 14.4562194651047 | 21.9892246806869 | 14.1968233297267 | 14.0689004138724 | 24.0890803080201 | 20.7923979452556 | 18.7394056300105 | 21.7105112933701 | Sap18b |
| **ENSMUSG00000061111** | -0.0965194284247637 | 0.63604066798682 | 0.932595977875587 | 772.504227666534 | 618.141538245975 | 670.0900611631 | 957.623154837583 | 811.364023101951 | 1051.85071958352 | 656.920275140922 | 707.762668163867 | Mcrip1 |
| **ENSMUSG00000061118** | 0.0355230274214919 | 0.69681740412918 | 0.944933386973863 | 739.977733870048 | 800.570661522785 | 815.844114014961 | 837.568537972538 | 806.984190318674 | 731.403174780168 | 777.685333645434 | 798.946815596022 | Dnajc30 |
| **ENSMUSG00000061119** | -0.42871417294913 | 0.140720614328661 | 0.779711473022872 | 140.044626068202 | 135.193010999779 | 104.110037751329 | 167.888878272211 | 234.321053905287 | 248.285693110993 | 107.231043327282 | 147.631476794917 | Prcp |
| **ENSMUSG00000061130** | 0.334384859641274 | 0.125006316156545 | 0.771949105062466 | 5069.61546366891 | 6052.73769729129 | 5714.69461765932 | 5330.23740346913 | 3595.84271506991 | 2962.30516608289 | 5704.06685815707 | 5317.98974131102 | Ppm1b |
| **ENSMUSG00000061132** | -0.562593166471322 | 0.180170253322845 | 0.802731406993634 | 11.7456783153976 | 11.4018202048006 | 31.2330113253987 | 25.3240207449704 | 31.7537876787538 | 37.9155491942896 | 27.0680303544596 | 20.6249857287016 | Blnk |
| **ENSMUSG00000061136** | 0.0991853492637672 | 0.390584078124632 | 0.877797322358001 | 733.653137854065 | 861.65184119136 | 731.609628925249 | 697.817460528072 | 686.538788778574 | 645.787418534998 | 735.001131932632 | 756.61131857395 | Prpf40a |
| **ENSMUSG00000061143** | 0.568770992447455 | 0.0414979977965468 | 0.683543358336265 | 61.4389327266951 | 101.801966114291 | 83.2880302010633 | 57.2135283497478 | 39.4184950494875 | 44.0309603546589 | 56.2182168900314 | 65.1315338801105 | Maml3 |
| **ENSMUSG00000061171** | -0.530235044799162 | 0.42059598253943 | 0.882568531013176 | 18.0702743313809 | 11.4018202048006 | 5.67872933189068 | 2.81378008277448 | 21.8991639163819 | 18.3462334811079 | 5.20539045278068 | 9.76973008201657 | Slc38a11 |
| **ENSMUSG00000061175** | -0.312803013810578 | 0.230027620325984 | 0.832048466795696 | 189.737880479499 | 260.613033252585 | 305.704929033448 | 288.881421831514 | 259.505092409126 | 225.04713070159 | 344.596847974081 | 467.861518372127 | Fnip2 |
| **ENSMUSG00000061207** | -0.259598994090495 | 0.233856232638316 | 0.832048466795696 | 105.711104838578 | 78.9983257046899 | 82.3415753124149 | 82.5375490947182 | 112.780694169367 | 129.646716599829 | 75.998700610598 | 100.953877514171 | Stk19 |
| **ENSMUSG00000061232** | -0.612720710061267 | 0.0954914802256219 | 0.757085157577544 | 1935.32638089089 | 1771.35421038866 | 2018.78827748714 | 1786.7503525618 | 4019.5915368519 | 4577.99679465246 | 1355.48367390409 | 1534.93314844127 | H2-K1 |
| **ENSMUSG00000061244** | 0.12496559948841 | 0.41916404176602 | 0.882550938324998 | 680.345828576491 | 609.182965227918 | 619.927952064733 | 635.914298707033 | 549.669014301187 | 450.09426140318 | 642.345181873136 | 690.394259129171 | Exoc5 |
| **ENSMUSG00000061273** | 0.0904342682336603 | 0.677611432689854 | 0.9442237961539 | 796.899098013898 | 918.660942215363 | 983.366629305736 | 741.900015158206 | 664.639624862192 | 566.287073450197 | 985.900951756661 | 1013.88087740039 | Mmgt1 |
| **ENSMUSG00000061286** | 0.156704334697813 | 0.423647327224911 | 0.884482395183227 | 343.335212296237 | 276.086932101957 | 302.865564367503 | 364.853484066425 | 314.253002200081 | 371.816998550453 | 241.530117009024 | 227.960368580387 | Exosc5 |
| **ENSMUSG00000061288** | 0.0205178851813139 | 0.883376581433682 | 0.97910597465636 | 216.843291976571 | 237.809392842984 | 180.772883731853 | 211.971432902344 | 213.516848184724 | 221.377884005369 | 193.640524843441 | 208.420908416353 | Taok3 |
| **ENSMUSG00000061306** | 0.0103874668050006 | 0.891347762579461 | 0.98237089758222 | 2318.41619671617 | 2407.41289467076 | 2481.60471803623 | 2263.21711324494 | 2221.67017931695 | 2356.87946120633 | 2415.30117009024 | 2408.78122799942 | Slc38a10 |
| **ENSMUSG00000061313** | 0.370508995435115 | 0.0945380200968587 | 0.757085157577544 | 317.133314515735 | 394.991628523449 | 389.93941412316 | 377.046531091781 | 236.510970296925 | 195.693157131818 | 334.18606706852 | 376.677370939972 | Ddhd2 |
| **ENSMUSG00000061315** | 0.210685127041956 | 0.0101026721068827 | 0.635862253526407 | 10541.294531211 | 10501.8908243503 | 10671.2788695112 | 11300.1408124223 | 9813.01535093075 | 8490.63685505673 | 9204.1713986068 | 9660.09199998505 | Naca |
| **ENSMUSG00000061322** | 0.00918563663488726 | 0.965999499525674 | 0.994284908685797 | 79.509207058076 | 67.5965054998893 | 54.8943835416099 | 77.8479156234274 | 58.0327843784121 | 72.1618516923577 | 66.6289977955927 | 81.4144173501381 | Dnaic1 |
| **ENSMUSG00000061353** | -0.436228519319523 | 0.128691032722936 | 0.77303252008843 | 1945.26503177315 | 2070.24478290022 | 2325.43966140923 | 2535.21585457981 | 3821.40410340865 | 4373.74206189612 | 1880.18703154438 | 1935.49208180395 | Cxcl12 |
| **ENSMUSG00000061360** | -0.0795758471309411 | 0.579152433029617 | 0.922450244597442 | 482.47632464787 | 416.980853204136 | 385.207139679918 | 482.094320848695 | 446.742943894192 | 555.279333361532 | 433.088485671353 | 433.124700302735 | Phf5a |
| **ENSMUSG00000061371** | -0.100340344842034 | 0.691647592668841 | 0.94469184542218 | 58.7283915769879 | 46.4216965481167 | 37.8581955459379 | 43.1446279358754 | 39.4184950494875 | 45.2540425867328 | 59.3414511616998 | 55.3618037980939 | Zfp873 |
| **ENSMUSG00000061374** | 0.087654075231477 | 0.465731031671407 | 0.895328112556867 | 423.747933070882 | 487.020605890769 | 463.762895437739 | 411.749818779333 | 446.742943894192 | 380.37857417497 | 453.910047482476 | 399.473407798011 | Fiz1 |
| **ENSMUSG00000061393** | -0.027395613135618 | 0.928332737834299 | 0.986901071486195 | 36.1405486627618 | 30.1333819698302 | 49.2156542097192 | 28.1378008277448 | 47.0832024202212 | 35.4693847301419 | 37.4788112600209 | 26.0526135520442 | Acvr2b |
| **ENSMUSG00000061410** | -0.160930650707565 | 0.335813509696055 | 0.858570960161243 | 508.678222428372 | 591.265819191803 | 567.872933189068 | 425.818719193205 | 644.930377337448 | 454.986590331476 | 623.605776243126 | 616.578520731712 | Zcchc14 |
| **ENSMUSG00000061411** | -0.648678195588001 | 0.0559096326970115 | 0.703396849069134 | 46.0791995450213 | 39.0919549878878 | 34.0723759913441 | 27.1998741334867 | 91.9764884488041 | 56.2617826753975 | 43.7252798033577 | 37.9933947633978 | Nol4l |
| **ENSMUSG00000061414** | 0.289032095648165 | 0.532791590165116 | 0.909082872831377 | 9.93865088225949 | 14.6594831204579 | 10.4110037751329 | 9.37926694258161 | 6.56974917491458 | 8.56157562451702 | 9.36970281500523 | 11.9407812113536 | Cracr2a |
| **ENSMUSG00000061436** | 0.244441678301376 | 0.539969864597291 | 0.913837378403038 | 833.03964667666 | 1822.66240131027 | 1498.23808873049 | 729.70696813285 | 607.701798679599 | 562.617826753975 | 1517.89185603085 | 1433.9792709271 | Hipk2 |
| **ENSMUSG00000061451** | 0.462784448516798 | 0.186180647899154 | 0.806376426354616 | 61.4389327266951 | 59.452348210746 | 34.0723759913441 | 30.0136542162612 | 33.943704070392 | 31.8001380339204 | 45.80743598447 | 22.7960368580387 | Tmem151a |
| **ENSMUSG00000061455** | 0.160462484008879 | 0.527068182438089 | 0.908217276872524 | 530.362551626029 | 630.35777417969 | 640.749959614998 | 567.445650026188 | 358.051330032845 | 371.816998550453 | 649.632728507029 | 739.242909539254 | Stx17 |
| **ENSMUSG00000061458** | -0.0608393287777208 | 0.734319771804049 | 0.954227437328031 | 152.693818100169 | 148.223662662408 | 138.182413742673 | 138.813150750208 | 181.76306050597 | 113.746647582869 | 141.586620315635 | 164.999885829613 | Nol10 |

| | | | | | | | | | | | | |
|---|---|---|---|---|---|---|---|---|---|---|---|---|
| **ENSMUSG00000061461** | -0.0264137874658179 | 0.871315751417751 | 0.975613767803763 | 1565.78927081416 | 1037.56563863685 | 1236.07008457487 | 1350.61443973175 | 1179.26997689717 | 1353.95203090576 | 1209.73274122623 | 1542.53182739395 | Smim20 |
| **ENSMUSG00000061462** | 0.380325278802596 | 0.0233910069097647 | 0.654124840968651 | 38351.4467272063 | 46619.5995702429 | 42815.7262526784 | 40422.7646691382 | 33309.7232750127 | 23152.9466531582 | 35763.1145667844 | 37002.3099228554 | Obscn |
| **ENSMUSG00000061474** | 0.0301949900182039 | 0.815767672257234 | 0.966177091475091 | 1417.61302129683 | 1084.80175091389 | 1118.70967838246 | 1334.66968592936 | 1209.9288063801 | 1336.82887965673 | 1169.13069569454 | 1137.6307917726 | Mrps36 |
| **ENSMUSG00000061477** | 0.0362761612803964 | 0.813448288625561 | 0.966039392015138 | 9348.65642533991 | 7024.33566188608 | 8147.08368148583 | 9882.93357739825 | 8428.98819141541 | 9889.84292854923 | 7336.47730414909 | 7893.94190626939 | Rps7 |
| **ENSMUSG00000061479** | -0.139136930734143 | 0.244086673198827 | 0.832048466795696 | 821.293968361262 | 728.08766164941 | 680.501064938233 | 846.94780491512 | 863.922016501267 | 905.080851734656 | 862.012658980481 | 757.696844138618 | Snrpa |
| **ENSMUSG00000061518** | 0.0280445549231375 | 0.894016652204218 | 0.983730237253949 | 24421.0722451447 | 17774.6232835552 | 18863.7923856522 | 23579.4770936502 | 20615.872910882 | 28818.2635521243 | 16623.9349500004 | 16952.6527434281 | Cox5b |
| **ENSMUSG00000061533** | -0.108930445512492 | 0.577208062420846 | 0.922450244597442 | 245.75573090678 | 195.459774939439 | 236.613722162112 | 159.447538023887 | 208.042057205628 | 184.685417043153 | 249.858741733473 | 259.440609955773 | Cep128 |
| **ENSMUSG00000061535** | -0.75505795235806 | 0.0533500236828322 | 0.703396849069134 | 77.7021796249379 | 66.7820897709749 | 36.9117406572894 | 68.4686486808458 | 155.484063806312 | 144.323703384715 | 51.0128264372507 | 71.6446872681215 | C1qtnf7 |
| **ENSMUSG00000061536** | -0.0049505059560665 | 0.979797388429374 | 0.996872525500296 | 196.062476495483 | 223.149909722526 | 194.96970706158 | 186.647412157374 | 140.154649064844 | 193.24699266767 | 224.872867560126 | 245.328777615083 | Sec22c |
| **ENSMUSG00000061544** | -0.133768645077438 | 0.527237677434039 | 0.908217276872524 | 49.6932544112975 | 55.3802695661743 | 46.3762895437739 | 51.5859681841989 | 56.937826182593 | 48.9232892829544 | 63.5057635239243 | 53.1907526687569 | Zfp229 |
| **ENSMUSG00000061559** | 0.0530853526447003 | 0.689264798736675 | 0.94469184542218 | 580.959319753896 | 464.216965481167 | 432.52988411234 | 468.963347129081 | 502.585811880965 | 480.671317205027 | 454.951125573032 | 438.552328126077 | Wdr61 |
| **ENSMUSG00000061589** | 0.137492558689512 | 0.613946004859694 | 0.928075610917191 | 188.83436676293 | 307.849145529616 | 361.545767463707 | 288.881421831514 | 190.522726072523 | 193.24699266767 | 318.569895710178 | 339.769501741243 | Dot1l |
| **ENSMUSG00000061601** | -0.119784931807633 | 0.786820172891046 | 0.961141387070613 | 12.6491920319666 | 21.1748089517725 | 10.4110037751329 | 8.44134024832345 | 13.1394983498292 | 17.123151249034 | 16.6572494488982 | 10.8552556466851 | Pclo |
| **ENSMUSG00000061603** | 0.188035890955062 | 0.525420026801335 | 0.908006595346551 | 3194.82450178814 | 4427.16390237829 | 2761.75536507617 | 2375.76831655592 | 2116.55419251831 | 1696.41505588644 | 3584.43186578478 | 3802.59605303378 | Akap6 |
| **ENSMUSG00000061607** | 0.0439305575914335 | 0.806875078536669 | 0.964026716319739 | 150.886790667031 | 201.160685041839 | 200.648436393471 | 153.819977858338 | 154.389105610493 | 165.116101329971 | 206.133461930115 | 159.572258006271 | Mdc1 |
| **ENSMUSG00000061613** | 0.0336908870320791 | 0.801724939531463 | 0.963461896828986 | 478.862269781594 | 455.25839246311 | 487.42426765395 | 550.562969529541 | 522.295059405709 | 533.263853184203 | 406.020455316893 | 465.69046724279 | U2af1 |
| **ENSMUSG00000061650** | 0.026349991057816 | 0.829674467366721 | 0.967523998969336 | 920.680477183857 | 853.507683902216 | 789.343377132805 | 908.850966736158 | 724.862325632242 | 836.58824673852 | 941.134593862747 | 906.413846498204 | Med9 |
| **ENSMUSG00000061665** | -0.136944373521757 | 0.399588090806956 | 0.880459952732461 | 343.335212296237 | 351.013179162076 | 390.885869011809 | 298.260688774095 | 467.547149614754 | 376.709327478749 | 361.254097422979 | 315.887939318536 | Cd2ap |
| **ENSMUSG00000061666** | 0.0532829785931763 | 0.719332880242431 | 0.951674101666722 | 291.834930451802 | 236.99497711407 | 245.131816159948 | 264.495327780801 | 267.16979977986 | 204.254732756335 | 274.84461590682 | 252.927456567762 | Gdpd1 |
| **ENSMUSG00000061689** | 0.0812461963612753 | 0.396403543723389 | 0.879359239049715 | 1404.96382926487 | 1290.0345146003 | 1583.41902870885 | 1431.27613543795 | 1390.59690869025 | 1325.82113956806 | 1323.21025309685 | 1355.82143027097 | Dlgap4 |
| **ENSMUSG00000061723** | -2.43913220743991 | 0.0192111100406721 | 0.642273034804458 | 6.32459601598331 | 2.44324718674299 | 1.89290977729689 | 5.62756016554897 | 4.37983278327639 | 73.3849339244316 | 1.04107809055614 | 9.76973008201657 | Tnnt3 |
| **ENSMUSG00000061731** | -0.35549005872318 | 0.011391315244773 | 0.635862253526407 | 451.756858284523 | 472.361122770311 | 460.923530771794 | 497.101147956825 | 508.060602860061 | 736.295503708463 | 572.592949805875 | 592.696958309005 | Ext1 |
| **ENSMUSG00000061740** | -0.594390774894953 | 0.055787987811592 | 0.703396849069134 | 187.027339329792 | 156.367819951551 | 159.950876181588 | 210.095579513828 | 329.582416941548 | 409.732547744743 | 153.038479311752 | 185.624871558315 | Cyp2d22 |
| **ENSMUSG00000061742** | -0.884788108662887 | 0.105673106595245 | 0.762094734041179 | 8.13162344912141 | 22.8036404096012 | 31.2330113253987 | 10.3171936368398 | 37.2285786578493 | 37.9155491942896 | 47.8895921655823 | 10.8552556466851 | Slc22a12 |
| **ENSMUSG00000061751** | 0.0626267788341567 | 0.753715242290012 | 0.956497786093635 | 383.089815825275 | 415.352021746308 | 522.443098533943 | 592.769670771158 | 459.882442244021 | 344.909189444828 | 531.990904274186 | 493.914131924171 | Kalrn |
| **ENSMUSG00000061755** | 0.239199283213748 | 0.0051672348231478 | 0.635862253526407 | 1198.05918817055 | 1270.48853710635 | 1201.99770858353 | 1174.28422121122 | 987.652292628825 | 967.458045570423 | 1081.68013608783 | 1067.07163006914 | Bod1l |
| **ENSMUSG00000061758** | 0.0704277079550939 | 0.563832934971754 | 0.918590066040684 | 1089.63754218227 | 965.897054492394 | 928.472245764126 | 999.8298560792 | 914.290093508946 | 809.680437632895 | 1031.70838774113 | 1036.67691425842 | Akr1b10 |
| **ENSMUSG00000061759** | 0.0394810964943856 | 0.797223399623088 | 0.962405094799643 | 160.82544154929 | 169.39847161418 | 208.220075502658 | 169.764731660727 | 169.71852035196 | 150.439114545085 | 187.394056300105 | 180.197243734972 | Armt1 |
| **ENSMUSG00000061762** | 0.2207479358834 | 0.523883397514255 | 0.907098029353404 | 47.8862269781594 | 33.3910448854875 | 67.1982970940397 | 30.9515809105193 | 42.7033696369448 | 28.1308913376988 | 52.0539045278068 | 30.3947158107182 | Tac1 |
| **ENSMUSG00000061778** | -0.135111378205688 | 0.396950295907593 | 0.879613019351548 | 169.86057871498 | 167.769640156352 | 159.950876181588 | 166.013024883695 | 225.561388338734 | 152.885279009232 | 173.860041122875 | 175.855141476298 | Mospd2 |
| **ENSMUSG00000061780** | -0.495161638579297 | 0.505639430327155 | 0.901494649536949 | 14.4562194651047 | 17.1027303072009 | 10.4110037751329 | 29.075727522003 | 19.7092475247437 | 63.6002760678407 | 2.08215618111227 | 15.1973579053591 | Cfd |
| **ENSMUSG00000061787** | 0.149075991746457 | 0.463943319149288 | 0.895328112556867 | 10937.0335390683 | 6951.85266201271 | 8814.33437798298 | 11229.796310353 | 8524.24955445167 | 10969.8245394704 | 7217.79440182569 | 7497.72507516538 | Rps17 |
| **ENSMUSG00000061808** | -0.148528558037631 | 0.598959686456353 | 0.924848033480823 | 887.250469670802 | 1459.43298621448 | 979.580809751142 | 1118.94654624999 | 951.518672166795 | 1089.76626877781 | 891.162845516053 | 1995.19598786072 | Ttr |
| **ENSMUSG00000061815** | 0.565991895831657 | 0.0410257621688766 | 0.683226273624098 | 59.631905293557 | 34.2054606144018 | 45.4298346551254 | 50.6480414899407 | 30.6588294829347 | 28.1308913376988 | 38.5198893505771 | 30.3947158107182 | Rufy4 |
| **ENSMUSG00000061816** | -0.491572564780138 | 0.293048819680734 | 0.851785577030345 | 678.538801143353 | 394.991628523449 | 392.778778789105 | 675.307219865876 | 362.431162816121 | 1733.10752284866 | 454.951125573032 | 461.348364984116 | Myl1 |
| **ENSMUSG00000061825** | -0.115925263197162 | 0.823525417343648 | 0.966637987288264 | 15.3597331816738 | 11.4018202048006 | 5.67872933189068 | 17.8206071909051 | 8.75966556655277 | 14.6769867848863 | 8.32862472444909 | 22.7960368580387 | Ces2c |
| **ENSMUSG00000061838** | 0.328098688554743 | 0.143336505320401 | 0.780607990324772 | 5119.30871808021 | 5295.33106940097 | 5877.48485850685 | 5014.15610750413 | 3719.57299119747 | 2650.41919690405 | 5193.93859378457 | 5407.00283761384 | Suclg2 |
| **ENSMUSG00000061848** | -0.0774288674929543 | 0.864408973879238 | 0.974868041734905 | 19.877301764519 | 13.8450673915436 | 6.62518422053913 | 13.1309737196143 | 16.4243729372865 | 13.4539045528125 | 9.36970281500523 | 17.3684090346961 | Gm5805 |
| **ENSMUSG00000061859** | 0.673682621747045 | 0.0077804816890078 | 0.635862253526407 | 700.22313034101 | 883.641065872047 | 1068.5475692841 | 984.823028971069 | 492.731188118594 | 352.247682837272 | 657.961353231478 | 776.150778737983 | Patj |
| **ENSMUSG00000061877** | 0.178171799837623 | 0.654543684529648 | 0.937269843859626 | 27.1054114970714 | 16.2883145782866 | 14.1968233297267 | 19.6964605794214 | 16.4243729372865 | 11.0077400886647 | 15.616171358342 | 24.9670879873757 | BC048679 |
| **ENSMUSG00000061878** | -0.385398535246046 | 0.514732225029011 | 0.904145806837306 | 19.877301764519 | 27.6901347830872 | 12.3039135524298 | 18.7585338851632 | 42.7033696369448 | 41.5847958905112 | 13.5340151772298 | 5.42762782334254 | Sphk1 |
| **ENSMUSG00000061882** | 0.0146464610027154 | 0.962644588416883 | 0.994140449160496 | 28.9124389302094 | 23.6180561385155 | 42.5904699891801 | 29.075727522003 | 31.7537876787538 | 37.9155491942896 | 21.8626399016789 | 31.4802413753867 | Ccdc62 |
| **ENSMUSG00000061887** | 0.00936646192006726 | 0.912912900685609 | 0.986226496507248 | 945.075347531221 | 892.599638890104 | 884.935320886298 | 888.216579462479 | 908.81530252985 | 920.980920751616 | 927.600578685518 | 830.427056971408 | Ssbp3 |
| **ENSMUSG00000061894** | 0.00875674868305492 | 0.975792787883277 | 0.99575536949131 | 280.992765852973 | 396.620459981278 | 241.345996605354 | 212.909359596603 | 250.745426842573 | 171.23151249034 | 396.650752501888 | 306.118209236519 | Zscan20 |
| **ENSMUSG00000061898** | 0.00408549567696355 | 0.985902873246853 | 0.997682566618009 | 41.5616309621761 | 50.4937751926884 | 62.4660226507975 | 57.2135283497478 | 56.937826182593 | 58.7079471395453 | 51.0128264372507 | 44.5065481514088 | Rbak |
| **ENSMUSG00000061904** | 0.0935829927276335 | 0.316138389976715 | 0.853356198600137 | 41209.2606127141 | 38442.0512362141 | 41136.7152802161 | 41698.3449733293 | 37947.9661925024 | 33609.0766551576 | 39962.8235840879 | 40758.2283766085 | Slc25a3 |
| **ENSMUSG00000061947** | -0.247538228610464 | 0.448601737303651 | 0.89108031640712 | 69.5705561758165 | 92.0289773673191 | 41.6440151005317 | 70.3445020693621 | 62.4126171616885 | 92.9542496376133 | 54.1360607089191 | 116.15123541953 | Serpina10 |
| **ENSMUSG00000061950** | 0.065564683601907 | 0.74653650837079 | 0.956085046483146 | 487.897406947284 | 592.894650649631 | 591.534305405279 | 401.432625142493 | 468.642107810573 | 364.47850515801 | 549.68923181364 | 598.124586132348 | Ppp4r1 |
| **ENSMUSG00000061959** | -0.313264812525031 | 0.396888528892968 | 0.879613019351548 | 26.2018977805023 | 40.7207864457164 | 29.3401015481019 | 33.7653609932938 | 27.3739548954774 | 37.9155491942896 | 27.0680303544596 | 69.4736361387845 | Ces1e |
| **ENSMUSG00000061979** | 0.0998450944285945 | 0.355257055308429 | 0.864215962387542 | 866.469654189714 | 813.601313185414 | 844.237760674415 | 1008.27119632752 | 825.598479647599 | 791.334204151787 | 872.423439886043 | 805.459968984033 | Rcc1l |
| **ENSMUSG00000061981** | 0.06753392057589 | 0.708029726927002 | 0.949174314815223 | 1706.73741059893 | 2000.20503021359 | 2024.46700681903 | 1622.61318106662 | 1565.79022002131 | 1317.25956394355 | 2063.41677548226 | 2070.09725182284 | Flot2 |
| **ENSMUSG00000061983** | 0.183869310539395 | 0.281697469488858 | 0.850048154552447 | 5588.23233697954 | 3845.67107193346 | 4396.28295777204 | 5725.10454175182 | 4069.95961385958 | 5149.17619703095 | 3926.94655757775 | 4069.63534194223 | Rps12 |
| **ENSMUSG00000062006** | 0.266465127767669 | 0.0960336787375395 | 0.757809915356977 | 4661.2272637797 | 3056.50223061548 | 3642.90486640787 | 4320.09035375309 | 3083.40227942658 | 3703.49299871965 | 2996.22274462056 | 3253.32011731152 | Rpl34 |
| **ENSMUSG00000062012** | -0.434950709961639 | 0.212666127693827 | 0.823370267313127 | 23.4913566307952 | 21.9892246806869 | 33.1259211026956 | 40.3308478531009 | 29.5638712871156 | 59.9310293716191 | 29.1501865355718 | 42.3354970220718 | Zfp13 |
| **ENSMUSG00000062014** | 0.0326877643843949 | 0.778088975579673 | 0.960126492111037 | 785.1534196985 | 780.210268299927 | 771.360734248484 | 744.71379524098 | 706.248036303317 | 653.125911927441 | 802.671207818781 | 848.880991570773 | Gmfb |
| **ENSMUSG00000062031** | -0.332928010946702 | 0.249687491915596 | 0.835314899646652 | 29.8159526467785 | 51.3081909216027 | 26.5007368821565 | 31.8895076047775 | 47.0832024202212 | 48.9232892829544 | 38.5198893505771 | 42.3354970220718 | Pgghg |
| **ENSMUSG00000062040** | 0.123511932615795 | 0.504069800898436 | 0.900927059989498 | 244.852217190211 | 225.593156909269 | 196.862616838877 | 225.102406621959 | 186.142893289246 | 152.885279009232 | 247.776585552361 | 231.216945274392 | Zfp27 |
| **ENSMUSG00000062054** | -0.202583891088264 | 0.405526554639129 | 0.881158207349486 | 344.238726012806 | 295.632909595901 | 196.862616838877 | 354.536290429585 | 300.018545654433 | 475.778988276731 | 298.789411989611 | 297.434004719171 | Iah1 |
| **ENSMUSG00000062070** | 0.0847909071114741 | 0.353130980467416 | 0.862728204353539 | 8924.00497855246 | 8586.38502994377 | 8493.48617073116 | 8762.11117775974 | 8395.04448734502 | 7175.82345557733 | 8606.59257462758 | 8601.70457443325 | Pgk1 |
| **ENSMUSG00000062075** | -0.283631549185728 | 0.128609539184409 | 0.77303252008843 | 85.8338030740593 | 72.4829998733752 | 69.0912068713366 | 64.7169419038131 | 77.7420319031559 | 95.400414101761 | 89.5327157878278 | 93.3551985614916 | Lmnb2 |
| **ENSMUSG00000062077** | -0.0517569343951592 | 0.75429202855876 | 0.956497786093635 | 17169.4711559616 | 14525.1045251871 | 14894.3605826606 | 18237.0466431557 | 18083.2346039524 | 20659.0819819596 | 14954.0456927483 | 13498.5103966529 | Trim54 |
| **ENSMUSG00000062078** | 0.122007142292537 | 0.577537015829447 | 0.922450244597442 | 5119.30871808021 | 6467.27530330868 | 6425.48223903431 | 5273.02387511938 | 4348.07899559763 | 3754.86245246675 | 6186.08601408456 | 7106.93587188472 | Qk |
| **ENSMUSG00000062081** | 0.472933720791381 | 0.143695323064158 | 0.780607990324772 | 25.2983840639333 | 28.5045505120015 | 41.6440151005317 | 30.9515809105193 | 21.8991639163819 | 22.0154801773295 | 16.6572494488982 | 30.3947158107182 | Gm6055 |
| **ENSMUSG00000062093** | 0.43062762833737 | 0.0307937567232274 | 0.667184591665619 | 112.035700854562 | 87.9568987227475 | 92.7525790875478 | 114.427056699496 | 75.5521155115177 | 66.0464405319884 | 67.6700758861489 | 92.2696729968231 | Gm10110 |
| **ENSMUSG00000062098** | -0.285246106324242 | 0.174523485356242 | 0.799889021902745 | 204.194099944604 | 221.521078264697 | 179.826428843205 | 211.971432902344 | 338.342082508101 | 265.408844360028 | 194.681602933998 | 198.651178334337 | Btbd3 |
| **ENSMUSG00000062101** | 0.255323639606755 | 0.476462518790234 | 0.897553735056661 | 80.412720774645 | 31.7622134276588 | 37.8581955459379 | 40.3308478531009 | 27.3739548954774 | 41.5847958905112 | 43.7252798033577 | 46.6775992807458 | Zfp119b |
| **ENSMUSG00000062110** | 0.200001740194019 | 0.242398130362805 | 0.832048466795696 | 262.922491521592 | 312.735639903102 | 260.275094378323 | 275.750448111899 | 275.929465346412 | 182.239252579005 | 231.119336103462 | 277.894544555138 | Scfd2 |
| **ENSMUSG00000062115** | 0.0722340260344009 | 0.653885042144644 | 0.937077409179068 | 297.256012751216 | 398.249291439107 | 475.12035410152 | 401.432625142493 | 381.045452145046 | 355.916929533493 | 381.034581143546 | 376.677370939972 | Rai1 |
| **ENSMUSG00000062116** | 0.045287454360439 | 0.855110516585037 | 0.973268281968411 | 66.8600150261093 | 83.8848200781758 | 67.1982970940397 | 106.92364314543 | 78.836990098975 | 103.961989726278 | 66.6289977955927 | 66.217059444779 | Zfp954 |
| **ENSMUSG00000062127** | -0.318199792778781 | 0.16674860926166 | 0.796745160868404 | 409.291713605777 | 535.885549625628 | 397.511053232348 | 357.350070512359 | 631.790878987619 | 667.802898712327 | 461.197594116369 | 360.394487469945 | Cttnbp2nl |
| **ENSMUSG00000062157** | -0.165422296539337 | 0.686247399199931 | 0.944452327903597 | 40.658117245607 | 37.4631235300591 | 50.1621090983677 | 65.6548685980713 | 33.943704070392 | 24.4616446414772 | 82.2451691539348 | 75.9867895267955 | Ifnlr1 |
| **ENSMUSG00000062169** | 0.0424590377101838 | 0.708845227891954 | 0.949267137065756 | 1400.44626068202 | 1278.6326943955 | 1318.41165988729 | 1303.71810501884 | 1202.26409900937 | 1144.80496922113 | 1312.79947219129 | 1486.08449803119 | Cnih4 |
| **ENSMUSG00000062170** | 0.489972847208846 | 0.384645206666245 | 0.875126835008975 | 22.5878429142261 | 13.0306516626293 | 17.9826428843205 | 20.6343872736795 | 2.18991639163819 | 24.4616446414772 | 16.6572494488982 | 9.76973008201657 | Fmr1nb |
| **ENSMUSG00000062181** | -0.144620016858516 | 0.674397347752471 | 0.944023556644042 | 41.5616309621761 | 60.2667639396603 | 29.3401015481019 | 60.0273084325223 | 41.6084114411257 | 50.1463715150282 | 38.5198893505771 | 81.4144173501381 | Ces3b |
| **ENSMUSG00000062190** | -0.258691185822874 | 0.100838371846263 | 0.759800953500741 | 181.606257030378 | 247.582381589956 | 255.542819935081 | 209.15765281957 | 271.549632563136 | 236.054870790255 | 274.84461590682 | 286.578749072486 | Lancl2 |
| **ENSMUSG00000062198** | 0.0995459337374137 | 0.550446082042671 | 0.914699131434573 | 120.167324303683 | 121.347943608235 | 125.878500190243 | 103.171936368398 | 114.970610561005 | 96.6234963338349 | 124.929370866736 | 102.03940307884 | 2700097O09 |
| **ENSMUSG00000062202** | 0.103744223244215 | 0.511336459694392 | 0.902394714365738 | 393.028466707535 | 447.114235173966 | 428.744064557746 | 417.377378944882 | 440.173194719277 | 289.870489001505 | 409.143689588562 | 428.78259804406 | Btbd9 |
| **ENSMUSG00000062203** | 0.188336159661383 | 0.366037662044659 | 0.869253398153585 | 1585.66657257867 | 1903.28955847279 | 1792.58555910016 | 1586.97196668481 | 1367.60278657805 | 994.365854676048 | 1777.12030057933 | 1887.72895695853 | Gspt1 |
| **ENSMUSG00000062209** | 0.297731216316672 | 0.3386749660295 | 0.858570960161243 | 110.228673421423 | 205.232763686411 | 183.612248397799 | 113.489130005238 | 102.926070406995 | 77.0541806206532 | 155.120635492864 | 162.828834700276 | Erbb4 |
| **ENSMUSG00000062210** | 0.126684296135522 | 0.503719861083257 | 0.900927059989498 | 1393.21815094947 | 1159.727997974 | 1095.04830616625 | 1154.5877606318 | 1094.9581958191 | 754.641737189571 | 1242.00616203347 | 1305.88725429621 | Tnfaip8 |
| **ENSMUSG00000062232** | 0.14889955567389 | 0.571079200805186 | 0.921126010531982 | 932.426155499254 | 1224.06684055824 | 753.378091364164 | 523.363095396054 | 985.462376237187 | 647.010500767072 | 783.931802188771 | 680.624529047154 | Rapgef2 |
| **ENSMUSG00000062234** | 0.0252330017342877 | 0.869398750009502 | 0.975613767803763 | 398.449549006949 | 494.350347450997 | 520.550188756646 | 405.184331919526 | 516.820268426614 | 425.632616761703 | 458.0743598447 | 386.447101021989 | Gak |
| **ENSMUSG00000062248** | -0.403375876794001 | 0.325157063066397 | 0.855977678491063 | 14.4562194651047 | 13.8450673915436 | 10.4110037751329 | 18.7585338851632 | 17.5193311331055 | 25.6847268735511 | 10.4107809055614 | 22.7960368580387 | Cks2 |
| **ENSMUSG00000062257** | -0.170237357994884 | 0.701524758386528 | 0.945991110533699 | 12.6491920319666 | 13.8450673915436 | 8.51809399783602 | 15.0068271081306 | 22.994122112201 | 11.0077400886647 | 11.4518589961175 | 10.8552556466851 | Opcml |
| **ENSMUSG00000062270** | 0.149829404305892 | 0.143923456938597 | 0.780607990324772 | 6697.74718092633 | 6183.04421391758 | 7177.91387550982 | 6981.92631205775 | 5933.57846314369 | 5430.48511040794 | 6518.18992497197 | 6488.18630002367 | Morf4l1 |
| **ENSMUSG00000062300** | -0.200881879633881 | 0.134599006389941 | 0.773512701181749 | 731.846110420926 | 721.572335818095 | 883.042411109001 | 774.727449457241 | 1046.78003520306 | 851.265233523406 | 896.368235968834 | 780.492880996657 | Nectin2 |
| **ENSMUSG00000062309** | -0.187957249840696 | 0.617558375343646 | 0.929314954729851 | 22.5878429142261 | 36.6487078011448 | 24.6078271048596 | 20.6343872736795 | 28.4689130912965 | 42.8078781225851 | 33.3144988977964 | 15.1973579053591 | Rpp25 |

| | | | | | | | | | | | | |
|---|---|---|---|---|---|---|---|---|---|---|---|---|
| **ENSMUSG00000062312** | -0.189067432745845 | 0.401166318354462 | 0.880924293718604 | 127.395434036235 | 114.018202048006 | 196.862616838877 | 136.937297361692 | 206.947099009809 | 140.654456688494 | 155.120635492864 | 151.973579053591 | Erbb2 |
| **ENSMUSG00000062328** | 0.117274958220529 | 0.50933565788319 | 0.902394714365738 | 7535.30439618584 | 4609.5930256551 | 5737.40953498688 | 7240.794079673 | 5610.56579537705 | 6488.45124115183 | 5212.67799941458 | 5849.89726799859 | Rpl17 |
| **ENSMUSG00000062329** | -0.57278614512536 | 0.244025099494587 | 0.832048466795696 | 201.483558794897 | 102.616381843205 | 134.396594188079 | 97.5443762028488 | 285.784089108784 | 333.901449356164 | 127.011527047849 | 51.0197015394199 | Cytl1 |
| **ENSMUSG00000062352** | 0.0863716387260974 | 0.544039484750382 | 0.913837378403038 | 923.391018333564 | 710.984931342209 | 699.430162711202 | 779.417082928532 | 664.639624862192 | 662.910569784032 | 805.79444209045 | 797.861290031353 | Itgb1bp1 |
| **ENSMUSG00000062353** | -0.109724645963151 | 0.826637735657555 | 0.967319591893432 | 28.9124389302094 | 8.14415728914329 | 17.036187995672 | 17.8206071909051 | 8.75966556655277 | 30.5770558018465 | 16.6572494488982 | 21.7105112933701 | Gm15772 |
| **ENSMUSG00000062373** | 0.121047029865782 | 0.455141735772561 | 0.892775943845882 | 4905.17596725335 | 4960.60620481718 | 4868.56394720761 | 4603.34421541906 | 3732.7124895473 | 3562.83854203115 | 5097.11833136285 | 5387.4633774498 | Tmem65 |
| **ENSMUSG00000062376** | 0.0379772607806027 | 0.837124089148529 | 0.968924788429433 | 371.344137509877 | 240.252640029727 | 276.364827485346 | 335.777756544422 | 264.979883388221 | 355.916929533493 | 298.789411989611 | 272.466916731795 | Borcs7 |
| **ENSMUSG00000062380** | -1.13356212786939 | 0.0231350263916407 | 0.651816463650892 | 14.4562194651047 | 15.4738988493722 | 9.46454888648447 | 11.2551203310979 | 16.4243729372865 | 57.4848649074714 | 20.8215618111227 | 17.3684090346961 | Tubb3 |
| **ENSMUSG00000062382** | -0.273234020032424 | 0.475748340099921 | 0.897549741595057 | 19.877301764519 | 31.7622134276588 | 25.5542819935081 | 37.5170677703264 | 32.8487458745729 | 58.7079471395453 | 22.903717992235 | 24.9670879873757 | Ftl1-ps1 |
| **ENSMUSG00000062397** | 0.126468589691708 | 0.111967181984915 | 0.764116514278315 | 3806.50328790539 | 3493.84347704247 | 3543.52710309978 | 3898.02334133692 | 3330.86283168169 | 3310.88360222394 | 3567.77461633588 | 3293.48456320425 | Zfp706 |
| **ENSMUSG00000062410** | -0.207661110202921 | 0.692676497316689 | 0.94469184542218 | 5.42108229941427 | 26.0613033252585 | 9.46454888648447 | 14.0689004138724 | 14.2344565456483 | 12.2308223207386 | 14.5750932677859 | 22.7960368580387 | Hsd3b3 |
| **ENSMUSG00000062421** | -0.0889013495692465 | 0.383473042222919 | 0.875126835008975 | 538.494175075151 | 508.195414842541 | 573.551662520959 | 534.618215727152 | 616.461464246152 | 591.971800323748 | 533.031982364742 | 550.361461286933 | Arf2 |
| **ENSMUSG00000062456** | 0.0451133997531897 | 0.853370180040119 | 0.973120292022893 | 67.7635287426784 | 41.5352021746308 | 65.3053873167428 | 60.9652351267805 | 49.2731188118594 | 52.592535979176 | 56.2182168900314 | 69.4736361387845 | Rpl9-ps6 |
| **ENSMUSG00000062480** | -0.565653213969178 | 0.144802998661513 | 0.780607990324772 | 27.1054114970714 | 37.4631235300591 | 37.8581955459379 | 28.1378008277448 | 36.1336204620302 | 37.9155491942896 | 30.191264626128 | 89.0130963028176 | Acat3 |
| **ENSMUSG00000062488** | -0.204163876011084 | 0.421186894538659 | 0.882568531013176 | 69.5705561758165 | 72.4829998733752 | 45.4298346551254 | 75.9720622349111 | 72.2672409240604 | 100.292743030056 | 58.3003730711437 | 73.8157383974585 | Ifit3b |
| **ENSMUSG00000062515** | -0.124288347614311 | 0.68329775547244 | 0.944452327903597 | 9615.19297172778 | 7139.98269539192 | 9846.91666149844 | 14565.063635135 | 14583.7482101146 | 15142.9811153065 | 6805.52747796547 | 8339.00738778347 | Fabp4 |
| **ENSMUSG00000062519** | 0.169623293798097 | 0.463940082563009 | 0.895328112556867 | 100.290022539164 | 167.769640156352 | 163.736695736181 | 146.316564304273 | 131.394983498292 | 91.7311674055395 | 144.709854587303 | 145.46042566558 | Zfp398 |
| **ENSMUSG00000062526** | -0.0216851103647727 | 0.882258949387258 | 0.978854883720278 | 271.054114970713 | 291.56083095133 | 248.917635714542 | 251.364354061187 | 233.226095709468 | 258.070350967584 | 322.734208072402 | 264.868237779116 | Mppe1 |
| **ENSMUSG00000062563** | -0.46776763192113 | 0.18465383093014 | 0.805963874499992 | 41.5616309621761 | 46.4216965481167 | 39.7511053232348 | 54.3997482669734 | 68.9823663366031 | 105.185071958352 | 39.5609674411332 | 39.0789203280663 | Cys1 |
| **ENSMUSG00000062580** | 0.141661296722514 | 0.145448169378971 | 0.780607990324772 | 4423.60315632204 | 3794.36288101186 | 3837.87457346945 | 4449.52423756072 | 3863.01251484977 | 3534.70765069345 | 3693.74506529317 | 3868.81311247856 | Timm17a |
| **ENSMUSG00000062590** | -0.174605378299437 | 0.475732048589629 | 0.897549741595057 | 57.8248778604189 | 45.6072808192024 | 52.0550188756646 | 52.523894878457 | 74.4571573156986 | 62.3771938357668 | 40.6020455316893 | 57.5328549274309 | Armc9 |
| **ENSMUSG00000062591** | -0.613054040941232 | 0.292679692382753 | 0.851785577030345 | 28.9124389302094 | 22.8036404096012 | 12.3039135524298 | 30.0136542162612 | 51.4630352034976 | 67.2695227640623 | 10.4107809055614 | 15.1973579053591 | Tubb4a |
| **ENSMUSG00000062593** | -0.240514719348324 | 0.472948139280891 | 0.89728597737823 | 14.4562194651047 | 22.8036404096012 | 24.6078271048596 | 15.9447538023887 | 19.7092475247437 | 25.6847268735511 | 22.903717992235 | 23.8815624227072 | Lilrb4a |
| **ENSMUSG00000062604** | -0.0405420784626603 | 0.839099975036999 | 0.96956148850045 | 1003.80373910821 | 1163.80007661858 | 697.537252933905 | 842.258171443829 | 789.464859185569 | 839.034411202668 | 1073.35151136338 | 1111.57817822055 | Srpk2 |
| **ENSMUSG00000062609** | -0.485824542275364 | 0.396909618900765 | 0.879613019351548 | 16.2632468982428 | 13.8450673915436 | 8.51809399783602 | 11.2551203310979 | 29.5638712871156 | 26.9078091056249 | 6.24646854333682 | 7.59867895267955 | Kcnj15 |
| **ENSMUSG00000062611** | -0.198160616452864 | 0.342640227820647 | 0.860161352689099 | 58.7283915769879 | 46.4216965481167 | 46.3762895437739 | 55.3376749612315 | 54.7479097909548 | 62.3771938357668 | 59.3414511616998 | 60.7894316214364 | Rps3a2 |
| **ENSMUSG00000062619** | -0.0815544938165379 | 0.704209630061172 | 0.946931219685582 | 343.335212296237 | 283.416673662186 | 275.418372596698 | 395.805064976944 | 378.855535753408 | 441.532685778663 | 300.871568170723 | 252.927456567762 | 2310039H08 |
| **ENSMUSG00000062624** | -0.309110330192338 | 0.317321372856357 | 0.854351922245337 | 33.4300075130547 | 43.9784493613737 | 20.8220075502658 | 30.0136542162612 | 32.8487458745729 | 40.3617136584374 | 35.3966550789086 | 51.0197015394199 | Cyp2c67 |
| **ENSMUSG00000062627** | 0.0311333561416288 | 0.863665500709192 | 0.974868041734905 | 113.8427282877 | 145.780415475665 | 149.539872406455 | 105.047789756914 | 125.920192519196 | 114.969729814943 | 138.463386043966 | 123.74991437221 | Mysm1 |
| **ENSMUSG00000062646** | 0.0627020679357995 | 0.796930582698185 | 0.962278883153809 | 234.010052591383 | 374.631235300591 | 282.043556817237 | 265.43325447506 | 226.656346534553 | 193.24699266767 | 342.514691792969 | 344.111603999917 | Ganc |
| **ENSMUSG00000062647** | -0.00474656946125084 | 0.976668127134284 | 0.996134981677394 | 8561.69597820827 | 6296.24800023667 | 7258.36254104494 | 8722.7182566009 | 7790.62756325288 | 9395.71770679139 | 6705.58398127208 | 7049.40301695729 | Rpl7a |
| **ENSMUSG00000062661** | 0.107988306380805 | 0.566819785914825 | 0.919931558693298 | 226.78194285883 | 222.335493993612 | 176.98706417726 | 229.79204009325 | 213.516848184724 | 140.654456688494 | 218.626399016789 | 220.361689627707 | Ncs1 |
| **ENSMUSG00000062683** | 0.163389036100416 | 0.0590950672823645 | 0.712406493939829 | 4998.23788005996 | 4274.0537453424 | 4767.29327412223 | 5048.85939519168 | 4279.09662926103 | 4374.9651441282 | 4150.77834704732 | 4238.97733003052 | Atp5g2 |
| **ENSMUSG00000062691** | 0.0848197396905988 | 0.556800559518279 | 0.916239900758826 | 1168.24323552378 | 816.044560372157 | 899.132144216024 | 1103.93971914186 | 894.580845984202 | 985.804279051531 | 918.230875870512 | 960.690124731629 | Cebpzos |
| **ENSMUSG00000062694** | -0.0678062532409809 | 0.643234256826578 | 0.934616627390525 | 1116.74295367934 | 913.774447841877 | 784.611102689562 | 1098.31215897631 | 1016.12120572012 | 1120.34332457966 | 1018.1743725639 | 947.663817955607 | Cav3 |
| **ENSMUSG00000062729** | 0.10794661824742 | 0.469116675818458 | 0.895328112556867 | 206.001127377742 | 202.789516499668 | 196.862616838877 | 189.461192240149 | 186.142893289246 | 199.362403828039 | 205.092383839559 | 147.631476794917 | Ppox |
| **ENSMUSG00000062743** | -0.0823435802062054 | 0.804280401976764 | 0.963461896828986 | 20.780815481088 | 21.9892246806869 | 21.7684624389143 | 21.5723139679377 | 25.1840385038392 | 13.4539045528125 | 29.1501865355718 | 22.7960368580387 | Zfp677 |
| **ENSMUSG00000062753** | 0.0539758429251735 | 0.814845298941733 | 0.966177091475091 | 174.378147297826 | 116.461449234749 | 116.413951303759 | 184.771558768858 | 150.009272827216 | 174.900759186562 | 116.600746142287 | 129.177542195552 | AI413582 |
| **ENSMUSG00000062760** | 0.561487465863757 | 0.159620865469102 | 0.790303302997254 | 30.7194663633475 | 19.5459774939439 | 12.3039135524298 | 20.6343872736795 | 13.1394983498292 | 13.4539045528125 | 13.5340151772298 | 16.2828834700276 | 1810041L15 |
| **ENSMUSG00000062761** | 0.247514115762788 | 0.105112837441051 | 0.762094734041179 | 738.17070643691 | 795.684167149299 | 840.451941119821 | 786.920496482597 | 640.550544554172 | 508.802208542726 | 746.45299092875 | 765.295523091298 | Zfp512 |
| **ENSMUSG00000062762** | 0.107350020849782 | 0.30476597123439 | 0.851785577030345 | 1767.27282960905 | 1690.72705322615 | 1848.42639753042 | 1683.5784161934 | 1478.19356435578 | 1521.51429669988 | 1654.2730858937 | 1833.45267872511 | Ei24 |
| **ENSMUSG00000062785** | -0.355878965592942 | 0.472127437082911 | 0.897171173241185 | 18.9737880479499 | 58.6379324818317 | 49.2156542097192 | 23.448167356454 | 22.994122112201 | 29.3539735697726 | 80.1630129728225 | 59.7039060567679 | Kcnc3 |
| **ENSMUSG00000062797** | 0.00663743553660326 | 0.945714052908291 | 0.990505437194639 | 730.942596704357 | 657.233493233863 | 661.571967165264 | 649.983199120906 | 720.482492848966 | 676.364474336844 | 629.852244786463 | 661.085068883121 | Hikeshi |
| **ENSMUSG00000062822** | -0.0810611588804811 | 0.619026280550198 | 0.929568738467614 | 228.588970291968 | 157.99665140938 | 173.201244622666 | 186.647412157374 | 177.383227722694 | 200.585486060113 | 211.338852382896 | 199.736703899005 | 4833420G17 |
| **ENSMUSG00000062825** | -0.657901221947355 | 0.0222978515885991 | 0.648762042745232 | 1784.43959022386 | 1614.1719747082 | 1874.92713441257 | 1826.14327372064 | 3879.43688778706 | 3829.47046862325 | 1695.91620951595 | 1797.63033509105 | Actg1 |
| **ENSMUSG00000062861** | -0.0491955638956402 | 0.807443538577263 | 0.964026716319739 | 84.0267756409212 | 89.5857301805761 | 70.9841166486335 | 80.6616957062019 | 64.6025335533267 | 81.9465095489486 | 99.9434966933891 | 90.0986218674861 | Zfp28 |
| **ENSMUSG00000062866** | 0.187880825540568 | 0.320407154113296 | 0.855228644171892 | 1287.50704611089 | 1571.00794107574 | 1680.90388223964 | 1261.51140377723 | 1108.09769416893 | 976.01962119494 | 1461.67363914082 | 1545.78840408795 | Phactr2 |
| **ENSMUSG00000062867** | 0.0581099375385984 | 0.738763809077486 | 0.954383574136061 | 996.575629375657 | 897.48613326359 | 1042.99328729059 | 1121.76032633276 | 1166.13047854734 | 1133.79722913247 | 833.903550535465 | 765.295523091298 | Impdh2 |
| **ENSMUSG00000062901** | 0.261464824691276 | 0.446172258026347 | 0.89108031640712 | 4215.79500151116 | 6737.66132530824 | 6712.25807029478 | 4211.29085721914 | 3627.59650274867 | 2065.78588997275 | 6399.50702264857 | 6157.10100279978 | Klhl24 |
| **ENSMUSG00000062906** | 0.17159796796857 | 0.260420126810861 | 0.840316047847749 | 161.728955265859 | 158.811067138294 | 181.719338620502 | 186.647412157374 | 158.768938393769 | 176.123841418636 | 141.586620315635 | 135.690695583563 | Hdac10 |
| **ENSMUSG00000062908** | 0.296554064427425 | 0.118780265806343 | 0.769868040577702 | 45786.4611008529 | 45156.0945053839 | 50345.7213467655 | 45347.8177406878 | 33466.3022970149 | 26646.0695079611 | 44976.6556682062 | 46867.5662545628 | Acadm |
| **ENSMUSG00000062929** | 0.158305680306648 | 0.240665577569809 | 0.832048466795696 | 8280.7032123553 | 7592.79784066829 | 7491.19044365246 | 8509.8088970043 | 6231.40709240648 | 6180.23451866921 | 7998.6029697428 | 8150.12593953115 | Cfl2 |
| **ENSMUSG00000062931** | 0.0606301645504329 | 0.747516640917785 | 0.956119556124081 | 233.106538874814 | 174.284965987666 | 178.879973954556 | 200.716312571247 | 164.243729372865 | 244.616446414772 | 165.531416398426 | 181.282769299641 | Zfp938 |
| **ENSMUSG00000062933** | 0.212631314760604 | 0.24456822895517 | 0.832048466795696 | 163.535982698997 | 130.306516626293 | 160.897331070236 | 173.51643843776 | 136.869774477387 | 167.562265794119 | 112.436433780063 | 125.920965501547 | Gm10123 |
| **ENSMUSG00000062937** | 0.0434528974029952 | 0.750634893076862 | 0.956197192463153 | 439.107666252556 | 495.164763179912 | 484.584902988005 | 492.411514485535 | 465.357233223116 | 373.040080782527 | 490.34778065194 | 524.308847734889 | Mtap |
| **ENSMUSG00000062944** | -0.0554619324656237 | 0.814942332079797 | 0.966177091475091 | 56.0178504272808 | 46.4216965481167 | 60.5731128735006 | 76.9099889291692 | 74.4571573156986 | 53.8156182112498 | 57.2592949805875 | 62.9604827507734 | 9130023H24 |
| **ENSMUSG00000062949** | -0.0436569963966434 | 0.826120230607367 | 0.967195718403264 | 139.141112351633 | 193.83094348161 | 199.701981504822 | 157.571684635371 | 225.561388338734 | 160.223772401676 | 149.915245040084 | 175.855141476298 | Atp11c |
| **ENSMUSG00000062960** | -0.410435323936001 | 0.0822973430990724 | 0.739855948845396 | 3086.40285579986 | 3000.30754532039 | 3825.57065991702 | 4029.33307853306 | 6369.37182507969 | 5254.3612689893 | 3078.4679137745 | 3827.56314102116 | Kdr |
| **ENSMUSG00000062963** | 0.112161124609904 | 0.424846668133183 | 0.885066977635295 | 1976.88801185307 | 1542.50339056374 | 1719.70853267423 | 2005.28727232395 | 1618.34821342063 | 1974.05472256721 | 1505.39891894417 | 1605.49231014472 | Ufc1 |
| **ENSMUSG00000062980** | -0.221333859363384 | 0.0953731042394985 | 0.757085157577544 | 383.089815825275 | 399.878122896935 | 422.118880337207 | 374.232751009006 | 513.535393839156 | 507.579126310652 | 390.404283958551 | 430.953649173397 | Cped1 |
| **ENSMUSG00000062981** | 0.114167451347644 | 0.266461177709794 | 0.843513595393324 | 7453.98816169462 | 6404.56529218228 | 6750.11626584072 | 7298.94553471701 | 5687.21286908439 | 6565.50542177248 | 6744.10387062265 | 6786.70583030751 | Mrpl42 |
| **ENSMUSG00000062995** | 0.0159255209998859 | 0.940025965020181 | 0.989269716906553 | 210.518695960587 | 204.418347957496 | 256.489274823729 | 226.978260010475 | 236.510970296925 | 280.085831144914 | 226.955023741238 | 145.46042566558 | Ica1 |
| **ENSMUSG00000062997** | 0.174446976564861 | 0.338428139293445 | 0.858570960161243 | 4639.54293458205 | 3025.55443291673 | 3649.53005062841 | 4450.46216425498 | 3454.59310780925 | 4310.14178582828 | 3089.91977277061 | 3115.45837059862 | Rpl35 |
| **ENSMUSG00000063015** | 0.109464550610974 | 0.306987472372413 | 0.851785577030345 | 4234.76878955911 | 4721.16798051636 | 5464.83052705613 | 4631.4820162468 | 4163.03106050421 | 4587.78145250905 | 4390.22630787523 | 4519.042925715 | Ccni |
| **ENSMUSG00000063019** | -0.227437513773609 | 0.209939041067712 | 0.82165491136391 | 282.799793286111 | 280.159010746529 | 268.793188376159 | 329.212269684615 | 278.119381738051 | 451.317343635254 | 322.734208072402 | 308.289260365856 | Manbal |
| **ENSMUSG00000063047** | 0.122138631849484 | 0.473218492647199 | 0.89728597737823 | 118.360296870545 | 136.007426728693 | 133.450139299431 | 124.744250336335 | 109.49581958191 | 97.8465785659088 | 121.806136595068 | 141.118323406906 | Zfp780b |
| **ENSMUSG00000063049** | -0.0133118944549936 | 0.943022185091413 | 0.990505437194639 | 89.4478579403355 | 84.6992358070902 | 107.895857305923 | 90.9788893430416 | 90.881530252985 | 113.746647582869 | 81.2040910633787 | 91.1841474321546 | Ing2 |
| **ENSMUSG00000063052** | -0.0872677882787286 | 0.632297285584008 | 0.931825800800958 | 109.325159704854 | 122.976775066064 | 122.09268063565 | 106.92364314543 | 108.400861386091 | 99.0696607979826 | 136.381229862854 | 145.46042566558 | Lrrc40 |
| **ENSMUSG00000063060** | -0.759663903501071 | 0.0105694424489345 | 0.635862253526407 | 299.966553900923 | 237.809392842984 | 298.133289924261 | 290.75727522003 | 705.153078107498 | 593.194882555822 | 266.515991182371 | 343.026078435248 | Sox7 |
| **ENSMUSG00000063065** | -0.327712842044396 | 0.200545100209772 | 0.812921911423739 | 992.961574509381 | 972.412380323708 | 1013.65318574249 | 1242.75286989206 | 1581.11963476278 | 1887.21588408997 | 895.327157878277 | 935.723036744253 | Mapk3 |
| **ENSMUSG00000063077** | 0.158092725887767 | 0.486459299585413 | 0.899236955296196 | 4185.97904886439 | 5723.7137428099 | 4811.7766538887 | 4016.20210481345 | 3345.09728822734 | 2980.651399564 | 4985.72297567334 | 5480.81857601129 | Kif1b |
| **ENSMUSG00000063108** | -0.00823605488932485 | 0.967846234615072 | 0.994406701590602 | 61.4389327266951 | 67.5965054998893 | 62.4660226507975 | 50.6480414899407 | 61.3176589658694 | 63.6002760678407 | 54.1360607089191 | 65.1315338801105 | Zfp26 |
| **ENSMUSG00000063145** | -0.00146405947954334 | 0.995897882539077 | 0.999262355603273 | 68.6670424592474 | 54.56585383726 | 87.0738497556571 | 43.1446279358754 | 75.5521155115177 | 46.4771248188067 | 57.2592949805875 | 73.8157383974585 | Bbs5 |
| **ENSMUSG00000063146** | -0.648442741685287 | 0.0110860240002081 | 0.635862253526407 | 120.167324303683 | 127.048853710635 | 104.110037751329 | 166.013024883695 | 227.751304730372 | 276.416584448692 | 140.545542225078 | 167.17093695895 | Clip2 |
| **ENSMUSG00000063160** | -0.62253259221459 | 0.0638100022349394 | 0.716991331547963 | 73.1846110420926 | 98.5443031986338 | 85.1809399783602 | 83.4754757889764 | 180.668102310151 | 188.354663739374 | 84.3273253350471 | 71.6446872681215 | Numbl |
| **ENSMUSG00000063171** | -0.351668286915375 | 0.157779071090567 | 0.789230661864335 | 467.116591466196 | 342.054606144018 | 388.046504345863 | 429.570425970238 | 551.858930692825 | 736.295503708463 | 317.528817619622 | 471.118095066132 | Rps4l |
| **ENSMUSG00000063172** | 0.106854174235141 | 0.62233183433021 | 0.93078489223231 | 121.070838020252 | 100.173134656462 | 76.6628459805242 | 109.737423228205 | 109.49581958191 | 105.185071958352 | 90.5737938783839 | 73.8157383974585 | Hspb11 |
| **ENSMUSG00000063179** | -0.0532658225184649 | 0.754850947548918 | 0.956497786093635 | 497.836057829544 | 377.074482487334 | 339.777305024792 | 453.018593326692 | 410.609323432161 | 491.679057293692 | 366.45948787576 | 462.433890548784 | Pstk |
| **ENSMUSG00000063193** | -0.309086958028788 | 0.465058553388641 | 0.895328112556867 | 9.03513716569045 | 10.5874044758863 | 22.7149173275627 | 18.7585338851632 | 13.1394983498292 | 19.5693157131818 | 19.7804837205666 | 22.7960368580387 | Cd300lb |
| **ENSMUSG00000063200** | 0.0518022009870137 | 0.646790671030009 | 0.935063240511793 | 1760.0447198765 | 1484.67987381082 | 1487.82708495536 | 1830.83290719193 | 1623.82300439972 | 1624.25320419409 | 1615.75319654312 | 1467.63056343182 | Nol7 |
| **ENSMUSG00000063229** | -0.14477201627977 | 0.245196158366713 | 0.832048466795696 | 21854.1897763721 | 18889.5584164389 | 19073.9053709321 | 18567.1968395346 | 22242.9807898691 | 25344.7100130345 | 19512.9266512937 | 19560.0851497618 | Ldha |
| **ENSMUSG00000063232** | -0.338958915380095 | 0.56135465396719 | 0.917031459819748 | 11.7456783153976 | 13.0306516626293 | 2.83936466594534 | 15.0068271081306 | 9.85462376237187 | 9.78465785659088 | 9.36970281500523 | 24.9670879873757 | Serpina11 |
| **ENSMUSG00000063235** | 0.0503963223670696 | 0.742497011494276 | 0.954383574136061 | 620.713923282934 | 500.051257553398 | 490.263632319895 | 659.362466063487 | 514.630352034976 | 633.556596214259 | 493.471014923609 | 551.446986851602 | Ptpmt1 |
| **ENSMUSG00000063236** | -0.0332021725038409 | 0.855548022626463 | 0.973387788552416 | 262.922491521592 | 231.294067011669 | 225.25626349833 | 271.998741334867 | 280.309298129689 | 316.77829810713 | 213.421008564008 | 205.164331722348 | 1110038F14 |
| **ENSMUSG00000063253** | 0.122086343912483 | 0.319411002948883 | 0.854670142081901 | 603.547162668122 | 532.627886709971 | 554.62256474799 | 584.328330522834 | 473.02194059385 | 470.886659348436 | 593.414511616998 | 551.446986851602 | Scoc |
| **ENSMUSG00000063268** | -0.134205028820354 | 0.555381745935062 | 0.916239900758826 | 105.711104838578 | 179.171460361152 | 118.306861081056 | 136.937297361692 | 188.332809680885 | 150.439114545085 | 130.134761319517 | 124.835439936878 | Parp10 |
| **ENSMUSG00000063273** | 0.0637676323245082 | 0.494878372495833 | 0.900200915538898 | 2377.14458829316 | 2260.00364773726 | 2164.542330339 | 2411.40953093773 | 1999.39366556567 | 2156.29397514621 | 2307.0290486724 | 2351.24837307199 | Naa15 |

| | | | | | | | | | | | | |
|---|---|---|---|---|---|---|---|---|---|---|---|---|
| **ENSMUSG00000063275** | -0.0134822774404534 | 0.921156937777252 | 0.986713918026868 | 625.231491865779 | 552.988279932829 | 479.852628544763 | 610.590277962063 | 590.182467546493 | 510.025290774799 | 547.607075632528 | 641.545608719088 | Hacd1 |
| **ENSMUSG00000063281** | 0.116134751344791 | 0.541228251235378 | 0.913837378403038 | 131.913002619081 | 118.904696421492 | 165.629605513478 | 152.88205116408 | 127.015150715015 | 112.523565350795 | 121.806136595068 | 162.828834700276 | Zfp35 |
| **ENSMUSG00000063286** | -0.0562691695162604 | 0.647144255473764 | 0.935063240511793 | 192.448421629207 | 184.872370463553 | 204.434255948065 | 185.709485463116 | 205.85214081399 | 204.254732756335 | 197.804837205666 | 189.966973816989 | Gm8995 |
| **ENSMUSG00000063296** | 0.0784222196690497 | 0.711230840636769 | 0.950020479830613 | 310.808718499751 | 279.344595017615 | 313.276568142636 | 299.198615468353 | 228.846262926191 | 201.808568292187 | 348.761160336306 | 358.223436340608 | Tmem117 |
| **ENSMUSG00000063316** | 0.0564674399650706 | 0.777755905569006 | 0.960041127374359 | 7849.72716955186 | 5347.45367605148 | 6241.86999063651 | 7629.09573109588 | 6527.04580527764 | 8617.83740719242 | 5351.14138545854 | 5534.00932868005 | Rpl27 |
| **ENSMUSG00000063317** | 0.210025606677227 | 0.22602024295796 | 0.832048466795696 | 287.317361868956 | 332.281617397046 | 323.687571917769 | 272.936668029125 | 267.16979977986 | 195.693157131818 | 270.680303544596 | 316.973464883204 | Usp31 |
| **ENSMUSG00000063320** | 0.187341530221231 | 0.273343350724087 | 0.844738346035895 | 294.545471601509 | 247.582381589956 | 249.86409060319 | 347.03287687552 | 266.074841584041 | 287.424324537357 | 229.03717992235 | 218.19063849837 | 1190007I07R |
| **ENSMUSG00000063334** | 0.050176644151497 | 0.737927742995197 | 0.954383574136061 | 533.976606492306 | 592.080234920717 | 510.139184981513 | 492.411514485535 | 489.446313531136 | 410.955629976817 | 596.537745888666 | 557.960140239613 | Krr1 |
| **ENSMUSG00000063358** | 0.0334900054461988 | 0.817858420681681 | 0.966315773308182 | 2140.42399455207 | 2422.07237779121 | 2303.67119897032 | 1902.11533595555 | 2084.80040483956 | 1728.21519392036 | 2348.67217229464 | 2404.43912574074 | Mapk1 |
| **ENSMUSG00000063362** | 0.0369473506359336 | 0.81711191336993 | 0.966315773308182 | 149.983276950461 | 160.439898596123 | 135.343049076728 | 166.013024883695 | 177.383227722694 | 138.208292224346 | 134.299073681742 | 146.545951230249 | Alg11 |
| **ENSMUSG00000063382** | -0.0718860053431938 | 0.759766084017939 | 0.957357521291862 | 735.460165287203 | 969.154717408051 | 830.987392233336 | 556.19052969509 | 865.016974697087 | 554.056251129458 | 1042.11916864669 | 788.091559949337 | Bcl9l |
| **ENSMUSG00000063388** | 1.12363906565761 | 0.0209456599564535 | 0.648762042745232 | 10.8421645988285 | 22.8036404096012 | 29.3401015481019 | 24.3860940507122 | 4.37983278327639 | 14.6769867848863 | 12.4929370866736 | 8.68420451734806 | BC023105 |
| **ENSMUSG00000063406** | -0.18913967873518 | 0.431016127466254 | 0.886278906070506 | 1438.39383677792 | 1086.43058237171 | 1312.7329305554 | 1510.06197775564 | 1920.5566754667 | 1981.39321595965 | 1082.72121417838 | 1112.66370378522 | Tmed5 |
| **ENSMUSG00000063410** | 0.0649670923177333 | 0.811968076541329 | 0.965224347191244 | 868.276681622852 | 1178.45955973903 | 1195.37252436299 | 709.07258085917 | 690.91862156185 | 650.679747463293 | 1189.95225750566 | 1245.09782267478 | Stk24 |
| **ENSMUSG00000063415** | 0.483928959962165 | 0.306541537412297 | 0.851785577030345 | 131.913002619081 | 94.4722245540621 | 520.550188756646 | 295.446908691321 | 234.321053905287 | 183.462334811079 | 114.518589961175 | 212.763010675027 | Cyp26b1 |
| **ENSMUSG00000063428** | 0.24556171241164 | 0.28965268892632 | 0.851785577030345 | 1094.15511076511 | 1205.33527879321 | 1026.90355418356 | 1107.69142591889 | 710.627869086594 | 629.887349518038 | 1204.52735077345 | 1194.07812113536 | Ddo |
| **ENSMUSG00000063439** | -0.109530720033274 | 0.635122367180652 | 0.932543559775713 | 170.764092431549 | 102.616381843205 | 132.503684410783 | 130.371810501884 | 140.154649064844 | 194.470074899744 | 119.723980413956 | 124.835439936878 | B9d2 |
| **ENSMUSG00000063442** | -0.422404452856667 | 0.160886766072574 | 0.792229941270026 | 32.5264937964856 | 22.8036404096012 | 21.7684624389143 | 37.5170677703264 | 40.5134532453066 | 29.3539735697726 | 36.4377331694648 | 46.6775992807458 | Gm7117 |
| **ENSMUSG00000063445** | 0.0369160687956349 | 0.896384525720045 | 0.984095879090375 | 32.5264937964856 | 26.0613033252585 | 38.8046504345863 | 31.8895076047775 | 37.2285786578493 | 23.2385624094033 | 34.3555769883525 | 30.3947158107182 | Nmral1 |
| **ENSMUSG00000063450** | 0.0560334328636402 | 0.652399840681819 | 0.936720465895231 | 489.704434380422 | 503.308920469055 | 519.603733867997 | 493.349441179793 | 560.618596259378 | 439.086521314516 | 426.842017128016 | 502.598336441519 | Syne2 |
| **ENSMUSG00000063455** | -0.253648590035952 | 0.441678307357409 | 0.89108031640712 | 24.3948703473642 | 30.1333819698302 | 25.5542819935081 | 13.1309737196143 | 28.4689130912965 | 24.4616446414772 | 29.1501865355718 | 29.3091902460497 | D630045J12I |
| **ENSMUSG00000063457** | 0.116569308390274 | 0.55074346396021 | 0.914699131434573 | 9202.28720325572 | 6190.37395547781 | 7558.3887407465 | 9464.61827175911 | 7883.6990098975 | 9409.1716113442 | 6281.86519841573 | 6325.35746532339 | Rps15 |
| **ENSMUSG00000063458** | -0.240421806760218 | 0.42099043258522 | 0.882568531013176 | 30.7194663633475 | 35.0198763433161 | 26.5007368821565 | 22.5102406621959 | 45.9882442244021 | 28.1308913376988 | 29.1501865355718 | 32.5657669400552 | Lrmda |
| **ENSMUSG00000063480** | 0.031159424947246 | 0.808300639212115 | 0.964026716319739 | 1152.8835023421 | 1033.49355999228 | 1023.11773462897 | 1296.21469146478 | 1253.72713421287 | 1100.77400886647 | 1022.33868492613 | 1032.33481199975 | Snu13 |
| **ENSMUSG00000063488** | 0.125283447665674 | 0.629819026998829 | 0.931825800800958 | 67.7635287426784 | 92.8433930962335 | 87.0738497556571 | 99.4202295913651 | 85.4067392738896 | 47.7002070508805 | 81.2040910633787 | 103.124928643508 | Zkscan7 |
| **ENSMUSG00000063506** | -0.755492652735743 | 0.186859936666495 | 0.807540361382744 | 7.22810973255236 | 6.51532583131463 | 8.51809399783602 | 11.2551203310979 | 20.8042057205628 | 23.2385624094033 | 6.24646854333682 | 6.51315338801105 | Arhgap22 |
| **ENSMUSG00000063511** | 0.00045684819171937 | 0.996679201220904 | 0.999262355603273 | 1030.00563688871 | 947.165492727364 | 963.491076644119 | 1072.05021153708 | 1023.78591309086 | 1117.89716011551 | 983.818795575549 | 886.874386334171 | Snrnp70 |
| **ENSMUSG00000063524** | -0.242125461604824 | 0.0628309771743928 | 0.715615581523459 | 3245.42126991601 | 3032.88417447696 | 3105.31848965555 | 3134.55101221077 | 3833.44864356266 | 4459.35781814129 | 3267.94412625571 | 3246.80696392351 | Eno1 |
| **ENSMUSG00000063550** | -0.00406819839201151 | 0.981991822691755 | 0.996940244108351 | 397.54603529038 | 601.038807938775 | 550.836745193396 | 442.701399689852 | 517.915226622433 | 401.170972120226 | 524.703357640293 | 553.618037980939 | Nup98 |
| **ENSMUSG00000063558** | 0.0688148967194461 | 0.687083489034212 | 0.944452327903597 | 1450.13951509332 | 1915.5057944065 | 1580.57966404291 | 1555.08245908003 | 1287.67083828326 | 1317.25956394355 | 1908.2961399894 | 1684.73567636552 | Aox1 |
| **ENSMUSG00000063564** | -0.534062818843086 | 0.102722904404774 | 0.759800953500741 | 28.0089252136404 | 32.5766291565731 | 17.036187995672 | 25.3240207449704 | 43.7983278327639 | 48.9232892829544 | 28.1091084450157 | 29.3091902460497 | Col23a1 |
| **ENSMUSG00000063568** | 0.244015991092037 | 0.47404229124309 | 0.897549741595057 | 43.3686583953142 | 39.0919549878878 | 25.5542819935081 | 42.2067012416173 | 47.0832024202212 | 35.4693847301419 | 20.8215618111227 | 23.8815624227072 | Jazf1 |
| **ENSMUSG00000063576** | 0.0474165796815575 | 0.679505559253366 | 0.944452327903597 | 1861.23825613223 | 1758.32355872604 | 1967.67971350012 | 2050.30775364834 | 2099.03486138521 | 1701.30738481474 | 1911.41937426107 | 1677.13699741284 | Klhdc3 |
| **ENSMUSG00000063605** | -0.435840084081567 | 0.229713965270009 | 0.832048466795696 | 99.3865088225949 | 74.1118313312039 | 99.3777633080869 | 132.247663890401 | 181.76306050597 | 207.923979452556 | 70.7933101578173 | 87.9275707381491 | Ccdc102a |
| **ENSMUSG00000063623** | -0.242151385933213 | 0.533767471814406 | 0.909585349466581 | 32.5264937964856 | 13.8450673915436 | 15.1432782183751 | 23.448167356454 | 19.7092475247437 | 34.2463024980681 | 26.0269522639034 | 20.6249857287016 | C230062I16F |
| **ENSMUSG00000063659** | 0.00737278727377829 | 0.979416817723325 | 0.996872525500296 | 683.056369726198 | 794.05533569147 | 803.540200462531 | 679.996853337167 | 522.295059405709 | 421.963370065482 | 945.298906224972 | 1055.13084885779 | Zbtb18 |
| **ENSMUSG00000063663** | 0.169674434659124 | 0.466190302165671 | 0.895328112556867 | 53.3073092775737 | 70.0397526866323 | 59.6266579848522 | 55.3376749612315 | 47.0832024202212 | 53.8156182112498 | 67.6700758861489 | 43.4210225867403 | Brwd3 |
| **ENSMUSG00000063681** | 0.0987945019192313 | 0.852190274971242 | 0.972951086581502 | 12.6491920319666 | 35.0198763433161 | 43.5369248778286 | 10.3171936368398 | 21.8991639163819 | 12.2308223207386 | 33.3144988977964 | 27.1381391167127 | Crb1 |
| **ENSMUSG00000063683** | -0.291473938517536 | 0.313487973797464 | 0.85295001595893 | 57.8248778604189 | 77.3694942468612 | 61.519567762149 | 76.9099889291692 | 58.0327843784121 | 75.8310983885793 | 69.7522320672611 | 131.348593324889 | Glyat |
| **ENSMUSG00000063684** | 0.178884323168544 | 0.47284112257967 | 0.89728597737823 | 459.888481733644 | 514.710740673856 | 556.515474525287 | 526.176875478828 | 408.419407040523 | 259.293433199658 | 534.073060455298 | 614.407469602375 | Gm13910 |
| **ENSMUSG00000063687** | 0.540814488364626 | 0.3108588006482 | 0.851785577030345 | 7.22810973255236 | 19.5459774939439 | 15.1432782183751 | 21.5723139679377 | 4.37983278327639 | 17.123151249034 | 8.32862472444909 | 14.1118323406906 | Pcdhb5 |
| **ENSMUSG00000063694** | 0.0611353829094091 | 0.532536310421815 | 0.909082872831377 | 26402.4778255806 | 21668.3448834946 | 21899.0732135478 | 24000.6061793721 | 21934.2025786481 | 23848.8804432082 | 22080.2252226051 | 22208.767527553 | Cycs |
| **ENSMUSG00000063696** | -0.353789070504677 | 0.19447842208856 | 0.809698611654466 | 55.1143367107117 | 59.452348210746 | 44.483379766477 | 68.4686486808458 | 65.6974917491458 | 84.3926740130963 | 46.8485140750261 | 94.4407241261602 | Gm8730 |
| **ENSMUSG00000063698** | 0.0633393602500621 | 0.685689537018548 | 0.944452327903597 | 132.81651633565 | 114.018202048006 | 134.396594188079 | 149.130344387048 | 120.445401540101 | 127.200552135681 | 136.381229862854 | 122.664388807541 | Sfxn4 |
| **ENSMUSG00000063714** | 0.34905915886131 | 0.107572655703247 | 0.762287160109383 | 97.5794813894569 | 88.7713144516618 | 86.1273948670087 | 125.682177030594 | 93.0714466446232 | 72.1618516923577 | 62.4646854333682 | 84.6709940441436 | Sp3os |
| **ENSMUSG00000063760** | 0.051913004146924 | 0.70249820335257 | 0.946451777392559 | 235.817080024521 | 249.211213047785 | 256.489274823729 | 229.79204009325 | 237.605928492744 | 195.693157131818 | 261.31060072959 | 240.986675356409 | Rnf217 |
| **ENSMUSG00000063765** | -0.333532222591055 | 0.350106445474185 | 0.861774532843376 | 13.5527057485357 | 18.7315617650296 | 26.5007368821565 | 30.0136542162612 | 29.5638712871156 | 36.6924669622158 | 22.903717992235 | 22.7960368580387 | Chadl |
| **ENSMUSG00000063785** | 0.140765076524253 | 0.238949114358801 | 0.832048466795696 | 459.888481733644 | 390.105134149963 | 393.725233677754 | 463.335786963532 | 399.65974147397 | 391.386314263635 | 401.856142954669 | 354.966859646602 | Utp14a |
| **ENSMUSG00000063787** | 0.156260580492076 | 0.268925320261291 | 0.843793744782845 | 1419.42004872997 | 1059.55486331754 | 1023.11773462897 | 1276.51823088536 | 1109.19265236475 | 1139.91264029284 | 974.449092760544 | 1064.90057893981 | Chchd1 |
| **ENSMUSG00000063800** | -0.0575065666822153 | 0.733953687787526 | 0.954182918936468 | 295.448985318078 | 229.665235553841 | 244.185361271299 | 333.901903155905 | 317.537876787538 | 305.770558018465 | 274.84461590682 | 249.670879873757 | Prpf38a |
| **ENSMUSG00000063801** | 0.142135670442499 | 0.152536791506619 | 0.783412422888014 | 791.478015714483 | 803.828324438442 | 853.702309560899 | 780.35500962279 | 686.538788778574 | 694.710707817952 | 812.040910633787 | 731.644230586574 | Ap3s2 |
| **ENSMUSG00000063802** | -0.120606020532873 | 0.353740643779695 | 0.86286441023679 | 688.477452025612 | 562.761268679801 | 565.033568523123 | 574.011136885995 | 675.589206820383 | 726.510845851873 | 629.852244786463 | 567.72987032163 | Hspbp1 |
| **ENSMUSG00000063808** | 0.15093722017921 | 0.212141445095526 | 0.823370267313127 | 383.089815825275 | 443.856572258309 | 428.744064557746 | 423.004939110431 | 433.603445544362 | 354.693847301419 | 368.541644056872 | 354.966859646602 | Gpatch1 |
| **ENSMUSG00000063810** | 0.617739388971095 | 0.0160705765899073 | 0.63777231095375 | 46.0791995450213 | 52.9370223794314 | 53.001473764313 | 44.0825546301336 | 26.2789966996583 | 25.6847268735511 | 39.5609674411332 | 35.8223436340608 | Alms1 |
| **ENSMUSG00000063838** | -0.836701000764855 | 0.0667953004324837 | 0.719725956118125 | 27.1054114970714 | 30.1333819698302 | 39.7511053232348 | 39.3929211588428 | 72.2672409240604 | 112.523565350795 | 35.3966550789086 | 23.8815624227072 | 1700027J19R |
| **ENSMUSG00000063849** | -0.0432679204789328 | 0.800126002602108 | 0.963461896828986 | 136.430571201926 | 134.378595270864 | 102.217127974032 | 134.123517278917 | 111.685735973548 | 138.208292224346 | 132.216917500629 | 141.118323406906 | Ppcdc |
| **ENSMUSG00000063856** | 0.100347458841759 | 0.697261501414009 | 0.944933386973863 | 2411.47810952278 | 2018.12217624971 | 3440.3635202371 | 2296.98247423824 | 1915.0818844876 | 3294.98353320698 | 1612.62996227146 | 2661.70868456718 | Gpx1 |
| **ENSMUSG00000063870** | 0.262306950263576 | 0.0957969417781097 | 0.757475844708466 | 4374.81341562732 | 4581.90289087201 | 4661.2903265936 | 4412.94509648465 | 3559.70909460788 | 2806.97372260951 | 4418.33541632024 | 4246.5760089832 | Chd4 |
| **ENSMUSG00000063873** | -0.51869671997109 | 0.175869973327174 | 0.799889021902745 | 37.9475760958999 | 43.1640336324594 | 46.3762895437739 | 28.1378008277448 | 68.9823663366031 | 88.0619207093179 | 39.5609674411332 | 27.1381391167127 | Slc24a3 |
| **ENSMUSG00000063882** | 0.146541908436862 | 0.30715397107723 | 0.851785577030345 | 12234.4792360614 | 9127.15707394288 | 10097.7272069903 | 12379.6944375135 | 9756.07752474815 | 11363.6570181982 | 8972.01098441279 | 9513.5460487548 | Uqcrh |
| **ENSMUSG00000063884** | 0.205734069213586 | 0.356381173448805 | 0.864965376583892 | 2555.13679045726 | 2772.27114122437 | 2665.21696643403 | 2432.98184490567 | 1927.12642464161 | 1444.46011607923 | 2674.52961463871 | 2992.79398179108 | Ptcd3 |
| **ENSMUSG00000063885** | -0.463004595181838 | 0.278724549315998 | 0.848075053184273 | 9.03513716569045 | 8.95857301805762 | 11.3574586637814 | 10.3171936368398 | 14.2344565456483 | 18.3462334811079 | 12.4929370866736 | 9.76973008201657 | Gm6498 |
| **ENSMUSG00000063888** | 0.0291805519115546 | 0.762852529112698 | 0.957842155335615 | 1426.64815846252 | 1179.27397546795 | 1197.26543414029 | 1253.0700635289 | 1228.54309570903 | 1231.64380769838 | 1251.37586484848 | 1242.92677154544 | Rpl7l1 |
| **ENSMUSG00000063889** | 0.0154040917255075 | 0.920974645195098 | 0.986713918026868 | 258.404922938747 | 268.757190541728 | 219.57753416644 | 275.750448111899 | 297.828629262794 | 216.485555077073 | 231.119336103462 | 265.953763343784 | Crem |
| **ENSMUSG00000063894** | 0.147309985375581 | 0.63265043346593 | 0.93206560467138 | 133.720030052219 | 205.232763686411 | 168.468970179424 | 173.51643843776 | 102.926070406995 | 91.7311674055395 | 216.544242835676 | 202.993280593011 | Zkscan8 |
| **ENSMUSG00000063895** | 0.135680087586095 | 0.715348225331359 | 0.951176382858794 | 215.036264543433 | 295.632909595901 | 242.292451494002 | 166.013024883695 | 174.098353135236 | 79.5003450848009 | 293.584021536831 | 288.749800201823 | Nupl1 |
| **ENSMUSG00000063897** | -0.47296536207221 | 0.0759590024611456 | 0.735399943172214 | 432.783070236573 | 411.279943101736 | 423.065335225856 | 482.094320848695 | 534.339599559719 | 987.027361283605 | 399.773986773556 | 508.025964264862 | CAAA011183 |
| **ENSMUSG00000063902** | 0.0699814303660015 | 0.647158330732158 | 0.935063240511793 | 128.298947752804 | 129.492100897378 | 143.861143074564 | 137.87522405595 | 122.635317931739 | 136.985209992272 | 112.436433780063 | 142.203848971575 | Gm7964 |
| **ENSMUSG00000063904** | 0.08090714335224 | 0.433804368107511 | 0.887868762931002 | 590.897970636155 | 575.79192034243 | 523.389553422591 | 543.997482669733 | 571.568178217569 | 535.71001764835 | 517.4158110064 | 488.486504100828 | Dpp3 |
| **ENSMUSG00000063919** | 0.0876676244847559 | 0.655382360454573 | 0.937625787683528 | 269.247087537575 | 315.178887089845 | 330.312756138308 | 264.495327780801 | 233.226095709468 | 210.370143916704 | 339.391457521301 | 325.657669400552 | Srrm4 |
| **ENSMUSG00000063931** | -0.0826800429242 | 0.416087242679694 | 0.882192126163777 | 524.037955610046 | 556.245942848486 | 534.747012086372 | 593.707597465416 | 605.511882287961 | 626.218102821816 | 556.976778447533 | 551.446986851602 | Pepd |
| **ENSMUSG00000063952** | 0.279283342912457 | 0.24618779385668 | 0.832048466795696 | 275.571683553559 | 359.157336451219 | 403.189782564238 | 256.053987532478 | 247.460552255116 | 177.34692365071 | 301.91264626128 | 338.683976176574 | Brpf3 |
| **ENSMUSG00000063953** | 0.058010254790568 | 0.801634815686346 | 0.963461896828986 | 65.0529875929712 | 63.5244268553176 | 49.2156542097192 | 43.1446279358754 | 47.0832024202212 | 56.2617826753975 | 54.1360607089191 | 55.3618037980939 | Amd2 |
| **ENSMUSG00000063972** | 0.396150153000835 | 0.411182823970864 | 0.882192126163777 | 11.7456783153976 | 30.9477976987445 | 17.036187995672 | 21.5723139679377 | 5.47479097909548 | 17.123151249034 | 18.7394056300105 | 20.6249857287016 | Nr6a1 |
| **ENSMUSG00000064023** | -0.444591089463536 | 0.228766305866586 | 0.832048466795696 | 133.720030052219 | 57.8235167529173 | 63.4124775394459 | 110.675349922463 | 154.389105610493 | 176.123841418636 | 70.7933101578173 | 96.6117752554972 | Klk8 |
| **ENSMUSG00000064030** | -0.201781068467391 | 0.388462000382919 | 0.876358159057779 | 87.6408305071974 | 61.895595397489 | 89.9132144216024 | 106.92364314543 | 105.115986798633 | 114.969729814943 | 73.9165444294857 | 104.210454208177 | Pym1 |
| **ENSMUSG00000064037** | 0.0989470066438975 | 0.474267711892015 | 0.897549741595057 | 452.660372001092 | 385.218639776477 | 402.24332767559 | 485.846027625728 | 382.140410340865 | 351.024600605198 | 443.499266576914 | 433.124700302735 | Gpn1 |
| **ENSMUSG00000064043** | -0.107798400209756 | 0.821078576773629 | 0.96650690096489 | 14.4562194651047 | 8.95857301805762 | 22.7149173275627 | 31.8895076047775 | 24.0890803080201 | 28.1308913376988 | 20.8215618111227 | 10.8552556466851 | Trerf1 |
| **ENSMUSG00000064061** | -0.0580354731113972 | 0.823963185198034 | 0.966637987288264 | 84.9302893574902 | 92.8433930962335 | 80.448665535118 | 66.5927952923295 | 56.937826182593 | 72.1618516923577 | 119.723980413956 | 89.0130963028176 | Dzip3 |
| **ENSMUSG00000064068** | -0.135706559899893 | 0.34953185850668 | 0.861774532843376 | 449.949830851384 | 392.548381336706 | 375.742590793433 | 353.598363735327 | 455.502609460744 | 501.463715150282 | 388.322127777439 | 383.190524327983 | Mtx1 |
| **ENSMUSG00000064080** | -1.15236045758491 | 0.0077210428239436 | 0.635862253526407 | 957.724539563188 | 958.567312932165 | 1011.76027596519 | 950.119741283517 | 3250.9308833869 | 3604.42333792166 | 953.627530949421 | 811.973122372044 | Fbln2 |
| **ENSMUSG00000064090** | -0.0043673005021577 | 0.984084101087645 | 0.996940244108351 | 131.009488902512 | 90.4001459094905 | 85.1809399783602 | 93.7926694258161 | 76.6470737073368 | 103.961989726278 | 114.518589961175 | 106.381505337514 | Vrk2 |
| **ENSMUSG00000064105** | -0.0927704268460749 | 0.700280740740912 | 0.94561797945347 | 138.237598635064 | 153.110157035894 | 177.933519065908 | 104.109863062656 | 134.679858085749 | 124.754387671534 | 211.338852382896 | 140.032797842237 | Cnnm2 |
| **ENSMUSG00000064120** | 0.00877106923635385 | 0.946149938488875 | 0.990505437194639 | 337.914129996823 | 351.013179162076 | 344.509579468035 | 416.439452250624 | 394.184950494875 | 362.032340693862 | 365.418409785204 | 319.144516012541 | Mocs1 |
| **ENSMUSG00000064127** | 0.202333002453052 | 0.371287723211077 | 0.873374202289323 | 449.949830851384 | 568.462178782201 | 556.515474525287 | 427.694572581722 | 374.475702970131 | 291.093571233579 | 548.648153723084 | 525.394373299558 | Med14 |
| **ENSMUSG00000064128** | -0.353523914488846 | 0.120065966957093 | 0.769868040577702 | 53.3073092775737 | 74.9262470601182 | 67.1982970940397 | 60.0273084325223 | 67.887408140784 | 70.9387694602839 | 81.2040910633787 | 106.381505337514 | Cenpj |
| **ENSMUSG00000064138** | 0.00304990846695277 | 0.985825517134259 | 0.997676971438605 | 210.518695960587 | 289.117583764587 | 318.955297474527 | 245.736793895638 | 269.359716171498 | 236.054870790255 | 299.830490080167 | 256.184033261768 | Fam172a |

| | | | | | | | | | | | | |
|---|---|---|---|---|---|---|---|---|---|---|---|---|
| **ENSMUSG00000064141** | -0.0370234544187371 | 0.954625805805641 | 0.992371505557225 | 11.7456783153976 | 24.4324718674299 | 0.946454888648447 | 18.7585338851632 | 17.5193311331055 | 13.4539045528125 | 12.4929370866736 | 14.1118323406906 | Zfp69 |
| **ENSMUSG00000064145** | 0.232714698694705 | 0.187698137386 | 0.808411512648538 | 1160.11161207465 | 1285.14802022681 | 1325.98329899647 | 1157.40154071457 | 1008.45649834939 | 743.633997100907 | 1232.63645921847 | 1208.18995347605 | Arih2 |
| **ENSMUSG00000064179** | -0.145350094780854 | 0.528663639350615 | 0.908295716206993 | 199.676531361759 | 123.791190794978 | 186.451613063744 | 211.033506208086 | 198.187433443257 | 261.739597663806 | 178.024353485099 | 159.572258006271 | Tnnt1 |
| **ENSMUSG00000064181** | 0.199306502511746 | 0.0762824283224355 | 0.736318922445191 | 3043.03419740454 | 2806.47660183878 | 2992.69035790639 | 3076.39955716677 | 2454.89627502642 | 2252.91747148005 | 2931.67590300608 | 2738.78099965864 | Rab3ip |
| **ENSMUSG00000064202** | -0.0563568158073466 | 0.674873042937293 | 0.944023556644042 | 420.133878204606 | 386.033055505392 | 374.796135904785 | 387.363724728621 | 360.241246424483 | 375.486245246675 | 407.061533407449 | 487.40097853616 | 4430402I18R |
| **ENSMUSG00000064210** | -0.631712843812967 | 0.0533741033648493 | 0.703396849069134 | 461.695509166782 | 447.928650902881 | 524.33600831124 | 457.708226797983 | 1067.58424092362 | 1031.05832163826 | 423.718782856348 | 409.243137880027 | Ano6 |
| **ENSMUSG00000064215** | -0.268727146623069 | 0.304896652157021 | 0.851785577030345 | 468.020105182765 | 361.600583637962 | 401.296872786941 | 436.135912830045 | 573.758094609207 | 739.964750404685 | 334.18606706852 | 361.480013034613 | Ifi27 |
| **ENSMUSG00000064225** | 0.125631487515287 | 0.501616206688561 | 0.900927059989498 | 1516.99953011943 | 1618.24405335277 | 1462.27280296185 | 1589.78574676758 | 1250.44225962541 | 1012.71208815716 | 1614.71211845257 | 1792.20270726771 | Paqr9 |
| **ENSMUSG00000064247** | -0.0488859119736161 | 0.900793731687344 | 0.984802943363903 | 18.9737880479499 | 10.5874044758863 | 15.1432782183751 | 23.448167356454 | 15.3294147414674 | 23.2385624094033 | 14.5750932677859 | 17.3684090346961 | Plcxd1 |
| **ENSMUSG00000064254** | -0.0112257117847345 | 0.935517298920887 | 0.98818087308472 | 842.07478384235 | 691.438953848265 | 733.502538702546 | 863.830485411767 | 889.106055005107 | 852.48831575548 | 723.549272936515 | 690.394259129171 | Ethe1 |
| **ENSMUSG00000064262** | -0.116205447914641 | 0.514495056642809 | 0.904135765061573 | 123.781379169959 | 128.677685168464 | 123.039135524298 | 151.944124469822 | 142.344565456483 | 178.570005882784 | 127.011527047849 | 124.835439936878 | Gimap8 |
| **ENSMUSG00000064264** | -0.214131744118379 | 0.307676042521296 | 0.851785577030345 | 341.528184863099 | 212.56250524664 | 228.095628164276 | 268.247034557834 | 313.158044004262 | 385.270903103266 | 261.31060072959 | 259.440609955773 | Zfp428 |
| **ENSMUSG00000064280** | -0.425228890028402 | 0.125582843893434 | 0.771949105062466 | 24.3948703473642 | 30.9477976987445 | 24.6078271048596 | 29.075727522003 | 30.6588294829347 | 30.5770558018465 | 43.7252798033577 | 41.2499714574033 | Ccdc146 |
| **ENSMUSG00000064284** | 0.138892637636016 | 0.328359023974284 | 0.855977678491063 | 257.501409222178 | 226.407572638183 | 269.739643264807 | 280.44008158319 | 204.757182618171 | 266.631926592101 | 226.955023741238 | 240.986675356409 | Cdpf1 |
| **ENSMUSG00000064289** | 0.00470098555665089 | 0.982461212071389 | 0.996940244108351 | 278.282224703266 | 299.704988240473 | 262.16800415562 | 247.612647284155 | 245.270635863478 | 179.793088114857 | 348.761160336306 | 309.374785930525 | Tank |
| **ENSMUSG00000064294** | -0.537608035369059 | 0.193160561649703 | 0.808863491502674 | 41.5616309621761 | 82.2559886203472 | 51.1085639870161 | 60.0273084325223 | 53.6529515951357 | 77.0541806206532 | 47.8895921655823 | 162.828834700276 | Aox3 |
| **ENSMUSG00000064302** | 0.0573234533938776 | 0.828106354730367 | 0.967319591893432 | 5735.5050727803 | 7903.09023338465 | 6345.03357349919 | 4671.8128640999 | 4387.49749064712 | 4020.27129682678 | 7260.4786035385 | 8026.37602515894 | Clasp1 |
| **ENSMUSG00000064307** | -0.00190536109508621 | 0.992829648253333 | 0.999111773824344 | 74.9916384752307 | 57.8235167529173 | 56.7872933189068 | 65.6548685980713 | 53.6529515951357 | 64.8233582999146 | 74.9576225200418 | 61.8749571861049 | Lrrc51 |
| **ENSMUSG00000064308** | 0.484382034821004 | 0.243471170822562 | 0.832048466795696 | 27.1054114970714 | 15.4738988493722 | 25.5542819935081 | 16.8826804966469 | 15.3294147414674 | 9.78465785659088 | 11.4518589961175 | 23.8815624227072 | 2410137M14 |
| **ENSMUSG00000064326** | 0.110428420934317 | 0.593134635464023 | 0.923935529355527 | 305.387636200337 | 231.294067011669 | 286.775831260479 | 295.446908691321 | 262.789966996583 | 353.470765069345 | 203.010227658447 | 218.19063849837 | Siva1 |
| **ENSMUSG00000064337** | 0.0306323156876743 | 0.854049332585373 | 0.9731539929935 | 237995.451595169 | 180295.354067054 | 190515.6903556 | 218694.491446787 | 217205.762346438 | 248590.24058678 | 170678.506478135 | 173642.840375504 | mt-Rnr1 |
| **ENSMUSG00000064339** | 0.149525190719891 | 0.305701635081158 | 0.851785577030345 | 698806.42083343 | 531870.480082081 | 528826.936757876 | 618229.690886796 | 542339.364138374 | 624107.062889257 | 458976.974549212 | 518216.87826597 | mt-Rnr2 |
| **ENSMUSG00000064340** | 0.361709426862897 | 0.0286341525938856 | 0.665956123973759 | 805.030721463019 | 867.35275129376 | 759.056820696054 | 621.845398293161 | 494.921104510232 | 713.05694129906 | 622.56469815257 | 547.104884592928 | mt-Tl1 |
| **ENSMUSG00000064341** | 0.0154501560345559 | 0.918850889673303 | 0.986713918026868 | 1080821.9588497 | 936104.912712979 | 986317.675648542 | 1086346.09021096 | 986401.8503692 | 1289227.74226665 | 877051.031998564 | 893349.546327418 | mt-Nd1 |
| **ENSMUSG00000064342** | 0.204566589632337 | 0.355261346390174 | 0.864215962387542 | 2584.04922938747 | 1829.17772714158 | 1932.66088262013 | 2120.6522557177 | 2101.22477777685 | 2443.71829968357 | 1359.64798626631 | 1443.74900100912 | mt-Ti |
| **ENSMUSG00000064343** | 0.270185115929574 | 0.107277188051645 | 0.762287160109383 | 1115.83943996277 | 1079.10084081149 | 908.596693102509 | 705.320874082137 | 751.1413223319 | 834.142082274372 | 733.960053842076 | 840.196787053425 | mt-Tq |
| **ENSMUSG00000064344** | 0.33073487191557 | 0.204141320714436 | 0.816447723790243 | 317.133314515735 | 443.856572258309 | 420.22597055991 | 245.736793895638 | 213.516848184724 | 221.377884005369 | 375.829190690765 | 323.486618271215 | mt-Tm |
| **ENSMUSG00000064345** | 0.134931399620975 | 0.242449093296274 | 0.832048466795696 | 710776.170550536 | 685132.932859281 | 705989.095089536 | 596189.351498423 | 532650.07906357 | 649526.380918448 | 601348.567745126 | 673659.797024241 | mt-Nd2 |
| **ENSMUSG00000064346** | 0.266380839570929 | 0.141035018912282 | 0.779711473022872 | 227.685456575399 | 213.376920975554 | 243.238906382651 | 255.11606083822 | 170.813478547779 | 168.785348026193 | 181.147587756768 | 259.440609955773 | mt-Tw |
| **ENSMUSG00000064347** | 0.356093989223677 | 0.0437845534764421 | 0.691603097094229 | 2010.31801936613 | 2141.91336704468 | 1796.37137865475 | 1778.30901231347 | 1257.01200880032 | 1280.56709698133 | 1492.9059818575 | 2006.0512435074 | mt-Ta |
| **ENSMUSG00000064348** | 0.577174330529733 | 0.0271962612127069 | 0.664353746352133 | 7367.25084490399 | 8564.39580526308 | 9207.11315677209 | 7191.08396487732 | 4108.28315071325 | 3340.23757579371 | 6456.76631762916 | 7763.67883850917 | mt-Tn |
| **ENSMUSG00000064349** | 0.746101999143634 | 5.74265682582098E-0 | 0.398827516553267 | 1378.76193148436 | 1463.50506485905 | 1593.83003248398 | 1442.53125576905 | 717.197618261509 | 665.35673424818 | 1045.24240291836 | 1075.75583458649 | mt-Tc |
| **ENSMUSG00000064350** | 0.662739812054761 | 0.00047463118814118 | 0.522942841846969 | 2070.85343837625 | 2458.72108559236 | 2501.48027069784 | 1595.41330693313 | 1235.11284488394 | 1136.24339359662 | 1427.31806215246 | 1649.99885829613 | mt-Ty |
| **ENSMUSG00000064351** | 0.19859495658012 | 0.0150847015371208 | 0.635862253526407 | 3542143.30606073 | 3443686.05554582 | 3393964.515776 | 3454829.53013258 | 3011863.18570274 | 2766417.53887617 | 2980070.41804558 | 3297122.15937146 | mt-Co1 |
| **ENSMUSG00000064354** | 0.116271138899361 | 0.397643230643169 | 0.88020150974241 | 1309.19137530855 | 1198.81995296189 | 1214.30162213596 | 1378.7522405595 | 1048.96995159469 | 1431.00621152642 | 1038.99593437502 | 1188.65049331202 | mt-Co2 |
| **ENSMUSG00000064356** | 0.121513537795858 | 0.50310153553182 | 0.900927059989498 | 271.054114970713 | 253.283291692356 | 229.988537941573 | 244.79886720138 | 213.516848184724 | 178.570005882784 | 224.872867560126 | 300.690581413177 | mt-Atp8 |
| **ENSMUSG00000064357** | 0.136769300123406 | 0.189048889390292 | 0.808677608936066 | 1542.29791418336 | 1430.11401997356 | 1423.46815252726 | 1537.26185188913 | 1298.62042024145 | 1544.75285910928 | 1252.41694293903 | 1302.63067760221 | mt-Atp6 |
| **ENSMUSG00000064358** | 0.0294344152041339 | 0.849957880488244 | 0.972640876584422 | 1938.94043575717 | 1466.76272777471 | 1603.29458137047 | 2093.45238158422 | 1725.6541166109 | 1925.13143328426 | 1481.45412286138 | 1826.9395253371 | mt-Co3 |
| **ENSMUSG00000064359** | 0.495130126362223 | 0.0826391877116191 | 0.739855948845396 | 45.1756858284523 | 30.1333819698302 | 46.3762895437739 | 54.3997482669734 | 28.4689130912965 | 37.9155491942896 | 27.0680303544596 | 31.4802413753867 | mt-Tg |
| **ENSMUSG00000064360** | -0.220693506628483 | 0.503974873492795 | 0.900927059989498 | 4311.56745546748 | 3207.98355619354 | 2308.40347341356 | 4024.64344506177 | 3158.95439493809 | 7140.35407084719 | 2599.57199211867 | 3244.63591279417 | mt-Nd3 |
| **ENSMUSG00000064361** | 0.759963384068498 | 0.0091621895334528 | 0.635862253526407 | 30.7194663633475 | 33.3910448854875 | 39.7511053232348 | 36.5791410760683 | 17.5193311331055 | 18.3462334811079 | 24.9858741733473 | 21.7105112933701 | mt-Tr |
| **ENSMUSG00000064363** | 0.0685895779466394 | 0.694386314697652 | 0.944771261777587 | 764630.105626634 | 588055.392388422 | 571257.455870875 | 741105.591248169 | 627286.220970019 | 798689.820695479 | 516673.522327833 | 598661.921286859 | mt-Nd4 |
| **ENSMUSG00000064364** | 0.804566303779148 | 0.0121483272581119 | 0.635862253526407 | 45.1756858284523 | 43.1640336324594 | 46.3762895437739 | 45.9584080186499 | 32.8487458745729 | 19.5693157131818 | 36.4377331694648 | 14.1118323406906 | mt-Th |
| **ENSMUSG00000064365** | 0.933609912537975 | 0.0326147697207533 | 0.671606027118506 | 27.1054114970714 | 20.3603932228582 | 16.0897331070236 | 12.1930470253561 | 7.66470737073368 | 8.56157562451702 | 11.4518589961175 | 11.9407812113536 | mt-Ts2 |
| **ENSMUSG00000064366** | 0.694027137721729 | 0.00014637226165386 | 0.399150048388892 | 243.948703473642 | 229.665235553841 | 242.292451494002 | 342.343243404229 | 147.819356435578 | 149.216032313011 | 172.818963032319 | 183.453820428978 | mt-Tl2 |
| **ENSMUSG00000064367** | 0.253284228871815 | 0.260055082753569 | 0.840316047847749 | 597943.570597961 | 718411.588374179 | 710129.835227373 | 522698.105369825 | 413020.421379355 | 386800.97897559 | 654750.668400203 | 684152.487132327 | mt-Nd5 |
| **ENSMUSG00000064368** | 0.229864467505469 | 0.00887105680759315 | 0.635862253526407 | 282640.774871995 | 242809.091002789 | 259046.595932857 | 272856.944260113 | 216561.926927297 | 221744.808674991 | 220158.865966087 | 243154.469909052 | mt-Nd6 |
| **ENSMUSG00000064369** | 0.608453753352199 | 7.89673470715512E-0 | 0.0109685645082385 | 14312.5607841702 | 11863.593923095 | 14408.829224784 | 12035.4753407207 | 8536.29409460568 | 7665.05634840688 | 9716.38181916042 | 8594.10589548057 | mt-Te |
| **ENSMUSG00000064370** | 0.0981628974087097 | 0.447998779868235 | 0.89108031640712 | 1708040.27737822 | 1407299.79215948 | 1435978.39324542 | 1583550.41010416 | 1393781.0471237 | 1704580.35286777 | 1282278.18581045 | 1350691.23645234 | mt-Cytb |
| **ENSMUSG00000064371** | 0.490446439081728 | 0.20393040633592 | 0.816074141171399 | 35.2370349461928 | 20.3603932228582 | 31.2330113253987 | 15.0068271081306 | 18.6142893289246 | 23.2385624094033 | 14.5750932677859 | 16.2828834700276 | mt-Tt |
| **ENSMUSG00000064372** | 0.352404848732712 | 0.00051952872932798 | 0.522942841846969 | 8985.44391127915 | 8500.87137840776 | 9300.81219074829 | 9866.98882359586 | 6778.88619031603 | 7255.32380066213 | 6648.32468629149 | 8027.46155072361 | mt-Tp |
| **ENSMUSG00000064373** | -0.0162203456310041 | 0.922242736288473 | 0.986713918026868 | 9956.72115659088 | 8059.4580533362 | 9735.23498463792 | 11090.0452329085 | 9491.09764135993 | 10426.7760284297 | 7649.84180940649 | 11712.8208427732 | Selenop |
| **ENSMUSG00000064493** | 0.782316680487366 | 0.146884816769851 | 0.780607990324772 | 29.8159526467785 | 46.4216965481167 | 53.001473764313 | 27.1998741334867 | 7.66470737073368 | 8.56157562451702 | 38.5198893505771 | 35.8223436340608 | Snora28 |
| **ENSMUSG00000065037** | 1.23345622608813 | 0.0174622458821677 | 0.642273034804458 | 33.4300075130547 | 33.3910448854875 | 60.5731128735006 | 92.854742731558 | 37.2285786578493 | 34.2463024980681 | 8.32862472444909 | 14.1118323406906 | Rn7sk |
| **ENSMUSG00000065126** | 0.248379623527259 | 0.514671414002076 | 0.904145806837306 | 20.780815481088 | 13.0306516626293 | 18.9290977729689 | 18.7585338851632 | 17.5193311331055 | 18.3462334811079 | 14.5750932677859 | 9.76973008201657 | Snord104 |
| **ENSMUSG00000065947** | 0.240293218066582 | 0.262649493127801 | 0.841957410465073 | 1005.61076654135 | 1088.87382955846 | 1076.11920839328 | 885.402799379704 | 613.176589658694 | 697.1568722821 | 952.586452858865 | 1170.19655871265 | mt-Nd4l |
| **ENSMUSG00000065954** | -0.0555711184494322 | 0.708906806910502 | 0.949267137065756 | 1013.74238999047 | 1100.27564976326 | 1097.8876708322 | 939.802547646678 | 1345.70362266167 | 956.450305481758 | 1074.39258945393 | 937.894087873591 | Tacc1 |
| **ENSMUSG00000065979** | -0.0213291675395291 | 0.893303198730204 | 0.983433576156181 | 1282.08596381147 | 1370.66167176282 | 1108.29867460733 | 1212.7392156758 | 1070.86911551108 | 1055.51996627974 | 1430.44129642413 | 1490.42660028986 | Cpped1 |
| **ENSMUSG00000065990** | -0.0236021125759675 | 0.895228503579157 | 0.983733637008786 | 2268.72294230487 | 1652.44951396717 | 1805.83592754124 | 2219.13455861481 | 2045.38190979007 | 2562.35727619474 | 1704.2448342404 | 1766.15009371566 | Aurkaip1 |
| **ENSMUSG00000065999** | -0.104322589467212 | 0.839096680381986 | 0.96956148850045 | 17.1667606148119 | 8.14415728914329 | 10.4110037751329 | 16.8826804966469 | 7.66470737073368 | 26.9078091056249 | 11.4518589961175 | 10.8552556466851 | Zfp985 |
| **ENSMUSG00000066000** | -0.0502148560996701 | 0.847623866686361 | 0.972455319662159 | 140.948139784771 | 65.9676740420606 | 101.270673085384 | 144.440710915757 | 122.635317931739 | 123.53130543946 | 115.559668051731 | 106.381505337514 | Zfp979 |
| **ENSMUSG00000066026** | -0.113116734844112 | 0.545133227767086 | 0.914211323944814 | 1486.28006375608 | 1435.00051434705 | 1843.69412308717 | 2112.21091546938 | 2295.03237843683 | 1964.27006471062 | 1575.15115101143 | 1603.32125901539 | Dhrs3 |
| **ENSMUSG00000066036** | 0.216009425627843 | 0.19812321679081 | 0.812921911423739 | 2965.33201777961 | 4141.30398152936 | 3503.77599777655 | 3127.98552535097 | 2999.09049834851 | 2306.7330896913 | 3441.80416737859 | 3079.63602696456 | Ubr4 |
| **ENSMUSG00000066037** | 0.128219681406331 | 0.469665829783035 | 0.895614823680169 | 807.741262612726 | 1012.31875104051 | 1025.01064440627 | 888.216579462479 | 840.927894389066 | 620.102691661447 | 1003.59927929612 | 949.834869084944 | Hnrnpr |
| **ENSMUSG00000066042** | -0.17790361743608 | 0.58314856386308 | 0.922450244597442 | 38.8510898124689 | 24.4324718674299 | 19.8755526616174 | 29.075727522003 | 35.0386622662111 | 24.4616446414772 | 27.0680303544596 | 40.1644458927348 | Med18 |
| **ENSMUSG00000066043** | -0.150896623121234 | 0.468093607734882 | 0.895328112556867 | 442.721721118832 | 473.989954228139 | 500.674636095028 | 268.247034557834 | 512.440435643337 | 495.348303989913 | 450.786813210807 | 413.585240138701 | Phactr4 |
| **ENSMUSG00000066057** | -0.0609123407706914 | 0.856383982279902 | 0.97363563840968 | 39.754603529038 | 30.1333819698302 | 25.5542819935081 | 31.8895076047775 | 20.8042057205628 | 24.4616446414772 | 44.7663578939139 | 42.3354970220718 | Gm1976 |
| **ENSMUSG00000066071** | 0.215135608015006 | 0.723670993368347 | 0.951674101666722 | 8.13162344912141 | 45.6072808192024 | 11.3574586637814 | 34.703287687552 | 19.7092475247437 | 15.9000690169602 | 11.4518589961175 | 39.0789203280663 | Cyp4a12a |
| **ENSMUSG00000066072** | -0.27583674436486 | 0.484936687844938 | 0.899236955296196 | 155.404359249876 | 342.054606144018 | 145.754052851861 | 186.647412157374 | 116.065568756824 | 188.354663739374 | 321.693129981846 | 378.848422069309 | Cyp4a10 |
| **ENSMUSG00000066113** | -0.353888167010984 | 0.468767958055927 | 0.895328112556867 | 26.2018977805023 | 43.9784493613737 | 26.5007368821565 | 19.6964605794214 | 42.7033696369448 | 68.4926049961361 | 13.5340151772298 | 24.9670879873757 | Adamtsl1 |
| **ENSMUSG00000066148** | 0.0325905882823372 | 0.845457441781354 | 0.972136081650911 | 338.817643713392 | 414.537606017393 | 386.153594568566 | 355.474217123843 | 377.760577557588 | 276.416584448692 | 439.33495421469 | 366.907640857956 | Prpf4 |
| **ENSMUSG00000066149** | 0.172737508825054 | 0.111975638524221 | 0.764116514278315 | 649.626362213143 | 551.359448475 | 588.694940739334 | 636.852225401292 | 526.674892188986 | 500.240632918209 | 542.401685179747 | 581.84170266232 | Cdc26 |
| **ENSMUSG00000066150** | -0.0541501453375231 | 0.774668195604874 | 0.959414024438738 | 479.765783498163 | 539.9576282702 | 535.693466975021 | 444.577253078368 | 527.769850384805 | 353.470765069345 | 585.085886892549 | 608.979841779033 | Slc31a1 |
| **ENSMUSG00000066151** | 0.307736532180186 | 0.062749948198481 | 0.715615581523459 | 372.247651226447 | 434.083583511337 | 493.102996985841 | 421.129085721914 | 346.006789878835 | 266.631926592101 | 387.281049686883 | 388.618152151326 | Fkbp15 |
| **ENSMUSG00000066152** | -0.0124790564953318 | 0.951445886846019 | 0.991915011884503 | 670.407177694231 | 493.535931722083 | 533.800557197724 | 568.383576720446 | 613.176589658694 | 450.09426140318 | 762.069162287092 | 459.177313854779 | Slc31a2 |
| **ENSMUSG00000066153** | -0.370596766191933 | 0.307822250958022 | 0.851785577030345 | 99.3865088225949 | 92.8433930962335 | 54.8943835416099 | 98.4823028971069 | 70.0773245324222 | 99.0696607979826 | 78.0808567917102 | 199.736703899005 | Mup21 |
| **ENSMUSG00000066154** | -0.436673499031858 | 0.143764828416162 | 0.780607990324772 | 4635.0253659992 | 4144.56164444502 | 3065.56738433232 | 4399.81412276503 | 3599.12758965737 | 5863.45622056208 | 3604.21234950535 | 8920.8490904458 | Mup3 |
| **ENSMUSG00000066170** | -0.382321196490606 | 0.379219483000684 | 0.87422797717331 | 20.780815481088 | 8.95857301805762 | 18.9290977729689 | 11.2551203310979 | 16.4243729372865 | 25.6847268735511 | 23.9447960827911 | 11.9407812113536 | E230001N04 |
| **ENSMUSG00000066175** | 0.386532043951401 | 0.452170733369821 | 0.891110666112841 | 15.3597331816738 | 4.88649437348597 | 11.3574586637814 | 15.0068271081306 | 8.75966556655277 | 9.78465785659088 | 10.4107809055614 | 6.51315338801105 | 2510046G10 |
| **ENSMUSG00000066191** | 0.0686673682952205 | 0.811808742055325 | 0.965224347191244 | 32.5264937964856 | 31.7622134276588 | 34.0723759913441 | 42.2067012416173 | 29.5638712871156 | 47.7002070508805 | 31.2323427166841 | 26.0526135520442 | Anks6 |
| **ENSMUSG00000066232** | 0.292971224729428 | 0.145519156317683 | 0.780607990324772 | 4137.18930816966 | 4491.50274496252 | 4356.5318524488 | 4044.33990564119 | 2989.23587458613 | 2374.00261245536 | 4249.68076565015 | 4285.65492931127 | Ipo7 |
| **ENSMUSG00000066233** | -0.00960127725864062 | 0.96576109287806 | 0.994284908685797 | 242.141676040504 | 168.584055885266 | 174.147699511314 | 218.536919762152 | 147.819356435578 | 267.855008824175 | 212.379930473452 | 181.282769299641 | Tmem42 |
| **ENSMUSG00000066235** | -0.205316634555387 | 0.319861528979885 | 0.855057089594033 | 328.878992831132 | 312.735639903102 | 327.473391472363 | 373.294824314748 | 455.502609460744 | 485.563646133322 | 342.514691792969 | 264.868237779116 | Pomgnt2 |
| **ENSMUSG00000066258** | 0.0380556256820923 | 0.820361710796397 | 0.966398681144067 | 163.535982698997 | 153.110157035894 | 150.486327295103 | 196.026679099956 | 178.478185918513 | 173.677676954488 | 161.367104036201 | 132.434118889558 | Trim12a |
| **ENSMUSG00000066278** | -0.0707967983717949 | 0.708525346104381 | 0.949174314815223 | 215.036264543433 | 231.294067011669 | 334.098575692902 | 307.639955716677 | 266.074841584041 | 253.178022039289 | 330.021754706295 | 292.006376895829 | Vps37b |
| **ENSMUSG00000066279** | 0.166447194789034 | 0.667880021058233 | 0.942024383071524 | 11.7456783153976 | 17.9171460361152 | 21.7684624389143 | 20.6343872736795 | 20.8042057205628 | 14.6769867848863 | 17.6983275394543 | 10.8552556466851 | Chrna10 |
| **ENSMUSG00000066306** | 0.066136776800207 | 0.658512412767712 | 0.938989601151841 | 1548.62251019934 | 2028.70958072559 | 1916.5711495131 | 1598.22708701591 | 1823.1053960388 | 1374.74442885102 | 1830.21528319769 | 1745.52510798696 | Numa1 |

| | | | | | | | | | | | | |
|---|---|---|---|---|---|---|---|---|---|---|---|---|
| **ENSMUSG00000066315** | 0.22669957199862 | 0.543669367631221 | 0.913837378403038 | 18.9737880479499 | 18.7315617650296 | 13.2503684410783 | 24.3860940507122 | 19.7092475247437 | 18.3462334811079 | 14.5750932677859 | 11.9407812113536 | Gm12918 |
| **ENSMUSG00000066319** | -0.176132772368921 | 0.622922458248753 | 0.931059568526633 | 50.5967681278665 | 65.9676740420606 | 83.2880302010633 | 85.3513291774927 | 87.5966556655277 | 134.539045528125 | 55.1771387994752 | 45.5920737160773 | Rtp3 |
| **ENSMUSG00000066324** | 0.0497312203841745 | 0.732834757473231 | 0.953733898884637 | 1330.8757045062 | 1413.82570539527 | 1148.04977993057 | 1365.62126683988 | 1054.44474257379 | 1154.58962707772 | 1435.64668687691 | 1435.06479649177 | Impad1 |
| **ENSMUSG00000066357** | -0.396183835339644 | 0.00304776631433464 | 0.635862253526407 | 256.597895505609 | 219.077831077954 | 270.686098153456 | 269.184961252092 | 365.716037403578 | 298.432064626022 | 347.72008224575 | 322.401092706547 | Wdr6 |
| **ENSMUSG00000066361** | -0.00763433580511395 | 0.988034072235721 | 0.998437672973644 | 5.42108229941427 | 19.5459774939439 | 18.9290977729689 | 18.7585338851632 | 16.4243729372865 | 25.6847268735511 | 9.36970281500523 | 11.9407812113536 | Serpina3c |
| **ENSMUSG00000066366** | -0.227381283257809 | 0.554579037931564 | 0.915945640531442 | 164.439496415566 | 405.579032999336 | 164.68315062483 | 235.419600258798 | 166.433645764503 | 220.154801773295 | 267.557069272927 | 481.973350712817 | Serpina1a |
| **ENSMUSG00000066406** | 0.12754900256139 | 0.550741757523708 | 0.914699131434573 | 1204.38378418654 | 1713.53069363575 | 1477.41608118023 | 1169.59458773993 | 1102.62290318983 | 942.996400928946 | 1661.56063252759 | 1386.21614608168 | Akap13 |
| **ENSMUSG00000066415** | -0.0496676345012975 | 0.678309998385752 | 0.944452327903597 | 580.959319753896 | 719.943504360267 | 718.359260484171 | 635.914298707033 | 689.823663366031 | 621.325773893521 | 733.960053842076 | 702.335040340524 | Msl2 |
| **ENSMUSG00000066440** | 0.09259373918306 | 0.643237153392047 | 0.934616627390525 | 224.974915425692 | 286.674336577844 | 266.900278598862 | 192.274972322923 | 210.231973597267 | 174.900759186562 | 268.598147363483 | 256.184033261768 | Zfyve26 |
| **ENSMUSG00000066441** | -0.445309988713069 | 0.145139799776551 | 0.780607990324772 | 58.7283915769879 | 57.8235167529173 | 87.0738497556571 | 94.7305961200743 | 128.110108910834 | 133.315963296051 | 62.4646854333682 | 82.4999429148066 | Rdh11 |
| **ENSMUSG00000066442** | 0.119439002367723 | 0.658929281390963 | 0.938989601151841 | 85.8338030740593 | 49.679359463774 | 67.1982970940397 | 68.4686486808458 | 49.2731188118594 | 88.0619207093179 | 53.094982618363 | 59.7039060567679 | Mthfs |
| **ENSMUSG00000066456** | -0.564679117293939 | 0.0472804901367078 | 0.703396849069134 | 23.4913566307952 | 33.3910448854875 | 32.1794662140472 | 36.5791410760683 | 54.7479097909548 | 56.2617826753975 | 42.6842017128016 | 32.5657669400552 | Hmgn3 |
| **ENSMUSG00000066491** | -0.222110286847589 | 0.614839426439058 | 0.928075610917191 | 20.780815481088 | 13.0306516626293 | 23.6613722162112 | 11.2551203310979 | 32.8487458745729 | 19.5693157131818 | 12.4929370866736 | 15.1973579053591 | Gm6265 |
| **ENSMUSG00000066538** | -0.00933432424974749 | 0.98015032978816 | 0.996872525500296 | 26.2018977805023 | 17.9171460361152 | 31.2330113253987 | 14.0689004138724 | 29.5638712871156 | 23.2385624094033 | 20.8215618111227 | 16.2828834700276 | Gm6254 |
| **ENSMUSG00000066551** | 0.0178981104654832 | 0.809619623806417 | 0.964412705343932 | 3670.97623042003 | 3524.79127474121 | 3640.06550174193 | 3840.80981298717 | 3692.19903630199 | 3773.20868594786 | 3406.40751229968 | 3624.56986042815 | Hmgb1 |
| **ENSMUSG00000066553** | 0.532289173597945 | 0.109296666865438 | 0.763154315676794 | 21.6843291976571 | 31.7622134276588 | 31.2330113253987 | 25.3240207449704 | 25.1840385038392 | 13.4539045528125 | 17.6983275394543 | 19.5394601640331 | Gm6969 |
| **ENSMUSG00000066554** | 0.158842435612272 | 0.681988231368691 | 0.944452327903597 | 32.5264937964856 | 24.4324718674299 | 28.3936466594534 | 20.6343872736795 | 21.8991639163819 | 11.0077400886647 | 37.4788112600209 | 23.8815624227072 | Gm10167 |
| **ENSMUSG00000066568** | 0.176866662077938 | 0.131373758063216 | 0.77303252008843 | 2137.71345340236 | 2188.3350635928 | 2225.1154432125 | 2047.49397356557 | 1727.84403300253 | 1674.39957570911 | 2172.72997499066 | 2029.93280593011 | Lsm14a |
| **ENSMUSG00000066571** | -0.102043145213489 | 0.611542457911053 | 0.928055903343287 | 306.291149916906 | 467.474628396825 | 441.047978110176 | 319.833002742033 | 512.440435643337 | 351.024600605198 | 407.061533407449 | 376.677370939972 | 4931406P16 |
| **ENSMUSG00000066595** | -0.179936806499841 | 0.553234135318783 | 0.915115571755578 | 30.7194663633475 | 28.5045505120015 | 28.3936466594534 | 21.5723139679377 | 25.1840385038392 | 42.8078781225851 | 24.9858741733473 | 31.4802413753867 | Flvcr1 |
| **ENSMUSG00000066607** | -0.0944039881997659 | 0.50285234484534 | 0.900927059989498 | 1304.6738067257 | 1166.24332380532 | 1065.70820461815 | 939.802547646678 | 1028.16574587413 | 1280.56709698133 | 1254.49909912014 | 1216.8741579934 | 6030419C18 |
| **ENSMUSG00000066613** | 0.317690028835304 | 0.176503392479477 | 0.799889021902745 | 84.0267756409212 | 73.2974156022896 | 52.0550188756646 | 57.2135283497478 | 56.937826182593 | 46.4771248188067 | 55.1771387994752 | 55.3618037980939 | Zfp932 |
| **ENSMUSG00000066621** | 0.252249140237528 | 0.0784460474660472 | 0.739855948845396 | 1669.6933482196 | 1369.8472560339 | 1874.92713441257 | 1890.86021562445 | 1609.58854785407 | 1297.69024823037 | 1351.31936154187 | 1453.51873109113 | Tecpr1 |
| **ENSMUSG00000066637** | 0.27464378156342 | 0.101603836673597 | 0.759800953500741 | 458.081454300506 | 311.921224174188 | 379.528410348027 | 438.011766218561 | 316.442918591719 | 341.239942748607 | 283.173240631269 | 371.24974311663 | Ttc32 |
| **ENSMUSG00000066640** | -0.0573534918238452 | 0.763689071528381 | 0.95785116771606 | 100.290022539164 | 118.090280692578 | 115.467496415111 | 108.799496533947 | 122.635317931739 | 86.838838477244 | 109.313199508394 | 141.118323406906 | Fbxl18 |
| **ENSMUSG00000066643** | 0.176077039438708 | 0.391819147356247 | 0.878285927324016 | 314.422773366028 | 321.69421292116 | 299.079744812909 | 271.998741334867 | 234.321053905287 | 183.462334811079 | 318.569895710178 | 331.085297223895 | Wdr35 |
| **ENSMUSG00000066647** | -0.0660130521695837 | 0.821452241261713 | 0.966637987288264 | 149.079763233892 | 171.027303072009 | 148.593417517806 | 151.006197775564 | 131.394983498292 | 84.3926740130963 | 200.928071477334 | 231.216945274392 | Gm5113 |
| **ENSMUSG00000066677** | 0.189547049612099 | 0.687671390216416 | 0.944452327903597 | 22.5878429142261 | 28.5045505120015 | 8.51809399783602 | 30.0136542162612 | 15.3294147414674 | 33.0232202659942 | 14.5750932677859 | 16.2828834700276 | Pydc3 |
| **ENSMUSG00000066687** | 0.166654393628228 | 0.619334995353808 | 0.929568738467614 | 2737.64656120421 | 4085.10929623427 | 3690.22761084029 | 1567.27550610539 | 2335.54583168213 | 1539.86053018099 | 3686.45751865928 | 3200.12936464276 | Zbtb16 |
| **ENSMUSG00000066705** | -0.381603146471042 | 0.100980587269916 | 0.759800953500741 | 786.056933415069 | 485.39177443294 | 481.745538322059 | 566.507723331929 | 738.001823982071 | 1038.39681503071 | 624.646854333682 | 622.006148555055 | Fxyd6 |
| **ENSMUSG00000066724** | -0.13543002978081 | 0.613516413508164 | 0.928075610917191 | 26.2018977805023 | 27.6901347830872 | 36.9117406572894 | 35.6412143818101 | 31.7537876787538 | 37.9155491942896 | 37.4788112600209 | 31.4802413753867 | Gm10175 |
| **ENSMUSG00000066735** | 0.0505331397020257 | 0.690102435824849 | 0.94469184542218 | 515.002818444356 | 545.6585383726 | 554.62256474799 | 513.045901759214 | 525.579933993166 | 418.29412336926 | 558.017856538089 | 551.446986851602 | Vkorc1l1 |
| **ENSMUSG00000066798** | 0.28827647800224 | 0.181130492335508 | 0.803056849297055 | 82.2197482077831 | 78.1839099757755 | 98.4313084194385 | 115.364983393754 | 91.9764884488041 | 78.277262852727 | 72.8754663389296 | 62.9604827507734 | Zbtb6 |
| **ENSMUSG00000066800** | 0.485733121009766 | 0.121186914456767 | 0.769868040577702 | 39.754603529038 | 42.3496179035451 | 81.3951204237664 | 66.5927952923295 | 45.9882442244021 | 30.5770558018465 | 47.8895921655823 | 39.0789203280663 | Rnasel |
| **ENSMUSG00000066809** | 0.854512637869386 | 0.105242564300145 | 0.762094734041179 | 22.5878429142261 | 5.7009101024003 | 16.0897331070236 | 17.8206071909051 | 7.66470737073368 | 8.56157562451702 | 11.4518589961175 | 6.51315338801105 | Gm10180 |
| **ENSMUSG00000066829** | -0.159050170606115 | 0.442172782795376 | 0.89108031640712 | 60.535419010126 | 47.2361122770311 | 56.7872933189068 | 61.9031618210386 | 58.0327843784121 | 61.154111603693 | 61.4236073428121 | 71.6446872681215 | Zfp810 |
| **ENSMUSG00000066838** | 0.0110729177320494 | 0.973278148036181 | 0.995495837718892 | 44.2721721118832 | 28.5045505120015 | 43.5369248778286 | 30.9515809105193 | 41.6084114411257 | 28.1308913376988 | 23.9447960827911 | 52.1052271040884 | Zfp772 |
| **ENSMUSG00000066839** | 0.079586947099972 | 0.472606532403995 | 0.89728597737823 | 3130.67502791174 | 2523.8743439055 | 2611.26903778106 | 2947.9036000534 | 2711.11649284808 | 2864.45858751698 | 2544.3948533192 | 2492.36669647889 | Ecsit |
| **ENSMUSG00000066842** | 0.259477956663002 | 0.334776018218287 | 0.858570960161243 | 87.6408305071974 | 162.068730053951 | 88.966759532954 | 90.9788893430416 | 78.836990098975 | 77.0541806206532 | 90.5737938783839 | 112.894658725525 | Hmcn1 |
| **ENSMUSG00000066877** | -0.0462758133282407 | 0.769335855576659 | 0.95879680569409 | 287.317361868956 | 242.69588721647 | 243.238906382651 | 287.943495137256 | 320.822751374995 | 277.639666680766 | 220.708555197901 | 276.809018990469 | Nck2 |
| **ENSMUSG00000066880** | 0.129610712706461 | 0.424259619829027 | 0.884586455610141 | 446.335775985108 | 386.033055505392 | 303.812019256151 | 356.412143818101 | 295.638712871156 | 340.016860516533 | 339.391457521301 | 389.703677715994 | Zfp617 |
| **ENSMUSG00000066892** | 0.084672327354573 | 0.711739323416411 | 0.950034518763592 | 82.2197482077831 | 48.8649437348597 | 72.8770264259304 | 81.59962240046 | 64.6025335533267 | 74.6080161565054 | 63.5057635239243 | 66.217059444779 | Fbxl12 |
| **ENSMUSG00000066894** | -0.0746446730223732 | 0.746771857427508 | 0.956093750545496 | 101.193536255733 | 106.688460487777 | 118.306861081056 | 142.564857527241 | 151.104231023035 | 150.439114545085 | 90.5737938783839 | 102.03940307884 | Vsig10 |
| **ENSMUSG00000066900** | 0.15708853828615 | 0.0704972363689285 | 0.72606739345142 | 3349.32534732145 | 2966.10208470598 | 3228.35762517985 | 3489.08730264036 | 2804.18793949271 | 3036.91318223939 | 2977.48333899055 | 2869.04406741887 | Suds3 |
| **ENSMUSG00000066975** | -0.101037753985367 | 0.643349140351109 | 0.934616627390525 | 143.658680934478 | 103.43079757212 | 118.306861081056 | 114.427056699496 | 131.394983498292 | 172.454594722414 | 99.9434966933891 | 111.809133160856 | Cryba4 |
| **ENSMUSG00000066979** | -0.158227816842372 | 0.269426014141651 | 0.843793744782845 | 795.995584297329 | 694.696616763922 | 711.734076263632 | 764.410255820401 | 958.08842134171 | 923.427085215764 | 699.604476853724 | 730.558705021906 | Bub3 |
| **ENSMUSG00000067071** | -0.170330978039036 | 0.414010354521284 | 0.882192126163777 | 403.870631306363 | 277.715763559786 | 243.238906382651 | 317.019222659259 | 324.107625962453 | 447.648096939033 | 295.666177717943 | 331.085297223895 | Hes6 |
| **ENSMUSG00000067081** | 0.0644428499607544 | 0.793307107136405 | 0.961804768755116 | 280.089252136404 | 356.714089264476 | 300.972654590206 | 254.178134143962 | 235.416012101106 | 182.239252579005 | 364.377331694648 | 357.137910775939 | Asb18 |
| **ENSMUSG00000067148** | 0.0204343730138679 | 0.903425410980987 | 0.985027469232262 | 558.37147683967 | 416.166437475222 | 427.797609669098 | 550.562969529541 | 522.295059405709 | 549.163922201163 | 457.033281754144 | 397.302356668674 | Polr1c |
| **ENSMUSG00000067150** | 0.332042995990357 | 0.215491255417804 | 0.824338622350122 | 365.923055210463 | 519.597235047342 | 512.03209475881 | 334.839829850164 | 342.721915291377 | 195.693157131818 | 403.938299135781 | 433.124700302735 | Xpo5 |
| **ENSMUSG00000067158** | -0.979796954848182 | 0.0515398599299254 | 0.703396849069134 | 28.9124389302094 | 28.5045505120015 | 22.7149173275627 | 33.7653609932938 | 96.3563212320805 | 83.1695917810225 | 27.0680303544596 | 18.4539345993646 | Col4a4 |
| **ENSMUSG00000067194** | 0.0947342771119721 | 0.458307289511382 | 0.893708865830843 | 2076.27452067567 | 1988.80321000879 | 1978.09071727525 | 1934.00484356033 | 1659.95662486175 | 1612.02238187335 | 2031.14335467502 | 2165.62350151367 | Eif1ax |
| **ENSMUSG00000067199** | -0.406325581318757 | 0.131698885843935 | 0.77303252008843 | 62.3424464432641 | 61.895595397489 | 57.7337482075553 | 101.296082979881 | 123.730276127558 | 96.6234963338349 | 84.3273253350471 | 70.559161703453 | Frat1 |
| **ENSMUSG00000067212** | -0.659631402987965 | 0.0705896664935821 | 0.72606739345142 | 477.055242348456 | 390.919549878878 | 501.621090983677 | 549.625042835282 | 982.17750164973 | 1253.65928787571 | 387.281049686883 | 409.243137880027 | H2-T23 |
| **ENSMUSG00000067225** | -0.343203837480754 | 0.343605720171661 | 0.861266047293794 | 66.8600150261093 | 65.9676740420606 | 24.6078271048596 | 52.523894878457 | 42.7033696369448 | 75.8310983885793 | 57.2592949805875 | 91.1841474321546 | Cyp2c54 |
| **ENSMUSG00000067235** | -0.228682070724953 | 0.455973915614606 | 0.892805623137304 | 355.984404328204 | 547.287369830429 | 406.029147230184 | 481.156394154437 | 430.318570956905 | 407.286383280595 | 351.884394607974 | 908.584897627541 | H2-Q10 |
| **ENSMUSG00000067242** | -0.256110066573069 | 0.398715723409224 | 0.880459952732461 | 84.0267756409212 | 58.6379324818317 | 32.1794662140472 | 50.6480414899407 | 72.2672409240604 | 52.592535979176 | 75.998700610598 | 68.388110574116 | Lgi1 |
| **ENSMUSG00000067274** | -0.0120784702205036 | 0.937795554142573 | 0.988970340396118 | 24871.9255897127 | 18479.907304795 | 21564.9746378549 | 26037.7829593008 | 23571.1650813977 | 26595.9231364461 | 19481.694308577 | 22070.9057808401 | Rplp0 |
| **ENSMUSG00000067279** | 0.0301166645275943 | 0.92121560242162 | 0.986713918026868 | 7426.88275019755 | 6284.03176430296 | 4193.74161160127 | 3233.97124180214 | 4224.34871947008 | 3433.19182543132 | 6905.47097465885 | 6138.64706820041 | Ppp1r3c |
| **ENSMUSG00000067288** | 0.22429412233472 | 0.318472186104546 | 0.854351922245337 | 2516.28570064479 | 1820.21915412352 | 2041.5031948147 | 2640.26364433672 | 2041.0020770068 | 2669.98851261724 | 1561.6171358342 | 1448.09110326779 | Rps28 |
| **ENSMUSG00000067297** | 0.134084334317604 | 0.619748209674209 | 0.929594515086012 | 53.3073092775737 | 66.7820897709749 | 63.4124775394459 | 78.7858423176855 | 70.0773245324222 | 34.2463024980681 | 70.7933101578173 | 62.9604827507734 | Ifit1bl2 |
| **ENSMUSG00000067321** | 0.116469864088117 | 0.818118704215044 | 0.966315773308182 | 14.4562194651047 | 6.51532583131463 | 9.46454888648447 | 18.7585338851632 | 10.949581958191 | 7.33849339244316 | 9.36970281500523 | 17.3684090346961 | Gm7931 |
| **ENSMUSG00000067336** | 0.0635784099854922 | 0.691294616697486 | 0.94469184542218 | 1549.52602391591 | 1948.08242356307 | 2007.43081882336 | 1502.55856420157 | 1587.68938393769 | 1384.52908670761 | 1872.89948491049 | 1859.50529227715 | Bmpr2 |
| **ENSMUSG00000067344** | 0.0740702067223748 | 0.853785835527727 | 0.9731539929935 | 10.8421645988285 | 13.0306516626293 | 14.1968233297267 | 17.8206071909051 | 10.949581958191 | 11.0077400886647 | 14.5750932677859 | 16.2828834700276 | Rps25-ps1 |
| **ENSMUSG00000067365** | -0.177079665747209 | 0.308458885024466 | 0.851785577030345 | 156.307872966445 | 120.533527879321 | 167.522515290775 | 143.502784221499 | 198.187433443257 | 151.662196777159 | 151.997401221196 | 161.743309135608 | Tmem128 |
| **ENSMUSG00000067367** | 0.202967710285437 | 0.0582277227334248 | 0.710862211775517 | 1163.72566694093 | 1053.03953748623 | 1127.2277723803 | 1279.33201096813 | 1024.88087128667 | 1088.54318654574 | 979.654483213325 | 923.7822555329 | Lyar |
| **ENSMUSG00000067369** | 0.0992894733619228 | 0.589531319317265 | 0.923594634030771 | 1249.55947001499 | 1338.89945833516 | 1337.34075766026 | 1157.40154071457 | 1043.4951606156 | 867.165302540366 | 1346.11397108908 | 1487.17002359586 | Trmt2b |
| **ENSMUSG00000067370** | -0.596460011246434 | 0.0710952346040001 | 0.726501344493452 | 23.4913566307952 | 24.4324718674299 | 27.447191770805 | 24.3860940507122 | 42.7033696369448 | 52.592535979176 | 35.3966550789086 | 20.6249857287016 | B3galt4 |
| **ENSMUSG00000067377** | -0.439598210306985 | 0.201190012079705 | 0.813072233862993 | 119.263810587114 | 73.2974156022896 | 62.4660226507975 | 73.1582821521366 | 180.668102310151 | 110.077400886647 | 65.5879197050366 | 89.0130963028176 | Tspan6 |
| **ENSMUSG00000067424** | 0.0482458544151098 | 0.864303872370352 | 0.974868041734905 | 31.6229800799166 | 34.2054606144018 | 38.8046504345863 | 50.6480414899407 | 40.5134532453066 | 26.9078091056249 | 36.4377331694648 | 45.5920737160773 | Zfp563 |
| **ENSMUSG00000067430** | -0.179789596957265 | 0.415967793381826 | 0.882192126163777 | 55.1143367107117 | 65.1532583131463 | 64.3589324280944 | 56.2756016554897 | 59.1277425742312 | 55.0387004433237 | 86.4094815161593 | 71.6446872681215 | Zfp763 |
| **ENSMUSG00000067547** | 0.0853746965034818 | 0.71068427180736 | 0.949886733585872 | 89.4478579403355 | 58.6379324818317 | 65.3053873167428 | 90.0409626487835 | 65.6974917491458 | 62.3771938357668 | 85.3684034256032 | 71.6446872681215 | Gm7666 |
| **ENSMUSG00000067567** | -0.233916846447798 | 0.268551325909761 | 0.843793744782845 | 61.4389327266951 | 59.452348210746 | 48.2691993210708 | 72.2203554578784 | 60.2227007700503 | 78.277262852727 | 71.8343882483734 | 73.8157383974585 | Hdac8 |
| **ENSMUSG00000067586** | -0.894278019805029 | 0.00729690630848684 | 0.635862253526407 | 112.035700854562 | 153.924572764808 | 133.450139299431 | 151.006197775564 | 371.190828382674 | 352.247682837272 | 158.243869764533 | 142.203848971575 | S1pr3 |
| **ENSMUSG00000067629** | 0.0805125084474709 | 0.784036493097416 | 0.961141387070613 | 38.8510898124689 | 49.679359463774 | 35.0188308799925 | 37.5170677703264 | 32.8487458745729 | 24.4616446414772 | 44.7663578939139 | 49.9341759747513 | Syngap1 |
| **ENSMUSG00000067653** | 0.309884625013802 | 0.37976823592544 | 0.87432013076003 | 6179.1303076157 | 6272.62994409816 | 4292.17292002071 | 5805.76623745802 | 2485.55510450935 | 2409.4719971855 | 6308.93322877019 | 6986.44253420652 | Ankrd23 |
| **ENSMUSG00000067713** | -0.112353933584659 | 0.340163114601439 | 0.858570960161243 | 1817.86959773692 | 1535.17364900351 | 1626.00949869803 | 1743.60572462592 | 1840.6247271719 | 2067.00897220482 | 1763.5862854021 | 1596.80810562737 | Prkag1 |
| **ENSMUSG00000067719** | 0.0176775195916323 | 0.929237401133213 | 0.987083779576348 | 158.114900399583 | 123.791190794978 | 106.949402417274 | 137.87522405595 | 137.964732673206 | 159.000690169602 | 110.35427759895 | 113.980184290193 | Gm10221 |
| **ENSMUSG00000067722** | -0.0571012427161768 | 0.680960551773622 | 0.944452327903597 | 833.03964667666 | 637.687515739919 | 624.660226507975 | 695.941607139556 | 682.158955995297 | 656.795158623663 | 795.383661184888 | 768.552099785303 | BC003965 |
| **ENSMUSG00000067736** | 0.124221198017885 | 0.624093850579136 | 0.931145988862144 | 108.421645988285 | 118.904696421492 | 81.3951204237664 | 71.2824287636203 | 86.5016974697087 | 59.9310293716191 | 98.902418602833 | 103.124928643508 | Gm10222 |
| **ENSMUSG00000067780** | -1.06224768848336 | 0.0717154161935949 | 0.728065810245271 | 9.03513716569045 | 19.5459774939439 | 10.4110037751329 | 17.8206071909051 | 36.1336204620302 | 59.9310293716191 | 13.5340151772298 | 9.76973008201657 | Pi15 |
| **ENSMUSG00000067787** | 0.0793156564540526 | 0.446140113970751 | 0.89108031640712 | 991.154547076242 | 1022.09173978748 | 1016.49255040843 | 965.126568391648 | 892.390929592564 | 842.703657898889 | 1032.74946583169 | 1011.70982627105 | Blcap |
| **ENSMUSG00000067818** | -0.684311580474845 | 0.133694965770824 | 0.77303252008843 | 492.41497553013 | 677.593886456721 | 530.961192531779 | 332.026049767389 | 918.669926292222 | 1637.7071087469 | 366.45948787576 | 345.197129564585 | Myl9 |
| **ENSMUSG00000067825** | 0.0270613531160503 | 0.909644421714538 | 0.985873987017395 | 75.8951521917998 | 96.1010560118908 | 111.681676860517 | 137.87522405595 | 85.4067392738896 | 89.2850029413918 | 116.600746142287 | 121.578863242873 | Pex26 |
| **ENSMUSG00000067847** | 0.0281049567562562 | 0.883777154068044 | 0.979250426961432 | 1530.55223586796 | 1180.90280692578 | 1231.33781013163 | 1574.77891965945 | 1320.51958415783 | 1801.6001278448 | 1204.52735077345 | 1085.52556466851 | Romo1 |
| **ENSMUSG00000067851** | 0.341646429500928 | 0.237334325407639 | 0.832048466795696 | 2642.77762096446 | 3225.90070222966 | 3454.56034356683 | 2484.56781308987 | 1851.57430913009 | 1238.98230109082 | 3154.46661438509 | 3072.03734801188 | Arfgef1 |
| **ENSMUSG00000067870** | 0.0403722047231973 | 0.917164997286966 | 0.986713918026868 | 46.9827132615903 | 21.9892246806869 | 22.7149173275627 | 41.2687745473591 | 26.2789966996583 | 51.3694537471021 | 27.0680303544596 | 24.9670879873757 | Rpl31-ps8 |

| | | | | | | | | | | | | |
|---|---|---|---|---|---|---|---|---|---|---|---|---|
| **ENSMUSG00000067873** | 0.198927209506242 | 0.0500595669065607 | 0.703396849069134 | 3260.78100309768 | 3031.25534301913 | 2961.45734658099 | 3365.28097899828 | 2612.57025522437 | 2508.54165798349 | 3029.53724351836 | 2840.82040273748 | Htatsf1 |
| **ENSMUSG00000067889** | -0.361342247694946 | 0.322934875504735 | 0.855977678491063 | 10.8421645988285 | 21.1748089517725 | 17.9826428843205 | 15.0068271081306 | 19.7092475247437 | 15.9000690169602 | 22.903717992235 | 24.9670879873757 | Sptbn2 |
| **ENSMUSG00000067916** | 0.0169865880114845 | 0.946051465261581 | 0.990505437194639 | 53.3073092775737 | 33.3910448854875 | 41.6440151005317 | 53.4618215727152 | 45.9882442244021 | 41.5847958905112 | 42.6842017128016 | 48.8486504100828 | Zfp991 |
| **ENSMUSG00000067924** | -0.398325938796189 | 0.0931533471788447 | 0.754058637722175 | 359.59845919448 | 291.56083095133 | 278.257737262643 | 392.99128489417 | 469.737066006393 | 610.318033804856 | 316.487739529066 | 347.368180693922 | Rtl8b |
| **ENSMUSG00000067925** | -0.507505932717873 | 0.0130827197142595 | 0.635862253526407 | 430.976042803434 | 365.672662282534 | 312.330113253987 | 413.625672167849 | 578.137927392483 | 715.503105763208 | 444.54034466747 | 427.697072479392 | Rtl8a |
| **ENSMUSG00000067928** | -0.25947916960043 | 0.572694615566232 | 0.921859799538181 | 19.877301764519 | 6.51532583131463 | 21.7684624389143 | 19.6964605794214 | 16.4243729372865 | 31.8001380339204 | 15.616171358342 | 17.3684090346961 | Zfp760 |
| **ENSMUSG00000067931** | -0.161467555222809 | 0.552877725229339 | 0.914977712327965 | 33.4300075130547 | 48.0505280059454 | 33.1259211026956 | 40.3308478531009 | 56.937826182593 | 40.3617136584374 | 42.6842017128016 | 33.6512925047237 | Zfp948 |
| **ENSMUSG00000067942** | -0.0865114773709474 | 0.736921076047372 | 0.954383574136061 | 65.0529875929712 | 60.2667639396603 | 53.9479286529615 | 72.2203554578784 | 49.2731188118594 | 50.1463715150282 | 84.3273253350471 | 82.4999429148066 | Zfp160 |
| **ENSMUSG00000067995** | 0.146877189031228 | 0.286629671199747 | 0.851785577030345 | 216.843291976571 | 218.26341534904 | 264.060913932917 | 248.550573978413 | 234.321053905287 | 210.370143916704 | 193.640524843441 | 217.105112933702 | Gtf2f2 |
| **ENSMUSG00000068011** | -0.105709580360208 | 0.756631995948769 | 0.956566265296272 | 45.1756858284523 | 25.2468875963442 | 22.7149173275627 | 26.2619474392285 | 27.3739548954774 | 30.5770558018465 | 26.0269522639034 | 44.5065481514088 | Mkrn2os |
| **ENSMUSG00000068015** | -0.00985119493315281 | 0.942763478446417 | 0.990470440696959 | 393.931980424104 | 377.888898216248 | 370.063861461543 | 340.467390015713 | 430.318570956905 | 335.124531588238 | 400.815064864113 | 325.657669400552 | Lrch1 |
| **ENSMUSG00000068036** | -0.211474748569413 | 0.125448719107857 | 0.771949105062466 | 1185.40999613859 | 1127.15136881743 | 1077.06566328193 | 1451.91052271163 | 1605.2087150708 | 1369.85209992272 | 1293.01898847072 | 1337.3674956716 | Afdn |
| **ENSMUSG00000068039** | 0.111156699182253 | 0.163043567457708 | 0.793968226938511 | 4944.93057078238 | 4680.44719407065 | 4453.07025109094 | 4993.52172023045 | 4434.58069306734 | 4179.27198699638 | 4420.41757250136 | 4622.1678543585 | Tcp1 |
| **ENSMUSG00000068040** | 0.00963015619533963 | 0.943956813333119 | 0.990505437194639 | 758.951521917998 | 926.805099504506 | 885.881775774946 | 738.148308381173 | 763.185862485911 | 753.418654957498 | 960.915077583314 | 809.802071242707 | Tm9sf4 |
| **ENSMUSG00000068079** | -0.730400305716341 | 0.0359277534978799 | 0.679566075569698 | 468.923618899334 | 311.921224174188 | 384.260684791269 | 596.521377548191 | 999.696832782835 | 1067.75078860048 | 428.924173309128 | 426.611546914723 | Tcf15 |
| **ENSMUSG00000068083** | 0.0322371173437304 | 0.952733338378925 | 0.992000114323903 | 33.4300075130547 | 29.3189662409158 | 16.0897331070236 | 23.448167356454 | 19.7092475247437 | 23.2385624094033 | 6.24646854333682 | 51.0197015394199 | Cyp2d40 |
| **ENSMUSG00000068086** | -0.422321967277106 | 0.298589282269977 | 0.851785577030345 | 198.77301764519 | 360.786167909048 | 180.772883731853 | 215.723139679377 | 198.187433443257 | 239.724117486476 | 198.845915296222 | 644.802185413094 | Cyp2d9 |
| **ENSMUSG00000068114** | 0.191204463415482 | 0.447470818254572 | 0.89108031640712 | 53.3073092775737 | 59.452348210746 | 52.0550188756646 | 59.0893817382642 | 33.943704070392 | 67.2695227640623 | 44.7663578939139 | 51.0197015394199 | Ccdc134 |
| **ENSMUSG00000068115** | -0.255792166546922 | 0.488674553644966 | 0.899236955296196 | 27.1054114970714 | 21.1748089517725 | 9.46454888648447 | 23.448167356454 | 25.1840385038392 | 20.7923979452556 | 26.0269522639034 | 24.9670879873757 | Ninl |
| **ENSMUSG00000068130** | 0.267207887207676 | 0.508821783040514 | 0.902394714365738 | 21.6843291976571 | 27.6901347830872 | 23.6613722162112 | 37.5170677703264 | 27.3739548954774 | 17.123151249034 | 11.4518589961175 | 35.8223436340608 | Zfp442 |
| **ENSMUSG00000068134** | -0.0235099607612994 | 0.875288273919815 | 0.976986380159756 | 191.544907912638 | 178.357044632238 | 211.059440168604 | 186.647412157374 | 159.863896589588 | 195.693157131818 | 215.50316474512 | 208.420908416353 | Zfp120 |
| **ENSMUSG00000068184** | 0.0694715779582714 | 0.652391953703139 | 0.936720465895231 | 780.635851115655 | 611.626212414661 | 588.694940739334 | 712.824287636203 | 618.65138063779 | 776.657217366901 | 595.49666779811 | 577.499600403646 | Ndufaf2 |
| **ENSMUSG00000068196** | -0.972565245611651 | 0.0381581804365442 | 0.679566075569698 | 135.527057485357 | 112.389370590177 | 93.6990339761962 | 173.51643843776 | 315.3479603959 | 485.563646133322 | 102.025652874501 | 108.552556466851 | Col8a1 |
| **ENSMUSG00000068205** | 0.0276441872759114 | 0.907745948703469 | 0.98504921011344 | 128.298947752804 | 101.801966114291 | 120.199770858353 | 127.55803041911 | 98.5462376237187 | 84.3926740130963 | 116.600746142287 | 168.256462523619 | Macrod2 |
| **ENSMUSG00000068206** | 0.0491908344903176 | 0.587427791431335 | 0.923058967588378 | 561.082017989377 | 560.318021493058 | 519.603733867997 | 517.735535230505 | 521.20010120989 | 512.471455238947 | 529.908748093074 | 523.223322170221 | Pick1 |
| **ENSMUSG00000068220** | -0.20058440245932 | 0.404012238856381 | 0.881158207349486 | 6138.47219037009 | 3995.5235660537 | 3570.97429487059 | 5236.44473404331 | 5440.84727502509 | 7568.43285207304 | 4260.09154655571 | 4498.4179399863 | Lgals1 |
| **ENSMUSG00000068240** | 0.0689790987164631 | 0.784376977476975 | 0.961141387070613 | 94.8689402397497 | 66.7820897709749 | 79.5022106464695 | 106.92364314543 | 63.5075753575076 | 113.746647582869 | 83.2862472444909 | 71.6446872681215 | Gm11808 |
| **ENSMUSG00000068245** | -0.0775976053682512 | 0.801955831852167 | 0.963461896828986 | 89.4478579403355 | 72.4829998733752 | 71.930571537282 | 117.24083678227 | 101.831112211176 | 139.43137445642 | 58.3003730711437 | 71.6446872681215 | Phf11d |
| **ENSMUSG00000068246** | -0.354710536376492 | 0.412998727501333 | 0.882192126163777 | 10.8421645988285 | 22.8036404096012 | 14.1968233297267 | 12.1930470253561 | 12.0445401540101 | 28.1308913376988 | 14.5750932677859 | 22.7960368580387 | Apol9b |
| **ENSMUSG00000068250** | 0.124102949626243 | 0.441963558742116 | 0.89108031640712 | 174.378147297826 | 136.821842457607 | 133.450139299431 | 139.751077444466 | 148.914314631397 | 133.315963296051 | 124.929370866736 | 129.177542195552 | Amn1 |
| **ENSMUSG00000068264** | -0.11043528089275 | 0.361821571844097 | 0.868897239438884 | 654.143930795989 | 714.242594257866 | 675.768790494991 | 755.03098887782 | 873.776640263639 | 763.203312814088 | 706.892023487617 | 678.453477917817 | Ap5s1 |
| **ENSMUSG00000068267** | 0.0395734784625952 | 0.609415874185903 | 0.927932112008768 | 7247.08352060031 | 7021.89241469934 | 7188.32487928495 | 8008.01811557618 | 7031.82153355024 | 7399.64750404685 | 7257.35536926683 | 6978.84385525384 | Cenpb |
| **ENSMUSG00000068270** | -0.336549218145528 | 0.135739895581848 | 0.776087404131513 | 345.142239729375 | 377.074482487334 | 377.63550057073 | 346.094950181261 | 648.215251924905 | 494.125221757839 | 323.775286162958 | 360.394487469945 | Shroom4 |
| **ENSMUSG00000068284** | 0.168045903924722 | 0.454073570516969 | 0.891964629399053 | 364.116027777325 | 491.09268453534 | 452.405436773958 | 357.350070512359 | 336.152166116463 | 245.839528646846 | 451.827891301363 | 447.236532643425 | Usf3 |
| **ENSMUSG00000068290** | -0.007706430557966 | 0.968344398180172 | 0.994697802893254 | 2755.71683553559 | 2247.78741180355 | 2437.12133826975 | 3206.77136766865 | 2798.71314851361 | 3494.34593703502 | 2195.63369298289 | 2216.64320305309 | Ddrgk1 |
| **ENSMUSG00000068299** | -0.419149487709243 | 0.318301630598147 | 0.854351922245337 | 16.2632468982428 | 11.4018202048006 | 10.4110037751329 | 11.2551203310979 | 15.3294147414674 | 9.78465785659088 | 16.6572494488982 | 23.8815624227072 | Nat8f4 |
| **ENSMUSG00000068328** | 0.0158078274081172 | 0.906616603341161 | 0.98504921011344 | 955.91751213005 | 895.857301805761 | 946.454888648447 | 1041.09863062656 | 989.842209020464 | 1130.12798243625 | 850.560799984364 | 828.256005842071 | Aup1 |
| **ENSMUSG00000068329** | 0.0880295951746378 | 0.356976235740106 | 0.865163598066054 | 561.985531705946 | 566.833347324373 | 554.62256474799 | 627.47295845871 | 540.909348734634 | 532.040770952129 | 569.469715534207 | 530.8220011229 | Htra2 |
| **ENSMUSG00000068335** | -0.833536616250891 | 0.0199159907808501 | 0.642273034804458 | 36.1405486627618 | 30.1333819698302 | 26.5007368821565 | 30.0136542162612 | 53.6529515951357 | 92.9542496376133 | 40.6020455316893 | 32.5657669400552 | Dok1 |
| **ENSMUSG00000068391** | -0.0504317511741652 | 0.741545778662729 | 0.954383574136061 | 680.345828576491 | 531.813470981057 | 565.033568523123 | 688.43819358549 | 695.298454345127 | 725.287763619799 | 587.168043073661 | 546.019359028259 | Chrac1 |
| **ENSMUSG00000068394** | 0.228030929685247 | 0.371482706056538 | 0.873374202289323 | 38.8510898124689 | 47.2361122770311 | 38.8046504345863 | 46.8963347129081 | 38.3235368536684 | 33.0232202659942 | 45.80743598447 | 29.3091902460497 | Cep152 |
| **ENSMUSG00000068457** | 0.0137669835536325 | 0.961673216898672 | 0.993822507448007 | 259.308436655316 | 343.683437601847 | 368.170951684246 | 260.743621003769 | 234.321053905287 | 179.793088114857 | 438.293876124134 | 366.907640857956 | Uty |
| **ENSMUSG00000068479** | 0.265581244742677 | 0.119502959295391 | 0.769868040577702 | 593.608511785863 | 627.100111264033 | 659.679057387967 | 560.88016316638 | 489.446313531136 | 369.370834086306 | 577.798340258656 | 592.696958309005 | Mfap1a |
| **ENSMUSG00000068523** | 0.0485642674844571 | 0.755205830939594 | 0.956566265296272 | 1621.80712124144 | 1404.86713237722 | 1430.0933367478 | 1846.77766099432 | 1575.64484378368 | 1837.06951257494 | 1328.41564354963 | 1354.7359047063 | Gng5 |
| **ENSMUSG00000068551** | -0.591388621132671 | 0.0364572157981224 | 0.679566075569698 | 52.4037955610046 | 65.1532583131463 | 58.6802030962037 | 62.8410885152968 | 99.6411958195378 | 133.315963296051 | 68.711153976705 | 59.7039060567679 | Zfp467 |
| **ENSMUSG00000068566** | -0.49355731938538 | 0.00566316772852029 | 0.635862253526407 | 1671.50037565273 | 1663.85133417197 | 1670.49287846451 | 1512.87575783841 | 2967.33671066975 | 2476.74151994957 | 2001.99316813945 | 1732.49880121094 | Myadm |
| **ENSMUSG00000068580** | 0.217461276890715 | 0.140635183556586 | 0.779711473022872 | 231.299511441676 | 215.820168162297 | 226.202718386979 | 272.936668029125 | 197.092475247437 | 236.054870790255 | 197.804837205666 | 183.453820428978 | Zfyve19 |
| **ENSMUSG00000068587** | -0.486999626830607 | 0.297294067827001 | 0.851785577030345 | 7.22810973255236 | 11.4018202048006 | 6.62518422053913 | 14.0689004138724 | 17.5193311331055 | 9.78465785659088 | 10.4107809055614 | 17.3684090346961 | Mgam |
| **ENSMUSG00000068614** | -0.0511270894414437 | 0.781994240753622 | 0.960742105865155 | 232738.80879217 | 218936.937156852 | 145560.029599688 | 217409.531875653 | 201109.876867898 | 171473.682772291 | 235676.134905827 | 235772.896069306 | Actc1 |
| **ENSMUSG00000068663** | 0.14816889990826 | 0.687773348775022 | 0.944452327903597 | 1335.39327308905 | 1770.53979465975 | 1551.2395624948 | 1129.26373988683 | 792.749733773026 | 588.302553627526 | 1792.73647193767 | 2047.30121496481 | Clec16a |
| **ENSMUSG00000068686** | -0.153740314008669 | 0.506014459451757 | 0.901494649536949 | 61.4389327266951 | 57.8235167529173 | 52.0550188756646 | 37.5170677703264 | 58.0327843784121 | 62.3771938357668 | 60.3825292522559 | 52.1052271040884 | Cd59b |
| **ENSMUSG00000068699** | 0.332872335453466 | 0.0706203778655772 | 0.72606739345142 | 12733.2188076076 | 14911.1375806924 | 12027.5487249445 | 13177.8700543272 | 10178.7313883343 | 7287.12393869606 | 12011.9590088367 | 12481.3729425585 | Flnc |
| **ENSMUSG00000068706** | 0.0482795348939786 | 0.804034969504497 | 0.963461896828986 | 163.535982698997 | 109.13170767452 | 138.182413742673 | 173.51643843776 | 151.104231023035 | 156.554525705454 | 127.011527047849 | 130.263067760221 | Gm10250 |
| **ENSMUSG00000068732** | 0.0751587762670042 | 0.594797718571825 | 0.924461858177432 | 657.757985662265 | 741.118313312039 | 821.522843346852 | 667.803806311811 | 637.265669966714 | 588.302553627526 | 725.631429117627 | 789.177085514005 | Tmem167b |
| **ENSMUSG00000068735** | -0.873495606911746 | 0.00486487030135009 | 0.635862253526407 | 630.652574165193 | 419.424100390879 | 362.492222352355 | 524.301022090312 | 1415.78094719409 | 956.450305481758 | 588.209121164217 | 588.354856050331 | Trp53i11 |
| **ENSMUSG00000068739** | 0.045827712809393 | 0.757317764540161 | 0.956693827734505 | 2471.11001481634 | 2166.34583891211 | 2412.51351116489 | 2478.00232623006 | 2515.11897579647 | 2772.72742011144 | 2042.59521367114 | 1900.75526373456 | Sars |
| **ENSMUSG00000068740** | 0.168930016263361 | 0.556564783555013 | 0.916239900758826 | 101.193536255733 | 162.068730053951 | 130.610774633486 | 78.7858423176855 | 100.736154015357 | 68.4926049961361 | 121.806136595068 | 129.177542195552 | Celsr2 |
| **ENSMUSG00000068742** | 0.0516946251756576 | 0.826679942611654 | 0.967319591893432 | 602.643648951553 | 786.725594131241 | 908.596693102509 | 718.451847801752 | 785.085026402292 | 459.878919259771 | 968.202624217207 | 695.821886952513 | Cry2 |
| **ENSMUSG00000068745** | -6.46578671530821 | NA | NA | 2.71054114970714 | 0.814415728914329 | 0 | 2.81378008277448 | 1.0949581958191 | 558.948580057754 | 0 | 0 | Mybphl |
| **ENSMUSG00000068747** | -0.0677248981415235 | 0.824124934856661 | 0.966637987288264 | 505.967681278665 | 1010.68991958268 | 718.359260484171 | 325.460562907582 | 671.209374037106 | 697.1568722821 | 730.836819570408 | 585.098279356326 | Sort1 |
| **ENSMUSG00000068749** | 0.0895880309898259 | 0.511544646894142 | 0.902394714365738 | 1966.94936097081 | 1579.15209836488 | 1481.20190073482 | 2047.49397356557 | 1717.98940924016 | 1708.64587820718 | 1621.99966508646 | 1600.06468232138 | Psma5 |
| **ENSMUSG00000068758** | -0.987738231749094 | 0.0215076446114617 | 0.648762042745232 | 61.4389327266951 | 52.122606650517 | 31.2330113253987 | 60.0273084325223 | 131.394983498292 | 170.008430258266 | 43.7252798033577 | 61.8749571861049 | Il3ra |
| **ENSMUSG00000068762** | -0.563027104682033 | 0.104958596395664 | 0.762094734041179 | 36.1405486627618 | 44.7928650902881 | 33.1259211026956 | 35.6412143818101 | 43.7983278327639 | 69.71568722821 | 28.1091084450157 | 80.3288917854696 | Gstm6 |
| **ENSMUSG00000068794** | -0.515188207005012 | 0.28516408507259 | 0.851275833092507 | 18.9737880479499 | 25.2468875963442 | 27.447191770805 | 20.6343872736795 | 52.5579933993167 | 50.1463715150282 | 16.6572494488982 | 13.0263067760221 | Col28a1 |
| **ENSMUSG00000068798** | 0.092695037366694 | 0.362110664231679 | 0.86924083181804 | 1927.19475744177 | 1718.41718800923 | 1968.62616838877 | 1904.92911603833 | 1688.42553795305 | 1607.13005294505 | 1814.59911183935 | 1938.74865849795 | Rap1a |
| **ENSMUSG00000068823** | 0.193200930136683 | 0.238581487647266 | 0.832048466795696 | 22735.1156500269 | 21118.6142664775 | 19500.7565257126 | 21375.3493621435 | 16079.4611056034 | 14500.8629434677 | 21309.8274355936 | 22218.537257635 | Csde1 |
| **ENSMUSG00000068854** | 0.244344529684505 | 0.254749111805422 | 0.837406774112203 | 232.203025158245 | 250.025628776699 | 267.84673348751 | 203.530092654021 | 154.389105610493 | 155.33144347338 | 241.530117009024 | 252.927456567762 | Hist2h2be |
| **ENSMUSG00000068856** | -0.294379490230195 | 0.233233109754725 | 0.832048466795696 | 252.080326922764 | 248.39679731887 | 246.078271048596 | 223.226553233442 | 373.380744774312 | 398.724807656078 | 219.667477107345 | 198.651178334337 | Sf3b4 |
| **ENSMUSG00000068874** | -0.139616398411276 | 0.325965423592428 | 0.855977678491063 | 2188.31022153023 | 2353.66145656241 | 2729.57589886212 | 2247.27235944255 | 2933.39300659936 | 2956.18975492252 | 2361.16510938132 | 2236.18266321713 | Selenbp1 |
| **ENSMUSG00000068877** | 0.102433000479306 | 0.793911151281735 | 0.961804768755116 | 16.2632468982428 | 36.6487078011448 | 20.8220075502658 | 26.2619474392285 | 27.3739548954774 | 13.4539045528125 | 20.8215618111227 | 31.4802413753867 | Selenbp2 |
| **ENSMUSG00000068882** | 0.0473417315219327 | 0.646094065584731 | 0.935063240511793 | 1506.1573655206 | 1285.14802022681 | 1341.12657721485 | 1499.7447841188 | 1373.07757755715 | 1429.78312929434 | 1406.49650034134 | 1240.7557204161 | Ssb |
| **ENSMUSG00000068917** | 0.260114241665055 | 0.0227800801135863 | 0.650119841310877 | 309.001691066613 | 302.96265115613 | 334.098575692902 | 318.895076047775 | 278.119381738051 | 248.285693110993 | 280.050006359601 | 248.585354309088 | Clk2 |
| **ENSMUSG00000068921** | 0.0353317025459082 | 0.637426259312998 | 0.932931968211785 | 2725.90088288881 | 2553.19331014642 | 2567.73211290324 | 2839.10410351945 | 2598.33579867872 | 2544.01104271363 | 2609.98277302423 | 2673.64946577853 | Dap3 |
| **ENSMUSG00000068922** | 0.0680578968419523 | 0.607422846249358 | 0.927459968605428 | 349.65980831222 | 346.941100517504 | 353.027673465871 | 434.260059441529 | 329.582416941548 | 381.601656407044 | 330.021754706295 | 374.506319810635 | Msto1 |
| **ENSMUSG00000068923** | -0.335693492882012 | 0.12534776182339 | 0.771949105062466 | 421.940905637744 | 410.465527372822 | 372.903226127488 | 403.308478531009 | 516.820268426614 | 733.849339244316 | 393.52751823022 | 387.532626586657 | Syt11 |
| **ENSMUSG00000068959** | 0.233509410089285 | 0.468226549088736 | 0.895328112556867 | 40.658117245607 | 54.56585383726 | 39.7511053232348 | 30.9515809105193 | 27.3739548954774 | 23.2385624094033 | 46.8485140750261 | 43.4210225867403 | Zfp619 |
| **ENSMUSG00000068966** | -0.0717959906453868 | 0.764464242679248 | 0.958110466593105 | 82.2197482077831 | 70.8541684155466 | 77.6093008691726 | 66.5927952923295 | 94.1664048404423 | 50.1463715150282 | 72.8754663389296 | 94.4407241261602 | Zbtb34 |
| **ENSMUSG00000069020** | -0.258000661248663 | 0.145935895505195 | 0.780607990324772 | 445.432262268539 | 340.425774686189 | 330.312756138308 | 399.556771753977 | 353.671497249568 | 551.610086665311 | 446.622500848583 | 461.348364984116 | Urm1 |
| **ENSMUSG00000069045** | 0.115837908931634 | 0.337791772915774 | 0.858570960161243 | 1553.14007878219 | 1656.52159261174 | 1637.36695736181 | 1645.12342172881 | 1463.95910781013 | 1316.03648171147 | 1788.57215957544 | 1420.95296415108 | Ddx3y |
| **ENSMUSG00000069049** | 0.0625367774256103 | 0.696983409156244 | 0.944933386973863 | 1133.00620057758 | 1073.39993070909 | 1106.40576483003 | 930.423280704096 | 845.307727172343 | 858.60372691585 | 1183.70578896233 | 1174.53866097133 | Eif2s3y |
| **ENSMUSG00000069089** | 0.113922567453791 | 0.305505544001929 | 0.851785577030345 | 424.651446787451 | 371.373572384934 | 398.457508120996 | 355.474217123843 | 362.431162816121 | 342.463024980681 | 372.705956419097 | 353.881334081933 | Cdk7 |
| **ENSMUSG00000069094** | 0.310333779649776 | 0.153781079221524 | 0.785320762994975 | 508.678222428372 | 617.327122517061 | 742.967087589031 | 526.176875478828 | 419.368988998714 | 346.132271676902 | 553.853544175865 | 611.15089290837 | Pde7a |
| **ENSMUSG00000069114** | -0.146965482769295 | 0.349724573314345 | 0.861774532843376 | 208.711668527449 | 210.933673788811 | 242.292451494002 | 225.102406621959 | 269.359716171498 | 190.800828203522 | 244.653351280692 | 275.723493425801 | Zbtb10 |
| **ENSMUSG00000069135** | 0.102047594568274 | 0.695250930620672 | 0.944933386973863 | 399.353062723518 | 372.187988113848 | 343.563124579386 | 341.405316709971 | 264.979883388221 | 204.254732756335 | 406.020455316893 | 480.887825148149 | Fgfr1op |
| **ENSMUSG00000069171** | -0.431562981143798 | 0.309450573314418 | 0.851785577030345 | 28.9124389302094 | 28.5045505120015 | 31.2330113253987 | 45.9584080186499 | 67.887408140784 | 66.0464405319884 | 20.8215618111227 | 27.1381391167127 | Nr2f1 |
| **ENSMUSG00000069184** | -0.0925525989835576 | 0.844471018439344 | 0.971806333564415 | 9.93865088225949 | 25.2468875963442 | 22.7149173275627 | 19.6964605794214 | 19.7092475247437 | 7.33849339244316 | 26.0269522639034 | 29.3091902460497 | Zfp72 |
| **ENSMUSG00000069206** | 0.00539337491142616 | 0.976066464303344 | 0.99575536949131 | 137.334084918495 | 109.946123403434 | 106.002947528626 | 121.930470253561 | 102.926070406995 | 103.961989726278 | 132.216917500629 | 133.519644454226 | Zfp874a |

| | | | | | | | | | | | | |
|---|---|---|---|---|---|---|---|---|---|---|---|---|
| **ENSMUSG00000069208** | -0.10312102893792 | 0.592085971589467 | 0.923935529355527 | 83.1232619243521 | 79.8127414336042 | 60.5731128735006 | 81.59962240046 | 72.2672409240604 | 86.838838477244 | 81.2040910633787 | 87.9275707381491 | Zfp825 |
| **ENSMUSG00000069237** | 0.159299661951388 | 0.243602526506819 | 0.832048466795696 | 965.856163012309 | 1129.59461600417 | 1258.78500190243 | 996.078149302167 | 946.0438811877 | 842.703657898889 | 1051.4888714617 | 1054.04532329312 | Fam8a1 |
| **ENSMUSG00000069255** | -0.205811500016084 | 0.188893659623749 | 0.808677608936066 | 266.536546387868 | 246.767965861042 | 268.793188376159 | 270.12288794635 | 261.695008800764 | 385.270903103266 | 275.885693997376 | 292.006376895829 | Dusp22 |
| **ENSMUSG00000069270** | 0.429980837979976 | 0.41298837057568 | 0.882192126163777 | 8.13162344912141 | 8.95857301805762 | 13.2503684410783 | 15.9447538023887 | 9.85462376237187 | 8.56157562451702 | 12.4929370866736 | 3.25657669400552 | Hist1h2ac |
| **ENSMUSG00000069456** | -0.212121010747178 | 0.646218887722501 | 0.935063240511793 | 20.780815481088 | 41.5352021746308 | 35.965285768641 | 41.2687745473591 | 25.1840385038392 | 25.6847268735511 | 27.0680303544596 | 83.5854684794751 | Rdh16 |
| **ENSMUSG00000069495** | 0.181049031590107 | 0.199499907477755 | 0.812921911423739 | 571.924182588206 | 663.748819065178 | 589.641395627982 | 512.107975064956 | 486.161438943679 | 441.532685778663 | 551.771387994752 | 581.84170266232 | Epc2 |
| **ENSMUSG00000069515** | -0.736722162054955 | 0.263459160880202 | 0.842510961779757 | 7.22810973255236 | 6.51532583131463 | 8.51809399783602 | 14.0689004138724 | 18.6142893289246 | 28.1308913376988 | 2.08215618111227 | 11.9407812113536 | Lyz1 |
| **ENSMUSG00000069516** | -0.152371199921419 | 0.466617048153744 | 0.895328112556867 | 1311.90191645825 | 1285.96243595572 | 1425.36106230456 | 1949.94959736272 | 1803.39614851405 | 1982.61629819173 | 1164.96638333232 | 1687.99225305953 | Lyz2 |
| **ENSMUSG00000069520** | -0.133940862557126 | 0.328242577766029 | 0.855977678491063 | 276.475197270128 | 296.447325324816 | 258.382184601026 | 281.378008277448 | 339.43704070392 | 338.793778284459 | 266.515991182371 | 277.894544555138 | Tmem19 |
| **ENSMUSG00000069539** | 0.0821440686288721 | 0.644902817481091 | 0.935041767725715 | 391.221439274397 | 493.535931722083 | 484.584902988005 | 425.818719193205 | 360.241246424483 | 338.793778284459 | 499.717483466946 | 496.085183053508 | Scyl2 |
| **ENSMUSG00000069565** | 0.0771912558305074 | 0.47621878497711 | 0.897553735056661 | 789.670988281345 | 899.929380450333 | 903.864418659267 | 880.713165908413 | 888.011096809288 | 875.726878164883 | 790.178270732108 | 740.328435103922 | Dazap1 |
| **ENSMUSG00000069601** | 0.102846830740184 | 0.561538937806052 | 0.917031459819748 | 1616.38603894202 | 1720.86043519598 | 1767.03127710665 | 1440.65540238054 | 1342.41874807421 | 1155.8127093098 | 1886.43350008772 | 1708.61723878823 | Ank3 |
| **ENSMUSG00000069631** | 0.0176111056377991 | 0.915887939600971 | 0.986713918026868 | 149.079763233892 | 171.027303072009 | 122.09268063565 | 158.509611329629 | 129.205067106653 | 149.216032313011 | 160.326025945645 | 155.230155747597 | Strada |
| **ENSMUSG00000069633** | -0.107220373415317 | 0.729866803220428 | 0.952921973914044 | 68.6670424592474 | 52.9370223794314 | 36.9117406572894 | 53.4618215727152 | 49.2731188118594 | 45.2540425867328 | 45.80743598447 | 87.9275707381491 | Pex11g |
| **ENSMUSG00000069662** | -0.487680792754062 | 0.173582396794003 | 0.799889021902745 | 496.029030396406 | 356.714089264476 | 446.726707442067 | 773.789522762983 | 1042.40020241978 | 991.9196902119 | 376.870268781321 | 496.085183053508 | Marcks |
| **ENSMUSG00000069678** | -0.150605507269789 | 0.382365035229099 | 0.875126835008975 | 108.421645988285 | 96.1010560118908 | 101.270673085384 | 118.178763476528 | 117.160526952643 | 130.869798831903 | 125.970448957293 | 96.6117752554972 | Pcgf1 |
| **ENSMUSG00000069682** | 0.10810682953048 | 0.68846290404242 | 0.944584424505182 | 56.0178504272808 | 43.1640336324594 | 38.8046504345863 | 63.779015209555 | 36.1336204620302 | 59.9310293716191 | 43.7252798033577 | 47.7631248454143 | Gm10275 |
| **ENSMUSG00000069713** | -0.035397555870387 | 0.949511806306294 | 0.991267322147698 | 22.5878429142261 | 19.5459774939439 | 6.62518422053913 | 7.50341355406529 | 20.8042057205628 | 6.1154111603693 | 17.6983275394543 | 13.0263067760221 | 4933406P04 |
| **ENSMUSG00000069727** | 0.214363100162062 | 0.511276710455147 | 0.902394714365738 | 74.9916384752307 | 60.2667639396603 | 44.483379766477 | 82.5375490947182 | 61.3176589658694 | 28.1308913376988 | 72.8754663389296 | 62.9604827507734 | Zfp975 |
| **ENSMUSG00000069729** | 0.103260331259227 | 0.547302106811549 | 0.914699131434573 | 790.574501997914 | 939.835751167135 | 975.794990196549 | 743.775868546722 | 785.085026402292 | 609.094951572782 | 908.861173055507 | 907.499372062872 | Arid1b |
| **ENSMUSG00000069743** | 0.261085964794412 | 0.570447134978602 | 0.920872458148558 | 11.7456783153976 | 14.6594831204579 | 14.1968233297267 | 10.3171936368398 | 7.66470737073368 | 18.3462334811079 | 8.32862472444909 | 8.68420451734806 | Zfp820 |
| **ENSMUSG00000069744** | -0.052375572975302 | 0.751635250959707 | 0.956197192463153 | 1984.11612158562 | 1575.89443544923 | 1495.39872406455 | 2000.59763885266 | 1954.50037953709 | 2174.64020862732 | 1564.74037010587 | 1623.94624474409 | Psmb3 |
| **ENSMUSG00000069755** | -0.0554581966678461 | 0.853859852212208 | 0.9731539929935 | 26.2018977805023 | 21.1748089517725 | 23.6613722162112 | 33.7653609932938 | 31.7537876787538 | 26.9078091056249 | 22.903717992235 | 27.1381391167127 | Zfp125 |
| **ENSMUSG00000069763** | -0.378492261058698 | 0.123258658623157 | 0.770251097497048 | 215.036264543433 | 272.8292691863 | 352.081218577222 | 232.605820176024 | 444.553027502553 | 359.586176229715 | 366.45948787576 | 223.618266321713 | Tmem100 |
| **ENSMUSG00000069769** | 0.202244257599385 | 0.332358769839289 | 0.856655692540538 | 2602.11950371885 | 2967.73091616381 | 2989.85099324044 | 2650.58083797356 | 2010.34324752386 | 1686.63039802985 | 2988.93519798667 | 3056.83999010652 | Msi2 |
| **ENSMUSG00000069792** | -0.223031608759392 | 0.49864426666743 | 0.900927059989498 | 31.6229800799166 | 30.9477976987445 | 24.6078271048596 | 46.8963347129081 | 35.0386622662111 | 37.9155491942896 | 27.0680303544596 | 56.4473293627624 | Wfdc17 |
| **ENSMUSG00000069793** | -0.181981632666014 | 0.615397407612965 | 0.928422072203769 | 25.2983840639333 | 30.1333819698302 | 23.6613722162112 | 15.0068271081306 | 26.2789966996583 | 33.0232202659942 | 15.616171358342 | 32.5657669400552 | Slfn9 |
| **ENSMUSG00000069805** | -0.375811684874791 | 0.353181811284597 | 0.862728204353539 | 120.167324303683 | 267.942774812814 | 114.521041526462 | 133.185590584659 | 112.780694169367 | 152.885279009232 | 186.352978209548 | 373.420794245967 | Fbp1 |
| **ENSMUSG00000069806** | -0.316477272356209 | 0.2580834967614 | 0.838310965110319 | 44.2721721118832 | 58.6379324818317 | 65.3053873167428 | 76.9099889291692 | 94.1664048404423 | 95.400414101761 | 52.0539045278068 | 64.046008315442 | Cacng7 |
| **ENSMUSG00000069808** | -0.481786124728832 | 0.200381499411908 | 0.812921911423739 | 37.9475760958999 | 46.4216965481167 | 34.0723759913441 | 47.8342614071662 | 74.4571573156986 | 91.7311674055395 | 35.3966550789086 | 31.4802413753867 | Fam57a |
| **ENSMUSG00000069833** | 0.238372597790517 | 0.146473828400993 | 0.780607990324772 | 12618.4725656033 | 15015.3827939935 | 13783.2225433873 | 11787.8626934366 | 12599.6839592903 | 8493.08301952088 | 10963.5933716467 | 13044.7607106215 | Ahnak |
| **ENSMUSG00000069835** | -0.0106343794301787 | 0.964674737080847 | 0.994284908685797 | 46.9827132615903 | 47.2361122770311 | 49.2156542097192 | 48.7721881014244 | 56.937826182593 | 40.3617136584374 | 37.4788112600209 | 58.6183804920994 | Sat2 |
| **ENSMUSG00000069844** | -0.0601155517543222 | 0.688667329094567 | 0.94469184542218 | 535.783633925444 | 471.546707041396 | 404.136237452887 | 456.770300103725 | 457.692525852383 | 409.732547744743 | 522.621201459181 | 556.874614674944 | Sco1 |
| **ENSMUSG00000069874** | -0.17361261624758 | 0.416684692793098 | 0.882192126163777 | 288.220875585525 | 340.425774686189 | 301.919109478855 | 259.805694309511 | 457.692525852383 | 344.909189444828 | 302.953724351836 | 237.730098662403 | Irgm2 |
| **ENSMUSG00000069892** | -0.267829989737947 | 0.143205926780815 | 0.780607990324772 | 84.9302893574902 | 87.1424829938332 | 114.521041526462 | 114.427056699496 | 114.970610561005 | 136.985209992272 | 115.559668051731 | 115.065709854862 | 9930111J21 |
| **ENSMUSG00000069893** | 0.250508126411992 | 0.538630362115096 | 0.913055373416973 | 15.3597331816738 | 35.8342920722305 | 28.3936466594534 | 24.3860940507122 | 21.8991639163819 | 19.5693157131818 | 12.4929370866736 | 33.6512925047237 | 9930111J21 |
| **ENSMUSG00000069895** | 0.0113735254834908 | 0.951826964603514 | 0.991915011884503 | 362.309000344187 | 445.485403716138 | 471.334534546927 | 347.03287687552 | 356.956371837026 | 315.555215875056 | 459.115437935256 | 480.887825148149 | Atxn1l |
| **ENSMUSG00000069899** | 0.247335161918318 | 0.385191658011763 | 0.875126835008975 | 37.0440623793308 | 32.5766291565731 | 32.1794662140472 | 44.0825546301336 | 41.6084114411257 | 26.9078091056249 | 29.1501865355718 | 24.9670879873757 | Gm12166 |
| **ENSMUSG00000069917** | 0.128969786313244 | 0.894751494258561 | 0.983733637008786 | 2026.58126626437 | 980.556537612852 | 6112.20567089167 | 759.720622349111 | 1097.14811221074 | 6401.61240267458 | 642.345181873136 | 893.387539722182 | Hba-a2 |
| **ENSMUSG00000069919** | -0.168745127524233 | 0.872872587970422 | 0.976162349063696 | 15769.0248952795 | 8147.41495205894 | 47367.2278121888 | 6491.39065096073 | 10611.2398756829 | 61327.7892806475 | 5876.88582118939 | 9610.1578240103 | Hba-a1 |
| **ENSMUSG00000069920** | -1.01890829558172 | 0.0218181178052686 | 0.648762042745232 | 27.1054114970714 | 30.9477976987445 | 18.9290977729689 | 34.703287687552 | 75.5521155115177 | 94.1773318696872 | 29.1501865355718 | 28.2236646813812 | B3gnt9 |
| **ENSMUSG00000069922** | -0.432537187369183 | 0.289401029793393 | 0.851785577030345 | 167.150037565273 | 376.26006675842 | 141.968233297267 | 234.48167356454 | 186.142893289246 | 239.724117486476 | 235.283648465687 | 580.756177097652 | Ces3a |
| **ENSMUSG00000069972** | -0.0481651748830508 | 0.902360699985034 | 0.984902747274457 | 21.6843291976571 | 20.3603932228582 | 13.2503684410783 | 36.5791410760683 | 20.8042057205628 | 31.8001380339204 | 17.6983275394543 | 24.9670879873757 | Rps13-ps2 |
| **ENSMUSG00000070002** | -0.0487380741326823 | 0.737773416255783 | 0.954383574136061 | 230.395997725106 | 302.148235427216 | 316.115932808581 | 304.826175633902 | 290.163921892061 | 280.085831144914 | 326.898520434627 | 295.262953589834 | Ell |
| **ENSMUSG00000070003** | -0.265749129136777 | 0.387314564786038 | 0.875903501282656 | 247.562758339918 | 307.034729800702 | 257.435729712378 | 301.07446885687 | 332.867291529005 | 567.510155682271 | 239.447960827911 | 199.736703899005 | Ssbp4 |
| **ENSMUSG00000070031** | -0.230342201560296 | 0.543719831811182 | 0.913837378403038 | 31.6229800799166 | 34.2054606144018 | 37.8581955459379 | 25.3240207449704 | 38.3235368536684 | 64.8233582999146 | 19.7804837205666 | 29.3091902460497 | Sp140 |
| **ENSMUSG00000070034** | -0.377839874941884 | 0.250834790682477 | 0.836703579022786 | 83.1232619243521 | 56.1946852950887 | 96.5383986421416 | 88.1651092602672 | 121.54035973592 | 162.669936865823 | 65.5879197050366 | 71.6446872681215 | Sp110 |
| **ENSMUSG00000070044** | -0.00952143159746177 | 0.965435100058339 | 0.994284908685797 | 93.9654265231807 | 96.9154717408051 | 138.182413742673 | 77.8479156234274 | 108.400861386091 | 86.838838477244 | 106.189965236726 | 107.467030902182 | Fam149a |
| **ENSMUSG00000070047** | -0.103184201887178 | 0.52313787142795 | 0.90648515894888 | 224.974915425692 | 257.355370336928 | 214.845259723197 | 234.48167356454 | 193.80760065998 | 247.06261087892 | 282.132162540713 | 277.894544555138 | Fat1 |
| **ENSMUSG00000070056** | -0.0084300223570009 | 0.963850217143107 | 0.994284908685797 | 1202.5767567534 | 1548.20430066614 | 1396.96741564511 | 1334.66968592936 | 1163.9405621557 | 1057.96613074389 | 1564.74037010587 | 1727.0711733876 | Mfhas1 |
| **ENSMUSG00000070283** | -0.0281873945535301 | 0.851369447342485 | 0.972692784625411 | 688.477452025612 | 532.627886709971 | 551.783200082044 | 659.362466063487 | 666.82954125383 | 703.272283442469 | 525.744435730849 | 585.098279356326 | Ndufaf3 |
| **ENSMUSG00000070284** | -0.40365743709373 | 0.0187084505674773 | 0.642273034804458 | 102.097049972302 | 100.987550385377 | 92.7525790875478 | 116.302910088012 | 125.920192519196 | 165.116101329971 | 124.929370866736 | 130.263067760221 | Gmppb |
| **ENSMUSG00000070287** | -0.464164892119408 | 0.0751355074072255 | 0.735399943172214 | 37.9475760958999 | 41.5352021746308 | 34.0723759913441 | 27.1998741334867 | 48.1781606160403 | 61.154111603693 | 46.8485140750261 | 39.0789203280663 | Slc35g2 |
| **ENSMUSG00000070304** | -0.121086430380276 | 0.794159181658809 | 0.961804768755116 | 12.6491920319666 | 13.8450673915436 | 17.036187995672 | 9.37926694258161 | 22.994122112201 | 13.4539045528125 | 14.5750932677859 | 6.51315338801105 | Scn2b |
| **ENSMUSG00000070305** | -0.244745341069293 | 0.375662289310444 | 0.873821423608136 | 39.754603529038 | 64.338842584232 | 42.5904699891801 | 63.779015209555 | 78.836990098975 | 50.1463715150282 | 49.9717483466946 | 70.559161703453 | Mpzl3 |
| **ENSMUSG00000070315** | 0.11003957044025 | 0.807380842864542 | 0.964026716319739 | 13.5527057485357 | 17.9171460361152 | 14.1968233297267 | 27.1998741334867 | 6.56974917491458 | 18.3462334811079 | 20.8215618111227 | 21.7105112933701 | 4930581F22 |
| **ENSMUSG00000070319** | 0.145150897485232 | 0.405264377681709 | 0.881158207349486 | 2196.44184497935 | 1830.80655859941 | 2386.01277428273 | 2370.14075639037 | 2239.18951045005 | 2409.4719971855 | 1676.13572579538 | 1618.51861692074 | Eif3g |
| **ENSMUSG00000070327** | -0.278608607760199 | 0.165446921772431 | 0.795216960550936 | 237.624107457659 | 310.292392716359 | 291.508105703722 | 253.240207449704 | 445.647985698372 | 336.347613820311 | 272.762459725708 | 271.381391167127 | Rnf213 |
| **ENSMUSG00000070343** | -0.138151715438615 | 0.703117457397853 | 0.946451777392559 | 45.1756858284523 | 17.1027303072009 | 29.3401015481019 | 26.2619474392285 | 36.1336204620302 | 41.5847958905112 | 26.0269522639034 | 26.0526135520442 | Gm10288 |
| **ENSMUSG00000070348** | -0.118384204665109 | 0.502966778525818 | 0.900927059989498 | 1164.6291806575 | 1014.76199822725 | 882.095956220352 | 1052.35375095766 | 1217.59351375084 | 793.780368615935 | 1227.43106876569 | 1225.55836251075 | Ccnd1 |
| **ENSMUSG00000070371** | 0.0212602426470607 | 0.885601849228893 | 0.979948700426504 | 220.457346842847 | 184.872370463553 | 192.130342395635 | 248.550573978413 | 222.276513751277 | 198.139321595965 | 208.215618111227 | 204.078806157679 | Prss36 |
| **ENSMUSG00000070372** | -0.149196202172453 | 0.144738094117878 | 0.780607990324772 | 546.625798524272 | 539.143212541285 | 560.301294079881 | 516.797608536247 | 579.232885588302 | 543.048511040794 | 628.811166695907 | 645.887710977762 | Capza1 |
| **ENSMUSG00000070394** | 0.130968960410589 | 0.501067164774534 | 0.900927059989498 | 1384.18301378378 | 773.694942468612 | 942.669069093853 | 1098.31215897631 | 918.669926292222 | 1114.22791341929 | 869.300205614374 | 932.466460050248 | Tmem256 |
| **ENSMUSG00000070407** | -0.812651020128947 | 0.0384396466515927 | 0.679566075569698 | 19.877301764519 | 15.4738988493722 | 15.1432782183751 | 20.6343872736795 | 45.9882442244021 | 33.0232202659942 | 14.5750932677859 | 31.4802413753867 | Hs3st3b1 |
| **ENSMUSG00000070420** | 0.0689514419949055 | 0.739443110900123 | 0.954383574136061 | 102.097049972302 | 117.275864963663 | 133.450139299431 | 78.7858423176855 | 106.210944994452 | 89.2850029413918 | 110.35427759895 | 105.295979772845 | Zscan25 |
| **ENSMUSG00000070423** | -0.357647765309057 | 0.224825755475331 | 0.832048466795696 | 71.3775836089546 | 66.7820897709749 | 53.001473764313 | 59.0893817382642 | 110.590777777729 | 95.400414101761 | 70.7933101578173 | 44.5065481514088 | Olfr558 |
| **ENSMUSG00000070424** | -0.0995547671298683 | 0.699083246475569 | 0.945173655771957 | 412.905768472054 | 450.371898089624 | 507.299820315568 | 515.859681841989 | 516.820268426614 | 771.764888438605 | 409.143689588562 | 324.572143835884 | Art5 |
| **ENSMUSG00000070426** | 0.0879230704997799 | 0.453440261339267 | 0.891651261195907 | 531.266065342598 | 472.361122770311 | 478.906173656114 | 523.363095396054 | 492.731188118594 | 483.117481669175 | 407.061533407449 | 504.769387570856 | Rnf121 |
| **ENSMUSG00000070427** | -0.0539216126884611 | 0.787323168671178 | 0.961224689823763 | 84.9302893574902 | 66.7820897709749 | 75.7163910918757 | 76.9099889291692 | 90.881530252985 | 89.2850029413918 | 66.6289977955927 | 69.4736361387845 | Il18bp |
| **ENSMUSG00000070436** | -0.348630477215659 | 0.246100509583703 | 0.832048466795696 | 4939.50948848297 | 4085.10929623427 | 3050.42410611394 | 3931.78870233021 | 5691.59270186767 | 8001.40396222719 | 2961.86716763221 | 3728.78031463632 | Serpinh1 |
| **ENSMUSG00000070461** | 0.200104173747492 | 0.355631133306433 | 0.864644922392151 | 123.781379169959 | 115.647033505835 | 114.521041526462 | 107.861569839689 | 98.5462376237187 | 66.0464405319884 | 107.231043327282 | 129.177542195552 | 9230112E08 |
| **ENSMUSG00000070462** | -0.649142348165859 | 0.0233837592392389 | 0.654124840968651 | 341.528184863099 | 314.364471360931 | 379.528410348027 | 454.894446715208 | 840.927894389066 | 727.733928083946 | 402.897221045225 | 365.822115293287 | Tlnrd1 |
| **ENSMUSG00000070469** | -0.51824657349293 | 0.189994568557043 | 0.808824684419004 | 52.4037955610046 | 45.6072808192024 | 46.3762895437739 | 44.0825546301336 | 116.065568756824 | 79.5003450848009 | 29.1501865355718 | 45.5920737160773 | Adamtsl3 |
| **ENSMUSG00000070473** | -0.542397293131354 | 0.268281314251922 | 0.843793744782845 | 16.2632468982428 | 13.0306516626293 | 8.51809399783602 | 16.8826804966469 | 12.0445401540101 | 17.123151249034 | 12.4929370866736 | 37.9933947633978 | Cldn3 |
| **ENSMUSG00000070476** | -0.101347665608794 | 0.41863231281298 | 0.882468496126562 | 381.282788392137 | 375.445651029505 | 402.24332767559 | 375.170677703265 | 487.256397139498 | 388.940149799487 | 393.52751823022 | 375.591845375304 | Fam217b |
| **ENSMUSG00000070493** | 0.011226336646798 | 0.941782606467629 | 0.989934163630124 | 15406.7158949354 | 12880.799168509 | 14359.6135705742 | 17163.1205782301 | 15645.8576600591 | 17833.7620258689 | 12921.8612599828 | 12945.9778842366 | Chchd2 |
| **ENSMUSG00000070509** | -0.428955617285552 | 0.0054049899918339 | 0.635862253526407 | 431.879556520004 | 426.753841951108 | 427.797609669098 | 432.384206053012 | 593.467342133951 | 688.595296657583 | 595.49666779811 | 437.466802561409 | Rgma |
| **ENSMUSG00000070520** | -0.22483840418675 | 0.336716055494415 | 0.858570960161243 | 296.352499034647 | 228.850819824926 | 248.917635714542 | 252.302280755445 | 367.905953795217 | 392.609396495709 | 222.790711379013 | 217.105112933702 | Nsmce3 |
| **ENSMUSG00000070526** | 0.118429997698697 | 0.789752915302608 | 0.961659329450003 | 12.6491920319666 | 14.6594831204579 | 16.0897331070236 | 15.0068271081306 | 17.5193311331055 | 20.7923979452556 | 8.32862472444909 | 7.59867895267955 | Peg12 |
| **ENSMUSG00000070544** | 0.02179017027945 | 0.863136828804377 | 0.974868041734905 | 2266.91591487173 | 1860.93994056924 | 1840.85475842123 | 1914.30838298091 | 1731.12890758999 | 2173.41712639525 | 1800.02401857156 | 2061.4130473055 | Top1 |
| **ENSMUSG00000070565** | -0.0665829549600585 | 0.708909144188847 | 0.949267137065756 | 687.573938309043 | 770.437279552955 | 788.396922244156 | 537.431995809926 | 849.687559955619 | 611.54111603693 | 809.958754452674 | 643.716659848425 | Rasal2 |
| **ENSMUSG00000070570** | -0.276093790659862 | 0.0796867735425614 | 0.739855948845396 | 1589.28062744495 | 1600.32690731666 | 1132.90650171219 | 1435.96576890924 | 1470.52885698505 | 1673.17649347704 | 1957.22681024554 | 1872.53159905318 | Slc17a7 |
| **ENSMUSG00000070574** | 0.425177643304162 | 0.0843727780657553 | 0.74314387275418 | 50.5967681278665 | 78.1839099757755 | 67.1982970940397 | 52.523894878457 | 52.5579933993167 | 39.1386314263635 | 51.0128264372507 | 42.3354970220718 | 2310016G11 |
| **ENSMUSG00000070576** | -0.0724131001275614 | 0.600609760966674 | 0.925755180289997 | 483.379838364439 | 469.917875583568 | 380.474865236676 | 444.577253078368 | 511.345477447518 | 512.471455238947 | 407.061533407449 | 440.723379255414 | Mn1 |
| **ENSMUSG00000070577** | 0.147067460239116 | 0.69786328371369 | 0.944933386973863 | 74.9916384752307 | 93.6578088251478 | 117.360406192407 | 60.9652351267805 | 73.3621991198795 | 30.5770558018465 | 93.6970281500523 | 115.065709854862 | Gm572 |
| **ENSMUSG00000070583** | -0.1585348597667 | 0.785307168007081 | 0.961141387070613 | 88.5443442237664 | 135.193010999779 | 123.985590412947 | 66.5927952923295 | 53.6529515951357 | 19.5693157131818 | 223.831789469569 | 164.999885829613 | Fv1 |
| **ENSMUSG00000070594** | -0.429839913749383 | 0.26115216111152 | 0.841039535784608 | 57.8248778604189 | 89.5857301805761 | 44.483379766477 | 53.4618215727152 | 48.1781606160403 | 85.6157562451702 | 55.1771387994752 | 142.203848971575 | Gm4788 |

| | | | | | | | | | | | | |
|---|---|---|---|---|---|---|---|---|---|---|---|---|
| **ENSMUSG00000070604** | 0.334307538644125 | 0.229807914325492 | 0.832048466795696 | 28.9124389302094 | 42.3496179035451 | 34.0723759913441 | 33.7653609932938 | 24.0890803080201 | 25.6847268735511 | 28.1091084450157 | 32.5657669400552 | Vsig10l |
| **ENSMUSG00000070605** | -0.345526970413845 | 0.27987134792863 | 0.848943557485723 | 20.780815481088 | 28.5045505120015 | 19.8755526616174 | 18.7585338851632 | 26.2789966996583 | 30.5770558018465 | 20.8215618111227 | 34.7368180693922 | Zfp992 |
| **ENSMUSG00000070610** | -0.198501805149567 | 0.437073457815477 | 0.888701139843674 | 33.4300075130547 | 35.8342920722305 | 29.3401015481019 | 35.6412143818101 | 35.0386622662111 | 41.5847958905112 | 32.2734208072402 | 45.5920737160773 | Gm13127 |
| **ENSMUSG00000070637** | -0.329068238906653 | 0.56996324556469 | 0.920872458148558 | 18.9737880479499 | 8.14415728914329 | 23.6613722162112 | 11.2551203310979 | 14.2344565456483 | 42.8078781225851 | 11.4518589961175 | 9.76973008201657 | Srarp |
| **ENSMUSG00000070639** | -0.292414517703241 | 0.291623188956844 | 0.851785577030345 | 45.1756858284523 | 65.1532583131463 | 84.2344850897118 | 66.5927952923295 | 107.305903190271 | 74.6080161565054 | 82.2451691539348 | 55.3618037980939 | Lrrc8b |
| **ENSMUSG00000070643** | -0.502192546711548 | 0.0406204826461296 | 0.68307324933988 | 245.75573090678 | 215.005752433383 | 225.25626349833 | 249.488500672671 | 453.312693069106 | 399.947889888152 | 251.940897914585 | 220.361689627707 | Sox13 |
| **ENSMUSG00000070644** | -0.117803741245925 | 0.770321533126303 | 0.958992744413128 | 47.8862269781594 | 109.946123403434 | 43.5369248778286 | 64.7169419038131 | 53.6529515951357 | 66.0464405319884 | 45.80743598447 | 123.74991437221 | Etnk2 |
| **ENSMUSG00000070691** | 0.294227950200652 | 0.555593838579606 | 0.916239900758826 | 17.1667606148119 | 20.3603932228582 | 10.4110037751329 | 6.56548685980713 | 10.949581958191 | 15.9000690169602 | 7.28754663389296 | 10.8552556466851 | Runx3 |
| **ENSMUSG00000070697** | 0.0920070661245621 | 0.269087805529116 | 0.843793744782845 | 862.855599323438 | 846.992358070902 | 879.256591554407 | 923.857793844289 | 814.648897689408 | 867.165302540366 | 804.753363999894 | 809.802071242707 | Utp3 |
| **ENSMUSG00000070699** | -0.112254564417424 | 0.42171399138477 | 0.882568531013176 | 208.711668527449 | 184.872370463553 | 216.738169500494 | 182.895705380341 | 211.326931793086 | 232.385624094033 | 225.913945650682 | 187.795922687652 | Sars2 |
| **ENSMUSG00000070704** | -0.352660091269096 | 0.237983696847135 | 0.832048466795696 | 243.948703473642 | 267.942774812814 | 247.024725937245 | 291.695201914288 | 227.751304730372 | 352.247682837272 | 209.256696201783 | 552.53251241627 | Ugt2b36 |
| **ENSMUSG00000070705** | 0.0364101521746678 | 0.832448470097551 | 0.967671708900743 | 246.659244623349 | 208.490426602068 | 206.327165725361 | 223.226553233442 | 162.053812981226 | 222.600966237442 | 221.749633288457 | 256.184033261768 | Eid2b |
| **ENSMUSG00000070709** | 0.0265531667828849 | 0.917795251961941 | 0.986713918026868 | 59.631905293557 | 48.8649437348597 | 37.8581955459379 | 47.8342614071662 | 38.3235368536684 | 40.3617136584374 | 53.094982618363 | 58.6183804920994 | Zfp974 |
| **ENSMUSG00000070713** | 0.182516636744115 | 0.243876162708405 | 0.832048466795696 | 149.983276950461 | 171.027303072009 | 182.66579350915 | 146.316564304273 | 134.679858085749 | 143.100621152642 | 163.449260217313 | 131.348593324889 | Gm10282 |
| **ENSMUSG00000070720** | -0.819392930174543 | 0.0732723351626309 | 0.730864975478592 | 131.009488902512 | 113.203786319092 | 135.343049076728 | 125.682177030594 | 296.733671066975 | 415.847958905112 | 89.5327157878278 | 90.0986218674861 | Tmem200b |
| **ENSMUSG00000070730** | -0.133480273756471 | 0.282975351012614 | 0.850587941933635 | 308.098177350044 | 282.602257933272 | 298.133289924261 | 325.460562907582 | 308.778211220985 | 321.670627035425 | 381.034581143546 | 319.144516012541 | Rmdn3 |
| **ENSMUSG00000070733** | 0.116231611511956 | 0.518701301235714 | 0.905689488434585 | 489.704434380422 | 672.707392083235 | 618.035042287436 | 468.963347129081 | 498.205979097689 | 415.847958905112 | 603.825292522559 | 556.874614674944 | Fryl |
| **ENSMUSG00000070737** | -0.240424151925658 | 0.4937623999892 | 0.900117970535229 | 30.7194663633475 | 31.7622134276588 | 24.6078271048596 | 14.0689004138724 | 39.4184950494875 | 31.8001380339204 | 26.0269522639034 | 22.7960368580387 | Tmem35b |
| **ENSMUSG00000070738** | 0.210828307070254 | 0.353289994698555 | 0.862728204353539 | 916.162908601012 | 1238.72632367869 | 1166.97887770353 | 1034.53314376675 | 892.390929592564 | 584.633306931305 | 1198.28088223011 | 1087.69661579784 | Dgkd |
| **ENSMUSG00000070780** | -0.457837704054106 | 0.367449972217034 | 0.870820698531752 | 9.03513716569045 | 28.5045505120015 | 8.51809399783602 | 8.44134024832345 | 21.8991639163819 | 19.5693157131818 | 13.5340151772298 | 20.6249857287016 | Rbm47 |
| **ENSMUSG00000070802** | -0.0595244789458382 | 0.75174034137492 | 0.956197192463153 | 92.1583990900426 | 85.5136515360045 | 95.5919437534931 | 100.358156285623 | 87.5966556655277 | 83.1695917810225 | 121.806136595068 | 95.5262496908287 | Pnmal2 |
| **ENSMUSG00000070803** | -0.32786082969177 | 0.230172319414573 | 0.832048466795696 | 196.965990212052 | 144.96599974675 | 187.398067952392 | 334.839829850164 | 221.181555555458 | 292.316653465652 | 237.365804646799 | 333.256348353232 | Cited4 |
| **ENSMUSG00000070808** | -0.00332641993335065 | 0.984044661986566 | 0.996940244108351 | 177.992202164102 | 191.387696294867 | 223.363353721033 | 207.281799431054 | 231.036179317829 | 168.785348026193 | 224.872867560126 | 175.855141476298 | Bicra |
| **ENSMUSG00000070814** | 0.191224712900095 | 0.16485363011843 | 0.795216960550936 | 492.41497553013 | 501.680089011226 | 493.102996985841 | 529.928582255861 | 421.558905390352 | 355.916929533493 | 491.388858742496 | 496.085183053508 | Zswim9 |
| **ENSMUSG00000070822** | 0.527013933958192 | 0.159123972207334 | 0.789393312599749 | 25.2983840639333 | 34.2054606144018 | 17.9826428843205 | 23.448167356454 | 14.2344565456483 | 12.2308223207386 | 24.9858741733473 | 18.4539345993646 | Zscan18 |
| **ENSMUSG00000070858** | -0.221061846113139 | 0.63390748177645 | 0.9325328237529 | 12.6491920319666 | 20.3603932228582 | 13.2503684410783 | 24.3860940507122 | 22.994122112201 | 35.4693847301419 | 10.4107809055614 | 14.1118323406906 | Gm1673 |
| **ENSMUSG00000070867** | -0.00265286492285209 | 0.988377477612673 | 0.998437672973644 | 4316.9885377669 | 3823.68184725277 | 3266.21582072579 | 3826.7409125733 | 2717.686242023 | 3785.4395082686 | 4133.08001950786 | 4625.42443105251 | Trabd2b |
| **ENSMUSG00000070871** | 0.0923739926635186 | 0.573326230274782 | 0.922207298536842 | 229.492484008537 | 275.272516373043 | 230.934992830221 | 323.584709519066 | 283.594172717146 | 236.054870790255 | 241.530117009024 | 232.302470839061 | Ccnyl1 |
| **ENSMUSG00000070873** | 0.42556992101076 | 0.315616152316284 | 0.853277176422981 | 13.5527057485357 | 14.6594831204579 | 32.1794662140472 | 24.3860940507122 | 13.1394983498292 | 12.2308223207386 | 15.616171358342 | 21.7105112933701 | Lilra5 |
| **ENSMUSG00000070883** | -0.388144314658354 | 0.33782106743807 | 0.858570960161243 | 19.877301764519 | 12.2162359337149 | 24.6078271048596 | 26.2619474392285 | 16.4243729372865 | 39.1386314263635 | 33.3144988977964 | 19.5394601640331 | Ccdc173 |
| **ENSMUSG00000070923** | -0.115790952868498 | 0.21923415051877 | 0.827535829146756 | 1111.32187137993 | 1095.38915538977 | 1011.76027596519 | 1185.53934154232 | 1184.74476787626 | 1113.00483118721 | 1211.81489740734 | 1261.38070614481 | Klhl9 |
| **ENSMUSG00000070934** | -0.0913106858633952 | 0.449557426050465 | 0.89108031640712 | 1118.54998111248 | 1158.09916651618 | 1293.80383278243 | 1135.82922674663 | 1350.08345544495 | 1093.43551547403 | 1398.16787561689 | 1170.19655871265 | Rraga |
| **ENSMUSG00000070939** | 0.147390441419266 | 0.427319354411005 | 0.885066977635295 | 369.537110076739 | 490.278268806426 | 474.173899212872 | 365.791410760683 | 352.576539053749 | 298.432064626022 | 448.704657029695 | 434.210225867403 | Tgfbrap1 |
| **ENSMUSG00000070942** | -0.727871806092139 | 0.113959016305573 | 0.767518180080285 | 6.32459601598331 | 13.0306516626293 | 11.3574586637814 | 12.1930470253561 | 28.4689130912965 | 18.3462334811079 | 11.4518589961175 | 13.0263067760221 | Il1rl2 |
| **ENSMUSG00000070953** | -0.0432016791668108 | 0.812772737130083 | 0.965402199310489 | 317.133314515735 | 240.252640029727 | 213.898804834549 | 292.633128608546 | 286.879047304603 | 324.116791499573 | 233.201492284575 | 252.927456567762 | Rabepk |
| **ENSMUSG00000070972** | 0.189497566785105 | 0.0835397666622009 | 0.742148948523533 | 843.881811275488 | 738.675066125296 | 707.948256709038 | 741.900015158206 | 650.405168316544 | 604.202622644487 | 690.234774038719 | 713.190295987209 | Dnajc25 |
| **ENSMUSG00000071001** | -0.470350365986367 | 0.197530286038556 | 0.812921911423739 | 74.0881247586617 | 64.338842584232 | 85.1809399783602 | 96.6064495085906 | 154.389105610493 | 165.116101329971 | 57.2592949805875 | 67.3025850094475 | Hrct1 |
| **ENSMUSG00000071014** | 0.0876559681496044 | 0.596254870580477 | 0.92448662081833 | 5469.872040109 | 3894.53601566832 | 4246.74308536558 | 5281.46521536771 | 4463.04960615864 | 5363.21558764387 | 3870.72834068772 | 4082.66164871826 | Ndufb6 |
| **ENSMUSG00000071037** | -0.336831454109537 | 0.385817882833061 | 0.875126835008975 | 14.4562194651047 | 13.8450673915436 | 17.036187995672 | 13.1309737196143 | 13.1394983498292 | 28.1308913376988 | 14.5750932677859 | 18.4539345993646 | Camkmt |
| **ENSMUSG00000071041** | -0.141018518009224 | 0.80274942762158 | 0.963461896828986 | 18.9737880479499 | 13.0306516626293 | 16.0897331070236 | 16.8826804966469 | 33.943704070392 | 13.4539045528125 | 20.8215618111227 | 3.25657669400552 | Impdh2-ps |
| **ENSMUSG00000071042** | -0.139235713480175 | 0.197168384116907 | 0.812565226112804 | 484.283352081008 | 509.82424630037 | 539.479286529615 | 515.859681841989 | 522.295059405709 | 535.71001764835 | 566.346481262538 | 631.775878637071 | Rasgrp3 |
| **ENSMUSG00000071054** | 0.023771153185267 | 0.831465846948864 | 0.967583158900775 | 1815.15905658721 | 1808.81733391872 | 1643.0456866937 | 1684.51634288766 | 1977.49450164929 | 1702.53046704681 | 1550.16527683809 | 1608.74888683873 | Safb |
| **ENSMUSG00000071064** | -0.10852793205093 | 0.667153391620541 | 0.942024383071524 | 271.054114970713 | 186.501201921381 | 258.382184601026 | 345.157023487003 | 271.549632563136 | 194.470074899744 | 378.952424962434 | 297.434004719171 | Zfp827 |
| **ENSMUSG00000071068** | 1.33371471113132 | 0.0682148667144316 | 0.722112580225498 | 12.6491920319666 | 10.5874044758863 | 77.6093008691726 | 12.1930470253561 | 6.56974917491458 | 14.6769867848863 | 6.24646854333682 | 17.3684090346961 | Treml2 |
| **ENSMUSG00000071072** | 0.07180787946504 | 0.492843834588415 | 0.899671555057574 | 2652.71627184672 | 2373.20743405635 | 2433.33551871516 | 2814.71800946874 | 2701.26186908571 | 2437.6028885232 | 2271.63239359349 | 2364.27467984801 | Ptges3 |
| **ENSMUSG00000071074** | -0.142203715088564 | 0.329061930056282 | 0.855977678491063 | 1278.4719089452 | 1250.94255961241 | 1334.50139299431 | 1311.22151857291 | 1650.10200109938 | 1632.8147798186 | 1273.23850475016 | 1156.08472637196 | Yipf3 |
| **ENSMUSG00000071076** | -0.2954465668945 | 0.150854095038256 | 0.783020695097675 | 2579.53166080462 | 2558.89422024882 | 2472.14016914974 | 2642.13949772524 | 3213.70230472905 | 4474.03480492618 | 2519.40897914585 | 2377.30098662403 | Jund |
| **ENSMUSG00000071078** | -0.300817732053309 | 0.49270853908285 | 0.899542798088958 | 19.877301764519 | 8.95857301805762 | 7.57163910918757 | 15.0068271081306 | 18.6142893289246 | 15.9000690169602 | 12.4929370866736 | 16.2828834700276 | Nr2c2ap |
| **ENSMUSG00000071172** | -0.0110898792199622 | 0.88407148655543 | 0.979384300778265 | 1607.35090177633 | 1624.75937918409 | 1504.86327295103 | 1536.32392519487 | 1662.14654125339 | 1563.09909259039 | 1538.71341784197 | 1558.81471086398 | Srsf3 |
| **ENSMUSG00000071176** | -0.0865779424670818 | 0.628742431830286 | 0.931825800800958 | 517.713359594063 | 452.815145276367 | 585.855576073389 | 436.135912830045 | 660.259792078915 | 560.171662289828 | 460.156516025812 | 435.295751432072 | Arhgef10 |
| **ENSMUSG00000071177** | -0.150250924076155 | 0.705722705559414 | 0.947760509100828 | 227.685456575399 | 552.173864203915 | 246.078271048596 | 311.39166249371 | 217.896680968 | 262.96267989588 | 318.569895710178 | 684.966631305828 | Serpina1d |
| **ENSMUSG00000071178** | -0.411977541293722 | 0.274007020077465 | 0.845204865395511 | 502.353626412389 | 1210.22177316669 | 547.997380527451 | 749.403428712271 | 640.550544554172 | 735.07242147639 | 828.698160082685 | 1800.88691178505 | Serpina1b |
| **ENSMUSG00000071180** | 0.0281145537538764 | 0.819996596646416 | 0.966398681144067 | 505.967681278665 | 423.496179035451 | 441.994432998825 | 513.983828453472 | 444.553027502553 | 518.586866399316 | 441.417110395802 | 445.065481514088 | Smim15 |
| **ENSMUSG00000071226** | 0.0603738888370586 | 0.802764513310654 | 0.963461896828986 | 140.044626068202 | 168.584055885266 | 220.523989055088 | 211.971432902344 | 145.62944004394 | 130.869798831903 | 232.160414194018 | 200.822229463674 | Cecr2 |
| **ENSMUSG00000071252** | 0.138899594814933 | 0.599053834213604 | 0.924848033480823 | 39.754603529038 | 39.0919549878878 | 49.2156542097192 | 50.6480414899407 | 32.8487458745729 | 31.8001380339204 | 44.7663578939139 | 52.1052271040884 | 2210408I21R |
| **ENSMUSG00000071253** | 0.286582291028118 | 0.241636991184133 | 0.832048466795696 | 583.669860903603 | 742.747144769868 | 756.217456030109 | 580.576623745802 | 404.039574257247 | 380.37857417497 | 660.043509412591 | 738.157383974585 | Slc25a16 |
| **ENSMUSG00000071256** | -0.228557735674312 | 0.3238907987774 | 0.855977678491063 | 67.7635287426784 | 93.6578088251478 | 105.056492639978 | 101.296082979881 | 116.065568756824 | 138.208292224346 | 92.6559500594962 | 84.6709940441436 | Zfp213 |
| **ENSMUSG00000071266** | -0.00864308134654658 | 0.973990333731202 | 0.995510898988495 | 68.6670424592474 | 83.8848200781758 | 63.4124775394459 | 42.2067012416173 | 65.6974917491458 | 79.5003450848009 | 64.5468416144805 | 51.0197015394199 | Zfp946 |
| **ENSMUSG00000071267** | -0.0684799717605039 | 0.69957233082213 | 0.945280238056015 | 88.5443442237664 | 99.3587189275481 | 93.6990339761962 | 105.047789756914 | 85.4067392738896 | 97.8465785659088 | 119.723980413956 | 102.03940307884 | Zfp942 |
| **ENSMUSG00000071281** | -0.175699149199153 | 0.414532155562611 | 0.882192126163777 | 75.8951521917998 | 93.6578088251478 | 63.4124775394459 | 90.9788893430416 | 74.4571573156986 | 85.6157562451702 | 110.35427759895 | 95.5262496908287 | Zfp65 |
| **ENSMUSG00000071291** | -0.197139402998186 | 0.527602966672743 | 0.908217276872524 | 45.1756858284523 | 21.9892246806869 | 31.2330113253987 | 46.8963347129081 | 49.2731188118594 | 37.9155491942896 | 34.3555769883525 | 44.5065481514088 | Zfp58 |
| **ENSMUSG00000071302** | -0.0133404829218634 | 0.955637105580467 | 0.992371505557225 | 206.904641094311 | 208.490426602068 | 159.950876181588 | 172.578511743502 | 154.389105610493 | 125.977469903608 | 242.57119509958 | 231.216945274392 | 2610044O15 |
| **ENSMUSG00000071317** | 0.0127050212842926 | 0.91754140481258 | 0.986713918026868 | 946.882374964359 | 1031.86472853445 | 907.65023821386 | 962.312788308873 | 867.206891088725 | 836.58824673852 | 1069.18719900115 | 1041.0190165171 | Bves |
| **ENSMUSG00000071324** | 0.0607935713138163 | 0.772607917238035 | 0.959045638244974 | 1181.79594127231 | 1260.71554835938 | 1255.94563723649 | 1116.13276616721 | 989.842209020464 | 769.318723974458 | 1435.64668687691 | 1419.86743858641 | Armc2 |
| **ENSMUSG00000071335** | -0.0257180755008429 | 0.905280889382666 | 0.98504921011344 | 184.316798180085 | 133.56417954195 | 115.467496415111 | 138.813150750208 | 107.305903190271 | 146.769867848863 | 167.613572579538 | 160.657783570939 | Mfsd4b3 |
| **ENSMUSG00000071337** | 0.0457045874516244 | 0.828984728951898 | 0.967483872587428 | 169.86057871498 | 173.470550258752 | 138.182413742673 | 160.385464718146 | 123.730276127558 | 124.754387671534 | 162.408182126757 | 210.59195954569 | Tia1 |
| **ENSMUSG00000071342** | 0.131870870147978 | 0.441009248341512 | 0.89108031640712 | 833.943160393229 | 571.719841697859 | 553.676109859341 | 619.031618210386 | 516.820268426614 | 598.087211484117 | 567.387559353095 | 670.854798965138 | Lsmem1 |
| **ENSMUSG00000071347** | -0.412411852161942 | 0.149511466657618 | 0.782641594375979 | 936.9437240821 | 818.4878075589 | 848.023580229008 | 1020.46424335288 | 1619.44317161644 | 1670.73032901289 | 715.220648212066 | 818.486275760055 | C1qtnf9 |
| **ENSMUSG00000071350** | -0.157955261874907 | 0.702059367703602 | 0.946390199670325 | 112.035700854562 | 139.26508964435 | 86.1273948670087 | 66.5927952923295 | 156.579022002131 | 40.3617136584374 | 163.449260217313 | 90.0986218674861 | Setdb2 |
| **ENSMUSG00000071359** | 0.0442020054423397 | 0.706028557752237 | 0.947877118420508 | 326.168451681425 | 308.663561258531 | 326.526936583714 | 360.163850595134 | 323.012667766634 | 287.424324537357 | 340.432535611857 | 328.914246094558 | Tbpl1 |
| **ENSMUSG00000071369** | 0.0724220972380806 | 0.669740545994341 | 0.942388853397102 | 1142.04133774327 | 1162.98566088966 | 1282.44637411865 | 1125.51203310979 | 1099.33802860237 | 803.565026472526 | 1257.62233339181 | 1319.99908663691 | Map3k5 |
| **ENSMUSG00000071379** | -0.57182201625341 | 0.157667623738551 | 0.789230661864335 | 300.870067617492 | 295.632909595901 | 320.848207251823 | 210.095579513828 | 610.986673267056 | 670.249063176475 | 197.804837205666 | 197.565652769668 | Hpcal1 |
| **ENSMUSG00000071415** | 0.0392497605101351 | 0.849830454855152 | 0.972640876584422 | 6420.36846993963 | 4914.18450826906 | 5917.23596383009 | 7122.61531619648 | 6212.79280307756 | 8012.41170231585 | 4837.88988681437 | 4657.99019799257 | Rpl23 |
| **ENSMUSG00000071451** | 0.219850327492688 | 0.302794170409502 | 0.851785577030345 | 237.624107457659 | 151.481325578065 | 158.057966404291 | 200.716312571247 | 155.484063806312 | 201.808568292187 | 154.079557402308 | 131.348593324889 | Psmg4 |
| **ENSMUSG00000071454** | -0.135966208650624 | 0.602679581980742 | 0.926448151609826 | 43.3686583953142 | 35.8342920722305 | 54.8943835416099 | 60.0273084325223 | 53.6529515951357 | 57.4848649074714 | 61.4236073428121 | 40.1644458927348 | Dtnb |
| **ENSMUSG00000071456** | 0.194320025100736 | 0.593682701157283 | 0.923935529355527 | 28.0089252136404 | 34.2054606144018 | 46.3762895437739 | 24.3860940507122 | 14.2344565456483 | 31.8001380339204 | 35.3966550789086 | 34.7368180693922 | 1110002L01R |
| **ENSMUSG00000071477** | -0.122796873574821 | 0.2988490777848 | 0.851785577030345 | 280.992765852973 | 306.220314071788 | 302.865564367503 | 256.053987532478 | 329.582416941548 | 313.109051410908 | 295.666177717943 | 310.460311495193 | Zfp777 |
| **ENSMUSG00000071497** | -0.287088300746244 | 0.509296613400246 | 0.902394714365738 | 41.5616309621761 | 26.0613033252585 | 12.3039135524298 | 27.1998741334867 | 29.5638712871156 | 20.7923979452556 | 27.0680303544596 | 53.1907526687569 | Nutf2-ps1 |
| **ENSMUSG00000071528** | 0.281725522017395 | 0.0512484824947 | 0.703396849069134 | 13555.4162896854 | 9162.1769502862 | 10592.7231137534 | 12634.8104983517 | 9391.45644554039 | 9876.38902399642 | 8849.16376972716 | 9677.46040901974 | Usmg5 |
| **ENSMUSG00000071533** | 0.0128186232836612 | 0.904533926270914 | 0.98504921011344 | 940.557778948376 | 969.154717408051 | 914.2754224344 | 966.064495085906 | 848.5926017598 | 914.865509591247 | 915.107641598844 | 1077.92688571583 | Pcnp |
| **ENSMUSG00000071540** | 0.387847075995307 | 0.131969239147456 | 0.77303252008843 | 3699.88866935024 | 3641.25272397596 | 3036.22728278422 | 3922.40943538763 | 1942.45583938308 | 1773.4692365071 | 3641.69116076537 | 3570.29358219472 | 3425401B19 |
| **ENSMUSG00000071547** | 0.0746681643156801 | 0.606543819252884 | 0.927110332696882 | 391.221439274397 | 346.12668478859 | 441.047978110176 | 431.446279358754 | 398.564783278151 | 349.801518373124 | 338.350379430744 | 440.723379255414 | Nt5dc2 |
| **ENSMUSG00000071551** | -0.181609636846923 | 0.621082055085426 | 0.93041735818988 | 19.877301764519 | 30.9477976987445 | 31.2330113253987 | 49.7101147956825 | 31.7537876787538 | 42.8078781225851 | 23.9447960827911 | 51.0197015394199 | Akr1c19 |
| **ENSMUSG00000071573** | -0.0777347489102265 | 0.672137849716893 | 0.943216235131458 | 102.097049972302 | 115.647033505835 | 104.110037751329 | 126.620103724852 | 93.0714466446232 | 116.192812047017 | 124.929370866736 | 138.947272277569 | Rnls |
| **ENSMUSG00000071604** | 0.182111675234913 | 0.102929185113923 | 0.759800953500741 | 1963.33530610453 | 2027.89516499668 | 2074.6291159174 | 2345.75466233966 | 2031.14745324442 | 1944.70074899744 | 1656.35524207481 | 1782.43297718569 | Fam189a2 |
| **ENSMUSG00000071632** | 0.0373533243115709 | 0.816968247507416 | 0.966315773308182 | 333.396561413978 | 282.602257933272 | 298.133289924261 | 374.232751009006 | 315.3479603959 | 365.701587390084 | 319.610973800734 | 255.098507697099 | 2510002D24 |
| **ENSMUSG00000071633** | -0.219878457500446 | 0.43573813370962 | 0.888687115594624 | 31.6229800799166 | 39.9063707168021 | 37.8581955459379 | 36.5791410760683 | 35.0386622662111 | 47.7002070508805 | 30.191264626128 | 57.5328549274309 | Gm4952 |

| | | | | | | | | | | | | |
|---|---|---|---|---|---|---|---|---|---|---|---|---|
| **ENSMUSG00000071637** | -0.685753042182514 | 0.324624565241724 | 0.855977678491063 | 276.475197270128 | 267.942774812814 | 214.845259723197 | 125.682177030594 | 469.737066006393 | 735.07242147639 | 104.107809055614 | 115.065709854862 | Cebpd |
| **ENSMUSG00000071644** | 0.00974794256862188 | 0.923091125056311 | 0.986713918026868 | 8011.45612481772 | 6638.30260638069 | 7231.86180416278 | 8137.45199938381 | 7346.07453575032 | 7947.58834401594 | 7171.98696584123 | 7351.17912393513 | Eef1g |
| **ENSMUSG00000071645** | -0.101354605235966 | 0.56932786077331 | 0.920872458148558 | 203.290586228035 | 272.8292691863 | 289.615195926425 | 249.488500672671 | 323.012667766634 | 291.093571233579 | 223.831789469569 | 251.841931003094 | Tut1 |
| **ENSMUSG00000071646** | -0.154692322581382 | 0.190510245537799 | 0.808824684419004 | 645.108793630298 | 642.574010113405 | 677.661700272288 | 582.452477134318 | 812.45898129777 | 714.280023531134 | 669.413212227596 | 640.46008315442 | Mta2 |
| **ENSMUSG00000071647** | -0.162325301568202 | 0.45200764726324 | 0.891110666112841 | 117.456783153976 | 155.553404222637 | 124.932045301595 | 150.068271081306 | 182.858018701789 | 188.354663739374 | 123.88829277618 | 119.407812113536 | Eml3 |
| **ENSMUSG00000071648** | -0.0560394798244466 | 0.751147928105023 | 0.956197192463153 | 576.441751171051 | 443.042156529395 | 353.027673465871 | 491.473587791276 | 489.446313531136 | 547.940839969089 | 423.718782856348 | 477.631248454143 | Rom1 |
| **ENSMUSG00000071649** | -0.0445469190556916 | 0.69193825668669 | 0.94469184542218 | 795.995584297329 | 833.961706408273 | 878.310136665759 | 788.796349871114 | 791.654775577207 | 971.127292266645 | 865.13589325215 | 773.979727608646 | B3gat3 |
| **ENSMUSG00000071650** | -0.114718699816209 | 0.395602672633921 | 0.879215782741314 | 832.13613296009 | 996.844852191138 | 965.383986421416 | 831.940977806989 | 1133.28173267277 | 1018.82749931753 | 846.396487622139 | 929.209883356242 | Ganab |
| **ENSMUSG00000071652** | -0.273205052437468 | 0.206637914726497 | 0.819960256425221 | 171.667606148119 | 192.202112023782 | 197.809071727525 | 198.84045918273 | 280.309298129689 | 291.093571233579 | 171.777884941763 | 176.940667040967 | Ints5 |
| **ENSMUSG00000071653** | -0.142925946261504 | 0.551595865027622 | 0.914873920695527 | 101.193536255733 | 64.338842584232 | 89.9132144216024 | 101.296082979881 | 100.736154015357 | 128.423634367755 | 74.9576225200418 | 90.0986218674861 | 1810009A15 |
| **ENSMUSG00000071654** | 0.045473714623598 | 0.837572999822678 | 0.968924788429433 | 985.733464776828 | 685.738043745865 | 738.234813145789 | 975.443762028488 | 839.832936193247 | 1127.6818179721 | 660.043509412591 | 653.486389930442 | Uqcc3 |
| **ENSMUSG00000071655** | 0.0573836687542551 | 0.738725742800628 | 0.954383574136061 | 2954.48985318078 | 2278.73520950229 | 2367.08367650977 | 3071.70992369548 | 2796.52323212197 | 3065.04407357709 | 2295.57718967628 | 2099.40644206889 | Ubxn1 |
| **ENSMUSG00000071656** | -1.19777359342002 | 0.00879928089136476 | 0.635862253526407 | 25.2983840639333 | 36.6487078011448 | 29.3401015481019 | 27.1998741334867 | 85.4067392738896 | 122.308223207386 | 27.0680303544596 | 37.9933947633978 | Lrrn4cl |
| **ENSMUSG00000071657** | -0.199653789926487 | 0.174263089766871 | 0.799889021902745 | 321.65088309858 | 306.220314071788 | 313.276568142636 | 314.205442576484 | 388.710159515779 | 431.748027922072 | 314.405583347953 | 308.289260365856 | Bscl2 |
| **ENSMUSG00000071659** | 0.191388856687529 | 0.202517600948839 | 0.815117205789446 | 3054.77987571994 | 3548.40933087973 | 3452.66743378953 | 2816.59386285726 | 2501.97947744664 | 2375.22569468744 | 3223.1777683618 | 3171.90569996138 | Hnrnpul2 |
| **ENSMUSG00000071660** | 0.249944906754594 | 0.188664415238674 | 0.808677608936066 | 261.115464088454 | 352.642010619904 | 372.903226127488 | 279.502154888932 | 271.549632563136 | 201.808568292187 | 307.11803671406 | 283.322172378481 | Ttc9c |
| **ENSMUSG00000071661** | -0.186228350470669 | 0.576699130912302 | 0.922450244597442 | 18.0702743313809 | 16.2883145782866 | 17.9826428843205 | 18.7585338851632 | 16.4243729372865 | 19.5693157131818 | 19.7804837205666 | 24.9670879873757 | Zbtb3 |
| **ENSMUSG00000071662** | 0.0745242305991735 | 0.582669073184577 | 0.922450244597442 | 882.732901087957 | 684.109212288036 | 737.28835825714 | 849.761584997894 | 776.32536083574 | 845.149822363037 | 688.152617857606 | 686.052156870497 | Polr2g |
| **ENSMUSG00000071669** | -0.0725449875881755 | 0.783331264929802 | 0.961051227381536 | 30.7194663633475 | 47.2361122770311 | 41.6440151005317 | 46.8963347129081 | 39.4184950494875 | 34.2463024980681 | 48.9306702561384 | 52.1052271040884 | Snx29 |
| **ENSMUSG00000071708** | 0.222063067032525 | 0.163732150489217 | 0.793968226938511 | 2138.61696711893 | 2043.36906384605 | 1937.39315706337 | 2208.81736497797 | 1449.72465126448 | 1520.29121446781 | 1996.78777768667 | 2172.13665490168 | Sms |
| **ENSMUSG00000071711** | -0.0524447907328452 | 0.697375266151698 | 0.944933386973863 | 865.566140473145 | 747.633639143354 | 766.628459805242 | 860.078778634734 | 943.853964796062 | 841.480575666815 | 889.080689334941 | 684.966631305828 | Mpst |
| **ENSMUSG00000071713** | -0.558748546940653 | 0.144447897665244 | 0.780607990324772 | 14.4562194651047 | 13.8450673915436 | 13.2503684410783 | 15.0068271081306 | 18.6142893289246 | 26.9078091056249 | 26.0269522639034 | 11.9407812113536 | Csf2rb |
| **ENSMUSG00000071714** | -0.0501078873324874 | 0.930533996119103 | 0.987630259501364 | 8.13162344912141 | 10.5874044758863 | 6.62518422053913 | 15.0068271081306 | 12.0445401540101 | 19.5693157131818 | 6.24646854333682 | 4.34210225867403 | Csf2rb2 |
| **ENSMUSG00000071715** | -0.141507385626479 | 0.817741660464039 | 0.966315773308182 | 6.32459601598331 | 7.32974156022896 | 8.51809399783602 | 16.8826804966469 | 14.2344565456483 | 18.3462334811079 | 3.12323427166841 | 7.59867895267955 | Ncf4 |
| **ENSMUSG00000071722** | -0.793717992036091 | 0.187872699399604 | 0.808661851459713 | 17.1667606148119 | 5.7009101024003 | 8.51809399783602 | 28.1378008277448 | 47.0832024202212 | 29.3539735697726 | 13.5340151772298 | 13.0263067760221 | Spin4 |
| **ENSMUSG00000071748** | 0.169821592621632 | 0.680529409950683 | 0.944452327903597 | 18.0702743313809 | 17.1027303072009 | 12.3039135524298 | 13.1309737196143 | 14.2344565456483 | 8.56157562451702 | 11.4518589961175 | 19.5394601640331 | Gm14698 |
| **ENSMUSG00000071757** | 0.184498417752059 | 0.327113716685933 | 0.855977678491063 | 744.495302452893 | 890.156391703361 | 834.77321178793 | 729.70696813285 | 652.595084708182 | 508.802208542726 | 748.535147109862 | 904.242795368867 | Zhx2 |
| **ENSMUSG00000071793** | 0.102211619227304 | 0.745310572383385 | 0.955985211044899 | 43.3686583953142 | 37.4631235300591 | 38.8046504345863 | 30.9515809105193 | 32.8487458745729 | 24.4616446414772 | 54.1360607089191 | 28.2236646813812 | 2610005L07 |
| **ENSMUSG00000071796** | 0.29111420973093 | 0.284545389646471 | 0.851125503777472 | 28.9124389302094 | 37.4631235300591 | 34.0723759913441 | 31.8895076047775 | 27.3739548954774 | 23.2385624094033 | 27.0680303544596 | 30.3947158107182 | 6820431F20 |
| **ENSMUSG00000071847** | -0.707491074928502 | 0.106322577776721 | 0.762094734041179 | 26.2018977805023 | 29.3189662409158 | 17.9826428843205 | 31.8895076047775 | 65.6974917491458 | 62.3771938357668 | 21.8626399016789 | 22.7960368580387 | Apcdd1 |
| **ENSMUSG00000071855** | -0.169316923277644 | 0.694159125767084 | 0.944725676749441 | 18.0702743313809 | 10.5874044758863 | 14.1968233297267 | 12.1930470253561 | 10.949581958191 | 15.9000690169602 | 23.9447960827911 | 10.8552556466851 | Ccdc112 |
| **ENSMUSG00000071856** | -0.0938468440711646 | 0.710581680580877 | 0.949886733585872 | 187.027339329792 | 174.284965987666 | 202.541346170768 | 321.708856130549 | 316.442918591719 | 189.577745971448 | 211.338852382896 | 226.874843015718 | Mcc |
| **ENSMUSG00000071866** | -0.0076266962832074 | 0.956619549449945 | 0.992371505557225 | 3037.61311510513 | 2535.27616411031 | 2860.18667349561 | 3238.66087527343 | 3029.74932783144 | 3440.53031882377 | 2480.88908979527 | 2783.28754781005 | Ppia |
| **ENSMUSG00000071984** | -0.847823200436699 | 0.0964834421661625 | 0.757809915356977 | 62.3424464432641 | 74.1118313312039 | 40.6975602118832 | 74.0962088463947 | 146.724398239759 | 216.485555077073 | 32.2734208072402 | 57.5328549274309 | Fndc1 |
| **ENSMUSG00000072082** | -0.0430992370962788 | 0.918340112714019 | 0.986713918026868 | 9.03513716569045 | 13.0306516626293 | 15.1432782183751 | 12.1930470253561 | 10.949581958191 | 13.4539045528125 | 16.6572494488982 | 9.76973008201657 | Ccnf |
| **ENSMUSG00000072115** | -0.579288528667507 | 0.0297825075580383 | 0.665956123973759 | 49.6932544112975 | 64.338842584232 | 49.2156542097192 | 59.0893817382642 | 95.2613630362614 | 113.746647582869 | 58.3003730711437 | 66.217059444779 | Ang |
| **ENSMUSG00000072214** | -0.652671260395529 | 0.053800574821808 | 0.703396849069134 | 48.7897406947284 | 61.0811796685746 | 64.3589324280944 | 70.3445020693621 | 105.115986798633 | 157.777607937528 | 55.1771387994752 | 67.3025850094475 | Sept5 |
| **ENSMUSG00000072235** | -0.631097478575689 | 0.0623937039985951 | 0.715615581523459 | 1212.51540763566 | 763.92195372164 | 777.985918469023 | 1307.46981179588 | 1999.39366556567 | 2361.77179013462 | 835.985706716578 | 1094.20976918586 | Tuba1a |
| **ENSMUSG00000072258** | 0.175058480027196 | 0.266115777418287 | 0.843513595393324 | 527.652010476322 | 573.348673155687 | 593.427215182576 | 562.756016554897 | 436.88832013182 | 401.170972120226 | 575.716184077544 | 584.012753791657 | Taf1a |
| **ENSMUSG00000072294** | 0.681175085214851 | 0.152063315277502 | 0.783412422888014 | 38.8510898124689 | 74.1118313312039 | 56.7872933189068 | 23.448167356454 | 29.5638712871156 | 9.78465785659088 | 38.5198893505771 | 42.3354970220718 | Klf12 |
| **ENSMUSG00000072494** | -0.159918454955061 | 0.597149550714225 | 0.924496535139492 | 24.3948703473642 | 48.8649437348597 | 42.5904699891801 | 35.6412143818101 | 50.3680770076785 | 39.1386314263635 | 31.2323427166841 | 48.8486504100828 | Ppp1r3e |
| **ENSMUSG00000072501** | 0.186230470004437 | 0.324613100181592 | 0.855977678491063 | 1015.54941742361 | 1015.57641395617 | 1086.53021216842 | 921.981940455772 | 729.242158415519 | 667.802898712327 | 1053.57102764281 | 1098.55187144453 | Phf20l1 |
| **ENSMUSG00000072568** | -0.239863126092173 | 0.551933584033006 | 0.91494897746968 | 97.5794813894569 | 75.7406627890326 | 163.736695736181 | 251.364354061187 | 228.846262926191 | 198.139321595965 | 88.4916376972716 | 179.111718170304 | Fam84b |
| **ENSMUSG00000072571** | 0.299806862901869 | 0.520191082095889 | 0.905962179970576 | 19.877301764519 | 24.4324718674299 | 17.9826428843205 | 24.3860940507122 | 4.37983278327639 | 17.123151249034 | 26.0269522639034 | 22.7960368580387 | Tmem253 |
| **ENSMUSG00000072582** | -0.0575053001986845 | 0.661797207014672 | 0.939611397743053 | 420.133878204606 | 354.270842077733 | 362.492222352355 | 370.481044231974 | 417.179072607076 | 437.863439082442 | 375.829190690765 | 338.683976176574 | Ptrh2 |
| **ENSMUSG00000072589** | 0.295037410989606 | 0.782432385039205 | 0.960742105865155 | 5.42108229941427 | 7.32974156022896 | 39.7511053232348 | 0 | 9.85462376237187 | 24.4616446414772 | 2.08215618111227 | 6.51315338801105 | Gm10371 |
| **ENSMUSG00000072594** | 0.237675761205353 | 0.52249041652862 | 0.90648515894888 | 19.877301764519 | 26.0613033252585 | 42.5904699891801 | 28.1378008277448 | 29.5638712871156 | 15.9000690169602 | 33.3144988977964 | 19.5394601640331 | Gm16439 |
| **ENSMUSG00000072612** | 0.57805620318743 | 0.225751388408831 | 0.832048466795696 | 16.2632468982428 | 17.1027303072009 | 19.8755526616174 | 18.7585338851632 | 7.66470737073368 | 4.89232892829544 | 12.4929370866736 | 22.7960368580387 | Gm10382 |
| **ENSMUSG00000072620** | -0.465665556133216 | 0.0682601740702721 | 0.722112580225498 | 155.404359249876 | 129.492100897378 | 119.253315969704 | 135.999370667433 | 225.561388338734 | 250.731857575141 | 117.641824232843 | 153.05910461826 | Slfn2 |
| **ENSMUSG00000072623** | 0.0451819925114068 | 0.837637717513859 | 0.968924788429433 | 81.316234491214 | 71.6685841444609 | 65.3053873167428 | 81.59962240046 | 86.5016974697087 | 50.1463715150282 | 74.9576225200418 | 78.1578406561325 | Zfp9 |
| **ENSMUSG00000072640** | 0.0555268413338843 | 0.783937647972178 | 0.961141387070613 | 751.723412185445 | 675.150639269978 | 533.800557197724 | 558.066383083606 | 473.02194059385 | 501.463715150282 | 749.576225200418 | 699.078463646519 | Lyrm9 |
| **ENSMUSG00000072664** | -0.163868835235711 | 0.700465326896815 | 0.94561797945347 | 11.7456783153976 | 24.4324718674299 | 18.9290977729689 | 24.3860940507122 | 21.8991639163819 | 13.4539045528125 | 16.6572494488982 | 36.9078691987293 | Ugt3a1 |
| **ENSMUSG00000072692** | -0.101539063395926 | 0.738490049493468 | 0.954383574136061 | 20.780815481088 | 22.8036404096012 | 28.3936466594534 | 25.3240207449704 | 20.8042057205628 | 30.5770558018465 | 30.191264626128 | 22.7960368580387 | Rpl37rt |
| **ENSMUSG00000072694** | -0.0814778483974762 | 0.861931648816203 | 0.974868041734905 | 9.03513716569045 | 11.4018202048006 | 22.7149173275627 | 16.8826804966469 | 25.1840385038392 | 14.6769867848863 | 9.36970281500523 | 14.1118323406906 | 1500011B03 |
| **ENSMUSG00000072704** | 0.0273337681333008 | 0.862704640541701 | 0.974868041734905 | 2355.4602590955 | 2168.78908609886 | 2029.19928126227 | 2170.36237051339 | 1687.33057975723 | 1971.60855810306 | 2223.74280142791 | 2676.90604247254 | Smim10l1 |
| **ENSMUSG00000072720** | 0.276865714358586 | 0.208358971859766 | 0.82055744801025 | 11505.3436667902 | 12638.1032812926 | 11521.1953595175 | 11388.3059216826 | 8332.63187018333 | 6188.79609429373 | 12372.1720281691 | 11941.8667369183 | Myo18b |
| **ENSMUSG00000072761** | 0.208936086674009 | 0.541349871432727 | 0.913837378403038 | 22.5878429142261 | 26.0613033252585 | 20.8220075502658 | 26.2619474392285 | 22.994122112201 | 15.9000690169602 | 14.5750932677859 | 29.3091902460497 | Gm6712 |
| **ENSMUSG00000072762** | -0.516875861915727 | 0.366436870749248 | 0.869754677013001 | 17.1667606148119 | 8.14415728914329 | 2.83936466594534 | 7.50341355406529 | 18.6142893289246 | 13.4539045528125 | 9.36970281500523 | 9.76973008201657 | 4930522L14 |
| **ENSMUSG00000072769** | -0.428880178826816 | 0.362435752847166 | 0.86924083181804 | 8.13162344912141 | 8.14415728914329 | 8.51809399783602 | 10.3171936368398 | 10.949581958191 | 6.1154111603693 | 14.5750932677859 | 15.1973579053591 | Gm10419 |
| **ENSMUSG00000072772** | -0.117635273142216 | 0.535821129100262 | 0.911073017897251 | 3980.88143520321 | 3092.33652268771 | 3588.95693775491 | 3978.68503704312 | 3820.30914521283 | 5418.2542880872 | 3265.8619700746 | 3381.4121339424 | Grcc10 |
| **ENSMUSG00000072774** | -0.929629474550566 | 0.105238459149153 | 0.762094734041179 | 2.71054114970714 | 11.4018202048006 | 11.3574586637814 | 21.5723139679377 | 10.949581958191 | 20.7923979452556 | 22.903717992235 | 34.7368180693922 | Zfp951 |
| **ENSMUSG00000072812** | -0.295748376111013 | 0.481045121161667 | 0.897732059433541 | 148.176249517323 | 188.13003337921 | 130.610774633486 | 248.550573978413 | 249.650468646754 | 408.509465512669 | 113.477511870619 | 107.467030902182 | Ahnak2 |
| **ENSMUSG00000072825** | 0.0385704193654381 | 0.779086808263113 | 0.960290688328569 | 299.063040184354 | 358.342920722305 | 313.276568142636 | 293.571055302804 | 327.39250054991 | 261.739597663806 | 305.035880532948 | 336.512925047237 | Cep170b |
| **ENSMUSG00000072847** | 0.149747897704176 | 0.47984447357198 | 0.897732059433541 | 72.2810973255236 | 57.8235167529173 | 56.7872933189068 | 55.3376749612315 | 53.6529515951357 | 59.9310293716191 | 57.2592949805875 | 47.7631248454143 | A530017D24 |
| **ENSMUSG00000072849** | -0.16535824852908 | 0.687449091079349 | 0.944452327903597 | 570.117155155067 | 1367.40400884716 | 384.260684791269 | 662.176246146262 | 555.143805280282 | 491.679057293692 | 1232.63645921847 | 1067.07163006914 | Serpina1e |
| **ENSMUSG00000072852** | 0.0610182932202214 | 0.875707946211701 | 0.977159268104838 | 57.8248778604189 | 114.83261777692 | 91.8061241988993 | 64.7169419038131 | 39.4184950494875 | 51.3694537471021 | 105.14888714617 | 119.407812113536 | 2310040G07 |
| **ENSMUSG00000072872** | 0.0376821805140363 | 0.855641595822048 | 0.973387788552416 | 136.430571201926 | 147.409246933493 | 182.66579350915 | 111.613276616721 | 166.433645764503 | 112.523565350795 | 146.792010768415 | 136.776221148232 | Rybp |
| **ENSMUSG00000072875** | 0.243768025418884 | 0.0604111430032425 | 0.71282992338528 | 654.143930795989 | 687.366875203693 | 758.110365807406 | 758.782695654852 | 608.796756875418 | 518.586866399316 | 701.686633034836 | 582.927228226989 | Gpr27 |
| **ENSMUSG00000072889** | 0.125203576947318 | 0.715020942730365 | 0.951176382858794 | 33.4300075130547 | 74.1118313312039 | 70.0376617599851 | 57.2135283497478 | 45.9882442244021 | 31.8001380339204 | 70.7933101578173 | 66.217059444779 | Nfxl1 |
| **ENSMUSG00000072902** | 0.12254762215742 | 0.79315660742036 | 0.961804768755116 | 25.2983840639333 | 33.3910448854875 | 29.3401015481019 | 23.448167356454 | 9.85462376237187 | 13.4539045528125 | 36.4377331694648 | 42.3354970220718 | Gm10435 |
| **ENSMUSG00000072941** | -0.896407791852005 | 0.0105446211018825 | 0.635862253526407 | 760.758549351136 | 720.757920089181 | 625.606681396623 | 722.203554578784 | 1309.57000219964 | 2447.38754637979 | 741.247600475969 | 769.637625349972 | Sod3 |
| **ENSMUSG00000072946** | 0.147459740044708 | 0.466781502407263 | 0.895328112556867 | 6824.239101246 | 6997.45994283191 | 6321.37220128298 | 5826.4006247317 | 4939.35642133995 | 4107.11013530402 | 7341.68269460188 | 7057.00169590997 | Ptgr2 |
| **ENSMUSG00000072949** | 0.581991519025496 | 0.0489923907167319 | 0.703396849069134 | 1741.07093182855 | 2067.80153571348 | 4723.7563492444 | 2528.65036772 | 2209.62563916294 | 1822.39252579005 | 1715.69669323651 | 1641.31465377878 | Acot1 |
| **ENSMUSG00000072964** | 0.222600329445579 | 0.257088858401575 | 0.837517407767513 | 347.852780879082 | 334.724864583789 | 411.707876562074 | 305.764102328161 | 275.929465346412 | 218.931719541221 | 322.734208072402 | 381.019473198646 | Bhlhb9 |
| **ENSMUSG00000072969** | 0.3063588966639 | 0.13912837768266 | 0.779711473022872 | 131.009488902512 | 175.099381716581 | 177.933519065908 | 126.620103724852 | 116.065568756824 | 95.400414101761 | 138.463386043966 | 143.289374536243 | Armcx5 |
| **ENSMUSG00000073002** | 0.140027752872115 | 0.494319000897073 | 0.900141680336539 | 1156.49755720838 | 803.828324438442 | 889.66759532954 | 1358.11785328582 | 1051.15986798633 | 1119.12024234758 | 816.205222996011 | 832.598108100745 | Vamp5 |
| **ENSMUSG00000073016** | 0.333460052344028 | 0.607609973142476 | 0.927510709653048 | 39.754603529038 | 87.1424829938332 | 74.7699362032273 | 21.5723139679377 | 20.8042057205628 | 7.33849339244316 | 74.9576225200418 | 73.8157383974585 | Uprt |
| **ENSMUSG00000073062** | -0.0126958844135126 | 0.963046671610793 | 0.994284908685797 | 37.9475760958999 | 29.3189662409158 | 45.4298346551254 | 55.3376749612315 | 41.6084114411257 | 48.9232892829544 | 37.4788112600209 | 41.2499714574033 | Zxdb |
| **ENSMUSG00000073096** | 0.0628078931835921 | 0.768415114953493 | 0.95879680569409 | 173.474633581257 | 229.665235553841 | 224.309808609682 | 187.585338851632 | 199.282391639076 | 128.423634367755 | 227.996101831794 | 223.618266321713 | Lrrc61 |
| **ENSMUSG00000073131** | 0.128473699602361 | 0.226050928773474 | 0.832048466795696 | 541.204716224858 | 496.79359463774 | 468.495169880981 | 500.852854733858 | 432.508487348543 | 473.332823812584 | 437.252798033577 | 493.914131924171 | Vma21 |
| **ENSMUSG00000073139** | 0.0138680891162526 | 0.916038334743812 | 0.986713918026868 | 449.046317134815 | 470.732291312482 | 499.72818120638 | 444.577253078368 | 405.134532453066 | 412.178712208891 | 516.374732915844 | 511.282540958867 | Tmem185a |
| **ENSMUSG00000073147** | -0.229309162574993 | 0.276814501780267 | 0.847279292579971 | 155.404359249876 | 115.647033505835 | 141.021778408619 | 180.081925297567 | 153.294147414674 | 225.04713070159 | 144.709854587303 | 171.513039217624 | 5031425E22 |
| **ENSMUSG00000073154** | -0.215082600828864 | 0.243731689331393 | 0.832048466795696 | 299.063040184354 | 274.458100644129 | 255.542819935081 | 259.805694309511 | 255.12525962585 | 425.632616761703 | 285.255396812381 | 299.605055848508 | 9330158H04 |
| **ENSMUSG00000073155** | 0.229242791026098 | 0.0684285691363867 | 0.72237439886039 | 2029.29180741408 | 1808.81733391872 | 2043.396104592 | 2211.63114506074 | 1633.67762816209 | 1991.17787381624 | 1505.39891894417 | 1774.83429823301 | 1810058I24R |
| **ENSMUSG00000073176** | 0.0773213184899206 | 0.774802044255966 | 0.959414024438738 | 168.053551281842 | 140.893921102179 | 145.754052851861 | 110.675349922463 | 94.1664048404423 | 94.1773318696872 | 164.49033830787 | 182.368294864309 | Zfp449 |
| **ENSMUSG00000073198** | 0.580417198012531 | 0.280000954362361 | 0.848943557485723 | 7.22810973255236 | 12.2162359337149 | 12.3039135524298 | 16.8826804966469 | 4.37983278327639 | 14.6769867848863 | 8.32862472444909 | 5.42762782334254 | Bnip3l-ps |
| **ENSMUSG00000073236** | 0.122189363083864 | 0.612129745768734 | 0.928055903343287 | 40.658117245607 | 55.3802695661743 | 46.3762895437739 | 41.2687745473591 | 38.3235368536684 | 41.5847958905112 | 49.9717483466946 | 39.0789203280663 | 2500004C02 |

| | | | | | | | | | | | | |
|---|---|---|---|---|---|---|---|---|---|---|---|---|
| **ENSMUSG00000073293** | -0.194980566396403 | 0.541910764060487 | 0.913837378403038 | 22.5878429142261 | 39.9063707168021 | 20.8220075502658 | 31.8895076047775 | 40.5134532453066 | 26.9078091056249 | 35.3966550789086 | 29.3091902460497 | Nudt10 |
| **ENSMUSG00000073295** | -0.33230802107887 | 0.316374635510358 | 0.85345575592132 | 53.3073092775737 | 70.0397526866323 | 33.1259211026956 | 45.0204813243917 | 49.2731188118594 | 41.5847958905112 | 73.9165444294857 | 89.0130963028176 | Nudt11 |
| **ENSMUSG00000073374** | 0.205888160801217 | 0.558286287933002 | 0.916574868225118 | 17.1667606148119 | 23.6180561385155 | 24.6078271048596 | 20.6343872736795 | 22.994122112201 | 23.2385624094033 | 12.4929370866736 | 16.2828834700276 | C030034I22 |
| **ENSMUSG00000073375** | -0.662534777096105 | 0.0313391397211035 | 0.669064667366769 | 23.4913566307952 | 16.2883145782866 | 22.7149173275627 | 17.8206071909051 | 30.6588294829347 | 24.4616446414772 | 33.3144988977964 | 37.9933947633978 | Lrrc30 |
| **ENSMUSG00000073380** | -0.23223061525716 | 0.668256379729016 | 0.942024383071524 | 4.51756858284523 | 12.2162359337149 | 14.1968233297267 | 10.3171936368398 | 5.47479097909548 | 12.2308223207386 | 19.7804837205666 | 10.8552556466851 | Arrdc5 |
| **ENSMUSG00000073400** | 0.826023483590556 | 0.529410439124082 | 0.908560090446144 | 31.6229800799166 | 8.95857301805762 | 162.790240847533 | 4.68963347129081 | 15.3294147414674 | 89.2850029413918 | 3.12323427166841 | 9.76973008201657 | Trim10 |
| **ENSMUSG00000073409** | 0.158089536081783 | 0.681231384483429 | 0.944452327903597 | 305.387636200337 | 182.42912327681 | 385.207139679918 | 215.723139679377 | 375.57066116595 | 327.786038195794 | 148.874166949528 | 123.74991437221 | H2-Q6 |
| **ENSMUSG00000073411** | -0.800010104318808 | 0.0582405469402835 | 0.710862211775517 | 3323.12344954095 | 2760.05490529066 | 3431.84542623927 | 3076.39955716677 | 7788.43764686124 | 9467.87955848375 | 2245.60544132959 | 2421.80753477544 | H2-D1 |
| **ENSMUSG00000073414** | 0.700156491499659 | 0.403799669188386 | 0.881158207349486 | 18.0702743313809 | 13.0306516626293 | 63.4124775394459 | 7.50341355406529 | 15.3294147414674 | 39.1386314263635 | 2.08215618111227 | 6.51315338801105 | Mpig6b |
| **ENSMUSG00000073418** | -1.1777637353815 | 0.00461235103463074 | 0.635862253526407 | 198.77301764519 | 316.807718547674 | 316.115932808581 | 223.226553233442 | 430.318570956905 | 1147.25113368528 | 222.790711379013 | 587.269330485663 | C4b |
| **ENSMUSG00000073421** | -0.686896971401756 | 0.0541822305827532 | 0.703396849069134 | 378.57224724243 | 452.815145276367 | 512.978549647458 | 526.176875478828 | 1086.19853025254 | 1122.7894890438 | 391.445362049107 | 411.414189009364 | H2-Ab1 |
| **ENSMUSG00000073422** | -0.0865762609424748 | 0.70138371208676 | 0.94593841740801 | 159.921927832721 | 99.3587189275481 | 106.949402417274 | 150.068271081306 | 120.445401540101 | 171.23151249034 | 138.463386043966 | 118.322286548867 | H2-Ke6 |
| **ENSMUSG00000073423** | -0.0600130138091594 | 0.7550066037963 | 0.956497786093635 | 107.518132271716 | 88.7713144516618 | 69.0912068713366 | 82.5375490947182 | 90.881530252985 | 89.2850029413918 | 90.5737938783839 | 92.2696729968231 | Zfp414 |
| **ENSMUSG00000073424** | -0.293349631237607 | 0.396059047145576 | 0.879251804756997 | 16.2632468982428 | 33.3910448854875 | 44.483379766477 | 32.8274342990356 | 36.1336204620302 | 30.5770558018465 | 37.4788112600209 | 51.0197015394199 | Cyp4f15 |
| **ENSMUSG00000073427** | 0.0902260212904106 | 0.869704096327893 | 0.975613767803763 | 16.2632468982428 | 8.14415728914329 | 13.2503684410783 | 6.56548685980713 | 6.56974917491458 | 4.89232892829544 | 16.6572494488982 | 13.0263067760221 | Gm4924 |
| **ENSMUSG00000073433** | 0.0756849679168804 | 0.872364736826922 | 0.976162349063696 | 16.2632468982428 | 38.2775392589734 | 37.8581955459379 | 42.2067012416173 | 42.7033696369448 | 51.3694537471021 | 13.5340151772298 | 20.6249857287016 | Arhgdig |
| **ENSMUSG00000073434** | -0.707230821095097 | 0.178023941628414 | 0.801696999069105 | 7.22810973255236 | 7.32974156022896 | 10.4110037751329 | 7.50341355406529 | 10.949581958191 | 24.4616446414772 | 11.4518589961175 | 6.51315338801105 | Wdr90 |
| **ENSMUSG00000073436** | 0.436597937465951 | 0.0188608808295563 | 0.642273034804458 | 155.404359249876 | 143.337168288922 | 144.807597963212 | 126.620103724852 | 99.6411958195378 | 79.5003450848009 | 125.970448957293 | 115.065709854862 | Eme2 |
| **ENSMUSG00000073468** | -0.0994914430123144 | 0.597790804095199 | 0.924496535139492 | 223.167887992554 | 150.666909849151 | 188.344522841041 | 205.405946042537 | 221.181555555458 | 234.831788558181 | 160.326025945645 | 206.249857287016 | Sft2d1 |
| **ENSMUSG00000073471** | -0.016322899107749 | 0.920471371573259 | 0.986713918026868 | 415.616309621761 | 458.516055378767 | 473.227444324223 | 538.369922504185 | 404.039574257247 | 591.971800323748 | 421.636626675235 | 490.657555230165 | Rsph3a |
| **ENSMUSG00000073478** | -0.721620688126504 | 0.0585633624358114 | 0.711712402279519 | 18.9737880479499 | 10.5874044758863 | 17.036187995672 | 13.1309737196143 | 25.1840385038392 | 30.5770558018465 | 14.5750932677859 | 28.2236646813812 | D730003I15 |
| **ENSMUSG00000073481** | -0.0438570324046085 | 0.795037344953322 | 0.962032824235414 | 956.821025846619 | 934.134841064735 | 893.453414884134 | 1085.18118525669 | 1099.33802860237 | 1220.63606760971 | 767.274552739873 | 903.157269804198 | Marc2 |
| **ENSMUSG00000073485** | 0.768446870130575 | 0.0985304313682231 | 0.759800953500741 | 15.3597331816738 | 31.7622134276588 | 40.6975602118832 | 27.1998741334867 | 15.3294147414674 | 6.1154111603693 | 26.0269522639034 | 19.5394601640331 | H3f3aos |
| **ENSMUSG00000073489** | -0.324484959949982 | 0.301143925863167 | 0.851785577030345 | 64.1494738764022 | 87.1424829938332 | 75.7163910918757 | 73.1582821521366 | 120.445401540101 | 134.539045528125 | 69.7522320672611 | 52.1052271040884 | Ifi204 |
| **ENSMUSG00000073490** | -0.203360011690609 | 0.488347719700814 | 0.899236955296196 | 148.176249517323 | 188.944449108124 | 183.612248397799 | 120.054616865045 | 247.460552255116 | 240.94719971855 | 137.42230795341 | 112.894658725525 | Ifi207 |
| **ENSMUSG00000073535** | 0.303455563981847 | 0.19205505388866 | 0.808824684419004 | 3352.93940218773 | 3155.04653381411 | 4820.29474788654 | 4288.20084614831 | 2512.92905940483 | 2296.94843183471 | 3942.56272893609 | 3900.29335385395 | Gm5532 |
| **ENSMUSG00000073542** | -0.169721019227137 | 0.371361506781436 | 0.873374202289323 | 167.150037565273 | 197.088606397268 | 187.398067952392 | 154.757904552597 | 194.902558855799 | 144.323703384715 | 238.406882737355 | 216.019587369033 | Cep76 |
| **ENSMUSG00000073555** | -0.441634449360413 | 0.227777606135935 | 0.832048466795696 | 15.3597331816738 | 24.4324718674299 | 46.3762895437739 | 38.4549944645846 | 47.0832024202212 | 33.0232202659942 | 38.5198893505771 | 49.9341759747513 | Gm4951 |
| **ENSMUSG00000073557** | 0.246200126749174 | 0.460540103816285 | 0.894774044236998 | 5487.03880072381 | 8083.89052520363 | 6752.95563050667 | 4538.62727351524 | 3740.37719691804 | 2635.74221011917 | 7548.85723462255 | 7036.37671018127 | Ppp1r12b |
| **ENSMUSG00000073563** | -0.0305249064632716 | 0.807986165026597 | 0.964026716319739 | 289.124389302094 | 292.375246680244 | 248.917635714542 | 307.639955716677 | 290.163921892061 | 266.631926592101 | 291.501865355718 | 313.716888189199 | Csnk1g3 |
| **ENSMUSG00000073565** | -0.638717137072562 | 0.348638849758374 | 0.861266047293794 | 9.93865088225949 | 12.2162359337149 | 2.83936466594534 | 8.44134024832345 | 9.85462376237187 | 12.2308223207386 | 3.12323427166841 | 27.1381391167127 | Prr16 |
| **ENSMUSG00000073590** | -0.121245555321919 | 0.679091522770259 | 0.944452327903597 | 30.7194663633475 | 36.6487078011448 | 35.0188308799925 | 27.1998741334867 | 25.1840385038392 | 29.3539735697726 | 39.5609674411332 | 46.6775992807458 | 3222401L13 |
| **ENSMUSG00000073591** | -0.339750947620117 | 0.178570715389402 | 0.801696999069105 | 60.535419010126 | 48.0505280059454 | 42.5904699891801 | 40.3308478531009 | 71.1722827282413 | 72.1618516923577 | 53.094982618363 | 46.6775992807458 | Pcdhb22 |
| **ENSMUSG00000073599** | -0.266262754353273 | 0.292863302326793 | 0.851785577030345 | 123.781379169959 | 101.801966114291 | 81.3951204237664 | 134.123517278917 | 144.534481848121 | 172.454594722414 | 89.5327157878278 | 124.835439936878 | Ecscr |
| **ENSMUSG00000073600** | 0.108881038103841 | 0.501313693186784 | 0.900927059989498 | 3727.89759456388 | 3787.84755518054 | 3355.18258025874 | 3397.17048660306 | 2823.89718701745 | 2650.41919690405 | 3981.08261828667 | 3774.3723883524 | Prob1 |
| **ENSMUSG00000073608** | -0.215622583474412 | 0.667754747710532 | 0.942024383071524 | 44.2721721118832 | 39.9063707168021 | 25.5542819935081 | 32.8274342990356 | 35.0386622662111 | 8.56157562451702 | 51.0128264372507 | 70.559161703453 | Gal3st2c |
| **ENSMUSG00000073609** | 0.253129018840943 | 0.325701637748621 | 0.855977678491063 | 641.494738764022 | 867.35275129376 | 883.988865997649 | 664.990026229036 | 500.395895489327 | 408.509465512669 | 826.616003901573 | 829.34153140674 | D2hgdh |
| **ENSMUSG00000073616** | 0.159626646309656 | 0.351985036921114 | 0.862728204353539 | 1303.77029300913 | 866.538335564846 | 1072.33338883869 | 1239.00116311503 | 1011.74137293685 | 1198.62058743238 | 908.861173055507 | 893.387539722182 | Cops9 |
| **ENSMUSG00000073627** | 0.270620352769369 | 0.394192659284541 | 0.878942305112115 | 22.5878429142261 | 22.8036404096012 | 31.2330113253987 | 22.5102406621959 | 20.8042057205628 | 20.7923979452556 | 16.6572494488982 | 23.8815624227072 | C130036L24 |
| **ENSMUSG00000073633** | -0.137503187647641 | 0.597894339464223 | 0.924496535139492 | 39.754603529038 | 35.0198763433161 | 39.7511053232348 | 43.1446279358754 | 54.7479097909548 | 44.0309603546589 | 30.191264626128 | 44.5065481514088 | Fbxo36 |
| **ENSMUSG00000073639** | 0.0740040860335592 | 0.51947258689126 | 0.905962179970576 | 2009.41450564956 | 2311.31183865886 | 2272.43818764492 | 1950.88752405698 | 1923.84155005415 | 1819.9463613259 | 2167.52458453788 | 2204.70242184174 | Rab18 |
| **ENSMUSG00000073643** | -0.120014307686038 | 0.603402183050948 | 0.926448151609826 | 84.9302893574902 | 63.5244268553176 | 68.1447519826882 | 68.4686486808458 | 59.1277425742312 | 102.738907494204 | 80.1630129728225 | 68.388110574116 | Wdfy1 |
| **ENSMUSG00000073647** | 0.0416987866154983 | 0.907648015299309 | 0.98504921011344 | 19.877301764519 | 26.8757190541728 | 20.8220075502658 | 15.9447538023887 | 15.3294147414674 | 15.9000690169602 | 28.1091084450157 | 21.7105112933701 | Gm10557 |
| **ENSMUSG00000073664** | 0.272159503006414 | 0.290228583619819 | 0.851785577030345 | 852.01343472461 | 896.671717534676 | 975.794990196549 | 751.279282100787 | 564.998429042654 | 434.19419238622 | 967.161546126651 | 910.755948756878 | Nbeal1 |
| **ENSMUSG00000073676** | 0.172052475398475 | 0.201647753190147 | 0.813395633194566 | 3489.36997338965 | 2663.95384927877 | 2650.07368821565 | 3469.39084206094 | 2800.90306490525 | 2542.78796048155 | 2741.15861243431 | 2807.16911023276 | Hspe1 |
| **ENSMUSG00000073678** | 0.226478835198768 | 0.357756817942016 | 0.865834559821921 | 66.8600150261093 | 85.5136515360045 | 109.78876708322 | 75.0341355406529 | 65.6974917491458 | 57.4848649074714 | 94.7381062406084 | 69.4736361387845 | Pgap1 |
| **ENSMUSG00000073680** | 0.716483485986528 | 0.230466162256944 | 0.832048466795696 | 10.8421645988285 | 17.9171460361152 | 12.3039135524298 | 12.1930470253561 | 7.66470737073368 | 1.22308223207386 | 16.6572494488982 | 6.51315338801105 | Tmem88b |
| **ENSMUSG00000073684** | -0.253439469230719 | 0.424897458194204 | 0.885066977635295 | 66.8600150261093 | 74.9262470601182 | 70.0376617599851 | 111.613276616721 | 95.2613630362614 | 154.108361241306 | 71.8343882483734 | 65.1315338801105 | Faap20 |
| **ENSMUSG00000073700** | 0.0979993569748134 | 0.58746157476467 | 0.923058967588378 | 2809.92765852973 | 3438.4632074763 | 3394.93368558198 | 3269.61245618395 | 3216.98717931651 | 2084.13212345386 | 3571.9389286981 | 3190.35963456074 | Klhl21 |
| **ENSMUSG00000073702** | -0.00558254853362215 | 0.979517295518279 | 0.996872525500296 | 5979.45377625394 | 3949.10186950558 | 4712.39889058062 | 5756.05612266234 | 5093.74552695044 | 7058.40756129824 | 4146.61403468509 | 4178.18789840909 | Rpl31 |
| **ENSMUSG00000073705** | -0.0912105580165542 | 0.827646583625587 | 0.967319591893432 | 41.5616309621761 | 18.7315617650296 | 11.3574586637814 | 19.6964605794214 | 26.2789966996583 | 23.2385624094033 | 23.9447960827911 | 23.8815624227072 | Cenps |
| **ENSMUSG00000073725** | 0.0812684981480423 | 0.488001593468267 | 0.899236955296196 | 1021.87401343959 | 1100.27564976326 | 1325.98329899647 | 1139.58093352367 | 1055.53970076961 | 1006.59667699679 | 1128.52865016285 | 1144.14394516061 | Lmbrd1 |
| **ENSMUSG00000073752** | -0.0688255071883904 | 0.88185402448969 | 0.978854883720278 | 18.0702743313809 | 16.2883145782866 | 12.3039135524298 | 19.6964605794214 | 14.2344565456483 | 25.6847268735511 | 6.24646854333682 | 23.8815624227072 | Gm10570 |
| **ENSMUSG00000073755** | 0.142537515058537 | 0.245551672175722 | 0.832048466795696 | 543.915257374565 | 513.081909216027 | 513.925004536107 | 541.183702586959 | 419.368988998714 | 468.440494884288 | 561.141090809758 | 463.519416113453 | 5730409E04 |
| **ENSMUSG00000073758** | -0.0891864350191512 | 0.727544228453478 | 0.952237115597464 | 103.000563688871 | 107.502876216691 | 112.628131749165 | 102.23400967414 | 151.104231023035 | 141.877538920568 | 85.3684034256032 | 74.901263962127 | Sh3d21 |
| **ENSMUSG00000073771** | -0.251453889454683 | 0.414374937826286 | 0.882192126163777 | 22.5878429142261 | 17.9171460361152 | 20.8220075502658 | 28.1378008277448 | 30.6588294829347 | 23.2385624094033 | 23.9447960827911 | 28.2236646813812 | Btbd19 |
| **ENSMUSG00000073775** | 0.0018148580337385 | 0.988404056723316 | 0.998437672973644 | 402.063603873225 | 349.384347704247 | 330.312756138308 | 400.494698448235 | 406.229490648885 | 347.355353908976 | 356.048706970199 | 370.164217551961 | Kti12 |
| **ENSMUSG00000073787** | -0.0584744850685699 | 0.800067116528222 | 0.963461896828986 | 54.2108229941427 | 43.9784493613737 | 42.5904699891801 | 61.9031618210386 | 58.0327843784121 | 56.2617826753975 | 46.8485140750261 | 49.9341759747513 | Gm10575 |
| **ENSMUSG00000073792** | 0.0409461852789765 | 0.864855263644181 | 0.974868041734905 | 94.8689402397497 | 119.719112150406 | 96.5383986421416 | 75.9720622349111 | 63.5075753575076 | 92.9542496376133 | 107.231043327282 | 112.894658725525 | Alg6 |
| **ENSMUSG00000073830** | -0.0741121554694028 | 0.866280396921591 | 0.974868041734905 | 88.5443442237664 | 122.162359337149 | 31.2330113253987 | 106.92364314543 | 68.9823663366031 | 89.2850029413918 | 54.1360607089191 | 155.230155747597 | Mup14 |
| **ENSMUSG00000073834** | -0.109264308729857 | 0.813719253428565 | 0.966177091475091 | 80.412720774645 | 91.2145616384048 | 28.3936466594534 | 86.2892558717508 | 41.6084114411257 | 70.9387694602839 | 49.9717483466946 | 146.545951230249 | Mup11 |
| **ENSMUSG00000073838** | 0.0757463098886175 | 0.390713418539521 | 0.877797322358001 | 6491.74605354859 | 6055.99536020695 | 5965.50516315116 | 6614.25904790855 | 6518.28613971108 | 5627.40134977183 | 5952.88452179999 | 5742.43023709641 | Tufm |
| **ENSMUSG00000073842** | -0.289359804699007 | 0.445713005810264 | 0.89108031640712 | 604.450676384691 | 626.285695535119 | 256.489274823729 | 631.224665235743 | 458.787484048202 | 648.233582999146 | 367.500565966316 | 1114.83475491456 | Mup7 |
| **ENSMUSG00000073888** | -0.132861873252127 | 0.634629882616762 | 0.932543559775713 | 93.9654265231807 | 62.7100111264033 | 71.930571537282 | 75.9720622349111 | 74.4571573156986 | 129.646716599829 | 64.5468416144805 | 66.217059444779 | Ccl27a |
| **ENSMUSG00000073889** | 0.041980561077185 | 0.733113861824346 | 0.953733898884637 | 1958.81773752169 | 1595.44041294317 | 1505.80972783968 | 1817.70193347232 | 1716.89445104434 | 1811.38478570139 | 1613.67104036201 | 1539.27525069994 | Il11ra1 |
| **ENSMUSG00000073910** | -0.928137112470959 | 0.0574897372593273 | 0.710862211775517 | 6.32459601598331 | 9.77298874697194 | 11.3574586637814 | 14.0689004138724 | 10.949581958191 | 35.4693847301419 | 16.6572494488982 | 16.2828834700276 | Mob3b |
| **ENSMUSG00000073940** | 0.556624548349867 | 0.532549072815565 | 0.909082872831377 | 3458.6505070263 | 1513.18442432282 | 11241.0447124776 | 1182.72556145954 | 1483.66835533488 | 6977.68413398137 | 1251.37586484848 | 2114.60379997425 | Hbb-bt |
| **ENSMUSG00000073968** | 0.180287882948639 | 0.239258336077669 | 0.832048466795696 | 179.79922959724 | 161.254314325037 | 191.183887506986 | 182.895705380341 | 129.205067106653 | 160.223772401676 | 174.901119213431 | 166.085411394282 | Trim68 |
| **ENSMUSG00000073982** | -0.576323413776976 | 0.0837546011428796 | 0.742148948523533 | 294.545471601509 | 228.036404096012 | 217.684624389143 | 252.302280755445 | 491.636229922775 | 555.279333361532 | 185.311900118992 | 248.585354309088 | Rhog |
| **ENSMUSG00000073985** | 0.0571328005303743 | 0.776924223519068 | 0.959738279197945 | 57.8248778604189 | 74.9262470601182 | 76.6628459805242 | 68.4686486808458 | 62.4126171616885 | 58.7079471395453 | 74.9576225200418 | 70.559161703453 | Gm10602 |
| **ENSMUSG00000073987** | -0.0232079552467361 | 0.906788356482035 | 0.98504921011344 | 177.992202164102 | 151.481325578065 | 145.754052851861 | 211.033506208086 | 178.478185918513 | 212.816308380852 | 136.381229862854 | 170.427513652956 | Ggh |
| **ENSMUSG00000073988** | -0.110585349582015 | 0.757503147402742 | 0.956693827734505 | 112.939214571131 | 246.767965861042 | 125.878500190243 | 158.509611329629 | 122.635317931739 | 178.570005882784 | 108.272121417838 | 286.578749072486 | Ttpa |
| **ENSMUSG00000074001** | -0.0932298166372856 | 0.705531269395353 | 0.947760509100828 | 474.344701198749 | 438.970077884823 | 271.632553042104 | 518.673461924763 | 460.97740043984 | 604.202622644487 | 410.184767679118 | 343.026078435248 | Klhl40 |
| **ENSMUSG00000074024** | -0.0271311477059224 | 0.96503964240341 | 0.994284908685797 | 12.6491920319666 | 11.4018202048006 | 2.83936466594534 | 15.0068271081306 | 4.37983278327639 | 19.5693157131818 | 10.4107809055614 | 8.68420451734806 | 4632427E13 |
| **ENSMUSG00000074030** | -0.132635555216715 | 0.326181491468889 | 0.855977678491063 | 321.65088309858 | 381.146561131906 | 420.22597055991 | 384.549944645846 | 431.413529152724 | 355.916929533493 | 435.170641852465 | 428.78259804406 | Exoc8 |
| **ENSMUSG00000074039** | -0.465730080280122 | 0.306610686716613 | 0.851785577030345 | 9.03513716569045 | 7.32974156022896 | 10.4110037751329 | 9.37926694258161 | 7.66470737073368 | 13.4539045528125 | 16.6572494488982 | 11.9407812113536 | 4930520O04 |
| **ENSMUSG00000074063** | -0.590783798811382 | 0.00328832819224563 | 0.635862253526407 | 127.395434036235 | 171.841718800923 | 122.09268063565 | 141.626930832982 | 217.896680968 | 271.524255520397 | 172.818963032319 | 187.795922687652 | Osgin1 |
| **ENSMUSG00000074064** | 0.127139148944521 | 0.265222344320277 | 0.842510961779757 | 4752.48214915318 | 4591.67587961899 | 5555.69019636638 | 5353.68557082558 | 5215.28588668636 | 4236.75685190385 | 4579.70252035645 | 4511.44424676232 | Mlycd |
| **ENSMUSG00000074088** | -0.0990767129846974 | 0.557073911084811 | 0.916253004732743 | 290.931416735233 | 254.912123150185 | 274.47191770805 | 281.378008277448 | 344.911831683016 | 346.132271676902 | 247.776585552361 | 242.072200921077 | Snrnp40 |
| **ENSMUSG00000074093** | 0.204570499697023 | 0.345487698101457 | 0.861266047293794 | 911.645340018166 | 1075.02876216691 | 1097.8876708322 | 1048.60204418062 | 709.532910890775 | 631.110431750112 | 1065.02288663893 | 1179.96628879467 | Svip |
| **ENSMUSG00000074102** | -0.0808347154468653 | 0.572239224238485 | 0.921574199900435 | 260.211950371885 | 323.323044378988 | 317.06238769723 | 264.495327780801 | 333.962249724825 | 336.347613820311 | 282.132162540713 | 281.151121249143 | Rbm15b |
| **ENSMUSG00000074115** | 0.683427907695138 | 0.172796161935992 | 0.799889021902745 | 56.9213641438498 | 277.715763559786 | 142.914688185915 | 47.8342614071662 | 90.881530252985 | 89.2850029413918 | 56.2182168900314 | 91.1841474321546 | Saa1 |
| **ENSMUSG00000074129** | -0.0176277432010097 | 0.924438719132635 | 0.986713918026868 | 5712.0137161495 | 3860.33055505392 | 4409.53332621311 | 5900.49683357809 | 4748.83369526742 | 6165.55753188433 | 4245.51645328793 | 4967.36498392309 | Rpl13a |
| **ENSMUSG00000074151** | 0.203671390622482 | 0.666473580020583 | 0.942024383071524 | 21.6843291976571 | 39.0919549878878 | 26.5007368821565 | 19.6964605794214 | 45.9882442244021 | 19.5693157131818 | 15.616171358342 | 11.9407812113536 | Nlrc5 |
| **ENSMUSG00000074158** | -0.218554065728316 | 0.4712088793869 | 0.897091462240771 | 46.9827132615903 | 33.3910448854875 | 33.1259211026956 | 24.3860940507122 | 44.893286028583 | 29.3539735697726 | 49.9717483466946 | 35.8223436340608 | Zfp976 |
| **ENSMUSG00000074165** | -0.0906202794665688 | 0.661097171439286 | 0.939611397743053 | 193.351935345776 | 130.306516626293 | 126.824955078892 | 151.944124469822 | 159.863896589588 | 143.100621152642 | 199.886993386778 | 137.8617467129 | Zfp788 |
| **ENSMUSG00000074166** | -0.511805472431745 | 0.0496416891669145 | 0.703396849069134 | 36.1405486627618 | 28.5045505120015 | 36.9117406572894 | 33.7653609932938 | 53.6529515951357 | 34.2463024980681 | 57.2592949805875 | 46.6775992807458 | AW146154 |

| | | | | | | | | | | | | |
|---|---|---|---|---|---|---|---|---|---|---|---|---|
| **ENSMUSG00000074170** | -0.354924092443417 | 0.144779990521454 | 0.780607990324772 | 441.818207402263 | 407.207864457164 | 294.347470369667 | 238.233380341573 | 469.737066006393 | 568.733237914345 | 367.500565966316 | 362.565538599282 | Plekhf1 |
| **ENSMUSG00000074171** | 0.121053497775667 | 0.815108591350785 | 0.966177091475091 | 25.2983840639333 | 15.4738988493722 | 27.447191770805 | 6.56548685980713 | 15.3294147414674 | 8.56157562451702 | 26.0269522639034 | 18.4539345993646 | Gm6658 |
| **ENSMUSG00000074178** | 0.0751712518272048 | 0.848684719476358 | 0.972640876584422 | 138.237598635064 | 198.717437855096 | 331.259211026956 | 249.488500672671 | 241.98576127602 | 351.024600605198 | 189.476212481217 | 89.0130963028176 | Gm10638 |
| **ENSMUSG00000074179** | -0.519397219852674 | 0.392098706647676 | 0.878285927324016 | 5.42108229941427 | 23.6180561385155 | 14.1968233297267 | 7.50341355406529 | 7.66470737073368 | 13.4539045528125 | 34.3555769883525 | 17.3684090346961 | Gm10639 |
| **ENSMUSG00000074182** | 0.282318406037785 | 0.0283098011049921 | 0.665956123973759 | 518.616873310632 | 482.134111517283 | 443.887342776122 | 600.273084325223 | 407.324448844704 | 437.863439082442 | 421.636626675235 | 414.67076570337 | Znhit6 |
| **ENSMUSG00000074183** | -0.46734188680223 | 0.151494479559964 | 0.783412422888014 | 75.8951521917998 | 74.9262470601182 | 36.9117406572894 | 82.5375490947182 | 68.9823663366031 | 143.100621152642 | 81.2040910633787 | 81.4144173501381 | Gsta1 |
| **ENSMUSG00000074194** | -0.341606216213583 | 0.378104929928243 | 0.87422797717331 | 36.1405486627618 | 30.1333819698302 | 19.8755526616174 | 31.8895076047775 | 26.2789966996583 | 19.5693157131818 | 46.8485140750261 | 56.4473293627624 | Zfp791 |
| **ENSMUSG00000074207** | -0.487794400266366 | 0.0577524479250511 | 0.710862211775517 | 486.993893230715 | 653.975830318206 | 484.584902988005 | 517.735535230505 | 858.447225522172 | 958.896469945906 | 408.102611498006 | 781.578406561325 | Adh1 |
| **ENSMUSG00000074211** | -0.0800590870566314 | 0.710876356776468 | 0.949886733585872 | 303.580608767199 | 210.933673788811 | 188.344522841041 | 294.508981997063 | 247.460552255116 | 319.224462571277 | 210.29777429234 | 277.894544555138 | Sdhaf1 |
| **ENSMUSG00000074212** | -0.0987475509112016 | 0.431541732409217 | 0.886559861988535 | 523.134441893477 | 564.39010013763 | 511.085639870161 | 489.59773440276 | 469.737066006393 | 565.063991218123 | 577.798340258656 | 624.177199684392 | Dnajb14 |
| **ENSMUSG00000074215** | -0.175002399782286 | 0.38796639423578 | 0.875951433019341 | 75.8951521917998 | 66.7820897709749 | 76.6628459805242 | 65.6548685980713 | 81.0269064906132 | 90.5080851734656 | 89.5327157878278 | 60.7894316214364 | Gm10643 |
| **ENSMUSG00000074217** | -0.124852218964951 | 0.575308690891557 | 0.922450244597442 | 70.4740698923855 | 72.4829998733752 | 55.8408384302584 | 95.6685228143324 | 87.5966556655277 | 89.2850029413918 | 70.7933101578173 | 73.8157383974585 | 2210011C24 |
| **ENSMUSG00000074218** | 0.209399637750914 | 0.206063077589109 | 0.819960256425221 | 26832.5503546675 | 16855.1479256109 | 20742.5053396194 | 22918.2387741982 | 19961.0879097821 | 21094.4992565779 | 16692.6461039771 | 17799.3626838695 | Cox7a1 |
| **ENSMUSG00000074219** | 0.0323589239007978 | 0.922607486201432 | 0.986713918026868 | 37.9475760958999 | 37.4631235300591 | 53.9479286529615 | 31.8895076047775 | 33.943704070392 | 25.6847268735511 | 61.4236073428121 | 35.8223436340608 | Gm10644 |
| **ENSMUSG00000074221** | 0.296806594956895 | 0.271104710731863 | 0.844129257012535 | 259.308436655316 | 408.836695914993 | 400.350417898293 | 283.253861665965 | 238.700886688563 | 168.785348026193 | 339.391457521301 | 352.795808517265 | Zfp568 |
| **ENSMUSG00000074227** | -0.16816231772593 | 0.329496435732594 | 0.855977678491063 | 721.907459538667 | 640.130762926662 | 679.554610049585 | 802.865250284986 | 821.218646864323 | 1026.16599270997 | 699.604476853724 | 650.229813236436 | Spint2 |
| **ENSMUSG00000074238** | 0.198713992350481 | 0.373273474469652 | 0.873700364743909 | 640.591225047453 | 676.779470727807 | 705.108892043093 | 647.169419038131 | 483.971522552041 | 376.709327478749 | 742.288678566525 | 721.874500504558 | Ap1ar |
| **ENSMUSG00000074240** | 0.556815536737968 | 0.163456800869851 | 0.793968226938511 | 24.3948703473642 | 11.4018202048006 | 23.6613722162112 | 27.1998741334867 | 13.1394983498292 | 13.4539045528125 | 13.5340151772298 | 18.4539345993646 | Cib3 |
| **ENSMUSG00000074247** | 0.0542966954553607 | 0.568232739908994 | 0.920544991525067 | 906.224257718752 | 837.21936932393 | 954.026527757634 | 839.444391361054 | 862.827058305448 | 852.48831575548 | 893.245001697165 | 796.775764466685 | Dda1 |
| **ENSMUSG00000074264** | 0.0229911178993151 | 0.872655950920801 | 0.976162349063696 | 709.2582675067 | 675.150639269978 | 514.871459424755 | 547.749189446766 | 566.093387238473 | 649.456665231219 | 577.798340258656 | 616.578520731712 | Amy1 |
| **ENSMUSG00000074272** | -0.454273883086683 | 0.120983356228997 | 0.769868040577702 | 299.063040184354 | 281.787842204358 | 308.544293699394 | 243.860940507122 | 469.737066006393 | 587.079471395453 | 260.269522639034 | 236.644573097735 | Ceacam1 |
| **ENSMUSG00000074282** | 0.337527343067068 | 0.320836796388344 | 0.855228644171892 | 27.1054114970714 | 28.5045505120015 | 42.5904699891801 | 37.5170677703264 | 26.2789966996583 | 28.1308913376988 | 37.4788112600209 | 15.1973579053591 | Zfp94 |
| **ENSMUSG00000074283** | 0.28626238208193 | 0.491685688725491 | 0.899236955296196 | 18.9737880479499 | 33.3910448854875 | 25.5542819935081 | 17.8206071909051 | 16.4243729372865 | 12.2308223207386 | 33.3144988977964 | 16.2828834700276 | Zfp109 |
| **ENSMUSG00000074305** | 0.0595616541999071 | 0.733324552922097 | 0.953733898884637 | 541.204716224858 | 642.574010113405 | 583.962666296092 | 515.859681841989 | 558.428679867739 | 392.609396495709 | 622.56469815257 | 616.578520731712 | Peak1 |
| **ENSMUSG00000074336** | -0.301131485339182 | 0.341529705829369 | 0.859283395807012 | 208.711668527449 | 255.726538879099 | 204.434255948065 | 239.171307035831 | 162.053812981226 | 250.731857575141 | 216.544242835676 | 489.572029665497 | Apoc4 |
| **ENSMUSG00000074346** | 0.352942248298676 | 0.398472559703861 | 0.880459952732461 | 16.2632468982428 | 14.6594831204579 | 20.8220075502658 | 14.0689004138724 | 12.0445401540101 | 20.7923979452556 | 10.4107809055614 | 8.68420451734806 | Kcnd3os |
| **ENSMUSG00000074358** | -0.1886521487146 | 0.38340079839005 | 0.875126835008975 | 177.088688447533 | 193.016527752696 | 149.539872406455 | 174.454365132018 | 245.270635863478 | 242.170281950624 | 154.079557402308 | 150.888053488923 | Ccdc61 |
| **ENSMUSG00000074361** | -0.285860969842744 | 0.370252358146313 | 0.872997029438264 | 21.6843291976571 | 17.1027303072009 | 26.5007368821565 | 24.3860940507122 | 32.8487458745729 | 31.8001380339204 | 23.9447960827911 | 20.6249857287016 | C5ar2 |
| **ENSMUSG00000074364** | -0.696894341895697 | 0.0131052115638264 | 0.635862253526407 | 1704.93038316579 | 1577.52326690705 | 1942.12543150661 | 2166.61066373635 | 4371.07311770984 | 3692.48525863098 | 1876.02271918216 | 2041.87358714146 | Ehd2 |
| **ENSMUSG00000074384** | 0.295886225141389 | 0.496091357250285 | 0.900927059989498 | 22.5878429142261 | 10.5874044758863 | 14.1968233297267 | 11.2551203310979 | 14.2344565456483 | 12.2308223207386 | 10.4107809055614 | 10.8552556466851 | AI429214 |
| **ENSMUSG00000074398** | 0.140096982243628 | 0.771994139147615 | 0.959045638244974 | 19.877301764519 | 8.95857301805762 | 20.8220075502658 | 8.44134024832345 | 9.85462376237187 | 19.5693157131818 | 10.4107809055614 | 13.0263067760221 | Gm15441 |
| **ENSMUSG00000074405** | 0.0195643778191744 | 0.874331008017688 | 0.97671366425653 | 257.501409222178 | 257.355370336928 | 303.812019256151 | 298.260688774095 | 283.594172717146 | 281.308913376988 | 280.050006359601 | 256.184033261768 | Zfp865 |
| **ENSMUSG00000074406** | 0.106994484864138 | 0.530718630266479 | 0.909082872831377 | 195.158962778914 | 191.387696294867 | 184.558703286447 | 167.888878272211 | 157.67398019795 | 212.816308380852 | 179.065431575656 | 137.8617467129 | Zfp628 |
| **ENSMUSG00000074415** | 0.242413920540016 | 0.426167885868624 | 0.885066977635295 | 25.2983840639333 | 27.6901347830872 | 22.7149173275627 | 30.0136542162612 | 19.7092475247437 | 18.3462334811079 | 23.9447960827911 | 27.1381391167127 | 2610203C20 |
| **ENSMUSG00000074457** | -0.274421896338418 | 0.249492447190711 | 0.835314899646652 | 925.198045766702 | 652.346998860377 | 645.482234058241 | 909.788893430416 | 1045.68507700724 | 1285.45942590963 | 662.125665593703 | 796.775764466685 | S100a16 |
| **ENSMUSG00000074461** | -0.0318533990142611 | 0.925807217149002 | 0.986713918026868 | 31.6229800799166 | 21.1748089517725 | 16.0897331070236 | 27.1998741334867 | 28.4689130912965 | 18.3462334811079 | 21.8626399016789 | 29.3091902460497 | Gm10699 |
| **ENSMUSG00000074466** | -0.014047959758216 | 0.913917299788182 | 0.986339811354287 | 449.949830851384 | 343.683437601847 | 380.474865236676 | 412.687745473591 | 376.665619361769 | 436.640356850368 | 383.116737324658 | 405.986561186022 | Gm15417 |
| **ENSMUSG00000074476** | -0.0747752719046959 | 0.794600807159207 | 0.961804768755116 | 179.79922959724 | 104.245213301034 | 65.3053873167428 | 110.675349922463 | 116.065568756824 | 121.085140975312 | 128.052605138405 | 119.407812113536 | Spc24 |
| **ENSMUSG00000074491** | -0.486654078526149 | 0.428650842269068 | 0.885378691567738 | 11.7456783153976 | 11.4018202048006 | 3.78581955459379 | 12.1930470253561 | 8.75966556655277 | 18.3462334811079 | 4.16431236222455 | 23.8815624227072 | Clec4g |
| **ENSMUSG00000074500** | 0.348255170062307 | 0.602322264991282 | 0.926448151609826 | 9.93865088225949 | 16.2883145782866 | 13.2503684410783 | 4.68963347129081 | 1.0949581958191 | 14.6769867848863 | 7.28754663389296 | 11.9407812113536 | Zfp558 |
| **ENSMUSG00000074513** | -0.259322779788246 | 0.130538744340531 | 0.77303252008843 | 121.974351736821 | 125.420022252807 | 123.039135524298 | 145.378637610015 | 148.914314631397 | 192.023910435596 | 134.299073681742 | 143.289374536243 | Arfip1 |
| **ENSMUSG00000074516** | -0.237750517743654 | 0.608657563639357 | 0.927932112008768 | 9.93865088225949 | 7.32974156022896 | 11.3574586637814 | 9.37926694258161 | 9.85462376237187 | 15.9000690169602 | 7.28754663389296 | 11.9407812113536 | Gm10709 |
| **ENSMUSG00000074519** | 0.154723930469821 | 0.734853986953391 | 0.954383574136061 | 129.202461469373 | 209.304842330982 | 135.343049076728 | 97.5443762028488 | 72.2672409240604 | 45.2540425867328 | 222.790711379013 | 172.598564782293 | Zfp971 |
| **ENSMUSG00000074521** | 0.208595530151057 | 0.591235714729331 | 0.923935529355527 | 138.237598635064 | 124.605606523892 | 159.004421292939 | 64.7169419038131 | 79.9319482947941 | 56.2617826753975 | 182.188665847324 | 102.03940307884 | Gm14327 |
| **ENSMUSG00000074576** | -0.151279760577159 | 0.494672094136402 | 0.900200915538898 | 180.702743313809 | 149.038078391322 | 153.325691961048 | 198.84045918273 | 197.092475247437 | 255.624186503437 | 169.69572876065 | 135.690695583563 | Mocs3 |
| **ENSMUSG00000074578** | 0.243827465748735 | 0.431812707679539 | 0.886559861988535 | 182.509770746947 | 85.5136515360045 | 132.503684410783 | 207.281799431054 | 117.160526952643 | 174.900759186562 | 130.134761319517 | 91.1841474321546 | Zfas1 |
| **ENSMUSG00000074579** | -0.146154543411689 | 0.663081040481907 | 0.939911792253667 | 32.5264937964856 | 20.3603932228582 | 27.447191770805 | 24.3860940507122 | 35.0386622662111 | 18.3462334811079 | 23.9447960827911 | 37.9933947633978 | Lekr1 |
| **ENSMUSG00000074580** | -0.134799695974095 | 0.846149484406204 | 0.97220748932105 | 4.51756858284523 | 20.3603932228582 | 19.8755526616174 | 1.87585338851632 | 9.85462376237187 | 18.3462334811079 | 14.5750932677859 | 8.68420451734806 | 4931440P22 |
| **ENSMUSG00000074582** | 0.253755507661923 | 0.339897696512227 | 0.858570960161243 | 1597.41225089407 | 2107.70790643028 | 2146.55968745468 | 1520.37917139248 | 1253.72713421287 | 924.650167447838 | 2051.96491648615 | 1951.77496527398 | Arfgef2 |
| **ENSMUSG00000074607** | -0.729890186514398 | 0.190324693925012 | 0.808824684419004 | 18.0702743313809 | 27.6901347830872 | 42.5904699891801 | 26.2619474392285 | 67.887408140784 | 85.6157562451702 | 24.9858741733473 | 11.9407812113536 | Tox2 |
| **ENSMUSG00000074622** | -0.198519164429932 | 0.360837992382054 | 0.868539734006824 | 385.800356974982 | 451.186313818538 | 346.402489245332 | 245.736793895638 | 418.274030802895 | 491.679057293692 | 410.184767679118 | 321.315567141878 | Mafb |
| **ENSMUSG00000074623** | 0.0311256824120988 | 0.87291477568845 | 0.976162349063696 | 157.211386683014 | 134.378595270864 | 172.254789734017 | 140.689004138724 | 111.685735973548 | 140.654456688494 | 185.311900118992 | 153.05910461826 | Gm826 |
| **ENSMUSG00000074634** | 0.226229886687382 | 0.643469588185736 | 0.934616627390525 | 33.4300075130547 | 70.0397526866323 | 72.8770264259304 | 29.075727522003 | 32.8487458745729 | 14.6769867848863 | 59.3414511616998 | 68.388110574116 | Tmem267 |
| **ENSMUSG00000074635** | -0.39037486684953 | 0.397798116656816 | 0.880282219458479 | 8.13162344912141 | 17.9171460361152 | 8.51809399783602 | 12.1930470253561 | 21.8991639163819 | 11.0077400886647 | 17.6983275394543 | 10.8552556466851 | 3110070M22 |
| **ENSMUSG00000074639** | -0.469086949756148 | 0.120865598634797 | 0.769868040577702 | 30.7194663633475 | 30.9477976987445 | 20.8220075502658 | 17.8206071909051 | 28.4689130912965 | 35.4693847301419 | 37.4788112600209 | 37.9933947633978 | Rdh16f2 |
| **ENSMUSG00000074649** | 0.108717387926341 | 0.405871979962744 | 0.881158207349486 | 805.934235179588 | 734.602987480724 | 641.696414503647 | 778.479156234274 | 598.942133113046 | 676.364474336844 | 780.808567917102 | 689.308733564502 | BC029722 |
| **ENSMUSG00000074652** | 0.558082208022349 | 0.032567256101014 | 0.671606027118506 | 645.108793630298 | 601.853223667689 | 447.673162330715 | 793.485983342404 | 341.626957095558 | 280.085831144914 | 527.826591911961 | 539.506205640248 | Myh7b |
| **ENSMUSG00000074656** | 0.00392932696955662 | 0.983201854971894 | 0.996940244108351 | 4141.7068767525 | 3001.1219610493 | 3300.28819671713 | 4359.48327491193 | 3546.56959625805 | 4729.65899142962 | 3080.55006995561 | 3406.37922192978 | Eif2s2 |
| **ENSMUSG00000074657** | 0.22361953562376 | 0.465884651222623 | 0.895328112556867 | 46.0791995450213 | 41.5352021746308 | 33.1259211026956 | 29.075727522003 | 27.3739548954774 | 23.2385624094033 | 44.7663578939139 | 32.5657669400552 | Kif5a |
| **ENSMUSG00000074671** | -0.283046658255073 | 0.586364697440069 | 0.923058967588378 | 6.32459601598331 | 16.2883145782866 | 18.9290977729689 | 20.6343872736795 | 16.4243729372865 | 30.5770558018465 | 7.28754663389296 | 21.7105112933701 | Tspyl3 |
| **ENSMUSG00000074676** | -1.14984176498626 | 0.0172154665001199 | 0.642273034804458 | 20.780815481088 | 13.8450673915436 | 23.6613722162112 | 24.3860940507122 | 56.937826182593 | 83.1695917810225 | 21.8626399016789 | 21.7105112933701 | Foxs1 |
| **ENSMUSG00000074682** | 0.0237730446258277 | 0.889284031757548 | 0.98126431530921 | 137.334084918495 | 126.234437981721 | 150.486327295103 | 170.702658354985 | 131.394983498292 | 129.646716599829 | 144.709854587303 | 168.256462523619 | Zcchc3 |
| **ENSMUSG00000074698** | 0.124998322281554 | 0.40003564466617 | 0.880487015130533 | 3066.52555403534 | 3294.31162345846 | 3039.06664745016 | 2900.06933864623 | 2518.40385038392 | 2268.81754049701 | 3247.12256444459 | 3243.5503872295 | Csnk2a1 |
| **ENSMUSG00000074733** | -0.0188805259369615 | 0.920257112518098 | 0.986713918026868 | 93.9654265231807 | 89.5857301805761 | 70.9841166486335 | 86.2892558717508 | 84.3117810780705 | 102.738907494204 | 80.1630129728225 | 79.2433662208011 | Zfp950 |
| **ENSMUSG00000074738** | -0.679572123461165 | 0.110121371571866 | 0.763647454385034 | 13.5527057485357 | 20.3603932228582 | 26.5007368821565 | 21.5723139679377 | 31.7537876787538 | 58.7079471395453 | 20.8215618111227 | 20.6249857287016 | Fndc10 |
| **ENSMUSG00000074743** | -0.749204544950285 | 0.0398430192963648 | 0.679566075569698 | 1151.07647490896 | 1012.31875104051 | 1073.27984372734 | 963.250715003132 | 2365.10970296925 | 2888.92023215846 | 940.093515772191 | 866.249400605469 | Thbd |
| **ENSMUSG00000074746** | 0.029866251294362 | 0.876949211525783 | 0.977515813184586 | 530.362551626029 | 667.006481980835 | 716.466350706874 | 516.797608536247 | 513.535393839156 | 485.563646133322 | 762.069162287092 | 618.749571861049 | Pdzd8 |
| **ENSMUSG00000074748** | -0.331994365064584 | 0.141160939646746 | 0.779711473022872 | 561.985531705946 | 504.937751926884 | 488.370722542599 | 545.87333605825 | 842.022852584886 | 835.365164506446 | 452.868969391919 | 515.624643217541 | Atxn7l3b |
| **ENSMUSG00000074749** | 0.229386997358696 | 0.0301765690121512 | 0.665956123973759 | 2465.68893251692 | 2450.57692830322 | 2839.36466594534 | 2715.29777987738 | 1979.68441804093 | 2404.57966825721 | 2313.27551721574 | 2234.01161208779 | Kiz |
| **ENSMUSG00000074754** | 0.289802275771262 | 0.0711446422095837 | 0.726501344493452 | 564.696072855653 | 402.321370083678 | 450.512526996661 | 606.83857118503 | 422.653863586171 | 426.855698993777 | 393.52751823022 | 412.499714574033 | Smim26 |
| **ENSMUSG00000074766** | -0.830154694338205 | 0.0153038475550648 | 0.635862253526407 | 27.1054114970714 | 18.7315617650296 | 16.0897331070236 | 17.8206071909051 | 36.1336204620302 | 50.1463715150282 | 32.2734208072402 | 23.8815624227072 | Ism1 |
| **ENSMUSG00000074768** | -0.290975443153941 | 0.351991538749643 | 0.862728204353539 | 587.283915769879 | 1208.59294170886 | 743.913542477679 | 656.548685980713 | 638.360628162534 | 763.203312814088 | 999.434966933891 | 1509.96606045389 | Bhmt |
| **ENSMUSG00000074781** | -0.0828291786375966 | 0.510028456121003 | 0.902394714365738 | 1457.36762482587 | 1351.93010999779 | 1199.15834391758 | 1207.11165551025 | 1182.55485148462 | 1362.51360653028 | 1494.98813803861 | 1483.91344690185 | Ube2n |
| **ENSMUSG00000074782** | -0.0697151745441196 | 0.897567792468164 | 0.984332015045905 | 6.32459601598331 | 17.9171460361152 | 14.1968233297267 | 10.3171936368398 | 16.4243729372865 | 3.66924669622158 | 13.5340151772298 | 17.3684090346961 | 4833422C13 |
| **ENSMUSG00000074785** | -0.313297040711141 | 0.331428885897344 | 0.856333478432716 | 21.6843291976571 | 26.8757190541728 | 28.3936466594534 | 21.5723139679377 | 40.5134532453066 | 35.4693847301419 | 19.7804837205666 | 27.1381391167127 | Plxnc1 |
| **ENSMUSG00000074793** | -0.463545726229925 | 0.085139535688035 | 0.743541556158628 | 781.539364832224 | 737.046234667467 | 926.579335986829 | 899.471699793577 | 1659.95662486175 | 1411.43689581323 | 799.547973547113 | 741.413960668591 | Hspa12b |
| **ENSMUSG00000074794** | -0.571363757954441 | 0.00018017721456127 | 0.399150048388892 | 528.555524192891 | 511.453077758198 | 418.333060782613 | 465.211640352048 | 707.342994499137 | 672.695227640623 | 870.34128370493 | 607.894316214364 | Arrdc3 |
| **ENSMUSG00000074797** | 0.0487923550320385 | 0.715135331670576 | 0.951176382858794 | 499.643085262682 | 416.980853204136 | 427.797609669098 | 423.004939110431 | 355.861413641207 | 456.20967256355 | 455.992203663588 | 440.723379255414 | Itpa |
| **ENSMUSG00000074800** | -0.064767061969085 | 0.827995120459207 | 0.967319591893432 | 69.5705561758165 | 43.1640336324594 | 63.4124775394459 | 58.151455044006 | 76.6470737073368 | 74.6080161565054 | 34.3555769883525 | 59.7039060567679 | Gm4149 |
| **ENSMUSG00000074802** | 0.19580028794928 | 0.647358231478298 | 0.935063240511793 | 36.1405486627618 | 54.56585383726 | 52.0550188756646 | 96.6064495085906 | 26.2789966996583 | 39.1386314263635 | 52.0539045278068 | 91.1841474321546 | Gas2l3 |
| **ENSMUSG00000074807** | -0.194310656186891 | 0.238195472506808 | 0.832048466795696 | 168.957064998411 | 152.295741306979 | 148.593417517806 | 190.399118934407 | 182.858018701789 | 214.039390612925 | 156.16171358342 | 202.993280593011 | Gm10762 |
| **ENSMUSG00000074811** | -0.604460752180383 | 0.00694541217902485 | 0.635862253526407 | 99.3865088225949 | 80.6271571625185 | 93.6990339761962 | 108.799496533947 | 167.528603960322 | 183.462334811079 | 105.14888714617 | 125.920965501547 | Hps6 |
| **ENSMUSG00000074813** | -0.252768185272708 | 0.504055740476563 | 0.900927059989498 | 27.1054114970714 | 11.4018202048006 | 18.9290977729689 | 22.5102406621959 | 25.1840385038392 | 31.8001380339204 | 17.6983275394543 | 20.6249857287016 | Gm14005 |
| **ENSMUSG00000074818** | 0.257311535135691 | 0.211744641999 | 0.823370267313127 | 120.167324303683 | 125.420022252807 | 157.111511515642 | 92.854742731558 | 108.400861386091 | 100.292743030056 | 115.559668051731 | 90.0986218674861 | Pdzd7 |
| **ENSMUSG00000074824** | 0.455092205811854 | 0.276036957793869 | 0.846578349250793 | 22.5878429142261 | 19.5459774939439 | 11.3574586637814 | 26.2619474392285 | 17.5193311331055 | 8.56157562451702 | 18.7394056300105 | 13.0263067760221 | Rslcan18 |
| **ENSMUSG00000074825** | -0.481001801661293 | 0.277776925986444 | 0.848075053184273 | 215.939778260002 | 201.160685041839 | 164.68315062483 | 135.999370667433 | 343.816873487196 | 441.532685778663 | 121.806136595068 | 95.5262496908287 | Itpripl1 |
| **ENSMUSG00000074829** | 0.125475329009057 | 0.526565180600778 | 0.908217276872524 | 102.097049972302 | 100.173134656462 | 106.949402417274 | 103.171936368398 | 77.7420319031559 | 84.3926740130963 | 89.5327157878278 | 125.920965501547 | 2010315B03 |

| | | | | | | | | | | | | |
|---|---|---|---|---|---|---|---|---|---|---|---|---|
| **ENSMUSG00000074862** | 0.0509598605344955 | 0.919843367502382 | 0.986713918026868 | 21.6843291976571 | 21.1748089517725 | 37.8581955459379 | 12.1930470253561 | 8.75966556655277 | 15.9000690169602 | 36.4377331694648 | 28.2236646813812 | BC025920 |
| **ENSMUSG00000074865** | 0.302677911727596 | 0.463684013505746 | 0.895328112556867 | 36.1405486627618 | 48.8649437348597 | 39.7511053232348 | 18.7585338851632 | 41.6084114411257 | 15.9000690169602 | 20.8215618111227 | 37.9933947633978 | Zfp934 |
| **ENSMUSG00000074867** | -0.489503175948081 | 0.29303607875833 | 0.851785577030345 | 13.5527057485357 | 10.5874044758863 | 10.4110037751329 | 12.1930470253561 | 9.85462376237187 | 8.56157562451702 | 23.9447960827911 | 22.7960368580387 | Zfp808 |
| **ENSMUSG00000074873** | 0.115423178482461 | 0.646466023285245 | 0.935063240511793 | 125.588406603097 | 67.5965054998893 | 86.1273948670087 | 125.682177030594 | 110.590777777729 | 92.9542496376133 | 89.5327157878278 | 80.3288917854696 | AI606181 |
| **ENSMUSG00000074874** | 0.0950996111774186 | 0.801934978688341 | 0.963461896828986 | 112.939214571131 | 70.0397526866323 | 175.094154399963 | 73.1582821521366 | 108.400861386091 | 149.216032313011 | 54.1360607089191 | 92.2696729968231 | Ctla2b |
| **ENSMUSG00000074876** | 0.136340244222716 | 0.615717051738328 | 0.928422072203769 | 58.7283915769879 | 96.9154717408051 | 70.0376617599851 | 87.227182566009 | 49.2731188118594 | 62.3771938357668 | 95.7791843311646 | 77.072315091464 | Spata5l1 |
| **ENSMUSG00000074882** | -0.287354208025304 | 0.289845473235263 | 0.851785577030345 | 29.8159526467785 | 30.9477976987445 | 25.5542819935081 | 36.5791410760683 | 32.8487458745729 | 30.5770558018465 | 41.6431236222455 | 44.5065481514088 | Cyp2c68 |
| **ENSMUSG00000074884** | -0.0403626188123103 | 0.804948199266911 | 0.963461896828986 | 7118.78457284751 | 5447.62681070794 | 5493.22417371559 | 6861.87169519271 | 5769.33473377082 | 8040.54259365355 | 5987.24009878834 | 5832.52885896389 | Serf2 |
| **ENSMUSG00000074886** | -0.0624176733306628 | 0.67757134028448 | 0.9442237961539 | 258.404922938747 | 307.034729800702 | 364.385132129652 | 274.812521417641 | 314.253002200081 | 288.647406769431 | 311.282349076285 | 343.026078435248 | Grk6 |
| **ENSMUSG00000074890** | -0.144405364591025 | 0.359062670148134 | 0.866767550983243 | 280.089252136404 | 303.777066885045 | 239.453086828057 | 328.274342990356 | 369.000911991036 | 321.670627035425 | 311.282349076285 | 271.381391167127 | Lcmt2 |
| **ENSMUSG00000074896** | -0.122861020511201 | 0.666449779126543 | 0.942024383071524 | 130.105975185942 | 157.182235680465 | 130.610774633486 | 178.206071909051 | 199.282391639076 | 218.931719541221 | 87.4505596067155 | 144.374900100912 | Ifit3 |
| **ENSMUSG00000074909** | -0.109884898735274 | 0.414098251548932 | 0.882192126163777 | 335.203588847116 | 353.456426348819 | 351.134763688574 | 333.901903155905 | 349.291664466292 | 308.216722482613 | 418.513392403567 | 404.901035621353 | Ranbp6 |
| **ENSMUSG00000074916** | -0.456184533465626 | 0.135420331983675 | 0.776010327117824 | 50.5967681278665 | 48.0505280059454 | 50.1621090983677 | 78.7858423176855 | 102.926070406995 | 99.0696607979826 | 56.2182168900314 | 54.2762782334254 | Chst14 |
| **ENSMUSG00000074918** | 0.0662163920652983 | 0.74665637176758 | 0.956085046483146 | 1320.03353990737 | 1278.6326943955 | 1009.86736618789 | 891.030359545253 | 1195.69434983445 | 1336.82887965673 | 952.586452858865 | 814.144173501381 | Inafm2 |
| **ENSMUSG00000074922** | 0.057596881119672 | 0.6321949331072 | 0.931825800800958 | 632.459601598332 | 548.101785559343 | 508.246275204216 | 532.742362338636 | 574.853052805026 | 558.948580057754 | 514.292576734731 | 487.40097853616 | Fam122a |
| **ENSMUSG00000074923** | 0.179866848303337 | 0.395123510881835 | 0.879215782741314 | 635.170142748039 | 697.95427967958 | 494.049451874489 | 606.83857118503 | 473.02194059385 | 386.49398533534 | 715.220648212066 | 573.157498144972 | Pak6 |
| **ENSMUSG00000074925** | 0.189530155465526 | 0.613645666876066 | 0.928075610917191 | 29.8159526467785 | 46.4216965481167 | 51.1085639870161 | 16.8826804966469 | 33.943704070392 | 25.6847268735511 | 37.4788112600209 | 29.3091902460497 | Ptar1 |
| **ENSMUSG00000074971** | -0.911970811474273 | 0.0129179124924518 | 0.635862253526407 | 109.325159704854 | 87.1424829938332 | 94.6454888648447 | 87.227182566009 | 209.137015401448 | 306.993640250539 | 97.8613405122768 | 98.7828263848342 | Fibin |
| **ENSMUSG00000074994** | 0.277270949303331 | 0.306413988553926 | 0.851785577030345 | 1256.78757974754 | 1584.85300846728 | 1480.25544584617 | 1225.87018939542 | 860.63714191381 | 675.14139210477 | 1538.71341784197 | 1502.36738150121 | Qser1 |
| **ENSMUSG00000075000** | 0.139977992263923 | 0.433896055267528 | 0.887868762931002 | 182.509770746947 | 176.728213174409 | 220.523989055088 | 271.998741334867 | 182.858018701789 | 187.131581507301 | 193.640524843441 | 208.420908416353 | Nrbf2 |
| **ENSMUSG00000075010** | -0.25683062416471 | 0.357090358807181 | 0.865163977643771 | 159.921927832721 | 114.018202048006 | 106.002947528626 | 198.84045918273 | 158.768938393769 | 243.393364182698 | 123.8882927618 | 166.085411394282 | AW112010 |
| **ENSMUSG00000075014** | 0.414432538840731 | 0.499325255000989 | 0.900927059989498 | 68.6670424592474 | 20.3603932228582 | 8.51809399783602 | 50.6480414899407 | 20.8042057205628 | 48.9232892829544 | 13.5340151772298 | 28.2236646813812 | Gm10800 |
| **ENSMUSG00000075027** | -0.0643357386052713 | 0.774545271004573 | 0.959414024438738 | 93.0619128066116 | 75.7406627890326 | 98.4313084194385 | 91.9168160372998 | 71.1722827282413 | 127.200552135681 | 82.2451691539348 | 95.5262496908287 | 4631405J19 |
| **ENSMUSG00000075028** | -0.022271899727415 | 0.934770541184023 | 0.98818087308472 | 81.316234491214 | 116.461449234749 | 176.040609288611 | 122.868396947819 | 146.724398239759 | 86.838838477244 | 136.381229862854 | 133.519644454226 | Prdm11 |
| **ENSMUSG00000075033** | 0.161541478251949 | 0.631966380896116 | 0.931825800800958 | 39.754603529038 | 33.3910448854875 | 34.0723759913441 | 36.5791410760683 | 44.893286028583 | 42.8078781225851 | 22.903717992235 | 18.4539345993646 | Nxpe3 |
| **ENSMUSG00000075040** | 0.408973414971061 | 0.0660960323847907 | 0.719725956118125 | 69.5705561758165 | 63.5244268553176 | 98.4313084194385 | 70.3445020693621 | 50.3680770076785 | 57.4848649074714 | 59.3414511616998 | 59.7039060567679 | Zfp408 |
| **ENSMUSG00000075053** | -0.0480387254937792 | 0.832093652488823 | 0.967583158900775 | 98.4829951060259 | 69.2253369577179 | 70.9841166486335 | 65.6548685980713 | 67.887408140784 | 78.277262852727 | 69.7522320672611 | 98.7828263848342 | Vdac3-ps1 |
| **ENSMUSG00000075054** | 0.1440681156361 | 0.491545775357699 | 0.899236955296196 | 1037.23374662126 | 1004.98900948028 | 994.724087969518 | 967.940348474422 | 741.286698569529 | 626.218102821816 | 1071.26935518226 | 1184.30839105334 | Yae1d1 |
| **ENSMUSG00000075227** | -0.231985022380424 | 0.390674828544337 | 0.877797322358001 | 279.185738419835 | 233.737314198412 | 231.881447718869 | 320.770929436291 | 354.766455445387 | 461.102001491845 | 225.913945650682 | 210.59195954569 | Znhit2 |
| **ENSMUSG00000075229** | 0.149659695716273 | 0.416873180144652 | 0.882192126163777 | 302.67709505063 | 253.283291692356 | 181.719338620502 | 279.502154888932 | 205.85214081399 | 236.054870790255 | 231.119336103462 | 244.243252050414 | Ccdc58 |
| **ENSMUSG00000075232** | 0.0954810408574912 | 0.568914835861221 | 0.920790849465435 | 4800.36837613134 | 5337.68068730451 | 4479.5709879731 | 3509.72168991404 | 3849.87301649994 | 3638.66964041973 | 4829.56126208992 | 4648.22046791055 | Amd1 |
| **ENSMUSG00000075254** | -0.0548576125483899 | 0.668418089121865 | 0.942024383071524 | 1833.22933091859 | 1993.68970438228 | 2157.91714611846 | 1894.61192240149 | 2412.19290538947 | 1752.67683856184 | 2005.11640241112 | 2013.64992246008 | Heg1 |
| **ENSMUSG00000075266** | 0.973182024493647 | 0.233993030345833 | 0.832048466795696 | 16.2632468982428 | 13.8450673915436 | 5.67872933189068 | 17.8206071909051 | 16.4243729372865 | 0 | 2.08215618111227 | 8.68420451734806 | Cenpw |
| **ENSMUSG00000075268** | 0.0146060063779694 | 0.967138653545674 | 0.994284908685797 | 20.780815481088 | 21.1748089517725 | 14.1968233297267 | 17.8206071909051 | 17.5193311331055 | 24.4616446414772 | 17.6983275394543 | 14.1118323406906 | Gm10819 |
| **ENSMUSG00000075270** | 0.248796217605612 | 0.364823552604833 | 0.86924083181804 | 234.913566307952 | 267.1283590839 | 226.202718386979 | 137.87522405595 | 112.780694169367 | 211.593226148778 | 225.913945650682 | 179.111718170304 | Pde11a |
| **ENSMUSG00000075271** | 0.261419068709636 | 0.273296830850937 | 0.844738346035895 | 92.1583990900426 | 103.43079757212 | 120.199770858353 | 148.192417692789 | 94.1664048404423 | 67.2695227640623 | 106.189965236726 | 118.322286548867 | Ttc30a1 |
| **ENSMUSG00000075273** | -0.117532529286884 | 0.636230589892553 | 0.932595977875587 | 225.878429142261 | 234.551729927327 | 242.292451494002 | 298.260688774095 | 318.632834983357 | 151.662196777159 | 333.144988977964 | 281.151121249143 | Ttc30b |
| **ENSMUSG00000075279** | -0.115918016399001 | 0.777451817670879 | 0.959808527904053 | 13.5527057485357 | 10.5874044758863 | 23.6613722162112 | 29.075727522003 | 21.8991639163819 | 17.123151249034 | 18.7394056300105 | 24.9670879873757 | Mrpl23-ps1 |
| **ENSMUSG00000075284** | -0.269376031604519 | 0.0166436231633223 | 0.642273034804458 | 292.738444168371 | 301.333819698302 | 314.223023031284 | 287.005568442997 | 373.380744774312 | 386.49398533534 | 357.089785060755 | 324.572143835884 | Wipf1 |
| **ENSMUSG00000075289** | 0.0859933368064185 | 0.628705047260414 | 0.931825800800958 | 282.799793286111 | 329.023954481389 | 254.596365046432 | 239.171307035831 | 231.036179317829 | 214.039390612925 | 296.707255808499 | 299.605055848508 | Carns1 |
| **ENSMUSG00000075307** | 0.207805978128156 | 0.0289426182342484 | 0.665956123973759 | 2783.72576074923 | 3159.9330281876 | 2681.30669954105 | 3030.44114914812 | 2341.02062266123 | 2523.21864476837 | 2634.96864719758 | 2593.32057399306 | Klhl41 |
| **ENSMUSG00000075324** | -0.189685648345999 | 0.347954789015143 | 0.861266047293794 | 145.465708367616 | 112.389370590177 | 113.574586637814 | 140.689004138724 | 112.780694169367 | 163.893019097897 | 176.983275394543 | 130.263067760221 | Fign |
| **ENSMUSG00000075327** | -0.284745536209574 | 0.218105524737266 | 0.827050433688403 | 128.298947752804 | 94.4722245540621 | 145.754052851861 | 155.695831246855 | 201.472308030714 | 172.454594722414 | 143.668776496747 | 120.493337678204 | Zbtb2 |
| **ENSMUSG00000075376** | -0.005655575714833406 | 0.967492463067294 | 0.994284908685797 | 624.32797814921 | 535.071133896714 | 446.726707442067 | 588.080037299867 | 563.903470846835 | 497.794468454061 | 567.387559353095 | 573.157498144972 | Rc3h2 |
| **ENSMUSG00000075389** | -0.311633896488799 | 0.18015859035432 | 0.802731406993634 | 70.4740698923855 | 46.4216965481167 | 59.6266579848522 | 52.523894878457 | 77.7420319031559 | 85.6157562451702 | 59.3414511616998 | 61.8749571861049 | 2810410L24 |
| **ENSMUSG00000075408** | -0.518212754379299 | 0.27443402203124 | 0.845770704684697 | 40.658117245607 | 25.2468875963442 | 18.9290977729689 | 28.1378008277448 | 54.7479097909548 | 61.154111603693 | 34.3555769883525 | 11.9407812113536 | 6030408B16 |
| **ENSMUSG00000075410** | 0.0351589775730822 | 0.938084891659234 | 0.989069314190585 | 21.6843291976571 | 10.5874044758863 | 20.8220075502658 | 19.6964605794214 | 19.7092475247437 | 26.9078091056249 | 7.28754663389296 | 17.3684090346961 | Prcd |
| **ENSMUSG00000075415** | 0.193770395169863 | 0.418216909916114 | 0.882468496126562 | 1061.62861696863 | 1296.54984043161 | 1427.25397208186 | 965.126568391648 | 886.916138613469 | 676.364474336844 | 1290.93683228961 | 1298.28857534354 | Fnbp1 |
| **ENSMUSG00000075419** | -0.206569813946685 | 0.150149620771169 | 0.782641594375979 | 433.686583953142 | 491.907100264254 | 428.744064557746 | 421.129085721914 | 523.390017601528 | 406.063301048521 | 577.798340258656 | 540.591731204917 | Dolk |
| **ENSMUSG00000075467** | -0.0725473180734993 | 0.724682664595154 | 0.951674101666722 | 1895.57177736186 | 1424.41310987116 | 1424.41460741591 | 1936.8186236431 | 1583.30955115441 | 2426.59514843454 | 1492.9059818575 | 1524.07789279458 | Dnlz |
| **ENSMUSG00000075470** | 0.153799677525748 | 0.71522984916766 | 0.951176382858794 | 168.053551281842 | 371.373572384934 | 363.438677241004 | 135.999370667433 | 185.047935093427 | 106.408154190426 | 345.637926064637 | 296.348479154503 | Deaf1 |
| **ENSMUSG00000075486** | 0.171416889120604 | 0.096905347373609 | 0.757809915356977 | 431.879556520004 | 403.135785812593 | 434.422793889637 | 438.011766218561 | 377.760577557588 | 406.063301048521 | 350.843316517418 | 382.104998763315 | Commd6 |
| **ENSMUSG00000075514** | -0.0518169932621695 | 0.645455017270465 | 0.935063240511793 | 589.090943203017 | 525.298145149742 | 498.781726317731 | 527.114802173087 | 534.339599559719 | 607.871869340708 | 571.551871715319 | 505.854913135525 | Gm13375 |
| **ENSMUSG00000075528** | -0.0699349494202802 | 0.665858592780342 | 0.942024383071524 | 132.81651633565 | 101.801966114291 | 121.146225747001 | 132.247663890401 | 128.110108910834 | 139.43137445642 | 120.765058504512 | 123.74991437221 | Aarsd1 |
| **ENSMUSG00000075585** | -0.0151343393013916 | 0.948218174465551 | 0.991217716191233 | 555.660935689963 | 668.635313438664 | 542.31865119556 | 418.31530563914 | 410.609323432161 | 428.078781225851 | 682.947227404826 | 686.052156870497 | 6330403L08 |
| **ENSMUSG00000075590** | 0.283547607123943 | 0.0551491217583674 | 0.703396849069134 | 309.001691066613 | 315.178887089845 | 334.098575692902 | 282.315934971707 | 213.516848184724 | 234.831788558181 | 271.721381635152 | 298.51953028384 | Nrbp2 |
| **ENSMUSG00000075591** | -0.504775363568951 | 0.258899588452639 | 0.838606660810377 | 7.22810973255236 | 8.95857301805762 | 9.46454888648447 | 14.0689004138724 | 13.1394983498292 | 9.78465785659088 | 13.5340151772298 | 19.5394601640331 | Gm10874 |
| **ENSMUSG00000075592** | -0.391515961339645 | 0.190531050512043 | 0.808824684419004 | 22.5878429142261 | 30.1333819698302 | 38.8046504345863 | 23.448167356454 | 48.1781606160403 | 36.6924669622158 | 31.2323427166841 | 34.7368180693922 | Nynrin |
| **ENSMUSG00000075595** | 0.165205816074036 | 0.394363910746332 | 0.878965776679486 | 1140.23431031013 | 1307.95166063641 | 1174.55051681272 | 1110.50520600166 | 1009.55145654521 | 715.503105763208 | 1158.71991478898 | 1336.28197010693 | Zfp652 |
| **ENSMUSG00000075600** | -0.0785842380972456 | 0.723275955345499 | 0.951674101666722 | 84.9302893574902 | 79.8127414336042 | 71.930571537282 | 100.358156285623 | 100.736154015357 | 105.185071958352 | 87.4505596067155 | 62.9604827507734 | Zc3h3 |
| **ENSMUSG00000075602** | -0.668253151452693 | 0.0926640669823076 | 0.754058637722175 | 1629.03523097399 | 1307.95166063641 | 1343.96594188079 | 1544.76526544319 | 2759.29465346412 | 4291.79555234717 | 1099.37846362728 | 1108.32160152655 | Ly6a |
| **ENSMUSG00000075700** | 0.0699513013725917 | 0.692177067728268 | 0.94469184542218 | 3769.45922552606 | 3847.29990339129 | 3719.5677123884 | 3315.5708642026 | 2933.39300659936 | 2680.9962527059 | 4129.95678523619 | 4212.92471647848 | Selenot |
| **ENSMUSG00000075701** | 0.0443637618991598 | 0.776214613006437 | 0.959731259983925 | 628.845546732055 | 533.442302438885 | 614.249222732842 | 624.659178375935 | 630.6959207918 | 705.718447906617 | 486.183468289716 | 506.940438700193 | Selenos |
| **ENSMUSG00000075702** | -0.556222246493953 | 0.111207041525386 | 0.764116514278315 | 399.353062723518 | 263.056280439328 | 268.793188376159 | 377.984457786039 | 497.11102090187 | 842.703657898889 | 256.10521027681 | 329.999771659226 | Selenom |
| **ENSMUSG00000075703** | -0.037193686166273 | 0.80484283828221 | 0.963461896828986 | 313.519259649459 | 357.52850499339 | 365.3315870183 | 329.212269684615 | 360.241246424483 | 272.747337752471 | 368.541644056872 | 398.387882233342 | Selenoi |
| **ENSMUSG00000075704** | 0.0937699541460533 | 0.327444497732859 | 0.855977678491063 | 1014.64590370704 | 1060.36927904646 | 1160.353693483 | 1175.22214790548 | 1097.14811221074 | 980.911950123235 | 1025.46191919779 | 1027.99270974108 | Txnrd2 |
| **ENSMUSG00000075705** | 0.0163513628896901 | 0.85991141048898 | 0.974868041734905 | 3638.44973662354 | 3269.87915159103 | 3026.76273389773 | 3316.50879089686 | 3139.24514741335 | 3308.43743775979 | 3474.07758818583 | 3180.58990447873 | Msrb1 |
| **ENSMUSG00000075706** | 0.0532700046732408 | 0.754657440668243 | 0.956497786093635 | 23290.7765857168 | 16133.5755897929 | 16062.2859152528 | 21067.7094064268 | 17742.7026050526 | 22019.1494240257 | 16325.1455380108 | 17692.981178532 | Gpx4 |
| **ENSMUSG00000076258** | 0.305424538864479 | 0.345330116591786 | 0.861266047293794 | 16686.9948313137 | 14700.2039069036 | 18785.2366298944 | 29531.5598954125 | 26534.1219592842 | 14575.4709596242 | 9599.78107301814 | 13787.2601968547 | Gm23935 |
| **ENSMUSG00000076281** | 0.198292503239691 | 0.549830010072622 | 0.914699131434573 | 2675.30411476094 | 2051.51322113519 | 2602.75094378323 | 4587.39946161667 | 4221.06384488262 | 2323.85624094033 | 1556.41174538142 | 2285.03131362721 | Gm24270 |
| **ENSMUSG00000076431** | -0.0389998047925034 | 0.908018821154431 | 0.98504921011344 | 147.272735800754 | 104.245213301034 | 292.45456059237 | 314.205442576484 | 217.896680968 | 215.262472844999 | 189.476212481217 | 258.355084391105 | Sox4 |
| **ENSMUSG00000076432** | -0.243752141001141 | 0.217264891827412 | 0.825996545635915 | 915.259394884443 | 735.417403209639 | 682.39397471553 | 790.67220325963 | 1047.87499339888 | 1148.47421591735 | 656.920275140922 | 846.709940441436 | Ywhaq |
| **ENSMUSG00000076435** | 0.373717864123227 | 0.0251030125369268 | 0.655303666217042 | 2300.34592238479 | 2475.00940017064 | 2616.00131222431 | 2719.04948665441 | 2022.38778767787 | 1429.78312929434 | 1964.51435687943 | 2384.89966557671 | Acsf2 |
| **ENSMUSG00000076437** | -0.0840506997128351 | 0.683872742504353 | 0.944452327903597 | 316.229800799166 | 235.366145656241 | 316.115932808581 | 366.729337454941 | 304.398378437709 | 435.417274618294 | 270.680303544596 | 298.51953028384 | Selenoh |
| **ENSMUSG00000076441** | -0.309202575268474 | 0.211834230441934 | 0.823370267313127 | 484.283352081008 | 826.631964848044 | 461.869985660442 | 561.818089860639 | 536.529515951357 | 688.595296657583 | 688.152617857606 | 980.229584895662 | Ass1 |
| **ENSMUSG00000076609** | -0.740571891391062 | 0.161756556314478 | 0.793081033253828 | 19.877301764519 | 8.14415728914329 | 10.4110037751329 | 9.37926694258161 | 27.3739548954774 | 30.5770558018465 | 14.5750932677859 | 7.59867895267955 | Igkc |
| **ENSMUSG00000076612** | 0.0447134338457289 | 0.828505131045086 | 0.967436464724085 | 383.089815825275 | 384.404224047563 | 214.845259723197 | 390.177504811395 | 337.247124312282 | 340.016860516533 | 345.637926064637 | 308.289260365856 | Ighg2c |
| **ENSMUSG00000076617** | -0.307156129890315 | 0.44164026653251 | 0.89108031640712 | 33.4300075130547 | 35.0198763433161 | 76.6628459805242 | 109.737423228205 | 81.0269064906132 | 83.1695917810225 | 56.2182168900314 | 94.4407241261602 | Ighm |
| **ENSMUSG00000077450** | 0.0923289072764898 | 0.302134979692673 | 0.851785577030345 | 3939.31980424104 | 4426.34948664938 | 4712.39889058062 | 4158.76696234069 | 3858.6326820665 | 4129.12561548135 | 4062.28670935004 | 4118.48399235232 | Rab11b |
| **ENSMUSG00000077928** | -0.0787239300810458 | 0.829360232979584 | 0.967483872587428 | 19.877301764519 | 15.4738988493722 | 18.9290977729689 | 18.7585338851632 | 22.994122112201 | 13.4539045528125 | 26.0269522639034 | 14.1118323406906 | Mir208b |
| **ENSMUSG00000078126** | -0.0733853340879251 | 0.857138740951101 | 0.974029039597056 | 12.6491920319666 | 18.7315617650296 | 11.3574586637814 | 19.6964605794214 | 14.2344565456483 | 22.0154801773295 | 10.4107809055614 | 19.5394601640331 | Rpl23a-ps3 |
| **ENSMUSG00000078137** | -0.709549717482044 | 0.0393513548138531 | 0.679566075569698 | 55.1143367107117 | 69.2253369577179 | 56.7872933189068 | 23.448167356454 | 81.0269064906132 | 118.638976511164 | 66.6289977955927 | 69.4736361387845 | Ankrd63 |
| **ENSMUSG00000078139** | 0.188061691580713 | 0.697430865910805 | 0.944933386973863 | 10.8421645988285 | 22.8036404096012 | 12.3039135524298 | 9.37926694258161 | 6.56974917491458 | 14.6769867848863 | 11.4518589961175 | 16.2828834700276 | AK157302 |
| **ENSMUSG00000078153** | 0.0458970094096024 | 0.846883601547258 | 0.972334438148346 | 121.070838020252 | 72.4829998733752 | 106.002947528626 | 113.489130005238 | 105.115986798633 | 128.423634367755 | 84.3273253350471 | 82.4999429148066 | Psme2b |
| **ENSMUSG00000078154** | 0.423335706484682 | 0.155048383407313 | 0.787275540343856 | 28.9124389302094 | 28.5045505120015 | 32.1794662140472 | 44.0825546301336 | 20.8042057205628 | 23.2385624094033 | 28.1091084450157 | 27.1381391167127 | Gm12184 |
| **ENSMUSG00000078185** | 0.138789682385002 | 0.54461775331883 | 0.913937916428821 | 121.974351736821 | 129.492100897378 | 139.128868631322 | 125.682177030594 | 85.4067392738896 | 96.6234963338349 | 166.572494488982 | 119.407812113536 | Chml |
| **ENSMUSG00000078193** | -0.0820403253445374 | 0.819074204381606 | 0.966315773308182 | 33.4300075130547 | 21.1748089517725 | 17.9826428843205 | 39.3929211588428 | 24.0890803080201 | 41.5847958905112 | 27.0680303544596 | 26.0526135520442 | Gm2000 |

| | | | | | | | | | | | | |
|---|---|---|---|---|---|---|---|---|---|---|---|---|
| ENSMUSG00000078201 | 0.0448237852287492 | 0.788238841777559 | 0.961444367900485 | 177.992202164102 | 156.367819951551 | 157.111511515642 | 166.013024883695 | 170.813478547779 | 181.016170346931 | 164.49033830787 | 121.578863242873 | Tmem203 |
| ENSMUSG00000078202 | -0.97952481229409 | 0.020518821514737 | 0.646348290773668 | 79.509207058076 | 38.2775392589734 | 41.6440151005317 | 94.7305961200743 | 224.466430142915 | 116.192812047017 | 88.4916376972716 | 71.6446872681215 | Nrarp |
| ENSMUSG00000078234 | -0.199423229794477 | 0.517386113090676 | 0.905197321821966 | 167.150037565273 | 161.254314325037 | 147.646962629158 | 103.171936368398 | 136.869774477387 | 102.738907494204 | 270.680303544596 | 154.144630182928 | Klhdc7a |
| ENSMUSG00000078247 | 0.315953005452665 | 0.092573121036134 | 0.754058637722175 | 75.8951521917998 | 68.4109212288036 | 79.5022106464695 | 79.7237690119437 | 55.8428679867739 | 61.154111603693 | 64.5468416144805 | 61.8749571861049 | Airn |
| ENSMUSG00000078300 | -1.11420605316225 | 0.0285803928984211 | 0.665956123973759 | 9.03513716569045 | 4.07207864457164 | 7.57163910918757 | 7.50341355406529 | 10.949581958191 | 25.6847268735511 | 11.4518589961175 | 13.0263067760221 | Gm2606 |
| ENSMUSG00000078317 | -0.54507900969016 | 0.0330631555705205 | 0.671606027118506 | 128.298947752804 | 127.048853710635 | 118.306861081056 | 155.695831246855 | 226.656346534553 | 269.078091056249 | 133.257995591185 | 144.374900100912 | F8a |
| ENSMUSG00000078348 | -0.157641798614301 | 0.370593811377184 | 0.873323500696802 | 725.521514404943 | 612.440628143575 | 629.392500951217 | 805.679030367761 | 779.610235423197 | 985.804279051531 | 630.893322877019 | 697.99293808185 | Sf3b5 |
| ENSMUSG00000078349 | 0.252865313681348 | 0.37565710365216 | 0.873821423608136 | 151.7903043836 | 168.584055885266 | 181.719338620502 | 131.309737196143 | 91.9764884488041 | 86.838838477244 | 198.845915296222 | 153.05910461826 | AW011738 |
| ENSMUSG00000078350 | -0.581312135773028 | 0.187640878304607 | 0.808411512648538 | 24.3948703473642 | 17.9171460361152 | 22.7149173275627 | 30.0136542162612 | 32.8487458745729 | 68.4926049961361 | 21.8626399016789 | 19.5394601640331 | Smim1 |
| ENSMUSG00000078427 | 0.0169045638224708 | 0.898439187391593 | 0.984462340635154 | 608.064731250967 | 489.463853077511 | 479.852628544763 | 602.14893771374 | 509.15556105588 | 582.187142467157 | 555.935700356977 | 506.940438700193 | Sarnp |
| ENSMUSG00000078429 | 0.00882657830957808 | 0.947034848234587 | 0.990684895464559 | 2126.87128880353 | 2488.85446756219 | 3026.76273389773 | 2636.51193755969 | 2758.19969526831 | 2452.27987530809 | 2605.81846066201 | 2399.0114979174 | Ctdsp2 |
| ENSMUSG00000078435 | 0.440888482161224 | 0.1734081764143 | 0.799889021902745 | 39.754603529038 | 31.7622134276588 | 38.8046504345863 | 30.0136542162612 | 33.943704070392 | 13.4539045528125 | 24.9858741733473 | 30.3947158107182 | AU041133 |
| ENSMUSG00000078439 | -0.384712735270267 | 0.224121070692914 | 0.832048466795696 | 22.5878429142261 | 18.7315617650296 | 30.2865564367503 | 22.5102406621959 | 36.1336204620302 | 36.6924669622158 | 21.8626399016789 | 28.2236646813812 | Smim24 |
| ENSMUSG00000078440 | -0.107732883328564 | 0.586841028009219 | 0.923058967588378 | 1133.90971429415 | 947.165492727364 | 994.724087969518 | 1300.90432493607 | 1267.96159075851 | 1543.52977687721 | 956.75076522109 | 948.749343520276 | Dohh |
| ENSMUSG00000078451 | 0.150732252187641 | 0.694350499406849 | 0.944771261777587 | 20.780815481088 | 26.0613033252585 | 18.9290977729689 | 11.2551203310979 | 13.1394983498292 | 18.3462334811079 | 20.8215618111227 | 17.3684090346961 | Ppil6 |
| ENSMUSG00000078453 | -0.0704456171012079 | 0.513789357137591 | 0.903358755777359 | 347.852780879082 | 378.703313945163 | 345.456034356683 | 386.425798034362 | 375.57066116595 | 406.063301048521 | 358.130863151311 | 392.96025441 | Abracl |
| ENSMUSG00000078484 | -0.06491685457601 | 0.661842381713478 | 0.939611397743053 | 193.351935345776 | 174.284965987666 | 187.398067952392 | 189.461192240149 | 234.321053905287 | 177.34692365071 | 191.558368662329 | 174.76961591163 | NA |
| ENSMUSG00000078486 | 0.125814008239911 | 0.6194011249009 | 0.929568738467614 | 9758.85165266226 | 10260.8237685916 | 10251.0528989513 | 8149.64504640916 | 6695.66936743378 | 5489.19305754748 | 11217.6164257424 | 11808.347092464 | Perm1 |
| ENSMUSG00000078490 | 0.635923834443958 | 0.175020516124001 | 0.799889021902745 | 10.8421645988285 | 8.95857301805762 | 17.9826428843205 | 14.0689004138724 | 9.85462376237187 | 7.33849339244316 | 7.28754663389296 | 8.68420451734806 | Cfap74 |
| ENSMUSG00000078495 | 0.0881828078061862 | 0.613618401292712 | 0.928075610917191 | 115.649755720838 | 83.8848200781758 | 99.3777633080869 | 103.171936368398 | 91.9764884488041 | 97.8465785659088 | 96.8202624217207 | 91.1841474321546 | Zfp984 |
| ENSMUSG00000078496 | 0.585275007400573 | 0.181014601345165 | 0.803056849297055 | 58.7283915769879 | 88.7713144516618 | 66.2518422053913 | 58.151455044006 | 30.6588294829347 | 14.6769867848863 | 56.2182168900314 | 79.2433662208011 | Zfp982 |
| ENSMUSG00000078502 | 0.0123115892059227 | 0.959928922242805 | 0.993625746680416 | 302.67709505063 | 290.746415222415 | 163.736695736181 | 193.212899017181 | 192.712642464161 | 292.316653465652 | 245.694429371248 | 212.763010675027 | Gm13212 |
| ENSMUSG00000078515 | 0.127860797584521 | 0.543834125365991 | 0.913837378403038 | 723.714486971805 | 1019.64849260074 | 950.240708203041 | 804.741103673502 | 752.23628052772 | 544.271593272868 | 935.929203409967 | 968.288803684309 | Ddi2 |
| ENSMUSG00000078517 | 0.0183359341401139 | 0.885688556674164 | 0.979948700426504 | 870.987222772559 | 1095.38915538977 | 1063.81529484085 | 933.23706078687 | 961.373295929167 | 858.60372691585 | 1075.43366754449 | 1017.13745409439 | Emc1 |
| ENSMUSG00000078532 | 0.156037414203886 | 0.545170824549042 | 0.914211323944814 | 354.177376895066 | 399.063707168021 | 283.936466594534 | 275.750448111899 | 228.846262926191 | 193.24699266767 | 376.870268781321 | 378.848422069309 | Nkain1 |
| ENSMUSG00000078566 | 0.447431195550644 | 0.058553272736212 | 0.711712402279519 | 7285.03109669621 | 6744.17665113956 | 13946.0127842349 | 9176.67477662185 | 6760.2719009871 | 6111.74191367308 | 7987.15111074668 | 6385.06137138016 | Bnip3 |
| ENSMUSG00000078570 | -0.208029297325018 | 0.392968091100888 | 0.878770935869648 | 342.431698579668 | 232.108482740584 | 241.345996605354 | 279.502154888932 | 256.220217821669 | 463.548165955993 | 240.489038918468 | 306.118209236519 | 1110065P20 |
| ENSMUSG00000078572 | -0.0498272869621868 | 0.805013915071207 | 0.963461896828986 | 571.020668871636 | 503.308920469055 | 434.422793889637 | 649.045272426648 | 583.612718371579 | 727.733928083946 | 466.402984569149 | 457.006262725442 | Ndufaf8 |
| ENSMUSG00000078578 | 0.192141638584484 | 0.136471225100946 | 0.776456211338551 | 11170.1400779431 | 9652.45521909262 | 10267.1426320584 | 11317.9614196132 | 8918.43450494654 | 7717.64888438605 | 9998.51398170114 | 10482.9203780038 | Ube2d3 |
| ENSMUSG00000078580 | -0.156604325707446 | 0.434351170670538 | 0.888121557461676 | 54.2108229941427 | 74.9262470601182 | 65.3053873167428 | 77.8479156234274 | 77.7420319031559 | 80.7234273168747 | 77.0397787011541 | 68.388110574116 | E430018J23 |
| ENSMUSG00000078584 | 0.128628687621422 | 0.507671992900356 | 0.901849850541751 | 622.520950716072 | 709.35609988438 | 646.428688946889 | 634.976372012775 | 506.965644664242 | 429.301863457925 | 724.590351027071 | 728.387653892569 | AU022252 |
| ENSMUSG00000078606 | -0.106372887646938 | 0.750985330089796 | 0.956197192463153 | 28.0089252136404 | 32.5766291565731 | 56.7872933189068 | 35.6412143818101 | 54.7479097909548 | 35.4693847301419 | 46.8485140750261 | 27.1381391167127 | Gm4070 |
| ENSMUSG00000078607 | 0.24711826032675 | 0.483189963905112 | 0.89863290032536 | 32.5264937964856 | 39.0919549878878 | 21.7684624389143 | 17.8206071909051 | 18.6142893289246 | 24.4616446414772 | 23.9447960827911 | 27.1381391167127 | 1810010H24 |
| ENSMUSG00000078619 | -0.333333176362454 | 0.123123194724908 | 0.770251097497048 | 298.159526467785 | 266.313943354985 | 329.36630124966 | 274.812521417641 | 445.647985698372 | 469.663577116362 | 281.091084450157 | 276.809018990469 | Smarcd2 |
| ENSMUSG00000078622 | 0.109093408095905 | 0.488633156881597 | 0.899236955296196 | 3137.90313764429 | 3357.02163458486 | 3375.05813292036 | 2765.00789467306 | 2673.88791419023 | 2322.63315870826 | 3489.69375954417 | 3227.26750375947 | Ccdc47 |
| ENSMUSG00000078640 | -0.875469039803686 | 0.11127917714894 | 0.764116514278315 | 13.5527057485357 | 7.32974156022896 | 7.57163910918757 | 8.44134024832345 | 16.4243729372865 | 30.5770558018465 | 15.616171358342 | 5.42762782334254 | Gm11627 |
| ENSMUSG00000078650 | -0.00032857870217666 | 0.999470437242158 | 0.999758344612816 | 21.6843291976571 | 52.9370223794314 | 39.7511053232348 | 25.3240207449704 | 22.994122112201 | 15.9000690169602 | 30.191264626128 | 70.559161703453 | G6pc |
| ENSMUSG00000078651 | -0.0315632747205006 | 0.919435067313071 | 0.986713918026868 | 19.877301764519 | 26.0613033252585 | 21.7684624389143 | 20.6343872736795 | 20.8042057205628 | 19.5693157131818 | 27.0680303544596 | 22.7960368580387 | Aoc2 |
| ENSMUSG00000078652 | 0.170714284810451 | 0.13900827871934 | 0.779711473022872 | 1519.71007126913 | 1580.78092982271 | 1663.86769424397 | 1449.09674262886 | 1204.45401540101 | 1303.80565939073 | 1482.49520095194 | 1528.41999505326 | Psme3 |
| ENSMUSG00000078656 | 0.117144799858694 | 0.639098646445261 | 0.933801135937988 | 77.7021796249379 | 52.122606650517 | 68.1447519826882 | 75.9720622349111 | 70.0773245324222 | 79.5003450848009 | 44.7663578939139 | 58.6183804920994 | Vps25 |
| ENSMUSG00000078670 | 0.142559154742198 | 0.248229859258636 | 0.834621660998471 | 7053.73158525453 | 6369.54541583896 | 6064.88292645925 | 6504.52162468035 | 5599.61621341886 | 5004.85249364623 | 6166.305530364 | 6774.76504909616 | Fam174b |
| ENSMUSG00000078671 | 0.0645168516410118 | 0.751239915708218 | 0.956197192463153 | 666.793122827955 | 753.334549245754 | 777.039463580375 | 622.783324987419 | 608.796756875418 | 464.771248188067 | 864.094815161593 | 757.696844138618 | Chd2 |
| ENSMUSG00000078672 | -0.442678771888971 | 0.139780658913019 | 0.779711473022872 | 3972.74981175409 | 3905.12342014421 | 3233.08989962309 | 4200.03573688805 | 3233.41155225379 | 5277.5998313987 | 3513.63855562696 | 8785.15839486223 | Mup20 |
| ENSMUSG00000078673 | -0.0673710359003879 | 0.928068034573993 | 0.986901071486195 | 11.7456783153976 | 11.4018202048006 | 9.46454888648447 | 6.56548685980713 | 5.47479097909548 | 8.56157562451702 | 1.04107809055614 | 26.0526135520442 | Mup19 |
| ENSMUSG00000078675 | -0.376680405339623 | 0.52932024872777 | 0.908560090446144 | 10.8421645988285 | 10.5874044758863 | 5.67872933189068 | 13.1309737196143 | 12.0445401540101 | 12.2308223207386 | 3.12323427166841 | 24.9670879873757 | Mup16 |
| ENSMUSG00000078676 | 0.0735332788247942 | 0.614051506850458 | 0.928075610917191 | 402.967117589794 | 460.144886836596 | 448.619617219364 | 372.35689762049 | 418.274030802895 | 321.670627035425 | 439.33495421469 | 420.098393526712 | Casc3 |
| ENSMUSG00000078681 | -0.130146622095044 | 0.511695725347447 | 0.902394714365738 | 261.115464088454 | 187.315617650296 | 247.024725937245 | 276.688374806158 | 318.632834983357 | 303.324393554317 | 225.913945650682 | 216.019587369033 | Tm2d3 |
| ENSMUSG00000078684 | 0.188489431963631 | 0.516102543519534 | 0.904444737050323 | 118.360296870545 | 194.645359210525 | 209.166530391307 | 142.564857527241 | 122.635317931739 | 89.2850029413918 | 191.558368662329 | 179.111718170304 | 5830417I10R |
| ENSMUSG00000078695 | -0.132019241103163 | 0.508430339321248 | 0.902394714365738 | 1576.63143541298 | 1272.11736856418 | 1226.60553568839 | 1480.04832353938 | 1557.03055445476 | 2074.34746559727 | 1190.99333559622 | 1266.80833396815 | Cisd3 |
| ENSMUSG00000078713 | 0.0939205584840814 | 0.509123053072308 | 0.902394714365738 | 2374.43404714345 | 1837.32188443073 | 1656.29605513478 | 2212.569071755 | 1831.86506160535 | 1772.24615427502 | 1984.2948406 | 1982.16968108469 | Tomm5 |
| ENSMUSG00000078716 | 0.0611251354049134 | 0.820123528349717 | 0.966398681144067 | 840.267756409212 | 1006.61784093811 | 834.77321178793 | 688.43819358549 | 593.467342133951 | 505.132961846504 | 1090.00876081228 | 1041.0190165171 | Tmem8b |
| ENSMUSG00000078762 | -0.571505473798425 | 0.139764580951891 | 0.779711473022872 | 14.4562194651047 | 22.8036404096012 | 19.8755526616174 | 14.0689004138724 | 26.2789966996583 | 33.0232202659942 | 13.5340151772298 | 33.6512925047237 | Haus5 |
| ENSMUSG00000078765 | 0.221118378397799 | 0.427725207927617 | 0.885066977635295 | 74.9916384752307 | 61.895595397489 | 50.1621090983677 | 60.9652351267805 | 54.7479097909548 | 78.277262852727 | 39.5609674411332 | 41.2499714574033 | U2af1l4 |
| ENSMUSG00000078768 | 0.0949240162816692 | 0.816496824285184 | 0.966315773308182 | 18.0702743313809 | 13.8450673915436 | 18.9290977729689 | 39.3929211588428 | 25.1840385038392 | 19.5693157131818 | 17.6983275394543 | 21.7105112933701 | Zfp566 |
| ENSMUSG00000078771 | 0.068282606929107 | 0.912879024989035 | 0.986226496507248 | 18.9737880479499 | 3.25766291565731 | 42.5904699891801 | 44.0825546301336 | 18.6142893289246 | 34.2463024980681 | 31.2323427166841 | 19.5394601640331 | Evi2a |
| ENSMUSG00000078779 | -0.116591348262797 | 0.653540210098718 | 0.936991241819063 | 72.2810973255236 | 75.7406627890326 | 68.1447519826882 | 83.4754757889764 | 75.5521155115177 | 48.9232892829544 | 111.395355689507 | 87.9275707381491 | Zfp59 |
| ENSMUSG00000078784 | 0.294957757351882 | 0.0676145633185125 | 0.722038584319139 | 319.843855665442 | 237.809392842984 | 247.024725937245 | 318.895076047775 | 204.757182618171 | 227.493295165738 | 229.03717992235 | 254.012982132431 | 1810022K09 |
| ENSMUSG00000078786 | 0.213453706677113 | 0.227572352523323 | 0.832048466795696 | 228.588970291968 | 213.376920975554 | 173.201244622666 | 189.461192240149 | 189.427767876704 | 134.539045528125 | 199.886993386778 | 169.341988088287 | BC024978 |
| ENSMUSG00000078789 | 0.0278495422217154 | 0.932336570269312 | 0.987982116552499 | 53.3073092775737 | 87.9568987227475 | 83.2880302010633 | 57.2135283497478 | 62.4126171616885 | 41.5847958905112 | 62.4646854333682 | 109.638082031519 | Dph1 |
| ENSMUSG00000078794 | -0.268275732422 | 0.329650528387374 | 0.855977678491063 | 340.62467114653 | 411.279943101736 | 166.576060402127 | 310.453735799451 | 307.683253025166 | 480.671317205027 | 342.514691792969 | 350.624757387928 | Dact3 |
| ENSMUSG00000078810 | -0.154851162142273 | 0.870952127410252 | 0.975613767803763 | 5.42108229941427 | 6.51532583131463 | 33.1259211026956 | 0 | 19.7092475247437 | 11.0077400886647 | 4.16431236222455 | 15.1973579053591 | Gp6 |
| ENSMUSG00000078812 | 0.0130591555523306 | 0.883792897940137 | 0.979250426961432 | 9779.63246814334 | 8882.01793953967 | 9379.36794650611 | 9702.85165210068 | 8957.85299999603 | 10407.2067127165 | 9006.36656140114 | 9033.74374917132 | Eif5a |
| ENSMUSG00000078813 | -0.0544444578737142 | 0.73288421680947 | 0.953733898884637 | 684.863397159336 | 607.554133770089 | 685.233339381476 | 778.479156234274 | 822.313605060142 | 821.911259953634 | 625.687932424238 | 592.696958309005 | Leng1 |
| ENSMUSG00000078815 | 0.824458336841369 | 0.0602724680302062 | 0.71282992338528 | 93.9654265231807 | 116.461449234749 | 168.468970179424 | 39.3929211588428 | 47.0832024202212 | 56.2617826753975 | 95.7791843311646 | 36.9078691987293 | Cacng6 |
| ENSMUSG00000078816 | 0.23334014897758 | 0.547573766051841 | 0.914699131434573 | 15.3597331816738 | 18.7315617650296 | 11.3574586637814 | 20.6343872736795 | 16.4243729372865 | 11.0077400886647 | 13.5340151772298 | 15.1973579053591 | Prkcg |
| ENSMUSG00000078817 | -0.200824892552762 | 0.537923034841842 | 0.912413109531467 | 36.1405486627618 | 49.679359463774 | 39.7511053232348 | 40.3308478531009 | 29.5638712871156 | 50.1463715150282 | 36.4377331694648 | 74.901263962127 | Nlrp12 |
| ENSMUSG00000078851 | -0.206075806073782 | 0.38596866721553 | 0.875126835008975 | 197.869503928621 | 142.522752560008 | 126.824955078892 | 199.778385876988 | 210.231973597267 | 243.393364182698 | 138.463386043966 | 178.026192605635 | Hist3h2a |
| ENSMUSG00000078853 | -0.21501538180561 | 0.396864137855372 | 0.879613019351548 | 172.571119864688 | 249.211213047785 | 190.237432618338 | 153.819977858338 | 308.778211220985 | 243.393364182698 | 172.818963032319 | 164.999885829613 | Igtp |
| ENSMUSG00000078861 | 0.00519103388497381 | 0.977427011986702 | 0.996179425433073 | 159.018414116152 | 121.347943608235 | 108.842312194571 | 130.371810501884 | 116.065568756824 | 116.192812047017 | 140.545542225078 | 144.374900100912 | Zfp931 |
| ENSMUSG00000078862 | 0.220892139245054 | 0.340787578345417 | 0.859081572271842 | 318.036828232304 | 268.757190541728 | 257.435729712378 | 258.867767615252 | 189.427767876704 | 160.223772401676 | 282.132162540713 | 313.716888189199 | Gm14326 |
| ENSMUSG00000078864 | -0.107071291145069 | 0.765789634171339 | 0.958345041873658 | 28.9124389302094 | 24.4324718674299 | 13.2503684410783 | 25.3240207449704 | 18.6142893289246 | 34.2463024980681 | 24.9858741733473 | 21.7105112933701 | Gm14322 |
| ENSMUSG00000078866 | 0.0701701317824184 | 0.805424186898842 | 0.963475137341661 | 622.520950716072 | 718.314672902438 | 709.841166486335 | 570.259430108962 | 488.351355335317 | 342.463024980681 | 926.559500594962 | 738.157383974585 | Zfp970 |
| ENSMUSG00000078867 | 0.462301777901619 | 0.257787024258335 | 0.838169889266919 | 18.0702743313809 | 13.8450673915436 | 17.036187995672 | 17.8206071909051 | 12.0445401540101 | 6.1154111603693 | 13.5340151772298 | 16.2828834700276 | Gm14418 |
| ENSMUSG00000078870 | -0.0859744044839767 | 0.757810075584571 | 0.95677573228595 | 36.1405486627618 | 40.7207864457164 | 27.447191770805 | 27.1998741334867 | 30.6588294829347 | 42.8078781225851 | 31.2323427166841 | 35.8223436340608 | Gm14410 |
| ENSMUSG00000078877 | 0.243731361605622 | 0.281954744918479 | 0.850271690603054 | 107.518132271716 | 93.6578088251478 | 77.6093008691726 | 75.0341355406529 | 55.8428679867739 | 69.71568722821 | 88.4916376972716 | 84.6709940441436 | Gm14295 |
| ENSMUSG00000078878 | 0.0785348683676991 | 0.74687529282735 | 0.95610346389357 | 74.0881247586617 | 51.3081909216027 | 53.001473764313 | 60.0273084325223 | 52.5579933993167 | 41.5847958905112 | 63.5057635239243 | 67.3025850094475 | Gm14305 |
| ENSMUSG00000078908 | -0.0710636624525194 | 0.583917518949783 | 0.922450244597442 | 186.123825613223 | 207.676010873154 | 188.344522841041 | 181.019851991825 | 211.326931793086 | 181.016170346931 | 203.010227658447 | 206.249857287016 | Mon1b |
| ENSMUSG00000078915 | -0.0905403742607667 | 0.824306291048253 | 0.966637987288264 | 30.7194663633475 | 25.2468875963442 | 31.2330113253987 | 62.8410885152968 | 50.3680770076785 | 57.4848649074714 | 26.0269522639034 | 26.0526135520442 | Hsp25-ps1 |
| ENSMUSG00000078919 | 0.186401299520942 | 0.274159722573205 | 0.845300454282312 | 105.711104838578 | 114.018202048006 | 117.360406192407 | 141.626930832982 | 114.970610561005 | 100.292743030056 | 108.272121417838 | 96.6117752554972 | Dpm1 |
| ENSMUSG00000078920 | -0.273392433377957 | 0.388965839291819 | 0.876536136761342 | 65.0529875929712 | 38.2775392589734 | 51.1085639870161 | 63.779015209555 | 81.0269064906132 | 91.7311674055395 | 47.8895921655823 | 43.4210225867403 | Ifi47 |
| ENSMUSG00000078921 | -0.18352282145149 | 0.594916423206015 | 0.924461858177432 | 104.807591122009 | 70.0397526866323 | 90.8596693102509 | 94.7305961200743 | 146.724398239759 | 136.985209992272 | 47.8895921655823 | 78.1578406561325 | Tgtp2 |
| ENSMUSG00000078922 | -0.127745141320695 | 0.772571793068889 | 0.959045638244974 | 41.5616309621761 | 30.1333819698302 | 64.3589324280944 | 53.4618215727152 | 74.4571573156986 | 79.5003450848009 | 21.8626399016789 | 31.4802413753867 | Tgtp1 |
| ENSMUSG00000078923 | -0.0195416467082873 | 0.931179768759756 | 0.987775860606096 | 93.9654265231807 | 119.719112150406 | 117.360406192407 | 102.23400967414 | 85.4067392738896 | 84.3926740130963 | 140.545542225078 | 128.092016630884 | Ube2v1 |
| ENSMUSG00000078931 | 0.347158988064789 | 0.446347554376906 | 0.89108031640712 | 9.03513716569045 | 13.0306516626293 | 15.1432782183751 | 15.9447538023887 | 12.0445401540101 | 4.89232892829544 | 12.4929370866736 | 11.9407812113536 | Pdf |
| ENSMUSG00000078932 | 0.305753936696458 | 0.403441871518441 | 0.881158207349486 | 27.1054114970714 | 40.7207864457164 | 35.0188308799925 | 46.8963347129081 | 24.0890803080201 | 15.9000690169602 | 46.8485140750261 | 33.6512925047237 | CN725425 |
| ENSMUSG00000078937 | 0.204824246283065 | 0.427107090645908 | 0.885066977635295 | 2577.72463337149 | 3081.74911821182 | 3106.2649445442 | 2281.97564713011 | 2116.55419251831 | 1363.73668876235 | 3095.12516322339 | 3009.0768652611 | Cpt1b |
| ENSMUSG00000078941 | 0.0819186858750365 | 0.480896203860341 | 0.897732059433541 | 894.478579403355 | 725.644414462667 | 794.075651576047 | 725.955261355817 | 690.91862156185 | 710.610776834912 | 788.096114550995 | 776.150778737983 | Ak6 |

| | | | | | | | | | | | | |
|---|---|---|---|---|---|---|---|---|---|---|---|---|
| **ENSMUSG00000078945** | -0.0548052823076166 | 0.873209983291048 | 0.976334877610406 | 22.5878429142261 | 33.3910448854875 | 34.0723759913441 | 27.1998741334867 | 37.2285786578493 | 42.8078781225851 | 23.9447960827911 | 18.4539345993646 | Naip2 |
| **ENSMUSG00000078954** | -0.0996616583671006 | 0.820004421879367 | 0.966398681144067 | 15.3597331816738 | 17.1027303072009 | 22.7149173275627 | 19.6964605794214 | 31.7537876787538 | 23.2385624094033 | 17.6983275394543 | 7.59867895267955 | Arhgap8 |
| **ENSMUSG00000078963** | 0.319700600575558 | 0.406119992702744 | 0.881158207349486 | 13.5527057485357 | 25.2468875963442 | 31.2330113253987 | 26.2619474392285 | 24.0890803080201 | 14.6769867848863 | 23.9447960827911 | 14.1118323406906 | Hsbp1l1 |
| **ENSMUSG00000078964** | -0.554647843272849 | 0.156221146579949 | 0.787275540343856 | 16.2632468982428 | 17.9171460361152 | 17.9826428843205 | 17.8206071909051 | 14.2344565456483 | 23.2385624094033 | 22.903717992235 | 42.3354970220718 | Ces1b |
| **ENSMUSG00000078965** | -0.606091145130885 | 0.0537761105099064 | 0.703396849069134 | 17.1667606148119 | 21.9892246806869 | 20.8220075502658 | 15.9447538023887 | 24.0890803080201 | 24.4616446414772 | 34.3555769883525 | 32.5657669400552 | Gm12033 |
| **ENSMUSG00000078970** | -0.0365338715166126 | 0.837642106512644 | 0.968924788429433 | 262.922491521592 | 229.665235553841 | 177.933519065908 | 188.52326554589 | 237.605928492744 | 185.908499275227 | 216.544242835676 | 240.986675356409 | Wdr92 |
| **ENSMUSG00000078974** | 0.327403203802207 | 0.0374463111254257 | 0.679566075569698 | 789.670988281345 | 541.586459728029 | 692.804978490663 | 702.507093999363 | 498.205979097689 | 634.779678446333 | 497.635327285833 | 542.762782334254 | Sec61g |
| **ENSMUSG00000078994** | -0.332834507115758 | 0.39467692408871 | 0.879215782741314 | 9.93865088225949 | 16.2883145782866 | 15.1432782183751 | 13.1309737196143 | 15.3294147414674 | 20.7923979452556 | 20.8215618111227 | 11.9407812113536 | Zfp429 |
| **ENSMUSG00000079003** | -0.119399286123185 | 0.561924533131205 | 0.917387372495585 | 483.379838364439 | 406.39344872825 | 433.476339000989 | 547.749189446766 | 548.574056105368 | 676.364474336844 | 413.308001950786 | 395.131305539337 | Samd1 |
| **ENSMUSG00000079012** | -0.317470430559543 | 0.356670907872266 | 0.864965376583892 | 68.6670424592474 | 103.43079757212 | 75.7163910918757 | 70.3445020693621 | 48.1781606160403 | 94.1773318696872 | 92.6559500594962 | 161.743309135608 | Serpina3m |
| **ENSMUSG00000079015** | -0.233071081813027 | 0.506661439492887 | 0.901698316485955 | 562.889045422515 | 1132.03786319092 | 478.906173656114 | 761.596475737627 | 621.936255225247 | 726.510845851873 | 621.523620062014 | 1480.65687020784 | Serpina1c |
| **ENSMUSG00000079017** | -0.0668953970676265 | 0.832318093913965 | 0.967671708900743 | 161.728955265859 | 107.502876216691 | 73.8234813145789 | 161.323391412404 | 118.255485148462 | 198.139321595965 | 104.107809055614 | 108.552556466851 | Ifi27l2a |
| **ENSMUSG00000079018** | -0.60765817377075 | 0.105663320520822 | 0.762094734041179 | 4005.27630555058 | 3074.41937665159 | 3076.9248429961 | 3801.41689182833 | 6973.78874917183 | 8940.73111645991 | 2481.93016788583 | 2873.38616967754 | Ly6c1 |
| **ENSMUSG00000079020** | -0.173525590013801 | 0.272388628110285 | 0.844738346035895 | 269.247087537575 | 278.5301792887 | 227.149173275627 | 229.79204009325 | 308.778211220985 | 225.04713070159 | 285.255396812381 | 313.716888189199 | Slc45a4 |
| **ENSMUSG00000079022** | 0.00749550629885208 | 0.989236807448761 | 0.998437672973644 | 15.3597331816738 | 31.7622134276588 | 7.57163910918757 | 32.8274342990356 | 14.2344565456483 | 15.9000690169602 | 41.6431236222455 | 15.1973579053591 | Col22a1 |
| **ENSMUSG00000079036** | 0.0751171354163912 | 0.724902334494622 | 0.951674101666722 | 89.4478579403355 | 97.7298874697194 | 78.5557557578211 | 64.7169419038131 | 70.0773245324222 | 68.4926049961361 | 93.6970281500523 | 81.4144173501381 | Alkbh1 |
| **ENSMUSG00000079037** | -0.0896120274312281 | 0.462941460788164 | 0.895328112556867 | 4015.21495643284 | 4431.23598102286 | 3676.97724239922 | 3627.90045339057 | 4101.71340153834 | 3896.73999138732 | 4856.62929244438 | 3905.72098167729 | Prnp |
| **ENSMUSG00000079043** | 0.0924658676410298 | 0.678331933222948 | 0.944452327903597 | 101.193536255733 | 87.1424829938332 | 74.7699362032273 | 90.9788893430416 | 84.3117810780705 | 108.854318654574 | 79.1219348822664 | 60.7894316214364 | Fastkd5 |
| **ENSMUSG00000079045** | 0.648830301970984 | 0.0913365484112715 | 0.753823363185192 | 19.877301764519 | 18.7315617650296 | 17.036187995672 | 29.075727522003 | 12.0445401540101 | 9.78465785659088 | 17.6983275394543 | 14.1118323406906 | Prox1os |
| **ENSMUSG00000079049** | 0.471355302941162 | 0.495926905769737 | 0.900927059989498 | 15.3597331816738 | 8.95857301805762 | 2.83936466594534 | 24.3860940507122 | 5.47479097909548 | 17.123151249034 | 10.4107809055614 | 4.34210225867403 | Serpinb1c |
| **ENSMUSG00000079056** | -1.14499360002231 | 0.0092606405231261 | 0.635862253526407 | 32.5264937964856 | 26.8757190541728 | 25.5542819935081 | 19.6964605794214 | 95.2613630362614 | 78.277262852727 | 29.1501865355718 | 29.3091902460497 | Kcnip3 |
| **ENSMUSG00000079057** | -0.684999558387388 | 0.0096940135105398 | 0.635862253526407 | 121.974351736821 | 122.976775066064 | 98.4313084194385 | 140.689004138724 | 198.187433443257 | 245.839528646846 | 108.272121417838 | 226.874843015718 | Cyp4v3 |
| **ENSMUSG00000079065** | -0.153994259019358 | 0.515727487541066 | 0.904239892322932 | 59.631905293557 | 65.9676740420606 | 58.6802030962037 | 79.7237690119437 | 63.5075753575076 | 99.0696607979826 | 58.3003730711437 | 73.8157383974585 | BC005561 |
| **ENSMUSG00000079083** | -0.357192453402199 | 0.103368041470455 | 0.759800953500741 | 124.684892886528 | 79.8127414336042 | 98.4313084194385 | 96.6064495085906 | 109.49581958191 | 105.185071958352 | 134.299073681742 | 161.743309135608 | Jrkl |
| **ENSMUSG00000079084** | 0.00293317533953189 | 0.982931380528979 | 0.996940244108351 | 231.299511441676 | 283.416673662186 | 297.186835035612 | 283.253861665965 | 289.068963696242 | 266.631926592101 | 291.501865355718 | 245.328777615083 | Ccdc82 |
| **ENSMUSG00000079104** | 0.00739723867358908 | 0.957821357944563 | 0.992623939554576 | 264.72951895473 | 272.014853457386 | 260.275094378323 | 210.095579513828 | 257.315176017488 | 222.600966237442 | 262.351678820146 | 259.440609955773 | Prps1l3 |
| **ENSMUSG00000079105** | -1.25093968346717 | 0.0371642116835885 | 0.679566075569698 | 39.754603529038 | 43.1640336324594 | 60.5731128735006 | 75.9720622349111 | 155.484063806312 | 291.093571233579 | 48.9306702561384 | 27.1381391167127 | C7 |
| **ENSMUSG00000079108** | 0.15300518273033 | 0.322315766821573 | 0.855977678491063 | 786.056933415069 | 803.828324438442 | 828.148027567391 | 725.955261355817 | 670.114415841287 | 569.956320146419 | 889.080689334941 | 696.907412517182 | Srp54b |
| **ENSMUSG00000079109** | 0.107500676454292 | 0.610556631368326 | 0.927932112008768 | 70.4740698923855 | 57.009101024003 | 81.3951204237664 | 71.2824287636203 | 60.2227007700503 | 73.3849339244316 | 70.7933101578173 | 55.3618037980939 | Pms2 |
| **ENSMUSG00000079110** | -0.0492523119194107 | 0.784953513793113 | 0.961141387070613 | 148.176249517323 | 121.347943608235 | 128.717864856189 | 119.116690170786 | 121.54035973592 | 107.6312364225 | 153.038479311752 | 151.973579053591 | Capn3 |
| **ENSMUSG00000079111** | -0.42206584877465 | 0.11829421965041 | 0.769868040577702 | 421.940905637744 | 354.270842077733 | 409.814966784777 | 470.839200517597 | 735.811907590433 | 743.633997100907 | 363.336253604092 | 377.762896504641 | Kdelr2 |
| **ENSMUSG00000079139** | 0.0356961729850057 | 0.792002431327757 | 0.961804768755116 | 213.229237110295 | 192.202112023782 | 192.130342395635 | 222.288626539184 | 199.282391639076 | 183.462334811079 | 192.599446752885 | 223.618266321713 | Gm4204 |
| **ENSMUSG00000079144** | -0.241716669547123 | 0.290233770461398 | 0.851785577030345 | 81.316234491214 | 87.9568987227475 | 76.6628459805242 | 82.5375490947182 | 77.7420319031559 | 72.1618516923577 | 128.052605138405 | 109.638082031519 | A130010J15 |
| **ENSMUSG00000079179** | -0.142030517648487 | 0.518641007807371 | 0.905689488434585 | 62.3424464432641 | 71.6685841444609 | 92.7525790875478 | 86.2892558717508 | 65.6974917491458 | 99.0696607979826 | 94.7381062406084 | 85.7565196088121 | Rab10os |
| **ENSMUSG00000079184** | 0.113025468429517 | 0.39188560075085 | 0.878285927324016 | 1013.74238999047 | 829.075212034787 | 930.365155541423 | 1030.78143698972 | 962.468254124986 | 967.458045570423 | 769.356708920985 | 818.486275760055 | Mphosph8 |
| **ENSMUSG00000079197** | -0.0805603962134057 | 0.650765962457826 | 0.936463929813731 | 1531.45574958453 | 1144.25409912463 | 1301.37547189161 | 1443.46918246331 | 1359.93807920732 | 1841.96184150323 | 1128.52865016285 | 1402.49902955171 | Psme2 |
| **ENSMUSG00000079215** | 0.213989707031549 | 0.419317716186779 | 0.882568531013176 | 909.838312585028 | 1237.09749222087 | 1212.40871235866 | 879.775239214155 | 754.426196919358 | 544.271593272868 | 1178.50039850955 | 1176.70971210066 | Zfp664 |
| **ENSMUSG00000079225** | -0.512850347992042 | 0.234414509255395 | 0.832048466795696 | 10.8421645988285 | 18.7315617650296 | 10.4110037751329 | 15.0068271081306 | 24.0890803080201 | 9.78465785659088 | 27.0680303544596 | 17.3684090346961 | Gm9531 |
| **ENSMUSG00000079227** | -0.22196981164068 | 0.581938546126386 | 0.922450244597442 | 12.6491920319666 | 30.9477976987445 | 33.1259211026956 | 50.6480414899407 | 40.5134532453066 | 25.6847268735511 | 39.5609674411332 | 42.3354970220718 | Ccr5 |
| **ENSMUSG00000079242** | 0.689688506663936 | 0.0227606529437256 | 0.650119841310877 | 98.4829951060259 | 44.7928650902881 | 65.3053873167428 | 70.3445020693621 | 50.3680770076785 | 31.8001380339204 | 51.0128264372507 | 39.0789203280663 | C730034F03 |
| **ENSMUSG00000079243** | 0.577313364339972 | 0.0154901344231921 | 0.635862253526407 | 4383.84855279301 | 6875.29758349476 | 8783.10136665759 | 7242.66993306152 | 3977.98312541078 | 3494.34593703502 | 5379.25049390356 | 5434.14097673055 | Xirp1 |
| **ENSMUSG00000079260** | -0.434937836224054 | 0.164900419137129 | 0.795216960550936 | 19.877301764519 | 30.1333819698302 | 35.0188308799925 | 19.6964605794214 | 43.7983278327639 | 34.2463024980681 | 35.3966550789086 | 28.2236646813812 | Tmppe |
| **ENSMUSG00000079283** | 0.0735625030550846 | 0.721392815094992 | 0.951674101666722 | 92.1583990900426 | 73.2974156022896 | 101.270673085384 | 113.489130005238 | 86.5016974697087 | 91.7311674055395 | 105.14888714617 | 77.072315091464 | 2310009B15 |
| **ENSMUSG00000079293** | 0.392390926866471 | 0.46615008856752 | 0.895328112556867 | 26.2018977805023 | 9.77298874697194 | 46.3762895437739 | 67.5307219865876 | 28.4689130912965 | 15.9000690169602 | 29.1501865355718 | 40.1644458927348 | Clec7a |
| **ENSMUSG00000079297** | -0.0539723246917864 | 0.928900223722146 | 0.986952046775844 | 7.22810973255236 | 8.14415728914329 | 17.9826428843205 | 6.56548685980713 | 9.85462376237187 | 2.44616446414772 | 14.5750932677859 | 14.1118323406906 | Gm2223 |
| **ENSMUSG00000079316** | 0.224518698003766 | 0.0758887307669161 | 0.735399943172214 | 803.223694029881 | 687.366875203693 | 937.936794650611 | 820.685857475891 | 706.248036303317 | 640.895089606702 | 710.015257759285 | 721.874500504558 | Rab9 |
| **ENSMUSG00000079317** | 0.0688041969429656 | 0.652316437249712 | 0.936720465895231 | 284.606820719249 | 232.108482740584 | 221.470443943737 | 238.233380341573 | 226.656346534553 | 196.916239363891 | 241.530117009024 | 264.868237779116 | Trappc2 |
| **ENSMUSG00000079334** | -0.117148217843308 | 0.326212192900394 | 0.855977678491063 | 449.046317134815 | 457.701639649853 | 429.690519446395 | 416.439452250624 | 529.959766776443 | 503.90987961443 | 452.868969391919 | 415.756291268038 | Nat6 |
| **ENSMUSG00000079339** | 0.301230768470966 | 0.244270536799002 | 0.832048466795696 | 66.8600150261093 | 86.3280672649188 | 56.7872933189068 | 59.0893817382642 | 50.3680770076785 | 44.0309603546589 | 74.9576225200418 | 48.8486504100828 | Ifit1bl1 |
| **ENSMUSG00000079355** | -0.663821527306329 | 0.213418318952426 | 0.824248286808591 | 31.6229800799166 | 42.3496179035451 | 12.3039135524298 | 30.9515809105193 | 49.2731188118594 | 88.0619207093179 | 14.5750932677859 | 34.7368180693922 | Ackr4 |
| **ENSMUSG00000079363** | 0.0780812667888242 | 0.740774393400942 | 0.954383574136061 | 131.913002619081 | 103.43079757212 | 143.861143074564 | 113.489130005238 | 163.148771177045 | 118.638976511164 | 93.6970281500523 | 91.1841474321546 | Gbp4 |
| **ENSMUSG00000079418** | -0.041332019768273 | 0.794147136865952 | 0.961804768755116 | 269.247087537575 | 280.159010746529 | 265.007368821565 | 231.667893481766 | 241.98576127602 | 221.377884005369 | 303.994802442392 | 308.289260365856 | Atg4a |
| **ENSMUSG00000079419** | -0.226219605373405 | 0.400988163292066 | 0.880899654809123 | 38.8510898124689 | 52.122606650517 | 65.3053873167428 | 44.0825546301336 | 70.0773245324222 | 58.7079471395453 | 40.6020455316893 | 65.1315338801105 | Ms4a6c |
| **ENSMUSG00000079426** | -0.1471559800832 | 0.417427987557741 | 0.882436023575407 | 870.987222772559 | 723.201167275924 | 795.022106464695 | 867.582192188799 | 1011.74137293685 | 1137.46647582869 | 725.631429117627 | 732.729756151243 | Arpc4 |
| **ENSMUSG00000079427** | 0.073302913145977 | 0.717424068447079 | 0.951530419413962 | 121.070838020252 | 97.7298874697194 | 105.056492639978 | 105.047789756914 | 119.350443344282 | 122.308223207386 | 87.4505596067155 | 79.2433662208011 | Mthfsl |
| **ENSMUSG00000079429** | 0.270216805619297 | 0.486638790165775 | 0.899236955296196 | 20.780815481088 | 21.1748089517725 | 15.1432782183751 | 16.8826804966469 | 13.1394983498292 | 24.4616446414772 | 12.4929370866736 | 11.9407812113536 | Mroh2a |
| **ENSMUSG00000079435** | 0.147550463392131 | 0.448557029859399 | 0.89108031640712 | 2263.30186000546 | 1579.9665140938 | 1921.30342395635 | 2232.26553233442 | 1831.86506160535 | 2365.44103683084 | 1557.45282347198 | 1465.45951230249 | Rpl36a |
| **ENSMUSG00000079442** | 0.0354011468153577 | 0.808217433260697 | 0.964026716319739 | 594.512025502432 | 561.946852950887 | 574.498117409607 | 721.265627884526 | 594.56230032977 | 709.387694602839 | 564.264325081426 | 525.394373299558 | St6galnac4 |
| **ENSMUSG00000079450** | -0.0152440633288853 | 0.971910739243098 | 0.995123114262614 | 9.93865088225949 | 12.2162359337149 | 14.1968233297267 | 25.3240207449704 | 10.949581958191 | 18.3462334811079 | 17.6983275394543 | 15.1973579053591 | Cldn34c1 |
| **ENSMUSG00000079457** | -0.105500541952268 | 0.792956831834096 | 0.961804768755116 | 22.5878429142261 | 17.9171460361152 | 19.8755526616174 | 32.8274342990356 | 25.1840385038392 | 39.1386314263635 | 12.4929370866736 | 23.8815624227072 | Gm7609 |
| **ENSMUSG00000079462** | 0.283070478761818 | 0.454047499670628 | 0.891964629399053 | 24.3948703473642 | 26.8757190541728 | 17.036187995672 | 22.5102406621959 | 10.949581958191 | 15.9000690169602 | 28.1091084450157 | 19.5394601640331 | Gm15737 |
| **ENSMUSG00000079465** | -0.56477422748209 | 0.227396341001975 | 0.832048466795696 | 27.1054114970714 | 11.4018202048006 | 8.51809399783602 | 17.8206071909051 | 36.1336204620302 | 26.9078091056249 | 15.616171358342 | 17.3684090346961 | Col4a3 |
| **ENSMUSG00000079469** | 0.469064411050309 | 0.0136555125275682 | 0.635862253526407 | 168.957064998411 | 175.099381716581 | 176.040609288611 | 186.647412157374 | 132.489941694111 | 86.838838477244 | 154.079557402308 | 135.690695583563 | Pigb |
| **ENSMUSG00000079470** | -0.40933010154964 | 0.341422780850745 | 0.859283395807012 | 13.5527057485357 | 12.2162359337149 | 23.6613722162112 | 15.0068271081306 | 17.5193311331055 | 13.4539045528125 | 35.3966550789086 | 18.4539345993646 | Utp14b |
| **ENSMUSG00000079477** | 0.0338582033005498 | 0.794262915436686 | 0.961804768755116 | 6240.56924034239 | 6595.13857274823 | 6849.49402914881 | 5812.33172431783 | 5583.19184048158 | 5406.02346576646 | 7115.76874895119 | 6799.73213708353 | Rab7 |
| **ENSMUSG00000079478** | -0.106637792963321 | 0.603038479864466 | 0.926448151609826 | 471.634160049041 | 374.631235300591 | 364.385132129652 | 468.963347129081 | 486.161438943679 | 588.302553627526 | 406.020455316893 | 328.914246094558 | Sssca1 |
| **ENSMUSG00000079480** | 0.0950636554065343 | 0.534729540641794 | 0.910706162399539 | 311.712232216321 | 241.067055758641 | 286.775831260479 | 326.39848960184 | 259.505092409126 | 313.109051410908 | 276.926772087932 | 242.072200921077 | Pin4 |
| **ENSMUSG00000079481** | -0.199823832362551 | 0.493100085045711 | 0.899931765958683 | 80.412720774645 | 87.1424829938332 | 114.521041526462 | 61.9031618210386 | 134.679858085749 | 117.415894279091 | 72.8754663389296 | 70.559161703453 | Nhsl2 |
| **ENSMUSG00000079484** | 0.203719131158972 | 0.142646498581402 | 0.780607990324772 | 359.59845919448 | 377.888898216248 | 433.476339000989 | 373.294824314748 | 286.879047304603 | 333.901449356164 | 328.980676615739 | 390.789203280663 | Phyhd1 |
| **ENSMUSG00000079487** | 0.37646581929829 | 0.082486519208167 | 0.739855948845396 | 302.67709505063 | 435.712414969166 | 479.852628544763 | 288.881421831514 | 313.158044004262 | 229.939459629886 | 284.214318721825 | 333.256348353232 | Med12 |
| **ENSMUSG00000079491** | -0.414330856232619 | 0.406798861625721 | 0.881296485258502 | 6.32459601598331 | 9.77298874697194 | 14.1968233297267 | 19.6964605794214 | 15.3294147414674 | 28.1308913376988 | 12.4929370866736 | 10.8552556466851 | H2-T10 |
| **ENSMUSG00000079499** | 0.0428098854373712 | 0.832816907006851 | 0.967854847250989 | 119.263810587114 | 179.171460361152 | 118.306861081056 | 132.247663890401 | 153.294147414674 | 108.854318654574 | 134.299073681742 | 136.776221148232 | 6530402F18 |
| **ENSMUSG00000079505** | -0.287223466046753 | 0.444654943351432 | 0.89108031640712 | 18.9737880479499 | 13.0306516626293 | 26.5007368821565 | 17.8206071909051 | 20.8042057205628 | 34.2463024980681 | 20.8215618111227 | 17.3684090346961 | Gm11131 |
| **ENSMUSG00000079508** | 0.165402710187239 | 0.11478228860518 | 0.767518180080285 | 3779.39787640832 | 3102.10951143468 | 3101.53267010096 | 3349.33622519589 | 2805.28289768853 | 2891.3663966226 | 2998.30490080167 | 3192.53068569008 | Apoo |
| **ENSMUSG00000079509** | -0.0963205474312734 | 0.49708688096461 | 0.900927059989498 | 291.834930451802 | 360.786167909048 | 361.545767463707 | 276.688374806158 | 369.000911991036 | 322.893709267499 | 357.089785060755 | 331.085297223895 | Zfx |
| **ENSMUSG00000079523** | -0.401885257564543 | 0.191005897686945 | 0.808824684419004 | 565.599586572222 | 390.105134149963 | 458.084166105848 | 564.631869943413 | 713.912743674051 | 1053.07380181559 | 381.034581143546 | 466.775992807458 | Tmsb10 |
| **ENSMUSG00000079547** | -0.663948953562293 | 0.0605005699563414 | 0.71282992338528 | 22.5878429142261 | 22.8036404096012 | 38.8046504345863 | 41.2687745473591 | 59.1277425742312 | 64.8233582999146 | 28.1091084450157 | 46.6775992807458 | H2-DMb1 |
| **ENSMUSG00000079553** | -0.500996056295259 | 0.338663813519536 | 0.858570960161243 | 10.8421645988285 | 8.14415728914329 | 12.3039135524298 | 11.2551203310979 | 27.3739548954774 | 15.9000690169602 | 11.4518589961175 | 5.42762782334254 | Kifc1 |
| **ENSMUSG00000079555** | -0.237629524314985 | 0.330805948858096 | 0.856303993370193 | 98.4829951060259 | 57.009101024003 | 57.7337482075553 | 62.8410885152968 | 90.881530252985 | 86.838838477244 | 68.711153976705 | 79.2433662208011 | Haus3 |
| **ENSMUSG00000079557** | 0.0725554736209085 | 0.471716575940168 | 0.897171173241185 | 2077.17803439223 | 2058.84296269542 | 2394.53086828057 | 2138.47286290861 | 1944.64575577472 | 2278.6021983536 | 1969.71974733221 | 2051.64331722348 | March2 |
| **ENSMUSG00000079562** | 0.0682807455686795 | 0.562738866918887 | 0.917961580916424 | 1687.76362255098 | 1821.03356985244 | 1794.47846887746 | 1571.02721288242 | 1598.63896589588 | 1400.42915572457 | 1754.21658258709 | 1801.97243734972 | Maea |
| **ENSMUSG00000079563** | -0.0238460167985028 | 0.96458021519897 | 0.994284908685797 | 8.13162344912141 | 35.8342920722305 | 13.2503684410783 | 22.5102406621959 | 14.2344565456483 | 13.4539045528125 | 17.6983275394543 | 35.8223436340608 | Pglyrp2 |
| **ENSMUSG00000079588** | 0.147413748168 | 0.242701346585205 | 0.832048466795696 | 4798.5613486982 | 5033.90362041947 | 5729.8378958777 | 5061.05244221704 | 4583.49500769874 | 3901.63232031561 | 5302.2107152024 | 4830.5887627486 | Tmem182 |
| **ENSMUSG00000079593** | -0.526925828518531 | 0.159185793275155 | 0.789393312599749 | 27.1054114970714 | 27.6901347830872 | 24.6078271048596 | 35.6412143818101 | 62.4126171616885 | 52.592535979176 | 21.8626399016789 | 29.3091902460497 | Gm14597 |
| **ENSMUSG00000079610** | 0.114621527096271 | 0.558885167081683 | 0.916574868225118 | 97.5794813894569 | 89.5857301805761 | 100.324218196735 | 96.6064495085906 | 78.836990098975 | 116.192812047017 | 73.9165444294857 | 86.8420451734806 | Ankrd39 |
| **ENSMUSG00000079614** | 0.19324689085602 | 0.102998508335845 | 0.759800953500741 | 608.064731250967 | 600.22439220986 | 676.71524538364 | 667.803806311811 | 618.65138063779 | 499.017550686135 | 595.49666779811 | 517.795694346878 | Seh1l |
| **ENSMUSG00000079641** | 0.228447720939497 | 0.220136805864117 | 0.828720352121776 | 2466.59244623349 | 1687.46939031049 | 1938.33961195202 | 2477.0643995358 | 1788.06673377259 | 2307.95617192337 | 1561.6171358342 | 1657.59753724881 | Rpl39 |

| | | | | | | | | | | | | |
|---|---|---|---|---|---|---|---|---|---|---|---|---|
| ENSMUSG00000079654 | -0.895402885595366 | 0.109225055389725 | 0.763154315676794 | 44.2721721118832 | 22.8036404096012 | 15.1432782183751 | 44.0825546301336 | 104.021028602814 | 84.3926740130963 | 23.9447960827911 | 22.7960368580387 | Prrt4 |
| ENSMUSG00000079658 | 0.0168351228709673 | 0.89703115889091 | 0.984332015045905 | 1625.42117610771 | 1317.72464938338 | 1301.37547189161 | 1565.39965271687 | 1271.24646534597 | 1601.01464178468 | 1487.70059140472 | 1382.95956938768 | Eloc |
| ENSMUSG00000079659 | 0.064766260875502 | 0.628316437708793 | 0.931825800800958 | 202.387072511466 | 242.69588721647 | 230.934992830221 | 266.371181169318 | 223.371471947096 | 221.377884005369 | 224.872867560126 | 231.216945274392 | Tmem243 |
| ENSMUSG00000079671 | 0.784571457252631 | 0.167144635343839 | 0.797814084166983 | 14.4562194651047 | 12.2162359337149 | 7.57163910918757 | 19.6964605794214 | 2.18991639163819 | 12.2308223207386 | 6.24646854333682 | 10.8552556466851 | 2610203C22I |
| ENSMUSG00000079737 | 0.113495814635512 | 0.797616221383315 | 0.9625446841889 | 14.4562194651047 | 37.4631235300591 | 25.5542819935081 | 15.9447538023887 | 32.8487458745729 | 20.7923979452556 | 12.4929370866736 | 20.6249857287016 | 3110001I22R |
| ENSMUSG00000079834 | 0.244897190426214 | 0.251346487998909 | 0.83723604018117 | 263.826005238161 | 320.065381463331 | 315.169477919933 | 234.48167356454 | 213.516848184724 | 168.785348026193 | 272.762459725708 | 300.690581413177 | Tmlhe |
| ENSMUSG00000079845 | 0.479309793724046 | 0.655038032751829 | 0.937504201434612 | 26.2018977805023 | 8.14415728914329 | 17.9826428843205 | 3.75170677703264 | 7.66470737073368 | 31.8001380339204 | 1.04107809055614 | 0 | Xlr4a |
| ENSMUSG00000079884 | -0.236226994819041 | 0.517071509221617 | 0.905197321821966 | 21.6843291976571 | 16.2883145782866 | 17.036187995672 | 13.1309737196143 | 17.5193311331055 | 14.6769867848863 | 27.0680303544596 | 20.6249857287016 | Gm10698 |
| ENSMUSG00000079941 | 0.118748072100337 | 0.587406942406522 | 0.923058967588378 | 112.939214571131 | 68.4109212288036 | 88.966759532954 | 92.854742731558 | 98.5462376237187 | 86.838838477244 | 71.8343882483734 | 77.072315091464 | Gm11273 |
| ENSMUSG00000080059 | -0.0422114137345688 | 0.925703203116676 | 0.986713918026868 | 26.2018977805023 | 12.2162359337149 | 14.1968233297267 | 10.3171936368398 | 19.7092475247437 | 22.0154801773295 | 12.4929370866736 | 10.8552556466851 | Rps19-ps3 |
| ENSMUSG00000080242 | -0.399179356836572 | 0.347523918998996 | 0.861266047293794 | 14.4562194651047 | 13.8450673915436 | 9.46454888648447 | 8.44134024832345 | 17.5193311331055 | 19.5693157131818 | 14.5750932677859 | 9.76973008201657 | Gm15487 |
| ENSMUSG00000080268 | 0.032370286632482 | 0.830437464879846 | 0.967583158900775 | 484.283352081008 | 398.249291439107 | 503.514000760974 | 507.418341593665 | 457.692525852383 | 556.502415593606 | 439.33495421469 | 398.387882233342 | Brms1 |
| ENSMUSG00000080316 | 0.595465925921358 | 0.172233381832052 | 0.799889021902745 | 29.8159526467785 | 13.8450673915436 | 14.1968233297267 | 28.1378008277448 | 9.85462376237187 | 18.3462334811079 | 11.4518589961175 | 17.3684090346961 | Spaca6 |
| ENSMUSG00000080717 | -0.0625572098233153 | 0.896321090518803 | 0.984095879090375 | 11.7456783153976 | 6.51532583131463 | 14.1968233297267 | 9.37926694258161 | 8.75966556655277 | 15.9000690169602 | 11.4518589961175 | 7.59867895267955 | B230307C23 |
| ENSMUSG00000080776 | -0.777525038798398 | 0.0384049442453554 | 0.679566075569698 | 13.5527057485357 | 8.95857301805762 | 19.8755526616174 | 15.9447538023887 | 32.8487458745729 | 19.5693157131818 | 23.9447960827911 | 22.7960368580387 | Gm12174 |
| ENSMUSG00000080836 | 0.293005252682593 | 0.544552231456555 | 0.913937916428821 | 17.1667606148119 | 7.32974156022896 | 14.1968233297267 | 17.8206071909051 | 13.1394983498292 | 17.123151249034 | 6.24646854333682 | 9.76973008201657 | Gm6444 |
| ENSMUSG00000080848 | -0.235472617576527 | 0.33168280158157 | 0.856333478432716 | 63.2459601598332 | 48.8649437348597 | 56.7872933189068 | 77.8479156234274 | 73.3621991198795 | 94.1773318696872 | 62.4646854333682 | 60.7894316214364 | Gm9385 |
| ENSMUSG00000080850 | 0.0502207436973798 | 0.794412415109308 | 0.961804768755116 | 307.194663633475 | 280.159010746529 | 222.416898832385 | 328.274342990356 | 231.036179317829 | 228.716377397812 | 324.816364253515 | 313.716888189199 | Gm12439 |
| ENSMUSG00000080875 | 0.296213194797521 | 0.412206381615287 | 0.882192126163777 | 16.2632468982428 | 20.3603932228582 | 18.9290977729689 | 17.8206071909051 | 19.7092475247437 | 14.6769867848863 | 13.5340151772298 | 11.9407812113536 | Gm7332 |
| ENSMUSG00000080893 | 0.167110576908804 | 0.69033847991482 | 0.94469184542218 | 45.1756858284523 | 30.9477976987445 | 28.3936466594534 | 42.2067012416173 | 27.3739548954774 | 57.4848649074714 | 33.3144988977964 | 13.0263067760221 | Gm15920 |
| ENSMUSG00000080921 | 0.277772516335173 | 0.479482881296766 | 0.897732059433541 | 28.0089252136404 | 13.8450673915436 | 40.6975602118832 | 35.6412143818101 | 20.8042057205628 | 31.8001380339204 | 18.7394056300105 | 26.0526135520442 | Rpl38-ps2 |
| ENSMUSG00000080935 | 0.142584207151368 | 0.78957279327156 | 0.961659329450003 | 13.5527057485357 | 25.2468875963442 | 12.3039135524298 | 15.0068271081306 | 8.75966556655277 | 6.1154111603693 | 15.616171358342 | 29.3091902460497 | Got2-ps1 |
| ENSMUSG00000081049 | 0.366350479462468 | 0.330722747876775 | 0.856303993370193 | 27.1054114970714 | 14.6594831204579 | 19.8755526616174 | 22.5102406621959 | 12.0445401540101 | 13.4539045528125 | 17.6983275394543 | 21.7105112933701 | Rps24-ps3 |
| ENSMUSG00000081051 | 0.0462011348996676 | 0.869568852181715 | 0.975613767803763 | 56.0178504272808 | 48.0505280059454 | 70.0376617599851 | 75.9720622349111 | 58.0327843784121 | 88.0619207093179 | 53.094982618363 | 43.4210225867403 | Gm15427 |
| ENSMUSG00000081058 | -0.489210249550424 | 0.0812021941960079 | 0.739855948845396 | 41.5616309621761 | 26.8757190541728 | 25.5542819935081 | 33.7653609932938 | 36.1336204620302 | 56.2617826753975 | 40.6020455316893 | 46.6775992807458 | Hist2h3c2 |
| ENSMUSG00000081067 | -0.619804828644051 | 0.206328183894635 | 0.819960256425221 | 14.4562194651047 | 14.6594831204579 | 11.3574586637814 | 9.37926694258161 | 6.56974917491458 | 15.9000690169602 | 30.191264626128 | 23.8815624227072 | Gm12912 |
| ENSMUSG00000081094 | -0.168340746513927 | 0.481324206444134 | 0.897732059433541 | 120.167324303683 | 72.4829998733752 | 73.8234813145789 | 90.0409626487835 | 102.926070406995 | 125.977469903608 | 83.2862472444909 | 89.0130963028176 | Rpl19-ps11 |
| ENSMUSG00000081214 | -0.328597079374293 | 0.47781080888216 | 0.897674307047416 | 16.2632468982428 | 8.95857301805762 | 10.4110037751329 | 19.6964605794214 | 9.85462376237187 | 26.9078091056249 | 19.7804837205666 | 13.0263067760221 | Rpl35a-ps2 |
| ENSMUSG00000081219 | -0.518484732533662 | 0.200585415723326 | 0.812921911423739 | 20.780815481088 | 17.9171460361152 | 46.3762895437739 | 58.151455044006 | 48.1781606160403 | 66.0464405319884 | 40.6020455316893 | 49.9341759747513 | Bambi-ps1 |
| ENSMUSG00000081221 | 0.621690992425402 | 0.151923781850334 | 0.783412422888014 | 15.3597331816738 | 16.2883145782866 | 11.3574586637814 | 19.6964605794214 | 10.949581958191 | 6.1154111603693 | 9.36970281500523 | 14.1118323406906 | Gm14760 |
| ENSMUSG00000081228 | -0.262656102492201 | 0.456451840800643 | 0.892805623137304 | 18.9737880479499 | 13.0306516626293 | 16.0897331070236 | 18.7585338851632 | 19.7092475247437 | 22.0154801773295 | 14.5750932677859 | 23.8815624227072 | Gm16089 |
| ENSMUSG00000081272 | -0.0278975062963297 | 0.901404919707929 | 0.984802943363903 | 52.4037955610046 | 48.8649437348597 | 77.6093008691726 | 57.2135283497478 | 62.4126171616885 | 57.4848649074714 | 60.3825292522559 | 59.7039060567679 | Gm13509 |
| ENSMUSG00000081344 | 0.161283972819445 | 0.75095380928811 | 0.956197192463153 | 41.5616309621761 | 23.6180561385155 | 27.447191770805 | 16.8826804966469 | 20.8042057205628 | 47.7002070508805 | 8.32862472444909 | 21.7105112933701 | Gm14303 |
| ENSMUSG00000081406 | -0.405195408813796 | 0.122166337049677 | 0.770251097497048 | 60.535419010126 | 49.679359463774 | 72.8770264259304 | 90.0409626487835 | 75.5521155115177 | 122.308223207386 | 77.0397787011541 | 86.8420451734806 | Rps6-ps4 |
| ENSMUSG00000081476 | 0.18827225960459 | 0.618893266048546 | 0.929568738467614 | 18.0702743313809 | 21.1748089517725 | 26.5007368821565 | 15.0068271081306 | 13.1394983498292 | 20.7923979452556 | 23.9447960827911 | 13.0263067760221 | Itpa-ps1 |
| ENSMUSG00000081485 | 0.049365322534529 | 0.791249698039234 | 0.961804768755116 | 174.378147297826 | 142.522752560008 | 141.021778408619 | 189.461192240149 | 155.484063806312 | 192.023910435596 | 129.093683228961 | 149.802527924254 | Gm12338 |
| ENSMUSG00000081534 | -0.00243378592246125 | 0.981482685125608 | 0.996940244108351 | 1960.62476495483 | 1858.4966933825 | 2008.377273712 | 2099.07994174977 | 1984.0642508242 | 2259.03288264042 | 1835.42067365047 | 1862.76186897116 | Slc48a1 |
| ENSMUSG00000081603 | -0.332984512985784 | 0.474627517913202 | 0.897549741595057 | 9.93865088225949 | 7.32974156022896 | 11.3574586637814 | 9.37926694258161 | 17.5193311331055 | 11.0077400886647 | 7.28754663389296 | 11.9407812113536 | Gm14681 |
| ENSMUSG00000081721 | 0.143657306919298 | 0.723874605466393 | 0.951674101666722 | 25.2983840639333 | 21.9892246806869 | 11.3574586637814 | 12.1930470253561 | 17.5193311331055 | 18.3462334811079 | 14.5750932677859 | 14.1118323406906 | Gm13882 |
| ENSMUSG00000081752 | 0.404711352453158 | 0.371584965670411 | 0.873433041763747 | 20.780815481088 | 13.0306516626293 | 16.0897331070236 | 36.5791410760683 | 9.85462376237187 | 14.6769867848863 | 17.6983275394543 | 22.7960368580387 | Sms-ps |
| ENSMUSG00000081824 | -0.119668159063234 | 0.738166210885528 | 0.954383574136061 | 45.1756858284523 | 31.7622134276588 | 56.7872933189068 | 67.5307219865876 | 60.2227007700503 | 84.3926740130963 | 39.5609674411332 | 34.7368180693922 | BC002163 |
| ENSMUSG00000081855 | -0.318818425955899 | 0.499884865536477 | 0.900927059989498 | 8.13162344912141 | 5.7009101024003 | 12.3039135524298 | 17.8206071909051 | 14.2344565456483 | 15.9000690169602 | 13.5340151772298 | 10.8552556466851 | Rpl17-ps5 |
| ENSMUSG00000081895 | 0.437129488746791 | 0.16307653677361 | 0.793968226938511 | 36.1405486627618 | 22.8036404096012 | 29.3401015481019 | 24.3860940507122 | 20.8042057205628 | 20.7923979452556 | 18.7394056300105 | 22.7960368580387 | Rpl17-ps10 |
| ENSMUSG00000081974 | -0.0357464116866517 | 0.912441771830062 | 0.986226496507248 | 21.6843291976571 | 26.8757190541728 | 30.2865564367503 | 18.7585338851632 | 18.6142893289246 | 33.0232202659942 | 24.9858741733473 | 23.8815624227072 | Gm11960 |
| ENSMUSG00000081999 | 0.217065197545933 | 0.629410571579266 | 0.931825800800958 | 12.6491920319666 | 13.8450673915436 | 12.3039135524298 | 7.50341355406529 | 9.85462376237187 | 12.2308223207386 | 10.4107809055614 | 7.59867895267955 | Gm13461 |
| ENSMUSG00000082044 | 0.520396033905041 | 0.15575718234946 | 0.787275540343856 | 40.658117245607 | 24.4324718674299 | 22.7149173275627 | 42.2067012416173 | 18.6142893289246 | 31.8001380339204 | 15.616171358342 | 24.9670879873757 | Snrpert |
| ENSMUSG00000082045 | -0.274790080163712 | 0.423214586937582 | 0.884281490205862 | 31.6229800799166 | 30.1333819698302 | 22.7149173275627 | 25.3240207449704 | 33.943704070392 | 42.8078781225851 | 16.6572494488982 | 40.1644458927348 | Gm11585 |
| ENSMUSG00000082127 | -0.214569975888325 | 0.337671644446847 | 0.858570960161243 | 77.7021796249379 | 75.7406627890326 | 69.0912068713366 | 73.1582821521366 | 82.1218646864323 | 118.638976511164 | 65.5879197050366 | 78.1578406561325 | Gm13577 |
| ENSMUSG00000082179 | -0.393997907736688 | 0.457853676251614 | 0.893021247686005 | 7.22810973255236 | 14.6594831204579 | 14.1968233297267 | 4.68963347129081 | 8.75966556655277 | 11.0077400886647 | 21.8626399016789 | 11.9407812113536 | Gm11407 |
| ENSMUSG00000082284 | -0.485298958276828 | 0.341029528438207 | 0.859283395807012 | 13.5527057485357 | 10.5874044758863 | 12.3039135524298 | 9.37926694258161 | 5.47479097909548 | 28.1308913376988 | 12.4929370866736 | 18.4539345993646 | H3f3a-ps1 |
| ENSMUSG00000082292 | -0.254108676830262 | 0.647062002930146 | 0.935063240511793 | 14.4562194651047 | 22.8036404096012 | 15.1432782183751 | 7.50341355406529 | 25.1840385038392 | 17.123151249034 | 24.9858741733473 | 4.34210225867403 | Gm12250 |
| ENSMUSG00000082319 | -0.581208498837926 | 0.214205069420278 | 0.824248286808591 | 15.3597331816738 | 7.32974156022896 | 7.57163910918757 | 6.56548685980713 | 15.3294147414674 | 9.78465785659088 | 14.5750932677859 | 15.1973579053591 | Gm8822 |
| ENSMUSG00000082361 | 0.00481879840461733 | 0.992263415405909 | 0.999024270802267 | 15.3597331816738 | 31.7622134276588 | 6.62518422053913 | 26.2619474392285 | 18.6142893289246 | 12.2308223207386 | 22.903717992235 | 26.0526135520442 | Btc |
| ENSMUSG00000082424 | 0.160456256288827 | 0.172771275603469 | 0.799889021902745 | 349.65980831222 | 304.591482613959 | 367.224496795597 | 329.212269684615 | 297.828629262794 | 305.770558018465 | 319.610973800734 | 284.407697943149 | Gm13292 |
| ENSMUSG00000082454 | -0.0616173313624344 | 0.890843289215324 | 0.982142775127356 | 7.22810973255236 | 9.77298874697194 | 13.2503684410783 | 12.1930470253561 | 12.0445401540101 | 9.78465785659088 | 13.5340151772298 | 8.68420451734806 | Gm12183 |
| ENSMUSG00000082536 | -0.695373592101378 | 0.00389784167497896 | 0.635862253526407 | 256.597895505609 | 208.490426602068 | 215.791714611846 | 198.84045918273 | 251.840385038392 | 331.455284892016 | 309.200192895173 | 531.907526687569 | Gm13456 |
| ENSMUSG00000082575 | 0.162773165644098 | 0.561574485041524 | 0.917031459819748 | 45.1756858284523 | 30.1333819698302 | 41.6440151005317 | 39.3929211588428 | 36.1336204620302 | 46.4771248188067 | 28.1091084450157 | 29.3091902460497 | Gm13050 |
| ENSMUSG00000082585 | -0.408460051625955 | 0.390932292932135 | 0.877896341705281 | 9.93865088225949 | 15.4738988493722 | 14.1968233297267 | 8.44134024832345 | 10.949581958191 | 24.4616446414772 | 8.32862472444909 | 20.6249857287016 | Gm15387 |
| ENSMUSG00000082644 | -0.58196005728436 | 0.167653272625481 | 0.798595321251004 | 16.2632468982428 | 21.9892246806869 | 23.6613722162112 | 18.7585338851632 | 18.6142893289246 | 39.1386314263635 | 15.616171358342 | 47.7631248454143 | Mup-ps19 |
| ENSMUSG00000082674 | 0.553916388255693 | 0.0511617421581821 | 0.703396849069134 | 39.754603529038 | 35.0198763433161 | 35.0188308799925 | 44.0825546301336 | 21.8991639163819 | 22.0154801773295 | 35.3966550789086 | 24.9670879873757 | Gm11914 |
| ENSMUSG00000082809 | -0.958517111795757 | 0.0517299303139695 | 0.703396849069134 | 9.93865088225949 | 8.14415728914329 | 6.62518422053913 | 8.44134024832345 | 10.949581958191 | 17.123151249034 | 9.36970281500523 | 27.1381391167127 | Gm14150 |
| ENSMUSG00000082816 | -0.0844458816923989 | 0.82007412228085 | 0.966398681144067 | 18.0702743313809 | 12.2162359337149 | 26.5007368821565 | 22.5102406621959 | 22.994122112201 | 14.6769867848863 | 24.9858741733473 | 20.6249857287016 | Gm11953 |
| ENSMUSG00000082884 | -0.118815734315785 | 0.58978481950239 | 0.92378339455212 | 43.3686583953142 | 42.3496179035451 | 41.6440151005317 | 44.0825546301336 | 42.7033696369448 | 50.1463715150282 | 48.9306702561384 | 44.5065481514088 | Gm13339 |
| ENSMUSG00000082896 | -0.838934697410619 | 0.0969552411429929 | 0.757809915356977 | 6.32459601598331 | 5.7009101024003 | 5.67872933189068 | 15.9447538023887 | 9.85462376237187 | 17.123151249034 | 16.6572494488982 | 16.2828834700276 | Gm5844 |
| ENSMUSG00000082901 | 0.0511262785577333 | 0.914622413790448 | 0.986675594624755 | 11.7456783153976 | 10.5874044758863 | 7.57163910918757 | 18.7585338851632 | 14.2344565456483 | 15.9000690169602 | 7.28754663389296 | 9.76973008201657 | Gm14323 |
| ENSMUSG00000082935 | 0.107300176407828 | 0.816068368407903 | 0.966177091475091 | 8.13162344912141 | 8.95857301805762 | 11.3574586637814 | 15.9447538023887 | 9.85462376237187 | 12.2308223207386 | 11.4518589961175 | 7.59867895267955 | Gm7658 |
| ENSMUSG00000083012 | 0.0917855838538608 | 0.449023658737989 | 0.89108031640712 | 2556.9438178904 | 2579.25461347168 | 3355.18258025874 | 2901.94519203475 | 2504.16939383827 | 2744.59652877374 | 2808.82868832046 | 2632.39949432113 | Fam220a |
| ENSMUSG00000083061 | 0.223926546333057 | 0.467538971432413 | 0.895328112556867 | 24.3948703473642 | 26.0613033252585 | 30.2865564367503 | 33.7653609932938 | 21.8991639163819 | 29.3539735697726 | 17.6983275394543 | 29.3091902460497 | Gm12191 |
| ENSMUSG00000083076 | 0.640294471468675 | 0.176287395086303 | 0.799889021902745 | 14.4562194651047 | 11.4018202048006 | 14.1968233297267 | 15.9447538023887 | 5.47479097909548 | 15.9000690169602 | 6.24646854333682 | 8.68420451734806 | Gm11408 |
| ENSMUSG00000083097 | -0.153808510072714 | 0.716205860585857 | 0.951348124093194 | 17.1667606148119 | 13.0306516626293 | 10.4110037751329 | 12.1930470253561 | 13.1394983498292 | 14.6769867848863 | 9.36970281500523 | 21.7105112933701 | Gm14494 |
| ENSMUSG00000083179 | 0.859180656576199 | 0.0430359193024433 | 0.687730553581215 | 21.6843291976571 | 12.2162359337149 | 17.9826428843205 | 16.8826804966469 | 9.85462376237187 | 12.2308223207386 | 8.32862472444909 | 7.59867895267955 | Gm12693 |
| ENSMUSG00000083282 | -0.0754020000843092 | 0.580746207094772 | 0.922450244597442 | 880.925873654819 | 1018.01966114291 | 1102.61994527544 | 909.788893430416 | 1102.62290318983 | 1143.58188698906 | 1012.96898211112 | 862.992823911464 | Ctsf |
| ENSMUSG00000083325 | 0.229203083708522 | 0.632247359034661 | 0.931825800800958 | 11.7456783153976 | 10.5874044758863 | 15.1432782183751 | 27.1998741334867 | 8.75966556655277 | 9.78465785659088 | 16.6572494488982 | 19.5394601640331 | Gm14121 |
| ENSMUSG00000083327 | 0.115410626689696 | 0.543980745968693 | 0.913837378403038 | 85.8338030740593 | 104.245213301034 | 123.039135524298 | 107.861569839689 | 95.2613630362614 | 80.7234273168747 | 100.984574783945 | 110.723607596188 | Vcp-rs |
| ENSMUSG00000083380 | 0.179249552281577 | 0.645364388261615 | 0.935063240511793 | 14.4562194651047 | 15.4738988493722 | 17.036187995672 | 12.1930470253561 | 12.0445401540101 | 17.123151249034 | 11.4518589961175 | 11.9407812113536 | Gm3244 |
| ENSMUSG00000083396 | -0.243865259551124 | 0.518513567418996 | 0.905589519860411 | 26.2018977805023 | 24.4324718674299 | 14.1968233297267 | 12.1930470253561 | 17.5193311331055 | 20.7923979452556 | 26.0269522639034 | 27.1381391167127 | Gm15542 |
| ENSMUSG00000083563 | 0.186972478480049 | 0.136930309843121 | 0.776456211338551 | 663.179067961679 | 692.253369577179 | 666.304241608507 | 600.273084325223 | 497.11102090187 | 550.387004433237 | 591.332355435886 | 664.341645577127 | Gm13340 |
| ENSMUSG00000083567 | -0.0445399463444781 | 0.887451619441211 | 0.980702840906172 | 32.5264937964856 | 20.3603932228582 | 31.2330113253987 | 32.8274342990356 | 26.2789966996583 | 23.2385624094033 | 29.1501865355718 | 41.2499714574033 | Gm11451 |
| ENSMUSG00000083619 | 0.0395046688149634 | 0.945672730433177 | 0.990505437194639 | 13.5527057485357 | 6.51532583131463 | 7.57163910918757 | 15.9447538023887 | 17.5193311331055 | 9.78465785659088 | 3.12323427166841 | 11.9407812113536 | Gm14414 |
| ENSMUSG00000083621 | 0.0331738745891792 | 0.899980449068138 | 0.984802943363903 | 50.5967681278665 | 44.7928650902881 | 53.001473764313 | 76.9099889291692 | 55.8428679867739 | 63.6002760678407 | 60.3825292522559 | 40.1644458927348 | Gm14586 |
| ENSMUSG00000083626 | 0.210912985720802 | 0.414528036497268 | 0.882192126163777 | 49.6932544112975 | 46.4216965481167 | 34.0723759913441 | 56.2756016554897 | 43.7983278327639 | 46.4771248188067 | 33.3144988977964 | 37.9933947633978 | Gm4459 |
| ENSMUSG00000083649 | 0.0902032183574684 | 0.557541858463982 | 0.916341990969984 | 110.228673421423 | 100.987550385377 | 109.78876708322 | 106.92364314543 | 102.926070406995 | 97.8465785659088 | 103.066730965058 | 97.6973008201657 | Rasl2-9 |
| ENSMUSG00000083674 | 0.217285032858271 | 0.529281783751355 | 0.908560090446144 | 24.3948703473642 | 26.0613033252585 | 16.0897331070236 | 28.1378008277448 | 16.4243729372865 | 15.9000690169602 | 22.903717992235 | 26.0526135520442 | Zfp133-ps |
| ENSMUSG00000083716 | 0.0186405648742592 | 0.952466057757882 | 0.992000114323903 | 46.9827132615903 | 43.1640336324594 | 31.2330113253987 | 47.8342614071662 | 39.4184950494875 | 62.3771938357668 | 26.0269522639034 | 40.1644458927348 | Gm13436 |
| ENSMUSG00000083718 | -0.128184239246766 | 0.760956516891933 | 0.957468401713057 | 15.3597331816738 | 8.14415728914329 | 11.3574586637814 | 14.0689004138724 | 10.949581958191 | 14.6769867848863 | 11.4518589961175 | 16.2828834700276 | Ccnb2-ps |
| ENSMUSG00000083773 | -0.0133794718094238 | 0.964307318346904 | 0.994284908685797 | 28.9124389302094 | 23.6180561385155 | 24.6078271048596 | 30.0136542162612 | 27.3739548954774 | 34.2463024980681 | 19.7804837205666 | 27.1381391167127 | Gm13394 |
| ENSMUSG00000083820 | 0.126461068300875 | 0.749172657623022 | 0.956197192463153 | 17.1667606148119 | 13.0306516626293 | 12.3039135524298 | 21.5723139679377 | 10.949581958191 | 14.6769867848863 | 18.7394056300105 | 14.1118323406906 | Gm6415 |

| | | | | | | | | | | | | |
|---|---|---|---|---|---|---|---|---|---|---|---|---|
| ENSMUSG00000083844 | 0.111465132284195 | 0.477965387660632 | 0.897674307047416 | 483.379838364439 | 379.517729674077 | 376.689045682082 | 487.721881014244 | 342.721915291377 | 407.286383280595 | 388.322127777439 | 460.262839419447 | Ube2d-ps |
| ENSMUSG00000083863 | -0.0088865540586009 | 0.963256124412306 | 0.994284908685797 | 209.615182244018 | 169.39847161418 | 170.36187995672 | 221.350699844926 | 187.237851485066 | 251.954939807215 | 159.284947855089 | 178.026192605635 | Gm13341 |
| ENSMUSG00000083899 | -0.0299470998508031 | 0.902044156376838 | 0.984871597499125 | 66.8600150261093 | 53.7514381083457 | 39.7511053232348 | 51.5859681841989 | 54.7479097909548 | 59.9310293716191 | 43.7252798033577 | 58.6183804920994 | Gm12346 |
| ENSMUSG00000083992 | -0.012650229897325 | 0.973191235754229 | 0.995480246308729 | 36.1405486627618 | 29.3189662409158 | 24.6078271048596 | 22.5102406621959 | 20.8042057205628 | 22.0154801773295 | 21.8626399016789 | 48.8486504100828 | Gm11478 |
| ENSMUSG00000084093 | 0.0144903012823082 | 0.969050541995048 | 0.994908125383341 | 30.7194663633475 | 25.2468875963442 | 23.6613722162112 | 18.7585338851632 | 22.994122112201 | 39.1386314263635 | 21.8626399016789 | 14.1118323406906 | Gm16418 |
| ENSMUSG00000084106 | -0.13905979192341 | 0.598761815818898 | 0.924848033480823 | 37.9475760958999 | 33.3910448854875 | 30.2865564367503 | 50.6480414899407 | 44.893286028583 | 42.8078781225851 | 35.3966550789086 | 44.5065481514088 | Gm6136 |
| ENSMUSG00000084128 | -0.343464912842634 | 0.586912124469477 | 0.923058967588378 | 10.8421645988285 | 20.3603932228582 | 8.51809399783602 | 6.56548685980713 | 7.66470737073368 | 6.1154111603693 | 12.4929370866736 | 32.5657669400552 | Esrp2 |
| ENSMUSG00000084159 | 0.323825156048867 | 0.240904935216354 | 0.832048466795696 | 40.658117245607 | 39.9063707168021 | 64.3589324280944 | 40.3308478531009 | 36.1336204620302 | 29.3539735697726 | 41.6431236222455 | 40.1644458927348 | Gm12696 |
| ENSMUSG00000084166 | -0.241657092235766 | 0.589598964475375 | 0.923596438092135 | 11.7456783153976 | 12.2162359337149 | 12.3039135524298 | 16.8826804966469 | 8.75966556655277 | 26.9078091056249 | 15.616171358342 | 11.9407812113536 | Gm6451 |
| ENSMUSG00000084168 | -0.332780486286089 | 0.450781725229337 | 0.89108031640712 | 15.3597331816738 | 8.14415728914329 | 8.51809399783602 | 10.3171936368398 | 12.0445401540101 | 15.9000690169602 | 15.616171358342 | 9.76973008201657 | Atp5l2-ps |
| ENSMUSG00000084260 | -0.233292713250626 | 0.591255440272568 | 0.923935529355527 | 12.6491920319666 | 7.32974156022896 | 10.4110037751329 | 11.2551203310979 | 10.949581958191 | 12.2308223207386 | 10.4107809055614 | 15.1973579053591 | Gm12844 |
| ENSMUSG00000084319 | -0.0050918414813185 | 0.981830454405031 | 0.996940244108351 | 473.44118748218 | 320.065381463331 | 381.421320125324 | 473.652980600371 | 396.374866886513 | 566.287073450197 | 305.035880532948 | 387.532626586657 | Tpt1-ps3 |
| ENSMUSG00000084347 | 0.371616705413761 | 0.401857437544697 | 0.881158207349486 | 18.0702743313809 | 15.4738988493722 | 14.1968233297267 | 13.1309737196143 | 8.75966556655277 | 6.1154111603693 | 18.7394056300105 | 13.0263067760221 | Akt2-ps |
| ENSMUSG00000084349 | -0.0583766426527434 | 0.881963433380747 | 0.978854883720278 | 35.2370349461928 | 19.5459774939439 | 37.8581955459379 | 43.1446279358754 | 28.4689130912965 | 59.9310293716191 | 21.8626399016789 | 31.4802413753867 | Rpl3-ps1 |
| ENSMUSG00000084350 | -0.018868342462101 | 0.946150190115081 | 0.990505437194639 | 65.0529875929712 | 43.9784493613737 | 51.1085639870161 | 69.4065753751039 | 60.2227007700503 | 50.1463715150282 | 80.1630129728225 | 41.2499714574033 | Znf41-ps |
| ENSMUSG00000084384 | -0.0196945675901715 | 0.940224384588678 | 0.989269716906553 | 37.0440623793308 | 35.8342920722305 | 35.965285768641 | 29.075727522003 | 32.8487458745729 | 42.8078781225851 | 36.4377331694648 | 28.2236646813812 | Gm12251 |
| ENSMUSG00000084407 | 0.0596445678874667 | 0.779205268340212 | 0.960307976179513 | 91.2548853734735 | 80.6271571625185 | 91.8061241988993 | 95.6685228143324 | 104.021028602814 | 58.7079471395453 | 85.3684034256032 | 95.5262496908287 | Gm14018 |
| ENSMUSG00000084416 | -0.272581270866276 | 0.395602127302728 | 0.879215782741314 | 57.8248778604189 | 35.8342920722305 | 32.1794662140472 | 57.2135283497478 | 45.9882442244021 | 81.9465095489486 | 41.6431236222455 | 52.1052271040884 | Rpl10a-ps1 |
| ENSMUSG00000084756 | 0.568528844853077 | 0.379150034546555 | 0.87422797717331 | 16.2632468982428 | 10.5874044758863 | 10.4110037751329 | 17.8206071909051 | 1.0949581958191 | 4.89232892829544 | 13.5340151772298 | 17.3684090346961 | Gm13881 |
| ENSMUSG00000084786 | -0.11236679973936 | 0.599682874938201 | 0.925099415025723 | 2601.21599000228 | 2084.90426602068 | 2126.68413479306 | 2583.05011598698 | 2447.23156765568 | 3654.56970943669 | 2063.41677548226 | 1993.02493673138 | Ubl5 |
| ENSMUSG00000084799 | 0.0321812354508514 | 0.946716149741414 | 0.990510945542179 | 13.5527057485357 | 19.5459774939439 | 8.51809399783602 | 7.50341355406529 | 9.85462376237187 | 8.56157562451702 | 14.5750932677859 | 15.1973579053591 | Ino80dos |
| ENSMUSG00000084803 | -0.0326650831743582 | 0.940095813047104 | 0.989269716906553 | 28.9124389302094 | 16.2883145782866 | 53.9479286529615 | 42.2067012416173 | 52.5579933993167 | 42.8078781225851 | 20.8215618111227 | 28.2236646813812 | 5830444B04 |
| ENSMUSG00000084808 | 0.232763108155639 | 0.530284807211346 | 0.908923862033581 | 166.246523848704 | 250.840044505613 | 196.862616838877 | 135.061443973175 | 131.394983498292 | 63.6002760678407 | 226.955023741238 | 214.934061804364 | 9430091E24 |
| ENSMUSG00000084817 | 0.0867978292527699 | 0.796151685739509 | 0.962200030474509 | 25.2983840639333 | 63.5244268553176 | 40.6975602118832 | 42.2067012416173 | 49.2731188118594 | 40.3617136584374 | 28.1091084450157 | 44.5065481514088 | Gm5526 |
| ENSMUSG00000084819 | 0.298506869758825 | 0.352509815061503 | 0.862728204353539 | 107.518132271716 | 57.8235167529173 | 39.7511053232348 | 64.7169419038131 | 61.3176589658694 | 44.0309603546589 | 54.1360607089191 | 59.7039060567679 | Gm11967 |
| ENSMUSG00000084830 | 0.284735468205048 | 0.437707259784885 | 0.888722970093854 | 83.1232619243521 | 33.3910448854875 | 61.519567762149 | 85.3513291774927 | 52.5579933993167 | 46.4771248188067 | 35.3966550789086 | 81.4144173501381 | Gm14539 |
| ENSMUSG00000084839 | -0.963056730515552 | 0.0070204358370625 | 0.635862253526407 | 11.7456783153976 | 15.4738988493722 | 22.7149173275627 | 16.8826804966469 | 26.2789966996583 | 29.3539735697726 | 46.8485140750261 | 27.1381391167127 | Gm14097 |
| ENSMUSG00000084843 | 0.91924155368342 | 0.0308257886662809 | 0.667184591665619 | 18.9737880479499 | 17.1027303072009 | 22.7149173275627 | 18.7585338851632 | 7.66470737073368 | 12.2308223207386 | 15.616171358342 | 5.42762782334254 | B230312C02 |
| ENSMUSG00000084850 | 0.169069901266983 | 0.670104654381861 | 0.94249140110542 | 40.658117245607 | 33.3910448854875 | 11.3574586637814 | 28.1378008277448 | 22.994122112201 | 30.5770558018465 | 22.903717992235 | 24.9670879873757 | Gm12092 |
| ENSMUSG00000084883 | 0.265998117306621 | 0.468339430011273 | 0.895328112556867 | 234.010052591383 | 445.485403716138 | 453.351891662606 | 230.729966787508 | 218.991639163819 | 140.654456688494 | 411.225845769674 | 362.565538599282 | Ccdc85c |
| ENSMUSG00000084929 | -0.191222150957894 | 0.645434733816695 | 0.935063240511793 | 37.9475760958999 | 32.5766291565731 | 14.1968233297267 | 23.448167356454 | 18.6142893289246 | 23.2385624094033 | 46.8485140750261 | 34.7368180693922 | Foxo6os |
| ENSMUSG00000084941 | 0.188698221066376 | 0.704703826211337 | 0.947228924036081 | 18.9737880479499 | 10.5874044758863 | 9.46454888648447 | 22.5102406621959 | 22.994122112201 | 7.33849339244316 | 12.4929370866736 | 10.8552556466851 | Gm11944 |
| ENSMUSG00000084957 | 0.118491482399637 | 0.476597104331266 | 0.897553735056661 | 453.563885717661 | 310.292392716359 | 365.3315870183 | 453.95652002095 | 330.677375137367 | 366.924669622158 | 409.143689588562 | 350.624757387928 | Bbip1 |
| ENSMUSG00000085022 | 0.154899893905559 | 0.369392782349129 | 0.872151240324563 | 228.588970291968 | 159.625482867208 | 176.040609288611 | 211.033506208086 | 167.528603960322 | 155.33144347338 | 173.860041122875 | 198.651178334337 | Gm5860 |
| ENSMUSG00000085024 | 0.305004260238968 | 0.377698390709341 | 0.87422797717331 | 35.2370349461928 | 23.6180561385155 | 18.9290977729689 | 34.703287687552 | 21.8991639163819 | 23.2385624094033 | 16.6572494488982 | 29.3091902460497 | C230035I16 |
| ENSMUSG00000085037 | 0.146142396283568 | 0.471465355305482 | 0.897171173241185 | 70.4740698923855 | 65.1532583131463 | 57.7337482075553 | 61.9031618210386 | 59.1277425742312 | 66.0464405319884 | 56.2182168900314 | 49.9341759747513 | 4933421O10 |
| ENSMUSG00000085042 | 0.35878665539893 | 0.365081064747059 | 0.86924083181804 | 31.6229800799166 | 26.8757190541728 | 12.3039135524298 | 26.2619474392285 | 15.3294147414674 | 13.4539045528125 | 23.9447960827911 | 22.7960368580387 | Abhd11os |
| ENSMUSG00000085069 | 0.354185011531799 | 0.314822522479615 | 0.853277176422981 | 109.325159704854 | 148.223662662408 | 196.862616838877 | 96.6064495085906 | 78.836990098975 | 58.7079471395453 | 159.284947855089 | 133.519644454226 | Gm13111 |
| ENSMUSG00000085087 | 0.0570463294614438 | 0.873781434297339 | 0.97671366425653 | 28.0089252136404 | 31.7622134276588 | 31.2330113253987 | 27.1998741334867 | 13.1394983498292 | 41.5847958905112 | 34.3555769883525 | 24.9670879873757 | Gm13528 |
| ENSMUSG00000085095 | -0.144997161307154 | 0.791235142900046 | 0.961804768755116 | 16.2632468982428 | 18.7315617650296 | 35.965285768641 | 20.6343872736795 | 53.6529515951357 | 23.2385624094033 | 15.616171358342 | 8.68420451734806 | Gm15635 |
| ENSMUSG00000085133 | -0.102937225367506 | 0.71623737933666 | 0.951348124093194 | 37.9475760958999 | 43.1640336324594 | 46.3762895437739 | 54.3997482669734 | 41.6084114411257 | 31.8001380339204 | 65.5879197050366 | 55.3618037980939 | B930095G15 |
| ENSMUSG00000085148 | -0.152923828236434 | 0.575681211609213 | 0.922450244597442 | 33.4300075130547 | 60.2667639396603 | 43.5369248778286 | 66.5927952923295 | 49.2731188118594 | 55.0387004433237 | 56.2182168900314 | 66.217059444779 | Mir22hg |
| ENSMUSG00000085151 | 0.10027805981214 | 0.579285985920304 | 0.922450244597442 | 152.693818100169 | 109.946123403434 | 112.628131749165 | 151.944124469822 | 129.205067106653 | 117.415894279091 | 117.641824232843 | 127.006491066215 | 1110018N20 |
| ENSMUSG00000085154 | -0.565924021724445 | 0.260119722847333 | 0.840316047847749 | 13.5527057485357 | 7.32974156022896 | 14.1968233297267 | 5.62756016554897 | 14.2344565456483 | 7.33849339244316 | 21.8626399016789 | 16.2828834700276 | C130046K22 |
| ENSMUSG00000085156 | 0.614074006864511 | 0.255853732670952 | 0.837517407767513 | 11.7456783153976 | 8.14415728914329 | 11.3574586637814 | 24.3860940507122 | 5.47479097909548 | 6.1154111603693 | 13.5340151772298 | 10.8552556466851 | Snhg15 |
| ENSMUSG00000085162 | 0.047670783509495 | 0.905749448943619 | 0.98504921011344 | 81.316234491214 | 81.4415728914329 | 57.7337482075553 | 75.9720622349111 | 22.994122112201 | 107.6312364225 | 94.7381062406084 | 61.8749571861049 | Gm12295 |
| ENSMUSG00000085175 | 0.637945550109489 | 0.230222515423428 | 0.832048466795696 | 18.0702743313809 | 12.2162359337149 | 18.9290977729689 | 23.448167356454 | 2.18991639163819 | 13.4539045528125 | 11.4518589961175 | 19.5394601640331 | Gm11423 |
| ENSMUSG00000085208 | -0.253045394125226 | 0.119991220068401 | 0.769868040577702 | 231.299511441676 | 230.479651282755 | 207.27362061401 | 232.605820176024 | 307.683253025166 | 313.109051410908 | 220.708555197901 | 234.473521968398 | Brip1os |
| ENSMUSG00000085218 | -0.0735985122768941 | 0.654486818414796 | 0.937269843859626 | 93.9654265231807 | 92.0289773673191 | 82.3415753124149 | 86.2892558717508 | 96.3563212320805 | 96.6234963338349 | 87.4505596067155 | 93.3551985614916 | BB218582 |
| ENSMUSG00000085241 | -0.135889985810868 | 0.646197188579322 | 0.935063240511793 | 48.7897406947284 | 36.6487078011448 | 35.0188308799925 | 62.8410885152968 | 66.7924499449649 | 53.8156182112498 | 40.6020455316893 | 40.1644458927348 | Snhg3 |
| ENSMUSG00000085260 | 0.181814321462123 | 0.724779626624434 | 0.951674101666722 | 9.03513716569045 | 12.2162359337149 | 11.3574586637814 | 10.3171936368398 | 7.66470737073368 | 14.6769867848863 | 12.4929370866736 | 3.25657669400552 | Med9os |
| ENSMUSG00000085272 | 0.205932302804851 | 0.385968009918631 | 0.875126835008975 | 205.097613661173 | 194.645359210525 | 324.634026806417 | 180.081925297567 | 194.902558855799 | 216.485555077073 | 224.872867560126 | 147.631476794917 | Sbk3 |
| ENSMUSG00000085277 | 0.261998830645431 | 0.622563573590241 | 0.930829713365818 | 65.9565013095403 | 76.5550785179469 | 40.6975602118832 | 37.5170677703264 | 18.6142893289246 | 15.9000690169602 | 89.5327157878278 | 59.7039060567679 | Gm13943 |
| ENSMUSG00000085282 | -0.105868603906233 | 0.815936771358512 | 0.966177091475091 | 15.3597331816738 | 10.5874044758863 | 17.9826428843205 | 25.3240207449704 | 17.5193311331055 | 31.8001380339204 | 14.5750932677859 | 10.8552556466851 | Gm15663 |
| ENSMUSG00000085287 | 0.130817042025057 | 0.7098656564823 | 0.949571160524938 | 23.4913566307952 | 31.7622134276588 | 25.5542819935081 | 30.0136542162612 | 13.1394983498292 | 22.0154801773295 | 33.3144988977964 | 32.5657669400552 | 4833418N02 |
| ENSMUSG00000085322 | 0.214133909750777 | 0.224516312606975 | 0.832048466795696 | 159.921927832721 | 119.719112150406 | 130.610774633486 | 125.682177030594 | 98.5462376237187 | 133.315963296051 | 117.641824232843 | 112.894658725525 | Gm14261 |
| ENSMUSG00000085334 | 0.302913922811663 | 0.572670846285219 | 0.921859799538181 | 22.5878429142261 | 8.95857301805762 | 10.4110037751329 | 21.5723139679377 | 10.949581958191 | 4.89232892829544 | 13.5340151772298 | 21.7105112933701 | Gm12940 |
| ENSMUSG00000085336 | -0.0213467428610264 | 0.965020913304239 | 0.994284908685797 | 11.7456783153976 | 17.1027303072009 | 6.62518422053913 | 14.0689004138724 | 7.66470737073368 | 8.56157562451702 | 16.6572494488982 | 17.3684090346961 | Gm11732 |
| ENSMUSG00000085342 | 0.671595870748953 | 0.245843436906605 | 0.832048466795696 | 9.93865088225949 | 10.5874044758863 | 15.1432782183751 | 13.1309737196143 | 7.66470737073368 | 15.9000690169602 | 2.08215618111227 | 5.42762782334254 | Gm12254 |
| ENSMUSG00000085379 | -0.248421425929732 | 0.513008112243041 | 0.903078589555164 | 15.3597331816738 | 16.2883145782866 | 11.3574586637814 | 14.0689004138724 | 17.5193311331055 | 23.2385624094033 | 13.5340151772298 | 14.1118323406906 | 2310058D17 |
| ENSMUSG00000085382 | -0.0948589346866265 | 0.837250117149545 | 0.968924788429433 | 18.0702743313809 | 12.2162359337149 | 10.4110037751329 | 8.44134024832345 | 8.75966556655277 | 20.7923979452556 | 11.4518589961175 | 11.9407812113536 | Gm13861 |
| ENSMUSG00000085385 | -0.260420296925825 | 0.589278997957056 | 0.92352104528132 | 7.22810973255236 | 12.2162359337149 | 12.3039135524298 | 8.44134024832345 | 9.85462376237187 | 6.1154111603693 | 15.616171358342 | 16.2828834700276 | Snhg17 |
| ENSMUSG00000085396 | 0.517329897670176 | 0.163832833019428 | 0.793968226938511 | 26.2018977805023 | 26.0613033252585 | 32.1794662140472 | 43.1446279358754 | 13.1394983498292 | 15.9000690169602 | 29.1501865355718 | 30.3947158107182 | Firre |
| ENSMUSG00000085408 | 0.148958802757564 | 0.581112067689407 | 0.922450244597442 | 41.5616309621761 | 58.6379324818317 | 62.4660226507975 | 36.5791410760683 | 39.4184950494875 | 41.5847958905112 | 55.1771387994752 | 43.4210225867403 | C530005A16 |
| ENSMUSG00000085412 | 0.177948826939689 | 0.751627146001528 | 0.956197192463153 | 6.32459601598331 | 13.8450673915436 | 12.3039135524298 | 25.3240207449704 | 6.56974917491458 | 17.123151249034 | 19.7804837205666 | 7.59867895267955 | Halr1 |
| ENSMUSG00000085414 | -0.468228528021257 | 0.356290317305594 | 0.864965376583892 | 14.4562194651047 | 13.0306516626293 | 7.57163910918757 | 14.0689004138724 | 9.85462376237187 | 9.78465785659088 | 15.616171358342 | 32.5657669400552 | Sspnos |
| ENSMUSG00000085427 | 0.772884038131887 | 0.0599581399211593 | 0.71282992338528 | 13.5527057485357 | 21.1748089517725 | 22.7149173275627 | 27.1998741334867 | 10.949581958191 | 7.33849339244316 | 14.5750932677859 | 16.2828834700276 | 6430710C18 |
| ENSMUSG00000085436 | -0.143549731606656 | 0.780664737585948 | 0.960742105865155 | 18.0702743313809 | 9.77298874697194 | 3.78581955459379 | 13.1309737196143 | 13.1394983498292 | 8.56157562451702 | 14.5750932677859 | 13.0263067760221 | Zfp335os |
| ENSMUSG00000085438 | 0.13082331945381 | 0.567975725641411 | 0.920432752255281 | 8275.28213005588 | 9044.90108532253 | 8225.63943724365 | 7116.04982933667 | 5808.75322882031 | 5036.65263168015 | 9320.77214474909 | 9663.34857667905 | 1700020I14R |
| ENSMUSG00000085457 | -0.76771251538806 | 0.029098058803034 | 0.665956123973759 | 19.877301764519 | 17.1027303072009 | 13.2503684410783 | 15.9447538023887 | 20.8042057205628 | 41.5847958905112 | 24.9858741733473 | 26.0526135520442 | 1110046J04 |
| ENSMUSG00000085492 | 0.131158302823958 | 0.680446143466443 | 0.944452327903597 | 45.1756858284523 | 35.8342920722305 | 49.2156542097192 | 51.5859681841989 | 29.5638712871156 | 26.9078091056249 | 46.8485140750261 | 61.8749571861049 | Trmt61b |
| ENSMUSG00000085498 | 0.147021043010517 | 0.639475623113185 | 0.933801135937988 | 23.4913566307952 | 28.5045505120015 | 34.0723759913441 | 31.8895076047775 | 29.5638712871156 | 24.4616446414772 | 17.6983275394543 | 34.7368180693922 | Gm14023 |
| ENSMUSG00000085527 | 0.269177771594861 | 0.493210370346586 | 0.899931765958683 | 16.2632468982428 | 23.6180561385155 | 20.8220075502658 | 24.3860940507122 | 12.0445401540101 | 29.3539735697726 | 12.4929370866736 | 17.3684090346961 | Gm15535 |
| ENSMUSG00000085558 | 0.300840712408427 | 0.507486881606456 | 0.90183410140733 | 21.6843291976571 | 22.8036404096012 | 29.3401015481019 | 24.3860940507122 | 10.949581958191 | 25.6847268735511 | 34.3555769883525 | 8.68420451734806 | 4930412C18 |
| ENSMUSG00000085566 | 0.214352585835753 | 0.552930146843255 | 0.914977712327965 | 26.2018977805023 | 42.3496179035451 | 52.0550188756646 | 30.9515809105193 | 25.1840385038392 | 20.7923979452556 | 35.3966550789086 | 48.8486504100828 | A730017L22 |
| ENSMUSG00000085586 | 0.15867761056238 | 0.499911360582736 | 0.900927059989498 | 114.746242004269 | 94.4722245540621 | 114.521041526462 | 138.813150750208 | 72.2672409240604 | 91.7311674055395 | 133.257995591185 | 116.15123541953 | Gm11613 |
| ENSMUSG00000085614 | -0.0871455459400032 | 0.683720328701137 | 0.944452327903597 | 60.535419010126 | 50.4937751926884 | 59.6266579848522 | 51.5859681841989 | 47.0832024202212 | 61.154111603693 | 61.4236073428121 | 66.217059444779 | 1700123M08 |
| ENSMUSG00000085615 | 0.173746956733028 | 0.746254108178771 | 0.956085046483146 | 9.93865088225949 | 10.5874044758863 | 15.1432782183751 | 14.0689004138724 | 4.37983278327639 | 7.33849339244316 | 10.4107809055614 | 21.7105112933701 | A330035P11 |
| ENSMUSG00000085622 | -0.0685976762888671 | 0.82098613691036 | 0.96650690096489 | 50.5967681278665 | 42.3496179035451 | 30.2865564367503 | 27.1998741334867 | 30.6588294829347 | 50.1463715150282 | 35.3966550789086 | 42.3354970220718 | 3110056K07 |
| ENSMUSG00000085632 | 0.34870327523169 | 0.25565108243973 | 0.837479467570303 | 69.5705561758165 | 46.4216965481167 | 107.895857305923 | 60.0273084325223 | 49.2731188118594 | 63.6002760678407 | 43.7252798033577 | 66.217059444779 | Gm12381 |
| ENSMUSG00000085642 | 0.187732507171223 | 0.741508116054606 | 0.954383574136061 | 10.8421645988285 | 9.77298874697194 | 13.2503684410783 | 12.1930470253561 | 16.4243729372865 | 1.22308223207386 | 9.36970281500523 | 13.0263067760221 | 3110053B16 |
| ENSMUSG00000085643 | -0.0044726782989583 | 0.98372604189869 | 0.996940244108351 | 68.6670424592474 | 61.0811796685746 | 84.2344850897118 | 60.0273084325223 | 67.887408140784 | 53.8156182112498 | 71.8343882483734 | 80.3288917854696 | Gm12519 |
| ENSMUSG00000085665 | 1.24055066611636 | 0.0159229878123956 | 0.63777231095375 | 18.0702743313809 | 14.6594831204579 | 17.9826428843205 | 9.37926694258161 | 5.47479097909548 | 4.89232892829544 | 4.16431236222455 | 10.8552556466851 | Gm12059 |
| ENSMUSG00000085704 | -0.0233803622176962 | 0.920981143985095 | 0.986713918026868 | 140.044626068202 | 112.389370590177 | 109.78876708322 | 173.51643843776 | 97.4512794278996 | 152.885279009232 | 128.052605138405 | 166.085411394282 | 4921531C22 |
| ENSMUSG00000085705 | -0.648933279012501 | 0.18965103516397 | 0.808824684419004 | 9.93865088225949 | 7.32974156022896 | 8.51809399783602 | 18.7585338851632 | 10.949581958191 | 26.9078091056249 | 11.4518589961175 | 20.6249857287016 | Gm16046 |
| ENSMUSG00000085711 | 0.018448294286251 | 0.960505973335411 | 0.993694458032251 | 31.6229800799166 | 23.6180561385155 | 28.3936466594534 | 27.1998741334867 | 27.3739548954774 | 11.0077400886647 | 31.2323427166841 | 39.0789203280663 | Gm15163 |
| ENSMUSG00000085741 | -0.539058463154657 | 0.315291848621256 | 0.853277176422981 | 9.93865088225949 | 13.0306516626293 | 6.62518422053913 | 16.8826804966469 | 21.8991639163819 | 25.6847268735511 | 5.20539045278068 | 15.1973579053591 | 5430405H02 |

| | | | | | | | | | | | | |
|---|---|---|---|---|---|---|---|---|---|---|---|---|
| **ENSMUSG00000085772** | 0.696217671589075 | 0.0559923574708103 | 0.703396849069134 | 31.6229800799166 | 34.2054606144018 | 29.3401015481019 | 19.6964605794214 | 12.0445401540101 | 23.2385624094033 | 22.903717992235 | 13.0263067760221 | D630024D03 |
| **ENSMUSG00000085779** | 0.413472793221824 | 0.00206560991700871 | 0.635862253526407 | 1062.5321306852 | 996.844852191138 | 813.951204237664 | 982.009248888295 | 655.879959295639 | 664.133652016106 | 757.904849924867 | 816.315224630718 | Atcayos |
| **ENSMUSG00000085783** | -0.134655526666869 | 0.787181023708805 | 0.961141387070613 | 7.22810973255236 | 14.6594831204579 | 9.46454888648447 | 10.3171936368398 | 9.85462376237187 | 6.1154111603693 | 18.7394056300105 | 10.8552556466851 | Gm9816 |
| **ENSMUSG00000085793** | -0.0698571734166511 | 0.684369867568281 | 0.944452327903597 | 257.501409222178 | 200.346269312925 | 240.399541716705 | 263.557401086543 | 239.795844884382 | 207.923979452556 | 310.241270985729 | 249.670879873757 | Lin52 |
| **ENSMUSG00000085795** | -0.43254175704859 | 0.0594954307393954 | 0.712406493939829 | 262.922491521592 | 324.951875836817 | 370.063861461543 | 379.860311174555 | 467.547149614754 | 620.102691661447 | 406.020455316893 | 312.63136262453 | Zfp703 |
| **ENSMUSG00000085829** | 0.246295597662655 | 0.289136172891171 | 0.851785577030345 | 58.7283915769879 | 51.3081909216027 | 53.9479286529615 | 72.2203554578784 | 49.2731188118594 | 59.9310293716191 | 49.9717483466946 | 40.1644458927348 | Gm4285 |
| **ENSMUSG00000085853** | 0.157237778170134 | 0.444929957611101 | 0.89108031640712 | 317.133314515735 | 195.459774939439 | 234.720812384815 | 250.426427366929 | 237.605928492744 | 178.570005882784 | 204.051305749003 | 273.552442296464 | Gm12167 |
| **ENSMUSG00000085881** | -0.736898266095577 | 0.0294980606573677 | 0.665956123973759 | 22.5878429142261 | 16.2883145782866 | 16.0897331070236 | 11.2551203310979 | 26.2789966996583 | 26.9078091056249 | 33.3144988977964 | 23.8815624227072 | Gm15912 |
| **ENSMUSG00000085882** | 0.118849913167107 | 0.769630150371346 | 0.95879680569409 | 166.246523848704 | 222.335493993612 | 148.593417517806 | 134.123517278917 | 107.305903190271 | 55.0387004433237 | 241.530117009024 | 213.848536239696 | 2610507I01R |
| **ENSMUSG00000085888** | 0.148647923004275 | 0.611101644650509 | 0.927932112008768 | 84.0267756409212 | 41.5352021746308 | 43.5369248778286 | 54.3997482669734 | 54.7479097909548 | 51.3694537471021 | 55.1771387994752 | 40.1644458927348 | Gm12224 |
| **ENSMUSG00000085939** | 0.325987651468196 | 0.322640024352335 | 0.855977678491063 | 26.2018977805023 | 25.2468875963442 | 18.9290977729689 | 36.5791410760683 | 25.1840385038392 | 20.7923979452556 | 19.7804837205666 | 19.5394601640331 | Cd63-ps |
| **ENSMUSG00000085945** | 0.112476532467745 | 0.803133398451883 | 0.963461896828986 | 23.4913566307952 | 18.7315617650296 | 17.036187995672 | 26.2619474392285 | 6.56974917491458 | 17.123151249034 | 31.2323427166841 | 23.8815624227072 | 2310014F06R |
| **ENSMUSG00000085995** | 0.0848950540022795 | 0.877278807717152 | 0.977569405470617 | 16.2632468982428 | 14.6594831204579 | 16.0897331070236 | 28.1378008277448 | 14.2344565456483 | 12.2308223207386 | 6.24646854333682 | 37.9933947633978 | Gm2788 |
| **ENSMUSG00000086013** | 0.111627482231674 | 0.807495895977269 | 0.964026716319739 | 13.5527057485357 | 7.32974156022896 | 14.1968233297267 | 22.5102406621959 | 14.2344565456483 | 12.2308223207386 | 16.6572494488982 | 9.76973008201657 | Gm15706 |
| **ENSMUSG00000086040** | 0.094310924138839 | 0.698189442081552 | 0.944933386973863 | 1013.74238999047 | 1220.80917764258 | 1166.97887770353 | 1170.53251443419 | 1032.54557865741 | 600.533375948265 | 1404.41434416023 | 1244.01229711011 | Wipf3 |
| **ENSMUSG00000086043** | -0.0108678358164856 | 0.978945641349901 | 0.996668984706452 | 41.5616309621761 | 75.7406627890326 | 70.0376617599851 | 55.3376749612315 | 48.1781606160403 | 24.4616446414772 | 107.231043327282 | 64.046008315442 | Gm12473 |
| **ENSMUSG00000086058** | 0.135187998303799 | 0.635129784242852 | 0.932543559775713 | 49.6932544112975 | 29.3189662409158 | 51.1085639870161 | 39.3929211588428 | 49.2731188118594 | 36.6924669622158 | 35.3966550789086 | 32.5657669400552 | Unc45bos |
| **ENSMUSG00000086095** | 0.518878619339641 | 0.36383574410241 | 0.86924083181804 | 10.8421645988285 | 17.9171460361152 | 20.8220075502658 | 20.6343872736795 | 5.47479097909548 | 3.66924669622158 | 15.616171358342 | 23.8815624227072 | Gm15328 |
| **ENSMUSG00000086108** | 0.534261779045573 | 0.267206911754317 | 0.843793744782845 | 18.0702743313809 | 14.6594831204579 | 21.7684624389143 | 7.50341355406529 | 12.0445401540101 | 6.1154111603693 | 14.5750932677859 | 9.76973008201657 | Gm5602 |
| **ENSMUSG00000086119** | -0.230702263161652 | 0.26033911361787 | 0.840316047847749 | 84.9302893574902 | 83.8848200781758 | 64.3589324280944 | 71.2824287636203 | 75.5521155115177 | 108.854318654574 | 94.7381062406084 | 79.2433662208011 | Gm2415 |
| **ENSMUSG00000086158** | 0.166535543426762 | 0.402145923965621 | 0.881158207349486 | 65.9565013095403 | 65.9676740420606 | 76.6628459805242 | 65.6548685980713 | 66.7924499449649 | 66.0464405319884 | 51.0128264372507 | 60.7894316214364 | Ccpg1os |
| **ENSMUSG00000086212** | -0.193737064450713 | 0.703127133975077 | 0.946451777392559 | 17.1667606148119 | 11.4018202048006 | 5.67872933189068 | 13.1309737196143 | 5.47479097909548 | 15.9000690169602 | 17.6983275394543 | 15.1973579053591 | Mkln1os |
| **ENSMUSG00000086233** | -0.0668097472084194 | 0.905766136701748 | 0.98504921011344 | 5.42108229941427 | 11.4018202048006 | 12.3039135524298 | 17.8206071909051 | 21.8991639163819 | 13.4539045528125 | 5.20539045278068 | 8.68420451734806 | Gm11816 |
| **ENSMUSG00000086236** | 0.272599126801629 | 0.526004526167148 | 0.90806673740897 | 11.7456783153976 | 17.1027303072009 | 11.3574586637814 | 10.3171936368398 | 12.0445401540101 | 9.78465785659088 | 10.4107809055614 | 9.76973008201657 | 5830418P13R |
| **ENSMUSG00000086240** | 0.148113933696004 | 0.740115681777768 | 0.954383574136061 | 15.3597331816738 | 10.5874044758863 | 11.3574586637814 | 15.9447538023887 | 16.4243729372865 | 15.9000690169602 | 8.32862472444909 | 7.59867895267955 | Gm7846 |
| **ENSMUSG00000086247** | 0.795230507353882 | 0.038579541260464 | 0.679566075569698 | 37.0440623793308 | 71.6685841444609 | 72.8770264259304 | 44.0825546301336 | 19.7092475247437 | 20.7923979452556 | 47.8895921655823 | 41.2499714574033 | Gm15787 |
| **ENSMUSG00000086287** | 0.457615230006682 | 0.0936924183702803 | 0.755303361092974 | 32.5264937964856 | 35.8342920722305 | 29.3401015481019 | 37.5170677703264 | 25.1840385038392 | 24.4616446414772 | 23.9447960827911 | 24.9670879873757 | Gm15972 |
| **ENSMUSG00000086290** | 0.267337666551289 | 0.30441686970383 | 0.851785577030345 | 91.2548853734735 | 56.1946852950887 | 80.448665535118 | 87.227182566009 | 72.2672409240604 | 85.6157562451702 | 47.8895921655823 | 56.4473293627624 | Snhg12 |
| **ENSMUSG00000086298** | 0.0855664429642371 | 0.608892252575159 | 0.927932112008768 | 292.738444168371 | 207.676010873154 | 274.47191770805 | 257.929840920994 | 221.181555555458 | 287.424324537357 | 248.817663642917 | 216.019587369033 | Gm11716 |
| **ENSMUSG00000086316** | 0.091197497434807 | 0.550323973829189 | 0.914699131434573 | 303.580608767199 | 244.324718674299 | 229.988537941573 | 302.950322245386 | 257.315176017488 | 282.531995609062 | 231.119336103462 | 244.243252050414 | Nbdy |
| **ENSMUSG00000086324** | 0.740482189796266 | 0.0740424938142767 | 0.73408296865118 | 15.3597331816738 | 15.4738988493722 | 29.3401015481019 | 32.8274342990356 | 16.4243729372865 | 14.6769867848863 | 13.5340151772298 | 10.8552556466851 | Gm15564 |
| **ENSMUSG00000086327** | 0.327944280086017 | 0.282521923069355 | 0.850587941933635 | 28.0089252136404 | 21.9892246806869 | 28.3936466594534 | 29.075727522003 | 18.6142893289246 | 23.2385624094033 | 18.7394056300105 | 24.9670879873757 | Slfn5os |
| **ENSMUSG00000086328** | 0.747708087353044 | 0.0423654577266636 | 0.685752963940267 | 41.5616309621761 | 25.2468875963442 | 30.2865564367503 | 30.9515809105193 | 19.7092475247437 | 30.5770558018465 | 14.5750932677859 | 11.9407812113536 | 2700033N17R |
| **ENSMUSG00000086350** | 0.26936429081985 | 0.3461688890111 | 0.861266047293794 | 162.632468982428 | 169.39847161418 | 165.629605513478 | 129.433883807626 | 94.1664048404423 | 78.277262852727 | 186.352978209548 | 160.657783570939 | B230369F24R |
| **ENSMUSG00000086353** | 0.0517148591754944 | 0.878648289004169 | 0.97807539143035 | 54.2108229941427 | 52.9370223794314 | 21.7684624389143 | 47.8342614071662 | 37.2285786578493 | 58.7079471395453 | 39.5609674411332 | 35.8223436340608 | Gm13481 |
| **ENSMUSG00000086361** | 0.396667517316485 | 0.528476560796153 | 0.908232038364594 | 8.13162344912141 | 13.8450673915436 | 10.4110037751329 | 16.8826804966469 | 21.8991639163819 | 4.89232892829544 | 7.28754663389296 | 3.25657669400552 | Gm6569 |
| **ENSMUSG00000086363** | 0.692915754769709 | 0.176506470131174 | 0.799889021902745 | 13.5527057485357 | 8.14415728914329 | 10.4110037751329 | 16.8826804966469 | 7.66470737073368 | 12.2308223207386 | 5.20539045278068 | 5.42762782334254 | A330102I10R |
| **ENSMUSG00000086369** | -0.228436508366287 | 0.404706594792852 | 0.881158207349486 | 51.5002818444356 | 67.5965054998893 | 53.9479286529615 | 86.2892558717508 | 54.7479097909548 | 103.961989726278 | 67.6700758861489 | 78.1578406561325 | E330017L17R |
| **ENSMUSG00000086384** | -0.381363647670817 | 0.338843066327024 | 0.858570960161243 | 13.5527057485357 | 13.0306516626293 | 19.8755526616174 | 10.3171936368398 | 25.1840385038392 | 12.2308223207386 | 17.6983275394543 | 18.4539345993646 | Banf2os |
| **ENSMUSG00000086392** | 0.0883130364439696 | 0.769660502770993 | 0.95879680569409 | 36.1405486627618 | 22.8036404096012 | 39.7511053232348 | 32.8274342990356 | 30.6588294829347 | 22.0154801773295 | 35.3966550789086 | 34.7368180693922 | Mccc1os |
| **ENSMUSG00000086417** | 0.594934633195787 | 0.108089706614152 | 0.762287160109383 | 33.4300075130547 | 21.9892246806869 | 18.9290977729689 | 21.5723139679377 | 13.1394983498292 | 22.0154801773295 | 12.4929370866736 | 16.2828834700276 | Gm12996 |
| **ENSMUSG00000086429** | -0.163358748675061 | 0.453823773093917 | 0.891964629399053 | 58.7283915769879 | 51.3081909216027 | 45.4298346551254 | 57.2135283497478 | 52.5579933993167 | 59.9310293716191 | 54.1360607089191 | 71.6446872681215 | Gt(ROSA)26S |
| **ENSMUSG00000086448** | 0.642815638083008 | 0.065819089282471 | 0.718732036268493 | 27.1054114970714 | 30.9477976987445 | 25.5542819935081 | 21.5723139679377 | 15.3294147414674 | 14.6769867848863 | 12.4929370866736 | 24.9670879873757 | 9330162012R |
| **ENSMUSG00000086451** | -0.0218381558794152 | 0.919906549276113 | 0.986713918026868 | 63.2459601598332 | 59.452348210746 | 75.7163910918757 | 75.9720622349111 | 55.8428679867739 | 64.8233582999146 | 86.4094815161593 | 70.559161703453 | 4933431K23R |
| **ENSMUSG00000086539** | 0.179336934487349 | 0.662300128993284 | 0.939611397743053 | 18.0702743313809 | 12.2162359337149 | 13.2503684410783 | 21.5723139679377 | 10.949581958191 | 13.4539045528125 | 20.8215618111227 | 11.9407812113536 | Gm16759 |
| **ENSMUSG00000086567** | 0.118516549068851 | 0.556838253722329 | 0.916239900758826 | 1219.74351736821 | 840.477032239587 | 1005.13509174465 | 1345.92480626046 | 1005.17162376193 | 1307.47490608696 | 855.766190437144 | 895.558590851519 | Gm2830 |
| **ENSMUSG00000086583** | -0.130980857782719 | 0.574310269266073 | 0.922280709351328 | 137.334084918495 | 109.13170767452 | 167.522515290775 | 182.895705380341 | 137.964732673206 | 218.931719541221 | 139.504464134522 | 157.401206876934 | Gm15500 |
| **ENSMUSG00000086584** | 0.0996609765822003 | 0.754701203037448 | 0.956497786093635 | 159.921927832721 | 136.007426728693 | 70.0376617599851 | 101.296082979881 | 124.825234323377 | 149.216032313011 | 70.7933101578173 | 92.2696729968231 | Gm12002 |
| **ENSMUSG00000086587** | -0.0629448569959044 | 0.868646614915598 | 0.975147618296101 | 19.877301764519 | 16.2883145782866 | 16.0897331070236 | 10.3171936368398 | 18.6142893289246 | 17.123151249034 | 12.4929370866736 | 17.3684090346961 | Gm11837 |
| **ENSMUSG00000086591** | -0.104078053356806 | 0.837737532073478 | 0.968954477516913 | 15.3597331816738 | 8.95857301805762 | 6.62518422053913 | 15.0068271081306 | 14.2344565456483 | 19.5693157131818 | 9.36970281500523 | 6.51315338801105 | Dnah2os |
| **ENSMUSG00000086596** | -0.750374764016514 | 0.124953364352401 | 0.771949105062466 | 19.877301764519 | 45.6072808192024 | 20.8220075502658 | 10.3171936368398 | 50.3680770076785 | 61.154111603693 | 23.9447960827911 | 28.2236646813812 | Susd5 |
| **ENSMUSG00000086604** | 0.398624438431784 | 0.415037058062291 | 0.882192126163777 | 11.7456783153976 | 5.7009101024003 | 16.0897331070236 | 15.0068271081306 | 7.66470737073368 | 9.78465785659088 | 9.36970281500523 | 9.76973008201657 | Gm15510 |
| **ENSMUSG00000086610** | -0.216972144840211 | 0.630877292419108 | 0.931825800800958 | 15.3597331816738 | 6.51532583131463 | 10.4110037751329 | 20.6343872736795 | 16.4243729372865 | 18.3462334811079 | 13.5340151772298 | 13.0263067760221 | Gm15408 |
| **ENSMUSG00000086629** | 0.91369262086958 | 0.013672046110436 | 0.635862253526407 | 30.7194663633475 | 31.7622134276588 | 21.7684624389143 | 23.448167356454 | 15.3294147414674 | 8.56157562451702 | 12.4929370866736 | 20.6249857287016 | 2810403D21R |
| **ENSMUSG00000086645** | -0.136877486651843 | 0.619336493967438 | 0.929568738467614 | 33.4300075130547 | 49.679359463774 | 32.1794662140472 | 37.5170677703264 | 50.3680770076785 | 31.8001380339204 | 47.8895921655823 | 37.9933947633978 | Gm15743 |
| **ENSMUSG00000086670** | 0.190142181066304 | 0.691618024175929 | 0.94469184542218 | 18.9737880479499 | 8.14415728914329 | 12.3039135524298 | 13.1309737196143 | 12.0445401540101 | 18.3462334811079 | 8.32862472444909 | 7.59867895267955 | Gm13194 |
| **ENSMUSG00000086688** | -0.205298057497218 | 0.47343261877783 | 0.89728597737823 | 28.0089252136404 | 26.8757190541728 | 37.8581955459379 | 30.0136542162612 | 43.7983278327639 | 40.3617136584374 | 27.0680303544596 | 30.3947158107182 | Gm11560 |
| **ENSMUSG00000086712** | 0.254981697422964 | 0.589401515670044 | 0.923594634030771 | 26.2018977805023 | 7.32974156022896 | 22.7149173275627 | 36.5791410760683 | 16.4243729372865 | 18.3462334811079 | 17.6983275394543 | 24.9670879873757 | AI427809 |
| **ENSMUSG00000086714** | -0.302312734270753 | 0.486045743304726 | 0.899236955296196 | 10.8421645988285 | 13.0306516626293 | 9.46454888648447 | 8.44134024832345 | 12.0445401540101 | 8.56157562451702 | 14.5750932677859 | 16.2828834700276 | 0610009E02R |
| **ENSMUSG00000086725** | -0.656393785016292 | 0.150152190822884 | 0.782641594375979 | 8.13162344912141 | 8.14415728914329 | 17.036187995672 | 9.37926694258161 | 10.949581958191 | 23.2385624094033 | 13.5340151772298 | 19.5394601640331 | A630052C17 |
| **ENSMUSG00000086740** | -0.0346064706348149 | 0.941999932019169 | 0.989965881497031 | 28.0089252136404 | 27.6901347830872 | 11.3574586637814 | 17.8206071909051 | 17.5193311331055 | 8.56157562451702 | 28.1091084450157 | 32.5657669400552 | Gm17029 |
| **ENSMUSG00000086741** | -0.289588658997907 | 0.347689320349318 | 0.861266047293794 | 56.0178504272808 | 54.56585383726 | 33.1259211026956 | 49.7101147956825 | 39.4184950494875 | 47.7002070508805 | 63.5057635239243 | 85.7565196088121 | Gm15816 |
| **ENSMUSG00000086765** | 0.0612889563779882 | 0.916837185420252 | 0.986713918026868 | 8.13162344912141 | 8.95857301805762 | 27.447191770805 | 7.50341355406529 | 5.47479097909548 | 12.2308223207386 | 14.5750932677859 | 17.3684090346961 | Gm11827 |
| **ENSMUSG00000086773** | 0.365824774430965 | 0.468851831752022 | 0.895328112556867 | 15.3597331816738 | 42.3496179035451 | 39.7511053232348 | 25.3240207449704 | 16.4243729372865 | 8.56157562451702 | 40.6020455316893 | 29.3091902460497 | Gm16192 |
| **ENSMUSG00000086784** | 0.17147385616468 | 0.117238445532333 | 0.769868040577702 | 1807.93094685466 | 1565.30703097334 | 1653.45669046884 | 1589.78574676758 | 1491.33306270561 | 1279.34401474926 | 1607.42457181867 | 1494.76870254853 | Isoc2a |
| **ENSMUSG00000086801** | -0.134616857127755 | 0.786778473439985 | 0.961141387070613 | 14.4562194651047 | 8.95857301805762 | 7.57163910918757 | 13.1309737196143 | 8.75966556655277 | 11.0077400886647 | 20.8215618111227 | 7.59867895267955 | Gm15943 |
| **ENSMUSG00000086806** | -2.84180972384731 | 0.00157749661066591 | 0.635862253526407 | 9.93865088225949 | 2.44324718674299 | 0.946454888648447 | 4.68963347129081 | 8.75966556655277 | 95.400414101761 | 16.6572494488982 | 8.68420451734806 | Gm13054 |
| **ENSMUSG00000086813** | 0.177085132983461 | 0.680330694158325 | 0.944452327903597 | 15.3597331816738 | 12.2162359337149 | 14.1968233297267 | 16.8826804966469 | 20.8042057205628 | 7.33849339244316 | 11.4518589961175 | 11.9407812113536 | Gm13657 |
| **ENSMUSG00000086841** | 0.285910383508186 | 0.300997640910583 | 0.851785577030345 | 261.115464088454 | 157.99665140938 | 302.865564367503 | 332.026049767389 | 262.789966996583 | 264.185762127954 | 153.038479311752 | 184.539345993646 | 2410006H16R |
| **ENSMUSG00000086859** | 0.0758491300322765 | 0.818973297545086 | 0.966315773308182 | 84.0267756409212 | 32.5766291565731 | 45.4298346551254 | 67.5307219865876 | 44.893286028583 | 70.9387694602839 | 51.0128264372507 | 51.0197015394199 | Snhg20 |
| **ENSMUSG00000086863** | 0.0796948483156023 | 0.485143034433962 | 0.899236955296196 | 430.072529086865 | 358.342920722305 | 371.95677123884 | 356.412143818101 | 371.190828382674 | 355.916929533493 | 349.802238426862 | 358.223436340608 | Gm15543 |
| **ENSMUSG00000086877** | 0.176964472326458 | 0.614675415799698 | 0.928075610917191 | 25.2983840639333 | 23.6180561385155 | 35.965285768641 | 23.448167356454 | 15.3294147414674 | 18.3462334811079 | 32.2734208072402 | 29.3091902460497 | A230072C01 |
| **ENSMUSG00000086881** | 0.178991991669159 | 0.54820194236649 | 0.914699131434573 | 48.7897406947284 | 41.5352021746308 | 43.5369248778286 | 69.4065753751039 | 56.937826182593 | 50.1463715150282 | 28.1091084450157 | 44.5065481514088 | Gm13594 |
| **ENSMUSG00000086914** | 0.259404106121256 | 0.437198783788077 | 0.888722970093854 | 18.9737880479499 | 20.3603932228582 | 28.3936466594534 | 21.5723139679377 | 17.5193311331055 | 23.2385624094033 | 17.6983275394543 | 16.2828834700276 | Gm16124 |
| **ENSMUSG00000086918** | 0.122176503567271 | 0.634588171791285 | 0.932543559775713 | 317.133314515735 | 340.425774686189 | 336.937940358847 | 441.763472995594 | 292.353838283699 | 192.023910435596 | 424.759860946904 | 409.243137880027 | 4930429F24R |
| **ENSMUSG00000086922** | 0.247946680782211 | 0.556787694419244 | 0.916239900758826 | 29.8159526467785 | 11.4018202048006 | 17.9826428843205 | 18.7585338851632 | 18.6142893289246 | 15.9000690169602 | 10.4107809055614 | 20.6249857287016 | Gm13835 |
| **ENSMUSG00000086932** | 0.498248798210923 | 0.418478200593678 | 0.882468496126562 | 13.5527057485357 | 5.7009101024003 | 10.4110037751329 | 22.5102406621959 | 4.37983278327639 | 14.6769867848863 | 13.5340151772298 | 4.34210225867403 | Gm22000 |
| **ENSMUSG00000086935** | 0.498954599633329 | 0.257622879220392 | 0.838048372687018 | 11.7456783153976 | 13.8450673915436 | 19.8755526616174 | 15.9447538023887 | 10.949581958191 | 14.6769867848863 | 12.4929370866736 | 5.42762782334254 | Gm15956 |
| **ENSMUSG00000086938** | -0.00492680407670142 | 0.965275947659265 | 0.994284908685797 | 613.485813550382 | 574.163088884602 | 709.841166486335 | 633.100518624259 | 594.56230032977 | 626.218102821816 | 624.646854333682 | 692.565310258508 | 4930481A15R |
| **ENSMUSG00000086947** | 0.908420191769271 | 0.0687819589244265 | 0.723457921755197 | 17.1667606148119 | 8.95857301805762 | 15.1432782183751 | 17.8206071909051 | 7.66470737073368 | 13.4539045528125 | 5.20539045278068 | 5.42762782334254 | 4930522O17R |
| **ENSMUSG00000086965** | 0.183213317174282 | 0.694874070514323 | 0.944772987416205 | 8.13162344912141 | 13.8450673915436 | 13.2503684410783 | 8.44134024832345 | 9.85462376237187 | 7.33849339244316 | 12.4929370866736 | 8.68420451734806 | Rtl10 |
| **ENSMUSG00000086968** | -0.254552000879772 | 0.493866022566521 | 0.900117970535229 | 26.2018977805023 | 35.8342920722305 | 17.036187995672 | 23.448167356454 | 40.5134532453066 | 15.9000690169602 | 33.3144988977964 | 32.5657669400552 | 4933431E20R |
| **ENSMUSG00000087006** | -0.817591923011311 | 0.0925732887704494 | 0.754058637722175 | 93.0619128066116 | 111.574954861263 | 43.5369248778286 | 82.5375490947182 | 211.326931793086 | 248.285693110993 | 65.5879197050366 | 58.6183804920994 | Gm13889 |
| **ENSMUSG00000087057** | -0.446678748547699 | 0.190928800703371 | 0.808824684419004 | 37.9475760958999 | 30.9477976987445 | 29.3401015481019 | 22.5102406621959 | 50.3680770076785 | 58.7079471395453 | 29.1501865355718 | 27.1381391167127 | Gm11730 |
| **ENSMUSG00000087107** | 0.197370022818031 | 0.622872138999384 | 0.931059568526633 | 22.5878429142261 | 18.7315617650296 | 50.1621090983677 | 26.2619474392285 | 22.994122112201 | 33.0232202659942 | 29.1501865355718 | 17.3684090346961 | AI662270 |
| **ENSMUSG00000087110** | 0.278133311675605 | 0.49024578151865 | 0.899236955296196 | 14.4562194651047 | 12.2162359337149 | 17.036187995672 | 22.5102406621959 | 9.85462376237187 | 17.123151249034 | 15.616171358342 | 11.9407812113536 | Gm11264 |
| **ENSMUSG00000087128** | 0.294059334136998 | 0.315227470080807 | 0.853277176422981 | 62.3424464432641 | 47.2361122770311 | 44.483379766477 | 54.3997482669734 | 38.3235368536684 | 30.5770558018465 | 64.5468416144805 | 35.8223436340608 | Gm12655 |

| | | | | | | | | | | | | |
|---|---|---|---|---|---|---|---|---|---|---|---|---|
| **ENSMUSG00000087129** | -0.094342875136119 | 0.885259374034821 | 0.979866999597423 | 10.8421645988285 | 15.4738988493722 | 23.6613722162112 | 1.87585338851632 | 5.47479097909548 | 15.9000690169602 | 16.6572494488982 | 17.3684090346961 | Gm16316 |
| **ENSMUSG00000087138** | 0.432200304647951 | 0.442284710904502 | 0.89108031640712 | 21.6843291976571 | 6.51532583131463 | 6.62518422053913 | 15.9447538023887 | 6.56974917491458 | 9.78465785659088 | 13.5340151772298 | 7.59867895267955 | Gm15545 |
| **ENSMUSG00000087141** | 0.115990542191025 | 0.785594344512827 | 0.961141387070613 | 18.0702743313809 | 25.2468875963442 | 11.3574586637814 | 15.0068271081306 | 14.2344565456483 | 19.5693157131818 | 9.36970281500523 | 21.7105112933701 | Plcxd2 |
| **ENSMUSG00000087165** | 0.0342017952021356 | 0.900250899194013 | 0.984802943363903 | 46.0791995450213 | 28.5045505120015 | 42.5904699891801 | 36.5791410760683 | 31.7537876787538 | 44.0309603546589 | 41.6431236222455 | 32.5657669400552 | 2010001A14 |
| **ENSMUSG00000087174** | 0.214627899752287 | 0.465219328108838 | 0.895328112556867 | 39.754603529038 | 34.2054606144018 | 29.3401015481019 | 44.0825546301336 | 37.2285786578493 | 37.9155491942896 | 20.8215618111227 | 31.4802413753867 | 5530601H04 |
| **ENSMUSG00000087177** | -0.114110988961278 | 0.562419542639523 | 0.917693310295102 | 103.90407740544 | 91.2145616384048 | 103.163582862681 | 76.9099889291692 | 83.2168228822514 | 110.077400886647 | 93.6970281500523 | 119.407812113536 | E130307A14 |
| **ENSMUSG00000087178** | 0.0713745903018306 | 0.804786309085018 | 0.963461896828986 | 38.8510898124689 | 26.0613033252585 | 29.3401015481019 | 43.1446279358754 | 39.4184950494875 | 29.3539735697726 | 33.3144988977964 | 28.2236646813812 | A230056P14 |
| **ENSMUSG00000087221** | 0.0260532019916874 | 0.958655196291763 | 0.993101678702062 | 58.7283915769879 | 66.7820897709749 | 50.1621090983677 | 34.703287687552 | 35.0386622662111 | 12.2308223207386 | 91.61487196894 | 67.3025850094475 | BC037032 |
| **ENSMUSG00000087222** | -0.0873782096123155 | 0.810399584425652 | 0.964412705343932 | 19.877301764519 | 30.1333819698302 | 22.7149173275627 | 16.8826804966469 | 17.5193311331055 | 17.123151249034 | 33.3144988977964 | 27.1381391167127 | E030042O20 |
| **ENSMUSG00000087236** | 0.518879377344331 | 0.358589073960091 | 0.866648170843829 | 18.0702743313809 | 13.0306516626293 | 6.62518422053913 | 12.1930470253561 | 9.85462376237187 | 3.66924669622158 | 15.616171358342 | 5.42762782334254 | Kif28 |
| **ENSMUSG00000087249** | 0.0923849694619095 | 0.770717639650413 | 0.958995611819782 | 28.9124389302094 | 21.9892246806869 | 22.7149173275627 | 24.3860940507122 | 29.5638712871156 | 19.5693157131818 | 17.6983275394543 | 24.9670879873757 | Gm16062 |
| **ENSMUSG00000087253** | 0.858703178275632 | 0.112037938032132 | 0.764116514278315 | 61.4389327266951 | 60.2667639396603 | 41.6440151005317 | 30.0136542162612 | 22.994122112201 | 3.66924669622158 | 43.7252798033577 | 35.8223436340608 | Gm12043 |
| **ENSMUSG00000087259** | 0.192515751470054 | 0.553512022978474 | 0.915271666567977 | 103.90407740544 | 74.1118313312039 | 106.002947528626 | 141.626930832982 | 150.009272827216 | 91.7311674055395 | 69.7522320672611 | 60.7894316214364 | 2610035D17 |
| **ENSMUSG00000087260** | -0.0918814376580989 | 0.557589813367768 | 0.916341990969984 | 1021.87401343959 | 816.858976101072 | 900.078599104673 | 1091.7466721165 | 1036.92541144068 | 1207.1821630569 | 834.944628626022 | 1004.11114731837 | Lamtor5 |
| **ENSMUSG00000087266** | 0.513261680524735 | 0.269451553098491 | 0.843793744782845 | 12.6491920319666 | 21.9892246806869 | 12.3039135524298 | 11.2551203310979 | 5.47479097909548 | 12.2308223207386 | 12.4929370866736 | 10.8552556466851 | Gm15991 |
| **ENSMUSG00000087269** | 0.0658894145329906 | 0.785618381967333 | 0.961141387070613 | 76.7986659083688 | 57.8235167529173 | 46.3762895437739 | 71.2824287636203 | 63.5075753575076 | 69.71568722821 | 60.3825292522559 | 47.7631248454143 | D330023K18 |
| **ENSMUSG00000087298** | 0.0756311273868293 | 0.864407150729051 | 0.974868041734905 | 12.6491920319666 | 10.5874044758863 | 9.46454888648447 | 9.37926694258161 | 8.75966556655277 | 12.2308223207386 | 9.36970281500523 | 9.76973008201657 | Gm9392 |
| **ENSMUSG00000087299** | -0.458715928949965 | 0.0674670696757225 | 0.722038584319139 | 41.5616309621761 | 44.7928650902881 | 40.6975602118832 | 56.2756016554897 | 53.6529515951357 | 84.3926740130963 | 63.5057635239243 | 51.0197015394199 | Gm12953 |
| **ENSMUSG00000087305** | 0.716089456832814 | 0.0432427493694051 | 0.687730553581215 | 20.780815481088 | 35.0198763433161 | 25.5542819935081 | 28.1378008277448 | 17.5193311331055 | 20.7923979452556 | 10.4107809055614 | 18.4539345993646 | A430035B10 |
| **ENSMUSG00000087307** | 0.00311521173515985 | 0.995039673145956 | 0.999262355603273 | 8.13162344912141 | 15.4738988493722 | 8.51809399783602 | 9.37926694258161 | 7.66470737073368 | 6.1154111603693 | 15.616171358342 | 11.9407812113536 | Gm12925 |
| **ENSMUSG00000087331** | 0.045554671276905 | 0.864056810645873 | 0.974868041734905 | 72.2810973255236 | 57.009101024003 | 44.483379766477 | 60.0273084325223 | 49.2731188118594 | 79.5003450848009 | 54.1360607089191 | 44.5065481514088 | 1810021B22 |
| **ENSMUSG00000087366** | 0.182341228516468 | 0.725972874295469 | 0.951837193124793 | 5.42108229941427 | 17.9171460361152 | 18.9290977729689 | 18.7585338851632 | 17.5193311331055 | 6.1154111603693 | 11.4518589961175 | 18.4539345993646 | Junos |
| **ENSMUSG00000087370** | 0.0515355772700336 | 0.831807336626263 | 0.967583158900775 | 392.124952990966 | 476.433201414882 | 419.279515671262 | 341.405316709971 | 363.52612101194 | 234.831788558181 | 476.813765474711 | 496.085183053508 | Tmem170b |
| **ENSMUSG00000087380** | 0.184352246796348 | 0.160357903761794 | 0.79125089991166 | 527.652010476322 | 469.917875583568 | 531.907647420427 | 567.445650026188 | 408.419407040523 | 532.040770952129 | 425.80093903746 | 479.80229958348 | 2210408F21 |
| **ENSMUSG00000087385** | 0.288123133649838 | 0.199703947788822 | 0.812921911423739 | 68.6670424592474 | 60.2667639396603 | 76.6628459805242 | 50.6480414899407 | 54.7479097909548 | 44.0309603546589 | 56.2182168900314 | 54.2762782334254 | Frg2f1 |
| **ENSMUSG00000087399** | 0.0720002794873797 | 0.788200740167803 | 0.961444367900485 | 56.0178504272808 | 57.009101024003 | 57.7337482075553 | 77.8479156234274 | 55.8428679867739 | 85.6157562451702 | 42.6842017128016 | 53.1907526687569 | Gm11899 |
| **ENSMUSG00000087400** | -0.16731958028173 | 0.539974268259342 | 0.913837378403038 | 38.8510898124689 | 35.0198763433161 | 49.2156542097192 | 57.2135283497478 | 54.7479097909548 | 57.4848649074714 | 55.1771387994752 | 34.7368180693922 | Gm15270 |
| **ENSMUSG00000087404** | 0.129197273507295 | 0.787761683760019 | 0.961258876168556 | 15.3597331816738 | 13.0306516626293 | 25.5542819935081 | 8.44134024832345 | 13.1394983498292 | 8.56157562451702 | 21.8626399016789 | 13.0263067760221 | Gm11752 |
| **ENSMUSG00000087412** | 0.381030335709582 | 0.238906685958304 | 0.832048466795696 | 28.9124389302094 | 26.8757190541728 | 19.8755526616174 | 30.0136542162612 | 16.4243729372865 | 26.9078091056249 | 18.7394056300105 | 19.5394601640331 | Gm15501 |
| **ENSMUSG00000087413** | -0.0281295271358325 | 0.932669372361985 | 0.988114361264825 | 28.0089252136404 | 35.8342920722305 | 51.1085639870161 | 21.5723139679377 | 27.3739548954774 | 34.2463024980681 | 42.6842017128016 | 34.7368180693922 | Gm11266 |
| **ENSMUSG00000087440** | 0.11404446617076 | 0.704798154994729 | 0.947232353447198 | 38.8510898124689 | 31.7622134276588 | 28.3936466594534 | 31.8895076047775 | 31.7537876787538 | 18.3462334811079 | 31.2323427166841 | 39.0789203280663 | 4930577N17 |
| **ENSMUSG00000087466** | -0.627636448484979 | 0.125085577154281 | 0.771949105062466 | 11.7456783153976 | 8.14415728914329 | 10.4110037751329 | 8.44134024832345 | 14.2344565456483 | 14.6769867848863 | 16.6572494488982 | 14.1118323406906 | A330041J22 |
| **ENSMUSG00000087484** | 0.881738344467407 | 0.0333455948573487 | 0.671606027118506 | 37.9475760958999 | 17.1027303072009 | 17.9826428843205 | 31.8895076047775 | 12.0445401540101 | 19.5693157131818 | 12.4929370866736 | 13.0263067760221 | 2900089D17 |
| **ENSMUSG00000087497** | 0.0142101275892053 | 0.937492313575625 | 0.988970340396118 | 133.720030052219 | 100.173134656462 | 98.4313084194385 | 127.55803041911 | 113.875652365186 | 100.292743030056 | 115.559668051731 | 124.835439936878 | 2810001G20 |
| **ENSMUSG00000087523** | 0.390891692085297 | 0.315755557006056 | 0.853277176422981 | 44.2721721118832 | 54.56585383726 | 28.3936466594534 | 27.1998741334867 | 15.3294147414674 | 26.9078091056249 | 31.2323427166841 | 44.5065481514088 | Gm12319 |
| **ENSMUSG00000087530** | 0.162338578051102 | 0.646600594699746 | 0.935063240511793 | 128.298947752804 | 172.656134529838 | 141.968233297267 | 118.178763476528 | 104.021028602814 | 50.1463715150282 | 170.736806851206 | 175.855141476298 | Gm15533 |
| **ENSMUSG00000087543** | 0.14582526060437 | 0.655591034522296 | 0.937625787683528 | 64.1494738764022 | 98.5443031986338 | 105.056492639978 | 46.8963347129081 | 62.4126171616885 | 47.7002070508805 | 96.8202624217207 | 77.072315091464 | Gm16576 |
| **ENSMUSG00000087574** | 0.0734282590012972 | 0.665814093719869 | 0.942024383071524 | 199.676531361759 | 188.13003337921 | 170.36187995672 | 207.281799431054 | 136.869774477387 | 194.470074899744 | 204.051305749003 | 192.138024946326 | C030037D09 |
| **ENSMUSG00000087578** | 0.335101823901257 | 0.536230490084148 | 0.911291453479697 | 15.3597331816738 | 13.8450673915436 | 13.2503684410783 | 17.8206071909051 | 7.66470737073368 | 2.44616446414772 | 18.7394056300105 | 18.4539345993646 | Ccdc142os |
| **ENSMUSG00000087579** | 0.0431339357810304 | 0.886227303213938 | 0.980068672363141 | 292.738444168371 | 210.933673788811 | 206.327165725361 | 255.11606083822 | 162.053812981226 | 139.43137445642 | 349.802238426862 | 284.407697943149 | Hectd2os |
| **ENSMUSG00000087589** | -0.970434573237151 | 0.0055556794187527 | 0.635862253526407 | 37.9475760958999 | 24.4324718674299 | 27.447191770805 | 37.5170677703264 | 86.5016974697087 | 79.5003450848009 | 49.9717483466946 | 33.6512925047237 | D430040D24 |
| **ENSMUSG00000087590** | 0.0112562097114817 | 0.971697040391185 | 0.995123114262614 | 64.1494738764022 | 48.8649437348597 | 87.0738497556571 | 90.0409626487835 | 82.1218646864323 | 103.961989726278 | 49.9717483466946 | 52.1052271040884 | Epb41l4aos |
| **ENSMUSG00000087595** | 0.272769934383762 | 0.545319035665244 | 0.914211323944814 | 18.0702743313809 | 24.4324718674299 | 20.8220075502658 | 19.6964605794214 | 7.66470737073368 | 9.78465785659088 | 28.1091084450157 | 22.7960368580387 | 1810012K08 |
| **ENSMUSG00000087598** | 0.121703197786288 | 0.690335842349048 | 0.94469184542218 | 34.3335212296237 | 35.8342920722305 | 35.0188308799925 | 64.7169419038131 | 39.4184950494875 | 47.7002070508805 | 32.2734208072402 | 36.9078691987293 | Zfp111 |
| **ENSMUSG00000087610** | 0.0397623679498976 | 0.900013768183327 | 0.984802943363903 | 33.4300075130547 | 21.9892246806869 | 31.2330113253987 | 36.5791410760683 | 28.4689130912965 | 42.8078781225851 | 23.9447960827911 | 24.9670879873757 | Gm16253 |
| **ENSMUSG00000087611** | 0.475270808804328 | 0.0926974688917113 | 0.754058637722175 | 38.8510898124689 | 70.0397526866323 | 51.1085639870161 | 71.2824287636203 | 35.0386622662111 | 35.4693847301419 | 46.8485140750261 | 48.8486504100828 | 4930458D05 |
| **ENSMUSG00000087644** | 0.452446696403634 | 0.375092411822262 | 0.873700364743909 | 17.1667606148119 | 7.32974156022896 | 15.1432782183751 | 15.0068271081306 | 8.75966556655277 | 11.0077400886647 | 15.616171358342 | 4.34210225867403 | Gm14703 |
| **ENSMUSG00000087651** | 0.312283775436155 | 0.342474737898728 | 0.860056790709335 | 94.8689402397497 | 61.0811796685746 | 39.7511053232348 | 102.23400967414 | 44.893286028583 | 62.3771938357668 | 73.9165444294857 | 58.6183804920994 | 1500009L16 |
| **ENSMUSG00000087679** | -0.0434801653920841 | 0.683437666581249 | 0.944452327903597 | 2052.78316404487 | 2112.59440080377 | 1711.19043867639 | 1839.27424744025 | 1850.47935093427 | 2027.87034077846 | 1980.13052823777 | 2095.06433981022 | Tmem250-ps |
| **ENSMUSG00000087684** | -0.552699846277682 | 0.24582903968345 | 0.832048466795696 | 16.2632468982428 | 9.77298874697194 | 14.1968233297267 | 7.50341355406529 | 14.2344565456483 | 11.0077400886647 | 30.191264626128 | 14.1118323406906 | 1200007C13 |
| **ENSMUSG00000087687** | 0.1531204080314 | 0.457121646687132 | 0.892805623137304 | 274.66816983699 | 189.758864837039 | 179.826428843205 | 229.79204009325 | 198.187433443257 | 249.508775343067 | 164.49033830787 | 174.76961591163 | Pet100 |
| **ENSMUSG00000087700** | -0.486554367611518 | 0.210730245530704 | 0.822896573073231 | 12.6491920319666 | 13.0306516626293 | 13.2503684410783 | 16.8826804966469 | 19.7092475247437 | 28.1308913376988 | 18.7394056300105 | 11.9407812113536 | Gm15283 |
| **ENSMUSG00000088185** | 0.0198295854772747 | 0.959227366301778 | 0.993265850449657 | 40.658117245607 | 28.5045505120015 | 40.6975602118832 | 55.3376749612315 | 73.3621991198795 | 31.8001380339204 | 27.0680303544596 | 30.3947158107182 | Scarna2 |
| **ENSMUSG00000088609** | 0.466759393477952 | 0.181784114399184 | 0.80356567265522 | 159.921927832721 | 120.533527879321 | 161.843785958884 | 232.605820176024 | 203.662224422352 | 130.869798831903 | 83.2862472444909 | 70.559161703453 | Gm24187 |
| **ENSMUSG00000089235** | 0.262780360703824 | 0.294216766520925 | 0.851785577030345 | 46.0791995450213 | 43.1640336324594 | 35.965285768641 | 36.5791410760683 | 35.0386622662111 | 31.8001380339204 | 30.191264626128 | 37.9933947633978 | Gm23119 |
| **ENSMUSG00000089417** | 0.821842640415652 | 0.0027267902659722 | 0.635862253526407 | 149.079763233892 | 122.162359337149 | 84.2344850897118 | 159.447538023887 | 97.4512794278996 | 55.0387004433237 | 60.3825292522559 | 78.1578406561325 | Gm22009 |
| **ENSMUSG00000089656** | 0.529311752687505 | 0.258724755419788 | 0.838606660810377 | 13.5527057485357 | 13.0306516626293 | 10.4110037751329 | 12.1930470253561 | 9.85462376237187 | 6.1154111603693 | 12.4929370866736 | 5.42762782334254 | Gm16271 |
| **ENSMUSG00000089662** | 0.484544752679816 | 0.112872369495912 | 0.767399516543425 | 33.4300075130547 | 26.8757190541728 | 25.5542819935081 | 30.0136542162612 | 25.1840385038392 | 15.9000690169602 | 21.8626399016789 | 19.5394601640331 | Gm14057 |
| **ENSMUSG00000089678** | -0.249505928654284 | 0.508783173472622 | 0.902394714365738 | 26.2018977805023 | 37.4631235300591 | 30.2865564367503 | 39.3929211588428 | 41.6084114411257 | 34.2463024980681 | 18.7394056300105 | 64.046008315442 | Agxt2 |
| **ENSMUSG00000089682** | 0.20344128547034 | 0.286341503648967 | 0.851785577030345 | 1492.60465977206 | 1547.38988493722 | 1496.34517895319 | 1427.52442866092 | 952.613630362614 | 1068.97387083255 | 1648.02661735036 | 1508.88053488923 | Bcl2l2 |
| **ENSMUSG00000089687** | 0.0460959937730041 | 0.929569510217248 | 0.987361053522794 | 9.93865088225949 | 16.2883145782866 | 10.4110037751329 | 5.62756016554897 | 7.66470737073368 | 12.2308223207386 | 6.24646854333682 | 15.1973579053591 | Rab42 |
| **ENSMUSG00000089698** | 0.0493241575351072 | 0.922087096605444 | 0.986713918026868 | 13.5527057485357 | 5.7009101024003 | 13.2503684410783 | 12.1930470253561 | 5.47479097909548 | 9.78465785659088 | 15.616171358342 | 11.9407812113536 | Gm2541 |
| **ENSMUSG00000089704** | 0.279868884816959 | 0.403650495944018 | 0.881158207349486 | 50.5967681278665 | 72.4829998733752 | 91.8061241988993 | 51.5859681841989 | 44.893286028583 | 31.8001380339204 | 79.1219348822664 | 62.9604827507734 | Galnt2 |
| **ENSMUSG00000089715** | -0.379049879501767 | 0.108362643916345 | 0.762287160109383 | 594.512025502432 | 511.453077758198 | 484.584902988005 | 679.996853337167 | 825.598479647599 | 1006.59667699679 | 632.975479058131 | 488.486504100828 | Cbx6 |
| **ENSMUSG00000089736** | 0.179110865673114 | 0.689076957149848 | 0.94469184542218 | 11.7456783153976 | 11.4018202048006 | 14.1968233297267 | 10.3171936368398 | 6.56974917491458 | 12.2308223207386 | 14.5750932677859 | 8.68420451734806 | Tgfbr3l |
| **ENSMUSG00000089762** | -0.873513829879842 | 0.0794664553107374 | 0.739855948845396 | 99.3865088225949 | 53.7514381083457 | 94.6454888648447 | 127.55803041911 | 234.321053905287 | 299.655146858096 | 109.313199508394 | 44.5065481514088 | Ier5l |
| **ENSMUSG00000089764** | -0.545921453002455 | 0.22754078542785 | 0.832048466795696 | 8.13162344912141 | 8.95857301805762 | 13.2503684410783 | 7.50341355406529 | 19.7092475247437 | 9.78465785659088 | 13.5340151772298 | 11.9407812113536 | Gm16580 |
| **ENSMUSG00000089774** | -0.336491541479622 | 0.306817317332752 | 0.851785577030345 | 42.4651446787451 | 40.7207864457164 | 39.7511053232348 | 38.4549944645846 | 42.7033696369448 | 86.838838477244 | 42.6842017128016 | 32.5657669400552 | Slc5a3 |
| **ENSMUSG00000089782** | -0.36484608735213 | 0.14726436425656 | 0.780607990324772 | 37.9475760958999 | 30.1333819698302 | 32.1794662140472 | 45.9584080186499 | 52.5579933993167 | 40.3617136584374 | 44.7663578939139 | 49.9341759747513 | Gm3531 |
| **ENSMUSG00000089809** | 0.0715001843762356 | 0.780064661837276 | 0.960643510321816 | 214.132750826864 | 303.777066885045 | 240.399541716705 | 154.757904552597 | 301.113503850252 | 179.793088114857 | 183.22974393788 | 205.164331722348 | Rasgef1b |
| **ENSMUSG00000089810** | 0.265416766080817 | 0.168244212915158 | 0.79907358295209 | 213.229237110295 | 178.357044632238 | 146.700507740509 | 243.860940507122 | 169.71852035196 | 173.677676954488 | 143.668776496747 | 163.914360264945 | Gm16536 |
| **ENSMUSG00000089817** | 1.20835583287715 | 0.01913856824272 | 0.642273034804458 | 13.5527057485357 | 22.8036404096012 | 18.9290977729689 | 28.1378008277448 | 14.2344565456483 | 3.66924669622158 | 13.5340151772298 | 4.34210225867403 | Gm7162 |
| **ENSMUSG00000089824** | -0.0317716267800234 | 0.861072676244423 | 0.974868041734905 | 155.404359249876 | 116.461449234749 | 116.413951303759 | 141.626930832982 | 135.774816281568 | 160.223772401676 | 131.175839410073 | 115.065709854862 | Rbm12 |
| **ENSMUSG00000089832** | -0.0868137903050653 | 0.738399916901236 | 0.954383574136061 | 109.325159704854 | 107.502876216691 | 122.09268063565 | 150.068271081306 | 166.433645764503 | 166.339183562045 | 93.6970281500523 | 93.3551985614916 | Shkbp1 |
| **ENSMUSG00000089838** | 0.168721209236423 | 0.625011211412153 | 0.931529302402539 | 41.5616309621761 | 21.9892246806869 | 33.1259211026956 | 40.3308478531009 | 18.6142893289246 | 41.5847958905112 | 27.0680303544596 | 34.7368180693922 | Gm2962 |
| **ENSMUSG00000089844** | -0.0185133512508165 | 0.968619757529554 | 0.994790165280964 | 11.7456783153976 | 8.95857301805762 | 20.8220075502658 | 15.9447538023887 | 15.3294147414674 | 6.1154111603693 | 19.7804837205666 | 16.2828834700276 | A530032D15 |
| **ENSMUSG00000089847** | -0.0403301540870005 | 0.857588661279541 | 0.974046056228403 | 277.378710986697 | 190.573280565953 | 263.114459044268 | 375.170677703265 | 293.448796479518 | 335.124531588238 | 234.242570375131 | 274.637967861132 | Timm10b |
| **ENSMUSG00000089855** | 0.248985888567998 | 0.662131285222463 | 0.939611397743053 | 10.8421645988285 | 13.8450673915436 | 8.51809399783602 | 23.448167356454 | 21.8991639163819 | 8.56157562451702 | 4.16431236222455 | 13.0263067760221 | NA |
| **ENSMUSG00000089857** | 0.00217787649288344 | 0.99524780934963 | 0.999262355603273 | 41.5616309621761 | 26.8757190541728 | 29.3401015481019 | 14.0689004138724 | 21.8991639163819 | 29.3539735697726 | 32.2734208072402 | 28.2236646813812 | Zfp882 |
| **ENSMUSG00000089862** | 0.20670746883858 | 0.237355785929744 | 0.832048466795696 | 224.071401709123 | 212.56250524664 | 227.149173275627 | 166.950951577953 | 181.76306050597 | 141.877538920568 | 200.928071477334 | 194.309076075663 | Umad1 |
| **ENSMUSG00000089872** | 0.0688646378551308 | 0.74768250217206 | 0.956119556124081 | 335.203588847116 | 384.404224047563 | 393.725233677754 | 331.088123073131 | 330.677375137367 | 221.377884005369 | 381.034581143546 | 442.894430384751 | Rps6kc1 |
| **ENSMUSG00000089875** | 0.165916551353895 | 0.534031893403658 | 0.909924312975565 | 71.3775836089546 | 73.2974156022896 | 111.681676860517 | 77.8479156234274 | 77.7420319031559 | 57.4848649074714 | 103.066730965058 | 58.6183804920994 | Etohd2 |
| **ENSMUSG00000089889** | -0.06728765818971 | 0.792827244453929 | 0.961804768755116 | 46.0791995450213 | 31.7622134276588 | 36.9117406572894 | 49.7101147956825 | 37.2285786578493 | 48.9232892829544 | 45.80743598447 | 40.1644458927348 | 0610040B10 |
| **ENSMUSG00000089901** | 0.147552963363526 | 0.661372625419809 | 0.939611397743053 | 54.2108229941427 | 41.5352021746308 | 37.8581955459379 | 30.9515809105193 | 39.4184950494875 | 18.3462334811079 | 48.9306702561384 | 41.2499714574033 | Gm8113 |
| **ENSMUSG00000089911** | 0.0375172894292364 | 0.780018035097064 | 0.960643510321816 | 320.747369382011 | 324.951875836817 | 335.04503058155 | 271.060814640609 | 338.342082508101 | 266.631926592101 | 314.405583347953 | 299.605055848508 | Mfsd14a |
| **ENSMUSG00000089917** | 0.242802895113603 | 0.0264717238896305 | 0.664353746352133 | 824.908023227538 | 746.819223414439 | 783.664647800914 | 847.885731609378 | 719.387534653147 | 594.417964787896 | 665.248899865371 | 726.216602763232 | Uckl1 |
| **ENSMUSG00000089945** | -0.113482546388355 | 0.855656220174586 | 0.973387788552416 | 23.4913566307952 | 58.6379324818317 | 29.3401015481019 | 17.8206071909051 | 16.4243729372865 | 7.33849339244316 | 54.1360607089191 | 61.8749571861049 | Pakap |

| | | | | | | | | | | | | |
|---|---|---|---|---|---|---|---|---|---|---|---|---|
| **ENSMUSG00000089960** | -0.586623499695164 | 0.177523783254967 | 0.800846167395741 | 19.877301764519 | 43.1640336324594 | 15.1432782183751 | 30.9515809105193 | 28.4689130912965 | 33.0232202659942 | 32.2734208072402 | 70.559161703453 | Ugt1a1 |
| **ENSMUSG00000090000** | 0.0136331951313775 | 0.923627659293363 | 0.986713918026868 | 985.733464776828 | 783.467931215584 | 783.664647800914 | 996.078149302167 | 851.877476347257 | 1016.38133485338 | 807.876598271562 | 840.196787053425 | Ier3ip1 |
| **ENSMUSG00000090015** | 0.215697572883355 | 0.498441186930188 | 0.900927059989498 | 42.4651446787451 | 40.7207864457164 | 40.6975602118832 | 51.5859681841989 | 29.5638712871156 | 20.7923979452556 | 49.9717483466946 | 49.9341759747513 | Gm15446 |
| **ENSMUSG00000090019** | -0.457219274260721 | 0.121924900687559 | 0.769868040577702 | 195.158962778914 | 150.666909849151 | 152.3792370724 | 166.950951577953 | 287.974005500422 | 332.67836712409 | 144.709854587303 | 148.717002359586 | Gimap1 |
| **ENSMUSG00000090035** | -0.470217158601432 | 0.133933869009072 | 0.77303252008843 | 80.412720774645 | 83.8848200781758 | 93.6990339761962 | 89.1030359545253 | 141.249607260664 | 188.354663739374 | 71.8343882483734 | 80.3288917854696 | Galnt4 |
| **ENSMUSG00000090038** | 0.138466018177112 | 0.683158607109097 | 0.944452327903597 | 23.4913566307952 | 24.4324718674299 | 25.5542819935081 | 21.5723139679377 | 18.6142893289246 | 19.5693157131818 | 15.616171358342 | 32.5657669400552 | Gm16573 |
| **ENSMUSG00000090053** | 0.13141382057366 | 0.865174129876769 | 0.974868041734905 | 10.8421645988285 | 37.4631235300591 | 19.8755526616174 | 10.3171936368398 | 4.37983278327639 | 2.44616446414772 | 35.3966550789086 | 29.3091902460497 | Palm2 |
| **ENSMUSG00000090066** | 0.227308420191823 | 0.467318746405885 | 0.895328112556867 | 1568.49981196386 | 2088.16192893634 | 1801.10365309799 | 1731.41267760057 | 1044.59011881142 | 799.895779776304 | 2123.79930473452 | 2172.13665490168 | 1110002E22 |
| **ENSMUSG00000090083** | -0.00574278723791747 | 0.98328260906355 | 0.996940244108351 | 101.193536255733 | 77.3694942468612 | 95.5919437534931 | 149.130344387048 | 123.730276127558 | 135.762127760198 | 86.4094815161593 | 79.2433662208011 | Rnf8 |
| **ENSMUSG00000090084** | -0.621759451141995 | 0.286523105541207 | 0.851785577030345 | 35.2370349461928 | 30.9477976987445 | 24.6078271048596 | 35.6412143818101 | 44.893286028583 | 117.415894279091 | 16.6572494488982 | 16.2828834700276 | Srpx |
| **ENSMUSG00000090086** | 0.00269279062550287 | 0.994463976259431 | 0.999262355603273 | 22.5878429142261 | 14.6594831204579 | 16.0897331070236 | 25.3240207449704 | 17.5193311331055 | 30.5770558018465 | 16.6572494488982 | 14.1118323406906 | AI480526 |
| **ENSMUSG00000090093** | -0.236884649787936 | 0.482032863166482 | 0.897732059433541 | 23.4913566307952 | 32.5766291565731 | 23.6613722162112 | 22.5102406621959 | 21.8991639163819 | 20.7923979452556 | 37.4788112600209 | 40.1644458927348 | Gm14399 |
| **ENSMUSG00000090100** | 0.252107684583863 | 0.185458367851454 | 0.805963874499992 | 219.553833126278 | 250.025628776699 | 176.98706417726 | 195.088752405698 | 136.869774477387 | 166.339183562045 | 207.174540020671 | 196.480127205 | Ttbk2 |
| **ENSMUSG00000090110** | 0.126535130884528 | 0.617354454973988 | 0.929242888988805 | 71.3775836089546 | 48.0505280059454 | 47.3227444324223 | 52.523894878457 | 40.5134532453066 | 46.4771248188067 | 63.5057635239243 | 49.9341759747513 | Cmc4 |
| **ENSMUSG00000090112** | 0.236601239048127 | 0.297903126381228 | 0.851785577030345 | 151.7903043836 | 216.634583891211 | 219.57753416644 | 149.130344387048 | 147.819356435578 | 112.523565350795 | 180.106509666212 | 184.539345993646 | Shprh |
| **ENSMUSG00000090115** | 0.0285634486914582 | 0.92463645489707 | 0.986713918026868 | 51.5002818444356 | 44.7928650902881 | 63.4124775394459 | 62.8410885152968 | 40.5134532453066 | 34.2463024980681 | 72.8754663389296 | 69.4736361387845 | Usp49 |
| **ENSMUSG00000090125** | 0.120466932013469 | 0.777291553639089 | 0.95974290923544 | 23.4913566307952 | 29.3189662409158 | 18.9290977729689 | 46.8963347129081 | 26.2789966996583 | 45.2540425867328 | 22.903717992235 | 15.1973579053591 | Pou3f1 |
| **ENSMUSG00000090137** | -0.048845656687572 | 0.82216465168681 | 0.966637987288264 | 5171.71251364121 | 4021.58486937895 | 4769.18618389952 | 5704.47015447814 | 5285.36321121878 | 7102.4385216529 | 3904.04283958551 | 4053.35245847221 | Uba52 |
| **ENSMUSG00000090150** | 0.226313923118273 | 0.278506901842935 | 0.848075053184273 | 4018.82901129911 | 4927.21515993169 | 5390.0605908529 | 3771.40323761207 | 3669.20491418979 | 2662.65001922479 | 4728.57668730597 | 4417.00352263616 | Acad11 |
| **ENSMUSG00000090157** | -0.299907078920242 | 0.368145201666483 | 0.871495599696183 | 63.2459601598332 | 36.6487078011448 | 27.447191770805 | 43.1446279358754 | 35.0386622662111 | 58.7079471395453 | 46.8485140750261 | 69.4736361387845 | Gm16534 |
| **ENSMUSG00000090213** | 0.0954057840253047 | 0.396767272392376 | 0.879613019351548 | 674.021232560508 | 579.863998987002 | 744.859997366328 | 649.045272426648 | 595.657258525589 | 631.110431750112 | 638.180869510912 | 612.236418473038 | Tmem189 |
| **ENSMUSG00000090222** | 0.170624156146032 | 0.66609502214824 | 0.942024383071524 | 20.780815481088 | 21.1748089517725 | 17.9826428843205 | 17.8206071909051 | 8.75966556655277 | 24.4616446414772 | 13.5340151772298 | 22.7960368580387 | Ifi203-ps |
| **ENSMUSG00000090231** | -0.201799732485481 | 0.571511861312305 | 0.921130164032017 | 23.4913566307952 | 31.7622134276588 | 24.6078271048596 | 36.5791410760683 | 28.4689130912965 | 29.3539735697726 | 21.8626399016789 | 54.2762782334254 | Cfb |
| **ENSMUSG00000090247** | -0.108155189231403 | 0.611915266317551 | 0.928055903343287 | 1215.22594878537 | 842.92027942633 | 1042.99328729059 | 1266.20103724852 | 1224.16326292575 | 1578.99916160735 | 920.313032051625 | 984.571687154336 | Bloc1s1 |
| **ENSMUSG00000090257** | -0.939126594467229 | 0.0132963254244602 | 0.635862253526407 | 11.7456783153976 | 11.4018202048006 | 10.4110037751329 | 12.1930470253561 | 16.4243729372865 | 28.1308913376988 | 26.0269522639034 | 17.3684090346961 | Gm4524 |
| **ENSMUSG00000090258** | 0.106212689651634 | 0.481149543051188 | 0.897732059433541 | 2125.06426137039 | 1477.35013225059 | 1766.084822218 | 2038.11470662298 | 1669.81124862412 | 1938.58533783707 | 1529.34371502696 | 1743.35405685762 | Churc1 |
| **ENSMUSG00000090266** | 0.0696328269886051 | 0.625236124689109 | 0.931573470007391 | 885.443442237664 | 660.491156149521 | 725.930899593359 | 831.940977806989 | 708.437952694956 | 689.818378889657 | 698.563398763168 | 859.736247217458 | Mettl23 |
| **ENSMUSG00000090272** | -0.503295438245835 | 0.087050342758412 | 0.747269672026173 | 117.456783153976 | 143.337168288922 | 123.039135524298 | 157.571684635371 | 215.706764576362 | 282.531995609062 | 102.025652874501 | 168.256462523619 | Mndal |
| **ENSMUSG00000090290** | 0.112784220465729 | 0.607718144948268 | 0.927510709653048 | 83.1232619243521 | 106.688460487777 | 70.9841166486335 | 76.9099889291692 | 67.887408140784 | 66.0464405319884 | 91.61487196894 | 86.8420451734806 | Tarbp1 |
| **ENSMUSG00000090330** | -0.215484218396485 | 0.33301719448895 | 0.856918526232803 | 55.1143367107117 | 79.8127414336042 | 72.8770264259304 | 60.9652351267805 | 90.881530252985 | 88.0619207093179 | 62.4646854333682 | 71.6446872681215 | 9130221H12 |
| **ENSMUSG00000090386** | 0.106433248446229 | 0.713065901231781 | 0.950526426881904 | 173.474633581257 | 81.4415728914329 | 73.8234813145789 | 136.937297361692 | 111.685735973548 | 110.077400886647 | 99.9434966933891 | 110.723607596188 | Mir99ahg |
| **ENSMUSG00000090394** | -0.457798508711234 | 0.0494783181839663 | 0.703396849069134 | 362.309000344187 | 245.139134403213 | 232.827902607518 | 256.053987532478 | 464.262275027297 | 445.201932474885 | 334.18606706852 | 262.697186649779 | 4930523C07 |
| **ENSMUSG00000090523** | -0.181652063052044 | 0.265520164580258 | 0.842603400964081 | 374.958192376154 | 364.043830824705 | 531.907647420427 | 418.31530563914 | 411.70428162798 | 492.902139525765 | 490.34778065194 | 519.966745476215 | Gypc |
| **ENSMUSG00000090553** | 0.0419764952601267 | 0.763589659788276 | 0.95785116771606 | 459.888481733644 | 369.744740927105 | 380.474865236676 | 392.99128489417 | 388.710159515779 | 456.20967256355 | 349.802238426862 | 363.65106416395 | Snrpe |
| **ENSMUSG00000090625** | -0.0423723641160136 | 0.876462094532415 | 0.977469019294424 | 61.4389327266951 | 47.2361122770311 | 30.2865564367503 | 43.1446279358754 | 45.9882442244021 | 55.0387004433237 | 43.7252798033577 | 43.4210225867403 | Gm20721 |
| **ENSMUSG00000090641** | 0.47652733352834 | 0.182159480505701 | 0.804505478598291 | 37.0440623793308 | 18.7315617650296 | 28.3936466594534 | 20.6343872736795 | 20.8042057205628 | 15.9000690169602 | 16.6572494488982 | 21.7105112933701 | Zfp712 |
| **ENSMUSG00000090659** | -0.548265542329086 | 0.200895296452392 | 0.812921911423739 | 17.1667606148119 | 10.5874044758863 | 5.67872933189068 | 11.2551203310979 | 15.3294147414674 | 18.3462334811079 | 15.616171358342 | 16.2828834700276 | Zfp493 |
| **ENSMUSG00000090673** | -0.243623432070889 | 0.642293481811524 | 0.934616627390525 | 12.6491920319666 | 13.8450673915436 | 22.7149173275627 | 17.8206071909051 | 8.75966556655277 | 8.56157562451702 | 35.3966550789086 | 26.0526135520442 | Gm340 |
| **ENSMUSG00000090674** | -0.0838323552055586 | 0.86130999978398 | 0.974868041734905 | 9.93865088225949 | 10.5874044758863 | 22.7149173275627 | 8.44134024832345 | 8.75966556655277 | 18.3462334811079 | 12.4929370866736 | 15.1973579053591 | Gm17082 |
| **ENSMUSG00000090698** | -0.496914582568704 | 0.107761349709668 | 0.762287160109383 | 481.572810931301 | 374.631235300591 | 245.131816159948 | 247.612647284155 | 700.773245324222 | 516.140701935169 | 373.747034509653 | 313.716888189199 | Apold1 |
| **ENSMUSG00000090733** | 0.225182622985112 | 0.174541463216255 | 0.799889021902745 | 3485.75591852338 | 2243.71533315898 | 3017.29818501125 | 3242.41258205046 | 2593.95596589544 | 2920.72037019238 | 2226.86603569958 | 2515.16273333693 | Rps27 |
| **ENSMUSG00000090761** | 0.0347757866483065 | 0.878103792539183 | 0.977934951033036 | 145.465708367616 | 128.677685168464 | 112.628131749165 | 198.84045918273 | 179.573144114332 | 135.762127760198 | 130.134761319517 | 125.920965501547 | Gm17201 |
| **ENSMUSG00000090778** | -0.044253450353224 | 0.903123316404018 | 0.985027469232262 | 26.2018977805023 | 16.2883145782866 | 23.6613722162112 | 31.8895076047775 | 14.2344565456483 | 26.9078091056249 | 27.0680303544596 | 32.5657669400552 | Gm3235 |
| **ENSMUSG00000090799** | 0.841020524973883 | 0.0401684128186942 | 0.679566075569698 | 84.0267756409212 | 116.461449234749 | 217.684624389143 | 74.0962088463947 | 71.1722827282413 | 31.8001380339204 | 82.2451691539348 | 89.0130963028176 | Klhl33 |
| **ENSMUSG00000090841** | -0.211063779293292 | 0.374399333600863 | 0.873700364743909 | 1529.64872215139 | 1341.3427055219 | 1451.86179918672 | 1489.42759048196 | 1786.97177557677 | 2490.19542450238 | 1146.22697770231 | 1305.88725429621 | Myl6 |
| **ENSMUSG00000090862** | 0.13325805854171 | 0.361234992322349 | 0.868539734006824 | 5228.63387778506 | 3578.54271284956 | 4560.01965350822 | 5090.12816973904 | 4124.70752365054 | 4646.48939964859 | 3950.89135366054 | 4106.54321114096 | Rps13 |
| **ENSMUSG00000090877** | 0.400050750932048 | 0.2566011476856 | 0.837517407767513 | 116.553269437407 | 206.047179415325 | 112.628131749165 | 75.0341355406529 | 78.836990098975 | 151.662196777159 | 77.0397787011541 | 80.3288917854696 | Hspa1b |
| **ENSMUSG00000090935** | 0.0830637162446152 | 0.56089483788857 | 0.916762264997884 | 1925.38773000863 | 1723.30368238272 | 1587.20484826345 | 1715.46792379818 | 1443.15490208957 | 1395.53682679627 | 1849.99576691825 | 1872.53159905318 | Synj2bp |
| **ENSMUSG00000090942** | 0.430227149817625 | 0.398134999321906 | 0.880459952732461 | 38.8510898124689 | 136.821842457607 | 101.270673085384 | 46.8963347129081 | 25.1840385038392 | 36.6924669622158 | 94.7381062406084 | 83.5854684794751 | F830016B08 |
| **ENSMUSG00000090946** | -0.108490486042223 | 0.315674285916258 | 0.853277176422981 | 320.747369382011 | 359.157336451219 | 326.526936583714 | 322.646782824807 | 362.431162816121 | 338.793778284459 | 381.034581143546 | 350.624757387928 | Ccdc71l |
| **ENSMUSG00000090952** | 0.241452589555594 | 0.243894435952522 | 0.832048466795696 | 96.6759676728878 | 105.059629029948 | 146.700507740509 | 127.55803041911 | 117.160526952643 | 108.854318654574 | 84.3273253350471 | 92.2696729968231 | Gm17251 |
| **ENSMUSG00000090958** | -1.06112405457671 | 0.00462701263446737 | 0.635862253526407 | 121.070838020252 | 84.6992358070902 | 140.07532351997 | 130.371810501884 | 374.475702970131 | 344.909189444828 | 161.367104036201 | 112.894658725525 | Lrrc32 |
| **ENSMUSG00000091086** | 0.0202556441541044 | 0.962360659553588 | 0.994128583764339 | 17.1667606148119 | 10.5874044758863 | 17.9826428843205 | 12.1930470253561 | 12.0445401540101 | 19.5693157131818 | 8.32862472444909 | 17.3684090346961 | Rpl6l |
| **ENSMUSG00000091089** | 1.04199998043141 | 0.0307307578401368 | 0.667184591665619 | 130.105975185942 | 189.758864837039 | 488.370722542599 | 180.081925297567 | 99.6411958195378 | 46.4771248188067 | 204.051305749003 | 129.177542195552 | NA |
| **ENSMUSG00000091119** | -0.199797569783886 | 0.625662422206667 | 0.931573470007391 | 19.877301764519 | 12.2162359337149 | 14.1968233297267 | 19.6964605794214 | 24.0890803080201 | 8.56157562451702 | 21.8626399016789 | 20.6249857287016 | Ccdc152 |
| **ENSMUSG00000091154** | 0.155719834776403 | 0.497821199526008 | 0.900927059989498 | 108.421645988285 | 102.616381843205 | 123.985590412947 | 156.633757941113 | 144.534481848121 | 112.523565350795 | 85.3684034256032 | 98.7828263848342 | Proscos |
| **ENSMUSG00000091243** | -0.39652606403924 | 0.305554178072447 | 0.851785577030345 | 15.3597331816738 | 18.7315617650296 | 41.6440151005317 | 41.2687745473591 | 45.9882442244021 | 42.8078781225851 | 33.3144988977964 | 31.4802413753867 | Vgll3 |
| **ENSMUSG00000091264** | 0.0959175242706671 | 0.70853743265795 | 0.949174314815223 | 884.539928521095 | 981.370953341766 | 973.902080419252 | 793.485983342404 | 659.164833883096 | 516.140701935169 | 1141.02158724953 | 1082.2689879745 | Smim13 |
| **ENSMUSG00000091277** | 0.200726574055291 | 0.346877149925132 | 0.861266047293794 | 65.9565013095403 | 70.8541684155466 | 63.4124775394459 | 76.9099889291692 | 61.3176589658694 | 44.0309603546589 | 68.711153976705 | 66.217059444779 | Gm1818 |
| **ENSMUSG00000091337** | -0.0370588424403313 | 0.670822946725318 | 0.942994710051074 | 988.444005926535 | 954.495234287593 | 966.330441310064 | 1072.05021153708 | 973.417836083177 | 1064.08154190426 | 1043.16024673725 | 1004.11114731837 | Eid1 |
| **ENSMUSG00000091421** | -0.437013939132474 | 0.317554869960448 | 0.854351922245337 | 9.03513716569045 | 12.2162359337149 | 16.0897331070236 | 7.50341355406529 | 19.7092475247437 | 11.0077400886647 | 12.4929370866736 | 17.3684090346961 | Gm4202 |
| **ENSMUSG00000091474** | -0.0571538593349504 | 0.786881966001621 | 0.961141387070613 | 85.8338030740593 | 94.4722245540621 | 81.3951204237664 | 95.6685228143324 | 63.5075753575076 | 97.8465785659088 | 100.984574783945 | 109.638082031519 | 2610021A01 |
| **ENSMUSG00000091478** | 0.0140968937341648 | 0.947962703446495 | 0.991192530276814 | 139.141112351633 | 92.0289773673191 | 91.8061241988993 | 144.440710915757 | 127.015150715015 | 118.638976511164 | 104.107809055614 | 112.894658725525 | Gm10039 |
| **ENSMUSG00000091498** | 0.170896596722761 | 0.172695107793967 | 0.799889021902745 | 1273.05082664578 | 1026.97823416097 | 1088.42312194571 | 1311.22151857291 | 966.848086908263 | 1028.61215717412 | 1023.37976301668 | 1154.99920080729 | Mpc1-ps |
| **ENSMUSG00000091509** | -0.428775920715113 | 0.265295176325092 | 0.842510961779757 | 51.5002818444356 | 25.2468875963442 | 30.2865564367503 | 52.523894878457 | 63.5075753575076 | 79.5003450848009 | 42.6842017128016 | 29.3091902460497 | Gm17066 |
| **ENSMUSG00000091512** | 0.106197527930565 | 0.463386559726366 | 0.895328112556867 | 1494.4116872052 | 1318.5390651123 | 1534.20337449913 | 1301.84225163033 | 1240.58763586304 | 1061.63537744011 | 1421.07159360913 | 1522.99236722992 | Lamtor3 |
| **ENSMUSG00000091514** | -0.261916745341433 | 0.654302802732225 | 0.937269843859626 | 11.7456783153976 | 19.5459774939439 | 12.3039135524298 | 2.81378008277448 | 15.3294147414674 | 11.0077400886647 | 21.8626399016789 | 7.59867895267955 | Gm17484 |
| **ENSMUSG00000091515** | -0.0894568970519639 | 0.779557431658782 | 0.960444627083598 | 32.5264937964856 | 50.4937751926884 | 36.9117406572894 | 27.1998741334867 | 30.6588294829347 | 28.1308913376988 | 51.0128264372507 | 46.6775992807458 | Zfp422-ps |
| **ENSMUSG00000091537** | 0.143135270371649 | 0.318848208622433 | 0.854670142081901 | 2511.76813206195 | 1891.07332253907 | 1968.62616838877 | 2529.58829441426 | 1965.44996149528 | 2233.34815576687 | 1814.59911183935 | 2047.30121496481 | Tma7 |
| **ENSMUSG00000091549** | -0.282570606609523 | 0.213588220150056 | 0.824248286808591 | 74.0881247586617 | 61.895595397489 | 62.4660226507975 | 71.2824287636203 | 111.685735973548 | 64.8233582999146 | 72.8754663389296 | 78.1578406561325 | Gm6548 |
| **ENSMUSG00000091562** | 0.437294230824581 | 0.568598780037004 | 0.92059532906293 | 93.0619128066116 | 302.96265115613 | 312.330113253987 | 81.59962240046 | 65.6974917491458 | 37.9155491942896 | 259.228444548478 | 220.361689627707 | Crybg3 |
| **ENSMUSG00000091586** | 0.588552613755782 | 0.0375078700681719 | 0.679566075569698 | 44.2721721118832 | 36.6487078011448 | 65.3053873167428 | 48.7721881014244 | 30.6588294829347 | 28.1308913376988 | 30.191264626128 | 40.1644458927348 | Cyp4f17 |
| **ENSMUSG00000091619** | -0.0478142504187343 | 0.896902857845647 | 0.984332015045905 | 13.5527057485357 | 18.7315617650296 | 16.0897331070236 | 16.8826804966469 | 18.6142893289246 | 18.3462334811079 | 19.7804837205666 | 10.8552556466851 | Gm3625 |
| **ENSMUSG00000091625** | 0.135488699791095 | 0.531395170563111 | 0.909082872831377 | 112.939214571131 | 96.9154717408051 | 76.6628459805242 | 107.861569839689 | 82.1218646864323 | 117.415894279091 | 78.0808567917102 | 82.4999429148066 | Lsm5 |
| **ENSMUSG00000091649** | -0.690973910451733 | 0.229850543732309 | 0.832048466795696 | 7.22810973255236 | 9.77298874697194 | 7.57163910918757 | 7.50341355406529 | 17.5193311331055 | 20.7923979452556 | 3.12323427166841 | 10.8552556466851 | Phf11b |
| **ENSMUSG00000091694** | 1.8497419765159 | 0.229718596365325 | 0.832048466795696 | 26.2018977805023 | 8.95857301805762 | 246.078271048596 | 1.87585338851632 | 12.0445401540101 | 62.3771938357668 | 2.08215618111227 | 2.17105112933702 | Apol11b |
| **ENSMUSG00000091764** | 0.00836626925388612 | 0.979757586272077 | 0.996872525500296 | 29.8159526467785 | 35.8342920722305 | 17.9826428843205 | 31.8895076047775 | 35.0386622662111 | 25.6847268735511 | 20.8215618111227 | 33.6512925047237 | Zfp964 |
| **ENSMUSG00000091803** | 0.0489612330641537 | 0.781918796767591 | 0.960742105865155 | 504.160653845527 | 355.899673535562 | 344.509579468035 | 462.397860269274 | 388.710159515779 | 443.978850242811 | 339.391457521301 | 439.637853690746 | Cox16 |
| **ENSMUSG00000091811** | -0.0113755453053664 | 0.939338168021024 | 0.989269716906553 | 505.064167562096 | 469.917875583568 | 517.7108240907 | 624.659178375935 | 589.087509350674 | 595.64104701997 | 488.265624470828 | 461.348364984116 | Inafm1 |
| **ENSMUSG00000091896** | 0.0778155302657562 | 0.337408668874674 | 0.858570960161243 | 1751.91309642738 | 1656.52159261174 | 1736.7447206699 | 1781.12279239625 | 1563.60030362967 | 1564.32217482247 | 1699.03944378762 | 1733.58432677561 | Ube2d2a |
| **ENSMUSG00000091898** | 0.0746991980798175 | 0.690742410797777 | 0.94469184542218 | 66328.7489607667 | 49453.7663068648 | 41424.4375663652 | 68354.2216241463 | 55708.1881286882 | 60845.8948812104 | 49933.228457344 | 47692.5656837109 | Tnnc1 |
| **ENSMUSG00000091931** | -0.12938408811204 | 0.566823908350014 | 0.919931558693298 | 119.263810587114 | 70.8541684155466 | 83.2880302010633 | 100.358156285623 | 93.0714466446232 | 125.977469903608 | 102.025652874501 | 87.9275707381491 | Gon7 |
| **ENSMUSG00000091937** | 0.494341965988578 | 0.0676913526032954 | 0.722112580225498 | 71.3775836089546 | 72.4829998733752 | 47.3227444324223 | 89.1030359545253 | 44.893286028583 | 37.9155491942896 | 62.4646854333682 | 53.1907526687569 | Gm3646 |
| **ENSMUSG00000091957** | -0.439963860852681 | 0.0811910233946544 | 0.739855948845396 | 28.9124389302094 | 29.3189662409158 | 27.447191770805 | 39.3929211588428 | 44.893286028583 | 42.8078781225851 | 41.6431236222455 | 40.1644458927348 | Rps2-ps10 |
| **ENSMUSG00000091971** | 0.546519246602579 | 0.101000603255066 | 0.759800953500741 | 335.203588847116 | 552.173864203915 | 352.081218577222 | 224.164479927701 | 248.555510450935 | 393.832478727783 | 164.49033830787 | 196.480127205 | Hspa1a |
| **ENSMUSG00000091985** | 0.22307936560256 | 0.380449783344894 | 0.87432013076003 | 46.0791995450213 | 53.7514381083457 | 43.5369248778286 | 60.9652351267805 | 37.2285786578493 | 44.0309603546589 | 36.4377331694648 | 57.5328549274309 | Gm17354 |
| **ENSMUSG00000091994** | 0.225475134336578 | 0.637388359799843 | 0.932931968211785 | 29.8159526467785 | 18.7315617650296 | 25.5542819935081 | 12.1930470253561 | 7.66470737073368 | 19.5693157131818 | 31.2323427166841 | 15.1973579053591 | E130317F20 |

| | | | | | | | | | | | | |
|---|---|---|---|---|---|---|---|---|---|---|---|---|
| **ENSMUSG00000092014** | -1.14900139847145 | 0.029461538989811 | 0.665956123973759 | 4.51756858284523 | 4.88649437348597 | 9.46454888648447 | 7.50341355406529 | 13.1394983498292 | 7.33849339244316 | 14.5750932677859 | 22.7960368580387 | Gm4468 |
| **ENSMUSG00000092035** | -0.429046187819229 | 0.310867571176213 | 0.851785577030345 | 19.877301764519 | 12.2162359337149 | 20.8220075502658 | 12.1930470253561 | 26.2789966996583 | 33.0232202659942 | 15.616171358342 | 13.0263067760221 | Peg10 |
| **ENSMUSG00000092060** | 0.115704298181569 | 0.743836457914251 | 0.954466085361166 | 27.1054114970714 | 24.4324718674299 | 25.5542819935081 | 46.8963347129081 | 35.0386622662111 | 18.3462334811079 | 35.3966550789086 | 24.9670879873757 | Bend4 |
| **ENSMUSG00000092071** | 0.142454855190431 | 0.764205674446578 | 0.958041307101859 | 32.5264937964856 | 14.6594831204579 | 10.4110037751329 | 17.8206071909051 | 10.949581958191 | 25.6847268735511 | 12.4929370866736 | 19.5394601640331 | A230065N10 |
| **ENSMUSG00000092118** | -0.923293594401249 | 0.0057751224462826 | 0.635862253526407 | 9.93865088225949 | 21.1748089517725 | 17.036187995672 | 17.8206071909051 | 36.1336204620302 | 34.2463024980681 | 27.0680303544596 | 28.2236646813812 | Fancf |
| **ENSMUSG00000092124** | 0.132984614839356 | 0.679089246726712 | 0.944452327903597 | 28.0089252136404 | 26.0613033252585 | 24.6078271048596 | 23.448167356454 | 16.4243729372865 | 29.3539735697726 | 29.1501865355718 | 18.4539345993646 | B930094E09 |
| **ENSMUSG00000092187** | 0.239315021074347 | 0.572227450711433 | 0.921574199900435 | 73.1846110420926 | 26.0613033252585 | 33.1259211026956 | 52.523894878457 | 24.0890803080201 | 30.5770558018465 | 64.5468416144805 | 36.9078691987293 | Gm20457 |
| **ENSMUSG00000092201** | -0.236143098388555 | 0.36902626014353 | 0.871495599696183 | 474.344701198749 | 420.238516119794 | 230.934992830221 | 400.494698448235 | 294.543754675337 | 478.225152740879 | 500.758561557502 | 524.308847734889 | A530058N18 |
| **ENSMUSG00000092203** | 0.106701371913082 | 0.774718440572176 | 0.959414024438738 | 29.8159526467785 | 28.5045505120015 | 29.3401015481019 | 61.9031618210386 | 41.6084114411257 | 45.2540425867328 | 22.903717992235 | 29.3091902460497 | 1110038B12 |
| **ENSMUSG00000092233** | 0.344579681662172 | 0.531988296624126 | 0.909082872831377 | 6.32459601598331 | 14.6594831204579 | 15.1432782183751 | 16.8826804966469 | 6.56974917491458 | 9.78465785659088 | 19.7804837205666 | 5.42762782334254 | Gm20403 |
| **ENSMUSG00000092241** | -0.271696470765145 | 0.384933926682173 | 0.875126835008975 | 91.2548853734735 | 34.2054606144018 | 62.4660226507975 | 52.523894878457 | 63.5075753575076 | 91.7311674055395 | 67.6700758861489 | 67.3025850094475 | Gm20522 |
| **ENSMUSG00000092260** | -0.441973707384013 | 0.157288515898923 | 0.788998730890588 | 20.780815481088 | 19.5459774939439 | 34.0723759913441 | 31.8895076047775 | 35.0386622662111 | 46.4771248188067 | 30.191264626128 | 32.5657669400552 | Zfp963 |
| **ENSMUSG00000092274** | 0.171589550189619 | 0.527851590871186 | 0.908232038364594 | 1082.40943244972 | 771.251695281869 | 856.541674226844 | 1379.69016725376 | 692.013579757669 | 746.080161565054 | 1387.75709471133 | 804.374443419364 | Neat1 |
| **ENSMUSG00000092335** | 0.542423422863161 | 0.26182778401652 | 0.841137531367976 | 13.5527057485357 | 9.77298874697194 | 15.1432782183751 | 22.5102406621959 | 6.56974917491458 | 7.33849339244316 | 16.6572494488982 | 10.8552556466851 | Zfp977 |
| **ENSMUSG00000092341** | 0.153540655463959 | 0.56363364857431 | 0.918571155207735 | 4054.96955996187 | 2651.73761334505 | 2348.1545787368 | 4570.51678112002 | 3152.38464576318 | 4180.49506922845 | 2336.17923520797 | 2581.37979278171 | Malat1 |
| **ENSMUSG00000092397** | -0.13146530439932 | 0.701132191866168 | 0.945920819861844 | 1126.6816045616 | 1445.58791882293 | 905.757328436564 | 891.968286239511 | 671.209374037106 | 699.603036746248 | 1711.53238087429 | 1704.27513652956 | C130080G10 |
| **ENSMUSG00000092416** | -0.281153906604025 | 0.29663832033636 | 0.851785577030345 | 34.3335212296237 | 47.2361122770311 | 56.7872933189068 | 49.7101147956825 | 41.6084114411257 | 75.8310983885793 | 58.3003730711437 | 53.1907526687569 | Zfp141 |
| **ENSMUSG00000092417** | -0.13197716218424 | 0.485216118799001 | 0.899236955296196 | 237.624107457659 | 219.892246806869 | 189.290977729689 | 246.674720589896 | 285.784089108784 | 291.093571233579 | 189.476212481217 | 213.848536239696 | Gpank1 |
| **ENSMUSG00000092545** | -0.486685131532854 | 0.439807386579088 | 0.890124522742755 | 14.4562194651047 | 13.0306516626293 | 3.78581955459379 | 18.7585338851632 | 6.56974917491458 | 19.5693157131818 | 9.36970281500523 | 34.7368180693922 | Gm20319 |
| **ENSMUSG00000092558** | 0.145364586162611 | 0.49449429450851 | 0.900199967329385 | 149.079763233892 | 153.924572764808 | 201.594891282119 | 220.412773150668 | 152.199189218854 | 123.53130543946 | 190.517290571773 | 187.795922687652 | Med20 |
| **ENSMUSG00000092564** | -0.00959443774561892 | 0.978284480482631 | 0.996580229842593 | 26.2018977805023 | 23.6180561385155 | 17.036187995672 | 21.5723139679377 | 26.2789966996583 | 26.9078091056249 | 12.4929370866736 | 23.8815624227072 | BC051226 |
| **ENSMUSG00000092569** | 0.0305592626981028 | 0.915906039392903 | 0.986713918026868 | 70.4740698923855 | 96.1010560118908 | 79.5022106464695 | 65.6548685980713 | 42.7033696369448 | 78.277262852727 | 107.231043327282 | 77.072315091464 | Gm20544 |
| **ENSMUSG00000092595** | 0.126275995244195 | 0.664888993044986 | 0.941513723457524 | 48.7897406947284 | 40.7207864457164 | 31.2330113253987 | 47.8342614071662 | 54.7479097909548 | 29.3539735697726 | 36.4377331694648 | 33.6512925047237 | Gm20427 |
| **ENSMUSG00000092607** | -0.0314324669216436 | 0.861799540770444 | 0.974868041734905 | 575.538237454482 | 404.764617270421 | 403.189782564238 | 493.349441179793 | 502.585811880965 | 580.964060235083 | 415.390158131898 | 420.098393526712 | Scnm1 |
| **ENSMUSG00000093402** | 0.415641761787936 | 0.287833430664933 | 0.851785577030345 | 15.3597331816738 | 21.1748089517725 | 22.7149173275627 | 14.0689004138724 | 14.2344565456483 | 8.56157562451702 | 15.616171358342 | 16.2828834700276 | Gm18588 |
| **ENSMUSG00000093424** | 0.398032490013502 | 0.34299326977851 | 0.860892034192898 | 14.4562194651047 | 15.4738988493722 | 11.3574586637814 | 21.5723139679377 | 14.2344565456483 | 13.4539045528125 | 12.4929370866736 | 7.59867895267955 | 6330562C20 |
| **ENSMUSG00000093482** | -0.131993614476744 | 0.497284385799097 | 0.900927059989498 | 115.649755720838 | 93.6578088251478 | 106.949402417274 | 75.9720622349111 | 95.2613630362614 | 100.292743030056 | 115.559668051731 | 118.322286548867 | Gm20619 |
| **ENSMUSG00000093483** | 0.213042212856348 | 0.6978885373486 | 0.944933386973863 | 13.5527057485357 | 11.4018202048006 | 8.51809399783602 | 12.1930470253561 | 6.56974917491458 | 3.66924669622158 | 9.36970281500523 | 19.5394601640331 | AA465934 |
| **ENSMUSG00000093577** | 0.164290166532843 | 0.538035647471302 | 0.912492691498947 | 83.1232619243521 | 101.801966114291 | 101.270673085384 | 90.0409626487835 | 74.4571573156986 | 57.4848649074714 | 129.093683228961 | 73.8157383974585 | Gm20632 |
| **ENSMUSG00000093661** | 0.144459245939986 | 0.171311596257086 | 0.799531027639052 | 2860.5244266576 | 2609.38799544151 | 2521.35582335946 | 2852.23507723907 | 2326.78616611558 | 2210.10959335746 | 2583.95582076033 | 2687.76129811922 | Eif4e3 |
| **ENSMUSG00000093674** | 0.234784929329587 | 0.148352706379096 | 0.781357001591751 | 5155.44926674297 | 3610.30492627722 | 4272.29736735909 | 4959.75635923716 | 3658.2553322316 | 4658.72022196933 | 3466.79004155193 | 3511.67520170262 | Rpl41 |
| **ENSMUSG00000093677** | -0.214966597665752 | 0.399418032301117 | 0.880459952732461 | 45.1756858284523 | 28.5045505120015 | 34.0723759913441 | 41.2687745473591 | 48.1781606160403 | 40.3617136584374 | 43.7252798033577 | 40.1644458927348 | Gm20712 |
| **ENSMUSG00000093798** | 0.16032566689217 | 0.477426895416587 | 0.897674307047416 | 67.7635287426784 | 67.5965054998893 | 80.448665535118 | 87.227182566009 | 63.5075753575076 | 84.3926740130963 | 49.9717483466946 | 73.8157383974585 | Gm8355 |
| **ENSMUSG00000093803** | 0.0511384343185107 | 0.667164006284449 | 0.942024383071524 | 554.757421973394 | 576.606336071345 | 537.586376752318 | 612.466131350579 | 597.847174917227 | 588.302553627526 | 545.524919451416 | 471.118095066132 | Ppp2r3d |
| **ENSMUSG00000093826** | -0.00418102042010809 | 0.989159126352852 | 0.998437672973644 | 21.6843291976571 | 26.0613033252585 | 26.5007368821565 | 19.6964605794214 | 27.3739548954774 | 25.6847268735511 | 19.7804837205666 | 21.7105112933701 | Gm6900 |
| **ENSMUSG00000093904** | 0.13474563629458 | 0.305922270463195 | 0.851785577030345 | 2557.84733160697 | 1990.43204146662 | 2393.58441339192 | 2630.88437739414 | 2196.48614081311 | 1894.55437748241 | 2250.81083178237 | 2375.12993549469 | Tomm20 |
| **ENSMUSG00000093930** | 0.0602565535472411 | 0.773956466257865 | 0.959203117135841 | 627.038519298917 | 733.78857175181 | 709.841166486335 | 528.990655561603 | 508.060602860061 | 465.99433042014 | 740.206522385413 | 778.32182986732 | Hmgcs1 |
| **ENSMUSG00000093954** | 0.167533910272333 | 0.772760171579391 | 0.959045638244974 | 20.780815481088 | 8.14415728914329 | 19.8755526616174 | 6.56548685980713 | 4.37983278327639 | 19.5693157131818 | 14.5750932677859 | 10.8552556466851 | Gm16867 |
| **ENSMUSG00000093962** | 0.372268665401809 | 0.503113153831458 | 0.900927059989498 | 15.3597331816738 | 8.14415728914329 | 26.5007368821565 | 20.6343872736795 | 17.5193311331055 | 4.89232892829544 | 22.903717992235 | 8.68420451734806 | Gm11756 |
| **ENSMUSG00000093989** | -0.054588809570292 | 0.77201606970636 | 0.959045638244974 | 87.6408305071974 | 92.0289773673191 | 94.6454888648447 | 119.116690170786 | 105.115986798633 | 113.746647582869 | 106.189965236726 | 83.5854684794751 | Rnasek |
| **ENSMUSG00000094114** | 0.161522096873962 | 0.339201416373344 | 0.858570960161243 | 178.895715880671 | 191.387696294867 | 222.416898832385 | 175.392291826276 | 210.231973597267 | 160.223772401676 | 146.792010768415 | 169.341988088287 | NA |
| **ENSMUSG00000094320** | 0.678953375353324 | 0.161586313243226 | 0.792786631432503 | 13.5527057485357 | 13.8450673915436 | 10.4110037751329 | 10.3171936368398 | 6.56974917491458 | 12.2308223207386 | 6.24646854333682 | 5.42762782334254 | Chchd2-ps |
| **ENSMUSG00000094410** | 0.0102572712030239 | 0.984696917115702 | 0.99711600049115 | 26.2018977805023 | 36.6487078011448 | 16.0897331070236 | 10.3171936368398 | 18.6142893289246 | 7.33849339244316 | 34.3555769883525 | 28.2236646813812 | Gm38394 |
| **ENSMUSG00000094441** | 0.238788484303698 | 0.579749803735992 | 0.922450244597442 | 91.2548853734735 | 101.801966114291 | 133.450139299431 | 82.5375490947182 | 61.3176589658694 | 26.9078091056249 | 142.627698406191 | 115.065709854862 | Zfp955a |
| **ENSMUSG00000094483** | 0.0569477140455281 | 0.441618865265318 | 0.89108031640712 | 2103.37993217274 | 2010.79243468948 | 1962.94743905688 | 2139.41078960287 | 2036.62224422352 | 1947.14691346158 | 1905.17290571773 | 2009.30782020141 | Purb |
| **ENSMUSG00000094518** | 0.113536458067347 | 0.74789990448135 | 0.956197192463153 | 18.0702743313809 | 19.5459774939439 | 17.9826428843205 | 22.5102406621959 | 13.1394983498292 | 20.7923979452556 | 14.5750932677859 | 23.8815624227072 | NA |
| **ENSMUSG00000094530** | -0.307370805490969 | 0.50087440474782 | 0.900927059989498 | 14.4562194651047 | 8.14415728914329 | 9.46454888648447 | 13.1309737196143 | 18.6142893289246 | 17.123151249034 | 7.28754663389296 | 13.0263067760221 | Gm21399 |
| **ENSMUSG00000094685** | -0.0656316381536465 | 0.864457737992129 | 0.974868041734905 | 16.2632468982428 | 10.5874044758863 | 13.2503684410783 | 15.0068271081306 | 13.1394983498292 | 13.4539045528125 | 15.616171358342 | 15.1973579053591 | Gm5900 |
| **ENSMUSG00000094724** | 0.148186463419611 | 0.144593733089733 | 0.780607990324772 | 871.890736489128 | 784.282346944498 | 936.043884873314 | 858.202925246218 | 812.45898129777 | 757.087901653719 | 727.71358529874 | 815.229699066049 | Rnaset2b |
| **ENSMUSG00000094786** | 0.240466715947653 | 0.434978709274326 | 0.888379878975956 | 200.580045078328 | 215.820168162297 | 175.094154399963 | 151.006197775564 | 135.774816281568 | 75.8310983885793 | 219.667477107345 | 196.480127205 | Gm14403 |
| **ENSMUSG00000094806** | -0.432853361486082 | 0.139432090488553 | 0.779711473022872 | 128.298947752804 | 198.717437855096 | 135.343049076728 | 163.19924480092 | 160.958854785407 | 176.123841418636 | 161.367104036201 | 346.282655129254 | Cyp2d10 |
| **ENSMUSG00000094870** | 0.120916072924288 | 0.516165113793946 | 0.904444737050323 | 475.248214915318 | 448.743066631795 | 395.618143455051 | 387.363724728621 | 359.146288228664 | 284.978160073209 | 465.361906478593 | 459.177313854779 | Zfp131 |
| **ENSMUSG00000094936** | 0.101850564132954 | 0.718418138141461 | 0.951674101666722 | 30.7194663633475 | 58.6379324818317 | 40.6975602118832 | 36.5791410760683 | 42.7033696369448 | 35.4693847301419 | 41.6431236222455 | 35.8223436340608 | Rbm4 |
| **ENSMUSG00000094942** | 0.060824046597089 | 0.902030206108642 | 0.984871597499125 | 17.1667606148119 | 26.0613033252585 | 9.46454888648447 | 12.1930470253561 | 25.1840385038392 | 8.56157562451702 | 17.6983275394543 | 10.8552556466851 | Gm3604 |
| **ENSMUSG00000095041** | -0.169529614161437 | 0.531962236865996 | 0.909082872831377 | 78.6056933415069 | 65.9676740420606 | 93.6990339761962 | 102.23400967414 | 74.4571573156986 | 139.43137445642 | 70.7933101578173 | 98.7828263848342 | AC149090.1 |
| **ENSMUSG00000095098** | -0.388565940494395 | 0.191996519078876 | 0.808824684419004 | 544.818771091134 | 400.69253862585 | 553.676109859341 | 738.148308381173 | 904.435469746574 | 1045.73530842315 | 511.169342463063 | 467.861518372127 | Ccdc85b |
| **ENSMUSG00000095115** | -0.239258240447784 | 0.0972505926660086 | 0.757809915356977 | 737.267192720341 | 888.527560245533 | 788.396922244156 | 700.631240610846 | 1100.43298679819 | 929.542496376133 | 807.876598271562 | 840.196787053425 | Itpripl2 |
| **ENSMUSG00000095180** | -0.222855606081485 | 0.709755506864633 | 0.949571160524938 | 11.7456783153976 | 6.51532583131463 | 16.0897331070236 | 21.5723139679377 | 30.6588294829347 | 12.2308223207386 | 17.6983275394543 | 4.34210225867403 | Rhox5 |
| **ENSMUSG00000095253** | -0.166645660381509 | 0.498409173023128 | 0.900927059989498 | 115.649755720838 | 105.874044758863 | 109.78876708322 | 109.737423228205 | 97.4512794278996 | 92.9542496376133 | 186.352978209548 | 117.236760984199 | Zfp799 |
| **ENSMUSG00000095325** | 0.00987779083075919 | 0.960571309431176 | 0.993694458032251 | 198.77301764519 | 160.439898596123 | 185.505158175096 | 136.937297361692 | 147.819356435578 | 135.762127760198 | 188.435134390661 | 204.078806157679 | Zfp870 |
| **ENSMUSG00000095332** | -0.00836180881805989 | 0.98777852731649 | 0.998437672973644 | 13.5527057485357 | 8.14415728914329 | 10.4110037751329 | 9.37926694258161 | 3.28487458745729 | 7.33849339244316 | 15.616171358342 | 15.1973579053591 | Gm9821 |
| **ENSMUSG00000095362** | 0.189398010560088 | 0.414653020817807 | 0.882192126163777 | 242.141676040504 | 230.479651282755 | 211.059440168604 | 245.736793895638 | 157.67398019795 | 139.43137445642 | 248.817663642917 | 268.124814473121 | Gm14325 |
| **ENSMUSG00000095403** | 0.387973885497406 | 0.408987690572555 | 0.882192126163777 | 9.93865088225949 | 20.3603932228582 | 12.3039135524298 | 20.6343872736795 | 14.2344565456483 | 9.78465785659088 | 17.6983275394543 | 6.51315338801105 | Gm21092 |
| **ENSMUSG00000095427** | 0.203299144367429 | 0.62907676094723 | 0.931825800800958 | 11.7456783153976 | 15.4738988493722 | 13.2503684410783 | 21.5723139679377 | 16.4243729372865 | 7.33849339244316 | 15.616171358342 | 14.1118323406906 | Rps2-ps6 |
| **ENSMUSG00000095432** | -0.23914200136486 | 0.293352430299692 | 0.851785577030345 | 58.7283915769879 | 54.56585383726 | 48.2691993210708 | 71.2824287636203 | 68.9823663366031 | 64.8233582999146 | 56.2182168900314 | 84.6709940441436 | Zfp748 |
| **ENSMUSG00000095438** | -0.0875289816008629 | 0.735873625319801 | 0.954383574136061 | 60.535419010126 | 57.8235167529173 | 34.0723759913441 | 55.3376749612315 | 44.893286028583 | 64.8233582999146 | 51.0128264372507 | 60.7894316214364 | Mir133a-1hg |
| **ENSMUSG00000095440** | 0.0838411520727398 | 0.831749648621435 | 0.967583158900775 | 24.3948703473642 | 26.0613033252585 | 23.6613722162112 | 22.5102406621959 | 39.4184950494875 | 23.2385624094033 | 12.4929370866736 | 16.2828834700276 | Fignl2 |
| **ENSMUSG00000095478** | -0.486411426878643 | 0.238427593389814 | 0.832048466795696 | 12.6491920319666 | 13.0306516626293 | 13.2503684410783 | 12.1930470253561 | 12.0445401540101 | 29.3539735697726 | 16.6572494488982 | 14.1118323406906 | Gm9824 |
| **ENSMUSG00000095567** | 0.156217368094328 | 0.0867885681550724 | 0.747269672026173 | 1061.62861696863 | 1083.98733518497 | 1118.70967838246 | 1200.54616865045 | 1028.16574587413 | 923.427085215764 | 1035.87270010336 | 1017.13745409439 | Noc2l |
| **ENSMUSG00000095588** | 0.14745346848901 | 0.636717916331248 | 0.932878870437987 | 32.5264937964856 | 26.0613033252585 | 27.447191770805 | 19.6964605794214 | 22.994122112201 | 26.9078091056249 | 18.7394056300105 | 27.1381391167127 | Gm12350 |
| **ENSMUSG00000095597** | -0.307916711006614 | 0.414916583687531 | 0.882192126163777 | 29.8159526467785 | 17.1027303072009 | 16.0897331070236 | 34.703287687552 | 35.0386622662111 | 40.3617136584374 | 21.8626399016789 | 23.8815624227072 | Gm6472 |
| **ENSMUSG00000095654** | 0.566470585753168 | 0.0256450499898981 | 0.663332857280604 | 38.8510898124689 | 43.1640336324594 | 47.3227444324223 | 41.2687745473591 | 33.943704070392 | 28.1308913376988 | 27.0680303544596 | 26.0526135520442 | Plscr5 |
| **ENSMUSG00000095677** | -0.284674109698473 | 0.474205784818559 | 0.897549741595057 | 27.1054114970714 | 14.6594831204579 | 29.3401015481019 | 24.3860940507122 | 29.5638712871156 | 47.7002070508805 | 18.7394056300105 | 20.6249857287016 | Dynlt1f |
| **ENSMUSG00000095687** | 0.122343270064982 | 0.249145308085358 | 0.835314899646652 | 910.741826301597 | 808.714818811928 | 986.205993971682 | 877.899385825639 | 875.966556655277 | 845.149822363037 | 764.151318468204 | 806.545494548701 | Rnaset2a |
| **ENSMUSG00000095690** | 0.276531788876118 | 0.419120058769722 | 0.882550938324998 | 25.2983840639333 | 30.1333819698302 | 42.5904699891801 | 24.3860940507122 | 22.994122112201 | 15.9000690169602 | 33.3144988977964 | 28.2236646813812 | Rab11b-ps2 |
| **ENSMUSG00000095742** | -0.388096474506278 | 0.241098561931519 | 0.832048466795696 | 121.974351736821 | 64.338842584232 | 81.3951204237664 | 104.109863062656 | 148.914314631397 | 174.900759186562 | 94.7381062406084 | 68.388110574116 | CAAA011473 |
| **ENSMUSG00000095762** | 0.0395025289407124 | 0.887732210524518 | 0.980702840906172 | 30.7194663633475 | 35.8342920722305 | 34.0723759913441 | 43.1446279358754 | 42.7033696369448 | 23.2385624094033 | 37.4788112600209 | 35.8223436340608 | Gm16378 |
| **ENSMUSG00000095930** | 0.14777104241006 | 0.399956213960946 | 0.880487015130533 | 114.746242004269 | 107.502876216691 | 140.07532351997 | 135.999370667433 | 131.394983498292 | 102.738907494204 | 105.14888714617 | 109.638082031519 | Nim1k |
| **ENSMUSG00000095990** | 0.0192314565047464 | 0.921656503987601 | 0.986713918026868 | 128.298947752804 | 109.946123403434 | 112.628131749165 | 110.675349922463 | 104.021028602814 | 84.3926740130963 | 129.093683228961 | 136.776221148232 | Zfp97 |
| **ENSMUSG00000096054** | 0.160097735059809 | 0.541760712022647 | 0.913837378403038 | 1168.24323552378 | 1642.6765252202 | 1599.50876181588 | 1092.68459881076 | 1064.29936633616 | 743.633997100907 | 1606.38349372812 | 1509.96606045389 | Syne1 |
| **ENSMUSG00000096145** | -0.216482053692333 | 0.431846962389863 | 0.886559861988535 | 209.615182244018 | 127.048853710635 | 138.182413742673 | 181.019851991825 | 210.231973597267 | 271.524255520397 | 130.134761319517 | 150.888053488923 | Vkorc1 |
| **ENSMUSG00000096146** | 0.200748272391682 | 0.157249667151953 | 0.788998730890588 | 4504.01587709669 | 4362.01064406514 | 4586.52039039037 | 4327.59376730716 | 3418.45948734722 | 3176.34455669581 | 4532.85400628142 | 4341.01673310936 | Kcnj11 |
| **ENSMUSG00000096188** | 0.207815046812888 | 0.464435856701425 | 0.895328112556867 | 457.177940583937 | 685.738043745865 | 628.446046062569 | 467.087493740564 | 423.748821781991 | 265.408844360028 | 586.126964983105 | 662.17059444779 | Cmtm4 |
| **ENSMUSG00000096199** | -0.119805362051279 | 0.508902870536281 | 0.902394714365738 | 146.369222084185 | 109.946123403434 | 103.163582862681 | 128.495957113368 | 137.964732673206 | 151.662196777159 | 122.847214685624 | 118.322286548867 | Ptrhd1 |
| **ENSMUSG00000096210** | 0.0673685697411334 | 0.618638602248178 | 0.929568738467614 | 2555.13679045726 | 2777.97205132677 | 3182.92779052473 | 2372.95453647315 | 2504.16939383827 | 2343.42555665351 | 2983.72980753389 | 2559.66928148834 | H1f0 |
| **ENSMUSG00000096255** | 0.0675454346927167 | 0.800030632432276 | 0.963461896828986 | 55.1143367107117 | 34.2054606144018 | 35.965285768641 | 37.5170677703264 | 42.7033696369448 | 41.5847958905112 | 37.4788112600209 | 33.6512925047237 | Dynlt1 |
| **ENSMUSG00000096401** | 0.295300801583013 | 0.515730976855781 | 0.904239892322932 | 9.03513716569045 | 19.5459774939439 | 14.1968233297267 | 13.1309737196143 | 9.85462376237187 | 17.123151249034 | 11.4518589961175 | 7.59867895267955 | Gm21811 |

| | | | | | | | | | | | | |
|---|---|---|---|---|---|---|---|---|---|---|---|---|
| **ENSMUSG00000096433** | -0.00271062104026535 | 0.990135045335116 | 0.998628446692557 | 72.2810973255236 | 65.9676740420606 | 45.4298346551254 | 65.6548685980713 | 67.887408140784 | 61.154111603693 | 63.5057635239243 | 57.5328549274309 | Zfp994 |
| **ENSMUSG00000096458** | -0.0331752841969761 | 0.943326615279069 | 0.990505437194639 | 17.1667606148119 | 33.3910448854875 | 24.6078271048596 | 15.9447538023887 | 27.3739548954774 | 7.33849339244316 | 34.3555769883525 | 23.8815624227072 | Moap1 |
| **ENSMUSG00000096472** | -0.118933334650549 | 0.481204424382931 | 0.897732059433541 | 345.142239729375 | 356.714089264476 | 341.670214802089 | 368.605190843457 | 469.737066006393 | 413.401794440965 | 365.418409785204 | 285.493223507818 | Cdkn2d |
| **ENSMUSG00000096544** | 0.0858857472762692 | 0.84276102574174 | 0.97128746259298 | 20.780815481088 | 18.7315617650296 | 7.57163910918757 | 22.5102406621959 | 20.8042057205628 | 19.5693157131818 | 12.4929370866736 | 13.0263067760221 | Gm4617 |
| **ENSMUSG00000096586** | 0.308497936321653 | 0.431695014581485 | 0.886559861988535 | 35.2370349461928 | 16.2883145782866 | 16.0897331070236 | 28.1378008277448 | 19.7092475247437 | 17.123151249034 | 24.9858741733473 | 15.1973579053591 | Gm22918 |
| **ENSMUSG00000096606** | -0.893064201722958 | 0.0432286687301516 | 0.687730553581215 | 24.3948703473642 | 29.3189662409158 | 46.3762895437739 | 50.6480414899407 | 79.9319482947941 | 123.53130543946 | 32.2734208072402 | 44.5065481514088 | Tpbgl |
| **ENSMUSG00000096674** | -0.287910850983289 | 0.492378284924233 | 0.899236955296196 | 124.684892886528 | 141.708336831093 | 51.1085639870161 | 106.92364314543 | 93.0714466446232 | 122.308223207386 | 67.6700758861489 | 235.559047533066 | Mup15 |
| **ENSMUSG00000096687** | -0.0514198947609859 | 0.857306105832848 | 0.974029039597056 | 65.9565013095403 | 67.5965054998893 | 56.7872933189068 | 68.4686486808458 | 58.0327843784121 | 37.9155491942896 | 75.998700610598 | 95.5262496908287 | Mfsd4b4 |
| **ENSMUSG00000096688** | -0.256395745222435 | 0.633467237391978 | 0.9325328237529 | 137.334084918495 | 167.769640156352 | 39.7511053232348 | 96.6064495085906 | 85.4067392738896 | 136.985209992272 | 42.6842017128016 | 262.697186649779 | Mup17 |
| **ENSMUSG00000096696** | 0.252144988888527 | 0.320521961081525 | 0.855228644171892 | 77.7021796249379 | 57.8235167529173 | 67.1982970940397 | 100.358156285623 | 70.0773245324222 | 45.2540425867328 | 70.7933101578173 | 67.3025850094475 | Zfp960 |
| **ENSMUSG00000096727** | -0.178750290251547 | 0.623387616076147 | 0.931059568526633 | 126.491920319666 | 96.9154717408051 | 99.3777633080869 | 146.316564304273 | 150.009272827216 | 228.716377397812 | 77.0397787011541 | 75.9867895267955 | Psmb9 |
| **ENSMUSG00000096740** | -0.497327198616811 | 0.115933624279409 | 0.769868040577702 | 39.754603529038 | 17.1027303072009 | 26.5007368821565 | 23.448167356454 | 41.6084114411257 | 34.2463024980681 | 37.4788112600209 | 36.9078691987293 | Lbhd1 |
| **ENSMUSG00000096768** | -0.0881792532244084 | 0.841582854193204 | 0.970640319105748 | 51.5002818444356 | 42.3496179035451 | 44.483379766477 | 28.1378008277448 | 60.2227007700503 | 29.3539735697726 | 70.7933101578173 | 16.2828834700276 | Gm47283 |
| **ENSMUSG00000096780** | 0.610546880316396 | 0.0773720761127835 | 0.739855948845396 | 28.0089252136404 | 21.9892246806869 | 27.447191770805 | 25.3240207449704 | 17.5193311331055 | 13.4539045528125 | 23.9447960827911 | 11.9407812113536 | Tmem181b-p |
| **ENSMUSG00000096795** | -0.161349138559296 | 0.721525190256519 | 0.951674101666722 | 20.780815481088 | 21.9892246806869 | 16.0897331070236 | 22.5102406621959 | 12.0445401540101 | 13.4539045528125 | 21.8626399016789 | 43.4210225867403 | Zfp433 |
| **ENSMUSG00000096842** | 0.331776224861822 | 0.0778370078473676 | 0.739855948845396 | 400.256576440087 | 309.477976987445 | 347.34894413398 | 334.839829850164 | 228.846262926191 | 221.377884005369 | 305.035880532948 | 349.539231823259 | Gm10736 |
| **ENSMUSG00000096910** | -0.125601579830754 | 0.760065927151106 | 0.957357521291862 | 56.0178504272808 | 92.8433930962335 | 94.6454888648447 | 60.0273084325223 | 63.5075753575076 | 30.5770558018465 | 131.175839410073 | 105.295979772845 | Zfp955b |
| **ENSMUSG00000096916** | -0.0782557849655665 | 0.827927770603863 | 0.967319591893432 | 24.3948703473642 | 21.1748089517725 | 14.1968233297267 | 21.5723139679377 | 19.7092475247437 | 31.8001380339204 | 18.7394056300105 | 16.2828834700276 | Zfp850 |
| **ENSMUSG00000096917** | 0.373019789987881 | 0.063250198568912 | 0.716594827179599 | 76.7986659083688 | 73.2974156022896 | 75.7163910918757 | 78.7858423176855 | 70.0773245324222 | 47.7002070508805 | 63.5057635239243 | 53.1907526687569 | 2500002B13I |
| **ENSMUSG00000096929** | 0.0139681960142953 | 0.980678117794148 | 0.996888334226108 | 14.4562194651047 | 4.88649437348597 | 13.2503684410783 | 12.1930470253561 | 17.5193311331055 | 9.78465785659088 | 13.5340151772298 | 3.25657669400552 | A330023F24I |
| **ENSMUSG00000096942** | -0.770814522886066 | 0.0017584751950615 | 0.635862253526407 | 48.7897406947284 | 39.9063707168021 | 53.9479286529615 | 34.703287687552 | 61.3176589658694 | 74.6080161565054 | 69.7522320672611 | 96.6117752554972 | Rps19-ps6 |
| **ENSMUSG00000096956** | -0.161097161271985 | 0.592841326438591 | 0.923935529355527 | 103.90407740544 | 69.2253369577179 | 62.4660226507975 | 106.92364314543 | 67.887408140784 | 145.546785616789 | 81.2040910633787 | 89.0130963028176 | Snhg18 |
| **ENSMUSG00000096957** | 0.386027282651146 | 0.27397655984425 | 0.845204865395511 | 182.509770746947 | 198.717437855096 | 206.327165725361 | 285.129715054481 | 94.1664048404423 | 92.9542496376133 | 252.981976005141 | 226.874843015718 | E230013L22I |
| **ENSMUSG00000096971** | -0.367856466097066 | 0.494139345675405 | 0.900141680336539 | 8.13162344912141 | 10.5874044758863 | 12.3039135524298 | 14.0689004138724 | 7.66470737073368 | 18.3462334811079 | 6.24646854333682 | 26.0526135520442 | 4930556M19 |
| **ENSMUSG00000096974** | -0.195133340151603 | 0.385794864623134 | 0.875126835008975 | 79.509207058076 | 64.338842584232 | 64.3589324280944 | 96.6064495085906 | 77.7420319031559 | 106.408154190426 | 72.8754663389296 | 92.2696729968231 | Gm26881 |
| **ENSMUSG00000096980** | -0.404459198608524 | 0.437120675839596 | 0.888701139843674 | 6.32459601598331 | 13.0306516626293 | 11.3574586637814 | 5.62756016554897 | 12.0445401540101 | 6.1154111603693 | 11.4518589961175 | 18.4539345993646 | Gm26526 |
| **ENSMUSG00000096983** | -0.0274581973545902 | 0.930368322034042 | 0.987529878729394 | 19.877301764519 | 24.4324718674299 | 29.3401015481019 | 22.5102406621959 | 26.2789966996583 | 30.5770558018465 | 18.7394056300105 | 22.7960368580387 | 2010015M23 |
| **ENSMUSG00000097006** | 0.00558632652624582 | 0.964264374834362 | 0.994284908685797 | 346.949267162513 | 334.724864583789 | 303.812019256151 | 383.612017951588 | 349.291664466292 | 362.032340693862 | 343.555769883525 | 309.374785930525 | 9530082P21I |
| **ENSMUSG00000097039** | 0.869690194347906 | 0.023836235507385 | 0.654124840968651 | 19.877301764519 | 24.4324718674299 | 30.2865564367503 | 42.2067012416173 | 19.7092475247437 | 9.78465785659088 | 18.7394056300105 | 15.1973579053591 | Pvt1 |
| **ENSMUSG00000097042** | 0.703849201577711 | 0.133547676056762 | 0.77303252008843 | 9.93865088225949 | 13.8450673915436 | 15.1432782183751 | 13.1309737196143 | 8.75966556655277 | 6.1154111603693 | 11.4518589961175 | 5.42762782334254 | Gm17491 |
| **ENSMUSG00000097047** | 0.440199130884676 | 0.244763940671396 | 0.832048466795696 | 20.780815481088 | 21.9892246806869 | 32.1794662140472 | 28.1378008277448 | 10.949581958191 | 14.6769867848863 | 28.1091084450157 | 21.7105112933701 | 1110020A21I |
| **ENSMUSG00000097048** | 0.0135852476201413 | 0.967327107638983 | 0.994284908685797 | 30.7194663633475 | 16.2883145782866 | 34.0723759913441 | 30.0136542162612 | 22.994122112201 | 29.3539735697726 | 33.3144988977964 | 23.8815624227072 | 1600020E01I |
| **ENSMUSG00000097057** | -0.374495535750403 | 0.364823595164059 | 0.86924083181804 | 16.2632468982428 | 12.2162359337149 | 19.8755526616174 | 10.3171936368398 | 16.4243729372865 | 28.1308913376988 | 19.7804837205666 | 11.9407812113536 | Gm17638 |
| **ENSMUSG00000097059** | 0.139959737719051 | 0.439131383407406 | 0.889583654898613 | 228.588970291968 | 153.924572764808 | 188.344522841041 | 219.47484645641 | 198.187433443257 | 194.470074899744 | 153.038479311752 | 171.513039217624 | Fam120aos |
| **ENSMUSG00000097061** | -0.0515804604471258 | 0.82877949333647 | 0.967483872587428 | 35.2370349461928 | 36.6487078011448 | 38.8046504345863 | 36.5791410760683 | 38.3235368536684 | 34.2463024980681 | 38.5198893505771 | 41.2499714574033 | 9330151L19I |
| **ENSMUSG00000097062** | -0.0367228808930594 | 0.874083254009195 | 0.97671366425653 | 78.6056933415069 | 67.5965054998893 | 57.7337482075553 | 60.0273084325223 | 84.3117810780705 | 74.6080161565054 | 61.4236073428121 | 51.0197015394199 | Gm17586 |
| **ENSMUSG00000097068** | 0.282644940153697 | 0.605514291568643 | 0.92658295801349 | 12.6491920319666 | 17.1027303072009 | 10.4110037751329 | 9.37926694258161 | 12.0445401540101 | 11.0077400886647 | 15.616171358342 | 2.17105112933702 | Gm26760 |
| **ENSMUSG00000097073** | 0.00319281153231 | 0.99315937672131 | 0.999133450854041 | 32.5264937964856 | 24.4324718674299 | 44.483379766477 | 26.2619474392285 | 16.4243729372865 | 36.6924669622158 | 45.80743598447 | 28.2236646813812 | 9430037G07I |
| **ENSMUSG00000097080** | 0.0230187431763075 | 0.961258626190415 | 0.993822507448007 | 9.03513716569045 | 6.51532583131463 | 13.2503684410783 | 18.7585338851632 | 10.949581958191 | 12.2308223207386 | 13.5340151772298 | 9.76973008201657 | 1700086O06I |
| **ENSMUSG00000097093** | -0.248888061546089 | 0.488803610552377 | 0.899236955296196 | 21.6843291976571 | 24.4324718674299 | 18.9290977729689 | 24.3860940507122 | 21.8991639163819 | 14.6769867848863 | 30.191264626128 | 39.0789203280663 | C330013E15I |
| **ENSMUSG00000097099** | -0.339508994474369 | 0.245576483027876 | 0.832048466795696 | 32.5264937964856 | 34.2054606144018 | 40.6975602118832 | 53.4618215727152 | 60.2227007700503 | 64.8233582999146 | 36.4377331694648 | 42.3354970220718 | Gm9917 |
| **ENSMUSG00000097101** | -0.136697627706037 | 0.654043824226813 | 0.937077409179068 | 121.974351736821 | 108.317291945606 | 49.2156542097192 | 83.4754757889764 | 77.7420319031559 | 89.2850029413918 | 100.984574783945 | 131.348593324889 | 1810034E14I |
| **ENSMUSG00000097102** | -0.0246990033368197 | 0.926268298629758 | 0.986713918026868 | 55.1143367107117 | 35.0198763433161 | 36.9117406572894 | 59.0893817382642 | 48.1781606160403 | 41.5847958905112 | 44.7663578939139 | 54.2762782334254 | NA |
| **ENSMUSG00000097119** | 0.210968161332042 | 0.0910185296456292 | 0.753823363185192 | 217.74680569314 | 231.294067011669 | 227.149173275627 | 253.240207449704 | 185.047935093427 | 205.477814988408 | 206.133461930115 | 206.249857287016 | B230354K17 |
| **ENSMUSG00000097121** | -0.0963469959721403 | 0.715621781758082 | 0.951176382858794 | 31.6229800799166 | 34.2054606144018 | 33.1259211026956 | 41.2687745473591 | 27.3739548954774 | 44.0309603546589 | 38.5198893505771 | 40.1644458927348 | D130020L05I |
| **ENSMUSG00000097125** | 0.0739178313394158 | 0.863964950701948 | 0.974868041734905 | 10.8421645988285 | 14.6594831204579 | 11.3574586637814 | 14.0689004138724 | 8.75966556655277 | 8.56157562451702 | 13.5340151772298 | 17.3684090346961 | Gm26885 |
| **ENSMUSG00000097131** | 0.116896258163031 | 0.698520184991589 | 0.945012697918882 | 24.3948703473642 | 28.5045505120015 | 29.3401015481019 | 26.2619474392285 | 18.6142893289246 | 31.8001380339204 | 21.8626399016789 | 28.2236646813812 | D230017M19 |
| **ENSMUSG00000097141** | 0.0616156638602381 | 0.910908707887836 | 0.986166948757759 | 46.9827132615903 | 159.625482867208 | 134.396594188079 | 43.1446279358754 | 43.7983278327639 | 41.5847958905112 | 171.777884941763 | 110.723607596188 | Gm10524 |
| **ENSMUSG00000097145** | 0.208450616038664 | 0.407446511032042 | 0.881573350402261 | 45.1756858284523 | 41.5352021746308 | 69.0912068713366 | 56.2756016554897 | 40.5134532453066 | 51.3694537471021 | 46.8485140750261 | 44.5065481514088 | 9230114K14I |
| **ENSMUSG00000097162** | 0.375829092129269 | 0.121090025178207 | 0.769868040577702 | 63.2459601598332 | 41.5352021746308 | 48.2691993210708 | 52.523894878457 | 36.1336204620302 | 39.1386314263635 | 43.7252798033577 | 39.0789203280663 | 2310010J17I |
| **ENSMUSG00000097164** | 0.276595368240928 | 0.233407475793043 | 0.832048466795696 | 49.6932544112975 | 51.3081909216027 | 65.3053873167428 | 63.779015209555 | 47.0832024202212 | 36.6924669622158 | 53.094982618363 | 52.1052271040884 | Cep83os |
| **ENSMUSG00000097167** | -0.261237395796181 | 0.546829007993871 | 0.914451747770672 | 9.03513716569045 | 13.8450673915436 | 15.1432782183751 | 9.37926694258161 | 16.4243729372865 | 19.5693157131818 | 11.4518589961175 | 9.76973008201657 | Gm16740 |
| **ENSMUSG00000097177** | 0.255640890998043 | 0.282417489261239 | 0.850587941933635 | 71.3775836089546 | 87.9568987227475 | 74.7699362032273 | 73.1582821521366 | 49.2731188118594 | 50.1463715150282 | 72.8754663389296 | 84.6709940441436 | 9330159M07 |
| **ENSMUSG00000097183** | -0.158806821414449 | 0.691677063756679 | 0.94469184542218 | 14.4562194651047 | 18.7315617650296 | 24.6078271048596 | 19.6964605794214 | 15.3294147414674 | 36.6924669622158 | 19.7804837205666 | 15.1973579053591 | Gm17501 |
| **ENSMUSG00000097184** | 0.21684366501423 | 0.358449832006666 | 0.866492893590776 | 109.325159704854 | 87.9568987227475 | 119.253315969704 | 119.116690170786 | 66.7924499449649 | 78.277262852727 | 107.231043327282 | 121.578863242873 | 4632428C04I |
| **ENSMUSG00000097187** | 0.857601837871115 | 0.0603314489948448 | 0.71282992338528 | 20.780815481088 | 22.8036404096012 | 21.7684624389143 | 22.5102406621959 | 7.66470737073368 | 23.2385624094033 | 5.20539045278068 | 13.0263067760221 | Gm19426 |
| **ENSMUSG00000097195** | -0.19242651242622 | 0.583899672107205 | 0.922450244597442 | 22.5878429142261 | 35.8342920722305 | 53.9479286529615 | 48.7721881014244 | 35.0386622662111 | 68.4926049961361 | 37.4788112600209 | 43.4210225867403 | Snhg5 |
| **ENSMUSG00000097204** | 0.123817068993535 | 0.784339292192089 | 0.961141387070613 | 28.0089252136404 | 35.0198763433161 | 31.2330113253987 | 16.8826804966469 | 18.6142893289246 | 13.4539045528125 | 48.9306702561384 | 20.6249857287016 | Gm17690 |
| **ENSMUSG00000097216** | 0.0896875246082413 | 0.716991904499897 | 0.95140862628518 | 69.5705561758165 | 105.874044758863 | 97.48485353079 | 83.4754757889764 | 64.6025335533267 | 67.2695227640623 | 111.395355689507 | 91.1841474321546 | 4932441J04I |
| **ENSMUSG00000097221** | 0.187062658856173 | 0.645698887951359 | 0.935063240511793 | 34.3335212296237 | 55.3802695661743 | 48.2691993210708 | 20.6343872736795 | 33.943704070392 | 17.123151249034 | 49.9717483466946 | 37.9933947633978 | 1810049J17I |
| **ENSMUSG00000097222** | 0.19873519470335 | 0.502545051358994 | 0.900927059989498 | 118.360296870545 | 109.946123403434 | 164.68315062483 | 83.4754757889764 | 87.5966556655277 | 79.5003450848009 | 158.243869764533 | 89.0130963028176 | 1010001N08I |
| **ENSMUSG00000097233** | -0.124221576803896 | 0.805562432790538 | 0.963475137341661 | 36.1405486627618 | 43.9784493613737 | 27.447191770805 | 34.703287687552 | 13.1394983498292 | 17.123151249034 | 58.3003730711437 | 66.217059444779 | Gm17552 |
| **ENSMUSG00000097243** | -0.417923588986426 | 0.383392344973356 | 0.875126835008975 | 15.3597331816738 | 7.32974156022896 | 9.46454888648447 | 6.56548685980713 | 10.949581958191 | 8.56157562451702 | 15.616171358342 | 16.2828834700276 | Gm26908 |
| **ENSMUSG00000097245** | 0.3434487388061 | 0.153115007525703 | 0.783412422888014 | 56.0178504272808 | 69.2253369577179 | 65.3053873167428 | 49.7101147956825 | 53.6529515951357 | 34.2463024980681 | 46.8485140750261 | 54.2762782334254 | Gm5421 |
| **ENSMUSG00000097251** | -0.294690136269924 | 0.480688218811645 | 0.897732059433541 | 11.7456783153976 | 10.5874044758863 | 14.1968233297267 | 8.44134024832345 | 12.0445401540101 | 11.0077400886647 | 17.6983275394543 | 14.1118323406906 | 5033417F24I |
| **ENSMUSG00000097252** | 0.736942332266774 | 0.329266301174496 | 0.855977678491063 | 30.7194663633475 | 43.9784493613737 | 113.574586637814 | 9.37926694258161 | 17.5193311331055 | 63.6002760678407 | 5.20539045278068 | 32.5657669400552 | Gm6634 |
| **ENSMUSG00000097271** | 0.404323323976406 | 0.44879459122459 | 0.89108031640712 | 9.93865088225949 | 10.5874044758863 | 19.8755526616174 | 7.50341355406529 | 7.66470737073368 | 4.89232892829544 | 14.5750932677859 | 8.68420451734806 | Gm9903 |
| **ENSMUSG00000097275** | 0.25165886394753 | 0.522649958948311 | 0.90648515894888 | 11.7456783153976 | 19.5459774939439 | 15.1432782183751 | 14.0689004138724 | 14.2344565456483 | 13.4539045528125 | 11.4518589961175 | 11.9407812113536 | Gm26648 |
| **ENSMUSG00000097277** | 0.220609710161949 | 0.315558563695937 | 0.853277176422981 | 572.827696304775 | 615.698291059232 | 510.139184981513 | 545.87333605825 | 428.128654565267 | 313.109051410908 | 538.237372817523 | 645.887710977762 | 2900076A07I |
| **ENSMUSG00000097284** | 0.22180780686761 | 0.707094818343145 | 0.948710970699398 | 34.3335212296237 | 21.9892246806869 | 25.5542819935081 | 17.8206071909051 | 2.18991639163819 | 18.3462334811079 | 33.3144988977964 | 31.4802413753867 | 4930480K23I |
| **ENSMUSG00000097287** | -0.109538935198748 | 0.742063936548516 | 0.954383574136061 | 37.9475760958999 | 70.8541684155466 | 53.001473764313 | 33.7653609932938 | 42.7033696369448 | 37.9155491942896 | 74.9576225200418 | 55.3618037980939 | D130017N08 |
| **ENSMUSG00000097290** | 0.0902398832998823 | 0.692935805408776 | 0.94469184542218 | 65.9565013095403 | 78.9983257046899 | 79.5022106464695 | 102.23400967414 | 61.3176589658694 | 84.3926740130963 | 93.6970281500523 | 67.3025850094475 | 1300002E11I |
| **ENSMUSG00000097316** | -0.0238621928506962 | 0.92553020607643 | 0.986713918026868 | 54.2108229941427 | 61.895595397489 | 53.9479286529615 | 45.9584080186499 | 38.3235368536684 | 51.3694537471021 | 56.2182168900314 | 73.8157383974585 | Gm10516 |
| **ENSMUSG00000097317** | 0.327812999411494 | 0.108513853220686 | 0.762287160109383 | 438.204152535987 | 395.806044252364 | 357.759947909113 | 399.556771753977 | 286.879047304603 | 215.262472844999 | 401.856142954669 | 362.565538599282 | Gm17281 |
| **ENSMUSG00000097318** | 0.12255212651824 | 0.708350339715689 | 0.949174314815223 | 53.3073092775737 | 29.3189662409158 | 25.5542819935081 | 37.5170677703264 | 37.2285786578493 | 41.5847958905112 | 24.9858741733473 | 30.3947158107182 | 1700007L15I |
| **ENSMUSG00000097320** | -0.841440468373858 | 0.0519811504035393 | 0.703396849069134 | 10.8421645988285 | 17.1027303072009 | 9.46454888648447 | 19.6964605794214 | 31.7537876787538 | 33.0232202659942 | 26.0269522639034 | 11.9407812113536 | Tmem147os |
| **ENSMUSG00000097321** | 0.478613271300193 | 0.345928495838597 | 0.861266047293794 | 14.4562194651047 | 17.1027303072009 | 21.7684624389143 | 6.56548685980713 | 17.5193311331055 | 7.33849339244316 | 9.36970281500523 | 8.68420451734806 | 1700028E10I |
| **ENSMUSG00000097328** | -0.725216571222527 | 0.0113287504425343 | 0.635862253526407 | 128.298947752804 | 109.13170767452 | 100.324218196735 | 134.123517278917 | 191.617684268342 | 308.216722482613 | 121.806136595068 | 159.572258006271 | Tnfsf12 |
| **ENSMUSG00000097330** | 0.386060257117026 | 0.461513183696321 | 0.895328112556867 | 19.877301764519 | 13.8450673915436 | 10.4110037751329 | 21.5723139679377 | 21.8991639163819 | 3.66924669622158 | 14.5750932677859 | 9.76973008201657 | Gm26672 |
| **ENSMUSG00000097333** | -0.340471420824082 | 0.105104004068659 | 0.762094734041179 | 49.6932544112975 | 58.6379324818317 | 62.4660226507975 | 59.0893817382642 | 62.4126171616885 | 88.0619207093179 | 77.0397787011541 | 64.046008315442 | Zfp87 |
| **ENSMUSG00000097340** | -0.162609030093926 | 0.611228243715211 | 0.927932112008768 | 43.3686583953142 | 25.2468875963442 | 47.3227444324223 | 57.2135283497478 | 53.6529515951357 | 63.6002760678407 | 39.5609674411332 | 36.9078691987293 | Gm26617 |
| **ENSMUSG00000097343** | -0.349083070207419 | 0.40293025446597 | 0.881158207349486 | 18.0702743313809 | 8.14415728914329 | 12.3039135524298 | 11.2551203310979 | 13.1394983498292 | 18.3462334811079 | 18.7394056300105 | 13.0263067760221 | 9030407P20I |
| **ENSMUSG00000097350** | -0.263668344374336 | 0.324156511668285 | 0.855977678491063 | 68.6670424592474 | 47.2361122770311 | 53.9479286529615 | 73.1582821521366 | 97.4512794278996 | 62.3771938357668 | 79.1219348822664 | 52.1052271040884 | 4732491K20I |
| **ENSMUSG00000097353** | 0.100911124961721 | 0.463026859431092 | 0.895328112556867 | 389.414411841258 | 364.858246553619 | 304.7584741448 | 355.474217123843 | 285.784089108784 | 368.147751854232 | 338.350379430744 | 327.828720529889 | A430046D13 |
| **ENSMUSG00000097354** | 0.193813458020974 | 0.483605538512028 | 0.89863290032536 | 66.8600150261093 | 37.4631235300591 | 70.9841166486335 | 76.9099889291692 | 64.6025335533267 | 50.1463715150282 | 58.3003730711437 | 46.6775992807458 | 2310001H17I |
| **ENSMUSG00000097357** | -0.217173564447542 | 0.560112830082244 | 0.916574868225118 | 72.2810973255236 | 87.1424829938332 | 55.8408384302584 | 51.5859681841989 | 54.7479097909548 | 40.3617136584374 | 132.216917500629 | 82.4999429148066 | Gm16793 |
| **ENSMUSG00000097383** | -0.13181568273909 | 0.758894639206598 | 0.957232704193574 | 18.0702743313809 | 22.8036404096012 | 11.3574586637814 | 9.37926694258161 | 12.0445401540101 | 23.2385624094033 | 17.6983275394543 | 15.1973579053591 | 1500026H17I |

| | | | | | | | | | | | | |
|---|---|---|---|---|---|---|---|---|---|---|---|---|
| **ENSMUSG00000097392** | -0.0904870501038753 | 0.743780143279781 | 0.954466085361166 | 46.9827132615903 | 32.5766291565731 | 39.7511053232348 | 65.6548685980713 | 50.3680770076785 | 42.8078781225851 | 55.1771387994752 | 47.7631248454143 | D930016D06 |
| **ENSMUSG00000097403** | -0.0457902303865197 | 0.933076006140178 | 0.988138588387242 | 7.22810973255236 | 19.5459774939439 | 9.46454888648447 | 8.44134024832345 | 20.8042057205628 | 8.56157562451702 | 8.32862472444909 | 8.68420451734806 | 9230116N13 |
| **ENSMUSG00000097412** | 0.283870105756592 | 0.178361526432107 | 0.801696999069105 | 150.886790667031 | 160.439898596123 | 176.98706417726 | 181.957778686083 | 152.199189218854 | 90.5080851734656 | 171.777884941763 | 134.605170018895 | 1810014B01 |
| **ENSMUSG00000097414** | -0.0327298663302847 | 0.927920739444355 | 0.986901071486195 | 25.2983840639333 | 17.9171460361152 | 12.3039135524298 | 17.8206071909051 | 16.4243729372865 | 17.123151249034 | 20.8215618111227 | 20.6249857287016 | B130046B21 |
| **ENSMUSG00000097415** | -0.26154865753813 | 0.446965611772064 | 0.89108031640712 | 112.939214571131 | 115.647033505835 | 110.735221971868 | 184.771558768858 | 195.997517051618 | 228.716377397812 | 74.9576225200418 | 129.177542195552 | AU020206 |
| **ENSMUSG00000097431** | 0.122479410793642 | 0.544045429587036 | 0.913837378403038 | 175.281661014395 | 188.13003337921 | 131.557229522134 | 142.564857527241 | 132.489941694111 | 114.969729814943 | 170.736806851206 | 167.17093695895 | Gm26782 |
| **ENSMUSG00000097439** | 0.168600984187997 | 0.739007892283186 | 0.954383574136061 | 8.13162344912141 | 14.6594831204579 | 10.4110037751329 | 15.0068271081306 | 15.3294147414674 | 8.56157562451702 | 14.5750932677859 | 4.34210225867403 | Gm16754 |
| **ENSMUSG00000097440** | 0.429443917797199 | 0.372646861112039 | 0.873700364743909 | 9.03513716569045 | 13.8450673915436 | 32.1794662140472 | 20.6343872736795 | 16.4243729372865 | 8.56157562451702 | 16.6572494488982 | 14.1118323406906 | Gm6277 |
| **ENSMUSG00000097445** | 0.0766673683474779 | 0.692626255462502 | 0.94469184542218 | 105.711104838578 | 79.8127414336042 | 84.2344850897118 | 107.861569839689 | 82.1218646864323 | 105.185071958352 | 88.4916376972716 | 82.4999429148066 | Gm26631 |
| **ENSMUSG00000097451** | -0.225792697877604 | 0.490224272654002 | 0.899236955296196 | 24.3948703473642 | 17.9171460361152 | 17.036187995672 | 20.6343872736795 | 19.7092475247437 | 30.5770558018465 | 20.8215618111227 | 22.7960368580387 | Rian |
| **ENSMUSG00000097462** | 0.537556922746594 | 0.0403208447872961 | 0.679680259824687 | 74.9916384752307 | 67.5965054998893 | 133.450139299431 | 103.171936368398 | 64.6025335533267 | 68.4926049961361 | 57.2592949805875 | 70.559161703453 | 9530026P05 |
| **ENSMUSG00000097487** | 0.0195723764661517 | 0.917856584975792 | 0.986713918026868 | 508.678222428372 | 374.631235300591 | 311.383658365339 | 483.970174237211 | 366.810995599397 | 406.063301048521 | 477.854843565267 | 404.901035621353 | Ptges3l |
| **ENSMUSG00000097488** | -0.0532900687023752 | 0.811215414071645 | 0.964955219787201 | 58.7283915769879 | 48.8649437348597 | 71.930571537282 | 66.5927952923295 | 64.6025335533267 | 75.8310983885793 | 53.094982618363 | 61.8749571861049 | 4732487G21 |
| **ENSMUSG00000097489** | 0.853094450788876 | 0.081909418462271 | 0.739855948845396 | 9.03513716569045 | 10.5874044758863 | 18.9290977729689 | 21.5723139679377 | 6.56974917491458 | 7.33849339244316 | 9.36970281500523 | 9.76973008201657 | Gm26808 |
| **ENSMUSG00000097493** | 0.0984916518241183 | 0.766993038116795 | 0.958655025595455 | 30.7194663633475 | 24.4324718674299 | 35.0188308799925 | 54.3997482669734 | 25.1840385038392 | 42.8078781225851 | 34.3555769883525 | 32.5657669400552 | 9930014A18 |
| **ENSMUSG00000097504** | 0.131746057389342 | 0.68633104629258 | 0.944452327903597 | 20.780815481088 | 40.7207864457164 | 43.5369248778286 | 45.9584080186499 | 43.7983278327639 | 36.6924669622158 | 27.0680303544596 | 30.3947158107182 | 4930516B21 |
| **ENSMUSG00000097509** | 0.506206155776718 | 0.302845901802387 | 0.851785577030345 | 18.0702743313809 | 13.8450673915436 | 9.46454888648447 | 8.44134024832345 | 10.949581958191 | 4.89232892829544 | 10.4107809055614 | 8.68420451734806 | B230322F03 |
| **ENSMUSG00000097518** | -0.472482941206859 | 0.181448280185796 | 0.803056849297055 | 29.8159526467785 | 30.1333819698302 | 17.036187995672 | 29.075727522003 | 50.3680770076785 | 46.4771248188067 | 24.9858741733473 | 26.0526135520442 | Gm26694 |
| **ENSMUSG00000097530** | 0.306099788192006 | 0.545813079734853 | 0.914211323944814 | 13.5527057485357 | 7.32974156022896 | 14.1968233297267 | 10.3171936368398 | 4.37983278327639 | 9.78465785659088 | 8.32862472444909 | 14.1118323406906 | Kansl2-ps |
| **ENSMUSG00000097547** | 0.254152413574176 | 0.38456722143681 | 0.875126835008975 | 35.2370349461928 | 48.0505280059454 | 45.4298346551254 | 30.9515809105193 | 21.8991639163819 | 37.9155491942896 | 37.4788112600209 | 36.9078691987293 | B230110C06 |
| **ENSMUSG00000097554** | -0.114032475755915 | 0.829601636747688 | 0.967523998969336 | 28.0089252136404 | 10.5874044758863 | 5.67872933189068 | 23.448167356454 | 14.2344565456483 | 18.3462334811079 | 13.5340151772298 | 27.1381391167127 | Gm26825 |
| **ENSMUSG00000097571** | 0.282406626241753 | 0.538280788706137 | 0.912708496930502 | 10.8421645988285 | 12.2162359337149 | 16.0897331070236 | 9.37926694258161 | 12.0445401540101 | 8.56157562451702 | 6.24646854333682 | 13.0263067760221 | Jpx |
| **ENSMUSG00000097573** | 0.437363315021306 | 0.191552458543839 | 0.808824684419004 | 39.754603529038 | 24.4324718674299 | 28.3936466594534 | 25.3240207449704 | 22.994122112201 | 29.3539735697726 | 15.616171358342 | 19.5394601640331 | G730003C15 |
| **ENSMUSG00000097574** | 0.94208457425661 | 0.0716008495175209 | 0.728064275108613 | 16.2632468982428 | 17.1027303072009 | 24.6078271048596 | 38.4549944645846 | 16.4243729372865 | 2.44616446414772 | 15.616171358342 | 15.1973579053591 | C920006O11 |
| **ENSMUSG00000097577** | 0.516594826392379 | 0.0847907528076489 | 0.743541556158628 | 56.0178504272808 | 58.6379324818317 | 56.7872933189068 | 86.2892558717508 | 45.9882442244021 | 56.2617826753975 | 53.094982618363 | 24.9670879873757 | 6230400D17 |
| **ENSMUSG00000097578** | 0.45260060426348 | 0.316833802489636 | 0.854196722938869 | 39.754603529038 | 63.5244268553176 | 36.9117406572894 | 42.2067012416173 | 16.4243729372865 | 14.6769867848863 | 56.2182168900314 | 45.5920737160773 | Gm26798 |
| **ENSMUSG00000097589** | 0.0388116401015945 | 0.867846508693593 | 0.974868041734905 | 170.764092431549 | 118.904696421492 | 123.039135524298 | 187.585338851632 | 168.623562156141 | 176.123841418636 | 127.011527047849 | 112.894658725525 | Dleu2 |
| **ENSMUSG00000097595** | 0.219075883340655 | 0.621857574331228 | 0.93078489223231 | 20.780815481088 | 8.95857301805762 | 17.036187995672 | 26.2619474392285 | 15.3294147414674 | 19.5693157131818 | 9.36970281500523 | 18.4539345993646 | 1500002F19 |
| **ENSMUSG00000097616** | 0.134995668693319 | 0.574136404870629 | 0.922280709351328 | 64.1494738764022 | 55.3802695661743 | 70.9841166486335 | 61.9031618210386 | 37.2285786578493 | 66.0464405319884 | 61.4236073428121 | 65.1315338801105 | 1110019D14 |
| **ENSMUSG00000097617** | -0.657868870448717 | 0.19492145998083 | 0.810131382146537 | 18.9737880479499 | 10.5874044758863 | 8.51809399783602 | 11.2551203310979 | 24.0890803080201 | 33.0232202659942 | 11.4518589961175 | 9.76973008201657 | Gm10687 |
| **ENSMUSG00000097626** | 0.166543928095195 | 0.443378633300474 | 0.89108031640712 | 60.535419010126 | 66.7820897709749 | 48.2691993210708 | 60.9652351267805 | 47.0832024202212 | 48.9232892829544 | 55.1771387994752 | 59.7039060567679 | 4921504A21 |
| **ENSMUSG00000097632** | 0.319040292806036 | 0.521223019074415 | 0.90609507420585 | 13.5527057485357 | 10.5874044758863 | 37.8581955459379 | 24.3860940507122 | 17.5193311331055 | 11.0077400886647 | 26.0269522639034 | 14.1118323406906 | 4930552P12 |
| **ENSMUSG00000097640** | 0.349419047571166 | 0.497692087617719 | 0.900927059989498 | 4.51756858284523 | 12.2162359337149 | 15.1432782183751 | 18.7585338851632 | 9.85462376237187 | 7.33849339244316 | 11.4518589961175 | 10.8552556466851 | Gm20033 |
| **ENSMUSG00000097644** | 0.0101304142176895 | 0.985289999958293 | 0.99735301701069 | 51.5002818444356 | 21.1748089517725 | 17.036187995672 | 30.0136542162612 | 15.3294147414674 | 11.0077400886647 | 55.1771387994752 | 36.9078691987293 | Gm26862 |
| **ENSMUSG00000097652** | -0.0730209444841779 | 0.604074422011682 | 0.926448151609826 | 450.853344567953 | 386.033055505392 | 335.04503058155 | 435.197986135787 | 375.57066116595 | 457.432754795624 | 418.513392403567 | 439.637853690746 | Mhrt |
| **ENSMUSG00000097666** | 0.0683546636452934 | 0.82631591120859 | 0.967261756842012 | 32.5264937964856 | 38.2775392589734 | 26.5007368821565 | 39.3929211588428 | 20.8042057205628 | 31.8001380339204 | 32.2734208072402 | 45.5920737160773 | A330094K24 |
| **ENSMUSG00000097675** | 0.15562099849276 | 0.643523545622574 | 0.934616627390525 | 28.9124389302094 | 40.7207864457164 | 31.2330113253987 | 29.075727522003 | 19.7092475247437 | 23.2385624094033 | 28.1091084450157 | 45.5920737160773 | 1700101I11R |
| **ENSMUSG00000097715** | -0.401272036408483 | 0.0780934312482663 | 0.739855948845396 | 38.8510898124689 | 42.3496179035451 | 45.4298346551254 | 42.2067012416173 | 66.7924499449649 | 58.7079471395453 | 52.0539045278068 | 45.5920737160773 | Gpr137b-ps |
| **ENSMUSG00000097718** | -0.0783463006608982 | 0.939981216418938 | 0.989269716906553 | 3.61405486627618 | 19.5459774939439 | 10.4110037751329 | 8.44134024832345 | 1.0949581958191 | 0 | 32.2734208072402 | 10.8552556466851 | Gm26896 |
| **ENSMUSG00000097729** | 0.133925257512179 | 0.378775741329154 | 0.87422797717331 | 404.774145022932 | 335.539280312703 | 379.528410348027 | 373.294824314748 | 308.778211220985 | 280.085831144914 | 396.650752501888 | 373.420794245967 | 2310015A10 |
| **ENSMUSG00000097743** | -0.0645963261346691 | 0.87149564671793 | 0.975613767803763 | 20.780815481088 | 11.4018202048006 | 22.7149173275627 | 18.7585338851632 | 18.6142893289246 | 11.0077400886647 | 27.0680303544596 | 19.5394601640331 | Gm16973 |
| **ENSMUSG00000097747** | 0.354279628950278 | 0.36237507726141 | 0.86924083181804 | 19.877301764519 | 17.1027303072009 | 15.1432782183751 | 13.1309737196143 | 10.949581958191 | 15.9000690169602 | 10.4107809055614 | 14.1118323406906 | Gm26534 |
| **ENSMUSG00000097750** | -0.25408525153204 | 0.407157894737712 | 0.881573350402261 | 74.9916384752307 | 75.7406627890326 | 121.146225747001 | 76.9099889291692 | 127.015150715015 | 143.100621152642 | 84.3273253350471 | 61.8749571861049 | Gm4673 |
| **ENSMUSG00000097756** | -0.504100256939133 | 0.310217672308389 | 0.851785577030345 | 9.93865088225949 | 4.07207864457164 | 8.51809399783602 | 15.9447538023887 | 16.4243729372865 | 11.0077400886647 | 11.4518589961175 | 15.1973579053591 | A730056A06 |
| **ENSMUSG00000097769** | 0.10311074720706 | 0.606836277886929 | 0.927177252274348 | 84.9302893574902 | 75.7406627890326 | 90.8596693102509 | 83.4754757889764 | 79.9319482947941 | 91.7311674055395 | 58.3003730711437 | 82.4999429148066 | Snhg4 |
| **ENSMUSG00000097776** | 0.137122910411173 | 0.697862912754391 | 0.944933386973863 | 83.1232619243521 | 105.059629029948 | 48.2691993210708 | 54.3997482669734 | 43.7983278327639 | 50.1463715150282 | 69.7522320672611 | 100.953877514171 | Gm26795 |
| **ENSMUSG00000097781** | 0.407633535140691 | 0.34335112424793 | 0.861167771001038 | 14.4562194651047 | 15.4738988493722 | 22.7149173275627 | 18.7585338851632 | 18.6142893289246 | 6.1154111603693 | 17.6983275394543 | 10.8552556466851 | 9330136K24 |
| **ENSMUSG00000097785** | 0.0522938527044552 | 0.901464761983722 | 0.984802943363903 | 14.4562194651047 | 10.5874044758863 | 9.46454888648447 | 15.9447538023887 | 10.949581958191 | 9.78465785659088 | 12.4929370866736 | 15.1973579053591 | B230217O12 |
| **ENSMUSG00000097787** | 0.192895242506121 | 0.453419442605853 | 0.891651261195907 | 37.0440623793308 | 59.452348210746 | 38.8046504345863 | 44.0825546301336 | 38.3235368536684 | 42.8078781225851 | 38.5198893505771 | 37.9933947633978 | 2700046G09 |
| **ENSMUSG00000097792** | -0.123406515309199 | 0.748710176410743 | 0.956197192463153 | 14.4562194651047 | 15.4738988493722 | 16.0897331070236 | 18.7585338851632 | 13.1394983498292 | 15.9000690169602 | 27.0680303544596 | 14.1118323406906 | Gm27042 |
| **ENSMUSG00000097793** | 0.91622503267331 | 0.115729207054917 | 0.769868040577702 | 23.4913566307952 | 6.51532583131463 | 19.8755526616174 | 12.1930470253561 | 8.75966556655277 | 2.44616446414772 | 9.36970281500523 | 11.9407812113536 | Gm17259 |
| **ENSMUSG00000097799** | 0.0959310935062419 | 0.78517221877002 | 0.961141387070613 | 28.9124389302094 | 40.7207864457164 | 27.447191770805 | 33.7653609932938 | 19.7092475247437 | 20.7923979452556 | 46.8485140750261 | 34.7368180693922 | Gm26899 |
| **ENSMUSG00000097804** | 0.71712194410062 | 0.153903617837782 | 0.785495071155314 | 18.9737880479499 | 48.8649437348597 | 14.1968233297267 | 22.5102406621959 | 8.75966556655277 | 25.6847268735511 | 15.616171358342 | 14.1118323406906 | Gm16685 |
| **ENSMUSG00000097815** | 0.154975870702313 | 0.582873933949523 | 0.922450244597442 | 252.080326922764 | 288.303168035672 | 313.276568142636 | 229.79204009325 | 187.237851485066 | 139.43137445642 | 340.432535611857 | 305.032683671851 | Gm26809 |
| **ENSMUSG00000097820** | -0.00864801645654792 | 0.97038821055901 | 0.9951105526746 | 92.1583990900426 | 83.8848200781758 | 65.3053873167428 | 60.0273084325223 | 77.7420319031559 | 56.2617826753975 | 87.4505596067155 | 81.4144173501381 | E530011L22 |
| **ENSMUSG00000097823** | -0.132814582666068 | 0.758291184147021 | 0.95677573228595 | 6.32459601598331 | 14.6594831204579 | 17.036187995672 | 16.8826804966469 | 14.2344565456483 | 19.5693157131818 | 13.5340151772298 | 13.0263067760221 | Gm16701 |
| **ENSMUSG00000097828** | -0.537248161652652 | 0.0542439129732887 | 0.703396849069134 | 21.6843291976571 | 25.2468875963442 | 29.3401015481019 | 33.7653609932938 | 49.2731188118594 | 35.4693847301419 | 34.3555769883525 | 40.1644458927348 | 6430562O15 |
| **ENSMUSG00000097835** | 0.0580453662837714 | 0.919429222476435 | 0.986713918026868 | 13.5527057485357 | 4.07207864457164 | 13.2503684410783 | 11.2551203310979 | 4.37983278327639 | 14.6769867848863 | 7.28754663389296 | 14.1118323406906 | Gm26910 |
| **ENSMUSG00000097837** | -0.423866296980569 | 0.552576659510124 | 0.914977712327965 | 14.4562194651047 | 84.6992358070902 | 36.9117406572894 | 9.37926694258161 | 16.4243729372865 | 19.5693157131818 | 83.2862472444909 | 75.9867895267955 | Gm26794 |
| **ENSMUSG00000097863** | 0.80176713767587 | 0.106495743612234 | 0.762094734041179 | 11.7456783153976 | 12.2162359337149 | 13.2503684410783 | 26.2619474392285 | 9.85462376237187 | 4.89232892829544 | 8.32862472444909 | 13.0263067760221 | 1010001B22 |
| **ENSMUSG00000097867** | 0.0377816380772654 | 0.842709415310681 | 0.97128746259298 | 224.071401709123 | 178.357044632238 | 163.736695736181 | 235.419600258798 | 205.85214081399 | 234.831788558181 | 178.024353485099 | 162.828834700276 | Lppos |
| **ENSMUSG00000097868** | -0.253748570483994 | 0.6493034899098 | 0.936461253686283 | 46.9827132615903 | 6.51532583131463 | 18.9290977729689 | 26.2619474392285 | 26.2789966996583 | 50.1463715150282 | 22.903717992235 | 18.4539345993646 | A330069E16 |
| **ENSMUSG00000097882** | 0.464461008208815 | 0.197914773897189 | 0.812921911423739 | 31.6229800799166 | 23.6180561385155 | 33.1259211026956 | 32.8274342990356 | 15.3294147414674 | 14.6769867848863 | 35.3966550789086 | 21.7105112933701 | 0610038B21 |
| **ENSMUSG00000097885** | 0.308023762656261 | 0.457201324505086 | 0.892805623137304 | 14.4562194651047 | 13.0306516626293 | 14.1968233297267 | 19.6964605794214 | 10.949581958191 | 7.33849339244316 | 15.616171358342 | 15.1973579053591 | 5031434O11 |
| **ENSMUSG00000097906** | -0.359195985941842 | 0.161286458966994 | 0.792786631432503 | 35.2370349461928 | 39.0919549878878 | 48.2691993210708 | 37.5170677703264 | 36.1336204620302 | 57.4848649074714 | 51.0128264372507 | 60.7894316214364 | Gm9625 |
| **ENSMUSG00000097908** | 0.0457298242769365 | 0.919476016083107 | 0.986713918026868 | 13.5527057485357 | 17.9171460361152 | 10.4110037751329 | 12.1930470253561 | 10.949581958191 | 8.56157562451702 | 21.8626399016789 | 10.8552556466851 | 4933404O12 |
| **ENSMUSG00000097910** | 0.0721388399773236 | 0.83660747363762 | 0.968924788429433 | 38.8510898124689 | 56.1946852950887 | 35.965285768641 | 25.3240207449704 | 47.0832024202212 | 45.2540425867328 | 35.3966550789086 | 21.7105112933701 | 5033428I22R |
| **ENSMUSG00000097915** | -0.298450302874485 | 0.310741073948673 | 0.851785577030345 | 36.1405486627618 | 32.5766291565731 | 29.3401015481019 | 32.8274342990356 | 58.0327843784121 | 40.3617136584374 | 28.1091084450157 | 34.7368180693922 | A330009N23 |
| **ENSMUSG00000097944** | -0.159628480018325 | 0.651272367844363 | 0.936463929813731 | 25.2983840639333 | 33.3910448854875 | 42.5904699891801 | 37.5170677703264 | 20.8042057205628 | 30.5770558018465 | 51.0128264372507 | 52.1052271040884 | A130014A01 |
| **ENSMUSG00000097971** | -1.16085244003939 | 0.0771234008333268 | 0.739855948845396 | 309.001691066613 | 195.459774939439 | 220.523989055088 | 364.853484066425 | 908.81530252985 | 1120.34332457966 | 160.326025945645 | 247.49982874442 | Gm26917 |
| **ENSMUSG00000097994** | 0.284292637962698 | 0.238094686169289 | 0.832048466795696 | 45.1756858284523 | 39.0919549878878 | 46.3762895437739 | 45.9584080186499 | 37.2285786578493 | 30.5770558018465 | 34.3555769883525 | 42.3354970220718 | Gm26982 |
| **ENSMUSG00000098040** | 1.00245292005308 | 0.0303690206190815 | 0.665956123973759 | 20.780815481088 | 12.2162359337149 | 12.3039135524298 | 23.448167356454 | 10.949581958191 | 6.1154111603693 | 7.28754663389296 | 9.76973008201657 | Gm20757 |
| **ENSMUSG00000098076** | -1.07596308371795 | 0.0154588445378836 | 0.635862253526407 | 7.22810973255236 | 11.4018202048006 | 6.62518422053913 | 5.62756016554897 | 16.4243729372865 | 20.7923979452556 | 13.5340151772298 | 15.1973579053591 | Gm6981 |
| **ENSMUSG00000098098** | -0.17891517514506 | 0.547926554731101 | 0.914699131434573 | 134.623543768788 | 70.8541684155466 | 106.949402417274 | 125.682177030594 | 133.58489988993 | 181.016170346931 | 90.5737938783839 | 91.1841474321546 | Bvht |
| **ENSMUSG00000098112** | -0.00696494298711341 | 0.988147594887256 | 0.998437672973644 | 14.4562194651047 | 15.4738988493722 | 47.3227444324223 | 22.5102406621959 | 24.0890803080201 | 36.6924669622158 | 15.616171358342 | 23.8815624227072 | Bin2 |
| **ENSMUSG00000098123** | -0.47904384200838 | 0.101647866995507 | 0.759800953500741 | 29.8159526467785 | 23.6180561385155 | 28.3936466594534 | 28.1378008277448 | 38.3235368536684 | 53.8156182112498 | 30.191264626128 | 31.4802413753867 | Gm4518 |
| **ENSMUSG00000098132** | -0.269165648364769 | 0.555807012314371 | 0.916239900758826 | 23.4913566307952 | 14.6594831204579 | 33.1259211026956 | 53.4618215727152 | 54.7479097909548 | 45.2540425867328 | 20.8215618111227 | 29.3091902460497 | Rassf10 |
| **ENSMUSG00000098134** | 0.169522599361572 | 0.297974874482769 | 0.851785577030345 | 278.282224703266 | 291.56083095133 | 256.489274823729 | 297.322762079837 | 293.448796479518 | 280.085831144914 | 230.078258012906 | 196.480127205 | Rnf113a2 |
| **ENSMUSG00000098156** | 1.27444805983755 | 0.0088265119452370 | 0.635862253526407 | 16.2632468982428 | 17.9171460361152 | 14.1968233297267 | 10.3171936368398 | 6.56974917491458 | 6.1154111603693 | 7.28754663389296 | 4.34210225867403 | Gm27023 |
| **ENSMUSG00000098176** | -0.544467755300033 | 0.228717750344468 | 0.832048466795696 | 25.2983840639333 | 16.2883145782866 | 15.1432782183751 | 16.8826804966469 | 29.5638712871156 | 47.7002070508805 | 17.6983275394543 | 13.0263067760221 | Ccdc166 |
| **ENSMUSG00000098178** | -0.796314257019661 | 0.0716772515491387 | 0.728065810245271 | 67.7635287426784 | 54.56585383726 | 39.7511053232348 | 79.7237690119437 | 96.3563212320805 | 211.593226148778 | 53.094982618363 | 59.7039060567679 | Gm42418 |
| **ENSMUSG00000098188** | -0.252835426436665 | 0.411886941703419 | 0.882192126163777 | 32.5264937964856 | 35.0198763433161 | 34.0723759913441 | 30.9515809105193 | 58.0327843784121 | 29.3539735697726 | 27.0680303544596 | 43.4210225867403 | Sowahc |
| **ENSMUSG00000098192** | 0.0364921553716959 | 0.89310164315126 | 0.983367564278319 | 45.1756858284523 | 31.7622134276588 | 51.1085639870161 | 35.6412143818101 | 38.3235368536684 | 50.1463715150282 | 33.3144988977964 | 37.9933947633978 | Gm3534 |
| **ENSMUSG00000098196** | -0.0570653245099693 | 0.930689450528881 | 0.987719779022475 | 1.80702743313809 | 19.5459774939439 | 13.2503684410783 | 7.50341355406529 | 6.56974917491458 | 8.56157562451702 | 12.4929370866736 | 16.2828834700276 | Gm26964 |
| **ENSMUSG00000098234** | 0.7161471362761 | 0.0036997897449620 | 0.635862253526407 | 151.7903043836 | 114.83261777692 | 159.950876181588 | 205.405946042537 | 114.970610561005 | 114.969729814943 | 83.2862472444909 | 71.6446872681215 | Snhg6 |
| **ENSMUSG00000098274** | 0.00437820836576825 | 0.982868854235437 | 0.996940244108351 | 2225.35428390956 | 1485.49428953974 | 1785.96037487962 | 2557.72609524201 | 2091.37015401448 | 2438.82597075528 | 1667.80710107093 | 1832.36715316044 | Rpl24 |
| **ENSMUSG00000098332** | 0.189691781496956 | 0.28597837481634 | 0.851593746788839 | 171.667606148119 | 150.666909849151 | 124.932045301595 | 162.261318106662 | 136.869774477387 | 156.554525705454 | 131.175839410073 | 110.723607596188 | 2310009A05 |

| | | | | | | | | | | | | |
|---|---|---|---|---|---|---|---|---|---|---|---|---|
| **ENSMUSG00000098374** | 0.0196340169411917 | 0.941695821677974 | 0.989934163630124 | 34.3335212296237 | 35.8342920722305 | 34.0723759913441 | 35.6412143818101 | 27.3739548954774 | 29.3539735697726 | 35.3966550789086 | 45.5920737160773 | Gm28043 |
| **ENSMUSG00000098424** | 0.444350927373334 | 0.32742752507331 | 0.855977678491063 | 8.13162344912141 | 21.1748089517725 | 17.036187995672 | 21.5723139679377 | 7.66470737073368 | 15.9000690169602 | 14.5750932677859 | 11.9407812113536 | Gm27202 |
| **ENSMUSG00000098557** | -0.350751698711713 | 0.0955768649869871 | 0.757085157577544 | 188.83436676293 | 162.883145782866 | 213.898804834549 | 210.095579513828 | 308.778211220985 | 276.416584448692 | 170.736806851206 | 233.387996403729 | Kctd12 |
| **ENSMUSG00000098679** | 0.688247641299031 | 0.264827287754584 | 0.842510961779757 | 26.2018977805023 | 6.51532583131463 | 8.51809399783602 | 17.8206071909051 | 3.28487458745729 | 9.78465785659088 | 8.32862472444909 | 15.1973579053591 | Mir7688 |
| **ENSMUSG00000098702** | 0.0583247512645004 | 0.827928928202457 | 0.967319591893432 | 32.5264937964856 | 35.0198763433161 | 39.7511053232348 | 41.2687745473591 | 29.5638712871156 | 35.4693847301419 | 46.8485140750261 | 30.3947158107182 | 1500015A07 |
| **ENSMUSG00000098708** | -0.216556304565195 | 0.444667597740241 | 0.89108031640712 | 39.754603529038 | 35.8342920722305 | 29.3401015481019 | 26.2619474392285 | 33.943704070392 | 41.5847958905112 | 47.8895921655823 | 29.3091902460497 | Gm27252 |
| **ENSMUSG00000098789** | 0.326064600993629 | 0.541159059967696 | 0.913837378403038 | 9.03513716569045 | 15.4738988493722 | 5.67872933189068 | 18.7585338851632 | 6.56974917491458 | 14.6769867848863 | 7.28754663389296 | 10.8552556466851 | Jmjd7 |
| **ENSMUSG00000098900** | 0.0704018096587941 | 0.923699355340328 | 0.986713918026868 | 7.22810973255236 | 10.5874044758863 | 32.1794662140472 | 9.37926694258161 | 2.18991639163819 | 7.33849339244316 | 24.9858741733473 | 21.7105112933701 | Gm18190 |
| **ENSMUSG00000098905** | -0.143941053551081 | 0.650680862288052 | 0.936463929813731 | 50.5967681278665 | 35.0198763433161 | 20.8220075502658 | 39.3929211588428 | 36.1336204620302 | 50.1463715150282 | 41.6431236222455 | 33.6512925047237 | Zfp953 |
| **ENSMUSG00000098912** | -0.529295827254579 | 0.17717468623577 | 0.800237299431217 | 9.93865088225949 | 13.8450673915436 | 13.2503684410783 | 13.1309737196143 | 21.8991639163819 | 23.2385624094033 | 12.4929370866736 | 15.1973579053591 | 1500004A13 |
| **ENSMUSG00000098923** | -0.106592610352408 | 0.506413055632979 | 0.901689186353298 | 219.553833126278 | 230.479651282755 | 210.112985279955 | 210.095579513828 | 247.460552255116 | 276.416584448692 | 234.242570375131 | 180.197243734972 | Tmem185b |
| **ENSMUSG00000098973** | 0.503478767235189 | 0.116613644626438 | 0.769868040577702 | 721.907459538667 | 612.440628143575 | 847.07712534036 | 1326.22834568104 | 1010.64641474103 | 513.694537471021 | 437.252798033577 | 512.368066523536 | Mir6236 |
| **ENSMUSG00000099034** | 0.0838511788596769 | 0.847744551869312 | 0.972455319662159 | 20.780815481088 | 26.0613033252585 | 9.46454888648447 | 13.1309737196143 | 15.3294147414674 | 11.0077400886647 | 20.8215618111227 | 18.4539345993646 | 2810039B14 |
| **ENSMUSG00000099041** | 0.179974645338045 | 0.472008069096267 | 0.897171173241185 | 46.0791995450213 | 44.7928650902881 | 54.8943835416099 | 59.0893817382642 | 41.6084114411257 | 35.4693847301419 | 59.3414511616998 | 43.4210225867403 | Gm28035 |
| **ENSMUSG00000099083** | 0.0144468206666995 | 0.923029965072456 | 0.986713918026868 | 402.063603873225 | 442.22774080048 | 493.102996985841 | 363.915557372167 | 370.095870186855 | 387.717067567413 | 459.115437935256 | 466.775992807458 | Atf7 |
| **ENSMUSG00000099137** | 0.505803173175545 | 0.0600740630053365 | 0.71282992338528 | 36.1405486627618 | 39.0919549878878 | 50.1621090983677 | 38.4549944645846 | 31.7537876787538 | 28.1308913376988 | 22.903717992235 | 32.5657669400552 | Gm10603 |
| **ENSMUSG00000099146** | -0.216667970301768 | 0.713645547989099 | 0.951025296130537 | 9.93865088225949 | 8.95857301805762 | 13.2503684410783 | 15.0068271081306 | 9.85462376237187 | 14.6769867848863 | 3.12323427166841 | 27.1381391167127 | 0610031O16 |
| **ENSMUSG00000099411** | 0.0606224070956081 | 0.921935744356675 | 0.986713918026868 | 14.4562194651047 | 14.6594831204579 | 8.51809399783602 | 10.3171936368398 | 2.18991639163819 | 18.3462334811079 | 6.24646854333682 | 19.5394601640331 | 2310015D24 |
| **ENSMUSG00000099478** | 0.0874531644497337 | 0.820367103734717 | 0.966398681144067 | 20.780815481088 | 16.2883145782866 | 19.8755526616174 | 10.3171936368398 | 13.1394983498292 | 19.5693157131818 | 15.616171358342 | 15.1973579053591 | Gm28370 |
| **ENSMUSG00000099481** | 0.00135586225805729 | 0.996336315583651 | 0.999262355603273 | 30.7194663633475 | 31.7622134276588 | 23.6613722162112 | 39.3929211588428 | 27.3739548954774 | 34.2463024980681 | 24.9858741733473 | 39.0789203280663 | Xndc1 |
| **ENSMUSG00000099583** | -0.679737832193757 | 0.125866834018386 | 0.771949105062466 | 11.7456783153976 | 8.14415728914329 | 7.57163910918757 | 8.44134024832345 | 13.1394983498292 | 12.2308223207386 | 11.4518589961175 | 20.6249857287016 | Hist1h3d |
| **ENSMUSG00000099597** | 0.0129495231345047 | 0.945853491638522 | 0.990505437194639 | 649.626362213143 | 507.380999113627 | 499.72818120638 | 742.837941852464 | 667.924499449649 | 665.35673424818 | 461.197594116369 | 584.012753791657 | NA |
| **ENSMUSG00000099622** | 0.273315972226269 | 0.620103810036127 | 0.929720349068849 | 11.7456783153976 | 10.5874044758863 | 16.0897331070236 | 6.56548685980713 | 6.56974917491458 | 18.3462334811079 | 7.28754663389296 | 5.42762782334254 | Gm29188 |
| **ENSMUSG00000099624** | -0.353797763682044 | 0.337248716747385 | 0.858570960161243 | 35.2370349461928 | 45.6072808192024 | 18.9290977729689 | 19.6964605794214 | 28.4689130912965 | 34.2463024980681 | 40.6020455316893 | 49.9341759747513 | Gm7889 |
| **ENSMUSG00000099632** | 0.0150264526342328 | 0.966228690024684 | 0.994284908685797 | 11.7456783153976 | 28.5045505120015 | 23.6613722162112 | 25.3240207449704 | 21.8991639163819 | 20.7923979452556 | 22.903717992235 | 22.7960368580387 | 2900093K20 |
| **ENSMUSG00000099647** | -0.401775851495862 | 0.425888358592871 | 0.885066977635295 | 15.3597331816738 | 7.32974156022896 | 10.4110037751329 | 11.2551203310979 | 13.1394983498292 | 25.6847268735511 | 13.5340151772298 | 6.51315338801105 | Gm5776 |
| **ENSMUSG00000099655** | 0.130135946062198 | 0.71151797388065 | 0.950020479830613 | 30.7194663633475 | 17.1027303072009 | 17.9826428843205 | 28.1378008277448 | 17.5193311331055 | 26.9078091056249 | 18.7394056300105 | 22.7960368580387 | 2310034G01 |
| **ENSMUSG00000099681** | -0.0458338198159106 | 0.820986781566463 | 0.96650690096489 | 84.9302893574902 | 72.4829998733752 | 63.4124775394459 | 74.0962088463947 | 82.1218646864323 | 79.5003450848009 | 83.2862472444909 | 59.7039060567679 | 1700052K11 |
| **ENSMUSG00000099689** | 0.113184976519694 | 0.518134081791479 | 0.905473752527583 | 100.290022539164 | 122.162359337149 | 96.5383986421416 | 92.854742731558 | 100.736154015357 | 101.51582526213 | 88.4916376972716 | 91.1841474321546 | Zfp383 |
| **ENSMUSG00000099696** | -0.0468121385071898 | 0.90392150981125 | 0.985039825636533 | 26.2018977805023 | 26.0613033252585 | 17.9826428843205 | 15.9447538023887 | 13.1394983498292 | 18.3462334811079 | 23.9447960827911 | 33.6512925047237 | 2900052N01 |
| **ENSMUSG00000099757** | -0.901033007851885 | 0.12150591592001 | 0.769868040577702 | 9.93865088225949 | 8.95857301805762 | 10.4110037751329 | 3.75170677703264 | 17.5193311331055 | 25.6847268735511 | 14.5750932677859 | 4.34210225867403 | BE692007 |
| **ENSMUSG00000099764** | 0.0457025379851974 | 0.862788060590478 | 0.974868041734905 | 39.754603529038 | 36.6487078011448 | 33.1259211026956 | 32.8274342990356 | 33.943704070392 | 44.0309603546589 | 27.0680303544596 | 33.6512925047237 | Rps10-ps2 |
| **ENSMUSG00000099835** | -0.0149454553752129 | 0.968850743551967 | 0.994850064902552 | 12.6491920319666 | 20.3603932228582 | 18.9290977729689 | 11.2551203310979 | 17.5193311331055 | 13.4539045528125 | 16.6572494488982 | 16.2828834700276 | Gm29668 |
| **ENSMUSG00000099843** | -0.0844237984559772 | 0.700927039900228 | 0.945920819861844 | 54.2108229941427 | 52.122606650517 | 61.519567762149 | 51.5859681841989 | 49.2731188118594 | 72.1618516923577 | 51.0128264372507 | 60.7894316214364 | Gm7160 |
| **ENSMUSG00000099858** | 0.360928231403333 | 0.199267491179247 | 0.812921911423739 | 58.7283915769879 | 61.895595397489 | 87.0738497556571 | 67.5307219865876 | 49.2731188118594 | 33.0232202659942 | 56.2182168900314 | 74.901263962127 | Gm6652 |
| **ENSMUSG00000099881** | 0.119515619248708 | 0.530382768735419 | 0.908923862033581 | 250.273299489625 | 209.304842330982 | 246.078271048596 | 268.247034557834 | 180.668102310151 | 183.462334811079 | 249.858741733473 | 281.151121249143 | 2810013P06 |
| **ENSMUSG00000099966** | -0.022288242291656 | 0.90257848715848 | 0.985027469232262 | 80.412720774645 | 81.4415728914329 | 87.0738497556571 | 83.4754757889764 | 88.6916138613469 | 68.4926049961361 | 96.8202624217207 | 82.4999429148066 | 2810402E24 |
| **ENSMUSG00000100017** | -0.224955095470744 | 0.272951337255583 | 0.844738346035895 | 84.0267756409212 | 74.9262470601182 | 65.3053873167428 | 62.8410885152968 | 72.2672409240604 | 103.961989726278 | 81.2040910633787 | 79.2433662208011 | 2410022M11 |
| **ENSMUSG00000100075** | -0.043203350687244 | 0.896151466441291 | 0.984073355116573 | 24.3948703473642 | 45.6072808192024 | 27.447191770805 | 27.1998741334867 | 22.994122112201 | 28.1308913376988 | 39.5609674411332 | 37.9933947633978 | 1700018L02 |
| **ENSMUSG00000100078** | -0.812801175923232 | 0.0659877286578476 | 0.719725956118125 | 15.3597331816738 | 13.0306516626293 | 11.3574586637814 | 12.1930470253561 | 15.3294147414674 | 25.6847268735511 | 12.4929370866736 | 37.9933947633978 | Gm6170 |
| **ENSMUSG00000100084** | 0.156618828144598 | 0.721975136649244 | 0.951674101666722 | 10.8421645988285 | 17.9171460361152 | 19.8755526616174 | 22.5102406621959 | 15.3294147414674 | 8.56157562451702 | 13.5340151772298 | 26.0526135520442 | 2310068J16 |
| **ENSMUSG00000100131** | 0.106269357943146 | 0.570025882116864 | 0.920872458148558 | 436.397125102849 | 419.424100390879 | 391.832323900457 | 458.646153492241 | 300.018545654433 | 344.909189444828 | 419.554470494123 | 519.966745476215 | Gm28439 |
| **ENSMUSG00000100141** | 0.221146549091845 | 0.533169499350457 | 0.909124029705112 | 27.1054114970714 | 25.2468875963442 | 54.8943835416099 | 30.0136542162612 | 30.6588294829347 | 26.9078091056249 | 21.8626399016789 | 37.9933947633978 | Gm6445 |
| **ENSMUSG00000100153** | -0.0640050290679233 | 0.875419185562874 | 0.976986380159756 | 23.4913566307952 | 15.4738988493722 | 29.3401015481019 | 21.5723139679377 | 17.5193311331055 | 28.1308913376988 | 12.4929370866736 | 35.8223436340608 | Gm5601 |
| **ENSMUSG00000100164** | 0.0782281204782037 | 0.645587205127189 | 0.935063240511793 | 115.649755720838 | 120.533527879321 | 105.056492639978 | 131.309737196143 | 98.5462376237187 | 123.53130543946 | 124.929370866736 | 100.953877514171 | 2610306M01 |
| **ENSMUSG00000100215** | 0.356725805911247 | 0.286010426837288 | 0.851593746788839 | 28.0089252136404 | 21.1748089517725 | 38.8046504345863 | 40.3308478531009 | 17.5193311331055 | 28.1308913376988 | 24.9858741733473 | 29.3091902460497 | Gm8292 |
| **ENSMUSG00000100235** | -0.454902073156005 | 0.308080598651287 | 0.851785577030345 | 11.7456783153976 | 8.95857301805762 | 11.3574586637814 | 11.2551203310979 | 13.1394983498292 | 7.33849339244316 | 16.6572494488982 | 21.7105112933701 | Gm28557 |
| **ENSMUSG00000100291** | -0.0402772148146259 | 0.929887462832967 | 0.987529878729394 | 14.4562194651047 | 18.7315617650296 | 16.0897331070236 | 6.56548685980713 | 14.2344565456483 | 13.4539045528125 | 9.36970281500523 | 20.6249857287016 | 2310069B03 |
| **ENSMUSG00000100347** | 0.610325471794332 | 0.131111020853841 | 0.77303252008843 | 21.6843291976571 | 26.8757190541728 | 26.5007368821565 | 23.448167356454 | 10.949581958191 | 7.33849339244316 | 21.8626399016789 | 23.8815624227072 | Gm7895 |
| **ENSMUSG00000100410** | 0.178684556232393 | 0.566194614033883 | 0.919931558693298 | 98.4829951060259 | 62.7100111264033 | 51.1085639870161 | 120.992543559303 | 65.6974917491458 | 59.9310293716191 | 73.9165444294857 | 94.4407241261602 | 2310020H05 |
| **ENSMUSG00000100455** | 0.210427163704855 | 0.540448548183992 | 0.913837378403038 | 60.535419010126 | 65.1532583131463 | 104.110037751329 | 46.8963347129081 | 39.4184950494875 | 48.9232892829544 | 91.61487196894 | 58.6183804920994 | Gm29170 |
| **ENSMUSG00000100457** | 0.0654798697737583 | 0.831968440948353 | 0.967583158900775 | 92.1583990900426 | 72.4829998733752 | 126.824955078892 | 92.854742731558 | 75.5521155115177 | 51.3694537471021 | 118.6829023234 | 120.493337678204 | D830032E09 |
| **ENSMUSG00000100510** | 0.158036237381132 | 0.268451453998556 | 0.843793744782845 | 736.363679003772 | 592.894650649631 | 696.590798045257 | 755.03098887782 | 605.511882287961 | 667.802898712327 | 699.604476853724 | 518.881219911547 | AV026068 |
| **ENSMUSG00000100550** | 0.139519786669578 | 0.582995449626243 | 0.922450244597442 | 55.1143367107117 | 43.1640336324594 | 52.0550188756646 | 40.3308478531009 | 55.8428679867739 | 44.0309603546589 | 38.5198893505771 | 34.7368180693922 | 2310039L15 |
| **ENSMUSG00000100594** | 0.321137909102995 | 0.166363972163852 | 0.796299608935682 | 60.535419010126 | 66.7820897709749 | 65.3053873167428 | 46.8963347129081 | 48.1781606160403 | 37.9155491942896 | 49.9717483466946 | 55.3618037980939 | 2810414N06 |
| **ENSMUSG00000100599** | -0.23373918937726 | 0.425525439230843 | 0.885066977635295 | 159.018414116152 | 100.987550385377 | 120.199770858353 | 120.054616865045 | 99.6411958195378 | 236.054870790255 | 115.559668051731 | 137.8617467129 | 1700120C14 |
| **ENSMUSG00000100680** | 0.249535290757722 | 0.271459471408147 | 0.844129257012535 | 72.2810973255236 | 64.338842584232 | 49.2156542097192 | 81.59962240046 | 55.8428679867739 | 59.9310293716191 | 58.3003730711437 | 51.0197015394199 | 1810044D09 |
| **ENSMUSG00000100691** | 0.049616782510901 | 0.837202486715299 | 0.968924788429433 | 219.553833126278 | 201.160685041839 | 156.165056626994 | 220.412773150668 | 141.249607260664 | 281.308913376988 | 163.449260217313 | 185.624871558315 | 2010320M18 |
| **ENSMUSG00000100707** | 0.265148470777957 | 0.596681565217594 | 0.924496535139492 | 18.9737880479499 | 10.5874044758863 | 21.7684624389143 | 24.3860940507122 | 10.949581958191 | 17.123151249034 | 28.1091084450157 | 6.51315338801105 | Gm28523 |
| **ENSMUSG00000100747** | 0.2562635095448 | 0.593973895912333 | 0.923935529355527 | 12.6491920319666 | 19.5459774939439 | 25.5542819935081 | 30.0136542162612 | 13.1394983498292 | 8.56157562451702 | 17.6983275394543 | 33.6512925047237 | 1700084E18 |
| **ENSMUSG00000100755** | -0.193950934282707 | 0.519553845695006 | 0.905962179970576 | 27.1054114970714 | 24.4324718674299 | 29.3401015481019 | 40.3308478531009 | 32.8487458745729 | 45.2540425867328 | 26.0269522639034 | 34.7368180693922 | Rps23-ps1 |
| **ENSMUSG00000100782** | -0.0315171268007482 | 0.951518454778235 | 0.991915011884503 | 18.0702743313809 | 9.77298874697194 | 5.67872933189068 | 20.6343872736795 | 13.1394983498292 | 11.0077400886647 | 9.36970281500523 | 21.7105112933701 | Gm28231 |
| **ENSMUSG00000100794** | 0.354247785951551 | 0.421706133491896 | 0.882568531013176 | 11.7456783153976 | 20.3603932228582 | 12.3039135524298 | 12.1930470253561 | 7.66470737073368 | 14.6769867848863 | 9.36970281500523 | 13.0263067760221 | Gm29667 |
| **ENSMUSG00000100801** | -0.174416711159894 | 0.573254912363403 | 0.922207298536842 | 22.5878429142261 | 26.8757190541728 | 23.6613722162112 | 18.7585338851632 | 27.3739548954774 | 22.0154801773295 | 21.8626399016789 | 32.5657669400552 | Gm15459 |
| **ENSMUSG00000100819** | 0.501299906732687 | 0.427866237776309 | 0.885066977635295 | 13.5527057485357 | 19.5459774939439 | 5.67872933189068 | 7.50341355406529 | 6.56974917491458 | 17.123151249034 | 4.16431236222455 | 5.42762782334254 | Gm2693 |
| **ENSMUSG00000100826** | 0.31704005233793 | 0.541437597699189 | 0.913837378403038 | 14.4562194651047 | 13.0306516626293 | 8.51809399783602 | 7.50341355406529 | 7.66470737073368 | 3.66924669622158 | 11.4518589961175 | 11.9407812113536 | Snhg14 |
| **ENSMUSG00000100862** | -0.129731471833671 | 0.44993805668683 | 0.89108031640712 | 805.934235179588 | 660.491156149521 | 618.981497176084 | 844.134024832345 | 765.375778877549 | 991.9196902119 | 665.248899865371 | 783.749457690662 | Gm10925 |
| **ENSMUSG00000100863** | 0.0927572219531232 | 0.762335744892702 | 0.957746336519503 | 24.3948703473642 | 28.5045505120015 | 28.3936466594534 | 36.5791410760683 | 30.6588294829347 | 20.7923979452556 | 22.903717992235 | 35.8223436340608 | Gm12669 |
| **ENSMUSG00000100954** | 0.0954470728069603 | 0.502417421521439 | 0.900927059989498 | 171.667606148119 | 151.481325578065 | 143.861143074564 | 155.695831246855 | 151.104231023035 | 134.539045528125 | 141.586620315635 | 155.230155747597 | Gm10138 |
| **ENSMUSG00000100975** | -0.417460051652495 | 0.192976602171676 | 0.808824684419004 | 18.9737880479499 | 17.9171460361152 | 25.5542819935081 | 25.3240207449704 | 36.1336204620302 | 34.2463024980681 | 21.8626399016789 | 24.9670879873757 | Gm28875 |
| **ENSMUSG00000101013** | -0.0924920135481424 | 0.825114967415598 | 0.96672635991852 | 18.9737880479499 | 15.4738988493722 | 9.46454888648447 | 14.0689004138724 | 18.6142893289246 | 15.9000690169602 | 18.7394056300105 | 8.68420451734806 | A630072M18 |
| **ENSMUSG00000101028** | 0.137216901577938 | 0.722406527497632 | 0.951674101666722 | 34.3335212296237 | 27.6901347830872 | 12.3039135524298 | 41.2687745473591 | 26.2789966996583 | 24.4616446414772 | 22.903717992235 | 31.4802413753867 | Gm28723 |
| **ENSMUSG00000101086** | 0.395050506557491 | 0.205585007907707 | 0.819960256425221 | 32.5264937964856 | 30.9477976987445 | 43.5369248778286 | 45.0204813243917 | 22.994122112201 | 19.5693157131818 | 37.4788112600209 | 34.7368180693922 | Gm28651 |
| **ENSMUSG00000101102** | 0.382088278400156 | 0.34765344950454 | 0.861266047293794 | 14.4562194651047 | 14.6594831204579 | 25.5542819935081 | 15.9447538023887 | 9.85462376237187 | 11.0077400886647 | 16.6572494488982 | 16.2828834700276 | Gm28609 |
| **ENSMUSG00000101111** | -0.134168201150612 | 0.459026531586822 | 0.894609025360033 | 973.987786461431 | 702.026358324151 | 736.341903368492 | 936.988767563903 | 924.144717271318 | 1133.79722913247 | 733.960053842076 | 884.703335204834 | Gm28437 |
| **ENSMUSG00000101188** | 0.0104430646712852 | 0.969698018880547 | 0.9951105526746 | 57.8248778604189 | 41.5352021746308 | 53.9479286529615 | 70.3445020693621 | 63.5075753575076 | 62.3771938357668 | 35.3966550789086 | 60.7894316214364 | Eif4a-ps4 |
| **ENSMUSG00000101225** | -0.206258626212452 | 0.480244033079745 | 0.897732059433541 | 35.2370349461928 | 27.6901347830872 | 42.5904699891801 | 48.7721881014244 | 36.1336204620302 | 35.4693847301419 | 46.8485140750261 | 58.6183804920994 | 1700008J07 |
| **ENSMUSG00000101249** | 0.109117165288628 | 0.615916541296708 | 0.928422072203769 | 611.678786117243 | 398.249291439107 | 441.047978110176 | 597.459304242449 | 545.28918151791 | 578.517895770936 | 361.254097422979 | 414.67076570337 | Gm29216 |
| **ENSMUSG00000101304** | 0.380299489403294 | 0.0134029813953648 | 0.635862253526407 | 454.46739943423 | 399.063707168021 | 336.937940358847 | 434.260059441529 | 331.772333333186 | 255.624186503437 | 335.227145159076 | 324.572143835884 | Plet1os |
| **ENSMUSG00000101320** | -0.332086116885411 | 0.372997253478813 | 0.873700364743909 | 14.4562194651047 | 21.9892246806869 | 22.7149173275627 | 35.6412143818101 | 32.8487458745729 | 37.9155491942896 | 30.191264626128 | 18.4539345993646 | Gm28529 |
| **ENSMUSG00000101386** | 0.0669533918233278 | 0.889167392653456 | 0.98126431530921 | 30.7194663633475 | 17.1027303072009 | 8.51809399783602 | 18.7585338851632 | 16.4243729372865 | 25.6847268735511 | 9.36970281500523 | 20.6249857287016 | Gm29346 |
| **ENSMUSG00000101389** | -0.643421530176694 | 0.0928138260375187 | 0.754058637722175 | 101.193536255733 | 122.162359337149 | 125.878500190243 | 108.799496533947 | 250.745426842573 | 283.755077841135 | 90.5737938783839 | 91.1841474321546 | Ms4a4a |
| **ENSMUSG00000101523** | -0.122906086107009 | 0.696568437321982 | 0.944933386973863 | 28.9124389302094 | 29.3189662409158 | 31.2330113253987 | 19.6964605794214 | 25.1840385038392 | 22.0154801773295 | 31.2323427166841 | 40.1644458927348 | Gm10031 |
| **ENSMUSG00000101609** | 0.229472463371136 | 0.556309083460949 | 0.916239900758826 | 39.754603529038 | 33.3910448854875 | 17.9826428843205 | 25.3240207449704 | 29.5638712871156 | 35.4693847301419 | 20.8215618111227 | 14.1118323406906 | Kcnq1ot1 |
| **ENSMUSG00000101634** | 0.477312320445568 | 0.279603041167374 | 0.848522228930484 | 17.1667606148119 | 10.5874044758863 | 16.0897331070236 | 10.3171936368398 | 7.66470737073368 | 11.0077400886647 | 9.36970281500523 | 10.8552556466851 | 1700066B17 |
| **ENSMUSG00000101655** | 0.191165155733413 | 0.265849297212274 | 0.843277879501733 | 241.238162323935 | 196.274190668353 | 198.755526616174 | 266.371181169318 | 187.237851485066 | 232.385624094033 | 169.69572876065 | 201.907755028342 | 2310040G24 |
| **ENSMUSG00000101660** | -0.641466099794253 | 0.428681676592615 | 0.885378691567738 | 9.93865088225949 | 11.4018202048006 | 13.2503684410783 | 12.1930470253561 | 3.28487458745729 | 1.22308223207386 | 30.191264626128 | 37.9933947633978 | Gm18666 |

| | | | | | | | | | | | | |
|---|---|---|---|---|---|---|---|---|---|---|---|---|
| ENSMUSG00000101693 | -0.112303271576419 | 0.773839654996797 | 0.959203117135841 | 84.9302893574902 | 141.708336831093 | 112.628131749165 | 54.3997482669734 | 63.5075753575076 | 66.0464405319884 | 165.531416398426 | 130.263067760221 | Gm19461 |
| ENSMUSG00000101795 | -0.384008038333273 | 0.28576145535319 | 0.85139996028653 | 22.5878429142261 | 14.6594831204579 | 11.3574586637814 | 17.8206071909051 | 18.6142893289246 | 25.6847268735511 | 18.7394056300105 | 23.8815624227072 | Gm5835 |
| ENSMUSG00000101856 | -0.0984718102247766 | 0.608703690824602 | 0.927932112008768 | 64.1494738764022 | 75.7406627890326 | 82.3415753124149 | 63.779015209555 | 70.0773245324222 | 74.6080161565054 | 85.3684034256032 | 75.9867895267955 | 1700096K18 |
| ENSMUSG00000101872 | 0.52837726042534 | 0.333438227121637 | 0.85715806744223 | 116.553269437407 | 175.099381716581 | 107.895857305923 | 105.047789756914 | 54.7479097909548 | 13.4539045528125 | 123.88829277618 | 157.401206876934 | Gm29237 |
| ENSMUSG00000101892 | -0.0112623072542654 | 0.956351627299681 | 0.992371505557225 | 233.106538874814 | 173.470550258752 | 172.254789734017 | 248.550573978413 | 223.371471947096 | 258.070350967584 | 171.777884941763 | 181.282769299641 | 9130401M01 |
| ENSMUSG00000101939 | -0.0440467969781871 | 0.853596771324793 | 0.9731539929935 | 82.2197482077831 | 57.009101024003 | 74.7699362032273 | 88.1651092602672 | 96.3563212320805 | 79.5003450848009 | 56.2182168900314 | 79.2433662208011 | Gm28438 |
| ENSMUSG00000101941 | -0.138046645512286 | 0.769140150562143 | 0.95879680569409 | 25.2983840639333 | 22.8036404096012 | 10.4110037751329 | 14.0689004138724 | 10.949581958191 | 13.4539045528125 | 22.903717992235 | 32.5657669400552 | Gm28979 |
| ENSMUSG00000101970 | 0.323260800305385 | 0.133509190531035 | 0.77303252008843 | 182.509770746947 | 156.367819951551 | 204.434255948065 | 271.060814640609 | 169.71852035196 | 163.893019097897 | 187.394056300105 | 129.177542195552 | 1810026B05 |
| ENSMUSG00000102014 | 0.382125561896055 | 0.321797736032233 | 0.85578605274511 | 19.877301764519 | 19.5459774939439 | 12.3039135524298 | 24.3860940507122 | 13.1394983498292 | 11.0077400886647 | 15.616171358342 | 18.4539345993646 | 2900009J06 |
| ENSMUSG00000102062 | 0.172018309105811 | 0.678135265801363 | 0.944452327903597 | 13.5527057485357 | 25.2468875963442 | 16.0897331070236 | 14.0689004138724 | 15.3294147414674 | 8.56157562451702 | 17.6983275394543 | 19.5394601640331 | Gm29246 |
| ENSMUSG00000102070 | 0.0158915038065085 | 0.928684990241831 | 0.986901071486195 | 725.521514404943 | 595.337897836374 | 612.356312955545 | 797.237690119437 | 623.031213421066 | 873.280713700736 | 575.716184077544 | 629.604827507734 | Gm28661 |
| ENSMUSG00000102091 | -0.0310498839377546 | 0.936351237697307 | 0.988592177836394 | 24.3948703473642 | 15.4738988493722 | 14.1968233297267 | 20.6343872736795 | 10.949581958191 | 18.3462334811079 | 20.8215618111227 | 26.0526135520442 | Olfr1034 |
| ENSMUSG00000102101 | 0.03397676715617 | 0.889573318916421 | 0.981427593308109 | 113.8427282877 | 108.317291945606 | 110.735221971868 | 137.87522405595 | 148.914314631397 | 140.654456688494 | 93.6970281500523 | 77.072315091464 | Zbtb11os1 |
| ENSMUSG00000102252 | -0.00108094894574676 | 0.992141591780111 | 0.998990770555763 | 2174.75751578169 | 2203.80896244217 | 2214.70443943737 | 1858.97070801968 | 1863.6188492841 | 2158.74013961036 | 2208.12663006957 | 2228.58398426445 | Snrpn |
| ENSMUSG00000102260 | 0.0772857639758819 | 0.730834054550901 | 0.952921973914044 | 170.764092431549 | 177.542628903324 | 204.434255948065 | 203.530092654021 | 163.148771177045 | 135.762127760198 | 260.269522639034 | 156.315681312265 | D330025C20 |
| ENSMUSG00000102426 | -0.192737573531834 | 0.72576910212567 | 0.951749700578319 | 10.8421645988285 | 7.32974156022896 | 15.1432782183751 | 11.2551203310979 | 17.5193311331055 | 20.7923979452556 | 4.16431236222455 | 8.68420451734806 | NA |
| ENSMUSG00000102428 | -0.563440303976302 | 0.20933322513894 | 0.821566820583092 | 14.4562194651047 | 24.4324718674299 | 17.9826428843205 | 21.5723139679377 | 25.1840385038392 | 11.0077400886647 | 49.9717483466946 | 29.3091902460497 | Pcdhga12 |
| ENSMUSG00000102531 | -0.272503504136047 | 0.58296378687998 | 0.922450244597442 | 16.2632468982428 | 12.2162359337149 | 4.73227444324223 | 10.3171936368398 | 16.4243729372865 | 7.33849339244316 | 12.4929370866736 | 16.2828834700276 | 1110002O04 |
| ENSMUSG00000102545 | -0.0986227257423991 | 0.792938189016002 | 0.961804768755116 | 19.877301764519 | 30.9477976987445 | 34.0723759913441 | 18.7585338851632 | 33.943704070392 | 33.0232202659942 | 30.191264626128 | 14.1118323406906 | 6430573P05 |
| ENSMUSG00000102644 | 0.0281331410340321 | 0.921039893777634 | 0.986713918026868 | 113.8427282877 | 114.83261777692 | 83.2880302010633 | 99.4202295913651 | 87.5966556655277 | 57.4848649074714 | 144.709854587303 | 112.894658725525 | Thap6 |
| ENSMUSG00000102742 | -0.358801117407861 | 0.273634882264987 | 0.845204865395511 | 15.3597331816738 | 17.9171460361152 | 22.7149173275627 | 18.7585338851632 | 24.0890803080201 | 30.5770558018465 | 20.8215618111227 | 20.6249857287016 | Pcdhga11 |
| ENSMUSG00000102748 | -0.199688484014547 | 0.502581171852935 | 0.900927059989498 | 20.780815481088 | 42.3496179035451 | 35.965285768641 | 43.1446279358754 | 39.4184950494875 | 35.4693847301419 | 39.5609674411332 | 48.8486504100828 | Pcdhgb2 |
| ENSMUSG00000102752 | 0.0735197293712817 | 0.81717853084482 | 0.966315773308182 | 263.826005238161 | 251.654460234528 | 364.385132129652 | 261.681547698027 | 197.092475247437 | 141.877538920568 | 404.979377226337 | 339.769501741243 | Gm7694 |
| ENSMUSG00000102805 | 0.0260733812986059 | 0.960369704896273 | 0.993694458032251 | 36.1405486627618 | 50.4937751926884 | 70.9841166486335 | 15.9447538023887 | 22.994122112201 | 19.5693157131818 | 63.5057635239243 | 64.046008315442 | Gm37240 |
| ENSMUSG00000102824 | -0.0201795653065286 | 0.944366243582602 | 0.990505437194639 | 55.1143367107117 | 41.5352021746308 | 60.5731128735006 | 66.5927952923295 | 68.9823663366031 | 73.3849339244316 | 48.9306702561384 | 35.8223436340608 | Pdcd5-ps |
| ENSMUSG00000102869 | 0.272315674089974 | 0.69286157058198 | 0.94469184542218 | 924.294532050133 | 1963.55632241245 | 1836.12248397799 | 725.017334661559 | 582.51776017576 | 260.516515431732 | 1870.81732872938 | 1797.63033509105 | 2900097C17 |
| ENSMUSG00000102918 | -0.76446919604064 | 0.00104120999316527 | 0.556246415579446 | 57.8248778604189 | 83.0704043492615 | 84.2344850897118 | 59.0893817382642 | 142.344565456483 | 141.877538920568 | 104.107809055614 | 95.5262496908287 | Pcdhgc3 |
| ENSMUSG00000102976 | 0.0947651693856058 | 0.772750904630985 | 0.959045638244974 | 67.7635287426784 | 91.2145616384048 | 88.0203046443056 | 61.9031618210386 | 53.6529515951357 | 39.1386314263635 | 89.5327157878278 | 106.381505337514 | Zc3h11a |
| ENSMUSG00000103009 | -0.0313406845095422 | 0.934185141678488 | 0.98818087308472 | 23.4913566307952 | 31.7622134276588 | 48.2691993210708 | 35.6412143818101 | 28.4689130912965 | 45.2540425867328 | 49.9717483466946 | 18.4539345993646 | Gm4430 |
| ENSMUSG00000103034 | -0.639825710707639 | 0.0380590069160511 | 0.679566075569698 | 46.9827132615903 | 43.9784493613737 | 52.0550188756646 | 73.1582821521366 | 83.2168228822514 | 129.646716599829 | 59.3414511616998 | 65.1315338801105 | Gm8797 |
| ENSMUSG00000103041 | 0.40758466709116 | 0.133754484329814 | 0.77303252008843 | 35.2370349461928 | 38.2775392589734 | 50.1621090983677 | 41.2687745473591 | 30.6588294829347 | 35.4693847301419 | 35.3966550789086 | 22.7960368580387 | Gm37305 |
| ENSMUSG00000103081 | -0.189363294434135 | 0.730056672377728 | 0.952921973914044 | 16.2632468982428 | 8.14415728914329 | 17.9826428843205 | 12.1930470253561 | 7.66470737073368 | 6.1154111603693 | 21.8626399016789 | 26.0526135520442 | Pcdhgb8 |
| ENSMUSG00000103088 | -0.602940877443579 | 0.0826678497198024 | 0.739855948845396 | 19.877301764519 | 31.7622134276588 | 24.6078271048596 | 13.1309737196143 | 44.893286028583 | 33.0232202659942 | 28.1091084450157 | 30.3947158107182 | Pcdhgb6 |
| ENSMUSG00000103283 | -0.65255849825989 | 0.214684603533605 | 0.824248286808591 | 9.03513716569045 | 4.88649437348597 | 12.3039135524298 | 10.3171936368398 | 18.6142893289246 | 18.3462334811079 | 5.20539045278068 | 15.1973579053591 | NA |
| ENSMUSG00000103332 | 0.122299784572678 | 0.800365733042439 | 0.963461896828986 | 20.780815481088 | 50.4937751926884 | 46.3762895437739 | 20.6343872736795 | 19.7092475247437 | 14.6769867848863 | 47.8895921655823 | 44.5065481514088 | Pcdhga2 |
| ENSMUSG00000103400 | -0.122648683565589 | 0.730352870208471 | 0.952921973914044 | 32.5264937964856 | 24.4324718674299 | 21.7684624389143 | 15.9447538023887 | 30.6588294829347 | 15.9000690169602 | 31.2323427166841 | 24.9670879873757 | Gm15853 |
| ENSMUSG00000103409 | 0.109381296972388 | 0.639756825916848 | 0.933917216183397 | 310.808718499751 | 310.292392716359 | 281.097101928589 | 303.888248939644 | 168.623562156141 | 250.731857575141 | 350.843316517418 | 347.368180693922 | Lsmem2 |
| ENSMUSG00000103472 | -0.444319557623363 | 0.118893621524727 | 0.769868040577702 | 23.4913566307952 | 22.8036404096012 | 22.7149173275627 | 25.3240207449704 | 27.3739548954774 | 33.0232202659942 | 39.5609674411332 | 28.2236646813812 | Pcdhga7 |
| ENSMUSG00000103476 | 0.126992036560404 | 0.57813799883344 | 0.922450244597442 | 515.002818444356 | 303.777066885045 | 360.599312575058 | 461.459933575015 | 331.772333333186 | 502.686797382356 | 302.953724351836 | 365.822115293287 | Gm34302 |
| ENSMUSG00000103567 | 0.335220870953355 | 0.318490586996858 | 0.854351922245337 | 19.877301764519 | 34.2054606144018 | 35.0188308799925 | 26.2619474392285 | 28.4689130912965 | 15.9000690169602 | 20.8215618111227 | 26.0526135520442 | Pcdhga5 |
| ENSMUSG00000103585 | -0.704093313246119 | 0.0556547721460352 | 0.703396849069134 | 11.7456783153976 | 21.9892246806869 | 19.8755526616174 | 11.2551203310979 | 28.4689130912965 | 25.6847268735511 | 32.2734208072402 | 19.5394601640331 | Pcdhgb4 |
| ENSMUSG00000103622 | 0.192471692440225 | 0.693281413247009 | 0.94469184542218 | 10.8421645988285 | 9.77298874697194 | 28.3936466594534 | 21.5723139679377 | 10.949581958191 | 12.2308223207386 | 23.9447960827911 | 14.1118323406906 | Gm38391 |
| ENSMUSG00000103649 | -0.805522079370658 | 0.0172901191961837 | 0.642273034804458 | 24.3948703473642 | 20.3603932228582 | 10.4110037751329 | 25.3240207449704 | 42.7033696369448 | 36.6924669622158 | 32.2734208072402 | 29.3091902460497 | Gm37768 |
| ENSMUSG00000103653 | -0.27524331657727 | 0.525052104296381 | 0.907989757056366 | 13.5527057485357 | 8.14415728914329 | 15.1432782183751 | 21.5723139679377 | 19.7092475247437 | 22.0154801773295 | 10.4107809055614 | 18.4539345993646 | Gstp-ps |
| ENSMUSG00000103703 | 0.0536210910769185 | 0.920160334176138 | 0.986713918026868 | 12.6491920319666 | 18.7315617650296 | 11.3574586637814 | 21.5723139679377 | 3.28487458745729 | 17.123151249034 | 16.6572494488982 | 24.9670879873757 | Gm42568 |
| ENSMUSG00000103711 | 0.141030900929846 | 0.77521751035063 | 0.959523366491736 | 8.13162344912141 | 10.5874044758863 | 15.1432782183751 | 13.1309737196143 | 13.1394983498292 | 14.6769867848863 | 4.16431236222455 | 10.8552556466851 | Tstd1 |
| ENSMUSG00000103715 | 0.00787319916778432 | 0.973129895595748 | 0.995480246308729 | 56.0178504272808 | 45.6072808192024 | 73.8234813145789 | 47.8342614071662 | 54.7479097909548 | 57.4848649074714 | 56.2182168900314 | 53.1907526687569 | 4933431K14 |
| ENSMUSG00000103749 | -0.168777701668247 | 0.759274754883555 | 0.957357521291862 | 9.93865088225949 | 7.32974156022896 | 15.1432782183751 | 9.37926694258161 | 14.2344565456483 | 2.44616446414772 | 15.616171358342 | 14.1118323406906 | Pcdhgb5 |
| ENSMUSG00000103789 | 0.201126839276277 | 0.788754240146888 | 0.961659329450003 | 5.42108229941427 | 32.5766291565731 | 22.7149173275627 | 13.1309737196143 | 6.56974917491458 | 2.44616446414772 | 33.3144988977964 | 21.7105112933701 | Gm10129 |
| ENSMUSG00000103793 | -0.435292286439672 | 0.249724109240291 | 0.835314899646652 | 13.5527057485357 | 21.9892246806869 | 18.9290977729689 | 11.2551203310979 | 25.1840385038392 | 29.3539735697726 | 17.6983275394543 | 17.3684090346961 | Pcdhga6 |
| ENSMUSG00000103906 | 0.0922487921015027 | 0.725189357588589 | 0.951674101666722 | 80.412720774645 | 66.7820897709749 | 88.966759532954 | 86.2892558717508 | 101.831112211176 | 90.5080851734656 | 58.3003730711437 | 52.1052271040884 | Tigd5 |
| ENSMUSG00000103922 | 0.290090700892589 | 0.311560848895454 | 0.851785577030345 | 25.2983840639333 | 30.9477976987445 | 33.1259211026956 | 28.1378008277448 | 22.994122112201 | 20.7923979452556 | 26.0269522639034 | 26.0526135520442 | Gm6123 |
| ENSMUSG00000104011 | 0.00567362513997135 | 0.986129691437642 | 0.997766711397789 | 32.5264937964856 | 32.5766291565731 | 24.6078271048596 | 30.9515809105193 | 20.8042057205628 | 31.8001380339204 | 44.7663578939139 | 22.7960368580387 | Gm32391 |
| ENSMUSG00000104063 | 0.0189390689763084 | 0.951850365035743 | 0.991915011884503 | 37.9475760958999 | 38.2775392589734 | 35.965285768641 | 35.6412143818101 | 42.7033696369448 | 25.6847268735511 | 53.094982618363 | 23.8815624227072 | Pcdhgb7 |
| ENSMUSG00000104168 | 0.625310063652357 | 0.192837276999092 | 0.808824684419004 | 15.3597331816738 | 17.1027303072009 | 13.2503684410783 | 10.3171936368398 | 4.37983278327639 | 9.78465785659088 | 14.5750932677859 | 7.59867895267955 | Gm38250 |
| ENSMUSG00000104195 | -0.270962423590053 | 0.614521876343778 | 0.928075610917191 | 7.22810973255236 | 5.7009101024003 | 17.036187995672 | 15.0068271081306 | 9.85462376237187 | 23.2385624094033 | 12.4929370866736 | 8.68420451734806 | B230377A18 |
| ENSMUSG00000104204 | -0.0563380741042973 | 0.885266449096333 | 0.979866999597423 | 18.0702743313809 | 12.2162359337149 | 12.3039135524298 | 21.5723139679377 | 12.0445401540101 | 18.3462334811079 | 15.616171358342 | 20.6249857287016 | Gm37397 |
| ENSMUSG00000104279 | 0.133909267963354 | 0.669442039418938 | 0.942334932906175 | 133.720030052219 | 93.6578088251478 | 47.3227444324223 | 83.4754757889764 | 88.6916138613469 | 99.0696607979826 | 67.6700758861489 | 71.6446872681215 | Gm38300 |
| ENSMUSG00000104346 | -0.152600013565485 | 0.757487505185119 | 0.956693827734505 | 13.5527057485357 | 16.2883145782866 | 6.62518422053913 | 7.50341355406529 | 13.1394983498292 | 18.3462334811079 | 9.36970281500523 | 8.68420451734806 | Pcdhga3 |
| ENSMUSG00000104348 | -0.0142722038755965 | 0.953123574000666 | 0.992049433957868 | 76.7986659083688 | 96.1010560118908 | 123.985590412947 | 88.1651092602672 | 79.9319482947941 | 77.0541806206532 | 103.066730965058 | 128.092016630884 | Gm37691 |
| ENSMUSG00000104394 | -0.0967266333251015 | 0.781628456297051 | 0.960742105865155 | 31.6229800799166 | 30.9477976987445 | 20.8220075502658 | 32.8274342990356 | 24.0890803080201 | 50.1463715150282 | 29.1501865355718 | 21.7105112933701 | Gm37254 |
| ENSMUSG00000104435 | 0.361438044840593 | 0.31277542109883 | 0.852613848149207 | 56.0178504272808 | 32.5766291565731 | 24.6078271048596 | 35.6412143818101 | 35.0386622662111 | 36.6924669622158 | 26.0269522639034 | 18.4539345993646 | Gm37422 |
| ENSMUSG00000104453 | 0.241299456233187 | 0.339763056759802 | 0.858570960161243 | 1964.2388198211 | 2222.5405242072 | 1865.46258552609 | 1702.33695007856 | 1275.62629812925 | 1015.1582526213 | 2050.92383839559 | 2217.72872861776 | Gm37829 |
| ENSMUSG00000104621 | -0.73538789535629 | 0.200903840192211 | 0.812921911423739 | 11.7456783153976 | 7.32974156022896 | 4.73227444324223 | 12.1930470253561 | 10.949581958191 | 25.6847268735511 | 5.20539045278068 | 18.4539345993646 | Gm43185 |
| ENSMUSG00000104682 | -0.0481172632108956 | 0.923154443089955 | 0.986713918026868 | 13.5527057485357 | 8.14415728914329 | 8.51809399783602 | 12.1930470253561 | 12.0445401540101 | 18.3462334811079 | 7.28754663389296 | 6.51315338801105 | Gm42636 |
| ENSMUSG00000104713 | 0.194110834820508 | 0.499765644791738 | 0.900927059989498 | 469.827132615903 | 489.463853077511 | 787.450467355508 | 382.67409125733 | 342.721915291377 | 347.355353908976 | 656.920275140922 | 513.453592088204 | Gbp6 |
| ENSMUSG00000104736 | -0.0339182415249001 | 0.908186622270561 | 0.98504921011344 | 109.325159704854 | 111.574954861263 | 74.7699362032273 | 141.626930832982 | 98.5462376237187 | 69.71568722821 | 160.326025945645 | 118.322286548867 | Gm33609 |
| ENSMUSG00000104784 | 0.375154579886797 | 0.285283038560196 | 0.851275833092507 | 23.4913566307952 | 52.122606650517 | 43.5369248778286 | 30.9515809105193 | 19.7092475247437 | 26.9078091056249 | 31.2323427166841 | 37.9933947633978 | Gm42984 |
| ENSMUSG00000104913 | -0.0537430389913241 | 0.791782109043732 | 0.961804768755116 | 90.3513716569045 | 79.8127414336042 | 56.7872933189068 | 82.5375490947182 | 81.0269064906132 | 84.3926740130963 | 74.9576225200418 | 81.4144173501381 | Gm6560 |
| ENSMUSG00000104950 | 0.123855879558737 | 0.794211669888219 | 0.961804768755116 | 19.877301764519 | 20.3603932228582 | 19.8755526616174 | 22.5102406621959 | 21.8991639163819 | 33.0232202659942 | 5.20539045278068 | 16.2828834700276 | 4833413G10 |
| ENSMUSG00000104960 | 0.0670223992599602 | 0.784884150368496 | 0.961141387070613 | 147.272735800754 | 109.13170767452 | 104.110037751329 | 174.454365132018 | 140.154649064844 | 159.000690169602 | 93.6970281500523 | 118.322286548867 | Snhg8 |
| ENSMUSG00000105004 | -0.0246442009238898 | 0.964914375021439 | 0.994284908685797 | 5.42108229941427 | 19.5459774939439 | 10.4110037751329 | 8.44134024832345 | 7.66470737073368 | 6.1154111603693 | 15.616171358342 | 15.1973579053591 | Gm43075 |
| ENSMUSG00000105095 | 0.0204926238830148 | 0.974450798237167 | 0.995510898988495 | 18.0702743313809 | 8.95857301805762 | 6.62518422053913 | 10.3171936368398 | 9.85462376237187 | 1.22308223207386 | 14.5750932677859 | 17.3684090346961 | 8430422M14 |
| ENSMUSG00000105096 | 0.248952922198405 | 0.676979298997701 | 0.94409858171788 | 27.1054114970714 | 39.0919549878878 | 106.949402417274 | 28.1378008277448 | 25.1840385038392 | 12.2308223207386 | 73.9165444294857 | 57.5328549274309 | Gbp10 |
| ENSMUSG00000105100 | 0.353326097088157 | 0.394140177505854 | 0.878942305112115 | 20.780815481088 | 30.1333819698302 | 26.5007368821565 | 18.7585338851632 | 12.0445401540101 | 9.78465785659088 | 28.1091084450157 | 24.9670879873757 | Gm44349 |
| ENSMUSG00000105135 | -0.338940003783866 | 0.555656267841991 | 0.916239900758826 | 19.877301764519 | 8.14415728914329 | 5.67872933189068 | 13.1309737196143 | 18.6142893289246 | 20.7923979452556 | 15.616171358342 | 4.34210225867403 | Gm43667 |
| ENSMUSG00000105161 | -1.10124171155965 | 0.0666274684236236 | 0.719725956118125 | 12.6491920319666 | 2.44324718674299 | 6.62518422053913 | 9.37926694258161 | 20.8042057205628 | 6.1154111603693 | 24.9858741733473 | 14.1118323406906 | Gm42595 |
| ENSMUSG00000105168 | -0.0678505737585833 | 0.87423558294495 | 0.97671366425653 | 42.4651446787451 | 39.0919549878878 | 33.1259211026956 | 85.3513291774927 | 40.5134532453066 | 90.5080851734656 | 28.1091084450157 | 51.0197015394199 | Gm30735 |
| ENSMUSG00000105176 | -0.144714751386739 | 0.629511936379317 | 0.931825800800958 | 55.1143367107117 | 44.7928650902881 | 23.6613722162112 | 46.8963347129081 | 40.5134532453066 | 41.5847958905112 | 51.0128264372507 | 55.3618037980939 | Gm43668 |
| ENSMUSG00000105245 | 0.393801531636485 | 0.303032025361766 | 0.851785577030345 | 77.7021796249379 | 92.0289773673191 | 92.7525790875478 | 79.7237690119437 | 39.4184950494875 | 28.1308913376988 | 99.9434966933891 | 92.2696729968231 | Gm31305 |
| ENSMUSG00000105383 | 0.33765645451313 | 0.0285124356330201 | 0.665956123973759 | 180.702743313809 | 240.252640029727 | 217.684624389143 | 195.088752405698 | 165.338687568684 | 149.216032313011 | 183.22974393788 | 161.743309135608 | Gm42641 |
| ENSMUSG00000105388 | 0.119350225687088 | 0.786243261830676 | 0.961141387070613 | 16.2632468982428 | 16.2883145782866 | 23.6613722162112 | 13.1309737196143 | 22.994122112201 | 22.0154801773295 | 9.36970281500523 | 9.76973008201657 | Rpl36a-ps2 |
| ENSMUSG00000105419 | -0.360962787409168 | 0.410769901117921 | 0.882192126163777 | 18.9737880479499 | 11.4018202048006 | 8.51809399783602 | 8.44134024832345 | 17.5193311331055 | 15.9000690169602 | 16.6572494488982 | 10.8552556466851 | Gm43205 |
| ENSMUSG00000105427 | 1.11207919169892 | 0.00047358642006091 | 0.522942841846969 | 48.7897406947284 | 56.1946852950887 | 37.8581955459379 | 75.9720622349111 | 25.1840385038392 | 17.123151249034 | 28.1091084450157 | 30.3947158107182 | Gm31678 |
| ENSMUSG00000105444 | 0.333918035213857 | 0.647408436573212 | 0.935063240511793 | 7.22810973255236 | 2.44324718674299 | 8.51809399783602 | 32.8274342990356 | 9.85462376237187 | 13.4539045528125 | 6.24646854333682 | 10.8552556466851 | Gm10727 |
| ENSMUSG00000105449 | 0.406756822904989 | 0.515501941473482 | 0.904239892322932 | 20.780815481088 | 8.14415728914329 | 12.3039135524298 | 5.62756016554897 | 12.0445401540101 | 8.56157562451702 | 12.4929370866736 | 2.17105112933702 | Gm43379 |
| ENSMUSG00000105504 | -0.107762515090638 | 0.623561253295639 | 0.931145988862144 | 136.430571201926 | 124.605606523892 | 147.646962629158 | 166.013024883695 | 204.757182618171 | 171.23151249034 | 120.765058504512 | 122.664388807541 | Gbp5 |

| | | | | | | | | | | | | |
|---|---|---|---|---|---|---|---|---|---|---|---|---|
| ENSMUSG00000105509 | 0.288847017892066 | 0.378263701365377 | 0.87422797717331 | 23.4913566307952 | 21.9892246806869 | 25.5542819935081 | 19.6964605794214 | 14.2344565456483 | 19.5693157131818 | 17.6983275394543 | 22.7960368580387 | C130013H08 |
| ENSMUSG00000105541 | 0.46992579154942 | 0.103877754549885 | 0.759800953500741 | 32.5264937964856 | 38.2775392589734 | 41.6440151005317 | 56.2756016554897 | 25.1840385038392 | 28.1308913376988 | 30.191264626128 | 37.9933947633978 | Gm43136 |
| ENSMUSG00000105692 | -0.216896253601207 | 0.459420722424355 | 0.894774044236998 | 30.7194663633475 | 31.7622134276588 | 36.9117406572894 | 33.7653609932938 | 49.2731188118594 | 46.4771248188067 | 35.3966550789086 | 23.8815624227072 | Gm43737 |
| ENSMUSG00000105703 | 0.190927618299519 | 0.648790226616388 | 0.936461253686283 | 51.5002818444356 | 165.326392969609 | 176.040609288611 | 283.253861665965 | 130.300025302473 | 194.470074899744 | 111.395355689507 | 156.315681312265 | Gm43305 |
| ENSMUSG00000105741 | -0.139250697301427 | 0.822726170931592 | 0.966637987288264 | 13.5527057485357 | 8.95857301805762 | 11.3574586637814 | 5.62756016554897 | 3.28487458745729 | 6.1154111603693 | 21.8626399016789 | 11.9407812113536 | C79130 |
| ENSMUSG00000105843 | -0.0121445404729332 | 0.970020814638407 | 0.9951105526746 | 46.0791995450213 | 30.1333819698302 | 49.2156542097192 | 61.9031618210386 | 40.5134532453066 | 33.0232202659942 | 68.711153976705 | 45.5920737160773 | Gm19439 |
| ENSMUSG00000105867 | 0.0372838160707711 | 0.880967946007506 | 0.97846486749867 | 42.4651446787451 | 43.1640336324594 | 38.8046504345863 | 44.0825546301336 | 43.7983278327639 | 34.2463024980681 | 52.0539045278068 | 33.6512925047237 | Gm42517 |
| ENSMUSG00000105879 | 0.0906267769024672 | 0.805547180999481 | 0.963475137341661 | 21.6843291976571 | 13.0306516626293 | 17.036187995672 | 21.5723139679377 | 12.0445401540101 | 18.3462334811079 | 19.7804837205666 | 18.4539345993646 | Gm6204 |
| ENSMUSG00000105957 | 0.053028260371427 | 0.87152020380915 | 0.975613767803763 | 28.0089252136404 | 26.8757190541728 | 35.0188308799925 | 45.0204813243917 | 36.1336204620302 | 45.2540425867328 | 21.8626399016789 | 27.1381391167127 | Gm43254 |
| ENSMUSG00000105960 | 0.0578222087867852 | 0.871277366597723 | 0.975613767803763 | 19.877301764519 | 15.4738988493722 | 15.1432782183751 | 17.8206071909051 | 14.2344565456483 | 20.7923979452556 | 14.5750932677859 | 16.2828834700276 | 6330410L21R |
| ENSMUSG00000105962 | -0.147324521493101 | 0.774852881652539 | 0.959414024438738 | 12.6491920319666 | 16.2883145782866 | 11.3574586637814 | 15.9447538023887 | 14.2344565456483 | 31.8001380339204 | 10.4107809055614 | 6.51315338801105 | Gm42432 |
| ENSMUSG00000105987 | 0.0497000381664462 | 0.6554255062057 | 0.937625787683528 | 293.64195788494 | 304.591482613959 | 279.204192151292 | 300.136542162612 | 271.549632563136 | 308.216722482613 | 272.762459725708 | 286.578749072486 | AI506816 |
| ENSMUSG00000106019 | 0.122075521352828 | 0.440716209355477 | 0.890970307297912 | 744.495302452893 | 590.451403462888 | 813.004749349016 | 606.83857118503 | 548.574056105368 | 588.302553627526 | 702.727711125392 | 690.394259129171 | Gm43672 |
| ENSMUSG00000106025 | 0.0979213590556933 | 0.654501372182755 | 0.937269843859626 | 96.6759676728878 | 95.2866402829764 | 72.8770264259304 | 88.1651092602672 | 62.4126171616885 | 84.3926740130963 | 77.0397787011541 | 106.381505337514 | Gm42940 |
| ENSMUSG00000106037 | 0.224340335233749 | 0.609184246192564 | 0.927932112008768 | 18.9737880479499 | 8.95857301805762 | 14.1968233297267 | 14.0689004138724 | 8.75966556655277 | 17.123151249034 | 10.4107809055614 | 11.9407812113536 | Gm4332 |
| ENSMUSG00000106106 | 0.714461955558407 | 0.0700421846382079 | 0.724952268721839 | 47298.0395486729 | 40077.3980198741 | 72026.1634810354 | 127161.287427439 | 70430.9960296718 | 38665.2986025509 | 24579.8537180304 | 40964.4782338955 | CT010467.1 |
| ENSMUSG00000106140 | 0.344081929911849 | 0.529830787235824 | 0.908620485539384 | 31.6229800799166 | 54.56585383726 | 45.4298346551254 | 11.2551203310979 | 18.6142893289246 | 11.0077400886647 | 45.80743598447 | 36.9078691987293 | Gm42208 |
| ENSMUSG00000106164 | 0.985495885858809 | 0.011307776032837 | 0.635862253526407 | 17.1667606148119 | 33.3910448854875 | 29.3401015481019 | 32.8274342990356 | 14.2344565456483 | 20.7923979452556 | 9.36970281500523 | 13.0263067760221 | 9430085M18 |
| ENSMUSG00000106209 | -0.360322381197465 | 0.350237981823153 | 0.861774532843376 | 23.4913566307952 | 17.1027303072009 | 28.3936466594534 | 29.075272522003 | 27.3739548954774 | 18.3462334811079 | 52.0539045278068 | 27.1381391167127 | Gm42918 |
| ENSMUSG00000106224 | 0.46021291449721 | 0.317878604796421 | 0.854351922245337 | 13.5527057485357 | 11.4018202048006 | 20.8220075502658 | 10.3171936368398 | 9.85462376237187 | 6.1154111603693 | 13.5340151772298 | 10.8552556466851 | Gm43823 |
| ENSMUSG00000106237 | 0.193495124811688 | 0.770641181822644 | 0.958995611819782 | 11.7456783153976 | 30.1333819698302 | 23.6613722162112 | 3.75170677703264 | 20.8042057205628 | 4.89232892829544 | 23.9447960827911 | 10.8552556466851 | Gm8066 |
| ENSMUSG00000106317 | 0.330056321947708 | 0.437469138931752 | 0.888722970093854 | 26.2018977805023 | 21.1748089517725 | 11.3574586637814 | 19.6964605794214 | 18.6142893289246 | 22.0154801773295 | 13.5340151772298 | 8.68420451734806 | Gm43848 |
| ENSMUSG00000106383 | 0.113373987193743 | 0.772984843939868 | 0.959121750920843 | 40.658117245607 | 18.7315617650296 | 17.9826428843205 | 48.7721881014244 | 27.3739548954774 | 29.3539735697726 | 26.0269522639034 | 33.6512925047237 | E330034L11R |
| ENSMUSG00000106464 | 0.192438699396944 | 0.601012261824151 | 0.925969784079517 | 57.8248778604189 | 60.2667639396603 | 63.4124775394459 | 38.4549944645846 | 39.4184950494875 | 20.7923979452556 | 69.7522320672611 | 61.8749571861049 | C130083M11 |
| ENSMUSG00000106498 | 0.281755496047551 | 0.56529873406358 | 0.919692942798738 | 30.7194663633475 | 32.5766291565731 | 29.3401015481019 | 18.7585338851632 | 5.47479097909548 | 33.0232202659942 | 18.7394056300105 | 34.7368180693922 | Gm43032 |
| ENSMUSG00000106662 | -0.0509670378412758 | 0.83868235242623 | 0.969320841670855 | 40.658117245607 | 45.6072808192024 | 31.2330113253987 | 34.703287687552 | 36.1336204620302 | 39.1386314263635 | 43.7252798033577 | 39.0789203280663 | Gm43034 |
| ENSMUSG00000106734 | -0.0110185431870498 | 0.976250366387177 | 0.9958226914238 | 45.1756858284523 | 22.8036404096012 | 29.3401015481019 | 32.8274342990356 | 47.0832024202212 | 29.3539735697726 | 17.6983275394543 | 36.9078691987293 | Gm20559 |
| ENSMUSG00000106735 | -0.350006411585294 | 0.469228220530229 | 0.895328112556867 | 7.22810973255236 | 8.14415728914329 | 8.51809399783602 | 14.0689004138724 | 12.0445401540101 | 6.1154111603693 | 15.616171358342 | 14.1118323406906 | A330058E17R |
| ENSMUSG00000106755 | -0.163249231614195 | 0.611203349224408 | 0.927932112008768 | 19.877301764519 | 17.1027303072009 | 20.8220075502658 | 26.2619474392285 | 26.2789966996583 | 18.3462334811079 | 22.903717992235 | 26.0526135520442 | Tpi-rs11 |
| ENSMUSG00000106783 | 0.566503304195486 | 0.218602833709993 | 0.827129763070499 | 24.3948703473642 | 30.9477976987445 | 46.3762895437739 | 18.7585338851632 | 7.66470737073368 | 17.123151249034 | 28.1091084450157 | 28.2236646813812 | Chaer1 |
| ENSMUSG00000106795 | -0.411454346298276 | 0.103784577971206 | 0.759800953500741 | 44.2721721118832 | 57.009101024003 | 47.3227444324223 | 44.0825546301336 | 42.7033696369448 | 77.0541806206532 | 62.4646854333682 | 74.901263962127 | Gm43050 |
| ENSMUSG00000106831 | 0.015902182912194 | 0.975457417195263 | 0.99575536949131 | 12.6491920319666 | 18.7315617650296 | 7.57163910918757 | 14.0689004138724 | 9.85462376237187 | 18.3462334811079 | 5.20539045278068 | 19.5394601640331 | Ube2n-ps1 |
| ENSMUSG00000106847 | -0.116053813691072 | 0.437649581838331 | 0.888722970093854 | 1128.48863199474 | 1409.7536267507 | 1611.81267536831 | 1199.60824195619 | 1567.98013641295 | 1424.89080036605 | 1532.46694929863 | 1272.23596179149 | Peg13 |
| ENSMUSG00000106864 | -0.0584700280861234 | 0.703262327401007 | 0.946542027868216 | 210.518695960587 | 232.108482740584 | 294.347470369667 | 236.357526953057 | 282.499214521327 | 249.508775343067 | 251.940897914585 | 229.045894145055 | Gtf3c2 |
| ENSMUSG00000106874 | 0.629738114941192 | 0.466754037528524 | 0.895328112556867 | 19.877301764519 | 13.8450673915436 | 23.6613722162112 | 4.68963347129081 | 0 | 3.66924669622158 | 14.5750932677859 | 21.7105112933701 | Gm20186 |
| ENSMUSG00000106918 | 0.0605132708204464 | 0.742102011406921 | 0.954383574136061 | 1740.16741811198 | 1262.34437981721 | 1314.62584033269 | 1801.75717966993 | 1303.00025302473 | 1849.30033489568 | 1296.14222274239 | 1419.86743858641 | Mrpl33 |
| ENSMUSG00000106951 | 0.344478893673609 | 0.331752348493621 | 0.856353859984463 | 18.0702743313809 | 26.8757190541728 | 22.7149173275627 | 24.3860940507122 | 14.2344565456483 | 12.2308223207386 | 22.903717992235 | 22.7960368580387 | 5930430L01R |
| ENSMUSG00000106978 | -0.476576245348994 | 0.0820458397879409 | 0.739855948845396 | 26.2018977805023 | 48.0505280059454 | 44.483379766477 | 37.5170677703264 | 65.6974917491458 | 59.9310293716191 | 47.8895921655823 | 44.5065481514088 | N4bp2os |
| ENSMUSG00000106981 | 0.0124091689729046 | 0.960355943035991 | 0.993694458032251 | 61.4389327266951 | 62.7100111264033 | 67.1982970940397 | 57.2135283497478 | 44.893286028583 | 79.5003450848009 | 47.8895921655823 | 74.901263962127 | Gm45495 |
| ENSMUSG00000106990 | -0.157914902664585 | 0.563090956598822 | 0.918253929144354 | 32.5264937964856 | 37.4631235300591 | 31.2330113253987 | 46.8963347129081 | 53.6529515951357 | 41.5847958905112 | 35.3966550789086 | 34.7368180693922 | Gm42547 |
| ENSMUSG00000107002 | -0.0727411645394454 | 0.660358366617886 | 0.939611397743053 | 2716.86574572312 | 2194.0359736952 | 2343.42230429355 | 2892.56592509217 | 2733.01565676447 | 3326.7836712409 | 2355.95971892854 | 2256.80764894583 | 0610012G03R |
| ENSMUSG00000107023 | 0.0461398683819203 | 0.928712184425426 | 0.986901071486195 | 12.6491920319666 | 12.2162359337149 | 13.2503684410783 | 31.8895076047775 | 30.6588294829347 | 14.6769867848863 | 11.4518589961175 | 10.8552556466851 | Gm42715 |
| ENSMUSG00000107035 | -0.180197369996693 | 0.680284095917178 | 0.944452327903597 | 13.5527057485357 | 10.5874044758863 | 14.1968233297267 | 10.3171936368398 | 9.85462376237187 | 12.2308223207386 | 21.8626399016789 | 10.8552556466851 | Ybx1-ps2 |
| ENSMUSG00000107096 | 0.295962863547619 | 0.229888839453401 | 0.832048466795696 | 77.7021796249379 | 60.2667639396603 | 72.8770264259304 | 77.8479156234274 | 54.7479097909548 | 37.9155491942896 | 72.8754663389296 | 68.388110574116 | Gm43597 |
| ENSMUSG00000107176 | -0.125946278586929 | 0.658225466294254 | 0.93897008594302 | 45.1756858284523 | 45.6072808192024 | 41.6440151005317 | 64.7169419038131 | 48.1781606160403 | 77.0541806206532 | 38.5198893505771 | 52.1052271040884 | Gm9794 |
| ENSMUSG00000107265 | -0.185669517710367 | 0.586818754855316 | 0.923058967588378 | 252.080326922764 | 191.387696294867 | 172.254789734017 | 255.11606083822 | 261.695008800764 | 413.401794440965 | 196.76375911511 | 119.407812113536 | Gm15469 |
| ENSMUSG00000107277 | 0.0932706684309477 | 0.794693314033327 | 0.961804768755116 | 41.5616309621761 | 22.8036404096012 | 26.5007368821565 | 37.5170677703264 | 16.4243729372865 | 28.1308913376988 | 39.5609674411332 | 35.8223436340608 | Gm10461 |
| ENSMUSG00000107283 | 0.0124033252612587 | 0.920590889960178 | 0.986713918026868 | 1295.63866956001 | 1022.9061555164 | 1184.01506569921 | 1311.22151857291 | 1100.43298679819 | 1301.35949492659 | 1121.24110352896 | 1249.43992493345 | Mpv17 |
| ENSMUSG00000107331 | 0.413363249589183 | 0.205290025448246 | 0.819390360194292 | 125.588406603097 | 121.347943608235 | 104.110037751329 | 113.489130005238 | 98.5462376237187 | 34.2463024980681 | 93.6970281500523 | 121.578863242873 | Gm42732 |
| ENSMUSG00000107352 | 0.859724355597678 | 0.129770249023088 | 0.77303252008843 | 12.6491920319666 | 8.14415728914329 | 17.9826428843205 | 20.6343872736795 | 5.47479097909548 | 15.9000690169602 | 8.32862472444909 | 3.25657669400552 | Gm43660 |
| ENSMUSG00000107355 | -0.240686021434069 | 0.446776315247924 | 0.89108031640712 | 30.7194663633475 | 26.8757190541728 | 20.8220075502658 | 47.8342614071662 | 35.0386622662111 | 44.0309603546589 | 34.3555769883525 | 35.8223436340608 | AI839979 |
| ENSMUSG00000107369 | -0.914549708615001 | 0.0500202769394646 | 0.703396849069134 | 10.8421645988285 | 8.14415728914329 | 4.73227444324223 | 11.2551203310979 | 13.1394983498292 | 13.4539045528125 | 24.9858741733473 | 14.1118323406906 | Gstm2-ps1 |
| ENSMUSG00000107374 | 0.152400231336231 | 0.721778985054017 | 0.951674101666722 | 13.5527057485357 | 9.77298874697194 | 10.4110037751329 | 16.8826804966469 | 8.75966556655277 | 12.2308223207386 | 12.4929370866736 | 11.9407812113536 | Gm43172 |
| ENSMUSG00000107383 | -0.0717083123870862 | 0.74211322969786 | 0.954383574136061 | 83.1232619243521 | 65.1532583131463 | 80.448665535118 | 95.6685228143324 | 68.9823663366031 | 78.277262852727 | 85.3684034256032 | 107.467030902182 | Gm4366 |
| ENSMUSG00000107440 | 0.127832175417958 | 0.782449031599989 | 0.960742105865155 | 17.1667606148119 | 11.4018202048006 | 11.3574586637814 | 9.37926694258161 | 14.2344565456483 | 6.1154111603693 | 9.36970281500523 | 15.1973579053591 | Gm40377 |
| ENSMUSG00000107451 | 0.0859666166363393 | 0.691778227611 | 0.94469184542218 | 54.2108229941427 | 52.9370223794314 | 57.7337482075553 | 55.3376749612315 | 55.8428679867739 | 39.1386314263635 | 54.1360607089191 | 57.5328549274309 | Gm44421 |
| ENSMUSG00000107470 | 0.509846265201563 | 0.270170710021148 | 0.843865788665109 | 10.8421645988285 | 12.2162359337149 | 9.46454888648447 | 19.6964605794214 | 8.75966556655277 | 11.0077400886647 | 9.36970281500523 | 7.59867895267955 | Gm3375 |
| ENSMUSG00000107476 | -0.448258015999609 | 0.17021261335549 | 0.799274239184501 | 24.3948703473642 | 31.7622134276588 | 41.6440151005317 | 36.5791410760683 | 67.887408140784 | 46.4771248188067 | 39.5609674411332 | 29.3091902460497 | Zfp862-ps |
| ENSMUSG00000107577 | -0.254264639434118 | 0.316214856313528 | 0.853356198600137 | 46.9827132615903 | 30.9477976987445 | 34.0723759913441 | 32.8274342990356 | 39.4184950494875 | 41.5847958905112 | 43.7252798033577 | 47.7631248454143 | Gm44103 |
| ENSMUSG00000107653 | -0.526468523783122 | 0.446754793760936 | 0.89108031640712 | 1.80702743313809 | 6.51532583131463 | 19.8755526616174 | 10.3171936368398 | 12.0445401540101 | 24.4616446414772 | 13.5340151772298 | 5.42762782334254 | Gm31520 |
| ENSMUSG00000107655 | 0.0870434718911237 | 0.740063142526822 | 0.954383574136061 | 68.6670424592474 | 74.1118313312039 | 99.3777633080869 | 87.227182566009 | 102.926070406995 | 91.7311674055395 | 61.4236073428121 | 54.2762782334254 | Gm44220 |
| ENSMUSG00000107689 | 0.26988554601966 | 0.12082793215449 | 0.769868040577702 | 1014.64590370704 | 701.211942595237 | 815.844114014961 | 936.050840869645 | 718.292576457328 | 889.180782717696 | 651.714884688142 | 617.664046296381 | Gm44386 |
| ENSMUSG00000107728 | -0.536566057782603 | 0.272250561033245 | 0.844663902781278 | 18.0702743313809 | 8.14415728914329 | 11.3574586637814 | 17.8206071909051 | 35.0386622662111 | 20.7923979452556 | 13.5340151772298 | 10.8552556466851 | Gm38910 |
| ENSMUSG00000107741 | 0.18791588120054 | 0.581194592442029 | 0.922450244597442 | 37.0440623793308 | 17.9171460361152 | 35.0188308799925 | 27.1998741334867 | 21.8991639163819 | 34.2463024980681 | 22.903717992235 | 23.8815624227072 | Gm2011 |
| ENSMUSG00000107742 | -0.507131879641459 | 0.247104691001784 | 0.833120697990749 | 19.877301764519 | 8.95857301805762 | 9.46454888648447 | 9.37926694258161 | 13.1394983498292 | 20.7923979452556 | 19.7804837205666 | 14.1118323406906 | 9530085L11R |
| ENSMUSG00000107810 | 0.384329067493783 | 0.635950223714119 | 0.932595977875587 | 10.8421645988285 | 40.7207864457164 | 32.1794662140472 | 14.0689004138724 | 10.949581958191 | 0 | 31.2323427166841 | 32.5657669400552 | Gm18609 |
| ENSMUSG00000107838 | 0.22931352884424 | 0.285976329249439 | 0.851593746788839 | 73.1846110420926 | 71.6685841444609 | 60.5731128735006 | 78.7858423176855 | 67.887408140784 | 72.1618516923577 | 48.9306702561384 | 54.2762782334254 | Gm45769 |
| ENSMUSG00000107868 | 0.346667057059844 | 0.313607565502354 | 0.85295001595893 | 34.3335212296237 | 30.9477976987445 | 35.0188308799925 | 33.7653609932938 | 29.5638712871156 | 12.2308223207386 | 23.9447960827911 | 39.0789203280663 | Gm5112 |
| ENSMUSG00000107879 | 0.408035775653039 | 0.185548090726541 | 0.805963874499992 | 100.290022539164 | 124.605606523892 | 105.056492639978 | 93.7926694258161 | 61.3176589658694 | 41.5847958905112 | 108.272121417838 | 107.467030902182 | 9330102E08R |
| ENSMUSG00000107881 | 0.0808450425157103 | 0.583702706050202 | 0.922450244597442 | 655.950958229127 | 760.664290805983 | 691.858523602015 | 633.100518624259 | 506.965644664242 | 660.464405319884 | 689.193695948162 | 735.986332845248 | Gm44250 |
| ENSMUSG00000107994 | -0.201016301886929 | 0.18240076080628 | 0.804505478598291 | 219.553833126278 | 200.346269312925 | 186.451613063744 | 182.895705380341 | 202.567266226533 | 272.747337752471 | 219.667477107345 | 213.848536239696 | D830050J10R |
| ENSMUSG00000108181 | -0.354917492318983 | 0.341587362008722 | 0.859283395807012 | 10.8421645988285 | 16.2883145782866 | 14.1968233297267 | 12.1930470253561 | 18.6142893289246 | 15.9000690169602 | 14.5750932677859 | 19.5394601640331 | C030015A19 |
| ENSMUSG00000108207 | -0.161658446753393 | 0.579459402997728 | 0.922450244597442 | 76.7986659083688 | 46.4216965481167 | 73.8234813145789 | 35.6412143818101 | 59.1277425742312 | 79.5003450848009 | 65.5879197050366 | 56.4473293627624 | 1810059H22 |
| ENSMUSG00000108218 | 0.541064977127283 | 0.190720886513614 | 0.808824684419004 | 18.9737880479499 | 15.4738988493722 | 29.3401015481019 | 15.0068271081306 | 17.5193311331055 | 8.56157562451702 | 12.4929370866736 | 15.1973579053591 | Olfr1372-ps1 |
| ENSMUSG00000108231 | 0.108963280681205 | 0.715434726426276 | 0.951176382858794 | 56.9213641438498 | 37.4631235300591 | 38.8046504345863 | 52.523894878457 | 49.2731188118594 | 58.7079471395453 | 35.3966550789086 | 29.3091902460497 | Gm4045 |
| ENSMUSG00000108298 | -0.0781917656408016 | 0.897396583999901 | 0.984332015045905 | 20.780815481088 | 14.6594831204579 | 10.4110037751329 | 12.1930470253561 | 20.8042057205628 | 29.3539735697726 | 8.32862472444909 | 3.25657669400552 | Gm44226 |
| ENSMUSG00000108322 | 0.645143582579731 | 0.0066561959133411 | 0.635862253526407 | 54.2108229941427 | 53.7514381083457 | 65.3053873167428 | 70.3445020693621 | 43.7983278327639 | 45.2540425867328 | 34.3555769883525 | 32.5657669400552 | 5430431A17R |
| ENSMUSG00000108324 | 0.171277070957823 | 0.662468406323993 | 0.939611397743053 | 46.0791995450213 | 27.6901347830872 | 33.1259211026956 | 45.0204813243917 | 40.5134532453066 | 24.4616446414772 | 53.094982618363 | 16.2828834700276 | Gm44608 |
| ENSMUSG00000108332 | -0.2723803033522 | 0.685854704197636 | 0.944452327903597 | 12.6491920319666 | 20.3603932228582 | 13.2503684410783 | 8.44134024832345 | 6.56974917491458 | 3.66924669622158 | 16.6572494486982 | 39.0789203280663 | D530033B14 |
| ENSMUSG00000108350 | -0.212850329283096 | 0.387031787877492 | 0.875697130512848 | 53.3073092775737 | 61.895595397489 | 43.5369248778286 | 58.151455044006 | 75.5521155115177 | 75.8310983885793 | 48.9306702561384 | 52.1052271040884 | Gm44950 |
| ENSMUSG00000108358 | -0.111787075679751 | 0.828320408683358 | 0.967406918070448 | 13.5527057485357 | 8.95857301805762 | 6.62518422053913 | 12.1930470253561 | 17.5193311331055 | 13.4539045528125 | 5.20539045278068 | 8.68420451734806 | Gm44509 |
| ENSMUSG00000108371 | 0.481245912308933 | 0.340162291064269 | 0.858570960161243 | 46.0791995450213 | 17.1027303072009 | 65.3053873167428 | 15.9447538023887 | 17.5193311331055 | 22.0154801773295 | 36.4377331694648 | 27.1381391167127 | Gm38832 |
| ENSMUSG00000108414 | 0.204140211130943 | 0.50883248244826 | 0.902394714365738 | 53.3073092775737 | 46.4216965481167 | 47.3227444324223 | 59.0893817382642 | 62.4126171616885 | 56.2617826753975 | 26.0269522639034 | 34.7368180693922 | Snhg1 |
| ENSMUSG00000108435 | -0.16782003769467 | 0.654058385678323 | 0.937077409179068 | 37.0440623793308 | 40.7207864457164 | 34.0723759913441 | 60.0273084325223 | 30.6588294829347 | 40.6020455316893 | 40.6020455316893 | 36.9078691987293 | Gm45051 |
| ENSMUSG00000108452 | 0.15562819787026 | 0.505084927141851 | 0.901404296286819 | 59.631905293557 | 79.8127414336042 | 78.5557557578211 | 86.2892558717508 | 61.3176589658694 | 72.1618516923577 | 87.4505596067155 | 52.1052271040884 | 4930413G21R |
| ENSMUSG00000108466 | -0.346133064219342 | 0.573284489194891 | 0.922207298536842 | 18.0702743313809 | 8.95857301805762 | 5.67872933189068 | 11.2551203310979 | 7.66470737073368 | 23.2385624094033 | 20.8215618111227 | 4.34210225867403 | Gm44771 |
| ENSMUSG00000108468 | 0.566243169125938 | 0.221224758055631 | 0.830349645871581 | 18.9737880479499 | 17.9171460361152 | 8.51809399783602 | 26.2619474392285 | 13.1394983498292 | 7.33849339244316 | 11.4518589961175 | 16.2828834700276 | Gm30459 |

| | | | | | | | | | | | | |
|---|---|---|---|---|---|---|---|---|---|---|---|---|
| **ENSMUSG00000108494** | 0.292547159356009 | 0.617834272375351 | 0.929461501493948 | 13.5527057485357 | 39.0919549878878 | 41.6440151005317 | 14.0689004138724 | 16.4243729372865 | 6.1154111603693 | 39.5609674411332 | 26.0526135520442 | Gm45203 |
| **ENSMUSG00000108591** | 0.0157909716561266 | 0.968249393652345 | 0.994697802893254 | 36.1405486627618 | 16.2883145782866 | 36.9117406572894 | 48.7721881014244 | 24.0890803080201 | 48.9232892829544 | 38.5198893505771 | 24.9670879873757 | Particl |
| **ENSMUSG00000108601** | -1.07089083159835 | 0.0478481661146513 | 0.703396849069134 | 12.6491920319666 | 3.25766291565731 | 8.51809399783602 | 9.37926694258161 | 25.1840385038392 | 23.2385624094033 | 8.32862472444909 | 14.1118323406906 | Gm44645 |
| **ENSMUSG00000108621** | 0.128485737330763 | 0.499750086580358 | 0.900927059989498 | 155.404359249876 | 188.944449108124 | 194.023252172932 | 176.330218520534 | 141.249607260664 | 128.423634367755 | 176.983275394543 | 206.249857287016 | Gm37494 |
| **ENSMUSG00000108626** | 0.239307442039276 | 0.535810256572246 | 0.911073017897251 | 91.2548853734735 | 55.3802695661743 | 42.5904699891801 | 60.0273084325223 | 44.893286028583 | 92.9542496376133 | 44.7663578939139 | 29.3091902460497 | Gm40457 |
| **ENSMUSG00000108658** | -0.298244014075864 | 0.292449627030638 | 0.851785577030345 | 79.509207058076 | 74.9262470601182 | 68.1447519826882 | 54.3997482669734 | 90.881530252985 | 124.754387671534 | 71.8343882483734 | 54.2762782334254 | NA |
| **ENSMUSG00000108701** | -0.730452393590909 | 0.142128732644222 | 0.780607990324772 | 7.22810973255236 | 9.77298874697194 | 3.78581955459379 | 11.2551203310979 | 8.75966556655277 | 15.9000690169602 | 11.4518589961175 | 17.3684090346961 | Gm44616 |
| **ENSMUSG00000108763** | 0.478667690889183 | 0.131665171388967 | 0.77303252008843 | 27.1054114970714 | 24.4324718674299 | 38.8046504345863 | 35.6412143818101 | 25.1840385038392 | 28.1308913376988 | 18.7394056300105 | 18.4539345993646 | Gm36028 |
| **ENSMUSG00000108994** | -0.275335535328205 | 0.400554470854397 | 0.880487015130533 | 75.8951521917998 | 72.4829998733752 | 53.001473764313 | 40.3308478531009 | 60.2227007700503 | 51.3694537471021 | 115.559668051731 | 65.1315338801105 | Gm31663 |
| **ENSMUSG00000109008** | 0.0263532039580713 | 0.895354740839629 | 0.983733637008786 | 79.509207058076 | 87.1424829938332 | 72.8770264259304 | 61.9031618210386 | 71.1722827282413 | 75.8310983885793 | 84.3273253350471 | 65.1315338801105 | Gm44899 |
| **ENSMUSG00000109029** | -0.0389676009999803 | 0.945203050690669 | 0.990505437194639 | 29.8159526467785 | 11.4018202048006 | 12.3039135524298 | 30.9515809105193 | 27.3739548954774 | 8.56157562451702 | 11.4518589961175 | 39.0789203280663 | Gm44717 |
| **ENSMUSG00000109052** | 0.335074479234854 | 0.298844383282257 | 0.851785577030345 | 96.6759676728878 | 122.976775066064 | 102.217127974032 | 45.9584080186499 | 53.6529515951357 | 63.6002760678407 | 84.3273253350471 | 90.0986218674861 | Gm45012 |
| **ENSMUSG00000109108** | -0.217571766404365 | 0.538315259564849 | 0.912708496930502 | 16.2632468982428 | 26.0613033252585 | 16.0897331070236 | 19.6964605794214 | 22.994122112201 | 29.3539735697726 | 23.9447960827911 | 15.1973579053591 | Gm36371 |
| **ENSMUSG00000109115** | 0.150514804127001 | 0.767407700100487 | 0.95879680569409 | 28.9124389302094 | 48.0505280059454 | 29.3401015481019 | 32.8274342990356 | 25.1840385038392 | 6.1154111603693 | 42.6842017128016 | 51.0197015394199 | Gm44669 |
| **ENSMUSG00000109118** | -0.0011740718412228 | 0.998104853607145 | 0.999262355603273 | 20.780815481088 | 10.5874044758863 | 14.1968233297267 | 9.37926694258161 | 20.8042057205628 | 6.1154111603693 | 10.4107809055614 | 17.3684090346961 | Gm32031 |
| **ENSMUSG00000109147** | 0.107208674312253 | 0.699591087199929 | 0.945280238056015 | 29.8159526467785 | 42.3496179035451 | 45.4298346551254 | 31.8895076047775 | 39.4184950494875 | 40.3617136584374 | 31.2323427166841 | 28.2236646813812 | 4930431P19 |
| **ENSMUSG00000109194** | 0.425142731633227 | 0.322589755232765 | 0.855977678491063 | 14.4562194651047 | 11.4018202048006 | 19.8755526616174 | 12.1930470253561 | 8.75966556655277 | 8.56157562451702 | 13.5340151772298 | 11.9407812113536 | Gm44570 |
| **ENSMUSG00000109222** | 0.3967858861197 | 0.246105320937022 | 0.832048466795696 | 26.2018977805023 | 26.8757190541728 | 30.2865564367503 | 18.7585338851632 | 21.8991639163819 | 22.0154801773295 | 21.8626399016789 | 11.9407812113536 | Gm10297 |
| **ENSMUSG00000109274** | 0.295672628923322 | 0.221826567148555 | 0.831327997735572 | 112.939214571131 | 120.533527879321 | 118.306861081056 | 81.59962240046 | 102.926070406995 | 56.2617826753975 | 103.066730965058 | 90.0986218674861 | Gm45133 |
| **ENSMUSG00000109324** | -0.0584533758890267 | 0.760178397171451 | 0.957357521291862 | 1643.49145043909 | 1179.27397546795 | 1088.42312194571 | 1677.01292933359 | 1544.98601430075 | 1696.41505588644 | 1213.89705358846 | 1364.50563478831 | Prmt1 |
| **ENSMUSG00000109336** | 0.00611989673988213 | 0.965880629165463 | 0.994284908685797 | 543.011743657996 | 616.512706788147 | 602.891764069061 | 482.094320848695 | 514.630352034976 | 660.464405319884 | 502.840717738614 | 559.045665804281 | Samd4b |
| **ENSMUSG00000109372** | 0.336330939243454 | 0.0791777401884874 | 0.739855948845396 | 98.4829951060259 | 100.173134656462 | 79.5022106464695 | 100.358156285623 | 72.2672409240604 | 61.154111603693 | 81.2040910633787 | 84.6709940441436 | Gm19410 |
| **ENSMUSG00000109401** | 0.087235931292216 | 0.705488249844431 | 0.947760509100828 | 56.9213641438498 | 66.7820897709749 | 52.0550188756646 | 78.7858423176855 | 49.2731188118594 | 55.0387004433237 | 65.5879197050366 | 69.4736361387845 | Cpeb1os1 |
| **ENSMUSG00000109419** | 0.206512718189149 | 0.657204439565902 | 0.938389079274206 | 24.3948703473642 | 13.0306516626293 | 12.3039135524298 | 12.1930470253561 | 10.949581958191 | 8.56157562451702 | 21.8626399016789 | 11.9407812113536 | Gm45163 |
| **ENSMUSG00000109455** | 0.353583427734945 | 0.398650114337614 | 0.880459952732461 | 28.0089252136404 | 39.9063707168021 | 44.483379766477 | 25.3240207449704 | 19.7092475247437 | 11.0077400886647 | 37.4788112600209 | 39.0789203280663 | Gm44710 |
| **ENSMUSG00000109482** | 0.300046662426378 | 0.596869921498187 | 0.924496535139492 | 5.42108229941427 | 14.6594831204579 | 14.1968233297267 | 18.7585338851632 | 5.47479097909548 | 8.56157562451702 | 8.32862472444909 | 20.6249857287016 | Gm4756 |
| **ENSMUSG00000109511** | -0.0416303773666995 | 0.688478957069148 | 0.944584424505182 | 505.967681278665 | 520.411650776256 | 508.246275204216 | 471.777127211855 | 564.998429042654 | 472.10974158051 | 520.539045278068 | 506.940438700193 | Nup62 |
| **ENSMUSG00000109536** | 0.39288438066851 | 0.275493998596969 | 0.846289746011912 | 25.2983840639333 | 19.5459774939439 | 15.1432782183751 | 22.5102406621959 | 14.2344565456483 | 12.2308223207386 | 18.7394056300105 | 17.3684090346961 | 9330162G02 |
| **ENSMUSG00000109587** | 0.356140602289297 | 0.135147037001791 | 0.775642208276055 | 69.5705561758165 | 69.2253369577179 | 56.7872933189068 | 77.8479156234274 | 58.0327843784121 | 36.6924669622158 | 52.0539045278068 | 66.217059444779 | Gm31105 |
| **ENSMUSG00000109628** | -0.254240812126623 | 0.55694562961255 | 0.916239900758826 | 23.4913566307952 | 10.5874044758863 | 17.9826428843205 | 27.1998741334867 | 30.6588294829347 | 19.5693157131818 | 12.4929370866736 | 31.4802413753867 | BC024386 |
| **ENSMUSG00000109635** | 0.0800928232026896 | 0.6953737097508 | 0.944933386973863 | 1197.15567445398 | 868.981582751589 | 598.159489625818 | 867.582192188799 | 829.978312430875 | 770.541806206532 | 851.60187807492 | 889.045437463508 | Gm45670 |
| **ENSMUSG00000109644** | 0.0884730664223804 | 0.771100817392134 | 0.959045638244974 | 49.6932544112975 | 65.9676740420606 | 42.5904699891801 | 49.7101147956825 | 40.5134532453066 | 44.0309603546589 | 35.3966550789086 | 75.9867895267955 | 0610005C13 |
| **ENSMUSG00000109695** | 0.083067939388401 | 0.632317002195139 | 0.931825800800958 | 472.537673765611 | 416.980853204136 | 304.7584741448 | 453.95652002095 | 331.772333333186 | 403.617136584374 | 426.842017128016 | 394.045779974668 | Gm31166 |
| **ENSMUSG00000109697** | -0.439336290394826 | 0.117533069978893 | 0.769868040577702 | 100.290022539164 | 89.5857301805761 | 91.8061241988993 | 98.4823028971069 | 130.300025302473 | 201.808568292187 | 104.107809055614 | 80.3288917854696 | Gm34030 |
| **ENSMUSG00000109715** | 0.664028350143029 | 0.188463455678965 | 0.808677608936066 | 8.13162344912141 | 9.77298874697194 | 19.8755526616174 | 13.1309737196143 | 6.56974917491458 | 6.1154111603693 | 8.32862472444909 | 10.8552556466851 | Gm45606 |
| **ENSMUSG00000109754** | 0.241073270740443 | 0.401204253568315 | 0.880924293718604 | 468.923618899334 | 355.085257806647 | 217.684624389143 | 494.287367874051 | 274.834507150593 | 234.831788558181 | 402.897221045225 | 386.447101021989 | Gm39214 |
| **ENSMUSG00000109764** | -0.0571505977195701 | 0.884467412138695 | 0.979450877350433 | 25.2983840639333 | 25.2468875963442 | 17.036187995672 | 27.1998741334867 | 13.1394983498292 | 25.6847268735511 | 19.7804837205666 | 40.1644458927348 | Klkb1 |
| **ENSMUSG00000109770** | -0.049884403492949 | 0.884959111981744 | 0.979839144314582 | 39.754603529038 | 37.4631235300591 | 18.9290977729689 | 48.7721881014244 | 27.3739548954774 | 44.0309603546589 | 32.2734208072402 | 46.6775992807458 | Gm30085 |
| **ENSMUSG00000109783** | 0.159683353282671 | 0.698083460501473 | 0.944933386973863 | 18.9737880479499 | 24.4324718674299 | 12.3039135524298 | 21.5723139679377 | 13.1394983498292 | 28.1308913376988 | 16.6572494488982 | 11.9407812113536 | Gm45338 |
| **ENSMUSG00000109812** | -0.272461131819326 | 0.543182048319278 | 0.913837378403038 | 138.237598635064 | 65.9676740420606 | 42.5904699891801 | 145.378637610015 | 178.478185918513 | 151.662196777159 | 65.5879197050366 | 78.1578406561325 | Gm45640 |
| **ENSMUSG00000109836** | 0.242983808655887 | 0.558840956217721 | 0.916574868225118 | 72.2810973255236 | 112.389370590177 | 122.09268063565 | 69.4065753751039 | 59.1277425742312 | 30.5770558018465 | 137.42230795341 | 90.0986218674861 | Gm45884 |
| **ENSMUSG00000109841** | 0.131445290235154 | 0.525807165142914 | 0.90806673740897 | 187.027339329792 | 127.86326943955 | 110.735221971868 | 145.378637610015 | 113.875652365186 | 121.085140975312 | 141.586620315635 | 144.374900100912 | E330011O21 |
| **ENSMUSG00000109865** | -0.140908087101124 | 0.252017858149785 | 0.83723604018117 | 427.361987937158 | 366.487078011448 | 364.385132129652 | 335.777756544422 | 432.508487348543 | 375.486245246675 | 416.431236222455 | 422.269444656049 | Hspa14 |
| **ENSMUSG00000109901** | -0.064131125534878 | 0.542464188159397 | 0.913837378403038 | 629.749060448624 | 682.480380830207 | 610.463403178248 | 601.211011019481 | 660.259792078915 | 589.5256358596 | 687.11153976705 | 701.249514775856 | Chmp1b |
| **ENSMUSG00000109904** | 0.231851803449533 | 0.487038243495686 | 0.899236955296196 | 140.044626068202 | 197.088606397268 | 136.289503965376 | 74.0962088463947 | 96.3563212320805 | 74.6080161565054 | 156.16171358342 | 138.947272277569 | Gm45819 |
| **ENSMUSG00000109946** | -0.47398143023339 | 0.146750044904926 | 0.780607990324772 | 28.9124389302094 | 35.0198763433161 | 23.6613722162112 | 29.075727522003 | 37.2285786578493 | 62.3771938357668 | 37.4788112600209 | 26.0526135520442 | Brd3os |
| **ENSMUSG00000109953** | 1.08069760705574 | 0.00015550846887759 | 0.399150048388892 | 48.7897406947284 | 51.3081909216027 | 68.1447519826882 | 82.5375490947182 | 26.2789966996583 | 26.9078091056249 | 27.0680303544596 | 37.9933947633978 | 5430430B14 |
| **ENSMUSG00000109975** | 0.293484951235093 | 0.265238490155045 | 0.842510961779757 | 78.6056933415069 | 51.3081909216027 | 83.2880302010633 | 104.109863062656 | 62.4126171616885 | 77.0541806206532 | 52.0539045278068 | 67.3025850094475 | Gm19196 |
| **ENSMUSG00000110057** | -0.0494467246601995 | 0.91380124569152 | 0.986300357654458 | 13.5527057485357 | 8.95857301805762 | 12.3039135524298 | 9.37926694258161 | 7.66470737073368 | 14.6769867848863 | 8.32862472444909 | 15.1973579053591 | Gm2225 |
| **ENSMUSG00000110062** | 0.0424950109791344 | 0.944867872685992 | 0.990505437194639 | 7.22810973255236 | 23.6180561385155 | 16.0897331070236 | 4.68963347129081 | 9.85462376237187 | 6.1154111603693 | 12.4929370866736 | 21.7105112933701 | B430319F04 |
| **ENSMUSG00000110105** | -0.530116131460421 | 0.183026308158681 | 0.805716353538343 | 18.9737880479499 | 8.95857301805762 | 16.0897331070236 | 13.1309737196143 | 13.1394983498292 | 24.4616446414772 | 19.7804837205666 | 24.9670879873757 | Gm45844 |
| **ENSMUSG00000110156** | -0.27675791744758 | 0.406128376123625 | 0.881158207349486 | 22.5878429142261 | 29.3189662409158 | 15.1432782183751 | 30.0136542162612 | 25.1840385038392 | 37.9155491942896 | 31.2323427166841 | 23.8815624227072 | Gm42067 |
| **ENSMUSG00000110161** | 0.203974666558039 | 0.526554652654711 | 0.908217276872524 | 50.5967681278665 | 63.5244268553176 | 49.2156542097192 | 31.8895076047775 | 43.7983278327639 | 28.1308913376988 | 60.3825292522559 | 36.9078691987293 | Gm45494 |
| **ENSMUSG00000110185** | 0.0364425624082012 | 0.8415606487785 | 0.970640319105748 | 355.080890611635 | 264.685111897157 | 228.095628164276 | 326.39848960184 | 324.107625962453 | 304.547475786391 | 255.064132186253 | 261.61166108511 | Igip |
| **ENSMUSG00000110195** | -0.187701514107348 | 0.403840771167055 | 0.881158207349486 | 117.456783153976 | 135.193010999779 | 150.486327295103 | 97.5443762028488 | 182.858018701789 | 150.439114545085 | 106.189965236726 | 131.348593324889 | Pde2a |
| **ENSMUSG00000110206** | -0.303757159066768 | 0.426823004013712 | 0.885066977635295 | 13.5527057485357 | 25.2468875963442 | 11.3574586637814 | 22.5102406621959 | 21.8991639163819 | 25.6847268735511 | 16.6572494488982 | 26.0526135520442 | Flt3l |
| **ENSMUSG00000110221** | 1.13742039815652 | 0.0481719261398243 | 0.703396849069134 | 8.13162344912141 | 26.8757190541728 | 16.0897331070236 | 15.9447538023887 | 5.47479097909548 | 2.44616446414772 | 10.4107809055614 | 11.9407812113536 | Gm36210 |
| **ENSMUSG00000110275** | 0.36511261140703 | 0.287658273514457 | 0.851785577030345 | 27.1054114970714 | 28.5045505120015 | 44.483379766477 | 29.075727522003 | 18.6142893289246 | 34.2463024980681 | 30.191264626128 | 17.3684090346961 | Gm5905 |
| **ENSMUSG00000110277** | -0.205855915353518 | 0.576349842228468 | 0.922450244597442 | 56.9213641438498 | 20.3603932228582 | 26.5007368821565 | 41.2687745473591 | 49.2731188118594 | 46.4771248188067 | 29.1501865355718 | 42.3354970220718 | Gm45871 |
| **ENSMUSG00000110279** | 0.206859511325999 | 0.596983970330657 | 0.924496535139492 | 19.877301764519 | 14.6594831204579 | 21.7684624389143 | 26.2619474392285 | 25.1840385038392 | 20.7923979452556 | 10.4107809055614 | 15.1973579053591 | Gm45552 |
| **ENSMUSG00000110289** | -0.0902022961178586 | 0.860671578228374 | 0.974868041734905 | 5.42108229941427 | 13.0306516626293 | 19.8755526616174 | 7.50341355406529 | 9.85462376237187 | 12.2308223207386 | 10.4107809055614 | 16.2828834700276 | 4930412F12 |
| **ENSMUSG00000110344** | 0.278972133856665 | 0.0249510453300751 | 0.655303666217042 | 478.862269781594 | 473.175538499225 | 450.512526996661 | 502.728708122374 | 400.754699669789 | 442.755768010737 | 331.062832796851 | 397.302356668674 | Gm45716 |
| **ENSMUSG00000110350** | -0.000663873658826666 | 0.998914573997356 | 0.999562238514752 | 13.5527057485357 | 15.4738988493722 | 6.62518422053913 | 14.0689004138724 | 20.8042057205628 | 11.0077400886647 | 7.28754663389296 | 10.8552556466851 | Gm10252 |
| **ENSMUSG00000110353** | -0.386836192154616 | 0.116359245946131 | 0.769868040577702 | 41.5616309621761 | 34.2054606144018 | 42.5904699891801 | 39.3929211588428 | 41.6084114411257 | 67.2695227640623 | 49.9717483466946 | 47.7631248454143 | Gm33543 |
| **ENSMUSG00000110357** | 0.27464325841796 | 0.416395771339014 | 0.882192126163777 | 56.0178504272808 | 61.895595397489 | 33.1259211026956 | 41.2687745473591 | 24.0890803080201 | 33.0232202659942 | 56.2182168900314 | 45.5920737160773 | A030001D20 |
| **ENSMUSG00000110360** | -0.050593627496745 | 0.87004133545003 | 0.975613767803763 | 51.5002818444356 | 54.56585383726 | 35.0188308799925 | 24.3860940507122 | 38.3235368536684 | 48.9232892829544 | 46.8485140750261 | 37.9933947633978 | Gm45380 |
| **ENSMUSG00000110439** | -0.397003570204891 | 0.298014310345011 | 0.851785577030345 | 19.877301764519 | 25.2468875963442 | 19.8755526616174 | 26.2619474392285 | 20.8042057205628 | 34.2463024980681 | 17.6983275394543 | 47.7631248454143 | Gm21320 |
| **ENSMUSG00000110444** | -0.0483115557098988 | 0.818951269246749 | 0.966315773308182 | 170.764092431549 | 163.69756151178 | 184.558703286447 | 133.185590584659 | 141.249607260664 | 130.869798831903 | 181.147587756768 | 220.361689627707 | Gm10033 |
| **ENSMUSG00000110453** | 0.0768442977509438 | 0.753130664331651 | 0.956497786093635 | 49.6932544112975 | 44.7928650902881 | 39.7511053232348 | 44.0825546301336 | 31.7537876787538 | 42.8078781225851 | 51.0128264372507 | 43.4210225867403 | Gm32352 |
| **ENSMUSG00000110489** | -0.40824131791507 | 0.329744297593414 | 0.855977678491063 | 65.9565013095403 | 43.9784493613737 | 38.8046504345863 | 77.8479156234274 | 48.1781606160403 | 143.100621152642 | 67.6700758861489 | 42.3354970220718 | Gm31659 |
| **ENSMUSG00000110528** | 0.051645204716639 | 0.849748202444279 | 0.972640876584422 | 54.2108229941427 | 37.4631235300591 | 55.8408384302584 | 62.8410885152968 | 44.893286028583 | 36.6924669622158 | 55.1771387994752 | 65.1315338801105 | Gm1943 |
| **ENSMUSG00000110545** | 0.849082163281437 | 0.0421872366742579 | 0.685406501171951 | 18.9737880479499 | 15.4738988493722 | 19.8755526616174 | 25.3240207449704 | 8.75966556655277 | 7.33849339244316 | 10.4107809055614 | 17.3684090346961 | Gm7730 |
| **ENSMUSG00000110580** | -0.12588933923126 | 0.575524510729044 | 0.922450244597442 | 390.317925557827 | 293.189662409158 | 217.684624389143 | 258.867767615252 | 270.454674367317 | 259.293433199658 | 388.322127777439 | 347.368180693922 | D830024N08 |
| **ENSMUSG00000110588** | -0.226155244931272 | 0.554969459765792 | 0.916239900758826 | 23.4913566307952 | 20.3603932228582 | 23.6613722162112 | 17.8206071909051 | 30.6588294829347 | 37.9155491942896 | 17.6983275394543 | 14.1118323406906 | Gm45774 |
| **ENSMUSG00000110618** | -0.814328060042416 | 0.0770756051605303 | 0.739855948845396 | 10.8421645988285 | 9.77298874697194 | 15.1432782183751 | 8.44134024832345 | 18.6142893289246 | 33.0232202659942 | 15.616171358342 | 10.8552556466851 | Gm39822 |
| **ENSMUSG00000110626** | -0.142764671491638 | 0.786136121185197 | 0.961141387070613 | 14.4562194651047 | 21.1748089517725 | 46.3762895437739 | 17.8206071909051 | 25.1840385038392 | 28.1308913376988 | 46.8485140750261 | 9.76973008201657 | Gm45805 |
| **ENSMUSG00000110630** | 0.181283552012044 | 0.54961656786331 | 0.914699131434573 | 33.4300075130547 | 35.0198763433161 | 28.3936466594534 | 30.9515809105193 | 20.8042057205628 | 31.8001380339204 | 38.5198893505771 | 21.7105112933701 | K230015D01 |
| **ENSMUSG00000110631** | -0.157538555812742 | 0.756733986141574 | 0.956566265296272 | 23.4913566307952 | 14.6594831204579 | 10.4110037751329 | 49.7101147956825 | 16.4243729372865 | 37.9155491942896 | 28.1091084450157 | 27.1381391167127 | Gm42047 |
| **ENSMUSG00000110649** | -0.017556251881407 | 0.968680379683212 | 0.994790165280964 | 16.2632468982428 | 11.4018202048006 | 9.46454888648447 | 7.50341355406529 | 12.0445401540101 | 9.78465785659088 | 12.4929370866736 | 10.8552556466851 | Gm40466 |
| **ENSMUSG00000110652** | 0.117455535361771 | 0.697529020211266 | 0.944933386973863 | 28.0089252136404 | 28.5045505120015 | 25.5542819935081 | 25.3240207449704 | 27.3739548954774 | 15.9000690169602 | 27.0680303544596 | 28.2236646813812 | Gm33023 |
| **ENSMUSG00000110702** | 0.0340287887276843 | 0.934131334088268 | 0.98818087308472 | 9.93865088225949 | 19.5459774939439 | 17.036187995672 | 15.0068271081306 | 13.1394983498292 | 22.0154801773295 | 14.5750932677859 | 10.8552556466851 | Gm45767 |
| **ENSMUSG00000110834** | 0.300388265138209 | 0.338496767372616 | 0.858570960161243 | 59.631905293557 | 59.452348210746 | 62.4660226507975 | 58.151455044006 | 32.8487458745729 | 34.2463024980681 | 81.2040910633787 | 45.5920737160773 | Gm39469 |
| **ENSMUSG00000110841** | -0.0679533455784711 | 0.740843581825997 | 0.954383574136061 | 1483.56952260637 | 1035.12239145011 | 991.884723303572 | 1543.82733874893 | 1304.09521122054 | 1660.9456711563 | 1095.21415126506 | 1238.58466928677 | Gpx4-ps2 |
| **ENSMUSG00000110922** | 0.242574897009357 | 0.577287759581232 | 0.922450244597442 | 18.9737880479499 | 12.2162359337149 | 11.3574586637814 | 12.1930470253561 | 8.75966556655277 | 9.78465785659088 | 16.6572494488982 | 10.8552556466851 | Gm35177 |
| **ENSMUSG00000110949** | 0.0671293087202244 | 0.610673741505919 | 0.927932112008768 | 224.071401709123 | 188.13003337921 | 193.076797284283 | 225.102406621959 | 200.377349834895 | 203.031650524261 | 195.722681024554 | 193.223550510994 | Nudt8 |
| **ENSMUSG00000111080** | 0.0482425398540223 | 0.808341058326017 | 0.964026716319739 | 782.442878548793 | 627.100111264033 | 533.800557197724 | 733.458674909882 | 521.20010120989 | 535.71001764835 | 726.672507208183 | 804.374443419364 | Gm20300 |
| **ENSMUSG00000111082** | 0.575408884340816 | 0.150840705732824 | 0.783020695097675 | 18.9737880479499 | 34.2054606144018 | 29.3401015481019 | 13.1309737196143 | 16.4243729372865 | 18.3462334811079 | 14.5750932677859 | 15.1973579053591 | Gm30934 |

| | | | | | | | | | | | | |
|---|---|---|---|---|---|---|---|---|---|---|---|---|
| ENSMUSG00000111128 | -0.0696680723618085 | 0.679884304712507 | 0.944452327903597 | 134.623543768788 | 136.007426728693 | 123.039135524298 | 123.806323642077 | 142.344565456483 | 105.185071958352 | 138.463386043966 | 156.315681312265 | AC126804.1 |
| ENSMUSG00000111291 | -0.207868496078695 | 0.589276818445544 | 0.92352104528132 | 20.780815481088 | 12.2162359337149 | 25.5542819935081 | 28.1378008277448 | 21.8991639163819 | 36.6924669622158 | 17.6983275394543 | 23.8815624227072 | Gm48604 |
| ENSMUSG00000111312 | 0.215718515097556 | 0.666517482108138 | 0.942024383071524 | 33.4300075130547 | 70.8541684155466 | 69.0912068713366 | 22.5102406621959 | 32.8487458745729 | 15.9000690169602 | 71.8343882483734 | 47.7631248454143 | Gm47205 |
| ENSMUSG00000111318 | -0.200473408533712 | 0.655787191425791 | 0.937803365479691 | 17.1667606148119 | 16.2883145782866 | 15.1432782183751 | 14.0689004138724 | 6.56974917491458 | 19.5693157131818 | 16.6572494488982 | 29.3091902460497 | Gm48483 |
| ENSMUSG00000111329 | 0.0911038306196159 | 0.804821960921938 | 0.963461896828986 | 14.4562194651047 | 24.4324718674299 | 31.2330113253987 | 15.9447538023887 | 18.6142893289246 | 19.5693157131818 | 18.7394056300105 | 23.8815624227072 | A830035O19 |
| ENSMUSG00000111394 | 0.311931308485976 | 0.486754177525653 | 0.899236955296196 | 30.7194663633475 | 17.1027303072009 | 11.3574586637814 | 18.7585338851632 | 13.1394983498292 | 11.0077400886647 | 13.5340151772298 | 24.9670879873757 | AC160637.1 |
| ENSMUSG00000111521 | -0.748153458899452 | 0.0699003051460246 | 0.724916894583216 | 14.4562194651047 | 14.6594831204579 | 16.0897331070236 | 15.0068271081306 | 32.8487458745729 | 37.9155491942896 | 13.5340151772298 | 17.3684090346961 | Gm48529 |
| ENSMUSG00000111594 | -0.0527783542322518 | 0.922905921439647 | 0.986713918026868 | 9.93865088225949 | 4.07207864457164 | 14.1968233297267 | 12.1930470253561 | 7.66470737073368 | 6.1154111603693 | 15.616171358342 | 11.9407812113536 | Gm3365 |
| ENSMUSG00000111709 | -0.56086640337133 | 0.101335044498511 | 0.759800953500741 | 81.316234491214 | 104.245213301034 | 51.1085639870161 | 137.87522405595 | 86.5016974697087 | 135.762127760198 | 198.845915296222 | 131.348593324889 | Gm3776 |
| ENSMUSG00000111761 | 0.38125017186033 | 0.0315909202822937 | 0.671337170953485 | 351.466835745359 | 355.085257806647 | 361.545767463707 | 352.660437041069 | 213.516848184724 | 226.270212933664 | 339.391457521301 | 310.460311495193 | 9230112J17[illegible] |
| ENSMUSG00000111765 | 0.377675075850017 | 0.126897544782711 | 0.772551378490123 | 72.2810973255236 | 70.0397526866323 | 63.4124775394459 | 66.5927952923295 | 35.0386622662111 | 46.4771248188067 | 57.2592949805875 | 70.559161703453 | Gm10635 |
| ENSMUSG00000111809 | 0.179876634434805 | 0.715640428601624 | 0.951176382858794 | 18.0702743313809 | 10.5874044758863 | 8.51809399783602 | 13.1309737196143 | 8.75966556655277 | 11.0077400886647 | 6.24646854333682 | 18.4539345993646 | AC112680.2 |
| ENSMUSG00000111815 | -0.148496109074007 | 0.750333331312076 | 0.956197192463153 | 56.0178504272808 | 38.2775392589734 | 66.2518422053913 | 42.2067012416173 | 38.3235368536684 | 17.123151249034 | 67.6700758861489 | 100.953877514171 | Gm6018 |
| ENSMUSG00000111828 | 0.507888327483479 | 0.236353851219998 | 0.832048466795696 | 20.780815481088 | 10.5874044758863 | 16.0897331070236 | 25.3240207449704 | 12.0445401540101 | 14.6769867848863 | 15.616171358342 | 8.68420451734806 | D830035M03 |
| ENSMUSG00000111867 | 0.198532565023419 | 0.322578281713252 | 0.855977678491063 | 293.64195788494 | 168.584055885266 | 204.434255948065 | 204.468019348279 | 170.813478547779 | 192.023910435596 | 183.22974393788 | 212.763010675027 | Gm31013 |
| ENSMUSG00000111875 | 0.244795727744243 | 0.604241236923465 | 0.926448151609826 | 13.5527057485357 | 8.95857301805762 | 14.1968233297267 | 20.6343872736795 | 13.1394983498292 | 18.3462334811079 | 6.24646854333682 | 10.8552556466851 | Gm5425 |
| ENSMUSG00000111877 | 0.369495928644433 | 0.355214792283563 | 0.864215962387542 | 38.8510898124689 | 18.7315617650296 | 16.0897331070236 | 27.1998741334867 | 17.5193311331055 | 26.9078091056249 | 19.7804837205666 | 14.1118323406906 | Gm6477 |
| ERCC-00002 | -0.401658008905493 | 0.211994220736682 | 0.823370267313127 | 104018.823647444 | 67765.0895557745 | 65388.6753469439 | 90811.0004647694 | 101539.853331088 | 188995.558828981 | 70547.615728446 | 72192.8776782791 | NA |
| ERCC-00003 | -0.329109977397219 | 0.292898391056073 | 0.851785577030345 | 9048.68987143899 | 6138.25134882729 | 5631.40658745826 | 8061.4799371489 | 8340.29657755406 | 15615.090856887 | 5935.18619426053 | 6390.4889992035 | NA |
| ERCC-00004 | -0.337217136435021 | 0.270554097273325 | 0.844129257012535 | 34022.712511124 | 23356.628689534 | 22345.7999209898 | 29751.9726685631 | 31909.2717425601 | 59281.5727063879 | 23095.2763608973 | 24019.4241694201 | NA |
| ERCC-00009 | -0.409939460348548 | 0.209499420566152 | 0.821566820583092 | 3514.66835745359 | 2322.71365886367 | 2189.15015744386 | 3044.51004956199 | 3443.64352585106 | 6473.77425436694 | 2355.95971892854 | 2437.0048926808 | NA |
| ERCC-00014 | -0.521482155831462 | 0.294160132903512 | 0.851785577030345 | 21.6843291976571 | 9.77298874697194 | 7.57163910918757 | 16.8826804966469 | 17.5193311331055 | 35.4693847301419 | 12.4929370866736 | 15.1973579053591 | NA |
| ERCC-00019 | -0.137975144915001 | 0.656089577574422 | 0.938034442101846 | 134.623543768788 | 114.83261777692 | 88.966759532954 | 81.59962240046 | 107.305903190271 | 188.354663739374 | 86.4094815161593 | 81.4144173501381 | NA |
| ERCC-00022 | -0.482612601484146 | 0.188280614624599 | 0.808677608936066 | 1436.58680934478 | 939.835751167135 | 844.237760674415 | 1281.20786435665 | 1458.48431683104 | 2954.96667269044 | 994.22957648111 | 883.617809640165 | NA |
| ERCC-00025 | -0.41987499862643 | 0.20712525795264 | 0.819960256425221 | 477.055242348456 | 312.735639903102 | 262.16800415562 | 407.9981120023 | 474.116898789669 | 836.58824673852 | 311.282349076285 | 332.170822788563 | NA |
| ERCC-00028 | -0.0734022368979499 | 0.862255696855118 | 0.974868041734905 | 15.3597331816738 | 23.6180561385155 | 18.9290977729689 | 34.703287687552 | 29.5638712871156 | 35.4693847301419 | 20.8215618111227 | 11.9407812113536 | NA |
| ERCC-00031 | -0.196489439567636 | 0.753071562098792 | 0.956497786093635 | 8.13162344912141 | 8.95857301805762 | 13.2503684410783 | 8.44134024832345 | 13.1394983498292 | 23.2385624094033 | 5.20539045278068 | 3.25657669400552 | NA |
| ERCC-00033 | 0.0416892634799186 | 0.946683237036751 | 0.990510945542179 | 15.3597331816738 | 21.1748089517725 | 3.78581955459379 | 6.56548685980713 | 9.85462376237187 | 14.6769867848863 | 5.20539045278068 | 16.2828834700276 | NA |
| ERCC-00034 | -0.410173404202163 | 0.299622256468022 | 0.851785577030345 | 121.070838020252 | 78.9983257046899 | 82.3415753124149 | 80.6616957062019 | 114.970610561005 | 228.716377397812 | 58.3003730711437 | 81.4144173501381 | NA |
| ERCC-00035 | -0.413536104660849 | 0.186385812012766 | 0.807013381813378 | 586.38040205331 | 364.043830824705 | 444.83379766477 | 509.294194982182 | 630.6959207918 | 1068.97387083255 | 408.102611498006 | 429.868123608729 | NA |
| ERCC-00039 | -0.481201939995698 | 0.294546570309267 | 0.851785577030345 | 12.6491920319666 | 25.2468875963442 | 13.2503684410783 | 18.7585338851632 | 30.6588294829347 | 39.1386314263635 | 16.6572494488982 | 11.9407812113536 | NA |
| ERCC-00042 | -0.362490812341237 | 0.238998268497862 | 0.832048466795696 | 2640.06707981475 | 1941.56709773176 | 1779.33519065908 | 2396.4027038296 | 2415.47777997693 | 4935.13680641802 | 1997.82885577723 | 1911.61051938124 | NA |
| ERCC-00043 | -0.324903013010139 | 0.279206190902233 | 0.848075053184273 | 4636.83239343234 | 3124.09873611536 | 3097.74685054637 | 4023.70551836751 | 4275.81175467357 | 7899.88813696506 | 3145.09691157009 | 3321.70822788563 | NA |
| ERCC-00044 | -0.38212510534629 | 0.249227574520147 | 0.835314899646652 | 574.634723737913 | 369.744740927105 | 310.437203476691 | 466.149567046306 | 545.28918151791 | 958.896469945906 | 370.623800237985 | 369.078691987293 | NA |
| ERCC-00046 | -0.340495884134918 | 0.292289139483221 | 0.851785577030345 | 21230.7653119394 | 15504.846647071 | 14855.555932226 | 18618.7828077188 | 19607.4164125326 | 40245.5208463903 | 14518.8750508959 | 14528.6741575233 | NA |
| ERCC-00051 | -0.147172679687615 | 0.712367257317995 | 0.950233477782287 | 81.316234491214 | 30.9477976987445 | 43.5369248778286 | 51.5859681841989 | 55.8428679867739 | 96.6234963338349 | 36.4377331694648 | 41.2499714574033 | NA |
| ERCC-00053 | -0.466178594586059 | 0.196679266343608 | 0.812065429583558 | 283.70330700268 | 145.780415475665 | 143.861143074564 | 225.102406621959 | 262.789966996583 | 480.671317205027 | 180.106509666212 | 180.197243734972 | NA |
| ERCC-00058 | -0.424473596921579 | 0.382192723521324 | 0.875126835008975 | 17.1667606148119 | 9.77298874697194 | 8.51809399783602 | 14.0689004138724 | 14.2344565456483 | 26.9078091056249 | 7.28754663389296 | 18.4539345993646 | NA |
| ERCC-00059 | -0.470725534459959 | 0.284294760326594 | 0.851125503777472 | 56.0178504272808 | 25.2468875963442 | 30.2865564367503 | 43.1446279358754 | 64.6025335533267 | 91.7311674055395 | 28.1091084450157 | 30.3947158107182 | NA |
| ERCC-00060 | -0.443979748673029 | 0.175137973386075 | 0.799889021902745 | 2661.75140901241 | 1825.92006422592 | 1767.03127710665 | 2266.96882002198 | 2715.49632563136 | 5163.85318381584 | 1866.65301636715 | 1847.5645110658 | NA |
| ERCC-00062 | -0.248135393069185 | 0.423632801549217 | 0.884482395183227 | 655.950958229127 | 436.52683069808 | 429.690519446395 | 543.059555975475 | 580.327843784121 | 1043.289143959 | 410.184767679118 | 420.098393526712 | NA |
| ERCC-00069 | -0.583165079714147 | 0.239012135406447 | 0.832048466795696 | 20.780815481088 | 10.5874044758863 | 15.1432782183751 | 14.0689004138724 | 27.3739548954774 | 40.3617136584374 | 10.4107809055614 | 13.0263067760221 | NA |
| ERCC-00071 | -0.43761501056279 | 0.198297920790295 | 0.812921911423739 | 343.335212296237 | 162.883145782866 | 171.308334845369 | 262.619474392285 | 316.442918591719 | 513.694537471021 | 207.174540020671 | 236.644573097735 | NA |
| ERCC-00074 | -0.306795960835284 | 0.324787296388853 | 0.855977678491063 | 64844.2759244438 | 45741.6594144733 | 44122.7804539019 | 56233.3949542481 | 61213.6379372666 | 113394.399900032 | 43559.7483869593 | 42759.9375178572 | NA |
| ERCC-00076 | -0.445560643245888 | 0.182447246766351 | 0.804505478598291 | 1014.64590370704 | 675.965054998893 | 660.625512276616 | 907.9130400419 | 993.127083607921 | 2003.40869613698 | 730.836819570408 | 712.104770422541 | NA |
| ERCC-00078 | -0.470442985970522 | 0.200746678675289 | 0.812921911423739 | 295.448985318078 | 220.706662535783 | 182.66579350915 | 297.322762079837 | 332.867291529005 | 634.779678446333 | 231.119336103462 | 182.368294864309 | NA |
| ERCC-00079 | -0.399696653625613 | 0.2242101919597 | 0.832048466795696 | 334.300075130547 | 226.407572638183 | 246.078271048596 | 260.743621003769 | 354.766455445387 | 611.54111603693 | 225.913945650682 | 217.105112933702 | NA |
| ERCC-00084 | -0.354536099259551 | 0.301889623709117 | 0.851785577030345 | 369.537110076739 | 261.427448981499 | 236.613722162112 | 296.384835385579 | 318.632834983357 | 687.372214425509 | 247.776585552361 | 235.559047533066 | NA |
| ERCC-00085 | -0.548858645531455 | 0.178829500275699 | 0.801755154770274 | 32.5264937964856 | 21.1748089517725 | 22.7149173275627 | 35.6412143818101 | 38.3235368536684 | 72.1618516923577 | 33.3144988977964 | 20.6249857287016 | NA |
| ERCC-00092 | -0.435314959826619 | 0.195617001692479 | 0.811094945724481 | 2339.19701219726 | 1549.01871639505 | 1365.73440431971 | 2038.11470662298 | 2274.22817271626 | 4398.2037065376 | 1608.46564990923 | 1580.52522215735 | NA |
| ERCC-00095 | -0.374520027710085 | 0.214579448854205 | 0.824248286808591 | 425.55496050402 | 307.034729800702 | 279.204192151292 | 391.115431505653 | 421.558905390352 | 759.534066117867 | 293.584021536831 | 345.197129564585 | NA |
| ERCC-00096 | -0.391776396803387 | 0.235816508960449 | 0.832048466795696 | 132353.917312766 | 82971.045630334 | 78870.925235741 | 110621.88810089 | 124581.058645709 | 233766.483934045 | 85907.6818765113 | 86870.268838162 | NA |
| ERCC-00099 | -0.567263284294447 | 0.144768765140617 | 0.780607990324772 | 158.114900399583 | 111.574954861263 | 101.270673085384 | 139.751077444466 | 181.76306050597 | 362.032340693862 | 100.984574783945 | 112.894658725525 | NA |
| ERCC-00108 | -0.432421993535483 | 0.18850752420516 | 0.808677608936066 | 5262.96739901469 | 3496.28672422921 | 3220.78598607066 | 4378.2418087971 | 5081.70098679643 | 9807.89641900028 | 3566.73353824532 | 3620.22775816947 | NA |
| ERCC-00111 | -0.35996453933239 | 0.269997961478383 | 0.843793744782845 | 3518.28241231986 | 2449.7625125743 | 2352.88685318004 | 3069.83407030696 | 3284.87458745729 | 6571.62083293285 | 2411.13685772801 | 2352.33389863666 | NA |
| ERCC-00112 | -0.391052151300304 | 0.26105635519494 | 0.840925967916911 | 852.916948441179 | 517.153987860599 | 498.781726317731 | 715.638067718977 | 733.621991198795 | 1550.86827026965 | 545.524919451416 | 560.13119136895 | NA |
| ERCC-00113 | -0.421921237941891 | 0.206138464794697 | 0.819960256425221 | 24408.4230531128 | 16800.5820717737 | 15902.3350390712 | 21426.9353303277 | 24663.9333608252 | 47363.8594370602 | 16379.2815987197 | 16812.6199455858 | NA |
| ERCC-00116 | -0.356693083641206 | 0.255531936361447 | 0.837479467570303 | 870.08370905599 | 543.215291185857 | 498.781726317731 | 740.962088463947 | 823.408563255961 | 1407.76764911701 | 558.017856538089 | 608.979841779033 | NA |
| ERCC-00126 | -0.0509043039689456 | 0.898293852727136 | 0.984462340635154 | 23.4913566307952 | 27.6901347830872 | 24.6078271048596 | 44.0825546301336 | 30.6588294829347 | 52.592535979176 | 18.7394056300105 | 22.7960368580387 | NA |
| ERCC-00130 | -0.370504129599376 | 0.251409906435139 | 0.83723604018117 | 212933.788124977 | 149112.190222653 | 143718.228386378 | 188868.422569377 | 206529.919937202 | 399952.78221708 | 146521.330464871 | 145021.873337454 | NA |
| ERCC-00131 | -0.336957421267369 | 0.336770940619862 | 0.858570960161243 | 847.495866141764 | 545.6585383726 | 576.391027186904 | 725.017334661559 | 846.402685368162 | 1542.30669464514 | 516.374732915844 | 499.341759747513 | NA |
| ERCC-00136 | -0.345593084613824 | 0.242889218952222 | 0.832048466795696 | 9687.4740690533 | 6542.2015503688 | 6075.29393023438 | 8380.37501319667 | 9383.79173816966 | 16046.838884809 | 6481.75219180251 | 7079.79773276801 | NA |
| ERCC-00143 | -0.545583750770759 | 0.318174837280426 | 0.854351922245337 | 21.6843291976571 | 16.2883145782866 | 13.2503684410783 | 11.2551203310979 | 25.1840385038392 | 46.4771248188067 | 9.36970281500523 | 10.8552556466851 | NA |
| ERCC-00144 | -0.582819504343985 | 0.177893308267621 | 0.801696999069105 | 89.4478579403355 | 45.6072808192024 | 61.519567762149 | 73.1582821521366 | 84.3117810780705 | 201.808568292187 | 45.80743598447 | 72.73021283279 | NA |
| ERCC-00145 | -0.389101026799727 | 0.235383777967272 | 0.832048466795696 | 4961.19381768063 | 3242.18901680794 | 3029.60209856368 | 4194.4081767225 | 4585.68492409038 | 8966.41584333347 | 3325.2034212363 | 3327.13585570898 | NA |
| ERCC-00148 | -0.461522189142769 | 0.387555696460279 | 0.875951433019341 | 36.1405486627618 | 23.6180561385155 | 33.1259211026956 | 29.075727522003 | 39.4184950494875 | 94.1773318696872 | 20.8215618111227 | 14.1118323406906 | NA |
| ERCC-00150 | -0.460513584105137 | 0.303809714215425 | 0.851785577030345 | 15.3597331816738 | 7.32974156022896 | 8.51809399783602 | 9.37926694258161 | 14.2344565456483 | 18.3462334811079 | 10.4107809055614 | 13.0263067760221 | NA |
| ERCC-00154 | -0.387313476290385 | 0.415387821945055 | 0.882192126163777 | 11.7456783153976 | 18.7315617650296 | 17.9826428843205 | 34.703287687552 | 21.8991639163819 | 50.1463715150282 | 19.7804837205666 | 17.3684090346961 | NA |
| ERCC-00157 | -0.327920603329675 | 0.387023911911417 | 0.875691730512848 | 91.2548853734735 | 54.56585383726 | 46.3762895437739 | 56.2756016554897 | 60.2227007700503 | 141.877538920568 | 51.0128264372507 | 59.7039060567679 | NA |
| ERCC-00160 | -0.15322540556926 | 0.702269050346765 | 0.946434671388391 | 120.167324303683 | 56.1946852950887 | 68.1447519826882 | 89.1030359545253 | 84.3117810780705 | 172.454594722414 | 54.1360607089191 | 60.7894316214364 | NA |
| ERCC-00162 | -0.277372149451712 | 0.430259762282879 | 0.886017087288711 | 263.826005238161 | 162.883145782866 | 187.398067952392 | 198.84045918273 | 202.567266226533 | 459.878919259771 | 170.736806851206 | 153.05910461826 | NA |
| ERCC-00163 | -0.558910618799631 | 0.201853705855521 | 0.813395633194566 | 59.631905293557 | 28.5045505120015 | 30.2865564367503 | 52.523894878457 | 76.6470737073368 | 106.408154190426 | 31.2323427166841 | 37.9933947633978 | NA |
| ERCC-00165 | -0.432549566324034 | 0.205757507790623 | 0.819960256425221 | 205.097613661173 | 132.749763813036 | 124.932045301595 | 172.578511743502 | 178.478185918513 | 386.49398533534 | 142.627698406191 | 150.888053488923 | NA |
| ERCC-00170 | -0.322140257290235 | 0.307909905902854 | 0.851785577030345 | 219.553833126278 | 153.110157035894 | 159.004421292939 | 150.068271081306 | 175.193311331055 | 371.816998550453 | 144.709854587303 | 161.743309135608 | NA |
| ERCC-00171 | -0.319715208349071 | 0.338323812256241 | 0.858570960161243 | 22659.2204978351 | 15140.8028162463 | 14895.3070375493 | 18598.1484204451 | 20794.3510968005 | 40085.2970739887 | 14755.1997774521 | 13346.5368175993 | NA |
