## Supplemental Table 3 for "Adropin protects against cardiac remodeling and metabolic dysfunction in a mouse model of HFpEF"

Sup Table 3

| | Pathway total | Hits.total | Hits.sig | Expected | P(Fisher) | P(EASE) | P(Gamma) | Emp.Hits | Empirical | AdjP.Fisher | AdjP.EASE | AdjP.Gamma | Pathway Number | cpd.hits |
|---|---|---|---|---|---|---|---|---|---|---|---|---|---|---|
| **UDP-<i>N</i>-acetyl-D-galactosamine biosynthesis II** | 13 | 4 | 3 | 0.83505 | 0.067528 | 0.39813 | 0.00042534 | 4 | 0.04 | 1 | 1 | 0.03062448 | P1 | EC000116;EC00093 |
| **putrescine degradation III** | 15 | 3 | 2 | 0.83505 | 0.067528 | 0.39813 | 0.00042534 | 1 | 0.01 | 1 | 1 | 0.03062448 | P2 | EC00099;EC00093 |
| **biosynthesis of serotonin and melatonin** | 14 | 1 | 1 | 0.27835 | 0.16071 | 1 | 0.00067237 | 5 | 0.05 | 1 | 1 | 0.0470659 | P3 | EC00093 |
| **serotonin and melatonin biosynthesis** | 14 | 1 | 1 | 0.27835 | 0.16071 | 1 | 0.00067237 | 5 | 0.05 | 1 | 1 | 0.0470659 | P4 | EC00093 |
| **wax esters biosynthesis I** | 8 | 1 | 1 | 0.27835 | 0.16071 | 1 | 0.00067237 | 5 | 0.05 | 1 | 1 | 0.0470659 | P5 | EC00093 |
| **wax esters biosynthesis II** | 7 | 1 | 1 | 0.27835 | 0.16071 | 1 | 0.00067237 | 5 | 0.05 | 1 | 1 | 0.0470659 | P6 | EC00093 |
| **fatty acid activation** | 6 | 1 | 1 | 0.27835 | 0.16071 | 1 | 0.00067237 | 5 | 0.05 | 1 | 1 | 0.0470659 | P7 | EC00093 |
| **acyl-CoA hydrolysis** | 5 | 1 | 1 | 0.27835 | 0.16071 | 1 | 0.00067237 | 5 | 0.05 | 1 | 1 | 0.0470659 | P8 | EC00093 |
| **fatty acid biosynthesis initiation II** | 7 | 1 | 1 | 0.27835 | 0.16071 | 1 | 0.00067237 | 5 | 0.05 | 1 | 1 | 0.0470659 | P9 | EC00093 |
| **very long chain fatty acid biosynthesis** | 11 | 1 | 1 | 0.27835 | 0.16071 | 1 | 0.00067237 | 5 | 0.05 | 1 | 1 | 0.0470659 | P10 | EC00093 |
| **palmitate biosynthesis I (animals)** | 38 | 1 | 1 | 0.27835 | 0.16071 | 1 | 0.00067237 | 5 | 0.05 | 1 | 1 | 0.0470659 | P11 | EC00093 |
| **2-ketoglutarate dehydrogenase complex** | 10 | 1 | 1 | 0.27835 | 0.16071 | 1 | 0.00067237 | 5 | 0.05 | 1 | 1 | 0.0470659 | P12 | EC00093 |
| **branched-chain α-keto acid dehydrogenase complex** | 10 | 1 | 1 | 0.27835 | 0.16071 | 1 | 0.00067237 | 5 | 0.05 | 1 | 1 | 0.0470659 | P13 | EC00093 |
| **acetyl-CoA biosynthesis (from pyruvate)** | 10 | 1 | 1 | 0.27835 | 0.16071 | 1 | 0.00067237 | 5 | 0.05 | 1 | 1 | 0.0470659 | P14 | EC00093 |
| **ethanol degradation II (cytosol)** | 12 | 1 | 1 | 0.27835 | 0.16071 | 1 | 0.00067237 | 5 | 0.05 | 1 | 1 | 0.0470659 | P15 | EC00093 |
| **<i>N</i>-acetylneuraminate and <i>N</i>-acetylmannosamine degradation** | 8 | 2 | 2 | 0.27835 | 0.16071 | 1 | 0.00067237 | 5 | 0.05 | 1 | 1 | 0.0470659 | P16 | EC000116 |
| **formaldehyde oxidation II (glutathione-dependent)** | 9 | 1 | 1 | 0.27835 | 0.16071 | 1 | 0.00067237 | 10 | 0.1 | 1 | 1 | 0.0470659 | P17 | EC000165 |
| **L-arabinose degradation II** | 9 | 1 | 1 | 0.27835 | 0.16071 | 1 | 0.00067237 | 9 | 0.09 | 1 | 1 | 0.0470659 | P18 | EC00063 |
| **acetate conversion to acetyl-CoA** | 6 | 1 | 1 | 0.27835 | 0.16071 | 1 | 0.00067237 | 5 | 0.05 | 1 | 1 | 0.0470659 | P19 | EC00093 |
| **2-amino-3-carboxymuconate semialdehyde degradation to glutaryl-CoA** | 12 | 1 | 1 | 0.27835 | 0.16071 | 1 | 0.00067237 | 5 | 0.05 | 1 | 1 | 0.0470659 | P20 | EC00093 |
| **glutaryl-CoA degradation** | 14 | 1 | 1 | 0.27835 | 0.16071 | 1 | 0.00067237 | 5 | 0.05 | 1 | 1 | 0.0470659 | P21 | EC00093 |
| **acetate formation from acetyl-CoA II** | 6 | 1 | 1 | 0.27835 | 0.16071 | 1 | 0.00067237 | 5 | 0.05 | 1 | 1 | 0.0470659 | P22 | EC00093 |
| **pyrimidine deoxyribonucleosides degradation** | 18 | 1 | 1 | 0.27835 | 0.16071 | 1 | 0.00067237 | 5 | 0.05 | 1 | 1 | 0.0470659 | P23 | EC00093 |
| **acetoacetate degradation (to acetyl CoA)** | 5 | 1 | 1 | 0.27835 | 0.16071 | 1 | 0.00067237 | 5 | 0.05 | 1 | 1 | 0.0470659 | P24 | EC00093 |
| **fatty acid β-oxidation I** | 18 | 1 | 1 | 0.27835 | 0.16071 | 1 | 0.00067237 | 5 | 0.05 | 1 | 1 | 0.0470659 | P25 | EC00093 |
| **fatty acid β-oxidation II (core pathway)** | 16 | 1 | 1 | 0.27835 | 0.16071 | 1 | 0.00067237 | 5 | 0.05 | 1 | 1 | 0.0470659 | P26 | EC00093 |
| **lysine degradation II** | 17 | 4 | 2 | 1.3918 | 0.18276 | 0.57335 | 0.00075017 | 25 | 0.25 | 1 | 1 | 0.0470659 | P27 | EC000148;EC00093 |
| **UDP-<i>N</i>-acetyl-D-glucosamine biosynthesis II** | 12 | 6 | 3 | 1.6701 | 0.24746 | 0.64141 | 0.0010374 | 38 | 0.38 | 1 | 1 | 0.0470659 | P28 | EC000116;EC00093 |
| **2-methylbutyrate biosynthesis** | 15 | 2 | 1 | 0.5567 | 0.29641 | 1 | 0.0013299 | 10 | 0.1 | 1 | 1 | 0.0585156 | P29 | EC00093 |
| **CMP-<i>N</i>-acetylneuraminate biosynthesis I (eukaryotes)** | 15 | 2 | 1 | 0.5567 | 0.29641 | 1 | 0.0013299 | 10 | 0.1 | 1 | 1 | 0.0585156 | P30 | EC000116 |
| **coenzyme A biosynthesis** | 13 | 2 | 1 | 0.5567 | 0.29641 | 1 | 0.0013299 | 6 | 0.06 | 1 | 1 | 0.0585156 | P31 | EC00093 |
| **tetrapyrrole biosynthesis II** | 11 | 2 | 1 | 0.5567 | 0.29641 | 1 | 0.0013299 | 11 | 0.11 | 1 | 1 | 0.0585156 | P32 | EC00093 |
| **ceramide biosynthesis** | 12 | 2 | 1 | 0.5567 | 0.29641 | 1 | 0.0013299 | 15 | 0.15 | 1 | 1 | 0.0585156 | P33 | EC00093 |
| **ketogenesis** | 12 | 2 | 1 | 0.5567 | 0.29641 | 1 | 0.0013299 | 19 | 0.19 | 1 | 1 | 0.0585156 | P34 | EC00093 |
| **bile acid biosynthesis, neutral pathway** | 48 | 2 | 1 | 0.5567 | 0.29641 | 1 | 0.0013299 | 11 | 0.11 | 1 | 1 | 0.0585156 | P35 | EC00093 |
| **oleate biosynthesis II (animals)** | 9 | 2 | 1 | 0.5567 | 0.29641 | 1 | 0.0013299 | 11 | 0.11 | 1 | 1 | 0.0585156 | P36 | EC00093 |
| **γ-linolenate biosynthesis II (animals)** | 11 | 2 | 1 | 0.5567 | 0.29641 | 1 | 0.0013299 | 5 | 0.05 | 1 | 1 | 0.0585156 | P37 | EC00093 |
| **ketogenesis** | 12 | 2 | 1 | 0.5567 | 0.29641 | 1 | 0.0013299 | 19 | 0.19 | 1 | 1 | 0.0585156 | P38 | EC00093 |
| **<i>N</i>-acetylglucosamine degradation I** | 7 | 2 | 1 | 0.5567 | 0.29641 | 1 | 0.0013299 | 18 | 0.18 | 1 | 1 | 0.0585156 | P39 | EC000116 |
| **ribose degradation** | 7 | 3 | 2 | 0.5567 | 0.29641 | 1 | 0.0013299 | 14 | 0.14 | 1 | 1 | 0.0585156 | P40 | EC00063 |
| **β-alanine degradation I** | 9 | 3 | 2 | 1.9485 | 0.31312 | 0.69896 | 0.0014486 | 24 | 0.24 | 1 | 1 | 0.0585156 | P41 | EC00093;EC00035 |
| **triacylglycerol biosynthesis** | 12 | 2 | 1 | 0.83505 | 0.41085 | 1 | 0.0024079 | 8 | 0.08 | 1 | 1 | 0.0746449 | P42 | EC00093 |
| **CDP-diacylglycerol biosynthesis I** | 12 | 2 | 1 | 0.83505 | 0.41085 | 1 | 0.0024079 | 8 | 0.08 | 1 | 1 | 0.0746449 | P43 | EC00093 |
| **acetyl-CoA biosynthesis (from citrate)** | 7 | 3 | 1 | 0.83505 | 0.41085 | 1 | 0.0024079 | 20 | 0.2 | 1 | 1 | 0.0746449 | P44 | EC00093 |
| **D-glucuronate degradation I** | 11 | 3 | 1 | 0.83505 | 0.41085 | 1 | 0.0024079 | 25 | 0.25 | 1 | 1 | 0.0746449 | P45 | EC00063 |
| **3-oxoadipate degradation** | 6 | 2 | 1 | 0.83505 | 0.41085 | 1 | 0.0024079 | 18 | 0.18 | 1 | 1 | 0.0746449 | P46 | EC00093 |
| **β-alanine biosynthesis I** | 10 | 1 | 1 | 1.1134 | 0.50725 | 1 | 0.0040422 | 27 | 0.27 | 1 | 1 | 0.1050972 | P47 | EC00035 |
| **mevalonate pathway I** | 17 | 3 | 1 | 1.1134 | 0.50725 | 1 | 0.0040422 | 16 | 0.16 | 1 | 1 | 0.1050972 | P48 | EC00093 |
| **molybdenum cofactor (sulfide) biosynthesis** | 6 | 1 | 1 | 1.1134 | 0.50725 | 1 | 0.0040422 | 27 | 0.27 | 1 | 1 | 0.1050972 | P49 | EC00035 |
| **lipoate biosynthesis and incorporation II** | 11 | 3 | 1 | 1.1134 | 0.50725 | 1 | 0.0040422 | 43 | 0.43 | 1 | 1 | 0.1050972 | P50 | EC00096 |
| **ketolysis** | 10 | 3 | 1 | 1.1134 | 0.50725 | 1 | 0.0040422 | 33 | 0.33 | 1 | 1 | 0.1050972 | P51 | EC00093 |
| **glutathione-mediated detoxification** | 15 | 3 | 1 | 1.1134 | 0.50725 | 1 | 0.0040422 | 43 | 0.43 | 1 | 1 | 0.1050972 | P52 | EC00093 |
| **tryptophan degradation to 2-amino-3-carboxymuconate semialdehyde** | 13 | 1 | 1 | 1.1134 | 0.50725 | 1 | 0.0040422 | 27 | 0.27 | 1 | 1 | 0.1050972 | P53 | EC00035 |
| **isoleucine degradation** | 28 | 3 | 1 | 1.1134 | 0.50725 | 1 | 0.0040422 | 57 | 0.57 | 1 | 1 | 0.1050972 | P54 | EC00093 |
| **Leucine Catabolism** | 25 | 3 | 1 | 1.1134 | 0.50725 | 1 | 0.0040422 | 57 | 0.57 | 1 | 1 | 0.1050972 | P55 | EC00093 |
| **leucine degradation I** | 22 | 3 | 1 | 1.1134 | 0.50725 | 1 | 0.0040422 | 57 | 0.57 | 1 | 1 | 0.1050972 | P56 | EC00093 |
| **ketone oxidation** | 10 | 3 | 1 | 1.1134 | 0.50725 | 1 | 0.0040422 | 33 | 0.33 | 1 | 1 | 0.1050972 | P57 | EC00093 |
| **glycine biosynthesis III** | 4 | 2 | 1 | 1.3918 | 0.58838 | 1 | 0.0063606 | 37 | 0.37 | 1 | 1 | 0.1050972 | P58 | EC00035 |
| **valine degradation** | 29 | 4 | 1 | 1.3918 | 0.58838 | 1 | 0.0063606 | 58 | 0.58 | 1 | 1 | 0.1050972 | P59 | EC00093 |
| **4-hydroxyproline degradation I** | 13 | 3 | 1 | 1.3918 | 0.58838 | 1 | 0.0063606 | 51 | 0.51 | 1 | 1 | 0.1050972 | P60 | EC000123 |
| **alanine biosynthesis II** | 4 | 2 | 1 | 1.6701 | 0.65656 | 1 | 0.0094689 | 59 | 0.59 | 1 | 1 | 0.1136268 | P61 | EC00035 |
| **tryptophan degradation I (via anthranilate)** | 9 | 2 | 1 | 1.6701 | 0.65656 | 1 | 0.0094689 | 41 | 0.41 | 1 | 1 | 0.1136268 | P62 | EC00035 |
| **valine degradation I** | 20 | 5 | 1 | 1.6701 | 0.65656 | 1 | 0.0094689 | 61 | 0.61 | 1 | 1 | 0.1136268 | P63 | EC00093 |
| **alanine degradation III** | 4 | 2 | 1 | 1.6701 | 0.65656 | 1 | 0.0094689 | 59 | 0.59 | 1 | 1 | 0.1136268 | P64 | EC00035 |
| **uracil degradation II (reductive)** | 10 | 3 | 1 | 1.9485 | 0.7138 | 1 | 0.013442 | 50 | 0.5 | 1 | 1 | 0.1136268 | P65 | EC00035 |
| **4-aminobutyrate degradation IV** | 9 | 3 | 1 | 1.9485 | 0.7138 | 1 | 0.013442 | 51 | 0.51 | 1 | 1 | 0.1136268 | P66 | EC00035 |
| **phenylalanine degradation III** | 12 | 3 | 1 | 1.9485 | 0.7138 | 1 | 0.013442 | 69 | 0.69 | 1 | 1 | 0.1136268 | P67 | EC00035 |
| **TCA cycle variation III (eukaryotic)** | 21 | 8 | 1 | 2.7835 | 0.83562 | 1 | 0.030767 | 51 | 0.51 | 1 | 1 | 0.153835 | P68 | EC00093 |
| **TCA cycle** | 23 | 8 | 1 | 2.7835 | 0.83562 | 1 | 0.030767 | 51 | 0.51 | 1 | 1 | 0.153835 | P69 | EC00093 |
| **glutamate degradation IV** | 15 | 5 | 1 | 2.7835 | 0.83562 | 1 | 0.030767 | 84 | 0.84 | 1 | 1 | 0.153835 | P70 | EC00035 |
| **glutamate degradation VII** | 19 | 7 | 1 | 3.3402 | 0.88715 | 1 | 0.04652 | 82 | 0.82 | 1 | 1 | 0.153835 | P71 | EC00093 |
| **tRNA charging pathway** | 64 | 17 | 2 | 6.6804 | 0.9318 | 0.98736 | 0.071346 | 97 | 0.97 | 1 | 1 | 0.153835 | P72 | EC000148;EC00035 |
